# Generalist temporal niche buffers against extinction risk among non-avian tetrapods

**DOI:** 10.1101/2025.10.06.680248

**Authors:** Jacinta Guirguis, Catherine Sheard, Daniel Pincheira-Donoso

**Author notes:** Corresponding author: Daniel Pincheira-Donoso.

## Abstract

Anthropocene extinctions of biodiversity vary dramatically through geographic and phylogenetic space, with certain regions and lineages concentrating high defaunation rates, while others remain nearly unchanged.This heterogeneity in biodiversity erosion results from (spatial and phylogenetic) variation in the interactions between species fitness-relevant traits and environmental threats that trigger extinctions. Evidence reveals that factors including life histories, ecophysiology, and resource-use generalism influence variation in species declines under rapidly changing environments. An emerging hypothesis predicts that species spread across different ‘locations’ of the 24h diel spectrum are exposed to different anthropogenic pressures on demographic stability (e.g. daytime contact with humans), which can trigger differential extinction risk among diurnal, nocturnal and cathemeral (day-night active) species. However, this hypothesis remains largely neglected. Only a single large-scale test exists, which documented higher declines among diurnal mammals. Here, we address this hypothesis across tetrapods globally, spanning >23,000 species across both endotherms and ectotherms for the first time. Our results reveal that cathemerality is significantly associated with a lower probability of undergoing population declines across tetrapods (except birds), while diurnality is associated with higher extinction risk category in amphibians. Collectively, our results suggest that the role of temporal niche in extinction is significant but variable across tetrapods.

## Introduction

Human-induced erosion of animal biodiversity is structured into ‘defaunation hotspots’ –disproportionate concentrations of threatened species– heterogeneously distributed across regions of the planet. Traditionally, the identification and driving factors shaping these hotspots have focused on their distribution across geographic (1–3) and phylogenetic (i.e. differential aggregation of threatened species across clades) space (4–6). However, whether defaunation hotspots are predictably distributed across ‘temporal space’ –which we define as the spread of phylogenetic diversity in the same geographic region over the 24h diel cycle– has rarely been considered. Therefore, the hypothesis that the likelihood for populations to undergo demographic collapses that lead to extinction is influenced by their temporal niche (i.e. nocturnal, diurnal, cathemeral) remains fundamentally untested.

Overlooking this temporal dimension can potentially eclipse the role of critical processes underlying the effects of human perturbations on species declines. First, the greater concentration of human activities during the daytime has led to the prediction that diurnal species may experience higher extinction risk (7,8). Second, the flexibility that cathemeral species –essentially, ‘temporal generalists’– require to remain active throughout day and night may facilitate avoidance of threats in certain parts of the 24h cycle (8–14).Incorporating temporal niche may, consequently, shed light on the role that the position of species along the generalist (cathemeral)-specialist (diurnal/nocturnal/crepuscular) continuum plays in their resilience under human perturbations and thus, in their risk of undergoing declines. In fact, a parallel can be made between species with small geographic range sizes (i.e. intrinsically susceptible to extinction) and day- or night-specialists, relative to species with large geographic ranges and cathemeral temporal generalists. From this parallel, we derive the prediction that cathemeral species have lower risk of extinction than species that are inflexibly restricted to either a diurnal or a nocturnal temporal niche.

Surprisingly, only one large-scale study, on mammals worldwide, has addressed the hypothesis that extinction risk varies across the 24h cycle (7). Cox et al (2023) observed that population declines are associated with diurnal activity. This effect, however, vanished when primates were removed from the analysis, suggesting that results were influenced by phylogenetic biases (7). In addition, the role of cathemerality in contributing to demographic resilience was not considered, and thus, the role of temporal generalism remains entirely unknown. Finally, no tests of any of these hypotheses exist for ectothermic animals (e.g., reptiles and amphibians). Therefore, the major components of this ‘temporal extinctions’ hypothesis remains untested.

Here, using a novel global scale dataset spanning >23,000 species from across the entire tetrapod tree of life (mammals, birds, reptiles and amphibians), we address the hypotheses that diurnality increases extinction risk and cathemerality decreases extinction risk using both IUCN Red List conservation categories and population trends as alternative proxies for extinction risk (3) under a phylogenetic Bayesian approach.

## Results

### Distribution of threatened and decreasing species across the 24h diel spectrum

Species decreasing in population sizes and threatened species are generally not uniformly distributed among temporal niches. For both IUCN Red List population trends and categories, our data reveal that the diurnal niche space accommodates the greatest proportion of species at risk of extinction among amphibians and mammals, with a disproportionally higher frequency for population trends (Fig 1). Specifically, 66.5% of diurnal amphibians and 67.5% of diurnal mammals are decreasing in population sizes, while 52% of diurnal amphibians and 34.4% of diurnal mammals are threatened based on IUCN Red List categories (Fig 1). When defining extinction risk as population declines, diurnal niche space across tetrapods consistently accommodates the highest proportions of species at risk (Fig 1). Yet, while these differences were pronounced in amphibians and mammals, they appear marginal in lizards and birds (Fig 1). When defining extinction risk using threat categories, diurnal niche space accommodates the highest proportions of species at risk among amphibians and mammals (see above), but not in lizards and birds (Fig 1). Among lizards, 22.6% of nocturnal species, 20.0% of diurnal species and 18.1% of cathemeral species are threatened with extinction (Fig 1). Among birds, 24.1% of cathemeral species, 11.0% of diurnal and 9.1% of nocturnal species were threatened with extinction.

**Fig 1.**
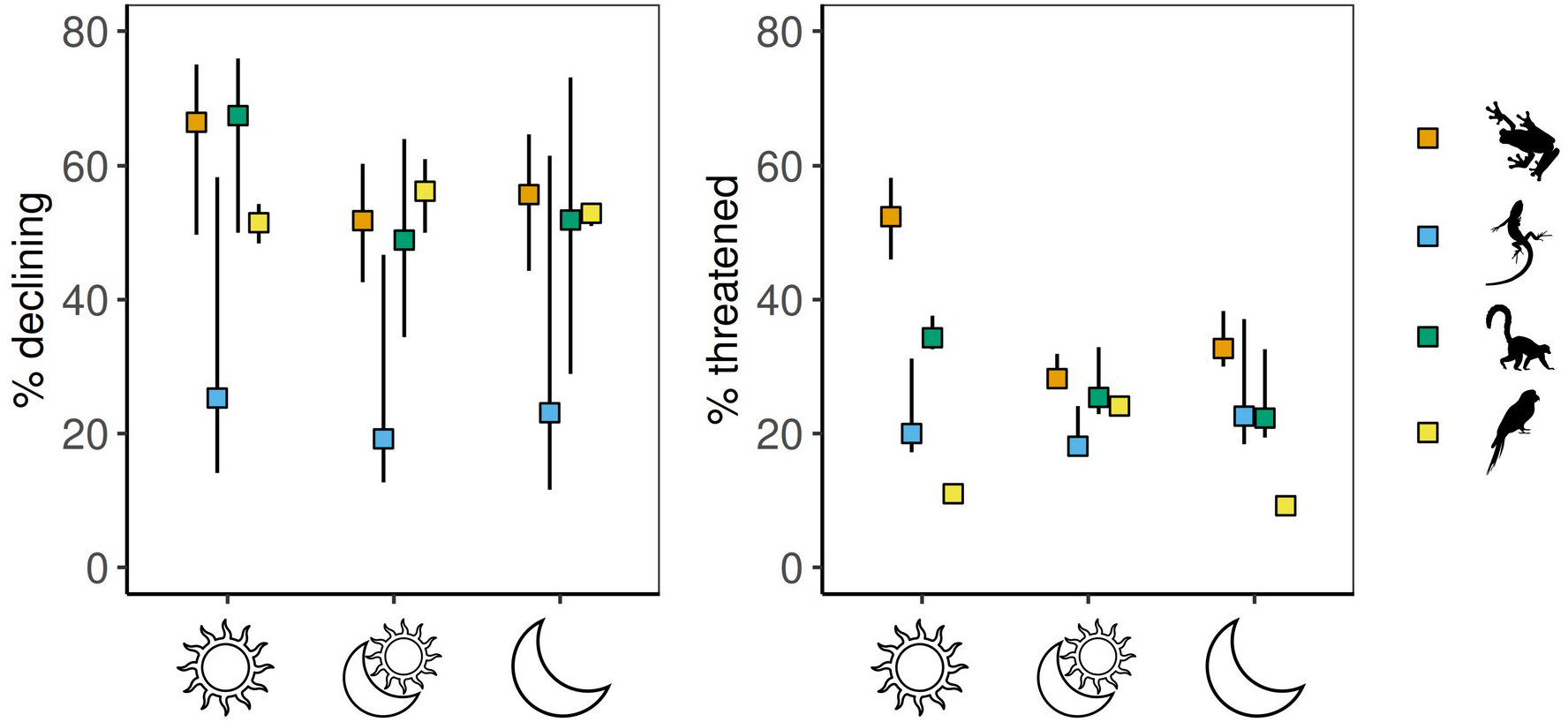
The percentage of species in each temporal niche that are considered at risk of extinction, which we defined as the as the probability of experiencing population declines (based on IUCN population trends) and the probability of being threatened (based on IUCN categories). Using population trends, we assigned species listed as ‘Decreasing’ as ‘declining’, whilst those listed as ‘Increasing’ and ‘Stable’ were assigned ‘not declining’. Similarly, using categories, we considered species listed as ‘Vulnerable’, ‘Endangered’, ‘Critically Engendered’, ‘Extinction in the Wild’ and ‘Extinct’ as ‘threatened’, whilst species listed as ‘Least Concern’ and ‘Near Threatened’ were assigned ‘not threatened’. We incorporated species assigned ‘Unknown’ population trends or ‘Data Deficient’ category using the formula in Böhm et al. (2013). The lower error bar represents a ‘best case scenario’, where all species assigned ‘Unknown’ or ‘Data Deficient’ are assumed to be not threatened/not declining, whereas the upper error bar represents a ‘worst case scenario’, where those species were instead assumed to be threatened/declining. Silhouettes representing taxonomic groups were obtained from phylopic.org.

### Effect of temporal niche on extinction risk

Our findings reveal variable support for the hypotheses that cathemerality and diurnality influence extinction risk among tetrapods. This variability expresses both between proxies for extinction risk (population trend or threat category) and across tetrapod classes. First, analyses addressing the cathemerality hypothesis reveal that the probability of experiencing population declines decreases among non-volant tetrapods with a cathemeral temporal niche (Table 1; Table S1). Second, analyses addressing the diurnality hypothesis reveal that the probability of being threatened increases among amphibians with a diurnal temporal niche (Table 1; Table S1).

**Table 1.** Predictor estimations from Bayesian phylogenetic single-response models, performed using the R package *MCMCglmm* to test the prediction that temporal niche (binary) influences extinction risk (binary) modelled using a threshold liability (probit) family function among tetrapods (amphibians, lizards, mammals and birds). The response variable was either (1) the probability of experiencing population declines or (2) the probability of being threatened threatened. For the probability of experiencing population declines, we assigned ‘1’ to species if their population trend was listed as ‘Decreasing’ and ‘0’ if their population trend was ‘Increasing’ or ‘Stable’ while removing species with a population trend of ‘Unknown’. For the probability of being threatened we assigned ‘1’ to species if their threat status was ‘Vulnerable’, ‘Endangered’ and ‘Critically Engendered’ and ‘0’ to species if their threat status was ‘Least Concern’ and ‘Near Threatened’ while excluding ‘Data Deficient’ and unassessed species. The explanatory variable was either (1) diurnal, (2) cathemeral or (3) nocturnal. For diurnal, we encoded diurnal species as ‘1’ and cathemeral, crepuscular and nocturnal species as ‘0’. For cathemeral, we encoded cathemeral species as ‘1’ and diurnal, crepuscular and nocturnal species as ‘0’. For nocturnal, we encoded nocturnal species as ‘1’ and diurnal, cathemeral and crepuscular species as ‘0’. We did not model the effect of crepuscularity on extinction risk due to low numbers of crepuscular species among vertebrates. Table also shows sample size (*n*), posterior mean with 95% credible interval (CI), and effective sample size (ESS). Predictors are considered significant when the 95% credible interval excludes include zero and is indicated by bold-face. Full model outputs are displayed in Table S1.

|  | *n* | Posterior mean<br>(95% CI) | ESS |
| --- | --- | --- | --- |
| **1 Population trends** |  |  |  |
| **1.1 Diurnal** |  |  |  |
| Amphibians | 2668 | 0.201 (-0.037, 0.448) | 50000 |
| Lizards | 2851 | 0.214 (-0.133, 0.574) | 30570 |
| Mammals | 2904 | 0.084 (-0.162, 0.31) | 49286 |
| Birds | 8896 | 0.090 (-0.143, 0.319) | 50000 |
| **1.2 Cathemeral** |  |  |  |
| Amphibians | 2668 | **-0.266 (-0.452, -0.082)** | 47452 |
| Lizards | 2851 | **-0.421 (-0.795, -0.06)** | 18074 |
| Mammals | 2904 | **-0.257 (-0.479, -0.049)** | 50000 |
| Birds | 8896 | 0.010 (-0.33, 0.336) | 50806 |
| **1.3 Nocturnal** |  |  |  |
| Amphibians | 2668 | 0.151 (-0.026, 0.32) | 48111 |
| Lizards | 2851 | 0.302 (-0.131, 0.734) | 33522 |
| Mammals | 2904 | 0.158 (-0.029, 0.353) | 50000 |
| Birds | 8896 | -0.173 (-0.51, 0.172) | 49301 |
| **2 Category** |  |  |  |
| **2.1 Diurnal** |  |  |  |
| Amphibians | 3098 | **0.441 (0.226, 0.655)** | 45045 |
| Lizards | 4392 | -0.049 (-0.305, 0.213) | 44132 |
| Mammals | 4232 | 0.021 (-0.199, 0.242) | 50000 |
| Birds | 9384 | **0.248 (0.006, 0.498)** | 42034 |
| **2.2 Cathemeral** |  |  |  |
| Amphibians | 3098 | **-0.206 (-0.381, -0.035)** | 48799 |
| Lizards | 4392 | -0.249 (-0.533, 0.022) | 35585 |
| Mammals | 4232 | -0.189 (-0.401, 0.027) | 42724 |
| Birds | 9384 | 0.079 (-0.254, 0.411) | 48138 |
| **2.3 Nocturnal** |  |  |  |
| Amphibians | 3098 | -0.061 (-0.221, 0.097) | 49190 |
| Lizards | 4392 | **0.412 (0.100, 0.732)** | 28401 |
| Mammals | 4232 | 0.136 (-0.047, 0.322) | 43050 |
| Birds | 9384 | -0.208 (-0.576, 0.156) | 46895 |

When analyses are performed for population trends, we observe that cathemerality is significantly associated with a lower probability of undergoing population declines among amphibians, lizards and mammals, but not birds (Table 1; Table S1). Importantly, we observe no correlation between the random effect and predictor value in the models where cathemerality was significant (Table S1), indicating that these findings are not impacted by Mundlak bias (see Methods). The effects of cathemerality are observed to be primarily concentrated between clades, as the interclass correlation coefficients of population declines in these models are 0.844 (amphibians), 0.898 (lizards) and 0.819 (mammals) (Table S1). Our exploratory data analysis of between- and within-clade patterns on the probability of population decline generally follow model findings, but also reveal some heterogeneity (Fig 2; Fig 3).

**Fig 2.**
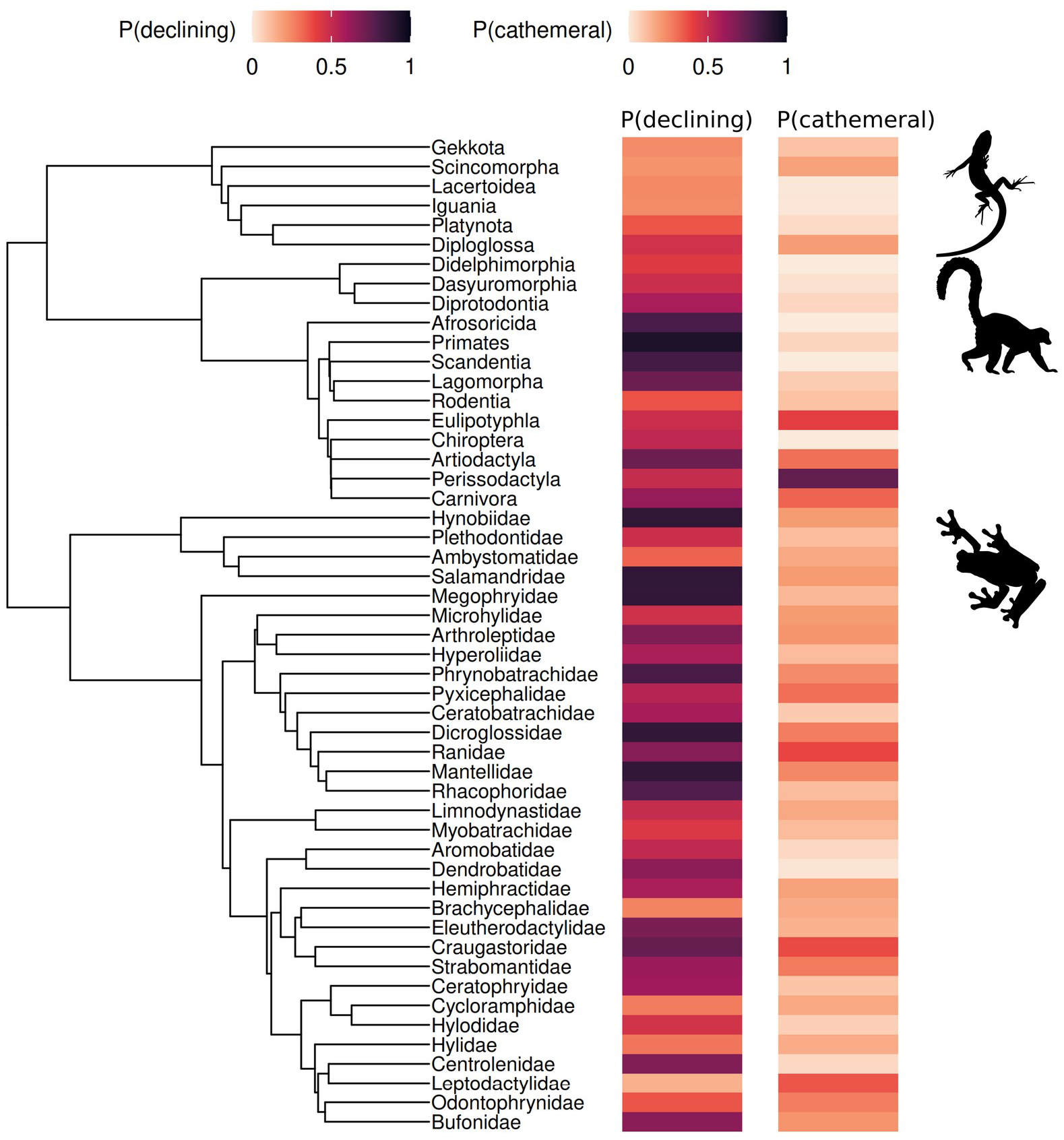
Phylogeny depicting between-clade association of the probability of experiencing population declines (based on IUCN population trend) and the probability of being cathemeral for amphibians, mammals and lizards. Tips represent families for amphibians, orders for mammals and superfamilies for lizards. Tips with less than 10 species were removed. Silhouettes representing taxonomic groups were obtained from phylopic.org.

**Fig 3.**
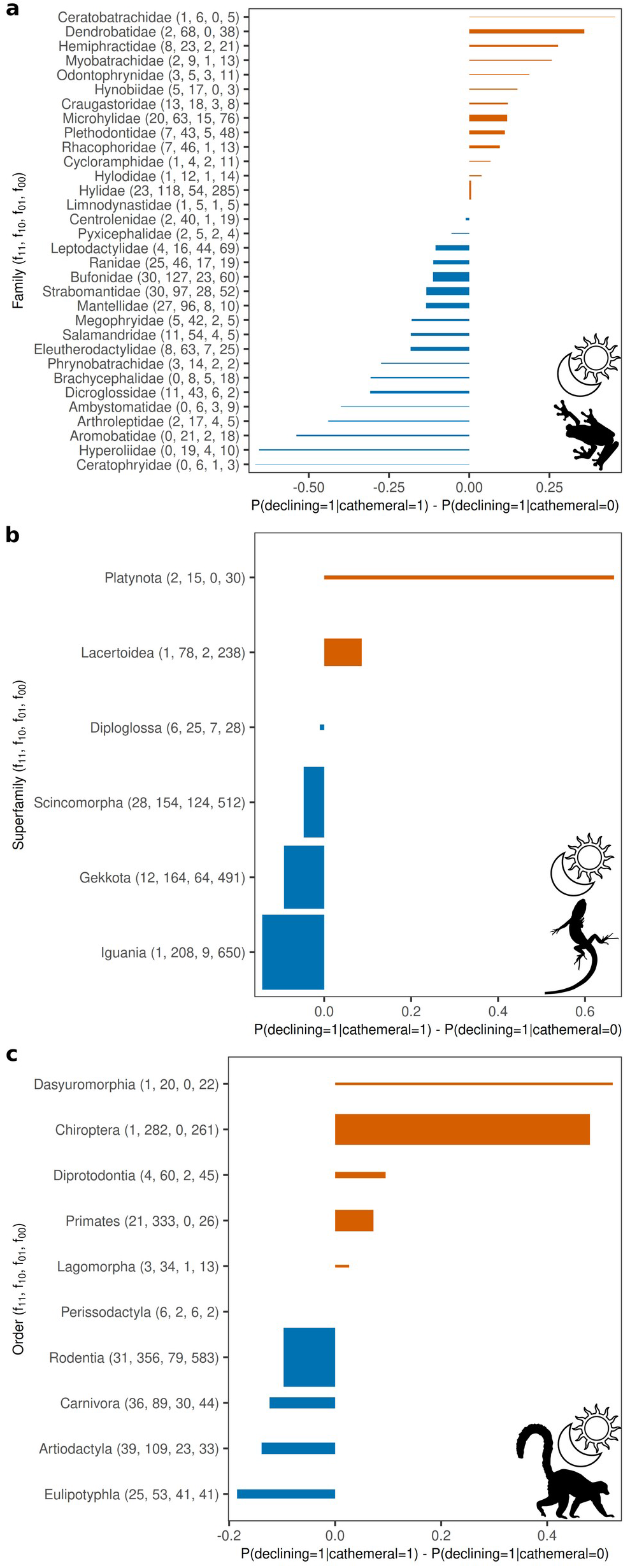
Bar plots showing within-clade association (represented by risk differences) of the probability of experiencing population declines (based on IUCN population trend) and the probability of being cathemeral for amphibians (**a**), lizards (**b**) and mammals (**c**). The risk difference (*x*-axis) compares the probability of experiencing population declines (declining=1) between cathemeral species (cathemeral=1) and non-cathemeral species (cathemeral=0) within clades (*y*-axis). Risk difference is calculated as P(declining=1| cathemeral=1) – P(declining=1|cathemeral=0). Positive values (coloured orange-brown in the plot) indicate higher risk in the cathemeral group, negative values (coloured blue in the plot) indicate lower risk in the cathemeral group, while a risk difference of 0 indicates no difference between cathemeral and non-cathemeral species is observed for that clade. Width of bar represents the total sample size of each clade.Numbers in brackets (*y*-axis) indicate observed frequencies in each cell of the 2×2 contingency table (f_11_: declining=1,cathemeral=1; f_10_: declining=1,cathemeral=0; f_01_: declining=0,cathemeral=1; f_00_: declining=0,cathemeral=0) for each clade. Clades represent families for amphibians, orders for mammals and superfamilies for lizards. Clades with less than 10 species were removed. Silhouettes representing taxonomic groups were obtained from phylopic.org.

When analyses are performed for threat categories, our findings support the hypothesis that temporal niche influences the probability of being threatened with extinction among amphibians only (Table 1; Table S1). Specifically, we observe higher extinction risk among diurnal amphibians, while extinction risk decreases among cathemeral amphibians (Table 1; Table S1). Importantly, we observe a significant, positive relationship between the phylogenetic deviation and the predictor in the diurnal model and a significant, negative relationship in the cathemeral model, revealing that the true effects are likely to be more pronounced or further from zero than what are estimated by the models (Table S1). The effect of both diurnality and cathemerality were concentrated primarily between clades, as ICC of the outcome for each model was 0.833 (diurnal) and 0.834 (cathemeral) (Table S1). As with analyses of population trends, our exploratory data analysis of between- and within-clade patterns on the probability of being threatened generally follow model findings, but also reveal some heterogeneity (Fig 4; Fig 5).

**Fig 4.**
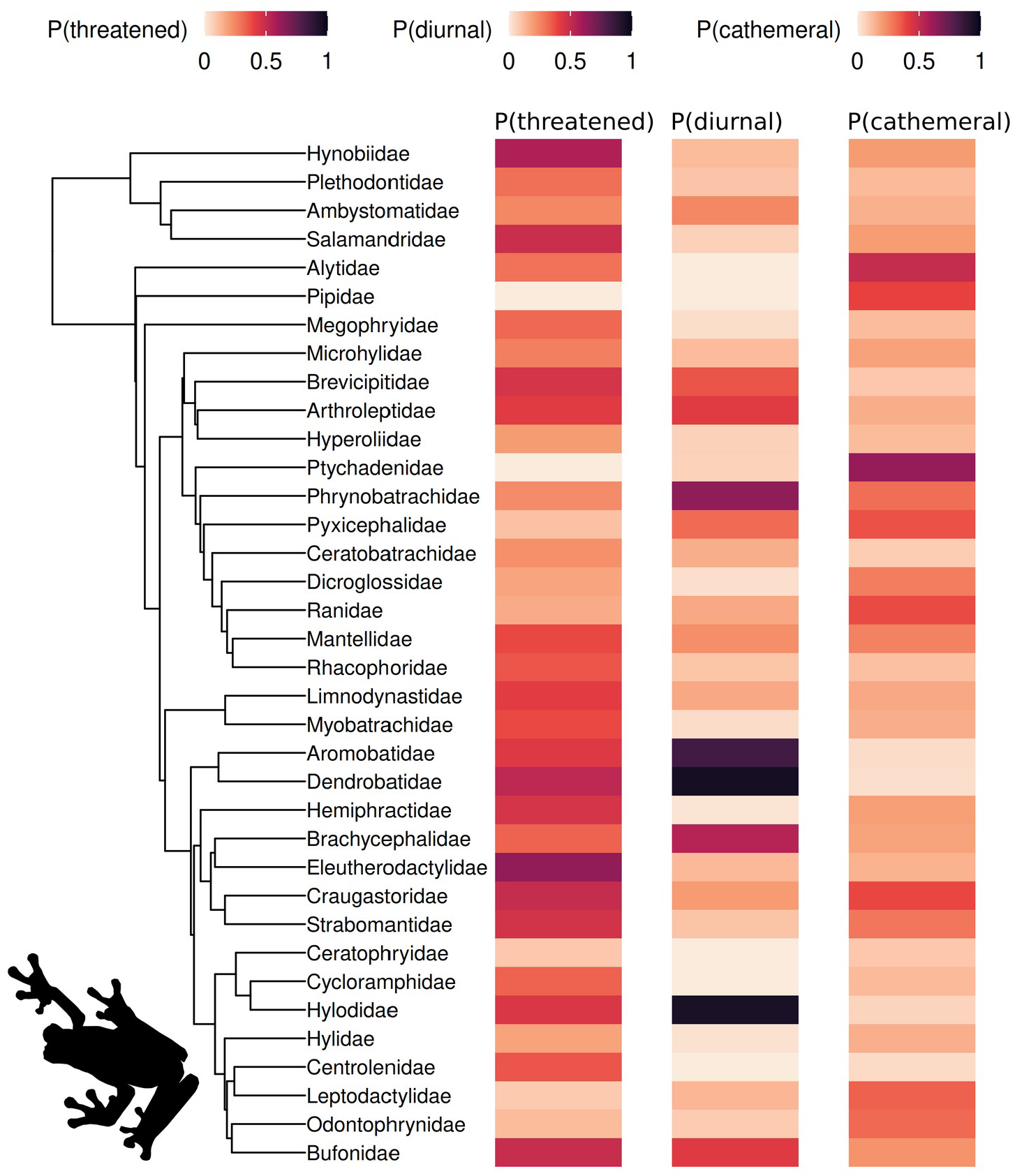
Phylogeny depicting between-clade association of the probability of being threatened (based on IUCN threat categories) the probabilities of being diurnal and cathemeral for amphibians. Tips represent families. Tips with less than 10 species were removed. Silhouettes representing taxonomic groups were obtained from phylopic.org.

**Fig 5.**
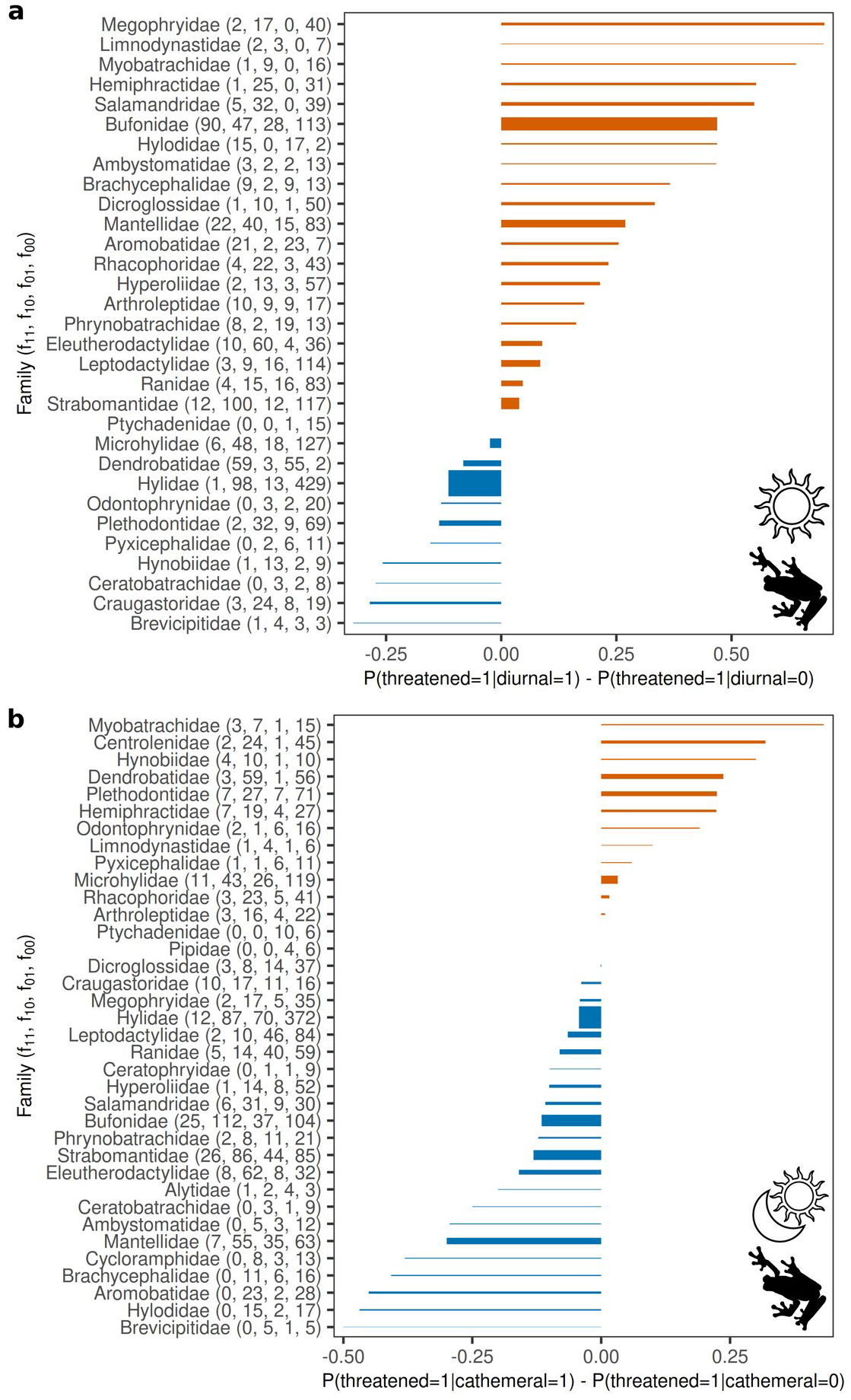
(**a**) Bar plot showing within-clade association (represented by risk differences) of the probability of being threatened (based on IUCN threat categories) and the probabilities of being diurnal (**a**) and cathemeral (**b**) for amphibians. The risk difference (*x*-axis) compares the probability of being threatened (threatened=1) between diurnal and cathemeral species (diurnal=1 in (**a**) and cathemeral=1 in (**b**)) and non-diurnal/non-cathemeral species (diurnal=0 in (**a**) and cathemeral=0 in (**b**)) within clades (*y*-axis). Risk difference is calculated as P(threatened=1|diurnal=1) – P(threatened=1|diurnal=0) in (**a**) and P(threatened=1| cathemeral=1) – P(threatened=1|cathemeral=0) in (**b**). Positive values (coloured orange-brown in the plot) indicate higher risk in the diurnal/cathemeral group, negative values (coloured blue in the plot) indicate lower risk in the diurnal/cathemeral group, while a risk difference of 0 indicates no difference between diurnal/cathemeral and non-diurnal/non-cathemeral species is observed for that clade. Width of bar represents the total sample size of each clade. Numbers in brackets (*y*-axis) indicate observed frequencies in each cell of the 2×2 contingency table (_11_: threatened=1,diurnal=1; f_10_: threatened=1,diurnal=0; f_01_: threatened=0,diurnal=1; f_00_: threatened=0,diurnal=0 in (**a**) and f_11_: threatened=1,cathemeral=1; f_10_: threatened=1,cathemeral=0; f_01_: threatened=0,cathemeral=1; f_00_: threatened=0,cathemeral=0 in (**b**)) for each clade. Clades represent families. Clades with less than 10 species were removed. Silhouettes representing taxonomic groups were obtained from phylopic.org.

Our models using threat categories additionally reveal that diurnal birds and nocturnal lizards have increased risk of extinction (Table 1; Table S1). However, we do not consider these effects to support the hypothesis that temporal niche influences extinction risk as we observe significant, negative relationships between the phylogenetic deviation and the temporal niche predictor, indicating that both models systematically over-predict extinction risk in these lineages (Table S1). Thus, the true effect of diurnality in birds and nocturnality in lizards is likely to be closer to zero than what each model estimates.

## Discussion

Our study provides the first tetrapod-wide, global-scale test of the ‘temporal-niche extinction hypothesis’ that inter-specific variation in extinction risk is influenced by variation in the temporal niche of species. As predicted by theory, our analyses showed that a cathemeral temporal niche is associated with a lower risk of undergoing population declines across all predominantly non-volant tetrapod groups. Why no signal was detected in cathemeral birds is unknown, but we suggest the possibility that a substantial proportion of cathemeral birds could be decoupled from the demographic pressures operating among other terrestrial tetrapods, given that cathemerality in birds is heavily concentrated among seabirds (Procellariiformes; 92.6% cathemeral) and penguins (Sphenisciformes; 100% cathemeral). These findings are thus consistent with the hypothesis that temporal generalism (cathemerality) buffers against the impacts of human activities and of human-induced changing environments. In contrast with the widespread role of temporal generalism on population trends, our analyses showed that temporal niche only influences the probability of extinction risk among amphibians. In these animals, higher risk of extinction is associated with the diurnal ‘temporal specialism’ niche. Finally, our analyses also reveal that model parameters must be interpreted with care when predictors have phylogenetic signal.

### Demographic mechanisms underpinning the temporal niche hypothesis

The mechanism underlying extinctions can be framed as the process when the onset of environmental pressures (i.e., the extrinsic trigger) exert stress on a population’s functional (fitness-relevant) traits beyond their capability to maintain viable levels of demographic stability (15–17). When this process persists, populations can enter trajectories of progressive decline until they reach a demographic ‘tipping point’, beyond which a demographic collapse is expected to be inevitable, thus consolidating risk of extinction, or extinction (3,15,18). Therefore, hypotheses aimed to identify signals of the role that an environmental trigger plays in shaping variation in extinction risk require a connection with some demographic feature or process within the population (i.e. intrinsic triggers). For instance, classic hypotheses on the role of body size and geographic range size as drivers of extinction risk are intrinsically linked with demography, as body size correlates with essentially most life history and metabolic traits and functions underpinning reproductive and ecological performance, while species with progressively smaller geographic ranges are progressively more vulnerable to demographic and genetic erosion caused by environmental factors (2,17,19–26).

Our aim to expand the range of traditionally used organismic predictors of extinction risk by addressing the role that variation in temporal niche –a largely neglected factor– plays as a driver of variation in population erosion across species reveals some promising findings. In particular, the observation that the cathemeral temporal niche buffers against risk of extinction across most tetrapods (except for birds). As suggested in our hypothesis, this association could be explained by the much higher ecological versatility that a cathemeral temporal niche (a form of generalism through the 24h diel cycle) offers to species compared with the narrower (‘temporally specialist’) versatility of species adapted to either a diurnal or to a nocturnal niche. Therefore, cathemeral species are expected to have the advantage to flexibly adjust their activity to concentrate over a region of the 24h diel cycle where mortality is lower, thus resulting in a positive effect on demographic stability that would delay a population erosion towards a tipping-point. If certain environmental threats are specific to the day or the night, temporal specialists are less likely to have the ecological versatility to adjust, or at least, if such adjustments are possible, they are likely to take much longer to realise, enhancing the probability of approaching a tipping point more rapidly. This is precisely the rationale underpinning the ‘diurnality hypothesis’ – given that environmental factors with potential to become threat are potentially more concentrated during the day, this hypothesis predicted that diurnal specialists are more likely to face extinction risk. However, our evidence only marginally supported this prediction, with amphibians being the only tetrapod class showing an association between diurnality and extinction risk. While the causes behind this observation remain an open question, the explanation could relate to the multiple exceptional features of these organisms – they are the only anamniotes, they are the only lineage that lacks some more specialised type of skin, they primarily rely on oviparous reproduction with a subsequent larval stage. Moreover, given that diurnal temporal niche in amphibians is associated with higher-elevation habitats (Table S2), this finding could relate to or reflect the amphibian-specific global-scale pattern of increased extinction risk in mountains (20). Overall, therefore, the cathemerality hypothesis offers a rationale that links ecological versatility as a pathway to demographic resilience in rapidly changing environments –thus offering a clear mechanism that transcends the trait profile idiosyncrasies of different lineages–, in contrast with the diurnality hypothesis, which makes is a promising avenue for future research.

### Conclusions

Our study addressed the hypotheses that cathemeral (generalist) temporal niche will decrease extinction risk due to the predisposition to flexibly alter temporal activity throughout the 24h diel cycle to where mortality is lower, and that in diurnal niche space, the aggregation of pressures involving human activities will increase extinction risk (7,8,27). While our global-scale evidence reveals that generalist temporal niche is associated with a decreasing probability of experiencing population declines among non-volant tetrapods, our analyses failed to identify evidence that extinction risk among tetrapods is concentrated in diurnal niche space – only an amphibian-specific pattern of increased extinction risk in diurnal niches was observed. In light of our findings, we call for action aimed towards understanding generalist temporal niche and the potential impacts of biotic homogenisation expressing across the day-night temporal niche spectrum (28,29). We emphasise that extinction risk is the (often synergistic) outcome of the interplay between extrinsic, anthropogenic pressures and intrinsic, evolutionary predispositions.

## Methods

### Data collection

We assembled a novel database spanning temporal niche and extinction risk (IUCN Red List conservation status) data for 23,277 species from across all four classes of tetrapod vertebrates – amphibians, ‘reptiles’ (restricted to lizards), mammals and birds. These data were obtained from the scientific literature (field guides, journal articles, books), existing class-specific databases (see details below), as well as field and museum records collected by the authors. First, for temporal niche data (defined as the range of time at which a species is active over the 24h diel spectrum), we employed the high-level category system (30), where each species was classed as either diurnal (active during the day hours), nocturnal (active during the night hours), crepuscular (active during twilight) or cathemeral (irregularly active throughout the 24h of the day). Temporal niche data were collated from existing databases for mammals (7,27) and lizards (31,32). For lizards, these sources of data included both cathemeral and crepuscular species within the same category.Therefore, these species were re-classified into the separate categories of cathemeral and crepuscular. Global-scale databases for birds and amphibians are not currently available, and thus, bird data were obtained primarily from the Handbook of the Birds of the World (33), whereas amphibian data were collated as part of the Global Amphibian Biodiversity Project (GABiP (17,26)). Second, we employed two alternative proxies for extinction risk from the IUCN Red List for all tetrapod species for which these assessments exist (3). Firstly, we used the classic conservation categories (the current estimated position of a species over an extinction risk spectrum). This proxy consists of two categories of non-threatened species (‘Least Concern’ or LC, and ‘Near Threatened’ or NT), and three categories of threatened species (‘Vulnerable’ or VU, ‘Endangered’ or EN, and ‘Critically Endangered’ or CR). Secondly, population trends (the trajectory of population sizes of a species through time), which assign species to either of the three categories of ‘Decreasing’, ‘Stable’ and ‘Increasing’ in population size (see 3). We downloaded IUCN Red List data using IUCN API 2024-1, accessed with the R package *rredlist* (34).

Finally, the taxonomic nomenclature used in our database follows a range of primary repositories of tetrapod taxonomy. For amphibians, we follow Frost (2023) (35). For lizards, Uetz et al. (2022) (36). For birds, we follow version 8.1 (January 2024) of the HBW Birdlife taxonomic checklist, maintained by Birdlife International. For mammals, we follow version 1.12.1 of the Mammal Diversity Database species checklist (37). We reviewed all data values corresponding to all species names that had been synonymised before assigning a single value to the valid species name in our taxonomy. When dealing with conflicting values, our decision-making depended specifically on each variable. For temporal niche, we assigned ‘cathemeral’ to species in each of the following cases: (1) where both nocturnal and diurnal were reported, (2) where cathemeral or diurnal were reported, and (3) where cathemeral and nocturnal were reported, and (4) where cathemeral and crepuscular were reported. Similarly, we categorised species as ‘nocturnal’ where both nocturnal and crepuscular were reported, and as ‘diurnal’ where both diurnal and crepuscular were reported.

For Red List categories and population trends, the majority of conflicting values did not affect our ability to assign species our binary proxy of extinction risk (e.g., synonyms listed as Endangered and Critically Endangered were nevertheless all considered ‘threatened’, see below). Where IUCN Red List assessments of synonyms revealed conflicting values for category or population trend that would affect the subsequent assignment of our binary extinction risk proxy, we used the value assigned to the valid species name only. We took values for category or population trend of a synonym in place of the respective values for the valid species name only when the valid species name had been assessed as Data Deficient for category or Unknown for population trend. We observed no situations where the value of category and population trends differed across synonyms in a way that would affect the assignment of our binary extinction risk proxy and where the IUCN assessment for the valid species name was absent. For our phylogenies (see below), where a valid species name was absent from a tree but corresponded to synonyms across multiple tip labels, we chose the most senior synonym. All data are available as supplementary material (Dataset 1).

### Phylogenetic trees

For phylogenetic analyses, we used consensus trees. For amphibians, we used Jetz and Pyron (2018) (6) phylogenetic supertree that contains ∼82% of known extant species diversity. For lizards, mammals and birds, we summarised entire posterior distributions of time-calibrated trees available from Tonini et al. (2016) (38), the Phylacine 1.2 database (39), and the Birdtree database (40), respectively, using median node heights with the TreeAnnotator app of Beast 2.5 (41). For birds, we used the Hackett backbone. These phylogenies represent ∼85% of squamate, ∼90% of mammal and ∼89% of bird diversity.

### Model specification

We performed our analyses using a Bayesian Markov chain Monte Carlo (MCMC) phylogenetic generalised mixed modelling approach, implemented in the package *MCMCglmm* version 2.36 (42) in R version 4.4.1 (43). We used MCMC to sample the posterior distributions of mixed models which included the phylogeny, in form of an inverse-relatedness matrix produced by the function ‘inverseA()’, as a random effect. We implemented a single-response model structure, with extinction risk (binary) as the response variable and temporal niche (binary) as the predictor, using a liability-threshold (probit) family function. In other words, we assume extinction risk is expressed by a latent, normally distributed continuous process which contains a threshold value at which species are separated into the binary outcome. We chose this approach because directly modelling the threshold or ‘tipping point’ (3,15,18) more accurately represents the mechanism of demographic decline to extinction.

We incorporated extinction risk into our models as the response variable using two proxies based on the IUCN Red List. Thus, each test ran twice. For our first proxy, we used the probability of experiencing population declines, based on the IUCN population trend. We assigned ‘1’ to species if their population trend was listed as ‘Decreasing’ and ‘0’ if their population trend was ‘Increasing’ or ‘Stable’ while removing species with a population trend of ‘Unknown’. For our second proxy, we used the probability of being threatened, based on the IUCN threat category. We assigned ‘1’ to species if their threat status was ‘Vulnerable’, ‘Endangered’ and ‘Critically Engendered’ and ‘0’ to species if their threat status was ‘Least Concern’ and ‘Near Threatened’ while excluding ‘Data Deficient’ and unassessed species.

We incorporated temporal niche into our models as the single predictor using binary contrasts, which compares a single group against all other groups combined. Thus, for each proxy of extinction risk, we ran three different models. For the first model, we encoded diurnal species as ‘1’ and cathemeral, crepuscular and nocturnal species as ‘0’. For the second model, we encoded cathemeral species as ‘1’ and diurnal, crepuscular and nocturnal species as ‘0’. For the third model, we encoded nocturnal species as ‘1’ and diurnal, cathemeral and crepuscular species as ‘0’. We did not model the effect of crepuscularity on extinction risk due to low numbers of crepuscular species among vertebrates.

While priors are less influential in analyses on large datasets, they may aid convergence and improve sampling efficiency by reducing the complexity of the parameter space. For the fixed effects priors, we implemented a normal distribution with variance of 9. The intercept prior mean was set to the probit-transformed baseline probability (calculated from the ‘0’ cases of the temporal niche predictor), while the temporal niche predictor prior was centred on 0. For the random effects phylogeny variance parameter, we specified a weakly informative inverse-Wishart distribution with a scale parameter of 1 with 1 degree of freedom. We employed the redundant working parameter ‘alpha.V’, using alpha.V=1,000 and alpha.mu=0, which provide a very diffuse prior (thus allowing the data to determine the random effects) while leveraging parameter expanded algorithms to aid MCMC in searching complex parameter space. Given that the outcome was binary, we fixed the residual variance parameter to 1.

We subsetted our data for vertebrates into amphibians, lizards, mammals and birds, and analysed each group separately. The package *MCMCglmm* employs a single chain to sample the posterior of each model, which we ran for 1,000,000 iterations with a thinning interval of 10, discarding the first 50% of iterations as burn-in. Convergence was determined by visually examining trace plots and we ensured effect sample size for each posterior was greater than 1,000. To aid interpretation of results, we calculated the intraclass correlation coefficient for each model, whereby values greater than 0.5 indicate that the predictor effect is concentrated more between clades whereas values less than 0.5 indicate that the predictor effect is concentrated within clades. Model outputs are displayed in Table S1.

### Hypothesis test

Generalised linear mixed models assume that random effects are independent of fixed effects. This assumption may be violated in single-response phylogenetic linear models when a predictor has high phylogenetic signal, resulting in Mundlak’s bias (44). When both predictor and outcome are binary, the impact of this bias on the model could be particularly problematic. This is because a binary variable with high phylogenetic signal expresses as severe class imbalance in certain regions of the phylogenetic space, which (regardless of the contingency table for the full dataset) could disproportionately influence the model estimates as the effect is actually based on very little information for that region of phylogenetic space. This bias is indicated to statistically by a correlation between the binary predictor value and the random intercept deviation from the outcome for each observation. In non-phylogenetic contexts, the observation of this correlation invalidates inferences as the effect of the random intercept may not be reliably separated from the effect of the predictor. However, phylogenetic linear models are inherently descriptive (see (45)), therefore this correlation does not necessarily invalidate inferences but its consideration is required as it reveals systematic under- or over-prediction for certain predictor values. Specifically, a positive correlation reveals that the model under-predicts at higher predictor values (i.e. observations at high predictor values tend to have higher outcomes than the model predicts), while a negative correlation reveals that the model over-predicts at higher predictor values (i.e. observations at high predictor values tend to have lower outcomes than the model predicts). As both extinction risk and temporal niche are binary predictors, our hypothesis test is vulnerable to such bias. Consequently, we implement an approach in which empirical support for our hypothesis is found when two conditions are satisfied. The first condition is the exclusion of zero from the 95% credible interval of the temporal niche predictor’s posterior distribution, while the second is either the absence of a significant relationship between random effect deviation and predictor value or the observation of such significant relationship but in a direction which does not bring closer to zero the predictor’s observed effect. We used the base R ‘glm()’ function with a probit link to test for significance in the relationship where the random effect deviation causes the predictor value. Model outputs are displayed in Table S1.

### Exploratory data analysis

To characterise the data features that could underlie the model findings and to examine variation in outcomes across different regions of phylogenetic space, we performed a descriptive exploratory data analysis on the datasets represented by the models which supported the hypothesis. This exploratory data analysis examined both between- and within-clade variation, defining ‘clade’ as families for amphibians, superfamilies for lizards and orders for mammals. To examine between-clade variation, we calculated the probability of extinction risk as well as the probability of the exhibiting the relevant temporal niche for each clade in the phylogeny with more than 10 lineages, and plotted these features using the R packages *ggtree* and *ggtreeExtra* (46,47). To examine within-clade variation, we calculated the risk differences for each clade in the phylogeny with more than 10 lineages. The we defined the risk difference as the difference in the probability of the outcome (extinction risk) between observations with and without the predictor (the specific temporal niche state), i.e. P(outcome=1|predictor=1) – P(outcome=1|predictor=0). The risk difference reveals the absolute percent point reduction or increase in the extinction risk attributed the specific temporal niche effect.

