## Supplementary material for "Generalist temporal niche buffers against extinction risk among non-avian tetrapods": Table S1 (from another version of this preprint: version 2, 2026-01-15)

**Table S1.** Numbers and percentages of species by temporal niche that are considered threatened/not threatened or declining/not declining based on IUCN threat status/population trend (respectively). Using threat status, we considered species listed as 'Vulnerable', 'Endangered', 'Critically Engendered', 'Extinction in the Wild' and 'Extinct' as 'threatened', whereas species listed as 'Least Concern' and 'Near Threatened' were assigned 'not threatened'. We excluded all 'Data Deficient' species from our analyses, but included them in the table below. Similarly, using population trends, we assigned species listed as 'Decreasing' as 'declining', whereas those listed as 'Increasing' and 'Stable' were assigned 'not declining'. Species with population trends listed as 'Unknown' were removed from all analyses, but are included in the table below. We accounted for Unknown/Data Deficient species in the table below by including error brackets on the percentage declining/threatened, as in Böhm et al. (2013). The lower number in brackets represents a 'best case scenario', where all species that are assigned 'Unknown' population trends or 'Data Deficient' threat category are assumed to be not declining/not threatened, whereas the higher number in brackets represents a 'worst case scenario', where those species were instead assumed to be declining/threatened.

| Taxonomic Group | Threat Status | | | | Population Trend | | | |
|---|---|---|---|---|---|---|---|---|
| | Threatened | Not Threatened | Data Deficient | % Threatened with Data Deficient species as errors | Declining | Not Declining | Unknown | % Declining with Unknown species as errors |
| **1 Amphibians** | | | | | | | | |
| Diurnal | 350 | 318 | 93 | 52.4 (46 - 58.2) | 378 | 190 | 193 | 66.5 (49.7 - 75) |
| Cathemeral | 183 | 465 | 35 | 28.2 (26.8 - 31.9) | 291 | 271 | 121 | 51.8 (42.6 - 60.3) |
| Nocturnal | 660 | 1359 | 182 | 32.7 (30 - 38.3) | 975 | 777 | 449 | 55.7 (44.3 - 64.7) |
| **2 Lizards** | | | | | | | | |
| Diurnal | 640 | 2558 | 522 | 20 (17.2 - 31.2) | 526 | 1553 | 1641 | 25.3 (14.1 - 58.3) |
| Cathemeral | 66 | 299 | 29 | 18.1 (16.8 - 24.1) | 50 | 210 | 134 | 19.2 (12.7 - 46.7) |
| Nocturnal | 224 | 767 | 228 | 22.6 (18.4 - 37.1) | 141 | 469 | 609 | 23.1 (11.6 - 61.5) |
| **3 Mammals** | | | | | | | | |
| Diurnal | 283 | 542 | 43 | 34.3 (32.6 - 37.6) | 434 | 209 | 225 | 67.5 (50 - 75.9) |
| Cathemeral | 117 | 343 | 51 | 25.4 (22.9 - 32.9) | 176 | 184 | 151 | 48.9 (34.4 - 64) |
| Nocturnal | 641 | 2229 | 438 | 22.3 (19.4 - 32.6) | 955 | 885 | 1468 | 51.9 (28.9 - 73.2) |
| **4 Birds** | | | | | | | | |
| Diurnal | 963 | 7822 | 78 | 11 (10.9 - 11.7) | 4292 | 4046 | 525 | 51.5 (48.4 - 54.3) |
| Cathemeral | 55 | 173 | 0 | 24.1 (24.1 - 24.1) | 114 | 89 | 25 | 56.2 (50 - 61) |
| Nocturnal | 26 | 258 | 2 | 9.2 (9.1 - 9.8) | 146 | 130 | 10 | 52.9 (51 - 54.5) |
