## Supplementary material for "Generalist temporal niche buffers against extinction risk among non-avian tetrapods": Table S2 (from another version of this preprint: version 2, 2026-01-15)

**Table S2.** Outputs from Bayesian phylogenetic models, where extinction risk (binary, based on either IUCN Category (1), or Population Trend (2)) was modelled under a liability-threshold specification, with the R package 'MCMCglmm', for each tetrapod group (amphibians (1), lizards (2), mammals (3) and birds (4)). Prior to analysis, body mass and geographic range size were log-transformed then *z*-transformed, while human footprint was *z*-transformed. We considered continuous predictors to be significant when the 95% Credible Interval did not include zero (indicated in bold-face text). We incorporated the phylogeny as an inverse-relatedness matrix as a random effect using the inverseA() function of the 'MCMCglmm' package, and estimated the phylogenetic variance, represented in the table as 'Phylo Name'. Note that the global intercept was removed, and thus each biome-temporal niche interaction parameter serves simultaneously as an absolute effect and an intercept. The removal of the global intercept impacts the interpretation of the credible intervals for these parameters in that the significance of credible intervals indicates significant difference from 0, which on the liability scale corresponds to a proportion of 0.5. In other words, if a credible interval for the biome-temporal niche interaction term excludes zero, this indicates that the percentage of species at risk of extinction attributed to that group is significantly greater or less than 50% after the effects of other predictors were considered, which is unrelated to our hypothesis. Due to *z*-transformation, the interpretation of the credible intervals for the continuous variables is unaffected by the removal of the global intercept. For each parameter, table shows the sample size (*n*), posterior mean, 95% credible interval, *p*-value (*pMCMC*) and effective sample size (*ESS*). While we report *pMCMC*, the decision for significance was based solely on the 95% Credible Interval. Convergence of the MCMC chain was determined by examination of trace plots and we ensured *ESS* > 1000.

| Extinction Risk Proxy | Taxonomic Group | *n* | Parameter | Posterior Mean | 95% Credible Interval | *pMCMC* | *ESS* |
|---|---|---|---|---|---|---|---|
| 1 Category | 1.1 Amphibians | 2924 | Tropical:Diurnal | -0.69 | -2.067, 0.616 | 0.308 | 24157 |
| | | | Temperate:Diurnal | -0.527 | -1.938, 0.851 | 0.452 | 22279 |
| | | | Tropical:Nocturnal | -0.905 | -2.246, 0.378 | 0.176 | 23785 |
| | | | Temperate:Nocturnal | -1.305 | -2.66, 0.041 | 0.06 | 22151 |
| | | | Tropical:Cathemeral | -1.058 | -2.393, 0.286 | 0.118 | 22126 |
| | | | Temperate:Cathemeral | -0.77 | -2.14, 0.631 | 0.271 | 23345 |
| | | | Tropical:Diurnal:Body Mass | 0.079 | -0.18, 0.337 | 0.545 | 22319 |
| | | | Temperate:Diurnal:Body Mass | 0.486 | -0.047, 1.04 | 0.07 | 19739 |
| | | | Tropical:Nocturnal:Body Mass | 0.055 | -0.106, 0.228 | 0.51 | 18052 |
| | | | Temperate:Nocturnal:Body Mass | **0.788** | **0.442, 1.14** | **<0.001** | **8988** |
| | | | Tropical:Cathemeral:Body Mass | **0.327** | **0.082, 0.587** | **0.011** | **15156** |
| | | | Temperate:Cathemeral:Body Mass | **0.565** | **0.122, 1.034** | **0.013** | **12824** |
| | | | Tropical:Diurnal:Geographic Range Size | **-1.646** | **-1.963, -1.341** | **<0.001** | **4842** |
| | | | Temperate:Diurnal:Geographic Range Size | **-1.723** | **-2.432, -1.032** | **<0.001** | **10176** |
| | | | Tropical:Nocturnal:Geographic Range Size | **-1.777** | **-2.005, -1.543** | **<0.001** | **2732** |
| | | | Temperate:Nocturnal:Geographic Range Size | **-2.36** | **-2.922, -1.839** | **<0.001** | **4377** |
| | | | Tropical:Cathemeral:Geographic Range Size | **-2.13** | **-2.55, -1.688** | **<0.001** | **2885** |
| | | | Temperate:Cathemeral:Geographic Range Size | **-2.084** | **-2.779, -1.412** | **<0.001** | **6641** |
| | | | Tropical:Diurnal:Human Footprint Mean | **0.53** | **0.294, 0.771** | **<0.001** | **10849** |
| | | | Temperate:Diurnal:Human Footprint Mean | -0.12 | -0.542, 0.341 | 0.57 | 15341 |
| | | | Tropical:Nocturnal:Human Footprint Mean | **0.244** | **0.112, 0.385** | **<0.001** | **12240** |
| | | | Temperate:Nocturnal:Human Footprint Mean | 0.265 | -0.034, 0.558 | 0.076 | 13967 |

| | | | | | | | |
|---|---|---|---|---|---|---|---|
| | | | Tropical:Cathemeral:Human Footprint Mean | 0.27 | -0.045, 0.59 | 0.089 | 11637 |
| | | | Temperate:Cathemeral:Human Footprint Mean | 0.376 | -0.034, 0.773 | 0.06 | 16575 |
| | | | Phylo Name | 3.279 | 1.817, 4.947 | NA | 15721 |
| 1 Category | 1.2 Lizards | 2972 | Tropical:Diurnal | -1.532 | -2.868, -0.211 | 0.026 | 91322 |
| | | | Temperate:Diurnal | -1.054 | -2.387, 0.289 | 0.123 | 71908 |
| | | | Tropical:Nocturnal | -0.997 | -2.389, 0.402 | 0.156 | 65442 |
| | | | Temperate:Nocturnal | -1.708 | -3.212, -0.234 | 0.025 | 61377 |
| | | | Tropical:Cathemeral | -1.481 | -2.916, -0.072 | 0.042 | 75738 |
| | | | Temperate:Cathemeral | -1.92 | -3.418, -0.488 | 0.011 | 60697 |
| | | | Tropical:Diurnal:Body Mass | 0.043 | -0.163, 0.245 | 0.674 | 65494 |
| | | | Temperate:Diurnal:Body Mass | 0.012 | -0.251, 0.28 | 0.922 | 64734 |
| | | | Tropical:Nocturnal:Body Mass | -0.16 | -0.537, 0.215 | 0.398 | 71053 |
| | | | Temperate:Nocturnal:Body Mass | -0.538 | -1.287, 0.217 | 0.156 | 44342 |
| | | | Tropical:Cathemeral:Body Mass | 0.386 | -0.154, 0.935 | 0.155 | 23571 |
| | | | Temperate:Cathemeral:Body Mass | **-0.841** | **-1.592, -0.09** | **0.021** | **27532** |
| | | | Tropical:Diurnal:Geographic Range Size | **-1.291** | **-1.594, -1.017** | **<0.001** | **2144** |
| | | | Temperate:Diurnal:Geographic Range Size | **-1.344** | **-1.675, -1.046** | **<0.001** | **3304** |
| | | | Tropical:Nocturnal:Geographic Range Size | **-0.848** | **-1.169, -0.543** | **<0.001** | **5622** |
| | | | Temperate:Nocturnal:Geographic Range Size | **-1.262** | **-1.84, -0.7** | **<0.001** | **5206** |
| | | | Tropical:Cathemeral:Geographic Range Size | **-1.38** | **-2.012, -0.793** | **<0.001** | **5630** |
| | | | Temperate:Cathemeral:Geographic Range Size | **-1.004** | **-1.634, -0.399** | **<0.001** | **15878** |
| | | | Tropical:Diurnal:Human Footprint Mean | **0.161** | **0.015, 0.309** | **0.028** | **38547** |
| | | | Temperate:Diurnal:Human Footprint Mean | **0.441** | **0.205, 0.671** | **<0.001** | **13776** |
| | | | Tropical:Nocturnal:Human Footprint Mean | 0.172 | -0.061, 0.41 | 0.144 | 39827 |
| | | | Temperate:Nocturnal:Human Footprint Mean | **0.659** | **0.128, 1.205** | **0.011** | **15831** |
| | | | Tropical:Cathemeral:Human Footprint Mean | -0.091 | -0.538, 0.366 | 0.695 | 66247 |
| | | | Temperate:Cathemeral:Human Footprint Mean | **-0.802** | **-1.549, -0.125** | **0.009** | **22450** |
| | | | Phylo Name | 5.694 | 2.306, 9.919 | NA | 1383 |
| 1 Category | 1.3 Mammals | 4112 | Tropical:Diurnal | -0.812 | -2.367, 0.735 | 0.305 | 19835 |
| | | | Temperate:Diurnal | -1.172 | -2.803, 0.407 | 0.153 | 20836 |
| | | | Tropical:Nocturnal | -0.549 | -2.078, 0.955 | 0.476 | 21118 |
| | | | Temperate:Nocturnal | -0.4 | -1.899, 1.144 | 0.601 | 21300 |
| | | | Tropical:Cathemeral | -0.414 | -1.982, 1.11 | 0.596 | 20813 |
| | | | Temperate:Cathemeral | -1.024 | -2.679, 0.476 | 0.195 | 20447 |
| | | | Tropical:Diurnal:Body Mass | **0.998** | **0.696, 1.297** | **<0.001** | **17518** |
| | | | Temperate:Diurnal:Body Mass | **0.765** | **0.376, 1.165** | **<0.001** | **15185** |
| | | | Tropical:Nocturnal:Body Mass | **0.671** | **0.471, 0.871** | **<0.001** | **13543** |

| | | | | | | | |
|---|---|---|---|---|---|---|---|
| | | | Temperate:Nocturnal:Body Mass | **0.625** | **0.381, 0.867** | **<0.001** | **18746** |
| | | | Tropical:Cathemeral:Body Mass | **0.831** | **0.532, 1.117** | **<0.001** | **19118** |
| | | | Temperate:Cathemeral:Body Mass | **0.88** | **0.541, 1.226** | **<0.001** | **8159** |
| | | | Tropical:Diurnal:Geographic Range Size | **-1.383** | **-1.638, -1.131** | **<0.001** | **5520** |
| | | | Temperate:Diurnal:Geographic Range Size | **-1.175** | **-1.569, -0.821** | **<0.001** | **13745** |
| | | | Tropical:Nocturnal:Geographic Range Size | **-1.376** | **-1.529, -1.231** | **<0.001** | **2743** |
| | | | Temperate:Nocturnal:Geographic Range Size | **-1.226** | **-1.435, -1.023** | **<0.001** | **4401** |
| | | | Tropical:Cathemeral:Geographic Range Size | **-1.275** | **-1.615, -0.943** | **<0.001** | **8951** |
| | | | Temperate:Cathemeral:Geographic Range Size | **-1.222** | **-1.678, -0.787** | **<0.001** | **7960** |
| | | | Tropical:Diurnal:Human Footprint Mean | **0.208** | **0.028, 0.385** | **0.022** | **21666** |
| | | | Temperate:Diurnal:Human Footprint Mean | -0.078 | -0.36, 0.206 | 0.595 | 23190 |
| | | | Tropical:Nocturnal:Human Footprint Mean | -0.02 | -0.117, 0.076 | 0.684 | 22216 |
| | | | Temperate:Nocturnal:Human Footprint Mean | **0.154** | **<0.001, 0.302** | **0.042** | **18203** |
| | | | Tropical:Cathemeral:Human Footprint Mean | -0.131 | -0.431, 0.179 | 0.401 | 24087 |
| | | | Temperate:Cathemeral:Human Footprint Mean | 0.194 | -0.058, 0.453 | 0.139 | 17825 |
| | | | Phylo Name | 3.445 | 1.673, 5.431 | NA | 1246 |
| 1 Category | 1.4 Birds | 8857 | Tropical:Diurnal | -1.76 | -2.561, -0.922 | <0.001 | 24368 |
| | | | Temperate:Diurnal | -1.335 | -2.178, -0.534 | 0.002 | 24495 |
| | | | Tropical:Nocturnal | -2.231 | -3.367, -1.146 | <0.001 | 17470 |
| | | | Temperate:Nocturnal | -2.642 | -4.227, -1.03 | 0.001 | 14992 |
| | | | Tropical:Cathemeral | -1.828 | -3.062, -0.638 | 0.003 | 21283 |
| | | | Temperate:Cathemeral | -0.802 | -1.933, 0.396 | 0.174 | 23419 |
| | | | Tropical:Diurnal:Body Mass | **0.495** | **0.378, 0.614** | **<0.001** | **16446** |
| | | | Temperate:Diurnal:Body Mass | **0.303** | **0.176, 0.437** | **<0.001** | **20088** |
| | | | Tropical:Nocturnal:Body Mass | 0.48 | -0.141, 1.099 | 0.125 | 17257 |
| | | | Temperate:Nocturnal:Body Mass | **0.898** | **0.026, 1.75** | **0.034** | **16169** |
| | | | Tropical:Cathemeral:Body Mass | 0.449 | -0.138, 1.03 | 0.126 | 21225 |
| | | | Temperate:Cathemeral:Body Mass | **0.741** | **0.292, 1.201** | **0.001** | **21022** |
| | | | Tropical:Diurnal:Geographic Range Size | **-0.879** | **-0.961, -0.794** | **<0.001** | **2800** |
| | | | Temperate:Diurnal:Geographic Range Size | **-0.787** | **-0.909, -0.663** | **<0.001** | **7783** |
| | | | Tropical:Nocturnal:Geographic Range Size | **-0.834** | **-1.213, -0.454** | **<0.001** | **11521** |
| | | | Temperate:Nocturnal:Geographic Range Size | -0.465 | -1.083, 0.177 | 0.145 | 21951 |
| | | | Tropical:Cathemeral:Geographic Range Size | -0.46 | -0.956, 0.06 | 0.074 | 17115 |
| | | | Temperate:Cathemeral:Geographic Range Size | **-1.493** | **-2.131, -0.876** | **<0.001** | **14464** |
| | | | Tropical:Diurnal:Human Footprint Mean | **0.193** | **0.139, 0.247** | **<0.001** | **17709** |
| | | | Temperate:Diurnal:Human Footprint Mean | **0.231** | **0.125, 0.335** | **<0.001** | **19018** |
| | | | Tropical:Nocturnal:Human Footprint Mean | 0.142 | -0.177, 0.463 | 0.379 | 19767 |

| | | | | | | | |
|---|---|---|---|---|---|---|---|
| | | | Temperate:Nocturnal:Human Footprint Mean | 0.068 | -0.644, 0.841 | 0.856 | 19685 |
| | | | Tropical:Cathemeral:Human Footprint Mean | -0.175 | -0.703, 0.341 | 0.513 | 20938 |
| | | | Temperate:Cathemeral:Human Footprint Mean | -0.092 | -0.483, 0.305 | 0.644 | 23128 |
| | | | Phylo Name | 1.36 | 0.809, 1.985 | NA | 1496 |
| 2 Population Trend | 2.1 Amphibians | 2520 | Tropical:Diurnal | 0.426 | -1.007, 1.93 | 0.566 | 25000 |
| | | | Temperate:Diurnal | -0.081 | -1.546, 1.455 | 0.908 | 25000 |
| | | | Tropical:Nocturnal | 0.429 | -1.014, 1.879 | 0.557 | 25000 |
| | | | Temperate:Nocturnal | 0.159 | -1.335, 1.588 | 0.836 | 24000 |
| | | | Tropical:Cathemeral | 0.35 | -1.075, 1.834 | 0.637 | 25000 |
| | | | Temperate:Cathemeral | -0.021 | -1.587, 1.411 | 0.978 | 25000 |
| | | | Tropical:Diurnal:Body Mass | -0.149 | -0.411, 0.117 | 0.27 | 24129 |
| | | | Temperate:Diurnal:Body Mass | 0.347 | -0.15, 0.839 | 0.163 | 24040 |
| | | | Tropical:Nocturnal:Body Mass | **0.166** | **0.01, 0.323** | **0.038** | **23752** |
| | | | Temperate:Nocturnal:Body Mass | **0.344** | **0.098, 0.594** | **0.007** | **21204** |
| | | | Tropical:Cathemeral:Body Mass | 0.153 | -0.058, 0.367 | 0.157 | 23722 |
| | | | Temperate:Cathemeral:Body Mass | 0.364 | 0.027, 0.711 | 0.033 | 22984 |
| | | | Tropical:Diurnal:Geographic Range Size | **-0.671** | **-0.906, -0.446** | **<0.001** | **15763** |
| | | | Temperate:Diurnal:Geographic Range Size | **-0.74** | **-1.257, -0.268** | **0.001** | **17463** |
| | | | Tropical:Nocturnal:Geographic Range Size | **-0.789** | **-0.95, -0.632** | **<0.001** | **9280** |
| | | | Temperate:Nocturnal:Geographic Range Size | **-0.502** | **-0.79, -0.203** | **<0.001** | **21663** |
| | | | Tropical:Cathemeral:Geographic Range Size | **-0.876** | **-1.115, -0.641** | **<0.001** | **13939** |
| | | | Temperate:Cathemeral:Geographic Range Size | **-0.811** | **-1.239, -0.386** | **<0.001** | **23015** |
| | | | Tropical:Diurnal:Human Footprint Mean | **0.329** | **0.099, 0.556** | **0.004** | **22361** |
| | | | Temperate:Diurnal:Human Footprint Mean | 0.286 | -0.107, 0.668 | 0.133 | 20554 |
| | | | Tropical:Nocturnal:Human Footprint Mean | 0.115 | -0.014, 0.243 | 0.076 | 23316 |
| | | | Temperate:Nocturnal:Human Footprint Mean | 0.087 | -0.177, 0.347 | 0.517 | 23732 |
| | | | Tropical:Cathemeral:Human Footprint Mean | 0.13 | -0.123, 0.39 | 0.318 | 23220 |
| | | | Temperate:Cathemeral:Human Footprint Mean | 0.153 | -0.188, 0.482 | 0.373 | 24083 |
| | | | Phylo Name | 4.572 | 2.753, 6.573 | NA | 2589 |
| 2 Population Trend | 2.2 Lizards | 2039 | Tropical:Diurnal | -0.908 | -2.455, 0.61 | 0.239 | 95186 |
| | | | Temperate:Diurnal | -0.547 | -2.065, 1.021 | 0.479 | 74529 |
| | | | Tropical:Nocturnal | -0.401 | -1.992, 1.241 | 0.619 | 66267 |
| | | | Temperate:Nocturnal | -0.895 | -2.541, 0.771 | 0.287 | 80180 |
| | | | Tropical:Cathemeral | -1.118 | -2.75, 0.508 | 0.173 | 84771 |
| | | | Temperate:Cathemeral | -1.204 | -2.798, 0.402 | 0.137 | 93133 |
| | | | Tropical:Diurnal:Body Mass | -0.032 | -0.265, 0.195 | 0.788 | 47864 |
| | | | Temperate:Diurnal:Body Mass | 0.011 | -0.255, 0.284 | 0.927 | 75055 |

| | | | | | | | |
|---|---|---|---|---|---|---|---|
| | | | Tropical:Nocturnal:Body Mass | -0.088 | -0.601, 0.408 | 0.731 | 65601 |
| | | | Temperate:Nocturnal:Body Mass | -0.575 | -1.385, 0.232 | 0.151 | 32090 |
| | | | Tropical:Cathemeral:Body Mass | 0.3 | -0.319, 0.936 | 0.336 | 51718 |
| | | | Temperate:Cathemeral:Body Mass | -0.151 | -0.765, 0.474 | 0.627 | 75968 |
| | | | Tropical:Diurnal:Geographic Range Size | **-0.67** | **-0.917, -0.435** | **<0.001** | **6108** |
| | | | Temperate:Diurnal:Geographic Range Size | **-0.843** | **-1.161, -0.554** | **<0.001** | **3789** |
| | | | Tropical:Nocturnal:Geographic Range Size | **-0.711** | **-1.169, -0.271** | **0.001** | **10467** |
| | | | Temperate:Nocturnal:Geographic Range Size | **-0.797** | **-1.39, -0.237** | **0.004** | **14659** |
| | | | Tropical:Cathemeral:Geographic Range Size | **-0.822** | **-1.455, -0.184** | **0.008** | **20709** |
| | | | Temperate:Cathemeral:Geographic Range Size | 0.097 | -0.447, 0.672 | 0.75 | 75222 |
| | | | Tropical:Diurnal:Human Footprint Mean | 0.169 | -0.031, 0.375 | 0.092 | 39081 |
| | | | Temperate:Diurnal:Human Footprint Mean | **0.751** | **0.466, 1.047** | **<0.001** | **5969** |
| | | | Tropical:Nocturnal:Human Footprint Mean | -0.031 | -0.426, 0.359 | 0.875 | 79958 |
| | | | Temperate:Nocturnal:Human Footprint Mean | **0.968** | **0.326, 1.644** | **0.002** | **13325** |
| | | | Tropical:Cathemeral:Human Footprint Mean | -0.337 | -0.946, 0.254 | 0.257 | 33184 |
| | | | Temperate:Cathemeral:Human Footprint Mean | 0.153 | -0.348, 0.65 | 0.536 | 85427 |
| | | | Phylo Name | 8.833 | 3.496, 15.478 | NA | 1594 |
| 2 Population Trend | 2.3 Mammals | 2812 | Tropical:Diurnal | 0.562 | -0.876, 1.982 | 0.445 | 23772 |
| | | | Temperate:Diurnal | 0.453 | -0.987, 1.925 | 0.548 | 23229 |
| | | | Tropical:Nocturnal | 0.719 | -0.687, 2.134 | 0.322 | 23685 |
| | | | Temperate:Nocturnal | 0.569 | -0.828, 2.011 | 0.434 | 23493 |
| | | | Tropical:Cathemeral | 0.707 | -0.718, 2.151 | 0.339 | 23492 |
| | | | Temperate:Cathemeral | 0.427 | -0.985, 1.917 | 0.568 | 23357 |
| | | | Tropical:Diurnal:Body Mass | **0.682** | **0.404, 0.97** | **<0.001** | **20034** |
| | | | Temperate:Diurnal:Body Mass | 0.164 | -0.166, 0.5 | 0.332 | 21998 |
| | | | Tropical:Nocturnal:Body Mass | **0.552** | **0.36, 0.752** | **<0.001** | **16004** |
| | | | Temperate:Nocturnal:Body Mass | **0.301** | **0.074, 0.527** | **0.008** | **24243** |
| | | | Tropical:Cathemeral:Body Mass | **0.526** | **0.253, 0.814** | **<0.001** | **16424** |
| | | | Temperate:Cathemeral:Body Mass | 0.212 | -0.048, 0.471 | 0.108 | 22423 |
| | | | Tropical:Diurnal:Geographic Range Size | **-0.415** | **-0.611, -0.226** | **<0.001** | **23397** |
| | | | Temperate:Diurnal:Geographic Range Size | **-0.596** | **-0.92, -0.277** | **<0.001** | **23287** |
| | | | Tropical:Nocturnal:Geographic Range Size | **-0.741** | **-0.85, -0.628** | **<0.001** | **12604** |
| | | | Temperate:Nocturnal:Geographic Range Size | **-0.645** | **-0.803, -0.487** | **<0.001** | **20684** |
| | | | Tropical:Cathemeral:Geographic Range Size | **-0.623** | **-0.932, -0.305** | **<0.001** | **17912** |
| | | | Temperate:Cathemeral:Geographic Range Size | **-0.764** | **-1.093, -0.425** | **<0.001** | **23747** |
| | | | Tropical:Diurnal:Human Footprint Mean | -0.04 | -0.229, 0.152 | 0.683 | 22464 |
| | | | Temperate:Diurnal:Human Footprint Mean | -0.021 | -0.269, 0.229 | 0.867 | 23357 |

| | | | | | | | |
|---|---|---|---|---|---|---|---|
| | | | Tropical:Nocturnal:Human Footprint Mean | **0.131** | **0.026, 0.237** | **0.013** | **24253** |
| | | | Temperate:Nocturnal:Human Footprint Mean | **0.199** | **0.055, 0.333** | **0.005** | **23402** |
| | | | Tropical:Cathemeral:Human Footprint Mean | -0.112 | -0.388, 0.172 | 0.439 | 24525 |
| | | | Temperate:Cathemeral:Human Footprint Mean | -0.015 | -0.23, 0.198 | 0.893 | 24347 |
| | | | Phylo Name | 2.891 | 1.409, 4.463 | NA | 1893 |
| 2 Population Trend | 2.4 Birds | 8403 | Tropical:Diurnal | 0.66 | -0.401, 1.661 | 0.208 | 25000 |
| | | | Temperate:Diurnal | 0.602 | -0.427, 1.632 | 0.252 | 25000 |
| | | | Tropical:Nocturnal | 0.145 | -1.001, 1.301 | 0.799 | 24452 |
| | | | Temperate:Nocturnal | 0.034 | -1.317, 1.405 | 0.962 | 25000 |
| | | | Tropical:Cathemeral | 0.696 | -0.587, 1.937 | 0.276 | 25000 |
| | | | Temperate:Cathemeral | 0.958 | -0.333, 2.234 | 0.147 | 25000 |
| | | | Tropical:Diurnal:Body Mass | **0.177** | **0.073, 0.282** | **0.001** | **25000** |
| | | | Temperate:Diurnal:Body Mass | -0.017 | -0.135, 0.091 | 0.77 | 25000 |
| | | | Tropical:Nocturnal:Body Mass | 0.378 | -0.094, 0.834 | 0.111 | 23237 |
| | | | Temperate:Nocturnal:Body Mass | 0.186 | -0.452, 0.799 | 0.564 | 25553 |
| | | | Tropical:Cathemeral:Body Mass | -0.228 | -0.765, 0.338 | 0.417 | 25000 |
| | | | Temperate:Cathemeral:Body Mass | -0.137 | -0.573, 0.285 | 0.531 | 25000 |
| | | | Tropical:Diurnal:Geographic Range Size | **-0.254** | **-0.302, -0.207** | **<0.001** | **20964** |
| | | | Temperate:Diurnal:Geographic Range Size | **-0.169** | **-0.257, -0.082** | **<0.001** | **23851** |
| | | | Tropical:Nocturnal:Geographic Range Size | -0.223 | -0.491, 0.055 | 0.109 | 25000 |
| | | | Temperate:Nocturnal:Geographic Range Size | 0.068 | -0.445, 0.585 | 0.803 | 25000 |
| | | | Tropical:Cathemeral:Geographic Range Size | -0.006 | -0.528, 0.497 | 0.984 | 25000 |
| | | | Temperate:Cathemeral:Geographic Range Size | -0.213 | -0.761, 0.301 | 0.434 | 24227 |
| | | | Tropical:Diurnal:Human Footprint Mean | 0.04 | -0.005, 0.081 | 0.072 | 24166 |
| | | | Temperate:Diurnal:Human Footprint Mean | 0.044 | -0.038, 0.124 | 0.297 | 25000 |
| | | | Tropical:Nocturnal:Human Footprint Mean | **-0.388** | **-0.64, -0.128** | **0.003** | **23845** |
| | | | Temperate:Nocturnal:Human Footprint Mean | 0.022 | -0.515, 0.557 | 0.941 | 25000 |
| | | | Tropical:Cathemeral:Human Footprint Mean | **0.516** | **0.024, 1.028** | **0.04** | **25000** |
| | | | Temperate:Cathemeral:Human Footprint Mean | 0.051 | -0.309, 0.429 | 0.789 | 24331 |
| | | | Phylo Name | 2.519 | 1.823, 3.296 | NA | 2510 |
