## Supplementary material for "Generalist temporal niche buffers against extinction risk among non-avian tetrapods": Table S3 (from another version of this preprint: version 2, 2026-01-15)

**Table S3.** Outputs from the test of posterior differences among posterior distributions of models. For each model component ('Absolute Effect', 'Body Mass', 'Geographic Range Size' and 'Human Footprint'), we performed pairwise tests between temporal niches ('Diurnal > Nocturnal', 'Diurnal > Cathemeral', as well as 'Nocturnal > Cathemeral'). We calculated the mean difference of two marginal posterior distributions accounting for the fact that model parameters are estimated jointly. Firstly, we determined the proportion (or probability, as odds) that the hypothesis is supported. For each iteration, we assigned '1' if the test was supported (e.g., for 'Diurnal > Nocturnal', if the value for diurnal was greater than the value for nocturnal), '0' if the test was not supported, and '0.5' if the parameter estimates were exactly equal. We then calculated the mean of these assigned values, which when converted to odds, represents the probability that the the hypothesis is supported. Secondly, we determined the average magnitude of differences by subtracting the values of one posterior from the other, and taking the mean, the 2.5% and 97.5% quantiles for the resulting distribution. We considered the difference between two posteriors to be significantly different from zero when the 95% credible interval did not include zero, which is indicated below in bold-face text. Note that for parameters that were negative in predicting extinction risk (e.g., geographic range size), a significant positive difference (e.g., in the test Diurnal > Cathemeral), indicates that the effect for diurnals is closer to zero (i.e., the effect is diminished by diurnality) whereas for parameters that were positive (e.g., body mass and most often time human footprint), a significant positive difference would indicate that the effect for diurnals is further from zero (i.e., the effect is intensified by diurnality). Also note that significance in the difference of posterior distributions is unrelated to whether those parameters significantly predicted extinction risk (see Table S2) – it may be that within a model component, one parameter may predict extinction risk significantly (e.g., human footprint for diurnality) while the other may not (e.g., human footprint for cathemerality), but these may not show significant a difference in posterior distributions due to the wide credible interval of the non-significant predictor. Therefore, we used the examination of differences in posterior distributions to test for temporal niche variation among parameters that significantly predict extinction risk, and also to examine differences in the absolute effects of temporal niches.

| Extinction Risk Proxy | Taxonomic Group | Model Component | Test | Mean Difference | 95% Credible Interval of Differences | Proportion | Odds |
|---|---|---|---|---|---|---|---|
| 1 Category | 1.1 Amphibians | Tropics: Absolute Effect | Diurnal > Nocturnal | 0.215 | -0.147, 0.569 | 0.879 | 7.232 |
| | | | Diurnal > Cathemeral | 0.367 | -0.048, 0.785 | 0.958 | 22.855 |
| | | | Nocturnal > Cathemeral | 0.153 | -0.163, 0.484 | 0.822 | 4.608 |
| | | Tropics: Body Mass | Diurnal > Nocturnal | 0.024 | -0.276, 0.324 | 0.562 | 1.282 |
| | | | Diurnal > Cathemeral | -0.248 | -0.599, 0.098 | 0.08 | 0.087 |
| | | | Nocturnal > Cathemeral | -0.271 | -0.558, 0.008 | 0.028 | 0.029 |
| | | Tropics: Geographic Range Size | Diurnal > Nocturnal | 0.131 | -0.206, 0.466 | 0.774 | 3.425 |
| | | | Diurnal > Cathemeral | **0.484** | **0.018, 0.979** | **0.979** | **47.263** |
| | | | Nocturnal > Cathemeral | 0.353 | -0.066, 0.802 | 0.948 | 18.231 |
| | | Tropics:Human Footprint | Diurnal > Nocturnal | **0.285** | **0.023, 0.553** | **0.983** | **58.102** |
| | | | Diurnal > Cathemeral | 0.259 | -0.122, 0.646 | 0.909 | 9.975 |
| | | | Nocturnal > Cathemeral | -0.026 | -0.361, 0.307 | 0.443 | 0.795 |
| | | Temperate:Absolute Effect | Diurnal > Nocturnal | **0.779** | **0.118, 1.439** | **0.99** | **98.602** |
| | | | Diurnal > Cathemeral | 0.243 | -0.49, 0.97 | 0.742 | 2.872 |
| | | | Nocturnal > Cathemeral | -0.535 | -1.15, 0.09 | 0.046 | 0.048 |
| | | Temperate:Body Mass | Diurnal > Nocturnal | -0.302 | -0.929, 0.341 | 0.174 | 0.21 |
| | | | Diurnal > Cathemeral | -0.079 | -0.763, 0.62 | 0.41 | 0.696 |
| | | | Nocturnal > Cathemeral | 0.223 | -0.324, 0.763 | 0.791 | 3.777 |
| | | Temperate:Geographic Range Size | Diurnal > Nocturnal | 0.637 | -0.249, 1.48 | 0.924 | 12.2 |
| | | | Diurnal > Cathemeral | 0.361 | -0.62, 1.319 | 0.772 | 3.387 |

| | | | | | | | |
|---|---|---|---|---|---|---|---|
| | | | Nocturnal > Cathemeral | -0.276 | -1.092, 0.561 | 0.255 | 0.341 |
| | | Temperate:Human Footprint | Diurnal > Nocturnal | -0.385 | -0.891, 0.162 | 0.081 | 0.089 |
| | | | Diurnal > Cathemeral | -0.496 | -1.088, 0.104 | 0.054 | 0.057 |
| | | | Nocturnal > Cathemeral | -0.111 | -0.612, 0.368 | 0.333 | 0.498 |
| 1 Category | 1.2 Lizards | Tropics: Absolute Effect | Diurnal > Nocturnal | -0.534 | -1.16, 0.068 | 0.041 | 0.042 |
| | | | Diurnal > Cathemeral | -0.051 | -0.687, 0.632 | 0.43 | 0.754 |
| | | | Nocturnal > Cathemeral | 0.484 | -0.237, 1.26 | 0.901 | 9.144 |
| | | Tropics: Body Mass | Diurnal > Nocturnal | 0.204 | -0.209, 0.63 | 0.834 | 5.037 |
| | | | Diurnal > Cathemeral | -0.343 | -0.933, 0.207 | 0.115 | 0.129 |
| | | | Nocturnal > Cathemeral | -0.546 | -1.228, 0.081 | 0.045 | 0.047 |
| | | Tropics: Geographic Range Size | Diurnal > Nocturnal | **-0.444** | **-0.792, -0.117** | **0.004** | **0.004** |
| | | | Diurnal > Cathemeral | 0.088 | -0.449, 0.675 | 0.612 | 1.575 |
| | | | Nocturnal > Cathemeral | 0.532 | -0.051, 1.18 | 0.963 | 25.667 |
| | | Tropics:Human Footprint | Diurnal > Nocturnal | -0.011 | -0.286, 0.259 | 0.47 | 0.888 |
| | | | Diurnal > Cathemeral | 0.252 | -0.21, 0.736 | 0.857 | 6.01 |
| | | | Nocturnal > Cathemeral | 0.263 | -0.227, 0.78 | 0.851 | 5.696 |
| | | Temperate:Absolute Effect | Diurnal > Nocturnal | 0.654 | -0.158, 1.54 | 0.942 | 16.129 |
| | | | **Diurnal > Cathemeral** | **0.866** | **0.141, 1.679** | **0.992** | **117.765** |
| | | | Nocturnal > Cathemeral | 0.212 | -0.773, 1.198 | 0.666 | 1.992 |
| | | Temperate:Body Mass | Diurnal > Nocturnal | 0.55 | -0.237, 1.345 | 0.915 | 10.705 |
| | | | Diurnal > Cathemeral | **0.853** | **0.099, 1.663** | **0.987** | **78.872** |
| | | | Nocturnal > Cathemeral | 0.303 | -0.691, 1.324 | 0.722 | 2.601 |
| | | Temperate:Geographic Range Size | Diurnal > Nocturnal | -0.082 | -0.635, 0.51 | 0.378 | 0.609 |
| | | | Diurnal > Cathemeral | -0.341 | -0.973, 0.326 | 0.151 | 0.177 |
| | | | Nocturnal > Cathemeral | -0.258 | -1.06, 0.534 | 0.261 | 0.353 |
| | | Temperate:Human Footprint | Diurnal > Nocturnal | -0.219 | -0.804, 0.338 | 0.225 | 0.291 |
| | | | Diurnal > Cathemeral | **1.243** | **0.567, 2.094** | **1** | **99999** |
| | | | Nocturnal > Cathemeral | **1.462** | **0.625, 2.456** | **1** | **11110.111** |
| 1 Category | 1.3 Mammals | Tropics: Absolute Effect | Diurnal > Nocturnal | -0.264 | -0.688, 0.148 | 0.104 | 0.115 |
| | | | Diurnal > Cathemeral | -0.398 | -0.914, 0.115 | 0.064 | 0.068 |
| | | | Nocturnal > Cathemeral | -0.134 | -0.501, 0.247 | 0.238 | 0.312 |
| | | Tropics: Body Mass | Diurnal > Nocturnal | **0.328** | **0.019, 0.641** | **0.981** | **51.854** |
| | | | Diurnal > Cathemeral | 0.167 | -0.205, 0.544 | 0.811 | 4.302 |
| | | | Nocturnal > Cathemeral | -0.16 | -0.469, 0.141 | 0.151 | 0.178 |
| | | Tropics: Geographic Range Size | Diurnal > Nocturnal | -0.007 | -0.272, 0.252 | 0.48 | 0.925 |
| | | | Diurnal > Cathemeral | -0.108 | -0.503, 0.296 | 0.297 | 0.423 |
| | | | Nocturnal > Cathemeral | -0.101 | -0.439, 0.244 | 0.277 | 0.384 |

| | | | | | | | |
|---|---|---|---|---|---|---|---|
| | | Tropics:Human Footprint | Diurnal > Nocturnal | **0.228** | **0.028, 0.431** | **0.987** | **74.988** |
| | | | Diurnal > Cathemeral | 0.339 | -0.013, 0.692 | 0.97 | 31.895 |
| | | | Nocturnal > Cathemeral | 0.111 | -0.208, 0.427 | 0.751 | 3.018 |
| | | Temperate:Absolute Effect | Diurnal > Nocturnal | **-0.772** | **-1.44, -0.143** | **0.008** | **0.008** |
| | | | Diurnal > Cathemeral | -0.148 | -0.934, 0.616 | 0.358 | 0.557 |
| | | | Nocturnal > Cathemeral | **0.625** | **0.112, 1.166** | **0.993** | **137.889** |
| | | Temperate:Body Mass | Diurnal > Nocturnal | 0.14 | -0.275, 0.569 | 0.742 | 2.872 |
| | | | Diurnal > Cathemeral | -0.115 | -0.575, 0.344 | 0.312 | 0.453 |
| | | | Nocturnal > Cathemeral | -0.255 | -0.637, 0.111 | 0.089 | 0.098 |
| | | Temperate:Geographic Range Size | Diurnal > Nocturnal | 0.051 | -0.372, 0.461 | 0.599 | 1.491 |
| | | | Diurnal > Cathemeral | 0.047 | -0.524, 0.636 | 0.562 | 1.282 |
| | | | Nocturnal > Cathemeral | -0.005 | -0.471, 0.48 | 0.486 | 0.944 |
| | | Temperate:Human Footprint | Diurnal > Nocturnal | -0.232 | -0.557, 0.089 | 0.078 | 0.085 |
| | | | Diurnal > Cathemeral | -0.272 | -0.647, 0.105 | 0.077 | 0.084 |
| | | | Nocturnal > Cathemeral | -0.04 | -0.336, 0.256 | 0.393 | 0.647 |
| 1 Category | 1.4 Birds | Tropics: Absolute Effect | Diurnal > Nocturnal | 0.471 | -0.35, 1.335 | 0.864 | 6.372 |
| | | | Diurnal > Cathemeral | 0.068 | -0.83, 1.054 | 0.542 | 1.185 |
| | | | Nocturnal > Cathemeral | -0.403 | -1.595, 0.841 | 0.254 | 0.341 |
| | | Tropics: Body Mass | Diurnal > Nocturnal | 0.015 | -0.616, 0.631 | 0.525 | 1.104 |
| | | | Diurnal > Cathemeral | 0.046 | -0.562, 0.623 | 0.566 | 1.305 |
| | | | Nocturnal > Cathemeral | 0.031 | -0.818, 0.867 | 0.527 | 1.113 |
| | | Tropics: Geographic Range Size | Diurnal > Nocturnal | -0.044 | -0.415, 0.35 | 0.401 | 0.671 |
| | | | Diurnal > Cathemeral | -0.419 | -0.924, 0.085 | 0.052 | 0.055 |
| | | | Nocturnal > Cathemeral | -0.374 | -0.994, 0.236 | 0.117 | 0.132 |
| | | Tropics:Human Footprint | Diurnal > Nocturnal | 0.051 | -0.274, 0.38 | 0.621 | 1.639 |
| | | | Diurnal > Cathemeral | 0.369 | -0.15, 0.901 | 0.917 | 11.095 |
| | | | Nocturnal > Cathemeral | 0.318 | -0.288, 0.943 | 0.844 | 5.412 |
| | | Temperate:Absolute Effect | Diurnal > Nocturnal | 1.307 | -0.059, 2.822 | 0.969 | 31.01 |
| | | | Diurnal > Cathemeral | -0.533 | -1.401, 0.328 | 0.113 | 0.127 |
| | | | Nocturnal > Cathemeral | **-1.84** | **-3.545, -0.247** | **0.011** | **0.011** |
| | | Temperate:Body Mass | Diurnal > Nocturnal | -0.595 | -1.488, 0.246 | 0.085 | 0.093 |
| | | | Diurnal > Cathemeral | -0.438 | -0.91, 0.021 | 0.03 | 0.031 |
| | | | Nocturnal > Cathemeral | 0.157 | -0.785, 1.132 | 0.619 | 1.624 |
| | | Temperate:Geographic Range Size | Diurnal > Nocturnal | -0.321 | -0.96, 0.323 | 0.159 | 0.189 |
| | | | Diurnal > Cathemeral | **0.706** | **0.089, 1.362** | **0.988** | **83.175** |
| | | | Nocturnal > Cathemeral | **1.028** | **0.152, 1.933** | **0.989** | **93.34** |
| | | Temperate:Human Footprint | Diurnal > Nocturnal | 0.163 | -0.579, 0.913 | 0.666 | 1.992 |

| | | | | | | | |
|---|---|---|---|---|---|---|---|
| | | | Diurnal > Cathemeral | 0.323 | -0.08, 0.731 | 0.939 | 15.469 |
| | | | Nocturnal > Cathemeral | 0.16 | -0.666, 0.982 | 0.648 | 1.837 |
| 2 Population Trend | 2.1 Amphibians | Tropics: Absolute Effect | Diurnal > Nocturnal | -0.004 | -0.34, 0.333 | 0.493 | 0.973 |
| | | | Diurnal > Cathemeral | 0.076 | -0.292, 0.44 | 0.66 | 1.942 |
| | | | Nocturnal > Cathemeral | 0.079 | -0.17, 0.33 | 0.732 | 2.725 |
| | | Tropics: Body Mass | Diurnal > Nocturnal | **-0.315** | **-0.614, -0.019** | **0.019** | **0.019** |
| | | | Diurnal > Cathemeral | -0.302 | -0.632, 0.026 | 0.036 | 0.037 |
| | | | Nocturnal > Cathemeral | 0.012 | -0.231, 0.255 | 0.539 | 1.167 |
| | | Tropics: Geographic Range Size | Diurnal > Nocturnal | 0.118 | -0.147, 0.383 | 0.809 | 4.229 |
| | | | Diurnal > Cathemeral | 0.206 | -0.112, 0.524 | 0.897 | 8.701 |
| | | | Nocturnal > Cathemeral | 0.087 | -0.174, 0.353 | 0.739 | 2.825 |
| | | Tropics:Human Footprint | Diurnal > Nocturnal | 0.213 | -0.045, 0.472 | 0.948 | 18.246 |
| | | | Diurnal > Cathemeral | 0.199 | -0.139, 0.534 | 0.875 | 6.987 |
| | | | Nocturnal > Cathemeral | -0.014 | -0.294, 0.264 | 0.465 | 0.868 |
| | | Temperate:Absolute Effect | Diurnal > Nocturnal | -0.241 | -0.784, 0.311 | 0.195 | 0.242 |
| | | | Diurnal > Cathemeral | -0.061 | -0.702, 0.583 | 0.426 | 0.741 |
| | | | Nocturnal > Cathemeral | 0.18 | -0.354, 0.716 | 0.743 | 2.895 |
| | | Temperate:Body Mass | Diurnal > Nocturnal | 0.003 | -0.53, 0.54 | 0.502 | 1.006 |
| | | | Diurnal > Cathemeral | -0.016 | -0.596, 0.562 | 0.479 | 0.92 |
| | | | Nocturnal > Cathemeral | -0.019 | -0.419, 0.371 | 0.46 | 0.85 |
| | | Temperate:Geographic Range Size | Diurnal > Nocturnal | -0.237 | -0.825, 0.315 | 0.21 | 0.265 |
| | | | Diurnal > Cathemeral | 0.071 | -0.586, 0.704 | 0.589 | 1.436 |
| | | | Nocturnal > Cathemeral | 0.309 | -0.204, 0.824 | 0.884 | 7.612 |
| | | Temperate:Human Footprint | Diurnal > Nocturnal | 0.199 | -0.237, 0.675 | 0.805 | 4.13 |
| | | | Diurnal > Cathemeral | 0.133 | -0.358, 0.641 | 0.694 | 2.267 |
| | | | Nocturnal > Cathemeral | -0.066 | -0.481, 0.345 | 0.378 | 0.609 |
| 2 Population Trend | 2.2 Lizards | Tropics: Absolute Effect | Diurnal > Nocturnal | -0.506 | -1.281, 0.244 | 0.089 | 0.098 |
| | | | Diurnal > Cathemeral | 0.21 | -0.505, 0.973 | 0.71 | 2.446 |
| | | | Nocturnal > Cathemeral | 0.717 | -0.16, 1.647 | 0.946 | 17.423 |
| | | Tropics: Body Mass | Diurnal > Nocturnal | 0.056 | -0.485, 0.601 | 0.581 | 1.387 |
| | | | Diurnal > Cathemeral | -0.332 | -1.012, 0.287 | 0.152 | 0.179 |
| | | | Nocturnal > Cathemeral | -0.388 | -1.205, 0.378 | 0.163 | 0.194 |
| | | Tropics: Geographic Range Size | Diurnal > Nocturnal | 0.041 | -0.419, 0.526 | 0.562 | 1.286 |
| | | | Diurnal > Cathemeral | 0.153 | -0.484, 0.809 | 0.679 | 2.116 |
| | | | Nocturnal > Cathemeral | 0.111 | -0.639, 0.861 | 0.615 | 1.599 |
| | | Tropics:Human Footprint | Diurnal > Nocturnal | 0.2 | -0.232, 0.645 | 0.819 | 4.512 |
| | | | Diurnal > Cathemeral | 0.506 | -0.094, 1.178 | 0.95 | 19.101 |

| | | | | | | | |
|---|---|---|---|---|---|---|---|
| | | | Nocturnal > Cathemeral | 0.306 | -0.369, 1.032 | 0.807 | 4.195 |
| | | Temperate:Absolute Effect | Diurnal > Nocturnal | 0.348 | -0.504, 1.254 | 0.785 | 3.647 |
| | | | Diurnal > Cathemeral | **0.657** | **0.013, 1.342** | **0.977** | **42.687** |
| | | | Nocturnal > Cathemeral | 0.309 | -0.602, 1.214 | 0.752 | 3.038 |
| | | Temperate:Body Mass | Diurnal > Nocturnal | 0.586 | -0.235, 1.454 | 0.919 | 11.364 |
| | | | Diurnal > Cathemeral | 0.162 | -0.484, 0.813 | 0.69 | 2.228 |
| | | | Nocturnal > Cathemeral | -0.424 | -1.417, 0.53 | 0.193 | 0.239 |
| | | Temperate:Geographic Range Size | Diurnal > Nocturnal | -0.046 | -0.644, 0.557 | 0.438 | 0.78 |
| | | | Diurnal > Cathemeral | **-0.94** | **-1.613, -0.357** | **0.001** | **0.001** |
| | | | Nocturnal > Cathemeral | **-0.894** | **-1.724, -0.147** | **0.009** | **0.009** |
| | | Temperate:Human Footprint | Diurnal > Nocturnal | -0.217 | -0.915, 0.446 | 0.265 | 0.36 |
| | | | Diurnal > Cathemeral | **0.597** | **0.062, 1.178** | **0.985** | **67.823** |
| | | | Nocturnal > Cathemeral | **0.815** | **0.034, 1.671** | **0.979** | **47.356** |
| 2 Population Trend | 2.3 Mammals | Tropics: Absolute Effect | Diurnal > Nocturnal | -0.157 | -0.471, 0.157 | 0.161 | 0.192 |
| | | | Diurnal > Cathemeral | -0.145 | -0.542, 0.245 | 0.236 | 0.308 |
| | | | Nocturnal > Cathemeral | 0.012 | -0.296, 0.32 | 0.531 | 1.133 |
| | | Tropics: Body Mass | Diurnal > Nocturnal | 0.129 | -0.157, 0.416 | 0.811 | 4.291 |
| | | | Diurnal > Cathemeral | 0.156 | -0.18, 0.489 | 0.818 | 4.502 |
| | | | Nocturnal > Cathemeral | 0.026 | -0.263, 0.308 | 0.576 | 1.356 |
| | | Tropics: Geographic Range Size | Diurnal > Nocturnal | **0.326** | **0.111, 0.539** | **0.998** | **624** |
| | | | Diurnal > Cathemeral | 0.208 | -0.157, 0.573 | 0.868 | 6.585 |
| | | | Nocturnal > Cathemeral | -0.118 | -0.441, 0.212 | 0.237 | 0.311 |
| | | Tropics:Human Footprint | Diurnal > Nocturnal | -0.171 | -0.387, 0.047 | 0.062 | 0.066 |
| | | | Diurnal > Cathemeral | 0.072 | -0.269, 0.414 | 0.66 | 1.938 |
| | | | Nocturnal > Cathemeral | 0.243 | -0.051, 0.546 | 0.946 | 17.477 |
| | | Temperate:Absolute Effect | Diurnal > Nocturnal | -0.116 | -0.534, 0.308 | 0.294 | 0.416 |
| | | | Diurnal > Cathemeral | 0.025 | -0.479, 0.536 | 0.537 | 1.161 |
| | | | Nocturnal > Cathemeral | 0.141 | -0.255, 0.534 | 0.76 | 3.167 |
| | | Temperate:Body Mass | Diurnal > Nocturnal | -0.137 | -0.496, 0.221 | 0.228 | 0.296 |
| | | | Diurnal > Cathemeral | -0.047 | -0.409, 0.316 | 0.396 | 0.656 |
| | | | Nocturnal > Cathemeral | 0.089 | -0.197, 0.374 | 0.729 | 2.686 |
| | | Temperate:Geographic Range Size | Diurnal > Nocturnal | 0.049 | -0.314, 0.397 | 0.615 | 1.594 |
| | | | Diurnal > Cathemeral | 0.168 | -0.29, 0.627 | 0.761 | 3.182 |
| | | | Nocturnal > Cathemeral | 0.119 | -0.239, 0.488 | 0.735 | 2.78 |
| | | Temperate:Human Footprint | Diurnal > Nocturnal | -0.22 | -0.505, 0.061 | 0.065 | 0.07 |
| | | | Diurnal > Cathemeral | -0.007 | -0.333, 0.317 | 0.485 | 0.94 |
| | | | Nocturnal > Cathemeral | 0.213 | -0.038, 0.467 | 0.95 | 19.194 |

| | | | | | | | |
|---|---|---|---|---|---|---|---|
| 2 Population Trend | 2.4 Birds | Tropics: Absolute Effect | Diurnal > Nocturnal | 0.515 | -0.087, 1.121 | 0.952 | 20.008 |
| | | | Diurnal > Cathemeral | -0.036 | -0.812, 0.737 | 0.465 | 0.869 |
| | | | Nocturnal > Cathemeral | -0.551 | -1.457, 0.349 | 0.119 | 0.135 |
| | | Tropics: Body Mass | Diurnal > Nocturnal | -0.201 | -0.67, 0.265 | 0.197 | 0.246 |
| | | | Diurnal > Cathemeral | 0.404 | -0.153, 0.959 | 0.925 | 12.405 |
| | | | Nocturnal > Cathemeral | 0.605 | -0.072, 1.29 | 0.961 | 24.381 |
| | | Tropics: Geographic Range Size | Diurnal > Nocturnal | -0.031 | -0.308, 0.247 | 0.416 | 0.712 |
| | | | Diurnal > Cathemeral | -0.248 | -0.758, 0.268 | 0.173 | 0.209 |
| | | | Nocturnal > Cathemeral | -0.217 | -0.788, 0.368 | 0.231 | 0.3 |
| | | Tropics:Human Footprint | Diurnal > Nocturnal | **0.427** | **0.17, 0.691** | **0.999** | **1135.364** |
| | | | Diurnal > Cathemeral | -0.477 | -0.99, 0.014 | 0.029 | 0.03 |
| | | | Nocturnal > Cathemeral | **-0.904** | **-1.481, -0.356** | **0.001** | **0.001** |
| | | Temperate:Absolute Effect | Diurnal > Nocturnal | 0.568 | -0.4, 1.535 | 0.876 | 7.046 |
| | | | Diurnal > Cathemeral | -0.356 | -1.192, 0.45 | 0.201 | 0.251 |
| | | | Nocturnal > Cathemeral | -0.924 | -2.129, 0.325 | 0.069 | 0.075 |
| | | Temperate:Body Mass | Diurnal > Nocturnal | -0.203 | -0.832, 0.427 | 0.264 | 0.358 |
| | | | Diurnal > Cathemeral | 0.12 | -0.319, 0.561 | 0.703 | 2.363 |
| | | | Nocturnal > Cathemeral | 0.323 | -0.401, 1.056 | 0.807 | 4.182 |
| | | Temperate:Geographic Range Size | Diurnal > Nocturnal | -0.236 | -0.76, 0.286 | 0.188 | 0.232 |
| | | | Diurnal > Cathemeral | 0.044 | -0.481, 0.598 | 0.554 | 1.243 |
| | | | Nocturnal > Cathemeral | 0.281 | -0.445, 1.038 | 0.771 | 3.36 |
| | | Temperate:Human Footprint | Diurnal > Nocturnal | 0.022 | -0.522, 0.566 | 0.535 | 1.15 |
| | | | Diurnal > Cathemeral | -0.008 | -0.385, 0.368 | 0.484 | 0.939 |
| | | | Nocturnal > Cathemeral | -0.029 | -0.679, 0.622 | 0.468 | 0.879 |
