## Supplementary material for "Generalist temporal niche buffers against extinction risk among non-avian tetrapods": Table S4 (from another version of this preprint: version 2, 2026-01-15)

### supp_table_s4_day_night_ext_risk

| group | scientific_name_updated | phylo_name | category | population_trend | biome | activity | body_mass | grs | hf_mean |
|---|---|---|---|---|---|---|---|---|---|
| amphibians | "Centrolene"_medemi | Centrolene_medemi | EN | Decreasing | tropical | nocturnal | 7.456 | 2031.581 | 10.054 |
| amphibians | "Cochranella"_duidaeana | NA | VU | Unknown | NA | NA | NA | NA | NA |
| amphibians | "Colostethus"_poecilonotus | NA | DD | Unknown | NA | NA | NA | NA | NA |
| amphibians | "Litoria"_castanea | NA | CR | Decreasing | NA | NA | NA | NA | NA |
| amphibians | Abavorana_luctuosa | Hylarana_luctuosa | LC | Unknown | tropical | nocturnal | 29.887 | 128378.792 | 7.542 |
| amphibians | Acanthixalus_sonjae | Acanthixalus_sonjae | VU | Decreasing | tropical | nocturnal | 4.934 | 6030.019 | 7.025 |
| amphibians | Acanthixalus_spinosus | Acanthixalus_spinosus | LC | Unknown | tropical | nocturnal | 4.934 | 1213834.976 | 5.904 |
| amphibians | Acris_blanchardi | Acris_blanchardi | LC | Stable | temperate | diurnal | 4.575 | 2345138.949 | 17.02 |
| amphibians | Acris_crepitans | Acris_crepitans | LC | Stable | temperate | cathemeral | 4.536 | 970974.661 | 17.785 |
| amphibians | Acris_gryllus | Acris_gryllus | LC | Stable | temperate | cathemeral | 2.872 | 667451.522 | 16.649 |
| amphibians | Adelastes_hylonomos | Adelastes_hylonomos | LC | Unknown | tropical | nocturnal | 1.889 | 17735.585 | 0.729 |
| amphibians | Adelophryne_adiastola | Adelophryne_adiastola | LC | Stable | tropical | nocturnal | 0.335 | 306381.381 | 1.093 |
| amphibians | Adelophryne_baturitensis | Adelophryne_baturitensis | LC | Stable | tropical | cathemeral | 0.335 | 1866.425 | 14.501 |
| amphibians | Adelophryne_gutturosa | Adelophryne_gutturosa | LC | Unknown | tropical | cathemeral | 0.275 | 161196.361 | 1.069 |
| amphibians | Adelophryne_maranguapensis | Adelophryne_maranguapensis | CR | Decreasing | tropical | diurnal | 0.361 | 1.291 | 14.194 |
| amphibians | Adelophryne_mucronata | Adelophryne_mucronatus | LC | Stable | tropical | nocturnal | 0.178 | 25444.524 | 9.128 |
| amphibians | Adelophryne_pachydactyla | Adelophryne_pachydactyla | LC | Stable | tropical | nocturnal | 0.178 | 4069.449 | 7.638 |
| amphibians | Adelophryne_patamona | NA | DD | Unknown | NA | NA | NA | NA | NA |
| amphibians | Adelphobates_castaneoticus | Adelphobates_castaneoticus | LC | Stable | tropical | diurnal | 1.17 | 630666.803 | 1.725 |
| amphibians | Adelphobates_galactonotus | Adelphobates_galactonotus | LC | Stable | tropical | diurnal | 5.576 | 996582.337 | 3.44 |
| amphibians | Adelphobates_quinquevittatus | Adelphobates_quinquevittatus | LC | Stable | tropical | diurnal | 0.585 | 777019.41 | 1.777 |
| amphibians | Adenomera_ajurauna | Adenomera_ajurauna | LC | Stable | temperate | nocturnal | 0.567 | 25993.333 | 9.672 |
| amphibians | Adenomera_andreae | Adenomera_andreae | LC | Stable | tropical | cathemeral | 1.868 | 5953365.832 | 2.044 |
| amphibians | Adenomera_araucaria | Adenomera_araucaria | LC | Stable | temperate | cathemeral | 0.567 | 57646.59 | 10.409 |
| amphibians | Adenomera_bokermanni | Adenomera_bokermanni | LC | Stable | temperate | nocturnal | 1.183 | 10978.036 | 15.554 |
| amphibians | Adenomera_chicomendesi | NA | LC | Stable | NA | NA | NA | NA | NA |
| amphibians | Adenomera_diptyx | Adenomera_diptyx | LC | Stable | tropical | diurnal | 1.168 | 568792.849 | 7.629 |
| amphibians | Adenomera_engelsi | Adenomera_engelsi | LC | Stable | temperate | cathemeral | 0.854 | 4825.172 | 13.239 |
| amphibians | Adenomera_heyeri | Adenomera_heyeri | LC | Unknown | tropical | diurnal | 1.326 | 306841.67 | 0.98 |
| amphibians | Adenomera_hylaedactyla | Adenomera_hylaedactyla | LC | Stable | tropical | cathemeral | 2.109 | 8309254.301 | 3.069 |
| amphibians | Adenomera_kweti | NA | EN | Decreasing | NA | NA | NA | NA | NA |
| amphibians | Adenomera_lutzi | Adenomera_lutzi | EN | Decreasing | tropical | cathemeral | 3.015 | 2985.451 | 0.307 |
| amphibians | Adenomera_marmorata | Adenomera_marmorata | LC | Stable | temperate | cathemeral | 2.109 | 52409.524 | 13.898 |

| | | | | | | | | | |
|---|---|---|---|---|---|---|---|---|---|
| amphibians | Adenomera_martinezi | Adenomera_martinezi | NT | Stable | tropical | diurnal | 1.168 | 5534.837 | 1.569 |
| amphibians | Adenomera_nana | Adenomera_nana | LC | Stable | temperate | nocturnal | 0.567 | 37230.271 | 11.313 |
| amphibians | Adenomera_saci | NA | LC | Stable | NA | NA | NA | NA | NA |
| amphibians | Adenomera_thomei | Adenomera_thomei | LC | Stable | tropical | nocturnal | 1.023 | 401569.877 | 13.068 |
| amphibians | Adenomus_kandianus | Adenomus_kandianus | EN | Decreasing | tropical | nocturnal | 2.652 | 267.435 | 14.991 |
| amphibians | Adenomus_kelaartii | Adenomus_kelaartii | VU | Decreasing | tropical | cathemeral | 11.036 | 11262.452 | 17.296 |
| amphibians | Afrixalus_aureus | Afrixalus_aureus | LC | Unknown | temperate | cathemeral | 1.023 | 100277.56 | 12.675 |
| amphibians | Afrixalus_brachycnemis | NA | LC | Decreasing | NA | NA | NA | NA | NA |
| amphibians | Afrixalus_clarkei | Afrixalus_clarkei | EN | Unknown | tropical | nocturnal | 0.892 | 1426.089 | 12.249 |
| amphibians | Afrixalus_delicatus | Afrixalus_delicatus | LC | Unknown | tropical | nocturnal | 1.499 | 748759.801 | 10.854 |
| amphibians | Afrixalus_dorsalis | Afrixalus_dorsalis | LC | Increasing | tropical | nocturnal | 4.536 | 609648.183 | 13.923 |
| amphibians | Afrixalus_dorsimaculatus | Afrixalus_dorsimaculatus | EN | Decreasing | tropical | diurnal | 1.023 | 3034.304 | 16.917 |
| amphibians | Afrixalus_equatorialis | Afrixalus_equatorialis | LC | Unknown | tropical | nocturnal | 4.536 | 553134.806 | 4.438 |
| amphibians | Afrixalus_fornasini | Afrixalus_fornasini | LC | Unknown | tropical | nocturnal | 5.356 | 944389.109 | 9.899 |
| amphibians | Afrixalus_lacteus | Afrixalus_lacteus | EN | Decreasing | tropical | nocturnal | 1.889 | 609.524 | 14.735 |
| amphibians | Afrixalus_manengubensis | NA | not_available | not_available | NA | NA | NA | NA | NA |
| amphibians | Afrixalus_nigeriensis | Afrixalus_nigeriensis | LC | Decreasing | tropical | nocturnal | 3.475 | 357705.721 | 16.489 |
| amphibians | Afrixalus_paradorsalis | Afrixalus_paradorsalis | LC | Unknown | tropical | nocturnal | 3.475 | 450758.517 | 6.671 |
| amphibians | Afrixalus_quadrivittatus | Afrixalus_quadrivittatus | LC | Unknown | tropical | nocturnal | 2.599 | 5187441.858 | 7.864 |
| amphibians | Afrixalus_uluguruensis | Afrixalus_uluguruensis | VU | Unknown | tropical | diurnal | 2.345 | 593.272 | 9.546 |
| amphibians | Afrixalus_vibekensis | Afrixalus_vibekensis | LC | Decreasing | tropical | nocturnal | 1.499 | 6059.557 | 6.645 |
| amphibians | Agalychnis_annae | Agalychnis_annae | VU | Stable | tropical | nocturnal | 59.726 | 2708.233 | 22.642 |
| amphibians | Agalychnis_buckleyi | Hylomantis_buckleyi | LC | Decreasing | tropical | nocturnal | 14.765 | 52017.815 | 6.703 |
| amphibians | Agalychnis_callidryas | Agalychnis_callidryas | LC | Decreasing | tropical | nocturnal | 45.085 | 501190.034 | 9.446 |
| amphibians | Agalychnis_dacnicolor | Pachymedusa_dacnicolor | LC | Decreasing | tropical | nocturnal | 116.928 | 150009.609 | 10.406 |
| amphibians | Agalychnis_lemur | Agalychnis_lemur | CR | Decreasing | tropical | nocturnal | 13.329 | 16484.301 | 7.555 |
| amphibians | Agalychnis_moreletii | Agalychnis_moreletii | LC | Decreasing | tropical | nocturnal | 61.584 | 59737.544 | 13.562 |
| amphibians | Agalychnis_psilopygion | Hylomantis_psilopygion | LC | Decreasing | tropical | nocturnal | 9.219 | 18937.266 | 5.983 |
| amphibians | Agalychnis_saltator | Agalychnis_saltator | LC | Decreasing | tropical | cathemeral | 27.131 | 11425.698 | 7.836 |
| amphibians | Agalychnis_spurrelli | Agalychnis_spurrelli | LC | Decreasing | tropical | nocturnal | 82.421 | 132657.119 | 6.849 |
| amphibians | Aglyptodactylus_chorus | NA | NT | Decreasing | NA | NA | NA | NA | NA |
| amphibians | Aglyptodactylus_inguinalis | NA | LC | Decreasing | NA | NA | NA | NA | NA |
| amphibians | Aglyptodactylus_madagascariensis | Aglyptodactylus_madagascariensis | LC | Decreasing | tropical | cathemeral | 31.334 | 110547.762 | 8.652 |
| amphibians | Aglyptodactylus_securifer | Aglyptodactylus_securifer | LC | Decreasing | tropical | cathemeral | 11.036 | 117095 | 7.707 |

| | | | | | | | | | |
|---|---|---|---|---|---|---|---|---|---|
| amphibians | Allobates_albiventris | NA | not_available | not_available | NA | NA | NA | NA | NA |
| amphibians | Allobates_algorei | Allobates_algorei | NT | Stable | tropical | cathemeral | 0.522 | 5220.101 | 12.956 |
| amphibians | Allobates_brunneus | Allobates_brunneus | EN | Decreasing | tropical | diurnal | 1.023 | 28521.751 | 3.411 |
| amphibians | Allobates_femoralis | Allobates_femoralis | LC | Stable | tropical | diurnal | 3.807 | 4353103.975 | 1.511 |
| amphibians | Allobates_goianus | NA | NT | Decreasing | NA | NA | NA | NA | NA |
| amphibians | Allobates_granti | Allobates_granti | LC | Unknown | tropical | diurnal | 0.433 | 33358.99 | 0.868 |
| amphibians | Allobates_hodli | Allobates_hodli | LC | Unknown | tropical | nocturnal | 1.706 | 76820.222 | 2.928 |
| amphibians | Allobates_insperatus | Allobates_insperatus | LC | Stable | tropical | diurnal | 0.433 | 74685.486 | 3.606 |
| amphibians | Allobates_juanii | Allobates_juanii | EN | Decreasing | tropical | diurnal | 1.065 | 945.253 | 12.325 |
| amphibians | Allobates_kingsburyi | Allobates_kingsburyi | LC | Decreasing | tropical | diurnal | 1.36 | 15817.206 | 6.624 |
| amphibians | Allobates_magnussoni | NA | LC | Unknown | NA | NA | NA | NA | NA |
| amphibians | Allobates_marchesianus | Allobates_marchesianus | LC | Stable | tropical | cathemeral | 0.568 | 2390953.861 | 1.045 |
| amphibians | Allobates_mcdiarmidi | Allobates_mcdiarmidi | CR | Decreasing | tropical | diurnal | 1.065 | 20649.025 | 5.381 |
| amphibians | Allobates_melanolaemus | Allobates_melanolaemus | LC | Unknown | tropical | diurnal | 0.969 | 27127.971 | 1.997 |
| amphibians | Allobates_myersi | Allobates_myersi | LC | Unknown | tropical | diurnal | 2.872 | 162443.401 | 0.91 |
| amphibians | Allobates_niputidea | Allobates_niputidea | LC | Unknown | tropical | diurnal | 0.403 | 10771.622 | 15.313 |
| amphibians | Allobates_nunciatus | NA | LC | Stable | NA | NA | NA | NA | NA |
| amphibians | Allobates_olfersioides | Allobates_olfersioides | LC | Decreasing | tropical | diurnal | 0.48 | 92525.877 | 13.044 |
| amphibians | Allobates_ornatus | NA | DD | Unknown | NA | NA | NA | NA | NA |
| amphibians | Allobates_pacaas | NA | VU | Stable | NA | NA | NA | NA | NA |
| amphibians | Allobates_ranoides | NA | CR | Decreasing | NA | NA | NA | NA | NA |
| amphibians | Allobates_subfolionidificans | Allobates_subfolionidificans | LC | Stable | tropical | diurnal | 0.456 | 152731.776 | 1.934 |
| amphibians | Allobates_talamancae | Allobates_talamancae | LC | Decreasing | tropical | diurnal | 1.168 | 117497.08 | 5.91 |
| amphibians | Allobates_tapajos | NA | LC | Stable | NA | NA | NA | NA | NA |
| amphibians | Allobates_trilineatus | Allobates_trilineatus | LC | Stable | tropical | diurnal | 0.505 | 1039919.362 | 2.608 |
| amphibians | Allobates_undulatus | Allobates_undulatus | VU | Unknown | tropical | diurnal | 1.168 | 244.289 | 0.377 |
| amphibians | Allobates_zaparo | Allobates_zaparo | LC | Stable | tropical | diurnal | 2.345 | 47502.846 | 1.981 |
| amphibians | Allophryne_relicta | Allophryne_relicta | EN | Decreasing | tropical | nocturnal | 0.761 | 176.806 | 10.106 |
| amphibians | Allophryne_ruthveni | Allophryne_ruthveni | LC | Unknown | tropical | cathemeral | 2.345 | 3055083.074 | 1.576 |
| amphibians | Alsodes_cantillanensis | NA | CR | Unknown | NA | NA | NA | NA | NA |
| amphibians | Alsodes_vanzolinii | Alsodes_vanzolinii | EN | Decreasing | temperate | nocturnal | 14.504 | 1065.623 | 8.007 |
| amphibians | Altiphrynoides_malcolmi | Altiphrynoides_malcolmi | EN | Decreasing | tropical | diurnal | 2.599 | 629.854 | 9.481 |
| amphibians | Alytes_cisternasii | Alytes_cisternasii | LC | Decreasing | temperate | nocturnal | 11.036 | 168715.797 | 14.937 |
| amphibians | Alytes_dickhilleni | Alytes_dickhilleni | EN | Decreasing | temperate | nocturnal | 16.873 | 31457.443 | 15.228 |

| | | | | | | | | | |
|---|---|---|---|---|---|---|---|---|---|
| amphibians | Alytes_maurus | Alytes_maurus | EN | Decreasing | temperate | cathemeral | 16.873 | 5263.822 | 14.417 |
| amphibians | Alytes_muletensis | Alytes_muletensis | EN | Increasing | temperate | nocturnal | 5.356 | 228.511 | 15.732 |
| amphibians | Alytes_obstetricans | Alytes_obstetricans | LC | Decreasing | temperate | nocturnal | 15.029 | 957570.842 | 21.049 |
| amphibians | Amazophrynella_bilinguis | NA | not_available | not_available | NA | NA | NA | NA | NA |
| amphibians | Amazophrynella_javierbustamantei | NA | LC | Stable | NA | NA | NA | NA | NA |
| amphibians | Amazophrynella_minuta | Amazophrynella_minuta | LC | Stable | tropical | cathemeral | 2.178 | 992304.103 | 1.624 |
| amphibians | Amazophrynella_siona | NA | LC | Decreasing | NA | NA | NA | NA | NA |
| amphibians | Amazophrynella_teko | NA | LC | Unknown | NA | NA | NA | NA | NA |
| amphibians | Ambystoma_amblycephalum | NA | CR | Decreasing | NA | NA | NA | NA | NA |
| amphibians | Ambystoma_annulatum | Ambystoma_annulatum | LC | Stable | temperate | nocturnal | 21.876 | 140918.309 | 13.995 |
| amphibians | Ambystoma_californiense | Ambystoma_californiense | VU | Decreasing | temperate | nocturnal | 56.865 | 38873.704 | 19.723 |
| amphibians | Ambystoma_cingulatum | Ambystoma_cingulatum | EN | Decreasing | temperate | diurnal | 3.492 | 89204.973 | 15.465 |
| amphibians | Ambystoma_gracile | Ambystoma_gracile | LC | Stable | temperate | cathemeral | 35.462 | 350902.189 | 7.587 |
| amphibians | Ambystoma_jeffersonianum | Ambystoma_jeffersonianum | LC | Stable | temperate | diurnal | 16.143 | 470696.231 | 19.654 |
| amphibians | Ambystoma_laterale | Ambystoma_laterale | LC | Stable | temperate | nocturnal | 6.729 | 1767652.417 | 7.959 |
| amphibians | Ambystoma_mabeei | Ambystoma_mabeei | LC | Unknown | temperate | diurnal | 2.604 | 112024.466 | 18.804 |
| amphibians | Ambystoma_macrodactylum | Ambystoma_macrodactylum | LC | Stable | temperate | nocturnal | 11.13 | 1443398.783 | 6.399 |
| amphibians | Ambystoma_maculatum | Ambystoma_maculatum | LC | Stable | temperate | cathemeral | 11.277 | 2992980.174 | 14.846 |
| amphibians | Ambystoma_mavortium | Ambystoma_mavortium | LC | Stable | temperate | nocturnal | 23.224 | 3659783.254 | 11.945 |
| amphibians | Ambystoma_mexicanum | Ambystoma_mexicanum | CR | Decreasing | tropical | diurnal | 32.024 | 32.181 | 27.316 |
| amphibians | Ambystoma_opacum | Ambystoma_opacum | LC | Stable | temperate | nocturnal | 6.729 | 1683648.272 | 17.654 |
| amphibians | Ambystoma_ordinarium | Ambystoma_ordinarium | EN | Decreasing | tropical | diurnal | 10.062 | 3394.332 | 11.451 |
| amphibians | Ambystoma_rosaceum | Ambystoma_rosaceum | LC | Unknown | temperate | nocturnal | 7.695 | 163030.963 | 4.553 |
| amphibians | Ambystoma_talpoideum | Ambystoma_talpoideum | LC | Stable | temperate | cathemeral | 3.315 | 678160.024 | 15.448 |
| amphibians | Ambystoma_texanum | Ambystoma_texanum | LC | Stable | temperate | nocturnal | 8.757 | 1389181.649 | 17.671 |
| amphibians | Ambystoma_tigrinum | Ambystoma_tigrinum | LC | Stable | temperate | nocturnal | 68.34 | 3989359.357 | 13.4 |
| amphibians | Ameerega_bassleri | Ameerega_bassleri | VU | Decreasing | tropical | diurnal | 7.844 | 17729.528 | 6.008 |
| amphibians | Ameerega_berohoka | NA | LC | Unknown | NA | NA | NA | NA | NA |
| amphibians | Ameerega_bilinguis | Ameerega_bilinguis | LC | Stable | tropical | diurnal | 0.854 | 69393.411 | 3.881 |
| amphibians | Ameerega_boehmei | NA | EN | Decreasing | NA | NA | NA | NA | NA |
| amphibians | Ameerega_boliviana | Ameerega_boliviana | NT | Stable | tropical | diurnal | 1.428 | 12764.906 | 6.394 |
| amphibians | Ameerega_flavopicta | Ameerega_flavopicta | LC | Stable | tropical | diurnal | 2.225 | 375628.255 | 11.238 |
| amphibians | Ameerega_hahneli | Ameerega_hahneli | LC | Stable | tropical | diurnal | 1.168 | 3909906.31 | 1.38 |
| amphibians | Ameerega_ignipedis | Ameerega_ignipedis | LC | Stable | tropical | diurnal | 1.051 | 197.047 | 2.052 |

| | | | | | | | | | |
|---|---|---|---|---|---|---|---|---|---|
| amphibians | Ameerega_ingeri | NA | DD | Unknown | NA | NA | NA | NA | NA |
| amphibians | Ameerega_macero | Ameerega_macero | LC | Stable | tropical | diurnal | 1.997 | 155114.811 | 0.802 |
| amphibians | Ameerega_panguana | NA | LC | Stable | NA | NA | NA | NA | NA |
| amphibians | Ameerega_parvula | Ameerega_parvula | LC | Stable | tropical | diurnal | 1.023 | 192116.22 | 1.935 |
| amphibians | Ameerega_pepperi | Ameerega_pepperi | VU | Decreasing | tropical | diurnal | 3.285 | 2962.194 | 5.462 |
| amphibians | Ameerega_petersi | Ameerega_petersi | LC | Stable | tropical | diurnal | 2.345 | 126813.337 | 2.483 |
| amphibians | Ameerega_picta | Ameerega_picta | LC | Stable | tropical | diurnal | 1.168 | 3960562.049 | 2.973 |
| amphibians | Ameerega_planipaleae | Ameerega_planipaleae | CR | Unknown | tropical | diurnal | 2.019 | 79.124 | 9.158 |
| amphibians | Ameerega_pongoensis | Ameerega_pongoensis | VU | Decreasing | tropical | diurnal | 1.326 | 10105.996 | 3.378 |
| amphibians | Ameerega_pulchripecta | Ameerega_pulchripecta | LC | Stable | tropical | diurnal | 1.499 | 8075.265 | 1.315 |
| amphibians | Ameerega_rubriventris | Ameerega_rubriventris | EN | Decreasing | tropical | diurnal | 1.094 | 753.3 | 3.559 |
| amphibians | Ameerega_silverstonei | Ameerega_silverstonei | EN | Decreasing | tropical | diurnal | 7.844 | 2764.054 | 3.731 |
| amphibians | Ameerega_simulans | Ameerega_simulans | LC | Stable | tropical | diurnal | 1.847 | 20899.892 | 1.888 |
| amphibians | Ameerega_trivittata | Ameerega_trivittata | LC | Stable | tropical | diurnal | 15.029 | 3357773.101 | 1.563 |
| amphibians | Ameerega_yoshina | Ameerega_yoshina | EN | Decreasing | tropical | diurnal | 3.411 | 753.523 | 2.256 |
| amphibians | Amerana_aurora | NA | LC | Decreasing | NA | NA | NA | NA | NA |
| amphibians | Amerana_boylii | NA | NT | Decreasing | NA | NA | NA | NA | NA |
| amphibians | Amerana_cascadae | NA | LC | Decreasing | NA | NA | NA | NA | NA |
| amphibians | Amerana_draytonii | NA | NT | Decreasing | NA | NA | NA | NA | NA |
| amphibians | Amerana_luteiventris | NA | LC | Decreasing | NA | NA | NA | NA | NA |
| amphibians | Amerana_muscosa | NA | EN | Decreasing | NA | NA | NA | NA | NA |
| amphibians | Amerana_pretiosa | NA | VU | Decreasing | NA | NA | NA | NA | NA |
| amphibians | Amerana_sierrae | NA | VU | Decreasing | NA | NA | NA | NA | NA |
| amphibians | Amietia_angolensis | Amietia_angolensis | LC | Decreasing | tropical | cathemeral | 141.992 | 343127.223 | 8.508 |
| amphibians | Amietia_delalandii | Amietia_quecketti | LC | Stable | temperate | cathemeral | 74.113 | 1731220.204 | 10.146 |
| amphibians | Amietia_fuscigula | Amietia_fuscigula | LC | Stable | temperate | diurnal | 214.852 | 333657.094 | 11.27 |
| amphibians | Amietia_tenuoplicata | Amietia_tenuoplicata | LC | Decreasing | tropical | diurnal | 82.421 | 6558.046 | 7.705 |
| amphibians | Amietia_vertebralis | Amietia_vertebralis | LC | Unknown | temperate | diurnal | 387.871 | 19409.501 | 10.673 |
| amphibians | Amietia_wittei | Amietia_wittei | LC | Stable | tropical | diurnal | 66.652 | 19491.811 | 13.142 |
| amphibians | Amolops_australis | NA | EN | Decreasing | NA | NA | NA | NA | NA |
| amphibians | Amolops_bellulus | Amolops_bellulus | NT | Decreasing | temperate | nocturnal | 24.557 | 1295.152 | 8.299 |
| amphibians | Amolops_cremnobatus | Amolops_cremnobatus | LC | Decreasing | tropical | nocturnal | 5.621 | 121959.738 | 6.745 |
| amphibians | Amolops_gerutu | NA | VU | Decreasing | NA | NA | NA | NA | NA |
| amphibians | Amolops_larutensis | Amolops_larutensis | LC | Stable | tropical | nocturnal | 41.053 | 51200.897 | 11.522 |

| | | | | | | | | | |
|---|---|---|---|---|---|---|---|---|---|
| amphibians | Amolops_lifanensis | Amolops_lifanensis | LC | Stable | temperate | nocturnal | 48.581 | 15793.416 | 7.589 |
| amphibians | Amolops_mantzorum | Amolops_mantzorum | LC | Unknown | temperate | nocturnal | 39.306 | 132411.256 | 8.753 |
| amphibians | Amolops_mengdingensis | NA | DD | Unknown | NA | NA | NA | NA | NA |
| amphibians | Amolops_minutus | Amolops_minutus | EN | Decreasing | tropical | nocturnal | 11.194 | 895.637 | 6.136 |
| amphibians | Amolops_pallasitatus | NA | not_available | not_available | NA | NA | NA | NA | NA |
| amphibians | Amolops_viridimaculatus | Amolops_viridimaculatus | LC | Decreasing | temperate | nocturnal | 141.992 | 39994.751 | 5.895 |
| amphibians | Amolops_wenshanensis | NA | DD | Unknown | NA | NA | NA | NA | NA |
| amphibians | Amphiuma_means | Amphiuma_means | LC | Unknown | temperate | nocturnal | 3026.268 | 491481.779 | 16.675 |
| amphibians | Anaxyrus_americanus | Anaxyrus_americanus | LC | Stable | temperate | nocturnal | 431.346 | 5652731.676 | 10.118 |
| amphibians | Anaxyrus_baxteri | NA | EW | not_available | NA | NA | NA | NA | NA |
| amphibians | Anaxyrus_canorus | Anaxyrus_canorus | VU | Decreasing | temperate | diurnal | 39.306 | 10686.359 | 3.25 |
| amphibians | Anaxyrus_cognatus | Anaxyrus_cognatus | LC | Stable | temperate | diurnal | 163.988 | 2795125.82 | 11.108 |
| amphibians | Anaxyrus_compactilis | Anaxyrus_compactilis | LC | Unknown | tropical | diurnal | 76.815 | 314894.543 | 11.873 |
| amphibians | Anaxyrus_debilis | Anaxyrus_debilis | LC | Stable | temperate | nocturnal | 14.161 | 1460610.639 | 10.408 |
| amphibians | Anaxyrus_exsul | Anaxyrus_exsul | VU | Stable | temperate | cathemeral | 48.581 | 31.951 | 5.359 |
| amphibians | Anaxyrus_fowleri | Anaxyrus_fowleri | LC | Stable | temperate | diurnal | 88.303 | 1991769.671 | 17.937 |
| amphibians | Anaxyrus_hemiophrys | Anaxyrus_hemiophrys | LC | Stable | temperate | cathemeral | 76.815 | 1401163.637 | 10.702 |
| amphibians | Anaxyrus_houstonensis | Anaxyrus_houstonensis | CR | Decreasing | temperate | cathemeral | 50.601 | 30066.839 | 19.323 |
| amphibians | Anaxyrus_microscaphus | Anaxyrus_microscaphus | LC | Decreasing | temperate | nocturnal | 76.815 | 242190.778 | 8.423 |
| amphibians | Anaxyrus_monfontanus | NA | CR | Unknown | NA | NA | NA | NA | NA |
| amphibians | Anaxyrus_nelsoni | Anaxyrus_nelsoni | CR | Decreasing | temperate | cathemeral | 198.59 | 76.394 | 9.498 |
| amphibians | Anaxyrus_nevadensis | NA | VU | Unknown | NA | NA | NA | NA | NA |
| amphibians | Anaxyrus_punctatus | Anaxyrus_punctatus | LC | Stable | temperate | nocturnal | 43.22 | 2436573.847 | 9.336 |
| amphibians | Anaxyrus_quercicus | Anaxyrus_quercicus | LC | Stable | temperate | cathemeral | 3.475 | 450929.867 | 16.653 |
| amphibians | Anaxyrus_retiformis | Anaxyrus_retiformis | LC | Unknown | temperate | nocturnal | 34.374 | 55041.857 | 6.603 |
| amphibians | Anaxyrus_speciosus | Anaxyrus_speciosus | LC | Stable | temperate | diurnal | 104.268 | 1063699.007 | 12.02 |
| amphibians | Anaxyrus_terrestris | Anaxyrus_terrestris | LC | Stable | temperate | nocturnal | 170.071 | 545693.552 | 16.231 |
| amphibians | Anaxyrus_williamsi | NA | CR | Unknown | NA | NA | NA | NA | NA |
| amphibians | Anaxyrus_woodhousii | Anaxyrus_woodhousii | LC | Stable | temperate | nocturnal | 226.191 | 3242461.375 | 12.325 |
| amphibians | Andinobates_abditus | Andinobates_abditus | CR | Decreasing | tropical | diurnal | 0.382 | 86.412 | 6.65 |
| amphibians | Andinobates_altobueyensis | Andinobates_altobueyensis | DD | Decreasing | tropical | diurnal | 0.335 | 89.699 | 3.741 |
| amphibians | Andinobates_bombetes | Andinobates_bombetes | VU | Decreasing | tropical | diurnal | 0.567 | 569.992 | 16.774 |
| amphibians | Andinobates_cassidyhornae | NA | VU | Stable | NA | NA | NA | NA | NA |
| amphibians | Andinobates_daleswansoni | Andinobates_daleswansoni | EN | Decreasing | tropical | diurnal | 0.54 | 641.422 | 13.152 |

| | | | | | | | | | |
|---|---|---|---|---|---|---|---|---|---|
| amphibians | Andinobates_dorisswansonae | Andinobates_dorisswansonae | VU | Stable | tropical | diurnal | 0.531 | 32.455 | 16.2 |
| amphibians | Andinobates_fulguritus | Andinobates_fulguritus | LC | Decreasing | tropical | diurnal | 0.304 | 39940.743 | 4.124 |
| amphibians | Andinobates_geminisae | NA | CR | Decreasing | NA | NA | NA | NA | NA |
| amphibians | Andinobates_minutus | Andinobates_minutus | LC | Decreasing | tropical | diurnal | 0.248 | 55926.215 | 4.763 |
| amphibians | Andinobates_opisthomelas | Andinobates_opisthomelas | VU | Stable | tropical | diurnal | 0.567 | 7404.466 | 15.252 |
| amphibians | Andinobates_tolimense | NA | VU | Unknown | NA | NA | NA | NA | NA |
| amphibians | Andinobates_victimatus | NA | EN | Decreasing | NA | NA | NA | NA | NA |
| amphibians | Andinobates_viridis | Andinobates_viridis | CR | Unknown | tropical | diurnal | 0.233 | 909.147 | 3.985 |
| amphibians | Andinobates_virolinensis | Andinobates_virolinensis | VU | Decreasing | tropical | diurnal | 0.472 | 3158.116 | 12.571 |
| amphibians | Andrias_davidianus | Andrias_davidianus | CR | Decreasing | temperate | nocturnal | 3845.762 | 330599.837 | 13.59 |
| amphibians | Andrias_japonicus | Andrias_japonicus | VU | Decreasing | temperate | nocturnal | 10792.735 | 57086.656 | 17.61 |
| amphibians | Aneides_aeneus | Aneides_aeneus | NT | Decreasing | temperate | nocturnal | 2.465 | 169792.054 | 16.009 |
| amphibians | Aneides_flavipunctatus | Aneides_flavipunctatus | LC | Stable | temperate | nocturnal | 13.483 | 15597.08 | 10.745 |
| amphibians | Aneides_lugubris | Aneides_lugubris | LC | Stable | temperate | nocturnal | 16.143 | 105848.838 | 16.121 |
| amphibians | Aneides_niger | NA | EN | Decreasing | NA | NA | NA | NA | NA |
| amphibians | Aneides_vagrans | Aneides_vagrans | LC | Decreasing | temperate | nocturnal | 6.996 | 52040.649 | 7.447 |
| amphibians | Anhydrophryne_ngongoniensis | Anhydrophryne_ngongoniensis | EN | Stable | temperate | cathemeral | 0.772 | 412.411 | 9.092 |
| amphibians | Anilany_helenae | Stumpffia_helenae | CR | Decreasing | tropical | diurnal | 0.223 | 20.32 | 11.203 |
| amphibians | Anodonthyla_boulengerii | Anodonthyla_boulengerii | NT | Decreasing | tropical | nocturnal | 0.893 | 80954.069 | 7.917 |
| amphibians | Anodonthyla_emilei | Anodonthyla_emilei | EN | Decreasing | tropical | nocturnal | 1.911 | 405.287 | 5.909 |
| amphibians | Anodonthyla_hutchisoni | Anodonthyla_hutchisoni | EN | Decreasing | tropical | nocturnal | 1.08 | 429.576 | 5.361 |
| amphibians | Anodonthyla_jeanbai | Anodonthyla_jeanbai | EN | Decreasing | temperate | nocturnal | 0.567 | 12.647 | 4.406 |
| amphibians | Anodonthyla_moramora | Anodonthyla_moramora | EN | Decreasing | tropical | nocturnal | 0.335 | 405.287 | 5.909 |
| amphibians | Anodonthyla_pollicaris | Anodonthyla_pollicaris | EN | Decreasing | tropical | nocturnal | 0.685 | 745.081 | 8.038 |
| amphibians | Anodonthyla_rouxae | Anodonthyla_rouxae | EN | Decreasing | temperate | nocturnal | 3.163 | 103.593 | 2.978 |
| amphibians | Anodonthyla_theoi | Anodonthyla_theoi | CR | Decreasing | tropical | nocturnal | 0.522 | 52.609 | 11.186 |
| amphibians | Anodonthyla_vallani | Anodonthyla_vallani | CR | Decreasing | tropical | nocturnal | 1.037 | 20.32 | 11.203 |
| amphibians | Anomaloglossus_baeobatrachus | Anomaloglossus_baeobatrachus | LC | Stable | tropical | diurnal | 0.514 | 490103.261 | 0.778 |
| amphibians | Anomaloglossus_blanci | NA | EN | Decreasing | NA | NA | NA | NA | NA |
| amphibians | Anomaloglossus_degranvillei | Anomaloglossus_degranvillei | CR | Decreasing | tropical | diurnal | 1.01 | 9.875 | 4.875 |
| amphibians | Anomaloglossus_dewynteri | NA | CR | Decreasing | NA | NA | NA | NA | NA |
| amphibians | Anomaloglossus_guanayensis | Anomaloglossus_guanayensis | NT | Unknown | tropical | diurnal | 0.956 | 76.534 | 0.44 |
| amphibians | Anomaloglossus_kaiei | Anomaloglossus_kaiei | EN | Stable | tropical | diurnal | 0.549 | 3399.36 | 0.895 |
| amphibians | Anomaloglossus_meansi | NA | not_available | not_available | NA | NA | NA | NA | NA |

| | | | | | | | | | |
|---|---|---|---|---|---|---|---|---|---|
| amphibians | Anomaloglossus_megacephalus | NA | DD | Unknown | NA | NA | NA | NA | NA |
| amphibians | Anomaloglossus_mitaraka | NA | not_available | not_available | NA | NA | NA | NA | NA |
| amphibians | Anomaloglossus_parkerae | NA | DD | Unknown | NA | NA | NA | NA | NA |
| amphibians | Anomaloglossus_praderioi | Anomaloglossus_praderioi | EN | Unknown | tropical | diurnal | 0.854 | 247.719 | 3.374 |
| amphibians | Anomaloglossus_roraima | Anomaloglossus_roraima | EN | Unknown | tropical | diurnal | 0.505 | 248.57 | 1.819 |
| amphibians | Anomaloglossus_rufulus | Anomaloglossus_rufulus | NT | Stable | tropical | diurnal | 0.943 | 478.724 | 0.661 |
| amphibians | Anomaloglossus_saramaka | NA | not_available | not_available | NA | NA | NA | NA | NA |
| amphibians | Anomaloglossus_stepheni | Anomaloglossus_stepheni | LC | Stable | tropical | diurnal | 0.421 | 388182.008 | 1.223 |
| amphibians | Anomaloglossus_tamacuarensis | Anomaloglossus_tamacuarensis | LC | Stable | tropical | diurnal | 1.168 | 11363.897 | 0.209 |
| amphibians | Anomaloglossus_tepequem | NA | CR | Unknown | NA | NA | NA | NA | NA |
| amphibians | Anomaloglossus_triunfo | NA | DD | Unknown | NA | NA | NA | NA | NA |
| amphibians | Anomaloglossus_vacheri | NA | not_available | not_available | NA | NA | NA | NA | NA |
| amphibians | Anomaloglossus_verbeeksnyderorum | Anomaloglossus_verbeeksnyderorum | LC | Unknown | tropical | diurnal | 0.807 | 11066.927 | 1.868 |
| amphibians | Anomaloglossus_wothuja | Anomaloglossus_wothuja | LC | Unknown | tropical | diurnal | 0.772 | 14001.304 | 0.932 |
| amphibians | Ansonia_endauensis | Ansonia_endauensis | NT | Unknown | tropical | nocturnal | 1.786 | 33.452 | 2.238 |
| amphibians | Ansonia_fuliginea | Ansonia_fuliginea | LC | Unknown | tropical | nocturnal | 7.294 | 290.493 | 5.289 |
| amphibians | Ansonia_hanitschi | Ansonia_hanitschi | LC | Decreasing | tropical | cathemeral | 4.16 | 21080.183 | 3.144 |
| amphibians | Ansonia_inthanon | Ansonia_inthanon | LC | Decreasing | tropical | nocturnal | 1.199 | 21404.452 | 6.218 |
| amphibians | Ansonia_jeetsukumarani | Ansonia_jeetsukumarani | VU | Decreasing | tropical | nocturnal | 1.214 | 3485.072 | 5.035 |
| amphibians | Ansonia_longidigita | Ansonia_longidigita | LC | Stable | tropical | nocturnal | 32.83 | 103142.789 | 4.079 |
| amphibians | Ansonia_phuketensis | NA | EN | Decreasing | NA | NA | NA | NA | NA |
| amphibians | Ansonia_pilokensis | NA | EN | Unknown | NA | NA | NA | NA | NA |
| amphibians | Ansonia_platysoma | Ansonia_platysoma | LC | Unknown | tropical | diurnal | 1.326 | 3497.691 | 6.026 |
| amphibians | Ansonia_smeagol | NA | VU | Unknown | NA | NA | NA | NA | NA |
| amphibians | Ansonia_spinulifer | Ansonia_spinulifer | LC | Decreasing | tropical | nocturnal | 11.036 | 53930.428 | 3.856 |
| amphibians | Ansonia_teneritas | NA | EN | Decreasing | NA | NA | NA | NA | NA |
| amphibians | Ansonia_tiomanica | Ansonia_tiomanica | LC | Stable | tropical | nocturnal | 4.692 | 45.924 | 6.088 |
| amphibians | Anstisia_lutea | NA | LC | Stable | NA | NA | NA | NA | NA |
| amphibians | Anstisia_vitellina | NA | VU | Stable | NA | NA | NA | NA | NA |
| amphibians | Aphantophryne_pansa | Aphantophryne_pansa | LC | Stable | tropical | diurnal | 2.495 | 848.608 | 2.998 |
| amphibians | Aplastodiscus_albofrenatus | Aplastodiscus_albofrenatus | LC | Stable | tropical | nocturnal | 6.082 | 19643.129 | 16.496 |
| amphibians | Aplastodiscus_albosignatus | Aplastodiscus_albosignatus | LC | Stable | tropical | nocturnal | 12.531 | 25307.53 | 13.237 |
| amphibians | Aplastodiscus_arildae | Aplastodiscus_arildae | LC | Stable | tropical | nocturnal | 6.082 | 166613.886 | 14.006 |
| amphibians | Aplastodiscus_cavicola | Aplastodiscus_cavicola | LC | Decreasing | tropical | nocturnal | 4.271 | 208748.254 | 10.318 |

| | | | | | | | | | |
|---|---|---|---|---|---|---|---|---|---|
| amphibians | Aplastodiscus_cochranae | Aplastodiscus_cochranae | LC | Decreasing | temperate | nocturnal | 11.252 | 31715.657 | 12.41 |
| amphibians | Aplastodiscus_ehrhardti | Aplastodiscus_ehrhardti | LC | Decreasing | temperate | nocturnal | 4.975 | 43757.831 | 10.195 |
| amphibians | Aplastodiscus_eugenioi | Aplastodiscus_eugenioi | LC | Stable | tropical | nocturnal | 4.934 | 16103.427 | 15.172 |
| amphibians | Aplastodiscus_flumineus | Aplastodiscus_flumineus | CR | Unknown | tropical | nocturnal | 11.325 | 69.718 | 5.878 |
| amphibians | Aplastodiscus_ibirapitanga | Aplastodiscus_ibirapitanga | LC | Unknown | tropical | nocturnal | 6.976 | 45298.128 | 7.567 |
| amphibians | Aplastodiscus_leucopygius | Aplastodiscus_leucopygius | LC | Stable | tropical | nocturnal | 9.031 | 116912.507 | 15.076 |
| amphibians | Aplastodiscus_lutzorum | NA | LC | Unknown | NA | NA | NA | NA | NA |
| amphibians | Aplastodiscus_perviridis | Aplastodiscus_perviridis | LC | Stable | temperate | nocturnal | 9.031 | 234555.883 | 12.668 |
| amphibians | Aplastodiscus_sibilatus | Aplastodiscus_sibilatus | LC | Unknown | tropical | nocturnal | 3.044 | 38340.726 | 8.952 |
| amphibians | Aplastodiscus_weygoldti | Aplastodiscus_weygoldti | LC | Decreasing | tropical | nocturnal | 6.129 | 116025.098 | 9.362 |
| amphibians | Aquarana_catesbeiana | Rana_catesbeiana | LC | Increasing | temperate | cathemeral | 1033.804 | 7019850.753 | 17.51 |
| amphibians | Aquarana_clamitans | Rana_clamitans | LC | Stable | temperate | cathemeral | 122.125 | 3829381.027 | 15.526 |
| amphibians | Aquarana_grylio | Rana_grylio | LC | Decreasing | temperate | nocturnal | 528.201 | 365850.877 | 15.945 |
| amphibians | Aquarana_heckscheri | Rana_heckscheri | LC | Stable | temperate | nocturnal | 431.346 | 253556.061 | 16.805 |
| amphibians | Aquarana_septentrionalis | Rana_septentrionalis | LC | Stable | temperate | cathemeral | 42.854 | 1981074.123 | 6.745 |
| amphibians | Aquarana_virgatipes | Rana_virgatipes | LC | Unknown | temperate | diurnal | 37.611 | 166516.816 | 17.726 |
| amphibians | Arcovomer_passarellii | Arcovomer_passarellii | LC | Decreasing | tropical | nocturnal | 1.326 | 42462.211 | 15.206 |
| amphibians | Arenophryne_rotunda | Arenophryne_rotunda | LC | Stable | temperate | nocturnal | 4.934 | 1871.974 | 7.81 |
| amphibians | Aromobates_cannatellai | Aromobates_cannatellai | CR | Stable | tropical | diurnal | 1.806 | 3.357 | 15.624 |
| amphibians | Aromobates_capurinensis | NA | DD | Unknown | NA | NA | NA | NA | NA |
| amphibians | Aromobates_duranti | Aromobates_duranti | CR | Decreasing | tropical | diurnal | 2.272 | 36.378 | 14.075 |
| amphibians | Aromobates_ericksonae | Aromobates_ericksonae | EN | Decreasing | tropical | diurnal | 1.277 | 356.777 | 15.445 |
| amphibians | Aromobates_leopardalis | NA | CR | Decreasing | NA | NA | NA | NA | NA |
| amphibians | Aromobates_mayorgai | Aromobates_mayorgai | EN | Decreasing | tropical | diurnal | 1.277 | 1126.477 | 17.545 |
| amphibians | Aromobates_molinarii | Aromobates_molinarii | CR | Decreasing | tropical | diurnal | 2.178 | 6.831 | 29.794 |
| amphibians | Aromobates_nocturnus | Aromobates_nocturnus | CR | Decreasing | tropical | nocturnal | 24.557 | 20.766 | 10.315 |
| amphibians | Aromobates_saltuensis | Aromobates_saltuensis | EN | Decreasing | tropical | diurnal | 1.481 | 191.723 | 27.305 |
| amphibians | Aromobates_serranus | Aromobates_serranus | CR | Decreasing | tropical | diurnal | 1.806 | 77.272 | 12.737 |
| amphibians | Arthroleptella_drewesii | Arthroleptella_drewesii | NT | Unknown | temperate | cathemeral | 0.664 | 44.04 | 7.949 |
| amphibians | Arthroleptides_martiensseni | Arthroleptides_martiensseni | EN | Unknown | tropical | diurnal | 39.306 | 635.452 | 13.38 |
| amphibians | Arthroleptis_adelphus | Arthroleptis_adelphus | LC | Unknown | tropical | diurnal | 4.16 | 471773.178 | 6.423 |
| amphibians | Arthroleptis_adolfifriederici | Arthroleptis_adolfifriederici | LC | Unknown | tropical | diurnal | 8.423 | 12500.444 | 13.342 |
| amphibians | Arthroleptis_bioko | Arthroleptis_bioko | EN | Decreasing | tropical | diurnal | 3.163 | 127.951 | 11.711 |
| amphibians | Arthroleptis_crusculum | Arthroleptis_crusculum | NT | Decreasing | tropical | diurnal | 0.664 | 1521.832 | 10.961 |

| | | | | | | | | | |
|---|---|---|---|---|---|---|---|---|---|
| amphibians | Arthroleptis_fichika | Arthroleptis_fichika | EN | Unknown | tropical | diurnal | 0.223 | 24.933 | 15.55 |
| amphibians | Arthroleptis_formosus | NA | DD | Unknown | NA | NA | NA | NA | NA |
| amphibians | Arthroleptis_kidogo | Arthroleptis_kidogo | CR | Unknown | tropical | diurnal | 0.223 | 69.194 | 7.486 |
| amphibians | Arthroleptis_langeri | Arthroleptis_langeri | EN | Decreasing | tropical | diurnal | 0.505 | 264.228 | 11.69 |
| amphibians | Arthroleptis_nimbaensis | NA | DD | Unknown | NA | NA | NA | NA | NA |
| amphibians | Arthroleptis_nlonakoensis | Arthroleptis_nlonakoensis | EN | Decreasing | tropical | nocturnal | 0.795 | 26.283 | 8.521 |
| amphibians | Arthroleptis_perreti | Arthroleptis_perreti | EN | Unknown | tropical | diurnal | 7.844 | 283.542 | 13.649 |
| amphibians | Arthroleptis_poecilonotus | Arthroleptis_poecilonotus | LC | Stable | tropical | diurnal | 3.163 | 2565497.12 | 8.483 |
| amphibians | Arthroleptis_stenodactylus | Arthroleptis_stenodactylus | LC | Stable | tropical | diurnal | 7.844 | 2386920.608 | 9.341 |
| amphibians | Arthroleptis_sylvaticus | Arthroleptis_sylvaticus | LC | Stable | tropical | diurnal | 1.023 | 1013604.902 | 5.673 |
| amphibians | Arthroleptis_taeniatus | Arthroleptis_taeniatus | LC | Unknown | tropical | diurnal | 1.109 | 723729.612 | 6.154 |
| amphibians | Arthroleptis_tanneri | Arthroleptis_tanneri | EN | Decreasing | tropical | diurnal | 15.029 | 647.981 | 19.249 |
| amphibians | Arthroleptis_troglodytes | Arthroleptis_troglodytes | CR | Unknown | tropical | nocturnal | 1.499 | 15.098 | 7.236 |
| amphibians | Arthroleptis_tuberosus | Arthroleptis_tuberosus | DD | Decreasing | tropical | diurnal | 1.726 | 1128791.568 | 5.612 |
| amphibians | Arthroleptis_variabilis | Arthroleptis_variabilis | LC | Decreasing | tropical | diurnal | 4.536 | 1636868.726 | 8.706 |
| amphibians | Arthroleptis_wahlbergii | Arthroleptis_wahlbergii | LC | Stable | temperate | cathemeral | 3.348 | 50856.125 | 15.068 |
| amphibians | Arthroleptis_xenodactyloides | Arthroleptis_xenodactyloides | LC | Stable | tropical | cathemeral | 1.168 | 1109401.587 | 9.883 |
| amphibians | Arthroleptis_xenodactylus | Arthroleptis_xenodactylus | EN | Decreasing | tropical | cathemeral | 0.772 | 599.661 | 12.789 |
| amphibians | Ascaphus_montanus | Ascaphus_montanus | LC | Stable | temperate | cathemeral | 15.932 | 239008.973 | 8.423 |
| amphibians | Ascaphus_truei | Ascaphus_truei | LC | Unknown | temperate | nocturnal | 11.767 | 304263.383 | 7.1 |
| amphibians | Assa_darlingtoni | Assa_darlingtoni | VU | Decreasing | temperate | cathemeral | 2.109 | 6062.968 | 7.697 |
| amphibians | Asterophrys_pullifer | NA | DD | Unknown | NA | NA | NA | NA | NA |
| amphibians | Asterophrys_senglaubi | NA | DD | Unknown | NA | NA | NA | NA | NA |
| amphibians | Astrobatrachus_kurichiyana | NA | DD | Unknown | NA | NA | NA | NA | NA |
| amphibians | Astylosternus_batesi | Astylosternus_batesi | LC | Decreasing | tropical | nocturnal | 39.306 | 652386.815 | 6.141 |
| amphibians | Astylosternus_diadematus | Astylosternus_diadematus | LC | Decreasing | tropical | nocturnal | 16.873 | 32334.278 | 9.472 |
| amphibians | Astylosternus_laticephalus | Astylosternus_laticephalus | NT | Unknown | tropical | nocturnal | 25.822 | 50739.636 | 17.012 |
| amphibians | Astylosternus_occidentalis | Astylosternus_occidentalis | LC | Unknown | tropical | nocturnal | 27.131 | 185806.719 | 14.31 |
| amphibians | Astylosternus_perreti | Astylosternus_perreti | EN | Decreasing | tropical | nocturnal | 29.887 | 416.348 | 15 |
| amphibians | Astylosternus_ranoides | Astylosternus_ranoides | EN | Decreasing | tropical | diurnal | 11.767 | 2055.427 | 13.918 |
| amphibians | Astylosternus_rheophilus | Astylosternus_rheophilus | NT | Decreasing | tropical | cathemeral | 29.887 | 9515.228 | 13.928 |
| amphibians | Astylosternus_schioetzi | Astylosternus_schioetzi | EN | Decreasing | tropical | nocturnal | 7.294 | 1018.106 | 12.662 |
| amphibians | Atelopus_andinus | Atelopus_andinus | EN | Unknown | tropical | diurnal | 4.16 | 2415.381 | 1.315 |
| amphibians | Atelopus_angelito | Atelopus_angelito | CR | Decreasing | tropical | diurnal | 10.474 | 100.498 | 11.125 |

| | | | | | | | | | |
|---|---|---|---|---|---|---|---|---|---|
| amphibians | Atelopus_ardila | NA | CR | Decreasing | NA | NA | NA | NA | NA |
| amphibians | Atelopus_arsyecue | Atelopus_arsyecue | CR | Decreasing | tropical | diurnal | 20.466 | 79.931 | 5.696 |
| amphibians | Atelopus_arthuri | Atelopus_arthuri | CR | Decreasing | tropical | diurnal | 8.969 | 129.529 | 10.609 |
| amphibians | Atelopus_balios | Atelopus_balios | CR | Decreasing | tropical | diurnal | 5.155 | 173.529 | 10.792 |
| amphibians | Atelopus_barbotini | NA | LC | Stable | NA | NA | NA | NA | NA |
| amphibians | Atelopus_bomolochos | Atelopus_bomolochos | CR | Decreasing | tropical | diurnal | 11.767 | 3922.282 | 18.858 |
| amphibians | Atelopus_boulengeri | Atelopus_boulengeri | CR | Decreasing | tropical | nocturnal | 26.999 | 4911.532 | 6.427 |
| amphibians | Atelopus_carauta | Atelopus_carauta | DD | Decreasing | tropical | diurnal | 8.846 | 115.054 | 8.324 |
| amphibians | Atelopus_carbonerensis | Atelopus_carbonerensis | CR | Decreasing | tropical | diurnal | 14.678 | 142.875 | 11.324 |
| amphibians | Atelopus_carrikeri | Atelopus_carrikeri | EN | Decreasing | tropical | diurnal | 22.272 | 627.246 | 5.044 |
| amphibians | Atelopus_certus | Atelopus_certus | CR | Decreasing | tropical | diurnal | 6.273 | 784.607 | 2.207 |
| amphibians | Atelopus_chiriquiensis | NA | EX | not_available | NA | NA | NA | NA | NA |
| amphibians | Atelopus_chirripoensis | NA | DD | Unknown | NA | NA | NA | NA | NA |
| amphibians | Atelopus_chocoensis | Atelopus_chocoensis | CR | Decreasing | tropical | diurnal | 9.866 | 233.1 | 2.059 |
| amphibians | Atelopus_chrysocorallus | Atelopus_chrysocorallus | CR | Decreasing | tropical | diurnal | 11.325 | 55.215 | 12.334 |
| amphibians | Atelopus_coynei | Atelopus_coynei | CR | Decreasing | tropical | diurnal | 3.946 | 50.677 | 10.223 |
| amphibians | Atelopus_cruciger | Atelopus_cruciger | CR | Stable | tropical | diurnal | 11.036 | 16489.973 | 15.364 |
| amphibians | Atelopus_dimorphus | Atelopus_dimorphus | DD | Decreasing | tropical | diurnal | 2.419 | 64.775 | 6.819 |
| amphibians | Atelopus_ebenoides | Atelopus_ebenoides | CR | Decreasing | tropical | diurnal | 9.669 | 807.903 | 13.383 |
| amphibians | Atelopus_elegans | Atelopus_elegans | EN | Decreasing | tropical | diurnal | 3.705 | 8362.61 | 9.402 |
| amphibians | Atelopus_epikeisthos | NA | EN | Decreasing | NA | NA | NA | NA | NA |
| amphibians | Atelopus_erythropus | Atelopus_erythropus | CR | Decreasing | tropical | diurnal | 1.686 | 12195.71 | 2.955 |
| amphibians | Atelopus_eusebianus | Atelopus_eusebianus | CR | Decreasing | tropical | diurnal | 8.073 | 824.86 | 10.887 |
| amphibians | Atelopus_eusebiodiazi | NA | CR | Decreasing | NA | NA | NA | NA | NA |
| amphibians | Atelopus_exiguus | Atelopus_exiguus | EN | Decreasing | tropical | diurnal | 3.605 | 410.87 | 10.535 |
| amphibians | Atelopus_famelicus | Atelopus_famelicus | CR | Decreasing | tropical | diurnal | 7.844 | 20.166 | 16.846 |
| amphibians | Atelopus_farci | Atelopus_farci | CR | Decreasing | tropical | diurnal | 3.946 | 1.616 | 31.049 |
| amphibians | Atelopus_flavescens | Atelopus_flavescens | LC | Unknown | tropical | diurnal | 5.356 | 135579.385 | 0.922 |
| amphibians | Atelopus_franciscus | Atelopus_franciscus | LC | Stable | tropical | diurnal | 1.411 | 35332.26 | 1.504 |
| amphibians | Atelopus_galactogaster | NA | DD | Unknown | NA | NA | NA | NA | NA |
| amphibians | Atelopus_gigas | NA | CR | Unknown | NA | NA | NA | NA | NA |
| amphibians | Atelopus_glyphus | Atelopus_glyphus | CR | Decreasing | tropical | diurnal | 9.669 | 363.977 | 2.067 |
| amphibians | Atelopus_guanujo | Atelopus_guanujo | CR | Decreasing | tropical | diurnal | 8.423 | 143.596 | 14.421 |
| amphibians | Atelopus_guitarraensis | NA | DD | Unknown | NA | NA | NA | NA | NA |

| | | | | | | | | | |
|---|---|---|---|---|---|---|---|---|---|
| amphibians | Atelopus_halihelos | Atelopus_halihelos | CR | Decreasing | tropical | diurnal | 3.841 | 58.852 | 5.41 |
| amphibians | Atelopus_hoogmoedi | Atelopus_hoogmoedi | LC | Stable | tropical | diurnal | 5.894 | 750741.451 | 1.069 |
| amphibians | Atelopus_ignescens | Atelopus_ignescens | CR | Decreasing | tropical | diurnal | 9.8 | 10227.639 | 12.461 |
| amphibians | Atelopus_laetissimus | Atelopus_laetissimus | EN | Stable | tropical | diurnal | 14.504 | 164.836 | 4.939 |
| amphibians | Atelopus_limosus | Atelopus_limosus | CR | Decreasing | tropical | diurnal | 5.443 | 1407.527 | 5.448 |
| amphibians | Atelopus_longibrachius | Atelopus_longibrachius | EN | Decreasing | tropical | diurnal | 4.016 | 5.046 | 12.103 |
| amphibians | Atelopus_longirostris | Atelopus_longirostris | CR | Decreasing | tropical | cathemeral | 9.094 | 1727.95 | 8.285 |
| amphibians | Atelopus_lozanoi | Atelopus_lozanoi | EN | Decreasing | tropical | diurnal | 4.536 | 4052.107 | 12.275 |
| amphibians | Atelopus_lynchi | Atelopus_lynchi | CR | Decreasing | tropical | cathemeral | 8.073 | 11.121 | 8.295 |
| amphibians | Atelopus_manauensis | NA | EN | Decreasing | NA | NA | NA | NA | NA |
| amphibians | Atelopus_mandingues | NA | DD | Unknown | NA | NA | NA | NA | NA |
| amphibians | Atelopus_marinkellei | Atelopus_marinkellei | EN | Decreasing | tropical | diurnal | 9.669 | 1787.79 | 14.772 |
| amphibians | Atelopus_mindoensis | Atelopus_mindoensis | CR | Decreasing | tropical | diurnal | 1.889 | 2029.815 | 6.051 |
| amphibians | Atelopus_minutulus | Atelopus_minutulus | CR | Decreasing | tropical | diurnal | 1.245 | 10.089 | 5.524 |
| amphibians | Atelopus_mittermeieri | NA | EN | Decreasing | NA | NA | NA | NA | NA |
| amphibians | Atelopus_monohernandezii | Atelopus_monohernandezii | CR | Decreasing | tropical | diurnal | 5.576 | 51.043 | 7.312 |
| amphibians | Atelopus_mucubajiensis | Atelopus_mucubajiensis | CR | Decreasing | tropical | diurnal | 7.294 | 10.699 | 11.975 |
| amphibians | Atelopus_muisca | Atelopus_muisca | CR | Decreasing | tropical | diurnal | 6.469 | 87.364 | 10.443 |
| amphibians | Atelopus_nahumae | Atelopus_nahumae | EN | Decreasing | tropical | diurnal | 11.767 | 3236.14 | 6.245 |
| amphibians | Atelopus_nanay | Atelopus_nanay | CR | Decreasing | tropical | diurnal | 10.371 | 76.895 | 11.095 |
| amphibians | Atelopus_nepiozomus | Atelopus_nepiozomus | EN | Unknown | tropical | diurnal | 2.706 | 2361.54 | 3.591 |
| amphibians | Atelopus_nicefori | Atelopus_nicefori | CR | Decreasing | tropical | diurnal | 2.706 | 90.393 | 4.426 |
| amphibians | Atelopus_onorei | Atelopus_onorei | CR | Decreasing | tropical | diurnal | 9.604 | 11.15 | 13.572 |
| amphibians | Atelopus_orcesi | Atelopus_orcesi | CR | Decreasing | tropical | diurnal | 6.322 | 4.23 | 8.735 |
| amphibians | Atelopus_oxyrhynchus | Atelopus_oxyrhynchus | CR | Decreasing | tropical | diurnal | 10.894 | 346.083 | 15.919 |
| amphibians | Atelopus_pachydermus | Atelopus_pachydermus | CR | Decreasing | tropical | diurnal | 21.697 | 4236.959 | 5.157 |
| amphibians | Atelopus_palmatus | Atelopus_palmatus | CR | Decreasing | tropical | cathemeral | 2.394 | 137.56 | 15.331 |
| amphibians | Atelopus_pastuso | Atelopus_pastuso | CR | Unknown | tropical | diurnal | 11.545 | 3666.513 | 14.504 |
| amphibians | Atelopus_patazensis | Atelopus_patazensis | CR | Decreasing | tropical | diurnal | 10.201 | 27.736 | 11.099 |
| amphibians | Atelopus_pedimarmoratus | Atelopus_pedimarmoratus | CR | Decreasing | tropical | diurnal | 2.064 | 10.098 | 3.882 |
| amphibians | Atelopus_peruensis | Atelopus_peruensis | CR | Decreasing | tropical | diurnal | 7.958 | 22730.303 | 9.447 |
| amphibians | Atelopus_petersi | Atelopus_petersi | CR | Decreasing | tropical | diurnal | 11.18 | 572.902 | 6.952 |
| amphibians | Atelopus_petriruizi | NA | CR | Decreasing | NA | NA | NA | NA | NA |
| amphibians | Atelopus_pictiventris | Atelopus_pictiventris | CR | Decreasing | tropical | diurnal | 4.692 | 11.458 | 13.575 |

| | | | | | | | | | |
|---|---|---|---|---|---|---|---|---|---|
| amphibians | Atelopus_pinangoi | Atelopus_pinangoi | CR | Decreasing | tropical | diurnal | 5.802 | 35.33 | 10.548 |
| amphibians | Atelopus_planispina | Atelopus_planispina | CR | Decreasing | tropical | diurnal | 3.981 | 272.283 | 5.867 |
| amphibians | Atelopus_podocarpus | Atelopus_podocarpus | CR | Decreasing | tropical | diurnal | 13.248 | 1732.359 | 7.816 |
| amphibians | Atelopus_pulcher | Atelopus_pulcher | VU | Decreasing | tropical | diurnal | 5.356 | 9255.545 | 5.307 |
| amphibians | Atelopus_pyrodactylus | NA | CR | Decreasing | NA | NA | NA | NA | NA |
| amphibians | Atelopus_quimbaya | Atelopus_quimbaya | CR | Unknown | tropical | diurnal | 2.957 | 20.207 | 29.722 |
| amphibians | Atelopus_reticulatus | Atelopus_reticulatus | DD | Decreasing | tropical | diurnal | 1.572 | 11.015 | 7.702 |
| amphibians | Atelopus_sanjosei | Atelopus_sanjosei | CR | Decreasing | tropical | diurnal | 2.788 | 21.052 | 11.595 |
| amphibians | Atelopus_seminiferus | Atelopus_seminiferus | EN | Unknown | tropical | diurnal | 5.356 | 2302.213 | 5.32 |
| amphibians | Atelopus_senex | NA | EX | not_available | NA | NA | NA | NA | NA |
| amphibians | Atelopus_sernai | Atelopus_sernai | CR | Decreasing | tropical | diurnal | 2.957 | 318.578 | 18.069 |
| amphibians | Atelopus_simulatus | Atelopus_simulatus | CR | Unknown | tropical | diurnal | 4.308 | 577.33 | 8.672 |
| amphibians | Atelopus_siranus | NA | DD | Unknown | NA | NA | NA | NA | NA |
| amphibians | Atelopus_sonsonensis | Atelopus_sonsonensis | CR | Unknown | tropical | diurnal | 2.345 | 359.221 | 12.95 |
| amphibians | Atelopus_sorianoi | Atelopus_sorianoi | CR | Decreasing | tropical | diurnal | 11.398 | 5.89 | 14.013 |
| amphibians | Atelopus_spumarius | Atelopus_spumarius | LC | Unknown | tropical | diurnal | 5.356 | 90169.925 | 2.728 |
| amphibians | Atelopus_spurrelli | Atelopus_spurrelli | NT | Stable | tropical | diurnal | 3.163 | 26789.532 | 4.12 |
| amphibians | Atelopus_subornatus | Atelopus_subornatus | CR | Decreasing | tropical | diurnal | 4.934 | 617.178 | 16.473 |
| amphibians | Atelopus_tamaense | Atelopus_tamaensis | CR | Decreasing | tropical | diurnal | 7.788 | 55.928 | 4.764 |
| amphibians | Atelopus_tricolor | Atelopus_tricolor | CR | Decreasing | tropical | diurnal | 3.807 | 43897.393 | 3.804 |
| amphibians | Atelopus_varius | Atelopus_varius | CR | Unknown | tropical | diurnal | 19.923 | 10807.36 | 11.508 |
| amphibians | Atelopus_vogli | NA | EX | not_available | NA | NA | NA | NA | NA |
| amphibians | Atelopus_walkeri | Atelopus_walkeri | DD | Decreasing | tropical | diurnal | 10.543 | 194.887 | 6.134 |
| amphibians | Atelopus_zeteki | Atelopus_zeteki | CR | Decreasing | tropical | diurnal | 23.335 | 580.72 | 13.12 |
| amphibians | Atlantihyla_spinipollex | Ptychohyla_spinipollex | EN | Decreasing | tropical | nocturnal | 8.483 | 445.948 | 4.921 |
| amphibians | Aubria_masako | Aubria_masako | LC | Unknown | tropical | nocturnal | 74.113 | 1137719.591 | 5.088 |
| amphibians | Aubria_subsigillata | Aubria_subsigillata | LC | Decreasing | tropical | nocturnal | 88.303 | 579127.973 | 14.097 |
| amphibians | Austrochaperina_alexanderi | NA | LC | Unknown | NA | NA | NA | NA | NA |
| amphibians | Austrochaperina_basipalmata | Austrochaperina_basipalmata | LC | Unknown | tropical | diurnal | 4.934 | 12356.166 | 6.574 |
| amphibians | Austrochaperina_beehleri | NA | not_available | not_available | NA | NA | NA | NA | NA |
| amphibians | Austrochaperina_brevipes | NA | DD | Unknown | NA | NA | NA | NA | NA |
| amphibians | Austrochaperina_fryi | Austrochaperina_fryi | LC | Stable | tropical | diurnal | 3.475 | 4191.132 | 4.346 |
| amphibians | Austrochaperina_fulva | NA | not_available | not_available | NA | NA | NA | NA | NA |
| amphibians | Austrochaperina_hooglandi | Austrochaperina_hooglandi | LC | Stable | tropical | diurnal | 8.015 | 112.837 | 2.872 |

| | | | | | | | | | |
|---|---|---|---|---|---|---|---|---|---|
| amphibians | Austrochaperina_laurae | NA | LC | Stable | NA | NA | NA | NA | NA |
| amphibians | Austrochaperina_palmipes | Austrochaperina_palmipes | LC | Stable | tropical | nocturnal | 10.337 | 29095.836 | 3.792 |
| amphibians | Austrochaperina_septentrionalis | Austrochaperina_septentrionalis | LC | Unknown | tropical | diurnal | 2.495 | 4223.573 | 3.446 |
| amphibians | Balebreviceps_hillmani | Balebreviceps_hillmani | CR | Decreasing | tropical | diurnal | 13.329 | 4.899 | 13.233 |
| amphibians | Barbarophryne_brongersmai | Barbarophryne_brongersmai | LC | Unknown | temperate | nocturnal | 12.688 | 131702.782 | 7.933 |
| amphibians | Barygenys_maculata | Barygenys_maculata | LC | Unknown | tropical | diurnal | 11.618 | 1413.89 | 2.615 |
| amphibians | Barygenys_parvula | Barygenys_parvula | NT | Unknown | tropical | nocturnal | 0.456 | 481.551 | 3.857 |
| amphibians | Batrachoseps_bramei | Batrachoseps_bramei | NT | Stable | temperate | nocturnal | 0.746 | 217.516 | 9.848 |
| amphibians | Batrachoseps_campi | Batrachoseps_campi | NT | Stable | temperate | nocturnal | 3.666 | 326.178 | 1.541 |
| amphibians | Batrachoseps_pacificus | Batrachoseps_pacificus | LC | Stable | temperate | nocturnal | 5.494 | 513.58 | 7.184 |
| amphibians | Batrachuperus_tibetanus | Batrachuperus_tibetanus | VU | Decreasing | temperate | cathemeral | 13.938 | 171618.723 | 9.574 |
| amphibians | Batrachyla_taeniata | Batrachyla_taeniata | LC | Decreasing | temperate | nocturnal | 7.844 | 145739.871 | 7.194 |
| amphibians | Beddomixalus_bijui | Beddomixalus_bijui | EN | Decreasing | tropical | nocturnal | 21.019 | 100.267 | 6.407 |
| amphibians | Blaira_ornata | NA | VU | Decreasing | NA | NA | NA | NA | NA |
| amphibians | Blommersia_angolafa | Blommersia_angolafa | LC | Decreasing | tropical | cathemeral | 0.664 | 23457.428 | 8.271 |
| amphibians | Blommersia_bara | NA | not_available | not_available | NA | NA | NA | NA | NA |
| amphibians | Blommersia_blommersae | Blommersia_blommersae | LC | Unknown | tropical | cathemeral | 0.664 | 17704.956 | 7.812 |
| amphibians | Blommersia_dejongi | Blommersia_dejongi | LC | Stable | tropical | nocturnal | 0.969 | 1226.149 | 14.267 |
| amphibians | Blommersia_domerguei | Blommersia_domerguei | LC | Unknown | tropical | cathemeral | 0.595 | 51578.289 | 11.175 |
| amphibians | Blommersia_galani | Blommersia_galani | LC | Stable | tropical | cathemeral | 1.051 | 1009.156 | 14.319 |
| amphibians | Blommersia_grandisonae | Blommersia_grandisonae | LC | Decreasing | tropical | cathemeral | 1.023 | 64654.362 | 7.21 |
| amphibians | Blommersia_kely | Blommersia_kely | LC | Unknown | tropical | diurnal | 0.275 | 9244.875 | 12.476 |
| amphibians | Blommersia_sarotra | Blommersia_sarotra | LC | Unknown | tropical | cathemeral | 0.335 | 6079.763 | 8.722 |
| amphibians | Blommersia_transmarina | NA | NT | Stable | NA | NA | NA | NA | NA |
| amphibians | Blommersia_variabilis | Blommersia_variabilis | LC | Stable | tropical | cathemeral | 1.889 | 1345.63 | 10.522 |
| amphibians | Blommersia_wittei | Blommersia_wittei | LC | Stable | tropical | cathemeral | 1.499 | 69134.077 | 8.63 |
| amphibians | Blythophryne_beryet | NA | VU | Stable | NA | NA | NA | NA | NA |
| amphibians | Boana_aguilari | Hypsiboas_aguilari | LC | Stable | tropical | nocturnal | 11.036 | 1040.681 | 6.653 |
| amphibians | Boana_albomarginata | Hypsiboas_albomarginatus | LC | Stable | tropical | nocturnal | 22.156 | 504714.802 | 12.438 |
| amphibians | Boana_albopunctata | Hypsiboas_albopunctatus | LC | Stable | tropical | nocturnal | 41.053 | 4885110.928 | 6.921 |
| amphibians | Boana_alfaroi | Hypsiboas_alfaroi | LC | Stable | tropical | nocturnal | 10.474 | 77059.352 | 2.869 |
| amphibians | Boana_almendarizae | Hypsiboas_almendarizae | NT | Decreasing | tropical | nocturnal | 12.454 | 14757.849 | 6.659 |
| amphibians | Boana_atlantica | Hypsiboas_atlanticus | LC | Stable | tropical | nocturnal | 7.028 | 42358.187 | 12.044 |
| amphibians | Boana_balzani | Hypsiboas_balzani | LC | Stable | tropical | nocturnal | 12.767 | 24585.413 | 4.911 |

| | | | | | | | | | |
|---|---|---|---|---|---|---|---|---|---|
| amphibians | Boana_beckeri | NA | DD | Unknown | NA | NA | NA | NA | NA |
| amphibians | Boana_benitezi | Hypsiboas_benitezi | LC | Stable | tropical | nocturnal | 12.069 | 5445.415 | 0.243 |
| amphibians | Boana_bischoffi | Hypsiboas_bischoffi | LC | Stable | temperate | nocturnal | 31.334 | 164695.227 | 12.044 |
| amphibians | Boana_boans | Hypsiboas_boans | LC | Stable | tropical | cathemeral | 256.337 | 7123580.791 | 2.549 |
| amphibians | Boana_botumirim | Hypsiboas_botumirim | LC | Decreasing | tropical | nocturnal | 3.044 | 54975.293 | 6.087 |
| amphibians | Boana_buriti | Hypsiboas_buriti | EN | Decreasing | tropical | nocturnal | 2.599 | 17759.239 | 11.513 |
| amphibians | Boana_caingua | Hypsiboas_caingua | LC | Stable | temperate | nocturnal | 4.934 | 275003.996 | 13.382 |
| amphibians | Boana_caipora | Hypsiboas_caipora | LC | Stable | temperate | nocturnal | 7.456 | 3405.128 | 7.522 |
| amphibians | Boana_calcarata | Hypsiboas_calcaratus | LC | Unknown | tropical | nocturnal | 22.156 | 4707380.255 | 1.395 |
| amphibians | Boana_callipleura | Hypsiboas_callipleura | LC | Stable | tropical | nocturnal | 12.145 | 15105.32 | 7.656 |
| amphibians | Boana_cinerascens | Hypsiboas_cinerascens | LC | Stable | tropical | nocturnal | 5.894 | 5780987.859 | 1.713 |
| amphibians | Boana_cipoensis | Hypsiboas_cipoensis | LC | Decreasing | tropical | nocturnal | 4.345 | 33311.84 | 6.782 |
| amphibians | Boana_crepitans | Hypsiboas_crepitans | LC | Stable | tropical | nocturnal | 41.053 | 935349.716 | 10.045 |
| amphibians | Boana_curupi | Hypsiboas_curupi | LC | Unknown | temperate | nocturnal | 9.031 | 76642.57 | 12.317 |
| amphibians | Boana_dentei | Hypsiboas_dentei | LC | Unknown | tropical | nocturnal | 14.161 | 186724.763 | 0.744 |
| amphibians | Boana_diabolica | NA | LC | Stable | NA | NA | NA | NA | NA |
| amphibians | Boana_ericae | Hypsiboas_ericae | EN | Decreasing | tropical | nocturnal | 4.16 | 9736.717 | 5.58 |
| amphibians | Boana_exastis | Hypsiboas_exastis | LC | Stable | tropical | nocturnal | 100.927 | 67340.913 | 13.121 |
| amphibians | Boana_faber | Hypsiboas_faber | LC | Stable | tropical | nocturnal | 118.397 | 1288354.879 | 13.063 |
| amphibians | Boana_fasciata | Hypsiboas_fasciatus | LC | Decreasing | tropical | nocturnal | 14.853 | 17312.193 | 5.021 |
| amphibians | Boana_freicanecae | Hypsiboas_freicanecae | EN | Decreasing | tropical | nocturnal | 6.37 | 2005.183 | 9.97 |
| amphibians | Boana_geographica | Hypsiboas_geographicus | LC | Stable | tropical | nocturnal | 61.584 | 6749152.751 | 2.312 |
| amphibians | Boana_goiana | Hypsiboas_goianus | LC | Decreasing | tropical | nocturnal | 4.653 | 51882.458 | 12.698 |
| amphibians | Boana_guentheri | Hypsiboas_guentheri | LC | Stable | temperate | nocturnal | 9.031 | 23644.053 | 17.076 |
| amphibians | Boana_heilprini | Hypsiboas_heilprini | VU | Decreasing | tropical | cathemeral | 14.418 | 37933.013 | 15.139 |
| amphibians | Boana_hobbsi | Hypsiboas_hobbsi | LC | Stable | tropical | nocturnal | 6.518 | 356380.433 | 0.735 |
| amphibians | Boana_hutchinsi | Hypsiboas_hutchinsi | LC | Unknown | tropical | nocturnal | 15.932 | 60245.041 | 0.89 |
| amphibians | Boana_jimenezi | Hypsiboas_jimenezi | LC | Unknown | tropical | nocturnal | 3.475 | 4233.41 | 2.951 |
| amphibians | Boana_joaquini | Hypsiboas_joaquini | LC | Stable | temperate | nocturnal | 16.304 | 11319.698 | 7.513 |
| amphibians | Boana_lanciformis | Hypsiboas_lanciformis | LC | Stable | tropical | nocturnal | 104.268 | 4670364.267 | 2.1 |
| amphibians | Boana_latistriata | NA | not_available | not_available | NA | NA | NA | NA | NA |
| amphibians | Boana_lemai | Hypsiboas_lemai | LC | Stable | tropical | nocturnal | 5.356 | 38398.082 | 1.508 |
| amphibians | Boana_leptolineata | Hypsiboas_leptolineatus | LC | Stable | temperate | nocturnal | 4.934 | 114377.644 | 10.783 |
| amphibians | Boana_lundii | Hypsiboas_lundii | LC | Stable | tropical | nocturnal | 42.854 | 688059.338 | 10.987 |

| | | | | | | | | | |
|---|---|---|---|---|---|---|---|---|---|
| amphibians | Boana_maculateralis | Hypsiboas_maculateralis | LC | Stable | tropical | nocturnal | 15.296 | 64262.39 | 3.355 |
| amphibians | Boana_marginata | Hypsiboas_marginatus | LC | Stable | temperate | nocturnal | 16.873 | 12500.968 | 12.082 |
| amphibians | Boana_marianitae | Hypsiboas_marianitae | LC | Stable | tropical | nocturnal | 16.682 | 70845.719 | 7.402 |
| amphibians | Boana_melanopleura | Hypsiboas_melanopleura | LC | Stable | tropical | nocturnal | 9.474 | 92.19 | 13.739 |
| amphibians | Boana_microderma | Hypsiboas_microderma | LC | Unknown | tropical | nocturnal | 3.163 | 1050477.994 | 0.888 |
| amphibians | Boana_multifasciata | Hypsiboas_multifasciatus | LC | Stable | tropical | nocturnal | 57.012 | 164135.101 | 4.882 |
| amphibians | Boana_nigra | NA | VU | Decreasing | NA | NA | NA | NA | NA |
| amphibians | Boana_nympha | Hypsiboas_nympha | LC | Unknown | tropical | nocturnal | 3.807 | 416259.317 | 1.582 |
| amphibians | Boana_ornatissima | Hypsiboas_ornatissimus | LC | Unknown | tropical | nocturnal | 6.273 | 838190.713 | 1.101 |
| amphibians | Boana_palaestes | Hypsiboas_palaestes | EN | Decreasing | tropical | nocturnal | 11.693 | 534.288 | 5.879 |
| amphibians | Boana_paranaiba | Hypsiboas_paranaiba | LC | Stable | tropical | nocturnal | 12.767 | 1355076.589 | 7.994 |
| amphibians | Boana_pardalis | Hypsiboas_pardalis | LC | Stable | tropical | nocturnal | 41.053 | 318984.469 | 12.815 |
| amphibians | Boana_pellucens | Hypsiboas_pellucens | LC | Unknown | tropical | nocturnal | 21.697 | 76138.717 | 11.097 |
| amphibians | Boana_picturata | Hypsiboas_picturatus | LC | Decreasing | tropical | nocturnal | 31.927 | 83028.812 | 6.789 |
| amphibians | Boana_poaju | Hypsiboas_poaju | LC | Stable | temperate | nocturnal | 8.188 | 9722.43 | 11.026 |
| amphibians | Boana_polytaenia | Hypsiboas_polytaenius | LC | Stable | tropical | nocturnal | 6.273 | 209403.864 | 13.256 |
| amphibians | Boana_pombali | Hypsiboas_pombali | LC | Stable | tropical | nocturnal | 26.734 | 95449.074 | 10.07 |
| amphibians | Boana_prasina | Hypsiboas_prasinus | LC | Stable | temperate | nocturnal | 15.029 | 80899.988 | 14.345 |
| amphibians | Boana_pugnax | Hypsiboas_pugnax | LC | Stable | tropical | nocturnal | 50.601 | 286727.002 | 12.703 |
| amphibians | Boana_pulchella | Hypsiboas_pulchellus | LC | Stable | temperate | nocturnal | 11.036 | 1267157.233 | 11.443 |
| amphibians | Boana_punctata | Hypsiboas_punctatus | LC | Stable | tropical | cathemeral | 6.129 | 8320786.592 | 3.671 |
| amphibians | Boana_raniceps | Hypsiboas_raniceps | LC | Stable | tropical | cathemeral | 54.816 | 7490502.562 | 5.935 |
| amphibians | Boana_rhythmica | Hypsiboas_rhythmicus | NT | Unknown | tropical | nocturnal | 3.224 | 796.558 | 0.779 |
| amphibians | Boana_riojana | Hypsiboas_riojanus | LC | Stable | tropical | cathemeral | 22.977 | 271454.513 | 7.105 |
| amphibians | Boana_roraima | Hypsiboas_roraima | LC | Stable | tropical | nocturnal | 9.156 | 2193.637 | 1.717 |
| amphibians | Boana_rosenbergi | Hypsiboas_rosenbergi | LC | Stable | tropical | nocturnal | 88.303 | 164358.874 | 8.641 |
| amphibians | Boana_rubracyla | Hypsiboas_rubracylus | LC | Decreasing | tropical | nocturnal | 18.867 | 25777.128 | 4.7 |
| amphibians | Boana_rufitela | Hypsiboas_rufitelus | LC | Stable | tropical | nocturnal | 15.029 | 39575.387 | 9.678 |
| amphibians | Boana_secedens | Hypsiboas_secedens | EN | Unknown | tropical | nocturnal | 16.873 | 3504.46 | 14.902 |
| amphibians | Boana_semiguttata | Hypsiboas_semiguttatus | LC | Stable | temperate | nocturnal | 9.669 | 84206.965 | 10.166 |
| amphibians | Boana_semilineata | Hypsiboas_semilineatus | LC | Stable | tropical | nocturnal | 15.932 | 444831.274 | 13.232 |
| amphibians | Boana_sibleszi | Hypsiboas_sibleszi | LC | Stable | tropical | nocturnal | 3.705 | 145943.683 | 1.193 |
| amphibians | Boana_stellae | Hypsiboas_stellae | LC | Stable | temperate | nocturnal | 15.932 | 10397.946 | 11.228 |
| amphibians | Boana_stenocephala | Hypsiboas_stenocephalus | LC | Unknown | tropical | nocturnal | 4.934 | 87320.485 | 13.276 |

| | | | | | | | | | |
|---|---|---|---|---|---|---|---|---|---|
| amphibians | Boana_tepuiana | Hypsiboas_tepuianus | LC | Stable | tropical | nocturnal | 9.474 | 127716.238 | 1.074 |
| amphibians | Boana_tetete | Hypsiboas_tetete | LC | Stable | tropical | nocturnal | 8.305 | 101701.69 | 1.523 |
| amphibians | Boana_ventrimaculata | NA | LC | Stable | NA | NA | NA | NA | NA |
| amphibians | Boehmantis_microtympanum | Boehmantis_microtympanum | VU | Decreasing | temperate | nocturnal | 104.268 | 4734.578 | 6.445 |
| amphibians | Bokermannohyla_ahenea | Bokermannohyla_ahenea | LC | Stable | tropical | nocturnal | 16.587 | 1479.381 | 5.178 |
| amphibians | Bokermannohyla_alvarengai | Bokermannohyla_alvarengai | LC | Stable | tropical | nocturnal | 316.682 | 60203.623 | 8.84 |
| amphibians | Bokermannohyla_astartea | Bokermannohyla_astartea | LC | Stable | tropical | nocturnal | 7.844 | 79225.159 | 14.206 |
| amphibians | Bokermannohyla_capra | Bokermannohyla_capra | LC | Stable | tropical | nocturnal | 24.681 | 46733.039 | 9.163 |
| amphibians | Bokermannohyla_caramaschii | Bokermannohyla_caramaschii | LC | Stable | tropical | nocturnal | 32.83 | 38584.119 | 12.92 |
| amphibians | Bokermannohyla_carvalhoi | Bokermannohyla_carvalhoi | NT | Stable | tropical | nocturnal | 28.486 | 729.543 | 12.091 |
| amphibians | Bokermannohyla_circumdata | Bokermannohyla_circumdata | LC | Stable | tropical | nocturnal | 34.374 | 948421.356 | 11.668 |
| amphibians | Bokermannohyla_diamantina | Bokermannohyla_diamantina | LC | Stable | tropical | nocturnal | 12.299 | 77943.098 | 7.686 |
| amphibians | Bokermannohyla_gouveai | Bokermannohyla_gouveai | LC | Stable | tropical | nocturnal | 31.334 | 647.209 | 5.664 |
| amphibians | Bokermannohyla_hylax | Bokermannohyla_hylax | LC | Stable | temperate | nocturnal | 27.131 | 44682.891 | 10.211 |
| amphibians | Bokermannohyla_ibitiguara | Bokermannohyla_ibitiguara | LC | Stable | tropical | nocturnal | 9.932 | 12112.506 | 12.659 |
| amphibians | Bokermannohyla_ibitipoca | Bokermannohyla_ibitipoca | LC | Stable | tropical | nocturnal | 6.419 | 26789.686 | 11.424 |
| amphibians | Bokermannohyla_itapoty | Bokermannohyla_itapoty | LC | Stable | tropical | nocturnal | 11.187 | 818.387 | 2.744 |
| amphibians | Bokermannohyla_izecksohni | Bokermannohyla_izecksohni | CR | Decreasing | tropical | nocturnal | 8.579 | 65.622 | 35.536 |
| amphibians | Bokermannohyla_lucianae | Bokermannohyla_lucianae | VU | Decreasing | tropical | nocturnal | 10.474 | 6483.59 | 6.354 |
| amphibians | Bokermannohyla_luctuosa | Bokermannohyla_luctuosa | LC | Decreasing | tropical | nocturnal | 32.83 | 42440.044 | 16.904 |
| amphibians | Bokermannohyla_martinsi | Bokermannohyla_martinsi | VU | Decreasing | tropical | nocturnal | 37.611 | 12544.282 | 13.086 |
| amphibians | Bokermannohyla_nanuzae | Bokermannohyla_nanuzae | LC | Decreasing | tropical | nocturnal | 7.294 | 27757.771 | 10.089 |
| amphibians | Bokermannohyla_oxente | Bokermannohyla_oxente | LC | Stable | tropical | nocturnal | 10.337 | 9126.444 | 6.596 |
| amphibians | Bokermannohyla_sapiranga | NA | LC | Decreasing | NA | NA | NA | NA | NA |
| amphibians | Bokermannohyla_saxicola | Bokermannohyla_saxicola | LC | Stable | tropical | nocturnal | 15.029 | 74214.509 | 6.754 |
| amphibians | Bokermannohyla_vulcaniae | Bokermannohyla_vulcaniae | EN | Decreasing | tropical | nocturnal | 15.386 | 934.066 | 16.342 |
| amphibians | Bolitoglossa_adspersa | Bolitoglossa_adspersa | NT | Decreasing | tropical | diurnal | 2.66 | 22488.825 | 16.864 |
| amphibians | Bolitoglossa_altamazonica | Bolitoglossa_altamazonica | NT | Decreasing | tropical | nocturnal | 2.941 | 1058929.165 | 2.364 |
| amphibians | Bolitoglossa_alvaradoi | Bolitoglossa_alvaradoi | VU | Decreasing | tropical | nocturnal | 7.84 | 2771.283 | 8.903 |
| amphibians | Bolitoglossa_aurae | NA | DD | Unknown | NA | NA | NA | NA | NA |
| amphibians | Bolitoglossa_biseriata | Bolitoglossa_biseriata | LC | Decreasing | tropical | nocturnal | 1.891 | 113714.41 | 5.286 |
| amphibians | Bolitoglossa_bramei | Bolitoglossa_bramei | LC | Stable | tropical | nocturnal | 1.14 | 238.869 | 2.753 |
| amphibians | Bolitoglossa_chica | Bolitoglossa_chica | CR | Decreasing | tropical | nocturnal | 1.084 | 11394.799 | 9.651 |
| amphibians | Bolitoglossa_coaxtlahuacana | NA | not_available | not_available | NA | NA | NA | NA | NA |

| | | | | | | | | | |
|---|---|---|---|---|---|---|---|---|---|
| amphibians | Bolitoglossa_colonnea | Bolitoglossa_colonnea | LC | Stable | tropical | nocturnal | 2.562 | 21089.341 | 9.236 |
| amphibians | Bolitoglossa_compacta | Bolitoglossa_compacta | EN | Decreasing | tropical | nocturnal | 6.52 | 562.39 | 5.796 |
| amphibians | Bolitoglossa_decora | Bolitoglossa_decora | CR | Decreasing | tropical | nocturnal | 3.863 | 54.586 | 2.431 |
| amphibians | Bolitoglossa_diaphora | Bolitoglossa_diaphora | EN | Decreasing | tropical | nocturnal | 2.28 | 80.9 | 9.213 |
| amphibians | Bolitoglossa_dunni | Bolitoglossa_dunni | EN | Decreasing | tropical | nocturnal | 6.265 | 277.022 | 9.248 |
| amphibians | Bolitoglossa_epimela | NA | DD | Unknown | NA | NA | NA | NA | NA |
| amphibians | Bolitoglossa_equatoriana | Bolitoglossa_equatoriana | LC | Unknown | tropical | nocturnal | 3.144 | 50146.926 | 4.135 |
| amphibians | Bolitoglossa_gomezi | Bolitoglossa_gomezi | EN | Decreasing | tropical | nocturnal | 2.632 | 74.321 | 10.798 |
| amphibians | Bolitoglossa_lignicolor | Bolitoglossa_lignicolor | LC | Decreasing | tropical | nocturnal | 8.499 | 19076.434 | 12.993 |
| amphibians | Bolitoglossa_longissima | Bolitoglossa_longissima | CR | Decreasing | tropical | nocturnal | 3.683 | 62.236 | 3.447 |
| amphibians | Bolitoglossa_medemi | Bolitoglossa_medemi | LC | Decreasing | tropical | cathemeral | 1.834 | 69439.377 | 4.297 |
| amphibians | Bolitoglossa_mexicana | Bolitoglossa_mexicana | LC | Decreasing | tropical | nocturnal | 8.748 | 205820.185 | 9.899 |
| amphibians | Bolitoglossa_minutula | Bolitoglossa_minutula | EN | Decreasing | tropical | nocturnal | 0.81 | 594.081 | 6.341 |
| amphibians | Bolitoglossa_muisca | NA | not_available | not_available | NA | NA | NA | NA | NA |
| amphibians | Bolitoglossa_nigrescens | NA | DD | Unknown | NA | NA | NA | NA | NA |
| amphibians | Bolitoglossa_occidentalis | Bolitoglossa_occidentalis | LC | Decreasing | tropical | nocturnal | 0.984 | 13229.503 | 8.734 |
| amphibians | Bolitoglossa_odonnelli | Bolitoglossa_odonnelli | NT | Unknown | tropical | nocturnal | 10.062 | 1600.969 | 12.931 |
| amphibians | Bolitoglossa_palmata | Bolitoglossa_palmata | LC | Unknown | tropical | nocturnal | 2.465 | 5733.576 | 3.377 |
| amphibians | Bolitoglossa_peruviana | NA | DD | Unknown | NA | NA | NA | NA | NA |
| amphibians | Bolitoglossa_pesrubra | Bolitoglossa_pesrubra | LC | Stable | tropical | nocturnal | 4.83 | 629.988 | 5.233 |
| amphibians | Bolitoglossa_platydactyla | Bolitoglossa_platydactyla | LC | Stable | tropical | nocturnal | 7.84 | 63430.276 | 15.435 |
| amphibians | Bolitoglossa_porrasorum | Bolitoglossa_porrasorum | EN | Decreasing | tropical | nocturnal | 4.559 | 301.907 | 4.415 |
| amphibians | Bolitoglossa_robusta | Bolitoglossa_robusta | VU | Decreasing | tropical | nocturnal | 36.66 | 7018.743 | 12.054 |
| amphibians | Bolitoglossa_rufescens | Bolitoglossa_rufescens | LC | Stable | tropical | nocturnal | 1.499 | 52145.183 | 11.36 |
| amphibians | Bolitoglossa_schizodactyla | Bolitoglossa_schizodactyla | LC | Decreasing | tropical | nocturnal | 3.845 | 14670.324 | 6.838 |
| amphibians | Bolitoglossa_sima | Bolitoglossa_sima | LC | Decreasing | tropical | nocturnal | 2.231 | 18232.435 | 8.651 |
| amphibians | Bolitoglossa_sooyorum | Bolitoglossa_sooyorum | EN | Decreasing | tropical | nocturnal | 5.968 | 60.705 | 7.918 |
| amphibians | Bolitoglossa_striatula | Bolitoglossa_striatula | LC | Decreasing | tropical | nocturnal | 4.6 | 122829.477 | 8.628 |
| amphibians | Bolitoglossa_stuarti | Bolitoglossa_stuarti | VU | Decreasing | tropical | nocturnal | 3.003 | 1839.889 | 11.128 |
| amphibians | Bolitoglossa_subpalmata | Bolitoglossa_subpalmata | LC | Stable | tropical | nocturnal | 3.492 | 1237.402 | 12.429 |
| amphibians | Bolitoglossa_synoria | Bolitoglossa_synoria | CR | Decreasing | tropical | nocturnal | 3.458 | 48.254 | 11.916 |
| amphibians | Bolitoglossa_yariguiensis | NA | EN | Decreasing | NA | NA | NA | NA | NA |
| amphibians | Bombina_bombina | Bombina_bombina | LC | Decreasing | temperate | diurnal | 19.923 | 3052238.788 | 17.422 |
| amphibians | Bombina_maxima | Bombina_maxima | LC | Stable | temperate | diurnal | 44.708 | 237704.912 | 12.447 |

| | | | | | | | | | |
|---|---|---|---|---|---|---|---|---|---|
| amphibians | Bombina_orientalis | Bombina_orientalis | LC | Decreasing | temperate | nocturnal | 25.822 | 942069.946 | 14.278 |
| amphibians | Bombina_variegata | Bombina_variegata | LC | Decreasing | temperate | diurnal | 15.932 | 1205594.997 | 20.19 |
| amphibians | Boophis_albilabris | Boophis_albilabris | LC | Decreasing | tropical | nocturnal | 94.785 | 82597.191 | 7 |
| amphibians | Boophis_albipunctatus | Boophis_albipunctatus | LC | Decreasing | tropical | nocturnal | 5.356 | 30209.27 | 7.175 |
| amphibians | Boophis_andohahela | Boophis_andohahela | VU | Decreasing | tropical | nocturnal | 2.109 | 7853.749 | 5.277 |
| amphibians | Boophis_andreonei | Boophis_andreonei | VU | Decreasing | tropical | nocturnal | 2.872 | 8190.078 | 4.924 |
| amphibians | Boophis_anjanaharibeensis | Boophis_anjanaharibeensis | EN | Decreasing | tropical | nocturnal | 13.658 | 2805.885 | 6.15 |
| amphibians | Boophis_ankaratra | Boophis_ankaratra | LC | Unknown | tropical | nocturnal | 1.686 | 31291.237 | 12.918 |
| amphibians | Boophis_axelmeyeri | Boophis_axelmeyeri | LC | Decreasing | tropical | nocturnal | 18.867 | 17940.21 | 5.678 |
| amphibians | Boophis_baetkei | Boophis_baetkei | CR | Decreasing | tropical | nocturnal | 2.296 | 30.354 | 6.482 |
| amphibians | Boophis_blommersae | Boophis_blommersae | VU | Decreasing | tropical | nocturnal | 1.499 | 2397.079 | 4.89 |
| amphibians | Boophis_boehmei | Boophis_boehmei | EN | Decreasing | tropical | nocturnal | 5.802 | 60222.881 | 8.487 |
| amphibians | Boophis_bottae | Boophis_bottae | LC | Decreasing | tropical | nocturnal | 3.507 | 17626.68 | 7.543 |
| amphibians | Boophis_brachychir | Boophis_brachychir | VU | Decreasing | tropical | nocturnal | 22.389 | 6080.003 | 5.729 |
| amphibians | Boophis_calcaratus | Boophis_calcaratus | LC | Decreasing | tropical | nocturnal | 5.356 | 38376.948 | 9.245 |
| amphibians | Boophis_doulioti | Boophis_doulioti | LC | Stable | tropical | nocturnal | 12.846 | 243894.944 | 8.564 |
| amphibians | Boophis_elenae | Boophis_elenae | NT | Unknown | tropical | nocturnal | 12.688 | 4512.279 | 8.278 |
| amphibians | Boophis_englaenderi | Boophis_englaenderi | VU | Decreasing | tropical | nocturnal | 8.724 | 5480.272 | 7.357 |
| amphibians | Boophis_entingae | Boophis_entingae | LC | Decreasing | tropical | nocturnal | 65.175 | 8087.146 | 6.678 |
| amphibians | Boophis_erythrodactylus | Boophis_erythrodactylus | LC | Decreasing | tropical | nocturnal | 2.872 | 14122.692 | 8.803 |
| amphibians | Boophis_feonnyala | Boophis_feonnyala | EN | Unknown | tropical | nocturnal | 1.168 | 2750.458 | 7.493 |
| amphibians | Boophis_goudotii | Boophis_goudotii | LC | Stable | tropical | nocturnal | 105.967 | 52329.98 | 11.074 |
| amphibians | Boophis_guibei | Boophis_guibei | LC | Decreasing | tropical | cathemeral | 5.356 | 12246.012 | 7.823 |
| amphibians | Boophis_idae | Boophis_idae | LC | Unknown | tropical | nocturnal | 3.807 | 32470.258 | 8.173 |
| amphibians | Boophis_jaegeri | Boophis_jaegeri | EN | Decreasing | tropical | nocturnal | 2.345 | 391.32 | 15.504 |
| amphibians | Boophis_liami | Boophis_liami | CR | Decreasing | tropical | nocturnal | 0.664 | 34.411 | 7.59 |
| amphibians | Boophis_lichenoides | Boophis_lichenoides | LC | Decreasing | tropical | nocturnal | 6.77 | 57689.12 | 7.052 |
| amphibians | Boophis_lilianae | NA | DD | Unknown | NA | NA | NA | NA | NA |
| amphibians | Boophis_luteus | Boophis_luteus | LC | Decreasing | tropical | nocturnal | 19.923 | 40589.22 | 7.862 |
| amphibians | Boophis_madagascariensis | Boophis_madagascariensis | LC | Decreasing | tropical | nocturnal | 114.748 | 84306.247 | 9.127 |
| amphibians | Boophis_marojezensis | Boophis_marojezensis | LC | Decreasing | tropical | nocturnal | 2.872 | 43462.675 | 7.003 |
| amphibians | Boophis_masoala | NA | not_available | not_available | NA | NA | NA | NA | NA |
| amphibians | Boophis_microtympanum | Boophis_microtympanum | LC | Decreasing | tropical | cathemeral | 6.273 | 30492.787 | 11.218 |
| amphibians | Boophis_miniatus | Boophis_miniatus | VU | Decreasing | tropical | nocturnal | 4.536 | 7853.749 | 5.277 |

| | | | | | | | | | |
|---|---|---|---|---|---|---|---|---|---|
| amphibians | Boophis_opisthodon | Boophis_opisthodon | LC | Stable | tropical | nocturnal | 61.584 | 41219.089 | 9.578 |
| amphibians | Boophis_picturatus | Boophis_picturatus | LC | Decreasing | tropical | nocturnal | 2.957 | 28021.676 | 7.503 |
| amphibians | Boophis_praedictus | Boophis_praedictus | LC | Decreasing | tropical | nocturnal | 72.262 | 6034.906 | 3.842 |
| amphibians | Boophis_pyrrhus | Boophis_pyrrhus | LC | Decreasing | tropical | nocturnal | 4.16 | 29571.016 | 7.949 |
| amphibians | Boophis_rappiodes | Boophis_rappiodes | LC | Decreasing | tropical | nocturnal | 3.163 | 31810.043 | 7.401 |
| amphibians | Boophis_reticulatus | Boophis_reticulatus | LC | Decreasing | tropical | nocturnal | 7.294 | 62357.647 | 7.689 |
| amphibians | Boophis_rhodoscelis | Boophis_rhodoscelis | EN | Decreasing | tropical | nocturnal | 4.088 | 19750.396 | 9.413 |
| amphibians | Boophis_roseipalmatus | Boophis_roseipalmatus | LC | Decreasing | tropical | nocturnal | 65.42 | 3429.056 | 4.472 |
| amphibians | Boophis_rufioculis | Boophis_rufioculis | NT | Decreasing | tropical | nocturnal | 9.031 | 15879.046 | 8.042 |
| amphibians | Boophis_sambirano | Boophis_sambirano | EN | Decreasing | tropical | nocturnal | 1.326 | 1201.713 | 6.427 |
| amphibians | Boophis_schuboeae | Boophis_schuboeae | EN | Decreasing | tropical | nocturnal | 1.326 | 405.287 | 5.909 |
| amphibians | Boophis_septentrionalis | Boophis_septentrionalis | LC | Decreasing | tropical | nocturnal | 12.531 | 18112.671 | 5.375 |
| amphibians | Boophis_sibilans | Boophis_sibilans | LC | Decreasing | tropical | nocturnal | 3.224 | 32359.643 | 5.369 |
| amphibians | Boophis_solomaso | Boophis_solomaso | EN | Decreasing | tropical | nocturnal | 0.772 | 71.148 | 8.09 |
| amphibians | Boophis_tampoka | Boophis_tampoka | LC | Decreasing | tropical | nocturnal | 8.423 | 42.882 | 4.621 |
| amphibians | Boophis_tasymena | Boophis_tasymena | LC | Decreasing | tropical | nocturnal | 2.599 | 60493.955 | 7.73 |
| amphibians | Boophis_tephraeomystax | Boophis_tephraeomystax | LC | Stable | tropical | nocturnal | 32.83 | 112253.606 | 8.344 |
| amphibians | Boophis_tsilomaro | Boophis_tsilomaro | CR | Decreasing | tropical | nocturnal | 61.584 | 89.932 | 6.347 |
| amphibians | Boophis_ulftunni | Boophis_ulftunni | VU | Decreasing | tropical | nocturnal | 4.536 | 2056.787 | 5.083 |
| amphibians | Boophis_viridis | Boophis_viridis | LC | Decreasing | tropical | nocturnal | 3.475 | 64241.127 | 7.696 |
| amphibians | Boophis_vittatus | Boophis_vittatus | VU | Decreasing | tropical | nocturnal | 1.23 | 11694.154 | 5.498 |
| amphibians | Boophis_williamsi | Boophis_williamsi | CR | Decreasing | tropical | nocturnal | 7.456 | 12.771 | 8.01 |
| amphibians | Boophis_xerophilus | Boophis_xerophilus | LC | Decreasing | tropical | nocturnal | 10.337 | 63546.651 | 10.416 |
| amphibians | Boreorana_sylvatica | Rana_sylvatica | LC | Stable | temperate | diurnal | 57.012 | 9135971.737 | 11.46 |
| amphibians | Boulengerula_taitana | Boulengerula_taitana | EN | Stable | tropical | cathemeral | 27.944 | 340.742 | 20.47 |
| amphibians | Boulenophrys_baolongensis | Xenophrys_baolongensis | CR | Decreasing | temperate | nocturnal | 7.844 | 10.879 | 12.448 |
| amphibians | Boulenophrys_binlingensis | Xenophrys_binlingensis | VU | Unknown | temperate | nocturnal | 11.767 | 35.053 | 40.541 |
| amphibians | Boulenophrys_boettgeri | Xenophrys_boettgeri | LC | Decreasing | temperate | nocturnal | 8.907 | 367630.191 | 5.193 |
| amphibians | Boulenophrys_caobangensis | NA | DD | Decreasing | NA | NA | NA | NA | NA |
| amphibians | Boulenophrys_fansipanensis | NA | EN | Decreasing | NA | NA | NA | NA | NA |
| amphibians | Boulenophrys_hoanglienensis | NA | EN | Decreasing | NA | NA | NA | NA | NA |
| amphibians | Boulenophrys_kuatunensis | Xenophrys_kuatunensis | LC | Decreasing | temperate | nocturnal | 4.308 | 231307.506 | 5.103 |
| amphibians | Boulenophrys_minor | NA | LC | Decreasing | NA | NA | NA | NA | NA |
| amphibians | Boulenophrys_ombrophila | NA | NT | Stable | NA | NA | NA | NA | NA |

| | | | | | | | | | |
|---|---|---|---|---|---|---|---|---|---|
| amphibians | Boulenophrys_omeimontis | NA | LC | Stable | NA | NA | NA | NA | NA |
| amphibians | Boulenophrys_palpebralespinosa | Xenophrys_palpebralespinosa | LC | Decreasing | tropical | cathemeral | 5.802 | 156994.733 | 9.408 |
| amphibians | Boulenophrys_parva | NA | DD | Unknown | NA | NA | NA | NA | NA |
| amphibians | Boulenophrys_shunhuangensis | NA | not_available | not_available | NA | NA | NA | NA | NA |
| amphibians | Boulenophrys_wuliangshanensis | NA | NT | Unknown | NA | NA | NA | NA | NA |
| amphibians | Brachycephalus_actaeus | NA | EN | Decreasing | NA | NA | NA | NA | NA |
| amphibians | Brachycephalus_alipioi | Brachycephalus_alipioi | EN | Decreasing | tropical | diurnal | 0.286 | 5754.455 | 12.188 |
| amphibians | Brachycephalus_brunneus | Brachycephalus_brunneus | VU | Stable | temperate | diurnal | 0.108 | 27.821 | 1.95 |
| amphibians | Brachycephalus_crispus | NA | VU | Stable | NA | NA | NA | NA | NA |
| amphibians | Brachycephalus_curupira | NA | EN | Decreasing | NA | NA | NA | NA | NA |
| amphibians | Brachycephalus_darkside | NA | EN | Decreasing | NA | NA | NA | NA | NA |
| amphibians | Brachycephalus_didactylus | Brachycephalus_didactylus | LC | Decreasing | tropical | diurnal | 0.075 | 12793.083 | 12.849 |
| amphibians | Brachycephalus_ephippium | Brachycephalus_ephippium | LC | Decreasing | tropical | diurnal | 0.567 | 57893.631 | 17.211 |
| amphibians | Brachycephalus_ferruginus | Brachycephalus_ferruginus | LC | Stable | temperate | diurnal | 0.2 | 113.606 | 5.038 |
| amphibians | Brachycephalus_garbeanus | Brachycephalus_garbeana | EN | Decreasing | tropical | diurnal | 0.375 | 912.898 | 10.5 |
| amphibians | Brachycephalus_guarani | Brachycephalus_guarani | LC | Stable | tropical | diurnal | 0.155 | 9.056 | 0.581 |
| amphibians | Brachycephalus_hermogenesi | Brachycephalus_hermogenesi | LC | Stable | temperate | diurnal | 0.223 | 6564.151 | 10.157 |
| amphibians | Brachycephalus_izecksohni | Brachycephalus_izecksohni | VU | Stable | temperate | diurnal | 0.144 | 34.877 | 2.264 |
| amphibians | Brachycephalus_leopardus | NA | VU | Stable | NA | NA | NA | NA | NA |
| amphibians | Brachycephalus_margaritatus | Brachycephalus_margaritatus | EN | Decreasing | tropical | diurnal | 0.472 | 1242.134 | 16.639 |
| amphibians | Brachycephalus_mirissimus | NA | CR | Decreasing | NA | NA | NA | NA | NA |
| amphibians | Brachycephalus_nodoterga | Brachycephalus_nodoterga | LC | Stable | temperate | diurnal | 0.155 | 2769.69 | 11.671 |
| amphibians | Brachycephalus_olivaceus | NA | EN | Decreasing | NA | NA | NA | NA | NA |
| amphibians | Brachycephalus_pernix | Brachycephalus_pernix | CR | Stable | temperate | diurnal | 0.264 | 0.469 | 8.17 |
| amphibians | Brachycephalus_pitanga | Brachycephalus_pitanga | LC | Stable | tropical | diurnal | 0.178 | 32.396 | 6.181 |
| amphibians | Brachycephalus_pombali | Brachycephalus_pombali | VU | Stable | temperate | diurnal | 0.238 | 38.239 | 1.149 |
| amphibians | Brachycephalus_sulfuratus | NA | NT | Decreasing | NA | NA | NA | NA | NA |
| amphibians | Brachycephalus_tabuleiro | NA | not_available | not_available | NA | NA | NA | NA | NA |
| amphibians | Brachycephalus_toby | Brachycephalus_toby | LC | Stable | temperate | diurnal | 0.187 | 7.164 | 2.281 |
| amphibians | Brachycephalus_tridactylus | Brachycephalus_tridactylus | EN | Decreasing | temperate | diurnal | 0.171 | 1031.999 | 3.432 |
| amphibians | Brachycephalus_vertebralis | Brachycephalus_vertebralis | EN | Decreasing | tropical | diurnal | 0.228 | 499.631 | 7.927 |
| amphibians | Brachytarsophrys_chuannanensis | NA | NT | Decreasing | NA | NA | NA | NA | NA |
| amphibians | Brachytarsophrys_platyparietus | Brachytarsophrys_carinense | LC | Decreasing | temperate | nocturnal | 559.955 | 941761.078 | 17.499 |
| amphibians | Breviceps_adspersus | Breviceps_adspersus | LC | Unknown | tropical | nocturnal | 19.923 | 2329054.014 | 8.241 |

| | | | | | | | | | |
|---|---|---|---|---|---|---|---|---|---|
| amphibians | Breviceps_bagginsi | Breviceps_bagginsi | NT | Unknown | temperate | diurnal | 1.889 | 2431.619 | 12.124 |
| amphibians | Breviceps_carruthersi | NA | not_available | not_available | NA | NA | NA | NA | NA |
| amphibians | Breviceps_fichus | Breviceps_fichus | LC | Unknown | tropical | diurnal | 7.187 | 17897.803 | 13.846 |
| amphibians | Breviceps_macrops | Breviceps_macrops | NT | Unknown | temperate | nocturnal | 14.161 | 5555.545 | 4.963 |
| amphibians | Breviceps_passmorei | NA | not_available | not_available | NA | NA | NA | NA | NA |
| amphibians | Breviceps_sopranus | Breviceps_sopranus | LC | Unknown | temperate | cathemeral | 1.36 | 12868.82 | 15.137 |
| amphibians | Bromeliohyla_bromeliacia | Bromeliohyla_bromeliacia | LC | Decreasing | tropical | nocturnal | 3.705 | 6976.633 | 12.552 |
| amphibians | Bryophryne_bakersfield | NA | LC | Stable | NA | NA | NA | NA | NA |
| amphibians | Buergeria_buergeri | Buergeria_buergeri | LC | Decreasing | temperate | cathemeral | 42.854 | 284874.192 | 21.11 |
| amphibians | Bufo_bankorensis | Bufo_bankorensis | LC | Decreasing | temperate | nocturnal | 152.716 | 21413.553 | 16.412 |
| amphibians | Bufo_bufo | Bufo_bufo | LC | Decreasing | temperate | nocturnal | 1341.531 | 9388939.921 | 12.716 |
| amphibians | Bufo_eichwaldi | Bufo_eichwaldi | VU | Decreasing | temperate | nocturnal | 47.005 | 18620.407 | 16.846 |
| amphibians | Bufo_gargarizans | Bufo_gargarizans | LC | Stable | temperate | cathemeral | 269.135 | 5233722.264 | 14.102 |
| amphibians | Bufo_praetextatus | NA | LC | Decreasing | NA | NA | NA | NA | NA |
| amphibians | Bufo_spinosus | Bufo_spinosus | LC | Decreasing | temperate | nocturnal | 1341.531 | 1155874.717 | 18.669 |
| amphibians | Bufo_stejnegeri | Bufo_stejnegeri | LC | Decreasing | temperate | nocturnal | 22.156 | 150839.178 | 14.166 |
| amphibians | Bufo_torrenticola | Bufo_torrenticola | LC | Stable | temperate | cathemeral | 559.955 | 21579.831 | 15.904 |
| amphibians | Bufo_tuberculatus | Bufo_tuberculatus | NT | Decreasing | temperate | nocturnal | 71.478 | 19308.474 | 6.806 |
| amphibians | Bufotes_boulengeri | Bufotes_boulengeri | LC | Unknown | temperate | nocturnal | 129.824 | 1966248.488 | 8.202 |
| amphibians | Bufotes_pewzowi | Bufotes_pewzowi | LC | Stable | temperate | nocturnal | 54.6 | 1286335.425 | 7.553 |
| amphibians | Bufotes_surdus | Bufotes_surdus | LC | Decreasing | temperate | nocturnal | 23.335 | 213988.755 | 6.059 |
| amphibians | Bufotes_viridis | Bufotes_viridis | LC | Decreasing | temperate | cathemeral | 310.174 | 10026528.871 | 16.587 |
| amphibians | Cacosternum_capense | Cacosternum_capense | NT | Decreasing | temperate | nocturnal | 4.934 | 6661.778 | 17.372 |
| amphibians | Cacosternum_namaquense | Cacosternum_namaquense | LC | Unknown | temperate | nocturnal | 1.868 | 45611.114 | 6.209 |
| amphibians | Cacosternum_thorini | NA | EN | Unknown | NA | NA | NA | NA | NA |
| amphibians | Caecilia_museugoeldi | NA | not_available | not_available | NA | NA | NA | NA | NA |
| amphibians | Caecilia_volcani | Caecilia_volcani | LC | Unknown | tropical | cathemeral | 17.105 | 8250.149 | 8.815 |
| amphibians | Callimedusa_atelopoides | Phyllomedusa_atelopoides | LC | Stable | tropical | nocturnal | 10.337 | 815329.15 | 0.902 |
| amphibians | Callimedusa_baltea | Phyllomedusa_baltea | EN | Unknown | tropical | nocturnal | 23.941 | 116.866 | 1.109 |
| amphibians | Callimedusa_duellmani | NA | DD | Unknown | NA | NA | NA | NA | NA |
| amphibians | Callimedusa_tomopterna | Phyllomedusa_tomopterna | LC | Stable | tropical | nocturnal | 53.102 | 5426896.299 | 1.71 |
| amphibians | Callixalus_pictus | Callixalus_pictus | VU | Decreasing | tropical | nocturnal | 6.77 | 6721.888 | 11.918 |
| amphibians | Callulina_dawida | Callulina_dawida | CR | Decreasing | tropical | nocturnal | 11.036 | 249.818 | 21.592 |
| amphibians | Callulina_laphami | Callulina_laphami | CR | Unknown | tropical | nocturnal | 7.844 | 21.985 | 16.791 |

| | | | | | | | | | |
|---|---|---|---|---|---|---|---|---|---|
| amphibians | Callulina_shengena | Callulina_shengena | CR | Unknown | tropical | nocturnal | 6.873 | 19.584 | 7.836 |
| amphibians | Callulina_stanleyi | Callulina_stanleyi | CR | Unknown | tropical | nocturnal | 6.77 | 6.92 | 13.637 |
| amphibians | Callulops_doriae | Callulops_doriae | LC | Stable | tropical | nocturnal | 104.268 | 75570.009 | 3.236 |
| amphibians | Callulops_stellatus | NA | LC | Stable | NA | NA | NA | NA | NA |
| amphibians | Callulops_wondiwoiensis | NA | LC | Unknown | NA | NA | NA | NA | NA |
| amphibians | Calotriton_arnoldi | Calotriton_arnoldi | CR | Decreasing | temperate | nocturnal | 3.315 | 289.548 | 14.37 |
| amphibians | Calotriton_asper | Calotriton_asper | LC | Decreasing | temperate | nocturnal | 9.722 | 27428.331 | 11.724 |
| amphibians | Calyptocephalella_gayi | Calyptocephalella_gayi | VU | Decreasing | temperate | cathemeral | 4516.866 | 137474.045 | 11.979 |
| amphibians | Cardioglossa_alsco | Cardioglossa_alsco | EN | Decreasing | tropical | nocturnal | 3.163 | 74.944 | 7.402 |
| amphibians | Cardioglossa_occidentalis | Cardioglossa_occidentalis | LC | Decreasing | tropical | nocturnal | 3.411 | 209290.825 | 14.473 |
| amphibians | Cardioglossa_oreas | Cardioglossa_oreas | EN | Unknown | tropical | diurnal | 2.394 | 1387.384 | 13.317 |
| amphibians | Cardioglossa_trifasciata | Cardioglossa_trifasciata | CR | Decreasing | tropical | diurnal | 5.313 | 14.429 | 13.067 |
| amphibians | Celsiella_revocata | Celsiella_revocata | VU | Decreasing | tropical | nocturnal | 1.199 | 684.415 | 17.319 |
| amphibians | Centrolene_ballux | Centrolene_ballux | EN | Decreasing | tropical | nocturnal | 0.93 | 481.542 | 6.181 |
| amphibians | Centrolene_buckleyi | Centrolene_buckleyi | CR | Decreasing | tropical | nocturnal | 3.379 | 74234 | 12.951 |
| amphibians | Centrolene_condor | Centrolene_condor | EN | Decreasing | tropical | nocturnal | 1.61 | 802.11 | 4.475 |
| amphibians | Centrolene_daidalea | Centrolene_daidaleum | LC | Decreasing | tropical | nocturnal | 1.245 | 5781.089 | 11.842 |
| amphibians | Centrolene_heloderma | Centrolene_heloderma | VU | Decreasing | tropical | cathemeral | 2.679 | 4737.584 | 12.61 |
| amphibians | Centrolene_hesperia | Centrolene_hesperium | EN | Decreasing | tropical | cathemeral | 1.847 | 31.114 | 10.424 |
| amphibians | Centrolene_huilensis | Centrolene_huilense | EN | Decreasing | tropical | nocturnal | 1.827 | 205.69 | 8.019 |
| amphibians | Centrolene_lemniscata | NA | DD | Unknown | NA | NA | NA | NA | NA |
| amphibians | Centrolene_lynchi | Centrolene_lynchi | EN | Decreasing | tropical | nocturnal | 1.394 | 672.681 | 7.211 |
| amphibians | Centrolene_muelleri | Centrolene_muelleri | DD | Decreasing | tropical | nocturnal | 0.956 | 140.046 | 6.798 |
| amphibians | Centrolene_notosticta | Centrolene_notostictum | LC | Stable | tropical | nocturnal | 1.153 | 5600.176 | 12.532 |
| amphibians | Centrolene_pipilata | Centrolene_pipilatum | CR | Decreasing | tropical | nocturnal | 0.904 | 35.328 | 8.405 |
| amphibians | Centrolene_sabini | Centrolene_sabini | VU | Unknown | tropical | nocturnal | 2.345 | 9.781 | 10.089 |
| amphibians | Centrolene_sanchezi | Centrolene_sanchezi | EN | Decreasing | tropical | nocturnal | 1.411 | 337.681 | 8.99 |
| amphibians | Centrolene_savagei | Centrolene_savagei | LC | Stable | tropical | nocturnal | 1.023 | 6382.967 | 17.349 |
| amphibians | Centrolene_venezuelense | Centrolene_venezuelense | LC | Decreasing | tropical | nocturnal | 3.103 | 7812.529 | 9.953 |
| amphibians | Centrolene_zarza | NA | not_available | not_available | NA | NA | NA | NA | NA |
| amphibians | Ceratophrys_aurita | Ceratophrys_aurita | LC | Unknown | tropical | nocturnal | 379.555 | 140904.158 | 11.221 |
| amphibians | Ceratophrys_calcarata | Ceratophrys_calcarata | LC | Stable | tropical | nocturnal | 74.113 | 134165.124 | 15.461 |
| amphibians | Ceratophrys_cornuta | Ceratophrys_cornuta | LC | Stable | tropical | cathemeral | 175.822 | 5479643.703 | 1.965 |
| amphibians | Ceratophrys_cranwelli | Ceratophrys_cranwelli | LC | Decreasing | temperate | nocturnal | 458.987 | 1124371.245 | 7.212 |

| | | | | | | | | | |
|---|---|---|---|---|---|---|---|---|---|
| amphibians | Ceratophrys_joazeirensis | Ceratophrys_joazeirensis | LC | Stable | tropical | nocturnal | 122.125 | 580713.411 | 8.807 |
| amphibians | Ceratophrys_ornata | Ceratophrys_ornata | NT | Decreasing | temperate | nocturnal | 528.201 | 513220.267 | 12.827 |
| amphibians | Ceratophrys_stolzmanni | Ceratophrys_stolzmanni | VU | Decreasing | tropical | nocturnal | 54.816 | 11075.087 | 12.391 |
| amphibians | Ceuthomantis_aracamuni | Ceuthomantis_aracamuni | VU | Unknown | tropical | diurnal | 0.418 | 3.748 | 0.03 |
| amphibians | Ceuthomantis_cavernibardus | Ceuthomantis_cavernibardus | LC | Unknown | tropical | diurnal | 2.599 | 16.586 | 0.757 |
| amphibians | Ceuthomantis_duellmani | Ceuthomantis_duellmani | NT | Unknown | tropical | diurnal | 1.168 | 1090.599 | 0.327 |
| amphibians | Ceuthomantis_smaragdinus | Ceuthomantis_smaragdinus | VU | Unknown | tropical | nocturnal | 0.567 | 28.443 | 1.591 |
| amphibians | Chacophrys_pierottii | Chacophrys_pierottii | LC | Decreasing | tropical | nocturnal | 15.029 | 349409.729 | 4.583 |
| amphibians | Chaperina_fusca | Chaperina_fusca | LC | Unknown | tropical | cathemeral | 1.326 | 169007.349 | 6.795 |
| amphibians | Charadrahyla_chaneque | Charadrahyla_chaneque | VU | Decreasing | tropical | nocturnal | 49.181 | 417.641 | 8.118 |
| amphibians | Charadrahyla_sakbah | NA | EN | Unknown | NA | NA | NA | NA | NA |
| amphibians | Charadrahyla_taeniopus | Charadrahyla_taeniopus | VU | Decreasing | tropical | nocturnal | 32.83 | 6562.931 | 17.318 |
| amphibians | Charadrahyla_tecuani | NA | DD | Unknown | NA | NA | NA | NA | NA |
| amphibians | Chiasmocleis_alagoana | Chiasmocleis_alagoanus | VU | Decreasing | tropical | diurnal | 1.648 | 18506.602 | 15.639 |
| amphibians | Chiasmocleis_albopunctata | Chiasmocleis_albopunctata | LC | Stable | tropical | nocturnal | 4.536 | 2319703.745 | 9.729 |
| amphibians | Chiasmocleis_anatipes | Chiasmocleis_anatipes | LC | Stable | tropical | nocturnal | 2.109 | 41482.562 | 3.649 |
| amphibians | Chiasmocleis_antenori | Syncope_antenori | LC | Stable | tropical | cathemeral | 0.17 | 609221.623 | 1.79 |
| amphibians | Chiasmocleis_atlantica | Chiasmocleis_atlantica | LC | Stable | tropical | nocturnal | 2.599 | 11602.141 | 21.492 |
| amphibians | Chiasmocleis_bassleri | Syncope_bassleri | LC | Stable | tropical | nocturnal | 2.086 | 3212224.954 | 1.678 |
| amphibians | Chiasmocleis_capixaba | Chiasmocleis_capixaba | LC | Stable | tropical | nocturnal | 0.585 | 28751.472 | 11.486 |
| amphibians | Chiasmocleis_carvalhoi | Chiasmocleis_carvalhoi | LC | Stable | tropical | nocturnal | 1.08 | 122088.123 | 1.663 |
| amphibians | Chiasmocleis_crucis | Chiasmocleis_crucis | LC | Stable | tropical | nocturnal | 0.624 | 48154.688 | 9.436 |
| amphibians | Chiasmocleis_gnoma | Chiasmocleis_gnoma | LC | Stable | tropical | nocturnal | 0.275 | 52083.544 | 8.473 |
| amphibians | Chiasmocleis_hudsoni | Syncope_hudsoni | LC | Stable | tropical | nocturnal | 1.766 | 2530095.986 | 1.075 |
| amphibians | Chiasmocleis_leucosticta | Chiasmocleis_leucosticta | LC | Stable | temperate | cathemeral | 1.326 | 167239.024 | 11.412 |
| amphibians | Chiasmocleis_mantiqueira | Chiasmocleis_mantiqueira | LC | Stable | tropical | nocturnal | 0.867 | 20325.947 | 10.129 |
| amphibians | Chiasmocleis_parkeri | NA | EN | Decreasing | NA | NA | NA | NA | NA |
| amphibians | Chiasmocleis_schubarti | Chiasmocleis_schubarti | LC | Unknown | tropical | nocturnal | 3.317 | 243150.516 | 9.712 |
| amphibians | Chiasmocleis_shudikarensis | Chiasmocleis_shudikarensis | LC | Stable | tropical | cathemeral | 2.599 | 2076572.054 | 1.077 |
| amphibians | Chiasmocleis_supercilialba | Syncope_supercilialba | LC | Unknown | tropical | nocturnal | 0.456 | 154803.93 | 2.532 |
| amphibians | Chiasmocleis_ventrimaculata | Chiasmocleis_ventrimaculata | LC | Stable | tropical | nocturnal | 3.163 | 1805618.132 | 1.366 |
| amphibians | Chimerella_mariaelenae | Chimerella_mariaelenae | LC | Unknown | tropical | nocturnal | 0.867 | 41571.78 | 4.516 |
| amphibians | Chioglossa_lusitanica | Chioglossa_lusitanica | NT | Decreasing | temperate | nocturnal | 2.14 | 54174.606 | 19.485 |
| amphibians | Chirixalus_dudhwaensis | Chiromantis_dudhwaensis | VU | Decreasing | temperate | nocturnal | 5.356 | 47.835 | 25.726 |

| | | | | | | | | | |
|---|---|---|---|---|---|---|---|---|---|
| amphibians | Chirixalus_nongkhorensis | Chiromantis_nongkhorensis | LC | Decreasing | tropical | cathemeral | 3.807 | 1344686.889 | 10.762 |
| amphibians | Chirixalus_simus | Chiromantis_simus | LC | Stable | temperate | nocturnal | 0.772 | 169100.981 | 18.215 |
| amphibians | Chiromantis_petersii | Chiromantis_petersii | LC | Unknown | tropical | nocturnal | 25.822 | 732830.404 | 11.282 |
| amphibians | Chiromantis_xerampelina | Chiromantis_xerampelina | LC | Unknown | tropical | nocturnal | 74.113 | 2783181.512 | 9.08 |
| amphibians | Chiropterotriton_perotensis | NA | CR | Decreasing | NA | NA | NA | NA | NA |
| amphibians | Choerophryne_alainduboisi | NA | LC | Stable | NA | NA | NA | NA | NA |
| amphibians | Choerophryne_arndtorum | NA | DD | Unknown | NA | NA | NA | NA | NA |
| amphibians | Choerophryne_bisyllaba | NA | LC | Stable | NA | NA | NA | NA | NA |
| amphibians | Choerophryne_burtoni | Choerophryne_burtoni | LC | Stable | tropical | nocturnal | 0.12 | 2225.153 | 7.712 |
| amphibians | Choerophryne_crucifer | NA | LC | Stable | NA | NA | NA | NA | NA |
| amphibians | Choerophryne_epirrhina | NA | LC | Unknown | NA | NA | NA | NA | NA |
| amphibians | Choerophryne_grylloides | NA | LC | Unknown | NA | NA | NA | NA | NA |
| amphibians | Choerophryne_microps | Choerophryne_microps | LC | Unknown | tropical | nocturnal | 0.567 | 9.893 | 1.579 |
| amphibians | Choerophryne_pipiens | NA | LC | Unknown | NA | NA | NA | NA | NA |
| amphibians | Choerophryne_siegfriedi | Albericus_siegfriedi | CR | Decreasing | tropical | nocturnal | 0.695 | 17.207 | 8.743 |
| amphibians | Chthonerpeton_noctinectes | Chthonerpeton_noctinectes | LC | Unknown | tropical | nocturnal | 15.427 | 2598.912 | 21.615 |
| amphibians | Clinotarsus_curtipes | Clinotarsus_curtipes | LC | Decreasing | tropical | nocturnal | 50.601 | 121934.475 | 18.367 |
| amphibians | Cochranella_euknemos | Cochranella_euknemos | LC | Decreasing | tropical | nocturnal | 2.599 | 25146.668 | 4.871 |
| amphibians | Cochranella_granulosa | Cochranella_granulosa | LC | Decreasing | tropical | nocturnal | 2.599 | 94988.315 | 10.385 |
| amphibians | Cochranella_mache | Cochranella_mache | NT | Decreasing | tropical | nocturnal | 2.986 | 29709.145 | 6.751 |
| amphibians | Colostethus_agilis | Colostethus_agilis | EN | Unknown | tropical | diurnal | 1.686 | 317.148 | 15.016 |
| amphibians | Colostethus_inguinalis | Colostethus_inguinalis | LC | Decreasing | tropical | diurnal | 1.61 | 25223.216 | 7.492 |
| amphibians | Colostethus_jacobuspetersi | Colostethus_jacobuspetersi | CR | Decreasing | tropical | diurnal | 1.997 | 19838.782 | 15.049 |
| amphibians | Colostethus_latinasus | Colostethus_latinasus | CR | Decreasing | tropical | diurnal | 3.163 | 281.082 | 2.364 |
| amphibians | Colostethus_panamansis | Colostethus_panamansis | LC | Decreasing | tropical | diurnal | 1.932 | 28472.762 | 4.956 |
| amphibians | Colostethus_pratti | Colostethus_pratti | LC | Decreasing | tropical | diurnal | 1.168 | 40070.037 | 5.644 |
| amphibians | Colostethus_ucumari | Colostethus_ucumari | EN | Unknown | tropical | cathemeral | 2.109 | 451.265 | 12.337 |
| amphibians | Conraua_alleni | Conraua_alleni | LC | Decreasing | tropical | cathemeral | 28.486 | 156455.676 | 13.586 |
| amphibians | Conraua_beccarii | Conraua_beccarii | LC | Unknown | tropical | cathemeral | 806.163 | 289875.467 | 13.208 |
| amphibians | Conraua_crassipes | Conraua_crassipes | LC | Decreasing | tropical | cathemeral | 52.68 | 557445.377 | 7.154 |
| amphibians | Conraua_derooi | Conraua_derooi | CR | Decreasing | tropical | cathemeral | 63.963 | 413.319 | 18.236 |
| amphibians | Conraua_goliath | Conraua_goliath | EN | Decreasing | tropical | cathemeral | 5497.218 | 69715.976 | 9.437 |
| amphibians | Conraua_robusta | Conraua_robusta | VU | Decreasing | tropical | cathemeral | 806.163 | 12792.211 | 10.383 |
| amphibians | Cophixalus_aenigma | Cophixalus_aenigma | EN | Stable | tropical | nocturnal | 0.892 | 360.474 | 0.877 |

| | | | | | | | | | |
|---|---|---|---|---|---|---|---|---|---|
| amphibians | Cophixalus_australis | Cophixalus_australis | LC | Stable | tropical | nocturnal | 1.889 | 5598.8 | 4.874 |
| amphibians | Cophixalus_balbus | Cophixalus_balbus | LC | Decreasing | tropical | nocturnal | 2.296 | 25794.32 | 2.576 |
| amphibians | Cophixalus_bombiens | Cophixalus_bombiens | LC | Stable | tropical | nocturnal | 0.335 | 2471.098 | 2.415 |
| amphibians | Cophixalus_caverniphilus | Cophixalus_caverniphilus | LC | Unknown | tropical | diurnal | 4.052 | 9.79 | 9.825 |
| amphibians | Cophixalus_cheesmanae | Cophixalus_cheesmanae | LC | Stable | tropical | nocturnal | 4.934 | 101120.247 | 4.226 |
| amphibians | Cophixalus_concinnus | Cophixalus_concinnus | CR | Stable | tropical | nocturnal | 1.411 | 11.834 | 2.841 |
| amphibians | Cophixalus_cryptotympanum | Cophixalus_cryptotympanum | LC | Stable | tropical | nocturnal | 1.766 | 992.876 | 2.765 |
| amphibians | Cophixalus_cupricarenus | Cophixalus_cupricarenus | LC | Unknown | tropical | nocturnal | 3.255 | 294.535 | 3.394 |
| amphibians | Cophixalus_desticans | Cophixalus_desticans | LC | Stable | tropical | nocturnal | 0.488 | 7230.22 | 5.114 |
| amphibians | Cophixalus_exiguus | Cophixalus_exiguus | LC | Stable | tropical | nocturnal | 0.48 | 99.512 | 2.794 |
| amphibians | Cophixalus_hannahae | NA | LC | Stable | NA | NA | NA | NA | NA |
| amphibians | Cophixalus_hosmeri | Cophixalus_hosmeri | EN | Stable | tropical | nocturnal | 0.335 | 224.646 | 0.492 |
| amphibians | Cophixalus_infacetus | Cophixalus_infacetus | LC | Stable | tropical | nocturnal | 0.403 | 4731.32 | 5.128 |
| amphibians | Cophixalus_kaindiensis | Cophixalus_kaindiensis | NT | Stable | tropical | nocturnal | 1.628 | 19.399 | 5.556 |
| amphibians | Cophixalus_mcdonaldi | Cophixalus_mcdonaldi | CR | Stable | tropical | nocturnal | 1.326 | 11.621 | 0.688 |
| amphibians | Cophixalus_misimae | Cophixalus_misimae | CR | Decreasing | tropical | nocturnal | 0.531 | 212.672 | 6.624 |
| amphibians | Cophixalus_monticola | Cophixalus_monticola | CR | Stable | tropical | nocturnal | 1.023 | 11.809 | 0.84 |
| amphibians | Cophixalus_neglectus | Cophixalus_neglectus | CR | Decreasing | tropical | nocturnal | 1.889 | 66.522 | 0.599 |
| amphibians | Cophixalus_ornatus | Cophixalus_ornatus | LC | Stable | tropical | nocturnal | 1.889 | 2804.621 | 6.709 |
| amphibians | Cophixalus_parkeri | Cophixalus_parkeri | LC | Stable | tropical | nocturnal | 2.9 | 14025.52 | 8.51 |
| amphibians | Cophixalus_rajampatensis | NA | DD | Unknown | NA | NA | NA | NA | NA |
| amphibians | Cophixalus_riparius | Cophixalus_riparius | LC | Stable | tropical | nocturnal | 11.036 | 4293.381 | 8.837 |
| amphibians | Cophixalus_salawatiensis | NA | DD | Unknown | NA | NA | NA | NA | NA |
| amphibians | Cophixalus_sphagnicola | Cophixalus_sphagnicola | EN | Decreasing | tropical | diurnal | 0.624 | 496.648 | 5.324 |
| amphibians | Cophixalus_tetzlaffi | NA | DD | Unknown | NA | NA | NA | NA | NA |
| amphibians | Cophixalus_timidus | Cophixalus_timidus | CR | Unknown | tropical | cathemeral | 0.674 | 5.055 | 3.962 |
| amphibians | Cophixalus_variabilis | Cophixalus_variabilis | LC | Stable | tropical | cathemeral | 0.54 | 21134.765 | 4.457 |
| amphibians | Cophixalus_verecundus | Cophixalus_verecundus | LC | Unknown | tropical | nocturnal | 0.48 | 145.51 | 4.088 |
| amphibians | Cophyla_berara | Cophyla_berara | EN | Decreasing | tropical | nocturnal | 1.326 | 44.989 | 6.193 |
| amphibians | Cophyla_fortuna | NA | not_available | not_available | NA | NA | NA | NA | NA |
| amphibians | Cophyla_maharipeo | NA | CR | Decreasing | NA | NA | NA | NA | NA |
| amphibians | Cophyla_occultans | Cophyla_occultans | VU | Decreasing | tropical | nocturnal | 0.664 | 3646.307 | 9.561 |
| amphibians | Cophyla_phyllodactyla | Cophyla_phyllodactyla | LC | Stable | tropical | nocturnal | 2.109 | 9136.551 | 8.497 |
| amphibians | Cophyla_puellarum | NA | CR | Decreasing | NA | NA | NA | NA | NA |

| | | | | | | | | | |
|---|---|---|---|---|---|---|---|---|---|
| amphibians | Copiula_annanoreenae | NA | LC | Stable | NA | NA | NA | NA | NA |
| amphibians | Copiula_bisyllaba | NA | not_available | not_available | NA | NA | NA | NA | NA |
| amphibians | Copiula_derongo | NA | LC | Stable | NA | NA | NA | NA | NA |
| amphibians | Copiula_guttata | Austrochaperina_guttata | LC | Stable | tropical | diurnal | 7.294 | 22479.91 | 2.575 |
| amphibians | Copiula_lennarti | NA | LC | Stable | NA | NA | NA | NA | NA |
| amphibians | Copiula_mosbyae | NA | not_available | not_available | NA | NA | NA | NA | NA |
| amphibians | Cornufer_akarithymus | Platymantis_akarithymus | LC | Decreasing | tropical | diurnal | 1.051 | 16228.942 | 5.02 |
| amphibians | Cornufer_browni | Platymantis_browni | LC | Stable | tropical | nocturnal | 1.446 | 903.894 | 2.998 |
| amphibians | Cornufer_exedrus | NA | LC | Stable | NA | NA | NA | NA | NA |
| amphibians | Cornufer_latro | Platymantis_latro | LC | Stable | tropical | nocturnal | 4.536 | 1985.564 | 6.343 |
| amphibians | Cornufer_magnus | Platymantis_magnus | LC | Stable | tropical | nocturnal | 23.335 | 7148.564 | 5.371 |
| amphibians | Cornufer_nakanaiorum | Platymantis_nakanaiorum | LC | Unknown | tropical | nocturnal | 4.536 | 3185.645 | 2.887 |
| amphibians | Cornufer_nexipus | Platymantis_nexipus | VU | Decreasing | tropical | nocturnal | 7.134 | 22622.32 | 3.348 |
| amphibians | Cornufer_paepkei | NA | DD | Unknown | NA | NA | NA | NA | NA |
| amphibians | Cornufer_pelewensis | Platymantis_pelewensis | LC | Stable | tropical | diurnal | 19.923 | 467.034 | 11.17 |
| amphibians | Cornufer_vitianus | Platymantis_vitianus | NT | Decreasing | tropical | cathemeral | 141.992 | 9910.783 | 8.511 |
| amphibians | Corythomantis_greeningi | Corythomantis_greeningi | LC | Stable | tropical | nocturnal | 66.404 | 1230069.466 | 8.117 |
| amphibians | Craugastor_alfredi | Craugastor_alfredi | LC | Decreasing | tropical | nocturnal | 7.844 | 53107.948 | 11.078 |
| amphibians | Craugastor_anciano | NA | EX | not_available | NA | NA | NA | NA | NA |
| amphibians | Craugastor_andi | Craugastor_andi | CR | Decreasing | tropical | nocturnal | 50.601 | 1988.722 | 14.343 |
| amphibians | Craugastor_angelicus | Craugastor_angelicus | CR | Decreasing | tropical | nocturnal | 41.053 | 1469.47 | 11.383 |
| amphibians | Craugastor_augusti | Craugastor_augusti | LC | Stable | temperate | nocturnal | 88.303 | 934920.214 | 10.619 |
| amphibians | Craugastor_aurilegulus | Craugastor_aurilegulus | VU | Decreasing | tropical | cathemeral | 51.633 | 1673.911 | 6.61 |
| amphibians | Craugastor_blairi | NA | EN | Decreasing | NA | NA | NA | NA | NA |
| amphibians | Craugastor_bransfordii | Craugastor_bransfordii | LC | Decreasing | tropical | cathemeral | 2.109 | 62031.136 | 9.687 |
| amphibians | Craugastor_chac | Craugastor_chac | LC | Decreasing | tropical | cathemeral | 4.934 | 11216.965 | 9.641 |
| amphibians | Craugastor_charadra | Craugastor_charadra | VU | Unknown | tropical | nocturnal | 28.486 | 3108.629 | 11.234 |
| amphibians | Craugastor_chrysozetetes | Craugastor_chrysozetetes | CR | Decreasing | tropical | cathemeral | 8.188 | 15.04 | 3.213 |
| amphibians | Craugastor_crassidigitus | Craugastor_crassidigitus | LC | Decreasing | tropical | cathemeral | 9.669 | 84725.121 | 8.803 |
| amphibians | Craugastor_cruzi | Craugastor_cruzi | CR | Decreasing | tropical | diurnal | 2.928 | 13.653 | 2.329 |
| amphibians | Craugastor_cuaquero | NA | DD | Unknown | NA | NA | NA | NA | NA |
| amphibians | Craugastor_emcelae | Craugastor_emcelae | CR | Decreasing | tropical | nocturnal | 33.596 | 1705.079 | 4.099 |
| amphibians | Craugastor_emleni | Craugastor_emleni | EN | Decreasing | tropical | cathemeral | 10.894 | 391.529 | 14.607 |
| amphibians | Craugastor_epochthidius | Craugastor_epochthidius | CR | Decreasing | tropical | cathemeral | 4.052 | 2095.963 | 3.236 |

| | | | | | | | | | |
|---|---|---|---|---|---|---|---|---|---|
| amphibians | Craugastor_escoces | Craugastor_escoces | CR | Decreasing | tropical | nocturnal | 35.967 | 586.854 | 11.274 |
| amphibians | Craugastor_fecundus | Craugastor_fecundus | CR | Decreasing | tropical | cathemeral | 4.271 | 128.047 | 5.338 |
| amphibians | Craugastor_fitzingeri | Craugastor_fitzingeri | LC | Stable | tropical | cathemeral | 13.329 | 282360.7 | 8.411 |
| amphibians | Craugastor_fleischmanni | Craugastor_fleischmanni | CR | Decreasing | tropical | nocturnal | 41.053 | 2626.052 | 21.553 |
| amphibians | Craugastor_gabbi | NA | LC | Stable | NA | NA | NA | NA | NA |
| amphibians | Craugastor_gollmeri | Craugastor_gollmeri | LC | Stable | tropical | diurnal | 14.161 | 32284.478 | 9.765 |
| amphibians | Craugastor_gutschei | NA | EN | Decreasing | NA | NA | NA | NA | NA |
| amphibians | Craugastor_laevissimus | Craugastor_laevissimus | EN | Decreasing | tropical | cathemeral | 20.908 | 36504.736 | 10.039 |
| amphibians | Craugastor_laticeps | Craugastor_laticeps | LC | Unknown | tropical | cathemeral | 50.601 | 19121.913 | 9.664 |
| amphibians | Craugastor_lauraster | Craugastor_lauraster | LC | Decreasing | tropical | cathemeral | 0.807 | 40686.568 | 5.525 |
| amphibians | Craugastor_longirostris | Craugastor_longirostris | LC | Stable | tropical | nocturnal | 19.496 | 111361.033 | 6.268 |
| amphibians | Craugastor_megacephalus | Craugastor_megacephalus | LC | Stable | tropical | nocturnal | 32.83 | 99890.717 | 8.417 |
| amphibians | Craugastor_melanostictus | Craugastor_melanostictus | LC | Unknown | tropical | nocturnal | 16.873 | 11076.846 | 6.949 |
| amphibians | Craugastor_merendonensis | Craugastor_merendonensis | CR | Unknown | tropical | nocturnal | 57.012 | 1.346 | 25.077 |
| amphibians | Craugastor_milesi | Craugastor_milesi | CR | Unknown | tropical | cathemeral | 4.197 | 87.832 | 8.134 |
| amphibians | Craugastor_mimus | Craugastor_mimus | LC | Unknown | tropical | cathemeral | 17.851 | 39493.603 | 8.225 |
| amphibians | Craugastor_noblei | Craugastor_noblei | LC | Unknown | tropical | cathemeral | 27.131 | 151238.919 | 8.009 |
| amphibians | Craugastor_obesus | Craugastor_obesus | CR | Unknown | tropical | nocturnal | 64.688 | 3447.634 | 5.202 |
| amphibians | Craugastor_occidentalis | Craugastor_occidentalis | LC | Stable | tropical | diurnal | 1.686 | 126686.226 | 9.726 |
| amphibians | Craugastor_olanchano | Craugastor_olanchano | CR | Decreasing | tropical | cathemeral | 2.064 | 597.719 | 5.312 |
| amphibians | Craugastor_omoaensis | NA | EX | not_available | NA | NA | NA | NA | NA |
| amphibians | Craugastor_pechorum | Craugastor_pechorum | EN | Decreasing | tropical | nocturnal | 22.272 | 4693.026 | 1.688 |
| amphibians | Craugastor_persimilis | Craugastor_persimilis | LC | Unknown | tropical | diurnal | 0.892 | 9670.587 | 14.413 |
| amphibians | Craugastor_podiciferus | Craugastor_podiciferus | LC | Stable | tropical | diurnal | 5.356 | 9474.103 | 10.05 |
| amphibians | Craugastor_polyptychus | Craugastor_polyptychus | LC | Stable | tropical | diurnal | 1.827 | 4777.937 | 9.73 |
| amphibians | Craugastor_psephosypharus | Craugastor_psephosypharus | NT | Decreasing | tropical | nocturnal | 26.21 | 5869.668 | 6.039 |
| amphibians | Craugastor_punctariolus | Craugastor_punctariolus | CR | Decreasing | tropical | nocturnal | 52.68 | 3793.471 | 7.513 |
| amphibians | Craugastor_pygmaeus | Craugastor_pygmaeus | LC | Unknown | tropical | diurnal | 0.403 | 49839.61 | 10.508 |
| amphibians | Craugastor_ranoides | Craugastor_ranoides | CR | Decreasing | tropical | nocturnal | 39.306 | 52873.425 | 11.322 |
| amphibians | Craugastor_rayo | Craugastor_rayo | EN | Decreasing | tropical | diurnal | 34.374 | 124.071 | 8.49 |
| amphibians | Craugastor_rhodopis | Craugastor_rhodopis | LC | Stable | tropical | cathemeral | 8.247 | 29284.826 | 14.69 |
| amphibians | Craugastor_rostralis | Craugastor_rostralis | VU | Decreasing | tropical | cathemeral | 18.867 | 1639.556 | 11.109 |
| amphibians | Craugastor_rugulosus | NA | LC | Unknown | NA | NA | NA | NA | NA |
| amphibians | Craugastor_saltuarius | Craugastor_saltuarius | CR | Decreasing | tropical | diurnal | 0.818 | 615.91 | 2.841 |

| | | | | | | | | | |
|---|---|---|---|---|---|---|---|---|---|
| amphibians | Craugastor_sandersoni | Craugastor_sandersoni | EN | Decreasing | tropical | nocturnal | 62.055 | 7194.886 | 8.282 |
| amphibians | Craugastor_stadelmani | Craugastor_stadelmani | CR | Decreasing | tropical | cathemeral | 9.283 | 762.318 | 4.483 |
| amphibians | Craugastor_stejnegerianus | Craugastor_stejnegerianus | LC | Stable | tropical | diurnal | 0.818 | 18354.381 | 16.258 |
| amphibians | Craugastor_talamancae | Craugastor_talamancae | LC | Decreasing | tropical | nocturnal | 11.036 | 49503.822 | 8.645 |
| amphibians | Craugastor_taurus | Craugastor_taurus | EN | Decreasing | tropical | nocturnal | 50.601 | 5821.841 | 13.153 |
| amphibians | Craugastor_underwoodi | Craugastor_underwoodi | LC | Unknown | tropical | diurnal | 2.109 | 5425.209 | 15.787 |
| amphibians | Craugastor_yucatanensis | Craugastor_yucatanensis | NT | Decreasing | tropical | nocturnal | 3.807 | 24234.273 | 6.308 |
| amphibians | Crinia_fimbriata | Crinia_fimbriata | LC | Stable | tropical | nocturnal | 0.403 | 7866.803 | 1.353 |
| amphibians | Crinia_georgiana | Crinia_georgiana | LC | Stable | temperate | nocturnal | 7.844 | 115254.344 | 9.92 |
| amphibians | Crinia_parinsignifera | Crinia_parinsignifera | LC | Stable | temperate | cathemeral | 0.892 | 1121430.763 | 8.922 |
| amphibians | Crinia_signifera | Crinia_signifera | LC | Stable | temperate | nocturnal | 3.475 | 1079554.515 | 9.741 |
| amphibians | Crossodactylodes_bokermanni | Crossodactylodes_bokermanni | LC | Stable | tropical | nocturnal | 0.335 | 6437.665 | 12.151 |
| amphibians | Crossodactylodes_izecksohni | Crossodactylodes_izecksohni | EN | Decreasing | tropical | nocturnal | 0.223 | 96.482 | 10.917 |
| amphibians | Crossodactylus_caramaschii | Crossodactylus_caramaschii | LC | Stable | temperate | diurnal | 1.535 | 110395.031 | 13.728 |
| amphibians | Crossodactylus_cyclospinus | Crossodactylus_cyclospinus | EN | Decreasing | tropical | diurnal | 1.094 | 187.31 | 4.14 |
| amphibians | Crossodactylus_dantei | Crossodactylus_dantei | CR | Stable | tropical | diurnal | 2.109 | 162.013 | 7.226 |
| amphibians | Crossodactylus_dispar | Crossodactylus_dispar | CR | Unknown | tropical | diurnal | 2.345 | 879.774 | 6.749 |
| amphibians | Crossodactylus_gaudichaudii | Crossodactylus_gaudichaudii | LC | Stable | tropical | cathemeral | 3.507 | 49389.165 | 15.059 |
| amphibians | Crossodactylus_grandis | Crossodactylus_grandis | CR | Unknown | tropical | diurnal | 6.273 | 3968.046 | 9.713 |
| amphibians | Crossodactylus_lutzorum | Crossodactylus_lutzorum | LC | Unknown | tropical | diurnal | 1.394 | 2234.926 | 8.023 |
| amphibians | Crossodactylus_schmidti | Crossodactylus_schmidti | LC | Decreasing | temperate | cathemeral | 2.957 | 44947.229 | 11.409 |
| amphibians | Crossodactylus_trachystomus | Crossodactylus_trachystomus | NT | Decreasing | tropical | diurnal | 1.499 | 13617.37 | 11.233 |
| amphibians | Cruziohyla_calcarifer | Cruziohyla_calcarifer | LC | Decreasing | tropical | cathemeral | 70.959 | 27099.783 | 5.099 |
| amphibians | Cruziohyla_craspedopus | Cruziohyla_craspedopus | LC | Unknown | tropical | nocturnal | 41.947 | 2096426.802 | 1.243 |
| amphibians | Cryptobatrachus_pedroruizi | Cryptobatrachus_pedroruizi | EN | Decreasing | tropical | nocturnal | 41.053 | 4803.16 | 8.317 |
| amphibians | Cryptobranchus_alleganiensis | Cryptobranchus_alleganiensis | VU | Decreasing | temperate | nocturnal | 923.27 | 435321.18 | 17.033 |
| amphibians | Cryptothylax_greshoffii | Cryptothylax_greshoffii | LC | Unknown | tropical | nocturnal | 19.923 | 1418395.847 | 5.79 |
| amphibians | Ctenophryne_aterrima | Ctenophryne_aterrima | LC | Decreasing | tropical | nocturnal | 32.83 | 153830.15 | 11.116 |
| amphibians | Ctenophryne_geayi | Ctenophryne_geayi | LC | Stable | tropical | nocturnal | 15.118 | 6199340.374 | 1.979 |
| amphibians | Cycloramphus_acangatan | Cycloramphus_acangatan | LC | Stable | temperate | nocturnal | 9.734 | 20804.336 | 12.062 |
| amphibians | Cycloramphus_bolitoglossus | Cycloramphus_bolitoglossus | LC | Decreasing | temperate | nocturnal | 5.27 | 17171.693 | 15.035 |
| amphibians | Cycloramphus_boraceiensis | Cycloramphus_boraceiensis | LC | Stable | tropical | nocturnal | 18.867 | 6189.129 | 9.597 |
| amphibians | Cycloramphus_brasiliensis | Cycloramphus_brasiliensis | EN | Decreasing | tropical | nocturnal | 46.23 | 3929.111 | 13.48 |
| amphibians | Cycloramphus_carvalhoi | Cycloramphus_carvalhoi | LC | Decreasing | tropical | cathemeral | 22.156 | 23506.041 | 12.819 |

| | | | | | | | | | |
|---|---|---|---|---|---|---|---|---|---|
| amphibians | Cycloramphus_catarinensis | Cycloramphus_catarinensis | CR | Unknown | temperate | nocturnal | 5.016 | 6.325 | 25.552 |
| amphibians | Cycloramphus_dubius | Cycloramphus_dubius | LC | Stable | temperate | nocturnal | 20.248 | 4376.418 | 16.51 |
| amphibians | Cycloramphus_eleutherodactylus | Cycloramphus_eleutherodactylus | LC | Stable | temperate | cathemeral | 18.152 | 110566.144 | 14.973 |
| amphibians | Cycloramphus_faustoi | NA | VU | Stable | NA | NA | NA | NA | NA |
| amphibians | Cycloramphus_fuliginosus | Cycloramphus_fuliginosus | LC | Stable | tropical | nocturnal | 24.932 | 88661.057 | 12.805 |
| amphibians | Cycloramphus_izecksohni | Cycloramphus_izecksohni | EN | Decreasing | temperate | nocturnal | 7.294 | 4162.782 | 15.861 |
| amphibians | Cycloramphus_juimirim | Cycloramphus_juimirim | LC | Unknown | temperate | nocturnal | 4.812 | 1434.784 | 4.908 |
| amphibians | Cycloramphus_lithomimeticus | Cycloramphus_lithomimeticus | NT | Stable | tropical | nocturnal | 2.345 | 1.892 | 18.457 |
| amphibians | Cycloramphus_lutzorum | Cycloramphus_lutzorum | LC | Stable | temperate | cathemeral | 17.851 | 24372.881 | 8.376 |
| amphibians | Cycloramphus_organensis | Cycloramphus_organensis | VU | Stable | tropical | nocturnal | 2.957 | 75.867 | 8.295 |
| amphibians | Cycloramphus_parvulus | Zachaenus_parvulus | LC | Decreasing | tropical | nocturnal | 2.019 | 21092.731 | 17.12 |
| amphibians | Cycloramphus_rhyakonastes | Cycloramphus_rhyakonastes | LC | Unknown | temperate | nocturnal | 16.304 | 1855.093 | 12.195 |
| amphibians | Cycloramphus_semipalmatus | Cycloramphus_semipalmatus | CR | Unknown | temperate | nocturnal | 11.693 | 3755.088 | 18.41 |
| amphibians | Cynops_ensicauda | Cynops_ensicauda | VU | Decreasing | temperate | nocturnal | 7.27 | 2048.685 | 23.352 |
| amphibians | Cynops_pyrrhogaster | Cynops_pyrrhogaster | NT | Decreasing | temperate | nocturnal | 6.729 | 287453.985 | 21.068 |
| amphibians | Dasypops_schirchi | Dasypops_schirchi | LC | Decreasing | tropical | nocturnal | 24.063 | 50800.681 | 9.741 |
| amphibians | Dendrobates_auratus | Dendrobates_auratus | LC | Decreasing | tropical | diurnal | 6.273 | 116892.069 | 8.924 |
| amphibians | Dendrobates_leucomelas | Dendrobates_leucomelas | LC | Stable | tropical | diurnal | 11.036 | 844723.624 | 1.663 |
| amphibians | Dendrobates_tinctorius | Dendrobates_tinctorius | LC | Stable | tropical | diurnal | 11.036 | 528727.862 | 0.645 |
| amphibians | Dendrobates_truncatus | Dendrobates_truncatus | LC | Stable | tropical | diurnal | 2.345 | 66001.847 | 14.186 |
| amphibians | Dendrophryniscus_berthalutzae | Dendrophryniscus_berthalutzae | LC | Stable | temperate | nocturnal | 1.023 | 45737.221 | 12.52 |
| amphibians | Dendrophryniscus_brevipollicatus | Dendrophryniscus_brevipollicatus | LC | Stable | tropical | cathemeral | 1.168 | 3288.221 | 16.026 |
| amphibians | Dendrophryniscus_leucomystax | Dendrophryniscus_leucomystax | LC | Stable | temperate | nocturnal | 1.168 | 44305.508 | 12.994 |
| amphibians | Dendrophryniscus_oreites | NA | DD | Unknown | NA | NA | NA | NA | NA |
| amphibians | Dendrophryniscus_proboscideus | Dendrophryniscus_proboscideus | LC | Decreasing | tropical | nocturnal | 8.663 | 64225.44 | 9.76 |
| amphibians | Dendropsophus_anceps | Dendropsophus_anceps | LC | Stable | tropical | nocturnal | 6.273 | 453887.219 | 13.194 |
| amphibians | Dendropsophus_arndti | NA | LC | Stable | NA | NA | NA | NA | NA |
| amphibians | Dendropsophus_berthalutzae | Dendropsophus_berthalutzae | LC | Stable | tropical | nocturnal | 1.023 | 122224.432 | 15.896 |
| amphibians | Dendropsophus_bifurcus | Dendropsophus_bifurcus | LC | Stable | tropical | nocturnal | 3.475 | 1059615.48 | 1.61 |
| amphibians | Dendropsophus_bipunctatus | Dendropsophus_bipunctatus | LC | Stable | tropical | nocturnal | 2.109 | 148090.392 | 12.037 |
| amphibians | Dendropsophus_bokermanni | Dendropsophus_bokermanni | LC | Stable | tropical | nocturnal | 1.686 | 1374636.812 | 1.901 |
| amphibians | Dendropsophus_branneri | Dendropsophus_branneri | LC | Stable | tropical | nocturnal | 1.168 | 1222358.23 | 9.341 |
| amphibians | Dendropsophus_brevifrons | Dendropsophus_brevifrons | LC | Stable | tropical | nocturnal | 1.168 | 1471694.421 | 1.447 |
| amphibians | Dendropsophus_bromeliaceus | NA | LC | Stable | NA | NA | NA | NA | NA |

| | | | | | | | | | |
|---|---|---|---|---|---|---|---|---|---|
| amphibians | Dendropsophus_carnifex | Dendropsophus_carnifex | LC | Stable | tropical | nocturnal | 2.872 | 3308.192 | 7.002 |
| amphibians | Dendropsophus_columbianus | Dendropsophus_columbianus | LC | Stable | tropical | nocturnal | 3.605 | 15247.809 | 20.939 |
| amphibians | Dendropsophus_decipiens | Dendropsophus_decipiens | LC | Stable | tropical | nocturnal | 0.664 | 929276.404 | 11.612 |
| amphibians | Dendropsophus_ebraccatus | Dendropsophus_ebraccatus | LC | Stable | tropical | nocturnal | 4.16 | 234634.685 | 8.464 |
| amphibians | Dendropsophus_elegans | Dendropsophus_elegans | LC | Stable | tropical | nocturnal | 3.705 | 545786.357 | 12.475 |
| amphibians | Dendropsophus_elianeae | Dendropsophus_elianeae | LC | Stable | tropical | nocturnal | 1.326 | 1138241.002 | 11.311 |
| amphibians | Dendropsophus_garagoensis | Dendropsophus_garagoensis | LC | Stable | tropical | nocturnal | 3.163 | 134.829 | 10.524 |
| amphibians | Dendropsophus_giesleri | Dendropsophus_giesleri | LC | Stable | tropical | nocturnal | 3.773 | 327238.307 | 12.609 |
| amphibians | Dendropsophus_haddadi | Dendropsophus_haddadi | LC | Stable | tropical | nocturnal | 1.023 | 112513.181 | 11.862 |
| amphibians | Dendropsophus_haraldschultzi | Dendropsophus_haraldschultzi | LC | Stable | tropical | nocturnal | 1.168 | 979393.962 | 1.221 |
| amphibians | Dendropsophus_jimi | Dendropsophus_jimi | LC | Stable | tropical | nocturnal | 0.807 | 683404.079 | 13.921 |
| amphibians | Dendropsophus_joannae | Dendropsophus_joannae | LC | Stable | tropical | nocturnal | 0.624 | 354126.837 | 1.996 |
| amphibians | Dendropsophus_juliani | Dendropsophus_juliani | LC | Unknown | tropical | nocturnal | 0.685 | 6003.933 | 5.457 |
| amphibians | Dendropsophus_kamagarini | NA | LC | Stable | NA | NA | NA | NA | NA |
| amphibians | Dendropsophus_kubricki | NA | LC | Stable | NA | NA | NA | NA | NA |
| amphibians | Dendropsophus_leali | Dendropsophus_leali | LC | Stable | tropical | nocturnal | 1.686 | 1207985.643 | 2.278 |
| amphibians | Dendropsophus_leucophyllatus | Dendropsophus_leucophyllatus | LC | Stable | tropical | nocturnal | 11.036 | 2574716.357 | 1.453 |
| amphibians | Dendropsophus_mapinguari | NA | LC | Stable | NA | NA | NA | NA | NA |
| amphibians | Dendropsophus_marmoratus | Dendropsophus_marmoratus | LC | Stable | tropical | nocturnal | 15.932 | 4020292.394 | 1.677 |
| amphibians | Dendropsophus_mathiassoni | Dendropsophus_mathiassoni | LC | Stable | tropical | nocturnal | 0.706 | 77538.269 | 9.223 |
| amphibians | Dendropsophus_meridensis | Dendropsophus_meridensis | EN | Decreasing | tropical | nocturnal | 11.036 | 882.886 | 14.738 |
| amphibians | Dendropsophus_meridianus | Dendropsophus_meridianus | LC | Stable | tropical | nocturnal | 1.023 | 23231.373 | 17.798 |
| amphibians | Dendropsophus_microcephalus | Dendropsophus_microcephalus | LC | Increasing | tropical | cathemeral | 2.599 | 4742856.852 | 3.868 |
| amphibians | Dendropsophus_microps | Dendropsophus_microps | LC | Stable | tropical | nocturnal | 2.872 | 350316.314 | 13.66 |
| amphibians | Dendropsophus_minusculus | Dendropsophus_minusculus | LC | Stable | tropical | nocturnal | 1.023 | 1240670.189 | 3.775 |
| amphibians | Dendropsophus_minutus | Dendropsophus_minutus | LC | Stable | tropical | nocturnal | 1.686 | 10714770.448 | 5.562 |
| amphibians | Dendropsophus_miyatai | Dendropsophus_miyatai | LC | Stable | tropical | nocturnal | 1.168 | 1186964.798 | 0.927 |
| amphibians | Dendropsophus_molitor | Dendropsophus_labialis | LC | Stable | tropical | nocturnal | 21.019 | 11470.241 | 18.87 |
| amphibians | Dendropsophus_nahdereri | Dendropsophus_nahdereri | LC | Stable | temperate | cathemeral | 11.398 | 77990.183 | 10.673 |
| amphibians | Dendropsophus_nanus | Dendropsophus_nanus | LC | Stable | tropical | cathemeral | 0.996 | 8670974.691 | 6.289 |
| amphibians | Dendropsophus_novaisi | Dendropsophus_novaisi | LC | Stable | tropical | nocturnal | 3.807 | 57821.881 | 8.633 |
| amphibians | Dendropsophus_oliveirai | Dendropsophus_oliveirai | LC | Stable | tropical | nocturnal | 0.567 | 572686.872 | 10.896 |
| amphibians | Dendropsophus_ozzyi | NA | LC | Stable | NA | NA | NA | NA | NA |
| amphibians | Dendropsophus_parviceps | Dendropsophus_parviceps | LC | Stable | tropical | nocturnal | 1.499 | 3216087.935 | 1.414 |

| | | | | | | | | | |
|---|---|---|---|---|---|---|---|---|---|
| amphibians | Dendropsophus_pauiniensis | NA | DD | Unknown | NA | NA | NA | NA | NA |
| amphibians | Dendropsophus_phlebodes | Dendropsophus_phlebodes | LC | Stable | tropical | nocturnal | 1.686 | 89516.897 | 7.243 |
| amphibians | Dendropsophus_praestans | Dendropsophus_praestans | LC | Stable | tropical | nocturnal | 2.47 | 314.213 | 15.713 |
| amphibians | Dendropsophus_pseudomeridianus | Dendropsophus_pseudomeridianus | LC | Stable | tropical | nocturnal | 0.854 | 30903.106 | 15.807 |
| amphibians | Dendropsophus_reichlei | Dendropsophus_reichlei | LC | Unknown | tropical | nocturnal | 0.717 | 13901.208 | 4.224 |
| amphibians | Dendropsophus_rhodopeplus | Dendropsophus_rhodopeplus | LC | Stable | tropical | nocturnal | 1.889 | 2517105.763 | 1.975 |
| amphibians | Dendropsophus_riveroi | Dendropsophus_riveroi | LC | Stable | tropical | nocturnal | 0.892 | 1957343.498 | 1.977 |
| amphibians | Dendropsophus_robertmertensi | Dendropsophus_robertmertensi | LC | Stable | tropical | diurnal | 1.706 | 19651.816 | 18.742 |
| amphibians | Dendropsophus_rossalleni | Dendropsophus_rossalleni | LC | Stable | tropical | nocturnal | 2.109 | 2083437.493 | 1.379 |
| amphibians | Dendropsophus_rubicundulus | Dendropsophus_rubicundulus | LC | Decreasing | tropical | nocturnal | 1.23 | 960542.159 | 8.901 |
| amphibians | Dendropsophus_ruschii | Dendropsophus_ruschii | LC | Stable | tropical | nocturnal | 1.889 | 26721.867 | 12.716 |
| amphibians | Dendropsophus_salli | Dendropsophus_salli | LC | Stable | tropical | nocturnal | 4.16 | 753071.83 | 4.377 |
| amphibians | Dendropsophus_sanborni | Dendropsophus_sanborni | LC | Stable | temperate | nocturnal | 0.674 | 1187849.234 | 12.164 |
| amphibians | Dendropsophus_sarayacuensis | Dendropsophus_sarayacuensis | LC | Stable | tropical | nocturnal | 7.294 | 3370986.482 | 1.465 |
| amphibians | Dendropsophus_sartori | Dendropsophus_sartori | LC | Stable | tropical | nocturnal | 1.806 | 55772.47 | 7.632 |
| amphibians | Dendropsophus_schubarti | Dendropsophus_schubarti | LC | Unknown | tropical | nocturnal | 1.245 | 1871193.775 | 2.131 |
| amphibians | Dendropsophus_seniculus | Dendropsophus_seniculus | LC | Stable | tropical | nocturnal | 6.77 | 338314.467 | 11.754 |
| amphibians | Dendropsophus_soaresi | Dendropsophus_soaresi | LC | Stable | tropical | nocturnal | 3.475 | 1428862.354 | 8.958 |
| amphibians | Dendropsophus_studerae | Dendropsophus_studerae | EN | Decreasing | tropical | nocturnal | 2.019 | 96.401 | 10.523 |
| amphibians | Dendropsophus_subocularis | Dendropsophus_subocularis | LC | Stable | tropical | nocturnal | 1.343 | 38979.152 | 9.252 |
| amphibians | Dendropsophus_timbeba | Dendropsophus_timbeba | LC | Stable | tropical | nocturnal | 1.36 | 486391.548 | 1.255 |
| amphibians | Dendropsophus_triangulum | Dendropsophus_triangulum | LC | Stable | tropical | nocturnal | 6.273 | 2397126.377 | 1.412 |
| amphibians | Dendropsophus_virolinensis | Dendropsophus_virolinensis | LC | Stable | tropical | nocturnal | 3.163 | 2636.296 | 9.179 |
| amphibians | Dendropsophus_vraemi | NA | LC | Stable | NA | NA | NA | NA | NA |
| amphibians | Dendropsophus_werneri | Dendropsophus_werneri | LC | Stable | temperate | nocturnal | 0.892 | 58187.862 | 13.063 |
| amphibians | Dermatonotus_muelleri | Dermatonotus_muelleri | LC | Stable | tropical | nocturnal | 39.565 | 3775072.171 | 8.175 |
| amphibians | Dermophis_costaricense | NA | DD | Unknown | NA | NA | NA | NA | NA |
| amphibians | Dermophis_mexicanus | Dermophis_mexicanus | LC | Decreasing | tropical | nocturnal | 82.051 | 78747.826 | 12.674 |
| amphibians | Dermophis_oaxacae | Dermophis_oaxacae | LC | Stable | tropical | nocturnal | 35.349 | 105295.251 | 8.278 |
| amphibians | Desmognathus_aeneus | Desmognathus_aeneus | LC | Stable | temperate | nocturnal | 0.194 | 45549.23 | 15.277 |
| amphibians | Desmognathus_apalachicolae | Desmognathus_apalachicolae | LC | Stable | temperate | cathemeral | 2.267 | 20334.694 | 13.643 |
| amphibians | Desmognathus_auriculatus | Desmognathus_auriculatus | LC | Decreasing | temperate | diurnal | 2.548 | 325799.609 | 17.347 |
| amphibians | Desmognathus_carolinensis | Desmognathus_carolinensis | LC | Stable | temperate | diurnal | 1.925 | 11431.372 | 15.914 |
| amphibians | Desmognathus_conanti | Desmognathus_conanti | LC | Stable | temperate | cathemeral | 1.47 | 622621.851 | 15.766 |

| | | | | | | | | | |
|---|---|---|---|---|---|---|---|---|---|
| amphibians | Desmognathus_fuscus | Desmognathus_fuscus | LC | Stable | temperate | nocturnal | 3.039 | 1007633.862 | 17.794 |
| amphibians | Desmognathus_imitator | Desmognathus_imitator | NT | Unknown | temperate | cathemeral | 1.605 | 3698.892 | 9.871 |
| amphibians | Desmognathus_marmoratus | Desmognathus_marmoratus | LC | Stable | temperate | nocturnal | 8.867 | 7803.442 | 16.433 |
| amphibians | Desmognathus_monticola | Desmognathus_monticola | LC | Stable | temperate | nocturnal | 3.809 | 324006.869 | 16.525 |
| amphibians | Desmognathus_ochrophaeus | Desmognathus_ochrophaeus | LC | Unknown | temperate | nocturnal | 1.092 | 287424.363 | 16.893 |
| amphibians | Desmognathus_ocoee | Desmognathus_ocoee | LC | Stable | temperate | nocturnal | 1.367 | 42012.809 | 17.488 |
| amphibians | Desmognathus_orestes | Desmognathus_orestes | LC | Stable | temperate | diurnal | 0.977 | 22135.663 | 16.018 |
| amphibians | Desmognathus_santeetlah | Desmognathus_santeetlah | NT | Unknown | temperate | cathemeral | 1.518 | 6447.54 | 9.876 |
| amphibians | Desmognathus_welteri | Desmognathus_welteri | LC | Decreasing | temperate | nocturnal | 6.215 | 57404.943 | 14.658 |
| amphibians | Diasporus_amirae | NA | LC | Stable | NA | NA | NA | NA | NA |
| amphibians | Diasporus_anthrax | Diasporus_anthrax | VU | Decreasing | tropical | cathemeral | 0.456 | 4982.698 | 11.609 |
| amphibians | Diasporus_darienensis | NA | NT | Stable | NA | NA | NA | NA | NA |
| amphibians | Diasporus_diastema | Diasporus_diastema | LC | Decreasing | tropical | cathemeral | 1.168 | 109484.477 | 9.278 |
| amphibians | Diasporus_gularis | Diasporus_gularis | LC | Stable | tropical | nocturnal | 1.138 | 57123.645 | 6.361 |
| amphibians | Diasporus_hylaeformis | Diasporus_hylaeformis | LC | Stable | tropical | nocturnal | 1.326 | 7513.159 | 10.449 |
| amphibians | Diasporus_majeensis | NA | CR | Unknown | NA | NA | NA | NA | NA |
| amphibians | Diasporus_sapo | NA | CR | Unknown | NA | NA | NA | NA | NA |
| amphibians | Diasporus_ventrimaculatus | Diasporus_ventrimaculatus | LC | Stable | tropical | nocturnal | 1.123 | 22.552 | 1.993 |
| amphibians | Diasporus_vocator | Diasporus_vocator | LC | Stable | tropical | nocturnal | 0.567 | 23566.486 | 10.128 |
| amphibians | Dicamptodon_ensatus | Dicamptodon_ensatus | NT | Decreasing | temperate | nocturnal | 78.547 | 16015.828 | 16.903 |
| amphibians | Dicamptodon_tenebrosus | Dicamptodon_tenebrosus | LC | Stable | temperate | nocturnal | 62.429 | 169824.687 | 10.435 |
| amphibians | Discoglossus_galganoi | Discoglossus_galganoi | LC | Decreasing | temperate | cathemeral | 50.601 | 488944.416 | 16.671 |
| amphibians | Discoglossus_montalentii | Discoglossus_montalentii | NT | Decreasing | temperate | cathemeral | 25.822 | 4154.892 | 9.52 |
| amphibians | Discoglossus_pictus | Discoglossus_pictus | LC | Stable | temperate | cathemeral | 50.601 | 356005.89 | 16.774 |
| amphibians | Discoglossus_sardus | Discoglossus_sardus | LC | Decreasing | temperate | cathemeral | 32.83 | 32995.846 | 16.518 |
| amphibians | Discoglossus_scovazzi | Discoglossus_scovazzi | LC | Unknown | temperate | nocturnal | 50.601 | 181223.136 | 14.531 |
| amphibians | Dryophytes_andersonii | Hyla_andersonii | LC | Stable | temperate | diurnal | 11.767 | 67727.631 | 20.566 |
| amphibians | Dryophytes_arenicolor | Hyla_arenicolor | LC | Stable | temperate | nocturnal | 16.969 | 1286044.855 | 9.156 |
| amphibians | Dryophytes_avivoca | Hyla_avivoca | LC | Stable | temperate | nocturnal | 13.329 | 446462.519 | 15.193 |
| amphibians | Dryophytes_chrysoscelis | Hyla_chrysoscelis | LC | Stable | temperate | cathemeral | 22.156 | 3693368.955 | 17.032 |
| amphibians | Dryophytes_cinereus | Hyla_cinerea | LC | Stable | temperate | cathemeral | 27.131 | 1290978.868 | 16.525 |
| amphibians | Dryophytes_eximius | Hyla_eximia | LC | Stable | tropical | nocturnal | 15.932 | 357544.209 | 12.385 |
| amphibians | Dryophytes_femoralis | Hyla_femoralis | LC | Stable | temperate | cathemeral | 7.294 | 482056.203 | 16.668 |
| amphibians | Dryophytes_gratiosus | Hyla_gratiosa | LC | Stable | temperate | nocturnal | 32.83 | 593264.039 | 16.598 |

| | | | | | | | | | |
|---|---|---|---|---|---|---|---|---|---|
| amphibians | Dryophytes_japonicus | Hyla_japonica | LC | Stable | temperate | cathemeral | 9.669 | 2185028.148 | 12.533 |
| amphibians | Dryophytes_squirellus | Hyla_squirella | LC | Stable | temperate | nocturnal | 7.844 | 747619.538 | 16.473 |
| amphibians | Dryophytes_suweonensis | Hyla_suweonensis | EN | Decreasing | temperate | cathemeral | 1.686 | 4430.27 | 23.937 |
| amphibians | Dryophytes_versicolor | Hyla_versicolor | LC | Stable | temperate | nocturnal | 19.923 | 2240480.061 | 16.982 |
| amphibians | Duellmanohyla_legleri | NA | EN | Unknown | NA | NA | NA | NA | NA |
| amphibians | Duellmanohyla_rufioculis | Duellmanohyla_rufioculis | LC | Stable | tropical | nocturnal | 5.356 | 20510.268 | 13.202 |
| amphibians | Duellmanohyla_soralia | Duellmanohyla_soralia | EN | Decreasing | tropical | nocturnal | 4.536 | 1194.233 | 8.478 |
| amphibians | Duellmanohyla_uranochroa | Duellmanohyla_uranochroa | VU | Increasing | tropical | nocturnal | 5.356 | 7897.406 | 10.915 |
| amphibians | Duttaphrynus_beddomii | Duttaphrynus_beddomii | VU | Decreasing | tropical | diurnal | 7.844 | 217.978 | 8.898 |
| amphibians | Duttaphrynus_himalayanus | Duttaphrynus_himalayanus | LC | Decreasing | temperate | nocturnal | 226.191 | 301662.309 | 9.303 |
| amphibians | Duttaphrynus_melanostictus | Duttaphrynus_melanostictus | LC | Increasing | tropical | nocturnal | 310.174 | 7410138.944 | 14.536 |
| amphibians | Duttaphrynus_microtympanum | Duttaphrynus_microtympanum | LC | Decreasing | tropical | nocturnal | 50.601 | 13413.317 | 12.485 |
| amphibians | Duttaphrynus_parietalis | Duttaphrynus_parietalis | LC | Decreasing | tropical | cathemeral | 79.584 | 22121.811 | 12.747 |
| amphibians | Duttaphrynus_scaber | Duttaphrynus_scaber | LC | Stable | tropical | nocturnal | 8.423 | 1359020.83 | 18.962 |
| amphibians | Dyscophus_antongilii | Dyscophus_antongilii | LC | Decreasing | tropical | cathemeral | 188.234 | 37130.041 | 8.241 |
| amphibians | Dyscophus_guineti | Dyscophus_guineti | LC | Decreasing | tropical | cathemeral | 152.277 | 6095.396 | 7.142 |
| amphibians | Dyscophus_insularis | Dyscophus_insularis | LC | Decreasing | tropical | cathemeral | 12.531 | 161545.532 | 8.246 |
| amphibians | Echinotriton_andersoni | Echinotriton_andersoni | VU | Decreasing | temperate | diurnal | 15.677 | 2193.919 | 23.375 |
| amphibians | Echinotriton_chinhaiensis | Echinotriton_chinhaiensis | CR | Decreasing | temperate | nocturnal | 6.709 | 114.649 | 16.998 |
| amphibians | Ecnomiohyla_fimbrimembra | Ecnomiohyla_fimbrimembra | VU | Unknown | tropical | nocturnal | 79.584 | 2127.435 | 9.239 |
| amphibians | Ecnomiohyla_miliaria | Ecnomiohyla_miliaria | LC | Decreasing | tropical | nocturnal | 154.926 | 13433.209 | 4.522 |
| amphibians | Ecnomiohyla_rabborum | NA | CR | Decreasing | NA | NA | NA | NA | NA |
| amphibians | Ecnomiohyla_salvaje | Ecnomiohyla_salvaje | EN | Decreasing | tropical | cathemeral | 63.963 | 106.795 | 6.938 |
| amphibians | Ecnomiohyla_thysanota | NA | DD | Unknown | NA | NA | NA | NA | NA |
| amphibians | Ecnomiohyla_valancifer | Ecnomiohyla_valancifer | CR | Unknown | tropical | nocturnal | 61.584 | 68.266 | 9.332 |
| amphibians | Ectopoglossus_atopoglossus | Anomaloglossus_atopoglossus | CR | Decreasing | tropical | cathemeral | 1.199 | 15.809 | 4.104 |
| amphibians | Ectopoglossus_confusus | Anomaloglossus_confusus | EN | Decreasing | tropical | diurnal | 1.326 | 430.827 | 8.035 |
| amphibians | Ectopoglossus_saxatilis | NA | DD | Unknown | NA | NA | NA | NA | NA |
| amphibians | Edalorhina_perezi | Edalorhina_perezi | LC | Stable | tropical | cathemeral | 5.356 | 1481341.994 | 1.321 |
| amphibians | Elachistocleis_bicolor | Elachistocleis_bicolor | LC | Stable | temperate | cathemeral | 11.036 | 2588357.628 | 9.711 |
| amphibians | Elachistocleis_cesarii | Elachistocleis_cesarii | LC | Stable | tropical | nocturnal | 6.618 | 2162380.35 | 10.27 |
| amphibians | Elachistocleis_erythrogaster | Elachistocleis_erythrogaster | EN | Decreasing | temperate | cathemeral | 4.536 | 5054.727 | 8.49 |
| amphibians | Elachistocleis_piauiensis | Elachistocleis_piauiensis | LC | Stable | tropical | diurnal | 3.163 | 367579.976 | 6.596 |
| amphibians | Elachistocleis_skotogaster | Elachistocleis_skotogaster | LC | Stable | tropical | nocturnal | 3.285 | 8322.222 | 4.379 |

| | | | | | | | | | |
|---|---|---|---|---|---|---|---|---|---|
| amphibians | Eleutherodactylus_abbotti | Eleutherodactylus_abbotti | LC | Stable | tropical | diurnal | 1.168 | 66828.194 | 16.108 |
| amphibians | Eleutherodactylus_adelus | Eleutherodactylus_adelus | EN | Decreasing | tropical | diurnal | 0.223 | 302.552 | 11.149 |
| amphibians | Eleutherodactylus_alcoae | Eleutherodactylus_alcoae | LC | Stable | tropical | nocturnal | 7.844 | 2848.043 | 10.159 |
| amphibians | Eleutherodactylus_amadeus | Eleutherodactylus_amadeus | CR | Decreasing | tropical | cathemeral | 1.168 | 500.761 | 12.977 |
| amphibians | Eleutherodactylus_amplinympha | Eleutherodactylus_amplinympha | EN | Decreasing | tropical | nocturnal | 11.036 | 217.365 | 9.168 |
| amphibians | Eleutherodactylus_angustidigitorum | Eleutherodactylus_angustidigitorum | LC | Stable | tropical | diurnal | 1.168 | 23214.173 | 11.676 |
| amphibians | Eleutherodactylus_antillensis | Eleutherodactylus_antillensis | LC | Stable | tropical | nocturnal | 9.031 | 9408.594 | 27.187 |
| amphibians | Eleutherodactylus_armstrongi | Eleutherodactylus_armstrongi | EN | Decreasing | tropical | nocturnal | 6.77 | 985.302 | 10.152 |
| amphibians | Eleutherodactylus_atkinsi | Eleutherodactylus_atkinsi | LC | Stable | tropical | nocturnal | 6.77 | 22235.382 | 13.702 |
| amphibians | Eleutherodactylus_audanti | Eleutherodactylus_audanti | VU | Decreasing | tropical | nocturnal | 1.168 | 2879.601 | 11.568 |
| amphibians | Eleutherodactylus_auriculatoides | Eleutherodactylus_auriculatoides | VU | Decreasing | tropical | nocturnal | 2.872 | 3107.083 | 9.751 |
| amphibians | Eleutherodactylus_auriculatus | Eleutherodactylus_auriculatus | LC | Stable | tropical | cathemeral | 1.023 | 16894.644 | 12.894 |
| amphibians | Eleutherodactylus_bakeri | Eleutherodactylus_bakeri | CR | Decreasing | tropical | nocturnal | 3.475 | 417.598 | 12.334 |
| amphibians | Eleutherodactylus_barlagnei | Eleutherodactylus_barlagnei | EN | Decreasing | tropical | nocturnal | 2.872 | 411.342 | 12.977 |
| amphibians | Eleutherodactylus_bartonsmithi | Eleutherodactylus_bartonsmithi | CR | Decreasing | tropical | nocturnal | 1.326 | 13.955 | 14.427 |
| amphibians | Eleutherodactylus_beguei | NA | CR | Decreasing | NA | NA | NA | NA | NA |
| amphibians | Eleutherodactylus_blairhedgesi | Eleutherodactylus_blairhedgesi | CR | Decreasing | tropical | nocturnal | 2.445 | 11.451 | 28.351 |
| amphibians | Eleutherodactylus_bresslerae | Eleutherodactylus_bresslerae | CR | Decreasing | tropical | nocturnal | 8.423 | 25.073 | 13.877 |
| amphibians | Eleutherodactylus_caribe | Eleutherodactylus_caribe | CR | Decreasing | tropical | nocturnal | 0.567 | 2.518 | 19.254 |
| amphibians | Eleutherodactylus_cattus | Eleutherodactylus_cavernicola | EN | Decreasing | tropical | nocturnal | 1.123 | 6.915 | 8.919 |
| amphibians | Eleutherodactylus_cochranae | Eleutherodactylus_cochranae | LC | Stable | tropical | nocturnal | 0.892 | 3963.48 | 25.577 |
| amphibians | Eleutherodactylus_cooki | Eleutherodactylus_cooki | EN | Decreasing | tropical | nocturnal | 14.161 | 238.775 | 25.903 |
| amphibians | Eleutherodactylus_coqui | Eleutherodactylus_coqui | LC | Increasing | tropical | nocturnal | 23.335 | 13154.859 | 27.561 |
| amphibians | Eleutherodactylus_corona | Eleutherodactylus_corona | CR | Decreasing | tropical | nocturnal | 0.488 | 31.539 | 13.643 |
| amphibians | Eleutherodactylus_cundalli | Eleutherodactylus_cundalli | VU | Decreasing | tropical | nocturnal | 19.923 | 6292.648 | 19.004 |
| amphibians | Eleutherodactylus_cuneatus | Eleutherodactylus_cuneatus | LC | Stable | tropical | cathemeral | 20.248 | 10968.078 | 13.991 |
| amphibians | Eleutherodactylus_cystignathoides | Eleutherodactylus_cystignathoides | LC | Stable | tropical | cathemeral | 1.294 | 107464.672 | 13.904 |
| amphibians | Eleutherodactylus_darlingtoni | Eleutherodactylus_darlingtoni | CR | Decreasing | tropical | diurnal | 5.802 | 317.585 | 12.968 |
| amphibians | Eleutherodactylus_dilatus | Eleutherodactylus_dilatus | LC | Stable | tropical | nocturnal | 2.109 | 511.23 | 5.516 |
| amphibians | Eleutherodactylus_dolomedes | Eleutherodactylus_dolomedes | CR | Decreasing | tropical | nocturnal | 0.727 | 9.995 | 12.154 |
| amphibians | Eleutherodactylus_eileenae | Eleutherodactylus_eileenae | LC | Decreasing | tropical | nocturnal | 2.872 | 7705.134 | 12.354 |
| amphibians | Eleutherodactylus_eneidae | Eleutherodactylus_eneidae | CR | Unknown | tropical | nocturnal | 2.109 | 1921.111 | 22.74 |
| amphibians | Eleutherodactylus_erythrochomus | NA | not_available | not_available | NA | NA | NA | NA | NA |
| amphibians | Eleutherodactylus_etheridgei | Eleutherodactylus_etheridgei | VU | Decreasing | tropical | nocturnal | 1.023 | 428.494 | 19.694 |

| | | | | | | | | | |
|---|---|---|---|---|---|---|---|---|---|
| amphibians | Eleutherodactylus_fowleri | Eleutherodactylus_fowleri | CR | Decreasing | tropical | nocturnal | 9.669 | 236.062 | 14.266 |
| amphibians | Eleutherodactylus_geitonos | NA | EN | Decreasing | NA | NA | NA | NA | NA |
| amphibians | Eleutherodactylus_glamyrus | Eleutherodactylus_glamyrus | EN | Decreasing | tropical | cathemeral | 1.535 | 860.279 | 8.313 |
| amphibians | Eleutherodactylus_glaphycompus | Eleutherodactylus_glaphycompus | EN | Decreasing | tropical | cathemeral | 3.163 | 1226.797 | 14.115 |
| amphibians | Eleutherodactylus_gossei | Eleutherodactylus_gossei | VU | Decreasing | tropical | nocturnal | 3.163 | 8661.193 | 18.52 |
| amphibians | Eleutherodactylus_grahami | Eleutherodactylus_grahami | EN | Decreasing | tropical | nocturnal | 1.168 | 1004.821 | 13.986 |
| amphibians | Eleutherodactylus_griphus | Eleutherodactylus_griphus | EN | Decreasing | tropical | nocturnal | 0.433 | 305.31 | 6.699 |
| amphibians | Eleutherodactylus_gryllus | Eleutherodactylus_gryllus | CR | Decreasing | tropical | nocturnal | 0.396 | 1092.052 | 19.751 |
| amphibians | Eleutherodactylus_guanahacabibes | Eleutherodactylus_guanahacabibes | LC | Unknown | tropical | nocturnal | 0.892 | 951.795 | 6.55 |
| amphibians | Eleutherodactylus_guantanamera | Eleutherodactylus_guantanamera | VU | Decreasing | tropical | nocturnal | 4.271 | 2108.339 | 10.719 |
| amphibians | Eleutherodactylus_gundlachi | Eleutherodactylus_gundlachi | VU | Decreasing | tropical | nocturnal | 1.326 | 1903.094 | 9.387 |
| amphibians | Eleutherodactylus_guttilatus | Eleutherodactylus_guttilatus | LC | Unknown | tropical | nocturnal | 2.599 | 72979.531 | 10.536 |
| amphibians | Eleutherodactylus_hedricki | Eleutherodactylus_hedricki | EN | Decreasing | tropical | nocturnal | 3.475 | 137.724 | 14.438 |
| amphibians | Eleutherodactylus_iberia | Eleutherodactylus_iberia | VU | Decreasing | tropical | cathemeral | 0.082 | 1867.959 | 11.169 |
| amphibians | Eleutherodactylus_intermedius | Eleutherodactylus_intermedius | EN | Decreasing | tropical | nocturnal | 0.567 | 793.516 | 8.165 |
| amphibians | Eleutherodactylus_interorbitalis | Eleutherodactylus_interorbitalis | LC | Stable | temperate | nocturnal | 1.499 | 56696.098 | 6.043 |
| amphibians | Eleutherodactylus_ionthus | Eleutherodactylus_ionthus | VU | Decreasing | tropical | nocturnal | 5.356 | 4006.264 | 14.387 |
| amphibians | Eleutherodactylus_jamaicensis | Eleutherodactylus_jamaicensis | CR | Decreasing | tropical | nocturnal | 2.109 | 3409.603 | 16.376 |
| amphibians | Eleutherodactylus_jasperi | Eleutherodactylus_jasperi | CR | Unknown | tropical | nocturnal | 1.168 | 12.642 | 22.187 |
| amphibians | Eleutherodactylus_johnstonei | Eleutherodactylus_johnstonei | LC | Increasing | tropical | nocturnal | 3.475 | 89710.273 | 18.424 |
| amphibians | Eleutherodactylus_juanariveroi | Eleutherodactylus_juanariveroi | CR | Decreasing | tropical | nocturnal | 0.354 | 2.469 | 34.023 |
| amphibians | Eleutherodactylus_klinikowskii | Eleutherodactylus_klinikowskii | EN | Decreasing | tropical | diurnal | 1.499 | 1497.706 | 13.391 |
| amphibians | Eleutherodactylus_lamprotes | Eleutherodactylus_lamprotes | CR | Decreasing | tropical | diurnal | 2.109 | 624.829 | 12.494 |
| amphibians | Eleutherodactylus_leberi | Eleutherodactylus_leberi | CR | Decreasing | tropical | nocturnal | 2.872 | 85.186 | 13.76 |
| amphibians | Eleutherodactylus_lentus | Eleutherodactylus_lentus | EN | Decreasing | tropical | nocturnal | 4.536 | 267.275 | 29.085 |
| amphibians | Eleutherodactylus_leoncei | Eleutherodactylus_leoncei | EN | Decreasing | tropical | diurnal | 7.844 | 1230.302 | 9.694 |
| amphibians | Eleutherodactylus_leprus | Eleutherodactylus_leprus | LC | Stable | tropical | diurnal | 1.889 | 105732.807 | 9.407 |
| amphibians | Eleutherodactylus_ligiae | NA | CR | Decreasing | NA | NA | NA | NA | NA |
| amphibians | Eleutherodactylus_limbatus | Eleutherodactylus_limbatus | LC | Stable | tropical | cathemeral | 0.335 | 8614.959 | 14.447 |
| amphibians | Eleutherodactylus_locustus | Eleutherodactylus_locustus | EN | Decreasing | tropical | nocturnal | 1.023 | 115.284 | 16.702 |
| amphibians | Eleutherodactylus_mariposa | Eleutherodactylus_mariposa | CR | Decreasing | tropical | nocturnal | 3.807 | 80.414 | 13.023 |
| amphibians | Eleutherodactylus_marnockii | Eleutherodactylus_marnockii | LC | Stable | temperate | nocturnal | 5.142 | 178842.93 | 11.347 |
| amphibians | Eleutherodactylus_martinicensis | Eleutherodactylus_martinicensis | NT | Stable | tropical | nocturnal | 9.031 | 3655.202 | 22.67 |
| amphibians | Eleutherodactylus_maurus | Eleutherodactylus_maurus | VU | Unknown | tropical | nocturnal | 1.499 | 4498.155 | 16.578 |

| | | | | | | | | | |
|---|---|---|---|---|---|---|---|---|---|
| amphibians | Eleutherodactylus_melacara | Eleutherodactylus_melacara | EN | Decreasing | tropical | nocturnal | 3.807 | 496.714 | 7.12 |
| amphibians | Eleutherodactylus_monensis | Eleutherodactylus_monensis | VU | Stable | tropical | cathemeral | 10.613 | 50.569 | 6.722 |
| amphibians | Eleutherodactylus_montanus | Eleutherodactylus_montanus | EN | Decreasing | tropical | nocturnal | 7.294 | 1772.631 | 8.678 |
| amphibians | Eleutherodactylus_nortoni | Eleutherodactylus_nortoni | CR | Decreasing | tropical | diurnal | 27.131 | 2687.182 | 14.305 |
| amphibians | Eleutherodactylus_nubicola | Eleutherodactylus_nubicola | EN | Decreasing | tropical | nocturnal | 10.337 | 195.162 | 12.528 |
| amphibians | Eleutherodactylus_orcutti | Eleutherodactylus_orcutti | CR | Decreasing | tropical | nocturnal | 4.934 | 381.153 | 11.244 |
| amphibians | Eleutherodactylus_parapelates | Eleutherodactylus_parapelates | CR | Decreasing | tropical | diurnal | 12.531 | 396.552 | 11.952 |
| amphibians | Eleutherodactylus_pictissimus | Eleutherodactylus_pictissimus | LC | Stable | tropical | diurnal | 6.77 | 18126.801 | 15.711 |
| amphibians | Eleutherodactylus_pinchoni | Eleutherodactylus_pinchoni | EN | Decreasing | tropical | diurnal | 0.567 | 467.384 | 14.633 |
| amphibians | Eleutherodactylus_pipilans | Eleutherodactylus_pipilans | LC | Stable | tropical | nocturnal | 2.345 | 63412.218 | 9.444 |
| amphibians | Eleutherodactylus_planirostris | Eleutherodactylus_planirostris | LC | Increasing | temperate | nocturnal | 11.036 | 330361.6 | 16.385 |
| amphibians | Eleutherodactylus_principalis | Eleutherodactylus_principalis | EN | Decreasing | tropical | nocturnal | 0.892 | 735.596 | 8.507 |
| amphibians | Eleutherodactylus_richmondi | Eleutherodactylus_richmondi | EN | Decreasing | tropical | nocturnal | 11.036 | 218.397 | 16.854 |
| amphibians | Eleutherodactylus_ricordii | Eleutherodactylus_ricordii | VU | Decreasing | tropical | nocturnal | 5.356 | 4518.7 | 13.372 |
| amphibians | Eleutherodactylus_rivularis | Eleutherodactylus_rivularis | CR | Decreasing | tropical | nocturnal | 2.345 | 56.931 | 13.137 |
| amphibians | Eleutherodactylus_ronaldi | Eleutherodactylus_ronaldi | NT | Decreasing | tropical | nocturnal | 1.686 | 5697.114 | 13.018 |
| amphibians | Eleutherodactylus_ruthae | Eleutherodactylus_ruthae | EN | Decreasing | tropical | cathemeral | 17.851 | 3292.896 | 16.296 |
| amphibians | Eleutherodactylus_schmidti | Eleutherodactylus_schmidti | CR | Decreasing | tropical | nocturnal | 17.851 | 10990.906 | 16.42 |
| amphibians | Eleutherodactylus_schwartzi | Eleutherodactylus_schwartzi | EN | Decreasing | tropical | nocturnal | 3.475 | 144.737 | 23.57 |
| amphibians | Eleutherodactylus_simulans | Eleutherodactylus_simulans | VU | Decreasing | tropical | nocturnal | 1.023 | 1893.859 | 11.273 |
| amphibians | Eleutherodactylus_sisyphodemus | Eleutherodactylus_sisyphodemus | CR | Decreasing | tropical | nocturnal | 0.403 | 55.178 | 3.991 |
| amphibians | Eleutherodactylus_tetajulia | Eleutherodactylus_tetajulia | EN | Decreasing | tropical | cathemeral | 0.178 | 444.168 | 8.007 |
| amphibians | Eleutherodactylus_thomasi | Eleutherodactylus_thomasi | NT | Decreasing | tropical | nocturnal | 6.77 | 2267.475 | 13.57 |
| amphibians | Eleutherodactylus_toa | Eleutherodactylus_toa | EN | Decreasing | tropical | nocturnal | 2.872 | 2451.137 | 10.127 |
| amphibians | Eleutherodactylus_tonyi | Eleutherodactylus_tonyi | VU | Decreasing | tropical | diurnal | 3.074 | 590.488 | 11.399 |
| amphibians | Eleutherodactylus_turquinensis | Eleutherodactylus_turquinensis | CR | Decreasing | tropical | nocturnal | 14.161 | 269.13 | 6.263 |
| amphibians | Eleutherodactylus_tychathrous | Eleutherodactylus_tychathrous | EN | Decreasing | tropical | nocturnal | 17.652 | 510.143 | 9.944 |
| amphibians | Eleutherodactylus_unicolor | Eleutherodactylus_unicolor | CR | Decreasing | tropical | nocturnal | 0.335 | 53.727 | 12.008 |
| amphibians | Eleutherodactylus_varians | Eleutherodactylus_varians | LC | Decreasing | tropical | nocturnal | 5.356 | 4268.577 | 13.092 |
| amphibians | Eleutherodactylus_varleyi | Eleutherodactylus_varleyi | LC | Unknown | tropical | nocturnal | 0.44 | 66329.936 | 16.206 |
| amphibians | Eleutherodactylus_weinlandi | Eleutherodactylus_weinlandi | LC | Stable | tropical | nocturnal | 5.356 | 33150.371 | 16.274 |
| amphibians | Eleutherodactylus_zugi | Eleutherodactylus_zugi | EN | Decreasing | tropical | nocturnal | 0.567 | 1073.881 | 12.68 |
| amphibians | Engystomops_coloradorum | Engystomops_coloradorum | EN | Decreasing | tropical | nocturnal | 1.277 | 2919.119 | 13.12 |
| amphibians | Engystomops_freibergi | Engystomops_freibergi | LC | Unknown | tropical | nocturnal | 4.934 | 1665882.707 | 2.535 |

| | | | | | | | | | |
|---|---|---|---|---|---|---|---|---|---|
| amphibians | Engystomops_montubio | Engystomops_montubio | LC | Stable | tropical | nocturnal | 0.867 | 18497.626 | 15.242 |
| amphibians | Engystomops_petersi | Engystomops_petersi | LC | Stable | tropical | cathemeral | 4.975 | 2015286.397 | 1.402 |
| amphibians | Engystomops_pustulatus | Engystomops_pustulatus | LC | Unknown | tropical | nocturnal | 3.981 | 23877.514 | 16.283 |
| amphibians | Engystomops_pustulosus | Engystomops_pustulosus | LC | Stable | tropical | nocturnal | 9.8 | 894865.275 | 11.2 |
| amphibians | Engystomops_randi | Engystomops_randi | LC | Stable | tropical | nocturnal | 1.023 | 8331.1 | 15.54 |
| amphibians | Ensatina_eschscholtzii | Ensatina_eschscholtzii | LC | Stable | temperate | nocturnal | 8.438 | 325428.35 | 12.415 |
| amphibians | Epicrionops_bicolor | Epicrionops_bicolor | LC | Unknown | tropical | nocturnal | 17.988 | 58008.429 | 2.668 |
| amphibians | Epidalea_calamita | Epidalea_calamita | LC | Decreasing | temperate | nocturnal | 104.268 | 2234800.717 | 21.073 |
| amphibians | Epipedobates_anthonyi | Epipedobates_anthonyi | NT | Decreasing | tropical | diurnal | 1.499 | 20770.836 | 9.867 |
| amphibians | Epipedobates_boulengeri | Epipedobates_boulengeri | LC | Decreasing | tropical | diurnal | 0.749 | 55338.229 | 7.456 |
| amphibians | Epipedobates_darwinwallacei | Epipedobates_darwinwallacei | EN | Decreasing | tropical | diurnal | 0.567 | 1809.593 | 7.444 |
| amphibians | Epipedobates_espinosai | Epipedobates_espinosai | DD | Decreasing | tropical | diurnal | 0.595 | 4668.424 | 12.218 |
| amphibians | Epipedobates_machalilla | Epipedobates_machalilla | LC | Decreasing | tropical | diurnal | 0.375 | 37054.527 | 15.899 |
| amphibians | Epipedobates_narinensis | NA | DD | Unknown | NA | NA | NA | NA | NA |
| amphibians | Epipedobates_tricolor | Epipedobates_tricolor | VU | Decreasing | tropical | diurnal | 1.411 | 4671.221 | 12.412 |
| amphibians | Espadarana_andina | Espadarana_andina | LC | Stable | tropical | nocturnal | 1.61 | 34642.717 | 11.477 |
| amphibians | Espadarana_callistomma | Espadarana_callistomma | LC | Decreasing | tropical | nocturnal | 2.547 | 30974.08 | 5.621 |
| amphibians | Espadarana_durrellorum | Centrolene_durrellorum | LC | Decreasing | tropical | nocturnal | 1.953 | 11952.006 | 6.713 |
| amphibians | Espadarana_prosoblepon | Espadarana_prosoblepon | LC | Stable | tropical | nocturnal | 2.345 | 346069.787 | 8.536 |
| amphibians | Euparkerella_robusta | Euparkerella_robusta | CR | Decreasing | tropical | cathemeral | 0.892 | 9.252 | 6.735 |
| amphibians | Euparkerella_tridactyla | Euparkerella_tridactyla | LC | Stable | tropical | cathemeral | 0.634 | 5677.968 | 12.086 |
| amphibians | Euphlyctis_cyanophlyctis | Euphlyctis_cyanophlyctis | LC | Stable | tropical | cathemeral | 79.584 | 4408170.133 | 15.489 |
| amphibians | Euproctus_montanus | Euproctus_montanus | LC | Stable | temperate | nocturnal | 3.16 | 7955.859 | 12.597 |
| amphibians | Euproctus_platycephalus | Euproctus_platycephalus | VU | Decreasing | temperate | diurnal | 5.54 | 5723.496 | 12.387 |
| amphibians | Eupsophus_altor | Eupsophus_altor | EN | Decreasing | temperate | nocturnal | 4.016 | 312.665 | 9.218 |
| amphibians | Eupsophus_contulmoensis | Eupsophus_contulmoensis | EN | Decreasing | temperate | nocturnal | 6.77 | 1008.94 | 7.619 |
| amphibians | Eupsophus_roseus | Eupsophus_roseus | LC | Decreasing | temperate | nocturnal | 28.486 | 39011.09 | 10.908 |
| amphibians | Eurycea_bislineata | Eurycea_bislineata | LC | Stable | temperate | nocturnal | 1.092 | 1198568.494 | 12.191 |
| amphibians | Eurycea_guttolineata | Eurycea_guttolineata | LC | Stable | temperate | nocturnal | 3.666 | 738309.305 | 16.972 |
| amphibians | Eurycea_longicauda | Eurycea_longicauda | LC | Stable | temperate | nocturnal | 3.34 | 848916.905 | 17.959 |
| amphibians | Eurycea_lucifuga | Eurycea_lucifuga | LC | Unknown | temperate | nocturnal | 3.509 | 449641.606 | 16.965 |
| amphibians | Eurycea_nana | Eurycea_nana | CR | Decreasing | temperate | diurnal | 0.352 | 0.088 | 46.456 |
| amphibians | Eurycea_quadridigitata | Eurycea_quadridigitata | LC | Stable | temperate | diurnal | 0.646 | 251090.108 | 16.826 |
| amphibians | Eurycea_spelaea | Eurycea_spelaea | LC | Stable | temperate | nocturnal | 5.494 | 32788.251 | 17.57 |

| | | | | | | | | | |
|---|---|---|---|---|---|---|---|---|---|
| amphibians | Eurycea_wilderae | Eurycea_wilderae | LC | Stable | temperate | nocturnal | 1.925 | 76390.174 | 16.934 |
| amphibians | Excidobates_captivus | Excidobates_captivus | VU | Decreasing | tropical | diurnal | 0.335 | 9146.584 | 3.382 |
| amphibians | Excidobates_condor | Excidobates_condor | EN | Stable | tropical | diurnal | 0.842 | 1025.709 | 4.244 |
| amphibians | Excidobates_mysteriosus | Excidobates_mysteriosus | EN | Decreasing | tropical | diurnal | 2.109 | 1306.16 | 7.224 |
| amphibians | Exerodonta_bivocata | Exerodonta_bivocata | EN | Unknown | tropical | nocturnal | 1.786 | 319.281 | 9.844 |
| amphibians | Exerodonta_catracha | Exerodonta_catracha | NT | Stable | tropical | cathemeral | 2.626 | 578.067 | 8.893 |
| amphibians | Exerodonta_melanomma | Exerodonta_melanomma | VU | Unknown | tropical | nocturnal | 2.47 | 1328.467 | 7.695 |
| amphibians | Exerodonta_perkinsi | Exerodonta_perkinsi | EN | Decreasing | tropical | nocturnal | 4.345 | 24.536 | 12.897 |
| amphibians | Exerodonta_xera | Exerodonta_xera | VU | Unknown | tropical | nocturnal | 3.475 | 7308.892 | 10.427 |
| amphibians | Feihyla_hansenae | NA | LC | Decreasing | NA | NA | NA | NA | NA |
| amphibians | Feihyla_kajau | Feihyla_kajau | LC | Decreasing | tropical | nocturnal | 0.567 | 152145.106 | 3.306 |
| amphibians | Feihyla_vittata | NA | LC | Stable | NA | NA | NA | NA | NA |
| amphibians | Fejervarya_cancrivora | Fejervarya_cancrivora | LC | Decreasing | tropical | nocturnal | 188.234 | 817338.524 | 11.937 |
| amphibians | Fejervarya_limnocharis | Fejervarya_limnocharis | LC | Stable | tropical | cathemeral | 42.854 | 6160012.435 | 14.861 |
| amphibians | Fejervarya_triora | Fejervarya_triora | LC | Decreasing | tropical | nocturnal | 20.797 | 10876.548 | 12.175 |
| amphibians | Fejervarya_vittigera | Fejervarya_vittigera | LC | Decreasing | tropical | nocturnal | 51.841 | 288730.345 | 13.672 |
| amphibians | Firouzophrynus_dhufarensis | Duttaphrynus_dhufarensis | LC | Stable | tropical | nocturnal | 25.822 | 479285.494 | 8.005 |
| amphibians | Firouzophrynus_stomaticus | Duttaphrynus_stomaticus | LC | Stable | tropical | cathemeral | 74.113 | 4282362.48 | 15.691 |
| amphibians | Flectonotus_fitzgeraldi | Flectonotus_fitzgeraldi | LC | Stable | tropical | nocturnal | 1.023 | 4192.852 | 12.757 |
| amphibians | Flectonotus_pygmaeus | Flectonotus_pygmaeus | LC | Stable | tropical | nocturnal | 2.706 | 41606.16 | 12.788 |
| amphibians | Fritziana_fissilis | Fritziana_fissilis | LC | Stable | tropical | nocturnal | 4.812 | 1963.944 | 14.807 |
| amphibians | Fritziana_goeldii | Fritziana_goeldii | LC | Stable | tropical | nocturnal | 4.16 | 31471.82 | 16.182 |
| amphibians | Fritziana_mitus | NA | LC | Unknown | NA | NA | NA | NA | NA |
| amphibians | Fritziana_ohausi | Fritziana_ohausi | LC | Decreasing | temperate | nocturnal | 3.379 | 38985.788 | 13.111 |
| amphibians | Frostius_erythrophthalmus | Frostius_erythrophthalmus | LC | Stable | tropical | nocturnal | 1.686 | 12221.613 | 7.942 |
| amphibians | Frostius_pernambucensis | Frostius_pernambucensis | LC | Stable | tropical | nocturnal | 0.664 | 18695.202 | 15.746 |
| amphibians | Gastrophryne_carolinensis | Gastrophryne_carolinensis | LC | Stable | temperate | nocturnal | 4.934 | 1576052.905 | 16.44 |
| amphibians | Gastrophryne_elegans | Gastrophryne_elegans | LC | Stable | tropical | nocturnal | 4.536 | 185653.566 | 10.758 |
| amphibians | Gastrophryne_olivacea | Gastrophryne_olivacea | LC | Stable | temperate | cathemeral | 6.273 | 1462157.645 | 13.027 |
| amphibians | Gastrotheca_albolineata | Gastrotheca_albolineata | LC | Stable | tropical | nocturnal | 19.923 | 18226.116 | 15.678 |
| amphibians | Gastrotheca_andaquiensis | Gastrotheca_andaquiensis | LC | Stable | tropical | nocturnal | 44.708 | 9079.865 | 5.316 |
| amphibians | Gastrotheca_angustifrons | Gastrotheca_angustifrons | CR | Decreasing | tropical | nocturnal | 79.024 | 1790.877 | 4.823 |
| amphibians | Gastrotheca_antomia | Gastrotheca_antomia | CR | Decreasing | tropical | nocturnal | 18.558 | 2115.983 | 8.215 |
| amphibians | Gastrotheca_antoniiochoai | NA | DD | Unknown | NA | NA | NA | NA | NA |

| | | | | | | | | | |
|---|---|---|---|---|---|---|---|---|---|
| amphibians | Gastrotheca_aratia | NA | LC | Stable | NA | NA | NA | NA | NA |
| amphibians | Gastrotheca_argenteovirens | Gastrotheca_argenteovirens | LC | Stable | tropical | nocturnal | 25.822 | 4237.163 | 13.522 |
| amphibians | Gastrotheca_aureomaculata | Gastrotheca_aureomaculata | EN | Decreasing | tropical | cathemeral | 42.854 | 1186.194 | 12.605 |
| amphibians | Gastrotheca_christiani | Gastrotheca_christiani | CR | Decreasing | tropical | nocturnal | 5.356 | 56.554 | 5.754 |
| amphibians | Gastrotheca_cornuta | Gastrotheca_cornuta | CR | Decreasing | tropical | nocturnal | 52.891 | 45654.312 | 4.98 |
| amphibians | Gastrotheca_cuencana | NA | EN | Decreasing | NA | NA | NA | NA | NA |
| amphibians | Gastrotheca_dendronastes | Gastrotheca_dendronastes | EN | Decreasing | tropical | nocturnal | 49.584 | 3405.576 | 12.429 |
| amphibians | Gastrotheca_dunni | Gastrotheca_dunni | LC | Stable | tropical | nocturnal | 15.932 | 2109.016 | 16.584 |
| amphibians | Gastrotheca_dysprosita | NA | DD | Unknown | NA | NA | NA | NA | NA |
| amphibians | Gastrotheca_elicioi | NA | not_available | not_available | NA | NA | NA | NA | NA |
| amphibians | Gastrotheca_ernestoi | Gastrotheca_ernestoi | LC | Stable | tropical | nocturnal | 41.053 | 55978.949 | 14.766 |
| amphibians | Gastrotheca_espeletia | Gastrotheca_espeletia | EN | Decreasing | tropical | cathemeral | 13.006 | 3877.469 | 16.424 |
| amphibians | Gastrotheca_fissipes | Gastrotheca_fissipes | LC | Decreasing | tropical | nocturnal | 75.997 | 45618.322 | 15.412 |
| amphibians | Gastrotheca_fulvorufa | Gastrotheca_fulvorufa | LC | Decreasing | tropical | nocturnal | 45.085 | 23229.113 | 16.458 |
| amphibians | Gastrotheca_galeata | NA | DD | Unknown | NA | NA | NA | NA | NA |
| amphibians | Gastrotheca_guentheri | Gastrotheca_guentheri | DD | Decreasing | tropical | nocturnal | 68.909 | 9010.522 | 7.407 |
| amphibians | Gastrotheca_helenae | Gastrotheca_helenae | EN | Stable | tropical | nocturnal | 27.131 | 446.866 | 5.672 |
| amphibians | Gastrotheca_litonedis | Gastrotheca_litonedis | CR | Decreasing | tropical | nocturnal | 22.623 | 81.001 | 26.701 |
| amphibians | Gastrotheca_lojana | NA | VU | Decreasing | NA | NA | NA | NA | NA |
| amphibians | Gastrotheca_longipes | Gastrotheca_longipes | LC | Decreasing | tropical | nocturnal | 69.673 | 54620.765 | 2.617 |
| amphibians | Gastrotheca_marsupiata | Gastrotheca_marsupiata | LC | Stable | tropical | nocturnal | 10.753 | 245721.672 | 9.645 |
| amphibians | Gastrotheca_megacephala | Gastrotheca_megacephala | LC | Decreasing | tropical | nocturnal | 141.992 | 72659.368 | 9.165 |
| amphibians | Gastrotheca_microdiscus | Gastrotheca_microdiscus | LC | Decreasing | temperate | nocturnal | 28.349 | 69344.745 | 11.966 |
| amphibians | Gastrotheca_monticola | Gastrotheca_monticola | LC | Stable | tropical | cathemeral | 44.708 | 30970.555 | 8.644 |
| amphibians | Gastrotheca_nicefori | Gastrotheca_nicefori | LC | Stable | tropical | nocturnal | 54.816 | 113015.127 | 10.578 |
| amphibians | Gastrotheca_orophylax | Gastrotheca_orophylax | VU | Decreasing | tropical | nocturnal | 39.306 | 3165.922 | 5.197 |
| amphibians | Gastrotheca_ossilaginis | NA | DD | Unknown | NA | NA | NA | NA | NA |
| amphibians | Gastrotheca_ovifera | Gastrotheca_ovifera | VU | Decreasing | tropical | cathemeral | 57.012 | 1413.662 | 10.601 |
| amphibians | Gastrotheca_pacchamama | Gastrotheca_pacchamama | EN | Unknown | tropical | diurnal | 4.536 | 224.06 | 8.54 |
| amphibians | Gastrotheca_pachachacae | Gastrotheca_pachachacae | VU | Unknown | tropical | nocturnal | 3.572 | 9.657 | 6.984 |
| amphibians | Gastrotheca_phalarosa | NA | DD | Unknown | NA | NA | NA | NA | NA |
| amphibians | Gastrotheca_plumbea | Gastrotheca_plumbea | VU | Decreasing | tropical | cathemeral | 37.611 | 7778.499 | 10.102 |
| amphibians | Gastrotheca_prasina | NA | DD | Unknown | NA | NA | NA | NA | NA |
| amphibians | Gastrotheca_pseustes | Gastrotheca_pseustes | NT | Decreasing | tropical | nocturnal | 24.186 | 34203.836 | 13.4 |

| | | | | | | | | | |
|---|---|---|---|---|---|---|---|---|---|
| amphibians | Gastrotheca_psychrophila | Gastrotheca_psychrophila | EN | Decreasing | tropical | cathemeral | 23.941 | 227.751 | 7.486 |
| amphibians | Gastrotheca_pulchra | Gastrotheca_pulchra | LC | Stable | tropical | nocturnal | 3.224 | 76462.292 | 12.364 |
| amphibians | Gastrotheca_rebeccae | Gastrotheca_rebeccae | EN | Unknown | tropical | nocturnal | 8.247 | 309.997 | 6.434 |
| amphibians | Gastrotheca_recava | Gastrotheca_recava | LC | Stable | tropical | nocturnal | 61.35 | 34642.798 | 7.394 |
| amphibians | Gastrotheca_riobambae | Gastrotheca_riobambae | VU | Decreasing | tropical | nocturnal | 27.668 | 9041.945 | 20.819 |
| amphibians | Gastrotheca_spectabilis | NA | DD | Unknown | NA | NA | NA | NA | NA |
| amphibians | Gastrotheca_splendens | NA | DD | Unknown | NA | NA | NA | NA | NA |
| amphibians | Gastrotheca_testudinea | Gastrotheca_testudinea | LC | Decreasing | tropical | nocturnal | 56.126 | 204436.356 | 4.972 |
| amphibians | Gastrotheca_trachyceps | Gastrotheca_trachyceps | EN | Decreasing | tropical | cathemeral | 29.887 | 99.924 | 8.913 |
| amphibians | Gastrotheca_turnerorum | NA | EN | Decreasing | NA | NA | NA | NA | NA |
| amphibians | Gastrotheca_walkeri | Gastrotheca_walkeri | VU | Unknown | tropical | nocturnal | 50.601 | 1181.874 | 8.958 |
| amphibians | Gastrotheca_weinlandii | Gastrotheca_weinlandii | LC | Decreasing | tropical | nocturnal | 63.963 | 25298.417 | 5.358 |
| amphibians | Gastrotheca_yacuri | NA | EN | Decreasing | NA | NA | NA | NA | NA |
| amphibians | Gastrotheca_zeugocystis | Gastrotheca_zeugocystis | DD | Decreasing | tropical | nocturnal | 4.345 | 10.005 | 9.375 |
| amphibians | Geobatrachus_walkeri | Geobatrachus_walkeri | EN | Stable | tropical | cathemeral | 0.772 | 478.406 | 5.218 |
| amphibians | Geotrypetes_seraphini | Geotrypetes_seraphini | LC | Decreasing | tropical | nocturnal | 24.115 | 1139615.211 | 10.533 |
| amphibians | Gephyromantis_ambohitra | Gephyromantis_ambohitra | VU | Decreasing | tropical | cathemeral | 5.665 | 4609.657 | 4.979 |
| amphibians | Gephyromantis_angano | NA | not_available | not_available | NA | NA | NA | NA | NA |
| amphibians | Gephyromantis_asper | Gephyromantis_asper | LC | Decreasing | tropical | cathemeral | 2.345 | 34830.199 | 8.006 |
| amphibians | Gephyromantis_blanci | Gephyromantis_blanci | NT | Decreasing | tropical | diurnal | 0.892 | 18076.393 | 8.273 |
| amphibians | Gephyromantis_boulengeri | Gephyromantis_boulengeri | LC | Decreasing | tropical | cathemeral | 3.475 | 27306.747 | 9.071 |
| amphibians | Gephyromantis_corvus | Gephyromantis_corvus | EN | Decreasing | tropical | cathemeral | 4.536 | 963.518 | 6.883 |
| amphibians | Gephyromantis_decaryi | Gephyromantis_decaryi | NT | Decreasing | tropical | cathemeral | 1.911 | 20003.247 | 8.32 |
| amphibians | Gephyromantis_eiselti | Gephyromantis_eiselti | EN | Decreasing | tropical | diurnal | 0.892 | 585.738 | 6.727 |
| amphibians | Gephyromantis_enki | Gephyromantis_enki | VU | Decreasing | tropical | diurnal | 0.664 | 7312.15 | 8.93 |
| amphibians | Gephyromantis_granulatus | Gephyromantis_granulatus | LC | Stable | tropical | nocturnal | 7.844 | 14033.981 | 6.027 |
| amphibians | Gephyromantis_grosjeani | NA | not_available | not_available | NA | NA | NA | NA | NA |
| amphibians | Gephyromantis_horridus | Gephyromantis_horridus | VU | Decreasing | tropical | nocturnal | 3.475 | 1003.383 | 7.911 |
| amphibians | Gephyromantis_kintana | NA | not_available | not_available | NA | NA | NA | NA | NA |
| amphibians | Gephyromantis_klemmeri | Gephyromantis_klemmeri | EN | Decreasing | tropical | diurnal | 1.326 | 1336.428 | 5.582 |
| amphibians | Gephyromantis_leucocephalus | Gephyromantis_leucocephalus | NT | Stable | temperate | diurnal | 2.872 | 10854.377 | 8.36 |
| amphibians | Gephyromantis_leucomaculatus | Gephyromantis_leucomaculatus | LC | Decreasing | tropical | nocturnal | 9.031 | 19419.018 | 7.278 |
| amphibians | Gephyromantis_luteus | Gephyromantis_luteus | LC | Decreasing | tropical | cathemeral | 19.923 | 73777.667 | 8.714 |
| amphibians | Gephyromantis_malagasius | Gephyromantis_malagasius | LC | Decreasing | tropical | nocturnal | 1.889 | 83503.159 | 8.727 |

| | | | | | | | | | |
|---|---|---|---|---|---|---|---|---|---|
| amphibians | Gephyromantis_mitsinjo | NA | not_available | not_available | NA | NA | NA | NA | NA |
| amphibians | Gephyromantis_moseri | Gephyromantis_moseri | LC | Decreasing | tropical | nocturnal | 5.356 | 38143.01 | 7.296 |
| amphibians | Gephyromantis_plicifer | Gephyromantis_plicifer | LC | Decreasing | tropical | cathemeral | 11.036 | 26307.335 | 5.678 |
| amphibians | Gephyromantis_pseudoasper | Gephyromantis_pseudoasper | LC | Stable | tropical | cathemeral | 3.163 | 20436.006 | 6.22 |
| amphibians | Gephyromantis_ranjomavo | Gephyromantis_ranjomavo | EN | Decreasing | tropical | nocturnal | 1.686 | 597.631 | 4.623 |
| amphibians | Gephyromantis_redimitus | Gephyromantis_redimitus | LC | Decreasing | tropical | cathemeral | 13.329 | 51901.96 | 7.86 |
| amphibians | Gephyromantis_rivicola | Gephyromantis_rivicola | VU | Decreasing | tropical | diurnal | 1.023 | 8412.826 | 7.012 |
| amphibians | Gephyromantis_runewsweeki | Gephyromantis_runewsweeki | VU | Unknown | tropical | diurnal | 1.023 | 1.419 | 3.508 |
| amphibians | Gephyromantis_salegy | Gephyromantis_salegy | VU | Decreasing | tropical | nocturnal | 11.036 | 4037.461 | 5.167 |
| amphibians | Gephyromantis_saturnini | NA | not_available | not_available | NA | NA | NA | NA | NA |
| amphibians | Gephyromantis_schilfi | Gephyromantis_schilfi | VU | Decreasing | tropical | nocturnal | 2.109 | 3639.508 | 4.2 |
| amphibians | Gephyromantis_silvanus | Gephyromantis_silvanus | VU | Decreasing | tropical | cathemeral | 2.345 | 3068.074 | 10.746 |
| amphibians | Gephyromantis_spiniferus | Gephyromantis_spiniferus | VU | Decreasing | temperate | diurnal | 4.536 | 7462.233 | 5.281 |
| amphibians | Gephyromantis_striatus | Gephyromantis_striatus | VU | Decreasing | tropical | nocturnal | 1.499 | 10455.972 | 5.906 |
| amphibians | Gephyromantis_tahotra | Gephyromantis_tahotra | VU | Decreasing | tropical | cathemeral | 3.807 | 4431.216 | 4.202 |
| amphibians | Gephyromantis_tandroka | Gephyromantis_tandroka | VU | Decreasing | tropical | nocturnal | 7.844 | 4431.216 | 4.202 |
| amphibians | Gephyromantis_thelenae | Gephyromantis_thelenae | EN | Decreasing | tropical | diurnal | 0.996 | 585.738 | 6.727 |
| amphibians | Gephyromantis_tohatra | NA | not_available | not_available | NA | NA | NA | NA | NA |
| amphibians | Gephyromantis_tschenki | Gephyromantis_tschenki | LC | Decreasing | tropical | nocturnal | 3.807 | 11082.666 | 6.425 |
| amphibians | Gephyromantis_webbi | Gephyromantis_webbi | EN | Decreasing | tropical | cathemeral | 2.872 | 2489.085 | 10.586 |
| amphibians | Gephyromantis_zavona | Gephyromantis_zavona | EN | Decreasing | tropical | nocturnal | 7.294 | 3012.957 | 4.773 |
| amphibians | Glandirana_rugosa | Rugosa_rugosa | LC | Decreasing | temperate | nocturnal | 19.923 | 310817.988 | 20.382 |
| amphibians | Glyphoglossus_guttulatus | Calluella_guttulata | LC | Decreasing | tropical | nocturnal | 14.161 | 372937.283 | 10.163 |
| amphibians | Glyphoglossus_molossus | Glyphoglossus_molossus | NT | Decreasing | tropical | nocturnal | 63.963 | 871247.256 | 15.76 |
| amphibians | Gracixalus_supercornutus | Gracixalus_supercornutus | NT | Decreasing | tropical | nocturnal | 2.626 | 2976.643 | 3.393 |
| amphibians | Guibemantis_bicalcaratus | Guibemantis_bicalcaratus | LC | Decreasing | tropical | diurnal | 1.889 | 112874.115 | 8.114 |
| amphibians | Guibemantis_depressiceps | Guibemantis_depressiceps | LC | Unknown | tropical | nocturnal | 7.844 | 134877.676 | 9.324 |
| amphibians | Guibemantis_diphonus | NA | CR | Decreasing | NA | NA | NA | NA | NA |
| amphibians | Guibemantis_kathrinae | Guibemantis_kathrinae | VU | Decreasing | tropical | nocturnal | 18.867 | 449.529 | 8.292 |
| amphibians | Guibemantis_liber | Guibemantis_liber | LC | Unknown | tropical | cathemeral | 1.889 | 143701.416 | 9.122 |
| amphibians | Guibemantis_pulcher | Guibemantis_pulcher | LC | Decreasing | tropical | cathemeral | 1.686 | 62804.354 | 7.53 |
| amphibians | Guibemantis_punctatus | Guibemantis_punctatus | CR | Decreasing | tropical | diurnal | 1.499 | 49.479 | 9.609 |
| amphibians | Guibemantis_timidus | Guibemantis_timidus | LC | Unknown | tropical | nocturnal | 15.029 | 23490.275 | 9.795 |
| amphibians | Guibemantis_tornieri | Guibemantis_tornieri | LC | Decreasing | tropical | nocturnal | 18.253 | 61565.49 | 7.666 |

| | | | | | | | | | |
|---|---|---|---|---|---|---|---|---|---|
| amphibians | Gymnopis_multiplicata | Gymnopis_multiplicata | LC | Decreasing | tropical | cathemeral | 41.823 | 182860.26 | 8.98 |
| amphibians | Gyrinophilus_palleucus | Gyrinophilus_palleucus | NT | Stable | temperate | nocturnal | 15.677 | 12512.721 | 18.757 |
| amphibians | Gyrinophilus_porphyriticus | Gyrinophilus_porphyriticus | LC | Stable | temperate | nocturnal | 26.796 | 764568.268 | 17.216 |
| amphibians | Haddadus_aramunha | Haddadus_aramunha | VU | Decreasing | tropical | nocturnal | 22.156 | 7731.241 | 7.888 |
| amphibians | Haddadus_binotatus | Haddadus_binotatus | LC | Stable | tropical | cathemeral | 24.557 | 496585.753 | 12.062 |
| amphibians | Hadromophryne_natalensis | Hadromophryne_natalensis | LC | Unknown | temperate | nocturnal | 25.822 | 121695.095 | 13.079 |
| amphibians | Hamptophryne_boliviana | Hamptophryne_boliviana | LC | Stable | tropical | cathemeral | 7.844 | 4439329.92 | 1.872 |
| amphibians | Heleioporus_australiacus | Heleioporus_australiacus | EN | Decreasing | temperate | nocturnal | 94.47 | 42343.838 | 9.487 |
| amphibians | Heleophryne_orientalis | Heleophryne_orientalis | LC | Unknown | temperate | cathemeral | 8.423 | 1191.272 | 7.29 |
| amphibians | Heleophryne_purcelli | Heleophryne_purcelli | LC | Unknown | temperate | diurnal | 15.932 | 14223.904 | 10.908 |
| amphibians | Hemidactylium_scutatum | Hemidactylium_scutatum | LC | Stable | temperate | diurnal | 2.293 | 1511888.541 | 17.88 |
| amphibians | Hemiphractus_bubalus | Hemiphractus_bubalus | VU | Decreasing | tropical | nocturnal | 27.533 | 32624.986 | 5.73 |
| amphibians | Hemiphractus_elioti | NA | CR | Decreasing | NA | NA | NA | NA | NA |
| amphibians | Hemiphractus_fasciatus | Hemiphractus_fasciatus | VU | Decreasing | tropical | cathemeral | 30.895 | 57066.597 | 6.135 |
| amphibians | Hemiphractus_helioi | Hemiphractus_helioi | LC | Decreasing | tropical | nocturnal | 41.053 | 400478.006 | 1.642 |
| amphibians | Hemiphractus_johnsoni | Hemiphractus_johnsoni | EN | Decreasing | tropical | nocturnal | 45.085 | 267.741 | 21.02 |
| amphibians | Hemiphractus_kaylockae | NA | EN | Decreasing | NA | NA | NA | NA | NA |
| amphibians | Hemiphractus_panamensis | NA | CR | Decreasing | NA | NA | NA | NA | NA |
| amphibians | Hemiphractus_proboscideus | Hemiphractus_proboscideus | LC | Decreasing | tropical | cathemeral | 28.486 | 338926.374 | 1.772 |
| amphibians | Hemiphractus_scutatus | Hemiphractus_scutatus | LC | Unknown | tropical | cathemeral | 52.68 | 2046879.868 | 1.36 |
| amphibians | Hemisus_guttatus | Hemisus_guttatus | NT | Decreasing | temperate | nocturnal | 50.601 | 8652.857 | 18.723 |
| amphibians | Hemisus_marmoratus | Hemisus_marmoratus | LC | Unknown | tropical | nocturnal | 19.923 | 9852655.862 | 9.716 |
| amphibians | Heterixalus_andrakata | Heterixalus_andrakata | LC | Stable | tropical | nocturnal | 2.599 | 1128.235 | 9.423 |
| amphibians | Heterixalus_betsileo | Heterixalus_betsileo | LC | Stable | tropical | nocturnal | 1.889 | 106400.593 | 10.467 |
| amphibians | Heterixalus_boettgeri | Heterixalus_boettgeri | LC | Unknown | temperate | cathemeral | 2.109 | 4328.294 | 15.533 |
| amphibians | Heterixalus_carbonei | Heterixalus_carbonei | LC | Decreasing | tropical | nocturnal | 1.326 | 3066.059 | 5.8 |
| amphibians | Heterixalus_luteostriatus | Heterixalus_luteostriatus | LC | Stable | tropical | nocturnal | 1.889 | 69382.224 | 8.545 |
| amphibians | Heterixalus_madagascariensis | Heterixalus_madagascariensis | LC | Stable | tropical | cathemeral | 5.356 | 29982.451 | 8.949 |
| amphibians | Heterixalus_punctatus | Heterixalus_punctatus | LC | Stable | tropical | cathemeral | 2.9 | 42198.26 | 8.498 |
| amphibians | Heterixalus_tricolor | Heterixalus_tricolor | LC | Stable | tropical | cathemeral | 2.109 | 26.632 | 7.745 |
| amphibians | Heterixalus_variabilis | Heterixalus_variabilis | LC | Stable | tropical | nocturnal | 3.163 | 3179.235 | 11.931 |
| amphibians | Hildebrandtia_macrotympanum | Hildebrandtia_macrotympanum | LC | Unknown | tropical | nocturnal | 11.767 | 738017.667 | 9.065 |
| amphibians | Hildebrandtia_ornata | Hildebrandtia_ornata | LC | Unknown | tropical | cathemeral | 32.83 | 6748590.698 | 9.244 |
| amphibians | Holoaden_luederwaldti | Holoaden_luederwaldti | NT | Stable | tropical | nocturnal | 9.669 | 2364.034 | 10.093 |

| | | | | | | | | | |
|---|---|---|---|---|---|---|---|---|---|
| amphibians | Holoaden_pholeter | NA | DD | Unknown | NA | NA | NA | NA | NA |
| amphibians | Hoplobatrachus_chinensis | NA | LC | Decreasing | NA | NA | NA | NA | NA |
| amphibians | Hoplobatrachus_crassus | Hoplobatrachus_crassus | LC | Decreasing | tropical | nocturnal | 275.697 | 2411778.567 | 18.511 |
| amphibians | Hoplobatrachus_occipitalis | Hoplobatrachus_occipitalis | LC | Stable | tropical | cathemeral | 478.079 | 8574915.608 | 10.488 |
| amphibians | Hoplobatrachus_tigerinus | Hoplobatrachus_tigerinus | LC | Stable | tropical | cathemeral | 675.324 | 3535067.892 | 17.623 |
| amphibians | Hoplophryne_rogersi | Hoplophryne_rogersi | EN | Unknown | tropical | diurnal | 2.599 | 1002.536 | 13.813 |
| amphibians | Huia_cavitympanum | Huia_cavitympanum | LC | Decreasing | tropical | nocturnal | 50.601 | 220248.906 | 3.166 |
| amphibians | Hyalinobatrachium_aureoguttatum | Hyalinobatrachium_aureoguttatum | LC | Stable | tropical | nocturnal | 1.023 | 25282.311 | 5.425 |
| amphibians | Hyalinobatrachium_cappellei | Hyalinobatrachium_cappellei | LC | Stable | tropical | nocturnal | 1.153 | 2321705.746 | 1.291 |
| amphibians | Hyalinobatrachium_carlesvilai | Hyalinobatrachium_carlesvilai | LC | Stable | tropical | nocturnal | 1.01 | 204335.595 | 3.351 |
| amphibians | Hyalinobatrachium_chirripoi | Hyalinobatrachium_chirripoi | LC | Stable | tropical | nocturnal | 2.109 | 57455.948 | 5.319 |
| amphibians | Hyalinobatrachium_colymbiphyllum | Hyalinobatrachium_colymbiphyllum | LC | Stable | tropical | nocturnal | 1.889 | 113097.155 | 10.91 |
| amphibians | Hyalinobatrachium_dianae | NA | LC | Stable | NA | NA | NA | NA | NA |
| amphibians | Hyalinobatrachium_fleischmanni | Hyalinobatrachium_fleischmanni | LC | Stable | tropical | nocturnal | 2.599 | 37182.307 | 10.96 |
| amphibians | Hyalinobatrachium_guairarepanense | Hyalinobatrachium_guairarepanense | EN | Decreasing | tropical | nocturnal | 0.969 | 1934.254 | 20.16 |
| amphibians | Hyalinobatrachium_mondolfii | Hyalinobatrachium_mondolfii | LC | Stable | tropical | cathemeral | 0.83 | 1129956.559 | 1.065 |
| amphibians | Hyalinobatrachium_munozorum | Hyalinobatrachium_munozorum | LC | Stable | tropical | nocturnal | 0.969 | 39779.407 | 5.089 |
| amphibians | Hyalinobatrachium_orientale | Hyalinobatrachium_orientale | VU | Decreasing | tropical | nocturnal | 0.969 | 2704.257 | 7.899 |
| amphibians | Hyalinobatrachium_orocostale | Hyalinobatrachium_orocostale | EN | Decreasing | tropical | nocturnal | 0.614 | 460.012 | 8.646 |
| amphibians | Hyalinobatrachium_pallidum | Hyalinobatrachium_pallidum | NT | Stable | tropical | nocturnal | 1.168 | 2564.451 | 5.935 |
| amphibians | Hyalinobatrachium_pellucidum | Hyalinobatrachium_pellucidum | NT | Decreasing | tropical | nocturnal | 0.772 | 9557.73 | 6.471 |
| amphibians | Hyalinobatrachium_talamancae | Hyalinobatrachium_talamancae | LC | Stable | tropical | nocturnal | 1.499 | 519.064 | 4.945 |
| amphibians | Hyalinobatrachium_tatayoi | Hyalinobatrachium_tatayoi | LC | Stable | tropical | nocturnal | 0.818 | 232033.577 | 8.734 |
| amphibians | Hyalinobatrachium_taylori | Hyalinobatrachium_taylori | LC | Stable | tropical | nocturnal | 0.772 | 463793.297 | 1.059 |
| amphibians | Hyalinobatrachium_tricolor | Hyalinobatrachium_tricolor | LC | Unknown | tropical | nocturnal | 0.664 | 37061.529 | 1.043 |
| amphibians | Hyalinobatrachium_valerioi | Hyalinobatrachium_valerioi | LC | Decreasing | tropical | nocturnal | 1.326 | 69861.615 | 9.052 |
| amphibians | Hydrolaetare_dantasi | Hydrolaetare_dantasi | LC | Stable | tropical | cathemeral | 101.59 | 36488.038 | 1.19 |
| amphibians | Hydrolaetare_schmidti | Hydrolaetare_schmidti | LC | Stable | tropical | nocturnal | 188.234 | 3669046.04 | 1.704 |
| amphibians | Hydromantes_samweli | NA | NT | Stable | NA | NA | NA | NA | NA |
| amphibians | Hydromantes_shastae | Hydromantes_shastae | NT | Stable | temperate | nocturnal | 4.144 | 747.514 | 10.145 |
| amphibians | Hydromantes_wintu | NA | VU | Stable | NA | NA | NA | NA | NA |
| amphibians | Hyla_annectans | Hyla_annectans | LC | Decreasing | temperate | nocturnal | 7.028 | 1002137.489 | 12.059 |
| amphibians | Hyla_arborea | Hyla_arborea | LC | Decreasing | temperate | cathemeral | 12.531 | 1771404.495 | 22.025 |
| amphibians | Hyla_chinensis | Hyla_chinensis | LC | Decreasing | temperate | nocturnal | 4.653 | 640031.699 | 15.864 |

| | | | | | | | | | |
|---|---|---|---|---|---|---|---|---|---|
| amphibians | Hyla_felixarabica | Hyla_felixarabica | LC | Stable | tropical | nocturnal | 7.028 | 94584.209 | 11.899 |
| amphibians | Hyla_intermedia | Hyla_intermedia | LC | Decreasing | temperate | cathemeral | 19.923 | 230703.125 | 23.431 |
| amphibians | Hyla_meridionalis | Hyla_meridionalis | LC | Decreasing | temperate | cathemeral | 25.822 | 643813.43 | 18.004 |
| amphibians | Hyla_orientalis | Hyla_orientalis | LC | Decreasing | temperate | cathemeral | 13.329 | 1852620.443 | 16.783 |
| amphibians | Hyla_sanchiangensis | Hyla_sanchiangensis | LC | Decreasing | temperate | nocturnal | 4.345 | 442581.777 | 12.785 |
| amphibians | Hyla_sarda | Hyla_sarda | LC | Stable | temperate | cathemeral | 11.036 | 31503.519 | 17.018 |
| amphibians | Hyla_savignyi | Hyla_savignyi | LC | Stable | temperate | cathemeral | 10.337 | 694154.181 | 16.429 |
| amphibians | Hylambates_boulengeri | NA | LC | Unknown | NA | NA | NA | NA | NA |
| amphibians | Hylambates_keithae | NA | EN | Decreasing | NA | NA | NA | NA | NA |
| amphibians | Hylambates_leonardi | NA | LC | Stable | NA | NA | NA | NA | NA |
| amphibians | Hylambates_maculatus | NA | LC | Unknown | NA | NA | NA | NA | NA |
| amphibians | Hylambates_verrucosus | NA | LC | Unknown | NA | NA | NA | NA | NA |
| amphibians | Hylarana_albolabris | NA | LC | Unknown | NA | NA | NA | NA | NA |
| amphibians | Hylarana_amnicola | NA | LC | Decreasing | NA | NA | NA | NA | NA |
| amphibians | Hylarana_aurantiaca | Hylarana_aurantiaca | EN | Decreasing | tropical | nocturnal | 6.77 | 2811.038 | 12.207 |
| amphibians | Hylarana_aurata | NA | DD | Unknown | NA | NA | NA | NA | NA |
| amphibians | Hylarana_chalconota | NA | LC | Stable | NA | NA | NA | NA | NA |
| amphibians | Hylarana_chitwanensis | NA | DD | Unknown | NA | NA | NA | NA | NA |
| amphibians | Hylarana_daemeli | NA | LC | Stable | NA | NA | NA | NA | NA |
| amphibians | Hylarana_darlingi | NA | LC | Unknown | NA | NA | NA | NA | NA |
| amphibians | Hylarana_erythraea | Hylarana_erythraea | LC | Decreasing | tropical | cathemeral | 46.616 | 2299374.729 | 13.174 |
| amphibians | Hylarana_fantastica | NA | not_available | not_available | NA | NA | NA | NA | NA |
| amphibians | Hylarana_galamensis | NA | LC | Unknown | NA | NA | NA | NA | NA |
| amphibians | Hylarana_garritor | NA | LC | Decreasing | NA | NA | NA | NA | NA |
| amphibians | Hylarana_gracilis | Hylarana_gracilis | LC | Decreasing | tropical | cathemeral | 23.941 | 41299.366 | 18.66 |
| amphibians | Hylarana_labialis | NA | LC | Stable | NA | NA | NA | NA | NA |
| amphibians | Hylarana_lateralis | NA | LC | Unknown | NA | NA | NA | NA | NA |
| amphibians | Hylarana_latouchii | Hylarana_latouchii | LC | Stable | temperate | diurnal | 10.648 | 875372.513 | 15.901 |
| amphibians | Hylarana_macrodactyla | Hylarana_macrodactyla | LC | Stable | tropical | nocturnal | 7.294 | 1762927.828 | 14.179 |
| amphibians | Hylarana_malabarica | NA | LC | Decreasing | NA | NA | NA | NA | NA |
| amphibians | Hylarana_megalonesa | Hylarana_megalonesa | LC | Unknown | tropical | nocturnal | 26.602 | 22774.667 | 1.9 |
| amphibians | Hylarana_milleti | NA | LC | Decreasing | NA | NA | NA | NA | NA |
| amphibians | Hylarana_milneana | NA | LC | Stable | NA | NA | NA | NA | NA |
| amphibians | Hylarana_nigrovittata | Hylarana_nigrovittata | LC | Decreasing | tropical | cathemeral | 32.83 | 783557.812 | 9.966 |

| | | | | | | | | | |
|---|---|---|---|---|---|---|---|---|---|
| amphibians | Hylarana_occidentalis | NA | LC | Decreasing | NA | NA | NA | NA | NA |
| amphibians | Hylarana_papua | NA | LC | Stable | NA | NA | NA | NA | NA |
| amphibians | Hylarana_picturata | NA | LC | Decreasing | NA | NA | NA | NA | NA |
| amphibians | Hylarana_serendipi | NA | VU | Decreasing | NA | NA | NA | NA | NA |
| amphibians | Hylarana_supragrisea | NA | LC | Stable | NA | NA | NA | NA | NA |
| amphibians | Hylarana_taipehensis | Hylarana_taipehensis | LC | Decreasing | tropical | cathemeral | 7.844 | 1177402.892 | 14.397 |
| amphibians | Hylarana_temporalis | Hylarana_temporalis | VU | Decreasing | tropical | nocturnal | 54.816 | 17658.528 | 17.203 |
| amphibians | Hylarana_volkerjane | NA | LC | Unknown | NA | NA | NA | NA | NA |
| amphibians | Hylodes_amnicola | Hylodes_amnicola | LC | Stable | tropical | diurnal | 2.321 | 23.196 | 4.783 |
| amphibians | Hylodes_asper | Hylodes_asper | LC | Stable | tropical | diurnal | 11.036 | 23535.623 | 15.93 |
| amphibians | Hylodes_babax | Hylodes_babax | NT | Stable | tropical | diurnal | 2.599 | 1244.95 | 6.523 |
| amphibians | Hylodes_caete | NA | LC | Stable | NA | NA | NA | NA | NA |
| amphibians | Hylodes_cardosoi | Hylodes_cardosoi | LC | Stable | temperate | diurnal | 8.724 | 11043.917 | 6.947 |
| amphibians | Hylodes_charadranaetes | Hylodes_charadranaetes | LC | Unknown | tropical | diurnal | 6.77 | 863.285 | 10.805 |
| amphibians | Hylodes_dactylocinus | Hylodes_dactylocinus | EN | Decreasing | temperate | diurnal | 2.345 | 7122.307 | 8.865 |
| amphibians | Hylodes_heyeri | Hylodes_heyeri | LC | Stable | temperate | diurnal | 8.247 | 45673.362 | 10.657 |
| amphibians | Hylodes_japi | NA | NT | Stable | NA | NA | NA | NA | NA |
| amphibians | Hylodes_lateristrigatus | Hylodes_lateristrigatus | LC | Stable | tropical | diurnal | 5.848 | 97067.258 | 14.341 |
| amphibians | Hylodes_magalhaesi | Hylodes_magalhaesi | EN | Decreasing | tropical | diurnal | 3.255 | 2304.986 | 10.415 |
| amphibians | Hylodes_meridionalis | Hylodes_meridionalis | LC | Stable | temperate | diurnal | 11.036 | 34414.972 | 10.951 |
| amphibians | Hylodes_nasus | Hylodes_nasus | EN | Decreasing | temperate | diurnal | 4.536 | 39520.358 | 16.649 |
| amphibians | Hylodes_ornatus | Hylodes_ornatus | VU | Unknown | tropical | diurnal | 1.245 | 11.694 | 8.496 |
| amphibians | Hylodes_otavioi | Hylodes_otavioi | VU | Decreasing | tropical | diurnal | 3.739 | 6194.52 | 6.921 |
| amphibians | Hylodes_perere | Hylodes_perere | EN | Decreasing | tropical | diurnal | 1.766 | 223.562 | 6.56 |
| amphibians | Hylodes_perplicatus | Hylodes_perplicatus | LC | Decreasing | temperate | diurnal | 8.188 | 35693.239 | 11.122 |
| amphibians | Hylodes_phyllodes | Hylodes_phyllodes | LC | Stable | tropical | diurnal | 2.652 | 22660.214 | 15.636 |
| amphibians | Hylodes_pipilans | Hylodes_pipilans | LC | Stable | tropical | diurnal | 1.686 | 3106.997 | 15.202 |
| amphibians | Hylodes_regius | Hylodes_regius | CR | Unknown | tropical | diurnal | 4.536 | 78.027 | 6.355 |
| amphibians | Hylodes_sazimai | Hylodes_sazimai | VU | Decreasing | tropical | diurnal | 2.178 | 5953.209 | 16.073 |
| amphibians | Hylodes_uai | Hylodes_uai | VU | Decreasing | tropical | diurnal | 4.016 | 9169.002 | 17.103 |
| amphibians | Hylomantis_aspera | Hylomantis_aspera | LC | Stable | tropical | nocturnal | 9.669 | 37306.786 | 9.495 |
| amphibians | Hylomantis_granulosa | Hylomantis_granulosa | LC | Stable | tropical | nocturnal | 9.669 | 110840.517 | 13.943 |
| amphibians | Hylophorbus_rufescens | NA | DD | Unknown | NA | NA | NA | NA | NA |
| amphibians | Hylophorbus_sigridae | NA | LC | Stable | NA | NA | NA | NA | NA |

| | | | | | | | | | |
|---|---|---|---|---|---|---|---|---|---|
| amphibians | Hylorina_sylvatica | Hylorina_sylvatica | LC | Decreasing | temperate | cathemeral | 41.053 | 91167.035 | 4.488 |
| amphibians | Hyloscirtus_albopunctulatus | Hyloscirtus_albopunctulatus | LC | Decreasing | tropical | nocturnal | 6.034 | 42987.24 | 3.688 |
| amphibians | Hyloscirtus_alytolylax | Hyloscirtus_alytolylax | LC | Decreasing | tropical | nocturnal | 8.423 | 36912.052 | 8.014 |
| amphibians | Hyloscirtus_colymba | Hyloscirtus_colymba | EN | Decreasing | tropical | nocturnal | 6.924 | 15862.067 | 3.107 |
| amphibians | Hyloscirtus_diabolus | NA | VU | Unknown | NA | NA | NA | NA | NA |
| amphibians | Hyloscirtus_japreria | NA | EN | Decreasing | NA | NA | NA | NA | NA |
| amphibians | Hyloscirtus_lascinius | Hyloscirtus_lascinius | LC | Stable | tropical | nocturnal | 9.031 | 2991.657 | 14.67 |
| amphibians | Hyloscirtus_lindae | Hyloscirtus_lindae | LC | Stable | tropical | nocturnal | 44.708 | 42055.517 | 5.945 |
| amphibians | Hyloscirtus_mashpi | NA | EN | Decreasing | NA | NA | NA | NA | NA |
| amphibians | Hyloscirtus_pacha | Hyloscirtus_pacha | EN | Decreasing | tropical | nocturnal | 30.417 | 70.835 | 8.05 |
| amphibians | Hyloscirtus_palmeri | Hyloscirtus_palmeri | LC | Decreasing | tropical | nocturnal | 11.036 | 91696.296 | 6.957 |
| amphibians | Hyloscirtus_pantostictus | Hyloscirtus_pantostictus | CR | Decreasing | tropical | nocturnal | 30.087 | 40.394 | 13.632 |
| amphibians | Hyloscirtus_phyllognathus | Hyloscirtus_phyllognathus | LC | Decreasing | tropical | nocturnal | 5.058 | 129430.389 | 4.525 |
| amphibians | Hyloscirtus_piceigularis | Hyloscirtus_piceigularis | EN | Decreasing | tropical | cathemeral | 5.802 | 219.372 | 6.945 |
| amphibians | Hyloscirtus_platydactylus | Hyloscirtus_platydactylus | VU | Decreasing | tropical | nocturnal | 6.419 | 5255.638 | 12.092 |
| amphibians | Hyloscirtus_princecharlesi | Hyloscirtus_princecharlesi | CR | Decreasing | tropical | nocturnal | 34.374 | 24.17 | 6.614 |
| amphibians | Hyloscirtus_psarolaimus | Hyloscirtus_psarolaimus | VU | Decreasing | tropical | nocturnal | 23.819 | 3852.128 | 3.309 |
| amphibians | Hyloscirtus_ptychodactylus | Hyloscirtus_ptychodactylus | EN | Decreasing | tropical | cathemeral | 45.274 | 487.427 | 6.424 |
| amphibians | Hyloscirtus_sarampiona | Hyloscirtus_sarampiona | EN | Unknown | tropical | nocturnal | 31.041 | 109.775 | 5.301 |
| amphibians | Hyloscirtus_staufferorum | Hyloscirtus_staufferorum | EN | Unknown | tropical | nocturnal | 22.882 | 476.968 | 5.55 |
| amphibians | Hyloscirtus_torrenticola | Hyloscirtus_torrenticola | VU | Decreasing | tropical | nocturnal | 3.638 | 5416.207 | 5.263 |
| amphibians | Hyloxalus_alessandroi | Allobates_alessandroi | EN | Decreasing | tropical | diurnal | 0.783 | 1113.515 | 1.928 |
| amphibians | Hyloxalus_anthracinus | Hyloxalus_anthracinus | CR | Decreasing | tropical | diurnal | 0.969 | 4418.938 | 15.626 |
| amphibians | Hyloxalus_arliensis | NA | not_available | not_available | NA | NA | NA | NA | NA |
| amphibians | Hyloxalus_awa | Hyloxalus_awa | LC | Stable | tropical | diurnal | 1.31 | 31202.067 | 10.336 |
| amphibians | Hyloxalus_bocagei | Hyloxalus_bocagei | VU | Decreasing | tropical | diurnal | 1.997 | 3003.421 | 5.135 |
| amphibians | Hyloxalus_breviquartus | Hyloxalus_breviquartus | LC | Decreasing | tropical | diurnal | 0.322 | 85489.658 | 11.931 |
| amphibians | Hyloxalus_craspedoceps | NA | DD | Unknown | NA | NA | NA | NA | NA |
| amphibians | Hyloxalus_delatorreae | Hyloxalus_delatorreae | CR | Decreasing | tropical | diurnal | 0.644 | 113.952 | 7.745 |
| amphibians | Hyloxalus_elachyhistus | Hyloxalus_elachyhistus | LC | Unknown | tropical | cathemeral | 1.277 | 24992.2 | 10.188 |
| amphibians | Hyloxalus_eleutherodactylus | NA | DD | Unknown | NA | NA | NA | NA | NA |
| amphibians | Hyloxalus_fascianigrus | Hyloxalus_fascianigrus | VU | Unknown | tropical | diurnal | 0.717 | 1057.691 | 9.652 |
| amphibians | Hyloxalus_felixcoperari | NA | not_available | not_available | NA | NA | NA | NA | NA |
| amphibians | Hyloxalus_fuliginosus | Hyloxalus_fuliginosus | EN | Decreasing | tropical | diurnal | 2.733 | 1407.725 | 5.752 |

| | | | | | | | | | |
|---|---|---|---|---|---|---|---|---|---|
| amphibians | Hyloxalus_infraguttatus | Hyloxalus_infraguttatus | VU | Decreasing | tropical | diurnal | 0.943 | 59007.583 | 15.226 |
| amphibians | Hyloxalus_insulatus | Hyloxalus_insulatus | VU | Unknown | tropical | diurnal | 1.326 | 15135.118 | 8.734 |
| amphibians | Hyloxalus_italoi | Hyloxalus_italoi | LC | Decreasing | tropical | diurnal | 2.132 | 36406.043 | 2.433 |
| amphibians | Hyloxalus_lehmanni | Hyloxalus_lehmanni | NT | Decreasing | tropical | diurnal | 0.867 | 64425.782 | 10.54 |
| amphibians | Hyloxalus_leucophaeus | NA | DD | Unknown | NA | NA | NA | NA | NA |
| amphibians | Hyloxalus_maculosus | Hyloxalus_maculosus | EN | Decreasing | tropical | diurnal | 1.889 | 2130.192 | 10.502 |
| amphibians | Hyloxalus_mittermeieri | NA | DD | Unknown | NA | NA | NA | NA | NA |
| amphibians | Hyloxalus_mystax | Hyloxalus_mystax | EN | Decreasing | tropical | diurnal | 0.749 | 2754.185 | 4.238 |
| amphibians | Hyloxalus_nexipus | Hyloxalus_nexipus | LC | Stable | tropical | diurnal | 3.475 | 31137.264 | 3.593 |
| amphibians | Hyloxalus_patitae | Hyloxalus_patitae | DD | Decreasing | tropical | diurnal | 0.917 | 5.9 | 1.09 |
| amphibians | Hyloxalus_picachos | Allobates_picachos | EN | Decreasing | tropical | diurnal | 0.464 | 358.9 | 7.537 |
| amphibians | Hyloxalus_pinguis | Hyloxalus_pinguis | EN | Decreasing | tropical | diurnal | 0.614 | 105.33 | 14.003 |
| amphibians | Hyloxalus_pulchellus | Hyloxalus_pulchellus | NT | Decreasing | tropical | diurnal | 1.08 | 13555.84 | 7.836 |
| amphibians | Hyloxalus_pulcherrimus | NA | DD | Unknown | NA | NA | NA | NA | NA |
| amphibians | Hyloxalus_ramosi | Hyloxalus_ramosi | EN | Decreasing | tropical | diurnal | 0.567 | 286.672 | 12.114 |
| amphibians | Hyloxalus_saltuarius | NA | DD | Unknown | NA | NA | NA | NA | NA |
| amphibians | Hyloxalus_sanctamariensis | NA | not_available | not_available | NA | NA | NA | NA | NA |
| amphibians | Hyloxalus_sauli | Hyloxalus_sauli | LC | Stable | tropical | diurnal | 1.827 | 56197.949 | 5.181 |
| amphibians | Hyloxalus_shuar | Hyloxalus_shuar | CR | Decreasing | tropical | cathemeral | 2.47 | 1218.124 | 7.101 |
| amphibians | Hyloxalus_sordidatus | NA | DD | Unknown | NA | NA | NA | NA | NA |
| amphibians | Hyloxalus_sylvaticus | Hyloxalus_sylvaticus | EN | Decreasing | tropical | diurnal | 2.109 | 784.382 | 10.626 |
| amphibians | Hyloxalus_toachi | Hyloxalus_toachi | VU | Decreasing | tropical | diurnal | 1.726 | 8072.704 | 5.482 |
| amphibians | Hyloxalus_vertebralis | Hyloxalus_vertebralis | VU | Decreasing | tropical | diurnal | 0.585 | 13409.498 | 11.86 |
| amphibians | Hyloxalus_yasuni | Hyloxalus_yasuni | LC | Decreasing | tropical | diurnal | 1.868 | 36611.134 | 5.573 |
| amphibians | Hymenochirus_boettgeri | Hymenochirus_boettgeri | LC | Unknown | tropical | nocturnal | 5.802 | 2034734.845 | 6.059 |
| amphibians | Hynobius_boulengeri | Hynobius_boulengeri | EN | Decreasing | temperate | nocturnal | 19.128 | 2837.244 | 9.814 |
| amphibians | Hynobius_dunni | Hynobius_dunni | VU | Decreasing | temperate | nocturnal | 8.135 | 4935.015 | 20.671 |
| amphibians | Hynobius_formosanus | Hynobius_formosanus | EN | Decreasing | temperate | nocturnal | 3.791 | 1985.854 | 7.233 |
| amphibians | Hynobius_glacialis | Hynobius_glacialis | CR | Decreasing | temperate | nocturnal | 4.915 | 99.631 | 5.973 |
| amphibians | Hynobius_kimurae | Hynobius_kimurae | LC | Decreasing | temperate | cathemeral | 14.334 | 33130.53 | 15.525 |
| amphibians | Hynobius_leechii | Hynobius_leechii | LC | Decreasing | temperate | nocturnal | 4.568 | 542724.125 | 14.678 |
| amphibians | Hynobius_naevius | Hynobius_naevius | EN | Decreasing | temperate | cathemeral | 13.358 | 2850.592 | 25.627 |
| amphibians | Hynobius_nebulosus | Hynobius_nebulosus | LC | Decreasing | temperate | nocturnal | 5.494 | 10929.533 | 28.496 |
| amphibians | Hynobius_nigrescens | Hynobius_nigrescens | LC | Decreasing | temperate | nocturnal | 9.386 | 82972.241 | 17.968 |

| | | | | | | | | | |
|---|---|---|---|---|---|---|---|---|---|
| amphibians | Hynobius_quelpaertensis | Hynobius_quelpaertensis | VU | Decreasing | temperate | diurnal | 5.775 | 7743.622 | 24.969 |
| amphibians | Hynobius_retardatus | Hynobius_retardatus | LC | Decreasing | temperate | nocturnal | 9.722 | 77446.256 | 15.163 |
| amphibians | Hynobius_sonani | Hynobius_sonani | EN | Unknown | temperate | nocturnal | 4.767 | 3090.833 | 3.297 |
| amphibians | Hynobius_tokyoensis | Hynobius_tokyoensis | VU | Decreasing | temperate | cathemeral | 9.39 | 8430.91 | 28.264 |
| amphibians | Hyperolius_castaneus | Hyperolius_castaneus | LC | Stable | tropical | nocturnal | 3.807 | 17288.833 | 18.298 |
| amphibians | Hyperolius_cinereus | Hyperolius_cinereus | LC | Stable | tropical | diurnal | 1.889 | 111826.256 | 10.15 |
| amphibians | Hyperolius_cinnamomeoventris | Hyperolius_cinnamomeoventris | LC | Unknown | tropical | nocturnal | 2.872 | 4028644.908 | 6.561 |
| amphibians | Hyperolius_cystocandicans | Hyperolius_cystocandicans | EN | Decreasing | tropical | nocturnal | 3.807 | 7790.335 | 14.501 |
| amphibians | Hyperolius_discodactylus | Hyperolius_discodactylus | LC | Unknown | tropical | diurnal | 4.16 | 12383.504 | 16.52 |
| amphibians | Hyperolius_drewesi | NA | not_available | not_available | NA | NA | NA | NA | NA |
| amphibians | Hyperolius_horstockii | Hyperolius_horstockii | LC | Unknown | temperate | diurnal | 6.77 | 18583.631 | 14.034 |
| amphibians | Hyperolius_kivuensis | Hyperolius_kivuensis | LC | Unknown | tropical | nocturnal | 4.934 | 2983645.538 | 9.001 |
| amphibians | Hyperolius_langi | Hyperolius_langi | LC | Unknown | tropical | nocturnal | 2.345 | 90318.334 | 8.107 |
| amphibians | Hyperolius_lateralis | Hyperolius_lateralis | LC | Unknown | tropical | nocturnal | 2.109 | 302571.604 | 15.15 |
| amphibians | Hyperolius_laurenti | Hyperolius_laurenti | NT | Decreasing | tropical | nocturnal | 4.536 | 3496.969 | 11.198 |
| amphibians | Hyperolius_marmoratus | Hyperolius_marmoratus | LC | Increasing | tropical | nocturnal | 6.77 | 1214080.419 | 10.641 |
| amphibians | Hyperolius_microps | NA | LC | Unknown | NA | NA | NA | NA | NA |
| amphibians | Hyperolius_montanus | Hyperolius_montanus | LC | Unknown | tropical | nocturnal | 3.163 | 20430.901 | 14.816 |
| amphibians | Hyperolius_mosaicus | Hyperolius_mosaicus | LC | Unknown | tropical | nocturnal | 1.499 | 145843.805 | 5.775 |
| amphibians | Hyperolius_nasutus | Hyperolius_nasutus | LC | Unknown | tropical | nocturnal | 1.326 | 1164689.559 | 6.351 |
| amphibians | Hyperolius_nitidulus | Hyperolius_nitidulus | LC | Stable | tropical | nocturnal | 3.475 | 3147892.171 | 12.528 |
| amphibians | Hyperolius_pardalis | Hyperolius_pardalis | LC | Unknown | tropical | nocturnal | 2.872 | 345372.373 | 5.423 |
| amphibians | Hyperolius_phantasticus | Hyperolius_phantasticus | LC | Unknown | tropical | nocturnal | 4.16 | 1203569.657 | 5.318 |
| amphibians | Hyperolius_puncticulatus | Hyperolius_puncticulatus | EN | Decreasing | tropical | nocturnal | 6.77 | 920.561 | 22.814 |
| amphibians | Hyperolius_pusillus | Hyperolius_pusillus | LC | Unknown | tropical | nocturnal | 1.168 | 802133.782 | 10.438 |
| amphibians | Hyperolius_spinigularis | Hyperolius_spinigularis | VU | Decreasing | tropical | nocturnal | 2.345 | 1022.346 | 8.295 |
| amphibians | Hyperolius_tuberculatus | Hyperolius_tuberculatus | LC | Decreasing | tropical | nocturnal | 3.807 | 1956619.434 | 5.762 |
| amphibians | Hyperolius_tuberilinguis | Hyperolius_tuberilinguis | LC | Unknown | tropical | nocturnal | 5.356 | 1134975.174 | 9.898 |
| amphibians | Hyperolius_veithi | Hyperolius_veithi | LC | Unknown | tropical | nocturnal | 4.16 | 7.072 | 1.046 |
| amphibians | Hyperolius_viridiflavus | Hyperolius_viridiflavus | LC | Unknown | tropical | cathemeral | 6.77 | 1960115.391 | 9.146 |
| amphibians | Hypopachus_barberi | Hypopachus_barberi | NT | Decreasing | tropical | nocturnal | 8.015 | 5846.654 | 15.921 |
| amphibians | Hypopachus_pictiventris | Hypopachus_pictiventris | LC | Stable | tropical | nocturnal | 4.536 | 20108.382 | 10.377 |
| amphibians | Hypopachus_variolosus | Hypopachus_variolosus | LC | Stable | tropical | nocturnal | 13.329 | 759251.162 | 10.979 |
| amphibians | Hypselotriton_fudingensis | Cynops_fudingensis | CR | Decreasing | temperate | diurnal | 1.834 | 7.022 | 11.963 |

| | | | | | | | | | |
|---|---|---|---|---|---|---|---|---|---|
| amphibians | Ichthyophis_beddomei | Ichthyophis_beddomei | LC | Unknown | tropical | nocturnal | 7.778 | 17913.693 | 13.617 |
| amphibians | Ichthyophis_glutinosus | Ichthyophis_glutinosus | VU | Decreasing | tropical | nocturnal | 34.417 | 8162.63 | 16.741 |
| amphibians | Ichthyophis_javanicus | NA | DD | Unknown | NA | NA | NA | NA | NA |
| amphibians | Ichthyophis_kohtaoensis | NA | DD | Unknown | NA | NA | NA | NA | NA |
| amphibians | Ichthyophis_sikkimensis | Ichthyophis_sikkimensis | LC | Unknown | temperate | nocturnal | 12.292 | 347.195 | 18.285 |
| amphibians | Ichthyophis_tricolor | Ichthyophis_tricolor | LC | Unknown | tropical | nocturnal | 13.489 | 15132.708 | 18.299 |
| amphibians | Ichthyosaura_alpestris | Ichthyosaura_alpestris | LC | Decreasing | temperate | cathemeral | 5.494 | 1366049.96 | 21.447 |
| amphibians | Incilius_alvarius | Incilius_alvarius | LC | Stable | temperate | nocturnal | 848.593 | 358467.986 | 8.265 |
| amphibians | Incilius_aucoinae | Incilius_aucoinae | LC | Stable | tropical | nocturnal | 120.251 | 5055.522 | 13.948 |
| amphibians | Incilius_campbelli | Incilius_campbelli | LC | Stable | tropical | diurnal | 74.113 | 4812.752 | 6.079 |
| amphibians | Incilius_coccifer | Incilius_coccifer | LC | Stable | tropical | cathemeral | 100.927 | 139093.147 | 14.57 |
| amphibians | Incilius_coniferus | Incilius_coniferus | LC | Stable | tropical | nocturnal | 88.303 | 120109.558 | 6.895 |
| amphibians | Incilius_epioticus | Incilius_epioticus | VU | Decreasing | tropical | diurnal | 5.443 | 959.75 | 6.021 |
| amphibians | Incilius_fastidiosus | Incilius_fastidiosus | CR | Decreasing | tropical | diurnal | 22.156 | 252.377 | 4.677 |
| amphibians | Incilius_holdridgei | Incilius_holdridgei | CR | Unknown | tropical | diurnal | 13.329 | 7.616 | 20.279 |
| amphibians | Incilius_luetkenii | Incilius_luetkenii | LC | Stable | tropical | nocturnal | 129.824 | 107380.941 | 15.328 |
| amphibians | Incilius_mazatlanensis | Incilius_mazatlanensis | LC | Stable | tropical | cathemeral | 21.926 | 46234.734 | 13.126 |
| amphibians | Incilius_melanochlorus | Incilius_melanochlorus | LC | Stable | tropical | nocturnal | 129.824 | 19579.372 | 12.564 |
| amphibians | Incilius_nebulifer | Incilius_nebulifer | LC | Stable | temperate | nocturnal | 243.965 | 668550.662 | 14.353 |
| amphibians | Incilius_periglenes | NA | EX | not_available | NA | NA | NA | NA | NA |
| amphibians | Incilius_valliceps | Incilius_valliceps | LC | Stable | tropical | cathemeral | 243.965 | 565201.886 | 9.406 |
| amphibians | Indirana_brachytarsus | Indirana_brachytarsus | LC | Decreasing | tropical | nocturnal | 17.851 | 15836.25 | 13.06 |
| amphibians | Indirana_gundia | Indirana_gundia | NT | Decreasing | tropical | cathemeral | 14.332 | 6899.848 | 12.178 |
| amphibians | Indirana_leithii | Indirana_leithii | LC | Decreasing | tropical | cathemeral | 4.536 | 24723.709 | 16.549 |
| amphibians | Indirana_semipalmata | Indirana_semipalmata | LC | Decreasing | tropical | cathemeral | 3.015 | 21471.759 | 14.008 |
| amphibians | Indirana_yadera | NA | VU | Decreasing | NA | NA | NA | NA | NA |
| amphibians | Ingerophrynus_biporcatus | Ingerophrynus_biporcatus | LC | Stable | tropical | nocturnal | 50.601 | 372621.868 | 14.543 |
| amphibians | Ingerophrynus_divergens | Ingerophrynus_divergens | LC | Decreasing | tropical | nocturnal | 25.822 | 330572.098 | 5.201 |
| amphibians | Ingerophrynus_galeatus | Ingerophrynus_galeatus | LC | Decreasing | tropical | nocturnal | 76.815 | 279353.294 | 9.077 |
| amphibians | Ingerophrynus_ledongensis | Ingerophrynus_ledongensis | NT | Stable | tropical | diurnal | 24.806 | 494.636 | 8.196 |
| amphibians | Ingerophrynus_parvus | Ingerophrynus_parvus | LC | Decreasing | tropical | cathemeral | 12.531 | 555777.267 | 10.228 |
| amphibians | Insuetophrynus_acarpicus | Insuetophrynus_acarpicus | EN | Decreasing | temperate | nocturnal | 16.211 | 975.731 | 10.938 |
| amphibians | Ischnocnema_bolbodactyla | Ischnocnema_bolbodactyla | LC | Stable | tropical | cathemeral | 0.48 | 4985.661 | 8.377 |
| amphibians | Ischnocnema_colibri | NA | DD | Unknown | NA | NA | NA | NA | NA |

| | | | | | | | | | |
|---|---|---|---|---|---|---|---|---|---|
| amphibians | Ischnocnema_concolor | Ischnocnema_concolor | CR | Stable | tropical | nocturnal | 0.567 | 9.937 | 10.323 |
| amphibians | Ischnocnema_guentheri | Ischnocnema_guentheri | NT | Stable | temperate | nocturnal | 8.423 | 110849.116 | 14.269 |
| amphibians | Ischnocnema_henselii | Ischnocnema_henselii | LC | Unknown | temperate | cathemeral | 4.934 | 223011.773 | 11.08 |
| amphibians | Ischnocnema_holti | Ischnocnema_holti | LC | Stable | tropical | cathemeral | 2.521 | 779.61 | 8.141 |
| amphibians | Ischnocnema_juipoca | Ischnocnema_juipoca | LC | Stable | tropical | cathemeral | 1.37 | 443008.053 | 13.784 |
| amphibians | Ischnocnema_manezinho | Ischnocnema_manezinho | EN | Decreasing | temperate | nocturnal | 3.411 | 15190.208 | 15.306 |
| amphibians | Ischnocnema_octavioi | Ischnocnema_octavioi | LC | Stable | tropical | nocturnal | 2.599 | 5192.704 | 18.2 |
| amphibians | Ischnocnema_oea | Ischnocnema_oea | LC | Stable | tropical | cathemeral | 1.168 | 4823.003 | 11.655 |
| amphibians | Ischnocnema_parnaso | NA | VU | Stable | NA | NA | NA | NA | NA |
| amphibians | Ischnocnema_parva | Ischnocnema_parva | LC | Stable | tropical | cathemeral | 1.245 | 298074.013 | 12.413 |
| amphibians | Ischnocnema_randorum | Ischnocnema_randorum | LC | Unknown | temperate | nocturnal | 0.418 | 166.569 | 3.396 |
| amphibians | Ischnocnema_sambaqui | Ischnocnema_sambaqui | LC | Stable | temperate | nocturnal | 5.356 | 15474.267 | 6.66 |
| amphibians | Ischnocnema_spanios | Ischnocnema_spanios | LC | Stable | temperate | diurnal | 0.706 | 9659.705 | 16.5 |
| amphibians | Ischnocnema_verrucosa | Ischnocnema_verrucosa | LC | Stable | tropical | nocturnal | 1.023 | 224958.036 | 10.437 |
| amphibians | Ischnocnema_vizottoi | Ischnocnema_vizottoi | LC | Stable | tropical | nocturnal | 0.585 | 4156.772 | 9.463 |
| amphibians | Isthmohyla_angustilineata | Isthmohyla_angustilineata | CR | Decreasing | tropical | nocturnal | 4.16 | 475.733 | 9.947 |
| amphibians | Isthmohyla_calypsa | Isthmohyla_calypsa | CR | Unknown | tropical | nocturnal | 5.802 | 323.317 | 7.181 |
| amphibians | Isthmohyla_infucata | Isthmohyla_infucata | EN | Unknown | tropical | nocturnal | 8.188 | 144.631 | 3.289 |
| amphibians | Isthmohyla_insolita | Isthmohyla_insolita | EN | Decreasing | tropical | nocturnal | 4.536 | 66.356 | 2.649 |
| amphibians | Isthmohyla_lancasteri | Isthmohyla_lancasteri | LC | Stable | tropical | nocturnal | 5.848 | 5171.358 | 8.041 |
| amphibians | Isthmohyla_picadoi | Isthmohyla_picadoi | LC | Stable | tropical | nocturnal | 3.54 | 4853.058 | 7.327 |
| amphibians | Isthmohyla_pictipes | Isthmohyla_pictipes | CR | Decreasing | tropical | nocturnal | 7.901 | 2795.174 | 7.041 |
| amphibians | Isthmohyla_pseudopuma | Isthmohyla_pseudopuma | LC | Decreasing | tropical | nocturnal | 12.531 | 5381.228 | 15.787 |
| amphibians | Isthmohyla_rivularis | Isthmohyla_rivularis | EN | Decreasing | tropical | nocturnal | 6.77 | 3827.419 | 11.757 |
| amphibians | Isthmohyla_xanthosticta | NA | DD | Unknown | NA | NA | NA | NA | NA |
| amphibians | Isthmohyla_zeteki | Isthmohyla_zeteki | VU | Decreasing | tropical | nocturnal | 3.54 | 1242.585 | 8.304 |
| amphibians | Isthmura_corrugata | NA | CR | Unknown | NA | NA | NA | NA | NA |
| amphibians | Itapotihyla_langsdorffii | Itapotihyla_langsdorffii | LC | Unknown | tropical | cathemeral | 150.528 | 729247.131 | 13.198 |
| amphibians | Julianus_fontanarrosai | NA | LC | Stable | NA | NA | NA | NA | NA |
| amphibians | Julianus_uruguayus | Scinax_uruguayus | LC | Stable | temperate | nocturnal | 2.109 | 122136.432 | 10.083 |
| amphibians | Kalophrynus_baluensis | Kalophrynus_baluensis | LC | Stable | tropical | cathemeral | 9.031 | 1487.148 | 6.877 |
| amphibians | Kalophrynus_calciphilus | Kalophrynus_calciphilus | LC | Stable | tropical | cathemeral | 4.934 | 9.857 | 4.279 |
| amphibians | Kalophrynus_cryptophonus | NA | EN | Decreasing | NA | NA | NA | NA | NA |
| amphibians | Kalophrynus_interlineatus | Kalophrynus_interlineatus | LC | Unknown | tropical | nocturnal | 9.474 | 2249570.467 | 11.99 |

| | | | | | | | | | |
|---|---|---|---|---|---|---|---|---|---|
| amphibians | Kalophrynus_kiewi | NA | LC | Decreasing | NA | NA | NA | NA | NA |
| amphibians | Kalophrynus_nubicola | Kalophrynus_nubicola | LC | Stable | tropical | diurnal | 1.08 | 1392.704 | 1.896 |
| amphibians | Kalophrynus_pleurostigma | Kalophrynus_pleurostigma | LC | Decreasing | tropical | cathemeral | 19.923 | 687690.043 | 9.614 |
| amphibians | Kalophrynus_sinensis | NA | not_available | not_available | NA | NA | NA | NA | NA |
| amphibians | Kalophrynus_yongi | Kalophrynus_yongi | CR | Unknown | tropical | nocturnal | 2.345 | 9.827 | 16.994 |
| amphibians | Kaloula_baleata | Kaloula_baleata | LC | Stable | tropical | cathemeral | 28.486 | 1372199.516 | 9.49 |
| amphibians | Kaloula_borealis | Kaloula_borealis | LC | Decreasing | temperate | nocturnal | 11.18 | 1369292.785 | 19.884 |
| amphibians | Kaloula_conjuncta | Kaloula_conjuncta | LC | Stable | tropical | nocturnal | 9.031 | 27864.973 | 11.56 |
| amphibians | Kaloula_indochinensis | Kaloula_indochinensis | LC | Decreasing | tropical | nocturnal | 14.161 | 144389.125 | 8.218 |
| amphibians | Kaloula_picta | Kaloula_picta | LC | Stable | tropical | nocturnal | 16.398 | 272030.778 | 13.883 |
| amphibians | Kaloula_pulchra | Kaloula_pulchra | LC | Stable | tropical | nocturnal | 57.012 | 2300321.309 | 13.282 |
| amphibians | Kaloula_walteri | Kaloula_walteri | VU | Decreasing | tropical | cathemeral | 2.679 | 641.272 | 10.877 |
| amphibians | Kassina_arboricola | Kassina_arboricola | VU | Decreasing | tropical | nocturnal | 5.356 | 153336.269 | 17.248 |
| amphibians | Kassina_cassinoides | Kassina_cassinoides | LC | Unknown | tropical | nocturnal | 11.036 | 1590492.976 | 13.691 |
| amphibians | Kassina_cochranae | Kassina_cochranae | LC | Decreasing | tropical | nocturnal | 6.273 | 174153.539 | 13.756 |
| amphibians | Kassina_decorata | Kassina_decorata | VU | Decreasing | tropical | nocturnal | 3.475 | 1892.536 | 13.066 |
| amphibians | Kassina_fusca | Kassina_fusca | LC | Unknown | tropical | nocturnal | 5.802 | 819826.686 | 12.576 |
| amphibians | Kassina_jozani | Kassina_jozani | EN | Decreasing | tropical | nocturnal | 6.37 | 55.491 | 16.529 |
| amphibians | Kassina_kuvangensis | Kassina_kuvangensis | LC | Unknown | tropical | nocturnal | 11.767 | 561513.544 | 5.594 |
| amphibians | Kassina_lamottei | Kassina_lamottei | LC | Decreasing | tropical | nocturnal | 9.031 | 72132.138 | 12.347 |
| amphibians | Kassina_maculifer | Kassina_maculifer | LC | Unknown | tropical | nocturnal | 7.294 | 160054.899 | 9.132 |
| amphibians | Kassina_maculosa | Kassina_maculosa | LC | Unknown | tropical | nocturnal | 4.536 | 706125.579 | 6.345 |
| amphibians | Kassina_schioetzi | Kassina_schioetzi | LC | Unknown | tropical | nocturnal | 5.356 | 181030.696 | 13.553 |
| amphibians | Kassina_senegalensis | Kassina_senegalensis | LC | Unknown | tropical | cathemeral | 12.531 | 12196935.289 | 9.751 |
| amphibians | Kassina_wazae | NA | DD | Unknown | NA | NA | NA | NA | NA |
| amphibians | Kurixalus_appendiculatus | Kurixalus_appendiculatus | LC | Decreasing | temperate | nocturnal | 11.036 | 119263.653 | 7.525 |
| amphibians | Kurixalus_banaensis | Kurixalus_banaensis | LC | Decreasing | tropical | nocturnal | 4.16 | 33846.688 | 7.396 |
| amphibians | Kurixalus_bisacculus | Kurixalus_bisacculus | LC | Unknown | tropical | nocturnal | 3.638 | 100778.35 | 10.481 |
| amphibians | Kurixalus_eiffingeri | Kurixalus_eiffingeri | LC | Decreasing | temperate | nocturnal | 7.294 | 12611.515 | 15.195 |
| amphibians | Kurixalus_gracilloides | NA | DD | Decreasing | NA | NA | NA | NA | NA |
| amphibians | Kurixalus_yangi | NA | LC | Decreasing | NA | NA | NA | NA | NA |
| amphibians | Laliostoma_labrosum | Laliostoma_labrosum | LC | Stable | tropical | nocturnal | 50.601 | 307700.357 | 8.623 |
| amphibians | Lankanectes_pera | NA | CR | Decreasing | NA | NA | NA | NA | NA |
| amphibians | Laotriton_laoensis | Laotriton_laoensis | EN | Decreasing | tropical | diurnal | 23.045 | 4559.116 | 6.355 |

| | | | | | | | | | |
|---|---|---|---|---|---|---|---|---|---|
| amphibians | Leiopelma_archeyi | Leiopelma_archeyi | CR | Stable | temperate | nocturnal | 4.536 | 1376.635 | 7.336 |
| amphibians | Leiopelma_hamiltoni | Leiopelma_hamiltoni | VU | Increasing | temperate | nocturnal | 12.531 | 9.649 | 2.836 |
| amphibians | Leiopelma_hochstetteri | Leiopelma_hochstetteri | LC | Decreasing | temperate | nocturnal | 9.031 | 10109.771 | 5.686 |
| amphibians | Lepidobatrachus_asper | Lepidobatrachus_asper | LC | Decreasing | temperate | nocturnal | 79.584 | 294120.193 | 7.163 |
| amphibians | Lepidobatrachus_laevis | Lepidobatrachus_laevis | LC | Stable | temperate | nocturnal | 190.787 | 465963.739 | 5.869 |
| amphibians | Lepidobatrachus_llanensis | Lepidobatrachus_llanensis | LC | Decreasing | tropical | nocturnal | 104.268 | 302269.196 | 4.515 |
| amphibians | Leptobrachella_aerea | Leptolalax_aereus | LC | Decreasing | tropical | nocturnal | 4.772 | 19064.165 | 8.447 |
| amphibians | Leptobrachella_arayai | Leptolalax_arayai | LC | Decreasing | tropical | nocturnal | 5.356 | 3267.555 | 5.33 |
| amphibians | Leptobrachella_baluensis | Leptobrachella_baluensis | LC | Decreasing | tropical | nocturnal | 0.892 | 7972.021 | 4.347 |
| amphibians | Leptobrachella_bijie | NA | DD | Unknown | NA | NA | NA | NA | NA |
| amphibians | Leptobrachella_crocea | NA | DD | Unknown | NA | NA | NA | NA | NA |
| amphibians | Leptobrachella_fuliginosa | Leptolalax_fuliginosus | LC | Unknown | tropical | nocturnal | 2.109 | 67999.716 | 8.004 |
| amphibians | Leptobrachella_isos | NA | VU | Decreasing | NA | NA | NA | NA | NA |
| amphibians | Leptobrachella_kajangensis | Leptolalax_kajangensis | LC | Unknown | tropical | nocturnal | 8.423 | 9.791 | 4.831 |
| amphibians | Leptobrachella_kecil | Leptolalax_kecil | CR | Decreasing | tropical | nocturnal | 1.168 | 9.842 | 29.745 |
| amphibians | Leptobrachella_liui | Leptolalax_liui | LC | Stable | temperate | nocturnal | 1.827 | 328958.896 | 12.542 |
| amphibians | Leptobrachella_macrops | NA | EN | Decreasing | NA | NA | NA | NA | NA |
| amphibians | Leptobrachella_melanoleuca | Leptolalax_melanoleucus | LC | Decreasing | tropical | nocturnal | 2.788 | 6645.811 | 2.973 |
| amphibians | Leptobrachella_melica | Leptolalax_melicus | EN | Decreasing | tropical | nocturnal | 0.854 | 1606.329 | 1.893 |
| amphibians | Leptobrachella_neangi | NA | not_available | not_available | NA | NA | NA | NA | NA |
| amphibians | Leptobrachella_petrops | NA | NT | Decreasing | NA | NA | NA | NA | NA |
| amphibians | Leptobrachella_puhoatensis | NA | EN | Decreasing | NA | NA | NA | NA | NA |
| amphibians | Leptobrachella_purpuraventra | NA | DD | Unknown | NA | NA | NA | NA | NA |
| amphibians | Leptobrachella_pyrrhops | NA | EN | Decreasing | NA | NA | NA | NA | NA |
| amphibians | Leptobrachella_shangsiensis | NA | NT | Unknown | NA | NA | NA | NA | NA |
| amphibians | Leptobrachella_sola | Leptolalax_solus | LC | Decreasing | tropical | nocturnal | 1.61 | 15204.136 | 3.626 |
| amphibians | Leptobrachella_ventripunctata | Leptolalax_ventripunctatus | LC | Decreasing | tropical | nocturnal | 1.686 | 178229.569 | 6.856 |
| amphibians | Leptobrachella_wulingensis | NA | not_available | not_available | NA | NA | NA | NA | NA |
| amphibians | Leptobrachium_abbotti | Leptobrachium_abbotti | LC | Decreasing | tropical | nocturnal | 88.303 | 68786.233 | 5.024 |
| amphibians | Leptobrachium_banae | Leptobrachium_banae | LC | Decreasing | tropical | nocturnal | 59.726 | 18358.072 | 5.868 |
| amphibians | Leptobrachium_boringii | Leptobrachium_boringii | EN | Decreasing | temperate | nocturnal | 82.421 | 65807.468 | 10.845 |
| amphibians | Leptobrachium_hainanense | Leptobrachium_hainanense | VU | Decreasing | tropical | nocturnal | 24.557 | 11818.714 | 10.993 |
| amphibians | Leptobrachium_hasseltii | Leptobrachium_hasseltii | LC | Decreasing | tropical | nocturnal | 45.085 | 165491.382 | 20.862 |
| amphibians | Leptobrachium_huashen | Leptobrachium_huashen | LC | Decreasing | temperate | nocturnal | 19.923 | 151491.042 | 11.033 |

| | | | | | | | | | |
|---|---|---|---|---|---|---|---|---|---|
| amphibians | Leptobrachium_liui | Leptobrachium_liui | LC | Decreasing | temperate | nocturnal | 75.456 | 269304.844 | 12.147 |
| amphibians | Leptobrachium_montanum | Leptobrachium_montanum | LC | Decreasing | tropical | cathemeral | 66.404 | 26539.324 | 4.354 |
| amphibians | Leptobrachium_mouhoti | Leptobrachium_mouhoti | LC | Decreasing | tropical | nocturnal | 49.786 | 47969.485 | 6.759 |
| amphibians | Leptobrachium_pullum | Leptobrachium_pullum | LC | Decreasing | tropical | nocturnal | 62.055 | 58196.739 | 7.609 |
| amphibians | Leptobrachium_smithi | Leptobrachium_smithi | LC | Decreasing | tropical | nocturnal | 46.616 | 223795.347 | 11.354 |
| amphibians | Leptodactylodon_ventrimarmoratus | Leptodactylodon_ventrimarmoratus | VU | Decreasing | tropical | diurnal | 7.844 | 17819.103 | 10.832 |
| amphibians | Leptodactylus_albilabris | Leptodactylus_albilabris | LC | Stable | tropical | diurnal | 10.337 | 10649.908 | 24.728 |
| amphibians | Leptodactylus_bolivianus | Leptodactylus_bolivianus | LC | Stable | tropical | cathemeral | 310.174 | 2780429.653 | 1.943 |
| amphibians | Leptodactylus_bufonius | Leptodactylus_bufonius | LC | Stable | temperate | nocturnal | 25.822 | 1569066.457 | 7.268 |
| amphibians | Leptodactylus_caatingae | Leptodactylus_caatingae | LC | Stable | tropical | cathemeral | 3.572 | 428232.182 | 8.597 |
| amphibians | Leptodactylus_camaquara | Leptodactylus_camaquara | NT | Decreasing | tropical | diurnal | 4.536 | 13092.869 | 4.851 |
| amphibians | Leptodactylus_cunicularius | Leptodactylus_cunicularius | LC | Stable | tropical | nocturnal | 8.423 | 179461.698 | 12.646 |
| amphibians | Leptodactylus_cupreus | Leptodactylus_cupreus | LC | Decreasing | tropical | nocturnal | 16.873 | 249138.741 | 9.798 |
| amphibians | Leptodactylus_didymus | Leptodactylus_didymus | LC | Stable | tropical | cathemeral | 18.456 | 304466.131 | 2.704 |
| amphibians | Leptodactylus_diedrus | Leptodactylus_diedrus | LC | Stable | tropical | cathemeral | 32.83 | 962549.603 | 0.748 |
| amphibians | Leptodactylus_discodactylus | Leptodactylus_discodactylus | LC | Stable | tropical | cathemeral | 3.475 | 1332439.425 | 1.452 |
| amphibians | Leptodactylus_fallax | Leptodactylus_fallax | CR | Decreasing | tropical | cathemeral | 1153.83 | 152.025 | 17.274 |
| amphibians | Leptodactylus_flavopictus | Leptodactylus_flavopictus | LC | Stable | temperate | nocturnal | 256.337 | 50910.81 | 11.049 |
| amphibians | Leptodactylus_fragilis | Leptodactylus_fragilis | LC | Stable | tropical | diurnal | 7.294 | 1557081.885 | 10.773 |
| amphibians | Leptodactylus_furnarius | Leptodactylus_furnarius | LC | Stable | tropical | cathemeral | 7.294 | 1429867.372 | 11.641 |
| amphibians | Leptodactylus_fuscus | Leptodactylus_fuscus | LC | Stable | tropical | cathemeral | 15.029 | 12048757.37 | 6.007 |
| amphibians | Leptodactylus_gracilis | Leptodactylus_gracilis | LC | Stable | temperate | cathemeral | 11.036 | 2227779.98 | 9.971 |
| amphibians | Leptodactylus_griseigularis | NA | LC | Stable | NA | NA | NA | NA | NA |
| amphibians | Leptodactylus_hylodes | NA | DD | Unknown | NA | NA | NA | NA | NA |
| amphibians | Leptodactylus_insularum | Leptodactylus_insularum | LC | Stable | tropical | nocturnal | 61.584 | 1051258.572 | 10.907 |
| amphibians | Leptodactylus_jolyi | Leptodactylus_jolyi | LC | Stable | tropical | nocturnal | 14.161 | 1113014.947 | 11.72 |
| amphibians | Leptodactylus_knudseni | Leptodactylus_knudseni | LC | Stable | tropical | nocturnal | 581.843 | 5858442.048 | 1.855 |
| amphibians | Leptodactylus_labrosus | Leptodactylus_labrosus | LC | Stable | tropical | cathemeral | 31.334 | 98372.577 | 10.581 |
| amphibians | Leptodactylus_labyrinthicus | Leptodactylus_labyrinthicus | LC | Stable | tropical | nocturnal | 765.222 | 3926794.408 | 8.122 |
| amphibians | Leptodactylus_laticeps | Leptodactylus_laticeps | NT | Decreasing | temperate | cathemeral | 188.234 | 433225.602 | 5.352 |
| amphibians | Leptodactylus_latinasus | Leptodactylus_latinasus | LC | Stable | temperate | diurnal | 5.356 | 1908729.087 | 8.742 |
| amphibians | Leptodactylus_latrans | Leptodactylus_latrans | LC | Stable | tropical | cathemeral | 310.174 | 179747.556 | 12.879 |
| amphibians | Leptodactylus_leptodactyloides | Leptodactylus_leptodactyloides | LC | Stable | tropical | cathemeral | 16.117 | 6143679.614 | 2.056 |
| amphibians | Leptodactylus_lithonaetes | Leptodactylus_lithonaetes | LC | Unknown | tropical | diurnal | 46.616 | 116702.863 | 1.723 |

| | | | | | | | | | |
|---|---|---|---|---|---|---|---|---|---|
| amphibians | Leptodactylus_longirostris | Leptodactylus_longirostris | LC | Stable | tropical | nocturnal | 12.222 | 2135853.296 | 1.574 |
| amphibians | Leptodactylus_macrosternum | Leptodactylus_macrosternum | LC | Stable | tropical | nocturnal | 74.113 | 9590666.544 | 5.765 |
| amphibians | Leptodactylus_melanonotus | Leptodactylus_melanonotus | LC | Stable | tropical | cathemeral | 17.851 | 833155.684 | 10.163 |
| amphibians | Leptodactylus_myersi | Leptodactylus_myersi | LC | Stable | tropical | diurnal | 190.787 | 989925.173 | 1.195 |
| amphibians | Leptodactylus_mystaceus | Leptodactylus_mystaceus | LC | Stable | tropical | cathemeral | 19.923 | 6802464.719 | 4.215 |
| amphibians | Leptodactylus_mystacinus | Leptodactylus_mystacinus | LC | Stable | temperate | cathemeral | 27.265 | 4307146.391 | 10.981 |
| amphibians | Leptodactylus_natalensis | Leptodactylus_natalensis | LC | Stable | tropical | nocturnal | 12.531 | 701823.022 | 10.275 |
| amphibians | Leptodactylus_notoaktites | Leptodactylus_notoaktites | LC | Stable | temperate | cathemeral | 11.036 | 136973.582 | 11.213 |
| amphibians | Leptodactylus_paraensis | Leptodactylus_paraensis | LC | Stable | tropical | diurnal | 458.987 | 770708.401 | 2.407 |
| amphibians | Leptodactylus_pascoensis | NA | DD | Unknown | NA | NA | NA | NA | NA |
| amphibians | Leptodactylus_pentadactylus | Leptodactylus_pentadactylus | LC | Stable | tropical | nocturnal | 907.535 | 6672526.947 | 2.16 |
| amphibians | Leptodactylus_peritoaktites | Leptodactylus_peritoaktites | EN | Decreasing | tropical | nocturnal | 357.72 | 1397.118 | 6.411 |
| amphibians | Leptodactylus_petersii | Leptodactylus_petersii | LC | Stable | tropical | cathemeral | 11.993 | 4865944.704 | 1.538 |
| amphibians | Leptodactylus_plaumanni | Leptodactylus_plaumanni | LC | Stable | temperate | cathemeral | 8.423 | 211860.662 | 11.666 |
| amphibians | Leptodactylus_podicipinus | Leptodactylus_podicipinus | LC | Stable | tropical | nocturnal | 14.161 | 6340303.569 | 6.12 |
| amphibians | Leptodactylus_poecilochilus | Leptodactylus_poecilochilus | LC | Stable | tropical | nocturnal | 11.036 | 239801.751 | 13.809 |
| amphibians | Leptodactylus_pustulatus | Leptodactylus_pustulatus | LC | Stable | tropical | diurnal | 5.356 | 849980.147 | 6.252 |
| amphibians | Leptodactylus_rhodomerus | Leptodactylus_rhodomerus | LC | Decreasing | tropical | nocturnal | 457.107 | 89873.964 | 5.122 |
| amphibians | Leptodactylus_rhodomystax | Leptodactylus_rhodomystax | LC | Stable | tropical | nocturnal | 74.113 | 4578521.004 | 1.594 |
| amphibians | Leptodactylus_rhodonotus | Leptodactylus_rhodonotus | LC | Stable | tropical | nocturnal | 74.113 | 886872.677 | 2.061 |
| amphibians | Leptodactylus_riveroi | Leptodactylus_riveroi | LC | Stable | tropical | cathemeral | 39.306 | 924607.472 | 0.67 |
| amphibians | Leptodactylus_rugosus | Leptodactylus_rugosus | LC | Stable | tropical | nocturnal | 30.173 | 157996.824 | 1.323 |
| amphibians | Leptodactylus_savagei | Leptodactylus_savagei | LC | Stable | tropical | nocturnal | 765.222 | 255617.77 | 9.125 |
| amphibians | Leptodactylus_sertanejo | Leptodactylus_sertanejo | LC | Stable | tropical | cathemeral | 15.029 | 1455034.963 | 10.47 |
| amphibians | Leptodactylus_silvanimbus | Leptodactylus_silvanimbus | CR | Decreasing | tropical | diurnal | 15.029 | 150.871 | 10.631 |
| amphibians | Leptodactylus_spixi | Leptodactylus_spixi | LC | Stable | tropical | cathemeral | 12.531 | 2550264.271 | 8.802 |
| amphibians | Leptodactylus_stenodema | Leptodactylus_stenodema | LC | Stable | tropical | cathemeral | 355.349 | 3288623.81 | 1.2 |
| amphibians | Leptodactylus_syphax | Leptodactylus_syphax | LC | Stable | tropical | nocturnal | 54.816 | 3088798.518 | 8.902 |
| amphibians | Leptodactylus_troglodytes | Leptodactylus_troglodytes | LC | Stable | tropical | nocturnal | 13.824 | 1873310.41 | 8.227 |
| amphibians | Leptodactylus_turimiquensis | Leptodactylus_turimiquensis | NT | Unknown | tropical | diurnal | 478.079 | 9455.8 | 10.746 |
| amphibians | Leptodactylus_validus | Leptodactylus_validus | LC | Stable | tropical | diurnal | 12.531 | 1375049.325 | 4.016 |
| amphibians | Leptodactylus_vastus | Leptodactylus_vastus | LC | Stable | tropical | cathemeral | 2029.91 | 2889018.504 | 6.066 |
| amphibians | Leptodactylus_ventrimaculatus | Leptodactylus_ventrimaculatus | LC | Stable | tropical | nocturnal | 19.18 | 123067.802 | 8.508 |
| amphibians | Leptodactylus_viridis | Leptodactylus_viridis | LC | Stable | tropical | nocturnal | 35.967 | 38774.517 | 7.86 |

| | | | | | | | | | |
|---|---|---|---|---|---|---|---|---|---|
| amphibians | Leptodactylus_wagneri | Leptodactylus_wagneri | LC | Stable | tropical | nocturnal | 54.169 | 1058311.353 | 1.924 |
| amphibians | Leptomantis_angulirostris | Rhacophorus_angulirostris | NT | Decreasing | tropical | nocturnal | 11.767 | 3176.124 | 7.631 |
| amphibians | Leptomantis_cyanopunctatus | Rhacophorus_cyanopunctatus | LC | Decreasing | tropical | nocturnal | 6.77 | 219171.556 | 9.376 |
| amphibians | Leptomantis_gauni | Rhacophorus_gauni | LC | Decreasing | tropical | nocturnal | 5.487 | 309679.878 | 3.845 |
| amphibians | Leptopelis_brevirostris | Leptopelis_brevirostris | LC | Unknown | tropical | diurnal | 24.557 | 543141.564 | 7.252 |
| amphibians | Leptopelis_bufonides | Leptopelis_bufonides | LC | Unknown | tropical | nocturnal | 5.802 | 1259399.792 | 13.959 |
| amphibians | Leptopelis_christyi | Leptopelis_christyi | LC | Unknown | tropical | nocturnal | 25.822 | 372533.448 | 11.825 |
| amphibians | Leptopelis_concolor | Leptopelis_concolor | LC | Stable | tropical | nocturnal | 11.767 | 113154.662 | 12.537 |
| amphibians | Leptopelis_flavomaculatus | Leptopelis_flavomaculatus | LC | Unknown | tropical | nocturnal | 32.83 | 858622.21 | 9.769 |
| amphibians | Leptopelis_karissimbensis | Leptopelis_karissimbensis | VU | Stable | tropical | nocturnal | 9.094 | 2750.825 | 16.139 |
| amphibians | Leptopelis_ragazzii | Leptopelis_ragazzii | VU | Stable | tropical | cathemeral | 11.036 | 1543.152 | 17.725 |
| amphibians | Leptopelis_spiritusnoctis | Leptopelis_spiritusnoctis | LC | Unknown | tropical | nocturnal | 12.531 | 521402.46 | 15.742 |
| amphibians | Leptopelis_susanae | Leptopelis_susanae | EN | Unknown | tropical | cathemeral | 13.329 | 829.094 | 15.669 |
| amphibians | Leptopelis_vannutellii | Leptopelis_vannutellii | LC | Stable | tropical | cathemeral | 8.423 | 41588.992 | 11.216 |
| amphibians | Leptopelis_viridis | Leptopelis_viridis | LC | Unknown | tropical | nocturnal | 11.036 | 3428870.055 | 11.171 |
| amphibians | Leucostethus_alacris | NA | DD | Unknown | NA | NA | NA | NA | NA |
| amphibians | Leucostethus_fraterdanieli | Colostethus_fraterdanieli | LC | Stable | tropical | diurnal | 1.499 | 42537.709 | 14.968 |
| amphibians | Leucostethus_ramirezi | Colostethus_ramirezi | EN | Decreasing | tropical | diurnal | 0.867 | 328.523 | 5.771 |
| amphibians | Limnodynastes_interioris | Limnodynastes_interioris | LC | Stable | temperate | nocturnal | 74.113 | 273943.641 | 10.229 |
| amphibians | Limnodynastes_peronii | Limnodynastes_peronii | LC | Increasing | temperate | cathemeral | 37.611 | 939844.672 | 8.323 |
| amphibians | Limnomedusa_macroglossa | Limnomedusa_macroglossa | LC | Decreasing | temperate | nocturnal | 25.822 | 398365.417 | 10.614 |
| amphibians | Limnonectes_arathooni | Limnonectes_arathooni | VU | Decreasing | tropical | nocturnal | 5.184 | 3387.386 | 10.607 |
| amphibians | Limnonectes_blythii | Limnonectes_blythii | LC | Decreasing | tropical | nocturnal | 2304.968 | 416463.009 | 10.81 |
| amphibians | Limnonectes_coffeatus | NA | DD | Decreasing | NA | NA | NA | NA | NA |
| amphibians | Limnonectes_dabanus | Limnonectes_dabanus | LC | Decreasing | tropical | nocturnal | 25.822 | 75275.039 | 8.489 |
| amphibians | Limnonectes_finchi | Limnonectes_finchi | LC | Decreasing | tropical | nocturnal | 8.015 | 16721.925 | 4.572 |
| amphibians | Limnonectes_hascheanus | Limnonectes_hascheanus | LC | Decreasing | tropical | cathemeral | 5.356 | 178238.036 | 9.38 |
| amphibians | Limnonectes_ibanorum | Limnonectes_ibanorum | LC | Decreasing | tropical | nocturnal | 275.697 | 105854.461 | 4.253 |
| amphibians | Limnonectes_kiziriani | NA | LC | Decreasing | NA | NA | NA | NA | NA |
| amphibians | Limnonectes_kuhlii | Limnonectes_kuhlii | LC | Decreasing | tropical | nocturnal | 66.404 | 2038423.887 | 10.3 |
| amphibians | Limnonectes_leporinus | Limnonectes_leporinus | LC | Decreasing | tropical | nocturnal | 639.138 | 346338.314 | 5.4 |
| amphibians | Limnonectes_leytensis | Limnonectes_leytensis | LC | Decreasing | tropical | nocturnal | 17.455 | 144027.558 | 13.06 |
| amphibians | Limnonectes_limborgi | Limnonectes_limborgi | LC | Decreasing | tropical | nocturnal | 4.732 | 432176.209 | 7.059 |
| amphibians | Limnonectes_macrodon | Limnonectes_macrodon | LC | Decreasing | tropical | nocturnal | 387.871 | 66067.283 | 25.343 |

| | | | | | | | | | |
|---|---|---|---|---|---|---|---|---|---|
| amphibians | Limnonectes_malesianus | Limnonectes_malesianus | LC | Decreasing | tropical | nocturnal | 387.871 | 194486.155 | 11.571 |
| amphibians | Limnonectes_megastomias | Limnonectes_megastomias | VU | Decreasing | tropical | cathemeral | 207.696 | 9084.399 | 10.19 |
| amphibians | Limnonectes_microtympanum | Limnonectes_microtympanum | EN | Decreasing | tropical | nocturnal | 31.927 | 429.649 | 12.446 |
| amphibians | Limnonectes_palavanensis | Limnonectes_palavanensis | LC | Decreasing | tropical | nocturnal | 5.356 | 126845.169 | 4.206 |
| amphibians | Limnonectes_paramacrodon | Limnonectes_paramacrodon | LC | Decreasing | tropical | nocturnal | 25.822 | 114392.095 | 10.427 |
| amphibians | Limnonectes_poilani | Limnonectes_poilani | LC | Decreasing | tropical | nocturnal | 102.588 | 109743.863 | 8.273 |
| amphibians | Limnonectes_quangninhensis | NA | VU | Unknown | NA | NA | NA | NA | NA |
| amphibians | Limnonectes_shompenorum | Limnonectes_shompenorum | LC | Decreasing | tropical | nocturnal | 48.581 | 83071.443 | 9.298 |
| amphibians | Limnonectes_timorensis | NA | not_available | not_available | NA | NA | NA | NA | NA |
| amphibians | Limnonectes_visayanus | Limnonectes_visayanus | NT | Decreasing | tropical | nocturnal | 85.916 | 5846.228 | 14.032 |
| amphibians | Lissotriton_boscai | Lissotriton_boscai | LC | Decreasing | temperate | cathemeral | 1.499 | 201777.076 | 15.688 |
| amphibians | Lissotriton_helveticus | Lissotriton_helveticus | LC | Decreasing | temperate | nocturnal | 1.703 | 1084367.76 | 22.719 |
| amphibians | Lissotriton_italicus | Lissotriton_italicus | LC | Decreasing | temperate | nocturnal | 1.499 | 75200.411 | 23.795 |
| amphibians | Lissotriton_montandoni | Lissotriton_montandoni | LC | Decreasing | temperate | cathemeral | 2.851 | 123049.566 | 16.754 |
| amphibians | Lissotriton_vulgaris | Lissotriton_vulgaris | LC | Stable | temperate | cathemeral | 3.16 | 898593.445 | 13.616 |
| amphibians | Lithobates_areolatus | Rana_areolata | LC | Decreasing | temperate | nocturnal | 198.59 | 417915.339 | 16.672 |
| amphibians | Lithobates_berlandieri | Rana_berlandieri | LC | Stable | temperate | cathemeral | 188.234 | 888950.827 | 10.351 |
| amphibians | Lithobates_bwana | Rana_bwana | LC | Stable | tropical | nocturnal | 88.303 | 7376.79 | 7.128 |
| amphibians | Lithobates_capito | Rana_capito | VU | Decreasing | temperate | nocturnal | 154.926 | 241845.506 | 16.781 |
| amphibians | Lithobates_chiricahuensis | Rana_chiricahuensis | VU | Decreasing | temperate | nocturnal | 289.152 | 228519.323 | 7.939 |
| amphibians | Lithobates_forreri | Rana_forreri | LC | Stable | tropical | cathemeral | 159.413 | 227810.276 | 12.547 |
| amphibians | Lithobates_juliani | Rana_juliani | LC | Stable | tropical | cathemeral | 111.177 | 3479.859 | 3.889 |
| amphibians | Lithobates_kauffeldi | NA | LC | Stable | NA | NA | NA | NA | NA |
| amphibians | Lithobates_maculatus | Rana_maculata | LC | Decreasing | tropical | nocturnal | 154.926 | 184811.313 | 12.024 |
| amphibians | Lithobates_palmipes | Rana_palmipes | LC | Stable | tropical | cathemeral | 220.471 | 9806600.501 | 4.029 |
| amphibians | Lithobates_palustris | Rana_palustris | LC | Stable | temperate | nocturnal | 66.404 | 2479424.186 | 17.184 |
| amphibians | Lithobates_pipiens | Rana_pipiens | LC | Decreasing | temperate | nocturnal | 146.217 | 6012762.574 | 10.727 |
| amphibians | Lithobates_sevosus | Rana_sevosa | CR | Stable | temperate | diurnal | 122.125 | 21134.376 | 17.134 |
| amphibians | Lithobates_sphenocephalus | Rana_sphenocephala | LC | Stable | temperate | nocturnal | 226.191 | 1727205.352 | 17.093 |
| amphibians | Lithobates_taylori | Rana_taylori | LC | Stable | tropical | cathemeral | 68.909 | 28326.391 | 11.708 |
| amphibians | Lithobates_vaillanti | Rana_vaillanti | LC | Stable | tropical | cathemeral | 214.852 | 507244.706 | 8.926 |
| amphibians | Lithobates_vibicarius | Rana_vibicaria | EN | Decreasing | tropical | cathemeral | 79.584 | 1356.169 | 8.198 |
| amphibians | Lithobates_warszewitschii | Rana_warszewitschii | LC | Increasing | tropical | cathemeral | 50.601 | 157320.661 | 9.158 |
| amphibians | Lithodytes_lineatus | Lithodytes_lineatus | LC | Stable | tropical | cathemeral | 15.932 | 7015371.83 | 2.361 |

| | | | | | | | | | |
|---|---|---|---|---|---|---|---|---|---|
| amphibians | Litoria_aurifera | Litoria_aurifera | LC | Stable | tropical | cathemeral | 0.772 | 12301.728 | 0.875 |
| amphibians | Litoria_bibonius | Litoria_bibonius | LC | Stable | tropical | nocturnal | 3.047 | 0.793 | 11.125 |
| amphibians | Litoria_bicolor | Litoria_bicolor | LC | Stable | tropical | diurnal | 2.109 | 841293.874 | 2.589 |
| amphibians | Litoria_christianbergmanni | Litoria_christianbergmanni | LC | Decreasing | tropical | nocturnal | 2.394 | 148.545 | 8.408 |
| amphibians | Litoria_contrastens | Litoria_contrastens | LC | Stable | tropical | nocturnal | 4.463 | 985.016 | 12.509 |
| amphibians | Litoria_coplandi | Litoria_coplandi | LC | Stable | tropical | nocturnal | 6.77 | 736072.339 | 2.25 |
| amphibians | Litoria_dentata | Litoria_dentata | LC | Stable | temperate | nocturnal | 8.13 | 69176.196 | 7.08 |
| amphibians | Litoria_ewingii | Litoria_ewingii | LC | Stable | temperate | diurnal | 11.036 | 614726.565 | 9.316 |
| amphibians | Litoria_fallax | Litoria_fallax | LC | Stable | temperate | cathemeral | 2.599 | 591734.541 | 7.139 |
| amphibians | Litoria_flavescens | Litoria_flavescens | LC | Stable | tropical | nocturnal | 8.785 | 781.54 | 5.249 |
| amphibians | Litoria_gasconi | Litoria_gasconi | LC | Unknown | tropical | nocturnal | 6.082 | 8516.743 | 4.283 |
| amphibians | Litoria_hilli | Litoria_hilli | LC | Stable | tropical | nocturnal | 14.765 | 211.073 | 3.704 |
| amphibians | Litoria_humboldtorum | Litoria_humboldtorum | LC | Stable | tropical | nocturnal | 8.724 | 21.437 | 6.263 |
| amphibians | Litoria_latopalmata | Litoria_latopalmata | LC | Stable | tropical | nocturnal | 6.273 | 1925902.001 | 6.14 |
| amphibians | Litoria_littlejohni | Litoria_littlejohni | EN | Decreasing | temperate | nocturnal | 35.967 | 6476.168 | 7.005 |
| amphibians | Litoria_longirostris | Litoria_longirostris | LC | Stable | tropical | nocturnal | 1.499 | 525.44 | 1.011 |
| amphibians | Litoria_megalops | NA | DD | Unknown | NA | NA | NA | NA | NA |
| amphibians | Litoria_meiriana | Litoria_meiriana | LC | Stable | tropical | cathemeral | 0.892 | 367981 | 2.11 |
| amphibians | Litoria_nasuta | Litoria_nasuta | LC | Stable | tropical | nocturnal | 15.932 | 1189277.513 | 3.762 |
| amphibians | Litoria_olongburensis | Litoria_olongburensis | VU | Decreasing | temperate | nocturnal | 3.194 | 9065.689 | 13.645 |
| amphibians | Litoria_pallida | Litoria_pallida | LC | Stable | tropical | nocturnal | 5.802 | 1442166.841 | 2.93 |
| amphibians | Litoria_peronii | Litoria_peronii | LC | Stable | temperate | nocturnal | 32.83 | 1260639.647 | 8.235 |
| amphibians | Litoria_personata | Litoria_personata | LC | Stable | tropical | nocturnal | 3.015 | 70212.11 | 0.998 |
| amphibians | Litoria_revelata | Litoria_revelata | LC | Stable | temperate | nocturnal | 3.807 | 66672.918 | 8.151 |
| amphibians | Litoria_staccato | Litoria_staccato | LC | Stable | tropical | nocturnal | 4.16 | 82088.359 | 1.963 |
| amphibians | Litoria_verreauxii | Litoria_verreauxii | LC | Decreasing | temperate | nocturnal | 3.807 | 362379.979 | 9.216 |
| amphibians | Litoria_vocivincens | Litoria_vocivincens | LC | Stable | tropical | nocturnal | 1.411 | 28902.423 | 4.813 |
| amphibians | Liua_tsinpaensis | Liua_tsinpaensis | VU | Decreasing | temperate | nocturnal | 5.728 | 432.81 | 10.863 |
| amphibians | Lyciasalamandra_atifi | Lyciasalamandra_atifi | EN | Decreasing | temperate | nocturnal | 11.13 | 2610.118 | 10.888 |
| amphibians | Lyciasalamandra_billae | NA | EN | Decreasing | NA | NA | NA | NA | NA |
| amphibians | Lyciasalamandra_fazilae | Lyciasalamandra_fazilae | EN | Decreasing | temperate | nocturnal | 5.728 | 552.424 | 15.972 |
| amphibians | Lyciasalamandra_flavimembris | Lyciasalamandra_flavimembris | EN | Decreasing | temperate | nocturnal | 43.05 | 595.119 | 16.802 |
| amphibians | Lyciasalamandra_helverseni | NA | VU | Decreasing | NA | NA | NA | NA | NA |
| amphibians | Lyciasalamandra_luschani | Lyciasalamandra_luschani | EN | Decreasing | temperate | cathemeral | 4.106 | 1801.912 | 14.302 |

| | | | | | | | | | |
|---|---|---|---|---|---|---|---|---|---|
| amphibians | Lynchius_nebulanastes | Lynchius_nebulanastes | EN | Decreasing | tropical | nocturnal | 6.322 | 66.339 | 14.703 |
| amphibians | Lynchius_oblitus | NA | LC | Unknown | NA | NA | NA | NA | NA |
| amphibians | Lynchius_simmonsi | Lynchius_simmonsi | EN | Decreasing | tropical | cathemeral | 5.399 | 1609.381 | 2.931 |
| amphibians | Lysapsus_caraya | Lysapsus_caraya | LC | Stable | tropical | cathemeral | 0.304 | 121340.78 | 5.188 |
| amphibians | Lysapsus_limellum | Lysapsus_limellum | LC | Stable | tropical | nocturnal | 1.138 | 1061961.661 | 8.837 |
| amphibians | Macrogenioglottus_alipioi | Macrogenioglottus_alipioi | LC | Stable | tropical | nocturnal | 158.508 | 557849.16 | 12.411 |
| amphibians | Mannophryne_caquetio | Mannophryne_caquetio | EN | Decreasing | tropical | diurnal | 2.296 | 592.243 | 11.562 |
| amphibians | Mannophryne_herminae | Mannophryne_herminae | NT | Stable | tropical | diurnal | 1.889 | 599.176 | 14.627 |
| amphibians | Mannophryne_lamarcai | Mannophryne_lamarcai | EN | Unknown | tropical | diurnal | 2.296 | 824.854 | 8.901 |
| amphibians | Mannophryne_molinai | NA | EN | Decreasing | NA | NA | NA | NA | NA |
| amphibians | Mannophryne_olmonae | Mannophryne_olmonae | VU | Stable | tropical | diurnal | 0.727 | 159.165 | 14.871 |
| amphibians | Mannophryne_orellana | Mannophryne_orellana | EN | Stable | tropical | diurnal | 2.844 | 769.655 | 12.785 |
| amphibians | Mannophryne_vulcano | Mannophryne_vulcano | NT | Decreasing | tropical | diurnal | 0.904 | 751.13 | 18.559 |
| amphibians | Mantella_aurantiaca | Mantella_aurantiaca | EN | Decreasing | tropical | cathemeral | 2.345 | 10.39 | 10.869 |
| amphibians | Mantella_baroni | Mantella_baroni | LC | Unknown | tropical | diurnal | 2.109 | 32997.279 | 8.416 |
| amphibians | Mantella_bernhardi | Mantella_bernhardi | VU | Decreasing | tropical | diurnal | 1.023 | 12373.088 | 10.136 |
| amphibians | Mantella_betsileo | Mantella_betsileo | LC | Stable | tropical | cathemeral | 1.326 | 84844.812 | 7.646 |
| amphibians | Mantella_cowanii | Mantella_cowanii | EN | Unknown | tropical | diurnal | 2.345 | 252.557 | 12.658 |
| amphibians | Mantella_crocea | Mantella_crocea | VU | Decreasing | tropical | diurnal | 1.023 | 535.937 | 7.62 |
| amphibians | Mantella_ebenaui | Mantella_ebenaui | LC | Stable | tropical | diurnal | 1.326 | 49853.175 | 8.088 |
| amphibians | Mantella_expectata | Mantella_expectata | EN | Decreasing | tropical | diurnal | 1.61 | 1112.304 | 7.029 |
| amphibians | Mantella_haraldmeieri | Mantella_haraldmeieri | EN | Decreasing | temperate | diurnal | 2.109 | 1976.389 | 5.796 |
| amphibians | Mantella_laevigata | Mantella_laevigata | LC | Decreasing | tropical | diurnal | 2.109 | 17602.27 | 8.083 |
| amphibians | Mantella_madagascariensis | Mantella_madagascariensis | VU | Decreasing | tropical | diurnal | 2.345 | 5208.683 | 7.31 |
| amphibians | Mantella_manery | Mantella_manery | VU | Decreasing | tropical | diurnal | 1.889 | 55.6 | 5.391 |
| amphibians | Mantella_milotympanum | Mantella_milotympanum | CR | Decreasing | tropical | diurnal | 2.109 | 65.312 | 8.828 |
| amphibians | Mantella_nigricans | Mantella_nigricans | LC | Decreasing | tropical | diurnal | 1.686 | 45585.713 | 7.052 |
| amphibians | Mantella_pulchra | Mantella_pulchra | NT | Decreasing | tropical | diurnal | 2.109 | 13262.94 | 6.326 |
| amphibians | Mantella_viridis | Mantella_viridis | EN | Decreasing | tropical | diurnal | 2.872 | 802.865 | 12.617 |
| amphibians | Mantidactylus_aerumnalis | Mantidactylus_aerumnalis | LC | Decreasing | tropical | diurnal | 2.345 | 18112.724 | 6.926 |
| amphibians | Mantidactylus_albofrenatus | Mantidactylus_albofrenatus | EN | Decreasing | tropical | diurnal | 1.686 | 1848.868 | 7.487 |
| amphibians | Mantidactylus_alutus | Mantidactylus_alutus | LC | Stable | tropical | cathemeral | 2.345 | 33784.163 | 13.058 |
| amphibians | Mantidactylus_ambreensis | Mantidactylus_ambreensis | LC | Decreasing | tropical | cathemeral | 4.536 | 9698.011 | 7.505 |
| amphibians | Mantidactylus_argenteus | Mantidactylus_argenteus | LC | Decreasing | tropical | diurnal | 2.345 | 25689.655 | 6.635 |

| | | | | | | | | | |
|---|---|---|---|---|---|---|---|---|---|
| amphibians | Mantidactylus_bellyi | Mantidactylus_bellyi | LC | Stable | tropical | cathemeral | 8.423 | 13701.072 | 9.033 |
| amphibians | Mantidactylus_betsileanus | Mantidactylus_betsileanus | LC | Stable | tropical | cathemeral | 6.273 | 87443.426 | 7.981 |
| amphibians | Mantidactylus_biporus | Mantidactylus_biporus | LC | Decreasing | tropical | diurnal | 3.163 | 90071.821 | 7.821 |
| amphibians | Mantidactylus_brevipalmatus | Mantidactylus_brevipalmatus | LC | Unknown | tropical | cathemeral | 7.844 | 39368.285 | 12.68 |
| amphibians | Mantidactylus_charlotteae | Mantidactylus_charlotteae | LC | Decreasing | tropical | diurnal | 2.599 | 65893.43 | 8.716 |
| amphibians | Mantidactylus_delormei | Mantidactylus_delormei | EN | Decreasing | tropical | diurnal | 4.934 | 2238.733 | 6.645 |
| amphibians | Mantidactylus_femoralis | Mantidactylus_femoralis | LC | Unknown | tropical | nocturnal | 15.029 | 189024.628 | 8.865 |
| amphibians | Mantidactylus_grandidieri | Mantidactylus_grandidieri | LC | Decreasing | tropical | cathemeral | 133.796 | 32649.526 | 6.522 |
| amphibians | Mantidactylus_guttulatus | Mantidactylus_guttulatus | LC | Decreasing | tropical | cathemeral | 188.234 | 40565.096 | 6.501 |
| amphibians | Mantidactylus_lugubris | Mantidactylus_lugubris | LC | Stable | tropical | nocturnal | 7.844 | 104217.404 | 8.817 |
| amphibians | Mantidactylus_majori | Mantidactylus_majori | LC | Decreasing | tropical | cathemeral | 9.031 | 34406.402 | 8.634 |
| amphibians | Mantidactylus_melanopleura | Mantidactylus_melanopleura | LC | Decreasing | tropical | diurnal | 5.802 | 95044.753 | 7.932 |
| amphibians | Mantidactylus_mocquardi | Mantidactylus_mocquardi | LC | Unknown | tropical | cathemeral | 28.486 | 131716.89 | 8.922 |
| amphibians | Mantidactylus_opiparis | Mantidactylus_opiparis | LC | Decreasing | tropical | cathemeral | 2.872 | 74906.32 | 7.658 |
| amphibians | Mantidactylus_pauliani | Mantidactylus_pauliani | CR | Decreasing | tropical | diurnal | 3.163 | 5.953 | 8.351 |
| amphibians | Mantidactylus_ulcerosus | Mantidactylus_ulcerosus | LC | Unknown | tropical | cathemeral | 7.844 | 50502.912 | 8.267 |
| amphibians | Mantidactylus_zipperi | Mantidactylus_zipperi | LC | Decreasing | tropical | cathemeral | 2.109 | 43952.615 | 9.332 |
| amphibians | Mantophryne_axanthogaster | Mantophryne_axanthogaster | LC | Stable | tropical | nocturnal | 12.299 | 455.362 | 3.785 |
| amphibians | Mantophryne_lateralis | Mantophryne_lateralis | LC | Stable | tropical | nocturnal | 11.036 | 364434.635 | 3.46 |
| amphibians | Megaelosia_goeldii | Megaelosia_goeldii | LC | Stable | tropical | diurnal | 193.364 | 1756.462 | 16.776 |
| amphibians | Megastomatohyla_mixe | Megastomatohyla_mixe | CR | Unknown | tropical | nocturnal | 3.133 | 32.58 | 5.695 |
| amphibians | Megastomatohyla_nubicola | Megastomatohyla_nubicola | CR | Unknown | tropical | nocturnal | 4.271 | 10.35 | 28.088 |
| amphibians | Megastomatohyla_pellita | Megastomatohyla_pellita | CR | Decreasing | tropical | nocturnal | 2.495 | 234.399 | 6.761 |
| amphibians | Megophrys_montana | Megophrys_montana | LC | Decreasing | tropical | nocturnal | 74.113 | 19685.222 | 15.114 |
| amphibians | Melanobatrachus_indicus | Melanobatrachus_indicus | VU | Decreasing | tropical | nocturnal | 3.163 | 2466.829 | 10.483 |
| amphibians | Melanophryniscus_admirabilis | Melanophryniscus_admirabilis | CR | Unknown | temperate | diurnal | 5.802 | 2402.143 | 11.919 |
| amphibians | Melanophryniscus_alipioi | Melanophryniscus_alipioi | VU | Decreasing | temperate | diurnal | 1.648 | 5279.827 | 10.839 |
| amphibians | Melanophryniscus_atroluteus | Melanophryniscus_atroluteus | LC | Stable | temperate | diurnal | 1.686 | 203412.136 | 10.85 |
| amphibians | Melanophryniscus_biancae | NA | EN | Decreasing | NA | NA | NA | NA | NA |
| amphibians | Melanophryniscus_cambaraensis | Melanophryniscus_cambaraensis | CR | Unknown | temperate | cathemeral | 4.536 | 633.799 | 6.643 |
| amphibians | Melanophryniscus_devincenzii | Melanophryniscus_devincenzii | LC | Stable | temperate | diurnal | 1.889 | 39108.024 | 11.333 |
| amphibians | Melanophryniscus_dorsalis | Melanophryniscus_dorsalis | LC | Decreasing | temperate | diurnal | 1.499 | 7855.651 | 16.086 |
| amphibians | Melanophryniscus_milanoi | NA | EN | Decreasing | NA | NA | NA | NA | NA |
| amphibians | Melanophryniscus_montevidensis | Melanophryniscus_montevidensis | NT | Decreasing | temperate | diurnal | 1.889 | 123386.175 | 12.875 |

| | | | | | | | | | |
|---|---|---|---|---|---|---|---|---|---|
| amphibians | Melanophryniscus_moreirae | Melanophryniscus_moreirae | NT | Stable | tropical | diurnal | 2.109 | 3184.71 | 9.317 |
| amphibians | Melanophryniscus_setiba | Melanophryniscus_setiba | CR | Decreasing | tropical | diurnal | 0.281 | 144.622 | 30.755 |
| amphibians | Melanophryniscus_simplex | Melanophryniscus_simplex | LC | Stable | temperate | nocturnal | 2.109 | 23087.917 | 9.293 |
| amphibians | Melanophryniscus_spectabilis | Melanophryniscus_spectabilis | EN | Decreasing | temperate | diurnal | 2.345 | 1425.336 | 11.561 |
| amphibians | Melanophryniscus_stelzneri | Melanophryniscus_stelzneri | LC | Stable | temperate | diurnal | 2.109 | 35188.738 | 10.13 |
| amphibians | Melanophryniscus_tumifrons | Melanophryniscus_tumifrons | LC | Stable | temperate | diurnal | 2.345 | 9942.278 | 17.658 |
| amphibians | Melanophryniscus_vilavelhensis | Melanophryniscus_vilavelhensis | EN | Decreasing | temperate | cathemeral | 0.348 | 402.472 | 9.02 |
| amphibians | Melanophryniscus_xanthostomus | NA | NT | Decreasing | NA | NA | NA | NA | NA |
| amphibians | Mercurana_myristicapalustris | Mercurana_myristicapalustris | EN | Decreasing | tropical | nocturnal | 25.822 | 141.566 | 14.839 |
| amphibians | Meristogenys_amoropalamus | Meristogenys_amoropalamus | LC | Decreasing | tropical | nocturnal | 29.041 | 12721.21 | 5.547 |
| amphibians | Meristogenys_jerboa | Meristogenys_jerboa | VU | Decreasing | tropical | cathemeral | 70.701 | 7986.388 | 10.699 |
| amphibians | Meristogenys_kinabaluensis | Meristogenys_kinabaluensis | LC | Stable | tropical | nocturnal | 82.421 | 51127.799 | 3.543 |
| amphibians | Meristogenys_maryatiae | Meristogenys_maryatiae | LC | Decreasing | tropical | nocturnal | 26.471 | 1404.631 | 7.718 |
| amphibians | Meristogenys_orphnocnemis | Meristogenys_orphnocnemis | LC | Decreasing | tropical | cathemeral | 32.83 | 42640.057 | 5.019 |
| amphibians | Meristogenys_whiteheadi | Meristogenys_whiteheadi | LC | Decreasing | tropical | nocturnal | 82.421 | 31449.001 | 4.607 |
| amphibians | Mertensiella_caucasica | Mertensiella_caucasica | NT | Decreasing | temperate | nocturnal | 7.16 | 14693.643 | 10.765 |
| amphibians | Mertensophryne_taitana | Mertensophryne_taitana | LC | Unknown | tropical | cathemeral | 3.807 | 1144764.241 | 10.123 |
| amphibians | Metaphrynella_sundana | Metaphrynella_sundana | LC | Decreasing | tropical | nocturnal | 2.109 | 85353.702 | 5.933 |
| amphibians | Micrixalus_adonis | NA | EN | Decreasing | NA | NA | NA | NA | NA |
| amphibians | Micrixalus_candidus | NA | EN | Decreasing | NA | NA | NA | NA | NA |
| amphibians | Micrixalus_elegans | Micrixalus_elegans | VU | Decreasing | tropical | diurnal | 0.664 | 5143.689 | 13.99 |
| amphibians | Micrixalus_frigidus | NA | EN | Decreasing | NA | NA | NA | NA | NA |
| amphibians | Micrixalus_fuscus | Micrixalus_fuscus | EN | Decreasing | tropical | diurnal | 2.9 | 1353.888 | 9.718 |
| amphibians | Micrixalus_gadgili | Micrixalus_gadgili | VU | Decreasing | tropical | diurnal | 0.418 | 2385.997 | 11.651 |
| amphibians | Micrixalus_herrei | NA | EN | Decreasing | NA | NA | NA | NA | NA |
| amphibians | Micrixalus_kodayari | NA | NT | Stable | NA | NA | NA | NA | NA |
| amphibians | Micrixalus_kottigeharensis | Micrixalus_kottigeharensis | VU | Decreasing | tropical | diurnal | 2.76 | 3522.631 | 15.381 |
| amphibians | Micrixalus_kurichiyari | NA | EN | Decreasing | NA | NA | NA | NA | NA |
| amphibians | Micrixalus_mallani | NA | EN | Decreasing | NA | NA | NA | NA | NA |
| amphibians | Micrixalus_nelliyampathi | NA | VU | Decreasing | NA | NA | NA | NA | NA |
| amphibians | Micrixalus_nigraventris | NA | EN | Decreasing | NA | NA | NA | NA | NA |
| amphibians | Micrixalus_niluvasei | NA | EN | Decreasing | NA | NA | NA | NA | NA |
| amphibians | Micrixalus_nudis | Micrixalus_nudis | EN | Decreasing | tropical | diurnal | 0.634 | 199.618 | 9.999 |
| amphibians | Micrixalus_phyllophilus | Micrixalus_phyllophilus | VU | Decreasing | tropical | diurnal | 2.733 | 1673.936 | 13.864 |

| | | | | | | | | | |
|---|---|---|---|---|---|---|---|---|---|
| amphibians | Micrixalus_sairandhri | NA | EN | Decreasing | NA | NA | NA | NA | NA |
| amphibians | Micrixalus_sali | NA | EN | Decreasing | NA | NA | NA | NA | NA |
| amphibians | Micrixalus_saxicola | Micrixalus_saxicola | LC | Decreasing | tropical | diurnal | 1.847 | 28601.601 | 15.776 |
| amphibians | Micrixalus_specca | NA | CR | Decreasing | NA | NA | NA | NA | NA |
| amphibians | Micrixalus_spelunca | NA | CR | Decreasing | NA | NA | NA | NA | NA |
| amphibians | Micrixalus_thampii | Micrixalus_thampii | EN | Decreasing | tropical | diurnal | 0.674 | 429.652 | 9.076 |
| amphibians | Micrixalus_uttaraghati | NA | EN | Decreasing | NA | NA | NA | NA | NA |
| amphibians | Microbatrachella_capensis | Microbatrachella_capensis | CR | Decreasing | temperate | nocturnal | 0.403 | 120.92 | 27.11 |
| amphibians | Microhyla_achatina | Microhyla_achatina | LC | Stable | tropical | nocturnal | 1.686 | 94800.653 | 23.321 |
| amphibians | Microhyla_aurantiventris | NA | DD | Unknown | NA | NA | NA | NA | NA |
| amphibians | Microhyla_berdmorei | Microhyla_berdmorei | LC | Decreasing | tropical | cathemeral | 8.188 | 1671204.759 | 11.869 |
| amphibians | Microhyla_borneensis | Microhyla_borneensis | LC | Decreasing | tropical | cathemeral | 1.168 | 4.563 | 15.978 |
| amphibians | Microhyla_butleri | Microhyla_butleri | LC | Decreasing | tropical | nocturnal | 1.326 | 2590639.865 | 14.011 |
| amphibians | Microhyla_fissipes | Microhyla_fissipes | LC | Stable | tropical | nocturnal | 1.686 | 4078682.042 | 14.681 |
| amphibians | Microhyla_fodiens | NA | DD | Unknown | NA | NA | NA | NA | NA |
| amphibians | Microhyla_heymonsi | Microhyla_heymonsi | LC | Stable | tropical | cathemeral | 2.201 | 3505075.566 | 13.267 |
| amphibians | Microhyla_irrawaddy | NA | DD | Unknown | NA | NA | NA | NA | NA |
| amphibians | Microhyla_mantheyi | Microhyla_mantheyi | LC | Unknown | tropical | nocturnal | 1.932 | 50835.819 | 8.359 |
| amphibians | Microhyla_mihintalei | NA | LC | Unknown | NA | NA | NA | NA | NA |
| amphibians | Microhyla_minuta | NA | DD | Decreasing | NA | NA | NA | NA | NA |
| amphibians | Microhyla_nilphamariensis | NA | LC | Decreasing | NA | NA | NA | NA | NA |
| amphibians | Microhyla_ornata | Microhyla_ornata | LC | Stable | tropical | nocturnal | 2.599 | 790904.301 | 20.566 |
| amphibians | Microhyla_pulchra | Microhyla_pulchra | LC | Stable | tropical | nocturnal | 3.981 | 2011158.497 | 13.79 |
| amphibians | Microhyla_rubra | Microhyla_rubra | LC | Decreasing | tropical | nocturnal | 2.345 | 1136655.498 | 20.194 |
| amphibians | Microkayla_ankohuma | Psychrophrynella_ankohuma | CR | Unknown | tropical | nocturnal | 1.554 | 31.847 | 8.987 |
| amphibians | Microkayla_chacaltaya | Psychrophrynella_chacaltaya | EN | Decreasing | tropical | cathemeral | 0.604 | 87.997 | 9.883 |
| amphibians | Microkayla_chapi | NA | not_available | not_available | NA | NA | NA | NA | NA |
| amphibians | Microkayla_huayna | NA | VU | Stable | NA | NA | NA | NA | NA |
| amphibians | Microkayla_iatamasi | Psychrophrynella_iatamasi | VU | Stable | tropical | diurnal | 0.996 | 90.902 | 7.88 |
| amphibians | Microkayla_wettsteini | Psychrophrynella_wettsteini | VU | Stable | tropical | cathemeral | 2.986 | 63.966 | 9.271 |
| amphibians | Micryletta_dissimulans | NA | not_available | not_available | NA | NA | NA | NA | NA |
| amphibians | Micryletta_erythropoda | Microhyla_erythropoda | LC | Decreasing | tropical | nocturnal | 2.109 | 178754.018 | 14.703 |
| amphibians | Micryletta_nigromaculata | NA | EN | Decreasing | NA | NA | NA | NA | NA |
| amphibians | Micryletta_steinegeri | Micryletta_steinegeri | VU | Decreasing | tropical | diurnal | 2.132 | 2491.982 | 13.505 |

| | | | | | | | | | |
|---|---|---|---|---|---|---|---|---|---|
| amphibians | Minervarya_brevipalmata | NA | DD | Unknown | NA | NA | NA | NA | NA |
| amphibians | Minervarya_chiangmaiensis | NA | LC | Unknown | NA | NA | NA | NA | NA |
| amphibians | Minervarya_greenii | Fejervarya_greenii | EN | Decreasing | tropical | diurnal | 11.036 | 764.222 | 15.425 |
| amphibians | Minervarya_kirtisinghei | Fejervarya_kirtisinghei | NT | Stable | tropical | diurnal | 7.294 | 9162.476 | 16.847 |
| amphibians | Minervarya_marathi | NA | DD | Unknown | NA | NA | NA | NA | NA |
| amphibians | Minervarya_nepalensis | Fejervarya_nepalensis | LC | Decreasing | temperate | nocturnal | 6.419 | 179285.144 | 15.887 |
| amphibians | Minervarya_nicobariensis | Fejervarya_nicobariensis | VU | Stable | tropical | cathemeral | 13.329 | 1411.474 | 5.482 |
| amphibians | Minervarya_pierrei | Fejervarya_pierrei | LC | Decreasing | temperate | nocturnal | 8.423 | 1724309.184 | 17.509 |
| amphibians | Minervarya_rufescens | Fejervarya_rufescens | NT | Decreasing | tropical | nocturnal | 6.77 | 20504.45 | 16.105 |
| amphibians | Minervarya_syhadrensis | Fejervarya_syhadrensis | LC | Decreasing | tropical | cathemeral | 6.518 | 2634149.582 | 18.111 |
| amphibians | Minervarya_teraiensis | Fejervarya_teraiensis | LC | Stable | temperate | nocturnal | 21.243 | 297104.056 | 13.564 |
| amphibians | Mixophyes_carbinensis | Mixophyes_carbinensis | LC | Stable | tropical | nocturnal | 46.616 | 746.86 | 1.151 |
| amphibians | Mixophyes_coggeri | Mixophyes_coggeri | LC | Stable | tropical | nocturnal | 118.397 | 14191.38 | 5.657 |
| amphibians | Mixophyes_fasciolatus | Mixophyes_fasciolatus | LC | Stable | temperate | nocturnal | 107.684 | 184187.487 | 8.658 |
| amphibians | Mixophyes_fleayi | Mixophyes_fleayi | EN | Decreasing | temperate | nocturnal | 71.478 | 4870.908 | 6.654 |
| amphibians | Mixophyes_iteratus | Mixophyes_iteratus | VU | Decreasing | temperate | nocturnal | 163.988 | 23086.442 | 7.783 |
| amphibians | Mixophyes_schevilli | Mixophyes_schevilli | LC | Stable | tropical | nocturnal | 79.584 | 17561.592 | 5.512 |
| amphibians | Morerella_cyanophthalma | Morerella_cyanophthalma | VU | Decreasing | tropical | nocturnal | 3.475 | 375.386 | 11.197 |
| amphibians | Myersiella_microps | Myersiella_microps | LC | Stable | tropical | cathemeral | 5.356 | 171653.056 | 13.912 |
| amphibians | Myersiohyla_chamaeleo | Myersiohyla_chamaeleo | NT | Unknown | tropical | nocturnal | 16.777 | 405.558 | 1.086 |
| amphibians | Myersiohyla_liliae | Hypsiboas_liliae | EN | Decreasing | tropical | nocturnal | 4.197 | 599.068 | 1.568 |
| amphibians | Myersiohyla_loveridgei | Myersiohyla_loveridgei | NT | Unknown | tropical | diurnal | 7.844 | 307.543 | 0.404 |
| amphibians | Myersiohyla_neblinaria | Myersiohyla_neblinaria | LC | Unknown | tropical | nocturnal | 21.697 | 757.055 | 0.938 |
| amphibians | Nannophryne_apolobambica | Nannophryne_apolobambica | CR | Decreasing | tropical | cathemeral | 39.652 | 76.31 | 3.059 |
| amphibians | Nannophryne_variegata | Nannophryne_variegata | LC | Decreasing | temperate | nocturnal | 11.036 | 135856.651 | 1.864 |
| amphibians | Nannophrys_ceylonensis | Nannophrys_ceylonensis | VU | Decreasing | tropical | nocturnal | 14.161 | 8067.268 | 18.344 |
| amphibians | Nannophrys_naeyakai | Nannophrys_naeyakai | EN | Decreasing | tropical | nocturnal | 7.081 | 73.047 | 10.388 |
| amphibians | Nanohyla_marmorata | Microhyla_marmorata | LC | Decreasing | tropical | nocturnal | 0.917 | 36576.326 | 5.492 |
| amphibians | Nanorana_annandalii | Nanorana_annandalii | LC | Decreasing | temperate | nocturnal | 12.145 | 37343.072 | 8.201 |
| amphibians | Nanorana_blanfordii | Nanorana_blanfordii | LC | Decreasing | temperate | nocturnal | 9.669 | 168900.018 | 10.834 |
| amphibians | Nanorana_kangxianensis | Nanorana_kangxianensis | EN | Decreasing | temperate | nocturnal | 108.03 | 912.142 | 11.68 |
| amphibians | Nanorana_liebigii | Nanorana_liebigii | LC | Decreasing | temperate | nocturnal | 570.827 | 261384.525 | 7.28 |
| amphibians | Nanorana_parkeri | Nanorana_parkeri | LC | Decreasing | temperate | cathemeral | 17.851 | 384079.59 | 6.135 |
| amphibians | Nanorana_pleskei | Nanorana_pleskei | LC | Decreasing | temperate | nocturnal | 6.273 | 142696.498 | 8.178 |

| | | | | | | | | | |
|---|---|---|---|---|---|---|---|---|---|
| amphibians | Nanorana_polunini | Nanorana_polunini | LC | Decreasing | temperate | nocturnal | 15.932 | 121833.7 | 10.023 |
| amphibians | Nanorana_rostandi | Nanorana_rostandi | LC | Decreasing | temperate | cathemeral | 25.822 | 76398.178 | 11.525 |
| amphibians | Nanorana_zhaoermii | NA | not_available | not_available | NA | NA | NA | NA | NA |
| amphibians | Nasikabatrachus_sahyadrensis | Nasikabatrachus_sahyadrensis | NT | Decreasing | tropical | nocturnal | 74.113 | 11357.474 | 14.018 |
| amphibians | Nectophryne_afra | Nectophryne_afra | LC | Unknown | tropical | cathemeral | 4.536 | 951359.521 | 7.321 |
| amphibians | Nectophryne_batesii | Nectophryne_batesii | LC | Unknown | tropical | nocturnal | 1.168 | 946911.719 | 5.825 |
| amphibians | Nectophrynoides_asperginis | NA | EW | not_available | NA | NA | NA | NA | NA |
| amphibians | Nectophrynoides_laticeps | Nectophrynoides_laticeps | CR | Decreasing | tropical | cathemeral | 1.023 | 17.175 | 10.714 |
| amphibians | Nectophrynoides_paulae | Nectophrynoides_paulae | CR | Decreasing | tropical | nocturnal | 1.214 | 17.951 | 10.662 |
| amphibians | Nectophrynoides_poyntoni | Nectophrynoides_poyntoni | CR | Decreasing | tropical | diurnal | 3.163 | 3.088 | 5.925 |
| amphibians | Nectophrynoides_pseudotornieri | Nectophrynoides_pseudotornieri | CR | Unknown | tropical | cathemeral | 1.889 | 38.147 | 10.247 |
| amphibians | Necturus_beyeri | Necturus_beyeri | LC | Stable | temperate | nocturnal | 94.154 | 303770.784 | 14.822 |
| amphibians | Necturus_lewisi | Necturus_lewisi | LC | Decreasing | temperate | nocturnal | 41.28 | 21188.52 | 23.376 |
| amphibians | Necturus_maculosus | Necturus_maculosus | LC | Stable | temperate | nocturnal | 533.849 | 1636355.166 | 18.063 |
| amphibians | Necturus_punctatus | Necturus_punctatus | LC | Stable | temperate | nocturnal | 31.001 | 177790.947 | 18.093 |
| amphibians | Neobatrachus_albipes | Neobatrachus_albipes | LC | Stable | temperate | nocturnal | 11.036 | 282547.959 | 9.634 |
| amphibians | Neobatrachus_kunapalari | Neobatrachus_kunapalari | LC | Stable | temperate | nocturnal | 22.389 | 412663.844 | 6.572 |
| amphibians | Neobatrachus_sudelli | Neobatrachus_sudelli | LC | Decreasing | temperate | nocturnal | 15.029 | 3558999.632 | 4.927 |
| amphibians | Neurergus_crocatus | Neurergus_crocatus | VU | Decreasing | temperate | nocturnal | 8.623 | 11081.366 | 13.12 |
| amphibians | Neurergus_kaiseri | Neurergus_kaiseri | VU | Decreasing | temperate | diurnal | 8.561 | 9198.41 | 14.848 |
| amphibians | Neurergus_strauchii | Neurergus_strauchii | LC | Decreasing | temperate | nocturnal | 8.016 | 20397.414 | 14.876 |
| amphibians | Niceforonia_babax | Hypodactylus_babax | LC | Decreasing | tropical | nocturnal | 10.133 | 10091.359 | 7.266 |
| amphibians | Niceforonia_elassodiscus | Hypodactylus_elassodiscus | LC | Unknown | tropical | diurnal | 4.124 | 9455.496 | 5.322 |
| amphibians | Niceforonia_nigrovittata | Hypodactylus_nigrovittatus | LC | Stable | tropical | cathemeral | 2.345 | 939846.04 | 2.008 |
| amphibians | Niceforonia_philippi | NA | VU | Decreasing | NA | NA | NA | NA | NA |
| amphibians | Nidirana_chapaensis | Babina_chapaensis | LC | Decreasing | tropical | nocturnal | 80.146 | 132769.031 | 7.03 |
| amphibians | Nidirana_daunchina | Babina_daunchina | LC | Stable | temperate | cathemeral | 13.329 | 81236.901 | 13.46 |
| amphibians | Nidirana_pleuraden | Babina_pleuraden | LC | Decreasing | temperate | diurnal | 16.873 | 357284.947 | 13.332 |
| amphibians | Nimbaphrynoides_occidentalis | Nimbaphrynoides_occidentalis | CR | Stable | tropical | cathemeral | 2.872 | 6.463 | 15.272 |
| amphibians | Noblella_coloma | Noblella_coloma | CR | Decreasing | tropical | cathemeral | 0.275 | 28.74 | 5.709 |
| amphibians | Noblella_duellmani | NA | DD | Unknown | NA | NA | NA | NA | NA |
| amphibians | Noblella_lochites | Noblella_lochites | EN | Decreasing | tropical | diurnal | 0.514 | 4229.501 | 5.945 |
| amphibians | Noblella_madreselva | NA | CR | Decreasing | NA | NA | NA | NA | NA |
| amphibians | Noblella_myrmecoides | Noblella_myrmecoides | LC | Decreasing | tropical | diurnal | 0.209 | 1148099.265 | 1.684 |

| | | | | | | | | | |
|---|---|---|---|---|---|---|---|---|---|
| amphibians | Noblella_naturetrekii | NA | EN | Decreasing | NA | NA | NA | NA | NA |
| amphibians | Noblella_personina | Noblella_personina | EN | Decreasing | tropical | diurnal | 0.396 | 983.696 | 4.46 |
| amphibians | Noblella_worleyae | NA | not_available | not_available | NA | NA | NA | NA | NA |
| amphibians | Notophthalmus_perstriatus | Notophthalmus_perstriatus | NT | Decreasing | temperate | nocturnal | 1.06 | 24419.514 | 16.106 |
| amphibians | Notophthalmus_viridescens | Notophthalmus_viridescens | LC | Stable | temperate | nocturnal | 2.562 | 3440358.26 | 15.726 |
| amphibians | Nototriton_abscondens | Nototriton_abscondens | LC | Stable | tropical | nocturnal | 0.56 | 902.952 | 19.402 |
| amphibians | Nototriton_barbouri | Nototriton_barbouri | EN | Decreasing | tropical | nocturnal | 1.052 | 36.004 | 5.548 |
| amphibians | Nototriton_brodiei | Nototriton_brodiei | EN | Decreasing | tropical | nocturnal | 0.686 | 93.897 | 6.312 |
| amphibians | Nototriton_limnospectator | Nototriton_limnospectator | EN | Decreasing | tropical | nocturnal | 0.926 | 310.018 | 8.859 |
| amphibians | Nototriton_nelsoni | NA | CR | Decreasing | NA | NA | NA | NA | NA |
| amphibians | Nyctibates_corrugatus | Nyctibates_corrugatus | LC | Unknown | tropical | nocturnal | 27.131 | 82370.015 | 9.54 |
| amphibians | Nyctibatrachus_aliciae | Nyctibatrachus_aliciae | VU | Decreasing | tropical | nocturnal | 3.044 | 5553.373 | 10.247 |
| amphibians | Nyctibatrachus_beddomii | Nyctibatrachus_beddomii | EN | Decreasing | tropical | cathemeral | 0.403 | 1399.145 | 9.735 |
| amphibians | Nyctibatrachus_dattatreyaensis | Nyctibatrachus_dattatreyaensis | EN | Decreasing | tropical | nocturnal | 6.419 | 572.632 | 14.683 |
| amphibians | Nyctibatrachus_karnatakaensis | Nyctibatrachus_karnatakaensis | EN | Decreasing | tropical | nocturnal | 61.584 | 852.749 | 10.046 |
| amphibians | Nyctibatrachus_kempholeyensis | Nyctibatrachus_kempholeyensis | LC | Decreasing | tropical | nocturnal | 1.08 | 35495.473 | 14.48 |
| amphibians | Nyctibatrachus_major | Nyctibatrachus_major | EN | Decreasing | tropical | nocturnal | 14.332 | 2635.665 | 12.031 |
| amphibians | Nyctibatrachus_manalari | NA | DD | Unknown | NA | NA | NA | NA | NA |
| amphibians | Nyctibatrachus_petraeus | Nyctibatrachus_petraeus | NT | Decreasing | tropical | nocturnal | 9.41 | 6905.853 | 12.427 |
| amphibians | Nyctibatrachus_pulivijayani | NA | EN | Decreasing | NA | NA | NA | NA | NA |
| amphibians | Nyctibatrachus_robinmoorei | NA | DD | Unknown | NA | NA | NA | NA | NA |
| amphibians | Nyctibatrachus_sabarimalai | NA | EN | Decreasing | NA | NA | NA | NA | NA |
| amphibians | Nyctibatrachus_webilla | NA | EN | Decreasing | NA | NA | NA | NA | NA |
| amphibians | Nyctimantis_arapapa | Aparasphenodon_arapapa | VU | Decreasing | tropical | nocturnal | 19.923 | 4273.649 | 7.696 |
| amphibians | Nyctimantis_bokermanni | Aparasphenodon_bokermanni | LC | Stable | temperate | nocturnal | 52.68 | 17346.888 | 14.557 |
| amphibians | Nyctimantis_brunoi | Aparasphenodon_brunoi | LC | Decreasing | tropical | nocturnal | 52.68 | 77451.512 | 11.992 |
| amphibians | Nyctimantis_rugiceps | Nyctimantis_rugiceps | LC | Stable | tropical | nocturnal | 29.887 | 45141.112 | 5.041 |
| amphibians | Nyctimantis_siemersi | Argenteohyla_siemersi | LC | Decreasing | temperate | nocturnal | 57.012 | 91004.379 | 8.085 |
| amphibians | Nyctimystes_infrafrenatus | NA | LC | Stable | NA | NA | NA | NA | NA |
| amphibians | Nyctimystes_kuduki | NA | DD | Unknown | NA | NA | NA | NA | NA |
| amphibians | Nyctimystes_multicolor | NA | DD | Unknown | NA | NA | NA | NA | NA |
| amphibians | Nyctixalus_pictus | Nyctixalus_pictus | LC | Decreasing | tropical | nocturnal | 4.536 | 231431.813 | 7.881 |
| amphibians | Nyctixalus_spinosus | Nyctixalus_spinosus | LC | Decreasing | tropical | nocturnal | 4.16 | 12654.062 | 10.775 |
| amphibians | Nymphargus_anomalus | Nymphargus_anomalus | EN | Decreasing | tropical | nocturnal | 1.499 | 1446.178 | 3.693 |

| | | | | | | | | | |
|---|---|---|---|---|---|---|---|---|---|
| amphibians | Nymphargus_balionotus | Cochranella_balionota | EN | Decreasing | tropical | nocturnal | 0.772 | 68.852 | 7.316 |
| amphibians | Nymphargus_caucanus | NA | EN | Decreasing | NA | NA | NA | NA | NA |
| amphibians | Nymphargus_chancas | Nymphargus_chancas | EN | Decreasing | tropical | nocturnal | 1.667 | 1292.402 | 7.678 |
| amphibians | Nymphargus_cochranae | Nymphargus_cochranae | LC | Decreasing | tropical | nocturnal | 2.178 | 10607.18 | 5.716 |
| amphibians | Nymphargus_colomai | NA | EN | Decreasing | NA | NA | NA | NA | NA |
| amphibians | Nymphargus_grandisonae | Nymphargus_grandisonae | LC | Decreasing | tropical | nocturnal | 2.771 | 21229.613 | 9.762 |
| amphibians | Nymphargus_griffithsi | Nymphargus_griffithsi | LC | Decreasing | tropical | nocturnal | 2.521 | 45955.224 | 11.596 |
| amphibians | Nymphargus_humboldti | NA | DD | Unknown | NA | NA | NA | NA | NA |
| amphibians | Nymphargus_ignotus | Nymphargus_ignotus | LC | Decreasing | tropical | nocturnal | 1.23 | 3565.855 | 8.194 |
| amphibians | Nymphargus_lasgralarias | Nymphargus_lasgralarias | EN | Decreasing | tropical | nocturnal | 1.517 | 760.773 | 7.38 |
| amphibians | Nymphargus_manduriacu | NA | CR | Unknown | NA | NA | NA | NA | NA |
| amphibians | Nymphargus_mariae | Nymphargus_mariae | LC | Decreasing | tropical | nocturnal | 2.109 | 21225.62 | 4.45 |
| amphibians | Nymphargus_megacheirus | Nymphargus_megacheirus | CR | Decreasing | tropical | nocturnal | 2.816 | 313.029 | 7.412 |
| amphibians | Nymphargus_mixomaculatus | Nymphargus_mixomaculatus | CR | Unknown | tropical | nocturnal | 1.377 | 1.391 | 7.048 |
| amphibians | Nymphargus_posadae | Nymphargus_posadae | LC | Decreasing | tropical | nocturnal | 3.194 | 28832.138 | 5.935 |
| amphibians | Nymphargus_rosada | Nymphargus_rosada | VU | Stable | tropical | nocturnal | 1.786 | 994.491 | 13.219 |
| amphibians | Nymphargus_sucre | Nymphargus_sucre | CR | Decreasing | tropical | nocturnal | 1.065 | 5.125 | 8.048 |
| amphibians | Nymphargus_vicenteruedai | NA | DD | Unknown | NA | NA | NA | NA | NA |
| amphibians | Nymphargus_wileyi | Nymphargus_wileyi | CR | Decreasing | tropical | nocturnal | 1.517 | 10.86 | 6.183 |
| amphibians | Occidozyga_baluensis | Occidozyga_baluensis | LC | Decreasing | tropical | cathemeral | 3.475 | 213555.664 | 4.114 |
| amphibians | Occidozyga_laevis | Occidozyga_laevis | LC | Decreasing | tropical | cathemeral | 22.506 | 356736.467 | 11.968 |
| amphibians | Occidozyga_lima | Occidozyga_lima | LC | Decreasing | tropical | cathemeral | 11.036 | 2814348.174 | 13.403 |
| amphibians | Occidozyga_martensii | Occidozyga_martensii | LC | Decreasing | tropical | nocturnal | 2.872 | 1761564.644 | 13.12 |
| amphibians | Occidozyga_sumatrana | Occidozyga_sumatrana | LC | Decreasing | tropical | nocturnal | 11.036 | 137030.46 | 10.882 |
| amphibians | Odontobatrachus_arndti | NA | NT | Decreasing | NA | NA | NA | NA | NA |
| amphibians | Odontobatrachus_fouta | NA | EN | Decreasing | NA | NA | NA | NA | NA |
| amphibians | Odontobatrachus_natator | Odontobatrachus_natator | LC | Decreasing | tropical | nocturnal | 23.335 | 162449.165 | 13.362 |
| amphibians | Odontobatrachus_smithi | NA | VU | Decreasing | NA | NA | NA | NA | NA |
| amphibians | Odontobatrachus_ziama | NA | VU | Decreasing | NA | NA | NA | NA | NA |
| amphibians | Odontophrynus_americanus | Odontophrynus_americanus | LC | Stable | temperate | nocturnal | 32.83 | 2881157.375 | 10.586 |
| amphibians | Odontophrynus_carvalhoi | Odontophrynus_carvalhoi | LC | Unknown | tropical | nocturnal | 32.83 | 917568.082 | 7.982 |
| amphibians | Odontophrynus_maisuma | Odontophrynus_maisuma | LC | Stable | temperate | nocturnal | 7.081 | 33564.851 | 13.463 |
| amphibians | Odorrana_andersonii | NA | LC | Decreasing | NA | NA | NA | NA | NA |
| amphibians | Odorrana_aureola | Odorrana_aureola | LC | Decreasing | tropical | nocturnal | 92.901 | 68449.282 | 7.384 |

| | | | | | | | | | |
|---|---|---|---|---|---|---|---|---|---|
| amphibians | Odorrana_banaorum | Odorrana_banaorum | LC | Decreasing | tropical | nocturnal | 136.627 | 121051.004 | 8.797 |
| amphibians | Odorrana_hosii | Odorrana_hosii | LC | Decreasing | tropical | cathemeral | 104.268 | 1116465.616 | 7.95 |
| amphibians | Odorrana_khalam | Odorrana_khalam | LC | Decreasing | tropical | nocturnal | 23.697 | 23823.438 | 5.398 |
| amphibians | Odorrana_kweichowensis | NA | VU | Decreasing | NA | NA | NA | NA | NA |
| amphibians | Odorrana_mutschmanni | NA | DD | Decreasing | NA | NA | NA | NA | NA |
| amphibians | Odorrana_narina | Odorrana_narina | EN | Stable | temperate | nocturnal | 100.927 | 311.628 | 14.726 |
| amphibians | Odorrana_tormota | Odorrana_tormota | LC | Decreasing | temperate | nocturnal | 19.923 | 28319.157 | 14.366 |
| amphibians | Odorrana_yentuensis | Odorrana_yentuensis | EN | Decreasing | tropical | nocturnal | 23.456 | 2007.161 | 9.709 |
| amphibians | Oedipina_berlini | NA | LC | Stable | NA | NA | NA | NA | NA |
| amphibians | Oedipina_collaris | NA | DD | Unknown | NA | NA | NA | NA | NA |
| amphibians | Oedipina_cyclocauda | Oedipina_cyclocauda | NT | Decreasing | tropical | nocturnal | 2.776 | 15448.678 | 9.301 |
| amphibians | Oedipina_gephyra | Oedipina_gephyra | CR | Decreasing | tropical | nocturnal | 3.177 | 19.984 | 7.316 |
| amphibians | Oedipina_grandis | Oedipina_grandis | EN | Decreasing | tropical | nocturnal | 5.968 | 189.186 | 8.534 |
| amphibians | Oedipina_poelzi | Oedipina_poelzi | EN | Decreasing | tropical | nocturnal | 4.221 | 647.912 | 20.764 |
| amphibians | Ololygon_agilis | Scinax_agilis | LC | Stable | tropical | cathemeral | 0.448 | 71774.631 | 11.385 |
| amphibians | Ololygon_albicans | Scinax_albicans | LC | Stable | tropical | nocturnal | 8.994 | 2601.538 | 20.013 |
| amphibians | Ololygon_alcatraz | NA | VU | Stable | NA | NA | NA | NA | NA |
| amphibians | Ololygon_angrensis | Scinax_angrensis | LC | Stable | tropical | cathemeral | 5.016 | 2483.057 | 9.686 |
| amphibians | Ololygon_arduoa | Scinax_arduous | LC | Stable | tropical | cathemeral | 1.36 | 4843.615 | 11.635 |
| amphibians | Ololygon_argyreornata | Scinax_argyreornatus | LC | Stable | tropical | cathemeral | 1.294 | 266745.587 | 12.714 |
| amphibians | Ololygon_ariadne | Scinax_ariadne | LC | Stable | tropical | nocturnal | 5.802 | 545.693 | 3.228 |
| amphibians | Ololygon_aromothyella | Scinax_aromothyella | LC | Stable | temperate | nocturnal | 1.499 | 377908.122 | 12.049 |
| amphibians | Ololygon_belloni | Scinax_belloni | EN | Decreasing | tropical | nocturnal | 4.934 | 1534.089 | 11.177 |
| amphibians | Ololygon_berthae | Scinax_berthae | LC | Stable | temperate | nocturnal | 1.168 | 875363.172 | 12.109 |
| amphibians | Ololygon_brieni | Scinax_brieni | LC | Stable | temperate | nocturnal | 5.356 | 17433.889 | 6.491 |
| amphibians | Ololygon_carnevallii | Scinax_carnevallii | LC | Stable | tropical | nocturnal | 2.599 | 119509.66 | 10.041 |
| amphibians | Ololygon_catharinae | Scinax_catharinae | LC | Stable | tropical | nocturnal | 9.031 | 1123002.094 | 11.79 |
| amphibians | Ololygon_cosenzai | Scinax_cosenzai | LC | Stable | tropical | nocturnal | 0.892 | 9703.664 | 11.715 |
| amphibians | Ololygon_faivovichi | NA | VU | Stable | NA | NA | NA | NA | NA |
| amphibians | Ololygon_flavoguttata | Scinax_flavoguttatus | LC | Stable | tropical | nocturnal | 8.073 | 140652.091 | 14.111 |
| amphibians | Ololygon_garibaldiae | NA | not_available | not_available | NA | NA | NA | NA | NA |
| amphibians | Ololygon_goya | NA | DD | Unknown | NA | NA | NA | NA | NA |
| amphibians | Ololygon_heyeri | Scinax_heyeri | EN | Decreasing | tropical | nocturnal | 4.934 | 6057.204 | 12.373 |
| amphibians | Ololygon_hiemalis | Scinax_hiemalis | LC | Stable | tropical | nocturnal | 3.475 | 44065.078 | 19.038 |

| | | | | | | | | | |
|---|---|---|---|---|---|---|---|---|---|
| amphibians | Ololygon_humilis | Scinax_humilis | LC | Stable | tropical | nocturnal | 5.756 | 60327.607 | 16.25 |
| amphibians | Ololygon_insperata | Scinax_insperatus | EN | Decreasing | tropical | nocturnal | 0.892 | 12.626 | 18.785 |
| amphibians | Ololygon_jureia | NA | DD | Unknown | NA | NA | NA | NA | NA |
| amphibians | Ololygon_littoralis | Scinax_littoralis | LC | Decreasing | temperate | nocturnal | 5.313 | 22250.255 | 8.728 |
| amphibians | Ololygon_littorea | Scinax_littoreus | EN | Decreasing | tropical | nocturnal | 1.245 | 1771.676 | 25.579 |
| amphibians | Ololygon_longilinea | Scinax_longilineus | LC | Decreasing | tropical | nocturnal | 9.669 | 105076.001 | 13.771 |
| amphibians | Ololygon_luizotavioi | Scinax_luizotavioi | LC | Decreasing | tropical | nocturnal | 2.109 | 36407.359 | 12.358 |
| amphibians | Ololygon_machadoi | Scinax_machadoi | LC | Stable | tropical | nocturnal | 1.023 | 44615.761 | 15.093 |
| amphibians | Ololygon_obtriangulata | Scinax_obtriangulatus | LC | Decreasing | tropical | nocturnal | 4.934 | 86195.502 | 15.63 |
| amphibians | Ololygon_peixotoi | NA | CR | Decreasing | NA | NA | NA | NA | NA |
| amphibians | Ololygon_perpusilla | Scinax_perpusillus | LC | Stable | tropical | nocturnal | 1.168 | 15916.158 | 12.442 |
| amphibians | Ololygon_ranki | Scinax_ranki | LC | Stable | tropical | nocturnal | 1.827 | 8625.648 | 14.134 |
| amphibians | Ololygon_rizibilis | Scinax_rizibilis | LC | Stable | temperate | cathemeral | 3.807 | 202433.453 | 12.702 |
| amphibians | Ololygon_skaios | Scinax_skaios | LC | Unknown | tropical | nocturnal | 3.841 | 230182.793 | 11.478 |
| amphibians | Ololygon_strigilata | Scinax_strigilatus | LC | Unknown | tropical | nocturnal | 4.653 | 28382.767 | 8.273 |
| amphibians | Ololygon_trapicheiroi | Scinax_trapicheiroi | LC | Stable | tropical | nocturnal | 5.356 | 47518.332 | 14.998 |
| amphibians | Ololygon_v-signata | Scinax_v-signatus | LC | Stable | tropical | nocturnal | 1.499 | 5662.036 | 12.522 |
| amphibians | Ombrana_sikimensis | Ombrana_sikimensis | LC | Decreasing | temperate | nocturnal | 68.909 | 75449.88 | 11.021 |
| amphibians | Ommatotriton_ophryticus | Ommatotriton_ophryticus | NT | Decreasing | temperate | nocturnal | 10.766 | 118354.751 | 11.882 |
| amphibians | Ommatotriton_vittatus | Ommatotriton_vittatus | LC | Decreasing | temperate | nocturnal | 3.543 | 57209.041 | 19.53 |
| amphibians | Onychodactylus_fischeri | Onychodactylus_fischeri | NT | Decreasing | temperate | nocturnal | 9.226 | 31338.509 | 5.711 |
| amphibians | Onychodactylus_japonicus | Onychodactylus_japonicus | LC | Stable | temperate | diurnal | 11.501 | 78734.7 | 16.408 |
| amphibians | Onychodactylus_nipponoborealis | Onychodactylus_nipponoborealis | LC | Stable | temperate | nocturnal | 6.969 | 37139.141 | 16.315 |
| amphibians | Onychodactylus_zhangyapingi | Onychodactylus_zhangyapingi | DD | Decreasing | temperate | nocturnal | 9.456 | 175944.418 | 9.521 |
| amphibians | Onychodactylus_zhaoermii | Onychodactylus_zhaoermii | NT | Decreasing | temperate | nocturnal | 8.499 | 21631.119 | 13.154 |
| amphibians | Oophaga_arborea | NA | CR | Decreasing | NA | NA | NA | NA | NA |
| amphibians | Oophaga_granulifera | Oophaga_granulifera | VU | Decreasing | tropical | diurnal | 1.168 | 6349.829 | 12.351 |
| amphibians | Oophaga_histrionica | NA | CR | Decreasing | NA | NA | NA | NA | NA |
| amphibians | Oophaga_lehmanni | NA | CR | Decreasing | NA | NA | NA | NA | NA |
| amphibians | Oophaga_occultator | Oophaga_occultator | CR | Decreasing | tropical | diurnal | 1.517 | 49.352 | 3.648 |
| amphibians | Oophaga_pumilio | Oophaga_pumilio | LC | Unknown | tropical | diurnal | 1.168 | 61975.534 | 9.876 |
| amphibians | Oophaga_speciosa | NA | EX | not_available | NA | NA | NA | NA | NA |
| amphibians | Oophaga_sylvatica | Oophaga_sylvatica | NT | Decreasing | tropical | diurnal | 4.536 | 48578.298 | 7.204 |
| amphibians | Oophaga_vicentei | Oophaga_vicentei | EN | Decreasing | tropical | diurnal | 0.664 | 2942.174 | 4.524 |

| | | | | | | | | | |
|---|---|---|---|---|---|---|---|---|---|
| amphibians | Ophryophryne_elfina | NA | LC | Decreasing | NA | NA | NA | NA | NA |
| amphibians | Ophryophryne_hansi | Ophryophryne_hansi | LC | Decreasing | tropical | nocturnal | 14.077 | 75873.011 | 7.082 |
| amphibians | Ophryophryne_microstoma | NA | LC | Decreasing | NA | NA | NA | NA | NA |
| amphibians | Ophryophryne_synoria | NA | VU | Decreasing | NA | NA | NA | NA | NA |
| amphibians | Opisthothylax_immaculatus | Opisthothylax_immaculatus | LC | Unknown | tropical | nocturnal | 3.475 | 816120.039 | 7.317 |
| amphibians | Oreobates_amarakaeri | Oreobates_amarakaeri | VU | Unknown | tropical | nocturnal | 2.296 | 16828.28 | 2.289 |
| amphibians | Oreobates_antrum | NA | LC | Stable | NA | NA | NA | NA | NA |
| amphibians | Oreobates_ayacucho | Oreobates_ayacucho | EN | Unknown | tropical | diurnal | 1.932 | 103.463 | 12.822 |
| amphibians | Oreobates_barituensis | Oreobates_barituensis | NT | Decreasing | tropical | nocturnal | 2.086 | 11974.815 | 5.184 |
| amphibians | Oreobates_cruralis | Oreobates_cruralis | LC | Stable | tropical | nocturnal | 3.044 | 146293.043 | 3.923 |
| amphibians | Oreobates_gemcare | Oreobates_gemcare | LC | Unknown | tropical | nocturnal | 5.313 | 81.263 | 8.403 |
| amphibians | Oreobates_machiguenga | NA | DD | Unknown | NA | NA | NA | NA | NA |
| amphibians | Oreobates_quixensis | Oreobates_quixensis | LC | Stable | tropical | cathemeral | 23.335 | 1916308.789 | 1.607 |
| amphibians | Oreobates_sanderi | Oreobates_sanderi | LC | Decreasing | tropical | diurnal | 4.536 | 25318.661 | 4.992 |
| amphibians | Oreolalax_chuanbeiensis | Oreolalax_chuanbeiensis | EN | Decreasing | temperate | nocturnal | 18.867 | 3829.552 | 4.896 |
| amphibians | Oreolalax_liangbeiensis | Oreolalax_liangbeiensis | CR | Decreasing | temperate | nocturnal | 26.734 | 6.749 | 10.858 |
| amphibians | Oreolalax_omeimontis | Oreolalax_omeimontis | EN | Decreasing | temperate | nocturnal | 18.253 | 2901.348 | 10.888 |
| amphibians | Oreolalax_popei | Oreolalax_popei | LC | Decreasing | temperate | nocturnal | 31.334 | 34208.9 | 8.627 |
| amphibians | Oreolalax_xiangchengensis | Oreolalax_xiangchengensis | LC | Stable | temperate | nocturnal | 21.469 | 28066.298 | 6.461 |
| amphibians | Oreophryne_albitympanum | NA | LC | Stable | NA | NA | NA | NA | NA |
| amphibians | Oreophryne_albomaculata | NA | LC | Stable | NA | NA | NA | NA | NA |
| amphibians | Oreophryne_anamiatoi | Oreophryne_anamiatoi | LC | Stable | tropical | nocturnal | 2.041 | 1877.905 | 8.361 |
| amphibians | Oreophryne_atrigularis | NA | DD | Unknown | NA | NA | NA | NA | NA |
| amphibians | Oreophryne_banshee | NA | LC | Stable | NA | NA | NA | NA | NA |
| amphibians | Oreophryne_brunnea | NA | LC | Stable | NA | NA | NA | NA | NA |
| amphibians | Oreophryne_cameroni | NA | EN | Decreasing | NA | NA | NA | NA | NA |
| amphibians | Oreophryne_choerophrynoides | NA | DD | Unknown | NA | NA | NA | NA | NA |
| amphibians | Oreophryne_clamata | NA | DD | Unknown | NA | NA | NA | NA | NA |
| amphibians | Oreophryne_curator | NA | LC | Stable | NA | NA | NA | NA | NA |
| amphibians | Oreophryne_equus | NA | LC | Stable | NA | NA | NA | NA | NA |
| amphibians | Oreophryne_flavomaculata | NA | LC | Stable | NA | NA | NA | NA | NA |
| amphibians | Oreophryne_gagneorum | NA | LC | Stable | NA | NA | NA | NA | NA |
| amphibians | Oreophryne_geislerorum | Oreophryne_geislerorum | LC | Stable | tropical | nocturnal | 1.975 | 20913.789 | 3.932 |
| amphibians | Oreophryne_loriae | NA | DD | Unknown | NA | NA | NA | NA | NA |

| | | | | | | | | | |
|---|---|---|---|---|---|---|---|---|---|
| amphibians | Oreophryne_matawan | NA | CR | Decreasing | NA | NA | NA | NA | NA |
| amphibians | Oreophryne_minuta | Oreophryne_minuta | LC | Stable | tropical | nocturnal | 0.094 | 9.879 | 2.34 |
| amphibians | Oreophryne_nicolasi | NA | LC | Stable | NA | NA | NA | NA | NA |
| amphibians | Oreophryne_notata | Oreophryne_notata | LC | Stable | tropical | nocturnal | 0.644 | 15346.96 | 5.69 |
| amphibians | Oreophryne_oviprotector | Oreophryne_oviprotector | LC | Stable | tropical | nocturnal | 1.517 | 53939.181 | 2.627 |
| amphibians | Oreophryne_parkopanorum | NA | LC | Stable | NA | NA | NA | NA | NA |
| amphibians | Oreophryne_philosylleptoris | NA | LC | Stable | NA | NA | NA | NA | NA |
| amphibians | Oreophryne_phoebe | NA | VU | Stable | NA | NA | NA | NA | NA |
| amphibians | Oreophryne_picticrus | NA | EN | Decreasing | NA | NA | NA | NA | NA |
| amphibians | Oreophryne_sibilans | NA | DD | Unknown | NA | NA | NA | NA | NA |
| amphibians | Oreophryne_unicolor | NA | DD | Unknown | NA | NA | NA | NA | NA |
| amphibians | Oreophryne_wapoga | NA | DD | Unknown | NA | NA | NA | NA | NA |
| amphibians | Oreophrynella_macconnelli | Oreophrynella_macconnelli | VU | Decreasing | tropical | nocturnal | 3.379 | 318.182 | 1.186 |
| amphibians | Oreophrynella_seegobini | Oreophrynella_seegobini | VU | Unknown | tropical | diurnal | 0.664 | 2.131 | 0.567 |
| amphibians | Osornophryne_antisana | Osornophryne_antisana | EN | Decreasing | tropical | cathemeral | 1.934 | 3121.606 | 5.75 |
| amphibians | Osornophryne_cofanorum | Osornophryne_cofanorum | CR | Decreasing | tropical | nocturnal | 3.285 | 46.84 | 5.214 |
| amphibians | Osornophryne_guacamayo | Osornophryne_guacamayo | VU | Decreasing | tropical | cathemeral | 5.643 | 1891.788 | 4.36 |
| amphibians | Osornophryne_occidentalis | Osornophryne_occidentalis | EN | Decreasing | tropical | cathemeral | 3.579 | 1412.445 | 9.67 |
| amphibians | Osornophryne_puruanta | Osornophryne_puruanta | EN | Decreasing | tropical | nocturnal | 9.094 | 40.436 | 3.293 |
| amphibians | Osornophryne_simpsoni | Osornophryne_simpsoni | EN | Decreasing | tropical | nocturnal | 2.872 | 812.812 | 2.716 |
| amphibians | Osornophryne_sumacoensis | Osornophryne_sumacoensis | VU | Unknown | tropical | cathemeral | 5.443 | 7.357 | 2.881 |
| amphibians | Osteocephalus_alboguttatus | Osteocephalus_alboguttatus | LC | Stable | tropical | nocturnal | 25.822 | 17609.51 | 8.62 |
| amphibians | Osteocephalus_buckleyi | Osteocephalus_buckleyi | LC | Stable | tropical | nocturnal | 31.334 | 816911.226 | 2.248 |
| amphibians | Osteocephalus_cabrerai | Osteocephalus_cabrerai | LC | Stable | tropical | nocturnal | 35.005 | 2206491.396 | 1.394 |
| amphibians | Osteocephalus_deridens | Osteocephalus_deridens | LC | Stable | tropical | nocturnal | 11.471 | 51880.85 | 2.536 |
| amphibians | Osteocephalus_festae | Osteocephalus_festae | LC | Unknown | tropical | nocturnal | 61.35 | 26630.893 | 4.299 |
| amphibians | Osteocephalus_fuscifacies | Osteocephalus_fuscifacies | LC | Decreasing | tropical | nocturnal | 17.357 | 53457.93 | 2.763 |
| amphibians | Osteocephalus_heyeri | Osteocephalus_heyeri | LC | Stable | tropical | nocturnal | 9.474 | 14254.186 | 1.223 |
| amphibians | Osteocephalus_leprieurii | Osteocephalus_leprieurii | LC | Stable | tropical | nocturnal | 41.053 | 4980892.49 | 1.517 |
| amphibians | Osteocephalus_planiceps | Osteocephalus_planiceps | LC | Stable | tropical | nocturnal | 72 | 645401.996 | 2.03 |
| amphibians | Osteocephalus_sangay | NA | DD | Unknown | NA | NA | NA | NA | NA |
| amphibians | Osteocephalus_subtilis | Osteocephalus_subtilis | LC | Stable | tropical | nocturnal | 11.918 | 151237.285 | 0.774 |
| amphibians | Osteocephalus_taurinus | Osteocephalus_taurinus | LC | Stable | tropical | nocturnal | 118.028 | 7474189.448 | 2.534 |
| amphibians | Osteocephalus_yasuni | Osteocephalus_yasuni | LC | Stable | tropical | cathemeral | 22.041 | 112322.909 | 1.418 |

| | | | | | | | | | |
|---|---|---|---|---|---|---|---|---|---|
| amphibians | Osteopilus_crucialis | Osteopilus_crucialis | VU | Decreasing | tropical | nocturnal | 198.59 | 3741.938 | 18.04 |
| amphibians | Osteopilus_dominicensis | Osteopilus_dominicensis | LC | Stable | tropical | nocturnal | 100.927 | 74765.187 | 15.762 |
| amphibians | Osteopilus_ocellatus | Osteopilus_ocellatus | NT | Stable | tropical | nocturnal | 42.854 | 8550.172 | 18.196 |
| amphibians | Osteopilus_septentrionalis | Osteopilus_septentrionalis | LC | Increasing | temperate | nocturnal | 528.201 | 279117.601 | 16.673 |
| amphibians | Otophryne_pyburni | Otophryne_pyburni | LC | Stable | tropical | diurnal | 21.019 | 568268.009 | 0.542 |
| amphibians | Otophryne_robusta | Otophryne_robusta | LC | Stable | tropical | diurnal | 15.029 | 28933.053 | 1.612 |
| amphibians | Otophryne_steyermarki | Otophryne_steyermarki | LC | Stable | tropical | diurnal | 9.669 | 23873.37 | 1.69 |
| amphibians | Pachytriton_archospotus | Pachytriton_archospotus | NT | Decreasing | temperate | nocturnal | 20.198 | 22128.579 | 9.64 |
| amphibians | Pachytriton_brevipes | Pachytriton_brevipes | LC | Decreasing | temperate | nocturnal | 15.562 | 78142.064 | 14.293 |
| amphibians | Pachytriton_changi | Pachytriton_changi | EN | Decreasing | temperate | nocturnal | 10.463 | 390.808 | 6.964 |
| amphibians | Pachytriton_feii | Pachytriton_feii | VU | Decreasing | temperate | nocturnal | 8.623 | 16412.309 | 10.754 |
| amphibians | Pachytriton_granulosus | Pachytriton_granulosus | LC | Decreasing | temperate | nocturnal | 8.623 | 114712.757 | 14.717 |
| amphibians | Pachytriton_inexpectatus | NA | LC | Stable | NA | NA | NA | NA | NA |
| amphibians | Pachytriton_moi | Pachytriton_moi | EN | Unknown | temperate | nocturnal | 15.77 | 1363.682 | 6.473 |
| amphibians | Pachytriton_wuguanfui | NA | EN | Decreasing | NA | NA | NA | NA | NA |
| amphibians | Pachytriton_xanthospilos | NA | CR | Decreasing | NA | NA | NA | NA | NA |
| amphibians | Paedophryne_amauensis | Paedophryne_amauensis | LC | Stable | tropical | nocturnal | 0.029 | 3013.913 | 4.061 |
| amphibians | Paracassina_kounhiensis | Paracassina_kounhiensis | VU | Stable | tropical | cathemeral | 10.337 | 455.972 | 15.444 |
| amphibians | Paracassina_obscura | Paracassina_obscura | LC | Unknown | tropical | cathemeral | 8.423 | 151622.709 | 13.349 |
| amphibians | Paradoxophyla_palmata | Paradoxophyla_palmata | LC | Decreasing | tropical | nocturnal | 1.326 | 33457.187 | 8.002 |
| amphibians | Paradoxophyla_tiarano | Paradoxophyla_tiarano | DD | Decreasing | tropical | nocturnal | 2.272 | 9.2 | 5.168 |
| amphibians | Paramesotriton_caudopunctatus | Paramesotriton_caudopunctatus | NT | Decreasing | temperate | nocturnal | 5.177 | 28929.919 | 11.479 |
| amphibians | Paramesotriton_chinensis | Paramesotriton_chinensis | LC | Decreasing | temperate | nocturnal | 10.41 | 107136.06 | 16.036 |
| amphibians | Paramesotriton_guangxiensis | NA | VU | Decreasing | NA | NA | NA | NA | NA |
| amphibians | Paramesotriton_hongkongensis | Paramesotriton_hongkongensis | NT | Decreasing | tropical | nocturnal | 6.017 | 6650.475 | 30.494 |
| amphibians | Paramesotriton_longliensis | Paramesotriton_longliensis | VU | Decreasing | temperate | nocturnal | 12.948 | 1397.478 | 15.432 |
| amphibians | Paramesotriton_wulingensis | NA | LC | Stable | NA | NA | NA | NA | NA |
| amphibians | Paramesotriton_yunwuensis | Paramesotriton_yunwuensis | EN | Decreasing | tropical | cathemeral | 18.197 | 1063.797 | 16.74 |
| amphibians | Paramesotriton_zhijinensis | NA | EN | Decreasing | NA | NA | NA | NA | NA |
| amphibians | Paratelmatobius_cardosoi | Paratelmatobius_cardosoi | LC | Stable | temperate | nocturnal | 0.41 | 2229.42 | 9.353 |
| amphibians | Paratelmatobius_gaigeae | Paratelmatobius_gaigeae | EN | Decreasing | tropical | nocturnal | 0.567 | 21.546 | 5.12 |
| amphibians | Paratelmatobius_lutzii | Paratelmatobius_lutzii | CR | Unknown | tropical | diurnal | 1.023 | 10.75 | 6.841 |
| amphibians | Paratelmatobius_mantiqueira | Paratelmatobius_mantiqueira | CR | Unknown | tropical | nocturnal | 0.505 | 324.644 | 6.482 |
| amphibians | Paratelmatobius_poecilogaster | Paratelmatobius_poecilogaster | LC | Stable | temperate | nocturnal | 2.132 | 5433.504 | 8.759 |

| | | | | | | | | | |
|---|---|---|---|---|---|---|---|---|---|
| amphibians | Paratelmatobius_segallai | NA | DD | Unknown | NA | NA | NA | NA | NA |
| amphibians | Paratelmatobius_yepiranga | NA | DD | Unknown | NA | NA | NA | NA | NA |
| amphibians | Paruwrobates_andinus | Ameerega_andina | CR | Decreasing | tropical | diurnal | 0.772 | 41.783 | 5.045 |
| amphibians | Paruwrobates_erythromos | Ameerega_erythromos | CR | Decreasing | tropical | diurnal | 1.199 | 18.7 | 10.426 |
| amphibians | Pedostibes_tuberculosus | Pedostibes_tuberculosus | LC | Decreasing | tropical | nocturnal | 4.732 | 55936.534 | 13.66 |
| amphibians | Pelobates_cultripes | Pelobates_cultripes | VU | Decreasing | temperate | nocturnal | 214.852 | 493091.327 | 17.768 |
| amphibians | Pelobates_fuscus | Pelobates_fuscus | LC | Decreasing | temperate | nocturnal | 50.601 | 2386089.746 | 19.016 |
| amphibians | Pelobates_syriacus | Pelobates_syriacus | LC | Decreasing | temperate | cathemeral | 74.113 | 372668.294 | 17.017 |
| amphibians | Pelobates_varaldii | Pelobates_varaldii | EN | Decreasing | temperate | nocturnal | 32.83 | 6672.948 | 23.628 |
| amphibians | Pelobatrachus_baluensis | NA | NT | Stable | NA | NA | NA | NA | NA |
| amphibians | Pelobatrachus_edwardinae | NA | LC | Decreasing | NA | NA | NA | NA | NA |
| amphibians | Pelobatrachus_kalimantanensis | NA | not_available | not_available | NA | NA | NA | NA | NA |
| amphibians | Pelobatrachus_kobayashii | NA | LC | Stable | NA | NA | NA | NA | NA |
| amphibians | Pelobatrachus_nasutus | NA | LC | Decreasing | NA | NA | NA | NA | NA |
| amphibians | Pelodytes_caucasicus | Pelodytes_caucasicus | LC | Decreasing | temperate | nocturnal | 15.029 | 111848.595 | 10.434 |
| amphibians | Pelodytes_ibericus | Pelodytes_ibericus | LC | Decreasing | temperate | cathemeral | 9.669 | 102533.406 | 16.925 |
| amphibians | Pelodytes_punctatus | Pelodytes_punctatus | LC | Decreasing | temperate | nocturnal | 11.767 | 707169.23 | 20.748 |
| amphibians | Pelophryne_linanitensis | Pelophryne_linanitensis | CR | Decreasing | tropical | nocturnal | 0.448 | 9.863 | 2.676 |
| amphibians | Pelophryne_misera | Pelophryne_misera | LC | Decreasing | tropical | diurnal | 0.892 | 2654.7 | 5.303 |
| amphibians | Pelophryne_signata | Pelophryne_signata | LC | Decreasing | tropical | cathemeral | 1.168 | 181875.489 | 9.177 |
| amphibians | Pelophylax_bedriagae | Pelophylax_bedriagae | LC | Stable | temperate | cathemeral | 221.835 | 1027992.456 | 13.969 |
| amphibians | Pelophylax_epeiroticus | Pelophylax_epeiroticus | NT | Decreasing | temperate | diurnal | 100.927 | 12315.977 | 19.234 |
| amphibians | Pelophylax_lessonae | Pelophylax_lessonae | LC | Decreasing | temperate | cathemeral | 188.234 | 3431615.932 | 18.847 |
| amphibians | Pelophylax_nigromaculatus | Pelophylax_nigromaculatus | LC | Decreasing | temperate | cathemeral | 85.327 | 4306604.691 | 15.856 |
| amphibians | Pelophylax_perezi | Pelophylax_perezi | LC | Decreasing | temperate | cathemeral | 141.992 | 650025.704 | 18.174 |
| amphibians | Pelophylax_plancyi | Pelophylax_plancyi | LC | Decreasing | temperate | diurnal | 43.037 | 814241.553 | 24.18 |
| amphibians | Pelophylax_porosus | Pelophylax_porosus | LC | Decreasing | temperate | diurnal | 66.404 | 96735.433 | 25.813 |
| amphibians | Pelophylax_ridibundus | Pelophylax_ridibundus | LC | Stable | temperate | cathemeral | 700.219 | 7792587.741 | 15.488 |
| amphibians | Pelophylax_saharicus | Pelophylax_saharicus | LC | Stable | temperate | cathemeral | 310.174 | 1403222.426 | 8.905 |
| amphibians | Peltophryne_armata | NA | CR | Decreasing | NA | NA | NA | NA | NA |
| amphibians | Peltophryne_fustiger | Peltophryne_fustiger | LC | Stable | tropical | nocturnal | 953.557 | 7303.868 | 17.812 |
| amphibians | Peltophryne_guentheri | Peltophryne_guentheri | LC | Stable | tropical | nocturnal | 107.684 | 24581.623 | 17.529 |
| amphibians | Peltophryne_gundlachi | Peltophryne_gundlachi | LC | Decreasing | tropical | nocturnal | 3.807 | 14026.823 | 15.667 |
| amphibians | Peltophryne_lemur | Peltophryne_lemur | EN | Decreasing | tropical | nocturnal | 188.234 | 407.912 | 21.872 |

| | | | | | | | | | |
|---|---|---|---|---|---|---|---|---|---|
| amphibians | Peltophryne_longinasus | Peltophryne_longinasus | CR | Decreasing | tropical | cathemeral | 3.807 | 695.869 | 12.445 |
| amphibians | Peltophryne_peltocephala | Peltophryne_peltocephala | LC | Stable | tropical | nocturnal | 507.737 | 28736.11 | 14.133 |
| amphibians | Peltophryne_taladai | Peltophryne_taladai | LC | Decreasing | tropical | nocturnal | 363.295 | 13997.387 | 14.552 |
| amphibians | Petropedetes_cameronensis | Petropedetes_cameronensis | LC | Decreasing | tropical | nocturnal | 10.337 | 40795.07 | 9.782 |
| amphibians | Petropedetes_euskircheni | Petropedetes_euskircheni | EN | Decreasing | tropical | nocturnal | 22.156 | 359.449 | 8.466 |
| amphibians | Petropedetes_palmipes | Petropedetes_palmipes | VU | Decreasing | tropical | nocturnal | 17.851 | 2078.997 | 5.985 |
| amphibians | Petropedetes_perreti | Petropedetes_perreti | CR | Decreasing | tropical | nocturnal | 10.337 | 550.32 | 10.098 |
| amphibians | Petropedetes_vulpiae | Petropedetes_vulpiae | LC | Decreasing | tropical | cathemeral | 9.031 | 190194.297 | 8.339 |
| amphibians | Phaeognathus_hubrichti | Phaeognathus_hubrichti | EN | Decreasing | temperate | nocturnal | 20.252 | 1166.989 | 8.626 |
| amphibians | Phantasmarana_apuana | Megaelosia_apuana | LC | Stable | tropical | diurnal | 95.102 | 6614.515 | 10.977 |
| amphibians | Phantasmarana_bocainensis | NA | DD | Unknown | NA | NA | NA | NA | NA |
| amphibians | Phantasmarana_boticariana | Megaelosia_boticariana | EN | Decreasing | tropical | diurnal | 89.21 | 2796.843 | 16.307 |
| amphibians | Phantasmarana_lutzae | Megaelosia_lutzae | LC | Stable | tropical | diurnal | 80.994 | 35.494 | 7.814 |
| amphibians | Phantasmarana_massarti | Megaelosia_massarti | EN | Decreasing | temperate | diurnal | 208.786 | 469.887 | 8.132 |
| amphibians | Phasmahyla_cochranae | Phasmahyla_cochranae | LC | Decreasing | tropical | nocturnal | 8.423 | 63311.807 | 16.025 |
| amphibians | Phasmahyla_exilis | Phasmahyla_exilis | LC | Stable | tropical | nocturnal | 3.317 | 1147.286 | 11.289 |
| amphibians | Phasmahyla_guttata | Phasmahyla_guttata | LC | Stable | tropical | nocturnal | 7.844 | 9871.988 | 15.666 |
| amphibians | Phasmahyla_jandaia | Phasmahyla_jandaia | LC | Stable | tropical | nocturnal | 3.807 | 18654.366 | 5.766 |
| amphibians | Phasmahyla_lisbella | NA | EN | Decreasing | NA | NA | NA | NA | NA |
| amphibians | Phasmahyla_spectabilis | Phasmahyla_spectabilis | LC | Stable | tropical | nocturnal | 10.066 | 2688.486 | 6.545 |
| amphibians | Phasmahyla_timbo | Phasmahyla_timbo | LC | Stable | tropical | nocturnal | 3.739 | 12509.16 | 8.853 |
| amphibians | Philautus_aurantium | Philautus_aurantium | VU | Decreasing | tropical | nocturnal | 2.109 | 4994.789 | 5.691 |
| amphibians | Philautus_bunitus | Philautus_bunitus | LC | Stable | tropical | cathemeral | 8.603 | 4837.695 | 5.816 |
| amphibians | Philautus_ingeri | Philautus_ingeri | VU | Decreasing | tropical | nocturnal | 11.036 | 5476.514 | 3.307 |
| amphibians | Philautus_longicrus | Philautus_longicrus | VU | Decreasing | tropical | nocturnal | 1.889 | 9200.675 | 4.518 |
| amphibians | Philautus_macroscelis | Philautus_macroscelis | LC | Stable | tropical | nocturnal | 15.029 | 13672.481 | 2.204 |
| amphibians | Philautus_mjobergi | Philautus_mjobergi | LC | Decreasing | tropical | cathemeral | 2.599 | 15067.316 | 3.414 |
| amphibians | Philautus_nephophilus | NA | LC | Stable | NA | NA | NA | NA | NA |
| amphibians | Philautus_petersi | NA | DD | Unknown | NA | NA | NA | NA | NA |
| amphibians | Philautus_saueri | Philautus_saueri | LC | Stable | tropical | diurnal | 0.706 | 22.275 | 7.043 |
| amphibians | Philoria_frosti | Philoria_frosti | CR | Decreasing | temperate | diurnal | 15.029 | 113.318 | 4.602 |
| amphibians | Philoria_kundagungan | Philoria_kundagungan | EN | Decreasing | temperate | nocturnal | 2.109 | 607.107 | 5.893 |
| amphibians | Philoria_richmondensis | Philoria_richmondensis | EN | Decreasing | temperate | cathemeral | 1.686 | 669.573 | 6.716 |
| amphibians | Philoria_sphagnicola | Philoria_sphagnicolus | VU | Decreasing | temperate | diurnal | 4.16 | 10171.243 | 5.542 |

| | | | | | | | | | |
|---|---|---|---|---|---|---|---|---|---|
| amphibians | Phrynobatrachus_acridoides | Phrynobatrachus_acridoides | LC | Unknown | tropical | cathemeral | 2.345 | 1759744.508 | 10.715 |
| amphibians | Phrynobatrachus_afiabirago | NA | CR | Decreasing | NA | NA | NA | NA | NA |
| amphibians | Phrynobatrachus_africanus | Phrynobatrachus_africanus | LC | Stable | tropical | diurnal | 2.345 | 715736.655 | 6.275 |
| amphibians | Phrynobatrachus_alleni | Phrynobatrachus_alleni | LC | Decreasing | tropical | diurnal | 1.686 | 389259.346 | 15.685 |
| amphibians | Phrynobatrachus_ambanguluensis | NA | not_available | not_available | NA | NA | NA | NA | NA |
| amphibians | Phrynobatrachus_annulatus | Phrynobatrachus_annulatus | LC | Decreasing | tropical | diurnal | 1.686 | 175610.191 | 13.909 |
| amphibians | Phrynobatrachus_arcanus | NA | not_available | not_available | NA | NA | NA | NA | NA |
| amphibians | Phrynobatrachus_auritus | Phrynobatrachus_auritus | LC | Stable | tropical | cathemeral | 5.356 | 1793524.134 | 6.203 |
| amphibians | Phrynobatrachus_batesii | Phrynobatrachus_batesii | LC | Unknown | tropical | diurnal | 2.345 | 146554.305 | 8.125 |
| amphibians | Phrynobatrachus_bibita | NA | not_available | not_available | NA | NA | NA | NA | NA |
| amphibians | Phrynobatrachus_breviceps | NA | DD | Unknown | NA | NA | NA | NA | NA |
| amphibians | Phrynobatrachus_bullans | Phrynobatrachus_bullans | LC | Unknown | tropical | cathemeral | 1.499 | 223139.571 | 13.095 |
| amphibians | Phrynobatrachus_calcaratus | Phrynobatrachus_calcaratus | LC | Decreasing | tropical | cathemeral | 1.168 | 1531152.419 | 11.991 |
| amphibians | Phrynobatrachus_chukuchuku | Phrynobatrachus_chukuchuku | CR | Decreasing | tropical | diurnal | 0.497 | 148.569 | 10.335 |
| amphibians | Phrynobatrachus_cornutus | Phrynobatrachus_cornutus | LC | Unknown | tropical | diurnal | 0.567 | 791166.01 | 6.138 |
| amphibians | Phrynobatrachus_danko | Phrynobatrachus_danko | EN | Decreasing | tropical | diurnal | 0.772 | 10.143 | 6.635 |
| amphibians | Phrynobatrachus_discogularis | NA | not_available | not_available | NA | NA | NA | NA | NA |
| amphibians | Phrynobatrachus_dispar | Phrynobatrachus_dispar | LC | Unknown | tropical | cathemeral | 1.123 | 70.002 | 15.903 |
| amphibians | Phrynobatrachus_francisci | Phrynobatrachus_francisci | LC | Unknown | tropical | cathemeral | 0.772 | 1825263.547 | 13.027 |
| amphibians | Phrynobatrachus_fraterculus | Phrynobatrachus_fraterculus | LC | Unknown | tropical | cathemeral | 0.892 | 193128.198 | 13.912 |
| amphibians | Phrynobatrachus_ghanensis | Phrynobatrachus_ghanensis | NT | Decreasing | tropical | diurnal | 0.664 | 58946.691 | 16.303 |
| amphibians | Phrynobatrachus_guineensis | Phrynobatrachus_guineensis | LC | Decreasing | tropical | diurnal | 0.772 | 187114.788 | 14.25 |
| amphibians | Phrynobatrachus_gutturosus | Phrynobatrachus_gutturosus | LC | Unknown | tropical | cathemeral | 1.023 | 733802.554 | 13.579 |
| amphibians | Phrynobatrachus_horsti | NA | NT | Unknown | NA | NA | NA | NA | NA |
| amphibians | Phrynobatrachus_hylaios | Phrynobatrachus_hylaios | LC | Unknown | tropical | diurnal | 0.772 | 626218.441 | 5.949 |
| amphibians | Phrynobatrachus_irangi | Phrynobatrachus_irangi | CR | Decreasing | tropical | diurnal | 11.767 | 3.691 | 20.97 |
| amphibians | Phrynobatrachus_jimzimkusi | Phrynobatrachus_jimzimkusi | CR | Decreasing | tropical | diurnal | 3.163 | 306.768 | 12.02 |
| amphibians | Phrynobatrachus_kakamikro | NA | DD | Unknown | NA | NA | NA | NA | NA |
| amphibians | Phrynobatrachus_keniensis | Phrynobatrachus_keniensis | LC | Unknown | tropical | diurnal | 1.326 | 34775.564 | 16.797 |
| amphibians | Phrynobatrachus_krefftii | Phrynobatrachus_krefftii | EN | Unknown | tropical | diurnal | 11.767 | 2119.876 | 17.718 |
| amphibians | Phrynobatrachus_latifrons | Phrynobatrachus_latifrons | LC | Stable | tropical | diurnal | 2.109 | 2626407.086 | 14.227 |
| amphibians | Phrynobatrachus_leveleve | Phrynobatrachus_leveleve | LC | Unknown | tropical | diurnal | 0.664 | 858.664 | 12.812 |
| amphibians | Phrynobatrachus_liberiensis | Phrynobatrachus_liberiensis | LC | Decreasing | tropical | diurnal | 3.54 | 258033.292 | 14.992 |
| amphibians | Phrynobatrachus_mababiensis | Phrynobatrachus_mababiensis | LC | Unknown | tropical | cathemeral | 0.772 | 4355476.063 | 8.482 |

| | | | | | | | | | |
|---|---|---|---|---|---|---|---|---|---|
| amphibians | Phrynobatrachus_maculiventris | NA | not_available | not_available | NA | NA | NA | NA | NA |
| amphibians | Phrynobatrachus_manengoubensis | Phrynobatrachus_manengoubensis | CR | Decreasing | tropical | diurnal | 0.567 | 9.819 | 10.194 |
| amphibians | Phrynobatrachus_mayokoensis | NA | DD | Unknown | NA | NA | NA | NA | NA |
| amphibians | Phrynobatrachus_mbabo | NA | not_available | not_available | NA | NA | NA | NA | NA |
| amphibians | Phrynobatrachus_natalensis | Phrynobatrachus_natalensis | LC | Stable | tropical | cathemeral | 5.356 | 10523648.243 | 9.581 |
| amphibians | Phrynobatrachus_njiomock | Phrynobatrachus_njiomock | CR | Decreasing | tropical | cathemeral | 3.475 | 10.332 | 11.792 |
| amphibians | Phrynobatrachus_ogoensis | NA | DD | Unknown | NA | NA | NA | NA | NA |
| amphibians | Phrynobatrachus_parvulus | Phrynobatrachus_parvulus | LC | Unknown | tropical | diurnal | 1.168 | 2022241.561 | 7.452 |
| amphibians | Phrynobatrachus_phyllophilus | Phrynobatrachus_phyllophilus | LC | Decreasing | tropical | diurnal | 0.892 | 258033.292 | 14.992 |
| amphibians | Phrynobatrachus_pintoi | Phrynobatrachus_pintoi | EN | Unknown | tropical | diurnal | 0.275 | 385.871 | 15.424 |
| amphibians | Phrynobatrachus_plicatus | Phrynobatrachus_plicatus | LC | Unknown | tropical | diurnal | 6.273 | 285268.63 | 15.958 |
| amphibians | Phrynobatrachus_rainerguentheri | NA | DD | Unknown | NA | NA | NA | NA | NA |
| amphibians | Phrynobatrachus_ruthbeateae | Phrynobatrachus_ruthbeateae | DD | Decreasing | tropical | diurnal | 0.567 | 59.317 | 4.377 |
| amphibians | Phrynobatrachus_sandersoni | Phrynobatrachus_sandersoni | LC | Unknown | tropical | cathemeral | 1.168 | 64991.23 | 8.966 |
| amphibians | Phrynobatrachus_scheffleri | Phrynobatrachus_scheffleri | LC | Unknown | tropical | nocturnal | 0.772 | 319642.544 | 13.056 |
| amphibians | Phrynobatrachus_schioetzi | Phrynobatrachus_schioetzi | EN | Decreasing | tropical | diurnal | 1.686 | 2509.803 | 9.356 |
| amphibians | Phrynobatrachus_steindachneri | Phrynobatrachus_steindachneri | CR | Decreasing | tropical | cathemeral | 3.163 | 12220.634 | 9.06 |
| amphibians | Phrynobatrachus_stewartae | Phrynobatrachus_stewartae | LC | Unknown | tropical | diurnal | 0.892 | 2812.637 | 7.254 |
| amphibians | Phrynobatrachus_tanoeensis | NA | not_available | not_available | NA | NA | NA | NA | NA |
| amphibians | Phrynobatrachus_tokba | Phrynobatrachus_tokba | LC | Unknown | tropical | diurnal | 0.614 | 187123.531 | 14.397 |
| amphibians | Phrynobatrachus_villiersi | Phrynobatrachus_villiersi | LC | Decreasing | tropical | diurnal | 0.275 | 95299.565 | 11.623 |
| amphibians | Phrynobatrachus_werneri | Phrynobatrachus_werneri | LC | Unknown | tropical | diurnal | 1.686 | 9174.431 | 11.368 |
| amphibians | Phrynoidis_asper | Phrynoidis_aspera | LC | Decreasing | tropical | cathemeral | 310.174 | 1614710.697 | 9.155 |
| amphibians | Phrynoidis_juxtasper | Phrynoidis_juxtaspera | LC | Decreasing | tropical | cathemeral | 1245.237 | 259019.277 | 4.654 |
| amphibians | Phrynomantis_bifasciatus | Phrynomantis_bifasciatus | LC | Unknown | tropical | nocturnal | 50.601 | 4714507.507 | 8.665 |
| amphibians | Phrynomantis_microps | Phrynomantis_microps | LC | Unknown | tropical | nocturnal | 23.335 | 3023303.7 | 11.435 |
| amphibians | Phrynomedusa_dryade | NA | LC | Stable | NA | NA | NA | NA | NA |
| amphibians | Phrynomedusa_marginata | Phrynomedusa_marginata | CR | Unknown | tropical | nocturnal | 2.345 | 12.073 | 20.362 |
| amphibians | Phrynopus_bracki | NA | DD | Unknown | NA | NA | NA | NA | NA |
| amphibians | Phrynopus_dagmarae | Phrynopus_dagmarae | EN | Decreasing | tropical | diurnal | 1.554 | 1190.862 | 9.122 |
| amphibians | Phrynopus_horstpauli | Phrynopus_horstpauli | EN | Decreasing | tropical | nocturnal | 5.227 | 257.091 | 10.677 |
| amphibians | Phrynopus_kauneorum | Phrynopus_kauneorum | EN | Decreasing | tropical | diurnal | 16.304 | 306.912 | 8.889 |
| amphibians | Phrynopus_mariellaleo | NA | not_available | not_available | NA | NA | NA | NA | NA |
| amphibians | Phrynopus_oblivius | NA | DD | Unknown | NA | NA | NA | NA | NA |

| | | | | | | | | | |
|---|---|---|---|---|---|---|---|---|---|
| amphibians | Phrynopus_tribulosus | Phrynopus_tribulosus | LC | Stable | tropical | diurnal | 0.685 | 33.242 | 5.34 |
| amphibians | Phyllobates_aurotaenia | Phyllobates_aurotaenia | LC | Stable | tropical | diurnal | 3.163 | 21407.506 | 4.392 |
| amphibians | Phyllobates_bicolor | Phyllobates_bicolor | EN | Decreasing | tropical | diurnal | 6.77 | 2879.297 | 4.285 |
| amphibians | Phyllobates_lugubris | Phyllobates_lugubris | LC | Unknown | tropical | diurnal | 1.023 | 19092.003 | 8.759 |
| amphibians | Phyllobates_terribilis | Phyllobates_terribilis | EN | Decreasing | tropical | diurnal | 9.031 | 1288.437 | 4.413 |
| amphibians | Phyllobates_vittatus | Phyllobates_vittatus | VU | Decreasing | tropical | diurnal | 2.345 | 4258.392 | 11.813 |
| amphibians | Phyllodytes_amadoi | NA | EN | Decreasing | NA | NA | NA | NA | NA |
| amphibians | Phyllodytes_kautskyi | Phyllodytes_kautskyi | LC | Decreasing | tropical | cathemeral | 5.356 | 74366.827 | 9.433 |
| amphibians | Phyllodytes_luteolus | Phyllodytes_luteolus | LC | Stable | tropical | cathemeral | 1.023 | 93868.563 | 10.228 |
| amphibians | Phyllodytes_melanomystax | Phyllodytes_melanomystax | LC | Decreasing | tropical | nocturnal | 1.428 | 41080.589 | 10.466 |
| amphibians | Phyllodytes_tuberculosus | Phyllodytes_tuberculosus | VU | Decreasing | tropical | nocturnal | 1.326 | 7259.85 | 8.139 |
| amphibians | Phyllodytes_wuchereri | Phyllodytes_wuchereri | LC | Stable | tropical | nocturnal | 1.517 | 69591.845 | 8.537 |
| amphibians | Phyllomedusa_bahiana | Phyllomedusa_bahiana | LC | Unknown | tropical | nocturnal | 61.584 | 225806.804 | 10.214 |
| amphibians | Phyllomedusa_bicolor | Phyllomedusa_bicolor | LC | Stable | tropical | nocturnal | 275.697 | 5032516.904 | 1.49 |
| amphibians | Phyllomedusa_burmeisteri | Phyllomedusa_burmeisteri | LC | Stable | tropical | nocturnal | 48.581 | 555418.705 | 11.977 |
| amphibians | Phyllomedusa_camba | Phyllomedusa_camba | LC | Stable | tropical | nocturnal | 59.267 | 1129163.049 | 3.631 |
| amphibians | Phyllomedusa_coelestis | Phyllomedusa_coelestis | LC | Unknown | tropical | nocturnal | 25.565 | 63207.328 | 1.828 |
| amphibians | Phyllomedusa_distincta | Phyllomedusa_distincta | LC | Decreasing | temperate | nocturnal | 34.374 | 120035.964 | 11.897 |
| amphibians | Phyllomedusa_iheringii | Phyllomedusa_iheringii | LC | Unknown | temperate | nocturnal | 41.053 | 157479.794 | 10.357 |
| amphibians | Phyllomedusa_neildi | NA | DD | Unknown | NA | NA | NA | NA | NA |
| amphibians | Phyllomedusa_sauvagii | Phyllomedusa_sauvagii | LC | Decreasing | temperate | nocturnal | 93.213 | 1025952.816 | 5.961 |
| amphibians | Phyllomedusa_tarsius | Phyllomedusa_tarsius | LC | Stable | tropical | nocturnal | 150.528 | 2240122.378 | 1.755 |
| amphibians | Phyllomedusa_tetraploidea | Phyllomedusa_tetraploidea | LC | Stable | temperate | nocturnal | 31.927 | 292985.405 | 13.11 |
| amphibians | Phyllomedusa_vaillantii | Phyllomedusa_vaillantii | LC | Stable | tropical | nocturnal | 59.267 | 5960632.717 | 2.125 |
| amphibians | Physalaemus_aguirrei | Physalaemus_aguirrei | LC | Stable | tropical | nocturnal | 1.726 | 82434.947 | 10.726 |
| amphibians | Physalaemus_albonotatus | Physalaemus_albonotatus | LC | Stable | temperate | cathemeral | 3.194 | 1492280.791 | 6.952 |
| amphibians | Physalaemus_angrensis | Physalaemus_angrensis | CR | Unknown | tropical | nocturnal | 0.567 | 97.486 | 17.771 |
| amphibians | Physalaemus_atim | NA | LC | Stable | NA | NA | NA | NA | NA |
| amphibians | Physalaemus_atlanticus | Physalaemus_atlanticus | LC | Stable | temperate | nocturnal | 1.463 | 2359.019 | 12.523 |
| amphibians | Physalaemus_barrioi | Physalaemus_barrioi | LC | Stable | tropical | nocturnal | 1.889 | 71.251 | 4.913 |
| amphibians | Physalaemus_biligonigerus | Physalaemus_biligonigerus | LC | Stable | temperate | nocturnal | 5.356 | 2026421.104 | 8.132 |
| amphibians | Physalaemus_bokermanni | Physalaemus_bokermanni | LC | Stable | temperate | nocturnal | 0.519 | 1569.173 | 15.895 |
| amphibians | Physalaemus_camacan | Physalaemus_camacan | LC | Stable | tropical | nocturnal | 1.023 | 14856.416 | 7.004 |
| amphibians | Physalaemus_centralis | Physalaemus_centralis | LC | Stable | tropical | nocturnal | 5.356 | 2405327.648 | 8.614 |

| | | | | | | | | | |
|---|---|---|---|---|---|---|---|---|---|
| amphibians | Physalaemus_crombiei | Physalaemus_crombiei | LC | Decreasing | tropical | nocturnal | 1.199 | 122259.193 | 11.535 |
| amphibians | Physalaemus_cuqui | Physalaemus_cuqui | LC | Stable | tropical | nocturnal | 2.111 | 102136.303 | 5.476 |
| amphibians | Physalaemus_cuvieri | Physalaemus_cuvieri | LC | Stable | tropical | nocturnal | 3.981 | 7110840.084 | 6.987 |
| amphibians | Physalaemus_deimaticus | Physalaemus_deimaticus | LC | Stable | tropical | nocturnal | 1.08 | 10314.875 | 4.56 |
| amphibians | Physalaemus_erikae | Physalaemus_erikae | LC | Stable | tropical | cathemeral | 1.517 | 132102.929 | 11.641 |
| amphibians | Physalaemus_erythros | Physalaemus_erythros | NT | Stable | tropical | nocturnal | 1.428 | 338.889 | 6.349 |
| amphibians | Physalaemus_evangelistai | Physalaemus_evangelistai | NT | Stable | tropical | nocturnal | 0.892 | 15052.148 | 7.104 |
| amphibians | Physalaemus_feioi | Physalaemus_feioi | LC | Decreasing | tropical | nocturnal | 2.394 | 43741.011 | 12.424 |
| amphibians | Physalaemus_fischeri | Physalaemus_fischeri | LC | Stable | tropical | nocturnal | 2.109 | 422301.987 | 7.474 |
| amphibians | Physalaemus_gracilis | Physalaemus_gracilis | LC | Stable | temperate | cathemeral | 5.356 | 505595.374 | 11.802 |
| amphibians | Physalaemus_jordanensis | Physalaemus_jordanensis | LC | Stable | tropical | nocturnal | 1.686 | 2301.399 | 11.798 |
| amphibians | Physalaemus_kroyeri | Physalaemus_kroyeri | LC | Stable | tropical | nocturnal | 3.475 | 492507.059 | 9.269 |
| amphibians | Physalaemus_lateristriga | Physalaemus_lateristriga | LC | Stable | temperate | nocturnal | 5.487 | 57930.258 | 12.403 |
| amphibians | Physalaemus_lisei | Physalaemus_lisei | LC | Stable | temperate | cathemeral | 2.872 | 33751.744 | 12.966 |
| amphibians | Physalaemus_maculiventris | Physalaemus_maculiventris | LC | Decreasing | tropical | nocturnal | 1.168 | 85909.926 | 13.757 |
| amphibians | Physalaemus_marmoratus | Physalaemus_marmoratus | LC | Stable | tropical | nocturnal | 4.934 | 1349605.191 | 11.982 |
| amphibians | Physalaemus_maximus | Physalaemus_maximus | NT | Decreasing | tropical | nocturnal | 9.669 | 5080.574 | 12.237 |
| amphibians | Physalaemus_moreirae | Physalaemus_moreirae | LC | Stable | temperate | nocturnal | 1.326 | 3781.972 | 16.886 |
| amphibians | Physalaemus_nanus | Physalaemus_nanus | LC | Stable | temperate | nocturnal | 0.772 | 165830.697 | 11.436 |
| amphibians | Physalaemus_nattereri | Eupemphix_nattereri | LC | Stable | tropical | cathemeral | 16.304 | 1921704.337 | 9.521 |
| amphibians | Physalaemus_obtectus | Physalaemus_obtectus | LC | Decreasing | tropical | nocturnal | 1.686 | 45873.339 | 12.387 |
| amphibians | Physalaemus_olfersii | Physalaemus_olfersii | LC | Decreasing | tropical | nocturnal | 5.802 | 124940.924 | 15.894 |
| amphibians | Physalaemus_riograndensis | Physalaemus_riograndensis | LC | Stable | temperate | cathemeral | 1.168 | 470374.732 | 10.021 |
| amphibians | Physalaemus_signifer | Physalaemus_signifer | LC | Decreasing | tropical | cathemeral | 1.572 | 873.482 | 13.091 |
| amphibians | Physalaemus_soaresi | Physalaemus_soaresi | CR | Stable | tropical | nocturnal | 0.867 | 831.918 | 31.367 |
| amphibians | Physalaemus_spiniger | Physalaemus_spiniger | LC | Stable | temperate | cathemeral | 0.854 | 15997.534 | 6.29 |
| amphibians | Phyzelaphryne_miriamae | Phyzelaphryne_miriamae | LC | Stable | tropical | cathemeral | 0.567 | 863352.893 | 0.961 |
| amphibians | Phyzelaphryne_nimio | NA | LC | Stable | NA | NA | NA | NA | NA |
| amphibians | Pipa_arrabali | Pipa_arrabali | LC | Stable | tropical | cathemeral | 16.873 | 2129877.405 | 1.995 |
| amphibians | Pipa_carvalhoi | Pipa_carvalhoi | LC | Stable | tropical | cathemeral | 29.887 | 238909.687 | 10.572 |
| amphibians | Pipa_pipa | Pipa_pipa | LC | Stable | tropical | cathemeral | 985.118 | 7019956.387 | 2.373 |
| amphibians | Pithecopus_ayeaye | Phyllomedusa_ayeaye | LC | Decreasing | tropical | nocturnal | 8.483 | 87959.347 | 13.353 |
| amphibians | Pithecopus_azureus | Phyllomedusa_azurea | LC | Stable | tropical | nocturnal | 7.511 | 1220516.476 | 6.241 |
| amphibians | Pithecopus_hypochondrialis | Phyllomedusa_hypochondrialis | LC | Stable | tropical | cathemeral | 11.767 | 4002445.599 | 4.305 |

| | | | | | | | | | |
|---|---|---|---|---|---|---|---|---|---|
| amphibians | Pithecopus_megacephalus | Phyllomedusa_megacephala | LC | Stable | tropical | nocturnal | 20.139 | 43611.336 | 7.34 |
| amphibians | Pithecopus_nordestinus | Phyllomedusa_nordestina | LC | Stable | tropical | nocturnal | 7.134 | 570936.129 | 8.757 |
| amphibians | Pithecopus_oreades | Phyllomedusa_oreades | LC | Decreasing | tropical | nocturnal | 10.823 | 108460.6 | 11.152 |
| amphibians | Pithecopus_palliatus | Phyllomedusa_palliata | LC | Stable | tropical | nocturnal | 22.506 | 1120855.584 | 2.604 |
| amphibians | Pithecopus_rohdei | Phyllomedusa_rohdei | LC | Stable | tropical | nocturnal | 8.423 | 294359.015 | 11.672 |
| amphibians | Platymantis_corrugatus | Platymantis_corrugatus | LC | Stable | tropical | nocturnal | 11.18 | 264394.753 | 13.854 |
| amphibians | Platymantis_dorsalis | Platymantis_dorsalis | LC | Decreasing | tropical | nocturnal | 7.844 | 62585.797 | 16.689 |
| amphibians | Platymantis_navjoti | NA | not_available | not_available | NA | NA | NA | NA | NA |
| amphibians | Platymantis_negrosensis | Platymantis_negrosensis | NT | Decreasing | tropical | nocturnal | 11.18 | 2035.698 | 12.127 |
| amphibians | Platymantis_paengi | Platymantis_paengi | EN | Decreasing | tropical | nocturnal | 9.669 | 369.368 | 8.459 |
| amphibians | Platymantis_panayensis | Platymantis_panayensis | EN | Decreasing | tropical | nocturnal | 2.872 | 1778.316 | 6.455 |
| amphibians | Platypelis_alticola | NA | EN | Decreasing | NA | NA | NA | NA | NA |
| amphibians | Platypelis_ando | NA | not_available | not_available | NA | NA | NA | NA | NA |
| amphibians | Platypelis_barbouri | NA | LC | Decreasing | NA | NA | NA | NA | NA |
| amphibians | Platypelis_cowanii | NA | DD | Unknown | NA | NA | NA | NA | NA |
| amphibians | Platypelis_grandis | NA | LC | Unknown | NA | NA | NA | NA | NA |
| amphibians | Platypelis_mavomavo | NA | EN | Decreasing | NA | NA | NA | NA | NA |
| amphibians | Platypelis_milloti | NA | EN | Decreasing | NA | NA | NA | NA | NA |
| amphibians | Platypelis_olgae | NA | EN | Decreasing | NA | NA | NA | NA | NA |
| amphibians | Platypelis_pollicaris | NA | LC | Decreasing | NA | NA | NA | NA | NA |
| amphibians | Platypelis_tetra | NA | EN | Decreasing | NA | NA | NA | NA | NA |
| amphibians | Platypelis_tsaratananaensis | NA | EN | Decreasing | NA | NA | NA | NA | NA |
| amphibians | Platypelis_tuberifera | NA | LC | Unknown | NA | NA | NA | NA | NA |
| amphibians | Platyplectrum_ornatum | Platyplectrum_ornatum | LC | Stable | tropical | nocturnal | 7.844 | 2936283.594 | 4.448 |
| amphibians | Platyplectrum_spenceri | Platyplectrum_spenceri | LC | Stable | temperate | nocturnal | 11.036 | 3182540.986 | 2.636 |
| amphibians | Plectrohyla_calvata | NA | CR | Unknown | NA | NA | NA | NA | NA |
| amphibians | Plectrohyla_chrysopleura | Plectrohyla_chrysopleura | CR | Decreasing | tropical | nocturnal | 27.131 | 561.008 | 2.041 |
| amphibians | Plectrohyla_dasypus | Plectrohyla_dasypus | CR | Decreasing | tropical | nocturnal | 9.866 | 44.514 | 6.246 |
| amphibians | Plectrohyla_exquisita | Plectrohyla_exquisita | CR | Decreasing | tropical | nocturnal | 52.05 | 75.396 | 6.405 |
| amphibians | Plectrohyla_glandulosa | Plectrohyla_glandulosa | CR | Decreasing | tropical | diurnal | 11.036 | 778.441 | 14.813 |
| amphibians | Plectrohyla_guatemalensis | Plectrohyla_guatemalensis | NT | Decreasing | tropical | cathemeral | 43.037 | 36439.68 | 13.219 |
| amphibians | Plectrohyla_hartwegi | Plectrohyla_hartwegi | EN | Decreasing | tropical | nocturnal | 44.708 | 8321.273 | 9.584 |
| amphibians | Plectrohyla_ixil | Plectrohyla_ixil | VU | Decreasing | tropical | cathemeral | 9.031 | 2924.833 | 11.433 |
| amphibians | Plectrohyla_matudai | Plectrohyla_matudai | LC | Stable | tropical | cathemeral | 10.337 | 7723.689 | 12.149 |

| | | | | | | | | | |
|---|---|---|---|---|---|---|---|---|---|
| amphibians | Plectrohyla_psiloderma | Plectrohyla_psiloderma | EN | Decreasing | tropical | nocturnal | 10.337 | 37.971 | 5.564 |
| amphibians | Plectrohyla_quecchi | Plectrohyla_quecchi | EN | Decreasing | tropical | nocturnal | 9.031 | 3677.768 | 14.804 |
| amphibians | Plectrohyla_sagorum | Plectrohyla_sagorum | VU | Unknown | tropical | cathemeral | 12.531 | 3741.32 | 12.217 |
| amphibians | Plethodon_caddoensis | Plethodon_caddoensis | NT | Stable | temperate | diurnal | 2.293 | 961.745 | 7.194 |
| amphibians | Plethodon_cinereus | Plethodon_cinereus | LC | Stable | temperate | nocturnal | 1.357 | 1830423.812 | 15.115 |
| amphibians | Plethodon_cylindraceus | Plethodon_cylindraceus | LC | Stable | temperate | diurnal | 7.586 | 198488.116 | 19.262 |
| amphibians | Plethodon_dunni | Plethodon_dunni | LC | Stable | temperate | nocturnal | 6.996 | 73586.869 | 12.162 |
| amphibians | Plethodon_elongatus | Plethodon_elongatus | LC | Decreasing | temperate | nocturnal | 6.996 | 24407.802 | 7.717 |
| amphibians | Plethodon_glutinosus | Plethodon_glutinosus | LC | Stable | temperate | nocturnal | 4.615 | 1260100.04 | 18.037 |
| amphibians | Plethodon_hubrichti | Plethodon_hubrichti | VU | Stable | temperate | nocturnal | 1.499 | 49.258 | 7.955 |
| amphibians | Plethodon_idahoensis | Plethodon_idahoensis | LC | Unknown | temperate | nocturnal | 4.621 | 82818.054 | 7.573 |
| amphibians | Plethodon_jordani | Plethodon_jordani | NT | Stable | temperate | nocturnal | 2.435 | 3524.737 | 8.361 |
| amphibians | Plethodon_kentucki | Plethodon_kentucki | LC | Stable | temperate | nocturnal | 2.941 | 29180.693 | 15.181 |
| amphibians | Plethodon_metcalfi | Plethodon_metcalfi | LC | Stable | temperate | cathemeral | 3.683 | 7930.864 | 17.749 |
| amphibians | Plethodon_neomexicanus | Plethodon_neomexicanus | EN | Decreasing | temperate | nocturnal | 4.83 | 626.407 | 9.02 |
| amphibians | Plethodon_nettingi | Plethodon_nettingi | NT | Decreasing | temperate | nocturnal | 1.703 | 3468.122 | 9.466 |
| amphibians | Plethodon_ocmulgee | NA | LC | Unknown | NA | NA | NA | NA | NA |
| amphibians | Plethodon_ouachitae | Plethodon_ouachitae | NT | Stable | temperate | diurnal | 3.241 | 3272.452 | 8.754 |
| amphibians | Plethodon_punctatus | Plethodon_punctatus | NT | Stable | temperate | nocturnal | 4.725 | 6730.986 | 12.08 |
| amphibians | Plethodon_richmondi | Plethodon_richmondi | LC | Stable | temperate | nocturnal | 2.053 | 100792.819 | 15.94 |
| amphibians | Plethodon_savannah | NA | DD | Unknown | NA | NA | NA | NA | NA |
| amphibians | Plethodon_serratus | Plethodon_serratus | LC | Stable | temperate | cathemeral | 0.947 | 192070.27 | 15.359 |
| amphibians | Plethodon_stormi | Plethodon_stormi | EN | Decreasing | temperate | nocturnal | 4.011 | 2274.872 | 8.313 |
| amphibians | Plethodon_teyahalee | Plethodon_teyahalee | LC | Stable | temperate | nocturnal | 6.417 | 13506.964 | 13.503 |
| amphibians | Plethodon_vehiculum | Plethodon_vehiculum | LC | Stable | temperate | nocturnal | 4.03 | 166117.488 | 10.871 |
| amphibians | Plethodon_wehrlei | Plethodon_wehrlei | LC | Stable | temperate | nocturnal | 2.857 | 96744.457 | 14.735 |
| amphibians | Plethodon_welleri | Plethodon_welleri | VU | Decreasing | temperate | nocturnal | 1.45 | 3540.242 | 12.715 |
| amphibians | Plethodon_yonahlossee | Plethodon_yonahlossee | LC | Stable | temperate | nocturnal | 5.968 | 12313.334 | 14.452 |
| amphibians | Plethodontohyla_alluaudi | Rhombophryne_alluaudi | LC | Unknown | tropical | diurnal | 19.923 | 107063.272 | 8.521 |
| amphibians | Plethodontohyla_bipunctata | Plethodontohyla_bipunctata | LC | Decreasing | tropical | diurnal | 2.599 | 80025.999 | 8.568 |
| amphibians | Plethodontohyla_guentheri | Plethodontohyla_guentheri | EN | Decreasing | tropical | cathemeral | 3.475 | 1336.428 | 5.582 |
| amphibians | Plethodontohyla_inguinalis | Plethodontohyla_inguinalis | LC | Decreasing | tropical | nocturnal | 104.268 | 71208.12 | 7.246 |
| amphibians | Plethodontohyla_mihanika | Plethodontohyla_mihanika | LC | Unknown | tropical | nocturnal | 3.285 | 34747.557 | 10.128 |
| amphibians | Plethodontohyla_notosticta | Plethodontohyla_notosticta | LC | Unknown | tropical | nocturnal | 6.273 | 102707.457 | 7.962 |

| | | | | | | | | | |
|---|---|---|---|---|---|---|---|---|---|
| amphibians | Plethodontohyla_ocellata | Plethodontohyla_ocellata | LC | Decreasing | tropical | nocturnal | 25.822 | 58001.498 | 7.602 |
| amphibians | Plethodontohyla_tuberata | Plethodontohyla_tuberata | NT | Decreasing | tropical | nocturnal | 7.844 | 21100.604 | 11.643 |
| amphibians | Pleurodeles_nebulosus | Pleurodeles_nebulosus | LC | Stable | temperate | nocturnal | 95.666 | 59414.744 | 22.36 |
| amphibians | Pleurodeles_poireti | Pleurodeles_poireti | EN | Decreasing | temperate | nocturnal | 3.323 | 1851.635 | 21.515 |
| amphibians | Pleurodeles_waltl | Pleurodeles_waltl | LC | Decreasing | temperate | nocturnal | 28.813 | 377028.154 | 17.654 |
| amphibians | Pleurodema_brachyops | Pleurodema_brachyops | LC | Stable | tropical | nocturnal | 11.767 | 890578.523 | 7.736 |
| amphibians | Pleurodema_cordobae | Pleurodema_cordobae | LC | Stable | temperate | nocturnal | 6.719 | 2851.593 | 6.593 |
| amphibians | Pleurodema_diplolister | Pleurodema_diplolister | LC | Stable | tropical | nocturnal | 3.317 | 969995.983 | 8.822 |
| amphibians | Pleurodema_kriegi | Pleurodema_kriegi | NT | Stable | temperate | cathemeral | 4.088 | 2778.172 | 5.734 |
| amphibians | Pleurodema_nebulosum | Pleurodema_nebulosum | LC | Stable | temperate | diurnal | 7.844 | 575751.622 | 5.305 |
| amphibians | Pleurodema_thaul | Pleurodema_thaul | LC | Stable | temperate | nocturnal | 17.851 | 274397.658 | 8.81 |
| amphibians | Polypedates_bengalensis | NA | DD | Unknown | NA | NA | NA | NA | NA |
| amphibians | Polypedates_leucomystax | Polypedates_leucomystax | LC | Stable | tropical | nocturnal | 61.584 | 3798236.019 | 11.837 |
| amphibians | Polypedates_macrotis | Polypedates_macrotis | LC | Stable | tropical | nocturnal | 74.113 | 1053469.955 | 7.61 |
| amphibians | Polypedates_maculatus | Polypedates_maculatus | LC | Unknown | tropical | nocturnal | 71.478 | 2581270.749 | 18.869 |
| amphibians | Polypedates_megacephalus | Polypedates_megacephalus | LC | Stable | temperate | nocturnal | 37.611 | 1324419.434 | 14.388 |
| amphibians | Polypedates_otilophus | Polypedates_otilophus | LC | Decreasing | tropical | nocturnal | 104.268 | 317169.428 | 4.398 |
| amphibians | Polypedates_pseudocruciger | Polypedates_pseudocruciger | LC | Decreasing | tropical | nocturnal | 52.68 | 25647.048 | 16.664 |
| amphibians | Polypedates_taeniatus | Polypedates_taeniatus | LC | Unknown | temperate | nocturnal | 7.844 | 122228.882 | 16.677 |
| amphibians | Potomotyphlus_kaupii | Potamotyphlus_kaupii | LC | Unknown | tropical | nocturnal | 127.898 | 4849495.113 | 2.232 |
| amphibians | Poyntonia_paludicola | Poyntonia_paludicola | NT | Stable | temperate | diurnal | 2.109 | 695.276 | 10.366 |
| amphibians | Poyntonophrynus_damaranus | NA | DD | Unknown | NA | NA | NA | NA | NA |
| amphibians | Pristimantis_acerus | Pristimantis_acerus | EN | Decreasing | tropical | cathemeral | 7.901 | 95.797 | 7.003 |
| amphibians | Pristimantis_achatinus | Pristimantis_achatinus | LC | Stable | tropical | nocturnal | 8.483 | 187731.778 | 8.937 |
| amphibians | Pristimantis_actites | Pristimantis_actites | VU | Decreasing | tropical | cathemeral | 24.806 | 6897.918 | 10.707 |
| amphibians | Pristimantis_acuminatus | Pristimantis_acuminatus | LC | Stable | tropical | nocturnal | 3.163 | 96581.522 | 1.546 |
| amphibians | Pristimantis_adiastolus | Pristimantis_adiastolus | LC | Stable | tropical | nocturnal | 1.686 | 624.952 | 3.81 |
| amphibians | Pristimantis_albujai | NA | EN | Decreasing | NA | NA | NA | NA | NA |
| amphibians | Pristimantis_altae | Pristimantis_altae | LC | Decreasing | tropical | cathemeral | 1.499 | 8323.225 | 7.987 |
| amphibians | Pristimantis_altamazonicus | Pristimantis_altamazonicus | LC | Stable | tropical | cathemeral | 3.133 | 1034298.597 | 1.7 |
| amphibians | Pristimantis_altamnis | Pristimantis_altamnis | LC | Stable | tropical | nocturnal | 1.61 | 75169.634 | 4.779 |
| amphibians | Pristimantis_andinogigas | NA | CR | Decreasing | NA | NA | NA | NA | NA |
| amphibians | Pristimantis_andinognomus | Pristimantis_andinognomus | LC | Stable | tropical | nocturnal | 0.396 | 1554.261 | 6.05 |
| amphibians | Pristimantis_anemerus | NA | DD | Unknown | NA | NA | NA | NA | NA |

| | | | | | | | | | |
|---|---|---|---|---|---|---|---|---|---|
| amphibians | Pristimantis_aniptopalmatus | Pristimantis_aniptopalmatus | LC | Stable | tropical | nocturnal | 0.917 | 86.142 | 4.847 |
| amphibians | Pristimantis_anolirex | Pristimantis_anolirex | VU | Unknown | tropical | nocturnal | 5.356 | 3876.301 | 7.942 |
| amphibians | Pristimantis_apiculatus | Pristimantis_apiculatus | EN | Decreasing | tropical | nocturnal | 1.377 | 1887.429 | 7.311 |
| amphibians | Pristimantis_appendiculatus | Pristimantis_appendiculatus | VU | Decreasing | tropical | nocturnal | 3.475 | 2551.938 | 7.62 |
| amphibians | Pristimantis_aquilonaris | Pristimantis_aquilonaris | LC | Stable | tropical | cathemeral | 0.892 | 1371.476 | 9.726 |
| amphibians | Pristimantis_ardalonychus | Pristimantis_ardalonychus | EN | Unknown | tropical | cathemeral | 1.572 | 2821.615 | 10.86 |
| amphibians | Pristimantis_ashaninka | NA | LC | Unknown | NA | NA | NA | NA | NA |
| amphibians | Pristimantis_atillo | NA | CR | Decreasing | NA | NA | NA | NA | NA |
| amphibians | Pristimantis_atrabracus | NA | DD | Unknown | NA | NA | NA | NA | NA |
| amphibians | Pristimantis_atratus | Pristimantis_atratus | VU | Decreasing | tropical | nocturnal | 1.932 | 5361.52 | 5.262 |
| amphibians | Pristimantis_aureolineatus | Pristimantis_aureolineatus | LC | Stable | tropical | cathemeral | 2.225 | 205337.424 | 1.984 |
| amphibians | Pristimantis_auricarens | Pristimantis_auricarens | NT | Stable | tropical | nocturnal | 0.969 | 92.453 | 1.214 |
| amphibians | Pristimantis_avius | NA | DD | Unknown | NA | NA | NA | NA | NA |
| amphibians | Pristimantis_balionotus | Pristimantis_balionotus | EN | Decreasing | tropical | diurnal | 1.911 | 76.059 | 9.667 |
| amphibians | Pristimantis_bambu | Pristimantis_bambu | EN | Decreasing | tropical | nocturnal | 1.394 | 119.53 | 9.15 |
| amphibians | Pristimantis_bellator | Pristimantis_bellator | LC | Stable | tropical | cathemeral | 1.065 | 1741.227 | 8.708 |
| amphibians | Pristimantis_bicantus | Pristimantis_bicantus | VU | Decreasing | tropical | nocturnal | 0.738 | 6792.429 | 4.496 |
| amphibians | Pristimantis_bicumulus | Pristimantis_bicumulus | VU | Decreasing | tropical | nocturnal | 4.16 | 1386.203 | 11.927 |
| amphibians | Pristimantis_bipunctatus | Pristimantis_bipunctatus | LC | Stable | tropical | nocturnal | 6.034 | 572.443 | 8.134 |
| amphibians | Pristimantis_bogotensis | NA | LC | Stable | NA | NA | NA | NA | NA |
| amphibians | Pristimantis_boucephalus | NA | DD | Unknown | NA | NA | NA | NA | NA |
| amphibians | Pristimantis_brevicrus | NA | LC | Stable | NA | NA | NA | NA | NA |
| amphibians | Pristimantis_bromeliaceus | Pristimantis_bromeliaceus | LC | Unknown | tropical | nocturnal | 2.816 | 15238.952 | 5.641 |
| amphibians | Pristimantis_buccinator | Pristimantis_buccinator | LC | Stable | tropical | cathemeral | 5.356 | 146985.81 | 1.014 |
| amphibians | Pristimantis_buenaventura | NA | EN | Decreasing | NA | NA | NA | NA | NA |
| amphibians | Pristimantis_caeruleonotus | NA | DD | Unknown | NA | NA | NA | NA | NA |
| amphibians | Pristimantis_cajamarcensis | Pristimantis_cajamarcensis | LC | Stable | tropical | cathemeral | 3.103 | 5366.084 | 8.984 |
| amphibians | Pristimantis_cajanuma | NA | VU | Stable | NA | NA | NA | NA | NA |
| amphibians | Pristimantis_calcarulatus | Pristimantis_calcarulatus | VU | Stable | tropical | nocturnal | 1.868 | 3865.012 | 8.959 |
| amphibians | Pristimantis_calima | NA | not_available | not_available | NA | NA | NA | NA | NA |
| amphibians | Pristimantis_campinarana | NA | not_available | not_available | NA | NA | NA | NA | NA |
| amphibians | Pristimantis_caniari | NA | CR | Decreasing | NA | NA | NA | NA | NA |
| amphibians | Pristimantis_cantitans | Pristimantis_cantitans | NT | Unknown | tropical | cathemeral | 7.958 | 66.347 | 0.413 |
| amphibians | Pristimantis_caprifer | Pristimantis_caprifer | CR | Decreasing | tropical | nocturnal | 7.187 | 14636.845 | 5.631 |

| | | | | | | | | | |
|---|---|---|---|---|---|---|---|---|---|
| amphibians | Pristimantis_carlosceroni | Pristimantis_carlosceroni | EN | Decreasing | tropical | nocturnal | 1.343 | 14.079 | 10.64 |
| amphibians | Pristimantis_carvalhoi | Pristimantis_carvalhoi | LC | Stable | tropical | nocturnal | 1.168 | 1555839.27 | 0.986 |
| amphibians | Pristimantis_cedros | NA | EN | Decreasing | NA | NA | NA | NA | NA |
| amphibians | Pristimantis_celator | Pristimantis_celator | VU | Decreasing | tropical | cathemeral | 1.094 | 3963.655 | 6.939 |
| amphibians | Pristimantis_cerasinus | Pristimantis_cerasinus | LC | Stable | tropical | cathemeral | 3.475 | 54745.891 | 8.781 |
| amphibians | Pristimantis_ceuthospilus | Pristimantis_ceuthospilus | VU | Decreasing | tropical | cathemeral | 1.446 | 2475.539 | 7.342 |
| amphibians | Pristimantis_chalceus | Pristimantis_chalceus | LC | Decreasing | tropical | nocturnal | 2.394 | 75471.816 | 6.165 |
| amphibians | Pristimantis_chiastonotus | Pristimantis_chiastonotus | LC | Stable | tropical | diurnal | 13.329 | 518535.985 | 0.811 |
| amphibians | Pristimantis_chloronotus | Pristimantis_chloronotus | LC | Stable | tropical | cathemeral | 4.893 | 11265.687 | 4.602 |
| amphibians | Pristimantis_chomskyi | NA | EN | Decreasing | NA | NA | NA | NA | NA |
| amphibians | Pristimantis_citriogaster | Pristimantis_citriogaster | EN | Decreasing | tropical | nocturnal | 11.767 | 9900.997 | 5.947 |
| amphibians | Pristimantis_colodactylus | Pristimantis_colodactylus | LC | Unknown | tropical | diurnal | 1.294 | 5159.434 | 4.303 |
| amphibians | Pristimantis_colomai | Pristimantis_colomai | VU | Decreasing | tropical | nocturnal | 0.389 | 9.913 | 7.092 |
| amphibians | Pristimantis_colonensis | Pristimantis_colonensis | VU | Unknown | tropical | nocturnal | 2.132 | 5219.139 | 6.449 |
| amphibians | Pristimantis_condor | Pristimantis_condor | LC | Stable | tropical | cathemeral | 23.576 | 5328.542 | 3.25 |
| amphibians | Pristimantis_conspicillatus | Pristimantis_conspicillatus | LC | Stable | tropical | cathemeral | 10.337 | 2709642.985 | 1.455 |
| amphibians | Pristimantis_coronatus | NA | DD | Unknown | NA | NA | NA | NA | NA |
| amphibians | Pristimantis_cosnipatae | Pristimantis_cosnipatae | CR | Decreasing | tropical | nocturnal | 2.733 | 28.619 | 7.414 |
| amphibians | Pristimantis_cremnobates | Pristimantis_cremnobates | EN | Decreasing | tropical | nocturnal | 12.299 | 1483.901 | 4.552 |
| amphibians | Pristimantis_crenunguis | Pristimantis_crenunguis | VU | Decreasing | tropical | nocturnal | 35.967 | 3552.897 | 7.122 |
| amphibians | Pristimantis_croceoinguinis | Pristimantis_croceoinguinis | LC | Unknown | tropical | cathemeral | 0.892 | 149848.42 | 3.949 |
| amphibians | Pristimantis_crucifer | Pristimantis_crucifer | NT | Decreasing | tropical | nocturnal | 3.475 | 14334.889 | 7.024 |
| amphibians | Pristimantis_cruciocularis | Pristimantis_cruciocularis | LC | Stable | tropical | nocturnal | 0.943 | 744.573 | 6.885 |
| amphibians | Pristimantis_cruentus | Pristimantis_cruentus | LC | Stable | tropical | nocturnal | 6.273 | 74499.611 | 8.73 |
| amphibians | Pristimantis_cryptomelas | Pristimantis_cryptomelas | NT | Stable | tropical | nocturnal | 5.142 | 11265.587 | 6.747 |
| amphibians | Pristimantis_cuneirostris | NA | DD | Unknown | NA | NA | NA | NA | NA |
| amphibians | Pristimantis_danae | Pristimantis_danae | LC | Stable | tropical | nocturnal | 8.305 | 50095.511 | 3.42 |
| amphibians | Pristimantis_degener | Pristimantis_degener | EN | Decreasing | tropical | nocturnal | 2.573 | 749.637 | 4.398 |
| amphibians | Pristimantis_dendrobatoides | Pristimantis_dendrobatoides | LC | Unknown | tropical | nocturnal | 2.345 | 186.349 | 1.166 |
| amphibians | Pristimantis_devillei | Pristimantis_devillei | EN | Decreasing | tropical | cathemeral | 7.676 | 808.172 | 5.125 |
| amphibians | Pristimantis_diadematus | Pristimantis_diadematus | LC | Stable | tropical | cathemeral | 7.566 | 548888.366 | 2.048 |
| amphibians | Pristimantis_dissimulatus | Pristimantis_dissimulatus | EN | Decreasing | tropical | nocturnal | 2.788 | 95.548 | 5.854 |
| amphibians | Pristimantis_dorado | NA | EN | Decreasing | NA | NA | NA | NA | NA |
| amphibians | Pristimantis_duellmani | Pristimantis_duellmani | CR | Decreasing | tropical | nocturnal | 8.305 | 5750.689 | 6.576 |

| | | | | | | | | | |
|---|---|---|---|---|---|---|---|---|---|
| amphibians | Pristimantis_enigmaticus | NA | LC | Stable | NA | NA | NA | NA | NA |
| amphibians | Pristimantis_eriphus | Pristimantis_eriphus | VU | Decreasing | tropical | nocturnal | 1.889 | 8988.147 | 3.827 |
| amphibians | Pristimantis_erythropleura | NA | LC | Stable | NA | NA | NA | NA | NA |
| amphibians | Pristimantis_erythros | NA | CR | Decreasing | NA | NA | NA | NA | NA |
| amphibians | Pristimantis_esmeraldas | Pristimantis_esmeraldas | LC | Decreasing | tropical | nocturnal | 0.795 | 29959.848 | 6.668 |
| amphibians | Pristimantis_exoristus | Pristimantis_exoristus | VU | Decreasing | tropical | nocturnal | 0.956 | 4460.567 | 6.421 |
| amphibians | Pristimantis_fenestratus | Pristimantis_fenestratus | LC | Stable | tropical | cathemeral | 17.065 | 2603658.33 | 2.27 |
| amphibians | Pristimantis_ferwerdai | NA | not_available | not_available | NA | NA | NA | NA | NA |
| amphibians | Pristimantis_festae | Pristimantis_festae | EN | Decreasing | tropical | cathemeral | 1.245 | 197.295 | 8.801 |
| amphibians | Pristimantis_gaigei | Pristimantis_gaigei | LC | Stable | tropical | nocturnal | 7.294 | 193182.537 | 9.532 |
| amphibians | Pristimantis_galdi | Pristimantis_galdi | LC | Decreasing | tropical | nocturnal | 3.163 | 20753.33 | 6.247 |
| amphibians | Pristimantis_ganonotus | Pristimantis_ganonotus | VU | Decreasing | tropical | diurnal | 0.233 | 4053.725 | 3.935 |
| amphibians | Pristimantis_gentryi | Pristimantis_gentryi | EN | Decreasing | tropical | nocturnal | 11.036 | 785.475 | 17.218 |
| amphibians | Pristimantis_gladiator | Pristimantis_gladiator | VU | Decreasing | tropical | cathemeral | 0.264 | 5845.548 | 4.462 |
| amphibians | Pristimantis_glandulosus | Pristimantis_glandulosus | EN | Decreasing | tropical | nocturnal | 5.941 | 561.886 | 7.208 |
| amphibians | Pristimantis_gloria | NA | EN | Decreasing | NA | NA | NA | NA | NA |
| amphibians | Pristimantis_gryllus | Pristimantis_gryllus | NT | Unknown | tropical | cathemeral | 1.377 | 8146.794 | 15.445 |
| amphibians | Pristimantis_guaiquinimensis | Pristimantis_guaiquinimensis | NT | Unknown | tropical | cathemeral | 2.986 | 259.355 | 0.415 |
| amphibians | Pristimantis_gualacenio | NA | EN | Decreasing | NA | NA | NA | NA | NA |
| amphibians | Pristimantis_gutturalis | Pristimantis_gutturalis | LC | Stable | tropical | diurnal | 5.802 | 146206.553 | 0.671 |
| amphibians | Pristimantis_hampatusami | NA | EN | Decreasing | NA | NA | NA | NA | NA |
| amphibians | Pristimantis_hectus | Pristimantis_hectus | VU | Decreasing | tropical | diurnal | 0.892 | 3892.622 | 5.527 |
| amphibians | Pristimantis_hoogmoedi | NA | CR | Decreasing | NA | NA | NA | NA | NA |
| amphibians | Pristimantis_humboldti | NA | DD | Unknown | NA | NA | NA | NA | NA |
| amphibians | Pristimantis_ignicolor | Pristimantis_ignicolor | EN | Decreasing | tropical | nocturnal | 1.394 | 939.616 | 4.649 |
| amphibians | Pristimantis_iiap | NA | DD | Unknown | NA | NA | NA | NA | NA |
| amphibians | Pristimantis_illotus | Pristimantis_illotus | NT | Decreasing | tropical | nocturnal | 7.621 | 2895.799 | 7.838 |
| amphibians | Pristimantis_imitatrix | Pristimantis_imitatrix | LC | Unknown | tropical | cathemeral | 0.585 | 54873.07 | 2.916 |
| amphibians | Pristimantis_incomptus | Pristimantis_incomptus | LC | Unknown | tropical | nocturnal | 1.31 | 10587.9 | 3.635 |
| amphibians | Pristimantis_infraguttatus | NA | DD | Unknown | NA | NA | NA | NA | NA |
| amphibians | Pristimantis_inusitatus | Pristimantis_inusitatus | EN | Decreasing | tropical | nocturnal | 1.261 | 855.6 | 6.552 |
| amphibians | Pristimantis_jimenezi | NA | CR | Decreasing | NA | NA | NA | NA | NA |
| amphibians | Pristimantis_jorgevelosai | Pristimantis_jorgevelosai | EN | Decreasing | tropical | nocturnal | 7.958 | 489.014 | 19.635 |
| amphibians | Pristimantis_jubatus | NA | NT | Stable | NA | NA | NA | NA | NA |

| | | | | | | | | | |
|---|---|---|---|---|---|---|---|---|---|
| amphibians | Pristimantis_katoptroides | Pristimantis_katoptroides | LC | Stable | tropical | nocturnal | 1.023 | 13493.93 | 3.563 |
| amphibians | Pristimantis_kichwarum | Pristimantis_kichwarum | LC | Stable | tropical | nocturnal | 1.499 | 51514.834 | 4.431 |
| amphibians | Pristimantis_labiosus | Pristimantis_labiosus | LC | Decreasing | tropical | nocturnal | 12.767 | 38057.775 | 6.785 |
| amphibians | Pristimantis_lacrimosus | Pristimantis_lacrimosus | LC | Stable | tropical | cathemeral | 3.163 | 1144976.571 | 1.502 |
| amphibians | Pristimantis_lanthanites | Pristimantis_lanthanites | LC | Stable | tropical | cathemeral | 8.423 | 692688.535 | 1.535 |
| amphibians | Pristimantis_laticlavius | Pristimantis_laticlavius | VU | Decreasing | tropical | nocturnal | 6.719 | 5431.396 | 5.974 |
| amphibians | Pristimantis_latidiscus | Pristimantis_latidiscus | LC | Decreasing | tropical | nocturnal | 13.658 | 90430.555 | 6.502 |
| amphibians | Pristimantis_leoni | Pristimantis_leoni | NT | Decreasing | tropical | cathemeral | 1.168 | 7439.417 | 11.536 |
| amphibians | Pristimantis_leopardus | NA | VU | Stable | NA | NA | NA | NA | NA |
| amphibians | Pristimantis_leucopus | Pristimantis_leucopus | EN | Decreasing | tropical | nocturnal | 7.294 | 136.647 | 9.236 |
| amphibians | Pristimantis_leucorrhinus | NA | DD | Unknown | NA | NA | NA | NA | NA |
| amphibians | Pristimantis_librarius | Pristimantis_librarius | LC | Decreasing | tropical | nocturnal | 2.272 | 45357.676 | 4.825 |
| amphibians | Pristimantis_limoncochensis | NA | NT | Unknown | NA | NA | NA | NA | NA |
| amphibians | Pristimantis_lindae | Pristimantis_lindae | LC | Stable | tropical | nocturnal | 5.016 | 64.534 | 7.39 |
| amphibians | Pristimantis_lirellus | Pristimantis_lirellus | LC | Unknown | tropical | nocturnal | 1.094 | 3075.248 | 8.357 |
| amphibians | Pristimantis_lividus | Pristimantis_lividus | EN | Decreasing | tropical | nocturnal | 3.475 | 119.56 | 6.286 |
| amphibians | Pristimantis_llanganati | NA | LC | Stable | NA | NA | NA | NA | NA |
| amphibians | Pristimantis_loustes | Pristimantis_loustes | EN | Decreasing | tropical | nocturnal | 8.846 | 4.963 | 12.911 |
| amphibians | Pristimantis_lucasi | Pristimantis_lucasi | LC | Stable | tropical | nocturnal | 0.93 | 36.817 | 4.164 |
| amphibians | Pristimantis_luscombei | Pristimantis_luscombei | LC | Stable | tropical | nocturnal | 2.109 | 524184.884 | 1.752 |
| amphibians | Pristimantis_luteolateralis | Pristimantis_luteolateralis | LC | Stable | tropical | nocturnal | 1.997 | 3471.705 | 7.384 |
| amphibians | Pristimantis_lutzae | NA | VU | Decreasing | NA | NA | NA | NA | NA |
| amphibians | Pristimantis_lymani | Pristimantis_lymani | LC | Stable | tropical | diurnal | 37.611 | 50808.874 | 8.664 |
| amphibians | Pristimantis_lythrodes | Pristimantis_lythrodes | LC | Stable | tropical | nocturnal | 2.599 | 6480.23 | 1.051 |
| amphibians | Pristimantis_macrummendozai | NA | not_available | not_available | NA | NA | NA | NA | NA |
| amphibians | Pristimantis_malkini | Pristimantis_malkini | LC | Stable | tropical | nocturnal | 9.604 | 818760.772 | 1.489 |
| amphibians | Pristimantis_marcoreyesi | NA | EN | Decreasing | NA | NA | NA | NA | NA |
| amphibians | Pristimantis_marmoratus | Pristimantis_marmoratus | LC | Stable | tropical | cathemeral | 0.83 | 891530.757 | 1.179 |
| amphibians | Pristimantis_martiae | Pristimantis_martiae | LC | Stable | tropical | cathemeral | 1.168 | 968361.267 | 1.367 |
| amphibians | Pristimantis_matidiktyo | Pristimantis_matidiktyo | LC | Unknown | tropical | nocturnal | 1.037 | 63637.052 | 3.688 |
| amphibians | Pristimantis_matildae | NA | not_available | not_available | NA | NA | NA | NA | NA |
| amphibians | Pristimantis_mazar | Pristimantis_mazar | EN | Decreasing | tropical | nocturnal | 0.983 | 86.384 | 12.095 |
| amphibians | Pristimantis_memorans | NA | DD | Unknown | NA | NA | NA | NA | NA |
| amphibians | Pristimantis_mendax | Pristimantis_mendax | LC | Stable | tropical | cathemeral | 1.706 | 176882.868 | 2.784 |

| | | | | | | | | | |
|---|---|---|---|---|---|---|---|---|---|
| amphibians | Pristimantis_miktos | NA | LC | Unknown | NA | NA | NA | NA | NA |
| amphibians | Pristimantis_minimus | Pristimantis_minimus | EN | Decreasing | tropical | nocturnal | 0.472 | 296.767 | 4.92 |
| amphibians | Pristimantis_minutulus | NA | DD | Unknown | NA | NA | NA | NA | NA |
| amphibians | Pristimantis_modipeplus | Pristimantis_modipeplus | VU | Decreasing | tropical | nocturnal | 3.379 | 1070.409 | 19.22 |
| amphibians | Pristimantis_moro | Pristimantis_moro | LC | Unknown | tropical | nocturnal | 1.168 | 1571.104 | 9.941 |
| amphibians | Pristimantis_multicolor | NA | VU | Stable | NA | NA | NA | NA | NA |
| amphibians | Pristimantis_munozi | NA | DD | Unknown | NA | NA | NA | NA | NA |
| amphibians | Pristimantis_muranunka | NA | EN | Decreasing | NA | NA | NA | NA | NA |
| amphibians | Pristimantis_muricatus | Pristimantis_muricatus | VU | Decreasing | tropical | nocturnal | 5.665 | 14023.927 | 12.357 |
| amphibians | Pristimantis_muscosus | Pristimantis_muscosus | NT | Decreasing | tropical | cathemeral | 8.483 | 4802.854 | 3.013 |
| amphibians | Pristimantis_museosus | Pristimantis_museosus | VU | Decreasing | tropical | nocturnal | 5.356 | 3836.675 | 4.998 |
| amphibians | Pristimantis_mutabilis | NA | EN | Decreasing | NA | NA | NA | NA | NA |
| amphibians | Pristimantis_myersi | Pristimantis_myersi | VU | Decreasing | tropical | nocturnal | 1.168 | 2698.938 | 6.184 |
| amphibians | Pristimantis_nangaritza | NA | CR | Decreasing | NA | NA | NA | NA | NA |
| amphibians | Pristimantis_nelsongalloi | NA | not_available | not_available | NA | NA | NA | NA | NA |
| amphibians | Pristimantis_nephophilus | Pristimantis_nephophilus | NT | Decreasing | tropical | nocturnal | 3.163 | 8146.011 | 2.981 |
| amphibians | Pristimantis_nietoi | NA | EN | Decreasing | NA | NA | NA | NA | NA |
| amphibians | Pristimantis_nigrogriseus | Pristimantis_nigrogriseus | VU | Decreasing | tropical | nocturnal | 2.155 | 7429.832 | 5.612 |
| amphibians | Pristimantis_nimbus | NA | EN | Decreasing | NA | NA | NA | NA | NA |
| amphibians | Pristimantis_nubisilva | NA | EN | Decreasing | NA | NA | NA | NA | NA |
| amphibians | Pristimantis_numbala | NA | not_available | not_available | NA | NA | NA | NA | NA |
| amphibians | Pristimantis_nyctophylax | Pristimantis_nyctophylax | LC | Decreasing | tropical | cathemeral | 4.459 | 12331.949 | 8.716 |
| amphibians | Pristimantis_ocellatus | Pristimantis_ocellatus | EN | Decreasing | tropical | nocturnal | 8.247 | 357.99 | 5.514 |
| amphibians | Pristimantis_ockendeni | Pristimantis_ockendeni | LC | Stable | tropical | cathemeral | 2.47 | 1624566.157 | 1.085 |
| amphibians | Pristimantis_omeviridis | NA | LC | Unknown | NA | NA | NA | NA | NA |
| amphibians | Pristimantis_ornatissimus | Pristimantis_ornatissimus | EN | Decreasing | tropical | nocturnal | 5.356 | 7300.237 | 9.507 |
| amphibians | Pristimantis_ornatus | Pristimantis_ornatus | EN | Decreasing | tropical | nocturnal | 1.554 | 510.824 | 10.119 |
| amphibians | Pristimantis_ortizi | Pristimantis_ortizi | EN | Decreasing | tropical | nocturnal | 1.932 | 37.054 | 5.515 |
| amphibians | Pristimantis_pahuma | NA | EN | Decreasing | NA | NA | NA | NA | NA |
| amphibians | Pristimantis_paladines | NA | not_available | not_available | NA | NA | NA | NA | NA |
| amphibians | Pristimantis_paquishae | NA | CR | Decreasing | NA | NA | NA | NA | NA |
| amphibians | Pristimantis_pardalinus | Pristimantis_pardalinus | EN | Decreasing | tropical | diurnal | 3.194 | 22.856 | 16.262 |
| amphibians | Pristimantis_pardalis | Pristimantis_pardalis | LC | Decreasing | tropical | nocturnal | 3.194 | 6411.762 | 7.946 |
| amphibians | Pristimantis_parvillus | Pristimantis_parvillus | VU | Decreasing | tropical | nocturnal | 1.31 | 37549.884 | 9.193 |

| | | | | | | | | | |
|---|---|---|---|---|---|---|---|---|---|
| amphibians | Pristimantis_paulodutrai | Pristimantis_paulodutrai | LC | Stable | tropical | nocturnal | 3.807 | 93095.275 | 9.117 |
| amphibians | Pristimantis_paululus | Pristimantis_paululus | LC | Unknown | tropical | cathemeral | 0.664 | 54443.268 | 4.669 |
| amphibians | Pristimantis_pecki | NA | DD | Unknown | NA | NA | NA | NA | NA |
| amphibians | Pristimantis_percnopterus | Pristimantis_percnopterus | LC | Decreasing | tropical | cathemeral | 1.31 | 12386.366 | 4.515 |
| amphibians | Pristimantis_percultus | Pristimantis_percultus | EN | Decreasing | tropical | cathemeral | 4.614 | 547.73 | 7.799 |
| amphibians | Pristimantis_peruvianus | Pristimantis_peruvianus | LC | Stable | tropical | cathemeral | 8.663 | 1084951.517 | 1.355 |
| amphibians | Pristimantis_petersi | Pristimantis_petersi | NT | Decreasing | tropical | nocturnal | 0.904 | 11580.797 | 5.362 |
| amphibians | Pristimantis_petrobardus | Pristimantis_petrobardus | EN | Decreasing | tropical | nocturnal | 6.77 | 65.162 | 8.703 |
| amphibians | Pristimantis_phalaroinguinis | NA | DD | Unknown | NA | NA | NA | NA | NA |
| amphibians | Pristimantis_philipi | Pristimantis_philipi | NT | Decreasing | tropical | diurnal | 3.074 | 169.019 | 10.389 |
| amphibians | Pristimantis_phoxocephalus | Pristimantis_phoxocephalus | CR | Decreasing | tropical | cathemeral | 5.313 | 65.58 | 11.95 |
| amphibians | Pristimantis_pichincha | NA | EN | Decreasing | NA | NA | NA | NA | NA |
| amphibians | Pristimantis_pinguis | Pristimantis_pinguis | EN | Decreasing | tropical | diurnal | 2.064 | 1415.95 | 9.227 |
| amphibians | Pristimantis_pirrensis | Pristimantis_pirrensis | NT | Stable | tropical | nocturnal | 2.109 | 127.976 | 1.446 |
| amphibians | Pristimantis_pramukae | NA | not_available | not_available | NA | NA | NA | NA | NA |
| amphibians | Pristimantis_prolatus | Pristimantis_prolatus | LC | Decreasing | tropical | nocturnal | 1.037 | 37414.84 | 4.91 |
| amphibians | Pristimantis_proserpens | Pristimantis_proserpens | VU | Unknown | tropical | nocturnal | 0.956 | 4522.547 | 4.688 |
| amphibians | Pristimantis_pseudoacuminatus | Pristimantis_pseudoacuminatus | LC | Unknown | tropical | cathemeral | 0.892 | 39677.584 | 5.607 |
| amphibians | Pristimantis_pteridophilus | Pristimantis_pteridophilus | VU | Decreasing | tropical | nocturnal | 3.163 | 6705.386 | 11.417 |
| amphibians | Pristimantis_pugnax | Pristimantis_pugnax | CR | Decreasing | tropical | cathemeral | 2.928 | 4673.444 | 5.447 |
| amphibians | Pristimantis_puipui | NA | DD | Unknown | NA | NA | NA | NA | NA |
| amphibians | Pristimantis_pulchridormientes | NA | EN | Unknown | NA | NA | NA | NA | NA |
| amphibians | Pristimantis_punzan | NA | CR | Decreasing | NA | NA | NA | NA | NA |
| amphibians | Pristimantis_puruscafeum | NA | VU | Stable | NA | NA | NA | NA | NA |
| amphibians | Pristimantis_pycnodermis | Pristimantis_pycnodermis | EN | Decreasing | tropical | nocturnal | 7.511 | 2562.857 | 7.228 |
| amphibians | Pristimantis_pyrrhomerus | Pristimantis_pyrrhomerus | CR | Decreasing | tropical | cathemeral | 1.023 | 2842.401 | 12.4 |
| amphibians | Pristimantis_quaquaversus | Pristimantis_quaquaversus | LC | Stable | tropical | nocturnal | 2.419 | 90970.774 | 3.768 |
| amphibians | Pristimantis_quinquagesimus | Pristimantis_quinquagesimus | VU | Decreasing | tropical | nocturnal | 5.399 | 4271.868 | 6.237 |
| amphibians | Pristimantis_quintanai | NA | not_available | not_available | NA | NA | NA | NA | NA |
| amphibians | Pristimantis_ramagii | Pristimantis_ramagii | LC | Stable | tropical | nocturnal | 2.109 | 71229.483 | 15.836 |
| amphibians | Pristimantis_reichlei | Pristimantis_reichlei | LC | Stable | tropical | nocturnal | 4.197 | 676994.86 | 3.134 |
| amphibians | Pristimantis_reticulatus | NA | DD | Unknown | NA | NA | NA | NA | NA |
| amphibians | Pristimantis_rhabdocnemus | Pristimantis_rhabdocnemus | LC | Stable | tropical | nocturnal | 1.997 | 190.418 | 9.417 |
| amphibians | Pristimantis_rhabdolaemus | Pristimantis_rhabdolaemus | LC | Stable | tropical | nocturnal | 2.521 | 6628.694 | 5.623 |

| | | | | | | | | | |
|---|---|---|---|---|---|---|---|---|---|
| amphibians | Pristimantis_rhodoplichus | Pristimantis_rhodoplichus | EN | Decreasing | tropical | nocturnal | 3.224 | 1738 | 5.952 |
| amphibians | Pristimantis_rhodostichus | Pristimantis_rhodostichus | LC | Decreasing | tropical | cathemeral | 0.54 | 8648.664 | 3.908 |
| amphibians | Pristimantis_ridens | Pristimantis_ridens | LC | Stable | tropical | nocturnal | 1.394 | 241689.957 | 7.666 |
| amphibians | Pristimantis_riveti | Pristimantis_riveti | CR | Decreasing | tropical | nocturnal | 3.224 | 27.901 | 10.434 |
| amphibians | Pristimantis_romanorum | Pristimantis_romanorum | VU | Decreasing | tropical | nocturnal | 5.802 | 1572.873 | 6.395 |
| amphibians | Pristimantis_rosadoi | Pristimantis_rosadoi | VU | Decreasing | tropical | nocturnal | 1.277 | 15668.021 | 7.444 |
| amphibians | Pristimantis_royi | NA | DD | Unknown | NA | NA | NA | NA | NA |
| amphibians | Pristimantis_rubicundus | Pristimantis_rubicundus | NT | Decreasing | tropical | nocturnal | 11.918 | 9830.144 | 6.003 |
| amphibians | Pristimantis_rufioculis | Pristimantis_rufioculis | VU | Decreasing | tropical | nocturnal | 1.23 | 6898.286 | 2.523 |
| amphibians | Pristimantis_rufoviridis | NA | EN | Decreasing | NA | NA | NA | NA | NA |
| amphibians | Pristimantis_sagedunneae | NA | not_available | not_available | NA | NA | NA | NA | NA |
| amphibians | Pristimantis_sagittulus | Pristimantis_sagittulus | LC | Stable | tropical | nocturnal | 2.086 | 67.155 | 10.386 |
| amphibians | Pristimantis_salaputium | Pristimantis_salaputium | LC | Unknown | tropical | nocturnal | 0.892 | 56.759 | 6.152 |
| amphibians | Pristimantis_samaniegoi | NA | not_available | not_available | NA | NA | NA | NA | NA |
| amphibians | Pristimantis_schultei | Pristimantis_schultei | VU | Unknown | tropical | cathemeral | 3.163 | 8509.01 | 5.645 |
| amphibians | Pristimantis_scolodiscus | Pristimantis_scolodiscus | VU | Decreasing | tropical | nocturnal | 0.807 | 3292.836 | 3.679 |
| amphibians | Pristimantis_serendipitus | Pristimantis_serendipitus | EN | Unknown | tropical | nocturnal | 0.685 | 5158.593 | 5.261 |
| amphibians | Pristimantis_simonbolivari | Pristimantis_simonbolivari | CR | Decreasing | tropical | cathemeral | 0.772 | 29.616 | 12.566 |
| amphibians | Pristimantis_sirnigeli | Pristimantis_sirnigeli | EN | Decreasing | tropical | nocturnal | 1.037 | 698.491 | 8.536 |
| amphibians | Pristimantis_skydmainos | Pristimantis_skydmainos | LC | Stable | tropical | cathemeral | 2.706 | 581754.515 | 2.279 |
| amphibians | Pristimantis_sobetes | Pristimantis_sobetes | VU | Decreasing | tropical | nocturnal | 5.941 | 4035.204 | 6.847 |
| amphibians | Pristimantis_spinosus | Pristimantis_spinosus | EN | Decreasing | tropical | nocturnal | 3.317 | 1187.067 | 5.863 |
| amphibians | Pristimantis_sternothylax | Pristimantis_sternothylax | LC | Stable | tropical | nocturnal | 4.052 | 5465.458 | 8.11 |
| amphibians | Pristimantis_stictogaster | Pristimantis_stictogaster | LC | Unknown | tropical | nocturnal | 1.01 | 19.839 | 3.695 |
| amphibians | Pristimantis_stictus | NA | NT | Stable | NA | NA | NA | NA | NA |
| amphibians | Pristimantis_subsigillatus | Pristimantis_subsigillatus | LC | Decreasing | tropical | nocturnal | 4.536 | 62732.241 | 12.829 |
| amphibians | Pristimantis_sulculus | Pristimantis_sulculus | VU | Unknown | tropical | cathemeral | 14.161 | 5.475 | 5.095 |
| amphibians | Pristimantis_supernatis | Pristimantis_supernatis | VU | Decreasing | tropical | nocturnal | 5.621 | 11625.683 | 10.151 |
| amphibians | Pristimantis_surdus | Pristimantis_surdus | EN | Decreasing | tropical | cathemeral | 14.678 | 3291.018 | 9.818 |
| amphibians | Pristimantis_tantanti | Pristimantis_tantanti | LC | Unknown | tropical | diurnal | 0.761 | 539.525 | 1.055 |
| amphibians | Pristimantis_tenebrionis | Pristimantis_tenebrionis | VU | Decreasing | tropical | nocturnal | 4.124 | 4938.104 | 8.682 |
| amphibians | Pristimantis_terrapacis | NA | not_available | not_available | NA | NA | NA | NA | NA |
| amphibians | Pristimantis_teslai | NA | CR | Decreasing | NA | NA | NA | NA | NA |
| amphibians | Pristimantis_thymalopsoides | Pristimantis_thymalopsoides | CR | Decreasing | tropical | cathemeral | 15.386 | 4.832 | 11.701 |

| | | | | | | | | | |
|---|---|---|---|---|---|---|---|---|---|
| amphibians | Pristimantis_thymelensis | Pristimantis_thymelensis | NT | Decreasing | tropical | nocturnal | 3.015 | 6020.408 | 7.319 |
| amphibians | Pristimantis_tiktik | NA | EN | Decreasing | NA | NA | NA | NA | NA |
| amphibians | Pristimantis_tinajillas | NA | VU | Stable | NA | NA | NA | NA | NA |
| amphibians | Pristimantis_tinguichaca | NA | VU | Decreasing | NA | NA | NA | NA | NA |
| amphibians | Pristimantis_toftae | Pristimantis_toftae | LC | Unknown | tropical | cathemeral | 1.572 | 233565.134 | 2.539 |
| amphibians | Pristimantis_totoroi | NA | EN | Decreasing | NA | NA | NA | NA | NA |
| amphibians | Pristimantis_truebae | Pristimantis_truebae | VU | Decreasing | tropical | diurnal | 8.305 | 2176.641 | 12.219 |
| amphibians | Pristimantis_tungurahua | Pristimantis_tungurahua | EN | Decreasing | tropical | nocturnal | 1.667 | 57.808 | 10.037 |
| amphibians | Pristimantis_ujucami | NA | not_available | not_available | NA | NA | NA | NA | NA |
| amphibians | Pristimantis_unistrigatus | Pristimantis_unistrigatus | LC | Stable | tropical | nocturnal | 2.345 | 21267.201 | 15.606 |
| amphibians | Pristimantis_uranobates | Pristimantis_uranobates | LC | Stable | tropical | nocturnal | 2.109 | 7193.473 | 8.512 |
| amphibians | Pristimantis_variabilis | Pristimantis_variabilis | LC | Stable | tropical | cathemeral | 1.463 | 605438.526 | 1.857 |
| amphibians | Pristimantis_ventrimarmoratus | Pristimantis_ventrimarmoratus | LC | Stable | tropical | nocturnal | 7.187 | 481735.975 | 2.743 |
| amphibians | Pristimantis_ventristellatus | NA | not_available | not_available | NA | NA | NA | NA | NA |
| amphibians | Pristimantis_verecundus | Pristimantis_verecundus | NT | Decreasing | tropical | nocturnal | 0.83 | 6585.101 | 6.819 |
| amphibians | Pristimantis_verrucolatus | NA | CR | Decreasing | NA | NA | NA | NA | NA |
| amphibians | Pristimantis_versicolor | Pristimantis_versicolor | LC | Stable | tropical | nocturnal | 2.706 | 6967.018 | 6.104 |
| amphibians | Pristimantis_vertebralis | Pristimantis_vertebralis | VU | Decreasing | tropical | nocturnal | 7.958 | 4072.463 | 8.411 |
| amphibians | Pristimantis_vilarsi | Pristimantis_vilarsi | LC | Stable | tropical | nocturnal | 10.337 | 1453803.987 | 1.526 |
| amphibians | Pristimantis_w-nigrum | Pristimantis_w-nigrum | LC | Decreasing | tropical | cathemeral | 35.164 | 192435.408 | 11.966 |
| amphibians | Pristimantis_wagteri | Pristimantis_wagteri | EN | Decreasing | tropical | nocturnal | 7.294 | 529.239 | 4.733 |
| amphibians | Pristimantis_walkeri | Pristimantis_walkeri | LC | Decreasing | tropical | cathemeral | 1.214 | 24178.391 | 11.918 |
| amphibians | Pristimantis_waoranii | Pristimantis_waoranii | LC | Stable | tropical | diurnal | 2.37 | 1664.054 | 5.085 |
| amphibians | Pristimantis_wiensi | NA | DD | Unknown | NA | NA | NA | NA | NA |
| amphibians | Pristimantis_yanezi | NA | NT | Unknown | NA | NA | NA | NA | NA |
| amphibians | Pristimantis_yantzaza | NA | EN | Decreasing | NA | NA | NA | NA | NA |
| amphibians | Pristimantis_yumbo | Pristimantis_yumbo | EN | Decreasing | tropical | nocturnal | 0.879 | 1404.166 | 7.522 |
| amphibians | Pristimantis_zeuctotylus | Pristimantis_zeuctotylus | LC | Stable | tropical | cathemeral | 11.036 | 1093634.377 | 1.074 |
| amphibians | Pristimantis_zimmermanae | Pristimantis_zimmermanae | LC | Stable | tropical | nocturnal | 1.294 | 730524.525 | 1.005 |
| amphibians | Pristimantis_zoilae | NA | EN | Decreasing | NA | NA | NA | NA | NA |
| amphibians | Pristimantis_zorro | NA | not_available | not_available | NA | NA | NA | NA | NA |
| amphibians | Proceratophrys_appendiculata | Proceratophrys_appendiculata | LC | Stable | tropical | cathemeral | 23.552 | 6804.884 | 12.522 |
| amphibians | Proceratophrys_ararype | NA | CR | Decreasing | NA | NA | NA | NA | NA |
| amphibians | Proceratophrys_avelinoi | Proceratophrys_avelinoi | LC | Unknown | temperate | cathemeral | 3.981 | 176620.422 | 12.613 |

| | | | | | | | | | |
|---|---|---|---|---|---|---|---|---|---|
| amphibians | Proceratophrys_bigibbosa | Proceratophrys_bigibbosa | LC | Decreasing | temperate | cathemeral | 15.029 | 121080.503 | 12.674 |
| amphibians | Proceratophrys_boiei | Proceratophrys_boiei | LC | Stable | tropical | nocturnal | 39.825 | 385929.035 | 11.818 |
| amphibians | Proceratophrys_brauni | Proceratophrys_brauni | LC | Stable | temperate | diurnal | 5.356 | 101738.098 | 10.2 |
| amphibians | Proceratophrys_concavitympanum | Proceratophrys_concavitympanum | LC | Decreasing | tropical | nocturnal | 18.867 | 127433.62 | 2.642 |
| amphibians | Proceratophrys_cristiceps | Proceratophrys_cristiceps | LC | Stable | tropical | nocturnal | 29.887 | 642715.937 | 8.985 |
| amphibians | Proceratophrys_cururu | Proceratophrys_cururu | VU | Decreasing | tropical | nocturnal | 14.161 | 9114.502 | 4.057 |
| amphibians | Proceratophrys_goyana | Proceratophrys_goyana | LC | Decreasing | tropical | nocturnal | 22.859 | 215006.723 | 12.556 |
| amphibians | Proceratophrys_laticeps | Proceratophrys_laticeps | LC | Stable | tropical | cathemeral | 56.347 | 95861.185 | 10.947 |
| amphibians | Proceratophrys_melanopogon | Proceratophrys_melanopogon | LC | Stable | tropical | nocturnal | 26.866 | 18819.424 | 13.819 |
| amphibians | Proceratophrys_minuta | Proceratophrys_minuta | LC | Stable | tropical | nocturnal | 2.109 | 868.448 | 4.993 |
| amphibians | Proceratophrys_moehringi | Proceratophrys_moehringi | LC | Stable | tropical | diurnal | 24.063 | 5536.322 | 12.218 |
| amphibians | Proceratophrys_moratoi | Proceratophrys_moratoi | LC | Decreasing | tropical | cathemeral | 3.807 | 71628.755 | 15.374 |
| amphibians | Proceratophrys_palustris | Proceratophrys_palustris | CR | Decreasing | tropical | cathemeral | 3.103 | 934.066 | 16.342 |
| amphibians | Proceratophrys_paviotii | Proceratophrys_paviotii | NT | Decreasing | tropical | nocturnal | 16.492 | 19474.151 | 12.39 |
| amphibians | Proceratophrys_renalis | Proceratophrys_renalis | LC | Stable | tropical | nocturnal | 35.806 | 155592.5 | 11.703 |
| amphibians | Proceratophrys_sanctaritae | Proceratophrys_sanctaritae | EN | Unknown | tropical | cathemeral | 8.13 | 54.994 | 7.132 |
| amphibians | Proceratophrys_schirchi | Proceratophrys_schirchi | LC | Decreasing | tropical | nocturnal | 11.036 | 234248.613 | 11.215 |
| amphibians | Proceratophrys_subguttata | Proceratophrys_subguttata | LC | Stable | temperate | nocturnal | 18.764 | 20077.002 | 12.229 |
| amphibians | Proceratophrys_tupinamba | Proceratophrys_tupinambis | LC | Stable | tropical | cathemeral | 37.278 | 184.388 | 5.071 |
| amphibians | Proteus_anguinus | Proteus_anguinus | VU | Decreasing | temperate | cathemeral | 74.48 | 16162.805 | 18.458 |
| amphibians | Pseudacris_brachyphona | Pseudacris_brachyphona | LC | Decreasing | temperate | nocturnal | 4.536 | 190497.809 | 16.238 |
| amphibians | Pseudacris_brimleyi | Pseudacris_brimleyi | LC | Stable | temperate | cathemeral | 3.475 | 135121.316 | 18.816 |
| amphibians | Pseudacris_cadaverina | Pseudacris_cadaverina | LC | Stable | temperate | cathemeral | 11.767 | 75369.281 | 13.026 |
| amphibians | Pseudacris_clarkii | Pseudacris_clarkii | LC | Stable | temperate | cathemeral | 2.599 | 556285.47 | 16.262 |
| amphibians | Pseudacris_crucifer | Pseudacris_crucifer | LC | Stable | temperate | cathemeral | 4.16 | 4040940.028 | 13.956 |
| amphibians | Pseudacris_feriarum | Pseudacris_feriarum | LC | Stable | temperate | cathemeral | 4.536 | 759360.331 | 17.891 |
| amphibians | Pseudacris_fouquettei | Pseudacris_fouquettei | LC | Stable | temperate | cathemeral | 2.109 | 604902.108 | 15.419 |
| amphibians | Pseudacris_illinoensis | Pseudacris_illinoensis | LC | Unknown | temperate | nocturnal | 9.669 | 29551.183 | 19.839 |
| amphibians | Pseudacris_kalmi | Pseudacris_kalmi | LC | Stable | temperate | cathemeral | 3.807 | 34400.226 | 26.586 |
| amphibians | Pseudacris_maculata | Pseudacris_maculata | LC | Decreasing | temperate | cathemeral | 12.531 | 5606940.29 | 9.648 |
| amphibians | Pseudacris_nigrita | Pseudacris_nigrita | LC | Stable | temperate | cathemeral | 2.872 | 384813.584 | 16.685 |
| amphibians | Pseudacris_ocularis | Pseudacris_ocularis | LC | Stable | temperate | cathemeral | 0.368 | 341591.463 | 17.17 |
| amphibians | Pseudacris_ornata | Pseudacris_ornata | LC | Stable | temperate | nocturnal | 5.356 | 347129.287 | 16.23 |
| amphibians | Pseudacris_regilla | Pseudacris_regilla | LC | Stable | temperate | nocturnal | 11.036 | 1621880.114 | 7.872 |

| | | | | | | | | | |
|---|---|---|---|---|---|---|---|---|---|
| amphibians | Pseudacris_streckeri | Pseudacris_streckeri | LC | Unknown | temperate | cathemeral | 9.669 | 519936.445 | 16.496 |
| amphibians | Pseudacris_triseriata | Pseudacris_triseriata | LC | Decreasing | temperate | nocturnal | 5.142 | 409563.538 | 22.733 |
| amphibians | Pseudis_bolbodactyla | Pseudis_bolbodactyla | LC | Stable | tropical | cathemeral | 17.851 | 592141.847 | 10.289 |
| amphibians | Pseudis_cardosoi | Pseudis_cardosoi | LC | Stable | temperate | cathemeral | 15.932 | 47382.876 | 9.52 |
| amphibians | Pseudis_fusca | Pseudis_fusca | LC | Stable | tropical | nocturnal | 13.329 | 108172.926 | 9.894 |
| amphibians | Pseudis_minuta | Pseudis_minuta | LC | Stable | temperate | nocturnal | 11.842 | 453415.643 | 11.191 |
| amphibians | Pseudis_paradoxa | Pseudis_paradoxa | LC | Stable | tropical | cathemeral | 104.268 | 5040866.18 | 2.486 |
| amphibians | Pseudis_tocantins | Pseudis_tocantins | LC | Stable | tropical | cathemeral | 26.999 | 197314.642 | 6.677 |
| amphibians | Pseudobranchus_striatus | Pseudobranchus_striatus | LC | Stable | temperate | nocturnal | 11.554 | 102698.039 | 15.159 |
| amphibians | Pseudoeurycea_brunnata | Pseudoeurycea_brunnata | CR | Decreasing | tropical | diurnal | 3.492 | 170.585 | 14.884 |
| amphibians | Pseudohynobius_flavomaculatus | Pseudohynobius_flavomaculatus | VU | Decreasing | temperate | nocturnal | 9.756 | 1587.047 | 12.587 |
| amphibians | Pseudohynobius_jinfo | Pseudohynobius_jinfo | EN | Decreasing | temperate | cathemeral | 10.097 | 11.148 | 16.934 |
| amphibians | Pseudohynobius_shuichengensis | NA | CR | Decreasing | NA | NA | NA | NA | NA |
| amphibians | Pseudopaludicola_ameghini | Pseudopaludicola_ameghini | LC | Decreasing | tropical | diurnal | 0.772 | 95378.55 | 7.979 |
| amphibians | Pseudopaludicola_boliviana | Pseudopaludicola_boliviana | LC | Stable | tropical | cathemeral | 0.223 | 3016227.571 | 3.093 |
| amphibians | Pseudopaludicola_falcipes | Pseudopaludicola_falcipes | LC | Stable | temperate | cathemeral | 0.567 | 729436.769 | 11.278 |
| amphibians | Pseudopaludicola_florencei | NA | LC | Stable | NA | NA | NA | NA | NA |
| amphibians | Pseudopaludicola_jaredi | NA | LC | Stable | NA | NA | NA | NA | NA |
| amphibians | Pseudopaludicola_jazmynmcdonaldae | NA | EN | Decreasing | NA | NA | NA | NA | NA |
| amphibians | Pseudopaludicola_murundu | Pseudopaludicola_murundu | LC | Unknown | tropical | nocturnal | 0.48 | 203289.426 | 13.439 |
| amphibians | Pseudopaludicola_restinga | NA | VU | Decreasing | NA | NA | NA | NA | NA |
| amphibians | Pseudopaludicola_saltica | Pseudopaludicola_saltica | LC | Stable | tropical | cathemeral | 0.772 | 1526283.822 | 10.736 |
| amphibians | Pseudophilautus_alto | Pseudophilautus_alto | EN | Decreasing | tropical | cathemeral | 1.591 | 1793.924 | 16.48 |
| amphibians | Pseudophilautus_asankai | Pseudophilautus_asankai | EN | Decreasing | tropical | cathemeral | 1.554 | 1288.529 | 14.797 |
| amphibians | Pseudophilautus_caeruleus | Pseudophilautus_caeruleus | EN | Decreasing | tropical | nocturnal | 0.48 | 485.591 | 13.171 |
| amphibians | Pseudophilautus_cuspis | Pseudophilautus_cuspis | EN | Decreasing | tropical | nocturnal | 1.868 | 3004.688 | 14.785 |
| amphibians | Pseudophilautus_decoris | Pseudophilautus_decoris | CR | Decreasing | tropical | nocturnal | 1.01 | 49.312 | 11.312 |
| amphibians | Pseudophilautus_femoralis | Pseudophilautus_femoralis | EN | Decreasing | tropical | nocturnal | 1.889 | 1592.175 | 17.019 |
| amphibians | Pseudophilautus_fulvus | Pseudophilautus_fulvus | EN | Decreasing | tropical | nocturnal | 8.907 | 4202.276 | 17.367 |
| amphibians | Pseudophilautus_hoffmanni | NA | not_available | not_available | NA | NA | NA | NA | NA |
| amphibians | Pseudophilautus_lunatus | Pseudophilautus_lunatus | CR | Decreasing | tropical | nocturnal | 5.756 | 24.644 | 10.161 |
| amphibians | Pseudophilautus_mittermeieri | Pseudophilautus_mittermeieri | VU | Decreasing | tropical | nocturnal | 0.433 | 2862.943 | 15.12 |
| amphibians | Pseudophilautus_procax | NA | not_available | not_available | NA | NA | NA | NA | NA |
| amphibians | Pseudophilautus_puranappu | Pseudophilautus_puranappu | EN | Decreasing | tropical | cathemeral | 12.145 | 31.81 | 9.511 |

| | | | | | | | | | |
|---|---|---|---|---|---|---|---|---|---|
| amphibians | Pseudophilautus_rus | Pseudophilautus_rus | NT | Stable | tropical | nocturnal | 1.037 | 7785.453 | 18.445 |
| amphibians | Pseudophilautus_sarasinorum | Pseudophilautus_sarasinorum | EN | Decreasing | tropical | nocturnal | 4.575 | 2025.011 | 16.01 |
| amphibians | Pseudophilautus_semiruber | Pseudophilautus_semiruber | EN | Decreasing | tropical | diurnal | 0.155 | 989.788 | 17.308 |
| amphibians | Pseudophilautus_silus | NA | not_available | not_available | NA | NA | NA | NA | NA |
| amphibians | Pseudophilautus_silvaticus | NA | not_available | not_available | NA | NA | NA | NA | NA |
| amphibians | Pseudophilautus_variabilis | NA | EX | not_available | NA | NA | NA | NA | NA |
| amphibians | Pseudophilautus_zorro | Pseudophilautus_zorro | VU | Decreasing | tropical | diurnal | 2.132 | 4105.537 | 17.72 |
| amphibians | Pseudophryne_bibronii | Pseudophryne_bibronii | LC | Decreasing | temperate | nocturnal | 3.807 | 630882.107 | 10.166 |
| amphibians | Pseudophryne_dendyi | Pseudophryne_dendyi | LC | Decreasing | temperate | nocturnal | 3.475 | 73886.955 | 7.038 |
| amphibians | Pseudophryne_pengilleyi | Pseudophryne_pengilleyi | CR | Decreasing | temperate | nocturnal | 2.109 | 980.519 | 6.086 |
| amphibians | Pseudophryne_semimarmorata | Pseudophryne_semimarmorata | LC | Stable | temperate | nocturnal | 3.475 | 172714.54 | 10.827 |
| amphibians | Pseudotriton_montanus | Pseudotriton_montanus | LC | Stable | temperate | nocturnal | 6.716 | 758984.358 | 17.385 |
| amphibians | Pseudotriton_ruber | Pseudotriton_ruber | LC | Stable | temperate | nocturnal | 8.985 | 1068841.545 | 17.693 |
| amphibians | Psychrophrynella_usurpator | Psychrophrynella_usurpator | NT | Stable | tropical | diurnal | 2.225 | 61.664 | 9.415 |
| amphibians | Ptychadena_"mascareniensis" | NA | not_available | not_available | NA | NA | NA | NA | NA |
| amphibians | Ptychadena_anchietae | Ptychadena_anchietae | LC | Unknown | tropical | cathemeral | 22.156 | 6632225.537 | 9.36 |
| amphibians | Ptychadena_beka | NA | not_available | not_available | NA | NA | NA | NA | NA |
| amphibians | Ptychadena_bibroni | Ptychadena_bibroni | LC | Unknown | tropical | cathemeral | 24.557 | 3611728.65 | 11.031 |
| amphibians | Ptychadena_erlangeri | Ptychadena_erlangeri | NT | Unknown | tropical | cathemeral | 14.161 | 9566.518 | 12.752 |
| amphibians | Ptychadena_mascareniensis | Ptychadena_mascareniensis | LC | Unknown | tropical | cathemeral | 35.967 | 6801836.235 | 8.315 |
| amphibians | Ptychadena_mossambica | Ptychadena_mossambica | LC | Unknown | tropical | nocturnal | 13.329 | 2865059.686 | 10.013 |
| amphibians | Ptychadena_neumanni | Ptychadena_neumanni | LC | Unknown | tropical | cathemeral | 9.031 | 212375.571 | 13.541 |
| amphibians | Ptychadena_nilotica | Ptychadena_nilotica | LC | Stable | tropical | cathemeral | 5.356 | 1968905.855 | 11.953 |
| amphibians | Ptychadena_oxyrhynchus | Ptychadena_oxyrhynchus | LC | Stable | tropical | nocturnal | 61.584 | 8134935.628 | 9.681 |
| amphibians | Ptychadena_porosissima | Ptychadena_porosissima | LC | Unknown | tropical | nocturnal | 12.61 | 2959490.263 | 8.831 |
| amphibians | Ptychadena_pumilio | Ptychadena_pumilio | LC | Unknown | tropical | cathemeral | 5.802 | 4805500.594 | 11.207 |
| amphibians | Ptychadena_schillukorum | Ptychadena_schillukorum | LC | Unknown | tropical | cathemeral | 15.029 | 5068749.536 | 10.332 |
| amphibians | Ptychadena_stenocephala | Ptychadena_stenocephala | LC | Unknown | tropical | diurnal | 9.669 | 108273.246 | 6.871 |
| amphibians | Ptychadena_taenioscelis | Ptychadena_taenioscelis | LC | Unknown | tropical | cathemeral | 5.356 | 3975471.576 | 8.004 |
| amphibians | Ptychadena_trinodis | Ptychadena_trinodis | LC | Unknown | tropical | nocturnal | 19.923 | 3145425.019 | 10.581 |
| amphibians | Ptychohyla_hypomykter | Ptychohyla_hypomykter | VU | Decreasing | tropical | nocturnal | 7.294 | 8078.531 | 10.698 |
| amphibians | Pyxicephalus_adspersus | Pyxicephalus_adspersus | LC | Decreasing | tropical | diurnal | 1901.294 | 3658402.384 | 8.549 |
| amphibians | Pyxicephalus_edulis | Pyxicephalus_edulis | LC | Decreasing | tropical | cathemeral | 347.523 | 2945288.235 | 10.669 |
| amphibians | Qosqophryne_flammiventris | NA | DD | Unknown | NA | NA | NA | NA | NA |

| | | | | | | | | | |
|---|---|---|---|---|---|---|---|---|---|
| amphibians | Quasipaa_acanthophora | Quasipaa_acanthophora | VU | Decreasing | tropical | nocturnal | 111.885 | 1659.517 | 9.075 |
| amphibians | Quasipaa_boulengeri | Quasipaa_boulengeri | VU | Decreasing | temperate | nocturnal | 269.135 | 990447.789 | 14.502 |
| amphibians | Quasipaa_exilispinosa | Quasipaa_exilispinosa | LC | Decreasing | temperate | nocturnal | 25.694 | 323280.941 | 13.508 |
| amphibians | Quasipaa_fasciculispina | Quasipaa_fasciculispina | LC | Decreasing | tropical | nocturnal | 135.287 | 6052.749 | 2.599 |
| amphibians | Quasipaa_spinosa | Quasipaa_spinosa | VU | Decreasing | temperate | cathemeral | 107.684 | 765191.444 | 14.836 |
| amphibians | Quasipaa_verrucospinosa | Quasipaa_verrucospinosa | LC | Decreasing | tropical | nocturnal | 359.307 | 136064.678 | 6.45 |
| amphibians | Quilticohyla_acrochorda | Ptychohyla_acrochorda | CR | Decreasing | tropical | nocturnal | 17.455 | 25.723 | 9.469 |
| amphibians | Quilticohyla_erythromma | Ptychohyla_erythromma | EN | Decreasing | tropical | nocturnal | 17.455 | 40.808 | 9.37 |
| amphibians | Quilticohyla_sanctaecrucis | Ptychohyla_sanctaecrucis | EN | Decreasing | tropical | nocturnal | 11.767 | 170.354 | 9.485 |
| amphibians | Rana_amurensis | Rana_amurensis | LC | Decreasing | temperate | nocturnal | 32.83 | 7752968.844 | 3.821 |
| amphibians | Rana_arvalis | Rana_arvalis | LC | Decreasing | temperate | cathemeral | 50.601 | 9971161.181 | 9.552 |
| amphibians | Rana_chensinensis | Rana_chensinensis | LC | Decreasing | temperate | diurnal | 27.131 | 2084914.761 | 14.42 |
| amphibians | Rana_chevronta | Rana_chevronta | CR | Unknown | temperate | nocturnal | 15.932 | 9.855 | 8.361 |
| amphibians | Rana_coreana | Rana_coreana | LC | Stable | temperate | diurnal | 7.732 | 149953.329 | 20.919 |
| amphibians | Rana_dabieshanensis | NA | DD | Stable | NA | NA | NA | NA | NA |
| amphibians | Rana_dalmatina | Rana_dalmatina | LC | Decreasing | temperate | cathemeral | 88.303 | 1950863.518 | 20.176 |
| amphibians | Rana_graeca | Rana_graeca | LC | Stable | temperate | cathemeral | 50.601 | 236208.944 | 15.784 |
| amphibians | Rana_iberica | Rana_iberica | VU | Decreasing | temperate | cathemeral | 32.83 | 106637.612 | 17.52 |
| amphibians | Rana_italica | Rana_italica | LC | Unknown | temperate | cathemeral | 25.822 | 115730.938 | 22.365 |
| amphibians | Rana_japonica | Rana_japonica | LC | Decreasing | temperate | diurnal | 35.967 | 236424.417 | 21.939 |
| amphibians | Rana_kukunoris | Rana_kukunoris | LC | Stable | temperate | diurnal | 22.041 | 733978.076 | 8.352 |
| amphibians | Rana_latastei | Rana_latastei | VU | Decreasing | temperate | cathemeral | 41.053 | 33894.542 | 31.898 |
| amphibians | Rana_macrocnemis | Rana_macrocnemis | LC | Decreasing | temperate | cathemeral | 89.21 | 1083970.702 | 14.203 |
| amphibians | Rana_pirica | Rana_pirica | LC | Decreasing | temperate | cathemeral | 35.967 | 158262.496 | 9.96 |
| amphibians | Rana_pyrenaica | Rana_pyrenaica | EN | Decreasing | temperate | cathemeral | 11.036 | 2237.018 | 9.099 |
| amphibians | Rana_temporaria | Rana_temporaria | LC | Stable | temperate | cathemeral | 141.992 | 7787374.871 | 13.142 |
| amphibians | Ranitomeya_amazonica | Ranitomeya_amazonica | LC | Stable | tropical | diurnal | 0.48 | 1301154.766 | 1.425 |
| amphibians | Ranitomeya_benedicta | Ranitomeya_benedicta | VU | Decreasing | tropical | diurnal | 0.664 | 10994.672 | 1.81 |
| amphibians | Ranitomeya_cyanovittata | Ranitomeya_cyanovittata | LC | Stable | tropical | diurnal | 0.354 | 66.983 | 0.495 |
| amphibians | Ranitomeya_defleri | Ranitomeya_defleri | LC | Unknown | tropical | diurnal | 0.382 | 29.441 | 2.434 |
| amphibians | Ranitomeya_fantastica | Ranitomeya_fantastica | VU | Unknown | tropical | diurnal | 0.892 | 7057.573 | 5.417 |
| amphibians | Ranitomeya_flavovittata | Ranitomeya_flavovittata | LC | Stable | tropical | diurnal | 0.48 | 10342.869 | 1.077 |
| amphibians | Ranitomeya_imitator | Ranitomeya_imitator | LC | Stable | tropical | diurnal | 0.772 | 9929.381 | 4.842 |
| amphibians | Ranitomeya_reticulata | Ranitomeya_reticulata | LC | Stable | tropical | diurnal | 0.567 | 26307.979 | 2.061 |

| | | | | | | | | | |
|---|---|---|---|---|---|---|---|---|---|
| amphibians | Ranitomeya_sirensis | Ranitomeya_sirensis | LC | Decreasing | tropical | diurnal | 3.133 | 309963.933 | 2.293 |
| amphibians | Ranitomeya_summersi | Ranitomeya_summersi | EN | Decreasing | tropical | diurnal | 0.604 | 242.866 | 8.506 |
| amphibians | Ranitomeya_toraro | Ranitomeya_toraro | LC | Stable | tropical | diurnal | 0.335 | 866697.706 | 1.026 |
| amphibians | Ranitomeya_uakarii | Ranitomeya_uakarii | LC | Stable | tropical | diurnal | 0.368 | 312456.021 | 1.367 |
| amphibians | Ranitomeya_vanzolinii | Ranitomeya_vanzolinii | LC | Stable | tropical | diurnal | 0.558 | 149685.292 | 0.972 |
| amphibians | Ranitomeya_variabilis | Ranitomeya_variabilis | LC | Stable | tropical | diurnal | 0.664 | 896931.182 | 1.955 |
| amphibians | Ranitomeya_ventrimaculata | Ranitomeya_ventrimaculata | LC | Stable | tropical | diurnal | 0.772 | 304522.45 | 1.631 |
| amphibians | Ranitomeya_yavaricola | NA | DD | Unknown | NA | NA | NA | NA | NA |
| amphibians | Ranodon_sibiricus | Ranodon_sibiricus | EN | Decreasing | temperate | nocturnal | 10.055 | 3060.712 | 5.1 |
| amphibians | Ranoidea_alboguttata | NA | LC | Stable | NA | NA | NA | NA | NA |
| amphibians | Ranoidea_auae | NA | LC | Unknown | NA | NA | NA | NA | NA |
| amphibians | Ranoidea_aurea | NA | NT | Decreasing | NA | NA | NA | NA | NA |
| amphibians | Ranoidea_australis | NA | LC | Stable | NA | NA | NA | NA | NA |
| amphibians | Ranoidea_booroolongensis | NA | EN | Decreasing | NA | NA | NA | NA | NA |
| amphibians | Ranoidea_caerulea | NA | LC | Stable | NA | NA | NA | NA | NA |
| amphibians | Ranoidea_cavernicola | NA | LC | Stable | NA | NA | NA | NA | NA |
| amphibians | Ranoidea_chloris | NA | LC | Stable | NA | NA | NA | NA | NA |
| amphibians | Ranoidea_citropa | NA | LC | Decreasing | NA | NA | NA | NA | NA |
| amphibians | Ranoidea_cyclorhynchus | NA | LC | Stable | NA | NA | NA | NA | NA |
| amphibians | Ranoidea_dayi | NA | VU | Stable | NA | NA | NA | NA | NA |
| amphibians | Ranoidea_eucnemis | NA | LC | Stable | NA | NA | NA | NA | NA |
| amphibians | Ranoidea_genimaculata | NA | LC | Stable | NA | NA | NA | NA | NA |
| amphibians | Ranoidea_lesueuri | NA | LC | Stable | NA | NA | NA | NA | NA |
| amphibians | Ranoidea_lorica | NA | CR | Stable | NA | NA | NA | NA | NA |
| amphibians | Ranoidea_moorei | NA | LC | Decreasing | NA | NA | NA | NA | NA |
| amphibians | Ranoidea_nannotis | NA | LC | Stable | NA | NA | NA | NA | NA |
| amphibians | Ranoidea_nyakalensis | NA | EX | not_available | NA | NA | NA | NA | NA |
| amphibians | Ranoidea_occidentalis | NA | LC | Stable | NA | NA | NA | NA | NA |
| amphibians | Ranoidea_pearsoniana | NA | LC | Decreasing | NA | NA | NA | NA | NA |
| amphibians | Ranoidea_raniformis | NA | VU | Decreasing | NA | NA | NA | NA | NA |
| amphibians | Ranoidea_rheocola | NA | NT | Stable | NA | NA | NA | NA | NA |
| amphibians | Ranoidea_spenceri | NA | CR | Decreasing | NA | NA | NA | NA | NA |
| amphibians | Ranoidea_spinifera | NA | LC | Stable | NA | NA | NA | NA | NA |
| amphibians | Ranoidea_splendida | NA | LC | Stable | NA | NA | NA | NA | NA |

| | | | | | | | | | |
|---|---|---|---|---|---|---|---|---|---|
| amphibians | Ranoidea_subglandulosa | NA | VU | Decreasing | NA | NA | NA | NA | NA |
| amphibians | Ranoidea_wilcoxii | NA | LC | Decreasing | NA | NA | NA | NA | NA |
| amphibians | Ranoidea_xanthomera | NA | LC | Stable | NA | NA | NA | NA | NA |
| amphibians | Raorchestes_annandalii | Pseudophilautus_annandalii | LC | Stable | temperate | nocturnal | 0.567 | 18391.919 | 11.106 |
| amphibians | Raorchestes_bombayensis | Raorchestes_bombayensis | LC | Decreasing | tropical | nocturnal | 2.109 | 21714.115 | 18.01 |
| amphibians | Raorchestes_chalazodes | Pseudophilautus_chalazodes | VU | Decreasing | tropical | nocturnal | 1.686 | 454.519 | 8.235 |
| amphibians | Raorchestes_coonoorensis | NA | EN | Decreasing | NA | NA | NA | NA | NA |
| amphibians | Raorchestes_flaviventris | Pseudophilautus_flaviventris | VU | Decreasing | tropical | diurnal | 1.975 | 800.157 | 9 |
| amphibians | Raorchestes_glandulosus | Raorchestes_glandulosus | VU | Decreasing | tropical | nocturnal | 4.536 | 10082.295 | 16.151 |
| amphibians | Raorchestes_hillisi | NA | not_available | not_available | NA | NA | NA | NA | NA |
| amphibians | Raorchestes_honnametti | NA | NT | Stable | NA | NA | NA | NA | NA |
| amphibians | Raorchestes_ponmudi | Raorchestes_ponmudi | LC | Decreasing | tropical | cathemeral | 4.893 | 18282.619 | 11.772 |
| amphibians | Raorchestes_rezakhani | NA | DD | Unknown | NA | NA | NA | NA | NA |
| amphibians | Raorchestes_signatus | Raorchestes_signatus | EN | Decreasing | tropical | diurnal | 2.37 | 976.428 | 16.322 |
| amphibians | Rentapia_everetti | Pedostibes_everetti | LC | Decreasing | tropical | nocturnal | 66.404 | 10579.738 | 5.637 |
| amphibians | Rhacophorus_annamensis | Rhacophorus_annamensis | LC | Decreasing | tropical | nocturnal | 74.113 | 178585.829 | 9.264 |
| amphibians | Rhacophorus_baluensis | Rhacophorus_baluensis | LC | Stable | tropical | nocturnal | 25.822 | 23297.464 | 3.827 |
| amphibians | Rhacophorus_helenae | Rhacophorus_helenae | EN | Decreasing | tropical | nocturnal | 75.997 | 917.087 | 9.256 |
| amphibians | Rhacophorus_hoabinhensis | NA | DD | Unknown | NA | NA | NA | NA | NA |
| amphibians | Rhacophorus_kio | Rhacophorus_kio | LC | Decreasing | tropical | diurnal | 53.954 | 883418.815 | 9.132 |
| amphibians | Rhacophorus_malabaricus | Rhacophorus_malabaricus | LC | Decreasing | tropical | nocturnal | 52.68 | 54438.935 | 14.687 |
| amphibians | Rhacophorus_pardalis | Rhacophorus_pardalis | LC | Decreasing | tropical | nocturnal | 41.053 | 157763.775 | 8.142 |
| amphibians | Rhacophorus_pseudomalabaricus | Rhacophorus_pseudomalabaricus | VU | Decreasing | tropical | nocturnal | 27.131 | 2339.145 | 10.675 |
| amphibians | Rhacophorus_rhodopus | Rhacophorus_rhodopus | LC | Stable | tropical | nocturnal | 13.329 | 1098451.281 | 11.32 |
| amphibians | Rhaebo_andinophrynoides | Rhaebo_andinophrynoides | VU | Decreasing | tropical | cathemeral | 11.325 | 10872.153 | 4.627 |
| amphibians | Rhaebo_blombergi | Rhaebo_blombergi | CR | Decreasing | tropical | cathemeral | 2029.91 | 18995.372 | 5.376 |
| amphibians | Rhaebo_caeruleostictus | Rhaebo_caeruleostictus | CR | Unknown | tropical | diurnal | 80.428 | 36909.872 | 11.667 |
| amphibians | Rhaebo_ceratophrys | Rhinella_ceratophrys | LC | Stable | tropical | cathemeral | 128.258 | 953465.339 | 1.277 |
| amphibians | Rhaebo_colomai | Andinophryne_colomai | EN | Decreasing | tropical | cathemeral | 18.971 | 1461.802 | 6.241 |
| amphibians | Rhaebo_ecuadorensis | Rhaebo_ecuadorensis | LC | Unknown | tropical | nocturnal | 446.863 | 944293.261 | 1.589 |
| amphibians | Rhaebo_glaberrimus | Rhaebo_glaberrimus | LC | Stable | tropical | nocturnal | 141.992 | 42940.72 | 9.551 |
| amphibians | Rhaebo_guttatus | Rhaebo_guttatus | LC | Decreasing | tropical | nocturnal | 630.892 | 7015845.748 | 2.347 |
| amphibians | Rhaebo_haematiticus | Rhaebo_haematiticus | LC | Decreasing | tropical | cathemeral | 147.501 | 316858.718 | 7.839 |
| amphibians | Rhaebo_hypomelas | Rhaebo_hypomelas | LC | Decreasing | tropical | diurnal | 36.129 | 13615.908 | 5.047 |

| | | | | | | | | | |
|---|---|---|---|---|---|---|---|---|---|
| amphibians | Rhaebo_olallai | Andinophryne_olallai | CR | Decreasing | tropical | nocturnal | 19.923 | 52.376 | 9.37 |
| amphibians | Rheobates_palmatus | Rheobates_palmatus | LC | Decreasing | tropical | diurnal | 4.934 | 49585.931 | 13.08 |
| amphibians | Rheobatrachus_silus | NA | EX | not_available | NA | NA | NA | NA | NA |
| amphibians | Rheobatrachus_vitellinus | NA | EX | not_available | NA | NA | NA | NA | NA |
| amphibians | Rheohyla_miotympanum | Ecnomiohyla_miotympanum | LC | Stable | tropical | nocturnal | 12.222 | 25544.43 | 13.848 |
| amphibians | Rhinella_achavali | Rhinella_achavali | LC | Stable | temperate | nocturnal | 188.234 | 246827.903 | 11.35 |
| amphibians | Rhinella_alata | Rhinella_alata | LC | Stable | tropical | diurnal | 15.685 | 161642.494 | 7.345 |
| amphibians | Rhinella_amabilis | Rhinella_amabilis | CR | Unknown | tropical | nocturnal | 93.84 | 6.645 | 23.252 |
| amphibians | Rhinella_arborescandens | Rhinella_arborescandens | EN | Decreasing | tropical | diurnal | 4.536 | 377.614 | 3.347 |
| amphibians | Rhinella_arenarum | Rhinella_arenarum | LC | Stable | temperate | nocturnal | 188.234 | 1798246.512 | 8.855 |
| amphibians | Rhinella_arunco | Rhinella_arunco | NT | Decreasing | temperate | cathemeral | 163.988 | 73509.701 | 13.935 |
| amphibians | Rhinella_atacamensis | Rhinella_atacamensis | VU | Decreasing | temperate | nocturnal | 77.363 | 13595.03 | 5.846 |
| amphibians | Rhinella_chavin | Rhinella_chavin | EN | Decreasing | tropical | diurnal | 25.694 | 75.731 | 6.467 |
| amphibians | Rhinella_chrysophora | Rhinella_chrysophora | CR | Decreasing | tropical | cathemeral | 19.496 | 1027.595 | 2.843 |
| amphibians | Rhinella_crucifer | Rhinella_crucifer | LC | Stable | tropical | nocturnal | 107.684 | 1232177.542 | 11.591 |
| amphibians | Rhinella_dapsilis | Rhinella_dapsilis | LC | Unknown | tropical | diurnal | 81.848 | 692612.778 | 2.164 |
| amphibians | Rhinella_diptycha | Rhinella_diptycha | LC | Stable | tropical | nocturnal | 2029.91 | 6282831.931 | 8.662 |
| amphibians | Rhinella_dorbignyi | Rhinella_dorbignyi | LC | Stable | temperate | cathemeral | 42.854 | 873585.255 | 11.674 |
| amphibians | Rhinella_festae | Rhinella_festae | LC | Decreasing | tropical | nocturnal | 5.576 | 41336.279 | 4.484 |
| amphibians | Rhinella_granulosa | Rhinella_granulosa | LC | Stable | tropical | nocturnal | 74.113 | 1204162.533 | 9.329 |
| amphibians | Rhinella_henseli | Rhinella_henseli | LC | Stable | temperate | nocturnal | 48.581 | 212610.866 | 11.966 |
| amphibians | Rhinella_hoogmoedi | Rhinella_hoogmoedi | LC | Stable | tropical | nocturnal | 15.932 | 176366.848 | 11.86 |
| amphibians | Rhinella_horribilis | NA | LC | Increasing | NA | NA | NA | NA | NA |
| amphibians | Rhinella_humboldti | Rhinella_humboldti | LC | Stable | tropical | nocturnal | 23.941 | 256711.456 | 13.771 |
| amphibians | Rhinella_icterica | Rhinella_icterica | LC | Stable | temperate | cathemeral | 834.282 | 661347.211 | 13.267 |
| amphibians | Rhinella_justinianoi | Rhinella_justinianoi | VU | Decreasing | tropical | diurnal | 13.824 | 5542.233 | 2.812 |
| amphibians | Rhinella_lescurei | Rhinella_lescurei | LC | Stable | tropical | diurnal | 7.134 | 61130.008 | 0.695 |
| amphibians | Rhinella_lilyrodriguezae | NA | not_available | not_available | NA | NA | NA | NA | NA |
| amphibians | Rhinella_magnussoni | Rhinella_magnussoni | LC | Stable | tropical | diurnal | 14.161 | 17207.446 | 2.863 |
| amphibians | Rhinella_manu | Rhinella_manu | LC | Stable | tropical | nocturnal | 6.177 | 9176.621 | 2.411 |
| amphibians | Rhinella_margaritifera | Rhinella_margaritifera | LC | Stable | tropical | cathemeral | 46.616 | 7506418.387 | 2.475 |
| amphibians | Rhinella_marina | Rhinella_marina | LC | Increasing | tropical | cathemeral | 2029.91 | 9335781.425 | 3.403 |
| amphibians | Rhinella_martyi | NA | not_available | not_available | NA | NA | NA | NA | NA |
| amphibians | Rhinella_nesiotes | Rhinella_nesiotes | VU | Unknown | tropical | nocturnal | 1.889 | 63.196 | 1.312 |

| | | | | | | | | | |
|---|---|---|---|---|---|---|---|---|---|
| amphibians | Rhinella_ornata | Rhinella_ornata | LC | Stable | temperate | cathemeral | 82.421 | 439232.472 | 14.025 |
| amphibians | Rhinella_proboscidea | Rhinella_proboscidea | LC | Stable | tropical | cathemeral | 15.029 | 436709.566 | 1.776 |
| amphibians | Rhinella_pygmaea | Rhinella_pygmaea | LC | Stable | tropical | nocturnal | 11.036 | 34412.58 | 16.858 |
| amphibians | Rhinella_roqueana | Rhinella_roqueana | LC | Stable | tropical | nocturnal | 52.68 | 571368.379 | 1.445 |
| amphibians | Rhinella_rubescens | Rhinella_rubescens | LC | Stable | tropical | cathemeral | 107.684 | 1144986.984 | 11.429 |
| amphibians | Rhinella_spinulosa | Rhinella_spinulosa | LC | Unknown | temperate | cathemeral | 82.421 | 1145779.986 | 5.923 |
| amphibians | Rhinella_unapensis | NA | not_available | not_available | NA | NA | NA | NA | NA |
| amphibians | Rhinella_yanachaga | Rhinella_yanachaga | EN | Stable | tropical | nocturnal | 8.247 | 160.218 | 4.297 |
| amphibians | Rhinoderma_darwinii | Rhinoderma_darwinii | EN | Decreasing | temperate | cathemeral | 4.345 | 88596.826 | 7.054 |
| amphibians | Rhinophrynus_dorsalis | Rhinophrynus_dorsalis | LC | Stable | tropical | cathemeral | 74.113 | 342090.366 | 11.025 |
| amphibians | Rhombophryne_botabota | NA | EN | Decreasing | NA | NA | NA | NA | NA |
| amphibians | Rhombophryne_coronata | Rhombophryne_coronata | LC | Decreasing | tropical | cathemeral | 0.892 | 12780.799 | 6.473 |
| amphibians | Rhombophryne_coudreaui | Rhombophryne_coudreaui | NT | Decreasing | tropical | cathemeral | 1.686 | 14705.183 | 8.094 |
| amphibians | Rhombophryne_diadema | NA | not_available | not_available | NA | NA | NA | NA | NA |
| amphibians | Rhombophryne_ellae | NA | not_available | not_available | NA | NA | NA | NA | NA |
| amphibians | Rhombophryne_guentherpetersi | Rhombophryne_guentherpetersi | EN | Decreasing | tropical | nocturnal | 3.705 | 1018.22 | 4.209 |
| amphibians | Rhombophryne_laevipes | Rhombophryne_laevipes | LC | Unknown | tropical | cathemeral | 15.932 | 21653.438 | 7.258 |
| amphibians | Rhombophryne_longicrus | NA | EN | Decreasing | NA | NA | NA | NA | NA |
| amphibians | Rhombophryne_mangabensis | Rhombophryne_mangabensis | VU | Stable | tropical | nocturnal | 0.917 | 5.438 | 7.205 |
| amphibians | Rhombophryne_matavy | Rhombophryne_matavy | CR | Decreasing | tropical | cathemeral | 10.613 | 29.963 | 6.421 |
| amphibians | Rhombophryne_minuta | Rhombophryne_minuta | EN | Decreasing | tropical | nocturnal | 0.772 | 597.631 | 4.623 |
| amphibians | Rhombophryne_nilevina | NA | not_available | not_available | NA | NA | NA | NA | NA |
| amphibians | Rhombophryne_ornata | NA | EN | Decreasing | NA | NA | NA | NA | NA |
| amphibians | Rhombophryne_regalis | NA | not_available | not_available | NA | NA | NA | NA | NA |
| amphibians | Rhombophryne_savaka | NA | EN | Decreasing | NA | NA | NA | NA | NA |
| amphibians | Rhombophryne_serratopalpebrosa | Rhombophryne_serratopalpebrosa | EN | Decreasing | tropical | nocturnal | 12.531 | 597.631 | 4.623 |
| amphibians | Rhombophryne_testudo | Rhombophryne_testudo | EN | Decreasing | tropical | cathemeral | 7.844 | 311.156 | 18.395 |
| amphibians | Rhyacotriton_variegatus | Rhyacotriton_variegatus | LC | Stable | temperate | nocturnal | 1.609 | 62030.558 | 9.678 |
| amphibians | Rulyrana_mcdiarmidi | Rulyrana_mcdiarmidi | NT | Decreasing | tropical | nocturnal | 2.162 | 23055.017 | 5.66 |
| amphibians | Rulyrana_spiculata | Rulyrana_spiculata | NT | Decreasing | tropical | nocturnal | 1.628 | 37106.561 | 4.233 |
| amphibians | Rupirana_cardosoi | Rupirana_cardosoi | VU | Decreasing | tropical | diurnal | 2029.91 | 3322.886 | 4.642 |
| amphibians | Sachatamia_albomaculata | Sachatamia_albomaculata | LC | Decreasing | tropical | nocturnal | 2.599 | 81367.951 | 8.821 |
| amphibians | Sachatamia_electrops | NA | EN | Decreasing | NA | NA | NA | NA | NA |
| amphibians | Sachatamia_ilex | Sachatamia_ilex | LC | Decreasing | tropical | nocturnal | 4.16 | 85352.739 | 5.312 |

| | | | | | | | | | |
|---|---|---|---|---|---|---|---|---|---|
| amphibians | Sachatamia_orejuela | Sachatamia_orejuela | LC | Decreasing | tropical | nocturnal | 3.103 | 12499.879 | 5.259 |
| amphibians | Salamandra_algira | Salamandra_algira | VU | Decreasing | temperate | nocturnal | 20.198 | 29255.662 | 19.37 |
| amphibians | Salamandra_atra | Salamandra_atra | LC | Decreasing | temperate | nocturnal | 6.996 | 111210.661 | 14.543 |
| amphibians | Salamandra_corsica | Salamandra_corsica | LC | Stable | temperate | nocturnal | 26.796 | 4417.806 | 9.943 |
| amphibians | Salamandra_infraimmaculata | Salamandra_infraimmaculata | LC | Decreasing | temperate | nocturnal | 50.136 | 90468.936 | 16.078 |
| amphibians | Salamandra_lanzai | Salamandra_lanzai | CR | Decreasing | temperate | cathemeral | 10.516 | 779.622 | 10.21 |
| amphibians | Salamandra_salamandra | Salamandra_salamandra | VU | Decreasing | temperate | cathemeral | 57.135 | 2161490.748 | 20.202 |
| amphibians | Salamandrella_keyserlingii | Salamandrella_keyserlingii | LC | Stable | temperate | diurnal | 5.266 | 12290772.129 | 2.84 |
| amphibians | Salamandrella_tridactyla | NA | LC | Decreasing | NA | NA | NA | NA | NA |
| amphibians | Salamandrina_perspicillata | Salamandrina_perspicillata | EN | Decreasing | temperate | nocturnal | 2.043 | 74001.369 | 22.17 |
| amphibians | Salamandrina_terdigitata | Salamandrina_terdigitata | LC | Stable | temperate | cathemeral | 0.843 | 28138.54 | 21.017 |
| amphibians | Sanguirana_acai | NA | NT | Decreasing | NA | NA | NA | NA | NA |
| amphibians | Sanguirana_aurantipunctata | Sanguirana_aurantipunctata | VU | Decreasing | tropical | nocturnal | 64.688 | 4060.085 | 7.348 |
| amphibians | Sarcohyla_arborescandens | Plectrohyla_arborescandens | NT | Decreasing | tropical | diurnal | 12.222 | 13656.189 | 17.029 |
| amphibians | Sarcohyla_bistincta | Plectrohyla_bistincta | LC | Decreasing | tropical | cathemeral | 29.321 | 36667.243 | 7.224 |
| amphibians | Sarcohyla_charadricola | Plectrohyla_charadricola | CR | Decreasing | tropical | nocturnal | 11.693 | 697.118 | 15.032 |
| amphibians | Sarcohyla_chryses | Plectrohyla_chryses | EN | Decreasing | tropical | nocturnal | 6.37 | 108.345 | 5.194 |
| amphibians | Sarcohyla_crassa | Plectrohyla_crassa | CR | Decreasing | tropical | nocturnal | 19.412 | 511.774 | 6.661 |
| amphibians | Sarcohyla_hapsa | NA | LC | Unknown | NA | NA | NA | NA | NA |
| amphibians | Sarcohyla_labeculata | Plectrohyla_calthula | EN | Unknown | tropical | nocturnal | 21.356 | 765.779 | 4.866 |
| amphibians | Sarcohyla_pentheter | Plectrohyla_pentheter | VU | Stable | tropical | cathemeral | 16.304 | 6286.238 | 5.644 |
| amphibians | Sarcohyla_robertsorum | Plectrohyla_robertsorum | VU | Decreasing | tropical | nocturnal | 11.618 | 560.15 | 12.844 |
| amphibians | Sarcohyla_siopela | Plectrohyla_siopela | CR | Decreasing | tropical | cathemeral | 12.926 | 107.968 | 11.192 |
| amphibians | Sarcohyla_thorectes | Plectrohyla_thorectes | EN | Decreasing | tropical | nocturnal | 4.459 | 136.607 | 5.941 |
| amphibians | Scaphiophryne_boribory | Scaphiophryne_boribory | VU | Decreasing | tropical | nocturnal | 19.923 | 1096.224 | 7.362 |
| amphibians | Scaphiophryne_calcarata | Scaphiophryne_calcarata | LC | Stable | tropical | nocturnal | 2.872 | 146996.998 | 9.107 |
| amphibians | Scaphiophryne_gottlebei | Scaphiophryne_gottlebei | EN | Decreasing | tropical | cathemeral | 5.356 | 619.484 | 7.491 |
| amphibians | Scaphiophryne_madagascariensis | Scaphiophryne_madagascariensis | NT | Decreasing | tropical | nocturnal | 15.932 | 20746.508 | 12.726 |
| amphibians | Scaphiophryne_marmorata | Scaphiophryne_marmorata | VU | Decreasing | tropical | nocturnal | 15.029 | 12136.843 | 9.039 |
| amphibians | Scaphiophryne_matsoko | NA | NT | Decreasing | NA | NA | NA | NA | NA |
| amphibians | Scaphiophryne_menabensis | Scaphiophryne_menabensis | LC | Decreasing | tropical | nocturnal | 7.844 | 41657.735 | 6.929 |
| amphibians | Scaphiophryne_spinosa | Scaphiophryne_spinosa | LC | Unknown | tropical | cathemeral | 9.866 | 91129.549 | 8.56 |
| amphibians | Scaphiopus_couchii | Scaphiopus_couchii | LC | Stable | temperate | nocturnal | 74.113 | 1960473.334 | 9.982 |
| amphibians | Scaphiopus_holbrookii | Scaphiopus_holbrookii | LC | Stable | temperate | nocturnal | 46.616 | 1122187.245 | 18.022 |

| | | | | | | | | | |
|---|---|---|---|---|---|---|---|---|---|
| amphibians | Scaphiopus_hurterii | Scaphiopus_hurterii | LC | Stable | temperate | nocturnal | 57.012 | 453628.45 | 16.351 |
| amphibians | Scarthyla_goinorum | Scarthyla_goinorum | LC | Stable | tropical | cathemeral | 1.168 | 1659883.963 | 1.51 |
| amphibians | Scarthyla_vigilans | Scarthyla_vigilans | LC | Unknown | tropical | cathemeral | 0.892 | 256027.589 | 13.912 |
| amphibians | Schismaderma_carens | Schismaderma_carens | LC | Unknown | tropical | cathemeral | 79.584 | 2960397.402 | 9.973 |
| amphibians | Scinax_acuminatus | Scinax_acuminatus | LC | Stable | temperate | nocturnal | 9.669 | 769187.371 | 6.642 |
| amphibians | Scinax_alter | Scinax_alter | LC | Stable | tropical | cathemeral | 2.599 | 207023.415 | 12.025 |
| amphibians | Scinax_baumgardneri | NA | DD | Unknown | NA | NA | NA | NA | NA |
| amphibians | Scinax_blairi | Scinax_blairi | LC | Stable | tropical | nocturnal | 2.733 | 121127.374 | 6.565 |
| amphibians | Scinax_boesemani | Scinax_boesemani | LC | Stable | tropical | nocturnal | 5.356 | 4415912.24 | 1.618 |
| amphibians | Scinax_boulengeri | Scinax_boulengeri | LC | Stable | tropical | nocturnal | 18.354 | 122080.661 | 8.54 |
| amphibians | Scinax_caldarum | Scinax_caldarum | EN | Decreasing | tropical | nocturnal | 2.162 | 4432.27 | 15.205 |
| amphibians | Scinax_caprarius | NA | not_available | not_available | NA | NA | NA | NA | NA |
| amphibians | Scinax_cretatus | Scinax_cretatus | LC | Stable | tropical | nocturnal | 2.733 | 42424.802 | 14.46 |
| amphibians | Scinax_crospedospilus | Scinax_crospedospilus | LC | Stable | tropical | nocturnal | 4.383 | 101142.95 | 14.084 |
| amphibians | Scinax_cuspidatus | Scinax_cuspidatus | LC | Stable | tropical | nocturnal | 2.733 | 291909.038 | 11.714 |
| amphibians | Scinax_danae | NA | DD | Unknown | NA | NA | NA | NA | NA |
| amphibians | Scinax_duartei | Scinax_duartei | LC | Stable | tropical | nocturnal | 4.16 | 3306.411 | 8.418 |
| amphibians | Scinax_elaeochroa | Scinax_elaeochroa | LC | Stable | tropical | nocturnal | 5.487 | 46071.227 | 11.23 |
| amphibians | Scinax_eurydice | Scinax_eurydice | LC | Stable | tropical | nocturnal | 13.992 | 586523.826 | 10.803 |
| amphibians | Scinax_exiguus | Scinax_exiguus | LC | Unknown | tropical | nocturnal | 1.094 | 15074.726 | 4.86 |
| amphibians | Scinax_funereus | Scinax_funereus | LC | Stable | tropical | nocturnal | 6.77 | 959399.494 | 1.505 |
| amphibians | Scinax_fuscomarginatus | Scinax_fuscomarginatus | LC | Stable | tropical | nocturnal | 1.446 | 5108321.19 | 7.602 |
| amphibians | Scinax_fuscovarius | Scinax_fuscovarius | LC | Stable | tropical | nocturnal | 15.44 | 5763739.251 | 8.472 |
| amphibians | Scinax_garbei | Scinax_garbei | LC | Stable | tropical | nocturnal | 10.405 | 4506038.845 | 1.754 |
| amphibians | Scinax_granulatus | Scinax_granulatus | LC | Stable | temperate | nocturnal | 6.77 | 888877.141 | 11.924 |
| amphibians | Scinax_hayii | Scinax_hayii | LC | Stable | tropical | nocturnal | 13.329 | 222916.658 | 15.173 |
| amphibians | Scinax_ictericus | Scinax_ictericus | LC | Unknown | tropical | cathemeral | 3.411 | 564135.356 | 1.511 |
| amphibians | Scinax_imbegue | Scinax_imbegue | LC | Stable | temperate | nocturnal | 4.536 | 15620.545 | 13.473 |
| amphibians | Scinax_iquitorum | Scinax_iquitorum | LC | Unknown | tropical | nocturnal | 4.732 | 676736.145 | 0.741 |
| amphibians | Scinax_juncae | Scinax_juncae | LC | Stable | tropical | nocturnal | 1.499 | 39360.949 | 8.591 |
| amphibians | Scinax_kennedyi | Scinax_kennedyi | LC | Stable | tropical | nocturnal | 4.271 | 51646.485 | 3.413 |
| amphibians | Scinax_nasicus | Scinax_nasicus | LC | Stable | temperate | nocturnal | 4.934 | 2861068.444 | 9.272 |
| amphibians | Scinax_nebulosus | Scinax_nebulosus | LC | Stable | tropical | nocturnal | 5.356 | 6018245.05 | 4.29 |
| amphibians | Scinax_pachycrus | Scinax_pachycrus | LC | Stable | tropical | nocturnal | 2.872 | 462642.974 | 10.516 |

| | | | | | | | | | |
|---|---|---|---|---|---|---|---|---|---|
| amphibians | Scinax_perereca | Scinax_perereca | LC | Stable | temperate | nocturnal | 8.423 | 314816.56 | 11.84 |
| amphibians | Scinax_quinquefasciatus | Scinax_quinquefasciatus | LC | Stable | tropical | cathemeral | 4.893 | 75342.506 | 13.519 |
| amphibians | Scinax_rogerioi | Scinax_rogerioi | LC | Decreasing | tropical | cathemeral | 3.317 | 17827.993 | 11.472 |
| amphibians | Scinax_ruber | Scinax_ruber | LC | Stable | tropical | cathemeral | 7.844 | 8019881.42 | 3.471 |
| amphibians | Scinax_ruberoculatus | NA | LC | Decreasing | NA | NA | NA | NA | NA |
| amphibians | Scinax_similis | Scinax_similis | LC | Stable | tropical | nocturnal | 5.802 | 1314635.387 | 12.02 |
| amphibians | Scinax_squalirostris | Scinax_squalirostris | LC | Stable | temperate | nocturnal | 1.889 | 1411664.874 | 11.703 |
| amphibians | Scinax_staufferi | Scinax_staufferi | LC | Stable | tropical | nocturnal | 2.599 | 569553.528 | 11.639 |
| amphibians | Scinax_sugillatus | Scinax_sugillatus | LC | Stable | tropical | nocturnal | 8.13 | 52970.424 | 7.463 |
| amphibians | Scinax_tigrinus | Scinax_tigrinus | LC | Unknown | tropical | nocturnal | 2.296 | 76882.249 | 11.179 |
| amphibians | Scinax_tymbamirim | Scinax_tymbamirim | LC | Stable | tropical | nocturnal | 1.326 | 62669.983 | 15.357 |
| amphibians | Scinax_wandae | Scinax_wandae | LC | Stable | tropical | nocturnal | 1.481 | 198166.946 | 5.239 |
| amphibians | Scinax_x-signatus | Scinax_x-signatus | LC | Stable | tropical | nocturnal | 9.669 | 4512856.201 | 8.953 |
| amphibians | Sclerophrys_capensis | Amietophrynus_rangeri | LC | Decreasing | temperate | nocturnal | 173.41 | 731803.493 | 12.019 |
| amphibians | Sclerophrys_channingi | Amietophrynus_channingi | LC | Unknown | tropical | nocturnal | 332.231 | 166636.394 | 6.815 |
| amphibians | Sclerophrys_dodsoni | Duttaphrynus_dodsoni | LC | Unknown | tropical | cathemeral | 24.932 | 863705.232 | 7.482 |
| amphibians | Sclerophrys_garmani | Amietophrynus_garmani | LC | Unknown | tropical | nocturnal | 163.988 | 1754416.465 | 9.952 |
| amphibians | Sclerophrys_gracilipes | Amietophrynus_gracilipes | LC | Decreasing | tropical | cathemeral | 5.802 | 1129749.949 | 5.959 |
| amphibians | Sclerophrys_gutturalis | Amietophrynus_gutturalis | LC | Increasing | tropical | cathemeral | 188.234 | 5933376.326 | 8.826 |
| amphibians | Sclerophrys_hadramautinus | NA | not_available | not_available | NA | NA | NA | NA | NA |
| amphibians | Sclerophrys_kassasii | Amietophrynus_kassasii | LC | Increasing | temperate | cathemeral | 4.934 | 40165.07 | 25.034 |
| amphibians | Sclerophrys_maculata | Amietophrynus_maculatus | LC | Stable | tropical | nocturnal | 61.584 | 2329563.013 | 13.121 |
| amphibians | Sclerophrys_mauritanica | Amietophrynus_mauritanicus | LC | Stable | temperate | cathemeral | 387.871 | 776785.9 | 12.665 |
| amphibians | Sclerophrys_pantherina | Amietophrynus_pantherinus | EN | Unknown | temperate | nocturnal | 363.295 | 679.817 | 23.731 |
| amphibians | Sclerophrys_pentoni | Bufo_pentoni | LC | Unknown | tropical | cathemeral | 88.303 | 3382939.441 | 11.716 |
| amphibians | Sclerophrys_perreti | Amietophrynus_perreti | CR | Decreasing | tropical | cathemeral | 25.822 | 22.496 | 21.418 |
| amphibians | Sclerophrys_regularis | Amietophrynus_regularis | LC | Stable | tropical | nocturnal | 243.965 | 7462640 | 10.993 |
| amphibians | Sclerophrys_superciliaris | Amietophrynus_superciliaris | LC | Unknown | tropical | nocturnal | 507.737 | 821127.386 | 9.208 |
| amphibians | Sclerophrys_tuberosa | Amietophrynus_tuberosus | LC | Unknown | tropical | diurnal | 39.306 | 864201.92 | 5.946 |
| amphibians | Sclerophrys_villiersi | Amietophrynus_villiersi | VU | Decreasing | tropical | cathemeral | 32.83 | 7517.948 | 13.464 |
| amphibians | Sclerophrys_xeros | Amietophrynus_xeros | LC | Unknown | tropical | nocturnal | 94.47 | 6286630.201 | 8.188 |
| amphibians | Scotobleps_gabonicus | Scotobleps_gabonicus | LC | Decreasing | tropical | nocturnal | 32.83 | 325321.092 | 7.652 |
| amphibians | Scutiger_boulengeri | Scutiger_boulengeri | LC | Stable | temperate | nocturnal | 29.462 | 704209.278 | 7.998 |
| amphibians | Scutiger_chintingensis | Scutiger_chintingensis | VU | Decreasing | temperate | nocturnal | 13.167 | 5827.964 | 11.198 |

| | | | | | | | | | |
|---|---|---|---|---|---|---|---|---|---|
| amphibians | Scutiger_liupanensis | Scutiger_liupanensis | EN | Decreasing | temperate | nocturnal | 19.39 | 2210.985 | 15.687 |
| amphibians | Scutiger_mammatus | Scutiger_mammatus | LC | Unknown | temperate | nocturnal | 51.841 | 288247.186 | 7.81 |
| amphibians | Scutiger_muliensis | Scutiger_muliensis | EN | Decreasing | temperate | cathemeral | 50.601 | 2859.784 | 6.028 |
| amphibians | Scutiger_pingwuensis | Scutiger_pingwuensis | EN | Decreasing | temperate | nocturnal | 45.655 | 4280.82 | 7.543 |
| amphibians | Scutiger_spinosus | NA | CR | Decreasing | NA | NA | NA | NA | NA |
| amphibians | Scutiger_tengchongensis | NA | DD | Unknown | NA | NA | NA | NA | NA |
| amphibians | Scutiger_wuguanfui | NA | DD | Unknown | NA | NA | NA | NA | NA |
| amphibians | Scythrophrys_sawayae | Scythrophrys_sawayae | LC | Stable | temperate | cathemeral | 0.664 | 5108.919 | 13.31 |
| amphibians | Sechellophryne_gardineri | NA | EN | Decreasing | NA | NA | NA | NA | NA |
| amphibians | Siamophryne_troglodytes | NA | CR | Decreasing | NA | NA | NA | NA | NA |
| amphibians | Silverstoneia_dalyi | Silverstoneia_dalyi | EN | Decreasing | tropical | diurnal | 0.48 | 2356.514 | 6.643 |
| amphibians | Silverstoneia_flotator | Silverstoneia_flotator | LC | Decreasing | tropical | diurnal | 0.425 | 57612.365 | 7.055 |
| amphibians | Silverstoneia_gutturalis | NA | DD | Unknown | NA | NA | NA | NA | NA |
| amphibians | Silverstoneia_minima | NA | DD | Unknown | NA | NA | NA | NA | NA |
| amphibians | Silverstoneia_minutissima | Silverstoneia_minutissima | NT | Unknown | tropical | diurnal | 0.335 | 995.971 | 5.089 |
| amphibians | Silverstoneia_nubicola | Silverstoneia_nubicola | VU | Decreasing | tropical | diurnal | 0.892 | 67282.356 | 4.946 |
| amphibians | Silverstoneia_punctiventris | Silverstoneia_punctiventris | EN | Decreasing | tropical | diurnal | 0.595 | 254.248 | 4.038 |
| amphibians | Siphonops_annulatus | Siphonops_annulatus | LC | Unknown | tropical | nocturnal | 59.356 | 4201157.22 | 3.728 |
| amphibians | Siren_intermedia | Siren_intermedia | LC | Stable | temperate | nocturnal | 205.633 | 1150933.988 | 16.467 |
| amphibians | Siren_lacertina | Siren_lacertina | LC | Stable | temperate | nocturnal | 2111.755 | 418248.927 | 17.096 |
| amphibians | Smilisca_baudinii | Smilisca_baudinii | LC | Stable | tropical | nocturnal | 74.113 | 968474.898 | 10.559 |
| amphibians | Smilisca_cyanosticta | Smilisca_cyanosticta | LC | Stable | tropical | cathemeral | 32.83 | 26226.252 | 9.634 |
| amphibians | Smilisca_dentata | Smilisca_dentata | EN | Decreasing | tropical | nocturnal | 22.272 | 387.229 | 20.492 |
| amphibians | Smilisca_fodiens | Smilisca_fodiens | LC | Stable | temperate | diurnal | 24.186 | 253884.905 | 9.165 |
| amphibians | Smilisca_phaeota | Smilisca_phaeota | LC | Increasing | tropical | cathemeral | 52.68 | 322059.058 | 8.416 |
| amphibians | Smilisca_puma | Smilisca_puma | LC | Stable | tropical | nocturnal | 8.423 | 18050.357 | 11.281 |
| amphibians | Smilisca_sila | Smilisca_sila | LC | Stable | tropical | nocturnal | 22.389 | 169790.177 | 11.381 |
| amphibians | Smilisca_sordida | Smilisca_sordida | LC | Stable | tropical | cathemeral | 24.557 | 50630.821 | 11.449 |
| amphibians | Sooglossus_thomasseti | NA | CR | Unknown | NA | NA | NA | NA | NA |
| amphibians | Spea_bombifrons | Spea_bombifrons | LC | Stable | temperate | nocturnal | 24.557 | 2525493.515 | 12.266 |
| amphibians | Spea_hammondii | Spea_hammondii | NT | Decreasing | temperate | nocturnal | 41.053 | 134297.682 | 18.206 |
| amphibians | Spea_intermontana | Spea_intermontana | LC | Stable | temperate | nocturnal | 25.822 | 1000665.472 | 9.32 |
| amphibians | Spea_multiplicata | Spea_multiplicata | LC | Stable | temperate | nocturnal | 25.822 | 1909224.032 | 9.847 |
| amphibians | Spelaeophryne_methneri | Spelaeophryne_methneri | LC | Unknown | tropical | diurnal | 13.329 | 153490.315 | 7.732 |

| | | | | | | | | | |
|---|---|---|---|---|---|---|---|---|---|
| amphibians | Speleomantes_ambrosii | Hydromantes_ambrosii | CR | Decreasing | temperate | cathemeral | 5.045 | 393.044 | 26.899 |
| amphibians | Speleomantes_flavus | Hydromantes_flavus | EN | Decreasing | temperate | cathemeral | 4.202 | 630.542 | 13.8 |
| amphibians | Speleomantes_genei | Hydromantes_genei | VU | Decreasing | temperate | cathemeral | 3.906 | 1144.784 | 14.658 |
| amphibians | Speleomantes_imperialis | Hydromantes_imperialis | NT | Decreasing | temperate | nocturnal | 5.847 | 2657.76 | 13.895 |
| amphibians | Speleomantes_italicus | Hydromantes_italicus | EN | Stable | temperate | cathemeral | 5.968 | 21061.666 | 20.208 |
| amphibians | Speleomantes_sarrabusensis | Hydromantes_sarrabusensis | CR | Decreasing | temperate | cathemeral | 7.382 | 103.132 | 7.975 |
| amphibians | Speleomantes_strinatii | Hydromantes_strinatii | EN | Decreasing | temperate | cathemeral | 5.968 | 10276.41 | 13.906 |
| amphibians | Speleomantes_supramontis | Hydromantes_supramontis | EN | Decreasing | temperate | cathemeral | 4.681 | 573.896 | 9.914 |
| amphibians | Sphaenorhynchus_botocudo | Sphaenorhynchus_botocudo | LC | Unknown | tropical | nocturnal | 1.953 | 46676.617 | 9.121 |
| amphibians | Sphaenorhynchus_cammaeus | NA | CR | Stable | NA | NA | NA | NA | NA |
| amphibians | Sphaenorhynchus_caramaschii | Sphaenorhynchus_caramaschii | LC | Stable | temperate | nocturnal | 1.953 | 119942.222 | 12.92 |
| amphibians | Sphaenorhynchus_carneus | Sphaenorhynchus_carneus | LC | Stable | tropical | nocturnal | 0.892 | 1628057.291 | 1.211 |
| amphibians | Sphaenorhynchus_dorisae | Sphaenorhynchus_dorisae | LC | Stable | tropical | nocturnal | 5.356 | 1408682.424 | 1.153 |
| amphibians | Sphaenorhynchus_lacteus | Sphaenorhynchus_lacteus | LC | Stable | tropical | nocturnal | 9.669 | 4964958.559 | 2.093 |
| amphibians | Sphaenorhynchus_mirim | Sphaenorhynchus_mirim | LC | Stable | tropical | nocturnal | 0.418 | 1686.5 | 6.52 |
| amphibians | Sphaenorhynchus_palustris | Sphaenorhynchus_palustris | DD | Stable | tropical | nocturnal | 3.807 | 278.589 | 2.855 |
| amphibians | Sphaenorhynchus_pauloalvini | Sphaenorhynchus_pauloalvini | LC | Stable | tropical | nocturnal | 1.023 | 35065.546 | 7.916 |
| amphibians | Sphaenorhynchus_planicola | Sphaenorhynchus_planicola | LC | Stable | tropical | nocturnal | 1.037 | 42453.117 | 15.051 |
| amphibians | Sphaenorhynchus_platycephalus | Sphaenorhynchus_platycephalus | LC | Stable | tropical | nocturnal | 2.872 | 28880.264 | 13.586 |
| amphibians | Sphaenorhynchus_prasinus | Sphaenorhynchus_prasinus | LC | Stable | tropical | nocturnal | 2.345 | 244135.813 | 11.297 |
| amphibians | Sphaenorhynchus_surdus | Sphaenorhynchus_surdus | LC | Stable | temperate | nocturnal | 2.109 | 84209.27 | 10.481 |
| amphibians | Sphaerotheca_breviceps | Sphaerotheca_breviceps | LC | Stable | tropical | cathemeral | 41.053 | 3248554.306 | 18.009 |
| amphibians | Sphaerotheca_dobsonii | Sphaerotheca_dobsonii | LC | Decreasing | tropical | cathemeral | 19.923 | 256436.083 | 17.998 |
| amphibians | Sphaerotheca_rolandae | Sphaerotheca_rolandae | LC | Decreasing | tropical | nocturnal | 11.036 | 707273.664 | 18.284 |
| amphibians | Sphenophryne_cornuta | Sphenophryne_cornuta | LC | Stable | tropical | nocturnal | 6.273 | 470007.827 | 3.049 |
| amphibians | Sphenophryne_magnitympanum | Liophryne_magnitympanum | LC | Stable | tropical | nocturnal | 1.411 | 1941.789 | 2.08 |
| amphibians | Sphenophryne_miniafia | NA | LC | Stable | NA | NA | NA | NA | NA |
| amphibians | Sphenophryne_schlaginhaufeni | Liophryne_schlaginhaufeni | LC | Stable | tropical | cathemeral | 6.77 | 179361.338 | 4.244 |
| amphibians | Sphenophryne_similis | NA | DD | Unknown | NA | NA | NA | NA | NA |
| amphibians | Spicospina_flammocaerulea | Spicospina_flammocaerulea | EN | Decreasing | temperate | nocturnal | 3.807 | 331.129 | 5.533 |
| amphibians | Spinomantis_aglavei | Spinomantis_aglavei | LC | Decreasing | tropical | nocturnal | 11.767 | 91736.524 | 8.949 |
| amphibians | Spinomantis_beckei | NA | not_available | not_available | NA | NA | NA | NA | NA |
| amphibians | Spinomantis_bertini | Spinomantis_bertini | NT | Decreasing | tropical | diurnal | 1.686 | 8490.274 | 5.23 |
| amphibians | Spinomantis_brunae | Spinomantis_brunae | EN | Decreasing | temperate | cathemeral | 3.475 | 1338.781 | 7.145 |

| | | | | | | | | | |
|---|---|---|---|---|---|---|---|---|---|
| amphibians | Spinomantis_elegans | Spinomantis_elegans | NT | Decreasing | tropical | cathemeral | 19.923 | 11045.02 | 5.818 |
| amphibians | Spinomantis_fimbriatus | Spinomantis_fimbriatus | LC | Decreasing | tropical | nocturnal | 4.934 | 35060.693 | 7.095 |
| amphibians | Spinomantis_guibei | Spinomantis_guibei | VU | Decreasing | temperate | diurnal | 5.356 | 1760.073 | 4.84 |
| amphibians | Spinomantis_microtis | Spinomantis_microtis | EN | Decreasing | temperate | diurnal | 9.669 | 1450.28 | 4.929 |
| amphibians | Spinomantis_mirus | NA | not_available | not_available | NA | NA | NA | NA | NA |
| amphibians | Spinomantis_nussbaumi | Spinomantis_nussbaumi | CR | Decreasing | tropical | nocturnal | 16.873 | 10.374 | 3.847 |
| amphibians | Spinomantis_peraccae | Spinomantis_peraccae | LC | Decreasing | tropical | nocturnal | 16.873 | 100390.667 | 9.062 |
| amphibians | Spinomantis_phantasticus | Spinomantis_phantasticus | LC | Decreasing | tropical | nocturnal | 4.536 | 31119.417 | 6.744 |
| amphibians | Spinomantis_tavaratra | Spinomantis_tavaratra | VU | Decreasing | tropical | nocturnal | 3.807 | 2933.933 | 4.02 |
| amphibians | Staurois_guttatus | Staurois_guttatus | LC | Decreasing | tropical | diurnal | 15.029 | 143821.918 | 4.683 |
| amphibians | Staurois_latopalmatus | Staurois_latopalmatus | LC | Decreasing | tropical | cathemeral | 32.83 | 138400.631 | 4.55 |
| amphibians | Staurois_natator | Staurois_natator | LC | Decreasing | tropical | cathemeral | 15.029 | 18396.614 | 9.581 |
| amphibians | Staurois_parvus | Staurois_parvus | VU | Decreasing | tropical | diurnal | 2.345 | 1492.214 | 3.244 |
| amphibians | Staurois_tuberilinguis | Staurois_tuberilinguis | LC | Decreasing | tropical | diurnal | 6.77 | 82444.037 | 4.301 |
| amphibians | Stefania_ackawaio | Stefania_ackawaio | VU | Stable | tropical | nocturnal | 27.131 | 53.487 | 0.45 |
| amphibians | Stefania_ayangannae | Stefania_ayangannae | VU | Stable | tropical | nocturnal | 11.036 | 53.487 | 0.45 |
| amphibians | Stefania_coxi | Stefania_coxi | VU | Stable | tropical | nocturnal | 32.83 | 26.6 | 0.452 |
| amphibians | Stefania_evansi | Stefania_evansi | DD | Stable | tropical | diurnal | 97.662 | 46467.23 | 1.23 |
| amphibians | Stefania_maccullochi | NA | not_available | not_available | NA | NA | NA | NA | NA |
| amphibians | Stefania_neblinae | Stefania_neblinae | LC | Stable | tropical | cathemeral | 7.844 | 174.912 | 0.678 |
| amphibians | Stefania_riae | Stefania_riae | NT | Unknown | tropical | nocturnal | 19.39 | 2371.678 | 0.506 |
| amphibians | Stefania_satelles | Stefania_satelles | NT | Stable | tropical | nocturnal | 19.39 | 615.867 | 1.134 |
| amphibians | Stefania_scalae | Stefania_scalae | LC | Stable | tropical | nocturnal | 24.063 | 14133.875 | 1.321 |
| amphibians | Stefania_schuberti | Stefania_schuberti | NT | Stable | tropical | nocturnal | 42.854 | 480.353 | 0.842 |
| amphibians | Stefania_tamacuarina | NA | DD | Unknown | NA | NA | NA | NA | NA |
| amphibians | Stereocyclops_histrio | Stereocyclops_histrio | LC | Decreasing | tropical | nocturnal | 9.669 | 27430.994 | 7.497 |
| amphibians | Stereocyclops_incrassatus | Stereocyclops_incrassatus | LC | Stable | tropical | nocturnal | 22.156 | 267810.126 | 12.16 |
| amphibians | Stereocyclops_parkeri | Stereocyclops_parkeri | LC | Stable | tropical | nocturnal | 19.923 | 36405.999 | 15.895 |
| amphibians | Strabomantis_anatipes | Strabomantis_anatipes | VU | Decreasing | tropical | nocturnal | 99.612 | 5707.866 | 4.62 |
| amphibians | Strabomantis_anomalus | Strabomantis_anomalus | CR | Decreasing | tropical | nocturnal | 80.71 | 86973.105 | 5.873 |
| amphibians | Strabomantis_biporcatus | Strabomantis_biporcatus | LC | Unknown | tropical | nocturnal | 104.268 | 4780.522 | 8.044 |
| amphibians | Strabomantis_bufoniformis | Strabomantis_bufoniformis | EN | Decreasing | tropical | nocturnal | 85.327 | 83335.512 | 5.257 |
| amphibians | Strabomantis_cerastes | Strabomantis_cerastes | CR | Decreasing | tropical | cathemeral | 15.749 | 23795.189 | 8.448 |
| amphibians | Strabomantis_cornutus | Strabomantis_cornutus | EN | Decreasing | tropical | cathemeral | 19.39 | 12293.028 | 7.057 |

| | | | | | | | | | |
|---|---|---|---|---|---|---|---|---|---|
| amphibians | Strabomantis_helonotus | Strabomantis_helonotus | CR | Unknown | tropical | nocturnal | 32.83 | 10.613 | 14.884 |
| amphibians | Strabomantis_laticorpus | NA | DD | Unknown | NA | NA | NA | NA | NA |
| amphibians | Strabomantis_necerus | Strabomantis_necerus | CR | Decreasing | tropical | cathemeral | 83.285 | 2721.848 | 10.191 |
| amphibians | Strabomantis_sulcatus | Strabomantis_sulcatus | LC | Stable | tropical | cathemeral | 19.602 | 1112766.692 | 1.572 |
| amphibians | Strauchbufo_raddei | Strauchbufo_raddei | LC | Decreasing | temperate | diurnal | 71.478 | 4773678.418 | 9.982 |
| amphibians | Strongylopus_fasciatus | Strongylopus_fasciatus | LC | Unknown | temperate | cathemeral | 11.036 | 779915.337 | 12.769 |
| amphibians | Strongylopus_grayii | Strongylopus_grayii | LC | Unknown | temperate | cathemeral | 24.557 | 589930.752 | 11.329 |
| amphibians | Stumpffia_achillei | NA | not_available | not_available | NA | NA | NA | NA | NA |
| amphibians | Stumpffia_analamaina | Stumpffia_analamaina | CR | Decreasing | tropical | cathemeral | 0.155 | 10.414 | 10.843 |
| amphibians | Stumpffia_be | Stumpffia_be | EN | Decreasing | tropical | nocturnal | 1.199 | 253.119 | 7.304 |
| amphibians | Stumpffia_betampona | NA | not_available | not_available | NA | NA | NA | NA | NA |
| amphibians | Stumpffia_contumelia | NA | not_available | not_available | NA | NA | NA | NA | NA |
| amphibians | Stumpffia_davidattenboroughi | NA | not_available | not_available | NA | NA | NA | NA | NA |
| amphibians | Stumpffia_diutissima | NA | not_available | not_available | NA | NA | NA | NA | NA |
| amphibians | Stumpffia_froschaueri | NA | not_available | not_available | NA | NA | NA | NA | NA |
| amphibians | Stumpffia_fusca | NA | not_available | not_available | NA | NA | NA | NA | NA |
| amphibians | Stumpffia_garraffoi | NA | not_available | not_available | NA | NA | NA | NA | NA |
| amphibians | Stumpffia_gimmeli | Stumpffia_gimmeli | LC | Stable | tropical | nocturnal | 0.41 | 8336.938 | 6.673 |
| amphibians | Stumpffia_grandis | Stumpffia_grandis | LC | Decreasing | tropical | nocturnal | 1.168 | 23178.808 | 7.513 |
| amphibians | Stumpffia_hara | Stumpffia_hara | CR | Decreasing | tropical | cathemeral | 1.109 | 2.933 | 3.305 |
| amphibians | Stumpffia_jeannoeli | NA | not_available | not_available | NA | NA | NA | NA | NA |
| amphibians | Stumpffia_kibomena | NA | EN | Decreasing | NA | NA | NA | NA | NA |
| amphibians | Stumpffia_larinki | NA | not_available | not_available | NA | NA | NA | NA | NA |
| amphibians | Stumpffia_makira | NA | not_available | not_available | NA | NA | NA | NA | NA |
| amphibians | Stumpffia_maledicta | NA | not_available | not_available | NA | NA | NA | NA | NA |
| amphibians | Stumpffia_mamitika | NA | not_available | not_available | NA | NA | NA | NA | NA |
| amphibians | Stumpffia_miery | Stumpffia_miery | EN | Decreasing | tropical | nocturnal | 0.204 | 1307.52 | 7.488 |
| amphibians | Stumpffia_nigrorubra | NA | not_available | not_available | NA | NA | NA | NA | NA |
| amphibians | Stumpffia_obscoena | NA | not_available | not_available | NA | NA | NA | NA | NA |
| amphibians | Stumpffia_pardus | NA | not_available | not_available | NA | NA | NA | NA | NA |
| amphibians | Stumpffia_psologlossa | Stumpffia_psologlossa | EN | Decreasing | tropical | cathemeral | 0.275 | 2003.713 | 9.486 |
| amphibians | Stumpffia_pygmaea | Stumpffia_pygmaea | EN | Decreasing | tropical | cathemeral | 0.124 | 311.205 | 18.395 |
| amphibians | Stumpffia_roseifemoralis | Stumpffia_roseifemoralis | EN | Decreasing | tropical | nocturnal | 0.772 | 597.631 | 4.623 |
| amphibians | Stumpffia_spandei | NA | not_available | not_available | NA | NA | NA | NA | NA |

| | | | | | | | | | |
|---|---|---|---|---|---|---|---|---|---|
| amphibians | Stumpffia_staffordi | Stumpffia_staffordi | VU | Unknown | tropical | diurnal | 1.667 | 47.693 | 10.23 |
| amphibians | Stumpffia_tetradactyla | Stumpffia_tetradactyla | DD | Decreasing | temperate | cathemeral | 0.223 | 427.239 | 14.594 |
| amphibians | Stumpffia_tridactyla | Stumpffia_tridactyla | DD | Decreasing | tropical | cathemeral | 0.082 | 1097.298 | 8.205 |
| amphibians | Stumpffia_yanniki | NA | not_available | not_available | NA | NA | NA | NA | NA |
| amphibians | Synapturanus_mirandaribeiroi | Synapturanus_mirandaribeiroi | LC | Stable | tropical | cathemeral | 6.273 | 1189452.179 | 0.808 |
| amphibians | Synapturanus_rabus | Synapturanus_rabus | LC | Stable | tropical | nocturnal | 0.48 | 309467.917 | 1.661 |
| amphibians | Synapturanus_salseri | Synapturanus_salseri | LC | Stable | tropical | cathemeral | 3.475 | 411294.707 | 0.828 |
| amphibians | Tachycnemis_seychellensis | NA | LC | Unknown | NA | NA | NA | NA | NA |
| amphibians | Taricha_granulosa | Taricha_granulosa | LC | Stable | temperate | cathemeral | 11.13 | 372204.666 | 8.015 |
| amphibians | Taricha_rivularis | Taricha_rivularis | VU | Decreasing | temperate | nocturnal | 11.13 | 13931.342 | 11.795 |
| amphibians | Taricha_torosa | Taricha_torosa | NT | Decreasing | temperate | nocturnal | 11.13 | 53483.456 | 15.619 |
| amphibians | Taudactylus_acutirostris | NA | EX | not_available | NA | NA | NA | NA | NA |
| amphibians | Taudactylus_diurnus | NA | EX | not_available | NA | NA | NA | NA | NA |
| amphibians | Taudactylus_eungellensis | Taudactylus_eungellensis | EN | Decreasing | tropical | cathemeral | 3.807 | 464.477 | 2.69 |
| amphibians | Taudactylus_pleione | Taudactylus_pleione | CR | Decreasing | temperate | diurnal | 2.345 | 19.198 | 6.133 |
| amphibians | Taudactylus_rheophilus | Taudactylus_rheophilus | CR | Unknown | tropical | cathemeral | 2.345 | 724.815 | 2.585 |
| amphibians | Telmatobius_atahualpai | Telmatobius_atahualpai | VU | Decreasing | tropical | cathemeral | 25.565 | 5896.856 | 5.319 |
| amphibians | Telmatobius_bolivianus | Telmatobius_bolivianus | CR | Decreasing | tropical | nocturnal | 13.086 | 6005.801 | 6.297 |
| amphibians | Telmatobius_carrillae | Telmatobius_carrillae | VU | Decreasing | tropical | nocturnal | 19.923 | 15063.389 | 9.55 |
| amphibians | Telmatobius_cirrhacelis | Telmatobius_cirrhacelis | CR | Decreasing | tropical | nocturnal | 30.895 | 208.909 | 3.478 |
| amphibians | Telmatobius_culeus | Telmatobius_culeus | EN | Decreasing | tropical | nocturnal | 355.349 | 8452.398 | 9.536 |
| amphibians | Telmatobius_niger | Telmatobius_niger | CR | Decreasing | tropical | diurnal | 45.846 | 12667.533 | 13.101 |
| amphibians | Telmatobius_thompsoni | Telmatobius_thompsoni | DD | Stable | tropical | diurnal | 45.274 | 10.342 | 10.648 |
| amphibians | Telmatobius_truebae | Telmatobius_truebae | VU | Decreasing | tropical | nocturnal | 54.816 | 3641.688 | 6.615 |
| amphibians | Tepuihyla_edelcae | Tepuihyla_edelcae | LC | Stable | tropical | nocturnal | 8.247 | 12192.491 | 1.3 |
| amphibians | Tepuihyla_exophthalma | Tepuihyla_exophthalma | LC | Unknown | tropical | nocturnal | 2.109 | 11079.606 | 1.107 |
| amphibians | Tepuihyla_rodriguezi | Tepuihyla_rodriguezi | NT | Unknown | tropical | nocturnal | 10.682 | 18368.899 | 1.332 |
| amphibians | Tepuihyla_tuberculosa | Ecnomiohyla_tuberculosa | LC | Stable | tropical | nocturnal | 74.113 | 190930.293 | 1.304 |
| amphibians | Teratohyla_amelie | Teratohyla_amelie | LC | Unknown | tropical | nocturnal | 0.505 | 102289.583 | 2.917 |
| amphibians | Teratohyla_midas | Teratohyla_midas | LC | Stable | tropical | nocturnal | 1.499 | 3562726.294 | 1.448 |
| amphibians | Teratohyla_pulverata | Teratohyla_pulverata | LC | Unknown | tropical | nocturnal | 2.872 | 117386.029 | 6.815 |
| amphibians | Teratohyla_spinosa | Teratohyla_spinosa | LC | Stable | tropical | nocturnal | 1.686 | 100062.164 | 6.47 |
| amphibians | Theloderma_annae | NA | DD | Decreasing | NA | NA | NA | NA | NA |
| amphibians | Theloderma_asperum | Theloderma_asperum | LC | Decreasing | tropical | nocturnal | 3.317 | 126695.868 | 11.424 |

| | | | | | | | | | |
|---|---|---|---|---|---|---|---|---|---|
| amphibians | Theloderma_auratum | NA | VU | Decreasing | NA | NA | NA | NA | NA |
| amphibians | Theloderma_horridum | Theloderma_horridum | LC | Decreasing | tropical | nocturnal | 10.337 | 152304.473 | 12.576 |
| amphibians | Theloderma_lacustrinum | NA | DD | Decreasing | NA | NA | NA | NA | NA |
| amphibians | Theloderma_laeve | Theloderma_laeve | LC | Decreasing | tropical | nocturnal | 1.827 | 66346.99 | 9.728 |
| amphibians | Theloderma_vietnamense | NA | LC | Decreasing | NA | NA | NA | NA | NA |
| amphibians | Thoropa_lutzi | Thoropa_lutzi | CR | Unknown | tropical | nocturnal | 2.109 | 1848.958 | 32.441 |
| amphibians | Thoropa_megatympanum | Thoropa_megatympanum | LC | Stable | tropical | nocturnal | 13.329 | 27762.428 | 5.463 |
| amphibians | Thoropa_miliaris | Thoropa_miliaris | LC | Stable | tropical | nocturnal | 52.68 | 485972.856 | 11.143 |
| amphibians | Thoropa_petropolitana | Thoropa_petropolitana | CR | Unknown | tropical | nocturnal | 1.499 | 1175.44 | 16.232 |
| amphibians | Thoropa_saxatilis | Thoropa_saxatilis | LC | Stable | temperate | nocturnal | 29.887 | 12301.23 | 12.65 |
| amphibians | Thoropa_taophora | Thoropa_taophora | LC | Stable | tropical | nocturnal | 111.531 | 7736.778 | 11.082 |
| amphibians | Tlalocohyla_loquax | Tlalocohyla_loquax | LC | Stable | tropical | nocturnal | 9.031 | 348675.79 | 8.954 |
| amphibians | Tlalocohyla_picta | Tlalocohyla_picta | LC | Increasing | tropical | nocturnal | 2.345 | 313539.973 | 10.359 |
| amphibians | Tlalocohyla_smithii | Tlalocohyla_smithii | LC | Stable | tropical | nocturnal | 2.321 | 260450.694 | 9.384 |
| amphibians | Tomopterna_cryptotis | Tomopterna_cryptotis | LC | Stable | tropical | nocturnal | 24.557 | 9242511.753 | 8.248 |
| amphibians | Trachycephalus_atlas | Trachycephalus_atlas | LC | Stable | tropical | nocturnal | 129.824 | 396055.513 | 9.272 |
| amphibians | Trachycephalus_coriaceus | Trachycephalus_coriaceus | LC | Stable | tropical | nocturnal | 29.887 | 3944543.131 | 1.375 |
| amphibians | Trachycephalus_cunauaru | Trachycephalus_cunauaru | LC | Unknown | tropical | nocturnal | 61.35 | 2892478.97 | 1.434 |
| amphibians | Trachycephalus_dibernardoi | Trachycephalus_dibernardoi | LC | Unknown | temperate | nocturnal | 61.584 | 88815.887 | 11.026 |
| amphibians | Trachycephalus_imitatrix | Trachycephalus_imitatrix | LC | Unknown | tropical | nocturnal | 34.374 | 53963.974 | 12.154 |
| amphibians | Trachycephalus_jordani | Trachycephalus_jordani | LC | Unknown | tropical | nocturnal | 103.258 | 63866.081 | 14.89 |
| amphibians | Trachycephalus_lepidus | Trachycephalus_lepidus | LC | Stable | temperate | nocturnal | 18.867 | 1059.656 | 1.771 |
| amphibians | Trachycephalus_mambaiensis | Trachycephalus_mambaiensis | LC | Decreasing | tropical | nocturnal | 65.665 | 97506.279 | 7.677 |
| amphibians | Trachycephalus_mesophaeus | Trachycephalus_mesophaeus | LC | Stable | tropical | cathemeral | 51.633 | 327726.913 | 11.811 |
| amphibians | Trachycephalus_nigromaculatus | Trachycephalus_nigromaculatus | LC | Stable | tropical | nocturnal | 79.584 | 936356.48 | 10.555 |
| amphibians | Trachycephalus_resinifictrix | Trachycephalus_resinifictrix | LC | Unknown | tropical | nocturnal | 104.268 | 1369861.561 | 1.791 |
| amphibians | Trachycephalus_typhonius | Trachycephalus_venulosus | LC | Stable | tropical | cathemeral | 159.413 | 9757480.349 | 4.915 |
| amphibians | Trachycephalus_venezolanus | Aparasphenodon_venezolanus | LC | Stable | tropical | nocturnal | 17.851 | 185908.063 | 0.522 |
| amphibians | Triprion_petasatus | Triprion_petasatus | LC | Stable | tropical | cathemeral | 41.409 | 153656.138 | 6.402 |
| amphibians | Triprion_spatulatus | Diaglena_spatulata | LC | Stable | tropical | nocturnal | 154.926 | 73095.045 | 9.342 |
| amphibians | Triprion_spinosus | Anotheca_spinosa | NT | Decreasing | tropical | cathemeral | 50.601 | 22780.672 | 11.077 |
| amphibians | Triturus_carnifex | Triturus_carnifex | VU | Decreasing | temperate | nocturnal | 11.13 | 501475.034 | 20.334 |
| amphibians | Triturus_cristatus | Triturus_cristatus | LC | Decreasing | temperate | cathemeral | 14.334 | 4456023.274 | 17.023 |
| amphibians | Triturus_dobrogicus | Triturus_dobrogicus | LC | Decreasing | temperate | nocturnal | 11.501 | 271908.807 | 21.086 |

| | | | | | | | | | |
|---|---|---|---|---|---|---|---|---|---|
| amphibians | Triturus_karelinii | Triturus_karelinii | LC | Decreasing | temperate | nocturnal | 11.501 | 242121.655 | 13.804 |
| amphibians | Triturus_marmoratus | Triturus_marmoratus | VU | Decreasing | temperate | cathemeral | 9.722 | 474835.687 | 20.25 |
| amphibians | Triturus_pygmaeus | Triturus_pygmaeus | NT | Decreasing | temperate | nocturnal | 3.683 | 174284.917 | 16.531 |
| amphibians | Tsingymantis_antitra | Tsingymantis_antitra | EN | Unknown | tropical | nocturnal | 28.486 | 253.119 | 7.304 |
| amphibians | Tylototriton_asperrimus | Tylototriton_asperrimus | NT | Decreasing | temperate | nocturnal | 4.673 | 111992.196 | 13.829 |
| amphibians | Tylototriton_kachinorum | NA | DD | Decreasing | NA | NA | NA | NA | NA |
| amphibians | Tylototriton_kweichowensis | Tylototriton_kweichowensis | VU | Decreasing | temperate | nocturnal | 14.773 | 51079.42 | 16.868 |
| amphibians | Tylototriton_lizhenchangi | Tylototriton_lizhenchangi | CR | Decreasing | temperate | nocturnal | 4.183 | 16.902 | 13.378 |
| amphibians | Tylototriton_phukhaensis | NA | DD | Unknown | NA | NA | NA | NA | NA |
| amphibians | Tylototriton_pseudoverrucosus | Tylototriton_pseudoverrucosus | EN | Decreasing | temperate | cathemeral | 6.942 | 389.038 | 13.931 |
| amphibians | Tylototriton_shanjing | Tylototriton_shanjing | VU | Decreasing | temperate | nocturnal | 11.177 | 125306.092 | 11.363 |
| amphibians | Tylototriton_sparreboomi | NA | DD | Decreasing | NA | NA | NA | NA | NA |
| amphibians | Tylototriton_taliangensis | Tylototriton_taliangensis | VU | Decreasing | temperate | nocturnal | 21.307 | 29860.176 | 11.394 |
| amphibians | Tylototriton_uyenoi | Tylototriton_uyenoi | LC | Unknown | tropical | nocturnal | 7.637 | 13094.234 | 6.881 |
| amphibians | Tylototriton_verrucosus | Tylototriton_verrucosus | NT | Decreasing | temperate | cathemeral | 15.677 | 25986.713 | 10.119 |
| amphibians | Tylototriton_wenxianensis | Tylototriton_wenxianensis | VU | Decreasing | temperate | nocturnal | 3.701 | 43334.526 | 14.819 |
| amphibians | Tylototriton_yangi | Tylototriton_yangi | EN | Decreasing | tropical | nocturnal | 13.819 | 10452.153 | 14.12 |
| amphibians | Tylototriton_zaimeng | NA | not_available | not_available | NA | NA | NA | NA | NA |
| amphibians | Typhlonectes_compressicauda | Typhlonectes_compressicauda | LC | Stable | tropical | nocturnal | 195.614 | 3464909.432 | 1.355 |
| amphibians | Typhlonectes_natans | Typhlonectes_natans | LC | Stable | tropical | nocturnal | 95.077 | 180513.758 | 14.904 |
| amphibians | Uperodon_globulosus | Uperodon_globulosus | LC | Decreasing | tropical | nocturnal | 59.267 | 1828141.707 | 18.817 |
| amphibians | Uperodon_nagaoi | Ramanella_nagaoi | EN | Decreasing | tropical | nocturnal | 2.47 | 3295.809 | 15.842 |
| amphibians | Uperodon_systoma | Uperodon_systoma | LC | Stable | tropical | nocturnal | 32.83 | 2471095.806 | 18.947 |
| amphibians | Uperodon_taprobanicus | Kaloula_taprobanica | LC | Decreasing | tropical | nocturnal | 41.053 | 2194606.957 | 18.513 |
| amphibians | Uperoleia_aspera | Uperoleia_aspera | LC | Stable | tropical | nocturnal | 3.163 | 77683.766 | 3.374 |
| amphibians | Uperoleia_daviesae | Uperoleia_daviesae | EN | Decreasing | tropical | nocturnal | 0.772 | 296.213 | 9.9 |
| amphibians | Uperoleia_laevigata | Uperoleia_laevigata | LC | Decreasing | temperate | nocturnal | 2.599 | 546414.686 | 8.031 |
| amphibians | Uperoleia_micra | Uperoleia_micra | LC | Stable | tropical | nocturnal | 0.772 | 18623.505 | 0.698 |
| amphibians | Uperoleia_orientalis | NA | DD | Unknown | NA | NA | NA | NA | NA |
| amphibians | Vitreorana_eurygnatha | Vitreorana_eurygnatha | LC | Unknown | tropical | nocturnal | 1.023 | 630688.309 | 12.127 |
| amphibians | Vitreorana_franciscana | NA | LC | Stable | NA | NA | NA | NA | NA |
| amphibians | Vitreorana_gorzulae | Vitreorana_gorzulae | LC | Unknown | tropical | nocturnal | 0.83 | 23059.109 | 1.42 |
| amphibians | Vitreorana_helenae | Vitreorana_helenae | VU | Unknown | tropical | nocturnal | 0.604 | 643.002 | 2.632 |
| amphibians | Vitreorana_ritae | Cochranella_ritae | LC | Stable | tropical | nocturnal | 1.023 | 3428488.123 | 1.34 |

| | | | | | | | | | |
|---|---|---|---|---|---|---|---|---|---|
| amphibians | Vitreorana_uranoscopa | Vitreorana_uranoscopa | LC | Unknown | tropical | nocturnal | 1.499 | 613709.591 | 13.11 |
| amphibians | Wakea_madinika | NA | DD | Unknown | NA | NA | NA | NA | NA |
| amphibians | Walkerana_leptodactyla | Indirana_leptodactyla | VU | Decreasing | tropical | nocturnal | 7.844 | 4472.588 | 9.271 |
| amphibians | Walkerana_phrynoderma | Indirana_phrynoderma | EN | Decreasing | tropical | diurnal | 3.475 | 342.905 | 8.692 |
| amphibians | Werneria_mertensiana | Werneria_mertensiana | CR | Decreasing | tropical | diurnal | 9.669 | 714.72 | 10.501 |
| amphibians | Werneria_preussi | Werneria_preussi | EN | Decreasing | tropical | cathemeral | 10.474 | 918.665 | 12.303 |
| amphibians | Werneria_submontana | Werneria_submontana | EN | Decreasing | tropical | diurnal | 7.844 | 107.951 | 11.01 |
| amphibians | Wijayarana_masonii | Huia_masonii | LC | Decreasing | tropical | nocturnal | 26.734 | 29549.58 | 17.874 |
| amphibians | Wolterstorffina_parvipalmata | Wolterstorffina_parvipalmata | CR | Decreasing | tropical | nocturnal | 4.934 | 2466.475 | 13.073 |
| amphibians | Xenohyla_truncata | Xenohyla_truncata | VU | Decreasing | tropical | nocturnal | 11.036 | 1925.555 | 26.316 |
| amphibians | Xenophrys_auralensis | NA | VU | Decreasing | NA | NA | NA | NA | NA |
| amphibians | Xenophrys_awuh | NA | VU | Decreasing | NA | NA | NA | NA | NA |
| amphibians | Xenophrys_damrei | NA | CR | Decreasing | NA | NA | NA | NA | NA |
| amphibians | Xenophrys_flavipunctata | NA | EN | Decreasing | NA | NA | NA | NA | NA |
| amphibians | Xenophrys_glandulosa | NA | LC | Decreasing | NA | NA | NA | NA | NA |
| amphibians | Xenophrys_himalayana | NA | EN | Decreasing | NA | NA | NA | NA | NA |
| amphibians | Xenophrys_major | NA | LC | Stable | NA | NA | NA | NA | NA |
| amphibians | Xenophrys_medogensis | NA | EN | Decreasing | NA | NA | NA | NA | NA |
| amphibians | Xenophrys_megacephala | Megophrys_megacephala | EN | Decreasing | temperate | diurnal | 25.058 | 34.411 | 13.921 |
| amphibians | Xenophrys_periosa | NA | EN | Decreasing | NA | NA | NA | NA | NA |
| amphibians | Xenophrys_robusta | NA | LC | Decreasing | NA | NA | NA | NA | NA |
| amphibians | Xenophrys_zunhebotoensis | Xenophrys_zunhebotoensis | EN | Decreasing | temperate | diurnal | 4.934 | 129.575 | 14.126 |
| amphibians | Xenopus_andrei | Xenopus_andrei | LC | Unknown | tropical | nocturnal | 13.329 | 251791.212 | 5.159 |
| amphibians | Xenopus_borealis | Xenopus_borealis | LC | Unknown | tropical | nocturnal | 88.303 | 149364.56 | 13.861 |
| amphibians | Xenopus_laevis | Xenopus_laevis | LC | Increasing | tropical | cathemeral | 363.295 | 3812401.029 | 8.467 |
| amphibians | Xenopus_muelleri | Xenopus_muelleri | LC | Stable | tropical | nocturnal | 74.113 | 3295891.82 | 9.552 |
| amphibians | Xenopus_tropicalis | Xenopus_tropicalis | LC | Stable | tropical | nocturnal | 18.867 | 1853559.298 | 13.772 |
| amphibians | Xenopus_wittei | Xenopus_wittei | LC | Unknown | tropical | nocturnal | 21.019 | 24772.99 | 18.307 |
| amphibians | Xenorhina_arboricola | Xenorhina_arboricola | LC | Stable | tropical | nocturnal | 8.423 | 1200.285 | 2.461 |
| amphibians | Xenorhina_rostrata | Xenorhina_rostrata | LC | Stable | tropical | diurnal | 14.161 | 69082.953 | 4.178 |
| amphibians | Xenorhina_salawati | NA | not_available | not_available | NA | NA | NA | NA | NA |
| amphibians | Xenorhina_varia | NA | DD | Unknown | NA | NA | NA | NA | NA |
| amphibians | Xenorhina_waigeo | NA | not_available | not_available | NA | NA | NA | NA | NA |
| amphibians | Xenorhina_zweifeli | Xenorhina_zweifeli | LC | Unknown | tropical | diurnal | 4.536 | 63.001 | 1.99 |

| | | | | | | | | | |
|---|---|---|---|---|---|---|---|---|---|
| amphibians | Yunganastes_bisignatus | Pristimantis_bisignatus | EN | Decreasing | tropical | nocturnal | 3.475 | 826.691 | 8.032 |
| amphibians | Yunganastes_mercedesae | Pristimantis_mercedesae | LC | Unknown | tropical | cathemeral | 23.697 | 37297.978 | 4.937 |
| amphibians | Yunganastes_pluvicanorus | Pristimantis_pluvicanorus | VU | Decreasing | tropical | cathemeral | 23.335 | 3833.481 | 3.336 |
| amphibians | Zhangixalus_feae | Rhacophorus_feae | LC | Decreasing | temperate | nocturnal | 146.217 | 1042361.557 | 10.258 |
| amphibians | Zhangixalus_franki | NA | DD | Unknown | NA | NA | NA | NA | NA |
| amphibians | Zhangixalus_jodiae | NA | not_available | not_available | NA | NA | NA | NA | NA |
| amphibians | Zhangixalus_omeimontis | Rhacophorus_omeimontis | LC | Stable | temperate | nocturnal | 50.601 | 167218.047 | 13.213 |
| amphibians | Zhangixalus_schlegelii | Rhacophorus_schlegelii | LC | Decreasing | temperate | diurnal | 23.576 | 286417.985 | 21.104 |
| birds | Abeillia_abeillei | Abeillia_abeillei | LC | Decreasing | tropical | diurnal | 2.7 | 144610.397 | 11.991 |
| birds | Abroscopus_albogularis | Abroscopus_albogularis | LC | Stable | tropical | diurnal | 4.8 | 1704650.599 | 11.606 |
| birds | Abroscopus_schisticeps | Abroscopus_schisticeps | LC | Stable | temperate | diurnal | 4.7 | 909217.045 | 8.31 |
| birds | Abroscopus_superciliaris | Abroscopus_superciliaris | LC | Stable | tropical | diurnal | 6.5 | 2493437.473 | 10.208 |
| birds | Aburria_aburri | Aburria_aburri | LC | Decreasing | tropical | diurnal | 1405.1 | 354555.416 | 8.929 |
| birds | Acanthagenys_rufogularis | Acanthagenys_rufogularis | LC | Decreasing | temperate | diurnal | 47.8 | 5976565.195 | 4.45 |
| birds | Acanthidops_bairdi | Acanthidops_bairdii | LC | Stable | tropical | diurnal | 16 | 4743.279 | 5.427 |
| birds | Acanthis_flammea | Carduelis_flammea | LC | Decreasing | temperate | diurnal | 13 | 42235480.516 | 6.388 |
| birds | Acanthisitta_chloris | Acanthisitta_chloris | LC | Decreasing | temperate | diurnal | 6.9 | 152503.845 | 6.456 |
| birds | Acanthiza_chrysorrhoa | Acanthiza_chrysorrhoa | LC | Decreasing | temperate | diurnal | 9.3 | 4590956.335 | 5.394 |
| birds | Acanthiza_cinerea | Gerygone_cinerea | LC | Stable | tropical | diurnal | 7.6 | 124509.89 | 5.044 |
| birds | Acanthiza_ewingii | Acanthiza_ewingii | LC | Decreasing | temperate | diurnal | 6.9 | 68050.509 | 5.966 |
| birds | Acanthiza_inornata | Acanthiza_inornata | LC | Decreasing | temperate | diurnal | 7 | 137406.16 | 11.325 |
| birds | Acanthiza_iredalei | Acanthiza_iredalei | LC | Decreasing | temperate | diurnal | 6 | 987131.547 | 5.861 |
| birds | Acanthiza_katherina | Acanthiza_katherina | VU | Decreasing | tropical | diurnal | 7.2 | 6563.027 | 2.915 |
| birds | Acanthiza_lineata | Acanthiza_lineata | LC | Decreasing | temperate | diurnal | 6.9 | 1043817.354 | 9.539 |
| birds | Acanthiza_murina | Acanthiza_murina | LC | Stable | tropical | diurnal | 8.4 | 84339.375 | 5.223 |
| birds | Acanthiza_nana | Acanthiza_nana | LC | Decreasing | temperate | diurnal | 6.2 | 2021416.131 | 7.73 |
| birds | Acanthiza_pusilla | Acanthiza_pusilla | LC | Stable | temperate | diurnal | 7.6 | 1131818.686 | 9.206 |
| birds | Acanthiza_reguloides | Acanthiza_reguloides | LC | Decreasing | temperate | diurnal | 7.7 | 1843422.3 | 7.897 |
| birds | Acanthiza_robustirostris | Acanthiza_robustirostris | LC | Stable | temperate | diurnal | 6.1 | 2061848.423 | 2.518 |
| birds | Acanthiza_uropygialis | Acanthiza_uropygialis | LC | Decreasing | temperate | diurnal | 6.4 | 4788831.671 | 4.729 |
| birds | Acanthorhynchus_superciliosus | Acanthorhynchus_superciliosus | LC | Decreasing | temperate | diurnal | 9.7 | 182218.537 | 10.398 |
| birds | Acanthorhynchus_tenuirostris | Acanthorhynchus_tenuirostris | LC | Decreasing | temperate | diurnal | 10.4 | 763502.742 | 9.082 |
| birds | Acanthornis_magna | Acanthornis_magna | LC | Decreasing | temperate | diurnal | 10.2 | 45032.529 | 5.767 |
| birds | Accipiter_albogularis | Accipiter_albogularis | LC | Stable | tropical | diurnal | 248.8 | 37461.21 | 4.678 |

| | | | | | | | | | |
|---|---|---|---|---|---|---|---|---|---|
| birds | Accipiter_badius | Accipiter_badius | LC | Stable | temperate | diurnal | 131.2 | 22853794.982 | 11.254 |
| birds | Accipiter_bicolor | Accipiter_bicolor | LC | Decreasing | tropical | diurnal | 287.5 | 14309701.269 | 5.772 |
| birds | Accipiter_brachyurus | Accipiter_brachyurus | VU | Decreasing | tropical | diurnal | 142 | 35580.708 | 4.161 |
| birds | Accipiter_brevipes | Accipiter_brevipes | LC | Stable | temperate | diurnal | 186.5 | 7225515.306 | 11.192 |
| birds | Accipiter_butleri | Accipiter_butleri | VU | Decreasing | tropical | diurnal | 122 | 478.819 | 9.702 |
| birds | Accipiter_castanilius | Accipiter_castanilius | LC | Decreasing | tropical | diurnal | 157.5 | 2096401.626 | 6.165 |
| birds | Accipiter_cirrocephalus | Accipiter_cirrocephalus | LC | Stable | temperate | diurnal | 165.7 | 5488344.154 | 5.28 |
| birds | Accipiter_collaris | Accipiter_collaris | LC | Stable | tropical | diurnal | 97.8 | 274415.489 | 8.105 |
| birds | Accipiter_cooperii | Accipiter_cooperii | LC | Increasing | temperate | diurnal | 429.7 | 10487652.985 | 12.797 |
| birds | Accipiter_erythrauchen | Accipiter_erythrauchen | NT | Decreasing | tropical | diurnal | 156 | 48150.322 | 6.55 |
| birds | Accipiter_erythropus | Accipiter_erythropus | LC | Decreasing | tropical | diurnal | 111.2 | 3904812.091 | 8.885 |
| birds | Accipiter_fasciatus | Accipiter_fasciatus | LC | Decreasing | temperate | diurnal | 358.7 | 8180492.011 | 4.349 |
| birds | Accipiter_francesiae | Accipiter_francesiae | LC | Stable | tropical | diurnal | 136.8 | 519468.805 | 8.536 |
| birds | Accipiter_gentilis | Accipiter_gentilis | LC | Unknown | temperate | diurnal | 866 | 36029138.677 | 7.818 |
| birds | Accipiter_griseiceps | Accipiter_griseiceps | LC | Decreasing | tropical | diurnal | 251.8 | 140604.936 | 7.923 |
| birds | Accipiter_gularis | Accipiter_gularis | LC | Stable | temperate | diurnal | 120.7 | 10101618.367 | 11.067 |
| birds | Accipiter_gundlachi | Accipiter_gundlachi | EN | Decreasing | tropical | diurnal | 675 | 106868.731 | 16.287 |
| birds | Accipiter_haplochrous | Accipiter_haplochrous | NT | Decreasing | tropical | diurnal | 198.4 | 16389.595 | 7.513 |
| birds | Accipiter_henicogrammus | NA | NT | Decreasing | NA | NA | NA | NA | NA |
| birds | Accipiter_henstii | Accipiter_henstii | VU | Decreasing | tropical | diurnal | 1050 | 434639.698 | 8.137 |
| birds | Accipiter_imitator | Accipiter_imitator | NT | Decreasing | tropical | diurnal | 222.5 | 10050.207 | 4.985 |
| birds | Accipiter_luteoschistaceus | Accipiter_luteoschistaceus | VU | Decreasing | tropical | diurnal | 214 | 31056.514 | 4.055 |
| birds | Accipiter_madagascariensis | Accipiter_madagascariensis | NT | Decreasing | tropical | diurnal | 180.4 | 476517.512 | 8.603 |
| birds | Accipiter_melanochlamys | Accipiter_melanochlamys | LC | Stable | tropical | diurnal | 265 | 115449.938 | 5.234 |
| birds | Accipiter_melanoleucus | Accipiter_melanoleucus | LC | Decreasing | tropical | diurnal | 699 | 9071924.49 | 9.542 |
| birds | Accipiter_meyerianus | Accipiter_meyerianus | LC | Decreasing | tropical | diurnal | 763.5 | 269144.413 | 4.951 |
| birds | Accipiter_minullus | Accipiter_minullus | LC | Stable | tropical | diurnal | 87.4 | 6794381.579 | 9.267 |
| birds | Accipiter_nanus | NA | LC | Stable | NA | NA | NA | NA | NA |
| birds | Accipiter_nisus | Accipiter_nisus | LC | Stable | temperate | diurnal | 220.8 | 36695930.299 | 10.269 |
| birds | Accipiter_novaehollandiae | Accipiter_novaehollandiae | LC | Decreasing | temperate | diurnal | 326.1 | 1807067.302 | 6.691 |
| birds | Accipiter_ovampensis | Accipiter_ovampensis | LC | Increasing | tropical | diurnal | 195 | 7295779.859 | 9.687 |
| birds | Accipiter_poliocephalus | Accipiter_poliocephalus | LC | Stable | tropical | diurnal | 214.8 | 671500.51 | 3.06 |
| birds | Accipiter_poliogaster | Accipiter_poliogaster | NT | Decreasing | tropical | diurnal | 315 | 9760988.496 | 4.821 |
| birds | Accipiter_princeps | NA | VU | Decreasing | NA | NA | NA | NA | NA |

| | | | | | | | | | |
|---|---|---|---|---|---|---|---|---|---|
| birds | Accipiter_rhodogaster | Accipiter_rhodogaster | LC | Decreasing | tropical | diurnal | 172.7 | 146036.851 | 7.556 |
| birds | Accipiter_rufitorques | Accipiter_rufitorques | LC | Stable | tropical | diurnal | 209 | 17581.072 | 9.386 |
| birds | Accipiter_rufiventris | Accipiter_rufiventris | LC | Increasing | tropical | diurnal | 198 | 756057.211 | 13.03 |
| birds | Accipiter_soloensis | Accipiter_soloensis | LC | Decreasing | temperate | diurnal | 164.8 | 4004394.764 | 15.235 |
| birds | Accipiter_striatus | Accipiter_striatus | LC | Increasing | temperate | diurnal | 130.6 | 21636790.051 | 9.561 |
| birds | Accipiter_superciliosus | Accipiter_superciliosus | LC | Decreasing | tropical | diurnal | 96.4 | 8560355.565 | 4.061 |
| birds | Accipiter_trinotatus | Accipiter_trinotatus | LC | Stable | tropical | diurnal | 146 | 120238.86 | 7.489 |
| birds | Accipiter_trivirgatus | Accipiter_trivirgatus | LC | Decreasing | tropical | diurnal | 265 | 3366430.691 | 10.504 |
| birds | Accipiter_virgatus | Accipiter_virgatus | LC | Decreasing | temperate | diurnal | 117 | 4398429.706 | 12.368 |
| birds | Aceros_nipalensis | Aceros_nipalensis | VU | Decreasing | tropical | diurnal | 2385 | 484788.186 | 8.043 |
| birds | Achaetops_pycnopygius | Achaetops_pycnopygius | LC | Stable | tropical | diurnal | 28.3 | 423641.183 | 6.986 |
| birds | Acridotheres_albocinctus | NA | LC | Unknown | NA | NA | NA | NA | NA |
| birds | Acridotheres_burmannicus | NA | LC | Increasing | NA | NA | NA | NA | NA |
| birds | Acridotheres_cinereus | NA | VU | Decreasing | NA | NA | NA | NA | NA |
| birds | Acridotheres_cristatellus | Acridotheres_cristatellus | LC | Stable | tropical | diurnal | 116.3 | 3225824.494 | 14.577 |
| birds | Acridotheres_fuscus | Acridotheres_fuscus | LC | Decreasing | tropical | diurnal | 82.8 | 2422718.947 | 16.287 |
| birds | Acridotheres_ginginianus | Acridotheres_ginginianus | LC | Increasing | temperate | diurnal | 72 | 1641344.881 | 19.152 |
| birds | Acridotheres_grandis | Acridotheres_grandis | LC | Stable | tropical | diurnal | 98.9 | 1707140.604 | 13.382 |
| birds | Acridotheres_melanopterus | Sturnus_melanopterus | EN | Decreasing | tropical | diurnal | 89 | 136022.663 | 22.522 |
| birds | Acridotheres_tristis | Acridotheres_tristis | LC | Increasing | temperate | diurnal | 116.4 | 10019063.907 | 13.001 |
| birds | Acritillas_indica | Iole_indica | LC | Stable | tropical | diurnal | 30.6 | 219041.392 | 16.687 |
| birds | Acrobatornis_fonsecai | Acrobatornis_fonsecai | VU | Decreasing | tropical | diurnal | 13.7 | 3689.543 | 8.529 |
| birds | Acrocephalus_aequinoctialis | Acrocephalus_aequinoctialis | EN | Decreasing | tropical | diurnal | 25 | 271.489 | 12.633 |
| birds | Acrocephalus_agricola | Acrocephalus_agricola | LC | Decreasing | temperate | diurnal | 9.6 | 7862832.834 | 8.35 |
| birds | Acrocephalus_arundinaceus | Acrocephalus_arundinaceus | LC | Decreasing | temperate | diurnal | 30 | 36259434.41 | 8.7 |
| birds | Acrocephalus_atyphus | Acrocephalus_atyphus | LC | Stable | tropical | diurnal | 30 | 611.157 | 3.761 |
| birds | Acrocephalus_australis | Acrocephalus_australis | LC | Stable | temperate | diurnal | 17.7 | 6368774.963 | 4.767 |
| birds | Acrocephalus_bistrigiceps | Acrocephalus_bistrigiceps | LC | Stable | temperate | diurnal | 8.7 | 6253044.538 | 13.064 |
| birds | Acrocephalus_brevipennis | Acrocephalus_brevipennis | VU | Unknown | tropical | diurnal | 16 | 1388.989 | 14.499 |
| birds | Acrocephalus_caffer | Acrocephalus_caffer | VU | Stable | tropical | diurnal | 43.8 | 266.98 | 11.494 |
| birds | Acrocephalus_concinens | Acrocephalus_concinens | LC | Stable | temperate | diurnal | 8.3 | 4244812.465 | 15.71 |
| birds | Acrocephalus_dumetorum | Acrocephalus_dumetorum | LC | Increasing | temperate | diurnal | 11.2 | 6977440.849 | 9.742 |
| birds | Acrocephalus_familiaris | Acrocephalus_familiaris | EN | Unknown | temperate | diurnal | 18.3 | 4.803 | 0 |
| birds | Acrocephalus_gracilirostris | Acrocephalus_gracilirostris | LC | Stable | tropical | diurnal | 14.4 | 4809980.002 | 10.371 |

| | | | | | | | | | |
|---|---|---|---|---|---|---|---|---|---|
| birds | Acrocephalus_griseldis | Acrocephalus_griseldis | EN | Stable | temperate | diurnal | 16.7 | 693358.246 | 12.814 |
| birds | Acrocephalus_kerearako | Acrocephalus_kerearako | NT | Decreasing | tropical | diurnal | 22.5 | 73.236 | 9.824 |
| birds | Acrocephalus_luscinius | NA | EX | not_available | NA | NA | NA | NA | NA |
| birds | Acrocephalus_melanopogon | Acrocephalus_melanopogon | LC | Stable | temperate | diurnal | 10.9 | 6177192.638 | 13.062 |
| birds | Acrocephalus_mendanae | Acrocephalus_mendanae | LC | Stable | tropical | diurnal | 36.3 | 593.467 | 7.099 |
| birds | Acrocephalus_newtoni | Acrocephalus_newtoni | LC | Stable | tropical | diurnal | 17 | 591120.506 | 8.749 |
| birds | Acrocephalus_paludicola | Acrocephalus_paludicola | VU | Decreasing | temperate | diurnal | 11.6 | 9180072.4 | 13.284 |
| birds | Acrocephalus_palustris | Acrocephalus_palustris | LC | Stable | tropical | diurnal | 11.5 | 18399649.063 | 11.566 |
| birds | Acrocephalus_rehsei | NA | NT | Unknown | NA | NA | NA | NA | NA |
| birds | Acrocephalus_rimitarae | NA | CR | Decreasing | NA | NA | NA | NA | NA |
| birds | Acrocephalus_rodericanus | NA | NT | Increasing | NA | NA | NA | NA | NA |
| birds | Acrocephalus_rufescens | Acrocephalus_rufescens | LC | Stable | tropical | diurnal | 22.7 | 1400095.073 | 11.291 |
| birds | Acrocephalus_schoenobaenus | Acrocephalus_schoenobaenus | LC | Stable | temperate | diurnal | 11.8 | 39149311.527 | 8.147 |
| birds | Acrocephalus_scirpaceus | Acrocephalus_scirpaceus | LC | Stable | tropical | diurnal | 12.3 | 41060299.185 | 8.984 |
| birds | Acrocephalus_sechellensis | NA | NT | Increasing | NA | NA | NA | NA | NA |
| birds | Acrocephalus_sorghophilus | Acrocephalus_sorghophilus | CR | Decreasing | temperate | diurnal | 8.2 | 206086.235 | 17.033 |
| birds | Acrocephalus_stentoreus | Acrocephalus_stentoreus | LC | Stable | temperate | diurnal | 24.2 | 8982425.753 | 13.181 |
| birds | Acrocephalus_syrinx | Acrocephalus_syrinx | LC | Stable | tropical | diurnal | 34 | 577.834 | 12.028 |
| birds | Acrocephalus_taiti | Acrocephalus_taiti | VU | Stable | temperate | diurnal | 24.3 | 44.552 | 7.723 |
| birds | Acrocephalus_tangorum | Acrocephalus_tangorum | VU | Decreasing | temperate | diurnal | 8 | 973843.457 | 8.078 |
| birds | Acrocephalus_vaughani | NA | EN | Decreasing | NA | NA | NA | NA | NA |
| birds | Acropternis_orthonyx | Acropternis_orthonyx | LC | Stable | tropical | diurnal | 89.9 | 180173.728 | 11.368 |
| birds | Acryllium_vulturinum | Acryllium_vulturinum | LC | Stable | tropical | diurnal | 1330 | 1226666.244 | 8.38 |
| birds | Actenoides_bougainvillei | Actenoides_bougainvillei | LC | Decreasing | tropical | diurnal | 188 | 3784.544 | 3.856 |
| birds | Actenoides_concretus | Actenoides_concretus | NT | Decreasing | tropical | diurnal | 73.5 | 1364096.242 | 8.27 |
| birds | Actenoides_hombroni | Actenoides_hombroni | VU | Decreasing | tropical | diurnal | 117 | 24714.25 | 11.191 |
| birds | Actenoides_lindsayi | Actenoides_lindsayi | LC | Decreasing | tropical | diurnal | 92.8 | 106620.851 | 13.704 |
| birds | Actenoides_monachus | Actenoides_monachus | NT | Decreasing | tropical | diurnal | 150 | 72249.862 | 8.528 |
| birds | Actenoides_princeps | Actenoides_princeps | LC | Decreasing | tropical | diurnal | 105 | 48342.796 | 5.527 |
| birds | Actinodura_egertoni | Actinodura_egertoni | LC | Decreasing | temperate | diurnal | 36 | 262348.958 | 8.264 |
| birds | Actinodura_ramsayi | Actinodura_ramsayi | LC | Decreasing | tropical | diurnal | 38.4 | 58116.07 | 8.727 |
| birds | Actitis_hypoleucos | Actitis_hypoleucos | LC | Decreasing | temperate | diurnal | 48 | 61178024.784 | 8.894 |
| birds | Actitis_macularius | Actitis_macularius | LC | Decreasing | tropical | diurnal | 40.4 | 34169893.459 | 7.388 |
| birds | Actophilornis_africanus | Actophilornis_africanus | LC | Stable | tropical | diurnal | 189.1 | 16988448.659 | 9.259 |

| | | | | | | | | | |
|---|---|---|---|---|---|---|---|---|---|
| birds | Actophilornis_albinucha | Actophilornis_albinucha | EN | Decreasing | tropical | diurnal | 239 | 140285.771 | 8.143 |
| birds | Adelomyia_melanogenys | Adelomyia_melanogenys | LC | Unknown | tropical | diurnal | 4.9 | 620219.308 | 8.204 |
| birds | Aechmophorus_clarkii | Aechmophorus_clarkii | LC | Decreasing | temperate | diurnal | 1090.9 | 3589069.346 | 10.556 |
| birds | Aechmophorus_occidentalis | Aechmophorus_occidentalis | LC | Stable | temperate | diurnal | 1137 | 6366407.789 | 10.636 |
| birds | Aegithalos_caudatus | Aegithalos_caudatus | LC | Stable | temperate | diurnal | 8.6 | 18425691.873 | 10.501 |
| birds | Aegithalos_concinnus | Aegithalos_concinnus | LC | Stable | temperate | diurnal | 6.1 | 2797505.093 | 13.493 |
| birds | Aegithalos_fuliginosus | Aegithalos_fuliginosus | LC | Stable | temperate | diurnal | 5.7 | 216742.865 | 9.367 |
| birds | Aegithalos_iouschistos | Aegithalos_iouschistos | LC | Stable | temperate | diurnal | 7 | 68201.139 | 5.49 |
| birds | Aegithalos_leucogenys | Aegithalos_leucogenys | LC | Stable | temperate | diurnal | 6.7 | 289366.029 | 11.556 |
| birds | Aegithalos_niveogularis | Aegithalos_niveogularis | LC | Stable | temperate | diurnal | 7.8 | 180594.846 | 10.786 |
| birds | Aegithina_lafresnayei | Aegithina_lafresnayei | LC | Unknown | tropical | diurnal | 12 | 1675913.709 | 12.95 |
| birds | Aegithina_nigrolutea | Aegithina_nigrolutea | LC | Unknown | temperate | diurnal | 11.8 | 714297.742 | 18.427 |
| birds | Aegithina_tiphia | Aegithina_tiphia | LC | Unknown | tropical | diurnal | 12 | 6356008.515 | 14.205 |
| birds | Aegithina_viridissima | Aegithina_viridissima | NT | Decreasing | tropical | diurnal | 13.4 | 1109607.12 | 7.892 |
| birds | Aegolius_acadicus | Aegolius_acadicus | LC | Decreasing | temperate | nocturnal | 100.7 | 10821011.005 | 11.838 |
| birds | Aegolius_funereus | Aegolius_funereus | LC | Stable | temperate | cathemeral | 137.2 | 23807935.254 | 4.576 |
| birds | Aegolius_harrisii | Aegolius_harrisii | LC | Stable | temperate | nocturnal | 120 | 1098550.772 | 10.498 |
| birds | Aegolius_ridgwayi | Aegolius_ridgwayi | LC | Stable | tropical | nocturnal | 80 | 43992.748 | 13.405 |
| birds | Aegotheles_albertisi | Aegotheles_albertisi | LC | Stable | tropical | nocturnal | 35.2 | 168432.172 | 4.71 |
| birds | Aegotheles_archboldi | Aegotheles_archboldi | LC | Stable | tropical | nocturnal | 30.6 | 62391.493 | 5.069 |
| birds | Aegotheles_bennettii | Aegotheles_bennettii | LC | Decreasing | tropical | nocturnal | 60 | 198612.166 | 4.099 |
| birds | Aegotheles_crinifrons | Aegotheles_crinifrons | LC | Decreasing | tropical | nocturnal | 136 | 20463.319 | 6.526 |
| birds | Aegotheles_cristatus | Aegotheles_cristatus | LC | Stable | temperate | nocturnal | 42.8 | 7702773.064 | 4.26 |
| birds | Aegotheles_insignis | Aegotheles_insignis | LC | Stable | tropical | nocturnal | 78.5 | 168011.219 | 4.661 |
| birds | Aegotheles_savesi | NA | CR | Decreasing | NA | NA | NA | NA | NA |
| birds | Aegotheles_tatei | Aegotheles_tatei | LC | Decreasing | tropical | nocturnal | 52 | 84978.497 | 2.149 |
| birds | Aegotheles_wallacii | Aegotheles_wallacii | LC | Decreasing | tropical | nocturnal | 50.7 | 40086.297 | 2.991 |
| birds | Aegypius_monachus | Aegypius_monachus | NT | Decreasing | temperate | diurnal | 9320.6 | 17944995.136 | 9.506 |
| birds | Aepypodius_arfakianus | Aepypodius_arfakianus | LC | Stable | tropical | diurnal | 1463.8 | 195287.344 | 4.407 |
| birds | Aepypodius_bruijnii | Aepypodius_bruijnii | EN | Decreasing | tropical | diurnal | 192 | 1101.785 | 2.576 |
| birds | Aerodramus_elaphrus | NA | VU | Stable | NA | NA | NA | NA | NA |
| birds | Aerodramus_francicus | Collocalia_francica | NT | Decreasing | tropical | diurnal | 8.9 | 4418.342 | 22.866 |
| birds | Aerodramus_hirundinaceus | Collocalia_hirundinacea | LC | Stable | tropical | diurnal | 10.1 | 583622.073 | 3.684 |
| birds | Aerodramus_infuscatus | Collocalia_infuscata | LC | Decreasing | tropical | diurnal | 13.4 | 182567.315 | 8.682 |

| | | | | | | | | | |
|---|---|---|---|---|---|---|---|---|---|
| birds | Aerodramus_inquietus | Collocalia_inquieta | LC | Stable | tropical | diurnal | 11 | 543.527 | 12.146 |
| birds | Aerodramus_leucophaeus | Collocalia_leucophaea | LC | Stable | tropical | diurnal | 13.4 | 1414.183 | 9.495 |
| birds | Aerodramus_maximus | Collocalia_maxima | LC | Decreasing | tropical | nocturnal | 17 | 1057235.003 | 7.988 |
| birds | Aerodramus_mearnsi | Collocalia_mearnsi | LC | Stable | tropical | diurnal | 11.3 | 254011.615 | 13.87 |
| birds | Aerodramus_papuensis | NA | DD | Unknown | NA | NA | NA | NA | NA |
| birds | Aerodramus_spodiopygius | Collocalia_spodiopygia | LC | Stable | tropical | diurnal | 8 | 136466.16 | 5.701 |
| birds | Aerodramus_terraereginae | Collocalia_terraereginae | LC | Stable | tropical | diurnal | 12 | 85667.061 | 3.914 |
| birds | Aerodramus_unicolor | Collocalia_unicolor | LC | Decreasing | tropical | cathemeral | 11 | 248834.493 | 19.288 |
| birds | Aerodramus_vanikorensis | Collocalia_vanikorensis | LC | Stable | tropical | diurnal | 10.7 | 1212501.772 | 6.141 |
| birds | Aeronautes_andecolus | Aeronautes_andecolus | LC | Stable | tropical | diurnal | 17.6 | 891310.572 | 6.623 |
| birds | Aeronautes_montivagus | Aeronautes_montivagus | LC | Stable | tropical | diurnal | 20.1 | 842701.638 | 6.407 |
| birds | Aeronautes_saxatalis | Aeronautes_saxatalis | LC | Decreasing | temperate | diurnal | 32.1 | 4032383.431 | 9.143 |
| birds | Aethia_cristatella | Aethia_cristatella | LC | Decreasing | temperate | diurnal | 259.4 | 8096883.711 | 2.131 |
| birds | Aethia_psittacula | Aethia_psittacula | LC | Decreasing | temperate | diurnal | 270 | 19779368.725 | 1.24 |
| birds | Aethia_pusilla | Aethia_pusilla | LC | Decreasing | temperate | diurnal | 84 | 4834897.864 | 4.647 |
| birds | Aethia_pygmaea | Aethia_pygmaea | LC | Decreasing | temperate | diurnal | 112 | 1883338.338 | 4.063 |
| birds | Aethomyias_arfakianus | Sericornis_arfakianus | LC | Stable | tropical | diurnal | 8.7 | 181310.736 | 3.73 |
| birds | Aethomyias_nigrorufus | Crateroscelis_nigrorufa | LC | Stable | tropical | diurnal | 15.9 | 105189.031 | 4.985 |
| birds | Aethomyias_papuensis | Sericornis_papuensis | LC | Stable | tropical | diurnal | 11.2 | 86586.534 | 4.935 |
| birds | Aethomyias_perspicillatus | Sericornis_perspicillatus | LC | Stable | tropical | diurnal | 9 | 122449.638 | 4.845 |
| birds | Aethomyias_rufescens | Sericornis_rufescens | LC | Stable | tropical | diurnal | 8.5 | 8367.91 | 2.486 |
| birds | Aethomyias_spilodera | Sericornis_spilodera | LC | Stable | tropical | diurnal | 11.7 | 283199.335 | 3.028 |
| birds | Aethopyga_boltoni | Aethopyga_boltoni | LC | Stable | tropical | diurnal | 7.1 | 7469.316 | 8.529 |
| birds | Aethopyga_christinae | Aethopyga_christinae | LC | Decreasing | tropical | diurnal | 5.2 | 33798.317 | 16.405 |
| birds | Aethopyga_duyvenbodei | Aethopyga_duyvenbodei | EN | Decreasing | tropical | diurnal | 6 | 671.719 | 12.328 |
| birds | Aethopyga_eximia | NA | LC | Stable | NA | NA | NA | NA | NA |
| birds | Aethopyga_flagrans | Aethopyga_flagrans | LC | Decreasing | tropical | diurnal | 6.7 | 103535.946 | 15.036 |
| birds | Aethopyga_gouldiae | Aethopyga_gouldiae | LC | Stable | tropical | diurnal | 5 | 3562401.811 | 10.726 |
| birds | Aethopyga_ignicauda | Aethopyga_ignicauda | LC | Stable | temperate | diurnal | 8.2 | 755258.738 | 8.174 |
| birds | Aethopyga_mystacalis | Aethopyga_mystacalis | LC | Decreasing | tropical | diurnal | 6 | 125331.132 | 22.42 |
| birds | Aethopyga_nipalensis | Aethopyga_nipalensis | LC | Stable | temperate | diurnal | 6.9 | 1011015.385 | 8.245 |
| birds | Aethopyga_primigenia | Aethopyga_primigenia | LC | Stable | tropical | diurnal | 7 | 9971.502 | 8.376 |
| birds | Aethopyga_pulcherrima | Aethopyga_pulcherrima | LC | Stable | tropical | diurnal | 6.2 | 210928.17 | 13.352 |
| birds | Aethopyga_saturata | Aethopyga_saturata | LC | Stable | tropical | diurnal | 5.3 | 1679748.824 | 10.077 |

| | | | | | | | | | |
|---|---|---|---|---|---|---|---|---|---|
| birds | Aethopyga_shelleyi | Aethopyga_shelleyi | LC | Stable | tropical | diurnal | 4 | 13152.407 | 9.3 |
| birds | Aethopyga_siparaja | Aethopyga_siparaja | LC | Stable | tropical | diurnal | 6.7 | 4911554.194 | 11.588 |
| birds | Afropavo_congensis | Afropavo_congensis | NT | Decreasing | tropical | diurnal | 1273.7 | 697505.341 | 4.656 |
| birds | Afrotis_afra | Eupodotis_afra | VU | Decreasing | temperate | diurnal | 690 | 173167.297 | 9.133 |
| birds | Afrotis_afraoides | Eupodotis_afraoides | LC | Stable | temperate | diurnal | 745 | 1934353.366 | 6.701 |
| birds | Agamia_agami | Agamia_agami | NT | Decreasing | tropical | diurnal | 567 | 8217123.754 | 3.379 |
| birds | Agapornis_canus | Agapornis_canus | LC | Stable | tropical | diurnal | 30 | 415982.932 | 8.477 |
| birds | Agapornis_fischeri | Agapornis_fischeri | NT | Decreasing | tropical | diurnal | 48.3 | 162335.086 | 13.211 |
| birds | Agapornis_lilianae | Agapornis_lilianae | NT | Decreasing | tropical | diurnal | 37.3 | 211626.338 | 9.038 |
| birds | Agapornis_nigrigenis | Agapornis_nigrigenis | VU | Decreasing | tropical | diurnal | 40.5 | 17094.248 | 5.473 |
| birds | Agapornis_personatus | Agapornis_personatus | LC | Stable | tropical | diurnal | 52.4 | 233268.943 | 11.449 |
| birds | Agapornis_pullarius | Agapornis_pullarius | LC | Decreasing | tropical | diurnal | 39.3 | 2650930.738 | 9.443 |
| birds | Agapornis_roseicollis | Agapornis_roseicollis | LC | Decreasing | tropical | diurnal | 53.8 | 457223.494 | 6.558 |
| birds | Agapornis_swindernianus | Agapornis_swindernianus | LC | Decreasing | tropical | diurnal | 40 | 1494870.123 | 5.822 |
| birds | Agapornis_taranta | Agapornis_taranta | LC | Increasing | tropical | diurnal | 57.5 | 407732.604 | 13.254 |
| birds | Agelaioides_badius | Agelaioides_badius | LC | Stable | temperate | diurnal | 45.3 | 3734880.19 | 8.714 |
| birds | Agelaius_assimilis | Agelaius_assimilis | LC | Stable | tropical | diurnal | 47.7 | 4077.173 | 6.303 |
| birds | Agelaius_humeralis | Agelaius_humeralis | LC | Stable | tropical | diurnal | 36.3 | 110595.487 | 16.183 |
| birds | Agelaius_phoeniceus | Agelaius_phoeniceus | LC | Decreasing | temperate | diurnal | 50.8 | 14132462.279 | 10.308 |
| birds | Agelaius_tricolor | Agelaius_tricolor | EN | Decreasing | temperate | diurnal | 57.9 | 113509.641 | 18.833 |
| birds | Agelaius_xanthomus | Agelaius_xanthomus | EN | Decreasing | tropical | diurnal | 38.4 | 8726.659 | 27.173 |
| birds | Agelastes_meleagrides | Agelastes_meleagrides | VU | Decreasing | tropical | diurnal | 815 | 52658.795 | 12.399 |
| birds | Agelastes_niger | Agelastes_niger | LC | Decreasing | tropical | diurnal | 700 | 764292.383 | 5.815 |
| birds | Agelasticus_cyanopus | Agelasticus_cyanopus | LC | Stable | tropical | diurnal | 37.2 | 2871356.079 | 8.834 |
| birds | Agelasticus_thilius | Agelasticus_thilius | LC | Stable | temperate | diurnal | 31.5 | 2816597.825 | 8.946 |
| birds | Agelasticus_xanthophthalmus | Agelasticus_xanthophthalmus | LC | Stable | tropical | diurnal | 45.1 | 9690.534 | 2.322 |
| birds | Aglaeactis_aliciae | Aglaeactis_aliciae | VU | Decreasing | tropical | diurnal | 7 | 342.361 | 10.697 |
| birds | Aglaeactis_castelnaudii | Aglaeactis_castelnaudii | NT | Decreasing | tropical | diurnal | 6.7 | 23294.943 | 8.044 |
| birds | Aglaeactis_cupripennis | Aglaeactis_cupripennis | LC | Stable | tropical | diurnal | 7.6 | 224852.989 | 9.594 |
| birds | Aglaeactis_pamela | Aglaeactis_pamela | LC | Stable | tropical | diurnal | 5.5 | 16558.41 | 10.493 |
| birds | Aglaiocercus_berlepschi | Aglaiocercus_berlepschi | VU | Decreasing | tropical | diurnal | 5 | 1391.538 | 8.213 |
| birds | Aglaiocercus_coelestis | Aglaiocercus_coelestis | LC | Stable | tropical | diurnal | 5.6 | 29539.307 | 6.972 |
| birds | Aglaiocercus_kingii | Aglaiocercus_kingi | LC | Decreasing | tropical | diurnal | 4.8 | 267748.933 | 8.462 |
| birds | Agraphospiza_rubescens | NA | LC | Stable | NA | NA | NA | NA | NA |

| | | | | | | | | | |
|---|---|---|---|---|---|---|---|---|---|
| birds | Agricola_infuscatus | Bradornis_infuscatus | LC | Stable | tropical | diurnal | 32.4 | 1446786.715 | 5.909 |
| birds | Agricola_pallidus | Bradornis_pallidus | LC | Stable | tropical | diurnal | 23.6 | 9110671.416 | 10.255 |
| birds | Agriornis_albicauda | Agriornis_albicauda | VU | Decreasing | tropical | diurnal | 53 | 652209.019 | 7.388 |
| birds | Agriornis_lividus | Agriornis_lividus | LC | Stable | temperate | diurnal | 99.1 | 592911.812 | 6.026 |
| birds | Agriornis_micropterus | Agriornis_micropterus | LC | Stable | temperate | diurnal | 67.8 | 2809307.822 | 7.386 |
| birds | Agriornis_montanus | Agriornis_montanus | LC | Stable | temperate | diurnal | 63.3 | 2168659.566 | 6.109 |
| birds | Agriornis_murinus | Agriornis_murinus | LC | Stable | temperate | diurnal | 29.9 | 2112940.186 | 7.557 |
| birds | Agropsar_philippensis | Sturnus_philippensis | LC | Unknown | tropical | diurnal | 28.8 | 568180.199 | 14.467 |
| birds | Agropsar_sturninus | Sturnus_sturninus | LC | Unknown | temperate | diurnal | 49.4 | 8519690.439 | 13.032 |
| birds | Ailuroedus_buccoides | Ailuroedus_buccoides | LC | Decreasing | tropical | diurnal | 138 | 374654.825 | 3.396 |
| birds | Ailuroedus_crassirostris | Ailuroedus_crassirostris | LC | Decreasing | temperate | diurnal | 205.3 | 208835.982 | 8.604 |
| birds | Ailuroedus_melanotis | Ailuroedus_melanotis | LC | Decreasing | tropical | diurnal | 176.9 | 167335.291 | 3.536 |
| birds | Aimophila_notosticta | Aimophila_notosticta | LC | Decreasing | tropical | diurnal | 27.1 | 29968.209 | 6.523 |
| birds | Aimophila_rufescens | Aimophila_rufescens | LC | Stable | tropical | diurnal | 34.3 | 616683.253 | 10.093 |
| birds | Aimophila_ruficeps | Aimophila_ruficeps | LC | Decreasing | temperate | diurnal | 18.7 | 1184786.969 | 10.913 |
| birds | Aix_galericulata | Aix_galericulata | LC | Decreasing | temperate | diurnal | 567 | 3446284.161 | 13.141 |
| birds | Aix_sponsa | Aix_sponsa | LC | Increasing | temperate | diurnal | 657.6 | 7693839.612 | 13.952 |
| birds | Akletos_goeldii | Myrmeciza_goeldii | LC | Stable | tropical | diurnal | 42 | 342175.604 | 1.674 |
| birds | Akletos_melanoceps | Myrmeciza_melanoceps | LC | Stable | tropical | diurnal | 38.5 | 1321301.009 | 1.611 |
| birds | Alaemon_alaudipes | Alaemon_alaudipes | LC | Decreasing | temperate | diurnal | 43.7 | 10471834.068 | 4.392 |
| birds | Alaemon_hamertoni | NA | LC | Stable | NA | NA | NA | NA | NA |
| birds | Alauda_arvensis | Alauda_arvensis | LC | Decreasing | temperate | diurnal | 37.3 | 37787455.66 | 9.386 |
| birds | Alauda_gulgula | Alauda_gulgula | LC | Decreasing | temperate | diurnal | 26.3 | 11900668.988 | 12.77 |
| birds | Alauda_leucoptera | Melanocorypha_leucoptera | LC | Decreasing | temperate | diurnal | 45.3 | 5089458.733 | 8.84 |
| birds | Alauda_razae | Alauda_razae | CR | Stable | tropical | diurnal | 21.9 | 1054.711 | 11.945 |
| birds | Alaudala_raytal | Calandrella_raytal | LC | Stable | temperate | diurnal | 18.5 | 1260034.403 | 17.805 |
| birds | Alaudala_rufescens | Calandrella_cheleensis | LC | Decreasing | temperate | diurnal | 24.1 | 2316407.929 | 8.624 |
| birds | Alaudala_somalica | NA | LC | Decreasing | NA | NA | NA | NA | NA |
| birds | Alca_torda | Alca_torda | LC | Increasing | temperate | diurnal | 726 | 10735652.971 | 12.944 |
| birds | Alcedo_atthis | Alcedo_atthis | LC | Unknown | temperate | cathemeral | 31.1 | 28202464.974 | 13.211 |
| birds | Alcedo_coerulescens | NA | LC | Stable | NA | NA | NA | NA | NA |
| birds | Alcedo_euryzona | Alcedo_euryzona | CR | Decreasing | tropical | diurnal | 43.1 | 125365.642 | 22.372 |
| birds | Alcedo_hercules | Alcedo_hercules | NT | Decreasing | tropical | diurnal | 62.5 | 727664.628 | 8.819 |
| birds | Alcedo_meninting | Alcedo_meninting | LC | Decreasing | tropical | diurnal | 20.4 | 3655192.839 | 10.906 |

| | | | | | | | | | |
|---|---|---|---|---|---|---|---|---|---|
| birds | Alcedo_quadribrachys | Alcedo_quadribrachys | LC | Stable | tropical | diurnal | 34.9 | 4674328.275 | 8.797 |
| birds | Alcedo_semitorquata | Alcedo_semitorquata | LC | Decreasing | tropical | diurnal | 47.3 | 3463736.196 | 9.116 |
| birds | Alcippe_brunneicauda | Alcippe_brunneicauda | NT | Decreasing | tropical | diurnal | 14.3 | 764716.698 | 7.201 |
| birds | Alcippe_morrisonia | Alcippe_morrisonia | LC | Unknown | temperate | diurnal | 15.3 | 3103870.071 | 12.935 |
| birds | Alcippe_nipalensis | Alcippe_nipalensis | LC | Decreasing | temperate | diurnal | 15.8 | 350859.192 | 8.363 |
| birds | Alcippe_peracensis | Alcippe_peracensis | LC | Decreasing | tropical | diurnal | 14.3 | 148125.107 | 8.267 |
| birds | Alcippe_poioicephala | Alcippe_poioicephala | LC | Decreasing | tropical | diurnal | 20.7 | 2023970.47 | 12.286 |
| birds | Alcippe_pyrrhoptera | Alcippe_pyrrhoptera | LC | Decreasing | tropical | diurnal | 16.3 | 24849.192 | 18.886 |
| birds | Alcurus_striatus | Pycnonotus_striatus | LC | Stable | tropical | diurnal | 52 | 1082921.859 | 9.07 |
| birds | Aleadryas_rufinucha | Aleadryas_rufinucha | LC | Stable | tropical | diurnal | 40.5 | 150684.676 | 4.785 |
| birds | Alectoris_barbara | Alectoris_barbara | LC | Decreasing | temperate | diurnal | 416.3 | 1391816.955 | 9.462 |
| birds | Alectoris_chukar | Alectoris_chukar | LC | Stable | temperate | diurnal | 502.1 | 11388432.947 | 8.802 |
| birds | Alectoris_graeca | Alectoris_graeca | NT | Decreasing | temperate | diurnal | 594.1 | 652935.308 | 15.158 |
| birds | Alectoris_magna | Alectoris_magna | LC | Decreasing | temperate | diurnal | 541.4 | 624646.348 | 7.807 |
| birds | Alectoris_philbyi | Alectoris_philbyi | LC | Stable | tropical | diurnal | 441 | 80634.578 | 12.29 |
| birds | Alectoris_rufa | Alectoris_rufa | NT | Decreasing | temperate | diurnal | 527.9 | 1107486.346 | 19.756 |
| birds | Alectroenas_madagascariensis | Alectroenas_madagascariensis | LC | Decreasing | tropical | diurnal | 173 | 311714.723 | 8.797 |
| birds | Alectroenas_pulcherrimus | NA | LC | Decreasing | NA | NA | NA | NA | NA |
| birds | Alectroenas_sganzini | Alectroenas_sganzini | NT | Decreasing | tropical | diurnal | 158 | 2217.001 | 17.332 |
| birds | Alectrurus_risora | Alectrurus_risora | VU | Decreasing | temperate | diurnal | 21 | 935059.477 | 11.355 |
| birds | Alectrurus_tricolor | Alectrurus_tricolor | VU | Decreasing | tropical | diurnal | 16 | 1250046.969 | 11.321 |
| birds | Alectura_lathami | Alectura_lathami | LC | Decreasing | tropical | diurnal | 2333.1 | 647248.176 | 5.726 |
| birds | Alethe_diademata | Alethe_diademata | LC | Decreasing | tropical | diurnal | 31.5 | 448296.69 | 15.299 |
| birds | Alexandrinus_eques | Psittacula_eques | VU | Increasing | tropical | diurnal | 163 | 1010.364 | 22.371 |
| birds | Alexandrinus_krameri | Psittacula_krameri | LC | Increasing | tropical | diurnal | 116.1 | 8372649.271 | 15.116 |
| birds | Alipiopsitta_xanthops | NA | NT | Decreasing | NA | NA | NA | NA | NA |
| birds | Alisterus_amboinensis | Alisterus_amboinensis | LC | Decreasing | tropical | diurnal | 153 | 148979.734 | 4.421 |
| birds | Alisterus_chloropterus | Alisterus_chloropterus | LC | Stable | tropical | diurnal | 156 | 323894.91 | 3.136 |
| birds | Alisterus_scapularis | Alisterus_scapularis | LC | Decreasing | temperate | diurnal | 232.3 | 811455.942 | 8.023 |
| birds | Alle_alle | Alle_alle | LC | Decreasing | temperate | diurnal | 180.7 | 14713464.854 | 6.362 |
| birds | Allenia_fusca | Margarops_fuscus | LC | Unknown | tropical | diurnal | 67.1 | 5913.663 | 22.547 |
| birds | Alophoixus_bres | Alophoixus_bres | EN | Decreasing | tropical | diurnal | 35.3 | 87148.919 | 19.981 |
| birds | Alophoixus_flaveolus | Alophoixus_flaveolus | LC | Stable | tropical | diurnal | 48.3 | 559779.468 | 8.273 |
| birds | Alophoixus_ochraceus | Alophoixus_ochraceus | LC | Stable | tropical | diurnal | 41.5 | 407080.061 | 10.682 |

| | | | | | | | | | |
|---|---|---|---|---|---|---|---|---|---|
| birds | Alophoixus_pallidus | Alophoixus_pallidus | LC | Stable | tropical | diurnal | 46 | 1388165.229 | 12.199 |
| birds | Alophoixus_phaeocephalus | Alophoixus_phaeocephalus | LC | Stable | tropical | diurnal | 32 | 1368840.146 | 7.821 |
| birds | Alopochelidon_fucata | Alopochelidon_fucata | LC | Stable | temperate | diurnal | 14 | 4411696.773 | 9.352 |
| birds | Alopochen_aegyptiaca | Alopochen_aegyptiaca | LC | Decreasing | temperate | diurnal | 1873 | 17932898.42 | 9.206 |
| birds | Amadina_erythrocephala | Amadina_erythrocephala | LC | Stable | tropical | diurnal | 22.5 | 1628356.185 | 7.696 |
| birds | Amadina_fasciata | Amadina_fasciata | LC | Stable | tropical | diurnal | 18 | 3347451.843 | 11.191 |
| birds | Amalocichla_incerta | Amalocichla_incerta | LC | Stable | tropical | diurnal | 32.7 | 161107.325 | 4.6 |
| birds | Amalocichla_sclateriana | Amalocichla_sclateriana | LC | Stable | tropical | diurnal | 54.7 | 15280.985 | 4.628 |
| birds | Amandava_amandava | Amandava_amandava | LC | Stable | tropical | diurnal | 9 | 3949205.984 | 16.035 |
| birds | Amandava_formosa | Amandava_formosa | VU | Decreasing | tropical | diurnal | 9 | 686385.245 | 17.114 |
| birds | Amandava_subflava | Amandava_subflava | LC | Stable | tropical | diurnal | 7.4 | 3352898.936 | 11.45 |
| birds | Amaurocichla_bocagii | Amaurocichla_bocagei | VU | Stable | tropical | diurnal | 7 | 340.894 | 7.576 |
| birds | Amaurolimnas_concolor | Amaurolimnas_concolor | LC | Decreasing | tropical | diurnal | 133 | 1725851.554 | 4.991 |
| birds | Amaurornis_cinerea | Porzana_cinerea | LC | Unknown | tropical | diurnal | 52.1 | 3636563.105 | 7.387 |
| birds | Amaurornis_isabellina | NA | LC | Unknown | NA | NA | NA | NA | NA |
| birds | Amaurornis_moluccana | Amaurornis_moluccana | LC | Unknown | tropical | diurnal | 186.8 | 1266108.022 | 3.962 |
| birds | Amaurornis_olivacea | Amaurornis_olivacea | LC | Unknown | tropical | diurnal | 292 | 271176.413 | 13.874 |
| birds | Amaurornis_phoenicurus | Amaurornis_phoenicurus | LC | Unknown | tropical | diurnal | 180 | 10030319.804 | 14.35 |
| birds | Amaurospiza_moesta | Amaurospiza_moesta | LC | Decreasing | temperate | diurnal | 13.6 | 864091.666 | 12.855 |
| birds | Amaurospiza_relicta | Amaurospiza_concolor | LC | Stable | tropical | diurnal | 13.1 | 112893.361 | 9.097 |
| birds | Amazilia_boucardi | Amazilia_boucardi | EN | Decreasing | tropical | diurnal | 4.5 | 1301.757 | 16.102 |
| birds | Amazilia_cyanifrons | Amazilia_cyanifrons | LC | Stable | tropical | diurnal | 5 | 54825.566 | 14.491 |
| birds | Amazilia_franciae | Amazilia_franciae | LC | Unknown | tropical | diurnal | 5 | 247257.855 | 10.539 |
| birds | Amazilia_lactea | NA | not_available | not_available | NA | NA | NA | NA | NA |
| birds | Amazilia_luciae | NA | VU | Decreasing | NA | NA | NA | NA | NA |
| birds | Amazilia_rutila | Amazilia_rutila | LC | Stable | tropical | diurnal | 5.2 | 397592.641 | 10.514 |
| birds | Amazilia_tzacatl | Amazilia_tzacatl | LC | Decreasing | tropical | diurnal | 4.8 | 1019992.014 | 10.812 |
| birds | Amazilia_viridifrons | NA | not_available | not_available | NA | NA | NA | NA | NA |
| birds | Amazilia_viridigaster | NA | not_available | not_available | NA | NA | NA | NA | NA |
| birds | Amazilia_yucatanensis | Amazilia_yucatanensis | LC | Unknown | temperate | diurnal | 3.9 | 582084.956 | 12.218 |
| birds | Amazilis_amazilia | Amazilia_amazilia | LC | Unknown | tropical | diurnal | 4.3 | 169380.819 | 9.983 |
| birds | Amazona_aestiva | Amazona_aestiva | NT | Decreasing | tropical | diurnal | 451 | 5857429.401 | 8.551 |
| birds | Amazona_agilis | Amazona_agilis | EN | Decreasing | tropical | diurnal | 213 | 8263.643 | 18.078 |
| birds | Amazona_albifrons | Amazona_albifrons | LC | Increasing | temperate | diurnal | 206 | 546295.399 | 9.864 |

| | | | | | | | | | |
|---|---|---|---|---|---|---|---|---|---|
| birds | Amazona_amazonica | Amazona_amazonica | LC | Decreasing | tropical | diurnal | 370 | 7382920.984 | 4.437 |
| birds | Amazona_arausiaca | Amazona_arausiaca | VU | Increasing | tropical | diurnal | 500 | 750.798 | 13.442 |
| birds | Amazona_autumnalis | Amazona_autumnalis | LC | Decreasing | tropical | diurnal | 416 | 977125.183 | 10.318 |
| birds | Amazona_barbadensis | Amazona_barbadensis | NT | Decreasing | tropical | diurnal | 270 | 42206.45 | 13.3 |
| birds | Amazona_brasiliensis | Amazona_brasiliensis | NT | Increasing | temperate | diurnal | 430 | 4747.023 | 8.455 |
| birds | Amazona_collaria | Amazona_collaria | VU | Decreasing | tropical | diurnal | 187 | 7226.416 | 17.698 |
| birds | Amazona_dufresniana | Amazona_dufresniana | NT | Decreasing | tropical | diurnal | 623 | 546076.569 | 1.336 |
| birds | Amazona_farinosa | Amazona_farinosa | LC | Decreasing | tropical | diurnal | 626 | 7457891.935 | 3.27 |
| birds | Amazona_festiva | Amazona_festiva | LC | Decreasing | tropical | diurnal | 430 | 689102.96 | 2.042 |
| birds | Amazona_finschi | Amazona_finschi | EN | Decreasing | tropical | diurnal | 302 | 112128.493 | 6.001 |
| birds | Amazona_guildingii | Amazona_guildingii | VU | Increasing | tropical | diurnal | 646 | 43.099 | 10.677 |
| birds | Amazona_imperialis | Amazona_imperialis | CR | Decreasing | tropical | diurnal | 529.5 | 221.981 | 9.275 |
| birds | Amazona_kawalli | NA | LC | Decreasing | NA | NA | NA | NA | NA |
| birds | Amazona_leucocephala | Amazona_leucocephala | NT | Decreasing | tropical | diurnal | 235.3 | 31494.263 | 12.3 |
| birds | Amazona_mercenarius | Amazona_mercenaria | LC | Decreasing | tropical | diurnal | 340 | 432626.422 | 8.815 |
| birds | Amazona_ochrocephala | Amazona_ochrocephala | LC | Decreasing | tropical | diurnal | 476.9 | 6635282.503 | 2.888 |
| birds | Amazona_pretrei | Amazona_pretrei | VU | Stable | temperate | diurnal | 272.8 | 181933.238 | 12.298 |
| birds | Amazona_rhodocorytha | Amazona_rhodocorytha | VU | Decreasing | tropical | diurnal | 474.3 | 26932.409 | 8.238 |
| birds | Amazona_tucumana | Amazona_tucumana | VU | Decreasing | tropical | diurnal | 260 | 129668.062 | 6.759 |
| birds | Amazona_ventralis | Amazona_ventralis | VU | Decreasing | tropical | diurnal | 214 | 37516.116 | 16.141 |
| birds | Amazona_versicolor | Amazona_versicolor | VU | Increasing | tropical | diurnal | 676 | 617.462 | 20.773 |
| birds | Amazona_vinacea | Amazona_vinacea | EN | Decreasing | temperate | diurnal | 254 | 534573.198 | 13.126 |
| birds | Amazona_viridigenalis | Amazona_viridigenalis | EN | Decreasing | temperate | diurnal | 316 | 219413.111 | 13.824 |
| birds | Amazona_vittata | Amazona_vittata | CR | Increasing | tropical | diurnal | 279.3 | 8952.432 | 27.063 |
| birds | Amazona_xantholora | Amazona_xantholora | LC | Decreasing | tropical | diurnal | 217 | 141207.228 | 6.347 |
| birds | Amazonetta_brasiliensis | Amazonetta_brasiliensis | LC | Decreasing | tropical | diurnal | 500 | 8747518.746 | 7.506 |
| birds | Amblycercus_holosericeus | Amblycercus_holosericeus | LC | Decreasing | tropical | diurnal | 70.4 | 1152587.022 | 9.586 |
| birds | Amblyornis_flavifrons | Amblyornis_flavifrons | LC | Stable | tropical | diurnal | 120 | 978.259 | 0.437 |
| birds | Amblyornis_inornata | Amblyornis_inornata | LC | Decreasing | tropical | diurnal | 124 | 16136.476 | 2.862 |
| birds | Amblyornis_macgregoriae | Amblyornis_macgregoriae | LC | Decreasing | tropical | diurnal | 117.5 | 139562.122 | 4.941 |
| birds | Amblyornis_subalaris | Amblyornis_subalaris | LC | Decreasing | tropical | diurnal | 108.9 | 18481.038 | 2.72 |
| birds | Amblyospiza_albifrons | Amblyospiza_albifrons | LC | Stable | tropical | diurnal | 41.6 | 3499946.835 | 10.529 |
| birds | Amblyramphus_holosericeus | Amblyramphus_holosericeus | LC | Stable | temperate | diurnal | 56.8 | 1580707.217 | 10.363 |
| birds | Ammodramus_aurifrons | Ammodramus_aurifrons | LC | Stable | tropical | diurnal | 17.8 | 3867543.928 | 2.457 |

| | | | | | | | | | |
|---|---|---|---|---|---|---|---|---|---|
| birds | Ammodramus_humeralis | Ammodramus_humeralis | LC | Stable | tropical | diurnal | 16 | 9115374.376 | 8.127 |
| birds | Ammodramus_savannarum | Ammodramus_savannarum | LC | Decreasing | temperate | diurnal | 17.6 | 7888248.743 | 14.283 |
| birds | Ammomanes_cinctura | Ammomanes_cinctura | LC | Decreasing | temperate | diurnal | 18.1 | 7020113.233 | 3.748 |
| birds | Ammomanes_deserti | Ammomanes_deserti | LC | Increasing | temperate | diurnal | 23.8 | 12034050.905 | 5.358 |
| birds | Ammomanes_phoenicura | Ammomanes_phoenicura | LC | Stable | tropical | diurnal | 25.6 | 1826048.111 | 18.586 |
| birds | Ammonastes_pelzelni | Myrmeciza_pelzelni | LC | Stable | tropical | diurnal | 17.7 | 58892.641 | 0.654 |
| birds | Ammoperdix_griseogularis | Ammoperdix_griseogularis | LC | Stable | temperate | diurnal | 201.9 | 2996830.793 | 8.975 |
| birds | Ammoperdix_heyi | Ammoperdix_heyi | LC | Stable | temperate | diurnal | 189.7 | 1664830.949 | 5.397 |
| birds | Ammospiza_caudacuta | Ammodramus_caudacutus | EN | Decreasing | temperate | diurnal | 19.2 | 215557.244 | 20.542 |
| birds | Ammospiza_leconteii | Ammodramus_leconteii | LC | Decreasing | temperate | diurnal | 13 | 5593741.343 | 11.428 |
| birds | Ammospiza_maritima | Ammodramus_maritimus | LC | Stable | temperate | diurnal | 22.2 | 211096.75 | 18.324 |
| birds | Ammospiza_nelsoni | Ammodramus_nelsoni | LC | Increasing | temperate | diurnal | 15.1 | 2904622.438 | 13.384 |
| birds | Ampeliceps_coronatus | Ampeliceps_coronatus | LC | Decreasing | tropical | diurnal | 87.9 | 1377766.105 | 11.036 |
| birds | Ampelioides_tschudii | Ampelioides_tschudii | LC | Decreasing | tropical | diurnal | 78.4 | 201035.433 | 5.516 |
| birds | Ampelion_rubrocristatus | Ampelion_rubrocristatus | LC | Stable | tropical | diurnal | 66.3 | 413439.186 | 9.602 |
| birds | Ampelion_rufaxilla | Ampelion_rufaxilla | LC | Stable | tropical | diurnal | 73.9 | 79754.203 | 8.544 |
| birds | Ampelornis_griseiceps | Myrmeciza_griseiceps | VU | Decreasing | tropical | diurnal | 15 | 18162.58 | 9.161 |
| birds | Amphispiza_bilineata | Amphispiza_bilineata | LC | Decreasing | temperate | diurnal | 13.5 | 2559845.859 | 8.335 |
| birds | Amphispiza_quinquestriata | Aimophila_quinquestriata | LC | Decreasing | temperate | diurnal | 19.5 | 151617.216 | 5.597 |
| birds | Amytornis_barbatus | Amytornis_barbatus | LC | Decreasing | temperate | diurnal | 18.4 | 55946.509 | 2.313 |
| birds | Amytornis_dorotheae | Amytornis_dorotheae | VU | Decreasing | tropical | diurnal | 23.2 | 47008 | 3.202 |
| birds | Amytornis_goyderi | Amytornis_goyderi | LC | Stable | temperate | diurnal | 16.7 | 227659.69 | 1.102 |
| birds | Amytornis_housei | Amytornis_housei | LC | Stable | tropical | diurnal | 28.9 | 13972.818 | 1.158 |
| birds | Amytornis_purnelli | Amytornis_purnelli | LC | Increasing | tropical | diurnal | 21 | 653027.467 | 1.151 |
| birds | Amytornis_striatus | Amytornis_striatus | LC | Decreasing | temperate | diurnal | 19.6 | 912126.848 | 2.876 |
| birds | Amytornis_textilis | Amytornis_textilis | LC | Decreasing | temperate | diurnal | 23 | 108980.136 | 5.494 |
| birds | Amytornis_woodwardi | Amytornis_woodwardi | EN | Decreasing | tropical | diurnal | 35.1 | 5520.041 | 2.142 |
| birds | Anabacerthia_amaurotis | Anabacerthia_amaurotis | NT | Decreasing | temperate | diurnal | 19.2 | 499233.475 | 13.171 |
| birds | Anabacerthia_lichtensteini | Philydor_lichtensteini | LC | Decreasing | tropical | diurnal | 21 | 1589284.847 | 12.923 |
| birds | Anabacerthia_ruficaudata | Philydor_ruficaudatum | LC | Decreasing | tropical | diurnal | 30.1 | 4312236.415 | 2.114 |
| birds | Anabacerthia_striaticollis | Anabacerthia_striaticollis | LC | Stable | tropical | diurnal | 24.8 | 335322.021 | 8.689 |
| birds | Anabacerthia_variegaticeps | Anabacerthia_variegaticeps | LC | Decreasing | tropical | diurnal | 27.8 | 215334.521 | 9.784 |
| birds | Anabathmis_hartlaubii | Nectarinia_hartlaubii | LC | Stable | tropical | diurnal | 10.5 | 146.654 | 14.707 |
| birds | Anabathmis_newtonii | Nectarinia_newtonii | LC | Stable | tropical | diurnal | 6.5 | 858.514 | 12.812 |

| | | | | | | | | | |
|---|---|---|---|---|---|---|---|---|---|
| birds | Anabathmis_reichenbachii | Nectarinia_reichenbachii | LC | Stable | tropical | diurnal | 11.4 | 2222320.407 | 6.188 |
| birds | Anabazenops_dorsalis | Anabazenops_dorsalis | LC | Stable | tropical | diurnal | 35.4 | 480171.496 | 2.882 |
| birds | Anabazenops_fuscus | Anabazenops_fuscus | LC | Decreasing | tropical | diurnal | 38.5 | 372722.516 | 12.458 |
| birds | Anairetes_alpinus | Anairetes_alpinus | EN | Decreasing | tropical | diurnal | 9 | 23428.249 | 7.956 |
| birds | Anairetes_fernandezianus | NA | EN | Decreasing | NA | NA | NA | NA | NA |
| birds | Anairetes_flavirostris | Anairetes_flavirostris | LC | Stable | temperate | diurnal | 6 | 2046579.468 | 8.257 |
| birds | Anairetes_nigrocristatus | Anairetes_nigrocristatus | LC | Stable | tropical | diurnal | 9.2 | 48678.073 | 9.427 |
| birds | Anairetes_parulus | Anairetes_parulus | LC | Stable | temperate | diurnal | 6.2 | 2638454.775 | 6.097 |
| birds | Anairetes_reguloides | Anairetes_reguloides | LC | Stable | tropical | diurnal | 5.9 | 103712.534 | 6.995 |
| birds | Anaplectes_rubriceps | Anaplectes_rubriceps | LC | Stable | tropical | diurnal | 22.6 | 3258277.526 | 8.559 |
| birds | Anarhynchus_frontalis | Anarhynchus_frontalis | VU | Decreasing | temperate | cathemeral | 57.6 | 34959.295 | 10.873 |
| birds | Anas_acuta | Anas_acuta | LC | Decreasing | temperate | diurnal | 944.6 | 52934942.782 | 8.436 |
| birds | Anas_aucklandica | NA | NT | Stable | NA | NA | NA | NA | NA |
| birds | Anas_bahamensis | Anas_bahamensis | LC | Decreasing | temperate | diurnal | 490 | 2991584.698 | 9.281 |
| birds | Anas_bernieri | Anas_bernieri | EN | Decreasing | tropical | diurnal | 378.5 | 53437.434 | 8.714 |
| birds | Anas_capensis | Anas_capensis | LC | Increasing | tropical | diurnal | 402 | 4317287.048 | 8.402 |
| birds | Anas_castanea | Anas_castanea | LC | Stable | temperate | diurnal | 624 | 3131751.241 | 6.537 |
| birds | Anas_crecca | Anas_crecca | LC | Unknown | tropical | diurnal | 341.9 | 39459336.835 | 9.735 |
| birds | Anas_erythrorhyncha | Anas_erythrorhyncha | LC | Decreasing | tropical | diurnal | 558.8 | 10763443.791 | 8.698 |
| birds | Anas_flavirostris | Anas_flavirostris | LC | Decreasing | temperate | diurnal | 434 | 3942276.348 | 7.328 |
| birds | Anas_georgica | Anas_georgica | LC | Decreasing | temperate | diurnal | 623.8 | 4196176.914 | 7.467 |
| birds | Anas_gibberifrons | Anas_gibberifrons | NT | Stable | tropical | diurnal | 488.1 | 787609.113 | 11.539 |
| birds | Anas_luzonica | Anas_luzonica | VU | Decreasing | tropical | diurnal | 832 | 279937.111 | 13.718 |
| birds | Anas_melleri | Anas_melleri | EN | Decreasing | tropical | diurnal | 959.2 | 210476.274 | 9.393 |
| birds | Anas_platyrhynchos | Anas_platyrhynchos | LC | Increasing | temperate | diurnal | 843.4 | 48513515.011 | 9.219 |
| birds | Anas_poecilorhyncha | Anas_poecilorhyncha | LC | Decreasing | tropical | diurnal | 1000 | 5263129.877 | 16.185 |
| birds | Anas_rubripes | Anas_rubripes | LC | Decreasing | temperate | diurnal | 1241 | 6750626.952 | 10.447 |
| birds | Anas_sparsa | Anas_sparsa | LC | Decreasing | tropical | diurnal | 981.5 | 9213622.594 | 9.055 |
| birds | Anas_superciliosa | Anas_superciliosa | LC | Unknown | temperate | diurnal | 882.8 | 8845309.601 | 5.162 |
| birds | Anas_undulata | Anas_undulata | LC | Stable | tropical | diurnal | 1008 | 8476793.264 | 8.764 |
| birds | Anastomus_lamelligerus | Anastomus_lamelligerus | LC | Stable | tropical | diurnal | 1081 | 13409712.688 | 9.22 |
| birds | Anastomus_oscitans | Anastomus_oscitans | LC | Unknown | tropical | diurnal | 1590 | 3412455.461 | 18.017 |
| birds | Ancistrops_strigilatus | Ancistrops_strigilatus | LC | Stable | tropical | diurnal | 35.9 | 2508167.129 | 1.558 |
| birds | Andigena_cucullata | Andigena_cucullata | LC | Stable | tropical | diurnal | 311.8 | 48713.293 | 6.052 |

| | | | | | | | | | |
|---|---|---|---|---|---|---|---|---|---|
| birds | Andigena_hypoglauca | Andigena_hypoglauca | LC | Decreasing | tropical | diurnal | 312.8 | 132720.009 | 7.944 |
| birds | Andigena_laminirostris | Andigena_laminirostris | NT | Decreasing | tropical | diurnal | 311 | 8893.386 | 8.516 |
| birds | Andigena_nigrirostris | Andigena_nigrirostris | LC | Decreasing | tropical | diurnal | 350 | 110031.995 | 10.966 |
| birds | Androdon_aequatorialis | Androdon_aequatorialis | LC | Decreasing | tropical | diurnal | 7.3 | 83164.782 | 7.578 |
| birds | Andropadus_importunus | Andropadus_importunus | LC | Stable | tropical | diurnal | 27.4 | 1338546.667 | 11.455 |
| birds | Androphobus_viridis | Androphobus_viridis | LC | Stable | tropical | diurnal | 30.5 | 30787.301 | 5.179 |
| birds | Anhima_cornuta | Anhima_cornuta | LC | Decreasing | tropical | diurnal | 3150 | 10980198.811 | 4.94 |
| birds | Anhinga_anhinga | Anhinga_anhinga | LC | Decreasing | tropical | diurnal | 1235 | 14921101.487 | 6.472 |
| birds | Anhinga_melanogaster | Anhinga_melanogaster | NT | Decreasing | tropical | diurnal | 1480.1 | 6866922.355 | 14.651 |
| birds | Anisognathus_igniventris | Anisognathus_igniventris | LC | Stable | tropical | diurnal | 34 | 96719.414 | 6.861 |
| birds | Anisognathus_lacrymosus | Anisognathus_lacrymosus | LC | Stable | tropical | diurnal | 31 | 173969.015 | 9.264 |
| birds | Anisognathus_melanogenys | Anisognathus_melanogenys | LC | Decreasing | tropical | diurnal | 41 | 7906.889 | 6.841 |
| birds | Anisognathus_notabilis | Anisognathus_notabilis | LC | Decreasing | tropical | diurnal | 57.3 | 18610.622 | 5.225 |
| birds | Anisognathus_somptuosus | Anisognathus_somptuosus | LC | Stable | tropical | diurnal | 44.5 | 319648.428 | 8.268 |
| birds | Anodorhynchus_hyacinthinus | Anodorhynchus_hyacinthinus | VU | Decreasing | tropical | diurnal | 1331 | 2832918.269 | 5.607 |
| birds | Anodorhynchus_leari | Anodorhynchus_leari | EN | Increasing | tropical | diurnal | 940 | 44706.576 | 6.157 |
| birds | Anomalospiza_imberbis | Anomalospiza_imberbis | LC | Stable | temperate | diurnal | 20.5 | 1822866.7 | 10.528 |
| birds | Anopetia_gounellei | Anopetia_gounellei | LC | Unknown | tropical | diurnal | 3.2 | 534925.908 | 7.423 |
| birds | Anorrhinus_austeni | Anorrhinus_austeni | NT | Decreasing | tropical | diurnal | 933 | 774356.595 | 8.502 |
| birds | Anorrhinus_galeritus | Anorrhinus_galeritus | NT | Decreasing | tropical | diurnal | 1172 | 1381919.988 | 8.051 |
| birds | Anorrhinus_tickelli | Anorrhinus_tickelli | NT | Decreasing | tropical | diurnal | 792.6 | 160242.6 | 10.506 |
| birds | Anous_albivittus | Procelsterna_albivitta | LC | Stable | temperate | diurnal | 72.2 | 14237944.952 | 10.474 |
| birds | Anous_ceruleus | Procelsterna_cerulea | LC | Stable | tropical | diurnal | 53 | 33816689.353 | 8.714 |
| birds | Anous_minutus | Anous_minutus | LC | Decreasing | tropical | diurnal | 112.4 | 84112718.178 | 7.285 |
| birds | Anous_stolidus | Anous_stolidus | LC | Stable | tropical | diurnal | 177.8 | 168965335.411 | 9.213 |
| birds | Anous_tenuirostris | Anous_tenuirostris | LC | Stable | tropical | diurnal | 112 | 5483959.416 | 18.808 |
| birds | Anser_albifrons | Anser_albifrons | LC | Unknown | temperate | diurnal | 2506.4 | 10495879.002 | 7.487 |
| birds | Anser_anser | Anser_anser | LC | Increasing | temperate | diurnal | 3302.4 | 19080676.338 | 10.232 |
| birds | Anser_brachyrhynchus | Anser_brachyrhynchus | LC | Increasing | temperate | diurnal | 2642 | 245976.492 | 6.588 |
| birds | Anser_caerulescens | Chen_caerulescens | LC | Increasing | temperate | diurnal | 2636.2 | 9421501.275 | 7.804 |
| birds | Anser_canagicus | Chen_canagica | LC | Increasing | temperate | diurnal | 2136.5 | 464902.485 | 0.986 |
| birds | Anser_cygnoid | Anser_cygnoides | EN | Decreasing | temperate | diurnal | 3511.9 | 7307006.36 | 10.5 |
| birds | Anser_erythropus | Anser_erythropus | VU | Decreasing | temperate | diurnal | 1755.5 | 29185526.078 | 7.757 |
| birds | Anser_fabalis | Anser_fabalis | LC | Decreasing | temperate | diurnal | 2754.7 | 17921622.229 | 5.615 |

| | | | | | | | | | |
|---|---|---|---|---|---|---|---|---|---|
| birds | Anser_indicus | Anser_indicus | LC | Decreasing | temperate | diurnal | 2212.6 | 4470193.833 | 11.021 |
| birds | Anser_rossii | Chen_rossii | LC | Increasing | temperate | diurnal | 1636 | 2899429.996 | 9.833 |
| birds | Anseranas_semipalmata | Anseranas_semipalmata | LC | Stable | tropical | diurnal | 2393.4 | 1308440.743 | 3.795 |
| birds | Anthipes_monileger | Ficedula_monileger | LC | Stable | tropical | diurnal | 11 | 941257.108 | 8.627 |
| birds | Anthipes_solitaris | Ficedula_solitaris | LC | Stable | tropical | diurnal | 13.1 | 157229.381 | 6.831 |
| birds | Anthobaphes_violacea | Nectarinia_violacea | LC | Decreasing | temperate | diurnal | 9.4 | 91921.14 | 10.695 |
| birds | Anthocephala_floriceps | Anthocephala_floriceps | VU | Decreasing | tropical | diurnal | 3.9 | 3065.919 | 8.464 |
| birds | Anthochaera_carunculata | Anthochaera_carunculata | LC | Stable | temperate | diurnal | 106.1 | 2173024.904 | 7.343 |
| birds | Anthochaera_chrysoptera | Anthochaera_chrysoptera | LC | Unknown | temperate | diurnal | 67.8 | 671226.201 | 9.712 |
| birds | Anthochaera_paradoxa | Anthochaera_paradoxa | LC | Decreasing | temperate | diurnal | 143.7 | 46805.045 | 7.322 |
| birds | Anthochaera_phrygia | Xanthomyza_phrygia | CR | Decreasing | temperate | diurnal | 40 | 281979.507 | 9.937 |
| birds | Anthornis_melanura | Anthornis_melanura | LC | Decreasing | temperate | diurnal | 26.9 | 212871.757 | 7.341 |
| birds | Anthoscopus_caroli | Anthoscopus_caroli | LC | Decreasing | tropical | diurnal | 6.9 | 4646761.756 | 9.397 |
| birds | Anthoscopus_flavifrons | Anthoscopus_flavifrons | LC | Stable | tropical | diurnal | 8 | 1195620.856 | 7.229 |
| birds | Anthoscopus_minutus | Anthoscopus_minutus | LC | Stable | temperate | diurnal | 6.6 | 2370698.923 | 7.153 |
| birds | Anthoscopus_musculus | Anthoscopus_musculus | LC | Stable | tropical | diurnal | 5.2 | 1141427.54 | 9.134 |
| birds | Anthoscopus_parvulus | Anthoscopus_parvulus | LC | Stable | tropical | diurnal | 5.8 | 1993198.486 | 12.624 |
| birds | Anthoscopus_punctifrons | Anthoscopus_punctifrons | LC | Stable | tropical | diurnal | 7 | 2414155.959 | 8.893 |
| birds | Anthracoceros_albirostris | Anthracoceros_albirostris | LC | Stable | tropical | diurnal | 892.9 | 4467706.591 | 12.217 |
| birds | Anthracoceros_coronatus | Anthracoceros_coronatus | NT | Decreasing | tropical | diurnal | 806.7 | 701610.859 | 16.345 |
| birds | Anthracoceros_malayanus | Anthracoceros_malayanus | VU | Decreasing | tropical | diurnal | 1050 | 798560.435 | 8.035 |
| birds | Anthracoceros_marchei | Anthracoceros_marchei | VU | Decreasing | tropical | diurnal | 750 | 12863.666 | 9.509 |
| birds | Anthracoceros_montani | Anthracoceros_montani | CR | Decreasing | tropical | diurnal | 741 | 1624.397 | 10.821 |
| birds | Anthracothorax_dominicus | Anthracothorax_dominicus | LC | Unknown | tropical | diurnal | 5.8 | 75252.822 | 15.761 |
| birds | Anthracothorax_mango | Anthracothorax_mango | LC | Unknown | tropical | diurnal | 8.3 | 11021.791 | 19.741 |
| birds | Anthracothorax_nigricollis | Anthracothorax_nigricollis | LC | Stable | tropical | diurnal | 7 | 9821751.586 | 5.837 |
| birds | Anthracothorax_prevostii | Anthracothorax_prevostii | LC | Decreasing | tropical | diurnal | 6.4 | 919148.979 | 10.687 |
| birds | Anthracothorax_veraguensis | Anthracothorax_veraguensis | LC | Stable | tropical | diurnal | 7 | 30443.514 | 13.399 |
| birds | Anthracothorax_viridigula | Anthracothorax_viridigula | LC | Decreasing | tropical | diurnal | 7.5 | 591465.009 | 3.237 |
| birds | Anthracothorax_viridis | Anthracothorax_viridis | LC | Stable | tropical | diurnal | 6.6 | 8774.421 | 27.252 |
| birds | Anthreptes_anchietae | Anthreptes_anchietae | LC | Stable | tropical | diurnal | 8 | 465325.067 | 8.664 |
| birds | Anthreptes_aurantius | Anthreptes_aurantium | LC | Decreasing | tropical | diurnal | 12.5 | 1740388.212 | 6.032 |
| birds | Anthreptes_gabonicus | NA | LC | Stable | NA | NA | NA | NA | NA |
| birds | Anthreptes_longuemarei | Anthreptes_longuemarei | LC | Stable | tropical | diurnal | 11.4 | 5262392.931 | 8.631 |

| | | | | | | | | | |
|---|---|---|---|---|---|---|---|---|---|
| birds | Anthreptes_malacensis | Anthreptes_malacensis | LC | Stable | tropical | diurnal | 11.3 | 2166604.348 | 11.238 |
| birds | Anthreptes_neglectus | Anthreptes_neglectus | LC | Decreasing | tropical | diurnal | 11.8 | 250651.547 | 10.256 |
| birds | Anthreptes_orientalis | Anthreptes_orientalis | LC | Stable | tropical | diurnal | 10.3 | 1935952.464 | 8.974 |
| birds | Anthreptes_rectirostris | Anthreptes_rectirostris | LC | Decreasing | tropical | diurnal | 10.6 | 448143.892 | 15.714 |
| birds | Anthreptes_reichenowi | Anthreptes_reichenowi | NT | Decreasing | tropical | diurnal | 7.3 | 438283.12 | 9.528 |
| birds | Anthreptes_rhodolaemus | Anthreptes_rhodolaemus | NT | Decreasing | tropical | diurnal | 13.1 | 442077.764 | 9.557 |
| birds | Anthreptes_rubritorques | Anthreptes_rubritorques | VU | Decreasing | tropical | diurnal | 10.7 | 7092.318 | 11.586 |
| birds | Anthreptes_seimundi | Nectarinia_seimundi | LC | Stable | tropical | diurnal | 6.6 | 2083351.491 | 8.794 |
| birds | Anthreptes_simplex | Anthreptes_simplex | LC | Decreasing | tropical | diurnal | 8.9 | 1062052.924 | 8.688 |
| birds | Anthropoides_paradiseus | Grus_paradisea | VU | Decreasing | temperate | diurnal | 4540.6 | 568469.111 | 10.841 |
| birds | Anthropoides_virgo | Grus_virgo | LC | Increasing | temperate | diurnal | 2417 | 20925462.355 | 8.923 |
| birds | Anthus_antarcticus | NA | LC | Stable | NA | NA | NA | NA | NA |
| birds | Anthus_berthelotii | Anthus_berthelotii | LC | Stable | temperate | diurnal | 15.9 | 8283.442 | 18.874 |
| birds | Anthus_bogotensis | Anthus_bogotensis | LC | Stable | tropical | diurnal | 25.1 | 336033.741 | 9.78 |
| birds | Anthus_brachyurus | Anthus_brachyurus | LC | Stable | tropical | diurnal | 15.3 | 1510202.74 | 7.637 |
| birds | Anthus_caffer | Anthus_caffer | LC | Stable | tropical | diurnal | 16.8 | 777646.716 | 9.896 |
| birds | Anthus_campestris | Anthus_campestris | LC | Stable | temperate | diurnal | 23 | 36400563.444 | 8.526 |
| birds | Anthus_cervinus | Anthus_cervinus | LC | Stable | tropical | diurnal | 20.9 | 15867275.101 | 8.91 |
| birds | Anthus_chacoensis | NA | LC | Stable | NA | NA | NA | NA | NA |
| birds | Anthus_correndera | Anthus_correndera | LC | Stable | temperate | diurnal | 20.9 | 3686700.656 | 7.324 |
| birds | Anthus_crenatus | Anthus_crenatus | NT | Decreasing | temperate | diurnal | 30.7 | 343685.27 | 9.78 |
| birds | Anthus_furcatus | Anthus_furcatus | LC | Stable | temperate | diurnal | 20.1 | 1397721.228 | 10.117 |
| birds | Anthus_godlewskii | Anthus_godlewskii | LC | Stable | temperate | diurnal | 25.3 | 9014937.365 | 9.978 |
| birds | Anthus_gustavi | Anthus_gustavi | LC | Stable | temperate | diurnal | 19.8 | 11759006.193 | 4.755 |
| birds | Anthus_gutturalis | Anthus_gutturalis | LC | Stable | tropical | diurnal | 36.3 | 34559.907 | 4.929 |
| birds | Anthus_hellmayri | Anthus_hellmayri | LC | Stable | temperate | diurnal | 18.6 | 1204389.515 | 9.611 |
| birds | Anthus_hodgsoni | Anthus_hodgsoni | LC | Stable | temperate | diurnal | 21.3 | 18925518.719 | 7.91 |
| birds | Anthus_hoeschi | Anthus_hoeschi | NT | Decreasing | tropical | diurnal | 27 | 1210644.431 | 7.134 |
| birds | Anthus_leucophrys | Anthus_leucophrys | LC | Stable | temperate | diurnal | 25 | 8950742.457 | 9.665 |
| birds | Anthus_lineiventris | Anthus_lineiventris | LC | Stable | tropical | diurnal | 35 | 1635119.6 | 11.267 |
| birds | Anthus_lutescens | Anthus_lutescens | LC | Decreasing | tropical | diurnal | 12.6 | 7241509.046 | 8.487 |
| birds | Anthus_melindae | Anthus_melindae | LC | Stable | tropical | diurnal | 22.5 | 108383.572 | 13.667 |
| birds | Anthus_nattereri | Anthus_nattereri | VU | Decreasing | temperate | diurnal | 19.5 | 753720.224 | 12.122 |
| birds | Anthus_nilghiriensis | NA | VU | Decreasing | NA | NA | NA | NA | NA |

| | | | | | | | | | |
|---|---|---|---|---|---|---|---|---|---|
| birds | Anthus_novaeseelandiae | Anthus_novaeseelandiae | LC | Stable | temperate | diurnal | 24.1 | 6479742.405 | 5.007 |
| birds | Anthus_pallidiventris | Anthus_pallidiventris | LC | Stable | tropical | diurnal | 31 | 942199.865 | 6.546 |
| birds | Anthus_petrosus | Anthus_petrosus | LC | Stable | temperate | diurnal | 22.4 | 497601.289 | 18.911 |
| birds | Anthus_pratensis | Anthus_pratensis | LC | Decreasing | temperate | diurnal | 18.4 | 16554757.401 | 12.339 |
| birds | Anthus_richardi | Anthus_richardi | LC | Stable | temperate | diurnal | 32.2 | 23084325.307 | 9.326 |
| birds | Anthus_roseatus | Anthus_roseatus | LC | Stable | temperate | diurnal | 20.6 | 5517844.356 | 8.154 |
| birds | Anthus_rubescens | Anthus_rubescens | LC | Decreasing | temperate | diurnal | 20.7 | 37048750.127 | 7.378 |
| birds | Anthus_rufulus | Anthus_rufulus | LC | Stable | tropical | diurnal | 20.4 | 7678484.066 | 14.518 |
| birds | Anthus_similis | Anthus_similis | LC | Stable | temperate | diurnal | 25.3 | 9112709.117 | 11.126 |
| birds | Anthus_sokokensis | Anthus_sokokensis | EN | Decreasing | tropical | diurnal | 15 | 2265.147 | 12.918 |
| birds | Anthus_spinoletta | Anthus_spinoletta | LC | Stable | temperate | diurnal | 23.9 | 17694009.144 | 11.839 |
| birds | Anthus_spragueii | Anthus_spragueii | VU | Decreasing | temperate | diurnal | 25.5 | 5658382.234 | 12.23 |
| birds | Anthus_sylvanus | Anthus_sylvanus | LC | Stable | temperate | diurnal | 31 | 465229.125 | 13.084 |
| birds | Anthus_trivialis | Anthus_trivialis | LC | Decreasing | tropical | diurnal | 23.3 | 56587106.052 | 8.462 |
| birds | Anthus_vaalensis | Anthus_vaalensis | LC | Increasing | tropical | diurnal | 27 | 4452532.196 | 8.036 |
| birds | Antilophia_galeata | Antilophia_galeata | LC | Decreasing | tropical | diurnal | 21.5 | 1960407.092 | 9.706 |
| birds | Antrostomus_badius | Caprimulgus_badius | LC | Decreasing | tropical | nocturnal | 59.8 | 139506.92 | 7.143 |
| birds | Antrostomus_carolinensis | Caprimulgus_carolinensis | NT | Decreasing | tropical | nocturnal | 109 | 3554721.021 | 14.396 |
| birds | Antrostomus_cubanensis | Caprimulgus_cubanensis | LC | Decreasing | tropical | nocturnal | 63.5 | 109234.541 | 16.178 |
| birds | Antrostomus_ekmani | Caprimulgus_ekmani | LC | Stable | tropical | nocturnal | 64 | 18166.194 | 12.352 |
| birds | Antrostomus_noctitherus | Caprimulgus_noctitherus | VU | Decreasing | tropical | nocturnal | 36.1 | 7754.144 | 28.147 |
| birds | Antrostomus_ridgwayi | Caprimulgus_ridgwayi | LC | Decreasing | tropical | nocturnal | 48 | 631238.259 | 8.707 |
| birds | Antrostomus_rufus | Caprimulgus_rufus | LC | Decreasing | tropical | nocturnal | 94.2 | 7445742.526 | 6.919 |
| birds | Antrostomus_salvini | Caprimulgus_salvini | LC | Decreasing | tropical | nocturnal | 56 | 115696.333 | 11.432 |
| birds | Antrostomus_saturatus | Caprimulgus_saturatus | LC | Stable | tropical | nocturnal | 52 | 5381.48 | 6.056 |
| birds | Antrostomus_sericocaudatus | Caprimulgus_sericocaudatus | LC | Decreasing | temperate | nocturnal | 83 | 1167113.564 | 9.649 |
| birds | Antrostomus_vociferus | Caprimulgus_vociferus | NT | Stable | temperate | nocturnal | 53.3 | 4881057.024 | 15.55 |
| birds | Anumara_forbesi | NA | VU | Decreasing | NA | NA | NA | NA | NA |
| birds | Anumbius_annumbi | Anumbius_annumbi | LC | Increasing | temperate | diurnal | 41.5 | 2928674.726 | 10.325 |
| birds | Anurophasis_monorthonyx | Anurophasis_monorthonyx | LC | Stable | tropical | diurnal | 401 | 8642.935 | 4.897 |
| birds | Apalharpactes_mackloti | Apalharpactes_mackloti | LC | Stable | tropical | diurnal | 118 | 95751.559 | 7.535 |
| birds | Apalharpactes_reinwardtii | NA | VU | Decreasing | NA | NA | NA | NA | NA |
| birds | Apalis_alticola | Apalis_alticola | LC | Stable | tropical | diurnal | 12.1 | 618220.777 | 7.746 |
| birds | Apalis_bamendae | Apalis_bamendae | LC | Stable | tropical | diurnal | 9.2 | 6480.952 | 10.97 |

| | | | | | | | | | |
|---|---|---|---|---|---|---|---|---|---|
| birds | Apalis_binotata | Apalis_binotata | LC | Stable | tropical | diurnal | 8.2 | 144782.972 | 10.762 |
| birds | Apalis_chapini | Apalis_chapini | LC | Stable | tropical | diurnal | 8.8 | 57820.785 | 13.632 |
| birds | Apalis_chariessa | Apalis_chariessa | NT | Decreasing | tropical | diurnal | 9.2 | 21433.887 | 14.322 |
| birds | Apalis_chirindensis | Apalis_chirindensis | VU | Decreasing | tropical | diurnal | 8.6 | 61830.351 | 10.334 |
| birds | Apalis_cinerea | Apalis_cinerea | LC | Stable | tropical | diurnal | 9.9 | 438312.455 | 13.72 |
| birds | Apalis_flavida | Apalis_flavida | LC | Increasing | tropical | diurnal | 8.2 | 4880471.306 | 10.329 |
| birds | Apalis_goslingi | NA | LC | Stable | NA | NA | NA | NA | NA |
| birds | Apalis_jacksoni | Apalis_jacksoni | LC | Stable | tropical | diurnal | 8.7 | 541008.097 | 11.46 |
| birds | Apalis_karamojae | Apalis_karamojae | VU | Decreasing | tropical | diurnal | 9.2 | 36056.116 | 9.383 |
| birds | Apalis_melanocephala | Apalis_melanocephala | LC | Stable | tropical | diurnal | 8.8 | 375838.951 | 11.703 |
| birds | Apalis_nigriceps | Apalis_nigriceps | LC | Stable | tropical | diurnal | 8.4 | 835771.191 | 11.185 |
| birds | Apalis_personata | Apalis_personata | LC | Stable | tropical | diurnal | 11 | 154837.25 | 15.88 |
| birds | Apalis_porphyrolaema | Apalis_porphyrolaema | LC | Stable | tropical | diurnal | 8.4 | 148453.307 | 15.012 |
| birds | Apalis_ruddi | Apalis_ruddi | LC | Stable | temperate | diurnal | 9.7 | 196688.516 | 8.973 |
| birds | Apalis_rufogularis | Apalis_rufogularis | LC | Stable | tropical | diurnal | 8.3 | 2747284.944 | 7.099 |
| birds | Apalis_sharpii | Apalis_sharpii | NT | Decreasing | tropical | diurnal | 8.7 | 386331.112 | 15.447 |
| birds | Apalis_thoracica | Apalis_thoracica | LC | Stable | temperate | diurnal | 11.1 | 1229497.395 | 12.107 |
| birds | Apaloderma_aequatoriale | NA | LC | Stable | NA | NA | NA | NA | NA |
| birds | Apaloderma_narina | Apaloderma_narina | LC | Stable | tropical | diurnal | 67.7 | 10932030.887 | 9.281 |
| birds | Apaloderma_vittatum | Apaloderma_vittatum | LC | Decreasing | tropical | diurnal | 55 | 485903.026 | 13.017 |
| birds | Apalopteron_familiare | NA | NT | Stable | NA | NA | NA | NA | NA |
| birds | Aphanotriccus_audax | Aphanotriccus_audax | NT | Decreasing | tropical | diurnal | 11.2 | 63616.204 | 12.015 |
| birds | Aphanotriccus_capitalis | Aphanotriccus_capitalis | VU | Decreasing | tropical | diurnal | 11.8 | 10429.541 | 9.975 |
| birds | Aphelocephala_leucopsis | Aphelocephala_leucopsis | VU | Decreasing | temperate | diurnal | 12.3 | 3873177.352 | 4.746 |
| birds | Aphelocephala_nigricincta | Aphelocephala_nigricincta | LC | Stable | temperate | diurnal | 10.4 | 1619312.312 | 1.707 |
| birds | Aphelocephala_pectoralis | Aphelocephala_pectoralis | NT | Decreasing | temperate | diurnal | 9 | 225402.653 | 3.236 |
| birds | Aphelocoma_californica | Aphelocoma_californica | LC | Stable | temperate | diurnal | 85.7 | 2323260.891 | 9.797 |
| birds | Aphelocoma_coerulescens | Aphelocoma_coerulescens | VU | Decreasing | temperate | diurnal | 77.1 | 65032.204 | 21.42 |
| birds | Aphelocoma_ultramarina | Aphelocoma_ultramarina | LC | Decreasing | tropical | diurnal | 113.3 | 87898.138 | 19.149 |
| birds | Aphelocoma_unicolor | Aphelocoma_unicolor | LC | Decreasing | tropical | diurnal | 124 | 149103.752 | 11.506 |
| birds | Aphrastura_masafuerae | NA | CR | Stable | NA | NA | NA | NA | NA |
| birds | Aphrastura_spinicauda | Aphrastura_spinicauda | LC | Stable | temperate | diurnal | 11.5 | 491332.802 | 5.541 |
| birds | Aphrodroma_brevirostris | Lugensa_brevirostris | LC | Decreasing | temperate | cathemeral | 357 | 60101965.874 | 9.67 |
| birds | Aplonis_atrifusca | NA | LC | Unknown | NA | NA | NA | NA | NA |

| | | | | | | | | | |
|---|---|---|---|---|---|---|---|---|---|
| birds | Aplonis_brunneicapillus | Aplonis_brunneicapillus | VU | Decreasing | tropical | diurnal | 66.7 | 25066.239 | 4.68 |
| birds | Aplonis_cantoroides | Aplonis_cantoroides | LC | Unknown | tropical | diurnal | 55.7 | 822753.89 | 3.404 |
| birds | Aplonis_cinerascens | Aplonis_cinerascens | NT | Stable | tropical | diurnal | 82.9 | 47.471 | 15.463 |
| birds | Aplonis_crassa | Aplonis_crassa | NT | Decreasing | tropical | diurnal | 59 | 3445.476 | 6.292 |
| birds | Aplonis_dichroa | Aplonis_dichroa | LC | Decreasing | tropical | diurnal | 84 | 3331.938 | 4.604 |
| birds | Aplonis_feadensis | Aplonis_feadensis | NT | Stable | tropical | diurnal | 72 | 96.84 | 5.41 |
| birds | Aplonis_grandis | Aplonis_grandis | LC | Unknown | tropical | diurnal | 134 | 31140.455 | 4.719 |
| birds | Aplonis_insularis | Aplonis_insularis | LC | Decreasing | tropical | diurnal | 69.9 | 539.039 | 3.385 |
| birds | Aplonis_magna | Aplonis_magna | LC | Unknown | tropical | diurnal | 190 | 2863.253 | 8.947 |
| birds | Aplonis_mavornata | NA | EX | not_available | NA | NA | NA | NA | NA |
| birds | Aplonis_metallica | Aplonis_metallica | LC | Decreasing | tropical | diurnal | 56.5 | 762829.934 | 3.663 |
| birds | Aplonis_minor | Aplonis_minor | LC | Decreasing | tropical | diurnal | 42 | 325906.875 | 16.321 |
| birds | Aplonis_mysolensis | Aplonis_mysolensis | LC | Unknown | tropical | diurnal | 62 | 64402.762 | 6.434 |
| birds | Aplonis_mystacea | Aplonis_mystacea | LC | Decreasing | tropical | diurnal | 68.7 | 84944.016 | 2.251 |
| birds | Aplonis_opaca | Aplonis_opaca | LC | Unknown | tropical | diurnal | 83.4 | 2018.985 | 16.726 |
| birds | Aplonis_panayensis | Aplonis_panayensis | LC | Unknown | tropical | diurnal | 56.2 | 2145170.846 | 10.343 |
| birds | Aplonis_pelzelni | Aplonis_pelzelni | CR | Decreasing | tropical | diurnal | 47 | 61.486 | 6.682 |
| birds | Aplonis_santovestris | Aplonis_santovestris | EN | Decreasing | tropical | diurnal | 68.7 | 59.602 | 2.779 |
| birds | Aplonis_striata | Aplonis_striata | LC | Unknown | tropical | diurnal | 54.2 | 17941.658 | 7.523 |
| birds | Aplonis_tabuensis | Aplonis_tabuensis | LC | Unknown | tropical | diurnal | 62.5 | 23355.895 | 9.109 |
| birds | Aplonis_zelandica | Aplonis_zelandica | LC | Decreasing | tropical | diurnal | 68.7 | 12296.11 | 5.723 |
| birds | Aplopelia_larvata | Aplopelia_larvata | LC | Stable | tropical | diurnal | 155 | 2621498.987 | 10.387 |
| birds | Aprositornis_disjuncta | Myrmeciza_disjuncta | LC | Unknown | tropical | diurnal | 15 | 42618.454 | 0.875 |
| birds | Aprosmictus_erythropterus | Aprosmictus_erythropterus | LC | Increasing | tropical | diurnal | 136 | 2743055.211 | 4.5 |
| birds | Aprosmictus_jonquillaceus | Aprosmictus_jonquillaceus | NT | Decreasing | tropical | diurnal | 119 | 28185.23 | 12.909 |
| birds | Aptenodytes_forsteri | NA | NT | Decreasing | NA | NA | NA | NA | NA |
| birds | Aptenodytes_patagonicus | Aptenodytes_patagonicus | LC | Increasing | temperate | cathemeral | 11731.1 | 21657131.922 | 3.302 |
| birds | Apteryx_australis | Apteryx_australis | VU | Decreasing | temperate | nocturnal | 2320.5 | 11025.49 | 1.721 |
| birds | Apteryx_haastii | Apteryx_haastii | VU | Decreasing | temperate | nocturnal | 2022.7 | 40104.056 | 4.31 |
| birds | Apteryx_owenii | Apteryx_owenii | NT | Increasing | temperate | nocturnal | 1238.3 | 84.061 | 8.373 |
| birds | Apus_affinis | Apus_affinis | LC | Increasing | temperate | diurnal | 22.6 | 22143832.392 | 10.676 |
| birds | Apus_alexandri | NA | LC | Stable | NA | NA | NA | NA | NA |
| birds | Apus_apus | Apus_apus | LC | Stable | temperate | diurnal | 37.6 | 42973934.897 | 9.848 |
| birds | Apus_barbatus | Apus_barbatus | LC | Decreasing | temperate | diurnal | 42.8 | 1200605.416 | 12.829 |

| | | | | | | | | | |
|---|---|---|---|---|---|---|---|---|---|
| birds | Apus_berliozi | Apus_berliozi | LC | Stable | tropical | diurnal | 38.2 | 214046.126 | 8.22 |
| birds | Apus_bradfieldi | Apus_bradfieldi | LC | Stable | tropical | diurnal | 42.4 | 882341.758 | 5.401 |
| birds | Apus_caffer | Apus_caffer | LC | Increasing | tropical | diurnal | 22.1 | 12490681.946 | 10.294 |
| birds | Apus_horus | Apus_horus | LC | Increasing | tropical | diurnal | 27.5 | 2043600.713 | 11.612 |
| birds | Apus_niansae | Apus_niansae | LC | Stable | tropical | diurnal | 32.5 | 381006.405 | 12.465 |
| birds | Apus_nipalensis | Apus_nipalensis | LC | Increasing | tropical | diurnal | 29.8 | 5768014.176 | 12.053 |
| birds | Apus_pacificus | Apus_pacificus | LC | Stable | temperate | diurnal | 39.4 | 28562076.188 | 7.884 |
| birds | Apus_pallidus | Apus_pallidus | LC | Stable | tropical | diurnal | 41.9 | 9749752.171 | 10.327 |
| birds | Apus_unicolor | Apus_unicolor | LC | Unknown | temperate | diurnal | 22.5 | 16132.272 | 15.424 |
| birds | Aquila_adalberti | Aquila_adalberti | VU | Increasing | temperate | diurnal | 2958 | 1072953.524 | 7.861 |
| birds | Aquila_africana | Aquila_africanus | LC | Decreasing | tropical | diurnal | 1047 | 1918245.902 | 8.213 |
| birds | Aquila_audax | Aquila_audax | LC | Increasing | temperate | diurnal | 3449.9 | 7679648.031 | 4.263 |
| birds | Aquila_chrysaetos | Aquila_chrysaetos | LC | Stable | temperate | diurnal | 4248 | 43360026.833 | 7.519 |
| birds | Aquila_fasciata | Aquila_fasciatus | LC | Decreasing | tropical | diurnal | 1936.5 | 5912536.574 | 14.497 |
| birds | Aquila_gurneyi | Aquila_gurneyi | NT | Decreasing | tropical | diurnal | 3060 | 601810.346 | 3.435 |
| birds | Aquila_heliaca | Aquila_heliaca | VU | Decreasing | temperate | diurnal | 3125.2 | 31736715.066 | 9.441 |
| birds | Aquila_nipalensis | Aquila_nipalensis | EN | Decreasing | tropical | diurnal | 2714.3 | 30326757.836 | 9.434 |
| birds | Aquila_rapax | Aquila_rapax | VU | Decreasing | temperate | diurnal | 2236.1 | 17572054.356 | 10.458 |
| birds | Aquila_spilogaster | Hieraaetus_spilogaster | LC | Decreasing | tropical | diurnal | 1466.2 | 12423292.008 | 9.656 |
| birds | Aquila_verreauxii | Aquila_verreauxii | LC | Stable | temperate | diurnal | 4195 | 4020161.708 | 9.347 |
| birds | Ara_ambiguus | Ara_ambiguus | CR | Decreasing | tropical | diurnal | 1300 | 151979.977 | 5.875 |
| birds | Ara_ararauna | Ara_ararauna | LC | Decreasing | tropical | diurnal | 1125 | 7836201.125 | 3.929 |
| birds | Ara_chloropterus | Ara_chloropterus | LC | Decreasing | tropical | diurnal | 1214 | 8101232.431 | 3.459 |
| birds | Ara_glaucogularis | Ara_glaucogularis | CR | Stable | tropical | diurnal | 784 | 30260.157 | 6.996 |
| birds | Ara_macao | Ara_macao | LC | Decreasing | tropical | diurnal | 1015 | 7046076.254 | 3.024 |
| birds | Ara_militaris | Ara_militaris | VU | Decreasing | tropical | diurnal | 1134 | 591218.424 | 7.248 |
| birds | Ara_rubrogenys | Ara_rubrogenys | CR | Decreasing | tropical | diurnal | 468 | 27916.969 | 9.682 |
| birds | Ara_severus | Ara_severus | LC | Decreasing | tropical | diurnal | 343 | 5828903.929 | 3.538 |
| birds | Arachnothera_affinis | Arachnothera_affinis | LC | Decreasing | tropical | diurnal | 26.4 | 349980.156 | 11.33 |
| birds | Arachnothera_chrysogenys | Arachnothera_chrysogenys | LC | Decreasing | tropical | diurnal | 23.2 | 1237350.362 | 8.662 |
| birds | Arachnothera_clarae | Arachnothera_clarae | LC | Stable | tropical | diurnal | 30.1 | 183271.357 | 13.581 |
| birds | Arachnothera_crassirostris | Arachnothera_crassirostris | LC | Decreasing | tropical | diurnal | 16 | 899703.482 | 7.921 |
| birds | Arachnothera_flavigaster | Arachnothera_flavigaster | LC | Decreasing | tropical | diurnal | 38.4 | 1166970.148 | 7.89 |
| birds | Arachnothera_juliae | NA | LC | Decreasing | NA | NA | NA | NA | NA |

| | | | | | | | | | |
|---|---|---|---|---|---|---|---|---|---|
| birds | Arachnothera_longirostra | Arachnothera_longirostra | LC | Stable | tropical | diurnal | 12.5 | 3514761.529 | 10.897 |
| birds | Arachnothera_magna | Arachnothera_magna | LC | Stable | tropical | diurnal | 30.7 | 1984624.624 | 11.122 |
| birds | Arachnothera_robusta | Arachnothera_robusta | LC | Decreasing | tropical | diurnal | 29.3 | 630859.749 | 12.726 |
| birds | Aramides_cajaneus | Aramides_cajanea | LC | Stable | tropical | nocturnal | 397 | 13038120.137 | 5.664 |
| birds | Aramidopsis_plateni | NA | VU | Decreasing | NA | NA | NA | NA | NA |
| birds | Aramus_guarauna | Aramus_guarauna | LC | Stable | tropical | cathemeral | 1080 | 12204575.672 | 7.293 |
| birds | Aratinga_auricapillus | Aratinga_auricapillus | LC | Decreasing | tropical | diurnal | 130 | 1320183.999 | 12.294 |
| birds | Aratinga_jandaya | Aratinga_jandaya | LC | Stable | tropical | diurnal | 89.4 | 1514385.78 | 6.856 |
| birds | Aratinga_nenday | Nandayus_nenday | LC | Increasing | temperate | diurnal | 128 | 373769.01 | 8.742 |
| birds | Aratinga_solstitialis | Aratinga_solstitialis | EN | Decreasing | tropical | diurnal | 119.6 | 86113.473 | 3.858 |
| birds | Aratinga_weddellii | Aratinga_weddellii | LC | Increasing | tropical | diurnal | 108 | 2260164.386 | 2.307 |
| birds | Arborophila_ardens | Arborophila_ardens | VU | Decreasing | tropical | diurnal | 266.6 | 13199.579 | 11.647 |
| birds | Arborophila_atrogularis | Arborophila_atrogularis | LC | Decreasing | temperate | diurnal | 237.3 | 355074.89 | 10.303 |
| birds | Arborophila_brunneopectus | Arborophila_brunneopectus | LC | Decreasing | tropical | diurnal | 291.5 | 486283.059 | 8.357 |
| birds | Arborophila_cambodiana | Arborophila_cambodiana | LC | Decreasing | tropical | diurnal | 285.9 | 17928.219 | 9.465 |
| birds | Arborophila_charltonii | NA | not_available | not_available | NA | NA | NA | NA | NA |
| birds | Arborophila_chloropus | NA | not_available | not_available | NA | NA | NA | NA | NA |
| birds | Arborophila_crudigularis | Arborophila_crudigularis | LC | Decreasing | temperate | diurnal | 256.8 | 22768.902 | 12.653 |
| birds | Arborophila_davidi | Arborophila_davidi | NT | Decreasing | tropical | diurnal | 241 | 4465.394 | 7.275 |
| birds | Arborophila_gingica | Arborophila_gingica | NT | Decreasing | temperate | diurnal | 253 | 396418.35 | 12.478 |
| birds | Arborophila_hyperythra | Arborophila_hyperythra | LC | Decreasing | tropical | diurnal | 270 | 161020.223 | 2.428 |
| birds | Arborophila_javanica | Arborophila_javanica | LC | Decreasing | tropical | diurnal | 272 | 36961.991 | 17.866 |
| birds | Arborophila_mandellii | Arborophila_mandellii | NT | Decreasing | temperate | diurnal | 268 | 74187.758 | 5.984 |
| birds | Arborophila_orientalis | Arborophila_orientalis | VU | Decreasing | tropical | diurnal | 336 | 10707.851 | 19.9 |
| birds | Arborophila_rubrirostris | Arborophila_rubrirostris | LC | Decreasing | tropical | diurnal | 225.4 | 69473.398 | 7.907 |
| birds | Arborophila_rufipectus | Arborophila_rufipectus | EN | Decreasing | temperate | diurnal | 400.1 | 7707.359 | 11.924 |
| birds | Arborophila_rufogularis | Arborophila_rufogularis | LC | Decreasing | tropical | diurnal | 344.5 | 1245539.712 | 8.472 |
| birds | Arborophila_torqueola | Arborophila_torqueola | LC | Stable | temperate | diurnal | 344.5 | 721402.086 | 7.706 |
| birds | Arcanator_orostruthus | Modulatrix_orostruthus | NT | Decreasing | tropical | diurnal | 32.7 | 8795.864 | 8.087 |
| birds | Archboldia_papuensis | Archboldia_papuensis | LC | Stable | tropical | diurnal | 177.6 | 80739.179 | 5.185 |
| birds | Archilochus_alexandri | Archilochus_alexandri | LC | Increasing | temperate | diurnal | 3.4 | 3841632.298 | 9.576 |
| birds | Archilochus_colubris | Archilochus_colubris | LC | Increasing | temperate | diurnal | 3.1 | 7444664.684 | 14.207 |
| birds | Ardea_alba | Casmerodius_albus | LC | Unknown | temperate | diurnal | 871.3 | 80817244.414 | 9.749 |
| birds | Ardea_cinerea | Ardea_cinerea | LC | Unknown | tropical | cathemeral | 1443 | 51699618.43 | 10.846 |

| | | | | | | | | | |
|---|---|---|---|---|---|---|---|---|---|
| birds | Ardea_cocoi | Ardea_cocoi | LC | Stable | tropical | cathemeral | 1752.4 | 14742289.158 | 6.017 |
| birds | Ardea_goliath | Ardea_goliath | LC | Stable | tropical | diurnal | 4468 | 17921027.03 | 9.294 |
| birds | Ardea_herodias | Ardea_herodias | LC | Increasing | temperate | cathemeral | 2523.4 | 14906902.55 | 10.846 |
| birds | Ardea_humbloti | NA | EN | Decreasing | NA | NA | NA | NA | NA |
| birds | Ardea_insignis | NA | CR | Decreasing | NA | NA | NA | NA | NA |
| birds | Ardea_intermedia | Mesophoyx_intermedia | LC | Decreasing | tropical | diurnal | 458.8 | 8898068.598 | 15.316 |
| birds | Ardea_melanocephala | Ardea_melanocephala | LC | Increasing | tropical | cathemeral | 1060 | 19067243.575 | 8.986 |
| birds | Ardea_pacifica | Ardea_pacifica | LC | Stable | temperate | diurnal | 881 | 6934005.006 | 4.543 |
| birds | Ardea_purpurea | Ardea_purpurea | LC | Decreasing | tropical | diurnal | 1064.5 | 42294330.035 | 11.075 |
| birds | Ardea_sumatrana | Ardea_sumatrana | LC | Decreasing | tropical | diurnal | 2024 | 1764501.955 | 7.139 |
| birds | Ardenna_bulleri | Puffinus_bulleri | VU | Stable | tropical | cathemeral | 407 | 109799796.006 | 6.668 |
| birds | Ardenna_carneipes | Puffinus_carneipes | NT | Decreasing | tropical | cathemeral | 609 | 117469913.741 | 10.061 |
| birds | Ardenna_creatopus | Puffinus_creatopus | VU | Unknown | tropical | cathemeral | 744 | 14595563.547 | 5.818 |
| birds | Ardenna_gravis | Puffinus_gravis | LC | Stable | tropical | cathemeral | 849 | 63607169.789 | 8.697 |
| birds | Ardenna_grisea | Puffinus_griseus | NT | Decreasing | tropical | cathemeral | 787 | 213006301.652 | 8.783 |
| birds | Ardenna_pacifica | Puffinus_pacificus | LC | Decreasing | tropical | cathemeral | 388 | 119963361.11 | 7.442 |
| birds | Ardenna_tenuirostris | Puffinus_tenuirostris | LC | Decreasing | tropical | cathemeral | 559 | 112949641.511 | 9.724 |
| birds | Ardeola_bacchus | Ardeola_bacchus | LC | Stable | temperate | diurnal | 304.9 | 7868535.624 | 14.232 |
| birds | Ardeola_grayii | Ardeola_grayii | LC | Unknown | tropical | cathemeral | 253 | 3991850.348 | 17.087 |
| birds | Ardeola_idae | NA | EN | Decreasing | NA | NA | NA | NA | NA |
| birds | Ardeola_ralloides | Ardeola_ralloides | LC | Unknown | temperate | diurnal | 287 | 34073703.041 | 8.043 |
| birds | Ardeola_rufiventris | NA | LC | Unknown | NA | NA | NA | NA | NA |
| birds | Ardeotis_arabs | Ardeotis_arabs | NT | Decreasing | tropical | diurnal | 5943.5 | 2950367.617 | 9.527 |
| birds | Ardeotis_australis | Ardeotis_australis | LC | Decreasing | temperate | diurnal | 4182.4 | 6717248.953 | 3.46 |
| birds | Ardeotis_kori | Ardeotis_kori | NT | Decreasing | temperate | diurnal | 7994.2 | 3565934.248 | 7.387 |
| birds | Ardeotis_nigriceps | Ardeotis_nigriceps | CR | Decreasing | temperate | diurnal | 7235.3 | 1064221.447 | 19.768 |
| birds | Arenaria_interpres | Arenaria_interpres | LC | Decreasing | temperate | diurnal | 136 | 13072863.93 | 8.86 |
| birds | Arenaria_melanocephala | Arenaria_melanocephala | LC | Stable | temperate | diurnal | 126.2 | 1073634.759 | 4.605 |
| birds | Argusianus_argus | Argusianus_argus | VU | Decreasing | tropical | diurnal | 1959.9 | 1480121.151 | 9.059 |
| birds | Argya_affinis | Turdoides_affinis | LC | Stable | tropical | diurnal | 63.3 | 745811.265 | 20.282 |
| birds | Argya_altirostris | Turdoides_altirostris | LC | Decreasing | temperate | diurnal | 33 | 149200.158 | 18.027 |
| birds | Argya_aylmeri | Turdoides_aylmeri | LC | Stable | tropical | diurnal | 35.3 | 1055408.062 | 9.776 |
| birds | Argya_caudata | Turdoides_caudata | LC | Stable | tropical | diurnal | 39.2 | 3191144.088 | 16.031 |
| birds | Argya_cinereifrons | Garrulax_cinereifrons | VU | Decreasing | tropical | diurnal | 70 | 11916.823 | 20.912 |

| | | | | | | | | | |
|---|---|---|---|---|---|---|---|---|---|
| birds | Argya_earlei | Turdoides_earlei | LC | Decreasing | temperate | diurnal | 47 | 1024770.226 | 18.334 |
| birds | Argya_fulva | Turdoides_fulva | LC | Stable | temperate | diurnal | 61.4 | 3666952.83 | 4.622 |
| birds | Argya_gularis | Turdoides_gularis | LC | Stable | tropical | diurnal | 45.5 | 216441.284 | 14.029 |
| birds | Argya_longirostris | Turdoides_longirostris | VU | Decreasing | temperate | diurnal | 35 | 28660.299 | 15.298 |
| birds | Argya_malcolmi | Turdoides_malcolmi | LC | Stable | tropical | diurnal | 75.7 | 1922153.024 | 19.282 |
| birds | Argya_rubiginosa | Turdoides_rubiginosa | LC | Stable | tropical | diurnal | 40.6 | 1410146.948 | 9.61 |
| birds | Argya_rufescens | Turdoides_rufescens | LC | Decreasing | tropical | diurnal | 64.5 | 12170.707 | 19.729 |
| birds | Argya_squamiceps | Turdoides_squamiceps | LC | Increasing | tropical | diurnal | 72.9 | 1273158.322 | 6.912 |
| birds | Argya_subrufa | Turdoides_subrufa | LC | Stable | tropical | diurnal | 66.7 | 178058.684 | 19.562 |
| birds | Arizelocichla_masukuensis | Andropadus_masukuensis | LC | Stable | tropical | diurnal | 26.8 | 163721.679 | 13.879 |
| birds | Arizelocichla_milanjensis | Andropadus_milanjensis | NT | Decreasing | tropical | diurnal | 37.4 | 42374.095 | 10.18 |
| birds | Arizelocichla_montana | Andropadus_montanus | LC | Decreasing | tropical | diurnal | 32.6 | 61327.265 | 9.061 |
| birds | Arizelocichla_tephrolaema | Andropadus_tephrolaemus | LC | Stable | tropical | diurnal | 38.3 | 59489.94 | 9.676 |
| birds | Arizelopsar_femoralis | NA | EN | Decreasing | NA | NA | NA | NA | NA |
| birds | Arremon_abeillei | Arremon_abeillei | LC | Decreasing | tropical | diurnal | 25.9 | 52316.766 | 11.692 |
| birds | Arremon_aurantiirostris | Arremon_aurantiirostris | LC | Decreasing | tropical | diurnal | 34.5 | 693974.169 | 7.971 |
| birds | Arremon_brunneinucha | Arremon_brunneinucha | LC | Decreasing | tropical | diurnal | 44 | 618730.155 | 9.698 |
| birds | Arremon_castaneiceps | Arremon_castaneiceps | NT | Decreasing | tropical | diurnal | 36.9 | 122751.416 | 5.802 |
| birds | Arremon_crassirostris | Arremon_crassirostris | LC | Decreasing | tropical | diurnal | 38.7 | 9497.109 | 6.908 |
| birds | Arremon_flavirostris | Arremon_flavirostris | LC | Stable | tropical | diurnal | 26.1 | 1940265.881 | 10.895 |
| birds | Arremon_franciscanus | Arremon_franciscanus | NT | Decreasing | tropical | diurnal | 24.9 | 53090.524 | 10.483 |
| birds | Arremon_schlegeli | Arremon_schlegeli | LC | Stable | tropical | diurnal | 27.7 | 187790.908 | 14.135 |
| birds | Arremon_semitorquatus | Arremon_semitorquatus | LC | Decreasing | tropical | diurnal | 25 | 215137.213 | 14.065 |
| birds | Arremon_taciturnus | Arremon_taciturnus | LC | Stable | tropical | diurnal | 24.8 | 6755484.618 | 3.949 |
| birds | Arremon_torquatus | Arremon_torquatus | LC | Decreasing | tropical | diurnal | 42 | 113027.409 | 7.168 |
| birds | Arremon_virenticeps | Arremon_virenticeps | LC | Decreasing | tropical | diurnal | 41 | 79786.326 | 9.221 |
| birds | Arremonops_chloronotus | Arremonops_chloronotus | LC | Decreasing | tropical | diurnal | 27.4 | 234300.884 | 7.921 |
| birds | Arremonops_conirostris | Arremonops_conirostris | LC | Stable | tropical | diurnal | 35.9 | 1132799.464 | 9.041 |
| birds | Arremonops_rufivirgatus | Arremonops_rufivirgatus | LC | Stable | tropical | diurnal | 22.4 | 553968.491 | 11.133 |
| birds | Arremonops_tocuyensis | Arremonops_tocuyensis | LC | Stable | tropical | diurnal | 29.4 | 72802.633 | 12.109 |
| birds | Arses_insularis | Arses_insularis | LC | Stable | tropical | diurnal | 19.3 | 171995.995 | 3.403 |
| birds | Arses_kaupi | Arses_kaupi | LC | Stable | tropical | diurnal | 13.5 | 27770.733 | 5.407 |
| birds | Arses_telescopthalmus | Arses_telescophthalmus | LC | Stable | tropical | diurnal | 15.3 | 551895.051 | 3.24 |
| birds | Artamella_viridis | Artamella_viridis | LC | Unknown | tropical | diurnal | 48.8 | 415338.92 | 8.361 |

| | | | | | | | | | |
|---|---|---|---|---|---|---|---|---|---|
| birds | Artamus_cinereus | Artamus_cinereus | LC | Increasing | temperate | diurnal | 35.6 | 7423572.802 | 4.033 |
| birds | Artamus_cyanopterus | Artamus_cyanopterus | LC | Decreasing | temperate | diurnal | 34.6 | 3897118.378 | 6.358 |
| birds | Artamus_fuscus | Artamus_fuscus | LC | Stable | tropical | diurnal | 39.8 | 4701116.551 | 15.467 |
| birds | Artamus_insignis | Artamus_insignis | LC | Stable | tropical | diurnal | 51.3 | 37843.106 | 4.519 |
| birds | Artamus_leucoryn | Artamus_leucorynchus | LC | Stable | temperate | diurnal | 41.7 | 7650935.5 | 5.777 |
| birds | Artamus_maximus | Artamus_maximus | LC | Stable | tropical | diurnal | 58.2 | 248677.848 | 4.101 |
| birds | Artamus_mentalis | Artamus_mentalis | LC | Decreasing | tropical | diurnal | 41.9 | 17666.455 | 9.362 |
| birds | Artamus_minor | Artamus_minor | LC | Stable | tropical | diurnal | 16 | 6485166.891 | 3.58 |
| birds | Artamus_monachus | NA | LC | Stable | NA | NA | NA | NA | NA |
| birds | Artamus_personatus | Artamus_personatus | LC | Stable | temperate | diurnal | 34.6 | 7500863.716 | 4.275 |
| birds | Artamus_superciliosus | Artamus_superciliosus | LC | Stable | temperate | diurnal | 35.5 | 7162672.293 | 4.123 |
| birds | Artemisiospiza_belli | Amphispiza_belli | LC | Stable | temperate | diurnal | 17 | 304827.375 | 8.005 |
| birds | Artisornis_metopias | Artisornis_metopias | LC | Decreasing | tropical | diurnal | 8.4 | 68472.652 | 12.389 |
| birds | Artisornis_moreaui | Artisornis_moreaui | CR | Decreasing | tropical | diurnal | 9.2 | 396.938 | 13.972 |
| birds | Artomyias_fuliginosa | Muscicapa_infuscata | LC | Stable | tropical | diurnal | 17.7 | 3058848.689 | 6.918 |
| birds | Artomyias_ussheri | Muscicapa_ussheri | LC | Stable | tropical | diurnal | 17.8 | 473137.786 | 15.491 |
| birds | Arundinax_aedon | Acrocephalus_aedon | LC | Decreasing | temperate | diurnal | 22.4 | 14630790.955 | 10.677 |
| birds | Arundinicola_leucocephala | Arundinicola_leucocephala | LC | Stable | tropical | diurnal | 13.8 | 9087776.957 | 6.643 |
| birds | Asarcornis_scutulata | Cairina_scutulata | EN | Decreasing | temperate | nocturnal | 2937.4 | 999723.979 | 10.02 |
| birds | Asemospiza_fuliginosa | Tiaris_fuliginosus | LC | Stable | tropical | diurnal | 13.3 | 951776.681 | 12.062 |
| birds | Asemospiza_obscura | Tiaris_obscurus | LC | Increasing | tropical | diurnal | 11.2 | 1522806.977 | 7.491 |
| birds | Ashbyia_lovensis | Ashbyia_lovensis | LC | Increasing | temperate | diurnal | 17.5 | 699489.419 | 3.037 |
| birds | Asio_abyssinicus | Asio_abyssinicus | LC | Stable | tropical | nocturnal | 322.5 | 153995.929 | 14.192 |
| birds | Asio_capensis | Asio_capensis | LC | Stable | tropical | nocturnal | 334 | 7713158.388 | 9.474 |
| birds | Asio_clamator | Pseudoscops_clamator | LC | Decreasing | tropical | nocturnal | 443.3 | 9101422.656 | 8.837 |
| birds | Asio_flammeus | Asio_flammeus | LC | Decreasing | temperate | cathemeral | 322.6 | 60054481.497 | 8.514 |
| birds | Asio_madagascariensis | NA | LC | Decreasing | NA | NA | NA | NA | NA |
| birds | Asio_otus | Asio_otus | LC | Decreasing | temperate | nocturnal | 296.6 | 35423274.659 | 10.489 |
| birds | Asio_stygius | Asio_stygius | LC | Decreasing | tropical | nocturnal | 565 | 5772064.129 | 9.039 |
| birds | Aspatha_gularis | Aspatha_gularis | LC | Decreasing | tropical | diurnal | 62.7 | 99079.025 | 12.712 |
| birds | Asthenes_anthoides | Asthenes_anthoides | LC | Stable | temperate | diurnal | 22.3 | 345958.92 | 7.51 |
| birds | Asthenes_arequipae | Asthenes_arequipae | LC | Decreasing | tropical | diurnal | 20 | 98013.952 | 7.19 |
| birds | Asthenes_baeri | Asthenes_baeri | LC | Decreasing | temperate | diurnal | 17.8 | 1726715.051 | 8.76 |
| birds | Asthenes_berlepschi | Asthenes_berlepschi | NT | Decreasing | tropical | diurnal | 67 | 1405.372 | 9.237 |

| | | | | | | | | | |
|---|---|---|---|---|---|---|---|---|---|
| birds | Asthenes_coryi | Schizoeaca_coryi | LC | Stable | tropical | diurnal | 16.8 | 9010.227 | 14.135 |
| birds | Asthenes_dorbignyi | Asthenes_dorbignyi | LC | Decreasing | temperate | diurnal | 15.9 | 252131.25 | 7.323 |
| birds | Asthenes_flammulata | Asthenes_flammulata | LC | Unknown | tropical | diurnal | 22.5 | 122945.373 | 10.647 |
| birds | Asthenes_fuliginosa | Schizoeaca_fuliginosa | LC | Stable | tropical | diurnal | 18.3 | 96182.508 | 10.28 |
| birds | Asthenes_griseomurina | Schizoeaca_griseomurina | LC | Decreasing | tropical | diurnal | 17.6 | 18912.895 | 9.441 |
| birds | Asthenes_harterti | Schizoeaca_harterti | LC | Decreasing | tropical | diurnal | 12.9 | 23849.1 | 5.307 |
| birds | Asthenes_helleri | Schizoeaca_helleri | LC | Stable | tropical | diurnal | 15.3 | 17004.804 | 6.742 |
| birds | Asthenes_heterura | Asthenes_heterura | LC | Decreasing | tropical | diurnal | 13.5 | 93072.436 | 8.997 |
| birds | Asthenes_huancavelicae | Asthenes_huancavelicae | LC | Decreasing | tropical | diurnal | 22.2 | 18528.088 | 12.091 |
| birds | Asthenes_hudsoni | Asthenes_hudsoni | NT | Decreasing | temperate | diurnal | 19 | 803230.146 | 10.591 |
| birds | Asthenes_humilis | Asthenes_humilis | LC | Stable | tropical | diurnal | 21 | 146012.523 | 8.513 |
| birds | Asthenes_luizae | Asthenes_luizae | NT | Decreasing | tropical | diurnal | 28 | 7704.865 | 4.586 |
| birds | Asthenes_maculicauda | Asthenes_maculicauda | LC | Decreasing | tropical | diurnal | 19 | 48997.23 | 6.76 |
| birds | Asthenes_modesta | Asthenes_modesta | LC | Decreasing | temperate | diurnal | 16.8 | 2015310.779 | 5.72 |
| birds | Asthenes_moreirae | Oreophylax_moreirae | LC | Decreasing | tropical | diurnal | 10.5 | 39562.85 | 12.905 |
| birds | Asthenes_ottonis | Asthenes_ottonis | LC | Stable | tropical | diurnal | 13 | 48172.577 | 10.338 |
| birds | Asthenes_palpebralis | Schizoeaca_palpebralis | LC | Decreasing | tropical | diurnal | 17 | 11496.434 | 6.657 |
| birds | Asthenes_perijana | Schizoeaca_perijana | EN | Decreasing | tropical | diurnal | 18.3 | 111.255 | 6.731 |
| birds | Asthenes_pudibunda | Asthenes_pudibunda | LC | Decreasing | tropical | diurnal | 14.9 | 89205.068 | 7.141 |
| birds | Asthenes_pyrrholeuca | Asthenes_pyrrholeuca | LC | Decreasing | temperate | diurnal | 13.2 | 2740647.457 | 7.633 |
| birds | Asthenes_urubambensis | Asthenes_urubambensis | NT | Decreasing | tropical | diurnal | 16 | 63948.662 | 5.854 |
| birds | Asthenes_vilcabambae | Schizoeaca_vilcabambae | LC | Stable | tropical | diurnal | 20.2 | 13148.349 | 3.237 |
| birds | Asthenes_virgata | Asthenes_virgata | LC | Unknown | tropical | diurnal | 22 | 46703.347 | 8.639 |
| birds | Asthenes_wyatti | Asthenes_wyatti | LC | Stable | tropical | diurnal | 15.2 | 467539.801 | 7.922 |
| birds | Astrapia_mayeri | Astrapia_mayeri | LC | Decreasing | tropical | diurnal | 139.3 | 11001.105 | 5.361 |
| birds | Astrapia_nigra | Astrapia_nigra | LC | Stable | tropical | diurnal | 190 | 3721.195 | 3.31 |
| birds | Astrapia_rothschildi | Astrapia_rothschildi | LC | Stable | tropical | diurnal | 181.4 | 6673.295 | 3.645 |
| birds | Astrapia_splendidissima | Astrapia_splendidissima | LC | Stable | tropical | diurnal | 131.3 | 42648.979 | 4.457 |
| birds | Astrapia_stephaniae | Astrapia_stephaniae | LC | Stable | tropical | diurnal | 141.8 | 55589.818 | 5.813 |
| birds | Atalotriccus_pilaris | Atalotriccus_pilaris | LC | Stable | tropical | diurnal | 6.8 | 813969.642 | 9.338 |
| birds | Atelornis_crossleyi | Atelornis_crossleyi | NT | Decreasing | tropical | cathemeral | 79 | 22418.548 | 5.372 |
| birds | Atelornis_pittoides | Atelornis_pittoides | LC | Stable | tropical | cathemeral | 91.5 | 45907.821 | 5.224 |
| birds | Athene_blewitti | Heteroglaux_blewitti | EN | Decreasing | tropical | diurnal | 241 | 5435.608 | 11.307 |
| birds | Athene_brama | Athene_brama | LC | Stable | tropical | nocturnal | 112 | 5213398.062 | 15.867 |

| | | | | | | | | | |
|---|---|---|---|---|---|---|---|---|---|
| birds | Athene_noctua | Athene_noctua | LC | Stable | temperate | nocturnal | 168.9 | 26928713.112 | 9.748 |
| birds | Athene_superciliaris | Ninox_superciliaris | LC | Decreasing | tropical | nocturnal | 231 | 415419.42 | 8.469 |
| birds | Atimastillas_flavicollis | Chlorocichla_flavicollis | LC | Stable | tropical | diurnal | 48.7 | 1683203.479 | 12.834 |
| birds | Atlapetes_albiceps | Atlapetes_albiceps | LC | Stable | tropical | diurnal | 37 | 26300.947 | 7.739 |
| birds | Atlapetes_albinucha | Atlapetes_albinucha | LC | Decreasing | tropical | diurnal | 33.8 | 275050.2 | 11.972 |
| birds | Atlapetes_albofrenatus | Atlapetes_albofrenatus | LC | Stable | tropical | diurnal | 30 | 31052.546 | 13.742 |
| birds | Atlapetes_canigenis | Atlapetes_canigenis | LC | Decreasing | tropical | diurnal | 26.6 | 35227.792 | 4.417 |
| birds | Atlapetes_citrinellus | Atlapetes_citrinellus | LC | Decreasing | temperate | diurnal | 28 | 13460.089 | 8.286 |
| birds | Atlapetes_flaviceps | Atlapetes_flaviceps | NT | Decreasing | tropical | diurnal | 33.7 | 38053.809 | 13.239 |
| birds | Atlapetes_forbesi | Atlapetes_forbesi | LC | Decreasing | tropical | diurnal | 32.2 | 18195.915 | 9.93 |
| birds | Atlapetes_fulviceps | Atlapetes_fulviceps | LC | Decreasing | tropical | diurnal | 28.2 | 77616.92 | 8.177 |
| birds | Atlapetes_fuscoolivaceus | Atlapetes_fuscoolivaceus | NT | Decreasing | tropical | diurnal | 32.5 | 7366.712 | 12.631 |
| birds | Atlapetes_latinuchus | Atlapetes_latinuchus | LC | Decreasing | tropical | diurnal | 23.6 | 185718.082 | 9.898 |
| birds | Atlapetes_leucopis | Atlapetes_leucopis | LC | Stable | tropical | diurnal | 43 | 12200.303 | 4.072 |
| birds | Atlapetes_leucopterus | Atlapetes_leucopterus | LC | Stable | tropical | diurnal | 20.8 | 65183.834 | 10.216 |
| birds | Atlapetes_melanocephalus | Atlapetes_melanocephalus | LC | Stable | tropical | diurnal | 25 | 4032.812 | 5.406 |
| birds | Atlapetes_melanolaemus | Atlapetes_melanolaemus | LC | Decreasing | tropical | diurnal | 28.9 | 61848.895 | 5.792 |
| birds | Atlapetes_melanopsis | Atlapetes_melanopsis | NT | Decreasing | tropical | diurnal | 29.6 | 4307.821 | 7.629 |
| birds | Atlapetes_nationi | Atlapetes_nationi | LC | Stable | tropical | diurnal | 41.6 | 33065.045 | 7.049 |
| birds | Atlapetes_pallidiceps | Atlapetes_pallidiceps | EN | Stable | tropical | diurnal | 27.5 | 915.004 | 12.356 |
| birds | Atlapetes_pallidinucha | Atlapetes_pallidinucha | LC | Stable | tropical | diurnal | 34.6 | 97974.809 | 10.011 |
| birds | Atlapetes_personatus | Atlapetes_personatus | LC | Stable | tropical | diurnal | 32.7 | 99663.065 | 0.945 |
| birds | Atlapetes_pileatus | Atlapetes_pileatus | LC | Decreasing | tropical | diurnal | 24 | 348950.664 | 10.378 |
| birds | Atlapetes_rufigenis | Atlapetes_rufigenis | LC | Stable | tropical | diurnal | 32.2 | 28964.256 | 8.809 |
| birds | Atlapetes_rufinucha | Atlapetes_rufinucha | LC | Decreasing | tropical | diurnal | 21.5 | 39008.499 | 5.274 |
| birds | Atlapetes_schistaceus | Atlapetes_schistaceus | LC | Stable | tropical | diurnal | 28 | 143739.861 | 10.743 |
| birds | Atlapetes_seebohmi | Atlapetes_seebohmi | LC | Stable | tropical | diurnal | 29.5 | 47364.951 | 8.271 |
| birds | Atlapetes_semirufus | Atlapetes_semirufus | LC | Stable | tropical | diurnal | 30.7 | 67791.564 | 11.772 |
| birds | Atlapetes_tricolor | Atlapetes_tricolor | LC | Decreasing | tropical | diurnal | 36.5 | 42442.445 | 6.863 |
| birds | Atrichornis_clamosus | Atrichornis_clamosus | EN | Stable | temperate | diurnal | 42.2 | 19384.962 | 9.352 |
| birds | Atrichornis_rufescens | Atrichornis_rufescens | EN | Decreasing | temperate | diurnal | 24 | 32957.628 | 9.599 |
| birds | Atronanus_fuliginosus | NA | LC | Stable | NA | NA | NA | NA | NA |
| birds | Attagis_gayi | Attagis_gayi | LC | Stable | temperate | diurnal | 311 | 1031574.139 | 5.377 |
| birds | Attagis_malouinus | Attagis_malouinus | LC | Stable | temperate | diurnal | 340 | 323646.863 | 5.173 |

| | | | | | | | | | |
|---|---|---|---|---|---|---|---|---|---|
| birds | Atticora_fasciata | Atticora_fasciata | LC | Decreasing | tropical | diurnal | 13 | 6512985.455 | 1.961 |
| birds | Atticora_pileata | Notiochelidon_pileata | LC | Decreasing | tropical | diurnal | 12.2 | 75275.036 | 13.167 |
| birds | Atticora_tibialis | Neochelidon_tibialis | LC | Decreasing | tropical | diurnal | 10.6 | 4182240.086 | 2.645 |
| birds | Attila_bolivianus | Attila_bolivianus | LC | Decreasing | tropical | diurnal | 39.5 | 2790241.245 | 2.545 |
| birds | Attila_cinnamomeus | Attila_cinnamomeus | LC | Decreasing | tropical | diurnal | 38.8 | 5275009.266 | 1.736 |
| birds | Attila_citriniventris | Attila_citriniventris | LC | Stable | tropical | diurnal | 33.7 | 1546685.495 | 0.974 |
| birds | Attila_phoenicurus | Attila_phoenicurus | LC | Decreasing | tropical | diurnal | 32.3 | 4105204.697 | 6.187 |
| birds | Attila_rufus | Attila_rufus | LC | Decreasing | tropical | diurnal | 42.6 | 342429.819 | 12.657 |
| birds | Attila_spadiceus | Attila_spadiceus | LC | Decreasing | tropical | diurnal | 39.1 | 8099963.715 | 3.622 |
| birds | Attila_torridus | Attila_torridus | VU | Decreasing | tropical | diurnal | 46 | 80270.827 | 12.99 |
| birds | Augastes_lumachella | Augastes_lumachella | NT | Decreasing | tropical | diurnal | 4 | 20687.462 | 6.641 |
| birds | Augastes_scutatus | Augastes_scutatus | LC | Decreasing | tropical | diurnal | 3 | 28653.283 | 6.4 |
| birds | Aulacorhynchus_coeruleicinctis | Aulacorhynchus_coeruleicinctis | LC | Decreasing | tropical | diurnal | 208 | 103161.859 | 7.171 |
| birds | Aulacorhynchus_derbianus | Aulacorhynchus_derbianus | LC | Decreasing | tropical | diurnal | 209.7 | 302414.838 | 4.503 |
| birds | Aulacorhynchus_haematopygus | Aulacorhynchus_haematopygus | LC | Stable | tropical | diurnal | 194.4 | 179236.714 | 11.139 |
| birds | Aulacorhynchus_huallagae | Aulacorhynchus_huallagae | NT | Stable | tropical | diurnal | 268.8 | 1305.726 | 0.759 |
| birds | Aulacorhynchus_prasinus | Aulacorhynchus_prasinus | LC | Decreasing | tropical | diurnal | 170.6 | 420156.222 | 10.636 |
| birds | Aulacorhynchus_sulcatus | Aulacorhynchus_sulcatus | LC | Decreasing | tropical | diurnal | 167 | 30220.861 | 13.915 |
| birds | Auriparus_flaviceps | Auriparus_flaviceps | LC | Decreasing | temperate | diurnal | 6.8 | 1879707.382 | 9.239 |
| birds | Automolus_infuscatus | Automolus_infuscatus | LC | Decreasing | tropical | diurnal | 32.9 | 4323531.204 | 1.346 |
| birds | Automolus_leucophthalmus | Automolus_leucophthalmus | LC | Decreasing | tropical | diurnal | 34.5 | 1800565.069 | 12.852 |
| birds | Automolus_melanopezus | Automolus_melanopezus | LC | Decreasing | tropical | diurnal | 31.4 | 670422.082 | 2.408 |
| birds | Automolus_ochrolaemus | Automolus_ochrolaemus | LC | Stable | tropical | diurnal | 40.2 | 6344367.797 | 2.367 |
| birds | Automolus_rufipileatus | Automolus_rufipileatus | LC | Decreasing | tropical | diurnal | 34.8 | 4360535.296 | 1.983 |
| birds | Automolus_subulatus | Hyloctistes_subulatus | LC | Decreasing | tropical | diurnal | 28.8 | 3284717.792 | 1.384 |
| birds | Automolus_virgatus | NA | LC | Decreasing | NA | NA | NA | NA | NA |
| birds | Aviceda_cuculoides | Aviceda_cuculoides | LC | Stable | tropical | diurnal | 277 | 10593852.381 | 9.096 |
| birds | Aviceda_jerdoni | Aviceda_jerdoni | LC | Decreasing | temperate | diurnal | 363 | 1597402.793 | 8.576 |
| birds | Aviceda_leuphotes | Aviceda_leuphotes | LC | Decreasing | tropical | diurnal | 194 | 4574587.217 | 12.965 |
| birds | Aviceda_madagascariensis | NA | LC | Decreasing | NA | NA | NA | NA | NA |
| birds | Aviceda_subcristata | Aviceda_subcristata | LC | Stable | tropical | diurnal | 322.7 | 1718124.834 | 4.881 |
| birds | Avocettula_recurvirostris | Avocettula_recurvirostris | LC | Decreasing | tropical | diurnal | 4.2 | 2531616.036 | 2.023 |
| birds | Aythya_affinis | Aythya_affinis | LC | Stable | temperate | diurnal | 819.5 | 16453053.342 | 9.946 |
| birds | Aythya_americana | Aythya_americana | LC | Increasing | temperate | diurnal | 1075.7 | 11782105.961 | 11.882 |

| | | | | | | | | | |
|---|---|---|---|---|---|---|---|---|---|
| birds | Aythya_australis | Aythya_australis | LC | Stable | temperate | diurnal | 869.4 | 6825013.889 | 4.628 |
| birds | Aythya_baeri | Aythya_baeri | CR | Decreasing | temperate | diurnal | 681.2 | 9009013.369 | 12.045 |
| birds | Aythya_collaris | Aythya_collaris | LC | Increasing | temperate | diurnal | 704.6 | 14291364.289 | 10.619 |
| birds | Aythya_ferina | Aythya_ferina | VU | Decreasing | temperate | diurnal | 823 | 27595007.891 | 10.678 |
| birds | Aythya_fuligula | Aythya_fuligula | LC | Stable | temperate | diurnal | 701.2 | 33316870.642 | 9.508 |
| birds | Aythya_innotata | Aythya_innotata | CR | Stable | tropical | diurnal | 685 | 1163.184 | 14.047 |
| birds | Aythya_marila | Aythya_marila | LC | Decreasing | temperate | diurnal | 1005.4 | 16793614.239 | 2.631 |
| birds | Aythya_novaeseelandiae | Aythya_novaeseelandiae | LC | Increasing | temperate | diurnal | 682 | 162640.633 | 8.102 |
| birds | Aythya_nyroca | Aythya_nyroca | NT | Decreasing | tropical | diurnal | 574 | 36110689.766 | 8.691 |
| birds | Aythya_valisineria | Aythya_valisineria | LC | Increasing | temperate | diurnal | 1202 | 10988439.079 | 10.941 |
| birds | Baeolophus_atricristatus | Baeolophus_atricristatus | LC | Stable | temperate | diurnal | 17.5 | 741786.535 | 11.797 |
| birds | Baeolophus_bicolor | Baeolophus_bicolor | LC | Increasing | temperate | diurnal | 21.6 | 2969712.505 | 18.161 |
| birds | Baeolophus_inornatus | Baeolophus_inornatus | LC | Stable | temperate | diurnal | 15.9 | 178257.642 | 14.872 |
| birds | Baeolophus_ridgwayi | Baeolophus_ridgwayi | LC | Stable | temperate | diurnal | 15.9 | 1056208.764 | 8.576 |
| birds | Baeolophus_wollweberi | Baeolophus_wollweberi | LC | Stable | temperate | diurnal | 10.4 | 532101.616 | 9.442 |
| birds | Baeopogon_clamans | NA | LC | Stable | NA | NA | NA | NA | NA |
| birds | Baeopogon_indicator | Baeopogon_indicator | LC | Stable | tropical | diurnal | 45.8 | 4280773.283 | 8.302 |
| birds | Balaeniceps_rex | Balaeniceps_rex | VU | Decreasing | tropical | diurnal | 5941 | 1291415.43 | 9.715 |
| birds | Balearica_pavonina | Balearica_pavonina | VU | Decreasing | tropical | diurnal | 3590 | 2976931.847 | 10.705 |
| birds | Balearica_regulorum | Balearica_regulorum | EN | Decreasing | tropical | diurnal | 3772 | 3528882.187 | 10.001 |
| birds | Bambusicola_fytchii | Bambusicola_fytchii | LC | Decreasing | tropical | diurnal | 313.7 | 550319.01 | 8.174 |
| birds | Bambusicola_thoracicus | Bambusicola_thoracicus | LC | Decreasing | temperate | diurnal | 264.8 | 1491544.116 | 16.229 |
| birds | Bangsia_arcaei | Bangsia_arcaei | NT | Decreasing | tropical | diurnal | 37.2 | 9434.497 | 6.973 |
| birds | Bangsia_aureocincta | Bangsia_aureocincta | VU | Decreasing | tropical | diurnal | 41.7 | 1451.272 | 6.286 |
| birds | Bangsia_edwardsi | Bangsia_edwardsi | LC | Decreasing | tropical | diurnal | 37 | 19030.856 | 6.371 |
| birds | Bangsia_flavovirens | Chlorospingus_flavovirens | VU | Decreasing | tropical | diurnal | 25 | 3777.74 | 7.675 |
| birds | Bangsia_melanochlamys | Bangsia_melanochlamys | VU | Decreasing | tropical | diurnal | 37.2 | 8226.112 | 7.997 |
| birds | Bangsia_rothschildi | Bangsia_rothschildi | LC | Decreasing | tropical | diurnal | 48.5 | 20015.604 | 5.351 |
| birds | Barnardius_zonarius | Barnardius_zonarius | LC | Increasing | temperate | diurnal | 133 | 5305452.042 | 4.579 |
| birds | Bartramia_longicauda | Bartramia_longicauda | LC | Increasing | tropical | diurnal | 158.9 | 21249057.488 | 8.711 |
| birds | Baryphthengus_martii | Baryphthengus_martii | LC | Decreasing | tropical | diurnal | 165 | 3234290.455 | 2.946 |
| birds | Baryphthengus_ruficapillus | Baryphthengus_ruficapillus | LC | Decreasing | tropical | diurnal | 141.7 | 1565635.848 | 12.559 |
| birds | Basileuterus_belli | Basileuterus_belli | LC | Decreasing | tropical | diurnal | 10.7 | 334156.382 | 9.943 |
| birds | Basileuterus_culicivorus | Basileuterus_culicivorus | LC | Decreasing | tropical | diurnal | 10.5 | 522685.981 | 9.553 |

| | | | | | | | | | |
|---|---|---|---|---|---|---|---|---|---|
| birds | Basileuterus_griseiceps | Basileuterus_griseiceps | EN | Decreasing | tropical | diurnal | 13 | 902.98 | 6.975 |
| birds | Basileuterus_hypoleucus | Basileuterus_hypoleucus | LC | Decreasing | tropical | diurnal | 11.4 | 1487777.701 | 10.52 |
| birds | Basileuterus_ignotus | Basileuterus_ignotus | VU | Stable | tropical | diurnal | 10.9 | 180.742 | 1.669 |
| birds | Basileuterus_lachrymosus | Euthlypis_lachrymosa | LC | Decreasing | tropical | diurnal | 25.6 | 340350.723 | 9.31 |
| birds | Basileuterus_melanogenys | Basileuterus_melanogenys | LC | Decreasing | tropical | diurnal | 11.8 | 4831.716 | 5.716 |
| birds | Basileuterus_rufifrons | Basileuterus_rufifrons | LC | Decreasing | tropical | diurnal | 10.4 | 967644.872 | 11.635 |
| birds | Basileuterus_trifasciatus | Basileuterus_trifasciatus | LC | Decreasing | tropical | diurnal | 9.7 | 29296.984 | 9.148 |
| birds | Basileuterus_tristriatus | Basileuterus_tristriatus | LC | Decreasing | tropical | diurnal | 12 | 432295.338 | 9.359 |
| birds | Basilinna_leucotis | Hylocharis_leucotis | LC | Decreasing | tropical | diurnal | 3.4 | 556989.383 | 10.206 |
| birds | Basilinna_xantusii | Hylocharis_xantusii | LC | Stable | temperate | diurnal | 3.4 | 41923.513 | 4.33 |
| birds | Basilornis_celebensis | NA | LC | Unknown | NA | NA | NA | NA | NA |
| birds | Basilornis_corythaix | Basilornis_corythaix | LC | Decreasing | tropical | diurnal | 126 | 16752.799 | 7.143 |
| birds | Basilornis_galeatus | NA | NT | Decreasing | NA | NA | NA | NA | NA |
| birds | Batara_cinerea | Batara_cinerea | LC | Decreasing | temperate | diurnal | 131 | 600001.374 | 12.56 |
| birds | Bathmocercus_cerviniventris | Bathmocercus_cerviniventris | DD | Decreasing | tropical | diurnal | 15.8 | 271073.858 | 14.979 |
| birds | Bathmocercus_rufus | Bathmocercus_rufus | LC | Decreasing | tropical | diurnal | 15.4 | 638060.538 | 7.434 |
| birds | Batis_capensis | Batis_capensis | LC | Stable | temperate | diurnal | 11.7 | 417033.496 | 11.864 |
| birds | Batis_diops | Batis_diops | LC | Stable | tropical | diurnal | 12.7 | 68443.813 | 14.752 |
| birds | Batis_fratrum | Batis_fratrum | LC | Decreasing | tropical | diurnal | 11.9 | 426278.311 | 10.065 |
| birds | Batis_ituriensis | Batis_ituriensis | LC | Stable | tropical | diurnal | 10.7 | 267495.158 | 7.579 |
| birds | Batis_margaritae | Batis_margaritae | LC | Stable | tropical | diurnal | 11.7 | 103855.363 | 4.924 |
| birds | Batis_minima | Batis_minima | LC | Decreasing | tropical | diurnal | 10 | 138511.079 | 5.664 |
| birds | Batis_minor | Batis_minor | LC | Decreasing | tropical | diurnal | 12.5 | 280465.173 | 11.583 |
| birds | Batis_minulla | Batis_minulla | LC | Stable | tropical | diurnal | 9.9 | 474658.689 | 7.809 |
| birds | Batis_mixta | Batis_mixta | LC | Stable | tropical | diurnal | 12.2 | 57484.872 | 12.827 |
| birds | Batis_molitor | Batis_molitor | LC | Stable | tropical | diurnal | 11.6 | 5233583.285 | 9.436 |
| birds | Batis_orientalis | Batis_orientalis | LC | Stable | tropical | diurnal | 11.1 | 2679950.477 | 9.701 |
| birds | Batis_perkeo | Batis_perkeo | LC | Stable | tropical | diurnal | 6.9 | 949410.686 | 8.589 |
| birds | Batis_poensis | Batis_poensis | LC | Stable | tropical | diurnal | 9.6 | 1554.431 | 12.036 |
| birds | Batis_pririt | Batis_pririt | LC | Stable | tropical | diurnal | 9.3 | 1547981.31 | 6.165 |
| birds | Batis_senegalensis | Batis_senegalensis | LC | Decreasing | tropical | diurnal | 9.6 | 3034470.606 | 12.63 |
| birds | Batis_soror | Batis_soror | LC | Stable | tropical | diurnal | 9.5 | 863131.728 | 9.376 |
| birds | Batrachostomus_auritus | Batrachostomus_auritus | VU | Decreasing | tropical | nocturnal | 206 | 894119.339 | 7.6 |
| birds | Batrachostomus_cornutus | NA | LC | Stable | NA | NA | NA | NA | NA |

| | | | | | | | | | |
|---|---|---|---|---|---|---|---|---|---|
| birds | Batrachostomus_hodgsoni | Batrachostomus_hodgsoni | LC | Decreasing | tropical | nocturnal | 51 | 119417.564 | 9.01 |
| birds | Batrachostomus_javensis | Batrachostomus_javensis | LC | Stable | tropical | nocturnal | 46.3 | 1413593.972 | 9.271 |
| birds | Batrachostomus_moniliger | Batrachostomus_moniliger | LC | Stable | tropical | nocturnal | 52.4 | 109433.829 | 17.814 |
| birds | Batrachostomus_poliolophus | NA | LC | Decreasing | NA | NA | NA | NA | NA |
| birds | Batrachostomus_septimus | Batrachostomus_septimus | LC | Decreasing | tropical | nocturnal | 86.8 | 251854.348 | 13.838 |
| birds | Batrachostomus_stellatus | Batrachostomus_stellatus | NT | Decreasing | tropical | nocturnal | 51.3 | 748621.011 | 6.737 |
| birds | Belocercus_longicaudus | Psittacula_longicauda | VU | Decreasing | tropical | diurnal | 107 | 877581.599 | 8.655 |
| birds | Berenicornis_comatus | Aceros_comatus | EN | Decreasing | tropical | diurnal | 1470 | 1179089.516 | 8.232 |
| birds | Berlepschia_rikeri | Berlepschia_rikeri | LC | Stable | tropical | diurnal | 37 | 5179268.804 | 2.525 |
| birds | Bernieria_madagascariensis | Bernieria_madagascariensis | LC | Decreasing | tropical | diurnal | 25.3 | 445930.234 | 8.2 |
| birds | Bias_musicus | Bias_musicus | LC | Decreasing | tropical | diurnal | 21.9 | 3741138.348 | 8.656 |
| birds | Biatas_nigropectus | Biatas_nigropectus | VU | Decreasing | temperate | diurnal | 25.5 | 157889.543 | 14.544 |
| birds | Biziura_lobata | Biziura_lobata | LC | Decreasing | temperate | diurnal | 1928.5 | 2736871.391 | 7.144 |
| birds | Bleda_canicapillus | Bleda_canicapillus | LC | Stable | tropical | diurnal | 41.3 | 1094053.694 | 14.228 |
| birds | Bleda_eximius | Bleda_eximius | NT | Decreasing | tropical | diurnal | 35.7 | 339296.543 | 15.473 |
| birds | Bleda_syndactylus | Bleda_syndactylus | LC | Stable | tropical | diurnal | 46.1 | 3527560.887 | 7.825 |
| birds | Blythipicus_pyrrhotis | Blythipicus_pyrrhotis | LC | Decreasing | tropical | diurnal | 132 | 1622803.087 | 10.56 |
| birds | Blythipicus_rubiginosus | Blythipicus_rubiginosus | LC | Decreasing | tropical | diurnal | 81.9 | 1344203.735 | 7.622 |
| birds | Bocagia_minuta | Tchagra_minutus | LC | Decreasing | tropical | diurnal | 34.4 | 3369836.919 | 10.491 |
| birds | Boissonneaua_flavescens | Boissonneaua_flavescens | LC | Unknown | tropical | diurnal | 8.2 | 128710.183 | 11.193 |
| birds | Boissonneaua_jardini | Boissonneaua_jardini | LC | Decreasing | tropical | diurnal | 9.1 | 30988.103 | 8.656 |
| birds | Boissonneaua_matthewsii | Boissonneaua_matthewsii | LC | Unknown | tropical | diurnal | 7.7 | 127445.481 | 5.927 |
| birds | Bolbopsittacus_lunulatus | Bolbopsittacus_lunulatus | LC | Stable | tropical | diurnal | 67.6 | 210517.7 | 13.583 |
| birds | Bolborhynchus_ferrugineifrons | Bolborhynchus_ferrugineifrons | VU | Decreasing | tropical | diurnal | 64 | 10889.615 | 7.638 |
| birds | Bolborhynchus_lineola | Bolborhynchus_lineola | LC | Stable | tropical | diurnal | 56.2 | 279157.1 | 10.219 |
| birds | Bolborhynchus_orbygnesius | Bolborhynchus_orbygnesius | LC | Stable | tropical | diurnal | 43.1 | 191118.756 | 8.74 |
| birds | Bolemoreus_frenatus | Lichenostomus_frenatus | LC | Stable | tropical | diurnal | 34.1 | 58164.639 | 4.576 |
| birds | Bolemoreus_hindwoodi | Lichenostomus_hindwoodi | NT | Stable | tropical | diurnal | 22.9 | 1242.503 | 4.608 |
| birds | Bombycilla_cedrorum | Bombycilla_cedrorum | LC | Increasing | temperate | diurnal | 31.6 | 13624903.872 | 11.074 |
| birds | Bombycilla_garrulus | Bombycilla_garrulus | LC | Increasing | temperate | diurnal | 54.4 | 32415917 | 7.48 |
| birds | Bombycilla_japonica | Bombycilla_japonica | NT | Decreasing | temperate | diurnal | 42.3 | 2963080.814 | 12.025 |
| birds | Bonasa_umbellus | Bonasa_umbellus | LC | Stable | temperate | diurnal | 530.9 | 8076355.641 | 6.712 |
| birds | Bostrychia_carunculata | Bostrychia_carunculata | LC | Unknown | tropical | diurnal | 1500 | 539792.684 | 11.85 |
| birds | Bostrychia_hagedash | Bostrychia_hagedash | LC | Increasing | tropical | diurnal | 1238 | 16241597.551 | 9.615 |

| | | | | | | | | | |
|---|---|---|---|---|---|---|---|---|---|
| birds | Bostrychia_olivacea | NA | LC | Decreasing | NA | NA | NA | NA | NA |
| birds | Bostrychia_rara | Bostrychia_rara | LC | Decreasing | tropical | diurnal | 866 | 2806304.389 | 6.581 |
| birds | Botaurus_lentiginosus | Botaurus_lentiginosus | LC | Decreasing | temperate | nocturnal | 706 | 14235010.419 | 10.383 |
| birds | Botaurus_pinnatus | Botaurus_pinnatus | LC | Stable | temperate | nocturnal | 812.2 | 2262647.754 | 9.331 |
| birds | Botaurus_poiciloptilus | Botaurus_poiciloptilus | VU | Decreasing | temperate | cathemeral | 1083.7 | 1425382.926 | 9.05 |
| birds | Botaurus_stellaris | Botaurus_stellaris | LC | Decreasing | temperate | cathemeral | 1319.5 | 24816284.223 | 10.182 |
| birds | Brachycope_anomala | Brachycope_anomala | LC | Stable | tropical | diurnal | 25.3 | 252363.812 | 5.668 |
| birds | Brachygalba_albogularis | Brachygalba_albogularis | LC | Decreasing | tropical | diurnal | 17.4 | 660078.821 | 1.293 |
| birds | Brachygalba_goeringi | Brachygalba_goeringi | LC | Stable | tropical | diurnal | 17 | 196562.334 | 10.422 |
| birds | Brachygalba_lugubris | Brachygalba_lugubris | LC | Decreasing | tropical | diurnal | 15.9 | 4798770.729 | 3.916 |
| birds | Brachygalba_salmoni | Brachygalba_salmoni | LC | Decreasing | tropical | diurnal | 17.2 | 50824.856 | 12.161 |
| birds | Brachypodius_atriceps | Pycnonotus_atriceps | LC | Stable | tropical | diurnal | 25.5 | 2968877.332 | 10.092 |
| birds | Brachypodius_priocephalus | NA | NT | Decreasing | NA | NA | NA | NA | NA |
| birds | Brachypteracias_leptosomus | Brachypteracias_leptosomus | VU | Decreasing | tropical | diurnal | 185 | 48822.546 | 5.576 |
| birds | Brachypteryx_hyperythra | NA | NT | Decreasing | NA | NA | NA | NA | NA |
| birds | Brachypteryx_leucophris | Brachypteryx_leucophrys | LC | Decreasing | tropical | diurnal | 15.8 | 2289506.948 | 10.21 |
| birds | Brachypteryx_montana | Brachypteryx_montana | LC | Decreasing | tropical | diurnal | 17.7 | 103057.903 | 21.055 |
| birds | Brachyramphus_brevirostris | Brachyramphus_brevirostris | NT | Decreasing | temperate | diurnal | 234 | 2263571.779 | 1.241 |
| birds | Brachyramphus_marmoratus | Brachyramphus_marmoratus | EN | Decreasing | temperate | diurnal | 220 | 2521326.401 | 2.08 |
| birds | Bradornis_comitatus | Muscicapa_comitata | LC | Stable | tropical | diurnal | 14.1 | 3634771.069 | 8.323 |
| birds | Bradornis_mariquensis | Bradornis_mariquensis | LC | Stable | tropical | diurnal | 25.2 | 1651176.931 | 6.921 |
| birds | Bradornis_microrhynchus | Bradornis_microrhynchus | LC | Stable | tropical | diurnal | 19.5 | 2477905.95 | 9.452 |
| birds | Bradypterus_baboecala | Bradypterus_baboecala | LC | Stable | tropical | diurnal | 12.9 | 3040370.234 | 9.615 |
| birds | Bradypterus_barratti | Bradypterus_barratti | LC | Stable | temperate | diurnal | 18.8 | 227225.238 | 12.736 |
| birds | Bradypterus_brunneus | Dromaeocercus_brunneus | LC | Decreasing | tropical | diurnal | 14.2 | 52379.86 | 5.393 |
| birds | Bradypterus_carpalis | Bradypterus_carpalis | LC | Decreasing | tropical | diurnal | 22.4 | 116124.146 | 16.197 |
| birds | Bradypterus_cinnamomeus | Bradypterus_cinnamomeus | LC | Stable | tropical | diurnal | 17.4 | 513979.165 | 13.1 |
| birds | Bradypterus_grandis | Bradypterus_grandis | NT | Stable | tropical | diurnal | 18.5 | 12449.447 | 4.177 |
| birds | Bradypterus_graueri | Bradypterus_graueri | VU | Decreasing | tropical | diurnal | 17 | 19795.128 | 15.388 |
| birds | Bradypterus_lopezi | Bradypterus_lopezi | LC | Decreasing | tropical | diurnal | 19.2 | 424179.16 | 11.39 |
| birds | Bradypterus_seebohmi | Amphilais_seebohmi | LC | Stable | tropical | diurnal | 12.3 | 290843.733 | 8.188 |
| birds | Bradypterus_sylvaticus | Bradypterus_sylvaticus | VU | Decreasing | temperate | diurnal | 21 | 8825.281 | 12.347 |
| birds | Branta_bernicla | Branta_bernicla | LC | Unknown | temperate | diurnal | 1277.9 | 4854965.922 | 6.122 |
| birds | Branta_canadensis | Branta_canadensis | LC | Increasing | temperate | diurnal | 2811.7 | 16352791.223 | 9.016 |

| | | | | | | | | | |
|---|---|---|---|---|---|---|---|---|---|
| birds | Branta_leucopsis | Branta_leucopsis | LC | Increasing | temperate | diurnal | 1684 | 259364.55 | 9.227 |
| birds | Branta_ruficollis | Branta_ruficollis | VU | Decreasing | temperate | diurnal | 1226.5 | 2994790.393 | 6.386 |
| birds | Branta_sandvicensis | Branta_sandvicensis | NT | Increasing | tropical | diurnal | 2044.1 | 1590.277 | 7.292 |
| birds | Brotogeris_chiriri | Brotogeris_chiriri | LC | Decreasing | tropical | diurnal | 61.6 | 3838812.722 | 8.422 |
| birds | Brotogeris_chrysoptera | Brotogeris_chrysoptera | LC | Decreasing | tropical | diurnal | 54.5 | 3112139.059 | 2.09 |
| birds | Brotogeris_cyanoptera | Brotogeris_cyanoptera | LC | Stable | tropical | diurnal | 56 | 2567592.628 | 2.453 |
| birds | Brotogeris_jugularis | Brotogeris_jugularis | LC | Decreasing | tropical | diurnal | 63.2 | 781311.172 | 12.029 |
| birds | Brotogeris_pyrrhoptera | Brotogeris_pyrrhoptera | VU | Decreasing | tropical | diurnal | 63.9 | 25806.376 | 14.817 |
| birds | Brotogeris_sanctithomae | Brotogeris_sanctithomae | LC | Decreasing | tropical | diurnal | 59 | 854340.868 | 2.31 |
| birds | Brotogeris_tirica | Brotogeris_tirica | LC | Stable | tropical | diurnal | 63 | 288572.864 | 12.702 |
| birds | Brotogeris_versicolurus | Brotogeris_versicolurus | LC | Stable | tropical | diurnal | 71.5 | 714999.048 | 4.17 |
| birds | Brunhilda_charmosyna | Estrilda_charmosyna | LC | Stable | tropical | diurnal | 7.4 | 748636.983 | 8.548 |
| birds | Brunhilda_erythronotos | Estrilda_erythronotos | LC | Stable | temperate | diurnal | 9.2 | 2061140.193 | 8.551 |
| birds | Bubalornis_albirostris | Bubalornis_albirostris | LC | Stable | tropical | diurnal | 61.9 | 2320967.625 | 12.656 |
| birds | Bubalornis_niger | Bubalornis_niger | LC | Stable | tropical | diurnal | 76.2 | 2708371.228 | 8.807 |
| birds | Bubo_africanus | Bubo_africanus | LC | Stable | tropical | nocturnal | 645 | 8655406.597 | 8.417 |
| birds | Bubo_ascalaphus | Bubo_ascalaphus | LC | Stable | temperate | nocturnal | 2100 | 3200038.616 | 5.274 |
| birds | Bubo_bengalensis | Bubo_bengalensis | LC | Stable | tropical | nocturnal | 1100 | 3750743.185 | 16.942 |
| birds | Bubo_blakistoni | Ketupa_blakistoni | EN | Decreasing | temperate | cathemeral | 3850 | 2289996.146 | 6.055 |
| birds | Bubo_bubo | Bubo_bubo | LC | Decreasing | temperate | cathemeral | 2668.5 | 31974373.191 | 8.093 |
| birds | Bubo_capensis | Bubo_capensis | LC | Stable | temperate | nocturnal | 1118.6 | 1905194.148 | 10.126 |
| birds | Bubo_cinerascens | Bubo_cinerascens | LC | Stable | tropical | nocturnal | 500 | 9679896.354 | 9.484 |
| birds | Bubo_coromandus | Bubo_coromandus | LC | Decreasing | tropical | diurnal | 1392.2 | 1428965.166 | 17.539 |
| birds | Bubo_lacteus | Bubo_lacteus | LC | Stable | tropical | nocturnal | 2114.9 | 11298632.021 | 9.251 |
| birds | Bubo_leucostictus | Bubo_leucostictus | LC | Decreasing | tropical | nocturnal | 532.5 | 2567896.426 | 7.954 |
| birds | Bubo_magellanicus | Bubo_magellanicus | LC | Stable | temperate | diurnal | 1166.4 | 2370543.592 | 5.862 |
| birds | Bubo_nipalensis | Bubo_nipalensis | LC | Decreasing | tropical | cathemeral | 1396.4 | 2384160.558 | 10.657 |
| birds | Bubo_poensis | Bubo_poensis | LC | Decreasing | tropical | nocturnal | 654.9 | 3809271.042 | 7.847 |
| birds | Bubo_scandiacus | Bubo_scandiaca | VU | Decreasing | temperate | cathemeral | 2028.8 | 27057680.291 | 2.993 |
| birds | Bubo_sumatranus | Bubo_sumatranus | NT | Decreasing | tropical | nocturnal | 1525 | 1415096.112 | 9.29 |
| birds | Bubo_virginianus | Bubo_virginianus | LC | Stable | temperate | nocturnal | 1354.5 | 24427057.983 | 8.678 |
| birds | Bubulcus_ibis | Bubulcus_ibis | LC | Increasing | tropical | diurnal | 366 | 63608925.749 | 9.739 |
| birds | Bucanetes_githagineus | Bucanetes_githagineus | LC | Stable | temperate | diurnal | 19.6 | 7293211.131 | 5.467 |
| birds | Bucanetes_mongolicus | Eremopsaltria_mongolicus | LC | Stable | temperate | diurnal | 21.5 | 4289647.076 | 6.03 |

| | | | | | | | | | |
|---|---|---|---|---|---|---|---|---|---|
| birds | Buccanodon_duchaillui | Buccanodon_duchaillui | LC | Decreasing | tropical | diurnal | 41.8 | 3014600.114 | 7.358 |
| birds | Bucco_capensis | Bucco_capensis | LC | Decreasing | tropical | diurnal | 54 | 5454915.037 | 1.696 |
| birds | Bucephala_albeola | Bucephala_albeola | LC | Increasing | temperate | diurnal | 397.5 | 15692279.646 | 9.518 |
| birds | Bucephala_clangula | Bucephala_clangula | LC | Stable | temperate | diurnal | 918.6 | 37078063.737 | 7.997 |
| birds | Bucephala_islandica | Bucephala_islandica | LC | Stable | temperate | diurnal | 921.2 | 3452110.558 | 4.78 |
| birds | Buceros_bicornis | Buceros_bicornis | VU | Decreasing | tropical | diurnal | 2790.8 | 2951578.247 | 12.749 |
| birds | Buceros_hydrocorax | Buceros_hydrocorax | VU | Decreasing | tropical | diurnal | 1409.6 | 78462.718 | 11.959 |
| birds | Buceros_rhinoceros | Buceros_rhinoceros | VU | Decreasing | tropical | diurnal | 2371.6 | 1455791.747 | 9.106 |
| birds | Bucorvus_abyssinicus | Bucorvus_abyssinicus | VU | Decreasing | tropical | diurnal | 4000 | 4482170.579 | 11.201 |
| birds | Bucorvus_leadbeateri | Bucorvus_cafer | VU | Decreasing | tropical | diurnal | 3743.6 | 4867849.88 | 9.005 |
| birds | Bugeranus_carunculatus | Grus_carunculatus | VU | Decreasing | tropical | diurnal | 8159 | 2289042.191 | 7.736 |
| birds | Bulweria_bulwerii | Bulweria_bulwerii | LC | Stable | tropical | cathemeral | 99 | 117388743.869 | 12.657 |
| birds | Buphagus_africanus | Buphagus_africanus | LC | Decreasing | tropical | diurnal | 58.5 | 3672712.872 | 11.374 |
| birds | Buphagus_erythrorynchus | Buphagus_erythrorhynchus | LC | Decreasing | tropical | diurnal | 50.7 | 2881050.228 | 9.583 |
| birds | Burhinus_bistriatus | Burhinus_bistriatus | LC | Stable | tropical | nocturnal | 787 | 787421.538 | 9.482 |
| birds | Burhinus_capensis | Burhinus_capensis | LC | Stable | tropical | nocturnal | 423 | 12421028.766 | 9.642 |
| birds | Burhinus_grallarius | Burhinus_grallarius | LC | Decreasing | temperate | nocturnal | 648.1 | 2586160.811 | 5.561 |
| birds | Burhinus_oedicnemus | Burhinus_oedicnemus | LC | Decreasing | temperate | nocturnal | 459 | 21340481.537 | 7.75 |
| birds | Burhinus_senegalensis | Burhinus_senegalensis | LC | Unknown | tropical | nocturnal | 326 | 7037522.757 | 10.265 |
| birds | Burhinus_superciliaris | Burhinus_superciliaris | LC | Unknown | tropical | nocturnal | 480 | 79703.928 | 8.541 |
| birds | Burhinus_vermiculatus | Burhinus_vermiculatus | LC | Unknown | tropical | nocturnal | 320 | 5802844.891 | 9.888 |
| birds | Busarellus_nigricollis | Busarellus_nigricollis | LC | Decreasing | tropical | diurnal | 766.1 | 12825637.868 | 6.275 |
| birds | Butastur_indicus | Butastur_indicus | LC | Decreasing | temperate | diurnal | 414.6 | 6892597.524 | 12.403 |
| birds | Butastur_liventer | Butastur_liventer | LC | Decreasing | tropical | diurnal | 338 | 1357262.801 | 12.292 |
| birds | Butastur_rufipennis | Butastur_rufipennis | LC | Decreasing | tropical | diurnal | 334.9 | 7779786.481 | 10.533 |
| birds | Butastur_teesa | Butastur_teesa | LC | Stable | tropical | diurnal | 325 | 3969936.204 | 16.526 |
| birds | Buteo_albigula | Buteo_albigula | LC | Unknown | tropical | diurnal | 525 | 2041826.718 | 6.717 |
| birds | Buteo_albonotatus | Buteo_albonotatus | LC | Stable | tropical | diurnal | 745.9 | 8230721.237 | 7.447 |
| birds | Buteo_augur | Buteo_augur | LC | Stable | tropical | diurnal | 1099.2 | 3919632.936 | 10.605 |
| birds | Buteo_auguralis | Buteo_auguralis | LC | Stable | tropical | diurnal | 663.9 | 7090619.928 | 10.486 |
| birds | Buteo_brachypterus | Buteo_brachypterus | LC | Stable | tropical | diurnal | 511 | 591120.44 | 8.749 |
| birds | Buteo_brachyurus | Buteo_brachyurus | LC | Stable | tropical | diurnal | 496 | 14372802.038 | 6.054 |
| birds | Buteo_buteo | Buteo_buteo | LC | Increasing | temperate | diurnal | 759.1 | 21382495.551 | 11.802 |
| birds | Buteo_galapagoensis | NA | VU | Stable | NA | NA | NA | NA | NA |

| | | | | | | | | | |
|---|---|---|---|---|---|---|---|---|---|
| birds | Buteo_hemilasius | Buteo_hemilasius | LC | Stable | temperate | diurnal | 1358.9 | 9027045.284 | 7.734 |
| birds | Buteo_jamaicensis | Buteo_jamaicensis | LC | Increasing | temperate | diurnal | 1101.2 | 15893934.472 | 9.716 |
| birds | Buteo_lagopus | Buteo_lagopus | LC | Stable | temperate | diurnal | 949.8 | 37406946.745 | 7.337 |
| birds | Buteo_lineatus | Buteo_lineatus | LC | Increasing | temperate | diurnal | 603.7 | 4090576.967 | 16.516 |
| birds | Buteo_nitidus | Buteo_nitidus | LC | Stable | tropical | diurnal | 519 | 10412658.231 | 5.079 |
| birds | Buteo_oreophilus | Buteo_oreophilus | NT | Decreasing | tropical | diurnal | 700 | 674707.007 | 13.821 |
| birds | Buteo_platypterus | Buteo_platypterus | LC | Increasing | tropical | diurnal | 453.7 | 12158210.009 | 9.34 |
| birds | Buteo_regalis | Buteo_regalis | LC | Increasing | temperate | diurnal | 1437.2 | 4975226.355 | 10.521 |
| birds | Buteo_ridgwayi | Buteo_ridgwayi | CR | Increasing | tropical | diurnal | 315 | 35683.586 | 14.282 |
| birds | Buteo_rufinus | Buteo_rufinus | LC | Stable | temperate | diurnal | 1166.2 | 20474801.171 | 8.852 |
| birds | Buteo_rufofuscus | Buteo_rufofuscus | LC | Stable | temperate | diurnal | 1530 | 1533534.657 | 8.609 |
| birds | Buteo_solitarius | Buteo_solitarius | NT | Stable | tropical | diurnal | 500.9 | 6957.484 | 8.064 |
| birds | Buteo_swainsoni | Buteo_swainsoni | LC | Stable | tropical | diurnal | 946.6 | 13792497.701 | 7.625 |
| birds | Buteo_ventralis | Buteo_ventralis | VU | Stable | temperate | diurnal | 1190.5 | 536992.068 | 4.507 |
| birds | Buteogallus_aequinoctialis | Buteogallus_aequinoctialis | NT | Decreasing | tropical | diurnal | 657.6 | 132570.119 | 9.707 |
| birds | Buteogallus_anthracinus | Buteogallus_anthracinus | LC | Decreasing | tropical | diurnal | 975.1 | 1898078.973 | 9.447 |
| birds | Buteogallus_coronatus | Harpyhaliaetus_coronatus | EN | Decreasing | temperate | diurnal | 2950 | 4233989.277 | 8.304 |
| birds | Buteogallus_lacernulatus | NA | VU | Decreasing | NA | NA | NA | NA | NA |
| birds | Buteogallus_meridionalis | Buteogallus_meridionalis | LC | Stable | tropical | diurnal | 808 | 10432463.325 | 7.307 |
| birds | Buteogallus_schistaceus | Leucopternis_schistaceus | LC | Decreasing | tropical | diurnal | 1000 | 3642728.144 | 2.154 |
| birds | Buteogallus_solitarius | Harpyhaliaetus_solitarius | NT | Decreasing | tropical | diurnal | 3000 | 1037709.983 | 7.384 |
| birds | Buteogallus_urubitinga | Buteogallus_urubitinga | LC | Decreasing | tropical | diurnal | 1152.9 | 13126253.65 | 5.715 |
| birds | Buthraupis_montana | Buthraupis_montana | LC | Stable | tropical | diurnal | 87.1 | 299466.643 | 9.383 |
| birds | Butorides_striata | Butorides_striata | LC | Stable | temperate | cathemeral | 201.5 | 53954317.585 | 10.066 |
| birds | Bycanistes_brevis | Bycanistes_brevis | LC | Decreasing | tropical | diurnal | 1232.8 | 834397.486 | 11.841 |
| birds | Bycanistes_bucinator | Bycanistes_bucinator | LC | Decreasing | tropical | diurnal | 639.4 | 4073212.841 | 9.162 |
| birds | Bycanistes_cylindricus | Bycanistes_cylindricus | VU | Decreasing | tropical | diurnal | 921 | 477009.11 | 14.486 |
| birds | Bycanistes_fistulator | Bycanistes_fistulator | LC | Decreasing | tropical | diurnal | 508.1 | 811024.817 | 15.093 |
| birds | Bycanistes_subcylindricus | Bycanistes_subcylindricus | LC | Unknown | tropical | diurnal | 1195.4 | 1147243.508 | 11.177 |
| birds | Cacatua_alba | Cacatua_alba | EN | Decreasing | temperate | diurnal | 570 | 62157.617 | 14.371 |
| birds | Cacatua_ducorpsii | Cacatua_ducorpsii | LC | Decreasing | tropical | diurnal | 394 | 32952.615 | 4.712 |
| birds | Cacatua_galerita | Cacatua_galerita | LC | Decreasing | temperate | diurnal | 720.4 | 3967177.315 | 5.55 |
| birds | Cacatua_goffiniana | Cacatua_goffiniana | NT | Decreasing | tropical | diurnal | 275 | 49974.026 | 20.441 |
| birds | Cacatua_haematuropygia | Cacatua_haematuropygia | CR | Decreasing | tropical | diurnal | 288 | 216118.702 | 13.516 |

| | | | | | | | | | |
|---|---|---|---|---|---|---|---|---|---|
| birds | Cacatua_leadbeateri | Cacatua_leadbeateri | LC | Stable | temperate | diurnal | 396.8 | 2876418.662 | 4.002 |
| birds | Cacatua_moluccensis | Cacatua_moluccensis | VU | Decreasing | tropical | diurnal | 835 | 18717.289 | 7.389 |
| birds | Cacatua_ophthalmica | Cacatua_ophthalmica | VU | Decreasing | tropical | diurnal | 492 | 34781.174 | 4.22 |
| birds | Cacatua_pastinator | Cacatua_pastinator | LC | Increasing | temperate | diurnal | 631.7 | 200538.756 | 10.57 |
| birds | Cacatua_sanguinea | Cacatua_sanguinea | LC | Increasing | temperate | diurnal | 523.7 | 5317399.11 | 5.057 |
| birds | Cacatua_sulphurea | Cacatua_sulphurea | CR | Decreasing | tropical | diurnal | 303.3 | 1917.543 | 40.751 |
| birds | Cacatua_tenuirostris | Cacatua_tenuirostris | LC | Increasing | temperate | diurnal | 566.6 | 562525.184 | 11.213 |
| birds | Cacicus_cela | Cacicus_cela | LC | Decreasing | tropical | diurnal | 85.5 | 8569297.307 | 3.301 |
| birds | Cacicus_chrysonotus | Cacicus_chrysonotus | LC | Decreasing | tropical | diurnal | 79.5 | 285871.715 | 8.848 |
| birds | Cacicus_chrysopterus | Cacicus_chrysopterus | LC | Stable | temperate | diurnal | 36.2 | 1634780.203 | 10.538 |
| birds | Cacicus_haemorrhous | Cacicus_haemorrhous | LC | Decreasing | tropical | diurnal | 83.7 | 8042084.836 | 4.709 |
| birds | Cacicus_koepckeae | Cacicus_koepckeae | NT | Decreasing | tropical | diurnal | 48 | 45403.994 | 0.914 |
| birds | Cacicus_latirostris | Ocyalus_latirostris | LC | Decreasing | tropical | diurnal | 96.1 | 418793.305 | 1.271 |
| birds | Cacicus_leucoramphus | NA | not_available | not_available | NA | NA | NA | NA | NA |
| birds | Cacicus_microrhynchus | Cacicus_microrhynchus | LC | Stable | tropical | diurnal | 80 | 132764.09 | 8.571 |
| birds | Cacicus_oseryi | Clypicterus_oseryi | LC | Stable | tropical | diurnal | 137.7 | 630368.422 | 1.741 |
| birds | Cacicus_sclateri | Cacicus_sclateri | LC | Stable | tropical | diurnal | 49 | 293827.205 | 2.553 |
| birds | Cacicus_solitarius | Cacicus_solitarius | LC | Stable | tropical | diurnal | 79.8 | 8791817.693 | 5.206 |
| birds | Cacicus_uropygialis | Cacicus_uropygialis | LC | Decreasing | tropical | diurnal | 61.2 | 137670.603 | 7.538 |
| birds | Cacomantis_aeruginosus | NA | LC | Decreasing | NA | NA | NA | NA | NA |
| birds | Cacomantis_castaneiventris | Cacomantis_castaneiventris | LC | Stable | tropical | diurnal | 33.2 | 790922.128 | 3.365 |
| birds | Cacomantis_flabelliformis | Cacomantis_flabelliformis | LC | Stable | temperate | diurnal | 49.8 | 2434269.438 | 7.669 |
| birds | Cacomantis_merulinus | Cacomantis_merulinus | LC | Stable | tropical | diurnal | 25.3 | 5252530.135 | 11.933 |
| birds | Cacomantis_sonneratii | Cacomantis_sonneratii | LC | Stable | tropical | diurnal | 33.8 | 4946192.251 | 12.875 |
| birds | Cacomantis_variolosus | Cacomantis_variolosus | LC | Stable | tropical | diurnal | 35.8 | 3750782.695 | 7.279 |
| birds | Cairina_moschata | Cairina_moschata | LC | Decreasing | tropical | diurnal | 2415.8 | 14998211.902 | 6.621 |
| birds | Calamanthus_campestris | Calamanthus_campestris | LC | Decreasing | temperate | diurnal | 13.9 | 1452514.488 | 4.265 |
| birds | Calamanthus_fuliginosus | Calamanthus_fuliginosus | LC | Decreasing | temperate | diurnal | 20.8 | 197682.928 | 9.208 |
| birds | Calamonastes_fasciolatus | Camaroptera_fasciolata | LC | Stable | tropical | diurnal | 13.1 | 1422725.532 | 6.562 |
| birds | Calamonastes_simplex | Camaroptera_simplex | LC | Stable | tropical | diurnal | 12 | 1535343.538 | 8.864 |
| birds | Calamonastes_undosus | Camaroptera_undosa | LC | Stable | tropical | diurnal | 13 | 1613961.155 | 8.453 |
| birds | Calamonastides_gracilirostris | Chloropeta_gracilirostris | VU | Decreasing | tropical | diurnal | 10.8 | 93067.217 | 17.012 |
| birds | Calamospiza_melanocorys | Calamospiza_melanocorys | LC | Decreasing | temperate | diurnal | 37.6 | 3695893.112 | 10.994 |
| birds | Calandrella_acutirostris | Calandrella_acutirostris | LC | Stable | temperate | diurnal | 20.2 | 5326192.971 | 7.809 |

| | | | | | | | | | |
|---|---|---|---|---|---|---|---|---|---|
| birds | Calandrella_brachydactyla | Calandrella_brachydactyla | LC | Unknown | temperate | diurnal | 21.1 | 27542314.9 | 7.017 |
| birds | Calandrella_cinerea | Calandrella_cinerea | LC | Increasing | tropical | diurnal | 23.7 | 5733287.569 | 8.244 |
| birds | Calcarius_lapponicus | Calcarius_lapponicus | LC | Increasing | temperate | diurnal | 27.8 | 27580104.551 | 6.869 |
| birds | Calcarius_ornatus | Calcarius_ornatus | VU | Decreasing | temperate | diurnal | 20.3 | 3253026.168 | 11.78 |
| birds | Calcarius_pictus | Calcarius_pictus | LC | Unknown | temperate | diurnal | 26.7 | 6213416.01 | 7.074 |
| birds | Calendulauda_africanoides | Mirafra_africanoides | LC | Stable | tropical | diurnal | 23.4 | 2859748.757 | 7.298 |
| birds | Calendulauda_albescens | Certhilauda_albescens | LC | Decreasing | temperate | diurnal | 30.4 | 283792.891 | 7.842 |
| birds | Calendulauda_burra | Certhilauda_burra | VU | Decreasing | temperate | diurnal | 37.4 | 72170.125 | 5.378 |
| birds | Calendulauda_poecilosterna | Mirafra_poecilosterna | LC | Decreasing | tropical | diurnal | 24.7 | 703693.953 | 9 |
| birds | Calendulauda_rufa | Mirafra_rufa | LC | Stable | tropical | diurnal | 29.2 | 513902.1 | 11.489 |
| birds | Calendulauda_sabota | Mirafra_sabota | LC | Increasing | tropical | diurnal | 24.7 | 2588939.444 | 7.346 |
| birds | Calherodius_leuconotus | Gorsachius_leuconotus | LC | Decreasing | tropical | nocturnal | 480.3 | 7860889.295 | 10.341 |
| birds | Calicalicus_madagascariensis | Calicalicus_madagascariensis | LC | Decreasing | tropical | diurnal | 16.4 | 363727.601 | 8.083 |
| birds | Calidris_acuminata | Calidris_acuminata | VU | Decreasing | temperate | diurnal | 66.1 | 12623150.227 | 5.439 |
| birds | Calidris_alba | Calidris_alba | LC | Unknown | temperate | cathemeral | 51.7 | 13217165.642 | 9.149 |
| birds | Calidris_alpina | Calidris_alpina | LC | Decreasing | temperate | cathemeral | 51.9 | 21936163.391 | 9.199 |
| birds | Calidris_bairdii | Calidris_bairdii | LC | Stable | temperate | diurnal | 41 | 12711846.236 | 6.905 |
| birds | Calidris_canutus | Calidris_canutus | NT | Decreasing | temperate | cathemeral | 141.9 | 6898573.861 | 7.924 |
| birds | Calidris_falcinellus | Limicola_falcinellus | LC | Decreasing | temperate | diurnal | 37.1 | 3247967.413 | 9.549 |
| birds | Calidris_ferruginea | Calidris_ferruginea | NT | Decreasing | temperate | cathemeral | 58.1 | 17924722.618 | 9.163 |
| birds | Calidris_fuscicollis | Calidris_fuscicollis | LC | Decreasing | temperate | diurnal | 43 | 27065084.743 | 7.009 |
| birds | Calidris_himantopus | Calidris_himantopus | LC | Increasing | temperate | diurnal | 57.2 | 21365273.09 | 7.929 |
| birds | Calidris_maritima | Calidris_maritima | LC | Decreasing | temperate | diurnal | 65.1 | 3299407.832 | 3.482 |
| birds | Calidris_mauri | Calidris_mauri | LC | Decreasing | temperate | diurnal | 27.8 | 4634457.923 | 10.293 |
| birds | Calidris_melanotos | Calidris_melanotos | LC | Stable | temperate | diurnal | 79.7 | 33471970.381 | 7.333 |
| birds | Calidris_minuta | Calidris_minuta | LC | Increasing | temperate | diurnal | 21.1 | 22084747.49 | 9.286 |
| birds | Calidris_minutilla | Calidris_minutilla | LC | Decreasing | temperate | cathemeral | 22.9 | 25807753.606 | 7.305 |
| birds | Calidris_ptilocnemis | Calidris_ptilocnemis | LC | Decreasing | temperate | diurnal | 81.3 | 977273.686 | 3.468 |
| birds | Calidris_pugnax | Philomachus_pugnax | LC | Decreasing | temperate | cathemeral | 131.7 | 28953892.128 | 8.536 |
| birds | Calidris_pusilla | Calidris_pusilla | NT | Decreasing | temperate | diurnal | 27.5 | 16118055.097 | 7.838 |
| birds | Calidris_pygmaea | Eurynorhynchus_pygmeus | CR | Decreasing | tropical | diurnal | 31.7 | 537601.015 | 16.276 |
| birds | Calidris_ruficollis | Calidris_ruficollis | NT | Decreasing | tropical | diurnal | 26.8 | 9990778.178 | 8.641 |
| birds | Calidris_subminuta | Calidris_subminuta | LC | Unknown | tropical | diurnal | 30.2 | 3687288.12 | 10.873 |
| birds | Calidris_subruficollis | Tryngites_subruficollis | NT | Decreasing | temperate | diurnal | 62.1 | 11646075.891 | 6.427 |

| | | | | | | | | | |
|---|---|---|---|---|---|---|---|---|---|
| birds | Calidris_temminckii | Calidris_temminckii | LC | Unknown | temperate | diurnal | 23 | 17581833.277 | 10.236 |
| birds | Calidris_tenuirostris | Calidris_tenuirostris | EN | Decreasing | temperate | cathemeral | 192 | 3461961.534 | 5.884 |
| birds | Calidris_virgata | Aphriza_virgata | LC | Decreasing | temperate | diurnal | 201 | 2291134.379 | 4.755 |
| birds | Caliechthrus_leucolophus | Caliechthrus_leucolophus | LC | Stable | tropical | diurnal | 107.2 | 498715.91 | 3.197 |
| birds | Caligavis_chrysops | Lichenostomus_chrysops | LC | Decreasing | temperate | diurnal | 17.3 | 1457474.715 | 8.4 |
| birds | Caligavis_obscura | Lichenostomus_obscurus | LC | Decreasing | tropical | diurnal | 26.4 | 173670.691 | 3.018 |
| birds | Caligavis_subfrenata | Lichenostomus_subfrenatus | LC | Stable | tropical | diurnal | 29.7 | 132670.149 | 5.134 |
| birds | Callacanthis_burtoni | NA | LC | Stable | NA | NA | NA | NA | NA |
| birds | Callaeas_cinereus | Callaeas_cinereus | CR | Decreasing | temperate | diurnal | 225.4 | 152922.998 | 6.861 |
| birds | Calliope_calliope | Luscinia_calliope | LC | Stable | temperate | diurnal | 18.5 | 23381103.544 | 7.26 |
| birds | Calliope_obscura | Luscinia_obscura | VU | Decreasing | temperate | diurnal | 19.6 | 291670.812 | 11.999 |
| birds | Calliope_pectardens | Luscinia_pectardens | NT | Decreasing | temperate | diurnal | 14.5 | 1171410.064 | 9.878 |
| birds | Calliope_pectoralis | Luscinia_pectoralis | LC | Stable | temperate | diurnal | 23.3 | 3670860.206 | 8.721 |
| birds | Callipepla_californica | Callipepla_californica | LC | Increasing | temperate | diurnal | 165.7 | 1691291.35 | 9.457 |
| birds | Callipepla_douglasii | Callipepla_douglasii | LC | Decreasing | temperate | diurnal | 177.1 | 167016.602 | 7.046 |
| birds | Callipepla_gambelii | Callipepla_gambelii | LC | Stable | temperate | diurnal | 166 | 558986.969 | 8.763 |
| birds | Callipepla_squamata | Callipepla_squamata | LC | Decreasing | temperate | diurnal | 183.9 | 1432736.299 | 9.377 |
| birds | Calliphlox_amethystina | Calliphlox_amethystina | LC | Decreasing | tropical | diurnal | 2.8 | 7988128.33 | 6.764 |
| birds | Callocephalon_fimbriatum | Callocephalon_fimbriatum | VU | Decreasing | temperate | diurnal | 256 | 214821.225 | 9.753 |
| birds | Callonetta_leucophrys | Callonetta_leucophrys | LC | Decreasing | temperate | diurnal | 362.8 | 3218752.754 | 8.994 |
| birds | Calochaetes_coccineus | Calochaetes_coccineus | LC | Decreasing | tropical | diurnal | 46.4 | 99150.856 | 5.203 |
| birds | Calonectris_diomedea | Calonectris_diomedea | LC | Decreasing | tropical | cathemeral | 535 | 46036415.895 | 18.112 |
| birds | Calonectris_leucomelas | Calonectris_leucomelas | NT | Decreasing | tropical | cathemeral | 580 | 32884751.33 | 10.641 |
| birds | Caloperdix_oculeus | Caloperdix_oculeus | NT | Decreasing | tropical | diurnal | 209 | 534135.665 | 7.243 |
| birds | Caloramphus_fuliginosus | Calorhamphus_fuliginosus | LC | Decreasing | tropical | diurnal | 42.6 | 729941.664 | 5.616 |
| birds | Calothorax_lucifer | Calothorax_lucifer | LC | Stable | tropical | diurnal | 3.3 | 428359.875 | 11.339 |
| birds | Calothorax_pulcher | Calothorax_pulcher | LC | Decreasing | tropical | diurnal | 2.7 | 64442.512 | 9.312 |
| birds | Calypte_anna | Calypte_anna | LC | Increasing | temperate | diurnal | 4.2 | 1270071.918 | 9.135 |
| birds | Calypte_costae | Calypte_costae | LC | Increasing | temperate | diurnal | 3.1 | 645531.133 | 8.186 |
| birds | Calyptocichla_serinus | Calyptocichla_serina | LC | Stable | tropical | diurnal | 38.7 | 1315030.407 | 9.445 |
| birds | Calyptomena_hosii | Calyptomena_hosii | LC | Decreasing | tropical | diurnal | 92 | 146528.719 | 2.963 |
| birds | Calyptomena_viridis | Calyptomena_viridis | NT | Decreasing | tropical | diurnal | 58.5 | 1457426.484 | 8.085 |
| birds | Calyptomena_whiteheadi | Calyptomena_whiteheadi | LC | Decreasing | tropical | diurnal | 156.1 | 95505.364 | 2.378 |
| birds | Calyptophilus_frugivorus | Calyptophilus_frugivorus | NT | Decreasing | tropical | diurnal | 32.1 | 8600.716 | 10.173 |

| | | | | | | | | | |
|---|---|---|---|---|---|---|---|---|---|
| birds | Calyptophilus_tertius | Calyptophilus_tertius | VU | Decreasing | tropical | diurnal | 49.2 | 6314.226 | 12.465 |
| birds | Calyptorhynchus_banksii | Calyptorhynchus_banksii | LC | Decreasing | tropical | diurnal | 721.9 | 2798975.204 | 4.537 |
| birds | Calyptorhynchus_lathami | Calyptorhynchus_lathami | VU | Decreasing | temperate | diurnal | 437 | 661191.562 | 8.1 |
| birds | Calyptura_cristata | Calyptura_cristata | CR | Decreasing | tropical | diurnal | 7.8 | 2918.487 | 14.523 |
| birds | Camaroptera_brachyura | Camaroptera_brachyura | LC | Increasing | tropical | diurnal | 9.7 | 15929835.831 | 9.475 |
| birds | Camaroptera_chloronota | Camaroptera_chloronota | LC | Unknown | tropical | diurnal | 11.4 | 1659026.002 | 10.689 |
| birds | Camaroptera_superciliaris | Camaroptera_superciliaris | LC | Stable | tropical | diurnal | 9.7 | 3154212.794 | 8.278 |
| birds | Campephaga_flava | Campephaga_flava | LC | Stable | tropical | diurnal | 32.3 | 5048097.557 | 9.495 |
| birds | Campephaga_petiti | Campephaga_petiti | LC | Decreasing | tropical | diurnal | 31.3 | 508880.649 | 9.308 |
| birds | Campephaga_phoenicea | Campephaga_phoenicea | LC | Stable | tropical | diurnal | 27.1 | 4733993.798 | 10.249 |
| birds | Campephaga_quiscalina | Campephaga_quiscalina | LC | Decreasing | tropical | diurnal | 34.5 | 1373914.126 | 9.021 |
| birds | Campephilus_gayaquilensis | Campephilus_gayaquilensis | LC | Decreasing | tropical | diurnal | 241.2 | 117915.418 | 11.183 |
| birds | Campephilus_guatemalensis | Campephilus_guatemalensis | LC | Decreasing | tropical | diurnal | 242 | 888852.351 | 9.806 |
| birds | Campephilus_haematogaster | Campephilus_haematogaster | LC | Decreasing | tropical | diurnal | 237 | 305393.431 | 5.396 |
| birds | Campephilus_imperialis | Campephilus_imperialis | CR | Unknown | temperate | diurnal | 700 | 366656.604 | 9.795 |
| birds | Campephilus_leucopogon | Campephilus_leucopogon | LC | Stable | temperate | diurnal | 249 | 1077553.565 | 6.696 |
| birds | Campephilus_magellanicus | Campephilus_magellanicus | LC | Decreasing | temperate | diurnal | 324 | 395600.759 | 4.237 |
| birds | Campephilus_melanoleucos | Campephilus_melanoleucos | LC | Decreasing | tropical | diurnal | 256 | 11099024.679 | 4.693 |
| birds | Campephilus_pollens | Campephilus_pollens | LC | Decreasing | tropical | diurnal | 187.9 | 195689.246 | 9.883 |
| birds | Campephilus_principalis | Campephilus_principalis | CR | Unknown | temperate | diurnal | 516 | 845150.889 | 15.742 |
| birds | Campephilus_robustus | Campephilus_robustus | LC | Increasing | tropical | diurnal | 200 | 1885815.697 | 12.37 |
| birds | Campephilus_rubricollis | Campephilus_rubricollis | LC | Decreasing | tropical | diurnal | 210.7 | 6911322.083 | 2.291 |
| birds | Campethera_abingoni | Campethera_abingoni | LC | Stable | tropical | diurnal | 67.5 | 7184316.957 | 8.871 |
| birds | Campethera_bennettii | Campethera_bennettii | LC | Stable | tropical | diurnal | 70.1 | 4469921.881 | 7.844 |
| birds | Campethera_cailliautii | Campethera_cailliautii | LC | Stable | tropical | diurnal | 41.1 | 3823971.241 | 8.793 |
| birds | Campethera_maculosa | Campethera_maculosa | LC | Increasing | tropical | diurnal | 54 | 4611843.267 | 7.765 |
| birds | Campethera_mombassica | Campethera_mombassica | LC | Decreasing | tropical | diurnal | 59.6 | 57677.626 | 11.192 |
| birds | Campethera_notata | Campethera_notata | NT | Stable | temperate | diurnal | 62 | 95428.777 | 11.211 |
| birds | Campethera_nubica | Campethera_nubica | LC | Stable | tropical | diurnal | 59.7 | 3241336.041 | 10.16 |
| birds | Campethera_punctuligera | Campethera_punctuligera | LC | Stable | tropical | diurnal | 71.8 | 4070381.326 | 10.95 |
| birds | Campethera_scriptoricauda | NA | LC | Stable | NA | NA | NA | NA | NA |
| birds | Campethera_tullbergi | Campethera_tullbergi | LC | Stable | tropical | diurnal | 50.5 | 62707.498 | 9.965 |
| birds | Campicoloides_bifasciatus | Oenanthe_bifasciata | LC | Stable | temperate | diurnal | 35 | 244640.94 | 12.004 |
| birds | Campochaera_sloetii | Campochaera_sloetii | LC | Stable | tropical | diurnal | 41 | 229861.835 | 2.885 |

| | | | | | | | | | |
|---|---|---|---|---|---|---|---|---|---|
| birds | Campocolinus_albogularis | Francolinus_albogularis | LC | Stable | tropical | diurnal | 276 | 1104886.457 | 11.795 |
| birds | Campocolinus_coqui | Francolinus_coqui | LC | Stable | tropical | diurnal | 262 | 4180135.389 | 9.41 |
| birds | Campocolinus_schlegelii | Francolinus_schlegelii | LC | Stable | tropical | diurnal | 236.6 | 553212.323 | 5.152 |
| birds | Camptostoma_imberbe | Camptostoma_imberbe | LC | Decreasing | tropical | diurnal | 7.4 | 1448668.841 | 10.427 |
| birds | Camptostoma_obsoletum | Camptostoma_obsoletum | LC | Stable | tropical | diurnal | 8.1 | 14428561.738 | 5.973 |
| birds | Campylopterus_curvipennis | NA | not_available | not_available | NA | NA | NA | NA | NA |
| birds | Campylopterus_duidae | Campylopterus_duidae | LC | Unknown | tropical | diurnal | 6.1 | 102776.856 | 0.44 |
| birds | Campylopterus_ensipennis | Campylopterus_ensipennis | NT | Decreasing | tropical | diurnal | 9.6 | 2431.083 | 9.011 |
| birds | Campylopterus_falcatus | Campylopterus_falcatus | LC | Unknown | tropical | diurnal | 7.2 | 88173.023 | 10.931 |
| birds | Campylopterus_hemileucurus | Campylopterus_hemileucurus | LC | Decreasing | tropical | diurnal | 9.7 | 314208.021 | 11.36 |
| birds | Campylopterus_hyperythrus | Campylopterus_hyperythrus | LC | Stable | tropical | diurnal | 6.4 | 40875.453 | 1.833 |
| birds | Campylopterus_largipennis | Campylopterus_largipennis | LC | Decreasing | tropical | diurnal | 8.3 | 7116669.401 | 2.289 |
| birds | Campylopterus_phainopeplus | Campylopterus_phainopeplus | CR | Decreasing | tropical | diurnal | 7.7 | 2692.601 | 6.111 |
| birds | Campylopterus_villaviscensio | Campylopterus_villaviscensio | LC | Decreasing | tropical | diurnal | 7.4 | 28838.443 | 5.313 |
| birds | Campylorhamphus_falcularius | Campylorhamphus_falcularius | LC | Decreasing | tropical | diurnal | 42.6 | 876884.838 | 12.097 |
| birds | Campylorhamphus_procurvoides | Campylorhamphus_procurvoides | LC | Decreasing | tropical | diurnal | 33.5 | 2626695.78 | 1.146 |
| birds | Campylorhamphus_pusillus | Campylorhamphus_pusillus | LC | Decreasing | tropical | diurnal | 40.5 | 177568.928 | 7.596 |
| birds | Campylorhamphus_trochilirostris | Campylorhamphus_trochilirostris | LC | Decreasing | tropical | diurnal | 40.8 | 7392722.432 | 6.345 |
| birds | Campylorhynchus_albobrunneus | Campylorhynchus_albobrunneus | LC | Stable | tropical | diurnal | 32.9 | 148353.974 | 6.763 |
| birds | Campylorhynchus_brunneicapillus | Campylorhynchus_brunneicapillus | LC | Decreasing | temperate | diurnal | 38.9 | 1847263.759 | 9.16 |
| birds | Campylorhynchus_chiapensis | Campylorhynchus_chiapensis | LC | Increasing | tropical | diurnal | 50.9 | 9934.361 | 14.207 |
| birds | Campylorhynchus_fasciatus | Campylorhynchus_fasciatus | LC | Stable | tropical | diurnal | 30 | 121819.886 | 10.749 |
| birds | Campylorhynchus_griseus | Campylorhynchus_griseus | LC | Stable | tropical | diurnal | 42.4 | 982980.091 | 9.26 |
| birds | Campylorhynchus_gularis | Campylorhynchus_gularis | LC | Decreasing | temperate | diurnal | 30.1 | 370562.242 | 10.007 |
| birds | Campylorhynchus_jocosus | Campylorhynchus_jocosus | LC | Decreasing | tropical | diurnal | 27.6 | 89805.268 | 9.461 |
| birds | Campylorhynchus_megalopterus | Campylorhynchus_megalopterus | LC | Stable | tropical | diurnal | 33.1 | 80148.813 | 15.154 |
| birds | Campylorhynchus_nuchalis | Campylorhynchus_nuchalis | LC | Stable | tropical | diurnal | 23.2 | 506629.358 | 9.827 |
| birds | Campylorhynchus_rufinucha | Campylorhynchus_rufinucha | LC | Stable | tropical | diurnal | 31 | 8261.175 | 17.649 |
| birds | Campylorhynchus_turdinus | Campylorhynchus_turdinus | LC | Decreasing | tropical | diurnal | 32.6 | 4503316.808 | 3.547 |
| birds | Campylorhynchus_yucatanicus | Campylorhynchus_yucatanicus | NT | Stable | tropical | diurnal | 35.5 | 9706.723 | 11.572 |
| birds | Campylorhynchus_zonatus | Campylorhynchus_zonatus | LC | Decreasing | tropical | diurnal | 34.6 | 579082.336 | 12.464 |
| birds | Canachites_canadensis | Dendragapus_canadensis | LC | Stable | temperate | diurnal | 473.7 | 6068215.693 | 2.158 |
| birds | Canirallus_oculeus | Canirallus_oculeus | LC | Decreasing | tropical | diurnal | 277 | 1616653.929 | 8.883 |
| birds | Cantorchilus_griseus | Thryothorus_griseus | LC | Decreasing | tropical | diurnal | 19.1 | 90775.326 | 0.673 |

| | | | | | | | | | |
|---|---|---|---|---|---|---|---|---|---|
| birds | Cantorchilus_guarayanus | Thryothorus_guarayanus | LC | Decreasing | tropical | diurnal | 13.5 | 696010.23 | 4.992 |
| birds | Cantorchilus_leucopogon | Thryothorus_leucopogon | LC | Decreasing | tropical | diurnal | 19.1 | 140102.267 | 7.176 |
| birds | Cantorchilus_leucotis | Thryothorus_leucotis | LC | Decreasing | tropical | diurnal | 19.4 | 8709608.574 | 4.209 |
| birds | Cantorchilus_longirostris | Thryothorus_longirostris | LC | Unknown | tropical | diurnal | 21.3 | 1511309.579 | 9.702 |
| birds | Cantorchilus_modestus | Thryothorus_modestus | LC | Stable | tropical | diurnal | 17.6 | 263277.445 | 13.289 |
| birds | Cantorchilus_nigricapillus | Thryothorus_nigricapillus | LC | Decreasing | tropical | diurnal | 23.9 | 300564.01 | 8.439 |
| birds | Cantorchilus_semibadius | Thryothorus_semibadius | LC | Stable | tropical | diurnal | 17 | 14829.633 | 14.785 |
| birds | Cantorchilus_superciliaris | Thryothorus_superciliaris | LC | Unknown | tropical | diurnal | 22.8 | 96843.254 | 11.228 |
| birds | Cantorchilus_thoracicus | Thryothorus_thoracicus | LC | Decreasing | tropical | diurnal | 17.6 | 90302.605 | 7.798 |
| birds | Capito_aurovirens | Capito_aurovirens | LC | Stable | tropical | diurnal | 56.2 | 1228941.985 | 1.424 |
| birds | Capito_brunneipectus | Capito_brunneipectus | LC | Decreasing | tropical | diurnal | 57.9 | 129171.282 | 1.354 |
| birds | Capito_dayi | Capito_dayi | NT | Decreasing | tropical | diurnal | 67.2 | 1168130.045 | 2.825 |
| birds | Capito_hypoleucus | Capito_hypoleucus | LC | Decreasing | tropical | diurnal | 70.9 | 32873.935 | 11.328 |
| birds | Capito_maculicoronatus | Capito_maculicoronatus | LC | Decreasing | tropical | diurnal | 52.8 | 133684.282 | 6.602 |
| birds | Capito_niger | Capito_niger | LC | Stable | tropical | diurnal | 57.4 | 1107267.206 | 1.4 |
| birds | Capito_quinticolor | Capito_quinticolor | NT | Decreasing | tropical | diurnal | 60 | 37003.719 | 5.336 |
| birds | Capito_squamatus | Capito_squamatus | LC | Decreasing | tropical | diurnal | 59.3 | 63414.19 | 11.108 |
| birds | Capito_wallacei | Capito_wallacei | VU | Decreasing | tropical | diurnal | 69.5 | 1216.526 | 1.191 |
| birds | Caprimulgus_aegyptius | Caprimulgus_aegyptius | LC | Decreasing | temperate | nocturnal | 77.3 | 12441335.297 | 5.298 |
| birds | Caprimulgus_affinis | Caprimulgus_affinis | LC | Stable | tropical | nocturnal | 58.7 | 5312716.765 | 15.456 |
| birds | Caprimulgus_asiaticus | Caprimulgus_asiaticus | LC | Stable | tropical | nocturnal | 42 | 4043903.954 | 17.113 |
| birds | Caprimulgus_atripennis | Caprimulgus_atripennis | LC | Stable | tropical | nocturnal | 61 | 487705.092 | 16.64 |
| birds | Caprimulgus_batesi | Caprimulgus_batesi | LC | Decreasing | tropical | nocturnal | 100 | 1859436.146 | 5.581 |
| birds | Caprimulgus_celebensis | Caprimulgus_celebensis | LC | Decreasing | tropical | nocturnal | 57.5 | 31758.246 | 10.25 |
| birds | Caprimulgus_clarus | Caprimulgus_clarus | LC | Stable | tropical | nocturnal | 42.5 | 1364940.554 | 10.774 |
| birds | Caprimulgus_climacurus | Caprimulgus_climacurus | LC | Stable | tropical | nocturnal | 45.8 | 8738621.751 | 9.851 |
| birds | Caprimulgus_concretus | Caprimulgus_concretus | VU | Decreasing | tropical | nocturnal | 57.5 | 394039.807 | 7.323 |
| birds | Caprimulgus_donaldsoni | Caprimulgus_donaldsoni | LC | Stable | tropical | nocturnal | 29 | 751577.809 | 8.728 |
| birds | Caprimulgus_europaeus | Caprimulgus_europaeus | LC | Decreasing | temperate | cathemeral | 67 | 29394935.221 | 10.444 |
| birds | Caprimulgus_fossii | Caprimulgus_fossii | LC | Stable | tropical | nocturnal | 58.3 | 6491736.248 | 8.123 |
| birds | Caprimulgus_fraenatus | Caprimulgus_fraenatus | LC | Stable | tropical | nocturnal | 59.7 | 543293.668 | 11.115 |
| birds | Caprimulgus_indicus | Caprimulgus_indicus | LC | Stable | tropical | nocturnal | 91.6 | 1651666.549 | 18.641 |
| birds | Caprimulgus_inornatus | Caprimulgus_inornatus | LC | Stable | tropical | nocturnal | 50.3 | 8530651.411 | 10.192 |
| birds | Caprimulgus_longipennis | Macrodipteryx_longipennis | LC | Stable | tropical | nocturnal | 48 | 7317278.336 | 10.986 |

| | | | | | | | | | |
|---|---|---|---|---|---|---|---|---|---|
| birds | Caprimulgus_macrurus | Caprimulgus_macrurus | LC | Stable | tropical | nocturnal | 66.2 | 5990101.348 | 10.358 |
| birds | Caprimulgus_madagascariensis | Caprimulgus_madagascariensis | LC | Stable | tropical | nocturnal | 41.1 | 591293.273 | 8.749 |
| birds | Caprimulgus_mahrattensis | Caprimulgus_mahrattensis | LC | Stable | temperate | nocturnal | 57 | 2105063.725 | 13.908 |
| birds | Caprimulgus_manillensis | Caprimulgus_manillensis | LC | Stable | tropical | nocturnal | 70.3 | 269038.468 | 13.935 |
| birds | Caprimulgus_natalensis | Caprimulgus_natalensis | LC | Decreasing | tropical | nocturnal | 77.7 | 2207800.388 | 8.304 |
| birds | Caprimulgus_nubicus | Caprimulgus_nubicus | LC | Decreasing | tropical | nocturnal | 49.3 | 713220.637 | 9.088 |
| birds | Caprimulgus_pectoralis | Caprimulgus_pectoralis | LC | Stable | tropical | nocturnal | 51.4 | 5725005.927 | 8.957 |
| birds | Caprimulgus_poliocephalus | Caprimulgus_poliocephalus | LC | Stable | tropical | nocturnal | 41.8 | 807034.95 | 13.062 |
| birds | Caprimulgus_pulchellus | Caprimulgus_pulchellus | LC | Decreasing | tropical | nocturnal | 57.5 | 70914.301 | 8.425 |
| birds | Caprimulgus_ruficollis | Caprimulgus_ruficollis | NT | Decreasing | tropical | nocturnal | 68.5 | 4900398.316 | 6.79 |
| birds | Caprimulgus_rufigena | Caprimulgus_rufigena | LC | Stable | tropical | nocturnal | 52 | 3243389.387 | 7.516 |
| birds | Caprimulgus_tristigma | Caprimulgus_tristigma | LC | Stable | tropical | nocturnal | 79 | 4557630.148 | 10.723 |
| birds | Caprimulgus_vexillarius | Macrodipteryx_vexillarius | LC | Stable | tropical | cathemeral | 72.2 | 6625232.434 | 7.949 |
| birds | Capsiempis_flaveola | Capsiempis_flaveola | LC | Decreasing | tropical | diurnal | 7.7 | 5724076.644 | 7.171 |
| birds | Caracara_plancus | Caracara_plancus | LC | Stable | tropical | diurnal | 1078.6 | 12754773.654 | 8.59 |
| birds | Cardellina_canadensis | Wilsonia_canadensis | LC | Decreasing | tropical | diurnal | 10 | 7865400.501 | 11.97 |
| birds | Cardellina_pusilla | Wilsonia_pusilla | LC | Decreasing | temperate | diurnal | 7 | 18354114.384 | 8.421 |
| birds | Cardellina_rubra | Ergaticus_ruber | LC | Decreasing | tropical | diurnal | 8.1 | 232459.315 | 9.883 |
| birds | Cardellina_rubrifrons | Cardellina_rubrifrons | LC | Decreasing | temperate | diurnal | 8.1 | 1056864.129 | 9.638 |
| birds | Cardellina_versicolor | Ergaticus_versicolor | LC | Decreasing | tropical | diurnal | 10 | 17025.105 | 14.648 |
| birds | Cardinalis_cardinalis | Cardinalis_cardinalis | LC | Stable | temperate | diurnal | 42.6 | 5854913.925 | 14.856 |
| birds | Cardinalis_phoeniceus | Cardinalis_phoeniceus | LC | Decreasing | tropical | diurnal | 35 | 69681.592 | 12.448 |
| birds | Cardinalis_sinuatus | Cardinalis_sinuatus | LC | Decreasing | temperate | diurnal | 35.2 | 1519386.565 | 9.275 |
| birds | Carduelis_carduelis | Carduelis_carduelis | LC | Decreasing | temperate | diurnal | 16 | 15546092.385 | 13.241 |
| birds | Carduelis_citrinella | Carduelis_citrinella | LC | Decreasing | temperate | diurnal | 12 | 241931.151 | 15.85 |
| birds | Cariama_cristata | Cariama_cristata | LC | Stable | tropical | diurnal | 1400 | 5932013.381 | 9.101 |
| birds | Caridonax_fulgidus | Caridonax_fulgidus | LC | Stable | tropical | diurnal | 171.8 | 33455.221 | 13.577 |
| birds | Carpococcyx_radiceus | Carpococcyx_radiatus | VU | Decreasing | tropical | diurnal | 495.7 | 259047.555 | 4.372 |
| birds | Carpococcyx_renauldi | Carpococcyx_renauldi | VU | Decreasing | tropical | diurnal | 400 | 457300.489 | 11.589 |
| birds | Carpococcyx_viridis | NA | CR | Decreasing | NA | NA | NA | NA | NA |
| birds | Carpodacus_edwardsii | Carpodacus_edwardsii | LC | Stable | temperate | diurnal | 26.5 | 553558.679 | 7.118 |
| birds | Carpodacus_erythrinus | Carpodacus_erythrinus | LC | Decreasing | temperate | diurnal | 24 | 28438262.754 | 8.062 |
| birds | Carpodacus_pulcherrimus | Carpodacus_pulcherrimus | LC | Stable | temperate | diurnal | 19.2 | 2405247.899 | 8.853 |
| birds | Carpodacus_puniceus | Carpodacus_puniceus | LC | Stable | temperate | diurnal | 46.6 | 3485769.613 | 4.484 |

| | | | | | | | | | |
|---|---|---|---|---|---|---|---|---|---|
| birds | Carpodacus_rhodochlamys | Carpodacus_rhodochlamys | LC | Stable | temperate | diurnal | 33.4 | 3087746.169 | 6.224 |
| birds | Carpodacus_roborowskii | NA | LC | Stable | NA | NA | NA | NA | NA |
| birds | Carpodacus_rodochroa | Carpodacus_rodochroa | LC | Stable | temperate | diurnal | 17.8 | 251585.02 | 9.399 |
| birds | Carpodacus_rodopeplus | Carpodacus_rodopeplus | LC | Stable | temperate | diurnal | 23 | 184670.722 | 11.263 |
| birds | Carpodacus_roseus | Carpodacus_roseus | LC | Stable | temperate | diurnal | 28.6 | 9266325.215 | 6.734 |
| birds | Carpodacus_rubicilla | Carpodacus_rubicilla | LC | Stable | temperate | diurnal | 43.2 | 3210935.497 | 5.028 |
| birds | Carpodacus_rubicilloides | Carpodacus_rubicilloides | LC | Stable | temperate | diurnal | 42 | 1872670.057 | 7.042 |
| birds | Carpodacus_sibiricus | Uragus_sibiricus | LC | Stable | temperate | diurnal | 17.4 | 7744795.24 | 6.978 |
| birds | Carpodacus_sipahi | Haematospiza_sipahi | LC | Stable | temperate | diurnal | 39.5 | 710638.761 | 9.024 |
| birds | Carpodacus_subhimachalus | Pinicola_subhimachala | LC | Stable | temperate | diurnal | 46.1 | 824298.843 | 7.773 |
| birds | Carpodacus_synoicus | Carpodacus_synoicus | LC | Stable | temperate | diurnal | 20.8 | 172774.388 | 4.735 |
| birds | Carpodacus_thura | Carpodacus_thura | LC | Stable | temperate | diurnal | 31 | 296993.997 | 7.96 |
| birds | Carpodacus_trifasciatus | NA | LC | Stable | NA | NA | NA | NA | NA |
| birds | Carpodacus_vinaceus | Carpodacus_vinaceus | LC | Stable | temperate | diurnal | 22.1 | 1267546.737 | 13.469 |
| birds | Carpodacus_waltoni | Carpodacus_eos | LC | Stable | temperate | diurnal | 27 | 970503.289 | 7.093 |
| birds | Carpodectes_antoniae | Carpodectes_antoniae | NT | Decreasing | tropical | diurnal | 98 | 3915.869 | 10.42 |
| birds | Carpodectes_hopkei | Carpodectes_hopkei | LC | Decreasing | tropical | diurnal | 102 | 96274.784 | 5.826 |
| birds | Carpodectes_nitidus | Carpodectes_nitidus | LC | Decreasing | tropical | diurnal | 101.6 | 103015.664 | 8.071 |
| birds | Carpornis_cucullata | Carpornis_cucullata | LC | Decreasing | tropical | diurnal | 74.2 | 82768.081 | 11.846 |
| birds | Carpornis_melanocephala | Carpornis_melanocephala | NT | Decreasing | tropical | diurnal | 64.2 | 236112.705 | 12.791 |
| birds | Carpospiza_brachydactyla | Petronia_brachydactyla | LC | Stable | temperate | diurnal | 22.3 | 5369770.17 | 7.249 |
| birds | Carterornis_castus | NA | NT | Decreasing | NA | NA | NA | NA | NA |
| birds | Carterornis_chrysomela | Monarcha_chrysomela | LC | Stable | tropical | diurnal | 15.6 | 651592.49 | 3.126 |
| birds | Carterornis_leucotis | Monarcha_leucotis | LC | Stable | tropical | diurnal | 11.4 | 312561.628 | 6.067 |
| birds | Carterornis_pileatus | Monarcha_pileatus | LC | Stable | tropical | diurnal | 14 | 20864.032 | 6.492 |
| birds | Caryothraustes_canadensis | Caryothraustes_canadensis | LC | Decreasing | tropical | diurnal | 34.5 | 2650244.991 | 3.041 |
| birds | Caryothraustes_celaeno | Rhodothraupis_celaeno | LC | Decreasing | tropical | diurnal | 60 | 127727.171 | 11.703 |
| birds | Caryothraustes_erythromelas | Periporphyrus_erythromelas | LC | Decreasing | tropical | diurnal | 48 | 1288168.813 | 2.168 |
| birds | Caryothraustes_poliogaster | Caryothraustes_poliogaster | LC | Decreasing | tropical | diurnal | 41.8 | 317013.578 | 9.126 |
| birds | Casiornis_fuscus | Casiornis_fuscus | LC | Decreasing | tropical | diurnal | 19.5 | 2107102.83 | 6.55 |
| birds | Casiornis_rufus | Casiornis_rufus | LC | Decreasing | tropical | diurnal | 24.8 | 5566772.384 | 6.126 |
| birds | Cassiculus_melanicterus | Cacicus_melanicterus | LC | Decreasing | tropical | diurnal | 81.3 | 231633.39 | 8.943 |
| birds | Castanozoster_thoracicus | Poospiza_thoracica | LC | Decreasing | tropical | diurnal | 11.9 | 77555.645 | 11.811 |
| birds | Casuarius_bennetti | Casuarius_bennetti | LC | Stable | tropical | diurnal | 35000 | 396486.378 | 3.868 |

| | | | | | | | | | |
|---|---|---|---|---|---|---|---|---|---|
| birds | Casuarius_casuarius | Casuarius_casuarius | LC | Decreasing | tropical | diurnal | 44000 | 397042.35 | 3.053 |
| birds | Casuarius_unappendiculatus | Casuarius_unappendiculatus | LC | Decreasing | tropical | diurnal | 46073.9 | 185350.582 | 3.372 |
| birds | Catamblyrhynchus_diadema | Catamblyrhynchus_diadema | LC | Decreasing | tropical | diurnal | 14.1 | 358573.963 | 8.698 |
| birds | Catamenia_analis | Catamenia_analis | LC | Increasing | tropical | diurnal | 12 | 1027720.641 | 7.022 |
| birds | Catamenia_homochroa | Catamenia_homochroa | LC | Stable | tropical | diurnal | 12.2 | 283918.399 | 6.907 |
| birds | Catamenia_inornata | Catamenia_inornata | LC | Increasing | tropical | diurnal | 13.4 | 801551.997 | 8.005 |
| birds | Cataponera_turdoides | Cataponera_turdoides | LC | Decreasing | tropical | diurnal | 89.7 | 23212.789 | 4.913 |
| birds | Catharacta_antarctica | Catharacta_antarctica | LC | Decreasing | temperate | nocturnal | 1650 | 87990607.821 | 5.246 |
| birds | Catharacta_chilensis | Catharacta_chilensis | LC | Stable | temperate | diurnal | 1367.8 | 1879625.66 | 3.628 |
| birds | Catharacta_maccormicki | Catharacta_maccormicki | LC | Stable | temperate | diurnal | 1347.1 | 17885700.619 | 1.778 |
| birds | Catharacta_skua | Catharacta_skua | LC | Stable | temperate | diurnal | 1338.2 | 20146857.185 | 10.158 |
| birds | Catharopeza_bishopi | Catharopeza_bishopi | EN | Decreasing | tropical | diurnal | 15.1 | 83.112 | 8.862 |
| birds | Cathartes_aura | Cathartes_aura | LC | Stable | tropical | diurnal | 1518.2 | 28211569.025 | 8.623 |
| birds | Cathartes_burrovianus | Cathartes_burrovianus | LC | Stable | tropical | diurnal | 935 | 7839868.539 | 7.993 |
| birds | Cathartes_melambrotus | Cathartes_melambrotus | LC | Decreasing | tropical | diurnal | 1373 | 6652618.685 | 2.065 |
| birds | Catharus_aurantiirostris | Catharus_aurantiirostris | LC | Decreasing | tropical | diurnal | 29.8 | 854579.169 | 11.174 |
| birds | Catharus_bicknelli | Catharus_bicknelli | VU | Decreasing | temperate | diurnal | 27.8 | 49188.476 | 7.068 |
| birds | Catharus_dryas | Catharus_dryas | LC | Decreasing | tropical | diurnal | 37.7 | 88086.158 | 12.665 |
| birds | Catharus_frantzii | Catharus_frantzii | LC | Decreasing | tropical | diurnal | 28.9 | 283300.457 | 10.835 |
| birds | Catharus_fuscater | Catharus_fuscater | LC | Decreasing | tropical | diurnal | 33.7 | 313318.356 | 8.223 |
| birds | Catharus_fuscescens | Catharus_fuscescens | LC | Decreasing | temperate | diurnal | 31.9 | 16366792.634 | 9.046 |
| birds | Catharus_gracilirostris | Catharus_gracilirostris | LC | Stable | tropical | diurnal | 21 | 3827.252 | 5.583 |
| birds | Catharus_guttatus | Catharus_guttatus | LC | Stable | temperate | diurnal | 30.1 | 15387664.79 | 9.419 |
| birds | Catharus_mexicanus | Catharus_mexicanus | LC | Decreasing | tropical | diurnal | 33 | 143279.41 | 10.49 |
| birds | Catharus_minimus | Catharus_minimus | LC | Decreasing | temperate | diurnal | 31.6 | 17837621.978 | 6.603 |
| birds | Catharus_occidentalis | Catharus_occidentalis | LC | Decreasing | tropical | diurnal | 26.2 | 417881.836 | 9.048 |
| birds | Catharus_ustulatus | Catharus_ustulatus | LC | Stable | temperate | diurnal | 30.3 | 11035380.862 | 12.647 |
| birds | Catherpes_mexicanus | Catherpes_mexicanus | LC | Stable | temperate | diurnal | 12 | 4298192.457 | 9.249 |
| birds | Catreus_wallichii | Catreus_wallichi | VU | Decreasing | temperate | diurnal | 1436.9 | 210158.104 | 12.476 |
| birds | Catriscus_brevirostris | Schoenicola_brevirostris | LC | Decreasing | tropical | diurnal | 15.7 | 2804794.872 | 9.191 |
| birds | Ceblepyris_caesius | Coracina_caesia | LC | Stable | tropical | diurnal | 47.3 | 905645.625 | 12.766 |
| birds | Ceblepyris_cinereus | Coracina_cinerea | LC | Decreasing | tropical | diurnal | 47.2 | 415959.231 | 8.379 |
| birds | Ceblepyris_graueri | NA | LC | Decreasing | NA | NA | NA | NA | NA |
| birds | Ceblepyris_pectoralis | Coracina_pectoralis | LC | Decreasing | tropical | diurnal | 54.4 | 6110219.708 | 9.145 |

| | | | | | | | | | |
|---|---|---|---|---|---|---|---|---|---|
| birds | Cecropis_abyssinica | Hirundo_abyssinica | LC | Increasing | tropical | diurnal | 13.5 | 13676430.323 | 9.535 |
| birds | Cecropis_cucullata | Hirundo_cucullata | LC | Increasing | tropical | diurnal | 27.1 | 5535896.414 | 7.635 |
| birds | Cecropis_daurica | Hirundo_daurica | LC | Stable | temperate | diurnal | 22.2 | 21306722.748 | 12.751 |
| birds | Cecropis_semirufa | Hirundo_semirufa | LC | Increasing | tropical | diurnal | 30.1 | 8911972.362 | 9.541 |
| birds | Cecropis_senegalensis | Hirundo_senegalensis | LC | Increasing | tropical | diurnal | 45.4 | 12912848.808 | 9.23 |
| birds | Celebesica_abbotti | Coracina_abbotti | LC | Decreasing | tropical | diurnal | 68 | 28181.388 | 5.024 |
| birds | Celeus_castaneus | Celeus_castaneus | LC | Decreasing | tropical | diurnal | 85.6 | 396905.383 | 8.571 |
| birds | Celeus_elegans | Celeus_elegans | LC | Decreasing | tropical | diurnal | 135 | 6510380.155 | 2.115 |
| birds | Celeus_flavescens | Celeus_flavescens | LC | Decreasing | tropical | diurnal | 139 | 2619318.97 | 10.696 |
| birds | Celeus_flavus | Celeus_flavus | LC | Decreasing | tropical | diurnal | 147.3 | 6199098.969 | 2.742 |
| birds | Celeus_galeatus | Dryocopus_galeatus | VU | Decreasing | temperate | diurnal | 124 | 267579.916 | 12.011 |
| birds | Celeus_loricatus | Celeus_loricatus | LC | Decreasing | tropical | diurnal | 76.6 | 297945.587 | 9.886 |
| birds | Celeus_lugubris | Celeus_lugubris | LC | Decreasing | tropical | diurnal | 137 | 841523.9 | 6.69 |
| birds | Celeus_spectabilis | Celeus_spectabilis | LC | Unknown | tropical | diurnal | 114 | 528390.731 | 2.358 |
| birds | Celeus_torquatus | Celeus_torquatus | NT | Decreasing | tropical | diurnal | 134 | 1895802.554 | 2.512 |
| birds | Celeus_undatus | Celeus_undatus | LC | Decreasing | tropical | diurnal | 64.5 | 5521579.18 | 1.755 |
| birds | Centrocercus_urophasianus | Centrocercus_urophasianus | NT | Decreasing | temperate | diurnal | 1902.5 | 1315889.139 | 10.041 |
| birds | Centropus_andamanensis | Centropus_andamanensis | LC | Stable | tropical | diurnal | 164 | 7211.897 | 6.666 |
| birds | Centropus_anselli | Centropus_anselli | LC | Stable | tropical | diurnal | 210 | 1058987.746 | 5.393 |
| birds | Centropus_ateralbus | Centropus_ateralbus | LC | Stable | tropical | diurnal | 339 | 40984.982 | 4.479 |
| birds | Centropus_bengalensis | Centropus_bengalensis | LC | Increasing | tropical | diurnal | 148.9 | 6030216.392 | 12.496 |
| birds | Centropus_bernsteini | Centropus_bernsteini | LC | Stable | tropical | diurnal | 162.1 | 516862.901 | 2.981 |
| birds | Centropus_celebensis | Centropus_celebensis | LC | Stable | tropical | diurnal | 244.4 | 134955.483 | 9.614 |
| birds | Centropus_chalybeus | Centropus_chalybeus | NT | Decreasing | tropical | diurnal | 244.4 | 2526.514 | 8.959 |
| birds | Centropus_chlororhynchos | Centropus_chlororhynchus | VU | Decreasing | tropical | diurnal | 244.4 | 11188.067 | 18.373 |
| birds | Centropus_cupreicaudus | Centropus_cupreicaudus | LC | Stable | tropical | diurnal | 285.2 | 1750330.636 | 7.239 |
| birds | Centropus_goliath | Centropus_goliath | LC | Stable | tropical | diurnal | 499.4 | 25406.987 | 6.288 |
| birds | Centropus_grillii | Centropus_grillii | LC | Stable | tropical | diurnal | 122.9 | 8530549.353 | 9.273 |
| birds | Centropus_leucogaster | Centropus_leucogaster | LC | Stable | tropical | diurnal | 307.8 | 1335483.706 | 12.134 |
| birds | Centropus_melanops | Centropus_melanops | LC | Decreasing | tropical | diurnal | 221.2 | 106106.005 | 11.781 |
| birds | Centropus_menbeki | Centropus_menbeki | LC | Stable | tropical | diurnal | 519 | 645860.008 | 3.188 |
| birds | Centropus_milo | Centropus_milo | LC | Stable | tropical | diurnal | 769 | 11026.514 | 4.473 |
| birds | Centropus_monachus | Centropus_monachus | LC | Stable | tropical | diurnal | 201.3 | 2920157.621 | 10.486 |
| birds | Centropus_nigrorufus | Centropus_nigrorufus | VU | Decreasing | tropical | diurnal | 244.4 | 28067.428 | 24.104 |

| | | | | | | | | | |
|---|---|---|---|---|---|---|---|---|---|
| birds | Centropus_phasianinus | Centropus_phasianinus | LC | Stable | tropical | diurnal | 382 | 2665520.199 | 4.107 |
| birds | Centropus_rectunguis | Centropus_rectunguis | VU | Decreasing | tropical | diurnal | 238 | 1057784.938 | 8.257 |
| birds | Centropus_senegalensis | Centropus_senegalensis | LC | Stable | tropical | diurnal | 169 | 9875812.368 | 10.036 |
| birds | Centropus_sinensis | Centropus_sinensis | LC | Stable | tropical | diurnal | 280.7 | 7931391.585 | 14.429 |
| birds | Centropus_spilopterus | Centropus_spilopterus | LC | Stable | tropical | diurnal | 244.4 | 1110.387 | 12.883 |
| birds | Centropus_steerii | Centropus_steerii | CR | Decreasing | tropical | diurnal | 198.4 | 699.116 | 14.23 |
| birds | Centropus_superciliosus | Centropus_superciliosus | LC | Stable | tropical | diurnal | 148.5 | 7292900.161 | 9.482 |
| birds | Centropus_toulou | Centropus_toulou | LC | Decreasing | tropical | diurnal | 166.4 | 590995.808 | 8.745 |
| birds | Centropus_unirufus | Centropus_unirufus | NT | Decreasing | tropical | diurnal | 183.8 | 80442.958 | 11.92 |
| birds | Centropus_violaceus | Centropus_violaceus | LC | Decreasing | tropical | diurnal | 500 | 38717.48 | 4.553 |
| birds | Centropus_viridis | Centropus_viridis | LC | Stable | tropical | diurnal | 142 | 273885.377 | 13.81 |
| birds | Cephalopterus_glabricollis | Cephalopterus_glabricollis | EN | Decreasing | tropical | diurnal | 379.5 | 26873.009 | 8.941 |
| birds | Cephalopterus_ornatus | Cephalopterus_ornatus | LC | Decreasing | tropical | diurnal | 380 | 5506246.904 | 2.25 |
| birds | Cephalopterus_penduliger | Cephalopterus_penduliger | VU | Decreasing | tropical | diurnal | 338 | 56545.394 | 9.071 |
| birds | Cephalopyrus_flammiceps | Cephalopyrus_flammiceps | LC | Unknown | temperate | diurnal | 7 | 498757.593 | 11.817 |
| birds | Cepphus_carbo | Cepphus_carbo | LC | Decreasing | temperate | diurnal | 490 | 2647371.712 | 3.627 |
| birds | Cepphus_columba | Cepphus_columba | LC | Stable | temperate | diurnal | 530 | 6817081.263 | 2.652 |
| birds | Cepphus_grylle | Cepphus_grylle | LC | Unknown | temperate | diurnal | 378 | 6808695.902 | 5.714 |
| birds | Ceratogymna_atrata | Ceratogymna_atrata | LC | Decreasing | tropical | diurnal | 1194.8 | 2282542.358 | 7.649 |
| birds | Ceratogymna_elata | Ceratogymna_elata | VU | Decreasing | tropical | diurnal | 1917 | 495247.104 | 14.341 |
| birds | Ceratopipra_chloromeros | Pipra_chloromeros | LC | Decreasing | tropical | diurnal | 16.5 | 472743.225 | 3.111 |
| birds | Ceratopipra_cornuta | Pipra_cornuta | LC | Stable | tropical | diurnal | 25.3 | 345161.435 | 1.036 |
| birds | Ceratopipra_erythrocephala | Pipra_erythrocephala | LC | Stable | tropical | diurnal | 13.4 | 3456038.861 | 2.791 |
| birds | Ceratopipra_mentalis | Pipra_mentalis | LC | Decreasing | tropical | diurnal | 15 | 558740.473 | 8.666 |
| birds | Ceratopipra_rubrocapilla | Pipra_rubrocapilla | LC | Decreasing | tropical | diurnal | 12 | 3391760.241 | 3.029 |
| birds | Cercibis_oxycerca | Cercibis_oxycerca | LC | Stable | tropical | diurnal | 1819 | 395881.11 | 6.28 |
| birds | Cercococcyx_mechowi | Cercococcyx_mechowi | LC | Decreasing | tropical | diurnal | 55.6 | 1462828.008 | 7.433 |
| birds | Cercococcyx_montanus | Cercococcyx_montanus | LC | Decreasing | tropical | diurnal | 58.2 | 739949.797 | 10.953 |
| birds | Cercococcyx_olivinus | Cercococcyx_olivinus | LC | Stable | tropical | diurnal | 65.9 | 1841106.004 | 7.865 |
| birds | Cercomacra_brasiliana | Cercomacra_brasiliana | NT | Decreasing | tropical | diurnal | 12 | 129178.513 | 11.085 |
| birds | Cercomacra_carbonaria | Cercomacra_carbonaria | CR | Decreasing | tropical | diurnal | 14.5 | 19133.111 | 4.457 |
| birds | Cercomacra_cinerascens | Cercomacra_cinerascens | LC | Decreasing | tropical | diurnal | 14.3 | 6701396.999 | 2.108 |
| birds | Cercomacra_ferdinandi | Cercomacra_ferdinandi | NT | Decreasing | tropical | diurnal | 15 | 154169.619 | 5.967 |
| birds | Cercomacra_manu | Cercomacra_manu | LC | Decreasing | tropical | diurnal | 18.1 | 484743.724 | 2.852 |

| | | | | | | | | | |
|---|---|---|---|---|---|---|---|---|---|
| birds | Cercomacra_melanaria | Cercomacra_melanaria | LC | Decreasing | tropical | diurnal | 19 | 445529.075 | 6.418 |
| birds | Cercomacra_nigricans | Cercomacra_nigricans | LC | Decreasing | tropical | diurnal | 16.6 | 546994.416 | 11.596 |
| birds | Cercomacroides_laeta | Cercomacra_laeta | LC | Stable | tropical | diurnal | 16 | 393677.481 | 3.238 |
| birds | Cercomacroides_nigrescens | Cercomacra_nigrescens | LC | Decreasing | tropical | diurnal | 16.5 | 3701417.627 | 2.65 |
| birds | Cercomacroides_parkeri | Cercomacra_parkeri | LC | Decreasing | tropical | diurnal | 17.2 | 60670.311 | 13.996 |
| birds | Cercomacroides_serva | Cercomacra_serva | LC | Stable | tropical | diurnal | 17 | 1344617.261 | 1.916 |
| birds | Cercomacroides_tyrannina | Cercomacra_tyrannina | LC | Decreasing | tropical | diurnal | 16.3 | 3170819.203 | 3.811 |
| birds | Cercotrichas_galactotes | Erythropygia_galactotes | LC | Stable | tropical | diurnal | 20.3 | 18679450.743 | 7.322 |
| birds | Cercotrichas_hartlaubi | Erythropygia_hartlaubi | LC | Stable | tropical | diurnal | 19.2 | 612316.425 | 11.166 |
| birds | Cercotrichas_leucophrys | Erythropygia_leucophrys | LC | Stable | tropical | diurnal | 16.5 | 8126492.582 | 8.776 |
| birds | Cercotrichas_paena | Erythropygia_paena | LC | Stable | tropical | diurnal | 19.7 | 1909273.596 | 6.424 |
| birds | Cercotrichas_podobe | Cercotrichas_podobe | LC | Stable | tropical | diurnal | 23.9 | 3942700.06 | 8.796 |
| birds | Cereopsis_novaehollandiae | Cereopsis_novaehollandiae | LC | Stable | temperate | diurnal | 3560 | 595801.467 | 10.418 |
| birds | Cerorhinca_monocerata | Cerorhinca_monocerata | LC | Decreasing | temperate | diurnal | 482.2 | 5458729.595 | 11.064 |
| birds | Certhia_americana | Certhia_americana | LC | Increasing | temperate | diurnal | 8.1 | 11412723.753 | 11.031 |
| birds | Certhia_brachydactyla | Certhia_brachydactyla | LC | Increasing | temperate | diurnal | 8.2 | 3199552.332 | 20.377 |
| birds | Certhia_discolor | Certhia_discolor | LC | Decreasing | temperate | diurnal | 10.4 | 173809.738 | 8.396 |
| birds | Certhia_familiaris | Certhia_familiaris | LC | Stable | temperate | diurnal | 9 | 15914424.481 | 10.005 |
| birds | Certhia_himalayana | Certhia_himalayana | LC | Decreasing | temperate | diurnal | 8.8 | 1333092.384 | 11.58 |
| birds | Certhia_nipalensis | Certhia_nipalensis | LC | Decreasing | temperate | diurnal | 11.1 | 163631.814 | 6.211 |
| birds | Certhiasomus_stictolaemus | Deconychura_stictolaema | LC | Decreasing | tropical | diurnal | 18.5 | 3212301.464 | 1.364 |
| birds | Certhiaxis_cinnamomeus | Certhiaxis_cinnamomeus | LC | Decreasing | tropical | diurnal | 15.2 | 8364757.314 | 8.15 |
| birds | Certhiaxis_mustelinus | Certhiaxis_mustelinus | LC | Stable | tropical | diurnal | 15 | 693874.134 | 2.032 |
| birds | Certhidea_olivacea | NA | VU | Decreasing | NA | NA | NA | NA | NA |
| birds | Certhilauda_chuana | Certhilauda_chuana | LC | Decreasing | temperate | diurnal | 44.1 | 127491.6 | 10.784 |
| birds | Certhilauda_curvirostris | Certhilauda_curvirostris | LC | Decreasing | temperate | diurnal | 40.4 | 63461.484 | 10.178 |
| birds | Certhionyx_variegatus | Certhionyx_variegatus | LC | Stable | temperate | diurnal | 26.3 | 5122091.983 | 3.79 |
| birds | Cettia_brunnifrons | Cettia_brunnifrons | LC | Stable | temperate | diurnal | 7.3 | 1378233.278 | 10.258 |
| birds | Cettia_castaneocoronata | Tesia_castaneocoronata | LC | Stable | temperate | diurnal | 8.9 | 1980546.826 | 10.362 |
| birds | Cettia_cetti | Cettia_cetti | LC | Increasing | temperate | diurnal | 13.2 | 5830320.199 | 14.166 |
| birds | Cettia_major | Cettia_major | LC | Decreasing | temperate | diurnal | 9.7 | 1701566.019 | 10.139 |
| birds | Ceuthmochares_aereus | Ceuthmochares_aereus | LC | Stable | tropical | diurnal | 65.8 | 4005421.098 | 8.35 |
| birds | Ceyx_argentatus | Alcedo_argentata | NT | Decreasing | tropical | diurnal | 17.8 | 89541.508 | 12.497 |
| birds | Ceyx_azureus | Alcedo_azurea | LC | Decreasing | temperate | diurnal | 34.9 | 3398489.468 | 5.235 |

| | | | | | | | | | |
|---|---|---|---|---|---|---|---|---|---|
| birds | Ceyx_cyanopectus | Alcedo_cyanopectus | LC | Decreasing | tropical | diurnal | 21.5 | 111263.194 | 15.291 |
| birds | Ceyx_erithaca | Ceyx_erithaca | NT | Decreasing | tropical | diurnal | 17.8 | 3178476.075 | 12.406 |
| birds | Ceyx_fallax | Ceyx_fallax | LC | Decreasing | tropical | diurnal | 18 | 162180.836 | 9.555 |
| birds | Ceyx_lepidus | Ceyx_lepidus | LC | Decreasing | tropical | diurnal | 21.3 | 43842.146 | 6.774 |
| birds | Ceyx_melanurus | Ceyx_melanurus | VU | Decreasing | tropical | diurnal | 22.8 | 127390.353 | 14.262 |
| birds | Ceyx_pusillus | Alcedo_pusilla | LC | Decreasing | tropical | diurnal | 13.3 | 910124.535 | 3.357 |
| birds | Ceyx_websteri | Alcedo_websteri | VU | Decreasing | tropical | diurnal | 61 | 13725.074 | 6.433 |
| birds | Chaetocercus_astreans | NA | LC | Stable | NA | NA | NA | NA | NA |
| birds | Chaetocercus_berlepschi | Chaetocercus_berlepschi | VU | Decreasing | tropical | diurnal | 2 | 5021.77 | 15.461 |
| birds | Chaetocercus_bombus | Chaetocercus_bombus | NT | Decreasing | tropical | diurnal | 6.5 | 175410.374 | 10.172 |
| birds | Chaetocercus_heliodor | Chaetocercus_heliodor | LC | Stable | tropical | diurnal | 2 | 113213.814 | 12.07 |
| birds | Chaetocercus_jourdanii | Chaetocercus_jourdanii | LC | Stable | tropical | diurnal | 3.3 | 46450.267 | 14.28 |
| birds | Chaetocercus_mulsant | Chaetocercus_mulsant | LC | Stable | tropical | diurnal | 3.9 | 263636.46 | 9.545 |
| birds | Chaetops_frenatus | Chaetops_frenatus | NT | Decreasing | temperate | diurnal | 55 | 59624.787 | 10.042 |
| birds | Chaetorhynchus_papuensis | Chaetorhynchus_papuensis | LC | Decreasing | tropical | diurnal | 37.2 | 305618.376 | 3.332 |
| birds | Chaetura_andrei | NA | VU | Decreasing | NA | NA | NA | NA | NA |
| birds | Chaetura_brachyura | Chaetura_brachyura | LC | Stable | tropical | diurnal | 18.3 | 7468188.252 | 3.176 |
| birds | Chaetura_chapmani | Chaetura_chapmani | LC | Decreasing | tropical | diurnal | 23.8 | 4701276.356 | 2.208 |
| birds | Chaetura_cinereiventris | Chaetura_cinereiventris | LC | Decreasing | tropical | diurnal | 13.9 | 6102705.507 | 4.26 |
| birds | Chaetura_egregia | Chaetura_egregia | LC | Decreasing | tropical | diurnal | 23.4 | 1756675.456 | 3.404 |
| birds | Chaetura_martinica | Chaetura_martinica | LC | Stable | tropical | diurnal | 12.5 | 4342.258 | 21.792 |
| birds | Chaetura_pelagica | Chaetura_pelagica | VU | Decreasing | tropical | diurnal | 23.6 | 11481187.632 | 10.333 |
| birds | Chaetura_vauxi | Chaetura_vauxi | LC | Decreasing | temperate | diurnal | 17.1 | 3704613.512 | 8.958 |
| birds | Chalcites_basalis | Chrysococcyx_basalis | LC | Stable | temperate | diurnal | 23.1 | 8599616.124 | 4.804 |
| birds | Chalcites_lucidus | Chrysococcyx_lucidus | LC | Stable | temperate | diurnal | 24.4 | 5267824.28 | 6.104 |
| birds | Chalcites_megarhynchus | Rhamphomantis_megarhynchus | LC | Stable | tropical | diurnal | 31.5 | 152360.637 | 3.699 |
| birds | Chalcites_meyerii | Chrysococcyx_meyeri | LC | Decreasing | tropical | diurnal | 19.4 | 513273.656 | 3.42 |
| birds | Chalcites_minutillus | Chrysococcyx_minutillus | LC | Stable | tropical | diurnal | 18.6 | 2901381.592 | 5.684 |
| birds | Chalcites_osculans | Chrysococcyx_osculans | LC | Stable | temperate | diurnal | 30.9 | 7147734.341 | 4.22 |
| birds | Chalcites_ruficollis | Chrysococcyx_ruficollis | LC | Stable | tropical | diurnal | 21 | 156338.366 | 4.766 |
| birds | Chalcomitra_adelberti | Nectarinia_adelberti | LC | Stable | tropical | diurnal | 9.8 | 915043.566 | 15.183 |
| birds | Chalcomitra_amethystina | Nectarinia_amethystina | LC | Stable | tropical | diurnal | 13.7 | 3590561.914 | 9.707 |
| birds | Chalcomitra_balfouri | Nectarinia_balfouri | LC | Decreasing | tropical | diurnal | 11 | 3622.403 | 11.03 |
| birds | Chalcomitra_fuliginosa | Nectarinia_fuliginosa | LC | Stable | tropical | diurnal | 11.9 | 263939.426 | 10.639 |

| | | | | | | | | | |
|---|---|---|---|---|---|---|---|---|---|
| birds | Chalcomitra_hunteri | Nectarinia_hunteri | LC | Stable | tropical | diurnal | 11.8 | 1422643.427 | 8.366 |
| birds | Chalcomitra_rubescens | Nectarinia_rubescens | LC | Stable | tropical | diurnal | 10.3 | 2910535.163 | 6.912 |
| birds | Chalcomitra_senegalensis | Nectarinia_senegalensis | LC | Stable | tropical | diurnal | 12.2 | 9424883.183 | 9.903 |
| birds | Chalcoparia_singalensis | Anthreptes_singalensis | LC | Stable | tropical | diurnal | 8.2 | 2829894.282 | 11.149 |
| birds | Chalcophaps_indica | Chalcophaps_indica | LC | Decreasing | tropical | diurnal | 136.5 | 4704607.553 | 11.204 |
| birds | Chalcophaps_stephani | Chalcophaps_stephani | LC | Stable | tropical | diurnal | 129 | 866814.343 | 4.453 |
| birds | Chalcopsitta_atra | Chalcopsitta_atra | LC | Stable | tropical | diurnal | 195 | 77414.23 | 3.454 |
| birds | Chalcopsitta_cardinalis | Chalcopsitta_cardinalis | LC | Stable | tropical | diurnal | 183 | 35346.625 | 4.693 |
| birds | Chalcopsitta_duivenbodei | Chalcopsitta_duivenbodei | LC | Decreasing | tropical | diurnal | 218 | 63546.371 | 4.162 |
| birds | Chalcopsitta_scintillata | Chalcopsitta_sintillata | LC | Stable | tropical | diurnal | 190 | 311563.344 | 2.69 |
| birds | Chalcostigma_herrani | Chalcostigma_herrani | LC | Decreasing | tropical | diurnal | 5.8 | 75956.84 | 11.135 |
| birds | Chalcostigma_heteropogon | Chalcostigma_heteropogon | LC | Decreasing | tropical | diurnal | 6.1 | 29042.28 | 13.598 |
| birds | Chalcostigma_olivaceum | Chalcostigma_olivaceum | LC | Stable | tropical | diurnal | 8.2 | 65218.726 | 9.686 |
| birds | Chalcostigma_ruficeps | Chalcostigma_ruficeps | LC | Stable | tropical | diurnal | 3.4 | 123503.393 | 7.335 |
| birds | Chalcostigma_stanleyi | Chalcostigma_stanleyi | LC | Decreasing | tropical | diurnal | 5.8 | 120440.27 | 7.743 |
| birds | Chalybura_buffonii | Chalybura_buffonii | LC | Decreasing | tropical | diurnal | 6.8 | 731730.835 | 12.273 |
| birds | Chalybura_urochrysia | Chalybura_urochrysia | LC | Decreasing | tropical | diurnal | 6.4 | 299760.166 | 8.264 |
| birds | Chamaea_fasciata | Chamaea_fasciata | LC | Decreasing | temperate | diurnal | 14.7 | 246063.253 | 14.639 |
| birds | Chamaepetes_unicolor | Chamaepetes_unicolor | LC | Decreasing | tropical | diurnal | 1135 | 15572.603 | 9.607 |
| birds | Chamaetylas_choloensis | Alethe_choloensis | VU | Decreasing | tropical | diurnal | 41.3 | 4661.033 | 15.337 |
| birds | Chamaetylas_fuelleborni | Alethe_fuelleborni | LC | Decreasing | tropical | diurnal | 52 | 189262.111 | 10.855 |
| birds | Chamaetylas_poliophrys | Alethe_poliophrys | LC | Decreasing | tropical | diurnal | 35.2 | 76094.568 | 16.677 |
| birds | Chamaeza_campanisona | Chamaeza_campanisona | LC | Decreasing | tropical | diurnal | 90.6 | 1801868.242 | 10.26 |
| birds | Chamaeza_meruloides | Chamaeza_meruloides | LC | Stable | tropical | diurnal | 69 | 103365.424 | 13.252 |
| birds | Chamaeza_mollissima | Chamaeza_mollissima | LC | Stable | tropical | diurnal | 72 | 103368.735 | 5.46 |
| birds | Chamaeza_nobilis | Chamaeza_nobilis | LC | Stable | tropical | diurnal | 123 | 2594439.355 | 1.517 |
| birds | Chamaeza_ruficauda | Chamaeza_ruficauda | LC | Decreasing | temperate | diurnal | 71.2 | 371454.368 | 12.489 |
| birds | Chamaeza_turdina | Chamaeza_turdina | LC | Stable | tropical | diurnal | 68.3 | 14682.687 | 11.343 |
| birds | Charadrius_alexandrinus | Charadrius_alexandrinus | LC | Decreasing | temperate | diurnal | 42.3 | 22446543.509 | 11.376 |
| birds | Charadrius_alticola | Charadrius_alticola | LC | Unknown | tropical | diurnal | 44.9 | 605273.296 | 6.785 |
| birds | Charadrius_asiaticus | Charadrius_asiaticus | LC | Decreasing | temperate | diurnal | 77.1 | 18276948.614 | 8.086 |
| birds | Charadrius_bicinctus | Charadrius_bicinctus | NT | Decreasing | temperate | cathemeral | 59.4 | 1655193.864 | 8.666 |
| birds | Charadrius_collaris | Charadrius_collaris | LC | Decreasing | tropical | diurnal | 28.3 | 15100843.491 | 6.167 |
| birds | Charadrius_dubius | Charadrius_dubius | LC | Stable | tropical | diurnal | 38.7 | 46401384.752 | 10.672 |

| | | | | | | | | | |
|---|---|---|---|---|---|---|---|---|---|
| birds | Charadrius_falklandicus | Charadrius_falklandicus | LC | Stable | temperate | diurnal | 65 | 1010983.202 | 7.205 |
| birds | Charadrius_forbesi | NA | LC | Unknown | NA | NA | NA | NA | NA |
| birds | Charadrius_hiaticula | Charadrius_hiaticula | LC | Decreasing | temperate | cathemeral | 64 | 28297283.645 | 8.009 |
| birds | Charadrius_javanicus | NA | LC | Stable | NA | NA | NA | NA | NA |
| birds | Charadrius_leschenaultii | Charadrius_leschenaultii | LC | Decreasing | temperate | diurnal | 74.8 | 8942028.548 | 7.551 |
| birds | Charadrius_marginatus | Charadrius_marginatus | LC | Decreasing | tropical | cathemeral | 48.3 | 4442943.797 | 10.272 |
| birds | Charadrius_melodus | Charadrius_melodus | NT | Increasing | temperate | cathemeral | 54.4 | 5351896.504 | 16.012 |
| birds | Charadrius_modestus | Charadrius_modestus | LC | Unknown | temperate | diurnal | 78 | 1462237.279 | 8.793 |
| birds | Charadrius_mongolus | Charadrius_mongolus | EN | Decreasing | tropical | cathemeral | 64 | 61116255.058 | 6.526 |
| birds | Charadrius_montanus | Charadrius_montanus | NT | Decreasing | temperate | diurnal | 95.7 | 2169361.819 | 9.564 |
| birds | Charadrius_obscurus | Charadrius_obscurus | CR | Decreasing | temperate | cathemeral | 146 | 311.437 | 15.221 |
| birds | Charadrius_pecuarius | Charadrius_pecuarius | LC | Unknown | tropical | cathemeral | 34 | 16351827.572 | 9.443 |
| birds | Charadrius_peronii | Charadrius_peronii | NT | Decreasing | tropical | diurnal | 42 | 1266559.929 | 11.578 |
| birds | Charadrius_placidus | Charadrius_placidus | LC | Decreasing | temperate | diurnal | 62.7 | 5396919.96 | 14.428 |
| birds | Charadrius_ruficapillus | Charadrius_ruficapillus | LC | Unknown | temperate | diurnal | 37.4 | 4811331.656 | 4.871 |
| birds | Charadrius_sanctaehelenae | NA | VU | Increasing | NA | NA | NA | NA | NA |
| birds | Charadrius_semipalmatus | Charadrius_semipalmatus | LC | Stable | temperate | cathemeral | 46.7 | 17838316.672 | 7.477 |
| birds | Charadrius_thoracicus | Charadrius_thoracicus | VU | Decreasing | tropical | diurnal | 11.5 | 14469.492 | 10.799 |
| birds | Charadrius_tricollaris | Charadrius_tricollaris | LC | Unknown | tropical | cathemeral | 31.2 | 9455723.891 | 8.886 |
| birds | Charadrius_veredus | Charadrius_veredus | LC | Unknown | temperate | cathemeral | 95 | 4627324.646 | 7.206 |
| birds | Charadrius_vociferus | Charadrius_vociferus | LC | Decreasing | temperate | cathemeral | 96.4 | 16664790.808 | 10.093 |
| birds | Charadrius_wilsonia | Charadrius_wilsonia | LC | Decreasing | tropical | cathemeral | 55.1 | 966205.604 | 11.816 |
| birds | Charitospiza_eucosma | Charitospiza_eucosma | NT | Decreasing | tropical | diurnal | 10.5 | 1639318.027 | 8.19 |
| birds | Charmosyna_amabilis | Charmosyna_amabilis | CR | Decreasing | tropical | diurnal | 8.5 | 16517.621 | 9.462 |
| birds | Charmosyna_diadema | Charmosyna_diadema | CR | Unknown | tropical | diurnal | 37 | 65.248 | 1.567 |
| birds | Charmosyna_josefinae | Charmosyna_josefinae | LC | Stable | tropical | diurnal | 74 | 159632.979 | 3.476 |
| birds | Charmosyna_margarethae | Charmosyna_margarethae | LC | Decreasing | tropical | diurnal | 49.4 | 27077.996 | 5.369 |
| birds | Charmosyna_meeki | Charmosyna_meeki | LC | Stable | tropical | diurnal | 23.8 | 6507.88 | 4.098 |
| birds | Charmosyna_multistriata | Charmosyna_multistriata | LC | Decreasing | tropical | diurnal | 37 | 82448.178 | 2.68 |
| birds | Charmosyna_palmarum | Charmosyna_palmarum | VU | Decreasing | tropical | diurnal | 37 | 12900.833 | 5.707 |
| birds | Charmosyna_papou | Charmosyna_papou | LC | Stable | tropical | diurnal | 82.9 | 9550.655 | 3.217 |
| birds | Charmosyna_placentis | Charmosyna_placentis | LC | Stable | tropical | diurnal | 33.6 | 821124.433 | 3.524 |
| birds | Charmosyna_pulchella | Charmosyna_pulchella | LC | Stable | tropical | diurnal | 34.1 | 381133.695 | 3.582 |
| birds | Charmosyna_rubrigularis | Charmosyna_rubrigularis | LC | Stable | tropical | diurnal | 37 | 43980.908 | 4.448 |

| | | | | | | | | | |
|---|---|---|---|---|---|---|---|---|---|
| birds | Charmosyna_rubronotata | Charmosyna_rubronotata | LC | Stable | tropical | diurnal | 32.5 | 241413.47 | 3.236 |
| birds | Charmosyna_toxopei | Charmosyna_toxopei | CR | Decreasing | tropical | diurnal | 37 | 8554.363 | 6.663 |
| birds | Charmosyna_wilhelminae | Charmosyna_wilhelminae | LC | Stable | tropical | diurnal | 18 | 289944.544 | 4.139 |
| birds | Chasiempis_sandwichensis | Chasiempis_sandwichensis | NT | Decreasing | tropical | diurnal | 13.7 | 10507.221 | 6.573 |
| birds | Chauna_chavaria | Chauna_chavaria | LC | Stable | tropical | diurnal | 2557.6 | 114400.073 | 16.488 |
| birds | Chauna_torquata | Chauna_torquata | LC | Stable | temperate | diurnal | 4400 | 3828524.394 | 8.474 |
| birds | Chelictinia_riocourii | Chelictinia_riocourii | VU | Decreasing | tropical | diurnal | 110 | 6292258.571 | 8.526 |
| birds | Chelidoptera_tenebrosa | Chelidoptera_tenebrosa | LC | Increasing | tropical | diurnal | 35.9 | 8690116.856 | 3.726 |
| birds | Chelidorhynx_hypoxanthus | Chelidorhynx_hypoxantha | LC | Stable | temperate | diurnal | 6 | 1909981.693 | 10.076 |
| birds | Chenonetta_jubata | Chenonetta_jubata | LC | Stable | temperate | diurnal | 807.5 | 5996685.102 | 5.004 |
| birds | Chenorhamphus_grayi | Malurus_grayi | LC | Decreasing | tropical | diurnal | 12.5 | 188415.623 | 2.451 |
| birds | Cheramoeca_leucosterna | Cheramoeca_leucosterna | LC | Increasing | temperate | diurnal | 14.1 | 5215727.853 | 4.532 |
| birds | Chersophilus_duponti | Chersophilus_duponti | VU | Decreasing | temperate | diurnal | 39.4 | 288089.018 | 9.049 |
| birds | Chionis_albus | Chionis_albus | LC | Stable | temperate | diurnal | 675.7 | 2171631.618 | 6.08 |
| birds | Chionis_minor | NA | LC | Decreasing | NA | NA | NA | NA | NA |
| birds | Chionodacryon_speculiferum | NA | LC | Stable | NA | NA | NA | NA | NA |
| birds | Chionomesa_fimbriata | Amazilia_fimbriata | LC | Unknown | tropical | diurnal | 4.9 | 10740118.893 | 4.826 |
| birds | Chiroxiphia_boliviana | Chiroxiphia_boliviana | LC | Decreasing | tropical | diurnal | 17.1 | 78997.543 | 4.369 |
| birds | Chiroxiphia_caudata | Chiroxiphia_caudata | LC | Stable | temperate | diurnal | 25.6 | 1459549.859 | 13.251 |
| birds | Chiroxiphia_lanceolata | Chiroxiphia_lanceolata | LC | Decreasing | tropical | diurnal | 18.5 | 420162.322 | 12.652 |
| birds | Chiroxiphia_linearis | Chiroxiphia_linearis | LC | Stable | tropical | diurnal | 17.4 | 101728.762 | 15.323 |
| birds | Chiroxiphia_pareola | Chiroxiphia_pareola | LC | Stable | tropical | diurnal | 16.8 | 5040298.893 | 2.715 |
| birds | Chlamydera_cerviniventris | Chlamydera_cerviniventris | LC | Stable | tropical | diurnal | 156.4 | 152782.31 | 4.039 |
| birds | Chlamydera_guttata | Chlamydera_guttata | LC | Decreasing | temperate | diurnal | 136 | 1491137.417 | 2.846 |
| birds | Chlamydera_lauterbachi | Chlamydera_lauterbachi | LC | Decreasing | tropical | diurnal | 123.2 | 124403.095 | 3.905 |
| birds | Chlamydera_maculata | Chlamydera_maculata | LC | Decreasing | temperate | diurnal | 140 | 1784471.922 | 5.911 |
| birds | Chlamydera_nuchalis | Chlamydera_nuchalis | LC | Stable | tropical | diurnal | 200.4 | 1285385.855 | 2.954 |
| birds | Chlamydochaera_jefferyi | Chlamydochaera_jefferyi | LC | Decreasing | tropical | diurnal | 65.1 | 63345.306 | 2.247 |
| birds | Chlamydotis_undulata | Chlamydotis_undulata | VU | Decreasing | temperate | diurnal | 1501.3 | 4448380.487 | 2.539 |
| birds | Chlidonias_albostriatus | Sterna_albostriata | EN | Decreasing | temperate | diurnal | 94.7 | 149045.866 | 9.026 |
| birds | Chlidonias_hybrida | Chlidonias_hybrida | LC | Stable | tropical | diurnal | 83.7 | 31615735.931 | 10.308 |
| birds | Chlidonias_leucopterus | Chlidonias_leucopterus | LC | Stable | temperate | diurnal | 54.2 | 40459557.15 | 9.033 |
| birds | Chlidonias_niger | Chlidonias_niger | LC | Decreasing | tropical | diurnal | 65.3 | 37715753.791 | 12.603 |
| birds | Chloebia_gouldiae | Erythrura_gouldiae | LC | Stable | tropical | diurnal | 13.8 | 1163550.377 | 2.652 |

| | | | | | | | | | |
|---|---|---|---|---|---|---|---|---|---|
| birds | Chloephaga_hybrida | Chloephaga_hybrida | LC | Stable | temperate | diurnal | 2281.8 | 219432.233 | 3.567 |
| birds | Chloephaga_melanoptera | Chloephaga_melanoptera | LC | Stable | tropical | diurnal | 2037 | 1159462.303 | 6.3 |
| birds | Chloephaga_picta | Chloephaga_picta | LC | Decreasing | temperate | diurnal | 2880.4 | 1582798.872 | 6.123 |
| birds | Chloephaga_poliocephala | Chloephaga_poliocephala | LC | Decreasing | temperate | diurnal | 2007.1 | 1643991.769 | 6.364 |
| birds | Chloephaga_rubidiceps | Chloephaga_rubidiceps | LC | Decreasing | temperate | diurnal | 2086 | 248383.806 | 6.758 |
| birds | Chlorestes_candida | Amazilia_candida | LC | Decreasing | tropical | diurnal | 3.8 | 564871.44 | 9.764 |
| birds | Chlorestes_cyanus | Hylocharis_cyanus | LC | Unknown | tropical | diurnal | 3.3 | 7031990.781 | 3.172 |
| birds | Chlorestes_eliciae | Hylocharis_eliciae | LC | Decreasing | tropical | diurnal | 3.6 | 306966.665 | 10.972 |
| birds | Chlorestes_julie | Damophila_julie | LC | Decreasing | tropical | diurnal | 3.3 | 289656.608 | 13.09 |
| birds | Chlorestes_notata | Chlorestes_notata | LC | Unknown | tropical | diurnal | 4.2 | 4700543.44 | 2.896 |
| birds | Chloris_ambigua | NA | LC | Stable | NA | NA | NA | NA | NA |
| birds | Chloris_chloris | Carduelis_chloris | LC | Stable | temperate | diurnal | 26 | 11398923.388 | 15.04 |
| birds | Chloris_monguilloti | Carduelis_monguilloti | LC | Unknown | tropical | diurnal | 15.5 | 7399.285 | 8.063 |
| birds | Chloris_sinica | Carduelis_sinica | LC | Stable | temperate | diurnal | 18.6 | 6355782.172 | 14.173 |
| birds | Chloris_spinoides | Carduelis_spinoides | LC | Stable | temperate | diurnal | 18.6 | 557286.265 | 9.18 |
| birds | Chloroceryle_aenea | Chloroceryle_aenea | LC | Decreasing | tropical | diurnal | 13.8 | 9332280.13 | 4.476 |
| birds | Chloroceryle_americana | Chloroceryle_americana | LC | Decreasing | tropical | diurnal | 33.7 | 16946715.06 | 6.595 |
| birds | Chloroceryle_inda | Chloroceryle_inda | LC | Decreasing | tropical | diurnal | 52.1 | 8573926.252 | 3.362 |
| birds | Chlorochrysa_calliparaea | Chlorochrysa_calliparaea | LC | Decreasing | tropical | diurnal | 17 | 147445.405 | 4.868 |
| birds | Chlorochrysa_nitidissima | Chlorochrysa_nitidissima | NT | Decreasing | tropical | diurnal | 18.6 | 17380.716 | 14.289 |
| birds | Chlorochrysa_phoenicotis | Chlorochrysa_phoenicotis | LC | Decreasing | tropical | diurnal | 21.8 | 33619.563 | 8.905 |
| birds | Chlorocichla_falkensteini | Chlorocichla_falkensteini | LC | Stable | tropical | diurnal | 31.7 | 978923.364 | 7.135 |
| birds | Chlorocichla_flaviventris | Chlorocichla_flaviventris | LC | Stable | tropical | diurnal | 39.4 | 3986239.887 | 8.737 |
| birds | Chlorocichla_laetissima | Chlorocichla_laetissima | LC | Stable | tropical | diurnal | 50.2 | 137013.098 | 13.022 |
| birds | Chlorocichla_prigoginei | Chlorocichla_prigoginei | EN | Decreasing | tropical | diurnal | 45.3 | 868.589 | 19.642 |
| birds | Chlorocichla_simplex | Chlorocichla_simplex | LC | Stable | tropical | diurnal | 46.4 | 4003835.484 | 8.258 |
| birds | Chlorodrepanis_flava | Hemignathus_flavus | NT | Unknown | tropical | diurnal | 12.6 | 1563.724 | 21.148 |
| birds | Chlorodrepanis_stejnegeri | Hemignathus_kauaiensis | EN | Decreasing | tropical | diurnal | 16.6 | 1368.25 | 9.376 |
| birds | Chlorodrepanis_virens | Hemignathus_virens | LC | Stable | tropical | diurnal | 13.7 | 6317.402 | 6.274 |
| birds | Chlorophanes_spiza | Chlorophanes_spiza | LC | Decreasing | tropical | diurnal | 19 | 7921066.066 | 3.603 |
| birds | Chlorophoneus_bocagei | Telophorus_bocagei | LC | Decreasing | tropical | diurnal | 27 | 785394.391 | 8.538 |
| birds | Chlorophoneus_kupeensis | NA | EN | Decreasing | NA | NA | NA | NA | NA |
| birds | Chlorophoneus_multicolor | Telophorus_multicolor | LC | Stable | tropical | diurnal | 50.9 | 1682374.806 | 10.403 |
| birds | Chlorophoneus_nigrifrons | Telophorus_nigrifrons | LC | Stable | tropical | diurnal | 34.8 | 540075.234 | 7.727 |

| | | | | | | | | | |
|---|---|---|---|---|---|---|---|---|---|
| birds | Chlorophoneus_olivaceus | Telophorus_olivaceus | LC | Decreasing | temperate | diurnal | 33.6 | 332431.69 | 12.572 |
| birds | Chlorophoneus_sulfureopectus | Telophorus_sulfureopectus | LC | Stable | tropical | diurnal | 27.1 | 8777553.103 | 9.953 |
| birds | Chlorophonia_callophrys | Chlorophonia_callophrys | LC | Decreasing | tropical | diurnal | 25.8 | 17430.09 | 7.832 |
| birds | Chlorophonia_cyanea | Chlorophonia_cyanea | LC | Stable | tropical | diurnal | 14 | 2010700.009 | 8.996 |
| birds | Chlorophonia_flavirostris | Chlorophonia_flavirostris | LC | Decreasing | tropical | diurnal | 11 | 21348.224 | 5.428 |
| birds | Chlorophonia_occipitalis | Chlorophonia_occipitalis | LC | Decreasing | tropical | diurnal | 25.8 | 160768.755 | 11.044 |
| birds | Chlorophonia_pyrrhophrys | Chlorophonia_pyrrhophrys | LC | Decreasing | tropical | diurnal | 17 | 124114.793 | 8.273 |
| birds | Chloropicoides_rafflesii | Dinopium_rafflesii | NT | Decreasing | tropical | diurnal | 102 | 1030245.342 | 8.387 |
| birds | Chloropipo_flavicapilla | Xenopipo_flavicapilla | VU | Decreasing | tropical | diurnal | 17.8 | 18720.277 | 13.855 |
| birds | Chloropipo_unicolor | Xenopipo_unicolor | LC | Decreasing | tropical | diurnal | 15.5 | 57212.374 | 4.982 |
| birds | Chloropsis_aurifrons | Chloropsis_aurifrons | LC | Stable | tropical | diurnal | 32.3 | 3471632.851 | 14.145 |
| birds | Chloropsis_cochinchinensis | NA | EN | Decreasing | NA | NA | NA | NA | NA |
| birds | Chloropsis_cyanopogon | Chloropsis_cyanopogon | NT | Decreasing | tropical | diurnal | 22 | 1261145.13 | 8.565 |
| birds | Chloropsis_flavipennis | Chloropsis_flavipennis | VU | Decreasing | tropical | diurnal | 35.4 | 118181.127 | 12.273 |
| birds | Chloropsis_hardwickii | Chloropsis_hardwickii | LC | Decreasing | tropical | diurnal | 32.3 | 902456.729 | 8.843 |
| birds | Chloropsis_palawanensis | Chloropsis_palawanensis | LC | Stable | tropical | diurnal | 30.3 | 13485.036 | 9.366 |
| birds | Chloropsis_sonnerati | Chloropsis_sonnerati | EN | Decreasing | tropical | diurnal | 45.8 | 1386747.464 | 8.751 |
| birds | Chloropsis_venusta | NA | NT | Decreasing | NA | NA | NA | NA | NA |
| birds | Chlorornis_riefferii | Chlorornis_riefferii | LC | Decreasing | tropical | diurnal | 53 | 307394.783 | 8.279 |
| birds | Chlorospingus_canigularis | Chlorospingus_canigularis | LC | Decreasing | tropical | diurnal | 18 | 165238.269 | 7.439 |
| birds | Chlorospingus_flavigularis | Chlorospingus_flavigularis | LC | Stable | tropical | diurnal | 25.6 | 200407.232 | 6.191 |
| birds | Chlorospingus_flavopectus | Chlorospingus_ophthalmicus | LC | Decreasing | tropical | diurnal | 20 | 587009.938 | 8.774 |
| birds | Chlorospingus_inornatus | Chlorospingus_inornatus | LC | Stable | tropical | diurnal | 28 | 1510.6 | 1.742 |
| birds | Chlorospingus_parvirostris | Chlorospingus_parvirostris | LC | Decreasing | tropical | diurnal | 24 | 134279.076 | 6.098 |
| birds | Chlorospingus_pileatus | Chlorospingus_pileatus | LC | Stable | tropical | diurnal | 21 | 3941.012 | 6.275 |
| birds | Chlorospingus_semifuscus | Chlorospingus_semifuscus | LC | Decreasing | tropical | diurnal | 19 | 50126.886 | 10.202 |
| birds | Chlorospingus_tacarcunae | Chlorospingus_tacarcunae | LC | Decreasing | tropical | diurnal | 18.9 | 1499.099 | 7.794 |
| birds | Chlorostilbon_lucidus | Chlorostilbon_lucidus | LC | Unknown | tropical | diurnal | 3.5 | 5788276.165 | 9.452 |
| birds | Chlorostilbon_mellisugus | Chlorostilbon_mellisugus | LC | Stable | tropical | diurnal | 2.6 | 4111804.659 | 3.806 |
| birds | Chlorostilbon_olivaresi | Chlorostilbon_olivaresi | LC | Unknown | tropical | diurnal | 3.7 | 6529.262 | 0.796 |
| birds | Chlorostilbon_poortmani | Chlorostilbon_poortmani | LC | Stable | tropical | diurnal | 3.3 | 60449.068 | 12.216 |
| birds | Chlorostilbon_russatus | Chlorostilbon_russatus | LC | Unknown | tropical | diurnal | 3.4 | 27765.982 | 11.18 |
| birds | Chlorostilbon_stenurus | Chlorostilbon_stenurus | LC | Unknown | tropical | diurnal | 3.4 | 47007.363 | 10.959 |
| birds | Chondestes_grammacus | Chondestes_grammacus | LC | Decreasing | temperate | diurnal | 29 | 7065667.034 | 11.922 |

| | | | | | | | | | |
|---|---|---|---|---|---|---|---|---|---|
| birds | Chondrohierax_uncinatus | Chondrohierax_uncinatus | LC | Decreasing | tropical | diurnal | 286.1 | 9787616.081 | 5.203 |
| birds | Chordeiles_acutipennis | Chordeiles_acutipennis | LC | Increasing | tropical | cathemeral | 48.4 | 10717818.683 | 7.448 |
| birds | Chordeiles_gundlachii | Chordeiles_gundlachii | LC | Stable | tropical | cathemeral | 50 | 228316.661 | 16.281 |
| birds | Chordeiles_minor | Chordeiles_minor | LC | Decreasing | tropical | nocturnal | 79.3 | 28501967.683 | 8.199 |
| birds | Chordeiles_nacunda | Podager_nacunda | LC | Stable | tropical | nocturnal | 159 | 12270759.096 | 5.793 |
| birds | Chordeiles_pusillus | Chordeiles_pusillus | LC | Stable | tropical | nocturnal | 30.6 | 4968284.381 | 5.514 |
| birds | Chordeiles_rupestris | Chordeiles_rupestris | LC | Decreasing | tropical | nocturnal | 52.3 | 1639560.087 | 1.713 |
| birds | Chrysococcyx_caprius | Chrysococcyx_caprius | LC | Stable | tropical | diurnal | 32.8 | 18736977.456 | 8.982 |
| birds | Chrysococcyx_cupreus | Chrysococcyx_cupreus | LC | Stable | tropical | diurnal | 37.7 | 11402508.038 | 9.602 |
| birds | Chrysococcyx_flavigularis | Chrysococcyx_flavigularis | LC | Decreasing | tropical | diurnal | 29.2 | 2131118.512 | 6.821 |
| birds | Chrysococcyx_klaas | Chrysococcyx_klaas | LC | Stable | tropical | diurnal | 27.4 | 14138608.994 | 9.497 |
| birds | Chrysococcyx_maculatus | Chrysococcyx_maculatus | LC | Decreasing | tropical | diurnal | 27.4 | 2481014.751 | 11.742 |
| birds | Chrysococcyx_xanthorhynchus | Chrysococcyx_xanthorhynchus | LC | Decreasing | tropical | diurnal | 22.4 | 2959317.44 | 10.407 |
| birds | Chrysocolaptes_festivus | Chrysocolaptes_festivus | LC | Stable | tropical | diurnal | 213 | 1703319.401 | 18.513 |
| birds | Chrysocolaptes_lucidus | Chrysocolaptes_lucidus | LC | Decreasing | tropical | diurnal | 142 | 119803.153 | 12.109 |
| birds | Chrysocolaptes_validus | Reinwardtipicus_validus | LC | Stable | tropical | diurnal | 149 | 1291734.803 | 8.754 |
| birds | Chrysocorypha_delatrii | Tachyphonus_delatrii | LC | Decreasing | tropical | diurnal | 18 | 196748.24 | 7.315 |
| birds | Chrysocorythus_estherae | NA | LC | Stable | NA | NA | NA | NA | NA |
| birds | Chrysoena_luteovirens | Ptilinopus_luteovirens | LC | Stable | tropical | diurnal | 46.6 | 9129.14 | 7.953 |
| birds | Chrysoena_victor | Ptilinopus_victor | LC | Stable | tropical | diurnal | 46.6 | 6108.649 | 9.839 |
| birds | Chrysoena_viridis | Ptilinopus_layardi | NT | Decreasing | tropical | diurnal | 46.6 | 464.937 | 6.031 |
| birds | Chrysolampis_mosquitus | Chrysolampis_mosquitus | LC | Unknown | tropical | diurnal | 3.9 | 7397775.668 | 6.338 |
| birds | Chrysolophus_amherstiae | Chrysolophus_amherstiae | LC | Decreasing | temperate | diurnal | 732.5 | 387085.425 | 10.361 |
| birds | Chrysolophus_pictus | Chrysolophus_pictus | LC | Decreasing | temperate | diurnal | 555.2 | 975613.445 | 13.014 |
| birds | Chrysominla_strigula | Minla_strigula | LC | Decreasing | temperate | diurnal | 19.2 | 814874.368 | 9.029 |
| birds | Chrysomma_altirostre | Chrysomma_altirostre | VU | Decreasing | tropical | diurnal | 20 | 47346.575 | 15.907 |
| birds | Chrysomma_sinense | Chrysomma_sinense | LC | Stable | tropical | diurnal | 17.2 | 4332183.624 | 15.763 |
| birds | Chrysomus_icterocephalus | Chrysomus_icterocephalus | LC | Increasing | tropical | diurnal | 30.7 | 1581439.98 | 6.842 |
| birds | Chrysophlegma_flavinucha | Picus_flavinucha | LC | Decreasing | tropical | diurnal | 176.8 | 2932266.055 | 10.934 |
| birds | Chrysophlegma_mentale | Picus_mentalis | NT | Decreasing | tropical | diurnal | 99.7 | 40422.478 | 19.397 |
| birds | Chrysophlegma_miniaceum | Picus_mineaceus | LC | Stable | tropical | diurnal | 97.3 | 1291605.288 | 8.041 |
| birds | Chrysothlypis_chrysomelas | Chrysothlypis_chrysomelas | LC | Decreasing | tropical | diurnal | 13 | 15703.462 | 5.941 |
| birds | Chrysothlypis_salmoni | Chrysothlypis_salmoni | LC | Stable | tropical | diurnal | 12.5 | 75321.247 | 5.162 |
| birds | Chrysuronia_brevirostris | Amazilia_brevirostris | LC | Stable | tropical | diurnal | 4.3 | 584285.398 | 3.658 |

| | | | | | | | | | |
|---|---|---|---|---|---|---|---|---|---|
| birds | Chrysuronia_coeruleogularis | Lepidopyga_coeruleogularis | LC | Stable | tropical | diurnal | 4.2 | 91772.235 | 13.114 |
| birds | Chrysuronia_goudoti | Lepidopyga_goudoti | LC | Increasing | tropical | diurnal | 4 | 333328.714 | 13.747 |
| birds | Chrysuronia_grayi | Hylocharis_grayi | LC | Stable | tropical | diurnal | 5.6 | 51024.741 | 14.682 |
| birds | Chrysuronia_leucogaster | Amazilia_leucogaster | LC | Stable | tropical | diurnal | 4.7 | 800744.997 | 6.921 |
| birds | Chrysuronia_lilliae | Lepidopyga_lilliae | EN | Decreasing | tropical | diurnal | 4.3 | 2292.904 | 11.208 |
| birds | Chrysuronia_oenone | Chrysuronia_oenone | LC | Stable | tropical | diurnal | 5 | 1336265.934 | 3.934 |
| birds | Chrysuronia_versicolor | Amazilia_versicolor | LC | Stable | tropical | diurnal | 4.1 | 6914568.209 | 6.56 |
| birds | Chunga_burmeisteri | Chunga_burmeisteri | LC | Stable | temperate | diurnal | 1298 | 736438.813 | 6.227 |
| birds | Ciccaba_albitarsis | Strix_albitarsis | LC | Decreasing | tropical | nocturnal | 307.5 | 325371.541 | 9.165 |
| birds | Ciccaba_huhula | Strix_huhula | LC | Decreasing | tropical | nocturnal | 370 | 7366712.066 | 3.219 |
| birds | Ciccaba_nigrolineata | Strix_nigrolineata | LC | Decreasing | tropical | nocturnal | 457.9 | 991396.189 | 10.433 |
| birds | Ciccaba_virgata | Strix_virgata | LC | Decreasing | tropical | nocturnal | 284 | 11001158.194 | 5.465 |
| birds | Cichladusa_arquata | Cichladusa_arquata | LC | Stable | tropical | diurnal | 34.2 | 2314814.622 | 9.215 |
| birds | Cichladusa_guttata | Cichladusa_guttata | LC | Stable | tropical | diurnal | 23.1 | 1373382.956 | 9.805 |
| birds | Cichladusa_ruficauda | Cichladusa_ruficauda | LC | Stable | tropical | diurnal | 28.5 | 699193.301 | 7.984 |
| birds | Cichlocolaptes_leucophrus | Cichlocolaptes_leucophrus | LC | Decreasing | tropical | diurnal | 50 | 148328.509 | 12.793 |
| birds | Cichlopsis_leucogenys | Cichlopsis_leucogenys | EN | Decreasing | tropical | diurnal | 53.8 | 102344.991 | 10.584 |
| birds | Cicinnurus_magnificus | Cicinnurus_magnificus | LC | Decreasing | tropical | diurnal | 89.8 | 116997.401 | 4.454 |
| birds | Cicinnurus_regius | Cicinnurus_regius | LC | Decreasing | tropical | diurnal | 51.6 | 479513.907 | 3.153 |
| birds | Cicinnurus_respublica | Cicinnurus_respublica | NT | Decreasing | tropical | diurnal | 58.4 | 3966.986 | 2.444 |
| birds | Ciconia_abdimii | Ciconia_abdimii | LC | Decreasing | tropical | diurnal | 1398 | 17648125.863 | 8.592 |
| birds | Ciconia_boyciana | Ciconia_boyciana | EN | Decreasing | temperate | diurnal | 4847.7 | 3630446.147 | 15.937 |
| birds | Ciconia_ciconia | Ciconia_ciconia | LC | Increasing | temperate | diurnal | 3445.8 | 30530843.125 | 10.172 |
| birds | Ciconia_episcopus | Ciconia_episcopus | NT | Decreasing | tropical | diurnal | 2061 | 3697795.592 | 16.812 |
| birds | Ciconia_maguari | Ciconia_maguari | LC | Stable | temperate | diurnal | 3995 | 5190878.296 | 7.397 |
| birds | Ciconia_nigra | Ciconia_nigra | LC | Unknown | temperate | diurnal | 2926 | 35630654.882 | 10.552 |
| birds | Ciconia_stormi | NA | EN | Decreasing | NA | NA | NA | NA | NA |
| birds | Cinclidium_frontale | Cinclidium_frontale | LC | Decreasing | tropical | diurnal | 25.4 | 228181.875 | 7.887 |
| birds | Cinclocerthia_gutturalis | Cinclocerthia_gutturalis | LC | Stable | tropical | diurnal | 69.2 | 1756.732 | 24.276 |
| birds | Cinclocerthia_ruficauda | Cinclocerthia_ruficauda | LC | Unknown | tropical | diurnal | 55 | 3295.063 | 20.22 |
| birds | Cinclodes_antarcticus | Cinclodes_antarcticus | NT | Decreasing | temperate | diurnal | 63.2 | 12926.554 | 3.247 |
| birds | Cinclodes_aricomae | Cinclodes_aricomae | CR | Decreasing | tropical | diurnal | 50 | 73490.42 | 9.242 |
| birds | Cinclodes_atacamensis | Cinclodes_atacamensis | LC | Decreasing | tropical | diurnal | 53 | 870994.254 | 6.285 |
| birds | Cinclodes_comechingonus | Cinclodes_comechingonus | LC | Decreasing | temperate | diurnal | 28.3 | 44865.651 | 7.901 |

| | | | | | | | | | |
|---|---|---|---|---|---|---|---|---|---|
| birds | Cinclodes_excelsior | Cinclodes_excelsior | LC | Stable | tropical | diurnal | 64 | 37973.194 | 11.265 |
| birds | Cinclodes_fuscus | Cinclodes_fuscus | LC | Stable | temperate | diurnal | 30 | 2507430.837 | 7.998 |
| birds | Cinclodes_nigrofumosus | Cinclodes_nigrofumosus | LC | Stable | temperate | diurnal | 65 | 74685.21 | 8.657 |
| birds | Cinclodes_oustaleti | Cinclodes_oustaleti | LC | Decreasing | temperate | diurnal | 30.5 | 734371.461 | 5.27 |
| birds | Cinclodes_pabsti | Cinclodes_pabsti | NT | Decreasing | temperate | diurnal | 53 | 21292.88 | 7.93 |
| birds | Cinclodes_palliatus | Cinclodes_palliatus | CR | Decreasing | tropical | diurnal | 103.9 | 21570.118 | 7.887 |
| birds | Cinclodes_patagonicus | Cinclodes_patagonicus | LC | Stable | temperate | diurnal | 30.7 | 661640.555 | 5.526 |
| birds | Cinclodes_taczanowskii | Cinclodes_taczanowskii | LC | Stable | tropical | diurnal | 64.5 | 24111.681 | 9.945 |
| birds | Cincloramphus_bivittatus | Buettikoferella_bivittata | LC | Increasing | tropical | diurnal | 30 | 25788.079 | 13.929 |
| birds | Cincloramphus_cruralis | Cincloramphus_cruralis | LC | Stable | temperate | diurnal | 49.2 | 6388749.628 | 4.579 |
| birds | Cincloramphus_grosvenori | Megalurulus_grosvenori | VU | Decreasing | tropical | diurnal | 54 | 2381.311 | 2.374 |
| birds | Cincloramphus_llaneae | Megalurulus_llaneae | NT | Decreasing | tropical | diurnal | 25.5 | 814.059 | 4.822 |
| birds | Cincloramphus_mariae | Megalurulus_mariei | LC | Decreasing | tropical | diurnal | 25 | 16389.766 | 7.513 |
| birds | Cincloramphus_mathewsi | Cincloramphus_mathewsi | LC | Stable | temperate | diurnal | 29.3 | 6668777.97 | 4.493 |
| birds | Cincloramphus_rubiginosus | Megalurulus_rubiginosus | LC | Decreasing | tropical | diurnal | 41.2 | 33761.384 | 4.146 |
| birds | Cincloramphus_rufus | Trichocichla_rufa | EN | Stable | tropical | diurnal | 26 | 14343.452 | 8.287 |
| birds | Cincloramphus_timoriensis | Megalurus_timoriensis | LC | Stable | tropical | diurnal | 21.6 | 2127266.893 | 5.806 |
| birds | Cinclosoma_ajax | Cinclosoma_ajax | LC | Stable | tropical | diurnal | 65.3 | 78596.6 | 3.298 |
| birds | Cinclosoma_castaneothorax | Cinclosoma_castaneothorax | LC | Decreasing | temperate | diurnal | 62.4 | 488478.66 | 5.073 |
| birds | Cinclosoma_castanotum | Cinclosoma_castanotum | LC | Decreasing | temperate | diurnal | 73.3 | 2823366.519 | 4.096 |
| birds | Cinclosoma_cinnamomeum | Cinclosoma_cinnamomeum | LC | Decreasing | temperate | diurnal | 56.1 | 764829.56 | 2.877 |
| birds | Cinclosoma_punctatum | Cinclosoma_punctatum | LC | Decreasing | temperate | diurnal | 113.3 | 567920.578 | 9.164 |
| birds | Cinclus_cinclus | Cinclus_cinclus | LC | Decreasing | temperate | diurnal | 61.5 | 10252132.107 | 10.21 |
| birds | Cinclus_leucocephalus | Cinclus_leucocephalus | LC | Stable | tropical | diurnal | 44 | 619987.755 | 8.549 |
| birds | Cinclus_mexicanus | Cinclus_mexicanus | LC | Decreasing | temperate | diurnal | 56.7 | 4972505.502 | 5.018 |
| birds | Cinclus_pallasii | Cinclus_pallasii | LC | Stable | temperate | diurnal | 76 | 9425465.166 | 10.85 |
| birds | Cinclus_schulzii | Cinclus_schulzi | VU | Decreasing | temperate | diurnal | 37.6 | 133757.165 | 6.398 |
| birds | Cinnycerthia_fulva | Cinnycerthia_fulva | LC | Decreasing | tropical | diurnal | 16.5 | 108129.229 | 5.139 |
| birds | Cinnycerthia_olivascens | Cinnycerthia_olivascens | LC | Decreasing | tropical | diurnal | 24.4 | 95207.431 | 10.176 |
| birds | Cinnycerthia_peruana | Cinnycerthia_peruana | LC | Decreasing | tropical | diurnal | 19.5 | 75819.14 | 5.38 |
| birds | Cinnycerthia_unirufa | Cinnycerthia_unirufa | LC | Decreasing | tropical | diurnal | 29.4 | 153814.026 | 11.214 |
| birds | Cinnyricinclus_leucogaster | Cinnyricinclus_leucogaster | LC | Decreasing | tropical | diurnal | 44 | 12754823.21 | 9.722 |
| birds | Cinnyris_afer | Nectarinia_afra | LC | Stable | temperate | diurnal | 11.1 | 365697.143 | 11.685 |
| birds | Cinnyris_asiaticus | Nectarinia_asiatica | LC | Stable | tropical | diurnal | 8.1 | 5816862.298 | 14.975 |

| | | | | | | | | | |
|---|---|---|---|---|---|---|---|---|---|
| birds | Cinnyris_batesi | Nectarinia_batesi | LC | Stable | tropical | diurnal | 6.3 | 854825.131 | 7.821 |
| birds | Cinnyris_bifasciatus | Nectarinia_bifasciata | LC | Stable | tropical | diurnal | 7.2 | 1927794.278 | 9.689 |
| birds | Cinnyris_bouvieri | Nectarinia_bouvieri | LC | Stable | tropical | diurnal | 8.7 | 360313.74 | 8.734 |
| birds | Cinnyris_buettikoferi | NA | LC | Stable | NA | NA | NA | NA | NA |
| birds | Cinnyris_chalcomelas | Nectarinia_chalcomelas | LC | Stable | tropical | diurnal | 8.2 | 63807.065 | 14.218 |
| birds | Cinnyris_chalybeus | Nectarinia_chalybea | LC | Stable | temperate | diurnal | 7.7 | 496199.25 | 9.565 |
| birds | Cinnyris_chloropygius | Nectarinia_chloropygia | LC | Stable | tropical | diurnal | 6.2 | 4297123.95 | 8.253 |
| birds | Cinnyris_coccinigastrus | Nectarinia_coccinigaster | LC | Stable | tropical | diurnal | 14.3 | 1740703.507 | 9.658 |
| birds | Cinnyris_comorensis | Nectarinia_comorensis | LC | Stable | tropical | diurnal | 7.8 | 357.547 | 13.818 |
| birds | Cinnyris_congensis | Nectarinia_congensis | LC | Decreasing | tropical | diurnal | 17.3 | 130504.854 | 6.25 |
| birds | Cinnyris_coquerellii | Nectarinia_coquerellii | LC | Stable | tropical | diurnal | 7.9 | 370.972 | 20.934 |
| birds | Cinnyris_cupreus | Nectarinia_cuprea | LC | Stable | tropical | diurnal | 8.9 | 7848077.34 | 8.712 |
| birds | Cinnyris_dussumieri | NA | LC | Stable | NA | NA | NA | NA | NA |
| birds | Cinnyris_erythrocercus | Nectarinia_erythrocerca | LC | Stable | tropical | diurnal | 8.8 | 438707.331 | 14.742 |
| birds | Cinnyris_fuscus | Nectarinia_fusca | LC | Stable | tropical | diurnal | 7.8 | 1120628.468 | 6.038 |
| birds | Cinnyris_habessinicus | Nectarinia_habessinica | LC | Stable | tropical | diurnal | 9.5 | 1107754.717 | 8.089 |
| birds | Cinnyris_humbloti | Nectarinia_humbloti | LC | Stable | tropical | diurnal | 6.4 | 1240.87 | 17.47 |
| birds | Cinnyris_johannae | Nectarinia_johannae | LC | Decreasing | tropical | diurnal | 13 | 1631281.91 | 7.16 |
| birds | Cinnyris_jugularis | Nectarinia_jugularis | LC | Stable | tropical | diurnal | 9 | 5063701.812 | 10.385 |
| birds | Cinnyris_lotenius | Nectarinia_lotenia | LC | Stable | tropical | diurnal | 8.8 | 292016.325 | 20.948 |
| birds | Cinnyris_loveridgei | Nectarinia_loveridgei | EN | Decreasing | tropical | diurnal | 9.8 | 273.218 | 10.01 |
| birds | Cinnyris_ludovicensis | Nectarinia_ludovicensis | LC | Stable | tropical | diurnal | 8.7 | 85632.036 | 10.646 |
| birds | Cinnyris_manoensis | Nectarinia_manoensis | LC | Stable | tropical | diurnal | 9.5 | 454948.662 | 13.049 |
| birds | Cinnyris_mariquensis | Nectarinia_mariquensis | LC | Stable | tropical | diurnal | 11.5 | 2606050.207 | 8.54 |
| birds | Cinnyris_mediocris | Nectarinia_mediocris | LC | Stable | tropical | diurnal | 8.2 | 195956.739 | 13.482 |
| birds | Cinnyris_minullus | Nectarinia_minulla | LC | Stable | tropical | diurnal | 5.7 | 1198202.189 | 8.516 |
| birds | Cinnyris_moreaui | Nectarinia_moreaui | NT | Decreasing | tropical | diurnal | 8.2 | 29656.257 | 10.983 |
| birds | Cinnyris_nectarinioides | Nectarinia_nectarinioides | LC | Stable | tropical | diurnal | 4.8 | 496613.105 | 8.532 |
| birds | Cinnyris_neergaardi | Nectarinia_neergardi | NT | Decreasing | temperate | diurnal | 6.1 | 138996.947 | 9.025 |
| birds | Cinnyris_notatus | Nectarinia_notata | LC | Stable | tropical | diurnal | 14.6 | 590045.876 | 8.742 |
| birds | Cinnyris_osea | Nectarinia_osea | LC | Stable | tropical | diurnal | 7.2 | 568902.395 | 8.795 |
| birds | Cinnyris_oustaleti | NA | LC | Stable | NA | NA | NA | NA | NA |
| birds | Cinnyris_pembae | NA | LC | Stable | NA | NA | NA | NA | NA |
| birds | Cinnyris_pulchellus | Nectarinia_pulchella | LC | Stable | tropical | diurnal | 7 | 5431378.759 | 10.702 |

| | | | | | | | | | |
|---|---|---|---|---|---|---|---|---|---|
| birds | Cinnyris_regius | Nectarinia_regia | LC | Decreasing | tropical | diurnal | 6.6 | 76601.79 | 17.058 |
| birds | Cinnyris_reichenowi | Nectarinia_preussi | LC | Stable | tropical | diurnal | 6.8 | 243088.546 | 12.31 |
| birds | Cinnyris_rockefelleri | Nectarinia_rockefelleri | VU | Decreasing | tropical | diurnal | 5.3 | 10464.769 | 14.043 |
| birds | Cinnyris_rufipennis | Nectarinia_rufipennis | VU | Decreasing | tropical | diurnal | 8.8 | 4406.472 | 7.415 |
| birds | Cinnyris_shelleyi | Nectarinia_shelleyi | LC | Stable | tropical | diurnal | 9 | 416193.988 | 8.987 |
| birds | Cinnyris_solaris | NA | LC | Stable | NA | NA | NA | NA | NA |
| birds | Cinnyris_sovimanga | Nectarinia_sovimanga | LC | Stable | tropical | diurnal | 7.2 | 591013.442 | 8.745 |
| birds | Cinnyris_stuhlmanni | Nectarinia_stuhlmanni | LC | Stable | tropical | diurnal | 8.8 | 12102.101 | 13.396 |
| birds | Cinnyris_superbus | Nectarinia_superba | LC | Decreasing | tropical | diurnal | 16.8 | 2854180.159 | 8.072 |
| birds | Cinnyris_talatala | Nectarinia_talatala | LC | Stable | tropical | diurnal | 8.1 | 2820942.589 | 9.166 |
| birds | Cinnyris_tsavoensis | Nectarinia_tsavoensis | LC | Stable | tropical | diurnal | 6.8 | 114686.091 | 9.023 |
| birds | Cinnyris_ursulae | Nectarinia_ursulae | LC | Decreasing | tropical | diurnal | 6.5 | 27769.05 | 11.03 |
| birds | Cinnyris_venustus | Nectarinia_venusta | LC | Stable | tropical | diurnal | 6.6 | 5912223.607 | 10.34 |
| birds | Circaetus_beaudouini | NA | VU | Decreasing | NA | NA | NA | NA | NA |
| birds | Circaetus_cinerascens | Circaetus_cinerascens | LC | Decreasing | tropical | diurnal | 1126 | 5762003.005 | 9.748 |
| birds | Circaetus_cinereus | Circaetus_cinereus | LC | Decreasing | tropical | diurnal | 2048 | 11541044.038 | 9.987 |
| birds | Circaetus_fasciolatus | Circaetus_fasciolatus | NT | Decreasing | tropical | diurnal | 1013.6 | 734455.966 | 10.012 |
| birds | Circaetus_gallicus | Circaetus_gallicus | LC | Stable | tropical | diurnal | 1699.1 | 24616860.096 | 11.221 |
| birds | Circaetus_pectoralis | Circaetus_pectoralis | LC | Stable | tropical | diurnal | 1500 | 10113444.05 | 8.507 |
| birds | Circus_aeruginosus | Circus_aeruginosus | LC | Stable | tropical | diurnal | 704.1 | 38081226.308 | 10.924 |
| birds | Circus_approximans | Circus_approximans | LC | Stable | temperate | diurnal | 746.2 | 5134943.646 | 5.734 |
| birds | Circus_assimilis | Circus_assimilis | LC | Stable | temperate | diurnal | 558.6 | 7817547.592 | 4.415 |
| birds | Circus_buffoni | Circus_buffoni | LC | Decreasing | temperate | diurnal | 507.4 | 6350653.748 | 8.803 |
| birds | Circus_cinereus | Circus_cinereus | LC | Decreasing | temperate | diurnal | 399.7 | 3705694.071 | 7.728 |
| birds | Circus_cyaneus | Circus_cyaneus | LC | Decreasing | temperate | diurnal | 393 | 36969892.737 | 9.881 |
| birds | Circus_macrourus | Circus_macrourus | NT | Decreasing | tropical | diurnal | 384.4 | 46915251.631 | 9.38 |
| birds | Circus_maillardi | Circus_maillardi | EN | Decreasing | tropical | diurnal | 663.6 | 1488.838 | 19.581 |
| birds | Circus_maurus | Circus_maurus | EN | Decreasing | temperate | diurnal | 509 | 1464708.152 | 7.904 |
| birds | Circus_melanoleucos | Circus_melanoleucos | LC | Decreasing | temperate | diurnal | 336.3 | 13367833.885 | 13.235 |
| birds | Circus_pygargus | Circus_pygargus | LC | Decreasing | tropical | diurnal | 310.8 | 30385902.218 | 10.867 |
| birds | Circus_ranivorus | Circus_ranivorus | LC | Decreasing | tropical | diurnal | 507 | 4426338.224 | 10.331 |
| birds | Circus_spilonotus | Circus_spilonotus | LC | Stable | temperate | diurnal | 558.8 | 11639585.512 | 9.767 |
| birds | Cissa_chinensis | Cissa_chinensis | LC | Decreasing | tropical | diurnal | 126.6 | 2108736.382 | 10.05 |
| birds | Cissa_hypoleuca | Cissa_hypoleuca | LC | Decreasing | temperate | diurnal | 125 | 890296.664 | 14.696 |

| | | | | | | | | | |
|---|---|---|---|---|---|---|---|---|---|
| birds | Cissa_thalassina | NA | CR | Decreasing | NA | NA | NA | NA | NA |
| birds | Cissomela_pectoralis | Certhionyx_pectoralis | LC | Stable | tropical | diurnal | 10.1 | 1086177.679 | 2.575 |
| birds | Cissopis_leverianus | Cissopis_leverianus | LC | Stable | tropical | diurnal | 76 | 5410821.018 | 5.02 |
| birds | Cisticola_aberdare | Cisticola_aberdare | VU | Decreasing | tropical | diurnal | 24 | 1272.48 | 12.763 |
| birds | Cisticola_aberrans | Cisticola_aberrans | LC | Stable | tropical | diurnal | 14.2 | 2482236.373 | 11.981 |
| birds | Cisticola_angusticauda | NA | LC | Stable | NA | NA | NA | NA | NA |
| birds | Cisticola_anonymus | Cisticola_anonymus | LC | Stable | tropical | diurnal | 13.9 | 2357531.943 | 6.773 |
| birds | Cisticola_aridulus | Cisticola_aridulus | LC | Increasing | tropical | diurnal | 9.2 | 4468382.379 | 9.33 |
| birds | Cisticola_ayresii | Cisticola_ayresii | LC | Decreasing | temperate | diurnal | 7 | 1258911.914 | 11.231 |
| birds | Cisticola_bodessa | Cisticola_bodessa | LC | Stable | tropical | diurnal | 17.1 | 437584.832 | 9.744 |
| birds | Cisticola_brachypterus | Cisticola_brachypterus | LC | Stable | tropical | diurnal | 7.8 | 7674053.245 | 10.12 |
| birds | Cisticola_brunnescens | Cisticola_brunnescens | LC | Stable | tropical | diurnal | 8 | 465384.188 | 10.182 |
| birds | Cisticola_bulliens | Cisticola_bulliens | LC | Stable | tropical | diurnal | 15.8 | 308376.906 | 9.334 |
| birds | Cisticola_cantans | Cisticola_cantans | LC | Stable | tropical | diurnal | 11.7 | 5670957.167 | 11.588 |
| birds | Cisticola_carruthersi | Cisticola_carruthersi | LC | Decreasing | tropical | diurnal | 11.2 | 174334.597 | 17.882 |
| birds | Cisticola_cherina | Cisticola_cherina | LC | Stable | tropical | diurnal | 10 | 591128.963 | 8.749 |
| birds | Cisticola_chiniana | Cisticola_chiniana | LC | Stable | tropical | diurnal | 16.1 | 6716168.112 | 8.535 |
| birds | Cisticola_chubbi | Cisticola_chubbi | LC | Stable | tropical | diurnal | 16 | 322235.685 | 15.326 |
| birds | Cisticola_cinereolus | Cisticola_cinereolus | LC | Stable | tropical | diurnal | 17 | 614939.777 | 8.712 |
| birds | Cisticola_dambo | Cisticola_dambo | LC | Stable | tropical | diurnal | 9 | 472758.505 | 6.134 |
| birds | Cisticola_erythrops | Cisticola_erythrops | LC | Stable | tropical | diurnal | 14.8 | 6407914.953 | 10.858 |
| birds | Cisticola_exilis | Cisticola_exilis | LC | Increasing | tropical | diurnal | 7 | 7177348.998 | 9.464 |
| birds | Cisticola_eximius | Cisticola_eximius | LC | Stable | tropical | diurnal | 12 | 674103.661 | 13.207 |
| birds | Cisticola_fulvicapilla | Cisticola_fulvicapilla | LC | Stable | tropical | diurnal | 8.2 | 4604333.833 | 8.743 |
| birds | Cisticola_galactotes | Cisticola_galactotes | LC | Stable | tropical | diurnal | 13.1 | 287146.869 | 11.901 |
| birds | Cisticola_guinea | Cisticola_guinea | LC | Stable | tropical | diurnal | 9.3 | 646786.749 | 13.449 |
| birds | Cisticola_haesitatus | Cisticola_haesitatus | LC | Stable | tropical | diurnal | 6.9 | 796.531 | 11.374 |
| birds | Cisticola_hunteri | Cisticola_hunteri | LC | Stable | tropical | diurnal | 15.5 | 156331.321 | 14.207 |
| birds | Cisticola_juncidis | Cisticola_juncidis | LC | Increasing | temperate | diurnal | 6.8 | 18929166.746 | 13.439 |
| birds | Cisticola_lais | Cisticola_lais | LC | Stable | temperate | diurnal | 14.8 | 881974.018 | 12.496 |
| birds | Cisticola_lateralis | Cisticola_lateralis | LC | Stable | tropical | diurnal | 16.8 | 3066342.447 | 9.373 |
| birds | Cisticola_melanurus | Cisticola_melanurus | DD | Stable | tropical | diurnal | 8.7 | 442014.73 | 5.815 |
| birds | Cisticola_nana | Cisticola_nanus | LC | Stable | tropical | diurnal | 5 | 685105.623 | 9.456 |
| birds | Cisticola_natalensis | Cisticola_natalensis | LC | Stable | tropical | diurnal | 16.4 | 7970636.773 | 10.316 |

| | | | | | | | | | |
|---|---|---|---|---|---|---|---|---|---|
| birds | Cisticola_nigriloris | Cisticola_nigriloris | LC | Stable | tropical | diurnal | 19.6 | 71914.394 | 13.568 |
| birds | Cisticola_njombe | Cisticola_njombe | LC | Stable | tropical | diurnal | 10.9 | 33269.219 | 15.353 |
| birds | Cisticola_pipiens | Cisticola_pipiens | LC | Stable | tropical | diurnal | 13.5 | 1127999.588 | 7.167 |
| birds | Cisticola_restrictus | NA | DD | Unknown | NA | NA | NA | NA | NA |
| birds | Cisticola_robustus | Cisticola_robustus | LC | Stable | tropical | diurnal | 18.5 | 1229121.479 | 10.608 |
| birds | Cisticola_ruficeps | Cisticola_ruficeps | LC | Stable | tropical | diurnal | 9.7 | 1253642.08 | 11.013 |
| birds | Cisticola_rufilatus | Cisticola_rufilatus | LC | Stable | tropical | diurnal | 11.3 | 2748861.044 | 7.113 |
| birds | Cisticola_rufus | Cisticola_rufus | LC | Stable | tropical | diurnal | 7.3 | 876800.152 | 13.799 |
| birds | Cisticola_subruficapilla | Cisticola_subruficapilla | LC | Decreasing | temperate | diurnal | 9.9 | 910756.54 | 6.783 |
| birds | Cisticola_textrix | Cisticola_textrix | LC | Decreasing | temperate | diurnal | 10.3 | 702791.556 | 11.233 |
| birds | Cisticola_tinniens | Cisticola_tinniens | LC | Stable | temperate | diurnal | 11.4 | 1754614.863 | 10.083 |
| birds | Cisticola_troglodytes | NA | LC | Stable | NA | NA | NA | NA | NA |
| birds | Cisticola_woosnami | Cisticola_woosnami | LC | Stable | tropical | diurnal | 13.5 | 1290884.311 | 10.3 |
| birds | Cistothorus_apolinari | Cistothorus_apolinari | EN | Decreasing | tropical | diurnal | 17.7 | 4937.623 | 16.528 |
| birds | Cistothorus_meridae | Cistothorus_meridae | LC | Stable | tropical | diurnal | 9.7 | 4041.711 | 14.1 |
| birds | Cistothorus_palustris | Cistothorus_palustris | LC | Increasing | temperate | diurnal | 10.8 | 10354699.518 | 12.962 |
| birds | Cistothorus_platensis | Cistothorus_platensis | LC | Increasing | temperate | diurnal | 9 | 4772915.829 | 9.519 |
| birds | Cittura_cyanotis | Cittura_cyanotis | LC | Decreasing | tropical | diurnal | 147 | 73612.744 | 8.053 |
| birds | Cladorhynchus_leucocephalus | Cladorhynchus_leucocephalus | LC | Stable | temperate | diurnal | 216 | 1291070.247 | 7.727 |
| birds | Clamator_coromandus | Clamator_coromandus | LC | Stable | tropical | diurnal | 74.6 | 9231925.559 | 14.66 |
| birds | Clamator_glandarius | Clamator_glandarius | LC | Stable | tropical | diurnal | 143 | 15293675.327 | 10.156 |
| birds | Clamator_jacobinus | Clamator_jacobinus | LC | Stable | tropical | diurnal | 79.4 | 18575025.926 | 11.211 |
| birds | Clamator_levaillantii | Clamator_levaillantii | LC | Stable | tropical | diurnal | 122 | 14092853.759 | 9.29 |
| birds | Clanga_clanga | Aquila_clanga | VU | Decreasing | temperate | diurnal | 2154.3 | 30259017.147 | 10.141 |
| birds | Clanga_pomarina | Aquila_pomarina | LC | Stable | tropical | diurnal | 1359.4 | 12328656.869 | 11.73 |
| birds | Clangula_hyemalis | Clangula_hyemalis | VU | Decreasing | temperate | diurnal | 871 | 14175641.321 | 2.103 |
| birds | Claravis_pretiosa | Claravis_pretiosa | LC | Decreasing | tropical | diurnal | 68.2 | 12977823.951 | 5.637 |
| birds | Cleptornis_marchei | Cleptornis_marchei | EN | Decreasing | tropical | diurnal | 20.2 | 137.13 | 28.047 |
| birds | Clibanornis_dendrocolaptoides | Clibanornis_dendrocolaptoides | LC | Decreasing | temperate | diurnal | 48.2 | 317493.202 | 11.944 |
| birds | Clibanornis_erythrocephalus | Hylocryptus_erythrocephalus | NT | Decreasing | tropical | diurnal | 46.5 | 18640 | 8.348 |
| birds | Clibanornis_rectirostris | Hylocryptus_rectirostris | LC | Decreasing | tropical | diurnal | 48 | 1421864.094 | 10.868 |
| birds | Clibanornis_rubiginosus | Automolus_rubiginosus | LC | Decreasing | tropical | diurnal | 39.8 | 2137031.201 | 3.112 |
| birds | Climacteris_affinis | Climacteris_affinis | LC | Decreasing | temperate | diurnal | 21.7 | 2630036.348 | 4.121 |
| birds | Climacteris_erythrops | Climacteris_erythrops | LC | Decreasing | temperate | diurnal | 23.3 | 317210.536 | 8.05 |

| | | | | | | | | | |
|---|---|---|---|---|---|---|---|---|---|
| birds | Climacteris_melanurus | Climacteris_melanurus | LC | Stable | tropical | diurnal | 31.9 | 1122025.404 | 2.615 |
| birds | Climacteris_picumnus | Climacteris_picumnus | LC | Decreasing | temperate | diurnal | 28.9 | 2269397.305 | 7.087 |
| birds | Climacteris_rufus | Climacteris_rufus | LC | Decreasing | temperate | diurnal | 32.6 | 452307.175 | 6.82 |
| birds | Clytoctantes_alixii | Clytoctantes_alixii | EN | Decreasing | tropical | diurnal | 32.5 | 79436.569 | 10.414 |
| birds | Clytoctantes_atrogularis | Clytoctantes_atrogularis | NT | Decreasing | tropical | diurnal | 31 | 299551.948 | 1.952 |
| birds | Clytolaema_rubricauda | Clytolaema_rubricauda | LC | Unknown | tropical | diurnal | 7.9 | 352748.434 | 13.003 |
| birds | Clytomyias_insignis | Clytomyias_insignis | LC | Stable | tropical | diurnal | 12.4 | 139418.96 | 4.898 |
| birds | Clytorhynchus_hamlini | Clytorhynchus_hamlini | NT | Decreasing | tropical | diurnal | 34.6 | 672.673 | 3.725 |
| birds | Clytorhynchus_nigrogularis | Clytorhynchus_nigrogularis | NT | Decreasing | tropical | diurnal | 55.7 | 16947.716 | 9.386 |
| birds | Clytorhynchus_pachycephaloides | Clytorhynchus_pachycephaloides | LC | Decreasing | tropical | diurnal | 26.3 | 27352.858 | 6.723 |
| birds | Clytorhynchus_vitiensis | Clytorhynchus_vitiensis | LC | Decreasing | tropical | diurnal | 29.2 | 19242.454 | 9.375 |
| birds | Clytospiza_monteiri | Clytospiza_monteiri | LC | Stable | tropical | diurnal | 14.9 | 1193530.917 | 7.092 |
| birds | Cnemarchus_erythropygius | Cnemarchus_erythropygius | LC | Stable | tropical | diurnal | 41 | 178441.619 | 8.965 |
| birds | Cnemathraupis_aureodorsalis | Buthraupis_aureodorsalis | EN | Decreasing | tropical | diurnal | 85 | 2445.846 | 4.924 |
| birds | Cnemathraupis_eximia | Buthraupis_eximia | LC | Stable | tropical | diurnal | 62.7 | 60098.847 | 10.744 |
| birds | Cnemophilus_loriae | Cnemophilus_loriae | LC | Stable | tropical | diurnal | 84 | 138396.013 | 4.983 |
| birds | Cnemophilus_macgregorii | Cnemophilus_macgregorii | LC | Unknown | tropical | diurnal | 97.9 | 6880.909 | 3.383 |
| birds | Cnemoscopus_rubrirostris | Cnemoscopus_rubrirostris | LC | Decreasing | tropical | diurnal | 18 | 61667.01 | 8.595 |
| birds | Cnemotriccus_fuscatus | Cnemotriccus_fuscatus | LC | Decreasing | tropical | diurnal | 13.6 | 12243851.433 | 5.448 |
| birds | Cnipodectes_subbrunneus | Cnipodectes_subbrunneus | LC | Decreasing | tropical | diurnal | 23.2 | 1950473.586 | 1.941 |
| birds | Coccopygia_melanotis | Estrilda_melanotis | LC | Stable | temperate | diurnal | 6.5 | 285539.095 | 12.64 |
| birds | Coccopygia_quartinia | Estrilda_quartinia | LC | Stable | tropical | diurnal | 6.7 | 676732.962 | 13.38 |
| birds | Coccothraustes_coccothraustes | Coccothraustes_coccothraustes | LC | Increasing | temperate | diurnal | 56.6 | 15375684.185 | 13.182 |
| birds | Coccycua_cinerea | Coccycua_cinerea | LC | Decreasing | tropical | diurnal | 41 | 5265483.414 | 6.615 |
| birds | Coccycua_minuta | Coccycua_minuta | LC | Decreasing | tropical | diurnal | 39.3 | 8754206.579 | 3.622 |
| birds | Coccycua_pumila | Coccycua_pumila | LC | Increasing | tropical | diurnal | 37.1 | 647667.457 | 8.85 |
| birds | Coccyzus_americanus | Coccyzus_americanus | LC | Decreasing | tropical | diurnal | 64 | 21067044.673 | 8.774 |
| birds | Coccyzus_erythropthalmus | Coccyzus_erythropthalmus | LC | Decreasing | tropical | diurnal | 50.9 | 10934644.021 | 12.003 |
| birds | Coccyzus_euleri | Coccyzus_euleri | LC | Stable | tropical | diurnal | 52.3 | 9036508.428 | 6.052 |
| birds | Coccyzus_ferrugineus | NA | VU | Stable | NA | NA | NA | NA | NA |
| birds | Coccyzus_lansbergi | Coccyzus_lansbergi | LC | Decreasing | tropical | diurnal | 50.3 | 326111.202 | 13.788 |
| birds | Coccyzus_longirostris | Coccyzus_longirostris | LC | Decreasing | tropical | diurnal | 100 | 75087.702 | 15.77 |
| birds | Coccyzus_melacoryphus | Coccyzus_melacoryphus | LC | Stable | tropical | diurnal | 49.7 | 14444342.034 | 6.013 |
| birds | Coccyzus_merlini | Coccyzus_merlini | LC | Stable | tropical | diurnal | 169.6 | 109791.831 | 16.137 |

| | | | | | | | | | |
|---|---|---|---|---|---|---|---|---|---|
| birds | Coccyzus_minor | Coccyzus_minor | LC | Decreasing | tropical | diurnal | 63.9 | 1105292.176 | 10.318 |
| birds | Coccyzus_pluvialis | Coccyzus_pluvialis | LC | Stable | tropical | diurnal | 163 | 11021.791 | 19.741 |
| birds | Coccyzus_rufigularis | Coccyzus_rufigularis | EN | Decreasing | tropical | diurnal | 128 | 17282.423 | 14.346 |
| birds | Coccyzus_vetula | Coccyzus_vetula | LC | Decreasing | tropical | diurnal | 96 | 11021.791 | 19.741 |
| birds | Coccyzus_vieilloti | Coccyzus_vieilloti | LC | Stable | tropical | diurnal | 81.9 | 8774.421 | 27.252 |
| birds | Cochlearius_cochlearius | Cochlearius_cochlearius | LC | Decreasing | tropical | cathemeral | 643.2 | 11698377.397 | 5.287 |
| birds | Cochoa_azurea | Cochoa_azurea | VU | Decreasing | tropical | diurnal | 111.5 | 4003.48 | 19.174 |
| birds | Cochoa_beccarii | Cochoa_beccarii | LC | Decreasing | tropical | diurnal | 111.5 | 64349.299 | 7.139 |
| birds | Cochoa_purpurea | Cochoa_purpurea | LC | Decreasing | tropical | diurnal | 103 | 623216.088 | 11.348 |
| birds | Cochoa_viridis | Cochoa_viridis | LC | Decreasing | tropical | diurnal | 120 | 1337235.033 | 9.773 |
| birds | Coeligena_bonapartei | Coeligena_bonapartei | LC | Stable | tropical | diurnal | 6.5 | 16909.144 | 17.875 |
| birds | Coeligena_coeligena | Coeligena_coeligena | LC | Stable | tropical | diurnal | 6.8 | 387643.169 | 8.644 |
| birds | Coeligena_helianthea | Coeligena_helianthea | LC | Unknown | tropical | diurnal | 6.6 | 42112.578 | 11.853 |
| birds | Coeligena_iris | Coeligena_iris | LC | Decreasing | tropical | diurnal | 6.9 | 80713.592 | 8.172 |
| birds | Coeligena_lutetiae | Coeligena_lutetiae | LC | Decreasing | tropical | diurnal | 7.3 | 63339.366 | 9.56 |
| birds | Coeligena_phalerata | Coeligena_phalerata | NT | Decreasing | tropical | diurnal | 6.1 | 5344.358 | 5.744 |
| birds | Coeligena_prunellei | Coeligena_prunellei | VU | Decreasing | tropical | diurnal | 7 | 9336.578 | 15.469 |
| birds | Coeligena_torquata | Coeligena_torquata | LC | Decreasing | tropical | diurnal | 6.7 | 302874.556 | 9.219 |
| birds | Coeligena_violifer | Coeligena_violifer | LC | Decreasing | tropical | diurnal | 7.4 | 43536.518 | 5.914 |
| birds | Coeligena_wilsoni | Coeligena_wilsoni | LC | Decreasing | tropical | diurnal | 6.7 | 34158.963 | 7.255 |
| birds | Coenocorypha_aucklandica | NA | NT | Stable | NA | NA | NA | NA | NA |
| birds | Coenocorypha_pusilla | NA | VU | Stable | NA | NA | NA | NA | NA |
| birds | Coereba_flaveola | Coereba_flaveola | LC | Decreasing | tropical | diurnal | 10 | 10807432.935 | 6.635 |
| birds | Colaptes_atricollis | Colaptes_atricollis | LC | Stable | tropical | diurnal | 76.1 | 200853.998 | 7.649 |
| birds | Colaptes_auratus | Colaptes_auratus | LC | Decreasing | temperate | diurnal | 131.5 | 11444924.478 | 9.228 |
| birds | Colaptes_auricularis | Colaptes_auricularis | LC | Decreasing | tropical | diurnal | 65.4 | 203098.619 | 5.965 |
| birds | Colaptes_campestris | Colaptes_campestris | LC | Increasing | tropical | diurnal | 158 | 4575239.788 | 9.139 |
| birds | Colaptes_chrysoides | Colaptes_chrysoides | LC | Decreasing | temperate | diurnal | 111 | 377623.21 | 6.625 |
| birds | Colaptes_fernandinae | Colaptes_fernandinae | EN | Decreasing | tropical | diurnal | 135 | 106634.82 | 16.283 |
| birds | Colaptes_melanochloros | Colaptes_melanochloros | LC | Stable | tropical | diurnal | 127.3 | 4723714.992 | 9.708 |
| birds | Colaptes_pitius | Colaptes_pitius | LC | Stable | temperate | diurnal | 159 | 428459.347 | 6.936 |
| birds | Colaptes_punctigula | Colaptes_punctigula | LC | Stable | tropical | diurnal | 62 | 3394047.542 | 4.262 |
| birds | Colaptes_rivolii | Colaptes_rivolii | LC | Stable | tropical | diurnal | 102 | 463744.407 | 10.153 |
| birds | Colaptes_rubiginosus | Colaptes_rubiginosus | LC | Decreasing | tropical | diurnal | 55.8 | 2243128.494 | 6.751 |

| | | | | | | | | | |
|---|---|---|---|---|---|---|---|---|---|
| birds | Colaptes_rupicola | Colaptes_rupicola | LC | Stable | tropical | diurnal | 174 | 785415.83 | 7.131 |
| birds | Colibri_coruscans | Colibri_coruscans | LC | Unknown | tropical | diurnal | 6.7 | 920400.096 | 8.422 |
| birds | Colibri_delphinae | Colibri_delphinae | LC | Decreasing | tropical | diurnal | 6.4 | 1201162.312 | 5.032 |
| birds | Colibri_serrirostris | Colibri_serrirostris | LC | Unknown | tropical | diurnal | 6.7 | 2370635.537 | 10.765 |
| birds | Colibri_thalassinus | Colibri_thalassinus | LC | Decreasing | tropical | diurnal | 5.9 | 792390.561 | 10.628 |
| birds | Colinus_cristatus | Colinus_cristatus | LC | Stable | tropical | diurnal | 134.5 | 1970617.036 | 7.044 |
| birds | Colinus_leucopogon | Colinus_leucopogon | LC | Stable | tropical | diurnal | 128.7 | 145250.929 | 14.054 |
| birds | Colinus_nigrogularis | Colinus_nigrogularis | LC | Decreasing | tropical | diurnal | 129.8 | 136499.43 | 7.167 |
| birds | Colinus_virginianus | Colinus_virginianus | NT | Decreasing | temperate | diurnal | 171.9 | 4364464.182 | 16.275 |
| birds | Colius_castanotus | Colius_castanotus | LC | Stable | tropical | diurnal | 56.6 | 289063.971 | 8.819 |
| birds | Colius_colius | Colius_colius | LC | Increasing | temperate | diurnal | 41.4 | 1611547.388 | 7.204 |
| birds | Colius_leucocephalus | Colius_leucocephalus | LC | Stable | tropical | diurnal | 35.8 | 533022.161 | 10.658 |
| birds | Colius_striatus | Colius_striatus | LC | Increasing | tropical | diurnal | 51.1 | 8057145.03 | 9.241 |
| birds | Collocalia_esculenta | Collocalia_esculenta | LC | Stable | tropical | diurnal | 6.3 | 2759413.587 | 7.603 |
| birds | Collocalia_linchi | Collocalia_linchi | LC | Decreasing | tropical | diurnal | 5.9 | 316140.075 | 14.639 |
| birds | Collocalia_troglodytes | Collocalia_troglodytes | LC | Decreasing | tropical | diurnal | 5.4 | 283053.111 | 13.72 |
| birds | Colluricincla_boweri | Colluricincla_boweri | VU | Decreasing | tropical | diurnal | 43.8 | 7631.069 | 3.555 |
| birds | Colluricincla_harmonica | Colluricincla_harmonica | LC | Decreasing | temperate | diurnal | 65.8 | 6919512.585 | 4.599 |
| birds | Colluricincla_megarhyncha | Colluricincla_megarhyncha | LC | Stable | tropical | diurnal | 32.3 | 1204726.416 | 3.583 |
| birds | Colluricincla_tenebrosa | Colluricincla_umbrina | LC | Stable | tropical | diurnal | 46 | 40629.609 | 3.936 |
| birds | Colluricincla_woodwardi | Colluricincla_woodwardi | LC | Stable | tropical | diurnal | 54.5 | 549567.851 | 2.358 |
| birds | Colonia_colonus | Colonia_colonus | LC | Stable | tropical | diurnal | 18.3 | 6767109.678 | 6.833 |
| birds | Colorhamphus_parvirostris | Colorhamphus_parvirostris | LC | Stable | temperate | diurnal | 10.6 | 536388.916 | 5.709 |
| birds | Columba_albinucha | Columba_albinucha | LC | Decreasing | tropical | diurnal | 325 | 142261.786 | 7.567 |
| birds | Columba_albitorques | Columba_albitorques | LC | Stable | tropical | diurnal | 277 | 137598.58 | 14.874 |
| birds | Columba_argentina | NA | CR | Decreasing | NA | NA | NA | NA | NA |
| birds | Columba_arquatrix | Columba_arquatrix | LC | Decreasing | temperate | diurnal | 400 | 1486805.138 | 12.724 |
| birds | Columba_bollii | Columba_bollii | LC | Increasing | temperate | diurnal | 308.5 | 2531.081 | 16.597 |
| birds | Columba_delegorguei | Columba_delegorguei | LC | Decreasing | tropical | diurnal | 133 | 232244.082 | 12.931 |
| birds | Columba_elphinstonii | Columba_elphinstonii | VU | Decreasing | tropical | diurnal | 379 | 116434.818 | 18.004 |
| birds | Columba_eversmanni | Columba_eversmanni | VU | Decreasing | temperate | diurnal | 202 | 5250966.243 | 9.048 |
| birds | Columba_guinea | Columba_guinea | LC | Stable | temperate | diurnal | 352 | 10557795.158 | 9.895 |
| birds | Columba_hodgsonii | Columba_hodgsonii | LC | Stable | temperate | diurnal | 308.5 | 2007983.93 | 9.56 |
| birds | Columba_iriditorques | Columba_iriditorques | LC | Stable | tropical | diurnal | 125.9 | 2757795.122 | 8.116 |

| | | | | | | | | | |
|---|---|---|---|---|---|---|---|---|---|
| birds | Columba_janthina | NA | LC | Stable | NA | NA | NA | NA | NA |
| birds | Columba_junoniae | NA | NT | Increasing | NA | NA | NA | NA | NA |
| birds | Columba_leucomela | Columba_leucomela | LC | Increasing | temperate | diurnal | 420 | 275007.141 | 7.529 |
| birds | Columba_leuconota | Columba_leuconota | LC | Stable | temperate | diurnal | 284 | 2656832.053 | 6.015 |
| birds | Columba_livia | Columba_livia | LC | Decreasing | temperate | diurnal | 354.2 | 27446747.696 | 11.326 |
| birds | Columba_malherbii | Columba_malherbii | NT | Decreasing | tropical | diurnal | 308.5 | 1017.047 | 12.941 |
| birds | Columba_oenas | Columba_oenas | LC | Increasing | temperate | diurnal | 290.8 | 9639015.462 | 15.244 |
| birds | Columba_oliviae | Columba_oliviae | DD | Stable | tropical | diurnal | 308.5 | 24568.317 | 8.158 |
| birds | Columba_pallidiceps | Columba_pallidiceps | VU | Decreasing | tropical | diurnal | 459 | 63785.654 | 4.431 |
| birds | Columba_palumboides | Columba_palumboides | NT | Decreasing | tropical | diurnal | 520 | 7618.236 | 6.774 |
| birds | Columba_palumbus | Columba_palumbus | LC | Increasing | temperate | diurnal | 490 | 12401484.341 | 14.524 |
| birds | Columba_pollenii | Columba_pollenii | NT | Decreasing | tropical | diurnal | 308.5 | 2057.516 | 17.332 |
| birds | Columba_pulchricollis | Columba_pulchricollis | LC | Decreasing | temperate | diurnal | 330 | 565151.186 | 7.818 |
| birds | Columba_punicea | Columba_punicea | VU | Decreasing | tropical | diurnal | 434.4 | 358510.168 | 11.987 |
| birds | Columba_rupestris | Columba_rupestris | LC | Decreasing | temperate | diurnal | 245.4 | 11908353.579 | 6.312 |
| birds | Columba_sjostedti | Columba_sjostedti | LC | Decreasing | tropical | diurnal | 308.5 | 44188.924 | 10.065 |
| birds | Columba_thomensis | Columba_thomensis | EN | Decreasing | tropical | diurnal | 428.7 | 366.992 | 7.486 |
| birds | Columba_torringtoniae | Columba_torringtoniae | VU | Decreasing | tropical | diurnal | 308.5 | 5001.872 | 16.525 |
| birds | Columba_trocaz | Columba_trocaz | LC | Stable | temperate | diurnal | 440 | 549.162 | 18.933 |
| birds | Columba_unicincta | Columba_unicincta | LC | Decreasing | tropical | diurnal | 389.1 | 2699737.731 | 7.637 |
| birds | Columba_vitiensis | Columba_vitiensis | LC | Stable | tropical | diurnal | 353 | 1128735.55 | 7.07 |
| birds | Columbina_buckleyi | Columbina_buckleyi | LC | Decreasing | tropical | diurnal | 57.5 | 82687.641 | 14.021 |
| birds | Columbina_cruziana | Columbina_cruziana | LC | Increasing | tropical | diurnal | 50.1 | 334307.936 | 8.982 |
| birds | Columbina_cyanopis | Columbina_cyanopis | CR | Decreasing | tropical | diurnal | 45.6 | 3140.316 | 10.347 |
| birds | Columbina_inca | Columbina_inca | LC | Increasing | tropical | diurnal | 47.5 | 2760667.854 | 10.641 |
| birds | Columbina_minuta | Columbina_minuta | LC | Stable | tropical | diurnal | 32.6 | 7050160.835 | 8.181 |
| birds | Columbina_passerina | Columbina_passerina | LC | Stable | tropical | diurnal | 35.4 | 7245312.584 | 7.888 |
| birds | Columbina_picui | Columbina_picui | LC | Stable | tropical | diurnal | 47 | 6427122.471 | 7.07 |
| birds | Columbina_squammata | Columbina_squammata | LC | Stable | tropical | diurnal | 52.9 | 4814670.861 | 9.41 |
| birds | Columbina_talpacoti | Columbina_talpacoti | LC | Increasing | tropical | diurnal | 46 | 14617238.714 | 6.132 |
| birds | Compsothraupis_loricata | Compsothraupis_loricata | LC | Stable | tropical | diurnal | 72.5 | 1679211.048 | 8.314 |
| birds | Conioptilon_mcilhennyi | Conioptilon_mcilhennyi | LC | Stable | tropical | diurnal | 90 | 146164.972 | 0.844 |
| birds | Conirostrum_albifrons | Conirostrum_albifrons | LC | Decreasing | tropical | diurnal | 14.9 | 298087.624 | 8.316 |
| birds | Conirostrum_bicolor | Conirostrum_bicolor | NT | Decreasing | tropical | diurnal | 10.5 | 566641.852 | 5.482 |

| | | | | | | | | | |
|---|---|---|---|---|---|---|---|---|---|
| birds | Conirostrum_binghami | Oreomanes_fraseri | NT | Decreasing | tropical | diurnal | 25 | 233433.106 | 9.305 |
| birds | Conirostrum_cinereum | Conirostrum_cinereum | LC | Stable | tropical | diurnal | 8.8 | 616914.781 | 8.421 |
| birds | Conirostrum_ferrugineiventre | Conirostrum_ferrugineiventre | LC | Stable | tropical | diurnal | 11.5 | 147474.331 | 6.22 |
| birds | Conirostrum_leucogenys | Conirostrum_leucogenys | LC | Decreasing | tropical | diurnal | 7 | 271279.348 | 13.674 |
| birds | Conirostrum_margaritae | Conirostrum_margaritae | VU | Decreasing | tropical | diurnal | 9.7 | 77630.04 | 3.375 |
| birds | Conirostrum_rufum | Conirostrum_rufum | LC | Stable | tropical | diurnal | 11 | 39407.058 | 13.078 |
| birds | Conirostrum_sitticolor | Conirostrum_sitticolor | LC | Decreasing | tropical | diurnal | 11.9 | 300002.234 | 9.007 |
| birds | Conirostrum_speciosum | Conirostrum_speciosum | LC | Decreasing | tropical | diurnal | 8.8 | 7564709.166 | 6.902 |
| birds | Conirostrum_tamarugense | Conirostrum_tamarugense | LC | Increasing | tropical | diurnal | 10.5 | 45044.68 | 4.936 |
| birds | Conopias_albovittatus | Conopias_albovittatus | LC | Decreasing | tropical | diurnal | 24.4 | 198320.462 | 6.198 |
| birds | Conopias_cinchoneti | Conopias_cinchoneti | VU | Stable | tropical | diurnal | 25 | 171399.95 | 6.4 |
| birds | Conopias_parvus | Conopias_parvus | LC | Stable | tropical | diurnal | 21 | 3906792.832 | 1.182 |
| birds | Conopias_trivirgatus | NA | LC | Stable | NA | NA | NA | NA | NA |
| birds | Conopophaga_ardesiaca | Conopophaga_ardesiaca | LC | Stable | tropical | diurnal | 26.3 | 100334.387 | 5.603 |
| birds | Conopophaga_aurita | Conopophaga_aurita | LC | Decreasing | tropical | diurnal | 26.3 | 2800393.444 | 1.256 |
| birds | Conopophaga_castaneiceps | Conopophaga_castaneiceps | LC | Increasing | tropical | diurnal | 27 | 160640.323 | 6.049 |
| birds | Conopophaga_cearae | Conopophaga_cearae | NT | Decreasing | tropical | diurnal | 22 | 44374.138 | 11.559 |
| birds | Conopophaga_lineata | Conopophaga_lineata | LC | Stable | tropical | diurnal | 25.2 | 1896972.302 | 12.565 |
| birds | Conopophaga_melanogaster | Conopophaga_melanogaster | LC | Stable | tropical | diurnal | 42.5 | 911253.952 | 2.343 |
| birds | Conopophaga_melanops | Conopophaga_melanops | LC | Stable | tropical | diurnal | 20.1 | 242500.405 | 12.661 |
| birds | Conopophaga_peruviana | Conopophaga_peruviana | LC | Stable | tropical | diurnal | 23 | 1098743.55 | 1.279 |
| birds | Conopophaga_roberti | Conopophaga_roberti | LC | Decreasing | tropical | diurnal | 20.8 | 686966.069 | 5.524 |
| birds | Conopophila_albogularis | Conopophila_albogularis | LC | Stable | tropical | diurnal | 11.8 | 496534.414 | 2.652 |
| birds | Conopophila_rufogularis | Conopophila_rufogularis | LC | Stable | tropical | diurnal | 10.8 | 1954135.965 | 3.017 |
| birds | Conopophila_whitei | Conopophila_whitei | LC | Decreasing | temperate | diurnal | 9.8 | 1437139.32 | 1.935 |
| birds | Conothraupis_mesoleuca | Conothraupis_mesoleuca | EN | Decreasing | tropical | diurnal | 15.3 | 212289.769 | 6.856 |
| birds | Conothraupis_speculigera | Conothraupis_speculigera | LC | Stable | tropical | diurnal | 25 | 489921.33 | 3.15 |
| birds | Contopus_albogularis | Contopus_albogularis | LC | Stable | tropical | diurnal | 9.6 | 114553.482 | 1.169 |
| birds | Contopus_caribaeus | Contopus_caribaeus | LC | Stable | tropical | diurnal | 10.5 | 119645.894 | 15.462 |
| birds | Contopus_cinereus | Contopus_cinereus | LC | Stable | tropical | diurnal | 11.6 | 3837797.833 | 9.89 |
| birds | Contopus_cooperi | Contopus_cooperi | NT | Decreasing | temperate | diurnal | 32.1 | 20278748.216 | 5.4 |
| birds | Contopus_fumigatus | Contopus_fumigatus | LC | Stable | tropical | diurnal | 19.9 | 1021094.076 | 6.746 |
| birds | Contopus_hispaniolensis | Contopus_hispaniolensis | LC | Decreasing | tropical | diurnal | 11.7 | 74799.651 | 15.781 |
| birds | Contopus_latirostris | Contopus_latirostris | LC | Decreasing | tropical | diurnal | 11.2 | 12727.556 | 25.711 |

| | | | | | | | | | |
|---|---|---|---|---|---|---|---|---|---|
| birds | Contopus_lugubris | Contopus_lugubris | LC | Stable | tropical | diurnal | 21.5 | 8200.085 | 7.356 |
| birds | Contopus_nigrescens | Contopus_nigrescens | LC | Decreasing | tropical | diurnal | 9.5 | 254170.676 | 4.557 |
| birds | Contopus_ochraceus | Contopus_ochraceus | LC | Stable | tropical | diurnal | 23 | 3144.578 | 5.471 |
| birds | Contopus_pallidus | Contopus_pallidus | LC | Decreasing | tropical | diurnal | 9.3 | 11021.791 | 19.741 |
| birds | Contopus_pertinax | Contopus_pertinax | LC | Decreasing | tropical | diurnal | 27.2 | 855800.292 | 10.207 |
| birds | Contopus_punensis | Contopus_punensis | LC | Stable | tropical | diurnal | 12.3 | 197413.595 | 11.182 |
| birds | Contopus_sordidulus | Contopus_sordidulus | LC | Decreasing | temperate | diurnal | 13.1 | 11524307.35 | 7.58 |
| birds | Contopus_virens | Contopus_virens | LC | Decreasing | tropical | diurnal | 13.9 | 10575580.524 | 9.863 |
| birds | Copsychus_albospecularis | Copsychus_albospecularis | LC | Decreasing | tropical | diurnal | 22.7 | 239689.824 | 9.309 |
| birds | Copsychus_cebuensis | NA | LC | Stable | NA | NA | NA | NA | NA |
| birds | Copsychus_fulicatus | Saxicoloides_fulicatus | LC | Stable | tropical | diurnal | 16.6 | 3270463.129 | 17.954 |
| birds | Copsychus_pyrropygus | Trichixos_pyrropygus | NT | Decreasing | tropical | diurnal | 40.9 | 1050012.651 | 7.827 |
| birds | Copsychus_saularis | Copsychus_saularis | LC | Stable | tropical | diurnal | 36 | 9009660.203 | 14.525 |
| birds | Copsychus_sechellarum | NA | EN | Increasing | NA | NA | NA | NA | NA |
| birds | Coracias_abyssinicus | Coracias_abyssinicus | LC | Increasing | tropical | diurnal | 102 | 7043827.445 | 10.716 |
| birds | Coracias_benghalensis | Coracias_benghalensis | LC | Increasing | tropical | cathemeral | 157.5 | 4430960.425 | 15.719 |
| birds | Coracias_caudatus | Coracias_caudatus | LC | Stable | tropical | diurnal | 110 | 8821260.607 | 8.823 |
| birds | Coracias_cyanogaster | Coracias_cyanogaster | LC | Decreasing | tropical | diurnal | 142 | 2031616.711 | 11.153 |
| birds | Coracias_garrulus | Coracias_garrulus | LC | Decreasing | tropical | diurnal | 146 | 34073962.642 | 9.233 |
| birds | Coracias_naevius | Coracias_naevia | LC | Decreasing | tropical | diurnal | 168 | 9355490.241 | 9.444 |
| birds | Coracias_spatulatus | Coracias_spatulatus | LC | Decreasing | tropical | diurnal | 90.4 | 2917947.702 | 8.938 |
| birds | Coracias_temminckii | Coracias_temminckii | LC | Decreasing | tropical | diurnal | 153.7 | 146790.505 | 10.179 |
| birds | Coracina_atriceps | Coracina_atriceps | LC | Stable | tropical | diurnal | 139 | 39084.458 | 6.971 |
| birds | Coracina_bicolor | NA | NT | Decreasing | NA | NA | NA | NA | NA |
| birds | Coracina_boyeri | Coracina_boyeri | LC | Stable | tropical | diurnal | 66.4 | 604477.417 | 3.049 |
| birds | Coracina_caeruleogrisea | Coracina_caeruleogrisea | LC | Stable | tropical | diurnal | 138.2 | 404850.402 | 3.148 |
| birds | Coracina_caledonica | Coracina_caledonica | LC | Decreasing | tropical | diurnal | 140 | 25323.805 | 6.883 |
| birds | Coracina_fortis | NA | NT | Decreasing | NA | NA | NA | NA | NA |
| birds | Coracina_javensis | Coracina_javensis | LC | Stable | tropical | diurnal | 94.5 | 3632403.85 | 13.395 |
| birds | Coracina_larvata | NA | LC | Decreasing | NA | NA | NA | NA | NA |
| birds | Coracina_leucopygia | NA | LC | Stable | NA | NA | NA | NA | NA |
| birds | Coracina_lineata | Coracina_lineata | LC | Stable | tropical | diurnal | 76 | 643371.297 | 5.286 |
| birds | Coracina_longicauda | Coracina_longicauda | LC | Stable | tropical | diurnal | 101 | 135269.827 | 5.105 |
| birds | Coracina_macei | Coracina_macei | LC | Decreasing | tropical | diurnal | 105 | 2249595.089 | 18.897 |

| | | | | | | | | | |
|---|---|---|---|---|---|---|---|---|---|
| birds | Coracina_maxima | Coracina_maxima | LC | Decreasing | temperate | diurnal | 127 | 6390802.079 | 4.329 |
| birds | Coracina_novaehollandiae | Coracina_novaehollandiae | LC | Decreasing | temperate | diurnal | 118 | 8239647.624 | 4.3 |
| birds | Coracina_papuensis | Coracina_papuensis | LC | Increasing | temperate | diurnal | 63.5 | 4454361.88 | 5.167 |
| birds | Coracina_parvula | Coracina_parvula | LC | Decreasing | tropical | diurnal | 68 | 16787.043 | 6.241 |
| birds | Coracina_personata | NA | LC | Stable | NA | NA | NA | NA | NA |
| birds | Coracina_schistacea | NA | LC | Stable | NA | NA | NA | NA | NA |
| birds | Coracina_striata | Coracina_striata | LC | Decreasing | tropical | diurnal | 111 | 1446124.69 | 9.519 |
| birds | Coracina_temminckii | NA | LC | Stable | NA | NA | NA | NA | NA |
| birds | Coracopsis_nigra | Coracopsis_nigra | LC | Stable | tropical | diurnal | 186.2 | 591110.011 | 8.747 |
| birds | Coracopsis_vasa | Coracopsis_vasa | LC | Decreasing | tropical | diurnal | 525 | 413210.959 | 8.42 |
| birds | Coracornis_raveni | Coracornis_raveni | LC | Stable | tropical | diurnal | 36 | 11498.504 | 4.665 |
| birds | Coragyps_atratus | Coragyps_atratus | LC | Increasing | tropical | diurnal | 1881.7 | 21175692.939 | 7.338 |
| birds | Corapipo_gutturalis | Corapipo_gutturalis | LC | Stable | tropical | diurnal | 8.2 | 1107394.301 | 1.141 |
| birds | Corapipo_leucorrhoa | Corapipo_leucorrhoa | LC | Decreasing | tropical | diurnal | 11.7 | 267106.87 | 9.061 |
| birds | Corcorax_melanorhamphos | Corcorax_melanorhamphos | LC | Stable | temperate | diurnal | 364 | 1979313.446 | 7.915 |
| birds | Cormobates_leucophaea | Cormobates_leucophaea | LC | Stable | temperate | diurnal | 22 | 969111.729 | 9.367 |
| birds | Cormobates_placens | Cormobates_placens | LC | Stable | tropical | diurnal | 15.8 | 139549.582 | 4.978 |
| birds | Corthylio_calendula | Regulus_calendula | LC | Increasing | temperate | diurnal | 6.2 | 16933121.983 | 8.557 |
| birds | Corvus_albicollis | Corvus_albicollis | LC | Decreasing | temperate | diurnal | 894 | 1615598.279 | 12.276 |
| birds | Corvus_albus | Corvus_albus | LC | Stable | tropical | diurnal | 524 | 19544750.218 | 8.932 |
| birds | Corvus_bennetti | Corvus_bennetti | LC | Decreasing | temperate | diurnal | 389 | 5681813.472 | 3.753 |
| birds | Corvus_brachyrhynchos | Corvus_brachyrhynchos | LC | Increasing | temperate | diurnal | 448.8 | 12113678.307 | 10.45 |
| birds | Corvus_capensis | Corvus_capensis | LC | Increasing | tropical | diurnal | 553 | 4036026.483 | 8.506 |
| birds | Corvus_caurinus | Corvus_caurinus | LC | Increasing | temperate | diurnal | 390.8 | 396973.078 | 3.285 |
| birds | Corvus_corax | Corvus_corax | LC | Increasing | temperate | diurnal | 928 | 52342379.533 | 6.106 |
| birds | Corvus_corone | Corvus_corone | LC | Increasing | temperate | diurnal | 570 | 34119496.896 | 8.589 |
| birds | Corvus_coronoides | Corvus_coronoides | LC | Increasing | temperate | diurnal | 644.3 | 4803477.781 | 5.6 |
| birds | Corvus_crassirostris | Corvus_crassirostris | LC | Stable | tropical | diurnal | 1135 | 759238.977 | 11.117 |
| birds | Corvus_cryptoleucus | Corvus_cryptoleucus | LC | Stable | temperate | diurnal | 533.5 | 1508239.498 | 10.024 |
| birds | Corvus_dauuricus | Corvus_dauuricus | LC | Stable | temperate | diurnal | 208 | 7425668.893 | 11.2 |
| birds | Corvus_enca | Corvus_enca | LC | Stable | tropical | diurnal | 251 | 1629657.055 | 9.658 |
| birds | Corvus_florensis | Corvus_florensis | EN | Decreasing | tropical | diurnal | 175 | 9095.62 | 12.47 |
| birds | Corvus_frugilegus | Corvus_frugilegus | LC | Decreasing | temperate | diurnal | 452.1 | 23167955.273 | 10.961 |
| birds | Corvus_fuscicapillus | NA | LC | Decreasing | NA | NA | NA | NA | NA |

| | | | | | | | | | |
|---|---|---|---|---|---|---|---|---|---|
| birds | Corvus_hawaiiensis | NA | EW | not_available | NA | NA | NA | NA | NA |
| birds | Corvus_imparatus | Corvus_imparatus | LC | Decreasing | temperate | diurnal | 221 | 114579.131 | 11.742 |
| birds | Corvus_jamaicensis | Corvus_jamaicensis | NT | Increasing | tropical | diurnal | 338 | 11021.791 | 19.741 |
| birds | Corvus_kubaryi | Corvus_kubaryi | CR | Decreasing | tropical | diurnal | 249.9 | 216.796 | 19.33 |
| birds | Corvus_leucognaphalus | NA | VU | Decreasing | NA | NA | NA | NA | NA |
| birds | Corvus_macrorhynchos | Corvus_macrorhynchos | LC | Stable | temperate | diurnal | 513.1 | 13313192.581 | 13.595 |
| birds | Corvus_meeki | NA | LC | Stable | NA | NA | NA | NA | NA |
| birds | Corvus_mellori | Corvus_mellori | LC | Increasing | temperate | diurnal | 540.4 | 1353458.96 | 8.455 |
| birds | Corvus_monedula | Corvus_monedula | LC | Stable | temperate | diurnal | 246 | 15225329.415 | 12.831 |
| birds | Corvus_moneduloides | Corvus_moneduloides | LC | Stable | tropical | diurnal | 277.4 | 17048.53 | 7.532 |
| birds | Corvus_nasicus | Corvus_nasicus | LC | Stable | tropical | diurnal | 401 | 28429.705 | 13.332 |
| birds | Corvus_orru | Corvus_orru | LC | Increasing | tropical | diurnal | 508 | 5394057.72 | 3.323 |
| birds | Corvus_ossifragus | Corvus_ossifragus | LC | Increasing | temperate | diurnal | 284.6 | 970986.53 | 17.527 |
| birds | Corvus_palmarum | Corvus_palmarum | LC | Decreasing | tropical | diurnal | 289 | 70693.53 | 15.465 |
| birds | Corvus_pectoralis | Corvus_torquatus | VU | Decreasing | temperate | diurnal | 470.5 | 2074732.63 | 18.971 |
| birds | Corvus_rhipidurus | Corvus_rhipidurus | LC | Decreasing | tropical | diurnal | 611 | 2756382.953 | 8.321 |
| birds | Corvus_ruficollis | Corvus_ruficollis | LC | Increasing | temperate | diurnal | 573.5 | 12915376.882 | 4.407 |
| birds | Corvus_sinaloae | Corvus_sinaloae | LC | Stable | temperate | diurnal | 244 | 95101.338 | 6.667 |
| birds | Corvus_splendens | Corvus_splendens | LC | Stable | tropical | diurnal | 292.6 | 4860994.177 | 15.788 |
| birds | Corvus_tasmanicus | Corvus_tasmanicus | LC | Decreasing | temperate | diurnal | 676.3 | 374479.028 | 9.638 |
| birds | Corvus_tristis | Corvus_tristis | LC | Stable | tropical | diurnal | 635 | 693202.247 | 3.207 |
| birds | Corvus_typicus | Corvus_typicus | LC | Decreasing | tropical | diurnal | 175 | 104370.922 | 9.219 |
| birds | Corvus_unicolor | Corvus_unicolor | CR | Decreasing | tropical | diurnal | 175 | 403.846 | 9.063 |
| birds | Corvus_validus | Corvus_validus | NT | Decreasing | tropical | diurnal | 300 | 24724.98 | 6.238 |
| birds | Corvus_woodfordi | Corvus_woodfordi | LC | Stable | tropical | diurnal | 467 | 13034.761 | 3.802 |
| birds | Corydon_sumatranus | Corydon_sumatranus | LC | Decreasing | tropical | diurnal | 140 | 2209826.905 | 9.268 |
| birds | Corydospiza_alaudina | Phrygilus_alaudinus | LC | Stable | tropical | diurnal | 23.9 | 687385.351 | 8.938 |
| birds | Corydospiza_carbonaria | Phrygilus_carbonarius | LC | Stable | temperate | diurnal | 17.9 | 901444.776 | 7.713 |
| birds | Coryphaspiza_melanotis | Coryphaspiza_melanotis | VU | Decreasing | tropical | diurnal | 15.7 | 2184668.562 | 10.339 |
| birds | Coryphistera_alaudina | Coryphistera_alaudina | LC | Stable | temperate | diurnal | 30 | 1447474.656 | 7.471 |
| birds | Coryphospingus_cucullatus | Coryphospingus_cucullatus | LC | Stable | temperate | diurnal | 14.3 | 4910505 | 8.779 |
| birds | Coryphospingus_pileatus | Coryphospingus_pileatus | LC | Stable | tropical | diurnal | 15.3 | 2758347.739 | 8.527 |
| birds | Corythaeola_cristata | Corythaeola_cristata | LC | Stable | tropical | diurnal | 965 | 3896093.974 | 8.151 |
| birds | Corythaixoides_concolor | Corythaixoides_concolor | LC | Stable | tropical | diurnal | 258 | 3808237.762 | 8.211 |

| | | | | | | | | | |
|---|---|---|---|---|---|---|---|---|---|
| birds | Corythaixoides_personatus | Corythaixoides_personatus | LC | Stable | tropical | diurnal | 250 | 52976.098 | 10.317 |
| birds | Corythopis_delalandi | Corythopis_delalandi | LC | Stable | tropical | diurnal | 15 | 2526735.172 | 10.805 |
| birds | Corythopis_torquatus | Corythopis_torquatus | LC | Stable | tropical | diurnal | 14.3 | 6636740.568 | 2.029 |
| birds | Corythornis_cristatus | Alcedo_cristata | LC | Stable | tropical | diurnal | 15.7 | 14363969.716 | 9.447 |
| birds | Corythornis_leucogaster | Alcedo_leucogaster | LC | Stable | tropical | diurnal | 14.5 | 2739779.011 | 7.447 |
| birds | Corythornis_madagascariensis | Ceyx_madagascariensis | LC | Decreasing | tropical | diurnal | 17.7 | 158670.298 | 6.923 |
| birds | Corythornis_vintsioides | Alcedo_vintsioides | LC | Stable | tropical | diurnal | 17.9 | 572332.969 | 8.716 |
| birds | Coscoroba_coscoroba | Coscoroba_coscoroba | LC | Stable | temperate | diurnal | 4294.8 | 4096479.607 | 8.877 |
| birds | Cossypha_albicapillus | Cossypha_albicapilla | LC | Stable | tropical | diurnal | 58.5 | 1269495.304 | 11.844 |
| birds | Cossypha_cyanocampter | Cossypha_cyanocampter | LC | Decreasing | tropical | diurnal | 26.7 | 1817635.07 | 10.581 |
| birds | Cossypha_heinrichi | Cossypha_heinrichi | LC | Decreasing | tropical | diurnal | 60.3 | 6203.477 | 10.365 |
| birds | Cossypha_heuglini | Cossypha_heuglini | LC | Stable | tropical | diurnal | 34.8 | 6366999.372 | 9.137 |
| birds | Cossypha_natalensis | Cossypha_natalensis | LC | Stable | tropical | diurnal | 34.1 | 3643626.525 | 9.767 |
| birds | Cossypha_niveicapilla | Cossypha_niveicapilla | LC | Stable | tropical | diurnal | 36.7 | 5176063.97 | 11.855 |
| birds | Cossypha_semirufa | Cossypha_semirufa | LC | Decreasing | tropical | diurnal | 27.3 | 865714.308 | 11.39 |
| birds | Cossyphicula_isabellae | Cossypha_isabellae | LC | Decreasing | tropical | diurnal | 24 | 51494.149 | 9.722 |
| birds | Cossyphicula_roberti | Cossyphicula_roberti | LC | Decreasing | tropical | diurnal | 34 | 100769.002 | 14.19 |
| birds | Cotinga_amabilis | Cotinga_amabilis | LC | Decreasing | tropical | diurnal | 71.5 | 297726.278 | 9.652 |
| birds | Cotinga_cayana | Cotinga_cayana | LC | Decreasing | tropical | diurnal | 61.8 | 6353282.74 | 1.957 |
| birds | Cotinga_cotinga | Cotinga_cotinga | LC | Decreasing | tropical | diurnal | 54 | 1911359.272 | 1.778 |
| birds | Cotinga_maculata | Cotinga_maculata | CR | Decreasing | tropical | diurnal | 65 | 205824.528 | 11.206 |
| birds | Cotinga_maynana | Cotinga_maynana | LC | Decreasing | tropical | diurnal | 69 | 2547258.17 | 1.608 |
| birds | Cotinga_nattererii | NA | LC | Decreasing | NA | NA | NA | NA | NA |
| birds | Cotinga_ridgwayi | Cotinga_ridgwayi | VU | Decreasing | tropical | diurnal | 57.2 | 14415.574 | 15.438 |
| birds | Coturnicops_exquisitus | NA | VU | Decreasing | NA | NA | NA | NA | NA |
| birds | Coturnicops_notatus | Coturnicops_notatus | LC | Decreasing | temperate | diurnal | 30 | 1592673.321 | 10.985 |
| birds | Coturnicops_noveboracensis | Coturnicops_noveboracensis | LC | Decreasing | temperate | diurnal | 60.9 | 6754791.169 | 11.88 |
| birds | Coturnix_coromandelica | Coturnix_coromandelica | LC | Stable | tropical | diurnal | 73.8 | 2899254.357 | 18.668 |
| birds | Coturnix_coturnix | Coturnix_coturnix | LC | Decreasing | temperate | diurnal | 96.3 | 34416948.306 | 10.929 |
| birds | Coturnix_delegorguei | Coturnix_delegorguei | LC | Stable | tropical | diurnal | 76 | 9153237.05 | 9.442 |
| birds | Coturnix_japonica | Coturnix_japonica | NT | Decreasing | temperate | diurnal | 94.8 | 9041141.353 | 11.98 |
| birds | Coturnix_pectoralis | Coturnix_pectoralis | LC | Increasing | temperate | diurnal | 101.9 | 6323957.075 | 4.706 |
| birds | Coua_caerulea | Coua_caerulea | LC | Decreasing | tropical | diurnal | 244 | 333697.609 | 8.695 |
| birds | Coua_coquereli | Coua_coquereli | LC | Decreasing | tropical | diurnal | 160 | 158107.384 | 8.006 |

| | | | | | | | | | |
|---|---|---|---|---|---|---|---|---|---|
| birds | Coua_cristata | Coua_cristata | LC | Decreasing | tropical | diurnal | 147 | 271871.292 | 8.118 |
| birds | Coua_cursor | Coua_cursor | LC | Decreasing | temperate | diurnal | 110 | 51235.248 | 10.971 |
| birds | Coua_gigas | Coua_gigas | LC | Decreasing | tropical | diurnal | 424 | 149356.189 | 8.078 |
| birds | Coua_reynaudii | Coua_reynaudii | LC | Decreasing | tropical | diurnal | 153 | 320733.342 | 8.726 |
| birds | Coua_ruficeps | Coua_ruficeps | LC | Stable | tropical | diurnal | 180 | 107746.816 | 8.035 |
| birds | Coua_serriana | Coua_serriana | LC | Decreasing | tropical | diurnal | 293 | 62730.5 | 7.016 |
| birds | Coua_verreauxi | Coua_verreauxi | LC | Decreasing | temperate | diurnal | 99 | 20300.519 | 10.454 |
| birds | Cracticus_cassicus | Cracticus_cassicus | LC | Stable | tropical | diurnal | 142 | 554775.589 | 3.195 |
| birds | Cracticus_louisiadensis | NA | NT | Decreasing | NA | NA | NA | NA | NA |
| birds | Cracticus_mentalis | Cracticus_mentalis | LC | Stable | tropical | diurnal | 77 | 218872.782 | 2.76 |
| birds | Cracticus_nigrogularis | Cracticus_nigrogularis | LC | Unknown | temperate | diurnal | 128 | 6347951.063 | 4.353 |
| birds | Cracticus_torquatus | Cracticus_torquatus | LC | Unknown | temperate | diurnal | 82.9 | 6032491.249 | 4.921 |
| birds | Cranioleuca_albicapilla | Cranioleuca_albicapilla | LC | Stable | tropical | diurnal | 21.3 | 18638.245 | 7.409 |
| birds | Cranioleuca_albiceps | Cranioleuca_albiceps | LC | Stable | tropical | diurnal | 17.6 | 42265.843 | 5.791 |
| birds | Cranioleuca_antisiensis | Cranioleuca_antisiensis | LC | Stable | tropical | diurnal | 16.9 | 156556.037 | 8.729 |
| birds | Cranioleuca_baroni | NA | not_available | not_available | NA | NA | NA | NA | NA |
| birds | Cranioleuca_berlepschi | Thripophaga_berlepschi | NT | Decreasing | tropical | diurnal | 25.2 | 5557.825 | 6.143 |
| birds | Cranioleuca_curtata | Cranioleuca_curtata | LC | Decreasing | tropical | diurnal | 17.5 | 362298.095 | 7.129 |
| birds | Cranioleuca_demissa | Cranioleuca_demissa | LC | Stable | tropical | diurnal | 15.2 | 77222.433 | 1.31 |
| birds | Cranioleuca_dissita | NA | LC | Increasing | NA | NA | NA | NA | NA |
| birds | Cranioleuca_erythrops | Cranioleuca_erythrops | LC | Decreasing | tropical | diurnal | 16.9 | 76469.87 | 8.142 |
| birds | Cranioleuca_hellmayri | Cranioleuca_hellmayri | LC | Decreasing | tropical | diurnal | 15 | 6775.393 | 7.214 |
| birds | Cranioleuca_henricae | NA | VU | Decreasing | NA | NA | NA | NA | NA |
| birds | Cranioleuca_marcapatae | Cranioleuca_marcapatae | LC | Stable | tropical | diurnal | 19.8 | 15353.766 | 5.268 |
| birds | Cranioleuca_muelleri | Cranioleuca_muelleri | LC | Decreasing | tropical | diurnal | 14.5 | 115725.425 | 3.107 |
| birds | Cranioleuca_obsoleta | Cranioleuca_obsoleta | LC | Decreasing | temperate | diurnal | 13.4 | 653975.038 | 12.387 |
| birds | Cranioleuca_pallida | Cranioleuca_pallida | LC | Decreasing | tropical | diurnal | 11.5 | 694100.166 | 12.605 |
| birds | Cranioleuca_pyrrhophia | Cranioleuca_pyrrhophia | LC | Stable | temperate | diurnal | 14.9 | 2026269.348 | 7.635 |
| birds | Cranioleuca_semicinerea | Cranioleuca_semicinerea | LC | Decreasing | tropical | diurnal | 15 | 987554.478 | 8.822 |
| birds | Cranioleuca_subcristata | Cranioleuca_subcristata | LC | Stable | tropical | diurnal | 14.7 | 182930.19 | 12.829 |
| birds | Cranioleuca_vulpecula | Cranioleuca_vulpecula | LC | Stable | tropical | diurnal | 18.9 | 340457.777 | 2.322 |
| birds | Cranioleuca_vulpina | Cranioleuca_vulpina | LC | Unknown | tropical | diurnal | 15.7 | 5731362.687 | 5.937 |
| birds | Crax_alberti | NA | CR | Decreasing | NA | NA | NA | NA | NA |
| birds | Crax_alector | Crax_alector | LC | Decreasing | tropical | diurnal | 2872 | 2099461.66 | 1.581 |

| | | | | | | | | | |
|---|---|---|---|---|---|---|---|---|---|
| birds | Crax_blumenbachii | Crax_blumenbachii | EN | Decreasing | tropical | diurnal | 3500 | 148947.316 | 11.247 |
| birds | Crax_daubentoni | Crax_daubentoni | NT | Decreasing | tropical | diurnal | 2499.9 | 342112.862 | 9.902 |
| birds | Crax_fasciolata | Crax_fasciolata | VU | Decreasing | tropical | diurnal | 2600 | 3381019.119 | 7.104 |
| birds | Crax_globulosa | Crax_globulosa | EN | Decreasing | tropical | diurnal | 2500 | 44465.661 | 3.513 |
| birds | Crax_rubra | Crax_rubra | VU | Decreasing | tropical | diurnal | 4133 | 662063.72 | 9.716 |
| birds | Creagrus_furcatus | NA | LC | Unknown | NA | NA | NA | NA | NA |
| birds | Creatophora_cinerea | Creatophora_cinerea | LC | Stable | tropical | diurnal | 73 | 5407365.7 | 8.883 |
| birds | Creurgops_dentatus | Creurgops_dentatus | LC | Decreasing | tropical | diurnal | 19 | 109941.342 | 4.634 |
| birds | Creurgops_verticalis | Creurgops_verticalis | LC | Decreasing | tropical | diurnal | 24 | 97022.082 | 6.913 |
| birds | Crinifer_piscator | Crinifer_piscator | LC | Stable | tropical | diurnal | 370 | 3640485.507 | 12.472 |
| birds | Crinifer_zonurus | Crinifer_zonurus | LC | Stable | tropical | diurnal | 527 | 1014236.613 | 11.045 |
| birds | Criniferoides_leucogaster | Corythaixoides_leucogaster | LC | Stable | tropical | diurnal | 216.2 | 2293621.909 | 9.124 |
| birds | Criniger_barbatus | Criniger_barbatus | LC | Stable | tropical | diurnal | 42.6 | 646336.43 | 15.417 |
| birds | Criniger_calurus | Criniger_calurus | LC | Stable | tropical | diurnal | 37.7 | 3299941.84 | 7.961 |
| birds | Criniger_chloronotus | Criniger_chloronotus | LC | Stable | tropical | diurnal | 45 | 1204957.834 | 5.845 |
| birds | Criniger_ndussumensis | Criniger_ndussumensis | LC | Stable | tropical | diurnal | 23 | 1455617.856 | 5.911 |
| birds | Criniger_olivaceus | Criniger_olivaceus | VU | Decreasing | tropical | diurnal | 28.1 | 239103.584 | 14.646 |
| birds | Crithagra_albogularis | Serinus_albogularis | LC | Stable | temperate | diurnal | 27.1 | 916328.358 | 7.148 |
| birds | Crithagra_ankoberensis | Serinus_ankoberensis | VU | Decreasing | tropical | diurnal | 14.9 | 50697.188 | 13.955 |
| birds | Crithagra_atrogularis | Serinus_atrogularis | LC | Stable | tropical | diurnal | 11.4 | 3009542.679 | 8.156 |
| birds | Crithagra_buchanani | NA | LC | Stable | NA | NA | NA | NA | NA |
| birds | Crithagra_burtoni | Serinus_burtoni | LC | Stable | tropical | diurnal | 30.5 | 400739.278 | 13.459 |
| birds | Crithagra_capistrata | NA | LC | Stable | NA | NA | NA | NA | NA |
| birds | Crithagra_citrinelloides | Serinus_citrinelloides | LC | Stable | tropical | diurnal | 13 | 555034.987 | 12.974 |
| birds | Crithagra_citrinipectus | Serinus_citrinipectus | LC | Decreasing | tropical | diurnal | 11.2 | 583770.244 | 10.343 |
| birds | Crithagra_concolor | Neospiza_concolor | CR | Decreasing | tropical | diurnal | 50 | 204.88 | 6.681 |
| birds | Crithagra_donaldsoni | Serinus_donaldsoni | LC | Stable | tropical | diurnal | 24 | 604511.84 | 7.907 |
| birds | Crithagra_dorsostriata | Serinus_dorsostriatus | LC | Stable | tropical | diurnal | 15.1 | 864663.904 | 11.189 |
| birds | Crithagra_flavigula | NA | EN | Decreasing | NA | NA | NA | NA | NA |
| birds | Crithagra_flaviventris | Serinus_flaviventris | LC | Stable | temperate | diurnal | 16.2 | 1693983.546 | 7.388 |
| birds | Crithagra_frontalis | Serinus_frontalis | LC | Stable | tropical | diurnal | 12.3 | 238675.246 | 15.765 |
| birds | Crithagra_gularis | Serinus_gularis | LC | Stable | temperate | diurnal | 16 | 1507366.086 | 11.063 |
| birds | Crithagra_hyposticta | Serinus_hypostictus | LC | Stable | tropical | diurnal | 13 | 160632.877 | 14.937 |
| birds | Crithagra_koliensis | Serinus_koliensis | LC | Decreasing | tropical | diurnal | 13.6 | 54614.042 | 20.143 |

| | | | | | | | | | |
|---|---|---|---|---|---|---|---|---|---|
| birds | Crithagra_leucoptera | Serinus_leucopterus | NT | Decreasing | temperate | diurnal | 22.1 | 82476.235 | 10.51 |
| birds | Crithagra_leucopygia | Serinus_leucopygius | LC | Stable | tropical | diurnal | 10.8 | 3039637.603 | 11.536 |
| birds | Crithagra_melanochroa | NA | LC | Decreasing | NA | NA | NA | NA | NA |
| birds | Crithagra_menachensis | NA | LC | Stable | NA | NA | NA | NA | NA |
| birds | Crithagra_mennelli | Serinus_mennelli | LC | Stable | tropical | diurnal | 15.4 | 1495142.191 | 9.36 |
| birds | Crithagra_mozambica | Serinus_mozambicus | LC | Decreasing | tropical | diurnal | 11.9 | 9549283.637 | 9.886 |
| birds | Crithagra_reichardi | Serinus_reichardi | LC | Stable | tropical | diurnal | 15.4 | 580348.354 | 9.29 |
| birds | Crithagra_reichenowi | NA | LC | Stable | NA | NA | NA | NA | NA |
| birds | Crithagra_rothschildi | Serinus_rothschildi | LC | Stable | temperate | diurnal | 14 | 129560.584 | 9.669 |
| birds | Crithagra_rufobrunnea | Serinus_rufobrunneus | LC | Stable | tropical | diurnal | 25.9 | 928.928 | 13.476 |
| birds | Crithagra_scotops | Serinus_scotops | LC | Stable | temperate | diurnal | 14.2 | 184665.553 | 12.431 |
| birds | Crithagra_striolata | Serinus_striolatus | LC | Stable | tropical | diurnal | 22.4 | 867600.667 | 13.524 |
| birds | Crithagra_sulphurata | Serinus_sulphuratus | LC | Stable | tropical | diurnal | 27.9 | 1869165.541 | 11.277 |
| birds | Crithagra_symonsi | Serinus_symonsi | LC | Stable | temperate | diurnal | 15.8 | 36985.628 | 11.815 |
| birds | Crithagra_totta | Serinus_totta | LC | Stable | temperate | diurnal | 13.4 | 85526.402 | 11.317 |
| birds | Crithagra_tristriata | Serinus_tristriatus | LC | Stable | tropical | diurnal | 15.5 | 299348.436 | 14.057 |
| birds | Crithagra_xantholaema | Serinus_xantholaemus | NT | Decreasing | tropical | diurnal | 11.8 | 54048.086 | 7.576 |
| birds | Crithagra_xanthopygia | Serinus_xanthopygius | LC | Stable | tropical | diurnal | 15.4 | 103232.062 | 13.662 |
| birds | Crossleyia_tenebrosa | Bernieria_tenebrosa | DD | Decreasing | tropical | diurnal | 21.4 | 15950.292 | 5.911 |
| birds | Crossleyia_xanthophrys | Crossleyia_xanthophrys | LC | Decreasing | tropical | diurnal | 19.9 | 22717.104 | 5.338 |
| birds | Crossoptilon_auritum | Crossoptilon_auritum | LC | Stable | temperate | diurnal | 1768.3 | 512906.047 | 9.256 |
| birds | Crossoptilon_crossoptilon | Crossoptilon_crossoptilon | NT | Decreasing | temperate | diurnal | 2075.3 | 171921.781 | 6.682 |
| birds | Crossoptilon_mantchuricum | Crossoptilon_mantchuricum | VU | Decreasing | temperate | diurnal | 1860.9 | 188621.536 | 14.144 |
| birds | Crotophaga_ani | Crotophaga_ani | LC | Stable | tropical | diurnal | 110.1 | 14119059.349 | 6.034 |
| birds | Crotophaga_major | Crotophaga_major | LC | Stable | tropical | diurnal | 148.3 | 12117795.166 | 5.475 |
| birds | Crotophaga_sulcirostris | Crotophaga_sulcirostris | LC | Increasing | tropical | diurnal | 82 | 2717920.777 | 10.435 |
| birds | Crypsirina_cucullata | Crypsirina_cucullata | NT | Decreasing | tropical | diurnal | 132.5 | 112222.325 | 16.467 |
| birds | Crypsirina_temia | Crypsirina_temia | LC | Stable | tropical | diurnal | 129.5 | 1282466.775 | 12.954 |
| birds | Cryptillas_victorini | Bradypterus_victorini | LC | Unknown | temperate | diurnal | 10 | 78210.995 | 11.134 |
| birds | Cryptoleucopteryx_plumbea | Leucopternis_plumbeus | NT | Decreasing | tropical | diurnal | 482 | 162116.727 | 6.663 |
| birds | Cryptolybia_olivacea | Stactolaema_olivacea | LC | Decreasing | tropical | diurnal | 49.4 | 155265.952 | 13.465 |
| birds | Cryptomicroeca_flaviventris | Eopsaltria_flaviventris | LC | Stable | tropical | diurnal | 13 | 16068.839 | 7.459 |
| birds | Cryptopezus_nattereri | Hylopezus_nattereri | LC | Stable | temperate | diurnal | 32 | 493986.995 | 12.774 |
| birds | Cryptophaps_poecilorrhoa | Cryptophaps_poecilorrhoa | LC | Decreasing | tropical | diurnal | 467 | 97468.244 | 6.691 |

| | | | | | | | | | |
|---|---|---|---|---|---|---|---|---|---|
| birds | Cryptopipo_holochlora | Xenopipo_holochlora | LC | Decreasing | tropical | diurnal | 14.9 | 336141.322 | 3.836 |
| birds | Cryptospiza_jacksoni | Cryptospiza_jacksoni | LC | Stable | tropical | diurnal | 13 | 75938.17 | 17.15 |
| birds | Cryptospiza_reichenovii | Cryptospiza_reichenovii | LC | Stable | tropical | diurnal | 12.4 | 435889.108 | 12.9 |
| birds | Cryptospiza_salvadorii | Cryptospiza_salvadorii | LC | Stable | tropical | diurnal | 12.2 | 258138.36 | 12.657 |
| birds | Cryptospiza_shelleyi | Cryptospiza_shelleyi | EN | Decreasing | tropical | diurnal | 17.6 | 46042.017 | 14.161 |
| birds | Cryptosylvicola_randrianasoloi | Cryptosylvicola_randrianasoloi | LC | Decreasing | tropical | diurnal | 8.6 | 116215.04 | 7.904 |
| birds | Crypturellus_bartletti | Crypturellus_bartletti | LC | Decreasing | tropical | diurnal | 241 | 1611496.597 | 1.69 |
| birds | Crypturellus_boucardi | Crypturellus_boucardi | VU | Decreasing | tropical | diurnal | 435.6 | 334426.961 | 9.247 |
| birds | Crypturellus_brevirostris | Crypturellus_brevirostris | LC | Decreasing | tropical | diurnal | 221 | 833118.737 | 0.784 |
| birds | Crypturellus_casiquiare | Crypturellus_casiquiare | LC | Decreasing | tropical | diurnal | 250 | 86537.957 | 1.114 |
| birds | Crypturellus_cinereus | Crypturellus_cinereus | LC | Decreasing | tropical | diurnal | 506.6 | 5924779.184 | 1.894 |
| birds | Crypturellus_cinnamomeus | Crypturellus_cinnamomeus | LC | Decreasing | tropical | diurnal | 422 | 695817.1 | 10.941 |
| birds | Crypturellus_kerriae | NA | NT | Decreasing | NA | NA | NA | NA | NA |
| birds | Crypturellus_noctivagus | Crypturellus_noctivagus | NT | Decreasing | tropical | diurnal | 562 | 1469564.579 | 10.064 |
| birds | Crypturellus_obsoletus | Crypturellus_obsoletus | LC | Decreasing | tropical | diurnal | 443.4 | 2652131.252 | 9.105 |
| birds | Crypturellus_parvirostris | Crypturellus_parvirostris | LC | Decreasing | tropical | diurnal | 199 | 6659535.622 | 7.254 |
| birds | Crypturellus_soui | Crypturellus_soui | LC | Decreasing | tropical | diurnal | 216.2 | 11151222.759 | 4.795 |
| birds | Crypturellus_tataupa | Crypturellus_tataupa | LC | Decreasing | tropical | diurnal | 218.8 | 4855861.195 | 9.007 |
| birds | Crypturellus_undulatus | Crypturellus_undulatus | LC | Decreasing | tropical | diurnal | 564.4 | 8533511.589 | 4.247 |
| birds | Crypturellus_variegatus | Crypturellus_variegatus | LC | Decreasing | tropical | diurnal | 378 | 5436849.917 | 1.797 |
| birds | Cuculus_canorus | Cuculus_canorus | LC | Decreasing | temperate | diurnal | 111.4 | 49981665.273 | 9.278 |
| birds | Cuculus_clamosus | Cuculus_clamosus | LC | Stable | tropical | diurnal | 78.8 | 13974312.781 | 9.259 |
| birds | Cuculus_crassirostris | NA | LC | Decreasing | NA | NA | NA | NA | NA |
| birds | Cuculus_gularis | Cuculus_gularis | LC | Stable | tropical | diurnal | 103 | 12452908.779 | 9.571 |
| birds | Cuculus_micropterus | Cuculus_micropterus | LC | Decreasing | temperate | diurnal | 88.8 | 10510479.755 | 14.169 |
| birds | Cuculus_poliocephalus | Cuculus_poliocephalus | LC | Stable | temperate | diurnal | 46.6 | 8851019.005 | 14.405 |
| birds | Cuculus_rochii | Cuculus_rochii | LC | Stable | tropical | diurnal | 62.7 | 2686934.396 | 10.537 |
| birds | Cuculus_saturatus | Cuculus_saturatus | LC | Decreasing | tropical | diurnal | 87 | 7972105.86 | 12.052 |
| birds | Cuculus_solitarius | Cuculus_solitarius | LC | Stable | tropical | diurnal | 76.7 | 12082464.266 | 9.236 |
| birds | Culicicapa_ceylonensis | Culicicapa_ceylonensis | LC | Stable | tropical | diurnal | 7.7 | 6901649.267 | 12.972 |
| birds | Culicicapa_helianthea | Culicicapa_helianthea | LC | Decreasing | tropical | diurnal | 8.4 | 376783.238 | 11.794 |
| birds | Culicivora_caudacuta | Culicivora_caudacuta | VU | Decreasing | tropical | diurnal | 6.1 | 2101399.614 | 10.648 |
| birds | Curaeus_curaeus | Curaeus_curaeus | LC | Stable | temperate | diurnal | 83.7 | 1036456.268 | 5.74 |
| birds | Curruca_boehmi | Sylvia_boehmi | LC | Stable | tropical | diurnal | 14 | 757783.639 | 10.595 |

| | | | | | | | | | |
|---|---|---|---|---|---|---|---|---|---|
| birds | Curruca_buryi | Sylvia_buryi | NT | Decreasing | tropical | diurnal | 22 | 62421.27 | 12.569 |
| birds | Curruca_communis | Sylvia_communis | LC | Increasing | tropical | diurnal | 15.1 | 42176814.383 | 8.689 |
| birds | Curruca_conspicillata | Sylvia_conspicillata | LC | Unknown | temperate | diurnal | 9 | 4545002.928 | 7.243 |
| birds | Curruca_deserticola | Sylvia_deserticola | LC | Stable | temperate | diurnal | 8.6 | 3004118.253 | 3.663 |
| birds | Curruca_layardi | Sylvia_layardi | LC | Stable | temperate | diurnal | 14.3 | 780211.284 | 7.221 |
| birds | Curruca_leucomelaena | Sylvia_leucomelaena | LC | Decreasing | tropical | diurnal | 13.9 | 547018.243 | 7.657 |
| birds | Curruca_lugens | Sylvia_lugens | LC | Decreasing | tropical | diurnal | 14.9 | 401532.968 | 11.943 |
| birds | Curruca_melanocephala | Sylvia_melanocephala | LC | Increasing | tropical | diurnal | 11.7 | 4224670.015 | 10.302 |
| birds | Curruca_melanothorax | Sylvia_melanothorax | LC | Decreasing | temperate | diurnal | 11.4 | 149419.083 | 8.694 |
| birds | Curruca_mystacea | Sylvia_mystacea | LC | Stable | temperate | diurnal | 9.9 | 7953271.299 | 7.734 |
| birds | Curruca_nisoria | Sylvia_nisoria | LC | Stable | temperate | diurnal | 22.5 | 12451191.898 | 10.932 |
| birds | Curruca_ruppeli | Sylvia_rueppelli | LC | Decreasing | temperate | diurnal | 12.8 | 5404863.934 | 4.464 |
| birds | Curruca_subcoerulea | Sylvia_subcaerulea | LC | Stable | tropical | diurnal | 14.6 | 2960791.883 | 7.487 |
| birds | Curruca_undata | Sylvia_undata | NT | Decreasing | temperate | diurnal | 10.8 | 1420763.346 | 16.723 |
| birds | Cursorius_coromandelicus | NA | LC | Stable | NA | NA | NA | NA | NA |
| birds | Cursorius_cursor | Cursorius_cursor | LC | Decreasing | tropical | diurnal | 138 | 10509601.04 | 6.569 |
| birds | Cursorius_rufus | Cursorius_rufus | LC | Decreasing | temperate | diurnal | 75 | 1392070.573 | 6.825 |
| birds | Cutia_nipalensis | Cutia_nipalensis | LC | Stable | tropical | diurnal | 47.2 | 848554.474 | 8.13 |
| birds | Cyanerpes_caeruleus | Cyanerpes_caeruleus | LC | Decreasing | tropical | diurnal | 12 | 6673757.38 | 2.563 |
| birds | Cyanerpes_cyaneus | Cyanerpes_cyaneus | LC | Decreasing | tropical | diurnal | 14 | 8492094.204 | 4.078 |
| birds | Cyanerpes_lucidus | Cyanerpes_lucidus | LC | Stable | tropical | diurnal | 11.4 | 234923.274 | 8.218 |
| birds | Cyanerpes_nitidus | Cyanerpes_nitidus | LC | Decreasing | tropical | diurnal | 9 | 2672995.371 | 1.473 |
| birds | Cyanicterus_cyanicterus | Cyanicterus_cyanicterus | LC | Decreasing | tropical | diurnal | 34 | 422685.667 | 1.53 |
| birds | Cyanistes_caeruleus | Parus_caeruleus | LC | Increasing | temperate | diurnal | 11.1 | 8770881.493 | 16.132 |
| birds | Cyanistes_cyanus | Parus_cyanus | LC | Decreasing | temperate | diurnal | 14.6 | 8153289.292 | 7.363 |
| birds | Cyanistes_teneriffae | Parus_teneriffae | LC | Stable | temperate | diurnal | 10 | 624090.327 | 14.289 |
| birds | Cyanochen_cyanoptera | Cyanochen_cyanoptera | NT | Decreasing | tropical | diurnal | 1520 | 361268.408 | 12.692 |
| birds | Cyanocitta_cristata | Cyanocitta_cristata | LC | Stable | temperate | diurnal | 88 | 6809850.434 | 14.044 |
| birds | Cyanocitta_stelleri | Cyanocitta_stelleri | LC | Increasing | temperate | diurnal | 128 | 3560822.245 | 7.702 |
| birds | Cyanocompsa_parellina | Cyanocompsa_parellina | LC | Decreasing | tropical | diurnal | 15.6 | 577372.056 | 10.073 |
| birds | Cyanocorax_affinis | Cyanocorax_affinis | LC | Stable | tropical | diurnal | 212 | 473163.738 | 11.954 |
| birds | Cyanocorax_beecheii | Cyanocorax_beecheii | LC | Decreasing | temperate | diurnal | 193 | 86050.248 | 6.099 |
| birds | Cyanocorax_cayanus | Cyanocorax_cayanus | LC | Stable | tropical | diurnal | 175 | 1160987.207 | 1.504 |
| birds | Cyanocorax_chrysops | Cyanocorax_chrysops | LC | Decreasing | tropical | diurnal | 166 | 2883955.963 | 8.512 |

| | | | | | | | | | |
|---|---|---|---|---|---|---|---|---|---|
| birds | Cyanocorax_coeruleus | Cyanocorax_caeruleus | NT | Decreasing | temperate | diurnal | 272 | 465283.81 | 12.248 |
| birds | Cyanocorax_colliei | Calocitta_colliei | LC | Decreasing | temperate | diurnal | 234 | 158921.03 | 7.289 |
| birds | Cyanocorax_cristatellus | Cyanocorax_cristatellus | LC | Stable | tropical | diurnal | 178 | 3037012.195 | 8.586 |
| birds | Cyanocorax_cyanomelas | Cyanocorax_cyanomelas | LC | Stable | tropical | diurnal | 207 | 1680479.72 | 7.45 |
| birds | Cyanocorax_cyanopogon | Cyanocorax_cyanopogon | LC | Decreasing | tropical | diurnal | 146 | 3068149.169 | 8.166 |
| birds | Cyanocorax_dickeyi | Cyanocorax_dickeyi | NT | Decreasing | tropical | diurnal | 181 | 13431.358 | 2.763 |
| birds | Cyanocorax_formosus | Calocitta_formosa | LC | Stable | tropical | diurnal | 210 | 293348.648 | 12.29 |
| birds | Cyanocorax_heilprini | Cyanocorax_heilprini | LC | Stable | tropical | diurnal | 172 | 140727.061 | 0.908 |
| birds | Cyanocorax_melanocyaneus | Cyanocorax_melanocyaneus | LC | Stable | tropical | diurnal | 101 | 123771.877 | 13.161 |
| birds | Cyanocorax_morio | Cyanocorax_morio | LC | Stable | tropical | diurnal | 204 | 670090.878 | 9.859 |
| birds | Cyanocorax_mystacalis | Cyanocorax_mystacalis | LC | Decreasing | tropical | diurnal | 154.4 | 81043.679 | 10.13 |
| birds | Cyanocorax_sanblasianus | Cyanocorax_sanblasianus | LC | Decreasing | tropical | diurnal | 109 | 67589.942 | 8.043 |
| birds | Cyanocorax_violaceus | Cyanocorax_violaceus | LC | Stable | tropical | diurnal | 262 | 3381230.339 | 2.252 |
| birds | Cyanocorax_yncas | Cyanocorax_yncas | LC | Increasing | tropical | diurnal | 78.5 | 1092392.739 | 9.407 |
| birds | Cyanocorax_yucatanicus | Cyanocorax_yucatanicus | LC | Stable | tropical | diurnal | 118 | 187835.71 | 7.356 |
| birds | Cyanoderma_chrysaeum | Stachyris_chrysaea | LC | Decreasing | tropical | diurnal | 9 | 1020720.191 | 7.942 |
| birds | Cyanoderma_erythropterum | Stachyris_erythroptera | LC | Decreasing | tropical | diurnal | 12.6 | 593405.278 | 10.853 |
| birds | Cyanoderma_melanothorax | Stachyris_melanothorax | LC | Decreasing | tropical | diurnal | 16 | 62512.286 | 18.584 |
| birds | Cyanoderma_pyrrhops | Stachyris_pyrrhops | LC | Stable | temperate | diurnal | 9.6 | 173970.871 | 13.301 |
| birds | Cyanoderma_ruficeps | Stachyris_ruficeps | LC | Decreasing | temperate | diurnal | 10.3 | 2205806.998 | 12.323 |
| birds | Cyanoderma_rufifrons | Stachyris_rufifrons | LC | Stable | tropical | diurnal | 10.6 | 2386114.606 | 9.439 |
| birds | Cyanograucalus_azureus | Coracina_azurea | LC | Decreasing | tropical | diurnal | 46.1 | 1644557.765 | 9.165 |
| birds | Cyanolanius_madagascarinus | Cyanolanius_madagascarinus | LC | Unknown | tropical | diurnal | 22.3 | 445373.261 | 8.099 |
| birds | Cyanoliseus_patagonus | Cyanoliseus_patagonus | LC | Decreasing | temperate | diurnal | 277.9 | 1217792.538 | 7.346 |
| birds | Cyanoloxia_brissonii | Cyanocompsa_brissonii | LC | Unknown | tropical | diurnal | 27.5 | 5851913.647 | 9.527 |
| birds | Cyanoloxia_cyanoides | Cyanocompsa_cyanoides | LC | Decreasing | tropical | diurnal | 32.5 | 877385.202 | 10.669 |
| birds | Cyanoloxia_glaucocaerulea | Cyanoloxia_glaucocaerulea | LC | Decreasing | temperate | diurnal | 18.2 | 1195239.944 | 12.298 |
| birds | Cyanolyca_argentigula | Cyanolyca_argentigula | LC | Stable | tropical | diurnal | 200 | 3924.603 | 5.019 |
| birds | Cyanolyca_armillata | Cyanolyca_armillata | LC | Stable | tropical | diurnal | 200 | 53723.976 | 13.468 |
| birds | Cyanolyca_cucullata | Cyanolyca_cucullata | LC | Decreasing | tropical | diurnal | 101.3 | 105285.426 | 11.292 |
| birds | Cyanolyca_mirabilis | Cyanolyca_mirabilis | VU | Decreasing | tropical | diurnal | 52.4 | 11137.84 | 5.373 |
| birds | Cyanolyca_nanus | Cyanolyca_nana | NT | Decreasing | tropical | diurnal | 41 | 4670.683 | 7.503 |
| birds | Cyanolyca_pulchra | NA | NT | Decreasing | NA | NA | NA | NA | NA |
| birds | Cyanolyca_pumilo | Cyanolyca_pumilo | LC | Decreasing | tropical | diurnal | 47 | 103163.244 | 12.148 |

| | | | | | | | | | |
|---|---|---|---|---|---|---|---|---|---|
| birds | Cyanolyca_turcosa | Cyanolyca_turcosa | LC | Stable | tropical | diurnal | 97.5 | 62126.944 | 10.345 |
| birds | Cyanolyca_viridicyanus | Cyanolyca_viridicyanus | LC | Stable | tropical | diurnal | 119.8 | 197888.285 | 5.349 |
| birds | Cyanomitra_alinae | Nectarinia_alinae | LC | Decreasing | tropical | diurnal | 12.6 | 114321.465 | 15.435 |
| birds | Cyanomitra_bannermani | Nectarinia_bannermani | LC | Stable | tropical | diurnal | 14 | 454416.957 | 6.081 |
| birds | Cyanomitra_cyanolaema | Nectarinia_cyanolaema | LC | Stable | tropical | diurnal | 16.8 | 3142385.637 | 7.608 |
| birds | Cyanomitra_olivacea | Nectarinia_olivacea | LC | Stable | tropical | diurnal | 9.7 | 5394958.819 | 9.311 |
| birds | Cyanomitra_oritis | Nectarinia_oritis | LC | Decreasing | tropical | diurnal | 11.2 | 65561.745 | 9.036 |
| birds | Cyanomitra_verreauxii | Nectarinia_veroxii | LC | Stable | temperate | diurnal | 11.5 | 155165.635 | 12.871 |
| birds | Cyanomitra_verticalis | Nectarinia_verticalis | LC | Stable | tropical | diurnal | 14.4 | 5373209.669 | 8.59 |
| birds | Cyanopica_cyanus | Cyanopica_cyanus | LC | Increasing | temperate | diurnal | 95.9 | 4748269.368 | 12.099 |
| birds | Cyanopsitta_spixii | NA | EW | not_available | NA | NA | NA | NA | NA |
| birds | Cyanoptila_cyanomelana | Cyanoptila_cyanomelana | LC | Stable | temperate | diurnal | 22.5 | 3159304.218 | 13.088 |
| birds | Cyanoramphus_auriceps | Cyanoramphus_auriceps | NT | Decreasing | temperate | diurnal | 45.6 | 65448.207 | 3.937 |
| birds | Cyanoramphus_forbesi | NA | VU | Stable | NA | NA | NA | NA | NA |
| birds | Cyanoramphus_malherbi | Cyanoramphus_malherbi | CR | Stable | temperate | diurnal | 46.7 | 46289.791 | 4.279 |
| birds | Cyanoramphus_novaezelandiae | Cyanoramphus_novaezelandiae | LC | Decreasing | temperate | diurnal | 90.2 | 285769.676 | 7.974 |
| birds | Cyanoramphus_unicolor | NA | VU | Stable | NA | NA | NA | NA | NA |
| birds | Cyclarhis_gujanensis | Cyclarhis_gujanensis | LC | Decreasing | tropical | diurnal | 28.8 | 13485715.12 | 6.445 |
| birds | Cyclarhis_nigrirostris | Cyclarhis_nigrirostris | LC | Decreasing | tropical | diurnal | 31.9 | 83596.158 | 10.557 |
| birds | Cyclopsitta_diophthalma | Cyclopsitta_diophthalma | LC | Stable | tropical | diurnal | 41.8 | 448013.861 | 3.405 |
| birds | Cyclopsitta_gulielmitertii | Cyclopsitta_gulielmitertii | LC | Stable | tropical | diurnal | 31 | 30177.417 | 4.517 |
| birds | Cygnus_atratus | Cygnus_atratus | LC | Stable | temperate | diurnal | 5623.2 | 7027995.144 | 5.171 |
| birds | Cygnus_buccinator | Cygnus_buccinator | LC | Increasing | temperate | diurnal | 11071.1 | 1616822.382 | 5.505 |
| birds | Cygnus_columbianus | Cygnus_columbianus | LC | Unknown | temperate | diurnal | 6378 | 13948485.914 | 7.782 |
| birds | Cygnus_cygnus | Cygnus_cygnus | LC | Unknown | temperate | diurnal | 9350 | 19300590.683 | 5.533 |
| birds | Cygnus_melancoryphus | Cygnus_melancoryphus | LC | Stable | temperate | diurnal | 4711.7 | 3619491.762 | 8.909 |
| birds | Cygnus_olor | Cygnus_olor | LC | Increasing | temperate | diurnal | 10682 | 6112278.315 | 14.573 |
| birds | Cymbilaimus_lineatus | Cymbilaimus_lineatus | LC | Decreasing | tropical | diurnal | 35.8 | 6643566.156 | 2.35 |
| birds | Cymbilaimus_sanctaemariae | Cymbilaimus_sanctaemariae | LC | Decreasing | tropical | diurnal | 28.7 | 494689.317 | 2.586 |
| birds | Cymbirhynchus_macrorhynchos | Cymbirhynchus_macrorhynchos | LC | Decreasing | tropical | diurnal | 59.4 | 1624283.494 | 8.47 |
| birds | Cynanthus_latirostris | Cynanthus_latirostris | LC | Stable | temperate | diurnal | 3.1 | 990017.277 | 8.924 |
| birds | Cyornis_banyumas | NA | CR | Decreasing | NA | NA | NA | NA | NA |
| birds | Cyornis_brunneatus | Rhinomyias_brunneatus | VU | Decreasing | temperate | diurnal | 18.1 | 677461.884 | 12.606 |
| birds | Cyornis_caerulatus | Cyornis_caerulatus | VU | Decreasing | tropical | diurnal | 17.8 | 963047.211 | 7.657 |

| | | | | | | | | | |
|---|---|---|---|---|---|---|---|---|---|
| birds | Cyornis_colonus | Rhinomyias_colonus | NT | Decreasing | tropical | diurnal | 23.1 | 3612.357 | 6.111 |
| birds | Cyornis_concretus | Cyornis_concretus | LC | Decreasing | tropical | diurnal | 24.5 | 866448.98 | 7.052 |
| birds | Cyornis_hainanus | Cyornis_hainanus | LC | Decreasing | tropical | diurnal | 14.4 | 1595419.542 | 12.237 |
| birds | Cyornis_herioti | Cyornis_herioti | NT | Decreasing | tropical | diurnal | 19.6 | 57832.702 | 10.976 |
| birds | Cyornis_lemprieri | Cyornis_lemprieri | NT | Decreasing | tropical | diurnal | 21.5 | 10987.029 | 8.925 |
| birds | Cyornis_olivaceus | Rhinomyias_olivaceus | LC | Decreasing | tropical | diurnal | 15.7 | 595916.663 | 12.472 |
| birds | Cyornis_omissus | Cyornis_omissus | LC | Stable | tropical | diurnal | 18 | 162644.61 | 9.507 |
| birds | Cyornis_oscillans | NA | not_available | not_available | NA | NA | NA | NA | NA |
| birds | Cyornis_pallidipes | Cyornis_pallipes | LC | Decreasing | tropical | diurnal | 19 | 52751.567 | 13.272 |
| birds | Cyornis_poliogenys | NA | LC | Decreasing | NA | NA | NA | NA | NA |
| birds | Cyornis_rubeculoides | Cyornis_rubeculoides | LC | Stable | tropical | diurnal | 14.2 | 2120972.34 | 14.965 |
| birds | Cyornis_ruckii | NA | CR | Unknown | NA | NA | NA | NA | NA |
| birds | Cyornis_ruficauda | Rhinomyias_ruficauda | LC | Decreasing | tropical | diurnal | 17.3 | 81701.796 | 10.348 |
| birds | Cyornis_rufigastra | Cyornis_rufigastra | LC | Decreasing | tropical | diurnal | 18.3 | 429957.486 | 12.787 |
| birds | Cyornis_superbus | NA | LC | Decreasing | NA | NA | NA | NA | NA |
| birds | Cyornis_tickelliae | Cyornis_tickelliae | LC | Stable | tropical | diurnal | 16.2 | 2366894.589 | 18.825 |
| birds | Cyornis_turcosus | NA | NT | Decreasing | NA | NA | NA | NA | NA |
| birds | Cyornis_umbratilis | Rhinomyias_umbratilis | NT | Decreasing | tropical | diurnal | 18.2 | 1066649.101 | 8.377 |
| birds | Cyornis_unicolor | Cyornis_unicolor | LC | Decreasing | tropical | diurnal | 21 | 2146298.644 | 10.108 |
| birds | Cyphorhinus_arada | Cyphorhinus_arada | LC | Decreasing | tropical | diurnal | 20.9 | 4458849.324 | 1.576 |
| birds | Cyphorhinus_phaeocephalus | Cyphorhinus_phaeocephalus | LC | Decreasing | tropical | diurnal | 24.6 | 318922.639 | 9.116 |
| birds | Cyphorhinus_thoracicus | Cyphorhinus_thoracicus | LC | Decreasing | tropical | diurnal | 31.3 | 58720.021 | 4.817 |
| birds | Cyphos_macrodactylus | Bucco_macrodactylus | LC | Decreasing | tropical | diurnal | 25 | 3329953.343 | 1.645 |
| birds | Cypseloides_cherriei | Cypseloides_cherriei | LC | Decreasing | tropical | cathemeral | 22.1 | 277365.449 | 9.84 |
| birds | Cypseloides_cryptus | Cypseloides_cryptus | LC | Decreasing | tropical | diurnal | 34.5 | 413123.998 | 6.367 |
| birds | Cypseloides_fumigatus | Cypseloides_fumigatus | LC | Decreasing | tropical | diurnal | 42 | 2325060.024 | 11.566 |
| birds | Cypseloides_lemosi | Cypseloides_lemosi | LC | Decreasing | tropical | diurnal | 28.1 | 3577.911 | 22.269 |
| birds | Cypseloides_niger | Cypseloides_niger | VU | Decreasing | temperate | diurnal | 39.7 | 2153809.267 | 9.645 |
| birds | Cypseloides_senex | Cypseloides_senex | LC | Stable | tropical | diurnal | 82.3 | 2015575.959 | 9.864 |
| birds | Cypsnagra_hirundinacea | Cypsnagra_hirundinacea | LC | Stable | tropical | diurnal | 31.4 | 3217340.784 | 8.109 |
| birds | Cyrtonyx_montezumae | Cyrtonyx_montezumae | LC | Decreasing | temperate | diurnal | 185.4 | 754065.67 | 8.885 |
| birds | Cyrtonyx_ocellatus | Cyrtonyx_ocellatus | VU | Decreasing | tropical | diurnal | 199.2 | 130826.394 | 11.648 |
| birds | Dacelo_gaudichaud | Dacelo_gaudichaud | LC | Stable | tropical | diurnal | 143 | 671238.413 | 3.124 |
| birds | Dacelo_leachii | Dacelo_leachii | LC | Unknown | tropical | diurnal | 308 | 2183194.4 | 3.693 |

| | | | | | | | | | |
|---|---|---|---|---|---|---|---|---|---|
| birds | Dacelo_novaeguineae | Dacelo_novaeguineae | LC | Stable | temperate | diurnal | 333.8 | 2766080.001 | 7.446 |
| birds | Dacelo_rex | Clytoceyx_rex | LC | Stable | tropical | cathemeral | 242 | 340625.845 | 3.689 |
| birds | Dacelo_tyro | Dacelo_tyro | LC | Stable | tropical | diurnal | 145.9 | 36878.494 | 3.721 |
| birds | Dacnis_albiventris | Dacnis_albiventris | LC | Stable | tropical | diurnal | 11.2 | 1044010.155 | 1.8 |
| birds | Dacnis_berlepschi | Dacnis_berlepschi | VU | Decreasing | tropical | diurnal | 12.8 | 19193.451 | 8.563 |
| birds | Dacnis_cayana | Dacnis_cayana | LC | Decreasing | tropical | diurnal | 13 | 12058770.432 | 5.416 |
| birds | Dacnis_egregia | NA | LC | Decreasing | NA | NA | NA | NA | NA |
| birds | Dacnis_flaviventer | Dacnis_flaviventer | LC | Decreasing | tropical | diurnal | 13 | 4118837.403 | 1.857 |
| birds | Dacnis_hartlaubi | NA | VU | Decreasing | NA | NA | NA | NA | NA |
| birds | Dacnis_lineata | Dacnis_lineata | LC | Decreasing | tropical | diurnal | 11 | 6051189.291 | 1.937 |
| birds | Dacnis_nigripes | Dacnis_nigripes | NT | Decreasing | tropical | diurnal | 14 | 51646.727 | 13.915 |
| birds | Dacnis_venusta | Dacnis_venusta | LC | Decreasing | tropical | diurnal | 16.1 | 203133.564 | 7.602 |
| birds | Dacnis_viguieri | NA | LC | Decreasing | NA | NA | NA | NA | NA |
| birds | Dactylortyx_thoracicus | Dactylortyx_thoracicus | LC | Decreasing | tropical | diurnal | 201.8 | 492692.141 | 9.727 |
| birds | Daphoenositta_chrysoptera | Daphoenositta_chrysoptera | LC | Decreasing | temperate | diurnal | 11.7 | 4739552.349 | 5.309 |
| birds | Daphoenositta_miranda | Daphoenositta_miranda | LC | Stable | tropical | diurnal | 16.8 | 42567.266 | 4.891 |
| birds | Daption_capense | Daption_capense | LC | Stable | temperate | cathemeral | 429.6 | 136304432.162 | 7.192 |
| birds | Daptrius_ater | Daptrius_ater | LC | Stable | tropical | diurnal | 351.8 | 7266225.059 | 2.45 |
| birds | Dasycrotapha_plateni | Stachyris_plateni | NT | Decreasing | tropical | diurnal | 8.7 | 49089.237 | 10.151 |
| birds | Dasycrotapha_speciosa | Dasycrotapha_speciosa | EN | Decreasing | tropical | diurnal | 16.4 | 23404.427 | 18.286 |
| birds | Dasylophus_superciliosus | Phaenicophaeus_superciliosus | LC | Decreasing | tropical | diurnal | 123 | 16303.411 | 9.327 |
| birds | Dasyornis_brachypterus | Dasyornis_brachypterus | VU | Decreasing | temperate | diurnal | 41.8 | 51744.962 | 10.693 |
| birds | Dasyornis_broadbenti | Dasyornis_broadbenti | LC | Decreasing | temperate | diurnal | 67.9 | 26759.889 | 11.415 |
| birds | Dasyornis_longirostris | Dasyornis_longirostris | EN | Decreasing | temperate | diurnal | 32.7 | 52040.522 | 10.663 |
| birds | Deconychura_longicauda | Deconychura_longicauda | LC | Decreasing | tropical | diurnal | 23.8 | 1094105.421 | 1.433 |
| birds | Delacourella_capistrata | Nesocharis_capistrata | LC | Stable | tropical | diurnal | 11.6 | 976089.609 | 8.062 |
| birds | Deleornis_fraseri | Anthreptes_fraseri | LC | Decreasing | tropical | diurnal | 11.3 | 1859228.624 | 10.143 |
| birds | Delichon_dasypus | Delichon_dasypus | LC | Increasing | temperate | diurnal | 18 | 11430696.77 | 12.634 |
| birds | Delichon_nipalense | Delichon_nipalense | LC | Stable | temperate | diurnal | 15 | 543149.358 | 8.849 |
| birds | Delichon_urbicum | Delichon_urbicum | LC | Decreasing | temperate | diurnal | 14.5 | 38136395.967 | 9.601 |
| birds | Deltarhynchus_flammulatus | Deltarhynchus_flammulatus | LC | Decreasing | tropical | diurnal | 17.2 | 103125.628 | 10.227 |
| birds | Dendrexetastes_rufigula | Dendrexetastes_rufigula | LC | Stable | tropical | diurnal | 69.6 | 3670233.996 | 2.129 |
| birds | Dendrocincla_anabatina | Dendrocincla_anabatina | LC | Decreasing | tropical | diurnal | 34.2 | 378885.001 | 8.502 |
| birds | Dendrocincla_fuliginosa | Dendrocincla_fuliginosa | LC | Decreasing | tropical | diurnal | 38.7 | 8483632.637 | 3.476 |

| | | | | | | | | | |
|---|---|---|---|---|---|---|---|---|---|
| birds | Dendrocincla_homochroa | Dendrocincla_homochroa | LC | Decreasing | tropical | diurnal | 39.4 | 492694.514 | 8.987 |
| birds | Dendrocincla_merula | Dendrocincla_merula | LC | Decreasing | tropical | diurnal | 50 | 5359264.175 | 1.685 |
| birds | Dendrocincla_turdina | Dendrocincla_turdina | LC | Decreasing | tropical | diurnal | 39 | 1193430.601 | 13.228 |
| birds | Dendrocincla_tyrannina | Dendrocincla_tyrannina | LC | Decreasing | tropical | diurnal | 54.9 | 307936.87 | 8.736 |
| birds | Dendrocitta_bayleii | Dendrocitta_bayleyi | VU | Decreasing | tropical | diurnal | 102 | 5706.316 | 6.555 |
| birds | Dendrocitta_formosae | Dendrocitta_formosae | LC | Decreasing | temperate | diurnal | 102.7 | 3328585.523 | 13.104 |
| birds | Dendrocitta_frontalis | Dendrocitta_frontalis | LC | Stable | temperate | diurnal | 100 | 200384.694 | 6.827 |
| birds | Dendrocitta_leucogastra | Dendrocitta_leucogastra | LC | Stable | tropical | diurnal | 99.2 | 69368.219 | 16.82 |
| birds | Dendrocitta_occipitalis | Dendrocitta_occipitalis | LC | Stable | tropical | diurnal | 95.3 | 165533.336 | 8.055 |
| birds | Dendrocitta_vagabunda | Dendrocitta_vagabunda | LC | Decreasing | tropical | diurnal | 100 | 4361512.676 | 16.321 |
| birds | Dendrocolaptes_certhia | Dendrocolaptes_certhia | LC | Stable | tropical | diurnal | 68.7 | 6659679.414 | 2.114 |
| birds | Dendrocolaptes_hoffmannsi | Dendrocolaptes_hoffmannsi | VU | Decreasing | tropical | diurnal | 83 | 862883.577 | 2.422 |
| birds | Dendrocolaptes_picumnus | Dendrocolaptes_picumnus | LC | Decreasing | tropical | diurnal | 76.6 | 5749616.395 | 2.817 |
| birds | Dendrocolaptes_platyrostris | Dendrocolaptes_platyrostris | LC | Stable | tropical | diurnal | 61.7 | 3885238.338 | 10.074 |
| birds | Dendrocolaptes_sanctithomae | Dendrocolaptes_sanctithomae | LC | Decreasing | tropical | diurnal | 66.9 | 591394.868 | 8.478 |
| birds | Dendrocopos_assimilis | Dendrocopos_assimilis | LC | Stable | temperate | diurnal | 51.8 | 604188.643 | 11.343 |
| birds | Dendrocopos_atratus | Dendrocopos_atratus | LC | Stable | tropical | diurnal | 46.7 | 309610.665 | 7.47 |
| birds | Dendrocopos_darjellensis | Dendrocopos_darjellensis | LC | Stable | temperate | diurnal | 70.7 | 301933.906 | 7.671 |
| birds | Dendrocopos_himalayensis | Dendrocopos_himalayensis | LC | Stable | temperate | diurnal | 67.7 | 115066.113 | 12.727 |
| birds | Dendrocopos_hyperythrus | Dendrocopos_hyperythrus | LC | Decreasing | temperate | diurnal | 46.5 | 1790193.927 | 9.46 |
| birds | Dendrocopos_leucopterus | Dendrocopos_leucopterus | LC | Stable | temperate | diurnal | 67 | 2454403.28 | 6.183 |
| birds | Dendrocopos_leucotos | Dendrocopos_leucotos | LC | Decreasing | temperate | diurnal | 116.8 | 11229537.308 | 9.484 |
| birds | Dendrocopos_macei | Dendrocopos_macei | LC | Stable | temperate | diurnal | 44.9 | 904703.866 | 13.991 |
| birds | Dendrocopos_major | Dendrocopos_major | LC | Increasing | temperate | diurnal | 74.9 | 24413724.287 | 9.997 |
| birds | Dendrocopos_noguchii | Dendrocopos_noguchii | CR | Decreasing | temperate | diurnal | 200 | 585.245 | 19.477 |
| birds | Dendrocopos_syriacus | Dendrocopos_syriacus | LC | Stable | temperate | diurnal | 76.7 | 2931365.507 | 16.329 |
| birds | Dendrocygna_arborea | Dendrocygna_arborea | NT | Decreasing | tropical | nocturnal | 1150 | 115603.829 | 16.275 |
| birds | Dendrocygna_arcuata | Dendrocygna_arcuata | LC | Decreasing | temperate | diurnal | 796.2 | 4892718.608 | 6.083 |
| birds | Dendrocygna_autumnalis | Dendrocygna_autumnalis | LC | Increasing | tropical | diurnal | 755.3 | 9474186.495 | 6.72 |
| birds | Dendrocygna_bicolor | Dendrocygna_bicolor | LC | Decreasing | tropical | nocturnal | 756.4 | 18682192.413 | 10.805 |
| birds | Dendrocygna_eytoni | Dendrocygna_eytoni | LC | Stable | temperate | nocturnal | 790 | 4778055.166 | 4.264 |
| birds | Dendrocygna_guttata | Dendrocygna_guttata | LC | Stable | tropical | nocturnal | 800 | 933148.328 | 4.952 |
| birds | Dendrocygna_javanica | Dendrocygna_javanica | LC | Decreasing | tropical | diurnal | 519.6 | 7165332.295 | 15.434 |
| birds | Dendrocygna_viduata | Dendrocygna_viduata | LC | Increasing | tropical | nocturnal | 690 | 27416318.336 | 9.088 |

| | | | | | | | | | |
|---|---|---|---|---|---|---|---|---|---|
| birds | Dendroma_erythroptera | Philydor_erythropterum | LC | Decreasing | tropical | diurnal | 30 | 3018856.229 | 2.161 |
| birds | Dendroma_rufa | Philydor_rufum | LC | Decreasing | tropical | diurnal | 25 | 2187683.25 | 11.22 |
| birds | Dendronanthus_indicus | Dendronanthus_indicus | LC | Stable | temperate | diurnal | 16.4 | 7547714.457 | 14.201 |
| birds | Dendropicos_abyssinicus | Dendropicos_abyssinicus | LC | Decreasing | tropical | diurnal | 24.5 | 360073.267 | 13.062 |
| birds | Dendropicos_dorae | Dendrocopos_dorae | NT | Decreasing | tropical | diurnal | 19.5 | 129511.083 | 9.399 |
| birds | Dendropicos_elachus | Dendropicos_elachus | LC | Unknown | tropical | diurnal | 18.4 | 1588913.65 | 8.285 |
| birds | Dendropicos_elliotii | Mesopicos_elliotii | LC | Stable | tropical | diurnal | 37.2 | 755495.268 | 8.153 |
| birds | Dendropicos_fuscescens | Dendropicos_fuscescens | LC | Stable | tropical | diurnal | 26 | 12708768.944 | 9.403 |
| birds | Dendropicos_gabonensis | Dendropicos_gabonensis | LC | Stable | tropical | diurnal | 25.9 | 1688352.484 | 6.485 |
| birds | Dendropicos_goertae | Mesopicos_goertae | LC | Stable | tropical | diurnal | 44.5 | 7460292.399 | 10.304 |
| birds | Dendropicos_griseocephalus | Mesopicos_griseocephalus | LC | Stable | tropical | diurnal | 40.9 | 2206119.905 | 8.595 |
| birds | Dendropicos_lugubris | NA | LC | Stable | NA | NA | NA | NA | NA |
| birds | Dendropicos_namaquus | Thripias_namaquus | LC | Stable | tropical | diurnal | 81.5 | 7980417.701 | 8.647 |
| birds | Dendropicos_obsoletus | Dendrocopos_obsoletus | LC | Stable | tropical | diurnal | 21.4 | 5967593.219 | 11.064 |
| birds | Dendropicos_poecilolaemus | Dendropicos_poecilolaemus | LC | Stable | tropical | diurnal | 28.1 | 723686.324 | 8.294 |
| birds | Dendropicos_pyrrhogaster | Thripias_pyrrhogaster | LC | Increasing | tropical | diurnal | 68.5 | 800529.521 | 14.841 |
| birds | Dendropicos_spodocephalus | Dendropicos_spodocephalus | LC | Stable | tropical | diurnal | 47.3 | 595112.594 | 10.89 |
| birds | Dendropicos_stierlingi | Dendropicos_stierlingi | NT | Stable | tropical | diurnal | 28 | 534905.478 | 9.326 |
| birds | Dendropicos_xantholophus | Thripias_xantholophus | LC | Stable | tropical | diurnal | 81.5 | 2776353.451 | 6.803 |
| birds | Dendroplex_kienerii | Dendroplex_kienerii | NT | Decreasing | tropical | diurnal | 45 | 237821.16 | 2.373 |
| birds | Dendroplex_picus | Dendroplex_picus | LC | Decreasing | tropical | diurnal | 41.3 | 9856567.731 | 4.047 |
| birds | Dendrortyx_barbatus | Dendrortyx_barbatus | VU | Decreasing | tropical | diurnal | 432 | 8177.783 | 12.333 |
| birds | Dendrortyx_leucophrys | Dendrortyx_leucophrys | LC | Decreasing | tropical | diurnal | 367.4 | 117127.032 | 12.358 |
| birds | Dendrortyx_macroura | Dendrortyx_macroura | LC | Decreasing | tropical | diurnal | 431 | 136253.422 | 10.408 |
| birds | Deroptyus_accipitrinus | Deroptyus_accipitrinus | LC | Decreasing | tropical | diurnal | 246 | 4158548.58 | 1.882 |
| birds | Dessonornis_anomalus | Cossypha_anomala | NT | Decreasing | tropical | diurnal | 24.5 | 8084.354 | 13.564 |
| birds | Dessonornis_archeri | Cossypha_archeri | LC | Decreasing | tropical | diurnal | 22.8 | 60103.14 | 17.054 |
| birds | Dessonornis_caffer | Cossypha_caffra | LC | Stable | temperate | diurnal | 28.5 | 1773198.123 | 11.252 |
| birds | Dessonornis_humeralis | Cossypha_humeralis | LC | Stable | tropical | diurnal | 22.8 | 644971.306 | 11.429 |
| birds | Devioeca_papuana | Microeca_papuana | LC | Stable | tropical | diurnal | 14.4 | 142595.352 | 4.8 |
| birds | Dicaeum_aeneum | Dicaeum_aeneum | LC | Stable | tropical | diurnal | 9.2 | 32482.616 | 4.717 |
| birds | Dicaeum_agile | Dicaeum_agile | LC | Stable | tropical | diurnal | 9 | 4429303.46 | 15.558 |
| birds | Dicaeum_annae | Dicaeum_annae | LC | Stable | tropical | diurnal | 10 | 28449.462 | 12.451 |
| birds | Dicaeum_anthonyi | Dicaeum_anthonyi | LC | Stable | tropical | diurnal | 11.6 | 5389.946 | 6.683 |

| | | | | | | | | | |
|---|---|---|---|---|---|---|---|---|---|
| birds | Dicaeum_aureolimbatum | Dicaeum_aureolimbatum | LC | Stable | tropical | diurnal | 8.1 | 175271.93 | 9.374 |
| birds | Dicaeum_australe | Dicaeum_australe | LC | Stable | tropical | diurnal | 8.7 | 237198.679 | 13.595 |
| birds | Dicaeum_bicolor | Dicaeum_bicolor | LC | Stable | tropical | diurnal | 9.1 | 259805.983 | 13.739 |
| birds | Dicaeum_celebicum | Dicaeum_celebicum | LC | Stable | tropical | diurnal | 6.5 | 184624.474 | 9.237 |
| birds | Dicaeum_chrysorrheum | Dicaeum_chrysorrheum | LC | Stable | temperate | diurnal | 9 | 3155552.175 | 10.31 |
| birds | Dicaeum_concolor | Dicaeum_concolor | LC | Stable | tropical | diurnal | 6.2 | 221339.977 | 20.145 |
| birds | Dicaeum_cruentatum | Dicaeum_cruentatum | LC | Stable | tropical | diurnal | 5.6 | 4479690.765 | 12.046 |
| birds | Dicaeum_erythrorhynchos | Dicaeum_erythrorhynchos | LC | Stable | tropical | diurnal | 6.3 | 2149808.709 | 17.734 |
| birds | Dicaeum_erythrothorax | Dicaeum_erythrothorax | LC | Stable | tropical | diurnal | 8.9 | 8554.414 | 6.663 |
| birds | Dicaeum_everetti | Dicaeum_everetti | NT | Decreasing | tropical | diurnal | 8.5 | 214852.281 | 6.483 |
| birds | Dicaeum_eximium | Dicaeum_eximium | LC | Stable | tropical | diurnal | 7.9 | 42347.944 | 4.423 |
| birds | Dicaeum_geelvinkianum | Dicaeum_geelvinkianum | LC | Stable | tropical | diurnal | 6.8 | 535209.767 | 3.077 |
| birds | Dicaeum_haematostictum | Dicaeum_haematostictum | LC | Stable | tropical | diurnal | 8.8 | 24928.065 | 17.883 |
| birds | Dicaeum_hirundinaceum | Dicaeum_hirundinaceum | LC | Stable | temperate | diurnal | 9 | 6464367.1 | 4.62 |
| birds | Dicaeum_hypoleucum | Dicaeum_hypoleucum | LC | Decreasing | tropical | diurnal | 8 | 229004.916 | 13.374 |
| birds | Dicaeum_igniferum | Dicaeum_igniferum | LC | Stable | tropical | diurnal | 8.5 | 29611.183 | 11.664 |
| birds | Dicaeum_ignipectus | Dicaeum_ignipectus | LC | Stable | tropical | diurnal | 5.7 | 4029931.666 | 11.961 |
| birds | Dicaeum_maugei | Dicaeum_maugei | LC | Stable | tropical | diurnal | 8.5 | 39542.568 | 14.071 |
| birds | Dicaeum_melanozanthum | Dicaeum_melanoxanthum | LC | Stable | temperate | diurnal | 8.5 | 1518409.57 | 8.994 |
| birds | Dicaeum_monticolum | Dicaeum_monticolum | LC | Decreasing | tropical | diurnal | 8.5 | 62934.251 | 2.633 |
| birds | Dicaeum_nehrkorni | Dicaeum_nehrkorni | LC | Decreasing | tropical | diurnal | 8.5 | 88003.64 | 6.159 |
| birds | Dicaeum_nigrilore | Dicaeum_nigrilore | LC | Stable | tropical | diurnal | 11 | 13373.777 | 6.834 |
| birds | Dicaeum_nitidum | Dicaeum_nitidum | LC | Stable | tropical | diurnal | 8.5 | 1321.841 | 4.241 |
| birds | Dicaeum_pectorale | Dicaeum_pectorale | LC | Decreasing | tropical | diurnal | 7.6 | 97817.395 | 3.16 |
| birds | Dicaeum_proprium | Dicaeum_proprium | LC | Decreasing | tropical | diurnal | 10.2 | 13567.452 | 7.072 |
| birds | Dicaeum_pygmaeum | Dicaeum_pygmaeum | LC | Decreasing | tropical | diurnal | 5 | 269443.513 | 13.549 |
| birds | Dicaeum_quadricolor | Dicaeum_quadricolor | CR | Decreasing | tropical | diurnal | 8.5 | 117.271 | 16.145 |
| birds | Dicaeum_retrocinctum | Dicaeum_retrocinctum | VU | Decreasing | tropical | diurnal | 6.5 | 9862.364 | 11.248 |
| birds | Dicaeum_sanguinolentum | Dicaeum_sanguinolentum | LC | Stable | tropical | diurnal | 8.5 | 26508.699 | 16.465 |
| birds | Dicaeum_trigonostigma | Dicaeum_trigonostigma | LC | Stable | tropical | diurnal | 7.1 | 2335989.48 | 10.26 |
| birds | Dicaeum_tristrami | Dicaeum_tristrami | LC | Stable | tropical | diurnal | 11.7 | 3331.874 | 4.604 |
| birds | Dicaeum_trochileum | Dicaeum_trochileum | LC | Stable | tropical | diurnal | 8.5 | 322091.433 | 14.704 |
| birds | Dicaeum_vincens | Dicaeum_vincens | NT | Decreasing | tropical | diurnal | 8.5 | 12166.572 | 21.992 |
| birds | Dicaeum_vulneratum | Dicaeum_vulneratum | LC | Decreasing | tropical | diurnal | 8.9 | 17812.667 | 7.271 |

| | | | | | | | | | |
|---|---|---|---|---|---|---|---|---|---|
| birds | Dichrozona_cincta | Dichrozona_cincta | LC | Decreasing | tropical | diurnal | 16 | 3112856.937 | 1.611 |
| birds | Dicrurus_adsimilis | Dicrurus_adsimilis | LC | Stable | tropical | diurnal | 40.3 | 14611485.32 | 9.525 |
| birds | Dicrurus_aeneus | Dicrurus_aeneus | LC | Unknown | tropical | diurnal | 26 | 4639369.765 | 12.191 |
| birds | Dicrurus_aldabranus | NA | NT | Stable | NA | NA | NA | NA | NA |
| birds | Dicrurus_andamanensis | Dicrurus_andamanensis | LC | Decreasing | tropical | diurnal | 50.5 | 5726.812 | 6.552 |
| birds | Dicrurus_annectens | Dicrurus_annectans | LC | Unknown | tropical | diurnal | 44 | 2725019.979 | 11.83 |
| birds | Dicrurus_atripennis | Dicrurus_atripennis | LC | Decreasing | tropical | diurnal | 38.9 | 1831426.112 | 7.597 |
| birds | Dicrurus_balicassius | Dicrurus_balicassius | LC | Unknown | tropical | diurnal | 72.3 | 151573.527 | 15.322 |
| birds | Dicrurus_bracteatus | Dicrurus_bracteatus | LC | Stable | temperate | diurnal | 85.5 | 1975520.316 | 4.885 |
| birds | Dicrurus_caerulescens | Dicrurus_caerulescens | LC | Unknown | tropical | diurnal | 40 | 2282100.813 | 18.864 |
| birds | Dicrurus_densus | NA | LC | Unknown | NA | NA | NA | NA | NA |
| birds | Dicrurus_forficatus | Dicrurus_forficatus | LC | Unknown | tropical | diurnal | 47.1 | 591728.753 | 8.755 |
| birds | Dicrurus_fuscipennis | Dicrurus_fuscipennis | EN | Stable | tropical | diurnal | 50.5 | 809.499 | 19.035 |
| birds | Dicrurus_hottentottus | Dicrurus_hottentottus | LC | Decreasing | temperate | diurnal | 79.2 | 7412372.407 | 13.586 |
| birds | Dicrurus_leucophaeus | Dicrurus_leucophaeus | LC | Unknown | tropical | diurnal | 37.6 | 9278672.073 | 15.369 |
| birds | Dicrurus_ludwigii | Dicrurus_ludwigii | LC | Stable | tropical | diurnal | 29.4 | 2047517.597 | 9.393 |
| birds | Dicrurus_megarhynchus | Dicrurus_megarhynchus | NT | Decreasing | tropical | diurnal | 130 | 7019.613 | 5.412 |
| birds | Dicrurus_modestus | Dicrurus_modestus | LC | Stable | tropical | nocturnal | 45.6 | 3490807.077 | 8.234 |
| birds | Dicrurus_montanus | Dicrurus_montanus | LC | Unknown | tropical | diurnal | 61 | 52683.33 | 5.3 |
| birds | Dicrurus_paradiseus | Dicrurus_paradiseus | LC | Decreasing | tropical | diurnal | 85.8 | 5998619.698 | 13.809 |
| birds | Dicrurus_remifer | Dicrurus_remifer | LC | Unknown | tropical | diurnal | 43.1 | 2438166.851 | 10.728 |
| birds | Dicrurus_sumatranus | Dicrurus_sumatranus | LC | Decreasing | tropical | diurnal | 50.5 | 97350.195 | 6.823 |
| birds | Dicrurus_waldenii | Dicrurus_waldenii | VU | Decreasing | tropical | diurnal | 50.5 | 316.353 | 21.537 |
| birds | Didunculus_strigirostris | NA | CR | Decreasing | NA | NA | NA | NA | NA |
| birds | Diglossa_albilatera | Diglossa_albilatera | LC | Stable | tropical | diurnal | 9.8 | 254619.681 | 9.835 |
| birds | Diglossa_baritula | Diglossa_baritula | LC | Stable | tropical | diurnal | 9.2 | 259874.78 | 12.864 |
| birds | Diglossa_brunneiventris | Diglossa_brunneiventris | LC | Stable | tropical | diurnal | 12 | 317862.931 | 8.287 |
| birds | Diglossa_caerulescens | Diglossa_caerulescens | LC | Stable | tropical | diurnal | 14.4 | 280727.035 | 8.799 |
| birds | Diglossa_carbonaria | Diglossa_carbonaria | LC | Stable | tropical | diurnal | 11 | 70106.559 | 9.974 |
| birds | Diglossa_cyanea | Diglossa_cyanea | LC | Stable | tropical | diurnal | 17 | 392147.643 | 8.904 |
| birds | Diglossa_duidae | Diglossa_duidae | LC | Decreasing | tropical | diurnal | 14.8 | 107045.145 | 0.471 |
| birds | Diglossa_glauca | Diglossa_glauca | LC | Decreasing | tropical | diurnal | 12 | 114824.309 | 5.5 |
| birds | Diglossa_gloriosa | Diglossa_gloriosa | LC | Stable | tropical | diurnal | 11 | 11423.966 | 14.352 |
| birds | Diglossa_gloriosissima | Diglossa_gloriosissima | NT | Unknown | tropical | diurnal | 14 | 1479.421 | 5.648 |

| | | | | | | | | | |
|---|---|---|---|---|---|---|---|---|---|
| birds | Diglossa_humeralis | Diglossa_humeralis | LC | Stable | tropical | diurnal | 13.1 | 155425.661 | 9.99 |
| birds | Diglossa_indigotica | Diglossa_indigotica | LC | Decreasing | tropical | diurnal | 13.7 | 19981.414 | 6.325 |
| birds | Diglossa_lafresnayii | Diglossa_lafresnayii | LC | Stable | tropical | diurnal | 16 | 109695.824 | 11.745 |
| birds | Diglossa_major | Diglossa_major | LC | Decreasing | tropical | diurnal | 22.2 | 33712.965 | 2.113 |
| birds | Diglossa_mystacalis | Diglossa_mystacalis | LC | Stable | tropical | diurnal | 16.2 | 121320.583 | 6.878 |
| birds | Diglossa_plumbea | Diglossa_plumbea | LC | Decreasing | tropical | diurnal | 10 | 8930.97 | 5.027 |
| birds | Diglossa_sittoides | Diglossa_sittoides | LC | Stable | tropical | diurnal | 9 | 593695.051 | 9.614 |
| birds | Diglossa_venezuelensis | Diglossa_venezuelensis | EN | Decreasing | tropical | diurnal | 12.6 | 795.992 | 7.079 |
| birds | Dinemellia_dinemelli | Dinemellia_dinemelli | LC | Stable | tropical | diurnal | 63.9 | 2361338.485 | 9.389 |
| birds | Dinopium_benghalense | Dinopium_benghalense | LC | Stable | tropical | diurnal | 100 | 3000600.757 | 18.545 |
| birds | Dinopium_javanense | Dinopium_javanense | LC | Decreasing | tropical | diurnal | 72.8 | 2707154.044 | 12.452 |
| birds | Dinopium_shorii | Dinopium_shorii | LC | Decreasing | tropical | diurnal | 101 | 382744.404 | 12.764 |
| birds | Diomedea_amsterdamensis | Diomedea_amsterdamensis | EN | Increasing | temperate | cathemeral | 6270 | 11232098.416 | 11.209 |
| birds | Diomedea_epomophora | Diomedea_epomophora | VU | Stable | temperate | cathemeral | 8905.6 | 56488366.655 | 8.997 |
| birds | Diomedea_exulans | Diomedea_exulans | VU | Decreasing | temperate | cathemeral | 6961.3 | 97771379.92 | 11.356 |
| birds | Diopsittaca_nobilis | Diopsittaca_nobilis | LC | Stable | tropical | diurnal | 150.9 | 495203.464 | 2.844 |
| birds | Discosura_conversii | Discosura_conversii | LC | Decreasing | tropical | diurnal | 3 | 132865.468 | 6.833 |
| birds | Discosura_langsdorffi | Discosura_langsdorffi | LC | Decreasing | tropical | diurnal | 2.8 | 1980012.6 | 2.04 |
| birds | Discosura_letitiae | NA | DD | Unknown | NA | NA | NA | NA | NA |
| birds | Discosura_longicaudus | Discosura_longicaudus | LC | Decreasing | tropical | diurnal | 3.4 | 2150114.71 | 2.689 |
| birds | Discosura_popelairii | Discosura_popelairii | LC | Decreasing | tropical | diurnal | 2.5 | 228570.944 | 4.58 |
| birds | Diuca_diuca | Diuca_diuca | LC | Stable | temperate | diurnal | 36.8 | 2498001.181 | 7.496 |
| birds | Dives_dives | Dives_dives | LC | Increasing | tropical | diurnal | 91.3 | 592160.799 | 11.127 |
| birds | Dives_warczewiczi | Dives_warszewiczi | LC | Increasing | tropical | diurnal | 73.3 | 238870.263 | 10.375 |
| birds | Dolichonyx_oryzivorus | Dolichonyx_oryzivorus | LC | Decreasing | temperate | diurnal | 31.5 | 12537285.928 | 9.564 |
| birds | Doliornis_remseni | Doliornis_remseni | NT | Decreasing | tropical | diurnal | 63.1 | 17881.707 | 7.313 |
| birds | Doliornis_sclateri | Doliornis_sclateri | NT | Decreasing | tropical | diurnal | 60.2 | 11866.561 | 4.096 |
| birds | Donacobius_atricapilla | Donacobius_atricapilla | LC | Decreasing | tropical | diurnal | 36.8 | 8620634.564 | 5.481 |
| birds | Donacospiza_albifrons | Donacospiza_albifrons | LC | Stable | temperate | diurnal | 16.5 | 1671293.501 | 11.863 |
| birds | Doricha_eliza | Doricha_eliza | NT | Decreasing | tropical | diurnal | 2.5 | 4054.807 | 13.381 |
| birds | Doricha_enicura | Doricha_enicura | LC | Decreasing | tropical | diurnal | 2.4 | 78586.848 | 13.48 |
| birds | Doryfera_johannae | Doryfera_johannae | LC | Decreasing | tropical | diurnal | 4.8 | 459646.349 | 2.731 |
| birds | Doryfera_ludovicae | Doryfera_ludovicae | LC | Stable | tropical | diurnal | 6 | 387149.162 | 8.383 |
| birds | Drepanis_coccinea | Vestiaria_coccinea | VU | Decreasing | tropical | diurnal | 18.2 | 2601.127 | 4.222 |

| | | | | | | | | | |
|---|---|---|---|---|---|---|---|---|---|
| birds | Drepanoptila_holosericea | Drepanoptila_holosericea | NT | Unknown | tropical | diurnal | 215 | 16533.091 | 7.541 |
| birds | Drepanorhynchus_reichenowi | Nectarinia_reichenowi | LC | Stable | tropical | diurnal | 14.7 | 217754.734 | 14.84 |
| birds | Drepanornis_albertisi | Epimachus_albertisi | LC | Stable | tropical | diurnal | 110 | 23525.736 | 3.088 |
| birds | Drepanornis_bruijnii | Epimachus_bruijnii | LC | Decreasing | tropical | diurnal | 153.3 | 48314.252 | 2.598 |
| birds | Dreptes_thomensis | Nectarinia_thomensis | VU | Decreasing | tropical | diurnal | 26.2 | 333.921 | 7.517 |
| birds | Dromaius_novaehollandiae | Dromaius_novaehollandiae | LC | Stable | temperate | diurnal | 34093.3 | 6731967.076 | 4.566 |
| birds | Dromas_ardeola | Dromas_ardeola | LC | Stable | temperate | nocturnal | 325 | 1344551.83 | 10.913 |
| birds | Dromococcyx_pavoninus | Dromococcyx_pavoninus | LC | Decreasing | tropical | diurnal | 46.4 | 3100662.233 | 5.409 |
| birds | Dromococcyx_phasianellus | Dromococcyx_phasianellus | LC | Decreasing | tropical | diurnal | 84.5 | 9600099.101 | 6.466 |
| birds | Drymocichla_incana | Drymocichla_incana | LC | Stable | tropical | diurnal | 10 | 1110783.225 | 5.265 |
| birds | Drymodes_brunneopygia | Drymodes_brunneopygia | LC | Decreasing | temperate | diurnal | 33 | 916861.693 | 8.131 |
| birds | Drymodes_superciliaris | Drymodes_superciliaris | LC | Stable | tropical | diurnal | 45.9 | 12382.108 | 1.737 |
| birds | Drymophila_caudata | Drymophila_caudata | LC | Decreasing | tropical | diurnal | 11.9 | 16339.115 | 10.271 |
| birds | Drymophila_devillei | Drymophila_devillei | LC | Decreasing | tropical | diurnal | 10.2 | 2101473.55 | 2.377 |
| birds | Drymophila_ferruginea | Drymophila_ferruginea | LC | Stable | tropical | diurnal | 10.6 | 506171.475 | 13.501 |
| birds | Drymophila_genei | Drymophila_genei | LC | Decreasing | tropical | diurnal | 11.4 | 24959.283 | 10.231 |
| birds | Drymophila_malura | Drymophila_malura | LC | Stable | temperate | diurnal | 13 | 769759.911 | 13.246 |
| birds | Drymophila_ochropyga | Drymophila_ochropyga | LC | Decreasing | tropical | diurnal | 10.5 | 338688.581 | 12.749 |
| birds | Drymophila_rubricollis | Drymophila_rubricollis | LC | Stable | temperate | diurnal | 10 | 329402.935 | 13.187 |
| birds | Drymophila_squamata | Drymophila_squamata | LC | Stable | tropical | diurnal | 10.8 | 340038.68 | 11.548 |
| birds | Drymornis_bridgesii | Drymornis_bridgesii | LC | Stable | temperate | diurnal | 92.6 | 1433240.563 | 7.838 |
| birds | Drymotoxeres_pucheranii | Campylorhamphus_pucherani | LC | Decreasing | tropical | diurnal | 70.5 | 140351.894 | 7.118 |
| birds | Dryobates_cathpharius | Dendrocopos_cathpharius | LC | Stable | temperate | diurnal | 30.7 | 125806.917 | 6.206 |
| birds | Dryobates_minor | Dendrocopos_minor | LC | Decreasing | temperate | diurnal | 19.8 | 18548908.455 | 9.122 |
| birds | Dryobates_nuttallii | Picoides_nuttallii | LC | Increasing | temperate | diurnal | 37.4 | 183833.95 | 15.565 |
| birds | Dryobates_pubescens | Picoides_pubescens | LC | Stable | temperate | diurnal | 25.6 | 12800223.819 | 9.64 |
| birds | Dryobates_scalaris | Picoides_scalaris | LC | Stable | temperate | diurnal | 31.6 | 2902465.04 | 9.895 |
| birds | Dryocopus_hodgei | Dryocopus_hodgei | VU | Decreasing | tropical | diurnal | 156 | 5706.316 | 6.555 |
| birds | Dryocopus_javensis | Dryocopus_javensis | LC | Decreasing | tropical | diurnal | 271.4 | 2693444.767 | 11.337 |
| birds | Dryocopus_martius | Dryocopus_martius | LC | Increasing | temperate | diurnal | 321 | 19634445.894 | 7.772 |
| birds | Dryolimnas_cuvieri | Dryolimnas_cuvieri | LC | Stable | tropical | diurnal | 182.4 | 590995.525 | 8.745 |
| birds | Dryoscopus_angolensis | Dryoscopus_angolensis | LC | Stable | tropical | diurnal | 36 | 372335.763 | 10.73 |
| birds | Dryoscopus_cubla | Dryoscopus_cubla | LC | Decreasing | tropical | diurnal | 26.2 | 5720470.905 | 8.899 |
| birds | Dryoscopus_gambensis | Dryoscopus_gambensis | LC | Stable | tropical | diurnal | 31.9 | 5608614.975 | 10.814 |

| | | | | | | | | | |
|---|---|---|---|---|---|---|---|---|---|
| birds | Dryoscopus_pringlii | Dryoscopus_pringlii | LC | Stable | tropical | diurnal | 19.7 | 448813.351 | 9.308 |
| birds | Dryoscopus_sabini | Dryoscopus_sabini | LC | Stable | tropical | diurnal | 37.1 | 1205106.958 | 10.647 |
| birds | Dryoscopus_senegalensis | Dryoscopus_senegalensis | LC | Stable | tropical | diurnal | 29.3 | 1438969.003 | 6.739 |
| birds | Dryotriorchis_spectabilis | Dryotriorchis_spectabilis | LC | Decreasing | tropical | diurnal | 697 | 2884763.004 | 7.537 |
| birds | Dubusia_castaneoventris | Delothraupis_castaneoventris | LC | Decreasing | tropical | diurnal | 28 | 131249.534 | 5.862 |
| birds | Dubusia_taeniata | Dubusia_taeniata | LC | Stable | tropical | diurnal | 37 | 212512.381 | 11.084 |
| birds | Ducula_aenea | Ducula_aenea | NT | Decreasing | tropical | diurnal | 545 | 4813912.017 | 12.226 |
| birds | Ducula_aurorae | Ducula_aurorae | EN | Increasing | tropical | diurnal | 507.2 | 299.519 | 14.667 |
| birds | Ducula_badia | Ducula_badia | LC | Decreasing | tropical | diurnal | 486 | 2980351.804 | 9.138 |
| birds | Ducula_bakeri | Ducula_bakeri | LC | Unknown | tropical | diurnal | 394 | 3924.432 | 3.974 |
| birds | Ducula_basilica | Ducula_basilica | LC | Stable | tropical | diurnal | 507.2 | 22924.769 | 6.468 |
| birds | Ducula_bicolor | Ducula_bicolor | LC | Decreasing | tropical | diurnal | 440 | 770004.59 | 11 |
| birds | Ducula_brenchleyi | Ducula_brenchleyi | NT | Decreasing | tropical | diurnal | 380 | 12795.623 | 5.266 |
| birds | Ducula_carola | Ducula_carola | VU | Decreasing | tropical | diurnal | 289 | 222701.314 | 13.705 |
| birds | Ducula_chalconota | Ducula_chalconota | LC | Stable | tropical | diurnal | 502 | 165349.324 | 4.553 |
| birds | Ducula_cineracea | Ducula_cineracea | NT | Decreasing | tropical | diurnal | 507.2 | 3484.285 | 11.873 |
| birds | Ducula_concinna | Ducula_concinna | LC | Decreasing | tropical | diurnal | 804.7 | 10909.951 | 8.631 |
| birds | Ducula_finschii | Ducula_finschii | NT | Decreasing | tropical | diurnal | 343.9 | 38730.115 | 4.13 |
| birds | Ducula_forsteni | Ducula_forsteni | LC | Decreasing | tropical | diurnal | 510 | 145694.294 | 7.777 |
| birds | Ducula_galeata | Ducula_galeata | EN | Increasing | tropical | diurnal | 605 | 434.56 | 8.377 |
| birds | Ducula_goliath | Ducula_goliath | NT | Unknown | tropical | diurnal | 600 | 16547.108 | 7.541 |
| birds | Ducula_lacernulata | Ducula_lacernulata | LC | Stable | tropical | diurnal | 150 | 68766.832 | 15.832 |
| birds | Ducula_latrans | Ducula_latrans | LC | Decreasing | tropical | diurnal | 365.1 | 18103.652 | 9.374 |
| birds | Ducula_luctuosa | Ducula_luctuosa | LC | Decreasing | tropical | diurnal | 410 | 145005.343 | 9.922 |
| birds | Ducula_melanochroa | Ducula_melanochroa | LC | Decreasing | tropical | diurnal | 665 | 36132.411 | 3.87 |
| birds | Ducula_mindorensis | Ducula_mindorensis | EN | Decreasing | tropical | diurnal | 507.2 | 2472.819 | 5.6 |
| birds | Ducula_mullerii | Ducula_mullerii | LC | Stable | tropical | diurnal | 574.9 | 281060.321 | 2.678 |
| birds | Ducula_myristicivora | Ducula_myristicivora | LC | Stable | tropical | diurnal | 535 | 7545.932 | 3.443 |
| birds | Ducula_oceanica | Ducula_oceanica | VU | Decreasing | tropical | diurnal | 371.5 | 1374.538 | 12.24 |
| birds | Ducula_pacifica | Ducula_pacifica | LC | Decreasing | tropical | diurnal | 394.2 | 29576.605 | 5.878 |
| birds | Ducula_perspicillata | Ducula_perspicillata | LC | Stable | tropical | diurnal | 667 | 34605.551 | 6.406 |
| birds | Ducula_pickeringii | Ducula_pickeringii | VU | Decreasing | tropical | diurnal | 377 | 3060.901 | 10.008 |
| birds | Ducula_pinon | Ducula_pinon | LC | Stable | tropical | diurnal | 756 | 639390.556 | 3.093 |
| birds | Ducula_pistrinaria | Ducula_pistrinaria | LC | Decreasing | tropical | diurnal | 515 | 85422.325 | 4.614 |

| | | | | | | | | | |
|---|---|---|---|---|---|---|---|---|---|
| birds | Ducula_poliocephala | Ducula_poliocephala | NT | Decreasing | tropical | diurnal | 555 | 266333.854 | 13.962 |
| birds | Ducula_radiata | Ducula_radiata | LC | Decreasing | tropical | diurnal | 295 | 154747.933 | 8.601 |
| birds | Ducula_rosacea | Ducula_rosacea | NT | Decreasing | tropical | diurnal | 385.6 | 64852.557 | 12.052 |
| birds | Ducula_rubricera | Ducula_rubricera | LC | Decreasing | tropical | diurnal | 652 | 72417.843 | 4.799 |
| birds | Ducula_rufigaster | Ducula_rufigaster | LC | Stable | tropical | diurnal | 436 | 670937.244 | 3.083 |
| birds | Ducula_spilorrhoa | Ducula_spilorrhoa | LC | Decreasing | tropical | diurnal | 464 | 518577.491 | 3.024 |
| birds | Ducula_whartoni | Ducula_whartoni | LC | Stable | tropical | diurnal | 571.2 | 137.178 | 12.523 |
| birds | Ducula_zoeae | Ducula_zoeae | LC | Stable | tropical | diurnal | 568 | 706706.055 | 3.15 |
| birds | Dulus_dominicus | Dulus_dominicus | LC | Stable | tropical | diurnal | 48.4 | 74909.056 | 15.767 |
| birds | Dumetella_carolinensis | Dumetella_carolinensis | LC | Stable | temperate | diurnal | 35.3 | 8544736.275 | 14.03 |
| birds | Dumetia_atriceps | Rhopocichla_atriceps | LC | Decreasing | tropical | diurnal | 16.3 | 229794.306 | 20.202 |
| birds | Dumetia_hyperythra | Dumetia_hyperythra | LC | Decreasing | tropical | diurnal | 12.9 | 2111330.266 | 18.757 |
| birds | Dyaphorophyia_blissetti | Platysteira_blissetti | LC | Decreasing | tropical | diurnal | 11 | 473029.12 | 15.722 |
| birds | Dyaphorophyia_castanea | Platysteira_castanea | LC | Stable | tropical | diurnal | 14.1 | 3596708.417 | 6.913 |
| birds | Dyaphorophyia_concreta | Platysteira_concreta | LC | Stable | tropical | diurnal | 11.4 | 175431.497 | 14.048 |
| birds | Dyaphorophyia_tonsa | Platysteira_tonsa | LC | Stable | tropical | diurnal | 11.5 | 1168679.833 | 5.731 |
| birds | Dysithamnus_leucostictus | Dysithamnus_leucostictus | LC | Decreasing | tropical | diurnal | 20.2 | 52236.899 | 7.465 |
| birds | Dysithamnus_mentalis | Dysithamnus_mentalis | LC | Decreasing | tropical | diurnal | 14.9 | 4153057.293 | 9.801 |
| birds | Dysithamnus_occidentalis | Dysithamnus_occidentalis | NT | Decreasing | tropical | diurnal | 26.4 | 14566.796 | 6.744 |
| birds | Dysithamnus_plumbeus | Dysithamnus_plumbeus | VU | Decreasing | tropical | diurnal | 20.5 | 144601.211 | 10.869 |
| birds | Dysithamnus_puncticeps | Dysithamnus_puncticeps | LC | Decreasing | tropical | diurnal | 14.9 | 127823.894 | 6.026 |
| birds | Dysithamnus_stictothorax | Dysithamnus_stictothorax | NT | Decreasing | tropical | diurnal | 15.5 | 415220.513 | 11.735 |
| birds | Dysithamnus_striaticeps | Dysithamnus_striaticeps | LC | Decreasing | tropical | diurnal | 12 | 88175.269 | 7.628 |
| birds | Dysithamnus_xanthopterus | Dysithamnus_xanthopterus | LC | Decreasing | tropical | diurnal | 16 | 46590.475 | 13.827 |
| birds | Eclectus_roratus | Eclectus_roratus | LC | Decreasing | tropical | nocturnal | 480.8 | 52021.224 | 6.761 |
| birds | Edolisoma_anale | Coracina_analis | NT | Stable | tropical | diurnal | 82.4 | 5155.812 | 4.211 |
| birds | Edolisoma_ceramense | Coracina_ceramensis | LC | Stable | tropical | diurnal | 47.9 | 28618.491 | 6.734 |
| birds | Edolisoma_coerulescens | Coracina_coerulescens | LC | Stable | tropical | diurnal | 62.8 | 88474.567 | 13.082 |
| birds | Edolisoma_dispar | Coracina_dispar | LC | Stable | tropical | diurnal | 68 | 4846.919 | 7.03 |
| birds | Edolisoma_dohertyi | Coracina_dohertyi | LC | Stable | tropical | diurnal | 68 | 39073.162 | 13.136 |
| birds | Edolisoma_holopolium | Coracina_holopolia | LC | Decreasing | tropical | diurnal | 48.9 | 24872.855 | 5.121 |
| birds | Edolisoma_incertum | Coracina_incerta | LC | Stable | tropical | diurnal | 58.9 | 348235.084 | 3.297 |
| birds | Edolisoma_melas | Coracina_melas | LC | Stable | tropical | diurnal | 59.4 | 592908.619 | 3.064 |
| birds | Edolisoma_mindanense | Coracina_mindanensis | VU | Decreasing | tropical | diurnal | 58.7 | 234173.679 | 13.279 |

| | | | | | | | | | |
|---|---|---|---|---|---|---|---|---|---|
| birds | Edolisoma_montanum | Coracina_montana | LC | Stable | tropical | diurnal | 79.2 | 289775.956 | 3.961 |
| birds | Edolisoma_morio | Coracina_morio | LC | Stable | tropical | diurnal | 58.5 | 172301.977 | 9.541 |
| birds | Edolisoma_ostentum | Coracina_ostenta | VU | Decreasing | tropical | diurnal | 68 | 23988.157 | 18.336 |
| birds | Edolisoma_schisticeps | Coracina_schisticeps | LC | Stable | tropical | diurnal | 52.8 | 166507.113 | 2.867 |
| birds | Edolisoma_sula | Coracina_sula | LC | Stable | tropical | diurnal | 68 | 4678.129 | 5.687 |
| birds | Edolisoma_tenuirostre | Coracina_tenuirostris | LC | Stable | tropical | diurnal | 60.9 | 2156165.11 | 4.925 |
| birds | Egretta_ardesiaca | Egretta_ardesiaca | LC | Stable | tropical | diurnal | 324 | 8505004.79 | 10.218 |
| birds | Egretta_caerulea | Egretta_caerulea | LC | Decreasing | tropical | diurnal | 338.6 | 8483901.27 | 10.07 |
| birds | Egretta_eulophotes | Egretta_eulophotes | VU | Decreasing | tropical | diurnal | 461.8 | 860382.307 | 17.444 |
| birds | Egretta_garzetta | Egretta_garzetta | LC | Increasing | temperate | diurnal | 312 | 38343874.822 | 10.691 |
| birds | Egretta_novaehollandiae | Egretta_novaehollandiae | LC | Unknown | temperate | diurnal | 558.6 | 7533663.443 | 4.763 |
| birds | Egretta_picata | Ardea_picata | LC | Stable | tropical | diurnal | 259 | 1218679.446 | 3.517 |
| birds | Egretta_rufescens | Egretta_rufescens | NT | Decreasing | tropical | diurnal | 614 | 580944.39 | 12.931 |
| birds | Egretta_sacra | Egretta_sacra | LC | Stable | temperate | cathemeral | 502 | 4099474.265 | 9.896 |
| birds | Egretta_thula | Egretta_thula | LC | Increasing | tropical | diurnal | 371 | 22001375.456 | 8.147 |
| birds | Egretta_tricolor | Egretta_tricolor | LC | Stable | tropical | diurnal | 372.3 | 2501648.229 | 12.004 |
| birds | Egretta_vinaceigula | Egretta_vinaceigula | VU | Decreasing | tropical | diurnal | 288 | 1053459.648 | 9.938 |
| birds | Elachura_formosa | Spelaeornis_formosus | LC | Decreasing | temperate | diurnal | 5 | 663267.732 | 9.401 |
| birds | Elaenia_albiceps | Elaenia_albiceps | LC | Stable | temperate | diurnal | 15.5 | 13323047.2 | 6.225 |
| birds | Elaenia_chiriquensis | Elaenia_chiriquensis | LC | Stable | tropical | diurnal | 15.4 | 8478302.486 | 6.77 |
| birds | Elaenia_cristata | Elaenia_cristata | LC | Decreasing | tropical | diurnal | 18.2 | 5128248.102 | 5.641 |
| birds | Elaenia_dayi | Elaenia_dayi | LC | Decreasing | tropical | diurnal | 34.5 | 26627.088 | 1.395 |
| birds | Elaenia_fallax | Elaenia_fallax | LC | Decreasing | tropical | diurnal | 14.4 | 6030.051 | 18.606 |
| birds | Elaenia_flavogaster | Elaenia_flavogaster | LC | Stable | tropical | diurnal | 24.8 | 10327940.877 | 7.276 |
| birds | Elaenia_frantzii | Elaenia_frantzii | LC | Decreasing | tropical | diurnal | 19.6 | 266786.471 | 11.876 |
| birds | Elaenia_gigas | Elaenia_gigas | LC | Increasing | tropical | diurnal | 31 | 474517.612 | 4.296 |
| birds | Elaenia_martinica | Elaenia_martinica | LC | Decreasing | tropical | diurnal | 20.8 | 26788.389 | 17.874 |
| birds | Elaenia_mesoleuca | Elaenia_mesoleuca | LC | Decreasing | temperate | diurnal | 17.6 | 2216525.016 | 12.025 |
| birds | Elaenia_obscura | Elaenia_obscura | LC | Stable | tropical | diurnal | 23.9 | 326587.088 | 6.841 |
| birds | Elaenia_pallatangae | Elaenia_pallatangae | LC | Stable | tropical | diurnal | 17.8 | 281873.777 | 7.985 |
| birds | Elaenia_parvirostris | Elaenia_parvirostris | LC | Stable | tropical | diurnal | 13.8 | 10962786.124 | 5.334 |
| birds | Elaenia_pelzelni | NA | LC | Decreasing | NA | NA | NA | NA | NA |
| birds | Elaenia_ridleyana | NA | VU | Stable | NA | NA | NA | NA | NA |
| birds | Elaenia_ruficeps | Elaenia_ruficeps | LC | Stable | tropical | diurnal | 18.7 | 709154.378 | 1.746 |

| | | | | | | | | | |
|---|---|---|---|---|---|---|---|---|---|
| birds | Elaenia_spectabilis | Elaenia_spectabilis | LC | Decreasing | tropical | diurnal | 27.3 | 8533057.472 | 4.964 |
| birds | Elaenia_strepera | Elaenia_strepera | LC | Decreasing | tropical | diurnal | 19.3 | 2575735.657 | 3.271 |
| birds | Elanoides_forficatus | Elanoides_forficatus | LC | Increasing | tropical | diurnal | 416.2 | 14031977.443 | 6.411 |
| birds | Elanus_axillaris | Elanus_axillaris | LC | Increasing | temperate | cathemeral | 270.1 | 7649038.175 | 4.265 |
| birds | Elanus_caeruleus | Elanus_caeruleus | LC | Stable | tropical | diurnal | 259.8 | 26925792.465 | 10.73 |
| birds | Elanus_leucurus | Elanus_leucurus | LC | Increasing | tropical | diurnal | 346 | 12522294.388 | 8.417 |
| birds | Elanus_scriptus | Elanus_scriptus | NT | Stable | temperate | nocturnal | 316 | 427962.131 | 6.577 |
| birds | Elaphrornis_palliseri | Bradypterus_palliseri | NT | Decreasing | tropical | diurnal | 9 | 3806.138 | 16.974 |
| birds | Electron_carinatum | Electron_carinatum | VU | Decreasing | tropical | diurnal | 64.9 | 43323.086 | 9.08 |
| birds | Electron_platyrhynchum | Electron_platyrhynchum | LC | Decreasing | tropical | diurnal | 73 | 2864115.609 | 2.957 |
| birds | Eleoscytalopus_indigoticus | Eleoscytalopus_indigoticus | LC | Decreasing | tropical | diurnal | 16.2 | 455549.551 | 12.945 |
| birds | Eleoscytalopus_psychopompus | Eleoscytalopus_psychopompus | EN | Decreasing | tropical | diurnal | 17.8 | 2626.389 | 6.82 |
| birds | Eleothreptus_anomalus | Eleothreptus_anomalus | VU | Decreasing | temperate | nocturnal | 43.7 | 1451335.701 | 10.56 |
| birds | Eleothreptus_candicans | Eleothreptus_candicans | VU | Decreasing | tropical | nocturnal | 49.3 | 1398605.144 | 10.614 |
| birds | Elliotomyia_chionogaster | Amazilia_chionogaster | LC | Unknown | tropical | diurnal | 5 | 488801.524 | 7.477 |
| birds | Elliotomyia_viridicauda | Amazilia_viridicauda | LC | Unknown | tropical | diurnal | 6.1 | 28878.126 | 6.134 |
| birds | Elminia_albicauda | Elminia_albicauda | LC | Stable | tropical | diurnal | 8.3 | 1971246.252 | 8.173 |
| birds | Elminia_albiventris | Elminia_albiventris | LC | Stable | tropical | diurnal | 8.9 | 224843.848 | 13.121 |
| birds | Elminia_albonotata | Elminia_albonotata | LC | Stable | tropical | diurnal | 9.2 | 905274.179 | 11.871 |
| birds | Elminia_longicauda | Elminia_longicauda | LC | Stable | tropical | diurnal | 9.9 | 3910894.162 | 8.914 |
| birds | Elminia_nigromitrata | Elminia_nigromitrata | LC | Decreasing | tropical | diurnal | 9 | 2433804.714 | 7.994 |
| birds | Elseyornis_melanops | Elseyornis_melanops | LC | Increasing | temperate | diurnal | 33 | 7354478.038 | 4.569 |
| birds | Emarginata_schlegelii | Cercomela_schlegelii | LC | Stable | tropical | diurnal | 23.3 | 1144740.062 | 6.453 |
| birds | Emarginata_sinuata | Cercomela_sinuata | LC | Stable | temperate | diurnal | 18.1 | 793361.298 | 8.65 |
| birds | Emarginata_tractrac | Cercomela_tractrac | LC | Stable | tropical | diurnal | 20.8 | 1018879.118 | 5.936 |
| birds | Emberiza_affinis | Emberiza_affinis | LC | Stable | tropical | diurnal | 15.4 | 1397601.223 | 11.487 |
| birds | Emberiza_aureola | Emberiza_aureola | CR | Decreasing | temperate | diurnal | 19.6 | 17060140.554 | 5.51 |
| birds | Emberiza_bruniceps | Emberiza_bruniceps | LC | Stable | temperate | diurnal | 24.8 | 5514147.764 | 10.539 |
| birds | Emberiza_buchanani | Emberiza_buchanani | LC | Stable | temperate | diurnal | 21.2 | 4008927.894 | 11.664 |
| birds | Emberiza_cabanisi | Emberiza_cabanisi | LC | Stable | tropical | diurnal | 24.9 | 2851364.025 | 9.136 |
| birds | Emberiza_caesia | Emberiza_caesia | LC | Stable | temperate | diurnal | 19.9 | 1505590.674 | 8.072 |
| birds | Emberiza_calandra | Miliaria_calandra | LC | Decreasing | temperate | diurnal | 48.5 | 8461055.907 | 16.448 |
| birds | Emberiza_capensis | Emberiza_capensis | LC | Stable | temperate | diurnal | 20.9 | 1320689.836 | 9.615 |
| birds | Emberiza_chrysophrys | Emberiza_chrysophrys | LC | Stable | temperate | diurnal | 18 | 3119549.038 | 4.778 |

| | | | | | | | | | |
|---|---|---|---|---|---|---|---|---|---|
| birds | Emberiza_cia | Emberiza_cia | LC | Increasing | temperate | diurnal | 23.4 | 5760543.738 | 13.574 |
| birds | Emberiza_cineracea | Emberiza_cineracea | NT | Decreasing | temperate | diurnal | 24.3 | 1304311.533 | 10.359 |
| birds | Emberiza_cioides | Emberiza_cioides | LC | Stable | temperate | diurnal | 21 | 10532097.134 | 9.328 |
| birds | Emberiza_cirlus | Emberiza_cirlus | LC | Increasing | temperate | diurnal | 25.6 | 2210307.259 | 18.588 |
| birds | Emberiza_citrinella | Emberiza_citrinella | LC | Decreasing | temperate | diurnal | 29.7 | 16427829.18 | 10.99 |
| birds | Emberiza_elegans | Emberiza_elegans | LC | Stable | temperate | diurnal | 16.8 | 3666177.192 | 12.529 |
| birds | Emberiza_flaviventris | Emberiza_flaviventris | LC | Stable | tropical | diurnal | 18.2 | 6969113.536 | 9.263 |
| birds | Emberiza_fucata | Emberiza_fucata | LC | Stable | temperate | diurnal | 19.8 | 8426706.697 | 12.117 |
| birds | Emberiza_godlewskii | Emberiza_godlewskii | LC | Stable | temperate | diurnal | 19 | 6786390.708 | 7.982 |
| birds | Emberiza_hortulana | Emberiza_hortulana | LC | Decreasing | temperate | diurnal | 19.9 | 10888484.284 | 12.206 |
| birds | Emberiza_impetuani | Emberiza_impetuani | LC | Stable | tropical | diurnal | 15.1 | 1657377.126 | 6.943 |
| birds | Emberiza_jankowskii | Emberiza_jankowskii | EN | Decreasing | temperate | diurnal | 22.5 | 288918.887 | 15.672 |
| birds | Emberiza_koslowi | Emberiza_koslowi | NT | Stable | temperate | diurnal | 25 | 132468.272 | 8.667 |
| birds | Emberiza_lathami | Melophus_lathami | LC | Stable | temperate | diurnal | 21.9 | 3887662.385 | 14.201 |
| birds | Emberiza_leucocephalos | Emberiza_leucocephalos | LC | Stable | temperate | diurnal | 28.5 | 14053729.432 | 5.862 |
| birds | Emberiza_melanocephala | Emberiza_melanocephala | LC | Unknown | temperate | diurnal | 28.3 | 2718177.531 | 16.296 |
| birds | Emberiza_pallasi | Emberiza_pallasi | LC | Stable | temperate | diurnal | 14 | 14847467.728 | 5.085 |
| birds | Emberiza_poliopleura | Emberiza_poliopleura | LC | Stable | tropical | diurnal | 15.8 | 1250196.229 | 8.68 |
| birds | Emberiza_pusilla | Emberiza_pusilla | LC | Stable | temperate | diurnal | 14.6 | 13018743.801 | 4.511 |
| birds | Emberiza_rustica | Emberiza_rustica | VU | Decreasing | temperate | diurnal | 20.5 | 22901225.248 | 6.411 |
| birds | Emberiza_rutila | Emberiza_rutila | LC | Stable | temperate | diurnal | 17.5 | 5656279.097 | 4.809 |
| birds | Emberiza_schoeniclus | Emberiza_schoeniclus | LC | Decreasing | temperate | diurnal | 18.4 | 36135699.037 | 9.575 |
| birds | Emberiza_siemsseni | Latoucheornis_siemsseni | LC | Stable | temperate | diurnal | 20 | 991372.514 | 14.13 |
| birds | Emberiza_socotrana | Emberiza_socotrana | NT | Stable | tropical | diurnal | 14.4 | 630.829 | 10.816 |
| birds | Emberiza_spodocephala | Emberiza_spodocephala | LC | Stable | temperate | diurnal | 18.5 | 6997496.162 | 7.157 |
| birds | Emberiza_stewarti | Emberiza_stewarti | LC | Stable | temperate | diurnal | 16 | 1817857.286 | 14.969 |
| birds | Emberiza_striolata | Emberiza_striolata | LC | Increasing | temperate | diurnal | 14.8 | 2669183.701 | 8.467 |
| birds | Emberiza_sulphurata | Emberiza_sulphurata | LC | Stable | temperate | diurnal | 14.9 | 542896.505 | 20.605 |
| birds | Emberiza_tahapisi | Emberiza_tahapisi | LC | Stable | tropical | diurnal | 13 | 4092995.915 | 9.958 |
| birds | Emberiza_tristrami | Emberiza_tristrami | LC | Stable | temperate | diurnal | 18 | 1229215.115 | 11.034 |
| birds | Emberiza_variabilis | Emberiza_variabilis | LC | Stable | temperate | diurnal | 23.8 | 409045.4 | 15.584 |
| birds | Emberiza_yessoensis | Emberiza_yessoensis | NT | Decreasing | temperate | diurnal | 13.5 | 1815524.152 | 17.776 |
| birds | Emberizoides_duidae | NA | DD | Unknown | NA | NA | NA | NA | NA |
| birds | Emberizoides_herbicola | Emberizoides_herbicola | LC | Decreasing | tropical | diurnal | 27.6 | 5561384.252 | 8.736 |

| | | | | | | | | | |
|---|---|---|---|---|---|---|---|---|---|
| birds | Emberizoides_ypiranganus | Emberizoides_ypiranganus | LC | Stable | temperate | diurnal | 20.1 | 649452.653 | 11.105 |
| birds | Embernagra_longicauda | Embernagra_longicauda | LC | Decreasing | tropical | diurnal | 42.2 | 86158.64 | 7.467 |
| birds | Embernagra_platensis | Embernagra_platensis | LC | Stable | temperate | diurnal | 45.5 | 2735501.784 | 10.306 |
| birds | Emblema_pictum | Emblema_pictum | LC | Stable | tropical | diurnal | 10.4 | 1295960.935 | 3.079 |
| birds | Eminia_lepida | Eminia_lepida | LC | Stable | tropical | diurnal | 20.2 | 631640.646 | 14.455 |
| birds | Empidonax_affinis | Empidonax_affinis | LC | Decreasing | tropical | diurnal | 11.3 | 480233.438 | 10.707 |
| birds | Empidonax_albigularis | Empidonax_albigularis | LC | Decreasing | tropical | diurnal | 9.6 | 1056434.018 | 10.325 |
| birds | Empidonax_alnorum | Empidonax_alnorum | LC | Decreasing | temperate | diurnal | 12.7 | 13408407.309 | 7.655 |
| birds | Empidonax_atriceps | Empidonax_atriceps | LC | Stable | tropical | diurnal | 9 | 3661.215 | 6.362 |
| birds | Empidonax_difficilis | Empidonax_difficilis | LC | Stable | temperate | diurnal | 10.7 | 1682980.973 | 7.832 |
| birds | Empidonax_flavescens | Empidonax_flavescens | LC | Decreasing | tropical | diurnal | 12.5 | 167716.78 | 11.501 |
| birds | Empidonax_flaviventris | Empidonax_flaviventris | LC | Increasing | temperate | diurnal | 11.8 | 8805558.819 | 9.476 |
| birds | Empidonax_fulvifrons | Empidonax_fulvifrons | LC | Decreasing | temperate | diurnal | 7.9 | 729421.792 | 9.844 |
| birds | Empidonax_hammondii | Empidonax_hammondii | LC | Increasing | temperate | diurnal | 10.4 | 5603866.933 | 7.554 |
| birds | Empidonax_minimus | Empidonax_minimus | LC | Decreasing | temperate | diurnal | 10 | 11595266.468 | 10.907 |
| birds | Empidonax_oberholseri | Empidonax_oberholseri | LC | Stable | temperate | diurnal | 10.4 | 4614600.726 | 8.466 |
| birds | Empidonax_occidentalis | Empidonax_occidentalis | LC | Stable | temperate | diurnal | 11.6 | 2964611.734 | 8.725 |
| birds | Empidonax_traillii | Empidonax_traillii | LC | Decreasing | temperate | diurnal | 13.4 | 11375149 | 11.915 |
| birds | Empidonax_virescens | Empidonax_virescens | LC | Stable | tropical | diurnal | 12.6 | 5003149.637 | 15.536 |
| birds | Empidonax_wrightii | Empidonax_wrightii | LC | Increasing | temperate | diurnal | 12.3 | 2898385.442 | 8.337 |
| birds | Empidonomus_varius | Empidonomus_varius | LC | Stable | tropical | diurnal | 27.1 | 13118402.849 | 5.507 |
| birds | Empidornis_semipartitus | Empidornis_semipartitus | LC | Stable | tropical | diurnal | 22.5 | 662620.708 | 10.573 |
| birds | Enicognathus_ferrugineus | Enicognathus_ferrugineus | LC | Stable | temperate | diurnal | 160 | 572089.61 | 5.281 |
| birds | Enicognathus_leptorhynchus | Enicognathus_leptorhynchus | LC | Stable | temperate | diurnal | 193 | 107787.938 | 9.937 |
| birds | Enicurus_immaculatus | Enicurus_immaculatus | LC | Stable | temperate | diurnal | 25.5 | 921267.824 | 9.736 |
| birds | Enicurus_leschenaulti | Enicurus_leschenaulti | LC | Stable | tropical | diurnal | 33.8 | 6166589.043 | 12.107 |
| birds | Enicurus_maculatus | Enicurus_maculatus | LC | Stable | temperate | diurnal | 40.4 | 1640498.355 | 12.404 |
| birds | Enicurus_ruficapillus | Enicurus_ruficapillus | NT | Decreasing | tropical | diurnal | 28.2 | 1105095.597 | 8.075 |
| birds | Enicurus_schistaceus | Enicurus_schistaceus | LC | Stable | tropical | diurnal | 31 | 2441930.117 | 10.949 |
| birds | Enicurus_scouleri | Enicurus_scouleri | LC | Stable | temperate | diurnal | 17.8 | 3341040.761 | 12.401 |
| birds | Enicurus_velatus | Enicurus_velatus | LC | Stable | tropical | diurnal | 33 | 103361.851 | 9.469 |
| birds | Enodes_erythrophris | Enodes_erythrophris | LC | Decreasing | tropical | diurnal | 130 | 77247.817 | 6.028 |
| birds | Ensifera_ensifera | Ensifera_ensifera | LC | Stable | tropical | diurnal | 11.2 | 398282.991 | 8.565 |
| birds | Entomodestes_coracinus | Entomodestes_coracinus | LC | Decreasing | tropical | diurnal | 56 | 21017.951 | 6.374 |

| | | | | | | | | | |
|---|---|---|---|---|---|---|---|---|---|
| birds | Entomodestes_leucotis | Entomodestes_leucotis | LC | Decreasing | tropical | diurnal | 61.7 | 206932.102 | 4.683 |
| birds | Entomyzon_cyanotis | Entomyzon_cyanotis | LC | Stable | temperate | diurnal | 105.3 | 2004063.991 | 7.057 |
| birds | Eolophus_roseicapilla | Cacatua_roseicapilla | LC | Increasing | temperate | diurnal | 328 | 7020540.496 | 4.54 |
| birds | Eophona_migratoria | Eophona_migratoria | LC | Stable | temperate | diurnal | 46.9 | 6201191.13 | 13.146 |
| birds | Eophona_personata | Eophona_personata | LC | Stable | temperate | diurnal | 80 | 2227615.207 | 11.152 |
| birds | Eopsaltria_australis | Eopsaltria_australis | LC | Increasing | temperate | diurnal | 20.2 | 1174338.84 | 8.308 |
| birds | Eopsaltria_griseogularis | Eopsaltria_griseogularis | LC | Decreasing | temperate | diurnal | 19.4 | 587762.324 | 7.32 |
| birds | Eos_bornea | Eos_bornea | LC | Decreasing | temperate | diurnal | 156 | 65314.797 | 14.712 |
| birds | Eos_cyanogenia | Eos_cyanogenia | NT | Decreasing | tropical | diurnal | 167 | 2964.621 | 8.812 |
| birds | Eos_histrio | Eos_histrio | EN | Decreasing | tropical | diurnal | 204 | 1756.242 | 10.513 |
| birds | Eos_reticulata | Eos_reticulata | NT | Decreasing | tropical | diurnal | 153 | 4779.507 | 7.205 |
| birds | Eos_semilarvata | Eos_semilarvata | NT | Decreasing | tropical | diurnal | 94.5 | 1353.855 | 6.085 |
| birds | Eos_squamata | Eos_squamata | LC | Stable | tropical | diurnal | 91 | 32014.972 | 5.67 |
| birds | Ephippiorhynchus_asiaticus | Ephippiorhynchus_asiaticus | NT | Decreasing | tropical | diurnal | 4100 | 5889190.753 | 10.654 |
| birds | Ephippiorhynchus_senegalensis | Ephippiorhynchus_senegalensis | LC | Decreasing | tropical | diurnal | 6158.7 | 11071552.529 | 9.332 |
| birds | Ephippiospingus_dorsalis | Phrygilus_dorsalis | LC | Stable | tropical | diurnal | 38.1 | 187023.923 | 3.957 |
| birds | Ephippiospingus_erythronotus | Phrygilus_erythronotus | LC | Stable | tropical | diurnal | 20.7 | 138756.512 | 6.461 |
| birds | Epimachus_fastosus | Epimachus_fastuosus | LC | Decreasing | tropical | diurnal | 232.9 | 80656.88 | 5.639 |
| birds | Epimachus_meyeri | Epimachus_meyeri | LC | Stable | tropical | diurnal | 220.5 | 135052.616 | 5.076 |
| birds | Epinecrophylla_erythrura | Epinecrophylla_erythrura | LC | Stable | tropical | diurnal | 11.4 | 1668234.157 | 1.603 |
| birds | Epinecrophylla_fulviventris | Epinecrophylla_fulviventris | LC | Decreasing | tropical | diurnal | 10.4 | 281139.808 | 7.599 |
| birds | Epinecrophylla_gutturalis | Myrmotherula_gutturalis | LC | Decreasing | tropical | diurnal | 9.2 | 1137766.701 | 1.491 |
| birds | Epinecrophylla_haematonota | Epinecrophylla_haematonota | LC | Stable | tropical | diurnal | 8.7 | 2201917.233 | 1.172 |
| birds | Epinecrophylla_leucophthalma | Epinecrophylla_leucophthalma | LC | Decreasing | tropical | diurnal | 9.4 | 2106545.327 | 1.853 |
| birds | Epinecrophylla_ornata | Epinecrophylla_ornata | LC | Stable | tropical | diurnal | 9.4 | 779701.15 | 3.398 |
| birds | Epinecrophylla_spodionota | Epinecrophylla_spodionota | LC | Decreasing | tropical | diurnal | 10.7 | 124365.619 | 4.425 |
| birds | Epthianura_albifrons | Epthianura_albifrons | LC | Increasing | temperate | diurnal | 13.3 | 2667021.556 | 6.769 |
| birds | Epthianura_aurifrons | Epthianura_aurifrons | LC | Stable | temperate | diurnal | 10.5 | 5609912.511 | 3.838 |
| birds | Epthianura_crocea | Epthianura_crocea | LC | Unknown | tropical | diurnal | 9.3 | 2179684.966 | 2.73 |
| birds | Epthianura_tricolor | Epthianura_tricolor | LC | Stable | temperate | diurnal | 10.7 | 5486066.869 | 3.901 |
| birds | Eremalauda_dunni | Eremalauda_dunni | LC | Stable | tropical | diurnal | 20.5 | 1825439.355 | 3.388 |
| birds | Eremomela_atricollis | Eremomela_atricollis | LC | Stable | tropical | diurnal | 10.2 | 1063082.232 | 6.349 |
| birds | Eremomela_badiceps | Eremomela_badiceps | LC | Stable | tropical | diurnal | 10.6 | 3290010.909 | 7.733 |
| birds | Eremomela_canescens | Eremomela_canescens | LC | Stable | tropical | diurnal | 6.9 | 2254734.869 | 8.524 |

| | | | | | | | | | |
|---|---|---|---|---|---|---|---|---|---|
| birds | Eremomela_flavicrissalis | Eremomela_flavicrissalis | LC | Decreasing | tropical | diurnal | 4.5 | 893727.776 | 7.186 |
| birds | Eremomela_gregalis | Eremomela_gregalis | LC | Decreasing | temperate | diurnal | 6.8 | 271515.778 | 5.87 |
| birds | Eremomela_icteropygialis | Eremomela_icteropygialis | LC | Stable | tropical | diurnal | 7.5 | 8262133.255 | 9.045 |
| birds | Eremomela_pusilla | Eremomela_pusilla | LC | Stable | tropical | diurnal | 6.2 | 2960274.143 | 13.256 |
| birds | Eremomela_scotops | Eremomela_scotops | LC | Stable | tropical | diurnal | 9 | 4568185.564 | 8.58 |
| birds | Eremomela_turneri | Eremomela_turneri | NT | Decreasing | tropical | diurnal | 7.3 | 1486.675 | 11.104 |
| birds | Eremomela_usticollis | Eremomela_usticollis | LC | Stable | tropical | diurnal | 8.4 | 1430953.148 | 9.041 |
| birds | Eremophila_alpestris | Eremophila_alpestris | LC | Decreasing | temperate | diurnal | 33.3 | 44034741.82 | 6.992 |
| birds | Eremophila_bilopha | Eremophila_bilopha | LC | Decreasing | temperate | diurnal | 38.5 | 1422315.018 | 5.331 |
| birds | Eremopterix_australis | Eremopterix_australis | LC | Decreasing | temperate | diurnal | 14.2 | 642788.971 | 5.703 |
| birds | Eremopterix_griseus | Eremopterix_griseus | LC | Stable | tropical | diurnal | 16 | 2966642.699 | 18.407 |
| birds | Eremopterix_hova | Mirafra_hova | LC | Increasing | tropical | diurnal | 18.9 | 591120.506 | 8.749 |
| birds | Eremopterix_leucopareia | Eremopterix_leucopareia | LC | Unknown | tropical | diurnal | 14.5 | 859362.701 | 11.633 |
| birds | Eremopterix_leucotis | Eremopterix_leucotis | LC | Stable | tropical | diurnal | 16.4 | 7621320.473 | 10.629 |
| birds | Eremopterix_signatus | Eremopterix_signatus | LC | Increasing | tropical | diurnal | 15.7 | 970819.659 | 8.408 |
| birds | Eriocnemis_aline | Eriocnemis_alinae | LC | Decreasing | tropical | diurnal | 4.2 | 68344.266 | 5.163 |
| birds | Eriocnemis_cupreoventris | Eriocnemis_cupreoventris | LC | Stable | tropical | diurnal | 5.3 | 27252.173 | 13.827 |
| birds | Eriocnemis_derbyi | Eriocnemis_derbyi | LC | Stable | tropical | diurnal | 5.5 | 31057.373 | 11.957 |
| birds | Eriocnemis_glaucopoides | Eriocnemis_glaucopoides | LC | Stable | tropical | diurnal | 3.7 | 57142.213 | 6.929 |
| birds | Eriocnemis_godini | Eriocnemis_godini | CR | Unknown | tropical | diurnal | 5.1 | 408.046 | 26.402 |
| birds | Eriocnemis_luciani | Eriocnemis_luciani | LC | Stable | tropical | diurnal | 6 | 39822.97 | 13.344 |
| birds | Eriocnemis_mirabilis | Eriocnemis_mirabilis | EN | Decreasing | tropical | diurnal | 5.1 | 33.952 | 6.891 |
| birds | Eriocnemis_mosquera | Eriocnemis_mosquera | LC | Stable | tropical | diurnal | 5.3 | 36125.455 | 11.107 |
| birds | Eriocnemis_nigrivestis | Eriocnemis_nigrivestis | EN | Decreasing | tropical | diurnal | 8.5 | 967.348 | 18.298 |
| birds | Eriocnemis_sapphiropygia | NA | LC | Stable | NA | NA | NA | NA | NA |
| birds | Eriocnemis_vestita | Eriocnemis_vestita | LC | Stable | tropical | diurnal | 4.5 | 97851.278 | 10.958 |
| birds | Erithacus_rubecula | Erithacus_rubecula | LC | Increasing | temperate | diurnal | 17.7 | 13266206.914 | 13.869 |
| birds | Erpornis_zantholeuca | Erpornis_zantholeuca | LC | Unknown | tropical | diurnal | 11.8 | 3629429.65 | 9.965 |
| birds | Erythrocercus_holochlorus | Erythrocercus_holochlorus | LC | Stable | tropical | diurnal | 6 | 185127.387 | 12.014 |
| birds | Erythrocercus_livingstonei | Erythrocercus_livingstonei | LC | Stable | tropical | diurnal | 5.9 | 1133267.592 | 9.527 |
| birds | Erythrocercus_mccallii | Erythrocercus_mccallii | LC | Decreasing | tropical | diurnal | 7.2 | 2166425.082 | 8.937 |
| birds | Erythrogenys_erythrocnemis | Pomatorhinus_erythrocnemis | LC | Increasing | temperate | diurnal | 58.2 | 25723.198 | 17.627 |
| birds | Erythrogenys_erythrogenys | Pomatorhinus_erythrogenys | LC | Stable | temperate | diurnal | 54.9 | 400892.954 | 10.12 |
| birds | Erythrogenys_hypoleucos | Pomatorhinus_hypoleucos | LC | Decreasing | temperate | diurnal | 75.6 | 788871.399 | 9.284 |

| | | | | | | | | | |
|---|---|---|---|---|---|---|---|---|---|
| birds | Erythrogonys_cinctus | Erythrogonys_cinctus | LC | Stable | temperate | diurnal | 51.4 | 5111119.632 | 5.069 |
| birds | Erythropitta_arquata | Pitta_arcuata | LC | Decreasing | tropical | diurnal | 54 | 214049.061 | 3.059 |
| birds | Erythropitta_dohertyi | NA | LC | Decreasing | NA | NA | NA | NA | NA |
| birds | Erythropitta_erythrogaster | Pitta_erythrogaster | LC | Decreasing | tropical | diurnal | 72.4 | 289376.937 | 13.679 |
| birds | Erythropitta_granatina | Pitta_granatina | NT | Decreasing | tropical | diurnal | 61.5 | 852985.142 | 8.354 |
| birds | Erythropitta_kochi | Pitta_kochi | LC | Decreasing | tropical | diurnal | 116 | 37303.879 | 7.755 |
| birds | Erythropitta_ussheri | Pitta_ussheri | LC | Decreasing | tropical | diurnal | 55.3 | 41122.345 | 8.267 |
| birds | Erythropitta_venusta | NA | LC | Decreasing | NA | NA | NA | NA | NA |
| birds | Erythrotriorchis_buergersi | Erythrotriorchis_buergersi | DD | Stable | tropical | diurnal | 575 | 152406.319 | 3.327 |
| birds | Erythrotriorchis_radiatus | Erythrotriorchis_radiatus | EN | Decreasing | tropical | diurnal | 887.4 | 1042091.379 | 3.505 |
| birds | Erythrura_coloria | Erythrura_coloria | LC | Stable | tropical | diurnal | 11.3 | 12662.234 | 8.334 |
| birds | Erythrura_cyaneovirens | Erythrura_cyaneovirens | NT | Decreasing | tropical | diurnal | 13 | 14573.351 | 5.935 |
| birds | Erythrura_hyperythra | Erythrura_hyperythra | LC | Stable | tropical | diurnal | 11 | 294579.35 | 9.5 |
| birds | Erythrura_kleinschmidti | Erythrura_kleinschmidti | VU | Decreasing | tropical | diurnal | 21 | 4811.607 | 7.775 |
| birds | Erythrura_papuana | Erythrura_papuana | LC | Stable | tropical | diurnal | 19.4 | 180233.842 | 4.079 |
| birds | Erythrura_pealii | Erythrura_pealii | LC | Stable | tropical | diurnal | 12 | 17012.666 | 9.369 |
| birds | Erythrura_prasina | Erythrura_prasina | LC | Stable | tropical | diurnal | 14.7 | 1585424.029 | 8.891 |
| birds | Erythrura_psittacea | Erythrura_psittacea | LC | Stable | tropical | diurnal | 11.5 | 17405.19 | 7.634 |
| birds | Erythrura_trichroa | Erythrura_trichroa | LC | Stable | tropical | diurnal | 14.4 | 858792.099 | 5.105 |
| birds | Erythrura_tricolor | Erythrura_tricolor | LC | Stable | tropical | diurnal | 11 | 36517.843 | 12.249 |
| birds | Erythrura_viridifacies | Erythrura_viridifacies | VU | Decreasing | tropical | diurnal | 12.6 | 106324.246 | 14.919 |
| birds | Esacus_magnirostris | Esacus_giganteus | NT | Decreasing | tropical | diurnal | 1015.9 | 2207239.517 | 8.62 |
| birds | Esacus_recurvirostris | Esacus_recurvirostris | NT | Decreasing | tropical | nocturnal | 790 | 2969603.168 | 15.887 |
| birds | Estrilda_astrild | Estrilda_astrild | LC | Stable | tropical | diurnal | 8.3 | 6015986.23 | 9.978 |
| birds | Estrilda_atricapilla | Estrilda_atricapilla | LC | Stable | tropical | diurnal | 7.8 | 665104.344 | 6.695 |
| birds | Estrilda_kandti | Estrilda_kandti | LC | Stable | tropical | diurnal | 8 | 113134.745 | 13.556 |
| birds | Estrilda_melpoda | Estrilda_melpoda | LC | Stable | tropical | diurnal | 7.6 | 3755904.769 | 10.102 |
| birds | Estrilda_nigriloris | Estrilda_nigriloris | DD | Stable | tropical | diurnal | 7.5 | 28361.751 | 6.533 |
| birds | Estrilda_nonnula | Estrilda_nonnula | LC | Stable | tropical | diurnal | 7.7 | 1216438.929 | 8.935 |
| birds | Estrilda_paludicola | Estrilda_paludicola | LC | Stable | tropical | diurnal | 7.6 | 1983230.355 | 9.045 |
| birds | Estrilda_poliopareia | Estrilda_poliopareia | NT | Stable | tropical | diurnal | 7.5 | 43938.466 | 16.74 |
| birds | Estrilda_rhodopyga | Estrilda_rhodopyga | LC | Stable | tropical | diurnal | 7.7 | 1239412.969 | 11.806 |
| birds | Estrilda_rufibarba | Estrilda_rufibarba | LC | Decreasing | tropical | diurnal | 8.5 | 86115.995 | 10.92 |
| birds | Estrilda_troglodytes | Estrilda_troglodytes | LC | Stable | tropical | diurnal | 7.6 | 2017180.163 | 12.548 |

| | | | | | | | | | |
|---|---|---|---|---|---|---|---|---|---|
| birds | Eubucco_bourcierii | Eubucco_bourcierii | LC | Decreasing | tropical | diurnal | 37 | 234502.702 | 8.757 |
| birds | Eubucco_richardsoni | Eubucco_richardsoni | LC | Decreasing | tropical | diurnal | 33.9 | 971591.17 | 2.494 |
| birds | Eubucco_tucinkae | Eubucco_tucinkae | LC | Stable | tropical | diurnal | 41.7 | 177646.678 | 1.341 |
| birds | Eubucco_versicolor | Eubucco_versicolor | LC | Unknown | tropical | diurnal | 32.8 | 125625.221 | 4.245 |
| birds | Euchrepomis_callinota | Terenura_callinota | LC | Decreasing | tropical | diurnal | 7 | 227242.245 | 6.4 |
| birds | Euchrepomis_humeralis | Terenura_humeralis | LC | Decreasing | tropical | diurnal | 7.3 | 1272005.553 | 1.632 |
| birds | Euchrepomis_sharpei | Terenura_sharpei | NT | Decreasing | tropical | diurnal | 7.5 | 35451.949 | 4.163 |
| birds | Euchrepomis_spodioptila | Terenura_spodioptila | LC | Decreasing | tropical | diurnal | 6.5 | 2162349.558 | 1.419 |
| birds | Eucometis_penicillata | Eucometis_penicillata | LC | Stable | tropical | diurnal | 27 | 5712449.557 | 5.297 |
| birds | Eudocimus_albus | Eudocimus_albus | LC | Stable | tropical | diurnal | 889.7 | 1931978.981 | 12.527 |
| birds | Eudocimus_ruber | Eudocimus_ruber | LC | Decreasing | tropical | diurnal | 660.1 | 816290.778 | 7.478 |
| birds | Eudromia_elegans | Eudromia_elegans | LC | Decreasing | temperate | diurnal | 702.9 | 1493624.395 | 6.838 |
| birds | Eudromia_formosa | Eudromia_formosa | LC | Decreasing | temperate | diurnal | 772.9 | 285672.468 | 5.666 |
| birds | Eudromias_morinellus | Eudromias_morinellus | LC | Decreasing | temperate | cathemeral | 108.2 | 4095968.093 | 3.083 |
| birds | Eudynamys_scolopaceus | Eudynamys_scolopaceus | LC | Stable | tropical | diurnal | 194.9 | 9699096.473 | 14.504 |
| birds | Eudyptes_chrysocome | Eudyptes_chrysocome | VU | Decreasing | temperate | cathemeral | 2327.9 | 11153673.098 | 2.55 |
| birds | Eudyptes_chrysolophus | Eudyptes_chrysolophus | VU | Decreasing | temperate | cathemeral | 4485.1 | 20777681.261 | 2.036 |
| birds | Eudyptes_pachyrhynchus | Eudyptes_pachyrhynchus | NT | Decreasing | temperate | cathemeral | 3904.9 | 4074057.852 | 10.161 |
| birds | Eudyptes_robustus | Eudyptes_robustus | VU | Stable | temperate | cathemeral | 3038 | 3482085.778 | 6.042 |
| birds | Eudyptes_schlegeli | NA | LC | Unknown | NA | NA | NA | NA | NA |
| birds | Eudyptes_sclateri | NA | EN | Decreasing | NA | NA | NA | NA | NA |
| birds | Eudyptula_minor | Eudyptula_minor | LC | Stable | temperate | cathemeral | 1107.8 | 853439.178 | 10.694 |
| birds | Eugenes_fulgens | Eugenes_fulgens | LC | Stable | temperate | diurnal | 8 | 1476036.284 | 8.971 |
| birds | Eugerygone_rubra | Eugerygone_rubra | LC | Stable | tropical | diurnal | 8.4 | 122286.483 | 5.089 |
| birds | Eugralla_paradoxa | Eugralla_paradoxa | LC | Decreasing | temperate | diurnal | 44 | 124178.004 | 9.08 |
| birds | Eulabeornis_castaneoventris | Eulabeornis_castaneoventris | LC | Unknown | tropical | cathemeral | 684.5 | 85822.054 | 2.315 |
| birds | Eulacestoma_nigropectus | Eulacestoma_nigropectus | LC | Stable | tropical | diurnal | 20 | 89484.92 | 4.998 |
| birds | Eulampis_holosericeus | Eulampis_holosericeus | LC | Unknown | tropical | diurnal | 5.7 | 8331.82 | 23.476 |
| birds | Eulampis_jugularis | Eulampis_jugularis | LC | Decreasing | tropical | diurnal | 8.9 | 4968.075 | 22.111 |
| birds | Eulidia_yarrellii | Eulidia_yarrellii | CR | Decreasing | tropical | diurnal | 2.5 | 35771.468 | 4.66 |
| birds | Eulipoa_wallacei | Eulipoa_wallacei | VU | Decreasing | tropical | diurnal | 505 | 41464.66 | 6.196 |
| birds | Eumomota_superciliosa | Eumomota_superciliosa | LC | Decreasing | tropical | diurnal | 62.5 | 242415.116 | 11.823 |
| birds | Eumyias_additus | Rhinomyias_additus | NT | Decreasing | tropical | diurnal | 22.5 | 4091.969 | 5.208 |
| birds | Eumyias_albicaudatus | Eumyias_albicaudatus | LC | Decreasing | tropical | diurnal | 16.4 | 32809.705 | 12.861 |

| | | | | | | | | | |
|---|---|---|---|---|---|---|---|---|---|
| birds | Eumyias_hoevelli | NA | LC | Decreasing | NA | NA | NA | NA | NA |
| birds | Eumyias_hyacinthinus | Cyornis_hyacinthinus | LC | Decreasing | tropical | diurnal | 25.8 | 32681.542 | 13.22 |
| birds | Eumyias_indigo | Eumyias_indigo | LC | Decreasing | tropical | diurnal | 23.1 | 270383.731 | 8.02 |
| birds | Eumyias_panayensis | Eumyias_panayensis | LC | Decreasing | tropical | diurnal | 20 | 173823.511 | 7.203 |
| birds | Eumyias_sanfordi | NA | LC | Stable | NA | NA | NA | NA | NA |
| birds | Eumyias_sordidus | Eumyias_sordidus | NT | Decreasing | tropical | diurnal | 25.1 | 5844.08 | 16.925 |
| birds | Eumyias_thalassinus | Eumyias_thalassinus | LC | Stable | tropical | diurnal | 18.1 | 7039599.106 | 12.669 |
| birds | Euneornis_campestris | Euneornis_campestris | LC | Decreasing | tropical | diurnal | 16 | 11021.791 | 19.741 |
| birds | Eunymphicus_cornutus | Eunymphicus_cornutus | VU | Increasing | tropical | diurnal | 130 | 3594.344 | 4.105 |
| birds | Euodice_cantans | Lonchura_cantans | LC | Stable | tropical | diurnal | 12 | 3802160.031 | 10.424 |
| birds | Euodice_malabarica | Lonchura_malabarica | LC | Stable | tropical | diurnal | 12 | 3376406.354 | 17.462 |
| birds | Eupetes_macrocerus | Eupetes_macrocerus | NT | Decreasing | tropical | diurnal | 69 | 1150485.5 | 8.48 |
| birds | Eupetomena_cirrochloris | Aphantochroa_cirrochloris | LC | Unknown | tropical | diurnal | 7.7 | 1438300.989 | 12.145 |
| birds | Eupetomena_macroura | Eupetomena_macroura | LC | Unknown | tropical | diurnal | 9 | 4594117.6 | 9.08 |
| birds | Euphagus_carolinus | Euphagus_carolinus | VU | Decreasing | temperate | diurnal | 59.6 | 12263114.381 | 8.083 |
| birds | Euphagus_cyanocephalus | Euphagus_cyanocephalus | LC | Decreasing | temperate | diurnal | 62.5 | 10214123.729 | 11.526 |
| birds | Eupherusa_cyanophrys | Eupherusa_cyanophrys | EN | Decreasing | tropical | diurnal | 4.8 | 4063.858 | 5.875 |
| birds | Eupherusa_eximia | Eupherusa_eximia | LC | Decreasing | tropical | diurnal | 4.3 | 174512.432 | 10.119 |
| birds | Eupherusa_nigriventris | Eupherusa_nigriventris | LC | Decreasing | tropical | diurnal | 3.5 | 14338.293 | 6.239 |
| birds | Eupherusa_poliocerca | Eupherusa_poliocerca | NT | Decreasing | tropical | diurnal | 4.8 | 15663.297 | 5.379 |
| birds | Eupherusa_ridgwayi | Thalurania_ridgwayi | VU | Decreasing | tropical | diurnal | 3.8 | 14528.864 | 9.963 |
| birds | Euphonia_affinis | Euphonia_affinis | LC | Decreasing | tropical | diurnal | 10 | 724298.203 | 10.149 |
| birds | Euphonia_anneae | Euphonia_anneae | LC | Decreasing | tropical | diurnal | 14.9 | 20146.513 | 5.566 |
| birds | Euphonia_cayennensis | Euphonia_cayennensis | LC | Stable | tropical | diurnal | 14.3 | 1272525.447 | 1.876 |
| birds | Euphonia_chalybea | Euphonia_chalybea | NT | Decreasing | temperate | diurnal | 19 | 745412.863 | 13.615 |
| birds | Euphonia_chlorotica | Euphonia_chlorotica | LC | Stable | tropical | diurnal | 11 | 9620331.066 | 6.736 |
| birds | Euphonia_chrysopasta | Euphonia_chrysopasta | LC | Decreasing | tropical | diurnal | 14 | 5478664.301 | 1.736 |
| birds | Euphonia_concinna | Euphonia_concinna | LC | Decreasing | tropical | diurnal | 10.3 | 26361.573 | 17.152 |
| birds | Euphonia_cyanocephala | Euphonia_cyanocephala | LC | Stable | tropical | diurnal | 14 | 2711347.809 | 10.554 |
| birds | Euphonia_elegantissima | Euphonia_elegantissima | LC | Decreasing | tropical | diurnal | 15 | 473565.766 | 11.496 |
| birds | Euphonia_finschi | Euphonia_finschi | LC | Stable | tropical | diurnal | 10.5 | 327330.525 | 2.355 |
| birds | Euphonia_fulvicrissa | Euphonia_fulvicrissa | LC | Decreasing | tropical | diurnal | 11 | 198414.878 | 8.057 |
| birds | Euphonia_gouldi | Euphonia_gouldi | LC | Decreasing | tropical | diurnal | 14 | 326598.755 | 9.184 |
| birds | Euphonia_hirundinacea | Euphonia_hirundinacea | LC | Decreasing | tropical | diurnal | 14 | 642669.455 | 10.263 |

| | | | | | | | | | |
|---|---|---|---|---|---|---|---|---|---|
| birds | Euphonia_imitans | Euphonia_imitans | LC | Decreasing | tropical | diurnal | 14 | 14869.651 | 14.541 |
| birds | Euphonia_jamaica | Euphonia_jamaica | LC | Stable | tropical | diurnal | 16.6 | 11021.791 | 19.741 |
| birds | Euphonia_laniirostris | Euphonia_laniirostris | LC | Stable | tropical | diurnal | 15 | 4464958.75 | 4.7 |
| birds | Euphonia_luteicapilla | Euphonia_luteicapilla | LC | Stable | tropical | diurnal | 13 | 111584.472 | 10.992 |
| birds | Euphonia_mesochrysa | Euphonia_mesochrysa | LC | Decreasing | tropical | diurnal | 13 | 193537.981 | 5.421 |
| birds | Euphonia_minuta | Euphonia_minuta | LC | Decreasing | tropical | diurnal | 10 | 5149944.301 | 2.397 |
| birds | Euphonia_musica | Euphonia_musica | LC | Stable | tropical | diurnal | 14.7 | 35327.884 | 16.889 |
| birds | Euphonia_pectoralis | Euphonia_pectoralis | LC | Decreasing | tropical | diurnal | 14.4 | 1361244.455 | 13.07 |
| birds | Euphonia_plumbea | Euphonia_plumbea | LC | Stable | tropical | diurnal | 8.9 | 913453.903 | 1.468 |
| birds | Euphonia_rufiventris | Euphonia_rufiventris | LC | Stable | tropical | diurnal | 14 | 4409554.137 | 1.976 |
| birds | Euphonia_saturata | Euphonia_saturata | LC | Stable | tropical | diurnal | 13.3 | 95078.534 | 11.754 |
| birds | Euphonia_trinitatis | Euphonia_trinitatis | LC | Stable | tropical | diurnal | 11 | 610172.655 | 9.623 |
| birds | Euphonia_violacea | Euphonia_violacea | LC | Decreasing | tropical | diurnal | 15 | 4921834.311 | 6.383 |
| birds | Euphonia_xanthogaster | Euphonia_xanthogaster | LC | Decreasing | tropical | diurnal | 13 | 3715684.497 | 3.228 |
| birds | Euplectes_afer | Euplectes_afer | LC | Stable | tropical | diurnal | 14.3 | 3118130.651 | 11.423 |
| birds | Euplectes_albonotatus | Euplectes_albonotatus | LC | Stable | tropical | diurnal | 21.1 | 2575317.408 | 11.432 |
| birds | Euplectes_ardens | Euplectes_ardens | LC | Stable | tropical | diurnal | 19 | 3182674.539 | 10.538 |
| birds | Euplectes_aureus | Euplectes_aureus | LC | Stable | tropical | diurnal | 25 | 44203.058 | 10.504 |
| birds | Euplectes_axillaris | Euplectes_axillaris | LC | Stable | tropical | diurnal | 25.1 | 2371547.671 | 10.769 |
| birds | Euplectes_capensis | Euplectes_capensis | LC | Stable | tropical | diurnal | 30.5 | 2639722.61 | 11.144 |
| birds | Euplectes_diadematus | Euplectes_diadematus | LC | Stable | tropical | diurnal | 13.9 | 233386.144 | 12.627 |
| birds | Euplectes_franciscanus | Euplectes_franciscanus | LC | Stable | tropical | diurnal | 16 | 3267667.669 | 12.301 |
| birds | Euplectes_gierowii | Euplectes_gierowii | LC | Stable | tropical | diurnal | 28.3 | 246606.931 | 11.932 |
| birds | Euplectes_hartlaubi | Euplectes_hartlaubi | LC | Stable | tropical | diurnal | 35.5 | 741261.112 | 7.505 |
| birds | Euplectes_hordeaceus | Euplectes_hordeaceus | LC | Stable | tropical | diurnal | 19.4 | 5103979.032 | 11.02 |
| birds | Euplectes_jacksoni | Euplectes_jacksoni | NT | Stable | tropical | diurnal | 37 | 107944.776 | 15.077 |
| birds | Euplectes_macroura | Euplectes_macroura | LC | Stable | tropical | diurnal | 21.7 | 3993290.032 | 10.717 |
| birds | Euplectes_nigroventris | Euplectes_nigroventris | LC | Stable | tropical | diurnal | 12.5 | 170115.235 | 10.888 |
| birds | Euplectes_orix | Euplectes_orix | LC | Stable | temperate | diurnal | 23 | 2969008.668 | 10.738 |
| birds | Euplectes_progne | Euplectes_progne | LC | Stable | temperate | diurnal | 35.1 | 697961.279 | 11.561 |
| birds | Euplectes_psammacromius | Euplectes_psammocromius | LC | Stable | tropical | diurnal | 35.5 | 34873.18 | 14.896 |
| birds | Eupodotis_caerulescens | Eupodotis_caerulescens | NT | Decreasing | temperate | diurnal | 1366 | 264843.991 | 11.341 |
| birds | Eupodotis_senegalensis | Eupodotis_senegalensis | LC | Decreasing | tropical | diurnal | 585 | 4886826.685 | 10.935 |
| birds | Eupsittula_aurea | Aratinga_aurea | LC | Stable | tropical | diurnal | 84.6 | 5764751.613 | 7.438 |

| | | | | | | | | | |
|---|---|---|---|---|---|---|---|---|---|
| birds | Eupsittula_cactorum | Aratinga_cactorum | LC | Stable | tropical | diurnal | 82.5 | 1195222.877 | 7.562 |
| birds | Eupsittula_canicularis | Aratinga_canicularis | VU | Decreasing | tropical | diurnal | 85 | 283885.271 | 12.632 |
| birds | Eupsittula_nana | Aratinga_nana | NT | Decreasing | tropical | diurnal | 76.9 | 11021.791 | 19.741 |
| birds | Eupsittula_pertinax | Aratinga_pertinax | LC | Stable | tropical | diurnal | 84 | 1480960.995 | 6.727 |
| birds | Euptilotis_neoxenus | Euptilotis_neoxenus | LC | Decreasing | temperate | diurnal | 122.3 | 338167.416 | 7.144 |
| birds | Euptilotus_eutilotus | Pycnonotus_eutilotus | NT | Decreasing | tropical | diurnal | 35.3 | 932677.681 | 6.772 |
| birds | Eurillas_ansorgei | Andropadus_ansorgei | LC | Stable | tropical | diurnal | 18.7 | 1858480.057 | 9.036 |
| birds | Eurillas_curvirostris | Andropadus_curvirostris | LC | Stable | tropical | diurnal | 24.9 | 3502649.293 | 8.493 |
| birds | Eurillas_gracilis | Andropadus_gracilis | LC | Stable | tropical | diurnal | 21.6 | 3552112.863 | 7.986 |
| birds | Eurillas_latirostris | Andropadus_latirostris | LC | Stable | tropical | diurnal | 26.2 | 3682285.719 | 8.791 |
| birds | Eurillas_virens | Andropadus_virens | LC | Stable | tropical | diurnal | 26.6 | 5766996.331 | 8.43 |
| birds | Eurocephalus_anguitimens | Eurocephalus_anguitimens | LC | Stable | tropical | diurnal | 69 | 1521047.708 | 7.876 |
| birds | Eurocephalus_ruppelli | Eurocephalus_rueppelli | LC | Stable | tropical | diurnal | 50.6 | 1835004.726 | 9.601 |
| birds | Eurochelidon_sirintarae | Eurochelidon_sirintarae | CR | Decreasing | tropical | diurnal | 26 | 895268.431 | 12.727 |
| birds | Eurostopodus_archboldi | Eurostopodus_archboldi | LC | Stable | tropical | nocturnal | 77 | 74591.658 | 4.829 |
| birds | Eurostopodus_argus | Eurostopodus_argus | LC | Stable | temperate | nocturnal | 93.4 | 7014029.222 | 3.926 |
| birds | Eurostopodus_diabolicus | Eurostopodus_diabolicus | LC | Decreasing | tropical | nocturnal | 104.4 | 18765.428 | 4.76 |
| birds | Eurostopodus_mystacalis | Eurostopodus_mystacalis | LC | Decreasing | temperate | nocturnal | 134.5 | 1216308.423 | 6.44 |
| birds | Eurostopodus_papuensis | Eurostopodus_papuensis | LC | Decreasing | tropical | nocturnal | 80.5 | 502860.656 | 3.126 |
| birds | Euryceros_prevostii | Euryceros_prevostii | EN | Decreasing | tropical | diurnal | 96.9 | 16245.717 | 5.851 |
| birds | Eurylaimus_javanicus | Eurylaimus_javanicus | NT | Decreasing | tropical | diurnal | 78.2 | 126074.832 | 22.349 |
| birds | Eurylaimus_ochromalus | Eurylaimus_ochromalus | NT | Decreasing | tropical | diurnal | 33.3 | 1129454.951 | 8.164 |
| birds | Euryptila_subcinnamomea | Euryptila_subcinnamomea | LC | Stable | temperate | diurnal | 11.5 | 274057.075 | 6.437 |
| birds | Eurypyga_helias | Eurypyga_helias | LC | Decreasing | tropical | diurnal | 210 | 8423208.451 | 3.184 |
| birds | Eurystomus_gularis | Eurystomus_gularis | LC | Decreasing | tropical | diurnal | 96.3 | 2827968.863 | 8.468 |
| birds | Euscarthmus_meloryphus | Euscarthmus_meloryphus | LC | Increasing | tropical | diurnal | 6.8 | 6211591.054 | 9.116 |
| birds | Euscarthmus_rufomarginatus | Euscarthmus_rufomarginatus | NT | Decreasing | tropical | diurnal | 6 | 2489423.839 | 9.08 |
| birds | Euschistospiza_cinereovinacea | Euschistospiza_cinereovinacea | LC | Stable | tropical | diurnal | 12 | 143514.592 | 12.283 |
| birds | Euschistospiza_dybowskii | Euschistospiza_dybowskii | LC | Stable | tropical | diurnal | 13.4 | 776837.641 | 8.371 |
| birds | Eutoxeres_aquila | Eutoxeres_aquila | LC | Decreasing | tropical | diurnal | 10.6 | 306610.24 | 6.905 |
| birds | Eutoxeres_condamini | Eutoxeres_condamini | LC | Decreasing | tropical | diurnal | 10.8 | 370060.389 | 3.962 |
| birds | Eutrichomyias_rowleyi | Eutrichomyias_rowleyi | CR | Decreasing | tropical | diurnal | 19 | 555.847 | 11.621 |
| birds | Eutriorchis_astur | Eutriorchis_astur | EN | Decreasing | tropical | diurnal | 700 | 34460.566 | 5.578 |
| birds | Falcipennis_falcipennis | Dendragapus_falcipennis | NT | Decreasing | temperate | diurnal | 685.6 | 692147.347 | 1.358 |

| | | | | | | | | | |
|---|---|---|---|---|---|---|---|---|---|
| birds | Falco_alopex | Falco_alopex | LC | Stable | tropical | diurnal | 273.9 | 5286034.086 | 11.066 |
| birds | Falco_araeus | NA | VU | Stable | NA | NA | NA | NA | NA |
| birds | Falco_ardosiaceus | Falco_ardosiaceus | LC | Stable | tropical | diurnal | 234.2 | 7345675.559 | 10.549 |
| birds | Falco_berigora | Falco_berigora | LC | Decreasing | temperate | diurnal | 587 | 8273511.567 | 4.21 |
| birds | Falco_biarmicus | Falco_biarmicus | LC | Decreasing | tropical | nocturnal | 606.1 | 18096388.228 | 9.14 |
| birds | Falco_cenchroides | Falco_cenchroides | LC | Increasing | temperate | diurnal | 170.5 | 8034280.028 | 4.21 |
| birds | Falco_cherrug | Falco_cherrug | EN | Decreasing | temperate | diurnal | 961.2 | 38026625.063 | 7.919 |
| birds | Falco_chicquera | Falco_chicquera | NT | Decreasing | tropical | diurnal | 208 | 3166581.198 | 17.241 |
| birds | Falco_columbarius | Falco_columbarius | LC | Stable | temperate | diurnal | 188.5 | 47918643.638 | 8.715 |
| birds | Falco_deiroleucus | Falco_deiroleucus | NT | Decreasing | tropical | diurnal | 459.6 | 3323144.473 | 5.172 |
| birds | Falco_dickinsoni | Falco_dickinsoni | LC | Stable | tropical | diurnal | 209 | 3765759.754 | 8.265 |
| birds | Falco_eleonorae | Falco_eleonorae | LC | Increasing | tropical | nocturnal | 390 | 5141958.02 | 10.984 |
| birds | Falco_fasciinucha | Falco_fasciinucha | VU | Decreasing | tropical | diurnal | 254.7 | 501059.822 | 12.473 |
| birds | Falco_femoralis | Falco_femoralis | LC | Decreasing | tropical | diurnal | 335.8 | 12113135.738 | 8.167 |
| birds | Falco_hypoleucos | Falco_hypoleucos | VU | Stable | tropical | diurnal | 457.2 | 3627330.536 | 2.566 |
| birds | Falco_jugger | Falco_jugger | NT | Decreasing | tropical | diurnal | 755 | 2912240.729 | 17.888 |
| birds | Falco_longipennis | Falco_longipennis | LC | Increasing | temperate | cathemeral | 249.8 | 8078573.631 | 4.328 |
| birds | Falco_mexicanus | Falco_mexicanus | LC | Increasing | temperate | diurnal | 713.1 | 5425089.082 | 10.108 |
| birds | Falco_moluccensis | Falco_moluccensis | LC | Increasing | tropical | diurnal | 162 | 466435.959 | 13.619 |
| birds | Falco_naumanni | Falco_naumanni | LC | Stable | tropical | diurnal | 152.1 | 43373581.242 | 8.372 |
| birds | Falco_newtoni | Falco_newtoni | LC | Increasing | tropical | diurnal | 123.4 | 590995.808 | 8.745 |
| birds | Falco_novaeseelandiae | Falco_novaeseelandiae | LC | Stable | temperate | diurnal | 418.6 | 242840.057 | 7.831 |
| birds | Falco_peregrinus | Falco_peregrinus | LC | Increasing | temperate | diurnal | 760 | 92354726.04 | 8.291 |
| birds | Falco_punctatus | Falco_punctatus | EN | Decreasing | tropical | diurnal | 167.4 | 773.932 | 19.699 |
| birds | Falco_rufigularis | Falco_rufigularis | LC | Decreasing | tropical | nocturnal | 163.6 | 13377698.051 | 5.758 |
| birds | Falco_rupicoloides | Falco_rupicoloides | LC | Stable | tropical | diurnal | 261 | 3191508.184 | 7.535 |
| birds | Falco_rusticolus | Falco_rusticolus | LC | Stable | temperate | diurnal | 1431.7 | 26161318.429 | 2.747 |
| birds | Falco_sparverius | Falco_sparverius | LC | Stable | temperate | diurnal | 114.6 | 27776585.208 | 9.101 |
| birds | Falco_subniger | Falco_subniger | LC | Stable | temperate | diurnal | 776 | 5910471.622 | 4.317 |
| birds | Falco_tinnunculus | Falco_tinnunculus | LC | Decreasing | temperate | diurnal | 183.2 | 58595922.698 | 10.142 |
| birds | Falco_zoniventris | Falco_zoniventris | LC | Stable | tropical | diurnal | 174 | 411893.802 | 8.353 |
| birds | Falculea_palliata | Falculea_palliata | LC | Decreasing | tropical | diurnal | 111 | 301726.392 | 8.531 |
| birds | Falcunculus_frontatus | Falcunculus_frontatus | LC | Decreasing | temperate | diurnal | 27.9 | 981576.388 | 9.825 |
| birds | Ferminia_cerverai | Ferminia_cerverai | EN | Decreasing | tropical | diurnal | 26 | 2102.528 | 5.493 |

| | | | | | | | | | |
|---|---|---|---|---|---|---|---|---|---|
| birds | Ficedula_albicollis | Ficedula_albicollis | LC | Increasing | tropical | diurnal | 12.7 | 11815429.538 | 12.438 |
| birds | Ficedula_basilanica | Ficedula_basilanica | VU | Decreasing | tropical | diurnal | 13.5 | 116046.89 | 11.91 |
| birds | Ficedula_bonthaina | NA | EN | Decreasing | NA | NA | NA | NA | NA |
| birds | Ficedula_buruensis | Ficedula_buruensis | LC | Decreasing | tropical | diurnal | 14.5 | 26661.655 | 6.981 |
| birds | Ficedula_crypta | Ficedula_crypta | LC | Decreasing | tropical | diurnal | 11.5 | 20661.05 | 8.691 |
| birds | Ficedula_disposita | NA | NT | Decreasing | NA | NA | NA | NA | NA |
| birds | Ficedula_dumetoria | Ficedula_dumetoria | LC | Decreasing | tropical | diurnal | 9.4 | 377877.311 | 4.17 |
| birds | Ficedula_erithacus | Ficedula_hodgsonii | LC | Stable | temperate | diurnal | 10 | 2131133.68 | 9.405 |
| birds | Ficedula_harterti | NA | LC | Decreasing | NA | NA | NA | NA | NA |
| birds | Ficedula_henrici | NA | NT | Decreasing | NA | NA | NA | NA | NA |
| birds | Ficedula_hodgsoni | Muscicapella_hodgsoni | LC | Stable | tropical | diurnal | 10 | 847137.812 | 7.696 |
| birds | Ficedula_hyperythra | Ficedula_hyperythra | LC | Decreasing | tropical | diurnal | 8.2 | 2932874.664 | 11.65 |
| birds | Ficedula_hypoleuca | Ficedula_hypoleuca | LC | Decreasing | temperate | diurnal | 13.8 | 21870324.643 | 7.934 |
| birds | Ficedula_mugimaki | Ficedula_mugimaki | LC | Stable | temperate | diurnal | 11.7 | 7731077.745 | 6.911 |
| birds | Ficedula_narcissina | Ficedula_narcissina | LC | Stable | temperate | diurnal | 13.1 | 871204.768 | 13.85 |
| birds | Ficedula_nigrorufa | Ficedula_nigrorufa | LC | Decreasing | tropical | diurnal | 9.4 | 21353.516 | 12.264 |
| birds | Ficedula_parva | Ficedula_parva | LC | Increasing | temperate | diurnal | 10.8 | 21638109.76 | 11.619 |
| birds | Ficedula_platenae | Ficedula_platenae | VU | Decreasing | tropical | diurnal | 12.1 | 11490.794 | 9.024 |
| birds | Ficedula_ruficauda | Muscicapa_ruficauda | LC | Stable | temperate | diurnal | 13.3 | 2570084.11 | 15.471 |
| birds | Ficedula_rufigula | Ficedula_rufigula | NT | Decreasing | tropical | diurnal | 11.6 | 114166.087 | 7.39 |
| birds | Ficedula_sapphira | Ficedula_sapphira | LC | Stable | temperate | diurnal | 7.8 | 1517380.397 | 11.503 |
| birds | Ficedula_semitorquata | Ficedula_semitorquata | LC | Decreasing | tropical | diurnal | 14.4 | 5511481.462 | 9.158 |
| birds | Ficedula_strophiata | Ficedula_strophiata | LC | Stable | temperate | diurnal | 12.8 | 2339628.403 | 11.077 |
| birds | Ficedula_subrubra | Ficedula_subrubra | VU | Decreasing | temperate | diurnal | 10.4 | 18887.266 | 13.327 |
| birds | Ficedula_superciliaris | Ficedula_superciliaris | LC | Stable | temperate | diurnal | 8 | 3996907.954 | 14.282 |
| birds | Ficedula_timorensis | NA | NT | Decreasing | NA | NA | NA | NA | NA |
| birds | Ficedula_tricolor | Ficedula_tricolor | LC | Stable | temperate | diurnal | 8.3 | 2148183.472 | 11.419 |
| birds | Ficedula_westermanni | Ficedula_westermanni | LC | Decreasing | tropical | diurnal | 7.8 | 4080480.379 | 11.479 |
| birds | Ficedula_zanthopygia | Ficedula_zanthopygia | LC | Stable | temperate | diurnal | 12.1 | 3306839.918 | 12.899 |
| birds | Florisuga_fusca | Florisuga_fusca | LC | Unknown | temperate | diurnal | 8 | 675214.038 | 12.127 |
| birds | Florisuga_mellivora | Florisuga_mellivora | LC | Decreasing | tropical | diurnal | 7.4 | 8105538.694 | 3.361 |
| birds | Fluvicola_albiventer | Fluvicola_albiventer | LC | Stable | tropical | diurnal | 11.6 | 8121417.599 | 5.571 |
| birds | Fluvicola_nengeta | Fluvicola_nengeta | LC | Increasing | tropical | diurnal | 21 | 3642098.173 | 9.91 |
| birds | Fluvicola_pica | Fluvicola_pica | LC | Decreasing | tropical | diurnal | 12.3 | 1883895.028 | 5.645 |

| | | | | | | | | | |
|---|---|---|---|---|---|---|---|---|---|
| birds | Formicarius_analis | Formicarius_analis | LC | Decreasing | tropical | diurnal | 62.2 | 5715479.199 | 2.993 |
| birds | Formicarius_colma | Formicarius_colma | LC | Decreasing | tropical | diurnal | 47 | 6982041.191 | 2.77 |
| birds | Formicarius_moniliger | Formicarius_moniliger | LC | Decreasing | tropical | diurnal | 61.9 | 316042.055 | 9.102 |
| birds | Formicarius_nigricapillus | Formicarius_nigricapillus | LC | Decreasing | tropical | diurnal | 56.2 | 108463.369 | 7.531 |
| birds | Formicarius_rufifrons | Formicarius_rufifrons | LC | Decreasing | tropical | diurnal | 55.5 | 17407.431 | 3.709 |
| birds | Formicarius_rufipectus | Formicarius_rufipectus | LC | Decreasing | tropical | diurnal | 74.4 | 206331.947 | 6.113 |
| birds | Formicivora_acutirostris | Stymphalornis_acutirostris | NT | Decreasing | temperate | diurnal | 10 | 880.769 | 11.785 |
| birds | Formicivora_erythronotos | Formicivora_erythronotos | EN | Decreasing | tropical | diurnal | 10.3 | 105.33 | 20.403 |
| birds | Formicivora_grisea | Formicivora_grisea | LC | Decreasing | tropical | diurnal | 10.4 | 5032873.87 | 4.619 |
| birds | Formicivora_iheringi | Formicivora_iheringi | LC | Decreasing | tropical | diurnal | 8.2 | 62390.869 | 7.689 |
| birds | Formicivora_intermedia | Formicivora_intermedia | LC | Decreasing | tropical | diurnal | 9.8 | 779878.052 | 10.628 |
| birds | Formicivora_melanogaster | Formicivora_melanogaster | LC | Decreasing | tropical | diurnal | 10.5 | 2649645.382 | 9.012 |
| birds | Formicivora_rufa | Formicivora_rufa | LC | Stable | tropical | diurnal | 10.8 | 4090899.874 | 8.206 |
| birds | Formicivora_serrana | Formicivora_serrana | LC | Decreasing | tropical | diurnal | 11.4 | 209781.994 | 11.266 |
| birds | Forpus_coelestis | Forpus_coelestis | LC | Stable | tropical | diurnal | 26.1 | 147204.187 | 11.299 |
| birds | Forpus_conspicillatus | Forpus_conspicillatus | LC | Decreasing | tropical | diurnal | 26.4 | 321378.788 | 11.094 |
| birds | Forpus_cyanopygius | Forpus_cyanopygius | NT | Decreasing | temperate | diurnal | 33.5 | 115539.409 | 7.651 |
| birds | Forpus_modestus | Forpus_modestus | LC | Stable | tropical | diurnal | 27.2 | 2865167.852 | 2.225 |
| birds | Forpus_passerinus | Forpus_passerinus | LC | Decreasing | tropical | diurnal | 23 | 1072147.569 | 6.459 |
| birds | Forpus_xanthops | Forpus_xanthops | VU | Stable | tropical | diurnal | 35 | 2888.583 | 8.074 |
| birds | Forpus_xanthopterygius | Forpus_xanthopterygius | LC | Stable | tropical | diurnal | 31 | 5781313.449 | 7.675 |
| birds | Foudia_eminentissima | Foudia_eminentissima | LC | Stable | tropical | diurnal | 24.5 | 1026.053 | 17.229 |
| birds | Foudia_flavicans | NA | NT | Increasing | NA | NA | NA | NA | NA |
| birds | Foudia_madagascariensis | Foudia_madagascariensis | LC | Stable | tropical | diurnal | 16.6 | 599015.322 | 8.899 |
| birds | Foudia_omissa | Foudia_omissa | LC | Decreasing | tropical | diurnal | 18.9 | 127433.644 | 7.543 |
| birds | Foudia_rubra | Foudia_rubra | EN | Stable | tropical | diurnal | 17.7 | 1873.252 | 25.919 |
| birds | Foudia_sechellarum | NA | NT | Increasing | NA | NA | NA | NA | NA |
| birds | Foulehaio_carunculatus | NA | LC | Decreasing | NA | NA | NA | NA | NA |
| birds | Francolinus_francolinus | Francolinus_francolinus | LC | Stable | temperate | diurnal | 452.1 | 3027937.657 | 14.589 |
| birds | Francolinus_pictus | Francolinus_pictus | LC | Decreasing | tropical | diurnal | 286.8 | 1554139.426 | 18.489 |
| birds | Francolinus_pintadeanus | Francolinus_pintadeanus | LC | Stable | tropical | diurnal | 310 | 2072227.726 | 13.13 |
| birds | Fraseria_caerulescens | Muscicapa_caerulescens | LC | Stable | tropical | diurnal | 17.6 | 7513617.503 | 8.357 |
| birds | Fraseria_cinerascens | Fraseria_cinerascens | LC | Stable | tropical | diurnal | 23.1 | 3468124.588 | 8.411 |
| birds | Fraseria_griseigularis | Myioparus_griseigularis | LC | Decreasing | tropical | diurnal | 12.6 | 2490848.704 | 7.981 |

| | | | | | | | | | |
|---|---|---|---|---|---|---|---|---|---|
| birds | Fraseria_lendu | Muscicapa_lendu | VU | Decreasing | tropical | diurnal | 17.7 | 37404.928 | 13.873 |
| birds | Fraseria_ocreata | Fraseria_ocreata | LC | Stable | tropical | diurnal | 35.5 | 3285812.964 | 8.031 |
| birds | Fraseria_olivascens | Muscicapa_olivascens | LC | Stable | tropical | diurnal | 15.4 | 1385356.111 | 7.784 |
| birds | Fraseria_plumbea | Myioparus_plumbeus | LC | Stable | tropical | diurnal | 12.8 | 7861104.005 | 9.013 |
| birds | Fraseria_tessmanni | NA | LC | Unknown | NA | NA | NA | NA | NA |
| birds | Fratercula_arctica | Fratercula_arctica | VU | Decreasing | temperate | diurnal | 593 | 19128225.31 | 3.725 |
| birds | Fratercula_cirrhata | Fratercula_cirrhata | LC | Stable | temperate | diurnal | 774.4 | 10848785.582 | 6.433 |
| birds | Fratercula_corniculata | Fratercula_corniculata | LC | Decreasing | temperate | diurnal | 536.1 | 14508008.876 | 2.935 |
| birds | Frederickena_unduliger | Frederickena_unduligera | LC | Decreasing | tropical | diurnal | 83 | 1735694.573 | 1.18 |
| birds | Frederickena_viridis | Frederickena_viridis | LC | Decreasing | tropical | diurnal | 76 | 1266903.393 | 1.468 |
| birds | Fregata_andrewsi | Fregata_andrewsi | VU | Decreasing | tropical | diurnal | 1473.1 | 4236549.838 | 10.594 |
| birds | Fregata_aquila | NA | VU | Stable | NA | NA | NA | NA | NA |
| birds | Fregata_ariel | Fregata_ariel | LC | Decreasing | tropical | diurnal | 804.3 | 94658392.519 | 8.722 |
| birds | Fregata_magnificens | Fregata_magnificens | LC | Decreasing | tropical | diurnal | 1490.5 | 11561729.286 | 10.815 |
| birds | Fregata_minor | Fregata_minor | LC | Decreasing | tropical | diurnal | 1274.7 | 108234403.191 | 8.913 |
| birds | Fregetta_grallaria | Fregetta_grallaria | LC | Decreasing | temperate | cathemeral | 50.6 | 75851984.341 | 9.291 |
| birds | Fregetta_tropica | Fregetta_tropica | LC | Decreasing | tropical | cathemeral | 52 | 191838588.41 | 9.989 |
| birds | Fringilla_coelebs | Fringilla_coelebs | LC | Increasing | temperate | diurnal | 23.8 | 18764781.258 | 11.836 |
| birds | Fringilla_montifringilla | Fringilla_montifringilla | LC | Decreasing | temperate | diurnal | 23.2 | 24958109.429 | 9.582 |
| birds | Fringilla_teydea | Fringilla_teydea | NT | Increasing | temperate | diurnal | 30.3 | 349.939 | 11.992 |
| birds | Fulica_alai | Fulica_alai | NT | Stable | tropical | diurnal | 459 | 3693.544 | 14.063 |
| birds | Fulica_americana | Fulica_americana | LC | Decreasing | temperate | diurnal | 636.7 | 13778114.985 | 11.241 |
| birds | Fulica_ardesiaca | Fulica_ardesiaca | LC | Unknown | tropical | diurnal | 988 | 594594.346 | 7.723 |
| birds | Fulica_armillata | Fulica_armillata | LC | Unknown | temperate | diurnal | 1040 | 2733323.639 | 8.213 |
| birds | Fulica_atra | Fulica_atra | LC | Increasing | temperate | diurnal | 717.1 | 40041045.872 | 9.962 |
| birds | Fulica_cornuta | Fulica_cornuta | NT | Decreasing | temperate | diurnal | 2110.4 | 223464.939 | 2.951 |
| birds | Fulica_cristata | Fulica_cristata | LC | Decreasing | tropical | diurnal | 826 | 5291448.697 | 9.344 |
| birds | Fulica_gigantea | Fulica_gigantea | LC | Stable | tropical | diurnal | 2442.7 | 534521.718 | 7.352 |
| birds | Fulica_leucoptera | Fulica_leucoptera | LC | Stable | temperate | diurnal | 656 | 3041454.414 | 8.195 |
| birds | Fulica_rufifrons | Fulica_rufifrons | LC | Stable | temperate | diurnal | 644 | 1255982.206 | 11.076 |
| birds | Fulmarus_glacialis | Fulmarus_glacialis | LC | Increasing | temperate | cathemeral | 611.9 | 29451058.089 | 4.921 |
| birds | Fulmarus_glacialoides | Fulmarus_glacialoides | LC | Stable | temperate | cathemeral | 793.4 | 97370037.043 | 6.618 |
| birds | Fulvetta_cinereiceps | Alcippe_cinereiceps | LC | Unknown | temperate | diurnal | 10 | 974328.456 | 12.248 |
| birds | Fulvetta_ludlowi | Alcippe_ludlowi | LC | Decreasing | temperate | diurnal | 10.5 | 36772.325 | 3.523 |

| | | | | | | | | | |
|---|---|---|---|---|---|---|---|---|---|
| birds | Fulvetta_ruficapilla | Alcippe_ruficapilla | LC | Increasing | temperate | diurnal | 9 | 552478.797 | 11.516 |
| birds | Fulvetta_striaticollis | Alcippe_striaticollis | LC | Unknown | temperate | diurnal | 13.2 | 427621.632 | 7.971 |
| birds | Fulvetta_vinipectus | Alcippe_vinipectus | LC | Decreasing | temperate | diurnal | 12 | 559498.479 | 8.667 |
| birds | Furnarius_cinnamomeus | Furnarius_cinnamomeus | LC | Increasing | tropical | diurnal | 41.2 | 170280.999 | 10.894 |
| birds | Furnarius_cristatus | Furnarius_cristatus | LC | Stable | temperate | diurnal | 25.5 | 679538.364 | 6.978 |
| birds | Furnarius_figulus | Furnarius_figulus | LC | Increasing | tropical | diurnal | 28 | 1971356.279 | 7.711 |
| birds | Furnarius_leucopus | Furnarius_leucopus | LC | Increasing | tropical | diurnal | 54.8 | 4783071.894 | 5.033 |
| birds | Furnarius_longirostris | Furnarius_longirostris | LC | Increasing | tropical | diurnal | 49 | 269335.855 | 13.623 |
| birds | Furnarius_minor | Furnarius_minor | LC | Decreasing | tropical | diurnal | 25.8 | 319149.091 | 2.532 |
| birds | Furnarius_rufus | Furnarius_rufus | LC | Increasing | temperate | diurnal | 46.4 | 6181562.763 | 9.262 |
| birds | Furnarius_torridus | Furnarius_torridus | LC | Decreasing | tropical | diurnal | 52.3 | 87570.318 | 2.54 |
| birds | Gactornis_enarratus | Caprimulgus_enarratus | LC | Decreasing | tropical | nocturnal | 52.8 | 226375.928 | 8.758 |
| birds | Galbalcyrhynchus_leucotis | Galbalcyrhynchus_leucotis | LC | Stable | tropical | diurnal | 46.9 | 479718.915 | 2.454 |
| birds | Galbalcyrhynchus_purusianus | Galbalcyrhynchus_purusianus | LC | Stable | tropical | diurnal | 50 | 821108.367 | 1.494 |
| birds | Galbula_albirostris | Galbula_albirostris | LC | Decreasing | tropical | diurnal | 22.1 | 1881365.042 | 1.218 |
| birds | Galbula_chalcothorax | Galbula_chalcothorax | LC | Decreasing | tropical | diurnal | 23.9 | 816560.673 | 1.992 |
| birds | Galbula_cyanescens | Galbula_cyanescens | LC | Stable | tropical | diurnal | 25.3 | 1188886.924 | 1.595 |
| birds | Galbula_cyanicollis | Galbula_cyanicollis | LC | Decreasing | tropical | diurnal | 23.2 | 2705041.54 | 2.231 |
| birds | Galbula_dea | Galbula_dea | LC | Decreasing | tropical | diurnal | 27.4 | 5923896.484 | 1.825 |
| birds | Galbula_galbula | Galbula_galbula | LC | Stable | tropical | diurnal | 23 | 1927640.438 | 1.453 |
| birds | Galbula_leucogastra | Galbula_leucogastra | LC | Decreasing | tropical | diurnal | 15.9 | 2991415.412 | 1.919 |
| birds | Galbula_pastazae | Galbula_pastazae | LC | Decreasing | tropical | diurnal | 31.8 | 31392.628 | 5.65 |
| birds | Galbula_ruficauda | Galbula_ruficauda | LC | Decreasing | tropical | diurnal | 26.5 | 7607043.145 | 7.216 |
| birds | Galbula_tombacea | Galbula_tombacea | LC | Decreasing | tropical | diurnal | 23.2 | 969424.778 | 1.769 |
| birds | Galerida_cristata | Galerida_cristata | LC | Decreasing | temperate | diurnal | 42.7 | 24389170.687 | 11.319 |
| birds | Galerida_deva | Galerida_deva | LC | Stable | tropical | diurnal | 19.9 | 832847.979 | 19.91 |
| birds | Galerida_magnirostris | Galerida_magnirostris | LC | Increasing | temperate | diurnal | 41.6 | 692806.516 | 9.106 |
| birds | Galerida_malabarica | NA | LC | Stable | NA | NA | NA | NA | NA |
| birds | Galerida_modesta | Galerida_modesta | LC | Stable | tropical | diurnal | 20.2 | 3249298.437 | 10.069 |
| birds | Galerida_theklae | Galerida_theklae | LC | Decreasing | temperate | diurnal | 38.2 | 2270448.107 | 11.272 |
| birds | Gallicolumba_crinigera | Gallicolumba_crinigera | VU | Decreasing | tropical | diurnal | 192.2 | 94609.549 | 13.131 |
| birds | Gallicolumba_keayi | Gallicolumba_keayi | CR | Decreasing | tropical | diurnal | 190.5 | 24352.057 | 17.824 |
| birds | Gallicolumba_luzonica | Gallicolumba_luzonica | NT | Decreasing | tropical | diurnal | 196 | 107374.523 | 14.793 |
| birds | Gallicolumba_menagei | Gallicolumba_menagei | CR | Unknown | tropical | diurnal | 196 | 795.015 | 7.052 |

| | | | | | | | | | |
|---|---|---|---|---|---|---|---|---|---|
| birds | Gallicolumba_platenae | Gallicolumba_platenae | CR | Decreasing | tropical | diurnal | 196 | 475.229 | 13.684 |
| birds | Gallicolumba_rufigula | Gallicolumba_rufigula | LC | Decreasing | tropical | diurnal | 130 | 731477.388 | 3.212 |
| birds | Gallicolumba_tristigmata | Gallicolumba_tristigmata | LC | Decreasing | tropical | diurnal | 275 | 171123.298 | 9.364 |
| birds | Gallinago_imperialis | NA | NT | Decreasing | NA | NA | NA | NA | NA |
| birds | Gallinago_macrodactyla | Gallinago_macrodactyla | VU | Decreasing | tropical | diurnal | 216 | 220000.712 | 9.246 |
| birds | Gallinago_media | Gallinago_media | NT | Decreasing | temperate | nocturnal | 170 | 39693519.257 | 8.809 |
| birds | Gallinago_megala | Gallinago_megala | LC | Unknown | temperate | nocturnal | 121 | 7562818.051 | 6.902 |
| birds | Gallinago_nemoricola | Gallinago_nemoricola | VU | Decreasing | tropical | diurnal | 167.7 | 1723347.527 | 11.268 |
| birds | Gallinago_nigripennis | Gallinago_nigripennis | LC | Unknown | tropical | nocturnal | 112 | 5609482.861 | 9.483 |
| birds | Gallinago_paraguaiae | Gallinago_paraguaiae | LC | Stable | tropical | diurnal | 109.4 | 13818662.999 | 5.806 |
| birds | Gallinago_solitaria | Gallinago_solitaria | LC | Stable | temperate | diurnal | 140.1 | 7600182.792 | 7.385 |
| birds | Gallinago_stenura | Gallinago_stenura | LC | Unknown | temperate | cathemeral | 113 | 15061284.14 | 6.977 |
| birds | Gallinago_stricklandii | Gallinago_stricklandii | NT | Decreasing | temperate | nocturnal | 252 | 340969.804 | 3.936 |
| birds | Gallinago_undulata | Gallinago_undulata | LC | Decreasing | tropical | nocturnal | 312.4 | 2585357.362 | 8.916 |
| birds | Gallinula_chloropus | Gallinula_chloropus | LC | Stable | tropical | diurnal | 339.6 | 38971687.191 | 12.38 |
| birds | Gallinula_nesiotis | NA | EX | not_available | NA | NA | NA | NA | NA |
| birds | Gallinula_tenebrosa | Gallinula_tenebrosa | LC | Unknown | temperate | diurnal | 530.1 | 3816495.379 | 6.56 |
| birds | Gallirallus_australis | Gallirallus_australis | VU | Decreasing | temperate | diurnal | 875.7 | 42528.432 | 4.63 |
| birds | Gallirex_johnstoni | Ruwenzorornis_johnstoni | LC | Stable | tropical | diurnal | 240 | 16706.271 | 14.222 |
| birds | Gallirex_porphyreolophus | Tauraco_porphyreolophus | LC | Decreasing | tropical | diurnal | 259 | 2166499.58 | 10.395 |
| birds | Galloperdix_bicalcarata | Galloperdix_bicalcarata | LC | Decreasing | tropical | diurnal | 290.9 | 37679.968 | 18.413 |
| birds | Galloperdix_lunulata | Galloperdix_lunulata | LC | Stable | tropical | diurnal | 254.4 | 1390384.267 | 18.188 |
| birds | Galloperdix_spadicea | Galloperdix_spadicea | LC | Stable | tropical | diurnal | 359.1 | 1175974.864 | 18.033 |
| birds | Gallus_gallus | Gallus_gallus | LC | Decreasing | tropical | diurnal | 751.7 | 5219583.043 | 13.144 |
| birds | Gallus_lafayettii | Gallus_lafayetii | LC | Stable | tropical | diurnal | 825 | 55601.24 | 16.566 |
| birds | Gallus_sonneratii | Gallus_sonneratii | LC | Decreasing | tropical | diurnal | 840.8 | 1030202.675 | 19.583 |
| birds | Gallus_varius | Gallus_varius | LC | Stable | tropical | diurnal | 839.3 | 187783.16 | 19.994 |
| birds | Gampsonyx_swainsonii | Gampsonyx_swainsonii | LC | Increasing | tropical | diurnal | 92.9 | 10266134.716 | 6.266 |
| birds | Gampsorhynchus_rufulus | Gampsorhynchus_rufulus | LC | Decreasing | tropical | diurnal | 37 | 453698.087 | 7.96 |
| birds | Garritornis_isidorei | Pomatostomus_isidorei | LC | Stable | tropical | diurnal | 70 | 561083.933 | 3.135 |
| birds | Garrodia_nereis | Garrodia_nereis | LC | Decreasing | temperate | diurnal | 38.2 | 28460926.477 | 4.125 |
| birds | Garrulax_albogularis | NA | not_available | not_available | NA | NA | NA | NA | NA |
| birds | Garrulax_canorus | Garrulax_canorus | LC | Unknown | temperate | diurnal | 62.8 | 1868020.761 | 14.768 |
| birds | Garrulax_cineraceus | NA | not_available | not_available | NA | NA | NA | NA | NA |

| | | | | | | | | | |
|---|---|---|---|---|---|---|---|---|---|
| birds | Garrulax_galbanus | NA | not_available | not_available | NA | NA | NA | NA | NA |
| birds | Garrulax_leucolophus | Garrulax_leucolophus | LC | Decreasing | tropical | diurnal | 123.4 | 1715209.641 | 9.756 |
| birds | Garrulax_maesi | Garrulax_maesi | LC | Decreasing | temperate | diurnal | 112.5 | 725718.269 | 13.868 |
| birds | Garrulax_merulinus | Garrulax_merulinus | LC | Decreasing | tropical | diurnal | 78.8 | 498651.876 | 9.327 |
| birds | Garrulax_milleti | NA | LC | Decreasing | NA | NA | NA | NA | NA |
| birds | Garrulax_mitratus | NA | not_available | not_available | NA | NA | NA | NA | NA |
| birds | Garrulax_monileger | Garrulax_monileger | LC | Decreasing | tropical | diurnal | 83.7 | 2496742.03 | 11.378 |
| birds | Garrulax_palliatus | NA | NT | Decreasing | NA | NA | NA | NA | NA |
| birds | Garrulax_poecilorhynchus | NA | not_available | not_available | NA | NA | NA | NA | NA |
| birds | Garrulax_rufifrons | Garrulax_rufifrons | CR | Decreasing | tropical | diurnal | 101.5 | 4771.679 | 13.748 |
| birds | Garrulax_strepitans | Garrulax_strepitans | LC | Decreasing | tropical | diurnal | 131.5 | 247234.991 | 9.471 |
| birds | Garrulus_glandarius | Garrulus_glandarius | LC | Stable | temperate | diurnal | 159.5 | 19458203.901 | 11.936 |
| birds | Garrulus_lanceolatus | Garrulus_lanceolatus | LC | Stable | temperate | diurnal | 97.7 | 322480.194 | 12.181 |
| birds | Garrulus_lidthi | Garrulus_lidthi | VU | Decreasing | temperate | diurnal | 125 | 1064.169 | 16.15 |
| birds | Gavia_adamsii | Gavia_adamsii | NT | Decreasing | temperate | diurnal | 4944.3 | 5436574.877 | 0.694 |
| birds | Gavia_arctica | Gavia_arctica | LC | Decreasing | temperate | diurnal | 2251.1 | 22818831.522 | 4.052 |
| birds | Gavia_immer | Gavia_immer | LC | Stable | temperate | diurnal | 4956.8 | 12626924.281 | 2.738 |
| birds | Gavia_stellata | Gavia_stellata | LC | Decreasing | temperate | diurnal | 1486 | 23359436.096 | 1.838 |
| birds | Gavicalis_fasciogularis | Lichenostomus_fasciogularis | LC | Increasing | tropical | diurnal | 26.8 | 43256.673 | 9.846 |
| birds | Gavicalis_versicolor | Lichenostomus_versicolor | LC | Stable | tropical | diurnal | 36.7 | 161900.972 | 4.043 |
| birds | Gavicalis_virescens | Lichenostomus_virescens | LC | Increasing | temperate | diurnal | 23.3 | 7306702.437 | 4.235 |
| birds | Gecinulus_grantia | Gecinulus_grantia | LC | Decreasing | temperate | diurnal | 76 | 750304.841 | 10.889 |
| birds | Gecinulus_viridis | Gecinulus_viridis | LC | Decreasing | tropical | diurnal | 69.2 | 187614.015 | 9.098 |
| birds | Gelochelidon_nilotica | Sterna_nilotica | LC | Decreasing | temperate | diurnal | 218.3 | 14873944.315 | 10.226 |
| birds | Gennaeodryas_placens | Poecilodryas_placens | LC | Decreasing | tropical | diurnal | 25.5 | 26389.228 | 3.187 |
| birds | Geobiastes_squamiger | Brachypteracias_squamiger | VU | Decreasing | tropical | cathemeral | 155 | 21195.483 | 5.866 |
| birds | Geocerthia_serrana | Upucerthia_serrana | LC | Stable | tropical | diurnal | 48.7 | 92999.643 | 9.234 |
| birds | Geococcyx_californianus | Geococcyx_californianus | LC | Increasing | temperate | diurnal | 376 | 3008009.53 | 10.895 |
| birds | Geococcyx_velox | Geococcyx_velox | LC | Stable | tropical | diurnal | 179.9 | 517104.331 | 9.971 |
| birds | Geoffroyus_geoffroyi | Geoffroyus_geoffroyi | LC | Decreasing | tropical | diurnal | 164 | 793407.63 | 4.477 |
| birds | Geoffroyus_heteroclitus | Geoffroyus_heteroclitus | LC | Stable | tropical | diurnal | 170 | 78773.928 | 4.569 |
| birds | Geoffroyus_simplex | Geoffroyus_simplex | LC | Stable | tropical | diurnal | 161 | 251222.712 | 3.748 |
| birds | Geokichla_camaronensis | Zoothera_camaronensis | LC | Decreasing | tropical | diurnal | 44 | 164369.198 | 8.583 |
| birds | Geokichla_cinerea | Zoothera_cinerea | VU | Decreasing | tropical | diurnal | 63 | 114921.743 | 14.578 |

| | | | | | | | | | |
|---|---|---|---|---|---|---|---|---|---|
| birds | Geokichla_citrina | Zoothera_citrina | LC | Decreasing | tropical | diurnal | 53.3 | 4482928.591 | 14.613 |
| birds | Geokichla_crossleyi | Zoothera_crossleyi | LC | Decreasing | tropical | diurnal | 72 | 52658.209 | 8.761 |
| birds | Geokichla_dohertyi | NA | NT | Decreasing | NA | NA | NA | NA | NA |
| birds | Geokichla_dumasi | Zoothera_dumasi | NT | Decreasing | tropical | diurnal | 49.5 | 8554.352 | 6.663 |
| birds | Geokichla_erythronota | Zoothera_erythronota | NT | Decreasing | tropical | diurnal | 56.3 | 157827.321 | 9.558 |
| birds | Geokichla_gurneyi | Zoothera_gurneyi | LC | Decreasing | tropical | diurnal | 57.9 | 421268.025 | 12.076 |
| birds | Geokichla_guttata | Zoothera_guttata | VU | Decreasing | tropical | diurnal | 57.5 | 84741.261 | 14.176 |
| birds | Geokichla_interpres | Zoothera_interpres | EN | Decreasing | tropical | diurnal | 81.9 | 670164.001 | 8.102 |
| birds | Geokichla_oberlaenderi | Zoothera_oberlaenderi | NT | Decreasing | tropical | diurnal | 43.4 | 82488.061 | 11.282 |
| birds | Geokichla_peronii | NA | NT | Decreasing | NA | NA | NA | NA | NA |
| birds | Geokichla_piaggiae | Zoothera_piaggiae | LC | Decreasing | tropical | diurnal | 51.1 | 328100.346 | 12.86 |
| birds | Geokichla_princei | Zoothera_princei | LC | Decreasing | tropical | diurnal | 65.4 | 453479.328 | 11.659 |
| birds | Geokichla_schistacea | NA | NT | Decreasing | NA | NA | NA | NA | NA |
| birds | Geokichla_sibirica | Zoothera_sibirica | LC | Decreasing | tropical | diurnal | 75.5 | 10702412.67 | 6.311 |
| birds | Geokichla_spiloptera | Zoothera_spiloptera | NT | Decreasing | tropical | diurnal | 70 | 11408.726 | 16.595 |
| birds | Geokichla_wardii | Zoothera_wardii | LC | Decreasing | temperate | diurnal | 58.6 | 194220.034 | 15.883 |
| birds | Geopelia_cuneata | Geopelia_cuneata | LC | Stable | temperate | diurnal | 31.6 | 5703540.333 | 3.677 |
| birds | Geopelia_humeralis | Geopelia_humeralis | LC | Stable | temperate | diurnal | 128.4 | 2758465.778 | 4.556 |
| birds | Geopelia_maugeus | NA | LC | Stable | NA | NA | NA | NA | NA |
| birds | Geopelia_placida | Geopelia_placida | LC | Stable | temperate | diurnal | 47.8 | 5580497.28 | 4.703 |
| birds | Geopelia_striata | Geopelia_striata | LC | Stable | tropical | diurnal | 56.6 | 1478473.744 | 13.913 |
| birds | Geophaps_plumifera | Geophaps_plumifera | LC | Stable | tropical | diurnal | 94.3 | 1860384.758 | 3.088 |
| birds | Geophaps_scripta | Geophaps_scripta | LC | Decreasing | tropical | diurnal | 221.4 | 953627.901 | 5.467 |
| birds | Geophaps_smithii | Geophaps_smithii | LC | Decreasing | tropical | diurnal | 194 | 288800.127 | 1.726 |
| birds | Geositta_antarctica | Geositta_antarctica | LC | Stable | temperate | diurnal | 36.9 | 548387.441 | 5.36 |
| birds | Geositta_crassirostris | Geositta_crassirostris | LC | Decreasing | tropical | diurnal | 55.8 | 51583.499 | 6.65 |
| birds | Geositta_cunicularia | Geositta_cunicularia | LC | Decreasing | temperate | diurnal | 28.5 | 2912290.536 | 7.716 |
| birds | Geositta_isabellina | Geositta_isabellina | LC | Decreasing | temperate | diurnal | 42.7 | 71409.446 | 3.055 |
| birds | Geositta_maritima | Geositta_maritima | LC | Stable | tropical | diurnal | 16.5 | 158400.301 | 5.531 |
| birds | Geositta_peruviana | Geositta_peruviana | LC | Stable | tropical | diurnal | 20.8 | 54662.981 | 8.952 |
| birds | Geositta_poeciloptera | Geositta_poeciloptera | VU | Decreasing | tropical | diurnal | 18 | 1456963.678 | 11.139 |
| birds | Geositta_punensis | Geositta_punensis | LC | Stable | tropical | diurnal | 25.8 | 382860.928 | 4.787 |
| birds | Geositta_rufipennis | Geositta_rufipennis | LC | Decreasing | temperate | diurnal | 32.4 | 757787.996 | 5.489 |
| birds | Geositta_saxicolina | Geositta_saxicolina | LC | Stable | tropical | diurnal | 32.9 | 44751.849 | 9.826 |

| | | | | | | | | | |
|---|---|---|---|---|---|---|---|---|---|
| birds | Geositta_tenuirostris | Geositta_tenuirostris | LC | Stable | tropical | diurnal | 32.2 | 622360.287 | 8.278 |
| birds | Geospiza_conirostris | NA | VU | Stable | NA | NA | NA | NA | NA |
| birds | Geospiza_difficilis | NA | LC | Stable | NA | NA | NA | NA | NA |
| birds | Geospiza_fortis | NA | LC | Stable | NA | NA | NA | NA | NA |
| birds | Geospiza_fuliginosa | NA | LC | Stable | NA | NA | NA | NA | NA |
| birds | Geospiza_heliobates | NA | CR | Decreasing | NA | NA | NA | NA | NA |
| birds | Geospiza_magnirostris | NA | LC | Stable | NA | NA | NA | NA | NA |
| birds | Geospiza_pallida | NA | NT | Decreasing | NA | NA | NA | NA | NA |
| birds | Geospiza_parvula | NA | LC | Stable | NA | NA | NA | NA | NA |
| birds | Geospiza_pauper | NA | CR | Decreasing | NA | NA | NA | NA | NA |
| birds | Geospiza_psittacula | NA | VU | Decreasing | NA | NA | NA | NA | NA |
| birds | Geospiza_scandens | NA | LC | Stable | NA | NA | NA | NA | NA |
| birds | Geospizopsis_plebejus | Phrygilus_plebejus | LC | Stable | tropical | diurnal | 14.7 | 950611.697 | 7.344 |
| birds | Geospizopsis_unicolor | Phrygilus_unicolor | LC | Stable | temperate | diurnal | 21.9 | 1358301.992 | 6.67 |
| birds | Geothlypis_aequinoctialis | Geothlypis_aequinoctialis | LC | Stable | tropical | diurnal | 13.1 | 7376725.602 | 8.232 |
| birds | Geothlypis_auricularis | Geothlypis_auricularis | LC | Stable | tropical | diurnal | 13.8 | 196093.057 | 9.322 |
| birds | Geothlypis_beldingi | Geothlypis_beldingi | VU | Decreasing | temperate | diurnal | 15.7 | 1747.193 | 13.157 |
| birds | Geothlypis_chiriquensis | Geothlypis_chiriquensis | LC | Increasing | tropical | diurnal | 13.8 | 1382.255 | 13.501 |
| birds | Geothlypis_flavovelata | Geothlypis_flavovelata | NT | Decreasing | tropical | diurnal | 10.9 | 48085.793 | 13.463 |
| birds | Geothlypis_formosa | Oporornis_formosus | LC | Decreasing | tropical | diurnal | 14 | 3787752.22 | 15.301 |
| birds | Geothlypis_nelsoni | Geothlypis_nelsoni | LC | Decreasing | tropical | diurnal | 11 | 117193.329 | 12.368 |
| birds | Geothlypis_philadelphia | Oporornis_philadelphia | LC | Decreasing | temperate | diurnal | 11.7 | 7176055.479 | 12.784 |
| birds | Geothlypis_poliocephala | Geothlypis_poliocephala | LC | Decreasing | tropical | diurnal | 14.6 | 859573.043 | 10.504 |
| birds | Geothlypis_rostrata | Geothlypis_rostrata | LC | Decreasing | temperate | diurnal | 16.1 | 8764.155 | 6.866 |
| birds | Geothlypis_semiflava | Geothlypis_semiflava | LC | Increasing | tropical | diurnal | 17.2 | 115622.866 | 9.552 |
| birds | Geothlypis_speciosa | Geothlypis_speciosa | VU | Decreasing | tropical | diurnal | 10.8 | 645.116 | 28.389 |
| birds | Geothlypis_tolmiei | Oporornis_tolmiei | LC | Decreasing | temperate | diurnal | 10.4 | 5855497.185 | 8.243 |
| birds | Geothlypis_trichas | Geothlypis_trichas | LC | Decreasing | temperate | diurnal | 9.5 | 15147170.247 | 10.341 |
| birds | Geotrygon_caniceps | Geotrygon_caniceps | VU | Decreasing | tropical | diurnal | 192 | 107566.318 | 16.225 |
| birds | Geotrygon_chrysia | Geotrygon_chrysia | LC | Stable | tropical | diurnal | 169 | 207769.784 | 15.951 |
| birds | Geotrygon_montana | Geotrygon_montana | LC | Decreasing | tropical | diurnal | 133.9 | 13251114.737 | 5.88 |
| birds | Geotrygon_mystacea | Geotrygon_mystacea | LC | Decreasing | tropical | diurnal | 211 | 14198.649 | 25.432 |
| birds | Geotrygon_saphirina | Geotrygon_saphirina | LC | Decreasing | tropical | diurnal | 203 | 569182.482 | 2.049 |
| birds | Geotrygon_versicolor | Geotrygon_versicolor | NT | Decreasing | tropical | diurnal | 225 | 11021.791 | 19.741 |

| | | | | | | | | | |
|---|---|---|---|---|---|---|---|---|---|
| birds | Geotrygon_violacea | Geotrygon_violacea | LC | Decreasing | tropical | diurnal | 95.2 | 1490404.571 | 11.841 |
| birds | Geranoaetus_albicaudatus | Buteo_albicaudatus | LC | Unknown | tropical | diurnal | 885 | 9466789.108 | 8.212 |
| birds | Geranoaetus_melanoleucus | Geranoaetus_melanoleucus | LC | Stable | temperate | diurnal | 2378.6 | 6807951.844 | 8.295 |
| birds | Geranoaetus_polyosoma | Buteo_polyosoma | LC | Increasing | temperate | diurnal | 782.6 | 4720061.527 | 7.207 |
| birds | Geranospiza_caerulescens | Geranospiza_caerulescens | LC | Decreasing | tropical | diurnal | 302.5 | 14333669.118 | 6.011 |
| birds | Geronticus_calvus | NA | VU | Decreasing | NA | NA | NA | NA | NA |
| birds | Geronticus_eremita | Geronticus_eremita | EN | Stable | temperate | diurnal | 1202 | 2452918.916 | 6.386 |
| birds | Gerygone_albofrontata | NA | LC | Decreasing | NA | NA | NA | NA | NA |
| birds | Gerygone_chloronota | Gerygone_chloronota | LC | Stable | tropical | diurnal | 6.6 | 1009439.841 | 2.672 |
| birds | Gerygone_chrysogaster | Gerygone_chrysogaster | LC | Stable | tropical | diurnal | 8 | 557753.518 | 3.124 |
| birds | Gerygone_dorsalis | Gerygone_dorsalis | LC | Stable | tropical | diurnal | 7.2 | 7004.776 | 8.139 |
| birds | Gerygone_flavolateralis | Gerygone_flavolateralis | LC | Stable | tropical | diurnal | 6.3 | 28357.671 | 6.839 |
| birds | Gerygone_fusca | Gerygone_fusca | LC | Decreasing | temperate | diurnal | 5.4 | 3748888.93 | 5.637 |
| birds | Gerygone_hypoxantha | Gerygone_hypoxantha | NT | Decreasing | tropical | diurnal | 7.2 | 2524.27 | 8.959 |
| birds | Gerygone_igata | Gerygone_igata | LC | Stable | temperate | diurnal | 6.4 | 268886.757 | 7.991 |
| birds | Gerygone_inornata | Gerygone_inornata | LC | Decreasing | tropical | diurnal | 7.2 | 33197.315 | 13.311 |
| birds | Gerygone_levigaster | Gerygone_levigaster | LC | Decreasing | tropical | diurnal | 6.2 | 342256.683 | 4.5 |
| birds | Gerygone_magnirostris | Gerygone_magnirostris | LC | Stable | tropical | diurnal | 6.8 | 1205051.147 | 2.832 |
| birds | Gerygone_modesta | NA | LC | Unknown | NA | NA | NA | NA | NA |
| birds | Gerygone_mouki | Gerygone_mouki | LC | Increasing | temperate | diurnal | 5.2 | 387637.869 | 7.331 |
| birds | Gerygone_olivacea | Gerygone_olivacea | LC | Decreasing | temperate | diurnal | 6.7 | 2358084.056 | 5.405 |
| birds | Gerygone_palpebrosa | Gerygone_palpebrosa | LC | Stable | tropical | diurnal | 7.4 | 969224.378 | 3.815 |
| birds | Gerygone_ruficollis | Gerygone_ruficollis | LC | Stable | tropical | diurnal | 7 | 107780.49 | 4.949 |
| birds | Gerygone_sulphurea | Gerygone_sulphurea | LC | Decreasing | tropical | diurnal | 5.6 | 1733224.343 | 10.323 |
| birds | Gerygone_tenebrosa | Gerygone_tenebrosa | LC | Stable | tropical | diurnal | 7.7 | 39187.749 | 5.482 |
| birds | Glaucestrilda_coerulescens | Estrilda_caerulescens | LC | Stable | tropical | diurnal | 9.4 | 1693485.933 | 13.032 |
| birds | Glaucestrilda_perreini | Estrilda_perreini | LC | Decreasing | tropical | diurnal | 7.4 | 1035934.151 | 8.224 |
| birds | Glaucestrilda_thomensis | Estrilda_thomensis | LC | Decreasing | tropical | diurnal | 7.6 | 210666.08 | 8.686 |
| birds | Glaucidium_albertinum | Glaucidium_albertinum | NT | Decreasing | tropical | nocturnal | 73 | 32391.819 | 13.36 |
| birds | Glaucidium_bolivianum | Glaucidium_bolivianum | LC | Decreasing | tropical | cathemeral | 63 | 121188.266 | 5.165 |
| birds | Glaucidium_brasilianum | Glaucidium_brasilianum | LC | Decreasing | tropical | diurnal | 63.3 | 16036298.782 | 6.336 |
| birds | Glaucidium_brodiei | Glaucidium_brodiei | LC | Decreasing | tropical | diurnal | 59.4 | 2339208.31 | 10.908 |
| birds | Glaucidium_californicum | NA | not_available | not_available | NA | NA | NA | NA | NA |
| birds | Glaucidium_capense | Glaucidium_capense | LC | Decreasing | tropical | diurnal | 122 | 4071921.706 | 8.616 |

| | | | | | | | | | |
|---|---|---|---|---|---|---|---|---|---|
| birds | Glaucidium_castaneum | Glaucidium_castaneum | LC | Decreasing | tropical | diurnal | 100 | 91747.231 | 11.015 |
| birds | Glaucidium_castanopterum | NA | LC | Stable | NA | NA | NA | NA | NA |
| birds | Glaucidium_castanotum | Glaucidium_castanonotum | NT | Decreasing | tropical | diurnal | 100 | 46677.712 | 15.202 |
| birds | Glaucidium_cuculoides | Glaucidium_cuculoides | LC | Increasing | tropical | diurnal | 198.2 | 4612393.635 | 13.742 |
| birds | Glaucidium_gnoma | Glaucidium_gnoma | LC | Increasing | temperate | diurnal | 61.3 | 3631571.534 | 7.979 |
| birds | Glaucidium_griseiceps | Glaucidium_griseiceps | LC | Stable | tropical | diurnal | 50.6 | 270382.473 | 9.087 |
| birds | Glaucidium_hardyi | Glaucidium_hardyi | LC | Decreasing | tropical | diurnal | 58.3 | 4691899.428 | 2.242 |
| birds | Glaucidium_jardinii | Glaucidium_jardinii | LC | Stable | tropical | cathemeral | 66.9 | 205500.737 | 11.301 |
| birds | Glaucidium_minutissimum | Glaucidium_minutissimum | LC | Decreasing | tropical | cathemeral | 50 | 1209018.269 | 12.616 |
| birds | Glaucidium_nana | Glaucidium_nanum | LC | Stable | temperate | cathemeral | 73.7 | 656548.539 | 5.438 |
| birds | Glaucidium_perlatum | Glaucidium_perlatum | LC | Stable | tropical | diurnal | 79.2 | 11923401.874 | 9.951 |
| birds | Glaucidium_sjostedti | Glaucidium_sjostedti | LC | Decreasing | tropical | nocturnal | 139 | 1369630.825 | 5.815 |
| birds | Glaucidium_tephronotum | Glaucidium_tephronotum | LC | Stable | tropical | nocturnal | 87.4 | 2054685.258 | 8.216 |
| birds | Glaucis_aeneus | Glaucis_aeneus | LC | Decreasing | tropical | diurnal | 4.8 | 177367.008 | 7.818 |
| birds | Glaucis_dohrnii | Glaucis_dohrnii | VU | Decreasing | tropical | diurnal | 5.8 | 60089.028 | 8.46 |
| birds | Glaucis_hirsutus | Glaucis_hirsutus | LC | Decreasing | tropical | diurnal | 6.8 | 7820159.949 | 3.668 |
| birds | Gliciphila_melanops | Phylidonyris_melanops | LC | Decreasing | temperate | diurnal | 18.2 | 1135955.29 | 8.866 |
| birds | Gliciphila_notabilis | Phylidonyris_notabilis | LC | Stable | tropical | diurnal | 27.1 | 8862.874 | 5.406 |
| birds | Gliciphila_undulata | Phylidonyris_undulatus | LC | Stable | tropical | diurnal | 16 | 16533.984 | 7.505 |
| birds | Glossopsitta_concinna | Glossopsitta_concinna | LC | Stable | temperate | diurnal | 70.5 | 663167.13 | 10.108 |
| birds | Glossopsitta_porphyrocephala | Glossopsitta_porphyrocephala | LC | Decreasing | temperate | diurnal | 44.7 | 944646.288 | 8.54 |
| birds | Glossopsitta_pusilla | Glossopsitta_pusilla | LC | Stable | temperate | diurnal | 39.4 | 776503.487 | 8.861 |
| birds | Glycichaera_fallax | Glycichaera_fallax | LC | Stable | tropical | diurnal | 11.2 | 604148.568 | 3.062 |
| birds | Glyphorynchus_spirurus | Glyphorynchus_spirurus | LC | Decreasing | tropical | diurnal | 14.6 | 7368285.067 | 2.955 |
| birds | Gnorimopsar_chopi | Gnorimopsar_chopi | LC | Stable | tropical | diurnal | 65.9 | 5930564.57 | 9.099 |
| birds | Goldmania_bella | Goethalsia_bella | NT | Unknown | tropical | diurnal | 3.4 | 1893.246 | 1.908 |
| birds | Goldmania_violiceps | Goldmania_violiceps | NT | Decreasing | tropical | diurnal | 4.1 | 2852.716 | 3.456 |
| birds | Goodfellowia_miranda | Basilornis_mirandus | NT | Decreasing | tropical | diurnal | 110 | 1528.358 | 6.186 |
| birds | Gorsachius_goisagi | Gorsachius_goisagi | VU | Decreasing | temperate | cathemeral | 527 | 823887.536 | 17.505 |
| birds | Gorsachius_melanolophus | Gorsachius_melanolophus | LC | Unknown | tropical | cathemeral | 412.3 | 1619076.855 | 11.345 |
| birds | Goura_cristata | Goura_cristata | VU | Decreasing | tropical | diurnal | 2000 | 97947.972 | 4.103 |
| birds | Goura_scheepmakeri | Goura_scheepmakeri | VU | Decreasing | tropical | diurnal | 2235 | 112181.704 | 2.53 |
| birds | Goura_victoria | Goura_victoria | NT | Decreasing | tropical | diurnal | 2384 | 116530.354 | 3.418 |
| birds | Gracula_ptilogenys | Gracula_ptilogenys | NT | Decreasing | tropical | diurnal | 192 | 15783.496 | 17.719 |

| | | | | | | | | | |
|---|---|---|---|---|---|---|---|---|---|
| birds | Gracula_religiosa | Gracula_religiosa | LC | Decreasing | tropical | diurnal | 192 | 3864550.112 | 10.954 |
| birds | Gracupica_contra | Sturnus_contra | LC | Increasing | tropical | diurnal | 84 | 3064197.419 | 16.679 |
| birds | Gracupica_nigricollis | Sturnus_nigricollis | LC | Increasing | tropical | diurnal | 156.9 | 2900662.508 | 13.302 |
| birds | Grafisia_torquata | Grafisia_torquata | LC | Decreasing | tropical | diurnal | 64 | 186750.61 | 7.148 |
| birds | Grallaria_albigula | Grallaria_albigula | LC | Stable | tropical | diurnal | 95 | 45655.476 | 6.527 |
| birds | Grallaria_alleni | Grallaria_alleni | VU | Decreasing | tropical | diurnal | 70.5 | 14285.268 | 7.687 |
| birds | Grallaria_andicolus | Grallaria_andicolus | LC | Stable | tropical | diurnal | 56.3 | 141026.374 | 8.236 |
| birds | Grallaria_bangsi | Grallaria_bangsi | VU | Decreasing | tropical | diurnal | 62 | 1985.531 | 5.903 |
| birds | Grallaria_blakei | Grallaria_blakei | LC | Decreasing | tropical | diurnal | 42.3 | 51490.337 | 5.449 |
| birds | Grallaria_capitalis | Grallaria_capitalis | LC | Stable | tropical | diurnal | 73.7 | 12133.267 | 5.613 |
| birds | Grallaria_carrikeri | Grallaria_carrikeri | LC | Decreasing | tropical | diurnal | 109.1 | 29971.083 | 3.135 |
| birds | Grallaria_chthonia | NA | CR | Decreasing | NA | NA | NA | NA | NA |
| birds | Grallaria_dignissima | Grallaria_dignissima | LC | Stable | tropical | diurnal | 110 | 386285.827 | 1.652 |
| birds | Grallaria_eludens | Grallaria_eludens | LC | Unknown | tropical | diurnal | 115 | 383522.836 | 1.012 |
| birds | Grallaria_erythroleuca | Grallaria_erythroleuca | LC | Stable | tropical | diurnal | 80 | 32694.029 | 3.858 |
| birds | Grallaria_erythrotis | Grallaria_erythrotis | LC | Stable | tropical | diurnal | 57 | 39088.657 | 5.199 |
| birds | Grallaria_excelsa | Grallaria_excelsa | NT | Decreasing | tropical | diurnal | 240.8 | 8310.449 | 11.411 |
| birds | Grallaria_flavotincta | Grallaria_flavotincta | LC | Stable | tropical | diurnal | 63 | 22776.327 | 7.181 |
| birds | Grallaria_gigantea | Grallaria_gigantea | VU | Decreasing | tropical | diurnal | 236 | 21095.752 | 6.426 |
| birds | Grallaria_griseonucha | NA | LC | Stable | NA | NA | NA | NA | NA |
| birds | Grallaria_guatimalensis | Grallaria_guatimalensis | LC | Decreasing | tropical | diurnal | 94.2 | 1176261.457 | 6.48 |
| birds | Grallaria_haplonota | Grallaria_haplonota | LC | Stable | tropical | diurnal | 85 | 87992.125 | 6.766 |
| birds | Grallaria_hypoleuca | Grallaria_hypoleuca | LC | Stable | tropical | diurnal | 73.7 | 62353.378 | 8.351 |
| birds | Grallaria_kaestneri | Grallaria_kaestneri | EN | Decreasing | tropical | diurnal | 47.5 | 1395.93 | 9.847 |
| birds | Grallaria_milleri | Grallaria_milleri | VU | Decreasing | tropical | diurnal | 52.5 | 15668.093 | 13.936 |
| birds | Grallaria_nuchalis | Grallaria_nuchalis | LC | Stable | tropical | diurnal | 117 | 63486.155 | 9.394 |
| birds | Grallaria_przewalskii | Grallaria_przewalskii | LC | Stable | tropical | diurnal | 65.2 | 19046.164 | 3.923 |
| birds | Grallaria_quitensis | Grallaria_quitensis | LC | Stable | tropical | diurnal | 65.7 | 74340.838 | 11.224 |
| birds | Grallaria_ruficapilla | Grallaria_ruficapilla | LC | Increasing | tropical | diurnal | 82.7 | 239637.785 | 10.96 |
| birds | Grallaria_rufocinerea | Grallaria_rufocinerea | LC | Decreasing | tropical | diurnal | 44.8 | 37568.66 | 9.865 |
| birds | Grallaria_rufula | Grallaria_rufula | LC | Stable | tropical | diurnal | 35.7 | 236386.273 | 8.234 |
| birds | Grallaria_squamigera | Grallaria_squamigera | LC | Stable | tropical | diurnal | 136 | 269461.885 | 8.056 |
| birds | Grallaria_varia | Grallaria_varia | LC | Decreasing | tropical | diurnal | 119 | 3247752.05 | 5.296 |
| birds | Grallaria_watkinsi | Grallaria_watkinsi | NT | Decreasing | tropical | diurnal | 69.8 | 16613.44 | 9.736 |

| | | | | | | | | | |
|---|---|---|---|---|---|---|---|---|---|
| birds | Grallaricula_cucullata | Grallaricula_cucullata | NT | Decreasing | tropical | diurnal | 19.3 | 15988.808 | 11.362 |
| birds | Grallaricula_ferrugineipectus | Grallaricula_ferrugineipectus | LC | Stable | tropical | diurnal | 17.2 | 70002.379 | 13.044 |
| birds | Grallaricula_flavirostris | Grallaricula_flavirostris | LC | Decreasing | tropical | diurnal | 17.7 | 271741.926 | 6.854 |
| birds | Grallaricula_leymebambae | NA | LC | Stable | NA | NA | NA | NA | NA |
| birds | Grallaricula_lineifrons | Grallaricula_lineifrons | LC | Stable | tropical | diurnal | 21.1 | 14344.407 | 8.368 |
| birds | Grallaricula_nana | Grallaricula_nana | LC | Stable | tropical | diurnal | 19.8 | 267727.744 | 9.22 |
| birds | Grallaricula_ochraceifrons | Grallaricula_ochraceifrons | VU | Decreasing | tropical | diurnal | 23.1 | 2574.014 | 3.37 |
| birds | Grallaricula_peruviana | Grallaricula_peruviana | NT | Decreasing | tropical | diurnal | 18.8 | 6450.421 | 4.737 |
| birds | Grallina_bruijnii | Grallina_bruijni | LC | Unknown | tropical | diurnal | 38.4 | 259817.039 | 4.03 |
| birds | Grallina_cyanoleuca | Grallina_cyanoleuca | LC | Increasing | temperate | diurnal | 82.6 | 6745943.052 | 4.739 |
| birds | Graminicola_bengalensis | Graminicola_bengalensis | NT | Decreasing | temperate | diurnal | 14.6 | 174532.671 | 17.102 |
| birds | Grammatoptila_striata | Garrulax_striatus | LC | Decreasing | temperate | diurnal | 92 | 317905.881 | 8.276 |
| birds | Granatellus_pelzelni | Granatellus_pelzelni | LC | Stable | tropical | diurnal | 11.2 | 3464920.452 | 1.963 |
| birds | Granatellus_sallaei | Granatellus_sallaei | LC | Decreasing | tropical | diurnal | 9.9 | 315515.726 | 8.799 |
| birds | Granatellus_venustus | Granatellus_venustus | LC | Decreasing | tropical | diurnal | 10.8 | 171793.058 | 9.083 |
| birds | Granatina_granatina | Uraeginthus_granatinus | LC | Stable | tropical | diurnal | 11.8 | 2368190.337 | 7.566 |
| birds | Granatina_ianthinogaster | Uraeginthus_ianthinogaster | LC | Stable | tropical | diurnal | 14.1 | 1450518.631 | 9.815 |
| birds | Grandala_coelicolor | Grandala_coelicolor | LC | Stable | temperate | diurnal | 45.6 | 1886746.699 | 7.965 |
| birds | Grantiella_picta | Grantiella_picta | LC | Stable | temperate | diurnal | 21.5 | 1455898.86 | 6.254 |
| birds | Graueria_vittata | Graueria_vittata | LC | Stable | tropical | diurnal | 15.5 | 44741.944 | 16.91 |
| birds | Graydidascalus_brachyurus | Graydidascalus_brachyurus | LC | Decreasing | tropical | diurnal | 159 | 658158.939 | 2.407 |
| birds | Griseotyrannus_aurantioatrocristatus | Empidonomus_aurantioatrocristatus | LC | Stable | tropical | diurnal | 33 | 9464416.149 | 4.774 |
| birds | Grus_americana | Grus_americana | EN | Increasing | temperate | diurnal | 5826 | 1543239.244 | 16.418 |
| birds | Grus_antigone | Grus_antigone | VU | Decreasing | temperate | diurnal | 5916.2 | 1882141.611 | 14.355 |
| birds | Grus_canadensis | Grus_canadensis | LC | Increasing | temperate | diurnal | 4296.2 | 14715653.061 | 6.588 |
| birds | Grus_grus | Grus_grus | LC | Increasing | temperate | diurnal | 5500 | 20795140.003 | 8.213 |
| birds | Grus_japonensis | Grus_japonensis | VU | Decreasing | temperate | diurnal | 8786 | 1639871.123 | 14.547 |
| birds | Grus_monacha | Grus_monacha | VU | Increasing | temperate | diurnal | 3729.9 | 1728477.414 | 2.696 |
| birds | Grus_nigricollis | Grus_nigricollis | NT | Stable | temperate | diurnal | 6000 | 886789.647 | 3.754 |
| birds | Grus_rubicunda | Grus_rubicunda | LC | Decreasing | temperate | diurnal | 6222.8 | 3559865.45 | 4.53 |
| birds | Grus_vipio | Grus_vipio | VU | Decreasing | temperate | diurnal | 4663 | 2152136.877 | 12.132 |
| birds | Guadalcanaria_inexpectata | Guadalcanaria_inexpectata | LC | Decreasing | tropical | diurnal | 43.3 | 624.026 | 3.054 |
| birds | Guaruba_guarouba | Guaruba_guarouba | VU | Decreasing | tropical | diurnal | 194 | 932156.376 | 2.597 |
| birds | Gubernatrix_cristata | Gubernatrix_cristata | EN | Decreasing | temperate | diurnal | 47.6 | 1303745.932 | 8.42 |

| | | | | | | | | | |
|---|---|---|---|---|---|---|---|---|---|
| birds | Gubernetes_yetapa | Gubernetes_yetapa | LC | Stable | tropical | diurnal | 68.4 | 2082131.948 | 10.572 |
| birds | Guira_guira | Guira_guira | LC | Increasing | tropical | diurnal | 141 | 8027232.35 | 8.577 |
| birds | Gulosus_aristotelis | Phalacrocorax_aristotelis | LC | Decreasing | temperate | diurnal | 1764.3 | 2002487.086 | 15.918 |
| birds | Guttera_plumifera | Guttera_plumifera | LC | Decreasing | tropical | diurnal | 866 | 834485.491 | 5.812 |
| birds | Guttera_pucherani | Guttera_pucherani | LC | Stable | tropical | diurnal | 1149 | 519919.073 | 10.877 |
| birds | Guyramemua_affinis | Suiriri_islerorum | NT | Decreasing | tropical | diurnal | 22.8 | 1304378.528 | 8.777 |
| birds | Gygis_alba | NA | LC | Stable | NA | NA | NA | NA | NA |
| birds | Gymnobucco_bonapartei | Gymnobucco_bonapartei | LC | Decreasing | tropical | diurnal | 63 | 1792433.713 | 5.872 |
| birds | Gymnobucco_calvus | Gymnobucco_calvus | LC | Decreasing | tropical | diurnal | 54.5 | 1685321.508 | 10.881 |
| birds | Gymnobucco_peli | Gymnobucco_peli | LC | Unknown | tropical | diurnal | 52 | 1350939.246 | 10.517 |
| birds | Gymnobucco_sladeni | Gymnobucco_sladeni | LC | Decreasing | tropical | diurnal | 45 | 1357987.407 | 5.722 |
| birds | Gymnocichla_nudiceps | Gymnocichla_nudiceps | LC | Decreasing | tropical | diurnal | 30.5 | 355899.135 | 10.364 |
| birds | Gymnocrex_plumbeiventris | Gymnocrex_plumbeiventris | LC | Unknown | tropical | diurnal | 288 | 694042.396 | 3.227 |
| birds | Gymnocrex_rosenbergii | NA | VU | Decreasing | NA | NA | NA | NA | NA |
| birds | Gymnoderus_foetidus | Gymnoderus_foetidus | LC | Decreasing | tropical | diurnal | 275.5 | 5252269.507 | 2.784 |
| birds | Gymnogyps_californianus | Gymnogyps_californianus | CR | Increasing | temperate | diurnal | 8442.7 | 388190.919 | 13.591 |
| birds | Gymnomystax_mexicanus | Gymnomystax_mexicanus | LC | Stable | tropical | diurnal | 98 | 1563861.271 | 4.464 |
| birds | Gymnomyza_aubryana | Gymnomyza_aubryana | CR | Decreasing | tropical | diurnal | 195.1 | 3182.985 | 3.845 |
| birds | Gymnomyza_samoensis | NA | EN | Decreasing | NA | NA | NA | NA | NA |
| birds | Gymnomyza_viridis | Gymnomyza_viridis | LC | Decreasing | tropical | diurnal | 87.8 | 5433.552 | 9.916 |
| birds | Gymnophaps_albertisii | Gymnophaps_albertisii | LC | Stable | tropical | diurnal | 259 | 535761.111 | 3.76 |
| birds | Gymnophaps_mada | Gymnophaps_mada | LC | Stable | tropical | diurnal | 334 | 8554.414 | 6.663 |
| birds | Gymnophaps_solomonensis | Gymnophaps_solomonensis | LC | Stable | tropical | diurnal | 335 | 6241.28 | 3.679 |
| birds | Gymnopithys_leucaspis | Gymnopithys_leucaspis | LC | Decreasing | tropical | diurnal | 27.2 | 1029456.746 | 1.588 |
| birds | Gymnopithys_rufigula | Gymnopithys_rufigula | LC | Stable | tropical | diurnal | 29.1 | 1727736.891 | 1.31 |
| birds | Gymnorhina_tibicen | Gymnorhina_tibicen | LC | Increasing | temperate | diurnal | 284.9 | 6989482.967 | 4.825 |
| birds | Gymnorhinus_cyanocephalus | Gymnorhinus_cyanocephalus | VU | Decreasing | temperate | diurnal | 104.8 | 1284048.893 | 8.751 |
| birds | Gymnoris_dentata | Petronia_dentata | LC | Stable | tropical | diurnal | 18.4 | 4184728.084 | 11.684 |
| birds | Gymnoris_pyrgita | Petronia_pyrgita | LC | Stable | tropical | diurnal | 24.1 | 1608726.474 | 9.434 |
| birds | Gymnoris_superciliaris | Petronia_superciliaris | LC | Stable | tropical | diurnal | 24 | 3946522.231 | 8.661 |
| birds | Gymnoris_xanthocollis | Petronia_xanthocollis | LC | Stable | tropical | diurnal | 18.1 | 4502531.204 | 15.172 |
| birds | Gypaetus_barbatus | Gypaetus_barbatus | NT | Decreasing | temperate | diurnal | 5695 | 10893249.16 | 8.451 |
| birds | Gypohierax_angolensis | Gypohierax_angolensis | LC | Stable | tropical | diurnal | 1600 | 11874949.009 | 8.944 |
| birds | Gyps_africanus | Gyps_africanus | CR | Decreasing | tropical | diurnal | 5433 | 15114222.689 | 9.567 |

| | | | | | | | | | |
|---|---|---|---|---|---|---|---|---|---|
| birds | Gyps_bengalensis | Gyps_bengalensis | CR | Decreasing | tropical | diurnal | 4385 | 5834018.965 | 15.1 |
| birds | Gyps_coprotheres | Gyps_coprotheres | VU | Decreasing | temperate | diurnal | 8177 | 3702609.83 | 7.611 |
| birds | Gyps_fulvus | Gyps_fulvus | LC | Increasing | temperate | diurnal | 7436 | 18194036.067 | 10.018 |
| birds | Gyps_himalayensis | Gyps_himalayensis | NT | Decreasing | temperate | diurnal | 9798 | 6434312.573 | 8.946 |
| birds | Gyps_indicus | Gyps_indicus | CR | Decreasing | tropical | diurnal | 5515 | 2380044.79 | 18.372 |
| birds | Gyps_rueppelli | Gyps_rueppellii | CR | Decreasing | tropical | diurnal | 7400 | 10082232.276 | 9.285 |
| birds | Gypsophila_brevicaudata | Napothera_brevicaudata | LC | Decreasing | tropical | diurnal | 19.5 | 1190917.892 | 9.294 |
| birds | Gypsophila_crassa | Napothera_crassa | LC | Decreasing | tropical | diurnal | 26.8 | 34993.361 | 3.28 |
| birds | Gypsophila_rufipecta | Turdinus_rufipectus | LC | Decreasing | tropical | diurnal | 37 | 73599.834 | 7.957 |
| birds | Habia_atrimaxillaris | Habia_atrimaxillaris | NT | Stable | tropical | diurnal | 48.9 | 1819.738 | 8.619 |
| birds | Habia_carmioli | Chlorothraupis_carmioli | LC | Decreasing | tropical | diurnal | 37.6 | 93291.501 | 7.979 |
| birds | Habia_cristata | Habia_cristata | LC | Decreasing | tropical | diurnal | 46.5 | 54054.456 | 12.109 |
| birds | Habia_frenata | Chlorothraupis_frenata | LC | Decreasing | tropical | diurnal | 37.8 | 108990.051 | 4.425 |
| birds | Habia_fuscicauda | Habia_fuscicauda | LC | Decreasing | tropical | diurnal | 36.8 | 581686.915 | 9.889 |
| birds | Habia_gutturalis | Habia_gutturalis | LC | Decreasing | tropical | diurnal | 36.4 | 95132.689 | 11.889 |
| birds | Habia_olivacea | Chlorothraupis_olivacea | LC | Decreasing | tropical | diurnal | 39 | 92811.915 | 5.696 |
| birds | Habia_rubica | Habia_rubica | LC | Decreasing | tropical | diurnal | 32.5 | 5706719.88 | 5.167 |
| birds | Habia_stolzmanni | Chlorothraupis_stolzmanni | LC | Stable | tropical | diurnal | 38.8 | 50027.821 | 6.318 |
| birds | Haematoderus_militaris | Haematoderus_militaris | LC | Decreasing | tropical | diurnal | 227 | 1043733.972 | 1.641 |
| birds | Haematopus_ater | Haematopus_ater | LC | Unknown | temperate | diurnal | 724 | 667536.241 | 5.751 |
| birds | Haematopus_chathamensis | NA | EN | Stable | NA | NA | NA | NA | NA |
| birds | Haematopus_fuliginosus | Haematopus_fuliginosus | LC | Stable | temperate | diurnal | 819 | 812087.46 | 7.643 |
| birds | Haematopus_leucopodus | Haematopus_leucopodus | LC | Stable | temperate | diurnal | 618 | 437048.977 | 4.12 |
| birds | Haematopus_longirostris | Haematopus_longirostris | LC | Unknown | tropical | diurnal | 613.9 | 867914.608 | 6.848 |
| birds | Haematopus_moquini | Haematopus_moquini | LC | Increasing | temperate | diurnal | 692 | 99382.151 | 13.274 |
| birds | Haematopus_ostralegus | Haematopus_ostralegus | NT | Decreasing | temperate | diurnal | 526 | 15759416.902 | 10.178 |
| birds | Haematopus_palliatus | Haematopus_palliatus | LC | Stable | tropical | diurnal | 601.5 | 1821882.106 | 11.942 |
| birds | Haematopus_unicolor | Haematopus_unicolor | LC | Increasing | temperate | diurnal | 671 | 73499.485 | 9.749 |
| birds | Haematortyx_sanguiniceps | Haematortyx_sanguiniceps | LC | Decreasing | tropical | diurnal | 300 | 73122.113 | 2.435 |
| birds | Haemorhous_cassinii | Carpodacus_cassinii | LC | Decreasing | temperate | diurnal | 26.5 | 2814033.132 | 8.492 |
| birds | Haemorhous_mexicanus | Carpodacus_mexicanus | LC | Increasing | temperate | diurnal | 21.4 | 10299459.91 | 12.856 |
| birds | Haemorhous_purpureus | Carpodacus_purpureus | LC | Decreasing | temperate | diurnal | 23.3 | 7952995.845 | 11.465 |
| birds | Hafferia_fortis | Myrmeciza_fortis | LC | Stable | tropical | diurnal | 45.9 | 1945728.917 | 1.501 |
| birds | Hafferia_immaculata | Myrmeciza_immaculata | LC | Decreasing | tropical | diurnal | 47.1 | 103061.805 | 11.631 |

| | | | | | | | | | |
|---|---|---|---|---|---|---|---|---|---|
| birds | Halcyon_albiventris | Halcyon_albiventris | LC | Stable | tropical | diurnal | 65.1 | 3765344.979 | 9.542 |
| birds | Halcyon_badia | Halcyon_badia | LC | Decreasing | tropical | diurnal | 57.9 | 3197177.134 | 7.491 |
| birds | Halcyon_chelicuti | Halcyon_chelicuti | LC | Stable | tropical | diurnal | 43.2 | 13120453.966 | 9.667 |
| birds | Halcyon_coromanda | Halcyon_coromanda | LC | Decreasing | tropical | diurnal | 77.5 | 4234824.051 | 11.891 |
| birds | Halcyon_cyanoventris | Halcyon_cyanoventris | LC | Decreasing | tropical | diurnal | 93 | 131597.724 | 22.341 |
| birds | Halcyon_leucocephala | Halcyon_leucocephala | LC | Stable | tropical | diurnal | 41.8 | 14797715.208 | 9.402 |
| birds | Halcyon_malimbica | Halcyon_malimbica | LC | Decreasing | tropical | diurnal | 91.8 | 4774587.942 | 9.27 |
| birds | Halcyon_pileata | Halcyon_pileata | VU | Decreasing | tropical | diurnal | 84 | 7076437.93 | 14.118 |
| birds | Halcyon_senegalensis | Halcyon_senegalensis | LC | Stable | tropical | diurnal | 73.4 | 14752529.256 | 9.467 |
| birds | Halcyon_senegaloides | Halcyon_senegaloides | LC | Decreasing | temperate | diurnal | 61.8 | 193724.208 | 13.517 |
| birds | Halcyon_smyrnensis | Halcyon_smyrnensis | LC | Increasing | tropical | diurnal | 91.4 | 6823580.455 | 15.509 |
| birds | Haliaeetus_albicilla | Haliaeetus_albicilla | LC | Increasing | temperate | diurnal | 4729.3 | 23280177.99 | 7.139 |
| birds | Haliaeetus_leucocephalus | Haliaeetus_leucocephalus | LC | Increasing | temperate | diurnal | 4700.6 | 15549096.956 | 8.469 |
| birds | Haliaeetus_leucogaster | Haliaeetus_leucogaster | LC | Decreasing | temperate | diurnal | 2827 | 6333774.53 | 8.672 |
| birds | Haliaeetus_leucoryphus | Haliaeetus_leucoryphus | EN | Decreasing | temperate | diurnal | 2885.9 | 14108255.233 | 8.405 |
| birds | Haliaeetus_pelagicus | Haliaeetus_pelagicus | VU | Decreasing | temperate | diurnal | 7757 | 1505403.984 | 2.548 |
| birds | Haliaeetus_sanfordi | Haliaeetus_sanfordi | VU | Decreasing | tropical | diurnal | 1980 | 36161.247 | 4.697 |
| birds | Haliaeetus_vocifer | Haliaeetus_vocifer | LC | Stable | tropical | diurnal | 3400 | 18331524.507 | 9.114 |
| birds | Haliaeetus_vociferoides | Haliaeetus_vociferoides | CR | Decreasing | tropical | diurnal | 3000 | 52761.993 | 9.082 |
| birds | Haliastur_indus | Haliastur_indus | LC | Decreasing | tropical | diurnal | 529 | 8840397.205 | 12.605 |
| birds | Haliastur_sphenurus | Haliastur_sphenurus | LC | Decreasing | temperate | diurnal | 753 | 5585453.97 | 5.358 |
| birds | Halobaena_caerulea | Halobaena_caerulea | LC | Decreasing | temperate | cathemeral | 202 | 76003094.552 | 5.679 |
| birds | Hamirostra_melanosternon | Hamirostra_melanosternon | LC | Decreasing | temperate | diurnal | 1196 | 5297941.743 | 3.414 |
| birds | Hapalopsittaca_amazonina | Hapalopsittaca_amazonina | NT | Decreasing | tropical | diurnal | 106 | 43328.387 | 10.304 |
| birds | Hapalopsittaca_fuertesi | Hapalopsittaca_fuertesi | EN | Increasing | tropical | diurnal | 124 | 2677.643 | 9.202 |
| birds | Hapalopsittaca_melanotis | Hapalopsittaca_melanotis | LC | Stable | tropical | diurnal | 158 | 38602.75 | 5.27 |
| birds | Hapalopsittaca_pyrrhops | Hapalopsittaca_pyrrhops | EN | Decreasing | tropical | diurnal | 103 | 3517.66 | 6.46 |
| birds | Hapaloptila_castanea | Hapaloptila_castanea | LC | Stable | tropical | diurnal | 80.6 | 31778.471 | 5.237 |
| birds | Haplophaedia_assimilis | Haplophaedia_assimilis | LC | Decreasing | tropical | diurnal | 4.8 | 59077.289 | 5.181 |
| birds | Haplophaedia_aureliae | Haplophaedia_aureliae | LC | Decreasing | tropical | diurnal | 5.2 | 222015.495 | 10.461 |
| birds | Haplophaedia_lugens | Haplophaedia_lugens | NT | Stable | tropical | diurnal | 6.9 | 6460.084 | 8.017 |
| birds | Haplospiza_unicolor | Haplospiza_unicolor | LC | Decreasing | temperate | diurnal | 15.3 | 977656.665 | 13.435 |
| birds | Harpactes_ardens | Harpactes_ardens | LC | Decreasing | tropical | diurnal | 96.3 | 211441.239 | 12.338 |
| birds | Harpactes_diardii | Harpactes_diardii | NT | Decreasing | tropical | diurnal | 98.3 | 1108720.015 | 8.725 |

| | | | | | | | | | |
|---|---|---|---|---|---|---|---|---|---|
| birds | Harpactes_duvaucelii | Harpactes_duvaucelii | NT | Decreasing | tropical | diurnal | 38.9 | 1253065.678 | 8.543 |
| birds | Harpactes_erythrocephalus | Harpactes_erythrocephalus | LC | Decreasing | tropical | diurnal | 80.3 | 2608367.684 | 11.131 |
| birds | Harpactes_kasumba | Harpactes_kasumba | NT | Decreasing | tropical | diurnal | 96.9 | 1093996.28 | 8.509 |
| birds | Harpactes_oreskios | Harpactes_oreskios | LC | Decreasing | tropical | diurnal | 57.3 | 1639985.91 | 10.95 |
| birds | Harpactes_orrhophaeus | Harpactes_orrhophaeus | VU | Decreasing | tropical | diurnal | 52.8 | 1075727.376 | 8.79 |
| birds | Harpactes_wardi | Harpactes_wardi | LC | Decreasing | temperate | diurnal | 119 | 137378.038 | 4.643 |
| birds | Harpactes_whiteheadi | Harpactes_whiteheadi | LC | Decreasing | tropical | diurnal | 73 | 21411.306 | 2.138 |
| birds | Harpagus_bidentatus | Harpagus_bidentatus | LC | Decreasing | tropical | diurnal | 215.3 | 8976331.271 | 3.961 |
| birds | Harpagus_diodon | Harpagus_diodon | LC | Decreasing | tropical | diurnal | 200 | 8031409.732 | 6.022 |
| birds | Harpia_harpyja | Harpia_harpyja | VU | Decreasing | tropical | diurnal | 4800 | 7838092.629 | 2.695 |
| birds | Harpyopsis_novaeguineae | Harpyopsis_novaeguineae | VU | Decreasing | tropical | diurnal | 1959.6 | 730591.983 | 3.31 |
| birds | Hartertula_flavoviridis | Neomixis_flavoviridis | NT | Decreasing | tropical | diurnal | 9.7 | 66817.981 | 7.137 |
| birds | Hartlaubius_auratus | Saroglossa_aurata | LC | Decreasing | tropical | diurnal | 40.5 | 409517.942 | 8.202 |
| birds | Hedydipna_collaris | Anthreptes_collaris | LC | Stable | tropical | diurnal | 7 | 9197435.7 | 9.073 |
| birds | Hedydipna_metallica | Anthreptes_metallicus | LC | Stable | tropical | diurnal | 7.2 | 1304366.516 | 8.511 |
| birds | Hedydipna_pallidigaster | Anthreptes_pallidigaster | EN | Decreasing | tropical | diurnal | 6.8 | 753.185 | 9.059 |
| birds | Hedydipna_platura | Anthreptes_platurus | LC | Stable | tropical | diurnal | 6.7 | 4094486.354 | 10.86 |
| birds | Heleia_crassirostris | Heleia_crassirostris | LC | Decreasing | tropical | diurnal | 19 | 28274.735 | 12.455 |
| birds | Heleia_dohertyi | NA | LC | Decreasing | NA | NA | NA | NA | NA |
| birds | Heleia_goodfellowi | Lophozosterops_goodfellowi | LC | Decreasing | tropical | diurnal | 20 | 3458.601 | 4.898 |
| birds | Heleia_javanica | NA | LC | Decreasing | NA | NA | NA | NA | NA |
| birds | Heleia_muelleri | Heleia_muelleri | NT | Decreasing | tropical | diurnal | 16.7 | 26735.011 | 13.911 |
| birds | Heleia_pinaiae | Lophozosterops_pinaiae | LC | Decreasing | tropical | diurnal | 27.4 | 497.761 | 6.057 |
| birds | Heleia_squamiceps | Lophozosterops_squamiceps | LC | Decreasing | tropical | diurnal | 14.5 | 27806.243 | 4.574 |
| birds | Heleia_squamifrons | Oculocincta_squamifrons | LC | Decreasing | tropical | diurnal | 6.9 | 81974.586 | 2.353 |
| birds | Heleia_superciliaris | NA | LC | Unknown | NA | NA | NA | NA | NA |
| birds | Heleia_wallacei | Zosterops_wallacei | LC | Unknown | tropical | diurnal | 11 | 37730.844 | 13.493 |
| birds | Heliactin_bilophus | Heliactin_bilophus | LC | Increasing | tropical | diurnal | 4.4 | 3879596.576 | 7.948 |
| birds | Heliangelus_amethysticollis | Heliangelus_amethysticollis | LC | Stable | tropical | diurnal | 5 | 146657.785 | 8.102 |
| birds | Heliangelus_clarisse | NA | LC | Stable | NA | NA | NA | NA | NA |
| birds | Heliangelus_exortis | Heliangelus_exortis | LC | Stable | tropical | diurnal | 4.6 | 64064.871 | 10.812 |
| birds | Heliangelus_mavors | Heliangelus_mavors | LC | Stable | tropical | diurnal | 4.2 | 38610.246 | 13.061 |
| birds | Heliangelus_micraster | Heliangelus_micraster | LC | Unknown | tropical | diurnal | 3.9 | 17855.932 | 7.77 |
| birds | Heliangelus_regalis | Heliangelus_regalis | NT | Decreasing | tropical | diurnal | 3.6 | 5218.905 | 4.348 |

| | | | | | | | | | |
|---|---|---|---|---|---|---|---|---|---|
| birds | Heliangelus_strophianus | Heliangelus_strophianus | LC | Decreasing | tropical | diurnal | 5.5 | 10032.035 | 10.449 |
| birds | Heliangelus_viola | Heliangelus_viola | LC | Stable | tropical | diurnal | 5.4 | 35976.795 | 11.192 |
| birds | Helicolestes_hamatus | Helicolestes_hamatus | LC | Decreasing | tropical | diurnal | 425.5 | 2374863.312 | 2.765 |
| birds | Heliobletus_contaminatus | Heliobletus_contaminatus | LC | Decreasing | temperate | diurnal | 14 | 815159.114 | 13.321 |
| birds | Heliodoxa_aurescens | Heliodoxa_aurescens | LC | Decreasing | tropical | diurnal | 6.4 | 3600288.829 | 1.454 |
| birds | Heliodoxa_branickii | Heliodoxa_branickii | LC | Decreasing | tropical | diurnal | 5.2 | 126780.84 | 3.63 |
| birds | Heliodoxa_gularis | Heliodoxa_gularis | LC | Decreasing | tropical | diurnal | 6.3 | 38003.017 | 6.69 |
| birds | Heliodoxa_imperatrix | Heliodoxa_imperatrix | LC | Unknown | tropical | diurnal | 8.8 | 22063.191 | 6.238 |
| birds | Heliodoxa_jacula | Heliodoxa_jacula | LC | Stable | tropical | diurnal | 8.1 | 90867.369 | 9.407 |
| birds | Heliodoxa_leadbeateri | Heliodoxa_leadbeateri | LC | Decreasing | tropical | diurnal | 7.4 | 409940.206 | 8.76 |
| birds | Heliodoxa_rubinoides | Heliodoxa_rubinoides | LC | Unknown | tropical | diurnal | 7.8 | 203573.737 | 7.532 |
| birds | Heliodoxa_schreibersii | Heliodoxa_schreibersii | LC | Decreasing | tropical | diurnal | 9.2 | 713038.856 | 1.802 |
| birds | Heliodoxa_xanthogonys | Heliodoxa_xanthogonys | LC | Decreasing | tropical | diurnal | 7 | 222443.103 | 0.827 |
| birds | Heliomaster_constantii | Heliomaster_constantii | LC | Decreasing | tropical | diurnal | 7.3 | 554868.14 | 10.227 |
| birds | Heliomaster_furcifer | Heliomaster_furcifer | LC | Decreasing | temperate | diurnal | 5.4 | 2967951.959 | 8.52 |
| birds | Heliomaster_longirostris | Heliomaster_longirostris | LC | Decreasing | tropical | diurnal | 6.6 | 8290213.633 | 3.638 |
| birds | Heliomaster_squamosus | Heliomaster_squamosus | LC | Stable | tropical | diurnal | 5.9 | 1519261.766 | 10.976 |
| birds | Heliopais_personatus | Heliopais_personatus | CR | Decreasing | tropical | diurnal | 510 | 766693.113 | 12.511 |
| birds | Heliornis_fulica | Heliornis_fulica | LC | Decreasing | tropical | diurnal | 132 | 8812012.443 | 3.856 |
| birds | Heliothryx_auritus | Heliothryx_auritus | LC | Decreasing | tropical | diurnal | 5.4 | 6893450.958 | 2.621 |
| birds | Heliothryx_barroti | Heliothryx_barroti | LC | Decreasing | tropical | diurnal | 5.5 | 491040.04 | 8.783 |
| birds | Hellmayrea_gularis | Hellmayrea_gularis | LC | Stable | tropical | diurnal | 12.4 | 170416.442 | 10.051 |
| birds | Helmitheros_vermivorum | Helmitheros_vermivorum | LC | Increasing | tropical | diurnal | 14.2 | 3326592.3 | 15.377 |
| birds | Helopsaltes_certhiola | Locustella_certhiola | LC | Decreasing | temperate | diurnal | 14.4 | 18568127.916 | 7.561 |
| birds | Helopsaltes_ochotensis | Locustella_ochotensis | LC | Decreasing | temperate | diurnal | 18.5 | 1896276.858 | 9.931 |
| birds | Helopsaltes_pleskei | Locustella_pleskei | VU | Decreasing | tropical | diurnal | 20 | 5556.544 | 25.112 |
| birds | Helopsaltes_pryeri | Locustella_pryeri | NT | Decreasing | temperate | diurnal | 14.2 | 227242.528 | 16.661 |
| birds | Hemicircus_canente | Hemicircus_canente | LC | Decreasing | tropical | diurnal | 43 | 665192.853 | 10.649 |
| birds | Hemicircus_concretus | Hemicircus_concretus | LC | Decreasing | tropical | diurnal | 29.4 | 102171.064 | 22.744 |
| birds | Hemignathus_lucidus | NA | EX | not_available | NA | NA | NA | NA | NA |
| birds | Hemignathus_wilsoni | Hemignathus_munroi | EN | Decreasing | tropical | diurnal | 28 | 8415.595 | 7.216 |
| birds | Hemimacronyx_chloris | Anthus_chloris | VU | Decreasing | temperate | diurnal | 25.1 | 82125.981 | 11 |
| birds | Hemiphaga_novaeseelandiae | Hemiphaga_novaeseelandiae | LC | Increasing | temperate | diurnal | 653 | 269913.169 | 7.99 |
| birds | Hemiprocne_comata | Hemiprocne_comata | LC | Decreasing | tropical | diurnal | 20.6 | 1482225.388 | 9.387 |

| | | | | | | | | | |
|---|---|---|---|---|---|---|---|---|---|
| birds | Hemiprocne_coronata | Hemiprocne_coronata | LC | Decreasing | tropical | diurnal | 29 | 3347103.258 | 13.781 |
| birds | Hemiprocne_longipennis | Hemiprocne_longipennis | LC | Decreasing | tropical | diurnal | 42.8 | 1504844.465 | 9.654 |
| birds | Hemiprocne_mystacea | Hemiprocne_mystacea | LC | Stable | tropical | nocturnal | 70.1 | 805813.235 | 3.504 |
| birds | Hemipus_hirundinaceus | Hemipus_hirundinaceus | LC | Decreasing | tropical | diurnal | 10.2 | 1391676.195 | 9.714 |
| birds | Hemipus_picatus | Hemipus_picatus | LC | Decreasing | tropical | diurnal | 9 | 4982703.19 | 12.486 |
| birds | Hemitesia_neumanni | Hemitesia_neumanni | LC | Stable | tropical | diurnal | 14.2 | 37085.54 | 14.013 |
| birds | Hemitesia_pallidipes | Cettia_pallidipes | LC | Stable | temperate | diurnal | 10 | 1229162.305 | 9.904 |
| birds | Hemithraupis_flavicollis | Hemithraupis_flavicollis | LC | Decreasing | tropical | diurnal | 12.8 | 5620134.106 | 2.539 |
| birds | Hemithraupis_guira | Hemithraupis_guira | LC | Decreasing | tropical | diurnal | 12 | 9380161.988 | 5.442 |
| birds | Hemithraupis_ruficapilla | Hemithraupis_ruficapilla | LC | Stable | tropical | diurnal | 11 | 648694.042 | 12.776 |
| birds | Hemitriccus_cinnamomeipectus | Hemitriccus_cinnamomeipectus | LC | Stable | tropical | diurnal | 7.5 | 6101.276 | 3.992 |
| birds | Hemitriccus_diops | Hemitriccus_diops | LC | Decreasing | tropical | diurnal | 10 | 737548.445 | 12.794 |
| birds | Hemitriccus_flammulatus | Hemitriccus_flammulatus | LC | Decreasing | tropical | diurnal | 10 | 806774.082 | 3.525 |
| birds | Hemitriccus_furcatus | NA | VU | Decreasing | NA | NA | NA | NA | NA |
| birds | Hemitriccus_granadensis | Hemitriccus_granadensis | LC | Stable | tropical | diurnal | 7.9 | 276758.68 | 9.506 |
| birds | Hemitriccus_griseipectus | Hemitriccus_griseipectus | LC | Stable | tropical | diurnal | 9 | 2659877.651 | 2.245 |
| birds | Hemitriccus_inornatus | Hemitriccus_inornatus | LC | Decreasing | tropical | diurnal | 8.3 | 34481.44 | 1.079 |
| birds | Hemitriccus_iohannis | Hemitriccus_iohannis | LC | Stable | tropical | diurnal | 10 | 1134921.913 | 1.498 |
| birds | Hemitriccus_josephinae | Hemitriccus_josephinae | LC | Decreasing | tropical | diurnal | 10.8 | 666063.425 | 0.956 |
| birds | Hemitriccus_kaempferi | Hemitriccus_kaempferi | VU | Decreasing | temperate | diurnal | 8.5 | 7847.608 | 17.696 |
| birds | Hemitriccus_margaritaceiventer | Hemitriccus_margaritaceiventer | LC | Stable | tropical | diurnal | 8.4 | 5809388.942 | 8.365 |
| birds | Hemitriccus_minimus | Hemitriccus_minimus | LC | Stable | tropical | diurnal | 7 | 2469713.046 | 2.641 |
| birds | Hemitriccus_minor | Hemitriccus_minor | LC | Decreasing | tropical | diurnal | 7.4 | 2416974.334 | 2.012 |
| birds | Hemitriccus_mirandae | NA | VU | Decreasing | NA | NA | NA | NA | NA |
| birds | Hemitriccus_nidipendulus | Hemitriccus_nidipendulus | LC | Decreasing | tropical | diurnal | 7.5 | 542148.006 | 12.968 |
| birds | Hemitriccus_obsoletus | Hemitriccus_obsoletus | LC | Decreasing | temperate | diurnal | 11.6 | 262970.93 | 12.18 |
| birds | Hemitriccus_orbitatus | Hemitriccus_orbitatus | NT | Decreasing | temperate | diurnal | 9.7 | 663643.419 | 14.34 |
| birds | Hemitriccus_rufigularis | Hemitriccus_rufigularis | LC | Decreasing | tropical | diurnal | 9.4 | 76788.806 | 4.286 |
| birds | Hemitriccus_spodiops | Hemitriccus_spodiops | LC | Decreasing | tropical | diurnal | 7.2 | 67550.211 | 4.582 |
| birds | Hemitriccus_striaticollis | Hemitriccus_striaticollis | LC | Decreasing | tropical | diurnal | 8.6 | 4016133.454 | 4.897 |
| birds | Hemitriccus_zosterops | Hemitriccus_zosterops | LC | Stable | tropical | diurnal | 8.7 | 2154385.693 | 1.276 |
| birds | Hemixos_castanonotus | Hemixos_castanonotus | LC | Increasing | temperate | diurnal | 35.3 | 979934.257 | 14.88 |
| birds | Hemixos_flavala | Hemixos_flavala | LC | Stable | tropical | diurnal | 32.5 | 1098549.778 | 9.098 |
| birds | Hemixos_leucogrammicus | Pycnonotus_leucogrammicus | LC | Stable | tropical | diurnal | 33.5 | 63047.145 | 7.437 |

| | | | | | | | | | |
|---|---|---|---|---|---|---|---|---|---|
| birds | Henicopernis_infuscatus | NA | VU | Decreasing | NA | NA | NA | NA | NA |
| birds | Henicopernis_longicauda | Henicopernis_longicauda | LC | Decreasing | tropical | diurnal | 571.2 | 750041.011 | 3.312 |
| birds | Henicophaps_albifrons | Henicophaps_albifrons | LC | Stable | tropical | diurnal | 247 | 769384.492 | 3.349 |
| birds | Henicophaps_foersteri | Henicophaps_foersteri | VU | Decreasing | tropical | diurnal | 247 | 31332.272 | 4.446 |
| birds | Henicorhina_leucophrys | Henicorhina_leucophrys | LC | Decreasing | tropical | diurnal | 16.2 | 722586.814 | 9.471 |
| birds | Henicorhina_leucoptera | Henicorhina_leucoptera | LC | Decreasing | tropical | diurnal | 13.6 | 10930.352 | 4.292 |
| birds | Henicorhina_leucosticta | Henicorhina_leucosticta | LC | Decreasing | tropical | diurnal | 14.2 | 2745877.944 | 3.548 |
| birds | Herpetotheres_cachinnans | Herpetotheres_cachinnans | LC | Decreasing | tropical | diurnal | 623.6 | 13330796.594 | 5.731 |
| birds | Herpsilochmus_atricapillus | Herpsilochmus_atricapillus | LC | Decreasing | tropical | diurnal | 9.9 | 3731834.004 | 8.973 |
| birds | Herpsilochmus_axillaris | Herpsilochmus_axillaris | LC | Decreasing | tropical | diurnal | 11.4 | 186689.553 | 5.701 |
| birds | Herpsilochmus_dorsimaculatus | Herpsilochmus_dorsimaculatus | LC | Decreasing | tropical | diurnal | 9.9 | 907516.408 | 0.899 |
| birds | Herpsilochmus_dugandi | Herpsilochmus_dugandi | LC | Decreasing | tropical | diurnal | 10.3 | 231956.507 | 2.55 |
| birds | Herpsilochmus_gentryi | Herpsilochmus_gentryi | LC | Decreasing | tropical | diurnal | 10.6 | 9533.659 | 1.411 |
| birds | Herpsilochmus_longirostris | Herpsilochmus_longirostris | LC | Decreasing | tropical | diurnal | 12.8 | 1991289.015 | 8.861 |
| birds | Herpsilochmus_motacilloides | Herpsilochmus_motacilloides | LC | Decreasing | tropical | diurnal | 12.5 | 40630.337 | 3.935 |
| birds | Herpsilochmus_parkeri | Herpsilochmus_parkeri | VU | Decreasing | tropical | diurnal | 11.5 | 5953.549 | 3.574 |
| birds | Herpsilochmus_pectoralis | Herpsilochmus_pectoralis | VU | Decreasing | tropical | diurnal | 11.1 | 314941.879 | 10.91 |
| birds | Herpsilochmus_pileatus | Herpsilochmus_pileatus | NT | Decreasing | tropical | diurnal | 8.9 | 7783.801 | 9.487 |
| birds | Herpsilochmus_roraimae | Herpsilochmus_roraimae | LC | Decreasing | tropical | diurnal | 11.5 | 283885.894 | 0.853 |
| birds | Herpsilochmus_rufimarginatus | Herpsilochmus_rufimarginatus | LC | Stable | temperate | diurnal | 10.6 | 680842.367 | 14.166 |
| birds | Herpsilochmus_sellowi | Herpsilochmus_sellowi | LC | Decreasing | tropical | diurnal | 7.5 | 995787.856 | 7.613 |
| birds | Herpsilochmus_stictocephalus | Herpsilochmus_stictocephalus | LC | Decreasing | tropical | diurnal | 8.8 | 527219.134 | 1.412 |
| birds | Herpsilochmus_sticturus | Herpsilochmus_sticturus | LC | Stable | tropical | diurnal | 9.5 | 590129.275 | 1.732 |
| birds | Hesperiphona_abeillei | Coccothraustes_abeillei | LC | Decreasing | tropical | diurnal | 48.5 | 251295.188 | 8.57 |
| birds | Hesperiphona_vespertina | Coccothraustes_vespertinus | VU | Decreasing | temperate | diurnal | 57.3 | 9649540.805 | 12.064 |
| birds | Heterocercus_aurantiivertex | Heterocercus_aurantiivertex | LC | Stable | tropical | diurnal | 21.5 | 233954.069 | 1.602 |
| birds | Heterocercus_flavivertex | Heterocercus_flavivertex | LC | Stable | tropical | diurnal | 21.3 | 827963.874 | 1.326 |
| birds | Heterocercus_linteatus | Heterocercus_linteatus | LC | Decreasing | tropical | diurnal | 21.2 | 2146985.536 | 2.136 |
| birds | Heteromirafra_archeri | Heteromirafra_archeri | CR | Decreasing | tropical | diurnal | 30 | 1009.963 | 14.738 |
| birds | Heteromirafra_ruddi | Heteromirafra_ruddi | EN | Decreasing | temperate | diurnal | 26.6 | 21832.331 | 10.167 |
| birds | Heteromunia_pectoralis | Heteromunia_pectoralis | LC | Decreasing | tropical | diurnal | 15.2 | 1302318.406 | 2.631 |
| birds | Heteromyias_albispecularis | Heteromyias_albispecularis | LC | Stable | tropical | diurnal | 36.3 | 12665.387 | 2.242 |
| birds | Heteronetta_atricapilla | Heteronetta_atricapilla | LC | Stable | temperate | diurnal | 538.4 | 1954633.056 | 8.646 |
| birds | Heterophasia_auricularis | Heterophasia_auricularis | LC | Decreasing | temperate | diurnal | 41.8 | 10366.184 | 5.409 |

| | | | | | | | | | |
|---|---|---|---|---|---|---|---|---|---|
| birds | Heterophasia_capistrata | Heterophasia_capistrata | LC | Unknown | temperate | diurnal | 39.4 | 257460.17 | 11.053 |
| birds | Heterophasia_gracilis | Heterophasia_gracilis | LC | Unknown | temperate | diurnal | 37.8 | 166618.91 | 6.783 |
| birds | Heterophasia_melanoleuca | Heterophasia_melanoleuca | LC | Decreasing | tropical | diurnal | 32.6 | 177773.391 | 8.479 |
| birds | Heterophasia_picaoides | Heterophasia_picaoides | LC | Decreasing | tropical | diurnal | 42.4 | 1517923.865 | 8.617 |
| birds | Heterophasia_pulchella | Heterophasia_pulchella | LC | Decreasing | temperate | diurnal | 40.6 | 179332.118 | 6.436 |
| birds | Heteroscenes_pallidus | Cuculus_pallidus | LC | Increasing | temperate | diurnal | 87.7 | 7756186.472 | 4.312 |
| birds | Heterospingus_rubrifrons | Heterospingus_rubrifrons | LC | Decreasing | tropical | diurnal | 38 | 22680.839 | 6.649 |
| birds | Heterospingus_xanthopygius | Heterospingus_xanthopygius | LC | Decreasing | tropical | diurnal | 38.8 | 153049.115 | 8.324 |
| birds | Heterotetrax_humilis | Eupodotis_humilis | NT | Decreasing | tropical | diurnal | 698 | 474259.61 | 7.411 |
| birds | Heterotetrax_rueppelii | Eupodotis_rueppellii | LC | Stable | tropical | diurnal | 1107 | 280978.336 | 5.581 |
| birds | Heterotetrax_vigorsii | Eupodotis_vigorsii | LC | Increasing | temperate | diurnal | 1700 | 691336.876 | 6.68 |
| birds | Heteroxenicus_stellatus | Brachypteryx_stellata | LC | Stable | temperate | diurnal | 23 | 237692.155 | 7.242 |
| birds | Hieraaetus_ayresii | Hieraaetus_ayresii | LC | Decreasing | tropical | diurnal | 806.1 | 7512090.852 | 9.639 |
| birds | Hieraaetus_morphnoides | Hieraaetus_morphnoides | LC | Stable | temperate | diurnal | 775.1 | 7265023.225 | 4.368 |
| birds | Hieraaetus_pennatus | Hieraaetus_pennatus | LC | Stable | temperate | diurnal | 823.8 | 29211510.176 | 10.238 |
| birds | Hieraaetus_wahlbergi | Aquila_wahlbergi | LC | Stable | tropical | diurnal | 640 | 12004193.208 | 10.248 |
| birds | Hierococcyx_fugax | Cuculus_fugax | LC | Decreasing | tropical | diurnal | 79.7 | 1412734.291 | 8.353 |
| birds | Hierococcyx_pectoralis | NA | LC | Decreasing | NA | NA | NA | NA | NA |
| birds | Hierococcyx_sparverioides | Cuculus_sparverioides | LC | Stable | temperate | diurnal | 151 | 7309805.018 | 13.45 |
| birds | Hierococcyx_vagans | Cuculus_vagans | NT | Decreasing | tropical | diurnal | 57.7 | 976640.03 | 8.124 |
| birds | Hierococcyx_varius | Cuculus_varius | LC | Stable | tropical | diurnal | 103 | 3245384.679 | 17.924 |
| birds | Himalayapsitta_cyanocephala | Psittacula_cyanocephala | LC | Decreasing | tropical | diurnal | 66 | 2783564.237 | 18.946 |
| birds | Himalayapsitta_finschii | Psittacula_finschii | NT | Decreasing | tropical | diurnal | 89.6 | 1639892.996 | 10.359 |
| birds | Himalayapsitta_himalayana | Psittacula_himalayana | LC | Stable | temperate | diurnal | 108.5 | 149148.55 | 14.084 |
| birds | Himalayapsitta_roseata | NA | NT | Decreasing | NA | NA | NA | NA | NA |
| birds | Himantopus_himantopus | Himantopus_himantopus | LC | Increasing | tropical | cathemeral | 176.8 | 65772153.442 | 9.742 |
| birds | Himantopus_novaezelandiae | Himantopus_novaezelandiae | CR | Increasing | temperate | diurnal | 195 | 130531.691 | 9.319 |
| birds | Himantornis_haematopus | Himantornis_haematopus | LC | Decreasing | tropical | diurnal | 585.7 | 2345297.5 | 7.339 |
| birds | Himatione_sanguinea | Himatione_sanguinea | LC | Stable | tropical | diurnal | 15.2 | 8566.614 | 6.436 |
| birds | Hippolais_icterina | Hippolais_icterina | LC | Decreasing | temperate | diurnal | 13.2 | 29586051.33 | 8.759 |
| birds | Hippolais_languida | Hippolais_languida | LC | Stable | tropical | diurnal | 12 | 10092888.182 | 8.479 |
| birds | Hippolais_olivetorum | Hippolais_olivetorum | LC | Stable | tropical | diurnal | 17.5 | 13353897.559 | 8.095 |
| birds | Hippolais_polyglotta | Hippolais_polyglotta | LC | Increasing | tropical | diurnal | 11 | 10548115.162 | 8.908 |
| birds | Hirundapus_caudacutus | Hirundapus_caudacutus | LC | Stable | temperate | diurnal | 95.5 | 16971508.016 | 8.573 |

| | | | | | | | | | |
|---|---|---|---|---|---|---|---|---|---|
| birds | Hirundapus_celebensis | Hirundapus_celebensis | LC | Stable | tropical | diurnal | 180 | 285896.076 | 13.364 |
| birds | Hirundapus_cochinchinensis | Hirundapus_cochinchinensis | LC | Stable | tropical | diurnal | 83.8 | 2870461.268 | 12.376 |
| birds | Hirundapus_giganteus | Hirundapus_giganteus | LC | Decreasing | tropical | diurnal | 127 | 2715789.434 | 10.98 |
| birds | Hirundinea_ferruginea | Hirundinea_ferruginea | LC | Increasing | tropical | diurnal | 30.6 | 916120.422 | 3.029 |
| birds | Hirundo_aethiopica | Hirundo_aethiopica | LC | Increasing | tropical | diurnal | 13.3 | 4764528.413 | 10.829 |
| birds | Hirundo_albigularis | Hirundo_albigularis | LC | Increasing | tropical | diurnal | 21.8 | 3893507.606 | 8.212 |
| birds | Hirundo_angolensis | Hirundo_angolensis | LC | Increasing | tropical | diurnal | 17.8 | 1666808.129 | 10.643 |
| birds | Hirundo_atrocaerulea | Hirundo_atrocaerulea | VU | Decreasing | tropical | diurnal | 13.8 | 1643896.125 | 12.01 |
| birds | Hirundo_dimidiata | Hirundo_dimidiata | LC | Stable | tropical | diurnal | 12.1 | 3681173.146 | 7.651 |
| birds | Hirundo_javanica | Hirundo_domicola | LC | Increasing | tropical | diurnal | 17.8 | 3415869.811 | 9.095 |
| birds | Hirundo_leucosoma | Hirundo_leucosoma | LC | Increasing | tropical | diurnal | 17.6 | 1390558.123 | 13.643 |
| birds | Hirundo_lucida | Hirundo_lucida | LC | Increasing | tropical | diurnal | 13 | 2258983.812 | 11.749 |
| birds | Hirundo_megaensis | Hirundo_megaensis | VU | Decreasing | tropical | diurnal | 11 | 11472.457 | 7.705 |
| birds | Hirundo_neoxena | Hirundo_neoxena | LC | Increasing | temperate | cathemeral | 14.7 | 6029453.139 | 5.326 |
| birds | Hirundo_nigrita | Hirundo_nigrita | LC | Increasing | tropical | diurnal | 18.3 | 4040980.583 | 7.72 |
| birds | Hirundo_nigrorufa | Hirundo_nigrorufa | LC | Stable | tropical | diurnal | 14.2 | 581101.752 | 6.976 |
| birds | Hirundo_rustica | Hirundo_rustica | LC | Decreasing | temperate | diurnal | 17.9 | 94643183.387 | 8.996 |
| birds | Hirundo_smithii | Hirundo_smithii | LC | Increasing | tropical | diurnal | 12.4 | 15137552.427 | 11.526 |
| birds | Hirundo_tahitica | Hirundo_tahitica | LC | Unknown | tropical | diurnal | 17.8 | 1208.547 | 15.131 |
| birds | Histrionicus_histrionicus | Histrionicus_histrionicus | LC | Increasing | temperate | diurnal | 561.6 | 11164771.887 | 1.963 |
| birds | Histurgops_ruficauda | Histurgops_ruficaudus | LC | Stable | tropical | diurnal | 69 | 98235.585 | 13.459 |
| birds | Hoploxypterus_cayanus | Vanellus_cayanus | LC | Unknown | tropical | diurnal | 75.8 | 11467799.978 | 5.07 |
| birds | Horizocerus_albocristatus | Tropicranus_albocristatus | LC | Decreasing | tropical | diurnal | 289.4 | 479716.633 | 15.608 |
| birds | Horizocerus_hartlaubi | Tockus_hartlaubi | LC | Decreasing | tropical | diurnal | 98.5 | 1163200.914 | 10.527 |
| birds | Horornis_acanthizoides | Cettia_acanthizoides | LC | Decreasing | temperate | diurnal | 5.3 | 311800.424 | 10.422 |
| birds | Horornis_annae | Cettia_annae | LC | Stable | tropical | diurnal | 18.5 | 442.791 | 11.2 |
| birds | Horornis_canturians | Cettia_canturians | LC | Stable | temperate | diurnal | 22.3 | 3459855.87 | 16.947 |
| birds | Horornis_carolinae | Cettia_carolinae | NT | Decreasing | tropical | diurnal | 17.1 | 3225.697 | 6.085 |
| birds | Horornis_diphone | Cettia_diphone | LC | Stable | temperate | diurnal | 13.3 | 430218.555 | 19.189 |
| birds | Horornis_flavolivaceus | Cettia_flavolivacea | LC | Stable | temperate | diurnal | 8 | 1896851.773 | 9.995 |
| birds | Horornis_fortipes | Cettia_fortipes | LC | Decreasing | temperate | diurnal | 9.6 | 2834443.671 | 13.624 |
| birds | Horornis_parens | Cettia_parens | LC | Decreasing | tropical | diurnal | 14 | 202.262 | 3.35 |
| birds | Horornis_ruficapilla | Cettia_ruficapilla | LC | Decreasing | tropical | diurnal | 12.7 | 16891.281 | 9.37 |
| birds | Horornis_seebohmi | Cettia_seebohmi | LC | Stable | tropical | diurnal | 12.2 | 14356.216 | 8.565 |

| | | | | | | | | | |
|---|---|---|---|---|---|---|---|---|---|
| birds | Houbaropsis_bengalensis | Houbaropsis_bengalensis | CR | Decreasing | temperate | diurnal | 2012.5 | 260519.849 | 16.915 |
| birds | Humblotia_flavirostris | NA | VU | Decreasing | NA | NA | NA | NA | NA |
| birds | Hydrobates_castro | Oceanodroma_castro | LC | Decreasing | tropical | cathemeral | 46.1 | 86645652.286 | 12.077 |
| birds | Hydrobates_furcatus | Oceanodroma_furcata | LC | Increasing | temperate | cathemeral | 59.7 | 19594359.951 | 2.947 |
| birds | Hydrobates_homochroa | Oceanodroma_homochroa | EN | Decreasing | temperate | diurnal | 36.9 | 347898.564 | 8.385 |
| birds | Hydrobates_leucorhous | Oceanodroma_leucorhoa | VU | Decreasing | temperate | cathemeral | 36.7 | 110025337.723 | 6.628 |
| birds | Hydrobates_matsudairae | Oceanodroma_matsudairae | VU | Unknown | tropical | diurnal | 62 | 22464440.648 | 9.85 |
| birds | Hydrobates_melania | Oceanodroma_melania | LC | Decreasing | tropical | diurnal | 61.5 | 7927238.233 | 7.684 |
| birds | Hydrobates_microsoma | Halocyptena_microsoma | LC | Stable | tropical | cathemeral | 20.5 | 2556397.261 | 8.035 |
| birds | Hydrobates_pelagicus | Hydrobates_pelagicus | LC | Unknown | tropical | cathemeral | 25.2 | 50310627.511 | 18.181 |
| birds | Hydrobates_tethys | Oceanodroma_tethys | LC | Decreasing | tropical | cathemeral | 22.6 | 11745531.825 | 10.088 |
| birds | Hydrobates_tristrami | Oceanodroma_tristrami | LC | Stable | temperate | cathemeral | 89.1 | 13675265.591 | 8.289 |
| birds | Hydrocoloeus_minutus | Larus_minutus | LC | Increasing | temperate | diurnal | 118 | 10822470.827 | 6.929 |
| birds | Hydrophasianus_chirurgus | Hydrophasianus_chirurgus | LC | Decreasing | tropical | diurnal | 160 | 8823045.221 | 14.641 |
| birds | Hydroprogne_caspia | Sterna_caspia | LC | Increasing | temperate | diurnal | 655 | 13929787.88 | 11.452 |
| birds | Hydropsalis_cayennensis | Caprimulgus_cayennensis | LC | Decreasing | tropical | nocturnal | 35.5 | 1530293.865 | 6.418 |
| birds | Hydropsalis_climacocerca | Hydropsalis_climacocerca | LC | Stable | tropical | nocturnal | 46 | 5856089.268 | 1.848 |
| birds | Hydropsalis_maculicaudus | Caprimulgus_maculicaudus | LC | Stable | tropical | nocturnal | 30.3 | 3375852.779 | 7.648 |
| birds | Hydropsalis_torquata | Hydropsalis_torquata | LC | Stable | tropical | nocturnal | 58.5 | 7882112.565 | 7.902 |
| birds | Hydrornis_baudii | Pitta_baudii | VU | Decreasing | tropical | diurnal | 66 | 604986.598 | 6.153 |
| birds | Hydrornis_caeruleus | Pitta_caerulea | NT | Decreasing | tropical | diurnal | 202 | 779802.961 | 8.733 |
| birds | Hydrornis_cyaneus | Pitta_cyanea | LC | Unknown | tropical | diurnal | 109 | 1019585.764 | 9.452 |
| birds | Hydrornis_elliotii | Pitta_elliotii | LC | Decreasing | tropical | diurnal | 91 | 605196.037 | 11.59 |
| birds | Hydrornis_guajanus | Pitta_guajana | LC | Decreasing | tropical | diurnal | 81.5 | 129669.157 | 22.474 |
| birds | Hydrornis_gurneyi | Pitta_gurneyi | CR | Decreasing | tropical | diurnal | 60.5 | 1891.849 | 6.858 |
| birds | Hydrornis_nipalensis | Pitta_nipalensis | LC | Decreasing | tropical | diurnal | 124 | 668283.531 | 9.746 |
| birds | Hydrornis_oatesi | Pitta_oatesi | LC | Decreasing | tropical | diurnal | 120 | 547817.02 | 7.829 |
| birds | Hydrornis_phayrei | Pitta_phayrei | LC | Decreasing | tropical | diurnal | 100 | 927700.653 | 10.636 |
| birds | Hydrornis_schneideri | NA | LC | Decreasing | NA | NA | NA | NA | NA |
| birds | Hydrornis_soror | Pitta_soror | LC | Unknown | tropical | diurnal | 114 | 579014.957 | 12.822 |
| birds | Hylacola_cauta | Hylacola_cauta | LC | Decreasing | temperate | diurnal | 15.2 | 971888.935 | 8.036 |
| birds | Hylacola_pyrrhopygia | Hylacola_pyrrhopygia | LC | Decreasing | temperate | diurnal | 16.8 | 767767.194 | 10.259 |
| birds | Hylatomus_lineatus | Dryocopus_lineatus | LC | Increasing | tropical | diurnal | 183.2 | 13770543.52 | 5.942 |
| birds | Hylatomus_pileatus | Dryocopus_pileatus | LC | Increasing | temperate | diurnal | 286.6 | 5925872.323 | 10.989 |

| | | | | | | | | | |
|---|---|---|---|---|---|---|---|---|---|
| birds | Hylatomus_schulzii | Dryocopus_schulzi | NT | Decreasing | temperate | diurnal | 200 | 388434.651 | 6.839 |
| birds | Hylexetastes_perrotii | Hylexetastes_perrotii | LC | Decreasing | tropical | diurnal | 103.1 | 1064719.633 | 1.496 |
| birds | Hylexetastes_stresemanni | Hylexetastes_stresemanni | LC | Decreasing | tropical | diurnal | 111 | 1449365.225 | 0.981 |
| birds | Hylia_prasina | Hylia_prasina | LC | Stable | tropical | diurnal | 13.1 | 4144582.435 | 8.629 |
| birds | Hyliota_australis | Hyliota_australis | LC | Decreasing | tropical | diurnal | 11.2 | 1342986.673 | 8.868 |
| birds | Hyliota_flavigaster | Hyliota_flavigaster | LC | Decreasing | tropical | diurnal | 12.5 | 4822595.381 | 9.307 |
| birds | Hyliota_violacea | Hyliota_violacea | LC | Decreasing | tropical | diurnal | 15.8 | 1001100.978 | 8.013 |
| birds | Hylocharis_chrysura | Hylocharis_chrysura | LC | Increasing | temperate | diurnal | 4.5 | 3128401.377 | 8.824 |
| birds | Hylocharis_sapphirina | Hylocharis_sapphirina | LC | Stable | tropical | diurnal | 4.4 | 5411214.464 | 3.371 |
| birds | Hylocichla_mustelina | Hylocichla_mustelina | LC | Decreasing | temperate | diurnal | 50.1 | 5235140.131 | 15.485 |
| birds | Hylocitrea_bonensis | Hylocitrea_bonensis | LC | Stable | tropical | diurnal | 27.5 | 29444.564 | 5.103 |
| birds | Hylomanes_momotula | Hylomanes_momotula | LC | Decreasing | tropical | diurnal | 29.3 | 343980.587 | 8.504 |
| birds | Hylonympha_macrocerca | Hylonympha_macrocerca | EN | Decreasing | tropical | diurnal | 7.3 | 308.708 | 5.295 |
| birds | Hylopezus_auricularis | Hylopezus_auricularis | VU | Stable | tropical | diurnal | 40.4 | 691.162 | 11.806 |
| birds | Hylopezus_macularius | Hylopezus_macularius | LC | Decreasing | tropical | diurnal | 44.2 | 3675311.614 | 1.574 |
| birds | Hylopezus_ochroleucus | Hylopezus_ochroleucus | LC | Decreasing | tropical | diurnal | 28 | 654341.084 | 7.806 |
| birds | Hylopezus_perspicillatus | Hylopezus_perspicillatus | LC | Decreasing | tropical | diurnal | 43 | 279909.566 | 7.958 |
| birds | Hylophilus_amaurocephalus | Hylophilus_amaurocephalus | LC | Stable | tropical | diurnal | 11.7 | 1515487.228 | 9.594 |
| birds | Hylophilus_brunneiceps | Hylophilus_brunneiceps | LC | Stable | tropical | diurnal | 9.9 | 528917.172 | 0.677 |
| birds | Hylophilus_flavipes | Hylophilus_flavipes | LC | Stable | tropical | diurnal | 11.7 | 939938.913 | 10.039 |
| birds | Hylophilus_olivaceus | Hylophilus_olivaceus | LC | Stable | tropical | diurnal | 12 | 61750.073 | 5.63 |
| birds | Hylophilus_pectoralis | Hylophilus_pectoralis | LC | Stable | tropical | diurnal | 11.6 | 3214859.357 | 5.742 |
| birds | Hylophilus_poicilotis | Hylophilus_poicilotis | LC | Stable | temperate | diurnal | 10.4 | 673378.959 | 13.482 |
| birds | Hylophilus_semicinereus | Hylophilus_semicinereus | LC | Stable | tropical | diurnal | 13 | 3561762.581 | 2.115 |
| birds | Hylophilus_thoracicus | Hylophilus_thoracicus | LC | Decreasing | tropical | diurnal | 13.8 | 141073.993 | 13.055 |
| birds | Hylophylax_naevioides | Hylophylax_naevioides | LC | Decreasing | tropical | diurnal | 17.8 | 372510.891 | 8.802 |
| birds | Hylophylax_naevius | Hylophylax_naevius | LC | Stable | tropical | diurnal | 14.2 | 5848723.886 | 1.719 |
| birds | Hylophylax_punctulatus | Hylophylax_punctulatus | LC | Decreasing | tropical | diurnal | 12.4 | 3128205.429 | 1.571 |
| birds | Hylopsar_cupreocauda | Lamprotornis_cupreocauda | NT | Decreasing | tropical | diurnal | 58 | 458895.758 | 15.06 |
| birds | Hylopsar_purpureiceps | Lamprotornis_purpureiceps | LC | Decreasing | tropical | diurnal | 62.8 | 1844762.065 | 7.58 |
| birds | Hylorchilus_navai | Hylorchilus_navai | VU | Decreasing | tropical | diurnal | 29.3 | 4383.869 | 6.387 |
| birds | Hylorchilus_sumichrasti | Hylorchilus_sumichrasti | NT | Decreasing | tropical | diurnal | 28.4 | 6551.838 | 11.947 |
| birds | Hymenolaimus_malacorhynchos | Hymenolaimus_malacorhynchos | EN | Decreasing | temperate | diurnal | 830 | 199757.756 | 6.922 |
| birds | Hymenops_perspicillatus | Hymenops_perspicillatus | LC | Stable | temperate | diurnal | 22.9 | 3807233.089 | 7.982 |

| | | | | | | | | | |
|---|---|---|---|---|---|---|---|---|---|
| birds | Hypargos_margaritatus | Hypargos_margaritatus | LC | Stable | tropical | diurnal | 12.8 | 155741.296 | 9.546 |
| birds | Hypargos_niveoguttatus | Hypargos_niveoguttatus | LC | Stable | tropical | diurnal | 14.6 | 1748989.028 | 9.71 |
| birds | Hypergerus_atriceps | Hypergerus_atriceps | LC | Stable | tropical | diurnal | 29.8 | 2301821.941 | 10.904 |
| birds | Hypnelus_bicinctus | NA | LC | Stable | NA | NA | NA | NA | NA |
| birds | Hypnelus_ruficollis | Hypnelus_ruficollis | LC | Stable | tropical | diurnal | 49.8 | 239856.985 | 14.512 |
| birds | Hypocnemis_cantator | Hypocnemis_cantator | LC | Stable | tropical | diurnal | 12.3 | 968489.412 | 1.314 |
| birds | Hypocnemis_hypoxantha | Hypocnemis_hypoxantha | LC | Stable | tropical | diurnal | 11.5 | 1409240.165 | 1.627 |
| birds | Hypocnemoides_maculicauda | Hypocnemoides_maculicauda | LC | Decreasing | tropical | diurnal | 11.8 | 3324431.814 | 3.251 |
| birds | Hypocnemoides_melanopogon | Hypocnemoides_melanopogon | LC | Stable | tropical | diurnal | 14.1 | 3494411.768 | 1.594 |
| birds | Hypocolius_ampelinus | Hypocolius_ampelinus | LC | Unknown | temperate | diurnal | 53.3 | 2239945.425 | 6.536 |
| birds | Hypocryptadius_cinnamomeus | Hypocryptadius_cinnamomeus | LC | Decreasing | tropical | diurnal | 30 | 6672.792 | 5.568 |
| birds | Hypoedaleus_guttatus | Hypoedaleus_guttatus | LC | Decreasing | tropical | diurnal | 38.8 | 829945.452 | 13.674 |
| birds | Hypopyrrhus_pyrohypogaster | Hypopyrrhus_pyrohypogaster | VU | Decreasing | tropical | diurnal | 101 | 31843.129 | 14.683 |
| birds | Hypositta_corallirostris | Hypositta_corallirostris | LC | Unknown | tropical | diurnal | 14.1 | 159578.631 | 8.13 |
| birds | Hypotaenidia_insignis | Gallirallus_insignis | NT | Decreasing | tropical | diurnal | 353.1 | 33841.036 | 4.252 |
| birds | Hypotaenidia_okinawae | Gallirallus_okinawae | EN | Decreasing | temperate | diurnal | 433 | 594.314 | 19.61 |
| birds | Hypotaenidia_owstoni | Gallirallus_owstoni | CR | Increasing | tropical | nocturnal | 226 | 543.4 | 22.995 |
| birds | Hypotaenidia_philippensis | Gallirallus_philippensis | LC | Stable | tropical | cathemeral | 205.9 | 2532556.07 | 7.748 |
| birds | Hypotaenidia_poeciloptera | NA | EX | not_available | NA | NA | NA | NA | NA |
| birds | Hypotaenidia_rovianae | Gallirallus_rovianae | LC | Stable | tropical | diurnal | 353.1 | 3354.631 | 2.791 |
| birds | Hypotaenidia_sylvestris | Gallirallus_sylvestris | EN | Increasing | temperate | diurnal | 494.4 | 15.075 | 6.721 |
| birds | Hypotaenidia_torquata | Gallirallus_torquatus | LC | Unknown | tropical | diurnal | 280 | 412241.216 | 12.682 |
| birds | Hypotaenidia_woodfordi | Nesoclopeus_woodfordi | LC | Unknown | tropical | diurnal | 553.2 | 7714.363 | 6.26 |
| birds | Hypothymis_azurea | Hypothymis_azurea | LC | Stable | tropical | diurnal | 11.1 | 7041185.043 | 13.551 |
| birds | Hypothymis_coelestis | Hypothymis_coelestis | VU | Decreasing | tropical | diurnal | 13.6 | 179521.375 | 15.096 |
| birds | Hypothymis_helenae | Hypothymis_helenae | NT | Decreasing | tropical | diurnal | 9.9 | 123419.68 | 11.071 |
| birds | Hypsipetes_affinis | Alophoixus_affinis | LC | Stable | tropical | diurnal | 40.5 | 18011.547 | 7.323 |
| birds | Hypsipetes_amaurotis | Ixos_amaurotis | LC | Increasing | temperate | diurnal | 70.9 | 715067.908 | 19.434 |
| birds | Hypsipetes_borbonicus | Hypsipetes_borbonicus | NT | Increasing | tropical | diurnal | 55.3 | 2541.136 | 20.642 |
| birds | Hypsipetes_crassirostris | NA | LC | Stable | NA | NA | NA | NA | NA |
| birds | Hypsipetes_everetti | Ixos_everetti | LC | Stable | tropical | diurnal | 58.3 | 115844.175 | 11.916 |
| birds | Hypsipetes_leucocephalus | Hypsipetes_leucocephalus | LC | Stable | tropical | diurnal | 51.8 | 4955880.132 | 12.938 |
| birds | Hypsipetes_madagascariensis | Hypsipetes_madagascariensis | LC | Stable | tropical | diurnal | 46 | 591386.986 | 8.769 |
| birds | Hypsipetes_olivaceus | Hypsipetes_olivaceus | VU | Stable | tropical | diurnal | 75 | 1872.303 | 25.919 |

| | | | | | | | | | |
|---|---|---|---|---|---|---|---|---|---|
| birds | Hypsipetes_parvirostris | Hypsipetes_parvirostris | NT | Decreasing | tropical | diurnal | 59.4 | 483.966 | 15.979 |
| birds | Hypsipetes_philippinus | Ixos_philippinus | LC | Stable | tropical | diurnal | 38.6 | 232032.85 | 13.043 |
| birds | Hypsipetes_rufigularis | Ixos_rufigularis | NT | Decreasing | tropical | diurnal | 58.7 | 21991.101 | 10.641 |
| birds | Hypsipetes_siquijorensis | NA | EN | Decreasing | NA | NA | NA | NA | NA |
| birds | Hypsipetes_thompsoni | Hypsipetes_thompsoni | LC | Stable | tropical | diurnal | 41.8 | 196437.348 | 8.586 |
| birds | Ianthocincla_bieti | Garrulax_bieti | VU | Decreasing | temperate | diurnal | 78.6 | 28399.812 | 7.227 |
| birds | Ianthocincla_lunulata | Garrulax_lunulatus | LC | Decreasing | temperate | diurnal | 82 | 214199.97 | 9.897 |
| birds | Ianthocincla_maxima | Garrulax_maximus | LC | Decreasing | temperate | diurnal | 120.5 | 530133.338 | 8.757 |
| birds | Ianthocincla_ocellata | Garrulax_ocellatus | LC | Decreasing | temperate | diurnal | 114 | 1005117.199 | 9.956 |
| birds | Ianthocincla_rufogularis | Garrulax_rufogularis | LC | Decreasing | temperate | diurnal | 62.8 | 260288.236 | 11.512 |
| birds | Ianthocincla_sukatschewi | Garrulax_sukatschewi | VU | Decreasing | temperate | diurnal | 67.8 | 28519.278 | 9.718 |
| birds | Ibidorhyncha_struthersii | Ibidorhyncha_struthersii | LC | Unknown | temperate | diurnal | 294 | 5219019.959 | 8.123 |
| birds | Ibycter_americanus | Ibycter_americanus | LC | Decreasing | tropical | diurnal | 624 | 8193442.502 | 3.145 |
| birds | Icteria_virens | Icteria_virens | LC | Decreasing | temperate | diurnal | 24.9 | 8893581.554 | 12.501 |
| birds | Icterus_abeillei | Icterus_abeillei | LC | Decreasing | tropical | diurnal | 33 | 394832.544 | 11.871 |
| birds | Icterus_auratus | Icterus_auratus | LC | Decreasing | tropical | diurnal | 32.1 | 115652.926 | 6.374 |
| birds | Icterus_auricapillus | Icterus_auricapillus | LC | Stable | tropical | diurnal | 32.2 | 405614.957 | 12.444 |
| birds | Icterus_bonana | Icterus_bonana | VU | Decreasing | tropical | diurnal | 29 | 1126.59 | 26.256 |
| birds | Icterus_bullockiorum | Icterus_bullockii | LC | Stable | temperate | diurnal | 37.9 | 5157597.513 | 9.872 |
| birds | Icterus_cayanensis | Icterus_cayanensis | LC | Stable | tropical | diurnal | 35.4 | 4104473.13 | 2.136 |
| birds | Icterus_chrysater | Icterus_chrysater | LC | Decreasing | tropical | diurnal | 53.4 | 777224.612 | 11.06 |
| birds | Icterus_chrysocephalus | NA | LC | Stable | NA | NA | NA | NA | NA |
| birds | Icterus_croconotus | Icterus_croconotus | LC | Decreasing | tropical | diurnal | 44.5 | 3433213.879 | 3.449 |
| birds | Icterus_cucullatus | Icterus_cucullatus | LC | Increasing | temperate | diurnal | 24.3 | 2334013.11 | 9.115 |
| birds | Icterus_dominicensis | Icterus_dominicensis | LC | Decreasing | tropical | diurnal | 35.7 | 75136.963 | 15.771 |
| birds | Icterus_galbula | Icterus_galbula | LC | Stable | temperate | diurnal | 32.8 | 7852695.637 | 14.537 |
| birds | Icterus_graceannae | Icterus_graceannae | LC | Stable | tropical | diurnal | 41.5 | 77582.359 | 9.697 |
| birds | Icterus_graduacauda | Icterus_graduacauda | LC | Decreasing | tropical | diurnal | 42.2 | 347160.285 | 11.322 |
| birds | Icterus_gularis | Icterus_gularis | LC | Stable | tropical | diurnal | 55.2 | 559678.921 | 11.306 |
| birds | Icterus_icterus | Icterus_icterus | LC | Stable | tropical | diurnal | 60.9 | 442639.316 | 9.965 |
| birds | Icterus_jamacaii | Icterus_jamacaii | LC | Stable | tropical | diurnal | 62.9 | 2653360.728 | 8.818 |
| birds | Icterus_laudabilis | Icterus_laudabilis | EN | Decreasing | tropical | diurnal | 36.9 | 581.481 | 20.28 |
| birds | Icterus_leucopteryx | Icterus_leucopteryx | LC | Stable | tropical | diurnal | 36.9 | 11260.283 | 19.821 |
| birds | Icterus_maculialatus | Icterus_maculialatus | LC | Decreasing | tropical | diurnal | 37.4 | 87262.548 | 13.469 |

| | | | | | | | | | |
|---|---|---|---|---|---|---|---|---|---|
| birds | Icterus_mesomelas | Icterus_mesomelas | LC | Decreasing | tropical | diurnal | 39.9 | 995703.978 | 10.394 |
| birds | Icterus_nigrogularis | Icterus_nigrogularis | LC | Stable | tropical | diurnal | 40.2 | 1368029.344 | 6.879 |
| birds | Icterus_oberi | Icterus_oberi | VU | Stable | tropical | diurnal | 35.8 | 13.19 | 6.459 |
| birds | Icterus_parisorum | Icterus_parisorum | LC | Decreasing | temperate | diurnal | 36.5 | 2080143.577 | 8.514 |
| birds | Icterus_pectoralis | Icterus_pectoralis | LC | Decreasing | tropical | diurnal | 46.5 | 206586.992 | 13.788 |
| birds | Icterus_prosthemelas | Icterus_prosthemelas | LC | Decreasing | tropical | diurnal | 28 | 440232.426 | 8.787 |
| birds | Icterus_pustulatus | Icterus_pustulatus | LC | Stable | tropical | diurnal | 36.8 | 553521.836 | 10.034 |
| birds | Icterus_spurius | Icterus_spurius | LC | Stable | temperate | diurnal | 19.4 | 6755981.458 | 14.515 |
| birds | Icterus_wagleri | Icterus_wagleri | LC | Decreasing | tropical | diurnal | 41.8 | 767331.586 | 10.239 |
| birds | Icthyophaga_humilis | Ichthyophaga_humilis | NT | Decreasing | tropical | diurnal | 782 | 2907438.807 | 11.151 |
| birds | Icthyophaga_ichthyaetus | Ichthyophaga_ichthyaetus | NT | Decreasing | tropical | diurnal | 1590 | 3707887.548 | 13.134 |
| birds | Ictinaetus_malaiensis | Ictinaetus_malayensis | LC | Decreasing | tropical | diurnal | 1028 | 3231811.389 | 11.431 |
| birds | Ictinia_mississippiensis | Ictinia_mississippiensis | LC | Increasing | tropical | diurnal | 276 | 4891699.341 | 8.507 |
| birds | Ictinia_plumbea | Ictinia_plumbea | LC | Decreasing | tropical | diurnal | 249.9 | 12792063.813 | 5.556 |
| birds | Idiopsar_brachyurus | Idiopsar_brachyurus | LC | Stable | tropical | diurnal | 43 | 76433.699 | 8.485 |
| birds | Iduna_caligata | Hippolais_caligata | LC | Increasing | temperate | diurnal | 8.9 | 10100495.933 | 9.412 |
| birds | Iduna_natalensis | Chloropeta_natalensis | LC | Stable | tropical | diurnal | 11.7 | 3742433.69 | 9.341 |
| birds | Iduna_pallida | Hippolais_pallida | LC | Stable | tropical | diurnal | 9 | 19827150.117 | 7.757 |
| birds | Iduna_rama | Hippolais_rama | LC | Stable | temperate | diurnal | 9.3 | 5878685.671 | 11.026 |
| birds | Iduna_similis | Chloropeta_similis | LC | Stable | tropical | diurnal | 11.1 | 283650.623 | 13.144 |
| birds | Ifrita_kowaldi | Ifrita_kowaldi | LC | Stable | tropical | diurnal | 32.1 | 91932.484 | 5.204 |
| birds | Ilicura_militaris | Ilicura_militaris | LC | Decreasing | tropical | diurnal | 12.7 | 840759.231 | 13.648 |
| birds | Illadopsis_albipectus | Illadopsis_albipectus | LC | Stable | tropical | diurnal | 30.6 | 1215586.753 | 5.794 |
| birds | Illadopsis_cleaveri | Illadopsis_cleaveri | LC | Decreasing | tropical | diurnal | 30.8 | 977280.762 | 10.382 |
| birds | Illadopsis_fulvescens | Illadopsis_fulvescens | LC | Stable | tropical | diurnal | 27.9 | 2855381.869 | 8.063 |
| birds | Illadopsis_puveli | Illadopsis_puveli | LC | Stable | tropical | diurnal | 48.4 | 631343.922 | 14.905 |
| birds | Illadopsis_pyrrhoptera | Illadopsis_pyrrhoptera | LC | Stable | tropical | diurnal | 24.8 | 145566.301 | 14.922 |
| birds | Illadopsis_rufescens | Illadopsis_rufescens | NT | Decreasing | tropical | diurnal | 37 | 264186.882 | 14.84 |
| birds | Illadopsis_rufipennis | Illadopsis_rufipennis | LC | Stable | tropical | diurnal | 22.2 | 2029123.6 | 8.772 |
| birds | Illadopsis_turdina | Ptyrticus_turdinus | LC | Decreasing | tropical | diurnal | 65.8 | 231174.569 | 7.11 |
| birds | Incana_incana | Incana_incana | LC | Stable | tropical | diurnal | 10.4 | 3614.005 | 11.038 |
| birds | Incaspiza_laeta | Incaspiza_laeta | LC | Stable | tropical | diurnal | 22 | 28938.709 | 9.735 |
| birds | Incaspiza_ortizi | Incaspiza_ortizi | LC | Stable | tropical | diurnal | 32.4 | 1463.279 | 10.138 |
| birds | Incaspiza_personata | Incaspiza_personata | LC | Stable | tropical | diurnal | 33.2 | 88039.252 | 8.491 |

| | | | | | | | | | |
|---|---|---|---|---|---|---|---|---|---|
| birds | Incaspiza_pulchra | Incaspiza_pulchra | LC | Stable | tropical | diurnal | 27.6 | 21927.77 | 7.965 |
| birds | Incaspiza_watkinsi | Incaspiza_watkinsi | VU | Decreasing | tropical | diurnal | 20.4 | 2308.737 | 12.304 |
| birds | Indicator_archipelagicus | Indicator_archipelagicus | NT | Decreasing | tropical | diurnal | 32.4 | 1098002.154 | 8.115 |
| birds | Indicator_exilis | Indicator_exilis | LC | Unknown | tropical | diurnal | 17.9 | 4132641.709 | 8.203 |
| birds | Indicator_indicator | Indicator_indicator | LC | Increasing | tropical | diurnal | 50.4 | 11206655.869 | 9.915 |
| birds | Indicator_maculatus | Indicator_maculatus | LC | Decreasing | tropical | diurnal | 46.8 | 4072953.886 | 9.166 |
| birds | Indicator_meliphilus | Indicator_meliphilus | LC | Decreasing | tropical | diurnal | 14 | 1489690.395 | 9.493 |
| birds | Indicator_minor | Indicator_minor | LC | Stable | tropical | diurnal | 28.2 | 13121952.735 | 9.869 |
| birds | Indicator_pumilio | Indicator_pumilio | LC | Decreasing | tropical | diurnal | 13.1 | 49549.754 | 16.349 |
| birds | Indicator_variegatus | Indicator_variegatus | LC | Stable | tropical | diurnal | 48.8 | 3473509.989 | 9.936 |
| birds | Indicator_willcocksi | Indicator_willcocksi | LC | Unknown | tropical | diurnal | 15.4 | 3313622.403 | 8.051 |
| birds | Indicator_xanthonotus | Indicator_xanthonotus | NT | Decreasing | temperate | diurnal | 30.4 | 97279.815 | 7.383 |
| birds | Inezia_caudata | Inezia_caudata | LC | Stable | tropical | diurnal | 7.5 | 597282.079 | 8.543 |
| birds | Inezia_inornata | Inezia_inornata | LC | Decreasing | tropical | diurnal | 12 | 2125032.215 | 4.052 |
| birds | Inezia_subflava | Inezia_subflava | LC | Stable | tropical | diurnal | 8.4 | 972309.297 | 2.318 |
| birds | Inezia_tenuirostris | Inezia_tenuirostris | LC | Decreasing | tropical | diurnal | 5 | 59966.6 | 11.954 |
| birds | Iodopleura_fusca | Iodopleura_fusca | LC | Decreasing | tropical | diurnal | 15.3 | 444576.697 | 1.984 |
| birds | Iodopleura_isabellae | Iodopleura_isabellae | LC | Decreasing | tropical | diurnal | 20 | 4495732.173 | 2.021 |
| birds | Iodopleura_pipra | Iodopleura_pipra | EN | Decreasing | tropical | diurnal | 10 | 277077.996 | 13.076 |
| birds | Iole_finschii | Alophoixus_finschii | NT | Decreasing | tropical | diurnal | 24.2 | 779808.762 | 6.499 |
| birds | Iole_palawanensis | Ixos_palawanensis | LC | Decreasing | tropical | diurnal | 26.8 | 11492.84 | 9.023 |
| birds | Iole_propinqua | Iole_propinqua | LC | Stable | tropical | diurnal | 25.9 | 1237485.989 | 10.612 |
| birds | Iole_viridescens | NA | LC | Stable | NA | NA | NA | NA | NA |
| birds | Irania_gutturalis | Irania_gutturalis | LC | Stable | temperate | diurnal | 21.6 | 2646506.329 | 11.747 |
| birds | Irediparra_gallinacea | Irediparra_gallinacea | LC | Unknown | tropical | diurnal | 98.8 | 2181164.522 | 5.328 |
| birds | Irena_cyanogastra | Irena_cyanogastra | NT | Decreasing | tropical | diurnal | 80.3 | 229101.267 | 13.372 |
| birds | Irena_puella | Irena_puella | LC | Decreasing | tropical | diurnal | 64.9 | 3709810.221 | 11.101 |
| birds | Iridophanes_pulcherrimus | Iridophanes_pulcherrimus | LC | Decreasing | tropical | diurnal | 15.3 | 115363.41 | 5.272 |
| birds | Iridosornis_analis | Iridosornis_analis | LC | Decreasing | tropical | diurnal | 26 | 97006.404 | 5.009 |
| birds | Iridosornis_jelskii | Iridosornis_jelskii | LC | Stable | tropical | diurnal | 20 | 98574.562 | 5.037 |
| birds | Iridosornis_porphyrocephalus | Iridosornis_porphyrocephalus | NT | Decreasing | tropical | diurnal | 21.7 | 23009.898 | 10.434 |
| birds | Iridosornis_reinhardti | Iridosornis_reinhardti | LC | Stable | tropical | diurnal | 24 | 66917.021 | 4.509 |
| birds | Iridosornis_rufivertex | Iridosornis_rufivertex | LC | Stable | tropical | diurnal | 23 | 82384.672 | 11.114 |
| birds | Isleria_guttata | Myrmotherula_guttata | LC | Stable | tropical | diurnal | 11 | 1694815.797 | 1.297 |

| | | | | | | | | | |
|---|---|---|---|---|---|---|---|---|---|
| birds | Isleria_hauxwelli | Myrmotherula_hauxwelli | LC | Decreasing | tropical | diurnal | 10.7 | 4248364.67 | 2.339 |
| birds | Islerothraupis_cristata | Tachyphonus_cristatus | LC | Stable | tropical | diurnal | 18.8 | 6338485.759 | 2.919 |
| birds | Islerothraupis_luctuosa | Tachyphonus_luctuosus | LC | Decreasing | tropical | diurnal | 13 | 7144190.037 | 3.25 |
| birds | Islerothraupis_rufiventer | Tachyphonus_rufiventer | LC | Stable | tropical | diurnal | 19 | 737324.121 | 1.676 |
| birds | Ispidina_lecontei | Ceyx_lecontei | LC | Stable | tropical | diurnal | 9.5 | 2195930.359 | 6.926 |
| birds | Ispidina_picta | Ceyx_pictus | LC | Stable | tropical | diurnal | 12.7 | 13894765.2 | 9.611 |
| birds | Ithaginis_cruentus | Ithaginis_cruentus | LC | Decreasing | temperate | diurnal | 530.7 | 657902.92 | 9.071 |
| birds | Ixidia_cyaniventris | Pycnonotus_cyaniventris | NT | Decreasing | tropical | diurnal | 22 | 926808.776 | 6.748 |
| birds | Ixidia_erythropthalmos | Pycnonotus_erythropthalmos | LC | Decreasing | tropical | diurnal | 19.2 | 1241474.047 | 8.523 |
| birds | Ixidia_squamata | Pycnonotus_squamatus | NT | Decreasing | tropical | diurnal | 22 | 298738.781 | 6.258 |
| birds | Ixobrychus_cinnamomeus | Ixobrychus_cinnamomeus | LC | Stable | temperate | diurnal | 126.5 | 8161986.561 | 14.634 |
| birds | Ixobrychus_eurhythmus | Ixobrychus_eurhythmus | LC | Decreasing | temperate | diurnal | 139 | 8438029.061 | 12.384 |
| birds | Ixobrychus_flavicollis | Ixobrychus_flavicollis | LC | Decreasing | tropical | cathemeral | 321 | 8168174.459 | 10.712 |
| birds | Ixobrychus_involucris | Ixobrychus_involucris | LC | Stable | temperate | nocturnal | 80.1 | 2399740.232 | 9.596 |
| birds | Ixobrychus_minutus | Ixobrychus_minutus | LC | Decreasing | temperate | diurnal | 99.5 | 27727320.232 | 10.571 |
| birds | Ixobrychus_sinensis | Ixobrychus_sinensis | LC | Unknown | temperate | cathemeral | 94.3 | 9358675.068 | 13.543 |
| birds | Ixobrychus_sturmii | Ixobrychus_sturmii | LC | Unknown | tropical | cathemeral | 140 | 12872490.534 | 9.419 |
| birds | Ixonotus_guttatus | Ixonotus_guttatus | LC | Stable | tropical | diurnal | 35.2 | 2790568.636 | 7.584 |
| birds | Ixoreus_naevius | Zoothera_naevia | LC | Decreasing | temperate | diurnal | 77.5 | 4018779.517 | 3.955 |
| birds | Ixos_malaccensis | Ixos_malaccensis | NT | Decreasing | tropical | diurnal | 37.3 | 976235.844 | 7.034 |
| birds | Ixos_mcclellandii | Hypsipetes_mcclellandii | LC | Stable | temperate | diurnal | 32.5 | 2391920.689 | 12.098 |
| birds | Ixos_nicobariensis | Hypsipetes_nicobariensis | NT | Decreasing | tropical | diurnal | 36 | 472.608 | 7.939 |
| birds | Ixos_virescens | NA | LC | Stable | NA | NA | NA | NA | NA |
| birds | Jabiru_mycteria | Jabiru_mycteria | LC | Unknown | tropical | diurnal | 5996.3 | 9048327.31 | 5.599 |
| birds | Jacamaralcyon_tridactyla | Jacamaralcyon_tridactyla | NT | Decreasing | tropical | diurnal | 18.3 | 196625.184 | 10.986 |
| birds | Jacamerops_aureus | Jacamerops_aureus | LC | Decreasing | tropical | diurnal | 62.9 | 5748994.39 | 1.896 |
| birds | Jacana_jacana | Jacana_jacana | LC | Stable | tropical | diurnal | 106.2 | 13881637.25 | 5.969 |
| birds | Jacana_spinosa | Jacana_spinosa | LC | Unknown | tropical | diurnal | 94 | 1048144.394 | 11.664 |
| birds | Jubula_lettii | Jubula_lettii | DD | Stable | tropical | nocturnal | 183 | 2256025.42 | 6.784 |
| birds | Junco_bairdi | NA | NT | Unknown | NA | NA | NA | NA | NA |
| birds | Junco_hyemalis | Junco_hyemalis | LC | Decreasing | temperate | diurnal | 19.5 | 16177050.96 | 8.455 |
| birds | Junco_insularis | NA | EN | Increasing | NA | NA | NA | NA | NA |
| birds | Junco_phaeonotus | Junco_phaeonotus | LC | Stable | tropical | diurnal | 20.4 | 532188.093 | 10.46 |
| birds | Junco_vulcani | Junco_vulcani | LC | Stable | tropical | diurnal | 28 | 1035.176 | 5.823 |

| | | | | | | | | | |
|---|---|---|---|---|---|---|---|---|---|
| birds | Jynx_ruficollis | Jynx_ruficollis | LC | Increasing | temperate | diurnal | 46.4 | 1609043.049 | 10.389 |
| birds | Jynx_torquilla | Jynx_torquilla | LC | Decreasing | temperate | diurnal | 35 | 30131695.781 | 11.003 |
| birds | Kakamega_poliothorax | Kakamega_poliothorax | LC | Decreasing | tropical | diurnal | 36.4 | 140625.846 | 13.662 |
| birds | Kaupifalco_monogrammicus | Kaupifalco_monogrammicus | LC | Stable | tropical | diurnal | 278.9 | 13079368.524 | 9.412 |
| birds | Kempiella_flavovirescens | Microeca_flavovirescens | LC | Stable | tropical | diurnal | 15 | 669110.722 | 3.112 |
| birds | Kempiella_griseoceps | Microeca_griseoceps | LC | Stable | tropical | diurnal | 10.6 | 184726.518 | 3.443 |
| birds | Kenopia_striata | Kenopia_striata | NT | Decreasing | tropical | diurnal | 19.9 | 1176589.951 | 8.117 |
| birds | Ketupa_flavipes | Ketupa_flavipes | LC | Decreasing | tropical | cathemeral | 2315 | 2648554.298 | 14.805 |
| birds | Ketupa_ketupu | Ketupa_ketupu | LC | Stable | tropical | nocturnal | 1293 | 2556769.486 | 10.48 |
| birds | Ketupa_zeylonensis | Ketupa_zeylonensis | LC | Decreasing | tropical | cathemeral | 1105 | 5480900.283 | 15.742 |
| birds | Klais_guimeti | Klais_guimeti | LC | Decreasing | tropical | diurnal | 2.6 | 472690.687 | 5.976 |
| birds | Kleinothraupis_atropileus | Hemispingus_atropileus | LC | Decreasing | tropical | diurnal | 22 | 129533.742 | 11.216 |
| birds | Kleinothraupis_auricularis | Hemispingus_auricularis | LC | Stable | tropical | diurnal | 22 | 61932.599 | 5.611 |
| birds | Kleinothraupis_calophrys | Hemispingus_calophrys | LC | Decreasing | tropical | diurnal | 19.3 | 23103.819 | 5.449 |
| birds | Kleinothraupis_parodii | Hemispingus_parodii | NT | Decreasing | tropical | diurnal | 21.8 | 28661.282 | 4.103 |
| birds | Kleinothraupis_reyi | Hemispingus_reyi | NT | Stable | tropical | diurnal | 19 | 5979.417 | 11.853 |
| birds | Knipolegus_aterrimus | Knipolegus_aterrimus | LC | Stable | temperate | diurnal | 20.2 | 2222826.22 | 7.331 |
| birds | Knipolegus_cyanirostris | Knipolegus_cyanirostris | LC | Decreasing | temperate | diurnal | 15.4 | 2153351.026 | 11.782 |
| birds | Knipolegus_franciscanus | Knipolegus_franciscanus | LC | Decreasing | tropical | diurnal | 20.5 | 170942.418 | 10.13 |
| birds | Knipolegus_hudsoni | Knipolegus_hudsoni | LC | Stable | temperate | diurnal | 13.8 | 2040051.933 | 6.768 |
| birds | Knipolegus_lophotes | Knipolegus_lophotes | LC | Stable | tropical | diurnal | 31.8 | 1319771.822 | 11.653 |
| birds | Knipolegus_nigerrimus | Knipolegus_nigerrimus | LC | Stable | tropical | diurnal | 20.3 | 789107.651 | 11.463 |
| birds | Knipolegus_orenocensis | Knipolegus_orenocensis | LC | Stable | tropical | diurnal | 19 | 255299.565 | 4.464 |
| birds | Knipolegus_poecilocercus | Knipolegus_poecilocercus | LC | Decreasing | tropical | diurnal | 14.3 | 1167038.38 | 2.732 |
| birds | Knipolegus_poecilurus | Knipolegus_poecilurus | LC | Stable | tropical | diurnal | 14.6 | 524563.473 | 5.254 |
| birds | Knipolegus_signatus | Knipolegus_signatus | LC | Stable | tropical | diurnal | 17.8 | 32221.106 | 4.173 |
| birds | Knipolegus_striaticeps | Knipolegus_striaticeps | LC | Decreasing | temperate | diurnal | 11 | 946873.165 | 5.6 |
| birds | Kupeornis_chapini | Kupeornis_chapini | LC | Decreasing | tropical | diurnal | 50 | 43946.987 | 10.468 |
| birds | Kupeornis_gilberti | Kupeornis_gilberti | VU | Decreasing | tropical | diurnal | 64 | 1661.423 | 11.08 |
| birds | Kupeornis_rufocinctus | Kupeornis_rufocinctus | EN | Decreasing | tropical | diurnal | 47 | 41401.324 | 14.586 |
| birds | Kurochkinegramma_hypogrammica | Hypogramma_hypogrammicum | LC | Stable | tropical | diurnal | 11.4 | 2037212.163 | 9.28 |
| birds | Lacedo_pulchella | Lacedo_pulchella | LC | Decreasing | tropical | diurnal | 47.3 | 1429865.782 | 11.167 |
| birds | Lafresnaya_lafresnayi | Lafresnaya_lafresnayi | LC | Stable | tropical | diurnal | 5.3 | 312941.283 | 9.529 |
| birds | Lagonosticta_larvata | Lagonosticta_larvata | LC | Stable | tropical | diurnal | 9.9 | 309178.843 | 9.313 |

| | | | | | | | | | |
|---|---|---|---|---|---|---|---|---|---|
| birds | Lagonosticta_nitidula | Lagonosticta_nitidula | LC | Stable | tropical | diurnal | 10.7 | 1509814.087 | 6.845 |
| birds | Lagonosticta_rara | Lagonosticta_rara | LC | Stable | tropical | diurnal | 10.4 | 2433099.838 | 9.993 |
| birds | Lagonosticta_rhodopareia | Lagonosticta_rhodopareia | LC | Stable | tropical | diurnal | 9.3 | 2706237.275 | 9.883 |
| birds | Lagonosticta_rubricata | Lagonosticta_rubricata | LC | Stable | tropical | diurnal | 10.2 | 6048001.079 | 10.144 |
| birds | Lagonosticta_rufopicta | Lagonosticta_rufopicta | LC | Stable | tropical | diurnal | 9.3 | 3308898.335 | 10.475 |
| birds | Lagonosticta_sanguinodorsalis | Lagonosticta_sanguinodorsalis | LC | Stable | tropical | diurnal | 11.4 | 28120.619 | 18.395 |
| birds | Lagonosticta_senegala | Lagonosticta_senegala | LC | Stable | tropical | diurnal | 9.2 | 10048316.395 | 11.102 |
| birds | Lagonosticta_umbrinodorsalis | Lagonosticta_umbrinodorsalis | LC | Stable | tropical | diurnal | 10.6 | 42139.717 | 8.9 |
| birds | Lagonosticta_virata | Lagonosticta_virata | LC | Stable | tropical | diurnal | 9.3 | 110944.972 | 13.008 |
| birds | Lagopus_lagopus | Lagopus_lagopus | LC | Decreasing | temperate | diurnal | 566.9 | 26280139.196 | 2.713 |
| birds | Lagopus_leucura | Lagopus_leucura | LC | Decreasing | temperate | diurnal | 355 | 1846729.067 | 2.444 |
| birds | Lagopus_muta | Lagopus_muta | LC | Decreasing | temperate | diurnal | 535.3 | 17597025.309 | 1.419 |
| birds | Lalage_atrovirens | Lalage_atrovirens | LC | Stable | tropical | diurnal | 33 | 296184.727 | 3.209 |
| birds | Lalage_aurea | Lalage_aurea | LC | Decreasing | tropical | diurnal | 32 | 25487.733 | 6.297 |
| birds | Lalage_fimbriata | Coracina_fimbriata | LC | Decreasing | tropical | diurnal | 29.3 | 1316199.589 | 9.792 |
| birds | Lalage_leucomela | Lalage_leucomela | LC | Decreasing | tropical | diurnal | 29.3 | 1258230.144 | 4.117 |
| birds | Lalage_leucopyga | Lalage_leucopyga | LC | Stable | tropical | diurnal | 24.3 | 34357.569 | 6.653 |
| birds | Lalage_leucopygialis | Lalage_leucopygialis | LC | Increasing | tropical | diurnal | 24.3 | 142638.671 | 10.221 |
| birds | Lalage_maculosa | Lalage_maculosa | LC | Stable | tropical | diurnal | 30.3 | 30893.859 | 8.161 |
| birds | Lalage_melanoleuca | Lalage_melanoleuca | LC | Stable | tropical | diurnal | 46.8 | 104068.686 | 15.19 |
| birds | Lalage_melanoptera | Coracina_melanoptera | LC | Stable | tropical | diurnal | 29.4 | 2480148.803 | 18.149 |
| birds | Lalage_melaschistos | Coracina_melaschistos | LC | Decreasing | temperate | diurnal | 43 | 6841809.691 | 14.79 |
| birds | Lalage_moesta | Lalage_moesta | LC | Stable | tropical | diurnal | 29.5 | 4371.571 | 6.744 |
| birds | Lalage_newtoni | NA | CR | Decreasing | NA | NA | NA | NA | NA |
| birds | Lalage_nigra | Lalage_nigra | LC | Decreasing | tropical | diurnal | 28.4 | 1519309.395 | 10.206 |
| birds | Lalage_polioptera | Coracina_polioptera | LC | Stable | tropical | diurnal | 35.8 | 932555.614 | 11.726 |
| birds | Lalage_sharpei | NA | NT | Decreasing | NA | NA | NA | NA | NA |
| birds | Lalage_sueurii | Lalage_sueurii | LC | Increasing | tropical | diurnal | 26 | 259989.369 | 12.156 |
| birds | Lalage_tricolor | Lalage_tricolor | LC | Decreasing | temperate | diurnal | 25 | 7633376.406 | 4.247 |
| birds | Lalage_typica | Coracina_typica | VU | Stable | tropical | diurnal | 42.9 | 1872.301 | 25.919 |
| birds | Lampornis_amethystinus | Lampornis_amethystinus | LC | Decreasing | tropical | diurnal | 5.2 | 154790.447 | 12.771 |
| birds | Lampornis_castaneoventris | Lampornis_castaneoventris | LC | Stable | tropical | diurnal | 5.1 | 1367.408 | 6.088 |
| birds | Lampornis_clemenciae | Lampornis_clemenciae | LC | Stable | tropical | diurnal | 7.6 | 547582.862 | 9.43 |
| birds | Lampornis_hemileucus | Lampornis_hemileucus | LC | Stable | tropical | diurnal | 5.5 | 8048.255 | 11.177 |

| | | | | | | | | | |
|---|---|---|---|---|---|---|---|---|---|
| birds | Lampornis_sybillae | Lampornis_sybillae | LC | Decreasing | tropical | diurnal | 6.3 | 45190.48 | 9.697 |
| birds | Lampornis_viridipallens | Lampornis_viridipallens | LC | Decreasing | tropical | diurnal | 5.3 | 91755.511 | 12.959 |
| birds | Lamprolaima_rhami | Lamprolaima_rhami | LC | Stable | tropical | diurnal | 7.8 | 138239.013 | 11.59 |
| birds | Lamprolia_victoriae | Lamprolia_victoriae | NT | Decreasing | tropical | diurnal | 14.7 | 442.187 | 12.351 |
| birds | Lampropsar_tanagrinus | Lampropsar_tanagrinus | LC | Stable | tropical | diurnal | 57.3 | 2416259.166 | 2.416 |
| birds | Lamprospiza_melanoleuca | Lamprospiza_melanoleuca | LC | Decreasing | tropical | diurnal | 34 | 3392867.326 | 1.859 |
| birds | Lamprotornis_acuticaudus | Lamprotornis_acuticaudus | LC | Stable | tropical | diurnal | 67.6 | 1542032.329 | 6.342 |
| birds | Lamprotornis_albicapillus | Spreo_albicapillus | LC | Stable | tropical | diurnal | 76.8 | 792878.494 | 7.52 |
| birds | Lamprotornis_australis | Lamprotornis_australis | LC | Unknown | tropical | diurnal | 101.1 | 1557885.954 | 6.241 |
| birds | Lamprotornis_bicolor | Spreo_bicolor | LC | Stable | temperate | diurnal | 104 | 787929.605 | 10.75 |
| birds | Lamprotornis_caudatus | Lamprotornis_caudatus | LC | Stable | tropical | diurnal | 121 | 2770669.045 | 12.759 |
| birds | Lamprotornis_chalcurus | Lamprotornis_chalcurus | LC | Stable | tropical | diurnal | 63 | 2454558.775 | 10.936 |
| birds | Lamprotornis_chalybaeus | Lamprotornis_chalybaeus | LC | Stable | tropical | diurnal | 86.2 | 8694433.536 | 10.318 |
| birds | Lamprotornis_chloropterus | Lamprotornis_chloropterus | LC | Stable | tropical | diurnal | 73.1 | 5690333.674 | 10.531 |
| birds | Lamprotornis_fischeri | Spreo_fischeri | LC | Stable | tropical | diurnal | 51 | 642683.488 | 9.964 |
| birds | Lamprotornis_hildebrandti | Lamprotornis_hildebrandti | LC | Stable | tropical | diurnal | 55.9 | 413819.205 | 12.531 |
| birds | Lamprotornis_iris | Coccycolius_iris | LC | Unknown | tropical | diurnal | 69 | 148739.048 | 12.375 |
| birds | Lamprotornis_mevesii | Lamprotornis_mevesii | LC | Unknown | tropical | diurnal | 64.2 | 639807.372 | 8.652 |
| birds | Lamprotornis_nitens | Lamprotornis_nitens | LC | Stable | tropical | diurnal | 85.6 | 3006322.47 | 7.906 |
| birds | Lamprotornis_ornatus | Lamprotornis_ornatus | LC | Unknown | tropical | diurnal | 130 | 146.654 | 14.707 |
| birds | Lamprotornis_pulcher | Lamprotornis_pulcher | LC | Stable | tropical | diurnal | 67.3 | 3948428.544 | 9.483 |
| birds | Lamprotornis_purpureus | Lamprotornis_purpureus | LC | Stable | tropical | diurnal | 112.9 | 3464722.792 | 12.432 |
| birds | Lamprotornis_purpuroptera | Lamprotornis_purpuroptera | LC | Unknown | tropical | diurnal | 77.4 | 1362512.114 | 11.574 |
| birds | Lamprotornis_regius | Cosmopsarus_regius | LC | Stable | tropical | diurnal | 53.8 | 1243437.565 | 8.619 |
| birds | Lamprotornis_shelleyi | Lamprotornis_shelleyi | LC | Stable | tropical | diurnal | 45 | 1132646.842 | 7.882 |
| birds | Lamprotornis_splendidus | Lamprotornis_splendidus | LC | Unknown | tropical | diurnal | 123.1 | 3906139.826 | 8.138 |
| birds | Lamprotornis_superbus | Lamprotornis_superbus | LC | Unknown | tropical | diurnal | 64.9 | 2087560.01 | 9.364 |
| birds | Lamprotornis_unicolor | Cosmopsarus_unicolor | LC | Unknown | tropical | diurnal | 60.2 | 309310.939 | 12.103 |
| birds | Laniarius_aethiopicus | Laniarius_aethiopicus | LC | Stable | tropical | diurnal | 49.4 | 7211323.508 | 9.421 |
| birds | Laniarius_amboimensis | Laniarius_amboimensis | EN | Decreasing | tropical | diurnal | 45.9 | 2924.075 | 11.266 |
| birds | Laniarius_atrococcineus | Laniarius_atrococcineus | LC | Increasing | tropical | diurnal | 48.8 | 1962868.617 | 6.844 |
| birds | Laniarius_atroflavus | Laniarius_atroflavus | LC | Stable | tropical | diurnal | 43 | 54064.781 | 9.571 |
| birds | Laniarius_barbarus | Laniarius_barbarus | LC | Stable | tropical | diurnal | 47.1 | 2828290.548 | 13.076 |
| birds | Laniarius_bicolor | Laniarius_bicolor | LC | Decreasing | tropical | diurnal | 54.3 | 459587.117 | 6.856 |

| | | | | | | | | | |
|---|---|---|---|---|---|---|---|---|---|
| birds | Laniarius_brauni | Laniarius_brauni | EN | Decreasing | tropical | diurnal | 54.3 | 3469.335 | 7.699 |
| birds | Laniarius_erythrogaster | Laniarius_erythrogaster | LC | Stable | tropical | diurnal | 48.8 | 1988843.909 | 10.063 |
| birds | Laniarius_ferrugineus | Laniarius_ferrugineus | LC | Stable | temperate | diurnal | 46.8 | 721209.793 | 11.037 |
| birds | Laniarius_fuelleborni | Laniarius_fuelleborni | LC | Stable | tropical | diurnal | 46.4 | 72493.56 | 12.701 |
| birds | Laniarius_funebris | Laniarius_funebris | LC | Stable | tropical | diurnal | 40.4 | 1152635.015 | 10.768 |
| birds | Laniarius_leucorhynchus | Laniarius_leucorhynchus | LC | Stable | tropical | diurnal | 50.1 | 2498729.012 | 7.577 |
| birds | Laniarius_luehderi | Laniarius_luehderi | LC | Stable | tropical | diurnal | 42.9 | 1379816.378 | 8.041 |
| birds | Laniarius_mufumbiri | Laniarius_mufumbiri | NT | Decreasing | tropical | diurnal | 41.5 | 98961.392 | 19.585 |
| birds | Laniarius_nigerrimus | NA | LC | Stable | NA | NA | NA | NA | NA |
| birds | Laniarius_poensis | Laniarius_poensis | LC | Stable | tropical | diurnal | 44.9 | 65722.138 | 10.091 |
| birds | Laniarius_ruficeps | Laniarius_ruficeps | LC | Stable | tropical | diurnal | 32.7 | 660049.389 | 8.529 |
| birds | Laniarius_turatii | Laniarius_turatii | LC | Stable | tropical | diurnal | 45.9 | 134854.059 | 14.328 |
| birds | Laniellus_albonotatus | Crocias_albonotatus | NT | Decreasing | tropical | diurnal | 45 | 2839.061 | 12.63 |
| birds | Laniellus_langbianis | Crocias_langbianis | EN | Decreasing | tropical | diurnal | 45 | 1727.561 | 12.678 |
| birds | Laniisoma_buckleyi | Laniisoma_buckleyi | LC | Decreasing | tropical | diurnal | 47.1 | 68589.423 | 5.946 |
| birds | Laniisoma_elegans | Laniisoma_elegans | NT | Decreasing | tropical | diurnal | 47.4 | 524194.002 | 14.289 |
| birds | Lanio_aurantius | Lanio_aurantius | LC | Decreasing | tropical | diurnal | 35 | 248230.792 | 9.87 |
| birds | Lanio_fulvus | Lanio_fulvus | LC | Stable | tropical | diurnal | 24 | 2285784.149 | 1.526 |
| birds | Lanio_leucothorax | Lanio_leucothorax | NT | Decreasing | tropical | diurnal | 40 | 91070.958 | 7.689 |
| birds | Lanio_versicolor | Lanio_versicolor | LC | Decreasing | tropical | diurnal | 17 | 2830210.839 | 1.988 |
| birds | Laniocera_hypopyrra | Laniocera_hypopyrra | LC | Decreasing | tropical | diurnal | 51 | 6630155.401 | 2.064 |
| birds | Laniocera_rufescens | Laniocera_rufescens | LC | Decreasing | tropical | diurnal | 48.1 | 394176.437 | 8.355 |
| birds | Lanioturdus_torquatus | Lanioturdus_torquatus | LC | Increasing | tropical | diurnal | 29 | 342747.576 | 7.526 |
| birds | Lanius_bucephalus | Lanius_bucephalus | LC | Decreasing | temperate | diurnal | 39.4 | 2462379.63 | 16.748 |
| birds | Lanius_cabanisi | Lanius_cabanisi | LC | Stable | tropical | diurnal | 73.8 | 548346.11 | 11.788 |
| birds | Lanius_collaris | Lanius_collaris | LC | Increasing | tropical | diurnal | 36.4 | 7220481.633 | 9.128 |
| birds | Lanius_collurio | Lanius_collurio | LC | Decreasing | temperate | diurnal | 28.4 | 27572252.522 | 9.693 |
| birds | Lanius_collurioides | Lanius_collurioides | LC | Stable | tropical | diurnal | 26 | 2126567.6 | 12.364 |
| birds | Lanius_corvinus | Corvinella_corvina | LC | Unknown | tropical | diurnal | 65.4 | 3469655.335 | 11.874 |
| birds | Lanius_cristatus | Lanius_cristatus | LC | Decreasing | temperate | diurnal | 32.2 | 21999940.843 | 8.843 |
| birds | Lanius_dorsalis | Lanius_dorsalis | LC | Stable | tropical | diurnal | 49.9 | 1312304.946 | 9.057 |
| birds | Lanius_excubitor | Lanius_excubitor | LC | Decreasing | temperate | diurnal | 63.4 | 36121373.627 | 8.909 |
| birds | Lanius_excubitoroides | Lanius_excubitoroides | LC | Stable | tropical | diurnal | 53.7 | 2533065.502 | 10.672 |
| birds | Lanius_gubernator | Lanius_gubernator | LC | Stable | tropical | diurnal | 23.5 | 427904.644 | 10.233 |

| | | | | | | | | | |
|---|---|---|---|---|---|---|---|---|---|
| birds | Lanius_isabellinus | Lanius_isabellinus | LC | Stable | tropical | diurnal | 25.2 | 25169824.707 | 7.471 |
| birds | Lanius_ludovicianus | Lanius_ludovicianus | NT | Decreasing | temperate | diurnal | 51.6 | 9319415.628 | 13.04 |
| birds | Lanius_mackinnoni | Lanius_mackinnoni | LC | Increasing | tropical | diurnal | 34.3 | 1193617.213 | 10.196 |
| birds | Lanius_melanoleucus | Urolestes_melanoleucus | LC | Decreasing | tropical | diurnal | 82.2 | 1707357.858 | 9.057 |
| birds | Lanius_meridionalis | Lanius_meridionalis | VU | Decreasing | temperate | diurnal | 60.4 | 678211.419 | 17.75 |
| birds | Lanius_minor | Lanius_minor | LC | Decreasing | temperate | diurnal | 46.5 | 10497294.932 | 12.204 |
| birds | Lanius_newtoni | Lanius_newtoni | CR | Decreasing | tropical | diurnal | 22.4 | 150.769 | 5.698 |
| birds | Lanius_nubicus | Lanius_nubicus | LC | Decreasing | temperate | diurnal | 19.7 | 8736712.433 | 7.43 |
| birds | Lanius_schach | Lanius_schach | LC | Unknown | tropical | diurnal | 51.5 | 11170512.096 | 14.005 |
| birds | Lanius_senator | Lanius_senator | NT | Decreasing | tropical | diurnal | 36 | 19072012.904 | 7.33 |
| birds | Lanius_somalicus | Lanius_somalicus | LC | Stable | tropical | diurnal | 45.3 | 858092.071 | 7.761 |
| birds | Lanius_souzae | Lanius_souzae | LC | Decreasing | tropical | diurnal | 26.3 | 1679903.363 | 7.538 |
| birds | Lanius_sphenocercus | Lanius_sphenocercus | LC | Stable | temperate | diurnal | 92.1 | 4022019.933 | 15.064 |
| birds | Lanius_tephronotus | Lanius_tephronotus | LC | Stable | temperate | diurnal | 46.4 | 3050531.35 | 10.505 |
| birds | Lanius_tigrinus | Lanius_tigrinus | LC | Decreasing | temperate | diurnal | 30.1 | 5955735.751 | 14.371 |
| birds | Lanius_validirostris | Lanius_validirostris | LC | Stable | tropical | diurnal | 37.5 | 18662.932 | 7.079 |
| birds | Lanius_vittatus | Lanius_vittatus | LC | Stable | tropical | diurnal | 20.8 | 4168923.157 | 15.276 |
| birds | Larosterna_inca | Larosterna_inca | NT | Decreasing | temperate | diurnal | 219 | 558060.926 | 11.085 |
| birds | Larus_argentatus | Larus_argentatus | LC | Decreasing | temperate | diurnal | 1091 | 3521562.852 | 15.323 |
| birds | Larus_armenicus | Larus_armenicus | LC | Increasing | temperate | diurnal | 867 | 1322509.775 | 15.542 |
| birds | Larus_atlanticus | Larus_atlanticus | NT | Stable | temperate | diurnal | 842.4 | 140744.803 | 8.879 |
| birds | Larus_atricilla | Larus_atricilla | LC | Increasing | temperate | diurnal | 307.4 | 3760946.604 | 11.489 |
| birds | Larus_audouinii | Larus_audouinii | VU | Decreasing | temperate | diurnal | 535 | 1457965.531 | 21.793 |
| birds | Larus_belcheri | Larus_belcheri | LC | Increasing | tropical | diurnal | 638.6 | 389151.943 | 10.405 |
| birds | Larus_brunnicephalus | Larus_brunnicephalus | LC | Stable | temperate | diurnal | 569.3 | 2442213.001 | 13.026 |
| birds | Larus_bulleri | Larus_bulleri | NT | Unknown | temperate | diurnal | 230.2 | 133703.017 | 9.488 |
| birds | Larus_cachinnans | Larus_cachinnans | LC | Increasing | temperate | diurnal | 1111.9 | 7445647.391 | 11.339 |
| birds | Larus_californicus | Larus_californicus | LC | Decreasing | temperate | diurnal | 683.4 | 6058232.192 | 9.121 |
| birds | Larus_canus | Larus_canus | LC | Unknown | temperate | diurnal | 412.5 | 45586444.285 | 7.776 |
| birds | Larus_cirrocephalus | Larus_cirrocephalus | LC | Stable | tropical | diurnal | 374.7 | 33691966.611 | 8.793 |
| birds | Larus_crassirostris | Larus_crassirostris | LC | Stable | temperate | diurnal | 561 | 2745922.28 | 16.716 |
| birds | Larus_delawarensis | Larus_delawarensis | LC | Increasing | temperate | diurnal | 516.3 | 13105607.4 | 11.212 |
| birds | Larus_dominicanus | Larus_dominicanus | LC | Increasing | temperate | diurnal | 952.4 | 4370314.504 | 8.411 |
| birds | Larus_fuliginosus | NA | VU | Stable | NA | NA | NA | NA | NA |

| | | | | | | | | | |
|---|---|---|---|---|---|---|---|---|---|
| birds | Larus_fuscus | Larus_fuscus | LC | Increasing | temperate | diurnal | 762.4 | 47994979.501 | 9.636 |
| birds | Larus_genei | Larus_genei | LC | Unknown | temperate | diurnal | 281 | 5030312.642 | 10.619 |
| birds | Larus_glaucescens | Larus_glaucescens | LC | Increasing | temperate | diurnal | 1056.5 | 6471916.141 | 2.025 |
| birds | Larus_glaucoides | Larus_glaucoides | LC | Stable | temperate | diurnal | 693.3 | 12843087.943 | 1.995 |
| birds | Larus_hartlaubii | Larus_hartlaubii | LC | Increasing | temperate | diurnal | 292 | 69935.229 | 7.588 |
| birds | Larus_heermanni | Larus_heermanni | NT | Unknown | temperate | diurnal | 500 | 1481289.517 | 7.152 |
| birds | Larus_hemprichii | Larus_hemprichii | LC | Decreasing | tropical | diurnal | 451.7 | 1840156.065 | 10.64 |
| birds | Larus_hyperboreus | Larus_hyperboreus | LC | Stable | temperate | diurnal | 1529 | 8106310.4 | 6.433 |
| birds | Larus_ichthyaetus | Larus_ichthyaetus | LC | Increasing | temperate | diurnal | 1378.8 | 5985921.731 | 12.793 |
| birds | Larus_leucophthalmus | Larus_leucophthalmus | LC | Stable | tropical | diurnal | 328.9 | 720417.092 | 12.065 |
| birds | Larus_livens | Larus_livens | LC | Unknown | temperate | diurnal | 1322 | 199173.629 | 6.708 |
| birds | Larus_maculipennis | Larus_maculipennis | LC | Increasing | temperate | diurnal | 339 | 2378141.143 | 8.345 |
| birds | Larus_marinus | Larus_marinus | LC | Unknown | temperate | diurnal | 1649.7 | 12819967.113 | 13.704 |
| birds | Larus_melanocephalus | Larus_melanocephalus | LC | Decreasing | temperate | diurnal | 256 | 3340143.418 | 19.249 |
| birds | Larus_modestus | Larus_modestus | LC | Decreasing | temperate | cathemeral | 344 | 567776.38 | 6.514 |
| birds | Larus_novaehollandiae | Larus_novaehollandiae | LC | Increasing | temperate | diurnal | 269 | 8538944.924 | 5.14 |
| birds | Larus_occidentalis | Larus_occidentalis | LC | Increasing | temperate | diurnal | 922.8 | 157463.923 | 14.209 |
| birds | Larus_pacificus | Larus_pacificus | LC | Stable | temperate | diurnal | 1292 | 392125.029 | 9.05 |
| birds | Larus_philadelphia | Larus_philadelphia | LC | Increasing | temperate | diurnal | 210.7 | 15309356.605 | 9.306 |
| birds | Larus_pipixcan | Larus_pipixcan | LC | Increasing | temperate | diurnal | 280 | 8494299.949 | 11.829 |
| birds | Larus_relictus | Larus_relictus | VU | Decreasing | temperate | diurnal | 490.2 | 1443958.552 | 5.785 |
| birds | Larus_ridibundus | Larus_ridibundus | LC | Unknown | temperate | diurnal | 284 | 44468354.719 | 10.267 |
| birds | Larus_schistisagus | Larus_schistisagus | LC | Unknown | temperate | diurnal | 1323 | 2043799.553 | 12.473 |
| birds | Larus_scoresbii | Leucophaeus_scoresbii | LC | Stable | temperate | diurnal | 523.5 | 498432.006 | 2.45 |
| birds | Larus_serranus | Larus_serranus | LC | Unknown | tropical | diurnal | 478 | 1115843.782 | 6.344 |
| birds | Larvivora_akahige | Erithacus_akahige | LC | Stable | temperate | diurnal | 17 | 416504.963 | 14.909 |
| birds | Larvivora_brunnea | Luscinia_brunnea | LC | Decreasing | tropical | diurnal | 17.5 | 3708882.519 | 14.927 |
| birds | Larvivora_cyane | Luscinia_cyane | LC | Decreasing | temperate | diurnal | 14.8 | 9963872.424 | 6.69 |
| birds | Larvivora_komadori | Erithacus_komadori | NT | Stable | temperate | diurnal | 23.9 | 2993.148 | 16.154 |
| birds | Larvivora_ruficeps | Luscinia_ruficeps | EN | Decreasing | temperate | diurnal | 18 | 46895.977 | 9.119 |
| birds | Larvivora_sibilans | Luscinia_sibilans | LC | Stable | temperate | diurnal | 16.7 | 6856830.681 | 4.543 |
| birds | Laterallus_albigularis | Laterallus_albigularis | LC | Stable | tropical | diurnal | 47.3 | 661410.857 | 11.47 |
| birds | Laterallus_exilis | Laterallus_exilis | LC | Decreasing | tropical | diurnal | 33.5 | 6585440.308 | 4.398 |
| birds | Laterallus_flaviventer | Porzana_flaviventer | LC | Unknown | temperate | diurnal | 24.9 | 2593534.549 | 8.839 |

| | | | | | | | | | |
|---|---|---|---|---|---|---|---|---|---|
| birds | Laterallus_jamaicensis | Laterallus_jamaicensis | EN | Decreasing | tropical | diurnal | 32.6 | 1137301.503 | 10.826 |
| birds | Laterallus_leucopyrrhus | Laterallus_leucopyrrhus | LC | Unknown | temperate | diurnal | 45.5 | 882070.866 | 11.358 |
| birds | Laterallus_levraudi | Laterallus_levraudi | VU | Decreasing | tropical | diurnal | 43.5 | 37022.494 | 15.107 |
| birds | Laterallus_melanophaius | Laterallus_melanophaius | LC | Unknown | tropical | diurnal | 52.1 | 10160298.316 | 6.195 |
| birds | Laterallus_rogersi | NA | VU | Stable | NA | NA | NA | NA | NA |
| birds | Laterallus_ruber | Laterallus_ruber | LC | Stable | tropical | diurnal | 44.4 | 557484.257 | 10.858 |
| birds | Laterallus_spilonota | NA | VU | Decreasing | NA | NA | NA | NA | NA |
| birds | Laterallus_spilopterus | NA | VU | Decreasing | NA | NA | NA | NA | NA |
| birds | Laterallus_xenopterus | Laterallus_xenopterus | VU | Decreasing | tropical | diurnal | 56.9 | 1703536.277 | 10.663 |
| birds | Lathamus_discolor | Lathamus_discolor | CR | Decreasing | temperate | diurnal | 64.7 | 767482.189 | 9.887 |
| birds | Lathrotriccus_euleri | Lathrotriccus_euleri | LC | Stable | tropical | diurnal | 11.3 | 10279275.783 | 6.316 |
| birds | Lathrotriccus_griseipectus | Lathrotriccus_griseipectus | VU | Decreasing | tropical | diurnal | 11 | 55790.482 | 11.588 |
| birds | Laticilla_burnesii | Prinia_burnesii | NT | Decreasing | temperate | diurnal | 19 | 332103.287 | 17.748 |
| birds | Legatus_leucophaius | Legatus_leucophaius | LC | Decreasing | tropical | diurnal | 22.2 | 11408794.56 | 5.447 |
| birds | Leiopicus_auriceps | Dendrocopos_auriceps | LC | Stable | temperate | diurnal | 39.9 | 79474.365 | 14.775 |
| birds | Leiopicus_mahrattensis | Dendrocopos_mahrattensis | LC | Stable | tropical | diurnal | 34.3 | 2815279.671 | 18.097 |
| birds | Leiopicus_medius | Dendrocopos_medius | LC | Increasing | temperate | diurnal | 59 | 3966649.977 | 17.973 |
| birds | Leioptila_annectens | Heterophasia_annectens | LC | Decreasing | temperate | diurnal | 24.5 | 693171.677 | 7.458 |
| birds | Leiothlypis_celata | Vermivora_celata | LC | Decreasing | temperate | diurnal | 9.2 | 15777236.339 | 8.857 |
| birds | Leiothlypis_crissalis | Vermivora_crissalis | LC | Stable | temperate | diurnal | 9.3 | 557643.61 | 8.577 |
| birds | Leiothlypis_luciae | Vermivora_luciae | LC | Increasing | temperate | diurnal | 6.6 | 682328.055 | 8.126 |
| birds | Leiothlypis_peregrina | Vermivora_peregrina | LC | Decreasing | temperate | diurnal | 8.9 | 12155394.321 | 10.476 |
| birds | Leiothlypis_ruficapilla | Vermivora_ruficapilla | LC | Stable | temperate | diurnal | 8.1 | 10359677.996 | 11.874 |
| birds | Leiothlypis_virginiae | Vermivora_virginiae | LC | Decreasing | temperate | diurnal | 8.2 | 1692005.667 | 8.816 |
| birds | Leiothrix_argentauris | Leiothrix_argentauris | LC | Decreasing | tropical | diurnal | 28.4 | 1292489.317 | 9.872 |
| birds | Leiothrix_lutea | Leiothrix_lutea | LC | Decreasing | temperate | diurnal | 21.4 | 3017583.349 | 15.013 |
| birds | Leipoa_ocellata | Leipoa_ocellata | VU | Decreasing | temperate | diurnal | 1913.9 | 1718483.341 | 5.496 |
| birds | Leistes_bellicosus | Sturnella_bellicosa | LC | Stable | tropical | diurnal | 58 | 351063.411 | 8.647 |
| birds | Leistes_loyca | Sturnella_loyca | LC | Stable | temperate | diurnal | 113 | 1809024.922 | 7.674 |
| birds | Leistes_militaris | Sturnella_militaris | LC | Increasing | tropical | diurnal | 48 | 3688152.764 | 4.288 |
| birds | Leistes_superciliaris | Sturnella_superciliaris | LC | Stable | tropical | diurnal | 45.3 | 6588682.447 | 9.192 |
| birds | Leonardina_woodi | Leonardina_woodi | LC | Stable | tropical | diurnal | 21 | 30087.223 | 9.133 |
| birds | Lepidocolaptes_affinis | Lepidocolaptes_affinis | LC | Decreasing | tropical | diurnal | 35 | 238243.154 | 10.346 |
| birds | Lepidocolaptes_albolineatus | Lepidocolaptes_albolineatus | LC | Decreasing | tropical | diurnal | 20.3 | 1239516.134 | 1.526 |

| | | | | | | | | | |
|---|---|---|---|---|---|---|---|---|---|
| birds | Lepidocolaptes_angustirostris | Lepidocolaptes_angustirostris | LC | Increasing | tropical | diurnal | 29.6 | 6576318.885 | 8.597 |
| birds | Lepidocolaptes_falcinellus | Lepidocolaptes_falcinellus | LC | Stable | temperate | diurnal | 27.9 | 497665.618 | 12.067 |
| birds | Lepidocolaptes_lacrymiger | Lepidocolaptes_lacrymiger | LC | Stable | tropical | diurnal | 30.6 | 360596.478 | 8.967 |
| birds | Lepidocolaptes_leucogaster | Lepidocolaptes_leucogaster | LC | Decreasing | tropical | diurnal | 36 | 291440.841 | 7.121 |
| birds | Lepidocolaptes_souleyetii | Lepidocolaptes_souleyetii | LC | Decreasing | tropical | diurnal | 25.7 | 1484080.312 | 10.23 |
| birds | Lepidocolaptes_squamatus | Lepidocolaptes_squamatus | LC | Decreasing | tropical | diurnal | 28.1 | 745438.764 | 11.661 |
| birds | Lepidogrammus_cumingi | Phaenicophaeus_cumingi | LC | Decreasing | tropical | diurnal | 170.5 | 40477.514 | 10.155 |
| birds | Lepidopygia_nana | Lonchura_nana | LC | Stable | tropical | diurnal | 8.2 | 590836.069 | 8.745 |
| birds | Lepidothrix_coeruleocapilla | Lepidothrix_coeruleocapilla | LC | Stable | tropical | diurnal | 9.7 | 140252.42 | 4.045 |
| birds | Lepidothrix_coronata | Lepidothrix_coronata | LC | Decreasing | tropical | diurnal | 8.3 | 3221211.218 | 1.89 |
| birds | Lepidothrix_iris | Lepidothrix_iris | VU | Decreasing | tropical | diurnal | 8 | 543503.12 | 3.923 |
| birds | Lepidothrix_isidorei | Lepidothrix_isidorei | LC | Decreasing | tropical | diurnal | 7.8 | 78737.069 | 5.636 |
| birds | Lepidothrix_nattereri | Lepidothrix_nattereri | LC | Decreasing | tropical | diurnal | 8.4 | 865198.456 | 2.876 |
| birds | Lepidothrix_serena | Lepidothrix_serena | LC | Stable | tropical | diurnal | 11.4 | 583711.439 | 0.995 |
| birds | Lepidothrix_suavissima | Lepidothrix_suavissima | LC | Stable | tropical | diurnal | 8.5 | 310206.619 | 1.136 |
| birds | Lepidothrix_vilasboasi | NA | LC | Decreasing | NA | NA | NA | NA | NA |
| birds | Leptasthenura_aegithaloides | Leptasthenura_aegithaloides | LC | Stable | temperate | diurnal | 10.7 | 250972.15 | 8.708 |
| birds | Leptasthenura_andicola | Leptasthenura_andicola | LC | Stable | tropical | diurnal | 15.5 | 256098.228 | 10.496 |
| birds | Leptasthenura_fuliginiceps | Leptasthenura_fuliginiceps | LC | Stable | temperate | diurnal | 12.6 | 369467.91 | 6.597 |
| birds | Leptasthenura_pileata | Leptasthenura_pileata | LC | Stable | tropical | diurnal | 10.3 | 93356.541 | 8.503 |
| birds | Leptasthenura_platensis | Leptasthenura_platensis | LC | Stable | temperate | diurnal | 10.6 | 2007502.949 | 8.502 |
| birds | Leptasthenura_setaria | Leptasthenura_setaria | NT | Decreasing | temperate | diurnal | 11 | 394948.895 | 12.947 |
| birds | Leptasthenura_striata | Leptasthenura_striata | LC | Decreasing | tropical | diurnal | 8.9 | 188336.203 | 7.337 |
| birds | Leptasthenura_striolata | Leptasthenura_striolata | LC | Stable | temperate | diurnal | 10.5 | 208971.088 | 11.457 |
| birds | Leptasthenura_xenothorax | Leptasthenura_xenothorax | EN | Decreasing | tropical | diurnal | 10.5 | 2565.16 | 7.168 |
| birds | Leptocoma_aspasia | Nectarinia_aspasia | LC | Stable | tropical | diurnal | 8 | 916101.698 | 4.653 |
| birds | Leptocoma_calcostetha | Nectarinia_calcostetha | LC | Stable | tropical | diurnal | 8.3 | 568344.61 | 12.448 |
| birds | Leptocoma_minima | Nectarinia_minima | LC | Stable | tropical | diurnal | 4.9 | 57005.418 | 12.329 |
| birds | Leptocoma_sperata | Nectarinia_sperata | LC | Stable | tropical | diurnal | 6.4 | 229948.472 | 13.857 |
| birds | Leptocoma_zeylonica | Nectarinia_zeylonica | LC | Stable | tropical | diurnal | 8.8 | 1817376.928 | 18.34 |
| birds | Leptodon_cayanensis | Leptodon_cayanensis | LC | Decreasing | tropical | diurnal | 474 | 12071802.075 | 5.445 |
| birds | Leptopoecile_elegans | Leptopoecile_elegans | LC | Stable | temperate | diurnal | 6.2 | 815155.612 | 6.423 |
| birds | Leptopoecile_sophiae | Leptopoecile_sophiae | LC | Stable | temperate | diurnal | 6.9 | 1449954.199 | 6.865 |
| birds | Leptopogon_amaurocephalus | Leptopogon_amaurocephalus | LC | Decreasing | tropical | diurnal | 11.7 | 8060136.759 | 6.93 |

| | | | | | | | | | |
|---|---|---|---|---|---|---|---|---|---|
| birds | Leptopogon_rufipectus | Leptopogon_rufipectus | LC | Decreasing | tropical | diurnal | 13.4 | 100544.646 | 10.903 |
| birds | Leptopogon_superciliaris | Leptopogon_superciliaris | LC | Stable | tropical | diurnal | 11.7 | 474222.709 | 9.379 |
| birds | Leptopogon_taczanowskii | Leptopogon_taczanowskii | LC | Decreasing | tropical | diurnal | 11.5 | 65070.378 | 4.215 |
| birds | Leptopterus_chabert | Leptopterus_chabert | LC | Unknown | tropical | diurnal | 19.6 | 577947.597 | 8.683 |
| birds | Leptoptilos_crumenifer | Leptoptilos_crumeniferus | LC | Increasing | tropical | diurnal | 6067.9 | 12361408.835 | 9.071 |
| birds | Leptoptilos_dubius | Leptoptilos_dubius | NT | Increasing | tropical | diurnal | 9050 | 2906151.219 | 15.864 |
| birds | Leptoptilos_javanicus | Leptoptilos_javanicus | NT | Unknown | tropical | diurnal | 4651 | 4920849.104 | 14.281 |
| birds | Leptosittaca_branickii | Leptosittaca_branickii | LC | Decreasing | tropical | diurnal | 155.7 | 46449.52 | 8.47 |
| birds | Leptosomus_discolor | Leptosomus_discolor | LC | Decreasing | tropical | diurnal | 255 | 592877.2 | 8.772 |
| birds | Leptotila_battyi | Leptotila_battyi | VU | Decreasing | tropical | diurnal | 160.6 | 1791.986 | 3.827 |
| birds | Leptotila_cassinii | Leptotila_cassini | LC | Decreasing | tropical | diurnal | 159 | 314249.819 | 8.955 |
| birds | Leptotila_conoveri | Leptotila_conoveri | NT | Decreasing | tropical | diurnal | 153.5 | 7847.083 | 13.155 |
| birds | Leptotila_jamaicensis | Leptotila_jamaicensis | LC | Decreasing | tropical | diurnal | 165 | 92373.414 | 6.798 |
| birds | Leptotila_megalura | Leptotila_megalura | LC | Stable | tropical | diurnal | 218 | 147285.79 | 6.688 |
| birds | Leptotila_ochraceiventris | Leptotila_ochraceiventris | VU | Decreasing | tropical | diurnal | 146 | 43290.897 | 13.676 |
| birds | Leptotila_pallida | Leptotila_pallida | LC | Decreasing | tropical | diurnal | 122.8 | 117295.446 | 10.388 |
| birds | Leptotila_plumbeiceps | Leptotila_plumbeiceps | LC | Decreasing | tropical | diurnal | 170 | 360918.877 | 9.936 |
| birds | Leptotila_rufaxilla | Leptotila_rufaxilla | LC | Decreasing | tropical | diurnal | 157 | 11668214.678 | 5.337 |
| birds | Leptotila_verreauxi | Leptotila_verreauxi | LC | Stable | tropical | diurnal | 146.9 | 15542979.807 | 6.279 |
| birds | Leptotila_wellsi | Leptotila_wellsi | CR | Decreasing | tropical | diurnal | 200 | 69.349 | 26.465 |
| birds | Leptotrygon_veraguensis | Geotrygon_veraguensis | LC | Decreasing | tropical | diurnal | 155 | 127743.926 | 6.559 |
| birds | Lerwa_lerwa | Lerwa_lerwa | LC | Decreasing | temperate | diurnal | 567.3 | 831638.529 | 5.936 |
| birds | Lesbia_nuna | Lesbia_nuna | LC | Stable | tropical | diurnal | 3.7 | 251932.098 | 9.303 |
| birds | Lesbia_victoriae | Lesbia_victoriae | LC | Stable | tropical | diurnal | 5 | 197489.261 | 10.511 |
| birds | Lessonia_oreas | Lessonia_oreas | LC | Stable | tropical | diurnal | 13.8 | 831919.208 | 6.44 |
| birds | Lessonia_rufa | Lessonia_rufa | LC | Stable | temperate | diurnal | 13.4 | 3918173.55 | 7.323 |
| birds | Leucippus_fallax | Leucippus_fallax | LC | Decreasing | tropical | diurnal | 6 | 93947.055 | 12.51 |
| birds | Leucocarbo_atriceps | Phalacrocorax_atriceps | LC | Unknown | temperate | diurnal | 2000 | 442340.253 | 4.988 |
| birds | Leucocarbo_bougainvilliorum | Phalacrocorax_bougainvillii | NT | Decreasing | tropical | diurnal | 2485 | 599089.916 | 7.809 |
| birds | Leucocarbo_campbelli | NA | VU | Stable | NA | NA | NA | NA | NA |
| birds | Leucocarbo_carunculatus | Phalacrocorax_carunculatus | VU | Stable | temperate | diurnal | 2576.3 | 1161.585 | 6.653 |
| birds | Leucocarbo_chalconotus | Phalacrocorax_chalconotus | VU | Decreasing | temperate | diurnal | 2220.1 | 13812.173 | 5.217 |
| birds | Leucocarbo_colensoi | NA | VU | Stable | NA | NA | NA | NA | NA |
| birds | Leucocarbo_magellanicus | Phalacrocorax_magellanicus | LC | Unknown | temperate | diurnal | 1483.4 | 710005.322 | 2.944 |

| | | | | | | | | | |
|---|---|---|---|---|---|---|---|---|---|
| birds | Leucocarbo_onslowi | NA | CR | Decreasing | NA | NA | NA | NA | NA |
| birds | Leucocarbo_ranfurlyi | NA | VU | Stable | NA | NA | NA | NA | NA |
| birds | Leucocarbo_verrucosus | NA | LC | Unknown | NA | NA | NA | NA | NA |
| birds | Leucochloris_albicollis | Leucochloris_albicollis | LC | Unknown | temperate | diurnal | 6.3 | 2176859.286 | 12.06 |
| birds | Leucogeranus_leucogeranus | Grus_leucogeranus | CR | Decreasing | temperate | diurnal | 5913.4 | 276450.451 | 4.278 |
| birds | Leucolia_violiceps | Amazilia_violiceps | LC | Stable | temperate | diurnal | 5.1 | 737817.46 | 8.985 |
| birds | Leuconotopicus_albolarvatus | Picoides_albolarvatus | LC | Stable | temperate | diurnal | 61.1 | 499656.93 | 9.492 |
| birds | Leuconotopicus_borealis | Picoides_borealis | NT | Decreasing | temperate | diurnal | 48 | 908307.875 | 16.054 |
| birds | Leuconotopicus_fumigatus | Picoides_fumigatus | LC | Decreasing | tropical | diurnal | 36.2 | 1109542.216 | 9.448 |
| birds | Leuconotopicus_stricklandi | Picoides_stricklandi | LC | Decreasing | tropical | diurnal | 40.9 | 18699.265 | 22.964 |
| birds | Leuconotopicus_villosus | Picoides_villosus | LC | Increasing | temperate | diurnal | 62.7 | 13332874.44 | 9.226 |
| birds | Leucopeza_semperi | Leucopeza_semperi | CR | Unknown | tropical | diurnal | 19.8 | 78.043 | 8.728 |
| birds | Leucopsar_rothschildi | NA | CR | Decreasing | NA | NA | NA | NA | NA |
| birds | Leucopternis_kuhli | Leucopternis_kuhli | LC | Decreasing | tropical | diurnal | 352.1 | 2375316.625 | 1.792 |
| birds | Leucopternis_melanops | Leucopternis_melanops | LC | Decreasing | tropical | diurnal | 312 | 2570861.924 | 1.358 |
| birds | Leucopternis_semiplumbeus | Leucopternis_semiplumbeus | LC | Decreasing | tropical | diurnal | 285 | 236308.058 | 7.636 |
| birds | Leucosarcia_melanoleuca | Leucosarcia_melanoleuca | LC | Decreasing | temperate | diurnal | 428.6 | 503204.108 | 8.02 |
| birds | Leucosticte_arctoa | Leucosticte_arctoa | LC | Decreasing | temperate | diurnal | 30.7 | 3906698.216 | 5.459 |
| birds | Leucosticte_atrata | Leucosticte_atrata | EN | Decreasing | temperate | diurnal | 25.2 | 970789.688 | 9.093 |
| birds | Leucosticte_australis | Leucosticte_australis | EN | Decreasing | temperate | diurnal | 26.6 | 178378.622 | 9.342 |
| birds | Leucosticte_brandti | Leucosticte_brandti | LC | Stable | temperate | diurnal | 30.4 | 3640952.709 | 4.42 |
| birds | Leucosticte_nemoricola | Leucosticte_nemoricola | LC | Stable | temperate | diurnal | 22.5 | 4294529.955 | 5.398 |
| birds | Leucosticte_tephrocotis | Leucosticte_tephrocotis | LC | Stable | temperate | diurnal | 37.8 | 4226170.676 | 5.162 |
| birds | Lewinia_mirifica | NA | DD | Unknown | NA | NA | NA | NA | NA |
| birds | Lewinia_pectoralis | Lewinia_pectoralis | LC | Decreasing | tropical | diurnal | 78.1 | 1002596.393 | 8.013 |
| birds | Lewinia_striata | Gallirallus_striatus | LC | Increasing | tropical | diurnal | 116 | 4893360.604 | 13.121 |
| birds | Lichenostomus_cratitius | Lichenostomus_cratitius | LC | Decreasing | temperate | diurnal | 20.3 | 479802.478 | 9.1 |
| birds | Lichenostomus_melanops | Lichenostomus_melanops | LC | Decreasing | temperate | diurnal | 32.3 | 732848.751 | 9.026 |
| birds | Lichmera_alboauricularis | Lichmera_alboauricularis | LC | Unknown | tropical | diurnal | 13.3 | 76151.285 | 3.587 |
| birds | Lichmera_argentauris | Lichmera_argentauris | LC | Unknown | tropical | diurnal | 14 | 8366.809 | 8.254 |
| birds | Lichmera_deningeri | Lichmera_deningeri | LC | Unknown | tropical | diurnal | 16.6 | 8554.414 | 6.663 |
| birds | Lichmera_flavicans | Lichmera_flavicans | LC | Stable | tropical | diurnal | 14.2 | 25012.065 | 13.573 |
| birds | Lichmera_incana | Lichmera_incana | LC | Stable | tropical | diurnal | 13.1 | 24110.299 | 7.117 |
| birds | Lichmera_indistincta | Lichmera_indistincta | LC | Decreasing | tropical | diurnal | 11.2 | 5351801.877 | 4.295 |

| | | | | | | | | | |
|---|---|---|---|---|---|---|---|---|---|
| birds | Lichmera_lombokia | Lichmera_lombokia | LC | Stable | tropical | diurnal | 15.3 | 8705.538 | 10.336 |
| birds | Lichmera_monticola | Lichmera_monticola | LC | Stable | tropical | diurnal | 6 | 3713.671 | 6.186 |
| birds | Lichmera_notabilis | Lichmera_notabilis | LC | Stable | tropical | diurnal | 11.5 | 2638.287 | 4.678 |
| birds | Lichmera_squamata | Lichmera_squamata | LC | Stable | tropical | diurnal | 14.5 | 9998.941 | 7.259 |
| birds | Limnoctites_rectirostris | Limnoctites_rectirostris | NT | Decreasing | temperate | diurnal | 18.5 | 179597.008 | 12.148 |
| birds | Limnoctites_sulphuriferus | Cranioleuca_sulphurifera | LC | Decreasing | temperate | diurnal | 13.5 | 711047.931 | 11.369 |
| birds | Limnodromus_griseus | Limnodromus_griseus | LC | Decreasing | temperate | cathemeral | 110.5 | 4634264.809 | 6.87 |
| birds | Limnodromus_scolopaceus | Limnodromus_scolopaceus | LC | Unknown | temperate | diurnal | 104.4 | 10469880.218 | 8.691 |
| birds | Limnodromus_semipalmatus | Limnodromus_semipalmatus | NT | Decreasing | temperate | diurnal | 212 | 3508614.021 | 8.77 |
| birds | Limnornis_curvirostris | Limnornis_curvirostris | LC | Stable | temperate | diurnal | 28.6 | 217435.322 | 11.893 |
| birds | Limnothlypis_swainsonii | Limnothlypis_swainsonii | LC | Increasing | temperate | diurnal | 18.9 | 1752740.74 | 14.789 |
| birds | Limosa_fedoa | Limosa_fedoa | LC | Decreasing | temperate | diurnal | 357 | 1926237.949 | 13.449 |
| birds | Limosa_haemastica | Limosa_haemastica | LC | Decreasing | temperate | diurnal | 253.3 | 3016657.158 | 9.306 |
| birds | Limosa_lapponica | Limosa_lapponica | NT | Decreasing | temperate | diurnal | 291.7 | 9184830.813 | 9.356 |
| birds | Limosa_limosa | Limosa_limosa | NT | Decreasing | temperate | diurnal | 288.4 | 55640629.683 | 9.441 |
| birds | Linaria_cannabina | Carduelis_cannabina | LC | Decreasing | temperate | diurnal | 19.5 | 13585958.382 | 14.178 |
| birds | Linaria_flavirostris | Carduelis_flavirostris | LC | Decreasing | temperate | diurnal | 15.4 | 9158045.257 | 7.931 |
| birds | Linaria_johannis | Carduelis_johannis | NT | Stable | tropical | diurnal | 13.4 | 2340.975 | 10.335 |
| birds | Linaria_yemenensis | Carduelis_yemenensis | LC | Stable | tropical | diurnal | 14.3 | 79054.219 | 11.491 |
| birds | Linurgus_olivaceus | Linurgus_olivaceus | LC | Stable | tropical | diurnal | 20.3 | 220044.208 | 12.661 |
| birds | Liocichla_omeiensis | Liocichla_omeiensis | VU | Decreasing | temperate | diurnal | 40.4 | 8458.495 | 11.65 |
| birds | Liocichla_phoenicea | Liocichla_phoenicea | LC | Decreasing | temperate | diurnal | 49 | 421118.542 | 6.61 |
| birds | Lioparus_chrysotis | Alcippe_chrysotis | LC | Decreasing | temperate | diurnal | 5.5 | 1322119.824 | 11.528 |
| birds | Liosceles_thoracicus | Liosceles_thoracicus | LC | Decreasing | tropical | diurnal | 40.5 | 2273901.193 | 1.581 |
| birds | Lipaugus_ater | Tijuca_atra | LC | Decreasing | tropical | diurnal | 80 | 12392.117 | 11.594 |
| birds | Lipaugus_conditus | Tijuca_condita | VU | Stable | tropical | diurnal | 80 | 253.736 | 10.922 |
| birds | Lipaugus_fuscocinereus | Lipaugus_fuscocinereus | LC | Decreasing | tropical | diurnal | 138 | 42613.145 | 9.095 |
| birds | Lipaugus_lanioides | Lipaugus_lanioides | LC | Decreasing | temperate | diurnal | 94.8 | 225795.925 | 12.501 |
| birds | Lipaugus_streptophorus | Lipaugus_streptophorus | LC | Stable | tropical | diurnal | 57.2 | 47371.379 | 1.872 |
| birds | Lipaugus_unirufus | Lipaugus_unirufus | LC | Decreasing | tropical | diurnal | 82.1 | 462431.536 | 8.246 |
| birds | Lipaugus_uropygialis | Lipaugus_uropygialis | VU | Decreasing | tropical | diurnal | 116 | 15254.669 | 4.979 |
| birds | Lipaugus_vociferans | Lipaugus_vociferans | LC | Decreasing | tropical | diurnal | 75.4 | 7063057.918 | 2.405 |
| birds | Lissotis_hartlaubii | Eupodotis_hartlaubii | LC | Stable | tropical | diurnal | 1190 | 606100.639 | 10.158 |
| birds | Lissotis_melanogaster | Eupodotis_melanogaster | LC | Decreasing | tropical | diurnal | 1151.5 | 10623922.486 | 10.099 |

| | | | | | | | | | |
|---|---|---|---|---|---|---|---|---|---|
| birds | Loboparadisea_sericea | Loboparadisea_sericea | LC | Stable | tropical | diurnal | 67.3 | 161569.22 | 4.522 |
| birds | Lobotos_lobatus | Campephaga_lobata | VU | Decreasing | tropical | diurnal | 33.1 | 343073.957 | 15.427 |
| birds | Lobotos_oriolinus | NA | DD | Stable | NA | NA | NA | NA | NA |
| birds | Lochmias_nematura | Lochmias_nematura | LC | Decreasing | temperate | diurnal | 22 | 2648897.907 | 10.988 |
| birds | Locustella_accentor | NA | LC | Stable | NA | NA | NA | NA | NA |
| birds | Locustella_alfredi | Bradypterus_alfredi | LC | Decreasing | tropical | diurnal | 17.5 | 91453.245 | 11.49 |
| birds | Locustella_castanea | Bradypterus_castaneus | LC | Stable | tropical | diurnal | 19 | 29814.466 | 5.088 |
| birds | Locustella_caudata | Bradypterus_caudatus | LC | Decreasing | tropical | diurnal | 22.9 | 24616.943 | 7.183 |
| birds | Locustella_fasciolata | NA | not_available | not_available | NA | NA | NA | NA | NA |
| birds | Locustella_fluviatilis | Locustella_fluviatilis | LC | Decreasing | tropical | diurnal | 16.1 | 29677457.723 | 9.225 |
| birds | Locustella_lanceolata | Locustella_lanceolata | LC | Stable | temperate | diurnal | 10.6 | 16637022.11 | 8.152 |
| birds | Locustella_luscinioides | Locustella_luscinioides | LC | Stable | temperate | diurnal | 13.9 | 6637595.401 | 12.819 |
| birds | Locustella_luteoventris | Bradypterus_luteoventris | LC | Stable | temperate | diurnal | 13.2 | 1710173.631 | 11.703 |
| birds | Locustella_major | NA | NT | Decreasing | NA | NA | NA | NA | NA |
| birds | Locustella_naevia | Locustella_naevia | LC | Stable | temperate | diurnal | 13.3 | 9444583.997 | 13.156 |
| birds | Locustella_seebohmi | Bradypterus_seebohmi | LC | Stable | tropical | diurnal | 11.3 | 7104.368 | 10.271 |
| birds | Locustella_tacsanowskia | Bradypterus_tacsanowskius | LC | Stable | temperate | diurnal | 14 | 8726490.531 | 11.346 |
| birds | Locustella_thoracica | Bradypterus_thoracicus | LC | Stable | temperate | diurnal | 10.8 | 1323689.33 | 10.264 |
| birds | Loddigesia_mirabilis | Loddigesia_mirabilis | NT | Decreasing | tropical | diurnal | 3 | 3663.404 | 6.957 |
| birds | Lonchura_caniceps | Lonchura_caniceps | LC | Stable | tropical | diurnal | 11.5 | 45326.558 | 4.218 |
| birds | Lonchura_castaneothorax | Lonchura_castaneothorax | LC | Stable | tropical | diurnal | 13.6 | 1876586.426 | 4.692 |
| birds | Lonchura_ferruginosa | Lonchura_ferruginosa | LC | Stable | tropical | diurnal | 13 | 136732.068 | 22.448 |
| birds | Lonchura_flaviprymna | Lonchura_flaviprymna | LC | Stable | tropical | diurnal | 11.8 | 117098.738 | 2.341 |
| birds | Lonchura_forbesi | Lonchura_forbesi | LC | Stable | tropical | diurnal | 13.3 | 6311.21 | 5.665 |
| birds | Lonchura_fuscans | Lonchura_fuscans | LC | Stable | tropical | diurnal | 9.5 | 737265.038 | 5.597 |
| birds | Lonchura_grandis | Lonchura_grandis | LC | Stable | tropical | diurnal | 14.3 | 256441.096 | 3.774 |
| birds | Lonchura_hunsteini | Lonchura_hunsteini | LC | Stable | tropical | diurnal | 13.3 | 4118.333 | 7.016 |
| birds | Lonchura_kelaarti | Lonchura_kelaarti | LC | Stable | tropical | diurnal | 14.1 | 139953.155 | 14.805 |
| birds | Lonchura_leucogastra | Lonchura_leucogastra | LC | Stable | tropical | diurnal | 11.4 | 1481477.601 | 9.989 |
| birds | Lonchura_leucogastroides | Lonchura_leucogastroides | LC | Stable | tropical | diurnal | 11.7 | 198857.571 | 19.363 |
| birds | Lonchura_leucosticta | Lonchura_leucosticta | LC | Stable | tropical | diurnal | 13.3 | 170094.55 | 2.353 |
| birds | Lonchura_maja | Lonchura_maja | LC | Stable | tropical | diurnal | 12.5 | 768204.303 | 13.166 |
| birds | Lonchura_malacca | Lonchura_malacca | LC | Stable | tropical | diurnal | 12.6 | 636026.55 | 16.985 |
| birds | Lonchura_melaena | Lonchura_melaena | LC | Stable | tropical | diurnal | 13.3 | 12280.03 | 7 |

| | | | | | | | | | |
|---|---|---|---|---|---|---|---|---|---|
| birds | Lonchura_molucca | Lonchura_molucca | LC | Stable | tropical | diurnal | 12 | 313163.122 | 10.192 |
| birds | Lonchura_montana | Lonchura_montana | LC | Stable | tropical | diurnal | 15.5 | 48855.681 | 5.259 |
| birds | Lonchura_monticola | Lonchura_monticola | LC | Stable | tropical | diurnal | 13.3 | 5354.895 | 2.94 |
| birds | Lonchura_nevermanni | Lonchura_nevermanni | LC | Stable | tropical | diurnal | 11 | 74957.626 | 2.846 |
| birds | Lonchura_pallida | Lonchura_pallida | LC | Stable | tropical | diurnal | 12 | 226245.629 | 10.952 |
| birds | Lonchura_punctulata | Lonchura_punctulata | LC | Stable | tropical | diurnal | 13.6 | 7671914.83 | 14.057 |
| birds | Lonchura_quinticolor | Lonchura_quinticolor | LC | Stable | tropical | diurnal | 13.3 | 86198.595 | 13.168 |
| birds | Lonchura_spectabilis | Lonchura_spectabilis | LC | Stable | tropical | diurnal | 11.5 | 214041.333 | 4.507 |
| birds | Lonchura_striata | Lonchura_striata | LC | Stable | tropical | diurnal | 12.3 | 5429668.122 | 14.016 |
| birds | Lonchura_stygia | Lonchura_stygia | LC | Decreasing | tropical | diurnal | 11.2 | 31370.72 | 3.527 |
| birds | Lonchura_teerinki | Lonchura_teerinki | LC | Stable | tropical | diurnal | 13.3 | 16481.572 | 4.616 |
| birds | Lonchura_tristissima | Lonchura_tristissima | LC | Stable | tropical | diurnal | 8.8 | 672961.503 | 3.209 |
| birds | Lonchura_vana | Lonchura_vana | LC | Stable | tropical | diurnal | 13.3 | 1040.816 | 4.613 |
| birds | Lophaetus_occipitalis | Lophaetus_occipitalis | LC | Increasing | tropical | diurnal | 1067 | 14031980.869 | 9.733 |
| birds | Lophoceros_alboterminatus | Tockus_alboterminatus | LC | Decreasing | tropical | diurnal | 223.7 | 3002290.123 | 10.263 |
| birds | Lophoceros_bradfieldi | Tockus_bradfieldi | LC | Decreasing | tropical | diurnal | 196.5 | 790509.69 | 5.53 |
| birds | Lophoceros_camurus | Tockus_camurus | LC | Decreasing | tropical | diurnal | 103.8 | 1703130.599 | 9.129 |
| birds | Lophoceros_fasciatus | Tockus_fasciatus | LC | Unknown | tropical | diurnal | 258.8 | 3868026.54 | 6.566 |
| birds | Lophoceros_hemprichii | Tockus_hemprichii | LC | Decreasing | tropical | diurnal | 297 | 546665.057 | 11.486 |
| birds | Lophoceros_nasutus | Tockus_nasutus | LC | Stable | tropical | diurnal | 179.6 | 9830291.233 | 10.602 |
| birds | Lophoceros_pallidirostris | Tockus_pallidirostris | LC | Decreasing | tropical | diurnal | 233 | 1957669.068 | 7.921 |
| birds | Lophodytes_cucullatus | Lophodytes_cucullatus | LC | Increasing | temperate | diurnal | 613.8 | 8843716.439 | 12.644 |
| birds | Lophoictinia_isura | Lophoictinia_isura | LC | Decreasing | temperate | diurnal | 579.4 | 5951770.473 | 4.953 |
| birds | Lopholaimus_antarcticus | Lopholaimus_antarcticus | LC | Decreasing | temperate | diurnal | 496.4 | 317014.629 | 7.435 |
| birds | Lophonetta_specularioides | Lophonetta_specularioides | LC | Stable | temperate | diurnal | 1019.9 | 1859148.867 | 5.583 |
| birds | Lophophanes_cristatus | Parus_cristatus | LC | Decreasing | temperate | diurnal | 11 | 5918314.938 | 14.813 |
| birds | Lophophanes_dichrous | Parus_dichrous | LC | Stable | temperate | diurnal | 13.8 | 968325.053 | 8.655 |
| birds | Lophophorus_impejanus | Lophophorus_impejanus | LC | Decreasing | temperate | diurnal | 2066.5 | 480828.137 | 6.712 |
| birds | Lophophorus_lhuysii | Lophophorus_lhuysii | VU | Decreasing | temperate | diurnal | 3008 | 175682.11 | 7.872 |
| birds | Lophophorus_sclateri | Lophophorus_sclateri | LC | Decreasing | temperate | diurnal | 2500 | 185023.027 | 4.236 |
| birds | Lophorina_intercedens | Ptiloris_intercedens | LC | Decreasing | tropical | diurnal | 158 | 104258.24 | 4.51 |
| birds | Lophorina_magnifica | Ptiloris_magnificus | LC | Decreasing | tropical | diurnal | 149.9 | 495422.812 | 2.835 |
| birds | Lophorina_paradisea | Ptiloris_paradiseus | LC | Decreasing | temperate | diurnal | 119.5 | 128251.121 | 8.429 |
| birds | Lophorina_superba | Lophorina_superba | LC | Decreasing | tropical | diurnal | 76.7 | 14774.576 | 2.852 |

| | | | | | | | | | |
|---|---|---|---|---|---|---|---|---|---|
| birds | Lophorina_victoriae | Ptiloris_victoriae | VU | Decreasing | tropical | diurnal | 95 | 11240.764 | 3.572 |
| birds | Lophornis_adorabilis | Lophornis_adorabilis | LC | Decreasing | tropical | diurnal | 2.7 | 16797.998 | 16.487 |
| birds | Lophornis_brachylophus | Lophornis_brachylophus | CR | Decreasing | tropical | diurnal | 2.7 | 1180.346 | 3.111 |
| birds | Lophornis_chalybeus | Lophornis_chalybeus | LC | Stable | temperate | diurnal | 3 | 192385.772 | 13.705 |
| birds | Lophornis_delattrei | Lophornis_delattrei | LC | Decreasing | tropical | diurnal | 2.8 | 227133.7 | 5.499 |
| birds | Lophornis_gouldii | Lophornis_gouldii | NT | Decreasing | tropical | diurnal | 2.4 | 1585965.489 | 4.452 |
| birds | Lophornis_helenae | Lophornis_helenae | LC | Decreasing | tropical | diurnal | 2.6 | 254041.649 | 10.434 |
| birds | Lophornis_magnificus | Lophornis_magnificus | LC | Unknown | tropical | diurnal | 2.7 | 2557177.007 | 11.068 |
| birds | Lophornis_ornatus | Lophornis_ornatus | LC | Stable | tropical | diurnal | 2.3 | 828805.981 | 1.962 |
| birds | Lophornis_pavoninus | Lophornis_pavoninus | LC | Unknown | tropical | diurnal | 3 | 206408.305 | 0.987 |
| birds | Lophornis_stictolophus | Lophornis_stictolophus | LC | Unknown | tropical | diurnal | 2.7 | 134543.997 | 10.647 |
| birds | Lophospingus_griseocristatus | Lophospingus_griseocristatus | LC | Stable | tropical | diurnal | 17.9 | 84852.785 | 8.494 |
| birds | Lophospingus_pusillus | Lophospingus_pusillus | LC | Stable | temperate | diurnal | 14.4 | 559474.539 | 5.434 |
| birds | Lophostrix_cristata | Lophostrix_cristata | LC | Decreasing | tropical | nocturnal | 538.7 | 5025098.507 | 3.078 |
| birds | Lophotibis_cristata | Lophotibis_cristata | NT | Decreasing | tropical | diurnal | 583 | 535452.804 | 8.582 |
| birds | Lophotis_gindiana | Eupodotis_gindiana | LC | Stable | tropical | diurnal | 782.3 | 1777669.918 | 8.893 |
| birds | Lophotis_ruficrista | Eupodotis_ruficrista | LC | Stable | tropical | diurnal | 712.1 | 2013859.392 | 7.315 |
| birds | Lophotis_savilei | NA | LC | Stable | NA | NA | NA | NA | NA |
| birds | Lophotriccus_eulophotes | Lophotriccus_eulophotes | LC | Decreasing | tropical | diurnal | 6 | 328683.826 | 1.515 |
| birds | Lophotriccus_galeatus | Lophotriccus_galeatus | LC | Decreasing | tropical | diurnal | 6.6 | 2884993.061 | 1.763 |
| birds | Lophotriccus_pileatus | Lophotriccus_pileatus | LC | Decreasing | tropical | diurnal | 7.5 | 475468.117 | 8.484 |
| birds | Lophotriccus_vitiosus | Lophotriccus_vitiosus | LC | Stable | tropical | diurnal | 6.7 | 2198953.696 | 1.478 |
| birds | Lophotriorchis_kienerii | Lophotriorchis_kienerii | NT | Decreasing | tropical | diurnal | 799 | 3758691.083 | 11.485 |
| birds | Lophura_bulweri | Lophura_bulweri | VU | Decreasing | tropical | diurnal | 1245.2 | 274609.441 | 2.979 |
| birds | Lophura_diardi | Lophura_diardi | LC | Decreasing | tropical | diurnal | 1088.9 | 510759.032 | 9.736 |
| birds | Lophura_edwardsi | Lophura_edwardsi | CR | Decreasing | tropical | diurnal | 1091 | 8246.946 | 11.221 |
| birds | Lophura_erythrophthalma | Lophura_erythrophthalma | CR | Decreasing | tropical | diurnal | 934.3 | 374809.688 | 11.566 |
| birds | Lophura_ignita | Lophura_ignita | VU | Decreasing | tropical | diurnal | 1768.1 | 486548.863 | 5.086 |
| birds | Lophura_inornata | Lophura_inornata | NT | Decreasing | tropical | diurnal | 685 | 71415.825 | 7.051 |
| birds | Lophura_leucomelanos | Lophura_leucomelanos | LC | Decreasing | temperate | diurnal | 868 | 1315909.309 | 9.43 |
| birds | Lophura_nycthemera | Lophura_nycthemera | LC | Decreasing | tropical | diurnal | 1230 | 1072525.933 | 10.018 |
| birds | Loriculus_amabilis | NA | LC | Stable | NA | NA | NA | NA | NA |
| birds | Loriculus_aurantiifrons | Loriculus_aurantiifrons | LC | Stable | tropical | diurnal | 15.5 | 573982.763 | 3.391 |
| birds | Loriculus_beryllinus | Loriculus_beryllinus | LC | Stable | tropical | diurnal | 30 | 16049.969 | 18.967 |

| | | | | | | | | | |
|---|---|---|---|---|---|---|---|---|---|
| birds | Loriculus_catamene | Loriculus_catamene | NT | Decreasing | tropical | diurnal | 34 | 556.531 | 11.584 |
| birds | Loriculus_exilis | Loriculus_exilis | LC | Decreasing | tropical | diurnal | 34 | 104090.106 | 8.131 |
| birds | Loriculus_flosculus | Loriculus_flosculus | VU | Decreasing | tropical | diurnal | 34 | 1417.69 | 10.765 |
| birds | Loriculus_galgulus | Loriculus_galgulus | LC | Stable | tropical | diurnal | 28 | 1413389.846 | 8.147 |
| birds | Loriculus_philippensis | Loriculus_philippensis | LC | Decreasing | tropical | diurnal | 34.9 | 276863.003 | 13.875 |
| birds | Loriculus_pusillus | NA | NT | Decreasing | NA | NA | NA | NA | NA |
| birds | Loriculus_sclateri | Loriculus_sclateri | LC | Stable | tropical | diurnal | 34 | 7519.863 | 6.941 |
| birds | Loriculus_stigmatus | NA | LC | Stable | NA | NA | NA | NA | NA |
| birds | Loriculus_tener | Loriculus_tener | LC | Decreasing | tropical | diurnal | 12 | 30581.568 | 5.024 |
| birds | Loriculus_vernalis | Loriculus_vernalis | LC | Stable | tropical | diurnal | 35.7 | 2278295.574 | 12.632 |
| birds | Lorius_albidinucha | Lorius_albidinucha | NT | Decreasing | tropical | diurnal | 132 | 2255.829 | 3.684 |
| birds | Lorius_chlorocercus | Lorius_chlorocercus | LC | Decreasing | tropical | diurnal | 173.5 | 13625.11 | 5.188 |
| birds | Lorius_domicella | Lorius_domicella | EN | Decreasing | tropical | diurnal | 182 | 6298.737 | 5.778 |
| birds | Lorius_garrulus | Lorius_garrulus | VU | Decreasing | tropical | diurnal | 143 | 25448.005 | 6.243 |
| birds | Lorius_hypoinochrous | Lorius_hypoinochrous | LC | Stable | tropical | diurnal | 187 | 61928.795 | 4.55 |
| birds | Lorius_lory | Lorius_lory | LC | Decreasing | tropical | diurnal | 176.9 | 663810.012 | 3.175 |
| birds | Loxia_curvirostra | Loxia_curvirostra | LC | Stable | temperate | diurnal | 38.3 | 29578527.672 | 9.146 |
| birds | Loxia_leucoptera | Loxia_leucoptera | LC | Decreasing | temperate | diurnal | 28.7 | 20422272.711 | 5.769 |
| birds | Loxia_pytyopsittacus | Loxia_pytyopsittacus | LC | Stable | temperate | diurnal | 53 | 3155285.386 | 8.42 |
| birds | Loxia_scotica | Loxia_scotica | LC | Stable | temperate | diurnal | 43.2 | 13734.015 | 9.673 |
| birds | Loxigilla_noctis | Loxigilla_noctis | LC | Stable | tropical | diurnal | 16.6 | 6167.157 | 22.091 |
| birds | Loxioides_bailleui | Loxioides_bailleui | CR | Decreasing | tropical | diurnal | 37.8 | 1374.08 | 3.529 |
| birds | Loxipasser_anoxanthus | Loxipasser_anoxanthus | LC | Decreasing | tropical | diurnal | 11.4 | 11021.791 | 19.741 |
| birds | Loxops_caeruleirostris | Loxops_caeruleirostris | CR | Decreasing | tropical | diurnal | 10.8 | 5.222 | 5.624 |
| birds | Loxops_coccineus | Loxops_coccineus | EN | Decreasing | tropical | diurnal | 10.5 | 8106.925 | 7.325 |
| birds | Lullula_arborea | Lullula_arborea | LC | Increasing | temperate | diurnal | 26.9 | 8808902.549 | 16.817 |
| birds | Lurocalis_rufiventris | Lurocalis_rufiventris | LC | Decreasing | tropical | nocturnal | 128.5 | 307432.645 | 8.2 |
| birds | Lurocalis_semitorquatus | Lurocalis_semitorquatus | LC | Decreasing | tropical | nocturnal | 75.9 | 8957931.233 | 4.614 |
| birds | Luscinia_luscinia | Luscinia_luscinia | LC | Stable | temperate | diurnal | 23.8 | 8587306.747 | 10.832 |
| birds | Luscinia_megarhynchos | Luscinia_megarhynchos | LC | Stable | tropical | diurnal | 19.6 | 11512328.434 | 13.475 |
| birds | Luscinia_phaenicuroides | Hodgsonius_phaenicuroides | LC | Stable | temperate | diurnal | 21.1 | 1681247.712 | 8.542 |
| birds | Luscinia_svecica | Luscinia_svecica | LC | Stable | temperate | diurnal | 17.2 | 61151327.316 | 8.055 |
| birds | Lybius_chaplini | Lybius_chaplini | VU | Decreasing | tropical | diurnal | 69.5 | 76477.214 | 9.7 |
| birds | Lybius_guifsobalito | Lybius_guifsobalito | LC | Increasing | tropical | diurnal | 42.2 | 1310708.12 | 11.43 |

| | | | | | | | | | |
|---|---|---|---|---|---|---|---|---|---|
| birds | Lybius_leucocephalus | Lybius_leucocephalus | LC | Stable | tropical | diurnal | 62.8 | 1924719.609 | 10.186 |
| birds | Lybius_rubrifacies | Lybius_rubrifacies | NT | Decreasing | tropical | diurnal | 60.1 | 82626.47 | 16.766 |
| birds | Lybius_torquatus | Lybius_torquatus | LC | Decreasing | tropical | diurnal | 51.7 | 4663900.428 | 8.914 |
| birds | Lybius_undatus | Lybius_undatus | LC | Unknown | tropical | diurnal | 38.5 | 580218.291 | 11.701 |
| birds | Lybius_vieilloti | Lybius_vieilloti | LC | Unknown | tropical | diurnal | 36 | 6618281.306 | 10.491 |
| birds | Lycocorax_pyrrhopterus | Lycocorax_pyrrhopterus | LC | Stable | tropical | diurnal | 283.9 | 22792.492 | 6.411 |
| birds | Lymnocryptes_minimus | Lymnocryptes_minimus | LC | Stable | temperate | nocturnal | 50.1 | 19801071.209 | 9.09 |
| birds | Lyncornis_macrotis | Eurostopodus_macrotis | LC | Stable | tropical | nocturnal | 140.6 | 1650527.298 | 10.174 |
| birds | Lyncornis_temminckii | Eurostopodus_temminckii | LC | Stable | tropical | nocturnal | 95.9 | 1296744.776 | 7.874 |
| birds | Lyrurus_tetrix | Tetrao_tetrix | LC | Decreasing | temperate | diurnal | 1068.7 | 14007422.539 | 6.635 |
| birds | Macgregoria_pulchra | Macgregoria_pulchra | VU | Decreasing | tropical | diurnal | 225.1 | 14172.406 | 4.529 |
| birds | Machaerirhynchus_flaviventer | Machaerirhynchus_flaviventer | LC | Stable | tropical | diurnal | 10.3 | 708932.931 | 3.193 |
| birds | Machaerirhynchus_nigripectus | Machaerirhynchus_nigripectus | LC | Unknown | tropical | diurnal | 12 | 223752.258 | 4.334 |
| birds | Machaeropterus_deliciosus | Machaeropterus_deliciosus | LC | Decreasing | tropical | diurnal | 12.7 | 40123.574 | 6.986 |
| birds | Machaeropterus_pyrocephalus | Machaeropterus_pyrocephalus | LC | Decreasing | tropical | diurnal | 9.5 | 2227740.971 | 3.417 |
| birds | Machaeropterus_regulus | Machaeropterus_regulus | LC | Decreasing | tropical | diurnal | 9.3 | 125343.348 | 11.859 |
| birds | Macheiramphus_alcinus | Macheiramphus_alcinus | LC | Stable | tropical | diurnal | 650 | 13573105.09 | 9.252 |
| birds | Machetornis_rixosa | Machetornis_rixosa | LC | Stable | tropical | diurnal | 29.6 | 6805756.6 | 9.154 |
| birds | Machlolophus_holsti | Parus_holsti | LC | Stable | temperate | diurnal | 16.7 | 12987.011 | 6.729 |
| birds | Machlolophus_nuchalis | Parus_nuchalis | VU | Decreasing | tropical | diurnal | 13.8 | 75702.744 | 18.656 |
| birds | Machlolophus_spilonotus | Parus_spilonotus | LC | Increasing | tropical | diurnal | 18.8 | 1250920.253 | 9.985 |
| birds | Machlolophus_xanthogenys | Parus_xanthogenys | LC | Stable | temperate | diurnal | 14.8 | 1118561.25 | 16.45 |
| birds | Mackenziaena_leachii | Mackenziaena_leachii | LC | Decreasing | temperate | diurnal | 70.2 | 828525.046 | 12.869 |
| birds | Mackenziaena_severa | Mackenziaena_severa | LC | Stable | tropical | diurnal | 51.8 | 955402.975 | 13.594 |
| birds | Macroagelaius_imthurni | Macroagelaius_imthurni | LC | Stable | tropical | diurnal | 77.8 | 58909.677 | 1.479 |
| birds | Macroagelaius_subalaris | Macroagelaius_subalaris | EN | Decreasing | tropical | diurnal | 62 | 40245.612 | 12.54 |
| birds | Macrocephalon_maleo | Macrocephalon_maleo | CR | Decreasing | tropical | diurnal | 1563.3 | 109982.377 | 7.254 |
| birds | Macronectes_giganteus | Macronectes_giganteus | LC | Increasing | temperate | cathemeral | 3591.7 | 100191496.873 | 8.786 |
| birds | Macronectes_halli | Macronectes_halli | LC | Increasing | temperate | cathemeral | 4206.3 | 91967378.632 | 6.278 |
| birds | Macronus_ptilosus | Macronous_ptilosus | NT | Decreasing | tropical | diurnal | 18 | 444346.102 | 9.65 |
| birds | Macronus_striaticeps | Macronous_striaticeps | LC | Decreasing | tropical | diurnal | 16.5 | 109828 | 11.49 |
| birds | Macronyx_ameliae | Macronyx_ameliae | LC | Decreasing | tropical | diurnal | 33.5 | 1902624.72 | 7.488 |
| birds | Macronyx_aurantiigula | Macronyx_aurantiigula | LC | Stable | tropical | diurnal | 39.4 | 375644.562 | 11.688 |
| birds | Macronyx_capensis | Macronyx_capensis | LC | Stable | temperate | diurnal | 50 | 815796.1 | 12.626 |

| | | | | | | | | | |
|---|---|---|---|---|---|---|---|---|---|
| birds | Macronyx_croceus | Macronyx_croceus | LC | Stable | tropical | diurnal | 47.1 | 6192075.067 | 11.838 |
| birds | Macronyx_flavicollis | Macronyx_flavicollis | NT | Decreasing | tropical | diurnal | 45.1 | 404989.726 | 12.243 |
| birds | Macronyx_fuelleborni | Macronyx_fuellebornii | LC | Stable | tropical | diurnal | 55.2 | 2083060.163 | 7.119 |
| birds | Macronyx_grimwoodi | Macronyx_grimwoodi | LC | Stable | tropical | diurnal | 45.1 | 671345.147 | 5.666 |
| birds | Macronyx_sharpei | Macronyx_sharpei | EN | Decreasing | tropical | diurnal | 29.9 | 2927.167 | 16.286 |
| birds | Macropsalis_forcipata | Macropsalis_forcipata | LC | Decreasing | temperate | nocturnal | 139 | 517277.527 | 12.553 |
| birds | Macropygia_amboinensis | Macropygia_amboinensis | LC | Stable | tropical | diurnal | 237 | 1059941.302 | 4.625 |
| birds | Macropygia_emiliana | Macropygia_emiliana | LC | Decreasing | tropical | diurnal | 161 | 186249.02 | 19.777 |
| birds | Macropygia_mackinlayi | Macropygia_mackinlayi | LC | Stable | tropical | diurnal | 85.9 | 54491.472 | 4.974 |
| birds | Macropygia_magna | NA | LC | Stable | NA | NA | NA | NA | NA |
| birds | Macropygia_nigrirostris | Macropygia_nigrirostris | LC | Stable | tropical | diurnal | 90.1 | 646983.573 | 3.826 |
| birds | Macropygia_phasianella | Macropygia_phasianella | LC | Stable | temperate | diurnal | 237 | 582030.418 | 6.546 |
| birds | Macropygia_ruficeps | Macropygia_ruficeps | LC | Stable | tropical | diurnal | 81.9 | 1783125.512 | 8.858 |
| birds | Macropygia_rufipennis | Macropygia_rufipennis | LC | Decreasing | tropical | diurnal | 256 | 7597.74 | 6.777 |
| birds | Macropygia_tenuirostris | Macropygia_tenuirostris | LC | Decreasing | tropical | diurnal | 177 | 563655.836 | 8.261 |
| birds | Macropygia_unchall | Macropygia_unchall | LC | Stable | tropical | diurnal | 168 | 2550794.478 | 10.499 |
| birds | Macrosphenus_concolor | Macrosphenus_concolor | LC | Stable | tropical | diurnal | 13.9 | 3119489.654 | 8.327 |
| birds | Macrosphenus_flavicans | Macrosphenus_flavicans | LC | Stable | tropical | diurnal | 13.4 | 1810288.683 | 5.863 |
| birds | Macrosphenus_kempi | Macrosphenus_kempi | LC | Stable | tropical | diurnal | 12.6 | 456072.284 | 15.623 |
| birds | Macrosphenus_kretschmeri | Macrosphenus_kretschmeri | LC | Decreasing | tropical | diurnal | 20 | 120112.339 | 11.848 |
| birds | Macrosphenus_pulitzeri | Macrosphenus_pulitzeri | EN | Decreasing | tropical | diurnal | 11.7 | 5725.087 | 9.892 |
| birds | Madanga_ruficollis | Madanga_ruficollis | EN | Decreasing | tropical | diurnal | 15 | 1781.705 | 4.056 |
| birds | Magumma_parva | Hemignathus_parvus | EN | Decreasing | tropical | diurnal | 9.3 | 1432.919 | 9.654 |
| birds | Malacocincla_abbotti | Malacocincla_abbotti | LC | Stable | tropical | diurnal | 26.8 | 2072332.429 | 8.925 |
| birds | Malacocincla_perspicillata | NA | DD | Decreasing | NA | NA | NA | NA | NA |
| birds | Malacocincla_sepiaria | Malacocincla_sepiaria | LC | Decreasing | tropical | diurnal | 25.5 | 969773.936 | 8.809 |
| birds | Malaconotus_alius | NA | EN | Stable | NA | NA | NA | NA | NA |
| birds | Malaconotus_blanchoti | Malaconotus_blanchoti | LC | Increasing | tropical | diurnal | 78.7 | 7329839.515 | 10.13 |
| birds | Malaconotus_gladiator | Malaconotus_gladiator | VU | Decreasing | tropical | diurnal | 99 | 3290.457 | 11.82 |
| birds | Malaconotus_lagdeni | Malaconotus_lagdeni | NT | Decreasing | tropical | diurnal | 92.3 | 338592.196 | 14.79 |
| birds | Malaconotus_monteiri | Malaconotus_monteiri | NT | Decreasing | tropical | diurnal | 81.9 | 27000.525 | 8.594 |
| birds | Malacopteron_affine | Malacopteron_affine | NT | Decreasing | tropical | diurnal | 17.1 | 387896.02 | 9.751 |
| birds | Malacopteron_albogulare | Malacopteron_albogulare | NT | Decreasing | tropical | diurnal | 16.5 | 597337.377 | 6.604 |
| birds | Malacopteron_cinereum | Malacopteron_cinereum | LC | Stable | tropical | diurnal | 18.1 | 1526343.714 | 9.177 |

| | | | | | | | | | |
|---|---|---|---|---|---|---|---|---|---|
| birds | Malacopteron_magnirostre | Malacopteron_magnirostre | LC | Stable | tropical | diurnal | 19.3 | 1312918.001 | 7.851 |
| birds | Malacopteron_magnum | Malacopteron_magnum | NT | Decreasing | tropical | diurnal | 27.2 | 666939.159 | 5.915 |
| birds | Malacopteron_palawanense | Malacopteron_palawanense | NT | Decreasing | tropical | diurnal | 32.1 | 3507.145 | 11.312 |
| birds | Malacoptila_fulvogularis | Malacoptila_fulvogularis | LC | Decreasing | tropical | diurnal | 57.4 | 120052.812 | 5.169 |
| birds | Malacoptila_fusca | Malacoptila_fusca | LC | Stable | tropical | diurnal | 45.4 | 1912315.395 | 1.86 |
| birds | Malacoptila_mystacalis | Malacoptila_mystacalis | LC | Stable | tropical | diurnal | 49 | 94080.956 | 10.156 |
| birds | Malacoptila_panamensis | Malacoptila_panamensis | LC | Decreasing | tropical | diurnal | 42.6 | 516560.602 | 9.208 |
| birds | Malacoptila_rufa | Malacoptila_rufa | LC | Decreasing | tropical | diurnal | 47.5 | 2602548.701 | 2.036 |
| birds | Malacoptila_semicincta | Malacoptila_semicincta | LC | Decreasing | tropical | diurnal | 44 | 649678.522 | 1.457 |
| birds | Malacoptila_striata | Malacoptila_striata | LC | Decreasing | tropical | diurnal | 44.1 | 706984.899 | 13.316 |
| birds | Malacorhynchus_membranaceus | Malacorhynchus_membranaceus | LC | Stable | temperate | diurnal | 372.8 | 6997669.669 | 4.521 |
| birds | Malcorus_pectoralis | Malcorus_pectoralis | LC | Stable | temperate | diurnal | 10.1 | 1273750.667 | 6.96 |
| birds | Malia_grata | Malia_grata | LC | Decreasing | tropical | diurnal | 94.5 | 43962.269 | 5.587 |
| birds | Malimbus_ballmanni | Malimbus_ballmanni | NT | Decreasing | tropical | diurnal | 63 | 38326.098 | 8.942 |
| birds | Malimbus_cassini | Malimbus_cassini | LC | Stable | tropical | diurnal | 35 | 940615.557 | 5.311 |
| birds | Malimbus_coronatus | Malimbus_coronatus | LC | Decreasing | tropical | diurnal | 32 | 395465.271 | 5.589 |
| birds | Malimbus_erythrogaster | Malimbus_erythrogaster | LC | Decreasing | tropical | diurnal | 45 | 1446184.322 | 6.153 |
| birds | Malimbus_ibadanensis | Malimbus_ibadanensis | EN | Decreasing | tropical | diurnal | 38.8 | 23474.801 | 18.789 |
| birds | Malimbus_malimbicus | Malimbus_malimbicus | LC | Stable | tropical | diurnal | 36.1 | 2471944.166 | 7.897 |
| birds | Malimbus_nitens | Malimbus_nitens | LC | Stable | tropical | diurnal | 37.8 | 2636859.51 | 8.103 |
| birds | Malimbus_racheliae | Malimbus_racheliae | LC | Decreasing | tropical | diurnal | 26 | 209703.783 | 7.505 |
| birds | Malimbus_rubricollis | Malimbus_rubricollis | LC | Stable | tropical | diurnal | 42.2 | 2607128.644 | 8.502 |
| birds | Malimbus_scutatus | Malimbus_scutatus | LC | Stable | tropical | diurnal | 30 | 857359.487 | 14.98 |
| birds | Malindangia_mcgregori | Coracina_mcgregori | LC | Decreasing | tropical | diurnal | 42.6 | 13707.887 | 7.552 |
| birds | Malurus_alboscapulatus | Malurus_alboscapulatus | LC | Stable | tropical | diurnal | 9.3 | 778806.382 | 3.43 |
| birds | Malurus_amabilis | Malurus_amabilis | LC | Stable | tropical | diurnal | 8.6 | 126910.612 | 3.109 |
| birds | Malurus_coronatus | Malurus_coronatus | LC | Decreasing | tropical | diurnal | 11.1 | 363760.078 | 2.877 |
| birds | Malurus_cyaneus | Malurus_cyaneus | LC | Stable | temperate | diurnal | 10.3 | 1239779.313 | 9.053 |
| birds | Malurus_cyanocephalus | Malurus_cyanocephalus | LC | Stable | tropical | diurnal | 12.4 | 491129.371 | 2.994 |
| birds | Malurus_elegans | Malurus_elegans | LC | Decreasing | temperate | diurnal | 9.6 | 71313.857 | 11.216 |
| birds | Malurus_lamberti | Malurus_lamberti | LC | Stable | temperate | diurnal | 8 | 212534.109 | 8.31 |
| birds | Malurus_leucopterus | Malurus_leucopterus | LC | Stable | temperate | diurnal | 7.4 | 5612678.07 | 3.977 |
| birds | Malurus_melanocephalus | Malurus_melanocephalus | LC | Stable | tropical | diurnal | 7.9 | 2025747.646 | 3.776 |
| birds | Malurus_pulcherrimus | Malurus_pulcherrimus | LC | Stable | temperate | diurnal | 9.2 | 379685.746 | 8.464 |

| | | | | | | | | | |
|---|---|---|---|---|---|---|---|---|---|
| birds | Malurus_splendens | Malurus_splendens | LC | Stable | temperate | diurnal | 9.4 | 2710216.864 | 4.73 |
| birds | Manacus_aurantiacus | Manacus_aurantiacus | LC | Decreasing | tropical | diurnal | 15.5 | 32298.088 | 13.205 |
| birds | Manacus_candei | Manacus_candei | LC | Decreasing | tropical | diurnal | 19.8 | 283521.134 | 8.221 |
| birds | Manacus_manacus | Manacus_manacus | LC | Stable | tropical | diurnal | 16.7 | 7524946.194 | 4.263 |
| birds | Manacus_vitellinus | Manacus_vitellinus | LC | Decreasing | tropical | diurnal | 18.2 | 118914.747 | 6.999 |
| birds | Mandingoa_nitidula | Mandingoa_nitidula | LC | Stable | tropical | diurnal | 9.6 | 1540214.12 | 11.277 |
| birds | Manorina_flavigula | Manorina_flavigula | LC | Stable | temperate | diurnal | 55.9 | 6815495.206 | 3.882 |
| birds | Manorina_melanocephala | Manorina_melanocephala | LC | Increasing | temperate | diurnal | 60.2 | 2065259.237 | 7.548 |
| birds | Manorina_melanophrys | Manorina_melanophrys | LC | Decreasing | temperate | diurnal | 30.4 | 317354.762 | 8.649 |
| birds | Manorina_melanotis | Manorina_melanotis | EN | Decreasing | temperate | diurnal | 54 | 66850.624 | 9.299 |
| birds | Manucerthia_mana | Oreomystis_mana | EN | Decreasing | tropical | diurnal | 14.6 | 763.594 | 3.343 |
| birds | Manucodia_ater | Manucodia_ater | LC | Decreasing | tropical | diurnal | 228 | 136828.138 | 4.028 |
| birds | Manucodia_chalybatus | Manucodia_chalybatus | LC | Decreasing | tropical | diurnal | 215.9 | 81016.844 | 3.727 |
| birds | Manucodia_comrii | Manucodia_comrii | LC | Decreasing | tropical | diurnal | 432.7 | 3220.809 | 5.999 |
| birds | Manucodia_jobiensis | Manucodia_jobiensis | LC | Decreasing | tropical | diurnal | 202.6 | 22487.645 | 3.542 |
| birds | Mareca_americana | Anas_americana | LC | Decreasing | temperate | diurnal | 754.6 | 15063927.377 | 7.757 |
| birds | Mareca_falcata | Anas_falcata | NT | Decreasing | temperate | diurnal | 645.8 | 12027658.536 | 7.981 |
| birds | Mareca_penelope | Anas_penelope | LC | Decreasing | temperate | diurnal | 770 | 29250559.935 | 8.1 |
| birds | Mareca_sibilatrix | Anas_sibilatrix | LC | Stable | temperate | diurnal | 923.2 | 3564873.368 | 8.017 |
| birds | Mareca_strepera | Anas_strepera | LC | Increasing | temperate | diurnal | 915.6 | 34553002.066 | 11.097 |
| birds | Margaroperdix_madagarensis | Margaroperdix_madagascariensis | LC | Decreasing | tropical | diurnal | 245 | 545800.129 | 8.574 |
| birds | Margarops_fuscatus | Margarops_fuscatus | LC | Stable | tropical | diurnal | 98.4 | 20593.033 | 20.947 |
| birds | Margarornis_bellulus | Margarornis_bellulus | NT | Stable | tropical | diurnal | 18.5 | 270.82 | 1.961 |
| birds | Margarornis_rubiginosus | Margarornis_rubiginosus | LC | Stable | tropical | diurnal | 17.8 | 5601.93 | 6.437 |
| birds | Margarornis_squamiger | Margarornis_squamiger | LC | Stable | tropical | diurnal | 17.3 | 469622.441 | 8.868 |
| birds | Margarornis_stellatus | Margarornis_stellatus | NT | Decreasing | tropical | diurnal | 21.5 | 21929.049 | 8.103 |
| birds | Marmaronetta_angustirostris | Marmaronetta_angustirostris | NT | Decreasing | temperate | diurnal | 477 | 12485474.466 | 7.854 |
| birds | Maschalethraupis_surinama | Tachyphonus_surinamus | LC | Decreasing | tropical | diurnal | 19 | 5574463.696 | 1.817 |
| birds | Masius_chrysopterus | Masius_chrysopterus | LC | Stable | tropical | diurnal | 11.5 | 176828.862 | 8.325 |
| birds | Mayrornis_lessoni | Mayrornis_lessoni | LC | Decreasing | tropical | diurnal | 11.2 | 17898.174 | 9.368 |
| birds | Mayrornis_schistaceus | NA | VU | Decreasing | NA | NA | NA | NA | NA |
| birds | Mayrornis_versicolor | NA | NT | Stable | NA | NA | NA | NA | NA |
| birds | Mazaria_propinqua | Synallaxis_propinqua | LC | Stable | tropical | diurnal | 19.9 | 531677.408 | 2.206 |
| birds | Mearnsia_picina | Mearnsia_picina | NT | Decreasing | tropical | diurnal | 42.5 | 112656.134 | 13.279 |

| | | | | | | | | | |
|---|---|---|---|---|---|---|---|---|---|
| birds | Mecocerculus_calopterus | Mecocerculus_calopterus | LC | Stable | tropical | diurnal | 7 | 58759.678 | 7.899 |
| birds | Mecocerculus_hellmayri | Mecocerculus_hellmayri | LC | Stable | tropical | diurnal | 8 | 69941.23 | 6.642 |
| birds | Mecocerculus_leucophrys | Mecocerculus_leucophrys | LC | Stable | tropical | diurnal | 11 | 885630.6 | 7.288 |
| birds | Mecocerculus_minor | Mecocerculus_minor | LC | Increasing | tropical | diurnal | 11 | 62120.594 | 5.377 |
| birds | Mecocerculus_poecilocercus | Mecocerculus_poecilocercus | LC | Stable | tropical | diurnal | 7.7 | 177426.015 | 8.263 |
| birds | Mecocerculus_stictopterus | Mecocerculus_stictopterus | LC | Stable | tropical | diurnal | 9.4 | 265969.643 | 9.072 |
| birds | Megabyas_flammulatus | Megabyas_flammulatus | LC | Decreasing | tropical | diurnal | 28 | 4083781.249 | 8.395 |
| birds | Megaceryle_alcyon | Megaceryle_alcyon | LC | Stable | temperate | diurnal | 148 | 18076084.923 | 8.897 |
| birds | Megaceryle_lugubris | Megaceryle_lugubris | LC | Decreasing | temperate | diurnal | 271 | 5024200.397 | 15.089 |
| birds | Megaceryle_maxima | Megaceryle_maxima | LC | Decreasing | tropical | diurnal | 325 | 13133154.898 | 9.443 |
| birds | Megaceryle_torquata | Megaceryle_torquata | LC | Stable | tropical | diurnal | 317 | 16295521.194 | 6.331 |
| birds | Megadyptes_antipodes | Megadyptes_antipodes | EN | Decreasing | temperate | cathemeral | 5326.2 | 80802.253 | 12.612 |
| birds | Megalampitta_gigantea | Melampitta_gigantea | LC | Stable | tropical | diurnal | 205 | 8372.654 | 2.922 |
| birds | Megaloprepia_formosa | Ptilinopus_bernsteinii | LC | Decreasing | tropical | diurnal | 118.5 | 21523.733 | 6.469 |
| birds | Megaloprepia_magnifica | Ptilinopus_magnificus | LC | Decreasing | tropical | diurnal | 185.4 | 979740.406 | 4.256 |
| birds | Megalurulus_whitneyi | NA | not_available | not_available | NA | NA | NA | NA | NA |
| birds | Megalurus_palustris | Megalurus_palustris | LC | Unknown | tropical | diurnal | 44.1 | 2258407.86 | 14.642 |
| birds | Megapodius_bernsteinii | Megapodius_bernsteinii | VU | Decreasing | tropical | diurnal | 768.6 | 4713.271 | 7.133 |
| birds | Megapodius_cumingii | Megapodius_cumingii | LC | Decreasing | tropical | diurnal | 1044 | 471034.038 | 11.128 |
| birds | Megapodius_decollatus | Megapodius_affinis | LC | Stable | tropical | diurnal | 666.3 | 239677.892 | 3.465 |
| birds | Megapodius_eremita | Megapodius_eremita | LC | Decreasing | tropical | diurnal | 676.7 | 84419.947 | 4.602 |
| birds | Megapodius_freycinet | Megapodius_freycinet | LC | Decreasing | tropical | diurnal | 642.5 | 44006.889 | 6.987 |
| birds | Megapodius_laperouse | Megapodius_laperouse | NT | Unknown | tropical | diurnal | 308 | 1505.274 | 17.558 |
| birds | Megapodius_layardi | Megapodius_layardi | VU | Decreasing | tropical | diurnal | 768.6 | 6809.734 | 7.25 |
| birds | Megapodius_nicobariensis | Megapodius_nicobariensis | VU | Stable | tropical | diurnal | 836.7 | 2301.928 | 5.075 |
| birds | Megapodius_pritchardii | Megapodius_pritchardii | VU | Increasing | tropical | diurnal | 365 | 346.171 | 8.984 |
| birds | Megapodius_reinwardt | Megapodius_reinwardt | LC | Stable | tropical | diurnal | 875.8 | 795651.652 | 3.657 |
| birds | Megarynchus_pitangua | Megarynchus_pitangua | LC | Stable | tropical | diurnal | 69.9 | 13243099.503 | 5.866 |
| birds | Megascops_albogularis | Megascops_albogularis | LC | Stable | tropical | nocturnal | 159 | 244881.909 | 9.162 |
| birds | Megascops_asio | Megascops_asio | LC | Decreasing | temperate | cathemeral | 180 | 4938942.172 | 16.532 |
| birds | Megascops_atricapilla | Megascops_atricapilla | LC | Stable | tropical | nocturnal | 118.9 | 990012.513 | 13.775 |
| birds | Megascops_choliba | Megascops_choliba | LC | Decreasing | tropical | nocturnal | 132 | 13065137.778 | 6.031 |
| birds | Megascops_clarkii | Megascops_clarkii | LC | Stable | tropical | nocturnal | 158.5 | 9927.9 | 7.824 |
| birds | Megascops_cooperi | Megascops_cooperi | LC | Decreasing | tropical | nocturnal | 149.9 | 78738.886 | 16.353 |

| | | | | | | | | | |
|---|---|---|---|---|---|---|---|---|---|
| birds | Megascops_guatemalae | Megascops_guatemalae | LC | Decreasing | tropical | nocturnal | 118 | 660793.493 | 9.421 |
| birds | Megascops_ingens | Megascops_ingens | LC | Stable | tropical | nocturnal | 165.3 | 314909.8 | 7.831 |
| birds | Megascops_kennicottii | Megascops_kennicottii | LC | Decreasing | temperate | nocturnal | 141.8 | 3582764.438 | 8.892 |
| birds | Megascops_marshalli | Megascops_marshalli | NT | Stable | tropical | nocturnal | 110 | 24727.091 | 3.571 |
| birds | Megascops_nudipes | Megascops_nudipes | LC | Stable | tropical | nocturnal | 137.3 | 9396.11 | 27.076 |
| birds | Megascops_roboratus | Megascops_roboratus | LC | Stable | tropical | nocturnal | 126.7 | 79303.5 | 9.619 |
| birds | Megascops_sanctaecatarinae | Megascops_sanctaecatarinae | LC | Decreasing | temperate | nocturnal | 179.7 | 418281.169 | 11.723 |
| birds | Megascops_trichopsis | Megascops_trichopsis | LC | Increasing | temperate | nocturnal | 88.3 | 660977.773 | 10.377 |
| birds | Megastictus_margaritatus | Megastictus_margaritatus | LC | Decreasing | tropical | diurnal | 20.2 | 1937447.717 | 1.335 |
| birds | Megatriorchis_doriae | Megatriorchis_doriae | LC | Decreasing | tropical | diurnal | 1000 | 599836.921 | 3.16 |
| birds | Megaxenops_parnaguae | Megaxenops_parnaguae | LC | Stable | tropical | diurnal | 25 | 669565.681 | 7.958 |
| birds | Megazosterops_palauensis | Megazosterops_palauensis | NT | Decreasing | tropical | diurnal | 31 | 413.295 | 11.016 |
| birds | Meiglyptes_jugularis | Meiglyptes_jugularis | LC | Stable | tropical | diurnal | 53.4 | 580432.677 | 9.587 |
| birds | Meiglyptes_tristis | Meiglyptes_tristis | EN | Decreasing | tropical | diurnal | 31.7 | 50016.694 | 21.35 |
| birds | Meiglyptes_tukki | Meiglyptes_tukki | NT | Decreasing | tropical | diurnal | 53.1 | 772219.58 | 7.348 |
| birds | Melaenornis_annamarulae | Melaenornis_annamarulae | VU | Decreasing | tropical | diurnal | 39.9 | 132494.244 | 15.066 |
| birds | Melaenornis_ardesiacus | Melaenornis_ardesiacus | LC | Decreasing | tropical | diurnal | 29 | 60515.808 | 16.8 |
| birds | Melaenornis_brunneus | Dioptrornis_brunneus | LC | Decreasing | tropical | diurnal | 26.8 | 68829.732 | 10.825 |
| birds | Melaenornis_chocolatinus | Dioptrornis_chocolatinus | LC | Decreasing | tropical | diurnal | 23.4 | 202824.29 | 15.214 |
| birds | Melaenornis_edolioides | Melaenornis_edolioides | LC | Stable | tropical | diurnal | 30.7 | 3825770.162 | 10.903 |
| birds | Melaenornis_pammelaina | Melaenornis_pammelaina | LC | Stable | tropical | diurnal | 30.2 | 4080335.722 | 8.686 |
| birds | Melampitta_lugubris | Melampitta_lugubris | LC | Stable | tropical | diurnal | 42.7 | 59306.618 | 4.908 |
| birds | Melamprosops_phaeosoma | NA | EX | not_available | NA | NA | NA | NA | NA |
| birds | Melanerpes_aurifrons | Melanerpes_aurifrons | LC | Stable | temperate | diurnal | 80.8 | 1414625.355 | 10.921 |
| birds | Melanerpes_cactorum | Melanerpes_cactorum | LC | Decreasing | temperate | diurnal | 34.9 | 942033.845 | 7.283 |
| birds | Melanerpes_candidus | Melanerpes_candidus | LC | Increasing | tropical | diurnal | 108 | 5535970.531 | 8.777 |
| birds | Melanerpes_carolinus | Melanerpes_carolinus | LC | Increasing | temperate | diurnal | 69.5 | 3074161.816 | 18.305 |
| birds | Melanerpes_chrysauchen | Melanerpes_chrysauchen | LC | Decreasing | tropical | diurnal | 52.5 | 15256.011 | 14.3 |
| birds | Melanerpes_chrysogenys | Melanerpes_chrysogenys | LC | Stable | tropical | diurnal | 67.7 | 161373.556 | 9.653 |
| birds | Melanerpes_cruentatus | Melanerpes_cruentatus | LC | Stable | tropical | diurnal | 58.1 | 7237147.708 | 2.409 |
| birds | Melanerpes_erythrocephalus | Melanerpes_erythrocephalus | LC | Decreasing | temperate | diurnal | 71.6 | 5461683.203 | 15.663 |
| birds | Melanerpes_flavifrons | Melanerpes_flavifrons | LC | Unknown | tropical | diurnal | 57.8 | 1557771.435 | 13.204 |
| birds | Melanerpes_formicivorus | Melanerpes_formicivorus | LC | Increasing | temperate | diurnal | 79.6 | 1502706.446 | 10.682 |
| birds | Melanerpes_herminieri | Melanerpes_herminieri | LC | Increasing | tropical | diurnal | 100 | 1468.623 | 24.334 |

| | | | | | | | | | |
|---|---|---|---|---|---|---|---|---|---|
| birds | Melanerpes_hoffmannii | Melanerpes_hoffmannii | LC | Increasing | tropical | diurnal | 72.2 | 51111.87 | 16.298 |
| birds | Melanerpes_hypopolius | Melanerpes_hypopolius | LC | Decreasing | tropical | diurnal | 52.4 | 81784.169 | 8.556 |
| birds | Melanerpes_lewis | Melanerpes_lewis | LC | Decreasing | temperate | diurnal | 105.8 | 3401486.03 | 9.267 |
| birds | Melanerpes_portoricensis | Melanerpes_portoricensis | LC | Stable | tropical | diurnal | 59.9 | 8923.083 | 27.081 |
| birds | Melanerpes_pucherani | Melanerpes_pucherani | LC | Decreasing | tropical | diurnal | 62.4 | 486437.573 | 9.545 |
| birds | Melanerpes_pygmaeus | Melanerpes_pygmaeus | LC | Decreasing | tropical | diurnal | 38.8 | 126112.502 | 6.567 |
| birds | Melanerpes_radiolatus | Melanerpes_radiolatus | LC | Decreasing | tropical | diurnal | 108 | 11021.791 | 19.741 |
| birds | Melanerpes_rubricapillus | Melanerpes_rubricapillus | LC | Stable | tropical | diurnal | 52.3 | 877834.62 | 10.186 |
| birds | Melanerpes_striatus | Melanerpes_striatus | LC | Decreasing | tropical | diurnal | 73.4 | 74161.47 | 15.781 |
| birds | Melanerpes_superciliaris | Melanerpes_superciliaris | LC | Decreasing | tropical | diurnal | 77.3 | 114069.089 | 15.95 |
| birds | Melanerpes_uropygialis | Melanerpes_uropygialis | LC | Stable | temperate | diurnal | 64.7 | 567324.492 | 7.202 |
| birds | Melaniparus_afer | Parus_afer | LC | Stable | temperate | diurnal | 20.2 | 427814.631 | 7.914 |
| birds | Melaniparus_albiventris | Parus_albiventris | LC | Decreasing | tropical | diurnal | 21.2 | 322293.631 | 13.241 |
| birds | Melaniparus_carpi | Parus_carpi | LC | Stable | tropical | diurnal | 18.6 | 305301.814 | 6.063 |
| birds | Melaniparus_cinerascens | Parus_cinerascens | LC | Stable | tropical | diurnal | 20 | 1795200.593 | 6.991 |
| birds | Melaniparus_fasciiventer | Parus_fasciiventer | LC | Decreasing | tropical | diurnal | 15.2 | 38778.021 | 16.463 |
| birds | Melaniparus_fringillinus | Parus_fringillinus | LC | Stable | tropical | diurnal | 13.8 | 390583.137 | 12.724 |
| birds | Melaniparus_funereus | Parus_funereus | LC | Decreasing | tropical | diurnal | 25.4 | 2252792.06 | 7.273 |
| birds | Melaniparus_griseiventris | Parus_griseiventris | LC | Stable | tropical | diurnal | 15.5 | 1535847.664 | 8.607 |
| birds | Melaniparus_leucomelas | Parus_leucomelas | LC | Stable | tropical | diurnal | 16.1 | 2587731.879 | 8.365 |
| birds | Melaniparus_leuconotus | Parus_leuconotus | LC | Stable | tropical | diurnal | 16.7 | 357687.797 | 13.617 |
| birds | Melaniparus_niger | Parus_niger | LC | Stable | tropical | diurnal | 21.5 | 2180649.324 | 9.558 |
| birds | Melaniparus_rufiventris | Parus_rufiventris | LC | Stable | tropical | diurnal | 18.2 | 2428644.321 | 6.704 |
| birds | Melaniparus_thruppi | Parus_thruppi | LC | Stable | tropical | diurnal | 12 | 869174.264 | 8.836 |
| birds | Melanitta_fusca | Melanitta_fusca | VU | Decreasing | temperate | diurnal | 1800.2 | 5830953.159 | 5.237 |
| birds | Melanitta_nigra | Melanitta_nigra | LC | Unknown | temperate | diurnal | 1050 | 5162178.682 | 2.146 |
| birds | Melanitta_perspicillata | Melanitta_perspicillata | LC | Decreasing | temperate | diurnal | 1096.3 | 6744994.035 | 3.177 |
| birds | Melanocharis_arfakiana | Melanocharis_arfakiana | LC | Unknown | tropical | diurnal | 11.5 | 9004.629 | 3.307 |
| birds | Melanocharis_longicauda | Melanocharis_longicauda | LC | Unknown | tropical | diurnal | 12 | 141891.577 | 4.05 |
| birds | Melanocharis_nigra | Melanocharis_nigra | LC | Unknown | tropical | diurnal | 14.2 | 458523.978 | 3.283 |
| birds | Melanocharis_striativentris | Melanocharis_striativentris | LC | Unknown | tropical | diurnal | 17.6 | 152632.535 | 4.293 |
| birds | Melanocharis_versteri | Melanocharis_versteri | LC | Unknown | tropical | diurnal | 14.7 | 148463.409 | 4.745 |
| birds | Melanochlora_sultanea | Melanochlora_sultanea | LC | Decreasing | tropical | diurnal | 37.6 | 1656290.696 | 10.098 |
| birds | Melanocichla_calva | Garrulax_calvus | LC | Decreasing | tropical | diurnal | 78.6 | 25342.436 | 3.773 |

| | | | | | | | | | |
|---|---|---|---|---|---|---|---|---|---|
| birds | Melanocichla_lugubris | Garrulax_lugubris | LC | Decreasing | tropical | diurnal | 78.6 | 41578.041 | 7.707 |
| birds | Melanocorypha_bimaculata | Melanocorypha_bimaculata | LC | Stable | temperate | diurnal | 54.1 | 4993981.898 | 9.847 |
| birds | Melanocorypha_calandra | Melanocorypha_calandra | LC | Decreasing | temperate | diurnal | 61.6 | 6304656.357 | 12.62 |
| birds | Melanocorypha_maxima | Melanocorypha_maxima | LC | Stable | temperate | diurnal | 75 | 1042885.902 | 6.596 |
| birds | Melanocorypha_mongolica | Melanocorypha_mongolica | LC | Stable | temperate | diurnal | 53.9 | 4731925.501 | 7.561 |
| birds | Melanocorypha_yeltoniensis | Melanocorypha_yeltoniensis | LC | Decreasing | temperate | diurnal | 59.9 | 3219779.336 | 10.621 |
| birds | Melanodera_melanodera | Melanodera_melanodera | LC | Decreasing | temperate | diurnal | 28.7 | 145980.071 | 5.579 |
| birds | Melanodera_xanthogramma | Melanodera_xanthogramma | LC | Stable | temperate | diurnal | 35.1 | 353522.864 | 3.191 |
| birds | Melanodryas_cucullata | Melanodryas_cucullata | LC | Decreasing | temperate | diurnal | 21.1 | 6360149.093 | 4.312 |
| birds | Melanodryas_vittata | Melanodryas_vittata | VU | Decreasing | temperate | diurnal | 26.6 | 68050.364 | 5.966 |
| birds | Melanopareia_elegans | Melanopareia_elegans | LC | Increasing | tropical | diurnal | 16 | 81037.63 | 11.708 |
| birds | Melanopareia_maranonica | Melanopareia_maranonica | LC | Decreasing | tropical | diurnal | 23 | 16464.381 | 7.923 |
| birds | Melanopareia_maximiliani | Melanopareia_maximiliani | LC | Stable | temperate | diurnal | 18 | 752374.154 | 6.467 |
| birds | Melanopareia_torquata | Melanopareia_torquata | LC | Stable | tropical | diurnal | 17.2 | 2770968.602 | 8.178 |
| birds | Melanoperdix_niger | Melanoperdix_niger | VU | Decreasing | tropical | diurnal | 280 | 1057322.279 | 8.263 |
| birds | Melanoptila_glabrirostris | Melanoptila_glabrirostris | LC | Decreasing | tropical | diurnal | 35.4 | 151379.515 | 6.633 |
| birds | Melanorectes_nigrescens | Pitohui_nigrescens | LC | Stable | tropical | diurnal | 76.9 | 131590.715 | 4.773 |
| birds | Melanospiza_bicolor | Tiaris_bicolor | LC | Increasing | tropical | diurnal | 9.8 | 269426.802 | 14.661 |
| birds | Melanospiza_richardsoni | Melanospiza_richardsoni | EN | Decreasing | tropical | diurnal | 20.9 | 435.297 | 19.179 |
| birds | Melanotis_caerulescens | Melanotis_caerulescens | LC | Decreasing | tropical | diurnal | 61.6 | 577099.946 | 10.47 |
| birds | Melanotis_hypoleucus | Melanotis_hypoleucus | LC | Decreasing | tropical | diurnal | 68 | 98583.115 | 12.712 |
| birds | Meleagris_gallopavo | Meleagris_gallopavo | LC | Increasing | temperate | diurnal | 5791.4 | 7375475.415 | 14.065 |
| birds | Meleagris_ocellata | Meleagris_ocellata | NT | Decreasing | tropical | diurnal | 5525 | 163840.219 | 6.534 |
| birds | Meliarchus_sclateri | Melidectes_sclateri | LC | Stable | tropical | diurnal | 66.8 | 3338.039 | 4.602 |
| birds | Melichneutes_robustus | Melichneutes_robustus | LC | Decreasing | tropical | diurnal | 54.2 | 2772579.96 | 6.598 |
| birds | Melidectes_belfordi | Melidectes_belfordi | LC | Decreasing | tropical | diurnal | 73.1 | 124314.412 | 5.23 |
| birds | Melidectes_foersteri | Melidectes_foersteri | LC | Stable | tropical | diurnal | 85 | 6152.622 | 3.658 |
| birds | Melidectes_leucostephes | Melidectes_leucostephes | LC | Stable | tropical | diurnal | 75.3 | 11363.023 | 2.319 |
| birds | Melidectes_ochromelas | Melidectes_ochromelas | LC | Stable | tropical | diurnal | 63 | 51249.231 | 3.133 |
| birds | Melidectes_rufocrissalis | Melidectes_rufocrissalis | LC | Decreasing | tropical | diurnal | 71.6 | 64711.258 | 5.898 |
| birds | Melidectes_torquatus | Melidectes_torquatus | LC | Increasing | tropical | diurnal | 44.9 | 95822.014 | 4.321 |
| birds | Melidora_macrorrhina | Melidora_macrorrhina | LC | Stable | tropical | cathemeral | 97 | 654694.186 | 3.153 |
| birds | Melierax_canorus | Melierax_canorus | LC | Stable | temperate | diurnal | 663.3 | 2246754.628 | 6.582 |
| birds | Melierax_metabates | Melierax_metabates | LC | Stable | tropical | diurnal | 674.5 | 12484209.957 | 9.113 |

| | | | | | | | | | |
|---|---|---|---|---|---|---|---|---|---|
| birds | Melierax_poliopterus | Melierax_poliopterus | LC | Stable | tropical | diurnal | 629.9 | 2071175.933 | 9.056 |
| birds | Melignomon_eisentrauti | Melignomon_eisentrauti | NT | Decreasing | tropical | diurnal | 24.2 | 81908.398 | 15.907 |
| birds | Melignomon_zenkeri | Melignomon_zenkeri | LC | Unknown | tropical | diurnal | 23.6 | 1324791.273 | 5.803 |
| birds | Melilestes_megarhynchus | Melilestes_megarhynchus | LC | Stable | tropical | diurnal | 43.8 | 565758.843 | 3.244 |
| birds | Melionyx_fuscus | Melidectes_fuscus | LC | Stable | tropical | diurnal | 45 | 70507.071 | 4.989 |
| birds | Melionyx_nouhuysi | Melidectes_nouhuysi | LC | Stable | tropical | diurnal | 62.3 | 9644.796 | 4.916 |
| birds | Melionyx_princeps | Melidectes_princeps | VU | Decreasing | tropical | diurnal | 42 | 5573.181 | 6.076 |
| birds | Meliphacator_provocator | Xanthotis_provocator | LC | Stable | tropical | diurnal | 33.1 | 432.99 | 6.14 |
| birds | Meliphaga_aruensis | Meliphaga_aruensis | LC | Decreasing | tropical | diurnal | 26.2 | 665428.404 | 3.116 |
| birds | Meliphaga_lewinii | Meliphaga_lewinii | LC | Decreasing | temperate | diurnal | 33.6 | 643140.808 | 7.185 |
| birds | Meliphaga_notata | Meliphaga_notata | LC | Stable | tropical | diurnal | 26.4 | 110238.594 | 3.226 |
| birds | Melipotes_ater | Melipotes_ater | LC | Stable | tropical | diurnal | 114 | 5540.005 | 3.831 |
| birds | Melipotes_fumigatus | Melipotes_fumigatus | LC | Unknown | tropical | diurnal | 58.6 | 146497.21 | 4.958 |
| birds | Melipotes_gymnops | Melipotes_gymnops | LC | Stable | tropical | diurnal | 54.8 | 7348.931 | 2.737 |
| birds | Melithreptus_affinis | Melithreptus_affinis | LC | Decreasing | temperate | diurnal | 15 | 68030.002 | 5.966 |
| birds | Melithreptus_albogularis | Melithreptus_albogularis | LC | Decreasing | tropical | diurnal | 11 | 1473520.943 | 3.846 |
| birds | Melithreptus_brevirostris | Melithreptus_brevirostris | LC | Stable | temperate | diurnal | 12.6 | 2003053.237 | 8.348 |
| birds | Melithreptus_gularis | Melithreptus_gularis | LC | Stable | temperate | diurnal | 19.4 | 1260779.547 | 8.455 |
| birds | Melithreptus_lunatus | Melithreptus_lunatus | LC | Stable | temperate | diurnal | 14 | 1242453.6 | 8.922 |
| birds | Melithreptus_validirostris | Melithreptus_validirostris | VU | Decreasing | temperate | diurnal | 24.8 | 66891.321 | 5.956 |
| birds | Melitograis_gilolensis | Melitograis_gilolensis | LC | Stable | tropical | diurnal | 54 | 22755.074 | 6.411 |
| birds | Mellisuga_helenae | Mellisuga_helenae | NT | Decreasing | tropical | diurnal | 2.2 | 108875.849 | 16.192 |
| birds | Mellisuga_minima | Mellisuga_minima | LC | Stable | tropical | diurnal | 2.4 | 86168.427 | 16.275 |
| birds | Melloria_quoyi | Cracticus_quoyi | LC | Stable | tropical | diurnal | 168 | 1145717.608 | 3.071 |
| birds | Melocichla_mentalis | Melocichla_mentalis | LC | Stable | tropical | diurnal | 33.2 | 6248614.058 | 9.361 |
| birds | Melopsittacus_undulatus | Melopsittacus_undulatus | LC | Increasing | temperate | diurnal | 29 | 6191813 | 4.856 |
| birds | Melopyrrha_portoricensis | Loxigilla_portoricensis | VU | Decreasing | tropical | diurnal | 32 | 8774.421 | 27.252 |
| birds | Melopyrrha_violacea | Loxigilla_violacea | LC | Stable | tropical | diurnal | 22.2 | 97899.28 | 15.256 |
| birds | Melospiza_georgiana | Melospiza_georgiana | LC | Increasing | temperate | diurnal | 16.1 | 10922479.134 | 10.379 |
| birds | Melospiza_lincolnii | Melospiza_lincolnii | LC | Stable | temperate | diurnal | 16.6 | 16098839.353 | 8.646 |
| birds | Melospiza_melodia | Melospiza_melodia | LC | Decreasing | temperate | diurnal | 21.9 | 13306731.28 | 10.087 |
| birds | Melozone_aberti | Pipilo_aberti | LC | Increasing | temperate | diurnal | 45.9 | 170236.93 | 9.684 |
| birds | Melozone_albicollis | Pipilo_albicollis | LC | Stable | tropical | diurnal | 46.4 | 64681.629 | 7.264 |
| birds | Melozone_biarcuata | Melozone_biarcuata | LC | Decreasing | tropical | diurnal | 30.6 | 85901.92 | 13.746 |

| | | | | | | | | | |
|---|---|---|---|---|---|---|---|---|---|
| birds | Melozone_cabanisi | Melozone_cabanisi | NT | Decreasing | tropical | diurnal | 28 | 3056.525 | 27.459 |
| birds | Melozone_crissalis | Pipilo_crissalis | LC | Decreasing | temperate | diurnal | 52.8 | 333326.695 | 10.621 |
| birds | Melozone_fusca | Pipilo_fuscus | LC | Decreasing | temperate | diurnal | 44.4 | 1684944.382 | 9.238 |
| birds | Melozone_kieneri | Melozone_kieneri | LC | Decreasing | tropical | diurnal | 38.7 | 330219.499 | 9.231 |
| birds | Melozone_leucotis | Melozone_leucotis | LC | Decreasing | tropical | diurnal | 39.9 | 11301.539 | 16.19 |
| birds | Mentocrex_kioloides | Canirallus_kioloides | LC | Decreasing | tropical | diurnal | 246 | 14988.068 | 5.192 |
| birds | Menura_alberti | Menura_alberti | LC | Stable | temperate | diurnal | 928 | 40026.782 | 10.9 |
| birds | Menura_novaehollandiae | Menura_novaehollandiae | LC | Decreasing | temperate | diurnal | 972.6 | 325624.393 | 8.736 |
| birds | Merganetta_armata | Merganetta_armata | LC | Decreasing | tropical | diurnal | 398.8 | 1090659.132 | 7.079 |
| birds | Mergellus_albellus | Mergellus_albellus | LC | Decreasing | temperate | diurnal | 608.6 | 13903839.294 | 5.891 |
| birds | Mergus_merganser | Mergus_merganser | LC | Unknown | temperate | diurnal | 1451 | 35370520.544 | 7.513 |
| birds | Mergus_octosetaceus | Mergus_octosetaceus | CR | Decreasing | tropical | diurnal | 983 | 1316243.474 | 13.052 |
| birds | Mergus_serrator | Mergus_serrator | LC | Stable | temperate | diurnal | 1015.2 | 37225500.743 | 5.859 |
| birds | Mergus_squamatus | Mergus_squamatus | EN | Decreasing | temperate | diurnal | 1234 | 2240002.661 | 13.196 |
| birds | Meropogon_forsteni | Meropogon_forsteni | LC | Decreasing | tropical | diurnal | 55 | 135924.863 | 7.596 |
| birds | Merops_albicollis | Merops_albicollis | LC | Stable | tropical | diurnal | 25.9 | 10467706.234 | 9.907 |
| birds | Merops_apiaster | Merops_apiaster | LC | Stable | temperate | diurnal | 56.6 | 13243490.573 | 11.379 |
| birds | Merops_boehmi | Merops_boehmi | LC | Stable | tropical | diurnal | 16.6 | 700033.602 | 8.515 |
| birds | Merops_breweri | Merops_breweri | LC | Decreasing | tropical | diurnal | 49.7 | 1632583.201 | 6.098 |
| birds | Merops_bullockoides | Merops_bullockoides | LC | Increasing | tropical | diurnal | 34.8 | 5313566.688 | 8.588 |
| birds | Merops_bulocki | Merops_bulocki | LC | Stable | tropical | diurnal | 23.1 | 2933973.615 | 11.518 |
| birds | Merops_gularis | Merops_gularis | LC | Stable | tropical | diurnal | 27.3 | 3039017.174 | 8.079 |
| birds | Merops_hirundineus | Merops_hirundineus | LC | Stable | tropical | diurnal | 21.7 | 8552948.686 | 8.722 |
| birds | Merops_leschenaulti | Merops_leschenaulti | LC | Increasing | tropical | diurnal | 27.2 | 2822822.271 | 13.953 |
| birds | Merops_malimbicus | Merops_malimbicus | LC | Unknown | tropical | diurnal | 50.4 | 1453484.123 | 10.102 |
| birds | Merops_muelleri | Merops_muelleri | LC | Decreasing | tropical | diurnal | 22.5 | 1597560.381 | 6.09 |
| birds | Merops_nubicus | Merops_nubicus | LC | Decreasing | tropical | diurnal | 42.4 | 8296122.079 | 10.107 |
| birds | Merops_oreobates | Merops_oreobates | LC | Unknown | tropical | diurnal | 24 | 360934.322 | 13.225 |
| birds | Merops_orientalis | Merops_orientalis | LC | Increasing | tropical | diurnal | 14.8 | 5717890.997 | 15.293 |
| birds | Merops_ornatus | Merops_ornatus | LC | Stable | temperate | diurnal | 29.5 | 7621196.563 | 4.695 |
| birds | Merops_persicus | Merops_persicus | LC | Stable | tropical | diurnal | 49.3 | 16160299.227 | 10.088 |
| birds | Merops_philippinus | Merops_philippinus | LC | Stable | tropical | diurnal | 34 | 6418828.75 | 14.058 |
| birds | Merops_pusillus | Merops_pusillus | LC | Decreasing | tropical | diurnal | 15.1 | 14410773.706 | 9.875 |
| birds | Merops_revoilii | Merops_revoilii | LC | Increasing | tropical | diurnal | 12.9 | 958341.747 | 7.577 |

| | | | | | | | | | |
|---|---|---|---|---|---|---|---|---|---|
| birds | Merops_superciliosus | Merops_superciliosus | LC | Stable | tropical | diurnal | 38.5 | 4269964.774 | 10.149 |
| birds | Merops_variegatus | Merops_variegatus | LC | Stable | tropical | diurnal | 22.5 | 3491407.889 | 7.021 |
| birds | Merops_viridis | Merops_viridis | LC | Stable | tropical | diurnal | 34.8 | 3204518.176 | 12.274 |
| birds | Merulaxis_ater | Merulaxis_ater | LC | Decreasing | temperate | diurnal | 35 | 105767.759 | 11.878 |
| birds | Merulaxis_stresemanni | NA | CR | Decreasing | NA | NA | NA | NA | NA |
| birds | Mesembrinibis_cayennensis | Mesembrinibis_cayennensis | LC | Stable | tropical | diurnal | 756 | 10576805.928 | 4.956 |
| birds | Mesitornis_unicolor | Mesitornis_unicolor | VU | Decreasing | tropical | diurnal | 148 | 37168.054 | 5.64 |
| birds | Mesitornis_variegatus | Mesitornis_variegatus | VU | Decreasing | tropical | diurnal | 110 | 39538.157 | 9.175 |
| birds | Metabolus_godeffroyi | NA | NT | Stable | NA | NA | NA | NA | NA |
| birds | Metabolus_rugensis | NA | EN | Decreasing | NA | NA | NA | NA | NA |
| birds | Metabolus_takatsukasae | NA | NT | Stable | NA | NA | NA | NA | NA |
| birds | Metallura_aeneocauda | Metallura_aeneocauda | LC | Decreasing | tropical | diurnal | 5.2 | 49696.48 | 5.708 |
| birds | Metallura_baroni | Metallura_baroni | VU | Decreasing | tropical | diurnal | 4.3 | 2281.79 | 9.862 |
| birds | Metallura_eupogon | Metallura_eupogon | LC | Decreasing | tropical | diurnal | 4.6 | 35465.529 | 7.376 |
| birds | Metallura_iracunda | Metallura_iracunda | NT | Stable | tropical | diurnal | 3.9 | 849.292 | 6.66 |
| birds | Metallura_odomae | Metallura_odomae | LC | Decreasing | tropical | diurnal | 5.1 | 8418.362 | 7.084 |
| birds | Metallura_phoebe | Metallura_phoebe | LC | Decreasing | tropical | diurnal | 5.4 | 142044.518 | 8.033 |
| birds | Metallura_theresiae | Metallura_theresiae | LC | Decreasing | tropical | diurnal | 4.9 | 34923.522 | 4.363 |
| birds | Metallura_tyrianthina | Metallura_tyrianthina | LC | Stable | tropical | diurnal | 3.8 | 422575.202 | 9.782 |
| birds | Metallura_williami | Metallura_williami | LC | Decreasing | tropical | diurnal | 4.5 | 51630.478 | 8.806 |
| birds | Metopidius_indicus | Metopidius_indicus | LC | Unknown | tropical | diurnal | 155 | 2649332.708 | 16.361 |
| birds | Metopothrix_aurantiaca | Metopothrix_aurantiaca | LC | Stable | tropical | diurnal | 11.2 | 458107.408 | 2.953 |
| birds | Metriopelia_aymara | Metriopelia_aymara | LC | Stable | tropical | diurnal | 61.8 | 832237.501 | 5.538 |
| birds | Metriopelia_ceciliae | Metriopelia_ceciliae | LC | Stable | tropical | diurnal | 55.1 | 490195.482 | 8.402 |
| birds | Metriopelia_melanoptera | Metriopelia_melanoptera | LC | Stable | temperate | diurnal | 100 | 1845275.521 | 6.103 |
| birds | Metriopelia_morenoi | Metriopelia_morenoi | LC | Stable | temperate | diurnal | 28.6 | 106199.058 | 3.598 |
| birds | Micrastur_buckleyi | Micrastur_buckleyi | LC | Decreasing | tropical | diurnal | 464 | 619557.033 | 1.775 |
| birds | Micrastur_gilvicollis | Micrastur_gilvicollis | LC | Decreasing | tropical | diurnal | 209 | 4251681.562 | 1.522 |
| birds | Micrastur_mirandollei | Micrastur_mirandollei | LC | Decreasing | tropical | diurnal | 549 | 5269885.686 | 1.694 |
| birds | Micrastur_plumbeus | Micrastur_plumbeus | VU | Decreasing | tropical | diurnal | 185.4 | 69294.606 | 6.798 |
| birds | Micrastur_ruficollis | Micrastur_ruficollis | LC | Decreasing | tropical | diurnal | 177.6 | 12486169.393 | 5.638 |
| birds | Micrastur_semitorquatus | Micrastur_semitorquatus | LC | Decreasing | tropical | diurnal | 621.7 | 13182553.541 | 5.652 |
| birds | Microbates_cinereiventris | Microbates_cinereiventris | LC | Decreasing | tropical | diurnal | 11.9 | 551552.186 | 5.283 |
| birds | Microbates_collaris | Microbates_collaris | LC | Decreasing | tropical | diurnal | 10.2 | 2367560.871 | 1.147 |

| | | | | | | | | | |
|---|---|---|---|---|---|---|---|---|---|
| birds | Microcarbo_africanus | Phalacrocorax_africanus | LC | Decreasing | tropical | diurnal | 545 | 17477574.054 | 8.973 |
| birds | Microcarbo_coronatus | Phalacrocorax_coronatus | LC | Stable | temperate | diurnal | 762.2 | 20853.119 | 8.801 |
| birds | Microcarbo_melanoleucos | Phalacrocorax_melanoleucos | LC | Unknown | temperate | diurnal | 727.4 | 5853894.375 | 5.778 |
| birds | Microcarbo_niger | Phalacrocorax_niger | LC | Unknown | tropical | diurnal | 427 | 4902890.616 | 16.155 |
| birds | Microcarbo_pygmaeus | Phalacrocorax_pygmeus | LC | Increasing | temperate | diurnal | 676 | 3509944.418 | 11.66 |
| birds | Microcerculus_bambla | Microcerculus_bambla | LC | Decreasing | tropical | diurnal | 19.1 | 1714246.726 | 1.26 |
| birds | Microcerculus_marginatus | Microcerculus_marginatus | LC | Decreasing | tropical | diurnal | 18.2 | 4781168.47 | 3.189 |
| birds | Microcerculus_philomela | Microcerculus_philomela | LC | Decreasing | tropical | diurnal | 17.8 | 177560.998 | 8.759 |
| birds | Microcerculus_ustulatus | Microcerculus_ustulatus | LC | Stable | tropical | diurnal | 21 | 265691.925 | 0.81 |
| birds | Microchera_albocoronata | Microchera_albocoronata | LC | Decreasing | tropical | diurnal | 2.7 | 76810.962 | 7.663 |
| birds | Microchera_chionura | Elvira_chionura | LC | Decreasing | tropical | diurnal | 3 | 9597.915 | 9.654 |
| birds | Microchera_cupreiceps | Elvira_cupreiceps | LC | Stable | tropical | diurnal | 3 | 5223.26 | 13.069 |
| birds | Microdynamis_parva | Microdynamis_parva | LC | Stable | tropical | diurnal | 47.2 | 607992.827 | 3.094 |
| birds | Microeca_fascinans | Microeca_fascinans | LC | Decreasing | temperate | diurnal | 14.5 | 5302401.744 | 5.1 |
| birds | Microeca_flavigaster | Microeca_flavigaster | LC | Stable | tropical | diurnal | 12.1 | 1175473.948 | 2.973 |
| birds | Microeca_hemixantha | Microeca_hemixantha | NT | Decreasing | tropical | diurnal | 13.7 | 2954.97 | 5.415 |
| birds | Microhierax_caerulescens | Microhierax_caerulescens | LC | Stable | temperate | diurnal | 38.7 | 1206450.704 | 11.331 |
| birds | Microhierax_erythrogenys | Microhierax_erythrogenys | LC | Decreasing | tropical | diurnal | 50 | 253762.695 | 13.704 |
| birds | Microhierax_fringillarius | Microhierax_fringillarius | LC | Stable | tropical | diurnal | 42.9 | 1402352.924 | 9.609 |
| birds | Microhierax_latifrons | Microhierax_latifrons | NT | Decreasing | tropical | diurnal | 41.2 | 60479.234 | 6.701 |
| birds | Microhierax_melanoleucos | Microhierax_melanoleucos | LC | Stable | temperate | diurnal | 64.2 | 2360698.644 | 13.936 |
| birds | Microligea_palustris | Microligea_palustris | LC | Decreasing | tropical | diurnal | 13.2 | 30480.023 | 12.586 |
| birds | Micromacronus_leytensis | Micromacronus_leytensis | NT | Decreasing | tropical | diurnal | 5.5 | 4721.623 | 6.243 |
| birds | Micromonacha_lanceolata | Micromonacha_lanceolata | LC | Decreasing | tropical | diurnal | 19.8 | 1651230.161 | 1.836 |
| birds | Micronisus_gabar | Melierax_gabar | LC | Stable | tropical | diurnal | 169.6 | 13910595.474 | 9.661 |
| birds | Microparra_capensis | Microparra_capensis | LC | Unknown | tropical | diurnal | 41.3 | 3244082.826 | 9.392 |
| birds | Micropsitta_bruijnii | Micropsitta_bruijnii | LC | Stable | tropical | diurnal | 13.4 | 221432.598 | 4.594 |
| birds | Micropsitta_finschii | Micropsitta_finschii | LC | Stable | tropical | diurnal | 15 | 42665.365 | 4.905 |
| birds | Micropsitta_geelvinkiana | Micropsitta_geelvinkiana | LC | Stable | tropical | diurnal | 12.2 | 2555.604 | 9.202 |
| birds | Micropsitta_keiensis | Micropsitta_keiensis | LC | Stable | tropical | diurnal | 12.1 | 181899.434 | 3.053 |
| birds | Micropsitta_meeki | Micropsitta_meeki | LC | Stable | tropical | diurnal | 13.8 | 2353.011 | 5.815 |
| birds | Micropsitta_pusio | Micropsitta_pusio | LC | Stable | tropical | diurnal | 14 | 290278.953 | 3.711 |
| birds | Micropternus_brachyurus | Celeus_brachyurus | LC | Decreasing | tropical | diurnal | 85.9 | 6131273.927 | 12.823 |
| birds | Microptilotis_albilineatus | Meliphaga_albilineata | LC | Stable | tropical | diurnal | 23.7 | 53764.239 | 1.212 |

| | | | | | | | | | |
|---|---|---|---|---|---|---|---|---|---|
| birds | Microptilotis_albonotatus | Meliphaga_albonotata | LC | Unknown | tropical | diurnal | 22.4 | 320661.12 | 3.294 |
| birds | Microptilotis_analogus | Meliphaga_analoga | LC | Stable | tropical | diurnal | 21.6 | 636132.655 | 3.091 |
| birds | Microptilotis_flavirictus | Meliphaga_flavirictus | LC | Stable | tropical | diurnal | 18.5 | 439941.456 | 3.006 |
| birds | Microptilotis_gracilis | Meliphaga_gracilis | LC | Stable | tropical | diurnal | 15 | 185643.548 | 2.647 |
| birds | Microptilotis_mimikae | Meliphaga_mimikae | LC | Stable | tropical | diurnal | 26.9 | 89115.846 | 3.025 |
| birds | Microptilotis_montanus | Meliphaga_montana | LC | Stable | tropical | diurnal | 28 | 121416.33 | 2.945 |
| birds | Microptilotis_orientalis | Meliphaga_orientalis | LC | Stable | tropical | diurnal | 18.1 | 193375.679 | 3.49 |
| birds | Microptilotis_reticulatus | Meliphaga_reticulata | LC | Unknown | tropical | diurnal | 23.6 | 27533.731 | 13.895 |
| birds | Microptilotis_vicina | Meliphaga_vicina | LC | Stable | tropical | diurnal | 23.6 | 890.181 | 4.136 |
| birds | Micropygia_schomburgkii | Micropygia_schomburgkii | LC | Decreasing | tropical | diurnal | 31.2 | 3010330.874 | 8.394 |
| birds | Microrhopias_quixensis | Microrhopias_quixensis | LC | Decreasing | tropical | diurnal | 7.9 | 4104536.418 | 2.784 |
| birds | Microspingus_alticola | Poospiza_alticola | EN | Decreasing | tropical | diurnal | 24 | 23500.969 | 8.728 |
| birds | Microspingus_cinereus | Poospiza_cinerea | LC | Stable | tropical | diurnal | 11.6 | 1642836.981 | 10.34 |
| birds | Microspingus_erythrophrys | Poospiza_erythrophrys | LC | Stable | tropical | diurnal | 14.1 | 80138.703 | 7.253 |
| birds | Microspingus_lateralis | Poospiza_lateralis | LC | Decreasing | tropical | diurnal | 19.2 | 57423.085 | 13.244 |
| birds | Microspingus_melanoleucus | Poospiza_melanoleuca | LC | Stable | temperate | diurnal | 13.1 | 1482632.758 | 7.582 |
| birds | Microspingus_torquatus | Poospiza_torquata | LC | Stable | tropical | diurnal | 11 | 121446.365 | 10.173 |
| birds | Microspingus_trifasciatus | Hemispingus_trifasciatus | LC | Decreasing | tropical | diurnal | 14 | 82213.807 | 5.876 |
| birds | Microstilbon_burmeisteri | Microstilbon_burmeisteri | LC | Stable | tropical | diurnal | 2.5 | 109902.823 | 7.794 |
| birds | Microtarsus_melanoleucos | Pycnonotus_melanoleucos | NT | Decreasing | tropical | diurnal | 31 | 1181602.117 | 7.571 |
| birds | Microxenops_milleri | Xenops_milleri | LC | Stable | tropical | diurnal | 12.2 | 3558978.041 | 1.228 |
| birds | Milvago_chimachima | Milvago_chimachima | LC | Increasing | tropical | diurnal | 315.2 | 13624390.701 | 5.796 |
| birds | Milvus_migrans | Milvus_migrans | LC | Stable | tropical | diurnal | 734.1 | 71149092.138 | 8.774 |
| birds | Milvus_milvus | Milvus_milvus | LC | Increasing | temperate | diurnal | 1071.8 | 3974917.684 | 19.972 |
| birds | Mimus_dorsalis | Mimus_dorsalis | LC | Stable | tropical | diurnal | 58.7 | 173822.653 | 8.422 |
| birds | Mimus_gilvus | Mimus_gilvus | LC | Increasing | tropical | diurnal | 52 | 2168760.732 | 8.403 |
| birds | Mimus_graysoni | NA | CR | Stable | NA | NA | NA | NA | NA |
| birds | Mimus_gundlachii | Mimus_gundlachii | LC | Decreasing | temperate | diurnal | 66.8 | 15471.497 | 7.452 |
| birds | Mimus_longicaudatus | Mimus_longicaudatus | LC | Unknown | tropical | diurnal | 66.6 | 181590.289 | 8.974 |
| birds | Mimus_macdonaldi | NA | VU | Unknown | NA | NA | NA | NA | NA |
| birds | Mimus_melanotis | NA | NT | Stable | NA | NA | NA | NA | NA |
| birds | Mimus_parvulus | NA | LC | Stable | NA | NA | NA | NA | NA |
| birds | Mimus_patagonicus | Mimus_patagonicus | LC | Stable | temperate | diurnal | 57.8 | 1884037.794 | 7.554 |
| birds | Mimus_polyglottos | Mimus_polyglottos | LC | Stable | temperate | diurnal | 48.5 | 10313735.122 | 13.005 |

| | | | | | | | | | |
|---|---|---|---|---|---|---|---|---|---|
| birds | Mimus_saturninus | Mimus_saturninus | LC | Stable | tropical | diurnal | 63.7 | 7198431.034 | 8.97 |
| birds | Mimus_thenca | Mimus_thenca | LC | Stable | temperate | diurnal | 66 | 229319.437 | 10.165 |
| birds | Mimus_trifasciatus | NA | EN | Stable | NA | NA | NA | NA | NA |
| birds | Mimus_triurus | Mimus_triurus | LC | Unknown | temperate | diurnal | 47.2 | 3284288.516 | 7.896 |
| birds | Minla_ignotincta | Minla_ignotincta | LC | Decreasing | temperate | diurnal | 14.3 | 1168696.97 | 10.388 |
| birds | Mino_anais | Mino_anais | LC | Decreasing | tropical | diurnal | 152 | 375373.573 | 3.076 |
| birds | Mino_dumontii | Mino_dumontii | LC | Unknown | tropical | diurnal | 217 | 640179.954 | 3.116 |
| birds | Mionectes_macconnelli | Mionectes_macconnelli | LC | Decreasing | tropical | diurnal | 13.5 | 2588557.723 | 1.867 |
| birds | Mionectes_oleagineus | Mionectes_oleagineus | LC | Decreasing | tropical | diurnal | 11.2 | 8947569.124 | 3.932 |
| birds | Mionectes_olivaceus | Mionectes_olivaceus | LC | Decreasing | tropical | diurnal | 15.3 | 39754.753 | 11.634 |
| birds | Mionectes_rufiventris | Mionectes_rufiventris | LC | Decreasing | temperate | diurnal | 13.3 | 1057487.402 | 13.737 |
| birds | Mionectes_striaticollis | Mionectes_striaticollis | LC | Stable | tropical | diurnal | 15 | 318010.85 | 7.146 |
| birds | Mirafra_africana | Mirafra_africana | LC | Decreasing | tropical | diurnal | 45 | 5074225.027 | 9.227 |
| birds | Mirafra_albicauda | Mirafra_albicauda | LC | Decreasing | tropical | diurnal | 22.2 | 516863.45 | 12.27 |
| birds | Mirafra_angolensis | Mirafra_angolensis | LC | Decreasing | tropical | diurnal | 36.2 | 763784.633 | 6.586 |
| birds | Mirafra_apiata | Mirafra_apiata | LC | Decreasing | temperate | diurnal | 26.9 | 258562.289 | 8.154 |
| birds | Mirafra_ashi | NA | not_available | not_available | NA | NA | NA | NA | NA |
| birds | Mirafra_assamica | Mirafra_assamica | LC | Stable | temperate | diurnal | 26.2 | 808359.62 | 17.026 |
| birds | Mirafra_cheniana | Mirafra_cheniana | LC | Decreasing | temperate | diurnal | 29.2 | 258493.247 | 12.847 |
| birds | Mirafra_collaris | Mirafra_collaris | LC | Stable | tropical | diurnal | 29.2 | 319160.758 | 7.193 |
| birds | Mirafra_cordofanica | Mirafra_cordofanica | LC | Decreasing | tropical | diurnal | 29.2 | 1714275.177 | 5.088 |
| birds | Mirafra_erythroptera | Mirafra_erythroptera | LC | Stable | tropical | diurnal | 21.3 | 2147813.665 | 18.591 |
| birds | Mirafra_gilletti | NA | not_available | not_available | NA | NA | NA | NA | NA |
| birds | Mirafra_hypermetra | Mirafra_hypermetra | LC | Stable | tropical | diurnal | 61.4 | 451245.369 | 9.976 |
| birds | Mirafra_javanica | Mirafra_javanica | LC | Stable | tropical | diurnal | 23 | 11882223.561 | 9.666 |
| birds | Mirafra_passerina | Mirafra_passerina | LC | Decreasing | tropical | diurnal | 24.7 | 1870815.459 | 6.903 |
| birds | Mirafra_pulpa | Mirafra_pulpa | DD | Decreasing | tropical | diurnal | 21.9 | 42119.969 | 9.813 |
| birds | Mirafra_rufocinnamomea | Mirafra_rufocinnamomea | LC | Decreasing | tropical | diurnal | 26.3 | 9716191.662 | 9.842 |
| birds | Mirafra_somalica | Mirafra_somalica | LC | Decreasing | tropical | diurnal | 46.6 | 124775.108 | 9.33 |
| birds | Mirafra_williamsi | Mirafra_williamsi | LC | Stable | tropical | diurnal | 29.2 | 30949.615 | 4.846 |
| birds | Mitrephanes_olivaceus | Mitrephanes_olivaceus | LC | Stable | tropical | diurnal | 10 | 91009.825 | 4.672 |
| birds | Mitrephanes_phaeocercus | Mitrephanes_phaeocercus | LC | Decreasing | tropical | diurnal | 8.6 | 882909.562 | 9.789 |
| birds | Mitrospingus_cassinii | Mitrospingus_cassinii | LC | Decreasing | tropical | diurnal | 40.4 | 167653.279 | 7.131 |
| birds | Mitrospingus_oleagineus | Mitrospingus_oleagineus | LC | Decreasing | tropical | diurnal | 41.3 | 38825.738 | 1.917 |

| | | | | | | | | | |
|---|---|---|---|---|---|---|---|---|---|
| birds | Mitu_mitu | NA | EW | not_available | NA | NA | NA | NA | NA |
| birds | Mitu_salvini | Mitu_salvini | LC | Decreasing | tropical | diurnal | 3100 | 490126.152 | 2.136 |
| birds | Mitu_tomentosum | Mitu_tomentosum | LC | Decreasing | tropical | diurnal | 2187.6 | 1438067.905 | 2.06 |
| birds | Mitu_tuberosum | Mitu_tuberosum | NT | Decreasing | tropical | diurnal | 2769.5 | 4359960.925 | 2.667 |
| birds | Mixornis_flavicollis | Macronous_flavicollis | LC | Decreasing | tropical | diurnal | 20 | 62071.473 | 18.106 |
| birds | Mixornis_gularis | Macronous_gularis | LC | Stable | tropical | diurnal | 11.8 | 3009284.287 | 11.751 |
| birds | Mixornis_kelleyi | NA | LC | Decreasing | NA | NA | NA | NA | NA |
| birds | Mniotilta_varia | Mniotilta_varia | LC | Stable | temperate | diurnal | 10.9 | 12084523.738 | 11.193 |
| birds | Modulatrix_stictigula | Modulatrix_stictigula | LC | Decreasing | tropical | diurnal | 30.5 | 70768.866 | 12.837 |
| birds | Mohoua_albicilla | Mohoua_albicilla | LC | Stable | temperate | diurnal | 17.6 | 36004.251 | 6.417 |
| birds | Mohoua_novaeseelandiae | Mohoua_novaeseelandiae | LC | Decreasing | temperate | diurnal | 12.1 | 76987.829 | 5.042 |
| birds | Mohoua_ochrocephala | Mohoua_ochrocephala | NT | Decreasing | temperate | diurnal | 27.7 | 29357.999 | 5.195 |
| birds | Molothrus_aeneus | Molothrus_aeneus | LC | Stable | tropical | diurnal | 62.6 | 2518193.731 | 10.05 |
| birds | Molothrus_armenti | Molothrus_armenti | NT | Decreasing | tropical | diurnal | 40.8 | 5020.703 | 22.684 |
| birds | Molothrus_ater | Molothrus_ater | LC | Decreasing | temperate | diurnal | 40.3 | 12432730.676 | 11.492 |
| birds | Molothrus_bonariensis | Molothrus_bonariensis | LC | Increasing | tropical | diurnal | 41.5 | 13210499.693 | 7.16 |
| birds | Molothrus_oryzivorus | Molothrus_oryzivorus | LC | Decreasing | tropical | diurnal | 181.9 | 10872496.45 | 5.196 |
| birds | Molothrus_rufoaxillaris | Molothrus_rufoaxillaris | LC | Stable | temperate | diurnal | 47.5 | 4719929.578 | 9.835 |
| birds | Momotus_aequatorialis | Momotus_aequatorialis | LC | Decreasing | tropical | diurnal | 158 | 97380.484 | 8.818 |
| birds | Momotus_mexicanus | Momotus_mexicanus | LC | Decreasing | tropical | diurnal | 75.7 | 322967.33 | 9.063 |
| birds | Momotus_momota | Momotus_momota | LC | Decreasing | tropical | diurnal | 115 | 8096018.252 | 3.233 |
| birds | Monachella_muelleriana | Monachella_muelleriana | LC | Stable | tropical | diurnal | 24.5 | 405960.844 | 3.361 |
| birds | Monarcha_castaneiventris | Monarcha_castaneiventris | LC | Decreasing | tropical | diurnal | 27.3 | 27842.78 | 5.045 |
| birds | Monarcha_cinerascens | Monarcha_cinerascens | LC | Stable | tropical | diurnal | 29.6 | 148476.263 | 10.579 |
| birds | Monarcha_frater | Monarcha_frater | LC | Stable | tropical | diurnal | 22 | 226581.078 | 3.142 |
| birds | Monarcha_melanopsis | Monarcha_melanopsis | LC | Stable | temperate | diurnal | 22.7 | 571217.799 | 6.46 |
| birds | Monarcha_richardsii | NA | NT | Decreasing | NA | NA | NA | NA | NA |
| birds | Monasa_atra | Monasa_atra | LC | Stable | tropical | diurnal | 90.4 | 1731260.776 | 1.405 |
| birds | Monasa_flavirostris | Monasa_flavirostris | LC | Stable | tropical | diurnal | 62.8 | 785439.522 | 2.779 |
| birds | Monasa_morphoeus | Monasa_morphoeus | LC | Decreasing | tropical | diurnal | 87.9 | 5210334.014 | 2.72 |
| birds | Monasa_nigrifrons | Monasa_nigrifrons | LC | Decreasing | tropical | diurnal | 80.7 | 7021917.734 | 4.477 |
| birds | Monias_benschi | Monias_benschi | VU | Decreasing | tropical | diurnal | 137.9 | 12321.802 | 9.216 |
| birds | Montecincla_cachinnans | Strophocincla_cachinnans | EN | Decreasing | tropical | diurnal | 40.3 | 1070.705 | 10.55 |
| birds | Montecincla_jerdoni | NA | EN | Decreasing | NA | NA | NA | NA | NA |

| | | | | | | | | | |
|---|---|---|---|---|---|---|---|---|---|
| birds | Monticola_angolensis | Monticola_angolensis | LC | Stable | tropical | diurnal | 44.4 | 2218343.4 | 8.731 |
| birds | Monticola_brevipes | Monticola_brevipes | LC | Stable | tropical | diurnal | 33 | 1043690.358 | 7.321 |
| birds | Monticola_cinclorhyncha | Monticola_cinclorhynchus | LC | Stable | tropical | diurnal | 36 | 2943767.325 | 17.094 |
| birds | Monticola_explorator | Monticola_explorator | NT | Decreasing | temperate | diurnal | 44.4 | 368644.242 | 12.373 |
| birds | Monticola_gularis | Monticola_gularis | LC | Stable | temperate | diurnal | 34.6 | 3922614.369 | 9.958 |
| birds | Monticola_imerina | Monticola_imerinus | LC | Stable | temperate | diurnal | 28.3 | 18846.501 | 13.459 |
| birds | Monticola_rufiventris | Monticola_rufiventris | LC | Stable | temperate | diurnal | 53.2 | 3015321.115 | 12.165 |
| birds | Monticola_rupestris | Monticola_rupestris | LC | Stable | temperate | diurnal | 61.4 | 557569.05 | 12.054 |
| birds | Monticola_saxatilis | Monticola_saxatilis | LC | Decreasing | temperate | diurnal | 50.3 | 14463681.441 | 8.885 |
| birds | Monticola_semirufus | Myrmecocichla_semirufa | LC | Stable | tropical | diurnal | 46.7 | 386134.072 | 12.86 |
| birds | Monticola_sharpei | Monticola_sharpei | LC | Decreasing | tropical | diurnal | 24.8 | 242865.006 | 8.475 |
| birds | Monticola_solitarius | Monticola_solitarius | LC | Stable | tropical | diurnal | 53.7 | 21338038.167 | 12.441 |
| birds | Montifringilla_adamsi | Montifringilla_adamsi | LC | Stable | temperate | diurnal | 31.3 | 980500.668 | 5.955 |
| birds | Montifringilla_nivalis | Montifringilla_nivalis | LC | Stable | temperate | diurnal | 36.9 | 4085451.28 | 6.496 |
| birds | Morococcyx_erythropygus | Morococcyx_erythropygus | LC | Decreasing | tropical | diurnal | 64.5 | 358833.182 | 10.922 |
| birds | Morphnarchus_princeps | Leucopternis_princeps | LC | Decreasing | tropical | diurnal | 1000 | 113390.895 | 6.779 |
| birds | Morphnus_guianensis | Morphnus_guianensis | NT | Decreasing | tropical | diurnal | 1750 | 11702782.034 | 5.1 |
| birds | Morus_bassanus | Morus_bassanus | LC | Increasing | temperate | diurnal | 2998.7 | 18270116.025 | 12.518 |
| birds | Morus_capensis | Morus_capensis | EN | Decreasing | tropical | diurnal | 2643.4 | 1102841.446 | 11.966 |
| birds | Morus_serrator | Morus_serrator | LC | Increasing | temperate | diurnal | 2350 | 7286304.015 | 9.499 |
| birds | Motacilla_aguimp | Motacilla_aguimp | LC | Stable | tropical | diurnal | 27 | 11321309.288 | 9.005 |
| birds | Motacilla_alba | Motacilla_alba | LC | Stable | temperate | diurnal | 23.9 | 59320722.256 | 8.825 |
| birds | Motacilla_capensis | Motacilla_capensis | LC | Stable | tropical | diurnal | 20.8 | 3891210.174 | 8.186 |
| birds | Motacilla_cinerea | Motacilla_cinerea | LC | Stable | temperate | diurnal | 17.2 | 32926989.106 | 10.497 |
| birds | Motacilla_citreola | Motacilla_citreola | LC | Increasing | temperate | diurnal | 19.1 | 33176540.874 | 8.848 |
| birds | Motacilla_clara | Motacilla_clara | LC | Stable | tropical | diurnal | 17.9 | 3122291.692 | 9.947 |
| birds | Motacilla_flava | Motacilla_flava | LC | Decreasing | tropical | diurnal | 17.7 | 58182475.752 | 8.613 |
| birds | Motacilla_flaviventris | Motacilla_flaviventris | LC | Stable | tropical | diurnal | 22.5 | 591120.506 | 8.749 |
| birds | Motacilla_grandis | Motacilla_grandis | LC | Stable | temperate | diurnal | 30.2 | 435186.173 | 20.121 |
| birds | Motacilla_maderaspatensis | Motacilla_madaraspatensis | LC | Stable | tropical | diurnal | 30.5 | 2844265.944 | 18.879 |
| birds | Moupinia_poecilotis | Chrysomma_poecilotis | LC | Decreasing | temperate | diurnal | 18.7 | 261236.646 | 8.201 |
| birds | Mulleripicus_fulvus | Mulleripicus_fulvus | LC | Decreasing | tropical | diurnal | 312.3 | 103818.081 | 7 |
| birds | Mulleripicus_funebris | Mulleripicus_funebris | NT | Decreasing | tropical | diurnal | 164 | 102097.412 | 15.205 |
| birds | Mulleripicus_pulverulentus | Mulleripicus_pulverulentus | VU | Decreasing | tropical | diurnal | 450.2 | 2625286.938 | 11.062 |

| | | | | | | | | | |
|---|---|---|---|---|---|---|---|---|---|
| birds | Muscicapa_adusta | Muscicapa_adusta | LC | Decreasing | tropical | diurnal | 10.4 | 2389660.854 | 11.389 |
| birds | Muscicapa_aquatica | Muscicapa_aquatica | LC | Stable | tropical | diurnal | 11.5 | 1932273.674 | 11.929 |
| birds | Muscicapa_cassini | Muscicapa_cassini | LC | Stable | tropical | diurnal | 17.3 | 3716280.129 | 7.668 |
| birds | Muscicapa_dauurica | Muscicapa_dauurica | LC | Stable | temperate | diurnal | 9.8 | 17791128.025 | 11.413 |
| birds | Muscicapa_epulata | Muscicapa_epulata | LC | Stable | tropical | diurnal | 9.6 | 903510.24 | 9.795 |
| birds | Muscicapa_ferruginea | Muscicapa_ferruginea | LC | Decreasing | temperate | diurnal | 12 | 2544221.997 | 11.674 |
| birds | Muscicapa_gambagae | Muscicapa_gambagae | LC | Stable | tropical | diurnal | 13 | 669433.068 | 10.296 |
| birds | Muscicapa_griseisticta | Muscicapa_griseisticta | LC | Stable | temperate | diurnal | 15.9 | 2648878.537 | 5.841 |
| birds | Muscicapa_muttui | Muscicapa_muttui | LC | Decreasing | temperate | diurnal | 12 | 1292048.538 | 12.002 |
| birds | Muscicapa_randi | Muscicapa_randi | VU | Decreasing | tropical | diurnal | 20.1 | 105159.839 | 14.884 |
| birds | Muscicapa_segregata | NA | NT | Decreasing | NA | NA | NA | NA | NA |
| birds | Muscicapa_sethsmithi | Muscicapa_sethsmithi | LC | Stable | tropical | diurnal | 9.1 | 1294819.353 | 6.378 |
| birds | Muscicapa_sibirica | Muscicapa_sibirica | LC | Stable | temperate | diurnal | 13.2 | 17841855.276 | 9.804 |
| birds | Muscicapa_striata | Muscicapa_striata | LC | Decreasing | tropical | diurnal | 15.9 | 47138089.09 | 9.447 |
| birds | Muscigralla_brevicauda | Muscigralla_brevicauda | LC | Stable | tropical | diurnal | 12.6 | 166922.918 | 8.26 |
| birds | Muscipipra_vetula | Muscipipra_vetula | LC | Stable | temperate | diurnal | 27 | 828419.997 | 13.12 |
| birds | Muscisaxicola_albifrons | Muscisaxicola_albifrons | LC | Stable | tropical | diurnal | 34 | 285962.861 | 8.695 |
| birds | Muscisaxicola_albilora | Muscisaxicola_albilora | LC | Stable | temperate | diurnal | 22.6 | 1237528.622 | 6.389 |
| birds | Muscisaxicola_alpinus | Muscisaxicola_alpinus | LC | Stable | tropical | diurnal | 26.1 | 42973.19 | 13.123 |
| birds | Muscisaxicola_capistratus | Muscisaxicola_capistratus | LC | Unknown | temperate | diurnal | 26.6 | 2111097.807 | 5.597 |
| birds | Muscisaxicola_cinereus | Muscisaxicola_cinereus | LC | Stable | temperate | diurnal | 18.6 | 770735.32 | 6.143 |
| birds | Muscisaxicola_flavinucha | Muscisaxicola_flavinucha | LC | Stable | temperate | diurnal | 36.2 | 1456441.881 | 5.694 |
| birds | Muscisaxicola_fluviatilis | Muscisaxicola_fluviatilis | LC | Stable | tropical | diurnal | 14.7 | 1234514.008 | 3.369 |
| birds | Muscisaxicola_frontalis | Muscisaxicola_frontalis | LC | Stable | temperate | diurnal | 28.9 | 807477.647 | 5.118 |
| birds | Muscisaxicola_griseus | Muscisaxicola_griseus | LC | Stable | tropical | diurnal | 20.2 | 313132.732 | 8.173 |
| birds | Muscisaxicola_juninensis | Muscisaxicola_juninensis | LC | Stable | tropical | diurnal | 22.2 | 514054.922 | 7.523 |
| birds | Muscisaxicola_maclovianus | Muscisaxicola_maclovianus | LC | Decreasing | temperate | diurnal | 23.8 | 2835656.756 | 7.169 |
| birds | Muscisaxicola_maculirostris | Muscisaxicola_maculirostris | LC | Stable | temperate | diurnal | 14.2 | 1996565.976 | 5.967 |
| birds | Muscisaxicola_rufivertex | Muscisaxicola_rufivertex | LC | Stable | tropical | diurnal | 20 | 696474.956 | 5.656 |
| birds | Musophaga_rossae | Musophaga_rossae | LC | Stable | tropical | diurnal | 361.4 | 3118175.853 | 8.017 |
| birds | Musophaga_violacea | Musophaga_violacea | LC | Stable | tropical | diurnal | 360 | 2077717.138 | 13.187 |
| birds | Myadestes_coloratus | Myadestes_coloratus | NT | Decreasing | tropical | diurnal | 28.9 | 2087.017 | 2.013 |
| birds | Myadestes_elisabeth | Myadestes_elisabeth | NT | Decreasing | tropical | diurnal | 30 | 8505.105 | 14.003 |
| birds | Myadestes_genibarbis | Myadestes_genibarbis | LC | Decreasing | tropical | diurnal | 29.2 | 88012.576 | 16.429 |

| | | | | | | | | | |
|---|---|---|---|---|---|---|---|---|---|
| birds | Myadestes_lanaiensis | Myadestes_lanaiensis | CR | Unknown | tropical | diurnal | 50 | 1508.294 | 8.916 |
| birds | Myadestes_melanops | Myadestes_melanops | LC | Decreasing | tropical | diurnal | 32.1 | 31920.032 | 9.574 |
| birds | Myadestes_myadestinus | NA | EX | not_available | NA | NA | NA | NA | NA |
| birds | Myadestes_obscurus | Myadestes_obscurus | NT | Unknown | tropical | diurnal | 50.4 | 6883.285 | 5.832 |
| birds | Myadestes_occidentalis | Myadestes_occidentalis | LC | Decreasing | tropical | diurnal | 36.4 | 528718.342 | 10.917 |
| birds | Myadestes_palmeri | Myadestes_palmeri | CR | Stable | tropical | diurnal | 39.6 | 33.079 | 1.124 |
| birds | Myadestes_ralloides | Myadestes_ralloides | LC | Decreasing | tropical | diurnal | 29.1 | 457703.182 | 8.694 |
| birds | Myadestes_townsendi | Myadestes_townsendi | LC | Stable | temperate | diurnal | 33.2 | 6168503.62 | 7.639 |
| birds | Myadestes_unicolor | Myadestes_unicolor | LC | Decreasing | tropical | diurnal | 37.9 | 184166.539 | 10.572 |
| birds | Mycerobas_affinis | Mycerobas_affinis | LC | Stable | temperate | diurnal | 83 | 1568902.726 | 8.319 |
| birds | Mycerobas_carnipes | Mycerobas_carnipes | LC | Stable | temperate | diurnal | 58.8 | 6584591.304 | 6.069 |
| birds | Mycerobas_icterioides | Mycerobas_icterioides | LC | Stable | temperate | diurnal | 67 | 406197.585 | 10.369 |
| birds | Mycerobas_melanozanthos | Mycerobas_melanozanthos | LC | Stable | temperate | diurnal | 62 | 1761447.644 | 8.835 |
| birds | Mycteria_americana | Mycteria_americana | LC | Decreasing | tropical | diurnal | 2554.5 | 14862621.423 | 6.375 |
| birds | Mycteria_cinerea | Mycteria_cinerea | EN | Decreasing | tropical | diurnal | 2393 | 155396.257 | 15.653 |
| birds | Mycteria_ibis | Mycteria_ibis | LC | Decreasing | tropical | diurnal | 2155.6 | 15368408.037 | 9.292 |
| birds | Mycteria_leucocephala | Mycteria_leucocephala | LC | Increasing | tropical | diurnal | 3180 | 3983383.997 | 17.793 |
| birds | Myiagra_albiventris | NA | NT | Unknown | NA | NA | NA | NA | NA |
| birds | Myiagra_alecto | Myiagra_alecto | LC | Stable | tropical | diurnal | 23.6 | 1376502.43 | 3.289 |
| birds | Myiagra_atra | NA | NT | Decreasing | NA | NA | NA | NA | NA |
| birds | Myiagra_azureocapilla | Myiagra_azureocapilla | NT | Decreasing | tropical | diurnal | 12.9 | 441.315 | 12.495 |
| birds | Myiagra_caledonica | Myiagra_caledonica | LC | Stable | tropical | diurnal | 10.8 | 31488.185 | 6.782 |
| birds | Myiagra_cervinicauda | Myiagra_cervinicauda | NT | Decreasing | tropical | diurnal | 11.5 | 3391.572 | 4.614 |
| birds | Myiagra_cyanoleuca | Myiagra_cyanoleuca | LC | Decreasing | temperate | diurnal | 17.5 | 1375898.132 | 7.156 |
| birds | Myiagra_erythrops | NA | LC | Stable | NA | NA | NA | NA | NA |
| birds | Myiagra_ferrocyanea | Myiagra_ferrocyanea | LC | Stable | tropical | diurnal | 12.8 | 32643.695 | 4.707 |
| birds | Myiagra_freycineti | NA | EX | not_available | NA | NA | NA | NA | NA |
| birds | Myiagra_galeata | Myiagra_galeata | LC | Decreasing | tropical | diurnal | 11 | 44624.814 | 7.109 |
| birds | Myiagra_hebetior | Myiagra_hebetior | VU | Decreasing | tropical | diurnal | 20.2 | 342.837 | 4.079 |
| birds | Myiagra_inquieta | Myiagra_inquieta | LC | Decreasing | temperate | diurnal | 20.9 | 2820032.992 | 7.225 |
| birds | Myiagra_oceanica | NA | LC | Stable | NA | NA | NA | NA | NA |
| birds | Myiagra_pluto | NA | LC | Stable | NA | NA | NA | NA | NA |
| birds | Myiagra_rubecula | Myiagra_rubecula | LC | Stable | temperate | diurnal | 12.1 | 2268381.697 | 5.437 |
| birds | Myiagra_ruficollis | Myiagra_ruficollis | LC | Decreasing | tropical | diurnal | 10.1 | 580888.401 | 3.33 |

| | | | | | | | | | |
|---|---|---|---|---|---|---|---|---|---|
| birds | Myiagra_vanikorensis | Myiagra_vanikorensis | LC | Stable | tropical | diurnal | 12.3 | 18452.635 | 9.292 |
| birds | Myiarchus_antillarum | Myiarchus_antillarum | LC | Decreasing | tropical | diurnal | 23.4 | 9177.012 | 27.041 |
| birds | Myiarchus_apicalis | Myiarchus_apicalis | LC | Increasing | tropical | diurnal | 29.2 | 41557.905 | 14.429 |
| birds | Myiarchus_barbirostris | Myiarchus_barbirostris | LC | Decreasing | tropical | diurnal | 13.4 | 11021.791 | 19.741 |
| birds | Myiarchus_cephalotes | Myiarchus_cephalotes | LC | Stable | tropical | diurnal | 26.3 | 292641.439 | 8.171 |
| birds | Myiarchus_cinerascens | Myiarchus_cinerascens | LC | Increasing | temperate | diurnal | 28.2 | 4277784.591 | 9.815 |
| birds | Myiarchus_crinitus | Myiarchus_crinitus | LC | Stable | temperate | diurnal | 32.1 | 6748704.123 | 14.829 |
| birds | Myiarchus_ferox | Myiarchus_ferox | LC | Stable | tropical | diurnal | 27.5 | 11854121.155 | 5.136 |
| birds | Myiarchus_magnirostris | NA | LC | Decreasing | NA | NA | NA | NA | NA |
| birds | Myiarchus_nugator | Myiarchus_nugator | LC | Decreasing | tropical | diurnal | 37 | 774.933 | 18.553 |
| birds | Myiarchus_nuttingi | Myiarchus_nuttingi | LC | Decreasing | tropical | diurnal | 23 | 634487.915 | 10.092 |
| birds | Myiarchus_oberi | Myiarchus_oberi | LC | Decreasing | tropical | diurnal | 34.7 | 4421.692 | 22.021 |
| birds | Myiarchus_panamensis | Myiarchus_panamensis | LC | Stable | tropical | diurnal | 31.7 | 370862.952 | 12.14 |
| birds | Myiarchus_phaeocephalus | Myiarchus_phaeocephalus | LC | Decreasing | tropical | diurnal | 26.3 | 99662.269 | 11.325 |
| birds | Myiarchus_sagrae | Myiarchus_sagrae | LC | Decreasing | tropical | diurnal | 19.3 | 122416.833 | 15.276 |
| birds | Myiarchus_semirufus | Myiarchus_semirufus | VU | Decreasing | tropical | diurnal | 22.5 | 55766.984 | 8.497 |
| birds | Myiarchus_stolidus | Myiarchus_stolidus | LC | Decreasing | tropical | diurnal | 20.8 | 86228.495 | 16.269 |
| birds | Myiarchus_swainsoni | Myiarchus_swainsoni | LC | Stable | tropical | diurnal | 25.1 | 13661126.222 | 5.674 |
| birds | Myiarchus_tuberculifer | Myiarchus_tuberculifer | LC | Decreasing | tropical | diurnal | 17.7 | 10588401.296 | 4.715 |
| birds | Myiarchus_tyrannulus | Myiarchus_tyrannulus | LC | Increasing | tropical | diurnal | 35.5 | 10591424.148 | 7.427 |
| birds | Myiarchus_validus | Myiarchus_validus | LC | Decreasing | tropical | diurnal | 40.6 | 11021.791 | 19.741 |
| birds | Myiarchus_venezuelensis | Myiarchus_venezuelensis | LC | Stable | tropical | diurnal | 29.8 | 234706.397 | 12.238 |
| birds | Myiarchus_yucatanensis | Myiarchus_yucatanensis | LC | Decreasing | tropical | diurnal | 19.6 | 165390.723 | 6.521 |
| birds | Myiobius_atricaudus | Myiobius_atricaudus | LC | Decreasing | tropical | diurnal | 10 | 4897288.434 | 5.152 |
| birds | Myiobius_barbatus | Myiobius_barbatus | LC | Stable | tropical | diurnal | 11 | 6231670.561 | 2.691 |
| birds | Myiobius_sulphureipygius | Myiobius_sulphureipygius | LC | Decreasing | tropical | diurnal | 11.6 | 640104.92 | 8.925 |
| birds | Myiobius_villosus | Myiobius_villosus | LC | Decreasing | tropical | diurnal | 14.1 | 270819.423 | 6.274 |
| birds | Myioborus_albifacies | Myioborus_albifacies | LC | Decreasing | tropical | diurnal | 15 | 17030.045 | 0.632 |
| birds | Myioborus_albifrons | NA | NT | Stable | NA | NA | NA | NA | NA |
| birds | Myioborus_brunniceps | Myioborus_brunniceps | LC | Decreasing | temperate | diurnal | 9.2 | 112336.705 | 7.832 |
| birds | Myioborus_cardonai | NA | NT | Decreasing | NA | NA | NA | NA | NA |
| birds | Myioborus_castaneocapilla | Myioborus_castaneocapilla | LC | Decreasing | tropical | diurnal | 10.9 | 95602.505 | 1.074 |
| birds | Myioborus_flavivertex | Myioborus_flavivertex | LC | Stable | tropical | diurnal | 15 | 3571.556 | 6.118 |
| birds | Myioborus_melanocephalus | Myioborus_melanocephalus | LC | Decreasing | tropical | diurnal | 11.3 | 185791.055 | 7.132 |

| | | | | | | | | | |
|---|---|---|---|---|---|---|---|---|---|
| birds | Myioborus_miniatus | Myioborus_miniatus | LC | Decreasing | tropical | diurnal | 9 | 1254327.4 | 8.608 |
| birds | Myioborus_ornatus | Myioborus_ornatus | LC | Stable | tropical | diurnal | 11.7 | 39629.134 | 13.332 |
| birds | Myioborus_pariae | NA | EN | Decreasing | NA | NA | NA | NA | NA |
| birds | Myioborus_pictus | Myioborus_pictus | LC | Decreasing | temperate | diurnal | 9.8 | 1124631.495 | 9.184 |
| birds | Myioborus_torquatus | Myioborus_torquatus | LC | Decreasing | tropical | diurnal | 10.5 | 5261.716 | 5.945 |
| birds | Myiodynastes_bairdii | Myiodynastes_bairdii | LC | Stable | tropical | diurnal | 45 | 96352.031 | 10.804 |
| birds | Myiodynastes_chrysocephalus | Myiodynastes_chrysocephalus | LC | Increasing | tropical | diurnal | 38.3 | 176225.835 | 5.905 |
| birds | Myiodynastes_hemichrysus | Myiodynastes_hemichrysus | LC | Decreasing | tropical | diurnal | 43.4 | 319266.742 | 10.304 |
| birds | Myiodynastes_luteiventris | Myiodynastes_luteiventris | LC | Stable | tropical | diurnal | 46.9 | 2795270.692 | 6.878 |
| birds | Myiodynastes_maculatus | Myiodynastes_maculatus | LC | Stable | tropical | diurnal | 43.2 | 4786616.481 | 4.348 |
| birds | Myiomela_diana | Cinclidium_diana | LC | Decreasing | tropical | diurnal | 26.2 | 91597.851 | 8.005 |
| birds | Myiomela_leucura | Cinclidium_leucurum | LC | Decreasing | tropical | diurnal | 26.8 | 2713438.904 | 11.611 |
| birds | Myiopagis_caniceps | Myiopagis_caniceps | LC | Decreasing | tropical | diurnal | 10.5 | 2761484.965 | 10.752 |
| birds | Myiopagis_cotta | Myiopagis_cotta | LC | Decreasing | tropical | diurnal | 11.6 | 11021.791 | 19.741 |
| birds | Myiopagis_flavivertex | Myiopagis_flavivertex | LC | Decreasing | tropical | diurnal | 11.8 | 1303223.78 | 2.113 |
| birds | Myiopagis_gaimardii | Myiopagis_gaimardii | LC | Stable | tropical | diurnal | 12 | 8673168.363 | 3.615 |
| birds | Myiopagis_olallai | Myiopagis_olallai | LC | Decreasing | tropical | diurnal | 12.1 | 17655.696 | 5.672 |
| birds | Myiopagis_subplacens | Myiopagis_subplacens | LC | Decreasing | tropical | diurnal | 15.9 | 109447.791 | 11.847 |
| birds | Myiopagis_viridicata | Myiopagis_viridicata | LC | Decreasing | tropical | diurnal | 11.5 | 7381273.665 | 8.228 |
| birds | Myiophobus_cryptoxanthus | Myiophobus_cryptoxanthus | LC | Increasing | tropical | diurnal | 9.8 | 70456.924 | 4.739 |
| birds | Myiophobus_fasciatus | Myiophobus_fasciatus | LC | Increasing | tropical | diurnal | 9.9 | 10017556.285 | 7.577 |
| birds | Myiophobus_flavicans | Myiophobus_flavicans | LC | Stable | tropical | diurnal | 12.7 | 272300.039 | 9.783 |
| birds | Myiophobus_inornatus | Myiophobus_inornatus | LC | Decreasing | tropical | diurnal | 11.2 | 63282.278 | 4.698 |
| birds | Myiophobus_phoenicomitra | Myiophobus_phoenicomitra | LC | Decreasing | tropical | diurnal | 11 | 45578.521 | 5.946 |
| birds | Myiophobus_roraimae | Myiophobus_roraimae | LC | Stable | tropical | diurnal | 13.6 | 247059.834 | 1.17 |
| birds | Myiopsitta_luchsi | NA | NT | Decreasing | NA | NA | NA | NA | NA |
| birds | Myiopsitta_monachus | Myiopsitta_monachus | LC | Increasing | temperate | diurnal | 120 | 3053903.303 | 9.626 |
| birds | Myiornis_albiventris | Myiornis_albiventris | LC | Decreasing | tropical | diurnal | 5.5 | 102039.107 | 5.158 |
| birds | Myiornis_atricapillus | Myiornis_atricapillus | LC | Decreasing | tropical | diurnal | 5.7 | 184278.946 | 7.685 |
| birds | Myiornis_auricularis | Myiornis_auricularis | LC | Decreasing | tropical | diurnal | 5.3 | 1267042.855 | 12.981 |
| birds | Myiornis_ecaudatus | Myiornis_ecaudatus | LC | Decreasing | tropical | diurnal | 5.3 | 6431271.735 | 2.242 |
| birds | Myiotheretes_fumigatus | Myiotheretes_fumigatus | LC | Stable | tropical | diurnal | 33.4 | 214456.428 | 10.488 |
| birds | Myiotheretes_fuscorufus | Myiotheretes_fuscorufus | LC | Stable | tropical | diurnal | 31.7 | 109817.509 | 5.912 |
| birds | Myiotheretes_pernix | Myiotheretes_pernix | EN | Decreasing | tropical | diurnal | 33 | 698.896 | 4.866 |

| | | | | | | | | | |
|---|---|---|---|---|---|---|---|---|---|
| birds | Myiotheretes_striaticollis | Myiotheretes_striaticollis | LC | Stable | tropical | diurnal | 61.2 | 632466.904 | 8.525 |
| birds | Myiothlypis_basilica | Basileuterus_basilicus | NT | Stable | tropical | diurnal | 16.2 | 1405.825 | 5.648 |
| birds | Myiothlypis_bivittata | Basileuterus_bivittatus | LC | Unknown | tropical | diurnal | 14.6 | 116028.344 | 5.229 |
| birds | Myiothlypis_chlorophrys | Basileuterus_chlorophrys | LC | Unknown | tropical | diurnal | 16.2 | 24162.634 | 6.638 |
| birds | Myiothlypis_chrysogaster | Basileuterus_chrysogaster | LC | Unknown | tropical | diurnal | 11.2 | 42961.207 | 3.382 |
| birds | Myiothlypis_cinereicollis | Basileuterus_cinereicollis | LC | Decreasing | tropical | diurnal | 17.2 | 52299.378 | 12.149 |
| birds | Myiothlypis_conspicillata | Basileuterus_conspicillatus | NT | Decreasing | tropical | diurnal | 13.7 | 7320.248 | 7.478 |
| birds | Myiothlypis_coronata | Basileuterus_coronatus | LC | Decreasing | tropical | diurnal | 16.3 | 417599.18 | 8.471 |
| birds | Myiothlypis_flaveola | Basileuterus_flaveolus | LC | Decreasing | tropical | diurnal | 13.2 | 4100375.803 | 8.502 |
| birds | Myiothlypis_fraseri | Basileuterus_fraseri | LC | Decreasing | tropical | diurnal | 11.6 | 60912.852 | 14.1 |
| birds | Myiothlypis_fulvicauda | Phaeothlypis_fulvicauda | LC | Decreasing | tropical | diurnal | 14.9 | 2473113.958 | 2.893 |
| birds | Myiothlypis_leucoblephara | Basileuterus_leucoblepharus | LC | Decreasing | temperate | diurnal | 15.1 | 1572872.225 | 12.272 |
| birds | Myiothlypis_leucophrys | Basileuterus_leucophrys | LC | Decreasing | tropical | diurnal | 18.1 | 808600.056 | 11.824 |
| birds | Myiothlypis_luteoviridis | Basileuterus_luteoviridis | LC | Decreasing | tropical | diurnal | 12.9 | 124562.365 | 10.286 |
| birds | Myiothlypis_nigrocristata | Basileuterus_nigrocristatus | LC | Decreasing | tropical | diurnal | 13.6 | 193235.07 | 10.961 |
| birds | Myiothlypis_rivularis | Phaeothlypis_rivularis | LC | Decreasing | tropical | diurnal | 13.5 | 845622.984 | 13.274 |
| birds | Myiothlypis_signata | Basileuterus_signatus | LC | Decreasing | tropical | diurnal | 11.3 | 162524.353 | 7.845 |
| birds | Myiotriccus_ornatus | Myiotriccus_ornatus | LC | Stable | tropical | diurnal | 9.9 | 149294.032 | 11.515 |
| birds | Myiozetetes_cayanensis | Myiozetetes_cayanensis | LC | Stable | tropical | diurnal | 25.9 | 7536061.767 | 5.067 |
| birds | Myiozetetes_granadensis | Myiozetetes_granadensis | LC | Stable | tropical | diurnal | 29.3 | 3322072.636 | 2.801 |
| birds | Myiozetetes_luteiventris | Myiozetetes_luteiventris | LC | Stable | tropical | diurnal | 19 | 3959233.083 | 1.357 |
| birds | Myiozetetes_similis | Myiozetetes_similis | LC | Stable | tropical | diurnal | 28 | 10242322.01 | 6.112 |
| birds | Myophonus_blighi | NA | EN | Decreasing | NA | NA | NA | NA | NA |
| birds | Myophonus_caeruleus | Myophonus_caeruleus | LC | Unknown | tropical | diurnal | 158 | 7139938.197 | 12.177 |
| birds | Myophonus_glaucinus | Myophonus_glaucinus | LC | Decreasing | tropical | diurnal | 89.7 | 24651.374 | 15.901 |
| birds | Myophonus_horsfieldii | Myophonus_horsfieldii | LC | Unknown | tropical | diurnal | 117 | 412482.187 | 17.812 |
| birds | Myophonus_insularis | NA | LC | Decreasing | NA | NA | NA | NA | NA |
| birds | Myophonus_melanurus | Myophonus_melanurus | LC | Decreasing | tropical | diurnal | 66.6 | 66916.959 | 7.458 |
| birds | Myophonus_robinsoni | Myophonus_robinsoni | NT | Decreasing | tropical | diurnal | 99.2 | 10816.585 | 5.105 |
| birds | Myopornis_boehmi | Muscicapa_boehmi | LC | Decreasing | tropical | diurnal | 19.3 | 863704.753 | 7.674 |
| birds | Myornis_senilis | Myornis_senilis | LC | Stable | tropical | diurnal | 21.2 | 118973.231 | 10.114 |
| birds | Myrmeciza_longipes | Myrmeciza_longipes | LC | Decreasing | tropical | diurnal | 27.9 | 1110901.772 | 7.118 |
| birds | Myrmecocichla_aethiops | Myrmecocichla_aethiops | LC | Stable | tropical | diurnal | 56.6 | 2195986.845 | 11.749 |
| birds | Myrmecocichla_arnotti | Myrmecocichla_arnoti | LC | Decreasing | tropical | diurnal | 39.6 | 3102386.997 | 8.257 |

| | | | | | | | | | |
|---|---|---|---|---|---|---|---|---|---|
| birds | Myrmecocichla_formicivora | Myrmecocichla_formicivora | LC | Stable | temperate | diurnal | 41.6 | 2080546.725 | 7.315 |
| birds | Myrmecocichla_melaena | NA | LC | Stable | NA | NA | NA | NA | NA |
| birds | Myrmecocichla_monticola | Oenanthe_monticola | LC | Stable | temperate | diurnal | 33.1 | 1391804.191 | 8.508 |
| birds | Myrmecocichla_nigra | Myrmecocichla_nigra | LC | Stable | tropical | diurnal | 40.3 | 3505287.839 | 7.984 |
| birds | Myrmecocichla_tholloni | NA | LC | Stable | NA | NA | NA | NA | NA |
| birds | Myrmelastes_caurensis | Schistocichla_caurensis | LC | Decreasing | tropical | diurnal | 39.4 | 240244 | 0.798 |
| birds | Myrmelastes_hyperythrus | Myrmeciza_hyperythra | LC | Stable | tropical | diurnal | 41 | 1642348.581 | 1.646 |
| birds | Myrmelastes_leucostigma | Schistocichla_leucostigma | LC | Stable | tropical | diurnal | 25.7 | 2969595.639 | 1.692 |
| birds | Myrmelastes_schistaceus | Schistocichla_schistacea | LC | Decreasing | tropical | diurnal | 23.8 | 529491.189 | 1.217 |
| birds | Myrmia_micrura | Myrmia_micrura | LC | Stable | tropical | diurnal | 2.3 | 72174.994 | 9.719 |
| birds | Myrmoborus_leucophrys | Myrmoborus_leucophrys | LC | Decreasing | tropical | diurnal | 21.5 | 5127312.125 | 2.498 |
| birds | Myrmoborus_lophotes | Percnostola_lophotes | LC | Decreasing | tropical | diurnal | 28 | 202927.683 | 1.618 |
| birds | Myrmoborus_lugubris | Myrmoborus_lugubris | LC | Decreasing | tropical | diurnal | 21.4 | 214631.65 | 2.316 |
| birds | Myrmoborus_melanurus | Myrmoborus_melanurus | NT | Decreasing | tropical | diurnal | 18.1 | 92005.739 | 2.757 |
| birds | Myrmoborus_myotherinus | Myrmoborus_myotherinus | LC | Stable | tropical | diurnal | 18.8 | 4778353.328 | 2.209 |
| birds | Myrmochanes_hemileucus | Myrmochanes_hemileucus | LC | Stable | tropical | diurnal | 12.5 | 261134.969 | 2.957 |
| birds | Myrmoderus_ferrugineus | Myrmeciza_ferruginea | LC | Stable | tropical | diurnal | 26.1 | 1290324.219 | 1.27 |
| birds | Myrmoderus_loricatus | Myrmeciza_loricata | LC | Decreasing | tropical | diurnal | 15.5 | 177950.427 | 12.411 |
| birds | Myrmoderus_ruficauda | Myrmeciza_ruficauda | EN | Decreasing | tropical | diurnal | 26 | 129646.485 | 10.6 |
| birds | Myrmoderus_squamosus | Myrmeciza_squamosa | LC | Decreasing | temperate | diurnal | 18.5 | 241730.196 | 14.153 |
| birds | Myrmophylax_atrothorax | Myrmeciza_atrothorax | LC | Stable | tropical | diurnal | 16.1 | 6800794.009 | 2.228 |
| birds | Myrmorchilus_strigilatus | Myrmorchilus_strigilatus | LC | Decreasing | tropical | diurnal | 23 | 1728828.969 | 6.894 |
| birds | Myrmornis_torquata | Myrmornis_torquata | LC | Decreasing | tropical | diurnal | 46.7 | 2996202.86 | 1.813 |
| birds | Myrmothera_berlepschi | Hylopezus_berlepschi | LC | Stable | tropical | diurnal | 48 | 1422657.013 | 2.475 |
| birds | Myrmothera_campanisona | Myrmothera_campanisona | LC | Decreasing | tropical | diurnal | 47 | 4089925.725 | 1.517 |
| birds | Myrmothera_dives | Hylopezus_dives | LC | Decreasing | tropical | diurnal | 44 | 179845.908 | 6.27 |
| birds | Myrmothera_fulviventris | Hylopezus_fulviventris | LC | Stable | tropical | diurnal | 41 | 252414.721 | 2.245 |
| birds | Myrmothera_simplex | Myrmothera_simplex | LC | Stable | tropical | diurnal | 52.4 | 113218.798 | 1.148 |
| birds | Myrmotherula_ambigua | Myrmotherula_ambigua | LC | Stable | tropical | diurnal | 6.7 | 257668.14 | 0.483 |
| birds | Myrmotherula_assimilis | Myrmotherula_assimilis | LC | Decreasing | tropical | diurnal | 9.3 | 412230.35 | 2.057 |
| birds | Myrmotherula_axillaris | Myrmotherula_axillaris | LC | Decreasing | tropical | diurnal | 8.1 | 7734412.169 | 2.887 |
| birds | Myrmotherula_behni | Myrmotherula_behni | LC | Decreasing | tropical | diurnal | 8 | 139424.09 | 1.189 |
| birds | Myrmotherula_brachyura | Myrmotherula_brachyura | LC | Stable | tropical | diurnal | 6.4 | 6190843.58 | 1.888 |
| birds | Myrmotherula_cherriei | Myrmotherula_cherriei | LC | Decreasing | tropical | diurnal | 8.2 | 676511.226 | 1.69 |

| | | | | | | | | | |
|---|---|---|---|---|---|---|---|---|---|
| birds | Myrmotherula_grisea | Myrmotherula_grisea | LC | Decreasing | tropical | diurnal | 8.4 | 67024.249 | 4.529 |
| birds | Myrmotherula_ignota | Myrmotherula_ignota | LC | Decreasing | tropical | diurnal | 7.1 | 1558811.941 | 2.183 |
| birds | Myrmotherula_iheringi | Myrmotherula_iheringi | LC | Decreasing | tropical | diurnal | 8 | 1149280.579 | 1.336 |
| birds | Myrmotherula_klagesi | Myrmotherula_klagesi | VU | Decreasing | tropical | diurnal | 7.8 | 61302.44 | 3.056 |
| birds | Myrmotherula_longicauda | Myrmotherula_longicauda | LC | Decreasing | tropical | diurnal | 8.3 | 153775.139 | 3.956 |
| birds | Myrmotherula_longipennis | Myrmotherula_longipennis | LC | Stable | tropical | diurnal | 9.4 | 5117183.47 | 1.575 |
| birds | Myrmotherula_luctuosa | Myrmotherula_luctuosa | LC | Decreasing | tropical | diurnal | 8.5 | 247649.4 | 12.255 |
| birds | Myrmotherula_menetriesii | Myrmotherula_menetriesii | LC | Decreasing | tropical | diurnal | 8.6 | 6331647.422 | 1.914 |
| birds | Myrmotherula_minor | Myrmotherula_minor | VU | Decreasing | tropical | diurnal | 6.4 | 40787.077 | 12.687 |
| birds | Myrmotherula_multostriata | Myrmotherula_multostriata | LC | Stable | tropical | diurnal | 8 | 4598823.752 | 2.175 |
| birds | Myrmotherula_pacifica | Myrmotherula_pacifica | LC | Decreasing | tropical | diurnal | 10.1 | 236644.015 | 8.808 |
| birds | Myrmotherula_schisticolor | Myrmotherula_schisticolor | LC | Decreasing | tropical | diurnal | 9.6 | 534175.268 | 8.666 |
| birds | Myrmotherula_sclateri | Myrmotherula_sclateri | LC | Decreasing | tropical | diurnal | 8.4 | 2651999.979 | 1.833 |
| birds | Myrmotherula_snowi | Myrmotherula_snowi | CR | Decreasing | tropical | diurnal | 9.6 | 285.42 | 8.012 |
| birds | Myrmotherula_sunensis | Myrmotherula_sunensis | LC | Decreasing | tropical | diurnal | 9.4 | 178742.397 | 2.423 |
| birds | Myrmotherula_surinamensis | Myrmotherula_surinamensis | LC | Stable | tropical | diurnal | 8.3 | 1574543.303 | 1.447 |
| birds | Myrmotherula_unicolor | Myrmotherula_unicolor | LC | Decreasing | temperate | diurnal | 11.5 | 73866.154 | 12.518 |
| birds | Myrmotherula_urosticta | Myrmotherula_urosticta | VU | Decreasing | tropical | diurnal | 8.4 | 118830.458 | 11.172 |
| birds | Myrtis_fanny | Myrtis_fanny | LC | Stable | tropical | diurnal | 2.3 | 256471.505 | 7.969 |
| birds | Mystacornis_crossleyi | Mystacornis_crossleyi | LC | Unknown | tropical | diurnal | 24.5 | 159923.75 | 7.997 |
| birds | Myza_celebensis | Myza_celebensis | LC | Decreasing | tropical | diurnal | 20 | 42076.191 | 5.52 |
| birds | Myza_sarasinorum | Myza_sarasinorum | LC | Decreasing | tropical | diurnal | 22.5 | 9804.189 | 4.524 |
| birds | Myzomela_adolphinae | Myzomela_adolphinae | LC | Stable | tropical | diurnal | 7.4 | 80091.129 | 5.031 |
| birds | Myzomela_albigula | NA | DD | Unknown | NA | NA | NA | NA | NA |
| birds | Myzomela_blasii | Myzomela_blasii | LC | Stable | tropical | diurnal | 14.9 | 11340.376 | 5.948 |
| birds | Myzomela_boiei | Myzomela_boiei | LC | Stable | tropical | diurnal | 10.8 | 5137.121 | 6.775 |
| birds | Myzomela_caledonica | Myzomela_caledonica | LC | Stable | tropical | diurnal | 7 | 16533.262 | 7.541 |
| birds | Myzomela_cardinalis | Myzomela_cardinalis | LC | Decreasing | tropical | diurnal | 13.8 | 22202.102 | 5.616 |
| birds | Myzomela_chermesina | NA | VU | Stable | NA | NA | NA | NA | NA |
| birds | Myzomela_chloroptera | Myzomela_chloroptera | LC | Unknown | tropical | diurnal | 10.8 | 113064.207 | 6.824 |
| birds | Myzomela_cineracea | Myzomela_cineracea | LC | Unknown | tropical | diurnal | 13.8 | 32686.464 | 4.377 |
| birds | Myzomela_cruentata | Myzomela_cruentata | LC | Unknown | tropical | diurnal | 8.2 | 199161.108 | 3.216 |
| birds | Myzomela_dammermani | NA | LC | Unknown | NA | NA | NA | NA | NA |
| birds | Myzomela_eichhorni | Myzomela_eichhorni | LC | Decreasing | tropical | diurnal | 10.8 | 5099.333 | 2.868 |

| | | | | | | | | | |
|---|---|---|---|---|---|---|---|---|---|
| birds | Myzomela_eques | Myzomela_eques | LC | Unknown | tropical | diurnal | 13.9 | 465709.311 | 3.112 |
| birds | Myzomela_erythrocephala | Myzomela_erythrocephala | LC | Stable | tropical | diurnal | 7.8 | 442219.316 | 2.624 |
| birds | Myzomela_erythromelas | Myzomela_erythromelas | LC | Stable | tropical | diurnal | 8 | 31100.964 | 4.23 |
| birds | Myzomela_jugularis | Myzomela_jugularis | LC | Stable | tropical | diurnal | 8.7 | 18250.741 | 9.375 |
| birds | Myzomela_kuehni | Myzomela_kuehni | LC | Unknown | tropical | diurnal | 10.8 | 2582.536 | 4.645 |
| birds | Myzomela_lafargei | Myzomela_lafargei | LC | Stable | tropical | diurnal | 12.8 | 17304.73 | 4.72 |
| birds | Myzomela_malaitae | Myzomela_malaitae | NT | Unknown | tropical | diurnal | 13.8 | 4343.26 | 5.3 |
| birds | Myzomela_melanocephala | Myzomela_melanocephala | LC | Stable | tropical | diurnal | 12 | 5908.769 | 5.795 |
| birds | Myzomela_nigrita | Myzomela_nigrita | LC | Stable | tropical | diurnal | 9.3 | 166127.838 | 2.921 |
| birds | Myzomela_obscura | Myzomela_obscura | LC | Stable | tropical | diurnal | 11.9 | 763713.2 | 3.287 |
| birds | Myzomela_pammelaena | Myzomela_pammelaena | LC | Unknown | tropical | diurnal | 15.8 | 1845.459 | 5.202 |
| birds | Myzomela_pulchella | Myzomela_pulchella | LC | Unknown | tropical | diurnal | 10 | 2425.632 | 3.838 |
| birds | Myzomela_rosenbergii | Myzomela_rosenbergii | LC | Unknown | tropical | diurnal | 10.3 | 176793.865 | 4.59 |
| birds | Myzomela_rubratra | Myzomela_rubratra | LC | Unknown | tropical | diurnal | 14 | 2167.328 | 15.992 |
| birds | Myzomela_sanguinolenta | Myzomela_sanguinolenta | LC | Stable | temperate | diurnal | 8.3 | 912973.163 | 7.317 |
| birds | Myzomela_sclateri | Myzomela_sclateri | LC | Unknown | tropical | diurnal | 10.5 | 1986.711 | 4.81 |
| birds | Myzomela_tristrami | Myzomela_tristrami | LC | Stable | tropical | diurnal | 13.3 | 3402.194 | 4.613 |
| birds | Myzomela_vulnerata | Myzomela_vulnerata | LC | Stable | tropical | diurnal | 6.9 | 23721.158 | 13.393 |
| birds | Myzomela_wakoloensis | Myzomela_wakoloensis | LC | Stable | tropical | diurnal | 7.9 | 5355.329 | 5.405 |
| birds | Myzornis_pyrrhoura | Myzornis_pyrrhoura | LC | Unknown | temperate | diurnal | 11.9 | 99938.476 | 6.777 |
| birds | Nannopsittaca_dachilleae | Nannopsittaca_dachilleae | LC | Decreasing | tropical | diurnal | 41.5 | 164020.674 | 1.867 |
| birds | Nannopsittaca_panychlora | Nannopsittaca_panychlora | LC | Stable | tropical | diurnal | 44.1 | 49125.107 | 1.524 |
| birds | Nannopterum_auritum | Phalacrocorax_auritus | LC | Increasing | temperate | diurnal | 1806.5 | 10596564.378 | 12.434 |
| birds | Nannopterum_brasilianum | Phalacrocorax_brasilianus | LC | Increasing | tropical | diurnal | 1239.3 | 18298756.617 | 6.6 |
| birds | Nannopterum_harrisi | NA | VU | Stable | NA | NA | NA | NA | NA |
| birds | Napothera_epilepidota | Napothera_epilepidota | LC | Decreasing | tropical | diurnal | 16 | 1772374.966 | 9.307 |
| birds | Nasica_longirostris | Nasica_longirostris | LC | Decreasing | tropical | diurnal | 92 | 5406855.929 | 2.301 |
| birds | Neafrapus_boehmi | Neafrapus_boehmi | LC | Decreasing | tropical | diurnal | 14.4 | 1674288.301 | 8.513 |
| birds | Neafrapus_cassini | Neafrapus_cassini | LC | Stable | tropical | diurnal | 40 | 2065483.423 | 7.617 |
| birds | Necrosyrtes_monachus | Necrosyrtes_monachus | CR | Decreasing | tropical | diurnal | 2043 | 14512300.574 | 9.602 |
| birds | Nectarinia_bocagii | Nectarinia_bocagii | LC | Stable | tropical | diurnal | 14.8 | 114260.416 | 9.14 |
| birds | Nectarinia_famosa | Nectarinia_famosa | LC | Stable | temperate | diurnal | 14.4 | 1109036.727 | 11.829 |
| birds | Nectarinia_johnstoni | Nectarinia_johnstoni | LC | Stable | tropical | diurnal | 15.2 | 21310.446 | 13.824 |
| birds | Nectarinia_kilimensis | Nectarinia_kilimensis | LC | Stable | tropical | diurnal | 15.7 | 511131.658 | 14.824 |

| | | | | | | | | | |
|---|---|---|---|---|---|---|---|---|---|
| birds | Nectarinia_purpureiventris | Nectarinia_purpureiventris | LC | Decreasing | tropical | diurnal | 11.5 | 70154.051 | 17.118 |
| birds | Nectarinia_tacazze | Nectarinia_tacazze | LC | Stable | tropical | diurnal | 14.9 | 285225.046 | 14.549 |
| birds | Nemosia_pileata | Nemosia_pileata | LC | Stable | tropical | diurnal | 16 | 8991826.243 | 6.272 |
| birds | Nemosia_rourei | Nemosia_rourei | CR | Decreasing | tropical | diurnal | 22 | 526.071 | 15.142 |
| birds | Neochen_jubata | Neochen_jubata | NT | Decreasing | tropical | diurnal | 1250 | 7934083.334 | 2.792 |
| birds | Neochmia_modesta | Neochmia_modesta | LC | Stable | temperate | diurnal | 12.7 | 1594602.223 | 6.137 |
| birds | Neochmia_phaeton | Neochmia_phaeton | LC | Stable | tropical | diurnal | 10 | 1056985.144 | 2.664 |
| birds | Neochmia_ruficauda | Neochmia_ruficauda | LC | Decreasing | tropical | diurnal | 11.1 | 1490192.669 | 3.569 |
| birds | Neochmia_temporalis | Neochmia_temporalis | LC | Stable | temperate | diurnal | 11.4 | 1100465.39 | 8.506 |
| birds | Neocichla_gutturalis | Neocichla_gutturalis | LC | Unknown | tropical | diurnal | 66.7 | 292415.081 | 8.264 |
| birds | Neocossyphus_poensis | Neocossyphus_poensis | LC | Unknown | tropical | diurnal | 51.8 | 2510054.875 | 8.071 |
| birds | Neocossyphus_rufus | Neocossyphus_rufus | LC | Decreasing | tropical | diurnal | 65.9 | 1123680.468 | 6.476 |
| birds | Neoctantes_niger | Neoctantes_niger | LC | Decreasing | tropical | diurnal | 32 | 1354044.106 | 1.214 |
| birds | Neodrepanis_coruscans | Neodrepanis_coruscans | LC | Decreasing | tropical | diurnal | 6.5 | 55129.608 | 5.759 |
| birds | Neodrepanis_hypoxantha | Neodrepanis_hypoxantha | VU | Decreasing | tropical | diurnal | 7.1 | 13059.505 | 5.32 |
| birds | Neohierax_insignis | Polihierax_insignis | NT | Decreasing | tropical | diurnal | 97 | 532936.57 | 13.03 |
| birds | Neolalage_banksiana | Neolalage_banksiana | LC | Stable | tropical | diurnal | 16.8 | 9879.995 | 5.836 |
| birds | Neolestes_torquatus | Neolestes_torquatus | LC | Stable | tropical | diurnal | 22.5 | 813954.886 | 6.981 |
| birds | Neomixis_striatigula | Neomixis_striatigula | LC | Unknown | tropical | diurnal | 8.3 | 162877.993 | 8.027 |
| birds | Neomixis_tenella | Neomixis_tenella | LC | Stable | tropical | diurnal | 6.9 | 586347.223 | 8.727 |
| birds | Neomixis_viridis | Neomixis_viridis | LC | Decreasing | tropical | diurnal | 7 | 173179.419 | 8.239 |
| birds | Neomorphus_geoffroyi | Neomorphus_geoffroyi | VU | Decreasing | tropical | diurnal | 355 | 3567739.22 | 3.95 |
| birds | Neomorphus_pucheranii | Neomorphus_pucheranii | LC | Stable | tropical | diurnal | 330 | 1098665.74 | 0.801 |
| birds | Neomorphus_radiolosus | Neomorphus_radiolosus | EN | Decreasing | tropical | diurnal | 399 | 36045.216 | 6.545 |
| birds | Neomorphus_rufipennis | Neomorphus_rufipennis | LC | Decreasing | tropical | diurnal | 377.7 | 594191.146 | 1.707 |
| birds | Neopelma_aurifrons | Neopelma_aurifrons | NT | Decreasing | tropical | diurnal | 14 | 502648.799 | 10.98 |
| birds | Neopelma_chrysocephalum | Neopelma_chrysocephalum | LC | Stable | tropical | diurnal | 15.5 | 704022.973 | 1.408 |
| birds | Neopelma_chrysolophum | Neopelma_chrysolophum | LC | Stable | tropical | diurnal | 14.2 | 159645.192 | 14.557 |
| birds | Neopelma_pallescens | Neopelma_pallescens | LC | Decreasing | tropical | diurnal | 18.2 | 2439572.761 | 7.758 |
| birds | Neopelma_sulphureiventer | Neopelma_sulphureiventer | LC | Decreasing | tropical | diurnal | 15.7 | 1158804.599 | 2.849 |
| birds | Neophedina_cincta | Riparia_cincta | LC | Increasing | tropical | diurnal | 23.4 | 11447865.226 | 8.192 |
| birds | Neophema_chrysogaster | Neophema_chrysogaster | CR | Decreasing | temperate | diurnal | 45.9 | 32591.232 | 8.125 |
| birds | Neophema_chrysostoma | Neophema_chrysostoma | VU | Decreasing | temperate | diurnal | 46.1 | 1211024.378 | 7.2 |
| birds | Neophema_elegans | Neophema_elegans | LC | Increasing | temperate | diurnal | 43.4 | 865735.951 | 8.595 |

| | | | | | | | | | |
|---|---|---|---|---|---|---|---|---|---|
| birds | Neophema_petrophila | Neophema_petrophila | LC | Decreasing | temperate | diurnal | 53.2 | 142134.666 | 10.215 |
| birds | Neophema_pulchella | Neophema_pulchella | LC | Stable | temperate | diurnal | 41.4 | 492396.136 | 9.866 |
| birds | Neophema_splendida | Neophema_splendida | LC | Stable | temperate | diurnal | 38.6 | 262252.219 | 3.179 |
| birds | Neophron_percnopterus | Neophron_percnopterus | EN | Decreasing | tropical | diurnal | 2082 | 28188514.183 | 8.832 |
| birds | Neopipo_cinnamomea | Neopipo_cinnamomea | LC | Decreasing | tropical | diurnal | 7.7 | 3575202.375 | 1.272 |
| birds | Neopsephotus_bourkii | Neopsephotus_bourkii | LC | Increasing | temperate | diurnal | 44.4 | 1717936.073 | 3.34 |
| birds | Neopsittacus_musschenbroekii | Neopsittacus_musschenbroekii | LC | Increasing | tropical | diurnal | 50.9 | 229336.722 | 4.324 |
| birds | Neopsittacus_pullicauda | Neopsittacus_pullicauda | LC | Stable | tropical | diurnal | 33.2 | 114873.515 | 4.677 |
| birds | Neosericornis_citreogularis | Sericornis_citreogularis | LC | Stable | temperate | diurnal | 17.6 | 169010.506 | 7.625 |
| birds | Neothraupis_fasciata | Neothraupis_fasciata | NT | Decreasing | tropical | diurnal | 26.8 | 2242410.572 | 9.522 |
| birds | Neotis_denhami | Neotis_denhami | NT | Decreasing | tropical | diurnal | 3970.2 | 7686065.236 | 9.927 |
| birds | Neotis_heuglinii | Neotis_heuglinii | LC | Stable | tropical | diurnal | 4098.8 | 989206.103 | 7.451 |
| birds | Neotis_ludwigii | Neotis_ludwigii | EN | Decreasing | temperate | diurnal | 3260.9 | 1006199.573 | 6.162 |
| birds | Neotis_nuba | Neotis_nuba | NT | Decreasing | tropical | diurnal | 5440 | 2385968.924 | 3.89 |
| birds | Neoxolmis_rufiventris | Neoxolmis_rufiventris | LC | Unknown | temperate | diurnal | 77 | 1290844.786 | 8.935 |
| birds | Nephelomyias_lintoni | Myiophobus_lintoni | LC | Stable | tropical | diurnal | 9.8 | 11695.68 | 8.12 |
| birds | Nephelomyias_ochraceiventris | Myiophobus_ochraceiventris | LC | Stable | tropical | diurnal | 10.5 | 92901.367 | 5.988 |
| birds | Nephelomyias_pulcher | Myiophobus_pulcher | LC | Stable | tropical | diurnal | 9.5 | 92358.667 | 9.715 |
| birds | Nephelornis_oneilli | Nephelornis_oneilli | LC | Stable | tropical | diurnal | 16.1 | 17432.093 | 4.79 |
| birds | Nesillas_aldabrana | NA | EX | not_available | NA | NA | NA | NA | NA |
| birds | Nesillas_brevicaudata | Nesillas_brevicaudata | LC | Stable | tropical | diurnal | 16.1 | 1031.434 | 17.43 |
| birds | Nesillas_mariae | Nesillas_mariae | EN | Decreasing | tropical | diurnal | 16 | 209.397 | 17.676 |
| birds | Nesillas_typica | Nesillas_typica | LC | Stable | tropical | diurnal | 19.7 | 520534.891 | 8.509 |
| birds | Nesocharis_ansorgei | Nesocharis_ansorgei | LC | Stable | tropical | diurnal | 8.4 | 157481.364 | 15.609 |
| birds | Nesocharis_shelleyi | Nesocharis_shelleyi | LC | Stable | tropical | diurnal | 7.6 | 31246.095 | 10.52 |
| birds | Nesoctites_micromegas | Nesoctites_micromegas | LC | Decreasing | tropical | diurnal | 32.2 | 74799.651 | 15.781 |
| birds | Nesoenas_mayeri | Nesoenas_mayeri | VU | Stable | tropical | diurnal | 302.8 | 1873.245 | 25.919 |
| birds | Nesoenas_picturatus | Nesoenas_picturata | LC | Stable | tropical | diurnal | 176.8 | 598530.915 | 8.881 |
| birds | Nesofregetta_fuliginosa | Nesofregetta_fuliginosa | EN | Decreasing | tropical | cathemeral | 70 | 29441980.124 | 8.018 |
| birds | Nesophlox_evelynae | Calliphlox_evelynae | LC | Stable | temperate | diurnal | 2.4 | 12175.656 | 7.167 |
| birds | Nesopsar_nigerrimus | Nesopsar_nigerrimus | EN | Decreasing | tropical | diurnal | 39 | 1727.911 | 11.442 |
| birds | Nesoptilotis_flavicollis | Lichenostomus_flavicollis | LC | Stable | temperate | diurnal | 28.7 | 66098.269 | 6.117 |
| birds | Nesoptilotis_leucotis | Lichenostomus_leucotis | LC | Decreasing | temperate | diurnal | 23.9 | 1974990.003 | 8.166 |
| birds | Nesospingus_speculiferus | Nesospingus_speculiferus | VU | Decreasing | tropical | diurnal | 35.2 | 3383.537 | 22.391 |

| | | | | | | | | | |
|---|---|---|---|---|---|---|---|---|---|
| birds | Nesospiza_acunhae | NA | VU | Stable | NA | NA | NA | NA | NA |
| birds | Nesospiza_wilkinsi | NA | CR | Decreasing | NA | NA | NA | NA | NA |
| birds | Nesotriccus_ridgwayi | NA | VU | Stable | NA | NA | NA | NA | NA |
| birds | Nestor_meridionalis | Nestor_meridionalis | VU | Decreasing | temperate | diurnal | 428 | 93990.14 | 5.195 |
| birds | Nestor_notabilis | Nestor_notabilis | EN | Decreasing | temperate | diurnal | 863 | 63273.752 | 3.905 |
| birds | Netta_erythrophthalma | Netta_erythrophthalma | LC | Decreasing | tropical | diurnal | 780.4 | 10612856.99 | 9.148 |
| birds | Netta_peposaca | Netta_peposaca | LC | Increasing | temperate | diurnal | 1000 | 5216896.732 | 8.497 |
| birds | Netta_rufina | Netta_rufina | LC | Unknown | temperate | diurnal | 1118 | 8920116.415 | 10.896 |
| birds | Nettapus_auritus | Nettapus_auritus | LC | Decreasing | tropical | diurnal | 266 | 17754493.4 | 9.261 |
| birds | Nettapus_coromandelianus | Nettapus_coromandelianus | LC | Stable | tropical | diurnal | 391.3 | 9230293.273 | 14.188 |
| birds | Nettapus_pulchellus | Nettapus_pulchellus | LC | Decreasing | tropical | diurnal | 307 | 1855537.355 | 4.057 |
| birds | Newtonia_amphichroa | Newtonia_amphichroa | LC | Unknown | tropical | diurnal | 12.3 | 471535.042 | 8.744 |
| birds | Newtonia_archboldi | Newtonia_archboldi | LC | Unknown | tropical | diurnal | 7.8 | 77835.206 | 10.264 |
| birds | Newtonia_brunneicauda | Newtonia_brunneicauda | LC | Unknown | tropical | diurnal | 10.1 | 550725.408 | 8.706 |
| birds | Newtonia_fanovanae | Newtonia_fanovanae | VU | Decreasing | tropical | diurnal | 12.7 | 84117.106 | 7.791 |
| birds | Nicator_chloris | Nicator_chloris | LC | Stable | tropical | diurnal | 46.1 | 3969589.788 | 8.829 |
| birds | Nicator_gularis | Nicator_gularis | LC | Stable | tropical | diurnal | 45.6 | 940421.619 | 10.226 |
| birds | Nicator_vireo | Nicator_vireo | LC | Stable | tropical | diurnal | 23.6 | 1466486.16 | 6.36 |
| birds | Nicopsitta_calthrapae | NA | LC | Decreasing | NA | NA | NA | NA | NA |
| birds | Nicopsitta_columboides | Psittacula_columboides | LC | Stable | tropical | diurnal | 87 | 121366.207 | 17.859 |
| birds | Nigrita_bicolor | Nigrita_bicolor | LC | Stable | tropical | diurnal | 10.5 | 3277545.302 | 8.312 |
| birds | Nigrita_canicapillus | Nigrita_canicapillus | LC | Stable | tropical | diurnal | 18.7 | 3831512.004 | 8.5 |
| birds | Nigrita_fusconotus | Nigrita_fusconotus | LC | Stable | tropical | diurnal | 9.1 | 2969480.856 | 7.61 |
| birds | Nigrita_luteifrons | Nigrita_luteifrons | LC | Stable | tropical | diurnal | 13.2 | 2645400.348 | 6.864 |
| birds | Nilaus_afer | Nilaus_afer | LC | Stable | tropical | diurnal | 19.1 | 10555427.154 | 9.966 |
| birds | Niltava_davidi | Niltava_davidi | LC | Stable | tropical | diurnal | 24.2 | 2058395.308 | 12.77 |
| birds | Niltava_grandis | Niltava_grandis | LC | Stable | tropical | diurnal | 30.3 | 1622470.964 | 9.025 |
| birds | Niltava_sumatrana | Niltava_sumatrana | LC | Decreasing | tropical | diurnal | 20.5 | 42459.467 | 7.238 |
| birds | Niltava_sundara | Niltava_sundara | LC | Stable | temperate | diurnal | 21.1 | 3184561.706 | 10.922 |
| birds | Niltava_vivida | Niltava_vivida | LC | Decreasing | temperate | diurnal | 33 | 15979.778 | 9.729 |
| birds | Ninox_affinis | Ninox_affinis | LC | Decreasing | tropical | nocturnal | 146 | 5655.232 | 7.159 |
| birds | Ninox_boobook | Ninox_boobook | LC | Decreasing | temperate | nocturnal | 282.2 | 7661995.204 | 4.247 |
| birds | Ninox_connivens | Ninox_connivens | LC | Decreasing | tropical | nocturnal | 578 | 2759461.251 | 6.071 |
| birds | Ninox_jacquinoti | Ninox_jacquinoti | LC | Decreasing | tropical | nocturnal | 200.8 | 20322.954 | 4.531 |

| | | | | | | | | | |
|---|---|---|---|---|---|---|---|---|---|
| birds | Ninox_natalis | Ninox_natalis | VU | Stable | tropical | nocturnal | 184 | 137.178 | 12.523 |
| birds | Ninox_novaeseelandiae | Ninox_novaeseelandiae | LC | Stable | temperate | nocturnal | 183.4 | 269367.33 | 7.992 |
| birds | Ninox_ochracea | NA | NT | Decreasing | NA | NA | NA | NA | NA |
| birds | Ninox_odiosa | Ninox_odiosa | VU | Decreasing | tropical | nocturnal | 209 | 33162.438 | 4.296 |
| birds | Ninox_philippensis | Ninox_philippensis | LC | Decreasing | tropical | nocturnal | 127.9 | 160922.428 | 15.021 |
| birds | Ninox_punctulata | Ninox_punctulata | LC | Stable | tropical | nocturnal | 151 | 152203.911 | 10.049 |
| birds | Ninox_rudolfi | Ninox_rudolfi | NT | Decreasing | tropical | nocturnal | 222 | 10704.53 | 14.92 |
| birds | Ninox_rufa | Ninox_rufa | LC | Decreasing | tropical | nocturnal | 931.7 | 824417.843 | 3.081 |
| birds | Ninox_scutulata | Ninox_scutulata | LC | Decreasing | tropical | nocturnal | 191 | 5215181.313 | 13.167 |
| birds | Ninox_strenua | Ninox_strenua | LC | Stable | temperate | nocturnal | 1346 | 573814.152 | 9.181 |
| birds | Ninox_theomacha | Ninox_theomacha | LC | Stable | tropical | nocturnal | 125 | 596413.954 | 3.515 |
| birds | Nipponia_nippon | Nipponia_nippon | EN | Increasing | temperate | diurnal | 1900 | 1119.51 | 16.672 |
| birds | Nisaetus_alboniger | Nisaetus_alboniger | LC | Decreasing | tropical | diurnal | 830 | 800448.348 | 7.948 |
| birds | Nisaetus_bartelsi | NA | EN | Decreasing | NA | NA | NA | NA | NA |
| birds | Nisaetus_cirrhatus | Nisaetus_cirrhatus | LC | Decreasing | tropical | diurnal | 1475.1 | 4419799.672 | 13.341 |
| birds | Nisaetus_lanceolatus | NA | LC | Decreasing | NA | NA | NA | NA | NA |
| birds | Nisaetus_nanus | Nisaetus_nanus | VU | Decreasing | tropical | diurnal | 557.8 | 1094809.568 | 8.234 |
| birds | Nisaetus_nipalensis | Nisaetus_nipalensis | NT | Decreasing | temperate | diurnal | 3005 | 2230341.255 | 11.717 |
| birds | Nisaetus_philippensis | Nisaetus_philippensis | EN | Decreasing | tropical | diurnal | 1168 | 126414.693 | 14.082 |
| birds | Nomonyx_dominicus | Nomonyx_dominicus | LC | Stable | tropical | diurnal | 365 | 8511481.944 | 9.075 |
| birds | Nonnula_amaurocephala | Nonnula_amaurocephala | LC | Decreasing | tropical | diurnal | 15.5 | 25182.482 | 1.08 |
| birds | Nonnula_brunnea | Nonnula_brunnea | LC | Decreasing | tropical | diurnal | 25 | 251222.792 | 3.403 |
| birds | Nonnula_frontalis | Nonnula_frontalis | LC | Decreasing | tropical | diurnal | 15.7 | 174551.736 | 12.009 |
| birds | Nonnula_rubecula | Nonnula_rubecula | LC | Decreasing | tropical | diurnal | 18.4 | 5438725.748 | 4.996 |
| birds | Nonnula_ruficapilla | Nonnula_ruficapilla | LC | Decreasing | tropical | diurnal | 22 | 2844029.843 | 2.335 |
| birds | Nonnula_sclateri | Nonnula_sclateri | LC | Decreasing | tropical | diurnal | 16 | 642854.118 | 1.28 |
| birds | Northiella_haematogaster | Northiella_haematogaster | LC | Decreasing | temperate | diurnal | 85.9 | 1732521.297 | 6.336 |
| birds | Notharchus_macrorhynchos | Notharchus_macrorhynchos | LC | Stable | tropical | diurnal | 95.9 | 871879.196 | 1.409 |
| birds | Notharchus_ordii | Notharchus_ordii | LC | Stable | tropical | diurnal | 52.8 | 524007.293 | 1.315 |
| birds | Notharchus_pectoralis | Notharchus_pectoralis | LC | Decreasing | tropical | diurnal | 69.1 | 197997.214 | 8.588 |
| birds | Notharchus_swainsoni | Notharchus_swainsoni | LC | Stable | tropical | diurnal | 73.8 | 927376.946 | 13.872 |
| birds | Notharchus_tectus | Notharchus_tectus | LC | Decreasing | tropical | diurnal | 26.8 | 5067701.358 | 2.176 |
| birds | Nothocercus_bonapartei | Nothocercus_bonapartei | LC | Decreasing | tropical | diurnal | 763 | 214864.073 | 10.751 |
| birds | Nothocercus_nigrocapillus | Nothocercus_nigrocapillus | LC | Decreasing | tropical | diurnal | 540 | 206296.781 | 5.122 |

| | | | | | | | | | |
|---|---|---|---|---|---|---|---|---|---|
| birds | Nothocrax_urumutum | Nothocrax_urumutum | LC | Decreasing | tropical | diurnal | 1700 | 2023323.503 | 1.253 |
| birds | Nothoprocta_cinerascens | Nothoprocta_cinerascens | LC | Decreasing | temperate | diurnal | 523.9 | 1208917.267 | 6.766 |
| birds | Nothoprocta_curvirostris | Nothoprocta_curvirostris | LC | Stable | tropical | diurnal | 376 | 52527.781 | 11.193 |
| birds | Nothoprocta_ornata | Nothoprocta_ornata | LC | Decreasing | tropical | diurnal | 619.3 | 765344.426 | 7.151 |
| birds | Nothoprocta_pentlandii | Nothoprocta_pentlandii | LC | Stable | temperate | diurnal | 300.7 | 612683.685 | 6.595 |
| birds | Nothoprocta_perdicaria | Nothoprocta_perdicaria | LC | Decreasing | temperate | diurnal | 458 | 118759.012 | 12.144 |
| birds | Nothoprocta_taczanowskii | NA | VU | Decreasing | NA | NA | NA | NA | NA |
| birds | Nothura_boraquira | Nothura_boraquira | LC | Decreasing | tropical | diurnal | 283 | 1348296.472 | 7.631 |
| birds | Nothura_darwinii | Nothura_darwinii | LC | Increasing | temperate | diurnal | 244.6 | 1098896.656 | 7.631 |
| birds | Nothura_maculosa | Nothura_maculosa | LC | Decreasing | temperate | diurnal | 257.5 | 5645223.734 | 9.782 |
| birds | Nothura_minor | Nothura_minor | VU | Decreasing | tropical | diurnal | 165.8 | 1088582.893 | 12.295 |
| birds | Notiomystis_cincta | Notiomystis_cincta | VU | Stable | temperate | diurnal | 93.4 | 388.025 | 7.237 |
| birds | Notopholia_corusca | Lamprotornis_corruscus | LC | Decreasing | tropical | diurnal | 56.1 | 495997.971 | 10.922 |
| birds | Nucifraga_caryocatactes | Nucifraga_caryocatactes | LC | Decreasing | temperate | diurnal | 182.5 | 15486182.218 | 5.302 |
| birds | Nucifraga_columbiana | Nucifraga_columbiana | LC | Stable | temperate | diurnal | 129.8 | 2287629.396 | 8.433 |
| birds | Numenius_americanus | Numenius_americanus | LC | Decreasing | temperate | diurnal | 583.9 | 3937491.93 | 9.993 |
| birds | Numenius_arquata | Numenius_arquata | NT | Decreasing | tropical | diurnal | 803 | 42784794.192 | 11.196 |
| birds | Numenius_borealis | Numenius_borealis | CR | Unknown | temperate | diurnal | 350.1 | 2235504.858 | 7.722 |
| birds | Numenius_madagascariensis | Numenius_madagascariensis | EN | Decreasing | temperate | cathemeral | 792 | 6598384.144 | 5.745 |
| birds | Numenius_phaeopus | Numenius_phaeopus | LC | Decreasing | temperate | diurnal | 364.6 | 17013389.569 | 8.477 |
| birds | Numenius_tahitiensis | Numenius_tahitiensis | NT | Decreasing | temperate | diurnal | 400.1 | 102179.876 | 3.794 |
| birds | Numenius_tenuirostris | Numenius_tenuirostris | CR | Decreasing | temperate | diurnal | 308 | 887016.765 | 14.418 |
| birds | Numida_meleagris | Numida_meleagris | LC | Stable | tropical | diurnal | 1299 | 13597695.41 | 9.375 |
| birds | Nyctanassa_violacea | Nyctanassa_violacea | LC | Stable | tropical | cathemeral | 681.7 | 5236164.311 | 14.194 |
| birds | Nyctibius_aethereus | Nyctibius_aethereus | LC | Decreasing | tropical | nocturnal | 440.5 | 4975131.515 | 3.204 |
| birds | Nyctibius_grandis | Nyctibius_grandis | LC | Decreasing | tropical | nocturnal | 547 | 8477906.134 | 3.891 |
| birds | Nyctibius_griseus | Nyctibius_griseus | LC | Decreasing | tropical | nocturnal | 172 | 13806169.858 | 5.759 |
| birds | Nyctibius_jamaicensis | Nyctibius_jamaicensis | LC | Decreasing | tropical | nocturnal | 266 | 782201.431 | 11.044 |
| birds | Nyctibius_leucopterus | Nyctibius_leucopterus | LC | Decreasing | tropical | nocturnal | 86.4 | 3324869.235 | 1.684 |
| birds | Nyctibius_maculosus | Nyctibius_maculosus | LC | Decreasing | tropical | nocturnal | 170.3 | 117205.57 | 10.028 |
| birds | Nycticorax_caledonicus | Nycticorax_caledonicus | LC | Stable | temperate | cathemeral | 856 | 7791277.404 | 5.586 |
| birds | Nycticorax_nycticorax | Nycticorax_nycticorax | LC | Decreasing | temperate | cathemeral | 810 | 52753396.92 | 11.287 |
| birds | Nyctidromus_albicollis | Nyctidromus_albicollis | LC | Stable | tropical | nocturnal | 57.8 | 13748689.236 | 6.045 |
| birds | Nyctidromus_anthonyi | Caprimulgus_anthonyi | LC | Increasing | tropical | nocturnal | 35.9 | 57328.237 | 10.654 |

| | | | | | | | | | |
|---|---|---|---|---|---|---|---|---|---|
| birds | Nyctiphrynus_mcleodii | Nyctiphrynus_mcleodii | LC | Decreasing | temperate | nocturnal | 35 | 188114.397 | 5.636 |
| birds | Nyctiphrynus_ocellatus | Nyctiphrynus_ocellatus | LC | Decreasing | tropical | nocturnal | 39.6 | 3398673.414 | 6.549 |
| birds | Nyctiphrynus_rosenbergi | Nyctiphrynus_rosenbergi | LC | Decreasing | tropical | nocturnal | 52 | 57920.209 | 5.165 |
| birds | Nyctiphrynus_yucatanicus | Nyctiphrynus_yucatanicus | LC | Decreasing | tropical | nocturnal | 24.5 | 145737.263 | 6.078 |
| birds | Nyctipolus_hirundinaceus | Caprimulgus_hirundinaceus | LC | Decreasing | tropical | nocturnal | 32 | 677679.002 | 8.535 |
| birds | Nyctipolus_nigrescens | Caprimulgus_nigrescens | LC | Stable | tropical | nocturnal | 35.7 | 4123782.877 | 1.874 |
| birds | Nyctiprogne_leucopyga | Nyctiprogne_leucopyga | LC | Decreasing | tropical | nocturnal | 27.3 | 5244008.149 | 2.657 |
| birds | Nyctyornis_amictus | Nyctyornis_amictus | LC | Decreasing | tropical | diurnal | 71.9 | 1492016.657 | 8.274 |
| birds | Nyctyornis_athertoni | Nyctyornis_athertoni | LC | Stable | tropical | diurnal | 84.2 | 2622222.408 | 12.333 |
| birds | Nymphicus_hollandicus | Nymphicus_hollandicus | LC | Stable | temperate | diurnal | 94.6 | 6462512.617 | 4.128 |
| birds | Nystactes_noanamae | NA | NT | Decreasing | NA | NA | NA | NA | NA |
| birds | Nystactes_tamatia | Bucco_tamatia | LC | Decreasing | tropical | diurnal | 35.5 | 4349445.919 | 1.854 |
| birds | Nystalus_chacuru | Nystalus_chacuru | LC | Stable | tropical | diurnal | 52.3 | 4329404.398 | 8.947 |
| birds | Nystalus_maculatus | Nystalus_maculatus | LC | Stable | tropical | diurnal | 42 | 3643937.663 | 7.924 |
| birds | Nystalus_radiatus | Nystalus_radiatus | LC | Decreasing | tropical | diurnal | 61 | 236336.294 | 8.976 |
| birds | Nystalus_striolatus | Nystalus_striolatus | LC | Decreasing | tropical | diurnal | 47 | 2922975.075 | 3.031 |
| birds | Oceanites_gracilis | NA | DD | Unknown | NA | NA | NA | NA | NA |
| birds | Oceanites_oceanicus | Oceanites_oceanicus | LC | Stable | tropical | diurnal | 30.4 | 227464044.199 | 9.284 |
| birds | Ochetorhynchus_andaecola | Ochetorhynchus_andaecola | LC | Decreasing | tropical | diurnal | 30.9 | 398226.845 | 6.384 |
| birds | Ochetorhynchus_melanurus | Ochetorhynchus_melanura | LC | Stable | temperate | diurnal | 40 | 39429.476 | 9.993 |
| birds | Ochetorhynchus_phoenicurus | Ochetorhynchus_phoenicurus | LC | Decreasing | temperate | diurnal | 30.2 | 525155.506 | 5.694 |
| birds | Ochetorhynchus_ruficaudus | Ochetorhynchus_ruficaudus | LC | Decreasing | temperate | diurnal | 36.3 | 1066359.628 | 4.929 |
| birds | Ochthoeca_cinnamomeiventris | Ochthoeca_cinnamomeiventris | LC | Stable | tropical | diurnal | 12 | 174511.373 | 11.377 |
| birds | Ochthoeca_fumicolor | Ochthoeca_fumicolor | LC | Stable | tropical | diurnal | 16.6 | 246238.91 | 9.765 |
| birds | Ochthoeca_leucophrys | Ochthoeca_leucophrys | LC | Stable | tropical | diurnal | 13.1 | 781892.762 | 7.343 |
| birds | Ochthoeca_nigrita | Ochthoeca_nigrita | LC | Stable | tropical | diurnal | 10.9 | 8206.928 | 14.305 |
| birds | Ochthoeca_oenanthoides | Ochthoeca_oenanthoides | LC | Stable | tropical | diurnal | 17.7 | 624271.788 | 8.088 |
| birds | Ochthoeca_piurae | Ochthoeca_piurae | LC | Decreasing | tropical | diurnal | 8.9 | 20907.643 | 8.902 |
| birds | Ochthoeca_rufipectoralis | Ochthoeca_rufipectoralis | LC | Stable | tropical | diurnal | 10 | 333647.743 | 9.005 |
| birds | Ochthoeca_salvini | Tumbezia_salvini | NT | Decreasing | tropical | diurnal | 12.2 | 25442.542 | 8.79 |
| birds | Ochthoeca_thoracica | Ochthoeca_thoracica | LC | Stable | tropical | diurnal | 10.8 | 124837.51 | 6.631 |
| birds | Ochthornis_littoralis | Ochthornis_littoralis | LC | Stable | tropical | diurnal | 13.4 | 4562555.946 | 1.727 |
| birds | Ocreatus_underwoodii | Ocreatus_underwoodii | LC | Stable | tropical | diurnal | 3 | 385599.414 | 8.74 |
| birds | Ocyceros_birostris | Ocyceros_birostris | LC | Stable | tropical | diurnal | 375 | 2062731.904 | 18.904 |

| | | | | | | | | | |
|---|---|---|---|---|---|---|---|---|---|
| birds | Ocyceros_gingalensis | Ocyceros_gingalensis | LC | Stable | tropical | diurnal | 238 | 64916.294 | 17.576 |
| birds | Ocyceros_griseus | Ocyceros_griseus | VU | Decreasing | tropical | diurnal | 292 | 230760.925 | 20.13 |
| birds | Ocyphaps_lophotes | Ocyphaps_lophotes | LC | Increasing | temperate | diurnal | 204 | 7382616.533 | 4.312 |
| birds | Odontophorus_atrifrons | Odontophorus_atrifrons | LC | Decreasing | tropical | diurnal | 304 | 34304.835 | 8.723 |
| birds | Odontophorus_balliviani | Odontophorus_balliviani | LC | Decreasing | tropical | diurnal | 317.4 | 90660.549 | 4.95 |
| birds | Odontophorus_capueira | Odontophorus_capueira | LC | Decreasing | tropical | diurnal | 425.4 | 1724256.704 | 12.684 |
| birds | Odontophorus_columbianus | Odontophorus_columbianus | NT | Decreasing | tropical | diurnal | 340 | 5025.305 | 18.403 |
| birds | Odontophorus_dialeucos | Odontophorus_dialeucos | LC | Stable | tropical | diurnal | 260 | 233.705 | 1.739 |
| birds | Odontophorus_erythrops | Odontophorus_erythrops | LC | Decreasing | tropical | diurnal | 335 | 164035.668 | 8.646 |
| birds | Odontophorus_gujanensis | Odontophorus_gujanensis | LC | Decreasing | tropical | diurnal | 314.1 | 7718738.668 | 2.891 |
| birds | Odontophorus_guttatus | Odontophorus_guttatus | LC | Decreasing | tropical | diurnal | 303.8 | 339131.07 | 10.028 |
| birds | Odontophorus_hyperythrus | Odontophorus_hyperythrus | LC | Decreasing | tropical | diurnal | 339.3 | 60489.279 | 11.636 |
| birds | Odontophorus_leucolaemus | Odontophorus_leucolaemus | LC | Decreasing | tropical | diurnal | 295 | 11852.566 | 7.554 |
| birds | Odontophorus_melanonotus | Odontophorus_melanonotus | LC | Decreasing | tropical | diurnal | 322 | 6392.09 | 6.462 |
| birds | Odontophorus_melanotis | Odontophorus_melanotis | NT | Decreasing | tropical | diurnal | 335 | 128032.297 | 7.576 |
| birds | Odontophorus_speciosus | Odontophorus_speciosus | LC | Decreasing | tropical | diurnal | 317 | 450538.919 | 5.26 |
| birds | Odontophorus_stellatus | Odontophorus_stellatus | LC | Decreasing | tropical | diurnal | 335.8 | 1723957.587 | 1.93 |
| birds | Odontophorus_strophium | Odontophorus_strophium | VU | Decreasing | tropical | diurnal | 302 | 3431.341 | 15.486 |
| birds | Odontorchilus_branickii | Odontorchilus_branickii | LC | Decreasing | tropical | diurnal | 9.6 | 155452.647 | 4.786 |
| birds | Odontorchilus_cinereus | Odontorchilus_cinereus | LC | Decreasing | tropical | diurnal | 11 | 1089993.93 | 2.894 |
| birds | Oedistoma_iliolophus | Toxorhamphus_iliolophus | LC | Unknown | tropical | diurnal | 11 | 624407.835 | 3.619 |
| birds | Oedistoma_pygmaeum | Oedistoma_pygmaeum | LC | Unknown | tropical | diurnal | 5.8 | 595536.968 | 3.08 |
| birds | Oena_capensis | Oena_capensis | LC | Increasing | tropical | diurnal | 40.6 | 17910218.785 | 8.908 |
| birds | Oenanthe_albifrons | Myrmecocichla_albifrons | LC | Stable | tropical | diurnal | 20.6 | 3294680.417 | 11.18 |
| birds | Oenanthe_albonigra | Oenanthe_albonigra | LC | Stable | temperate | diurnal | 25.2 | 1325675.775 | 8.782 |
| birds | Oenanthe_bottae | Oenanthe_bottae | LC | Stable | tropical | diurnal | 19.9 | 116654.275 | 12.366 |
| birds | Oenanthe_cypriaca | Oenanthe_cypriaca | LC | Stable | tropical | diurnal | 16.6 | 1608130.429 | 8.313 |
| birds | Oenanthe_deserti | Oenanthe_deserti | LC | Stable | temperate | diurnal | 19.8 | 18380455.151 | 5.767 |
| birds | Oenanthe_familiaris | Cercomela_familiaris | LC | Stable | tropical | diurnal | 19.5 | 7293437.667 | 9.593 |
| birds | Oenanthe_finschii | Oenanthe_finschii | LC | Stable | temperate | diurnal | 27.4 | 3778500.051 | 9.584 |
| birds | Oenanthe_fusca | Cercomela_fusca | LC | Stable | temperate | diurnal | 12.7 | 1062229.72 | 18.565 |
| birds | Oenanthe_hispanica | Oenanthe_hispanica | LC | Decreasing | temperate | diurnal | 15 | 18516204.752 | 6.337 |
| birds | Oenanthe_isabellina | Oenanthe_isabellina | LC | Stable | tropical | diurnal | 29.4 | 23687917.958 | 8.048 |
| birds | Oenanthe_leucopyga | Oenanthe_leucopyga | LC | Stable | temperate | diurnal | 27.9 | 5398249.866 | 3.192 |

| | | | | | | | | | |
|---|---|---|---|---|---|---|---|---|---|
| birds | Oenanthe_leucura | Oenanthe_leucura | LC | Decreasing | temperate | diurnal | 41 | 1288299.839 | 10.887 |
| birds | Oenanthe_lugens | Oenanthe_lugens | LC | Stable | temperate | diurnal | 22.4 | 3562658.081 | 6.588 |
| birds | Oenanthe_melanura | Cercomela_melanura | LC | Stable | tropical | diurnal | 15.2 | 2174337.344 | 5.683 |
| birds | Oenanthe_moesta | Oenanthe_moesta | LC | Stable | temperate | diurnal | 30.2 | 987108.26 | 7.832 |
| birds | Oenanthe_monacha | Oenanthe_monacha | LC | Stable | temperate | diurnal | 21.6 | 938292.159 | 6.263 |
| birds | Oenanthe_oenanthe | Oenanthe_oenanthe | LC | Decreasing | temperate | diurnal | 25.4 | 39231050.358 | 7.464 |
| birds | Oenanthe_phillipsi | Oenanthe_phillipsi | LC | Stable | tropical | diurnal | 19.6 | 557674.641 | 7.51 |
| birds | Oenanthe_picata | Oenanthe_picata | LC | Stable | temperate | diurnal | 21.6 | 4008049.739 | 10.322 |
| birds | Oenanthe_pileata | Oenanthe_pileata | LC | Stable | tropical | diurnal | 26.8 | 5997462.333 | 7.933 |
| birds | Oenanthe_pleschanka | Oenanthe_pleschanka | LC | Stable | temperate | diurnal | 19.4 | 12225171.531 | 9.224 |
| birds | Oenanthe_scotocerca | Cercomela_scotocerca | LC | Stable | tropical | diurnal | 16 | 554755.074 | 9.073 |
| birds | Oenanthe_xanthoprymna | Oenanthe_xanthoprymna | LC | Stable | temperate | diurnal | 22.7 | 3804462.026 | 6.057 |
| birds | Ognorhynchus_icterotis | Ognorhynchus_icterotis | VU | Increasing | tropical | diurnal | 285 | 146924.812 | 12.081 |
| birds | Oncostoma_cinereigulare | Oncostoma_cinereigulare | LC | Decreasing | tropical | diurnal | 6.4 | 588153.898 | 9.954 |
| birds | Oncostoma_olivaceum | Oncostoma_olivaceum | LC | Decreasing | tropical | diurnal | 6.6 | 226322.397 | 11.766 |
| birds | Oneillornis_lunulatus | Gymnopithys_lunulatus | LC | Decreasing | tropical | diurnal | 23.9 | 297195.905 | 2.062 |
| birds | Oneillornis_salvini | Gymnopithys_salvini | LC | Decreasing | tropical | diurnal | 25.9 | 1355133.039 | 1.546 |
| birds | Onychognathus_albirostris | Onychognathus_albirostris | LC | Stable | tropical | diurnal | 137.3 | 162747.003 | 14.15 |
| birds | Onychognathus_blythii | Onychognathus_blythii | LC | Stable | tropical | diurnal | 100 | 440734.83 | 10.369 |
| birds | Onychognathus_frater | Onychognathus_frater | LC | Stable | tropical | diurnal | 100 | 3622.403 | 11.03 |
| birds | Onychognathus_fulgidus | Onychognathus_fulgidus | LC | Decreasing | tropical | diurnal | 109 | 2367641.442 | 8.407 |
| birds | Onychognathus_morio | Onychognathus_morio | LC | Increasing | tropical | cathemeral | 132.9 | 3660528.378 | 11.14 |
| birds | Onychognathus_nabouroup | Onychognathus_nabouroup | LC | Stable | temperate | diurnal | 102.4 | 984106.614 | 5.892 |
| birds | Onychognathus_neumanni | Onychognathus_neumanni | LC | Increasing | tropical | diurnal | 39.8 | 467656.081 | 11.927 |
| birds | Onychognathus_salvadorii | Onychognathus_salvadorii | LC | Stable | tropical | diurnal | 160 | 646688.673 | 7.583 |
| birds | Onychognathus_tenuirostris | Onychognathus_tenuirostris | LC | Decreasing | tropical | diurnal | 118 | 405677.732 | 13.471 |
| birds | Onychognathus_tristramii | Onychognathus_tristramii | LC | Stable | tropical | diurnal | 122.7 | 564513.208 | 7.014 |
| birds | Onychognathus_walleri | Onychognathus_walleri | LC | Decreasing | tropical | diurnal | 93 | 308237.317 | 13.489 |
| birds | Onychoprion_aleuticus | Sterna_aleutica | VU | Decreasing | temperate | diurnal | 120 | 1447946.406 | 2.98 |
| birds | Onychoprion_anaethetus | Sterna_anaethetus | LC | Unknown | tropical | diurnal | 119.4 | 43608241.991 | 9.2 |
| birds | Onychoprion_fuscatus | Sterna_fuscata | LC | Unknown | tropical | diurnal | 185.7 | 173979002.67 | 10.442 |
| birds | Onychoprion_lunatus | Sterna_lunata | LC | Decreasing | tropical | diurnal | 120 | 36453698.886 | 8.872 |
| birds | Onychorhynchus_coronatus | Onychorhynchus_coronatus | LC | Decreasing | tropical | diurnal | 14 | 5973721.862 | 1.857 |
| birds | Onychorhynchus_mexicanus | Onychorhynchus_mexicanus | LC | Decreasing | tropical | diurnal | 19 | 888001.001 | 10.245 |

| | | | | | | | | | |
|---|---|---|---|---|---|---|---|---|---|
| birds | Onychorhynchus_occidentalis | Onychorhynchus_occidentalis | VU | Decreasing | tropical | diurnal | 14.8 | 34880.48 | 15.81 |
| birds | Onychorhynchus_swainsoni | Onychorhynchus_swainsoni | VU | Decreasing | tropical | diurnal | 17 | 198858.894 | 13.23 |
| birds | Onychostruthus_taczanowskii | Montifringilla_taczanowskii | LC | Stable | temperate | diurnal | 32.5 | 1477066.845 | 6.266 |
| birds | Opisthocomus_hoazin | Opisthocomus_hoazin | LC | Decreasing | tropical | nocturnal | 696 | 7559878.736 | 2.649 |
| birds | Opisthoprora_euryptera | Opisthoprora_euryptera | LC | Stable | tropical | diurnal | 6.2 | 26992.637 | 4.945 |
| birds | Oporornis_agilis | Oporornis_agilis | LC | Decreasing | tropical | diurnal | 13.3 | 6641240.306 | 7.579 |
| birds | Orchesticus_abeillei | Orchesticus_abeillei | NT | Decreasing | temperate | diurnal | 31.5 | 163525.702 | 13.875 |
| birds | Oreocharis_arfaki | Oreocharis_arfaki | LC | Unknown | tropical | diurnal | 19.7 | 192214.004 | 4.535 |
| birds | Oreoica_gutturalis | Oreoica_gutturalis | LC | Decreasing | temperate | diurnal | 62.8 | 4957954.292 | 4.281 |
| birds | Oreolais_pulcher | Oreolais_pulchra | LC | Stable | tropical | diurnal | 8.4 | 100450.197 | 12.754 |
| birds | Oreolais_ruwenzorii | Oreolais_ruwenzorii | LC | Stable | tropical | diurnal | 9.9 | 123324.794 | 16.659 |
| birds | Oreomystis_bairdi | Oreomystis_bairdi | CR | Decreasing | tropical | diurnal | 14.2 | 1343.132 | 9.429 |
| birds | Oreonympha_nobilis | Oreonympha_nobilis | LC | Stable | tropical | diurnal | 7.7 | 36217.73 | 9.436 |
| birds | Oreophasis_derbianus | Oreophasis_derbianus | EN | Decreasing | tropical | diurnal | 2071.5 | 31132.276 | 14.159 |
| birds | Oreophilais_robertsi | Prinia_robertsi | LC | Stable | tropical | diurnal | 10 | 9712.822 | 11.243 |
| birds | Oreopholus_ruficollis | Oreopholus_ruficollis | LC | Decreasing | temperate | diurnal | 131 | 2680827.364 | 6.947 |
| birds | Oreopsar_bolivianus | Oreopsar_bolivianus | LC | Stable | tropical | diurnal | 72.9 | 77402.168 | 9.554 |
| birds | Oreopsittacus_arfaki | Oreopsittacus_arfaki | LC | Stable | tropical | diurnal | 20.2 | 108008.259 | 4.778 |
| birds | Oreornis_chrysogenys | Oreornis_chrysogenys | LC | Stable | tropical | diurnal | 69.5 | 22304.254 | 4.574 |
| birds | Oreortyx_pictus | Oreortyx_pictus | LC | Decreasing | temperate | diurnal | 233 | 386114.497 | 10.316 |
| birds | Oreoscoptes_montanus | Oreoscoptes_montanus | LC | Decreasing | temperate | diurnal | 44.2 | 3989086.927 | 9.641 |
| birds | Oreoscopus_gutturalis | Oreoscopus_gutturalis | EN | Decreasing | tropical | diurnal | 20 | 5869.254 | 2.662 |
| birds | Oreostruthus_fuliginosus | Oreostruthus_fuliginosus | LC | Stable | tropical | diurnal | 18.3 | 51165.483 | 5.003 |
| birds | Oreothlypis_gutturalis | Parula_gutturalis | LC | Stable | tropical | diurnal | 9.5 | 3913.068 | 6.044 |
| birds | Oreothlypis_superciliosa | Parula_superciliosa | LC | Decreasing | tropical | diurnal | 9 | 402973.465 | 10.927 |
| birds | Oreothraupis_arremonops | Oreothraupis_arremonops | LC | Decreasing | tropical | diurnal | 48 | 22305.925 | 8.925 |
| birds | Oreotrochilus_adela | Oreotrochilus_adela | LC | Decreasing | tropical | diurnal | 7.9 | 62744.185 | 10.089 |
| birds | Oreotrochilus_chimborazo | Oreotrochilus_chimborazo | LC | Stable | tropical | diurnal | 8.2 | 36894.509 | 11.912 |
| birds | Oreotrochilus_estella | Oreotrochilus_estella | LC | Stable | tropical | diurnal | 8.1 | 595797.116 | 7.128 |
| birds | Oreotrochilus_leucopleurus | Oreotrochilus_leucopleurus | LC | Stable | temperate | diurnal | 8.2 | 404047.636 | 5.139 |
| birds | Oreotrochilus_melanogaster | Oreotrochilus_melanogaster | LC | Stable | tropical | diurnal | 9.1 | 55336.69 | 9.587 |
| birds | Oreotrochilus_stolzmanni | NA | LC | Stable | NA | NA | NA | NA | NA |
| birds | Origma_murina | Crateroscelis_murina | LC | Stable | tropical | diurnal | 15 | 531992.386 | 3.074 |
| birds | Origma_robusta | Crateroscelis_robusta | LC | Stable | tropical | diurnal | 18.6 | 164313.706 | 4.699 |

| | | | | | | | | | |
|---|---|---|---|---|---|---|---|---|---|
| birds | Origma_solitaria | Origma_solitaria | LC | Stable | temperate | diurnal | 14.3 | 56963.22 | 8.296 |
| birds | Oriolia_bernieri | Oriolia_bernieri | EN | Decreasing | tropical | diurnal | 52 | 20172.623 | 5.708 |
| birds | Oriolus_albiloris | Oriolus_albiloris | LC | Decreasing | tropical | diurnal | 38.6 | 22182.672 | 6.084 |
| birds | Oriolus_auratus | Oriolus_auratus | LC | Decreasing | tropical | diurnal | 66.2 | 10077113.977 | 9.708 |
| birds | Oriolus_bouroensis | NA | LC | Unknown | NA | NA | NA | NA | NA |
| birds | Oriolus_brachyrynchus | Oriolus_brachyrhynchus | LC | Stable | tropical | diurnal | 49 | 2640549.203 | 9.135 |
| birds | Oriolus_chinensis | Oriolus_chinensis | LC | Decreasing | temperate | diurnal | 82.6 | 8169313.157 | 14.811 |
| birds | Oriolus_chlorocephalus | NA | LC | Decreasing | NA | NA | NA | NA | NA |
| birds | Oriolus_crassirostris | Oriolus_crassirostris | VU | Decreasing | tropical | diurnal | 50 | 481.689 | 8.663 |
| birds | Oriolus_cruentus | Oriolus_cruentus | DD | Decreasing | tropical | diurnal | 64 | 3882.047 | 9.85 |
| birds | Oriolus_flavocinctus | Oriolus_flavocinctus | LC | Stable | tropical | diurnal | 109 | 535100.567 | 2.461 |
| birds | Oriolus_forsteni | Oriolus_forsteni | LC | Stable | tropical | diurnal | 109 | 15001.228 | 6.802 |
| birds | Oriolus_hosii | NA | NT | Decreasing | NA | NA | NA | NA | NA |
| birds | Oriolus_isabellae | Oriolus_isabellae | CR | Decreasing | tropical | diurnal | 46.1 | 1158.852 | 12.162 |
| birds | Oriolus_larvatus | Oriolus_larvatus | LC | Increasing | tropical | diurnal | 66.1 | 6081428.22 | 9.546 |
| birds | Oriolus_melanotis | NA | LC | Unknown | NA | NA | NA | NA | NA |
| birds | Oriolus_mellianus | Oriolus_mellianus | EN | Decreasing | temperate | diurnal | 93 | 166532.371 | 14.212 |
| birds | Oriolus_monacha | Oriolus_monacha | LC | Stable | tropical | diurnal | 68.5 | 337144.379 | 13.227 |
| birds | Oriolus_nigripennis | Oriolus_nigripennis | LC | Stable | tropical | diurnal | 56.7 | 2050735.039 | 8.758 |
| birds | Oriolus_oriolus | Oriolus_oriolus | LC | Stable | temperate | diurnal | 79 | 33861421.139 | 9.217 |
| birds | Oriolus_percivali | Oriolus_percivali | LC | Stable | tropical | diurnal | 61 | 164128.384 | 17.036 |
| birds | Oriolus_phaeochromus | NA | LC | Stable | NA | NA | NA | NA | NA |
| birds | Oriolus_sagittatus | Oriolus_sagittatus | LC | Decreasing | temperate | diurnal | 95.2 | 3411405.869 | 5.598 |
| birds | Oriolus_steerii | Oriolus_steerii | LC | Stable | tropical | diurnal | 46.5 | 140076.591 | 13.281 |
| birds | Oriolus_szalayi | Oriolus_szalayi | LC | Stable | tropical | diurnal | 95.2 | 680368.63 | 3.158 |
| birds | Oriolus_tenuirostris | Oriolus_tenuirostris | LC | Decreasing | tropical | diurnal | 81.4 | 1127097.862 | 10.029 |
| birds | Oriolus_traillii | Oriolus_traillii | LC | Stable | tropical | diurnal | 73.7 | 1982675.499 | 10.574 |
| birds | Oriolus_xanthonotus | Oriolus_xanthonotus | LC | Decreasing | tropical | diurnal | 45 | 1443272.167 | 9.465 |
| birds | Oriolus_xanthornus | Oriolus_xanthornus | LC | Stable | tropical | diurnal | 56.3 | 4355542.077 | 15.501 |
| birds | Oriturus_superciliosus | Oriturus_superciliosus | LC | Decreasing | temperate | diurnal | 41.4 | 295047.675 | 11.523 |
| birds | Ornithion_brunneicapillus | Ornithion_brunneicapillus | LC | Decreasing | tropical | diurnal | 7.1 | 282994.635 | 9.326 |
| birds | Ornithion_inerme | Ornithion_inerme | LC | Decreasing | tropical | diurnal | 7 | 6701180.645 | 2.227 |
| birds | Ornithion_semiflavum | Ornithion_semiflavum | LC | Decreasing | tropical | diurnal | 6.6 | 257158.919 | 9.65 |
| birds | Ornorectes_cristatus | Pitohui_cristatus | LC | Stable | tropical | diurnal | 79 | 192704.933 | 2.962 |

| | | | | | | | | | |
|---|---|---|---|---|---|---|---|---|---|
| birds | Oroanassa_magnifica | Gorsachius_magnificus | EN | Decreasing | temperate | nocturnal | 480.3 | 1647055.734 | 14.871 |
| birds | Orochelidon_andecola | Haplochelidon_andecola | LC | Stable | tropical | diurnal | 17 | 506072.712 | 7.995 |
| birds | Orochelidon_flavipes | Notiochelidon_flavipes | LC | Stable | tropical | diurnal | 9.8 | 127853.29 | 6.491 |
| birds | Orochelidon_murina | Notiochelidon_murina | LC | Stable | tropical | diurnal | 11.7 | 425662.431 | 9.495 |
| birds | Ortalis_canicollis | Ortalis_canicollis | LC | Decreasing | temperate | diurnal | 539 | 1171806.839 | 6.052 |
| birds | Ortalis_cinereiceps | Ortalis_cinereiceps | LC | Stable | tropical | diurnal | 493 | 236334.315 | 8.254 |
| birds | Ortalis_erythroptera | Ortalis_erythroptera | VU | Decreasing | tropical | diurnal | 632 | 74565.606 | 13.346 |
| birds | Ortalis_garrula | Ortalis_garrula | LC | Stable | tropical | diurnal | 532 | 109708.022 | 14.096 |
| birds | Ortalis_guttata | Ortalis_guttata | LC | Unknown | tropical | diurnal | 547.7 | 4434552.3 | 2.818 |
| birds | Ortalis_leucogastra | Ortalis_leucogastra | LC | Stable | tropical | diurnal | 495.8 | 57398.424 | 17.671 |
| birds | Ortalis_motmot | Ortalis_motmot | LC | Decreasing | tropical | diurnal | 520 | 2134685.165 | 1.594 |
| birds | Ortalis_poliocephala | Ortalis_poliocephala | LC | Decreasing | tropical | diurnal | 760 | 200072.196 | 9.538 |
| birds | Ortalis_ruficauda | Ortalis_ruficauda | LC | Stable | tropical | diurnal | 586.2 | 662199.339 | 10.38 |
| birds | Ortalis_superciliaris | NA | LC | Decreasing | NA | NA | NA | NA | NA |
| birds | Ortalis_vetula | Ortalis_vetula | LC | Stable | tropical | diurnal | 562.6 | 603428.429 | 10.436 |
| birds | Ortalis_wagleri | Ortalis_wagleri | LC | Decreasing | temperate | diurnal | 834 | 114061.433 | 6.484 |
| birds | Orthogonys_chloricterus | Orthogonys_chloricterus | LC | Decreasing | temperate | diurnal | 39 | 121968.516 | 13.967 |
| birds | Orthonyx_spaldingii | Orthonyx_spaldingii | LC | Stable | tropical | diurnal | 149.9 | 23436.787 | 4.813 |
| birds | Orthonyx_temminckii | Orthonyx_temminckii | LC | Decreasing | temperate | diurnal | 58.1 | 166548.281 | 8.984 |
| birds | Orthopsittaca_manilatus | Orthopsittaca_manilata | LC | Stable | tropical | diurnal | 372 | 7148717.743 | 3.56 |
| birds | Orthorhyncus_cristatus | Orthorhyncus_cristatus | LC | Unknown | tropical | diurnal | 3 | 7736.301 | 22.95 |
| birds | Orthotomus_atrogularis | Orthotomus_atrogularis | LC | Stable | tropical | diurnal | 7.7 | 2752593.837 | 11.158 |
| birds | Orthotomus_castaneiceps | Orthotomus_castaneiceps | LC | Stable | tropical | diurnal | 12.8 | 33106.016 | 18.38 |
| birds | Orthotomus_cinereiceps | Orthotomus_cinereiceps | LC | Stable | tropical | diurnal | 9 | 20650.364 | 11.909 |
| birds | Orthotomus_derbianus | Orthotomus_derbianus | LC | Stable | tropical | diurnal | 10.7 | 72219.626 | 15.732 |
| birds | Orthotomus_frontalis | Orthotomus_frontalis | LC | Stable | tropical | diurnal | 8 | 120551.539 | 12.089 |
| birds | Orthotomus_nigriceps | Orthotomus_nigriceps | LC | Stable | tropical | diurnal | 9.8 | 24461.655 | 10.861 |
| birds | Orthotomus_ruficeps | Orthotomus_ruficeps | LC | Stable | tropical | diurnal | 7 | 887360.639 | 10.093 |
| birds | Orthotomus_samarensis | Orthotomus_samarensis | NT | Decreasing | tropical | diurnal | 8.9 | 18296.836 | 11.454 |
| birds | Orthotomus_sepium | Orthotomus_sepium | LC | Stable | tropical | diurnal | 8.4 | 140271.594 | 22.489 |
| birds | Orthotomus_sericeus | Orthotomus_sericeus | LC | Stable | tropical | diurnal | 10.8 | 1068502.74 | 8.977 |
| birds | Orthotomus_sutorius | Orthotomus_sutorius | LC | Stable | tropical | diurnal | 7.5 | 6849946.152 | 15.439 |
| birds | Ortygornis_pondicerianus | Francolinus_pondicerianus | LC | Stable | tropical | diurnal | 249.9 | 3601349.63 | 16.404 |
| birds | Ortygornis_sephaena | Francolinus_sephaena | LC | Stable | tropical | diurnal | 340.5 | 3350686.796 | 9.214 |

| | | | | | | | | | |
|---|---|---|---|---|---|---|---|---|---|
| birds | Ortygospiza_atricollis | Ortygospiza_atricollis | LC | Stable | tropical | diurnal | 10.6 | 3231575.556 | 11.523 |
| birds | Otidiphaps_nobilis | Otidiphaps_nobilis | LC | Decreasing | tropical | diurnal | 500 | 260128.691 | 2.888 |
| birds | Otis_tarda | Otis_tarda | EN | Decreasing | temperate | diurnal | 6759.9 | 10008660.6 | 10.839 |
| birds | Otocichla_mupinensis | Turdus_mupinensis | LC | Stable | temperate | diurnal | 62.5 | 1082838.831 | 12.919 |
| birds | Otus_angelinae | Otus_angelinae | VU | Decreasing | tropical | nocturnal | 82.2 | 16745.86 | 14.132 |
| birds | Otus_bakkamoena | Otus_bakkamoena | LC | Stable | tropical | nocturnal | 138.5 | 2947848.958 | 18.201 |
| birds | Otus_balli | NA | LC | Stable | NA | NA | NA | NA | NA |
| birds | Otus_beccarii | Otus_beccarii | NT | Decreasing | tropical | nocturnal | 114 | 2524.27 | 8.959 |
| birds | Otus_brookii | NA | LC | Decreasing | NA | NA | NA | NA | NA |
| birds | Otus_brucei | Otus_brucei | LC | Stable | temperate | cathemeral | 84.4 | 1794970.432 | 11.548 |
| birds | Otus_elegans | Otus_elegans | NT | Decreasing | temperate | nocturnal | 124 | 4014.395 | 20.585 |
| birds | Otus_hartlaubi | Otus_hartlaubi | VU | Decreasing | tropical | diurnal | 79 | 388.973 | 7.875 |
| birds | Otus_insularis | NA | CR | Decreasing | NA | NA | NA | NA | NA |
| birds | Otus_ireneae | Otus_ireneae | EN | Decreasing | tropical | nocturnal | 50.3 | 634.785 | 9.88 |
| birds | Otus_lempiji | Otus_lempiji | LC | Stable | tropical | nocturnal | 115 | 1486775.343 | 9.482 |
| birds | Otus_lettia | Otus_lettia | LC | Stable | tropical | nocturnal | 139 | 4204562.896 | 13.834 |
| birds | Otus_magicus | Otus_magicus | LC | Decreasing | tropical | nocturnal | 165 | 84643.893 | 8.888 |
| birds | Otus_manadensis | Otus_manadensis | LC | Decreasing | tropical | nocturnal | 88 | 177304.105 | 9.363 |
| birds | Otus_mentawi | NA | LC | Stable | NA | NA | NA | NA | NA |
| birds | Otus_mindorensis | NA | NT | Decreasing | NA | NA | NA | NA | NA |
| birds | Otus_pembaensis | NA | VU | Decreasing | NA | NA | NA | NA | NA |
| birds | Otus_sagittatus | Otus_sagittatus | VU | Decreasing | tropical | nocturnal | 122.6 | 148393.297 | 9.811 |
| birds | Otus_scops | Otus_scops | LC | Decreasing | tropical | cathemeral | 92 | 26192272.395 | 8.788 |
| birds | Otus_semitorques | Otus_semitorques | LC | Stable | temperate | nocturnal | 121.5 | 1982105.073 | 14.75 |
| birds | Otus_senegalensis | Otus_senegalensis | LC | Stable | tropical | nocturnal | 67 | 13485667.888 | 9.668 |
| birds | Otus_spilocephalus | Otus_spilocephalus | LC | Stable | tropical | nocturnal | 67.5 | 3350281.146 | 10.818 |
| birds | Otus_sunia | Otus_sunia | LC | Stable | temperate | nocturnal | 78.1 | 8012147.551 | 14.526 |
| birds | Otus_umbra | Otus_umbra | LC | Stable | tropical | nocturnal | 95 | 1736.07 | 9.36 |
| birds | Oxylabes_madagascariensis | Oxylabes_madagascariensis | LC | Decreasing | tropical | diurnal | 23.3 | 172738.737 | 8.301 |
| birds | Oxypogon_guerinii | Oxypogon_guerinii | LC | Decreasing | tropical | diurnal | 4.8 | 19693.678 | 13.524 |
| birds | Oxyruncus_cristatus | Oxyruncus_cristatus | LC | Decreasing | tropical | diurnal | 42 | 1872045.952 | 7.44 |
| birds | Oxyura_australis | Oxyura_australis | LC | Stable | temperate | diurnal | 798 | 3286126.167 | 6.605 |
| birds | Oxyura_jamaicensis | Oxyura_jamaicensis | LC | Decreasing | temperate | diurnal | 608.2 | 12454394.743 | 11.922 |
| birds | Oxyura_leucocephala | Oxyura_leucocephala | EN | Decreasing | temperate | diurnal | 661.1 | 10904918.604 | 9.554 |

| | | | | | | | | | |
|---|---|---|---|---|---|---|---|---|---|
| birds | Oxyura_maccoa | Oxyura_maccoa | EN | Decreasing | temperate | diurnal | 674 | 1133393.565 | 9.984 |
| birds | Oxyura_vittata | Oxyura_vittata | LC | Stable | temperate | diurnal | 642.7 | 3138365.749 | 8.273 |
| birds | Pachycare_flavogriseum | Pachycare_flavogriseum | LC | Stable | tropical | diurnal | 16.2 | 171110.445 | 3.536 |
| birds | Pachycephala_albiventris | Pachycephala_albiventris | LC | Decreasing | tropical | diurnal | 21.1 | 104394.317 | 13.228 |
| birds | Pachycephala_arctitorquis | NA | LC | Stable | NA | NA | NA | NA | NA |
| birds | Pachycephala_aurea | NA | LC | Stable | NA | NA | NA | NA | NA |
| birds | Pachycephala_caledonica | Pachycephala_caledonica | LC | Stable | tropical | diurnal | 22.7 | 16533.262 | 7.541 |
| birds | Pachycephala_cinerea | Pachycephala_grisola | LC | Stable | tropical | diurnal | 21 | 1077022.849 | 13.251 |
| birds | Pachycephala_flavifrons | NA | LC | Stable | NA | NA | NA | NA | NA |
| birds | Pachycephala_griseiceps | NA | LC | Stable | NA | NA | NA | NA | NA |
| birds | Pachycephala_griseonota | Pachycephala_griseonota | LC | Stable | tropical | diurnal | 22.4 | 43720.568 | 6.073 |
| birds | Pachycephala_homeyeri | Pachycephala_homeyeri | LC | Stable | tropical | diurnal | 25.8 | 125156.591 | 13.123 |
| birds | Pachycephala_hyperythra | Pachycephala_hyperythra | LC | Stable | tropical | diurnal | 27.7 | 103362.374 | 2.814 |
| birds | Pachycephala_hypoxantha | NA | LC | Decreasing | NA | NA | NA | NA | NA |
| birds | Pachycephala_implicata | Pachycephala_implicata | LC | Stable | tropical | diurnal | 34.5 | 260.24 | 2.801 |
| birds | Pachycephala_inornata | Pachycephala_inornata | LC | Decreasing | temperate | diurnal | 30.8 | 1593660.359 | 6.433 |
| birds | Pachycephala_jacquinoti | NA | NT | Decreasing | NA | NA | NA | NA | NA |
| birds | Pachycephala_lanioides | Pachycephala_lanioides | LC | Stable | tropical | diurnal | 38.7 | 255698.626 | 3.264 |
| birds | Pachycephala_leucogastra | Pachycephala_leucogastra | LC | Stable | tropical | diurnal | 18.5 | 3984.197 | 5.706 |
| birds | Pachycephala_lorentzi | Pachycephala_lorentzi | LC | Stable | tropical | diurnal | 19.4 | 47469.457 | 4.51 |
| birds | Pachycephala_melanura | Pachycephala_melanura | LC | Stable | tropical | diurnal | 23.8 | 524241.757 | 3.536 |
| birds | Pachycephala_meyeri | Pachycephala_meyeri | LC | Stable | tropical | diurnal | 19 | 8657.874 | 2.425 |
| birds | Pachycephala_modesta | Pachycephala_modesta | LC | Stable | tropical | diurnal | 18.3 | 68146.904 | 5.267 |
| birds | Pachycephala_monacha | Pachycephala_monacha | LC | Increasing | tropical | diurnal | 26 | 33190.897 | 5.315 |
| birds | Pachycephala_nudigula | NA | LC | Decreasing | NA | NA | NA | NA | NA |
| birds | Pachycephala_olivacea | Pachycephala_olivacea | LC | Decreasing | temperate | diurnal | 41.2 | 298358.813 | 9.057 |
| birds | Pachycephala_orpheus | Pachycephala_orpheus | LC | Stable | tropical | diurnal | 15.5 | 21909.217 | 11.87 |
| birds | Pachycephala_pectoralis | Pachycephala_pectoralis | LC | Stable | temperate | diurnal | 30.4 | 1963546.275 | 8.558 |
| birds | Pachycephala_phaionota | Pachycephala_phaionota | LC | Stable | tropical | diurnal | 24.8 | 9616.801 | 4.156 |
| birds | Pachycephala_philippinensis | Pachycephala_philippinensis | LC | Decreasing | tropical | diurnal | 22.3 | 193262.302 | 12.145 |
| birds | Pachycephala_rufiventris | Pachycephala_rufiventris | LC | Decreasing | temperate | diurnal | 22.5 | 7227395.749 | 4.422 |
| birds | Pachycephala_rufogularis | Pachycephala_rufogularis | VU | Decreasing | temperate | diurnal | 36.6 | 133566.321 | 9.542 |
| birds | Pachycephala_schlegelii | Pachycephala_schlegelii | LC | Stable | tropical | diurnal | 22.7 | 132622.242 | 5.059 |
| birds | Pachycephala_simplex | Pachycephala_simplex | LC | Stable | tropical | diurnal | 17.4 | 93200.995 | 2.417 |

| | | | | | | | | | |
|---|---|---|---|---|---|---|---|---|---|
| birds | Pachycephala_soror | Pachycephala_soror | LC | Stable | tropical | diurnal | 24.8 | 221930.328 | 4.109 |
| birds | Pachycephala_sulfuriventer | Pachycephala_sulfuriventer | LC | Decreasing | tropical | diurnal | 21.4 | 49473.027 | 5.74 |
| birds | Pachycephala_tenebrosa | Colluricincla_tenebrosa | LC | Decreasing | tropical | diurnal | 46 | 456.668 | 11.182 |
| birds | Pachycephalopsis_hattamensis | Pachycephalopsis_hattamensis | LC | Stable | tropical | diurnal | 34.6 | 75632.902 | 3.545 |
| birds | Pachycephalopsis_poliosoma | Pachycephalopsis_poliosoma | LC | Stable | tropical | diurnal | 38 | 184655.431 | 4.156 |
| birds | Pachycoccyx_audeberti | Pachycoccyx_audeberti | LC | Decreasing | tropical | diurnal | 102.9 | 4407880.924 | 9.225 |
| birds | Pachyptila_belcheri | Pachyptila_belcheri | LC | Stable | temperate | cathemeral | 145 | 56286348.413 | 5.432 |
| birds | Pachyptila_crassirostris | Pachyptila_crassirostris | LC | Stable | temperate | cathemeral | 138.3 | 6979661.914 | 10.471 |
| birds | Pachyptila_desolata | Pachyptila_desolata | LC | Decreasing | temperate | cathemeral | 147 | 84155477.564 | 7.053 |
| birds | Pachyptila_salvini | Pachyptila_salvini | LC | Stable | temperate | cathemeral | 159 | 29593329.822 | 11.803 |
| birds | Pachyptila_turtur | Pachyptila_turtur | LC | Stable | temperate | cathemeral | 137 | 21293222.128 | 10.495 |
| birds | Pachyptila_vittata | Pachyptila_vittata | LC | Decreasing | temperate | cathemeral | 196 | 50803001.588 | 12.363 |
| birds | Pachyramphus_aglaiae | Pachyramphus_aglaiae | LC | Decreasing | tropical | diurnal | 29.7 | 1345543.813 | 9.965 |
| birds | Pachyramphus_albogriseus | Pachyramphus_albogriseus | LC | Decreasing | tropical | diurnal | 17.3 | 341646.914 | 9.547 |
| birds | Pachyramphus_castaneus | Pachyramphus_castaneus | LC | Stable | tropical | diurnal | 19.5 | 5788367.532 | 5.498 |
| birds | Pachyramphus_cinnamomeus | Pachyramphus_cinnamomeus | LC | Decreasing | tropical | diurnal | 20.3 | 701563.571 | 10.421 |
| birds | Pachyramphus_homochrous | Pachyramphus_homochrous | LC | Decreasing | tropical | diurnal | 37.3 | 370203.78 | 11.354 |
| birds | Pachyramphus_major | Pachyramphus_major | LC | Decreasing | tropical | diurnal | 24.8 | 468168.148 | 9.047 |
| birds | Pachyramphus_marginatus | Pachyramphus_marginatus | LC | Decreasing | tropical | diurnal | 18.4 | 6344951.425 | 2.437 |
| birds | Pachyramphus_minor | Pachyramphus_minor | LC | Decreasing | tropical | diurnal | 36.6 | 6895984.62 | 2.275 |
| birds | Pachyramphus_niger | Pachyramphus_niger | LC | Stable | tropical | diurnal | 40.2 | 11021.791 | 19.741 |
| birds | Pachyramphus_polychopterus | Pachyramphus_polychopterus | LC | Stable | tropical | diurnal | 20.8 | 13935213.896 | 5.837 |
| birds | Pachyramphus_rufus | Pachyramphus_rufus | LC | Decreasing | tropical | diurnal | 18.8 | 3994359.028 | 3.354 |
| birds | Pachyramphus_spodiurus | Pachyramphus_spodiurus | VU | Decreasing | tropical | diurnal | 18.2 | 40652.05 | 11.36 |
| birds | Pachyramphus_surinamus | Pachyramphus_surinamus | LC | Stable | tropical | diurnal | 19.7 | 707437.361 | 1.024 |
| birds | Pachyramphus_validus | Pachyramphus_validus | LC | Decreasing | tropical | diurnal | 43 | 6063334.02 | 8.537 |
| birds | Pachyramphus_versicolor | Pachyramphus_versicolor | LC | Decreasing | tropical | diurnal | 15.5 | 302157.644 | 8.174 |
| birds | Pachyramphus_viridis | Pachyramphus_viridis | LC | Stable | tropical | diurnal | 21 | 4959688.094 | 9.577 |
| birds | Pachyramphus_xanthogenys | NA | LC | Stable | NA | NA | NA | NA | NA |
| birds | Pachysylvia_aurantiifrons | Hylophilus_aurantiifrons | LC | Decreasing | tropical | diurnal | 9.5 | 611619.822 | 11.382 |
| birds | Pachysylvia_decurtata | Hylophilus_decurtatus | LC | Decreasing | tropical | diurnal | 8.5 | 856443.061 | 10.133 |
| birds | Pachysylvia_hypochrysea | Vireo_hypochryseus | LC | Decreasing | tropical | diurnal | 12.2 | 356386.016 | 8.224 |
| birds | Pachysylvia_hypoxantha | Hylophilus_hypoxanthus | LC | Decreasing | tropical | diurnal | 17 | 3940574.211 | 1.78 |
| birds | Pachysylvia_muscicapina | Hylophilus_muscicapinus | LC | Decreasing | tropical | diurnal | 11.7 | 2386727.389 | 2.489 |

| | | | | | | | | | |
|---|---|---|---|---|---|---|---|---|---|
| birds | Pachysylvia_semibrunnea | Hylophilus_semibrunneus | LC | Stable | tropical | diurnal | 11.4 | 42187.561 | 9.69 |
| birds | Padda_fuscata | Padda_fuscata | NT | Decreasing | tropical | diurnal | 20.7 | 24603.859 | 14.521 |
| birds | Padda_oryzivora | Padda_oryzivora | EN | Decreasing | tropical | diurnal | 24.8 | 960325.856 | 18.039 |
| birds | Pagodroma_nivea | NA | LC | Stable | NA | NA | NA | NA | NA |
| birds | Pagophila_eburnea | Pagophila_eburnea | NT | Decreasing | temperate | diurnal | 550 | 17367389.82 | 0.2 |
| birds | Palaeornis_eupatria | Psittacula_eupatria | NT | Decreasing | tropical | diurnal | 214 | 3231032.199 | 15.314 |
| birds | Palmeria_dolei | Palmeria_dolei | CR | Decreasing | tropical | diurnal | 26.1 | 1012.579 | 5.954 |
| birds | Paludipasser_locustella | NA | LC | Stable | NA | NA | NA | NA | NA |
| birds | Pampa_rufa | Campylopterus_rufus | LC | Decreasing | tropical | diurnal | 7.1 | 31219.993 | 16.547 |
| birds | Pampusana_beccarii | Gallicolumba_beccarii | LC | Stable | tropical | diurnal | 78.7 | 167246.955 | 4.686 |
| birds | Pampusana_canifrons | NA | EN | Decreasing | NA | NA | NA | NA | NA |
| birds | Pampusana_erythroptera | Gallicolumba_erythroptera | CR | Decreasing | tropical | diurnal | 113.2 | 2477.53 | 13.57 |
| birds | Pampusana_hoedtii | NA | EN | Decreasing | NA | NA | NA | NA | NA |
| birds | Pampusana_jobiensis | Gallicolumba_jobiensis | LC | Stable | tropical | diurnal | 135 | 647409.704 | 3.582 |
| birds | Pampusana_kubaryi | Gallicolumba_kubaryi | EN | Decreasing | tropical | diurnal | 162 | 424.306 | 12.519 |
| birds | Pampusana_rubescens | Gallicolumba_rubescens | EN | Stable | tropical | diurnal | 60 | 350.26 | 7.697 |
| birds | Pampusana_salamonis | NA | EX | not_available | NA | NA | NA | NA | NA |
| birds | Pampusana_sanctaecrucis | NA | EN | Decreasing | NA | NA | NA | NA | NA |
| birds | Pampusana_stairi | Gallicolumba_stairi | VU | Decreasing | tropical | diurnal | 171 | 21209.625 | 9.375 |
| birds | Pampusana_xanthonura | Gallicolumba_xanthonura | NT | Unknown | tropical | diurnal | 110.6 | 1148.685 | 19.878 |
| birds | Pandion_haliaetus | Pandion_haliaetus | LC | Increasing | tropical | diurnal | 1483.2 | 91738111.448 | 8.967 |
| birds | Panterpe_insignis | Panterpe_insignis | LC | Stable | tropical | diurnal | 5.7 | 8271.569 | 7.328 |
| birds | Panurus_biarmicus | Panurus_biarmicus | LC | Unknown | temperate | diurnal | 13.8 | 12154631.847 | 8.131 |
| birds | Panyptila_cayennensis | Panyptila_cayennensis | LC | Decreasing | tropical | diurnal | 21.1 | 8548025.435 | 3.636 |
| birds | Panyptila_sanctihieronymi | Panyptila_sanctihieronymi | LC | Decreasing | tropical | diurnal | 48.1 | 366914.496 | 10.462 |
| birds | Papasula_abbotti | Papasula_abbotti | EN | Stable | tropical | diurnal | 1550.7 | 677593.38 | 12.551 |
| birds | Parabuteo_leucorrhous | Buteo_leucorrhous | LC | Decreasing | temperate | diurnal | 290 | 1189248.195 | 11.437 |
| birds | Parabuteo_unicinctus | Parabuteo_unicinctus | LC | Decreasing | tropical | diurnal | 850.3 | 9015595.351 | 9.253 |
| birds | Paraclaravis_geoffroyi | Claravis_godefrida | CR | Decreasing | tropical | diurnal | 43.5 | 612044.339 | 12.658 |
| birds | Paraclaravis_mondetoura | Claravis_mondetoura | LC | Decreasing | tropical | diurnal | 89.7 | 342167.22 | 9.45 |
| birds | Paradigalla_brevicauda | Paradigalla_brevicauda | LC | Stable | tropical | diurnal | 167.9 | 91714.964 | 5.454 |
| birds | Paradigalla_carunculata | Paradigalla_carunculata | NT | Decreasing | tropical | diurnal | 170 | 3319.374 | 3.798 |
| birds | Paradisaea_apoda | Paradisaea_apoda | LC | Decreasing | tropical | diurnal | 172 | 99261.897 | 2.54 |
| birds | Paradisaea_decora | Paradisaea_decora | VU | Decreasing | tropical | diurnal | 237 | 834.458 | 4.511 |

| | | | | | | | | | |
|---|---|---|---|---|---|---|---|---|---|
| birds | Paradisaea_guilielmi | Paradisaea_guilielmi | NT | Decreasing | tropical | diurnal | 256 | 10730.797 | 4.632 |
| birds | Paradisaea_minor | Paradisaea_minor | LC | Decreasing | tropical | diurnal | 207.9 | 273509.568 | 3.227 |
| birds | Paradisaea_raggiana | Paradisaea_raggiana | LC | Decreasing | tropical | diurnal | 212.6 | 202567.577 | 3.603 |
| birds | Paradisaea_rubra | Paradisaea_rubra | NT | Decreasing | tropical | diurnal | 178.2 | 3877.441 | 2.458 |
| birds | Paradisornis_rudolphi | Paradisaea_rudolphi | NT | Decreasing | tropical | diurnal | 164.1 | 51027.787 | 6.03 |
| birds | Paradoxornis_aemodium | Conostoma_oemodium | LC | Decreasing | temperate | diurnal | 84 | 747028.87 | 8.868 |
| birds | Paradoxornis_flavirostris | Paradoxornis_flavirostris | VU | Decreasing | temperate | diurnal | 26 | 36993.819 | 15.827 |
| birds | Paradoxornis_gularis | Paradoxornis_gularis | LC | Unknown | temperate | diurnal | 29 | 1985394.87 | 11.753 |
| birds | Paradoxornis_guttaticollis | Paradoxornis_guttaticollis | LC | Unknown | temperate | diurnal | 26 | 1409894.644 | 10.682 |
| birds | Paradoxornis_heudei | Paradoxornis_heudei | NT | Decreasing | temperate | diurnal | 32 | 452378.336 | 14.204 |
| birds | Paradoxornis_paradoxus | Paradoxornis_paradoxus | LC | Stable | temperate | diurnal | 35 | 137179.648 | 7.824 |
| birds | Paradoxornis_unicolor | Paradoxornis_unicolor | LC | Decreasing | temperate | diurnal | 34 | 150398.919 | 6.158 |
| birds | Paragallinula_angulata | Gallinula_angulata | LC | Unknown | tropical | diurnal | 130 | 11738770.118 | 9.559 |
| birds | Paramythia_montium | Paramythia_montium | LC | Unknown | tropical | diurnal | 41.8 | 43513.484 | 5.541 |
| birds | Parayuhina_diademata | Yuhina_diademata | LC | Unknown | temperate | diurnal | 22 | 1124238.68 | 11.991 |
| birds | Pardaliparus_amabilis | Parus_amabilis | NT | Decreasing | tropical | diurnal | 12.7 | 14312.452 | 9.25 |
| birds | Pardaliparus_elegans | Parus_elegans | LC | Stable | tropical | diurnal | 12.8 | 272069.254 | 13.817 |
| birds | Pardaliparus_venustulus | Parus_venustulus | LC | Decreasing | temperate | diurnal | 10.6 | 1942840.091 | 16.068 |
| birds | Pardalotus_punctatus | Pardalotus_punctatus | LC | Decreasing | temperate | diurnal | 8.7 | 2170812.264 | 7.992 |
| birds | Pardalotus_quadragintus | Pardalotus_quadragintus | EN | Decreasing | temperate | diurnal | 10.7 | 1433.298 | 11.412 |
| birds | Pardalotus_rubricatus | Pardalotus_rubricatus | LC | Stable | tropical | diurnal | 10.9 | 5191015.827 | 3.045 |
| birds | Pardipicus_caroli | Campethera_caroli | LC | Decreasing | tropical | diurnal | 44.2 | 3849726.472 | 7.422 |
| birds | Pardipicus_nivosus | Campethera_nivosa | LC | Stable | tropical | diurnal | 37.8 | 3825842.411 | 8.371 |
| birds | Pardirallus_nigricans | Pardirallus_nigricans | LC | Unknown | tropical | diurnal | 217 | 3254810.225 | 10.027 |
| birds | Pardirallus_sanguinolentus | Pardirallus_sanguinolentus | LC | Stable | temperate | cathemeral | 232 | 4579906.012 | 8.368 |
| birds | Pareudiastes_silvestris | Gallinula_silvestris | CR | Unknown | tropical | diurnal | 450 | 0.985 | 1.833 |
| birds | Parkerthraustes_humeralis | Parkerthraustes_humeralis | LC | Decreasing | tropical | diurnal | 37 | 1929725.376 | 2.08 |
| birds | Parkesia_motacilla | Seiurus_motacilla | LC | Increasing | tropical | diurnal | 19.9 | 4813246.081 | 15.123 |
| birds | Parkesia_noveboracensis | Seiurus_noveboracensis | LC | Increasing | temperate | diurnal | 16.3 | 17656885.931 | 8.751 |
| birds | Parmoptila_jamesoni | Parmoptila_jamesoni | LC | Stable | tropical | diurnal | 9.5 | 365703.462 | 7.19 |
| birds | Parmoptila_rubrifrons | Parmoptila_rubrifrons | NT | Decreasing | tropical | diurnal | 9.5 | 570085.791 | 14.074 |
| birds | Parmoptila_woodhousei | Parmoptila_woodhousei | LC | Stable | tropical | diurnal | 9.4 | 716141.127 | 6.342 |
| birds | Paroaria_baeri | Paroaria_baeri | LC | Stable | tropical | diurnal | 25.8 | 107338.324 | 4.992 |
| birds | Paroaria_capitata | Paroaria_capitata | LC | Stable | temperate | diurnal | 22.3 | 1522354.912 | 8.712 |

| | | | | | | | | | |
|---|---|---|---|---|---|---|---|---|---|
| birds | Paroaria_coronata | Paroaria_coronata | LC | Stable | temperate | diurnal | 37.8 | 2412937.447 | 8.491 |
| birds | Paroaria_dominicana | Paroaria_dominicana | LC | Stable | tropical | diurnal | 33.2 | 1212763.443 | 8.647 |
| birds | Paroaria_gularis | Paroaria_gularis | LC | Stable | tropical | diurnal | 23.6 | 4448336.299 | 2.434 |
| birds | Paroreomyza_maculata | Paroreomyza_maculata | CR | Unknown | tropical | diurnal | 11.3 | 1560.381 | 21.133 |
| birds | Paroreomyza_montana | Paroreomyza_montana | EN | Decreasing | tropical | diurnal | 13.1 | 1598.666 | 9.598 |
| birds | Parotia_carolae | Parotia_carolae | LC | Stable | tropical | diurnal | 163.2 | 23107.828 | 6.106 |
| birds | Parotia_lawesii | Parotia_lawesii | LC | Stable | tropical | diurnal | 161.4 | 96297.666 | 4.926 |
| birds | Parotia_sefilata | Parotia_sefilata | LC | Stable | tropical | diurnal | 181.7 | 7209.382 | 2.311 |
| birds | Parotia_wahnesi | Parotia_wahnesi | NT | Decreasing | tropical | diurnal | 158 | 4260.369 | 3.85 |
| birds | Parus_major | Parus_major | LC | Increasing | temperate | diurnal | 16.3 | 32577984.276 | 11.567 |
| birds | Parus_monticolus | Parus_monticolus | LC | Stable | temperate | diurnal | 14 | 1684621.77 | 12.658 |
| birds | Passer_ammodendri | Passer_ammodendri | LC | Stable | temperate | diurnal | 28.9 | 1302317.474 | 3.925 |
| birds | Passer_castanopterus | Passer_castanopterus | LC | Stable | tropical | diurnal | 18 | 712666.844 | 7.475 |
| birds | Passer_cinnamomeus | Passer_rutilans | LC | Stable | temperate | diurnal | 18.5 | 4964243.405 | 14.563 |
| birds | Passer_cordofanicus | Passer_cordofanicus | LC | Stable | tropical | diurnal | 22 | 143587.997 | 12.101 |
| birds | Passer_diffusus | Passer_diffusus | LC | Stable | tropical | diurnal | 24.2 | 3365122.435 | 8.92 |
| birds | Passer_domesticus | Passer_domesticus | LC | Decreasing | temperate | diurnal | 26.5 | 64177419.85 | 9.764 |
| birds | Passer_eminibey | Passer_eminibey | LC | Stable | tropical | diurnal | 13.4 | 549271.111 | 11.331 |
| birds | Passer_euchlorus | Passer_euchlorus | LC | Stable | tropical | diurnal | 14 | 163824.346 | 11.432 |
| birds | Passer_flaveolus | Passer_flaveolus | LC | Stable | tropical | diurnal | 19.4 | 1319875.05 | 13.783 |
| birds | Passer_gongonensis | Passer_gongonensis | LC | Stable | tropical | diurnal | 39.9 | 1125295.411 | 9.776 |
| birds | Passer_griseus | Passer_griseus | LC | Stable | tropical | diurnal | 27.8 | 10513534.527 | 9.564 |
| birds | Passer_hispaniolensis | Passer_hispaniolensis | LC | Decreasing | temperate | diurnal | 24.2 | 8060752.71 | 10.472 |
| birds | Passer_iagoensis | Passer_iagoensis | LC | Stable | tropical | diurnal | 31.5 | 3585.234 | 12.672 |
| birds | Passer_insularis | Passer_insularis | LC | Stable | tropical | diurnal | 27.3 | 3671.443 | 11.027 |
| birds | Passer_luteus | Passer_luteus | LC | Stable | tropical | diurnal | 11.2 | 2874074.904 | 7.791 |
| birds | Passer_melanurus | Passer_melanurus | LC | Stable | temperate | diurnal | 29.5 | 1765360.169 | 8.345 |
| birds | Passer_moabiticus | Passer_moabiticus | LC | Decreasing | temperate | diurnal | 16.5 | 565850.917 | 13.855 |
| birds | Passer_montanus | Passer_montanus | LC | Decreasing | temperate | diurnal | 21.4 | 39493440.331 | 9.259 |
| birds | Passer_motitensis | Passer_motitensis | LC | Stable | tropical | diurnal | 30.2 | 1648538.382 | 6.381 |
| birds | Passer_pyrrhonotus | Passer_pyrrhonotus | LC | Stable | temperate | diurnal | 22 | 524876.721 | 13.683 |
| birds | Passer_rufocinctus | Passer_rufocinctus | LC | Stable | tropical | diurnal | 27.8 | 90120.457 | 12.189 |
| birds | Passer_shelleyi | Passer_shelleyi | LC | Stable | tropical | diurnal | 22 | 201266.837 | 8.914 |
| birds | Passer_simplex | Passer_simplex | LC | Decreasing | temperate | diurnal | 19.5 | 660184.35 | 2.869 |

| | | | | | | | | | |
|---|---|---|---|---|---|---|---|---|---|
| birds | Passer_suahelicus | Passer_suahelicus | LC | Stable | tropical | diurnal | 25 | 282661.661 | 14.161 |
| birds | Passer_swainsonii | Passer_swainsonii | LC | Stable | tropical | diurnal | 31.6 | 752104.104 | 10.84 |
| birds | Passerculus_bairdii | Ammodramus_bairdii | LC | Stable | temperate | diurnal | 18.1 | 2235509.58 | 11.905 |
| birds | Passerculus_henslowii | Ammodramus_henslowii | LC | Increasing | temperate | diurnal | 12.8 | 2912755.702 | 18.128 |
| birds | Passerculus_sandwichensis | Passerculus_sandwichensis | LC | Decreasing | temperate | diurnal | 20 | 19177797.532 | 7.875 |
| birds | Passerella_arborea | Spizella_arborea | LC | Decreasing | temperate | diurnal | 17.8 | 14358243.446 | 7.194 |
| birds | Passerella_iliaca | Passerella_iliaca | LC | Decreasing | temperate | diurnal | 33.3 | 12314409.985 | 8.176 |
| birds | Passerina_amoena | Passerina_amoena | LC | Increasing | temperate | diurnal | 15.5 | 4666546.614 | 9.361 |
| birds | Passerina_caerulea | Passerina_caerulea | LC | Increasing | temperate | diurnal | 27.4 | 6742464.863 | 12.303 |
| birds | Passerina_ciris | Passerina_ciris | LC | Stable | tropical | diurnal | 15.5 | 3706093.071 | 11.618 |
| birds | Passerina_cyanea | Passerina_cyanea | LC | Decreasing | temperate | diurnal | 14.7 | 7436442.545 | 14.613 |
| birds | Passerina_leclancherii | Passerina_leclancherii | LC | Decreasing | tropical | diurnal | 14 | 140628.187 | 9.782 |
| birds | Passerina_rositae | Passerina_rositae | NT | Stable | tropical | diurnal | 20 | 6979.796 | 6.171 |
| birds | Passerina_versicolor | Passerina_versicolor | LC | Stable | temperate | diurnal | 12.9 | 1540631.554 | 9.04 |
| birds | Pastor_roseus | Sturnus_roseus | LC | Unknown | temperate | diurnal | 73 | 7512915.799 | 11.265 |
| birds | Patagioenas_araucana | Patagioenas_araucana | LC | Increasing | temperate | diurnal | 270 | 289225.923 | 7.954 |
| birds | Patagioenas_caribaea | Patagioenas_caribaea | VU | Decreasing | tropical | diurnal | 250 | 7105.931 | 17.37 |
| birds | Patagioenas_cayennensis | Patagioenas_cayennensis | LC | Decreasing | tropical | diurnal | 229 | 13847694.767 | 5.869 |
| birds | Patagioenas_corensis | Patagioenas_corensis | LC | Stable | tropical | diurnal | 272.3 | 129079.857 | 12.428 |
| birds | Patagioenas_fasciata | Patagioenas_fasciata | LC | Decreasing | temperate | diurnal | 366.3 | 2745784.923 | 9.396 |
| birds | Patagioenas_flavirostris | Patagioenas_flavirostris | LC | Decreasing | tropical | diurnal | 270.8 | 928061.872 | 10.468 |
| birds | Patagioenas_goodsoni | NA | LC | Decreasing | NA | NA | NA | NA | NA |
| birds | Patagioenas_inornata | Patagioenas_inornata | NT | Decreasing | tropical | diurnal | 365 | 205258.361 | 16.67 |
| birds | Patagioenas_leucocephala | Patagioenas_leucocephala | NT | Decreasing | tropical | diurnal | 240.1 | 243969.098 | 15.781 |
| birds | Patagioenas_maculosa | Patagioenas_maculosa | LC | Increasing | temperate | diurnal | 347 | 3659053.309 | 8.575 |
| birds | Patagioenas_nigrirostris | Patagioenas_nigrirostris | LC | Decreasing | tropical | diurnal | 176 | 352468.534 | 8.703 |
| birds | Patagioenas_oenops | Patagioenas_oenops | NT | Decreasing | tropical | diurnal | 264.6 | 12881.115 | 8.306 |
| birds | Patagioenas_picazuro | Patagioenas_picazuro | LC | Increasing | tropical | diurnal | 279 | 6181623.946 | 9.256 |
| birds | Patagioenas_plumbea | Patagioenas_plumbea | LC | Decreasing | tropical | diurnal | 178.8 | 6564806.47 | 4.241 |
| birds | Patagioenas_speciosa | Patagioenas_speciosa | LC | Decreasing | tropical | diurnal | 258.5 | 9715779.78 | 4.525 |
| birds | Patagioenas_squamosa | Patagioenas_squamosa | LC | Decreasing | tropical | diurnal | 312 | 202802.195 | 16.669 |
| birds | Patagioenas_subvinacea | Patagioenas_subvinacea | LC | Decreasing | tropical | diurnal | 162.5 | 8499137.272 | 3.478 |
| birds | Patagona_gigas | Patagona_gigas | LC | Stable | tropical | diurnal | 20.2 | 1270749.201 | 7.091 |
| birds | Pauxi_pauxi | Pauxi_pauxi | VU | Decreasing | tropical | diurnal | 3117.7 | 79524.862 | 11.805 |

| | | | | | | | | | |
|---|---|---|---|---|---|---|---|---|---|
| birds | Pauxi_unicornis | Pauxi_unicornis | CR | Decreasing | tropical | nocturnal | 3743.2 | 11846.327 | 3.204 |
| birds | Pavo_cristatus | Pavo_cristatus | LC | Stable | tropical | diurnal | 4154.8 | 3442849.758 | 17.029 |
| birds | Pavo_muticus | Pavo_muticus | EN | Decreasing | tropical | diurnal | 2205.7 | 1851214.378 | 13.022 |
| birds | Pedionomus_torquatus | Pedionomus_torquatus | EN | Decreasing | temperate | diurnal | 62.5 | 2237290.612 | 5.989 |
| birds | Pelagodroma_marina | Pelagodroma_marina | LC | Decreasing | tropical | cathemeral | 47.2 | 104837176.204 | 9.951 |
| birds | Pelargopsis_amauroptera | Pelargopsis_amauroptera | NT | Decreasing | tropical | diurnal | 162 | 41842.356 | 13.043 |
| birds | Pelargopsis_capensis | Pelargopsis_capensis | LC | Decreasing | tropical | diurnal | 178.7 | 5091973.15 | 12.91 |
| birds | Pelargopsis_melanorhyncha | Pelargopsis_melanorhyncha | LC | Decreasing | tropical | diurnal | 193.3 | 168034.982 | 9.747 |
| birds | Pelecanoides_garnotii | Pelecanoides_garnotii | NT | Increasing | tropical | cathemeral | 219.7 | 469387.157 | 11.351 |
| birds | Pelecanoides_georgicus | Pelecanoides_georgicus | LC | Decreasing | temperate | cathemeral | 121 | 8731179.275 | 0.201 |
| birds | Pelecanoides_magellani | Pelecanoides_magellani | LC | Decreasing | temperate | cathemeral | 134 | 1187555.986 | 2.092 |
| birds | Pelecanoides_urinatrix | Pelecanoides_urinatrix | LC | Decreasing | temperate | cathemeral | 152.1 | 21746036.601 | 5.399 |
| birds | Pelecanus_conspicillatus | Pelecanus_conspicillatus | LC | Stable | temperate | diurnal | 5505 | 6565542.055 | 5.223 |
| birds | Pelecanus_crispus | Pelecanus_crispus | NT | Decreasing | temperate | diurnal | 9512.1 | 12289083.634 | 10.608 |
| birds | Pelecanus_erythrorhynchos | Pelecanus_erythrorhynchos | LC | Increasing | temperate | diurnal | 5608.5 | 8104341.009 | 11.449 |
| birds | Pelecanus_occidentalis | Pelecanus_occidentalis | LC | Increasing | tropical | diurnal | 3427.8 | 6500402.255 | 13.319 |
| birds | Pelecanus_onocrotalus | Pelecanus_onocrotalus | LC | Unknown | temperate | diurnal | 9322.3 | 16249898.892 | 9.32 |
| birds | Pelecanus_philippensis | Pelecanus_philippensis | NT | Decreasing | temperate | diurnal | 5011 | 3052519.602 | 15.93 |
| birds | Pelecanus_rufescens | Pelecanus_rufescens | LC | Stable | tropical | diurnal | 5419.6 | 14141427.797 | 8.872 |
| birds | Peliperdix_lathami | Francolinus_lathami | LC | Decreasing | tropical | diurnal | 269 | 2655327.382 | 8.486 |
| birds | Pellorneum_albiventre | Pellorneum_albiventre | LC | Stable | tropical | diurnal | 21.5 | 461554.393 | 9.555 |
| birds | Pellorneum_bicolor | Trichastoma_bicolor | LC | Stable | tropical | diurnal | 27.2 | 785296.582 | 9.157 |
| birds | Pellorneum_capistratum | Pellorneum_capistratum | LC | Decreasing | tropical | diurnal | 25.1 | 121654.546 | 22.663 |
| birds | Pellorneum_celebense | Trichastoma_celebense | LC | Stable | tropical | diurnal | 23.6 | 179475.524 | 9.34 |
| birds | Pellorneum_cinereiceps | Malacocincla_cinereiceps | LC | Decreasing | tropical | diurnal | 23.9 | 12143.56 | 9.075 |
| birds | Pellorneum_fuscocapillus | Pellorneum_fuscocapillus | LC | Stable | tropical | diurnal | 30 | 53280.693 | 17.621 |
| birds | Pellorneum_malaccense | Malacocincla_malaccensis | NT | Decreasing | tropical | diurnal | 21.4 | 1073369.635 | 7.216 |
| birds | Pellorneum_palustre | NA | VU | Decreasing | NA | NA | NA | NA | NA |
| birds | Pellorneum_pyrrogenys | Trichastoma_pyrrogenys | LC | Decreasing | tropical | diurnal | 23.2 | 193222.644 | 4.625 |
| birds | Pellorneum_rostratum | Trichastoma_rostratum | NT | Decreasing | tropical | diurnal | 21 | 789435.459 | 7.23 |
| birds | Pellorneum_ruficeps | Pellorneum_ruficeps | LC | Stable | tropical | diurnal | 26 | 2044549.112 | 12.333 |
| birds | Pellorneum_tickelli | Trichastoma_tickelli | LC | Stable | temperate | diurnal | 17.1 | 851979.332 | 8.898 |
| birds | Peltohyas_australis | Charadrius_australis | LC | Unknown | temperate | nocturnal | 83.5 | 2556918.743 | 4.941 |
| birds | Peltops_blainvillii | Peltops_blainvillii | LC | Decreasing | tropical | diurnal | 30 | 529218 | 3.115 |

| | | | | | | | | | |
|---|---|---|---|---|---|---|---|---|---|
| birds | Peltops_montanus | Peltops_montanus | LC | Stable | tropical | diurnal | 31.6 | 324099.399 | 4.02 |
| birds | Penelope_albipennis | Penelope_albipennis | EN | Stable | tropical | diurnal | 1750 | 18595.674 | 7.957 |
| birds | Penelope_argyrotis | Penelope_argyrotis | LC | Decreasing | tropical | diurnal | 806.7 | 90114.257 | 11.499 |
| birds | Penelope_barbata | Penelope_barbata | NT | Decreasing | tropical | diurnal | 665 | 24010.993 | 8.643 |
| birds | Penelope_dabbenei | Penelope_dabbenei | LC | Decreasing | tropical | diurnal | 1230 | 31904.432 | 7.196 |
| birds | Penelope_jacquacu | Penelope_jacquacu | LC | Decreasing | tropical | diurnal | 1487.9 | 5007343.09 | 2.195 |
| birds | Penelope_jacucaca | Penelope_jacucaca | VU | Decreasing | tropical | diurnal | 1400 | 599768.772 | 7.935 |
| birds | Penelope_marail | Penelope_marail | LC | Decreasing | tropical | diurnal | 879.7 | 1217966.933 | 1.577 |
| birds | Penelope_montagnii | Penelope_montagnii | LC | Decreasing | tropical | diurnal | 706 | 368616.684 | 8.891 |
| birds | Penelope_obscura | Penelope_obscura | LC | Decreasing | temperate | diurnal | 1770 | 1012773.034 | 11.182 |
| birds | Penelope_ochrogaster | NA | VU | Decreasing | NA | NA | NA | NA | NA |
| birds | Penelope_ortoni | Penelope_ortoni | EN | Decreasing | tropical | diurnal | 872 | 77623.513 | 5.763 |
| birds | Penelope_perspicax | Penelope_perspicax | VU | Decreasing | tropical | diurnal | 1197 | 16807.618 | 17.514 |
| birds | Penelope_pileata | Penelope_pileata | VU | Decreasing | tropical | diurnal | 1249.8 | 903532.552 | 3.027 |
| birds | Penelope_purpurascens | Penelope_purpurascens | NT | Decreasing | tropical | diurnal | 2060 | 1277537.175 | 10.528 |
| birds | Penelope_superciliaris | Penelope_superciliaris | NT | Decreasing | tropical | diurnal | 895 | 6014652.883 | 7.584 |
| birds | Penelopides_affinis | Penelopides_affinis | LC | Decreasing | tropical | diurnal | 485.1 | 117669.514 | 12.31 |
| birds | Penelopides_manillae | Penelopides_manillae | LC | Decreasing | tropical | diurnal | 460.9 | 104660.897 | 15.015 |
| birds | Penelopides_mindorensis | Penelopides_mindorensis | EN | Decreasing | tropical | diurnal | 462 | 8535.866 | 12.232 |
| birds | Penelopides_panini | Penelopides_panini | EN | Decreasing | tropical | diurnal | 470.5 | 27587.92 | 18.216 |
| birds | Penelopina_nigra | Penelopina_nigra | VU | Decreasing | tropical | diurnal | 890 | 103251.957 | 11.928 |
| birds | Peneoenanthe_pulverulenta | Eopsaltria_pulverulenta | LC | Stable | tropical | diurnal | 19.2 | 435422.985 | 3.393 |
| birds | Peneothello_bimaculata | Peneothello_bimaculata | LC | Stable | tropical | diurnal | 23.8 | 51621.469 | 3.333 |
| birds | Peneothello_cryptoleuca | Peneothello_cryptoleuca | LC | Stable | tropical | diurnal | 18.9 | 45602.008 | 4.008 |
| birds | Peneothello_cyanus | Peneothello_cyanus | LC | Stable | tropical | diurnal | 25 | 167864.705 | 4.543 |
| birds | Peneothello_sigillata | Peneothello_sigillatus | LC | Stable | tropical | diurnal | 22.8 | 77436.989 | 5.099 |
| birds | Percnostola_rufifrons | Percnostola_rufifrons | LC | Stable | tropical | diurnal | 23.9 | 1139425.122 | 1.24 |
| birds | Perdicula_argoondah | Perdicula_argoondah | LC | Decreasing | tropical | diurnal | 60.5 | 1182863.734 | 19.86 |
| birds | Perdicula_asiatica | Perdicula_asiatica | LC | Stable | tropical | diurnal | 68.4 | 2187881.266 | 18.938 |
| birds | Perdicula_erythrorhyncha | Perdicula_erythrorhyncha | LC | Stable | tropical | diurnal | 67.6 | 502809.332 | 16.63 |
| birds | Perdicula_manipurensis | Perdicula_manipurensis | EN | Decreasing | temperate | diurnal | 70.7 | 137525.139 | 14.211 |
| birds | Perdix_dauurica | Perdix_dauurica | LC | Decreasing | temperate | diurnal | 263.4 | 7725915.517 | 6.939 |
| birds | Perdix_hodgsoniae | Perdix_hodgsoniae | LC | Stable | temperate | diurnal | 363.7 | 1000982.756 | 5.916 |
| birds | Perdix_perdix | Perdix_perdix | LC | Decreasing | temperate | diurnal | 405.3 | 12799976.795 | 13.73 |

| | | | | | | | | | |
|---|---|---|---|---|---|---|---|---|---|
| birds | Pericrocotus_brevirostris | Pericrocotus_brevirostris | LC | Decreasing | tropical | diurnal | 16.5 | 2416801.638 | 9.991 |
| birds | Pericrocotus_cantonensis | Pericrocotus_cantonensis | LC | Decreasing | tropical | diurnal | 16.3 | 3495894.366 | 14.331 |
| birds | Pericrocotus_cinnamomeus | Pericrocotus_cinnamomeus | LC | Stable | tropical | diurnal | 8.6 | 4793425.28 | 16.695 |
| birds | Pericrocotus_divaricatus | Pericrocotus_divaricatus | LC | Decreasing | temperate | diurnal | 24.6 | 5823422.319 | 14.918 |
| birds | Pericrocotus_erythropygius | Pericrocotus_erythropygius | LC | Stable | tropical | diurnal | 10.8 | 1571440.411 | 18.692 |
| birds | Pericrocotus_ethologus | Pericrocotus_ethologus | LC | Decreasing | temperate | diurnal | 19 | 6237997.801 | 13.639 |
| birds | Pericrocotus_flammeus | Pericrocotus_flammeus | LC | Decreasing | tropical | diurnal | 23.3 | 6801040.969 | 13.044 |
| birds | Pericrocotus_igneus | Pericrocotus_igneus | NT | Decreasing | tropical | diurnal | 15 | 1266299.035 | 8.63 |
| birds | Pericrocotus_lansbergei | Pericrocotus_lansbergei | LC | Stable | tropical | diurnal | 16.3 | 30021.145 | 12.377 |
| birds | Pericrocotus_miniatus | Pericrocotus_miniatus | LC | Stable | tropical | diurnal | 16.3 | 155206.179 | 11.3 |
| birds | Pericrocotus_roseus | Pericrocotus_roseus | LC | Decreasing | tropical | diurnal | 19.2 | 3379728.881 | 12.952 |
| birds | Pericrocotus_solaris | Pericrocotus_solaris | LC | Stable | tropical | diurnal | 14.5 | 2947693.855 | 12.296 |
| birds | Pericrocotus_tegimae | Pericrocotus_tegimae | LC | Increasing | temperate | diurnal | 21 | 58093.737 | 20.222 |
| birds | Periparus_ater | Parus_ater | LC | Decreasing | temperate | diurnal | 9.2 | 22198675.929 | 10.083 |
| birds | Periparus_rubidiventris | Parus_rubidiventris | LC | Stable | temperate | diurnal | 11.8 | 949945.598 | 7.259 |
| birds | Periparus_rufonuchalis | Parus_rufonuchalis | LC | Stable | temperate | diurnal | 12.6 | 530583.683 | 10.981 |
| birds | Perisoreus_canadensis | Perisoreus_canadensis | LC | Decreasing | temperate | diurnal | 71.6 | 7301159.582 | 2.615 |
| birds | Perisoreus_infaustus | Perisoreus_infaustus | LC | Decreasing | temperate | diurnal | 84.4 | 11840129.849 | 3.182 |
| birds | Perisoreus_internigrans | Perisoreus_internigrans | NT | Decreasing | temperate | diurnal | 102 | 90049.413 | 8.113 |
| birds | Perissocephalus_tricolor | Perissocephalus_tricolor | LC | Decreasing | tropical | diurnal | 338.9 | 1655180.773 | 1.302 |
| birds | Pernis_apivorus | Pernis_apivorus | LC | Stable | tropical | diurnal | 754.4 | 32679909.865 | 10.368 |
| birds | Pernis_celebensis | Pernis_celebensis | LC | Decreasing | tropical | diurnal | 722 | 155679.508 | 9.965 |
| birds | Pernis_ptilorhynchus | Pernis_ptilorhyncus | LC | Decreasing | temperate | diurnal | 1141.1 | 16484980.319 | 12.495 |
| birds | Petrochelidon_ariel | Hirundo_ariel | LC | Increasing | temperate | diurnal | 10.8 | 5371208.208 | 5.161 |
| birds | Petrochelidon_fluvicola | Hirundo_fluvicola | LC | Increasing | tropical | diurnal | 9.7 | 2700681.599 | 18.562 |
| birds | Petrochelidon_fulva | Petrochelidon_fulva | LC | Increasing | temperate | diurnal | 19 | 1053297.208 | 11.075 |
| birds | Petrochelidon_nigricans | Hirundo_nigricans | LC | Increasing | temperate | diurnal | 15.4 | 8047366.996 | 4.46 |
| birds | Petrochelidon_perdita | NA | DD | Stable | NA | NA | NA | NA | NA |
| birds | Petrochelidon_preussi | Hirundo_preussi | LC | Increasing | tropical | diurnal | 13.3 | 2226117.984 | 12.501 |
| birds | Petrochelidon_pyrrhonota | Petrochelidon_pyrrhonota | LC | Increasing | tropical | diurnal | 21.6 | 26655865.334 | 8.217 |
| birds | Petrochelidon_rufigula | Hirundo_rufigula | LC | Increasing | tropical | diurnal | 15.9 | 1298041.093 | 6.909 |
| birds | Petrochelidon_rufocollaris | Petrochelidon_rufocollaris | LC | Increasing | tropical | diurnal | 15.9 | 109832.355 | 10.757 |
| birds | Petrochelidon_spilodera | Hirundo_spilodera | LC | Increasing | tropical | diurnal | 20.6 | 3660516.717 | 6.875 |
| birds | Petroica_archboldi | Petroica_archboldi | NT | Unknown | tropical | diurnal | 12.2 | 581.389 | 6.94 |

| | | | | | | | | | |
|---|---|---|---|---|---|---|---|---|---|
| birds | Petroica_australis | Petroica_australis | LC | Decreasing | temperate | diurnal | 31.3 | 43012.79 | 4.633 |
| birds | Petroica_bivittata | Petroica_bivittata | LC | Stable | tropical | diurnal | 9 | 16203.377 | 4.63 |
| birds | Petroica_goodenovii | Petroica_goodenovii | LC | Stable | temperate | diurnal | 8.9 | 6551523.568 | 4.527 |
| birds | Petroica_macrocephala | Petroica_macrocephala | LC | Decreasing | temperate | diurnal | 11.8 | 204209.107 | 7.113 |
| birds | Petroica_multicolor | NA | EN | Decreasing | NA | NA | NA | NA | NA |
| birds | Petroica_phoenicea | Petroica_phoenicea | LC | Decreasing | temperate | diurnal | 13 | 667819.799 | 9.969 |
| birds | Petroica_rodinogaster | Petroica_rodinogaster | LC | Decreasing | temperate | diurnal | 10.2 | 273895.479 | 9.755 |
| birds | Petroica_rosea | Petroica_rosea | LC | Decreasing | temperate | diurnal | 8.2 | 694633.379 | 9.378 |
| birds | Petroica_traversi | NA | VU | Stable | NA | NA | NA | NA | NA |
| birds | Petronia_petronia | Petronia_petronia | LC | Increasing | temperate | diurnal | 30.2 | 8115554.533 | 9.46 |
| birds | Petrophassa_albipennis | Petrophassa_albipennis | LC | Decreasing | tropical | diurnal | 124.3 | 235719.787 | 2.421 |
| birds | Petrophassa_rufipennis | Petrophassa_rufipennis | LC | Stable | tropical | diurnal | 159 | 55159.815 | 1.409 |
| birds | Peucaea_aestivalis | Aimophila_aestivalis | NT | Decreasing | temperate | diurnal | 19.6 | 761850.699 | 15.377 |
| birds | Peucaea_botterii | Aimophila_botterii | LC | Stable | temperate | diurnal | 19.9 | 998316.536 | 10.694 |
| birds | Peucaea_carpalis | Aimophila_carpalis | LC | Decreasing | temperate | diurnal | 15.3 | 175983.965 | 8.172 |
| birds | Peucaea_cassinii | Aimophila_cassinii | LC | Decreasing | temperate | diurnal | 18.9 | 2273142.062 | 10.394 |
| birds | Peucaea_humeralis | Aimophila_humeralis | LC | Decreasing | tropical | diurnal | 23.9 | 83912.611 | 9.088 |
| birds | Peucaea_mystacalis | Aimophila_mystacalis | LC | Decreasing | tropical | diurnal | 22.2 | 52598.295 | 9.258 |
| birds | Peucaea_ruficauda | Aimophila_ruficauda | LC | Stable | tropical | diurnal | 28.2 | 263649.932 | 11.464 |
| birds | Peucaea_sumichrasti | Aimophila_sumichrasti | NT | Stable | tropical | diurnal | 29.4 | 7147.339 | 11.341 |
| birds | Peucedramus_taeniatus | Peucedramus_taeniatus | LC | Decreasing | tropical | diurnal | 11 | 692877.181 | 9.006 |
| birds | Pezopetes_capitalis | Pezopetes_capitalis | LC | Stable | tropical | diurnal | 55.8 | 3401.861 | 5.856 |
| birds | Pezoporus_occidentalis | Pezoporus_occidentalis | CR | Decreasing | temperate | diurnal | 130 | 2740521.515 | 2.893 |
| birds | Pezoporus_wallicus | Pezoporus_wallicus | LC | Stable | temperate | diurnal | 71.2 | 195370.721 | 8.52 |
| birds | Phacellodomus_dorsalis | Phacellodomus_dorsalis | NT | Decreasing | tropical | diurnal | 35.9 | 2363.2 | 8.766 |
| birds | Phacellodomus_erythrophthalmus | Phacellodomus_erythrophthalmus | LC | Decreasing | tropical | diurnal | 24.5 | 394570.108 | 11.061 |
| birds | Phacellodomus_inornatus | Phacellodomus_inornatus | LC | Stable | tropical | diurnal | 24 | 378209.087 | 8.984 |
| birds | Phacellodomus_maculipectus | Phacellodomus_maculipectus | LC | Stable | temperate | diurnal | 21.9 | 119584.525 | 6.657 |
| birds | Phacellodomus_ruber | Phacellodomus_ruber | LC | Decreasing | tropical | diurnal | 41 | 2766219.769 | 7.91 |
| birds | Phacellodomus_rufifrons | Phacellodomus_rufifrons | LC | Stable | tropical | diurnal | 24.6 | 2793273.525 | 8.531 |
| birds | Phacellodomus_sibilatrix | Phacellodomus_sibilatrix | LC | Decreasing | temperate | diurnal | 15.5 | 1150059.834 | 8.093 |
| birds | Phacellodomus_striaticeps | Phacellodomus_striaticeps | LC | Stable | tropical | diurnal | 29.1 | 304772.682 | 8.694 |
| birds | Phacellodomus_striaticollis | Phacellodomus_striaticollis | LC | Stable | temperate | diurnal | 25.7 | 651178.34 | 11.83 |
| birds | Phaenicophaeus_curvirostris | Phaenicophaeus_curvirostris | LC | Stable | tropical | diurnal | 154 | 1217500.23 | 10.304 |

| | | | | | | | | | |
|---|---|---|---|---|---|---|---|---|---|
| birds | Phaenicophaeus_diardi | Phaenicophaeus_diardi | NT | Decreasing | tropical | diurnal | 62.1 | 1255042.985 | 8.366 |
| birds | Phaenicophaeus_pyrrhocephalus | NA | VU | Decreasing | NA | NA | NA | NA | NA |
| birds | Phaenicophaeus_sumatranus | Phaenicophaeus_sumatranus | NT | Decreasing | tropical | diurnal | 91.9 | 1280940.392 | 8.445 |
| birds | Phaenicophaeus_tristis | Phaenicophaeus_tristis | LC | Stable | tropical | diurnal | 117 | 4373319.662 | 12.955 |
| birds | Phaenicophaeus_viridirostris | Phaenicophaeus_viridirostris | LC | Stable | tropical | diurnal | 67 | 785946.331 | 19.556 |
| birds | Phaenicophilus_palmarum | Phaenicophilus_palmarum | LC | Stable | tropical | diurnal | 30.5 | 66937.13 | 15.622 |
| birds | Phaenicophilus_poliocephalus | Phaenicophilus_poliocephalus | NT | Decreasing | tropical | diurnal | 31 | 9268.82 | 16.89 |
| birds | Phaenostictus_mcleannani | Phaenostictus_mcleannani | LC | Decreasing | tropical | diurnal | 51.1 | 254363.891 | 6.938 |
| birds | Phaeochroa_cuvierii | Phaeochroa_cuvierii | LC | Stable | tropical | diurnal | 8.9 | 238479.539 | 10.599 |
| birds | Phaeomyias_murina | Phaeomyias_murina | LC | Increasing | tropical | diurnal | 10 | 10817801.085 | 5.35 |
| birds | Phaeomyias_tumbezana | Phaeomyias_tumbezana | LC | Increasing | tropical | diurnal | 11.5 | 123816.236 | 9.989 |
| birds | Phaeoptila_sordida | Cynanthus_sordidus | LC | Stable | tropical | diurnal | 4 | 137024.776 | 9.851 |
| birds | Phaethornis_anthophilus | Phaethornis_anthophilus | LC | Decreasing | tropical | diurnal | 5.9 | 426150.71 | 13.489 |
| birds | Phaethornis_atrimentalis | Phaethornis_atrimentalis | LC | Decreasing | tropical | diurnal | 2.6 | 829137.725 | 3.111 |
| birds | Phaethornis_augusti | Phaethornis_augusti | LC | Stable | tropical | diurnal | 6 | 557848.617 | 6.187 |
| birds | Phaethornis_bourcieri | Phaethornis_bourcieri | LC | Stable | tropical | diurnal | 4.3 | 2982755.215 | 1.35 |
| birds | Phaethornis_eurynome | Phaethornis_eurynome | LC | Decreasing | tropical | diurnal | 5.3 | 934246.248 | 13.142 |
| birds | Phaethornis_griseogularis | Phaethornis_griseogularis | LC | Unknown | tropical | diurnal | 2.3 | 363135.551 | 4.31 |
| birds | Phaethornis_guy | Phaethornis_guy | LC | Decreasing | tropical | diurnal | 6.3 | 555160.159 | 7.467 |
| birds | Phaethornis_hispidus | Phaethornis_hispidus | LC | Decreasing | tropical | diurnal | 5 | 4008483.064 | 2.695 |
| birds | Phaethornis_idaliae | Phaethornis_idaliae | LC | Decreasing | tropical | diurnal | 2.6 | 154965.577 | 12.352 |
| birds | Phaethornis_koepckeae | Phaethornis_koepckeae | LC | Decreasing | tropical | diurnal | 5 | 96811.648 | 3.671 |
| birds | Phaethornis_longirostris | Phaethornis_longirostris | LC | Decreasing | tropical | diurnal | 6 | 533601.88 | 10.775 |
| birds | Phaethornis_longuemareus | Phaethornis_longuemareus | LC | Stable | tropical | diurnal | 3 | 281727.997 | 3.056 |
| birds | Phaethornis_malaris | Phaethornis_malaris | LC | Decreasing | tropical | diurnal | 5.9 | 3801993.443 | 2.48 |
| birds | Phaethornis_nattereri | Phaethornis_nattereri | LC | Decreasing | tropical | diurnal | 3.1 | 1411591.704 | 5.538 |
| birds | Phaethornis_philippii | Phaethornis_philippii | LC | Unknown | tropical | diurnal | 5 | 1596505.836 | 1.263 |
| birds | Phaethornis_pretrei | Phaethornis_pretrei | LC | Unknown | tropical | diurnal | 5.6 | 3736605.659 | 8.993 |
| birds | Phaethornis_ruber | Phaethornis_ruber | LC | Decreasing | tropical | diurnal | 2.4 | 9729125.657 | 4.398 |
| birds | Phaethornis_rupurumii | Phaethornis_rupurumii | LC | Unknown | tropical | diurnal | 3.7 | 570013.043 | 1.893 |
| birds | Phaethornis_squalidus | Phaethornis_squalidus | LC | Unknown | tropical | diurnal | 3.4 | 272378.961 | 13.529 |
| birds | Phaethornis_striigularis | Phaethornis_striigularis | LC | Decreasing | tropical | diurnal | 3 | 764879.777 | 10.583 |
| birds | Phaethornis_stuarti | Phaethornis_stuarti | LC | Decreasing | tropical | diurnal | 2.9 | 147916.919 | 4.452 |
| birds | Phaethornis_subochraceus | Phaethornis_subochraceus | LC | Decreasing | tropical | diurnal | 3.8 | 338446.166 | 6.006 |

| | | | | | | | | | |
|---|---|---|---|---|---|---|---|---|---|
| birds | Phaethornis_superciliosus | Phaethornis_superciliosus | LC | Decreasing | tropical | diurnal | 6.3 | 2562623.831 | 1.896 |
| birds | Phaethornis_syrmatophorus | Phaethornis_syrmatophorus | LC | Unknown | tropical | diurnal | 5.8 | 159638.354 | 8.137 |
| birds | Phaethornis_yaruqui | Phaethornis_yaruqui | LC | Decreasing | tropical | diurnal | 5.4 | 119931.581 | 7.729 |
| birds | Phaetusa_simplex | Phaetusa_simplex | LC | Stable | tropical | diurnal | 235 | 13316076.041 | 5.698 |
| birds | Phainopepla_nitens | Phainopepla_nitens | LC | Stable | temperate | diurnal | 22.1 | 1423921.856 | 9.144 |
| birds | Phainoptila_melanoxantha | Phainoptila_melanoxantha | LC | Decreasing | tropical | diurnal | 56 | 6565.629 | 5.876 |
| birds | Phalacrocorax_capensis | Phalacrocorax_capensis | EN | Decreasing | tropical | diurnal | 1211.7 | 503893.889 | 7.962 |
| birds | Phalacrocorax_capillatus | Phalacrocorax_capillatus | LC | Unknown | temperate | diurnal | 2801.8 | 2090157.797 | 23.136 |
| birds | Phalacrocorax_carbo | Phalacrocorax_carbo | LC | Increasing | temperate | diurnal | 2529 | 32279804.491 | 10.598 |
| birds | Phalacrocorax_featherstoni | NA | EN | Decreasing | NA | NA | NA | NA | NA |
| birds | Phalacrocorax_fuscescens | Phalacrocorax_fuscescens | LC | Unknown | temperate | diurnal | 1515 | 203014.548 | 10.338 |
| birds | Phalacrocorax_fuscicollis | Phalacrocorax_fuscicollis | LC | Unknown | temperate | diurnal | 706 | 1187570.011 | 18.345 |
| birds | Phalacrocorax_neglectus | Phalacrocorax_neglectus | EN | Decreasing | temperate | diurnal | 1944.2 | 107865.707 | 10.042 |
| birds | Phalacrocorax_nigrogularis | Phalacrocorax_nigrogularis | VU | Decreasing | temperate | diurnal | 1560 | 776333.059 | 14.526 |
| birds | Phalacrocorax_punctatus | Phalacrocorax_punctatus | LC | Unknown | temperate | diurnal | 1184.7 | 85166.45 | 11.371 |
| birds | Phalacrocorax_sulcirostris | Phalacrocorax_sulcirostris | LC | Unknown | tropical | diurnal | 995 | 4510039.584 | 6.214 |
| birds | Phalacrocorax_varius | Phalacrocorax_varius | LC | Unknown | temperate | diurnal | 1940.7 | 4379236.116 | 5.646 |
| birds | Phalaenoptilus_nuttallii | Phalaenoptilus_nuttallii | LC | Stable | temperate | nocturnal | 48.3 | 4441424.234 | 9.645 |
| birds | Phalaropus_fulicarius | Phalaropus_fulicarius | LC | Unknown | temperate | diurnal | 55.6 | 15687867.412 | 0.435 |
| birds | Phalaropus_lobatus | Phalaropus_lobatus | LC | Decreasing | temperate | diurnal | 36.7 | 18813928.957 | 0.961 |
| birds | Phalcoboenus_albogularis | Phalcoboenus_albogularis | LC | Stable | temperate | diurnal | 809 | 341178.689 | 2.819 |
| birds | Phalcoboenus_australis | Phalcoboenus_australis | NT | Stable | temperate | diurnal | 1187 | 23812.364 | 2.543 |
| birds | Phalcoboenus_carunculatus | NA | LC | Stable | NA | NA | NA | NA | NA |
| birds | Phalcoboenus_chimango | Milvago_chimango | LC | Increasing | temperate | diurnal | 296 | 4070781.952 | 8.288 |
| birds | Phalcoboenus_megalopterus | Phalcoboenus_megalopterus | LC | Stable | tropical | diurnal | 788 | 1011292.224 | 6.424 |
| birds | Phapitreron_amethystinus | Phapitreron_amethystinus | LC | Stable | tropical | diurnal | 145 | 227781.898 | 13.379 |
| birds | Phapitreron_cinereiceps | Phapitreron_cinereiceps | EN | Decreasing | tropical | diurnal | 146 | 667.844 | 6.998 |
| birds | Phapitreron_leucotis | Phapitreron_leucotis | LC | Stable | tropical | diurnal | 108 | 118342.177 | 14.478 |
| birds | Phaps_chalcoptera | Phaps_chalcoptera | LC | Stable | temperate | diurnal | 330.7 | 5889502.174 | 5.107 |
| birds | Phaps_elegans | Phaps_elegans | LC | Decreasing | temperate | diurnal | 211.4 | 694565.107 | 10.175 |
| birds | Phaps_histrionica | Phaps_histrionica | LC | Decreasing | tropical | diurnal | 289 | 3734100.704 | 2.729 |
| birds | Pharomachrus_antisianus | Pharomachrus_antisianus | LC | Stable | tropical | diurnal | 152.1 | 427099.999 | 7.831 |
| birds | Pharomachrus_auriceps | Pharomachrus_auriceps | LC | Decreasing | tropical | diurnal | 180 | 494429.381 | 7.971 |
| birds | Pharomachrus_fulgidus | Pharomachrus_fulgidus | LC | Decreasing | tropical | diurnal | 160 | 16767.161 | 11.105 |

| | | | | | | | | | |
|---|---|---|---|---|---|---|---|---|---|
| birds | Pharomachrus_mocinno | Pharomachrus_mocinno | NT | Decreasing | tropical | diurnal | 202.5 | 155797.199 | 11.214 |
| birds | Pharomachrus_pavoninus | Pharomachrus_pavoninus | LC | Decreasing | tropical | diurnal | 163 | 3760744.349 | 1.418 |
| birds | Phasianus_colchicus | Phasianus_colchicus | LC | Decreasing | temperate | diurnal | 1120.3 | 16757729.727 | 14.158 |
| birds | Phasianus_versicolor | Phasianus_versicolor | LC | Decreasing | temperate | diurnal | 918.8 | 306118.98 | 20.41 |
| birds | Phedina_borbonica | Phedina_borbonica | LC | Stable | tropical | diurnal | 22.2 | 624323.077 | 8.941 |
| birds | Phedinopsis_brazzae | Phedina_brazzae | LC | Stable | tropical | diurnal | 13 | 403109.287 | 7.588 |
| birds | Phegornis_mitchellii | Phegornis_mitchellii | NT | Decreasing | tropical | diurnal | 35.5 | 761982.956 | 6.068 |
| birds | Phelpsia_inornata | Phelpsia_inornatus | LC | Stable | tropical | diurnal | 29.4 | 278027.72 | 7.957 |
| birds | Pheucticus_aureoventris | Pheucticus_aureoventris | LC | Unknown | tropical | diurnal | 49 | 1918971.638 | 6.539 |
| birds | Pheucticus_chrysogaster | Pheucticus_chrysogaster | LC | Stable | tropical | diurnal | 55.8 | 537171.299 | 8.556 |
| birds | Pheucticus_chrysopeplus | Pheucticus_chrysopeplus | LC | Decreasing | tropical | diurnal | 77.6 | 321297.585 | 8.176 |
| birds | Pheucticus_ludovicianus | Pheucticus_ludovicianus | LC | Decreasing | temperate | diurnal | 42 | 9302746.811 | 13.401 |
| birds | Pheucticus_melanocephalus | Pheucticus_melanocephalus | LC | Increasing | temperate | diurnal | 47.1 | 5971695.748 | 10.067 |
| birds | Pheucticus_tibialis | Pheucticus_tibialis | LC | Decreasing | tropical | diurnal | 62.3 | 19605.628 | 8.301 |
| birds | Pheugopedius_atrogularis | Thryothorus_atrogularis | LC | Decreasing | tropical | diurnal | 24.2 | 34566.039 | 9.798 |
| birds | Pheugopedius_coraya | Thryothorus_coraya | LC | Unknown | tropical | diurnal | 17.2 | 3811040.455 | 1.792 |
| birds | Pheugopedius_eisenmanni | Thryothorus_eisenmanni | LC | Increasing | tropical | diurnal | 23.1 | 17967.707 | 5.673 |
| birds | Pheugopedius_euophrys | Thryothorus_euophrys | LC | Unknown | tropical | diurnal | 30.3 | 50266.615 | 10.538 |
| birds | Pheugopedius_fasciatoventris | Thryothorus_fasciatoventris | LC | Decreasing | tropical | diurnal | 24 | 264878.845 | 11.878 |
| birds | Pheugopedius_felix | Thryothorus_felix | LC | Decreasing | tropical | diurnal | 12.8 | 364236.123 | 7.944 |
| birds | Pheugopedius_genibarbis | Thryothorus_genibarbis | LC | Unknown | tropical | diurnal | 19.2 | 6326924.93 | 4.972 |
| birds | Pheugopedius_maculipectus | Thryothorus_maculipectus | LC | Decreasing | tropical | diurnal | 15 | 604088.843 | 9.889 |
| birds | Pheugopedius_mystacalis | Thryothorus_mystacalis | LC | Unknown | tropical | diurnal | 29 | 271375.884 | 12.623 |
| birds | Pheugopedius_paucimaculatus | NA | LC | Decreasing | NA | NA | NA | NA | NA |
| birds | Pheugopedius_rutilus | Thryothorus_rutilus | LC | Decreasing | tropical | diurnal | 16.5 | 206155.133 | 12.738 |
| birds | Pheugopedius_sclateri | Thryothorus_sclateri | LC | Decreasing | tropical | diurnal | 20 | 15611.593 | 7.362 |
| birds | Pheugopedius_spadix | Thryothorus_spadix | LC | Decreasing | tropical | diurnal | 19.1 | 51078.732 | 8.515 |
| birds | Phibalura_flavirostris | Phibalura_flavirostris | LC | Decreasing | tropical | diurnal | 46.5 | 1154826.618 | 13.24 |
| birds | Phigys_solitarius | Phigys_solitarius | LC | Stable | tropical | diurnal | 83.3 | 18131.757 | 9.375 |
| birds | Philemon_albitorques | Philemon_albitorques | LC | Unknown | tropical | diurnal | 85.7 | 1887.905 | 6.45 |
| birds | Philemon_argenticeps | Philemon_argenticeps | LC | Stable | tropical | diurnal | 80.4 | 885758.869 | 2.479 |
| birds | Philemon_brassi | Philemon_brassi | LC | Stable | tropical | diurnal | 85.7 | 15799.409 | 1.764 |
| birds | Philemon_buceroides | Philemon_buceroides | LC | Stable | tropical | diurnal | 108.6 | 1092195.197 | 3.89 |
| birds | Philemon_citreogularis | Philemon_citreogularis | LC | Stable | temperate | diurnal | 60.6 | 3161053.216 | 5.331 |

| | | | | | | | | | |
|---|---|---|---|---|---|---|---|---|---|
| birds | Philemon_cockerelli | Philemon_cockerelli | LC | Stable | tropical | diurnal | 154.1 | 35335.215 | 4.184 |
| birds | Philemon_corniculatus | Philemon_corniculatus | LC | Stable | temperate | diurnal | 100.7 | 1758116.368 | 7.089 |
| birds | Philemon_diemenensis | Philemon_diemenensis | LC | Stable | tropical | diurnal | 65.9 | 17820.103 | 7.501 |
| birds | Philemon_eichhorni | Philemon_eichhorni | LC | Unknown | tropical | diurnal | 101 | 1714.737 | 3.735 |
| birds | Philemon_fuscicapillus | Philemon_fuscicapillus | NT | Decreasing | tropical | diurnal | 85.7 | 2036.37 | 5.7 |
| birds | Philemon_inornatus | Philemon_inornatus | LC | Stable | tropical | diurnal | 85.7 | 28799.068 | 13.847 |
| birds | Philemon_kisserensis | Philemon_kisserensis | LC | Stable | tropical | diurnal | 62.8 | 534.056 | 15.115 |
| birds | Philemon_meyeri | Philemon_meyeri | LC | Stable | tropical | diurnal | 52 | 466260.954 | 3.09 |
| birds | Philemon_moluccensis | Philemon_moluccensis | LC | Unknown | tropical | diurnal | 85.7 | 8456.571 | 6.715 |
| birds | Philemon_subcorniculatus | Philemon_subcorniculatus | LC | Stable | tropical | diurnal | 194 | 17074.166 | 7.08 |
| birds | Philentoma_pyrhoptera | Philentoma_pyrhoptera | LC | Decreasing | tropical | diurnal | 16.9 | 1224085.412 | 7.643 |
| birds | Philentoma_velata | Philentoma_velata | NT | Decreasing | tropical | diurnal | 26.1 | 1151168.58 | 8.173 |
| birds | Philepitta_castanea | Philepitta_castanea | LC | Decreasing | tropical | diurnal | 35.4 | 46337.011 | 5.16 |
| birds | Philepitta_schlegeli | Philepitta_schlegeli | NT | Decreasing | tropical | diurnal | 27.9 | 75629.577 | 7.566 |
| birds | Philesturnus_carunculatus | Philesturnus_carunculatus | LC | Increasing | temperate | diurnal | 74.7 | 45.368 | 0.632 |
| birds | Philetairus_socius | Philetairus_socius | LC | Stable | temperate | diurnal | 27.4 | 658355.718 | 5.829 |
| birds | Philodice_bryantae | Calliphlox_bryantae | LC | Decreasing | tropical | diurnal | 3.3 | 9185.275 | 12.228 |
| birds | Philodice_mitchellii | Calliphlox_mitchellii | LC | Decreasing | tropical | diurnal | 3.1 | 59801.221 | 9.206 |
| birds | Philohydor_lictor | Pitangus_lictor | LC | Decreasing | tropical | diurnal | 25.5 | 9185994.706 | 3.946 |
| birds | Philortyx_fasciatus | Philortyx_fasciatus | LC | Stable | tropical | diurnal | 130 | 133413.974 | 9.867 |
| birds | Philydor_atricapillus | Philydor_atricapillus | LC | Decreasing | tropical | diurnal | 22.2 | 1129776.92 | 13.15 |
| birds | Philydor_erythrocercum | Philydor_erythrocercum | LC | Stable | tropical | diurnal | 25.3 | 5418434.134 | 2.22 |
| birds | Philydor_fuscipenne | Philydor_fuscipenne | LC | Decreasing | tropical | diurnal | 26.5 | 111453.838 | 9.094 |
| birds | Philydor_novaesi | NA | EX | not_available | NA | NA | NA | NA | NA |
| birds | Philydor_pyrrhodes | Philydor_pyrrhodes | LC | Stable | tropical | diurnal | 30.5 | 5868049.15 | 1.758 |
| birds | Phimosus_infuscatus | Phimosus_infuscatus | LC | Stable | tropical | diurnal | 559 | 6585514.175 | 8.662 |
| birds | Phlegopsis_borbae | Skutchia_borbae | LC | Decreasing | tropical | diurnal | 50 | 234463.476 | 1.135 |
| birds | Phlegopsis_erythroptera | Phlegopsis_erythroptera | LC | Decreasing | tropical | diurnal | 54.6 | 2093169.386 | 1.209 |
| birds | Phlegopsis_nigromaculata | Phlegopsis_nigromaculata | LC | Decreasing | tropical | diurnal | 45.2 | 3887704.806 | 2.356 |
| birds | Phleocryptes_melanops | Phleocryptes_melanops | LC | Decreasing | temperate | diurnal | 14.6 | 3823377.463 | 7.852 |
| birds | Phlogophilus_harterti | Phlogophilus_harterti | LC | Stable | tropical | diurnal | 2.7 | 41157.244 | 3.783 |
| birds | Phlogophilus_hemileucurus | Phlogophilus_hemileucurus | LC | Decreasing | tropical | diurnal | 2.8 | 93348.287 | 6.371 |
| birds | Phodilus_badius | Phodilus_badius | LC | Stable | tropical | nocturnal | 280.2 | 3497562.234 | 10.988 |
| birds | Phoebastria_albatrus | Phoebastria_albatrus | VU | Increasing | temperate | cathemeral | 4411 | 48012504.021 | 5.429 |

| | | | | | | | | | |
|---|---|---|---|---|---|---|---|---|---|
| birds | Phoebastria_immutabilis | Phoebastria_immutabilis | NT | Stable | temperate | cathemeral | 3145.9 | 45350845.456 | 7.721 |
| birds | Phoebastria_irrorata | Phoebastria_irrorata | CR | Decreasing | tropical | cathemeral | 3376.4 | 1736846.608 | 10.169 |
| birds | Phoebastria_nigripes | Phoebastria_nigripes | NT | Increasing | tropical | cathemeral | 3188.4 | 72160530.94 | 6.185 |
| birds | Phoebetria_fusca | Phoebetria_fusca | EN | Decreasing | temperate | cathemeral | 2553.7 | 61330907.011 | 11.848 |
| birds | Phoebetria_palpebrata | Phoebetria_palpebrata | NT | Decreasing | temperate | cathemeral | 3150 | 74642407.908 | 5.32 |
| birds | Phoenicircus_carnifex | Phoenicircus_carnifex | LC | Decreasing | tropical | diurnal | 82.9 | 1387247.24 | 1.867 |
| birds | Phoenicircus_nigricollis | Phoenicircus_nigricollis | LC | Decreasing | tropical | diurnal | 96.3 | 2636332.672 | 1.369 |
| birds | Phoeniconaias_minor | Phoeniconaias_minor | NT | Decreasing | tropical | diurnal | 1500 | 5490439.67 | 9.357 |
| birds | Phoenicoparrus_andinus | Phoenicoparrus_andinus | VU | Decreasing | tropical | diurnal | 4900 | 354440.55 | 4.526 |
| birds | Phoenicoparrus_jamesi | Phoenicoparrus_jamesi | NT | Stable | tropical | diurnal | 2000 | 345049.405 | 4.594 |
| birds | Phoenicopterus_chilensis | Phoenicopterus_chilensis | NT | Decreasing | temperate | diurnal | 2277 | 3679115.35 | 7.285 |
| birds | Phoenicopterus_ruber | Phoenicopterus_ruber | LC | Increasing | tropical | diurnal | 3031.6 | 490729.153 | 13.996 |
| birds | Phoeniculus_bollei | Phoeniculus_bollei | LC | Decreasing | tropical | diurnal | 54.6 | 1297397.491 | 12.678 |
| birds | Phoeniculus_damarensis | Phoeniculus_damarensis | LC | Decreasing | tropical | diurnal | 81.8 | 828686.305 | 7.264 |
| birds | Phoeniculus_purpureus | Phoeniculus_purpureus | LC | Decreasing | tropical | diurnal | 73.2 | 12280558.029 | 9.804 |
| birds | Phoeniculus_somaliensis | Phoeniculus_somaliensis | LC | Decreasing | tropical | diurnal | 26.5 | 1065209.4 | 9.586 |
| birds | Phoenicurus_alaschanicus | Phoenicurus_alaschanicus | NT | Decreasing | temperate | diurnal | 16.8 | 925983.453 | 9.496 |
| birds | Phoenicurus_auroreus | Phoenicurus_auroreus | LC | Stable | temperate | diurnal | 16.2 | 8472998.968 | 11.477 |
| birds | Phoenicurus_bicolor | Rhyacornis_bicolor | NT | Decreasing | tropical | diurnal | 23 | 31763.671 | 7.604 |
| birds | Phoenicurus_coeruleocephala | Phoenicurus_caeruleocephala | LC | Stable | temperate | diurnal | 14.9 | 818739.248 | 7.281 |
| birds | Phoenicurus_erythrogastrus | Phoenicurus_erythrogastrus | LC | Stable | temperate | diurnal | 24.5 | 4461790.549 | 6.875 |
| birds | Phoenicurus_erythronotus | Phoenicurus_erythronotus | LC | Stable | temperate | diurnal | 18.5 | 4478602.326 | 8.366 |
| birds | Phoenicurus_frontalis | Phoenicurus_frontalis | LC | Stable | temperate | diurnal | 15.8 | 3065153.65 | 9.208 |
| birds | Phoenicurus_fuliginosus | Rhyacornis_fuliginosa | LC | Stable | temperate | diurnal | 20.3 | 5292160.317 | 13.92 |
| birds | Phoenicurus_hodgsoni | Phoenicurus_hodgsoni | LC | Stable | temperate | diurnal | 17 | 2029614.174 | 9.468 |
| birds | Phoenicurus_leucocephalus | Chaimarrornis_leucocephalus | LC | Stable | temperate | diurnal | 30 | 4372940.271 | 10.73 |
| birds | Phoenicurus_moussieri | Phoenicurus_moussieri | LC | Stable | temperate | diurnal | 14.9 | 1093552.501 | 10.516 |
| birds | Phoenicurus_ochruros | Phoenicurus_ochruros | LC | Increasing | temperate | diurnal | 16.5 | 29007225.13 | 10.511 |
| birds | Phoenicurus_phoenicurus | Phoenicurus_phoenicurus | LC | Increasing | temperate | diurnal | 14.6 | 42756527.378 | 8.367 |
| birds | Phoenicurus_schisticeps | Phoenicurus_schisticeps | LC | Stable | temperate | diurnal | 18.1 | 754510.631 | 8.087 |
| birds | Pholia_sharpii | Cinnyricinclus_sharpii | LC | Decreasing | tropical | diurnal | 41 | 195406.838 | 12.666 |
| birds | Pholidornis_rushiae | Pholidornis_rushiae | LC | Stable | tropical | diurnal | 6 | 3139298.251 | 7.83 |
| birds | Phonipara_canora | Tiaris_canorus | LC | Decreasing | tropical | diurnal | 8.1 | 93007.108 | 16.133 |
| birds | Phonygammus_keraudrenii | Manucodia_keraudrenii | LC | Decreasing | tropical | diurnal | 154.7 | 767082.804 | 3.196 |

| | | | | | | | | | |
|---|---|---|---|---|---|---|---|---|---|
| birds | Phragmacia_substriata | Phragmacia_substriata | LC | Increasing | temperate | diurnal | 10.6 | 466617.529 | 7.342 |
| birds | Phrygilus_atriceps | Phrygilus_atriceps | LC | Stable | tropical | diurnal | 24.3 | 526947.203 | 5.63 |
| birds | Phrygilus_gayi | Phrygilus_gayi | LC | Stable | temperate | diurnal | 25.6 | 1649829.111 | 6.155 |
| birds | Phrygilus_patagonicus | Phrygilus_patagonicus | LC | Stable | temperate | diurnal | 22.6 | 569692.026 | 5.731 |
| birds | Phrygilus_punensis | Phrygilus_punensis | LC | Stable | tropical | diurnal | 37.2 | 309713.42 | 8.748 |
| birds | Phylidonyris_niger | Phylidonyris_niger | LC | Stable | temperate | diurnal | 18.2 | 1047225.931 | 7.793 |
| birds | Phylidonyris_novaehollandiae | Phylidonyris_novaehollandiae | LC | Stable | temperate | diurnal | 20.2 | 851798.64 | 9.918 |
| birds | Phylidonyris_pyrrhopterus | Phylidonyris_pyrrhopterus | LC | Stable | temperate | diurnal | 16.3 | 357930.02 | 9.743 |
| birds | Phyllaemulor_bracteatus | Nyctibius_bracteatus | LC | Decreasing | tropical | nocturnal | 51.7 | 5116065.28 | 1.834 |
| birds | Phyllanthus_atripennis | Phyllanthus_atripennis | VU | Decreasing | tropical | diurnal | 85.6 | 230594.303 | 15.459 |
| birds | Phyllastrephus_albigularis | Phyllastrephus_albigularis | LC | Stable | tropical | diurnal | 24.5 | 2136746.716 | 8.291 |
| birds | Phyllastrephus_baumanni | Phyllastrephus_baumanni | LC | Stable | tropical | diurnal | 27.5 | 446395.595 | 15.351 |
| birds | Phyllastrephus_cabanisi | Phyllastrephus_cabanisi | LC | Stable | tropical | diurnal | 34.4 | 1995009.618 | 8.898 |
| birds | Phyllastrephus_cerviniventris | Phyllastrephus_cerviniventris | LC | Stable | tropical | diurnal | 27 | 1170058.225 | 9.389 |
| birds | Phyllastrephus_debilis | Phyllastrephus_debilis | LC | Decreasing | tropical | diurnal | 14.3 | 199654.823 | 10.647 |
| birds | Phyllastrephus_fischeri | Phyllastrephus_fischeri | LC | Decreasing | tropical | diurnal | 27 | 270855.04 | 9.975 |
| birds | Phyllastrephus_flavostriatus | Phyllastrephus_flavostriatus | LC | Stable | tropical | diurnal | 28.3 | 444379.943 | 12.989 |
| birds | Phyllastrephus_fulviventris | Phyllastrephus_fulviventris | LC | Stable | tropical | diurnal | 33.3 | 152087.251 | 9.25 |
| birds | Phyllastrephus_hypochloris | Phyllastrephus_hypochloris | LC | Stable | tropical | diurnal | 23.1 | 134659.571 | 15.01 |
| birds | Phyllastrephus_icterinus | Phyllastrephus_icterinus | LC | Stable | tropical | diurnal | 18.9 | 2473054.921 | 7.953 |
| birds | Phyllastrephus_lorenzi | Phyllastrephus_lorenzi | LC | Decreasing | tropical | diurnal | 19 | 39316.139 | 14.551 |
| birds | Phyllastrephus_poensis | Phyllastrephus_poensis | LC | Stable | tropical | diurnal | 28.9 | 44370.69 | 10.035 |
| birds | Phyllastrephus_poliocephalus | Phyllastrephus_poliocephalus | LC | Decreasing | tropical | diurnal | 53 | 32217.094 | 9.954 |
| birds | Phyllastrephus_strepitans | Phyllastrephus_strepitans | LC | Decreasing | tropical | diurnal | 27.1 | 1347241.745 | 9.572 |
| birds | Phyllastrephus_terrestris | Phyllastrephus_terrestris | LC | Stable | tropical | diurnal | 31.1 | 2434586.591 | 10.032 |
| birds | Phyllastrephus_xavieri | Phyllastrephus_xavieri | LC | Stable | tropical | diurnal | 22.6 | 968575.231 | 5.903 |
| birds | Phyllergates_cucullatus | Orthotomus_cuculatus | LC | Stable | tropical | diurnal | 5.9 | 1549112.241 | 9.824 |
| birds | Phyllergates_heterolaemus | Orthotomus_heterolaemus | LC | Decreasing | tropical | diurnal | 8.3 | 7657.176 | 7.673 |
| birds | Phyllolais_pulchella | Phyllolais_pulchella | LC | Stable | tropical | diurnal | 6 | 2078702.064 | 11.875 |
| birds | Phyllomyias_burmeisteri | Phyllomyias_burmeisteri | LC | Decreasing | tropical | diurnal | 11.1 | 971005.03 | 12.179 |
| birds | Phyllomyias_cinereiceps | Phyllomyias_cinereiceps | LC | Stable | tropical | diurnal | 9.5 | 144525.034 | 7.851 |
| birds | Phyllomyias_fasciatus | Phyllomyias_fasciatus | LC | Decreasing | tropical | diurnal | 10.3 | 3909215.377 | 10.033 |
| birds | Phyllomyias_griseiceps | Phyllomyias_griseiceps | LC | Stable | tropical | diurnal | 7.2 | 707633.088 | 6.698 |
| birds | Phyllomyias_griseocapilla | Phyllomyias_griseocapilla | NT | Decreasing | tropical | diurnal | 8 | 190093.197 | 13.259 |

| | | | | | | | | | |
|---|---|---|---|---|---|---|---|---|---|
| birds | Phyllomyias_nigrocapillus | Phyllomyias_nigrocapillus | LC | Stable | tropical | diurnal | 9.5 | 254835.698 | 9.865 |
| birds | Phyllomyias_plumbeiceps | Phyllomyias_plumbeiceps | LC | Stable | tropical | diurnal | 10.3 | 71922.726 | 5.909 |
| birds | Phyllomyias_reiseri | Phyllomyias_reiseri | LC | Decreasing | tropical | diurnal | 7.5 | 650161.746 | 10.255 |
| birds | Phyllomyias_sclateri | Phyllomyias_sclateri | LC | Stable | tropical | diurnal | 9.2 | 87000.009 | 7.005 |
| birds | Phyllomyias_urichi | Phyllomyias_urichi | EN | Decreasing | tropical | diurnal | 9.1 | 1078.78 | 9.132 |
| birds | Phyllomyias_uropygialis | Phyllomyias_uropygialis | LC | Stable | tropical | diurnal | 9 | 232128.958 | 8.674 |
| birds | Phyllomyias_virescens | Phyllomyias_virescens | LC | Decreasing | temperate | diurnal | 8.2 | 1029199.707 | 13.483 |
| birds | Phyllomyias_zeledoni | Phyllomyias_zeledoni | LC | Decreasing | tropical | diurnal | 10.3 | 92249.438 | 7.743 |
| birds | Phylloscartes_beckeri | Phylloscartes_beckeri | EN | Decreasing | tropical | diurnal | 8.7 | 2304.781 | 7.846 |
| birds | Phylloscartes_ceciliae | Phylloscartes_ceciliae | CR | Decreasing | tropical | diurnal | 7.6 | 11223.535 | 14.118 |
| birds | Phylloscartes_difficilis | Phylloscartes_difficilis | LC | Decreasing | temperate | diurnal | 6.9 | 150332.53 | 13.16 |
| birds | Phylloscartes_flaviventris | Phylloscartes_flaviventris | LC | Decreasing | tropical | diurnal | 8 | 5225.565 | 20.055 |
| birds | Phylloscartes_flavovirens | Phylloscartes_flavovirens | LC | Stable | tropical | diurnal | 8.8 | 17272.593 | 7.1 |
| birds | Phylloscartes_gualaquizae | Phylloscartes_gualaquizae | LC | Decreasing | tropical | diurnal | 8 | 21301.676 | 7.409 |
| birds | Phylloscartes_kronei | Phylloscartes_kronei | LC | Decreasing | temperate | diurnal | 8.3 | 5479.393 | 17.345 |
| birds | Phylloscartes_nigrifrons | Phylloscartes_nigrifrons | LC | Decreasing | tropical | diurnal | 9.1 | 172912.875 | 0.797 |
| birds | Phylloscartes_oustaleti | Phylloscartes_oustaleti | LC | Decreasing | temperate | diurnal | 10 | 97272.795 | 12.364 |
| birds | Phylloscartes_parkeri | Phylloscartes_parkeri | LC | Decreasing | tropical | diurnal | 8.1 | 94588.996 | 3.903 |
| birds | Phylloscartes_paulista | Phylloscartes_paulista | NT | Decreasing | temperate | diurnal | 7.5 | 369792.802 | 14.274 |
| birds | Phylloscartes_roquettei | Phylloscartes_roquettei | EN | Decreasing | tropical | diurnal | 6 | 261686.728 | 9.759 |
| birds | Phylloscartes_superciliaris | Phylloscartes_superciliaris | LC | Decreasing | tropical | diurnal | 7.7 | 21396.152 | 6.664 |
| birds | Phylloscartes_sylviolus | Phylloscartes_sylviolus | LC | Decreasing | tropical | diurnal | 8 | 524513.855 | 12.141 |
| birds | Phylloscartes_ventralis | Phylloscartes_ventralis | LC | Decreasing | temperate | diurnal | 8.3 | 1586787.276 | 11.925 |
| birds | Phylloscartes_virescens | Phylloscartes_virescens | LC | Stable | tropical | diurnal | 8.5 | 798656.796 | 1.868 |
| birds | Phylloscopus_affinis | Phylloscopus_affinis | LC | Stable | tropical | diurnal | 7 | 6032455.644 | 14.19 |
| birds | Phylloscopus_amoenus | Phylloscopus_amoenus | NT | Stable | tropical | diurnal | 10.5 | 37.602 | 1.119 |
| birds | Phylloscopus_armandii | Phylloscopus_armandii | LC | Stable | temperate | diurnal | 9.4 | 4630323.06 | 13.454 |
| birds | Phylloscopus_bonelli | Phylloscopus_bonelli | LC | Stable | temperate | diurnal | 8.7 | 10369392.474 | 6.551 |
| birds | Phylloscopus_borealis | Phylloscopus_borealis | LC | Increasing | temperate | diurnal | 10.9 | 15296466.246 | 3.158 |
| birds | Phylloscopus_borealoides | Phylloscopus_borealoides | LC | Decreasing | tropical | diurnal | 9.8 | 2985602.647 | 15.462 |
| birds | Phylloscopus_budongoensis | Phylloscopus_budongoensis | LC | Decreasing | tropical | diurnal | 8.2 | 176695.471 | 9.981 |
| birds | Phylloscopus_burkii | Seicercus_burkii | LC | Stable | temperate | diurnal | 7.3 | 1414029.025 | 16.035 |
| birds | Phylloscopus_cantator | Phylloscopus_cantator | LC | Stable | tropical | diurnal | 6 | 788069.432 | 9.222 |
| birds | Phylloscopus_castaniceps | Seicercus_castaniceps | LC | Stable | tropical | diurnal | 5.3 | 3275730.695 | 12.003 |

| | | | | | | | | | |
|---|---|---|---|---|---|---|---|---|---|
| birds | Phylloscopus_cebuensis | Phylloscopus_cebuensis | LC | Stable | tropical | diurnal | 9 | 121172.882 | 15.551 |
| birds | Phylloscopus_chloronotus | Phylloscopus_chloronotus | LC | Stable | temperate | diurnal | 4.9 | 452141.095 | 10.738 |
| birds | Phylloscopus_collybita | Phylloscopus_collybita | LC | Increasing | temperate | diurnal | 8.3 | 27400112.713 | 9.092 |
| birds | Phylloscopus_coronatus | Phylloscopus_coronatus | LC | Stable | temperate | diurnal | 10.1 | 6469893.738 | 13.897 |
| birds | Phylloscopus_emeiensis | Phylloscopus_emeiensis | LC | Stable | temperate | diurnal | 10.5 | 21846.579 | 9.886 |
| birds | Phylloscopus_fuligiventer | Phylloscopus_fuligiventer | LC | Stable | temperate | diurnal | 8.8 | 1181471.605 | 7.714 |
| birds | Phylloscopus_fuscatus | Phylloscopus_fuscatus | LC | Stable | temperate | diurnal | 8.7 | 20202368.256 | 7.598 |
| birds | Phylloscopus_grammiceps | Seicercus_grammiceps | LC | Stable | tropical | diurnal | 5 | 17394.874 | 16.296 |
| birds | Phylloscopus_griseolus | Phylloscopus_griseolus | LC | Stable | temperate | diurnal | 7.6 | 7386291.381 | 8.571 |
| birds | Phylloscopus_hainanus | Phylloscopus_hainanus | VU | Decreasing | tropical | diurnal | 9.5 | 13199.579 | 11.647 |
| birds | Phylloscopus_herberti | Phylloscopus_herberti | LC | Stable | tropical | diurnal | 8.9 | 50664.198 | 9.113 |
| birds | Phylloscopus_humei | Phylloscopus_humei | LC | Stable | temperate | diurnal | 6 | 6707584.554 | 11.387 |
| birds | Phylloscopus_ijimae | Phylloscopus_ijimae | VU | Decreasing | tropical | diurnal | 8.7 | 159185.737 | 17.122 |
| birds | Phylloscopus_inornatus | Phylloscopus_inornatus | LC | Stable | temperate | diurnal | 6.6 | 25153244.428 | 6.949 |
| birds | Phylloscopus_intensior | Phylloscopus_davisoni | LC | Stable | tropical | diurnal | 6.4 | 873724.181 | 10.03 |
| birds | Phylloscopus_intermedius | Seicercus_affinis | LC | Stable | temperate | diurnal | 6.9 | 658460.766 | 10.265 |
| birds | Phylloscopus_kansuensis | Phylloscopus_kansuensis | LC | Stable | temperate | diurnal | 6.6 | 157431.45 | 10.365 |
| birds | Phylloscopus_laetus | Phylloscopus_laetus | LC | Stable | tropical | diurnal | 9.5 | 81632.113 | 17.025 |
| birds | Phylloscopus_laurae | Phylloscopus_laurae | LC | Stable | tropical | diurnal | 8.5 | 364414.575 | 7.729 |
| birds | Phylloscopus_maculipennis | Phylloscopus_maculipennis | LC | Stable | temperate | diurnal | 5.1 | 972285.742 | 9.648 |
| birds | Phylloscopus_magnirostris | Phylloscopus_magnirostris | LC | Stable | tropical | diurnal | 11.6 | 2364522.293 | 14.677 |
| birds | Phylloscopus_montis | Seicercus_montis | LC | Stable | tropical | diurnal | 6.4 | 369636.968 | 6.243 |
| birds | Phylloscopus_neglectus | Phylloscopus_neglectus | LC | Stable | temperate | diurnal | 5.3 | 3222380.78 | 9.643 |
| birds | Phylloscopus_nitidus | Phylloscopus_nitidus | LC | Stable | temperate | diurnal | 7.8 | 4625690.047 | 12.596 |
| birds | Phylloscopus_occipitalis | Phylloscopus_occipitalis | LC | Stable | temperate | diurnal | 9 | 1246331.273 | 17.105 |
| birds | Phylloscopus_olivaceus | Phylloscopus_olivaceus | LC | Stable | tropical | diurnal | 9.7 | 133609.086 | 12.724 |
| birds | Phylloscopus_orientalis | Phylloscopus_orientalis | LC | Increasing | tropical | diurnal | 8.5 | 3917108.949 | 4.902 |
| birds | Phylloscopus_poliocephalus | Phylloscopus_poliocephalus | LC | Stable | tropical | diurnal | 10.8 | 474523.859 | 4.049 |
| birds | Phylloscopus_poliogenys | Seicercus_poliogenys | LC | Stable | tropical | diurnal | 6.3 | 1049263.635 | 9.167 |
| birds | Phylloscopus_presbytes | Phylloscopus_presbytes | LC | Stable | tropical | diurnal | 9.5 | 34990.033 | 13.486 |
| birds | Phylloscopus_proregulus | Phylloscopus_proregulus | LC | Stable | temperate | diurnal | 6 | 9985414.535 | 9.303 |
| birds | Phylloscopus_pulcher | Phylloscopus_pulcher | LC | Stable | temperate | diurnal | 6.8 | 1521950.816 | 9.492 |
| birds | Phylloscopus_reguloides | Phylloscopus_reguloides | LC | Stable | tropical | diurnal | 7.6 | 3194582.199 | 12.36 |
| birds | Phylloscopus_ricketti | Phylloscopus_ricketti | LC | Stable | tropical | diurnal | 6.3 | 3059069.625 | 14.024 |

| | | | | | | | | | |
|---|---|---|---|---|---|---|---|---|---|
| birds | Phylloscopus_ruficapilla | Phylloscopus_ruficapilla | LC | Decreasing | temperate | diurnal | 7.7 | 423407.958 | 12.497 |
| birds | Phylloscopus_sarasinorum | Phylloscopus_sarasinorum | NT | Decreasing | tropical | diurnal | 10 | 2393.882 | 13.588 |
| birds | Phylloscopus_schwarzi | Phylloscopus_schwarzi | LC | Stable | temperate | diurnal | 11.4 | 8407744.291 | 11.493 |
| birds | Phylloscopus_sibilatrix | Phylloscopus_sibilatrix | LC | Decreasing | tropical | diurnal | 9.2 | 23976906.186 | 8.682 |
| birds | Phylloscopus_sindianus | Phylloscopus_sindianus | LC | Stable | temperate | diurnal | 7.5 | 4635151.234 | 9.605 |
| birds | Phylloscopus_subaffinis | Phylloscopus_subaffinis | LC | Stable | temperate | diurnal | 6.2 | 3400992.32 | 12.889 |
| birds | Phylloscopus_subviridis | Phylloscopus_subviridis | LC | Stable | temperate | diurnal | 5.5 | 523915.027 | 18.412 |
| birds | Phylloscopus_tenellipes | Phylloscopus_tenellipes | LC | Stable | temperate | diurnal | 11.8 | 4607603.525 | 14.009 |
| birds | Phylloscopus_trivirgatus | Phylloscopus_trivirgatus | LC | Stable | tropical | diurnal | 9 | 539476.749 | 10.358 |
| birds | Phylloscopus_trochiloides | Phylloscopus_trochiloides | LC | Increasing | temperate | diurnal | 8.1 | 22292683.075 | 9.876 |
| birds | Phylloscopus_trochilus | Phylloscopus_trochilus | LC | Decreasing | tropical | diurnal | 8.7 | 59842121.382 | 7.606 |
| birds | Phylloscopus_tytleri | Phylloscopus_tytleri | LC | Unknown | tropical | diurnal | 7.2 | 1418121.01 | 17.833 |
| birds | Phylloscopus_umbrovirens | Phylloscopus_umbrovirens | LC | Stable | tropical | diurnal | 8.6 | 611974.74 | 11.849 |
| birds | Phylloscopus_xanthoschistos | Phylloscopus_xanthoschistos | LC | Stable | temperate | diurnal | 6.9 | 269573.706 | 10.147 |
| birds | Phylloscopus_yunnanensis | Phylloscopus_yunnanensis | LC | Stable | tropical | diurnal | 6 | 467272.955 | 12.354 |
| birds | Phytotoma_raimondii | Phytotoma_raimondii | VU | Decreasing | tropical | diurnal | 39.8 | 58073.47 | 8.033 |
| birds | Phytotoma_rara | Phytotoma_rara | LC | Stable | temperate | diurnal | 47 | 478137.958 | 6.856 |
| birds | Phytotoma_rutila | Phytotoma_rutila | LC | Decreasing | temperate | diurnal | 40.5 | 2362046.179 | 8.491 |
| birds | Piaya_cayana | Piaya_cayana | LC | Decreasing | tropical | diurnal | 102 | 13448425.195 | 5.854 |
| birds | Piaya_melanogaster | Piaya_melanogaster | LC | Decreasing | tropical | diurnal | 102.8 | 4840731.882 | 1.568 |
| birds | Pica_hudsonia | Pica_hudsonia | LC | Stable | temperate | diurnal | 177.1 | 6726173.66 | 8.602 |
| birds | Pica_nutalli | Pica_nuttalli | VU | Decreasing | temperate | diurnal | 158.3 | 95191.282 | 16.922 |
| birds | Pica_pica | Pica_pica | LC | Stable | temperate | diurnal | 217.5 | 31870223.471 | 10.189 |
| birds | Picathartes_gymnocephalus | Picathartes_gymnocephalus | VU | Decreasing | tropical | diurnal | 196 | 388878.275 | 15.552 |
| birds | Picathartes_oreas | Picathartes_oreas | NT | Decreasing | tropical | diurnal | 220 | 381185.415 | 6.443 |
| birds | Picoides_arcticus | Picoides_arcticus | LC | Stable | temperate | diurnal | 69.2 | 6771211.075 | 4.326 |
| birds | Picoides_canicapillus | Dendrocopos_canicapillus | LC | Stable | temperate | diurnal | 25.8 | 8010810.884 | 12.701 |
| birds | Picoides_kizuki | Dendrocopos_kizuki | LC | Stable | temperate | diurnal | 19.2 | 1603020.854 | 13.679 |
| birds | Picoides_maculatus | Dendrocopos_maculatus | LC | Stable | tropical | diurnal | 24.5 | 267253.511 | 13.856 |
| birds | Picoides_moluccensis | Dendrocopos_moluccensis | LC | Increasing | tropical | diurnal | 16.2 | 742649.604 | 12.146 |
| birds | Picoides_temminckii | Dendrocopos_temminckii | LC | Stable | tropical | diurnal | 66.7 | 173473.636 | 9.289 |
| birds | Picoides_tridactylus | Picoides_tridactylus | LC | Stable | temperate | diurnal | 65.5 | 23486930.713 | 4.166 |
| birds | Piculus_aurulentus | Piculus_aurulentus | NT | Decreasing | temperate | diurnal | 75 | 748842.173 | 12.662 |
| birds | Piculus_callopterus | Piculus_callopterus | LC | Decreasing | tropical | diurnal | 51 | 5548.219 | 4.003 |

| | | | | | | | | | |
|---|---|---|---|---|---|---|---|---|---|
| birds | Piculus_chrysochloros | Piculus_chrysochloros | LC | Decreasing | tropical | diurnal | 88 | 8623251.859 | 3.552 |
| birds | Piculus_flavigula | Piculus_flavigula | LC | Decreasing | tropical | diurnal | 55 | 6464140.534 | 2.718 |
| birds | Piculus_leucolaemus | Piculus_leucolaemus | LC | Decreasing | tropical | diurnal | 69 | 885342.239 | 2.891 |
| birds | Piculus_litae | NA | LC | Decreasing | NA | NA | NA | NA | NA |
| birds | Piculus_simplex | Piculus_simplex | LC | Decreasing | tropical | diurnal | 55 | 92589.277 | 7.352 |
| birds | Picumnus_albosquamatus | Picumnus_albosquamatus | LC | Decreasing | tropical | diurnal | 11.9 | 2007279.992 | 9.119 |
| birds | Picumnus_aurifrons | Picumnus_aurifrons | LC | Decreasing | tropical | diurnal | 8.9 | 2989266.488 | 2.301 |
| birds | Picumnus_castelnau | Picumnus_castelnau | LC | Decreasing | tropical | diurnal | 11.4 | 57699.815 | 3.622 |
| birds | Picumnus_cinnamomeus | Picumnus_cinnamomeus | LC | Unknown | tropical | diurnal | 11.3 | 130928.203 | 15.134 |
| birds | Picumnus_cirratus | Picumnus_cirratus | LC | Decreasing | tropical | diurnal | 10.2 | 2348460.612 | 9.096 |
| birds | Picumnus_dorbignyanus | Picumnus_dorbignyanus | LC | Stable | tropical | diurnal | 9.6 | 100469.477 | 8.56 |
| birds | Picumnus_exilis | Picumnus_exilis | LC | Decreasing | tropical | diurnal | 9.3 | 1264006.827 | 2.759 |
| birds | Picumnus_fulvescens | Picumnus_fulvescens | NT | Decreasing | tropical | diurnal | 11.3 | 119592.592 | 11.532 |
| birds | Picumnus_fuscus | Picumnus_fuscus | LC | Decreasing | tropical | diurnal | 11.3 | 11590.987 | 1.657 |
| birds | Picumnus_granadensis | Picumnus_granadensis | LC | Decreasing | tropical | diurnal | 12.2 | 27384.642 | 15.857 |
| birds | Picumnus_innominatus | Picumnus_innominatus | LC | Decreasing | tropical | diurnal | 10.2 | 4531626.734 | 12.669 |
| birds | Picumnus_lafresnayi | Picumnus_lafresnayi | LC | Decreasing | tropical | diurnal | 9.5 | 779374.899 | 2.954 |
| birds | Picumnus_limae | Picumnus_limae | LC | Stable | tropical | diurnal | 11.3 | 303100.712 | 10.911 |
| birds | Picumnus_minutissimus | Picumnus_minutissimus | LC | Stable | tropical | diurnal | 13.3 | 36687.972 | 4.176 |
| birds | Picumnus_nebulosus | Picumnus_nebulosus | NT | Decreasing | temperate | diurnal | 11.6 | 517928.065 | 11.311 |
| birds | Picumnus_nigropunctatus | Picumnus_nigropunctatus | LC | Decreasing | tropical | diurnal | 11.3 | 24797.174 | 2.68 |
| birds | Picumnus_olivaceus | Picumnus_olivaceus | LC | Decreasing | tropical | diurnal | 10.6 | 493974.891 | 11.468 |
| birds | Picumnus_pumilus | Picumnus_pumilus | LC | Unknown | tropical | diurnal | 9.5 | 210423.826 | 0.871 |
| birds | Picumnus_pygmaeus | Picumnus_pygmaeus | LC | Unknown | tropical | diurnal | 11 | 837474.167 | 7.564 |
| birds | Picumnus_rufiventris | Picumnus_rufiventris | LC | Stable | tropical | diurnal | 21 | 1172719.045 | 2.226 |
| birds | Picumnus_sclateri | Picumnus_sclateri | LC | Stable | tropical | diurnal | 11 | 44945.62 | 12.431 |
| birds | Picumnus_spilogaster | Picumnus_spilogaster | VU | Decreasing | tropical | diurnal | 13.2 | 280875.097 | 4.043 |
| birds | Picumnus_squamulatus | Picumnus_squamulatus | LC | Stable | tropical | diurnal | 10.2 | 437386.968 | 9.934 |
| birds | Picumnus_steindachneri | Picumnus_steindachneri | NT | Decreasing | tropical | diurnal | 9.9 | 7043.951 | 6.842 |
| birds | Picumnus_subtilis | Picumnus_subtilis | LC | Stable | tropical | diurnal | 10.4 | 39544.256 | 2.454 |
| birds | Picumnus_temminckii | Picumnus_temminckii | LC | Unknown | temperate | diurnal | 11.5 | 514581.531 | 13.09 |
| birds | Picumnus_varzeae | Picumnus_varzeae | NT | Decreasing | tropical | diurnal | 11.5 | 189643.391 | 2.232 |
| birds | Picus_awokera | Picus_awokera | LC | Stable | temperate | diurnal | 143 | 281098.752 | 20.834 |
| birds | Picus_canus | Picus_canus | LC | Increasing | temperate | diurnal | 137 | 11476968.697 | 11.558 |

| | | | | | | | | | |
|---|---|---|---|---|---|---|---|---|---|
| birds | Picus_chlorolophus | Picus_chlorolophus | LC | Stable | tropical | diurnal | 65.4 | 3444974.536 | 12.597 |
| birds | Picus_erythropygius | Picus_erythropygius | LC | Decreasing | tropical | diurnal | 116.2 | 465293.359 | 12.455 |
| birds | Picus_puniceus | Picus_puniceus | LC | Decreasing | tropical | diurnal | 79.1 | 1380604.208 | 8.67 |
| birds | Picus_rabieri | NA | NT | Decreasing | NA | NA | NA | NA | NA |
| birds | Picus_squamatus | Picus_squamatus | LC | Stable | temperate | diurnal | 170 | 781014.453 | 8.748 |
| birds | Picus_viridanus | Picus_viridanus | LC | Decreasing | tropical | diurnal | 111.4 | 390821.745 | 12.272 |
| birds | Picus_viridis | Picus_viridis | LC | Increasing | temperate | diurnal | 176 | 5981746.045 | 17.74 |
| birds | Picus_vittatus | Picus_vittatus | LC | Decreasing | tropical | diurnal | 108.1 | 1352502.995 | 12.742 |
| birds | Picus_xanthopygaeus | Picus_xanthopygaeus | LC | Unknown | temperate | diurnal | 99.7 | 2078020.449 | 16.059 |
| birds | Piezorina_cinerea | Piezorhina_cinerea | LC | Stable | tropical | diurnal | 32.2 | 41880.069 | 8.237 |
| birds | Pilherodius_pileatus | Pilherodius_pileatus | LC | Decreasing | tropical | diurnal | 568.6 | 10271266.316 | 4.403 |
| birds | Pinarochroa_sordida | Cercomela_sordida | LC | Stable | tropical | diurnal | 21 | 188372.16 | 14.544 |
| birds | Pinarocorys_erythropygia | Pinarocorys_erythropygia | LC | Stable | tropical | diurnal | 30.1 | 5722826.261 | 9.162 |
| birds | Pinarocorys_nigricans | Pinarocorys_nigricans | LC | Stable | tropical | diurnal | 38.1 | 3553770.125 | 7.294 |
| birds | Pinaroloxias_inornata | NA | LC | Stable | NA | NA | NA | NA | NA |
| birds | Pinarornis_plumosus | Pinarornis_plumosus | LC | Stable | tropical | diurnal | 65.8 | 213455.193 | 12.934 |
| birds | Pinicola_enucleator | Pinicola_enucleator | LC | Decreasing | temperate | diurnal | 56.4 | 24288557.5 | 4.785 |
| birds | Pionites_leucogaster | Pionites_leucogaster | VU | Decreasing | tropical | diurnal | 155 | 1982282.572 | 2.865 |
| birds | Pionites_melanocephalus | Pionites_melanocephalus | LC | Stable | tropical | diurnal | 148.8 | 2920163.541 | 1.624 |
| birds | Pionopsitta_pileata | Pionopsitta_pileata | LC | Decreasing | tropical | diurnal | 119 | 728878.227 | 12.682 |
| birds | Pionus_chalcopterus | Pionus_chalcopterus | LC | Decreasing | tropical | diurnal | 210 | 158502.651 | 13.03 |
| birds | Pionus_fuscus | Pionus_fuscus | LC | Decreasing | tropical | diurnal | 206 | 1865113.688 | 2.121 |
| birds | Pionus_maximiliani | Pionus_maximiliani | LC | Decreasing | tropical | diurnal | 293 | 4524942.978 | 9.587 |
| birds | Pionus_menstruus | Pionus_menstruus | LC | Stable | tropical | diurnal | 251 | 8375262.878 | 3.341 |
| birds | Pionus_senilis | Pionus_senilis | LC | Decreasing | tropical | diurnal | 212 | 458536.209 | 9.923 |
| birds | Pionus_sordidus | Pionus_sordidus | LC | Decreasing | tropical | diurnal | 266 | 177970.605 | 7.754 |
| birds | Pionus_tumultuosus | Pionus_tumultuosus | LC | Decreasing | tropical | diurnal | 239 | 150467.551 | 5.972 |
| birds | Pipile_cujubi | Pipile_cujubi | VU | Decreasing | tropical | diurnal | 1195.8 | 2530765.848 | 4.02 |
| birds | Pipile_cumanensis | Pipile_cumanensis | LC | Decreasing | tropical | diurnal | 1239.2 | 3138935.129 | 1.996 |
| birds | Pipile_jacutinga | Pipile_jacutinga | EN | Decreasing | tropical | diurnal | 1241 | 1063311.161 | 13.524 |
| birds | Pipile_pipile | Pipile_pipile | CR | Increasing | tropical | nocturnal | 1816.6 | 3403.877 | 18.099 |
| birds | Pipilo_chlorurus | Pipilo_chlorurus | LC | Stable | temperate | diurnal | 29.4 | 4121928.586 | 9.486 |
| birds | Pipilo_erythrophthalmus | Pipilo_erythrophthalmus | LC | Decreasing | temperate | diurnal | 40 | 3619655.583 | 17.602 |
| birds | Pipilo_maculatus | Pipilo_maculatus | LC | Stable | temperate | diurnal | 39.3 | 5605136.849 | 11.127 |

| | | | | | | | | | |
|---|---|---|---|---|---|---|---|---|---|
| birds | Pipilo_ocai | Pipilo_ocai | LC | Decreasing | tropical | diurnal | 61 | 114057.776 | 10.192 |
| birds | Pipra_aureola | Pipra_aureola | LC | Decreasing | tropical | diurnal | 16.1 | 671432.518 | 2.836 |
| birds | Pipra_fasciicauda | Pipra_fasciicauda | LC | Decreasing | tropical | diurnal | 15.9 | 4779358.798 | 6.012 |
| birds | Pipra_filicauda | Pipra_filicauda | LC | Stable | tropical | diurnal | 15.4 | 2614150.068 | 2.383 |
| birds | Pipraeidea_bonariensis | Thraupis_bonariensis | LC | Stable | temperate | diurnal | 36 | 3316978.369 | 9.235 |
| birds | Pipraeidea_melanonota | Pipraeidea_melanonota | LC | Stable | tropical | diurnal | 21 | 2111740.744 | 11.344 |
| birds | Pipreola_arcuata | Pipreola_arcuata | LC | Stable | tropical | diurnal | 118 | 259332.068 | 9.272 |
| birds | Pipreola_aureopectus | Pipreola_aureopectus | LC | Decreasing | tropical | diurnal | 46 | 83804.892 | 11.55 |
| birds | Pipreola_chlorolepidota | Pipreola_chlorolepidota | LC | Decreasing | tropical | diurnal | 29.5 | 136102.383 | 5.442 |
| birds | Pipreola_formosa | Pipreola_formosa | LC | Decreasing | tropical | diurnal | 47.2 | 5381.637 | 13.614 |
| birds | Pipreola_frontalis | Pipreola_frontalis | LC | Decreasing | tropical | diurnal | 42.4 | 113721.277 | 4.962 |
| birds | Pipreola_intermedia | Pipreola_intermedia | LC | Stable | tropical | diurnal | 51.2 | 121136.093 | 5.133 |
| birds | Pipreola_jucunda | NA | LC | Decreasing | NA | NA | NA | NA | NA |
| birds | Pipreola_lubomirskii | Pipreola_lubomirskii | LC | Decreasing | tropical | diurnal | 57 | 28329.189 | 5.261 |
| birds | Pipreola_pulchra | Pipreola_pulchra | LC | Decreasing | tropical | diurnal | 58.2 | 78215.762 | 4.332 |
| birds | Pipreola_riefferii | Pipreola_riefferii | LC | Stable | tropical | diurnal | 49.3 | 257255.048 | 9.556 |
| birds | Pipreola_whitelyi | Pipreola_whitelyi | LC | Stable | tropical | diurnal | 50 | 24788.578 | 1.638 |
| birds | Piprites_chloris | Piprites_chloris | LC | Decreasing | tropical | diurnal | 16 | 7073219.6 | 3.3 |
| birds | Piprites_griseiceps | Piprites_griseiceps | LC | Decreasing | tropical | diurnal | 16 | 119822.907 | 8.164 |
| birds | Piprites_pileata | Piprites_pileata | NT | Decreasing | temperate | diurnal | 15 | 134629.412 | 12.392 |
| birds | Piranga_bidentata | Piranga_bidentata | LC | Decreasing | temperate | diurnal | 36.1 | 417994.015 | 9.701 |
| birds | Piranga_erythrocephala | Piranga_erythrocephala | LC | Decreasing | tropical | diurnal | 21.8 | 256230.431 | 7.394 |
| birds | Piranga_flava | Piranga_flava | LC | Unknown | tropical | diurnal | 37.7 | 5776535.166 | 8.783 |
| birds | Piranga_hepatica | Piranga_hepatica | LC | Increasing | temperate | diurnal | 32.3 | 3539367.178 | 8.517 |
| birds | Piranga_leucoptera | Piranga_leucoptera | LC | Decreasing | tropical | diurnal | 16 | 567150.477 | 8.58 |
| birds | Piranga_ludoviciana | Piranga_ludoviciana | LC | Increasing | temperate | diurnal | 28.1 | 7784975.199 | 9.131 |
| birds | Piranga_olivacea | Piranga_olivacea | LC | Stable | temperate | diurnal | 28.2 | 6581671.34 | 12.955 |
| birds | Piranga_roseogularis | Piranga_roseogularis | LC | Decreasing | tropical | diurnal | 24 | 105691.579 | 4.955 |
| birds | Piranga_rubra | Piranga_rubra | LC | Stable | tropical | diurnal | 29.1 | 8740188.438 | 9.793 |
| birds | Piranga_rubriceps | Piranga_rubriceps | LC | Decreasing | tropical | diurnal | 35.9 | 80398.643 | 7.514 |
| birds | Pitangus_sulphuratus | Pitangus_sulphuratus | LC | Increasing | tropical | diurnal | 62.9 | 16134345.057 | 6.295 |
| birds | Pithecophaga_jefferyi | Pithecophaga_jefferyi | CR | Decreasing | tropical | diurnal | 5175.3 | 142256.552 | 11.456 |
| birds | Pithys_albifrons | Pithys_albifrons | LC | Stable | tropical | diurnal | 19.5 | 3102848.345 | 1.785 |
| birds | Pithys_castaneus | Pithys_castaneus | NT | Decreasing | tropical | diurnal | 29.8 | 3885.126 | 2.084 |

| | | | | | | | | | |
|---|---|---|---|---|---|---|---|---|---|
| birds | Pitohui_dichrous | Pitohui_dichrous | LC | Stable | tropical | diurnal | 92.9 | 224365.726 | 3.372 |
| birds | Pitohui_kirhocephalus | Pitohui_kirhocephalus | LC | Unknown | tropical | diurnal | 72 | 243662.143 | 3.087 |
| birds | Pitta_anerythra | Pitta_anerythra | NT | Decreasing | tropical | diurnal | 86 | 14211.153 | 4.595 |
| birds | Pitta_angolensis | Pitta_angolensis | LC | Decreasing | tropical | diurnal | 62.7 | 3968093.861 | 10.287 |
| birds | Pitta_brachyura | Pitta_brachyura | LC | Decreasing | tropical | diurnal | 55.5 | 2625350.305 | 19.022 |
| birds | Pitta_elegans | Pitta_elegans | LC | Decreasing | tropical | diurnal | 56.6 | 92181.315 | 11.009 |
| birds | Pitta_iris | Pitta_iris | LC | Decreasing | tropical | diurnal | 62.2 | 258796.968 | 1.874 |
| birds | Pitta_maxima | Pitta_maxima | LC | Stable | tropical | diurnal | 190 | 23409.647 | 6.33 |
| birds | Pitta_megarhyncha | Pitta_megarhyncha | NT | Decreasing | tropical | diurnal | 106 | 56694.661 | 10.149 |
| birds | Pitta_moluccensis | Pitta_moluccensis | LC | Decreasing | tropical | diurnal | 76 | 2249725.199 | 8.998 |
| birds | Pitta_nympha | Pitta_nympha | VU | Decreasing | temperate | diurnal | 88.3 | 1374640.888 | 12.949 |
| birds | Pitta_reichenowi | NA | LC | Decreasing | NA | NA | NA | NA | NA |
| birds | Pitta_sordida | Pitta_sordida | LC | Decreasing | tropical | diurnal | 64.5 | 3188235.868 | 10.237 |
| birds | Pitta_steerii | Pitta_steerii | VU | Decreasing | tropical | diurnal | 94.8 | 117483.708 | 12.097 |
| birds | Pitta_superba | Pitta_superba | EN | Decreasing | tropical | diurnal | 115 | 1830.09 | 6.316 |
| birds | Pitta_versicolor | Pitta_versicolor | LC | Decreasing | tropical | diurnal | 88.9 | 565329.427 | 5.523 |
| birds | Pittasoma_michleri | Pittasoma_michleri | LC | Decreasing | tropical | diurnal | 109 | 49173.734 | 4.891 |
| birds | Pittasoma_rufopileatum | Pittasoma_rufopileatum | LC | Decreasing | tropical | diurnal | 96.5 | 75066.712 | 5.25 |
| birds | Pityriasis_gymnocephala | Pityriasis_gymnocephala | VU | Decreasing | tropical | diurnal | 128.5 | 620878.32 | 6.189 |
| birds | Platalea_ajaja | Platalea_ajaja | LC | Stable | tropical | diurnal | 1490 | 8415237.441 | 8.516 |
| birds | Platalea_alba | Platalea_alba | LC | Stable | tropical | diurnal | 1521 | 14157612.5 | 10.036 |
| birds | Platalea_flavipes | Platalea_flavipes | LC | Stable | temperate | diurnal | 1741.3 | 4698362.966 | 5.704 |
| birds | Platalea_leucorodia | Platalea_leucorodia | LC | Unknown | temperate | cathemeral | 1868 | 9201391.344 | 12.456 |
| birds | Platalea_minor | Platalea_minor | EN | Increasing | tropical | diurnal | 1228 | 210695.085 | 24.344 |
| birds | Platalea_regia | Platalea_regia | LC | Stable | temperate | cathemeral | 1723.5 | 6262402.868 | 5.588 |
| birds | Platycercus_adscitus | Platycercus_adscitus | LC | Increasing | tropical | diurnal | 111 | 1465853.969 | 5.821 |
| birds | Platycercus_caledonicus | Platycercus_caledonicus | LC | Decreasing | temperate | diurnal | 132.6 | 68050.364 | 5.966 |
| birds | Platycercus_elegans | Platycercus_elegans | LC | Decreasing | temperate | diurnal | 123.9 | 743673.241 | 9.746 |
| birds | Platycercus_eximius | Platycercus_eximius | LC | Increasing | temperate | diurnal | 104 | 1189978.195 | 9.484 |
| birds | Platycercus_icterotis | Platycercus_icterotis | LC | Decreasing | temperate | diurnal | 63.3 | 214002.525 | 8.613 |
| birds | Platycercus_venustus | Platycercus_venustus | LC | Stable | tropical | diurnal | 89.4 | 785415.826 | 2.126 |
| birds | Platylophus_galericulatus | Platylophus_galericulatus | NT | Decreasing | tropical | diurnal | 77.9 | 1559387.648 | 9.278 |
| birds | Platyrinchus_cancrominus | Platyrinchus_cancrominus | LC | Decreasing | tropical | diurnal | 9.2 | 443828.033 | 9.551 |
| birds | Platyrinchus_coronatus | Platyrinchus_coronatus | LC | Decreasing | tropical | diurnal | 9.1 | 5183010.815 | 1.885 |

| | | | | | | | | | |
|---|---|---|---|---|---|---|---|---|---|
| birds | Platyrinchus_flavigularis | Platyrinchus_flavigularis | LC | Decreasing | tropical | diurnal | 10 | 114162.948 | 6.216 |
| birds | Platyrinchus_leucoryphus | Platyrinchus_leucoryphus | VU | Decreasing | temperate | diurnal | 16 | 481741.402 | 13.589 |
| birds | Platyrinchus_mystaceus | Platyrinchus_mystaceus | LC | Decreasing | tropical | diurnal | 9.7 | 5068239.901 | 9.002 |
| birds | Platyrinchus_platyrhynchos | Platyrinchus_platyrhynchos | LC | Decreasing | tropical | diurnal | 12 | 5529652.291 | 1.6 |
| birds | Platyrinchus_saturatus | Platyrinchus_saturatus | LC | Stable | tropical | diurnal | 10.7 | 3011725.404 | 1.62 |
| birds | Platysmurus_leucopterus | Platysmurus_leucopterus | LC | Decreasing | tropical | diurnal | 180 | 624071.229 | 11.122 |
| birds | Platyspiza_crassirostris | NA | LC | Stable | NA | NA | NA | NA | NA |
| birds | Platysteira_albifrons | Platysteira_albifrons | NT | Decreasing | tropical | diurnal | 12.2 | 155796.964 | 8.872 |
| birds | Platysteira_cyanea | Platysteira_cyanea | LC | Stable | tropical | diurnal | 14.6 | 5598291.675 | 9.727 |
| birds | Platysteira_peltata | Platysteira_peltata | LC | Decreasing | tropical | diurnal | 13.7 | 3620454.797 | 9.774 |
| birds | Plectorhyncha_lanceolata | Plectorhyncha_lanceolata | LC | Stable | temperate | diurnal | 40.6 | 2020455.086 | 7.222 |
| birds | Plectrophenax_hyperboreus | Plectrophenax_hyperboreus | LC | Stable | temperate | diurnal | 54.5 | 77792.428 | 1.408 |
| birds | Plectrophenax_nivalis | Plectrophenax_nivalis | LC | Decreasing | temperate | diurnal | 42.2 | 29206717.754 | 5.863 |
| birds | Plectropterus_gambensis | Plectropterus_gambensis | LC | Increasing | tropical | diurnal | 3869 | 15772780.822 | 9.508 |
| birds | Plegadis_chihi | Plegadis_chihi | LC | Increasing | temperate | diurnal | 616.9 | 9579778.672 | 10.357 |
| birds | Plegadis_falcinellus | Plegadis_falcinellus | LC | Decreasing | tropical | diurnal | 626.6 | 28264905.805 | 9.24 |
| birds | Plegadis_ridgwayi | Plegadis_ridgwayi | LC | Decreasing | tropical | diurnal | 579.4 | 495452.93 | 8.298 |
| birds | Plesiodryas_albonotata | Poecilodryas_albonotata | LC | Stable | tropical | diurnal | 38.6 | 117306.589 | 4.925 |
| birds | Plocepasser_donaldsoni | Plocepasser_donaldsoni | LC | Stable | tropical | diurnal | 39.9 | 190025.35 | 6.205 |
| birds | Plocepasser_mahali | Plocepasser_mahali | LC | Stable | tropical | diurnal | 41.5 | 3098861.258 | 8.552 |
| birds | Plocepasser_rufoscapulatus | Plocepasser_rufoscapulatus | LC | Stable | tropical | diurnal | 44.6 | 403630.329 | 7.067 |
| birds | Plocepasser_superciliosus | Plocepasser_superciliosus | LC | Stable | tropical | diurnal | 37.1 | 2062715.245 | 11.757 |
| birds | Ploceus_albinucha | Ploceus_albinucha | LC | Stable | tropical | diurnal | 27.2 | 745212.437 | 8.651 |
| birds | Ploceus_alienus | Ploceus_alienus | LC | Decreasing | tropical | diurnal | 22.2 | 61881.532 | 15.894 |
| birds | Ploceus_angolensis | Ploceus_angolensis | LC | Stable | tropical | diurnal | 26.6 | 260942.79 | 7.38 |
| birds | Ploceus_aurantius | Ploceus_aurantius | LC | Stable | tropical | diurnal | 23 | 441030.16 | 9.736 |
| birds | Ploceus_aureonucha | Ploceus_aureonucha | EN | Decreasing | tropical | diurnal | 26.6 | 16614.788 | 8.164 |
| birds | Ploceus_badius | Ploceus_badius | LC | Stable | tropical | diurnal | 22 | 365559.995 | 10.601 |
| birds | Ploceus_baglafecht | Ploceus_baglafecht | LC | Stable | tropical | diurnal | 31.6 | 1651063.488 | 11.618 |
| birds | Ploceus_bannermani | Ploceus_bannermani | VU | Decreasing | tropical | diurnal | 30.8 | 4569.345 | 11.904 |
| birds | Ploceus_batesi | Ploceus_batesi | EN | Decreasing | tropical | diurnal | 26.6 | 53494.778 | 6.81 |
| birds | Ploceus_benghalensis | Ploceus_benghalensis | LC | Stable | temperate | diurnal | 19.8 | 1001878.116 | 19.448 |
| birds | Ploceus_bertrandi | Ploceus_bertrandi | LC | Stable | tropical | diurnal | 39 | 185221.004 | 13.343 |
| birds | Ploceus_bicolor | Ploceus_bicolor | LC | Stable | tropical | diurnal | 34.5 | 1377098.306 | 9.823 |

| | | | | | | | | | |
|---|---|---|---|---|---|---|---|---|---|
| birds | Ploceus_bojeri | Ploceus_bojeri | LC | Stable | tropical | diurnal | 22.4 | 223498.083 | 11.414 |
| birds | Ploceus_burnieri | Ploceus_burnieri | VU | Decreasing | tropical | diurnal | 18 | 1464.962 | 5.49 |
| birds | Ploceus_capensis | Ploceus_capensis | LC | Stable | temperate | diurnal | 44 | 666169.707 | 11.275 |
| birds | Ploceus_castaneiceps | Ploceus_castaneiceps | LC | Stable | tropical | diurnal | 18.5 | 45796.602 | 11.295 |
| birds | Ploceus_castanops | Ploceus_castanops | LC | Stable | tropical | diurnal | 21.4 | 193380.609 | 17.093 |
| birds | Ploceus_cucullatus | Ploceus_cucullatus | LC | Stable | tropical | diurnal | 36.1 | 13754902.578 | 9.611 |
| birds | Ploceus_dichrocephalus | Ploceus_dichrocephalus | LC | Stable | tropical | diurnal | 18.5 | 262572.094 | 10.818 |
| birds | Ploceus_dorsomaculatus | Ploceus_dorsomaculatus | LC | Stable | tropical | diurnal | 26.6 | 334876.309 | 5.274 |
| birds | Ploceus_flavipes | Ploceus_flavipes | VU | Decreasing | tropical | diurnal | 46.3 | 18694.159 | 5.438 |
| birds | Ploceus_galbula | Ploceus_galbula | LC | Stable | tropical | diurnal | 21.7 | 943847.182 | 10.198 |
| birds | Ploceus_golandi | Ploceus_golandi | EN | Decreasing | tropical | diurnal | 23.5 | 837.187 | 9.482 |
| birds | Ploceus_grandis | Ploceus_grandis | NT | Decreasing | tropical | diurnal | 63.5 | 858.514 | 12.812 |
| birds | Ploceus_heuglini | Ploceus_heuglini | LC | Stable | tropical | diurnal | 22.8 | 2515613.143 | 13.27 |
| birds | Ploceus_hypoxanthus | Ploceus_hypoxanthus | NT | Decreasing | tropical | diurnal | 18.6 | 861169.037 | 14.789 |
| birds | Ploceus_insignis | Ploceus_insignis | LC | Stable | tropical | diurnal | 27.5 | 247177.256 | 13.373 |
| birds | Ploceus_intermedius | Ploceus_intermedius | LC | Stable | tropical | diurnal | 20.6 | 3847715.308 | 10.094 |
| birds | Ploceus_jacksoni | Ploceus_jacksoni | LC | Stable | tropical | diurnal | 23.4 | 525034.954 | 14.623 |
| birds | Ploceus_katangae | Ploceus_katangae | LC | Unknown | tropical | diurnal | 18.5 | 61902.355 | 7.611 |
| birds | Ploceus_luteolus | Ploceus_luteolus | LC | Stable | tropical | diurnal | 15.1 | 3085847.315 | 12.024 |
| birds | Ploceus_manyar | Ploceus_manyar | LC | Stable | temperate | diurnal | 17.4 | 3778604.755 | 16.092 |
| birds | Ploceus_megarhynchus | Ploceus_megarhynchus | EN | Decreasing | temperate | diurnal | 34.3 | 125839.04 | 19.067 |
| birds | Ploceus_melanocephalus | Ploceus_melanocephalus | LC | Stable | tropical | diurnal | 23 | 1189276.856 | 11.812 |
| birds | Ploceus_melanogaster | Ploceus_melanogaster | LC | Decreasing | tropical | diurnal | 22.4 | 221733.168 | 16.011 |
| birds | Ploceus_nelicourvi | Ploceus_nelicourvi | LC | Stable | tropical | diurnal | 23.8 | 139470.306 | 7.629 |
| birds | Ploceus_nicolli | Ploceus_nicolli | NT | Decreasing | tropical | diurnal | 29.8 | 5984.297 | 9.02 |
| birds | Ploceus_nigerrimus | Ploceus_nigerrimus | LC | Stable | tropical | diurnal | 34.3 | 1840056.418 | 7.903 |
| birds | Ploceus_nigricollis | Ploceus_nigricollis | LC | Stable | tropical | diurnal | 27.1 | 2474612.529 | 8.863 |
| birds | Ploceus_nigrimentus | Ploceus_nigrimentus | LC | Stable | tropical | diurnal | 26.6 | 84397.42 | 7.303 |
| birds | Ploceus_ocularis | Ploceus_ocularis | LC | Stable | tropical | diurnal | 25.2 | 2998767.223 | 10.383 |
| birds | Ploceus_olivaceiceps | Ploceus_olivaceiceps | NT | Decreasing | tropical | diurnal | 20.8 | 233911.572 | 12.037 |
| birds | Ploceus_pelzelni | Ploceus_pelzelni | LC | Stable | tropical | diurnal | 12.7 | 559286.207 | 13.892 |
| birds | Ploceus_philippinus | Ploceus_philippinus | LC | Stable | tropical | diurnal | 28.2 | 6341236.33 | 15.258 |
| birds | Ploceus_preussi | Ploceus_preussi | LC | Stable | tropical | diurnal | 31.6 | 596680.152 | 7.637 |
| birds | Ploceus_princeps | Ploceus_princeps | LC | Increasing | tropical | diurnal | 31.2 | 146.654 | 14.707 |

| | | | | | | | | | |
|---|---|---|---|---|---|---|---|---|---|
| birds | Ploceus_reichardi | Ploceus_reichardi | LC | Decreasing | tropical | diurnal | 26.6 | 34740.2 | 7.513 |
| birds | Ploceus_rubiginosus | Ploceus_rubiginosus | LC | Stable | tropical | diurnal | 30 | 1228691.951 | 8.928 |
| birds | Ploceus_ruweti | Ploceus_ruweti | LC | Stable | tropical | diurnal | 17 | 2607.316 | 9.099 |
| birds | Ploceus_sakalava | Ploceus_sakalava | LC | Stable | tropical | diurnal | 23.8 | 268618.92 | 8.657 |
| birds | Ploceus_sanctithomae | Ploceus_sanctithomae | LC | Stable | tropical | diurnal | 22 | 858.514 | 12.812 |
| birds | Ploceus_spekei | Ploceus_spekei | LC | Stable | tropical | diurnal | 35.1 | 369188.997 | 11.411 |
| birds | Ploceus_spekeoides | Ploceus_spekeoides | NT | Decreasing | tropical | diurnal | 26.6 | 12674.616 | 14.525 |
| birds | Ploceus_subaureus | Ploceus_subaureus | LC | Stable | tropical | diurnal | 30.4 | 493658.92 | 11.471 |
| birds | Ploceus_subpersonatus | Ploceus_subpersonatus | VU | Decreasing | tropical | diurnal | 22.5 | 3562.655 | 11.601 |
| birds | Ploceus_superciliosus | Ploceus_superciliosus | LC | Stable | tropical | diurnal | 21.4 | 1002709.092 | 10.92 |
| birds | Ploceus_taeniopterus | Ploceus_taeniopterus | LC | Stable | tropical | diurnal | 17.6 | 187720.655 | 10.526 |
| birds | Ploceus_temporalis | Ploceus_temporalis | LC | Stable | tropical | diurnal | 35.8 | 77920.105 | 7.18 |
| birds | Ploceus_tricolor | Ploceus_tricolor | LC | Stable | tropical | diurnal | 36.8 | 1061576.05 | 10.512 |
| birds | Ploceus_velatus | Ploceus_velatus | LC | Stable | tropical | diurnal | 30.9 | 3539457.497 | 8.461 |
| birds | Ploceus_vitellinus | Ploceus_vitellinus | LC | Stable | tropical | diurnal | 20 | 4213194.469 | 11.107 |
| birds | Ploceus_weynsi | Ploceus_weynsi | LC | Stable | tropical | diurnal | 29 | 56949.544 | 18.877 |
| birds | Ploceus_xanthops | Ploceus_xanthops | LC | Stable | tropical | diurnal | 40.6 | 2485649.705 | 10.603 |
| birds | Ploceus_xanthopterus | Ploceus_xanthopterus | LC | Decreasing | tropical | diurnal | 22 | 299643.547 | 9.914 |
| birds | Pluvialis_apricaria | Pluvialis_apricaria | LC | Increasing | temperate | cathemeral | 214 | 4417307.887 | 9.227 |
| birds | Pluvialis_dominica | Pluvialis_dominica | LC | Decreasing | temperate | diurnal | 151.5 | 28684530.209 | 7.089 |
| birds | Pluvialis_fulva | Pluvialis_fulva | LC | Decreasing | temperate | diurnal | 134.9 | 8070887.146 | 7.583 |
| birds | Pluvialis_squatarola | Pluvialis_squatarola | LC | Decreasing | temperate | cathemeral | 250 | 15201972.98 | 8.403 |
| birds | Pluvianellus_socialis | Pluvianellus_socialis | VU | Stable | temperate | diurnal | 84.1 | 172032.985 | 7.491 |
| birds | Pluvianus_aegyptius | Pluvianus_aegyptius | LC | Decreasing | tropical | diurnal | 82 | 5977374.573 | 10.698 |
| birds | Pnoepyga_albiventer | Pnoepyga_albiventer | LC | Unknown | temperate | diurnal | 20.9 | 588072.588 | 8.539 |
| birds | Pnoepyga_immaculata | Pnoepyga_immaculata | LC | Stable | temperate | diurnal | 20.5 | 97921.23 | 13.072 |
| birds | Pnoepyga_pusilla | Pnoepyga_pusilla | LC | Decreasing | temperate | diurnal | 12 | 3686790.812 | 12.038 |
| birds | Podargus_ocellatus | Podargus_ocellatus | LC | Decreasing | tropical | nocturnal | 175.4 | 761242.934 | 3.544 |
| birds | Podargus_papuensis | Podargus_papuensis | LC | Stable | tropical | cathemeral | 375.7 | 958302.867 | 3.292 |
| birds | Podargus_strigoides | Podargus_strigoides | LC | Stable | temperate | nocturnal | 308 | 7674636.859 | 4.261 |
| birds | Podica_senegalensis | Podica_senegalensis | LC | Decreasing | tropical | diurnal | 599 | 11384786.124 | 9.256 |
| birds | Podiceps_auritus | Podiceps_auritus | VU | Decreasing | temperate | diurnal | 453 | 23714565.82 | 7.477 |
| birds | Podiceps_cristatus | Podiceps_cristatus | LC | Unknown | temperate | diurnal | 731 | 35889933.129 | 11.407 |
| birds | Podiceps_gallardoi | Podiceps_gallardoi | CR | Stable | temperate | diurnal | 575 | 106791.025 | 6.211 |

| | | | | | | | | | |
|---|---|---|---|---|---|---|---|---|---|
| birds | Podiceps_grisegena | Podiceps_grisegena | LC | Decreasing | temperate | diurnal | 1023 | 23965746.916 | 6.698 |
| birds | Podiceps_major | Podiceps_major | LC | Stable | temperate | diurnal | 1646 | 2909991.501 | 8.226 |
| birds | Podiceps_nigricollis | Podiceps_nigricollis | LC | Unknown | temperate | diurnal | 412.4 | 23548498.593 | 11.178 |
| birds | Podiceps_occipitalis | Podiceps_occipitalis | LC | Decreasing | temperate | diurnal | 320.7 | 2692417.519 | 7.668 |
| birds | Podiceps_taczanowskii | Podiceps_taczanowskii | EN | Decreasing | tropical | diurnal | 390.2 | 145.944 | 2.348 |
| birds | Podilymbus_podiceps | Podilymbus_podiceps | LC | Increasing | tropical | diurnal | 411.9 | 21861728.489 | 10.288 |
| birds | Podoces_biddulphi | Podoces_biddulphi | NT | Decreasing | temperate | diurnal | 130.5 | 420108.258 | 2.554 |
| birds | Podoces_hendersoni | Podoces_hendersoni | LC | Decreasing | temperate | diurnal | 120.5 | 1783211.453 | 3.376 |
| birds | Podoces_panderi | Podoces_panderi | LC | Decreasing | temperate | diurnal | 91.1 | 291415.991 | 5.895 |
| birds | Podoces_pleskei | Podoces_pleskei | LC | Stable | temperate | diurnal | 87.5 | 219917.486 | 5.009 |
| birds | Poecile_atricapillus | Parus_atricapillus | LC | Increasing | temperate | diurnal | 10.8 | 9078514.177 | 8.699 |
| birds | Poecile_carolinensis | Parus_carolinensis | LC | Stable | temperate | diurnal | 10 | 2312694.524 | 17.157 |
| birds | Poecile_cinctus | Parus_cinctus | LC | Decreasing | temperate | diurnal | 11.4 | 10800997.614 | 1.677 |
| birds | Poecile_davidi | Parus_davidi | LC | Decreasing | temperate | diurnal | 10.3 | 56804.656 | 6.254 |
| birds | Poecile_gambeli | Parus_gambeli | LC | Stable | temperate | diurnal | 11.2 | 2711124.484 | 7.933 |
| birds | Poecile_hudsonicus | Parus_hudsonicus | LC | Decreasing | temperate | diurnal | 9.8 | 7102487.578 | 2.04 |
| birds | Poecile_lugubris | Parus_lugubris | LC | Stable | temperate | diurnal | 15.8 | 1281185.716 | 15.705 |
| birds | Poecile_montanus | Parus_montanus | LC | Decreasing | temperate | diurnal | 11.1 | 21263364.575 | 8.326 |
| birds | Poecile_palustris | Parus_palustris | LC | Decreasing | temperate | diurnal | 11.1 | 9072135.09 | 14.027 |
| birds | Poecile_rufescens | Parus_rufescens | LC | Increasing | temperate | diurnal | 9.4 | 958071.951 | 6.665 |
| birds | Poecile_sclateri | Parus_sclateri | LC | Decreasing | tropical | diurnal | 10.1 | 201847.75 | 7.578 |
| birds | Poecile_superciliosus | Parus_superciliosus | LC | Stable | temperate | diurnal | 10.7 | 546757.409 | 9.854 |
| birds | Poecilodryas_brachyura | Poecilodryas_brachyura | LC | Stable | tropical | diurnal | 25.1 | 113208.308 | 4.257 |
| birds | Poecilodryas_hypoleuca | Poecilodryas_hypoleuca | LC | Stable | tropical | diurnal | 18 | 419659.376 | 2.985 |
| birds | Poecilodryas_superciliosa | Poecilodryas_superciliosa | LC | Increasing | tropical | diurnal | 19.8 | 238285.191 | 3.452 |
| birds | Poecilotriccus_albifacies | Poecilotriccus_albifacies | LC | Decreasing | tropical | diurnal | 8 | 4050.694 | 2.684 |
| birds | Poecilotriccus_calopterus | Poecilotriccus_calopterus | LC | Decreasing | tropical | diurnal | 7.5 | 208588.827 | 2.915 |
| birds | Poecilotriccus_capitalis | Poecilotriccus_capitalis | LC | Decreasing | tropical | diurnal | 8 | 525289.167 | 2.737 |
| birds | Poecilotriccus_fumifrons | Poecilotriccus_fumifrons | LC | Decreasing | tropical | diurnal | 6.6 | 2055264.876 | 3.494 |
| birds | Poecilotriccus_latirostris | Poecilotriccus_latirostris | LC | Stable | tropical | diurnal | 8.1 | 4661103.333 | 4.03 |
| birds | Poecilotriccus_luluae | Poecilotriccus_luluae | EN | Decreasing | tropical | diurnal | 7.3 | 361.84 | 3.138 |
| birds | Poecilotriccus_plumbeiceps | Poecilotriccus_plumbeiceps | LC | Stable | tropical | diurnal | 5.7 | 1475066.736 | 11.921 |
| birds | Poecilotriccus_pulchellus | Poecilotriccus_pulchellus | LC | Decreasing | tropical | diurnal | 8 | 6877.805 | 2.031 |
| birds | Poecilotriccus_ruficeps | Poecilotriccus_ruficeps | LC | Decreasing | tropical | diurnal | 8.8 | 129577.782 | 10.754 |

| | | | | | | | | | |
|---|---|---|---|---|---|---|---|---|---|
| birds | Poecilotriccus_russatus | Poecilotriccus_russatus | LC | Stable | tropical | diurnal | 8 | 28836.767 | 2.269 |
| birds | Poecilotriccus_senex | NA | LC | Stable | NA | NA | NA | NA | NA |
| birds | Poecilotriccus_sylvia | Poecilotriccus_sylvia | LC | Decreasing | tropical | diurnal | 7.1 | 1909899.709 | 7.759 |
| birds | Poeoptera_kenricki | Poeoptera_kenricki | LC | Decreasing | tropical | diurnal | 46.4 | 203009.077 | 12.708 |
| birds | Poeoptera_lugubris | Poeoptera_lugubris | LC | Decreasing | tropical | diurnal | 38.8 | 611884.694 | 9.921 |
| birds | Poeoptera_stuhlmanni | Poeoptera_stuhlmanni | LC | Decreasing | tropical | diurnal | 39.3 | 175255.699 | 14.79 |
| birds | Poephila_acuticauda | Poephila_acuticauda | LC | Stable | tropical | diurnal | 14 | 844828.161 | 2.284 |
| birds | Poephila_cincta | Poephila_cincta | LC | Decreasing | tropical | diurnal | 14.9 | 826709.363 | 5.091 |
| birds | Poephila_personata | Poephila_personata | LC | Stable | tropical | diurnal | 13 | 972681.315 | 2.501 |
| birds | Pogoniulus_atroflavus | Pogoniulus_atroflavus | LC | Stable | tropical | diurnal | 18.2 | 3231917.567 | 7.932 |
| birds | Pogoniulus_bilineatus | Pogoniulus_bilineatus | LC | Stable | tropical | diurnal | 13.1 | 7153853.604 | 9.007 |
| birds | Pogoniulus_chrysoconus | Pogoniulus_chrysoconus | LC | Stable | tropical | diurnal | 10.8 | 10202745.337 | 9.851 |
| birds | Pogoniulus_coryphaea | Pogoniulus_coryphaeus | LC | Decreasing | tropical | diurnal | 10.7 | 307301.792 | 12.293 |
| birds | Pogoniulus_leucomystax | Pogoniulus_leucomystax | LC | Stable | tropical | diurnal | 11.8 | 552561.746 | 12.317 |
| birds | Pogoniulus_pusillus | Pogoniulus_pusillus | LC | Stable | tropical | diurnal | 9.6 | 2446279.621 | 10.944 |
| birds | Pogoniulus_scolopaceus | Pogoniulus_scolopaceus | LC | Decreasing | tropical | diurnal | 14.1 | 3952077.919 | 8.301 |
| birds | Pogoniulus_simplex | Pogoniulus_simplex | LC | Decreasing | tropical | diurnal | 8.8 | 519676.857 | 10.207 |
| birds | Pogoniulus_subsulphureus | Pogoniulus_subsulphureus | LC | Stable | tropical | diurnal | 10.5 | 3512045.742 | 8.388 |
| birds | Pogonocichla_stellata | Pogonocichla_stellata | LC | Decreasing | temperate | diurnal | 18.6 | 1563462.547 | 12.279 |
| birds | Pogonornis_bidentatus | Lybius_bidentatus | LC | Stable | tropical | diurnal | 79.3 | 2585581.976 | 10.964 |
| birds | Pogonornis_dubius | Lybius_dubius | LC | Unknown | tropical | diurnal | 91 | 2333237.283 | 12.79 |
| birds | Pogonornis_melanopterus | Lybius_melanopterus | LC | Unknown | tropical | diurnal | 52.4 | 809569.533 | 10.149 |
| birds | Pogonornis_minor | Lybius_minor | LC | Stable | tropical | diurnal | 46.7 | 750053.05 | 7.465 |
| birds | Pogonornis_rolleti | Lybius_rolleti | LC | Decreasing | tropical | diurnal | 100.4 | 1106455.935 | 9.668 |
| birds | Pogonotriccus_chapmani | Phylloscartes_chapmani | LC | Decreasing | tropical | diurnal | 8 | 175661.162 | 0.783 |
| birds | Pogonotriccus_eximius | Phylloscartes_eximius | NT | Decreasing | temperate | diurnal | 7.5 | 1008457.467 | 13.51 |
| birds | Pogonotriccus_lanyoni | Phylloscartes_lanyoni | EN | Decreasing | tropical | diurnal | 8 | 7102.551 | 11.208 |
| birds | Pogonotriccus_ophthalmicus | Phylloscartes_ophthalmicus | LC | Stable | tropical | diurnal | 9.2 | 179706.353 | 7.665 |
| birds | Pogonotriccus_orbitalis | Phylloscartes_orbitalis | LC | Decreasing | tropical | diurnal | 7.1 | 121722.451 | 4.835 |
| birds | Pogonotriccus_poecilotis | Phylloscartes_poecilotis | LC | Stable | tropical | diurnal | 8 | 213006.739 | 7.561 |
| birds | Pogonotriccus_venezuelanus | Phylloscartes_venezuelanus | NT | Decreasing | tropical | diurnal | 9 | 10622.116 | 19.991 |
| birds | Poicephalus_crassus | NA | LC | Stable | NA | NA | NA | NA | NA |
| birds | Poicephalus_cryptoxanthus | Poicephalus_cryptoxanthus | LC | Decreasing | tropical | diurnal | 140 | 1182606.673 | 9.699 |
| birds | Poicephalus_flavifrons | Poicephalus_flavifrons | LC | Stable | tropical | diurnal | 174 | 205738.935 | 13.387 |

| | | | | | | | | | |
|---|---|---|---|---|---|---|---|---|---|
| birds | Poicephalus_gulielmi | Poicephalus_gulielmi | LC | Decreasing | tropical | diurnal | 274.6 | 1483974.657 | 7.492 |
| birds | Poicephalus_meyeri | Poicephalus_meyeri | LC | Stable | tropical | diurnal | 117.5 | 5835819.268 | 9.028 |
| birds | Poicephalus_robustus | Poicephalus_robustus | VU | Stable | temperate | diurnal | 343 | 95197.084 | 13.744 |
| birds | Poicephalus_rueppellii | Poicephalus_rueppellii | LC | Decreasing | tropical | diurnal | 116 | 597941.08 | 6.737 |
| birds | Poicephalus_rufiventris | Poicephalus_rufiventris | LC | Stable | tropical | diurnal | 120 | 1544074.334 | 9.141 |
| birds | Poicephalus_senegalus | Poicephalus_senegalus | LC | Decreasing | tropical | diurnal | 147 | 2714553.583 | 13.456 |
| birds | Poikilocarbo_gaimardi | Phalacrocorax_gaimardi | NT | Decreasing | temperate | diurnal | 1358.3 | 529277.902 | 6.423 |
| birds | Polemaetus_bellicosus | Polemaetus_bellicosus | EN | Decreasing | tropical | diurnal | 3964.5 | 14888019.222 | 9.473 |
| birds | Polihierax_semitorquatus | Polihierax_semitorquatus | LC | Stable | tropical | diurnal | 61.5 | 3103663.96 | 8.766 |
| birds | Poliocephalus_poliocephalus | Poliocephalus_poliocephalus | LC | Stable | temperate | diurnal | 239.9 | 7046975.371 | 4.486 |
| birds | Poliocephalus_rufopectus | Poliocephalus_rufopectus | LC | Increasing | temperate | diurnal | 249 | 173213.589 | 7.66 |
| birds | Poliocrania_exsul | Myrmeciza_exsul | LC | Decreasing | tropical | diurnal | 26.5 | 180127.532 | 8.32 |
| birds | Poliolais_lopezi | Poliolais_lopezi | LC | Decreasing | tropical | diurnal | 12 | 23329.928 | 9.196 |
| birds | Poliolophus_urostictus | Pycnonotus_urostictus | LC | Stable | tropical | diurnal | 24.8 | 207679.007 | 12.762 |
| birds | Polioptila_albiloris | Polioptila_albiloris | LC | Decreasing | tropical | diurnal | 6.3 | 221828.648 | 11.928 |
| birds | Polioptila_caerulea | Polioptila_caerulea | LC | Increasing | temperate | diurnal | 5.8 | 7262670.826 | 13.388 |
| birds | Polioptila_californica | Polioptila_californica | LC | Decreasing | temperate | diurnal | 5.8 | 135910.747 | 6.192 |
| birds | Polioptila_dumicola | Polioptila_dumicola | LC | Decreasing | tropical | diurnal | 7 | 4483982.866 | 8.482 |
| birds | Polioptila_guianensis | Polioptila_guianensis | LC | Stable | tropical | diurnal | 6 | 3402937.826 | 1.802 |
| birds | Polioptila_lactea | Polioptila_lactea | LC | Decreasing | temperate | diurnal | 6.5 | 294003.08 | 12.564 |
| birds | Polioptila_lembeyei | Polioptila_lembeyei | LC | Decreasing | tropical | diurnal | 5 | 13553.509 | 14.274 |
| birds | Polioptila_maior | NA | LC | Decreasing | NA | NA | NA | NA | NA |
| birds | Polioptila_nigriceps | Polioptila_nigriceps | LC | Decreasing | temperate | diurnal | 5.6 | 222914.087 | 8.413 |
| birds | Polioptila_plumbea | Polioptila_plumbea | LC | Decreasing | tropical | diurnal | 6.4 | 7562341.501 | 4.953 |
| birds | Polioptila_schistaceigula | Polioptila_schistaceigula | LC | Decreasing | tropical | diurnal | 6 | 144511.456 | 7.714 |
| birds | Polioxolmis_rufipennis | Polioxolmis_rufipennis | LC | Decreasing | tropical | diurnal | 34.5 | 402534.915 | 9.112 |
| birds | Polyboroides_radiatus | NA | LC | Stable | NA | NA | NA | NA | NA |
| birds | Polyboroides_typus | Polyboroides_typus | LC | Stable | tropical | diurnal | 633.2 | 14893143.158 | 9.201 |
| birds | Polyerata_amabilis | Amazilia_amabilis | LC | Decreasing | tropical | diurnal | 4.3 | 298612.116 | 8.972 |
| birds | Polyerata_rosenbergi | Amazilia_rosenbergi | LC | Stable | tropical | diurnal | 3.7 | 76061.183 | 5.962 |
| birds | Polyonymus_caroli | Polyonymus_caroli | LC | Stable | tropical | diurnal | 4.8 | 159552.964 | 8.068 |
| birds | Polyplectron_bicalcaratum | Polyplectron_bicalcaratum | LC | Decreasing | tropical | diurnal | 467.8 | 1095333.727 | 8.326 |
| birds | Polyplectron_chalcurum | Polyplectron_chalcurum | LC | Decreasing | tropical | diurnal | 508 | 83208.603 | 8.09 |
| birds | Polyplectron_germaini | Polyplectron_germaini | VU | Decreasing | tropical | diurnal | 450 | 117609.349 | 13.85 |

| | | | | | | | | | |
|---|---|---|---|---|---|---|---|---|---|
| birds | Polyplectron_inopinatum | Polyplectron_inopinatum | NT | Decreasing | tropical | diurnal | 497.5 | 10661.169 | 4.533 |
| birds | Polyplectron_malacense | Polyplectron_malacense | EN | Decreasing | tropical | diurnal | 608 | 47675.752 | 5.496 |
| birds | Polyplectron_napoleonis | Polyplectron_napoleonis | VU | Decreasing | tropical | diurnal | 374.7 | 11490.413 | 9.023 |
| birds | Polyplectron_schleiermacheri | NA | EN | Decreasing | NA | NA | NA | NA | NA |
| birds | Polysticta_stelleri | Polysticta_stelleri | VU | Decreasing | temperate | diurnal | 806.8 | 3544014.159 | 0.908 |
| birds | Polystictus_pectoralis | Polystictus_pectoralis | NT | Decreasing | temperate | diurnal | 6.2 | 4115700.934 | 10.03 |
| birds | Polystictus_superciliaris | Polystictus_superciliaris | LC | Decreasing | tropical | diurnal | 6 | 313983.578 | 10.767 |
| birds | Polytelis_alexandrae | Polytelis_alexandrae | NT | Stable | temperate | diurnal | 103.7 | 1852628.362 | 1.47 |
| birds | Polytelis_anthopeplus | Polytelis_anthopeplus | LC | Decreasing | temperate | diurnal | 170 | 387356.366 | 9.278 |
| birds | Polytelis_swainsonii | Polytelis_swainsonii | LC | Increasing | temperate | diurnal | 153 | 229296.888 | 10.885 |
| birds | Polytmus_guainumbi | Polytmus_guainumbi | LC | Unknown | tropical | diurnal | 5.2 | 4835885.747 | 7.789 |
| birds | Polytmus_milleri | Polytmus_milleri | LC | Unknown | tropical | diurnal | 4.1 | 74967.653 | 1.13 |
| birds | Polytmus_theresiae | Polytmus_theresiae | LC | Unknown | tropical | diurnal | 3.8 | 1942723.666 | 1.414 |
| birds | Pomarea_dimidiata | Pomarea_dimidiata | VU | Increasing | tropical | diurnal | 23 | 98.98 | 15.392 |
| birds | Pomarea_iphis | Pomarea_iphis | CR | Stable | tropical | diurnal | 23.9 | 126.477 | 6.389 |
| birds | Pomarea_mendozae | Pomarea_mendozae | EN | Stable | tropical | diurnal | 29.6 | 849.802 | 7.712 |
| birds | Pomarea_nigra | Pomarea_nigra | CR | Increasing | tropical | diurnal | 25.4 | 57.951 | 14.86 |
| birds | Pomarea_whitneyi | Pomarea_whitneyi | CR | Decreasing | tropical | diurnal | 42.5 | 4.029 | 8.258 |
| birds | Pomatorhinus_ferruginosus | Pomatorhinus_ferruginosus | LC | Decreasing | temperate | diurnal | 40 | 130932.122 | 5.88 |
| birds | Pomatorhinus_horsfieldii | Pomatorhinus_horsfieldii | LC | Increasing | tropical | diurnal | 43 | 1024241.881 | 17.386 |
| birds | Pomatorhinus_montanus | Pomatorhinus_montanus | VU | Decreasing | tropical | diurnal | 28.6 | 12471.672 | 13.826 |
| birds | Pomatorhinus_ochraceiceps | Pomatorhinus_ochraceiceps | LC | Decreasing | temperate | diurnal | 34 | 662845.401 | 7.947 |
| birds | Pomatorhinus_ruficollis | Pomatorhinus_ruficollis | LC | Stable | temperate | diurnal | 31.6 | 2706746.888 | 12.341 |
| birds | Pomatorhinus_schisticeps | Pomatorhinus_schisticeps | LC | Decreasing | tropical | diurnal | 43 | 1222844.413 | 8.938 |
| birds | Pomatorhinus_superciliaris | Xiphirhynchus_superciliaris | LC | Stable | temperate | diurnal | 28 | 664803.498 | 7.986 |
| birds | Pomatostomus_halli | Pomatostomus_halli | LC | Decreasing | temperate | diurnal | 41.8 | 487442.038 | 4.814 |
| birds | Pomatostomus_ruficeps | Pomatostomus_ruficeps | LC | Stable | temperate | diurnal | 57.8 | 1074854.311 | 6.296 |
| birds | Pomatostomus_superciliosus | Pomatostomus_superciliosus | LC | Decreasing | temperate | diurnal | 41.5 | 4621673.341 | 4.769 |
| birds | Pomatostomus_temporalis | Pomatostomus_temporalis | LC | Decreasing | temperate | diurnal | 75 | 3774038.587 | 4.491 |
| birds | Poodytes_albolimbatus | Megalurus_albolimbatus | VU | Decreasing | tropical | diurnal | 14 | 2389.389 | 4.036 |
| birds | Poodytes_carteri | Eremiornis_carteri | LC | Decreasing | tropical | diurnal | 12.2 | 1881674.614 | 2.248 |
| birds | Poodytes_gramineus | Megalurus_gramineus | LC | Increasing | temperate | diurnal | 14.1 | 3277909.054 | 6.868 |
| birds | Poodytes_punctatus | Bowdleria_punctata | LC | Decreasing | temperate | diurnal | 25.2 | 196470.834 | 7.522 |
| birds | Pooecetes_gramineus | Pooecetes_gramineus | LC | Decreasing | temperate | diurnal | 25.7 | 11093552.212 | 12.209 |

| | | | | | | | | | |
|---|---|---|---|---|---|---|---|---|---|
| birds | Poospiza_baeri | Compsospiza_baeri | VU | Decreasing | temperate | diurnal | 32.5 | 24723.162 | 4.454 |
| birds | Poospiza_boliviana | Poospiza_boliviana | LC | Stable | tropical | diurnal | 28.6 | 65161.27 | 9.443 |
| birds | Poospiza_garleppi | NA | NT | Decreasing | NA | NA | NA | NA | NA |
| birds | Poospiza_goeringi | Hemispingus_goeringi | NT | Stable | tropical | diurnal | 17.1 | 3842.505 | 10.428 |
| birds | Poospiza_hispaniolensis | Poospiza_hispaniolensis | LC | Stable | tropical | diurnal | 16.5 | 115454.538 | 8.573 |
| birds | Poospiza_nigrorufa | Poospiza_nigrorufa | LC | Stable | temperate | diurnal | 17.4 | 952615.997 | 11.09 |
| birds | Poospiza_ornata | Poospiza_ornata | LC | Stable | temperate | diurnal | 11.8 | 708515.274 | 7.205 |
| birds | Poospiza_rubecula | Poospiza_rubecula | EN | Decreasing | tropical | diurnal | 24.4 | 16057.935 | 11.472 |
| birds | Poospiza_rufosuperciliaris | Hemispingus_rufosuperciliaris | LC | Unknown | tropical | diurnal | 29 | 8368.583 | 4.923 |
| birds | Poospiza_whitii | Poospiza_whitii | LC | Stable | tropical | diurnal | 17 | 294938.889 | 7.136 |
| birds | Poospizopsis_caesar | Poospiza_caesar | LC | Stable | tropical | diurnal | 51.2 | 32343.462 | 9.224 |
| birds | Poospizopsis_hypocondria | Poospiza_hypochondria | LC | Stable | temperate | diurnal | 21.2 | 286396.55 | 7.504 |
| birds | Porphyrio_alleni | Porphyrio_alleni | LC | Decreasing | tropical | cathemeral | 138.7 | 16360428.259 | 9.21 |
| birds | Porphyrio_flavirostris | Porphyrio_flavirostris | LC | Unknown | tropical | diurnal | 92.6 | 5270055.286 | 4.631 |
| birds | Porphyrio_mantelli | NA | EX | not_available | NA | NA | NA | NA | NA |
| birds | Porphyrio_martinicus | Porphyrio_martinica | LC | Decreasing | tropical | diurnal | 235.1 | 16405883.334 | 6.808 |
| birds | Porphyrio_porphyrio | Porphyrio_porphyrio | LC | Unknown | tropical | nocturnal | 773.9 | 18686249.252 | 10.991 |
| birds | Porphyriops_melanops | Gallinula_melanops | LC | Stable | tropical | diurnal | 191 | 4136401.213 | 10.223 |
| birds | Porphyrolaema_porphyrolaema | Porphyrolaema_porphyrolaema | LC | Decreasing | tropical | diurnal | 60 | 2187394.02 | 1.339 |
| birds | Porphyrospiza_caerulescens | Porphyrospiza_caerulescens | NT | Decreasing | tropical | diurnal | 13 | 1671290.255 | 8.639 |
| birds | Porzana_albicollis | Porzana_albicollis | LC | Unknown | tropical | diurnal | 100 | 6152059.487 | 8.338 |
| birds | Porzana_carolina | Porzana_carolina | LC | Increasing | temperate | diurnal | 74.8 | 16460975.591 | 10.147 |
| birds | Porzana_fasciata | Anurolimnas_fasciatus | LC | Unknown | tropical | diurnal | 69 | 1297085.555 | 1.261 |
| birds | Porzana_fluminea | Porzana_fluminea | LC | Unknown | temperate | diurnal | 60.8 | 1975889.794 | 6.652 |
| birds | Premnoplex_brunnescens | Premnoplex_brunnescens | LC | Decreasing | tropical | diurnal | 16.3 | 406258.31 | 8.138 |
| birds | Premnoplex_tatei | NA | EN | Decreasing | NA | NA | NA | NA | NA |
| birds | Premnornis_guttuliger | Premnornis_guttuligera | LC | Stable | tropical | diurnal | 15.9 | 289937.61 | 8.032 |
| birds | Primolius_auricollis | Primolius_auricollis | LC | Increasing | tropical | diurnal | 245 | 838451.741 | 5.418 |
| birds | Primolius_couloni | Primolius_couloni | VU | Decreasing | tropical | diurnal | 250 | 729954.854 | 2.792 |
| birds | Primolius_maracana | Primolius_maracana | NT | Decreasing | tropical | diurnal | 256 | 5423813.074 | 8.024 |
| birds | Prinia_atrogularis | Prinia_atrogularis | LC | Stable | temperate | diurnal | 11.8 | 90149.057 | 6.524 |
| birds | Prinia_bairdii | Prinia_bairdii | LC | Stable | tropical | diurnal | 13.4 | 1110524.41 | 7.821 |
| birds | Prinia_buchanani | Prinia_buchanani | LC | Stable | temperate | diurnal | 6.7 | 1899856.568 | 17.292 |
| birds | Prinia_cinereocapilla | Prinia_cinereocapilla | VU | Decreasing | temperate | diurnal | 7.3 | 33032.598 | 14.401 |

| | | | | | | | | | |
|---|---|---|---|---|---|---|---|---|---|
| birds | Prinia_crinigera | Prinia_crinigera | LC | Stable | temperate | diurnal | 13.7 | 2843218.329 | 13.275 |
| birds | Prinia_erythroptera | Heliolais_erythropterus | LC | Stable | tropical | diurnal | 11.5 | 3486114.044 | 11.609 |
| birds | Prinia_familiaris | Prinia_familiaris | NT | Decreasing | tropical | diurnal | 9 | 497167.738 | 13.182 |
| birds | Prinia_flavicans | Prinia_flavicans | LC | Stable | tropical | diurnal | 8.7 | 2717287.687 | 6.859 |
| birds | Prinia_flaviventris | Prinia_flaviventris | LC | Decreasing | tropical | diurnal | 7 | 4037497.014 | 11.401 |
| birds | Prinia_fluviatilis | Prinia_fluviatilis | LC | Stable | tropical | diurnal | 8 | 98560.021 | 9.287 |
| birds | Prinia_gracilis | Prinia_gracilis | LC | Stable | temperate | diurnal | 7.1 | 4034571.259 | 12.895 |
| birds | Prinia_hodgsonii | Prinia_hodgsonii | LC | Stable | tropical | diurnal | 6.4 | 4680228.234 | 16.12 |
| birds | Prinia_inornata | Prinia_inornata | LC | Stable | tropical | diurnal | 7 | 7403210.361 | 15.805 |
| birds | Prinia_maculosa | Prinia_maculosa | LC | Decreasing | temperate | diurnal | 9 | 581622.97 | 8.952 |
| birds | Prinia_molleri | Prinia_molleri | LC | Increasing | tropical | diurnal | 8.8 | 858.514 | 12.812 |
| birds | Prinia_polychroa | Prinia_polychroa | LC | Stable | tropical | diurnal | 11 | 734233.978 | 11.659 |
| birds | Prinia_rufescens | Prinia_rufescens | LC | Stable | tropical | diurnal | 6.5 | 2213362.616 | 11.648 |
| birds | Prinia_rufifrons | Spiloptila_rufifrons | LC | Stable | tropical | diurnal | 6.5 | 1242957.363 | 8.215 |
| birds | Prinia_socialis | Prinia_socialis | LC | Stable | tropical | diurnal | 8 | 2758579.476 | 18.74 |
| birds | Prinia_somalica | Prinia_somalica | LC | Stable | tropical | diurnal | 7.5 | 1273080.064 | 7.977 |
| birds | Prinia_subflava | Prinia_subflava | LC | Stable | tropical | diurnal | 8.6 | 14611740.022 | 9.991 |
| birds | Prinia_sylvatica | Prinia_sylvatica | LC | Decreasing | tropical | diurnal | 16.1 | 2267095.605 | 18.737 |
| birds | Prioniturus_discurus | Prioniturus_discurus | LC | Stable | tropical | diurnal | 148.1 | 171737.972 | 13.472 |
| birds | Prioniturus_flavicans | Prioniturus_flavicans | NT | Decreasing | tropical | diurnal | 116.8 | 19945.559 | 9.401 |
| birds | Prioniturus_luconensis | Prioniturus_luconensis | EN | Decreasing | tropical | diurnal | 108 | 93519.259 | 14.928 |
| birds | Prioniturus_mada | Prioniturus_mada | LC | Stable | tropical | diurnal | 116.8 | 6285.537 | 5.988 |
| birds | Prioniturus_montanus | Prioniturus_montanus | NT | Decreasing | tropical | diurnal | 118.3 | 10411.713 | 5.379 |
| birds | Prioniturus_platenae | Prioniturus_platenae | VU | Decreasing | tropical | diurnal | 112 | 13583.923 | 9.357 |
| birds | Prioniturus_platurus | Prioniturus_platurus | LC | Decreasing | tropical | cathemeral | 66.3 | 184161.11 | 9.27 |
| birds | Prioniturus_verticalis | Prioniturus_verticalis | CR | Decreasing | tropical | diurnal | 45 | 813.767 | 7.651 |
| birds | Prioniturus_waterstradti | Prioniturus_waterstradti | NT | Decreasing | tropical | diurnal | 119 | 11282.822 | 6.872 |
| birds | Prionochilus_maculatus | Prionochilus_maculatus | LC | Stable | tropical | diurnal | 8 | 1401405.5 | 7.972 |
| birds | Prionochilus_olivaceus | Prionochilus_olivaceus | LC | Stable | tropical | diurnal | 9.4 | 226152.004 | 13.379 |
| birds | Prionochilus_percussus | Prionochilus_percussus | LC | Stable | tropical | diurnal | 8.5 | 1530525.146 | 9.261 |
| birds | Prionochilus_plateni | Prionochilus_plateni | LC | Decreasing | tropical | diurnal | 8.5 | 13498.605 | 9.258 |
| birds | Prionochilus_thoracicus | Prionochilus_thoracicus | NT | Decreasing | tropical | diurnal | 8.9 | 1166737.855 | 8.03 |
| birds | Prionochilus_xanthopygius | Prionochilus_xanthopygius | LC | Stable | tropical | diurnal | 7.6 | 383880.169 | 4.963 |
| birds | Prionodura_newtoniana | Prionodura_newtoniana | NT | Decreasing | tropical | diurnal | 78.3 | 5474.012 | 3.373 |

| | | | | | | | | | |
|---|---|---|---|---|---|---|---|---|---|
| birds | Prionops_alberti | Prionops_alberti | VU | Decreasing | tropical | diurnal | 61.3 | 33733.367 | 13.197 |
| birds | Prionops_caniceps | Prionops_caniceps | LC | Decreasing | tropical | diurnal | 48.2 | 892308.986 | 14.83 |
| birds | Prionops_gabela | NA | EN | Decreasing | NA | NA | NA | NA | NA |
| birds | Prionops_plumatus | Prionops_plumatus | LC | Stable | tropical | diurnal | 33.8 | 10670489.609 | 9.625 |
| birds | Prionops_poliolophus | Prionops_poliolophus | NT | Decreasing | tropical | diurnal | 49 | 55433.142 | 17.234 |
| birds | Prionops_retzii | Prionops_retzii | LC | Decreasing | tropical | diurnal | 50.1 | 4451615.159 | 8.343 |
| birds | Prionops_rufiventris | Prionops_rufiventris | LC | Decreasing | tropical | diurnal | 50 | 1903359.865 | 5.823 |
| birds | Prionops_scopifrons | Prionops_scopifrons | LC | Decreasing | tropical | diurnal | 29.4 | 827650.544 | 9.843 |
| birds | Priotelus_temnurus | Priotelus_temnurus | LC | Decreasing | tropical | diurnal | 53.4 | 108974.518 | 16.183 |
| birds | Probosciger_aterrimus | Probosciger_aterrimus | NT | Decreasing | tropical | diurnal | 841 | 715363.09 | 3.07 |
| birds | Procarduelis_nipalensis | Carpodacus_nipalensis | LC | Stable | temperate | diurnal | 22.1 | 713091.482 | 8.759 |
| birds | Procellaria_aequinoctialis | Procellaria_aequinoctialis | VU | Decreasing | temperate | cathemeral | 1213 | 107346797.963 | 7.925 |
| birds | Procellaria_cinerea | Procellaria_cinerea | NT | Decreasing | temperate | cathemeral | 1131 | 83049738.822 | 5.957 |
| birds | Procellaria_parkinsoni | Procellaria_parkinsoni | VU | Stable | tropical | cathemeral | 702.2 | 39453259.779 | 9.489 |
| birds | Procellaria_westlandica | Procellaria_westlandica | EN | Unknown | temperate | cathemeral | 1199 | 18867087.614 | 3.45 |
| birds | Procnias_albus | Procnias_albus | LC | Decreasing | tropical | diurnal | 210 | 527310.402 | 1.467 |
| birds | Procnias_averano | Procnias_averano | LC | Decreasing | tropical | diurnal | 146.1 | 1123871.614 | 7.171 |
| birds | Procnias_nudicollis | Procnias_nudicollis | NT | Decreasing | tropical | diurnal | 172 | 1131224.33 | 12.353 |
| birds | Procnias_tricarunculatus | Procnias_tricarunculatus | VU | Decreasing | tropical | diurnal | 210 | 155461.676 | 10.287 |
| birds | Prodotiscus_insignis | Prodotiscus_insignis | LC | Decreasing | tropical | diurnal | 10.4 | 3022105.251 | 9.245 |
| birds | Prodotiscus_regulus | Prodotiscus_regulus | LC | Increasing | tropical | diurnal | 14.1 | 3186750.811 | 10.311 |
| birds | Prodotiscus_zambesiae | Prodotiscus_zambesiae | LC | Stable | tropical | diurnal | 10.2 | 3997564.298 | 8.795 |
| birds | Progne_chalybea | Progne_chalybea | LC | Stable | tropical | diurnal | 50 | 15629572.136 | 6.315 |
| birds | Progne_cryptoleuca | Progne_cryptoleuca | LC | Stable | tropical | diurnal | 46 | 109839.23 | 16.136 |
| birds | Progne_dominicensis | Progne_dominicensis | LC | Increasing | tropical | diurnal | 39.6 | 102598.204 | 17.621 |
| birds | Progne_elegans | Progne_elegans | LC | Stable | tropical | diurnal | 50.6 | 5048767.485 | 4.573 |
| birds | Progne_modesta | NA | EN | Decreasing | NA | NA | NA | NA | NA |
| birds | Progne_murphyi | Progne_murphyi | NT | Decreasing | tropical | diurnal | 41.4 | 134702.657 | 7.336 |
| birds | Progne_sinaloae | Progne_sinaloae | VU | Decreasing | temperate | diurnal | 42.6 | 197014.514 | 7.11 |
| birds | Progne_subis | Progne_subis | LC | Stable | tropical | diurnal | 53.8 | 21193891.3 | 9.17 |
| birds | Progne_tapera | Progne_tapera | LC | Decreasing | tropical | diurnal | 32 | 14549505.198 | 6.065 |
| birds | Promerops_cafer | Promerops_cafer | LC | Stable | temperate | diurnal | 34.7 | 114533.352 | 10.664 |
| birds | Promerops_gurneyi | Promerops_gurneyi | NT | Decreasing | temperate | diurnal | 34.9 | 103702.815 | 12.488 |
| birds | Prosobonia_cancellata | NA | EX | not_available | NA | NA | NA | NA | NA |

| | | | | | | | | | |
|---|---|---|---|---|---|---|---|---|---|
| birds | Prosopeia_personata | Prosopeia_personata | NT | Decreasing | tropical | diurnal | 40.3 | 10553.159 | 9.269 |
| birds | Prosopeia_splendens | NA | NT | Unknown | NA | NA | NA | NA | NA |
| birds | Prosopeia_tabuensis | Prosopeia_tabuensis | LC | Decreasing | tropical | diurnal | 232 | 6621.965 | 9.841 |
| birds | Prosthemadera_novaeseelandiae | Prosthemadera_novaeseelandiae | LC | Decreasing | temperate | diurnal | 105.8 | 125476.533 | 9.676 |
| birds | Protonotaria_citrea | Protonotaria_citrea | LC | Decreasing | tropical | diurnal | 14.3 | 4050752.012 | 15.338 |
| birds | Prunella_atrogularis | Prunella_atrogularis | LC | Stable | temperate | diurnal | 18.5 | 2939605.659 | 8.74 |
| birds | Prunella_collaris | Prunella_collaris | LC | Stable | temperate | diurnal | 38.6 | 11553059.39 | 8.042 |
| birds | Prunella_fulvescens | Prunella_fulvescens | LC | Stable | temperate | diurnal | 18.9 | 3210424.28 | 5.895 |
| birds | Prunella_himalayana | Prunella_himalayana | LC | Stable | temperate | diurnal | 25.7 | 2117170.288 | 4.863 |
| birds | Prunella_immaculata | Prunella_immaculata | LC | Stable | temperate | diurnal | 20.7 | 859718.291 | 8.405 |
| birds | Prunella_koslowi | Prunella_koslowi | LC | Stable | temperate | diurnal | 17.5 | 721915.952 | 4.043 |
| birds | Prunella_modularis | Prunella_modularis | LC | Decreasing | temperate | diurnal | 20.2 | 9380149.326 | 14.431 |
| birds | Prunella_montanella | Prunella_montanella | LC | Stable | temperate | diurnal | 17 | 14996026.233 | 4.365 |
| birds | Prunella_ocularis | Prunella_ocularis | LC | Stable | temperate | diurnal | 22.5 | 753696.609 | 13.78 |
| birds | Prunella_rubeculoides | Prunella_rubeculoides | LC | Stable | temperate | diurnal | 23.9 | 1417873.848 | 6.046 |
| birds | Prunella_rubida | Prunella_rubida | LC | Stable | temperate | diurnal | 16.7 | 395971.399 | 17.648 |
| birds | Prunella_strophiata | Prunella_strophiata | LC | Stable | temperate | diurnal | 18.1 | 1390796.425 | 8.924 |
| birds | Psalidoprocne_albiceps | Psalidoprocne_albiceps | LC | Stable | tropical | diurnal | 11.1 | 1147033.422 | 12.103 |
| birds | Psalidoprocne_fuliginosa | Psalidoprocne_fuliginosa | LC | Stable | tropical | diurnal | 12.2 | 22148.509 | 8.98 |
| birds | Psalidoprocne_nitens | Psalidoprocne_nitens | LC | Decreasing | tropical | diurnal | 9.8 | 2890566.405 | 8.036 |
| birds | Psalidoprocne_obscura | Psalidoprocne_obscura | LC | Stable | tropical | diurnal | 9.4 | 1468905.411 | 14.214 |
| birds | Psaltria_exilis | Psaltria_exilis | LC | Decreasing | tropical | diurnal | 4 | 11281.753 | 16.124 |
| birds | Psaltriparus_minimus | Psaltriparus_minimus | LC | Decreasing | temperate | diurnal | 5.3 | 2692275.517 | 9.867 |
| birds | Psarisomus_dalhousiae | Psarisomus_dalhousiae | LC | Decreasing | tropical | diurnal | 67 | 1318582.998 | 9.125 |
| birds | Psarocolius_angustifrons | Psarocolius_angustifrons | LC | Decreasing | tropical | diurnal | 271.5 | 2224154.866 | 2.921 |
| birds | Psarocolius_atrovirens | Psarocolius_atrovirens | LC | Stable | tropical | diurnal | 152 | 139269.139 | 5.256 |
| birds | Psarocolius_bifasciatus | Psarocolius_bifasciatus | LC | Decreasing | tropical | diurnal | 335.7 | 5789777.685 | 2.438 |
| birds | Psarocolius_cassini | Psarocolius_cassini | VU | Decreasing | tropical | diurnal | 383.9 | 9315.659 | 4.414 |
| birds | Psarocolius_decumanus | Psarocolius_decumanus | LC | Decreasing | tropical | diurnal | 206.3 | 10475235.797 | 4.913 |
| birds | Psarocolius_guatimozinus | Psarocolius_guatimozinus | LC | Decreasing | tropical | diurnal | 405.3 | 107561.074 | 10.79 |
| birds | Psarocolius_montezuma | Psarocolius_montezuma | LC | Decreasing | tropical | diurnal | 309.9 | 497553.812 | 10.237 |
| birds | Psarocolius_viridis | Psarocolius_viridis | LC | Stable | tropical | diurnal | 287 | 5050471.199 | 1.729 |
| birds | Psarocolius_wagleri | Psarocolius_wagleri | LC | Decreasing | tropical | diurnal | 155.5 | 481772.49 | 9.13 |
| birds | Psarocolius_yuracares | NA | not_available | not_available | NA | NA | NA | NA | NA |

| | | | | | | | | | |
|---|---|---|---|---|---|---|---|---|---|
| birds | Pselliophorus_luteoviridis | NA | VU | Decreasing | NA | NA | NA | NA | NA |
| birds | Pselliophorus_tibialis | Pselliophorus_tibialis | LC | Stable | tropical | diurnal | 30 | 6558.822 | 6.561 |
| birds | Psephotellus_chrysopterygius | Psephotus_chrysopterygius | EN | Decreasing | tropical | diurnal | 46.8 | 155156.228 | 2.592 |
| birds | Psephotellus_dissimilis | Psephotus_dissimilis | LC | Stable | tropical | diurnal | 45.8 | 212031.581 | 1.51 |
| birds | Psephotellus_varius | Psephotus_varius | LC | Decreasing | temperate | diurnal | 61.5 | 3329382.064 | 4.524 |
| birds | Psephotus_haematonotus | Psephotus_haematonotus | LC | Increasing | temperate | diurnal | 61.4 | 1548738.738 | 8.229 |
| birds | Pseudasthenes_cactorum | Asthenes_cactorum | LC | Decreasing | tropical | diurnal | 17.4 | 64992.804 | 6.69 |
| birds | Pseudasthenes_humicola | Asthenes_humicola | LC | Decreasing | temperate | diurnal | 20.8 | 118680.456 | 9.61 |
| birds | Pseudasthenes_patagonica | Asthenes_patagonica | LC | Decreasing | temperate | diurnal | 16.4 | 585683.996 | 5.327 |
| birds | Pseudasthenes_steinbachi | Asthenes_steinbachi | LC | Stable | temperate | diurnal | 19.5 | 273597.842 | 4.804 |
| birds | Pseudastur_albicollis | Leucopternis_albicollis | LC | Decreasing | tropical | diurnal | 703.7 | 8478535.509 | 3.347 |
| birds | Pseudastur_occidentalis | Leucopternis_occidentalis | EN | Decreasing | tropical | diurnal | 660 | 49655.96 | 14.477 |
| birds | Pseudastur_polionotus | NA | NT | Decreasing | NA | NA | NA | NA | NA |
| birds | Pseudelaenia_leucospodia | Pseudelaenia_leucospodia | LC | Decreasing | tropical | diurnal | 11.9 | 51916.21 | 8.362 |
| birds | Pseudeos_fuscata | Pseudeos_fuscata | LC | Stable | tropical | diurnal | 149 | 766350.357 | 3.378 |
| birds | Pseudhirundo_griseopyga | Pseudhirundo_griseopyga | LC | Stable | tropical | diurnal | 10.2 | 5583454.867 | 9.958 |
| birds | Pseudibis_davisoni | Pseudibis_davisoni | CR | Decreasing | tropical | diurnal | 1588 | 61742.397 | 10.763 |
| birds | Pseudibis_papillosa | Pseudibis_papillosa | LC | Decreasing | temperate | diurnal | 823 | 1740563.754 | 19.077 |
| birds | Pseudobias_wardi | Pseudobias_wardi | LC | Unknown | tropical | diurnal | 12.7 | 172356.276 | 8.355 |
| birds | Pseudocalyptomena_graueri | Pseudocalyptomena_graueri | VU | Decreasing | tropical | diurnal | 31.8 | 18563.545 | 12.47 |
| birds | Pseudochelidon_eurystomina | Pseudochelidon_eurystomina | DD | Decreasing | tropical | diurnal | 26 | 557559.877 | 5.892 |
| birds | Pseudocolaptes_boissonneauii | Pseudocolaptes_boissonneautii | LC | Stable | tropical | diurnal | 43.9 | 395920.181 | 9.526 |
| birds | Pseudocolaptes_johnsoni | NA | LC | Stable | NA | NA | NA | NA | NA |
| birds | Pseudocolaptes_lawrencii | Pseudocolaptes_lawrencii | LC | Stable | tropical | diurnal | 51 | 6603.07 | 6.308 |
| birds | Pseudocolopteryx_acutipennis | Pseudocolopteryx_acutipennis | LC | Decreasing | tropical | diurnal | 6.9 | 1594435.282 | 6.521 |
| birds | Pseudocolopteryx_dinelliana | NA | NT | Decreasing | NA | NA | NA | NA | NA |
| birds | Pseudocolopteryx_flaviventris | Pseudocolopteryx_flaviventris | LC | Decreasing | temperate | diurnal | 7.5 | 1685669.254 | 10.079 |
| birds | Pseudocolopteryx_sclateri | Pseudocolopteryx_sclateri | LC | Stable | temperate | diurnal | 8 | 816154.8 | 9.921 |
| birds | Pseudoleistes_guirahuro | Pseudoleistes_guirahuro | LC | Stable | temperate | diurnal | 86.4 | 2138341.082 | 11.957 |
| birds | Pseudoleistes_virescens | Pseudoleistes_virescens | LC | Stable | temperate | diurnal | 79.9 | 1218337.213 | 10.851 |
| birds | Pseudonestor_xanthophrys | Pseudonestor_xanthophrys | CR | Decreasing | tropical | diurnal | 22.1 | 34.887 | 1.441 |
| birds | Pseudonigrita_arnaudi | Pseudonigrita_arnaudi | LC | Stable | tropical | diurnal | 20 | 462385.542 | 12.5 |
| birds | Pseudonigrita_cabanisi | Pseudonigrita_cabanisi | LC | Stable | tropical | diurnal | 20.6 | 266415.079 | 8.435 |
| birds | Pseudopipra_pipra | Pipra_pipra | LC | Stable | tropical | diurnal | 11.1 | 4953783.656 | 2.107 |

| | | | | | | | | | |
|---|---|---|---|---|---|---|---|---|---|
| birds | Pseudopodoces_humilis | Pseudopodoces_humilis | LC | Stable | temperate | diurnal | 45.4 | 2316477.816 | 4.033 |
| birds | Pseudorectes_ferrugineus | Pitohui_ferrugineus | LC | Stable | tropical | diurnal | 93.9 | 614817.292 | 3.11 |
| birds | Pseudorectes_incertus | Pitohui_incertus | LC | Stable | tropical | diurnal | 75 | 110302.726 | 2.222 |
| birds | Pseudosaltator_rufiventris | Saltator_rufiventris | LC | Decreasing | tropical | diurnal | 74.2 | 42777.657 | 9.387 |
| birds | Pseudoscops_grammicus | Pseudoscops_grammicus | LC | Decreasing | tropical | nocturnal | 335 | 11021.788 | 19.741 |
| birds | Pseudoseisura_cristata | Pseudoseisura_cristata | LC | Increasing | tropical | diurnal | 53.6 | 909279.078 | 8.514 |
| birds | Pseudoseisura_gutturalis | Pseudoseisura_gutturalis | LC | Stable | temperate | diurnal | 70.4 | 779615.045 | 5.371 |
| birds | Pseudoseisura_lophotes | Pseudoseisura_lophotes | LC | Decreasing | temperate | diurnal | 72.2 | 1313910.999 | 7.866 |
| birds | Pseudoseisura_unirufa | Pseudoseisura_unirufa | LC | Stable | tropical | diurnal | 44.9 | 435035.551 | 5.955 |
| birds | Pseudospingus_verticalis | Hemispingus_verticalis | LC | Decreasing | tropical | diurnal | 14 | 67457.698 | 10.472 |
| birds | Pseudospingus_xanthophthalmus | Hemispingus_xanthophthalmus | LC | Decreasing | tropical | diurnal | 12 | 92237.269 | 4.873 |
| birds | Pseudotriccus_pelzelni | Pseudotriccus_pelzelni | LC | Decreasing | tropical | diurnal | 10.9 | 148376.196 | 7.207 |
| birds | Pseudotriccus_ruficeps | Pseudotriccus_ruficeps | LC | Decreasing | tropical | diurnal | 11.3 | 194101.794 | 7.283 |
| birds | Pseudotriccus_simplex | Pseudotriccus_simplex | LC | Decreasing | tropical | diurnal | 9.5 | 68986.063 | 4.164 |
| birds | Psilopogon_armillaris | Megalaima_armillaris | LC | Stable | tropical | diurnal | 69.4 | 136023.895 | 22.477 |
| birds | Psilopogon_asiaticus | Megalaima_asiatica | LC | Stable | temperate | diurnal | 90.5 | 1991863.598 | 11.873 |
| birds | Psilopogon_australis | Megalaima_australis | LC | Decreasing | tropical | diurnal | 33.3 | 78190.522 | 19.608 |
| birds | Psilopogon_chrysopogon | Megalaima_chrysopogon | LC | Decreasing | tropical | diurnal | 151 | 924419.876 | 7.439 |
| birds | Psilopogon_corvinus | Megalaima_corvina | LC | Unknown | tropical | diurnal | 130 | 6748.314 | 15.063 |
| birds | Psilopogon_eximius | NA | LC | Decreasing | NA | NA | NA | NA | NA |
| birds | Psilopogon_faiostrictus | Megalaima_faiostricta | LC | Decreasing | tropical | diurnal | 93 | 709761.142 | 12.118 |
| birds | Psilopogon_flavifrons | Megalaima_flavifrons | LC | Stable | tropical | diurnal | 58.5 | 39998.464 | 18.2 |
| birds | Psilopogon_franklinii | Megalaima_franklinii | LC | Stable | tropical | diurnal | 63.5 | 747848.447 | 8.51 |
| birds | Psilopogon_haemacephalus | Megalaima_haemacephala | LC | Increasing | tropical | diurnal | 44.5 | 5975609.967 | 15.544 |
| birds | Psilopogon_henricii | Megalaima_henricii | NT | Decreasing | tropical | diurnal | 74.1 | 715928.635 | 7.189 |
| birds | Psilopogon_incognitus | Megalaima_incognita | LC | Decreasing | tropical | diurnal | 93 | 709194.712 | 11.475 |
| birds | Psilopogon_javensis | Megalaima_javensis | LC | Decreasing | tropical | diurnal | 115 | 10739.019 | 12.134 |
| birds | Psilopogon_lagrandieri | Megalaima_lagrandieri | LC | Decreasing | tropical | diurnal | 93 | 353200.77 | 12.543 |
| birds | Psilopogon_lineatus | Megalaima_lineata | LC | Stable | tropical | diurnal | 149 | 1966842.259 | 12.728 |
| birds | Psilopogon_malabaricus | NA | LC | Decreasing | NA | NA | NA | NA | NA |
| birds | Psilopogon_monticola | Megalaima_monticola | LC | Unknown | tropical | diurnal | 93.4 | 53959.572 | 2.32 |
| birds | Psilopogon_mystacophanos | Megalaima_mystacophanos | NT | Decreasing | tropical | diurnal | 77.4 | 1239604.691 | 8.463 |
| birds | Psilopogon_oorti | Megalaima_oorti | LC | Decreasing | tropical | diurnal | 87.7 | 78643.098 | 6.524 |
| birds | Psilopogon_pulcherrimus | Megalaima_pulcherrima | LC | Decreasing | tropical | diurnal | 65.4 | 5208.539 | 3.268 |

| | | | | | | | | | |
|---|---|---|---|---|---|---|---|---|---|
| birds | Psilopogon_pyrolophus | Psilopogon_pyrolophus | LC | Decreasing | tropical | diurnal | 129 | 474947.384 | 10.225 |
| birds | Psilopogon_rafflesii | Megalaima_rafflesii | NT | Decreasing | tropical | diurnal | 118 | 1091543.703 | 8.581 |
| birds | Psilopogon_virens | Megalaima_virens | LC | Decreasing | temperate | diurnal | 202 | 3256344.791 | 12.304 |
| birds | Psilopogon_viridis | Megalaima_viridis | LC | Stable | tropical | diurnal | 80.5 | 263685.813 | 20.476 |
| birds | Psilopogon_zeylanicus | Megalaima_zeylanica | LC | Stable | tropical | diurnal | 119 | 2194453.733 | 18.645 |
| birds | Psilopsiagon_aurifrons | Psilopsiagon_aurifrons | LC | Stable | tropical | diurnal | 33.6 | 947209.798 | 6.428 |
| birds | Psilopsiagon_aymara | Psilopsiagon_aymara | LC | Stable | temperate | diurnal | 30 | 382901.126 | 6.656 |
| birds | Psilorhamphus_guttatus | Psilorhamphus_guttatus | LC | Decreasing | temperate | diurnal | 11.3 | 183478.246 | 12.713 |
| birds | Psiloscops_flammeolus | Otus_flammeolus | LC | Decreasing | temperate | nocturnal | 59.9 | 1654758.998 | 9.055 |
| birds | Psittacara_acuticaudatus | Aratinga_acuticaudata | LC | Decreasing | temperate | diurnal | 171 | 3363879.676 | 7.818 |
| birds | Psittacara_chloropterus | Aratinga_chloroptera | VU | Decreasing | tropical | diurnal | 146 | 74178.221 | 15.783 |
| birds | Psittacara_erythrogenys | Aratinga_erythrogenys | NT | Decreasing | tropical | diurnal | 151 | 164717.144 | 14.291 |
| birds | Psittacara_euops | Aratinga_euops | VU | Decreasing | tropical | diurnal | 87 | 108504.916 | 16.218 |
| birds | Psittacara_finschi | Aratinga_finschi | LC | Stable | tropical | diurnal | 150 | 92560.877 | 11.497 |
| birds | Psittacara_holochlorus | Aratinga_holochlora | LC | Decreasing | tropical | diurnal | 138 | 299273.469 | 12.064 |
| birds | Psittacara_leucophthalmus | Aratinga_leucophthalma | LC | Decreasing | tropical | diurnal | 158 | 8731069.107 | 5.641 |
| birds | Psittacara_mitratus | Aratinga_mitrata | LC | Stable | tropical | diurnal | 249 | 291306.045 | 8.784 |
| birds | Psittacara_rubritorquis | Aratinga_rubritorquis | LC | Stable | tropical | diurnal | 212 | 108828.415 | 10.785 |
| birds | Psittacara_wagleri | Aratinga_wagleri | NT | Decreasing | tropical | diurnal | 194 | 216336.616 | 12.409 |
| birds | Psittacella_brehmii | Psittacella_brehmii | LC | Stable | tropical | diurnal | 107 | 123546.512 | 5.036 |
| birds | Psittacella_madaraszi | Psittacella_madaraszi | LC | Stable | tropical | diurnal | 38.8 | 193283.602 | 4.412 |
| birds | Psittacella_modesta | Psittacella_modesta | LC | Stable | tropical | diurnal | 44.7 | 98065.584 | 4.796 |
| birds | Psittacella_picta | Psittacella_picta | LC | Stable | tropical | diurnal | 62.9 | 56415.702 | 6.089 |
| birds | Psittacula_alexandri | Psittacula_alexandri | NT | Decreasing | tropical | diurnal | 147 | 2602483.038 | 12.122 |
| birds | Psittacula_caniceps | Psittacula_caniceps | NT | Decreasing | tropical | diurnal | 224 | 1111.02 | 4.108 |
| birds | Psittacula_derbiana | Psittacula_derbiana | NT | Decreasing | temperate | diurnal | 251 | 332466.751 | 7.29 |
| birds | Psittaculirostris_desmarestii | Psittaculirostris_desmarestii | LC | Decreasing | tropical | diurnal | 117 | 82418.216 | 3.337 |
| birds | Psittaculirostris_edwardsii | Psittaculirostris_edwardsii | LC | Stable | tropical | diurnal | 118 | 99151.911 | 4.841 |
| birds | Psittaculirostris_salvadorii | Psittaculirostris_salvadorii | LC | Decreasing | tropical | diurnal | 118 | 50674.665 | 2.183 |
| birds | Psittacus_erithacus | Psittacus_erithacus | EN | Decreasing | tropical | diurnal | 333 | 2775743.777 | 7.762 |
| birds | Psitteuteles_goldiei | Psitteuteles_goldiei | LC | Decreasing | tropical | diurnal | 58.7 | 307384.239 | 4.006 |
| birds | Psitteuteles_versicolor | Psitteuteles_versicolor | LC | Stable | tropical | diurnal | 53.5 | 1214241.532 | 2.899 |
| birds | Psittinus_cyanurus | Psittinus_cyanurus | NT | Decreasing | tropical | diurnal | 60.5 | 1398599.343 | 8.094 |
| birds | Psittiparus_ruficeps | NA | not_available | not_available | NA | NA | NA | NA | NA |

| | | | | | | | | | |
|---|---|---|---|---|---|---|---|---|---|
| birds | Psittirostra_psittacea | Psittirostra_psittacea | CR | Unknown | tropical | diurnal | 28.6 | 16385.915 | 8.655 |
| birds | Psittrichas_fulgidus | Psittrichas_fulgidus | VU | Decreasing | tropical | diurnal | 865 | 355263.53 | 3.705 |
| birds | Psophia_crepitans | Psophia_crepitans | LC | Decreasing | tropical | diurnal | 1026 | 2708717.398 | 1.481 |
| birds | Psophia_leucoptera | Psophia_leucoptera | LC | Decreasing | tropical | diurnal | 1315.6 | 1556201.474 | 1.888 |
| birds | Psophia_viridis | Psophia_viridis | VU | Decreasing | tropical | diurnal | 1071 | 641583.01 | 2.136 |
| birds | Psophocichla_litsitsirupa | Psophocichla_litsitsirupa | LC | Unknown | tropical | diurnal | 74 | 3146030.544 | 7.838 |
| birds | Psophodes_cristatus | Psophodes_cristatus | LC | Stable | temperate | diurnal | 40.7 | 990722.543 | 4.481 |
| birds | Psophodes_nigrogularis | Psophodes_nigrogularis | LC | Unknown | temperate | diurnal | 44.5 | 209988.362 | 11.41 |
| birds | Psophodes_occidentalis | Psophodes_occidentalis | LC | Decreasing | temperate | diurnal | 39.6 | 2326926.646 | 2.234 |
| birds | Psophodes_olivaceus | Psophodes_olivaceus | LC | Decreasing | temperate | diurnal | 62.4 | 428340.702 | 8.325 |
| birds | Pteridophora_alberti | Pteridophora_alberti | LC | Decreasing | tropical | diurnal | 81.8 | 109282.789 | 5.287 |
| birds | Pternistis_adspersus | Francolinus_adspersus | LC | Stable | tropical | diurnal | 428 | 1150093.127 | 5.628 |
| birds | Pternistis_afer | Francolinus_afer | LC | Decreasing | tropical | diurnal | 454.8 | 4561470.056 | 9.043 |
| birds | Pternistis_ahantensis | Francolinus_ahantensis | LC | Decreasing | tropical | diurnal | 544.1 | 683993.474 | 15.299 |
| birds | Pternistis_bicalcaratus | Francolinus_bicalcaratus | LC | Decreasing | tropical | diurnal | 439.5 | 3597861.299 | 12.599 |
| birds | Pternistis_camerunensis | Francolinus_camerunensis | EN | Decreasing | tropical | diurnal | 549.4 | 193.348 | 13.343 |
| birds | Pternistis_capensis | Francolinus_capensis | LC | Stable | temperate | diurnal | 629.8 | 150806.37 | 8.993 |
| birds | Pternistis_castaneicollis | Francolinus_castaneicollis | LC | Stable | tropical | diurnal | 791.5 | 264511.059 | 12.693 |
| birds | Pternistis_clappertoni | Francolinus_clappertoni | LC | Decreasing | tropical | diurnal | 469.6 | 2323019.725 | 10.289 |
| birds | Pternistis_erckelii | Francolinus_erckelii | LC | Stable | tropical | diurnal | 1197.8 | 162711.963 | 13.186 |
| birds | Pternistis_hartlaubi | Francolinus_hartlaubi | LC | Stable | tropical | diurnal | 241.5 | 353895.578 | 6.778 |
| birds | Pternistis_harwoodi | Francolinus_harwoodi | NT | Decreasing | tropical | diurnal | 493 | 49847.915 | 12.603 |
| birds | Pternistis_hildebrandti | Francolinus_hildebrandti | LC | Stable | tropical | diurnal | 532 | 900428.973 | 12.328 |
| birds | Pternistis_icterorhynchus | Francolinus_icterorhynchus | LC | Stable | tropical | diurnal | 571 | 1444290.429 | 5.972 |
| birds | Pternistis_jacksoni | Francolinus_jacksoni | LC | Decreasing | tropical | diurnal | 1097 | 16544.058 | 14.084 |
| birds | Pternistis_leucoscepus | Francolinus_leucoscepus | LC | Decreasing | tropical | diurnal | 640.6 | 2071108.37 | 9.168 |
| birds | Pternistis_natalensis | Francolinus_natalensis | LC | Stable | tropical | diurnal | 508.1 | 1136104.662 | 11.235 |
| birds | Pternistis_ochropectus | Francolinus_ochropectus | CR | Decreasing | tropical | diurnal | 699.6 | 607.573 | 11.292 |
| birds | Pternistis_rufopictus | Francolinus_rufopictus | LC | Decreasing | tropical | diurnal | 706.1 | 77700.064 | 14.361 |
| birds | Pternistis_squamatus | Francolinus_squamatus | LC | Decreasing | tropical | diurnal | 449.5 | 2593426.096 | 9.128 |
| birds | Pternistis_swierstrai | Francolinus_swierstrai | VU | Decreasing | tropical | diurnal | 460.3 | 17202.419 | 12.071 |
| birds | Pterocles_bicinctus | Pterocles_bicinctus | LC | Decreasing | tropical | nocturnal | 237 | 1654904.572 | 7.378 |
| birds | Pterocles_lichtensteinii | Pterocles_lichtensteinii | LC | Stable | temperate | nocturnal | 209.1 | 3826174.114 | 5.174 |
| birds | Pterocles_quadricinctus | Pterocles_quadricinctus | LC | Stable | tropical | nocturnal | 178.9 | 3917295.509 | 11.672 |

| | | | | | | | | | |
|---|---|---|---|---|---|---|---|---|---|
| birds | Pterodroma_alba | Pterodroma_alba | VU | Decreasing | tropical | cathemeral | 259 | 22290948.301 | 7.553 |
| birds | Pterodroma_arminjoniana | Pterodroma_arminjoniana | VU | Stable | tropical | cathemeral | 394 | 47409362.676 | 8.656 |
| birds | Pterodroma_axillaris | Pterodroma_axillaris | VU | Increasing | temperate | cathemeral | 165 | 30397121.728 | 12.045 |
| birds | Pterodroma_cahow | Pterodroma_cahow | EN | Increasing | temperate | cathemeral | 246 | 15874369.539 | 14.111 |
| birds | Pterodroma_cervicalis | Pterodroma_cervicalis | VU | Increasing | tropical | cathemeral | 445 | 67284605.113 | 7.986 |
| birds | Pterodroma_cookii | Pterodroma_cookii | VU | Increasing | tropical | nocturnal | 177.9 | 120250809.723 | 7.394 |
| birds | Pterodroma_defilippiana | NA | VU | Stable | NA | NA | NA | NA | NA |
| birds | Pterodroma_externa | Pterodroma_externa | VU | Stable | tropical | cathemeral | 428 | 44060886.108 | 4.512 |
| birds | Pterodroma_feae | Pterodroma_feae | NT | Unknown | temperate | cathemeral | 311 | 12674661.91 | 12.945 |
| birds | Pterodroma_hasitata | Pterodroma_hasitata | EN | Decreasing | tropical | cathemeral | 278 | 7451677.196 | 11.389 |
| birds | Pterodroma_hypoleuca | Pterodroma_hypoleuca | LC | Decreasing | temperate | cathemeral | 176 | 18787810.665 | 15.959 |
| birds | Pterodroma_incerta | Pterodroma_incerta | EN | Decreasing | temperate | cathemeral | 520 | 19854482.975 | 9.827 |
| birds | Pterodroma_inexpectata | Pterodroma_inexpectata | NT | Decreasing | tropical | cathemeral | 316 | 119601815.08 | 6.678 |
| birds | Pterodroma_lessonii | Pterodroma_lessonii | LC | Decreasing | temperate | cathemeral | 698 | 74962974.708 | 5.939 |
| birds | Pterodroma_leucoptera | Pterodroma_leucoptera | VU | Decreasing | tropical | cathemeral | 147.1 | 122022847.776 | 7.445 |
| birds | Pterodroma_macroptera | Pterodroma_macroptera | LC | Decreasing | temperate | cathemeral | 531.8 | 47147490.288 | 12.546 |
| birds | Pterodroma_mollis | Pterodroma_mollis | LC | Stable | temperate | cathemeral | 312 | 52198187.39 | 10.207 |
| birds | Pterodroma_neglecta | Pterodroma_neglecta | LC | Decreasing | tropical | cathemeral | 501 | 87299058.054 | 7.514 |
| birds | Pterodroma_nigripennis | Pterodroma_nigripennis | LC | Decreasing | tropical | cathemeral | 173.7 | 118358781.118 | 5.166 |
| birds | Pterodroma_phaeopygia | Pterodroma_phaeopygia | CR | Decreasing | tropical | cathemeral | 400.8 | 15494657.472 | 9.679 |
| birds | Pterodroma_pycrofti | Pterodroma_pycrofti | VU | Increasing | tropical | cathemeral | 159 | 75938807.194 | 8.159 |
| birds | Pterodroma_solandri | Pterodroma_solandri | LC | Stable | tropical | cathemeral | 468.6 | 59746398.172 | 11.155 |
| birds | Pterodroma_ultima | Pterodroma_ultima | LC | Unknown | tropical | cathemeral | 360 | 66311108.884 | 10.23 |
| birds | Pteroglossus_aracari | Pteroglossus_aracari | LC | Decreasing | tropical | diurnal | 250.2 | 3319977.819 | 4.719 |
| birds | Pteroglossus_azara | Pteroglossus_azara | LC | Stable | tropical | diurnal | 144.5 | 3418746.757 | 1.726 |
| birds | Pteroglossus_bailloni | Pteroglossus_bailloni | NT | Decreasing | temperate | diurnal | 146 | 457341.043 | 12.809 |
| birds | Pteroglossus_beauharnaisii | Pteroglossus_beauharnaesii | LC | Decreasing | tropical | diurnal | 215.9 | 1824148.667 | 1.981 |
| birds | Pteroglossus_bitorquatus | Pteroglossus_bitorquatus | NT | Decreasing | tropical | diurnal | 142 | 936672.384 | 2.943 |
| birds | Pteroglossus_castanotis | Pteroglossus_castanotis | LC | Decreasing | tropical | diurnal | 258.1 | 5071451.427 | 4.894 |
| birds | Pteroglossus_frantzii | Pteroglossus_frantzii | LC | Decreasing | tropical | diurnal | 253 | 19662.61 | 14.804 |
| birds | Pteroglossus_inscriptus | Pteroglossus_inscriptus | LC | Decreasing | tropical | diurnal | 125.8 | 2978379.814 | 3.798 |
| birds | Pteroglossus_pluricinctus | Pteroglossus_pluricinctus | LC | Decreasing | tropical | diurnal | 261.7 | 1636986.613 | 1.681 |
| birds | Pteroglossus_torquatus | Pteroglossus_torquatus | LC | Decreasing | tropical | diurnal | 219.1 | 1018314.607 | 11.016 |
| birds | Pteroglossus_viridis | Pteroglossus_viridis | LC | Stable | tropical | diurnal | 135 | 1558794.854 | 1.369 |

| | | | | | | | | | |
|---|---|---|---|---|---|---|---|---|---|
| birds | Pteronetta_hartlaubii | Pteronetta_hartlaubii | LC | Decreasing | tropical | diurnal | 790 | 4283729.601 | 8.103 |
| birds | Pterophanes_cyanopterus | Pterophanes_cyanopterus | LC | Stable | tropical | diurnal | 9.3 | 232571.392 | 9.547 |
| birds | Pteroptochos_castaneus | Pteroptochos_castaneus | LC | Decreasing | temperate | diurnal | 160.6 | 53074.045 | 9.844 |
| birds | Pteroptochos_megapodius | Pteroptochos_megapodius | LC | Stable | temperate | diurnal | 114 | 132289.256 | 9.252 |
| birds | Pteroptochos_tarnii | Pteroptochos_tarnii | LC | Decreasing | temperate | diurnal | 147 | 275064.613 | 4.923 |
| birds | Pterorhinus_caerulatus | Garrulax_caerulatus | LC | Decreasing | temperate | diurnal | 85.7 | 219784.763 | 6.859 |
| birds | Pterorhinus_chinensis | Garrulax_chinensis | LC | Decreasing | tropical | diurnal | 88.5 | 1498163.608 | 11.68 |
| birds | Pterorhinus_davidi | Garrulax_davidi | LC | Stable | temperate | diurnal | 56.2 | 1085948.476 | 12.252 |
| birds | Pterorhinus_delesserti | Garrulax_delesserti | LC | Decreasing | tropical | diurnal | 78 | 125058.65 | 19.225 |
| birds | Pterorhinus_gularis | Garrulax_gularis | LC | Decreasing | temperate | diurnal | 92 | 307618.651 | 7.333 |
| birds | Pterorhinus_koslowi | Babax_koslowi | NT | Stable | temperate | diurnal | 112.5 | 44443.217 | 8.732 |
| birds | Pterorhinus_lanceolatus | Babax_lanceolatus | LC | Stable | temperate | diurnal | 80.5 | 1946905.495 | 11.913 |
| birds | Pterorhinus_nuchalis | Garrulax_nuchalis | NT | Stable | temperate | diurnal | 78 | 60702.135 | 6.228 |
| birds | Pterorhinus_pectoralis | Garrulax_pectoralis | LC | Decreasing | temperate | diurnal | 145.1 | 2170571.374 | 12.103 |
| birds | Pterorhinus_perspicillatus | Garrulax_perspicillatus | LC | Stable | temperate | diurnal | 117.8 | 1367037.218 | 16.44 |
| birds | Pterorhinus_ruficollis | Garrulax_ruficollis | LC | Stable | temperate | diurnal | 66.2 | 296349.203 | 8.503 |
| birds | Pterorhinus_sannio | Garrulax_sannio | LC | Stable | temperate | diurnal | 67.8 | 2031254.052 | 12.232 |
| birds | Pterorhinus_vassali | NA | LC | Stable | NA | NA | NA | NA | NA |
| birds | Pterorhinus_waddelli | Babax_waddelli | NT | Decreasing | temperate | diurnal | 46 | 59928.984 | 8.899 |
| birds | Pteruthius_aenobarbus | Pteruthius_aenobarbus | LC | Decreasing | tropical | diurnal | 11.8 | 996369.256 | 9.536 |
| birds | Pteruthius_flaviscapis | Pteruthius_flaviscapis | LC | Decreasing | tropical | diurnal | 38.7 | 63531.489 | 21.044 |
| birds | Pteruthius_melanotis | Pteruthius_melanotis | LC | Decreasing | temperate | diurnal | 13.3 | 841041.72 | 8.233 |
| birds | Pteruthius_rufiventer | Pteruthius_rufiventer | LC | Decreasing | temperate | diurnal | 43 | 251147.364 | 7.038 |
| birds | Pteruthius_xanthochlorus | Pteruthius_xanthochlorus | LC | Decreasing | temperate | diurnal | 14.3 | 786930.437 | 8.319 |
| birds | Ptilinopus_alligator | NA | LC | Stable | NA | NA | NA | NA | NA |
| birds | Ptilinopus_arcanus | NA | CR | Decreasing | NA | NA | NA | NA | NA |
| birds | Ptilinopus_aurantiifrons | Ptilinopus_aurantiifrons | LC | Stable | tropical | diurnal | 152 | 479656.997 | 3.039 |
| birds | Ptilinopus_chalcurus | NA | VU | Stable | NA | NA | NA | NA | NA |
| birds | Ptilinopus_cinctus | Ptilinopus_cinctus | LC | Decreasing | tropical | diurnal | 243 | 74691.524 | 13.785 |
| birds | Ptilinopus_coralensis | Ptilinopus_coralensis | NT | Decreasing | tropical | diurnal | 83.2 | 624.423 | 7.062 |
| birds | Ptilinopus_coronulatus | Ptilinopus_coronulatus | LC | Stable | tropical | diurnal | 76.6 | 670308.782 | 3.097 |
| birds | Ptilinopus_dohertyi | NA | VU | Decreasing | NA | NA | NA | NA | NA |
| birds | Ptilinopus_dupetithouarsii | Ptilinopus_dupetithouarsii | LC | Decreasing | tropical | diurnal | 99.5 | 1018.266 | 7.344 |
| birds | Ptilinopus_eugeniae | Ptilinopus_eugeniae | LC | Stable | tropical | diurnal | 114 | 3382.95 | 4.498 |

| | | | | | | | | | |
|---|---|---|---|---|---|---|---|---|---|
| birds | Ptilinopus_granulifrons | NA | VU | Decreasing | NA | NA | NA | NA | NA |
| birds | Ptilinopus_greyi | Ptilinopus_greyii | LC | Stable | tropical | diurnal | 82.7 | 32025.735 | 6.809 |
| birds | Ptilinopus_huttoni | NA | CR | Decreasing | NA | NA | NA | NA | NA |
| birds | Ptilinopus_hyogastrus | NA | LC | Stable | NA | NA | NA | NA | NA |
| birds | Ptilinopus_insolitus | Ptilinopus_insolitus | LC | Stable | tropical | diurnal | 136.5 | 42750.018 | 4.398 |
| birds | Ptilinopus_insularis | Ptilinopus_insularis | VU | Stable | temperate | diurnal | 104 | 41.112 | 6.407 |
| birds | Ptilinopus_iozonus | Ptilinopus_iozonus | LC | Stable | tropical | diurnal | 114 | 674305.793 | 3.098 |
| birds | Ptilinopus_melanospilus | Ptilinopus_melanospilus | LC | Stable | tropical | diurnal | 92 | 498198.967 | 13.845 |
| birds | Ptilinopus_mercierii | NA | EX | not_available | NA | NA | NA | NA | NA |
| birds | Ptilinopus_monacha | Ptilinopus_monacha | LC | Decreasing | tropical | diurnal | 53.9 | 26140.071 | 6.399 |
| birds | Ptilinopus_nainus | Ptilinopus_naina | LC | Stable | tropical | diurnal | 49 | 269374.328 | 3.081 |
| birds | Ptilinopus_ornatus | Ptilinopus_ornatus | LC | Stable | tropical | diurnal | 163 | 29156.11 | 3.455 |
| birds | Ptilinopus_pelewensis | Ptilinopus_pelewensis | LC | Stable | tropical | diurnal | 93 | 454.469 | 11.182 |
| birds | Ptilinopus_perlatus | Ptilinopus_perlatus | LC | Stable | tropical | diurnal | 210 | 673225.357 | 3.097 |
| birds | Ptilinopus_perousii | Ptilinopus_perousii | LC | Decreasing | tropical | diurnal | 93.5 | 22003.57 | 9.375 |
| birds | Ptilinopus_porphyraceus | Ptilinopus_porphyraceus | LC | Stable | tropical | diurnal | 105 | 3346.875 | 8.32 |
| birds | Ptilinopus_porphyreus | Ptilinopus_porphyreus | LC | Decreasing | tropical | diurnal | 168 | 27641.476 | 10.395 |
| birds | Ptilinopus_pulchellus | Ptilinopus_pulchellus | LC | Stable | tropical | diurnal | 70 | 339387.213 | 3.092 |
| birds | Ptilinopus_purpuratus | Ptilinopus_purpuratus | LC | Decreasing | tropical | diurnal | 97.6 | 1194.954 | 15.144 |
| birds | Ptilinopus_rarotongensis | NA | NT | Stable | NA | NA | NA | NA | NA |
| birds | Ptilinopus_regina | Ptilinopus_regina | LC | Decreasing | tropical | diurnal | 101.6 | 553574.471 | 5.682 |
| birds | Ptilinopus_richardsii | Ptilinopus_richardsii | LC | Stable | tropical | diurnal | 103.9 | 769.1 | 4.011 |
| birds | Ptilinopus_rivoli | Ptilinopus_rivoli | LC | Stable | tropical | diurnal | 149 | 294245.103 | 4.508 |
| birds | Ptilinopus_roseicapilla | Ptilinopus_roseicapilla | NT | Decreasing | tropical | diurnal | 90.4 | 913.667 | 22.335 |
| birds | Ptilinopus_solomonensis | Ptilinopus_solomonensis | LC | Stable | tropical | diurnal | 108.5 | 63045.626 | 4.465 |
| birds | Ptilinopus_superbus | Ptilinopus_superbus | LC | Decreasing | tropical | diurnal | 118 | 1524991.573 | 5.12 |
| birds | Ptilinopus_tannensis | Ptilinopus_tannensis | LC | Decreasing | tropical | diurnal | 208 | 12046.286 | 5.836 |
| birds | Ptilinopus_viridis | Ptilinopus_viridis | LC | Stable | tropical | diurnal | 121 | 95441.027 | 5.291 |
| birds | Ptilinopus_wallacii | NA | LC | Stable | NA | NA | NA | NA | NA |
| birds | Ptiliogonys_caudatus | Ptilogonys_caudatus | LC | Stable | tropical | diurnal | 36.2 | 9544.171 | 8.823 |
| birds | Ptiliogonys_cinereus | Ptilogonys_cinereus | LC | Decreasing | tropical | diurnal | 33.6 | 524061.636 | 10.684 |
| birds | Ptilocichla_falcata | NA | VU | Decreasing | NA | NA | NA | NA | NA |
| birds | Ptilocichla_leucogrammica | Ptilocichla_leucogrammica | VU | Decreasing | tropical | diurnal | 39.5 | 579272.51 | 6.271 |
| birds | Ptilocichla_mindanensis | Ptilocichla_mindanensis | LC | Decreasing | tropical | diurnal | 29.2 | 52312.865 | 9.671 |

| | | | | | | | | | |
|---|---|---|---|---|---|---|---|---|---|
| birds | Ptilonorhynchus_violaceus | Ptilonorhynchus_violaceus | LC | Decreasing | temperate | diurnal | 218.9 | 523221.884 | 8.365 |
| birds | Ptilopachus_nahani | Francolinus_nahani | VU | Decreasing | tropical | diurnal | 310 | 99521.542 | 6.126 |
| birds | Ptilopachus_petrosus | Ptilopachus_petrosus | LC | Stable | tropical | diurnal | 193 | 4691145.788 | 11.243 |
| birds | Ptiloprora_erythropleura | Ptiloprora_erythropleura | LC | Stable | tropical | diurnal | 21.1 | 60498.522 | 3.97 |
| birds | Ptiloprora_guisei | Ptiloprora_guisei | LC | Stable | tropical | diurnal | 25.8 | 61859.614 | 6 |
| birds | Ptiloprora_mayri | Ptiloprora_mayri | LC | Stable | tropical | diurnal | 30.5 | 1102.689 | 1.056 |
| birds | Ptiloprora_meekiana | Ptiloprora_meekiana | LC | Stable | tropical | diurnal | 18.5 | 139400.206 | 4.998 |
| birds | Ptiloprora_perstriata | Ptiloprora_perstriata | LC | Stable | tropical | diurnal | 26.6 | 102361.907 | 4.958 |
| birds | Ptiloprora_plumbea | Ptiloprora_plumbea | LC | Stable | tropical | diurnal | 14.1 | 136682.135 | 4.713 |
| birds | Ptilopsis_granti | Ptilopsis_granti | LC | Stable | tropical | nocturnal | 214.4 | 7822704.31 | 8.472 |
| birds | Ptilopsis_leucotis | Otus_leucotis | LC | Stable | tropical | nocturnal | 204 | 8309644.168 | 9.574 |
| birds | Ptilorrhoa_caerulescens | Ptilorrhoa_caerulescens | LC | Stable | tropical | diurnal | 49 | 399425.346 | 3.138 |
| birds | Ptilorrhoa_castanonota | Ptilorrhoa_castanonota | LC | Stable | tropical | diurnal | 69.7 | 246242.694 | 3.508 |
| birds | Ptilorrhoa_leucosticta | Ptilorrhoa_leucosticta | LC | Stable | tropical | diurnal | 49.2 | 235330.781 | 4.241 |
| birds | Ptilotula_flavescens | Lichenostomus_flavescens | LC | Stable | tropical | diurnal | 12.4 | 1038738.886 | 2.599 |
| birds | Ptilotula_fusca | Lichenostomus_fuscus | LC | Decreasing | temperate | diurnal | 17.7 | 949120.713 | 8.636 |
| birds | Ptilotula_keartlandi | Lichenostomus_keartlandi | LC | Stable | tropical | diurnal | 15.2 | 2753396.761 | 2.561 |
| birds | Ptilotula_ornata | Lichenostomus_ornatus | LC | Decreasing | temperate | diurnal | 17.6 | 1105315.888 | 8.112 |
| birds | Ptilotula_penicillata | Lichenostomus_penicillatus | LC | Stable | temperate | diurnal | 19.4 | 4922061.086 | 4.836 |
| birds | Ptilotula_plumula | Lichenostomus_plumulus | LC | Unknown | temperate | diurnal | 16.5 | 5368167.697 | 3.599 |
| birds | Ptiloxena_atroviolacea | Dives_atroviolaceus | LC | Stable | tropical | diurnal | 83.6 | 106633.354 | 16.283 |
| birds | Ptychoramphus_aleuticus | Ptychoramphus_aleuticus | NT | Decreasing | temperate | cathemeral | 184 | 4083740.369 | 6.001 |
| birds | Ptyonoprogne_concolor | Hirundo_concolor | LC | Increasing | tropical | diurnal | 13 | 2727893.22 | 16.764 |
| birds | Ptyonoprogne_obsoleta | Hirundo_obsoleta | LC | Increasing | temperate | diurnal | 10.2 | 7338605.45 | 5.178 |
| birds | Ptyonoprogne_rupestris | Hirundo_rupestris | LC | Stable | temperate | diurnal | 20.4 | 12884287.967 | 11.331 |
| birds | Pucrasia_macrolopha | Pucrasia_macrolopha | LC | Decreasing | temperate | diurnal | 1050.5 | 1384088.505 | 11.68 |
| birds | Puffinus_assimilis | Puffinus_assimilis | LC | Decreasing | temperate | cathemeral | 226 | 21735982.042 | 12.226 |
| birds | Puffinus_gavia | Puffinus_gavia | LC | Decreasing | temperate | cathemeral | 228.9 | 10247529.714 | 11.653 |
| birds | Puffinus_huttoni | Puffinus_huttoni | EN | Stable | temperate | cathemeral | 364 | 4657116.88 | 6.259 |
| birds | Puffinus_lherminieri | Puffinus_lherminieri | LC | Decreasing | temperate | cathemeral | 192.8 | 24673109.086 | 14.314 |
| birds | Puffinus_nativitatis | Puffinus_nativitatis | LC | Stable | tropical | cathemeral | 356 | 50917561.974 | 5.918 |
| birds | Puffinus_puffinus | Puffinus_puffinus | LC | Unknown | tropical | cathemeral | 453.3 | 69101740.745 | 11.25 |
| birds | Puffinus_yelkouan | Puffinus_yelkouan | VU | Decreasing | temperate | cathemeral | 497 | 2989515.985 | 19.334 |
| birds | Pulsatrix_koeniswaldiana | Pulsatrix_koeniswaldiana | LC | Decreasing | tropical | nocturnal | 481 | 651781.62 | 13.418 |

| | | | | | | | | | |
|---|---|---|---|---|---|---|---|---|---|
| birds | Pulsatrix_melanota | Pulsatrix_melanota | LC | Decreasing | tropical | nocturnal | 82 | 156909.182 | 4.217 |
| birds | Pulsatrix_perspicillata | Pulsatrix_perspicillata | LC | Decreasing | tropical | nocturnal | 983.6 | 12679502.975 | 5.681 |
| birds | Purnella_albifrons | Phylidonyris_albifrons | LC | Stable | temperate | diurnal | 17.4 | 4027281.707 | 3.858 |
| birds | Purpureicephalus_spurius | Purpureicephalus_spurius | LC | Increasing | temperate | diurnal | 116 | 126268.754 | 10.688 |
| birds | Pycnonotus_aurigaster | Pycnonotus_aurigaster | LC | Decreasing | tropical | diurnal | 44.3 | 3748833.483 | 12.694 |
| birds | Pycnonotus_barbatus | Pycnonotus_barbatus | LC | Increasing | tropical | diurnal | 32.1 | 19572184.189 | 9.152 |
| birds | Pycnonotus_bimaculatus | NA | NT | Decreasing | NA | NA | NA | NA | NA |
| birds | Pycnonotus_blanfordi | Pycnonotus_blanfordi | LC | Stable | tropical | diurnal | 33.4 | 1240819.119 | 13.981 |
| birds | Pycnonotus_brunneus | Pycnonotus_brunneus | LC | Decreasing | tropical | diurnal | 28.8 | 1138188.08 | 8.569 |
| birds | Pycnonotus_cafer | Pycnonotus_cafer | LC | Increasing | tropical | diurnal | 42.9 | 4241761.352 | 16.681 |
| birds | Pycnonotus_capensis | Pycnonotus_capensis | LC | Stable | temperate | diurnal | 38.6 | 151666.39 | 9.756 |
| birds | Pycnonotus_finlaysoni | Pycnonotus_finlaysoni | LC | Stable | tropical | diurnal | 26.3 | 1253924.24 | 11.456 |
| birds | Pycnonotus_flavescens | Pycnonotus_flavescens | LC | Stable | tropical | diurnal | 28.9 | 917016.171 | 8.227 |
| birds | Pycnonotus_goiavier | Pycnonotus_goiavier | LC | Increasing | tropical | diurnal | 27.8 | 2142578.357 | 11.059 |
| birds | Pycnonotus_jocosus | Pycnonotus_jocosus | LC | Decreasing | tropical | diurnal | 29.5 | 5257404.41 | 14.947 |
| birds | Pycnonotus_leucogenys | Pycnonotus_leucogenys | LC | Increasing | temperate | diurnal | 35.9 | 378023.582 | 10.895 |
| birds | Pycnonotus_leucotis | Pycnonotus_leucotis | LC | Decreasing | temperate | diurnal | 27.1 | 2206861.393 | 11.633 |
| birds | Pycnonotus_luteolus | Pycnonotus_luteolus | LC | Stable | tropical | diurnal | 34.7 | 1386882.2 | 19.38 |
| birds | Pycnonotus_nieuwenhuisii | NA | DD | Unknown | NA | NA | NA | NA | NA |
| birds | Pycnonotus_nigricans | Pycnonotus_nigricans | LC | Increasing | tropical | diurnal | 30.8 | 2193980.338 | 6.775 |
| birds | Pycnonotus_penicillatus | Pycnonotus_penicillatus | LC | Stable | tropical | diurnal | 36.5 | 4919.71 | 16.646 |
| birds | Pycnonotus_plumosus | Pycnonotus_plumosus | LC | Stable | tropical | diurnal | 34.9 | 1506344.747 | 9.327 |
| birds | Pycnonotus_simplex | Pycnonotus_simplex | LC | Decreasing | tropical | diurnal | 25.1 | 1340820.454 | 8.653 |
| birds | Pycnonotus_sinensis | Pycnonotus_sinensis | LC | Increasing | temperate | diurnal | 34.2 | 3460936.244 | 16.85 |
| birds | Pycnonotus_taivanus | Pycnonotus_taivanus | VU | Decreasing | tropical | diurnal | 26.1 | 7274.279 | 11.852 |
| birds | Pycnonotus_tympanistrigus | Pycnonotus_tympanistrigus | LC | Decreasing | tropical | diurnal | 33.5 | 97157.16 | 7.735 |
| birds | Pycnonotus_xantholaemus | Pycnonotus_xantholaemus | VU | Decreasing | tropical | diurnal | 28.1 | 362753.3 | 20.761 |
| birds | Pycnonotus_xanthopygos | Pycnonotus_xanthopygos | LC | Increasing | temperate | diurnal | 37.6 | 1313216.503 | 7.883 |
| birds | Pycnonotus_xanthorrhous | Pycnonotus_xanthorrhous | LC | Stable | temperate | diurnal | 26.9 | 1741022.34 | 11.07 |
| birds | Pycnonotus_zeylanicus | NA | CR | Decreasing | NA | NA | NA | NA | NA |
| birds | Pycnoptilus_floccosus | Pycnoptilus_floccosus | VU | Decreasing | temperate | diurnal | 28.9 | 79638.344 | 6.829 |
| birds | Pycnopygius_cinereus | Pycnopygius_cinereus | LC | Stable | tropical | diurnal | 44.8 | 106235.796 | 4.279 |
| birds | Pycnopygius_ixoides | Pycnopygius_ixoides | LC | Stable | tropical | diurnal | 27.6 | 459537.017 | 3.278 |
| birds | Pycnopygius_stictocephalus | Pycnopygius_stictocephalus | LC | Unknown | tropical | diurnal | 38.5 | 629677.658 | 3.083 |

| | | | | | | | | | |
|---|---|---|---|---|---|---|---|---|---|
| birds | Pygarrhichas_albogularis | Pygarrhichas_albogularis | LC | Decreasing | temperate | diurnal | 24 | 470695.913 | 5.28 |
| birds | Pygiptila_stellaris | Pygiptila_stellaris | LC | Stable | tropical | diurnal | 24.1 | 4691696.055 | 1.71 |
| birds | Pygochelidon_cyanoleuca | Pygochelidon_cyanoleuca | LC | Decreasing | tropical | diurnal | 9.7 | 10370376.427 | 8.365 |
| birds | Pygochelidon_melanoleuca | Atticora_melanoleuca | LC | Stable | tropical | diurnal | 10.8 | 2130687.813 | 2.367 |
| birds | Pygoscelis_adeliae | NA | LC | Increasing | NA | NA | NA | NA | NA |
| birds | Pygoscelis_antarcticus | Pygoscelis_antarcticus | LC | Decreasing | temperate | cathemeral | 4146.1 | 33818559.546 | 2.774 |
| birds | Pygoscelis_papua | Pygoscelis_papua | LC | Stable | temperate | cathemeral | 5933 | 26559452.173 | 3.159 |
| birds | Pyrenestes_minor | Pyrenestes_minor | LC | Stable | tropical | diurnal | 14 | 437153.576 | 9.789 |
| birds | Pyrenestes_ostrinus | Pyrenestes_ostrinus | LC | Stable | tropical | diurnal | 22.4 | 4722415.431 | 7.528 |
| birds | Pyrenestes_sanguineus | Pyrenestes_sanguineus | LC | Stable | tropical | diurnal | 19.2 | 420440.403 | 14.665 |
| birds | Pyrgilauda_blanfordi | NA | LC | Stable | NA | NA | NA | NA | NA |
| birds | Pyrgilauda_davidiana | Montifringilla_davidiana | LC | Stable | temperate | diurnal | 21.2 | 2260394.819 | 5.037 |
| birds | Pyrgilauda_ruficollis | Montifringilla_ruficollis | LC | Stable | temperate | diurnal | 22.8 | 1758725.414 | 4.436 |
| birds | Pyrgilauda_theresae | Montifringilla_theresae | LC | Stable | temperate | diurnal | 24.3 | 133286.61 | 9.328 |
| birds | Pyriglena_atra | Pyriglena_atra | EN | Decreasing | tropical | diurnal | 32 | 5169.733 | 12.071 |
| birds | Pyriglena_leuconota | Pyriglena_leuconota | LC | Unknown | tropical | diurnal | 32.3 | 1982603.486 | 5.41 |
| birds | Pyriglena_leucoptera | Pyriglena_leucoptera | LC | Decreasing | tropical | diurnal | 28.8 | 1228571.496 | 12.804 |
| birds | Pyrilia_barrabandi | Pyrilia_barrabandi | LC | Decreasing | tropical | diurnal | 140 | 3875903.966 | 1.69 |
| birds | Pyrilia_caica | Pyrilia_caica | LC | Decreasing | tropical | diurnal | 132 | 1290463.273 | 1.505 |
| birds | Pyrilia_haematotis | Pyrilia_haematotis | LC | Decreasing | tropical | diurnal | 149 | 387845.619 | 8.779 |
| birds | Pyrilia_pulchra | Pyrilia_pulchra | LC | Stable | tropical | diurnal | 150 | 92646.186 | 7.935 |
| birds | Pyrilia_pyrilia | NA | NT | Decreasing | NA | NA | NA | NA | NA |
| birds | Pyrilia_vulturina | Pyrilia_vulturina | LC | Decreasing | tropical | diurnal | 151.5 | 886478.711 | 2.093 |
| birds | Pyrocephalus_nanus | NA | NT | Decreasing | NA | NA | NA | NA | NA |
| birds | Pyrocephalus_rubinus | Pyrocephalus_rubinus | LC | Increasing | tropical | diurnal | 14.4 | 17320256.156 | 7.846 |
| birds | Pyroderus_scutatus | Pyroderus_scutatus | LC | Decreasing | tropical | diurnal | 357 | 2116677.795 | 12 |
| birds | Pyrrhocorax_graculus | Pyrrhocorax_graculus | LC | Stable | temperate | diurnal | 211 | 3556954.704 | 9.324 |
| birds | Pyrrhocorax_pyrrhocorax | Pyrrhocorax_pyrrhocorax | LC | Decreasing | temperate | diurnal | 276.7 | 12981520.531 | 7.898 |
| birds | Pyrrholaemus_brunneus | Pyrrholaemus_brunneus | LC | Decreasing | temperate | diurnal | 12.5 | 3285337.242 | 3.812 |
| birds | Pyrrholaemus_sagittatus | Chthonicola_sagittatus | LC | Decreasing | temperate | diurnal | 13.3 | 1269134.495 | 8.475 |
| birds | Pyrrhomyias_cinnamomeus | Pyrrhomyias_cinnamomeus | LC | Stable | tropical | diurnal | 9.9 | 539963.443 | 8.828 |
| birds | Pyrrhoplectes_epauletta | Pyrrhoplectes_epauletta | LC | Stable | temperate | diurnal | 19 | 284503.615 | 7.526 |
| birds | Pyrrhula_aurantiaca | Pyrrhula_aurantiaca | LC | Stable | temperate | diurnal | 19.3 | 115576.221 | 14.429 |
| birds | Pyrrhula_erythaca | Pyrrhula_erythaca | LC | Stable | temperate | diurnal | 19 | 2408569.638 | 11.115 |

| | | | | | | | | | |
|---|---|---|---|---|---|---|---|---|---|
| birds | Pyrrhula_erythrocephala | Pyrrhula_erythrocephala | LC | Stable | temperate | diurnal | 22.3 | 283133.571 | 8.997 |
| birds | Pyrrhula_leucogenis | Pyrrhula_leucogenis | LC | Stable | tropical | diurnal | 19 | 16930.874 | 7.835 |
| birds | Pyrrhula_murina | Pyrrhula_murina | VU | Stable | temperate | diurnal | 30 | 144.435 | 13.563 |
| birds | Pyrrhula_nipalensis | Pyrrhula_nipalensis | LC | Stable | temperate | diurnal | 24.6 | 2202759.211 | 11.421 |
| birds | Pyrrhula_pyrrhula | Pyrrhula_pyrrhula | LC | Decreasing | temperate | diurnal | 24.3 | 24467876.149 | 9.132 |
| birds | Pyrrhulagra_nigra | NA | not_available | not_available | NA | NA | NA | NA | NA |
| birds | Pyrrhura_albipectus | Pyrrhura_albipectus | VU | Decreasing | tropical | diurnal | 103.8 | 7670.992 | 6.158 |
| birds | Pyrrhura_calliptera | NA | VU | Decreasing | NA | NA | NA | NA | NA |
| birds | Pyrrhura_cruentata | Pyrrhura_cruentata | VU | Decreasing | tropical | diurnal | 75.9 | 233537.341 | 11.47 |
| birds | Pyrrhura_devillei | NA | LC | Decreasing | NA | NA | NA | NA | NA |
| birds | Pyrrhura_egregia | Pyrrhura_egregia | LC | Decreasing | tropical | diurnal | 66.3 | 51342.199 | 1.644 |
| birds | Pyrrhura_frontalis | Pyrrhura_frontalis | LC | Stable | tropical | diurnal | 72 | 1704534.467 | 12.169 |
| birds | Pyrrhura_hoematotis | Pyrrhura_hoematotis | LC | Decreasing | tropical | diurnal | 68.7 | 6303.913 | 17.06 |
| birds | Pyrrhura_hoffmanni | Pyrrhura_hoffmanni | LC | Stable | tropical | diurnal | 82.2 | 8001.112 | 5.135 |
| birds | Pyrrhura_lepida | Pyrrhura_lepida | VU | Decreasing | tropical | diurnal | 75 | 897166.386 | 3.469 |
| birds | Pyrrhura_leucotis | Pyrrhura_leucotis | VU | Decreasing | tropical | diurnal | 75.9 | 273322.173 | 11.618 |
| birds | Pyrrhura_melanura | Pyrrhura_melanura | LC | Decreasing | tropical | diurnal | 70.5 | 1490802.925 | 1.528 |
| birds | Pyrrhura_molinae | Pyrrhura_molinae | LC | Decreasing | tropical | diurnal | 77.1 | 675243.925 | 5.976 |
| birds | Pyrrhura_orcesi | Pyrrhura_orcesi | EN | Decreasing | tropical | diurnal | 70.8 | 1002.905 | 8.495 |
| birds | Pyrrhura_perlata | Pyrrhura_perlata | LC | Decreasing | tropical | diurnal | 75 | 1396027.274 | 2.942 |
| birds | Pyrrhura_picta | Pyrrhura_picta | LC | Decreasing | tropical | diurnal | 62.1 | 1473164.348 | 1.303 |
| birds | Pyrrhura_rhodocephala | NA | LC | Stable | NA | NA | NA | NA | NA |
| birds | Pyrrhura_rupicola | Pyrrhura_rupicola | LC | Stable | tropical | diurnal | 75 | 545590.252 | 2.547 |
| birds | Pyrrhura_viridicata | Pyrrhura_viridicata | EN | Decreasing | tropical | diurnal | 87.5 | 2159.899 | 5.776 |
| birds | Pyrrhurus_scandens | Pyrrhurus_scandens | LC | Stable | tropical | diurnal | 43.8 | 3269576.147 | 8.91 |
| birds | Pyrroglaux_podargina | NA | LC | Stable | NA | NA | NA | NA | NA |
| birds | Pytilia_afra | Pytilia_afra | LC | Stable | tropical | diurnal | 16.6 | 3386528.175 | 8.708 |
| birds | Pytilia_hypogrammica | Pytilia_hypogrammica | LC | Stable | tropical | diurnal | 14.8 | 818060.767 | 12.121 |
| birds | Pytilia_lineata | Pytilia_lineata | LC | Stable | tropical | diurnal | 14.3 | 222730.635 | 10.942 |
| birds | Pytilia_melba | Pytilia_melba | LC | Stable | tropical | diurnal | 15.4 | 6033515.562 | 10.343 |
| birds | Pytilia_phoenicoptera | Pytilia_phoenicoptera | LC | Stable | tropical | diurnal | 14.5 | 1507011.59 | 10.5 |
| birds | Quelea_cardinalis | Quelea_cardinalis | LC | Stable | tropical | diurnal | 12.7 | 675544.271 | 13.246 |
| birds | Quelea_erythrops | Quelea_erythrops | LC | Stable | tropical | diurnal | 19.3 | 2572026.918 | 11.251 |
| birds | Quelea_quelea | Quelea_quelea | LC | Stable | tropical | diurnal | 18.6 | 10621873.644 | 9.779 |

| | | | | | | | | | |
|---|---|---|---|---|---|---|---|---|---|
| birds | Querula_purpurata | Querula_purpurata | LC | Decreasing | tropical | diurnal | 107.4 | 5636816.574 | 2.581 |
| birds | Quiscalus_lugubris | Quiscalus_lugubris | LC | Stable | tropical | diurnal | 63 | 918524.953 | 8.473 |
| birds | Quiscalus_major | Quiscalus_major | LC | Increasing | temperate | diurnal | 151.2 | 207614.151 | 17.554 |
| birds | Quiscalus_mexicanus | Quiscalus_mexicanus | LC | Stable | tropical | diurnal | 160.5 | 5121812.211 | 11.068 |
| birds | Quiscalus_nicaraguensis | Quiscalus_nicaraguensis | LC | Decreasing | tropical | diurnal | 63.4 | 7099.203 | 16.091 |
| birds | Quiscalus_niger | Quiscalus_niger | LC | Stable | tropical | diurnal | 72.2 | 207006.006 | 16.64 |
| birds | Quiscalus_quiscula | Quiscalus_quiscula | NT | Decreasing | temperate | diurnal | 105.2 | 8475642.149 | 12.515 |
| birds | Quoyornis_georgianus | Eopsaltria_georgiana | LC | Stable | temperate | diurnal | 19.1 | 45266.45 | 10.053 |
| birds | Radjah_radjah | Tadorna_radjah | LC | Decreasing | tropical | diurnal | 885.2 | 2173785.404 | 3.444 |
| birds | Rallicula_forbesi | Rallina_forbesi | LC | Stable | tropical | diurnal | 85.1 | 196864.345 | 4.593 |
| birds | Rallicula_leucospila | Rallina_leucospila | LC | Decreasing | tropical | diurnal | 120 | 4386.054 | 2.216 |
| birds | Rallicula_mayri | Rallina_mayri | LC | Stable | tropical | diurnal | 122.9 | 19594.627 | 6.231 |
| birds | Rallicula_rubra | Rallina_rubra | LC | Stable | tropical | diurnal | 81 | 114047.121 | 4.558 |
| birds | Rallina_canningi | NA | LC | Decreasing | NA | NA | NA | NA | NA |
| birds | Rallina_eurizonoides | Rallina_eurizonoides | LC | Decreasing | tropical | cathemeral | 109 | 3016592.948 | 11.968 |
| birds | Rallina_fasciata | Rallina_fasciata | LC | Unknown | tropical | diurnal | 100 | 1965465.703 | 10.745 |
| birds | Rallina_tricolor | Rallina_tricolor | LC | Unknown | tropical | cathemeral | 178 | 727507.098 | 3.221 |
| birds | Rallus_antarcticus | Rallus_antarcticus | VU | Decreasing | temperate | cathemeral | 60.4 | 1079988.151 | 7.033 |
| birds | Rallus_aquaticus | Rallus_aquaticus | LC | Decreasing | temperate | diurnal | 110.7 | 13788813.363 | 12.804 |
| birds | Rallus_elegans | Rallus_elegans | NT | Decreasing | temperate | diurnal | 313.3 | 3180995.761 | 18.151 |
| birds | Rallus_longirostris | Rallus_longirostris | LC | Decreasing | tropical | cathemeral | 259.4 | 359148.154 | 8.861 |
| birds | Rallus_madagascariensis | Rallus_madagascariensis | VU | Decreasing | tropical | diurnal | 114 | 167662.322 | 8.214 |
| birds | Rallus_semiplumbeus | NA | VU | Decreasing | NA | NA | NA | NA | NA |
| birds | Rallus_wetmorei | NA | EN | Decreasing | NA | NA | NA | NA | NA |
| birds | Ramphastos_ambiguus | Ramphastos_ambiguus | LC | Decreasing | tropical | diurnal | 651.7 | 684339.23 | 9 |
| birds | Ramphastos_brevis | Ramphastos_brevis | LC | Decreasing | tropical | diurnal | 412 | 105684.496 | 7.081 |
| birds | Ramphastos_dicolorus | Ramphastos_dicolorus | LC | Decreasing | temperate | diurnal | 331 | 1394133.062 | 13.273 |
| birds | Ramphastos_sulfuratus | Ramphastos_sulfuratus | NT | Decreasing | tropical | diurnal | 409.7 | 717685.394 | 10.291 |
| birds | Ramphastos_toco | Ramphastos_toco | LC | Decreasing | tropical | diurnal | 618 | 4573489.807 | 8.606 |
| birds | Ramphastos_tucanus | Ramphastos_tucanus | LC | Decreasing | tropical | diurnal | 659.6 | 2365965.204 | 2.313 |
| birds | Ramphastos_vitellinus | Ramphastos_vitellinus | LC | Decreasing | tropical | diurnal | 360.4 | 1748066.749 | 1.511 |
| birds | Ramphiculus_fischeri | Ptilinopus_fischeri | LC | Decreasing | tropical | diurnal | 177 | 41322.27 | 5.401 |
| birds | Ramphiculus_jambu | Ptilinopus_jambu | NT | Decreasing | tropical | diurnal | 135 | 1433397.98 | 8.376 |
| birds | Ramphiculus_leclancheri | Ptilinopus_leclancheri | LC | Decreasing | tropical | diurnal | 164 | 288493.562 | 13.914 |

| | | | | | | | | | |
|---|---|---|---|---|---|---|---|---|---|
| birds | Ramphiculus_marchei | Ptilinopus_marchei | VU | Decreasing | tropical | diurnal | 326 | 53418.747 | 10.127 |
| birds | Ramphiculus_merrilli | Ptilinopus_merrilli | NT | Decreasing | tropical | diurnal | 263 | 63334.69 | 12.786 |
| birds | Ramphiculus_occipitalis | Ptilinopus_occipitalis | LC | Decreasing | tropical | diurnal | 238 | 267780.834 | 13.839 |
| birds | Ramphiculus_subgularis | Ptilinopus_subgularis | VU | Decreasing | tropical | diurnal | 142 | 2670.994 | 9.251 |
| birds | Ramphocaenus_melanurus | Ramphocaenus_melanurus | LC | Decreasing | tropical | diurnal | 9.7 | 9179813.285 | 4.045 |
| birds | Ramphocelus_bresilius | Ramphocelus_bresilius | LC | Stable | tropical | diurnal | 32.9 | 377620.268 | 12.36 |
| birds | Ramphocelus_carbo | Ramphocelus_carbo | LC | Stable | tropical | diurnal | 25.9 | 10498944.337 | 4.544 |
| birds | Ramphocelus_costaricensis | NA | not_available | not_available | NA | NA | NA | NA | NA |
| birds | Ramphocelus_dimidiatus | Ramphocelus_dimidiatus | LC | Stable | tropical | diurnal | 28 | 475288.523 | 12.413 |
| birds | Ramphocelus_flammigerus | Ramphocelus_flammigerus | LC | Increasing | tropical | diurnal | 33 | 347421.545 | 11.482 |
| birds | Ramphocelus_icteronotus | NA | not_available | not_available | NA | NA | NA | NA | NA |
| birds | Ramphocelus_melanogaster | Ramphocelus_melanogaster | LC | Stable | tropical | diurnal | 25 | 76666.78 | 5.097 |
| birds | Ramphocelus_nigrogularis | Ramphocelus_nigrogularis | LC | Stable | tropical | diurnal | 31 | 2107052.176 | 1.614 |
| birds | Ramphocelus_passerinii | Ramphocelus_passerinii | LC | Increasing | tropical | diurnal | 32 | 302241.047 | 9.588 |
| birds | Ramphocelus_sanguinolentus | Ramphocelus_sanguinolentus | LC | Decreasing | tropical | diurnal | 41 | 277681.383 | 9.541 |
| birds | Ramphocinclus_brachyurus | Ramphocinclus_brachyurus | EN | Decreasing | tropical | diurnal | 50 | 1382.846 | 26.154 |
| birds | Ramphocoris_clotbey | Rhamphocoris_clotbey | LC | Stable | temperate | diurnal | 49.1 | 934337.779 | 4.663 |
| birds | Ramphodon_naevius | Ramphodon_naevius | LC | Decreasing | temperate | diurnal | 7.9 | 143609.058 | 15.069 |
| birds | Ramphomicron_dorsale | Ramphomicron_dorsale | NT | Decreasing | tropical | diurnal | 3.5 | 5513.32 | 6.111 |
| birds | Ramphomicron_microrhynchum | Ramphomicron_microrhynchum | LC | Decreasing | tropical | diurnal | 3.6 | 203534.205 | 10.845 |
| birds | Ramphotrigon_fuscicauda | Ramphotrigon_fuscicauda | LC | Decreasing | tropical | diurnal | 18.6 | 472095.304 | 2.461 |
| birds | Ramphotrigon_megacephalum | Ramphotrigon_megacephalum | LC | Decreasing | tropical | diurnal | 14.2 | 2269306.342 | 3.694 |
| birds | Ramphotrigon_ruficauda | Ramphotrigon_ruficauda | LC | Decreasing | tropical | diurnal | 18.4 | 6736020.91 | 2.138 |
| birds | Ramsayornis_fasciatus | Ramsayornis_fasciatus | LC | Stable | tropical | diurnal | 12.7 | 931104.022 | 2.743 |
| birds | Ramsayornis_modestus | Ramsayornis_modestus | LC | Stable | tropical | diurnal | 12.1 | 290171.245 | 3.202 |
| birds | Randia_pseudozosterops | Randia_pseudozosterops | LC | Decreasing | tropical | diurnal | 11 | 149492.808 | 8.059 |
| birds | Recurvirostra_americana | Recurvirostra_americana | LC | Stable | temperate | diurnal | 304.5 | 6045989.844 | 10.961 |
| birds | Recurvirostra_andina | Recurvirostra_andina | LC | Unknown | tropical | diurnal | 361 | 708582.58 | 6.179 |
| birds | Recurvirostra_avosetta | Recurvirostra_avosetta | LC | Unknown | tropical | diurnal | 304 | 33474924.456 | 9.598 |
| birds | Recurvirostra_novaehollandiae | Recurvirostra_novaehollandiae | LC | Stable | temperate | diurnal | 325 | 3538477.508 | 5.976 |
| birds | Regulus_goodfellowi | Regulus_goodfellowi | LC | Stable | temperate | diurnal | 4.1 | 22144.911 | 11.11 |
| birds | Regulus_ignicapilla | Regulus_ignicapilla | LC | Stable | temperate | diurnal | 5.6 | 3980020.751 | 19.723 |
| birds | Regulus_regulus | Regulus_regulus | LC | Decreasing | temperate | diurnal | 5.5 | 16994335.778 | 12.097 |
| birds | Regulus_satrapa | Regulus_satrapa | LC | Stable | temperate | diurnal | 6.2 | 11944577.152 | 10.333 |

| | | | | | | | | | |
|---|---|---|---|---|---|---|---|---|---|
| birds | Reinwardtoena_browni | Reinwardtoena_browni | NT | Decreasing | tropical | diurnal | 304 | 45117.985 | 4.608 |
| birds | Reinwardtoena_crassirostris | Reinwardtoena_crassirostris | LC | Decreasing | tropical | diurnal | 259 | 34295.639 | 4.581 |
| birds | Reinwardtoena_reinwardti | Reinwardtoena_reinwardtsi | LC | Stable | tropical | diurnal | 275 | 656098.158 | 4.054 |
| birds | Remiz_consobrinus | Remiz_consobrinus | LC | Increasing | temperate | diurnal | 9.1 | 1693475.347 | 12.241 |
| birds | Remiz_coronatus | Remiz_coronatus | LC | Decreasing | temperate | diurnal | 6.9 | 3914348.344 | 8.063 |
| birds | Remiz_pendulinus | Remiz_pendulinus | LC | Increasing | temperate | diurnal | 9.3 | 12487369.779 | 12.569 |
| birds | Rhabdornis_grandis | Rhabdornis_grandis | LC | Decreasing | tropical | diurnal | 45.5 | 14055.762 | 7.244 |
| birds | Rhabdornis_inornatus | Rhabdornis_inornatus | LC | Decreasing | tropical | diurnal | 39.1 | 114902.722 | 11.934 |
| birds | Rhabdornis_mystacalis | Rhabdornis_mystacalis | LC | Decreasing | tropical | diurnal | 25.6 | 247778.102 | 14.012 |
| birds | Rhabdotorrhinus_corrugatus | Aceros_corrugatus | EN | Decreasing | tropical | diurnal | 1590 | 892566.532 | 9.241 |
| birds | Rhabdotorrhinus_exarhatus | Penelopides_exarhatus | VU | Decreasing | tropical | diurnal | 370 | 119932.785 | 8.815 |
| birds | Rhabdotorrhinus_leucocephalus | Aceros_leucocephalus | NT | Decreasing | tropical | diurnal | 1086 | 89999.667 | 12.332 |
| birds | Rhabdotorrhinus_waldeni | NA | CR | Decreasing | NA | NA | NA | NA | NA |
| birds | Rhagologus_leucostigma | Rhagologus_leucostigma | LC | Stable | tropical | diurnal | 26.8 | 150755.686 | 4.674 |
| birds | Rhamphocharis_crassirostris | Melanocharis_crassirostris | LC | Unknown | tropical | diurnal | 18 | 34344.011 | 2.543 |
| birds | Rhamphococcyx_calyorhynchus | NA | LC | Decreasing | NA | NA | NA | NA | NA |
| birds | Rhaphidura_leucopygialis | Rhaphidura_leucopygialis | LC | Stable | tropical | diurnal | 13.6 | 1203937.954 | 9.808 |
| birds | Rhaphidura_sabini | Rhaphidura_sabini | LC | Stable | tropical | diurnal | 16 | 1778431.253 | 9.333 |
| birds | Rhea_americana | Rhea_americana | NT | Decreasing | tropical | diurnal | 23000 | 6949731.032 | 9.037 |
| birds | Rhea_pennata | Rhea_pennata | LC | Decreasing | temperate | diurnal | 23900 | 1309452.501 | 4.832 |
| birds | Rhegmatorhina_berlepschi | Rhegmatorhina_berlepschi | LC | Decreasing | tropical | diurnal | 31.3 | 68143.221 | 1.654 |
| birds | Rhegmatorhina_cristata | Rhegmatorhina_cristata | LC | Stable | tropical | diurnal | 31.2 | 267428.697 | 0.618 |
| birds | Rhegmatorhina_gymnops | Rhegmatorhina_gymnops | VU | Decreasing | tropical | diurnal | 28.5 | 320154.806 | 2.327 |
| birds | Rhegmatorhina_hoffmannsi | Rhegmatorhina_hoffmannsi | LC | Decreasing | tropical | diurnal | 31.4 | 777485.332 | 2.584 |
| birds | Rhegmatorhina_melanosticta | Rhegmatorhina_melanosticta | LC | Decreasing | tropical | diurnal | 31 | 1909667.49 | 1.665 |
| birds | Rheinardia_ocellata | Rheinardia_ocellata | CR | Decreasing | tropical | diurnal | 1500 | 45112.576 | 6.247 |
| birds | Rhinocrypta_lanceolata | Rhinocrypta_lanceolata | LC | Stable | temperate | diurnal | 61.9 | 1006274.619 | 5.742 |
| birds | Rhinoplax_vigil | NA | CR | Decreasing | NA | NA | NA | NA | NA |
| birds | Rhinopomastus_aterrimus | Rhinopomastus_aterrimus | LC | Decreasing | tropical | diurnal | 27.1 | 7792738 | 9.421 |
| birds | Rhinopomastus_castaneiceps | Phoeniculus_castaneiceps | LC | Decreasing | tropical | diurnal | 24.4 | 622400.006 | 10.927 |
| birds | Rhinopomastus_cyanomelas | Rhinopomastus_cyanomelas | LC | Decreasing | tropical | diurnal | 27.1 | 6396418.321 | 8.682 |
| birds | Rhinopomastus_minor | Rhinopomastus_minor | LC | Decreasing | tropical | diurnal | 21.7 | 2064431.667 | 9.043 |
| birds | Rhinoptilus_bitorquatus | NA | CR | Decreasing | NA | NA | NA | NA | NA |
| birds | Rhinoptilus_chalcopterus | Rhinoptilus_chalcopterus | LC | Stable | tropical | nocturnal | 150 | 10165054.177 | 9.752 |

| | | | | | | | | | |
|---|---|---|---|---|---|---|---|---|---|
| birds | Rhinoptilus_cinctus | Rhinoptilus_cinctus | LC | Stable | tropical | nocturnal | 105 | 1957707.604 | 9.867 |
| birds | Rhinortha_chlorophaea | Phaenicophaeus_chlorophaeus | LC | Stable | tropical | diurnal | 51.2 | 1422260.564 | 8.033 |
| birds | Rhipidura_albicollis | Rhipidura_albicollis | LC | Stable | tropical | diurnal | 12.9 | 4734174.117 | 12.275 |
| birds | Rhipidura_albolimbata | Rhipidura_albolimbata | LC | Stable | tropical | diurnal | 10.9 | 148493.841 | 4.877 |
| birds | Rhipidura_atra | Rhipidura_atra | LC | Stable | tropical | diurnal | 12.2 | 179053.161 | 4.211 |
| birds | Rhipidura_aureola | Rhipidura_aureola | LC | Stable | tropical | diurnal | 11.2 | 4306980.922 | 16.244 |
| birds | Rhipidura_brachyrhyncha | Rhipidura_brachyrhyncha | LC | Stable | tropical | diurnal | 10.1 | 131180.684 | 4.794 |
| birds | Rhipidura_cockerelli | Rhipidura_cockerelli | LC | Decreasing | tropical | diurnal | 17.5 | 27207.093 | 4.655 |
| birds | Rhipidura_cyaniceps | Rhipidura_cyaniceps | LC | Stable | tropical | diurnal | 13.6 | 83397.442 | 12.258 |
| birds | Rhipidura_dahli | Rhipidura_dahli | LC | Stable | tropical | diurnal | 9.9 | 43231.522 | 4.394 |
| birds | Rhipidura_dedemi | Rhipidura_dedemi | LC | Stable | tropical | diurnal | 11 | 3375.296 | 5.823 |
| birds | Rhipidura_diluta | Rhipidura_diluta | LC | Stable | tropical | diurnal | 15 | 30545.409 | 12.504 |
| birds | Rhipidura_drownei | Rhipidura_drownei | LC | Stable | tropical | diurnal | 11.1 | 1905.469 | 4.876 |
| birds | Rhipidura_euryura | Rhipidura_euryura | LC | Decreasing | tropical | diurnal | 15 | 8512.961 | 12.458 |
| birds | Rhipidura_fuliginosa | Rhipidura_fuliginosa | LC | Unknown | temperate | diurnal | 7.5 | 267492.072 | 7.985 |
| birds | Rhipidura_fuscorufa | Rhipidura_fuscorufa | LC | Decreasing | tropical | diurnal | 15 | 4912.735 | 6.743 |
| birds | Rhipidura_hyperythra | Rhipidura_hyperythra | LC | Stable | tropical | diurnal | 11.2 | 465792.132 | 3.241 |
| birds | Rhipidura_javanica | Rhipidura_javanica | LC | Stable | tropical | diurnal | 12.5 | 1594085.243 | 13.044 |
| birds | Rhipidura_kubaryi | Rhipidura_kubaryi | LC | Stable | tropical | diurnal | 15 | 351.183 | 12.237 |
| birds | Rhipidura_lepida | Rhipidura_lepida | LC | Increasing | tropical | diurnal | 15 | 442.777 | 11.201 |
| birds | Rhipidura_leucophrys | Rhipidura_leucophrys | LC | Increasing | temperate | diurnal | 27.4 | 8400289.568 | 4.204 |
| birds | Rhipidura_leucothorax | Rhipidura_leucothorax | LC | Stable | tropical | diurnal | 17.6 | 565343.927 | 3.241 |
| birds | Rhipidura_maculipectus | Rhipidura_maculipectus | LC | Stable | tropical | diurnal | 18 | 137530.75 | 3.261 |
| birds | Rhipidura_malaitae | Rhipidura_malaitae | NT | Unknown | tropical | diurnal | 12 | 615.589 | 2.933 |
| birds | Rhipidura_matthiae | Rhipidura_matthiae | NT | Unknown | tropical | diurnal | 15 | 355.494 | 3.98 |
| birds | Rhipidura_nebulosa | NA | LC | Stable | NA | NA | NA | NA | NA |
| birds | Rhipidura_nigrocinnamomea | Rhipidura_nigrocinnamomea | LC | Decreasing | tropical | diurnal | 13 | 9000.73 | 6.707 |
| birds | Rhipidura_opistherythra | Rhipidura_opistherythra | NT | Decreasing | tropical | diurnal | 15 | 3122.041 | 5.622 |
| birds | Rhipidura_perlata | Rhipidura_perlata | LC | Decreasing | tropical | diurnal | 13.4 | 1287003.576 | 7.622 |
| birds | Rhipidura_personata | Rhipidura_personata | NT | Decreasing | tropical | diurnal | 35.7 | 462.602 | 6.033 |
| birds | Rhipidura_phasiana | Rhipidura_phasiana | LC | Stable | tropical | diurnal | 6.9 | 319575.426 | 2.98 |
| birds | Rhipidura_phoenicura | Rhipidura_phoenicura | LC | Decreasing | tropical | diurnal | 15 | 8270.501 | 12.44 |
| birds | Rhipidura_rennelliana | Rhipidura_rennelliana | LC | Stable | tropical | diurnal | 14.5 | 675.464 | 3.166 |
| birds | Rhipidura_rufidorsa | Rhipidura_rufidorsa | LC | Decreasing | tropical | diurnal | 9.8 | 489139.146 | 3.138 |

| | | | | | | | | | |
|---|---|---|---|---|---|---|---|---|---|
| birds | Rhipidura_rufifrons | Rhipidura_rufifrons | LC | Decreasing | temperate | diurnal | 10.1 | 918089.741 | 7.091 |
| birds | Rhipidura_rufiventris | Rhipidura_rufiventris | LC | Stable | tropical | diurnal | 12 | 31219.37 | 13.074 |
| birds | Rhipidura_semirubra | Rhipidura_semirubra | LC | Stable | tropical | diurnal | 8 | 2012.606 | 6.325 |
| birds | Rhipidura_spilodera | Rhipidura_verreauxi | LC | Stable | tropical | diurnal | 12.9 | 9510.788 | 5.757 |
| birds | Rhipidura_superciliaris | Rhipidura_superciliaris | LC | Stable | tropical | diurnal | 13.1 | 92445.848 | 12.233 |
| birds | Rhipidura_superflua | Rhipidura_superflua | LC | Stable | tropical | diurnal | 15 | 2601.615 | 4.781 |
| birds | Rhipidura_tenebrosa | Rhipidura_tenebrosa | NT | Decreasing | tropical | diurnal | 17.3 | 3338.039 | 4.602 |
| birds | Rhipidura_teysmanni | Rhipidura_teysmanni | LC | Decreasing | tropical | diurnal | 10.7 | 66641.374 | 5.739 |
| birds | Rhipidura_threnothorax | Rhipidura_threnothorax | LC | Decreasing | tropical | diurnal | 17.6 | 594363.366 | 3.145 |
| birds | Rhizothera_longirostris | Rhizothera_longirostris | NT | Decreasing | tropical | diurnal | 746.7 | 1292604.793 | 8.588 |
| birds | Rhodinocichla_rosea | Rhodinocichla_rosea | LC | Decreasing | tropical | diurnal | 48 | 163331.806 | 11.66 |
| birds | Rhodopechys_sanguineus | Rhodopechys_sanguineus | LC | Stable | temperate | diurnal | 38.1 | 1852767.153 | 11.456 |
| birds | Rhodophoneus_cruentus | Rhodophoneus_cruentus | LC | Stable | tropical | diurnal | 53.3 | 1845043.079 | 8.277 |
| birds | Rhodopis_vesper | Rhodopis_vesper | LC | Stable | tropical | diurnal | 3.8 | 305602.243 | 5.787 |
| birds | Rhodospingus_cruentus | Rhodospingus_cruentus | LC | Stable | tropical | diurnal | 11.6 | 95985.12 | 12.533 |
| birds | Rhodospiza_obsoleta | Rhodopechys_obsoletus | LC | Stable | temperate | diurnal | 25.5 | 3747421.266 | 7.375 |
| birds | Rhodostethia_rosea | Rhodostethia_rosea | LC | Unknown | temperate | diurnal | 187 | 3404341.738 | 0.429 |
| birds | Rhopias_gularis | Myrmotherula_gularis | LC | Decreasing | temperate | diurnal | 11.3 | 500928.793 | 13.858 |
| birds | Rhopophilus_pekinensis | Rhopophilus_pekinensis | LC | Decreasing | temperate | diurnal | 18 | 1353087.16 | 14.654 |
| birds | Rhopornis_ardesiacus | Rhopornis_ardesiacus | EN | Decreasing | tropical | diurnal | 26.3 | 2749.531 | 9.629 |
| birds | Rhopospina_fruticeti | Phrygilus_fruticeti | LC | Stable | temperate | diurnal | 38.8 | 2085092.664 | 6.209 |
| birds | Rhynchocyclus_brevirostris | Rhynchocyclus_brevirostris | LC | Decreasing | tropical | diurnal | 24.3 | 579326.26 | 9.098 |
| birds | Rhynchocyclus_fulvipectus | Rhynchocyclus_fulvipectus | LC | Stable | tropical | diurnal | 23.1 | 196891.42 | 6.014 |
| birds | Rhynchocyclus_olivaceus | Rhynchocyclus_olivaceus | LC | Stable | tropical | diurnal | 21.3 | 2806251.132 | 2.794 |
| birds | Rhynchocyclus_pacificus | Rhynchocyclus_pacificus | LC | Stable | tropical | diurnal | 25 | 105882.399 | 6.449 |
| birds | Rhynchophanes_mccownii | Calcarius_mccownii | LC | Stable | temperate | diurnal | 25.7 | 2569799.103 | 10.98 |
| birds | Rhynchopsitta_pachyrhyncha | Rhynchopsitta_pachyrhyncha | EN | Decreasing | temperate | diurnal | 324.8 | 173654.148 | 7.811 |
| birds | Rhynchopsitta_terrisi | Rhynchopsitta_terrisi | EN | Decreasing | temperate | diurnal | 442 | 12870.534 | 5.784 |
| birds | Rhynchortyx_cinctus | Rhynchortyx_cinctus | NT | Decreasing | tropical | diurnal | 150 | 221334.002 | 6.257 |
| birds | Rhynchospiza_stolzmanni | Aimophila_stolzmanni | LC | Stable | tropical | diurnal | 25.6 | 48995.911 | 8.319 |
| birds | Rhynchospiza_strigiceps | Aimophila_strigiceps | LC | Stable | temperate | diurnal | 22.2 | 666629.129 | 7.944 |
| birds | Rhynchostruthus_socotranus | Rhynchostruthus_socotranus | LC | Stable | tropical | diurnal | 28.3 | 3622.403 | 11.03 |
| birds | Rhynchotus_rufescens | Rhynchotus_rufescens | LC | Decreasing | tropical | diurnal | 843.5 | 5674200.175 | 9.685 |
| birds | Rhynochetos_jubatus | Rhynochetos_jubatus | EN | Decreasing | tropical | diurnal | 860 | 3143.338 | 3.56 |

| | | | | | | | | | |
|---|---|---|---|---|---|---|---|---|---|
| birds | Rhyticeros_cassidix | Aceros_cassidix | VU | Decreasing | tropical | diurnal | 2360 | 164525.735 | 9.686 |
| birds | Rhyticeros_everetti | NA | EN | Decreasing | NA | NA | NA | NA | NA |
| birds | Rhyticeros_narcondami | Aceros_narcondami | VU | Stable | tropical | diurnal | 697.2 | 6.511 | 0.06 |
| birds | Rhyticeros_plicatus | Aceros_plicatus | LC | Decreasing | tropical | diurnal | 1703.8 | 754091.955 | 3.427 |
| birds | Rhyticeros_subruficollis | Aceros_subruficollis | VU | Decreasing | tropical | diurnal | 2042 | 72279.839 | 7.136 |
| birds | Rhyticeros_undulatus | Aceros_undulatus | VU | Decreasing | tropical | diurnal | 2214.6 | 3225027.077 | 10.978 |
| birds | Rhytipterna_holerythra | Rhytipterna_holerythra | LC | Decreasing | tropical | diurnal | 36.8 | 469909.568 | 8.444 |
| birds | Rhytipterna_immunda | Rhytipterna_immunda | LC | Stable | tropical | diurnal | 27.4 | 1435282.352 | 2.875 |
| birds | Rhytipterna_simplex | Rhytipterna_simplex | LC | Decreasing | tropical | diurnal | 31.8 | 7376423.256 | 2.667 |
| birds | Riccordia_bicolor | Cyanophaia_bicolor | LC | Decreasing | tropical | diurnal | 4.5 | 1906.024 | 21.117 |
| birds | Riccordia_maugaeus | Chlorostilbon_maugaeus | LC | Stable | tropical | diurnal | 3.1 | 8728.542 | 27.268 |
| birds | Riccordia_ricordii | Chlorostilbon_ricordii | LC | Unknown | tropical | diurnal | 3.2 | 119014.468 | 15.422 |
| birds | Riccordia_swainsonii | Chlorostilbon_swainsonii | LC | Unknown | tropical | diurnal | 4.3 | 74117.206 | 15.787 |
| birds | Ridgwayia_pinicola | Zoothera_pinicola | LC | Decreasing | tropical | diurnal | 88 | 379299.808 | 8.498 |
| birds | Rimator_danjoui | NA | not_available | not_available | NA | NA | NA | NA | NA |
| birds | Rimator_malacoptilus | NA | not_available | not_available | NA | NA | NA | NA | NA |
| birds | Riparia_congica | Riparia_congica | LC | Stable | tropical | diurnal | 16.5 | 164353.549 | 6.64 |
| birds | Riparia_riparia | Riparia_riparia | LC | Decreasing | temperate | diurnal | 12.7 | 71794265.121 | 8.137 |
| birds | Rissa_brevirostris | Rissa_brevirostris | VU | Decreasing | temperate | diurnal | 377 | 12249906.819 | 1.831 |
| birds | Rissa_tridactyla | Rissa_tridactyla | VU | Decreasing | temperate | diurnal | 416 | 44680386.753 | 2.413 |
| birds | Robsonius_rabori | Robsonius_rabori | VU | Decreasing | tropical | diurnal | 61 | 5360.944 | 6.207 |
| birds | Rollandia_microptera | Rollandia_microptera | EN | Decreasing | tropical | diurnal | 706 | 48882.25 | 9.65 |
| birds | Rollandia_rolland | Rollandia_rolland | LC | Decreasing | temperate | diurnal | 324.3 | 4837890.416 | 7.565 |
| birds | Rollulus_rouloul | Rollulus_rouloul | VU | Decreasing | tropical | diurnal | 216.5 | 1418421.489 | 8.073 |
| birds | Roraimia_adusta | Roraimia_adusta | LC | Stable | tropical | diurnal | 16.2 | 71553.865 | 1.251 |
| birds | Rostrhamus_sociabilis | Rostrhamus_sociabilis | LC | Stable | tropical | diurnal | 366.9 | 11603405.026 | 7.257 |
| birds | Rougetius_rougetii | Rougetius_rougetii | NT | Decreasing | tropical | diurnal | 193.4 | 344840.952 | 13.09 |
| birds | Rowettia_goughensis | NA | CR | Decreasing | NA | NA | NA | NA | NA |
| birds | Rubigula_melanictera | Pycnonotus_melanicterus | LC | Decreasing | tropical | diurnal | 29.3 | 62208.809 | 17.267 |
| birds | Rufirallus_castaneiceps | Anurolimnas_castaneiceps | LC | Decreasing | tropical | diurnal | 126 | 607612.538 | 2.164 |
| birds | Rufirallus_viridis | Anurolimnas_viridis | LC | Unknown | tropical | diurnal | 64.3 | 6139478.63 | 4.057 |
| birds | Rukia_longirostra | Rukia_longirostra | NT | Decreasing | tropical | diurnal | 17.9 | 351.152 | 12.237 |
| birds | Rukia_ruki | Rukia_ruki | EN | Decreasing | tropical | diurnal | 21 | 35.569 | 10.543 |
| birds | Rupicola_peruvianus | Rupicola_peruvianus | LC | Decreasing | tropical | diurnal | 242.5 | 257511.056 | 7.49 |

| | | | | | | | | | |
|---|---|---|---|---|---|---|---|---|---|
| birds | Rupicola_rupicola | Rupicola_rupicola | LC | Stable | tropical | diurnal | 194.6 | 2067586.185 | 1.392 |
| birds | Rupornis_magnirostris | Buteo_magnirostris | LC | Increasing | tropical | diurnal | 269 | 15043360.16 | 6.218 |
| birds | Rynchops_albicollis | NA | EN | Decreasing | NA | NA | NA | NA | NA |
| birds | Rynchops_flavirostris | Rynchops_flavirostris | LC | Decreasing | tropical | cathemeral | 164 | 11670589.884 | 9.221 |
| birds | Rynchops_niger | Rynchops_niger | LC | Decreasing | tropical | cathemeral | 297.7 | 14191827.261 | 6.235 |
| birds | Sagittarius_serpentarius | Sagittarius_serpentarius | EN | Decreasing | tropical | diurnal | 4017 | 14023594.923 | 9.111 |
| birds | Sakesphorus_canadensis | Sakesphorus_canadensis | LC | Decreasing | tropical | diurnal | 24.4 | 1626726.492 | 3.752 |
| birds | Sakesphorus_cristatus | Sakesphorus_cristatus | LC | Decreasing | tropical | diurnal | 17.7 | 1141026.652 | 8.887 |
| birds | Sakesphorus_luctuosus | Sakesphorus_luctuosus | LC | Decreasing | tropical | diurnal | 31.4 | 1618420.864 | 2.363 |
| birds | Salpinctes_obsoletus | Salpinctes_obsoletus | LC | Decreasing | temperate | diurnal | 16.4 | 5577879.407 | 9.914 |
| birds | Salpornis_spilonota | Salpornis_spilonotus | LC | Decreasing | tropical | diurnal | 14.8 | 952342.709 | 18.273 |
| birds | Saltator_albicollis | Saltator_albicollis | LC | Unknown | tropical | diurnal | 41.3 | 3992.013 | 22.23 |
| birds | Saltator_atriceps | Saltator_atriceps | LC | Decreasing | tropical | diurnal | 83.8 | 710915.845 | 10.053 |
| birds | Saltator_atricollis | Saltator_atricollis | LC | Unknown | tropical | diurnal | 53.8 | 3347116.202 | 9.041 |
| birds | Saltator_atripennis | Saltator_atripennis | LC | Unknown | tropical | diurnal | 55 | 110254.944 | 10.297 |
| birds | Saltator_aurantiirostris | Saltator_aurantiirostris | LC | Unknown | temperate | diurnal | 41.9 | 2474154.664 | 7.677 |
| birds | Saltator_cinctus | Saltator_cinctus | LC | Decreasing | tropical | diurnal | 48.6 | 75858.599 | 9.229 |
| birds | Saltator_coerulescens | Saltator_coerulescens | LC | Stable | tropical | diurnal | 54.9 | 9192151.569 | 4.722 |
| birds | Saltator_fuliginosus | Saltator_fuliginosus | LC | Unknown | tropical | diurnal | 44 | 811739.376 | 13.404 |
| birds | Saltator_grossus | Saltator_grossus | LC | Decreasing | tropical | diurnal | 44.2 | 6243128.127 | 2.073 |
| birds | Saltator_maxillosus | Saltator_maxillosus | LC | Unknown | temperate | diurnal | 50.8 | 436886.303 | 12.64 |
| birds | Saltator_maximus | Saltator_maximus | LC | Decreasing | tropical | diurnal | 47.6 | 9544602.66 | 4.235 |
| birds | Saltator_multicolor | Saltatricula_multicolor | LC | Decreasing | temperate | diurnal | 22.2 | 1113148.292 | 6.678 |
| birds | Saltator_nigriceps | Saltator_nigriceps | LC | Unknown | tropical | diurnal | 71 | 26525.419 | 9.042 |
| birds | Saltator_orenocensis | Saltator_orenocensis | LC | Unknown | tropical | diurnal | 33.7 | 410839.036 | 9.318 |
| birds | Saltator_similis | Saltator_similis | LC | Decreasing | tropical | diurnal | 43.3 | 3782501.252 | 9.926 |
| birds | Saltator_striatipectus | Saltator_striatipectus | LC | Stable | tropical | diurnal | 39 | 889950.272 | 12.314 |
| birds | Salvadorina_waigiuensis | Salvadorina_waigiuensis | LC | Unknown | tropical | diurnal | 324 | 339423.008 | 3.927 |
| birds | Sapayoa_aenigma | Sapayoa_aenigma | LC | Decreasing | tropical | diurnal | 20.8 | 103347.576 | 5.346 |
| birds | Sappho_sparganurus | Sappho_sparganura | LC | Stable | temperate | diurnal | 5.3 | 409731.949 | 6.944 |
| birds | Sarcogyps_calvus | Sarcogyps_calvus | CR | Decreasing | tropical | diurnal | 4469.9 | 4444974.887 | 15.684 |
| birds | Sarcophanops_samarensis | Eurylaimus_samarensis | VU | Decreasing | tropical | diurnal | 27.1 | 22689.043 | 12.522 |
| birds | Sarcophanops_steerii | Eurylaimus_steerii | VU | Decreasing | tropical | diurnal | 38.7 | 96675.604 | 12.021 |
| birds | Sarcops_calvus | Sarcops_calvus | LC | Unknown | tropical | diurnal | 142 | 253849.176 | 14.179 |

| | | | | | | | | | |
|---|---|---|---|---|---|---|---|---|---|
| birds | Sarcoramphus_papa | Sarcoramphus_papa | LC | Decreasing | tropical | diurnal | 3400 | 14224410.709 | 5.916 |
| birds | Sarkidiornis_melanotos | Sarkidiornis_melanotos | LC | Decreasing | tropical | diurnal | 2610 | 18695511.284 | 9.928 |
| birds | Saroglossa_spilopterus | Saroglossa_spiloptera | LC | Decreasing | temperate | diurnal | 47.5 | 728867.555 | 13.354 |
| birds | Sarothrura_affinis | Sarothrura_affinis | LC | Decreasing | temperate | diurnal | 27.8 | 524225.878 | 12.54 |
| birds | Sarothrura_ayresi | Sarothrura_ayresi | CR | Decreasing | temperate | diurnal | 31.8 | 344822.653 | 13.343 |
| birds | Sarothrura_boehmi | Sarothrura_boehmi | LC | Decreasing | tropical | diurnal | 35 | 1772998.93 | 8.378 |
| birds | Sarothrura_elegans | Sarothrura_elegans | LC | Stable | tropical | diurnal | 46 | 7047799.89 | 8.693 |
| birds | Sarothrura_insularis | Sarothrura_insularis | LC | Stable | tropical | diurnal | 26.8 | 289629.495 | 9.111 |
| birds | Sarothrura_lugens | Sarothrura_lugens | LC | Decreasing | tropical | diurnal | 30.8 | 189277.403 | 8.177 |
| birds | Sarothrura_pulchra | Sarothrura_pulchra | LC | Decreasing | tropical | diurnal | 45.2 | 4635423.451 | 8.401 |
| birds | Sarothrura_rufa | Sarothrura_rufa | LC | Decreasing | temperate | diurnal | 37.7 | 4402554.999 | 9.194 |
| birds | Sarothrura_watersi | Sarothrura_watersi | NT | Decreasing | tropical | diurnal | 26.5 | 13260.929 | 10.63 |
| birds | Sasia_abnormis | Sasia_abnormis | LC | Stable | tropical | diurnal | 9.2 | 1159649.152 | 9.576 |
| birds | Sasia_ochracea | Sasia_ochracea | LC | Stable | tropical | diurnal | 9.7 | 1515543.877 | 10.253 |
| birds | Satrapa_icterophrys | Satrapa_icterophrys | LC | Stable | tropical | diurnal | 21.5 | 8028682.879 | 7.606 |
| birds | Saucerottia_beryllina | Amazilia_beryllina | LC | Stable | tropical | diurnal | 4.5 | 683715.078 | 10.737 |
| birds | Saucerottia_castaneiventris | Amazilia_castaneiventris | NT | Decreasing | tropical | diurnal | 5 | 4916.587 | 14.751 |
| birds | Saucerottia_cupreicauda | NA | LC | Decreasing | NA | NA | NA | NA | NA |
| birds | Saucerottia_cyanocephala | Amazilia_cyanocephala | LC | Decreasing | tropical | diurnal | 5.1 | 304040.65 | 11.135 |
| birds | Saucerottia_cyanura | Amazilia_cyanura | LC | Decreasing | tropical | diurnal | 3.9 | 89777.59 | 15.158 |
| birds | Saucerottia_edward | Amazilia_edward | LC | Stable | tropical | diurnal | 4.7 | 42606.222 | 11.933 |
| birds | Saucerottia_saucerottei | Amazilia_saucerrottei | LC | Decreasing | tropical | diurnal | 4.5 | 245059.299 | 14.932 |
| birds | Saucerottia_tobaci | Amazilia_tobaci | LC | Unknown | tropical | diurnal | 4.7 | 434771.479 | 7.253 |
| birds | Saundersilarus_saundersi | Larus_saundersi | VU | Decreasing | tropical | diurnal | 198 | 276484.039 | 23.58 |
| birds | Saxicola_caprata | Saxicola_caprata | LC | Stable | temperate | diurnal | 15.2 | 9769943 | 12.574 |
| birds | Saxicola_dacotiae | Saxicola_dacotiae | NT | Decreasing | temperate | diurnal | 12.2 | 1260.313 | 14.909 |
| birds | Saxicola_ferreus | Saxicola_ferreus | LC | Stable | temperate | diurnal | 14.7 | 4791183.216 | 13.703 |
| birds | Saxicola_gutturalis | Saxicola_gutturalis | LC | Decreasing | tropical | diurnal | 15.3 | 30091.323 | 13.957 |
| birds | Saxicola_insignis | Saxicola_insignis | VU | Decreasing | temperate | diurnal | 15.3 | 377390.337 | 9.663 |
| birds | Saxicola_jerdoni | Saxicola_jerdoni | LC | Stable | temperate | diurnal | 15.3 | 398091.359 | 11.301 |
| birds | Saxicola_leucurus | NA | LC | Stable | NA | NA | NA | NA | NA |
| birds | Saxicola_macrorhynchus | Saxicola_macrorhynchus | VU | Decreasing | temperate | diurnal | 15.3 | 715078.004 | 17.176 |
| birds | Saxicola_rubetra | Saxicola_rubetra | LC | Decreasing | temperate | diurnal | 16.6 | 18264973.039 | 11.728 |
| birds | Saxicola_tectes | Saxicola_tectes | LC | Stable | tropical | diurnal | 12.6 | 2541.136 | 20.642 |

| | | | | | | | | | |
|---|---|---|---|---|---|---|---|---|---|
| birds | Saxicola_torquatus | Saxicola_torquatus | LC | Stable | temperate | diurnal | 14.1 | 54335876.948 | 9.044 |
| birds | Sayornis_nigricans | Sayornis_nigricans | LC | Increasing | temperate | diurnal | 18.6 | 3793824.655 | 8.912 |
| birds | Sayornis_phoebe | Sayornis_phoebe | LC | Increasing | temperate | diurnal | 19.7 | 9059279.46 | 12.324 |
| birds | Sayornis_saya | Sayornis_saya | LC | Increasing | temperate | diurnal | 20.9 | 10053515.709 | 7.018 |
| birds | Scelorchilus_albicollis | Scelorchilus_albicollis | LC | Stable | temperate | diurnal | 47.3 | 112769.271 | 9.406 |
| birds | Scelorchilus_rubecula | Scelorchilus_rubecula | LC | Decreasing | temperate | diurnal | 56.7 | 271566.989 | 6.274 |
| birds | Scenopoeetes_dentirostris | Scenopoeetes_dentirostris | NT | Decreasing | tropical | diurnal | 163.4 | 4997.732 | 3.542 |
| birds | Scepomycter_winifredae | Bathmocercus_winifredae | NT | Decreasing | tropical | diurnal | 14.8 | 3652.206 | 13.311 |
| birds | Schetba_rufa | Schetba_rufa | LC | Unknown | tropical | diurnal | 37.3 | 408087.532 | 7.998 |
| birds | Schiffornis_major | Schiffornis_major | LC | Decreasing | tropical | diurnal | 31 | 3141459.064 | 1.466 |
| birds | Schiffornis_turdina | Schiffornis_turdina | LC | Decreasing | tropical | diurnal | 31.7 | 5587673.674 | 2.605 |
| birds | Schiffornis_virescens | Schiffornis_virescens | LC | Decreasing | tropical | diurnal | 25.6 | 1662556.585 | 12.985 |
| birds | Schistes_geoffroyi | Schistes_geoffroyi | LC | Stable | tropical | diurnal | 3.6 | 546183.795 | 6.653 |
| birds | Schistochlamys_melanopis | Schistochlamys_melanopis | LC | Stable | tropical | diurnal | 33 | 6472149.559 | 6.255 |
| birds | Schistochlamys_ruficapillus | Schistochlamys_ruficapillus | LC | Stable | tropical | diurnal | 31.2 | 2412337.048 | 9.286 |
| birds | Schistolais_leontica | Prinia_leontica | EN | Decreasing | tropical | diurnal | 12.4 | 17375.437 | 16.029 |
| birds | Schistolais_leucopogon | Prinia_leucopogon | LC | Stable | tropical | diurnal | 11.7 | 2377144.233 | 7.693 |
| birds | Schoenicola_platyurus | Schoenicola_platyurus | VU | Decreasing | tropical | diurnal | 20 | 39801.455 | 16.848 |
| birds | Schoenicola_striatus | Chaetornis_striata | VU | Decreasing | temperate | diurnal | 31.5 | 1189979.434 | 17.924 |
| birds | Schoeniophylax_phryganophilus | Schoeniophylax_phryganophilus | LC | Stable | temperate | diurnal | 18.6 | 2565832.928 | 8.591 |
| birds | Schoeniparus_brunneus | Alcippe_brunnea | LC | Decreasing | temperate | diurnal | 17.4 | 1327800.287 | 14.614 |
| birds | Schoeniparus_castaneceps | Alcippe_castaneceps | LC | Decreasing | tropical | diurnal | 12.5 | 1180063.302 | 8.348 |
| birds | Schoeniparus_cinereus | Alcippe_cinerea | LC | Decreasing | temperate | diurnal | 11 | 208968.529 | 6.104 |
| birds | Schoeniparus_dubius | Alcippe_dubia | LC | Decreasing | temperate | diurnal | 18 | 1489085.654 | 10.431 |
| birds | Schoeniparus_rufogularis | Alcippe_rufogularis | LC | Unknown | temperate | diurnal | 13.2 | 444866.631 | 7.831 |
| birds | Schoeniparus_variegaticeps | Alcippe_variegaticeps | VU | Decreasing | temperate | diurnal | 13.2 | 155236.215 | 14.113 |
| birds | Schoutedenapus_myoptilus | Schoutedenapus_myoptilus | LC | Stable | tropical | diurnal | 23.5 | 384804.791 | 13.572 |
| birds | Sciaphylax_hemimelaena | Myrmeciza_hemimelaena | LC | Stable | tropical | diurnal | 16 | 2503854.83 | 2.433 |
| birds | Scissirostrum_dubium | Scissirostrum_dubium | LC | Decreasing | tropical | diurnal | 50 | 141970.519 | 10.18 |
| birds | Sclateria_naevia | Sclateria_naevia | LC | Stable | tropical | diurnal | 24.6 | 6179033.304 | 1.921 |
| birds | Scleroptila_finschi | Francolinus_finschi | LC | Unknown | tropical | diurnal | 569.2 | 304435.846 | 8.324 |
| birds | Scleroptila_gutturalis | Francolinus_levaillantoides | LC | Stable | tropical | diurnal | 376.7 | 1483139.562 | 7.998 |
| birds | Scleroptila_levaillantii | Francolinus_levaillantii | LC | Stable | tropical | diurnal | 430.9 | 832216.058 | 12.351 |
| birds | Scleroptila_psilolaema | Francolinus_psilolaemus | NT | Decreasing | tropical | diurnal | 478.3 | 47075.9 | 12.725 |

| | | | | | | | | | |
|---|---|---|---|---|---|---|---|---|---|
| birds | Scleroptila_shelleyi | Francolinus_shelleyi | LC | Decreasing | tropical | diurnal | 488 | 1271472.682 | 11.137 |
| birds | Scleroptila_streptophora | Francolinus_streptophorus | NT | Decreasing | tropical | diurnal | 385 | 216468.87 | 13.594 |
| birds | Sclerurus_albigularis | Sclerurus_albigularis | LC | Decreasing | tropical | diurnal | 34.8 | 655407.302 | 4.99 |
| birds | Sclerurus_caudacutus | Sclerurus_caudacutus | LC | Decreasing | tropical | diurnal | 36.2 | 5508699.648 | 1.772 |
| birds | Sclerurus_guatemalensis | Sclerurus_guatemalensis | LC | Decreasing | tropical | diurnal | 34.7 | 417195.638 | 8.789 |
| birds | Sclerurus_mexicanus | Sclerurus_mexicanus | LC | Decreasing | tropical | diurnal | 25.1 | 6362030.4 | 2.988 |
| birds | Sclerurus_rufigularis | Sclerurus_rufigularis | LC | Decreasing | tropical | diurnal | 21.6 | 5320850.733 | 1.662 |
| birds | Sclerurus_scansor | Sclerurus_scansor | LC | Decreasing | tropical | diurnal | 36.9 | 2117654.842 | 12.152 |
| birds | Scolopax_minor | Scolopax_minor | LC | Decreasing | temperate | diurnal | 196.3 | 4661375.092 | 15.551 |
| birds | Scolopax_rusticola | Scolopax_rusticola | LC | Stable | temperate | nocturnal | 308.3 | 20716119.642 | 10.361 |
| birds | Scolopax_saturata | Scolopax_saturata | NT | Decreasing | tropical | nocturnal | 205 | 19141.404 | 5.823 |
| birds | Scopus_umbretta | Scopus_umbretta | LC | Stable | tropical | diurnal | 472 | 19881333.448 | 9.058 |
| birds | Scotocerca_inquieta | Scotocerca_inquieta | LC | Decreasing | temperate | diurnal | 8.2 | 4724610.245 | 7.134 |
| birds | Scotopelia_bouvieri | Scotopelia_bouvieri | LC | Decreasing | tropical | nocturnal | 637 | 3377961.537 | 6.521 |
| birds | Scotopelia_peli | Scotopelia_peli | LC | Decreasing | tropical | nocturnal | 1942.8 | 4649310.205 | 8.74 |
| birds | Scotopelia_ussheri | Scotopelia_ussheri | VU | Decreasing | tropical | nocturnal | 787.2 | 358370.995 | 15.561 |
| birds | Scytalopus_acutirostris | Scytalopus_acutirostris | LC | Stable | tropical | diurnal | 18.1 | 21326.436 | 5.22 |
| birds | Scytalopus_affinis | Scytalopus_affinis | LC | Stable | tropical | diurnal | 13.1 | 15372.719 | 10.329 |
| birds | Scytalopus_altirostris | Scytalopus_altirostris | LC | Stable | tropical | diurnal | 19.5 | 24016.162 | 6.053 |
| birds | Scytalopus_argentifrons | Scytalopus_argentifrons | LC | Stable | tropical | diurnal | 17 | 15549.461 | 6.384 |
| birds | Scytalopus_atratus | Scytalopus_atratus | LC | Stable | tropical | diurnal | 25.7 | 187476.084 | 8.559 |
| birds | Scytalopus_bolivianus | Scytalopus_bolivianus | LC | Decreasing | tropical | diurnal | 20.8 | 64274.245 | 6.373 |
| birds | Scytalopus_canus | Scytalopus_canus | NT | Decreasing | tropical | diurnal | 18.2 | 3607.179 | 6.645 |
| birds | Scytalopus_caracae | Scytalopus_caracae | LC | Stable | tropical | diurnal | 24 | 5235.877 | 15.626 |
| birds | Scytalopus_chocoensis | Scytalopus_chocoensis | LC | Stable | tropical | diurnal | 20 | 25911.283 | 4.878 |
| birds | Scytalopus_femoralis | Scytalopus_femoralis | LC | Stable | tropical | diurnal | 23.2 | 83386.491 | 5.161 |
| birds | Scytalopus_fuscus | NA | LC | Decreasing | NA | NA | NA | NA | NA |
| birds | Scytalopus_griseicollis | Scytalopus_griseicollis | LC | Stable | tropical | diurnal | 18.1 | 61348.219 | 11.299 |
| birds | Scytalopus_iraiensis | Scytalopus_iraiensis | VU | Decreasing | tropical | diurnal | 12.4 | 4509.83 | 7.76 |
| birds | Scytalopus_latebricola | NA | NT | Stable | NA | NA | NA | NA | NA |
| birds | Scytalopus_latrans | Scytalopus_latrans | LC | Stable | tropical | diurnal | 17.6 | 239676.207 | 10.577 |
| birds | Scytalopus_macropus | Scytalopus_macropus | LC | Stable | tropical | diurnal | 37.1 | 40202.468 | 4.786 |
| birds | Scytalopus_magellanicus | Scytalopus_magellanicus | LC | Unknown | temperate | diurnal | 13.7 | 454183.191 | 4.848 |
| birds | Scytalopus_meridanus | Scytalopus_meridanus | LC | Stable | tropical | diurnal | 20 | 20695.728 | 13.999 |

| | | | | | | | | | |
|---|---|---|---|---|---|---|---|---|---|
| birds | Scytalopus_micropterus | Scytalopus_micropterus | LC | Stable | tropical | diurnal | 29.7 | 45400.685 | 6.287 |
| birds | Scytalopus_novacapitalis | Scytalopus_novacapitalis | EN | Decreasing | tropical | diurnal | 18.9 | 118540.073 | 12.244 |
| birds | Scytalopus_panamensis | NA | NT | Stable | NA | NA | NA | NA | NA |
| birds | Scytalopus_parkeri | Scytalopus_parkeri | LC | Stable | tropical | diurnal | 21.3 | 14790.201 | 7.732 |
| birds | Scytalopus_parvirostris | Scytalopus_parvirostris | LC | Decreasing | tropical | diurnal | 17.8 | 124858.926 | 6.147 |
| birds | Scytalopus_robbinsi | Scytalopus_robbinsi | EN | Decreasing | tropical | diurnal | 19.3 | 1135.628 | 8.433 |
| birds | Scytalopus_sanctaemartae | NA | NT | Decreasing | NA | NA | NA | NA | NA |
| birds | Scytalopus_schulenbergi | Scytalopus_schulenbergi | LC | Stable | tropical | diurnal | 16.1 | 38231.313 | 5.52 |
| birds | Scytalopus_simonsi | Scytalopus_simonsi | LC | Stable | tropical | diurnal | 16.6 | 29912.191 | 4.401 |
| birds | Scytalopus_speluncae | Scytalopus_speluncae | LC | Decreasing | temperate | diurnal | 13 | 249767.517 | 12.178 |
| birds | Scytalopus_spillmanni | Scytalopus_spillmanni | LC | Stable | tropical | diurnal | 23.5 | 132293.773 | 10.861 |
| birds | Scytalopus_superciliaris | Scytalopus_superciliaris | LC | Stable | temperate | diurnal | 17.8 | 53108.46 | 5.791 |
| birds | Scytalopus_unicolor | Scytalopus_unicolor | NT | Decreasing | tropical | diurnal | 17.6 | 8500.648 | 10.575 |
| birds | Scytalopus_urubambae | Scytalopus_urubambae | LC | Stable | tropical | diurnal | 13 | 3465.393 | 6.989 |
| birds | Scytalopus_vicinior | Scytalopus_vicinior | LC | Stable | tropical | diurnal | 23 | 16812.484 | 5.737 |
| birds | Scytalopus_zimmeri | Scytalopus_zimmeri | LC | Stable | tropical | diurnal | 15.9 | 53350.144 | 8.341 |
| birds | Scythrops_novaehollandiae | Scythrops_novaehollandiae | LC | Stable | tropical | diurnal | 684 | 2988558.234 | 4.599 |
| birds | Seiurus_aurocapilla | Seiurus_aurocapilla | LC | Stable | temperate | diurnal | 18.8 | 8879216.957 | 12.449 |
| birds | Selasphorus_ardens | Selasphorus_ardens | EN | Decreasing | tropical | diurnal | 3.1 | 3166.343 | 4.725 |
| birds | Selasphorus_calliope | Stellula_calliope | LC | Increasing | temperate | diurnal | 2.6 | 3088048.773 | 8.458 |
| birds | Selasphorus_ellioti | Atthis_ellioti | LC | Decreasing | tropical | diurnal | 2.9 | 98380.355 | 13.182 |
| birds | Selasphorus_flammula | Selasphorus_flammula | LC | Stable | tropical | diurnal | 2.7 | 4598.215 | 5.596 |
| birds | Selasphorus_heloisa | Atthis_heloisa | LC | Stable | tropical | diurnal | 2.2 | 247558.029 | 9.656 |
| birds | Selasphorus_platycercus | Selasphorus_platycercus | LC | Decreasing | temperate | diurnal | 3.5 | 976207.06 | 9.005 |
| birds | Selasphorus_rufus | Selasphorus_rufus | NT | Decreasing | temperate | diurnal | 3.5 | 2817202.817 | 7.935 |
| birds | Selasphorus_sasin | Selasphorus_sasin | LC | Increasing | temperate | diurnal | 3.4 | 1128506.961 | 9.584 |
| birds | Selasphorus_scintilla | Selasphorus_scintilla | LC | Decreasing | tropical | diurnal | 2.2 | 9422.595 | 11.805 |
| birds | Selenidera_gouldii | Selenidera_gouldii | LC | Decreasing | tropical | diurnal | 180.9 | 2070480.105 | 2.997 |
| birds | Selenidera_maculirostris | Selenidera_maculirostris | LC | Decreasing | tropical | diurnal | 164 | 930797.494 | 13.058 |
| birds | Selenidera_nattereri | Selenidera_nattereri | LC | Unknown | tropical | diurnal | 158.4 | 879238.298 | 0.789 |
| birds | Selenidera_piperivora | Selenidera_piperivora | LC | Stable | tropical | diurnal | 147 | 1090580.683 | 1.399 |
| birds | Selenidera_reinwardtii | Selenidera_reinwardtii | LC | Unknown | tropical | diurnal | 161.8 | 512669.413 | 2.569 |
| birds | Selenidera_spectabilis | Selenidera_spectabilis | LC | Decreasing | tropical | diurnal | 219 | 153224.717 | 6.471 |
| birds | Seleucidis_melanoleucus | Seleucidis_melanoleucus | LC | Decreasing | tropical | diurnal | 174.4 | 334716.182 | 2.893 |

| | | | | | | | | | |
|---|---|---|---|---|---|---|---|---|---|
| birds | Semioptera_wallacii | Semioptera_wallacii | LC | Decreasing | tropical | diurnal | 148.3 | 13322.44 | 5.459 |
| birds | Semnornis_frantzii | Semnornis_frantzii | LC | Stable | tropical | diurnal | 57.3 | 15874.286 | 7.174 |
| birds | Semnornis_ramphastinus | Semnornis_ramphastinus | NT | Decreasing | tropical | diurnal | 97.7 | 36256.06 | 10.822 |
| birds | Sephanoides_fernandensis | NA | CR | Decreasing | NA | NA | NA | NA | NA |
| birds | Sephanoides_sephaniodes | Sephanoides_sephaniodes | LC | Decreasing | temperate | diurnal | 5.2 | 1104451.074 | 5.711 |
| birds | Sericornis_beccarii | Sericornis_beccarii | LC | Stable | tropical | diurnal | 11.9 | 246723.36 | 2.63 |
| birds | Sericornis_frontalis | Sericornis_frontalis | LC | Decreasing | temperate | diurnal | 13.2 | 1682487.01 | 8.396 |
| birds | Sericornis_humilis | Sericornis_humilis | LC | Decreasing | temperate | diurnal | 18.3 | 41698.308 | 6.668 |
| birds | Sericornis_keri | Sericornis_keri | VU | Decreasing | tropical | diurnal | 11.4 | 24710.745 | 4.804 |
| birds | Sericornis_magnirostra | Sericornis_magnirostra | LC | Stable | temperate | diurnal | 9.8 | 270681.408 | 7.776 |
| birds | Sericornis_nouhuysi | Sericornis_nouhuysi | LC | Stable | tropical | diurnal | 17.1 | 132591.21 | 4.529 |
| birds | Sericossypha_albocristata | Sericossypha_albocristata | VU | Decreasing | tropical | diurnal | 114 | 103262.328 | 7.728 |
| birds | Sericulus_aureus | Sericulus_aureus | LC | Decreasing | tropical | diurnal | 172 | 35679.743 | 2.793 |
| birds | Sericulus_bakeri | Sericulus_bakeri | NT | Stable | tropical | diurnal | 176.5 | 564.145 | 3.839 |
| birds | Sericulus_chrysocephalus | Sericulus_chrysocephalus | LC | Decreasing | temperate | diurnal | 99.5 | 89618.775 | 7.653 |
| birds | Serilophus_lunatus | Serilophus_lunatus | LC | Decreasing | tropical | diurnal | 34 | 1235508.382 | 9.462 |
| birds | Serinus_alario | Serinus_alario | LC | Stable | temperate | diurnal | 12.2 | 508153.435 | 7.366 |
| birds | Serinus_canaria | Serinus_canaria | LC | Stable | tropical | diurnal | 24.3 | 19528.293 | 22.874 |
| birds | Serinus_canicollis | Serinus_canicollis | LC | Stable | temperate | diurnal | 15.2 | 618013.335 | 11.409 |
| birds | Serinus_flavivertex | Serinus_flavivertex | LC | Stable | tropical | diurnal | 14 | 323625.704 | 13.886 |
| birds | Serinus_nigriceps | Serinus_nigriceps | LC | Stable | tropical | diurnal | 15.4 | 114938.98 | 15.235 |
| birds | Serinus_pusillus | Serinus_pusillus | LC | Stable | temperate | diurnal | 11.5 | 2537686.866 | 9.68 |
| birds | Serinus_serinus | Serinus_serinus | LC | Decreasing | temperate | diurnal | 11.2 | 5141384.689 | 18.645 |
| birds | Serinus_syriacus | Serinus_syriacus | VU | Decreasing | temperate | diurnal | 11.7 | 78766.521 | 14.926 |
| birds | Serpophaga_cinerea | Serpophaga_cinerea | LC | Decreasing | tropical | diurnal | 8.3 | 537348.407 | 8.631 |
| birds | Serpophaga_hypoleuca | Serpophaga_hypoleuca | LC | Stable | tropical | diurnal | 6.1 | 557853.943 | 3.626 |
| birds | Serpophaga_nigricans | Serpophaga_nigricans | LC | Stable | temperate | diurnal | 8.5 | 3661570.487 | 10.923 |
| birds | Serpophaga_subcristata | Serpophaga_subcristata | LC | Stable | temperate | diurnal | 6.6 | 5786710.077 | 9.536 |
| birds | Setopagis_parvula | Caprimulgus_parvulus | LC | Stable | tropical | nocturnal | 37 | 6789301.911 | 8.407 |
| birds | Setopagis_whitelyi | Caprimulgus_whitelyi | LC | Stable | tropical | nocturnal | 47 | 45695.271 | 1.13 |
| birds | Setophaga_adelaidae | Dendroica_adelaidae | LC | Stable | tropical | diurnal | 6.9 | 8923.083 | 27.081 |
| birds | Setophaga_americana | Parula_americana | LC | Increasing | temperate | diurnal | 7.8 | 5430698.009 | 15.349 |
| birds | Setophaga_angelae | Dendroica_angelae | EN | Decreasing | tropical | diurnal | 7.7 | 163.582 | 15.686 |
| birds | Setophaga_caerulescens | Dendroica_caerulescens | LC | Increasing | temperate | diurnal | 10.1 | 2978266.677 | 15.83 |

| | | | | | | | | | |
|---|---|---|---|---|---|---|---|---|---|
| birds | Setophaga_castanea | Dendroica_castanea | LC | Increasing | temperate | diurnal | 11.8 | 8432233.605 | 11.969 |
| birds | Setophaga_cerulea | Dendroica_cerulea | NT | Decreasing | tropical | diurnal | 9 | 4658662.601 | 14.834 |
| birds | Setophaga_chrysoparia | Dendroica_chrysoparia | EN | Decreasing | tropical | diurnal | 9.9 | 434557.34 | 11.922 |
| birds | Setophaga_citrina | Wilsonia_citrina | LC | Increasing | tropical | diurnal | 10.5 | 3115320.703 | 15.806 |
| birds | Setophaga_coronata | Dendroica_coronata | LC | Stable | temperate | diurnal | 11.9 | 15842705.296 | 8.87 |
| birds | Setophaga_delicata | Dendroica_delicata | LC | Stable | tropical | diurnal | 7.4 | 617.37 | 20.77 |
| birds | Setophaga_discolor | Dendroica_discolor | LC | Decreasing | temperate | diurnal | 7.6 | 2180545.91 | 16.976 |
| birds | Setophaga_dominica | Dendroica_dominica | LC | Increasing | tropical | diurnal | 9.7 | 3207549.903 | 15.356 |
| birds | Setophaga_fusca | Dendroica_fusca | LC | Increasing | tropical | diurnal | 9.7 | 8748176.98 | 12.655 |
| birds | Setophaga_graciae | Dendroica_graciae | LC | Decreasing | tropical | diurnal | 8.1 | 1217041.457 | 8.893 |
| birds | Setophaga_kirtlandii | Dendroica_kirtlandii | NT | Increasing | temperate | diurnal | 13.9 | 53047.643 | 10.292 |
| birds | Setophaga_magnolia | Dendroica_magnolia | LC | Increasing | temperate | diurnal | 8.1 | 8830411.551 | 11.697 |
| birds | Setophaga_nigrescens | Dendroica_nigrescens | LC | Decreasing | temperate | diurnal | 8.7 | 3560024.468 | 8.766 |
| birds | Setophaga_occidentalis | Dendroica_occidentalis | LC | Stable | temperate | diurnal | 10.4 | 2474138.363 | 9.22 |
| birds | Setophaga_palmarum | Dendroica_palmarum | LC | Increasing | temperate | diurnal | 10.3 | 7890903.867 | 11.053 |
| birds | Setophaga_pensylvanica | Dendroica_pensylvanica | LC | Decreasing | tropical | diurnal | 9.3 | 7337980.567 | 13.851 |
| birds | Setophaga_petechia | Dendroica_petechia | LC | Decreasing | temperate | diurnal | 10.2 | 21086713.837 | 8.204 |
| birds | Setophaga_pharetra | Dendroica_pharetra | LC | Decreasing | tropical | diurnal | 9.9 | 11021.791 | 19.741 |
| birds | Setophaga_pinus | Dendroica_pinus | LC | Increasing | temperate | diurnal | 11.8 | 3767948.835 | 16.825 |
| birds | Setophaga_pitiayumi | Parula_pitiayumi | LC | Decreasing | tropical | diurnal | 6.8 | 8364955.267 | 8.341 |
| birds | Setophaga_pityophila | Dendroica_pityophila | LC | Decreasing | tropical | diurnal | 8 | 7445.485 | 12.44 |
| birds | Setophaga_plumbea | Dendroica_plumbea | LC | Decreasing | tropical | diurnal | 10.1 | 2397.364 | 20.541 |
| birds | Setophaga_ruticilla | Setophaga_ruticilla | LC | Increasing | temperate | diurnal | 8.2 | 15136310.839 | 10.107 |
| birds | Setophaga_striata | Dendroica_striata | NT | Decreasing | temperate | diurnal | 11.8 | 16034144.171 | 6.658 |
| birds | Setophaga_subita | Dendroica_subita | VU | Decreasing | tropical | diurnal | 7.6 | 153.929 | 11.334 |
| birds | Setophaga_tigrina | Dendroica_tigrina | LC | Increasing | temperate | diurnal | 10 | 5707680.077 | 11.889 |
| birds | Setophaga_townsendi | Dendroica_townsendi | LC | Decreasing | temperate | diurnal | 8.8 | 5348431.125 | 7.687 |
| birds | Setophaga_virens | Dendroica_virens | LC | Stable | temperate | diurnal | 8.7 | 8653032.872 | 12.143 |
| birds | Setophaga_vitellina | Dendroica_vitellina | NT | Decreasing | tropical | diurnal | 6.7 | 279.505 | 22.085 |
| birds | Setornis_criniger | Setornis_criniger | VU | Decreasing | tropical | diurnal | 29.5 | 544495.929 | 6.918 |
| birds | Sheppardia_aequatorialis | Sheppardia_aequatorialis | LC | Decreasing | tropical | diurnal | 16.8 | 138552.17 | 17.921 |
| birds | Sheppardia_bocagei | Sheppardia_bocagei | LC | Decreasing | tropical | diurnal | 18.3 | 589392.079 | 7.992 |
| birds | Sheppardia_cyornithopsis | Sheppardia_cyornithopsis | LC | Decreasing | tropical | diurnal | 18.4 | 1077673.352 | 7.859 |
| birds | Sheppardia_gabela | Sheppardia_gabela | EN | Decreasing | tropical | diurnal | 11 | 8397.098 | 8.751 |

| | | | | | | | | | |
|---|---|---|---|---|---|---|---|---|---|
| birds | Sheppardia_gunningi | Sheppardia_gunningi | NT | Decreasing | tropical | diurnal | 17.3 | 58833.067 | 9.983 |
| birds | Sheppardia_lowei | Sheppardia_lowei | VU | Decreasing | tropical | diurnal | 18.4 | 6701.617 | 11.615 |
| birds | Sheppardia_montana | Sheppardia_montana | EN | Decreasing | tropical | diurnal | 21.8 | 367.996 | 15.542 |
| birds | Sheppardia_polioptera | Cossypha_polioptera | LC | Decreasing | tropical | diurnal | 18.7 | 644019.502 | 12.43 |
| birds | Sheppardia_sharpei | Sheppardia_sharpei | LC | Decreasing | tropical | diurnal | 14.7 | 92949.689 | 12.213 |
| birds | Sholicola_major | Myiomela_major | EN | Decreasing | tropical | diurnal | 38.5 | 1160.849 | 10.086 |
| birds | Sialia_currucoides | Sialia_currucoides | LC | Increasing | temperate | diurnal | 29.6 | 7775403.646 | 9.115 |
| birds | Sialia_mexicana | Sialia_mexicana | LC | Increasing | temperate | diurnal | 26.4 | 2595315.963 | 9.446 |
| birds | Sialia_sialis | Sialia_sialis | LC | Increasing | temperate | diurnal | 27.5 | 6065720.04 | 14.713 |
| birds | Sibia_morrisoniana | Actinodura_morrisoniana | LC | Decreasing | temperate | diurnal | 24 | 11919.305 | 5.611 |
| birds | Sibia_nipalensis | Actinodura_nipalensis | LC | Decreasing | temperate | diurnal | 43.9 | 108962.948 | 8.95 |
| birds | Sibia_souliei | Actinodura_souliei | LC | Decreasing | temperate | diurnal | 39.3 | 161149.523 | 11.353 |
| birds | Sibia_waldeni | Actinodura_waldeni | LC | Decreasing | temperate | diurnal | 46.7 | 344520.78 | 7.366 |
| birds | Sibirionetta_formosa | Anas_formosa | LC | Increasing | temperate | diurnal | 434 | 13029180.96 | 7.513 |
| birds | Sicalis_auriventris | Sicalis_auriventris | LC | Stable | temperate | diurnal | 55 | 135742.055 | 2.85 |
| birds | Sicalis_citrina | Sicalis_citrina | LC | Stable | tropical | diurnal | 11.6 | 1533817.56 | 8.925 |
| birds | Sicalis_columbiana | Sicalis_columbiana | LC | Stable | tropical | diurnal | 13.4 | 1844478.678 | 5.808 |
| birds | Sicalis_flaveola | Sicalis_flaveola | LC | Stable | tropical | diurnal | 16.9 | 8114458.537 | 9.281 |
| birds | Sicalis_lebruni | Sicalis_lebruni | LC | Stable | temperate | diurnal | 25.3 | 418583.419 | 5.86 |
| birds | Sicalis_lutea | Sicalis_lutea | LC | Stable | tropical | diurnal | 16.3 | 333221.128 | 7.255 |
| birds | Sicalis_luteocephala | Sicalis_luteocephala | LC | Stable | tropical | diurnal | 16.3 | 79079.453 | 9.591 |
| birds | Sicalis_luteola | Sicalis_luteola | LC | Increasing | temperate | diurnal | 15.9 | 6593159.399 | 9.136 |
| birds | Sicalis_olivascens | Sicalis_olivascens | LC | Stable | tropical | diurnal | 21.8 | 769970.913 | 6.026 |
| birds | Sicalis_raimondii | Sicalis_raimondii | LC | Stable | tropical | diurnal | 15.8 | 67931.75 | 7.412 |
| birds | Sicalis_taczanowskii | Sicalis_taczanowskii | LC | Stable | tropical | diurnal | 8 | 41438.321 | 8.153 |
| birds | Sicalis_uropigyalis | Sicalis_uropygialis | LC | Stable | tropical | diurnal | 22.5 | 705565.484 | 7.111 |
| birds | Sigelus_silens | Sigelus_silens | LC | Stable | temperate | diurnal | 25.7 | 982178.671 | 10.8 |
| birds | Silvicultrix_diadema | Ochthoeca_diadema | LC | Stable | tropical | diurnal | 11.5 | 130712.81 | 10.687 |
| birds | Silvicultrix_frontalis | Ochthoeca_frontalis | LC | Stable | tropical | diurnal | 10 | 229755.723 | 9.735 |
| birds | Silvicultrix_jelskii | Ochthoeca_jelskii | LC | Stable | tropical | diurnal | 11.6 | 73769.989 | 8.588 |
| birds | Silvicultrix_pulchella | Ochthoeca_pulchella | LC | Stable | tropical | diurnal | 12 | 116794.53 | 6.753 |
| birds | Sinosuthora_brunnea | NA | not_available | not_available | NA | NA | NA | NA | NA |
| birds | Siphonorhis_brewsteri | Siphonorhis_brewsteri | NT | Decreasing | tropical | nocturnal | 37 | 20562.36 | 13.942 |
| birds | Sipia_berlepschi | Myrmeciza_berlepschi | LC | Stable | tropical | diurnal | 25.6 | 55751.319 | 5.371 |

| | | | | | | | | | |
|---|---|---|---|---|---|---|---|---|---|
| birds | Sipia_laemosticta | Myrmeciza_laemosticta | LC | Decreasing | tropical | diurnal | 25 | 36688.837 | 6.525 |
| birds | Sipia_nigricauda | Myrmeciza_nigricauda | LC | Decreasing | tropical | diurnal | 24 | 64675.015 | 7.056 |
| birds | Sipodotus_wallacii | Sipodotus_wallacii | LC | Decreasing | tropical | diurnal | 7.7 | 489818.272 | 3.321 |
| birds | Siptornis_striaticollis | Siptornis_striaticollis | LC | Stable | tropical | diurnal | 13 | 20659.128 | 5.635 |
| birds | Sirystes_albogriseus | Sirystes_albogriseus | LC | Decreasing | tropical | diurnal | 34 | 174213.099 | 7.113 |
| birds | Sirystes_sibilator | Sirystes_sibilator | LC | Decreasing | tropical | diurnal | 32.3 | 4179503.46 | 7.023 |
| birds | Sitta_azurea | Sitta_azurea | LC | Decreasing | tropical | diurnal | 16.3 | 361034.344 | 12.843 |
| birds | Sitta_canadensis | Sitta_canadensis | LC | Increasing | temperate | diurnal | 9.8 | 12270262.343 | 10.483 |
| birds | Sitta_carolinensis | Sitta_carolinensis | LC | Increasing | temperate | diurnal | 21 | 9154405.083 | 12.777 |
| birds | Sitta_cashmirensis | Sitta_cashmirensis | LC | Decreasing | temperate | diurnal | 19 | 140130.444 | 9.804 |
| birds | Sitta_castanea | Sitta_castanea | LC | Unknown | temperate | diurnal | 19.6 | 1252045.842 | 17.474 |
| birds | Sitta_europaea | Sitta_europaea | LC | Stable | temperate | diurnal | 20.4 | 21575986.153 | 10 |
| birds | Sitta_formosa | NA | VU | Decreasing | NA | NA | NA | NA | NA |
| birds | Sitta_frontalis | Sitta_frontalis | LC | Decreasing | tropical | diurnal | 16.5 | 5214819.553 | 12.434 |
| birds | Sitta_himalayensis | Sitta_himalayensis | LC | Decreasing | temperate | diurnal | 14.3 | 421140.599 | 8.35 |
| birds | Sitta_krueperi | Sitta_krueperi | LC | Decreasing | temperate | diurnal | 12.4 | 282938.846 | 12.785 |
| birds | Sitta_ledanti | Sitta_ledanti | EN | Decreasing | temperate | diurnal | 17.2 | 324.46 | 15.877 |
| birds | Sitta_leucopsis | Sitta_leucopsis | LC | Decreasing | temperate | diurnal | 14.8 | 258577.792 | 10.337 |
| birds | Sitta_magna | Sitta_magna | EN | Decreasing | tropical | diurnal | 41.1 | 500173.512 | 11.242 |
| birds | Sitta_nagaensis | Sitta_nagaensis | LC | Decreasing | temperate | diurnal | 14.7 | 1053108.557 | 9.483 |
| birds | Sitta_neumayer | Sitta_neumayer | LC | Stable | temperate | diurnal | 26.5 | 1244088.586 | 14.998 |
| birds | Sitta_oenochlamys | Sitta_oenochlamys | LC | Decreasing | tropical | diurnal | 16.4 | 249953.707 | 13.931 |
| birds | Sitta_pusilla | Sitta_pusilla | LC | Decreasing | temperate | diurnal | 10.2 | 891316.405 | 16.461 |
| birds | Sitta_pygmaea | Sitta_pygmaea | LC | Increasing | temperate | diurnal | 10.6 | 1953869.524 | 9.143 |
| birds | Sitta_solangiae | Sitta_solangiae | NT | Decreasing | tropical | diurnal | 15 | 24723.097 | 6.291 |
| birds | Sitta_tephronota | Sitta_tephronota | LC | Stable | temperate | diurnal | 35.2 | 2326077.815 | 10.619 |
| birds | Sitta_victoriae | NA | EN | Decreasing | NA | NA | NA | NA | NA |
| birds | Sitta_villosa | Sitta_villosa | LC | Decreasing | temperate | diurnal | 9.4 | 1824265.591 | 12.453 |
| birds | Sitta_whiteheadi | Sitta_whiteheadi | VU | Decreasing | temperate | diurnal | 11.8 | 2318.9 | 8.329 |
| birds | Sitta_yunnanensis | Sitta_yunnanensis | LC | Decreasing | temperate | diurnal | 9.9 | 218133.913 | 10.591 |
| birds | Sittasomus_griseicapillus | Sittasomus_griseicapillus | LC | Decreasing | tropical | diurnal | 13.1 | 10911731.741 | 5.493 |
| birds | Sittiparus_semilarvatus | Parus_semilarvatus | NT | Decreasing | tropical | diurnal | 18 | 55106.291 | 12.567 |
| birds | Sittiparus_varius | Parus_varius | LC | Unknown | temperate | diurnal | 17 | 692847.172 | 18.207 |
| birds | Siva_cyanouroptera | Minla_cyanouroptera | LC | Decreasing | tropical | diurnal | 17 | 2841737.412 | 11.306 |

| | | | | | | | | | |
|---|---|---|---|---|---|---|---|---|---|
| birds | Smicrornis_brevirostris | Smicrornis_brevirostris | LC | Decreasing | temperate | diurnal | 5.3 | 7230924.882 | 4.376 |
| birds | Smithornis_capensis | Smithornis_capensis | LC | Decreasing | tropical | diurnal | 23.2 | 3751914.002 | 9.849 |
| birds | Smithornis_rufolateralis | Smithornis_rufolateralis | LC | Decreasing | tropical | diurnal | 19 | 1880114.415 | 8.279 |
| birds | Smithornis_sharpei | Smithornis_sharpei | LC | Decreasing | tropical | diurnal | 37.5 | 498936.687 | 7.169 |
| birds | Smutsornis_africanus | Rhinoptilus_africanus | LC | Stable | temperate | nocturnal | 87.5 | 2999057.706 | 7.511 |
| birds | Snowornis_cryptolophus | Snowornis_cryptolophus | LC | Decreasing | tropical | diurnal | 73.6 | 109184.972 | 4.974 |
| birds | Snowornis_subalaris | Snowornis_subalaris | LC | Decreasing | tropical | diurnal | 81.2 | 59762.195 | 4.125 |
| birds | Somateria_fischeri | Somateria_fischeri | NT | Decreasing | temperate | diurnal | 1557.2 | 969744.294 | 0.965 |
| birds | Somateria_mollissima | Somateria_mollissima | NT | Unknown | temperate | diurnal | 2060.9 | 16485948.204 | 4.825 |
| birds | Somateria_spectabilis | Somateria_spectabilis | LC | Decreasing | temperate | diurnal | 1616.7 | 7316509.599 | 0.847 |
| birds | Spartonoica_maluroides | Spartonoica_maluroides | LC | Decreasing | temperate | diurnal | 11 | 527135.992 | 12.671 |
| birds | Spatula_clypeata | Anas_clypeata | LC | Decreasing | temperate | diurnal | 612.6 | 50160551.832 | 9.797 |
| birds | Spatula_cyanoptera | Anas_cyanoptera | LC | Decreasing | temperate | diurnal | 377.5 | 8842984.669 | 9.734 |
| birds | Spatula_discors | Anas_discors | LC | Increasing | temperate | diurnal | 359.4 | 15868657.31 | 10.377 |
| birds | Spatula_hottentota | Anas_hottentota | LC | Decreasing | tropical | diurnal | 276.4 | 7754871.013 | 9.77 |
| birds | Spatula_platalea | Anas_platalea | LC | Stable | temperate | diurnal | 465.4 | 2947935.326 | 7.489 |
| birds | Spatula_querquedula | Anas_querquedula | LC | Decreasing | temperate | diurnal | 325.6 | 34165137.376 | 10.08 |
| birds | Spatula_rhynchotis | Anas_rhynchotis | LC | Stable | temperate | diurnal | 639 | 4454068.383 | 6.018 |
| birds | Spatula_smithii | Anas_smithii | LC | Increasing | temperate | diurnal | 591.5 | 2430691.949 | 8.192 |
| birds | Spatula_versicolor | Anas_versicolor | LC | Stable | temperate | diurnal | 595.2 | 3843988.901 | 8.666 |
| birds | Speculanas_specularis | Speculanas_specularis | NT | Stable | temperate | diurnal | 1496.1 | 1108774.137 | 5.074 |
| birds | Speculipastor_bicolor | Speculipastor_bicolor | LC | Unknown | tropical | diurnal | 64.2 | 1308255.702 | 8.508 |
| birds | Spelaeornis_badeigularis | Spelaeornis_badeigularis | NT | Decreasing | temperate | diurnal | 6 | 3777.321 | 4.421 |
| birds | Spelaeornis_caudatus | Spelaeornis_caudatus | NT | Decreasing | temperate | diurnal | 11 | 56850.866 | 6.558 |
| birds | Spelaeornis_chocolatinus | Spelaeornis_chocolatinus | VU | Decreasing | temperate | diurnal | 11.8 | 11101.972 | 12.483 |
| birds | Spelaeornis_longicaudatus | Spelaeornis_longicaudatus | NT | Decreasing | temperate | diurnal | 12 | 6234.116 | 10.429 |
| birds | Spelaeornis_troglodytoides | Spelaeornis_troglodytoides | LC | Decreasing | temperate | diurnal | 7 | 498405.283 | 8.705 |
| birds | Spermestes_bicolor | Lonchura_bicolor | LC | Stable | tropical | diurnal | 9.9 | 5169258.147 | 9.22 |
| birds | Spermestes_cucullata | Lonchura_cucullata | LC | Stable | tropical | diurnal | 9 | 8677298.347 | 10.306 |
| birds | Spermestes_fringilloides | Lonchura_fringilloides | LC | Stable | tropical | diurnal | 16.8 | 1503708.265 | 9.501 |
| birds | Spermestes_griseicapilla | Lonchura_griseicapilla | LC | Stable | tropical | diurnal | 15 | 431259.262 | 10.903 |
| birds | Spermophaga_haematina | Spermophaga_haematina | LC | Stable | tropical | diurnal | 23 | 1910654.247 | 9.281 |
| birds | Spermophaga_poliogenys | Spermophaga_poliogenys | LC | Stable | tropical | diurnal | 19.8 | 366771.57 | 6.105 |
| birds | Spermophaga_ruficapilla | Spermophaga_ruficapilla | LC | Stable | tropical | diurnal | 23.6 | 562817.835 | 10.042 |

| | | | | | | | | | |
|---|---|---|---|---|---|---|---|---|---|
| birds | Sphecotheres_hypoleucus | Sphecotheres_hypoleucus | LC | Stable | tropical | diurnal | 77.5 | 2638.287 | 4.678 |
| birds | Sphecotheres_vieilloti | Sphecotheres_vieilloti | LC | Stable | tropical | diurnal | 121.5 | 1005032.838 | 5.022 |
| birds | Sphecotheres_viridis | Sphecotheres_viridis | LC | Decreasing | tropical | diurnal | 121 | 30023.142 | 13.965 |
| birds | Spheniscus_demersus | Spheniscus_demersus | EN | Decreasing | temperate | cathemeral | 3130.1 | 330067.229 | 11.221 |
| birds | Spheniscus_humboldti | Spheniscus_humboldti | VU | Decreasing | tropical | cathemeral | 4366.4 | 885813.19 | 18.195 |
| birds | Spheniscus_magellanicus | Spheniscus_magellanicus | LC | Decreasing | temperate | cathemeral | 4105.1 | 2637292.608 | 4.56 |
| birds | Spheniscus_mendiculus | NA | EN | Decreasing | NA | NA | NA | NA | NA |
| birds | Sphenoeacus_afer | Sphenoeacus_afer | LC | Decreasing | temperate | diurnal | 30.4 | 430218.369 | 13.144 |
| birds | Sphenopsis_frontalis | Hemispingus_frontalis | LC | Decreasing | tropical | diurnal | 17 | 200867.546 | 8.9 |
| birds | Sphenopsis_melanotis | Hemispingus_melanotis | LC | Stable | tropical | diurnal | 16 | 183679.4 | 7.35 |
| birds | Sphenopsis_ochracea | NA | LC | Stable | NA | NA | NA | NA | NA |
| birds | Sphenopsis_piurae | NA | LC | Unknown | NA | NA | NA | NA | NA |
| birds | Sphyrapicus_nuchalis | Sphyrapicus_nuchalis | LC | Decreasing | temperate | diurnal | 48.5 | 3614392.641 | 8.22 |
| birds | Sphyrapicus_ruber | Sphyrapicus_ruber | LC | Decreasing | temperate | diurnal | 54.8 | 1189115.638 | 6.609 |
| birds | Sphyrapicus_thyroideus | Sphyrapicus_thyroideus | LC | Stable | temperate | diurnal | 47.6 | 1248737.494 | 8.564 |
| birds | Sphyrapicus_varius | Sphyrapicus_varius | LC | Decreasing | temperate | diurnal | 50.3 | 9946724.596 | 10.974 |
| birds | Spilopelia_chinensis | Stigmatopelia_chinensis | LC | Increasing | tropical | diurnal | 159 | 8414112.051 | 12.996 |
| birds | Spilopelia_senegalensis | Stigmatopelia_senegalensis | LC | Stable | tropical | diurnal | 82.1 | 29236937.738 | 9.431 |
| birds | Spiloptila_clamans | Spiloptila_clamans | LC | Stable | tropical | diurnal | 7 | 2719529.349 | 8.372 |
| birds | Spilornis_cheela | Spilornis_cheela | LC | Stable | tropical | diurnal | 597.7 | 7009544.381 | 13.584 |
| birds | Spilornis_elgini | Spilornis_elgini | VU | Decreasing | tropical | diurnal | 895 | 5798.894 | 7.27 |
| birds | Spilornis_holospilus | Spilornis_holospilus | LC | Decreasing | tropical | diurnal | 705 | 276025.538 | 13.883 |
| birds | Spilornis_kinabaluensis | NA | NT | Decreasing | NA | NA | NA | NA | NA |
| birds | Spilornis_rufipectus | NA | LC | Stable | NA | NA | NA | NA | NA |
| birds | Spindalis_dominicensis | Spindalis_dominicensis | LC | Stable | tropical | diurnal | 29.4 | 74799.651 | 15.781 |
| birds | Spindalis_nigricephala | Spindalis_nigricephala | LC | Stable | tropical | diurnal | 45.1 | 11021.791 | 19.741 |
| birds | Spindalis_portoricensis | Spindalis_portoricensis | LC | Stable | tropical | diurnal | 32.1 | 8774.421 | 27.252 |
| birds | Spindalis_zena | Spindalis_zena | LC | Stable | tropical | diurnal | 25.4 | 126553.5 | 15.227 |
| birds | Spinus_atratus | Carduelis_atrata | LC | Stable | tropical | diurnal | 16 | 839801.137 | 5.805 |
| birds | Spinus_atriceps | Carduelis_atriceps | LC | Decreasing | tropical | diurnal | 13.5 | 50051.076 | 13.365 |
| birds | Spinus_barbatus | Carduelis_barbata | LC | Stable | temperate | diurnal | 15.5 | 1234638.372 | 5.416 |
| birds | Spinus_crassirostris | Carduelis_crassirostris | LC | Stable | tropical | diurnal | 19 | 373239.18 | 6.151 |
| birds | Spinus_cucullatus | Carduelis_cucullata | EN | Decreasing | tropical | diurnal | 9.8 | 108994.244 | 13.052 |
| birds | Spinus_dominicensis | Carduelis_dominicensis | LC | Stable | tropical | diurnal | 12.6 | 9702.057 | 11.454 |

| | | | | | | | | | |
|---|---|---|---|---|---|---|---|---|---|
| birds | Spinus_lawrencei | Carduelis_lawrencei | LC | Decreasing | temperate | diurnal | 10.9 | 556776.949 | 8.972 |
| birds | Spinus_magellanicus | Carduelis_magellanica | LC | Stable | tropical | diurnal | 13.6 | 6077584.713 | 9.171 |
| birds | Spinus_notatus | Carduelis_notata | LC | Decreasing | tropical | diurnal | 10.9 | 488138.476 | 9.061 |
| birds | Spinus_olivaceus | NA | LC | Decreasing | NA | NA | NA | NA | NA |
| birds | Spinus_pinus | Carduelis_pinus | LC | Stable | temperate | diurnal | 12.7 | 14061824.422 | 9.996 |
| birds | Spinus_psaltria | Carduelis_psaltria | LC | Decreasing | temperate | diurnal | 8.8 | 4186163.772 | 9.541 |
| birds | Spinus_siemiradzkii | NA | LC | Decreasing | NA | NA | NA | NA | NA |
| birds | Spinus_spinescens | Carduelis_spinescens | LC | Stable | tropical | diurnal | 11.3 | 117953.766 | 11.84 |
| birds | Spinus_spinus | Carduelis_spinus | LC | Decreasing | temperate | diurnal | 13.2 | 15585639.538 | 13.202 |
| birds | Spinus_thibetanus | Serinus_thibetanus | LC | Stable | temperate | diurnal | 10.5 | 948405.552 | 6.8 |
| birds | Spinus_tristis | Carduelis_tristis | LC | Increasing | temperate | diurnal | 12.8 | 10571095.976 | 12.533 |
| birds | Spinus_uropygialis | Carduelis_uropygialis | LC | Stable | temperate | diurnal | 14 | 494166.635 | 6.156 |
| birds | Spinus_xanthogastrus | Carduelis_xanthogastra | LC | Decreasing | tropical | diurnal | 12.7 | 161327.428 | 9.047 |
| birds | Spinus_yarrellii | Carduelis_yarrellii | VU | Decreasing | tropical | diurnal | 10.5 | 496176.13 | 8.326 |
| birds | Spiza_americana | Spiza_americana | LC | Stable | temperate | diurnal | 26.2 | 6183307.842 | 13.644 |
| birds | Spizaetus_isidori | NA | EN | Decreasing | NA | NA | NA | NA | NA |
| birds | Spizaetus_melanoleucus | Spizaetus_melanoleucus | LC | Decreasing | tropical | diurnal | 850 | 8710494.019 | 4.346 |
| birds | Spizaetus_ornatus | Spizaetus_ornatus | NT | Decreasing | tropical | diurnal | 1197.4 | 10597697.765 | 5.19 |
| birds | Spizaetus_tyrannus | Spizaetus_tyrannus | LC | Decreasing | tropical | diurnal | 1007.1 | 8304333.121 | 4.113 |
| birds | Spizella_atrogularis | Spizella_atrogularis | LC | Decreasing | temperate | diurnal | 11.3 | 1397656.801 | 8.428 |
| birds | Spizella_breweri | Spizella_breweri | LC | Decreasing | temperate | diurnal | 10.9 | 5119315.916 | 8.603 |
| birds | Spizella_pallida | Spizella_pallida | LC | Decreasing | temperate | diurnal | 11.2 | 7311087.961 | 11.19 |
| birds | Spizella_passerina | Spizella_passerina | LC | Decreasing | temperate | diurnal | 12.2 | 14987835.946 | 9.683 |
| birds | Spizella_pusilla | Spizella_pusilla | LC | Decreasing | temperate | diurnal | 12.5 | 4818402.43 | 16.416 |
| birds | Spizella_wortheni | Spizella_wortheni | EN | Decreasing | temperate | diurnal | 12.4 | 60094.131 | 6.399 |
| birds | Spiziapteryx_circumcincta | Spiziapteryx_circumcincta | LC | Stable | temperate | diurnal | 166.2 | 1416372.049 | 7.688 |
| birds | Spizixos_canifrons | Spizixos_canifrons | LC | Increasing | tropical | diurnal | 44 | 1162572.346 | 10.364 |
| birds | Spizixos_semitorques | Spizixos_semitorques | LC | Stable | temperate | diurnal | 39 | 2328094.864 | 15.545 |
| birds | Spizocorys_conirostris | Spizocorys_conirostris | LC | Decreasing | temperate | diurnal | 14.3 | 1128198.555 | 8.218 |
| birds | Spizocorys_fringillaris | Spizocorys_fringillaris | EN | Decreasing | temperate | diurnal | 19.2 | 3270.794 | 9.946 |
| birds | Spizocorys_obbiensis | Spizocorys_obbiensis | DD | Decreasing | tropical | diurnal | 13.5 | 55209.01 | 11.767 |
| birds | Spizocorys_personata | Spizocorys_personata | LC | Stable | tropical | diurnal | 19.7 | 73283.544 | 5.432 |
| birds | Spizocorys_sclateri | Spizocorys_sclateri | NT | Unknown | temperate | diurnal | 19 | 226405.691 | 5.514 |
| birds | Spodiopsar_cineraceus | Sturnus_cineraceus | LC | Unknown | temperate | diurnal | 83 | 7910712.352 | 11.857 |

| | | | | | | | | | |
|---|---|---|---|---|---|---|---|---|---|
| birds | Spodiopsar_sericeus | Sturnus_sericeus | LC | Stable | temperate | diurnal | 65 | 1771850.626 | 15.711 |
| birds | Spodiornis_rusticus | Haplospiza_rustica | LC | Decreasing | tropical | diurnal | 15.6 | 480755.349 | 9.484 |
| birds | Sporathraupis_cyanocephala | Thraupis_cyanocephala | LC | Stable | tropical | diurnal | 36 | 429823.392 | 9.18 |
| birds | Sporophila_albogularis | Sporophila_albogularis | LC | Stable | tropical | diurnal | 9.7 | 910904.264 | 8.946 |
| birds | Sporophila_americana | Sporophila_americana | LC | Decreasing | tropical | diurnal | 12 | 1491401.162 | 2.638 |
| birds | Sporophila_angolensis | Oryzoborus_angolensis | LC | Increasing | tropical | diurnal | 13 | 11883533.04 | 5.299 |
| birds | Sporophila_atrirostris | Oryzoborus_atrirostris | LC | Stable | tropical | diurnal | 26 | 201164.716 | 3.767 |
| birds | Sporophila_bouvreuil | Sporophila_bouvreuil | LC | Decreasing | tropical | diurnal | 8.6 | 3349344.108 | 9.012 |
| birds | Sporophila_bouvronides | Sporophila_bouvronides | LC | Stable | tropical | diurnal | 9.1 | 4740689.394 | 2.496 |
| birds | Sporophila_caerulescens | Sporophila_caerulescens | LC | Increasing | tropical | diurnal | 9.7 | 7990264.015 | 7.005 |
| birds | Sporophila_castaneiventris | Sporophila_castaneiventris | LC | Stable | tropical | diurnal | 7.8 | 3489887.595 | 1.899 |
| birds | Sporophila_cinnamomea | Sporophila_cinnamomea | VU | Decreasing | tropical | diurnal | 8.9 | 2747258.158 | 10.658 |
| birds | Sporophila_collaris | Sporophila_collaris | LC | Stable | tropical | diurnal | 13.5 | 3264556.931 | 9.263 |
| birds | Sporophila_corvina | Sporophila_corvina | LC | Stable | tropical | diurnal | 10.6 | 288447.382 | 8.855 |
| birds | Sporophila_crassirostris | Oryzoborus_crassirostris | LC | Increasing | tropical | diurnal | 20.9 | 2533030.548 | 3.504 |
| birds | Sporophila_falcirostris | Sporophila_falcirostris | VU | Decreasing | tropical | diurnal | 13.5 | 175217.061 | 11.646 |
| birds | Sporophila_fringilloides | Dolospingus_fringilloides | LC | Stable | tropical | diurnal | 12.7 | 330728.089 | 0.756 |
| birds | Sporophila_frontalis | Sporophila_frontalis | VU | Decreasing | tropical | diurnal | 19.3 | 280054.212 | 12.716 |
| birds | Sporophila_hypochroma | Sporophila_hypochroma | NT | Decreasing | temperate | diurnal | 9.7 | 1603648.935 | 8.678 |
| birds | Sporophila_hypoxantha | Sporophila_hypoxantha | LC | Stable | tropical | diurnal | 10.5 | 3190597.65 | 9.905 |
| birds | Sporophila_intermedia | Sporophila_intermedia | LC | Stable | tropical | diurnal | 12.1 | 1593110.492 | 7.617 |
| birds | Sporophila_leucoptera | Sporophila_leucoptera | LC | Stable | tropical | diurnal | 15.5 | 3778100.992 | 9.19 |
| birds | Sporophila_lineola | Sporophila_lineola | LC | Stable | tropical | diurnal | 9.8 | 12050519.36 | 5.072 |
| birds | Sporophila_luctuosa | Sporophila_luctuosa | LC | Stable | tropical | diurnal | 12.5 | 1127825.555 | 5.343 |
| birds | Sporophila_maximiliani | Oryzoborus_maximiliani | EN | Decreasing | tropical | diurnal | 24.6 | 1854793.49 | 9.692 |
| birds | Sporophila_melanogaster | Sporophila_melanogaster | NT | Decreasing | tropical | diurnal | 9.3 | 837030.155 | 13.418 |
| birds | Sporophila_minuta | Sporophila_minuta | LC | Decreasing | tropical | diurnal | 7.8 | 2243040.212 | 6.509 |
| birds | Sporophila_murallae | Sporophila_murallae | LC | Increasing | tropical | diurnal | 10.8 | 973993.274 | 1.873 |
| birds | Sporophila_nigricollis | Sporophila_nigricollis | LC | Increasing | tropical | diurnal | 9.6 | 7194514.705 | 7.616 |
| birds | Sporophila_nigrorufa | Sporophila_nigrorufa | VU | Decreasing | tropical | diurnal | 10.8 | 283022.624 | 4.4 |
| birds | Sporophila_nuttingi | Oryzoborus_nuttingi | LC | Decreasing | tropical | diurnal | 24 | 52780.902 | 9.216 |
| birds | Sporophila_palustris | Sporophila_palustris | EN | Decreasing | tropical | diurnal | 8.2 | 2517089.605 | 10.798 |
| birds | Sporophila_peruviana | Sporophila_peruviana | LC | Increasing | tropical | diurnal | 12.6 | 102408.25 | 10.709 |
| birds | Sporophila_plumbea | Sporophila_plumbea | LC | Stable | tropical | diurnal | 9.7 | 4740053.967 | 7.625 |

| | | | | | | | | | |
|---|---|---|---|---|---|---|---|---|---|
| birds | Sporophila_ruficollis | Sporophila_ruficollis | NT | Decreasing | temperate | diurnal | 8.6 | 3429263.765 | 8.904 |
| birds | Sporophila_schistacea | Sporophila_schistacea | LC | Decreasing | tropical | diurnal | 12.7 | 1554851.819 | 3.814 |
| birds | Sporophila_simplex | Sporophila_simplex | LC | Increasing | tropical | diurnal | 28 | 95371.898 | 8.583 |
| birds | Sporophila_telasco | Sporophila_telasco | LC | Increasing | tropical | diurnal | 9.6 | 240640.682 | 9.629 |
| birds | Sporophila_torqueola | Sporophila_torqueola | LC | Increasing | tropical | diurnal | 8.7 | 543612.607 | 11.41 |
| birds | Sporopipes_frontalis | Sporopipes_frontalis | LC | Stable | tropical | diurnal | 17.3 | 3005541.444 | 11.937 |
| birds | Sporopipes_squamifrons | Sporopipes_squamifrons | LC | Stable | tropical | diurnal | 12.3 | 1945507.155 | 6.948 |
| birds | Stachyris_grammiceps | Stachyris_grammiceps | NT | Decreasing | tropical | diurnal | 13 | 28092.099 | 17.18 |
| birds | Stachyris_herberti | Stachyris_herberti | LC | Decreasing | tropical | diurnal | 29 | 21761.3 | 7.886 |
| birds | Stachyris_humei | Sphenocichla_humei | NT | Decreasing | temperate | diurnal | 34.1 | 92705.578 | 6.003 |
| birds | Stachyris_leucotis | Stachyris_leucotis | NT | Decreasing | tropical | diurnal | 23.7 | 1065906.973 | 8.215 |
| birds | Stachyris_maculata | Stachyris_maculata | NT | Decreasing | tropical | diurnal | 29.2 | 650885.412 | 7.332 |
| birds | Stachyris_nigriceps | Stachyris_nigriceps | LC | Decreasing | tropical | diurnal | 15.8 | 1916677.612 | 8.372 |
| birds | Stachyris_nigricollis | Stachyris_nigricollis | NT | Decreasing | tropical | diurnal | 26.2 | 238222.037 | 9.766 |
| birds | Stachyris_oglei | Stachyris_oglei | NT | Decreasing | temperate | diurnal | 33 | 11204.112 | 4.299 |
| birds | Stachyris_poliocephala | Stachyris_poliocephala | LC | Decreasing | tropical | diurnal | 23.3 | 715193.379 | 9.132 |
| birds | Stachyris_strialata | Stachyris_striolata | LC | Decreasing | tropical | diurnal | 28 | 593976.724 | 8.143 |
| birds | Stachyris_thoracica | Stachyris_thoracica | LC | Decreasing | tropical | diurnal | 12 | 63832.943 | 18.242 |
| birds | Stactolaema_anchietae | Stactolaema_anchietae | LC | Stable | tropical | diurnal | 47.3 | 1482309.278 | 6.616 |
| birds | Stactolaema_leucotis | Stactolaema_leucotis | LC | Decreasing | tropical | diurnal | 54.9 | 862758.909 | 11.135 |
| birds | Stactolaema_whytii | Stactolaema_whytii | LC | Decreasing | tropical | diurnal | 55.6 | 456236.863 | 11.498 |
| birds | Stagonopleura_bella | Stagonopleura_bella | LC | Stable | temperate | diurnal | 14 | 276741.258 | 9.58 |
| birds | Stagonopleura_guttata | Stagonopleura_guttata | VU | Decreasing | temperate | diurnal | 17.6 | 1192427.366 | 8.774 |
| birds | Stagonopleura_oculata | Stagonopleura_oculata | LC | Stable | temperate | diurnal | 12.9 | 82318.018 | 9.288 |
| birds | Staphida_everetti | Yuhina_everetti | LC | Decreasing | tropical | diurnal | 12.1 | 183515.143 | 2.811 |
| birds | Starnoenas_cyanocephala | Starnoenas_cyanocephala | EN | Decreasing | tropical | diurnal | 219 | 110641.899 | 16.093 |
| birds | Steatornis_caripensis | Steatornis_caripensis | LC | Stable | tropical | nocturnal | 408 | 1061013.513 | 5.414 |
| birds | Steganopus_tricolor | Steganopus_tricolor | LC | Increasing | temperate | diurnal | 59.4 | 15117456.652 | 9.589 |
| birds | Stelgidillas_gracilirostris | Andropadus_gracilirostris | LC | Stable | tropical | diurnal | 31.2 | 3807444.868 | 8.611 |
| birds | Stelgidopteryx_ridgwayi | NA | LC | Unknown | NA | NA | NA | NA | NA |
| birds | Stelgidopteryx_ruficollis | Stelgidopteryx_ruficollis | LC | Stable | tropical | diurnal | 16.1 | 13831495.263 | 5.851 |
| birds | Stelgidopteryx_serripennis | Stelgidopteryx_serripennis | LC | Decreasing | temperate | diurnal | 15.7 | 11481237.669 | 12.617 |
| birds | Stenostira_scita | Stenostira_scita | LC | Stable | temperate | diurnal | 5.9 | 1116923.254 | 9.054 |
| birds | Stephanoaetus_coronatus | Stephanoaetus_coronatus | NT | Decreasing | tropical | diurnal | 3640 | 6608300.515 | 9.617 |

| | | | | | | | | | |
|---|---|---|---|---|---|---|---|---|---|
| birds | Stephanophorus_diadematus | Stephanophorus_diadematus | LC | Stable | temperate | diurnal | 35.4 | 1088996.674 | 11.845 |
| birds | Stephanoxis_lalandi | Stephanoxis_lalandi | LC | Unknown | tropical | diurnal | 4 | 300448.067 | 15.002 |
| birds | Stercorarius_longicaudus | Stercorarius_longicaudus | LC | Stable | temperate | diurnal | 287.9 | 52268431.472 | 0.631 |
| birds | Stercorarius_parasiticus | Stercorarius_parasiticus | LC | Stable | temperate | diurnal | 444.3 | 21986115.797 | 0.842 |
| birds | Stercorarius_pomarinus | Stercorarius_pomarinus | LC | Stable | temperate | diurnal | 692.5 | 84401558.045 | 0.8 |
| birds | Sterna_acuticauda | NA | EN | Decreasing | NA | NA | NA | NA | NA |
| birds | Sterna_aurantia | NA | VU | Decreasing | NA | NA | NA | NA | NA |
| birds | Sterna_dougallii | Sterna_dougallii | LC | Unknown | tropical | diurnal | 112 | 34638646.076 | 10.828 |
| birds | Sterna_forsteri | Sterna_forsteri | LC | Increasing | temperate | diurnal | 149 | 10875479.249 | 12.677 |
| birds | Sterna_hirundinacea | Sterna_hirundinacea | LC | Decreasing | temperate | diurnal | 188 | 1717673.54 | 9.614 |
| birds | Sterna_hirundo | Sterna_hirundo | LC | Unknown | temperate | diurnal | 129.2 | 51361894.706 | 8.772 |
| birds | Sterna_paradisaea | Sterna_paradisaea | LC | Decreasing | temperate | diurnal | 110 | 61272189.474 | 1.421 |
| birds | Sterna_repressa | Sterna_repressa | LC | Decreasing | tropical | diurnal | 90 | 1914237.793 | 10.489 |
| birds | Sterna_striata | Sterna_striata | NT | Decreasing | temperate | diurnal | 129 | 4439208.047 | 10.578 |
| birds | Sterna_sumatrana | Sterna_sumatrana | LC | Unknown | tropical | diurnal | 106 | 41179216.023 | 9.209 |
| birds | Sterna_trudeaui | Sterna_trudeaui | LC | Stable | temperate | diurnal | 153 | 539946.351 | 12.844 |
| birds | Sterna_virgata | NA | NT | Stable | NA | NA | NA | NA | NA |
| birds | Sterna_vittata | Sterna_vittata | LC | Unknown | temperate | diurnal | 139 | 10872535.388 | 9.301 |
| birds | Sternoclyta_cyanopectus | Sternoclyta_cyanopectus | LC | Unknown | tropical | diurnal | 8 | 40400.064 | 13.534 |
| birds | Sternula_albifrons | Sterna_albifrons | LC | Decreasing | temperate | diurnal | 57 | 24714096.445 | 13.145 |
| birds | Sternula_antillarum | Sterna_antillarum | LC | Decreasing | tropical | diurnal | 45.9 | 3520113.384 | 14.605 |
| birds | Sternula_balaenarum | Sterna_balaenarum | LC | Decreasing | tropical | diurnal | 60 | 739942.626 | 5.848 |
| birds | Sternula_lorata | Sterna_lorata | EN | Decreasing | tropical | diurnal | 53.8 | 201942.778 | 11.205 |
| birds | Sternula_nereis | Sterna_nereis | VU | Decreasing | temperate | diurnal | 72.3 | 816751.806 | 9.72 |
| birds | Sternula_saundersi | Sterna_saundersi | LC | Decreasing | tropical | diurnal | 42.5 | 4572313.974 | 12.578 |
| birds | Sternula_superciliaris | Sterna_superciliaris | LC | Stable | tropical | diurnal | 46.4 | 11385573.195 | 5.199 |
| birds | Sterrhoptilus_capitalis | Stachyris_capitalis | LC | Decreasing | tropical | diurnal | 14.9 | 37149.261 | 9.466 |
| birds | Sterrhoptilus_dennistouni | Stachyris_dennistouni | NT | Decreasing | tropical | diurnal | 15.7 | 15801.313 | 4.879 |
| birds | Sterrhoptilus_nigrocapitatus | Stachyris_nigrocapitata | LC | Decreasing | tropical | diurnal | 14.1 | 30644.434 | 12.213 |
| birds | Stictonetta_naevosa | Stictonetta_naevosa | LC | Stable | temperate | diurnal | 903.3 | 5897864.382 | 4.753 |
| birds | Stigmatura_bahiae | NA | LC | Stable | NA | NA | NA | NA | NA |
| birds | Stigmatura_budytoides | Stigmatura_budytoides | LC | Stable | temperate | diurnal | 11.1 | 1866889.927 | 7.321 |
| birds | Stigmatura_napensis | Stigmatura_napensis | LC | Stable | tropical | diurnal | 9.8 | 322846.588 | 2.319 |
| birds | Stiltia_isabella | Stiltia_isabella | LC | Stable | tropical | diurnal | 65.5 | 5216126.15 | 4.769 |

| | | | | | | | | | |
|---|---|---|---|---|---|---|---|---|---|
| birds | Stiphrornis_erythrothorax | Stiphrornis_erythrothorax | LC | Decreasing | tropical | diurnal | 15.6 | 812936.307 | 13.741 |
| birds | Stipiturus_malachurus | Stipiturus_malachurus | LC | Decreasing | temperate | diurnal | 7.3 | 687011.192 | 9.489 |
| birds | Stipiturus_mallee | Stipiturus_mallee | EN | Decreasing | temperate | diurnal | 5.5 | 30237.259 | 7.858 |
| birds | Stipiturus_ruficeps | Stipiturus_ruficeps | LC | Decreasing | tropical | diurnal | 5 | 568075.871 | 1.986 |
| birds | Stizorhina_finschi | Stizorhina_finschi | LC | Decreasing | tropical | diurnal | 37 | 397355.963 | 15.761 |
| birds | Stizorhina_fraseri | Stizorhina_fraseri | LC | Decreasing | tropical | diurnal | 35.7 | 3553586.078 | 6.863 |
| birds | Stomiopera_flava | Lichenostomus_flavus | LC | Stable | tropical | diurnal | 21.4 | 415063.543 | 3.837 |
| birds | Stomiopera_unicolor | Lichenostomus_unicolor | LC | Stable | tropical | diurnal | 30.7 | 1189196.614 | 2.823 |
| birds | Strepera_fuliginosa | Strepera_fuliginosa | LC | Stable | temperate | diurnal | 375.5 | 68050.364 | 5.966 |
| birds | Strepera_graculina | Strepera_graculina | LC | Increasing | temperate | diurnal | 298.9 | 1580095.102 | 7.409 |
| birds | Strepera_versicolor | Strepera_versicolor | LC | Stable | temperate | diurnal | 395.5 | 2079178.273 | 6.6 |
| birds | Streptocitta_albertinae | Streptocitta_albertinae | NT | Decreasing | tropical | diurnal | 248 | 2737.236 | 5.753 |
| birds | Streptocitta_albicollis | Streptocitta_albicollis | LC | Decreasing | tropical | diurnal | 248 | 55724.428 | 10.919 |
| birds | Streptopelia_bitorquata | Streptopelia_bitorquata | LC | Decreasing | tropical | diurnal | 153 | 186292.822 | 20.01 |
| birds | Streptopelia_capicola | Streptopelia_capicola | LC | Increasing | tropical | diurnal | 142 | 9534302.211 | 8.608 |
| birds | Streptopelia_decaocto | Streptopelia_decaocto | LC | Increasing | temperate | diurnal | 149 | 19314230.802 | 14.002 |
| birds | Streptopelia_decipiens | Streptopelia_decipiens | LC | Stable | tropical | diurnal | 134 | 7210175.108 | 10.065 |
| birds | Streptopelia_hypopyrrha | Streptopelia_hypopyrrha | LC | Stable | tropical | diurnal | 162 | 529885.839 | 11.37 |
| birds | Streptopelia_lugens | Streptopelia_lugens | LC | Stable | tropical | diurnal | 155 | 945579.475 | 12.966 |
| birds | Streptopelia_orientalis | Streptopelia_orientalis | LC | Stable | temperate | diurnal | 232.9 | 20050458.531 | 9.702 |
| birds | Streptopelia_reichenowi | Streptopelia_reichenowi | NT | Stable | tropical | diurnal | 119 | 153525.54 | 9.748 |
| birds | Streptopelia_roseogrisea | Streptopelia_roseogrisea | LC | Stable | tropical | diurnal | 155 | 6669492.639 | 7.2 |
| birds | Streptopelia_semitorquata | Streptopelia_semitorquata | LC | Increasing | tropical | diurnal | 176 | 13753343.715 | 9.599 |
| birds | Streptopelia_tranquebarica | Streptopelia_tranquebarica | LC | Decreasing | temperate | diurnal | 103 | 9537403.558 | 14.941 |
| birds | Streptopelia_turtur | Streptopelia_turtur | VU | Decreasing | tropical | diurnal | 132 | 22086231.077 | 11.687 |
| birds | Streptopelia_vinacea | Streptopelia_vinacea | LC | Stable | tropical | diurnal | 110 | 6146574.468 | 10.691 |
| birds | Streptoprocne_biscutata | Streptoprocne_biscutata | LC | Decreasing | tropical | diurnal | 95.8 | 1741670.394 | 10.841 |
| birds | Streptoprocne_rutila | Streptoprocne_rutila | LC | Decreasing | tropical | diurnal | 20.2 | 1392664.2 | 9.961 |
| birds | Streptoprocne_semicollaris | Streptoprocne_semicollaris | LC | Decreasing | tropical | diurnal | 175 | 324675.194 | 8.72 |
| birds | Streptoprocne_zonaris | Streptoprocne_zonaris | LC | Decreasing | tropical | diurnal | 86.8 | 8315596.308 | 7.625 |
| birds | Stresemannia_bougainvillei | Stresemannia_bougainvillei | LC | Unknown | tropical | diurnal | 24.7 | 2008.771 | 4.886 |
| birds | Strigops_habroptila | Strigops_habroptila | CR | Increasing | temperate | nocturnal | 1732.1 | 17744.129 | 2.736 |
| birds | Strix_aluco | Strix_aluco | LC | Stable | temperate | cathemeral | 472.5 | 9594068.114 | 15.899 |
| birds | Strix_butleri | Strix_butleri | LC | Stable | tropical | nocturnal | 219 | 5226.95 | 14.906 |

| | | | | | | | | | |
|---|---|---|---|---|---|---|---|---|---|
| birds | Strix_chacoensis | Strix_chacoensis | NT | Decreasing | temperate | nocturnal | 330 | 766166.489 | 6.836 |
| birds | Strix_fulvescens | NA | LC | Decreasing | NA | NA | NA | NA | NA |
| birds | Strix_hylophila | Strix_hylophila | LC | Decreasing | temperate | nocturnal | 345.4 | 851827.979 | 13.146 |
| birds | Strix_leptogrammica | Strix_leptogrammica | LC | Decreasing | tropical | nocturnal | 970 | 5428569.659 | 11.748 |
| birds | Strix_nebulosa | Strix_nebulosa | LC | Increasing | temperate | cathemeral | 1061.9 | 19866544.632 | 3.662 |
| birds | Strix_occidentalis | Strix_occidentalis | NT | Decreasing | temperate | nocturnal | 586.4 | 1222682.339 | 8.715 |
| birds | Strix_rufipes | Strix_rufipes | LC | Decreasing | temperate | nocturnal | 380 | 639116.126 | 5.43 |
| birds | Strix_seloputo | Strix_seloputo | LC | Stable | tropical | nocturnal | 1011 | 610935.048 | 15.732 |
| birds | Strix_uralensis | Strix_uralensis | LC | Stable | temperate | cathemeral | 780.6 | 12224942.843 | 6.17 |
| birds | Strix_varia | Strix_varia | LC | Increasing | temperate | nocturnal | 711.5 | 6628563.517 | 12.534 |
| birds | Strix_woodfordii | Strix_woodfordii | LC | Stable | tropical | nocturnal | 350 | 9861983.738 | 9.039 |
| birds | Struthidea_cinerea | Struthidea_cinerea | LC | Stable | temperate | diurnal | 132 | 2500913.565 | 5.943 |
| birds | Struthio_camelus | Struthio_camelus | LC | Decreasing | temperate | diurnal | 111000 | 9266924.362 | 7.471 |
| birds | Sturnella_magna | Sturnella_magna | NT | Decreasing | temperate | diurnal | 91.8 | 7869636.907 | 12.857 |
| birds | Sturnella_neglecta | Sturnella_neglecta | LC | Decreasing | temperate | diurnal | 100.1 | 8668509.022 | 11.693 |
| birds | Sturnia_erythropygia | NA | LC | Unknown | NA | NA | NA | NA | NA |
| birds | Sturnia_malabarica | Sturnus_malabaricus | LC | Unknown | tropical | diurnal | 39.6 | 5278207.02 | 15.339 |
| birds | Sturnia_pagodarum | Sturnus_pagodarum | LC | Unknown | tropical | diurnal | 48.9 | 3846594.049 | 16.392 |
| birds | Sturnia_sinensis | Sturnus_sinensis | LC | Stable | tropical | diurnal | 61 | 1582507.906 | 14.881 |
| birds | Sturnornis_albofrontatus | NA | VU | Decreasing | NA | NA | NA | NA | NA |
| birds | Sturnus_unicolor | Sturnus_unicolor | LC | Increasing | temperate | diurnal | 83.7 | 826392.837 | 17.141 |
| birds | Sturnus_vulgaris | Sturnus_vulgaris | LC | Decreasing | temperate | diurnal | 77.1 | 40394100.724 | 10.463 |
| birds | Sublegatus_arenarum | Sublegatus_arenarum | LC | Stable | tropical | diurnal | 12.3 | 940862.165 | 8.41 |
| birds | Sublegatus_modestus | Sublegatus_modestus | LC | Increasing | temperate | diurnal | 14 | 6470238.679 | 7.776 |
| birds | Sublegatus_obscurior | Sublegatus_obscurior | LC | Stable | tropical | diurnal | 16 | 5373017.822 | 1.775 |
| birds | Sugomel_nigrum | Certhionyx_niger | LC | Decreasing | temperate | diurnal | 9.9 | 5440427.414 | 3.944 |
| birds | Suiriri_suiriri | Suiriri_suiriri | LC | Decreasing | tropical | diurnal | 14.5 | 6428767.243 | 8.752 |
| birds | Sula_dactylatra | Sula_dactylatra | LC | Decreasing | tropical | diurnal | 1732.1 | 130196830.591 | 9.488 |
| birds | Sula_leucogaster | Sula_leucogaster | LC | Decreasing | tropical | diurnal | 1288.8 | 136904984.896 | 7.948 |
| birds | Sula_nebouxii | Sula_nebouxii | LC | Decreasing | tropical | diurnal | 1520.1 | 4287194.459 | 5.985 |
| birds | Sula_sula | Sula_sula | LC | Decreasing | tropical | nocturnal | 1031.1 | 122461347.45 | 8.901 |
| birds | Sula_variegata | Sula_variegata | LC | Stable | tropical | diurnal | 1432.5 | 912459.658 | 10.709 |
| birds | Surnia_ulula | Surnia_ulula | LC | Stable | temperate | cathemeral | 319.9 | 21292793.546 | 3.117 |
| birds | Surniculus_lugubris | Surniculus_lugubris | LC | Decreasing | tropical | diurnal | 29.7 | 1813552.507 | 9.131 |

| | | | | | | | | | |
|---|---|---|---|---|---|---|---|---|---|
| birds | Surniculus_velutinus | Surniculus_velutinus | LC | Decreasing | tropical | diurnal | 35.6 | 259937.536 | 13.729 |
| birds | Suthora_alphonsiana | Paradoxornis_alphonsianus | LC | Stable | temperate | diurnal | 10.5 | 186111.218 | 15.082 |
| birds | Suthora_atrosuperciliaris | Paradoxornis_atrosuperciliaris | LC | Decreasing | tropical | diurnal | 16.8 | 204421.847 | 6.737 |
| birds | Suthora_conspicillata | Paradoxornis_conspicillatus | LC | Unknown | temperate | diurnal | 8.5 | 276903.925 | 10.389 |
| birds | Suthora_davidiana | Paradoxornis_davidianus | LC | Decreasing | tropical | diurnal | 8.5 | 172440.84 | 11.006 |
| birds | Suthora_fulvifrons | Paradoxornis_fulvifrons | LC | Decreasing | temperate | diurnal | 6.4 | 287751.919 | 7.187 |
| birds | Suthora_nipalensis | Paradoxornis_nipalensis | LC | Unknown | temperate | diurnal | 5.5 | 409687.66 | 7.334 |
| birds | Suthora_przewalskii | NA | LC | Stable | NA | NA | NA | NA | NA |
| birds | Suthora_verreauxi | Paradoxornis_verreauxi | LC | Decreasing | tropical | diurnal | 6 | 430054.136 | 11.511 |
| birds | Suthora_webbiana | Paradoxornis_webbianus | LC | Stable | temperate | diurnal | 10.9 | 4347110.321 | 15.874 |
| birds | Suthora_zappeyi | Paradoxornis_zappeyi | VU | Decreasing | temperate | diurnal | 9.5 | 41291.234 | 12.975 |
| birds | Swynnertonia_swynnertoni | Swynnertonia_swynnertoni | VU | Decreasing | tropical | diurnal | 16 | 7065.433 | 9.149 |
| birds | Sylvia_abyssinica | Pseudoalcippe_abyssinica | LC | Decreasing | tropical | diurnal | 18.8 | 387143.468 | 12.063 |
| birds | Sylvia_atricapilla | Sylvia_atricapilla | LC | Increasing | temperate | diurnal | 16.7 | 21049205.579 | 10.803 |
| birds | Sylvia_borin | Sylvia_borin | LC | Decreasing | tropical | diurnal | 18.2 | 48022839.559 | 8.741 |
| birds | Sylvia_cantillans | NA | not_available | not_available | NA | NA | NA | NA | NA |
| birds | Sylvia_curruca | NA | not_available | not_available | NA | NA | NA | NA | NA |
| birds | Sylvia_dohrni | Horizorhinus_dohrni | LC | Stable | tropical | diurnal | 19.3 | 150.944 | 14.677 |
| birds | Sylvia_galinieri | Parophasma_galinieri | LC | Decreasing | tropical | diurnal | 21.5 | 78935.61 | 14.774 |
| birds | Sylvia_hortensis | NA | not_available | not_available | NA | NA | NA | NA | NA |
| birds | Sylvia_nana | NA | not_available | not_available | NA | NA | NA | NA | NA |
| birds | Sylvia_nigricapillus | Lioptilus_nigricapillus | VU | Decreasing | temperate | diurnal | 29.6 | 170916.755 | 12.144 |
| birds | Sylvia_sarda | NA | not_available | not_available | NA | NA | NA | NA | NA |
| birds | Sylvietta_brachyura | Sylvietta_brachyura | LC | Stable | tropical | diurnal | 8.1 | 5897093.521 | 10.922 |
| birds | Sylvietta_denti | Sylvietta_denti | LC | Stable | tropical | diurnal | 8.4 | 1130212.974 | 8.975 |
| birds | Sylvietta_isabellina | Sylvietta_isabellina | LC | Stable | tropical | diurnal | 10 | 901911.547 | 7.627 |
| birds | Sylvietta_leucophrys | Sylvietta_leucophrys | LC | Stable | tropical | diurnal | 11.3 | 156344.043 | 15.889 |
| birds | Sylvietta_philippae | Sylvietta_philippae | DD | Stable | tropical | diurnal | 8.6 | 206306.366 | 8.014 |
| birds | Sylvietta_rufescens | Sylvietta_rufescens | LC | Stable | tropical | diurnal | 11.6 | 4186905.874 | 8.298 |
| birds | Sylvietta_ruficapilla | Sylvietta_ruficapilla | LC | Stable | tropical | diurnal | 11.1 | 2557018.327 | 7.479 |
| birds | Sylvietta_virens | Sylvietta_virens | LC | Stable | tropical | diurnal | 8.4 | 3738336.184 | 8.519 |
| birds | Sylvietta_whytii | Sylvietta_whytii | LC | Decreasing | tropical | diurnal | 9.9 | 1855770.987 | 12.177 |
| birds | Sylviorthorhynchus_desmurii | Sylviorthorhynchus_desmursii | LC | Decreasing | temperate | diurnal | 10.9 | 386264.464 | 5.065 |
| birds | Sylviorthorhynchus_yanacensis | Leptasthenura_yanacensis | LC | Decreasing | tropical | diurnal | 8.8 | 148145.611 | 8.66 |

| | | | | | | | | | |
|---|---|---|---|---|---|---|---|---|---|
| birds | Sylviparus_modestus | Sylviparus_modestus | LC | Stable | temperate | diurnal | 7.1 | 837783.043 | 13.049 |
| birds | Syma_megarhyncha | Syma_megarhyncha | LC | Stable | tropical | diurnal | 56 | 157417.529 | 4.255 |
| birds | Syma_torotoro | Syma_torotoro | LC | Stable | tropical | diurnal | 37.7 | 650715.396 | 3.03 |
| birds | Symposiachrus_axillaris | Monarcha_axillaris | LC | Stable | tropical | diurnal | 15.6 | 118942.749 | 4.356 |
| birds | Symposiachrus_barbatus | Monarcha_barbatus | LC | Decreasing | tropical | diurnal | 21 | 21960.623 | 5.076 |
| birds | Symposiachrus_boanensis | Monarcha_boanensis | CR | Decreasing | tropical | diurnal | 17.5 | 13.873 | 5.452 |
| birds | Symposiachrus_brehmii | Monarcha_brehmii | NT | Decreasing | tropical | diurnal | 41 | 2524.27 | 8.959 |
| birds | Symposiachrus_browni | NA | NT | Decreasing | NA | NA | NA | NA | NA |
| birds | Symposiachrus_everetti | NA | EN | Decreasing | NA | NA | NA | NA | NA |
| birds | Symposiachrus_guttula | Monarcha_guttula | LC | Stable | tropical | diurnal | 16.4 | 657660.08 | 3.11 |
| birds | Symposiachrus_infelix | NA | NT | Decreasing | NA | NA | NA | NA | NA |
| birds | Symposiachrus_julianae | NA | NT | Decreasing | NA | NA | NA | NA | NA |
| birds | Symposiachrus_leucurus | NA | NT | Decreasing | NA | NA | NA | NA | NA |
| birds | Symposiachrus_loricatus | NA | LC | Stable | NA | NA | NA | NA | NA |
| birds | Symposiachrus_manadensis | Monarcha_manadensis | LC | Stable | tropical | diurnal | 22.5 | 445281.042 | 3.132 |
| birds | Symposiachrus_menckei | NA | NT | Unknown | NA | NA | NA | NA | NA |
| birds | Symposiachrus_mundus | NA | LC | Stable | NA | NA | NA | NA | NA |
| birds | Symposiachrus_rubiensis | NA | LC | Stable | NA | NA | NA | NA | NA |
| birds | Symposiachrus_sacerdotum | Monarcha_sacerdotum | EN | Decreasing | tropical | diurnal | 37 | 696.218 | 10.956 |
| birds | Symposiachrus_trivirgatus | Monarcha_trivirgatus | LC | Stable | tropical | diurnal | 12.7 | 612420.232 | 5.882 |
| birds | Symposiachrus_verticalis | Monarcha_verticalis | LC | Stable | tropical | diurnal | 21.7 | 42001.393 | 4.36 |
| birds | Symposiachrus_vidua | Monarcha_viduus | NT | Decreasing | tropical | diurnal | 17.1 | 3376.333 | 4.61 |
| birds | Synallaxis_albescens | Synallaxis_albescens | LC | Stable | tropical | diurnal | 11.2 | 9891841.773 | 7.243 |
| birds | Synallaxis_albigularis | Synallaxis_albigularis | LC | Increasing | tropical | diurnal | 15.4 | 1133087.176 | 1.984 |
| birds | Synallaxis_albilora | Synallaxis_albilora | LC | Stable | tropical | diurnal | 14.9 | 218221.061 | 5.892 |
| birds | Synallaxis_azarae | Synallaxis_azarae | LC | Stable | tropical | diurnal | 16.9 | 477890.567 | 8.873 |
| birds | Synallaxis_brachyura | Synallaxis_brachyura | LC | Stable | tropical | diurnal | 18.3 | 403853.13 | 9.628 |
| birds | Synallaxis_cabanisi | Synallaxis_cabanisi | LC | Decreasing | tropical | diurnal | 22.7 | 197400.659 | 4.257 |
| birds | Synallaxis_candei | Synallaxis_candei | LC | Stable | tropical | diurnal | 15.3 | 156033.913 | 13.486 |
| birds | Synallaxis_castanea | Synallaxis_castanea | LC | Stable | tropical | diurnal | 15 | 1609.916 | 18.029 |
| birds | Synallaxis_cherriei | Synallaxis_cherriei | LC | Decreasing | tropical | diurnal | 16 | 792427.571 | 2.287 |
| birds | Synallaxis_chinchipensis | NA | LC | Stable | NA | NA | NA | NA | NA |
| birds | Synallaxis_cinerascens | Synallaxis_cinerascens | LC | Decreasing | temperate | diurnal | 13.1 | 1223916.17 | 12.97 |
| birds | Synallaxis_cinerea | Synallaxis_whitneyi | NT | Decreasing | tropical | diurnal | 16.2 | 24736.239 | 7.146 |

| | | | | | | | | | |
|---|---|---|---|---|---|---|---|---|---|
| birds | Synallaxis_cinnamomea | Synallaxis_cinnamomea | LC | Stable | tropical | diurnal | 18.4 | 85347.41 | 12.434 |
| birds | Synallaxis_courseni | Synallaxis_courseni | VU | Stable | tropical | diurnal | 16.8 | 184.946 | 10.851 |
| birds | Synallaxis_erythrothorax | Synallaxis_erythrothorax | LC | Stable | tropical | diurnal | 17.1 | 355771.381 | 10.022 |
| birds | Synallaxis_frontalis | Synallaxis_frontalis | LC | Increasing | tropical | diurnal | 14 | 5745641.866 | 8.527 |
| birds | Synallaxis_fuscorufa | Synallaxis_fuscorufa | NT | Stable | tropical | diurnal | 16 | 2113.742 | 5.713 |
| birds | Synallaxis_gujanensis | Synallaxis_gujanensis | LC | Stable | tropical | diurnal | 18.1 | 4462466.661 | 2.51 |
| birds | Synallaxis_hellmayri | Gyalophylax_hellmayri | LC | Decreasing | tropical | diurnal | 25.5 | 339711.967 | 7.411 |
| birds | Synallaxis_hypochondriaca | Siptornopsis_hypochondriaca | NT | Decreasing | tropical | diurnal | 24.5 | 8608.39 | 9.554 |
| birds | Synallaxis_hypospodia | Synallaxis_hypospodia | LC | Stable | tropical | diurnal | 16.9 | 3530294.959 | 7.36 |
| birds | Synallaxis_infuscata | Synallaxis_infuscata | EN | Decreasing | tropical | diurnal | 18.2 | 25054.162 | 15.634 |
| birds | Synallaxis_kollari | Synallaxis_kollari | CR | Decreasing | tropical | diurnal | 16 | 4101.242 | 7.094 |
| birds | Synallaxis_macconnelli | Synallaxis_macconnelli | LC | Stable | tropical | diurnal | 19.5 | 247511.893 | 1.193 |
| birds | Synallaxis_maranonica | Synallaxis_maranonica | CR | Decreasing | tropical | diurnal | 13 | 4053.009 | 10.466 |
| birds | Synallaxis_moesta | Synallaxis_moesta | LC | Decreasing | tropical | diurnal | 22.4 | 172710.818 | 6.186 |
| birds | Synallaxis_ruficapilla | Synallaxis_ruficapilla | LC | Stable | temperate | diurnal | 13.8 | 1174516.013 | 13.472 |
| birds | Synallaxis_rutilans | Synallaxis_rutilans | LC | Decreasing | tropical | diurnal | 16.7 | 5856806.856 | 1.833 |
| birds | Synallaxis_scutata | Synallaxis_scutata | LC | Decreasing | tropical | diurnal | 15.4 | 3789577.698 | 8.417 |
| birds | Synallaxis_simoni | NA | LC | Decreasing | NA | NA | NA | NA | NA |
| birds | Synallaxis_spixi | Synallaxis_spixi | LC | Increasing | temperate | diurnal | 12.6 | 1343787.353 | 12.118 |
| birds | Synallaxis_stictothorax | Synallaxis_stictothorax | LC | Stable | tropical | diurnal | 10.2 | 49674.421 | 9.181 |
| birds | Synallaxis_subpudica | NA | LC | Stable | NA | NA | NA | NA | NA |
| birds | Synallaxis_tithys | Synallaxis_tithys | VU | Decreasing | tropical | diurnal | 15.8 | 8430.302 | 12.911 |
| birds | Synallaxis_unirufa | Synallaxis_unirufa | LC | Stable | tropical | diurnal | 17.9 | 210448.016 | 8.169 |
| birds | Synallaxis_zimmeri | Synallaxis_zimmeri | VU | Decreasing | tropical | diurnal | 13 | 4318.409 | 8.975 |
| birds | Syndactyla_dimidiata | Syndactyla_dimidiata | LC | Decreasing | tropical | diurnal | 29.4 | 1723823.38 | 11.046 |
| birds | Syndactyla_guttulata | Syndactyla_guttulata | LC | Stable | tropical | diurnal | 36 | 12401.489 | 15.387 |
| birds | Syndactyla_roraimae | Syndactyla_roraimae | LC | Stable | tropical | diurnal | 27.5 | 99093.77 | 1.108 |
| birds | Syndactyla_ruficollis | Syndactyla_ruficollis | VU | Decreasing | tropical | diurnal | 31 | 19699.527 | 7.934 |
| birds | Syndactyla_rufosuperciliata | Syndactyla_rufosuperciliata | LC | Stable | temperate | diurnal | 25.6 | 1726834.003 | 11.076 |
| birds | Syndactyla_striata | Simoxenops_striatus | LC | Decreasing | tropical | diurnal | 41.6 | 81400.962 | 4.221 |
| birds | Syndactyla_subalaris | Syndactyla_subalaris | LC | Decreasing | tropical | diurnal | 28.9 | 258113.591 | 8.996 |
| birds | Syndactyla_ucayalae | Simoxenops_ucayalae | NT | Decreasing | tropical | diurnal | 49.8 | 286032.696 | 1.559 |
| birds | Synoicus_adansonii | NA | LC | Stable | NA | NA | NA | NA | NA |
| birds | Synoicus_chinensis | Coturnix_chinensis | LC | Stable | tropical | diurnal | 40.4 | 8165093.608 | 11.47 |

| | | | | | | | | | |
|---|---|---|---|---|---|---|---|---|---|
| birds | Synoicus_ypsilophorus | Coturnix_ypsilophora | LC | Decreasing | tropical | diurnal | 107.7 | 5840602.826 | 4.658 |
| birds | Synthliboramphus_antiquus | Synthliboramphus_antiquus | LC | Decreasing | temperate | diurnal | 217.7 | 5455056.64 | 10.342 |
| birds | Synthliboramphus_craveri | Synthliboramphus_craveri | VU | Decreasing | temperate | diurnal | 151 | 744430.32 | 4.654 |
| birds | Synthliboramphus_hypoleucus | Synthliboramphus_hypoleucus | EN | Decreasing | temperate | diurnal | 167 | 1155481.814 | 7.616 |
| birds | Synthliboramphus_wumizusume | Synthliboramphus_wumizusume | VU | Decreasing | temperate | diurnal | 198 | 919606.853 | 23.261 |
| birds | Sypheotides_indicus | Sypheotides_indicus | CR | Decreasing | tropical | diurnal | 614.3 | 2471338.063 | 18.41 |
| birds | Syrigma_sibilatrix | Syrigma_sibilatrix | LC | Unknown | temperate | diurnal | 463 | 4235220.188 | 9.043 |
| birds | Syrmaticus_ellioti | Syrmaticus_ellioti | LC | Decreasing | temperate | diurnal | 1007.5 | 496586.997 | 12.182 |
| birds | Syrmaticus_humiae | Syrmaticus_humiae | VU | Decreasing | tropical | diurnal | 873.4 | 327611.249 | 11.066 |
| birds | Syrmaticus_mikado | Syrmaticus_mikado | LC | Stable | temperate | diurnal | 1148.7 | 8435.298 | 4.428 |
| birds | Syrmaticus_reevesii | Syrmaticus_reevesii | VU | Decreasing | temperate | diurnal | 1204.6 | 722698.122 | 18.082 |
| birds | Syrmaticus_soemmerringii | Syrmaticus_soemmerringii | NT | Decreasing | temperate | diurnal | 1081.7 | 174734.057 | 18.937 |
| birds | Syrrhaptes_tibetanus | Syrrhaptes_tibetanus | LC | Decreasing | temperate | diurnal | 343.7 | 1238571.565 | 4.618 |
| birds | Systellura_longirostris | Caprimulgus_longirostris | LC | Stable | tropical | nocturnal | 45.6 | 5080733.587 | 7.728 |
| birds | Taccocua_leschenaultii | Phaenicophaeus_leschenaultii | LC | Stable | tropical | diurnal | 174.7 | 2673797.889 | 18.727 |
| birds | Tachornis_furcata | Tachornis_furcata | LC | Decreasing | tropical | diurnal | 6.5 | 32700.659 | 17.244 |
| birds | Tachornis_phoenicobia | Tachornis_phoenicobia | LC | Stable | tropical | diurnal | 10.2 | 194217.826 | 16.229 |
| birds | Tachornis_squamata | Tachornis_squamata | LC | Stable | tropical | diurnal | 11 | 9486634.444 | 3.799 |
| birds | Tachuris_rubrigastra | Tachuris_rubrigastra | LC | Stable | temperate | diurnal | 7.8 | 2891313.768 | 9.238 |
| birds | Tachybaptus_dominicus | Tachybaptus_dominicus | LC | Stable | tropical | diurnal | 132.6 | 13082260.665 | 7.309 |
| birds | Tachybaptus_novaehollandiae | Tachybaptus_novaehollandiae | LC | Increasing | temperate | diurnal | 219 | 8308868.822 | 4.822 |
| birds | Tachybaptus_pelzelnii | Tachybaptus_pelzelnii | EN | Decreasing | tropical | diurnal | 184 | 590930.764 | 8.745 |
| birds | Tachybaptus_ruficollis | Tachybaptus_ruficollis | LC | Decreasing | temperate | diurnal | 169.4 | 37429277.13 | 12.685 |
| birds | Tachybaptus_rufolavatus | NA | EX | not_available | NA | NA | NA | NA | NA |
| birds | Tachycineta_albiventer | Tachycineta_albiventer | LC | Stable | tropical | diurnal | 17.7 | 11742614.015 | 5.299 |
| birds | Tachycineta_bicolor | Tachycineta_bicolor | LC | Stable | temperate | diurnal | 21.2 | 17478048.629 | 8.851 |
| birds | Tachycineta_cyaneoviridis | Tachycineta_cyaneoviridis | EN | Decreasing | temperate | diurnal | 17.5 | 14730.665 | 7.692 |
| birds | Tachycineta_euchrysea | Tachycineta_euchrysea | VU | Decreasing | tropical | diurnal | 17 | 14021.594 | 11.544 |
| birds | Tachycineta_leucorrhoa | Tachycineta_leucorrhoa | LC | Increasing | tropical | diurnal | 15.3 | 6386014.55 | 8.061 |
| birds | Tachycineta_meyeni | Tachycineta_meyeni | LC | Increasing | temperate | diurnal | 14.9 | 4066377.87 | 6.969 |
| birds | Tachycineta_stolzmanni | Tachycineta_stolzmanni | LC | Stable | tropical | diurnal | 11.3 | 43489.157 | 7.954 |
| birds | Tachycineta_thalassina | Tachycineta_thalassina | LC | Increasing | temperate | diurnal | 14.1 | 6579514.327 | 7.33 |
| birds | Tachyeres_brachypterus | Tachyeres_brachypterus | LC | Stable | temperate | diurnal | 3450 | 12033.796 | 3.247 |
| birds | Tachyeres_leucocephalus | Tachyeres_leucocephalus | VU | Decreasing | temperate | diurnal | 3013 | 49163.716 | 8.019 |

| | | | | | | | | | |
|---|---|---|---|---|---|---|---|---|---|
| birds | Tachyeres_patachonicus | Tachyeres_patachonicus | LC | Decreasing | temperate | diurnal | 2821.6 | 1006665.135 | 5.234 |
| birds | Tachyeres_pteneres | Tachyeres_pteneres | LC | Stable | temperate | diurnal | 4737.6 | 350010.456 | 2.167 |
| birds | Tachymarptis_aequatorialis | Tachymarptis_aequatorialis | LC | Stable | tropical | diurnal | 92.8 | 1399846.867 | 12.265 |
| birds | Tachymarptis_melba | Tachymarptis_melba | LC | Stable | tropical | nocturnal | 100.4 | 12945294.109 | 12.599 |
| birds | Tachyphonus_coronatus | Tachyphonus_coronatus | LC | Stable | tropical | diurnal | 29.3 | 1492450.617 | 12.682 |
| birds | Tachyphonus_phoenicius | Tachyphonus_phoenicius | LC | Stable | tropical | diurnal | 21 | 2229357.676 | 1.939 |
| birds | Tachyphonus_rufus | Tachyphonus_rufus | LC | Stable | tropical | diurnal | 34.4 | 5942265.974 | 7.628 |
| birds | Tadorna_cana | Tadorna_cana | LC | Increasing | temperate | diurnal | 1182 | 1631456.112 | 8.196 |
| birds | Tadorna_ferruginea | Tadorna_ferruginea | LC | Unknown | temperate | diurnal | 1235 | 24406375.914 | 9.296 |
| birds | Tadorna_tadorna | Tadorna_tadorna | LC | Increasing | temperate | diurnal | 1146.8 | 11207269.885 | 10.754 |
| birds | Tadorna_tadornoides | Tadorna_tadornoides | LC | Increasing | temperate | diurnal | 1418.7 | 3300063.223 | 6.176 |
| birds | Tadorna_variegata | Tadorna_variegata | LC | Stable | temperate | diurnal | 1541 | 268823.502 | 7.991 |
| birds | Taeniopygia_bichenovii | Taeniopygia_bichenovii | LC | Stable | temperate | diurnal | 9.6 | 3108066.097 | 4.972 |
| birds | Taeniopygia_guttata | Taeniopygia_guttata | LC | Stable | tropical | diurnal | 12 | 83399.732 | 13.518 |
| birds | Taeniotriccus_andrei | Taeniotriccus_andrei | LC | Decreasing | tropical | diurnal | 9.5 | 857576.239 | 2.482 |
| birds | Talaphorus_chlorocercus | Leucippus_chlorocercus | LC | Stable | tropical | diurnal | 4.6 | 221320.451 | 2.647 |
| birds | Talegalla_cuvieri | Talegalla_cuvieri | LC | Decreasing | tropical | diurnal | 1785 | 97152.1 | 3.223 |
| birds | Talegalla_fuscirostris | Talegalla_fuscirostris | LC | Decreasing | tropical | diurnal | 1314.5 | 284369.793 | 2.764 |
| birds | Talegalla_jobiensis | Talegalla_jobiensis | LC | Decreasing | tropical | diurnal | 1478.8 | 222871.352 | 3.597 |
| birds | Tangara_abbas | Thraupis_abbas | LC | Increasing | tropical | diurnal | 45 | 677166.093 | 10.245 |
| birds | Tangara_argentea | Tangara_cyanoptera | LC | Decreasing | tropical | diurnal | 21.5 | 93683.155 | 10.828 |
| birds | Tangara_argyrofenges | Tangara_argyrofenges | VU | Decreasing | tropical | diurnal | 19 | 52084.231 | 5.581 |
| birds | Tangara_arthus | Tangara_arthus | LC | Decreasing | tropical | diurnal | 22 | 38919.719 | 13.968 |
| birds | Tangara_brasiliensis | NA | LC | Decreasing | NA | NA | NA | NA | NA |
| birds | Tangara_cabanisi | Tangara_cabanisi | VU | Decreasing | tropical | diurnal | 20 | 3574.173 | 10.42 |
| birds | Tangara_callophrys | Tangara_callophrys | LC | Decreasing | tropical | diurnal | 22.9 | 1481423.436 | 1.698 |
| birds | Tangara_cayana | Tangara_cayana | LC | Stable | tropical | diurnal | 18 | 1467704.42 | 4.426 |
| birds | Tangara_chilensis | Tangara_chilensis | LC | Decreasing | tropical | diurnal | 19.8 | 4594777.457 | 2.061 |
| birds | Tangara_chrysotis | Tangara_chrysotis | LC | Decreasing | tropical | diurnal | 24 | 99793.388 | 5.143 |
| birds | Tangara_cucullata | Tangara_cucullata | LC | Stable | tropical | diurnal | 27.9 | 318.731 | 20.469 |
| birds | Tangara_cyanicollis | Tangara_cyanicollis | LC | Stable | tropical | diurnal | 17 | 1248379.883 | 5.906 |
| birds | Tangara_cyanocephala | Tangara_cyanocephala | LC | Stable | tropical | diurnal | 18 | 444844.918 | 12.799 |
| birds | Tangara_cyanoptera | Thraupis_cyanoptera | NT | Decreasing | tropical | diurnal | 43.3 | 224069.568 | 13.234 |
| birds | Tangara_cyanotis | Tangara_cyanotis | LC | Decreasing | tropical | diurnal | 15 | 88910.48 | 5.933 |

| | | | | | | | | | |
|---|---|---|---|---|---|---|---|---|---|
| birds | Tangara_cyanoventris | Tangara_cyanoventris | LC | Stable | tropical | diurnal | 16.5 | 475625.934 | 12.673 |
| birds | Tangara_desmaresti | Tangara_desmaresti | LC | Stable | temperate | diurnal | 20.4 | 260465.016 | 13.457 |
| birds | Tangara_dowii | Tangara_dowii | LC | Decreasing | tropical | diurnal | 15 | 12083.47 | 6.767 |
| birds | Tangara_episcopus | Thraupis_episcopus | LC | Increasing | tropical | diurnal | 35 | 7925982.834 | 3.863 |
| birds | Tangara_fastuosa | Tangara_fastuosa | VU | Decreasing | tropical | diurnal | 25.5 | 15694.935 | 16.014 |
| birds | Tangara_florida | Tangara_florida | LC | Decreasing | tropical | diurnal | 19.3 | 37066.943 | 5.283 |
| birds | Tangara_fucosa | Tangara_fucosa | LC | Stable | tropical | diurnal | 21 | 1086.105 | 1.512 |
| birds | Tangara_glaucocolpa | Thraupis_glaucocolpa | LC | Stable | tropical | diurnal | 31.6 | 236044.282 | 11.264 |
| birds | Tangara_guttata | Tangara_guttata | LC | Decreasing | tropical | diurnal | 18.4 | 376030.938 | 4.104 |
| birds | Tangara_gyrola | Tangara_gyrola | LC | Decreasing | tropical | diurnal | 21 | 4181131.618 | 3.213 |
| birds | Tangara_heinei | Tangara_heinei | LC | Stable | tropical | diurnal | 20.2 | 133743.397 | 10.225 |
| birds | Tangara_icterocephala | Tangara_icterocephala | LC | Decreasing | tropical | diurnal | 22 | 75163.998 | 7.051 |
| birds | Tangara_inornata | Tangara_inornata | LC | Stable | tropical | diurnal | 18 | 180601.403 | 8.708 |
| birds | Tangara_johannae | Tangara_johannae | LC | Decreasing | tropical | diurnal | 20 | 78323.968 | 5.561 |
| birds | Tangara_labradorides | Tangara_labradorides | LC | Stable | tropical | diurnal | 15 | 104444.947 | 11.62 |
| birds | Tangara_larvata | Tangara_larvata | LC | Decreasing | tropical | diurnal | 20 | 480330.116 | 8.565 |
| birds | Tangara_lavinia | Tangara_lavinia | LC | Decreasing | tropical | diurnal | 24 | 135531.532 | 5.668 |
| birds | Tangara_mexicana | Tangara_mexicana | LC | Decreasing | tropical | diurnal | 20.5 | 7040026.361 | 2.278 |
| birds | Tangara_meyerdeschauenseei | Tangara_meyerdeschauenseei | NT | Increasing | tropical | diurnal | 26 | 11014.625 | 4.335 |
| birds | Tangara_nigrocincta | Tangara_nigrocincta | LC | Decreasing | tropical | diurnal | 17 | 3416902.143 | 2.247 |
| birds | Tangara_nigroviridis | Tangara_nigroviridis | LC | Stable | tropical | diurnal | 17 | 348796.622 | 8.005 |
| birds | Tangara_ornata | Thraupis_ornata | LC | Stable | tropical | diurnal | 33 | 392822.736 | 12.618 |
| birds | Tangara_palmarum | Thraupis_palmarum | LC | Stable | tropical | diurnal | 39 | 12189499.365 | 5.512 |
| birds | Tangara_palmeri | Tangara_palmeri | LC | Decreasing | tropical | diurnal | 32.3 | 44181.15 | 5.063 |
| birds | Tangara_parzudakii | Tangara_parzudakii | LC | Decreasing | tropical | diurnal | 28 | 196526.949 | 7.597 |
| birds | Tangara_peruviana | Tangara_peruviana | VU | Decreasing | temperate | diurnal | 22.2 | 125652.77 | 13.702 |
| birds | Tangara_phillipsi | Tangara_phillipsi | LC | Stable | tropical | diurnal | 19.9 | 3651.09 | 2.405 |
| birds | Tangara_preciosa | Tangara_preciosa | LC | Stable | temperate | diurnal | 22.8 | 1104627.469 | 11.546 |
| birds | Tangara_punctata | Tangara_punctata | LC | Decreasing | tropical | diurnal | 15 | 2558212.032 | 1.711 |
| birds | Tangara_ruficervix | Tangara_ruficervix | LC | Stable | tropical | diurnal | 19 | 134578.1 | 9.626 |
| birds | Tangara_rufigenis | Tangara_rufigenis | LC | Decreasing | tropical | diurnal | 16.5 | 6837.305 | 18.512 |
| birds | Tangara_rufigula | Tangara_rufigula | LC | Decreasing | tropical | diurnal | 19.1 | 23087.114 | 6.365 |
| birds | Tangara_sayaca | Thraupis_sayaca | LC | Stable | tropical | diurnal | 32.5 | 6747180.968 | 8.851 |
| birds | Tangara_schrankii | Tangara_schrankii | LC | Decreasing | tropical | diurnal | 19 | 2751315.134 | 1.503 |

| | | | | | | | | | |
|---|---|---|---|---|---|---|---|---|---|
| birds | Tangara_seledon | Tangara_seledon | LC | Stable | tropical | diurnal | 18.7 | 672563.84 | 13.005 |
| birds | Tangara_varia | Tangara_varia | LC | Stable | tropical | diurnal | 10 | 1477680.374 | 1.585 |
| birds | Tangara_vassorii | Tangara_vassorii | LC | Decreasing | tropical | diurnal | 18 | 206740.237 | 10.645 |
| birds | Tangara_velia | Tangara_velia | LC | Decreasing | tropical | diurnal | 21 | 4923995.698 | 1.658 |
| birds | Tangara_viridicollis | Tangara_viridicollis | LC | Stable | tropical | diurnal | 21 | 102254.127 | 6.233 |
| birds | Tangara_vitriolina | Tangara_vitriolina | LC | Stable | tropical | diurnal | 23 | 123913.111 | 12.599 |
| birds | Tangara_xanthocephala | Tangara_xanthocephala | LC | Decreasing | tropical | diurnal | 19 | 356998.99 | 7.973 |
| birds | Tangara_xanthogastra | Tangara_xanthogastra | LC | Decreasing | tropical | diurnal | 15 | 2799036.322 | 1.616 |
| birds | Tanygnathus_gramineus | Tanygnathus_gramineus | VU | Decreasing | tropical | diurnal | 237.3 | 1781.705 | 4.056 |
| birds | Tanygnathus_lucionensis | Tanygnathus_lucionensis | NT | Decreasing | tropical | diurnal | 215 | 311136.643 | 13.281 |
| birds | Tanygnathus_megalorynchos | Tanygnathus_megalorynchos | LC | Decreasing | tropical | diurnal | 325 | 96725.712 | 8.885 |
| birds | Tanygnathus_sumatranus | Tanygnathus_sumatranus | LC | Decreasing | tropical | cathemeral | 269.6 | 154743.611 | 10.037 |
| birds | Tanysiptera_carolinae | Tanysiptera_carolinae | NT | Decreasing | tropical | diurnal | 50.2 | 336.016 | 8.855 |
| birds | Tanysiptera_danae | Tanysiptera_danae | LC | Stable | tropical | diurnal | 43.5 | 27138.999 | 3.907 |
| birds | Tanysiptera_ellioti | Tanysiptera_ellioti | NT | Decreasing | tropical | diurnal | 50.2 | 176.044 | 1.655 |
| birds | Tanysiptera_galatea | Tanysiptera_galatea | LC | Decreasing | tropical | diurnal | 50 | 702146.788 | 3.35 |
| birds | Tanysiptera_hydrocharis | NA | DD | Unknown | NA | NA | NA | NA | NA |
| birds | Tanysiptera_nympha | Tanysiptera_nympha | LC | Decreasing | tropical | diurnal | 57 | 104476.663 | 3.496 |
| birds | Tanysiptera_riedelii | Tanysiptera_riedelii | NT | Decreasing | tropical | diurnal | 63.5 | 2526.514 | 8.959 |
| birds | Tanysiptera_sylvia | Tanysiptera_sylvia | LC | Decreasing | tropical | diurnal | 49.6 | 395148.654 | 3.072 |
| birds | Taoniscus_nanus | Taoniscus_nanus | EN | Decreasing | tropical | diurnal | 43 | 1083212.525 | 11.047 |
| birds | Tapera_naevia | Tapera_naevia | LC | Stable | tropical | diurnal | 48.4 | 12346242.134 | 6.374 |
| birds | Taphrolesbia_griseiventris | Taphrolesbia_griseiventris | EN | Decreasing | tropical | diurnal | 8.9 | 5547.215 | 10.967 |
| birds | Taphrospilus_hypostictus | Taphrospilus_hypostictus | LC | Decreasing | tropical | diurnal | 7 | 110112.14 | 4.991 |
| birds | Taraba_major | Taraba_major | LC | Decreasing | tropical | diurnal | 59.2 | 12899627.836 | 5.256 |
| birds | Tarphonomus_certhioides | Tarphonomus_certhioides | LC | Stable | temperate | diurnal | 22.1 | 1267589.78 | 6.716 |
| birds | Tarphonomus_harterti | Tarphonomus_harterti | LC | Decreasing | tropical | diurnal | 23.9 | 70447.433 | 8.95 |
| birds | Tarsiger_chrysaeus | Tarsiger_chrysaeus | LC | Stable | temperate | diurnal | 13.8 | 1061369.997 | 8.495 |
| birds | Tarsiger_cyanurus | Tarsiger_cyanurus | LC | Stable | temperate | diurnal | 13.4 | 13872524.623 | 5.617 |
| birds | Tarsiger_hyperythrus | Tarsiger_hyperythrus | LC | Stable | temperate | diurnal | 12 | 267035.259 | 6.765 |
| birds | Tarsiger_indicus | Tarsiger_indicus | LC | Stable | temperate | diurnal | 14.6 | 533602.164 | 8.048 |
| birds | Tarsiger_johnstoniae | Tarsiger_johnstoniae | LC | Stable | temperate | diurnal | 10.5 | 23533.291 | 12.125 |
| birds | Tauraco_bannermani | Tauraco_bannermani | EN | Decreasing | tropical | diurnal | 224 | 1978.6 | 12.927 |
| birds | Tauraco_corythaix | Tauraco_corythaix | LC | Decreasing | temperate | diurnal | 308 | 124836.473 | 11.871 |

| | | | | | | | | | |
|---|---|---|---|---|---|---|---|---|---|
| birds | Tauraco_erythrolophus | Tauraco_erythrolophus | LC | Decreasing | tropical | diurnal | 261.2 | 189792.611 | 8.097 |
| birds | Tauraco_fischeri | Tauraco_fischeri | NT | Decreasing | tropical | diurnal | 250 | 48655.561 | 13.169 |
| birds | Tauraco_hartlaubi | Tauraco_hartlaubi | LC | Decreasing | tropical | diurnal | 224 | 264805.146 | 12.197 |
| birds | Tauraco_leucolophus | Tauraco_leucolophus | LC | Stable | tropical | diurnal | 219 | 1433202.567 | 7.593 |
| birds | Tauraco_leucotis | Tauraco_leucotis | LC | Stable | tropical | diurnal | 265 | 807016.239 | 10.428 |
| birds | Tauraco_livingstonii | Tauraco_livingstonii | LC | Decreasing | tropical | diurnal | 249 | 974104.703 | 9.835 |
| birds | Tauraco_macrorhynchus | Tauraco_macrorhynchus | LC | Decreasing | tropical | diurnal | 243.7 | 1273570.691 | 11.152 |
| birds | Tauraco_persa | Tauraco_persa | LC | Stable | tropical | diurnal | 306 | 3260169.314 | 9.932 |
| birds | Tauraco_ruspolii | Tauraco_ruspolii | NT | Decreasing | tropical | diurnal | 263 | 24072.841 | 6.103 |
| birds | Tauraco_schalowi | Tauraco_schalowi | LC | Stable | tropical | diurnal | 206.4 | 1899419.962 | 7.948 |
| birds | Tauraco_schuettii | Tauraco_schuetti | LC | Stable | tropical | diurnal | 235 | 2085207.967 | 7.036 |
| birds | Tchagra_australis | Tchagra_australis | LC | Stable | tropical | diurnal | 32.3 | 8305150.13 | 8.9 |
| birds | Tchagra_jamesi | Tchagra_jamesi | LC | Stable | tropical | diurnal | 27.4 | 1196428.504 | 8.307 |
| birds | Tchagra_senegalus | Tchagra_senegalus | LC | Stable | tropical | diurnal | 53.5 | 14953687.107 | 9.943 |
| birds | Tchagra_tchagra | Tchagra_tchagra | LC | Increasing | temperate | diurnal | 50.3 | 251680.904 | 11.718 |
| birds | Telacanthura_melanopygia | Telacanthura_melanopygia | LC | Stable | tropical | diurnal | 52 | 588050.823 | 8.945 |
| birds | Telacanthura_ussheri | Telacanthura_ussheri | LC | Decreasing | tropical | diurnal | 32.6 | 5188546 | 9.879 |
| birds | Teledromas_fuscus | Teledromas_fuscus | LC | Stable | temperate | diurnal | 37.9 | 521735.986 | 4.598 |
| birds | Telespiza_cantans | NA | VU | Stable | NA | NA | NA | NA | NA |
| birds | Telespiza_ultima | Telespiza_ultima | CR | Stable | tropical | diurnal | 24.2 | 1.554 | 0 |
| birds | Telophorus_dohertyi | Telophorus_dohertyi | LC | Stable | tropical | diurnal | 35 | 113100.736 | 17.008 |
| birds | Telophorus_viridis | Telophorus_viridis | LC | Stable | temperate | diurnal | 37.1 | 1025110.515 | 8.818 |
| birds | Telophorus_zeylonus | Telophorus_zeylonus | LC | Stable | temperate | diurnal | 65 | 1314512.433 | 8.942 |
| birds | Temnotrogon_roseigaster | Priotelus_roseigaster | LC | Decreasing | tropical | diurnal | 74.2 | 15254.535 | 12.209 |
| birds | Temnurus_temnurus | Temnurus_temnurus | LC | Decreasing | tropical | diurnal | 138 | 266868.951 | 12.5 |
| birds | Tephrodornis_pondicerianus | Tephrodornis_pondicerianus | LC | Stable | tropical | diurnal | 20.2 | 4411317.051 | 16.28 |
| birds | Tephrodornis_sylvicola | Tephrodornis_gularis | LC | Stable | tropical | diurnal | 37.8 | 128127.363 | 19.074 |
| birds | Tephrophilus_wetmorei | Buthraupis_wetmorei | VU | Decreasing | tropical | diurnal | 62.5 | 12901.207 | 7.038 |
| birds | Tephrozosterops_stalkeri | Tephrozosterops_stalkeri | LC | Decreasing | tropical | diurnal | 17.7 | 3178.996 | 6.168 |
| birds | Terathopius_ecaudatus | Terathopius_ecaudatus | EN | Decreasing | tropical | diurnal | 2200 | 14023477.786 | 9.524 |
| birds | Terenotriccus_erythrurus | Terenotriccus_erythrurus | LC | Decreasing | tropical | diurnal | 7.4 | 7494686.858 | 2.96 |
| birds | Terenura_maculata | Terenura_maculata | LC | Decreasing | tropical | diurnal | 6.5 | 783959.127 | 13.246 |
| birds | Terenura_sicki | Terenura_sicki | CR | Decreasing | tropical | diurnal | 6.4 | 2173.238 | 12.019 |
| birds | Teretistris_fernandinae | Teretistris_fernandinae | LC | Stable | tropical | diurnal | 9.9 | 25700.55 | 15.018 |

| | | | | | | | | | |
|---|---|---|---|---|---|---|---|---|---|
| birds | Teretistris_fornsi | Teretistris_fornsi | LC | Stable | tropical | diurnal | 9.7 | 38336.733 | 15.206 |
| birds | Terpsiphone_atrocaudata | Terpsiphone_atrocaudata | LC | Stable | tropical | diurnal | 18.7 | 2897084.957 | 15.45 |
| birds | Terpsiphone_atrochalybeia | Terpsiphone_atrochalybeia | LC | Stable | tropical | diurnal | 12.2 | 858.514 | 12.812 |
| birds | Terpsiphone_bedfordi | Terpsiphone_bedfordi | LC | Decreasing | tropical | diurnal | 15.4 | 42552.862 | 11.204 |
| birds | Terpsiphone_bourbonnensis | Terpsiphone_bourbonnensis | LC | Stable | tropical | diurnal | 10.8 | 3297.843 | 21.831 |
| birds | Terpsiphone_cinnamomea | Terpsiphone_cinnamomea | LC | Stable | tropical | diurnal | 20.4 | 109426.267 | 11.878 |
| birds | Terpsiphone_corvina | NA | VU | Increasing | NA | NA | NA | NA | NA |
| birds | Terpsiphone_cyanescens | Terpsiphone_cyanescens | LC | Decreasing | tropical | diurnal | 24.6 | 11452.594 | 10.09 |
| birds | Terpsiphone_mutata | Terpsiphone_mutata | LC | Decreasing | tropical | diurnal | 13.2 | 592893.864 | 8.772 |
| birds | Terpsiphone_paradisi | Terpsiphone_paradisi | LC | Stable | temperate | diurnal | 19.2 | 4205670.728 | 16.481 |
| birds | Terpsiphone_rufiventer | Terpsiphone_rufiventer | LC | Decreasing | tropical | diurnal | 15.1 | 4275150.967 | 8.677 |
| birds | Terpsiphone_rufocinerea | Terpsiphone_rufocinerea | LC | Stable | tropical | diurnal | 15.4 | 444393.519 | 8.416 |
| birds | Terpsiphone_viridis | Terpsiphone_viridis | LC | Stable | tropical | diurnal | 13.2 | 14483913.965 | 9.535 |
| birds | Tersina_viridis | Tersina_viridis | LC | Stable | tropical | diurnal | 29 | 8757523.605 | 5.919 |
| birds | Tesia_cyaniventer | Tesia_cyaniventer | LC | Stable | temperate | diurnal | 9.7 | 2098380.828 | 10.836 |
| birds | Tesia_everetti | Tesia_everetti | LC | Stable | tropical | diurnal | 6.3 | 29083.296 | 12.592 |
| birds | Tesia_olivea | Tesia_olivea | LC | Stable | temperate | diurnal | 7 | 1813260.117 | 11.008 |
| birds | Tesia_superciliaris | NA | LC | Stable | NA | NA | NA | NA | NA |
| birds | Tetrao_urogalloides | Tetrao_parvirostris | LC | Decreasing | temperate | diurnal | 2752.3 | 7003788.429 | 1.854 |
| birds | Tetrao_urogallus | Tetrao_urogallus | LC | Decreasing | temperate | diurnal | 2716.6 | 11330230.843 | 6.008 |
| birds | Tetraogallus_altaicus | Tetraogallus_altaicus | LC | Decreasing | temperate | diurnal | 2760.4 | 819469.256 | 4.174 |
| birds | Tetraogallus_caspius | Tetraogallus_caspius | LC | Decreasing | temperate | diurnal | 2508.2 | 313937.424 | 12.686 |
| birds | Tetraogallus_caucasicus | Tetraogallus_caucasicus | LC | Stable | temperate | diurnal | 1830.8 | 23392.942 | 5.543 |
| birds | Tetraogallus_himalayensis | Tetraogallus_himalayensis | LC | Stable | temperate | diurnal | 2428 | 1912197.695 | 5.766 |
| birds | Tetraogallus_tibetanus | Tetraogallus_tibetanus | LC | Stable | temperate | diurnal | 1488.9 | 1066992.694 | 4.129 |
| birds | Tetraophasis_obscurus | Tetraophasis_obscurus | LC | Decreasing | temperate | diurnal | 855.4 | 118338.722 | 8.713 |
| birds | Tetraophasis_szechenyii | Tetraophasis_szechenyii | LC | Unknown | temperate | diurnal | 1053 | 524228.723 | 6.799 |
| birds | Tetrastes_bonasia | Bonasa_bonasia | LC | Decreasing | temperate | diurnal | 429 | 15669355.229 | 6.044 |
| birds | Tetrastes_sewerzowi | Bonasa_sewerzowi | NT | Decreasing | temperate | diurnal | 267.3 | 394309.683 | 8.249 |
| birds | Tetrax_tetrax | Tetrax_tetrax | NT | Decreasing | temperate | diurnal | 833.5 | 10243202.486 | 10.523 |
| birds | Thalassarche_bulleri | Thalassarche_bulleri | NT | Stable | temperate | cathemeral | 2781.5 | 29134433.318 | 10.989 |
| birds | Thalassarche_cauta | Thalassarche_cauta | NT | Stable | temperate | cathemeral | 3897.7 | 23868551.627 | 13.743 |
| birds | Thalassarche_chlororhynchos | Thalassarche_chlororhynchos | EN | Decreasing | temperate | cathemeral | 2128.8 | 21606227.427 | 13.288 |
| birds | Thalassarche_chrysostoma | Thalassarche_chrysostoma | EN | Decreasing | temperate | cathemeral | 3499 | 83986959.447 | 6.009 |

| | | | | | | | | | |
|---|---|---|---|---|---|---|---|---|---|
| birds | Thalassarche_melanophris | Thalassarche_melanophrys | LC | Increasing | temperate | cathemeral | 3202.9 | 109981586.658 | 4.622 |
| birds | Thalasseus_bengalensis | Sterna_bengalensis | LC | Stable | tropical | diurnal | 209 | 24346020.886 | 10.09 |
| birds | Thalasseus_bergii | Sterna_bergii | LC | Stable | tropical | diurnal | 328 | 47809209.072 | 11.332 |
| birds | Thalasseus_bernsteini | Sterna_bernsteini | CR | Decreasing | temperate | diurnal | 284 | 59.404 | 24.977 |
| birds | Thalasseus_elegans | Sterna_elegans | NT | Stable | tropical | diurnal | 260 | 2159166.976 | 8.777 |
| birds | Thalasseus_maximus | Sterna_maxima | LC | Stable | tropical | diurnal | 415.3 | 5211221.664 | 12.384 |
| birds | Thalasseus_sandvicensis | Sterna_sandvicensis | LC | Stable | tropical | diurnal | 199.8 | 8765162.127 | 13.354 |
| birds | Thalassoica_antarctica | Thalassoica_antarctica | LC | Stable | temperate | cathemeral | 708.4 | 58576534.09 | 1.986 |
| birds | Thalassornis_leuconotus | Thalassornis_leuconotus | LC | Decreasing | tropical | diurnal | 707.5 | 10032374.01 | 9.05 |
| birds | Thalurania_colombica | Thalurania_colombica | LC | Decreasing | tropical | diurnal | 4.5 | 705818.087 | 11.035 |
| birds | Thalurania_furcata | Thalurania_furcata | LC | Unknown | tropical | diurnal | 4.2 | 11225568.611 | 4.603 |
| birds | Thalurania_glaucopis | Thalurania_glaucopis | LC | Unknown | temperate | diurnal | 4.8 | 1415794.42 | 12.863 |
| birds | Thalurania_watertonii | Thalurania_watertonii | EN | Decreasing | tropical | diurnal | 4.6 | 44734.573 | 15.1 |
| birds | Thamnistes_anabatinus | Thamnistes_anabatinus | LC | Decreasing | tropical | diurnal | 20.7 | 398111.816 | 8.394 |
| birds | Thamnolaea_cinnamomeiventris | Myrmecocichla_cinnamomeiventris | LC | Stable | tropical | diurnal | 39.8 | 2978092.642 | 12.612 |
| birds | Thamnomanes_ardesiacus | Thamnomanes_ardesiacus | LC | Decreasing | tropical | diurnal | 17.7 | 3380827.139 | 1.478 |
| birds | Thamnomanes_caesius | Thamnomanes_caesius | LC | Decreasing | tropical | diurnal | 15.7 | 6049355.842 | 2.768 |
| birds | Thamnomanes_saturninus | Thamnomanes_saturninus | LC | Stable | tropical | diurnal | 20.4 | 1364638.562 | 1.385 |
| birds | Thamnomanes_schistogynus | Thamnomanes_schistogynus | LC | Decreasing | tropical | diurnal | 17.2 | 1226196.42 | 1.99 |
| birds | Thamnophilus_aethiops | Thamnophilus_aethiops | LC | Decreasing | tropical | diurnal | 25.7 | 4946417.179 | 2.246 |
| birds | Thamnophilus_amazonicus | Thamnophilus_amazonicus | LC | Stable | tropical | diurnal | 18.7 | 6192801.734 | 2.292 |
| birds | Thamnophilus_ambiguus | Thamnophilus_ambiguus | LC | Stable | tropical | diurnal | 23.4 | 209875.652 | 12.505 |
| birds | Thamnophilus_aroyae | Thamnophilus_aroyae | LC | Decreasing | tropical | diurnal | 20 | 62450.077 | 4.542 |
| birds | Thamnophilus_atrinucha | Thamnophilus_atrinucha | LC | Stable | tropical | diurnal | 23.6 | 583507.071 | 10.909 |
| birds | Thamnophilus_bernardi | Sakesphorus_bernardi | LC | Stable | tropical | diurnal | 34.4 | 102383.976 | 10.48 |
| birds | Thamnophilus_bridgesi | Thamnophilus_bridgesi | LC | Decreasing | tropical | diurnal | 27 | 28247.58 | 13.862 |
| birds | Thamnophilus_caerulescens | Thamnophilus_caerulescens | LC | Decreasing | tropical | diurnal | 21.1 | 3969673.889 | 10.487 |
| birds | Thamnophilus_cryptoleucus | Thamnophilus_cryptoleucus | LC | Decreasing | tropical | diurnal | 27.5 | 177320.51 | 2.903 |
| birds | Thamnophilus_doliatus | Thamnophilus_doliatus | LC | Stable | tropical | diurnal | 27 | 10725968.397 | 5.907 |
| birds | Thamnophilus_insignis | Thamnophilus_insignis | LC | Decreasing | tropical | diurnal | 24.9 | 143229.398 | 0.79 |
| birds | Thamnophilus_melanonotus | Sakesphorus_melanonotus | LC | Decreasing | tropical | diurnal | 25.5 | 113947.776 | 14.369 |
| birds | Thamnophilus_melanothorax | Sakesphorus_melanothorax | LC | Decreasing | tropical | diurnal | 31.8 | 304787.248 | 1.414 |
| birds | Thamnophilus_multistriatus | Thamnophilus_multistriatus | LC | Stable | tropical | diurnal | 27.5 | 121617.786 | 13.339 |
| birds | Thamnophilus_murinus | Thamnophilus_murinus | LC | Decreasing | tropical | diurnal | 19.3 | 4198741.885 | 1.362 |

| | | | | | | | | | |
|---|---|---|---|---|---|---|---|---|---|
| birds | Thamnophilus_nigriceps | Thamnophilus_nigriceps | LC | Decreasing | tropical | diurnal | 22.9 | 210463.798 | 13.054 |
| birds | Thamnophilus_nigrocinereus | Thamnophilus_nigrocinereus | LC | Decreasing | tropical | diurnal | 30 | 1299292.688 | 2.552 |
| birds | Thamnophilus_palliatus | Thamnophilus_palliatus | LC | Stable | tropical | diurnal | 27 | 2542297.141 | 4.991 |
| birds | Thamnophilus_pelzelni | Thamnophilus_pelzelni | LC | Decreasing | tropical | diurnal | 20.9 | 2622762.3 | 9.279 |
| birds | Thamnophilus_praecox | Thamnophilus_praecox | NT | Decreasing | tropical | diurnal | 29.5 | 25769.524 | 4.394 |
| birds | Thamnophilus_punctatus | Thamnophilus_punctatus | LC | Unknown | tropical | diurnal | 19.4 | 1569761.657 | 2.161 |
| birds | Thamnophilus_ruficapillus | Thamnophilus_ruficapillus | LC | Decreasing | temperate | diurnal | 20.4 | 1391729.072 | 11.45 |
| birds | Thamnophilus_schistaceus | Thamnophilus_schistaceus | LC | Decreasing | tropical | diurnal | 20.3 | 3991785.912 | 2.395 |
| birds | Thamnophilus_stictocephalus | Thamnophilus_stictocephalus | LC | Decreasing | tropical | diurnal | 21.5 | 1068500.362 | 2.275 |
| birds | Thamnophilus_sticturus | Thamnophilus_sticturus | LC | Unknown | tropical | diurnal | 18.2 | 424021.105 | 5.113 |
| birds | Thamnophilus_tenuepunctatus | Thamnophilus_tenuepunctatus | LC | Stable | tropical | diurnal | 23.3 | 136115.507 | 5.981 |
| birds | Thamnophilus_torquatus | Thamnophilus_torquatus | LC | Decreasing | tropical | diurnal | 21.1 | 3732484.235 | 8.412 |
| birds | Thamnophilus_unicolor | Thamnophilus_unicolor | LC | Decreasing | tropical | diurnal | 24.2 | 168168.058 | 8.377 |
| birds | Thamnophilus_zarumae | Thamnophilus_zarumae | LC | Decreasing | tropical | diurnal | 22.8 | 27875.191 | 9.158 |
| birds | Thamnornis_chloropetoides | Thamnornis_chloropetoides | LC | Decreasing | tropical | diurnal | 14.3 | 69624.439 | 8.747 |
| birds | Thaumasius_baeri | Leucippus_baeri | LC | Stable | tropical | diurnal | 4.5 | 29962.881 | 8.526 |
| birds | Thaumasius_taczanowskii | Leucippus_taczanowskii | LC | Stable | tropical | diurnal | 6.4 | 70512.134 | 8.799 |
| birds | Thaumastura_cora | Thaumastura_cora | LC | Stable | tropical | diurnal | 1.9 | 203316.511 | 7.153 |
| birds | Thaumatibis_gigantea | Thaumatibis_gigantea | CR | Decreasing | tropical | diurnal | 3515 | 50313.058 | 6.64 |
| birds | Theristicus_caerulescens | Theristicus_caerulescens | LC | Stable | temperate | diurnal | 1500 | 1749653.629 | 8.051 |
| birds | Theristicus_caudatus | Theristicus_caudatus | LC | Stable | tropical | diurnal | 1726 | 5146136.121 | 8.317 |
| birds | Theristicus_melanopis | Theristicus_melanopis | LC | Stable | temperate | diurnal | 1250 | 2020059.975 | 7.472 |
| birds | Thescelocichla_leucopleura | Thescelocichla_leucopleura | LC | Stable | tropical | diurnal | 62.1 | 3240183.587 | 8.455 |
| birds | Thinocorus_orbignyianus | Thinocorus_orbignyianus | LC | Stable | temperate | diurnal | 115 | 1770605.522 | 5.956 |
| birds | Thinocorus_rumicivorus | Thinocorus_rumicivorus | LC | Stable | temperate | diurnal | 53.1 | 2241860.15 | 7.175 |
| birds | Thinornis_cucullatus | Thinornis_rubricollis | VU | Decreasing | temperate | cathemeral | 100 | 608290.866 | 8.498 |
| birds | Thinornis_novaeseelandiae | NA | EN | Increasing | NA | NA | NA | NA | NA |
| birds | Thlypopsis_fulviceps | Thlypopsis_fulviceps | LC | Stable | tropical | diurnal | 12 | 40001.465 | 12.055 |
| birds | Thlypopsis_inornata | Thlypopsis_inornata | LC | Stable | tropical | diurnal | 15 | 20048.294 | 8.893 |
| birds | Thlypopsis_ornata | Thlypopsis_ornata | LC | Stable | tropical | diurnal | 12 | 181105.809 | 9.749 |
| birds | Thlypopsis_pectoralis | Thlypopsis_pectoralis | LC | Stable | tropical | diurnal | 15 | 12520.915 | 9.469 |
| birds | Thlypopsis_pyrrhocoma | Pyrrhocoma_ruficeps | LC | Stable | temperate | diurnal | 15.6 | 611568.556 | 12.9 |
| birds | Thlypopsis_ruficeps | Thlypopsis_ruficeps | LC | Stable | tropical | diurnal | 11 | 213849.229 | 7.696 |
| birds | Thlypopsis_sordida | Thlypopsis_sordida | LC | Stable | tropical | diurnal | 17 | 6836052.487 | 7.048 |

| | | | | | | | | | |
|---|---|---|---|---|---|---|---|---|---|
| birds | Thlypopsis_superciliaris | Hemispingus_superciliaris | LC | Decreasing | tropical | diurnal | 14 | 271703.986 | 9.032 |
| birds | Threnetes_niger | Threnetes_niger | LC | Stable | tropical | diurnal | 5.6 | 313115.482 | 1.289 |
| birds | Threnetes_ruckeri | Threnetes_ruckeri | LC | Decreasing | tropical | diurnal | 5.2 | 482644.974 | 9.988 |
| birds | Threskiornis_aethiopicus | Threskiornis_aethiopicus | LC | Stable | tropical | diurnal | 1493.2 | 17731713.856 | 8.912 |
| birds | Threskiornis_melanocephalus | Threskiornis_melanocephalus | NT | Decreasing | temperate | diurnal | 1303 | 2840531.68 | 16.931 |
| birds | Threskiornis_moluccus | Threskiornis_molucca | LC | Stable | temperate | diurnal | 1794.9 | 5697443.52 | 5.137 |
| birds | Threskiornis_spinicollis | Threskiornis_spinicollis | LC | Decreasing | temperate | diurnal | 1346.7 | 6556820.353 | 4.693 |
| birds | Thripadectes_flammulatus | Thripadectes_flammulatus | LC | Stable | tropical | diurnal | 55.9 | 57126.723 | 10.055 |
| birds | Thripadectes_holostictus | Thripadectes_holostictus | LC | Stable | tropical | diurnal | 40.7 | 214735.935 | 7.098 |
| birds | Thripadectes_ignobilis | Thripadectes_ignobilis | LC | Stable | tropical | diurnal | 47.2 | 41009.217 | 6.92 |
| birds | Thripadectes_melanorhynchus | Thripadectes_melanorhynchus | LC | Stable | tropical | diurnal | 44.3 | 121002.941 | 5.039 |
| birds | Thripadectes_rufobrunneus | Thripadectes_rufobrunneus | LC | Stable | tropical | diurnal | 56.2 | 17593.386 | 6.596 |
| birds | Thripadectes_scrutator | Thripadectes_scrutator | LC | Stable | tropical | diurnal | 66.3 | 116443.325 | 5.347 |
| birds | Thripadectes_virgaticeps | Thripadectes_virgaticeps | LC | Stable | tropical | diurnal | 65.8 | 62554.862 | 10.764 |
| birds | Thripophaga_cherriei | Thripophaga_cherriei | VU | Decreasing | tropical | diurnal | 12.8 | 9.867 | 3.175 |
| birds | Thripophaga_fusciceps | Thripophaga_fusciceps | LC | Stable | tropical | diurnal | 23 | 514581.935 | 3.142 |
| birds | Thripophaga_gutturata | Cranioleuca_gutturata | LC | Stable | tropical | diurnal | 14.9 | 4168734.036 | 1.807 |
| birds | Thripophaga_macroura | NA | VU | Decreasing | NA | NA | NA | NA | NA |
| birds | Thryomanes_bewickii | Thryomanes_bewickii | LC | Decreasing | temperate | diurnal | 9.9 | 3814727.236 | 10.874 |
| birds | Thryophilus_nicefori | NA | CR | Decreasing | NA | NA | NA | NA | NA |
| birds | Thryophilus_pleurostictus | Thryothorus_pleurostictus | LC | Decreasing | tropical | diurnal | 17.7 | 185486.447 | 13.07 |
| birds | Thryophilus_rufalbus | Thryothorus_rufalbus | LC | Decreasing | tropical | diurnal | 24.9 | 821636.361 | 10.345 |
| birds | Thryophilus_sinaloa | Thryothorus_sinaloa | LC | Decreasing | temperate | diurnal | 15.1 | 304927.467 | 7.746 |
| birds | Thryorchilus_browni | Thryorchilus_browni | LC | Stable | tropical | diurnal | 14 | 1835.353 | 5.69 |
| birds | Thryothorus_ludovicianus | Thryothorus_ludovicianus | LC | Increasing | temperate | diurnal | 19 | 3040027.221 | 17.386 |
| birds | Tiaris_olivaceus | Tiaris_olivaceus | LC | Increasing | tropical | diurnal | 8.5 | 968065.521 | 11.925 |
| birds | Tichodroma_muraria | Tichodroma_muraria | LC | Stable | temperate | diurnal | 17.6 | 9663228.796 | 11.029 |
| birds | Tickellia_hodgsoni | Tickellia_hodgsoni | LC | Decreasing | tropical | diurnal | 4.5 | 616057.052 | 7.86 |
| birds | Tigrisoma_fasciatum | Tigrisoma_fasciatum | LC | Decreasing | temperate | nocturnal | 850 | 1533469.083 | 9.633 |
| birds | Tigrisoma_lineatum | Tigrisoma_lineatum | LC | Stable | tropical | cathemeral | 813 | 13239678.284 | 5.682 |
| birds | Tigrisoma_mexicanum | Tigrisoma_mexicanum | LC | Decreasing | tropical | nocturnal | 1154.4 | 928353.013 | 9.671 |
| birds | Tilmatura_dupontii | Tilmatura_dupontii | LC | Stable | tropical | diurnal | 2.2 | 307275.413 | 10.616 |
| birds | Timalia_pileata | Timalia_pileata | LC | Decreasing | tropical | diurnal | 16 | 2665313.174 | 12.544 |
| birds | Timeliopsis_fulvigula | Timeliopsis_fulvigula | LC | Stable | tropical | diurnal | 18.3 | 137140.891 | 4.707 |

| | | | | | | | | | |
|---|---|---|---|---|---|---|---|---|---|
| birds | Timeliopsis_griseigula | Timeliopsis_griseigula | LC | Stable | tropical | diurnal | 32 | 170363.663 | 3.669 |
| birds | Tinamotis_ingoufi | Tinamotis_ingoufi | LC | Stable | temperate | diurnal | 950 | 403733.454 | 5.652 |
| birds | Tinamotis_pentlandii | Tinamotis_pentlandii | LC | Stable | tropical | diurnal | 650 | 632710.157 | 6.746 |
| birds | Tinamus_guttatus | Tinamus_guttatus | NT | Decreasing | tropical | diurnal | 686.2 | 3971186.795 | 1.902 |
| birds | Tinamus_major | Tinamus_major | LC | Decreasing | tropical | diurnal | 1026.2 | 8123168.476 | 3.402 |
| birds | Tinamus_solitarius | Tinamus_solitarius | NT | Decreasing | tropical | diurnal | 1386.4 | 1069608.942 | 13.205 |
| birds | Tinamus_tao | Tinamus_tao | VU | Decreasing | tropical | diurnal | 1600.1 | 3810484.552 | 3.609 |
| birds | Tityra_cayana | Tityra_cayana | LC | Decreasing | tropical | diurnal | 68.1 | 5115788.259 | 2.291 |
| birds | Tityra_inquisitor | Tityra_inquisitor | LC | Decreasing | tropical | diurnal | 43.1 | 11190798.84 | 6.33 |
| birds | Tityra_semifasciata | Tityra_semifasciata | LC | Decreasing | tropical | diurnal | 79.3 | 6821138.583 | 5.137 |
| birds | Tmetothylacus_tenellus | Tmetothylacus_tenellus | LC | Stable | tropical | diurnal | 20.1 | 1285795.149 | 9.593 |
| birds | Tockus_deckeni | Tockus_deckeni | LC | Decreasing | tropical | diurnal | 167.7 | 1628868.177 | 9.502 |
| birds | Tockus_erythrorhynchus | Tockus_erythrorhynchus | LC | Stable | tropical | diurnal | 138.6 | 9150146.923 | 10.164 |
| birds | Tockus_flavirostris | Tockus_flavirostris | LC | Decreasing | tropical | diurnal | 216.7 | 1449740.425 | 7.905 |
| birds | Tockus_leucomelas | Tockus_leucomelas | LC | Decreasing | tropical | diurnal | 188.3 | 1820119.324 | 7.414 |
| birds | Tockus_monteiri | Tockus_monteiri | LC | Stable | tropical | diurnal | 289.8 | 399497.556 | 5.889 |
| birds | Todiramphus_albonotatus | Todiramphus_albonotatus | NT | Decreasing | tropical | diurnal | 32 | 30015.576 | 3.973 |
| birds | Todiramphus_australasia | Todiramphus_australasia | LC | Decreasing | tropical | diurnal | 43 | 44763.907 | 14.066 |
| birds | Todiramphus_chloris | Todiramphus_chloris | LC | Decreasing | tropical | diurnal | 66.1 | 3828153.103 | 9.557 |
| birds | Todiramphus_cinnamominus | NA | EW | not_available | NA | NA | NA | NA | NA |
| birds | Todiramphus_diops | Todiramphus_diops | LC | Stable | tropical | diurnal | 47.4 | 25186.567 | 6.338 |
| birds | Todiramphus_enigma | NA | NT | Stable | NA | NA | NA | NA | NA |
| birds | Todiramphus_farquhari | Todiramphus_farquhari | NT | Decreasing | tropical | diurnal | 38.8 | 5637.518 | 5.747 |
| birds | Todiramphus_funebris | NA | LC | Decreasing | NA | NA | NA | NA | NA |
| birds | Todiramphus_gambieri | Todiramphus_gambieri | CR | Increasing | tropical | diurnal | 39 | 41.081 | 10.788 |
| birds | Todiramphus_godeffroyi | Todiramphus_godeffroyi | CR | Decreasing | tropical | diurnal | 63.4 | 382.097 | 6.991 |
| birds | Todiramphus_lazuli | NA | LC | Stable | NA | NA | NA | NA | NA |
| birds | Todiramphus_leucopygius | Todiramphus_leucopygius | LC | Stable | tropical | diurnal | 47.9 | 23194.377 | 4.985 |
| birds | Todiramphus_macleayii | Todiramphus_macleayii | LC | Decreasing | tropical | diurnal | 37.3 | 1316178.398 | 4.298 |
| birds | Todiramphus_nigrocyaneus | Todiramphus_nigrocyaneus | NT | Decreasing | tropical | diurnal | 54.2 | 463339.634 | 2.981 |
| birds | Todiramphus_pyrrhopygius | Todiramphus_pyrrhopygius | LC | Increasing | temperate | diurnal | 51.7 | 7435153.362 | 4.144 |
| birds | Todiramphus_sanctus | Todiramphus_sanctus | LC | Increasing | temperate | diurnal | 53 | 8193818.919 | 5.893 |
| birds | Todiramphus_saurophagus | Todiramphus_saurophaga | LC | Stable | tropical | diurnal | 126 | 134276.077 | 5.388 |
| birds | Todiramphus_tutus | Todiramphus_tutus | NT | Decreasing | tropical | diurnal | 42.3 | 447.024 | 16.671 |

| | | | | | | | | | |
|---|---|---|---|---|---|---|---|---|---|
| birds | Todiramphus_veneratus | Todiramphus_veneratus | NT | Decreasing | tropical | diurnal | 44.5 | 1061.309 | 14.626 |
| birds | Todiramphus_winchelli | Todiramphus_winchelli | VU | Decreasing | tropical | diurnal | 67.4 | 133080.355 | 13.553 |
| birds | Todirostrum_chrysocrotaphum | Todirostrum_chrysocrotaphum | LC | Stable | tropical | diurnal | 7 | 4205502.108 | 2.199 |
| birds | Todirostrum_cinereum | Todirostrum_cinereum | LC | Stable | tropical | diurnal | 6.3 | 8243668.88 | 7.564 |
| birds | Todirostrum_maculatum | Todirostrum_maculatum | LC | Stable | tropical | diurnal | 7.3 | 4363496.679 | 1.893 |
| birds | Todirostrum_nigriceps | Todirostrum_nigriceps | LC | Decreasing | tropical | diurnal | 6.5 | 423081.579 | 11.189 |
| birds | Todirostrum_pictum | Todirostrum_pictum | LC | Stable | tropical | diurnal | 6.8 | 1704255.676 | 1.302 |
| birds | Todirostrum_poliocephalum | Todirostrum_poliocephalum | LC | Stable | tropical | diurnal | 5.6 | 465051.923 | 13.69 |
| birds | Todirostrum_viridanum | Todirostrum_viridanum | LC | Decreasing | tropical | diurnal | 6.5 | 24015.081 | 13.623 |
| birds | Todus_angustirostris | Todus_angustirostris | LC | Decreasing | tropical | diurnal | 7.5 | 74117.206 | 15.787 |
| birds | Todus_mexicanus | Todus_mexicanus | LC | Decreasing | tropical | diurnal | 5.9 | 8774.421 | 27.252 |
| birds | Todus_multicolor | Todus_multicolor | LC | Decreasing | tropical | diurnal | 5.9 | 109693.074 | 16.145 |
| birds | Todus_subulatus | Todus_subulatus | LC | Decreasing | tropical | diurnal | 8.7 | 74799.651 | 15.781 |
| birds | Todus_todus | Todus_todus | LC | Decreasing | tropical | diurnal | 6.4 | 11021.788 | 19.741 |
| birds | Tolmomyias_assimilis | Tolmomyias_assimilis | LC | Decreasing | tropical | diurnal | 17 | 5871080.895 | 1.88 |
| birds | Tolmomyias_flaviventris | Tolmomyias_flaviventris | LC | Stable | tropical | diurnal | 12.2 | 7836566.046 | 5.401 |
| birds | Tolmomyias_flavotectus | Tolmomyias_flavotectus | LC | Decreasing | tropical | diurnal | 14.9 | 212437.983 | 8.238 |
| birds | Tolmomyias_poliocephalus | Tolmomyias_poliocephalus | LC | Stable | tropical | diurnal | 10.8 | 6112774.744 | 1.993 |
| birds | Tolmomyias_sulphurescens | Tolmomyias_sulphurescens | LC | Decreasing | tropical | diurnal | 14.3 | 12413249.737 | 5.916 |
| birds | Tolmomyias_traylori | Tolmomyias_traylori | LC | Decreasing | tropical | diurnal | 11.5 | 270037.117 | 1.786 |
| birds | Tolmomyias_viridiceps | Tolmomyias_viridiceps | LC | Stable | tropical | diurnal | 11.3 | 2382276.786 | 1.638 |
| birds | Topaza_pella | Topaza_pella | LC | Decreasing | tropical | diurnal | 12.1 | 1678442.908 | 1.81 |
| birds | Torgos_tracheliotos | Torgos_tracheliotos | EN | Decreasing | tropical | diurnal | 6969 | 14737739.362 | 8.264 |
| birds | Torreornis_inexpectata | Torreornis_inexpectata | NT | Decreasing | tropical | diurnal | 25.5 | 4568.064 | 5.567 |
| birds | Touit_costaricensis | Touit_costaricensis | NT | Decreasing | tropical | diurnal | 80 | 19135.588 | 7.508 |
| birds | Touit_dilectissimus | Touit_dilectissimus | LC | Decreasing | tropical | diurnal | 65 | 132570.471 | 7.641 |
| birds | Touit_huetii | Touit_huetii | LC | Decreasing | tropical | diurnal | 60 | 3874966.143 | 2.471 |
| birds | Touit_melanonotus | Touit_melanonotus | NT | Decreasing | temperate | diurnal | 66.5 | 51978.023 | 9.848 |
| birds | Touit_purpuratus | Touit_purpuratus | LC | Stable | tropical | diurnal | 59.7 | 3821297.271 | 1.461 |
| birds | Touit_stictopterus | Touit_stictopterus | NT | Decreasing | tropical | diurnal | 78.8 | 59769.965 | 6.846 |
| birds | Touit_surdus | Touit_surdus | VU | Decreasing | temperate | diurnal | 51 | 67159.176 | 11.301 |
| birds | Toxorhamphus_novaeguineae | Toxorhamphus_novaeguineae | LC | Unknown | tropical | diurnal | 11.8 | 463594.806 | 2.888 |
| birds | Toxorhamphus_poliopterus | Toxorhamphus_poliopterus | LC | Unknown | tropical | diurnal | 11.4 | 179355.789 | 4.083 |
| birds | Toxostoma_bendirei | Toxostoma_bendirei | VU | Decreasing | temperate | diurnal | 62.2 | 822797.041 | 8.426 |

| | | | | | | | | | |
|---|---|---|---|---|---|---|---|---|---|
| birds | Toxostoma_cinereum | Toxostoma_cinereum | LC | Stable | temperate | diurnal | 59 | 99643.399 | 3.687 |
| birds | Toxostoma_crissale | Toxostoma_crissale | LC | Stable | temperate | diurnal | 62.7 | 1010689.858 | 7.961 |
| birds | Toxostoma_curvirostre | Toxostoma_curvirostre | LC | Decreasing | temperate | diurnal | 80.5 | 2044910.563 | 9.771 |
| birds | Toxostoma_guttatum | Toxostoma_guttatum | CR | Decreasing | tropical | diurnal | 52.8 | 488.302 | 10.21 |
| birds | Toxostoma_lecontei | Toxostoma_lecontei | LC | Stable | temperate | diurnal | 61.9 | 251262.301 | 7.395 |
| birds | Toxostoma_longirostre | Toxostoma_longirostre | LC | Increasing | temperate | diurnal | 67.6 | 300961.71 | 12.166 |
| birds | Toxostoma_ocellatum | Toxostoma_ocellatum | LC | Decreasing | tropical | diurnal | 84.3 | 88725.786 | 17.429 |
| birds | Toxostoma_redivivum | Toxostoma_redivivum | LC | Decreasing | temperate | diurnal | 84.4 | 167320.21 | 13.937 |
| birds | Toxostoma_rufum | Toxostoma_rufum | LC | Decreasing | temperate | diurnal | 68.8 | 5859124.466 | 15.734 |
| birds | Trachylaemus_purpuratus | Trachyphonus_purpuratus | LC | Decreasing | tropical | diurnal | 76.1 | 3484026.334 | 7.345 |
| birds | Trachyphonus_darnaudii | Trachyphonus_darnaudii | LC | Stable | tropical | diurnal | 35.8 | 1278810.544 | 8.801 |
| birds | Trachyphonus_erythrocephalus | Trachyphonus_erythrocephalus | LC | Stable | tropical | diurnal | 63.5 | 1574108.268 | 8.361 |
| birds | Trachyphonus_margaritatus | Trachyphonus_margaritatus | LC | Stable | tropical | diurnal | 64 | 2214289.929 | 8.879 |
| birds | Trachyphonus_vaillantii | Trachyphonus_vaillantii | LC | Decreasing | tropical | diurnal | 69.4 | 3120428.043 | 8.939 |
| birds | Tragopan_blythii | Tragopan_blythii | VU | Decreasing | temperate | diurnal | 1930 | 120944.169 | 5.63 |
| birds | Tragopan_caboti | Tragopan_caboti | VU | Decreasing | temperate | diurnal | 1122.5 | 396418.35 | 12.478 |
| birds | Tragopan_melanocephalus | Tragopan_melanocephalus | VU | Decreasing | temperate | diurnal | 1613.2 | 21593.776 | 8.57 |
| birds | Tragopan_satyra | Tragopan_satyra | NT | Decreasing | temperate | diurnal | 1417 | 99176.64 | 7.658 |
| birds | Tragopan_temminckii | Tragopan_temminckii | LC | Decreasing | temperate | diurnal | 1163.4 | 1261852.145 | 12.028 |
| birds | Tregellasia_capito | Tregellasia_capito | LC | Decreasing | temperate | diurnal | 13.9 | 155466.502 | 7.297 |
| birds | Tregellasia_leucops | Tregellasia_leucops | LC | Stable | tropical | diurnal | 15.6 | 201214.414 | 3.885 |
| birds | Treron_apicauda | Treron_apicauda | LC | Decreasing | tropical | diurnal | 217.2 | 1636420.934 | 9.655 |
| birds | Treron_australis | Treron_australis | LC | Decreasing | tropical | diurnal | 229 | 505554.319 | 8.441 |
| birds | Treron_bicinctus | Treron_bicinctus | LC | Decreasing | tropical | diurnal | 174 | 2448171.733 | 14.084 |
| birds | Treron_calvus | Treron_calvus | LC | Decreasing | tropical | diurnal | 197.5 | 10739326.332 | 9.148 |
| birds | Treron_capellei | Treron_capellei | VU | Decreasing | tropical | diurnal | 401.3 | 1258129.64 | 8.368 |
| birds | Treron_curvirostra | Treron_curvirostra | LC | Decreasing | tropical | diurnal | 134.5 | 3273169.542 | 10.649 |
| birds | Treron_floris | Treron_floris | VU | Decreasing | tropical | diurnal | 205.7 | 37564.775 | 13.54 |
| birds | Treron_formosae | Treron_formosae | NT | Decreasing | temperate | diurnal | 140 | 36703.571 | 19.882 |
| birds | Treron_fulvicollis | Treron_fulvicollis | VU | Decreasing | tropical | diurnal | 167 | 932712.225 | 9.788 |
| birds | Treron_griseicauda | Treron_griseicauda | LC | Stable | tropical | diurnal | 205.7 | 245187.838 | 15.734 |
| birds | Treron_olax | Treron_olax | LC | Decreasing | tropical | diurnal | 77 | 1359140.895 | 8.221 |
| birds | Treron_oxyurus | Treron_oxyurus | NT | Decreasing | tropical | diurnal | 205.7 | 95078.115 | 7.841 |
| birds | Treron_pembaensis | Treron_pembaensis | VU | Decreasing | tropical | diurnal | 205.7 | 908.294 | 17.669 |

| | | | | | | | | | |
|---|---|---|---|---|---|---|---|---|---|
| birds | Treron_phoenicopterus | Treron_phoenicopterus | LC | Increasing | tropical | diurnal | 235 | 4843995.111 | 15.709 |
| birds | Treron_pompadora | Treron_pompadora | LC | Decreasing | tropical | diurnal | 234 | 65443.569 | 17.565 |
| birds | Treron_psittaceus | Treron_psittaceus | EN | Decreasing | tropical | diurnal | 205.7 | 19598.188 | 14.735 |
| birds | Treron_sanctithomae | Treron_sanctithomae | EN | Decreasing | tropical | diurnal | 205.7 | 643.431 | 9.869 |
| birds | Treron_seimundi | Treron_seimundi | LC | Decreasing | tropical | diurnal | 157.8 | 219840.389 | 10.962 |
| birds | Treron_sieboldii | Treron_sieboldii | LC | Decreasing | tropical | diurnal | 245 | 2159328.238 | 15.463 |
| birds | Treron_sphenurus | Treron_sphenurus | LC | Stable | temperate | diurnal | 210 | 1739910.382 | 9.622 |
| birds | Treron_teysmannii | Treron_teysmannii | NT | Decreasing | tropical | diurnal | 205.7 | 6075.208 | 14.332 |
| birds | Treron_vernans | Treron_vernans | LC | Stable | tropical | diurnal | 132 | 2260914.585 | 10.807 |
| birds | Treron_waalia | Treron_waalia | LC | Decreasing | tropical | diurnal | 254 | 5454383.589 | 11.477 |
| birds | Tribonyx_mortierii | Gallinula_mortierii | LC | Unknown | temperate | diurnal | 1291.8 | 42670.168 | 7.674 |
| birds | Tribonyx_ventralis | Gallinula_ventralis | LC | Stable | temperate | diurnal | 386.3 | 4599237.178 | 5.101 |
| birds | Trichodere_cockerelli | Trichodere_cockerelli | LC | Stable | tropical | diurnal | 14.7 | 62694.097 | 2.391 |
| birds | Trichoglossus_chlorolepidotus | Trichoglossus_chlorolepidotus | LC | Stable | temperate | diurnal | 87.1 | 1109928.467 | 6.424 |
| birds | Trichoglossus_euteles | Trichoglossus_euteles | LC | Stable | tropical | diurnal | 73.7 | 37893.112 | 12.796 |
| birds | Trichoglossus_flavoviridis | Trichoglossus_flavoviridis | LC | Stable | tropical | diurnal | 90.5 | 4102.714 | 5.221 |
| birds | Trichoglossus_haematodus | Trichoglossus_haematodus | LC | Decreasing | tropical | diurnal | 113.3 | 876908.035 | 3.661 |
| birds | Trichoglossus_iris | NA | NT | Decreasing | NA | NA | NA | NA | NA |
| birds | Trichoglossus_johnstoniae | Trichoglossus_johnstoniae | NT | Decreasing | tropical | diurnal | 55.2 | 8694.551 | 7.014 |
| birds | Trichoglossus_ornatus | Trichoglossus_ornatus | LC | Decreasing | tropical | diurnal | 120 | 172993.285 | 9.533 |
| birds | Trichoglossus_rubiginosus | Trichoglossus_rubiginosus | NT | Decreasing | tropical | diurnal | 77.5 | 355.651 | 12.208 |
| birds | Tricholaema_diademata | Tricholaema_diademata | LC | Stable | tropical | diurnal | 30.6 | 578757.822 | 10.621 |
| birds | Tricholaema_frontata | Tricholaema_frontata | LC | Decreasing | tropical | diurnal | 26.8 | 1375938.257 | 7.07 |
| birds | Tricholaema_hirsuta | Tricholaema_hirsuta | LC | Decreasing | tropical | diurnal | 55.2 | 3751171.457 | 8.413 |
| birds | Tricholaema_lacrymosa | Tricholaema_lacrymosa | LC | Stable | tropical | diurnal | 22.9 | 1269448.612 | 12.31 |
| birds | Tricholaema_leucomelas | Tricholaema_leucomelas | LC | Increasing | tropical | diurnal | 32.2 | 3123368.529 | 7.922 |
| birds | Tricholaema_melanocephala | Tricholaema_melanocephala | LC | Stable | tropical | diurnal | 20.5 | 1969018.583 | 8.96 |
| birds | Tricholestes_criniger | Tricholestes_criniger | LC | Decreasing | tropical | diurnal | 17.1 | 1220380.835 | 8.136 |
| birds | Trichothraupis_melanops | Trichothraupis_melanops | LC | Stable | tropical | diurnal | 22.6 | 2473090.346 | 11.53 |
| birds | Triclaria_malachitacea | Triclaria_malachitacea | LC | Decreasing | temperate | diurnal | 90 | 99479.822 | 11.58 |
| birds | Trigonoceps_occipitalis | Trigonoceps_occipitalis | CR | Decreasing | tropical | diurnal | 3016 | 11472482.168 | 9.895 |
| birds | Tringa_brevipes | Heteroscelus_brevipes | NT | Decreasing | temperate | diurnal | 127 | 4918509.31 | 4.646 |
| birds | Tringa_erythropus | Tringa_erythropus | LC | Stable | temperate | cathemeral | 158 | 17465707.916 | 9.698 |
| birds | Tringa_flavipes | Tringa_flavipes | LC | Decreasing | tropical | cathemeral | 77.5 | 35216181.962 | 7.539 |

| | | | | | | | | | |
|---|---|---|---|---|---|---|---|---|---|
| birds | Tringa_glareola | Tringa_glareola | LC | Stable | temperate | diurnal | 62.1 | 47087120.536 | 8.321 |
| birds | Tringa_guttifer | Tringa_guttifer | EN | Decreasing | temperate | diurnal | 158 | 413621.642 | 7.817 |
| birds | Tringa_incana | Heteroscelus_incanus | LC | Stable | temperate | diurnal | 108.2 | 2865961.573 | 4.121 |
| birds | Tringa_melanoleuca | Tringa_melanoleuca | LC | Stable | tropical | cathemeral | 161.7 | 34094264.542 | 7.769 |
| birds | Tringa_nebularia | Tringa_nebularia | LC | Stable | temperate | cathemeral | 187 | 46520227.983 | 8.683 |
| birds | Tringa_ochropus | Tringa_ochropus | LC | Increasing | temperate | diurnal | 71.4 | 43983761.472 | 9.479 |
| birds | Tringa_semipalmata | Catoptrophorus_semipalmatus | LC | Stable | temperate | cathemeral | 245.7 | 3922835.391 | 12.396 |
| birds | Tringa_solitaria | Tringa_solitaria | LC | Decreasing | tropical | diurnal | 48.4 | 29415391.205 | 7.548 |
| birds | Tringa_stagnatilis | Tringa_stagnatilis | LC | Decreasing | temperate | diurnal | 77.5 | 33164442.822 | 9.164 |
| birds | Tringa_totanus | Tringa_totanus | LC | Unknown | temperate | cathemeral | 129 | 46612790.942 | 11.448 |
| birds | Trochalopteron_affine | Garrulax_affinis | LC | Decreasing | temperate | diurnal | 72.7 | 740183.994 | 9.604 |
| birds | Trochalopteron_austeni | Garrulax_austeni | LC | Decreasing | temperate | diurnal | 59 | 54762.061 | 8.227 |
| birds | Trochalopteron_elliotii | Garrulax_elliotii | LC | Decreasing | temperate | diurnal | 45 | 1030829.657 | 10.764 |
| birds | Trochalopteron_erythrocephalum | Garrulax_erythrocephalus | LC | Increasing | temperate | diurnal | 71.7 | 334402.424 | 8.766 |
| birds | Trochalopteron_formosum | Garrulax_formosus | LC | Decreasing | tropical | diurnal | 94.5 | 136653.998 | 11.333 |
| birds | Trochalopteron_henrici | Garrulax_henrici | LC | Unknown | temperate | diurnal | 65 | 151106.187 | 5.006 |
| birds | Trochalopteron_lineatum | Garrulax_lineatus | LC | Stable | temperate | diurnal | 40.7 | 395193.544 | 11.151 |
| birds | Trochalopteron_milnei | Garrulax_milnei | LC | Decreasing | tropical | diurnal | 79.5 | 1489338.517 | 11.38 |
| birds | Trochalopteron_morrisonianum | Garrulax_morrisonianus | LC | Decreasing | temperate | diurnal | 73.1 | 13167.813 | 6.245 |
| birds | Trochalopteron_squamatum | Garrulax_squamatus | LC | Decreasing | temperate | diurnal | 84 | 639106.377 | 9.055 |
| birds | Trochalopteron_subunicolor | Garrulax_subunicolor | LC | Decreasing | temperate | diurnal | 65.9 | 340826.36 | 8.485 |
| birds | Trochalopteron_variegatum | Garrulax_variegatus | LC | Decreasing | temperate | diurnal | 64.5 | 281662.125 | 10.86 |
| birds | Trochalopteron_virgatum | Garrulax_virgatus | LC | Decreasing | temperate | diurnal | 48 | 112099.79 | 7.636 |
| birds | Trochalopteron_yersini | Garrulax_yersini | EN | Decreasing | tropical | diurnal | 78.6 | 719.172 | 6.488 |
| birds | Trochilus_polytmus | Trochilus_polytmus | LC | Unknown | tropical | diurnal | 4.8 | 10224.989 | 20.197 |
| birds | Trochilus_scitulus | Trochilus_scitulus | LC | Unknown | tropical | diurnal | 4 | 796.787 | 13.887 |
| birds | Trochocercus_cyanomelas | Trochocercus_cyanomelas | LC | Decreasing | temperate | diurnal | 10.2 | 179874.791 | 12.398 |
| birds | Trochocercus_nitens | Trochocercus_nitens | LC | Decreasing | tropical | diurnal | 11.8 | 3680877.595 | 8.209 |
| birds | Troglodytes_aedon | Troglodytes_aedon | LC | Increasing | tropical | diurnal | 10.9 | 28997428.72 | 8.66 |
| birds | Troglodytes_beani | NA | LC | Stable | NA | NA | NA | NA | NA |
| birds | Troglodytes_cobbi | Troglodytes_cobbi | LC | Stable | temperate | diurnal | 19.1 | 283.46 | 2.33 |
| birds | Troglodytes_monticola | Troglodytes_monticola | CR | Decreasing | tropical | diurnal | 9 | 1330.6 | 5.103 |
| birds | Troglodytes_ochraceus | Troglodytes_ochraceus | LC | Stable | tropical | diurnal | 9.4 | 17334.672 | 6.617 |
| birds | Troglodytes_rufociliatus | Troglodytes_rufociliatus | LC | Decreasing | tropical | diurnal | 10 | 135313.919 | 11.602 |

| | | | | | | | | | |
|---|---|---|---|---|---|---|---|---|---|
| birds | Troglodytes_rufulus | Troglodytes_rufulus | LC | Decreasing | tropical | diurnal | 16.1 | 67695.091 | 1.421 |
| birds | Troglodytes_sissonii | NA | NT | Unknown | NA | NA | NA | NA | NA |
| birds | Troglodytes_solstitialis | Troglodytes_solstitialis | LC | Decreasing | tropical | diurnal | 11.4 | 553583.655 | 8.612 |
| birds | Troglodytes_tanneri | NA | VU | Stable | NA | NA | NA | NA | NA |
| birds | Troglodytes_troglodytes | Troglodytes_troglodytes | LC | Increasing | temperate | diurnal | 9.7 | 18137186.27 | 13.464 |
| birds | Trogon_bairdii | Trogon_bairdii | NT | Decreasing | tropical | diurnal | 94.4 | 12068.833 | 13.274 |
| birds | Trogon_citreolus | Trogon_citreolus | LC | Stable | tropical | diurnal | 79 | 124015.578 | 8.704 |
| birds | Trogon_clathratus | Trogon_clathratus | LC | Decreasing | tropical | diurnal | 130 | 16280.048 | 9.229 |
| birds | Trogon_collaris | Trogon_collaris | LC | Decreasing | tropical | diurnal | 65.4 | 6799060.772 | 3.376 |
| birds | Trogon_comptus | Trogon_comptus | LC | Decreasing | tropical | diurnal | 114 | 79435.539 | 5.812 |
| birds | Trogon_curucui | Trogon_curucui | LC | Decreasing | tropical | diurnal | 54 | 7396318.686 | 4.407 |
| birds | Trogon_elegans | Trogon_elegans | LC | Decreasing | tropical | diurnal | 70.9 | 205850.355 | 13.063 |
| birds | Trogon_massena | Trogon_massena | LC | Decreasing | tropical | diurnal | 141 | 493358.7 | 8.468 |
| birds | Trogon_melanocephalus | Trogon_melanocephalus | LC | Decreasing | tropical | diurnal | 85.1 | 443923.306 | 10.106 |
| birds | Trogon_melanurus | Trogon_melanurus | LC | Decreasing | tropical | diurnal | 114 | 6755571.077 | 2.445 |
| birds | Trogon_mexicanus | Trogon_mexicanus | LC | Decreasing | tropical | diurnal | 71 | 375078.565 | 9.739 |
| birds | Trogon_personatus | Trogon_personatus | LC | Stable | tropical | diurnal | 63.4 | 745904.621 | 7.2 |
| birds | Trogon_rufus | Trogon_rufus | LC | Decreasing | tropical | diurnal | 53.8 | 7239946.908 | 3.99 |
| birds | Trogon_surrucura | Trogon_surrucura | LC | Decreasing | tropical | diurnal | 73.3 | 1366930.995 | 13.111 |
| birds | Trogon_violaceus | Trogon_violaceus | LC | Increasing | tropical | diurnal | 46.5 | 7284998.294 | 3.433 |
| birds | Trogon_viridis | Trogon_viridis | LC | Stable | tropical | diurnal | 89.7 | 7907953.755 | 3.132 |
| birds | Trugon_terrestris | Trugon_terrestris | LC | Stable | tropical | diurnal | 400 | 651761.908 | 3.075 |
| birds | Tunchiornis_ochraceiceps | Hylophilus_ochraceiceps | LC | Decreasing | tropical | diurnal | 11.6 | 3677162.236 | 2.582 |
| birds | Turacoena_manadensis | Turacoena_manadensis | LC | Decreasing | tropical | diurnal | 208.6 | 157651.986 | 9.935 |
| birds | Turacoena_modesta | Turacoena_modesta | NT | Decreasing | tropical | diurnal | 340 | 31798.328 | 13.079 |
| birds | Turdinus_atrigularis | Turdinus_atrigularis | NT | Decreasing | tropical | diurnal | 66.3 | 77387.709 | 3.763 |
| birds | Turdinus_crispifrons | NA | not_available | not_available | NA | NA | NA | NA | NA |
| birds | Turdinus_macrodactylus | Turdinus_macrodactylus | NT | Decreasing | tropical | diurnal | 48.7 | 129852.779 | 9.872 |
| birds | Turdinus_marmoratus | Turdinus_marmorata | LC | Decreasing | tropical | diurnal | 41 | 87991.715 | 6.986 |
| birds | Turdoides_bicolor | Turdoides_bicolor | LC | Decreasing | tropical | diurnal | 78.3 | 1132138.947 | 6.469 |
| birds | Turdoides_gymnogenys | Turdoides_gymnogenys | LC | Stable | tropical | diurnal | 77 | 411624.932 | 6.605 |
| birds | Turdoides_hartlaubii | Turdoides_hartlaubii | LC | Stable | tropical | diurnal | 86.2 | 1716629.628 | 6.738 |
| birds | Turdoides_hindei | Turdoides_hindei | VU | Decreasing | tropical | diurnal | 67.6 | 24091.463 | 18.312 |
| birds | Turdoides_hypoleuca | Turdoides_hypoleuca | LC | Stable | tropical | diurnal | 74 | 89042.464 | 13.788 |

| | | | | | | | | | |
|---|---|---|---|---|---|---|---|---|---|
| birds | Turdoides_jardineii | Turdoides_jardineii | LC | Stable | tropical | diurnal | 78.2 | 5822352.498 | 8.84 |
| birds | Turdoides_leucocephala | Turdoides_leucocephala | LC | Stable | tropical | diurnal | 63.3 | 265511.503 | 10.639 |
| birds | Turdoides_leucopygia | Turdoides_leucopygia | LC | Stable | tropical | diurnal | 82.4 | 693637.916 | 11.322 |
| birds | Turdoides_melanops | Turdoides_melanops | LC | Decreasing | tropical | diurnal | 77 | 224027.26 | 6.551 |
| birds | Turdoides_plebejus | Turdoides_plebejus | LC | Stable | tropical | diurnal | 62.5 | 5173060.806 | 11.201 |
| birds | Turdoides_reinwardtii | Turdoides_reinwardii | LC | Decreasing | tropical | diurnal | 78.8 | 2152069.641 | 12.464 |
| birds | Turdoides_sharpei | Turdoides_sharpei | LC | Decreasing | tropical | diurnal | 79.3 | 532508.01 | 14.615 |
| birds | Turdoides_squamulata | Turdoides_squamulata | LC | Decreasing | tropical | diurnal | 69.4 | 64347.573 | 10.741 |
| birds | Turdoides_striata | NA | not_available | not_available | NA | NA | NA | NA | NA |
| birds | Turdoides_tenebrosa | Turdoides_tenebrosa | LC | Decreasing | tropical | diurnal | 76 | 294743.093 | 8.884 |
| birds | Turdus_albicollis | Turdus_albicollis | LC | Decreasing | tropical | diurnal | 54 | 8533196.814 | 4.34 |
| birds | Turdus_albocinctus | Turdus_albocinctus | LC | Unknown | temperate | diurnal | 99.4 | 438247.5 | 8.794 |
| birds | Turdus_amaurochalinus | Turdus_amaurochalinus | LC | Stable | tropical | diurnal | 57.9 | 8248771.078 | 7.934 |
| birds | Turdus_assimilis | Turdus_assimilis | LC | Decreasing | tropical | diurnal | 70.2 | 548279.872 | 9.663 |
| birds | Turdus_aurantius | Turdus_aurantius | LC | Stable | tropical | diurnal | 81.9 | 11021.791 | 19.741 |
| birds | Turdus_bewsheri | Turdus_bewsheri | NT | Decreasing | tropical | diurnal | 63.3 | 1671.227 | 16.55 |
| birds | Turdus_boulboul | Turdus_boulboul | LC | Decreasing | temperate | diurnal | 97.1 | 1054926.295 | 9.966 |
| birds | Turdus_cardis | Turdus_cardis | LC | Unknown | temperate | diurnal | 58.9 | 1354131.507 | 16.491 |
| birds | Turdus_celaenops | Turdus_celaenops | VU | Decreasing | temperate | diurnal | 84.5 | 883.507 | 12.695 |
| birds | Turdus_chiguanco | Turdus_chiguanco | LC | Stable | tropical | diurnal | 93.3 | 478403.612 | 8.152 |
| birds | Turdus_chrysolaus | Turdus_chrysolaus | LC | Unknown | temperate | diurnal | 65.4 | 824574.826 | 18.149 |
| birds | Turdus_dissimilis | Turdus_dissimilis | LC | Decreasing | tropical | diurnal | 75 | 820644.32 | 10.413 |
| birds | Turdus_eremita | NA | NT | Stable | NA | NA | NA | NA | NA |
| birds | Turdus_falcklandii | Turdus_falcklandii | LC | Stable | temperate | diurnal | 93.9 | 987597.347 | 5.734 |
| birds | Turdus_feae | Turdus_feae | VU | Decreasing | temperate | diurnal | 71 | 2133337.983 | 13.797 |
| birds | Turdus_flavipes | Turdus_flavipes | LC | Stable | tropical | diurnal | 65.1 | 940921.437 | 10.811 |
| birds | Turdus_fulviventris | Turdus_fulviventris | LC | Decreasing | tropical | diurnal | 70 | 71579.244 | 7.153 |
| birds | Turdus_fumigatus | Turdus_fumigatus | LC | Decreasing | tropical | diurnal | 75.7 | 4070825.395 | 3.976 |
| birds | Turdus_fuscater | Turdus_fuscater | LC | Stable | tropical | diurnal | 143 | 411278.64 | 9.74 |
| birds | Turdus_grayi | Turdus_grayi | LC | Stable | tropical | diurnal | 79.5 | 1125160.826 | 11.341 |
| birds | Turdus_haplochrous | Turdus_haplochrous | NT | Decreasing | tropical | diurnal | 84 | 150807.053 | 7.512 |
| birds | Turdus_hauxwelli | Turdus_hauxwelli | LC | Decreasing | tropical | diurnal | 69 | 3257190.89 | 2.041 |
| birds | Turdus_helleri | Turdus_helleri | EN | Decreasing | tropical | diurnal | 68.7 | 5.375 | 21.794 |
| birds | Turdus_hortulorum | Turdus_hortulorum | LC | Unknown | temperate | diurnal | 66.7 | 4088520.404 | 12.599 |

| | | | | | | | | | |
|---|---|---|---|---|---|---|---|---|---|
| birds | Turdus_ignobilis | Turdus_ignobilis | LC | Stable | tropical | diurnal | 63.8 | 272969.908 | 12.413 |
| birds | Turdus_iliacus | Turdus_iliacus | NT | Decreasing | temperate | diurnal | 61.2 | 23032956.131 | 9.369 |
| birds | Turdus_infuscatus | Turdus_infuscatus | LC | Decreasing | tropical | diurnal | 72.4 | 198275.608 | 11.229 |
| birds | Turdus_jamaicensis | Turdus_jamaicensis | LC | Decreasing | tropical | diurnal | 59.1 | 11021.791 | 19.741 |
| birds | Turdus_kessleri | Turdus_kessleri | LC | Stable | temperate | diurnal | 76.3 | 1084912.877 | 7.213 |
| birds | Turdus_lawrencii | Turdus_lawrencii | LC | Decreasing | tropical | diurnal | 72.6 | 3085560.279 | 1.405 |
| birds | Turdus_leucomelas | Turdus_leucomelas | LC | Stable | tropical | diurnal | 69.1 | 7690463.382 | 7.456 |
| birds | Turdus_leucops | Turdus_leucops | LC | Decreasing | tropical | diurnal | 62.4 | 490113.807 | 6.669 |
| birds | Turdus_lherminieri | Turdus_lherminieri | NT | Unknown | tropical | diurnal | 105 | 1592.179 | 18.582 |
| birds | Turdus_libonyana | Turdus_libonyanus | LC | Unknown | tropical | diurnal | 60.6 | 4390032.162 | 8.785 |
| birds | Turdus_ludoviciae | Turdus_ludoviciae | LC | Stable | tropical | diurnal | 62 | 12100.208 | 10.576 |
| birds | Turdus_maculirostris | Turdus_maculirostris | LC | Stable | tropical | diurnal | 69.6 | 88017.543 | 13.076 |
| birds | Turdus_maranonicus | Turdus_maranonicus | LC | Decreasing | tropical | diurnal | 70 | 39621.902 | 8.226 |
| birds | Turdus_menachensis | Turdus_menachensis | NT | Decreasing | tropical | diurnal | 75.4 | 66735.6 | 12.157 |
| birds | Turdus_merula | Turdus_merula | LC | Increasing | temperate | diurnal | 102.7 | 14326330.936 | 13.917 |
| birds | Turdus_migratorius | Turdus_migratorius | LC | Stable | temperate | diurnal | 78.5 | 17922995.713 | 8.104 |
| birds | Turdus_naumanni | Turdus_naumanni | LC | Unknown | temperate | diurnal | 73.2 | 4484419.712 | 9.908 |
| birds | Turdus_nigrescens | Turdus_nigrescens | LC | Stable | tropical | diurnal | 96 | 2072.87 | 5.763 |
| birds | Turdus_nigriceps | Turdus_nigriceps | LC | Unknown | tropical | diurnal | 52.7 | 655026.184 | 5.297 |
| birds | Turdus_nudigenis | Turdus_nudigenis | LC | Stable | tropical | diurnal | 63.9 | 1554169.638 | 5.794 |
| birds | Turdus_obscurus | Turdus_obscurus | LC | Unknown | temperate | diurnal | 62.6 | 13237348.697 | 7.989 |
| birds | Turdus_obsoletus | Turdus_obsoletus | LC | Decreasing | tropical | diurnal | 74.8 | 247412.356 | 10.387 |
| birds | Turdus_olivaceofuscus | Turdus_olivaceofuscus | LC | Decreasing | tropical | diurnal | 87 | 856.749 | 12.812 |
| birds | Turdus_olivaceus | Turdus_olivaceus | LC | Unknown | temperate | diurnal | 67.3 | 399268.726 | 12.246 |
| birds | Turdus_olivater | Turdus_olivater | LC | Decreasing | tropical | diurnal | 86.6 | 227974.735 | 4.662 |
| birds | Turdus_pallidus | Turdus_pallidus | LC | Unknown | temperate | diurnal | 72.1 | 3500844.869 | 15.725 |
| birds | Turdus_philomelos | Turdus_philomelos | LC | Increasing | temperate | diurnal | 67.7 | 17749014.34 | 11.361 |
| birds | Turdus_pilaris | Turdus_pilaris | LC | Stable | temperate | diurnal | 106 | 20398397.008 | 9.278 |
| birds | Turdus_plebejus | Turdus_plebejus | LC | Decreasing | tropical | diurnal | 86.3 | 160186.827 | 11.432 |
| birds | Turdus_plumbeus | Turdus_plumbeus | LC | Stable | temperate | diurnal | 75.3 | 7073.934 | 7.776 |
| birds | Turdus_poliocephalus | Turdus_poliocephalus | LC | Decreasing | tropical | diurnal | 61.6 | 246956.22 | 6.555 |
| birds | Turdus_reevei | Turdus_reevei | LC | Decreasing | tropical | diurnal | 61 | 77222.874 | 12.924 |
| birds | Turdus_rubrocanus | Turdus_rubrocanus | LC | Unknown | temperate | diurnal | 92.2 | 1637844.189 | 10.586 |
| birds | Turdus_ruficollis | Turdus_ruficollis | LC | Unknown | temperate | diurnal | 80.6 | 7170455.486 | 4.239 |

| | | | | | | | | | |
|---|---|---|---|---|---|---|---|---|---|
| birds | Turdus_rufitorques | Turdus_rufitorques | LC | Decreasing | tropical | diurnal | 72 | 76537.366 | 13.449 |
| birds | Turdus_rufiventris | Turdus_rufiventris | LC | Stable | tropical | diurnal | 69.4 | 5028683.784 | 9.866 |
| birds | Turdus_rufopalliatus | Turdus_rufopalliatus | LC | Stable | tropical | diurnal | 75 | 478243.897 | 10.921 |
| birds | Turdus_serranus | Turdus_serranus | LC | Decreasing | tropical | diurnal | 84.9 | 408915.41 | 9.054 |
| birds | Turdus_subalaris | Turdus_subalaris | LC | Unknown | tropical | diurnal | 49.5 | 2163914.094 | 11.202 |
| birds | Turdus_swalesi | Turdus_swalesi | VU | Decreasing | tropical | diurnal | 75 | 4953.985 | 8.337 |
| birds | Turdus_tephronotus | Turdus_tephronotus | LC | Unknown | tropical | diurnal | 49.4 | 998012.214 | 9.794 |
| birds | Turdus_torquatus | Turdus_torquatus | LC | Stable | temperate | diurnal | 109 | 7426107.991 | 15.091 |
| birds | Turdus_unicolor | Turdus_unicolor | LC | Unknown | temperate | diurnal | 64.2 | 1400167.387 | 16.014 |
| birds | Turdus_viscivorus | Turdus_viscivorus | LC | Decreasing | temperate | diurnal | 117.4 | 15469229.78 | 12.082 |
| birds | Turnix_castanotus | Turnix_castanotus | LC | Stable | tropical | nocturnal | 82.7 | 432722.674 | 1.721 |
| birds | Turnix_hottentottus | Turnix_hottentottus | LC | Decreasing | temperate | diurnal | 50.6 | 58126.33 | 11.92 |
| birds | Turnix_maculosus | Turnix_maculosus | LC | Decreasing | tropical | nocturnal | 39.1 | 1110140.422 | 4.593 |
| birds | Turnix_melanogaster | Turnix_melanogaster | VU | Decreasing | temperate | cathemeral | 87.1 | 109323.746 | 7.862 |
| birds | Turnix_nigricollis | Turnix_nigricollis | LC | Stable | tropical | diurnal | 61.3 | 595286.916 | 8.847 |
| birds | Turnix_olivii | Turnix_olivii | CR | Unknown | tropical | diurnal | 56.4 | 2066.237 | 2.265 |
| birds | Turnix_pyrrhothorax | Turnix_pyrrhothorax | LC | Decreasing | temperate | cathemeral | 45.7 | 4287341.806 | 4.977 |
| birds | Turnix_suscitator | Turnix_suscitator | LC | Increasing | tropical | diurnal | 49.1 | 5511938.059 | 14.895 |
| birds | Turnix_sylvaticus | Turnix_sylvaticus | LC | Decreasing | tropical | nocturnal | 52.7 | 17314162.232 | 11.684 |
| birds | Turnix_tanki | Turnix_tanki | LC | Stable | temperate | diurnal | 52.8 | 9318008.546 | 15.767 |
| birds | Turnix_varius | Turnix_varius | LC | Decreasing | temperate | cathemeral | 90.2 | 2276694.741 | 8.056 |
| birds | Turnix_velox | Turnix_velox | LC | Decreasing | temperate | cathemeral | 44.3 | 6979116.249 | 4.322 |
| birds | Turtur_abyssinicus | Turtur_abyssinicus | LC | Stable | tropical | diurnal | 62.7 | 4644147.821 | 12.123 |
| birds | Turtur_afer | Turtur_afer | LC | Stable | tropical | diurnal | 65.6 | 10011361.738 | 9.39 |
| birds | Turtur_brehmeri | Turtur_brehmeri | LC | Decreasing | tropical | diurnal | 116 | 2760838.426 | 8.359 |
| birds | Turtur_chalcospilos | Turtur_chalcospilos | LC | Stable | tropical | diurnal | 60.6 | 5888722.681 | 9.361 |
| birds | Turtur_tympanistria | Turtur_tympanistria | LC | Stable | tropical | diurnal | 70.6 | 8134665.65 | 9.271 |
| birds | Tychaedon_barbata | Erythropygia_barbata | LC | Stable | tropical | diurnal | 25.6 | 1773141.403 | 7.054 |
| birds | Tychaedon_coryphoeus | Erythropygia_coryphaeus | LC | Stable | temperate | diurnal | 20.4 | 875738.692 | 7.211 |
| birds | Tychaedon_leucosticta | Erythropygia_leucosticta | LC | Decreasing | tropical | diurnal | 25.4 | 888229.898 | 10.168 |
| birds | Tychaedon_quadrivirgata | Erythropygia_quadrivirgata | LC | Stable | tropical | diurnal | 26.1 | 1616296.197 | 10.366 |
| birds | Tychaedon_signata | Erythropygia_signata | LC | Stable | temperate | diurnal | 38.2 | 167351.003 | 13.219 |
| birds | Tylas_eduardi | Tylas_eduardi | LC | Unknown | tropical | diurnal | 46.8 | 231862.91 | 8.199 |
| birds | Tympanuchus_pallidicinctus | Tympanuchus_pallidicinctus | VU | Decreasing | temperate | diurnal | 745.8 | 453431.175 | 13.371 |

| | | | | | | | | | |
|---|---|---|---|---|---|---|---|---|---|
| birds | Tympanuchus_phasianellus | Tympanuchus_phasianellus | LC | Increasing | temperate | diurnal | 882.4 | 6080574.603 | 4.972 |
| birds | Tyranneutes_stolzmanni | Tyranneutes_stolzmanni | LC | Decreasing | tropical | diurnal | 7.2 | 5293137.561 | 2.095 |
| birds | Tyranneutes_virescens | Tyranneutes_virescens | LC | Decreasing | tropical | diurnal | 7.2 | 1189030.515 | 1.533 |
| birds | Tyrannopsis_sulphurea | Tyrannopsis_sulphurea | LC | Stable | tropical | diurnal | 53.6 | 6496396.059 | 2.276 |
| birds | Tyrannulus_elatus | Tyrannulus_elatus | LC | Stable | tropical | diurnal | 7 | 6406376.291 | 2.666 |
| birds | Tyrannus_albogularis | Tyrannus_albogularis | LC | Stable | tropical | diurnal | 37.1 | 6990635.263 | 3.571 |
| birds | Tyrannus_caudifasciatus | Tyrannus_caudifasciatus | LC | Stable | tropical | diurnal | 42.4 | 211089.906 | 16.369 |
| birds | Tyrannus_couchii | Tyrannus_couchii | LC | Stable | tropical | diurnal | 39 | 551401.867 | 11.04 |
| birds | Tyrannus_crassirostris | Tyrannus_crassirostris | LC | Unknown | temperate | diurnal | 55.9 | 689580.18 | 7.95 |
| birds | Tyrannus_cubensis | Tyrannus_cubensis | EN | Decreasing | tropical | diurnal | 93.6 | 110766.904 | 16.023 |
| birds | Tyrannus_dominicensis | Tyrannus_dominicensis | LC | Stable | tropical | diurnal | 46.5 | 1743599.427 | 8.792 |
| birds | Tyrannus_forficatus | Tyrannus_forficatus | LC | Decreasing | temperate | diurnal | 39.3 | 2047593.807 | 13.78 |
| birds | Tyrannus_melancholicus | Tyrannus_melancholicus | LC | Increasing | tropical | diurnal | 37.4 | 16507028.161 | 6.415 |
| birds | Tyrannus_niveigularis | Tyrannus_niveigularis | LC | Stable | tropical | diurnal | 34.4 | 113185.585 | 11.679 |
| birds | Tyrannus_savana | Tyrannus_savana | LC | Stable | tropical | diurnal | 31.9 | 13572472.847 | 5.92 |
| birds | Tyrannus_tyrannus | Tyrannus_tyrannus | LC | Decreasing | tropical | diurnal | 39.9 | 16650317.212 | 9.107 |
| birds | Tyrannus_verticalis | Tyrannus_verticalis | LC | Decreasing | temperate | diurnal | 39.6 | 6509951.477 | 10.535 |
| birds | Tyrannus_vociferans | Tyrannus_vociferans | LC | Stable | temperate | diurnal | 45.6 | 2291219.895 | 8.926 |
| birds | Tyto_alba | Tyto_alba | LC | Stable | temperate | cathemeral | 403.3 | 63140770.909 | 9.559 |
| birds | Tyto_capensis | Tyto_capensis | LC | Decreasing | tropical | nocturnal | 364.7 | 2692718.843 | 9.881 |
| birds | Tyto_glaucops | Tyto_glaucops | LC | Stable | tropical | nocturnal | 397.5 | 75807.64 | 15.794 |
| birds | Tyto_longimembris | Tyto_longimembris | LC | Decreasing | tropical | nocturnal | 423.5 | 4282168.163 | 12.942 |
| birds | Tyto_multipunctata | NA | LC | Stable | NA | NA | NA | NA | NA |
| birds | Tyto_nigrobrunnea | NA | VU | Unknown | NA | NA | NA | NA | NA |
| birds | Tyto_novaehollandiae | Tyto_novaehollandiae | LC | Decreasing | temperate | nocturnal | 631.9 | 3178834.967 | 6.093 |
| birds | Tyto_rosenbergii | NA | LC | Stable | NA | NA | NA | NA | NA |
| birds | Tyto_soumagnei | Tyto_soumagnei | VU | Decreasing | tropical | nocturnal | 284 | 64190.801 | 6.055 |
| birds | Tyto_tenebricosa | Tyto_tenebricosa | LC | Decreasing | tropical | nocturnal | 660.6 | 1125489.71 | 4.928 |
| birds | Upucerthia_albigula | Upucerthia_albigula | LC | Decreasing | tropical | diurnal | 39.9 | 48222.851 | 5.637 |
| birds | Upucerthia_dumetaria | Upucerthia_dumetaria | LC | Decreasing | temperate | diurnal | 48.7 | 2678611.869 | 7.136 |
| birds | Upucerthia_validirostris | Upucerthia_validirostris | LC | Decreasing | tropical | diurnal | 33.6 | 731195.966 | 7.141 |
| birds | Upupa_epops | Upupa_epops | LC | Decreasing | tropical | diurnal | 66.9 | 61808940.043 | 9.172 |
| birds | Uraeginthus_angolensis | Uraeginthus_angolensis | LC | Stable | tropical | diurnal | 9.9 | 3460240.46 | 9.36 |
| birds | Uraeginthus_bengalus | Uraeginthus_bengalus | LC | Stable | tropical | diurnal | 9.9 | 7659842.25 | 10.612 |

| | | | | | | | | | |
|---|---|---|---|---|---|---|---|---|---|
| birds | Uraeginthus_cyanocephalus | Uraeginthus_cyanocephalus | LC | Stable | tropical | diurnal | 9.8 | 756367.533 | 11.11 |
| birds | Uratelornis_chimaera | Uratelornis_chimaera | VU | Decreasing | tropical | cathemeral | 226 | 12247.665 | 8.934 |
| birds | Uria_aalge | Uria_aalge | LC | Increasing | temperate | diurnal | 991.9 | 20362473.188 | 9.774 |
| birds | Uria_lomvia | Uria_lomvia | LC | Increasing | temperate | diurnal | 964 | 22158382.917 | 1.489 |
| birds | Urile_pelagicus | Phalacrocorax_pelagicus | LC | Decreasing | temperate | diurnal | 1844 | 6583940.552 | 2.168 |
| birds | Urile_penicillatus | Phalacrocorax_penicillatus | LC | Decreasing | temperate | diurnal | 2224.2 | 702773.834 | 4.967 |
| birds | Urile_urile | Phalacrocorax_urile | LC | Decreasing | temperate | diurnal | 2117.7 | 1348091.153 | 2.715 |
| birds | Urochroa_bougueri | Urochroa_bougueri | LC | Unknown | tropical | diurnal | 11.4 | 40202.28 | 9.042 |
| birds | Urocissa_caerulea | Urocissa_caerulea | LC | Decreasing | temperate | diurnal | 257 | 20500.707 | 17.715 |
| birds | Urocissa_erythroryncha | Urocissa_erythrorhyncha | LC | Stable | temperate | diurnal | 151.5 | 4461983.112 | 15.162 |
| birds | Urocissa_flavirostris | Urocissa_flavirostris | LC | Stable | temperate | diurnal | 149.2 | 304129.158 | 8.88 |
| birds | Urocissa_ornata | Urocissa_ornata | VU | Decreasing | tropical | diurnal | 196 | 15761.875 | 20.79 |
| birds | Urocissa_whiteheadi | Urocissa_whiteheadi | EN | Decreasing | tropical | diurnal | 200 | 33794.257 | 16.404 |
| birds | Urocolius_indicus | Urocolius_indicus | LC | Unknown | tropical | diurnal | 52.7 | 3462430.498 | 8.619 |
| birds | Urocolius_macrourus | Urocolius_macrourus | LC | Decreasing | tropical | diurnal | 45.5 | 5779732.9 | 10.054 |
| birds | Urocynchramus_pylzowi | Urocynchramus_pylzowi | LC | Stable | temperate | diurnal | 28.6 | 627499.683 | 7.909 |
| birds | Urodynamis_taitensis | Eudynamys_taitensis | LC | Decreasing | temperate | diurnal | 117 | 234604.065 | 5.822 |
| birds | Urolais_epichlorus | Urolais_epichlorus | LC | Decreasing | tropical | diurnal | 11.3 | 48907.515 | 9.861 |
| birds | Uromyias_agilis | Anairetes_agilis | LC | Stable | tropical | diurnal | 10.1 | 53360.05 | 11.573 |
| birds | Uromyias_agraphia | Anairetes_agraphia | LC | Stable | tropical | diurnal | 9.9 | 76899.648 | 4.77 |
| birds | Uropelia_campestris | Uropelia_campestris | LC | Stable | tropical | diurnal | 35.2 | 1791696.144 | 7.539 |
| birds | Uropsalis_lyra | Uropsalis_lyra | LC | Stable | tropical | nocturnal | 71.2 | 231615.962 | 8.775 |
| birds | Uropsalis_segmentata | Uropsalis_segmentata | LC | Stable | tropical | nocturnal | 42.9 | 221680.804 | 8.112 |
| birds | Uropsila_leucogastra | Uropsila_leucogastra | LC | Decreasing | tropical | diurnal | 9 | 409367.84 | 9.696 |
| birds | Urosphena_squameiceps | Urosphena_squameiceps | LC | Stable | tropical | diurnal | 9.3 | 2720076.548 | 15.312 |
| birds | Urosphena_subulata | Urosphena_subulata | LC | Stable | tropical | diurnal | 10.3 | 35526.192 | 12.892 |
| birds | Urosphena_whiteheadi | Urosphena_whiteheadi | LC | Stable | tropical | diurnal | 9 | 131729.062 | 2.853 |
| birds | Urosticte_benjamini | Urosticte_benjamini | LC | Decreasing | tropical | diurnal | 3.9 | 17377.94 | 7.571 |
| birds | Urosticte_ruficrissa | Urosticte_ruficrissa | LC | Decreasing | tropical | diurnal | 4.3 | 31247.82 | 5.45 |
| birds | Urothraupis_stolzmanni | Urothraupis_stolzmanni | LC | Stable | tropical | diurnal | 23.1 | 36296.607 | 8.648 |
| birds | Urotriorchis_macrourus | Urotriorchis_macrourus | LC | Decreasing | tropical | diurnal | 492 | 3073602.249 | 7.715 |
| birds | Vanellus_albiceps | Vanellus_albiceps | LC | Stable | tropical | diurnal | 192 | 7550710.295 | 9.137 |
| birds | Vanellus_armatus | Vanellus_armatus | LC | Increasing | tropical | diurnal | 156 | 5662037.204 | 8.225 |
| birds | Vanellus_chilensis | Vanellus_chilensis | LC | Increasing | tropical | diurnal | 327 | 13321164.291 | 7.102 |

| | | | | | | | | | |
|---|---|---|---|---|---|---|---|---|---|
| birds | Vanellus_cinereus | Vanellus_cinereus | LC | Decreasing | temperate | diurnal | 270 | 5700503.931 | 15.394 |
| birds | Vanellus_coronatus | Vanellus_coronatus | LC | Increasing | tropical | nocturnal | 165 | 6790147.371 | 8.556 |
| birds | Vanellus_crassirostris | Vanellus_crassirostris | LC | Unknown | tropical | diurnal | 170 | 3461973.987 | 9.299 |
| birds | Vanellus_duvaucelii | Vanellus_duvaucelii | NT | Decreasing | temperate | diurnal | 143 | 1666558.666 | 14.921 |
| birds | Vanellus_gregarius | Vanellus_gregarius | CR | Decreasing | temperate | diurnal | 224.5 | 11070005.966 | 11.002 |
| birds | Vanellus_leucurus | Vanellus_leucurus | LC | Unknown | temperate | diurnal | 132 | 6537401.048 | 10.656 |
| birds | Vanellus_lugubris | Vanellus_lugubris | LC | Unknown | tropical | cathemeral | 122 | 3561388.305 | 10.23 |
| birds | Vanellus_malabaricus | Vanellus_malarbaricus | LC | Stable | tropical | nocturnal | 140 | 2548296.229 | 18.619 |
| birds | Vanellus_melanocephalus | Vanellus_melanocephalus | LC | Unknown | tropical | diurnal | 214 | 179801.062 | 13.776 |
| birds | Vanellus_melanopterus | Vanellus_melanopterus | LC | Decreasing | tropical | diurnal | 184.9 | 959143.34 | 12.09 |
| birds | Vanellus_miles | Vanellus_miles | LC | Increasing | tropical | cathemeral | 387 | 3058008.891 | 2.484 |
| birds | Vanellus_senegallus | Vanellus_senegallus | LC | Stable | tropical | diurnal | 248 | 11259086.327 | 10.006 |
| birds | Vanellus_spinosus | Vanellus_spinosus | LC | Increasing | tropical | diurnal | 188 | 8392007.785 | 11.177 |
| birds | Vanellus_superciliosus | Vanellus_superciliosus | LC | Unknown | tropical | cathemeral | 150 | 3364640.176 | 9.281 |
| birds | Vanellus_tectus | Vanellus_tectus | LC | Unknown | tropical | nocturnal | 100 | 5265920.675 | 10.439 |
| birds | Vanellus_vanellus | Vanellus_vanellus | NT | Decreasing | temperate | cathemeral | 218.4 | 24038094.917 | 11.353 |
| birds | Vanga_curvirostris | Vanga_curvirostris | LC | Unknown | tropical | diurnal | 64.2 | 499102.364 | 8.5 |
| birds | Vauriella_albigularis | Rhinomyias_albigularis | EN | Decreasing | tropical | diurnal | 35.7 | 23988.062 | 18.336 |
| birds | Vauriella_goodfellowi | Rhinomyias_goodfellowi | NT | Decreasing | tropical | diurnal | 31.7 | 7732.347 | 6.863 |
| birds | Vauriella_gularis | Rhinomyias_gularis | LC | Decreasing | tropical | diurnal | 25.1 | 34379.012 | 3.209 |
| birds | Vauriella_insignis | Rhinomyias_insignis | VU | Decreasing | tropical | diurnal | 28.7 | 9756.909 | 7.518 |
| birds | Veles_binotatus | Caprimulgus_binotatus | LC | Decreasing | tropical | nocturnal | 63 | 701719.54 | 8.741 |
| birds | Veniliornis_affinis | Veniliornis_affinis | LC | Stable | tropical | diurnal | 34.8 | 5386514.863 | 2.628 |
| birds | Veniliornis_callonotus | Veniliornis_callonotus | LC | Increasing | tropical | diurnal | 26.5 | 136187.779 | 11.187 |
| birds | Veniliornis_cassini | Veniliornis_cassini | LC | Stable | tropical | diurnal | 34.6 | 1226065.002 | 1.578 |
| birds | Veniliornis_chocoensis | Veniliornis_chocoensis | NT | Decreasing | tropical | diurnal | 30 | 79496.042 | 5.259 |
| birds | Veniliornis_dignus | Veniliornis_dignus | LC | Stable | tropical | diurnal | 37.6 | 95384.146 | 8.951 |
| birds | Veniliornis_frontalis | Veniliornis_frontalis | LC | Decreasing | tropical | diurnal | 36.7 | 102721.217 | 6.904 |
| birds | Veniliornis_kirkii | Veniliornis_kirkii | LC | Decreasing | tropical | diurnal | 28.8 | 846131.892 | 11.057 |
| birds | Veniliornis_lignarius | Veniliornis_lignarius | LC | Stable | temperate | diurnal | 34 | 597481.814 | 7.394 |
| birds | Veniliornis_maculifrons | Veniliornis_maculifrons | LC | Stable | tropical | diurnal | 38.7 | 253876.592 | 12.119 |
| birds | Veniliornis_mixtus | Veniliornis_mixtus | LC | Stable | temperate | diurnal | 28.7 | 3656621.358 | 8.541 |
| birds | Veniliornis_nigriceps | Veniliornis_nigriceps | LC | Stable | tropical | diurnal | 44 | 181315.524 | 8.161 |
| birds | Veniliornis_passerinus | Veniliornis_passerinus | LC | Stable | tropical | diurnal | 32.1 | 11102862.121 | 4.777 |

| | | | | | | | | | |
|---|---|---|---|---|---|---|---|---|---|
| birds | Veniliornis_sanguineus | Veniliornis_sanguineus | LC | Stable | tropical | diurnal | 26.5 | 36223.603 | 5.425 |
| birds | Veniliornis_spilogaster | Veniliornis_spilogaster | LC | Stable | temperate | diurnal | 40.3 | 1540012.26 | 12.721 |
| birds | Vermivora_chrysoptera | Vermivora_chrysoptera | NT | Decreasing | tropical | diurnal | 8.7 | 4556072.355 | 15.772 |
| birds | Vermivora_cyanoptera | Vermivora_pinus | LC | Decreasing | tropical | diurnal | 8.9 | 3546336.022 | 16.571 |
| birds | Verreauxia_africana | Sasia_africana | LC | Stable | tropical | diurnal | 9.4 | 2527784.986 | 6.152 |
| birds | Vidua_camerunensis | Vidua_camerunensis | LC | Stable | tropical | diurnal | 12.1 | 614412.819 | 8.217 |
| birds | Vidua_chalybeata | Vidua_chalybeata | LC | Stable | tropical | diurnal | 12.5 | 6962831.565 | 11.964 |
| birds | Vidua_codringtoni | Vidua_codringtoni | LC | Stable | tropical | diurnal | 12.8 | 219030.785 | 10.982 |
| birds | Vidua_fischeri | Vidua_fischeri | LC | Stable | tropical | diurnal | 12.2 | 1423681.716 | 9.604 |
| birds | Vidua_funerea | Vidua_funerea | LC | Stable | tropical | diurnal | 14.3 | 1390191.122 | 10.459 |
| birds | Vidua_hypocherina | Vidua_hypocherina | LC | Stable | tropical | diurnal | 11.9 | 782303.109 | 11.063 |
| birds | Vidua_interjecta | Vidua_interjecta | LC | Stable | tropical | diurnal | 20.6 | 1775828.322 | 9.324 |
| birds | Vidua_larvaticola | Vidua_larvaticola | LC | Stable | tropical | diurnal | 13.1 | 572282.443 | 9.59 |
| birds | Vidua_macroura | Vidua_macroura | LC | Stable | tropical | diurnal | 15.7 | 13175399.572 | 10.126 |
| birds | Vidua_maryae | Vidua_maryae | LC | Decreasing | tropical | diurnal | 12.8 | 12038.874 | 18.185 |
| birds | Vidua_nigeriae | Vidua_nigeriae | LC | Stable | tropical | diurnal | 12 | 77109.515 | 15.26 |
| birds | Vidua_obtusa | Vidua_obtusa | LC | Stable | tropical | diurnal | 20.7 | 3242417.306 | 8.668 |
| birds | Vidua_orientalis | Vidua_orientalis | LC | Stable | tropical | diurnal | 18.5 | 2652424.775 | 12.478 |
| birds | Vidua_paradisaea | Vidua_paradisaea | LC | Stable | tropical | diurnal | 20.4 | 4674400.638 | 9.935 |
| birds | Vidua_purpurascens | Vidua_purpurascens | LC | Stable | tropical | diurnal | 13.1 | 995520.208 | 11.467 |
| birds | Vidua_raricola | Vidua_raricola | LC | Stable | tropical | diurnal | 11.9 | 236927.7 | 11.387 |
| birds | Vidua_regia | Vidua_regia | LC | Stable | tropical | diurnal | 14.4 | 1955547.715 | 7.414 |
| birds | Vidua_togoensis | Vidua_togoensis | LC | Stable | tropical | diurnal | 15.6 | 418624.533 | 12.104 |
| birds | Vidua_wilsoni | Vidua_wilsoni | LC | Stable | tropical | diurnal | 13.1 | 2114085.296 | 10.411 |
| birds | Vini_australis | Vini_australis | LC | Decreasing | tropical | diurnal | 49.5 | 4099.961 | 8.987 |
| birds | Vini_kuhlii | Vini_kuhlii | EN | Stable | tropical | diurnal | 40 | 824.365 | 11.909 |
| birds | Vini_peruviana | Vini_peruviana | VU | Decreasing | tropical | diurnal | 29 | 1833.444 | 15.198 |
| birds | Vini_stepheni | Vini_stepheni | VU | Stable | temperate | diurnal | 55 | 41.112 | 6.407 |
| birds | Vini_ultramarina | NA | CR | Stable | NA | NA | NA | NA | NA |
| birds | Vireo_altiloquus | Vireo_altiloquus | LC | Unknown | tropical | diurnal | 19 | 4891227.029 | 3.283 |
| birds | Vireo_atricapilla | Vireo_atricapilla | NT | Decreasing | temperate | diurnal | 9 | 1513757.313 | 10.984 |
| birds | Vireo_bairdi | Vireo_bairdi | NT | Unknown | tropical | diurnal | 12 | 488.334 | 10.21 |
| birds | Vireo_bellii | Vireo_bellii | LC | Increasing | temperate | diurnal | 8.5 | 3853184.168 | 12.013 |
| birds | Vireo_brevipennis | Vireo_brevipennis | LC | Decreasing | tropical | diurnal | 11.8 | 162984.256 | 9.341 |

| | | | | | | | | | |
|---|---|---|---|---|---|---|---|---|---|
| birds | Vireo_caribaeus | Vireo_caribaeus | VU | Stable | tropical | diurnal | 9.3 | 16.905 | 30.251 |
| birds | Vireo_carmioli | Vireo_carmioli | LC | Stable | tropical | diurnal | 13.1 | 4187.032 | 6.112 |
| birds | Vireo_cassinii | Vireo_cassinii | LC | Stable | temperate | diurnal | 14.7 | 4111688.479 | 8.646 |
| birds | Vireo_crassirostris | Vireo_crassirostris | LC | Stable | temperate | diurnal | 13 | 19800.804 | 8.64 |
| birds | Vireo_flavifrons | Vireo_flavifrons | LC | Increasing | temperate | diurnal | 18 | 5029981.242 | 15.807 |
| birds | Vireo_flavoviridis | Vireo_flavoviridis | LC | Stable | tropical | diurnal | 17.6 | 3010055.814 | 7.113 |
| birds | Vireo_gilvus | Vireo_gilvus | LC | Increasing | temperate | diurnal | 12.7 | 11735657.604 | 11.12 |
| birds | Vireo_gracilirostris | NA | NT | Unknown | NA | NA | NA | NA | NA |
| birds | Vireo_griseus | Vireo_griseus | LC | Increasing | temperate | diurnal | 11.4 | 3615420.04 | 16.317 |
| birds | Vireo_gundlachii | Vireo_gundlachii | LC | Stable | tropical | diurnal | 12.9 | 109844.769 | 16.133 |
| birds | Vireo_huttoni | Vireo_huttoni | LC | Increasing | temperate | diurnal | 11.3 | 1214843.427 | 9.993 |
| birds | Vireo_latimeri | Vireo_latimeri | LC | Decreasing | tropical | diurnal | 12 | 6995.483 | 26.913 |
| birds | Vireo_leucophrys | Vireo_leucophrys | LC | Decreasing | tropical | diurnal | 12.3 | 592873.421 | 9.459 |
| birds | Vireo_magister | Vireo_magister | LC | Decreasing | tropical | diurnal | 15.4 | 8592.847 | 6.888 |
| birds | Vireo_masteri | Vireo_masteri | NT | Decreasing | tropical | diurnal | 11.2 | 8292.656 | 6.066 |
| birds | Vireo_modestus | Vireo_modestus | LC | Stable | tropical | diurnal | 9.6 | 11021.791 | 19.741 |
| birds | Vireo_nanus | Vireo_nanus | LC | Unknown | tropical | diurnal | 10.5 | 13196.28 | 15.2 |
| birds | Vireo_nelsoni | Vireo_nelsoni | LC | Decreasing | tropical | diurnal | 9.4 | 190119.25 | 10.171 |
| birds | Vireo_olivaceus | Vireo_olivaceus | LC | Increasing | temperate | diurnal | 16.1 | 21662377.649 | 8.76 |
| birds | Vireo_osburni | Vireo_osburni | NT | Decreasing | tropical | diurnal | 19.9 | 11021.791 | 19.741 |
| birds | Vireo_pallens | Vireo_pallens | LC | Decreasing | tropical | diurnal | 11 | 264054.346 | 9.161 |
| birds | Vireo_philadelphicus | Vireo_philadelphicus | LC | Increasing | temperate | diurnal | 11.5 | 6279191.556 | 12.026 |
| birds | Vireo_plumbeus | Vireo_plumbeus | LC | Decreasing | tropical | diurnal | 16.4 | 2446894.167 | 9.06 |
| birds | Vireo_sclateri | Hylophilus_sclateri | LC | Stable | tropical | diurnal | 11.2 | 292061.957 | 0.86 |
| birds | Vireo_solitarius | Vireo_solitarius | LC | Increasing | temperate | diurnal | 15.3 | 8397663.45 | 12.199 |
| birds | Vireo_vicinior | Vireo_vicinior | LC | Increasing | temperate | diurnal | 12.8 | 966622.865 | 8.048 |
| birds | Vireolanius_eximius | NA | LC | Decreasing | NA | NA | NA | NA | NA |
| birds | Vireolanius_leucotis | Vireolanius_leucotis | LC | Decreasing | tropical | diurnal | 26 | 4216599.518 | 1.802 |
| birds | Vireolanius_melitophrys | Vireolanius_melitophrys | LC | Stable | tropical | diurnal | 34.7 | 170969.253 | 10.194 |
| birds | Vireolanius_pulchellus | Vireolanius_pulchellus | LC | Decreasing | tropical | diurnal | 24 | 372658.609 | 9.535 |
| birds | Volatinia_jacarina | Volatinia_jacarina | LC | Increasing | tropical | diurnal | 9.9 | 15248241.408 | 6.154 |
| birds | Vosea_whitemanensis | Melidectes_whitemanensis | NT | Decreasing | tropical | diurnal | 48 | 3884.625 | 2.607 |
| birds | Vultur_gryphus | Vultur_gryphus | VU | Decreasing | temperate | diurnal | 11236.1 | 2543864.992 | 6.103 |
| birds | Wetmorethraupis_sterrhopteron | Wetmorethraupis_sterrhopteron | VU | Decreasing | tropical | diurnal | 50.2 | 9638.829 | 2.744 |

| | | | | | | | | | |
|---|---|---|---|---|---|---|---|---|---|
| birds | Willisornis_poecilinotus | Willisornis_poecilinotus | LC | Stable | tropical | diurnal | 18.4 | 5210711.185 | 1.857 |
| birds | Xanthocephalus_xanthocephalus | Xanthocephalus_xanthocephalus | LC | Increasing | temperate | diurnal | 62.7 | 8088084.365 | 11.35 |
| birds | Xanthomixis_apperti | Bernieria_apperti | VU | Decreasing | tropical | diurnal | 14.6 | 1525.763 | 7.349 |
| birds | Xanthomixis_cinereiceps | Bernieria_cinereiceps | NT | Decreasing | tropical | diurnal | 18.2 | 163803.273 | 8.85 |
| birds | Xanthomixis_zosterops | Bernieria_zosterops | LC | Decreasing | tropical | diurnal | 10.9 | 210073.516 | 8.354 |
| birds | Xanthopsar_flavus | Xanthopsar_flavus | EN | Decreasing | temperate | diurnal | 43 | 732681.647 | 11.411 |
| birds | Xanthotis_flaviventer | Xanthotis_flaviventer | LC | Stable | tropical | diurnal | 40 | 759089.469 | 3.076 |
| birds | Xanthotis_macleayanus | Xanthotis_macleayanus | LC | Stable | tropical | diurnal | 31 | 27328.196 | 4.319 |
| birds | Xanthotis_polygrammus | Xanthotis_polygrammus | LC | Stable | tropical | diurnal | 20 | 420905.849 | 3.373 |
| birds | Xema_sabini | Xema_sabini | LC | Stable | temperate | diurnal | 198 | 60417075.337 | 1.709 |
| birds | Xenerpestes_minlosi | Xenerpestes_minlosi | LC | Decreasing | tropical | diurnal | 11 | 108320.343 | 8.169 |
| birds | Xenerpestes_singularis | Xenerpestes_singularis | LC | Decreasing | tropical | diurnal | 11.6 | 35692.786 | 5.134 |
| birds | Xenicus_gilviventris | Xenicus_gilviventris | EN | Decreasing | temperate | diurnal | 17.6 | 11251.47 | 2.277 |
| birds | Xenodacnis_parina | Xenodacnis_parina | LC | Decreasing | tropical | diurnal | 11.5 | 55452.637 | 9.916 |
| birds | Xenoligea_montana | Xenoligea_montana | VU | Decreasing | tropical | diurnal | 12.5 | 6517.463 | 8.967 |
| birds | Xenoperdix_udzungwensis | Xenoperdix_udzungwensis | EN | Decreasing | tropical | diurnal | 224.9 | 1098.162 | 4.332 |
| birds | Xenopipo_atronitens | Xenopipo_atronitens | LC | Decreasing | tropical | diurnal | 15.5 | 1985208.121 | 1.845 |
| birds | Xenopipo_uniformis | Xenopipo_uniformis | LC | Stable | tropical | diurnal | 17.9 | 70300.47 | 1.178 |
| birds | Xenopirostris_damii | NA | EN | Decreasing | NA | NA | NA | NA | NA |
| birds | Xenopirostris_polleni | Xenopirostris_polleni | NT | Unknown | tropical | diurnal | 61.5 | 61347.271 | 7.165 |
| birds | Xenopirostris_xenopirostris | Xenopirostris_xenopirostris | NT | Decreasing | temperate | diurnal | 59.5 | 40589.826 | 11.056 |
| birds | Xenops_minutus | Xenops_minutus | LC | Decreasing | tropical | diurnal | 10.6 | 945529.292 | 12.721 |
| birds | Xenops_rutilus | Xenops_rutilans | LC | Stable | tropical | diurnal | 11.2 | 6929954.15 | 7.535 |
| birds | Xenops_tenuirostris | Xenops_tenuirostris | LC | Stable | tropical | diurnal | 9.7 | 3583718.567 | 1.846 |
| birds | Xenopsaris_albinucha | Xenopsaris_albinucha | LC | Stable | tropical | diurnal | 10 | 4453736.322 | 7.41 |
| birds | Xenornis_setifrons | Xenornis_setifrons | NT | Decreasing | tropical | diurnal | 24.6 | 17510.593 | 3.507 |
| birds | Xenospingus_concolor | Xenospingus_concolor | NT | Decreasing | tropical | diurnal | 21 | 34562.576 | 7.968 |
| birds | Xenospiza_baileyi | Xenospiza_baileyi | EN | Decreasing | temperate | diurnal | 17.4 | 583.448 | 17.524 |
| birds | Xenotriccus_callizonus | Xenotriccus_callizonus | LC | Decreasing | tropical | diurnal | 11.2 | 53967.147 | 11.96 |
| birds | Xenotriccus_mexicanus | Xenotriccus_mexicanus | LC | Decreasing | tropical | diurnal | 13.8 | 112766.327 | 8.567 |
| birds | Xenus_cinereus | Xenus_cinereus | LC | Decreasing | temperate | diurnal | 78.8 | 14264708.236 | 5.487 |
| birds | Xiphidiopicus_percussus | Xiphidiopicus_percussus | LC | Stable | tropical | diurnal | 60.9 | 110277.337 | 16.107 |
| birds | Xiphocolaptes_albicollis | Xiphocolaptes_albicollis | LC | Stable | tropical | diurnal | 118 | 2097683.154 | 12.368 |
| birds | Xiphocolaptes_falcirostris | Xiphocolaptes_falcirostris | VU | Decreasing | tropical | diurnal | 110 | 507320.554 | 6.927 |

| | | | | | | | | | |
|---|---|---|---|---|---|---|---|---|---|
| birds | Xiphocolaptes_major | Xiphocolaptes_major | LC | Decreasing | tropical | diurnal | 156 | 1461863.846 | 6.309 |
| birds | Xiphocolaptes_promeropirhynchus | Xiphocolaptes_promeropirhynchus | LC | Decreasing | tropical | diurnal | 93 | 4853747.848 | 3.502 |
| birds | Xipholena_atropurpurea | Xipholena_atropurpurea | VU | Decreasing | tropical | diurnal | 61.3 | 214393.098 | 12.399 |
| birds | Xipholena_lamellipennis | Xipholena_lamellipennis | NT | Decreasing | tropical | diurnal | 53.8 | 1084022.355 | 2.631 |
| birds | Xipholena_punicea | Xipholena_punicea | LC | Decreasing | tropical | diurnal | 68.1 | 3624635.856 | 1.744 |
| birds | Xiphorhynchus_erythropygius | Xiphorhynchus_erythropygius | LC | Decreasing | tropical | diurnal | 46.8 | 175302.576 | 11.824 |
| birds | Xiphorhynchus_flavigaster | Xiphorhynchus_flavigaster | LC | Decreasing | tropical | diurnal | 46.2 | 767413.136 | 10.417 |
| birds | Xiphorhynchus_fuscus | Xiphorhynchus_fuscus | LC | Decreasing | tropical | diurnal | 21.8 | 1948346.792 | 11.707 |
| birds | Xiphorhynchus_guttatus | Xiphorhynchus_guttatus | LC | Decreasing | tropical | diurnal | 59.7 | 2005419.734 | 3.304 |
| birds | Xiphorhynchus_lachrymosus | Xiphorhynchus_lachrymosus | LC | Decreasing | tropical | diurnal | 56.4 | 271654.211 | 8.466 |
| birds | Xiphorhynchus_obsoletus | Xiphorhynchus_obsoletus | LC | Decreasing | tropical | diurnal | 32.6 | 6665026.213 | 2.098 |
| birds | Xiphorhynchus_ocellatus | Xiphorhynchus_ocellatus | LC | Decreasing | tropical | diurnal | 36.7 | 1260065.923 | 0.973 |
| birds | Xiphorhynchus_pardalotus | Xiphorhynchus_pardalotus | LC | Stable | tropical | diurnal | 34.6 | 1710329.556 | 1.326 |
| birds | Xiphorhynchus_spixii | Xiphorhynchus_spixii | LC | Decreasing | tropical | diurnal | 31.2 | 993444.469 | 2.989 |
| birds | Xiphorhynchus_susurrans | Xiphorhynchus_susurrans | LC | Decreasing | tropical | diurnal | 46.7 | 640926.308 | 12.258 |
| birds | Xiphorhynchus_triangularis | Xiphorhynchus_triangularis | LC | Decreasing | tropical | diurnal | 46.3 | 286355.727 | 8.307 |
| birds | Xolmis_cinereus | Xolmis_cinereus | LC | Decreasing | tropical | diurnal | 57.1 | 6894893.412 | 7.631 |
| birds | Xolmis_coronatus | Xolmis_coronatus | LC | Stable | temperate | diurnal | 46.8 | 2279881.376 | 8.129 |
| birds | Xolmis_dominicanus | Xolmis_dominicanus | VU | Decreasing | temperate | diurnal | 42.8 | 344144.447 | 10.908 |
| birds | Xolmis_irupero | Xolmis_irupero | LC | Stable | temperate | diurnal | 28.7 | 3570599.317 | 8.298 |
| birds | Xolmis_pyrope | Xolmis_pyrope | LC | Stable | temperate | diurnal | 35.3 | 747754.311 | 5.612 |
| birds | Xolmis_rubetra | Xolmis_rubetra | LC | Decreasing | temperate | diurnal | 36.3 | 1640531.884 | 8.475 |
| birds | Xolmis_salinarum | Xolmis_salinarum | LC | Stable | temperate | diurnal | 25.1 | 40541.42 | 3.504 |
| birds | Xolmis_velatus | NA | LC | Stable | NA | NA | NA | NA | NA |
| birds | Yuhina_bakeri | Yuhina_bakeri | LC | Decreasing | temperate | diurnal | 17.1 | 200853.535 | 6.81 |
| birds | Yuhina_brunneiceps | Yuhina_brunneiceps | LC | Decreasing | temperate | diurnal | 12.5 | 13673.615 | 6.278 |
| birds | Yuhina_castaniceps | NA | not_available | not_available | NA | NA | NA | NA | NA |
| birds | Yuhina_flavicollis | Yuhina_flavicollis | LC | Unknown | temperate | diurnal | 16.9 | 622981.086 | 8.458 |
| birds | Yuhina_gularis | Yuhina_gularis | LC | Decreasing | temperate | diurnal | 20.8 | 515717.007 | 8.755 |
| birds | Yuhina_humilis | Yuhina_humilis | LC | Decreasing | tropical | diurnal | 14 | 57796.561 | 8.514 |
| birds | Yuhina_nigrimenta | Yuhina_nigrimenta | LC | Decreasing | temperate | diurnal | 9.5 | 2065413.286 | 11.366 |
| birds | Yuhina_occipitalis | Yuhina_occipitalis | LC | Decreasing | temperate | diurnal | 13 | 355046.541 | 8.832 |
| birds | Zanclostomus_javanicus | Phaenicophaeus_javanicus | LC | Decreasing | tropical | diurnal | 109 | 1534805.959 | 9.192 |
| birds | Zanda_baudinii | Calyptorhynchus_baudinii | CR | Decreasing | temperate | diurnal | 620 | 56186.022 | 11.039 |

| | | | | | | | | | |
|---|---|---|---|---|---|---|---|---|---|
| birds | Zanda_funerea | Calyptorhynchus_funereus | LC | Stable | temperate | diurnal | 674.7 | 1045925.975 | 8.588 |
| birds | Zanda_latirostris | Calyptorhynchus_latirostris | EN | Decreasing | temperate | diurnal | 612 | 235747.476 | 11.304 |
| birds | Zapornia_atra | Porzana_atra | VU | Stable | temperate | diurnal | 77.4 | 41.112 | 6.407 |
| birds | Zapornia_bicolor | NA | LC | Decreasing | NA | NA | NA | NA | NA |
| birds | Zapornia_flavirostra | Amaurornis_flavirostra | LC | Unknown | tropical | diurnal | 86.2 | 16710583.488 | 9.415 |
| birds | Zapornia_olivieri | NA | EN | Decreasing | NA | NA | NA | NA | NA |
| birds | Zapornia_parva | Porzana_parva | LC | Stable | temperate | diurnal | 49.7 | 12207181.678 | 10.377 |
| birds | Zapornia_paykullii | Porzana_paykullii | NT | Decreasing | temperate | nocturnal | 111 | 2630845.245 | 13.179 |
| birds | Zapornia_tabuensis | Porzana_tabuensis | LC | Unknown | temperate | diurnal | 42.2 | 2288129.044 | 7.129 |
| birds | Zaratornis_stresemanni | Zaratornis_stresemanni | NT | Decreasing | tropical | diurnal | 52.1 | 61015.408 | 8.065 |
| birds | Zavattariornis_stresemanni | Zavattariornis_stresemanni | EN | Decreasing | tropical | diurnal | 130 | 4600.825 | 8.745 |
| birds | Zebrilus_undulatus | Zebrilus_undulatus | LC | Decreasing | tropical | diurnal | 123 | 6925813.734 | 2.252 |
| birds | Zeledonia_coronata | Zeledonia_coronata | LC | Stable | tropical | diurnal | 21 | 5169.989 | 5.872 |
| birds | Zenaida_asiatica | Zenaida_asiatica | LC | Increasing | tropical | diurnal | 153 | 3221953.539 | 10.271 |
| birds | Zenaida_auriculata | Zenaida_auriculata | LC | Increasing | tropical | diurnal | 110.2 | 11530888.212 | 8.217 |
| birds | Zenaida_aurita | Zenaida_aurita | LC | Stable | tropical | diurnal | 156 | 230814.876 | 16.157 |
| birds | Zenaida_galapagoensis | NA | NT | Decreasing | NA | NA | NA | NA | NA |
| birds | Zenaida_graysoni | NA | EW | not_available | NA | NA | NA | NA | NA |
| birds | Zenaida_macroura | Zenaida_macroura | LC | Increasing | temperate | diurnal | 118.9 | 12173012.557 | 12.477 |
| birds | Zenaida_meloda | Zenaida_meloda | LC | Increasing | tropical | diurnal | 209.2 | 265815.101 | 6.662 |
| birds | Zentrygon_albifacies | Geotrygon_albifacies | LC | Decreasing | tropical | diurnal | 238 | 192106.95 | 11.093 |
| birds | Zentrygon_carrikeri | Geotrygon_carrikeri | EN | Decreasing | tropical | diurnal | 267 | 127.541 | 4.563 |
| birds | Zentrygon_chiriquensis | Geotrygon_chiriquensis | LC | Decreasing | tropical | diurnal | 308 | 16384.722 | 9.153 |
| birds | Zentrygon_costaricensis | Geotrygon_costaricensis | LC | Decreasing | tropical | diurnal | 285.1 | 16523.838 | 8.009 |
| birds | Zentrygon_frenata | Geotrygon_frenata | LC | Stable | tropical | diurnal | 307 | 442995.434 | 7.241 |
| birds | Zentrygon_goldmani | Geotrygon_goldmani | NT | Stable | tropical | diurnal | 258 | 2089.727 | 2.366 |
| birds | Zentrygon_lawrencii | Geotrygon_lawrencii | LC | Decreasing | tropical | diurnal | 220 | 7970.026 | 7.473 |
| birds | Zentrygon_linearis | Geotrygon_linearis | LC | Decreasing | tropical | diurnal | 261 | 162092.826 | 11.68 |
| birds | Zimmerius_bolivianus | Zimmerius_bolivianus | LC | Stable | tropical | diurnal | 11 | 111971.31 | 5.29 |
| birds | Zimmerius_chrysops | Zimmerius_chrysops | LC | Stable | tropical | diurnal | 10.4 | 387308.825 | 9.658 |
| birds | Zimmerius_cinereicapilla | Zimmerius_cinereicapilla | VU | Decreasing | tropical | diurnal | 11.8 | 75707.756 | 4.388 |
| birds | Zimmerius_gracilipes | Zimmerius_gracilipes | LC | Increasing | tropical | diurnal | 7.1 | 3855220.6 | 1.632 |
| birds | Zimmerius_improbus | Zimmerius_improbus | LC | Stable | tropical | diurnal | 10.9 | 24090.834 | 10.251 |
| birds | Zimmerius_vilissimus | Zimmerius_vilissimus | LC | Decreasing | tropical | diurnal | 9.7 | 79107.682 | 12.723 |

| | | | | | | | | | |
|---|---|---|---|---|---|---|---|---|---|
| birds | Zimmerius_viridiflavus | Zimmerius_viridiflavus | LC | Decreasing | tropical | diurnal | 9.6 | 89059.906 | 5.627 |
| birds | Zonotrichia_albicollis | Zonotrichia_albicollis | LC | Decreasing | temperate | diurnal | 24.4 | 11733130.059 | 9.96 |
| birds | Zonotrichia_atricapilla | Zonotrichia_atricapilla | LC | Decreasing | temperate | diurnal | 32 | 2344274.019 | 3.736 |
| birds | Zonotrichia_capensis | Zonotrichia_capensis | LC | Increasing | tropical | diurnal | 20.3 | 12703362.392 | 7.251 |
| birds | Zonotrichia_leucophrys | Zonotrichia_leucophrys | LC | Stable | temperate | diurnal | 28 | 17967442.743 | 7.787 |
| birds | Zonotrichia_querula | Zonotrichia_querula | NT | Decreasing | temperate | diurnal | 35.5 | 6150624.353 | 7.755 |
| birds | Zoonavena_grandidieri | NA | LC | Decreasing | NA | NA | NA | NA | NA |
| birds | Zoonavena_sylvatica | NA | LC | Stable | NA | NA | NA | NA | NA |
| birds | Zoonavena_thomensis | Zoonavena_thomensis | LC | Stable | tropical | diurnal | 8 | 1008.509 | 13.068 |
| birds | Zoothera_andromedae | Zoothera_andromedae | LC | Decreasing | tropical | diurnal | 90.5 | 132967.794 | 8.48 |
| birds | Zoothera_dixoni | Zoothera_dixoni | LC | Unknown | temperate | diurnal | 90 | 1131286.289 | 9.407 |
| birds | Zoothera_everetti | Zoothera_everetti | NT | Decreasing | tropical | diurnal | 81.9 | 3523.431 | 4.034 |
| birds | Zoothera_heinei | Zoothera_heinei | LC | Decreasing | temperate | diurnal | 82.8 | 469278.966 | 6.129 |
| birds | Zoothera_heinrichi | Geomalia_heinrichi | LC | Stable | tropical | diurnal | 73.2 | 27407.533 | 4.996 |
| birds | Zoothera_lunulata | Zoothera_lunulata | LC | Decreasing | temperate | diurnal | 130 | 578456.612 | 9.335 |
| birds | Zoothera_machiki | Zoothera_machiki | NT | Decreasing | tropical | diurnal | 81.9 | 3241.381 | 5.813 |
| birds | Zoothera_major | Zoothera_major | NT | Increasing | temperate | diurnal | 172 | 723.01 | 16.316 |
| birds | Zoothera_margaretae | Zoothera_margaretae | NT | Decreasing | tropical | diurnal | 65.3 | 3338.039 | 4.602 |
| birds | Zoothera_marginata | Zoothera_marginata | LC | Decreasing | tropical | diurnal | 81 | 939864.555 | 8.357 |
| birds | Zoothera_mollissima | Zoothera_mollissima | LC | Stable | temperate | diurnal | 98.2 | 422988.163 | 7.771 |
| birds | Zoothera_monticola | Zoothera_monticola | LC | Decreasing | temperate | diurnal | 126 | 323451.379 | 8.936 |
| birds | Zoothera_talaseae | NA | LC | Stable | NA | NA | NA | NA | NA |
| birds | Zosterops_abyssinicus | Zosterops_abyssinicus | LC | Unknown | tropical | diurnal | 9.4 | 665331.72 | 10.609 |
| birds | Zosterops_albogularis | NA | CR | Decreasing | NA | NA | NA | NA | NA |
| birds | Zosterops_anomalus | Zosterops_anomalus | LC | Unknown | tropical | diurnal | 12 | 31315.061 | 16.569 |
| birds | Zosterops_atricapilla | Zosterops_atricapilla | LC | Decreasing | tropical | diurnal | 9.2 | 116342.05 | 5.459 |
| birds | Zosterops_atriceps | Zosterops_atriceps | LC | Decreasing | tropical | diurnal | 12 | 20724.436 | 6.419 |
| birds | Zosterops_atrifrons | Zosterops_atrifrons | LC | Unknown | tropical | diurnal | 10 | 51931.655 | 8.451 |
| birds | Zosterops_borbonicus | Zosterops_borbonicus | LC | Unknown | tropical | diurnal | 8.2 | 2541.136 | 20.642 |
| birds | Zosterops_brunneus | Speirops_brunneus | VU | Stable | tropical | diurnal | 16.1 | 63.217 | 12.071 |
| birds | Zosterops_buruensis | Zosterops_buruensis | LC | Unknown | tropical | diurnal | 12 | 8245.733 | 6.825 |
| birds | Zosterops_ceylonensis | Zosterops_ceylonensis | LC | Decreasing | tropical | diurnal | 12.6 | 8009.586 | 16.298 |
| birds | Zosterops_chloris | Zosterops_chloris | LC | Decreasing | tropical | diurnal | 11.5 | 167122.055 | 10.674 |
| birds | Zosterops_chloronothos | Zosterops_chloronothus | CR | Decreasing | tropical | diurnal | 8.1 | 1860.389 | 26.013 |

| | | | | | | | | | |
|---|---|---|---|---|---|---|---|---|---|
| birds | Zosterops_cinereus | Zosterops_cinereus | LC | Stable | tropical | diurnal | 11.2 | 102.517 | 10.477 |
| birds | Zosterops_citrinella | Zosterops_citrinella | LC | Decreasing | tropical | diurnal | 11.5 | 70904.803 | 11.981 |
| birds | Zosterops_consobrinorum | Zosterops_consobrinorum | LC | Unknown | tropical | diurnal | 10.1 | 42944.114 | 8.279 |
| birds | Zosterops_conspicillatus | NA | EX | not_available | NA | NA | NA | NA | NA |
| birds | Zosterops_emiliae | Chlorocharis_emiliae | LC | Decreasing | tropical | diurnal | 13.9 | 206417.301 | 3.864 |
| birds | Zosterops_erythropleurus | Zosterops_erythropleurus | LC | Decreasing | temperate | diurnal | 11 | 5256262.275 | 13.554 |
| birds | Zosterops_everetti | Zosterops_everetti | LC | Decreasing | tropical | diurnal | 10 | 122580.156 | 12.25 |
| birds | Zosterops_explorator | Zosterops_explorator | LC | Unknown | tropical | diurnal | 10.5 | 16653.441 | 9.386 |
| birds | Zosterops_ficedulinus | Zosterops_ficedulinus | EN | Decreasing | tropical | diurnal | 9.4 | 40.889 | 10.193 |
| birds | Zosterops_finschii | Zosterops_finschii | LC | Stable | tropical | diurnal | 10.6 | 445.062 | 11.176 |
| birds | Zosterops_flavifrons | Zosterops_flavifrons | LC | Unknown | tropical | diurnal | 10.7 | 12171.101 | 5.807 |
| birds | Zosterops_flavus | NA | EN | Decreasing | NA | NA | NA | NA | NA |
| birds | Zosterops_fuscicapilla | Zosterops_fuscicapilla | LC | Unknown | tropical | diurnal | 11.2 | 102101.025 | 4.114 |
| birds | Zosterops_grayi | Zosterops_grayi | LC | Stable | tropical | diurnal | 16 | 556.547 | 10.332 |
| birds | Zosterops_griseotinctus | Zosterops_griseotinctus | LC | Unknown | tropical | diurnal | 14 | 1046.448 | 3.804 |
| birds | Zosterops_griseovirescens | NA | LC | Stable | NA | NA | NA | NA | NA |
| birds | Zosterops_hypolais | Zosterops_hypolais | NT | Stable | tropical | diurnal | 10.6 | 97.968 | 16.751 |
| birds | Zosterops_hypoxanthus | Zosterops_hypoxanthus | LC | Decreasing | tropical | diurnal | 12.5 | 45547.011 | 4.499 |
| birds | Zosterops_inornatus | Zosterops_inornatus | LC | Unknown | tropical | diurnal | 22.9 | 1159.539 | 7.797 |
| birds | Zosterops_japonicus | Zosterops_japonicus | LC | Unknown | temperate | diurnal | 11.3 | 621194.124 | 15.216 |
| birds | Zosterops_kuehni | Zosterops_kuehni | LC | Stable | tropical | diurnal | 12 | 794.699 | 13.578 |
| birds | Zosterops_kulambangrae | Zosterops_kulambangrae | LC | Stable | tropical | diurnal | 10.6 | 3708.662 | 2.853 |
| birds | Zosterops_lacertosus | Woodfordia_lacertosa | NT | Unknown | tropical | diurnal | 28 | 537.136 | 4.291 |
| birds | Zosterops_lateralis | Zosterops_lateralis | LC | Stable | temperate | diurnal | 12.7 | 2702683.6 | 7.868 |
| birds | Zosterops_leucophaeus | Speirops_leucophoeus | LC | Unknown | tropical | diurnal | 14.8 | 146.64 | 14.708 |
| birds | Zosterops_lugubris | Speirops_lugubris | LC | Unknown | tropical | diurnal | 14.8 | 858.528 | 12.812 |
| birds | Zosterops_luteirostris | Zosterops_luteirostris | VU | Stable | tropical | diurnal | 10.6 | 35.157 | 11.529 |
| birds | Zosterops_luteus | Zosterops_luteus | LC | Stable | tropical | diurnal | 9.7 | 672778.056 | 3.025 |
| birds | Zosterops_maderaspatanus | Zosterops_maderaspatanus | LC | Decreasing | tropical | diurnal | 8.9 | 592006.089 | 8.755 |
| birds | Zosterops_mayottensis | Zosterops_mayottensis | LC | Decreasing | tropical | diurnal | 9.9 | 362.323 | 20.903 |
| birds | Zosterops_meeki | Zosterops_meeki | NT | Decreasing | tropical | diurnal | 9 | 812.836 | 4.012 |
| birds | Zosterops_melanocephalus | Speirops_melanocephalus | VU | Stable | tropical | diurnal | 10.9 | 148.501 | 9.385 |
| birds | Zosterops_metcalfii | Zosterops_metcalfii | LC | Decreasing | tropical | diurnal | 14 | 16905.997 | 4.811 |
| birds | Zosterops_meyeni | Zosterops_meyeni | LC | Stable | tropical | diurnal | 10 | 105626.257 | 15.274 |

| | | | | | | | | | |
|---|---|---|---|---|---|---|---|---|---|
| birds | Zosterops_minor | Zosterops_minor | LC | Unknown | tropical | diurnal | 11.1 | 82133.234 | 3.047 |
| birds | Zosterops_minutus | Zosterops_minutus | LC | Unknown | tropical | diurnal | 8.3 | 1159.539 | 7.797 |
| birds | Zosterops_modestus | NA | VU | Stable | NA | NA | NA | NA | NA |
| birds | Zosterops_mouroniensis | Zosterops_mouroniensis | VU | Decreasing | tropical | diurnal | 9.9 | 71.701 | 10.943 |
| birds | Zosterops_murphyi | Zosterops_murphyi | LC | Decreasing | tropical | diurnal | 10.6 | 227.7 | 1.951 |
| birds | Zosterops_mysorensis | Zosterops_mysorensis | NT | Decreasing | tropical | diurnal | 15 | 2311.918 | 8.484 |
| birds | Zosterops_natalis | Zosterops_natalis | LC | Stable | tropical | diurnal | 10 | 155.177 | 12.508 |
| birds | Zosterops_nigrorum | Zosterops_nigrorum | LC | Unknown | tropical | diurnal | 7.3 | 141165.471 | 15.281 |
| birds | Zosterops_novaeguineae | Zosterops_novaeguineae | LC | Unknown | tropical | diurnal | 12.5 | 103133.525 | 4.127 |
| birds | Zosterops_oleagineus | Zosterops_oleagineus | NT | Decreasing | tropical | diurnal | 10.6 | 97.968 | 16.751 |
| birds | Zosterops_olivaceus | Zosterops_olivaceus | LC | Unknown | tropical | diurnal | 9.1 | 2512.652 | 20.815 |
| birds | Zosterops_pallidus | Zosterops_pallidus | LC | Unknown | temperate | diurnal | 11.1 | 420000.758 | 8.102 |
| birds | Zosterops_palpebrosus | Zosterops_palpebrosus | LC | Decreasing | tropical | diurnal | 8.6 | 6382482.99 | 15.653 |
| birds | Zosterops_poliogastrus | Zosterops_poliogastrus | LC | Unknown | tropical | diurnal | 11 | 324865.264 | 12.11 |
| birds | Zosterops_rendovae | Zosterops_rendovae | LC | Decreasing | tropical | diurnal | 17.2 | 3011.814 | 5.769 |
| birds | Zosterops_rennellianus | Zosterops_rennellianus | LC | Decreasing | tropical | diurnal | 14.5 | 669.686 | 3.726 |
| birds | Zosterops_rotensis | Zosterops_rotensis | CR | Increasing | tropical | diurnal | 10 | 8.177 | 8.296 |
| birds | Zosterops_samoensis | NA | NT | Stable | NA | NA | NA | NA | NA |
| birds | Zosterops_sanctaecrucis | Zosterops_sanctaecrucis | LC | Unknown | tropical | diurnal | 14 | 501.65 | 4.434 |
| birds | Zosterops_semperi | Zosterops_semperi | LC | Unknown | tropical | diurnal | 10.6 | 847.824 | 11.902 |
| birds | Zosterops_senegalensis | Zosterops_senegalensis | LC | Stable | tropical | diurnal | 10.9 | 8737492.268 | 9.838 |
| birds | Zosterops_splendidus | Zosterops_splendidus | LC | Stable | tropical | diurnal | 10.6 | 145.066 | 3.9 |
| birds | Zosterops_stresemanni | Zosterops_stresemanni | LC | Stable | tropical | diurnal | 22 | 4360.573 | 5.293 |
| birds | Zosterops_superciliosus | Woodfordia_superciliosa | LC | Unknown | tropical | diurnal | 29.8 | 675.483 | 3.166 |
| birds | Zosterops_tenuirostris | NA | VU | Decreasing | NA | NA | NA | NA | NA |
| birds | Zosterops_uropygialis | Zosterops_uropygialis | NT | Decreasing | tropical | diurnal | 14 | 493.119 | 16.138 |
| birds | Zosterops_vaughani | NA | LC | Stable | NA | NA | NA | NA | NA |
| birds | Zosterops_vellalavella | Zosterops_vellalavella | NT | Unknown | tropical | diurnal | 10.6 | 673.738 | 3.679 |
| birds | Zosterops_winifredae | NA | NT | Decreasing | NA | NA | NA | NA | NA |
| birds | Zosterops_xanthochroa | Zosterops_xanthochroa | LC | Decreasing | tropical | diurnal | 10.3 | 17202.114 | 7.56 |
| birds | Zosterornis_hypogrammicus | NA | LC | Stable | NA | NA | NA | NA | NA |
| birds | Zosterornis_latistriatus | Stachyris_latistriata | LC | Stable | tropical | diurnal | 29.5 | 316.789 | 3.957 |
| birds | Zosterornis_nigrorum | Stachyris_nigrorum | EN | Decreasing | tropical | diurnal | 21.6 | 468.619 | 11.307 |
| birds | Zosterornis_striatus | Stachyris_striata | LC | Decreasing | tropical | diurnal | 13.9 | 20400.667 | 7.62 |

| | | | | | | | | | |
|---|---|---|---|---|---|---|---|---|---|
| birds | Zosterornis_whiteheadi | Stachyris_whiteheadi | LC | Stable | tropical | diurnal | 21.9 | 90503.908 | 15.018 |
| lizards | Ablepharus_alaicus | NA | LC | Unknown | NA | NA | NA | NA | NA |
| lizards | Ablepharus_bivittatus | Ablepharus_bivittatus | LC | Decreasing | temperate | diurnal | 4.365 | 1124383.734 | 11.313 |
| lizards | Ablepharus_budaki | Ablepharus_budaki | LC | Stable | temperate | diurnal | 1.995 | 121185.752 | 16.489 |
| lizards | Ablepharus_chernovi | Ablepharus_chernovi | LC | Stable | temperate | diurnal | 2.951 | 296009.761 | 14.304 |
| lizards | Ablepharus_darvazi | NA | DD | Unknown | NA | NA | NA | NA | NA |
| lizards | Ablepharus_deserti | Ablepharus_deserti | LC | Decreasing | temperate | diurnal | 3.89 | 889020.937 | 9.73 |
| lizards | Ablepharus_eremchenkoi | NA | LC | Stable | NA | NA | NA | NA | NA |
| lizards | Ablepharus_grayanus | Ablepharus_grayanus | LC | Stable | temperate | diurnal | 0.724 | 230787.607 | 10.792 |
| lizards | Ablepharus_himalayanus | NA | LC | Stable | NA | NA | NA | NA | NA |
| lizards | Ablepharus_kitaibelii | Ablepharus_kitaibelii | LC | Stable | temperate | cathemeral | 4.786 | 713068.086 | 16.782 |
| lizards | Ablepharus_ladacensis | NA | LC | Stable | NA | NA | NA | NA | NA |
| lizards | Ablepharus_lindbergi | Ablepharus_lindbergi | LC | Unknown | temperate | diurnal | 2.291 | 165909.252 | 9.413 |
| lizards | Ablepharus_mahabharatus | NA | DD | Unknown | NA | NA | NA | NA | NA |
| lizards | Ablepharus_nepalensis | NA | DD | Unknown | NA | NA | NA | NA | NA |
| lizards | Ablepharus_pannonicus | Ablepharus_pannonicus | LC | Unknown | temperate | diurnal | 3.09 | 2664129.081 | 9.941 |
| lizards | Ablepharus_rueppellii | Ablepharus_rueppellii | LC | Unknown | temperate | diurnal | 2.63 | 46688.691 | 20.961 |
| lizards | Ablepharus_sikimmensis | NA | LC | Stable | NA | NA | NA | NA | NA |
| lizards | Ablepharus_tragbulensis | NA | DD | Unknown | NA | NA | NA | NA | NA |
| lizards | Abronia_antauges | NA | DD | Unknown | NA | NA | NA | NA | NA |
| lizards | Abronia_anzuetoi | NA | VU | Unknown | NA | NA | NA | NA | NA |
| lizards | Abronia_aurita | Abronia_aurita | EN | Unknown | tropical | diurnal | 33.884 | 4835.247 | 20.711 |
| lizards | Abronia_bogerti | NA | DD | Unknown | NA | NA | NA | NA | NA |
| lizards | Abronia_campbelli | Abronia_campbelli | CR | Decreasing | tropical | diurnal | 36.308 | 124.207 | 12.932 |
| lizards | Abronia_chiszari | NA | EN | Decreasing | NA | NA | NA | NA | NA |
| lizards | Abronia_cuetzpali | NA | not_available | not_available | NA | NA | NA | NA | NA |
| lizards | Abronia_deppii | Abronia_deppii | EN | Decreasing | tropical | diurnal | 30.2 | 4814.261 | 16.625 |
| lizards | Abronia_fimbriata | Abronia_fimbriata | EN | Unknown | tropical | diurnal | 38.905 | 3088.495 | 13.381 |
| lizards | Abronia_frosti | NA | CR | Decreasing | NA | NA | NA | NA | NA |
| lizards | Abronia_gaiophantasma | NA | EN | Unknown | NA | NA | NA | NA | NA |
| lizards | Abronia_graminea | Abronia_graminea | EN | Decreasing | tropical | diurnal | 22.387 | 8507.048 | 15.917 |
| lizards | Abronia_juarezi | NA | EN | Decreasing | NA | NA | NA | NA | NA |
| lizards | Abronia_lythrochila | Abronia_lythrochila | LC | Stable | tropical | diurnal | 29.512 | 8425.632 | 13.104 |
| lizards | Abronia_matudai | NA | EN | Unknown | NA | NA | NA | NA | NA |

| | | | | | | | | | |
|---|---|---|---|---|---|---|---|---|---|
| lizards | Abronia_meledona | Abronia_meledona | EN | Unknown | tropical | diurnal | 29.512 | 19.982 | 14.001 |
| lizards | Abronia_mixteca | Abronia_mixteca | VU | Decreasing | tropical | diurnal | 61.66 | 7624.967 | 7.651 |
| lizards | Abronia_monticola | NA | LC | Stable | NA | NA | NA | NA | NA |
| lizards | Abronia_moreletii | NA | LC | Stable | NA | NA | NA | NA | NA |
| lizards | Abronia_oaxacae | Abronia_oaxacae | VU | Decreasing | tropical | diurnal | 26.915 | 6695.114 | 8.437 |
| lizards | Abronia_ochoterenai | NA | DD | Unknown | NA | NA | NA | NA | NA |
| lizards | Abronia_ornelasi | NA | DD | Unknown | NA | NA | NA | NA | NA |
| lizards | Abronia_ramirezi | NA | DD | Unknown | NA | NA | NA | NA | NA |
| lizards | Abronia_reidi | NA | DD | Unknown | NA | NA | NA | NA | NA |
| lizards | Abronia_salvadorensis | NA | EN | Decreasing | NA | NA | NA | NA | NA |
| lizards | Abronia_smithi | Abronia_smithi | LC | Stable | tropical | diurnal | 21.878 | 1998.075 | 4.774 |
| lizards | Abronia_vasconcelosii | NA | VU | Decreasing | NA | NA | NA | NA | NA |
| lizards | Acanthocercus_adramitanus | Acanthocercus_adramitanus | LC | Stable | tropical | diurnal | 134.896 | 236389.563 | 8.844 |
| lizards | Acanthocercus_annectans | NA | LC | Stable | NA | NA | NA | NA | NA |
| lizards | Acanthocercus_atricollis | Acanthocercus_atricollis | LC | Stable | tropical | diurnal | 177.828 | 6072021.842 | 9.416 |
| lizards | Acanthocercus_branchi | NA | LC | Stable | NA | NA | NA | NA | NA |
| lizards | Acanthocercus_cyanogaster | Acanthocercus_cyanogaster | LC | Unknown | tropical | diurnal | 165.959 | 2095557.485 | 9.956 |
| lizards | Acanthocercus_yemensis | NA | LC | Stable | NA | NA | NA | NA | NA |
| lizards | Acanthodactylus_aegyptius | NA | not_available | not_available | NA | NA | NA | NA | NA |
| lizards | Acanthodactylus_ahmaddisii | NA | EN | Decreasing | NA | NA | NA | NA | NA |
| lizards | Acanthodactylus_arabicus | NA | LC | Stable | NA | NA | NA | NA | NA |
| lizards | Acanthodactylus_aureus | NA | LC | Stable | NA | NA | NA | NA | NA |
| lizards | Acanthodactylus_bedriagai | Acanthodactylus_bedriagai | NT | Unknown | temperate | diurnal | 10.471 | 192655.867 | 11.376 |
| lizards | Acanthodactylus_beershebensis | NA | CR | Decreasing | NA | NA | NA | NA | NA |
| lizards | Acanthodactylus_blanci | Acanthodactylus_blanci | EN | Decreasing | temperate | diurnal | 19.498 | 129571.9 | 14.973 |
| lizards | Acanthodactylus_blanfordii | Acanthodactylus_blanfordii | LC | Stable | temperate | diurnal | 9.772 | 934334.519 | 6.317 |
| lizards | Acanthodactylus_boskianus | Acanthodactylus_boskianus | LC | Stable | temperate | diurnal | 19.498 | 12634604.797 | 4.47 |
| lizards | Acanthodactylus_busacki | NA | LC | Stable | NA | NA | NA | NA | NA |
| lizards | Acanthodactylus_cantoris | Acanthodactylus_cantoris | LC | Unknown | temperate | diurnal | 19.055 | 1416266.46 | 13.941 |
| lizards | Acanthodactylus_dumerilii | Acanthodactylus_dumerilii | LC | Stable | temperate | diurnal | 7.244 | 2618778.627 | 4.094 |
| lizards | Acanthodactylus_erythrurus | Acanthodactylus_erythrurus | LC | Stable | temperate | diurnal | 13.804 | 1030722.73 | 16.154 |
| lizards | Acanthodactylus_felicis | NA | VU | Unknown | NA | NA | NA | NA | NA |
| lizards | Acanthodactylus_gongrorhynchatus | NA | DD | Unknown | NA | NA | NA | NA | NA |
| lizards | Acanthodactylus_grandis | NA | LC | Stable | NA | NA | NA | NA | NA |

| | | | | | | | | | |
|---|---|---|---|---|---|---|---|---|---|
| lizards | Acanthodactylus_guineensis | NA | LC | Stable | NA | NA | NA | NA | NA |
| lizards | Acanthodactylus_haasi | NA | LC | Unknown | NA | NA | NA | NA | NA |
| lizards | Acanthodactylus_hardyi | NA | not_available | not_available | NA | NA | NA | NA | NA |
| lizards | Acanthodactylus_harranensis | NA | CR | Decreasing | NA | NA | NA | NA | NA |
| lizards | Acanthodactylus_khamirensis | NA | not_available | not_available | NA | NA | NA | NA | NA |
| lizards | Acanthodactylus_longipes | Acanthodactylus_longipes | LC | Stable | temperate | diurnal | 6.457 | 7831763.024 | 1.659 |
| lizards | Acanthodactylus_maculatus | Acanthodactylus_maculatus | LC | Unknown | temperate | diurnal | 5.623 | 1193367.151 | 7.595 |
| lizards | Acanthodactylus_margaritae | NA | not_available | not_available | NA | NA | NA | NA | NA |
| lizards | Acanthodactylus_masirae | NA | LC | Stable | NA | NA | NA | NA | NA |
| lizards | Acanthodactylus_micropholis | NA | LC | Stable | NA | NA | NA | NA | NA |
| lizards | Acanthodactylus_nilsoni | NA | DD | Unknown | NA | NA | NA | NA | NA |
| lizards | Acanthodactylus_opheodurus | Acanthodactylus_opheodurus | LC | Stable | temperate | diurnal | 5.623 | 1708910.228 | 5.383 |
| lizards | Acanthodactylus_orientalis | NA | LC | Stable | NA | NA | NA | NA | NA |
| lizards | Acanthodactylus_pardalis | Acanthodactylus_pardalis | VU | Decreasing | temperate | diurnal | 10.471 | 178962.186 | 7.298 |
| lizards | Acanthodactylus_robustus | NA | LC | Unknown | NA | NA | NA | NA | NA |
| lizards | Acanthodactylus_savignyi | Acanthodactylus_savignyi | NT | Unknown | temperate | diurnal | 9.772 | 515134.892 | 9.057 |
| lizards | Acanthodactylus_schmidti | Acanthodactylus_schmidti | LC | Stable | temperate | diurnal | 26.303 | 2034614.222 | 4.584 |
| lizards | Acanthodactylus_schreiberi | Acanthodactylus_schreiberi | EN | Decreasing | temperate | diurnal | 18.621 | 9272.66 | 19.898 |
| lizards | Acanthodactylus_scutellatus | Acanthodactylus_scutellatus | LC | Stable | temperate | diurnal | 14.125 | 8730741.196 | 2.193 |
| lizards | Acanthodactylus_spinicauda | Acanthodactylus_spinicauda | CR | Decreasing | temperate | diurnal | 4.169 | 365.624 | 11.08 |
| lizards | Acanthodactylus_taghitensis | NA | DD | Unknown | NA | NA | NA | NA | NA |
| lizards | Acanthodactylus_tilburyi | NA | LC | Stable | NA | NA | NA | NA | NA |
| lizards | Acanthodactylus_tristrami | Acanthodactylus_tristrami | LC | Unknown | temperate | diurnal | 19.498 | 76447.331 | 14.358 |
| lizards | Acanthodactylus_yemenicus | NA | LC | Unknown | NA | NA | NA | NA | NA |
| lizards | Acanthosaura_armata | Acanthosaura_armata | LC | Stable | tropical | diurnal | 95.499 | 194795.932 | 13.358 |
| lizards | Acanthosaura_bintangensis | NA | DD | Unknown | NA | NA | NA | NA | NA |
| lizards | Acanthosaura_brachypoda | NA | DD | Decreasing | NA | NA | NA | NA | NA |
| lizards | Acanthosaura_capra | Acanthosaura_capra | NT | Unknown | tropical | diurnal | 95.499 | 11351.153 | 8.756 |
| lizards | Acanthosaura_cardamomensis | Acanthosaura_cardamomensis | LC | Stable | tropical | cathemeral | 117.49 | 10121.348 | 4.145 |
| lizards | Acanthosaura_coronata | NA | LC | Unknown | NA | NA | NA | NA | NA |
| lizards | Acanthosaura_crucigera | Acanthosaura_crucigera | LC | Unknown | tropical | diurnal | 128.825 | 234223.495 | 12.729 |
| lizards | Acanthosaura_lepidogaster | Acanthosaura_lepidogaster | LC | Unknown | tropical | diurnal | 45.709 | 2548799.74 | 13.056 |
| lizards | Acanthosaura_nataliae | Acanthosaura_nataliae | LC | Unknown | tropical | diurnal | 138.038 | 56149.662 | 8.745 |
| lizards | Acanthosaura_phuketensis | NA | LC | Stable | NA | NA | NA | NA | NA |

| | | | | | | | | | |
|---|---|---|---|---|---|---|---|---|---|
| lizards | Acanthosaura_titiwangsaensis | NA | EN | Unknown | NA | NA | NA | NA | NA |
| lizards | Acontias_gariepensis | Acontias_gariepensis | LC | Stable | temperate | nocturnal | 5.37 | 51588.12 | 5.565 |
| lizards | Acontias_gracilicauda | Acontias_gracilicauda | LC | Stable | temperate | cathemeral | 22.387 | 314474.122 | 11.723 |
| lizards | Acontias_jappi | Acontias_jappi | LC | Stable | tropical | cathemeral | 10.471 | 26592.965 | 5.283 |
| lizards | Acontias_kgalagadi | Acontias_kgalagadi | LC | Stable | tropical | nocturnal | 10.233 | 643796.403 | 4.57 |
| lizards | Acontias_meleagris | Acontias_meleagris | LC | Stable | temperate | cathemeral | 29.512 | 147662.817 | 9.806 |
| lizards | Acontias_orientalis | Acontias_orientalis | LC | Stable | temperate | nocturnal | 12.882 | 54785.573 | 10.277 |
| lizards | Acontias_plumbeus | Acontias_plumbeus | LC | Stable | temperate | nocturnal | 100 | 316954.366 | 11.815 |
| lizards | Acratosaura_mentalis | Acratosaura_mentalis | LC | Unknown | tropical | diurnal | 4.677 | 603401.121 | 9.716 |
| lizards | Acritoscincus_duperreyi | Bassiana_duperreyi | LC | Unknown | temperate | diurnal | 10.471 | 413900.118 | 10.378 |
| lizards | Acritoscincus_platynotus | Bassiana_platynota | LC | Unknown | temperate | diurnal | 10.471 | 336960.157 | 8.703 |
| lizards | Acritoscincus_trilineatus | Bassiana_trilineata | LC | Stable | temperate | diurnal | 10.471 | 319903.678 | 8.821 |
| lizards | Adolfus_africanus | NA | LC | Stable | NA | NA | NA | NA | NA |
| lizards | Adolfus_alleni | Adolfus_alleni | NT | Stable | tropical | diurnal | 7.943 | 13638.534 | 12.676 |
| lizards | Adolfus_jacksoni | Adolfus_jacksoni | LC | Unknown | tropical | diurnal | 16.596 | 389063.454 | 12.98 |
| lizards | Adolfus_masavaensis | NA | NT | Stable | NA | NA | NA | NA | NA |
| lizards | Advenus_montisilvestris | NA | DD | Unknown | NA | NA | NA | NA | NA |
| lizards | Aeluroscalabotes_felinus | Aeluroscalabotes_felinus | LC | Unknown | tropical | nocturnal | 38.905 | 367109.593 | 5.178 |
| lizards | Afroedura_africana | Afroedura_africana | LC | Stable | tropical | diurnal | 5.495 | 57106.403 | 5.185 |
| lizards | Afroedura_bogerti | Afroedura_bogerti | LC | Stable | tropical | nocturnal | 3.311 | 67804.633 | 7.437 |
| lizards | Afroedura_broadleyi | Afroedura_broadleyi | LC | Stable | tropical | nocturnal | 3.802 | 325.437 | 5.848 |
| lizards | Afroedura_halli | Afroedura_halli | LC | Stable | temperate | nocturnal | 5.623 | 27850.976 | 11.634 |
| lizards | Afroedura_hawequensis | Afroedura_hawequensis | LC | Stable | temperate | diurnal | 11.482 | 12444.669 | 13.134 |
| lizards | Afroedura_karroica | Afroedura_karroica | LC | Stable | temperate | nocturnal | 5.495 | 21238.282 | 8.345 |
| lizards | Afroedura_major | Afroedura_major | LC | Decreasing | temperate | diurnal | 8.913 | 5165.431 | 18.909 |
| lizards | Afroedura_pienaari | Afroedura_pienaari | LC | Stable | tropical | nocturnal | 3.311 | 678.378 | 6.397 |
| lizards | Afroedura_pondolia | Afroedura_pondolia | LC | Stable | temperate | diurnal | 4.266 | 42402.427 | 16.035 |
| lizards | Afrogecko_porphyreus | Afrogecko_porphyreus | LC | Stable | temperate | nocturnal | 2.951 | 86912.029 | 11.478 |
| lizards | Agama_aculeata | Agama_aculeata | LC | Stable | tropical | diurnal | 56.234 | 4075015.476 | 6.906 |
| lizards | Agama_africana | Agama_africana | LC | Stable | tropical | diurnal | 58.884 | 1181582.683 | 12.616 |
| lizards | Agama_agama | Agama_agama | LC | Stable | tropical | diurnal | 117.49 | 3768296.525 | 9.695 |
| lizards | Agama_anchietae | Agama_anchietae | LC | Stable | tropical | diurnal | 95.499 | 875209.894 | 5.294 |
| lizards | Agama_armata | Agama_armata | LC | Stable | tropical | diurnal | 27.542 | 2843147.343 | 9.841 |
| lizards | Agama_atra | Agama_atra | LC | Stable | temperate | diurnal | 95.499 | 1292509.145 | 9.43 |

| | | | | | | | | | |
|---|---|---|---|---|---|---|---|---|---|
| lizards | Agama_bibronii | NA | LC | Stable | NA | NA | NA | NA | NA |
| lizards | Agama_boensis | Agama_boensis | LC | Unknown | tropical | diurnal | 17.378 | 135764.883 | 10.029 |
| lizards | Agama_boueti | Agama_boueti | LC | Stable | tropical | diurnal | 44.668 | 2366175.354 | 5.854 |
| lizards | Agama_boulengeri | NA | LC | Stable | NA | NA | NA | NA | NA |
| lizards | Agama_caudospinosa | Agama_caudospinosa | LC | Unknown | tropical | diurnal | 158.489 | 47508.755 | 14.092 |
| lizards | Agama_cristata | NA | LC | Unknown | NA | NA | NA | NA | NA |
| lizards | Agama_doriae | Agama_doriae | LC | Stable | tropical | diurnal | 51.286 | 2796459.645 | 10.769 |
| lizards | Agama_etoshae | Agama_etoshae | LC | Decreasing | tropical | diurnal | 19.953 | 70851.947 | 6.492 |
| lizards | Agama_finchi | NA | LC | Unknown | NA | NA | NA | NA | NA |
| lizards | Agama_gracilimembris | NA | LC | Stable | NA | NA | NA | NA | NA |
| lizards | Agama_hartmanni | NA | LC | Unknown | NA | NA | NA | NA | NA |
| lizards | Agama_hispida | Agama_hispida | LC | Stable | temperate | diurnal | 83.176 | 286034.832 | 6.905 |
| lizards | Agama_hulbertorum | NA | not_available | not_available | NA | NA | NA | NA | NA |
| lizards | Agama_kaimosae | Agama_kaimosae | LC | Unknown | tropical | diurnal | 104.713 | 32480.06 | 12.704 |
| lizards | Agama_kirkii | Agama_kirkii | LC | Stable | tropical | diurnal | 51.286 | 948984.735 | 11.599 |
| lizards | Agama_knobeli | NA | not_available | not_available | NA | NA | NA | NA | NA |
| lizards | Agama_lanzai | NA | LC | Unknown | NA | NA | NA | NA | NA |
| lizards | Agama_lebretoni | NA | LC | Stable | NA | NA | NA | NA | NA |
| lizards | Agama_lionotus | Agama_lionotus | LC | Stable | tropical | diurnal | 107.152 | 1013804.822 | 9.107 |
| lizards | Agama_montana | Agama_montana | VU | Unknown | tropical | diurnal | 27.542 | 14081.054 | 13.353 |
| lizards | Agama_mossambica | Agama_mossambica | LC | Unknown | tropical | diurnal | 58.884 | 658704.461 | 9.099 |
| lizards | Agama_mwanzae | Agama_mwanzae | LC | Stable | tropical | diurnal | 144.544 | 118000.274 | 16.182 |
| lizards | Agama_parafricana | NA | LC | Unknown | NA | NA | NA | NA | NA |
| lizards | Agama_paragama | NA | LC | Unknown | NA | NA | NA | NA | NA |
| lizards | Agama_persimilis | NA | LC | Unknown | NA | NA | NA | NA | NA |
| lizards | Agama_planiceps | Agama_planiceps | LC | Stable | tropical | diurnal | 112.202 | 778036.552 | 6.569 |
| lizards | Agama_robecchii | NA | LC | Unknown | NA | NA | NA | NA | NA |
| lizards | Agama_rueppelli | NA | LC | Stable | NA | NA | NA | NA | NA |
| lizards | Agama_sankaranica | NA | LC | Unknown | NA | NA | NA | NA | NA |
| lizards | Agama_somalica | NA | LC | Unknown | NA | NA | NA | NA | NA |
| lizards | Agama_spinosa | Agama_spinosa | LC | Stable | tropical | diurnal | 69.183 | 1244916.303 | 9.511 |
| lizards | Agama_tassiliensis | NA | LC | Stable | NA | NA | NA | NA | NA |
| lizards | Agama_weidholzi | Agama_weidholzi | LC | Stable | tropical | diurnal | 8.71 | 437786.783 | 11.616 |
| lizards | Agamura_persica | Agamura_persica | LC | Stable | temperate | nocturnal | 9.333 | 1509180.461 | 6.415 |

| | | | | | | | | | |
|---|---|---|---|---|---|---|---|---|---|
| lizards | Agasthyagama_beddomei | NA | EN | Stable | NA | NA | NA | NA | NA |
| lizards | Ailuronyx_seychellensis | NA | LC | Stable | NA | NA | NA | NA | NA |
| lizards | Ailuronyx_tachyscopaeus | NA | NT | Stable | NA | NA | NA | NA | NA |
| lizards | Ailuronyx_trachygaster | NA | CR | Decreasing | NA | NA | NA | NA | NA |
| lizards | Algyroides_fitzingeri | Algyroides_fitzingeri | LC | Stable | temperate | diurnal | 2.188 | 32877.901 | 16.516 |
| lizards | Algyroides_hidalgoi | NA | EN | Decreasing | NA | NA | NA | NA | NA |
| lizards | Algyroides_moreoticus | Algyroides_moreoticus | NT | Decreasing | temperate | diurnal | 3.02 | 22700.814 | 16.821 |
| lizards | Algyroides_nigropunctatus | Algyroides_nigropunctatus | LC | Unknown | temperate | diurnal | 7.943 | 85186.667 | 17.259 |
| lizards | Alopoglossus_angulatus | Alopoglossus_angulatus | LC | Unknown | tropical | diurnal | 6.166 | 4274943.145 | 1.466 |
| lizards | Alopoglossus_atriventris | Alopoglossus_atriventris | LC | Stable | tropical | diurnal | 3.715 | 1315927.394 | 1.358 |
| lizards | Alopoglossus_bicolor | NA | LC | Unknown | NA | NA | NA | NA | NA |
| lizards | Alopoglossus_brevifrontalis | NA | LC | Unknown | NA | NA | NA | NA | NA |
| lizards | Alopoglossus_buckleyi | Alopoglossus_buckleyi | LC | Unknown | tropical | diurnal | 5.623 | 812923.338 | 1.893 |
| lizards | Alopoglossus_copii | Alopoglossus_copii | LC | Unknown | tropical | diurnal | 12.882 | 144233.427 | 2.13 |
| lizards | Alopoglossus_eurylepis | NA | NT | Unknown | NA | NA | NA | NA | NA |
| lizards | Alopoglossus_festae | NA | LC | Unknown | NA | NA | NA | NA | NA |
| lizards | Alopoglossus_gorgonae | NA | DD | Unknown | NA | NA | NA | NA | NA |
| lizards | Alopoglossus_grandisquamatus | NA | DD | Unknown | NA | NA | NA | NA | NA |
| lizards | Alopoglossus_kugleri | NA | LC | Unknown | NA | NA | NA | NA | NA |
| lizards | Alopoglossus_lehmanni | NA | CR | Decreasing | NA | NA | NA | NA | NA |
| lizards | Alopoglossus_myersi | NA | LC | Stable | NA | NA | NA | NA | NA |
| lizards | Alopoglossus_plicatus | NA | LC | Stable | NA | NA | NA | NA | NA |
| lizards | Alopoglossus_romaleos | NA | DD | Unknown | NA | NA | NA | NA | NA |
| lizards | Alopoglossus_viridiceps | NA | not_available | not_available | NA | NA | NA | NA | NA |
| lizards | Alpinoscincus_alpinus | NA | LC | Unknown | NA | NA | NA | NA | NA |
| lizards | Alpinoscincus_subalpinus | NA | LC | Unknown | NA | NA | NA | NA | NA |
| lizards | Alsophylax_laevis | Alsophylax_laevis | CR | Decreasing | temperate | nocturnal | 1.288 | 193212.437 | 7.627 |
| lizards | Alsophylax_loricatus | Alsophylax_loricatus | VU | Decreasing | temperate | nocturnal | 0.871 | 128811.347 | 12.649 |
| lizards | Alsophylax_pipiens | Alsophylax_pipiens | LC | Stable | temperate | nocturnal | 1.698 | 2852234.856 | 5.11 |
| lizards | Alsophylax_przewalskii | Alsophylax_przewalskii | LC | Stable | temperate | nocturnal | 1.096 | 692562.462 | 2.566 |
| lizards | Alsophylax_szczerbaki | Alsophylax_szczerbaki | VU | Decreasing | temperate | cathemeral | 0.871 | 64182.067 | 8.915 |
| lizards | Alsophylax_tadjikiensis | Alsophylax_tadjikiensis | CR | Unknown | temperate | nocturnal | 0.759 | 68343.237 | 12.543 |
| lizards | Altiphylax_baturensis | NA | DD | Unknown | NA | NA | NA | NA | NA |
| lizards | Altiphylax_levitoni | Altiphylax_levitoni | LC | Unknown | temperate | nocturnal | 2.089 | 25938.062 | 10.727 |

| | | | | | | | | | |
|---|---|---|---|---|---|---|---|---|---|
| lizards | Altiphylax_mintoni | NA | DD | Unknown | NA | NA | NA | NA | NA |
| lizards | Altiphylax_stoliczkai | NA | LC | Stable | NA | NA | NA | NA | NA |
| lizards | Altiphylax_tokobajevi | Altiphylax_tokobajevi | LC | Stable | temperate | nocturnal | 2.754 | 6465.806 | 8.368 |
| lizards | Amalosia_jacovae | Amalosia_jacovae | LC | Unknown | temperate | nocturnal | 4.677 | 72413.465 | 8.421 |
| lizards | Amalosia_lesueurii | Amalosia_lesueurii | LC | Unknown | temperate | nocturnal | 10.233 | 52067.634 | 9.137 |
| lizards | Amalosia_obscura | Amalosia_obscura | LC | Stable | tropical | nocturnal | 4.786 | 28246.582 | 1.131 |
| lizards | Amalosia_rhombifer | Amalosia_rhombifer | LC | Unknown | tropical | nocturnal | 10.233 | 1984790.552 | 3.542 |
| lizards | Amalosia_robusta | NA | LC | Stable | NA | NA | NA | NA | NA |
| lizards | Amapasaurus_tetradactylus | Amapasaurus_tetradactylus | LC | Stable | tropical | diurnal | 0.912 | 72339.455 | 0.598 |
| lizards | Amblyrhynchus_cristatus | NA | VU | Decreasing | NA | NA | NA | NA | NA |
| lizards | Ameiva_aggerecusans | NA | NT | Stable | NA | NA | NA | NA | NA |
| lizards | Ameiva_ameiva | Ameiva_ameiva | LC | Stable | tropical | diurnal | 301.995 | 12706264.26 | 5.409 |
| lizards | Ameiva_atrigularis | Ameiva_atrigularis | LC | Stable | tropical | diurnal | 218.776 | 15371.231 | 14.13 |
| lizards | Ameiva_bifrontata | Ameiva_bifrontata | LC | Stable | tropical | diurnal | 64.565 | 1615919.57 | 5.613 |
| lizards | Ameiva_concolor | Ameiva_concolor | LC | Stable | tropical | diurnal | 64.565 | 565.233 | 9.421 |
| lizards | Ameiva_fuliginosa | NA | not_available | not_available | NA | NA | NA | NA | NA |
| lizards | Ameiva_nodam | NA | NT | Stable | NA | NA | NA | NA | NA |
| lizards | Ameiva_pantherina | NA | DD | Stable | NA | NA | NA | NA | NA |
| lizards | Ameiva_parecis | Ameiva_parecis | VU | Unknown | tropical | diurnal | 21.38 | 1505.572 | 5.406 |
| lizards | Ameiva_praesignis | Ameiva_praesignis | LC | Stable | tropical | diurnal | 467.735 | 240603.028 | 11.214 |
| lizards | Ameiva_provitaae | NA | EN | Decreasing | NA | NA | NA | NA | NA |
| lizards | Ameiva_reticulata | NA | not_available | not_available | NA | NA | NA | NA | NA |
| lizards | Ameivula_apipensis | NA | not_available | not_available | NA | NA | NA | NA | NA |
| lizards | Ameivula_cipoensis | NA | not_available | not_available | NA | NA | NA | NA | NA |
| lizards | Ameivula_confusioniba | NA | LC | Unknown | NA | NA | NA | NA | NA |
| lizards | Ameivula_jalapensis | Ameivula_jalapensis | LC | Unknown | tropical | diurnal | 5.495 | 11017.85 | 6.481 |
| lizards | Ameivula_mumbuca | Ameivula_mumbuca | LC | Unknown | tropical | diurnal | 5.754 | 49042.116 | 6.093 |
| lizards | Ameivula_nigrigula | NA | DD | Unknown | NA | NA | NA | NA | NA |
| lizards | Ameivula_ocellifera | Ameivula_ocellifera | LC | Stable | tropical | diurnal | 52.481 | 7006546.865 | 7.339 |
| lizards | Ameivula_pyrrhogularis | NA | LC | Unknown | NA | NA | NA | NA | NA |
| lizards | Amphibolurus_burnsi | Lophognathus_burnsi | LC | Stable | temperate | diurnal | 85.114 | 816579.804 | 6.428 |
| lizards | Amphibolurus_centralis | NA | not_available | not_available | NA | NA | NA | NA | NA |
| lizards | Amphibolurus_muricatus | Amphibolurus_muricatus | LC | Unknown | temperate | diurnal | 66.069 | 678727.049 | 8.88 |
| lizards | Amphibolurus_norrisi | Amphibolurus_norrisi | LC | Unknown | temperate | diurnal | 54.954 | 264247.622 | 7.148 |

| | | | | | | | | | |
|---|---|---|---|---|---|---|---|---|---|
| lizards | Amphiglossus_astrolabi | Amphiglossus_astrolabi | LC | Decreasing | tropical | diurnal | 301.995 | 128986.634 | 8.812 |
| lizards | Amphiglossus_reticulatus | Amphiglossus_reticulatus | LC | Decreasing | tropical | cathemeral | 275.423 | 75187.822 | 8.392 |
| lizards | Amphisbaena_alba | Amphisbaena_alba | LC | Stable | tropical | cathemeral | 1023.293 | 13781841.241 | 5.796 |
| lizards | Amphisbaena_anomala | NA | LC | Unknown | NA | NA | NA | NA | NA |
| lizards | Amphisbaena_camura | NA | LC | Stable | NA | NA | NA | NA | NA |
| lizards | Amphisbaena_dubia | Amphisbaena_dubia | LC | Unknown | temperate | nocturnal | 13.49 | 150218.379 | 13.728 |
| lizards | Amphisbaena_frontalis | NA | EN | Unknown | NA | NA | NA | NA | NA |
| lizards | Amphisbaena_fuliginosa | NA | LC | Stable | NA | NA | NA | NA | NA |
| lizards | Amphisbaena_gonavensis | NA | CR | Decreasing | NA | NA | NA | NA | NA |
| lizards | Amphisbaena_hastata | NA | LC | Unknown | NA | NA | NA | NA | NA |
| lizards | Amphisbaena_hyporissor | NA | EN | Decreasing | NA | NA | NA | NA | NA |
| lizards | Amphisbaena_manni | NA | LC | Unknown | NA | NA | NA | NA | NA |
| lizards | Amphisbaena_medemi | NA | DD | Unknown | NA | NA | NA | NA | NA |
| lizards | Amphisbaena_pericensis | NA | LC | Unknown | NA | NA | NA | NA | NA |
| lizards | Amphisbaena_slevini | NA | LC | Unknown | NA | NA | NA | NA | NA |
| lizards | Anadia_altaserrania | NA | LC | Unknown | NA | NA | NA | NA | NA |
| lizards | Anadia_bitaeniata | NA | LC | Stable | NA | NA | NA | NA | NA |
| lizards | Anadia_brevifrontalis | Anadia_brevifrontalis | EN | Decreasing | tropical | diurnal | 25.704 | 2502.953 | 13.223 |
| lizards | Anadia_bumanguesa | NA | DD | Unknown | NA | NA | NA | NA | NA |
| lizards | Anadia_escalerae | NA | LC | Unknown | NA | NA | NA | NA | NA |
| lizards | Anadia_hobarti | Anadia_hobarti | LC | Unknown | tropical | diurnal | 16.596 | 9.991 | 13.37 |
| lizards | Anadia_marmorata | NA | LC | Stable | NA | NA | NA | NA | NA |
| lizards | Anadia_ocellata | NA | LC | Unknown | NA | NA | NA | NA | NA |
| lizards | Anadia_rhombifera | Anadia_rhombifera | LC | Unknown | tropical | diurnal | 8.128 | 148126.681 | 14.537 |
| lizards | Anadia_steyeri | NA | LC | Unknown | NA | NA | NA | NA | NA |
| lizards | Anadia_vittata | NA | LC | Unknown | NA | NA | NA | NA | NA |
| lizards | Anatololacerta_anatolica | Anatololacerta_anatolica | LC | Stable | temperate | diurnal | 9.772 | 78409.314 | 16.363 |
| lizards | Anatololacerta_danfordi | Anatololacerta_danfordi | LC | Stable | temperate | diurnal | 9.772 | 56425.42 | 12.371 |
| lizards | Anatololacerta_finikensis | NA | not_available | not_available | NA | NA | NA | NA | NA |
| lizards | Anatololacerta_pelasgiana | NA | not_available | not_available | NA | NA | NA | NA | NA |
| lizards | Ancylodactylus_africanus | NA | LC | Unknown | NA | NA | NA | NA | NA |
| lizards | Ancylodactylus_dickersonae | NA | LC | Unknown | NA | NA | NA | NA | NA |
| lizards | Ancylodactylus_elgonensis | NA | VU | Unknown | NA | NA | NA | NA | NA |
| lizards | Ancylodactylus_gigas | NA | DD | Unknown | NA | NA | NA | NA | NA |

| | | | | | | | | | |
|---|---|---|---|---|---|---|---|---|---|
| lizards | Ancylodactylus_occidentalis | NA | EN | Unknown | NA | NA | NA | NA | NA |
| lizards | Ancylodactylus_petrodroma | NA | DD | Unknown | NA | NA | NA | NA | NA |
| lizards | Ancylodactylus_quattuorseriatus | NA | NT | Unknown | NA | NA | NA | NA | NA |
| lizards | Ancylodactylus_spinicollis | NA | LC | Unknown | NA | NA | NA | NA | NA |
| lizards | Ancylodactylus_uzungwae | NA | DD | Unknown | NA | NA | NA | NA | NA |
| lizards | Andinosaura_afrania | NA | DD | Unknown | NA | NA | NA | NA | NA |
| lizards | Andinosaura_aurea | NA | VU | Unknown | NA | NA | NA | NA | NA |
| lizards | Andinosaura_kiziriani | NA | EN | Unknown | NA | NA | NA | NA | NA |
| lizards | Andinosaura_oculata | Riama_oculata | EN | Unknown | tropical | diurnal | 17.378 | 4971.344 | 9.81 |
| lizards | Andinosaura_stellae | NA | DD | Unknown | NA | NA | NA | NA | NA |
| lizards | Andinosaura_vespertina | Riama_vespertina | VU | Unknown | tropical | diurnal | 7.413 | 9387.896 | 10.288 |
| lizards | Anepischetosia_maccoyi | Anepischetosia_maccoyi | LC | Unknown | temperate | cathemeral | 4.169 | 179855.275 | 10.313 |
| lizards | Anguis_cephallonica | Anguis_cephallonica | NT | Decreasing | temperate | diurnal | 75.858 | 22293.59 | 16.75 |
| lizards | Anguis_colchica | NA | not_available | not_available | NA | NA | NA | NA | NA |
| lizards | Anguis_fragilis | Anguis_fragilis | LC | Stable | temperate | diurnal | 79.433 | 2517204.348 | 20.848 |
| lizards | Anguis_graeca | NA | not_available | not_available | NA | NA | NA | NA | NA |
| lizards | Anguis_veronensis | NA | not_available | not_available | NA | NA | NA | NA | NA |
| lizards | Anniella_geronimensis | Anniella_geronimensis | EN | Decreasing | temperate | diurnal | 7.762 | 2929.912 | 5.745 |
| lizards | Anniella_pulchra | Anniella_pulchra | LC | Decreasing | temperate | cathemeral | 9.55 | 135562.361 | 15.114 |
| lizards | Anolis_acutus | Anolis_acutus | LC | Stable | tropical | diurnal | 6.31 | 193.129 | 29.374 |
| lizards | Anolis_aeneus | Anolis_aeneus | LC | Unknown | tropical | diurnal | 10.471 | 421.598 | 20.177 |
| lizards | Anolis_aequatorialis | Anolis_aequatorialis | LC | Stable | tropical | diurnal | 34.674 | 22401.358 | 6.721 |
| lizards | Anolis_agassizi | NA | LC | Stable | NA | NA | NA | NA | NA |
| lizards | Anolis_agueroi | NA | EN | Unknown | NA | NA | NA | NA | NA |
| lizards | Anolis_ahli | NA | EN | Unknown | NA | NA | NA | NA | NA |
| lizards | Anolis_alayoni | NA | LC | Stable | NA | NA | NA | NA | NA |
| lizards | Anolis_alfaroi | NA | DD | Unknown | NA | NA | NA | NA | NA |
| lizards | Anolis_aliniger | Anolis_aliniger | LC | Unknown | tropical | diurnal | 4.571 | 23061.525 | 13.442 |
| lizards | Anolis_allisoni | Anolis_allisoni | LC | Increasing | tropical | diurnal | 20.417 | 64972.75 | 16.727 |
| lizards | Anolis_allogus | Anolis_allogus | LC | Stable | tropical | diurnal | 5.248 | 77077.827 | 16.497 |
| lizards | Anolis_alocomyos | NA | not_available | not_available | NA | NA | NA | NA | NA |
| lizards | Anolis_altae | NA | LC | Stable | NA | NA | NA | NA | NA |
| lizards | Anolis_altavelensis | NA | CR | Unknown | NA | NA | NA | NA | NA |
| lizards | Anolis_altitudinalis | Anolis_altitudinalis | NT | Unknown | tropical | diurnal | 2.951 | 16.608 | 6.199 |

| | | | | | | | | | |
|---|---|---|---|---|---|---|---|---|---|
| lizards | Anolis_alumina | Anolis_alumina | NT | Stable | tropical | diurnal | 1.38 | 3921.83 | 10.853 |
| lizards | Anolis_alutaceus | Anolis_alutaceus | LC | Unknown | tropical | diurnal | 2.344 | 57368.135 | 16.611 |
| lizards | Anolis_alvarezdeltoroi | NA | DD | Unknown | NA | NA | NA | NA | NA |
| lizards | Anolis_amplisquamosus | NA | CR | Decreasing | NA | NA | NA | NA | NA |
| lizards | Anolis_anatoloros | NA | LC | Unknown | NA | NA | NA | NA | NA |
| lizards | Anolis_anchicayae | NA | LC | Unknown | NA | NA | NA | NA | NA |
| lizards | Anolis_anfiloquioi | NA | LC | Stable | NA | NA | NA | NA | NA |
| lizards | Anolis_angusticeps | Anolis_angusticeps | LC | Stable | tropical | diurnal | 3.162 | 100327.618 | 15.879 |
| lizards | Anolis_anisolepis | Anolis_anisolepis | LC | Stable | tropical | diurnal | 3.715 | 4345.909 | 11.644 |
| lizards | Anolis_annectens | NA | LC | Stable | NA | NA | NA | NA | NA |
| lizards | Anolis_antonii | NA | LC | Stable | NA | NA | NA | NA | NA |
| lizards | Anolis_apletolepis | NA | not_available | not_available | NA | NA | NA | NA | NA |
| lizards | Anolis_apletophallus | NA | LC | Stable | NA | NA | NA | NA | NA |
| lizards | Anolis_aquaticus | Anolis_aquaticus | LC | Stable | tropical | diurnal | 9.55 | 11011.775 | 14.364 |
| lizards | Anolis_argenteolus | Anolis_argenteolus | LC | Stable | tropical | diurnal | 5.754 | 32510.032 | 16.632 |
| lizards | Anolis_argillaceus | NA | LC | Stable | NA | NA | NA | NA | NA |
| lizards | Anolis_armouri | Anolis_armouri | EN | Unknown | tropical | diurnal | 8.511 | 4997.504 | 12.168 |
| lizards | Anolis_auratus | Anolis_auratus | LC | Stable | tropical | diurnal | 3.89 | 3453866.791 | 4.576 |
| lizards | Anolis_aurifer | NA | not_available | not_available | NA | NA | NA | NA | NA |
| lizards | Anolis_bahorucoensis | Anolis_bahorucoensis | VU | Decreasing | tropical | diurnal | 2.818 | 2842.226 | 11.158 |
| lizards | Anolis_baleatus | Anolis_baleatus | LC | Stable | tropical | diurnal | 114.815 | 23168.303 | 17.094 |
| lizards | Anolis_baracoae | Anolis_baracoae | NT | Stable | tropical | diurnal | 100 | 1052.442 | 12.422 |
| lizards | Anolis_barahonae | Anolis_barahonae | EN | Unknown | tropical | diurnal | 100 | 1713.683 | 9.155 |
| lizards | Anolis_barbatus | Anolis_barbatus | NT | Unknown | tropical | diurnal | 97.724 | 537.65 | 12.62 |
| lizards | Anolis_barbouri | Anolis_barbouri | VU | Unknown | tropical | diurnal | 3.548 | 7797.017 | 12.404 |
| lizards | Anolis_barkeri | Anolis_barkeri | VU | Decreasing | tropical | diurnal | 20.893 | 36541.563 | 11.68 |
| lizards | Anolis_bartschi | Anolis_bartschi | LC | Stable | tropical | diurnal | 10.471 | 1318.988 | 13.925 |
| lizards | Anolis_beckeri | Anolis_beckeri | LC | Stable | tropical | diurnal | 6.607 | 180350.65 | 9.001 |
| lizards | Anolis_bellipeniculus | NA | LC | Unknown | NA | NA | NA | NA | NA |
| lizards | Anolis_benedikti | NA | EN | Decreasing | NA | NA | NA | NA | NA |
| lizards | Anolis_bicaorum | Anolis_bicaorum | CR | Stable | tropical | diurnal | 13.183 | 42.084 | 9.659 |
| lizards | Anolis_bimaculatus | Anolis_bimaculatus | LC | Stable | tropical | diurnal | 39.811 | 298.224 | 24.198 |
| lizards | Anolis_biporcatus | Anolis_biporcatus | LC | Stable | tropical | diurnal | 30.903 | 650952.07 | 10.066 |
| lizards | Anolis_birama | NA | EN | Unknown | NA | NA | NA | NA | NA |

| | | | | | | | | | |
|---|---|---|---|---|---|---|---|---|---|
| lizards | Anolis_bitectus | Anolis_bitectus | EN | Decreasing | tropical | diurnal | 3.802 | 15453.422 | 16.205 |
| lizards | Anolis_blanquillanus | NA | LC | Stable | NA | NA | NA | NA | NA |
| lizards | Anolis_bombiceps | Anolis_bombiceps | LC | Unknown | tropical | diurnal | 8.511 | 371682.206 | 1.204 |
| lizards | Anolis_bonairensis | Anolis_bonairensis | VU | Stable | tropical | diurnal | 8.71 | 244.262 | 18.344 |
| lizards | Anolis_boulengerianus | NA | DD | Unknown | NA | NA | NA | NA | NA |
| lizards | Anolis_brasiliensis | Anolis_brasiliensis | LC | Unknown | tropical | diurnal | 8.71 | 1239579.558 | 7.069 |
| lizards | Anolis_bremeri | Anolis_bremeri | LC | Unknown | tropical | diurnal | 7.762 | 5260.159 | 16.254 |
| lizards | Anolis_breslini | NA | EN | Decreasing | NA | NA | NA | NA | NA |
| lizards | Anolis_brevirostris | Anolis_brevirostris | LC | Stable | tropical | diurnal | 3.162 | 15142.285 | 13.765 |
| lizards | Anolis_brooksi | NA | not_available | not_available | NA | NA | NA | NA | NA |
| lizards | Anolis_campbelli | NA | EN | Unknown | NA | NA | NA | NA | NA |
| lizards | Anolis_capito | Anolis_capito | LC | Stable | tropical | diurnal | 21.878 | 278977.933 | 9.72 |
| lizards | Anolis_caquetae | NA | DD | Unknown | NA | NA | NA | NA | NA |
| lizards | Anolis_carlliebi | NA | not_available | not_available | NA | NA | NA | NA | NA |
| lizards | Anolis_carlostoddi | NA | LC | Unknown | NA | NA | NA | NA | NA |
| lizards | Anolis_carolinensis | Anolis_carolinensis | LC | Stable | temperate | diurnal | 8.71 | 1275289.21 | 16.331 |
| lizards | Anolis_carpenteri | Anolis_carpenteri | LC | Unknown | tropical | diurnal | 2.239 | 39659.491 | 11.096 |
| lizards | Anolis_casildae | NA | EN | Unknown | NA | NA | NA | NA | NA |
| lizards | Anolis_caudalis | NA | LC | Stable | NA | NA | NA | NA | NA |
| lizards | Anolis_centralis | Anolis_centralis | LC | Stable | tropical | diurnal | 2.344 | 40134.292 | 16.592 |
| lizards | Anolis_chamaeleonides | Anolis_chamaeleonides | LC | Unknown | tropical | diurnal | 109.648 | 18057.055 | 16.725 |
| lizards | Anolis_chloris | Anolis_chloris | LC | Unknown | tropical | diurnal | 5.012 | 99568.492 | 6.798 |
| lizards | Anolis_chlorocyanus | Anolis_chlorocyanus | LC | Unknown | tropical | diurnal | 10.471 | 61926.908 | 15.638 |
| lizards | Anolis_chlorodius | NA | not_available | not_available | NA | NA | NA | NA | NA |
| lizards | Anolis_christophei | Anolis_christophei | NT | Decreasing | tropical | diurnal | 2.63 | 9856.012 | 17.016 |
| lizards | Anolis_chrysolepis | Anolis_chrysolepis | LC | Stable | tropical | diurnal | 8.511 | 5296831.931 | 3.031 |
| lizards | Anolis_chrysops | Anolis_chrysops | VU | Unknown | tropical | diurnal | 8.128 | 1.116 | 0.251 |
| lizards | Anolis_clivicola | Anolis_clivicola | NT | Unknown | tropical | diurnal | 2.57 | 1153.761 | 9.539 |
| lizards | Anolis_compressicauda | NA | LC | Stable | NA | NA | NA | NA | NA |
| lizards | Anolis_concolor | Anolis_concolor | LC | Stable | tropical | diurnal | 10.471 | 51.485 | 27.482 |
| lizards | Anolis_confusus | NA | EN | Stable | NA | NA | NA | NA | NA |
| lizards | Anolis_conspersus | Anolis_conspersus | LC | Stable | tropical | diurnal | 9.12 | 203.033 | 23.367 |
| lizards | Anolis_cooki | Anolis_cooki | EN | Decreasing | tropical | diurnal | 8.128 | 258.4 | 23.547 |
| lizards | Anolis_crassulus | Anolis_crassulus | LC | Stable | tropical | diurnal | 4.365 | 51762.53 | 15.041 |

| | | | | | | | | | |
|---|---|---|---|---|---|---|---|---|---|
| lizards | Anolis_cristatellus | Anolis_cristatellus | LC | Stable | tropical | diurnal | 10.471 | 9147.172 | 26.916 |
| lizards | Anolis_cryptolimifrons | NA | LC | Stable | NA | NA | NA | NA | NA |
| lizards | Anolis_cupeyalensis | NA | LC | Stable | NA | NA | NA | NA | NA |
| lizards | Anolis_cupreus | Anolis_cupreus | LC | Stable | tropical | diurnal | 3.89 | 150610.189 | 9.997 |
| lizards | Anolis_cuprinus | Anolis_cuprinus | LC | Stable | tropical | diurnal | 6.918 | 7965.569 | 9.707 |
| lizards | Anolis_cusuco | NA | CR | Decreasing | NA | NA | NA | NA | NA |
| lizards | Anolis_cuvieri | Anolis_cuvieri | LC | Stable | tropical | diurnal | 114.815 | 3890.573 | 26.657 |
| lizards | Anolis_cyanopleurus | NA | LC | Stable | NA | NA | NA | NA | NA |
| lizards | Anolis_cyanostictus | NA | not_available | not_available | NA | NA | NA | NA | NA |
| lizards | Anolis_cybotes | Anolis_cybotes | LC | Stable | tropical | diurnal | 10.965 | 70110.556 | 15.736 |
| lizards | Anolis_darlingtoni | NA | CR | Decreasing | NA | NA | NA | NA | NA |
| lizards | Anolis_delafuentei | NA | DD | Unknown | NA | NA | NA | NA | NA |
| lizards | Anolis_deltae | NA | DD | Unknown | NA | NA | NA | NA | NA |
| lizards | Anolis_desechensis | NA | LC | Unknown | NA | NA | NA | NA | NA |
| lizards | Anolis_desiradei | Anolis_desiradei | LC | Unknown | tropical | diurnal | 10.471 | 23.039 | 15.263 |
| lizards | Anolis_distichus | Anolis_distichus | LC | Stable | temperate | diurnal | 3.162 | 8963.129 | 8.091 |
| lizards | Anolis_divius | NA | not_available | not_available | NA | NA | NA | NA | NA |
| lizards | Anolis_dolichocephalus | NA | EN | Decreasing | NA | NA | NA | NA | NA |
| lizards | Anolis_dollfusianus | Anolis_dollfusianus | LC | Stable | tropical | diurnal | 1.82 | 2134.476 | 15.768 |
| lizards | Anolis_dominicensis | NA | not_available | not_available | NA | NA | NA | NA | NA |
| lizards | Anolis_duellmani | NA | DD | Unknown | NA | NA | NA | NA | NA |
| lizards | Anolis_dunni | Anolis_dunni | LC | Unknown | tropical | diurnal | 4.169 | 26633.779 | 8.856 |
| lizards | Anolis_eladioi | NA | not_available | not_available | NA | NA | NA | NA | NA |
| lizards | Anolis_elcopeensis | NA | not_available | not_available | NA | NA | NA | NA | NA |
| lizards | Anolis_equestris | Anolis_equestris | LC | Unknown | tropical | diurnal | 134.896 | 60238.858 | 17.462 |
| lizards | Anolis_ernestwilliamsi | Anolis_ernestwilliamsi | CR | Unknown | tropical | diurnal | 11.482 | 5.888 | 5.203 |
| lizards | Anolis_etheridgei | Anolis_etheridgei | VU | Unknown | tropical | diurnal | 1.82 | 1555.785 | 12.152 |
| lizards | Anolis_eugenegrahami | Anolis_eugenegrahami | CR | Decreasing | tropical | diurnal | 7.762 | 66.21 | 12.531 |
| lizards | Anolis_euskalerriari | NA | NT | Decreasing | NA | NA | NA | NA | NA |
| lizards | Anolis_evermanni | Anolis_evermanni | LC | Unknown | tropical | diurnal | 9.772 | 3511.443 | 25.809 |
| lizards | Anolis_extremus | Anolis_extremus | LC | Unknown | tropical | diurnal | 12.589 | 439.348 | 31.806 |
| lizards | Anolis_fasciatus | Anolis_fasciatus | LC | Stable | tropical | diurnal | 7.762 | 39215.845 | 13.908 |
| lizards | Anolis_favillarum | NA | not_available | not_available | NA | NA | NA | NA | NA |
| lizards | Anolis_ferreus | NA | LC | Stable | NA | NA | NA | NA | NA |

| | | | | | | | | | |
|---|---|---|---|---|---|---|---|---|---|
| lizards | Anolis_festae | Anolis_festae | NT | Decreasing | tropical | diurnal | 4.786 | 46452.957 | 15.665 |
| lizards | Anolis_fitchi | NA | LC | Unknown | NA | NA | NA | NA | NA |
| lizards | Anolis_forresti | NA | not_available | not_available | NA | NA | NA | NA | NA |
| lizards | Anolis_fowleri | Anolis_fowleri | EN | Unknown | tropical | diurnal | 9.772 | 148.314 | 12.046 |
| lizards | Anolis_fraseri | NA | LC | Unknown | NA | NA | NA | NA | NA |
| lizards | Anolis_frenatus | Anolis_frenatus | LC | Unknown | tropical | diurnal | 69.183 | 175594.615 | 9.631 |
| lizards | Anolis_fugitivus | NA | NT | Stable | NA | NA | NA | NA | NA |
| lizards | Anolis_fungosus | NA | NT | Unknown | NA | NA | NA | NA | NA |
| lizards | Anolis_fuscoauratus | Anolis_fuscoauratus | LC | Stable | tropical | diurnal | 3.162 | 9192893.898 | 3.798 |
| lizards | Anolis_gadovii | Anolis_gadovii | LC | Stable | tropical | diurnal | 10.471 | 3092.48 | 13.493 |
| lizards | Anolis_gaigei | Anolis_gaigei | LC | Stable | tropical | diurnal | 3.09 | 63716.664 | 13.173 |
| lizards | Anolis_garmani | Anolis_garmani | LC | Stable | tropical | diurnal | 52.481 | 9880.466 | 19.578 |
| lizards | Anolis_garridoi | NA | EN | Unknown | NA | NA | NA | NA | NA |
| lizards | Anolis_gemmosus | Anolis_gemmosus | LC | Unknown | tropical | diurnal | 6.026 | 21456.828 | 12.402 |
| lizards | Anolis_ginaelisae | NA | not_available | not_available | NA | NA | NA | NA | NA |
| lizards | Anolis_gingivinus | Anolis_gingivinus | LC | Unknown | tropical | diurnal | 7.762 | 200.332 | 28.678 |
| lizards | Anolis_gonavensis | NA | not_available | not_available | NA | NA | NA | NA | NA |
| lizards | Anolis_gorgonae | NA | NT | Decreasing | NA | NA | NA | NA | NA |
| lizards | Anolis_gracilipes | NA | LC | Stable | NA | NA | NA | NA | NA |
| lizards | Anolis_grahami | Anolis_grahami | LC | Stable | tropical | diurnal | 8.71 | 11021.537 | 19.74 |
| lizards | Anolis_granuliceps | NA | LC | Unknown | NA | NA | NA | NA | NA |
| lizards | Anolis_griseus | Anolis_griseus | LC | Stable | tropical | diurnal | 50.119 | 324.73 | 16.473 |
| lizards | Anolis_gruuo | NA | EN | Unknown | NA | NA | NA | NA | NA |
| lizards | Anolis_guafe | NA | EN | Unknown | NA | NA | NA | NA | NA |
| lizards | Anolis_guamuhaya | NA | EN | Unknown | NA | NA | NA | NA | NA |
| lizards | Anolis_guazuma | NA | LC | Unknown | NA | NA | NA | NA | NA |
| lizards | Anolis_gundlachi | Anolis_gundlachi | LC | Unknown | tropical | diurnal | 9.12 | 3942.499 | 24.962 |
| lizards | Anolis_haguei | NA | NT | Unknown | NA | NA | NA | NA | NA |
| lizards | Anolis_hendersoni | NA | VU | Decreasing | NA | NA | NA | NA | NA |
| lizards | Anolis_heterodermus | Anolis_heterodermus | LC | Stable | tropical | diurnal | 13.183 | 94840.934 | 14.208 |
| lizards | Anolis_heteropholidotus | NA | LC | Stable | NA | NA | NA | NA | NA |
| lizards | Anolis_hobartsmithi | NA | EN | Decreasing | NA | NA | NA | NA | NA |
| lizards | Anolis_homolechis | Anolis_homolechis | LC | Stable | tropical | diurnal | 7.079 | 88747.995 | 16.695 |
| lizards | Anolis_humilis | Anolis_humilis | LC | Stable | tropical | diurnal | 2.63 | 72019.78 | 11.796 |

| | | | | | | | | | |
|---|---|---|---|---|---|---|---|---|---|
| lizards | Anolis_hyacinthogularis | NA | not_available | not_available | NA | NA | NA | NA | NA |
| lizards | Anolis_ibague | NA | DD | Unknown | NA | NA | NA | NA | NA |
| lizards | Anolis_ignigularis | NA | not_available | not_available | NA | NA | NA | NA | NA |
| lizards | Anolis_imias | NA | LC | Unknown | NA | NA | NA | NA | NA |
| lizards | Anolis_immaculogularis | NA | not_available | not_available | NA | NA | NA | NA | NA |
| lizards | Anolis_inexpectatus | NA | EN | Unknown | NA | NA | NA | NA | NA |
| lizards | Anolis_insignis | NA | LC | Unknown | NA | NA | NA | NA | NA |
| lizards | Anolis_insolitus | Anolis_insolitus | EN | Unknown | tropical | diurnal | 2.239 | 677.752 | 11.94 |
| lizards | Anolis_isolepis | Anolis_isolepis | LC | Unknown | tropical | diurnal | 1.585 | 13567.019 | 16.23 |
| lizards | Anolis_jacare | Anolis_jacare | LC | Stable | tropical | diurnal | 8.511 | 31227.575 | 12.379 |
| lizards | Anolis_johnmeyeri | Anolis_johnmeyeri | EN | Decreasing | tropical | diurnal | 8.128 | 300.869 | 8.769 |
| lizards | Anolis_juangundlachi | NA | CR | Unknown | NA | NA | NA | NA | NA |
| lizards | Anolis_jubar | Anolis_jubar | LC | Stable | tropical | diurnal | 5.012 | 44811.352 | 16.103 |
| lizards | Anolis_kahouannensis | NA | NT | Unknown | NA | NA | NA | NA | NA |
| lizards | Anolis_kathydayae | NA | not_available | not_available | NA | NA | NA | NA | NA |
| lizards | Anolis_kemptoni | Anolis_kemptoni | LC | Stable | tropical | diurnal | 4.571 | 743.196 | 8.932 |
| lizards | Anolis_koopmani | NA | CR | Decreasing | NA | NA | NA | NA | NA |
| lizards | Anolis_kreutzi | NA | CR | Decreasing | NA | NA | NA | NA | NA |
| lizards | Anolis_krugi | Anolis_krugi | LC | Stable | tropical | diurnal | 3.548 | 5077.087 | 24.949 |
| lizards | Anolis_laeviventris | Anolis_laeviventris | LC | Stable | tropical | diurnal | 6.026 | 166442.109 | 11.709 |
| lizards | Anolis_lamari | NA | DD | Unknown | NA | NA | NA | NA | NA |
| lizards | Anolis_landestoyi | NA | not_available | not_available | NA | NA | NA | NA | NA |
| lizards | Anolis_latifrons | NA | LC | Stable | NA | NA | NA | NA | NA |
| lizards | Anolis_leachii | Anolis_leachii | LC | Unknown | tropical | diurnal | 37.154 | 322.468 | 27.18 |
| lizards | Anolis_leditzigorum | NA | not_available | not_available | NA | NA | NA | NA | NA |
| lizards | Anolis_lemurinus | Anolis_lemurinus | LC | Stable | tropical | diurnal | 10.233 | 569991.687 | 8.577 |
| lizards | Anolis_leucodera | NA | not_available | not_available | NA | NA | NA | NA | NA |
| lizards | Anolis_limifrons | Anolis_limifrons | LC | Unknown | tropical | diurnal | 2.818 | 208940.287 | 9.213 |
| lizards | Anolis_limon | NA | not_available | not_available | NA | NA | NA | NA | NA |
| lizards | Anolis_lineatopus | Anolis_lineatopus | LC | Stable | tropical | diurnal | 8.128 | 10925.386 | 19.669 |
| lizards | Anolis_lineatus | NA | NT | Decreasing | NA | NA | NA | NA | NA |
| lizards | Anolis_liogaster | Anolis_liogaster | LC | Stable | tropical | diurnal | 6.026 | 4209.27 | 5.028 |
| lizards | Anolis_lionotus | Anolis_lionotus | LC | Stable | tropical | diurnal | 9.772 | 55747.178 | 11.593 |
| lizards | Anolis_litoralis | Anolis_litoralis | LC | Stable | tropical | diurnal | 2.344 | 7433.593 | 18.532 |

| | | | | | | | | | |
|---|---|---|---|---|---|---|---|---|---|
| lizards | Anolis_lividus | Anolis_lividus | NT | Decreasing | tropical | diurnal | 11.482 | 96.566 | 12.512 |
| lizards | Anolis_longiceps | NA | LC | Stable | NA | NA | NA | NA | NA |
| lizards | Anolis_longitibialis | Anolis_longitibialis | EN | Unknown | tropical | diurnal | 7.762 | 1259.238 | 9.776 |
| lizards | Anolis_lososi | NA | not_available | not_available | NA | NA | NA | NA | NA |
| lizards | Anolis_loveridgei | Anolis_loveridgei | EN | Decreasing | tropical | diurnal | 33.113 | 3203.255 | 5.644 |
| lizards | Anolis_loysiana | Anolis_loysiana | LC | Unknown | tropical | diurnal | 2.951 | 5524.16 | 21.236 |
| lizards | Anolis_luciae | Anolis_luciae | EN | Decreasing | tropical | diurnal | 15.488 | 617.396 | 20.77 |
| lizards | Anolis_lucius | Anolis_lucius | LC | Stable | tropical | diurnal | 7.079 | 36088.103 | 18.111 |
| lizards | Anolis_luteogularis | Anolis_luteogularis | LC | Unknown | tropical | diurnal | 138.038 | 20864.645 | 15.467 |
| lizards | Anolis_lyra | Anolis_lyra | LC | Unknown | tropical | diurnal | 9.333 | 86778.798 | 5.415 |
| lizards | Anolis_macilentus | NA | DD | Unknown | NA | NA | NA | NA | NA |
| lizards | Anolis_macrinii | Anolis_macrinii | LC | Stable | tropical | diurnal | 18.197 | 166.262 | 10.232 |
| lizards | Anolis_macrolepis | NA | NT | Unknown | NA | NA | NA | NA | NA |
| lizards | Anolis_macrophallus | NA | LC | Stable | NA | NA | NA | NA | NA |
| lizards | Anolis_maculiventris | Anolis_maculiventris | LC | Stable | tropical | diurnal | 2.63 | 166970.536 | 10.085 |
| lizards | Anolis_magnaphallus | NA | LC | Stable | NA | NA | NA | NA | NA |
| lizards | Anolis_marcanoi | NA | NT | Unknown | NA | NA | NA | NA | NA |
| lizards | Anolis_mariarum | Anolis_mariarum | LC | Stable | tropical | diurnal | 3.715 | 34597.681 | 13.412 |
| lizards | Anolis_marmoratus | Anolis_marmoratus | LC | Stable | tropical | diurnal | 12.589 | 1510.931 | 24.241 |
| lizards | Anolis_marron | Anolis_marron | LC | Stable | tropical | diurnal | 9.12 | 354.79 | 17.547 |
| lizards | Anolis_marsupialis | NA | not_available | not_available | NA | NA | NA | NA | NA |
| lizards | Anolis_matudai | Anolis_matudai | LC | Stable | tropical | cathemeral | 3.548 | 11656.571 | 7.524 |
| lizards | Anolis_maynardii | NA | LC | Stable | NA | NA | NA | NA | NA |
| lizards | Anolis_mccraniei | NA | not_available | not_available | NA | NA | NA | NA | NA |
| lizards | Anolis_medemi | NA | LC | Stable | NA | NA | NA | NA | NA |
| lizards | Anolis_megalopithecus | NA | LC | Unknown | NA | NA | NA | NA | NA |
| lizards | Anolis_megapholidotus | Anolis_megapholidotus | LC | Stable | tropical | diurnal | 3.162 | 6213.572 | 9.27 |
| lizards | Anolis_menta | Anolis_menta | NT | Unknown | tropical | diurnal | 3.715 | 9.991 | 8.666 |
| lizards | Anolis_meridionalis | Anolis_meridionalis | LC | Stable | tropical | diurnal | 4.365 | 3299660.692 | 7.093 |
| lizards | Anolis_mestrei | Anolis_mestrei | LC | Unknown | tropical | diurnal | 4.571 | 3354.04 | 17.647 |
| lizards | Anolis_microlepidotus | Anolis_microlepidotus | LC | Stable | tropical | diurnal | 3.311 | 17303.266 | 9.09 |
| lizards | Anolis_milleri | NA | DD | Unknown | NA | NA | NA | NA | NA |
| lizards | Anolis_monensis | NA | LC | Stable | NA | NA | NA | NA | NA |
| lizards | Anolis_monteverde | NA | CR | Decreasing | NA | NA | NA | NA | NA |

| | | | | | | | | | |
|---|---|---|---|---|---|---|---|---|---|
| lizards | Anolis_monticola | NA | VU | Decreasing | NA | NA | NA | NA | NA |
| lizards | Anolis_morazani | NA | CR | Unknown | NA | NA | NA | NA | NA |
| lizards | Anolis_muralla | NA | CR | Decreasing | NA | NA | NA | NA | NA |
| lizards | Anolis_nasofrontalis | NA | EN | Unknown | NA | NA | NA | NA | NA |
| lizards | Anolis_naufragus | Anolis_naufragus | VU | Decreasing | tropical | diurnal | 4.571 | 8292.829 | 13.387 |
| lizards | Anolis_nebuloides | Anolis_nebuloides | LC | Stable | tropical | diurnal | 3.631 | 36060.502 | 7.344 |
| lizards | Anolis_nebulosus | Anolis_nebulosus | LC | Stable | tropical | diurnal | 4.365 | 575815.859 | 10.219 |
| lizards | Anolis_nelsoni | NA | CR | Decreasing | NA | NA | NA | NA | NA |
| lizards | Anolis_nietoi | NA | not_available | not_available | NA | NA | NA | NA | NA |
| lizards | Anolis_nigrolineatus | NA | DD | Unknown | NA | NA | NA | NA | NA |
| lizards | Anolis_noblei | Anolis_noblei | NT | Unknown | tropical | diurnal | 134.896 | 11591.647 | 15.367 |
| lizards | Anolis_notopholis | NA | LC | Stable | NA | NA | NA | NA | NA |
| lizards | Anolis_nubilus | NA | CR | Unknown | NA | NA | NA | NA | NA |
| lizards | Anolis_occultus | NA | LC | Stable | NA | NA | NA | NA | NA |
| lizards | Anolis_ocelloscapularis | NA | VU | Unknown | NA | NA | NA | NA | NA |
| lizards | Anolis_oculatus | Anolis_oculatus | NT | Decreasing | tropical | diurnal | 19.055 | 766.69 | 13.545 |
| lizards | Anolis_olssoni | Anolis_olssoni | LC | Unknown | tropical | diurnal | 3.162 | 46744.662 | 14.505 |
| lizards | Anolis_omiltemanus | Anolis_omiltemanus | LC | Stable | tropical | diurnal | 2.344 | 2889.196 | 6.904 |
| lizards | Anolis_onca | Anolis_onca | LC | Stable | tropical | diurnal | 15.849 | 25740.675 | 13.014 |
| lizards | Anolis_opalinus | Anolis_opalinus | LC | Stable | tropical | diurnal | 3.715 | 10293.324 | 19.499 |
| lizards | Anolis_ophiolepis | Anolis_ophiolepis | LC | Unknown | tropical | diurnal | 1.349 | 14915.435 | 18.071 |
| lizards | Anolis_oporinus | NA | DD | Unknown | NA | NA | NA | NA | NA |
| lizards | Anolis_ortonii | Anolis_ortonii | LC | Stable | tropical | diurnal | 4.571 | 5357221.04 | 1.948 |
| lizards | Anolis_osa | NA | LC | Stable | NA | NA | NA | NA | NA |
| lizards | Anolis_otongae | Anolis_otongae | VU | Unknown | tropical | diurnal | 6.31 | 214.851 | 7.895 |
| lizards | Anolis_oxylophus | Anolis_oxylophus | LC | Stable | tropical | diurnal | 12.589 | 118926.098 | 8.643 |
| lizards | Anolis_pachypus | Anolis_pachypus | LC | Unknown | tropical | diurnal | 3.311 | 11879.357 | 10.215 |
| lizards | Anolis_parvauritus | NA | not_available | not_available | NA | NA | NA | NA | NA |
| lizards | Anolis_parvicirculatus | Anolis_parvicirculatus | LC | Unknown | tropical | diurnal | 3.02 | 1246.575 | 9.302 |
| lizards | Anolis_paternus | NA | LC | Unknown | NA | NA | NA | NA | NA |
| lizards | Anolis_pecuarius | NA | not_available | not_available | NA | NA | NA | NA | NA |
| lizards | Anolis_pentaprion | Anolis_pentaprion | LC | Stable | tropical | diurnal | 10.471 | 355371.957 | 9.075 |
| lizards | Anolis_peraccae | Anolis_peraccae | LC | Unknown | tropical | diurnal | 5.248 | 44318.891 | 11.004 |
| lizards | Anolis_petersii | Anolis_petersii | NT | Decreasing | tropical | diurnal | 54.954 | 34767.382 | 13.739 |

| | | | | | | | | | |
|---|---|---|---|---|---|---|---|---|---|
| lizards | Anolis_peynadoi | NA | not_available | not_available | NA | NA | NA | NA | NA |
| lizards | Anolis_phyllorhinus | NA | LC | Unknown | NA | NA | NA | NA | NA |
| lizards | Anolis_pigmaequestris | Anolis_pigmaequestris | EN | Unknown | tropical | diurnal | 54.954 | 32.137 | 14.39 |
| lizards | Anolis_pijolense | NA | EN | Decreasing | NA | NA | NA | NA | NA |
| lizards | Anolis_placidus | Anolis_placidus | EN | Decreasing | tropical | diurnal | 2.089 | 265.899 | 13.402 |
| lizards | Anolis_planiceps | Anolis_planiceps | LC | Stable | tropical | diurnal | 9.12 | 1206913.454 | 3.341 |
| lizards | Anolis_podocarpus | Anolis_podocarpus | VU | Unknown | tropical | diurnal | 18.197 | 321.64 | 5.845 |
| lizards | Anolis_poecilopus | Anolis_poecilopus | LC | Unknown | tropical | diurnal | 8.511 | 89834.99 | 8.396 |
| lizards | Anolis_pogus | Anolis_pogus | NT | Increasing | tropical | diurnal | 4.074 | 192.762 | 28.853 |
| lizards | Anolis_polylepis | Anolis_polylepis | LC | Stable | tropical | diurnal | 4.365 | 12841.873 | 14.416 |
| lizards | Anolis_poncensis | NA | LC | Unknown | NA | NA | NA | NA | NA |
| lizards | Anolis_porcatus | Anolis_porcatus | LC | Stable | tropical | diurnal | 17.783 | 94983.288 | 16.484 |
| lizards | Anolis_porcus | Anolis_porcus | LC | Unknown | tropical | diurnal | 100 | 13537.828 | 16.947 |
| lizards | Anolis_prasinorius | NA | not_available | not_available | NA | NA | NA | NA | NA |
| lizards | Anolis_princeps | NA | LC | Unknown | NA | NA | NA | NA | NA |
| lizards | Anolis_proboscis | Anolis_proboscis | EN | Unknown | tropical | diurnal | 12.023 | 107.013 | 8.734 |
| lizards | Anolis_properus | NA | not_available | not_available | NA | NA | NA | NA | NA |
| lizards | Anolis_propinquus | NA | DD | Unknown | NA | NA | NA | NA | NA |
| lizards | Anolis_pseudokemptoni | NA | EN | Unknown | NA | NA | NA | NA | NA |
| lizards | Anolis_pseudopachypus | NA | LC | Stable | NA | NA | NA | NA | NA |
| lizards | Anolis_pseudotigrinus | NA | EN | Unknown | NA | NA | NA | NA | NA |
| lizards | Anolis_pulchellus | Anolis_pulchellus | LC | Stable | tropical | diurnal | 2.884 | 8714.061 | 26.804 |
| lizards | Anolis_pumilus | NA | LC | Unknown | NA | NA | NA | NA | NA |
| lizards | Anolis_punctatus | Anolis_punctatus | LC | Unknown | tropical | diurnal | 15.849 | 7465476.199 | 2.935 |
| lizards | Anolis_purpurescens | NA | LC | Unknown | NA | NA | NA | NA | NA |
| lizards | Anolis_purpurgularis | NA | EN | Decreasing | NA | NA | NA | NA | NA |
| lizards | Anolis_purpuronectes | NA | not_available | not_available | NA | NA | NA | NA | NA |
| lizards | Anolis_quadriocellifer | NA | LC | Unknown | NA | NA | NA | NA | NA |
| lizards | Anolis_quaggulus | Anolis_quaggulus | LC | Stable | tropical | diurnal | 1.778 | 130421.106 | 8.586 |
| lizards | Anolis_quercorum | NA | LC | Stable | NA | NA | NA | NA | NA |
| lizards | Anolis_ravitergum | NA | not_available | not_available | NA | NA | NA | NA | NA |
| lizards | Anolis_reconditus | Anolis_reconditus | EN | Decreasing | tropical | diurnal | 20.417 | 120.636 | 14.161 |
| lizards | Anolis_rejectus | Anolis_rejectus | EN | Unknown | tropical | diurnal | 1.096 | 358.438 | 23.106 |
| lizards | Anolis_richardii | Anolis_richardii | LC | Stable | tropical | diurnal | 54.954 | 377.093 | 20.407 |

| | | | | | | | | | |
|---|---|---|---|---|---|---|---|---|---|
| lizards | Anolis_ricordii | NA | NT | Decreasing | NA | NA | NA | NA | NA |
| lizards | Anolis_rimarum | NA | CR | Decreasing | NA | NA | NA | NA | NA |
| lizards | Anolis_roatanensis | NA | EN | Unknown | NA | NA | NA | NA | NA |
| lizards | Anolis_rodriguezii | Anolis_rodriguezi | LC | Stable | tropical | diurnal | 2.63 | 272090.601 | 8.608 |
| lizards | Anolis_roosevelti | NA | CR | Unknown | NA | NA | NA | NA | NA |
| lizards | Anolis_roquet | Anolis_roquet | LC | Stable | tropical | diurnal | 13.183 | 1141.645 | 26.204 |
| lizards | Anolis_rubribarbaris | Anolis_rubribarbaris | VU | Unknown | tropical | diurnal | 2.951 | 112.844 | 9.617 |
| lizards | Anolis_rubribarbus | NA | EN | Unknown | NA | NA | NA | NA | NA |
| lizards | Anolis_ruibali | NA | EN | Unknown | NA | NA | NA | NA | NA |
| lizards | Anolis_rupinae | NA | EN | Decreasing | NA | NA | NA | NA | NA |
| lizards | Anolis_sabanus | Anolis_sabanus | NT | Stable | tropical | diurnal | 6.918 | 12.488 | 18.219 |
| lizards | Anolis_sacamecatensis | NA | not_available | not_available | NA | NA | NA | NA | NA |
| lizards | Anolis_sagrei | Anolis_sagrei | LC | Stable | tropical | diurnal | 8.71 | 225473.674 | 12.168 |
| lizards | Anolis_santamartae | NA | DD | Unknown | NA | NA | NA | NA | NA |
| lizards | Anolis_savagei | NA | not_available | not_available | NA | NA | NA | NA | NA |
| lizards | Anolis_saxatilis | Anolis_whitemani | LC | Unknown | tropical | diurnal | 6.31 | 19236.284 | 14.671 |
| lizards | Anolis_schwartzi | Anolis_schwartzi | LC | Stable | tropical | diurnal | 2.512 | 230.587 | 22.45 |
| lizards | Anolis_scriptus | NA | LC | Stable | NA | NA | NA | NA | NA |
| lizards | Anolis_scypheus | Anolis_scypheus | LC | Stable | tropical | diurnal | 13.183 | 843487.135 | 3.259 |
| lizards | Anolis_semilineatus | Anolis_semilineatus | LC | Stable | tropical | diurnal | 2.239 | 72703.318 | 15.862 |
| lizards | Anolis_sericeus | Anolis_sericeus | LC | Stable | tropical | diurnal | 3.162 | 655922.233 | 11.322 |
| lizards | Anolis_serranoi | NA | LC | Stable | NA | NA | NA | NA | NA |
| lizards | Anolis_sheplani | Anolis_sheplani | EN | Decreasing | tropical | diurnal | 1.479 | 216.97 | 10.315 |
| lizards | Anolis_shrevei | Anolis_shrevei | NT | Unknown | tropical | diurnal | 4.571 | 1408.127 | 8.736 |
| lizards | Anolis_sierramaestrae | NA | DD | Unknown | NA | NA | NA | NA | NA |
| lizards | Anolis_singularis | Anolis_singularis | CR | Unknown | tropical | diurnal | 2.951 | 9.991 | 18.371 |
| lizards | Anolis_smallwoodi | Anolis_smallwoodi | NT | Unknown | tropical | diurnal | 134.896 | 4402.324 | 13.998 |
| lizards | Anolis_smaragdinus | Anolis_smaragdinus | LC | Stable | temperate | diurnal | 5.495 | 2713.054 | 9.232 |
| lizards | Anolis_sminthus | Anolis_sminthus | EN | Decreasing | tropical | diurnal | 4.365 | 13721.816 | 11.401 |
| lizards | Anolis_soinii | NA | LC | Unknown | NA | NA | NA | NA | NA |
| lizards | Anolis_solitarius | NA | NT | Unknown | NA | NA | NA | NA | NA |
| lizards | Anolis_spectrum | Anolis_spectrum | VU | Unknown | tropical | diurnal | 1.585 | 11960.202 | 19.59 |
| lizards | Anolis_squamulatus | NA | LC | Stable | NA | NA | NA | NA | NA |
| lizards | Anolis_stevepoei | NA | not_available | not_available | NA | NA | NA | NA | NA |

| | | | | | | | | | |
|---|---|---|---|---|---|---|---|---|---|
| lizards | Anolis_strahmi | Anolis_strahmi | EN | Decreasing | tropical | diurnal | 10.471 | 709.04 | 7.955 |
| lizards | Anolis_stratulus | NA | LC | Stable | NA | NA | NA | NA | NA |
| lizards | Anolis_subocularis | NA | DD | Unknown | NA | NA | NA | NA | NA |
| lizards | Anolis_tandai | Anolis_tandai | LC | Unknown | tropical | diurnal | 8.511 | 1400788.684 | 1.355 |
| lizards | Anolis_taylori | Anolis_taylori | LC | Stable | tropical | diurnal | 9.772 | 80.237 | 37.857 |
| lizards | Anolis_terraealtae | Anolis_terraealtae | LC | Unknown | tropical | diurnal | 10.471 | 16.94 | 21.836 |
| lizards | Anolis_terueli | NA | LC | Stable | NA | NA | NA | NA | NA |
| lizards | Anolis_tigrinus | NA | LC | Stable | NA | NA | NA | NA | NA |
| lizards | Anolis_toldo | NA | VU | Unknown | NA | NA | NA | NA | NA |
| lizards | Anolis_tolimensis | Anolis_tolimensis | LC | Unknown | tropical | diurnal | 4.571 | 48671.289 | 14.281 |
| lizards | Anolis_townsendi | NA | LC | Stable | NA | NA | NA | NA | NA |
| lizards | Anolis_trachyderma | Anolis_trachyderma | LC | Stable | tropical | diurnal | 4.786 | 1001940.694 | 1.414 |
| lizards | Anolis_transversalis | Anolis_transversalis | LC | Unknown | tropical | diurnal | 19.055 | 2234134.979 | 1.373 |
| lizards | Anolis_trinitatis | Anolis_trinitatis | LC | Stable | tropical | diurnal | 10.471 | 348.239 | 16.756 |
| lizards | Anolis_triumphalis | NA | not_available | not_available | NA | NA | NA | NA | NA |
| lizards | Anolis_tropidogaster | Anolis_tropidogaster | LC | Stable | tropical | diurnal | 5.248 | 422645.866 | 9.343 |
| lizards | Anolis_tropidolepis | Anolis_tropidolepis | LC | Stable | tropical | diurnal | 4.365 | 3965.11 | 17.659 |
| lizards | Anolis_tropidonotus | Anolis_tropidonotus | LC | Stable | tropical | diurnal | 5.754 | 359474.766 | 9.412 |
| lizards | Anolis_uniformis | Anolis_uniformis | LC | Decreasing | tropical | diurnal | 1.479 | 84815.201 | 10.449 |
| lizards | Anolis_unilobatus | NA | LC | Stable | NA | NA | NA | NA | NA |
| lizards | Anolis_urraoi | NA | not_available | not_available | NA | NA | NA | NA | NA |
| lizards | Anolis_ustus | NA | not_available | not_available | NA | NA | NA | NA | NA |
| lizards | Anolis_utilensis | Anolis_utilensis | CR | Decreasing | tropical | diurnal | 5.012 | 42.084 | 9.659 |
| lizards | Anolis_valencienni | Anolis_valencienni | LC | Stable | tropical | diurnal | 13.183 | 11021.537 | 19.74 |
| lizards | Anolis_vanidicus | NA | LC | Stable | NA | NA | NA | NA | NA |
| lizards | Anolis_vanzolinii | NA | CR | Unknown | NA | NA | NA | NA | NA |
| lizards | Anolis_vaupesianus | NA | LC | Unknown | NA | NA | NA | NA | NA |
| lizards | Anolis_vermiculatus | Anolis_vermiculatus | LC | Stable | tropical | diurnal | 48.978 | 2375.371 | 14.435 |
| lizards | Anolis_vescus | NA | DD | Unknown | NA | NA | NA | NA | NA |
| lizards | Anolis_vicarius | NA | DD | Unknown | NA | NA | NA | NA | NA |
| lizards | Anolis_villai | NA | CR | Decreasing | NA | NA | NA | NA | NA |
| lizards | Anolis_vinosus | NA | not_available | not_available | NA | NA | NA | NA | NA |
| lizards | Anolis_viridius | NA | not_available | not_available | NA | NA | NA | NA | NA |
| lizards | Anolis_vittigerus | Anolis_vittigerus | LC | Stable | tropical | diurnal | 7.762 | 83506.202 | 9.035 |

| | | | | | | | | | |
|---|---|---|---|---|---|---|---|---|---|
| lizards | Anolis_wampuensis | NA | DD | Unknown | NA | NA | NA | NA | NA |
| lizards | Anolis_wattsii | Anolis_wattsi | LC | Unknown | tropical | diurnal | 4.074 | 716.38 | 23.762 |
| lizards | Anolis_websteri | NA | LC | Stable | NA | NA | NA | NA | NA |
| lizards | Anolis_wellbornae | NA | LC | Stable | NA | NA | NA | NA | NA |
| lizards | Anolis_williamsmittermeierorum | NA | DD | Unknown | NA | NA | NA | NA | NA |
| lizards | Anolis_wilsoni | NA | not_available | not_available | NA | NA | NA | NA | NA |
| lizards | Anolis_woodi | Anolis_woodi | LC | Stable | tropical | diurnal | 20.417 | 11087.036 | 14.454 |
| lizards | Anolis_yoroensis | NA | EN | Decreasing | NA | NA | NA | NA | NA |
| lizards | Anolis_zapotecorum | NA | not_available | not_available | NA | NA | NA | NA | NA |
| lizards | Anolis_zeus | NA | VU | Decreasing | NA | NA | NA | NA | NA |
| lizards | Anomalopus_leuckartii | Anomalopus_leuckartii | LC | Stable | temperate | cathemeral | 11.749 | 397667.281 | 7.229 |
| lizards | Anomalopus_swansoni | Anomalopus_swansoni | LC | Unknown | temperate | diurnal | 2.884 | 18131.023 | 10.334 |
| lizards | Anomalopus_verreauxii | Anomalopus_verreauxi | LC | Stable | temperate | cathemeral | 25.119 | 344288.5 | 7.131 |
| lizards | Anotosaura_collaris | NA | DD | Unknown | NA | NA | NA | NA | NA |
| lizards | Anotosaura_vanzolinia | Anotosaura_vanzolinia | LC | Unknown | tropical | diurnal | 2.692 | 45489.943 | 12.071 |
| lizards | Apathya_cappadocica | Apathya_cappadocica | LC | Stable | temperate | diurnal | 14.454 | 336629.587 | 16.593 |
| lizards | Apathya_yassujica | NA | LC | Stable | NA | NA | NA | NA | NA |
| lizards | Aphaniotis_acutirostris | Aphaniotis_acutirostris | LC | Stable | tropical | diurnal | 12.023 | 63335.475 | 8.628 |
| lizards | Aphaniotis_fusca | Aphaniotis_fusca | LC | Stable | tropical | diurnal | 12.589 | 1322523.813 | 7.728 |
| lizards | Aphaniotis_ornata | Aphaniotis_ornata | LC | Unknown | tropical | diurnal | 5.888 | 26163.785 | 6.263 |
| lizards | Aprasia_aurita | Aprasia_aurita | NT | Stable | temperate | cathemeral | 6.761 | 72135.188 | 11.343 |
| lizards | Aprasia_haroldi | Aprasia_haroldi | LC | Unknown | temperate | nocturnal | 5.754 | 1629.499 | 8.078 |
| lizards | Aprasia_inaurita | Aprasia_inaurita | LC | Stable | temperate | cathemeral | 7.762 | 386166.528 | 6.054 |
| lizards | Aprasia_parapulchella | Aprasia_parapulchella | LC | Unknown | temperate | nocturnal | 8.318 | 152289.553 | 10.949 |
| lizards | Aprasia_picturata | NA | DD | Unknown | NA | NA | NA | NA | NA |
| lizards | Aprasia_pseudopulchella | Aprasia_pseudopulchella | LC | Stable | temperate | nocturnal | 9.12 | 40407.355 | 10.096 |
| lizards | Aprasia_pulchella | Aprasia_pulchella | LC | Unknown | temperate | cathemeral | 7.413 | 65114.576 | 12.027 |
| lizards | Aprasia_repens | Aprasia_repens | LC | Unknown | temperate | nocturnal | 6.918 | 156382.456 | 11.613 |
| lizards | Aprasia_rostrata | Aprasia_rostrata | VU | Decreasing | tropical | nocturnal | 6.918 | 12573.47 | 6.51 |
| lizards | Aprasia_smithi | NA | LC | Stable | NA | NA | NA | NA | NA |
| lizards | Aprasia_striolata | Aprasia_striolata | LC | Stable | temperate | cathemeral | 8.128 | 207815.333 | 9.75 |
| lizards | Archaeolacerta_bedriagae | Archaeolacerta_bedriagae | NT | Decreasing | temperate | diurnal | 14.125 | 12496.029 | 13.488 |
| lizards | Archaius_tigris | NA | EN | Unknown | NA | NA | NA | NA | NA |
| lizards | Aristelliger_barbouri | NA | DD | Unknown | NA | NA | NA | NA | NA |

| | | | | | | | | | |
|---|---|---|---|---|---|---|---|---|---|
| lizards | Aristelliger_cochranae | Aristelliger_cochranae | LC | Stable | tropical | cathemeral | 8.71 | 35091.313 | 14.675 |
| lizards | Aristelliger_expectatus | Aristelliger_expectatus | LC | Unknown | tropical | nocturnal | 4.074 | 32623.993 | 14.642 |
| lizards | Aristelliger_georgeensis | Aristelliger_georgeensis | LC | Decreasing | tropical | cathemeral | 36.308 | 2198.567 | 4.804 |
| lizards | Aristelliger_hechti | Aristelliger_hechti | VU | Unknown | tropical | nocturnal | 17.783 | 152.336 | 10.337 |
| lizards | Aristelliger_lar | Aristelliger_lar | NT | Unknown | tropical | nocturnal | 79.433 | 44003.84 | 16.353 |
| lizards | Aristelliger_praesignis | Aristelliger_praesignis | LC | Stable | tropical | cathemeral | 24.547 | 10815.102 | 19.681 |
| lizards | Aristelliger_reyesi | Aristelliger_reyesi | CR | Decreasing | tropical | cathemeral | 6.31 | 17.712 | 18.758 |
| lizards | Arthrosaura_kockii | Arthrosaura_kockii | LC | Stable | tropical | diurnal | 8.511 | 780816.699 | 1.875 |
| lizards | Arthrosaura_montigena | NA | LC | Unknown | NA | NA | NA | NA | NA |
| lizards | Arthrosaura_reticulata | Arthrosaura_reticulata | LC | Unknown | tropical | diurnal | 8.511 | 4621507.22 | 1.777 |
| lizards | Arthrosaura_synaptolepis | NA | LC | Unknown | NA | NA | NA | NA | NA |
| lizards | Arthrosaura_versteegii | NA | LC | Unknown | NA | NA | NA | NA | NA |
| lizards | Asaccus_andersoni | NA | not_available | not_available | NA | NA | NA | NA | NA |
| lizards | Asaccus_arnoldi | NA | LC | Stable | NA | NA | NA | NA | NA |
| lizards | Asaccus_barani | NA | not_available | not_available | NA | NA | NA | NA | NA |
| lizards | Asaccus_caudivolvulus | Asaccus_caudivolvulus | CR | Decreasing | temperate | nocturnal | 6.607 | 125.162 | 22.435 |
| lizards | Asaccus_elisae | Asaccus_elisae | LC | Stable | temperate | nocturnal | 7.244 | 329259.552 | 15.624 |
| lizards | Asaccus_gallagheri | Asaccus_gallagheri | LC | Stable | tropical | nocturnal | 1.738 | 14296.83 | 11.419 |
| lizards | Asaccus_gardneri | NA | LC | Stable | NA | NA | NA | NA | NA |
| lizards | Asaccus_granularis | NA | not_available | not_available | NA | NA | NA | NA | NA |
| lizards | Asaccus_griseonotus | NA | LC | Stable | NA | NA | NA | NA | NA |
| lizards | Asaccus_iranicus | NA | not_available | not_available | NA | NA | NA | NA | NA |
| lizards | Asaccus_kermanshahensis | NA | LC | Unknown | NA | NA | NA | NA | NA |
| lizards | Asaccus_kurdistanensis | NA | LC | Stable | NA | NA | NA | NA | NA |
| lizards | Asaccus_margaritae | NA | LC | Stable | NA | NA | NA | NA | NA |
| lizards | Asaccus_montanus | NA | LC | Stable | NA | NA | NA | NA | NA |
| lizards | Asaccus_nasrullahi | Asaccus_nasrullahi | LC | Stable | temperate | cathemeral | 8.913 | 5601.902 | 14.458 |
| lizards | Asaccus_platyrhynchus | Asaccus_platyrhynchus | LC | Stable | tropical | nocturnal | 6.607 | 5911.07 | 10.064 |
| lizards | Asaccus_saffinae | NA | not_available | not_available | NA | NA | NA | NA | NA |
| lizards | Asaccus_tangestanensis | NA | not_available | not_available | NA | NA | NA | NA | NA |
| lizards | Asaccus_zagrosicus | NA | not_available | not_available | NA | NA | NA | NA | NA |
| lizards | Aspidoscelis_angusticeps | NA | LC | Stable | NA | NA | NA | NA | NA |
| lizards | Aspidoscelis_burti | Aspidoscelis_burti | LC | Stable | temperate | diurnal | 85.114 | 165629.668 | 7.828 |
| lizards | Aspidoscelis_calidipes | Aspidoscelis_calidipes | LC | Unknown | tropical | diurnal | 14.125 | 8198.387 | 9.922 |

| | | | | | | | | | |
|---|---|---|---|---|---|---|---|---|---|
| lizards | Aspidoscelis_carmenensis | NA | LC | Stable | NA | NA | NA | NA | NA |
| lizards | Aspidoscelis_ceralbensis | Aspidoscelis_ceralbensis | LC | Stable | temperate | diurnal | 25.119 | 142.906 | 2.171 |
| lizards | Aspidoscelis_communis | Aspidoscelis_communis | LC | Stable | tropical | diurnal | 109.648 | 99380.575 | 10.354 |
| lizards | Aspidoscelis_costatus | Aspidoscelis_costata | LC | Stable | temperate | diurnal | 70.795 | 331734.788 | 10.353 |
| lizards | Aspidoscelis_cozumela | Aspidoscelis_cozumelae | LC | Stable | tropical | diurnal | 16.596 | 488.275 | 10.21 |
| lizards | Aspidoscelis_danheimae | Aspidoscelis_danheimae | LC | Stable | temperate | diurnal | 8.128 | 169.308 | 1.536 |
| lizards | Aspidoscelis_deppii | Aspidoscelis_deppei | LC | Stable | tropical | diurnal | 23.442 | 609448.246 | 11.137 |
| lizards | Aspidoscelis_espiritensis | NA | LC | Stable | NA | NA | NA | NA | NA |
| lizards | Aspidoscelis_exsanguis | Aspidoscelis_exsanguis | LC | Stable | temperate | diurnal | 36.308 | 409465.473 | 8.675 |
| lizards | Aspidoscelis_franciscensis | NA | LC | Stable | NA | NA | NA | NA | NA |
| lizards | Aspidoscelis_gularis | Aspidoscelis_gularis | LC | Stable | temperate | diurnal | 45.709 | 1417047.366 | 11.428 |
| lizards | Aspidoscelis_guttatus | Aspidoscelis_guttata | LC | Stable | tropical | diurnal | 93.325 | 169406.917 | 10.333 |
| lizards | Aspidoscelis_hyperythrus | Aspidoscelis_hyperythra | LC | Decreasing | temperate | diurnal | 10.715 | 140729.203 | 6.03 |
| lizards | Aspidoscelis_inornatus | Aspidoscelis_inornata | LC | Decreasing | temperate | diurnal | 18.621 | 659625.448 | 7.98 |
| lizards | Aspidoscelis_labialis | Aspidoscelis_labialis | VU | Decreasing | temperate | diurnal | 7.079 | 13910.992 | 3.995 |
| lizards | Aspidoscelis_laredoensis | Aspidoscelis_laredoensis | LC | Stable | temperate | diurnal | 21.38 | 40046.349 | 15.116 |
| lizards | Aspidoscelis_lineattissimus | Aspidoscelis_lineattissima | LC | Stable | tropical | diurnal | 42.658 | 65053.395 | 9.406 |
| lizards | Aspidoscelis_marmoratus | NA | not_available | not_available | NA | NA | NA | NA | NA |
| lizards | Aspidoscelis_martyris | Cnemidophorus_martyris | VU | Stable | temperate | diurnal | 14.791 | 1.223 | 2.107 |
| lizards | Aspidoscelis_maslini | NA | LC | Stable | NA | NA | NA | NA | NA |
| lizards | Aspidoscelis_maximus | NA | not_available | not_available | NA | NA | NA | NA | NA |
| lizards | Aspidoscelis_mexicanus | NA | LC | Stable | NA | NA | NA | NA | NA |
| lizards | Aspidoscelis_motaguae | Aspidoscelis_motaguae | LC | Stable | tropical | diurnal | 93.325 | 42636.38 | 9.574 |
| lizards | Aspidoscelis_neomexicanus | Aspidoscelis_neomexicana | LC | Stable | temperate | diurnal | 18.621 | 91254.626 | 10.377 |
| lizards | Aspidoscelis_neotesselatus | Aspidoscelis_neotesselata | NT | Decreasing | temperate | diurnal | 36.308 | 124729.527 | 9.974 |
| lizards | Aspidoscelis_opatae | NA | DD | Unknown | NA | NA | NA | NA | NA |
| lizards | Aspidoscelis_pai | NA | not_available | not_available | NA | NA | NA | NA | NA |
| lizards | Aspidoscelis_parvisocius | Aspidoscelis_parvisocia | LC | Stable | tropical | diurnal | 14.125 | 3556.21 | 8.994 |
| lizards | Aspidoscelis_pictus | Aspidoscelis_picta | LC | Stable | temperate | diurnal | 10.233 | 18.099 | 0.891 |
| lizards | Aspidoscelis_rodecki | Aspidoscelis_rodecki | NT | Unknown | tropical | diurnal | 12.589 | 223.591 | 9.06 |
| lizards | Aspidoscelis_sackii | Aspidoscelis_sackii | LC | Stable | tropical | diurnal | 112.202 | 46103.86 | 11.626 |
| lizards | Aspidoscelis_scalaris | NA | not_available | not_available | NA | NA | NA | NA | NA |
| lizards | Aspidoscelis_sexlineatus | Aspidoscelis_sexlineata | LC | Stable | temperate | diurnal | 22.387 | 2757748.862 | 15.952 |
| lizards | Aspidoscelis_sonorae | Aspidoscelis_sonorae | LC | Stable | temperate | diurnal | 30.903 | 183626.985 | 9.022 |

| | | | | | | | | | |
|---|---|---|---|---|---|---|---|---|---|
| lizards | Aspidoscelis_stictogrammus | Aspidoscelis_stictogramma | LC | Stable | temperate | diurnal | 85.114 | 85889.545 | 6.987 |
| lizards | Aspidoscelis_tesselatus | Aspidoscelis_tesselata | LC | Stable | temperate | diurnal | 36.308 | 339352.99 | 10.352 |
| lizards | Aspidoscelis_tigris | Aspidoscelis_tigris | LC | Stable | temperate | diurnal | 79.433 | 2084184.452 | 8.473 |
| lizards | Aspidoscelis_uniparens | Aspidoscelis_uniparens | LC | Stable | temperate | diurnal | 18.621 | 218191.048 | 9.262 |
| lizards | Aspidoscelis_velox | Aspidoscelis_velox | LC | Stable | temperate | diurnal | 20.893 | 369953.79 | 9.19 |
| lizards | Aspidoscelis_xanthonotus | Aspidoscelis_xanthonota | NT | Unknown | temperate | diurnal | 44.668 | 17223.337 | 7.553 |
| lizards | Aspronema_cochabambae | Aspronema_cochabambae | VU | Decreasing | tropical | diurnal | 11.482 | 3676.241 | 6.161 |
| lizards | Aspronema_dorsivittatum | Aspronema_dorsivittatum | LC | Stable | tropical | diurnal | 15.488 | 3779794.066 | 10.163 |
| lizards | Ateuchosaurus_chinensis | Ateuchosaurus_chinensis | LC | Stable | temperate | cathemeral | 21.38 | 822446.791 | 15.175 |
| lizards | Ateuchosaurus_pellopleurus | Ateuchosaurus_pellopleurus | LC | Stable | temperate | cathemeral | 6.457 | 2708.818 | 23.33 |
| lizards | Atlantolacerta_andreanskyi | Atlantolacerta_andreanskyi | NT | Stable | temperate | diurnal | 3.89 | 11816.941 | 11.306 |
| lizards | Aurivela_longicauda | NA | LC | Stable | NA | NA | NA | NA | NA |
| lizards | Australolacerta_australis | Australolacerta_australis | LC | Stable | temperate | diurnal | 7.943 | 20866.441 | 10.959 |
| lizards | Austroablepharus_kinghorni | NA | LC | Stable | NA | NA | NA | NA | NA |
| lizards | Austroablepharus_naranjicauda | NA | LC | Stable | NA | NA | NA | NA | NA |
| lizards | Bachia_bicolor | Bachia_bicolor | LC | Unknown | tropical | diurnal | 2.951 | 156298.955 | 13.474 |
| lizards | Bachia_bresslaui | Bachia_bresslaui | VU | Decreasing | tropical | diurnal | 6.026 | 1123654.727 | 9.853 |
| lizards | Bachia_flavescens | Bachia_flavescens | LC | Unknown | tropical | diurnal | 3.162 | 3587470.01 | 1.668 |
| lizards | Bachia_geralista | NA | DD | Unknown | NA | NA | NA | NA | NA |
| lizards | Bachia_heteropa | Bachia_heteropa | LC | Unknown | tropical | diurnal | 1.82 | 83731.216 | 13.44 |
| lizards | Bachia_micromela | NA | DD | Unknown | NA | NA | NA | NA | NA |
| lizards | Bachia_panoplia | NA | LC | Unknown | NA | NA | NA | NA | NA |
| lizards | Bachia_peruana | NA | LC | Unknown | NA | NA | NA | NA | NA |
| lizards | Bachia_psamophila | NA | CR | Decreasing | NA | NA | NA | NA | NA |
| lizards | Bachia_pyburni | NA | LC | Unknown | NA | NA | NA | NA | NA |
| lizards | Bachia_remota | NA | not_available | not_available | NA | NA | NA | NA | NA |
| lizards | Bachia_scaea | NA | DD | Unknown | NA | NA | NA | NA | NA |
| lizards | Bachia_trisanale | Bachia_trisanale | LC | Unknown | tropical | diurnal | 1.445 | 1489602.663 | 1.404 |
| lizards | Barisia_ciliaris | NA | not_available | not_available | NA | NA | NA | NA | NA |
| lizards | Barisia_imbricata | Barisia_imbricata | LC | Unknown | tropical | diurnal | 77.625 | 485129.42 | 10.447 |
| lizards | Barisia_jonesi | NA | not_available | not_available | NA | NA | NA | NA | NA |
| lizards | Barisia_planifrons | NA | not_available | not_available | NA | NA | NA | NA | NA |
| lizards | Barkudia_insularis | NA | CR | Unknown | NA | NA | NA | NA | NA |
| lizards | Barkudia_melanosticta | NA | DD | Unknown | NA | NA | NA | NA | NA |

| | | | | | | | | | |
|---|---|---|---|---|---|---|---|---|---|
| lizards | Basiliscus_basiliscus | Basiliscus_basiliscus | LC | Stable | tropical | diurnal | 549.541 | 432691.006 | 12.785 |
| lizards | Basiliscus_galeritus | Basiliscus_galeritus | LC | Unknown | tropical | diurnal | 489.779 | 323161.635 | 8.158 |
| lizards | Basiliscus_plumifrons | Basiliscus_plumifrons | LC | Stable | tropical | diurnal | 549.541 | 107491.885 | 8.602 |
| lizards | Basiliscus_vittatus | Basiliscus_vittatus | LC | Stable | tropical | diurnal | 398.107 | 880749.99 | 10.33 |
| lizards | Bauerius_ansorgii | NA | LC | Stable | NA | NA | NA | NA | NA |
| lizards | Bavayia_crassicollis | Bavayia_crassicollis | VU | Unknown | tropical | nocturnal | 12.882 | 1979.641 | 8.084 |
| lizards | Bavayia_cyclura | Bavayia_cyclura | EN | Decreasing | tropical | nocturnal | 7.413 | 17439.168 | 7.532 |
| lizards | Bavayia_exsuccida | Bavayia_exsuccida | CR | Decreasing | tropical | nocturnal | 2.042 | 1000.099 | 9.506 |
| lizards | Bavayia_geitaina | Bavayia_geitaina | NT | Decreasing | tropical | nocturnal | 7.413 | 3641.36 | 6.348 |
| lizards | Bavayia_goroensis | Bavayia_goroensis | EN | Decreasing | tropical | nocturnal | 2.239 | 146.484 | 9.232 |
| lizards | Bavayia_montana | Bavayia_montana | EN | Decreasing | tropical | nocturnal | 8.913 | 10593.66 | 7.075 |
| lizards | Bavayia_nubila | Bavayia_nubila | EN | Decreasing | tropical | nocturnal | 6.026 | 95.381 | 4.722 |
| lizards | Bavayia_ornata | Bavayia_ornata | CR | Decreasing | tropical | nocturnal | 6.607 | 270.036 | 3.113 |
| lizards | Bavayia_pulchella | Bavayia_pulchella | EN | Decreasing | tropical | nocturnal | 2.344 | 254.099 | 7.286 |
| lizards | Bavayia_robusta | Bavayia_robusta | NT | Stable | tropical | nocturnal | 11.482 | 3095.745 | 9.704 |
| lizards | Bavayia_sauvagii | Bavayia_sauvagii | EN | Decreasing | tropical | nocturnal | 4.786 | 16396.165 | 7.459 |
| lizards | Bavayia_septuiclavis | Bavayia_septuiclavis | NT | Unknown | tropical | nocturnal | 2.455 | 2941.22 | 9.642 |
| lizards | Bellatorias_frerei | Bellatorias_frerei | LC | Stable | tropical | cathemeral | 234.423 | 460068.122 | 5.461 |
| lizards | Bellatorias_major | Bellatorias_major | LC | Decreasing | temperate | diurnal | 1778.279 | 72332.577 | 10.121 |
| lizards | Bellatorias_obiri | NA | CR | Decreasing | NA | NA | NA | NA | NA |
| lizards | Bipes_biporus | Bipes_biporus | LC | Stable | temperate | diurnal | 14.125 | 42660.687 | 4.17 |
| lizards | Bipes_canaliculatus | Bipes_canaliculatus | LC | Stable | tropical | diurnal | 14.125 | 9816.829 | 9.527 |
| lizards | Blaesodactylus_ambonihazo | NA | DD | Unknown | NA | NA | NA | NA | NA |
| lizards | Blaesodactylus_antongilensis | Blaesodactylus_antongilensis | LC | Decreasing | tropical | nocturnal | 17.378 | 33912.215 | 8.28 |
| lizards | Blaesodactylus_boivini | Blaesodactylus_boivini | VU | Decreasing | tropical | cathemeral | 64.565 | 9379.962 | 8.973 |
| lizards | Blaesodactylus_microtuberculatus | NA | not_available | not_available | NA | NA | NA | NA | NA |
| lizards | Blaesodactylus_sakalava | NA | LC | Decreasing | NA | NA | NA | NA | NA |
| lizards | Blaesodactylus_victori | NA | not_available | not_available | NA | NA | NA | NA | NA |
| lizards | Blanus_cinereus | Blanus_cinereus | LC | Stable | temperate | cathemeral | 16.596 | 47115.081 | 15.512 |
| lizards | Blanus_mettetali | Blanus_mettetali | LC | Unknown | temperate | nocturnal | 9.55 | 111004.251 | 16.218 |
| lizards | Blanus_vandellii | NA | not_available | not_available | NA | NA | NA | NA | NA |
| lizards | Brachylophus_bulabula | Brachylophus_bulabula | EN | Decreasing | tropical | diurnal | 676.083 | 15793.112 | 8.754 |
| lizards | Brachylophus_fasciatus | Brachylophus_fasciatus | EN | Decreasing | tropical | diurnal | 676.083 | 12789.307 | 8.916 |
| lizards | Brachylophus_gau | NA | EN | Decreasing | NA | NA | NA | NA | NA |

| | | | | | | | | | |
|---|---|---|---|---|---|---|---|---|---|
| lizards | Brachylophus_vitiensis | Brachylophus_vitiensis | CR | Decreasing | tropical | diurnal | 707.946 | 1562.706 | 16.049 |
| lizards | Brachymeles_apus | Brachymeles_apus | LC | Unknown | tropical | cathemeral | 4.571 | 166.532 | 17.342 |
| lizards | Brachymeles_bicolor | NA | LC | Decreasing | NA | NA | NA | NA | NA |
| lizards | Brachymeles_bonitae | Brachymeles_bonitae | LC | Stable | tropical | diurnal | 4.677 | 13683.249 | 11.493 |
| lizards | Brachymeles_boulengeri | Brachymeles_boulengeri | LC | Stable | tropical | diurnal | 26.915 | 156558.458 | 15.355 |
| lizards | Brachymeles_gracilis | Brachymeles_gracilis | LC | Stable | tropical | diurnal | 31.623 | 122838.104 | 12.416 |
| lizards | Brachymeles_miriamae | NA | LC | Unknown | NA | NA | NA | NA | NA |
| lizards | Brachymeles_orientalis | Brachymeles_orientalis | LC | Unknown | tropical | diurnal | 33.884 | 10861.213 | 11.103 |
| lizards | Brachymeles_samarensis | Brachymeles_samarensis | NT | Unknown | tropical | diurnal | 1.95 | 24628.393 | 11.85 |
| lizards | Brachymeles_schadenbergi | Brachymeles_schadenbergi | LC | Stable | tropical | diurnal | 38.019 | 118956.831 | 12.097 |
| lizards | Brachymeles_talinis | Brachymeles_talinis | LC | Stable | tropical | nocturnal | 79.433 | 39075.482 | 16.903 |
| lizards | Brachymeles_taylori | Brachymeles_taylori | LC | Unknown | tropical | nocturnal | 25.119 | 17449.002 | 19.12 |
| lizards | Brachymeles_tridactylus | Brachymeles_tridactylus | LC | Stable | tropical | nocturnal | 3.548 | 24396.928 | 17.822 |
| lizards | Brachyseps_anosyensis | NA | VU | Decreasing | NA | NA | NA | NA | NA |
| lizards | Brachyseps_frontoparietalis | Amphiglossus_frontoparietalis | LC | Unknown | tropical | nocturnal | 8.913 | 58116.39 | 8.338 |
| lizards | Brachyseps_gastrostictus | NA | DD | Unknown | NA | NA | NA | NA | NA |
| lizards | Brachyseps_mandady | NA | DD | Unknown | NA | NA | NA | NA | NA |
| lizards | Brachyseps_punctatus | Amphiglossus_punctatus | LC | Unknown | tropical | diurnal | 7.762 | 54731.145 | 8.775 |
| lizards | Brachyseps_spilostichus | NA | DD | Unknown | NA | NA | NA | NA | NA |
| lizards | Bradypodion_damaranum | Bradypodion_damaranum | LC | Stable | temperate | diurnal | 13.804 | 2059.781 | 10.747 |
| lizards | Bradypodion_melanocephalum | Bradypodion_melanocephalum | NT | Decreasing | temperate | diurnal | 10.965 | 21790.406 | 17.245 |
| lizards | Bradypodion_occidentale | Bradypodion_occidentale | LC | Stable | temperate | diurnal | 20.893 | 34988.038 | 9.256 |
| lizards | Bradypodion_pumilum | Bradypodion_pumilum | NT | Decreasing | temperate | diurnal | 25.119 | 27711.518 | 14.809 |
| lizards | Bradypodion_setaroi | Bradypodion_setaroi | NT | Stable | temperate | diurnal | 9.55 | 8036.756 | 11.815 |
| lizards | Bradypodion_taeniabronchum | Bradypodion_taeniabronchum | LC | Stable | temperate | diurnal | 9.12 | 6644.588 | 13.542 |
| lizards | Bradypodion_thamnobates | Bradypodion_thamnobates | EN | Decreasing | temperate | diurnal | 25.119 | 17251.726 | 13.147 |
| lizards | Bradypodion_transvaalense | Bradypodion_transvaalense | LC | Stable | temperate | diurnal | 15.488 | 45421.132 | 11.542 |
| lizards | Bradypodion_ventrale | Bradypodion_ventrale | LC | Stable | temperate | diurnal | 24.547 | 242128.836 | 9.182 |
| lizards | Brasiliscincus_agilis | Brasiliscincus_agilis | LC | Unknown | tropical | diurnal | 19.055 | 479715.01 | 10.43 |
| lizards | Brasiliscincus_caissara | Brasiliscincus_caissara | EN | Unknown | temperate | diurnal | 15.488 | 694.127 | 16.384 |
| lizards | Brasiliscincus_heathi | Brasiliscincus_heathi | LC | Unknown | tropical | diurnal | 16.596 | 1824206.95 | 8.017 |
| lizards | Broadleysaurus_major | Broadleysaurus_major | LC | Unknown | tropical | diurnal | 389.045 | 3762454.512 | 10.72 |
| lizards | Bronchocela_burmana | NA | LC | Unknown | NA | NA | NA | NA | NA |
| lizards | Bronchocela_cristatella | Bronchocela_cristatella | LC | Stable | tropical | diurnal | 75.858 | 1963684.213 | 9.697 |

| | | | | | | | | | |
|---|---|---|---|---|---|---|---|---|---|
| lizards | Bronchocela_danieli | NA | LC | Unknown | NA | NA | NA | NA | NA |
| lizards | Bronchocela_hayeki | Bronchocela_hayeki | LC | Stable | tropical | diurnal | 58.884 | 2470.989 | 8.524 |
| lizards | Bronchocela_jubata | Bronchocela_jubata | LC | Stable | tropical | diurnal | 117.49 | 618236.176 | 11.72 |
| lizards | Bronchocela_marmorata | Bronchocela_marmorata | LC | Unknown | tropical | diurnal | 66.069 | 276676.405 | 13.909 |
| lizards | Bronchocela_orlovi | NA | DD | Unknown | NA | NA | NA | NA | NA |
| lizards | Bronchocela_rayaensis | NA | LC | Unknown | NA | NA | NA | NA | NA |
| lizards | Bronchocela_rubrigularis | NA | DD | Unknown | NA | NA | NA | NA | NA |
| lizards | Bronchocela_shenlong | NA | NT | Stable | NA | NA | NA | NA | NA |
| lizards | Bronchocela_smaragdina | Bronchocela_smaragdina | LC | Unknown | tropical | diurnal | 58.884 | 31053.283 | 11.55 |
| lizards | Bronchocela_vietnamensis | NA | VU | Unknown | NA | NA | NA | NA | NA |
| lizards | Brookesia_antakarana | Brookesia_antakarana | NT | Stable | tropical | diurnal | 6.026 | 45.968 | 6.673 |
| lizards | Brookesia_bekolosy | NA | EN | Decreasing | NA | NA | NA | NA | NA |
| lizards | Brookesia_bonsi | NA | CR | Decreasing | NA | NA | NA | NA | NA |
| lizards | Brookesia_brunoi | NA | NT | Unknown | NA | NA | NA | NA | NA |
| lizards | Brookesia_brygooi | Brookesia_brygooi | LC | Unknown | tropical | diurnal | 3.981 | 146282.968 | 7.987 |
| lizards | Brookesia_confidens | NA | NT | Unknown | NA | NA | NA | NA | NA |
| lizards | Brookesia_decaryi | Brookesia_decaryi | EN | Decreasing | tropical | diurnal | 4.169 | 1967.449 | 9.386 |
| lizards | Brookesia_dentata | NA | EN | Decreasing | NA | NA | NA | NA | NA |
| lizards | Brookesia_desperata | Brookesia_desperata | CR | Decreasing | tropical | diurnal | 0.912 | 50.054 | 7.263 |
| lizards | Brookesia_ebenaui | NA | VU | Decreasing | NA | NA | NA | NA | NA |
| lizards | Brookesia_exarmata | Brookesia_exarmata | EN | Decreasing | tropical | diurnal | 0.661 | 1727.581 | 3.778 |
| lizards | Brookesia_griveaudi | Brookesia_griveaudi | NT | Decreasing | tropical | diurnal | 6.918 | 21829.164 | 7.375 |
| lizards | Brookesia_lambertoni | NA | DD | Unknown | NA | NA | NA | NA | NA |
| lizards | Brookesia_micra | NA | NT | Unknown | NA | NA | NA | NA | NA |
| lizards | Brookesia_minima | Brookesia_minima | EN | Decreasing | tropical | diurnal | 0.562 | 7342.827 | 6.182 |
| lizards | Brookesia_perarmata | NA | EN | Decreasing | NA | NA | NA | NA | NA |
| lizards | Brookesia_peyrierasi | NA | EN | Decreasing | NA | NA | NA | NA | NA |
| lizards | Brookesia_stumpffi | Brookesia_stumpffi | LC | Stable | tropical | diurnal | 5.129 | 61920.454 | 8.525 |
| lizards | Brookesia_superciliaris | Brookesia_superciliaris | LC | Decreasing | tropical | diurnal | 4.467 | 90360.091 | 7.901 |
| lizards | Brookesia_thieli | Brookesia_thieli | LC | Decreasing | tropical | diurnal | 2.692 | 46977.037 | 8.055 |
| lizards | Brookesia_tristis | Brookesia_tristis | EN | Decreasing | tropical | diurnal | 0.49 | 215.305 | 12.229 |
| lizards | Brookesia_tuberculata | NA | VU | Stable | NA | NA | NA | NA | NA |
| lizards | Brookesia_valerieae | NA | EN | Unknown | NA | NA | NA | NA | NA |
| lizards | Bufoniceps_laungwalaensis | NA | LC | Unknown | NA | NA | NA | NA | NA |

| | | | | | | | | | |
|---|---|---|---|---|---|---|---|---|---|
| lizards | Bunopus_blanfordii | NA | not_available | not_available | NA | NA | NA | NA | NA |
| lizards | Bunopus_crassicauda | NA | DD | Unknown | NA | NA | NA | NA | NA |
| lizards | Bunopus_tuberculatus | Bunopus_tuberculatus | LC | Stable | temperate | cathemeral | 5.129 | 3918133.925 | 6.337 |
| lizards | Cachryx_defensor | Ctenosaura_defensor | VU | Unknown | tropical | diurnal | 162.181 | 45697.637 | 7.976 |
| lizards | Caesoris_novaecaledoniae | Lioscincus_novaecaledoniae | LC | Unknown | tropical | diurnal | 6.166 | 11307.158 | 7.131 |
| lizards | Caledoniscincus_aquilonius | Caledoniscincus_aquilonius | NT | Unknown | tropical | diurnal | 2.138 | 4378.346 | 5.927 |
| lizards | Caledoniscincus_atropunctatus | Caledoniscincus_atropunctatus | LC | Stable | tropical | diurnal | 3.09 | 20414.732 | 7.479 |
| lizards | Caledoniscincus_auratus | Caledoniscincus_auratus | EN | Decreasing | tropical | diurnal | 2.455 | 709.309 | 11.958 |
| lizards | Caledoniscincus_austrocaledonicus | Caledoniscincus_austrocaledonicus | LC | Stable | tropical | diurnal | 3.467 | 18804.953 | 7.571 |
| lizards | Caledoniscincus_chazeaui | Caledoniscincus_chazeaui | EN | Decreasing | tropical | diurnal | 1.413 | 1552.383 | 4.796 |
| lizards | Caledoniscincus_constellatus | NA | CR | Decreasing | NA | NA | NA | NA | NA |
| lizards | Caledoniscincus_cryptos | NA | DD | Unknown | NA | NA | NA | NA | NA |
| lizards | Caledoniscincus_festivus | Caledoniscincus_festivus | LC | Unknown | tropical | diurnal | 7.413 | 16395.487 | 7.512 |
| lizards | Caledoniscincus_haplorhinus | NA | LC | Stable | NA | NA | NA | NA | NA |
| lizards | Caledoniscincus_notialis | NA | NT | Unknown | NA | NA | NA | NA | NA |
| lizards | Caledoniscincus_orestes | Caledoniscincus_orestes | VU | Decreasing | tropical | diurnal | 5.37 | 1981.227 | 5.036 |
| lizards | Caledoniscincus_pelletieri | Caledoniscincus_pelletieri | EN | Decreasing | tropical | diurnal | 3.802 | 46.543 | 12.75 |
| lizards | Caledoniscincus_renevieri | Caledoniscincus_renevieri | EN | Decreasing | tropical | diurnal | 2.455 | 730.444 | 4.735 |
| lizards | Caledoniscincus_terma | NA | DD | Unknown | NA | NA | NA | NA | NA |
| lizards | Callisaurus_draconoides | Callisaurus_draconoides | LC | Stable | temperate | diurnal | 39.811 | 798757.742 | 7.177 |
| lizards | Callopistes_flavipunctatus | NA | NT | Decreasing | NA | NA | NA | NA | NA |
| lizards | Callopistes_maculatus | Callopistes_maculatus | LC | Decreasing | temperate | diurnal | 162.181 | 101772.523 | 7.886 |
| lizards | Calodactylodes_aureus | Calodactylodes_aureus | LC | not_available | tropical | cathemeral | 13.804 | 121948.801 | 15.705 |
| lizards | Calodactylodes_illingworthorum | NA | VU | Decreasing | NA | NA | NA | NA | NA |
| lizards | Calotes_bachae | Calotes_bachae | LC | Stable | tropical | diurnal | 30.2 | 103754.129 | 12.074 |
| lizards | Calotes_bhutanensis | NA | DD | Unknown | NA | NA | NA | NA | NA |
| lizards | Calotes_calotes | Calotes_calotes | LC | Stable | tropical | diurnal | 95.499 | 153325.473 | 20.441 |
| lizards | Calotes_ceylonensis | Calotes_ceylonensis | LC | Unknown | tropical | diurnal | 18.621 | 31880.401 | 15.769 |
| lizards | Calotes_chincollium | NA | LC | Stable | NA | NA | NA | NA | NA |
| lizards | Calotes_desilvai | Calotes_desilvai | CR | Unknown | tropical | diurnal | 16.596 | 9.991 | 13.276 |
| lizards | Calotes_emma | Calotes_emma | LC | Stable | tropical | diurnal | 66.069 | 2135284.533 | 12.75 |
| lizards | Calotes_grandisquamis | Calotes_grandisquamis | LC | Unknown | tropical | diurnal | 117.49 | 32181.388 | 14.55 |
| lizards | Calotes_htunwini | NA | LC | Unknown | NA | NA | NA | NA | NA |
| lizards | Calotes_irawadi | NA | LC | Unknown | NA | NA | NA | NA | NA |

| | | | | | | | | | |
|---|---|---|---|---|---|---|---|---|---|
| lizards | Calotes_jerdoni | Calotes_jerdoni | LC | Unknown | temperate | diurnal | 75.858 | 613086.384 | 9.953 |
| lizards | Calotes_liocephalus | Calotes_liocephalus | EN | Decreasing | tropical | diurnal | 33.113 | 2402.124 | 15.467 |
| lizards | Calotes_liolepis | Calotes_liolepis | LC | Unknown | tropical | diurnal | 33.113 | 29660.608 | 17.878 |
| lizards | Calotes_manamendrai | NA | EN | Unknown | NA | NA | NA | NA | NA |
| lizards | Calotes_maria | NA | LC | Unknown | NA | NA | NA | NA | NA |
| lizards | Calotes_medogensis | NA | LC | Stable | NA | NA | NA | NA | NA |
| lizards | Calotes_minor | Brachysaura_minor | LC | Decreasing | temperate | diurnal | 30.2 | 1862266.059 | 17.509 |
| lizards | Calotes_mystaceus | Calotes_mystaceus | LC | Stable | tropical | diurnal | 138.038 | 1617569.774 | 12.745 |
| lizards | Calotes_nemoricola | Calotes_nemoricola | LC | Unknown | tropical | diurnal | 117.49 | 20157.297 | 13.933 |
| lizards | Calotes_nigrilabris | Calotes_nigrilabris | EN | Decreasing | tropical | diurnal | 95.499 | 2047.695 | 17.483 |
| lizards | Calotes_paulus | Oriocalotes_paulus | EN | Unknown | temperate | diurnal | 12.023 | 49343.844 | 7.054 |
| lizards | Calotes_pethiyagodai | Calotes_pethiyagodai | EN | Unknown | tropical | diurnal | 25.704 | 240.345 | 8.674 |
| lizards | Calotes_versicolor | Calotes_versicolor | LC | Stable | tropical | diurnal | 117.49 | 6740870.497 | 15.142 |
| lizards | Calumma_amber | NA | NT | Stable | NA | NA | NA | NA | NA |
| lizards | Calumma_ambreense | NA | NT | Stable | NA | NA | NA | NA | NA |
| lizards | Calumma_brevicorne | Calumma_brevicorne | LC | Decreasing | tropical | diurnal | 95.499 | 89712.197 | 7.98 |
| lizards | Calumma_cucullatum | NA | VU | Decreasing | NA | NA | NA | NA | NA |
| lizards | Calumma_gallus | NA | EN | Decreasing | NA | NA | NA | NA | NA |
| lizards | Calumma_gastrotaenia | NA | LC | Unknown | NA | NA | NA | NA | NA |
| lizards | Calumma_gehringi | NA | not_available | not_available | NA | NA | NA | NA | NA |
| lizards | Calumma_hilleniusi | Calumma_hilleniusi | EN | Decreasing | tropical | diurnal | 10 | 820.952 | 6.161 |
| lizards | Calumma_jejy | NA | VU | Stable | NA | NA | NA | NA | NA |
| lizards | Calumma_nasutum | Calumma_nasutum | LC | Decreasing | tropical | diurnal | 3.631 | 203379.202 | 9.076 |
| lizards | Calumma_oshaughnessyi | NA | VU | Decreasing | NA | NA | NA | NA | NA |
| lizards | Calumma_parsonii | Calumma_parsonii | NT | Decreasing | tropical | diurnal | 436.516 | 63315.071 | 7.592 |
| lizards | Calumma_peyrierasi | NA | VU | Unknown | NA | NA | NA | NA | NA |
| lizards | Calumma_vohibola | NA | EN | Decreasing | NA | NA | NA | NA | NA |
| lizards | Calyptommatus_confusionibus | NA | LC | Unknown | NA | NA | NA | NA | NA |
| lizards | Calyptommatus_leiolepis | Calyptommatus_leiolepis | EN | Unknown | tropical | nocturnal | 3.02 | 28595.987 | 5.66 |
| lizards | Calyptommatus_nicterus | NA | EN | Unknown | NA | NA | NA | NA | NA |
| lizards | Calyptommatus_sinebrachiatus | Calyptommatus_sinebrachiatus | EN | Unknown | tropical | nocturnal | 2.344 | 106.289 | 5.938 |
| lizards | Calyptotis_lepidorostrum | Calyptotis_lepidorostrum | LC | Stable | temperate | cathemeral | 3.09 | 76721.46 | 7.318 |
| lizards | Calyptotis_ruficauda | Calyptotis_ruficauda | LC | Unknown | temperate | cathemeral | 3.09 | 36946.693 | 6.947 |
| lizards | Calyptotis_scutirostrum | Calyptotis_scutirostrum | LC | Stable | temperate | cathemeral | 3.89 | 145402.795 | 7.795 |

| | | | | | | | | | |
|---|---|---|---|---|---|---|---|---|---|
| lizards | Caparaonia_itaiquara | Caparaonia_itaiquara | LC | Unknown | tropical | diurnal | 4.898 | 193.443 | 5.548 |
| lizards | Caribicus_anelpistus | NA | CR | Unknown | NA | NA | NA | NA | NA |
| lizards | Caribicus_darlingtoni | NA | EN | Unknown | NA | NA | NA | NA | NA |
| lizards | Caribicus_warreni | NA | VU | Decreasing | NA | NA | NA | NA | NA |
| lizards | Carinascincus_coventryi | Niveoscincus_coventryi | LC | Unknown | temperate | diurnal | 2.951 | 133421.326 | 9.692 |
| lizards | Carinascincus_greeni | Niveoscincus_greeni | VU | Stable | temperate | diurnal | 8.511 | 15462.578 | 4.452 |
| lizards | Carinascincus_metallicus | Niveoscincus_metallicus | LC | Stable | temperate | diurnal | 7.079 | 96536.609 | 8.616 |
| lizards | Carinascincus_microlepidotus | Niveoscincus_microlepidotus | VU | Unknown | temperate | diurnal | 6.761 | 19919.415 | 3.969 |
| lizards | Carinascincus_ocellatus | Niveoscincus_ocellatus | LC | Stable | temperate | diurnal | 12.882 | 47312.972 | 6.746 |
| lizards | Carinascincus_orocryptus | Niveoscincus_orocryptus | VU | Unknown | temperate | diurnal | 5.754 | 13931.473 | 2.454 |
| lizards | Carinascincus_palfreymani | NA | VU | Unknown | NA | NA | NA | NA | NA |
| lizards | Carinascincus_pretiosus | Niveoscincus_pretiosus | LC | Stable | temperate | diurnal | 6.761 | 67928.473 | 5.969 |
| lizards | Carlia_aenigma | NA | LC | Unknown | NA | NA | NA | NA | NA |
| lizards | Carlia_ailanpalai | Carlia_ailanpalai | LC | Unknown | tropical | diurnal | 5.888 | 1977.083 | 6.377 |
| lizards | Carlia_amax | Carlia_amax | LC | Stable | tropical | diurnal | 1.202 | 753606.382 | 2.093 |
| lizards | Carlia_aramia | NA | LC | Unknown | NA | NA | NA | NA | NA |
| lizards | Carlia_babarensis | NA | LC | Stable | NA | NA | NA | NA | NA |
| lizards | Carlia_beccarii | NA | LC | Stable | NA | NA | NA | NA | NA |
| lizards | Carlia_bicarinata | Carlia_bicarinata | LC | Unknown | tropical | diurnal | 2.042 | 8061.148 | 7.038 |
| lizards | Carlia_bomberai | Carlia_bomberai | LC | Unknown | tropical | diurnal | 2.951 | 24.826 | 13.429 |
| lizards | Carlia_caesius | Carlia_caesius | LC | Unknown | tropical | diurnal | 5.248 | 60131.516 | 2.405 |
| lizards | Carlia_crypta | NA | not_available | not_available | NA | NA | NA | NA | NA |
| lizards | Carlia_decora | NA | LC | Unknown | NA | NA | NA | NA | NA |
| lizards | Carlia_diguliensis | NA | LC | Unknown | NA | NA | NA | NA | NA |
| lizards | Carlia_dogare | NA | LC | Stable | NA | NA | NA | NA | NA |
| lizards | Carlia_eothen | Carlia_eothen | LC | Unknown | tropical | diurnal | 6.31 | 7207.16 | 5.436 |
| lizards | Carlia_fusca | NA | LC | Unknown | NA | NA | NA | NA | NA |
| lizards | Carlia_gracilis | Carlia_gracilis | LC | Stable | tropical | diurnal | 1.202 | 280253.165 | 1.678 |
| lizards | Carlia_inconnexa | NA | LC | Unknown | NA | NA | NA | NA | NA |
| lizards | Carlia_isostriacantha | NA | not_available | not_available | NA | NA | NA | NA | NA |
| lizards | Carlia_jarnoldae | Carlia_jarnoldae | LC | Unknown | tropical | diurnal | 2.138 | 239066.046 | 3.653 |
| lizards | Carlia_johnstonei | NA | LC | Stable | NA | NA | NA | NA | NA |
| lizards | Carlia_longipes | Carlia_longipes | LC | Stable | tropical | diurnal | 7.762 | 60361.701 | 3.078 |
| lizards | Carlia_luctuosa | NA | LC | Unknown | NA | NA | NA | NA | NA |

| | | | | | | | | | |
|---|---|---|---|---|---|---|---|---|---|
| lizards | Carlia_munda | Carlia_munda | LC | Stable | tropical | diurnal | 1.514 | 2703686.596 | 3.298 |
| lizards | Carlia_mysi | Carlia_mysi | LC | Stable | tropical | diurnal | 4.898 | 71129.446 | 5.427 |
| lizards | Carlia_pectoralis | Carlia_pectoralis | LC | Stable | temperate | diurnal | 5.129 | 218228.788 | 6.614 |
| lizards | Carlia_peronii | Carlia_peronii | LC | Unknown | tropical | diurnal | 1.413 | 7243.731 | 14.796 |
| lizards | Carlia_pulla | NA | LC | Unknown | NA | NA | NA | NA | NA |
| lizards | Carlia_quinquecarinata | NA | DD | Unknown | NA | NA | NA | NA | NA |
| lizards | Carlia_rhomboidalis | Carlia_rhomboidalis | LC | Unknown | tropical | diurnal | 4.365 | 11002.224 | 6.815 |
| lizards | Carlia_rostralis | Carlia_rostralis | LC | Stable | tropical | diurnal | 6.761 | 41863.247 | 3.969 |
| lizards | Carlia_rubigo | NA | LC | Stable | NA | NA | NA | NA | NA |
| lizards | Carlia_rubrigularis | Carlia_rubrigularis | LC | Stable | tropical | diurnal | 4.365 | 21875.898 | 5.035 |
| lizards | Carlia_rufilatus | Carlia_rufilatus | LC | Stable | tropical | diurnal | 1.318 | 299086.368 | 2.207 |
| lizards | Carlia_schmeltzii | Carlia_schmeltzii | LC | Stable | tropical | diurnal | 6.457 | 690904.071 | 5.268 |
| lizards | Carlia_sexdentata | NA | LC | Stable | NA | NA | NA | NA | NA |
| lizards | Carlia_storri | Carlia_storri | LC | Unknown | tropical | diurnal | 1.738 | 147749.918 | 2.761 |
| lizards | Carlia_tetradactyla | Carlia_tetradactyla | LC | Unknown | temperate | diurnal | 5.129 | 242145.604 | 8.979 |
| lizards | Carlia_triacantha | Carlia_triacantha | LC | Stable | tropical | diurnal | 2.754 | 1878749.832 | 1.878 |
| lizards | Carlia_tutela | Carlia_tutela | LC | Stable | tropical | diurnal | 2.884 | 1773.801 | 9.825 |
| lizards | Carlia_vivax | Carlia_vivax | LC | Stable | tropical | diurnal | 2.291 | 745884.014 | 5.617 |
| lizards | Carlia_wundalthini | NA | VU | Unknown | NA | NA | NA | NA | NA |
| lizards | Carphodactylus_laevis | Carphodactylus_laevis | LC | Stable | tropical | nocturnal | 33.884 | 11029.809 | 5.255 |
| lizards | Celatiscincus_euryotis | Celatiscincus_euryotis | EN | Decreasing | tropical | diurnal | 1.318 | 143.523 | 10.853 |
| lizards | Celatiscincus_similis | Celatiscincus_similis | EN | Decreasing | tropical | diurnal | 1.698 | 451.676 | 5.178 |
| lizards | Celestus_fowleri | NA | VU | Stable | NA | NA | NA | NA | NA |
| lizards | Celestus_hewardi | NA | EN | Decreasing | NA | NA | NA | NA | NA |
| lizards | Centrosaura_apodema | NA | LC | Stable | NA | NA | NA | NA | NA |
| lizards | Ceratophora_aspera | Ceratophora_aspera | EN | Unknown | tropical | diurnal | 2.692 | 10376.647 | 20.002 |
| lizards | Ceratophora_erdeleni | Ceratophora_erdeleni | CR | Decreasing | tropical | diurnal | 19.498 | 85.304 | 14.675 |
| lizards | Ceratophora_karu | Ceratophora_karu | CR | Unknown | tropical | diurnal | 1.95 | 19.982 | 13.875 |
| lizards | Ceratophora_stoddartii | Ceratophora_stoddartii | EN | Unknown | tropical | diurnal | 22.909 | 3144.987 | 17.406 |
| lizards | Ceratophora_tennentii | Ceratophora_tennentii | EN | Decreasing | tropical | diurnal | 23.988 | 751.854 | 11.549 |
| lizards | Cercosaura_argulus | Cercosaura_argulus | LC | Unknown | tropical | diurnal | 2.692 | 4709101.625 | 2.307 |
| lizards | Cercosaura_bassleri | NA | not_available | not_available | NA | NA | NA | NA | NA |
| lizards | Cercosaura_doanae | NA | not_available | not_available | NA | NA | NA | NA | NA |
| lizards | Cercosaura_eigenmanni | Cercosaura_eigenmanni | LC | Stable | tropical | diurnal | 3.09 | 1229760.579 | 3.988 |

| | | | | | | | | | |
|---|---|---|---|---|---|---|---|---|---|
| lizards | Cercosaura_hypnoides | NA | DD | Unknown | NA | NA | NA | NA | NA |
| lizards | Cercosaura_manicata | Cercosaura_manicata | LC | Unknown | tropical | diurnal | 9.333 | 42492.95 | 8.054 |
| lizards | Cercosaura_ocellata | Cercosaura_ocellata | LC | Stable | tropical | diurnal | 6.457 | 8056581.455 | 4.715 |
| lizards | Cercosaura_oshaughnessyi | NA | not_available | not_available | NA | NA | NA | NA | NA |
| lizards | Cercosaura_parkeri | Cercosaura_parkeri | LC | Stable | tropical | diurnal | 2.344 | 769015.043 | 5.335 |
| lizards | Cercosaura_schreibersii | Cercosaura_schreibersii | LC | Stable | temperate | diurnal | 3.715 | 5106870.971 | 8.86 |
| lizards | Chalarodon_madagascariensis | NA | LC | Stable | NA | NA | NA | NA | NA |
| lizards | Chalarodon_steinkampi | NA | not_available | not_available | NA | NA | NA | NA | NA |
| lizards | Chalcides_bedriagai | Chalcides_bedriagai | NT | Decreasing | temperate | diurnal | 19.953 | 364193.696 | 17.082 |
| lizards | Chalcides_boulengeri | NA | LC | Stable | NA | NA | NA | NA | NA |
| lizards | Chalcides_chalcides | Chalcides_chalcides | LC | Decreasing | temperate | cathemeral | 33.884 | 241759.964 | 22.757 |
| lizards | Chalcides_coeruleopunctatus | NA | not_available | not_available | NA | NA | NA | NA | NA |
| lizards | Chalcides_colosii | Chalcides_colosii | LC | Decreasing | temperate | diurnal | 33.113 | 16857.227 | 17.679 |
| lizards | Chalcides_delislei | NA | LC | Stable | NA | NA | NA | NA | NA |
| lizards | Chalcides_ebneri | NA | CR | Decreasing | NA | NA | NA | NA | NA |
| lizards | Chalcides_guentheri | Chalcides_guentheri | VU | Decreasing | temperate | diurnal | 10.715 | 19774.001 | 29.587 |
| lizards | Chalcides_lanzai | Chalcides_lanzai | NT | Unknown | temperate | diurnal | 38.905 | 13339.026 | 14.484 |
| lizards | Chalcides_levitoni | NA | DD | not_available | NA | NA | NA | NA | NA |
| lizards | Chalcides_manueli | Chalcides_manueli | VU | Decreasing | temperate | diurnal | 8.913 | 24838.642 | 17.337 |
| lizards | Chalcides_mauritanicus | NA | EN | Decreasing | NA | NA | NA | NA | NA |
| lizards | Chalcides_mertensi | Chalcides_mertensi | LC | Unknown | temperate | diurnal | 28.184 | 234082.954 | 16.442 |
| lizards | Chalcides_minutus | Chalcides_minutus | VU | Decreasing | temperate | diurnal | 7.586 | 24432.161 | 13.228 |
| lizards | Chalcides_mionecton | Chalcides_mionecton | LC | Unknown | temperate | cathemeral | 6.457 | 166737.182 | 15.099 |
| lizards | Chalcides_montanus | Chalcides_montanus | NT | Stable | temperate | diurnal | 20.893 | 64863.929 | 9.889 |
| lizards | Chalcides_ocellatus | Chalcides_ocellatus | LC | Stable | tropical | cathemeral | 97.724 | 9309659.675 | 6.859 |
| lizards | Chalcides_parallelus | Chalcides_parallelus | EN | Decreasing | temperate | diurnal | 16.218 | 1479.52 | 24.768 |
| lizards | Chalcides_polylepis | Chalcides_polylepis | LC | Decreasing | temperate | cathemeral | 97.724 | 169524.698 | 14.566 |
| lizards | Chalcides_pseudostriatus | Chalcides_pseudostriatus | NT | Decreasing | temperate | diurnal | 30.903 | 101093.957 | 17.317 |
| lizards | Chalcides_ragazzii | Chalcides_ragazzii | LC | Unknown | tropical | diurnal | 46.774 | 941595.6 | 7.406 |
| lizards | Chalcides_sepsoides | Chalcides_sepsoides | LC | Stable | temperate | cathemeral | 7.762 | 893517.286 | 3.951 |
| lizards | Chalcides_sexlineatus | Chalcides_sexlineatus | LC | Stable | temperate | diurnal | 20.893 | 1538.293 | 21.897 |
| lizards | Chalcides_sphenopsiformis | NA | LC | Stable | NA | NA | NA | NA | NA |
| lizards | Chalcides_striatus | Chalcides_striatus | LC | Decreasing | temperate | diurnal | 33.884 | 530079.993 | 18.206 |
| lizards | Chalcides_viridanus | Chalcides_viridanus | LC | Stable | temperate | diurnal | 48.978 | 2033.771 | 20.379 |

| | | | | | | | | | |
|---|---|---|---|---|---|---|---|---|---|
| lizards | Chalcidoseps_thwaitesi | Chalcidoseps_thwaitesi | EN | Unknown | tropical | diurnal | 16.982 | 1290.761 | 16.382 |
| lizards | Chamaeleo_africanus | Chamaeleo_africanus | LC | Stable | tropical | diurnal | 147.911 | 4788798.972 | 10.404 |
| lizards | Chamaeleo_arabicus | NA | LC | not_available | NA | NA | NA | NA | NA |
| lizards | Chamaeleo_calyptratus | Chamaeleo_calyptratus | LC | Stable | tropical | diurnal | 239.883 | 96601.517 | 13.221 |
| lizards | Chamaeleo_chamaeleon | Chamaeleo_chamaeleon | LC | Stable | temperate | diurnal | 95.499 | 1963765.503 | 10.969 |
| lizards | Chamaeleo_dilepis | Chamaeleo_dilepis | LC | Stable | tropical | diurnal | 138.038 | 11094336.994 | 8.024 |
| lizards | Chamaeleo_gracilis | Chamaeleo_gracilis | LC | Stable | tropical | diurnal | 128.825 | 11136998.652 | 9.224 |
| lizards | Chamaeleo_laevigatus | Chamaeleo_laevigatus | LC | Stable | tropical | diurnal | 67.608 | 2382716.521 | 8.998 |
| lizards | Chamaeleo_monachus | Chamaeleo_monachus | NT | Unknown | tropical | diurnal | 102.329 | 2898.393 | 11.033 |
| lizards | Chamaeleo_namaquensis | Chamaeleo_namaquensis | LC | Unknown | tropical | diurnal | 81.283 | 875059.99 | 5.347 |
| lizards | Chamaeleo_senegalensis | Chamaeleo_senegalensis | LC | Unknown | tropical | diurnal | 70.795 | 3770462.909 | 11.189 |
| lizards | Chamaeleo_zeylanicus | Chamaeleo_zeylanicus | LC | Unknown | tropical | diurnal | 229.087 | 2090346.115 | 18.572 |
| lizards | Chamaesaura_aenea | Chamaesaura_aenea | NT | Decreasing | temperate | diurnal | 11.749 | 94654.009 | 13.066 |
| lizards | Chamaesaura_anguina | Chamaesaura_anguina | LC | Stable | tropical | diurnal | 15.488 | 334093.384 | 14.153 |
| lizards | Chamaesaura_macrolepis | Chamaesaura_macrolepis | LC | Stable | temperate | diurnal | 19.055 | 15302.753 | 14.767 |
| lizards | Chamaesaura_miopropus | Chamaesaura_miopropus | LC | Stable | tropical | diurnal | 8.511 | 218446.37 | 11.011 |
| lizards | Chamaesaura_tenuior | Chamaesaura_tenuior | LC | Unknown | tropical | diurnal | 14.791 | 105658.748 | 17.978 |
| lizards | Chatogekko_amazonicus | Chatogekko_amazonicus | LC | Stable | tropical | diurnal | 0.617 | 5812690.696 | 1.888 |
| lizards | Chelosania_brunnea | NA | DD | Unknown | NA | NA | NA | NA | NA |
| lizards | Chioninia_coctei | NA | EX | not_available | NA | NA | NA | NA | NA |
| lizards | Chioninia_delalandii | Chioninia_delalandii | LC | Stable | tropical | diurnal | 16.596 | 1522.711 | 14.105 |
| lizards | Chioninia_fogoensis | NA | LC | Unknown | NA | NA | NA | NA | NA |
| lizards | Chioninia_nicolauensis | NA | LC | Stable | NA | NA | NA | NA | NA |
| lizards | Chioninia_spinalis | NA | LC | Stable | NA | NA | NA | NA | NA |
| lizards | Chioninia_stangeri | NA | NT | Unknown | NA | NA | NA | NA | NA |
| lizards | Chioninia_vaillantii | Chioninia_vaillantii | EN | Decreasing | tropical | diurnal | 363.078 | 1212.235 | 14.735 |
| lizards | Chlamydosaurus_kingii | Chlamydosaurus_kingii | LC | Unknown | tropical | diurnal | 912.011 | 1508834.779 | 3.319 |
| lizards | Chondrodactylus_angulifer | Chondrodactylus_angulifer | LC | Stable | temperate | nocturnal | 26.915 | 886144.598 | 5.338 |
| lizards | Chondrodactylus_bibronii | Chondrodactylus_bibronii | LC | Stable | tropical | nocturnal | 19.055 | 1160964.314 | 6.649 |
| lizards | Chondrodactylus_fitzsimonsi | Chondrodactylus_fitzsimonsi | LC | Stable | tropical | nocturnal | 14.125 | 210931.635 | 5.725 |
| lizards | Chondrodactylus_laevigatus | Pachydactylus_laevigatus | LC | Stable | tropical | cathemeral | 14.791 | 803157.118 | 5.106 |
| lizards | Chondrodactylus_turneri | Chondrodactylus_turneri | LC | Stable | tropical | nocturnal | 16.596 | 2890005.347 | 7.607 |
| lizards | Christinus_alexanderi | Christinus_alexanderi | LC | Stable | temperate | nocturnal | 7.079 | 97691.6 | 3.192 |
| lizards | Christinus_guentheri | Christinus_guentheri | VU | Decreasing | temperate | nocturnal | 19.953 | 51.336 | 6.722 |

| | | | | | | | | | |
|---|---|---|---|---|---|---|---|---|---|
| lizards | Christinus_marmoratus | Christinus_marmoratus | LC | Stable | temperate | nocturnal | 7.079 | 1169407.926 | 8.308 |
| lizards | Cnemaspis_adii | NA | CR | Decreasing | NA | NA | NA | NA | NA |
| lizards | Cnemaspis_affinis | Cnemaspis_affinis | VU | Unknown | tropical | diurnal | 2.951 | 95.393 | 32.282 |
| lizards | Cnemaspis_alwisi | NA | EN | Unknown | NA | NA | NA | NA | NA |
| lizards | Cnemaspis_andalas | NA | LC | Unknown | NA | NA | NA | NA | NA |
| lizards | Cnemaspis_argus | Cnemaspis_argus | LC | Stable | tropical | diurnal | 6.026 | 487.24 | 1.228 |
| lizards | Cnemaspis_assamensis | Cnemaspis_assamensis | VU | Unknown | temperate | diurnal | 0.912 | 12358.349 | 11.58 |
| lizards | Cnemaspis_aurantiacopes | NA | DD | Unknown | NA | NA | NA | NA | NA |
| lizards | Cnemaspis_baueri | Cnemaspis_baueri | LC | Stable | tropical | cathemeral | 6.457 | 15.583 | 2.669 |
| lizards | Cnemaspis_bayuensis | Cnemaspis_bayuensis | VU | Unknown | tropical | diurnal | 2.239 | 9.991 | 13.785 |
| lizards | Cnemaspis_beddomei | NA | DD | Unknown | NA | NA | NA | NA | NA |
| lizards | Cnemaspis_bidongensis | Cnemaspis_bidongensis | LC | Stable | tropical | cathemeral | 4.266 | 2.196 | 2.821 |
| lizards | Cnemaspis_biocellata | Cnemaspis_biocellata | LC | Stable | tropical | cathemeral | 1.549 | 181.302 | 24.909 |
| lizards | Cnemaspis_boulengeri | Cnemaspis_boulengerii | NT | Stable | tropical | diurnal | 6.918 | 73.206 | 19.291 |
| lizards | Cnemaspis_caudanivea | Cnemaspis_caudanivea | VU | Stable | tropical | cathemeral | 2.399 | 3.072 | 17.665 |
| lizards | Cnemaspis_chanardi | Cnemaspis_chanardi | LC | Stable | tropical | diurnal | 1.514 | 6519.914 | 11.047 |
| lizards | Cnemaspis_chanthaburiensis | Cnemaspis_chanthaburiensis | LC | Stable | tropical | cathemeral | 1.778 | 12507.987 | 15.376 |
| lizards | Cnemaspis_dezwaani | NA | DD | Unknown | NA | NA | NA | NA | NA |
| lizards | Cnemaspis_flavigaster | Cnemaspis_flavigaster | LC | Stable | tropical | cathemeral | 2.818 | 172.871 | 26.396 |
| lizards | Cnemaspis_flaviventralis | NA | EN | Unknown | NA | NA | NA | NA | NA |
| lizards | Cnemaspis_flavolineata | NA | DD | Unknown | NA | NA | NA | NA | NA |
| lizards | Cnemaspis_gemunu | Cnemaspis_gemunu | VU | Unknown | tropical | diurnal | 0.977 | 379.735 | 16.021 |
| lizards | Cnemaspis_goaensis | NA | EN | Unknown | NA | NA | NA | NA | NA |
| lizards | Cnemaspis_gracilis | Cnemaspis_gracilis | LC | Unknown | tropical | nocturnal | 1.514 | 28544.644 | 20.065 |
| lizards | Cnemaspis_grismeri | Cnemaspis_grismeri | LC | Stable | tropical | diurnal | 2.884 | 10.951 | 20.482 |
| lizards | Cnemaspis_hangus | Cnemaspis_hangus | NT | Unknown | tropical | diurnal | 2.884 | 9.991 | 1.861 |
| lizards | Cnemaspis_harimau | Cnemaspis_harimau | VU | Unknown | tropical | nocturnal | 1.585 | 9.991 | 29.537 |
| lizards | Cnemaspis_heteropholis | Cnemaspis_heteropholis | NT | Unknown | tropical | cathemeral | 2.138 | 14604.049 | 11.99 |
| lizards | Cnemaspis_huaseesom | Cnemaspis_huaseesom | LC | Stable | tropical | nocturnal | 1.905 | 3078.8 | 12.478 |
| lizards | Cnemaspis_indica | NA | VU | Unknown | NA | NA | NA | NA | NA |
| lizards | Cnemaspis_jerdonii | Cnemaspis_jerdonii | VU | Unknown | tropical | cathemeral | 1.862 | 13830.96 | 12.078 |
| lizards | Cnemaspis_kallima | NA | EN | Unknown | NA | NA | NA | NA | NA |
| lizards | Cnemaspis_kamolnorranathi | NA | LC | Stable | NA | NA | NA | NA | NA |
| lizards | Cnemaspis_kandambyi | NA | DD | Unknown | NA | NA | NA | NA | NA |

| | | | | | | | | | |
|---|---|---|---|---|---|---|---|---|---|
| lizards | Cnemaspis_kandiana | Cnemaspis_kandiana | EN | Unknown | tropical | diurnal | 1.122 | 2342.12 | 16.088 |
| lizards | Cnemaspis_karsticola | NA | VU | Unknown | NA | NA | NA | NA | NA |
| lizards | Cnemaspis_kendallii | Cnemaspis_kendallii | LC | Unknown | tropical | diurnal | 4.898 | 45212.33 | 11.967 |
| lizards | Cnemaspis_kolhapurensis | NA | DD | Unknown | NA | NA | NA | NA | NA |
| lizards | Cnemaspis_kottiyoorensis | NA | CR | Unknown | NA | NA | NA | NA | NA |
| lizards | Cnemaspis_kumarasinghei | Cnemaspis_kumarasinghei | EN | Decreasing | tropical | nocturnal | 0.794 | 1042.508 | 9.211 |
| lizards | Cnemaspis_kumpoli | Cnemaspis_kumpoli | LC | Stable | tropical | nocturnal | 5.37 | 20135.986 | 9.862 |
| lizards | Cnemaspis_laoensis | NA | DD | Unknown | NA | NA | NA | NA | NA |
| lizards | Cnemaspis_leucura | NA | DD | Unknown | NA | NA | NA | NA | NA |
| lizards | Cnemaspis_limi | Cnemaspis_limi | LC | Stable | tropical | cathemeral | 14.125 | 144.804 | 8.052 |
| lizards | Cnemaspis_lineogularis | NA | LC | Unknown | NA | NA | NA | NA | NA |
| lizards | Cnemaspis_littoralis | NA | DD | Unknown | NA | NA | NA | NA | NA |
| lizards | Cnemaspis_mahsuriae | NA | LC | Stable | NA | NA | NA | NA | NA |
| lizards | Cnemaspis_mcguirei | Cnemaspis_mcguirei | LC | Unknown | tropical | cathemeral | 5.888 | 359.967 | 12.075 |
| lizards | Cnemaspis_molligodai | NA | EN | Unknown | NA | NA | NA | NA | NA |
| lizards | Cnemaspis_monachorum | Cnemaspis_monachorum | LC | Unknown | tropical | diurnal | 1.047 | 9.991 | 26.941 |
| lizards | Cnemaspis_mumpuniae | Cnemaspis_mumpuniae | LC | Unknown | tropical | nocturnal | 4.898 | 393.563 | 8.726 |
| lizards | Cnemaspis_mysoriensis | NA | LC | Unknown | NA | NA | NA | NA | NA |
| lizards | Cnemaspis_nairi | Cnemaspis_nairi | NT | Unknown | tropical | diurnal | 1.995 | 5454.061 | 13.761 |
| lizards | Cnemaspis_narathiwatensis | NA | LC | Unknown | NA | NA | NA | NA | NA |
| lizards | Cnemaspis_neangthyi | NA | EN | Unknown | NA | NA | NA | NA | NA |
| lizards | Cnemaspis_nigridia | Cnemaspis_nigridia | LC | Unknown | tropical | diurnal | 13.804 | 390.955 | 7.355 |
| lizards | Cnemaspis_niyomwanae | NA | EN | Decreasing | NA | NA | NA | NA | NA |
| lizards | Cnemaspis_nuicamensis | Cnemaspis_nuicamensis | VU | Decreasing | tropical | diurnal | 2.512 | 173.509 | 20.049 |
| lizards | Cnemaspis_omari | Cnemaspis_omari | LC | Stable | tropical | diurnal | 1.66 | 19.982 | 11.717 |
| lizards | Cnemaspis_ornata | Cnemaspis_ornata | NT | Decreasing | tropical | diurnal | 3.802 | 5890.179 | 15.26 |
| lizards | Cnemaspis_otai | Cnemaspis_otai | VU | Unknown | tropical | diurnal | 0.832 | 474.979 | 29.901 |
| lizards | Cnemaspis_paripari | NA | CR | Unknown | NA | NA | NA | NA | NA |
| lizards | Cnemaspis_pava | NA | NT | Unknown | NA | NA | NA | NA | NA |
| lizards | Cnemaspis_pemanggilensis | Cnemaspis_pemanggilensis | LC | Stable | tropical | cathemeral | 8.913 | 9.333 | 1.879 |
| lizards | Cnemaspis_peninsularis | Cnemaspis_peninsularis | LC | Unknown | tropical | diurnal | 4.677 | 20411.098 | 9.522 |
| lizards | Cnemaspis_perhentianensis | Cnemaspis_perhentianensis | NT | Unknown | tropical | cathemeral | 2.399 | 7.847 | 17.115 |
| lizards | Cnemaspis_phangngaensis | NA | LC | Stable | NA | NA | NA | NA | NA |
| lizards | Cnemaspis_phillipsi | NA | CR | Unknown | NA | NA | NA | NA | NA |

| | | | | | | | | | |
|---|---|---|---|---|---|---|---|---|---|
| lizards | Cnemaspis_phuketensis | NA | LC | Stable | NA | NA | NA | NA | NA |
| lizards | Cnemaspis_podihuna | Cnemaspis_podihuna | VU | Decreasing | tropical | diurnal | 0.49 | 7456.339 | 15.903 |
| lizards | Cnemaspis_pseudomcguirei | Cnemaspis_pseudomcguirei | NT | Unknown | tropical | cathemeral | 1.778 | 19.982 | 9.064 |
| lizards | Cnemaspis_psychedelica | Cnemaspis_psychedelica | EN | Unknown | tropical | diurnal | 8.71 | 6.292 | 0.795 |
| lizards | Cnemaspis_punctata | NA | EN | Decreasing | NA | NA | NA | NA | NA |
| lizards | Cnemaspis_punctatonuchalis | Cnemaspis_punctatonuchalis | LC | Unknown | tropical | nocturnal | 2.754 | 9.991 | 12.443 |
| lizards | Cnemaspis_purnamai | NA | DD | Unknown | NA | NA | NA | NA | NA |
| lizards | Cnemaspis_rajabasa | NA | VU | Unknown | NA | NA | NA | NA | NA |
| lizards | Cnemaspis_rammalensis | Cnemaspis_rammalensis | CR | Unknown | tropical | diurnal | 3.467 | 9.991 | 13.417 |
| lizards | Cnemaspis_retigalensis | NA | CR | Unknown | NA | NA | NA | NA | NA |
| lizards | Cnemaspis_roticanai | Cnemaspis_roticanai | LC | Unknown | tropical | cathemeral | 2.344 | 9.991 | 19.581 |
| lizards | Cnemaspis_samanalensis | NA | CR | Unknown | NA | NA | NA | NA | NA |
| lizards | Cnemaspis_scalpensis | Cnemaspis_scalpensis | CR | Decreasing | tropical | diurnal | 1.23 | 478.362 | 22.099 |
| lizards | Cnemaspis_selamatkanmerapoh | Cnemaspis_selamatkanmerapoh | VU | Unknown | tropical | cathemeral | 1.905 | 9.991 | 6.921 |
| lizards | Cnemaspis_shahruli | Cnemaspis_shahruli | LC | Unknown | tropical | cathemeral | 1.175 | 2842.217 | 23.546 |
| lizards | Cnemaspis_siamensis | Cnemaspis_siamensis | LC | Stable | tropical | cathemeral | 1.738 | 34842.31 | 11.079 |
| lizards | Cnemaspis_silvula | Cnemaspis_silvula | LC | Unknown | tropical | diurnal | 1.047 | 2004.101 | 16.103 |
| lizards | Cnemaspis_sisparensis | NA | NT | Unknown | NA | NA | NA | NA | NA |
| lizards | Cnemaspis_stongensis | Cnemaspis_stongensis | LC | Unknown | tropical | nocturnal | 2.692 | 9.991 | 10.724 |
| lizards | Cnemaspis_sundagekko | NA | DD | Unknown | NA | NA | NA | NA | NA |
| lizards | Cnemaspis_sundainsula | Cnemaspis_sundainsula | LC | Stable | tropical | diurnal | 12.023 | 41.401 | 9.576 |
| lizards | Cnemaspis_tapanuli | NA | LC | Unknown | NA | NA | NA | NA | NA |
| lizards | Cnemaspis_temiah | Cnemaspis_temiah | CR | Decreasing | tropical | diurnal | 1.38 | 9.991 | 36.232 |
| lizards | Cnemaspis_thachanaensis | NA | NT | Unknown | NA | NA | NA | NA | NA |
| lizards | Cnemaspis_tropidogaster | Cnemaspis_tropidogaster | CR | Unknown | tropical | diurnal | 1.047 | 4356.179 | 19.446 |
| lizards | Cnemaspis_tucdupensis | Cnemaspis_tucdupensis | VU | Stable | tropical | cathemeral | 2.951 | 13.603 | 17.959 |
| lizards | Cnemaspis_upendrai | NA | VU | Unknown | NA | NA | NA | NA | NA |
| lizards | Cnemaspis_vandeventeri | NA | LC | Unknown | NA | NA | NA | NA | NA |
| lizards | Cnemaspis_whittenorum | NA | LC | Unknown | NA | NA | NA | NA | NA |
| lizards | Cnemaspis_wynadensis | NA | EN | Unknown | NA | NA | NA | NA | NA |
| lizards | Cnemaspis_yercaudensis | Cnemaspis_yercaudensis | EN | Unknown | tropical | nocturnal | 0.692 | 311.747 | 13.835 |
| lizards | Cnemidophorus_arenivagus | Cnemidophorus_arenivagus | LC | Unknown | tropical | diurnal | 17.783 | 9012.899 | 10.276 |
| lizards | Cnemidophorus_arubensis | Cnemidophorus_arubensis | LC | Stable | tropical | diurnal | 63.096 | 181.607 | 31.444 |
| lizards | Cnemidophorus_cryptus | Cnemidophorus_cryptus | LC | Unknown | tropical | diurnal | 10.715 | 851696.147 | 2.537 |

| | | | | | | | | | |
|---|---|---|---|---|---|---|---|---|---|
| lizards | Cnemidophorus_duellmani | NA | not_available | not_available | NA | NA | NA | NA | NA |
| lizards | Cnemidophorus_espeuti | NA | not_available | not_available | NA | NA | NA | NA | NA |
| lizards | Cnemidophorus_flavissimus | NA | NT | Unknown | NA | NA | NA | NA | NA |
| lizards | Cnemidophorus_gaigei | NA | not_available | not_available | NA | NA | NA | NA | NA |
| lizards | Cnemidophorus_gramivagus | Cnemidophorus_gramivagus | LC | Stable | tropical | diurnal | 46.774 | 446489.476 | 4.92 |
| lizards | Cnemidophorus_lemniscatus | Cnemidophorus_lemniscatus | LC | Stable | tropical | diurnal | 43.652 | 3115711.906 | 4.628 |
| lizards | Cnemidophorus_leucopsammus | NA | LC | Unknown | NA | NA | NA | NA | NA |
| lizards | Cnemidophorus_murinus | Cnemidophorus_murinus | LC | Stable | tropical | diurnal | 125.893 | 840.112 | 17.532 |
| lizards | Cnemidophorus_rostralis | NA | NT | Stable | NA | NA | NA | NA | NA |
| lizards | Cnemidophorus_ruatanus | NA | not_available | not_available | NA | NA | NA | NA | NA |
| lizards | Cnemidophorus_ruthveni | Cnemidophorus_ruthveni | LC | Stable | tropical | diurnal | 107.152 | 17.023 | 18.945 |
| lizards | Cnemidophorus_senectus | NA | LC | Stable | NA | NA | NA | NA | NA |
| lizards | Cnemidophorus_splendidus | NA | not_available | not_available | NA | NA | NA | NA | NA |
| lizards | Cnemidophorus_vanzoi | NA | CR | Increasing | NA | NA | NA | NA | NA |
| lizards | Coeranoscincus_frontalis | NA | LC | Stable | NA | NA | NA | NA | NA |
| lizards | Coeranoscincus_reticulatus | Coeranoscincus_reticulatus | LC | Stable | temperate | cathemeral | 28.184 | 37562.284 | 10.25 |
| lizards | Coggeria_naufragus | NA | LC | Stable | NA | NA | NA | NA | NA |
| lizards | Coleodactylus_brachystoma | NA | LC | Unknown | NA | NA | NA | NA | NA |
| lizards | Coleodactylus_elizae | NA | DD | Unknown | NA | NA | NA | NA | NA |
| lizards | Coleodactylus_meridionalis | Coleodactylus_meridionalis | LC | Unknown | tropical | diurnal | 0.708 | 1486866.644 | 8.086 |
| lizards | Coleodactylus_natalensis | NA | EN | Unknown | NA | NA | NA | NA | NA |
| lizards | Coleodactylus_septentrionalis | Coleodactylus_septentrionalis | LC | Stable | tropical | diurnal | 0.813 | 399484.314 | 3.415 |
| lizards | Coleonyx_brevis | Coleonyx_brevis | LC | Stable | temperate | nocturnal | 5.37 | 572250.262 | 9.109 |
| lizards | Coleonyx_elegans | Coleonyx_elegans | LC | Stable | tropical | nocturnal | 37.154 | 504670.552 | 9.162 |
| lizards | Coleonyx_fasciatus | NA | LC | Decreasing | NA | NA | NA | NA | NA |
| lizards | Coleonyx_gypsicolus | Coleonyx_gypsicolus | LC | Stable | temperate | nocturnal | 13.804 | 28.412 | 8.929 |
| lizards | Coleonyx_mitratus | Coleonyx_mitratus | LC | Stable | tropical | nocturnal | 18.621 | 98543.902 | 15.476 |
| lizards | Coleonyx_reticulatus | Coleonyx_reticulatus | LC | Stable | temperate | nocturnal | 16.596 | 13245.73 | 5.927 |
| lizards | Coleonyx_switaki | Coleonyx_switaki | LC | Unknown | temperate | nocturnal | 12.589 | 35273.063 | 2.507 |
| lizards | Coleonyx_variegatus | Coleonyx_variegatus | LC | Stable | temperate | nocturnal | 8.71 | 624429.513 | 7.501 |
| lizards | Colobosaura_modesta | Colobosaura_modesta | LC | Unknown | tropical | diurnal | 3.715 | 3349736.173 | 7.26 |
| lizards | Colobosauroides_carvalhoi | NA | DD | Unknown | NA | NA | NA | NA | NA |
| lizards | Colobosauroides_cearensis | NA | LC | Unknown | NA | NA | NA | NA | NA |
| lizards | Complicitus_nigrigularis | NA | DD | Unknown | NA | NA | NA | NA | NA |

| | | | | | | | | | |
|---|---|---|---|---|---|---|---|---|---|
| lizards | Comptus_badius | NA | LC | Stable | NA | NA | NA | NA | NA |
| lizards | Comptus_stenurus | NA | LC | Unknown | NA | NA | NA | NA | NA |
| lizards | Concinnia_ampla | NA | LC | Unknown | NA | NA | NA | NA | NA |
| lizards | Concinnia_brachysoma | Concinnia_brachyosoma | LC | Stable | tropical | diurnal | 8.913 | 434526.519 | 5.333 |
| lizards | Concinnia_frerei | NA | VU | Unknown | NA | NA | NA | NA | NA |
| lizards | Concinnia_martini | Concinnia_martini | LC | Stable | temperate | diurnal | 7.079 | 281952.629 | 7.062 |
| lizards | Concinnia_sokosoma | Concinnia_sokosoma | LC | Stable | tropical | diurnal | 10 | 295686.266 | 5.855 |
| lizards | Concinnia_tenuis | Concinnia_tenuis | LC | Stable | temperate | cathemeral | 12.882 | 502803.818 | 7.414 |
| lizards | Concinnia_tigrina | NA | LC | Stable | NA | NA | NA | NA | NA |
| lizards | Congolacerta_asukului | NA | EN | Stable | NA | NA | NA | NA | NA |
| lizards | Congolacerta_vauereselli | Congolacerta_vauereselli | LC | Unknown | tropical | diurnal | 5.623 | 130221.862 | 14.722 |
| lizards | Conolophus_marthae | NA | CR | Unknown | NA | NA | NA | NA | NA |
| lizards | Conolophus_pallidus | NA | VU | Unknown | NA | NA | NA | NA | NA |
| lizards | Conolophus_subcristatus | NA | VU | Decreasing | NA | NA | NA | NA | NA |
| lizards | Contomastix_lacertoides | Contomastix_lacertoides | LC | Unknown | temperate | diurnal | 13.183 | 199535.483 | 10.155 |
| lizards | Contomastix_vacariensis | Contomastix_vacariensis | VU | Decreasing | temperate | diurnal | 15.849 | 84279.674 | 11.851 |
| lizards | Copeoglossum_arajara | Copeoglossum_arajara | LC | Unknown | tropical | diurnal | 33.113 | 60516.733 | 7.355 |
| lizards | Copeoglossum_aurae | Copeoglossum_aurae | LC | Unknown | tropical | diurnal | 28.184 | 4117.968 | 19.318 |
| lizards | Copeoglossum_margaritae | NA | DD | Unknown | NA | NA | NA | NA | NA |
| lizards | Copeoglossum_nigropunctatum | Copeoglossum_nigropunctatum | LC | Decreasing | tropical | diurnal | 31.623 | 10876578.194 | 4.548 |
| lizards | Cophosaurus_texanus | Cophosaurus_texanus | LC | Stable | temperate | diurnal | 22.909 | 1124004.216 | 9.741 |
| lizards | Cophoscincopus_durus | Cophoscincopus_durus | LC | Unknown | tropical | diurnal | 3.09 | 164915.903 | 14.069 |
| lizards | Cophoscincopus_greeri | Cophoscincopus_greeri | LC | Stable | tropical | diurnal | 5.623 | 146191.845 | 13.629 |
| lizards | Cophoscincopus_simulans | Cophoscincopus_simulans | LC | Unknown | tropical | diurnal | 4.365 | 357521.671 | 15.181 |
| lizards | Cophotis_ceylanica | Cophotis_ceylanica | EN | Unknown | tropical | diurnal | 9.55 | 2093.308 | 15.629 |
| lizards | Cophotis_dumbara | NA | CR | Unknown | NA | NA | NA | NA | NA |
| lizards | Cordylosaurus_subtessellatus | NA | LC | Stable | NA | NA | NA | NA | NA |
| lizards | Cordylus_beraduccii | NA | LC | Unknown | NA | NA | NA | NA | NA |
| lizards | Cordylus_cordylus | Cordylus_cordylus | LC | Stable | temperate | diurnal | 26.915 | 286245.434 | 10.484 |
| lizards | Cordylus_imkeae | Cordylus_imkeae | LC | Stable | temperate | diurnal | 6.761 | 11.287 | 7.31 |
| lizards | Cordylus_jonesii | Cordylus_jonesii | LC | Stable | temperate | diurnal | 19.953 | 683213.394 | 10.38 |
| lizards | Cordylus_macropholis | Cordylus_macropholis | NT | Stable | temperate | diurnal | 10.471 | 24268.673 | 9.695 |
| lizards | Cordylus_mclachlani | Cordylus_mclachlani | LC | Stable | temperate | diurnal | 8.71 | 10908.023 | 8.365 |
| lizards | Cordylus_meculae | Cordylus_meculae | LC | Unknown | tropical | diurnal | 21.38 | 27.343 | 5.55 |

| | | | | | | | | | |
|---|---|---|---|---|---|---|---|---|---|
| lizards | Cordylus_minor | Cordylus_minor | LC | Stable | temperate | diurnal | 8.318 | 10977.51 | 6.882 |
| lizards | Cordylus_namakuiyus | NA | LC | Stable | NA | NA | NA | NA | NA |
| lizards | Cordylus_niger | Cordylus_niger | LC | Stable | temperate | diurnal | 19.953 | 4029.396 | 25.375 |
| lizards | Cordylus_nyikae | Cordylus_nyikae | LC | Stable | tropical | diurnal | 22.387 | 10062.904 | 10.782 |
| lizards | Cordylus_oelofseni | Cordylus_oelofseni | LC | Stable | temperate | diurnal | 7.079 | 7056.995 | 15.664 |
| lizards | Cordylus_rhodesianus | Cordylus_rhodesianus | LC | Unknown | tropical | diurnal | 19.055 | 86524.117 | 13.512 |
| lizards | Cordylus_tropidosternum | Cordylus_tropidosternum | LC | Unknown | tropical | diurnal | 34.674 | 1784084.284 | 9.862 |
| lizards | Cordylus_ukingensis | NA | DD | Unknown | NA | NA | NA | NA | NA |
| lizards | Cordylus_vittifer | Cordylus_vittifer | LC | Stable | temperate | diurnal | 22.387 | 364265.968 | 12.912 |
| lizards | Correlophus_belepensis | Correlophus_belepensis | CR | Unknown | tropical | nocturnal | 20.417 | 51.746 | 6.73 |
| lizards | Correlophus_ciliatus | Correlophus_ciliatus | VU | Decreasing | tropical | nocturnal | 45.709 | 2529.783 | 9.914 |
| lizards | Correlophus_sarasinorum | Correlophus_sarasinorum | VU | Decreasing | tropical | nocturnal | 51.286 | 1473.558 | 9.423 |
| lizards | Corucia_zebrata | Corucia_zebrata | NT | Decreasing | tropical | nocturnal | 1230.269 | 30522.202 | 4.708 |
| lizards | Coryphophylax_brevicauda | NA | VU | Unknown | NA | NA | NA | NA | NA |
| lizards | Coryphophylax_subcristatus | Coryphophylax_subcristatus | LC | Stable | tropical | diurnal | 46.774 | 6111.469 | 7.393 |
| lizards | Corytophanes_cristatus | Corytophanes_cristatus | LC | Stable | tropical | diurnal | 58.884 | 568337.171 | 8.782 |
| lizards | Corytophanes_hernandesii | Corytophanes_hernandesii | LC | Stable | tropical | diurnal | 51.286 | 161179.62 | 10.384 |
| lizards | Corytophanes_percarinatus | Corytophanes_percarinatus | LC | Stable | tropical | diurnal | 38.905 | 45357.744 | 16.423 |
| lizards | Crenadactylus_horni | NA | LC | Stable | NA | NA | NA | NA | NA |
| lizards | Crenadactylus_naso | NA | LC | Stable | NA | NA | NA | NA | NA |
| lizards | Crenadactylus_occidentalis | NA | LC | Unknown | NA | NA | NA | NA | NA |
| lizards | Crenadactylus_ocellatus | Crenadactylus_ocellatus | LC | Unknown | temperate | nocturnal | 0.891 | 423447.531 | 7.418 |
| lizards | Crenadactylus_pilbarensis | NA | LC | Stable | NA | NA | NA | NA | NA |
| lizards | Crenadactylus_rostralis | NA | LC | Unknown | NA | NA | NA | NA | NA |
| lizards | Crenadactylus_tuberculatus | NA | LC | Stable | NA | NA | NA | NA | NA |
| lizards | Cricosaura_typica | Cricosaura_typica | NT | Stable | tropical | cathemeral | 1.23 | 4205.364 | 14.896 |
| lizards | Cristidorsa_otai | NA | DD | Unknown | NA | NA | NA | NA | NA |
| lizards | Cristidorsa_planidorsata | NA | LC | Decreasing | NA | NA | NA | NA | NA |
| lizards | Crocodilurus_amazonicus | Crocodilurus_amazonicus | LC | Unknown | tropical | diurnal | 1096.478 | 3319210.352 | 1.205 |
| lizards | Crossobamon_eversmanni | Crossobamon_eversmanni | LC | Stable | temperate | nocturnal | 7.079 | 1699213.715 | 7.033 |
| lizards | Crossobamon_orientalis | Crossobamon_orientalis | LC | Unknown | temperate | nocturnal | 3.981 | 300169.032 | 12.858 |
| lizards | Crotaphytus_antiquus | Crotaphytus_antiquus | EN | Decreasing | temperate | diurnal | 46.774 | 207.367 | 7.678 |
| lizards | Crotaphytus_bicinctores | Crotaphytus_bicinctores | LC | Stable | temperate | diurnal | 52.481 | 556345.849 | 7.821 |
| lizards | Crotaphytus_collaris | Crotaphytus_collaris | LC | Stable | temperate | diurnal | 61.66 | 1956785.655 | 11.087 |

| | | | | | | | | | |
|---|---|---|---|---|---|---|---|---|---|
| lizards | Crotaphytus_dickersonae | Crotaphytus_dickersonae | LC | Stable | temperate | diurnal | 51.286 | 2619.961 | 6.159 |
| lizards | Crotaphytus_grismeri | Crotaphytus_grismeri | LC | Stable | temperate | diurnal | 39.811 | 874.488 | 4.872 |
| lizards | Crotaphytus_insularis | Crotaphytus_insularis | LC | Stable | temperate | diurnal | 54.954 | 931.872 | 0.447 |
| lizards | Crotaphytus_nebrius | Crotaphytus_nebrius | LC | Stable | temperate | diurnal | 48.978 | 101563.801 | 6.391 |
| lizards | Crotaphytus_reticulatus | Crotaphytus_reticulatus | VU | Decreasing | temperate | diurnal | 66.069 | 95194.013 | 12.483 |
| lizards | Crotaphytus_vestigium | Crotaphytus_vestigium | LC | Stable | temperate | diurnal | 58.884 | 66129.09 | 3.669 |
| lizards | Cryptactites_peringueyi | Cryptactites_peringueyi | LC | Stable | temperate | nocturnal | 0.589 | 3963.458 | 16.524 |
| lizards | Cryptagama_aurita | NA | DD | Unknown | NA | NA | NA | NA | NA |
| lizards | Cryptoblepharus_adamsi | NA | LC | Stable | NA | NA | NA | NA | NA |
| lizards | Cryptoblepharus_africanus | Cryptoblepharus_africanus | LC | Unknown | temperate | diurnal | 2.344 | 43203.326 | 12.41 |
| lizards | Cryptoblepharus_aldabrae | NA | VU | Stable | NA | NA | NA | NA | NA |
| lizards | Cryptoblepharus_australis | Cryptoblepharus_australis | LC | Stable | temperate | diurnal | 1.778 | 3408736.82 | 4.296 |
| lizards | Cryptoblepharus_balinensis | NA | LC | Unknown | NA | NA | NA | NA | NA |
| lizards | Cryptoblepharus_bitaeniatus | NA | NT | Decreasing | NA | NA | NA | NA | NA |
| lizards | Cryptoblepharus_boutonii | Cryptoblepharus_boutonii | NT | Stable | tropical | diurnal | 1.318 | 2143.066 | 24.822 |
| lizards | Cryptoblepharus_buchananii | Cryptoblepharus_buchananii | LC | Stable | temperate | diurnal | 2.188 | 1126517.287 | 5.072 |
| lizards | Cryptoblepharus_burdeni | Cryptoblepharus_burdeni | LC | Stable | tropical | diurnal | 2.291 | 470.275 | 11.129 |
| lizards | Cryptoblepharus_caudatus | NA | CR | Decreasing | NA | NA | NA | NA | NA |
| lizards | Cryptoblepharus_cognatus | NA | DD | Unknown | NA | NA | NA | NA | NA |
| lizards | Cryptoblepharus_cursor | NA | LC | Stable | NA | NA | NA | NA | NA |
| lizards | Cryptoblepharus_cygnatus | Cryptoblepharus_cygnatus | LC | Stable | tropical | diurnal | 1.585 | 99121.234 | 1.989 |
| lizards | Cryptoblepharus_daedalos | NA | LC | Stable | NA | NA | NA | NA | NA |
| lizards | Cryptoblepharus_egeriae | NA | EW | not_available | NA | NA | NA | NA | NA |
| lizards | Cryptoblepharus_eximius | Cryptoblepharus_eximius | LC | Unknown | tropical | diurnal | 1.148 | 13854.643 | 9.298 |
| lizards | Cryptoblepharus_exochus | NA | LC | Stable | NA | NA | NA | NA | NA |
| lizards | Cryptoblepharus_fuhni | NA | LC | Stable | NA | NA | NA | NA | NA |
| lizards | Cryptoblepharus_furvus | Cryptoblepharus_furvus | LC | Unknown | tropical | diurnal | 1.862 | 886.795 | 5.243 |
| lizards | Cryptoblepharus_gloriosus | Cryptoblepharus_gloriosus | VU | Unknown | tropical | diurnal | 1.479 | 362.257 | 19.302 |
| lizards | Cryptoblepharus_gurrmul | NA | DD | Stable | NA | NA | NA | NA | NA |
| lizards | Cryptoblepharus_juno | NA | LC | Stable | NA | NA | NA | NA | NA |
| lizards | Cryptoblepharus_keiensis | NA | LC | Stable | NA | NA | NA | NA | NA |
| lizards | Cryptoblepharus_leschenault | NA | LC | Stable | NA | NA | NA | NA | NA |
| lizards | Cryptoblepharus_litoralis | Cryptoblepharus_litoralis | LC | Stable | tropical | diurnal | 3.09 | 19997.871 | 5.246 |
| lizards | Cryptoblepharus_megastictus | Cryptoblepharus_megastictus | LC | Stable | tropical | diurnal | 1.175 | 22543.991 | 1.584 |

| | | | | | | | | | |
|---|---|---|---|---|---|---|---|---|---|
| lizards | Cryptoblepharus_mertensi | NA | LC | Stable | NA | NA | NA | NA | NA |
| lizards | Cryptoblepharus_metallicus | Cryptoblepharus_metallicus | LC | Stable | tropical | diurnal | 1.995 | 2091715.059 | 3.182 |
| lizards | Cryptoblepharus_nigropunctatus | NA | NT | Decreasing | NA | NA | NA | NA | NA |
| lizards | Cryptoblepharus_novaeguineae | NA | LC | Stable | NA | NA | NA | NA | NA |
| lizards | Cryptoblepharus_novocaledonicus | Cryptoblepharus_novocaledonicus | LC | Stable | tropical | diurnal | 1.413 | 16548.094 | 7.409 |
| lizards | Cryptoblepharus_novohebridicus | Cryptoblepharus_novohebridicus | LC | Stable | tropical | diurnal | 0.871 | 10623.937 | 5.999 |
| lizards | Cryptoblepharus_ochrus | NA | LC | Stable | NA | NA | NA | NA | NA |
| lizards | Cryptoblepharus_pannosus | Cryptoblepharus_pannosus | LC | Stable | temperate | diurnal | 1.514 | 2570391.548 | 5.518 |
| lizards | Cryptoblepharus_plagiocephalus | Cryptoblepharus_plagiocephalus | LC | Stable | temperate | diurnal | 1.862 | 508873.63 | 4.228 |
| lizards | Cryptoblepharus_poecilopleurus | Cryptoblepharus_poecilopleurus | LC | Stable | tropical | diurnal | 2.63 | 44707.714 | 5.742 |
| lizards | Cryptoblepharus_pulcher | Cryptoblepharus_pulcher | LC | Stable | temperate | diurnal | 1.288 | 756469.164 | 7.279 |
| lizards | Cryptoblepharus_renschi | Cryptoblepharus_renschi | LC | Stable | tropical | diurnal | 2.291 | 17157.348 | 17.094 |
| lizards | Cryptoblepharus_richardsi | NA | DD | Unknown | NA | NA | NA | NA | NA |
| lizards | Cryptoblepharus_ruber | Cryptoblepharus_ruber | LC | Stable | tropical | diurnal | 1.905 | 497359.102 | 2.345 |
| lizards | Cryptoblepharus_rutilus | NA | LC | Stable | NA | NA | NA | NA | NA |
| lizards | Cryptoblepharus_schlegelianus | NA | LC | Unknown | NA | NA | NA | NA | NA |
| lizards | Cryptoblepharus_tytthos | NA | LC | Stable | NA | NA | NA | NA | NA |
| lizards | Cryptoblepharus_ustulatus | NA | LC | Stable | NA | NA | NA | NA | NA |
| lizards | Cryptoblepharus_virgatus | Cryptoblepharus_virgatus | LC | Stable | tropical | diurnal | 1.202 | 188197.606 | 2.887 |
| lizards | Cryptoblepharus_wulbu | NA | LC | Unknown | NA | NA | NA | NA | NA |
| lizards | Cryptoblepharus_xenikos | NA | DD | Unknown | NA | NA | NA | NA | NA |
| lizards | Cryptoblepharus_yulensis | Cryptoblepharus_yulensis | LC | Stable | tropical | diurnal | 1.288 | 2825.426 | 9.368 |
| lizards | Cryptoblepharus_zoticus | NA | LC | Stable | NA | NA | NA | NA | NA |
| lizards | Ctenoblepharys_adspersa | NA | VU | Unknown | NA | NA | NA | NA | NA |
| lizards | Ctenophorus_adelaidensis | Ctenophorus_adelaidensis | LC | Unknown | temperate | diurnal | 4.677 | 45133.672 | 10.899 |
| lizards | Ctenophorus_butlerorum | Ctenophorus_butlerorum | LC | Stable | temperate | diurnal | 2.818 | 3219.909 | 6.429 |
| lizards | Ctenophorus_caudicinctus | Ctenophorus_caudicinctus | LC | Stable | temperate | diurnal | 38.905 | 818951.302 | 3.763 |
| lizards | Ctenophorus_chapmani | NA | LC | Stable | NA | NA | NA | NA | NA |
| lizards | Ctenophorus_clayi | Ctenophorus_clayi | LC | Unknown | temperate | diurnal | 6.166 | 2030499.966 | 1.806 |
| lizards | Ctenophorus_cristatus | Ctenophorus_cristatus | LC | Stable | temperate | diurnal | 47.863 | 816178.422 | 3.843 |
| lizards | Ctenophorus_decresii | Ctenophorus_decresii | LC | Decreasing | temperate | diurnal | 23.988 | 122891.22 | 8.484 |
| lizards | Ctenophorus_femoralis | Ctenophorus_femoralis | LC | Stable | tropical | diurnal | 5.888 | 62060.05 | 5.653 |
| lizards | Ctenophorus_fionni | Ctenophorus_fionni | LC | Unknown | temperate | diurnal | 29.512 | 140174.982 | 6.287 |
| lizards | Ctenophorus_fordi | Ctenophorus_fordi | LC | Unknown | temperate | diurnal | 6.166 | 1229820.209 | 4.45 |

| | | | | | | | | | |
|---|---|---|---|---|---|---|---|---|---|
| lizards | Ctenophorus_gibba | NA | LC | Stable | NA | NA | NA | NA | NA |
| lizards | Ctenophorus_isolepis | Ctenophorus_isolepis | LC | Unknown | temperate | diurnal | 18.621 | 3313601.451 | 2.02 |
| lizards | Ctenophorus_maculatus | Ctenophorus_maculatus | LC | Stable | temperate | diurnal | 9.55 | 511180.998 | 6.411 |
| lizards | Ctenophorus_maculosus | Ctenophorus_maculosus | LC | Unknown | temperate | diurnal | 10.965 | 82125.216 | 2.9 |
| lizards | Ctenophorus_mckenziei | NA | LC | Stable | NA | NA | NA | NA | NA |
| lizards | Ctenophorus_mirrityana | Ctenophorus_mirrityana | NT | Unknown | temperate | diurnal | 24.547 | 2043.317 | 7.76 |
| lizards | Ctenophorus_nguyarna | Ctenophorus_nguyarna | VU | Unknown | tropical | diurnal | 15.849 | 3130.805 | 1.074 |
| lizards | Ctenophorus_nuchalis | Ctenophorus_nuchalis | LC | Unknown | temperate | diurnal | 70.795 | 4690120.933 | 2.978 |
| lizards | Ctenophorus_ornatus | Ctenophorus_ornatus | LC | Unknown | temperate | diurnal | 28.184 | 360252.675 | 7.786 |
| lizards | Ctenophorus_parviceps | Ctenophorus_parviceps | LC | Stable | temperate | diurnal | 4.365 | 38678.166 | 5.982 |
| lizards | Ctenophorus_pictus | Ctenophorus_pictus | LC | Stable | temperate | diurnal | 13.49 | 3023219.88 | 3.4 |
| lizards | Ctenophorus_reticulatus | Ctenophorus_reticulatus | LC | Unknown | temperate | diurnal | 42.658 | 2333990.522 | 2.937 |
| lizards | Ctenophorus_rubens | NA | LC | Stable | NA | NA | NA | NA | NA |
| lizards | Ctenophorus_salinarum | Ctenophorus_salinarum | LC | Stable | temperate | diurnal | 15.488 | 607776.366 | 4.516 |
| lizards | Ctenophorus_scutulatus | Ctenophorus_scutulatus | LC | Stable | temperate | diurnal | 51.286 | 755547.534 | 4.717 |
| lizards | Ctenophorus_tjantjalka | NA | LC | Unknown | NA | NA | NA | NA | NA |
| lizards | Ctenophorus_vadnappa | Ctenophorus_vadnappa | LC | Unknown | temperate | diurnal | 23.988 | 31990.408 | 5.542 |
| lizards | Ctenophorus_yinnietharra | NA | LC | Stable | NA | NA | NA | NA | NA |
| lizards | Ctenosaura_acanthura | Ctenosaura_acanthura | LC | Decreasing | tropical | diurnal | 3890.451 | 63252.752 | 15.469 |
| lizards | Ctenosaura_bakeri | Ctenosaura_bakeri | CR | Decreasing | tropical | diurnal | 1348.963 | 42.084 | 9.659 |
| lizards | Ctenosaura_clarki | Ctenosaura_clarki | VU | Unknown | tropical | diurnal | 676.083 | 9608.118 | 9.33 |
| lizards | Ctenosaura_conspicuosa | NA | VU | Unknown | NA | NA | NA | NA | NA |
| lizards | Ctenosaura_hemilopha | Ctenosaura_hemilopha | LC | Unknown | temperate | diurnal | 2754.229 | 141254.583 | 7.677 |
| lizards | Ctenosaura_macrolopha | Ctenosaura_macrolopha | LC | Decreasing | temperate | diurnal | 416.869 | 42785.87 | 5.388 |
| lizards | Ctenosaura_melanosterna | Ctenosaura_melanosterna | EN | Decreasing | tropical | diurnal | 1445.44 | 580.582 | 15.309 |
| lizards | Ctenosaura_nolascensis | NA | VU | Stable | NA | NA | NA | NA | NA |
| lizards | Ctenosaura_oaxacana | NA | CR | Decreasing | NA | NA | NA | NA | NA |
| lizards | Ctenosaura_oedirhina | Ctenosaura_oedirhina | EN | Decreasing | tropical | diurnal | 1348.963 | 160.302 | 19.805 |
| lizards | Ctenosaura_palearis | Ctenosaura_palearis | EN | Decreasing | tropical | diurnal | 1288.25 | 940.357 | 20.676 |
| lizards | Ctenosaura_pectinata | Ctenosaura_pectinata | LC | Decreasing | tropical | diurnal | 2691.535 | 358773.116 | 9.23 |
| lizards | Ctenosaura_quinquecarinata | Ctenosaura_quinquecarinata | DD | Decreasing | tropical | diurnal | 346.737 | 19897.348 | 12.667 |
| lizards | Ctenosaura_similis | Ctenosaura_similis | LC | Stable | tropical | diurnal | 5011.872 | 893356.472 | 10.139 |
| lizards | Ctenotus_agrestis | NA | LC | Unknown | NA | NA | NA | NA | NA |
| lizards | Ctenotus_alacer | Ctenotus_alacer | LC | Stable | tropical | diurnal | 6.026 | 722622.961 | 1.633 |

| | | | | | | | | | |
|---|---|---|---|---|---|---|---|---|---|
| lizards | Ctenotus_alleni | Ctenotus_alleni | NT | Unknown | temperate | diurnal | 16.982 | 13512.05 | 6.127 |
| lizards | Ctenotus_allotropis | NA | LC | Stable | NA | NA | NA | NA | NA |
| lizards | Ctenotus_angusticeps | NA | LC | Stable | NA | NA | NA | NA | NA |
| lizards | Ctenotus_aphrodite | NA | not_available | not_available | NA | NA | NA | NA | NA |
| lizards | Ctenotus_arcanus | NA | LC | Stable | NA | NA | NA | NA | NA |
| lizards | Ctenotus_ariadnae | Ctenotus_ariadnae | LC | Stable | temperate | diurnal | 5.129 | 1189930.553 | 1.485 |
| lizards | Ctenotus_arnhemensis | Ctenotus_arnhemensis | LC | Stable | tropical | diurnal | 3.162 | 10830.53 | 2.558 |
| lizards | Ctenotus_astarte | NA | LC | Stable | NA | NA | NA | NA | NA |
| lizards | Ctenotus_astictus | NA | LC | Stable | NA | NA | NA | NA | NA |
| lizards | Ctenotus_atlas | Ctenotus_atlas | LC | Stable | temperate | diurnal | 8.511 | 985391.664 | 4.214 |
| lizards | Ctenotus_australis | Ctenotus_australis | LC | Stable | temperate | diurnal | 29.512 | 72199.848 | 10.16 |
| lizards | Ctenotus_borealis | NA | LC | Stable | NA | NA | NA | NA | NA |
| lizards | Ctenotus_brevipes | Ctenotus_brevipes | LC | Stable | tropical | diurnal | 4.169 | 252239.383 | 3.227 |
| lizards | Ctenotus_brooksi | Ctenotus_brooksi | LC | Stable | temperate | diurnal | 3.467 | 1745904.081 | 1.492 |
| lizards | Ctenotus_burbidgei | NA | LC | Stable | NA | NA | NA | NA | NA |
| lizards | Ctenotus_calurus | Ctenotus_calurus | LC | Stable | temperate | diurnal | 2.291 | 1489493.443 | 1.282 |
| lizards | Ctenotus_capricorni | NA | LC | Stable | NA | NA | NA | NA | NA |
| lizards | Ctenotus_catenifer | NA | LC | Unknown | NA | NA | NA | NA | NA |
| lizards | Ctenotus_coggeri | Ctenotus_coggeri | LC | Stable | tropical | cathemeral | 13.804 | 50864.809 | 1.122 |
| lizards | Ctenotus_colletti | Ctenotus_colletti | LC | Stable | tropical | diurnal | 1.622 | 31414.945 | 2.846 |
| lizards | Ctenotus_decaneurus | NA | LC | Stable | NA | NA | NA | NA | NA |
| lizards | Ctenotus_delli | NA | LC | Stable | NA | NA | NA | NA | NA |
| lizards | Ctenotus_duricola | NA | LC | Stable | NA | NA | NA | NA | NA |
| lizards | Ctenotus_dux | Ctenotus_dux | LC | Stable | temperate | diurnal | 5.754 | 1196447.385 | 1.171 |
| lizards | Ctenotus_ehmanni | NA | LC | Stable | NA | NA | NA | NA | NA |
| lizards | Ctenotus_essingtonii | Ctenotus_essingtonii | LC | Stable | tropical | diurnal | 7.762 | 230922.522 | 1.73 |
| lizards | Ctenotus_euclae | NA | LC | Stable | NA | NA | NA | NA | NA |
| lizards | Ctenotus_eurydice | NA | LC | Unknown | NA | NA | NA | NA | NA |
| lizards | Ctenotus_eutaenius | NA | LC | Stable | NA | NA | NA | NA | NA |
| lizards | Ctenotus_gagudju | Ctenotus_gagudju | LC | Stable | tropical | diurnal | 2.951 | 4592.283 | 2.16 |
| lizards | Ctenotus_gemmula | Ctenotus_gemmula | LC | Stable | temperate | diurnal | 4.169 | 177921.285 | 8.502 |
| lizards | Ctenotus_grandis | Ctenotus_grandis | LC | Stable | tropical | cathemeral | 40.738 | 1509619.037 | 1.64 |
| lizards | Ctenotus_greeri | NA | LC | Stable | NA | NA | NA | NA | NA |
| lizards | Ctenotus_halysis | NA | LC | Unknown | NA | NA | NA | NA | NA |

| | | | | | | | | | |
|---|---|---|---|---|---|---|---|---|---|
| lizards | Ctenotus_hanloni | Ctenotus_hanloni | LC | Stable | temperate | diurnal | 7.943 | 1498577.694 | 2.146 |
| lizards | Ctenotus_hebetior | Ctenotus_hebetior | LC | Stable | tropical | diurnal | 4.169 | 580090.804 | 4.83 |
| lizards | Ctenotus_hilli | NA | LC | Stable | NA | NA | NA | NA | NA |
| lizards | Ctenotus_iapetus | NA | LC | Stable | NA | NA | NA | NA | NA |
| lizards | Ctenotus_impar | Ctenotus_impar | LC | Unknown | temperate | diurnal | 5.623 | 305520.158 | 8.949 |
| lizards | Ctenotus_ingrami | NA | LC | Stable | NA | NA | NA | NA | NA |
| lizards | Ctenotus_inornatus | Ctenotus_inornatus | LC | Stable | tropical | cathemeral | 18.197 | 4465089.519 | 2.824 |
| lizards | Ctenotus_joanae | NA | LC | Unknown | NA | NA | NA | NA | NA |
| lizards | Ctenotus_kurnbudj | Ctenotus_kurnbudj | EN | Unknown | tropical | diurnal | 3.311 | 3260.86 | 2.675 |
| lizards | Ctenotus_labillardieri | Ctenotus_labillardieri | LC | Unknown | temperate | diurnal | 8.913 | 118665.431 | 10.102 |
| lizards | Ctenotus_lancelini | Ctenotus_lancelini | CR | Unknown | temperate | diurnal | 13.804 | 12.102 | 20.464 |
| lizards | Ctenotus_lateralis | NA | LC | Stable | NA | NA | NA | NA | NA |
| lizards | Ctenotus_leae | Ctenotus_leae | LC | Stable | temperate | diurnal | 4.571 | 1951798.113 | 1.894 |
| lizards | Ctenotus_leonhardii | Ctenotus_leonhardii | LC | Stable | temperate | diurnal | 10 | 4712048.742 | 2.795 |
| lizards | Ctenotus_maryani | NA | LC | Stable | NA | NA | NA | NA | NA |
| lizards | Ctenotus_mastigura | NA | LC | Stable | NA | NA | NA | NA | NA |
| lizards | Ctenotus_mesotes | NA | LC | Unknown | NA | NA | NA | NA | NA |
| lizards | Ctenotus_militaris | NA | LC | Stable | NA | NA | NA | NA | NA |
| lizards | Ctenotus_mimetes | Ctenotus_mimetes | LC | Stable | temperate | diurnal | 11.22 | 182002.247 | 6.397 |
| lizards | Ctenotus_monticola | NA | LC | Unknown | NA | NA | NA | NA | NA |
| lizards | Ctenotus_nasutus | Ctenotus_nasutus | LC | Stable | temperate | diurnal | 1.738 | 936346.321 | 0.987 |
| lizards | Ctenotus_nigrilineatus | NA | LC | Unknown | NA | NA | NA | NA | NA |
| lizards | Ctenotus_nullum | NA | LC | Unknown | NA | NA | NA | NA | NA |
| lizards | Ctenotus_olympicus | Ctenotus_olympicus | LC | Stable | temperate | diurnal | 8.511 | 705214.885 | 4.296 |
| lizards | Ctenotus_ora | NA | VU | Decreasing | NA | NA | NA | NA | NA |
| lizards | Ctenotus_orientalis | Ctenotus_orientalis | LC | Stable | temperate | diurnal | 11.22 | 1008363.009 | 6.702 |
| lizards | Ctenotus_pallasotus | NA | not_available | not_available | NA | NA | NA | NA | NA |
| lizards | Ctenotus_pallescens | Ctenotus_pallescens | LC | Unknown | tropical | diurnal | 1.622 | 405369.892 | 1.653 |
| lizards | Ctenotus_pantherinus | Ctenotus_pantherinus | LC | Stable | tropical | cathemeral | 45.709 | 5203160.872 | 2.927 |
| lizards | Ctenotus_piankai | Ctenotus_piankai | LC | Stable | tropical | diurnal | 4.169 | 2013068.234 | 1.516 |
| lizards | Ctenotus_pulchellus | NA | LC | Stable | NA | NA | NA | NA | NA |
| lizards | Ctenotus_quattuordecimlineatus | Ctenotus_quattuordecimlineatus | LC | Unknown | temperate | diurnal | 7.079 | 1773965.61 | 1.447 |
| lizards | Ctenotus_quinkan | NA | LC | Stable | NA | NA | NA | NA | NA |
| lizards | Ctenotus_rawlinsoni | NA | LC | Unknown | NA | NA | NA | NA | NA |

| | | | | | | | | | |
|---|---|---|---|---|---|---|---|---|---|
| lizards | Ctenotus_regius | Ctenotus_regius | LC | Stable | temperate | diurnal | 11.749 | 1705313.275 | 3.778 |
| lizards | Ctenotus_rhabdotus | NA | not_available | not_available | NA | NA | NA | NA | NA |
| lizards | Ctenotus_rimacolus | NA | LC | Stable | NA | NA | NA | NA | NA |
| lizards | Ctenotus_robustus | Ctenotus_robustus | LC | Stable | temperate | cathemeral | 44.668 | 3394362.134 | 5.633 |
| lizards | Ctenotus_rosarium | NA | LC | Unknown | NA | NA | NA | NA | NA |
| lizards | Ctenotus_rubicundus | NA | LC | Stable | NA | NA | NA | NA | NA |
| lizards | Ctenotus_rufescens | Ctenotus_rufescens | LC | Unknown | tropical | cathemeral | 1.622 | 49511.453 | 5.86 |
| lizards | Ctenotus_rutilans | Ctenotus_rutilans | LC | Stable | tropical | diurnal | 3.981 | 119782.695 | 4.34 |
| lizards | Ctenotus_schevilli | NA | LC | Unknown | NA | NA | NA | NA | NA |
| lizards | Ctenotus_schomburgkii | Ctenotus_schomburgkii | LC | Unknown | temperate | diurnal | 3.467 | 4312611.49 | 3.498 |
| lizards | Ctenotus_septenarius | Ctenotus_septenarius | LC | Stable | temperate | diurnal | 7.413 | 651429.865 | 2.498 |
| lizards | Ctenotus_serotinus | NA | CR | Decreasing | NA | NA | NA | NA | NA |
| lizards | Ctenotus_serventyi | NA | LC | Stable | NA | NA | NA | NA | NA |
| lizards | Ctenotus_spaldingi | Ctenotus_spaldingi | LC | Stable | temperate | diurnal | 29.512 | 2891125.546 | 6.093 |
| lizards | Ctenotus_storri | Ctenotus_storri | LC | Stable | tropical | diurnal | 1.122 | 117064.7 | 1.997 |
| lizards | Ctenotus_strauchii | Ctenotus_strauchii | LC | Stable | temperate | diurnal | 4.898 | 2788871.033 | 4.648 |
| lizards | Ctenotus_striaticeps | NA | LC | Stable | NA | NA | NA | NA | NA |
| lizards | Ctenotus_stuarti | NA | EN | Decreasing | NA | NA | NA | NA | NA |
| lizards | Ctenotus_superciliaris | NA | LC | Stable | NA | NA | NA | NA | NA |
| lizards | Ctenotus_taeniatus | NA | LC | Stable | NA | NA | NA | NA | NA |
| lizards | Ctenotus_taeniolatus | Ctenotus_taeniolatus | LC | Stable | temperate | diurnal | 14.791 | 1203988.057 | 7.056 |
| lizards | Ctenotus_tanamiensis | NA | LC | Unknown | NA | NA | NA | NA | NA |
| lizards | Ctenotus_tantillus | NA | LC | Stable | NA | NA | NA | NA | NA |
| lizards | Ctenotus_terrareginae | NA | LC | Unknown | NA | NA | NA | NA | NA |
| lizards | Ctenotus_uber | Ctenotus_uber | LC | Stable | temperate | diurnal | 11.22 | 1062714.223 | 3.517 |
| lizards | Ctenotus_vagus | NA | DD | Unknown | NA | NA | NA | NA | NA |
| lizards | Ctenotus_vertebralis | Ctenotus_vertebralis | LC | Stable | tropical | diurnal | 3.09 | 179335.693 | 1.54 |
| lizards | Ctenotus_xenopleura | NA | LC | Stable | NA | NA | NA | NA | NA |
| lizards | Ctenotus_youngsoni | NA | LC | Stable | NA | NA | NA | NA | NA |
| lizards | Ctenotus_zastictus | NA | NT | Unknown | NA | NA | NA | NA | NA |
| lizards | Ctenotus_zebrilla | NA | LC | Stable | NA | NA | NA | NA | NA |
| lizards | Cyclodomorphus_branchialis | Cyclodomorphus_branchialis | NT | Decreasing | temperate | cathemeral | 38.905 | 56971.334 | 8.196 |
| lizards | Cyclodomorphus_casuarinae | Cyclodomorphus_casuarinae | LC | Stable | temperate | cathemeral | 128.825 | 67107.961 | 5.939 |
| lizards | Cyclodomorphus_celatus | Cyclodomorphus_celatus | LC | Stable | temperate | nocturnal | 39.811 | 72346.055 | 8.273 |

| | | | | | | | | | |
|---|---|---|---|---|---|---|---|---|---|
| lizards | Cyclodomorphus_gerrardii | Hemisphaeriodon_gerrardii | LC | Stable | temperate | cathemeral | 446.684 | 419636.984 | 6.903 |
| lizards | Cyclodomorphus_maximus | Cyclodomorphus_maxima | LC | Unknown | tropical | cathemeral | 323.594 | 82497.224 | 1.72 |
| lizards | Cyclodomorphus_melanops | Cyclodomorphus_melanops | LC | Stable | temperate | cathemeral | 52.481 | 3536483.12 | 2.621 |
| lizards | Cyclodomorphus_michaeli | Cyclodomorphus_michaeli | LC | Unknown | temperate | cathemeral | 128.825 | 37214.243 | 11.109 |
| lizards | Cyclodomorphus_praealtus | Cyclodomorphus_praealtus | EN | Decreasing | temperate | diurnal | 45.709 | 8620.451 | 6.77 |
| lizards | Cyclodomorphus_venustus | Cyclodomorphus_venustus | LC | Stable | temperate | nocturnal | 22.387 | 576717.488 | 3.916 |
| lizards | Cyclura_carinata | Cyclura_carinata | EN | Stable | tropical | diurnal | 5623.413 | 153.243 | 14.717 |
| lizards | Cyclura_collei | Cyclura_collei | CR | Increasing | tropical | diurnal | 3311.311 | 565.94 | 23.999 |
| lizards | Cyclura_cornuta | Cyclura_cornuta | EN | Decreasing | tropical | diurnal | 10964.782 | 29980.874 | 14.451 |
| lizards | Cyclura_cychlura | Cyclura_cychlura | VU | Decreasing | temperate | diurnal | 11748.976 | 154.761 | 4.612 |
| lizards | Cyclura_lewisi | Cyclura_lewisi | EN | Increasing | tropical | diurnal | 7079.458 | 210.62 | 23.235 |
| lizards | Cyclura_nubila | Cyclura_nubila | VU | not_available | tropical | diurnal | 17782.794 | 86522.678 | 16.304 |
| lizards | Cyclura_pinguis | Cyclura_pinguis | CR | Increasing | tropical | diurnal | 7413.102 | 41.59 | 11.412 |
| lizards | Cyclura_ricordi | Cyclura_ricordi | EN | Decreasing | tropical | diurnal | 4786.301 | 452.684 | 10.933 |
| lizards | Cyclura_rileyi | Cyclura_rileyi | EN | not_available | temperate | diurnal | 2630.268 | 32.584 | 10.292 |
| lizards | Cyclura_stejnegeri | NA | CR | Increasing | NA | NA | NA | NA | NA |
| lizards | Cyrtodactylus_aaroni | NA | DD | Unknown | NA | NA | NA | NA | NA |
| lizards | Cyrtodactylus_adleri | NA | EN | Decreasing | NA | NA | NA | NA | NA |
| lizards | Cyrtodactylus_aequalis | NA | DD | Unknown | NA | NA | NA | NA | NA |
| lizards | Cyrtodactylus_agamensis | NA | LC | Unknown | NA | NA | NA | NA | NA |
| lizards | Cyrtodactylus_agusanensis | Cyrtodactylus_agusanensis | LC | Unknown | tropical | cathemeral | 22.387 | 115002.596 | 11.912 |
| lizards | Cyrtodactylus_albofasciatus | Cyrtodactylus_albofasciatus | LC | Unknown | tropical | nocturnal | 8.913 | 22867.659 | 14.713 |
| lizards | Cyrtodactylus_angularis | Cyrtodactylus_angularis | LC | Unknown | tropical | nocturnal | 16.596 | 1568.175 | 18.731 |
| lizards | Cyrtodactylus_annandalei | Cyrtodactylus_annandalei | LC | Unknown | tropical | nocturnal | 3.715 | 12943.257 | 16.35 |
| lizards | Cyrtodactylus_annulatus | NA | LC | Stable | NA | NA | NA | NA | NA |
| lizards | Cyrtodactylus_astrum | Cyrtodactylus_astrum | LC | Stable | tropical | nocturnal | 23.988 | 2047.801 | 17.317 |
| lizards | Cyrtodactylus_aurensis | Cyrtodactylus_aurensis | LC | Unknown | tropical | nocturnal | 18.621 | 15.583 | 2.669 |
| lizards | Cyrtodactylus_auribalteatus | NA | VU | Stable | NA | NA | NA | NA | NA |
| lizards | Cyrtodactylus_australotitiwangsaensis | Cyrtodactylus_australotitiwangsaensis | LC | Unknown | tropical | nocturnal | 31.623 | 1668.784 | 10.545 |
| lizards | Cyrtodactylus_ayeyarwadyensis | Cyrtodactylus_ayeyarwadyensis | NT | Unknown | tropical | nocturnal | 9.55 | 27844.628 | 11.481 |
| lizards | Cyrtodactylus_badenensis | NA | VU | Unknown | NA | NA | NA | NA | NA |
| lizards | Cyrtodactylus_baluensis | Cyrtodactylus_baluensis | LC | Unknown | tropical | nocturnal | 18.197 | 39485.081 | 4.737 |
| lizards | Cyrtodactylus_bansocensis | NA | DD | Unknown | NA | NA | NA | NA | NA |
| lizards | Cyrtodactylus_battalensis | NA | DD | Unknown | NA | NA | NA | NA | NA |

| | | | | | | | | | |
|---|---|---|---|---|---|---|---|---|---|
| lizards | Cyrtodactylus_batucolus | Cyrtodactylus_batucolus | LC | Unknown | tropical | nocturnal | 8.71 | 1.325 | 12.337 |
| lizards | Cyrtodactylus_bichnganae | NA | VU | Unknown | NA | NA | NA | NA | NA |
| lizards | Cyrtodactylus_bidoupimontis | NA | LC | Stable | NA | NA | NA | NA | NA |
| lizards | Cyrtodactylus_bintangrendah | Cyrtodactylus_bintangrendah | LC | Unknown | tropical | nocturnal | 28.84 | 2649.875 | 10.838 |
| lizards | Cyrtodactylus_bintangtinggi | Cyrtodactylus_bintangtinggi | NT | Unknown | tropical | nocturnal | 25.704 | 134.088 | 13.944 |
| lizards | Cyrtodactylus_biordinis | Cyrtodactylus_biordinis | LC | Stable | tropical | nocturnal | 16.218 | 5461.348 | 5.671 |
| lizards | Cyrtodactylus_bobrovi | NA | LC | Unknown | NA | NA | NA | NA | NA |
| lizards | Cyrtodactylus_boreoclivus | Cyrtodactylus_boreoclivus | LC | Unknown | tropical | nocturnal | 23.988 | 4892.423 | 2.031 |
| lizards | Cyrtodactylus_brevidactylus | Cyrtodactylus_brevidactylus | EN | Unknown | tropical | nocturnal | 13.49 | 230.572 | 13.092 |
| lizards | Cyrtodactylus_brevipalmatus | Cyrtodactylus_brevipalmatus | LC | Unknown | tropical | nocturnal | 7.943 | 98073.656 | 9.104 |
| lizards | Cyrtodactylus_buchardi | NA | DD | Unknown | NA | NA | NA | NA | NA |
| lizards | Cyrtodactylus_bugiamapensis | NA | LC | Stable | NA | NA | NA | NA | NA |
| lizards | Cyrtodactylus_calamei | NA | LC | Stable | NA | NA | NA | NA | NA |
| lizards | Cyrtodactylus_caovansungi | NA | EN | Decreasing | NA | NA | NA | NA | NA |
| lizards | Cyrtodactylus_capreoloides | Cyrtodactylus_capreoloides | LC | Stable | tropical | nocturnal | 11.749 | 10629.385 | 4.357 |
| lizards | Cyrtodactylus_cattienensis | Cyrtodactylus_cattienensis | LC | Stable | tropical | nocturnal | 6.918 | 10730.452 | 19.529 |
| lizards | Cyrtodactylus_cavernicolus | Cyrtodactylus_cavernicolus | LC | Unknown | tropical | nocturnal | 10.715 | 986.352 | 5.348 |
| lizards | Cyrtodactylus_chanhomeae | Cyrtodactylus_chanhomeae | CR | Decreasing | tropical | nocturnal | 9.772 | 41.655 | 38.374 |
| lizards | Cyrtodactylus_chauquangensis | NA | LC | Unknown | NA | NA | NA | NA | NA |
| lizards | Cyrtodactylus_chrysopylos | Cyrtodactylus_chrysopylos | VU | Unknown | tropical | nocturnal | 11.749 | 120.105 | 5.619 |
| lizards | Cyrtodactylus_collegalensis | Cyrtodactylus_collegalensis | LC | Unknown | tropical | nocturnal | 3.631 | 260686.128 | 20.248 |
| lizards | Cyrtodactylus_condorensis | Cyrtodactylus_condorensis | LC | Stable | tropical | nocturnal | 14.791 | 2941.732 | 14.853 |
| lizards | Cyrtodactylus_consobrinoides | NA | DD | Unknown | NA | NA | NA | NA | NA |
| lizards | Cyrtodactylus_consobrinus | Cyrtodactylus_consobrinus | LC | Stable | tropical | nocturnal | 35.481 | 285419.7 | 8.454 |
| lizards | Cyrtodactylus_cracens | NA | NT | Unknown | NA | NA | NA | NA | NA |
| lizards | Cyrtodactylus_cryptus | Cyrtodactylus_cryptus | LC | Stable | tropical | nocturnal | 15.488 | 956.471 | 7.525 |
| lizards | Cyrtodactylus_cucdongensis | NA | LC | Unknown | NA | NA | NA | NA | NA |
| lizards | Cyrtodactylus_cucphuongensis | NA | LC | Unknown | NA | NA | NA | NA | NA |
| lizards | Cyrtodactylus_dammathetensis | NA | not_available | not_available | NA | NA | NA | NA | NA |
| lizards | Cyrtodactylus_darevskii | NA | DD | Unknown | NA | NA | NA | NA | NA |
| lizards | Cyrtodactylus_darmandvillei | NA | LC | Unknown | NA | NA | NA | NA | NA |
| lizards | Cyrtodactylus_dati | NA | DD | Unknown | NA | NA | NA | NA | NA |
| lizards | Cyrtodactylus_dattanensis | NA | DD | Unknown | NA | NA | NA | NA | NA |
| lizards | Cyrtodactylus_deccanensis | Cyrtodactylus_deccanensis | LC | Unknown | tropical | nocturnal | 12.303 | 71091.354 | 19.317 |

| | | | | | | | | | |
|---|---|---|---|---|---|---|---|---|---|
| lizards | Cyrtodactylus_deveti | NA | VU | Decreasing | NA | NA | NA | NA | NA |
| lizards | Cyrtodactylus_doisuthep | NA | LC | Unknown | NA | NA | NA | NA | NA |
| lizards | Cyrtodactylus_dumnuii | Cyrtodactylus_dumnuii | LC | Unknown | tropical | nocturnal | 11.749 | 10965.486 | 13.723 |
| lizards | Cyrtodactylus_durio | NA | DD | Unknown | NA | NA | NA | NA | NA |
| lizards | Cyrtodactylus_edwardtaylori | NA | EN | Unknown | NA | NA | NA | NA | NA |
| lizards | Cyrtodactylus_eisenmanae | NA | LC | Stable | NA | NA | NA | NA | NA |
| lizards | Cyrtodactylus_elok | Cyrtodactylus_elok | LC | Stable | tropical | nocturnal | 10.965 | 4193.565 | 8.609 |
| lizards | Cyrtodactylus_epiroticus | Cyrtodactylus_epiroticus | LC | Unknown | tropical | nocturnal | 42.658 | 28614.093 | 5.355 |
| lizards | Cyrtodactylus_erythrops | Cyrtodactylus_erythrops | LC | Unknown | tropical | cathemeral | 10.233 | 70.23 | 8.27 |
| lizards | Cyrtodactylus_fasciolatus | NA | NT | Stable | NA | NA | NA | NA | NA |
| lizards | Cyrtodactylus_feae | NA | DD | Unknown | NA | NA | NA | NA | NA |
| lizards | Cyrtodactylus_fraenatus | Cyrtodactylus_fraenatus | EN | Unknown | tropical | nocturnal | 19.055 | 728.33 | 23.592 |
| lizards | Cyrtodactylus_fumosus | NA | DD | Unknown | NA | NA | NA | NA | NA |
| lizards | Cyrtodactylus_gansi | Cyrtodactylus_gansi | LC | Unknown | tropical | nocturnal | 5.37 | 436.271 | 7.431 |
| lizards | Cyrtodactylus_gialaiensis | NA | CR | Unknown | NA | NA | NA | NA | NA |
| lizards | Cyrtodactylus_grismeri | NA | VU | Stable | NA | NA | NA | NA | NA |
| lizards | Cyrtodactylus_guakanthanensis | NA | CR | Unknown | NA | NA | NA | NA | NA |
| lizards | Cyrtodactylus_gubaot | NA | LC | Stable | NA | NA | NA | NA | NA |
| lizards | Cyrtodactylus_gubernatoris | NA | DD | Unknown | NA | NA | NA | NA | NA |
| lizards | Cyrtodactylus_gunungsenyumensis | NA | LC | Unknown | NA | NA | NA | NA | NA |
| lizards | Cyrtodactylus_halmahericus | NA | LC | Unknown | NA | NA | NA | NA | NA |
| lizards | Cyrtodactylus_hidupselamanya | NA | VU | Stable | NA | NA | NA | NA | NA |
| lizards | Cyrtodactylus_hikidai | NA | LC | Stable | NA | NA | NA | NA | NA |
| lizards | Cyrtodactylus_himalayanus | NA | DD | Unknown | NA | NA | NA | NA | NA |
| lizards | Cyrtodactylus_hinnamnoensis | NA | LC | Unknown | NA | NA | NA | NA | NA |
| lizards | Cyrtodactylus_hontreensis | NA | LC | Unknown | NA | NA | NA | NA | NA |
| lizards | Cyrtodactylus_huongsonensis | NA | VU | Decreasing | NA | NA | NA | NA | NA |
| lizards | Cyrtodactylus_huynhi | Cyrtodactylus_huynhi | VU | Stable | tropical | nocturnal | 10.233 | 19.982 | 23.605 |
| lizards | Cyrtodactylus_ingeri | Cyrtodactylus_ingeri | LC | Unknown | tropical | nocturnal | 10.471 | 24505.713 | 5.924 |
| lizards | Cyrtodactylus_interdigitalis | Cyrtodactylus_interdigitalis | LC | Unknown | tropical | nocturnal | 10.233 | 88252.151 | 8.202 |
| lizards | Cyrtodactylus_intermedius | Cyrtodactylus_intermedius | LC | Stable | tropical | nocturnal | 15.136 | 42675.807 | 13.021 |
| lizards | Cyrtodactylus_inthanon | NA | LC | Unknown | NA | NA | NA | NA | NA |
| lizards | Cyrtodactylus_irregularis | NA | DD | Unknown | NA | NA | NA | NA | NA |
| lizards | Cyrtodactylus_jaegeri | NA | CR | Unknown | NA | NA | NA | NA | NA |

| | | | | | | | | | |
|---|---|---|---|---|---|---|---|---|---|
| lizards | Cyrtodactylus_jambangan | NA | LC | Stable | NA | NA | NA | NA | NA |
| lizards | Cyrtodactylus_jarakensis | NA | CR | Unknown | NA | NA | NA | NA | NA |
| lizards | Cyrtodactylus_jarujini | NA | LC | Stable | NA | NA | NA | NA | NA |
| lizards | Cyrtodactylus_jelawangensis | NA | DD | Unknown | NA | NA | NA | NA | NA |
| lizards | Cyrtodactylus_jellesmae | Cyrtodactylus_jellesmae | LC | Decreasing | tropical | nocturnal | 10 | 138356.929 | 9.05 |
| lizards | Cyrtodactylus_jeyporensis | NA | EN | Unknown | NA | NA | NA | NA | NA |
| lizards | Cyrtodactylus_khammouanensis | NA | VU | Unknown | NA | NA | NA | NA | NA |
| lizards | Cyrtodactylus_khasiensis | NA | DD | Unknown | NA | NA | NA | NA | NA |
| lizards | Cyrtodactylus_khelangensis | NA | EN | Decreasing | NA | NA | NA | NA | NA |
| lizards | Cyrtodactylus_kingsadai | NA | LC | Unknown | NA | NA | NA | NA | NA |
| lizards | Cyrtodactylus_klugei | NA | LC | Unknown | NA | NA | NA | NA | NA |
| lizards | Cyrtodactylus_kunyai | NA | LC | Unknown | NA | NA | NA | NA | NA |
| lizards | Cyrtodactylus_laevigatus | Cyrtodactylus_laevigatus | LC | Stable | tropical | nocturnal | 2.344 | 1084.997 | 14.817 |
| lizards | Cyrtodactylus_langkawiensis | Cyrtodactylus_langkawiensis | LC | Unknown | tropical | nocturnal | 19.055 | 9.991 | 26.948 |
| lizards | Cyrtodactylus_lateralis | Cyrtodactylus_lateralis | LC | Unknown | tropical | nocturnal | 16.982 | 2186.892 | 13.359 |
| lizards | Cyrtodactylus_lawderanus | NA | DD | Unknown | NA | NA | NA | NA | NA |
| lizards | Cyrtodactylus_leegrismeri | Cyrtodactylus_leegrismeri | LC | Unknown | tropical | nocturnal | 15.136 | 4.039 | 1.876 |
| lizards | Cyrtodactylus_lekaguli | Cyrtodactylus_lekaguli | LC | Stable | tropical | nocturnal | 20.893 | 14886.837 | 13.9 |
| lizards | Cyrtodactylus_lenggongensis | NA | LC | Stable | NA | NA | NA | NA | NA |
| lizards | Cyrtodactylus_linnoensis | NA | not_available | not_available | NA | NA | NA | NA | NA |
| lizards | Cyrtodactylus_linnwayensis | NA | not_available | not_available | NA | NA | NA | NA | NA |
| lizards | Cyrtodactylus_lomyenensis | NA | VU | Unknown | NA | NA | NA | NA | NA |
| lizards | Cyrtodactylus_loriae | Cyrtodactylus_loriae | LC | Unknown | tropical | nocturnal | 64.565 | 16920.854 | 2.64 |
| lizards | Cyrtodactylus_louisiadensis | Cyrtodactylus_louisiadensis | LC | Stable | tropical | nocturnal | 41.687 | 6258.193 | 5.367 |
| lizards | Cyrtodactylus_macrotuberculatus | Cyrtodactylus_macrotuberculatus | LC | Unknown | tropical | nocturnal | 31.623 | 10334.433 | 19.517 |
| lizards | Cyrtodactylus_majulah | Cyrtodactylus_majulah | LC | Unknown | tropical | nocturnal | 6.761 | 19.982 | 29.978 |
| lizards | Cyrtodactylus_malayanus | Cyrtodactylus_malayanus | LC | Unknown | tropical | nocturnal | 30.2 | 13679.757 | 6.024 |
| lizards | Cyrtodactylus_mamanwa | NA | LC | Stable | NA | NA | NA | NA | NA |
| lizards | Cyrtodactylus_mandalayensis | NA | DD | Unknown | NA | NA | NA | NA | NA |
| lizards | Cyrtodactylus_markuscombaii | NA | DD | Unknown | NA | NA | NA | NA | NA |
| lizards | Cyrtodactylus_marmoratus | Cyrtodactylus_marmoratus | LC | Unknown | tropical | nocturnal | 12.303 | 7798 | 10.657 |
| lizards | Cyrtodactylus_martini | NA | DD | Unknown | NA | NA | NA | NA | NA |
| lizards | Cyrtodactylus_martinstolli | NA | DD | Unknown | NA | NA | NA | NA | NA |
| lizards | Cyrtodactylus_matsuii | NA | LC | Unknown | NA | NA | NA | NA | NA |

| | | | | | | | | | |
|---|---|---|---|---|---|---|---|---|---|
| lizards | Cyrtodactylus_mcdonaldi | Cyrtodactylus_mcdonaldi | LC | Stable | tropical | nocturnal | 21.878 | 17782.502 | 2.471 |
| lizards | Cyrtodactylus_metropolis | NA | LC | Unknown | NA | NA | NA | NA | NA |
| lizards | Cyrtodactylus_minor | NA | DD | Unknown | NA | NA | NA | NA | NA |
| lizards | Cyrtodactylus_multiporus | NA | LC | Stable | NA | NA | NA | NA | NA |
| lizards | Cyrtodactylus_nebulosus | NA | LC | Unknown | NA | NA | NA | NA | NA |
| lizards | Cyrtodactylus_nepalensis | NA | DD | Unknown | NA | NA | NA | NA | NA |
| lizards | Cyrtodactylus_nigriocularis | NA | CR | Unknown | NA | NA | NA | NA | NA |
| lizards | Cyrtodactylus_novaeguineae | NA | LC | Unknown | NA | NA | NA | NA | NA |
| lizards | Cyrtodactylus_nuaulu | NA | NT | Stable | NA | NA | NA | NA | NA |
| lizards | Cyrtodactylus_oldhami | Cyrtodactylus_oldhami | LC | Stable | tropical | nocturnal | 9.333 | 186494.617 | 11.407 |
| lizards | Cyrtodactylus_otai | NA | EN | Unknown | NA | NA | NA | NA | NA |
| lizards | Cyrtodactylus_pageli | Cyrtodactylus_pageli | LC | Stable | tropical | nocturnal | 13.804 | 24.338 | 14.666 |
| lizards | Cyrtodactylus_pantiensis | Cyrtodactylus_pantiensis | VU | Unknown | tropical | nocturnal | 9.333 | 13096.402 | 12.135 |
| lizards | Cyrtodactylus_papilionoides | Cyrtodactylus_papilionoides | LC | Stable | tropical | nocturnal | 15.488 | 6130.927 | 13.978 |
| lizards | Cyrtodactylus_paradoxus | NA | not_available | not_available | NA | NA | NA | NA | NA |
| lizards | Cyrtodactylus_payacola | Cyrtodactylus_payacola | DD | Stable | tropical | nocturnal | 6.457 | 1168.978 | 17.639 |
| lizards | Cyrtodactylus_peguensis | Cyrtodactylus_peguensis | LC | Unknown | tropical | nocturnal | 12.303 | 50780.145 | 12.35 |
| lizards | Cyrtodactylus_petani | NA | DD | Stable | NA | NA | NA | NA | NA |
| lizards | Cyrtodactylus_phetchaburiensis | NA | LC | Unknown | NA | NA | NA | NA | NA |
| lizards | Cyrtodactylus_philippinicus | Cyrtodactylus_philippinicus | LC | Stable | tropical | nocturnal | 17.783 | 165885.824 | 14.854 |
| lizards | Cyrtodactylus_phongnhakebangensis | Cyrtodactylus_phongnhakebangensis | LC | Stable | tropical | nocturnal | 21.38 | 267.062 | 6.99 |
| lizards | Cyrtodactylus_phuocbinhensis | NA | LC | Stable | NA | NA | NA | NA | NA |
| lizards | Cyrtodactylus_phuquocensis | NA | EN | Stable | NA | NA | NA | NA | NA |
| lizards | Cyrtodactylus_psarops | NA | EN | Unknown | NA | NA | NA | NA | NA |
| lizards | Cyrtodactylus_pseudoquadrivirgatus | Cyrtodactylus_pseudoquadrivirgatus | LC | Unknown | tropical | nocturnal | 13.183 | 43794.899 | 6.324 |
| lizards | Cyrtodactylus_pubisulcus | Cyrtodactylus_pubisulcus | LC | Unknown | tropical | nocturnal | 9.333 | 54306.811 | 6.628 |
| lizards | Cyrtodactylus_puhuensis | NA | LC | Unknown | NA | NA | NA | NA | NA |
| lizards | Cyrtodactylus_pulchellus | Cyrtodactylus_pulchellus | EN | Unknown | tropical | nocturnal | 35.481 | 319.177 | 29.572 |
| lizards | Cyrtodactylus_pyinyaungensis | NA | not_available | not_available | NA | NA | NA | NA | NA |
| lizards | Cyrtodactylus_quadrivirgatus | NA | LC | Unknown | NA | NA | NA | NA | NA |
| lizards | Cyrtodactylus_ranongensis | NA | LC | Unknown | NA | NA | NA | NA | NA |
| lizards | Cyrtodactylus_redimiculus | Cyrtodactylus_redimiculus | NT | Decreasing | tropical | nocturnal | 16.218 | 3158.921 | 5.064 |
| lizards | Cyrtodactylus_rishivalleyensis | NA | EN | Unknown | NA | NA | NA | NA | NA |
| lizards | Cyrtodactylus_robustus | NA | LC | Unknown | NA | NA | NA | NA | NA |

| | | | | | | | | | |
|---|---|---|---|---|---|---|---|---|---|
| lizards | Cyrtodactylus_roesleri | NA | LC | Unknown | NA | NA | NA | NA | NA |
| lizards | Cyrtodactylus_rosichonarieforum | NA | LC | Unknown | NA | NA | NA | NA | NA |
| lizards | Cyrtodactylus_rubidus | Cyrtodactylus_rubidus | LC | Stable | tropical | nocturnal | 10.233 | 1467.541 | 7.912 |
| lizards | Cyrtodactylus_rufford | NA | DD | Unknown | NA | NA | NA | NA | NA |
| lizards | Cyrtodactylus_russelli | NA | DD | Unknown | NA | NA | NA | NA | NA |
| lizards | Cyrtodactylus_sadanensis | NA | not_available | not_available | NA | NA | NA | NA | NA |
| lizards | Cyrtodactylus_sadansinensis | NA | not_available | not_available | NA | NA | NA | NA | NA |
| lizards | Cyrtodactylus_sadleiri | Cyrtodactylus_sadleiri | EN | Decreasing | tropical | nocturnal | 13.49 | 137.177 | 12.527 |
| lizards | Cyrtodactylus_saiyok | NA | LC | Stable | NA | NA | NA | NA | NA |
| lizards | Cyrtodactylus_salomonensis | NA | NT | Stable | NA | NA | NA | NA | NA |
| lizards | Cyrtodactylus_samroiyot | Cyrtodactylus_samroiyot | LC | Stable | tropical | nocturnal | 6.31 | 17.892 | 26.114 |
| lizards | Cyrtodactylus_sangi | NA | not_available | not_available | NA | NA | NA | NA | NA |
| lizards | Cyrtodactylus_sanook | NA | LC | Unknown | NA | NA | NA | NA | NA |
| lizards | Cyrtodactylus_sanpelensis | NA | not_available | not_available | NA | NA | NA | NA | NA |
| lizards | Cyrtodactylus_semenanjungensis | Cyrtodactylus_semenanjungensis | NT | Stable | tropical | nocturnal | 7.413 | 277.95 | 6.457 |
| lizards | Cyrtodactylus_semiadii | Cyrtodactylus_semiadii | LC | Stable | tropical | nocturnal | 2.399 | 873.763 | 27.203 |
| lizards | Cyrtodactylus_semicinctus | NA | LC | Unknown | NA | NA | NA | NA | NA |
| lizards | Cyrtodactylus_seribuatensis | Cyrtodactylus_seribuatensis | VU | Unknown | tropical | nocturnal | 8.71 | 28.395 | 3.771 |
| lizards | Cyrtodactylus_serratus | NA | LC | Unknown | NA | NA | NA | NA | NA |
| lizards | Cyrtodactylus_sharkari | NA | VU | Unknown | NA | NA | NA | NA | NA |
| lizards | Cyrtodactylus_shwetaungorum | NA | not_available | not_available | NA | NA | NA | NA | NA |
| lizards | Cyrtodactylus_sinyineensis | NA | not_available | not_available | NA | NA | NA | NA | NA |
| lizards | Cyrtodactylus_slowinskii | NA | LC | Unknown | NA | NA | NA | NA | NA |
| lizards | Cyrtodactylus_soba | Cyrtodactylus_soba | VU | Unknown | tropical | nocturnal | 22.387 | 83.873 | 14.391 |
| lizards | Cyrtodactylus_sommerladi | NA | LC | Unknown | NA | NA | NA | NA | NA |
| lizards | Cyrtodactylus_soni | NA | LC | Unknown | NA | NA | NA | NA | NA |
| lizards | Cyrtodactylus_sonlaensis | NA | not_available | not_available | NA | NA | NA | NA | NA |
| lizards | Cyrtodactylus_soudthichaki | NA | LC | Unknown | NA | NA | NA | NA | NA |
| lizards | Cyrtodactylus_speciosus | NA | EN | Decreasing | NA | NA | NA | NA | NA |
| lizards | Cyrtodactylus_spelaeus | NA | DD | Unknown | NA | NA | NA | NA | NA |
| lizards | Cyrtodactylus_spinosus | NA | DD | Unknown | NA | NA | NA | NA | NA |
| lizards | Cyrtodactylus_srilekhae | NA | NT | Unknown | NA | NA | NA | NA | NA |
| lizards | Cyrtodactylus_stresemanni | NA | DD | Unknown | NA | NA | NA | NA | NA |
| lizards | Cyrtodactylus_sumonthai | Cyrtodactylus_sumonthai | LC | Unknown | tropical | nocturnal | 10.471 | 9.991 | 24.022 |

| | | | | | | | | | |
|---|---|---|---|---|---|---|---|---|---|
| lizards | Cyrtodactylus_sumuroi | NA | LC | Stable | NA | NA | NA | NA | NA |
| lizards | Cyrtodactylus_surin | NA | LC | Unknown | NA | NA | NA | NA | NA |
| lizards | Cyrtodactylus_sworderi | Cyrtodactylus_sworderi | EN | Unknown | tropical | nocturnal | 10.233 | 116326.577 | 13.787 |
| lizards | Cyrtodactylus_tahuna | NA | LC | Unknown | NA | NA | NA | NA | NA |
| lizards | Cyrtodactylus_takouensis | Cyrtodactylus_takouensis | CR | Unknown | tropical | nocturnal | 10.715 | 9.991 | 28.577 |
| lizards | Cyrtodactylus_tamaiensis | NA | DD | Unknown | NA | NA | NA | NA | NA |
| lizards | Cyrtodactylus_tambora | NA | NT | Unknown | NA | NA | NA | NA | NA |
| lizards | Cyrtodactylus_tanim | NA | not_available | not_available | NA | NA | NA | NA | NA |
| lizards | Cyrtodactylus_tautbatorum | NA | LC | Unknown | NA | NA | NA | NA | NA |
| lizards | Cyrtodactylus_taynguyenensis | NA | LC | Stable | NA | NA | NA | NA | NA |
| lizards | Cyrtodactylus_tebuensis | Cyrtodactylus_tebuensis | LC | Unknown | tropical | nocturnal | 12.303 | 14.928 | 3.64 |
| lizards | Cyrtodactylus_teyniei | NA | DD | Unknown | NA | NA | NA | NA | NA |
| lizards | Cyrtodactylus_thirakhupti | NA | NT | Unknown | NA | NA | NA | NA | NA |
| lizards | Cyrtodactylus_thochuensis | NA | not_available | not_available | NA | NA | NA | NA | NA |
| lizards | Cyrtodactylus_thuongae | NA | VU | Unknown | NA | NA | NA | NA | NA |
| lizards | Cyrtodactylus_tibetanus | NA | LC | Stable | NA | NA | NA | NA | NA |
| lizards | Cyrtodactylus_tigroides | Cyrtodactylus_tigroides | LC | Unknown | tropical | nocturnal | 12.303 | 19.982 | 24.906 |
| lizards | Cyrtodactylus_timur | Cyrtodactylus_timur | LC | Unknown | tropical | nocturnal | 31.623 | 14.927 | 3.64 |
| lizards | Cyrtodactylus_tiomanensis | Cyrtodactylus_tiomanensis | NT | Unknown | tropical | nocturnal | 12.882 | 96.764 | 6.984 |
| lizards | Cyrtodactylus_triedrus | Cyrtodactylus_triedrus | NT | Stable | tropical | nocturnal | 5.129 | 23243.055 | 16.65 |
| lizards | Cyrtodactylus_trilatofasciatus | NA | EN | Unknown | NA | NA | NA | NA | NA |
| lizards | Cyrtodactylus_tripartitus | NA | LC | Unknown | NA | NA | NA | NA | NA |
| lizards | Cyrtodactylus_tuberculatus | Cyrtodactylus_tuberculatus | LC | Stable | tropical | nocturnal | 47.863 | 20487.105 | 2.185 |
| lizards | Cyrtodactylus_varadgirii | NA | LC | Unknown | NA | NA | NA | NA | NA |
| lizards | Cyrtodactylus_variegatus | NA | DD | Unknown | NA | NA | NA | NA | NA |
| lizards | Cyrtodactylus_vilaphongi | NA | DD | Unknown | NA | NA | NA | NA | NA |
| lizards | Cyrtodactylus_wakeorum | Cyrtodactylus_wakeorum | EN | Unknown | tropical | nocturnal | 5.495 | 5150.322 | 8.27 |
| lizards | Cyrtodactylus_wangkulangkulae | NA | LC | Unknown | NA | NA | NA | NA | NA |
| lizards | Cyrtodactylus_wayakonei | Cyrtodactylus_wayakonei | NT | Unknown | tropical | nocturnal | 15.488 | 19.982 | 6.976 |
| lizards | Cyrtodactylus_welpyanensis | NA | not_available | not_available | NA | NA | NA | NA | NA |
| lizards | Cyrtodactylus_yakhuna | Cyrtodactylus_yakhuna | VU | Unknown | tropical | nocturnal | 3.162 | 21228.655 | 17.346 |
| lizards | Cyrtodactylus_yangbayensis | Cyrtodactylus_yangbayensis | LC | Stable | tropical | nocturnal | 15.136 | 64.752 | 7.182 |
| lizards | Cyrtodactylus_yathepyanensis | NA | not_available | not_available | NA | NA | NA | NA | NA |
| lizards | Cyrtodactylus_yoshii | Cyrtodactylus_yoshii | LC | Unknown | tropical | nocturnal | 20.893 | 10162.524 | 6.281 |

| | | | | | | | | | |
|---|---|---|---|---|---|---|---|---|---|
| lizards | Cyrtodactylus_zhaoermii | NA | LC | Stable | NA | NA | NA | NA | NA |
| lizards | Cyrtodactylus_ziegleri | NA | LC | Stable | NA | NA | NA | NA | NA |
| lizards | Cyrtodactylus_zugi | NA | LC | Unknown | NA | NA | NA | NA | NA |
| lizards | Cyrtopodion_agamuroides | Cyrtopodion_agamuroides | LC | Unknown | temperate | cathemeral | 2.818 | 546163.082 | 6.141 |
| lizards | Cyrtopodion_aravallensis | NA | VU | Unknown | NA | NA | NA | NA | NA |
| lizards | Cyrtopodion_baigii | NA | DD | Unknown | NA | NA | NA | NA | NA |
| lizards | Cyrtopodion_belaense | NA | DD | Unknown | NA | NA | NA | NA | NA |
| lizards | Cyrtopodion_brevipes | NA | LC | Stable | NA | NA | NA | NA | NA |
| lizards | Cyrtopodion_fortmunroi | Cyrtopodion_fortmunroi | LC | Unknown | temperate | nocturnal | 2.884 | 14.998 | 18.79 |
| lizards | Cyrtopodion_gastrophole | NA | DD | Unknown | NA | NA | NA | NA | NA |
| lizards | Cyrtopodion_golubevi | NA | not_available | not_available | NA | NA | NA | NA | NA |
| lizards | Cyrtopodion_hormozganum | NA | not_available | not_available | NA | NA | NA | NA | NA |
| lizards | Cyrtopodion_indusoani | Cyrtopodion_indusoani | LC | Stable | temperate | cathemeral | 3.467 | 4704.025 | 14.322 |
| lizards | Cyrtopodion_kachhense | Cyrtopodion_kachhense | LC | Unknown | temperate | nocturnal | 3.631 | 317223.416 | 11.217 |
| lizards | Cyrtopodion_kiabii | NA | not_available | not_available | NA | NA | NA | NA | NA |
| lizards | Cyrtopodion_kirmanense | NA | LC | Stable | NA | NA | NA | NA | NA |
| lizards | Cyrtopodion_kohsulaimanai | Cyrtopodion_kohsulaimanai | LC | Unknown | temperate | nocturnal | 4.467 | 65691.78 | 7.368 |
| lizards | Cyrtopodion_mansarulus | NA | DD | Unknown | NA | NA | NA | NA | NA |
| lizards | Cyrtopodion_medogense | NA | LC | Unknown | NA | NA | NA | NA | NA |
| lizards | Cyrtopodion_montiumsalsorum | NA | LC | Stable | NA | NA | NA | NA | NA |
| lizards | Cyrtopodion_persepolense | NA | not_available | not_available | NA | NA | NA | NA | NA |
| lizards | Cyrtopodion_potoharense | Cyrtopodion_potoharense | LC | Unknown | temperate | nocturnal | 3.162 | 76284.19 | 16.423 |
| lizards | Cyrtopodion_rhodocauda | NA | LC | Unknown | NA | NA | NA | NA | NA |
| lizards | Cyrtopodion_rohtasfortai | Cyrtopodion_rohtasfortai | LC | Stable | temperate | nocturnal | 4.074 | 6935.431 | 21.264 |
| lizards | Cyrtopodion_scabrum | Cyrtopodion_scabrum | LC | Stable | temperate | nocturnal | 5.754 | 4041175.664 | 8.964 |
| lizards | Cyrtopodion_sistanense | NA | LC | Unknown | NA | NA | NA | NA | NA |
| lizards | Cyrtopodion_watsoni | Cyrtopodion_watsoni | LC | Unknown | temperate | cathemeral | 3.467 | 52928.676 | 13.439 |
| lizards | Dactylocnemis_pacificus | Dactylocnemis_pacificus | LC | Stable | temperate | nocturnal | 20.417 | 84526.141 | 9.473 |
| lizards | Dalmatolacerta_oxycephala | Dalmatolacerta_oxycephala | LC | Unknown | temperate | diurnal | 8.128 | 30034.602 | 16.66 |
| lizards | Darevskia_alpina | Darevskia_alpina | VU | Decreasing | temperate | diurnal | 7.244 | 6118.551 | 6.077 |
| lizards | Darevskia_armeniaca | Darevskia_armeniaca | LC | Stable | temperate | diurnal | 9.12 | 57387.049 | 14.143 |
| lizards | Darevskia_bendimahiensis | Darevskia_bendimahiensis | EN | Unknown | temperate | diurnal | 6.166 | 4282.666 | 12.502 |
| lizards | Darevskia_bithynica | NA | not_available | not_available | NA | NA | NA | NA | NA |
| lizards | Darevskia_brauneri | Darevskia_brauneri | LC | Stable | temperate | diurnal | 8.318 | 47237.888 | 9.946 |

| | | | | | | | | | |
|---|---|---|---|---|---|---|---|---|---|
| lizards | Darevskia_caspica | NA | not_available | not_available | NA | NA | NA | NA | NA |
| lizards | Darevskia_caucasica | Darevskia_caucasica | LC | Stable | temperate | diurnal | 7.079 | 142840.962 | 12.004 |
| lizards | Darevskia_chlorogaster | Darevskia_chlorogaster | LC | Decreasing | temperate | diurnal | 8.71 | 54120.582 | 18.179 |
| lizards | Darevskia_clarkorum | NA | EN | Decreasing | NA | NA | NA | NA | NA |
| lizards | Darevskia_daghestanica | Darevskia_daghestanica | LC | Stable | temperate | diurnal | 4.571 | 27783.677 | 10.106 |
| lizards | Darevskia_dahli | Darevskia_dahli | NT | Decreasing | temperate | diurnal | 6.166 | 22647.899 | 13.497 |
| lizards | Darevskia_defilippii | Darevskia_defilippii | LC | Stable | temperate | diurnal | 4.571 | 76748.357 | 16.642 |
| lizards | Darevskia_derjugini | Darevskia_derjugini | NT | Decreasing | temperate | diurnal | 6.457 | 51277.367 | 10.606 |
| lizards | Darevskia_kamii | NA | not_available | not_available | NA | NA | NA | NA | NA |
| lizards | Darevskia_kopetdaghica | NA | EN | Decreasing | NA | NA | NA | NA | NA |
| lizards | Darevskia_lindholmi | Darevskia_lindholmi | LC | Stable | temperate | diurnal | 9.772 | 25224.527 | 18.682 |
| lizards | Darevskia_mixta | Darevskia_mixta | NT | Decreasing | temperate | diurnal | 5.888 | 6446.269 | 8.931 |
| lizards | Darevskia_parvula | Darevskia_parvula | LC | Stable | temperate | diurnal | 5.012 | 39377.424 | 11.201 |
| lizards | Darevskia_portschinskii | Darevskia_portschinskii | LC | Decreasing | temperate | diurnal | 7.079 | 24669.687 | 15.434 |
| lizards | Darevskia_praticola | Darevskia_praticola | NT | Decreasing | temperate | diurnal | 6.761 | 548280.824 | 16.112 |
| lizards | Darevskia_raddei | Darevskia_raddei | LC | Stable | temperate | diurnal | 10.233 | 198744.706 | 14.176 |
| lizards | Darevskia_rostombekowi | Darevskia_rostombekovi | EN | Decreasing | temperate | diurnal | 5.012 | 6860.747 | 14.575 |
| lizards | Darevskia_rudis | Darevskia_rudis | LC | Stable | temperate | diurnal | 15.849 | 238514.976 | 13.281 |
| lizards | Darevskia_sapphirina | Darevskia_sapphirina | LC | Unknown | temperate | diurnal | 4.786 | 53.09 | 22.842 |
| lizards | Darevskia_saxicola | Darevskia_saxicola | LC | Stable | temperate | diurnal | 15.849 | 19671.502 | 12.195 |
| lizards | Darevskia_schaekeli | NA | not_available | not_available | NA | NA | NA | NA | NA |
| lizards | Darevskia_steineri | NA | DD | Unknown | NA | NA | NA | NA | NA |
| lizards | Darevskia_unisexualis | Darevskia_unisexualis | NT | Decreasing | temperate | diurnal | 9.12 | 79925.585 | 12.751 |
| lizards | Darevskia_uzzelli | NA | EN | Decreasing | NA | NA | NA | NA | NA |
| lizards | Darevskia_valentini | Darevskia_valentini | LC | Stable | temperate | diurnal | 11.749 | 259966.14 | 12.845 |
| lizards | Dasia_griffini | NA | NT | Stable | NA | NA | NA | NA | NA |
| lizards | Dasia_grisea | Dasia_grisea | LC | Unknown | tropical | diurnal | 50.119 | 166998.032 | 13.107 |
| lizards | Dasia_haliana | Dasia_haliana | VU | Unknown | tropical | diurnal | 15.488 | 34469.852 | 16.707 |
| lizards | Dasia_johnsinghi | NA | DD | Unknown | NA | NA | NA | NA | NA |
| lizards | Dasia_nicobarensis | NA | DD | Stable | NA | NA | NA | NA | NA |
| lizards | Dasia_olivacea | Dasia_olivacea | LC | Stable | tropical | diurnal | 79.433 | 1953300.126 | 10.236 |
| lizards | Dasia_semicincta | NA | DD | Unknown | NA | NA | NA | NA | NA |
| lizards | Dasia_subcaerulea | NA | EN | Unknown | NA | NA | NA | NA | NA |
| lizards | Dasia_vittata | Dasia_vittata | LC | Unknown | tropical | diurnal | 19.055 | 434667.819 | 4.425 |

| | | | | | | | | | |
|---|---|---|---|---|---|---|---|---|---|
| lizards | Dasia_vyneri | NA | LC | Unknown | NA | NA | NA | NA | NA |
| lizards | Delma_australis | Delma_australis | LC | Stable | temperate | diurnal | 4.571 | 1472695.692 | 5.101 |
| lizards | Delma_borea | Delma_borea | LC | Stable | tropical | nocturnal | 4.898 | 2244028.148 | 1.904 |
| lizards | Delma_butleri | Delma_butleri | LC | Stable | temperate | cathemeral | 5.754 | 3244974.974 | 2.873 |
| lizards | Delma_concinna | Delma_concinna | LC | Unknown | temperate | cathemeral | 5.888 | 12765.479 | 8.339 |
| lizards | Delma_desmosa | NA | LC | Stable | NA | NA | NA | NA | NA |
| lizards | Delma_elegans | NA | LC | Stable | NA | NA | NA | NA | NA |
| lizards | Delma_fraseri | Delma_fraseri | LC | Stable | temperate | cathemeral | 7.943 | 591977.388 | 6.581 |
| lizards | Delma_grayii | Delma_grayii | LC | Stable | temperate | diurnal | 6.918 | 115357.603 | 12.184 |
| lizards | Delma_impar | Delma_impar | EN | Decreasing | temperate | diurnal | 6.166 | 205008.694 | 11.65 |
| lizards | Delma_inornata | Delma_inornata | LC | Decreasing | temperate | cathemeral | 7.413 | 656642.124 | 9.625 |
| lizards | Delma_labialis | Delma_labialis | LC | Unknown | tropical | diurnal | 6.918 | 62756.533 | 5.224 |
| lizards | Delma_molleri | Delma_molleri | LC | Stable | temperate | cathemeral | 5.888 | 52247.841 | 13.398 |
| lizards | Delma_nasuta | Delma_nasuta | LC | Stable | tropical | cathemeral | 5.888 | 3222078.03 | 2.095 |
| lizards | Delma_pax | NA | LC | Stable | NA | NA | NA | NA | NA |
| lizards | Delma_plebeia | Delma_plebeia | LC | Stable | temperate | diurnal | 6.607 | 281255.728 | 8.149 |
| lizards | Delma_tincta | Delma_tincta | LC | Stable | tropical | cathemeral | 4.467 | 4730812.559 | 3.534 |
| lizards | Dendragama_australis | NA | LC | Unknown | NA | NA | NA | NA | NA |
| lizards | Dendragama_boulengeri | Dendragama_boulengeri | LC | Unknown | tropical | diurnal | 16.218 | 12700.222 | 11.311 |
| lizards | Dendragama_dioidema | NA | LC | Unknown | NA | NA | NA | NA | NA |
| lizards | Dibamus_bogadeki | NA | EN | Unknown | NA | NA | NA | NA | NA |
| lizards | Dibamus_booliati | NA | DD | Unknown | NA | NA | NA | NA | NA |
| lizards | Dibamus_celebensis | NA | NT | Unknown | NA | NA | NA | NA | NA |
| lizards | Dibamus_ingeri | NA | DD | Unknown | NA | NA | NA | NA | NA |
| lizards | Dibamus_leucurus | NA | LC | Unknown | NA | NA | NA | NA | NA |
| lizards | Dibamus_nicobaricum | NA | EN | Unknown | NA | NA | NA | NA | NA |
| lizards | Dibamus_tiomanensis | NA | EN | Stable | NA | NA | NA | NA | NA |
| lizards | Dibamus_vorisi | NA | DD | Unknown | NA | NA | NA | NA | NA |
| lizards | Dicrodon_guttulatum | NA | LC | Decreasing | NA | NA | NA | NA | NA |
| lizards | Dierogekko_baaba | NA | DD | Unknown | NA | NA | NA | NA | NA |
| lizards | Dierogekko_inexpectatus | Dierogekko_inexpectatus | CR | Decreasing | tropical | nocturnal | 1.202 | 29.764 | 8.251 |
| lizards | Dierogekko_insularis | Dierogekko_insularis | EN | Decreasing | tropical | nocturnal | 1.38 | 79.371 | 5.106 |
| lizards | Dierogekko_kaalaensis | Dierogekko_kaalaensis | CR | Decreasing | tropical | nocturnal | 1.82 | 54.654 | 7.107 |
| lizards | Dierogekko_koniambo | Dierogekko_koniambo | CR | Decreasing | tropical | nocturnal | 1.549 | 232.687 | 10.282 |

| | | | | | | | | | |
|---|---|---|---|---|---|---|---|---|---|
| lizards | Dierogekko_nehoueensis | Dierogekko_nehoueensis | EN | Decreasing | tropical | nocturnal | 1.259 | 179.514 | 10.892 |
| lizards | Dierogekko_poumensis | Dierogekko_poumensis | CR | Decreasing | tropical | nocturnal | 1.096 | 30.988 | 8.554 |
| lizards | Dierogekko_thomaswhitei | Dierogekko_thomaswhitei | CR | Decreasing | tropical | nocturnal | 1.778 | 62.17 | 4.365 |
| lizards | Dierogekko_validiclavis | Dierogekko_validiclavis | EN | Decreasing | tropical | nocturnal | 1.778 | 615.453 | 4.398 |
| lizards | Dinarolacerta_montenegrina | NA | LC | Unknown | NA | NA | NA | NA | NA |
| lizards | Dinarolacerta_mosorensis | Dinarolacerta_mosorensis | VU | Decreasing | temperate | diurnal | 11.749 | 17451.529 | 14.985 |
| lizards | Diplodactylus_barraganae | NA | LC | Unknown | NA | NA | NA | NA | NA |
| lizards | Diplodactylus_bilybara | NA | LC | Stable | NA | NA | NA | NA | NA |
| lizards | Diplodactylus_calcicolus | Diplodactylus_calcicolus | LC | Stable | temperate | nocturnal | 3.89 | 346267.468 | 6.235 |
| lizards | Diplodactylus_capensis | Diplodactylus_capensis | NT | Stable | tropical | nocturnal | 5.888 | 1436.859 | 6.804 |
| lizards | Diplodactylus_conspicillatus | Diplodactylus_conspicillatus | LC | Stable | temperate | nocturnal | 5.495 | 1922008.312 | 1.891 |
| lizards | Diplodactylus_custos | NA | LC | Stable | NA | NA | NA | NA | NA |
| lizards | Diplodactylus_fulleri | NA | VU | Unknown | NA | NA | NA | NA | NA |
| lizards | Diplodactylus_furcosus | NA | LC | Unknown | NA | NA | NA | NA | NA |
| lizards | Diplodactylus_galeatus | Diplodactylus_galeatus | LC | Stable | temperate | nocturnal | 4.266 | 154489.23 | 2.845 |
| lizards | Diplodactylus_granariensis | Diplodactylus_granariensis | LC | Stable | temperate | nocturnal | 7.413 | 578743.022 | 5.74 |
| lizards | Diplodactylus_hillii | NA | LC | Stable | NA | NA | NA | NA | NA |
| lizards | Diplodactylus_kenneallyi | NA | DD | Unknown | NA | NA | NA | NA | NA |
| lizards | Diplodactylus_klugei | Diplodactylus_klugei | LC | Stable | temperate | nocturnal | 3.89 | 48287.792 | 5.621 |
| lizards | Diplodactylus_laevis | NA | LC | Stable | NA | NA | NA | NA | NA |
| lizards | Diplodactylus_lateroides | Diplodactylus_lateroides | LC | Stable | temperate | nocturnal | 2.63 | 16169.892 | 10.995 |
| lizards | Diplodactylus_mitchelli | Diplodactylus_mitchelli | LC | Stable | tropical | nocturnal | 9.12 | 47363.124 | 4.006 |
| lizards | Diplodactylus_nebulosus | NA | LC | Unknown | NA | NA | NA | NA | NA |
| lizards | Diplodactylus_ornatus | Diplodactylus_ornatus | LC | Stable | temperate | nocturnal | 3.89 | 63632.697 | 7.86 |
| lizards | Diplodactylus_platyurus | NA | LC | Unknown | NA | NA | NA | NA | NA |
| lizards | Diplodactylus_polyophthalmus | NA | VU | Decreasing | NA | NA | NA | NA | NA |
| lizards | Diplodactylus_pulcher | Diplodactylus_pulcher | LC | Stable | temperate | nocturnal | 4.786 | 1324956.296 | 4.033 |
| lizards | Diplodactylus_savagei | Diplodactylus_savagei | LC | Stable | tropical | nocturnal | 2.455 | 137750.773 | 4.309 |
| lizards | Diplodactylus_tessellatus | Diplodactylus_tessellatus | LC | Stable | temperate | nocturnal | 4.074 | 2252041.593 | 4.524 |
| lizards | Diplodactylus_vittatus | Diplodactylus_vittatus | LC | Unknown | temperate | nocturnal | 4.169 | 1495846.668 | 7.942 |
| lizards | Diplodactylus_wiru | NA | LC | Stable | NA | NA | NA | NA | NA |
| lizards | Diploderma_batangense | NA | LC | Stable | NA | NA | NA | NA | NA |
| lizards | Diploderma_brevicauda | NA | LC | Stable | NA | NA | NA | NA | NA |
| lizards | Diploderma_brevipes | Japalura_brevipes | VU | Decreasing | temperate | diurnal | 12.589 | 4305.631 | 6.653 |

| | | | | | | | | | |
|---|---|---|---|---|---|---|---|---|---|
| lizards | Diploderma_chapaense | NA | DD | Unknown | NA | NA | NA | NA | NA |
| lizards | Diploderma_dymondi | Japalura_dymondi | LC | Decreasing | temperate | diurnal | 31.623 | 72735.079 | 10.944 |
| lizards | Diploderma_fasciatum | NA | LC | Stable | NA | NA | NA | NA | NA |
| lizards | Diploderma_flaviceps | Japalura_flaviceps | LC | Unknown | temperate | diurnal | 20.893 | 1519307.91 | 12.81 |
| lizards | Diploderma_grahami | NA | DD | Unknown | NA | NA | NA | NA | NA |
| lizards | Diploderma_hamptoni | NA | DD | Unknown | NA | NA | NA | NA | NA |
| lizards | Diploderma_iadinum | NA | NT | Stable | NA | NA | NA | NA | NA |
| lizards | Diploderma_laeviventre | NA | LC | Stable | NA | NA | NA | NA | NA |
| lizards | Diploderma_luei | Japalura_luei | EN | Decreasing | temperate | diurnal | 13.49 | 1403.705 | 4.126 |
| lizards | Diploderma_makii | Japalura_makii | VU | Decreasing | tropical | diurnal | 19.498 | 2111.319 | 8.818 |
| lizards | Diploderma_micangshanense | NA | LC | Stable | NA | NA | NA | NA | NA |
| lizards | Diploderma_polygonatum | Japalura_polygonata | NT | Decreasing | temperate | diurnal | 16.982 | 16072.943 | 20.533 |
| lizards | Diploderma_slowinskii | NA | DD | Unknown | NA | NA | NA | NA | NA |
| lizards | Diploderma_splendidum | Japalura_splendida | LC | Decreasing | temperate | diurnal | 33.113 | 2005622.259 | 13.453 |
| lizards | Diploderma_swinhonis | Japalura_swinhonis | LC | Stable | temperate | diurnal | 21.878 | 29434.903 | 22.918 |
| lizards | Diploderma_varcoae | Japalura_varcoae | LC | Stable | temperate | diurnal | 19.953 | 53387.638 | 14.356 |
| lizards | Diploderma_vela | NA | LC | Stable | NA | NA | NA | NA | NA |
| lizards | Diploderma_yulongense | NA | LC | Stable | NA | NA | NA | NA | NA |
| lizards | Diploderma_yunnanense | Japalura_yunnanensis | LC | Stable | temperate | diurnal | 22.387 | 120457.718 | 11.884 |
| lizards | Diploderma_zhaoermii | NA | LC | Stable | NA | NA | NA | NA | NA |
| lizards | Diploglossus_delasagra | Diploglossus_delasagra | LC | Unknown | tropical | diurnal | 30.2 | 16122.328 | 18.887 |
| lizards | Diploglossus_fasciatus | Diploglossus_fasciatus | LC | Unknown | tropical | diurnal | 100 | 751798.097 | 8.81 |
| lizards | Diploglossus_lessonae | Diploglossus_lessonae | LC | Unknown | tropical | diurnal | 83.176 | 323282.894 | 10.588 |
| lizards | Diploglossus_millepunctatus | NA | LC | Stable | NA | NA | NA | NA | NA |
| lizards | Diploglossus_monotropis | Diploglossus_monotropis | LC | Decreasing | tropical | diurnal | 223.872 | 329718.237 | 9.902 |
| lizards | Diploglossus_montisserrati | NA | CR | Unknown | NA | NA | NA | NA | NA |
| lizards | Diplolaemus_bibronii | Diplolaemus_bibronii | LC | Stable | temperate | diurnal | 56.234 | 295294.095 | 5.644 |
| lizards | Diplolaemus_darwinii | NA | LC | Stable | NA | NA | NA | NA | NA |
| lizards | Diplolaemus_leopardinus | NA | LC | Stable | NA | NA | NA | NA | NA |
| lizards | Diplolaemus_sexcinctus | NA | LC | Stable | NA | NA | NA | NA | NA |
| lizards | Diplometopon_zarudnyi | NA | LC | Stable | NA | NA | NA | NA | NA |
| lizards | Diporiphora_albilabris | Diporiphora_albilabris | LC | Stable | tropical | diurnal | 6.607 | 532682.43 | 2.015 |
| lizards | Diporiphora_amphiboluroides | Diporiphora_amphiboluroides | LC | Unknown | temperate | diurnal | 32.359 | 920616.43 | 3.393 |
| lizards | Diporiphora_australis | Diporiphora_australis | LC | Unknown | tropical | diurnal | 14.791 | 935824.653 | 5.28 |

| | | | | | | | | | |
|---|---|---|---|---|---|---|---|---|---|
| lizards | Diporiphora_bennettii | Diporiphora_bennettii | LC | Unknown | tropical | diurnal | 16.596 | 406968.263 | 1.874 |
| lizards | Diporiphora_bilineata | Diporiphora_bilineata | LC | Unknown | tropical | diurnal | 8.318 | 2132931.319 | 2.915 |
| lizards | Diporiphora_lalliae | NA | LC | Stable | NA | NA | NA | NA | NA |
| lizards | Diporiphora_linga | NA | LC | Unknown | NA | NA | NA | NA | NA |
| lizards | Diporiphora_magna | NA | LC | Stable | NA | NA | NA | NA | NA |
| lizards | Diporiphora_margaretae | NA | not_available | not_available | NA | NA | NA | NA | NA |
| lizards | Diporiphora_nobbi | Diporiphora_nobbi | LC | Unknown | temperate | diurnal | 19.498 | 1964903.236 | 6.991 |
| lizards | Diporiphora_paraconvergens | Diporiphora_paraconvergens | LC | Unknown | temperate | diurnal | 6.918 | 852657.997 | 1.06 |
| lizards | Diporiphora_phaeospinosa | NA | LC | Unknown | NA | NA | NA | NA | NA |
| lizards | Diporiphora_pindan | NA | LC | Stable | NA | NA | NA | NA | NA |
| lizards | Diporiphora_reginae | NA | LC | Unknown | NA | NA | NA | NA | NA |
| lizards | Diporiphora_superba | Diporiphora_superba | LC | Stable | tropical | diurnal | 28.184 | 70055.108 | 2.056 |
| lizards | Diporiphora_valens | NA | LC | Stable | NA | NA | NA | NA | NA |
| lizards | Diporiphora_vescus | NA | VU | Unknown | NA | NA | NA | NA | NA |
| lizards | Diporiphora_winneckei | Diporiphora_winneckei | LC | Unknown | temperate | diurnal | 10.965 | 653992.829 | 2.406 |
| lizards | Dipsosaurus_catalinensis | NA | not_available | not_available | NA | NA | NA | NA | NA |
| lizards | Dipsosaurus_dorsalis | Dipsosaurus_dorsalis | LC | Unknown | temperate | diurnal | 158.489 | 456965.137 | 6.982 |
| lizards | Dixonius_aaronbaueri | NA | LC | Stable | NA | NA | NA | NA | NA |
| lizards | Dixonius_hangseesom | Dixonius_hangseesom | LC | Stable | tropical | nocturnal | 1.738 | 6426.516 | 13.237 |
| lizards | Dixonius_kaweesaki | NA | CR | Decreasing | NA | NA | NA | NA | NA |
| lizards | Dixonius_melanostictus | NA | LC | Stable | NA | NA | NA | NA | NA |
| lizards | Dixonius_minhlei | NA | LC | Stable | NA | NA | NA | NA | NA |
| lizards | Dixonius_siamensis | Dixonius_siamensis | LC | Stable | tropical | nocturnal | 4.074 | 788329.94 | 14.866 |
| lizards | Dixonius_taoi | NA | VU | Stable | NA | NA | NA | NA | NA |
| lizards | Dixonius_vietnamensis | Dixonius_vietnamensis | LC | Unknown | tropical | nocturnal | 2.818 | 103532.319 | 10.465 |
| lizards | Dopasia_buettikoferi | NA | LC | Unknown | NA | NA | NA | NA | NA |
| lizards | Dopasia_gracilis | Dopasia_gracilis | LC | Decreasing | tropical | cathemeral | 134.896 | 1725322.864 | 11.638 |
| lizards | Dopasia_hainanensis | Dopasia_hainanensis | LC | Stable | tropical | diurnal | 75.858 | 5958.469 | 13.108 |
| lizards | Dopasia_harti | Dopasia_harti | LC | Unknown | temperate | diurnal | 75.858 | 1636253.615 | 14.886 |
| lizards | Dopasia_ludovici | NA | LC | Stable | NA | NA | NA | NA | NA |
| lizards | Dopasia_sokolovi | Dopasia_sokolovi | LC | Unknown | tropical | diurnal | 18.621 | 8994.149 | 12.338 |
| lizards | Dopasia_wegneri | NA | LC | Unknown | NA | NA | NA | NA | NA |
| lizards | Dracaena_guianensis | Dracaena_guianensis | LC | Unknown | tropical | diurnal | 2454.709 | 3151888.227 | 1.44 |
| lizards | Dracaena_paraguayensis | NA | LC | Stable | NA | NA | NA | NA | NA |

| | | | | | | | | | |
|---|---|---|---|---|---|---|---|---|---|
| lizards | Draco_abbreviatus | NA | not_available | not_available | NA | NA | NA | NA | NA |
| lizards | Draco_beccarii | Draco_beccarii | LC | Stable | tropical | diurnal | 5.37 | 31194.774 | 7.371 |
| lizards | Draco_biaro | Draco_biaro | LC | Unknown | tropical | diurnal | 4.571 | 26.306 | 7.739 |
| lizards | Draco_bimaculatus | Draco_bimaculatus | LC | Stable | tropical | diurnal | 4.074 | 62043.041 | 10.616 |
| lizards | Draco_blanfordii | Draco_blanfordii | LC | Stable | tropical | diurnal | 20.893 | 1393371.651 | 12.213 |
| lizards | Draco_caerulhians | NA | LC | Unknown | NA | NA | NA | NA | NA |
| lizards | Draco_cornutus | Draco_cornutus | LC | Unknown | tropical | diurnal | 7.586 | 734529.435 | 5.566 |
| lizards | Draco_cristatellus | NA | DD | Unknown | NA | NA | NA | NA | NA |
| lizards | Draco_cyanopterus | Draco_cyanopterus | LC | Stable | tropical | diurnal | 8.318 | 85923.128 | 12.049 |
| lizards | Draco_dussumieri | Draco_dussumieri | LC | Stable | tropical | diurnal | 9.333 | 178190.895 | 20.159 |
| lizards | Draco_fimbriatus | Draco_fimbriatus | LC | Stable | tropical | diurnal | 19.498 | 378214.493 | 8.853 |
| lizards | Draco_formosus | Draco_formosus | LC | Stable | tropical | diurnal | 12.589 | 99721.479 | 9.98 |
| lizards | Draco_guentheri | NA | LC | Stable | NA | NA | NA | NA | NA |
| lizards | Draco_haematopogon | Draco_haematopogon | LC | Stable | tropical | diurnal | 10.471 | 1810765.357 | 10.87 |
| lizards | Draco_indochinensis | Draco_indochinensis | LC | Unknown | tropical | diurnal | 14.125 | 56391.81 | 12.522 |
| lizards | Draco_iskandari | NA | LC | Unknown | NA | NA | NA | NA | NA |
| lizards | Draco_jareckii | NA | LC | Stable | NA | NA | NA | NA | NA |
| lizards | Draco_lineatus | Draco_lineatus | LC | Stable | tropical | diurnal | 7.413 | 16010.164 | 7.825 |
| lizards | Draco_maculatus | Draco_maculatus | LC | Decreasing | tropical | diurnal | 8.511 | 2591726.251 | 12.372 |
| lizards | Draco_maximus | Draco_maximus | LC | Unknown | tropical | diurnal | 28.84 | 427565.289 | 6.819 |
| lizards | Draco_melanopogon | Draco_melanopogon | LC | Stable | tropical | diurnal | 7.762 | 473941.954 | 8.135 |
| lizards | Draco_mindanensis | Draco_mindanensis | NT | Unknown | tropical | diurnal | 12.023 | 112729.234 | 11.824 |
| lizards | Draco_modiglianii | Draco_modiglianii | LC | Unknown | tropical | diurnal | 4.266 | 400.056 | 8.693 |
| lizards | Draco_norvillii | NA | NT | Unknown | NA | NA | NA | NA | NA |
| lizards | Draco_obscurus | Draco_obscurus | LC | Unknown | tropical | diurnal | 13.183 | 248262.76 | 8.66 |
| lizards | Draco_ornatus | NA | LC | Stable | NA | NA | NA | NA | NA |
| lizards | Draco_palawanensis | Draco_palawanensis | LC | Stable | tropical | diurnal | 6.166 | 11489.367 | 9.024 |
| lizards | Draco_quadrasi | NA | LC | Unknown | NA | NA | NA | NA | NA |
| lizards | Draco_quinquefasciatus | Draco_quinquefasciatus | LC | Stable | tropical | diurnal | 12.023 | 725920.546 | 7.157 |
| lizards | Draco_reticulatus | Draco_reticulatus | LC | Unknown | tropical | diurnal | 19.498 | 23324.445 | 12.375 |
| lizards | Draco_spilonotus | Draco_spilonotus | LC | Stable | tropical | diurnal | 5.248 | 25739.593 | 8.524 |
| lizards | Draco_spilopterus | Draco_spilopterus | LC | Unknown | tropical | diurnal | 10.233 | 142930.732 | 15.601 |
| lizards | Draco_sumatranus | Draco_sumatranus | LC | Unknown | tropical | diurnal | 7.413 | 1021895.798 | 8.151 |
| lizards | Draco_supriatnai | NA | LC | Stable | NA | NA | NA | NA | NA |

| | | | | | | | | | |
|---|---|---|---|---|---|---|---|---|---|
| lizards | Draco_taeniopterus | Draco_taeniopterus | LC | Unknown | tropical | diurnal | 7.079 | 402687.722 | 14.903 |
| lizards | Draco_timoriensis | Draco_timorensis | LC | Unknown | tropical | diurnal | 7.762 | 35281.877 | 13.048 |
| lizards | Draco_volans | Draco_volans | LC | Unknown | tropical | diurnal | 8.511 | 115530.677 | 22.437 |
| lizards | Draco_walkeri | NA | LC | Stable | NA | NA | NA | NA | NA |
| lizards | Dravidogecko_anamallensis | Hemidactylus_anamallensis | NT | Unknown | tropical | nocturnal | 8.913 | 11511.337 | 12.993 |
| lizards | Dravidoseps_goaensis | NA | DD | Unknown | NA | NA | NA | NA | NA |
| lizards | Dravidoseps_pruthi | NA | DD | Unknown | NA | NA | NA | NA | NA |
| lizards | Dryadosaura_nordestina | Dryadosaura_nordestina | LC | Unknown | tropical | diurnal | 4.169 | 108269.618 | 14.773 |
| lizards | Ebenavia_inunguis | Ebenavia_inunguis | LC | Unknown | tropical | cathemeral | 1.23 | 2096.976 | 7.591 |
| lizards | Ebenavia_maintimainty | Ebenavia_maintimainty | EN | Decreasing | temperate | nocturnal | 0.372 | 2613.741 | 7.926 |
| lizards | Echinosaura_brachycephala | NA | EN | Unknown | NA | NA | NA | NA | NA |
| lizards | Echinosaura_horrida | Echinosaura_horrida | LC | Unknown | tropical | cathemeral | 16.218 | 178484.412 | 8.635 |
| lizards | Echinosaura_keyi | NA | VU | Decreasing | NA | NA | NA | NA | NA |
| lizards | Egernia_cunninghami | Egernia_cunninghami | LC | Unknown | temperate | diurnal | 446.684 | 439015.887 | 9.984 |
| lizards | Egernia_cygnitos | NA | LC | Stable | NA | NA | NA | NA | NA |
| lizards | Egernia_depressa | Egernia_depressa | LC | Stable | temperate | diurnal | 35.481 | 450583.582 | 4.676 |
| lizards | Egernia_douglasi | NA | DD | Unknown | NA | NA | NA | NA | NA |
| lizards | Egernia_eos | NA | LC | Stable | NA | NA | NA | NA | NA |
| lizards | Egernia_epsisolus | NA | LC | Stable | NA | NA | NA | NA | NA |
| lizards | Egernia_formosa | Egernia_formosa | LC | Stable | temperate | diurnal | 26.915 | 316321.235 | 4.049 |
| lizards | Egernia_hosmeri | Egernia_hosmeri | LC | Unknown | tropical | diurnal | 144.544 | 215411.999 | 3.847 |
| lizards | Egernia_kingii | Egernia_kingii | LC | Stable | temperate | diurnal | 416.869 | 96409.899 | 10.949 |
| lizards | Egernia_mcpheei | Egernia_mcpheei | LC | Unknown | temperate | diurnal | 67.608 | 74857.14 | 7.616 |
| lizards | Egernia_napoleonis | Egernia_napoleonis | LC | Stable | temperate | diurnal | 53.703 | 206538.1 | 9.355 |
| lizards | Egernia_pilbarensis | NA | LC | Stable | NA | NA | NA | NA | NA |
| lizards | Egernia_richardi | NA | LC | Stable | NA | NA | NA | NA | NA |
| lizards | Egernia_roomi | NA | not_available | not_available | NA | NA | NA | NA | NA |
| lizards | Egernia_rugosa | Egernia_rugosa | LC | Decreasing | tropical | diurnal | 323.594 | 789989.539 | 5.101 |
| lizards | Egernia_saxatilis | Egernia_saxatilis | LC | Unknown | temperate | diurnal | 56.234 | 250861.566 | 10.209 |
| lizards | Egernia_stokesii | Egernia_stokesii | LC | Decreasing | temperate | diurnal | 223.872 | 1248794.26 | 4.752 |
| lizards | Egernia_striolata | Egernia_striolata | LC | Stable | temperate | diurnal | 38.019 | 1971703.667 | 7.086 |
| lizards | Elasmodactylus_tetensis | Elasmodactylus_tetensis | LC | Stable | tropical | nocturnal | 19.055 | 86757.046 | 11.219 |
| lizards | Elasmodactylus_tuberculosus | Elasmodactylus_tuberculosus | LC | Unknown | tropical | nocturnal | 10 | 862531.254 | 9.592 |
| lizards | Elgaria_cedrosensis | NA | LC | Stable | NA | NA | NA | NA | NA |

| | | | | | | | | | |
|---|---|---|---|---|---|---|---|---|---|
| lizards | Elgaria_coerulea | Elgaria_coerulea | LC | Stable | temperate | diurnal | 45.709 | 551499.256 | 10.532 |
| lizards | Elgaria_kingii | Elgaria_kingii | LC | Stable | temperate | diurnal | 50.119 | 305414.675 | 6.742 |
| lizards | Elgaria_multicarinata | Elgaria_multicarinata | LC | Decreasing | temperate | cathemeral | 117.49 | 217760.633 | 14.163 |
| lizards | Elgaria_panamintina | Elgaria_panamintina | VU | Decreasing | temperate | cathemeral | 67.608 | 8268.708 | 4.722 |
| lizards | Elgaria_paucicarinata | Elgaria_paucicarinata | LC | Stable | temperate | diurnal | 29.512 | 9952.259 | 7.468 |
| lizards | Elgaria_velazquezi | NA | LC | Stable | NA | NA | NA | NA | NA |
| lizards | Emoia_adspersa | Emoia_adspersa | EN | Decreasing | tropical | diurnal | 16.982 | 3024.587 | 1.07 |
| lizards | Emoia_aenea | Emoia_aenea | LC | Unknown | tropical | diurnal | 7.413 | 48412.518 | 2.78 |
| lizards | Emoia_arnoensis | Emoia_arnoensis | LC | Unknown | tropical | diurnal | 22.387 | 1195.757 | 12.19 |
| lizards | Emoia_atrocostata | Emoia_atrocostata | LC | Stable | tropical | diurnal | 21.38 | 2079573.146 | 7.886 |
| lizards | Emoia_aurulenta | Emoia_aurulenta | LC | Unknown | tropical | diurnal | 2.239 | 69136.425 | 2.012 |
| lizards | Emoia_baudini | Emoia_baudini | LC | Unknown | tropical | diurnal | 6.166 | 3102.972 | 6.15 |
| lizards | Emoia_beryllion | NA | not_available | not_available | NA | NA | NA | NA | NA |
| lizards | Emoia_bismarckensis | Emoia_bismarckensis | LC | Stable | tropical | diurnal | 7.413 | 19861.706 | 4.507 |
| lizards | Emoia_boettgeri | Emoia_boettgeri | EN | Decreasing | tropical | diurnal | 9.333 | 734.386 | 12.041 |
| lizards | Emoia_bogerti | Emoia_bogerti | LC | Unknown | tropical | diurnal | 3.89 | 44810.709 | 3.809 |
| lizards | Emoia_brongersmai | Emoia_brongersmai | LC | Unknown | tropical | diurnal | 6.761 | 3036.328 | 3.705 |
| lizards | Emoia_caeruleocauda | Emoia_caeruleocauda | LC | Stable | tropical | diurnal | 5.623 | 1089921.566 | 4.786 |
| lizards | Emoia_campbelli | Emoia_campbelli | EN | Decreasing | tropical | diurnal | 19.953 | 9.991 | 3.917 |
| lizards | Emoia_coggeri | NA | DD | Unknown | NA | NA | NA | NA | NA |
| lizards | Emoia_concolor | Emoia_concolor | NT | Unknown | tropical | diurnal | 18.197 | 9087.882 | 6.958 |
| lizards | Emoia_cyanogaster | Emoia_cyanogaster | LC | Stable | tropical | diurnal | 20.893 | 43311.808 | 4.666 |
| lizards | Emoia_cyanura | Emoia_cyanura | LC | Decreasing | tropical | diurnal | 5.37 | 93259.442 | 6.746 |
| lizards | Emoia_erronan | NA | VU | Stable | NA | NA | NA | NA | NA |
| lizards | Emoia_flavigularis | NA | LC | Stable | NA | NA | NA | NA | NA |
| lizards | Emoia_guttata | Emoia_guttata | LC | Unknown | tropical | diurnal | 7.762 | 405.658 | 9.717 |
| lizards | Emoia_impar | Emoia_impar | LC | Stable | tropical | diurnal | 2.291 | 53986.919 | 7.341 |
| lizards | Emoia_isolata | Emoia_isolata | LC | Stable | tropical | diurnal | 4.169 | 20.606 | 8.907 |
| lizards | Emoia_jakati | Emoia_jakati | LC | Stable | tropical | diurnal | 2.818 | 479940.329 | 3.858 |
| lizards | Emoia_jamur | NA | DD | Unknown | NA | NA | NA | NA | NA |
| lizards | Emoia_kordoana | Emoia_kordoana | LC | Unknown | tropical | diurnal | 4.365 | 398786.022 | 3.514 |
| lizards | Emoia_laobaoensis | NA | DD | Unknown | NA | NA | NA | NA | NA |
| lizards | Emoia_lawesii | NA | EN | Unknown | NA | NA | NA | NA | NA |
| lizards | Emoia_longicauda | Emoia_longicauda | LC | Stable | tropical | diurnal | 21.38 | 612690.268 | 3.636 |

| | | | | | | | | | |
|---|---|---|---|---|---|---|---|---|---|
| lizards | Emoia_loyaltiensis | Emoia_loyaltiensis | VU | Decreasing | tropical | diurnal | 12.023 | 1845.675 | 7.884 |
| lizards | Emoia_maculata | Emoia_maculata | LC | Stable | tropical | diurnal | 6.761 | 3100.328 | 4.476 |
| lizards | Emoia_mivarti | Emoia_mivarti | DD | Stable | tropical | diurnal | 3.388 | 1893.335 | 6.405 |
| lizards | Emoia_mokolahi | NA | not_available | not_available | NA | NA | NA | NA | NA |
| lizards | Emoia_mokosariniveikau | NA | EN | Decreasing | NA | NA | NA | NA | NA |
| lizards | Emoia_nativitatis | NA | EX | not_available | NA | NA | NA | NA | NA |
| lizards | Emoia_nigra | Emoia_nigra | LC | Stable | tropical | diurnal | 47.863 | 64085.694 | 6.071 |
| lizards | Emoia_nigromarginata | Emoia_nigromarginata | LC | Stable | tropical | diurnal | 10.471 | 8045.272 | 5.921 |
| lizards | Emoia_obscura | Emoia_obscura | LC | Unknown | tropical | diurnal | 5.012 | 272171.725 | 3.736 |
| lizards | Emoia_oriva | NA | not_available | not_available | NA | NA | NA | NA | NA |
| lizards | Emoia_pallidiceps | Emoia_pallidiceps | LC | Unknown | tropical | diurnal | 4.467 | 331355.804 | 4.134 |
| lizards | Emoia_parkeri | Emoia_parkeri | VU | Decreasing | tropical | diurnal | 2.884 | 11496.216 | 9.177 |
| lizards | Emoia_physicae | Emoia_physicae | LC | Unknown | tropical | diurnal | 9.55 | 717251.465 | 3.336 |
| lizards | Emoia_physicina | Emoia_physicina | LC | Unknown | tropical | diurnal | 2.399 | 109472.059 | 2.571 |
| lizards | Emoia_ponapea | Emoia_ponapea | EN | Decreasing | tropical | diurnal | 2.399 | 342.824 | 12.238 |
| lizards | Emoia_popei | Emoia_popei | LC | Unknown | tropical | diurnal | 5.37 | 38743.34 | 5.392 |
| lizards | Emoia_pseudocyanura | Emoia_pseudocyanura | LC | Stable | tropical | diurnal | 6.761 | 21644.989 | 5.331 |
| lizards | Emoia_reimschiisseli | Emoia_reimschiisseli | LC | Stable | tropical | diurnal | 9.55 | 192719.705 | 8.898 |
| lizards | Emoia_rennellensis | Emoia_rennellensis | LC | Stable | tropical | diurnal | 1.622 | 675.7 | 3.166 |
| lizards | Emoia_ruficauda | Emoia_ruficauda | LC | Unknown | tropical | diurnal | 2.951 | 9745.734 | 12.807 |
| lizards | Emoia_rufilabialis | NA | LC | Stable | NA | NA | NA | NA | NA |
| lizards | Emoia_samoensis | NA | EN | Decreasing | NA | NA | NA | NA | NA |
| lizards | Emoia_sanfordi | Emoia_sanfordi | LC | Stable | tropical | diurnal | 34.674 | 9879.166 | 5.592 |
| lizards | Emoia_schmidti | Emoia_schmidti | LC | Stable | tropical | diurnal | 4.898 | 2051.969 | 2.034 |
| lizards | Emoia_slevini | Emoia_slevini | CR | Decreasing | tropical | diurnal | 12.882 | 956.861 | 20.338 |
| lizards | Emoia_submetallica | Emoia_submetallica | LC | Unknown | tropical | diurnal | 5.248 | 14091.82 | 5.151 |
| lizards | Emoia_taumakoensis | NA | LC | Unknown | NA | NA | NA | NA | NA |
| lizards | Emoia_tetrataenia | Emoia_tetrataenia | LC | Unknown | tropical | diurnal | 5.012 | 2502.49 | 4.619 |
| lizards | Emoia_tongana | NA | LC | Unknown | NA | NA | NA | NA | NA |
| lizards | Emoia_tropidolepis | Emoia_tropidolepis | LC | Unknown | tropical | diurnal | 7.943 | 94680.786 | 2.382 |
| lizards | Emoia_trossula | Emoia_trossula | EN | Decreasing | tropical | diurnal | 28.184 | 13927.558 | 9.19 |
| lizards | Emoia_tuitarere | NA | VU | Unknown | NA | NA | NA | NA | NA |
| lizards | Emoia_veracunda | Emoia_veracunda | LC | Unknown | tropical | diurnal | 2.692 | 138722.05 | 4.576 |
| lizards | Enyalioides_altotambo | NA | EN | Unknown | NA | NA | NA | NA | NA |

| | | | | | | | | | |
|---|---|---|---|---|---|---|---|---|---|
| lizards | Enyalioides_anisolepis | NA | not_available | not_available | NA | NA | NA | NA | NA |
| lizards | Enyalioides_annularis | NA | VU | Decreasing | NA | NA | NA | NA | NA |
| lizards | Enyalioides_azulae | NA | LC | Stable | NA | NA | NA | NA | NA |
| lizards | Enyalioides_cofanorum | Enyalioides_cofanorum | LC | Unknown | tropical | diurnal | 85.114 | 358695.668 | 1.666 |
| lizards | Enyalioides_groi | NA | EN | Unknown | NA | NA | NA | NA | NA |
| lizards | Enyalioides_heterolepis | Enyalioides_heterolepis | LC | Stable | tropical | diurnal | 81.283 | 165895.816 | 7.106 |
| lizards | Enyalioides_laticeps | Enyalioides_laticeps | LC | Stable | tropical | diurnal | 123.027 | 1134082.42 | 2.307 |
| lizards | Enyalioides_microlepis | NA | LC | Unknown | NA | NA | NA | NA | NA |
| lizards | Enyalioides_oshaughnessyi | NA | VU | Unknown | NA | NA | NA | NA | NA |
| lizards | Enyalioides_palpebralis | Enyalioides_palpebralis | LC | Unknown | tropical | diurnal | 53.703 | 243180.164 | 1.475 |
| lizards | Enyalioides_peruvianus | NA | LC | Unknown | NA | NA | NA | NA | NA |
| lizards | Enyalioides_praestabilis | Enyalioides_praestabilis | LC | Unknown | tropical | diurnal | 63.096 | 137697.657 | 3.95 |
| lizards | Enyalioides_rubrigularis | Enyalioides_rubrigularis | VU | Decreasing | tropical | diurnal | 66.069 | 1002.191 | 8.885 |
| lizards | Enyalioides_rudolfarndti | NA | LC | Stable | NA | NA | NA | NA | NA |
| lizards | Enyalioides_sophiarothschildae | NA | not_available | not_available | NA | NA | NA | NA | NA |
| lizards | Enyalioides_touzeti | NA | VU | Unknown | NA | NA | NA | NA | NA |
| lizards | Enyalius_bibronii | Enyalius_bibronii | LC | Unknown | tropical | diurnal | 56.234 | 982903.851 | 9.745 |
| lizards | Enyalius_bilineatus | Enyalius_bilineatus | LC | Unknown | tropical | diurnal | 35.481 | 548759.194 | 11.324 |
| lizards | Enyalius_brasiliensis | Enyalius_brasiliensis | LC | Unknown | tropical | cathemeral | 51.286 | 44732.728 | 14.646 |
| lizards | Enyalius_catenatus | Enyalius_catenatus | LC | Unknown | tropical | diurnal | 61.66 | 133671.261 | 10.94 |
| lizards | Enyalius_erythroceneus | NA | CR | Unknown | NA | NA | NA | NA | NA |
| lizards | Enyalius_iheringii | Enyalius_iheringii | LC | Unknown | temperate | diurnal | 63.096 | 221445.831 | 13.893 |
| lizards | Enyalius_leechii | Enyalius_leechii | LC | Unknown | tropical | diurnal | 48.978 | 1438882.263 | 2.796 |
| lizards | Enyalius_perditus | Enyalius_perditus | LC | Unknown | tropical | diurnal | 30.2 | 450100.05 | 14.629 |
| lizards | Epibator_greeri | Lioscincus_greeri | EN | Decreasing | tropical | diurnal | 4.365 | 77.052 | 9.704 |
| lizards | Epibator_insularis | NA | not_available | not_available | NA | NA | NA | NA | NA |
| lizards | Epibator_nigrofasciolatus | Lioscincus_nigrofasciolatum | LC | Stable | tropical | diurnal | 30.903 | 18806.715 | 7.571 |
| lizards | Eremias_acutirostris | Eremias_acutirostris | LC | Stable | temperate | diurnal | 11.749 | 213043.527 | 5.493 |
| lizards | Eremias_afghanistanica | NA | LC | Unknown | NA | NA | NA | NA | NA |
| lizards | Eremias_andersoni | NA | LC | Stable | NA | NA | NA | NA | NA |
| lizards | Eremias_argus | Eremias_argus | LC | Stable | temperate | diurnal | 7.244 | 4271177.412 | 11.949 |
| lizards | Eremias_arguta | Eremias_arguta | LC | Stable | temperate | diurnal | 22.909 | 4578499.404 | 9.819 |
| lizards | Eremias_aria | NA | LC | Unknown | NA | NA | NA | NA | NA |
| lizards | Eremias_brenchleyi | Eremias_brenchleyi | LC | Stable | temperate | diurnal | 7.586 | 600869.288 | 21.706 |

| | | | | | | | | | |
|---|---|---|---|---|---|---|---|---|---|
| lizards | Eremias_buechneri | Eremias_buechneri | LC | Stable | temperate | diurnal | 6.761 | 14695.402 | 4.097 |
| lizards | Eremias_cholistanica | NA | DD | Unknown | NA | NA | NA | NA | NA |
| lizards | Eremias_dzungarica | NA | not_available | not_available | NA | NA | NA | NA | NA |
| lizards | Eremias_fasciata | NA | LC | Stable | NA | NA | NA | NA | NA |
| lizards | Eremias_grammica | Eremias_grammica | LC | Stable | temperate | diurnal | 22.909 | 1823617.275 | 6.685 |
| lizards | Eremias_intermedia | Eremias_intermedia | LC | Stable | temperate | diurnal | 7.586 | 2024194.595 | 6.52 |
| lizards | Eremias_isfahanica | NA | not_available | not_available | NA | NA | NA | NA | NA |
| lizards | Eremias_kavirensis | NA | LC | Stable | NA | NA | NA | NA | NA |
| lizards | Eremias_kokshaaliensis | NA | DD | Unknown | NA | NA | NA | NA | NA |
| lizards | Eremias_kopetdaghica | NA | not_available | not_available | NA | NA | NA | NA | NA |
| lizards | Eremias_lalezharica | NA | LC | Stable | NA | NA | NA | NA | NA |
| lizards | Eremias_lineolata | Eremias_lineolata | LC | Decreasing | temperate | diurnal | 3.981 | 1173018.608 | 7.463 |
| lizards | Eremias_montana | NA | LC | Stable | NA | NA | NA | NA | NA |
| lizards | Eremias_multiocellata | Eremias_multiocellata | LC | Stable | temperate | diurnal | 10.965 | 4762088.328 | 4.985 |
| lizards | Eremias_nigrocellata | Eremias_nigrocellata | LC | Decreasing | temperate | diurnal | 13.183 | 322777.745 | 9.619 |
| lizards | Eremias_nikolskii | Eremias_nikolskii | LC | Stable | temperate | diurnal | 9.772 | 115485.938 | 12.152 |
| lizards | Eremias_papenfussi | NA | not_available | not_available | NA | NA | NA | NA | NA |
| lizards | Eremias_persica | Eremias_persica | LC | Stable | temperate | diurnal | 21.38 | 1686829.414 | 8.472 |
| lizards | Eremias_pleskei | Eremias_pleskei | CR | Decreasing | temperate | diurnal | 5.012 | 47164.584 | 15.214 |
| lizards | Eremias_przewalskii | Eremias_przewalskii | LC | Stable | temperate | diurnal | 21.38 | 2448607.516 | 3.022 |
| lizards | Eremias_quadrifrons | NA | DD | Unknown | NA | NA | NA | NA | NA |
| lizards | Eremias_regeli | Eremias_regeli | LC | Stable | temperate | diurnal | 7.943 | 375839.707 | 11.12 |
| lizards | Eremias_scripta | Eremias_scripta | LC | Stable | temperate | diurnal | 6.761 | 1835545.751 | 7.286 |
| lizards | Eremias_strauchi | Eremias_strauchi | LC | Decreasing | temperate | diurnal | 11.749 | 309010.54 | 13.242 |
| lizards | Eremias_stummeri | Eremias_stummeri | LC | Stable | temperate | diurnal | 7.079 | 17321.549 | 8.402 |
| lizards | Eremias_suphani | Eremias_suphani | LC | Stable | temperate | diurnal | 8.318 | 10976.032 | 12.948 |
| lizards | Eremias_szczerbaki | NA | DD | Unknown | NA | NA | NA | NA | NA |
| lizards | Eremias_velox | Eremias_velox | LC | Stable | temperate | diurnal | 19.055 | 4796912.231 | 7.5 |
| lizards | Eremias_vermiculata | Eremias_vermiculata | LC | Stable | temperate | diurnal | 8.511 | 2114242.455 | 3.309 |
| lizards | Eremias_yarkandensis | Eremias_yarkandensis | LC | Unknown | temperate | diurnal | 5.248 | 173393.286 | 4.434 |
| lizards | Eremiascincus_brongersmai | NA | LC | Stable | NA | NA | NA | NA | NA |
| lizards | Eremiascincus_douglasi | Eremiascincus_douglasi | LC | Stable | tropical | cathemeral | 11.22 | 203711.029 | 1.712 |
| lizards | Eremiascincus_fasciolatus | Eremiascincus_fasciolatus | LC | Stable | tropical | nocturnal | 42.658 | 435748.856 | 5.902 |
| lizards | Eremiascincus_intermedius | Eremiascincus_intermedius | LC | Stable | tropical | nocturnal | 14.454 | 778700.396 | 1.385 |

| | | | | | | | | | |
|---|---|---|---|---|---|---|---|---|---|
| lizards | Eremiascincus_isolepis | Eremiascincus_isolepis | LC | Stable | tropical | cathemeral | 8.511 | 1417339.731 | 2.637 |
| lizards | Eremiascincus_musivus | Eremiascincus_musivus | LC | Stable | tropical | nocturnal | 3.981 | 155315.136 | 2.714 |
| lizards | Eremiascincus_pallidus | Eremiascincus_pallidus | LC | Stable | temperate | nocturnal | 9.333 | 1858227.786 | 2.303 |
| lizards | Eremiascincus_pardalis | Eremiascincus_pardalis | LC | Stable | tropical | cathemeral | 8.511 | 226382.767 | 2.917 |
| lizards | Eremiascincus_phantasmus | Eremiascincus_phantasmus | LC | Stable | temperate | nocturnal | 18.197 | 950164.669 | 3.115 |
| lizards | Eremiascincus_richardsonii | Eremiascincus_richardsonii | LC | Stable | temperate | nocturnal | 46.774 | 5310086.508 | 3.663 |
| lizards | Eremiascincus_rubiginosus | NA | not_available | not_available | NA | NA | NA | NA | NA |
| lizards | Eroticoscincus_graciloides | Eroticoscincus_graciloides | LC | Stable | temperate | cathemeral | 0.724 | 26757.303 | 7.988 |
| lizards | Eublepharis_angramainyu | Eublepharis_angramainyu | DD | Decreasing | temperate | nocturnal | 125.893 | 99296.006 | 15.394 |
| lizards | Eublepharis_fuscus | Eublepharis_fuscus | LC | Unknown | tropical | nocturnal | 416.869 | 76785.136 | 19.665 |
| lizards | Eublepharis_hardwickii | Eublepharis_hardwickii | LC | Unknown | tropical | nocturnal | 29.512 | 284089.622 | 15.398 |
| lizards | Eublepharis_macularius | Eublepharis_macularius | LC | Stable | temperate | nocturnal | 114.815 | 1631437.081 | 13.803 |
| lizards | Eublepharis_satpuraensis | NA | LC | Stable | NA | NA | NA | NA | NA |
| lizards | Eublepharis_turcmenicus | Eublepharis_turcmenicus | LC | Unknown | temperate | nocturnal | 64.565 | 70844.552 | 11.342 |
| lizards | Eugongylus_albofasciolatus | Eugongylus_albofasciolatus | LC | Stable | tropical | cathemeral | 245.471 | 88884.305 | 4.638 |
| lizards | Eugongylus_rufescens | Eugongylus_rufescens | LC | Stable | tropical | cathemeral | 117.49 | 830810.752 | 3.828 |
| lizards | Eulamprus_heatwolei | Eulamprus_heatwolei | LC | Unknown | temperate | diurnal | 22.387 | 283848.251 | 9.135 |
| lizards | Eulamprus_kosciuskoi | Eulamprus_kosciuskoi | LC | Unknown | temperate | diurnal | 12.882 | 34679.698 | 5.554 |
| lizards | Eulamprus_leuraensis | Eulamprus_leuraensis | EN | Decreasing | temperate | diurnal | 12.882 | 1217.176 | 10.652 |
| lizards | Eulamprus_quoyii | Eulamprus_quoyii | LC | Stable | temperate | diurnal | 46.774 | 1004641.982 | 7.693 |
| lizards | Eulamprus_tympanum | Eulamprus_tympanum | LC | Stable | temperate | diurnal | 21.38 | 243512.557 | 10.506 |
| lizards | Euleptes_europaea | Euleptes_europaea | NT | Unknown | temperate | nocturnal | 2.089 | 33808.587 | 16.778 |
| lizards | Eumeces_algeriensis | Eumeces_algeriensis | LC | Stable | temperate | diurnal | 234.423 | 294432.773 | 13.275 |
| lizards | Eumeces_blythianus | Eumeces_blythianus | LC | Unknown | temperate | diurnal | 30.2 | 1068638.79 | 8.35 |
| lizards | Eumeces_cholistanensis | NA | DD | Unknown | NA | NA | NA | NA | NA |
| lizards | Eumeces_indothalensis | NA | DD | Unknown | NA | NA | NA | NA | NA |
| lizards | Eumeces_persicus | NA | not_available | not_available | NA | NA | NA | NA | NA |
| lizards | Eumeces_schneiderii | Eumeces_schneideri | LC | Stable | temperate | diurnal | 158.489 | 2484338.484 | 10.959 |
| lizards | Eumecia_anchietae | Eumecia_anchietae | LC | Unknown | tropical | diurnal | 81.283 | 1405431.328 | 6.881 |
| lizards | Eurolophosaurus_amathites | Eurolophosaurus_amathites | EN | Unknown | tropical | diurnal | 12.589 | 1499.315 | 3.695 |
| lizards | Eurolophosaurus_divaricatus | NA | LC | Unknown | NA | NA | NA | NA | NA |
| lizards | Eurolophosaurus_nanuzae | Eurolophosaurus_nanuzae | LC | Unknown | tropical | diurnal | 9.772 | 49868.494 | 6.282 |
| lizards | Eurydactylodes_agricolae | Eurydactylodes_agricolae | NT | Stable | tropical | diurnal | 4.266 | 3524.975 | 6.779 |
| lizards | Eurydactylodes_occidentalis | Eurydactylodes_occidentalis | EN | Decreasing | tropical | diurnal | 3.02 | 279.85 | 11.793 |

| | | | | | | | | | |
|---|---|---|---|---|---|---|---|---|---|
| lizards | Eurydactylodes_symmetricus | Eurydactylodes_symmetricus | NT | Decreasing | tropical | cathemeral | 4.365 | 3621.015 | 6.026 |
| lizards | Eurydactylodes_vieillardi | Eurydactylodes_vieillardi | NT | Decreasing | tropical | diurnal | 4.365 | 12846.358 | 7.657 |
| lizards | Eurylepis_poonaensis | NA | EN | Unknown | NA | NA | NA | NA | NA |
| lizards | Eurylepis_taeniolata | Eurylepis_taeniolata | LC | Stable | temperate | cathemeral | 138.038 | 1826116.584 | 13.091 |
| lizards | Euspondylus_acutirostris | Euspondylus_acutirostris | LC | Unknown | tropical | diurnal | 3.89 | 5017.043 | 16.726 |
| lizards | Euspondylus_auyanensis | NA | LC | Stable | NA | NA | NA | NA | NA |
| lizards | Euspondylus_excelsum | NA | not_available | not_available | NA | NA | NA | NA | NA |
| lizards | Euspondylus_paxcorpus | NA | not_available | not_available | NA | NA | NA | NA | NA |
| lizards | Eutropis_allapallensis | NA | LC | Stable | NA | NA | NA | NA | NA |
| lizards | Eutropis_andamanensis | NA | LC | Stable | NA | NA | NA | NA | NA |
| lizards | Eutropis_ashwamedhi | NA | EN | Unknown | NA | NA | NA | NA | NA |
| lizards | Eutropis_austini | NA | DD | Unknown | NA | NA | NA | NA | NA |
| lizards | Eutropis_beddomei | Eutropis_beddomii | LC | Unknown | tropical | diurnal | 4.169 | 269543.691 | 20.51 |
| lizards | Eutropis_bibronii | NA | NT | Unknown | NA | NA | NA | NA | NA |
| lizards | Eutropis_bontocensis | NA | LC | Stable | NA | NA | NA | NA | NA |
| lizards | Eutropis_borealis | NA | LC | Stable | NA | NA | NA | NA | NA |
| lizards | Eutropis_caraga | NA | LC | Stable | NA | NA | NA | NA | NA |
| lizards | Eutropis_carinata | Eutropis_carinata | LC | Unknown | tropical | diurnal | 97.724 | 3483962.346 | 17.033 |
| lizards | Eutropis_chapaensis | NA | LC | Unknown | NA | NA | NA | NA | NA |
| lizards | Eutropis_clivicola | Eutropis_clivicola | EN | Unknown | tropical | diurnal | 4.898 | 5640.202 | 15.152 |
| lizards | Eutropis_cumingi | Eutropis_cumingi | LC | Stable | tropical | diurnal | 2.951 | 89632.003 | 14.917 |
| lizards | Eutropis_cuprea | NA | LC | Unknown | NA | NA | NA | NA | NA |
| lizards | Eutropis_darevskii | NA | DD | Unknown | NA | NA | NA | NA | NA |
| lizards | Eutropis_dattaroyi | NA | not_available | not_available | NA | NA | NA | NA | NA |
| lizards | Eutropis_dawsoni | NA | DD | Unknown | NA | NA | NA | NA | NA |
| lizards | Eutropis_englei | NA | LC | Unknown | NA | NA | NA | NA | NA |
| lizards | Eutropis_floweri | NA | DD | Unknown | NA | NA | NA | NA | NA |
| lizards | Eutropis_greeri | NA | DD | Unknown | NA | NA | NA | NA | NA |
| lizards | Eutropis_gubataas | NA | LC | Unknown | NA | NA | NA | NA | NA |
| lizards | Eutropis_indeprensa | Eutropis_indeprensa | LC | Unknown | tropical | diurnal | 5.888 | 169835.348 | 12.801 |
| lizards | Eutropis_innotata | NA | DD | Unknown | NA | NA | NA | NA | NA |
| lizards | Eutropis_islamaliit | NA | DD | Unknown | NA | NA | NA | NA | NA |
| lizards | Eutropis_lankae | NA | not_available | not_available | NA | NA | NA | NA | NA |
| lizards | Eutropis_lapulapu | NA | LC | Stable | NA | NA | NA | NA | NA |

| | | | | | | | | | |
|---|---|---|---|---|---|---|---|---|---|
| lizards | Eutropis_lewisi | NA | not_available | not_available | NA | NA | NA | NA | NA |
| lizards | Eutropis_longicaudata | Eutropis_longicaudata | LC | Stable | tropical | diurnal | 63.096 | 1965017.291 | 13.303 |
| lizards | Eutropis_macrophthalma | NA | VU | Decreasing | NA | NA | NA | NA | NA |
| lizards | Eutropis_macularia | Eutropis_macularia | LC | Stable | tropical | cathemeral | 9.333 | 4896988.014 | 15.994 |
| lizards | Eutropis_madaraszi | NA | LC | Stable | NA | NA | NA | NA | NA |
| lizards | Eutropis_multicarinata | Eutropis_multicarinata | LC | Stable | tropical | diurnal | 19.498 | 247749.894 | 14.177 |
| lizards | Eutropis_multifasciata | Eutropis_multifasciata | LC | Stable | tropical | diurnal | 63.096 | 6105593.017 | 11.907 |
| lizards | Eutropis_nagarjuni | NA | NT | Unknown | NA | NA | NA | NA | NA |
| lizards | Eutropis_palauensis | NA | not_available | not_available | NA | NA | NA | NA | NA |
| lizards | Eutropis_quadratilobus | NA | EN | Unknown | NA | NA | NA | NA | NA |
| lizards | Eutropis_quadricarinata | Eutropis_quadricarinata | LC | Unknown | temperate | diurnal | 2.399 | 33567.753 | 8.643 |
| lizards | Eutropis_resetarii | NA | not_available | not_available | NA | NA | NA | NA | NA |
| lizards | Eutropis_rudis | Eutropis_rudis | LC | Stable | tropical | diurnal | 60.256 | 571636.044 | 5.663 |
| lizards | Eutropis_rugifera | NA | LC | Unknown | NA | NA | NA | NA | NA |
| lizards | Eutropis_sahulinghangganan | NA | LC | Stable | NA | NA | NA | NA | NA |
| lizards | Eutropis_sibalom | NA | DD | Unknown | NA | NA | NA | NA | NA |
| lizards | Eutropis_tammanna | Eutropis_tammanna | LC | Stable | tropical | diurnal | 3.467 | 8845.863 | 18.53 |
| lizards | Eutropis_trivittata | NA | LC | Unknown | NA | NA | NA | NA | NA |
| lizards | Eutropis_tytleri | NA | LC | Stable | NA | NA | NA | NA | NA |
| lizards | Exila_nigropalmata | NA | LC | Unknown | NA | NA | NA | NA | NA |
| lizards | Feylinia_boulengeri | Chabanaudia_boulengeri | LC | Unknown | tropical | cathemeral | 2.455 | 37431.439 | 6.509 |
| lizards | Feylinia_currori | NA | LC | Stable | NA | NA | NA | NA | NA |
| lizards | Flexiseps_andranovahensis | NA | DD | Unknown | NA | NA | NA | NA | NA |
| lizards | Flexiseps_ardouini | NA | VU | Unknown | NA | NA | NA | NA | NA |
| lizards | Flexiseps_melanurus | Amphiglossus_melanurus | LC | Unknown | tropical | cathemeral | 30.903 | 35290.436 | 7.865 |
| lizards | Flexiseps_ornaticeps | NA | LC | Decreasing | NA | NA | NA | NA | NA |
| lizards | Flexiseps_stylus | NA | DD | Unknown | NA | NA | NA | NA | NA |
| lizards | Flexiseps_tanysoma | NA | LC | Unknown | NA | NA | NA | NA | NA |
| lizards | Fojia_bumui | NA | DD | Unknown | NA | NA | NA | NA | NA |
| lizards | Furcifer_antimena | Furcifer_antimena | VU | Decreasing | tropical | cathemeral | 95.499 | 6350.713 | 8.294 |
| lizards | Furcifer_balteatus | NA | EN | Decreasing | NA | NA | NA | NA | NA |
| lizards | Furcifer_belalandaensis | NA | CR | Decreasing | NA | NA | NA | NA | NA |
| lizards | Furcifer_cephalolepis | Furcifer_cephalolepis | LC | Stable | tropical | diurnal | 11.482 | 1031.546 | 17.428 |
| lizards | Furcifer_labordi | Furcifer_labordi | VU | Decreasing | tropical | diurnal | 64.565 | 16997.627 | 8.324 |

| | | | | | | | | | |
|---|---|---|---|---|---|---|---|---|---|
| lizards | Furcifer_lateralis | Furcifer_lateralis | LC | Stable | tropical | diurnal | 56.234 | 467150.957 | 8.93 |
| lizards | Furcifer_minor | Furcifer_minor | EN | Decreasing | tropical | diurnal | 22.909 | 7309.422 | 10.723 |
| lizards | Furcifer_nicosiai | NA | EN | Decreasing | NA | NA | NA | NA | NA |
| lizards | Furcifer_oustaleti | Furcifer_oustaleti | LC | Stable | tropical | diurnal | 436.516 | 591142.808 | 8.749 |
| lizards | Furcifer_pardalis | Furcifer_pardalis | LC | Stable | tropical | diurnal | 269.153 | 109567.079 | 7.745 |
| lizards | Furcifer_petteri | Furcifer_petteri | VU | Decreasing | tropical | diurnal | 17.378 | 11024.039 | 9.554 |
| lizards | Furcifer_rhinoceratus | NA | VU | Decreasing | NA | NA | NA | NA | NA |
| lizards | Furcifer_timoni | Furcifer_timoni | NT | Stable | tropical | diurnal | 20.893 | 1226.494 | 6.443 |
| lizards | Furcifer_tuzetae | NA | DD | Decreasing | NA | NA | NA | NA | NA |
| lizards | Furcifer_verrucosus | Furcifer_verrucosus | LC | Stable | tropical | diurnal | 316.228 | 352674.699 | 8.717 |
| lizards | Gallotia_atlantica | Gallotia_atlantica | LC | Stable | temperate | diurnal | 28.84 | 2586.702 | 17.521 |
| lizards | Gallotia_auaritae | NA | not_available | not_available | NA | NA | NA | NA | NA |
| lizards | Gallotia_bravoana | Gallotia_bravoana | CR | Increasing | temperate | diurnal | 208.93 | 108.503 | 11.745 |
| lizards | Gallotia_caesaris | Gallotia_caesaris | LC | Stable | temperate | diurnal | 30.903 | 660.383 | 15.041 |
| lizards | Gallotia_galloti | Gallotia_galloti | LC | Stable | temperate | diurnal | 69.183 | 2744.687 | 19.154 |
| lizards | Gallotia_intermedia | Gallotia_intermedia | CR | Increasing | temperate | diurnal | 117.49 | 161.343 | 19.503 |
| lizards | Gallotia_simonyi | Gallotia_simonyi | CR | Stable | temperate | diurnal | 2691.535 | 190.641 | 14.058 |
| lizards | Gallotia_stehlini | Gallotia_stehlini | LC | Stable | temperate | diurnal | 1096.478 | 1538.293 | 21.897 |
| lizards | Gambelia_copeii | Gambelia_copeii | LC | Decreasing | temperate | diurnal | 69.183 | 101483.761 | 3.593 |
| lizards | Gambelia_sila | Gambelia_sila | EN | Decreasing | temperate | diurnal | 57.544 | 37816.298 | 16.684 |
| lizards | Gambelia_wislizenii | Gambelia_wislizenii | LC | Stable | temperate | diurnal | 74.131 | 1387973.491 | 8.205 |
| lizards | Garthia_gaudichaudii | Garthia_gaudichaudii | LC | Stable | temperate | nocturnal | 2.754 | 69245.949 | 7.122 |
| lizards | Garthia_penai | NA | not_available | not_available | NA | NA | NA | NA | NA |
| lizards | Gastropholis_echinata | NA | LC | Unknown | NA | NA | NA | NA | NA |
| lizards | Gastropholis_prasina | Gastropholis_prasina | NT | Unknown | tropical | diurnal | 30.2 | 10592.745 | 14.338 |
| lizards | Gastropholis_tropidopholis | NA | LC | Stable | NA | NA | NA | NA | NA |
| lizards | Gastropholis_vittata | NA | LC | Unknown | NA | NA | NA | NA | NA |
| lizards | Geckolepis_humbloti | NA | not_available | not_available | NA | NA | NA | NA | NA |
| lizards | Geckolepis_maculata | Geckolepis_maculata | LC | Unknown | tropical | nocturnal | 10 | 314848.147 | 8.529 |
| lizards | Geckolepis_megalepis | NA | not_available | not_available | NA | NA | NA | NA | NA |
| lizards | Geckolepis_polylepis | NA | DD | Unknown | NA | NA | NA | NA | NA |
| lizards | Geckolepis_typica | Geckolepis_typica | LC | Unknown | tropical | nocturnal | 5.37 | 186421.545 | 8.946 |
| lizards | Gehyra_angusticaudata | NA | LC | Stable | NA | NA | NA | NA | NA |
| lizards | Gehyra_australis | Gehyra_australis | LC | Stable | tropical | nocturnal | 10.715 | 926589.681 | 2.067 |

| | | | | | | | | | |
|---|---|---|---|---|---|---|---|---|---|
| lizards | Gehyra_baliola | Gehyra_baliola | LC | Unknown | tropical | nocturnal | 21.878 | 197757.633 | 2.506 |
| lizards | Gehyra_borroloola | Gehyra_borroloola | LC | Stable | tropical | nocturnal | 6.918 | 79273.853 | 1.156 |
| lizards | Gehyra_catenata | Gehyra_catenata | LC | Decreasing | temperate | nocturnal | 3.631 | 355339.282 | 5.62 |
| lizards | Gehyra_dubia | Gehyra_dubia | LC | Unknown | tropical | nocturnal | 10.233 | 1500034.118 | 5.58 |
| lizards | Gehyra_einasleighensis | NA | LC | Stable | NA | NA | NA | NA | NA |
| lizards | Gehyra_fehlmanni | NA | LC | Unknown | NA | NA | NA | NA | NA |
| lizards | Gehyra_georgpotthasti | Gehyra_georgpotthasti | VU | Decreasing | tropical | nocturnal | 50.119 | 13053.854 | 6.113 |
| lizards | Gehyra_girloorloo | NA | LC | Stable | NA | NA | NA | NA | NA |
| lizards | Gehyra_insulensis | NA | LC | Stable | NA | NA | NA | NA | NA |
| lizards | Gehyra_koira | Gehyra_koira | LC | Stable | tropical | nocturnal | 16.982 | 314185.044 | 2.408 |
| lizards | Gehyra_lacerata | Gehyra_lacerata | LC | Stable | tropical | nocturnal | 4.467 | 14777.369 | 14.09 |
| lizards | Gehyra_lazelli | NA | LC | Stable | NA | NA | NA | NA | NA |
| lizards | Gehyra_minuta | Gehyra_minuta | LC | Stable | tropical | nocturnal | 2.089 | 119008.305 | 1.638 |
| lizards | Gehyra_montium | Gehyra_montium | LC | Unknown | tropical | nocturnal | 2.951 | 245462.192 | 3.162 |
| lizards | Gehyra_moritzi | Gehyra_moritzi | LC | Stable | tropical | nocturnal | 2.63 | 187148.156 | 1.889 |
| lizards | Gehyra_mutilata | Gehyra_mutilata | LC | Stable | tropical | nocturnal | 5.495 | 3740611.781 | 11.766 |
| lizards | Gehyra_nana | Gehyra_nana | LC | Stable | tropical | nocturnal | 3.467 | 570543.744 | 2.062 |
| lizards | Gehyra_occidentalis | NA | LC | Stable | NA | NA | NA | NA | NA |
| lizards | Gehyra_oceanica | Gehyra_oceanica | LC | Stable | tropical | nocturnal | 19.953 | 84829.699 | 4.645 |
| lizards | Gehyra_pamela | NA | LC | Stable | NA | NA | NA | NA | NA |
| lizards | Gehyra_pilbara | Gehyra_pilbara | LC | Stable | tropical | nocturnal | 6.026 | 1782077.767 | 2.104 |
| lizards | Gehyra_punctata | Gehyra_punctata | LC | Stable | temperate | nocturnal | 5.754 | 600961.027 | 4.262 |
| lizards | Gehyra_purpurascens | Gehyra_purpurascens | LC | Unknown | temperate | nocturnal | 6.31 | 3319362.414 | 2.048 |
| lizards | Gehyra_robusta | Gehyra_robusta | LC | Stable | tropical | nocturnal | 8.71 | 163324.912 | 4.738 |
| lizards | Gehyra_rohan | NA | not_available | not_available | NA | NA | NA | NA | NA |
| lizards | Gehyra_variegata | Gehyra_variegata | LC | Stable | temperate | nocturnal | 7.413 | 1717652.017 | 3.926 |
| lizards | Gehyra_versicolor | Gehyra_versicolor | LC | Unknown | temperate | cathemeral | 4.266 | 2543791.137 | 5.051 |
| lizards | Gehyra_vorax | Gehyra_vorax | NT | Unknown | tropical | cathemeral | 109.648 | 17009.28 | 9.379 |
| lizards | Gehyra_xenopus | Gehyra_xenopus | LC | Stable | tropical | nocturnal | 10 | 35702.295 | 1.258 |
| lizards | Gekko_aaronbaueri | NA | VU | Stable | NA | NA | NA | NA | NA |
| lizards | Gekko_adleri | NA | LC | Stable | NA | NA | NA | NA | NA |
| lizards | Gekko_albofasciolatus | NA | not_available | not_available | NA | NA | NA | NA | NA |
| lizards | Gekko_athymus | NA | NT | Unknown | NA | NA | NA | NA | NA |
| lizards | Gekko_auriverrucosus | NA | DD | Unknown | NA | NA | NA | NA | NA |

| | | | | | | | | | |
|---|---|---|---|---|---|---|---|---|---|
| lizards | Gekko_badenii | Gekko_badenii | EN | Decreasing | tropical | nocturnal | 47.863 | 25425.094 | 12.477 |
| lizards | Gekko_bannaense | NA | DD | Unknown | NA | NA | NA | NA | NA |
| lizards | Gekko_boehmei | NA | VU | Unknown | NA | NA | NA | NA | NA |
| lizards | Gekko_bonkowskii | NA | VU | Unknown | NA | NA | NA | NA | NA |
| lizards | Gekko_brooksii | NA | NT | Unknown | NA | NA | NA | NA | NA |
| lizards | Gekko_browni | NA | LC | Unknown | NA | NA | NA | NA | NA |
| lizards | Gekko_canaensis | Gekko_canaensis | LC | Stable | tropical | nocturnal | 28.184 | 5.694 | 35.496 |
| lizards | Gekko_canhi | NA | LC | Stable | NA | NA | NA | NA | NA |
| lizards | Gekko_carusadensis | NA | DD | Unknown | NA | NA | NA | NA | NA |
| lizards | Gekko_chinensis | Gekko_chinensis | LC | Stable | temperate | nocturnal | 10.233 | 842949.142 | 14.397 |
| lizards | Gekko_coi | NA | LC | Unknown | NA | NA | NA | NA | NA |
| lizards | Gekko_crombota | NA | LC | Stable | NA | NA | NA | NA | NA |
| lizards | Gekko_ernstkelleri | Gekko_ernstkelleri | LC | Stable | tropical | nocturnal | 15.136 | 540.436 | 12.598 |
| lizards | Gekko_gecko | Gekko_gecko | LC | Unknown | tropical | nocturnal | 169.824 | 4656309.145 | 12.433 |
| lizards | Gekko_gigante | NA | VU | Unknown | NA | NA | NA | NA | NA |
| lizards | Gekko_grossmanni | Gekko_grossmanni | DD | Stable | tropical | nocturnal | 14.125 | 6080.395 | 14.202 |
| lizards | Gekko_guishanicus | NA | DD | Unknown | NA | NA | NA | NA | NA |
| lizards | Gekko_gulat | NA | DD | Unknown | NA | NA | NA | NA | NA |
| lizards | Gekko_hokouensis | Gekko_hokouensis | LC | Stable | temperate | nocturnal | 12.303 | 651203.401 | 15.561 |
| lizards | Gekko_horsfieldii | Ptychozoon_horsfieldii | LC | Unknown | tropical | nocturnal | 16.982 | 38301.296 | 5.564 |
| lizards | Gekko_intermedium | Ptychozoon_intermedium | LC | Decreasing | tropical | nocturnal | 19.055 | 54309.976 | 15.288 |
| lizards | Gekko_iskandari | NA | DD | Unknown | NA | NA | NA | NA | NA |
| lizards | Gekko_japonicus | Gekko_japonicus | LC | Stable | temperate | nocturnal | 12.023 | 2304586.026 | 17.386 |
| lizards | Gekko_kaengkrachanense | NA | LC | Unknown | NA | NA | NA | NA | NA |
| lizards | Gekko_kikuchii | Gekko_kikuchii | LC | Stable | tropical | nocturnal | 10.233 | 9799.19 | 12.844 |
| lizards | Gekko_kuhli | Ptychozoon_kuhli | LC | Stable | tropical | nocturnal | 23.442 | 1423689.164 | 9.01 |
| lizards | Gekko_kwangsiensis | NA | DD | Decreasing | NA | NA | NA | NA | NA |
| lizards | Gekko_lauhachindai | NA | CR | Decreasing | NA | NA | NA | NA | NA |
| lizards | Gekko_liboensis | NA | DD | Unknown | NA | NA | NA | NA | NA |
| lizards | Gekko_lionotum | Ptychozoon_lionotum | LC | Stable | tropical | nocturnal | 20.893 | 1406078.456 | 13.13 |
| lizards | Gekko_melli | NA | LC | Stable | NA | NA | NA | NA | NA |
| lizards | Gekko_mindorensis | Gekko_mindorensis | LC | Stable | tropical | nocturnal | 13.49 | 55609.922 | 17.604 |
| lizards | Gekko_monarchus | Gekko_monarchus | LC | Stable | tropical | nocturnal | 19.953 | 1740240.708 | 9.233 |
| lizards | Gekko_nadenensis | NA | DD | Unknown | NA | NA | NA | NA | NA |

| | | | | | | | | | |
|---|---|---|---|---|---|---|---|---|---|
| lizards | Gekko_nicobarensis | Ptychozoon_nicobarensis | EN | Decreasing | tropical | nocturnal | 19.055 | 620.15 | 9.933 |
| lizards | Gekko_nutaphandi | Gekko_nutaphandi | LC | Stable | tropical | nocturnal | 29.512 | 19.982 | 16.126 |
| lizards | Gekko_palawanensis | NA | NT | Stable | NA | NA | NA | NA | NA |
| lizards | Gekko_palmatus | Gekko_palmatus | LC | Stable | tropical | nocturnal | 10.233 | 69777.324 | 15.789 |
| lizards | Gekko_petricolus | Gekko_petricolus | LC | Stable | tropical | cathemeral | 20.893 | 16575.654 | 13.448 |
| lizards | Gekko_porosus | NA | LC | Stable | NA | NA | NA | NA | NA |
| lizards | Gekko_reevesii | NA | LC | Decreasing | NA | NA | NA | NA | NA |
| lizards | Gekko_remotus | NA | not_available | not_available | NA | NA | NA | NA | NA |
| lizards | Gekko_rhacophorus | NA | DD | Unknown | NA | NA | NA | NA | NA |
| lizards | Gekko_romblon | NA | LC | Stable | NA | NA | NA | NA | NA |
| lizards | Gekko_rossi | NA | LC | Stable | NA | NA | NA | NA | NA |
| lizards | Gekko_russelltraini | Gekko_russelltraini | VU | Stable | tropical | nocturnal | 11.482 | 23.254 | 19.555 |
| lizards | Gekko_scabridus | NA | LC | Stable | NA | NA | NA | NA | NA |
| lizards | Gekko_scientiadventura | Gekko_scientiadventura | LC | Stable | tropical | nocturnal | 7.943 | 2736.162 | 5.695 |
| lizards | Gekko_sengchanthavongi | NA | VU | Unknown | NA | NA | NA | NA | NA |
| lizards | Gekko_shibatai | Gekko_shibatai | VU | Decreasing | temperate | nocturnal | 8.511 | 6.483 | 15.541 |
| lizards | Gekko_siamensis | Gekko_siamensis | LC | Stable | tropical | nocturnal | 58.884 | 5157.351 | 19.976 |
| lizards | Gekko_similignum | NA | LC | Stable | NA | NA | NA | NA | NA |
| lizards | Gekko_smithii | Gekko_smithii | LC | Stable | tropical | nocturnal | 114.815 | 1477995.914 | 9.197 |
| lizards | Gekko_sorok | NA | DD | Unknown | NA | NA | NA | NA | NA |
| lizards | Gekko_subpalmatus | Gekko_subpalmatus | LC | Stable | temperate | nocturnal | 10.965 | 1126594.906 | 15.847 |
| lizards | Gekko_swinhonis | Gekko_swinhonis | VU | Decreasing | temperate | nocturnal | 8.318 | 1375479.65 | 20.037 |
| lizards | Gekko_taibaiensis | NA | DD | Unknown | NA | NA | NA | NA | NA |
| lizards | Gekko_takouensis | Gekko_takouensis | VU | Stable | tropical | nocturnal | 22.909 | 9.991 | 19.293 |
| lizards | Gekko_tawaensis | NA | LC | Decreasing | NA | NA | NA | NA | NA |
| lizards | Gekko_thakhekensis | NA | VU | Unknown | NA | NA | NA | NA | NA |
| lizards | Gekko_trinotaterra | NA | LC | Stable | NA | NA | NA | NA | NA |
| lizards | Gekko_truongi | NA | DD | Unknown | NA | NA | NA | NA | NA |
| lizards | Gekko_verreauxi | NA | LC | Stable | NA | NA | NA | NA | NA |
| lizards | Gekko_vertebralis | NA | LC | Unknown | NA | NA | NA | NA | NA |
| lizards | Gekko_vietnamensis | NA | VU | Stable | NA | NA | NA | NA | NA |
| lizards | Gekko_vittatus | Gekko_vittatus | LC | Stable | tropical | nocturnal | 47.863 | 729666.633 | 5.159 |
| lizards | Gekko_wenxianensis | NA | DD | Unknown | NA | NA | NA | NA | NA |
| lizards | Gekko_yakuensis | Gekko_yakuensis | NT | Decreasing | temperate | nocturnal | 7.762 | 8949.277 | 20.869 |

| | | | | | | | | | |
|---|---|---|---|---|---|---|---|---|---|
| lizards | Geomyersia_glabra | Geomyersia_glabra | NT | Decreasing | tropical | nocturnal | 0.794 | 8778.262 | 6.34 |
| lizards | Gerrhonotus_farri | NA | not_available | not_available | NA | NA | NA | NA | NA |
| lizards | Gerrhonotus_infernalis | Gerrhonotus_infernalis | LC | Stable | temperate | cathemeral | 186.209 | 333221.594 | 7.263 |
| lizards | Gerrhonotus_liocephalus | Gerrhonotus_liocephalus | LC | Stable | tropical | diurnal | 186.209 | 365465.399 | 12.281 |
| lizards | Gerrhonotus_lugoi | Gerrhonotus_lugoi | LC | Unknown | temperate | nocturnal | 17.783 | 1486.106 | 2.477 |
| lizards | Gerrhonotus_ophiurus | Gerrhonotus_ophiurus | LC | Unknown | tropical | diurnal | 173.78 | 84300.562 | 15.216 |
| lizards | Gerrhonotus_parvus | Gerrhonotus_parvus | EN | Decreasing | temperate | diurnal | 8.71 | 6276.423 | 6.81 |
| lizards | Gerrhonotus_rhombifer | NA | DD | Unknown | NA | NA | NA | NA | NA |
| lizards | Gerrhosaurus_auritus | Gerrhosaurus_auritus | LC | Stable | tropical | diurnal | 257.04 | 612172.211 | 5.017 |
| lizards | Gerrhosaurus_bulsi | Gerrhosaurus_bulsi | LC | Stable | tropical | diurnal | 208.93 | 839027.36 | 6.42 |
| lizards | Gerrhosaurus_flavigularis | Gerrhosaurus_flavigularis | LC | Unknown | tropical | diurnal | 79.433 | 2867547.103 | 10.696 |
| lizards | Gerrhosaurus_multilineatus | Gerrhosaurus_multilineatus | LC | Stable | tropical | diurnal | 257.04 | 879482.256 | 6.083 |
| lizards | Gerrhosaurus_nigrolineatus | Gerrhosaurus_nigrolineatus | LC | Stable | tropical | diurnal | 158.489 | 4359388.444 | 8.707 |
| lizards | Gerrhosaurus_skoogi | Gerrhosaurus_skoogi | LC | Stable | tropical | diurnal | 104.713 | 33736.49 | 2.871 |
| lizards | Gerrhosaurus_typicus | Gerrhosaurus_typicus | LC | Stable | temperate | cathemeral | 69.183 | 122707.098 | 7.51 |
| lizards | Gigarcanum_delcourti | NA | EX | not_available | NA | NA | NA | NA | NA |
| lizards | Glaphyromorphus_cracens | NA | LC | Stable | NA | NA | NA | NA | NA |
| lizards | Glaphyromorphus_darwiniensis | NA | LC | Stable | NA | NA | NA | NA | NA |
| lizards | Glaphyromorphus_fuscicaudis | Glaphyromorphus_fuscicaudis | LC | Stable | tropical | cathemeral | 15.849 | 18803.061 | 5.243 |
| lizards | Glaphyromorphus_mjobergi | NA | LC | Unknown | NA | NA | NA | NA | NA |
| lizards | Glaphyromorphus_nigricaudis | NA | LC | Stable | NA | NA | NA | NA | NA |
| lizards | Glaphyromorphus_othelarrni | NA | LC | Unknown | NA | NA | NA | NA | NA |
| lizards | Glaphyromorphus_pumilus | Glaphyromorphus_pumilus | LC | Unknown | tropical | cathemeral | 3.09 | 137312.004 | 2.589 |
| lizards | Glaphyromorphus_punctulatus | Glaphyromorphus_punctulatus | LC | Stable | tropical | cathemeral | 6.761 | 201931.595 | 6.147 |
| lizards | Glaucomastix_abaetensis | Ameivula_abaetensis | EN | Decreasing | tropical | diurnal | 10.715 | 249.559 | 35.678 |
| lizards | Glaucomastix_littoralis | Ameivula_littoralis | EN | Unknown | tropical | diurnal | 15.849 | 5242.458 | 22.978 |
| lizards | Glaucomastix_venetacauda | NA | LC | Unknown | NA | NA | NA | NA | NA |
| lizards | Gnypetoscincus_queenslandiae | NA | LC | Stable | NA | NA | NA | NA | NA |
| lizards | Goggia_braacki | Goggia_braacki | LC | Stable | temperate | nocturnal | 1.023 | 1929.783 | 8.207 |
| lizards | Goggia_essexi | NA | LC | Stable | NA | NA | NA | NA | NA |
| lizards | Goggia_gemmula | Goggia_gemmula | LC | Stable | temperate | nocturnal | 0.692 | 2921.457 | 5.663 |
| lizards | Goggia_hewitti | Goggia_hewitti | LC | Stable | temperate | nocturnal | 1.259 | 10782.185 | 7.76 |
| lizards | Goggia_hexapora | NA | LC | Stable | NA | NA | NA | NA | NA |
| lizards | Goggia_incognita | NA | LC | Stable | NA | NA | NA | NA | NA |

| | | | | | | | | | |
|---|---|---|---|---|---|---|---|---|---|
| lizards | Goggia_lineata | Goggia_lineata | LC | Stable | temperate | nocturnal | 0.813 | 58711.373 | 5.459 |
| lizards | Goggia_microlepidota | NA | LC | Stable | NA | NA | NA | NA | NA |
| lizards | Goggia_rupicola | NA | LC | Stable | NA | NA | NA | NA | NA |
| lizards | Gonatodes_albogularis | Gonatodes_albogularis | LC | Stable | tropical | diurnal | 2.754 | 873793.037 | 12.493 |
| lizards | Gonatodes_alexandermendesi | Gonatodes_alexandermendesi | LC | Unknown | tropical | diurnal | 2.884 | 40483.843 | 1.877 |
| lizards | Gonatodes_annularis | Gonatodes_annularis | LC | Unknown | tropical | diurnal | 4.074 | 1140558.821 | 1.515 |
| lizards | Gonatodes_antillensis | Gonatodes_antillensis | LC | Stable | tropical | nocturnal | 1.349 | 99.471 | 18.988 |
| lizards | Gonatodes_astralis | NA | VU | Decreasing | NA | NA | NA | NA | NA |
| lizards | Gonatodes_atricucullaris | NA | NT | Stable | NA | NA | NA | NA | NA |
| lizards | Gonatodes_caudiscutatus | Gonatodes_caudiscutatus | LC | Unknown | tropical | diurnal | 2.291 | 106052.218 | 12.095 |
| lizards | Gonatodes_ceciliae | Gonatodes_ceciliae | LC | Stable | tropical | diurnal | 7.413 | 5164.753 | 17.042 |
| lizards | Gonatodes_concinnatus | Gonatodes_concinnatus | LC | Stable | tropical | diurnal | 3.548 | 851434.055 | 2.878 |
| lizards | Gonatodes_daudini | Gonatodes_daudini | CR | Stable | tropical | diurnal | 0.759 | 10.874 | 19.377 |
| lizards | Gonatodes_eladioi | NA | LC | Unknown | NA | NA | NA | NA | NA |
| lizards | Gonatodes_falconensis | NA | LC | Stable | NA | NA | NA | NA | NA |
| lizards | Gonatodes_hasemani | Gonatodes_hasemani | LC | Stable | tropical | diurnal | 2.399 | 2613126.805 | 2.349 |
| lizards | Gonatodes_humeralis | Gonatodes_humeralis | LC | Stable | tropical | diurnal | 2.57 | 7917317.285 | 2.769 |
| lizards | Gonatodes_infernalis | NA | CR | Decreasing | NA | NA | NA | NA | NA |
| lizards | Gonatodes_lichenosus | NA | DD | Unknown | NA | NA | NA | NA | NA |
| lizards | Gonatodes_ocellatus | Gonatodes_ocellatus | LC | Stable | tropical | diurnal | 3.09 | 86.285 | 16.673 |
| lizards | Gonatodes_petersi | Gonatodes_petersi | LC | Stable | tropical | diurnal | 1.862 | 3100.383 | 7.499 |
| lizards | Gonatodes_purpurogularis | NA | EN | Decreasing | NA | NA | NA | NA | NA |
| lizards | Gonatodes_rayito | NA | not_available | not_available | NA | NA | NA | NA | NA |
| lizards | Gonatodes_rozei | NA | LC | Stable | NA | NA | NA | NA | NA |
| lizards | Gonatodes_seigliei | NA | EN | Decreasing | NA | NA | NA | NA | NA |
| lizards | Gonatodes_superciliaris | Gonatodes_superciliaris | LC | Stable | tropical | diurnal | 2.884 | 14.306 | 0.326 |
| lizards | Gonatodes_taniae | Gonatodes_taniae | LC | Stable | tropical | diurnal | 3.467 | 840.391 | 14.523 |
| lizards | Gonatodes_tapajonicus | NA | VU | Unknown | NA | NA | NA | NA | NA |
| lizards | Gonatodes_timidus | NA | LC | Unknown | NA | NA | NA | NA | NA |
| lizards | Gonatodes_vittatus | Gonatodes_vittatus | LC | Stable | tropical | diurnal | 1.072 | 278124.207 | 12.542 |
| lizards | Gongylomorphus_bojerii | Gongylomorphus_bojerii | CR | Stable | tropical | diurnal | 7.413 | 1872.513 | 25.918 |
| lizards | Gongylomorphus_fontenayi | NA | EN | Decreasing | NA | NA | NA | NA | NA |
| lizards | Goniurosaurus_araneus | Goniurosaurus_araneus | EN | Unknown | tropical | nocturnal | 47.863 | 588.192 | 13.766 |
| lizards | Goniurosaurus_bawanglingensis | NA | EN | Decreasing | NA | NA | NA | NA | NA |

| | | | | | | | | | |
|---|---|---|---|---|---|---|---|---|---|
| lizards | Goniurosaurus_catbaensis | NA | EN | Unknown | NA | NA | NA | NA | NA |
| lizards | Goniurosaurus_hainanensis | Goniurosaurus_hainanensis | NT | Stable | tropical | nocturnal | 19.498 | 8319.642 | 13.84 |
| lizards | Goniurosaurus_huuliensis | NA | CR | Unknown | NA | NA | NA | NA | NA |
| lizards | Goniurosaurus_kadoorieorum | NA | EN | Decreasing | NA | NA | NA | NA | NA |
| lizards | Goniurosaurus_kuroiwae | Goniurosaurus_kuroiwae | VU | Decreasing | temperate | nocturnal | 20.417 | 1255.45 | 27.529 |
| lizards | Goniurosaurus_kwangsiensis | NA | EN | Decreasing | NA | NA | NA | NA | NA |
| lizards | Goniurosaurus_liboensis | NA | EN | Decreasing | NA | NA | NA | NA | NA |
| lizards | Goniurosaurus_lichtenfelderi | NA | VU | Decreasing | NA | NA | NA | NA | NA |
| lizards | Goniurosaurus_luii | Goniurosaurus_luii | VU | Decreasing | tropical | nocturnal | 39.811 | 10484.792 | 15.228 |
| lizards | Goniurosaurus_orientalis | Goniurosaurus_orientalis | EN | Decreasing | temperate | nocturnal | 16.982 | 3.475 | 19.175 |
| lizards | Goniurosaurus_sengokui | NA | not_available | not_available | NA | NA | NA | NA | NA |
| lizards | Goniurosaurus_splendens | NA | EN | Decreasing | NA | NA | NA | NA | NA |
| lizards | Goniurosaurus_toyamai | Goniurosaurus_toyamai | CR | Unknown | temperate | nocturnal | 11.749 | 18.933 | 19.792 |
| lizards | Goniurosaurus_yamashinae | Goniurosaurus_yamashinae | CR | Decreasing | temperate | nocturnal | 13.49 | 58.117 | 26.376 |
| lizards | Goniurosaurus_yingdeensis | Goniurosaurus_yingdeensis | CR | Decreasing | temperate | nocturnal | 17.783 | 9.991 | 5.152 |
| lizards | Goniurosaurus_zhelongi | NA | EN | Decreasing | NA | NA | NA | NA | NA |
| lizards | Goniurosaurus_zhoui | NA | DD | Unknown | NA | NA | NA | NA | NA |
| lizards | Gonocephalus_bellii | Gonocephalus_bellii | LC | Unknown | tropical | diurnal | 123.027 | 62519.174 | 9.405 |
| lizards | Gonocephalus_beyschlagi | NA | NT | Unknown | NA | NA | NA | NA | NA |
| lizards | Gonocephalus_bornensis | Gonocephalus_bornensis | LC | Unknown | tropical | diurnal | 91.201 | 308153.192 | 4.368 |
| lizards | Gonocephalus_chamaeleontinus | Gonocephalus_chamaeleontinus | LC | Stable | tropical | diurnal | 173.78 | 233857.854 | 9.932 |
| lizards | Gonocephalus_doriae | Gonocephalus_doriae | LC | Unknown | tropical | diurnal | 151.356 | 31728.302 | 7.541 |
| lizards | Gonocephalus_grandis | Gonocephalus_grandis | LC | Stable | tropical | diurnal | 144.544 | 1315206.165 | 7.681 |
| lizards | Gonocephalus_interruptus | NA | LC | Stable | NA | NA | NA | NA | NA |
| lizards | Gonocephalus_klossi | NA | NT | Unknown | NA | NA | NA | NA | NA |
| lizards | Gonocephalus_kuhlii | Gonocephalus_kuhlii | VU | Unknown | tropical | diurnal | 223.872 | 12094.161 | 20.508 |
| lizards | Gonocephalus_lacunosus | Gonocephalus_lacunosus | LC | Unknown | tropical | diurnal | 117.49 | 9.991 | 21.452 |
| lizards | Gonocephalus_liogaster | Gonocephalus_liogaster | LC | Unknown | tropical | diurnal | 107.152 | 658393.542 | 5.843 |
| lizards | Gonocephalus_megalepis | NA | LC | Unknown | NA | NA | NA | NA | NA |
| lizards | Gonocephalus_mjobergi | NA | DD | Unknown | NA | NA | NA | NA | NA |
| lizards | Gonocephalus_semperi | Gonocephalus_semperi | LC | Unknown | tropical | diurnal | 50.119 | 273776.479 | 13.672 |
| lizards | Gonocephalus_sophiae | Gonocephalus_sophiae | LC | Unknown | tropical | diurnal | 117.49 | 274059.4 | 13.673 |
| lizards | Gowidon_longirostris | Lophognathus_longirostris | LC | Stable | temperate | diurnal | 50.119 | 2537718.584 | 2.111 |
| lizards | Graciliscincus_shonae | NA | VU | Unknown | NA | NA | NA | NA | NA |

| | | | | | | | | | |
|---|---|---|---|---|---|---|---|---|---|
| lizards | Grandidierina_fierinensis | NA | LC | Stable | NA | NA | NA | NA | NA |
| lizards | Grandidierina_lineata | NA | LC | Stable | NA | NA | NA | NA | NA |
| lizards | Gymnodactylus_amarali | Gymnodactylus_amarali | LC | Stable | tropical | nocturnal | 4.365 | 1124711.459 | 8.027 |
| lizards | Gymnodactylus_darwinii | Gymnodactylus_darwinii | LC | Unknown | tropical | nocturnal | 5.495 | 399674.464 | 11.027 |
| lizards | Gymnodactylus_geckoides | Gymnodactylus_geckoides | LC | Unknown | tropical | cathemeral | 4.365 | 539398.376 | 9.034 |
| lizards | Gymnodactylus_vanzolinii | NA | DD | Unknown | NA | NA | NA | NA | NA |
| lizards | Gymnophthalmus_cryptus | NA | LC | Stable | NA | NA | NA | NA | NA |
| lizards | Gymnophthalmus_leucomystax | NA | DD | Unknown | NA | NA | NA | NA | NA |
| lizards | Gymnophthalmus_lineatus | NA | LC | Unknown | NA | NA | NA | NA | NA |
| lizards | Gymnophthalmus_marconaterai | NA | not_available | not_available | NA | NA | NA | NA | NA |
| lizards | Gymnophthalmus_pleii | NA | NT | Unknown | NA | NA | NA | NA | NA |
| lizards | Gymnophthalmus_speciosus | Gymnophthalmus_speciosus | LC | Unknown | tropical | diurnal | 2.344 | 1096550.609 | 10.411 |
| lizards | Gymnophthalmus_underwoodi | Gymnophthalmus_underwoodi | LC | Unknown | tropical | diurnal | 2.344 | 1215624.718 | 1.488 |
| lizards | Gymnophthalmus_vanzoi | Gymnophthalmus_vanzoi | LC | Unknown | tropical | diurnal | 1.66 | 27581.727 | 6.037 |
| lizards | Haemodracon_riebeckii | Haemodracon_riebeckii | LC | Unknown | tropical | cathemeral | 69.183 | 3465.903 | 10.956 |
| lizards | Haemodracon_trachyrhinus | Haemodracon_trachyrhinus | LC | Unknown | tropical | nocturnal | 4.365 | 3228.849 | 10.872 |
| lizards | Hakaria_simonyi | NA | NT | Unknown | NA | NA | NA | NA | NA |
| lizards | Harpesaurus_beccarii | NA | DD | Unknown | NA | NA | NA | NA | NA |
| lizards | Harpesaurus_borneensis | NA | DD | Unknown | NA | NA | NA | NA | NA |
| lizards | Harpesaurus_brooksi | NA | LC | Unknown | NA | NA | NA | NA | NA |
| lizards | Harpesaurus_ensicauda | NA | DD | Unknown | NA | NA | NA | NA | NA |
| lizards | Harpesaurus_modiglianii | NA | EN | Unknown | NA | NA | NA | NA | NA |
| lizards | Harpesaurus_tricinctus | NA | DD | Unknown | NA | NA | NA | NA | NA |
| lizards | Harrisoniascincus_zia | Harrisoniascincus_zia | LC | Stable | temperate | diurnal | 3.89 | 13695.321 | 5.918 |
| lizards | Heliobolus_lugubris | Heliobolus_lugubris | LC | Stable | tropical | diurnal | 6.457 | 2200737.129 | 6.392 |
| lizards | Heliobolus_nitidus | NA | LC | Unknown | NA | NA | NA | NA | NA |
| lizards | Heliobolus_spekii | Heliobolus_spekii | LC | Unknown | tropical | diurnal | 5.012 | 2240391.229 | 9.149 |
| lizards | Hellenolacerta_graeca | Hellenolacerta_graeca | NT | Decreasing | temperate | diurnal | 14.125 | 21404.315 | 16.603 |
| lizards | Heloderma_horridum | Heloderma_horridum | LC | Decreasing | tropical | cathemeral | 3019.952 | 171205.411 | 9.202 |
| lizards | Heloderma_suspectum | Heloderma_suspectum | NT | Decreasing | temperate | cathemeral | 977.237 | 345298.437 | 8.44 |
| lizards | Hemicordylus_capensis | Hemicordylus_capensis | LC | Stable | temperate | diurnal | 36.308 | 72370.851 | 9.251 |
| lizards | Hemicordylus_nebulosus | Hemicordylus_nebulosus | VU | Stable | temperate | diurnal | 12.589 | 10.551 | 8.434 |
| lizards | Hemidactylus_aaronbaueri | NA | LC | Unknown | NA | NA | NA | NA | NA |
| lizards | Hemidactylus_acanthopholis | NA | DD | Stable | NA | NA | NA | NA | NA |

| | | | | | | | | | |
|---|---|---|---|---|---|---|---|---|---|
| lizards | Hemidactylus_adensis | NA | not_available | not_available | NA | NA | NA | NA | NA |
| lizards | Hemidactylus_agrius | NA | LC | Unknown | NA | NA | NA | NA | NA |
| lizards | Hemidactylus_albituberculatus | NA | LC | Unknown | NA | NA | NA | NA | NA |
| lizards | Hemidactylus_albivertebralis | NA | DD | Unknown | NA | NA | NA | NA | NA |
| lizards | Hemidactylus_albofasciatus | Hemidactylus_albofasciatus | VU | Unknown | tropical | nocturnal | 1.122 | 1814.635 | 19.6 |
| lizards | Hemidactylus_alfarraji | NA | not_available | not_available | NA | NA | NA | NA | NA |
| lizards | Hemidactylus_alkiyumii | NA | not_available | not_available | NA | NA | NA | NA | NA |
| lizards | Hemidactylus_angulatus | Hemidactylus_angulatus | LC | Stable | tropical | nocturnal | 7.413 | 9223834.718 | 10.262 |
| lizards | Hemidactylus_ansorgii | NA | LC | Unknown | NA | NA | NA | NA | NA |
| lizards | Hemidactylus_aquilonius | Hemidactylus_aquilonius | LC | Unknown | temperate | nocturnal | 3.311 | 1034050.983 | 10.699 |
| lizards | Hemidactylus_asirensis | NA | not_available | not_available | NA | NA | NA | NA | NA |
| lizards | Hemidactylus_awashensis | NA | not_available | not_available | NA | NA | NA | NA | NA |
| lizards | Hemidactylus_barbierii | NA | DD | Unknown | NA | NA | NA | NA | NA |
| lizards | Hemidactylus_bavazzanoi | NA | DD | Unknown | NA | NA | NA | NA | NA |
| lizards | Hemidactylus_beninensis | Hemidactylus_beninensis | LC | Unknown | tropical | nocturnal | 10.233 | 19.982 | 20.558 |
| lizards | Hemidactylus_bouvieri | NA | CR | Unknown | NA | NA | NA | NA | NA |
| lizards | Hemidactylus_bowringii | Hemidactylus_bowringii | LC | Stable | temperate | nocturnal | 10.715 | 402160.744 | 17.667 |
| lizards | Hemidactylus_brasilianus | Hemidactylus_brasilianus | LC | Unknown | tropical | nocturnal | 5.495 | 925038.727 | 8.003 |
| lizards | Hemidactylus_brookii | Hemidactylus_brookii | LC | Unknown | tropical | nocturnal | 12.303 | 6273555.679 | 14.429 |
| lizards | Hemidactylus_chipkali | NA | LC | Unknown | NA | NA | NA | NA | NA |
| lizards | Hemidactylus_coalescens | Hemidactylus_coalescens | LC | Decreasing | tropical | nocturnal | 9.55 | 910124.808 | 5.803 |
| lizards | Hemidactylus_craspedotus | Hemidactylus_craspedotus | LC | Unknown | tropical | nocturnal | 5.754 | 125107.139 | 10.016 |
| lizards | Hemidactylus_dawudazraqi | NA | not_available | not_available | NA | NA | NA | NA | NA |
| lizards | Hemidactylus_depressus | Hemidactylus_depressus | LC | Stable | tropical | nocturnal | 12.303 | 56020.995 | 18.019 |
| lizards | Hemidactylus_dracaenacolus | Hemidactylus_dracaenacolus | CR | Decreasing | tropical | nocturnal | 8.71 | 94.138 | 11.22 |
| lizards | Hemidactylus_echinus | Hemidactylus_echinus | LC | Unknown | tropical | nocturnal | 7.079 | 29843.049 | 10.867 |
| lizards | Hemidactylus_eniangii | NA | LC | Decreasing | NA | NA | NA | NA | NA |
| lizards | Hemidactylus_fasciatus | Hemidactylus_fasciatus | LC | Unknown | tropical | nocturnal | 16.596 | 482648.755 | 15.488 |
| lizards | Hemidactylus_festivus | NA | not_available | not_available | NA | NA | NA | NA | NA |
| lizards | Hemidactylus_flaviviridis | Hemidactylus_flaviviridis | LC | Increasing | temperate | cathemeral | 16.596 | 3803737.441 | 15.643 |
| lizards | Hemidactylus_forbesii | NA | LC | Unknown | NA | NA | NA | NA | NA |
| lizards | Hemidactylus_foudaii | NA | LC | Stable | NA | NA | NA | NA | NA |
| lizards | Hemidactylus_frenatus | Hemidactylus_frenatus | LC | Stable | tropical | nocturnal | 6.31 | 7347932.06 | 12.505 |
| lizards | Hemidactylus_garnotii | Hemidactylus_garnotii | LC | Stable | tropical | nocturnal | 6.607 | 4533420.358 | 11.55 |

| | | | | | | | | | |
|---|---|---|---|---|---|---|---|---|---|
| lizards | Hemidactylus_giganteus | NA | LC | Unknown | NA | NA | NA | NA | NA |
| lizards | Hemidactylus_gleadowi | NA | DD | Unknown | NA | NA | NA | NA | NA |
| lizards | Hemidactylus_gracilis | NA | LC | Unknown | NA | NA | NA | NA | NA |
| lizards | Hemidactylus_graniticolus | NA | LC | Stable | NA | NA | NA | NA | NA |
| lizards | Hemidactylus_granosus | NA | not_available | not_available | NA | NA | NA | NA | NA |
| lizards | Hemidactylus_granti | NA | NT | Unknown | NA | NA | NA | NA | NA |
| lizards | Hemidactylus_gujaratensis | NA | VU | Unknown | NA | NA | NA | NA | NA |
| lizards | Hemidactylus_hajarensis | Hemidactylus_hajarensis | LC | Stable | tropical | nocturnal | 6.31 | 12343.192 | 9.465 |
| lizards | Hemidactylus_hemchandrai | NA | LC | Unknown | NA | NA | NA | NA | NA |
| lizards | Hemidactylus_homoeolepis | Hemidactylus_homoeolepis | LC | Stable | tropical | nocturnal | 2.692 | 97776.817 | 4.325 |
| lizards | Hemidactylus_hunae | Hemidactylus_hunae | EN | Unknown | tropical | nocturnal | 38.019 | 7821.032 | 11.577 |
| lizards | Hemidactylus_imbricatus | Hemidactylus_imbricatus | LC | Unknown | temperate | nocturnal | 3.802 | 211884.843 | 10.866 |
| lizards | Hemidactylus_inexpectatus | Hemidactylus_inexpectatus | LC | Stable | tropical | nocturnal | 1.995 | 15.96 | 7.382 |
| lizards | Hemidactylus_inintellectus | Hemidactylus_inintellectus | LC | Unknown | tropical | nocturnal | 5.495 | 2704.167 | 10.625 |
| lizards | Hemidactylus_isolepis | Hemidactylus_isolepis | LC | Unknown | tropical | nocturnal | 2.951 | 527262.518 | 7.684 |
| lizards | Hemidactylus_jumailiae | NA | not_available | not_available | NA | NA | NA | NA | NA |
| lizards | Hemidactylus_kamdemtohami | Hemidactylus_kamdemtohami | LC | Stable | tropical | diurnal | 7.413 | 132645.304 | 7.622 |
| lizards | Hemidactylus_karenorum | NA | LC | Stable | NA | NA | NA | NA | NA |
| lizards | Hemidactylus_kundaensis | NA | CR | Unknown | NA | NA | NA | NA | NA |
| lizards | Hemidactylus_kushmorensis | NA | DD | Unknown | NA | NA | NA | NA | NA |
| lizards | Hemidactylus_kyaboboensis | NA | not_available | not_available | NA | NA | NA | NA | NA |
| lizards | Hemidactylus_laevis | NA | DD | Unknown | NA | NA | NA | NA | NA |
| lizards | Hemidactylus_lankae | NA | not_available | not_available | NA | NA | NA | NA | NA |
| lizards | Hemidactylus_laticaudatus | NA | LC | Unknown | NA | NA | NA | NA | NA |
| lizards | Hemidactylus_lavadeserticus | NA | not_available | not_available | NA | NA | NA | NA | NA |
| lizards | Hemidactylus_lemurinus | NA | DD | Unknown | NA | NA | NA | NA | NA |
| lizards | Hemidactylus_leschenaultii | Hemidactylus_leschenaultii | LC | Stable | tropical | cathemeral | 15.136 | 1578465.794 | 18.763 |
| lizards | Hemidactylus_longicephalus | Hemidactylus_longicephalus | LC | Stable | tropical | nocturnal | 9.55 | 1525636.148 | 6.735 |
| lizards | Hemidactylus_luqueorum | NA | LC | Stable | NA | NA | NA | NA | NA |
| lizards | Hemidactylus_mabouia | Hemidactylus_mabouia | LC | Stable | tropical | cathemeral | 14.125 | 9561088.711 | 8.961 |
| lizards | Hemidactylus_macropholis | Hemidactylus_macropholis | LC | Unknown | tropical | nocturnal | 14.791 | 955988.769 | 8.752 |
| lizards | Hemidactylus_maculatus | Hemidactylus_maculatus | LC | Unknown | tropical | nocturnal | 33.113 | 438618.321 | 18.872 |
| lizards | Hemidactylus_malcolmsmithi | NA | DD | Unknown | NA | NA | NA | NA | NA |
| lizards | Hemidactylus_mandebensis | NA | not_available | not_available | NA | NA | NA | NA | NA |

| | | | | | | | | | |
|---|---|---|---|---|---|---|---|---|---|
| lizards | Hemidactylus_masirahensis | NA | LC | Stable | NA | NA | NA | NA | NA |
| lizards | Hemidactylus_matschiei | NA | LC | Unknown | NA | NA | NA | NA | NA |
| lizards | Hemidactylus_mercatorius | Hemidactylus_mercatorius | LC | Increasing | tropical | nocturnal | 4.467 | 647004.545 | 9.229 |
| lizards | Hemidactylus_mindiae | NA | LC | Increasing | NA | NA | NA | NA | NA |
| lizards | Hemidactylus_minutus | NA | not_available | not_available | NA | NA | NA | NA | NA |
| lizards | Hemidactylus_mrimaensis | NA | not_available | not_available | NA | NA | NA | NA | NA |
| lizards | Hemidactylus_muriceus | Hemidactylus_muriceus | LC | Unknown | tropical | cathemeral | 4.266 | 1805438.065 | 9.459 |
| lizards | Hemidactylus_ophiolepis | NA | LC | Unknown | NA | NA | NA | NA | NA |
| lizards | Hemidactylus_oxyrhinus | NA | LC | Unknown | NA | NA | NA | NA | NA |
| lizards | Hemidactylus_palaichthus | Hemidactylus_palaichthus | LC | Stable | tropical | nocturnal | 7.413 | 1432246.753 | 1.968 |
| lizards | Hemidactylus_parvimaculatus | Hemidactylus_parvimaculatus | LC | Unknown | tropical | nocturnal | 3.162 | 215161.375 | 20.311 |
| lizards | Hemidactylus_paucituberculatus | NA | LC | Stable | NA | NA | NA | NA | NA |
| lizards | Hemidactylus_persicus | Hemidactylus_persicus | LC | Unknown | temperate | nocturnal | 9.12 | 1174736.644 | 10.967 |
| lizards | Hemidactylus_pieresii | NA | EN | Decreasing | NA | NA | NA | NA | NA |
| lizards | Hemidactylus_platycephalus | Hemidactylus_platycephalus | LC | Unknown | tropical | nocturnal | 16.218 | 2633418.315 | 10.14 |
| lizards | Hemidactylus_platyurus | Hemidactylus_platyurus | LC | Increasing | tropical | cathemeral | 6.918 | 6316105.575 | 12.911 |
| lizards | Hemidactylus_prashadi | NA | LC | Unknown | NA | NA | NA | NA | NA |
| lizards | Hemidactylus_pseudomuriceus | NA | DD | Unknown | NA | NA | NA | NA | NA |
| lizards | Hemidactylus_pumilio | NA | LC | Unknown | NA | NA | NA | NA | NA |
| lizards | Hemidactylus_reticulatus | NA | LC | Unknown | NA | NA | NA | NA | NA |
| lizards | Hemidactylus_richardsonii | Hemidactylus_richardsonii | LC | Stable | tropical | nocturnal | 10.233 | 530792.47 | 6.284 |
| lizards | Hemidactylus_robustus | Hemidactylus_robustus | LC | Increasing | tropical | nocturnal | 4.677 | 2651796.373 | 7.627 |
| lizards | Hemidactylus_romeshkanicus | NA | not_available | not_available | NA | NA | NA | NA | NA |
| lizards | Hemidactylus_ruspolii | Hemidactylus_ruspolii | LC | Unknown | tropical | nocturnal | 3.467 | 574666.458 | 8.716 |
| lizards | Hemidactylus_saba | NA | not_available | not_available | NA | NA | NA | NA | NA |
| lizards | Hemidactylus_sataraensis | Hemidactylus_sataraensis | CR | Unknown | tropical | nocturnal | 2.291 | 59.648 | 13.177 |
| lizards | Hemidactylus_scabriceps | Hemidactylus_scabriceps | EN | Unknown | tropical | nocturnal | 2.399 | 49055.999 | 25.499 |
| lizards | Hemidactylus_shihraensis | NA | not_available | not_available | NA | NA | NA | NA | NA |
| lizards | Hemidactylus_sinaitus | NA | LC | Stable | NA | NA | NA | NA | NA |
| lizards | Hemidactylus_somalicus | NA | LC | Unknown | NA | NA | NA | NA | NA |
| lizards | Hemidactylus_squamulatus | Hemidactylus_squamulatus | LC | Unknown | tropical | nocturnal | 2.512 | 543958.5 | 10.615 |
| lizards | Hemidactylus_stejnegeri | Hemidactylus_stejnegeri | LC | Stable | tropical | cathemeral | 4.571 | 19726.88 | 19.784 |
| lizards | Hemidactylus_sushilduttai | NA | LC | Stable | NA | NA | NA | NA | NA |
| lizards | Hemidactylus_tanganicus | NA | DD | Unknown | NA | NA | NA | NA | NA |

| | | | | | | | | | |
|---|---|---|---|---|---|---|---|---|---|
| lizards | Hemidactylus_tenkatei | NA | LC | Unknown | NA | NA | NA | NA | NA |
| lizards | Hemidactylus_thayene | NA | LC | Stable | NA | NA | NA | NA | NA |
| lizards | Hemidactylus_treutleri | Hemidactylus_treutleri | LC | Unknown | tropical | nocturnal | 7.244 | 419611.506 | 17.551 |
| lizards | Hemidactylus_triedrus | Hemidactylus_triedrus | LC | Unknown | tropical | nocturnal | 16.218 | 253557.231 | 18.082 |
| lizards | Hemidactylus_tropidolepis | Hemidactylus_tropidolepis | LC | Unknown | tropical | nocturnal | 3.236 | 322141.25 | 9.973 |
| lizards | Hemidactylus_turcicus | Hemidactylus_turcicus | LC | Increasing | temperate | nocturnal | 5.754 | 2226414.004 | 13.821 |
| lizards | Hemidactylus_ulii | NA | not_available | not_available | NA | NA | NA | NA | NA |
| lizards | Hemidactylus_vanam | NA | DD | Unknown | NA | NA | NA | NA | NA |
| lizards | Hemidactylus_vietnamensis | NA | LC | Unknown | NA | NA | NA | NA | NA |
| lizards | Hemidactylus_yajurvedi | NA | EN | Stable | NA | NA | NA | NA | NA |
| lizards | Hemidactylus_yerburii | NA | LC | Unknown | NA | NA | NA | NA | NA |
| lizards | Hemiergis_decresiensis | Hemiergis_decresiensis | LC | Stable | temperate | diurnal | 3.02 | 141899 | 10.391 |
| lizards | Hemiergis_gracilipes | Hemiergis_gracilipes | LC | Stable | temperate | diurnal | 14.791 | 30507.046 | 9.565 |
| lizards | Hemiergis_initialis | Hemiergis_initialis | LC | Stable | temperate | cathemeral | 0.977 | 418696.124 | 6.18 |
| lizards | Hemiergis_millewae | Hemiergis_millewae | LC | Stable | temperate | cathemeral | 3.715 | 279036.18 | 6.625 |
| lizards | Hemiergis_peronii | Hemiergis_peronii | LC | Stable | temperate | cathemeral | 3.02 | 643819.971 | 8.104 |
| lizards | Hemiergis_quadrilineatus | NA | LC | Stable | NA | NA | NA | NA | NA |
| lizards | Hemiergis_talbingoensis | Hemiergis_talbingoensis | LC | Unknown | temperate | diurnal | 1.862 | 282908.791 | 8.421 |
| lizards | Hemiphyllodactylus_aurantiacus | Hemiphyllodactylus_aurantiacus | CR | Unknown | tropical | nocturnal | 1.622 | 68026.075 | 18.008 |
| lizards | Hemiphyllodactylus_banaensis | Hemiphyllodactylus_banaensis | LC | Unknown | tropical | nocturnal | 2.951 | 9.991 | 11.932 |
| lizards | Hemiphyllodactylus_bintik | NA | LC | Unknown | NA | NA | NA | NA | NA |
| lizards | Hemiphyllodactylus_changningensis | NA | LC | Stable | NA | NA | NA | NA | NA |
| lizards | Hemiphyllodactylus_chiangmaiensis | Hemiphyllodactylus_chiangmaiensis | LC | Stable | tropical | nocturnal | 1.66 | 19.982 | 32.341 |
| lizards | Hemiphyllodactylus_cicak | NA | VU | Unknown | NA | NA | NA | NA | NA |
| lizards | Hemiphyllodactylus_engganoensis | NA | LC | Unknown | NA | NA | NA | NA | NA |
| lizards | Hemiphyllodactylus_flaviventris | NA | not_available | not_available | NA | NA | NA | NA | NA |
| lizards | Hemiphyllodactylus_ganoklonis | Hemiphyllodactylus_ganoklonis | LC | Stable | tropical | nocturnal | 0.977 | 424.34 | 11.227 |
| lizards | Hemiphyllodactylus_harterti | NA | NT | Unknown | NA | NA | NA | NA | NA |
| lizards | Hemiphyllodactylus_huishuiensis | NA | DD | Stable | NA | NA | NA | NA | NA |
| lizards | Hemiphyllodactylus_insularis | NA | DD | Unknown | NA | NA | NA | NA | NA |
| lizards | Hemiphyllodactylus_khlonglanensis | NA | not_available | not_available | NA | NA | NA | NA | NA |
| lizards | Hemiphyllodactylus_kiziriani | Hemiphyllodactylus_kiziriani | LC | Stable | tropical | nocturnal | 1.622 | 12.512 | 13.814 |
| lizards | Hemiphyllodactylus_larutensis | NA | DD | Unknown | NA | NA | NA | NA | NA |
| lizards | Hemiphyllodactylus_margarethae | NA | DD | Unknown | NA | NA | NA | NA | NA |

| | | | | | | | | | |
|---|---|---|---|---|---|---|---|---|---|
| lizards | Hemiphyllodactylus_montawaensis | NA | not_available | not_available | NA | NA | NA | NA | NA |
| lizards | Hemiphyllodactylus_tehtarik | Hemiphyllodactylus_tehtarik | LC | Unknown | tropical | nocturnal | 1.549 | 9.991 | 3.869 |
| lizards | Hemiphyllodactylus_titiwangsaensis | NA | EN | Unknown | NA | NA | NA | NA | NA |
| lizards | Hemiphyllodactylus_typus | Hemiphyllodactylus_typus | LC | Stable | tropical | nocturnal | 4.677 | 2660225.563 | 10.017 |
| lizards | Hemiphyllodactylus_yunnanensis | NA | LC | Unknown | NA | NA | NA | NA | NA |
| lizards | Hemiphyllodactylus_zugi | NA | DD | Stable | NA | NA | NA | NA | NA |
| lizards | Hemitheconyx_caudicinctus | Hemitheconyx_caudicinctus | LC | Unknown | tropical | nocturnal | 93.325 | 1712974.329 | 13.382 |
| lizards | Hemitheconyx_taylori | Hemitheconyx_taylori | LC | Unknown | tropical | nocturnal | 56.234 | 79546.174 | 8.681 |
| lizards | Heremites_auratus | Trachylepis_aurata | LC | Stable | temperate | diurnal | 33.884 | 554428.114 | 13.984 |
| lizards | Heremites_septemtaeniatus | Trachylepis_septemtaeniata | LC | Stable | temperate | diurnal | 25.119 | 1364784.598 | 12.156 |
| lizards | Heremites_vittatus | Trachylepis_vittata | LC | Stable | temperate | cathemeral | 20.893 | 1163294.204 | 14.633 |
| lizards | Hesperoedura_reticulata | Hesperoedura_reticulata | LC | Stable | temperate | nocturnal | 7.244 | 291832.366 | 6.894 |
| lizards | Heteronotia_atra | Heteronotia_atra | LC | Unknown | tropical | nocturnal | 5.248 | 2236.829 | 3.783 |
| lizards | Heteronotia_binoei | Heteronotia_binoei | LC | Stable | tropical | nocturnal | 3.631 | 6898166.198 | 3.593 |
| lizards | Heteronotia_fasciolata | NA | LC | Stable | NA | NA | NA | NA | NA |
| lizards | Heteronotia_planiceps | Heteronotia_planiceps | LC | Stable | tropical | nocturnal | 3.631 | 226325.883 | 2.777 |
| lizards | Heteronotia_spelea | Heteronotia_spelea | LC | Stable | tropical | nocturnal | 3.802 | 196701.972 | 4.452 |
| lizards | Holaspis_guentheri | Holaspis_guentheri | LC | Unknown | tropical | diurnal | 3.548 | 2080975.373 | 8.623 |
| lizards | Holaspis_laevis | Holaspis_laevis | LC | Unknown | tropical | diurnal | 2.754 | 546758.093 | 10.223 |
| lizards | Holbrookia_approximans | NA | not_available | not_available | NA | NA | NA | NA | NA |
| lizards | Holbrookia_elegans | Holbrookia_elegans | LC | Stable | temperate | diurnal | 14.454 | 188396.537 | 7.751 |
| lizards | Holbrookia_lacerata | Holbrookia_lacerata | NT | Decreasing | temperate | diurnal | 12.589 | 267337.579 | 12.806 |
| lizards | Holbrookia_maculata | Holbrookia_maculata | LC | Stable | temperate | diurnal | 14.454 | 1870599.631 | 10.154 |
| lizards | Holbrookia_propinqua | Holbrookia_propinqua | LC | Stable | temperate | diurnal | 8.71 | 124685.745 | 16.281 |
| lizards | Holcosus_anomalus | NA | LC | Unknown | NA | NA | NA | NA | NA |
| lizards | Holcosus_bridgesii | NA | LC | Unknown | NA | NA | NA | NA | NA |
| lizards | Holcosus_chaitzami | NA | DD | Unknown | NA | NA | NA | NA | NA |
| lizards | Holcosus_festivus | Holcosus_festivus | LC | Stable | tropical | diurnal | 93.325 | 455421.274 | 9.682 |
| lizards | Holcosus_leptophrys | NA | LC | Stable | NA | NA | NA | NA | NA |
| lizards | Holcosus_niceforoi | Holcosus_niceforoi | LC | Unknown | tropical | diurnal | 15.849 | 4698.205 | 14.781 |
| lizards | Holcosus_orcesi | NA | CR | Unknown | NA | NA | NA | NA | NA |
| lizards | Holcosus_quadrilineatus | Holcosus_quadrilineatus | LC | Unknown | tropical | diurnal | 19.953 | 41564.399 | 10.977 |
| lizards | Holcosus_septemlineatus | NA | LC | Stable | NA | NA | NA | NA | NA |
| lizards | Holcosus_undulatus | Holcosus_undulatus | LC | Stable | tropical | diurnal | 81.283 | 909409.794 | 11.45 |

| | | | | | | | | | |
|---|---|---|---|---|---|---|---|---|---|
| lizards | Holodactylus_africanus | Holodactylus_africanus | LC | Unknown | tropical | nocturnal | 12.023 | 325908.444 | 8.996 |
| lizards | Holodactylus_cornii | NA | DD | Unknown | NA | NA | NA | NA | NA |
| lizards | Homonota_andicola | NA | LC | Unknown | NA | NA | NA | NA | NA |
| lizards | Homonota_borellii | Homonota_borellii | LC | Stable | temperate | nocturnal | 1.995 | 475244.793 | 6.889 |
| lizards | Homonota_darwinii | Homonota_darwinii | LC | Stable | temperate | nocturnal | 4.365 | 656478.512 | 5.333 |
| lizards | Homonota_fasciata | NA | LC | Unknown | NA | NA | NA | NA | NA |
| lizards | Homonota_horrida | NA | not_available | not_available | NA | NA | NA | NA | NA |
| lizards | Homonota_rupicola | NA | CR | Unknown | NA | NA | NA | NA | NA |
| lizards | Homonota_septentrionalis | NA | not_available | not_available | NA | NA | NA | NA | NA |
| lizards | Homonota_taragui | Homonota_taragui | CR | Unknown | temperate | nocturnal | 1.95 | 101.464 | 7.099 |
| lizards | Homonota_underwoodi | Homonota_underwoodi | LC | Stable | temperate | nocturnal | 4.266 | 395255.541 | 4.686 |
| lizards | Homonota_uruguayensis | Homonota_uruguayensis | EN | Decreasing | temperate | cathemeral | 3.311 | 167811.586 | 9.958 |
| lizards | Homonota_whitii | Homonota_whitii | LC | Stable | temperate | nocturnal | 2.138 | 84524.977 | 7.891 |
| lizards | Homonota_williamsii | NA | LC | Stable | NA | NA | NA | NA | NA |
| lizards | Homopholis_arnoldi | NA | LC | Stable | NA | NA | NA | NA | NA |
| lizards | Homopholis_fasciata | Homopholis_fasciata | LC | Unknown | tropical | nocturnal | 10.965 | 1188680.912 | 10.354 |
| lizards | Homopholis_mulleri | Homopholis_mulleri | NT | Stable | tropical | nocturnal | 10.715 | 15811.905 | 9.054 |
| lizards | Homopholis_walbergii | Homopholis_walbergii | LC | Stable | temperate | cathemeral | 33.884 | 330104.022 | 10.404 |
| lizards | Hoplocercus_spinosus | Hoplocercus_spinosus | LC | Unknown | tropical | diurnal | 33.113 | 2139549.623 | 6.215 |
| lizards | Hoplodactylus_duvaucelii | Hoplodactylus_duvaucelii | NT | Stable | temperate | nocturnal | 93.325 | 86164.498 | 9.074 |
| lizards | Hyalosaurus_koellikeri | NA | LC | Unknown | NA | NA | NA | NA | NA |
| lizards | Hydrosaurus_amboinensis | Hydrosaurus_amboinensis | LC | Unknown | tropical | diurnal | 1621.81 | 165540.29 | 4.519 |
| lizards | Hydrosaurus_pustulatus | Hydrosaurus_pustulatus | LC | Decreasing | tropical | diurnal | 812.831 | 160747.093 | 15.142 |
| lizards | Hydrosaurus_weberi | NA | VU | Decreasing | NA | NA | NA | NA | NA |
| lizards | Hypsicalotes_kinabaluensis | NA | DD | Unknown | NA | NA | NA | NA | NA |
| lizards | Hypsilurus_auritus | NA | LC | Unknown | NA | NA | NA | NA | NA |
| lizards | Hypsilurus_binotatus | NA | LC | Unknown | NA | NA | NA | NA | NA |
| lizards | Hypsilurus_godeffroyi | NA | DD | Unknown | NA | NA | NA | NA | NA |
| lizards | Hypsilurus_hikidanus | NA | LC | Unknown | NA | NA | NA | NA | NA |
| lizards | Hypsilurus_macrolepis | Hypsilurus_macrolepis | NT | Stable | tropical | diurnal | 95.499 | 15623.516 | 5.088 |
| lizards | Hypsilurus_modestus | Hypsilurus_modestus | LC | Stable | tropical | diurnal | 41.687 | 653605.492 | 3.183 |
| lizards | Hypsilurus_schoedei | Hypsilurus_schoedei | LC | Unknown | tropical | diurnal | 72.444 | 451.144 | 4.03 |
| lizards | Hypsilurus_schultzewestrumi | NA | LC | Unknown | NA | NA | NA | NA | NA |
| lizards | Hypsilurus_tenuicephalus | NA | DD | Unknown | NA | NA | NA | NA | NA |

| | | | | | | | | | |
|---|---|---|---|---|---|---|---|---|---|
| lizards | Iberolacerta_aranica | Iberolacerta_aranica | EN | Decreasing | temperate | diurnal | 6.918 | 59.947 | 7.81 |
| lizards | Iberolacerta_aurelioi | Iberolacerta_aurelioi | EN | Decreasing | temperate | diurnal | 6.457 | 122.446 | 6.762 |
| lizards | Iberolacerta_bonnali | Iberolacerta_bonnali | NT | Stable | temperate | diurnal | 6.457 | 1583.09 | 7.583 |
| lizards | Iberolacerta_cyreni | Iberolacerta_cyreni | EN | Decreasing | temperate | diurnal | 17.378 | 6606.261 | 14.512 |
| lizards | Iberolacerta_galani | Iberolacerta_galani | NT | Unknown | temperate | diurnal | 14.125 | 858.908 | 6.973 |
| lizards | Iberolacerta_horvathi | Iberolacerta_horvathi | NT | Stable | temperate | diurnal | 6.457 | 27101.731 | 14.889 |
| lizards | Iberolacerta_martinezricai | Iberolacerta_martinezricai | CR | Decreasing | temperate | diurnal | 7.762 | 248.026 | 10.479 |
| lizards | Iberolacerta_monticola | Iberolacerta_monticola | VU | Decreasing | temperate | diurnal | 13.804 | 31589.402 | 12.078 |
| lizards | Ichnotropis_bivittata | NA | LC | Stable | NA | NA | NA | NA | NA |
| lizards | Ichnotropis_capensis | Ichnotropis_capensis | LC | Stable | tropical | diurnal | 7.244 | 4125084.817 | 7.418 |
| lizards | Ichnotropis_grandiceps | Ichnotropis_grandiceps | LC | Unknown | tropical | diurnal | 7.943 | 93129.93 | 4.798 |
| lizards | Iguana_delicatissima | Iguana_delicatissima | CR | Decreasing | tropical | diurnal | 3467.369 | 4509.858 | 22.77 |
| lizards | Iguana_iguana | Iguana_iguana | LC | Unknown | tropical | diurnal | 8128.305 | 12236032.357 | 5.27 |
| lizards | Insulasaurus_arborens | Insulasaurus_arborens | LC | Unknown | tropical | diurnal | 5.754 | 28606.123 | 17.642 |
| lizards | Intellagama_lesueurii | Intellagama_lesueurii | LC | Stable | temperate | diurnal | 912.011 | 834282.101 | 8.036 |
| lizards | Iphisa_elegans | Iphisa_elegans | LC | Stable | tropical | diurnal | 5.754 | 6181116.342 | 1.989 |
| lizards | Iranolacerta_brandtii | NA | DD | Unknown | NA | NA | NA | NA | NA |
| lizards | Iranolacerta_zagrosica | NA | LC | Stable | NA | NA | NA | NA | NA |
| lizards | Isopachys_anguinoides | NA | LC | Unknown | NA | NA | NA | NA | NA |
| lizards | Isopachys_gyldenstolpei | Isopachys_gyldenstolpei | LC | Stable | tropical | cathemeral | 15.136 | 8617.875 | 19.251 |
| lizards | Janetaescincus_braueri | NA | EN | Decreasing | NA | NA | NA | NA | NA |
| lizards | Janetaescincus_veseyfitzgeraldi | NA | EN | Decreasing | NA | NA | NA | NA | NA |
| lizards | Japalura_andersoniana | Japalura_andersoniana | LC | Stable | temperate | diurnal | 13.804 | 66509.273 | 4.308 |
| lizards | Japalura_austeniana | NA | LC | Unknown | NA | NA | NA | NA | NA |
| lizards | Japalura_dasi | NA | VU | Unknown | NA | NA | NA | NA | NA |
| lizards | Japalura_kumaonensis | Japalura_kumaonensis | NT | Unknown | temperate | diurnal | 11.22 | 136935.076 | 13.394 |
| lizards | Japalura_major | Japalura_major | LC | Unknown | temperate | diurnal | 27.542 | 102211.666 | 8.772 |
| lizards | Japalura_sagittifera | NA | DD | Unknown | NA | NA | NA | NA | NA |
| lizards | Japalura_tricarinata | NA | LC | Stable | NA | NA | NA | NA | NA |
| lizards | Japalura_variegata | Japalura_variegata | LC | Unknown | temperate | diurnal | 61.66 | 39870.609 | 15.153 |
| lizards | Kaestlea_beddomii | NA | LC | Unknown | NA | NA | NA | NA | NA |
| lizards | Kaestlea_bilineata | NA | LC | Unknown | NA | NA | NA | NA | NA |
| lizards | Kaestlea_laterimaculata | NA | VU | Unknown | NA | NA | NA | NA | NA |
| lizards | Kaestlea_palnica | NA | DD | Unknown | NA | NA | NA | NA | NA |

| | | | | | | | | | |
|---|---|---|---|---|---|---|---|---|---|
| lizards | Kaestlea_travancorica | NA | LC | Unknown | NA | NA | NA | NA | NA |
| lizards | Kanakysaurus_viviparus | Kanakysaurus_viviparus | EN | Decreasing | tropical | cathemeral | 11.749 | 296.515 | 9.395 |
| lizards | Kanakysaurus_zebratus | NA | EN | Decreasing | NA | NA | NA | NA | NA |
| lizards | Karusasaurus_jordani | Karusasaurus_jordani | LC | Stable | temperate | diurnal | 63.096 | 213618.208 | 5.081 |
| lizards | Karusasaurus_polyzonus | Karusasaurus_polyzonus | LC | Stable | temperate | diurnal | 45.709 | 754298.666 | 7.495 |
| lizards | Kentropyx_altamazonica | Kentropyx_altamazonica | LC | Stable | tropical | diurnal | 44.668 | 3922337.694 | 2.065 |
| lizards | Kentropyx_borckiana | Kentropyx_borckiana | LC | Unknown | tropical | diurnal | 30.903 | 211722.645 | 2.398 |
| lizards | Kentropyx_calcarata | Kentropyx_calcarata | LC | Stable | tropical | diurnal | 51.286 | 6454200.008 | 3.311 |
| lizards | Kentropyx_lagartija | Kentropyx_lagartija | LC | Stable | temperate | diurnal | 19.055 | 27136.294 | 7.375 |
| lizards | Kentropyx_paulensis | Kentropyx_paulensis | LC | Unknown | tropical | diurnal | 13.183 | 1157731.095 | 11.445 |
| lizards | Kentropyx_pelviceps | Kentropyx_pelviceps | LC | Stable | tropical | diurnal | 67.608 | 2576988.516 | 1.842 |
| lizards | Kentropyx_striata | Kentropyx_striata | LC | Stable | tropical | diurnal | 63.096 | 2096781.336 | 2.291 |
| lizards | Kentropyx_vanzoi | Kentropyx_vanzoi | NT | Unknown | tropical | diurnal | 17.783 | 1399763.633 | 5.864 |
| lizards | Kentropyx_viridistriga | Kentropyx_viridistriga | LC | Unknown | tropical | diurnal | 36.308 | 1073309.004 | 6.548 |
| lizards | Kinyongia_adolfifriderici | Kinyongia_adolfifriderici | LC | Unknown | tropical | diurnal | 7.586 | 124018.616 | 12.003 |
| lizards | Kinyongia_asheorum | NA | NT | Unknown | NA | NA | NA | NA | NA |
| lizards | Kinyongia_boehmei | Kinyongia_boehmei | NT | Unknown | tropical | diurnal | 21.38 | 438.391 | 18.979 |
| lizards | Kinyongia_excubitor | Kinyongia_excubitor | VU | Stable | tropical | diurnal | 11.482 | 5340.724 | 11.763 |
| lizards | Kinyongia_fischeri | Kinyongia_fischeri | NT | Unknown | tropical | diurnal | 147.911 | 256.658 | 7.588 |
| lizards | Kinyongia_itombwensis | NA | not_available | not_available | NA | NA | NA | NA | NA |
| lizards | Kinyongia_oxyrhina | Kinyongia_oxyrhina | NT | Stable | tropical | diurnal | 9.55 | 12394.663 | 9.929 |
| lizards | Kinyongia_rugegensis | NA | not_available | not_available | NA | NA | NA | NA | NA |
| lizards | Kinyongia_tavetana | Kinyongia_tavetana | NT | Decreasing | tropical | diurnal | 21.38 | 10491.706 | 16.316 |
| lizards | Kinyongia_tenuis | Kinyongia_tenuis | EN | Unknown | tropical | diurnal | 8.913 | 28982.47 | 12.05 |
| lizards | Kinyongia_tolleyae | NA | not_available | not_available | NA | NA | NA | NA | NA |
| lizards | Kinyongia_uluguruensis | Kinyongia_uluguruensis | LC | Stable | tropical | diurnal | 19.055 | 878.744 | 12.253 |
| lizards | Kinyongia_uthmoelleri | Kinyongia_uthmoelleri | LC | Stable | tropical | diurnal | 19.055 | 23853.503 | 11.812 |
| lizards | Kolekanos_plumicauda | NA | LC | Stable | NA | NA | NA | NA | NA |
| lizards | Lacerta_agilis | Lacerta_agilis | LC | Stable | temperate | diurnal | 33.884 | 11148919.361 | 11.623 |
| lizards | Lacerta_bilineata | Lacerta_bilineata | LC | Decreasing | temperate | diurnal | 50.119 | 856006.706 | 21.745 |
| lizards | Lacerta_media | Lacerta_media | LC | Decreasing | temperate | diurnal | 91.201 | 927506.602 | 15.644 |
| lizards | Lacerta_pamphylica | Lacerta_pamphylica | LC | Stable | temperate | diurnal | 38.905 | 9661.298 | 13.028 |
| lizards | Lacerta_schreiberi | Lacerta_schreiberi | NT | Decreasing | temperate | diurnal | 75.858 | 145400.973 | 17.089 |
| lizards | Lacerta_strigata | Lacerta_strigata | LC | Decreasing | temperate | diurnal | 91.201 | 415809.47 | 14.44 |

| | | | | | | | | | |
|---|---|---|---|---|---|---|---|---|---|
| lizards | Lacerta_trilineata | Lacerta_trilineata | LC | Stable | temperate | diurnal | 117.49 | 596692.931 | 16.397 |
| lizards | Lacerta_viridis | Lacerta_viridis | LC | Decreasing | temperate | diurnal | 91.201 | 1369557.619 | 18.507 |
| lizards | Lacertaspis_chriswildi | NA | NT | Decreasing | NA | NA | NA | NA | NA |
| lizards | Lacertaspis_gemmiventris | NA | EN | Decreasing | NA | NA | NA | NA | NA |
| lizards | Lacertaspis_lepesmei | Lacertaspis_lepesmei | CR | Stable | tropical | diurnal | 3.715 | 146.501 | 14.446 |
| lizards | Lacertaspis_reichenowii | NA | LC | Stable | NA | NA | NA | NA | NA |
| lizards | Lacertaspis_rohdei | NA | LC | Stable | NA | NA | NA | NA | NA |
| lizards | Lacertoides_pardalis | Lacertoides_pardalis | VU | Decreasing | tropical | diurnal | 54.954 | 375.068 | 9.054 |
| lizards | Laemanctus_longipes | Laemanctus_longipes | LC | Unknown | tropical | diurnal | 104.713 | 177694.711 | 11.365 |
| lizards | Laemanctus_serratus | Laemanctus_serratus | LC | Stable | tropical | cathemeral | 229.087 | 208774.874 | 10.383 |
| lizards | Lamprolepis_leucosticta | NA | DD | Unknown | NA | NA | NA | NA | NA |
| lizards | Lamprolepis_nieuwenhuisii | NA | LC | Unknown | NA | NA | NA | NA | NA |
| lizards | Lamprolepis_smaragdina | Lamprolepis_smaragdina | LC | Stable | tropical | diurnal | 40.738 | 892245.134 | 9.349 |
| lizards | Lampropholis_adonis | Lampropholis_adonis | LC | Stable | temperate | diurnal | 2.754 | 43556.934 | 6.315 |
| lizards | Lampropholis_amicula | Lampropholis_amicula | LC | Stable | temperate | diurnal | 0.724 | 105352.544 | 8.011 |
| lizards | Lampropholis_bellendenkerensis | NA | not_available | not_available | NA | NA | NA | NA | NA |
| lizards | Lampropholis_caligula | Lampropholis_caligula | LC | Unknown | temperate | diurnal | 2.951 | 3501.025 | 4.873 |
| lizards | Lampropholis_coggeri | Lampropholis_coggeri | LC | Stable | tropical | diurnal | 1.622 | 5206.969 | 4.909 |
| lizards | Lampropholis_colossus | NA | DD | Unknown | NA | NA | NA | NA | NA |
| lizards | Lampropholis_couperi | NA | LC | Stable | NA | NA | NA | NA | NA |
| lizards | Lampropholis_delicata | Lampropholis_delicata | LC | Stable | temperate | diurnal | 3.09 | 636433.848 | 7.82 |
| lizards | Lampropholis_elliotensis | NA | not_available | not_available | NA | NA | NA | NA | NA |
| lizards | Lampropholis_elongata | NA | DD | Unknown | NA | NA | NA | NA | NA |
| lizards | Lampropholis_guichenoti | Lampropholis_guichenoti | LC | Unknown | temperate | diurnal | 2.63 | 688423.805 | 10.499 |
| lizards | Lampropholis_mirabilis | Lampropholis_mirabilis | LC | Unknown | tropical | diurnal | 2.63 | 9.991 | 0.39 |
| lizards | Lampropholis_robertsi | Lampropholis_robertsi | LC | Unknown | tropical | diurnal | 3.09 | 5902.826 | 5.58 |
| lizards | Lampropholis_similis | NA | not_available | not_available | NA | NA | NA | NA | NA |
| lizards | Lankascincus_deignani | Lankascincus_deignani | CR | Unknown | tropical | diurnal | 3.715 | 13434.623 | 16.982 |
| lizards | Lankascincus_dorsicatenatus | NA | LC | Stable | NA | NA | NA | NA | NA |
| lizards | Lankascincus_fallax | Lankascincus_fallax | LC | Stable | tropical | diurnal | 4.169 | 65499.276 | 17.597 |
| lizards | Lankascincus_gansi | Lankascincus_gansi | LC | Stable | tropical | diurnal | 3.09 | 20573.044 | 19.622 |
| lizards | Lankascincus_merrill | NA | not_available | not_available | NA | NA | NA | NA | NA |
| lizards | Lankascincus_sameerai | NA | not_available | not_available | NA | NA | NA | NA | NA |
| lizards | Lankascincus_sripadensis | Lankascincus_sripadensis | EN | Unknown | tropical | diurnal | 3.802 | 9.991 | 9.523 |

| | | | | | | | | | |
|---|---|---|---|---|---|---|---|---|---|
| lizards | Lankascincus_taprobanensis | Lankascincus_taprobanensis | EN | Decreasing | tropical | diurnal | 3.715 | 13064.495 | 16.997 |
| lizards | Lankascincus_taylori | Lankascincus_taylori | VU | Unknown | tropical | diurnal | 2.63 | 19983.826 | 19.638 |
| lizards | Lanthanotus_borneensis | Lanthanotus_borneensis | EN | Decreasing | tropical | nocturnal | 204.174 | 45789.228 | 5.978 |
| lizards | Larutia_larutensis | Larutia_larutensis | VU | Unknown | tropical | diurnal | 26.915 | 19.982 | 4.331 |
| lizards | Larutia_miodactyla | Larutia_miodactyla | VU | Unknown | tropical | diurnal | 15.136 | 6793 | 9.261 |
| lizards | Larutia_nubisilvicola | NA | LC | Unknown | NA | NA | NA | NA | NA |
| lizards | Larutia_seribuatensis | NA | NT | Stable | NA | NA | NA | NA | NA |
| lizards | Larutia_trifasciata | Larutia_trifasciata | NT | Stable | tropical | diurnal | 52.481 | 3203.049 | 7.808 |
| lizards | Latastia_doriai | NA | LC | Unknown | NA | NA | NA | NA | NA |
| lizards | Latastia_johnstonii | Latastia_johnstonii | LC | Unknown | tropical | diurnal | 6.166 | 411744.379 | 9.579 |
| lizards | Latastia_longicaudata | Latastia_longicaudata | LC | Stable | tropical | diurnal | 30.2 | 6995048.782 | 8.314 |
| lizards | Laudakia_agrorensis | Laudakia_agrorensis | LC | Unknown | temperate | diurnal | 48.978 | 377167.679 | 13.603 |
| lizards | Laudakia_dayana | NA | DD | Unknown | NA | NA | NA | NA | NA |
| lizards | Laudakia_melanura | Laudakia_melanura | LC | Stable | temperate | diurnal | 107.152 | 893887.523 | 9.626 |
| lizards | Laudakia_nupta | Laudakia_nupta | LC | Stable | temperate | diurnal | 181.97 | 1524079.157 | 8.7 |
| lizards | Laudakia_nuristanica | NA | DD | Unknown | NA | NA | NA | NA | NA |
| lizards | Laudakia_pakistanica | NA | LC | Stable | NA | NA | NA | NA | NA |
| lizards | Laudakia_papenfussi | NA | LC | Unknown | NA | NA | NA | NA | NA |
| lizards | Laudakia_sacra | NA | LC | Stable | NA | NA | NA | NA | NA |
| lizards | Laudakia_stellio | NA | LC | Stable | NA | NA | NA | NA | NA |
| lizards | Laudakia_tuberculata | Laudakia_tuberculata | LC | Stable | temperate | diurnal | 117.49 | 459162.023 | 9.836 |
| lizards | Laudakia_wui | NA | LC | Stable | NA | NA | NA | NA | NA |
| lizards | Leiocephalus_barahonensis | Leiocephalus_barahonensis | LC | Stable | tropical | diurnal | 16.218 | 4140.885 | 10.421 |
| lizards | Leiocephalus_carinatus | Leiocephalus_carinatus | LC | Stable | tropical | diurnal | 72.444 | 124368.487 | 15.194 |
| lizards | Leiocephalus_cubensis | Leiocephalus_cubensis | LC | Stable | tropical | diurnal | 52.481 | 55970.769 | 16.522 |
| lizards | Leiocephalus_cuneus | NA | EX | not_available | NA | NA | NA | NA | NA |
| lizards | Leiocephalus_greenwayi | NA | VU | Unknown | NA | NA | NA | NA | NA |
| lizards | Leiocephalus_inaguae | Leiocephalus_inaguae | LC | Stable | tropical | diurnal | 19.953 | 1581.958 | 5.317 |
| lizards | Leiocephalus_loxogrammus | NA | LC | Stable | NA | NA | NA | NA | NA |
| lizards | Leiocephalus_lunatus | Leiocephalus_lunatus | LC | Stable | tropical | diurnal | 7.762 | 943.969 | 20.013 |
| lizards | Leiocephalus_macropus | Leiocephalus_macropus | LC | Stable | tropical | diurnal | 23.988 | 69312.199 | 16.182 |
| lizards | Leiocephalus_melanochlorus | Leiocephalus_melanochlorus | NT | Unknown | tropical | diurnal | 66.069 | 6510.147 | 18.225 |
| lizards | Leiocephalus_onaneyi | Leiocephalus_onaneyi | CR | Unknown | tropical | diurnal | 10.233 | 14.697 | 13.964 |
| lizards | Leiocephalus_personatus | Leiocephalus_personatus | LC | Stable | tropical | diurnal | 36.308 | 50792.402 | 16.349 |

| | | | | | | | | | |
|---|---|---|---|---|---|---|---|---|---|
| lizards | Leiocephalus_psammodromus | Leiocephalus_psammodromus | VU | Decreasing | tropical | diurnal | 38.905 | 305.164 | 15.66 |
| lizards | Leiocephalus_raviceps | Leiocephalus_raviceps | LC | Stable | tropical | diurnal | 10.715 | 12074.383 | 17.343 |
| lizards | Leiocephalus_schreibersii | Leiocephalus_schreibersii | LC | Unknown | tropical | diurnal | 35.481 | 29792.503 | 14.916 |
| lizards | Leiocephalus_semilineatus | Leiocephalus_semilineatus | LC | Unknown | tropical | diurnal | 3.631 | 6069.81 | 14.138 |
| lizards | Leiocephalus_sixtoi | NA | not_available | not_available | NA | NA | NA | NA | NA |
| lizards | Leiocephalus_stictigaster | Leiocephalus_stictigaster | LC | Stable | tropical | diurnal | 28.184 | 44480.29 | 16.807 |
| lizards | Leiocephalus_varius | NA | EN | Decreasing | NA | NA | NA | NA | NA |
| lizards | Leiocephalus_vinculum | NA | EN | Unknown | NA | NA | NA | NA | NA |
| lizards | Leiolepis_belliana | Leiolepis_belliana | LC | Stable | tropical | diurnal | 199.526 | 819785.379 | 13.676 |
| lizards | Leiolepis_boehmei | NA | VU | Decreasing | NA | NA | NA | NA | NA |
| lizards | Leiolepis_guentherpetersi | Leiolepis_guentherpetersi | EN | Decreasing | tropical | diurnal | 131.826 | 4446.029 | 21.684 |
| lizards | Leiolepis_guttata | Leiolepis_guttata | DD | Decreasing | tropical | diurnal | 575.44 | 40549.257 | 15.958 |
| lizards | Leiolepis_ngovantrii | NA | VU | Unknown | NA | NA | NA | NA | NA |
| lizards | Leiolepis_peguensis | NA | LC | Unknown | NA | NA | NA | NA | NA |
| lizards | Leiolepis_reevesii | Leiolepis_reevesii | LC | Decreasing | tropical | diurnal | 245.471 | 446691.991 | 16.512 |
| lizards | Leiolepis_rubritaeniata | NA | LC | Stable | NA | NA | NA | NA | NA |
| lizards | Leiolepis_triploida | Leiolepis_triploida | DD | Increasing | tropical | diurnal | 112.202 | 9365.465 | 23.356 |
| lizards | Leiolopisma_alazon | NA | CR | Decreasing | NA | NA | NA | NA | NA |
| lizards | Leiolopisma_telfairii | NA | VU | Increasing | NA | NA | NA | NA | NA |
| lizards | Leiosaurus_bellii | Leiosaurus_bellii | LC | Stable | temperate | diurnal | 41.687 | 628194.606 | 5.375 |
| lizards | Leiosaurus_catamarcensis | Leiosaurus_catamarcensis | LC | Unknown | temperate | diurnal | 56.234 | 179019.493 | 4.596 |
| lizards | Leiosaurus_jaguaris | NA | LC | Stable | NA | NA | NA | NA | NA |
| lizards | Leiosaurus_paronae | NA | DD | Unknown | NA | NA | NA | NA | NA |
| lizards | Lepidoblepharis_buchwaldi | NA | LC | Stable | NA | NA | NA | NA | NA |
| lizards | Lepidoblepharis_conolepis | NA | EN | Unknown | NA | NA | NA | NA | NA |
| lizards | Lepidoblepharis_festae | NA | LC | Unknown | NA | NA | NA | NA | NA |
| lizards | Lepidoblepharis_grandis | NA | VU | Unknown | NA | NA | NA | NA | NA |
| lizards | Lepidoblepharis_heyerorum | Lepidoblepharis_heyerorum | LC | Unknown | tropical | cathemeral | 1.072 | 2547613.74 | 2.128 |
| lizards | Lepidoblepharis_intermedius | NA | LC | Stable | NA | NA | NA | NA | NA |
| lizards | Lepidoblepharis_miyatai | NA | CR | Unknown | NA | NA | NA | NA | NA |
| lizards | Lepidoblepharis_nukak | NA | not_available | not_available | NA | NA | NA | NA | NA |
| lizards | Lepidoblepharis_peraccae | NA | LC | Stable | NA | NA | NA | NA | NA |
| lizards | Lepidoblepharis_ruthveni | NA | LC | Unknown | NA | NA | NA | NA | NA |
| lizards | Lepidoblepharis_sanctaemartae | Lepidoblepharis_sanctaemartae | LC | Unknown | tropical | diurnal | 1.072 | 103968.006 | 14.414 |

| | | | | | | | | | |
|---|---|---|---|---|---|---|---|---|---|
| lizards | Lepidoblepharis_xanthostigma | Lepidoblepharis_xanthostigma | LC | Stable | tropical | diurnal | 2.239 | 192457.926 | 11.204 |
| lizards | Lepidodactylus_aureolineatus | NA | LC | Stable | NA | NA | NA | NA | NA |
| lizards | Lepidodactylus_balioburius | NA | LC | Stable | NA | NA | NA | NA | NA |
| lizards | Lepidodactylus_buleli | NA | DD | Unknown | NA | NA | NA | NA | NA |
| lizards | Lepidodactylus_christiani | NA | LC | Stable | NA | NA | NA | NA | NA |
| lizards | Lepidodactylus_euaensis | NA | CR | Stable | NA | NA | NA | NA | NA |
| lizards | Lepidodactylus_gardineri | NA | CR | Decreasing | NA | NA | NA | NA | NA |
| lizards | Lepidodactylus_guppyi | Lepidodactylus_guppyi | LC | Unknown | tropical | nocturnal | 3.548 | 36047.369 | 4.992 |
| lizards | Lepidodactylus_herrei | NA | LC | Stable | NA | NA | NA | NA | NA |
| lizards | Lepidodactylus_labialis | NA | DD | Unknown | NA | NA | NA | NA | NA |
| lizards | Lepidodactylus_listeri | NA | EW | not_available | NA | NA | NA | NA | NA |
| lizards | Lepidodactylus_lombocensis | NA | DD | Unknown | NA | NA | NA | NA | NA |
| lizards | Lepidodactylus_lugubris | Lepidodactylus_lugubris | LC | Unknown | tropical | nocturnal | 2.951 | 7666582.838 | 12.51 |
| lizards | Lepidodactylus_magnus | Lepidodactylus_magnus | LC | Unknown | tropical | nocturnal | 7.413 | 30113.204 | 5.574 |
| lizards | Lepidodactylus_manni | Lepidodactylus_manni | VU | Unknown | tropical | nocturnal | 2.512 | 16606.327 | 9.364 |
| lizards | Lepidodactylus_moestus | Lepidodactylus_moestus | LC | Stable | tropical | nocturnal | 2.089 | 893.465 | 11.606 |
| lizards | Lepidodactylus_oligoporus | NA | CR | Unknown | NA | NA | NA | NA | NA |
| lizards | Lepidodactylus_pantai | NA | DD | Unknown | NA | NA | NA | NA | NA |
| lizards | Lepidodactylus_paurolepis | NA | VU | Unknown | NA | NA | NA | NA | NA |
| lizards | Lepidodactylus_planicauda | NA | LC | Stable | NA | NA | NA | NA | NA |
| lizards | Lepidodactylus_pumilus | Lepidodactylus_pumilus | LC | Unknown | tropical | nocturnal | 2.512 | 17402.975 | 2.95 |
| lizards | Lepidodactylus_ranauensis | NA | DD | Unknown | NA | NA | NA | NA | NA |
| lizards | Lepidodactylus_vanuatuensis | Lepidodactylus_vanuatuensis | LC | Stable | tropical | nocturnal | 2.399 | 4769.355 | 6.249 |
| lizards | Lepidodactylus_woodfordi | NA | not_available | not_available | NA | NA | NA | NA | NA |
| lizards | Lepidodactylus_yami | Lepidodactylus_yami | NT | Stable | tropical | nocturnal | 1.738 | 49.72 | 13.386 |
| lizards | Lepidophyma_flavimaculatum | Lepidophyma_flavimaculatum | LC | Unknown | tropical | cathemeral | 72.444 | 311582.871 | 10.059 |
| lizards | Lepidophyma_gaigeae | Lepidophyma_gaigeae | VU | Decreasing | tropical | nocturnal | 5.754 | 6382.132 | 8.83 |
| lizards | Lepidophyma_mayae | NA | NT | Unknown | NA | NA | NA | NA | NA |
| lizards | Lepidophyma_micropholis | Lepidophyma_micropholis | VU | Unknown | tropical | cathemeral | 27.542 | 4050.182 | 12.886 |
| lizards | Lepidophyma_occulor | NA | LC | Stable | NA | NA | NA | NA | NA |
| lizards | Lepidophyma_pajapanensis | Lepidophyma_pajapanensis | LC | Stable | tropical | nocturnal | 16.982 | 1855.656 | 11.759 |
| lizards | Lepidophyma_smithii | Lepidophyma_smithii | LC | Stable | tropical | cathemeral | 30.903 | 52342.598 | 13.331 |
| lizards | Lepidophyma_tarascae | NA | DD | Unknown | NA | NA | NA | NA | NA |
| lizards | Lepidophyma_tuxtlae | Lepidophyma_tuxtlae | DD | Decreasing | tropical | nocturnal | 18.197 | 19919.385 | 14.252 |

| | | | | | | | | | |
|---|---|---|---|---|---|---|---|---|---|
| lizards | Leposoma_puk | NA | EN | Unknown | NA | NA | NA | NA | NA |
| lizards | Leposternon_polystegum | Amphisbaena_polystegum | LC | Unknown | tropical | diurnal | 114.815 | 1986716.106 | 5.263 |
| lizards | Leptosiaphos_aloysiisabaudiae | NA | LC | Unknown | NA | NA | NA | NA | NA |
| lizards | Leptosiaphos_blochmanni | NA | DD | Unknown | NA | NA | NA | NA | NA |
| lizards | Leptosiaphos_graueri | Leptosiaphos_graueri | EN | Unknown | tropical | diurnal | 2.692 | 80181.431 | 15.699 |
| lizards | Leptosiaphos_hackarsi | NA | EN | Unknown | NA | NA | NA | NA | NA |
| lizards | Leptosiaphos_kilimensis | Leptosiaphos_kilimensis | LC | Unknown | tropical | cathemeral | 7.762 | 225182.312 | 11.491 |
| lizards | Leptosiaphos_meleagris | Leptosiaphos_meleagris | VU | Decreasing | tropical | diurnal | 2.692 | 27533.291 | 15.1 |
| lizards | Leptosiaphos_rhomboidalis | NA | DD | Unknown | NA | NA | NA | NA | NA |
| lizards | Lerista_aericeps | NA | LC | Stable | NA | NA | NA | NA | NA |
| lizards | Lerista_allanae | Lerista_allanae | CR | Decreasing | tropical | nocturnal | 4.467 | 597.072 | 9.125 |
| lizards | Lerista_allochira | NA | LC | Unknown | NA | NA | NA | NA | NA |
| lizards | Lerista_ameles | NA | EN | Decreasing | NA | NA | NA | NA | NA |
| lizards | Lerista_amicorum | NA | LC | Stable | NA | NA | NA | NA | NA |
| lizards | Lerista_arenicola | Lerista_arenicola | LC | Stable | temperate | diurnal | 5.623 | 27969.537 | 6.171 |
| lizards | Lerista_axillaris | NA | DD | Unknown | NA | NA | NA | NA | NA |
| lizards | Lerista_baynesi | Lerista_baynesi | LC | Stable | temperate | cathemeral | 4.266 | 230.764 | 7.732 |
| lizards | Lerista_bipes | Lerista_bipes | LC | Stable | tropical | cathemeral | 2.042 | 1374937.738 | 1.267 |
| lizards | Lerista_borealis | Lerista_borealis | LC | Stable | tropical | cathemeral | 1.738 | 34070.917 | 1.679 |
| lizards | Lerista_bougainvillii | Lerista_bougainvillii | LC | Unknown | temperate | cathemeral | 8.128 | 694518.308 | 10.391 |
| lizards | Lerista_bunglebungle | NA | DD | Unknown | NA | NA | NA | NA | NA |
| lizards | Lerista_carpentariae | NA | LC | Unknown | NA | NA | NA | NA | NA |
| lizards | Lerista_chalybura | NA | LC | Stable | NA | NA | NA | NA | NA |
| lizards | Lerista_chordae | NA | VU | Decreasing | NA | NA | NA | NA | NA |
| lizards | Lerista_christinae | Lerista_christinae | VU | Unknown | temperate | cathemeral | 0.562 | 14329.851 | 10.881 |
| lizards | Lerista_cinerea | Lerista_cinerea | NT | Decreasing | tropical | nocturnal | 2.399 | 9926.016 | 5.95 |
| lizards | Lerista_connivens | Lerista_connivens | LC | Stable | temperate | cathemeral | 3.715 | 54227.559 | 5.339 |
| lizards | Lerista_desertorum | Lerista_desertorum | LC | Stable | temperate | cathemeral | 4.677 | 1491472.972 | 1.809 |
| lizards | Lerista_distinguenda | Lerista_distinguenda | LC | Stable | temperate | cathemeral | 1.023 | 405698.677 | 8.449 |
| lizards | Lerista_dorsalis | NA | LC | Stable | NA | NA | NA | NA | NA |
| lizards | Lerista_edwardsae | Lerista_edwardsae | LC | Stable | temperate | cathemeral | 4.786 | 174485.979 | 6.802 |
| lizards | Lerista_elegans | Lerista_elegans | LC | Stable | temperate | cathemeral | 0.759 | 199087.288 | 7.27 |
| lizards | Lerista_emmotti | Lerista_emmotti | LC | Decreasing | tropical | cathemeral | 5.888 | 259402.328 | 4.894 |
| lizards | Lerista_eupoda | NA | LC | Stable | NA | NA | NA | NA | NA |

| | | | | | | | | | |
|---|---|---|---|---|---|---|---|---|---|
| lizards | Lerista_flammicauda | NA | LC | Stable | NA | NA | NA | NA | NA |
| lizards | Lerista_fragilis | Lerista_fragilis | LC | Stable | temperate | cathemeral | 1.549 | 520820.909 | 6.043 |
| lizards | Lerista_frosti | Lerista_frosti | LC | Stable | tropical | diurnal | 2.089 | 65790.147 | 2.361 |
| lizards | Lerista_gascoynensis | NA | LC | Stable | NA | NA | NA | NA | NA |
| lizards | Lerista_gerrardii | Lerista_gerrardii | LC | Unknown | temperate | cathemeral | 4.266 | 174869.241 | 7.767 |
| lizards | Lerista_greeri | NA | LC | Stable | NA | NA | NA | NA | NA |
| lizards | Lerista_griffini | Lerista_griffini | LC | Stable | tropical | cathemeral | 2.042 | 175113.678 | 2.023 |
| lizards | Lerista_haroldi | NA | EN | Unknown | NA | NA | NA | NA | NA |
| lizards | Lerista_humphriesi | Lerista_humphriesi | LC | Stable | temperate | nocturnal | 0.912 | 3354.449 | 4.237 |
| lizards | Lerista_ingrami | NA | DD | Stable | NA | NA | NA | NA | NA |
| lizards | Lerista_ips | Lerista_ips | LC | Unknown | tropical | cathemeral | 2.399 | 434663.314 | 0.706 |
| lizards | Lerista_kalumburu | NA | LC | Stable | NA | NA | NA | NA | NA |
| lizards | Lerista_karlschmidti | Lerista_karlschmidti | LC | Unknown | tropical | nocturnal | 2.344 | 125442.089 | 1.694 |
| lizards | Lerista_kendricki | NA | LC | Stable | NA | NA | NA | NA | NA |
| lizards | Lerista_kennedyensis | NA | LC | Stable | NA | NA | NA | NA | NA |
| lizards | Lerista_kingi | NA | LC | Stable | NA | NA | NA | NA | NA |
| lizards | Lerista_labialis | Lerista_labialis | LC | Stable | temperate | cathemeral | 1.549 | 2260861.2 | 1.96 |
| lizards | Lerista_lineata | Lerista_lineata | EN | Unknown | temperate | cathemeral | 1.66 | 25103.439 | 9.465 |
| lizards | Lerista_lineopunctulata | Lerista_lineopunctulata | LC | Stable | temperate | cathemeral | 7.413 | 75800.402 | 7.808 |
| lizards | Lerista_macropisthopus | Lerista_macropisthopus | LC | Stable | temperate | cathemeral | 5.495 | 632433.739 | 5.21 |
| lizards | Lerista_micra | NA | LC | Stable | NA | NA | NA | NA | NA |
| lizards | Lerista_microtis | Lerista_microtis | LC | Stable | temperate | diurnal | 4.365 | 102837.815 | 9.709 |
| lizards | Lerista_miopus | NA | not_available | not_available | NA | NA | NA | NA | NA |
| lizards | Lerista_muelleri | Lerista_muelleri | LC | Stable | temperate | diurnal | 0.977 | 2180748.391 | 5.511 |
| lizards | Lerista_neander | NA | LC | Stable | NA | NA | NA | NA | NA |
| lizards | Lerista_nichollsi | NA | LC | Stable | NA | NA | NA | NA | NA |
| lizards | Lerista_onsloviana | NA | LC | Stable | NA | NA | NA | NA | NA |
| lizards | Lerista_orientalis | NA | LC | Stable | NA | NA | NA | NA | NA |
| lizards | Lerista_petersoni | Lerista_petersoni | LC | Stable | temperate | cathemeral | 2.344 | 53512.095 | 5.434 |
| lizards | Lerista_picturata | Lerista_picturata | LC | Stable | temperate | cathemeral | 4.467 | 172756.647 | 2.326 |
| lizards | Lerista_planiventralis | Lerista_planiventralis | LC | Stable | temperate | cathemeral | 2.399 | 111772.364 | 6.994 |
| lizards | Lerista_praefrontalis | NA | DD | Unknown | NA | NA | NA | NA | NA |
| lizards | Lerista_praepedita | Lerista_praepedita | LC | Stable | temperate | cathemeral | 0.955 | 106472.829 | 8.161 |
| lizards | Lerista_punctatovittata | Lerista_punctatovittata | LC | Stable | temperate | cathemeral | 6.166 | 1426418.07 | 6.519 |

| | | | | | | | | | |
|---|---|---|---|---|---|---|---|---|---|
| lizards | Lerista_robusta | NA | DD | Unknown | NA | NA | NA | NA | NA |
| lizards | Lerista_rochfordensis | NA | VU | Unknown | NA | NA | NA | NA | NA |
| lizards | Lerista_rolfei | NA | LC | Stable | NA | NA | NA | NA | NA |
| lizards | Lerista_simillima | NA | LC | Stable | NA | NA | NA | NA | NA |
| lizards | Lerista_storri | Lerista_storri | NT | Decreasing | tropical | nocturnal | 2.512 | 14805.459 | 3.949 |
| lizards | Lerista_stylis | Lerista_stylis | LC | Unknown | tropical | nocturnal | 1.318 | 11149.961 | 2.2 |
| lizards | Lerista_terdigitata | NA | LC | Stable | NA | NA | NA | NA | NA |
| lizards | Lerista_timida | Lerista_rhodonoides | LC | Stable | temperate | diurnal | 1.122 | 3880600.665 | 4.685 |
| lizards | Lerista_tridactyla | NA | LC | Stable | NA | NA | NA | NA | NA |
| lizards | Lerista_uniduo | Lerista_uniduo | LC | Stable | temperate | cathemeral | 1.585 | 137380.851 | 5.264 |
| lizards | Lerista_varia | NA | LC | Stable | NA | NA | NA | NA | NA |
| lizards | Lerista_verhmens | NA | LC | Stable | NA | NA | NA | NA | NA |
| lizards | Lerista_vermicularis | NA | LC | Unknown | NA | NA | NA | NA | NA |
| lizards | Lerista_viduata | NA | DD | Unknown | NA | NA | NA | NA | NA |
| lizards | Lerista_vittata | Lerista_vittata | CR | Decreasing | tropical | nocturnal | 2.754 | 518.343 | 5.011 |
| lizards | Lerista_walkeri | Lerista_walkeri | LC | Stable | tropical | cathemeral | 2.188 | 23437.026 | 1.067 |
| lizards | Lerista_wilkinsi | Lerista_wilkinsi | LC | Stable | tropical | nocturnal | 2.692 | 8726.063 | 4.022 |
| lizards | Lerista_xanthura | Lerista_xanthura | LC | Stable | tropical | cathemeral | 1.413 | 491754.568 | 0.768 |
| lizards | Lerista_yuna | NA | NT | Unknown | NA | NA | NA | NA | NA |
| lizards | Lerista_zonulata | NA | LC | Stable | NA | NA | NA | NA | NA |
| lizards | Lialis_burtonis | Lialis_burtonis | LC | Stable | temperate | cathemeral | 22.909 | 7207219.23 | 4.069 |
| lizards | Lialis_jicari | Lialis_jicari | LC | Stable | tropical | diurnal | 23.988 | 186471.096 | 3.989 |
| lizards | Liburnascincus_artemis | NA | LC | Unknown | NA | NA | NA | NA | NA |
| lizards | Liburnascincus_coensis | NA | LC | Unknown | NA | NA | NA | NA | NA |
| lizards | Liburnascincus_mundivensis | Liburnascincus_mundivensis | LC | Unknown | tropical | diurnal | 4.169 | 195037.916 | 5.293 |
| lizards | Liburnascincus_scirtetis | Liburnascincus_scirtetis | LC | Unknown | tropical | diurnal | 6.761 | 26.037 | 7.3 |
| lizards | Liolaemus_abaucan | NA | LC | Stable | NA | NA | NA | NA | NA |
| lizards | Liolaemus_abdalai | Liolaemus_abdalai | LC | Stable | temperate | diurnal | 3.89 | 5916.851 | 3.577 |
| lizards | Liolaemus_absconditus | NA | not_available | not_available | NA | NA | NA | NA | NA |
| lizards | Liolaemus_acostai | NA | LC | Unknown | NA | NA | NA | NA | NA |
| lizards | Liolaemus_albiceps | NA | LC | Stable | NA | NA | NA | NA | NA |
| lizards | Liolaemus_alticolor | Liolaemus_alticolor | LC | Stable | tropical | diurnal | 5.248 | 45889.032 | 6.493 |
| lizards | Liolaemus_andinus | Liolaemus_andinus | LC | Stable | temperate | diurnal | 24.547 | 63469.145 | 2.87 |
| lizards | Liolaemus_annectens | NA | LC | Unknown | NA | NA | NA | NA | NA |

| | | | | | | | | | |
|---|---|---|---|---|---|---|---|---|---|
| lizards | Liolaemus_anomalus | Liolaemus_anomalus | LC | Unknown | temperate | diurnal | 28.184 | 44677.069 | 4.527 |
| lizards | Liolaemus_antumalguen | NA | LC | Unknown | NA | NA | NA | NA | NA |
| lizards | Liolaemus_arambarensis | Liolaemus_arambarensis | EN | Decreasing | temperate | diurnal | 6.761 | 1239.656 | 13.716 |
| lizards | Liolaemus_araucaniensis | NA | LC | Stable | NA | NA | NA | NA | NA |
| lizards | Liolaemus_archeforus | Liolaemus_archeforus | LC | Stable | temperate | diurnal | 25.119 | 22380.616 | 3.494 |
| lizards | Liolaemus_atacamensis | Liolaemus_atacamensis | LC | Stable | temperate | diurnal | 14.454 | 119428.951 | 5.217 |
| lizards | Liolaemus_audituvelatus | Liolaemus_audituvelatus | VU | Unknown | temperate | diurnal | 6.918 | 39887.978 | 3.756 |
| lizards | Liolaemus_austromendocinus | NA | LC | Stable | NA | NA | NA | NA | NA |
| lizards | Liolaemus_azarai | NA | CR | Unknown | NA | NA | NA | NA | NA |
| lizards | Liolaemus_baguali | Liolaemus_baguali | LC | Stable | temperate | diurnal | 33.884 | 6012.625 | 5.268 |
| lizards | Liolaemus_bellii | Liolaemus_bellii | LC | Stable | temperate | diurnal | 26.915 | 10201.496 | 7.288 |
| lizards | Liolaemus_bibronii | Liolaemus_bibronii | LC | Stable | temperate | diurnal | 10.965 | 605195.196 | 5.282 |
| lizards | Liolaemus_bitaeniatus | Liolaemus_bitaeniatus | LC | Stable | temperate | diurnal | 7.413 | 47641.754 | 5.443 |
| lizards | Liolaemus_boulengeri | Liolaemus_boulengeri | LC | Stable | temperate | diurnal | 15.136 | 557576.762 | 5.355 |
| lizards | Liolaemus_buergeri | NA | LC | Stable | NA | NA | NA | NA | NA |
| lizards | Liolaemus_burmeisteri | NA | DD | Unknown | NA | NA | NA | NA | NA |
| lizards | Liolaemus_calchaqui | NA | LC | Stable | NA | NA | NA | NA | NA |
| lizards | Liolaemus_calliston | NA | not_available | not_available | NA | NA | NA | NA | NA |
| lizards | Liolaemus_canqueli | NA | LC | Stable | NA | NA | NA | NA | NA |
| lizards | Liolaemus_caparensis | NA | LC | Stable | NA | NA | NA | NA | NA |
| lizards | Liolaemus_capillitas | NA | LC | Unknown | NA | NA | NA | NA | NA |
| lizards | Liolaemus_carlosgarini | NA | VU | Stable | NA | NA | NA | NA | NA |
| lizards | Liolaemus_casamiquelai | NA | LC | Stable | NA | NA | NA | NA | NA |
| lizards | Liolaemus_cazianiae | Liolaemus_cazianiae | LC | Unknown | temperate | diurnal | 14.125 | 4758.887 | 3.326 |
| lizards | Liolaemus_chacabucoense | NA | LC | Unknown | NA | NA | NA | NA | NA |
| lizards | Liolaemus_chacoensis | Liolaemus_chacoensis | LC | Decreasing | temperate | diurnal | 5.129 | 484670.632 | 5.88 |
| lizards | Liolaemus_chaltin | Liolaemus_chaltin | LC | Stable | tropical | diurnal | 6.31 | 11870.571 | 6.97 |
| lizards | Liolaemus_chavin | Liolaemus_chavin | NT | Unknown | tropical | diurnal | 9.333 | 5142.544 | 8.031 |
| lizards | Liolaemus_chehuachekenk | NA | LC | Unknown | NA | NA | NA | NA | NA |
| lizards | Liolaemus_chiliensis | Liolaemus_chiliensis | LC | Stable | temperate | diurnal | 38.905 | 215500.982 | 9.568 |
| lizards | Liolaemus_chillanensis | Liolaemus_chillanensis | LC | Unknown | temperate | diurnal | 22.909 | 11442.453 | 4.63 |
| lizards | Liolaemus_chlorostictus | NA | LC | Stable | NA | NA | NA | NA | NA |
| lizards | Liolaemus_chungara | NA | LC | Stable | NA | NA | NA | NA | NA |
| lizards | Liolaemus_coeruleus | Liolaemus_coeruleus | LC | Stable | temperate | diurnal | 10.965 | 12446.755 | 4.814 |

| | | | | | | | | | |
|---|---|---|---|---|---|---|---|---|---|
| lizards | Liolaemus_confusus | Liolaemus_confusus | DD | Decreasing | temperate | diurnal | 8.71 | 33.908 | 9.962 |
| lizards | Liolaemus_constanzae | Liolaemus_constanzae | LC | Stable | temperate | diurnal | 14.454 | 33922.412 | 3.152 |
| lizards | Liolaemus_crandalli | NA | not_available | not_available | NA | NA | NA | NA | NA |
| lizards | Liolaemus_crepuscularis | Liolaemus_crepuscularis | LC | Stable | temperate | diurnal | 8.318 | 1298.029 | 4.235 |
| lizards | Liolaemus_cristiani | NA | LC | Stable | NA | NA | NA | NA | NA |
| lizards | Liolaemus_curicensis | NA | LC | Stable | NA | NA | NA | NA | NA |
| lizards | Liolaemus_curis | Liolaemus_curis | CR | Decreasing | temperate | diurnal | 36.308 | 105.601 | 4.449 |
| lizards | Liolaemus_cuyanus | NA | LC | Stable | NA | NA | NA | NA | NA |
| lizards | Liolaemus_cuyumhue | NA | CR | Unknown | NA | NA | NA | NA | NA |
| lizards | Liolaemus_cyaneinotatus | NA | DD | Unknown | NA | NA | NA | NA | NA |
| lizards | Liolaemus_cyanogaster | Liolaemus_cyanogaster | LC | Stable | temperate | diurnal | 11.482 | 140369.786 | 8.363 |
| lizards | Liolaemus_darwinii | Liolaemus_darwinii | LC | Stable | temperate | diurnal | 10.471 | 779703.748 | 5.135 |
| lizards | Liolaemus_diaguita | NA | LC | Stable | NA | NA | NA | NA | NA |
| lizards | Liolaemus_dicktracyi | NA | VU | Unknown | NA | NA | NA | NA | NA |
| lizards | Liolaemus_disjunctus | NA | DD | Unknown | NA | NA | NA | NA | NA |
| lizards | Liolaemus_ditadai | NA | DD | Unknown | NA | NA | NA | NA | NA |
| lizards | Liolaemus_donosobarrosi | NA | LC | Unknown | NA | NA | NA | NA | NA |
| lizards | Liolaemus_dorbignyi | Liolaemus_dorbignyi | LC | Stable | temperate | diurnal | 34.674 | 64763.059 | 3.228 |
| lizards | Liolaemus_dumerili | NA | LC | Stable | NA | NA | NA | NA | NA |
| lizards | Liolaemus_eleodori | NA | LC | Unknown | NA | NA | NA | NA | NA |
| lizards | Liolaemus_elongatus | Liolaemus_elongatus | LC | Stable | temperate | diurnal | 27.542 | 270073.045 | 4.51 |
| lizards | Liolaemus_erguetae | NA | LC | Unknown | NA | NA | NA | NA | NA |
| lizards | Liolaemus_espinozai | Liolaemus_espinozai | LC | Unknown | temperate | diurnal | 7.762 | 1216.307 | 4.54 |
| lizards | Liolaemus_etheridgei | NA | LC | Unknown | NA | NA | NA | NA | NA |
| lizards | Liolaemus_fabiani | Liolaemus_fabiani | EN | Unknown | tropical | diurnal | 23.988 | 1918.352 | 4.682 |
| lizards | Liolaemus_famatinae | NA | VU | Stable | NA | NA | NA | NA | NA |
| lizards | Liolaemus_fittkaui | NA | VU | Decreasing | NA | NA | NA | NA | NA |
| lizards | Liolaemus_fitzgeraldi | Liolaemus_fitzgeraldi | LC | Stable | temperate | diurnal | 8.71 | 54255.717 | 3.142 |
| lizards | Liolaemus_fitzingerii | NA | LC | Stable | NA | NA | NA | NA | NA |
| lizards | Liolaemus_flavipiceus | NA | LC | Unknown | NA | NA | NA | NA | NA |
| lizards | Liolaemus_foxi | Liolaemus_foxi | LC | Stable | tropical | diurnal | 18.197 | 771.085 | 3.557 |
| lizards | Liolaemus_frassinettii | NA | VU | Stable | NA | NA | NA | NA | NA |
| lizards | Liolaemus_fuscus | Liolaemus_fuscus | LC | Stable | temperate | diurnal | 4.677 | 2464.966 | 23.315 |
| lizards | Liolaemus_gallardoi | Liolaemus_gallardoi | LC | Unknown | temperate | diurnal | 25.119 | 34575.938 | 4.455 |

| | | | | | | | | | |
|---|---|---|---|---|---|---|---|---|---|
| lizards | Liolaemus_gardeli | NA | not_available | not_available | NA | NA | NA | NA | NA |
| lizards | Liolaemus_goetschi | Liolaemus_goetschi | LC | Stable | temperate | diurnal | 13.183 | 122943.802 | 5.405 |
| lizards | Liolaemus_gracielae | NA | LC | Unknown | NA | NA | NA | NA | NA |
| lizards | Liolaemus_gracilis | Liolaemus_gracilis | LC | Stable | temperate | diurnal | 6.166 | 577965.521 | 5.083 |
| lizards | Liolaemus_gravenhorstii | Liolaemus_gravenhorstii | EN | Decreasing | temperate | diurnal | 10.471 | 6065.403 | 18.144 |
| lizards | Liolaemus_griseus | NA | DD | Unknown | NA | NA | NA | NA | NA |
| lizards | Liolaemus_grosseorum | NA | LC | Unknown | NA | NA | NA | NA | NA |
| lizards | Liolaemus_gununakuna | NA | LC | Unknown | NA | NA | NA | NA | NA |
| lizards | Liolaemus_hajeki | NA | LC | Stable | NA | NA | NA | NA | NA |
| lizards | Liolaemus_halonastes | Liolaemus_halonastes | VU | Unknown | temperate | diurnal | 11.749 | 69.145 | 6.832 |
| lizards | Liolaemus_hatcheri | Liolaemus_hatcheri | LC | Stable | temperate | diurnal | 12.303 | 16719.838 | 4.159 |
| lizards | Liolaemus_heliodermis | NA | LC | Unknown | NA | NA | NA | NA | NA |
| lizards | Liolaemus_hellmichi | NA | VU | Stable | NA | NA | NA | NA | NA |
| lizards | Liolaemus_hermannunezi | NA | DD | Unknown | NA | NA | NA | NA | NA |
| lizards | Liolaemus_huacahuasicus | Liolaemus_huacahuasicus | LC | Stable | temperate | diurnal | 14.125 | 841.884 | 3.666 |
| lizards | Liolaemus_huayra | NA | LC | Stable | NA | NA | NA | NA | NA |
| lizards | Liolaemus_igneus | NA | not_available | not_available | NA | NA | NA | NA | NA |
| lizards | Liolaemus_inacayali | NA | LC | Stable | NA | NA | NA | NA | NA |
| lizards | Liolaemus_incaicus | Liolaemus_incaicus | LC | Unknown | tropical | diurnal | 6.026 | 1122.17 | 16.667 |
| lizards | Liolaemus_inti | NA | LC | Stable | NA | NA | NA | NA | NA |
| lizards | Liolaemus_irregularis | Liolaemus_irregularis | LC | Stable | temperate | diurnal | 23.442 | 7925.074 | 3.864 |
| lizards | Liolaemus_isabelae | NA | EN | Decreasing | NA | NA | NA | NA | NA |
| lizards | Liolaemus_islugensis | NA | LC | Stable | NA | NA | NA | NA | NA |
| lizards | Liolaemus_jamesi | Liolaemus_jamesi | LC | Stable | tropical | diurnal | 32.359 | 24376.308 | 4.138 |
| lizards | Liolaemus_janequeoae | NA | not_available | not_available | NA | NA | NA | NA | NA |
| lizards | Liolaemus_josei | NA | LC | Unknown | NA | NA | NA | NA | NA |
| lizards | Liolaemus_juanortizi | NA | LC | Stable | NA | NA | NA | NA | NA |
| lizards | Liolaemus_kingii | Liolaemus_kingii | LC | Stable | temperate | diurnal | 33.113 | 273140.787 | 5.876 |
| lizards | Liolaemus_kolengh | Liolaemus_kolengh | LC | Stable | temperate | diurnal | 7.586 | 1023.717 | 3.414 |
| lizards | Liolaemus_koslowskyi | Liolaemus_koslowskyi | LC | Unknown | temperate | diurnal | 17.783 | 33902.59 | 3.393 |
| lizards | Liolaemus_kriegi | Liolaemus_kriegi | LC | Stable | temperate | diurnal | 50.119 | 112291.95 | 4.876 |
| lizards | Liolaemus_laurenti | NA | LC | Stable | NA | NA | NA | NA | NA |
| lizards | Liolaemus_lavillai | NA | LC | Stable | NA | NA | NA | NA | NA |
| lizards | Liolaemus_leftrarui | NA | not_available | not_available | NA | NA | NA | NA | NA |

| | | | | | | | | | |
|---|---|---|---|---|---|---|---|---|---|
| lizards | Liolaemus_lemniscatus | Liolaemus_lemniscatus | LC | Stable | temperate | diurnal | 5.495 | 176579.803 | 9.905 |
| lizards | Liolaemus_lentus | NA | DD | Unknown | NA | NA | NA | NA | NA |
| lizards | Liolaemus_leopardinus | Liolaemus_leopardinus | EN | Decreasing | temperate | diurnal | 34.674 | 644.197 | 10.98 |
| lizards | Liolaemus_lineomaculatus | Liolaemus_lineomaculatus | LC | Unknown | temperate | diurnal | 10 | 435633.653 | 5.69 |
| lizards | Liolaemus_loboi | NA | EN | Stable | NA | NA | NA | NA | NA |
| lizards | Liolaemus_lonquimayensis | NA | not_available | not_available | NA | NA | NA | NA | NA |
| lizards | Liolaemus_lopezi | NA | LC | Stable | NA | NA | NA | NA | NA |
| lizards | Liolaemus_lorenzmuelleri | NA | EN | Decreasing | NA | NA | NA | NA | NA |
| lizards | Liolaemus_lutzae | Liolaemus_lutzae | CR | Decreasing | tropical | diurnal | 22.909 | 1707.581 | 28.203 |
| lizards | Liolaemus_magellanicus | Liolaemus_magellanicus | LC | Unknown | temperate | diurnal | 11.482 | 94721.804 | 6.02 |
| lizards | Liolaemus_maldonadae | NA | LC | Stable | NA | NA | NA | NA | NA |
| lizards | Liolaemus_mapuche | NA | LC | Stable | NA | NA | NA | NA | NA |
| lizards | Liolaemus_martorii | NA | VU | Decreasing | NA | NA | NA | NA | NA |
| lizards | Liolaemus_melaniceps | NA | LC | Unknown | NA | NA | NA | NA | NA |
| lizards | Liolaemus_melanopleurus | NA | DD | Unknown | NA | NA | NA | NA | NA |
| lizards | Liolaemus_melanops | Liolaemus_melanops | LC | Stable | temperate | diurnal | 31.623 | 127566.141 | 5.257 |
| lizards | Liolaemus_millcayac | Liolaemus_millcayac | LC | Unknown | temperate | diurnal | 15.488 | 21008.107 | 6.507 |
| lizards | Liolaemus_molinai | NA | LC | Stable | NA | NA | NA | NA | NA |
| lizards | Liolaemus_montanezi | NA | DD | Unknown | NA | NA | NA | NA | NA |
| lizards | Liolaemus_montanus | NA | LC | Unknown | NA | NA | NA | NA | NA |
| lizards | Liolaemus_monticola | Liolaemus_monticola | LC | Stable | temperate | diurnal | 22.909 | 31123.616 | 10.552 |
| lizards | Liolaemus_moradoensis | NA | LC | Stable | NA | NA | NA | NA | NA |
| lizards | Liolaemus_morenoi | Liolaemus_morenoi | VU | Unknown | temperate | diurnal | 22.387 | 22101.859 | 4.351 |
| lizards | Liolaemus_multicolor | Liolaemus_multicolor | LC | Stable | tropical | diurnal | 19.055 | 38154.879 | 5.091 |
| lizards | Liolaemus_multimaculatus | Liolaemus_multimaculatus | EN | Decreasing | temperate | diurnal | 12.023 | 6252.344 | 11.935 |
| lizards | Liolaemus_neuquensis | NA | DD | Unknown | NA | NA | NA | NA | NA |
| lizards | Liolaemus_nigriceps | NA | LC | Unknown | NA | NA | NA | NA | NA |
| lizards | Liolaemus_nigromaculatus | Liolaemus_nigromaculatus | LC | Stable | temperate | diurnal | 26.303 | 80035.075 | 6.584 |
| lizards | Liolaemus_nigroviridis | Liolaemus_nigroviridis | LC | Stable | temperate | diurnal | 22.387 | 17486.218 | 12.366 |
| lizards | Liolaemus_nitidus | Liolaemus_nitidus | LC | Stable | temperate | diurnal | 38.905 | 100923.325 | 10.254 |
| lizards | Liolaemus_occipitalis | NA | VU | Decreasing | NA | NA | NA | NA | NA |
| lizards | Liolaemus_olongasta | Liolaemus_olongasta | LC | Stable | temperate | diurnal | 9.55 | 52531.164 | 3.518 |
| lizards | Liolaemus_omorfi | NA | not_available | not_available | NA | NA | NA | NA | NA |
| lizards | Liolaemus_ornatus | Liolaemus_ornatus | LC | Stable | tropical | diurnal | 11.482 | 130150.229 | 4.945 |

| | | | | | | | | | |
|---|---|---|---|---|---|---|---|---|---|
| lizards | Liolaemus_pacha | NA | LC | Stable | NA | NA | NA | NA | NA |
| lizards | Liolaemus_pachacutec | Liolaemus_pachacutec | LC | Unknown | tropical | diurnal | 4.365 | 9627.033 | 10.669 |
| lizards | Liolaemus_pachecoi | NA | LC | Unknown | NA | NA | NA | NA | NA |
| lizards | Liolaemus_pagaburoi | Liolaemus_pagaburoi | LC | Stable | temperate | diurnal | 5.754 | 4627.174 | 4.143 |
| lizards | Liolaemus_pantherinus | NA | LC | Stable | NA | NA | NA | NA | NA |
| lizards | Liolaemus_parthenos | NA | not_available | not_available | NA | NA | NA | NA | NA |
| lizards | Liolaemus_parvus | Liolaemus_parvus | LC | Stable | temperate | diurnal | 11.482 | 19702.158 | 2.681 |
| lizards | Liolaemus_patriciaiturrae | Liolaemus_patriciaiturrae | NT | Stable | temperate | diurnal | 29.512 | 4065.763 | 3.288 |
| lizards | Liolaemus_paulinae | Liolaemus_paulinae | CR | Decreasing | tropical | diurnal | 7.079 | 215.258 | 18.904 |
| lizards | Liolaemus_petrophilus | Liolaemus_petrophilus | LC | Stable | temperate | diurnal | 33.113 | 187888.486 | 5.184 |
| lizards | Liolaemus_pictus | Liolaemus_pictus | LC | Decreasing | temperate | diurnal | 37.154 | 172561.914 | 8.096 |
| lizards | Liolaemus_pipanaco | NA | LC | Unknown | NA | NA | NA | NA | NA |
| lizards | Liolaemus_platei | Liolaemus_platei | LC | Stable | temperate | diurnal | 11.482 | 99102.123 | 5.707 |
| lizards | Liolaemus_pleopholis | Liolaemus_pleopholis | LC | Stable | tropical | diurnal | 16.982 | 4961.099 | 4.709 |
| lizards | Liolaemus_poconchilensis | NA | EN | Decreasing | NA | NA | NA | NA | NA |
| lizards | Liolaemus_poecilochromus | Liolaemus_poecilochromus | LC | Stable | temperate | diurnal | 14.125 | 5440.977 | 2.564 |
| lizards | Liolaemus_polystictus | NA | LC | Unknown | NA | NA | NA | NA | NA |
| lizards | Liolaemus_porosus | NA | LC | Stable | NA | NA | NA | NA | NA |
| lizards | Liolaemus_pseudoanomalus | Liolaemus_pseudoanomalus | LC | Stable | temperate | diurnal | 10 | 59606.585 | 2.713 |
| lizards | Liolaemus_pseudolemniscatus | NA | LC | Stable | NA | NA | NA | NA | NA |
| lizards | Liolaemus_puelche | NA | LC | Stable | NA | NA | NA | NA | NA |
| lizards | Liolaemus_pulcherrimus | NA | LC | Stable | NA | NA | NA | NA | NA |
| lizards | Liolaemus_puna | Liolaemus_puna | LC | Stable | tropical | diurnal | 5.495 | 278719.851 | 5.866 |
| lizards | Liolaemus_punmahuida | NA | LC | Stable | NA | NA | NA | NA | NA |
| lizards | Liolaemus_puritamensis | Liolaemus_puritamensis | LC | Stable | tropical | diurnal | 33.884 | 24529.919 | 3.317 |
| lizards | Liolaemus_purul | NA | LC | Stable | NA | NA | NA | NA | NA |
| lizards | Liolaemus_quilmes | NA | NT | Stable | NA | NA | NA | NA | NA |
| lizards | Liolaemus_rabinoi | NA | CR | Decreasing | NA | NA | NA | NA | NA |
| lizards | Liolaemus_ramirezae | Liolaemus_ramirezae | LC | Stable | temperate | diurnal | 5.888 | 19191.903 | 4.278 |
| lizards | Liolaemus_riojanus | Liolaemus_riojanus | LC | Stable | temperate | diurnal | 7.762 | 20256.936 | 4.07 |
| lizards | Liolaemus_robertmertensi | NA | LC | Stable | NA | NA | NA | NA | NA |
| lizards | Liolaemus_robertoi | Liolaemus_robertoi | EN | Unknown | temperate | diurnal | 11.749 | 778.306 | 3.996 |
| lizards | Liolaemus_rosenmanni | Liolaemus_rosenmanni | NT | Stable | temperate | diurnal | 15.136 | 10786.242 | 2.564 |
| lizards | Liolaemus_rothi | Liolaemus_rothi | LC | Stable | temperate | diurnal | 37.154 | 138928.597 | 4.923 |

| | | | | | | | | | |
|---|---|---|---|---|---|---|---|---|---|
| lizards | Liolaemus_ruibali | Liolaemus_ruibali | LC | Stable | temperate | diurnal | 9.333 | 19123.255 | 2.834 |
| lizards | Liolaemus_sagei | Liolaemus_sagei | LC | Stable | temperate | diurnal | 33.113 | 15129.838 | 3.792 |
| lizards | Liolaemus_salinicola | NA | EN | Decreasing | NA | NA | NA | NA | NA |
| lizards | Liolaemus_sanjuanensis | Liolaemus_sanjuanensis | LC | Stable | temperate | diurnal | 8.71 | 4084.61 | 2.428 |
| lizards | Liolaemus_sarmientoi | Liolaemus_sarmientoi | LC | Stable | temperate | diurnal | 30.903 | 82944.7 | 6.426 |
| lizards | Liolaemus_saxatilis | Liolaemus_saxatilis | LC | Stable | temperate | diurnal | 6.761 | 17738.856 | 8.568 |
| lizards | Liolaemus_scapularis | Liolaemus_scapularis | EN | Decreasing | temperate | diurnal | 14.454 | 13758.41 | 4.57 |
| lizards | Liolaemus_schmidti | NA | LC | Stable | NA | NA | NA | NA | NA |
| lizards | Liolaemus_schroederi | Liolaemus_schroederi | LC | Unknown | temperate | diurnal | 9.772 | 77024.669 | 11.823 |
| lizards | Liolaemus_scorialis | NA | not_available | not_available | NA | NA | NA | NA | NA |
| lizards | Liolaemus_scrocchii | NA | LC | Stable | NA | NA | NA | NA | NA |
| lizards | Liolaemus_senguer | NA | LC | Stable | NA | NA | NA | NA | NA |
| lizards | Liolaemus_septentrionalis | Liolaemus_septentrionalis | LC | Stable | temperate | diurnal | 6.607 | 3081.929 | 10.101 |
| lizards | Liolaemus_shehuen | NA | LC | Stable | NA | NA | NA | NA | NA |
| lizards | Liolaemus_shitan | NA | LC | Stable | NA | NA | NA | NA | NA |
| lizards | Liolaemus_signifer | Liolaemus_signifer | NT | Decreasing | tropical | diurnal | 41.687 | 373214.762 | 6.669 |
| lizards | Liolaemus_silvai | NA | NT | Stable | NA | NA | NA | NA | NA |
| lizards | Liolaemus_silvanae | Liolaemus_silvanae | LC | Stable | temperate | diurnal | 15.136 | 3113.521 | 2.473 |
| lizards | Liolaemus_sitesi | NA | LC | Stable | NA | NA | NA | NA | NA |
| lizards | Liolaemus_smaug | NA | LC | Stable | NA | NA | NA | NA | NA |
| lizards | Liolaemus_somuncurae | NA | LC | Stable | NA | NA | NA | NA | NA |
| lizards | Liolaemus_stolzmanni | Liolaemus_stolzmanni | EN | Decreasing | tropical | diurnal | 7.244 | 1268.651 | 7.319 |
| lizards | Liolaemus_tacnae | NA | NT | Unknown | NA | NA | NA | NA | NA |
| lizards | Liolaemus_tacora | NA | not_available | not_available | NA | NA | NA | NA | NA |
| lizards | Liolaemus_talampaya | NA | LC | Unknown | NA | NA | NA | NA | NA |
| lizards | Liolaemus_tandiliensis | Liolaemus_tandiliensis | VU | Decreasing | temperate | diurnal | 6.166 | 11868.176 | 12.571 |
| lizards | Liolaemus_tari | Liolaemus_tari | LC | Stable | temperate | diurnal | 34.674 | 4127.892 | 5.387 |
| lizards | Liolaemus_tehuelche | NA | LC | Stable | NA | NA | NA | NA | NA |
| lizards | Liolaemus_telsen | NA | LC | Stable | NA | NA | NA | NA | NA |
| lizards | Liolaemus_tenuis | Liolaemus_tenuis | LC | Stable | temperate | diurnal | 8.511 | 226075.717 | 9.6 |
| lizards | Liolaemus_thermarum | NA | VU | Unknown | NA | NA | NA | NA | NA |
| lizards | Liolaemus_tirantii | NA | not_available | not_available | NA | NA | NA | NA | NA |
| lizards | Liolaemus_tolhuaca | NA | not_available | not_available | NA | NA | NA | NA | NA |
| lizards | Liolaemus_torresi | Liolaemus_torresi | EN | Decreasing | tropical | diurnal | 8.71 | 3344.319 | 3.847 |

| | | | | | | | | | |
|---|---|---|---|---|---|---|---|---|---|
| lizards | Liolaemus_tregenzai | NA | VU | Unknown | NA | NA | NA | NA | NA |
| lizards | Liolaemus_tristis | Liolaemus_tristis | LC | Unknown | temperate | diurnal | 20.417 | 4459.623 | 4.191 |
| lizards | Liolaemus_tromen | NA | LC | Stable | NA | NA | NA | NA | NA |
| lizards | Liolaemus_tulkas | NA | LC | Stable | NA | NA | NA | NA | NA |
| lizards | Liolaemus_ubaghsi | NA | DD | Unknown | NA | NA | NA | NA | NA |
| lizards | Liolaemus_umbrifer | NA | LC | Stable | NA | NA | NA | NA | NA |
| lizards | Liolaemus_uniformis | NA | not_available | not_available | NA | NA | NA | NA | NA |
| lizards | Liolaemus_uspallatensis | Liolaemus_uspallatensis | LC | Stable | temperate | diurnal | 8.71 | 18567.895 | 3.04 |
| lizards | Liolaemus_valdesianus | Liolaemus_valdesianus | NT | Stable | temperate | diurnal | 28.184 | 481.535 | 3.323 |
| lizards | Liolaemus_vallecurensis | Liolaemus_vallecurensis | NT | Unknown | temperate | diurnal | 14.791 | 71303.661 | 5.485 |
| lizards | Liolaemus_variegatus | NA | VU | Decreasing | NA | NA | NA | NA | NA |
| lizards | Liolaemus_velosoi | Liolaemus_velosoi | LC | Stable | temperate | diurnal | 6.457 | 153.125 | 12.86 |
| lizards | Liolaemus_villaricensis | NA | VU | Unknown | NA | NA | NA | NA | NA |
| lizards | Liolaemus_vulcanus | NA | LC | Stable | NA | NA | NA | NA | NA |
| lizards | Liolaemus_walkeri | NA | NT | Unknown | NA | NA | NA | NA | NA |
| lizards | Liolaemus_wari | Liolaemus_wari | LC | Unknown | tropical | diurnal | 7.244 | 9364.507 | 10.106 |
| lizards | Liolaemus_wiegmannii | Liolaemus_wiegmannii | LC | Stable | temperate | diurnal | 9.12 | 1481143.349 | 9.473 |
| lizards | Liolaemus_williamsi | NA | LC | Unknown | NA | NA | NA | NA | NA |
| lizards | Liolaemus_xanthoviridis | Liolaemus_xanthoviridis | LC | Stable | temperate | diurnal | 33.884 | 15719.971 | 6.426 |
| lizards | Liolaemus_yalguaraz | NA | not_available | not_available | NA | NA | NA | NA | NA |
| lizards | Liolaemus_yanalcu | Liolaemus_yanalcu | LC | Stable | temperate | diurnal | 7.244 | 1339.475 | 2.781 |
| lizards | Liolaemus_yatel | NA | LC | Stable | NA | NA | NA | NA | NA |
| lizards | Liolaemus_zabalai | NA | not_available | not_available | NA | NA | NA | NA | NA |
| lizards | Liolaemus_zapallarensis | Liolaemus_zapallarensis | LC | Stable | temperate | diurnal | 43.652 | 1996.25 | 14.459 |
| lizards | Liolaemus_zullyae | Liolaemus_zullyae | LC | Stable | temperate | diurnal | 22.909 | 2053.643 | 3.965 |
| lizards | Liopholis_guthega | Liopholis_guthega | EN | Decreasing | temperate | diurnal | 32.359 | 1108.353 | 6.351 |
| lizards | Liopholis_inornata | Liopholis_inornata | LC | Stable | temperate | cathemeral | 12.882 | 2541556.984 | 3.306 |
| lizards | Liopholis_kintorei | Liopholis_kintorei | VU | Decreasing | tropical | cathemeral | 223.872 | 1336754.786 | 1.202 |
| lizards | Liopholis_margaretae | NA | LC | Stable | NA | NA | NA | NA | NA |
| lizards | Liopholis_modesta | Liopholis_modesta | LC | Stable | temperate | diurnal | 30.903 | 299350.966 | 7.8 |
| lizards | Liopholis_montana | Liopholis_montana | NT | Unknown | temperate | diurnal | 30.2 | 40855.75 | 6.802 |
| lizards | Liopholis_multiscutata | Liopholis_multiscutata | LC | Stable | temperate | diurnal | 19.055 | 568847.563 | 7.402 |
| lizards | Liopholis_personata | NA | not_available | not_available | NA | NA | NA | NA | NA |
| lizards | Liopholis_pulchra | Liopholis_pulchra | LC | Stable | temperate | diurnal | 29.512 | 82244.658 | 10.924 |

| | | | | | | | | | |
|---|---|---|---|---|---|---|---|---|---|
| lizards | Liopholis_slateri | Liopholis_slateri | VU | Decreasing | temperate | cathemeral | 19.498 | 73334.731 | 2.667 |
| lizards | Liopholis_striata | Liopholis_striata | LC | Stable | temperate | cathemeral | 35.481 | 2261385.718 | 1.88 |
| lizards | Liopholis_whitii | Liopholis_whitii | LC | Unknown | temperate | diurnal | 31.623 | 831367.907 | 10.166 |
| lizards | Lioscincus_steindachneri | Lioscincus_steindachneri | EN | Decreasing | tropical | diurnal | 31.623 | 1594.226 | 4.182 |
| lizards | Lioscincus_vivae | NA | CR | Decreasing | NA | NA | NA | NA | NA |
| lizards | Lipinia_auriculata | Lipinia_auriculata | LC | Unknown | tropical | diurnal | 2.455 | 40497.005 | 14.649 |
| lizards | Lipinia_inexpectata | Lipinia_inexpectata | LC | Unknown | tropical | cathemeral | 1.175 | 62144.219 | 6.849 |
| lizards | Lipinia_leptosoma | Lipinia_leptosoma | NT | Stable | tropical | diurnal | 1.514 | 455.637 | 11.185 |
| lizards | Lipinia_macrotympanum | NA | DD | Unknown | NA | NA | NA | NA | NA |
| lizards | Lipinia_microcerca | NA | not_available | not_available | NA | NA | NA | NA | NA |
| lizards | Lipinia_nitens | NA | DD | Unknown | NA | NA | NA | NA | NA |
| lizards | Lipinia_pulchella | Lipinia_pulchella | LC | Stable | tropical | diurnal | 2.291 | 186999.272 | 13.222 |
| lizards | Lipinia_quadrivittata | Lipinia_quadrivittata | LC | Stable | tropical | diurnal | 1.738 | 38346.877 | 13.567 |
| lizards | Lipinia_rabori | NA | DD | Unknown | NA | NA | NA | NA | NA |
| lizards | Lipinia_rouxi | Lipinia_rouxi | LC | Unknown | tropical | diurnal | 1.202 | 6878.303 | 5.423 |
| lizards | Lipinia_sekayuensis | NA | LC | Stable | NA | NA | NA | NA | NA |
| lizards | Lipinia_semperi | NA | DD | Unknown | NA | NA | NA | NA | NA |
| lizards | Lipinia_surda | NA | DD | Unknown | NA | NA | NA | NA | NA |
| lizards | Lipinia_trivittata | NA | not_available | not_available | NA | NA | NA | NA | NA |
| lizards | Lipinia_vassilievi | NA | not_available | not_available | NA | NA | NA | NA | NA |
| lizards | Lipinia_vittigera | Lipinia_vittigera | LC | Unknown | tropical | diurnal | 1.622 | 457593.082 | 12.796 |
| lizards | Lipinia_vulcania | NA | DD | Unknown | NA | NA | NA | NA | NA |
| lizards | Lissolepis_coventryi | Lissolepis_coventryi | EN | Decreasing | temperate | diurnal | 79.433 | 109117.348 | 11.837 |
| lizards | Lissolepis_luctuosa | NA | LC | Unknown | NA | NA | NA | NA | NA |
| lizards | Lobulia_brongersmai | Lobulia_brongersmai | LC | Unknown | tropical | diurnal | 3.09 | 3306.866 | 3.188 |
| lizards | Lobulia_elegans | Lobulia_elegans | LC | Stable | tropical | diurnal | 5.623 | 92798.493 | 5.33 |
| lizards | Lobulia_lobulus | NA | not_available | not_available | NA | NA | NA | NA | NA |
| lizards | Lophocalotes_achlios | NA | LC | Unknown | NA | NA | NA | NA | NA |
| lizards | Lophocalotes_ludekingi | NA | LC | Unknown | NA | NA | NA | NA | NA |
| lizards | Lophognathus_gilberti | Lophognathus_gilberti | LC | Stable | tropical | diurnal | 85.114 | 258890.606 | 1.761 |
| lizards | Lophosaurus_boydii | Hypsilurus_boydii | LC | Stable | tropical | diurnal | 190.546 | 28110.475 | 4.73 |
| lizards | Lophosaurus_dilophus | Hypsilurus_dilophus | LC | Unknown | tropical | diurnal | 389.045 | 772102.211 | 3.471 |
| lizards | Lophosaurus_spinipes | Hypsilurus_spinipes | LC | Unknown | temperate | diurnal | 87.096 | 78462.357 | 9.802 |
| lizards | Loxopholis_ferreirai | NA | LC | Unknown | NA | NA | NA | NA | NA |

| | | | | | | | | | |
|---|---|---|---|---|---|---|---|---|---|
| lizards | Loxopholis_guianense | Leposoma_guianense | LC | Stable | tropical | cathemeral | 1.202 | 1480769.955 | 2.032 |
| lizards | Loxopholis_hexalepis | Leposoma_hexalepis | LC | Unknown | tropical | diurnal | 1.413 | 67952.088 | 2.785 |
| lizards | Loxopholis_osvaldoi | Leposoma_osvaldoi | LC | Unknown | tropical | diurnal | 1 | 1602331.615 | 1.95 |
| lizards | Loxopholis_parietalis | Leposoma_parietale | LC | Stable | tropical | diurnal | 1.905 | 632209.615 | 2.423 |
| lizards | Loxopholis_percarinatum | Leposoma_percarinatum | LC | Unknown | tropical | diurnal | 1 | 5730663.91 | 2.015 |
| lizards | Loxopholis_rugiceps | Leposoma_rugiceps | LC | Unknown | tropical | diurnal | 1.778 | 465129.473 | 12.029 |
| lizards | Loxopholis_snethlageae | Leposoma_snethlageae | LC | Unknown | tropical | diurnal | 0.692 | 675574.208 | 0.711 |
| lizards | Loxopholis_southi | Leposoma_southi | LC | Stable | tropical | diurnal | 3.715 | 303496.731 | 10.004 |
| lizards | Lubuya_ivensii | Trachylepis_ivensii | LC | Stable | tropical | diurnal | 75.858 | 479094.026 | 6.102 |
| lizards | Lucasium_alboguttatum | Lucasium_alboguttatum | LC | Unknown | temperate | nocturnal | 3.631 | 80258.931 | 8.987 |
| lizards | Lucasium_bungabinna | Lucasium_bungabinna | LC | Stable | temperate | nocturnal | 3.89 | 515832.853 | 2.296 |
| lizards | Lucasium_byrnei | Lucasium_byrnei | LC | Stable | temperate | nocturnal | 3.548 | 873971.169 | 4.586 |
| lizards | Lucasium_damaeum | Lucasium_damaeum | LC | Stable | temperate | nocturnal | 4.266 | 2068398.97 | 3.127 |
| lizards | Lucasium_immaculatum | NA | LC | Stable | NA | NA | NA | NA | NA |
| lizards | Lucasium_maini | Lucasium_maini | LC | Unknown | temperate | nocturnal | 3.09 | 350641.045 | 5.471 |
| lizards | Lucasium_occultum | Lucasium_occultum | EN | Unknown | tropical | nocturnal | 1.349 | 2381.578 | 2.75 |
| lizards | Lucasium_squarrosum | Lucasium_squarrosum | LC | Stable | temperate | nocturnal | 3.631 | 505496.954 | 4.679 |
| lizards | Lucasium_steindachneri | Lucasium_steindachneri | LC | Stable | tropical | nocturnal | 4.074 | 1570837.404 | 5.348 |
| lizards | Lucasium_stenodactylus | NA | LC | Stable | NA | NA | NA | NA | NA |
| lizards | Lucasium_wombeyi | Lucasium_wombeyi | LC | Stable | tropical | nocturnal | 3.09 | 114715.144 | 4.051 |
| lizards | Luperosaurus_angliit | NA | DD | Decreasing | NA | NA | NA | NA | NA |
| lizards | Luperosaurus_corfieldi | NA | DD | Unknown | NA | NA | NA | NA | NA |
| lizards | Luperosaurus_cumingii | NA | DD | Unknown | NA | NA | NA | NA | NA |
| lizards | Luperosaurus_joloensis | NA | DD | Decreasing | NA | NA | NA | NA | NA |
| lizards | Luperosaurus_kubli | NA | LC | Unknown | NA | NA | NA | NA | NA |
| lizards | Luperosaurus_macgregori | NA | NT | Unknown | NA | NA | NA | NA | NA |
| lizards | Luperosaurus_palawanensis | NA | DD | Unknown | NA | NA | NA | NA | NA |
| lizards | Luperosaurus_yasumai | NA | DD | Unknown | NA | NA | NA | NA | NA |
| lizards | Lygisaurus_absconditus | NA | DD | Unknown | NA | NA | NA | NA | NA |
| lizards | Lygisaurus_aeratus | NA | LC | Stable | NA | NA | NA | NA | NA |
| lizards | Lygisaurus_curtus | Lygisaurus_curtus | LC | Unknown | tropical | diurnal | 1.445 | 39987.113 | 3.757 |
| lizards | Lygisaurus_foliorum | Lygisaurus_foliorum | LC | Stable | temperate | diurnal | 1.514 | 842999.212 | 6.779 |
| lizards | Lygisaurus_laevis | Lygisaurus_laevis | LC | Unknown | tropical | diurnal | 0.871 | 11367.902 | 6.148 |
| lizards | Lygisaurus_macfarlani | Lygisaurus_macfarlani | LC | Stable | tropical | diurnal | 1.072 | 94494.782 | 2.675 |

| | | | | | | | | | |
|---|---|---|---|---|---|---|---|---|---|
| lizards | Lygisaurus_malleolus | NA | LC | Unknown | NA | NA | NA | NA | NA |
| lizards | Lygisaurus_novaeguineae | NA | LC | Unknown | NA | NA | NA | NA | NA |
| lizards | Lygisaurus_parrhasius | NA | LC | Unknown | NA | NA | NA | NA | NA |
| lizards | Lygisaurus_rimula | NA | LC | Unknown | NA | NA | NA | NA | NA |
| lizards | Lygisaurus_rococo | Lygisaurus_rococo | LC | Unknown | tropical | diurnal | 1.622 | 950.587 | 3.99 |
| lizards | Lygisaurus_sesbrauna | NA | LC | Unknown | NA | NA | NA | NA | NA |
| lizards | Lygisaurus_tanneri | NA | LC | Unknown | NA | NA | NA | NA | NA |
| lizards | Lygisaurus_zuma | Lygisaurus_zuma | LC | Stable | tropical | diurnal | 0.708 | 14787.684 | 5.934 |
| lizards | Lygodactylus_angolensis | Lygodactylus_angolensis | LC | Stable | tropical | diurnal | 1.122 | 1958377.775 | 8.706 |
| lizards | Lygodactylus_angularis | Lygodactylus_angularis | LC | Unknown | tropical | cathemeral | 2.239 | 441775.809 | 10.198 |
| lizards | Lygodactylus_arnoulti | NA | LC | Stable | NA | NA | NA | NA | NA |
| lizards | Lygodactylus_bivittis | Lygodactylus_bivittis | VU | Decreasing | tropical | diurnal | 1.148 | 20306.383 | 6.725 |
| lizards | Lygodactylus_blancae | NA | NT | Stable | NA | NA | NA | NA | NA |
| lizards | Lygodactylus_blanci | NA | VU | Unknown | NA | NA | NA | NA | NA |
| lizards | Lygodactylus_bradfieldi | Lygodactylus_bradfieldi | LC | Stable | tropical | diurnal | 0.692 | 1028863.935 | 5.55 |
| lizards | Lygodactylus_capensis | Lygodactylus_capensis | LC | Stable | tropical | diurnal | 1.862 | 4100565.08 | 8.984 |
| lizards | Lygodactylus_conradti | NA | DD | Unknown | NA | NA | NA | NA | NA |
| lizards | Lygodactylus_conraui | Lygodactylus_conraui | LC | Unknown | tropical | diurnal | 1.514 | 560180.616 | 15.243 |
| lizards | Lygodactylus_decaryi | NA | DD | Unknown | NA | NA | NA | NA | NA |
| lizards | Lygodactylus_depressus | Lygodactylus_depressus | LC | Stable | tropical | diurnal | 1.318 | 645324.891 | 5.368 |
| lizards | Lygodactylus_expectatus | NA | NT | Stable | NA | NA | NA | NA | NA |
| lizards | Lygodactylus_fischeri | NA | LC | Unknown | NA | NA | NA | NA | NA |
| lizards | Lygodactylus_gravis | Lygodactylus_gravis | VU | Unknown | tropical | diurnal | 1.738 | 12320.678 | 14.145 |
| lizards | Lygodactylus_grotei | Lygodactylus_grotei | LC | Stable | tropical | diurnal | 0.832 | 379608.537 | 8.536 |
| lizards | Lygodactylus_guibei | NA | NT | Decreasing | NA | NA | NA | NA | NA |
| lizards | Lygodactylus_gutturalis | Lygodactylus_gutturalis | LC | Stable | tropical | diurnal | 2.089 | 6305237.655 | 9.847 |
| lizards | Lygodactylus_heterurus | NA | LC | Decreasing | NA | NA | NA | NA | NA |
| lizards | Lygodactylus_inexpectatus | NA | DD | Unknown | NA | NA | NA | NA | NA |
| lizards | Lygodactylus_insularis | NA | CR | Decreasing | NA | NA | NA | NA | NA |
| lizards | Lygodactylus_intermedius | NA | EN | Decreasing | NA | NA | NA | NA | NA |
| lizards | Lygodactylus_keniensis | Lygodactylus_keniensis | LC | Unknown | tropical | diurnal | 1.738 | 262812.177 | 7.436 |
| lizards | Lygodactylus_kimhowelli | Lygodactylus_kimhowelli | LC | Stable | tropical | diurnal | 1.096 | 8703.89 | 14.87 |
| lizards | Lygodactylus_klemmeri | NA | NT | Unknown | NA | NA | NA | NA | NA |
| lizards | Lygodactylus_klugei | Lygodactylus_klugei | LC | Unknown | tropical | diurnal | 0.977 | 247442.92 | 7.513 |

| | | | | | | | | | |
|---|---|---|---|---|---|---|---|---|---|
| lizards | Lygodactylus_laterimaculatus | NA | LC | Unknown | NA | NA | NA | NA | NA |
| lizards | Lygodactylus_lawrencei | Lygodactylus_lawrencei | LC | Stable | tropical | diurnal | 0.977 | 102734.88 | 5.785 |
| lizards | Lygodactylus_madagascariensis | NA | VU | Decreasing | NA | NA | NA | NA | NA |
| lizards | Lygodactylus_manni | Lygodactylus_manni | LC | Unknown | tropical | diurnal | 1.738 | 46936.309 | 19.41 |
| lizards | Lygodactylus_miops | NA | LC | Decreasing | NA | NA | NA | NA | NA |
| lizards | Lygodactylus_mirabilis | Lygodactylus_mirabilis | CR | Unknown | tropical | diurnal | 0.646 | 23.154 | 8.232 |
| lizards | Lygodactylus_mombasicus | Lygodactylus_mombasicus | LC | Stable | tropical | diurnal | 1.413 | 7241.407 | 14.784 |
| lizards | Lygodactylus_montanus | NA | LC | Stable | NA | NA | NA | NA | NA |
| lizards | Lygodactylus_ornatus | NA | EN | Unknown | NA | NA | NA | NA | NA |
| lizards | Lygodactylus_pauliani | NA | DD | Unknown | NA | NA | NA | NA | NA |
| lizards | Lygodactylus_picturatus | Lygodactylus_picturatus | LC | Unknown | tropical | diurnal | 1.862 | 1218978.755 | 10.231 |
| lizards | Lygodactylus_pictus | Lygodactylus_pictus | LC | Unknown | tropical | diurnal | 1.622 | 29084.424 | 10.562 |
| lizards | Lygodactylus_rarus | NA | NT | Unknown | NA | NA | NA | NA | NA |
| lizards | Lygodactylus_regulus | NA | NT | Decreasing | NA | NA | NA | NA | NA |
| lizards | Lygodactylus_rex | NA | NT | Decreasing | NA | NA | NA | NA | NA |
| lizards | Lygodactylus_roavolana | NA | EN | Decreasing | NA | NA | NA | NA | NA |
| lizards | Lygodactylus_scheffleri | NA | DD | Unknown | NA | NA | NA | NA | NA |
| lizards | Lygodactylus_scorteccii | NA | DD | Unknown | NA | NA | NA | NA | NA |
| lizards | Lygodactylus_somalicus | Lygodactylus_somalicus | LC | Unknown | tropical | diurnal | 0.871 | 153499.587 | 6.432 |
| lizards | Lygodactylus_thomensis | Lygodactylus_thomensis | LC | Stable | tropical | diurnal | 1.202 | 108.186 | 15.68 |
| lizards | Lygodactylus_tolampyae | NA | LC | Decreasing | NA | NA | NA | NA | NA |
| lizards | Lygodactylus_tuberosus | Lygodactylus_tuberosus | LC | Stable | temperate | diurnal | 1.349 | 50959.9 | 11.206 |
| lizards | Lygodactylus_verticillatus | Lygodactylus_verticillatus | LC | Stable | temperate | diurnal | 1.047 | 66079.371 | 10.58 |
| lizards | Lygodactylus_viscatus | NA | NT | Unknown | NA | NA | NA | NA | NA |
| lizards | Lygodactylus_williamsi | Lygodactylus_williamsi | CR | Decreasing | tropical | diurnal | 0.977 | 50.944 | 16.09 |
| lizards | Lygodactylus_wojnowskii | NA | not_available | not_available | NA | NA | NA | NA | NA |
| lizards | Lygosoma_angeli | Lygosoma_angeli | LC | Stable | tropical | nocturnal | 79.433 | 3681.571 | 12.698 |
| lizards | Lygosoma_boehmei | NA | DD | Unknown | NA | NA | NA | NA | NA |
| lizards | Lygosoma_corpulentum | Lygosoma_corpulentum | LC | Stable | tropical | cathemeral | 186.209 | 75138.056 | 9.889 |
| lizards | Lygosoma_haroldyoungi | NA | LC | Unknown | NA | NA | NA | NA | NA |
| lizards | Lygosoma_kinabatanganense | NA | not_available | not_available | NA | NA | NA | NA | NA |
| lizards | Lygosoma_koratense | Lygosoma_koratense | LC | Unknown | tropical | nocturnal | 38.905 | 2802.458 | 17.255 |
| lizards | Lygosoma_quadrupes | Lygosoma_quadrupes | LC | Stable | tropical | cathemeral | 4.898 | 110.905 | 29.035 |
| lizards | Lygosoma_siamense | NA | LC | Unknown | NA | NA | NA | NA | NA |

| | | | | | | | | | |
|---|---|---|---|---|---|---|---|---|---|
| lizards | Lygosoma_veunsaiense | NA | LC | Stable | NA | NA | NA | NA | NA |
| lizards | Lyriocephalus_scutatus | Lyriocephalus_scutatus | VU | Unknown | tropical | diurnal | 223.872 | 13983.564 | 19.11 |
| lizards | Mabuya_cochonae | NA | CR | Unknown | NA | NA | NA | NA | NA |
| lizards | Mabuya_desiradae | NA | CR | Decreasing | NA | NA | NA | NA | NA |
| lizards | Mabuya_dominicana | Mabuya_dominicana | LC | Stable | tropical | diurnal | 22.387 | 624.108 | 12.98 |
| lizards | Mabuya_mabouya | Mabuya_mabouya | CR | Unknown | tropical | diurnal | 22.387 | 1142.264 | 26.199 |
| lizards | Mabuya_parviterrae | NA | not_available | not_available | NA | NA | NA | NA | NA |
| lizards | Macropholidus_annectens | Macropholidus_annectens | EN | Unknown | tropical | diurnal | 4.898 | 3607.906 | 11.604 |
| lizards | Macropholidus_ataktolepis | NA | DD | Unknown | NA | NA | NA | NA | NA |
| lizards | Macropholidus_ruthveni | Macropholidus_ruthveni | LC | Unknown | tropical | diurnal | 2.042 | 24877.005 | 9.39 |
| lizards | Madascincus_arenicola | NA | CR | Unknown | NA | NA | NA | NA | NA |
| lizards | Madascincus_igneocaudatus | NA | LC | Decreasing | NA | NA | NA | NA | NA |
| lizards | Madascincus_macrolepis | NA | EN | Decreasing | NA | NA | NA | NA | NA |
| lizards | Madascincus_melanopleura | Madascincus_melanopleura | LC | Decreasing | tropical | diurnal | 3.467 | 141035.308 | 7.717 |
| lizards | Madascincus_miafina | NA | not_available | not_available | NA | NA | NA | NA | NA |
| lizards | Madascincus_minutus | NA | LC | Decreasing | NA | NA | NA | NA | NA |
| lizards | Madascincus_mouroundavae | Madascincus_mouroundavae | LC | Decreasing | tropical | diurnal | 7.413 | 69093.273 | 7.239 |
| lizards | Madascincus_polleni | NA | LC | Stable | NA | NA | NA | NA | NA |
| lizards | Madascincus_stumpffi | Madascincus_stumpffi | NT | Decreasing | tropical | nocturnal | 33.113 | 15269.165 | 8.755 |
| lizards | Malayodracon_robinsonii | Gonocephalus_robinsonii | NT | Stable | tropical | diurnal | 138.038 | 6669.145 | 7.162 |
| lizards | Manciola_guaporicola | Manciola_guaporicola | LC | Decreasing | tropical | diurnal | 19.953 | 1493212.379 | 7.002 |
| lizards | Mantheyus_phuwuanensis | Mantheyus_phuwuanensis | NT | Stable | tropical | cathemeral | 23.988 | 1887.291 | 17.342 |
| lizards | Maracaiba_meridensis | Maracaiba_meridensis | LC | Stable | tropical | diurnal | 9.12 | 11816.637 | 17.094 |
| lizards | Maracaiba_zuliae | NA | LC | Stable | NA | NA | NA | NA | NA |
| lizards | Marinussaurus_curupira | NA | EN | Unknown | NA | NA | NA | NA | NA |
| lizards | Marisora_alliacea | Marisora_alliacea | LC | Stable | tropical | diurnal | 14.125 | 7785.384 | 11.086 |
| lizards | Marisora_aquilonaria | NA | not_available | not_available | NA | NA | NA | NA | NA |
| lizards | Marisora_aurulae | NA | VU | Decreasing | NA | NA | NA | NA | NA |
| lizards | Marisora_berengerae | NA | LC | Stable | NA | NA | NA | NA | NA |
| lizards | Marisora_brachypoda | Marisora_brachypoda | LC | Stable | tropical | diurnal | 15.488 | 865336.97 | 10.081 |
| lizards | Marisora_falconensis | NA | LC | Stable | NA | NA | NA | NA | NA |
| lizards | Marisora_pergravis | Alinea_pergravis | LC | Stable | tropical | diurnal | 15.849 | 23.528 | 22.202 |
| lizards | Marisora_roatanae | NA | CR | Decreasing | NA | NA | NA | NA | NA |
| lizards | Marisora_unimarginata | Marisora_unimarginata | LC | Stable | tropical | diurnal | 16.596 | 938239.036 | 10.004 |

| | | | | | | | | | |
|---|---|---|---|---|---|---|---|---|---|
| lizards | Marisora_urtica | NA | not_available | not_available | NA | NA | NA | NA | NA |
| lizards | Marmorosphax_boulinda | Marmorosphax_boulinda | VU | Unknown | tropical | cathemeral | 2.754 | 99.943 | 2.5 |
| lizards | Marmorosphax_kaala | NA | CR | Decreasing | NA | NA | NA | NA | NA |
| lizards | Marmorosphax_montana | Marmorosphax_montana | EN | Decreasing | tropical | cathemeral | 5.888 | 461.722 | 4.682 |
| lizards | Marmorosphax_taom | Marmorosphax_taom | EN | Decreasing | tropical | cathemeral | 3.02 | 155.74 | 10.536 |
| lizards | Marmorosphax_tricolor | Marmorosphax_tricolor | LC | Stable | tropical | cathemeral | 5.623 | 14859.606 | 7.319 |
| lizards | Matoatoa_brevipes | NA | VU | Decreasing | NA | NA | NA | NA | NA |
| lizards | Matoatoa_spannringi | Matoatoa_spannringi | DD | Decreasing | tropical | nocturnal | 4.266 | 134.555 | 8.738 |
| lizards | Matobosaurus_maltzahni | Matobosaurus_maltzahni | LC | Stable | tropical | diurnal | 407.38 | 324748.28 | 7.254 |
| lizards | Matobosaurus_validus | Matobosaurus_validus | LC | Stable | tropical | diurnal | 616.595 | 820594.417 | 12.24 |
| lizards | Mediodactylus_amictopholis | Mediodactylus_amictopholis | EN | Decreasing | temperate | diurnal | 1.122 | 253.573 | 23.081 |
| lizards | Mediodactylus_aspratilis | NA | DD | Unknown | NA | NA | NA | NA | NA |
| lizards | Mediodactylus_brachykolon | NA | DD | Unknown | NA | NA | NA | NA | NA |
| lizards | Mediodactylus_heterocercus | Mediodactylus_heterocercus | LC | Stable | temperate | nocturnal | 3.162 | 163494.2 | 17.497 |
| lizards | Mediodactylus_heteropholis | NA | DD | Unknown | NA | NA | NA | NA | NA |
| lizards | Mediodactylus_ilamensis | NA | not_available | not_available | NA | NA | NA | NA | NA |
| lizards | Mediodactylus_kotschyi | Mediodactylus_kotschyi | LC | Unknown | temperate | cathemeral | 3.802 | 122055.923 | 17.35 |
| lizards | Mediodactylus_narynensis | NA | LC | Stable | NA | NA | NA | NA | NA |
| lizards | Mediodactylus_russowii | Mediodactylus_russowii | LC | Stable | temperate | cathemeral | 3.311 | 2115396.523 | 7.422 |
| lizards | Mediodactylus_sagittifer | NA | DD | Unknown | NA | NA | NA | NA | NA |
| lizards | Mediodactylus_spinicauda | Mediodactylus_spinicauda | LC | Stable | temperate | nocturnal | 2.512 | 70327.02 | 10.193 |
| lizards | Mediodactylus_stevenandersoni | NA | not_available | not_available | NA | NA | NA | NA | NA |
| lizards | Mediodactylus_walli | NA | LC | Unknown | NA | NA | NA | NA | NA |
| lizards | Medopheos_edracanthus | NA | LC | Stable | NA | NA | NA | NA | NA |
| lizards | Melanoseps_ater | Melanoseps_ater | LC | Unknown | tropical | diurnal | 10.233 | 85203.574 | 13.689 |
| lizards | Melanoseps_emmrichi | Melanoseps_emmrichi | EN | Unknown | tropical | cathemeral | 9.12 | 282.947 | 12.911 |
| lizards | Melanoseps_loveridgei | Melanoseps_loveridgei | LC | Unknown | tropical | diurnal | 5.754 | 350996.054 | 10.03 |
| lizards | Melanoseps_pygmaeus | Melanoseps_pygmaeus | LC | Unknown | tropical | diurnal | 1.778 | 8008.98 | 15.694 |
| lizards | Melanoseps_rondoensis | NA | DD | Unknown | NA | NA | NA | NA | NA |
| lizards | Menetia_alanae | Menetia_alanae | LC | Stable | tropical | diurnal | 0.437 | 138055.468 | 1.853 |
| lizards | Menetia_concinna | NA | LC | Unknown | NA | NA | NA | NA | NA |
| lizards | Menetia_greyii | Menetia_greyii | LC | Stable | temperate | diurnal | 1.122 | 7074442.807 | 4.039 |
| lizards | Menetia_maini | Menetia_maini | LC | Stable | tropical | diurnal | 0.55 | 1319458.927 | 2.84 |
| lizards | Menetia_surda | NA | LC | Stable | NA | NA | NA | NA | NA |

| | | | | | | | | | |
|---|---|---|---|---|---|---|---|---|---|
| lizards | Meroles_anchietae | Meroles_anchietae | LC | Stable | tropical | diurnal | 4.786 | 200330.97 | 4.006 |
| lizards | Meroles_ctenodactylus | Meroles_ctenodactylus | LC | Stable | temperate | diurnal | 20.893 | 58210.975 | 3.86 |
| lizards | Meroles_cuneirostris | Meroles_cuneirostris | LC | Stable | temperate | diurnal | 4.571 | 138082.052 | 3.684 |
| lizards | Meroles_knoxii | Meroles_knoxii | LC | Stable | temperate | diurnal | 7.244 | 165150.607 | 6.69 |
| lizards | Meroles_micropholidotus | NA | LC | Stable | NA | NA | NA | NA | NA |
| lizards | Meroles_reticulatus | Meroles_reticulatus | LC | Stable | tropical | diurnal | 3.89 | 187250.75 | 4.711 |
| lizards | Meroles_squamulosus | Meroles_squamulosa | LC | Stable | tropical | diurnal | 11.482 | 3461354.19 | 8.193 |
| lizards | Meroles_suborbitalis | Meroles_suborbitalis | LC | Stable | temperate | diurnal | 8.318 | 701307.964 | 5.133 |
| lizards | Mesalina_adramitana | Mesalina_adramitana | LC | not_available | temperate | diurnal | 2.291 | 742653.619 | 4.193 |
| lizards | Mesalina_ayunensis | NA | DD | Unknown | NA | NA | NA | NA | NA |
| lizards | Mesalina_bahaeldini | Mesalina_bahaeldini | LC | Stable | temperate | diurnal | 4.169 | 647041.204 | 5.564 |
| lizards | Mesalina_balfouri | Mesalina_balfouri | LC | Unknown | tropical | diurnal | 4.571 | 3631.199 | 11 |
| lizards | Mesalina_bernoullii | NA | not_available | not_available | NA | NA | NA | NA | NA |
| lizards | Mesalina_brevirostris | Mesalina_brevirostris | LC | Stable | temperate | diurnal | 5.012 | 1193015.604 | 7.775 |
| lizards | Mesalina_guttulata | Mesalina_guttulata | LC | Stable | temperate | diurnal | 5.012 | 9617162.617 | 2.459 |
| lizards | Mesalina_kuri | NA | LC | Unknown | NA | NA | NA | NA | NA |
| lizards | Mesalina_martini | Mesalina_martini | LC | Stable | tropical | diurnal | 2.951 | 100772.916 | 7.967 |
| lizards | Mesalina_microlepis | NA | not_available | not_available | NA | NA | NA | NA | NA |
| lizards | Mesalina_olivieri | Mesalina_olivieri | LC | Stable | temperate | diurnal | 3.311 | 2601525.96 | 6.164 |
| lizards | Mesalina_pasteuri | Mesalina_pasteuri | LC | Unknown | temperate | diurnal | 2.951 | 5154182.544 | 1.361 |
| lizards | Mesalina_rubropunctata | Mesalina_rubropunctata | LC | Stable | temperate | diurnal | 7.079 | 5592089.707 | 1.868 |
| lizards | Mesalina_saudiarabica | NA | not_available | not_available | NA | NA | NA | NA | NA |
| lizards | Mesalina_simoni | Mesalina_simoni | LC | Stable | temperate | diurnal | 2.951 | 55851.966 | 16.315 |
| lizards | Mesalina_watsonana | Mesalina_watsonana | LC | Stable | temperate | diurnal | 5.012 | 2510586.892 | 8.795 |
| lizards | Mesoamericus_bilobatus | NA | LC | Unknown | NA | NA | NA | NA | NA |
| lizards | Mesoscincus_altamirani | NA | DD | Unknown | NA | NA | NA | NA | NA |
| lizards | Mesoscincus_managuae | NA | LC | Unknown | NA | NA | NA | NA | NA |
| lizards | Mesoscincus_schwartzei | NA | LC | Stable | NA | NA | NA | NA | NA |
| lizards | Micrablepharus_atticolus | Micrablepharus_atticolus | LC | Unknown | tropical | diurnal | 2.344 | 1787612.212 | 8.512 |
| lizards | Micrablepharus_maximiliani | Micrablepharus_maximiliani | LC | Unknown | tropical | diurnal | 1.862 | 4221760.121 | 7.325 |
| lizards | Microauris_aurantolabium | Calotes_aurantolabium | EN | Unknown | tropical | diurnal | 16.218 | 621.174 | 9.144 |
| lizards | Microgecko_chabaharensis | NA | not_available | not_available | NA | NA | NA | NA | NA |
| lizards | Microgecko_depressus | NA | LC | Unknown | NA | NA | NA | NA | NA |
| lizards | Microgecko_helenae | NA | DD | Unknown | NA | NA | NA | NA | NA |

| | | | | | | | | | |
|---|---|---|---|---|---|---|---|---|---|
| lizards | Microgecko_latifi | NA | LC | Stable | NA | NA | NA | NA | NA |
| lizards | Microgecko_persicus | Microgecko_persicus | LC | Unknown | temperate | nocturnal | 1.122 | 182508.696 | 8.949 |
| lizards | Microlophus_albemarlensis | NA | LC | Stable | NA | NA | NA | NA | NA |
| lizards | Microlophus_arenarius | NA | DD | Unknown | NA | NA | NA | NA | NA |
| lizards | Microlophus_atacamensis | Microlophus_atacamensis | LC | Stable | temperate | diurnal | 72.444 | 18794.232 | 4.712 |
| lizards | Microlophus_barringtonensis | NA | not_available | not_available | NA | NA | NA | NA | NA |
| lizards | Microlophus_bivittatus | NA | NT | Unknown | NA | NA | NA | NA | NA |
| lizards | Microlophus_delanonis | NA | LC | Unknown | NA | NA | NA | NA | NA |
| lizards | Microlophus_duncanensis | NA | NT | Stable | NA | NA | NA | NA | NA |
| lizards | Microlophus_grayii | NA | NT | Stable | NA | NA | NA | NA | NA |
| lizards | Microlophus_habelii | NA | LC | Stable | NA | NA | NA | NA | NA |
| lizards | Microlophus_heterolepis | NA | DD | Unknown | NA | NA | NA | NA | NA |
| lizards | Microlophus_indefatigabilis | NA | LC | Stable | NA | NA | NA | NA | NA |
| lizards | Microlophus_jacobii | NA | LC | Stable | NA | NA | NA | NA | NA |
| lizards | Microlophus_occipitalis | Microlophus_occipitalis | LC | Stable | tropical | diurnal | 19.498 | 97167.951 | 9.879 |
| lizards | Microlophus_pacificus | NA | LC | Increasing | NA | NA | NA | NA | NA |
| lizards | Microlophus_peruvianus | Microlophus_peruvianus | LC | Unknown | tropical | diurnal | 89.125 | 60949.557 | 9.817 |
| lizards | Microlophus_quadrivittatus | Microlophus_quadrivittatus | LC | Stable | tropical | diurnal | 72.444 | 34776.741 | 5.172 |
| lizards | Microlophus_tarapacensis | NA | DD | Unknown | NA | NA | NA | NA | NA |
| lizards | Microlophus_theresiae | Microlophus_theresiae | LC | Unknown | tropical | diurnal | 42.658 | 11277.626 | 11.328 |
| lizards | Microlophus_theresioides | Microlophus_theresioides | LC | Stable | tropical | diurnal | 52.481 | 48894.593 | 3.516 |
| lizards | Microlophus_thoracicus | Microlophus_thoracicus | LC | Stable | tropical | diurnal | 30.2 | 54452.811 | 8.44 |
| lizards | Mniarogekko_chahoua | Mniarogekko_chahoua | VU | Decreasing | tropical | nocturnal | 66.069 | 3821.268 | 9.076 |
| lizards | Mniarogekko_jalu | Mniarogekko_jalu | EN | Unknown | tropical | nocturnal | 56.234 | 391.569 | 8.423 |
| lizards | Mochlus_brevicaudis | Mochlus_brevicaudis | LC | Unknown | tropical | diurnal | 9.55 | 358660.483 | 13.961 |
| lizards | Mochlus_fernandi | Lepidothyris_fernandi | LC | Unknown | tropical | cathemeral | 144.544 | 2980336.329 | 8.242 |
| lizards | Mochlus_guineensis | Mochlus_guineensis | LC | Unknown | tropical | diurnal | 10.471 | 2959270.674 | 9.137 |
| lizards | Mochlus_hinkeli | Lepidothyris_hinkeli | LC | Unknown | tropical | diurnal | 72.444 | 828947.62 | 7.314 |
| lizards | Mochlus_mabuiiformis | NA | DD | Unknown | NA | NA | NA | NA | NA |
| lizards | Mochlus_mafianus | NA | LC | Unknown | NA | NA | NA | NA | NA |
| lizards | Mochlus_pembanus | Lygosoma_pembanum | LC | Unknown | tropical | nocturnal | 16.596 | 3527.902 | 19.87 |
| lizards | Mochlus_simonettai | NA | DD | Unknown | NA | NA | NA | NA | NA |
| lizards | Mochlus_striatus | Lepidothyris_striatus | LC | Stable | tropical | cathemeral | 112.202 | 425436.302 | 5.382 |
| lizards | Mochlus_sundevallii | Mochlus_sundevalli | LC | Stable | tropical | diurnal | 63.096 | 7876204.793 | 8.501 |

| | | | | | | | | | |
|---|---|---|---|---|---|---|---|---|---|
| lizards | Mokopirirakau_cryptozoicus | Mokopirirakau_cryptozoicus | EN | Decreasing | temperate | nocturnal | 13.49 | 3963.552 | 6.202 |
| lizards | Mokopirirakau_granulatus | Mokopirirakau_granulatus | VU | Decreasing | temperate | cathemeral | 19.055 | 118586.854 | 8.681 |
| lizards | Mokopirirakau_kahutarae | Mokopirirakau_kahutarae | VU | Decreasing | temperate | nocturnal | 15.136 | 5639.707 | 4.442 |
| lizards | Mokopirirakau_nebulosus | Mokopirirakau_nebulosus | NT | Increasing | temperate | nocturnal | 14.791 | 995.26 | 3.533 |
| lizards | Moloch_horridus | Moloch_horridus | LC | Stable | temperate | diurnal | 61.66 | 3199451.13 | 2.66 |
| lizards | Monilesaurus_ellioti | Calotes_ellioti | LC | Unknown | tropical | diurnal | 14.454 | 85602.924 | 19.257 |
| lizards | Monilesaurus_rouxii | Calotes_rouxii | LC | Stable | tropical | diurnal | 18.197 | 280435.788 | 19.805 |
| lizards | Monopeltis_capensis | Monopeltis_capensis | LC | Stable | temperate | diurnal | 31.623 | 821772.489 | 10.188 |
| lizards | Morethia_adelaidensis | Morethia_adelaidensis | LC | Stable | temperate | diurnal | 4.169 | 1707088.541 | 4.51 |
| lizards | Morethia_boulengeri | Morethia_boulengeri | LC | Stable | temperate | diurnal | 3.467 | 3302096.32 | 5.151 |
| lizards | Morethia_butleri | Morethia_butleri | LC | Stable | temperate | diurnal | 3.467 | 1275905.508 | 3.367 |
| lizards | Morethia_lineoocellata | Morethia_lineoocellata | LC | Stable | temperate | diurnal | 3.467 | 259510.214 | 8.446 |
| lizards | Morethia_obscura | Morethia_obscura | LC | Unknown | temperate | diurnal | 3.311 | 1568218.055 | 6.142 |
| lizards | Morethia_ruficauda | Morethia_ruficauda | LC | Stable | tropical | diurnal | 1.738 | 3133787.402 | 1.978 |
| lizards | Morethia_storri | Morethia_storri | LC | Stable | tropical | diurnal | 0.955 | 910508.765 | 1.933 |
| lizards | Morethia_taeniopleura | Morethia_taeniopleura | LC | Stable | tropical | diurnal | 1.514 | 739225.908 | 5.129 |
| lizards | Nactus_acutus | NA | DD | Unknown | NA | NA | NA | NA | NA |
| lizards | Nactus_cheverti | NA | LC | Stable | NA | NA | NA | NA | NA |
| lizards | Nactus_coindemirensis | NA | VU | Stable | NA | NA | NA | NA | NA |
| lizards | Nactus_eboracensis | Nactus_eboracensis | LC | Stable | tropical | nocturnal | 4.266 | 48422.381 | 2.377 |
| lizards | Nactus_galgajuga | NA | LC | Unknown | NA | NA | NA | NA | NA |
| lizards | Nactus_kunan | NA | DD | Unknown | NA | NA | NA | NA | NA |
| lizards | Nactus_multicarinatus | Nactus_multicarinatus | LC | Stable | tropical | nocturnal | 5.37 | 92954.719 | 4.562 |
| lizards | Nactus_pelagicus | Nactus_pelagicus | LC | Stable | tropical | nocturnal | 10.233 | 266115.247 | 5.411 |
| lizards | Nactus_serpensinsula | NA | VU | Stable | NA | NA | NA | NA | NA |
| lizards | Nactus_soniae | NA | EX | not_available | NA | NA | NA | NA | NA |
| lizards | Nactus_sphaerodactylodes | Nactus_sphaerodactylodes | LC | Unknown | tropical | nocturnal | 0.457 | 1921.004 | 4.281 |
| lizards | Nactus_vankampeni | NA | LC | Unknown | NA | NA | NA | NA | NA |
| lizards | Namazonurus_campbelli | Namazonurus_campbelli | LC | Stable | temperate | diurnal | 11.482 | 6755.86 | 5.15 |
| lizards | Namazonurus_lawrenci | Namazonurus_lawrenci | LC | Stable | temperate | diurnal | 9.55 | 5891.686 | 5.162 |
| lizards | Namazonurus_namaquensis | Namazonurus_namaquensis | LC | Stable | temperate | diurnal | 13.183 | 5499.795 | 4.18 |
| lizards | Namazonurus_peersi | Namazonurus_peersi | LC | Stable | temperate | diurnal | 15.136 | 24004.951 | 6.913 |
| lizards | Namazonurus_pustulatus | Namazonurus_pustulatus | LC | Stable | tropical | diurnal | 13.183 | 17393.537 | 6.11 |
| lizards | Nangura_spinosa | NA | EN | Decreasing | NA | NA | NA | NA | NA |

| | | | | | | | | | |
|---|---|---|---|---|---|---|---|---|---|
| lizards | Nannoscincus_exos | NA | CR | Decreasing | NA | NA | NA | NA | NA |
| lizards | Nannoscincus_garrulus | Nannoscincus_garrulus | EN | Decreasing | tropical | cathemeral | 2.63 | 135.614 | 5.314 |
| lizards | Nannoscincus_gracilis | Nannoscincus_gracilis | VU | Unknown | tropical | cathemeral | 0.933 | 6301.309 | 6.718 |
| lizards | Nannoscincus_greeri | NA | EN | Decreasing | NA | NA | NA | NA | NA |
| lizards | Nannoscincus_hanchisteus | NA | CR | Decreasing | NA | NA | NA | NA | NA |
| lizards | Nannoscincus_humectus | NA | EN | Decreasing | NA | NA | NA | NA | NA |
| lizards | Nannoscincus_mariei | Nannoscincus_mariei | VU | Decreasing | tropical | cathemeral | 1.738 | 3111.79 | 9.269 |
| lizards | Nannoscincus_rankini | NA | CR | Decreasing | NA | NA | NA | NA | NA |
| lizards | Nannoscincus_slevini | NA | EN | Decreasing | NA | NA | NA | NA | NA |
| lizards | Narudasia_festiva | Narudasia_festiva | LC | Stable | temperate | diurnal | 0.759 | 184589.511 | 4.716 |
| lizards | Naultinus_elegans | Naultinus_elegans | VU | Decreasing | temperate | diurnal | 6.918 | 42897.183 | 11.561 |
| lizards | Naultinus_gemmeus | Naultinus_gemmeus | EN | Decreasing | temperate | diurnal | 10.233 | 72027.274 | 9.032 |
| lizards | Naultinus_grayii | Naultinus_grayii | NT | Decreasing | temperate | diurnal | 17.378 | 1952.653 | 9.628 |
| lizards | Naultinus_manukanus | Naultinus_manukanus | EN | Unknown | temperate | diurnal | 10.715 | 2842.957 | 6.546 |
| lizards | Naultinus_punctatus | Naultinus_punctatus | VU | Decreasing | temperate | diurnal | 17.378 | 56072.889 | 8.361 |
| lizards | Naultinus_rudis | Naultinus_rudis | EN | Decreasing | temperate | diurnal | 7.413 | 4131.881 | 6.153 |
| lizards | Naultinus_stellatus | Naultinus_stellatus | VU | Decreasing | temperate | diurnal | 10.233 | 11292.103 | 4.857 |
| lizards | Naultinus_tuberculatus | Naultinus_tuberculatus | VU | Decreasing | temperate | diurnal | 12.303 | 5594.208 | 3.768 |
| lizards | Nephrurus_amyae | Nephrurus_amyae | LC | Stable | tropical | nocturnal | 39.811 | 111723.021 | 2.172 |
| lizards | Nephrurus_asper | Nephrurus_asper | LC | Unknown | tropical | nocturnal | 26.303 | 878623.088 | 4.689 |
| lizards | Nephrurus_deleani | Nephrurus_deleani | LC | Stable | temperate | nocturnal | 18.197 | 8983.915 | 5.042 |
| lizards | Nephrurus_laevissimus | Nephrurus_laevissimus | LC | Stable | temperate | nocturnal | 18.197 | 1629449.613 | 1.381 |
| lizards | Nephrurus_levis | Nephrurus_levis | LC | Stable | temperate | nocturnal | 23.988 | 3537661.708 | 2.562 |
| lizards | Nephrurus_sheai | Nephrurus_sheai | LC | Stable | tropical | nocturnal | 28.84 | 320983.271 | 2.454 |
| lizards | Nephrurus_stellatus | Nephrurus_stellatus | LC | Stable | temperate | nocturnal | 16.596 | 180306.519 | 4.862 |
| lizards | Nephrurus_vertebralis | Nephrurus_vertebralis | LC | Stable | temperate | nocturnal | 15.488 | 555857.82 | 3.641 |
| lizards | Nephrurus_wheeleri | Nephrurus_wheeleri | LC | Unknown | temperate | nocturnal | 18.621 | 508326.239 | 4.239 |
| lizards | Nessia_bipes | Nessia_bipes | EN | Unknown | tropical | cathemeral | 1.479 | 716.523 | 12.773 |
| lizards | Nessia_burtonii | Nessia_burtonii | NT | Stable | tropical | nocturnal | 2.754 | 12090.069 | 18.752 |
| lizards | Nessia_deraniyagalai | NA | DD | Unknown | NA | NA | NA | NA | NA |
| lizards | Nessia_didactyla | NA | EN | Unknown | NA | NA | NA | NA | NA |
| lizards | Nessia_hickanala | NA | EN | Unknown | NA | NA | NA | NA | NA |
| lizards | Nessia_layardi | Nessia_layardi | CR | Decreasing | tropical | cathemeral | 2.138 | 3836.194 | 24.299 |
| lizards | Nessia_monodactyla | Nessia_monodactyla | EN | Unknown | tropical | nocturnal | 1.995 | 13848.97 | 20.507 |

| | | | | | | | | | |
|---|---|---|---|---|---|---|---|---|---|
| lizards | Nessia_sarasinorum | NA | VU | Unknown | NA | NA | NA | NA | NA |
| lizards | Neusticurus_bicarinatus | Neusticurus_bicarinatus | LC | Unknown | tropical | diurnal | 44.668 | 4356982.202 | 1.844 |
| lizards | Neusticurus_medemi | NA | LC | Unknown | NA | NA | NA | NA | NA |
| lizards | Neusticurus_racenisi | NA | LC | Stable | NA | NA | NA | NA | NA |
| lizards | Neusticurus_rudis | Neusticurus_rudis | LC | Decreasing | tropical | diurnal | 21.878 | 809214.026 | 1.155 |
| lizards | Neusticurus_tatei | Neusticurus_tatei | LC | Unknown | tropical | diurnal | 30.2 | 317553.835 | 0.593 |
| lizards | Ninurta_coeruleopunctatus | Ninurta_coeruleopunctatus | LC | Stable | temperate | diurnal | 13.183 | 15088.563 | 9.082 |
| lizards | Nothobachia_ablephara | Nothobachia_ablephara | LC | Unknown | tropical | cathemeral | 1.698 | 59192.358 | 6.744 |
| lizards | Notomabuya_frenata | Notomabuya_frenata | LC | Stable | tropical | diurnal | 15.849 | 4419369.536 | 8.176 |
| lizards | Notoscincus_butleri | NA | LC | Unknown | NA | NA | NA | NA | NA |
| lizards | Notoscincus_ornatus | Notoscincus_ornatus | LC | Stable | tropical | diurnal | 1.175 | 3561381.473 | 2.457 |
| lizards | Nubeoscincus_glacialis | NA | DD | Unknown | NA | NA | NA | NA | NA |
| lizards | Nubeoscincus_stellaris | NA | DD | Unknown | NA | NA | NA | NA | NA |
| lizards | Nucras_boulengeri | Nucras_boulengeri | LC | Unknown | tropical | diurnal | 6.457 | 523266.744 | 12.367 |
| lizards | Nucras_holubi | Nucras_holubi | LC | Stable | temperate | diurnal | 3.162 | 791737.586 | 10.428 |
| lizards | Nucras_intertexta | Nucras_intertexta | LC | Stable | temperate | diurnal | 19.055 | 1010673.719 | 7.624 |
| lizards | Nucras_lalandii | Nucras_lalandii | LC | Stable | temperate | diurnal | 38.905 | 303549.885 | 13.032 |
| lizards | Nucras_ornata | Nucras_ornata | LC | Stable | tropical | diurnal | 30.903 | 1027910.451 | 11.3 |
| lizards | Nucras_taeniolata | Nucras_taeniolata | LC | Stable | temperate | diurnal | 30.903 | 58948.726 | 8.836 |
| lizards | Nucras_tessellata | Nucras_tessellata | LC | Stable | temperate | diurnal | 19.055 | 697346.95 | 6.424 |
| lizards | Oedodera_marmorata | Oedodera_marmorata | EN | Decreasing | tropical | nocturnal | 5.495 | 121.1 | 11.584 |
| lizards | Oedura_castelnaui | Oedura_castelnaui | LC | Stable | tropical | nocturnal | 18.621 | 383086.6 | 3.753 |
| lizards | Oedura_coggeri | Oedura_coggeri | LC | Stable | tropical | nocturnal | 10.715 | 98443.023 | 3.676 |
| lizards | Oedura_filicipoda | Oedura_filicipoda | LC | Unknown | tropical | nocturnal | 24.547 | 21336.446 | 0.895 |
| lizards | Oedura_gemmata | Oedura_gemmata | LC | Stable | tropical | nocturnal | 27.542 | 38325.095 | 0.922 |
| lizards | Oedura_gracilis | Oedura_gracilis | LC | Stable | tropical | nocturnal | 12.303 | 189248.186 | 2.624 |
| lizards | Oedura_jowalbinna | Oedura_jowalbinna | NT | Stable | tropical | nocturnal | 6.607 | 66.233 | 2.301 |
| lizards | Oedura_luritja | NA | LC | Stable | NA | NA | NA | NA | NA |
| lizards | Oedura_marmorata | Oedura_marmorata | LC | Stable | tropical | nocturnal | 27.542 | 221682.197 | 1.741 |
| lizards | Oedura_monilis | Oedura_monilis | LC | Unknown | temperate | nocturnal | 19.055 | 574184.236 | 6.837 |
| lizards | Oedura_tryoni | Oedura_tryoni | LC | Stable | temperate | nocturnal | 22.387 | 250672.481 | 7.257 |
| lizards | Oligosoma_acrinasum | Oligosoma_acrinasum | NT | Increasing | temperate | diurnal | 14.125 | 1579.706 | 0.356 |
| lizards | Oligosoma_aeneum | Oligosoma_aeneum | LC | Stable | temperate | diurnal | 8.913 | 100139.804 | 9.747 |
| lizards | Oligosoma_alani | Oligosoma_alani | VU | Increasing | temperate | nocturnal | 79.433 | 103.057 | 9.511 |

| | | | | | | | | | |
|---|---|---|---|---|---|---|---|---|---|
| lizards | Oligosoma_albornense | NA | CR | Decreasing | NA | NA | NA | NA | NA |
| lizards | Oligosoma_auroraense | NA | VU | Stable | NA | NA | NA | NA | NA |
| lizards | Oligosoma_awakopaka | NA | CR | Decreasing | NA | NA | NA | NA | NA |
| lizards | Oligosoma_burganae | Oligosoma_burganae | CR | Decreasing | temperate | diurnal | 6.761 | 19.982 | 8.397 |
| lizards | Oligosoma_chloronoton | Oligosoma_chloronoton | EN | Decreasing | temperate | diurnal | 44.668 | 37946.172 | 8.737 |
| lizards | Oligosoma_elium | NA | EN | Decreasing | NA | NA | NA | NA | NA |
| lizards | Oligosoma_fallai | Oligosoma_fallai | VU | Stable | temperate | cathemeral | 70.795 | 4.572 | 1.169 |
| lizards | Oligosoma_grande | Oligosoma_grande | EN | Stable | temperate | diurnal | 37.154 | 9040.371 | 8.466 |
| lizards | Oligosoma_hardyi | Oligosoma_hardyi | VU | Unknown | temperate | cathemeral | 4.571 | 2.008 | 0.691 |
| lizards | Oligosoma_homalonotum | Oligosoma_homalonotum | EN | Stable | temperate | diurnal | 72.444 | 313.854 | 7.1 |
| lizards | Oligosoma_hoparatea | NA | CR | Decreasing | NA | NA | NA | NA | NA |
| lizards | Oligosoma_inconspicuum | Oligosoma_inconspicuum | VU | Decreasing | temperate | diurnal | 8.511 | 42485.119 | 7.633 |
| lizards | Oligosoma_infrapunctatum | Oligosoma_infrapunctatum | CR | Unknown | temperate | diurnal | 33.884 | 44858.158 | 7.843 |
| lizards | Oligosoma_judgei | Oligosoma_judgei | CR | Stable | temperate | diurnal | 15.849 | 795.613 | 4.835 |
| lizards | Oligosoma_kokowai | NA | LC | Increasing | NA | NA | NA | NA | NA |
| lizards | Oligosoma_levidensum | Oligosoma_levidensum | EN | Decreasing | temperate | diurnal | 2.399 | 509.573 | 6.777 |
| lizards | Oligosoma_lichenigerum | NA | VU | Unknown | NA | NA | NA | NA | NA |
| lizards | Oligosoma_lineoocellatum | Oligosoma_lineoocellatum | EN | Decreasing | temperate | diurnal | 30.2 | 10139.384 | 11.979 |
| lizards | Oligosoma_longipes | Oligosoma_longipes | EN | Decreasing | temperate | diurnal | 8.511 | 30550.547 | 5.908 |
| lizards | Oligosoma_maccanni | Oligosoma_maccanni | LC | Stable | temperate | diurnal | 9.55 | 57177.209 | 9.064 |
| lizards | Oligosoma_macgregori | Oligosoma_macgregori | VU | Increasing | temperate | cathemeral | 35.481 | 20.338 | 5.982 |
| lizards | Oligosoma_microlepis | Oligosoma_microlepis | EN | Decreasing | temperate | diurnal | 7.762 | 7616.178 | 5.653 |
| lizards | Oligosoma_moco | Oligosoma_moco | LC | Stable | temperate | diurnal | 10.965 | 4289.082 | 11.941 |
| lizards | Oligosoma_newmani | NA | VU | Decreasing | NA | NA | NA | NA | NA |
| lizards | Oligosoma_nigriplantare | NA | NT | Stable | NA | NA | NA | NA | NA |
| lizards | Oligosoma_notosaurus | Oligosoma_notosaurus | LC | Unknown | temperate | diurnal | 15.488 | 1288.428 | 3.159 |
| lizards | Oligosoma_oliveri | Oligosoma_oliveri | NT | Stable | temperate | cathemeral | 34.674 | 47.966 | 9.176 |
| lizards | Oligosoma_ornatum | Oligosoma_ornatum | LC | Decreasing | temperate | cathemeral | 13.804 | 94721.432 | 9.425 |
| lizards | Oligosoma_otagense | Oligosoma_otagense | EN | Decreasing | temperate | diurnal | 117.49 | 8968.577 | 8.606 |
| lizards | Oligosoma_pachysomaticum | NA | not_available | not_available | NA | NA | NA | NA | NA |
| lizards | Oligosoma_pikitanga | NA | CR | Decreasing | NA | NA | NA | NA | NA |
| lizards | Oligosoma_polychroma | Oligosoma_polychroma | LC | Stable | temperate | diurnal | 10.471 | 177591.856 | 8.104 |
| lizards | Oligosoma_prasinum | NA | EN | Decreasing | NA | NA | NA | NA | NA |
| lizards | Oligosoma_repens | NA | VU | Decreasing | NA | NA | NA | NA | NA |

| | | | | | | | | | |
|---|---|---|---|---|---|---|---|---|---|
| lizards | Oligosoma_robinsoni | NA | VU | Decreasing | NA | NA | NA | NA | NA |
| lizards | Oligosoma_roimata | NA | VU | Unknown | NA | NA | NA | NA | NA |
| lizards | Oligosoma_salmo | NA | CR | Decreasing | NA | NA | NA | NA | NA |
| lizards | Oligosoma_smithi | Oligosoma_smithi | LC | Unknown | temperate | diurnal | 11.22 | 14624.132 | 8.088 |
| lizards | Oligosoma_stenotis | Oligosoma_stenotis | EN | Decreasing | temperate | diurnal | 8.511 | 820.071 | 3.024 |
| lizards | Oligosoma_striatum | Oligosoma_striatum | VU | Decreasing | temperate | diurnal | 8.913 | 23867.617 | 11.886 |
| lizards | Oligosoma_suteri | Oligosoma_suteri | LC | Stable | temperate | cathemeral | 45.709 | 13451.259 | 10.008 |
| lizards | Oligosoma_taumakae | NA | EN | Decreasing | NA | NA | NA | NA | NA |
| lizards | Oligosoma_tekakahu | NA | VU | Increasing | NA | NA | NA | NA | NA |
| lizards | Oligosoma_toka | NA | EN | Decreasing | NA | NA | NA | NA | NA |
| lizards | Oligosoma_townsi | NA | VU | Increasing | NA | NA | NA | NA | NA |
| lizards | Oligosoma_waimatense | NA | EN | Decreasing | NA | NA | NA | NA | NA |
| lizards | Oligosoma_whitakeri | Oligosoma_whitakeri | EN | Stable | temperate | cathemeral | 31.623 | 136.676 | 12.927 |
| lizards | Oligosoma_zelandicum | Oligosoma_zelandicum | LC | Decreasing | temperate | diurnal | 8.511 | 13845.237 | 10.011 |
| lizards | Omanosaura_cyanura | Omanosaura_cyanura | LC | Stable | temperate | diurnal | 5.012 | 23800.099 | 11.449 |
| lizards | Omanosaura_jayakari | Omanosaura_jayakari | LC | Stable | temperate | diurnal | 177.828 | 35859.697 | 11.803 |
| lizards | Ophidiocephalus_taeniatus | NA | LC | Unknown | NA | NA | NA | NA | NA |
| lizards | Ophiodes_intermedius | Ophiodes_intermedius | LC | Stable | temperate | cathemeral | 63.096 | 1342019.387 | 8.413 |
| lizards | Ophiodes_striatus | Ophiodes_striatus | LC | Unknown | tropical | diurnal | 102.329 | 978405.101 | 12.915 |
| lizards | Ophiodes_vertebralis | Ophiodes_vertebralis | LC | Unknown | temperate | cathemeral | 33.884 | 243664.343 | 12.041 |
| lizards | Ophiomorus_blanfordii | NA | LC | Unknown | NA | NA | NA | NA | NA |
| lizards | Ophiomorus_brevipes | NA | LC | Stable | NA | NA | NA | NA | NA |
| lizards | Ophiomorus_chernovi | NA | LC | Unknown | NA | NA | NA | NA | NA |
| lizards | Ophiomorus_latastii | NA | DD | Unknown | NA | NA | NA | NA | NA |
| lizards | Ophiomorus_maranjabensis | NA | not_available | not_available | NA | NA | NA | NA | NA |
| lizards | Ophiomorus_nuchalis | Ophiomorus_nuchalis | LC | Stable | temperate | nocturnal | 7.413 | 10407.946 | 6.044 |
| lizards | Ophiomorus_persicus | NA | LC | Unknown | NA | NA | NA | NA | NA |
| lizards | Ophiomorus_punctatissimus | Ophiomorus_punctatissimus | LC | Unknown | temperate | diurnal | 2.455 | 19823.911 | 15.629 |
| lizards | Ophiomorus_raithmai | NA | LC | Unknown | NA | NA | NA | NA | NA |
| lizards | Ophiomorus_streeti | NA | LC | Stable | NA | NA | NA | NA | NA |
| lizards | Ophiomorus_tridactylus | Ophiomorus_tridactylus | LC | Stable | temperate | cathemeral | 6.166 | 385438.011 | 7.161 |
| lizards | Ophioscincus_ophioscincus | Ophioscincus_ophioscincus | LC | Stable | temperate | cathemeral | 2.291 | 41069.912 | 7.65 |
| lizards | Ophioscincus_truncatus | Ophioscincus_truncatus | LC | Stable | temperate | nocturnal | 1.445 | 37032.493 | 11.01 |
| lizards | Ophisaurus_attenuatus | Ophisaurus_attenuatus | LC | Stable | temperate | diurnal | 147.911 | 1936754.713 | 16.707 |

| | | | | | | | | | |
|---|---|---|---|---|---|---|---|---|---|
| lizards | Ophisaurus_ceroni | NA | EN | Decreasing | NA | NA | NA | NA | NA |
| lizards | Ophisaurus_compressus | Ophisaurus_compressus | LC | Decreasing | temperate | diurnal | 25.119 | 149110.699 | 16.774 |
| lizards | Ophisaurus_mimicus | Ophisaurus_mimicus | LC | Decreasing | temperate | cathemeral | 20.893 | 105948.081 | 15.349 |
| lizards | Ophisaurus_ventralis | Ophisaurus_ventralis | LC | Stable | temperate | cathemeral | 91.201 | 472335.083 | 16.556 |
| lizards | Ophisops_beddomei | NA | LC | Unknown | NA | NA | NA | NA | NA |
| lizards | Ophisops_elbaensis | NA | DD | Unknown | NA | NA | NA | NA | NA |
| lizards | Ophisops_elegans | Ophisops_elegans | LC | Stable | temperate | diurnal | 7.943 | 2393067.586 | 13.473 |
| lizards | Ophisops_jerdonii | Ophisops_jerdonii | LC | Stable | tropical | diurnal | 2.754 | 2891076.11 | 16.881 |
| lizards | Ophisops_leschenaultii | Ophisops_leschenaultii | LC | Unknown | tropical | diurnal | 4.365 | 1300623.338 | 19.329 |
| lizards | Ophisops_microlepis | Ophisops_microlepis | LC | Unknown | temperate | diurnal | 6.457 | 664093.061 | 17.248 |
| lizards | Ophisops_nictans | NA | LC | Unknown | NA | NA | NA | NA | NA |
| lizards | Ophisops_occidentalis | Ophisops_occidentalis | LC | Decreasing | temperate | diurnal | 2.63 | 333927.522 | 13.397 |
| lizards | Oplurus_cuvieri | Oplurus_cuvieri | LC | Stable | tropical | diurnal | 177.828 | 207380.258 | 7.895 |
| lizards | Oplurus_cyclurus | Oplurus_cyclurus | LC | Decreasing | tropical | diurnal | 177.828 | 230746.915 | 8.678 |
| lizards | Oplurus_fierinensis | Oplurus_fierinensis | LC | Stable | tropical | diurnal | 27.542 | 9875.942 | 9.184 |
| lizards | Oplurus_grandidieri | Oplurus_grandidieri | LC | Stable | tropical | diurnal | 52.481 | 16124.108 | 9.021 |
| lizards | Oplurus_quadrimaculatus | Oplurus_quadrimaculatus | LC | Unknown | tropical | diurnal | 120.226 | 149236.185 | 9.731 |
| lizards | Oplurus_saxicola | Oplurus_saxicola | LC | Stable | temperate | diurnal | 38.905 | 67455.675 | 10.65 |
| lizards | Oreosaurus_achlyens | Riama_achlyens | LC | Stable | tropical | diurnal | 17.378 | 4024.024 | 13.788 |
| lizards | Oreosaurus_mcdiarmidi | NA | LC | Stable | NA | NA | NA | NA | NA |
| lizards | Ornithuroscincus_noctua | NA | LC | Stable | NA | NA | NA | NA | NA |
| lizards | Ornithuroscincus_sabini | NA | not_available | not_available | NA | NA | NA | NA | NA |
| lizards | Orosaura_nebulosylvestris | NA | LC | Stable | NA | NA | NA | NA | NA |
| lizards | Orraya_occultus | Orraya_occultus | VU | Unknown | tropical | nocturnal | 21.878 | 617.734 | 0.633 |
| lizards | Otocryptis_nigristigma | Otocryptis_nigristigma | LC | Unknown | tropical | diurnal | 16.596 | 26990.697 | 15.184 |
| lizards | Otocryptis_wiegmanni | Otocryptis_wiegmanni | LC | Decreasing | tropical | diurnal | 16.596 | 17038.49 | 19.239 |
| lizards | Otosaurus_cumingii | NA | LC | Unknown | NA | NA | NA | NA | NA |
| lizards | Ouroborus_cataphractus | Ouroborus_cataphractus | NT | Unknown | temperate | cathemeral | 58.884 | 95329.769 | 7.293 |
| lizards | Pachydactylus_acuminatus | Pachydactylus_acuminatus | LC | Stable | temperate | nocturnal | 2.818 | 25198.33 | 3.964 |
| lizards | Pachydactylus_angolensis | NA | LC | Stable | NA | NA | NA | NA | NA |
| lizards | Pachydactylus_atorquatus | Pachydactylus_atorquatus | LC | Stable | temperate | nocturnal | 3.548 | 21400.862 | 5.398 |
| lizards | Pachydactylus_austeni | Pachydactylus_austeni | LC | Stable | temperate | nocturnal | 2.344 | 42139.856 | 10.076 |
| lizards | Pachydactylus_barnardi | Pachydactylus_barnardi | LC | Stable | temperate | nocturnal | 5.248 | 34338.917 | 6.653 |
| lizards | Pachydactylus_bicolor | Pachydactylus_bicolor | LC | Stable | tropical | nocturnal | 1.862 | 231661.678 | 5.336 |

| | | | | | | | | | |
|---|---|---|---|---|---|---|---|---|---|
| lizards | Pachydactylus_boehmei | Pachydactylus_boehmei | LC | Stable | tropical | nocturnal | 2.042 | 2040.264 | 6.13 |
| lizards | Pachydactylus_capensis | Pachydactylus_capensis | LC | Stable | temperate | nocturnal | 6.607 | 1796090.866 | 6.841 |
| lizards | Pachydactylus_etultra | NA | LC | Stable | NA | NA | NA | NA | NA |
| lizards | Pachydactylus_fasciatus | Pachydactylus_fasciatus | LC | Stable | tropical | nocturnal | 3.802 | 124432.485 | 4.985 |
| lizards | Pachydactylus_gaiasensis | NA | LC | Stable | NA | NA | NA | NA | NA |
| lizards | Pachydactylus_geitje | Pachydactylus_geitje | LC | Stable | temperate | nocturnal | 2.089 | 132809.23 | 9.659 |
| lizards | Pachydactylus_haackei | Pachydactylus_haackei | LC | Stable | temperate | cathemeral | 12.303 | 99726.083 | 3.938 |
| lizards | Pachydactylus_kladaroderma | NA | LC | Stable | NA | NA | NA | NA | NA |
| lizards | Pachydactylus_kobosensis | Pachydactylus_kobosensis | LC | Stable | tropical | nocturnal | 3.388 | 9742.769 | 6.553 |
| lizards | Pachydactylus_kochii | Colopus_kochii | LC | Stable | tropical | nocturnal | 4.677 | 96713.395 | 4.556 |
| lizards | Pachydactylus_latirostris | NA | LC | Stable | NA | NA | NA | NA | NA |
| lizards | Pachydactylus_maculatus | Pachydactylus_maculatus | LC | Stable | temperate | nocturnal | 4.266 | 338309.26 | 11.376 |
| lizards | Pachydactylus_maraisi | Pachydactylus_maraisi | NT | Stable | tropical | nocturnal | 1.622 | 1202.655 | 3.218 |
| lizards | Pachydactylus_mariquensis | Pachydactylus_mariquensis | LC | Stable | temperate | nocturnal | 4.266 | 351788.388 | 8.394 |
| lizards | Pachydactylus_namaquensis | Pachydactylus_namaquensis | LC | Stable | temperate | cathemeral | 12.303 | 27679.317 | 5.979 |
| lizards | Pachydactylus_oculatus | Pachydactylus_oculatus | LC | Stable | temperate | nocturnal | 3.311 | 122997.76 | 8.054 |
| lizards | Pachydactylus_oreophilus | NA | LC | Stable | NA | NA | NA | NA | NA |
| lizards | Pachydactylus_parascutatus | Pachydactylus_parascutatus | LC | Stable | tropical | nocturnal | 1.349 | 9094.749 | 4.262 |
| lizards | Pachydactylus_punctatus | Pachydactylus_punctatus | LC | Stable | tropical | nocturnal | 1.738 | 3247237.48 | 7.556 |
| lizards | Pachydactylus_purcelli | Pachydactylus_purcelli | LC | Stable | temperate | nocturnal | 2.951 | 217357.639 | 5.479 |
| lizards | Pachydactylus_rangei | Pachydactylus_rangei | LC | Stable | tropical | nocturnal | 10.233 | 229410.867 | 3.757 |
| lizards | Pachydactylus_rugosus | Pachydactylus_rugosus | LC | Stable | temperate | nocturnal | 6.026 | 476375.145 | 4.485 |
| lizards | Pachydactylus_sansteynae | Pachydactylus_sansteynae | LC | Stable | tropical | nocturnal | 2.512 | 47434.145 | 3.465 |
| lizards | Pachydactylus_scutatus | Pachydactylus_scutatus | LC | Stable | tropical | nocturnal | 2.344 | 217542.834 | 6.492 |
| lizards | Pachydactylus_tsodiloensis | Pachydactylus_tsodiloensis | LC | Unknown | tropical | nocturnal | 4.677 | 9.991 | 2.769 |
| lizards | Pachydactylus_vansoni | Pachydactylus_vansoni | LC | Stable | temperate | nocturnal | 4.467 | 346112.433 | 11.392 |
| lizards | Pachydactylus_vanzyli | NA | LC | Stable | NA | NA | NA | NA | NA |
| lizards | Pachydactylus_wahlbergii | Colopus_wahlbergii | LC | Stable | tropical | nocturnal | 4.898 | 834161.504 | 5.074 |
| lizards | Pachydactylus_weberi | Pachydactylus_weberi | LC | Stable | temperate | nocturnal | 2.818 | 264086.269 | 5.396 |
| lizards | Pachydactylus_werneri | Pachydactylus_werneri | LC | Stable | tropical | nocturnal | 3.236 | 23199.681 | 3.776 |
| lizards | Palleon_lolontany | Palleon_lolontany | NT | Stable | tropical | diurnal | 1.096 | 800.243 | 4.617 |
| lizards | Palleon_nasus | NA | VU | Decreasing | NA | NA | NA | NA | NA |
| lizards | Pamelaescincus_gardineri | NA | NT | Unknown | NA | NA | NA | NA | NA |
| lizards | Panaspis_africana | NA | LC | Stable | NA | NA | NA | NA | NA |

| | | | | | | | | | |
|---|---|---|---|---|---|---|---|---|---|
| lizards | Panaspis_breviceps | Panaspis_breviceps | LC | Stable | tropical | diurnal | 6.761 | 930288.505 | 5.779 |
| lizards | Panaspis_cabindae | Panaspis_cabindae | LC | Stable | tropical | cathemeral | 1.318 | 584514.861 | 7.576 |
| lizards | Panaspis_maculicollis | Afroablepharus_maculicollis | LC | Stable | tropical | diurnal | 1.622 | 863948.324 | 10.252 |
| lizards | Panaspis_massaiensis | NA | not_available | not_available | NA | NA | NA | NA | NA |
| lizards | Panaspis_megalurus | NA | LC | Unknown | NA | NA | NA | NA | NA |
| lizards | Panaspis_mocamedensis | NA | not_available | not_available | NA | NA | NA | NA | NA |
| lizards | Panaspis_namibiana | NA | LC | Stable | NA | NA | NA | NA | NA |
| lizards | Panaspis_tancredii | NA | DD | Unknown | NA | NA | NA | NA | NA |
| lizards | Panaspis_thomensis | NA | LC | Stable | NA | NA | NA | NA | NA |
| lizards | Panaspis_togoensis | Panaspis_togoensis | LC | Unknown | tropical | diurnal | 3.09 | 2676662.934 | 11.457 |
| lizards | Panaspis_tristaoi | Panaspis_nimbaensis | LC | Unknown | tropical | diurnal | 6.166 | 533551.175 | 13.328 |
| lizards | Panaspis_tsavoensis | NA | not_available | not_available | NA | NA | NA | NA | NA |
| lizards | Panaspis_wahlbergii | NA | LC | Unknown | NA | NA | NA | NA | NA |
| lizards | Panaspis_wilsoni | NA | DD | Unknown | NA | NA | NA | NA | NA |
| lizards | Paniegekko_madjo | Paniegekko_madjo | EN | Decreasing | tropical | nocturnal | 8.511 | 388.536 | 3.209 |
| lizards | Panolopus_costatus | NA | LC | Stable | NA | NA | NA | NA | NA |
| lizards | Panolopus_marcanoi | NA | LC | Unknown | NA | NA | NA | NA | NA |
| lizards | Panopa_carvalhoi | NA | LC | Unknown | NA | NA | NA | NA | NA |
| lizards | Panopa_croizati | NA | CR | Decreasing | NA | NA | NA | NA | NA |
| lizards | Pantepuisaurus_rodriguesi | NA | LC | Unknown | NA | NA | NA | NA | NA |
| lizards | Papuascincus_morokanus | Papuascincus_morokanus | LC | Unknown | tropical | diurnal | 2.63 | 112736.515 | 5.123 |
| lizards | Papuascincus_stanleyanus | Papuascincus_stanleyanus | LC | Stable | tropical | diurnal | 4.169 | 184604.107 | 4.53 |
| lizards | Paracontias_hafa | NA | NT | Decreasing | NA | NA | NA | NA | NA |
| lizards | Paracontias_hildebrandti | NA | LC | Decreasing | NA | NA | NA | NA | NA |
| lizards | Paracontias_minimus | Paracontias_minimus | CR | Decreasing | tropical | diurnal | 1.259 | 89.345 | 13.142 |
| lizards | Paracontias_rothschildi | NA | CR | Unknown | NA | NA | NA | NA | NA |
| lizards | Paracontias_tsararano | NA | DD | Unknown | NA | NA | NA | NA | NA |
| lizards | Paradelma_orientalis | Paradelma_orientalis | LC | Decreasing | temperate | cathemeral | 12.882 | 407132.732 | 6.127 |
| lizards | Paragehyra_austini | NA | not_available | not_available | NA | NA | NA | NA | NA |
| lizards | Paragehyra_felicitae | NA | not_available | not_available | NA | NA | NA | NA | NA |
| lizards | Paragehyra_gabriellae | Paragehyra_gabriellae | EN | Decreasing | temperate | nocturnal | 8.71 | 1348.876 | 7.838 |
| lizards | Paragehyra_petiti | NA | VU | Stable | NA | NA | NA | NA | NA |
| lizards | Paralaudakia_badakhshana | NA | LC | Stable | NA | NA | NA | NA | NA |
| lizards | Paralaudakia_bochariensis | Paralaudakia_bochariensis | LC | Stable | temperate | diurnal | 58.884 | 100406.896 | 11.331 |

| | | | | | | | | | |
|---|---|---|---|---|---|---|---|---|---|
| lizards | Paralaudakia_caucasia | Paralaudakia_caucasia | LC | Stable | temperate | diurnal | 186.209 | 1711153.775 | 10.621 |
| lizards | Paralaudakia_erythrogaster | Paralaudakia_erythrogaster | LC | Stable | temperate | diurnal | 128.825 | 237044.873 | 10.077 |
| lizards | Paralaudakia_himalayana | Paralaudakia_himalayana | LC | Stable | temperate | diurnal | 107.152 | 549740.627 | 6.334 |
| lizards | Paralaudakia_lehmanni | Paralaudakia_lehmanni | LC | Stable | temperate | diurnal | 117.49 | 386332.21 | 10.347 |
| lizards | Paralaudakia_microlepis | NA | LC | Stable | NA | NA | NA | NA | NA |
| lizards | Paralaudakia_stoliczkana | Paralaudakia_stoliczkana | LC | Stable | temperate | diurnal | 208.93 | 508336.607 | 2.517 |
| lizards | Paroedura_androyensis | Paroedura_androyensis | VU | Decreasing | temperate | nocturnal | 2.344 | 19244.037 | 11.791 |
| lizards | Paroedura_bastardi | Paroedura_bastardi | LC | Stable | tropical | nocturnal | 11.749 | 172114.553 | 8.893 |
| lizards | Paroedura_gracilis | Paroedura_gracilis | LC | Decreasing | tropical | nocturnal | 7.762 | 38038.337 | 7.303 |
| lizards | Paroedura_homalorhina | NA | LC | Stable | NA | NA | NA | NA | NA |
| lizards | Paroedura_hordiesi | NA | not_available | not_available | NA | NA | NA | NA | NA |
| lizards | Paroedura_ibityensis | Paroedura_ibityensis | NT | Stable | tropical | cathemeral | 4.898 | 2999.548 | 10.23 |
| lizards | Paroedura_karstophila | Paroedura_karstophila | LC | Stable | tropical | nocturnal | 3.631 | 4776.62 | 6.994 |
| lizards | Paroedura_lohatsara | Paroedura_lohatsara | CR | Decreasing | tropical | nocturnal | 10.471 | 288.1 | 12.739 |
| lizards | Paroedura_maingoka | NA | NT | Stable | NA | NA | NA | NA | NA |
| lizards | Paroedura_masobe | Paroedura_masobe | EN | Decreasing | tropical | nocturnal | 25.704 | 121.668 | 6.964 |
| lizards | Paroedura_oviceps | Paroedura_oviceps | NT | Decreasing | tropical | nocturnal | 7.413 | 30337.09 | 8.409 |
| lizards | Paroedura_picta | Paroedura_picta | LC | Stable | tropical | nocturnal | 25.704 | 136497.281 | 9.433 |
| lizards | Paroedura_sanctijohannis | NA | VU | Unknown | NA | NA | NA | NA | NA |
| lizards | Paroedura_stellata | Paroedura_stellata | VU | Unknown | tropical | nocturnal | 5.012 | 332.627 | 20.796 |
| lizards | Paroedura_stumpffi | Paroedura_stumpffi | LC | Stable | tropical | nocturnal | 8.511 | 59105.426 | 8.66 |
| lizards | Paroedura_tanjaka | Paroedura_tanjaka | EN | Unknown | tropical | cathemeral | 19.953 | 1302.97 | 6.444 |
| lizards | Paroedura_vahiny | Paroedura_vahiny | LC | Unknown | tropical | nocturnal | 1.738 | 55037.056 | 8.218 |
| lizards | Paroedura_vazimba | Paroedura_vazimba | VU | Decreasing | tropical | nocturnal | 2.818 | 8442.636 | 9.636 |
| lizards | Parsigecko_ziaiei | NA | not_available | not_available | NA | NA | NA | NA | NA |
| lizards | Parvilacerta_fraasii | Parvilacerta_fraasii | EN | Decreasing | temperate | diurnal | 5.012 | 1418.905 | 13.755 |
| lizards | Parvilacerta_parva | Parvilacerta_parva | LC | Stable | temperate | diurnal | 5.623 | 293339.169 | 13.93 |
| lizards | Parvoscincus_beyeri | Parvoscincus_beyeri | VU | Unknown | tropical | diurnal | 7.762 | 17317.818 | 20.055 |
| lizards | Parvoscincus_boyingi | NA | DD | Unknown | NA | NA | NA | NA | NA |
| lizards | Parvoscincus_decipiens | NA | LC | Stable | NA | NA | NA | NA | NA |
| lizards | Parvoscincus_duwendorum | NA | DD | Unknown | NA | NA | NA | NA | NA |
| lizards | Parvoscincus_leucospilos | NA | LC | Unknown | NA | NA | NA | NA | NA |
| lizards | Parvoscincus_luzonensis | NA | DD | Unknown | NA | NA | NA | NA | NA |
| lizards | Parvoscincus_manananggalae | NA | NT | Stable | NA | NA | NA | NA | NA |

| | | | | | | | | | |
|---|---|---|---|---|---|---|---|---|---|
| lizards | Parvoscincus_sisoni | Parvoscincus_sisoni | EN | Unknown | tropical | diurnal | 0.832 | 9.991 | 3.891 |
| lizards | Parvoscincus_steerei | Parvoscincus_steerei | LC | Unknown | tropical | diurnal | 1.318 | 268427.767 | 13.907 |
| lizards | Parvoscincus_tagapayo | Parvoscincus_tagapayo | NT | Decreasing | tropical | diurnal | 0.55 | 2455.429 | 7.73 |
| lizards | Pedioplanis_breviceps | Pedioplanis_breviceps | LC | Stable | tropical | diurnal | 2.291 | 149069.041 | 4.516 |
| lizards | Pedioplanis_burchelli | Pedioplanis_burchelli | LC | Stable | temperate | diurnal | 5.754 | 371177.278 | 9.876 |
| lizards | Pedioplanis_husabensis | Pedioplanis_husabensis | LC | Stable | tropical | diurnal | 5.37 | 19159.896 | 4.532 |
| lizards | Pedioplanis_inornata | Pedioplanis_inornata | LC | Stable | temperate | diurnal | 3.89 | 295832.052 | 4.268 |
| lizards | Pedioplanis_lineoocellata | Pedioplanis_lineoocellata | LC | Stable | temperate | diurnal | 7.079 | 1985439.31 | 7.12 |
| lizards | Pedioplanis_namaquensis | Pedioplanis_namaquensis | LC | Stable | tropical | diurnal | 4.169 | 1464925.101 | 5.466 |
| lizards | Pedioplanis_rubens | Pedioplanis_rubens | LC | Stable | tropical | diurnal | 2.951 | 879.548 | 6.881 |
| lizards | Pedioplanis_undata | Pedioplanis_undata | LC | Stable | tropical | diurnal | 5.623 | 611403.523 | 5.641 |
| lizards | Pelturagonia_borneensis | NA | LC | Unknown | NA | NA | NA | NA | NA |
| lizards | Pelturagonia_cephalum | NA | LC | Unknown | NA | NA | NA | NA | NA |
| lizards | Pelturagonia_nigrilabris | NA | LC | Unknown | NA | NA | NA | NA | NA |
| lizards | Pelturagonia_spiniceps | NA | LC | Unknown | NA | NA | NA | NA | NA |
| lizards | Perochirus_ateles | Perochirus_ateles | VU | Decreasing | tropical | nocturnal | 14.125 | 1550.827 | 17.506 |
| lizards | Perochirus_guentheri | NA | DD | Unknown | NA | NA | NA | NA | NA |
| lizards | Perochirus_scutellatus | NA | LC | Stable | NA | NA | NA | NA | NA |
| lizards | Petrosaurus_mearnsi | Petrosaurus_mearnsi | LC | Stable | temperate | diurnal | 37.154 | 24622.519 | 3.428 |
| lizards | Petrosaurus_repens | NA | LC | Stable | NA | NA | NA | NA | NA |
| lizards | Petrosaurus_slevini | NA | LC | Stable | NA | NA | NA | NA | NA |
| lizards | Petrosaurus_thalassinus | Petrosaurus_thalassinus | LC | Stable | temperate | diurnal | 141.254 | 7545.616 | 6.886 |
| lizards | Phasmasaurus_maruia | Lioscincus_maruia | VU | Decreasing | tropical | diurnal | 4.365 | 905.262 | 5.225 |
| lizards | Phasmasaurus_tillieri | Lioscincus_tillieri | NT | Unknown | tropical | diurnal | 5.129 | 3316.011 | 8.731 |
| lizards | Phelsuma_abbotti | Phelsuma_abbotti | LC | Stable | tropical | diurnal | 5.754 | 33813.532 | 8.059 |
| lizards | Phelsuma_andamanensis | Phelsuma_andamanense | LC | Stable | tropical | diurnal | 5.495 | 4566.085 | 7.702 |
| lizards | Phelsuma_antanosy | Phelsuma_antanosy | CR | Decreasing | temperate | diurnal | 2.512 | 524.92 | 14.059 |
| lizards | Phelsuma_astriata | NA | LC | Stable | NA | NA | NA | NA | NA |
| lizards | Phelsuma_barbouri | Phelsuma_barbouri | LC | Unknown | tropical | diurnal | 5.495 | 573.166 | 8.489 |
| lizards | Phelsuma_berghofi | Phelsuma_berghofi | NT | Unknown | temperate | diurnal | 4.266 | 1982.534 | 12.194 |
| lizards | Phelsuma_borbonica | Phelsuma_borbonica | EN | Decreasing | tropical | diurnal | 7.943 | 34.377 | 21.095 |
| lizards | Phelsuma_breviceps | Phelsuma_breviceps | VU | Decreasing | tropical | diurnal | 3.467 | 9278.483 | 9.011 |
| lizards | Phelsuma_cepediana | Phelsuma_cepediana | LC | Unknown | tropical | diurnal | 4.677 | 1872.583 | 25.918 |
| lizards | Phelsuma_comorensis | Phelsuma_comorensis | NT | Unknown | tropical | diurnal | 3.802 | 1031.546 | 17.428 |

| | | | | | | | | | |
|---|---|---|---|---|---|---|---|---|---|
| lizards | Phelsuma_dorsivittata | Phelsuma_dorsivittata | NT | Unknown | tropical | diurnal | 4.074 | 975.472 | 6.585 |
| lizards | Phelsuma_dubia | Phelsuma_dubia | LC | Stable | tropical | diurnal | 7.079 | 25456.112 | 10.955 |
| lizards | Phelsuma_edwardnewtoni | NA | EX | not_available | NA | NA | NA | NA | NA |
| lizards | Phelsuma_flavigularis | NA | EN | Decreasing | NA | NA | NA | NA | NA |
| lizards | Phelsuma_gigas | NA | EX | not_available | NA | NA | NA | NA | NA |
| lizards | Phelsuma_gouldi | NA | DD | Unknown | NA | NA | NA | NA | NA |
| lizards | Phelsuma_grandis | Phelsuma_grandis | LC | Unknown | tropical | diurnal | 30.2 | 9173.231 | 8.317 |
| lizards | Phelsuma_guentheri | Phelsuma_guentheri | VU | Increasing | tropical | cathemeral | 69.183 | 1872.605 | 25.918 |
| lizards | Phelsuma_guimbeaui | Phelsuma_guimbeaui | EN | Decreasing | tropical | diurnal | 7.079 | 1877.187 | 25.915 |
| lizards | Phelsuma_guttata | Phelsuma_guttata | LC | Decreasing | tropical | diurnal | 6.607 | 33638.221 | 7.652 |
| lizards | Phelsuma_hielscheri | NA | VU | Decreasing | NA | NA | NA | NA | NA |
| lizards | Phelsuma_hoeschi | NA | DD | Unknown | NA | NA | NA | NA | NA |
| lizards | Phelsuma_inexpectata | Phelsuma_inexpectata | CR | Decreasing | tropical | diurnal | 4.266 | 383.254 | 33.002 |
| lizards | Phelsuma_kely | NA | DD | not_available | NA | NA | NA | NA | NA |
| lizards | Phelsuma_klemmeri | Phelsuma_klemmeri | EN | Unknown | tropical | diurnal | 1.862 | 958.943 | 4.85 |
| lizards | Phelsuma_kochi | Phelsuma_kochi | LC | Decreasing | tropical | diurnal | 28.184 | 66294.699 | 8.016 |
| lizards | Phelsuma_laticauda | Phelsuma_laticauda | LC | Stable | tropical | diurnal | 5.754 | 39094.213 | 7.732 |
| lizards | Phelsuma_lineata | Phelsuma_lineata | LC | Stable | tropical | diurnal | 5.754 | 224353.229 | 9.26 |
| lizards | Phelsuma_madagascariensis | Phelsuma_madagascariensis | LC | Decreasing | tropical | diurnal | 31.623 | 79710.515 | 8.552 |
| lizards | Phelsuma_malamakibo | Phelsuma_malamakibo | NT | Unknown | temperate | diurnal | 4.898 | 836.708 | 6.304 |
| lizards | Phelsuma_masohoala | Phelsuma_masohoala | CR | Unknown | tropical | diurnal | 2.344 | 841.308 | 10.352 |
| lizards | Phelsuma_modesta | Phelsuma_modesta | LC | Stable | temperate | diurnal | 2.239 | 25546.69 | 13.224 |
| lizards | Phelsuma_mutabilis | Phelsuma_mutabilis | LC | Unknown | tropical | diurnal | 2.818 | 103490.069 | 9.925 |
| lizards | Phelsuma_nigristriata | Phelsuma_nigristriata | VU | Stable | tropical | diurnal | 2.089 | 386.174 | 20.933 |
| lizards | Phelsuma_ornata | Phelsuma_ornata | LC | Stable | tropical | diurnal | 4.467 | 882.071 | 28.095 |
| lizards | Phelsuma_parkeri | Phelsuma_parkeri | LC | Unknown | tropical | diurnal | 6.918 | 908.253 | 17.67 |
| lizards | Phelsuma_parva | Phelsuma_parva | LC | Decreasing | tropical | diurnal | 1.122 | 39161.317 | 9.286 |
| lizards | Phelsuma_pasteuri | NA | NT | Stable | NA | NA | NA | NA | NA |
| lizards | Phelsuma_pronki | Phelsuma_pronki | CR | Decreasing | tropical | diurnal | 2.818 | 3452.377 | 13.659 |
| lizards | Phelsuma_pusilla | Phelsuma_pusilla | LC | Stable | tropical | diurnal | 1.23 | 75550.235 | 8.034 |
| lizards | Phelsuma_quadriocellata | Phelsuma_quadriocellata | LC | Stable | tropical | diurnal | 4.898 | 96380.241 | 8.092 |
| lizards | Phelsuma_ravenala | NA | LC | Stable | NA | NA | NA | NA | NA |
| lizards | Phelsuma_robertmertensi | Phelsuma_robertmertensi | NT | Stable | tropical | diurnal | 2.818 | 386.174 | 20.933 |
| lizards | Phelsuma_roesleri | NA | EN | Unknown | NA | NA | NA | NA | NA |

| | | | | | | | | | |
|---|---|---|---|---|---|---|---|---|---|
| lizards | Phelsuma_rosagularis | Phelsuma_rosagularis | EN | Decreasing | tropical | diurnal | 6.918 | 1872.582 | 25.918 |
| lizards | Phelsuma_seippi | Phelsuma_seippi | EN | Unknown | tropical | diurnal | 3.631 | 3751.592 | 6.852 |
| lizards | Phelsuma_serraticauda | Phelsuma_serraticauda | EN | Decreasing | tropical | diurnal | 6.026 | 6783.578 | 10.973 |
| lizards | Phelsuma_standingi | Phelsuma_standingi | VU | Unknown | tropical | diurnal | 15.488 | 17234.111 | 8.399 |
| lizards | Phelsuma_sundbergi | NA | LC | Stable | NA | NA | NA | NA | NA |
| lizards | Phelsuma_v-nigra | Phelsuma_v-nigra | LC | Unknown | tropical | diurnal | 3.311 | 2039.613 | 17.331 |
| lizards | Phelsuma_vanheygeni | NA | EN | Unknown | NA | NA | NA | NA | NA |
| lizards | Philochortus_intermedius | NA | LC | Unknown | NA | NA | NA | NA | NA |
| lizards | Philochortus_neumanni | NA | LC | Unknown | NA | NA | NA | NA | NA |
| lizards | Philochortus_zolii | NA | EN | Decreasing | NA | NA | NA | NA | NA |
| lizards | Phoboscincus_bocourti | NA | CR | Decreasing | NA | NA | NA | NA | NA |
| lizards | Phoboscincus_garnieri | NA | LC | Unknown | NA | NA | NA | NA | NA |
| lizards | Phoenicolacerta_cyanisparsa | NA | LC | Stable | NA | NA | NA | NA | NA |
| lizards | Phoenicolacerta_kulzeri | Phoenicolacerta_kulzeri | EN | Decreasing | temperate | diurnal | 15.136 | 14652.333 | 20.981 |
| lizards | Phoenicolacerta_laevis | Phoenicolacerta_laevis | LC | Stable | temperate | diurnal | 15.136 | 70636.46 | 21.374 |
| lizards | Phoenicolacerta_troodica | Phoenicolacerta_troodica | LC | Stable | temperate | diurnal | 7.943 | 9272.651 | 19.898 |
| lizards | Pholidobolus_affinis | Pholidobolus_affinis | NT | Unknown | tropical | diurnal | 6.457 | 1731.867 | 23.185 |
| lizards | Pholidobolus_hillisi | NA | LC | Stable | NA | NA | NA | NA | NA |
| lizards | Pholidobolus_macbrydei | Pholidobolus_macbrydei | LC | Stable | tropical | diurnal | 4.898 | 29668.986 | 10.819 |
| lizards | Pholidobolus_montium | Pholidobolus_montium | NT | Decreasing | tropical | diurnal | 6.761 | 5905.262 | 18.81 |
| lizards | Pholidobolus_prefrontalis | Pholidobolus_prefrontalis | LC | Unknown | tropical | diurnal | 5.754 | 1269.338 | 18.073 |
| lizards | Pholidobolus_ulisesi | NA | not_available | not_available | NA | NA | NA | NA | NA |
| lizards | Pholidobolus_vertebralis | Cercosaura_vertebralis | LC | Unknown | tropical | diurnal | 7.413 | 149316.924 | 13.813 |
| lizards | Pholidoscelis_atratus | NA | CR | Decreasing | NA | NA | NA | NA | NA |
| lizards | Pholidoscelis_auberi | Ameiva_auberi | LC | Stable | tropical | diurnal | 77.625 | 111513.759 | 15.675 |
| lizards | Pholidoscelis_chrysolaemus | Ameiva_chrysolaema | LC | Stable | tropical | diurnal | 128.825 | 52162.876 | 15.075 |
| lizards | Pholidoscelis_cineraceus | NA | EX | not_available | NA | NA | NA | NA | NA |
| lizards | Pholidoscelis_corax | NA | EN | Unknown | NA | NA | NA | NA | NA |
| lizards | Pholidoscelis_corvinus | NA | CR | Unknown | NA | NA | NA | NA | NA |
| lizards | Pholidoscelis_dorsalis | NA | EN | Decreasing | NA | NA | NA | NA | NA |
| lizards | Pholidoscelis_erythrocephalus | NA | NT | Unknown | NA | NA | NA | NA | NA |
| lizards | Pholidoscelis_exsul | Ameiva_exsul | LC | Stable | tropical | diurnal | 263.027 | 9254.626 | 26.85 |
| lizards | Pholidoscelis_fuscatus | Ameiva_fuscata | LC | Stable | tropical | diurnal | 512.861 | 660.067 | 13.189 |
| lizards | Pholidoscelis_griswoldi | NA | NT | Decreasing | NA | NA | NA | NA | NA |

| | | | | | | | | | |
|---|---|---|---|---|---|---|---|---|---|
| lizards | Pholidoscelis_lineolatus | NA | LC | Unknown | NA | NA | NA | NA | NA |
| lizards | Pholidoscelis_maynardi | NA | LC | Unknown | NA | NA | NA | NA | NA |
| lizards | Pholidoscelis_plei | Ameiva_plei | LC | Unknown | tropical | diurnal | 186.209 | 199.358 | 28.678 |
| lizards | Pholidoscelis_pluvianotatus | NA | NT | Decreasing | NA | NA | NA | NA | NA |
| lizards | Pholidoscelis_polops | NA | EN | Unknown | NA | NA | NA | NA | NA |
| lizards | Pholidoscelis_taeniurus | Ameiva_taeniura | LC | Unknown | tropical | diurnal | 34.674 | 62690.911 | 15.566 |
| lizards | Pholidoscelis_wetmorei | Ameiva_wetmorei | VU | Decreasing | tropical | diurnal | 3.981 | 495.873 | 27.412 |
| lizards | Phoxophrys_tuberculata | NA | LC | Unknown | NA | NA | NA | NA | NA |
| lizards | Phrynocephalus_ananjevae | NA | not_available | not_available | NA | NA | NA | NA | NA |
| lizards | Phrynocephalus_arabicus | Phrynocephalus_arabicus | LC | Stable | temperate | diurnal | 6.918 | 1958315.278 | 4.289 |
| lizards | Phrynocephalus_axillaris | Phrynocephalus_axillaris | LC | Unknown | temperate | diurnal | 8.128 | 1029671.947 | 2.401 |
| lizards | Phrynocephalus_clarkorum | Phrynocephalus_clarkorum | LC | Unknown | temperate | diurnal | 2.818 | 130981.091 | 4.394 |
| lizards | Phrynocephalus_erythrurus | Phrynocephalus_erythrurus | LC | Stable | temperate | diurnal | 5.623 | 371161.081 | 2.256 |
| lizards | Phrynocephalus_euptilopus | NA | LC | Unknown | NA | NA | NA | NA | NA |
| lizards | Phrynocephalus_forsythii | Phrynocephalus_forsythii | LC | Stable | temperate | diurnal | 7.079 | 343495.005 | 3.295 |
| lizards | Phrynocephalus_frontalis | NA | LC | Stable | NA | NA | NA | NA | NA |
| lizards | Phrynocephalus_golubewii | Phrynocephalus_golubewii | CR | Decreasing | temperate | diurnal | 10.965 | 193.252 | 4.98 |
| lizards | Phrynocephalus_guttatus | Phrynocephalus_guttatus | LC | Decreasing | temperate | diurnal | 8.71 | 2125606.434 | 6.945 |
| lizards | Phrynocephalus_helioscopus | Phrynocephalus_helioscopus | LC | Unknown | temperate | diurnal | 10.965 | 2586056.742 | 7.379 |
| lizards | Phrynocephalus_interscapularis | Phrynocephalus_interscapularis | LC | Stable | temperate | diurnal | 3.236 | 1167045.406 | 7.09 |
| lizards | Phrynocephalus_lutensis | NA | not_available | not_available | NA | NA | NA | NA | NA |
| lizards | Phrynocephalus_luteoguttatus | Phrynocephalus_luteoguttatus | LC | Stable | temperate | diurnal | 3.236 | 291725.81 | 4.907 |
| lizards | Phrynocephalus_maculatus | Phrynocephalus_maculatus | LC | Stable | temperate | diurnal | 25.119 | 2828625.894 | 5.695 |
| lizards | Phrynocephalus_mystaceus | Phrynocephalus_mystaceus | LC | Unknown | temperate | diurnal | 63.096 | 2820240.64 | 7.334 |
| lizards | Phrynocephalus_ornatus | Phrynocephalus_ornatus | LC | Stable | temperate | diurnal | 3.236 | 364906.62 | 5.367 |
| lizards | Phrynocephalus_persicus | Phrynocephalus_persicus | VU | Stable | temperate | diurnal | 6.918 | 355797.135 | 13.728 |
| lizards | Phrynocephalus_przewalskii | Phrynocephalus_przewalskii | LC | Stable | temperate | diurnal | 23.988 | 3274656.581 | 7.585 |
| lizards | Phrynocephalus_putjatai | Phrynocephalus_putjatai | LC | Stable | temperate | diurnal | 19.498 | 11425.196 | 8.043 |
| lizards | Phrynocephalus_raddei | Phrynocephalus_raddei | LC | Unknown | temperate | diurnal | 6.166 | 315414.944 | 8.414 |
| lizards | Phrynocephalus_reticulatus | NA | LC | Unknown | NA | NA | NA | NA | NA |
| lizards | Phrynocephalus_roborowskii | NA | not_available | not_available | NA | NA | NA | NA | NA |
| lizards | Phrynocephalus_rossikowi | Phrynocephalus_rossikowi | EN | Decreasing | temperate | diurnal | 3.89 | 88378.17 | 6.37 |
| lizards | Phrynocephalus_sakoi | NA | LC | Stable | NA | NA | NA | NA | NA |
| lizards | Phrynocephalus_scutellatus | Phrynocephalus_scutellatus | LC | Stable | temperate | diurnal | 5.495 | 1614986.164 | 7.552 |

| | | | | | | | | | |
|---|---|---|---|---|---|---|---|---|---|
| lizards | Phrynocephalus_strauchi | Phrynocephalus_strauchi | VU | Decreasing | temperate | diurnal | 5.248 | 24633.995 | 18.107 |
| lizards | Phrynocephalus_theobaldi | Phrynocephalus_theobaldi | LC | Stable | temperate | diurnal | 8.128 | 1631165.54 | 3.299 |
| lizards | Phrynocephalus_versicolor | Phrynocephalus_versicolor | LC | Stable | temperate | diurnal | 9.55 | 4002944.428 | 5.166 |
| lizards | Phrynocephalus_vlangalii | Phrynocephalus_vlangalii | LC | Stable | temperate | diurnal | 18.621 | 1144045.765 | 2.93 |
| lizards | Phrynosoma_asio | Phrynosoma_asio | LC | Stable | tropical | diurnal | 56.234 | 115543.614 | 8.835 |
| lizards | Phrynosoma_bauri | NA | not_available | not_available | NA | NA | NA | NA | NA |
| lizards | Phrynosoma_blainvillii | Phrynosoma_blainvillii | LC | Decreasing | temperate | diurnal | 44.668 | 216760.096 | 13.898 |
| lizards | Phrynosoma_braconnieri | Phrynosoma_braconnieri | LC | Stable | tropical | diurnal | 16.596 | 12632.632 | 13.474 |
| lizards | Phrynosoma_cerroense | NA | not_available | not_available | NA | NA | NA | NA | NA |
| lizards | Phrynosoma_cornutum | Phrynosoma_cornutum | LC | Stable | temperate | diurnal | 63.096 | 1798114.315 | 11.869 |
| lizards | Phrynosoma_coronatum | Phrynosoma_coronatum | LC | Decreasing | temperate | diurnal | 44.668 | 287997.753 | 11.307 |
| lizards | Phrynosoma_ditmarsi | NA | DD | Unknown | NA | NA | NA | NA | NA |
| lizards | Phrynosoma_douglasii | Phrynosoma_douglasii | LC | Stable | temperate | diurnal | 17.378 | 209725.162 | 11.632 |
| lizards | Phrynosoma_goodei | Phrynosoma_goodei | LC | Unknown | temperate | diurnal | 23.988 | 72654.196 | 6.791 |
| lizards | Phrynosoma_hernandesi | Phrynosoma_hernandesi | LC | Stable | temperate | diurnal | 57.544 | 1210130.555 | 9.743 |
| lizards | Phrynosoma_mcallii | Phrynosoma_mcallii | NT | Decreasing | temperate | diurnal | 39.811 | 31422.87 | 5.203 |
| lizards | Phrynosoma_modestum | Phrynosoma_modestum | LC | Stable | temperate | cathemeral | 12.589 | 811535.925 | 8.64 |
| lizards | Phrynosoma_orbiculare | Phrynosoma_orbiculare | LC | Stable | temperate | diurnal | 28.84 | 448940.884 | 9.535 |
| lizards | Phrynosoma_ornatissimum | NA | not_available | not_available | NA | NA | NA | NA | NA |
| lizards | Phrynosoma_platyrhinos | Phrynosoma_platyrhinos | LC | Stable | temperate | diurnal | 27.542 | 836125.636 | 8.296 |
| lizards | Phrynosoma_sherbrookei | NA | not_available | not_available | NA | NA | NA | NA | NA |
| lizards | Phrynosoma_solare | Phrynosoma_solare | LC | Stable | temperate | diurnal | 47.863 | 330439.412 | 7.986 |
| lizards | Phrynosoma_taurus | Phrynosoma_taurus | LC | Stable | tropical | diurnal | 23.988 | 26989.027 | 11.183 |
| lizards | Phyllodactylus_angustidigitus | Phyllodactylus_angustidigitus | EN | Decreasing | tropical | nocturnal | 4.898 | 1017.358 | 5.43 |
| lizards | Phyllodactylus_barringtonensis | NA | LC | Unknown | NA | NA | NA | NA | NA |
| lizards | Phyllodactylus_baurii | NA | DD | Unknown | NA | NA | NA | NA | NA |
| lizards | Phyllodactylus_bugastrolepis | Phyllodactylus_bugastrolepis | LC | Stable | temperate | nocturnal | 6.607 | 27.728 | 1.218 |
| lizards | Phyllodactylus_clinatus | NA | DD | Unknown | NA | NA | NA | NA | NA |
| lizards | Phyllodactylus_darwini | NA | NT | Decreasing | NA | NA | NA | NA | NA |
| lizards | Phyllodactylus_delsolari | Phyllodactylus_delsolari | EN | Stable | tropical | nocturnal | 18.197 | 214.695 | 8.372 |
| lizards | Phyllodactylus_dixoni | NA | LC | Unknown | NA | NA | NA | NA | NA |
| lizards | Phyllodactylus_duellmani | Phyllodactylus_duellmani | LC | Unknown | tropical | nocturnal | 2.138 | 1646.19 | 8.863 |
| lizards | Phyllodactylus_duncanensis | NA | not_available | not_available | NA | NA | NA | NA | NA |
| lizards | Phyllodactylus_galapagensis | NA | NT | Decreasing | NA | NA | NA | NA | NA |

| | | | | | | | | | |
|---|---|---|---|---|---|---|---|---|---|
| lizards | Phyllodactylus_gerrhopygus | Phyllodactylus_gerrhopygus | LC | Stable | tropical | cathemeral | 4.677 | 125801.386 | 4.912 |
| lizards | Phyllodactylus_gilberti | NA | DD | Unknown | NA | NA | NA | NA | NA |
| lizards | Phyllodactylus_gorii | NA | not_available | not_available | NA | NA | NA | NA | NA |
| lizards | Phyllodactylus_homolepidurus | NA | LC | Stable | NA | NA | NA | NA | NA |
| lizards | Phyllodactylus_insularis | Phyllodactylus_insularis | VU | Decreasing | tropical | nocturnal | 8.511 | 11.469 | 16.929 |
| lizards | Phyllodactylus_interandinus | Phyllodactylus_interandinus | LC | Unknown | tropical | nocturnal | 3.162 | 23866.098 | 9.242 |
| lizards | Phyllodactylus_johnwrighti | Phyllodactylus_johnwrighti | EN | Unknown | tropical | nocturnal | 3.311 | 983.136 | 8.178 |
| lizards | Phyllodactylus_kofordi | Phyllodactylus_kofordi | LC | Stable | tropical | nocturnal | 2.57 | 43204.696 | 7.836 |
| lizards | Phyllodactylus_lanei | Phyllodactylus_lanei | LC | Stable | tropical | nocturnal | 13.804 | 124168.381 | 9.076 |
| lizards | Phyllodactylus_leei | NA | NT | Unknown | NA | NA | NA | NA | NA |
| lizards | Phyllodactylus_leoni | Phyllodactylus_leoni | VU | Unknown | tropical | nocturnal | 4.365 | 320.566 | 9.988 |
| lizards | Phyllodactylus_lepidopygus | Phyllodactylus_lepidopygus | VU | Unknown | tropical | nocturnal | 4.365 | 44015.771 | 8.85 |
| lizards | Phyllodactylus_magister | NA | not_available | not_available | NA | NA | NA | NA | NA |
| lizards | Phyllodactylus_martini | NA | LC | Decreasing | NA | NA | NA | NA | NA |
| lizards | Phyllodactylus_microphyllus | Phyllodactylus_microphyllus | LC | Decreasing | tropical | nocturnal | 5.129 | 56050.389 | 8.784 |
| lizards | Phyllodactylus_nocticolus | Phyllodactylus_nocticolus | LC | Stable | temperate | nocturnal | 6.607 | 92187.002 | 3.208 |
| lizards | Phyllodactylus_pachamama | NA | not_available | not_available | NA | NA | NA | NA | NA |
| lizards | Phyllodactylus_palmeus | Phyllodactylus_palmeus | LC | Stable | tropical | nocturnal | 12.882 | 246.88 | 15.798 |
| lizards | Phyllodactylus_paralepis | NA | not_available | not_available | NA | NA | NA | NA | NA |
| lizards | Phyllodactylus_partidus | NA | LC | Stable | NA | NA | NA | NA | NA |
| lizards | Phyllodactylus_paucituberculatus | NA | DD | Unknown | NA | NA | NA | NA | NA |
| lizards | Phyllodactylus_pulcher | Phyllodactylus_pulcher | CR | Decreasing | tropical | nocturnal | 7.943 | 179.098 | 30.231 |
| lizards | Phyllodactylus_reissii | Phyllodactylus_reissii | LC | Unknown | tropical | nocturnal | 13.183 | 228951.321 | 10.415 |
| lizards | Phyllodactylus_sentosus | NA | CR | Decreasing | NA | NA | NA | NA | NA |
| lizards | Phyllodactylus_thompsoni | Phyllodactylus_thompsoni | EN | Stable | tropical | nocturnal | 2.57 | 898.527 | 7.918 |
| lizards | Phyllodactylus_transversalis | NA | LC | Stable | NA | NA | NA | NA | NA |
| lizards | Phyllodactylus_tuberculosus | Phyllodactylus_tuberculosus | LC | Stable | tropical | nocturnal | 25.704 | 340776.315 | 11.305 |
| lizards | Phyllodactylus_ventralis | NA | LC | Stable | NA | NA | NA | NA | NA |
| lizards | Phyllodactylus_wirshingi | Phyllodactylus_wirshingi | NT | Unknown | tropical | nocturnal | 7.586 | 40.437 | 23.827 |
| lizards | Phyllodactylus_xanti | Phyllodactylus_xanti | LC | Stable | temperate | nocturnal | 11.482 | 126932.844 | 3.865 |
| lizards | Phyllopezus_lutzae | NA | LC | Stable | NA | NA | NA | NA | NA |
| lizards | Phyllopezus_maranjonensis | Phyllopezus_maranjonensis | NT | Unknown | tropical | nocturnal | 52.481 | 34376.79 | 5.378 |
| lizards | Phyllopezus_periosus | Phyllopezus_periosus | LC | Unknown | tropical | nocturnal | 51.286 | 278407.36 | 11.546 |
| lizards | Phyllopezus_pollicaris | Phyllopezus_pollicaris | LC | Unknown | tropical | cathemeral | 21.878 | 1857384.691 | 8.036 |

| | | | | | | | | | |
|---|---|---|---|---|---|---|---|---|---|
| lizards | Phyllurus_amnicola | Phyllurus_amnicola | NT | Unknown | tropical | nocturnal | 24.547 | 224.053 | 1.541 |
| lizards | Phyllurus_caudiannulatus | Phyllurus_caudiannulatus | NT | Stable | temperate | nocturnal | 23.988 | 239.07 | 3.681 |
| lizards | Phyllurus_championae | Phyllurus_championae | NT | Unknown | tropical | nocturnal | 10.965 | 192.739 | 4.615 |
| lizards | Phyllurus_gulbaru | Phyllurus_gulbaru | EN | Unknown | tropical | nocturnal | 17.378 | 513.712 | 1.756 |
| lizards | Phyllurus_isis | NA | VU | Unknown | NA | NA | NA | NA | NA |
| lizards | Phyllurus_kabikabi | NA | CR | Unknown | NA | NA | NA | NA | NA |
| lizards | Phyllurus_nepthys | Phyllurus_nepthys | LC | Unknown | tropical | nocturnal | 19.498 | 840.32 | 4.289 |
| lizards | Phyllurus_ossa | NA | LC | Unknown | NA | NA | NA | NA | NA |
| lizards | Phyllurus_platurus | Phyllurus_platurus | LC | Unknown | temperate | nocturnal | 22.909 | 28511.114 | 10.07 |
| lizards | Phymaturus_aguanegra | Phymaturus_aguanegra | LC | Unknown | temperate | diurnal | 46.774 | 10.062 | 4.882 |
| lizards | Phymaturus_aguedae | NA | DD | Unknown | NA | NA | NA | NA | NA |
| lizards | Phymaturus_alicahuense | NA | LC | Stable | NA | NA | NA | NA | NA |
| lizards | Phymaturus_antofagastensis | Phymaturus_antofagastensis | LC | Stable | temperate | diurnal | 40.738 | 1725.099 | 2.404 |
| lizards | Phymaturus_bibronii | NA | EN | Unknown | NA | NA | NA | NA | NA |
| lizards | Phymaturus_camilae | Phymaturus_camilae | LC | Stable | temperate | diurnal | 42.658 | 8558.51 | 6.41 |
| lizards | Phymaturus_castillensis | Phymaturus_castillensis | LC | Stable | temperate | diurnal | 33.113 | 50.661 | 7.093 |
| lizards | Phymaturus_curivilcun | NA | not_available | not_available | NA | NA | NA | NA | NA |
| lizards | Phymaturus_darwini | Phymaturus_darwini | EN | Unknown | temperate | diurnal | 48.978 | 370.102 | 6.878 |
| lizards | Phymaturus_delheyi | Phymaturus_delheyi | LC | Unknown | temperate | diurnal | 33.884 | 15.47 | 5.214 |
| lizards | Phymaturus_denotatus | NA | LC | Stable | NA | NA | NA | NA | NA |
| lizards | Phymaturus_dorsimaculatus | NA | LC | Stable | NA | NA | NA | NA | NA |
| lizards | Phymaturus_etheridgei | NA | LC | Unknown | NA | NA | NA | NA | NA |
| lizards | Phymaturus_extrilidus | NA | LC | Stable | NA | NA | NA | NA | NA |
| lizards | Phymaturus_maulense | NA | LC | Stable | NA | NA | NA | NA | NA |
| lizards | Phymaturus_palluma | Phymaturus_palluma | LC | Stable | temperate | diurnal | 56.234 | 180821.426 | 4.164 |
| lizards | Phymaturus_patagonicus | Phymaturus_patagonicus | LC | Stable | temperate | diurnal | 53.703 | 67493.558 | 5.629 |
| lizards | Phymaturus_payuniae | Phymaturus_payuniae | NT | Unknown | temperate | diurnal | 28.184 | 4022.972 | 1.31 |
| lizards | Phymaturus_punae | Phymaturus_punae | LC | Stable | temperate | diurnal | 63.096 | 840.196 | 2.466 |
| lizards | Phymaturus_querque | NA | LC | Stable | NA | NA | NA | NA | NA |
| lizards | Phymaturus_rahuensis | NA | not_available | not_available | NA | NA | NA | NA | NA |
| lizards | Phymaturus_roigorum | Phymaturus_roigorum | LC | Stable | temperate | diurnal | 51.286 | 5174.84 | 1.756 |
| lizards | Phymaturus_sinervoi | Phymaturus_sinervoi | LC | Unknown | temperate | diurnal | 45.709 | 86.28 | 7.564 |
| lizards | Phymaturus_sitesi | NA | LC | Stable | NA | NA | NA | NA | NA |
| lizards | Phymaturus_somuncurensis | Phymaturus_somuncurensis | LC | Stable | temperate | diurnal | 40.738 | 21525.136 | 4.28 |

| | | | | | | | | | |
|---|---|---|---|---|---|---|---|---|---|
| lizards | Phymaturus_spurcus | Phymaturus_spurcus | LC | Stable | temperate | diurnal | 33.113 | 11937.235 | 5.289 |
| lizards | Phymaturus_tenebrosus | Phymaturus_tenebrosus | EN | Decreasing | temperate | diurnal | 56.234 | 2691.936 | 4.924 |
| lizards | Phymaturus_verdugo | NA | LC | Decreasing | NA | NA | NA | NA | NA |
| lizards | Phymaturus_videlai | Phymaturus_videlai | LC | Unknown | temperate | diurnal | 33.884 | 124.73 | 7.906 |
| lizards | Phymaturus_vociferator | Phymaturus_vociferator | VU | Stable | temperate | diurnal | 44.668 | 14135.798 | 3.176 |
| lizards | Phymaturus_williamsi | Phymaturus_williamsi | LC | Stable | temperate | diurnal | 48.978 | 570.911 | 2.124 |
| lizards | Phymaturus_yachanana | NA | LC | Unknown | NA | NA | NA | NA | NA |
| lizards | Phymaturus_zapalensis | Phymaturus_zapalensis | LC | Stable | temperate | diurnal | 48.978 | 10818.321 | 4.149 |
| lizards | Physignathus_cocincinus | Physignathus_cocincinus | VU | Decreasing | tropical | diurnal | 575.44 | 1243408.496 | 13.78 |
| lizards | Pinoyscincus_abdictus | NA | LC | Stable | NA | NA | NA | NA | NA |
| lizards | Pinoyscincus_coxi | Pinoyscincus_coxi | LC | Unknown | tropical | diurnal | 15.488 | 134924.056 | 12.557 |
| lizards | Pinoyscincus_jagori | Pinoyscincus_jagori | LC | Stable | tropical | diurnal | 29.512 | 273098.444 | 13.89 |
| lizards | Pinoyscincus_mindanensis | Pinoyscincus_mindanensis | LC | Decreasing | tropical | diurnal | 5.248 | 29535.744 | 12.246 |
| lizards | Placosoma_limaverdorum | NA | not_available | not_available | NA | NA | NA | NA | NA |
| lizards | Platysaurus_attenboroughi | NA | LC | Stable | NA | NA | NA | NA | NA |
| lizards | Platysaurus_broadleyi | Platysaurus_broadleyi | LC | Stable | temperate | diurnal | 15.849 | 5396.49 | 7.03 |
| lizards | Platysaurus_capensis | Platysaurus_capensis | LC | Stable | temperate | diurnal | 16.982 | 54943.567 | 3.847 |
| lizards | Platysaurus_imperator | Platysaurus_imperator | NT | Unknown | tropical | diurnal | 104.713 | 34203.091 | 12.968 |
| lizards | Platysaurus_intermedius | Platysaurus_intermedius | LC | Stable | tropical | diurnal | 67.608 | 405952.402 | 12.39 |
| lizards | Platysaurus_lebomboensis | Platysaurus_lebomboensis | LC | Stable | temperate | diurnal | 9.55 | 8411.917 | 13.597 |
| lizards | Platysaurus_maculatus | Platysaurus_maculatus | LC | Stable | tropical | diurnal | 13.183 | 93.226 | 13.656 |
| lizards | Platysaurus_minor | Platysaurus_minor | LC | Stable | temperate | diurnal | 8.71 | 19496.333 | 6.815 |
| lizards | Platysaurus_mitchelli | NA | LC | Stable | NA | NA | NA | NA | NA |
| lizards | Platysaurus_monotropis | Platysaurus_monotropis | NT | Stable | tropical | diurnal | 10.471 | 2219.342 | 10.298 |
| lizards | Platysaurus_ocellatus | Platysaurus_ocellatus | LC | Stable | tropical | diurnal | 21.38 | 23479.505 | 13.004 |
| lizards | Platysaurus_orientalis | Platysaurus_orientalis | LC | Stable | temperate | diurnal | 19.055 | 28276.45 | 11.825 |
| lizards | Platysaurus_pungweensis | Platysaurus_pungweensis | LC | Stable | tropical | diurnal | 19.055 | 38507.696 | 9.512 |
| lizards | Platysaurus_relictus | NA | NT | Decreasing | NA | NA | NA | NA | NA |
| lizards | Plestiodon_anthracinus | Plestiodon_anthracinus | LC | Stable | temperate | diurnal | 6.761 | 1099280.691 | 15.658 |
| lizards | Plestiodon_barbouri | NA | VU | Decreasing | NA | NA | NA | NA | NA |
| lizards | Plestiodon_bilineatus | NA | not_available | not_available | NA | NA | NA | NA | NA |
| lizards | Plestiodon_brevirostris | Plestiodon_brevirostris | LC | Stable | tropical | diurnal | 7.079 | 88662.24 | 11.971 |
| lizards | Plestiodon_callicephalus | Plestiodon_callicephalus | LC | Stable | temperate | diurnal | 8.913 | 174367.883 | 7.175 |
| lizards | Plestiodon_capito | Plestiodon_capito | LC | Stable | temperate | diurnal | 10.471 | 1059338.261 | 15.708 |

| | | | | | | | | | |
|---|---|---|---|---|---|---|---|---|---|
| lizards | Plestiodon_chinensis | Plestiodon_chinensis | LC | Stable | temperate | diurnal | 54.954 | 1767825.959 | 15.738 |
| lizards | Plestiodon_colimensis | NA | DD | Unknown | NA | NA | NA | NA | NA |
| lizards | Plestiodon_copei | Plestiodon_copei | LC | Stable | tropical | diurnal | 8.913 | 9117.402 | 25.022 |
| lizards | Plestiodon_coreensis | NA | DD | Unknown | NA | NA | NA | NA | NA |
| lizards | Plestiodon_dicei | NA | not_available | not_available | NA | NA | NA | NA | NA |
| lizards | Plestiodon_dugesii | Plestiodon_dugesii | VU | Decreasing | tropical | diurnal | 6.457 | 19119.847 | 15.513 |
| lizards | Plestiodon_egregius | Plestiodon_egregius | LC | Decreasing | temperate | diurnal | 5.129 | 251532.979 | 16.788 |
| lizards | Plestiodon_elegans | Plestiodon_elegans | LC | Unknown | temperate | diurnal | 25.119 | 2780942.553 | 15.461 |
| lizards | Plestiodon_fasciatus | Plestiodon_fasciatus | LC | Stable | temperate | diurnal | 13.804 | 2540293.283 | 17.897 |
| lizards | Plestiodon_finitimus | NA | LC | Unknown | NA | NA | NA | NA | NA |
| lizards | Plestiodon_gilberti | Plestiodon_gilberti | LC | Stable | temperate | diurnal | 35.481 | 217185.413 | 12.051 |
| lizards | Plestiodon_indubitus | NA | not_available | not_available | NA | NA | NA | NA | NA |
| lizards | Plestiodon_inexpectatus | Plestiodon_inexpectatus | LC | Stable | temperate | diurnal | 14.791 | 862871.06 | 16.934 |
| lizards | Plestiodon_japonicus | Plestiodon_japonicus | LC | Stable | temperate | diurnal | 19.055 | 106243.356 | 21.364 |
| lizards | Plestiodon_kishinouyei | Plestiodon_kishinouyei | VU | Decreasing | temperate | diurnal | 123.027 | 796.552 | 17.703 |
| lizards | Plestiodon_kuchinoshimensis | NA | VU | Unknown | NA | NA | NA | NA | NA |
| lizards | Plestiodon_lagunensis | Plestiodon_lagunensis | LC | Stable | temperate | diurnal | 4.169 | 10333.943 | 2.9 |
| lizards | Plestiodon_laticeps | Plestiodon_laticeps | LC | Stable | temperate | diurnal | 67.608 | 1777565.655 | 17.003 |
| lizards | Plestiodon_latiscutatus | Plestiodon_latiscutatus | LC | Stable | temperate | diurnal | 19.055 | 2989.859 | 24.129 |
| lizards | Plestiodon_leucostictus | NA | NT | Unknown | NA | NA | NA | NA | NA |
| lizards | Plestiodon_liui | NA | DD | Unknown | NA | NA | NA | NA | NA |
| lizards | Plestiodon_longirostris | Plestiodon_longirostris | CR | Decreasing | temperate | diurnal | 17.783 | 63.196 | 32.618 |
| lizards | Plestiodon_lynxe | Plestiodon_lynxe | LC | Stable | tropical | diurnal | 8.511 | 252363.334 | 12.302 |
| lizards | Plestiodon_marginatus | NA | NT | Decreasing | NA | NA | NA | NA | NA |
| lizards | Plestiodon_multilineatus | NA | DD | Unknown | NA | NA | NA | NA | NA |
| lizards | Plestiodon_multivirgatus | Plestiodon_multivirgatus | LC | Stable | temperate | diurnal | 8.913 | 609662.31 | 10.458 |
| lizards | Plestiodon_obsoletus | Plestiodon_obsoletus | LC | Stable | temperate | cathemeral | 67.608 | 1800483.913 | 11.608 |
| lizards | Plestiodon_ochoterenae | NA | LC | Stable | NA | NA | NA | NA | NA |
| lizards | Plestiodon_oshimensis | NA | NT | Decreasing | NA | NA | NA | NA | NA |
| lizards | Plestiodon_parviauriculatus | NA | DD | Unknown | NA | NA | NA | NA | NA |
| lizards | Plestiodon_parvulus | NA | DD | Unknown | NA | NA | NA | NA | NA |
| lizards | Plestiodon_quadrilineatus | Plestiodon_quadrilineatus | LC | Stable | tropical | diurnal | 30.2 | 412691.726 | 17.369 |
| lizards | Plestiodon_reynoldsi | Plestiodon_reynoldsi | VU | Decreasing | temperate | diurnal | 1.862 | 8146.16 | 21.335 |
| lizards | Plestiodon_septentrionalis | Plestiodon_septentrionalis | LC | Stable | temperate | diurnal | 15.488 | 1851681.789 | 14.916 |

| | | | | | | | | | |
|---|---|---|---|---|---|---|---|---|---|
| lizards | Plestiodon_skiltonianus | Plestiodon_skiltonianus | LC | Stable | temperate | diurnal | 13.183 | 1164304.832 | 9.822 |
| lizards | Plestiodon_stimpsonii | Plestiodon_stimpsonii | NT | Decreasing | temperate | diurnal | 10.471 | 554.755 | 13.623 |
| lizards | Plestiodon_sumichrasti | Plestiodon_sumichrasti | LC | Decreasing | tropical | diurnal | 21.38 | 135378.519 | 11.546 |
| lizards | Plestiodon_takarai | NA | DD | Unknown | NA | NA | NA | NA | NA |
| lizards | Plestiodon_tamdaoensis | NA | LC | Unknown | NA | NA | NA | NA | NA |
| lizards | Plestiodon_tetragrammus | Plestiodon_tetragrammus | LC | Stable | temperate | diurnal | 8.913 | 584850.076 | 11.792 |
| lizards | Plestiodon_tunganus | NA | NT | Decreasing | NA | NA | NA | NA | NA |
| lizards | Pletholax_gracilis | Pletholax_gracilis | LC | Stable | temperate | diurnal | 4.365 | 40485.452 | 9.433 |
| lizards | Plica_caribeana | NA | not_available | not_available | NA | NA | NA | NA | NA |
| lizards | Plica_kathleenae | NA | not_available | not_available | NA | NA | NA | NA | NA |
| lizards | Plica_lumaria | NA | LC | Stable | NA | NA | NA | NA | NA |
| lizards | Plica_medemi | NA | not_available | not_available | NA | NA | NA | NA | NA |
| lizards | Plica_pansticta | NA | LC | Stable | NA | NA | NA | NA | NA |
| lizards | Plica_plica | Plica_plica | LC | Stable | tropical | diurnal | 169.824 | 6864445.872 | 2.4 |
| lizards | Plica_rayi | NA | not_available | not_available | NA | NA | NA | NA | NA |
| lizards | Plica_umbra | Plica_umbra | LC | Stable | tropical | diurnal | 35.481 | 5802933.827 | 1.809 |
| lizards | Podarcis_bocagei | Podarcis_bocagei | LC | Stable | temperate | diurnal | 7.943 | 59240.892 | 17.336 |
| lizards | Podarcis_carbonelli | Podarcis_carbonelli | EN | Decreasing | temperate | diurnal | 6.457 | 15441.882 | 23.24 |
| lizards | Podarcis_cretensis | Podarcis_cretensis | EN | Decreasing | temperate | diurnal | 8.511 | 6298.076 | 17.844 |
| lizards | Podarcis_erhardii | Podarcis_erhardii | LC | Stable | temperate | diurnal | 13.804 | 153937.828 | 16.254 |
| lizards | Podarcis_filfolensis | NA | LC | Stable | NA | NA | NA | NA | NA |
| lizards | Podarcis_gaigeae | Podarcis_gaigeae | VU | Unknown | temperate | diurnal | 22.909 | 227.05 | 13.907 |
| lizards | Podarcis_guadarramae | NA | not_available | not_available | NA | NA | NA | NA | NA |
| lizards | Podarcis_hispanicus | Podarcis_hispanicus | LC | Unknown | temperate | diurnal | 9.333 | 62282.823 | 17.585 |
| lizards | Podarcis_levendis | NA | VU | Stable | NA | NA | NA | NA | NA |
| lizards | Podarcis_lilfordi | Podarcis_lilfordi | EN | Decreasing | temperate | diurnal | 12.303 | 4346.499 | 24.171 |
| lizards | Podarcis_liolepis | NA | not_available | not_available | NA | NA | NA | NA | NA |
| lizards | Podarcis_melisellensis | Podarcis_melisellensis | LC | Stable | temperate | diurnal | 9.333 | 31611.339 | 20.023 |
| lizards | Podarcis_milensis | Podarcis_milensis | VU | Stable | temperate | diurnal | 9.772 | 242.86 | 15.29 |
| lizards | Podarcis_muralis | Podarcis_muralis | LC | Stable | temperate | diurnal | 10.965 | 1837171.642 | 19.691 |
| lizards | Podarcis_peloponnesiacus | Podarcis_peloponnesiacus | LC | Stable | temperate | diurnal | 14.125 | 21531.399 | 16.63 |
| lizards | Podarcis_pityusensis | Podarcis_pityusensis | NT | Stable | temperate | diurnal | 12.589 | 660.626 | 24.169 |
| lizards | Podarcis_raffonei | Podarcis_raffoneae | CR | Decreasing | temperate | diurnal | 14.125 | 71.512 | 14.111 |
| lizards | Podarcis_siculus | Podarcis_siculus | LC | Increasing | temperate | diurnal | 16.596 | 276524.864 | 23.986 |

| | | | | | | | | | |
|---|---|---|---|---|---|---|---|---|---|
| lizards | Podarcis_tauricus | Podarcis_tauricus | LC | Stable | temperate | diurnal | 17.378 | 543683.3 | 17.642 |
| lizards | Podarcis_tiliguerta | Podarcis_tiliguerta | LC | Unknown | temperate | diurnal | 15.136 | 32891.797 | 16.515 |
| lizards | Podarcis_vaucheri | Podarcis_vaucheri | LC | Stable | temperate | diurnal | 5.012 | 390655.226 | 16.778 |
| lizards | Podarcis_virescens | NA | not_available | not_available | NA | NA | NA | NA | NA |
| lizards | Podarcis_waglerianus | Podarcis_waglerianus | LC | Decreasing | temperate | diurnal | 10.233 | 23913.304 | 23.232 |
| lizards | Pogona_barbata | Pogona_barbata | LC | Stable | temperate | diurnal | 691.831 | 1641296.362 | 8.129 |
| lizards | Pogona_henrylawsoni | Pogona_henrylawsoni | LC | Unknown | tropical | diurnal | 123.027 | 167036.479 | 4.903 |
| lizards | Pogona_microlepidota | Pogona_microlepidota | LC | Stable | tropical | diurnal | 208.93 | 38139.888 | 1.795 |
| lizards | Pogona_minor | Pogona_minor | LC | Stable | temperate | diurnal | 173.78 | 3027728.338 | 3.26 |
| lizards | Pogona_vitticeps | Pogona_vitticeps | LC | Unknown | temperate | diurnal | 575.44 | 2466684.34 | 4.582 |
| lizards | Polychrus_acutirostris | Polychrus_acutirostris | LC | Stable | tropical | diurnal | 104.713 | 6733865.568 | 7.052 |
| lizards | Polychrus_auduboni | NA | not_available | not_available | NA | NA | NA | NA | NA |
| lizards | Polychrus_femoralis | Polychrus_femoralis | LC | Stable | tropical | diurnal | 36.308 | 56600.86 | 12.124 |
| lizards | Polychrus_gutturosus | Polychrus_gutturosus | LC | Unknown | tropical | diurnal | 165.959 | 422163.602 | 9.498 |
| lizards | Polychrus_jacquelinae | NA | DD | Unknown | NA | NA | NA | NA | NA |
| lizards | Polychrus_liogaster | Polychrus_liogaster | LC | Unknown | tropical | diurnal | 109.648 | 963491.217 | 3.261 |
| lizards | Polychrus_marmoratus | Polychrus_marmoratus | LC | Stable | tropical | diurnal | 102.329 | 8528534.28 | 3.613 |
| lizards | Polychrus_peruvianus | Polychrus_peruvianus | VU | Unknown | tropical | diurnal | 109.648 | 63098.51 | 6.254 |
| lizards | Poromera_fordii | NA | LC | Stable | NA | NA | NA | NA | NA |
| lizards | Potamites_ecpleopus | Potamites_ecpleopus | LC | Unknown | tropical | diurnal | 15.136 | 2258840.429 | 1.935 |
| lizards | Potamites_erythrocularis | NA | not_available | not_available | NA | NA | NA | NA | NA |
| lizards | Potamites_juruazensis | Potamites_juruazensis | LC | Unknown | tropical | diurnal | 4.365 | 47076.454 | 2.685 |
| lizards | Praeteropus_brevicollis | NA | LC | Decreasing | NA | NA | NA | NA | NA |
| lizards | Prasinohaema_flavipes | Prasinohaema_flavipes | LC | Unknown | tropical | diurnal | 23.442 | 49779.927 | 5.023 |
| lizards | Prasinohaema_prehensicauda | NA | LC | Unknown | NA | NA | NA | NA | NA |
| lizards | Prasinohaema_semoni | Prasinohaema_semoni | LC | Unknown | tropical | diurnal | 8.128 | 66690.409 | 4.391 |
| lizards | Prasinohaema_virens | Prasinohaema_virens | LC | Stable | tropical | diurnal | 5.37 | 103574.876 | 5.408 |
| lizards | Pristidactylus_achalensis | Pristidactylus_achalensis | NT | Stable | temperate | diurnal | 56.234 | 1357.228 | 7.661 |
| lizards | Pristidactylus_alvaroi | NA | EN | Unknown | NA | NA | NA | NA | NA |
| lizards | Pristidactylus_nigroiugulus | NA | LC | Stable | NA | NA | NA | NA | NA |
| lizards | Pristidactylus_scapulatus | Pristidactylus_scapulatus | LC | Stable | temperate | diurnal | 41.687 | 102259.227 | 4.01 |
| lizards | Pristidactylus_torquatus | Pristidactylus_torquatus | VU | Unknown | temperate | diurnal | 41.687 | 111288.843 | 10.875 |
| lizards | Pristidactylus_valeriae | NA | EN | Unknown | NA | NA | NA | NA | NA |
| lizards | Pristidactylus_volcanensis | Pristidactylus_volcanensis | EN | Decreasing | temperate | diurnal | 27.542 | 749.241 | 10.176 |

| | | | | | | | | | |
|---|---|---|---|---|---|---|---|---|---|
| lizards | Pristurus_abdelkuri | Pristurus_abdelkuri | LC | Increasing | tropical | diurnal | 1.259 | 1987.75 | 10.013 |
| lizards | Pristurus_adrarensis | NA | DD | Unknown | NA | NA | NA | NA | NA |
| lizards | Pristurus_carteri | Pristurus_carteri | LC | not_available | tropical | cathemeral | 12.303 | 217855.548 | 5.028 |
| lizards | Pristurus_celerrimus | Pristurus_celerrimus | LC | Stable | temperate | diurnal | 1.585 | 26382.493 | 17.044 |
| lizards | Pristurus_collaris | Pristurus_collaris | LC | Stable | tropical | cathemeral | 5.248 | 77478.832 | 6.532 |
| lizards | Pristurus_crucifer | Pristurus_crucifer | LC | Unknown | tropical | diurnal | 1.585 | 774631.521 | 8.108 |
| lizards | Pristurus_flavipunctatus | Pristurus_flavipunctatus | LC | Unknown | tropical | diurnal | 1.585 | 2017062.79 | 8.688 |
| lizards | Pristurus_gallagheri | Pristurus_gallagheri | LC | Stable | tropical | diurnal | 1.585 | 6567.876 | 11.646 |
| lizards | Pristurus_guichardi | Pristurus_guichardi | LC | Unknown | tropical | diurnal | 1.288 | 347.601 | 11.481 |
| lizards | Pristurus_insignis | Pristurus_insignis | LC | Stable | tropical | diurnal | 5.248 | 2940.826 | 10.574 |
| lizards | Pristurus_insignoides | Pristurus_insignoides | LC | Unknown | tropical | diurnal | 3.631 | 66.735 | 8.206 |
| lizards | Pristurus_longipes | NA | DD | not_available | NA | NA | NA | NA | NA |
| lizards | Pristurus_mazbah | NA | DD | Unknown | NA | NA | NA | NA | NA |
| lizards | Pristurus_minimus | Pristurus_minimus | LC | Stable | tropical | cathemeral | 0.676 | 702202.611 | 3.355 |
| lizards | Pristurus_obsti | Pristurus_obsti | LC | Stable | tropical | diurnal | 1.318 | 1033.699 | 9.549 |
| lizards | Pristurus_ornithocephalus | Pristurus_ornithocephalus | LC | Unknown | tropical | diurnal | 5.248 | 55646.253 | 9.641 |
| lizards | Pristurus_phillipsii | Pristurus_phillipsii | LC | Unknown | tropical | diurnal | 0.676 | 80172.126 | 7.327 |
| lizards | Pristurus_popovi | Pristurus_popovi | LC | not_available | tropical | diurnal | 1.318 | 28560.03 | 12.66 |
| lizards | Pristurus_rupestris | Pristurus_rupestris | LC | Stable | tropical | diurnal | 0.813 | 884378.486 | 8.685 |
| lizards | Pristurus_saada | Pristurus_saada | LC | Unknown | tropical | diurnal | 1.698 | 27459.209 | 11.878 |
| lizards | Pristurus_samhaensis | Pristurus_samhaensis | LC | Stable | tropical | diurnal | 1.413 | 49.462 | 2.95 |
| lizards | Pristurus_schneideri | NA | LC | Unknown | NA | NA | NA | NA | NA |
| lizards | Pristurus_simonettai | NA | LC | Stable | NA | NA | NA | NA | NA |
| lizards | Pristurus_sokotranus | Pristurus_sokotranus | LC | Stable | tropical | diurnal | 1.585 | 3551.629 | 10.997 |
| lizards | Pristurus_somalicus | Pristurus_somalicus | LC | Unknown | tropical | diurnal | 1.479 | 3640.306 | 8.919 |
| lizards | Proablepharus_reginae | Proablepharus_reginae | LC | Stable | tropical | diurnal | 1.202 | 2311047.714 | 1.771 |
| lizards | Proablepharus_tenuis | Proablepharus_tenuis | LC | Stable | tropical | diurnal | 0.55 | 1534087.383 | 3.196 |
| lizards | Procellosaurinus_erythrocercus | NA | LC | Unknown | NA | NA | NA | NA | NA |
| lizards | Procellosaurinus_tetradactylus | NA | EN | Unknown | NA | NA | NA | NA | NA |
| lizards | Proctoporus_chasqui | Proctoporus_chasqui | LC | Unknown | tropical | diurnal | 9.772 | 21044.16 | 4.89 |
| lizards | Proctoporus_laudahnae | NA | DD | Unknown | NA | NA | NA | NA | NA |
| lizards | Proctoporus_oreades | NA | DD | Unknown | NA | NA | NA | NA | NA |
| lizards | Proctoporus_sucullucu | Proctoporus_sucullucu | LC | Unknown | tropical | diurnal | 3.89 | 30309.595 | 9.334 |
| lizards | Proctoporus_unsaacae | Proctoporus_unsaacae | LC | Unknown | tropical | diurnal | 2.951 | 11651.601 | 10.012 |

| | | | | | | | | | |
|---|---|---|---|---|---|---|---|---|---|
| lizards | Proscelotes_arnoldi | Proscelotes_arnoldi | LC | Stable | tropical | diurnal | 4.786 | 49591.761 | 11.199 |
| lizards | Proscelotes_eggeli | Proscelotes_eggeli | NT | Unknown | tropical | diurnal | 5.754 | 5991.458 | 15.779 |
| lizards | Psammodromus_algirus | Psammodromus_algirus | LC | Stable | temperate | diurnal | 19.498 | 1118062.967 | 15.751 |
| lizards | Psammodromus_blanci | Psammodromus_blanci | NT | Decreasing | temperate | diurnal | 2.455 | 155249.47 | 19.698 |
| lizards | Psammodromus_edwarsianus | NA | not_available | not_available | NA | NA | NA | NA | NA |
| lizards | Psammodromus_hispanicus | Psammodromus_hispanicus | LC | Stable | temperate | diurnal | 4.169 | 149004.202 | 16.433 |
| lizards | Psammodromus_microdactylus | NA | EN | Decreasing | NA | NA | NA | NA | NA |
| lizards | Psammodromus_occidentalis | NA | not_available | not_available | NA | NA | NA | NA | NA |
| lizards | Psammophilus_blanfordanus | Psammophilus_blanfordanus | LC | Stable | tropical | diurnal | 38.019 | 889843.53 | 18.39 |
| lizards | Psammophilus_dorsalis | Psammophilus_dorsalis | LC | Stable | tropical | diurnal | 95.499 | 369569.722 | 20.918 |
| lizards | Pseudemoia_baudini | Pseudemoia_baudini | LC | Stable | temperate | diurnal | 2.754 | 89633.058 | 5.945 |
| lizards | Pseudemoia_cryodroma | Pseudemoia_cryodroma | EN | Unknown | temperate | diurnal | 4.169 | 22949.373 | 6.278 |
| lizards | Pseudemoia_entrecasteauxii | Pseudemoia_entrecasteauxii | LC | Stable | temperate | diurnal | 5.37 | 581993.391 | 10.358 |
| lizards | Pseudemoia_pagenstecheri | Pseudemoia_pagenstecheri | LC | Decreasing | temperate | diurnal | 6.457 | 250162.807 | 9.375 |
| lizards | Pseudemoia_rawlinsoni | NA | DD | Unknown | NA | NA | NA | NA | NA |
| lizards | Pseudemoia_spenceri | Pseudemoia_spenceri | LC | Unknown | temperate | diurnal | 5.37 | 191785.546 | 10.386 |
| lizards | Pseuderemias_mucronata | NA | LC | Unknown | NA | NA | NA | NA | NA |
| lizards | Pseudocalotes_andamanensis | NA | VU | Decreasing | NA | NA | NA | NA | NA |
| lizards | Pseudocalotes_baliomus | NA | DD | Unknown | NA | NA | NA | NA | NA |
| lizards | Pseudocalotes_brevipes | Pseudocalotes_brevipes | LC | Unknown | tropical | diurnal | 15.136 | 216284.453 | 15.607 |
| lizards | Pseudocalotes_cybelidermus | Pseudocalotes_cybelidermus | LC | Unknown | tropical | diurnal | 27.542 | 268.058 | 8.99 |
| lizards | Pseudocalotes_dringi | NA | LC | Stable | NA | NA | NA | NA | NA |
| lizards | Pseudocalotes_drogon | NA | DD | Unknown | NA | NA | NA | NA | NA |
| lizards | Pseudocalotes_flavigula | Pseudocalotes_flavigula | CR | Stable | tropical | diurnal | 16.982 | 9.991 | 2.97 |
| lizards | Pseudocalotes_floweri | Pseudocalotes_floweri | VU | Unknown | tropical | diurnal | 32.359 | 115266.662 | 10.386 |
| lizards | Pseudocalotes_guttalineatus | Pseudocalotes_guttalineatus | LC | Unknown | tropical | diurnal | 28.84 | 255.997 | 9.013 |
| lizards | Pseudocalotes_kakhienensis | Pseudocalotes_kakhienensis | LC | Unknown | tropical | diurnal | 70.795 | 123991.863 | 10.352 |
| lizards | Pseudocalotes_khaonanensis | NA | DD | Unknown | NA | NA | NA | NA | NA |
| lizards | Pseudocalotes_kingdonwardi | NA | LC | Unknown | NA | NA | NA | NA | NA |
| lizards | Pseudocalotes_larutensis | Pseudocalotes_larutensis | VU | Stable | tropical | diurnal | 15.136 | 9.991 | 5.605 |
| lizards | Pseudocalotes_microlepis | NA | LC | Stable | NA | NA | NA | NA | NA |
| lizards | Pseudocalotes_poilani | NA | EN | Decreasing | NA | NA | NA | NA | NA |
| lizards | Pseudocalotes_rhaegal | NA | CR | Unknown | NA | NA | NA | NA | NA |
| lizards | Pseudocalotes_rhammanotus | NA | DD | Unknown | NA | NA | NA | NA | NA |

| | | | | | | | | | |
|---|---|---|---|---|---|---|---|---|---|
| lizards | Pseudocalotes_saravacensis | NA | DD | Unknown | NA | NA | NA | NA | NA |
| lizards | Pseudocalotes_tympanistriga | Pseudocalotes_tympanistriga | LC | Stable | tropical | diurnal | 17.378 | 5078.068 | 21.371 |
| lizards | Pseudocalotes_viserion | NA | VU | Stable | NA | NA | NA | NA | NA |
| lizards | Pseudocalotes_ziegleri | NA | DD | Unknown | NA | NA | NA | NA | NA |
| lizards | Pseudoceramodactylus_khobarensis | Pseudoceramodactylus_khobarensis | LC | Decreasing | temperate | nocturnal | 6.918 | 99923.848 | 9.368 |
| lizards | Pseudocophotis_kontumensis | NA | DD | Unknown | NA | NA | NA | NA | NA |
| lizards | Pseudocophotis_sumatrana | NA | DD | Unknown | NA | NA | NA | NA | NA |
| lizards | Pseudocordylus_langi | Pseudocordylus_langi | LC | Stable | temperate | diurnal | 33.113 | 30620.212 | 11.914 |
| lizards | Pseudocordylus_melanotus | Pseudocordylus_melanotus | LC | Stable | temperate | diurnal | 117.49 | 165377.004 | 12.637 |
| lizards | Pseudocordylus_microlepidotus | Pseudocordylus_microlepidotus | LC | Stable | temperate | diurnal | 162.181 | 216570.517 | 9.766 |
| lizards | Pseudocordylus_spinosus | Pseudocordylus_spinosus | LC | Decreasing | temperate | diurnal | 17.783 | 15084.257 | 12.14 |
| lizards | Pseudocordylus_subviridis | NA | not_available | not_available | NA | NA | NA | NA | NA |
| lizards | Pseudocordylus_transvaalensis | Pseudocordylus_transvaalensis | LC | Stable | temperate | diurnal | 117.49 | 25028.988 | 10.633 |
| lizards | Pseudogekko_atiorum | NA | not_available | not_available | NA | NA | NA | NA | NA |
| lizards | Pseudogekko_brevipes | Pseudogekko_brevipes | VU | Decreasing | tropical | nocturnal | 3.467 | 29759.211 | 13.406 |
| lizards | Pseudogekko_chavacano | NA | not_available | not_available | NA | NA | NA | NA | NA |
| lizards | Pseudogekko_compresicorpus | Pseudogekko_compressicorpus | LC | Unknown | tropical | nocturnal | 5.129 | 86089.023 | 12.627 |
| lizards | Pseudogekko_ditoy | NA | DD | Unknown | NA | NA | NA | NA | NA |
| lizards | Pseudogekko_pungkaypinit | NA | LC | Unknown | NA | NA | NA | NA | NA |
| lizards | Pseudogekko_smaragdinus | Pseudogekko_smaragdinus | LC | Unknown | tropical | nocturnal | 5.623 | 920.814 | 8.992 |
| lizards | Pseudogekko_sumiklab | NA | EN | Unknown | NA | NA | NA | NA | NA |
| lizards | Pseudogonatodes_barbouri | NA | NT | Unknown | NA | NA | NA | NA | NA |
| lizards | Pseudogonatodes_furvus | NA | DD | Unknown | NA | NA | NA | NA | NA |
| lizards | Pseudogonatodes_gasconi | NA | LC | Unknown | NA | NA | NA | NA | NA |
| lizards | Pseudogonatodes_guianensis | Pseudogonatodes_guianensis | LC | Unknown | tropical | diurnal | 0.676 | 3849398.494 | 1.351 |
| lizards | Pseudogonatodes_lunulatus | Pseudogonatodes_lunulatus | LC | Unknown | tropical | cathemeral | 0.661 | 48421.532 | 11.733 |
| lizards | Pseudogonatodes_manessi | NA | LC | Unknown | NA | NA | NA | NA | NA |
| lizards | Pseudogonatodes_peruvianus | NA | NT | Unknown | NA | NA | NA | NA | NA |
| lizards | Pseudopus_apodus | Pseudopus_apodus | LC | Stable | temperate | diurnal | 588.844 | 1920903.703 | 14.212 |
| lizards | Pseudothecadactylus_australis | NA | LC | Stable | NA | NA | NA | NA | NA |
| lizards | Pseudothecadactylus_cavaticus | NA | LC | Unknown | NA | NA | NA | NA | NA |
| lizards | Pseudothecadactylus_lindneri | NA | LC | Unknown | NA | NA | NA | NA | NA |
| lizards | Pseudotrapelus_aqabensis | NA | not_available | not_available | NA | NA | NA | NA | NA |
| lizards | Pseudotrapelus_chlodnickii | NA | LC | Unknown | NA | NA | NA | NA | NA |

| | | | | | | | | | |
|---|---|---|---|---|---|---|---|---|---|
| lizards | Pseudotrapelus_dhofarensis | NA | LC | Unknown | NA | NA | NA | NA | NA |
| lizards | Pseudotrapelus_jensvindumi | NA | not_available | not_available | NA | NA | NA | NA | NA |
| lizards | Pseudotrapelus_neumanni | NA | not_available | not_available | NA | NA | NA | NA | NA |
| lizards | Pseudotrapelus_sinaitus | Pseudotrapelus_sinaitus | LC | Stable | temperate | diurnal | 44.668 | 255754.008 | 6.241 |
| lizards | Psilops_mucugensis | NA | not_available | not_available | NA | NA | NA | NA | NA |
| lizards | Psilops_paeminosus | Psilophthalmus_paeminosus | NT | Unknown | tropical | diurnal | 1.288 | 1706.715 | 4.954 |
| lizards | Psilops_seductus | NA | not_available | not_available | NA | NA | NA | NA | NA |
| lizards | Psychosaura_agmosticha | Psychosaura_agmosticha | LC | Unknown | tropical | diurnal | 19.055 | 24199.946 | 13.114 |
| lizards | Psychosaura_macrorhyncha | Psychosaura_macrorhyncha | LC | Unknown | tropical | diurnal | 12.882 | 889124.185 | 11.157 |
| lizards | Ptenopus_carpi | Ptenopus_carpi | LC | Stable | tropical | nocturnal | 4.786 | 38353.562 | 3.033 |
| lizards | Ptenopus_garrulus | Ptenopus_garrulus | LC | Stable | temperate | nocturnal | 5.37 | 1314010.357 | 5.032 |
| lizards | Ptenopus_kochi | NA | LC | Stable | NA | NA | NA | NA | NA |
| lizards | Ptyctolaemus_collicristatus | NA | LC | Unknown | NA | NA | NA | NA | NA |
| lizards | Ptyctolaemus_gularis | Ptyctolaemus_gularis | LC | Stable | temperate | cathemeral | 21.878 | 173499.409 | 9.917 |
| lizards | Ptyodactylus_ananjevae | NA | not_available | not_available | NA | NA | NA | NA | NA |
| lizards | Ptyodactylus_dhofarensis | NA | not_available | not_available | NA | NA | NA | NA | NA |
| lizards | Ptyodactylus_guttatus | Ptyodactylus_guttatus | LC | Stable | temperate | cathemeral | 21.38 | 525182.429 | 5.821 |
| lizards | Ptyodactylus_hasselquistii | Ptyodactylus_hasselquistii | LC | Stable | tropical | cathemeral | 29.512 | 2220688.308 | 5.567 |
| lizards | Ptyodactylus_homolepis | NA | DD | Unknown | NA | NA | NA | NA | NA |
| lizards | Ptyodactylus_orlovi | NA | LC | Stable | NA | NA | NA | NA | NA |
| lizards | Ptyodactylus_oudrii | Ptyodactylus_oudrii | LC | Stable | temperate | nocturnal | 6.31 | 509271.985 | 7.275 |
| lizards | Ptyodactylus_puiseuxi | Ptyodactylus_puiseuxi | LC | Stable | temperate | diurnal | 18.621 | 145812.085 | 12.906 |
| lizards | Ptyodactylus_ragazzii | Ptyodactylus_ragazzii | LC | Stable | tropical | cathemeral | 25.704 | 4056885.104 | 6.739 |
| lizards | Ptyodactylus_rivapadiali | NA | not_available | not_available | NA | NA | NA | NA | NA |
| lizards | Ptyodactylus_ruusaljibalicus | NA | LC | Stable | NA | NA | NA | NA | NA |
| lizards | Pygmaeascincus_koshlandae | NA | LC | Unknown | NA | NA | NA | NA | NA |
| lizards | Pygmaeascincus_sadlieri | Pygmaeascincus_sadlieri | LC | Stable | tropical | diurnal | 0.437 | 54.047 | 12.928 |
| lizards | Pygmaeascincus_timlowi | Pygmaeascincus_timlowi | LC | Unknown | tropical | diurnal | 0.398 | 530697.426 | 5.78 |
| lizards | Pygomeles_braconnieri | NA | LC | Decreasing | NA | NA | NA | NA | NA |
| lizards | Pygomeles_petteri | NA | EN | Decreasing | NA | NA | NA | NA | NA |
| lizards | Pygopus_lepidopodus | Pygopus_lepidopodus | LC | Stable | temperate | cathemeral | 19.953 | 3927238.309 | 5.656 |
| lizards | Pygopus_nigriceps | Pygopus_nigriceps | LC | Stable | temperate | nocturnal | 15.488 | 3851746.997 | 2.464 |
| lizards | Pygopus_schraderi | Pygopus_schraderi | LC | Stable | temperate | nocturnal | 12.882 | 2696954.242 | 5.141 |
| lizards | Pygopus_steelescotti | Pygopus_steelescotti | LC | Stable | tropical | cathemeral | 11.749 | 877195.403 | 2.324 |

| | | | | | | | | | |
|---|---|---|---|---|---|---|---|---|---|
| lizards | Quedenfeldtia_moerens | Quedenfeldtia_moerens | LC | Stable | temperate | diurnal | 1.585 | 60974.656 | 10.326 |
| lizards | Quedenfeldtia_trachyblepharus | Quedenfeldtia_trachyblepharus | NT | Stable | temperate | diurnal | 2.455 | 9521.657 | 12.711 |
| lizards | Ramigekko_swartbergensis | Ramigekko_swartbergensis | LC | Stable | temperate | nocturnal | 9.333 | 9427.102 | 7.85 |
| lizards | Rankinia_diemensis | Rankinia_diemensis | LC | Unknown | temperate | diurnal | 19.498 | 361951.848 | 9.125 |
| lizards | Rhacodactylus_auriculatus | Rhacodactylus_auriculatus | LC | Stable | tropical | nocturnal | 40.738 | 9897.768 | 8.246 |
| lizards | Rhacodactylus_leachianus | Rhacodactylus_leachianus | LC | Stable | tropical | nocturnal | 478.63 | 12858.559 | 7.027 |
| lizards | Rhacodactylus_trachycephalus | NA | CR | Decreasing | NA | NA | NA | NA | NA |
| lizards | Rhacodactylus_trachyrhynchus | NA | VU | Decreasing | NA | NA | NA | NA | NA |
| lizards | Rhampholeon_boulengeri | Rhampholeon_boulengeri | LC | Decreasing | tropical | diurnal | 6.457 | 465606.985 | 8.018 |
| lizards | Rhampholeon_marshalli | Rhampholeon_marshalli | VU | Decreasing | tropical | diurnal | 10 | 7879.209 | 11.381 |
| lizards | Rhampholeon_maspictus | Rhampholeon_maspictus | NT | Stable | tropical | diurnal | 7.586 | 13.967 | 4.327 |
| lizards | Rhampholeon_moyeri | Rhampholeon_moyeri | LC | Stable | tropical | diurnal | 3.89 | 7517.584 | 9.183 |
| lizards | Rhampholeon_platyceps | Rhampholeon_platyceps | EN | Decreasing | tropical | diurnal | 8.913 | 2018.291 | 13.048 |
| lizards | Rhampholeon_spectrum | Rhampholeon_spectrum | LC | Unknown | tropical | diurnal | 6.166 | 1599030.121 | 7.323 |
| lizards | Rhampholeon_spinosus | Rhampholeon_spinosus | EN | Decreasing | tropical | diurnal | 3.802 | 3516.748 | 17.753 |
| lizards | Rhampholeon_temporalis | Rhampholeon_temporalis | EN | Decreasing | tropical | diurnal | 3.631 | 731.365 | 13.103 |
| lizards | Rhampholeon_tilburyi | Rhampholeon_tilburyi | EN | Decreasing | tropical | diurnal | 8.71 | 61.012 | 10.595 |
| lizards | Rhampholeon_uluguruensis | Rhampholeon_uluguruensis | LC | Stable | tropical | diurnal | 3.631 | 412.664 | 10.768 |
| lizards | Rhinogekko_femoralis | NA | LC | Unknown | NA | NA | NA | NA | NA |
| lizards | Rhinogekko_misonnei | NA | LC | Stable | NA | NA | NA | NA | NA |
| lizards | Rhoptropella_ocellata | Rhoptropella_ocellata | LC | Stable | temperate | diurnal | 2.239 | 37666.958 | 5.502 |
| lizards | Rhoptropus_afer | Rhoptropus_afer | LC | Stable | tropical | diurnal | 3.631 | 148366.281 | 4.451 |
| lizards | Rhoptropus_barnardi | Rhoptropus_barnardi | LC | Stable | tropical | diurnal | 2.63 | 289977.887 | 5.481 |
| lizards | Rhoptropus_biporosus | NA | LC | Stable | NA | NA | NA | NA | NA |
| lizards | Rhoptropus_boultoni | Rhoptropus_boultoni | LC | Stable | tropical | diurnal | 8.318 | 248492.668 | 5.978 |
| lizards | Rhoptropus_bradfieldi | Rhoptropus_bradfieldi | LC | Stable | tropical | diurnal | 7.079 | 71747.538 | 4.074 |
| lizards | Rhoptropus_diporus | NA | LC | Stable | NA | NA | NA | NA | NA |
| lizards | Rhoptropus_taeniostictus | NA | LC | Stable | NA | NA | NA | NA | NA |
| lizards | Rhynchoedura_angusta | Rhynchoedura_angusta | LC | Unknown | temperate | nocturnal | 2.951 | 173564.141 | 4.658 |
| lizards | Rhynchoedura_eyrensis | NA | LC | Stable | NA | NA | NA | NA | NA |
| lizards | Rhynchoedura_mentalis | NA | LC | Unknown | NA | NA | NA | NA | NA |
| lizards | Rhynchoedura_ormsbyi | Rhynchoedura_ormsbyi | LC | Stable | temperate | nocturnal | 2.951 | 723260.834 | 6.247 |
| lizards | Rhynchoedura_ornata | Rhynchoedura_ornata | LC | Stable | temperate | nocturnal | 3.631 | 2671543.627 | 2.28 |
| lizards | Rhynchoedura_sexapora | Rhynchoedura_sexapora | LC | Stable | tropical | nocturnal | 2.239 | 229008.351 | 2.466 |

| | | | | | | | | | |
|---|---|---|---|---|---|---|---|---|---|
| lizards | Riama_columbiana | Riama_columbiana | EN | Unknown | tropical | diurnal | 13.804 | 15553.121 | 15.072 |
| lizards | Riama_unicolor | NA | VU | Decreasing | NA | NA | NA | NA | NA |
| lizards | Rieppeleon_brachyurus | Rieppeleon_brachyurus | LC | Unknown | tropical | diurnal | 3.631 | 542075.656 | 9.038 |
| lizards | Rieppeleon_brevicaudatus | Rieppeleon_brevicaudatus | LC | Unknown | tropical | diurnal | 10.715 | 178303.605 | 9.405 |
| lizards | Rieppeleon_kerstenii | Rieppeleon_kerstenii | LC | Unknown | tropical | diurnal | 9.12 | 1280227.059 | 9.161 |
| lizards | Riolama_inopinata | NA | not_available | not_available | NA | NA | NA | NA | NA |
| lizards | Riolama_luridiventris | NA | LC | Stable | NA | NA | NA | NA | NA |
| lizards | Riopa_albopunctata | NA | LC | Unknown | NA | NA | NA | NA | NA |
| lizards | Riopa_guentheri | NA | LC | Unknown | NA | NA | NA | NA | NA |
| lizards | Riopa_lineata | NA | LC | Unknown | NA | NA | NA | NA | NA |
| lizards | Riopa_lineolata | NA | LC | Stable | NA | NA | NA | NA | NA |
| lizards | Riopa_punctata | NA | LC | Unknown | NA | NA | NA | NA | NA |
| lizards | Ristella_beddomii | Ristella_beddomii | LC | Unknown | tropical | diurnal | 1.122 | 11463.764 | 14.698 |
| lizards | Ristella_guentheri | NA | DD | Unknown | NA | NA | NA | NA | NA |
| lizards | Ristella_rurkii | Ristella_rurkii | NT | Unknown | tropical | diurnal | 1.862 | 10912.546 | 19.328 |
| lizards | Ristella_travancorica | NA | DD | Unknown | NA | NA | NA | NA | NA |
| lizards | Rondonops_biscutatus | NA | not_available | not_available | NA | NA | NA | NA | NA |
| lizards | Rondonops_xanthomystax | NA | not_available | not_available | NA | NA | NA | NA | NA |
| lizards | Saara_asmussi | NA | LC | Unknown | NA | NA | NA | NA | NA |
| lizards | Saara_hardwickii | Saara_hardwickii | VU | Decreasing | temperate | diurnal | 575.44 | 1111718.164 | 16.474 |
| lizards | Saara_loricata | Saara_loricata | LC | Unknown | temperate | diurnal | 912.011 | 139510.088 | 16.402 |
| lizards | Saiphos_equalis | Saiphos_equalis | LC | Unknown | temperate | cathemeral | 4.467 | 245681.423 | 8.407 |
| lizards | Salea_anamallayana | Salea_anamallayana | LC | Stable | tropical | diurnal | 45.709 | 6379.907 | 14.122 |
| lizards | Salea_gularis | NA | DD | Unknown | NA | NA | NA | NA | NA |
| lizards | Salea_horsfieldii | Salea_horsfieldii | LC | Stable | tropical | diurnal | 28.184 | 4619.344 | 13.725 |
| lizards | Saltuarius_cornutus | Saltuarius_cornutus | LC | Unknown | tropical | nocturnal | 54.954 | 19859.74 | 5.265 |
| lizards | Saltuarius_eximius | NA | EN | Unknown | NA | NA | NA | NA | NA |
| lizards | Saltuarius_kateae | Saltuarius_kateae | LC | Stable | temperate | nocturnal | 22.909 | 257.686 | 1.286 |
| lizards | Saltuarius_moritzi | Saltuarius_moritzi | LC | Stable | temperate | nocturnal | 22.387 | 30847.591 | 6.833 |
| lizards | Saltuarius_salebrosus | Saltuarius_salebrosus | LC | Stable | temperate | nocturnal | 42.658 | 89491.311 | 5.196 |
| lizards | Saltuarius_swaini | Saltuarius_swaini | LC | Unknown | temperate | nocturnal | 37.154 | 11797.295 | 9.012 |
| lizards | Saltuarius_wyberba | Saltuarius_wyberba | LC | Stable | temperate | nocturnal | 22.387 | 6886.313 | 6.435 |
| lizards | Salvator_duseni | NA | LC | Unknown | NA | NA | NA | NA | NA |
| lizards | Salvator_merianae | Salvator_merianae | LC | Stable | tropical | diurnal | 4466.836 | 8025045.464 | 7.965 |

| | | | | | | | | | |
|---|---|---|---|---|---|---|---|---|---|
| lizards | Salvator_rufescens | Salvator_rufescens | LC | Unknown | temperate | diurnal | 8317.638 | 1371652.754 | 6.34 |
| lizards | Saproscincus_basiliscus | Saproscincus_basiliscus | LC | Stable | tropical | diurnal | 2.291 | 22520.228 | 5.825 |
| lizards | Saproscincus_challengeri | Saproscincus_challengeri | LC | Stable | temperate | diurnal | 3.89 | 19134.331 | 11.714 |
| lizards | Saproscincus_czechurai | Saproscincus_czechurai | LC | Stable | tropical | diurnal | 1.122 | 13189.238 | 5.318 |
| lizards | Saproscincus_eungellensis | NA | DD | Unknown | NA | NA | NA | NA | NA |
| lizards | Saproscincus_hannahae | NA | LC | Stable | NA | NA | NA | NA | NA |
| lizards | Saproscincus_lewisi | NA | LC | Stable | NA | NA | NA | NA | NA |
| lizards | Saproscincus_mustelinus | Saproscincus_mustelinus | LC | Unknown | temperate | cathemeral | 5.129 | 212712.415 | 9.477 |
| lizards | Saproscincus_oriarus | Saproscincus_oriarus | LC | Unknown | temperate | diurnal | 1.413 | 4899.017 | 11.638 |
| lizards | Saproscincus_rosei | Saproscincus_rosei | LC | Stable | temperate | diurnal | 5.37 | 65680.532 | 7.76 |
| lizards | Saproscincus_saltus | NA | VU | Unknown | NA | NA | NA | NA | NA |
| lizards | Saproscincus_spectabilis | Saproscincus_spectabilis | LC | Unknown | temperate | diurnal | 4.169 | 66641.798 | 9.283 |
| lizards | Saproscincus_tetradactylus | Saproscincus_tetradactylus | LC | Unknown | tropical | diurnal | 0.603 | 14781.593 | 5.692 |
| lizards | Sarada_darwini | NA | LC | Unknown | NA | NA | NA | NA | NA |
| lizards | Sarada_deccanensis | NA | LC | Unknown | NA | NA | NA | NA | NA |
| lizards | Sarada_superba | NA | CR | Stable | NA | NA | NA | NA | NA |
| lizards | Saurodactylus_brosseti | NA | LC | Stable | NA | NA | NA | NA | NA |
| lizards | Saurodactylus_fasciatus | Saurodactylus_fasciatus | VU | Decreasing | temperate | nocturnal | 1.349 | 42698.669 | 16.705 |
| lizards | Saurodactylus_mauritanicus | Saurodactylus_mauritanicus | LC | Stable | temperate | diurnal | 0.977 | 235990.434 | 12.51 |
| lizards | Sauromalus_ater | Sauromalus_ater | LC | Stable | temperate | diurnal | 524.807 | 387687.401 | 6.8 |
| lizards | Sauromalus_hispidus | Sauromalus_hispidus | EN | Decreasing | temperate | diurnal | 1380.384 | 991.442 | 0.452 |
| lizards | Sauromalus_klauberi | NA | VU | Unknown | NA | NA | NA | NA | NA |
| lizards | Sauromalus_slevini | NA | NT | Unknown | NA | NA | NA | NA | NA |
| lizards | Sauromalus_varius | Sauromalus_varius | VU | Unknown | temperate | diurnal | 1659.587 | 41.202 | 1.574 |
| lizards | Scelarcis_perspicillata | Scelarcis_perspicillata | LC | Stable | temperate | diurnal | 7.586 | 133305.076 | 13.925 |
| lizards | Sceloporus_acanthinus | Sceloporus_acanthinus | LC | Stable | tropical | diurnal | 30.903 | 13458.613 | 14.091 |
| lizards | Sceloporus_adleri | Sceloporus_adleri | LC | Stable | tropical | diurnal | 16.596 | 4633.155 | 4.601 |
| lizards | Sceloporus_aeneus | Sceloporus_aeneus | LC | Stable | tropical | diurnal | 9.12 | 52636.259 | 19.285 |
| lizards | Sceloporus_albiventris | NA | not_available | not_available | NA | NA | NA | NA | NA |
| lizards | Sceloporus_anahuacus | NA | LC | Stable | NA | NA | NA | NA | NA |
| lizards | Sceloporus_angustus | Sceloporus_angustus | LC | Stable | temperate | cathemeral | 32.359 | 45.047 | 1.325 |
| lizards | Sceloporus_arenicolus | Sceloporus_arenicolus | VU | Decreasing | temperate | diurnal | 12.023 | 5622.153 | 11.198 |
| lizards | Sceloporus_asper | NA | LC | Decreasing | NA | NA | NA | NA | NA |
| lizards | Sceloporus_aurantius | NA | not_available | not_available | NA | NA | NA | NA | NA |

| | | | | | | | | | |
|---|---|---|---|---|---|---|---|---|---|
| lizards | Sceloporus_becki | NA | not_available | not_available | NA | NA | NA | NA | NA |
| lizards | Sceloporus_bicanthalis | Sceloporus_bicanthalis | LC | Stable | tropical | diurnal | 8.71 | 24407.254 | 22.295 |
| lizards | Sceloporus_bimaculosus | NA | not_available | not_available | NA | NA | NA | NA | NA |
| lizards | Sceloporus_brownorum | NA | not_available | not_available | NA | NA | NA | NA | NA |
| lizards | Sceloporus_bulleri | Sceloporus_bulleri | LC | Stable | tropical | diurnal | 46.774 | 29304.513 | 6.262 |
| lizards | Sceloporus_caeruleus | NA | not_available | not_available | NA | NA | NA | NA | NA |
| lizards | Sceloporus_cautus | NA | LC | Decreasing | NA | NA | NA | NA | NA |
| lizards | Sceloporus_chaneyi | Sceloporus_chaneyi | EN | Decreasing | temperate | diurnal | 7.079 | 3018.858 | 5.075 |
| lizards | Sceloporus_chrysostictus | Sceloporus_chrysostictus | LC | Stable | tropical | diurnal | 8.71 | 156256.05 | 6.653 |
| lizards | Sceloporus_clarkii | Sceloporus_clarkii | LC | Stable | temperate | diurnal | 83.176 | 457350.442 | 7.577 |
| lizards | Sceloporus_consobrinus | NA | not_available | not_available | NA | NA | NA | NA | NA |
| lizards | Sceloporus_couchii | Sceloporus_couchii | LC | Stable | temperate | diurnal | 9.55 | 48041.725 | 7.359 |
| lizards | Sceloporus_cowlesi | NA | not_available | not_available | NA | NA | NA | NA | NA |
| lizards | Sceloporus_cozumelae | Sceloporus_cozumelae | LC | Stable | tropical | diurnal | 8.128 | 20751.442 | 8.042 |
| lizards | Sceloporus_cryptus | Sceloporus_cryptus | LC | Stable | tropical | diurnal | 15.488 | 1680.899 | 8.061 |
| lizards | Sceloporus_cyanogenys | NA | not_available | not_available | NA | NA | NA | NA | NA |
| lizards | Sceloporus_cyanostictus | Sceloporus_cyanostictus | EN | Decreasing | temperate | diurnal | 17.783 | 3325.919 | 5.096 |
| lizards | Sceloporus_druckercolini | NA | not_available | not_available | NA | NA | NA | NA | NA |
| lizards | Sceloporus_dugesii | Sceloporus_dugesii | LC | Stable | tropical | diurnal | 22.387 | 147176.865 | 13.157 |
| lizards | Sceloporus_edbelli | NA | not_available | not_available | NA | NA | NA | NA | NA |
| lizards | Sceloporus_edwardtaylori | Sceloporus_edwardtaylori | LC | Stable | tropical | diurnal | 38.019 | 18437.354 | 6.566 |
| lizards | Sceloporus_exsul | Sceloporus_exsul | CR | Decreasing | tropical | diurnal | 10 | 323.539 | 10.495 |
| lizards | Sceloporus_formosus | Sceloporus_formosus | LC | Stable | tropical | diurnal | 22.387 | 97765.324 | 10.594 |
| lizards | Sceloporus_gadoviae | Sceloporus_gadoviae | LC | Stable | tropical | diurnal | 15.136 | 22582.32 | 9.54 |
| lizards | Sceloporus_gadsdeni | NA | not_available | not_available | NA | NA | NA | NA | NA |
| lizards | Sceloporus_goldmani | Sceloporus_goldmani | EN | Decreasing | temperate | diurnal | 8.128 | 9208.607 | 7.38 |
| lizards | Sceloporus_graciosus | Sceloporus_graciosus | LC | Stable | temperate | diurnal | 22.909 | 1358846.804 | 9.587 |
| lizards | Sceloporus_grammicus | Sceloporus_grammicus | LC | Stable | tropical | diurnal | 17.783 | 981930.664 | 9.276 |
| lizards | Sceloporus_grandaevus | Sceloporus_grandaevus | LC | Stable | temperate | diurnal | 17.783 | 142.907 | 2.171 |
| lizards | Sceloporus_halli | NA | DD | Unknown | NA | NA | NA | NA | NA |
| lizards | Sceloporus_heterolepis | NA | LC | Unknown | NA | NA | NA | NA | NA |
| lizards | Sceloporus_horridus | Sceloporus_horridus | LC | Stable | tropical | diurnal | 48.978 | 330779.989 | 9.165 |
| lizards | Sceloporus_hunsakeri | Sceloporus_hunsakeri | LC | Stable | temperate | cathemeral | 21.878 | 9946.615 | 7.361 |
| lizards | Sceloporus_internasalis | Sceloporus_internasalis | LC | Decreasing | tropical | diurnal | 31.623 | 34714.514 | 11.034 |

| | | | | | | | | | |
|---|---|---|---|---|---|---|---|---|---|
| lizards | Sceloporus_jalapae | Sceloporus_jalapae | LC | Stable | tropical | diurnal | 8.71 | 34104.499 | 12.571 |
| lizards | Sceloporus_jarrovii | Sceloporus_jarrovii | LC | Stable | temperate | diurnal | 37.154 | 532172.569 | 7.335 |
| lizards | Sceloporus_lemosespinali | NA | DD | Unknown | NA | NA | NA | NA | NA |
| lizards | Sceloporus_licki | Sceloporus_licki | LC | Stable | temperate | diurnal | 26.915 | 4628.555 | 6.627 |
| lizards | Sceloporus_lunae | NA | LC | Unknown | NA | NA | NA | NA | NA |
| lizards | Sceloporus_lundelli | NA | LC | Stable | NA | NA | NA | NA | NA |
| lizards | Sceloporus_macdougalli | Sceloporus_macdougalli | LC | Stable | tropical | diurnal | 27.542 | 8909.215 | 9.887 |
| lizards | Sceloporus_magister | Sceloporus_magister | LC | Stable | temperate | diurnal | 81.283 | 1395390.781 | 7.487 |
| lizards | Sceloporus_malachiticus | Sceloporus_malachiticus | LC | Stable | tropical | diurnal | 30.2 | 50271.058 | 14.731 |
| lizards | Sceloporus_megalepidurus | Sceloporus_megalepidurus | VU | Decreasing | tropical | diurnal | 6.31 | 10818.615 | 18.181 |
| lizards | Sceloporus_melanorhinus | Sceloporus_melanorhinus | LC | Stable | tropical | diurnal | 36.308 | 164283.9 | 9.644 |
| lizards | Sceloporus_merriami | Sceloporus_merriami | LC | Stable | temperate | diurnal | 10.471 | 208660.182 | 7.261 |
| lizards | Sceloporus_minor | Sceloporus_minor | LC | Stable | tropical | diurnal | 21.878 | 133511.213 | 8.139 |
| lizards | Sceloporus_mucronatus | Sceloporus_mucronatus | LC | Stable | tropical | diurnal | 44.668 | 98787.588 | 15.354 |
| lizards | Sceloporus_nelsoni | Sceloporus_nelsoni | LC | Stable | temperate | diurnal | 11.22 | 117808.957 | 6.797 |
| lizards | Sceloporus_oberon | NA | VU | Decreasing | NA | NA | NA | NA | NA |
| lizards | Sceloporus_occidentalis | Sceloporus_occidentalis | LC | Stable | temperate | diurnal | 30.903 | 1010683.226 | 10.104 |
| lizards | Sceloporus_ochoterenae | Sceloporus_ochoterenae | LC | Stable | tropical | diurnal | 8.71 | 14081.843 | 12.529 |
| lizards | Sceloporus_olivaceus | Sceloporus_olivaceus | LC | Stable | temperate | diurnal | 56.234 | 646821.003 | 12.616 |
| lizards | Sceloporus_omiltemanus | NA | not_available | not_available | NA | NA | NA | NA | NA |
| lizards | Sceloporus_orcutti | Sceloporus_orcutti | LC | Stable | temperate | diurnal | 47.863 | 111356.738 | 4.932 |
| lizards | Sceloporus_ornatus | Sceloporus_ornatus | NT | Unknown | temperate | diurnal | 30.2 | 56330.395 | 6.108 |
| lizards | Sceloporus_palaciosi | Sceloporus_palaciosi | LC | Stable | tropical | diurnal | 14.454 | 985.532 | 14.881 |
| lizards | Sceloporus_parvus | Sceloporus_parvus | LC | Stable | temperate | diurnal | 5.248 | 163557.95 | 7.178 |
| lizards | Sceloporus_poinsettii | Sceloporus_poinsettii | LC | Stable | temperate | diurnal | 74.131 | 842198.089 | 7.572 |
| lizards | Sceloporus_pyrocephalus | Sceloporus_pyrocephalus | LC | Stable | tropical | diurnal | 14.454 | 62214.862 | 8.656 |
| lizards | Sceloporus_salvini | NA | DD | Unknown | NA | NA | NA | NA | NA |
| lizards | Sceloporus_samcolemani | Sceloporus_samcolemani | LC | Stable | temperate | diurnal | 8.318 | 25507.135 | 9.101 |
| lizards | Sceloporus_scalaris | Sceloporus_scalaris | LC | Stable | tropical | diurnal | 16.218 | 227283.399 | 14.025 |
| lizards | Sceloporus_serrifer | Sceloporus_serrifer | LC | Stable | temperate | diurnal | 43.652 | 225665.133 | 9.584 |
| lizards | Sceloporus_shannonorum | NA | DD | Unknown | NA | NA | NA | NA | NA |
| lizards | Sceloporus_siniferus | Sceloporus_siniferus | LC | Stable | tropical | diurnal | 12.589 | 156926.035 | 8.671 |
| lizards | Sceloporus_slevini | Sceloporus_slevini | LC | Decreasing | temperate | diurnal | 12.023 | 151398.696 | 5.347 |
| lizards | Sceloporus_smaragdinus | Sceloporus_smaragdinus | LC | Stable | tropical | diurnal | 20.893 | 24225.805 | 15.601 |

| | | | | | | | | | |
|---|---|---|---|---|---|---|---|---|---|
| lizards | Sceloporus_smithi | Sceloporus_smithi | LC | Stable | tropical | diurnal | 12.589 | 20200.242 | 7.506 |
| lizards | Sceloporus_spinosus | Sceloporus_spinosus | LC | Stable | tropical | diurnal | 58.884 | 304248 | 11.905 |
| lizards | Sceloporus_squamosus | Sceloporus_squamosus | LC | Stable | tropical | diurnal | 7.762 | 95046.167 | 16.441 |
| lizards | Sceloporus_stejnegeri | NA | LC | Stable | NA | NA | NA | NA | NA |
| lizards | Sceloporus_subpictus | NA | DD | Unknown | NA | NA | NA | NA | NA |
| lizards | Sceloporus_sugillatus | Sceloporus_sugillatus | LC | Stable | tropical | diurnal | 21.878 | 1224.515 | 18.024 |
| lizards | Sceloporus_taeniocnemis | Sceloporus_taeniocnemis | LC | Stable | tropical | diurnal | 18.621 | 19174.847 | 12.501 |
| lizards | Sceloporus_teapensis | Sceloporus_teapensis | LC | Stable | tropical | diurnal | 12.023 | 179968.887 | 10.909 |
| lizards | Sceloporus_torquatus | Sceloporus_torquatus | LC | Stable | tropical | diurnal | 79.433 | 321352.381 | 12.274 |
| lizards | Sceloporus_tristichus | NA | not_available | not_available | NA | NA | NA | NA | NA |
| lizards | Sceloporus_undulatus | Sceloporus_undulatus | LC | Stable | temperate | diurnal | 24.547 | 4098325.292 | 13.622 |
| lizards | Sceloporus_unicanthalis | NA | not_available | not_available | NA | NA | NA | NA | NA |
| lizards | Sceloporus_uniformis | NA | not_available | not_available | NA | NA | NA | NA | NA |
| lizards | Sceloporus_utiformis | Sceloporus_utiformis | LC | Stable | tropical | diurnal | 19.953 | 162522.64 | 9.034 |
| lizards | Sceloporus_variabilis | Sceloporus_variabilis | LC | Stable | tropical | diurnal | 15.849 | 953656.305 | 10.76 |
| lizards | Sceloporus_virgatus | Sceloporus_virgatus | LC | Stable | temperate | diurnal | 12.589 | 78813.04 | 5.667 |
| lizards | Sceloporus_woodi | Sceloporus_woodi | NT | Decreasing | temperate | diurnal | 10 | 13827.788 | 25.406 |
| lizards | Sceloporus_zosteromus | Sceloporus_zosteromus | LC | Stable | temperate | cathemeral | 64.565 | 126799.423 | 3.964 |
| lizards | Scelotes_anguinus | NA | LC | Stable | NA | NA | NA | NA | NA |
| lizards | Scelotes_caffer | Scelotes_caffer | LC | Stable | temperate | diurnal | 1.23 | 26702.136 | 10.094 |
| lizards | Scelotes_gronovii | Scelotes_gronovii | NT | Decreasing | temperate | diurnal | 2.239 | 25154.957 | 14.849 |
| lizards | Scelotes_limpopoensis | Scelotes_limpopoensis | LC | Stable | tropical | diurnal | 3.631 | 29130.268 | 8.435 |
| lizards | Scelotes_montispectus | NA | NT | Unknown | NA | NA | NA | NA | NA |
| lizards | Scincella_apraefrontalis | NA | DD | Unknown | NA | NA | NA | NA | NA |
| lizards | Scincella_assata | Scincella_assatus | LC | Stable | tropical | diurnal | 3.09 | 195617.958 | 11.621 |
| lizards | Scincella_badenensis | NA | not_available | not_available | NA | NA | NA | NA | NA |
| lizards | Scincella_baraensis | NA | not_available | not_available | NA | NA | NA | NA | NA |
| lizards | Scincella_boettgeri | Scincella_boettgeri | LC | Unknown | temperate | cathemeral | 3.311 | 798.67 | 17.703 |
| lizards | Scincella_capitanea | NA | LC | Unknown | NA | NA | NA | NA | NA |
| lizards | Scincella_caudaequinae | NA | not_available | not_available | NA | NA | NA | NA | NA |
| lizards | Scincella_cherriei | Scincella_cherriei | LC | Stable | tropical | diurnal | 6.166 | 482005.374 | 9.477 |
| lizards | Scincella_darevskii | NA | DD | Unknown | NA | NA | NA | NA | NA |
| lizards | Scincella_devorator | NA | DD | Unknown | NA | NA | NA | NA | NA |
| lizards | Scincella_doriae | NA | LC | Unknown | NA | NA | NA | NA | NA |

| | | | | | | | | | |
|---|---|---|---|---|---|---|---|---|---|
| lizards | Scincella_formosensis | NA | LC | Stable | NA | NA | NA | NA | NA |
| lizards | Scincella_gemmingeri | Scincella_gemmingeri | LC | Stable | tropical | diurnal | 5.37 | 146449.241 | 14.438 |
| lizards | Scincella_huanrenensis | Scincella_huanrenensis | CR | Decreasing | temperate | diurnal | 4.571 | 15069.213 | 19.559 |
| lizards | Scincella_incerta | NA | LC | Stable | NA | NA | NA | NA | NA |
| lizards | Scincella_lateralis | Scincella_lateralis | LC | Stable | temperate | cathemeral | 3.89 | 2107754.698 | 16.425 |
| lizards | Scincella_macrotis | NA | DD | Unknown | NA | NA | NA | NA | NA |
| lizards | Scincella_melanosticta | Scincella_melanosticta | LC | Stable | tropical | diurnal | 5.37 | 483325.97 | 13.123 |
| lizards | Scincella_modesta | Scincella_modesta | LC | Stable | temperate | diurnal | 5.754 | 1279283.811 | 16.121 |
| lizards | Scincella_monticola | Scincella_monticola | LC | Unknown | temperate | cathemeral | 3.89 | 908302.353 | 12.506 |
| lizards | Scincella_nigrofasciata | NA | not_available | not_available | NA | NA | NA | NA | NA |
| lizards | Scincella_ochracea | Scincella_ochracea | LC | Unknown | tropical | diurnal | 2.455 | 39206.702 | 10.972 |
| lizards | Scincella_potanini | NA | LC | Stable | NA | NA | NA | NA | NA |
| lizards | Scincella_przewalskii | NA | DD | Unknown | NA | NA | NA | NA | NA |
| lizards | Scincella_punctatolineata | NA | LC | Unknown | NA | NA | NA | NA | NA |
| lizards | Scincella_rara | NA | DD | Unknown | NA | NA | NA | NA | NA |
| lizards | Scincella_reevesii | Scincella_reevesii | LC | Stable | tropical | cathemeral | 4.169 | 2762379.807 | 13.346 |
| lizards | Scincella_rufocaudata | Scincella_rufocaudatus | LC | Stable | tropical | diurnal | 2.455 | 111699.274 | 9.416 |
| lizards | Scincella_rupicola | NA | LC | Unknown | NA | NA | NA | NA | NA |
| lizards | Scincella_silvicola | Scincella_silvicola | LC | Stable | tropical | diurnal | 3.715 | 18296.274 | 9.598 |
| lizards | Scincella_tsinlingensis | Scincella_tsinlingensis | LC | Unknown | temperate | diurnal | 6.761 | 282019.324 | 13.673 |
| lizards | Scincella_vandenburghi | Scincella_vandenburghi | LC | Unknown | temperate | diurnal | 2.884 | 57898.86 | 20.831 |
| lizards | Scincella_victoriana | Scincella_victoriana | LC | Stable | tropical | diurnal | 9.12 | 5227.434 | 5.141 |
| lizards | Scincopus_fasciatus | NA | LC | Unknown | NA | NA | NA | NA | NA |
| lizards | Scincus_albifasciatus | Scincus_albifasciatus | LC | Stable | tropical | diurnal | 51.286 | 2073161.134 | 2.076 |
| lizards | Scincus_conirostris | NA | not_available | not_available | NA | NA | NA | NA | NA |
| lizards | Scincus_hemprichii | NA | LC | not_available | NA | NA | NA | NA | NA |
| lizards | Scincus_mitranus | Scincus_mitranus | LC | Stable | temperate | diurnal | 54.954 | 1455446.249 | 4.112 |
| lizards | Scincus_scincus | Scincus_scincus | LC | Stable | temperate | cathemeral | 74.131 | 10611472.07 | 3.111 |
| lizards | Scriptosaura_catimbau | NA | LC | Unknown | NA | NA | NA | NA | NA |
| lizards | Sepsina_alberti | Sepsina_alberti | LC | Stable | tropical | diurnal | 1.23 | 41674.024 | 6.124 |
| lizards | Sepsina_angolensis | NA | LC | Stable | NA | NA | NA | NA | NA |
| lizards | Sepsophis_punctatus | NA | LC | Unknown | NA | NA | NA | NA | NA |
| lizards | Shinisaurus_crocodilurus | Shinisaurus_crocodilurus | EN | Decreasing | temperate | diurnal | 83.176 | 90732.085 | 15.821 |
| lizards | Siderolamprus_atitlanensis | NA | DD | Unknown | NA | NA | NA | NA | NA |

| | | | | | | | | | |
|---|---|---|---|---|---|---|---|---|---|
| lizards | Siderolamprus_cyanochloris | NA | LC | Unknown | NA | NA | NA | NA | NA |
| lizards | Siderolamprus_hylaius | NA | NT | Stable | NA | NA | NA | NA | NA |
| lizards | Siderolamprus_laf | NA | not_available | not_available | NA | NA | NA | NA | NA |
| lizards | Siderolamprus_montanus | NA | EN | Decreasing | NA | NA | NA | NA | NA |
| lizards | Siderolamprus_rozellae | NA | LC | Decreasing | NA | NA | NA | NA | NA |
| lizards | Siderolamprus_scansorius | NA | EN | Decreasing | NA | NA | NA | NA | NA |
| lizards | Sigaloseps_balios | NA | VU | Stable | NA | NA | NA | NA | NA |
| lizards | Sigaloseps_conditus | NA | EN | Decreasing | NA | NA | NA | NA | NA |
| lizards | Sigaloseps_deplanchei | Sigaloseps_deplanchei | LC | Unknown | tropical | cathemeral | 1.738 | 4672.604 | 7.904 |
| lizards | Sigaloseps_pisinnus | NA | EN | Decreasing | NA | NA | NA | NA | NA |
| lizards | Sigaloseps_ruficauda | Sigaloseps_ruficauda | CR | Decreasing | tropical | cathemeral | 4.169 | 193.792 | 4.195 |
| lizards | Silvascincus_murrayi | Silvascincus_murrayi | LC | Stable | temperate | diurnal | 27.542 | 75202.705 | 8.399 |
| lizards | Silvascincus_tryoni | Silvascincus_tryoni | LC | Stable | temperate | diurnal | 24.547 | 629.052 | 5.417 |
| lizards | Simiscincus_aurantiacus | Simiscincus_aurantiacus | VU | Decreasing | tropical | nocturnal | 12.882 | 1293.422 | 10.303 |
| lizards | Sitana_devakai | NA | VU | Unknown | NA | NA | NA | NA | NA |
| lizards | Sitana_fusca | Sitana_fusca | CR | Unknown | temperate | diurnal | 3.388 | 19.982 | 13.985 |
| lizards | Sitana_laticeps | NA | LC | Unknown | NA | NA | NA | NA | NA |
| lizards | Sitana_marudhamneydhal | NA | EN | Decreasing | NA | NA | NA | NA | NA |
| lizards | Sitana_ponticeriana | Sitana_ponticeriana | LC | Stable | tropical | diurnal | 17.378 | 173089.057 | 21.602 |
| lizards | Sitana_schleichi | NA | EN | Unknown | NA | NA | NA | NA | NA |
| lizards | Sitana_sivalensis | Sitana_sivalensis | LC | Stable | temperate | diurnal | 2.692 | 14229.842 | 13.469 |
| lizards | Sitana_spinaecephalus | NA | LC | Stable | NA | NA | NA | NA | NA |
| lizards | Sitana_visiri | NA | VU | Unknown | NA | NA | NA | NA | NA |
| lizards | Smaug_barbertonensis | Smaug_barbertonensis | LC | Stable | temperate | diurnal | 95.499 | 10140.02 | 16.223 |
| lizards | Smaug_breyeri | Smaug_breyeri | LC | Stable | temperate | diurnal | 102.329 | 17344.437 | 6.65 |
| lizards | Smaug_depressus | Smaug_depressus | LC | Stable | tropical | diurnal | 53.703 | 7008.352 | 10.122 |
| lizards | Smaug_giganteus | Smaug_giganteus | VU | Decreasing | temperate | diurnal | 457.088 | 119790.504 | 12.566 |
| lizards | Smaug_mossambicus | Smaug_mossambicus | LC | Stable | tropical | diurnal | 61.66 | 21926.194 | 9.522 |
| lizards | Smaug_regius | Smaug_regius | LC | Stable | tropical | diurnal | 102.329 | 26058.507 | 11.846 |
| lizards | Smaug_vandami | Smaug_vandami | LC | Stable | temperate | diurnal | 102.329 | 43320.671 | 10.982 |
| lizards | Smaug_warreni | Smaug_warreni | LC | Stable | temperate | diurnal | 128.825 | 6984.526 | 12.259 |
| lizards | Sphaerodactylus_altavelensis | Sphaerodactylus_altavelensis | LC | Stable | tropical | diurnal | 0.617 | 30077.36 | 14.527 |
| lizards | Sphaerodactylus_argus | Sphaerodactylus_argus | LC | Stable | tropical | diurnal | 0.891 | 55662.42 | 17.009 |
| lizards | Sphaerodactylus_becki | NA | LC | Stable | NA | NA | NA | NA | NA |

| | | | | | | | | | |
|---|---|---|---|---|---|---|---|---|---|
| lizards | Sphaerodactylus_cinereus | Sphaerodactylus_cinereus | VU | Decreasing | tropical | nocturnal | 1.259 | 5532.655 | 16.346 |
| lizards | Sphaerodactylus_clenchi | Sphaerodactylus_clenchi | NT | Unknown | tropical | diurnal | 0.977 | 1846.563 | 12.823 |
| lizards | Sphaerodactylus_cochranae | NA | CR | Unknown | NA | NA | NA | NA | NA |
| lizards | Sphaerodactylus_corticola | Sphaerodactylus_corticola | LC | Unknown | temperate | nocturnal | 1.479 | 1120.039 | 5.832 |
| lizards | Sphaerodactylus_cryphius | NA | EN | Decreasing | NA | NA | NA | NA | NA |
| lizards | Sphaerodactylus_darlingtoni | Sphaerodactylus_darlingtoni | NT | Decreasing | tropical | diurnal | 0.617 | 24299.547 | 15.431 |
| lizards | Sphaerodactylus_difficilis | Sphaerodactylus_difficilis | LC | Unknown | tropical | diurnal | 0.977 | 49802.776 | 16.146 |
| lizards | Sphaerodactylus_elegans | Sphaerodactylus_elegans | LC | Stable | tropical | nocturnal | 1.549 | 127682.273 | 16.247 |
| lizards | Sphaerodactylus_elegantulus | NA | DD | Unknown | NA | NA | NA | NA | NA |
| lizards | Sphaerodactylus_fantasticus | Sphaerodactylus_fantasticus | LC | Stable | tropical | diurnal | 0.617 | 1509.483 | 18.644 |
| lizards | Sphaerodactylus_gaigeae | NA | LC | Stable | NA | NA | NA | NA | NA |
| lizards | Sphaerodactylus_glaucus | Sphaerodactylus_glaucus | LC | Stable | tropical | cathemeral | 0.741 | 322192.514 | 9.194 |
| lizards | Sphaerodactylus_graptolaemus | NA | LC | Stable | NA | NA | NA | NA | NA |
| lizards | Sphaerodactylus_guanajae | NA | not_available | not_available | NA | NA | NA | NA | NA |
| lizards | Sphaerodactylus_homolepis | Sphaerodactylus_homolepis | LC | Stable | tropical | diurnal | 0.891 | 15779.127 | 8.953 |
| lizards | Sphaerodactylus_inaguae | Sphaerodactylus_inaguae | EN | Unknown | tropical | nocturnal | 0.617 | 436.104 | 7.19 |
| lizards | Sphaerodactylus_kirbyi | Sphaerodactylus_kirbyi | VU | Unknown | tropical | diurnal | 0.617 | 78.835 | 20.066 |
| lizards | Sphaerodactylus_klauberi | Sphaerodactylus_klauberi | LC | Unknown | tropical | diurnal | 1.259 | 4438.209 | 24.998 |
| lizards | Sphaerodactylus_ladae | NA | EN | Unknown | NA | NA | NA | NA | NA |
| lizards | Sphaerodactylus_leonardovaldesi | NA | not_available | not_available | NA | NA | NA | NA | NA |
| lizards | Sphaerodactylus_leucaster | Sphaerodactylus_leucaster | LC | Unknown | tropical | nocturnal | 0.741 | 2105.33 | 15.546 |
| lizards | Sphaerodactylus_levinsi | Sphaerodactylus_levinsi | LC | Increasing | tropical | diurnal | 0.661 | 1.298 | 1.067 |
| lizards | Sphaerodactylus_lineolatus | NA | LC | Stable | NA | NA | NA | NA | NA |
| lizards | Sphaerodactylus_macrolepis | Sphaerodactylus_macrolepis | LC | Stable | tropical | diurnal | 0.525 | 9157.782 | 27.062 |
| lizards | Sphaerodactylus_millepunctatus | Sphaerodactylus_millepunctatus | LC | Stable | tropical | cathemeral | 1.096 | 251986.701 | 9.553 |
| lizards | Sphaerodactylus_molei | Sphaerodactylus_molei | LC | Unknown | tropical | diurnal | 0.55 | 501975.055 | 8.825 |
| lizards | Sphaerodactylus_monensis | NA | LC | Stable | NA | NA | NA | NA | NA |
| lizards | Sphaerodactylus_nicholsi | Sphaerodactylus_nicholsi | LC | Stable | tropical | cathemeral | 0.389 | 5917.329 | 25.619 |
| lizards | Sphaerodactylus_notatus | Sphaerodactylus_notatus | LC | Stable | tropical | diurnal | 0.977 | 127588.643 | 15.774 |
| lizards | Sphaerodactylus_oliveri | Sphaerodactylus_oliveri | EN | Unknown | tropical | diurnal | 0.977 | 2033.262 | 16.615 |
| lizards | Sphaerodactylus_pacificus | NA | LC | Stable | NA | NA | NA | NA | NA |
| lizards | Sphaerodactylus_parkeri | Sphaerodactylus_parkeri | EN | Decreasing | tropical | nocturnal | 1.072 | 664.815 | 22.627 |
| lizards | Sphaerodactylus_parthenopion | NA | EN | Unknown | NA | NA | NA | NA | NA |
| lizards | Sphaerodactylus_parvus | Sphaerodactylus_parvus | LC | Unknown | tropical | nocturnal | 0.347 | 195.943 | 28.819 |

| | | | | | | | | | |
|---|---|---|---|---|---|---|---|---|---|
| lizards | Sphaerodactylus_plummeri | NA | EN | Unknown | NA | NA | NA | NA | NA |
| lizards | Sphaerodactylus_richardsonii | Sphaerodactylus_richardsonii | EN | Decreasing | tropical | diurnal | 1.585 | 569.39 | 25.054 |
| lizards | Sphaerodactylus_roosevelti | Sphaerodactylus_roosevelti | NT | Stable | tropical | nocturnal | 1.479 | 168.561 | 26.028 |
| lizards | Sphaerodactylus_rosaurae | NA | LC | Stable | NA | NA | NA | NA | NA |
| lizards | Sphaerodactylus_sabanus | Sphaerodactylus_sabanus | LC | Stable | tropical | cathemeral | 0.676 | 310.325 | 23.955 |
| lizards | Sphaerodactylus_savagei | Sphaerodactylus_savagei | NT | Unknown | tropical | cathemeral | 0.891 | 2896.712 | 18.167 |
| lizards | Sphaerodactylus_scapularis | NA | EN | Unknown | NA | NA | NA | NA | NA |
| lizards | Sphaerodactylus_sommeri | NA | CR | Decreasing | NA | NA | NA | NA | NA |
| lizards | Sphaerodactylus_sputator | Sphaerodactylus_sputator | LC | Unknown | tropical | cathemeral | 1.585 | 324.721 | 26.189 |
| lizards | Sphaerodactylus_thompsoni | NA | EN | Unknown | NA | NA | NA | NA | NA |
| lizards | Sphaerodactylus_torrei | Sphaerodactylus_torrei | EN | Stable | tropical | diurnal | 1.479 | 790.997 | 21.629 |
| lizards | Sphaerodactylus_townsendi | Sphaerodactylus_townsendi | LC | Stable | tropical | diurnal | 0.676 | 1539.524 | 25.579 |
| lizards | Sphaerodactylus_vincenti | Sphaerodactylus_vincenti | LC | Unknown | tropical | cathemeral | 1.585 | 2876.612 | 20.516 |
| lizards | Sphaerodactylus_williamsi | NA | CR | Decreasing | NA | NA | NA | NA | NA |
| lizards | Sphenomorphus_acutus | NA | LC | Unknown | NA | NA | NA | NA | NA |
| lizards | Sphenomorphus_aignanus | Sphenomorphus_aignanus | LC | Unknown | tropical | diurnal | 10.471 | 29903.329 | 4.116 |
| lizards | Sphenomorphus_annamiticus | NA | not_available | not_available | NA | NA | NA | NA | NA |
| lizards | Sphenomorphus_apalpebratus | NA | NT | Unknown | NA | NA | NA | NA | NA |
| lizards | Sphenomorphus_bacboensis | NA | DD | Unknown | NA | NA | NA | NA | NA |
| lizards | Sphenomorphus_bignelli | Sphenomorphus_bignelli | LC | Stable | tropical | diurnal | 0.724 | 12970.579 | 4.824 |
| lizards | Sphenomorphus_brunneus | Sphenomorphus_brunneus | LC | Stable | tropical | nocturnal | 13.804 | 7957.212 | 3.05 |
| lizards | Sphenomorphus_buenloicus | Sphenomorphus_buenloicus | LC | Stable | tropical | diurnal | 3.467 | 14831.681 | 11.749 |
| lizards | Sphenomorphus_cameronicus | NA | EN | Unknown | NA | NA | NA | NA | NA |
| lizards | Sphenomorphus_concinnatus | Sphenomorphus_concinnatus | LC | Stable | tropical | diurnal | 6.761 | 34463.945 | 4.753 |
| lizards | Sphenomorphus_courcyanus | NA | LC | Stable | NA | NA | NA | NA | NA |
| lizards | Sphenomorphus_cranei | Sphenomorphus_cranei | LC | Stable | tropical | diurnal | 10 | 28487.911 | 4.696 |
| lizards | Sphenomorphus_cryptotis | NA | LC | Unknown | NA | NA | NA | NA | NA |
| lizards | Sphenomorphus_cyanolaemus | Sphenomorphus_cyanolaemus | LC | Unknown | tropical | diurnal | 4.169 | 232642.675 | 6.281 |
| lizards | Sphenomorphus_darlingtoni | Sphenomorphus_darlingtoni | LC | Unknown | tropical | nocturnal | 5.129 | 37540.507 | 6.244 |
| lizards | Sphenomorphus_dussumieri | Sphenomorphus_dussumieri | LC | Stable | tropical | diurnal | 5.129 | 121173.563 | 17.918 |
| lizards | Sphenomorphus_fasciatus | Sphenomorphus_fasciatus | LC | Unknown | tropical | diurnal | 39.811 | 118957.086 | 12.097 |
| lizards | Sphenomorphus_forbesi | Sphenomorphus_forbesi | LC | Unknown | tropical | nocturnal | 3.89 | 302.126 | 4.79 |
| lizards | Sphenomorphus_fragilis | Sphenomorphus_fragilis | LC | Stable | tropical | nocturnal | 2.951 | 5816.936 | 7.771 |
| lizards | Sphenomorphus_granulatus | Sphenomorphus_granulatus | LC | Unknown | tropical | nocturnal | 6.761 | 29844.638 | 4.493 |

| | | | | | | | | | |
|---|---|---|---|---|---|---|---|---|---|
| lizards | Sphenomorphus_haasi | NA | DD | Unknown | NA | NA | NA | NA | NA |
| lizards | Sphenomorphus_incognitus | Sphenomorphus_incognitus | LC | Stable | temperate | diurnal | 28.84 | 1013128.303 | 14.437 |
| lizards | Sphenomorphus_indicus | Sphenomorphus_indicus | LC | Stable | tropical | diurnal | 25.119 | 5514148.775 | 13.369 |
| lizards | Sphenomorphus_jobiensis | Sphenomorphus_jobiensis | LC | Stable | tropical | cathemeral | 34.674 | 322943.895 | 3.769 |
| lizards | Sphenomorphus_kinabaluensis | NA | LC | Unknown | NA | NA | NA | NA | NA |
| lizards | Sphenomorphus_lineopunctulatus | NA | LC | Stable | NA | NA | NA | NA | NA |
| lizards | Sphenomorphus_louisiadensis | Sphenomorphus_louisiadensis | LC | Unknown | tropical | diurnal | 2.138 | 294.493 | 3.17 |
| lizards | Sphenomorphus_maculatus | Sphenomorphus_maculatus | LC | Stable | tropical | diurnal | 6.761 | 2700117.516 | 11.487 |
| lizards | Sphenomorphus_maculicollus | NA | DD | Unknown | NA | NA | NA | NA | NA |
| lizards | Sphenomorphus_maindroni | Sphenomorphus_maindroni | LC | Unknown | tropical | nocturnal | 9.333 | 72003.768 | 5.647 |
| lizards | Sphenomorphus_melanopogon | Sphenomorphus_melanopogon | LC | Unknown | tropical | cathemeral | 15.488 | 32867.408 | 12.734 |
| lizards | Sphenomorphus_meyeri | NA | LC | Unknown | NA | NA | NA | NA | NA |
| lizards | Sphenomorphus_mimicus | NA | DD | Unknown | NA | NA | NA | NA | NA |
| lizards | Sphenomorphus_minutus | Sphenomorphus_minutus | LC | Unknown | tropical | nocturnal | 1.202 | 78131.173 | 5.564 |
| lizards | Sphenomorphus_muelleri | NA | LC | Unknown | NA | NA | NA | NA | NA |
| lizards | Sphenomorphus_multisquamatus | Sphenomorphus_multisquamatus | LC | Unknown | tropical | diurnal | 7.762 | 174704.782 | 5.443 |
| lizards | Sphenomorphus_murudensis | NA | DD | Unknown | NA | NA | NA | NA | NA |
| lizards | Sphenomorphus_necopinatus | NA | DD | Unknown | NA | NA | NA | NA | NA |
| lizards | Sphenomorphus_nigriventris | Sphenomorphus_nigriventris | LC | Unknown | tropical | nocturnal | 15.488 | 42917.395 | 4.69 |
| lizards | Sphenomorphus_nigrolineatus | Sphenomorphus_nigrolineatus | LC | Unknown | tropical | cathemeral | 8.511 | 13453.898 | 2.742 |
| lizards | Sphenomorphus_papuae | Sphenomorphus_papuae | LC | Unknown | tropical | nocturnal | 10.471 | 12742.54 | 3.238 |
| lizards | Sphenomorphus_praesignis | Sphenomorphus_praesignis | LC | Unknown | tropical | diurnal | 39.811 | 14193.995 | 9.114 |
| lizards | Sphenomorphus_pratti | Sphenomorphus_pratti | LC | Stable | tropical | nocturnal | 18.621 | 252218.772 | 3.755 |
| lizards | Sphenomorphus_preylangensis | NA | not_available | not_available | NA | NA | NA | NA | NA |
| lizards | Sphenomorphus_puncticentralis | NA | LC | Stable | NA | NA | NA | NA | NA |
| lizards | Sphenomorphus_sabanus | Sphenomorphus_sabanus | LC | Unknown | tropical | diurnal | 4.169 | 36610.253 | 6.277 |
| lizards | Sphenomorphus_sanctus | NA | LC | Stable | NA | NA | NA | NA | NA |
| lizards | Sphenomorphus_schlegeli | Sphenomorphus_schlegeli | LC | Unknown | tropical | diurnal | 0.603 | 333.653 | 11.302 |
| lizards | Sphenomorphus_schultzei | Sphenomorphus_schultzei | LC | Unknown | tropical | nocturnal | 1.862 | 102107.234 | 4.852 |
| lizards | Sphenomorphus_scotophilus | Sphenomorphus_scotophilus | LC | Stable | tropical | diurnal | 6.761 | 17431.781 | 16.027 |
| lizards | Sphenomorphus_scutatus | Sphenomorphus_scutatus | LC | Stable | tropical | diurnal | 1.514 | 415.884 | 11.199 |
| lizards | Sphenomorphus_shelfordi | NA | DD | Unknown | NA | NA | NA | NA | NA |
| lizards | Sphenomorphus_simus | Sphenomorphus_simus | LC | Stable | tropical | diurnal | 3.715 | 693689.116 | 3.138 |
| lizards | Sphenomorphus_solomonis | Sphenomorphus_solomonis | LC | Stable | tropical | cathemeral | 10 | 203096.628 | 5.104 |

| | | | | | | | | | |
|---|---|---|---|---|---|---|---|---|---|
| lizards | Sphenomorphus_stellatus | NA | LC | Unknown | NA | NA | NA | NA | NA |
| lizards | Sphenomorphus_sungaicolus | NA | LC | Unknown | NA | NA | NA | NA | NA |
| lizards | Sphenomorphus_taiwanensis | Sphenomorphus_taiwanensis | LC | Stable | temperate | diurnal | 6.457 | 8542.109 | 10.568 |
| lizards | Sphenomorphus_tanahtinggi | NA | DD | Unknown | NA | NA | NA | NA | NA |
| lizards | Sphenomorphus_tenuiculus | NA | DD | Unknown | NA | NA | NA | NA | NA |
| lizards | Sphenomorphus_tersus | NA | LC | Unknown | NA | NA | NA | NA | NA |
| lizards | Sphenomorphus_tetradactylus | NA | LC | Unknown | NA | NA | NA | NA | NA |
| lizards | Sphenomorphus_tridigitus | NA | NT | Unknown | NA | NA | NA | NA | NA |
| lizards | Sphenomorphus_tritaeniatus | NA | NT | Unknown | NA | NA | NA | NA | NA |
| lizards | Sphenomorphus_vanheurni | Sphenomorphus_vanheurni | LC | Unknown | tropical | diurnal | 5.129 | 19.982 | 11.649 |
| lizards | Sphenomorphus_variegatus | NA | LC | Decreasing | NA | NA | NA | NA | NA |
| lizards | Sphenomorphus_woodfordi | NA | LC | Stable | NA | NA | NA | NA | NA |
| lizards | Sphenomorphus_yersini | NA | not_available | not_available | NA | NA | NA | NA | NA |
| lizards | Spondylurus_anegadae | Spondylurus_anegadae | CR | Unknown | tropical | diurnal | 6.918 | 41.59 | 11.412 |
| lizards | Spondylurus_caicosae | Spondylurus_caicosae | NT | Decreasing | tropical | diurnal | 9.55 | 513.485 | 12.237 |
| lizards | Spondylurus_fulgidus | Spondylurus_fulgidus | EN | Decreasing | tropical | diurnal | 12.589 | 1215.823 | 28.855 |
| lizards | Spondylurus_macleani | NA | CR | Decreasing | NA | NA | NA | NA | NA |
| lizards | Spondylurus_magnacruzae | NA | CR | Decreasing | NA | NA | NA | NA | NA |
| lizards | Spondylurus_monae | Spondylurus_monae | CR | Unknown | tropical | diurnal | 13.804 | 58.342 | 6.697 |
| lizards | Spondylurus_monitae | NA | CR | Unknown | NA | NA | NA | NA | NA |
| lizards | Spondylurus_nitidus | Spondylurus_nitidus | EN | Decreasing | tropical | diurnal | 18.621 | 752.93 | 31.319 |
| lizards | Spondylurus_powelli | Spondylurus_powelli | EN | Decreasing | tropical | diurnal | 7.413 | 104.302 | 26.119 |
| lizards | Spondylurus_semitaeniatus | NA | CR | Decreasing | NA | NA | NA | NA | NA |
| lizards | Spondylurus_sloanii | Spondylurus_sloanii | CR | Decreasing | tropical | diurnal | 20.893 | 244.26 | 25.828 |
| lizards | Spondylurus_turksae | Spondylurus_turksae | CR | Decreasing | tropical | diurnal | 10.233 | 31.991 | 24.38 |
| lizards | Stenocercus_aculeatus | NA | LC | Unknown | NA | NA | NA | NA | NA |
| lizards | Stenocercus_albolineatus | NA | not_available | not_available | NA | NA | NA | NA | NA |
| lizards | Stenocercus_amydrorhytus | NA | not_available | not_available | NA | NA | NA | NA | NA |
| lizards | Stenocercus_angel | Stenocercus_angel | NT | Unknown | tropical | diurnal | 24.547 | 5639.936 | 16.064 |
| lizards | Stenocercus_arndti | NA | EN | Unknown | NA | NA | NA | NA | NA |
| lizards | Stenocercus_azureus | Stenocercus_azureus | LC | Unknown | temperate | diurnal | 21.38 | 219015.232 | 10.917 |
| lizards | Stenocercus_bolivarensis | NA | DD | Unknown | NA | NA | NA | NA | NA |
| lizards | Stenocercus_cadlei | Stenocercus_cadlei | LC | Stable | tropical | diurnal | 22.909 | 5181.797 | 19.656 |
| lizards | Stenocercus_caducus | Stenocercus_caducus | LC | Stable | tropical | diurnal | 29.512 | 1582548.446 | 6.644 |

| | | | | | | | | | |
|---|---|---|---|---|---|---|---|---|---|
| lizards | Stenocercus_chinchaoensis | Stenocercus_chinchaoensis | VU | Unknown | tropical | diurnal | 23.988 | 19.982 | 9.815 |
| lizards | Stenocercus_cupreus | Stenocercus_cupreus | LC | Unknown | tropical | diurnal | 18.197 | 3610.494 | 10.384 |
| lizards | Stenocercus_dumerilii | Stenocercus_dumerilii | VU | Unknown | tropical | diurnal | 48.978 | 268590.035 | 5.065 |
| lizards | Stenocercus_erythrogaster | NA | LC | Unknown | NA | NA | NA | NA | NA |
| lizards | Stenocercus_eunetopsis | Stenocercus_eunetopsis | LC | Unknown | tropical | diurnal | 21.38 | 9.991 | 11.069 |
| lizards | Stenocercus_guentheri | Stenocercus_guentheri | LC | Stable | tropical | diurnal | 32.359 | 24528.27 | 15.014 |
| lizards | Stenocercus_huancabambae | Stenocercus_huancabambae | LC | Unknown | tropical | diurnal | 34.674 | 2558.805 | 11.82 |
| lizards | Stenocercus_humeralis | Stenocercus_humeralis | LC | Unknown | tropical | diurnal | 48.978 | 452.299 | 13.962 |
| lizards | Stenocercus_ivitus | NA | DD | Unknown | NA | NA | NA | NA | NA |
| lizards | Stenocercus_johaberfellneri | NA | not_available | not_available | NA | NA | NA | NA | NA |
| lizards | Stenocercus_latebrosus | NA | LC | Unknown | NA | NA | NA | NA | NA |
| lizards | Stenocercus_limitaris | NA | VU | Unknown | NA | NA | NA | NA | NA |
| lizards | Stenocercus_marmoratus | NA | LC | Unknown | NA | NA | NA | NA | NA |
| lizards | Stenocercus_nigromaculatus | NA | DD | Unknown | NA | NA | NA | NA | NA |
| lizards | Stenocercus_nubicola | NA | DD | Unknown | NA | NA | NA | NA | NA |
| lizards | Stenocercus_ochoai | NA | LC | Unknown | NA | NA | NA | NA | NA |
| lizards | Stenocercus_omari | NA | not_available | not_available | NA | NA | NA | NA | NA |
| lizards | Stenocercus_orientalis | NA | LC | Stable | NA | NA | NA | NA | NA |
| lizards | Stenocercus_percultus | Stenocercus_percultus | LC | Stable | tropical | diurnal | 42.658 | 3864.383 | 7.442 |
| lizards | Stenocercus_prionotus | Stenocercus_prionotus | LC | Unknown | tropical | diurnal | 29.512 | 289499.7 | 3.317 |
| lizards | Stenocercus_puyango | Stenocercus_puyango | LC | Stable | tropical | diurnal | 52.481 | 17869.229 | 8.249 |
| lizards | Stenocercus_roseiventris | Stenocercus_roseiventris | LC | Stable | tropical | diurnal | 37.154 | 1215734.902 | 4.442 |
| lizards | Stenocercus_squarrosus | NA | LC | Unknown | NA | NA | NA | NA | NA |
| lizards | Stenocercus_stigmosus | NA | VU | Decreasing | NA | NA | NA | NA | NA |
| lizards | Stenocercus_varius | Stenocercus_varius | EN | Unknown | tropical | diurnal | 22.909 | 998.812 | 6.69 |
| lizards | Stenodactylus_affinis | NA | LC | Unknown | NA | NA | NA | NA | NA |
| lizards | Stenodactylus_doriae | Stenodactylus_doriae | LC | Stable | temperate | nocturnal | 11.482 | 2966808.13 | 4.846 |
| lizards | Stenodactylus_grandiceps | Stenodactylus_grandiceps | LC | Stable | temperate | nocturnal | 4.898 | 468036.752 | 7.103 |
| lizards | Stenodactylus_leptocosymbotes | NA | LC | Stable | NA | NA | NA | NA | NA |
| lizards | Stenodactylus_mauritanicus | NA | not_available | not_available | NA | NA | NA | NA | NA |
| lizards | Stenodactylus_petrii | Stenodactylus_petrii | LC | Stable | temperate | nocturnal | 7.413 | 7334237.655 | 2.899 |
| lizards | Stenodactylus_slevini | Stenodactylus_slevini | LC | Stable | temperate | nocturnal | 13.49 | 1192838.56 | 5.847 |
| lizards | Stenodactylus_stenurus | NA | not_available | not_available | NA | NA | NA | NA | NA |
| lizards | Stenodactylus_sthenodactylus | NA | LC | Stable | NA | NA | NA | NA | NA |

| | | | | | | | | | |
|---|---|---|---|---|---|---|---|---|---|
| lizards | Stenodactylus_yemenensis | NA | LC | Unknown | NA | NA | NA | NA | NA |
| lizards | Strobilurus_torquatus | Strobilurus_torquatus | LC | Unknown | tropical | diurnal | 42.658 | 261835.172 | 9.926 |
| lizards | Strophurus_assimilis | Strophurus_assimilis | LC | Stable | temperate | nocturnal | 9.55 | 320714.056 | 4.292 |
| lizards | Strophurus_ciliaris | Strophurus_ciliaris | LC | Stable | tropical | nocturnal | 14.791 | 3961905.441 | 2.411 |
| lizards | Strophurus_congoo | NA | DD | Unknown | NA | NA | NA | NA | NA |
| lizards | Strophurus_elderi | Strophurus_elderi | LC | Stable | temperate | nocturnal | 2.63 | 2487114.839 | 2.374 |
| lizards | Strophurus_horneri | NA | VU | Unknown | NA | NA | NA | NA | NA |
| lizards | Strophurus_intermedius | Strophurus_intermedius | LC | Stable | temperate | nocturnal | 10.233 | 2115405.265 | 5.231 |
| lizards | Strophurus_jeanae | Strophurus_jeanae | LC | Stable | tropical | nocturnal | 2.455 | 1368479.807 | 2.247 |
| lizards | Strophurus_krisalys | Strophurus_krisalys | LC | Stable | tropical | nocturnal | 8.913 | 615003.324 | 4.842 |
| lizards | Strophurus_mcmillani | Strophurus_mcmillani | LC | Unknown | tropical | nocturnal | 2.951 | 59894.774 | 1.614 |
| lizards | Strophurus_michaelseni | Strophurus_michaelseni | LC | Unknown | temperate | nocturnal | 5.754 | 36254.144 | 8.771 |
| lizards | Strophurus_rankini | Strophurus_rankini | LC | Stable | tropical | nocturnal | 5.012 | 14827.05 | 5.904 |
| lizards | Strophurus_robinsoni | NA | LC | Unknown | NA | NA | NA | NA | NA |
| lizards | Strophurus_spinigerus | Strophurus_spinigerus | LC | Stable | temperate | nocturnal | 10 | 322149.952 | 9.018 |
| lizards | Strophurus_strophurus | Strophurus_strophurus | LC | Stable | temperate | nocturnal | 8.913 | 919280.074 | 3.451 |
| lizards | Strophurus_taeniatus | Strophurus_taeniatus | LC | Unknown | tropical | nocturnal | 2.455 | 229984.795 | 3.01 |
| lizards | Strophurus_taenicauda | Strophurus_taenicauda | LC | Decreasing | temperate | nocturnal | 7.943 | 183820.189 | 6.365 |
| lizards | Strophurus_trux | NA | not_available | not_available | NA | NA | NA | NA | NA |
| lizards | Strophurus_wellingtonae | NA | LC | Stable | NA | NA | NA | NA | NA |
| lizards | Strophurus_williamsi | Strophurus_williamsi | LC | Stable | temperate | nocturnal | 6.918 | 1142513.221 | 6.329 |
| lizards | Strophurus_wilsoni | Strophurus_wilsoni | LC | Unknown | temperate | nocturnal | 4.266 | 31963.976 | 3.961 |
| lizards | Subdoluseps_bowringii | NA | LC | Stable | NA | NA | NA | NA | NA |
| lizards | Subdoluseps_herberti | NA | LC | Unknown | NA | NA | NA | NA | NA |
| lizards | Takydromus_albomaculosus | NA | DD | Unknown | NA | NA | NA | NA | NA |
| lizards | Takydromus_amurensis | Takydromus_amurensis | LC | Stable | temperate | diurnal | 11.749 | 1082455.111 | 13.052 |
| lizards | Takydromus_dorsalis | Takydromus_dorsalis | EN | Decreasing | temperate | diurnal | 7.943 | 528.954 | 13.498 |
| lizards | Takydromus_formosanus | Takydromus_formosanus | LC | Stable | temperate | diurnal | 6.166 | 17564.48 | 25.145 |
| lizards | Takydromus_hani | NA | LC | Unknown | NA | NA | NA | NA | NA |
| lizards | Takydromus_haughtonianus | NA | DD | Unknown | NA | NA | NA | NA | NA |
| lizards | Takydromus_hsuehshanensis | Takydromus_hsuehshanensis | LC | Stable | temperate | diurnal | 8.71 | 3517.443 | 10.418 |
| lizards | Takydromus_intermedius | NA | LC | Stable | NA | NA | NA | NA | NA |
| lizards | Takydromus_khasiensis | Takydromus_khasiensis | LC | Unknown | temperate | diurnal | 9.772 | 189038.792 | 14.073 |
| lizards | Takydromus_kuehnei | Takydromus_kuehnei | LC | Stable | temperate | diurnal | 6.607 | 924777.089 | 15.567 |

| | | | | | | | | | |
|---|---|---|---|---|---|---|---|---|---|
| lizards | Takydromus_luyeanus | Takydromus_luyeanus | LC | Stable | tropical | diurnal | 3.715 | 4584.889 | 16.476 |
| lizards | Takydromus_madaensis | NA | DD | Unknown | NA | NA | NA | NA | NA |
| lizards | Takydromus_sauteri | Takydromus_sauteri | NT | Stable | tropical | diurnal | 10.471 | 6749.742 | 23.149 |
| lizards | Takydromus_septentrionalis | Takydromus_septentrionalis | LC | Stable | temperate | diurnal | 11.749 | 2465519.623 | 16.847 |
| lizards | Takydromus_sexlineatus | Takydromus_sexlineatus | LC | Stable | tropical | diurnal | 7.943 | 4405213.071 | 11.746 |
| lizards | Takydromus_sikkimensis | NA | EN | Unknown | NA | NA | NA | NA | NA |
| lizards | Takydromus_smaragdinus | Takydromus_smaragdinus | NT | Decreasing | temperate | diurnal | 6.457 | 2664.378 | 23.5 |
| lizards | Takydromus_stejnegeri | Takydromus_stejnegeri | LC | Stable | temperate | diurnal | 5.623 | 19041.827 | 28.745 |
| lizards | Takydromus_sylvaticus | Takydromus_sylvaticus | LC | Unknown | temperate | diurnal | 4.571 | 11695.308 | 12.104 |
| lizards | Takydromus_tachydromoides | Takydromus_tachydromoides | LC | Stable | temperate | diurnal | 7.943 | 368570.007 | 19.826 |
| lizards | Takydromus_toyamai | Takydromus_toyamai | EN | Decreasing | temperate | diurnal | 3.89 | 193.471 | 28.919 |
| lizards | Takydromus_viridipunctatus | Takydromus_viridipunctatus | LC | Stable | temperate | diurnal | 4.898 | 5773.19 | 26.79 |
| lizards | Takydromus_wolteri | Takydromus_wolteri | LC | Stable | temperate | diurnal | 6.761 | 1893523.426 | 14.371 |
| lizards | Tarentola_angustimentalis | Tarentola_angustimentalis | LC | Stable | temperate | nocturnal | 13.183 | 2521.198 | 17.132 |
| lizards | Tarentola_annularis | Tarentola_annularis | LC | Stable | tropical | nocturnal | 69.183 | 5606088.155 | 5.895 |
| lizards | Tarentola_boehmei | Tarentola_boehmei | LC | Stable | temperate | nocturnal | 14.125 | 74734.545 | 6.669 |
| lizards | Tarentola_boettgeri | Tarentola_boettgeri | LC | Stable | temperate | nocturnal | 10.715 | 1755.316 | 21.157 |
| lizards | Tarentola_chazaliae | Tarentola_chazaliae | VU | Decreasing | temperate | nocturnal | 10.471 | 31281.167 | 6.211 |
| lizards | Tarentola_crombiei | Tarentola_crombiei | LC | Stable | tropical | nocturnal | 5.129 | 212.841 | 16.57 |
| lizards | Tarentola_delalandii | Tarentola_delalandii | LC | Stable | temperate | cathemeral | 15.488 | 2744.124 | 19.154 |
| lizards | Tarentola_deserti | Tarentola_deserti | LC | Stable | temperate | nocturnal | 28.184 | 353807.283 | 5.22 |
| lizards | Tarentola_ephippiata | Tarentola_ephippiata | LC | Stable | tropical | nocturnal | 54.954 | 7434256.943 | 7.632 |
| lizards | Tarentola_fascicularis | NA | not_available | not_available | NA | NA | NA | NA | NA |
| lizards | Tarentola_gigas | Tarentola_gigas | EN | Stable | tropical | nocturnal | 93.325 | 10.719 | 0.103 |
| lizards | Tarentola_gomerensis | Tarentola_gomerensis | LC | Stable | temperate | nocturnal | 11.482 | 365.532 | 15.045 |
| lizards | Tarentola_mauritanica | Tarentola_mauritanica | LC | Stable | temperate | cathemeral | 16.596 | 1852529.318 | 15.621 |
| lizards | Tarentola_mindiae | Tarentola_mindiae | LC | Stable | temperate | nocturnal | 13.804 | 72616.341 | 2.343 |
| lizards | Tarentola_neglecta | Tarentola_neglecta | LC | Stable | temperate | nocturnal | 10.233 | 1321275.592 | 2.702 |
| lizards | Tarentola_parvicarinata | NA | LC | Stable | NA | NA | NA | NA | NA |
| lizards | Tarentola_pastoria | NA | LC | Unknown | NA | NA | NA | NA | NA |
| lizards | Tarentola_substituta | NA | LC | Unknown | NA | NA | NA | NA | NA |
| lizards | Techmarscincus_jigurru | Techmarscincus_jigurru | VU | Unknown | tropical | diurnal | 10.471 | 30.297 | 0.515 |
| lizards | Teira_dugesii | Teira_dugesii | LC | Stable | temperate | diurnal | 12.303 | 815.078 | 20.491 |
| lizards | Teius_oculatus | Teius_oculatus | LC | Unknown | temperate | diurnal | 52.481 | 1753401.041 | 11.194 |

| | | | | | | | | | |
|---|---|---|---|---|---|---|---|---|---|
| lizards | Teius_suquiensis | Teius_suquiensis | LC | Stable | temperate | diurnal | 58.884 | 74034.561 | 10.908 |
| lizards | Teius_teyou | Teius_teyou | LC | Unknown | temperate | diurnal | 104.713 | 1622772.271 | 6.556 |
| lizards | Tenuidactylus_bogdanovi | Tenuidactylus_bogdanovi | LC | Unknown | temperate | nocturnal | 7.762 | 40289.055 | 13.074 |
| lizards | Tenuidactylus_caspius | Tenuidactylus_caspius | LC | Stable | temperate | cathemeral | 10.233 | 2125031.923 | 8.648 |
| lizards | Tenuidactylus_dadunensis | NA | DD | Unknown | NA | NA | NA | NA | NA |
| lizards | Tenuidactylus_elongatus | Tenuidactylus_elongatus | LC | Unknown | temperate | nocturnal | 3.981 | 823802.865 | 2.897 |
| lizards | Tenuidactylus_fedtschenkoi | Tenuidactylus_fedtschenkoi | LC | Stable | temperate | cathemeral | 9.333 | 292971.551 | 11.248 |
| lizards | Tenuidactylus_longipes | Tenuidactylus_longipes | LC | Stable | temperate | cathemeral | 6.761 | 343865.492 | 7.596 |
| lizards | Tenuidactylus_turcmenicus | Tenuidactylus_turcmenicus | LC | Stable | temperate | nocturnal | 10.233 | 127397.069 | 12.281 |
| lizards | Tenuidactylus_voraginosus | NA | LC | Unknown | NA | NA | NA | NA | NA |
| lizards | Teratoscincus_bedriagai | Teratoscincus_bedriagai | LC | Stable | temperate | nocturnal | 9.55 | 309286.918 | 4.433 |
| lizards | Teratoscincus_keyserlingii | NA | LC | Stable | NA | NA | NA | NA | NA |
| lizards | Teratoscincus_microlepis | Teratoscincus_microlepis | LC | Stable | temperate | nocturnal | 18.197 | 365555.138 | 4.689 |
| lizards | Teratoscincus_przewalskii | Teratoscincus_przewalskii | LC | Stable | temperate | nocturnal | 21.878 | 1506810.984 | 2.625 |
| lizards | Teratoscincus_roborowskii | Teratoscincus_roborowskii | LC | Unknown | temperate | nocturnal | 20.893 | 33584.644 | 1.502 |
| lizards | Teratoscincus_scincus | Teratoscincus_scincus | LC | Stable | temperate | nocturnal | 41.687 | 1871823.423 | 7.476 |
| lizards | Teratoscincus_sistanense | NA | not_available | not_available | NA | NA | NA | NA | NA |
| lizards | Tetradactylus_africanus | Tetradactylus_africanus | LC | Stable | temperate | diurnal | 3.89 | 62541.746 | 14.815 |
| lizards | Tetradactylus_breyeri | Tetradactylus_breyeri | NT | Decreasing | temperate | diurnal | 2.399 | 18726.459 | 11.231 |
| lizards | Tetradactylus_ellenbergeri | Tetradactylus_ellenbergeri | LC | Unknown | tropical | diurnal | 2.399 | 470440.276 | 6.344 |
| lizards | Tetradactylus_seps | Tetradactylus_seps | LC | Stable | temperate | diurnal | 7.413 | 78876.683 | 10.689 |
| lizards | Tetradactylus_tetradactylus | Tetradactylus_tetradactylus | LC | Stable | temperate | diurnal | 2.512 | 84034.524 | 9.722 |
| lizards | Tetradactylus_udzungwensis | Tetradactylus_udzungwensis | EN | Unknown | tropical | diurnal | 1.995 | 19.982 | 10.395 |
| lizards | Thecadactylus_oskrobapreinorum | NA | DD | Unknown | NA | NA | NA | NA | NA |
| lizards | Thecadactylus_rapicauda | Thecadactylus_rapicauda | LC | Stable | tropical | nocturnal | 50.119 | 7382390.201 | 3.693 |
| lizards | Thecadactylus_solimoensis | Thecadactylus_solimoensis | LC | Stable | tropical | nocturnal | 50.119 | 2751783.832 | 1.818 |
| lizards | Tiliqua_adelaidensis | Tiliqua_adelaidensis | EN | Decreasing | temperate | diurnal | 26.915 | 11850.053 | 15.689 |
| lizards | Tiliqua_gigas | Tiliqua_gigas | LC | Unknown | tropical | diurnal | 1148.154 | 441063.24 | 3.789 |
| lizards | Tiliqua_multifasciata | Tiliqua_multifasciata | LC | Stable | tropical | cathemeral | 741.31 | 3512589.307 | 2.186 |
| lizards | Tiliqua_nigrolutea | Tiliqua_nigrolutea | LC | Stable | temperate | diurnal | 1445.44 | 362450.69 | 9.552 |
| lizards | Tiliqua_occipitalis | Tiliqua_occipitalis | LC | Stable | temperate | diurnal | 912.011 | 2510711.016 | 4.482 |
| lizards | Tiliqua_rugosa | Tiliqua_rugosa | LC | Stable | temperate | diurnal | 1230.269 | 3235976.761 | 6.477 |
| lizards | Tiliqua_scincoides | Tiliqua_scincoides | LC | Stable | temperate | cathemeral | 1621.81 | 3476688.889 | 5.774 |
| lizards | Timon_kurdistanicus | NA | not_available | not_available | NA | NA | NA | NA | NA |

| | | | | | | | | | |
|---|---|---|---|---|---|---|---|---|---|
| lizards | Timon_lepidus | Timon_lepidus | NT | Decreasing | temperate | diurnal | 380.189 | 715237.909 | 17.915 |
| lizards | Timon_nevadensis | NA | not_available | not_available | NA | NA | NA | NA | NA |
| lizards | Timon_pater | Timon_pater | LC | Decreasing | temperate | diurnal | 109.648 | 235602.927 | 17.633 |
| lizards | Timon_princeps | Timon_princeps | LC | Unknown | temperate | diurnal | 72.444 | 64199.218 | 14.26 |
| lizards | Timon_tangitanus | Timon_tangitanus | LC | Decreasing | temperate | diurnal | 57.544 | 341580.134 | 16.373 |
| lizards | Toenayar_novemcarinata | NA | LC | Stable | NA | NA | NA | NA | NA |
| lizards | Toropuku_stephensi | Toropuku_stephensi | EN | Unknown | temperate | nocturnal | 12.303 | 294.544 | 8.803 |
| lizards | Tracheloptychus_madagascariensis | Tracheloptychus_madagascariensis | LC | Decreasing | tropical | diurnal | 17.783 | 97239.921 | 10.11 |
| lizards | Tracheloptychus_petersi | NA | VU | Decreasing | NA | NA | NA | NA | NA |
| lizards | Trachydactylus_hajarensis | NA | LC | Stable | NA | NA | NA | NA | NA |
| lizards | Trachydactylus_spatalurus | Bunopus_spatalurus | LC | Stable | tropical | nocturnal | 7.079 | 446738.217 | 6.047 |
| lizards | Trachylepis_acutilabris | Trachylepis_acutilabris | LC | Stable | tropical | diurnal | 4.169 | 370708.954 | 6.105 |
| lizards | Trachylepis_adamastor | NA | VU | Stable | NA | NA | NA | NA | NA |
| lizards | Trachylepis_affinis | Trachylepis_affinis | LC | Stable | tropical | diurnal | 11.749 | 5568536.906 | 9.073 |
| lizards | Trachylepis_albilabris | Trachylepis_albilabris | LC | Stable | tropical | diurnal | 8.511 | 484550.495 | 6.682 |
| lizards | Trachylepis_albotaeniata | NA | LC | Unknown | NA | NA | NA | NA | NA |
| lizards | Trachylepis_atlantica | NA | LC | Stable | NA | NA | NA | NA | NA |
| lizards | Trachylepis_aureogularis | NA | LC | Unknown | NA | NA | NA | NA | NA |
| lizards | Trachylepis_aureopunctata | NA | LC | Decreasing | NA | NA | NA | NA | NA |
| lizards | Trachylepis_bayonii | Trachylepis_bayonii | LC | Stable | tropical | diurnal | 11.22 | 3822391.938 | 6.948 |
| lizards | Trachylepis_bensonii | NA | LC | Unknown | NA | NA | NA | NA | NA |
| lizards | Trachylepis_binotata | NA | LC | Stable | NA | NA | NA | NA | NA |
| lizards | Trachylepis_bocagii | Trachylepis_bocagii | LC | Stable | tropical | diurnal | 7.762 | 237104.925 | 7.74 |
| lizards | Trachylepis_boehmei | NA | not_available | not_available | NA | NA | NA | NA | NA |
| lizards | Trachylepis_boettgeri | Trachylepis_boettgeri | LC | Decreasing | tropical | diurnal | 3.09 | 43162.172 | 11.79 |
| lizards | Trachylepis_boulengeri | Trachylepis_boulengeri | LC | Unknown | tropical | diurnal | 30.903 | 766840.779 | 9.501 |
| lizards | Trachylepis_brauni | Trachylepis_brauni | NT | Unknown | tropical | diurnal | 11.22 | 3578.389 | 12.684 |
| lizards | Trachylepis_brevicollis | Trachylepis_brevicollis | LC | Stable | tropical | cathemeral | 144.544 | 3076857.861 | 7.734 |
| lizards | Trachylepis_buettneri | Trachylepis_buettneri | LC | Unknown | tropical | diurnal | 15.488 | 780397.386 | 8.026 |
| lizards | Trachylepis_capensis | Trachylepis_capensis | LC | Stable | temperate | diurnal | 56.234 | 1821762.395 | 8.356 |
| lizards | Trachylepis_chimbana | NA | LC | Stable | NA | NA | NA | NA | NA |
| lizards | Trachylepis_comorensis | Trachylepis_comorensis | LC | Stable | tropical | diurnal | 30.903 | 1964.522 | 17.312 |
| lizards | Trachylepis_damarana | NA | LC | Stable | NA | NA | NA | NA | NA |
| lizards | Trachylepis_depressa | Trachylepis_depressa | LC | Stable | temperate | diurnal | 13.804 | 170914.577 | 8.709 |

| | | | | | | | | | |
|---|---|---|---|---|---|---|---|---|---|
| lizards | Trachylepis_dichroma | Trachylepis_dichroma | LC | Unknown | tropical | diurnal | 158.489 | 105146.459 | 12.805 |
| lizards | Trachylepis_dumasi | NA | VU | Decreasing | NA | NA | NA | NA | NA |
| lizards | Trachylepis_elegans | NA | LC | Stable | NA | NA | NA | NA | NA |
| lizards | Trachylepis_gonwouoi | NA | LC | Stable | NA | NA | NA | NA | NA |
| lizards | Trachylepis_gravenhorstii | Trachylepis_gravenhorstii | LC | Stable | tropical | diurnal | 17.783 | 583710.133 | 8.736 |
| lizards | Trachylepis_hoeschi | NA | LC | Stable | NA | NA | NA | NA | NA |
| lizards | Trachylepis_homalocephala | Trachylepis_homalocephala | LC | Stable | temperate | diurnal | 19.953 | 246655.212 | 11.117 |
| lizards | Trachylepis_infralineata | NA | not_available | not_available | NA | NA | NA | NA | NA |
| lizards | Trachylepis_irregularis | Trachylepis_irregularis | NT | Stable | tropical | diurnal | 8.511 | 59335.264 | 13.592 |
| lizards | Trachylepis_keroanensis | NA | DD | Unknown | NA | NA | NA | NA | NA |
| lizards | Trachylepis_laevigata | NA | LC | Stable | NA | NA | NA | NA | NA |
| lizards | Trachylepis_lavarambo | Trachylepis_lavarambo | VU | Unknown | tropical | diurnal | 4.365 | 291.322 | 19.202 |
| lizards | Trachylepis_loluiensis | NA | VU | Unknown | NA | NA | NA | NA | NA |
| lizards | Trachylepis_maculilabris | Trachylepis_maculilabris | LC | Stable | tropical | diurnal | 19.953 | 6788234.543 | 8.412 |
| lizards | Trachylepis_madagascariensis | Trachylepis_madagascariensis | LC | Unknown | tropical | diurnal | 9.55 | 32941.709 | 11.655 |
| lizards | Trachylepis_margaritifera | Trachylepis_margaritifera | LC | Unknown | tropical | diurnal | 38.905 | 1806100.15 | 10.671 |
| lizards | Trachylepis_megalura | Trachylepis_megalura | LC | Unknown | tropical | diurnal | 12.882 | 1992826.734 | 11.027 |
| lizards | Trachylepis_mlanjensis | NA | LC | Stable | NA | NA | NA | NA | NA |
| lizards | Trachylepis_nganghae | Trachylepis_nganghae | CR | Decreasing | tropical | diurnal | 4.169 | 19.982 | 5.902 |
| lizards | Trachylepis_occidentalis | Trachylepis_occidentalis | LC | Stable | tropical | diurnal | 33.884 | 939225.654 | 5.294 |
| lizards | Trachylepis_ozorii | NA | LC | Stable | NA | NA | NA | NA | NA |
| lizards | Trachylepis_paucisquamis | NA | LC | Unknown | NA | NA | NA | NA | NA |
| lizards | Trachylepis_pendeana | NA | DD | Unknown | NA | NA | NA | NA | NA |
| lizards | Trachylepis_perrotetii | Trachylepis_perrotetii | LC | Stable | tropical | diurnal | 144.544 | 520829.887 | 10.554 |
| lizards | Trachylepis_planifrons | Trachylepis_planifrons | LC | Unknown | tropical | diurnal | 34.674 | 1456491.393 | 8.607 |
| lizards | Trachylepis_polytropis | Trachylepis_polytropis | LC | Stable | tropical | diurnal | 33.113 | 1592079.714 | 8.205 |
| lizards | Trachylepis_pulcherrima | Trachylepis_pulcherrima | LC | Stable | tropical | diurnal | 39.811 | 18494.22 | 6.566 |
| lizards | Trachylepis_punctatissima | Trachylepis_punctatissima | LC | Stable | temperate | diurnal | 26.915 | 1021857.202 | 9.738 |
| lizards | Trachylepis_punctulata | NA | LC | Stable | NA | NA | NA | NA | NA |
| lizards | Trachylepis_quinquetaeniata | NA | LC | Stable | NA | NA | NA | NA | NA |
| lizards | Trachylepis_raymondlaurenti | NA | LC | Unknown | NA | NA | NA | NA | NA |
| lizards | Trachylepis_rodenburgi | NA | LC | Stable | NA | NA | NA | NA | NA |
| lizards | Trachylepis_sechellensis | NA | LC | Stable | NA | NA | NA | NA | NA |
| lizards | Trachylepis_socotrana | NA | LC | Unknown | NA | NA | NA | NA | NA |

| | | | | | | | | | |
|---|---|---|---|---|---|---|---|---|---|
| lizards | Trachylepis_sparsa | Trachylepis_sparsa | LC | Stable | temperate | diurnal | 27.542 | 180145.355 | 4.958 |
| lizards | Trachylepis_spilogaster | Trachylepis_spilogaster | LC | Stable | tropical | diurnal | 16.982 | 892599.191 | 5.284 |
| lizards | Trachylepis_striata | Trachylepis_striata | LC | Unknown | tropical | diurnal | 31.623 | 5399481.889 | 9.767 |
| lizards | Trachylepis_sulcata | Trachylepis_sulcata | LC | Stable | tropical | diurnal | 29.512 | 1242359.535 | 6.13 |
| lizards | Trachylepis_tavaratra | Trachylepis_tavaratra | VU | Unknown | tropical | diurnal | 4.571 | 10529.287 | 9.632 |
| lizards | Trachylepis_tessellata | Trachylepis_tessellata | LC | Unknown | tropical | diurnal | 21.38 | 356472.452 | 8.259 |
| lizards | Trachylepis_thomensis | NA | LC | Stable | NA | NA | NA | NA | NA |
| lizards | Trachylepis_varia | Trachylepis_varia | LC | Unknown | tropical | diurnal | 7.079 | 7946224.685 | 8.85 |
| lizards | Trachylepis_variegata | Trachylepis_variegata | LC | Stable | temperate | diurnal | 3.467 | 906913.776 | 5.671 |
| lizards | Trachylepis_vato | Trachylepis_vato | LC | Stable | tropical | diurnal | 3.09 | 93023.691 | 9.876 |
| lizards | Trachylepis_vezo | NA | DD | Unknown | NA | NA | NA | NA | NA |
| lizards | Trachylepis_wahlbergii | NA | DD | Unknown | NA | NA | NA | NA | NA |
| lizards | Trachylepis_wingati | NA | LC | Unknown | NA | NA | NA | NA | NA |
| lizards | Trachylepis_wrightii | NA | VU | Stable | NA | NA | NA | NA | NA |
| lizards | Trapelus_agilis | Trapelus_agilis | LC | Stable | temperate | diurnal | 52.481 | 3146170.542 | 9.179 |
| lizards | Trapelus_agnetae | Trapelus_agnetae | LC | Stable | temperate | diurnal | 26.915 | 1353248.279 | 5.45 |
| lizards | Trapelus_boehmei | NA | LC | Stable | NA | NA | NA | NA | NA |
| lizards | Trapelus_flavimaculatus | Trapelus_flavimaculatus | LC | Stable | tropical | diurnal | 75.858 | 2148031.312 | 4.347 |
| lizards | Trapelus_megalonyx | Trapelus_megalonyx | LC | Stable | temperate | diurnal | 13.804 | 808903.298 | 7.875 |
| lizards | Trapelus_mutabilis | Trapelus_mutabilis | LC | Stable | temperate | diurnal | 32.359 | 4067499.626 | 2.206 |
| lizards | Trapelus_rubrigularis | NA | DD | Unknown | NA | NA | NA | NA | NA |
| lizards | Trapelus_ruderatus | Trapelus_ruderatus | LC | Stable | temperate | diurnal | 85.114 | 2535454.795 | 10.327 |
| lizards | Trapelus_sanguinolentus | Trapelus_sanguinolentus | LC | Stable | temperate | diurnal | 239.883 | 2563982.646 | 7.527 |
| lizards | Trapelus_savignii | Trapelus_savignii | VU | Decreasing | temperate | diurnal | 63.096 | 35892.019 | 10.186 |
| lizards | Trapelus_schmitzi | NA | DD | Unknown | NA | NA | NA | NA | NA |
| lizards | Trapelus_tournevillei | Trapelus_tournevillei | LC | Stable | temperate | diurnal | 33.113 | 487687.696 | 2.314 |
| lizards | Tretioscincus_agilis | NA | LC | Stable | NA | NA | NA | NA | NA |
| lizards | Tretioscincus_bifasciatus | NA | LC | Stable | NA | NA | NA | NA | NA |
| lizards | Tretioscincus_oriximinensis | NA | LC | Unknown | NA | NA | NA | NA | NA |
| lizards | Tribolonotus_blanchardi | Tribolonotus_blanchardi | LC | Decreasing | tropical | diurnal | 1.122 | 8572.391 | 6.34 |
| lizards | Tribolonotus_brongersmai | NA | DD | Unknown | NA | NA | NA | NA | NA |
| lizards | Tribolonotus_gracilis | Tribolonotus_gracilis | LC | Unknown | tropical | diurnal | 23.442 | 117110.41 | 3.898 |
| lizards | Tribolonotus_novaeguineae | Tribolonotus_novaeguineae | LC | Unknown | tropical | diurnal | 23.442 | 137845.662 | 2.951 |
| lizards | Tribolonotus_ponceleti | NA | DD | Unknown | NA | NA | NA | NA | NA |

| | | | | | | | | | |
|---|---|---|---|---|---|---|---|---|---|
| lizards | Tribolonotus_pseudoponceleti | Tribolonotus_pseudoponceleti | LC | Stable | tropical | diurnal | 8.913 | 12171.079 | 5.477 |
| lizards | Tribolonotus_schmidti | Tribolonotus_schmidti | LC | Stable | tropical | diurnal | 1.202 | 5461.348 | 5.671 |
| lizards | Trigonodactylus_arabicus | Stenodactylus_arabicus | LC | Stable | tropical | nocturnal | 1.738 | 839451.322 | 4.529 |
| lizards | Trigonodactylus_pulcher | NA | LC | not_available | NA | NA | NA | NA | NA |
| lizards | Trigonodactylus_sharqiyahensis | NA | LC | Stable | NA | NA | NA | NA | NA |
| lizards | Trioceros_affinis | Trioceros_affinis | LC | Stable | tropical | diurnal | 13.183 | 343971.01 | 12.725 |
| lizards | Trioceros_balebicornutus | Trioceros_balebicornutus | NT | Decreasing | tropical | diurnal | 10.965 | 1291.195 | 16.035 |
| lizards | Trioceros_bitaeniatus | Trioceros_bitaeniatus | LC | Stable | tropical | diurnal | 18.621 | 1350654.572 | 10.736 |
| lizards | Trioceros_conirostratus | Trioceros_conirostratus | LC | Unknown | tropical | diurnal | 14.125 | 21851.053 | 8.531 |
| lizards | Trioceros_cristatus | Trioceros_cristatus | LC | Unknown | tropical | diurnal | 81.283 | 1322406.386 | 7.62 |
| lizards | Trioceros_ellioti | Trioceros_ellioti | LC | Unknown | tropical | diurnal | 21.38 | 1007953.107 | 9.952 |
| lizards | Trioceros_hanangensis | Trioceros_hanangensis | NT | Unknown | tropical | diurnal | 10.965 | 98.005 | 15.911 |
| lizards | Trioceros_harennae | Trioceros_harennae | LC | Unknown | tropical | diurnal | 10.471 | 5204.66 | 13.689 |
| lizards | Trioceros_hoehnelii | Trioceros_hoehnelii | LC | Stable | tropical | diurnal | 29.512 | 56737.316 | 16.245 |
| lizards | Trioceros_ituriensis | Trioceros_ituriensis | LC | Unknown | tropical | diurnal | 46.774 | 459565.932 | 7.192 |
| lizards | Trioceros_jacksonii | Trioceros_jacksonii | LC | Stable | tropical | diurnal | 91.201 | 31172.363 | 15.325 |
| lizards | Trioceros_kinangopensis | NA | NT | Unknown | NA | NA | NA | NA | NA |
| lizards | Trioceros_laterispinis | Trioceros_laterispinis | EN | Decreasing | tropical | diurnal | 12.303 | 11538.536 | 15.73 |
| lizards | Trioceros_marsabitensis | NA | NT | Stable | NA | NA | NA | NA | NA |
| lizards | Trioceros_melleri | Trioceros_melleri | LC | Unknown | tropical | diurnal | 389.045 | 379655.572 | 10.923 |
| lizards | Trioceros_montium | Trioceros_montium | NT | Decreasing | tropical | diurnal | 38.905 | 21680.702 | 10.377 |
| lizards | Trioceros_narraioca | NA | NT | Unknown | NA | NA | NA | NA | NA |
| lizards | Trioceros_ntunte | NA | DD | Unknown | NA | NA | NA | NA | NA |
| lizards | Trioceros_nyirit | NA | LC | Stable | NA | NA | NA | NA | NA |
| lizards | Trioceros_oweni | Trioceros_oweni | LC | Unknown | tropical | diurnal | 66.069 | 2129943.347 | 5.883 |
| lizards | Trioceros_quadricornis | Trioceros_quadricornis | VU | Decreasing | tropical | diurnal | 93.325 | 13912.411 | 10.265 |
| lizards | Trioceros_rudis | Trioceros_rudis | LC | Stable | tropical | diurnal | 10.715 | 327813.057 | 10.58 |
| lizards | Trioceros_schubotzi | Trioceros_schubotzi | NT | Stable | tropical | diurnal | 14.125 | 516.091 | 3.461 |
| lizards | Trioceros_sternfeldi | Trioceros_sternfeldi | LC | Stable | tropical | diurnal | 16.218 | 22624.655 | 13.173 |
| lizards | Trogonophis_wiegmanni | Trogonophis_wiegmanni | LC | Unknown | temperate | cathemeral | 14.125 | 505845.319 | 14.512 |
| lizards | Tropicagama_temporalis | Lophognathus_temporalis | LC | Unknown | tropical | diurnal | 123.027 | 501763.586 | 2.441 |
| lizards | Tropidophorus_assamensis | NA | VU | Unknown | NA | NA | NA | NA | NA |
| lizards | Tropidophorus_baviensis | Tropidophorus_baviensis | LC | Stable | tropical | cathemeral | 16.596 | 120341.719 | 9.929 |
| lizards | Tropidophorus_beccarii | Tropidophorus_beccarii | LC | Unknown | tropical | diurnal | 19.953 | 298445.109 | 6.134 |

| | | | | | | | | | |
|---|---|---|---|---|---|---|---|---|---|
| lizards | Tropidophorus_berdmorei | Tropidophorus_berdmorei | LC | Stable | tropical | diurnal | 19.498 | 348334.766 | 10.953 |
| lizards | Tropidophorus_brookei | Tropidophorus_brookei | LC | Unknown | tropical | diurnal | 29.512 | 338300.752 | 5.377 |
| lizards | Tropidophorus_cocincinensis | Tropidophorus_cocincinensis | LC | Stable | tropical | cathemeral | 22.387 | 92976.812 | 8.588 |
| lizards | Tropidophorus_davaoensis | Tropidophorus_davaoensis | LC | Unknown | tropical | diurnal | 52.481 | 49600.775 | 11.69 |
| lizards | Tropidophorus_grayi | Tropidophorus_grayi | LC | Stable | tropical | diurnal | 38.019 | 71193.131 | 15.266 |
| lizards | Tropidophorus_hainanus | Tropidophorus_hainanus | LC | Unknown | tropical | diurnal | 3.631 | 287009.472 | 16.079 |
| lizards | Tropidophorus_hangnam | NA | DD | Unknown | NA | NA | NA | NA | NA |
| lizards | Tropidophorus_laotus | Tropidophorus_laotus | EN | Stable | tropical | cathemeral | 15.488 | 41996.528 | 11.593 |
| lizards | Tropidophorus_latiscutatus | NA | DD | Unknown | NA | NA | NA | NA | NA |
| lizards | Tropidophorus_microlepis | Tropidophorus_microlepis | LC | Stable | tropical | cathemeral | 11.749 | 72503.973 | 8.984 |
| lizards | Tropidophorus_micropus | NA | DD | Unknown | NA | NA | NA | NA | NA |
| lizards | Tropidophorus_misaminius | Tropidophorus_misaminius | LC | Stable | tropical | cathemeral | 30.903 | 52148.558 | 11.2 |
| lizards | Tropidophorus_mocquardii | NA | not_available | not_available | NA | NA | NA | NA | NA |
| lizards | Tropidophorus_murphyi | Tropidophorus_murphyi | LC | Unknown | tropical | nocturnal | 19.055 | 19.315 | 16.687 |
| lizards | Tropidophorus_noggei | Tropidophorus_noggei | LC | Unknown | tropical | nocturnal | 29.512 | 1222.769 | 5.809 |
| lizards | Tropidophorus_partelloi | Tropidophorus_partelloi | LC | Unknown | tropical | cathemeral | 46.774 | 94145.238 | 12.032 |
| lizards | Tropidophorus_perplexus | NA | LC | Unknown | NA | NA | NA | NA | NA |
| lizards | Tropidophorus_robinsoni | Tropidophorus_robinsoni | LC | Stable | tropical | diurnal | 8.511 | 48939.741 | 7.528 |
| lizards | Tropidophorus_sinicus | Tropidophorus_sinicus | LC | Stable | tropical | nocturnal | 7.079 | 281145.254 | 17.631 |
| lizards | Tropidophorus_thai | NA | LC | Unknown | NA | NA | NA | NA | NA |
| lizards | Tropidosaura_gularis | Tropidosaura_gularis | LC | Stable | temperate | diurnal | 5.623 | 72028.113 | 10.998 |
| lizards | Tropidosaura_montana | Tropidosaura_montana | LC | Stable | temperate | diurnal | 6.761 | 257256.863 | 9.794 |
| lizards | Tropidoscincus_aubrianus | NA | VU | Decreasing | NA | NA | NA | NA | NA |
| lizards | Tropidoscincus_boreus | Tropidoscincus_boreus | LC | Unknown | tropical | diurnal | 18.197 | 11545.475 | 7.179 |
| lizards | Tropidoscincus_variabilis | Tropidoscincus_variabilis | LC | Stable | tropical | diurnal | 18.197 | 4548.797 | 7.466 |
| lizards | Tropidurus_bogerti | Tropidurus_bogerti | LC | Stable | tropical | diurnal | 28.84 | 1525.26 | 1.592 |
| lizards | Tropidurus_callathelys | Tropidurus_callathelys | LC | Unknown | tropical | diurnal | 33.884 | 19.982 | 19.533 |
| lizards | Tropidurus_catalanensis | Tropidurus_catalanensis | LC | Unknown | temperate | diurnal | 77.625 | 544715.378 | 11.914 |
| lizards | Tropidurus_chromatops | Tropidurus_chromatops | LC | Decreasing | tropical | diurnal | 45.709 | 5886.913 | 1.963 |
| lizards | Tropidurus_cocorobensis | Tropidurus_cocorobensis | LC | Unknown | tropical | diurnal | 16.218 | 72805.583 | 7.845 |
| lizards | Tropidurus_erythrocephalus | Tropidurus_erythrocephalus | VU | Unknown | tropical | diurnal | 22.387 | 26701.211 | 8.726 |
| lizards | Tropidurus_etheridgei | Tropidurus_etheridgei | LC | Unknown | tropical | diurnal | 52.481 | 1548687.107 | 5.904 |
| lizards | Tropidurus_guarani | Tropidurus_guarani | LC | Unknown | tropical | diurnal | 52.481 | 438039.888 | 8.097 |
| lizards | Tropidurus_helenae | NA | LC | Unknown | NA | NA | NA | NA | NA |

| | | | | | | | | | |
|---|---|---|---|---|---|---|---|---|---|
| lizards | Tropidurus_hispidus | Tropidurus_hispidus | LC | Stable | tropical | diurnal | 95.499 | 3583749.069 | 5.086 |
| lizards | Tropidurus_hygomi | Tropidurus_hygomi | VU | Decreasing | tropical | diurnal | 19.498 | 15763.287 | 13.327 |
| lizards | Tropidurus_imbituba | NA | CR | Unknown | NA | NA | NA | NA | NA |
| lizards | Tropidurus_itambere | Tropidurus_itambere | LC | Unknown | tropical | diurnal | 30.903 | 1650871.849 | 11.014 |
| lizards | Tropidurus_jaguaribanus | NA | DD | Unknown | NA | NA | NA | NA | NA |
| lizards | Tropidurus_lagunablanca | NA | not_available | not_available | NA | NA | NA | NA | NA |
| lizards | Tropidurus_montanus | Tropidurus_montanus | LC | Unknown | tropical | diurnal | 34.674 | 133739.512 | 8.812 |
| lizards | Tropidurus_oreadicus | Tropidurus_oreadicus | LC | Unknown | tropical | diurnal | 54.954 | 4046111.739 | 5.135 |
| lizards | Tropidurus_pinima | Tropidurus_pinima | LC | Unknown | tropical | diurnal | 22.909 | 302.651 | 4.769 |
| lizards | Tropidurus_psammonastes | Tropidurus_psammonastes | EN | Unknown | tropical | diurnal | 34.674 | 5722.808 | 3.93 |
| lizards | Tropidurus_semitaeniatus | Tropidurus_semitaeniatus | LC | Unknown | tropical | diurnal | 35.481 | 964683.246 | 8.562 |
| lizards | Tropidurus_sertanejo | NA | not_available | not_available | NA | NA | NA | NA | NA |
| lizards | Tropidurus_spinulosus | Tropidurus_spinulosus | LC | Unknown | tropical | diurnal | 89.125 | 1027962.335 | 5.885 |
| lizards | Tropidurus_torquatus | Tropidurus_torquatus | LC | Unknown | tropical | diurnal | 79.433 | 4145465.985 | 9.547 |
| lizards | Tropidurus_xanthochilus | Tropidurus_xanthochilus | VU | Stable | tropical | diurnal | 66.069 | 9.991 | 5.669 |
| lizards | Tropiocolotes_algericus | Tropiocolotes_algericus | LC | Stable | temperate | nocturnal | 1.23 | 455120.052 | 4.529 |
| lizards | Tropiocolotes_bisharicus | NA | LC | Stable | NA | NA | NA | NA | NA |
| lizards | Tropiocolotes_hormozganensis | NA | not_available | not_available | NA | NA | NA | NA | NA |
| lizards | Tropiocolotes_nattereri | Tropiocolotes_nattereri | LC | Stable | temperate | nocturnal | 0.692 | 84669.694 | 6.259 |
| lizards | Tropiocolotes_naybandensis | NA | not_available | not_available | NA | NA | NA | NA | NA |
| lizards | Tropiocolotes_nubicus | NA | DD | Unknown | NA | NA | NA | NA | NA |
| lizards | Tropiocolotes_scorteccii | NA | LC | Stable | NA | NA | NA | NA | NA |
| lizards | Tropiocolotes_somalicus | Tropiocolotes_somalicus | LC | Unknown | tropical | nocturnal | 0.562 | 8022.589 | 8.72 |
| lizards | Tropiocolotes_steudneri | Tropiocolotes_steudneri | LC | Stable | tropical | nocturnal | 1.023 | 2758118.766 | 2.897 |
| lizards | Tropiocolotes_tripolitanus | Tropiocolotes_tripolitanus | LC | Stable | temperate | nocturnal | 1.23 | 5063135.148 | 3.791 |
| lizards | Tropiocolotes_wolfgangboehmei | NA | DD | Unknown | NA | NA | NA | NA | NA |
| lizards | Tukutuku_rakiurae | Tukutuku_rakiurae | EN | Decreasing | temperate | cathemeral | 7.244 | 1692.312 | 3.168 |
| lizards | Tumbunascincus_luteilateralis | Tumbunascincus_luteilateralis | DD | Stable | tropical | diurnal | 30.903 | 180.022 | 1.872 |
| lizards | Tupinambis_cryptus | NA | not_available | not_available | NA | NA | NA | NA | NA |
| lizards | Tupinambis_cuzcoensis | NA | not_available | not_available | NA | NA | NA | NA | NA |
| lizards | Tupinambis_longilineus | NA | LC | Unknown | NA | NA | NA | NA | NA |
| lizards | Tupinambis_matipu | NA | not_available | not_available | NA | NA | NA | NA | NA |
| lizards | Tupinambis_quadrilineatus | NA | LC | Unknown | NA | NA | NA | NA | NA |
| lizards | Tupinambis_teguixin | Tupinambis_teguixin | LC | Stable | tropical | diurnal | 4466.836 | 9385193.531 | 4.029 |

| | | | | | | | | | |
|---|---|---|---|---|---|---|---|---|---|
| lizards | Tympanocryptis_centralis | Tympanocryptis_centralis | LC | Stable | tropical | diurnal | 7.244 | 552386.112 | 1.63 |
| lizards | Tympanocryptis_cephalus | NA | DD | Unknown | NA | NA | NA | NA | NA |
| lizards | Tympanocryptis_condaminensis | NA | EN | Decreasing | NA | NA | NA | NA | NA |
| lizards | Tympanocryptis_diabolicus | NA | LC | Unknown | NA | NA | NA | NA | NA |
| lizards | Tympanocryptis_fortescuensis | NA | LC | Stable | NA | NA | NA | NA | NA |
| lizards | Tympanocryptis_gigas | NA | LC | Unknown | NA | NA | NA | NA | NA |
| lizards | Tympanocryptis_houstoni | NA | LC | Stable | NA | NA | NA | NA | NA |
| lizards | Tympanocryptis_intima | Tympanocryptis_intima | LC | Unknown | temperate | diurnal | 7.244 | 1159092.277 | 3.776 |
| lizards | Tympanocryptis_lineata | Tympanocryptis_lineata | LC | Unknown | temperate | diurnal | 12.023 | 981181.573 | 4.6 |
| lizards | Tympanocryptis_pentalineata | NA | DD | Unknown | NA | NA | NA | NA | NA |
| lizards | Tympanocryptis_pinguicolla | Tympanocryptis_pinguicolla | EN | Decreasing | temperate | diurnal | 8.71 | 8173.168 | 18.407 |
| lizards | Tympanocryptis_pseudopsephos | NA | LC | Stable | NA | NA | NA | NA | NA |
| lizards | Tympanocryptis_tetraporophora | Tympanocryptis_tetraporophora | LC | Unknown | temperate | diurnal | 13.183 | 2308571.203 | 4.412 |
| lizards | Tympanocryptis_wilsoni | NA | EN | Unknown | NA | NA | NA | NA | NA |
| lizards | Typhlacontias_brevipes | Typhlacontias_brevipes | LC | Stable | tropical | cathemeral | 4.786 | 57199.885 | 2.481 |
| lizards | Typhlacontias_johnsonii | NA | LC | Stable | NA | NA | NA | NA | NA |
| lizards | Typhlacontias_punctatissimus | NA | LC | Stable | NA | NA | NA | NA | NA |
| lizards | Typhlacontias_rohani | Typhlacontias_rohani | LC | Stable | tropical | cathemeral | 1.95 | 595828.278 | 5.268 |
| lizards | Typhlosaurus_braini | NA | LC | Stable | NA | NA | NA | NA | NA |
| lizards | Typhlosaurus_lomiae | NA | NT | Stable | NA | NA | NA | NA | NA |
| lizards | Tytthoscincus_aesculeticola | Tytthoscincus_aesculeticola | LC | Unknown | tropical | diurnal | 1.413 | 1433.871 | 8.451 |
| lizards | Tytthoscincus_atrigularis | NA | LC | Unknown | NA | NA | NA | NA | NA |
| lizards | Tytthoscincus_batupanggah | NA | DD | Unknown | NA | NA | NA | NA | NA |
| lizards | Tytthoscincus_biparietalis | NA | NT | Stable | NA | NA | NA | NA | NA |
| lizards | Tytthoscincus_bukitensis | Sphenomorphus_bukitensis | NT | Unknown | tropical | diurnal | 1.514 | 866.579 | 5.653 |
| lizards | Tytthoscincus_butleri | NA | LC | Stable | NA | NA | NA | NA | NA |
| lizards | Tytthoscincus_ishaki | NA | LC | Stable | NA | NA | NA | NA | NA |
| lizards | Tytthoscincus_jaripendek | NA | not_available | not_available | NA | NA | NA | NA | NA |
| lizards | Tytthoscincus_kakikecil | NA | not_available | not_available | NA | NA | NA | NA | NA |
| lizards | Tytthoscincus_leproauricularis | NA | DD | Unknown | NA | NA | NA | NA | NA |
| lizards | Tytthoscincus_perhentianensis | NA | LC | Unknown | NA | NA | NA | NA | NA |
| lizards | Tytthoscincus_sibuensis | NA | DD | Unknown | NA | NA | NA | NA | NA |
| lizards | Tytthoscincus_temasekensis | NA | not_available | not_available | NA | NA | NA | NA | NA |
| lizards | Tytthoscincus_temengorensis | NA | NT | Stable | NA | NA | NA | NA | NA |

| | | | | | | | | | |
|---|---|---|---|---|---|---|---|---|---|
| lizards | Tytthoscincus_temmincki | Sphenomorphus_temmincki | LC | Unknown | tropical | diurnal | 3.311 | 12431.976 | 18.841 |
| lizards | Uma_cowlesi | NA | not_available | not_available | NA | NA | NA | NA | NA |
| lizards | Uma_exsul | Uma_exsul | EN | Decreasing | temperate | diurnal | 31.623 | 10958.169 | 11.627 |
| lizards | Uma_inornata | Uma_inornata | EN | Decreasing | temperate | diurnal | 56.234 | 734.75 | 27.403 |
| lizards | Uma_notata | Uma_notata | NT | Decreasing | temperate | diurnal | 53.703 | 20568.982 | 6.751 |
| lizards | Uma_paraphygas | Uma_paraphygas | NT | Unknown | temperate | diurnal | 22.909 | 1945.977 | 5.019 |
| lizards | Uma_scoparia | Uma_scoparia | LC | Stable | temperate | diurnal | 46.774 | 39209.625 | 5.932 |
| lizards | Underwoodisaurus_milii | Underwoodisaurus_milii | LC | Stable | temperate | nocturnal | 22.909 | 2778149.505 | 6.152 |
| lizards | Underwoodisaurus_seorsus | Underwoodisaurus_seorsus | LC | Unknown | tropical | nocturnal | 18.197 | 14234.851 | 4.326 |
| lizards | Uracentron_azureum | Uracentron_azureum | LC | Unknown | tropical | diurnal | 25.119 | 3804464.194 | 1.546 |
| lizards | Uracentron_flaviceps | Uracentron_flaviceps | LC | Unknown | tropical | diurnal | 72.444 | 1189332.15 | 1.536 |
| lizards | Uranoscodon_superciliosus | Uranoscodon_superciliosus | LC | Stable | tropical | diurnal | 141.254 | 4847371.975 | 1.691 |
| lizards | Urocotyledon_inexpectata | NA | LC | Stable | NA | NA | NA | NA | NA |
| lizards | Urocotyledon_palmata | NA | DD | Unknown | NA | NA | NA | NA | NA |
| lizards | Urocotyledon_rasmusseni | NA | DD | Unknown | NA | NA | NA | NA | NA |
| lizards | Urocotyledon_weileri | NA | EN | Unknown | NA | NA | NA | NA | NA |
| lizards | Urocotyledon_wolterstorffi | NA | DD | Unknown | NA | NA | NA | NA | NA |
| lizards | Uromastyx_acanthinura | Uromastyx_acanthinura | NT | Decreasing | temperate | diurnal | 2454.709 | 1472842.994 | 4.341 |
| lizards | Uromastyx_aegyptia | Uromastyx_aegyptia | VU | Decreasing | temperate | diurnal | 2818.383 | 3447574.29 | 5.184 |
| lizards | Uromastyx_alfredschmidti | Uromastyx_alfredschmidti | NT | Stable | temperate | diurnal | 446.684 | 49623.26 | 2.085 |
| lizards | Uromastyx_benti | Uromastyx_benti | LC | Unknown | tropical | diurnal | 269.153 | 293988.35 | 6.624 |
| lizards | Uromastyx_dispar | Uromastyx_dispar | LC | Decreasing | tropical | diurnal | 457.088 | 4715456.514 | 1.683 |
| lizards | Uromastyx_geyri | Uromastyx_geyri | NT | Decreasing | tropical | diurnal | 275.423 | 920201.602 | 1.523 |
| lizards | Uromastyx_macfadyeni | Uromastyx_macfadyeni | LC | Unknown | tropical | diurnal | 61.66 | 115699.722 | 7.973 |
| lizards | Uromastyx_nigriventris | NA | not_available | not_available | NA | NA | NA | NA | NA |
| lizards | Uromastyx_occidentalis | NA | DD | Unknown | NA | NA | NA | NA | NA |
| lizards | Uromastyx_ocellata | Uromastyx_ocellata | LC | Decreasing | tropical | diurnal | 190.546 | 1527514.508 | 7.324 |
| lizards | Uromastyx_ornata | Uromastyx_ornata | LC | Decreasing | tropical | diurnal | 331.131 | 261938.167 | 8.227 |
| lizards | Uromastyx_princeps | NA | LC | Unknown | NA | NA | NA | NA | NA |
| lizards | Uromastyx_shobraki | NA | NT | Unknown | NA | NA | NA | NA | NA |
| lizards | Uromastyx_thomasi | Uromastyx_thomasi | EN | Decreasing | tropical | diurnal | 245.471 | 99513.078 | 4.134 |
| lizards | Uromastyx_yemenensis | NA | NT | Unknown | NA | NA | NA | NA | NA |
| lizards | Uroplatus_alluaudi | Uroplatus_alluaudi | NT | Unknown | tropical | nocturnal | 10 | 2811.333 | 6.355 |
| lizards | Uroplatus_ebenaui | Uroplatus_ebenaui | VU | Decreasing | tropical | nocturnal | 7.244 | 28776.818 | 7.646 |

| | | | | | | | | | |
|---|---|---|---|---|---|---|---|---|---|
| lizards | Uroplatus_fiera | NA | not_available | not_available | NA | NA | NA | NA | NA |
| lizards | Uroplatus_fimbriatus | Uroplatus_fimbriatus | LC | Decreasing | tropical | nocturnal | 120.226 | 94198.143 | 8.523 |
| lizards | Uroplatus_finiavana | NA | NT | Stable | NA | NA | NA | NA | NA |
| lizards | Uroplatus_fotsivava | NA | not_available | not_available | NA | NA | NA | NA | NA |
| lizards | Uroplatus_giganteus | NA | VU | Decreasing | NA | NA | NA | NA | NA |
| lizards | Uroplatus_guentheri | Uroplatus_guentheri | EN | Decreasing | tropical | nocturnal | 10 | 5471.967 | 9.108 |
| lizards | Uroplatus_henkeli | Uroplatus_henkeli | VU | Unknown | tropical | nocturnal | 69.183 | 12389.283 | 10.373 |
| lizards | Uroplatus_kelirambo | NA | not_available | not_available | NA | NA | NA | NA | NA |
| lizards | Uroplatus_lineatus | Uroplatus_lineatus | LC | Decreasing | tropical | nocturnal | 83.176 | 43159.485 | 7.258 |
| lizards | Uroplatus_malahelo | Uroplatus_malahelo | EN | Decreasing | temperate | nocturnal | 10 | 1088.333 | 5.257 |
| lizards | Uroplatus_malama | NA | VU | Decreasing | NA | NA | NA | NA | NA |
| lizards | Uroplatus_phantasticus | Uroplatus_phantasticus | LC | Decreasing | tropical | nocturnal | 12.589 | 44946.066 | 8.869 |
| lizards | Uroplatus_pietschmanni | Uroplatus_pietschmanni | EN | Unknown | tropical | nocturnal | 10.715 | 1264.585 | 7.866 |
| lizards | Uroplatus_sameiti | Uroplatus_sameiti | LC | Unknown | tropical | nocturnal | 28.184 | 52955.609 | 9.378 |
| lizards | Uroplatus_sikorae | Uroplatus_sikorae | LC | Decreasing | tropical | nocturnal | 33.884 | 88289.283 | 7.785 |
| lizards | Urosaurus_auriculatus | NA | EN | Decreasing | NA | NA | NA | NA | NA |
| lizards | Urosaurus_bicarinatus | Urosaurus_bicarinatus | LC | Stable | tropical | diurnal | 10 | 364544.766 | 8.471 |
| lizards | Urosaurus_clarionensis | NA | VU | Unknown | NA | NA | NA | NA | NA |
| lizards | Urosaurus_gadovi | NA | LC | Unknown | NA | NA | NA | NA | NA |
| lizards | Urosaurus_graciosus | Urosaurus_graciosus | LC | Stable | temperate | diurnal | 11.22 | 183159.458 | 8.276 |
| lizards | Urosaurus_lahtelai | NA | LC | Stable | NA | NA | NA | NA | NA |
| lizards | Urosaurus_nigricauda | NA | LC | Stable | NA | NA | NA | NA | NA |
| lizards | Urosaurus_ornatus | Urosaurus_ornatus | LC | Stable | temperate | diurnal | 11.749 | 1685548.261 | 8.623 |
| lizards | Urostrophus_gallardoi | Urostrophus_gallardoi | DD | Decreasing | temperate | diurnal | 13.183 | 214160.579 | 6.798 |
| lizards | Urostrophus_undulatus | NA | DD | Unknown | NA | NA | NA | NA | NA |
| lizards | Urostrophus_vautieri | Urostrophus_vautieri | LC | Unknown | temperate | diurnal | 39.811 | 506846.517 | 13.842 |
| lizards | Uta_encantadae | Uta_encantadae | VU | Stable | temperate | diurnal | 11.749 | 0.617 | 0.348 |
| lizards | Uta_lowei | NA | VU | Unknown | NA | NA | NA | NA | NA |
| lizards | Uta_nolascensis | Uta_nolascensis | LC | Stable | temperate | diurnal | 6.31 | 2.968 | 1.549 |
| lizards | Uta_palmeri | Uta_palmeri | VU | Stable | temperate | diurnal | 19.055 | 1.223 | 2.107 |
| lizards | Uta_squamata | Uta_squamata | LC | Stable | temperate | diurnal | 7.079 | 27.728 | 1.218 |
| lizards | Uta_stansburiana | Uta_stansburiana | LC | Stable | temperate | diurnal | 15.849 | 2345521.987 | 8.57 |
| lizards | Uta_tumidarostra | NA | VU | Unknown | NA | NA | NA | NA | NA |
| lizards | Uvidicolus_sphyrurus | Uvidicolus_sphyrurus | LC | Unknown | temperate | nocturnal | 9.333 | 59393.837 | 7.929 |

| | | | | | | | | | |
|---|---|---|---|---|---|---|---|---|---|
| lizards | Vanzosaura_multiscutata | Vanzosaura_multiscutatus | LC | Unknown | tropical | diurnal | 1.096 | 761798.832 | 9.197 |
| lizards | Vanzosaura_rubricauda | Vanzosaura_rubricauda | LC | Unknown | tropical | diurnal | 1.905 | 5109646.326 | 8.001 |
| lizards | Vanzosaura_savanicola | Vanzosaura_savanicola | LC | Unknown | tropical | diurnal | 0.871 | 152213.439 | 7.786 |
| lizards | Varanus_acanthurus | Varanus_acanthurus | LC | Stable | tropical | diurnal | 288.403 | 3077096.875 | 2.522 |
| lizards | Varanus_albigularis | Varanus_albigularis | LC | Stable | tropical | diurnal | 15135.612 | 8253607.15 | 8.162 |
| lizards | Varanus_auffenbergi | Varanus_auffenbergi | EN | Decreasing | tropical | diurnal | 151.356 | 1221.116 | 16.938 |
| lizards | Varanus_bangonorum | NA | LC | Unknown | NA | NA | NA | NA | NA |
| lizards | Varanus_beccarii | Varanus_beccarii | DD | Decreasing | tropical | diurnal | 776.247 | 7854.143 | 4.203 |
| lizards | Varanus_bengalensis | Varanus_bengalensis | NT | Decreasing | tropical | diurnal | 18197.009 | 4839323.295 | 15.376 |
| lizards | Varanus_bitatawa | NA | NT | Unknown | NA | NA | NA | NA | NA |
| lizards | Varanus_boehmei | NA | DD | Unknown | NA | NA | NA | NA | NA |
| lizards | Varanus_bogerti | Varanus_bogerti | LC | Unknown | tropical | diurnal | 446.684 | 1686.127 | 5.157 |
| lizards | Varanus_brevicauda | Varanus_brevicauda | LC | Stable | tropical | diurnal | 30.903 | 2296478.736 | 2.338 |
| lizards | Varanus_bushi | NA | LC | Stable | NA | NA | NA | NA | NA |
| lizards | Varanus_caerulivirens | Varanus_caerulivirens | LC | Unknown | tropical | diurnal | 1318.257 | 14590.354 | 7.206 |
| lizards | Varanus_caudolineatus | Varanus_caudolineatus | LC | Stable | temperate | diurnal | 37.154 | 1179363.035 | 3.968 |
| lizards | Varanus_cumingi | Varanus_cumingi | LC | Increasing | tropical | diurnal | 4897.788 | 96858.026 | 12.035 |
| lizards | Varanus_dalubhasa | NA | LC | Unknown | NA | NA | NA | NA | NA |
| lizards | Varanus_doreanus | Varanus_doreanus | LC | Unknown | tropical | diurnal | 14125.375 | 647811.208 | 3.119 |
| lizards | Varanus_douarrha | NA | not_available | not_available | NA | NA | NA | NA | NA |
| lizards | Varanus_dumerilii | NA | DD | Unknown | NA | NA | NA | NA | NA |
| lizards | Varanus_eremius | Varanus_eremius | LC | Stable | temperate | diurnal | 144.544 | 2912674.981 | 2.186 |
| lizards | Varanus_exanthematicus | Varanus_exanthematicus | LC | Unknown | tropical | diurnal | 10000 | 7554393.89 | 10.328 |
| lizards | Varanus_finschi | Varanus_finschi | LC | Unknown | tropical | diurnal | 1023.293 | 42667.276 | 4.613 |
| lizards | Varanus_flavescens | Varanus_flavescens | EN | Decreasing | temperate | diurnal | 2951.209 | 883802.277 | 18.547 |
| lizards | Varanus_giganteus | Varanus_giganteus | LC | Stable | temperate | diurnal | 21379.621 | 2464635.027 | 2.585 |
| lizards | Varanus_gilleni | Varanus_gilleni | LC | Stable | temperate | diurnal | 125.893 | 2155331.713 | 1.537 |
| lizards | Varanus_glauerti | Varanus_glauerti | LC | Stable | tropical | diurnal | 288.403 | 153277.132 | 2.411 |
| lizards | Varanus_glebopalma | Varanus_glebopalma | LC | Stable | tropical | diurnal | 1288.25 | 656187.114 | 2.118 |
| lizards | Varanus_gouldii | Varanus_gouldii | LC | Stable | temperate | diurnal | 6918.31 | 6329091.519 | 3.891 |
| lizards | Varanus_griseus | Varanus_griseus | LC | Decreasing | temperate | diurnal | 5495.409 | 17979964.401 | 4.981 |
| lizards | Varanus_hamersleyensis | NA | LC | Stable | NA | NA | NA | NA | NA |
| lizards | Varanus_indicus | Varanus_indicus | LC | Unknown | tropical | diurnal | 4365.158 | 651554.201 | 3.444 |
| lizards | Varanus_insulanicus | NA | LC | Stable | NA | NA | NA | NA | NA |

| | | | | | | | | | |
|---|---|---|---|---|---|---|---|---|---|
| lizards | Varanus_jobiensis | Varanus_jobiensis | LC | Unknown | tropical | diurnal | 1949.845 | 494063.002 | 3.159 |
| lizards | Varanus_juxtindicus | Varanus_juxtindicus | LC | Stable | tropical | diurnal | 2754.229 | 675.7 | 3.166 |
| lizards | Varanus_keithhornei | Varanus_keithhornei | LC | Stable | tropical | diurnal | 436.516 | 10996.049 | 1.987 |
| lizards | Varanus_kingorum | Varanus_kingorum | LC | Stable | tropical | diurnal | 26.915 | 40086.063 | 2.278 |
| lizards | Varanus_komodoensis | Varanus_komodoensis | EN | Stable | tropical | diurnal | 109647.82 | 1113.101 | 13.106 |
| lizards | Varanus_kordensis | NA | DD | Unknown | NA | NA | NA | NA | NA |
| lizards | Varanus_lirungensis | Varanus_lirungensis | EN | Decreasing | tropical | diurnal | 891.251 | 93.233 | 13.796 |
| lizards | Varanus_mabitang | Varanus_mabitang | EN | Decreasing | tropical | diurnal | 7943.282 | 336.892 | 4.929 |
| lizards | Varanus_macraei | Varanus_macraei | EN | Unknown | tropical | diurnal | 933.254 | 453.712 | 1.38 |
| lizards | Varanus_marmoratus | Varanus_marmoratus | LC | Unknown | tropical | diurnal | 11748.976 | 89087.084 | 14.934 |
| lizards | Varanus_melinus | Varanus_melinus | EN | Decreasing | tropical | diurnal | 2884.032 | 4741.454 | 5.677 |
| lizards | Varanus_mertensi | Varanus_mertensi | EN | Decreasing | tropical | diurnal | 3890.451 | 1201720.634 | 2.889 |
| lizards | Varanus_mitchelli | Varanus_mitchelli | CR | Decreasing | tropical | diurnal | 812.831 | 449926.026 | 1.782 |
| lizards | Varanus_nebulosus | NA | not_available | not_available | NA | NA | NA | NA | NA |
| lizards | Varanus_nesterovi | NA | not_available | not_available | NA | NA | NA | NA | NA |
| lizards | Varanus_niloticus | Varanus_niloticus | LC | Stable | tropical | diurnal | 23988.329 | 15915291.187 | 9.454 |
| lizards | Varanus_nuchalis | Varanus_nuchalis | LC | Increasing | tropical | diurnal | 5011.872 | 34950.569 | 18.206 |
| lizards | Varanus_obor | NA | EN | Decreasing | NA | NA | NA | NA | NA |
| lizards | Varanus_olivaceus | Varanus_olivaceus | VU | Decreasing | tropical | diurnal | 9120.108 | 26652.265 | 7.819 |
| lizards | Varanus_palawanensis | NA | LC | Stable | NA | NA | NA | NA | NA |
| lizards | Varanus_panoptes | Varanus_panoptes | LC | Decreasing | tropical | diurnal | 9549.926 | 3341093.981 | 3.068 |
| lizards | Varanus_pilbarensis | Varanus_pilbarensis | LC | Stable | tropical | diurnal | 107.152 | 31460.328 | 4.178 |
| lizards | Varanus_prasinus | Varanus_prasinus | LC | Stable | tropical | diurnal | 575.44 | 632368.528 | 3.139 |
| lizards | Varanus_primordius | Varanus_primordius | LC | Unknown | tropical | diurnal | 26.915 | 102040.888 | 1.648 |
| lizards | Varanus_rainerguentheri | NA | LC | Stable | NA | NA | NA | NA | NA |
| lizards | Varanus_rasmusseni | NA | LC | Unknown | NA | NA | NA | NA | NA |
| lizards | Varanus_reisingeri | NA | DD | Unknown | NA | NA | NA | NA | NA |
| lizards | Varanus_rosenbergi | Varanus_rosenbergi | LC | Unknown | temperate | diurnal | 3019.952 | 413035.439 | 7.944 |
| lizards | Varanus_rudicollis | NA | DD | Unknown | NA | NA | NA | NA | NA |
| lizards | Varanus_salvadorii | Varanus_salvadorii | LC | Unknown | tropical | diurnal | 15135.612 | 505211.348 | 2.803 |
| lizards | Varanus_salvator | Varanus_salvator | LC | Unknown | tropical | diurnal | 41686.938 | 4677934.788 | 12.098 |
| lizards | Varanus_samarensis | NA | LC | Unknown | NA | NA | NA | NA | NA |
| lizards | Varanus_scalaris | Varanus_scalaris | LC | Unknown | tropical | diurnal | 354.813 | 1203625.356 | 2.742 |
| lizards | Varanus_semiremex | Varanus_semiremex | LC | Unknown | tropical | diurnal | 426.58 | 202421.376 | 4.433 |

| | | | | | | | | | |
|---|---|---|---|---|---|---|---|---|---|
| lizards | Varanus_semotus | NA | not_available | not_available | NA | NA | NA | NA | NA |
| lizards | Varanus_similis | NA | not_available | not_available | NA | NA | NA | NA | NA |
| lizards | Varanus_sparnus | NA | DD | Unknown | NA | NA | NA | NA | NA |
| lizards | Varanus_spenceri | Varanus_spenceri | LC | Stable | tropical | diurnal | 4365.158 | 700991.097 | 3.985 |
| lizards | Varanus_spinulosus | Varanus_spinulosus | LC | Stable | tropical | diurnal | 630.957 | 1594.65 | 3.9 |
| lizards | Varanus_storri | Varanus_storri | LC | Stable | tropical | diurnal | 42.658 | 518428.556 | 3.133 |
| lizards | Varanus_telenesetes | NA | DD | Unknown | NA | NA | NA | NA | NA |
| lizards | Varanus_timorensis | Varanus_timorensis | LC | Decreasing | tropical | diurnal | 436.516 | 28857.863 | 13.927 |
| lizards | Varanus_togianus | Varanus_togianus | LC | Stable | tropical | diurnal | 3981.072 | 184110.681 | 9.344 |
| lizards | Varanus_tristis | Varanus_tristis | LC | Stable | tropical | diurnal | 549.541 | 5950251.606 | 3.208 |
| lizards | Varanus_varius | Varanus_varius | LC | Stable | temperate | diurnal | 10715.193 | 1219347.692 | 8.545 |
| lizards | Varanus_yemenensis | NA | DD | Unknown | NA | NA | NA | NA | NA |
| lizards | Varanus_yuwonoi | NA | VU | Unknown | NA | NA | NA | NA | NA |
| lizards | Varanus_zugorum | NA | DD | Unknown | NA | NA | NA | NA | NA |
| lizards | Varzea_altamazonica | NA | LC | Unknown | NA | NA | NA | NA | NA |
| lizards | Varzea_bistriata | Varzea_bistriata | LC | Unknown | tropical | diurnal | 28.184 | 8162647.765 | 3.941 |
| lizards | Vhembelacerta_rupicola | Vhembelacerta_rupicola | LC | Decreasing | tropical | diurnal | 3.236 | 15435.29 | 9.707 |
| lizards | Vietnascincus_rugosus | NA | DD | Unknown | NA | NA | NA | NA | NA |
| lizards | Voeltzkowia_mobydick | NA | not_available | not_available | NA | NA | NA | NA | NA |
| lizards | Voeltzkowia_yamagishii | NA | EN | Decreasing | NA | NA | NA | NA | NA |
| lizards | Woodworthia_brunnea | Woodworthia_brunneus | EN | Decreasing | temperate | nocturnal | 10.233 | 9177.537 | 10.269 |
| lizards | Woodworthia_chrysosiretica | Woodworthia_chrysosireticus | NT | Stable | temperate | nocturnal | 10.233 | 6515.379 | 9.271 |
| lizards | Woodworthia_maculata | Woodworthia_maculatus | LC | Stable | temperate | cathemeral | 14.454 | 7.113 | 41.003 |
| lizards | Xantusia_arizonae | Xantusia_arizonae | LC | Stable | temperate | cathemeral | 4.169 | 783.958 | 6.351 |
| lizards | Xantusia_bezyi | Xantusia_bezyi | LC | Unknown | temperate | diurnal | 6.761 | 4891.335 | 10.382 |
| lizards | Xantusia_extorris | Xantusia_extorris | LC | Unknown | temperate | nocturnal | 1.514 | 1353.53 | 8.222 |
| lizards | Xantusia_gracilis | Xantusia_gracilis | VU | Unknown | temperate | nocturnal | 7.079 | 44.937 | 2.648 |
| lizards | Xantusia_henshawi | Xantusia_henshawi | LC | Stable | temperate | cathemeral | 7.079 | 27942.41 | 8.417 |
| lizards | Xantusia_jaycolei | NA | not_available | not_available | NA | NA | NA | NA | NA |
| lizards | Xantusia_riversiana | Xantusia_riversiana | LC | Stable | temperate | diurnal | 32.359 | 208.079 | 9.731 |
| lizards | Xantusia_sanchezi | Xantusia_sanchezi | LC | Unknown | tropical | cathemeral | 3.236 | 1707.918 | 10.035 |
| lizards | Xantusia_vigilis | Xantusia_vigilis | LC | Stable | temperate | cathemeral | 6.761 | 261352.528 | 6.542 |
| lizards | Xenagama_batillifera | Xenagama_batillifera | LC | Unknown | tropical | diurnal | 33.884 | 40067.033 | 10.921 |
| lizards | Xenagama_taylori | Xenagama_taylori | LC | Unknown | tropical | diurnal | 20.893 | 51741.436 | 9.18 |

| | | | | | | | | | |
|---|---|---|---|---|---|---|---|---|---|
| lizards | Xenosaurus_agrenon | NA | not_available | not_available | NA | NA | NA | NA | NA |
| lizards | Xenosaurus_arboreus | NA | not_available | not_available | NA | NA | NA | NA | NA |
| lizards | Xenosaurus_grandis | Xenosaurus_grandis | VU | Decreasing | tropical | diurnal | 56.234 | 5386.36 | 11.66 |
| lizards | Xenosaurus_mendozai | NA | not_available | not_available | NA | NA | NA | NA | NA |
| lizards | Xenosaurus_newmanorum | Xenosaurus_newmanorum | EN | Decreasing | tropical | diurnal | 54.954 | 3894.728 | 12.168 |
| lizards | Xenosaurus_phalaroanthereon | Xenosaurus_phalaroanthereon | DD | Stable | tropical | cathemeral | 52.481 | 1363.398 | 4.686 |
| lizards | Xenosaurus_platyceps | Xenosaurus_platyceps | EN | Decreasing | tropical | diurnal | 42.658 | 4161.093 | 4.99 |
| lizards | Xenosaurus_rectocollaris | Xenosaurus_rectocollaris | LC | Stable | tropical | diurnal | 29.512 | 1263.723 | 13.098 |
| lizards | Xenosaurus_tzacualtipantecus | NA | not_available | not_available | NA | NA | NA | NA | NA |
| lizards | Yanomamia_guianensis | NA | LC | Unknown | NA | NA | NA | NA | NA |
| lizards | Yanomamia_hoogmoedi | NA | LC | Stable | NA | NA | NA | NA | NA |
| lizards | Zonosaurus_aeneus | NA | LC | Decreasing | NA | NA | NA | NA | NA |
| lizards | Zonosaurus_anelanelany | Zonosaurus_anelanelany | VU | Decreasing | temperate | diurnal | 19.498 | 6448.394 | 7.724 |
| lizards | Zonosaurus_bemaraha | Zonosaurus_bemaraha | LC | Stable | tropical | diurnal | 10 | 1604.399 | 3.522 |
| lizards | Zonosaurus_boettgeri | NA | VU | Decreasing | NA | NA | NA | NA | NA |
| lizards | Zonosaurus_brygooi | NA | LC | Decreasing | NA | NA | NA | NA | NA |
| lizards | Zonosaurus_haraldmeieri | Zonosaurus_haraldmeieri | NT | Unknown | tropical | diurnal | 69.183 | 1956.626 | 7.967 |
| lizards | Zonosaurus_karsteni | Zonosaurus_karsteni | LC | Unknown | tropical | diurnal | 58.884 | 28718.013 | 8.937 |
| lizards | Zonosaurus_laticaudatus | NA | LC | Unknown | NA | NA | NA | NA | NA |
| lizards | Zonosaurus_madagascariensis | NA | LC | Stable | NA | NA | NA | NA | NA |
| lizards | Zonosaurus_maximus | NA | VU | Decreasing | NA | NA | NA | NA | NA |
| lizards | Zonosaurus_ornatus | NA | LC | Unknown | NA | NA | NA | NA | NA |
| lizards | Zonosaurus_quadrilineatus | NA | VU | Unknown | NA | NA | NA | NA | NA |
| lizards | Zonosaurus_rufipes | Zonosaurus_rufipes | NT | Decreasing | tropical | diurnal | 16.596 | 23315.277 | 9.221 |
| lizards | Zonosaurus_subunicolor | NA | EN | Decreasing | NA | NA | NA | NA | NA |
| lizards | Zonosaurus_trilineatus | NA | LC | Stable | NA | NA | NA | NA | NA |
| lizards | Zonosaurus_tsingy | NA | LC | Stable | NA | NA | NA | NA | NA |
| lizards | Zootoca_vivipara | Zootoca_vivipara | LC | Unknown | temperate | diurnal | 10.715 | 17595247.879 | 8.426 |
| lizards | Zygaspis_dolichomenta | NA | LC | Stable | NA | NA | NA | NA | NA |
| lizards | Zygaspis_quadrifrons | Zygaspis_quadrifrons | LC | Stable | tropical | diurnal | 8.511 | 3374594.089 | 7.128 |
| mammals | Abditomys_latidens | NA | DD | Unknown | NA | NA | NA | NA | NA |
| mammals | Abeomelomys_sevia | Abeomelomys_sevia | LC | Unknown | tropical | nocturnal | 52 | 9683.812 | 8.606 |
| mammals | Abrawayaomys_ruschii | Abrawayaomys_ruschii | LC | Unknown | tropical | nocturnal | 63 | 103564.627 | 12.487 |
| mammals | Abrocoma_bennettii | Abrocoma_bennettii | LC | Unknown | temperate | nocturnal | 250 | 124895.992 | 7.656 |

| | | | | | | | | | |
|---|---|---|---|---|---|---|---|---|---|
| mammals | Abrocoma_boliviensis | Abrocoma_boliviensis | CR | Decreasing | tropical | diurnal | 158 | 64.739 | 14.011 |
| mammals | Abrocoma_budini | NA | DD | Unknown | NA | NA | NA | NA | NA |
| mammals | Abrocoma_cinerea | Abrocoma_cinerea | LC | Stable | tropical | diurnal | 250 | 529152.754 | 6.449 |
| mammals | Abrocoma_famatina | NA | DD | Unknown | NA | NA | NA | NA | NA |
| mammals | Abrocoma_schistacea | Abrocoma_shistacea | LC | Unknown | temperate | diurnal | 276.3 | 5496.476 | 4.252 |
| mammals | Abrocoma_uspallata | NA | DD | Unknown | NA | NA | NA | NA | NA |
| mammals | Abrocoma_vaccarum | NA | DD | Unknown | NA | NA | NA | NA | NA |
| mammals | Abrothrix_andina | Abrothrix_andinus | LC | Unknown | temperate | diurnal | 18 | 612688.887 | 5.231 |
| mammals | Abrothrix_illutea | Abrothrix_illutea | LC | Decreasing | temperate | cathemeral | 47.8 | 56345.628 | 7.389 |
| mammals | Abrothrix_jelskii | Abrothrix_jelskii | LC | Stable | tropical | cathemeral | 34.5 | 498022.482 | 7.901 |
| mammals | Abrothrix_lanosa | Abrothrix_lanosus | LC | Stable | temperate | nocturnal | 32.5 | 40587.894 | 4.015 |
| mammals | Abrothrix_longipilis | Abrothrix_longipilis | LC | Decreasing | temperate | cathemeral | 37.6 | 717949.899 | 6.355 |
| mammals | Abrothrix_olivacea | Abrothrix_olivaceus | LC | Stable | temperate | cathemeral | 27 | 1083345.055 | 5.845 |
| mammals | Abrothrix_sanborni | Abrothrix_sanborni | NT | Decreasing | temperate | cathemeral | 24.7 | 4530.172 | 4.523 |
| mammals | Acerodon_celebensis | Acerodon_celebensis | VU | Decreasing | tropical | nocturnal | 390 | 176922.809 | 9.266 |
| mammals | Acerodon_humilis | Acerodon_humilis | EN | Decreasing | tropical | nocturnal | 599.6 | 929.157 | 9.342 |
| mammals | Acerodon_jubatus | Acerodon_jubatus | EN | Decreasing | tropical | diurnal | 1075 | 155342.189 | 13.963 |
| mammals | Acerodon_leucotis | Acerodon_leucotis | VU | Decreasing | tropical | nocturnal | 513.2 | 12751.181 | 9.253 |
| mammals | Acerodon_macklotii | Acerodon_mackloti | VU | Decreasing | tropical | nocturnal | 470.3 | 74622.811 | 13.76 |
| mammals | Acinonyx_jubatus | Acinonyx_jubatus | VU | Decreasing | temperate | cathemeral | 46700 | 32853534.645 | 7.708 |
| mammals | Acomys_cahirinus | Acomys_cahirinus | LC | Stable | tropical | nocturnal | 52.5 | 4017285.727 | 2.943 |
| mammals | Acomys_chudeaui | Acomys_airensis | LC | Stable | tropical | nocturnal | 48.3 | 1825638.342 | 2.418 |
| mammals | Acomys_cilicicus | NA | DD | Unknown | NA | NA | NA | NA | NA |
| mammals | Acomys_cineraceus | Acomys_cineraceus | LC | Stable | tropical | nocturnal | 20.3 | 1049545.755 | 10.303 |
| mammals | Acomys_dimidiatus | Acomys_dimidiatus | LC | Unknown | temperate | nocturnal | 37.5 | 1558726.498 | 6.947 |
| mammals | Acomys_ignitus | Acomys_ignitus | LC | Stable | tropical | nocturnal | 24 | 105257.36 | 11.128 |
| mammals | Acomys_johannis | Acomys_johannis | LC | Stable | tropical | nocturnal | 18.5 | 426085.265 | 13.638 |
| mammals | Acomys_kempi | Acomys_kempi | LC | Stable | tropical | nocturnal | 22 | 569643.336 | 8.691 |
| mammals | Acomys_louisae | Acomys_louisae | LC | Stable | tropical | nocturnal | 20.2 | 688439.915 | 9.566 |
| mammals | Acomys_minous | NA | DD | Unknown | NA | NA | NA | NA | NA |
| mammals | Acomys_mullah | Acomys_mullah | LC | Unknown | tropical | nocturnal | 28.4 | 160420.635 | 8.059 |
| mammals | Acomys_nesiotes | NA | DD | Unknown | NA | NA | NA | NA | NA |
| mammals | Acomys_percivali | Acomys_percivali | LC | Stable | tropical | nocturnal | 22 | 222958.299 | 7.552 |
| mammals | Acomys_russatus | Acomys_russatus | LC | Unknown | tropical | diurnal | 49.8 | 702024.705 | 7.132 |

| | | | | | | | | | |
|---|---|---|---|---|---|---|---|---|---|
| mammals | Acomys_seurati | Acomys_seurati | LC | Stable | temperate | nocturnal | 71.2 | 298887.042 | 1.484 |
| mammals | Acomys_spinosissimus | Acomys_spinosissimus | LC | Stable | tropical | nocturnal | 26.5 | 101250.459 | 12.498 |
| mammals | Acomys_subspinosus | Acomys_subspinosus | LC | Stable | temperate | nocturnal | 21 | 94357.703 | 10.503 |
| mammals | Acomys_wilsoni | Acomys_wilsoni | LC | Stable | tropical | nocturnal | 18.5 | 1372812.729 | 10.051 |
| mammals | Aconaemys_fuscus | Aconaemys_fuscus | LC | Unknown | temperate | nocturnal | 133.5 | 31935.754 | 5.717 |
| mammals | Aconaemys_porteri | NA | DD | Unknown | NA | NA | NA | NA | NA |
| mammals | Acrobates_pygmaeus | Acrobates_pygmaeus | LC | Stable | temperate | nocturnal | 12 | 1200801.195 | 7.744 |
| mammals | Addax_nasomaculatus | Addax_nasomaculatus | CR | Decreasing | tropical | diurnal | 70000.3 | 49488.369 | 0.27 |
| mammals | Aegialomys_xanthaeolus | Aegialomys_xanthaeolus | LC | Stable | tropical | nocturnal | 79.8 | 258878.321 | 9.149 |
| mammals | Aepeomys_lugens | Aepeomys_lugens | LC | Unknown | tropical | nocturnal | 37 | 9497.265 | 11.838 |
| mammals | Aepeomys_reigi | Aepeomys_reigi | VU | Decreasing | tropical | nocturnal | 40.6 | 13016.499 | 11.651 |
| mammals | Aepyceros_melampus | Aepyceros_melampus | LC | Stable | tropical | cathemeral | 52500.1 | 3498111.537 | 9.521 |
| mammals | Aepyprymnus_rufescens | Aepyprymnus_rufescens | LC | Unknown | temperate | nocturnal | 3250 | 566664.643 | 6.456 |
| mammals | Aeretes_melanopterus | Aeretes_melanopterus | NT | Decreasing | temperate | nocturnal | 732.4 | 56180.82 | 7.035 |
| mammals | Aeromys_tephromelas | NA | DD | Unknown | NA | NA | NA | NA | NA |
| mammals | Aeromys_thomasi | Aeromys_thomasi | LC | Unknown | tropical | nocturnal | 1435 | 446207.387 | 5.012 |
| mammals | Aethalops_aequalis | Aethalops_aequalis | LC | Unknown | tropical | nocturnal | 11.6 | 93008.852 | 4.112 |
| mammals | Aethalops_alecto | Aethalops_alecto | LC | Decreasing | tropical | nocturnal | 15 | 79531.469 | 9.747 |
| mammals | Aethomys_bocagei | Aethomys_bocagei | LC | Unknown | tropical | nocturnal | 107.6 | 253323.277 | 8.309 |
| mammals | Aethomys_chrysophilus | Aethomys_chrysophilus | LC | Unknown | tropical | nocturnal | 74.9 | 4154330.279 | 8.422 |
| mammals | Aethomys_hindei | Aethomys_hindei | LC | Unknown | tropical | cathemeral | 103.5 | 2482352.209 | 7.759 |
| mammals | Aethomys_ineptus | Aethomys_ineptus | LC | Unknown | temperate | nocturnal | 74.9 | 592259.589 | 12.145 |
| mammals | Aethomys_kaiseri | Aethomys_kaiseri | LC | Unknown | tropical | nocturnal | 90 | 1693329.175 | 9.163 |
| mammals | Aethomys_nyikae | Aethomys_nyikae | LC | Unknown | tropical | nocturnal | 77 | 897965.406 | 7.081 |
| mammals | Aethomys_silindensis | NA | DD | Unknown | NA | NA | NA | NA | NA |
| mammals | Aethomys_stannarius | NA | DD | Unknown | NA | NA | NA | NA | NA |
| mammals | Aethomys_thomasi | Aethomys_thomasi | LC | Unknown | tropical | nocturnal | 107.6 | 270711.032 | 8.626 |
| mammals | Afronycteris_helios | NA | DD | Unknown | NA | NA | NA | NA | NA |
| mammals | Afronycteris_nanus | Neoromicia_nana | LC | Unknown | tropical | nocturnal | 3.9 | 10443728.671 | 9.027 |
| mammals | Ailuropoda_melanoleuca | Ailuropoda_melanoleuca | VU | Increasing | temperate | cathemeral | 108400 | 34603.985 | 4.266 |
| mammals | Ailurops_melanotis | Ailurops_melanotis | CR | Decreasing | tropical | nocturnal | 6669 | 274.514 | 11.149 |
| mammals | Ailurops_ursinus | Ailurops_ursinus | VU | Decreasing | tropical | diurnal | 10000 | 179858.765 | 9.32 |
| mammals | Ailurus_fulgens | Ailurus_fulgens | EN | Decreasing | temperate | cathemeral | 4900 | 338830.995 | 6.483 |
| mammals | Akodon_aerosus | Akodon_aerosus | LC | Stable | tropical | nocturnal | 60 | 114165.459 | 4.514 |

| | | | | | | | | | |
|---|---|---|---|---|---|---|---|---|---|
| mammals | Akodon_affinis | Akodon_affinis | LC | Stable | tropical | diurnal | 24.9 | 42197.044 | 10.998 |
| mammals | Akodon_albiventer | Akodon_albiventer | LC | Stable | tropical | diurnal | 21.8 | 402049.086 | 6.282 |
| mammals | Akodon_azarae | Akodon_azarae | LC | Stable | temperate | cathemeral | 25 | 1227616.004 | 10.896 |
| mammals | Akodon_boliviensis | Akodon_boliviensis | LC | Stable | tropical | cathemeral | 27.5 | 217533.181 | 9.362 |
| mammals | Akodon_budini | Akodon_budini | LC | Stable | tropical | nocturnal | 26.9 | 19876.491 | 5.832 |
| mammals | Akodon_caenosus | NA | DD | Unknown | NA | NA | NA | NA | NA |
| mammals | Akodon_cursor | Akodon_cursor | LC | Unknown | tropical | nocturnal | 39.9 | 824145.087 | 11.989 |
| mammals | Akodon_dayi | Akodon_dayi | LC | Stable | tropical | cathemeral | 32.5 | 289331.634 | 7.056 |
| mammals | Akodon_dolores | Akodon_dolores | LC | Stable | temperate | nocturnal | 50.5 | 885883.778 | 6.543 |
| mammals | Akodon_fumeus | Akodon_fumeus | LC | Stable | tropical | cathemeral | 22.7 | 151082.142 | 7.422 |
| mammals | Akodon_iniscatus | Akodon_iniscatus | LC | Stable | temperate | nocturnal | 28.7 | 573427.583 | 5.481 |
| mammals | Akodon_juninensis | Akodon_juninensis | LC | Stable | tropical | cathemeral | 39 | 115454.096 | 8.533 |
| mammals | Akodon_kofordi | Akodon_kofordi | LC | Unknown | tropical | cathemeral | 29.5 | 13610.994 | 5.009 |
| mammals | Akodon_lindberghi | NA | DD | Unknown | NA | NA | NA | NA | NA |
| mammals | Akodon_lutescens | Akodon_lutescens | LC | Stable | tropical | cathemeral | 20.7 | 118603.69 | 10.6 |
| mammals | Akodon_mimus | Akodon_mimus | LC | Stable | tropical | cathemeral | 24 | 57439.278 | 9.642 |
| mammals | Akodon_mollis | Akodon_mollis | LC | Stable | tropical | nocturnal | 30.4 | 265351.806 | 8.217 |
| mammals | Akodon_montensis | Akodon_montensis | LC | Decreasing | temperate | nocturnal | 44.1 | 1050409.371 | 13.599 |
| mammals | Akodon_mystax | NA | DD | Unknown | NA | NA | NA | NA | NA |
| mammals | Akodon_oenos | NA | not_available | not_available | NA | NA | NA | NA | NA |
| mammals | Akodon_orophilus | Akodon_orophilus | LC | Stable | tropical | nocturnal | 39 | 33882.521 | 4.661 |
| mammals | Akodon_paranaensis | Akodon_paranaensis | LC | Unknown | temperate | cathemeral | 40.3 | 482669.072 | 12.57 |
| mammals | Akodon_pervalens | NA | DD | Unknown | NA | NA | NA | NA | NA |
| mammals | Akodon_philipmyersi | NA | DD | Unknown | NA | NA | NA | NA | NA |
| mammals | Akodon_reigi | Akodon_reigi | LC | Stable | temperate | nocturnal | 41.3 | 65868.071 | 8.926 |
| mammals | Akodon_sanctipaulensis | NA | DD | Unknown | NA | NA | NA | NA | NA |
| mammals | Akodon_siberiae | Akodon_siberiae | NT | Decreasing | tropical | cathemeral | 34.6 | 25253.904 | 8.779 |
| mammals | Akodon_simulator | Akodon_simulator | LC | Stable | temperate | nocturnal | 42.5 | 156808.587 | 7.042 |
| mammals | Akodon_spegazzinii | Akodon_spegazzinii | LC | Stable | temperate | nocturnal | 28.6 | 219496.235 | 5.389 |
| mammals | Akodon_subfuscus | Akodon_subfuscus | LC | Stable | tropical | cathemeral | 30.4 | 171092.783 | 8.34 |
| mammals | Akodon_surdus | Akodon_surdus | VU | Unknown | tropical | cathemeral | 39 | 16434.639 | 6.894 |
| mammals | Akodon_sylvanus | Akodon_sylvanus | LC | Stable | tropical | cathemeral | 39 | 39317.405 | 7.078 |
| mammals | Akodon_toba | Akodon_toba | LC | Stable | tropical | cathemeral | 51.2 | 443710.584 | 5.145 |
| mammals | Akodon_torques | Akodon_torques | LC | Stable | tropical | cathemeral | 39 | 66658.263 | 6.019 |

| | | | | | | | | | |
|---|---|---|---|---|---|---|---|---|---|
| mammals | Akodon_varius | NA | DD | Unknown | NA | NA | NA | NA | NA |
| mammals | Alcelaphus_buselaphus | Alcelaphus_buselaphus | LC | Decreasing | tropical | diurnal | 171001.5 | 6856359.005 | 8.882 |
| mammals | Alces_alces | Alces_alces | LC | Increasing | temperate | cathemeral | 356998 | 23096272.63 | 4.301 |
| mammals | Alexandromys_evoronensis | NA | DD | Unknown | NA | NA | NA | NA | NA |
| mammals | Alexandromys_fortis | Microtus_fortis | LC | Unknown | temperate | cathemeral | 63 | 1970776.53 | 12.049 |
| mammals | Alexandromys_limnophilus | Microtus_limnophilus | LC | Unknown | temperate | cathemeral | 19.1 | 933449.663 | 6.366 |
| mammals | Alexandromys_maximowiczii | Microtus_maximowiczii | LC | Unknown | temperate | cathemeral | 41.7 | 1644298.677 | 6.965 |
| mammals | Alexandromys_middendorffii | Microtus_middendorffii | LC | Unknown | temperate | cathemeral | 56.6 | 3237474.784 | 0.87 |
| mammals | Alexandromys_mongolicus | Microtus_mongolicus | LC | Stable | temperate | cathemeral | 27.5 | 1040262.608 | 7.761 |
| mammals | Alexandromys_montebelli | Microtus_montebelli | LC | Stable | temperate | nocturnal | 39.9 | 235353.969 | 21.062 |
| mammals | Alexandromys_mujanensis | NA | DD | Unknown | NA | NA | NA | NA | NA |
| mammals | Alexandromys_oeconomus | Microtus_oeconomus | LC | Stable | temperate | cathemeral | 34.4 | 18612352.019 | 4.284 |
| mammals | Alexandromys_sachalinensis | Microtus_sachalinensis | LC | Unknown | temperate | cathemeral | 54.1 | 45125.985 | 4.286 |
| mammals | Alionycteris_paucidentata | Alionycteris_paucidentata | LC | Stable | tropical | nocturnal | 16.2 | 6417.838 | 9.278 |
| mammals | Allactaga_major | Allactaga_major | LC | Decreasing | temperate | nocturnal | 350 | 4297326.136 | 10.711 |
| mammals | Allactaga_severtzovi | Allactaga_severtzovi | LC | Unknown | temperate | nocturnal | 159 | 706422.241 | 6.727 |
| mammals | Allactodipus_bobrinskii | Allactodipus_bobrinskii | LC | Unknown | temperate | nocturnal | 70 | 275856.829 | 5.734 |
| mammals | Allenopithecus_nigroviridis | Allenopithecus_nigroviridis | LC | Decreasing | tropical | diurnal | 4750 | 526623.002 | 4.846 |
| mammals | Allocebus_trichotis | Allocebus_trichotis | EN | Decreasing | tropical | nocturnal | 77.5 | 7303.538 | 4.955 |
| mammals | Allochrocebus_lhoesti | Allochrocebus_lhoesti | VU | Decreasing | tropical | diurnal | 5375 | 216021.539 | 4.858 |
| mammals | Allochrocebus_preussi | Allochrocebus_preussi | EN | Decreasing | tropical | diurnal | 5250 | 17526.762 | 9.06 |
| mammals | Allochrocebus_solatus | Allochrocebus_solatus | NT | Unknown | tropical | diurnal | 6500 | 18941.854 | 6.086 |
| mammals | Allocricetulus_curtatus | Allocricetulus_curtatus | LC | Unknown | temperate | nocturnal | 47.8 | 1332299.367 | 5.455 |
| mammals | Allocricetulus_eversmanni | Allocricetulus_eversmanni | LC | Unknown | temperate | nocturnal | 115.6 | 1776745.67 | 8.489 |
| mammals | Alouatta_arctoidea | Alouatta_arctoidea | LC | Unknown | tropical | diurnal | 5390 | 406536.152 | 9.16 |
| mammals | Alouatta_belzebul | Alouatta_belzebul | VU | Decreasing | tropical | diurnal | 6400 | 864468.57 | 4.296 |
| mammals | Alouatta_caraya | Alouatta_caraya | NT | Decreasing | tropical | diurnal | 5862.5 | 3064214.8 | 8.544 |
| mammals | Alouatta_discolor | Alouatta_discolor | VU | Decreasing | tropical | diurnal | 6262.8 | 375751.384 | 1.858 |
| mammals | Alouatta_guariba | Alouatta_guariba | VU | Decreasing | tropical | diurnal | 5589.2 | 940355.902 | 12.805 |
| mammals | Alouatta_macconnelli | Alouatta_macconnelli | LC | Stable | tropical | diurnal | 7554.8 | 1755234.563 | 1.378 |
| mammals | Alouatta_nigerrima | Alouatta_nigerrima | LC | Decreasing | tropical | diurnal | 7662 | 236116.233 | 1.224 |
| mammals | Alouatta_palliata | Alouatta_palliata | VU | Decreasing | tropical | diurnal | 7275 | 549356.886 | 10.606 |
| mammals | Alouatta_pigra | Alouatta_pigra | EN | Decreasing | tropical | diurnal | 7000 | 246704.216 | 7.736 |
| mammals | Alouatta_sara | Alouatta_sara | NT | Decreasing | tropical | diurnal | 6611 | 341403.365 | 5.471 |

| | | | | | | | | | |
|---|---|---|---|---|---|---|---|---|---|
| mammals | Alticola_albicauda | NA | DD | Unknown | NA | NA | NA | NA | NA |
| mammals | Alticola_argentatus | Alticola_argentatus | LC | Unknown | temperate | diurnal | 37.7 | 1380073.109 | 7.404 |
| mammals | Alticola_barakshin | Alticola_barakshin | LC | Unknown | temperate | nocturnal | 35 | 225997.116 | 4.06 |
| mammals | Alticola_lemminus | Alticola_lemminus | LC | Unknown | temperate | cathemeral | 15.5 | 2068169.332 | 0.888 |
| mammals | Alticola_macrotis | Alticola_macrotis | LC | Unknown | temperate | cathemeral | 36 | 379222.679 | 2.124 |
| mammals | Alticola_montosus | Alticola_montosa | VU | Decreasing | temperate | diurnal | 19.7 | 19533.933 | 7.836 |
| mammals | Alticola_olchonensis | Alticola_olchonensis | EN | Unknown | temperate | cathemeral | 33.5 | 1348.368 | 7.495 |
| mammals | Alticola_roylei | Alticola_roylei | NT | Decreasing | temperate | diurnal | 37 | 38435.414 | 7.445 |
| mammals | Alticola_semicanus | Alticola_semicanus | LC | Unknown | temperate | nocturnal | 53.2 | 671213.975 | 6.433 |
| mammals | Alticola_stoliczkanus | Alticola_stoliczkanus | LC | Unknown | temperate | nocturnal | 27.5 | 2306359.627 | 3.252 |
| mammals | Alticola_strelzovi | Alticola_strelzowi | LC | Unknown | temperate | cathemeral | 50 | 441285.295 | 6.031 |
| mammals | Alticola_tuvinicus | Alticola_tuvinicus | LC | Unknown | temperate | cathemeral | 41 | 134541.195 | 3.535 |
| mammals | Ametrida_centurio | Ametrida_centurio | LC | Unknown | tropical | nocturnal | 10.3 | 5766469.12 | 3.586 |
| mammals | Ammodillus_imbellis | Ammodillus_imbellis | DD | Stable | tropical | nocturnal | 50 | 210233.71 | 8.049 |
| mammals | Ammodorcas_clarkei | Ammodorcas_clarkei | VU | Decreasing | tropical | diurnal | 28049.8 | 93566.481 | 6.593 |
| mammals | Ammospermophilus_harrisii | Ammospermophilus_harrisii | LC | Unknown | temperate | diurnal | 122 | 269488.41 | 8.326 |
| mammals | Ammospermophilus_interpres | Ammospermophilus_interpres | LC | Unknown | temperate | diurnal | 110.3 | 312449.218 | 7.745 |
| mammals | Ammospermophilus_leucurus | Ammospermophilus_leucurus | LC | Stable | temperate | diurnal | 120.5 | 990844.097 | 7.978 |
| mammals | Ammospermophilus_nelsoni | Ammospermophilus_nelsoni | EN | Decreasing | temperate | diurnal | 160.4 | 16430.668 | 14.867 |
| mammals | Ammotragus_lervia | Ammotragus_lervia | VU | Decreasing | temperate | diurnal | 48000 | 1167864.69 | 2.319 |
| mammals | Amorphochilus_schnablii | Amorphochilus_schnablii | VU | Stable | tropical | nocturnal | 3.3 | 178408.336 | 7.792 |
| mammals | Anathana_ellioti | Anathana_ellioti | LC | Decreasing | tropical | diurnal | 160 | 1366073.824 | 19.004 |
| mammals | Andalgalomys_olrogi | Andalgalomys_olrogi | LC | Unknown | temperate | nocturnal | 32.9 | 57114.934 | 3.206 |
| mammals | Andalgalomys_pearsoni | Andalgalomys_pearsoni | LC | Stable | tropical | nocturnal | 25.4 | 139008.134 | 3.88 |
| mammals | Andinomys_edax | Andinomys_edax | LC | Stable | tropical | nocturnal | 69.7 | 228613.141 | 7.722 |
| mammals | Anisomys_imitator | Anisomys_imitator | LC | Stable | tropical | nocturnal | 510.5 | 181682.711 | 4.712 |
| mammals | Anomalurus_beecrofti | Anomalurus_beecrofti | LC | Unknown | tropical | nocturnal | 389 | 3245229.516 | 8.26 |
| mammals | Anomalurus_derbianus | Anomalurus_derbianus | LC | Unknown | tropical | nocturnal | 650 | 5752727.174 | 8.508 |
| mammals | Anomalurus_pelii | Anomalurus_pelii | LC | Decreasing | tropical | nocturnal | 1770 | 104894.753 | 16.989 |
| mammals | Anomalurus_pusillus | Anomalurus_pusillus | LC | Unknown | tropical | nocturnal | 250 | 573139.914 | 6.371 |
| mammals | Anonymomys_mindorensis | NA | DD | Unknown | NA | NA | NA | NA | NA |
| mammals | Anotomys_leander | Anotomys_leander | EN | Decreasing | tropical | nocturnal | 66.4 | 12508.223 | 13.009 |
| mammals | Anoura_caudifer | Anoura_caudifer | LC | Unknown | tropical | nocturnal | 10.1 | 5940468.235 | 6.463 |
| mammals | Anoura_cultrata | Anoura_cultrata | LC | Decreasing | tropical | nocturnal | 16.9 | 757500.042 | 9.025 |

| | | | | | | | | | |
|---|---|---|---|---|---|---|---|---|---|
| mammals | Anoura_fistulata | NA | DD | Unknown | NA | NA | NA | NA | NA |
| mammals | Anoura_geoffroyi | Anoura_geoffroyi | LC | Stable | tropical | nocturnal | 14.6 | 8203646.832 | 8.193 |
| mammals | Anoura_latidens | Anoura_latidens | LC | Stable | tropical | nocturnal | 14.7 | 710044.889 | 7.294 |
| mammals | Anoura_luismanueli | Anoura_luismanueli | LC | Stable | tropical | nocturnal | 8.5 | 80298.605 | 11.516 |
| mammals | Anourosorex_assamensis | Anourosorex_assamensis | LC | Unknown | temperate | nocturnal | 25.2 | 17603.52 | 10.139 |
| mammals | Anourosorex_squamipes | Anourosorex_squamipes | LC | Unknown | temperate | nocturnal | 20 | 1189018.887 | 11.319 |
| mammals | Anourosorex_yamashinai | Anourosorex_yamashinai | LC | Unknown | temperate | nocturnal | 11.4 | 34157.023 | 20.929 |
| mammals | Antechinomys_laniger | Antechinomys_laniger | LC | Decreasing | temperate | nocturnal | 25 | 2438423.419 | 2.877 |
| mammals | Antechinomys_longicaudatus | Sminthopsis_longicaudata | LC | Unknown | temperate | nocturnal | 18 | 747801.205 | 2.956 |
| mammals | Antechinus_adustus | Antechinus_adustus | LC | Decreasing | tropical | diurnal | 28.2 | 4920.268 | 3.462 |
| mammals | Antechinus_agilis | Antechinus_agilis | LC | Unknown | temperate | nocturnal | 24 | 163654.965 | 9.831 |
| mammals | Antechinus_bellus | Antechinus_bellus | VU | Decreasing | tropical | nocturnal | 44.5 | 20577.834 | 2.44 |
| mammals | Antechinus_flavipes | Antechinus_flavipes | LC | Stable | temperate | nocturnal | 45 | 1008801.549 | 8.403 |
| mammals | Antechinus_godmani | Antechinus_godmani | LC | Stable | tropical | diurnal | 76.5 | 3599.522 | 3.71 |
| mammals | Antechinus_leo | Antechinus_leo | LC | Stable | tropical | nocturnal | 74 | 4393.221 | 1.296 |
| mammals | Antechinus_stuartii | Antechinus_stuartii | LC | Stable | temperate | nocturnal | 27.5 | 118336.847 | 8.32 |
| mammals | Antechinus_subtropicus | Antechinus_subtropicus | LC | Decreasing | temperate | nocturnal | 33.1 | 9618.052 | 13.608 |
| mammals | Antechinus_swainsonii | Antechinus_swainsonii | LC | Unknown | temperate | cathemeral | 53 | 314283.027 | 7.903 |
| mammals | Anteliomys_chinensis | Eothenomys_chinensis | LC | Unknown | temperate | cathemeral | 45.3 | 8999.065 | 8.471 |
| mammals | Anteliomys_custos | Eothenomys_custos | LC | Unknown | temperate | cathemeral | 30.8 | 115771.46 | 8.635 |
| mammals | Anteliomys_olitor | Eothenomys_olitor | LC | Unknown | temperate | cathemeral | 22 | 177906.741 | 12.367 |
| mammals | Anteliomys_proditor | NA | DD | Unknown | NA | NA | NA | NA | NA |
| mammals | Anteliomys_wardi | Eothenomys_wardi | NT | Unknown | temperate | cathemeral | 30.5 | 8331.727 | 8.005 |
| mammals | Anthops_ornatus | Anthops_ornatus | VU | Decreasing | tropical | nocturnal | 8 | 21294.852 | 4.929 |
| mammals | Antidorcas_marsupialis | Antidorcas_marsupialis | LC | Increasing | tropical | cathemeral | 31500 | 1950012.855 | 6.523 |
| mammals | Antilocapra_americana | Antilocapra_americana | LC | Stable | temperate | cathemeral | 46082.9 | 2007736.603 | 10.985 |
| mammals | Antilope_cervicapra | Antilope_cervicapra | LC | Unknown | tropical | diurnal | 37500 | 1754050.908 | 19.144 |
| mammals | Antrozous_pallidus | Antrozous_pallidus | LC | Stable | temperate | nocturnal | 20.8 | 4367598.171 | 9.792 |
| mammals | Aotus_azarae | Aotus_azarae | LC | Decreasing | tropical | cathemeral | 963 | 3156514.711 | 4.519 |
| mammals | Aotus_brumbacki | Aotus_brumbacki | VU | Decreasing | tropical | nocturnal | 800 | 123963.783 | 7.729 |
| mammals | Aotus_griseimembra | Aotus_griseimembra | VU | Decreasing | tropical | nocturnal | 966 | 234087.454 | 14.33 |
| mammals | Aotus_jorgehernandezi | NA | DD | Unknown | NA | NA | NA | NA | NA |
| mammals | Aotus_lemurinus | Aotus_lemurinus | VU | Decreasing | tropical | nocturnal | 873 | 177191.552 | 10.749 |
| mammals | Aotus_miconax | Aotus_miconax | EN | Decreasing | tropical | nocturnal | 800 | 27618.651 | 3.706 |

| | | | | | | | | | |
|---|---|---|---|---|---|---|---|---|---|
| mammals | Aotus_nancymai | Aotus_nancymaae | VU | Decreasing | tropical | nocturnal | 788 | 509597.724 | 1.441 |
| mammals | Aotus_nigriceps | Aotus_nigriceps | LC | Unknown | tropical | nocturnal | 1060 | 1569879.018 | 1.67 |
| mammals | Aotus_trivirgatus | Aotus_trivirgatus | LC | Decreasing | tropical | cathemeral | 900 | 752044.959 | 1.11 |
| mammals | Aotus_vociferans | Aotus_vociferans | LC | Unknown | tropical | nocturnal | 873 | 1054332.874 | 1.606 |
| mammals | Aotus_zonalis | Aotus_zonalis | NT | Unknown | tropical | nocturnal | 902.5 | 131878.681 | 6.795 |
| mammals | Aplodontia_rufa | Aplodontia_rufa | LC | Stable | temperate | cathemeral | 1004 | 245471.87 | 10.049 |
| mammals | Apodemus_agrarius | Apodemus_agrarius | LC | Stable | temperate | diurnal | 23.3 | 11531272.514 | 13.253 |
| mammals | Apodemus_argenteus | Apodemus_argenteus | LC | Stable | temperate | nocturnal | 21 | 366528.075 | 19.809 |
| mammals | Apodemus_chevrieri | Apodemus_chevrieri | LC | Unknown | temperate | diurnal | 32.9 | 720401.613 | 12.064 |
| mammals | Apodemus_draco | Apodemus_draco | LC | Stable | temperate | nocturnal | 22.3 | 1798566.219 | 12.962 |
| mammals | Apodemus_epimelas | Apodemus_epimelas | LC | Stable | temperate | nocturnal | 42 | 174107.252 | 16.921 |
| mammals | Apodemus_hyrcanicus | Apodemus_hyrcanicus | NT | Decreasing | temperate | nocturnal | 28.5 | 48562.597 | 18.38 |
| mammals | Apodemus_latronum | Apodemus_latronum | LC | Unknown | temperate | nocturnal | 35.2 | 536889.579 | 7.882 |
| mammals | Apodemus_mystacinus | Apodemus_mystacinus | LC | Stable | temperate | nocturnal | 45.6 | 804373.39 | 15.247 |
| mammals | Apodemus_peninsulae | Apodemus_peninsulae | LC | Stable | temperate | nocturnal | 32.9 | 8904776.966 | 7.945 |
| mammals | Apodemus_ponticus | NA | LC | Unknown | NA | NA | NA | NA | NA |
| mammals | Apodemus_rusiges | Apodemus_rusiges | LC | Unknown | temperate | nocturnal | 37 | 112760.373 | 9.047 |
| mammals | Apodemus_semotus | Apodemus_semotus | LC | Unknown | temperate | nocturnal | 28.8 | 33914.895 | 21.053 |
| mammals | Apodemus_speciosus | Apodemus_speciosus | LC | Stable | temperate | nocturnal | 33.8 | 368742.872 | 19.748 |
| mammals | Apodemus_witherbyi | Apodemus_witherbyi | LC | Unknown | temperate | nocturnal | 30 | 1581288.62 | 14.644 |
| mammals | Apomys_abrae | Apomys_abrae | LC | Unknown | tropical | nocturnal | 56.6 | 12463.631 | 9.097 |
| mammals | Apomys_datae | Apomys_datae | LC | Unknown | tropical | nocturnal | 106 | 11591.319 | 8.95 |
| mammals | Apomys_gracilirostris | Apomys_gracilirostris | LC | Unknown | tropical | nocturnal | 88.1 | 571.34 | 5.622 |
| mammals | Apomys_hylocoetes | Apomys_hylocetes | LC | Stable | tropical | nocturnal | 45.1 | 4399.293 | 9.558 |
| mammals | Apomys_insignis | Apomys_insignis | LC | Decreasing | tropical | nocturnal | 29.9 | 94959.732 | 12.024 |
| mammals | Apomys_littoralis | NA | DD | Unknown | NA | NA | NA | NA | NA |
| mammals | Apomys_microdon | Apomys_microdon | LC | Unknown | tropical | nocturnal | 35.1 | 61956.394 | 10.282 |
| mammals | Apomys_musculus | Apomys_musculus | LC | Stable | tropical | nocturnal | 21.2 | 47774.032 | 10.79 |
| mammals | Apomys_sacobianus | Apomys_sacobianus | LC | Unknown | tropical | nocturnal | 82.4 | 951.477 | 7.713 |
| mammals | Aproteles_bulmerae | Aproteles_bulmerae | CR | Decreasing | tropical | nocturnal | 600 | 48.155 | 3.225 |
| mammals | Arabitragus_jayakari | Arabitragus_jayakari | EN | Decreasing | temperate | cathemeral | 23000 | 28012.017 | 8.975 |
| mammals | Arborimus_albipes | Arborimus_albipes | LC | Unknown | temperate | nocturnal | 23 | 77753.369 | 11.488 |
| mammals | Arborimus_longicaudus | Arborimus_longicaudus | NT | Decreasing | temperate | nocturnal | 21.8 | 54110.767 | 12.091 |
| mammals | Arborimus_pomo | Arborimus_pomo | NT | Decreasing | temperate | nocturnal | 32.5 | 22003.978 | 9.866 |

| | | | | | | | | | |
|---|---|---|---|---|---|---|---|---|---|
| mammals | Archboldomys_luzonensis | Archboldomys_luzonensis | VU | Stable | tropical | diurnal | 35.4 | 22.211 | 7.741 |
| mammals | Arctictis_binturong | Arctictis_binturong | VU | Decreasing | tropical | cathemeral | 9875 | 3049767.958 | 9.694 |
| mammals | Arctocebus_aureus | Arctocebus_aureus | LC | Unknown | tropical | nocturnal | 235 | 667484.142 | 5.537 |
| mammals | Arctocebus_calabarensis | Arctocebus_calabarensis | NT | Decreasing | tropical | nocturnal | 210 | 148440.251 | 13.752 |
| mammals | Arctogalidia_trivirgata | Arctogalidia_trivirgata | LC | Decreasing | tropical | nocturnal | 2250 | 2539704.359 | 8.431 |
| mammals | Arctonyx_albogularis | Arctonyx_albogularis | LC | Decreasing | temperate | cathemeral | 9000 | 4210609.604 | 14.738 |
| mammals | Arctonyx_collaris | Arctonyx_collaris | VU | Decreasing | tropical | cathemeral | 10.5 | 1878579.031 | 11.081 |
| mammals | Arctonyx_hoevenii | Arctonyx_hoevenii | LC | Stable | tropical | cathemeral | 7000 | 98102.533 | 7.765 |
| mammals | Ardops_nichollsi | Ardops_nichollsi | LC | Unknown | tropical | nocturnal | 17.2 | 4742.707 | 21.748 |
| mammals | Arielulus_circumdatus | Arielulus_circumdatus | LC | Unknown | temperate | nocturnal | 10.4 | 170888.042 | 12.83 |
| mammals | Arielulus_cuprosus | Arielulus_cuprosus | VU | Unknown | tropical | nocturnal | 5.5 | 2630.005 | 7.341 |
| mammals | Arielulus_societatis | Arielulus_societatis | LC | Decreasing | tropical | nocturnal | 4.9 | 92389.411 | 8.621 |
| mammals | Ariteus_flavescens | Ariteus_flavescens | LC | Unknown | tropical | nocturnal | 11.9 | 11021.007 | 19.74 |
| mammals | Artibeus_amplus | Artibeus_amplus | LC | Unknown | tropical | nocturnal | 60 | 1459798.334 | 7.349 |
| mammals | Artibeus_concolor | Artibeus_concolor | LC | Stable | tropical | nocturnal | 19.4 | 6437731.246 | 2.767 |
| mammals | Artibeus_fimbriatus | Artibeus_fimbriatus | LC | Stable | tropical | nocturnal | 63.9 | 1480430.38 | 12.54 |
| mammals | Artibeus_fraterculus | Artibeus_fraterculus | LC | Decreasing | tropical | nocturnal | 39.5 | 302070.729 | 9.251 |
| mammals | Artibeus_hirsutus | Artibeus_hirsutus | LC | Stable | tropical | nocturnal | 38.7 | 228810.602 | 8.157 |
| mammals | Artibeus_inopinatus | NA | DD | Unknown | NA | NA | NA | NA | NA |
| mammals | Artibeus_jamaicensis | Artibeus_jamaicensis | LC | Stable | tropical | nocturnal | 42.2 | 1559896.27 | 11.323 |
| mammals | Artibeus_lituratus | Artibeus_lituratus | LC | Stable | tropical | nocturnal | 60.5 | 14235952.081 | 6.143 |
| mammals | Artibeus_obscurus | Artibeus_obscurus | LC | Unknown | tropical | nocturnal | 35.2 | 11122956.74 | 4.898 |
| mammals | Artibeus_planirostris | Artibeus_planirostris | LC | Stable | tropical | nocturnal | 47 | 7906983.194 | 4.591 |
| mammals | Artibeus_schwartzi | NA | DD | Unknown | NA | NA | NA | NA | NA |
| mammals | Arvicanthis_abyssinicus | Arvicanthis_abyssinicus | LC | Stable | tropical | diurnal | 77.1 | 296092.685 | 13.432 |
| mammals | Arvicanthis_ansorgei | Arvicanthis_ansorgei | LC | Unknown | tropical | diurnal | 124.5 | 651274.694 | 12.793 |
| mammals | Arvicanthis_blicki | Arvicanthis_blicki | NT | Unknown | tropical | diurnal | 128 | 17893.747 | 13.55 |
| mammals | Arvicanthis_nairobae | Arvicanthis_nairobae | LC | Stable | tropical | diurnal | 104 | 151138.937 | 12.274 |
| mammals | Arvicanthis_neumanni | Arvicanthis_neumanni | LC | Unknown | tropical | diurnal | 60 | 1218452.16 | 8.944 |
| mammals | Arvicanthis_niloticus | Arvicanthis_niloticus | LC | Unknown | tropical | diurnal | 106.9 | 6185272.895 | 10.919 |
| mammals | Arvicanthis_rufinus | Arvicanthis_rufinus | LC | Unknown | tropical | diurnal | 144.1 | 1154420.169 | 13.684 |
| mammals | Arvicola_amphibius | Arvicola_amphibius | LC | Unknown | temperate | cathemeral | 225 | 15657066.94 | 9.973 |
| mammals | Arvicola_sapidus | Arvicola_sapidus | VU | Decreasing | temperate | cathemeral | 220 | 882587.177 | 19.845 |
| mammals | Asellia_patrizii | Asellia_patrizii | LC | Unknown | tropical | nocturnal | 6 | 227346.991 | 11.039 |

| | | | | | | | | | |
|---|---|---|---|---|---|---|---|---|---|
| mammals | Asellia_tridens | Asellia_tridens | LC | Stable | tropical | nocturnal | 11.9 | 16925970.618 | 4.585 |
| mammals | Aselliscus_stoliczkanus | Aselliscus_stoliczkanus | LC | Decreasing | tropical | nocturnal | 4.9 | 1218727.088 | 11.351 |
| mammals | Aselliscus_tricuspidatus | Aselliscus_tricuspidatus | LC | Stable | tropical | nocturnal | 4.1 | 274415.136 | 4.706 |
| mammals | Atelerix_albiventris | Atelerix_albiventris | LC | Stable | tropical | nocturnal | 600 | 7028364.294 | 11.181 |
| mammals | Atelerix_algirus | Atelerix_algirus | LC | Unknown | temperate | nocturnal | 1250 | 734429.776 | 15.276 |
| mammals | Atelerix_frontalis | Atelerix_frontalis | LC | Stable | temperate | nocturnal | 309.5 | 1047880.248 | 9.251 |
| mammals | Atelerix_sclateri | Atelerix_sclateri | LC | Unknown | tropical | nocturnal | 436.5 | 309001.284 | 9.186 |
| mammals | Ateles_belzebuth | Ateles_belzebuth | EN | Decreasing | tropical | diurnal | 5000 | 815705.065 | 1.668 |
| mammals | Ateles_chamek | Ateles_chamek | EN | Decreasing | tropical | diurnal | 6000 | 2602958.433 | 2.319 |
| mammals | Ateles_fusciceps | Ateles_fusciceps | EN | Decreasing | tropical | diurnal | 9100 | 212235.693 | 8.913 |
| mammals | Ateles_geoffroyi | Ateles_geoffroyi | EN | Decreasing | tropical | diurnal | 5284.9 | 901722.852 | 10.645 |
| mammals | Ateles_hybridus | Ateles_hybridus | CR | Decreasing | tropical | diurnal | 8675 | 165736.604 | 13.009 |
| mammals | Ateles_marginatus | Ateles_marginatus | EN | Decreasing | tropical | diurnal | 6000 | 524099.558 | 2.102 |
| mammals | Ateles_paniscus | Ateles_paniscus | VU | Decreasing | tropical | diurnal | 7900.1 | 918126.827 | 1.203 |
| mammals | Atelocynus_microtis | Atelocynus_microtis | NT | Decreasing | tropical | cathemeral | 7750 | 2852893.505 | 1.494 |
| mammals | Atherurus_africanus | Atherurus_africanus | LC | Unknown | tropical | nocturnal | 1934 | 3544244.62 | 9.153 |
| mammals | Atherurus_macrourus | Atherurus_macrourus | LC | Decreasing | tropical | nocturnal | 2000 | 2672201.971 | 12.856 |
| mammals | Atilax_paludinosus | Atilax_paludinosus | LC | Decreasing | tropical | nocturnal | 3300 | 14049574.29 | 9.5 |
| mammals | Atlantoxerus_getulus | Atlantoxerus_getulus | LC | Stable | temperate | diurnal | 251 | 303645.313 | 8.352 |
| mammals | Atopogale_cubana | Solenodon_cubanus | EN | Decreasing | tropical | nocturnal | 650 | 1996.729 | 10.278 |
| mammals | Auliscomys_boliviensis | Auliscomys_boliviensis | LC | Stable | tropical | diurnal | 65.7 | 210614.075 | 8.859 |
| mammals | Auliscomys_pictus | Auliscomys_pictus | LC | Stable | tropical | cathemeral | 54.7 | 309598.291 | 8.635 |
| mammals | Auliscomys_sublimis | Auliscomys_sublimis | LC | Stable | tropical | nocturnal | 38 | 393512.532 | 6.734 |
| mammals | Austronomus_australis | Austronomus_australis | LC | Decreasing | temperate | nocturnal | 31.8 | 6025987.525 | 4.57 |
| mammals | Austronomus_kuboriensis | Austronomus_kuboriensis | LC | Stable | tropical | nocturnal | 25.9 | 107966.743 | 5.374 |
| mammals | Avahi_betsileo | Avahi_betsileo | EN | Decreasing | tropical | nocturnal | 1050 | 970.994 | 7.717 |
| mammals | Avahi_cleesei | Avahi_cleesei | CR | Decreasing | tropical | nocturnal | 830 | 459.838 | 4.978 |
| mammals | Avahi_laniger | Avahi_laniger | VU | Decreasing | tropical | nocturnal | 1300 | 51921.14 | 7.252 |
| mammals | Avahi_meridionalis | Avahi_meridionalis | EN | Decreasing | temperate | nocturnal | 1060 | 1559.768 | 6.051 |
| mammals | Avahi_mooreorum | Avahi_mooreorum | EN | Decreasing | tropical | nocturnal | 920 | 1992.272 | 3.647 |
| mammals | Avahi_occidentalis | Avahi_occidentalis | VU | Decreasing | tropical | nocturnal | 850 | 3050.161 | 8.771 |
| mammals | Avahi_peyrierasi | Avahi_peyrierasi | VU | Decreasing | tropical | nocturnal | 920 | 4374.325 | 6.721 |
| mammals | Avahi_ramanantsoavani | Avahi_ramanantsoavanai | VU | Decreasing | temperate | nocturnal | 980 | 5832.064 | 5.445 |
| mammals | Avahi_unicolor | Avahi_unicolor | CR | Decreasing | tropical | nocturnal | 850 | 581.333 | 4.507 |

| | | | | | | | | | |
|---|---|---|---|---|---|---|---|---|---|
| mammals | Axis_calamianensis | Axis_calamianensis | EN | Decreasing | tropical | diurnal | 40500 | 1345.322 | 12.081 |
| mammals | Axis_kuhlii | NA | CR | Stable | NA | NA | NA | NA | NA |
| mammals | Babyrousa_babyrussa | Babyrousa_babyrussa | VU | Decreasing | tropical | diurnal | 100000 | 13191.945 | 6.329 |
| mammals | Babyrousa_celebensis | Babyrousa_celebensis | VU | Decreasing | tropical | diurnal | 84327.5 | 135977.785 | 7.486 |
| mammals | Babyrousa_togeanensis | Babyrousa_togeanensis | EN | Decreasing | tropical | diurnal | 113762.8 | 533.084 | 6.448 |
| mammals | Baeodon_alleni | Rhogeessa_alleni | LC | Stable | tropical | nocturnal | 7.2 | 261934.25 | 13.993 |
| mammals | Baeodon_gracilis | Rhogeessa_gracilis | LC | Unknown | tropical | nocturnal | 6.1 | 178177.406 | 8.029 |
| mammals | Baiomys_musculus | Baiomys_musculus | LC | Stable | tropical | diurnal | 9 | 450251.276 | 11.431 |
| mammals | Baiomys_taylori | Baiomys_taylori | LC | Stable | temperate | nocturnal | 8 | 1275181.162 | 12.516 |
| mammals | Baiyankamys_habbema | NA | DD | Unknown | NA | NA | NA | NA | NA |
| mammals | Baiyankamys_shawmayeri | Baiyankamys_shawmayeri | LC | Stable | tropical | nocturnal | 78.4 | 65462.775 | 6.362 |
| mammals | Balantiopteryx_infusca | Balantiopteryx_infusca | VU | Decreasing | tropical | nocturnal | 4.4 | 2841.698 | 14.357 |
| mammals | Balantiopteryx_io | Balantiopteryx_io | VU | Decreasing | tropical | nocturnal | 4.3 | 170568.449 | 10.394 |
| mammals | Balantiopteryx_plicata | Balantiopteryx_plicata | LC | Unknown | tropical | nocturnal | 6.4 | 825040.693 | 12.324 |
| mammals | Balionycteris_maculata | Balionycteris_maculata | LC | Unknown | tropical | nocturnal | 13.4 | 439669.601 | 5.166 |
| mammals | Bandicota_bengalensis | Bandicota_bengalensis | LC | Increasing | tropical | nocturnal | 188.5 | 4165486.078 | 16.969 |
| mammals | Barbastella_barbastellus | Barbastella_barbastellus | NT | Decreasing | temperate | nocturnal | 8 | 3811256.039 | 20.062 |
| mammals | Barbastella_leucomelas | Barbastella_leucomelas | LC | Unknown | temperate | nocturnal | 15.1 | 3928085.941 | 10.564 |
| mammals | Bassaricyon_alleni | Bassaricyon_alleni | LC | Decreasing | tropical | nocturnal | 1235 | 4449618.019 | 2.138 |
| mammals | Bassaricyon_gabbii | Bassaricyon_gabbii | LC | Decreasing | tropical | nocturnal | 1250 | 156420.124 | 9.926 |
| mammals | Bassaricyon_medius | Bassaricyon_medius | LC | Decreasing | tropical | nocturnal | 1076 | 181889.986 | 9.297 |
| mammals | Bassaricyon_neblina | Bassaricyon_neblina | NT | Decreasing | tropical | nocturnal | 872 | 120399.355 | 12.378 |
| mammals | Bassariscus_astutus | Bassariscus_astutus | LC | Unknown | temperate | nocturnal | 1129.5 | 3903858.562 | 10.45 |
| mammals | Bassariscus_sumichrasti | Bassariscus_sumichrasti | LC | Unknown | tropical | nocturnal | 900 | 622168.546 | 10.139 |
| mammals | Batomys_dentatus | NA | DD | Unknown | NA | NA | NA | NA | NA |
| mammals | Batomys_grantii | Batomys_granti | LC | Stable | tropical | nocturnal | 177.8 | 5211.943 | 8.896 |
| mammals | Batomys_russatus | Batomys_russatus | EN | Decreasing | tropical | nocturnal | 115 | 826.357 | 11.016 |
| mammals | Batomys_salomonseni | Batomys_salomonseni | LC | Stable | tropical | nocturnal | 187 | 16015.509 | 14.31 |
| mammals | Bauerus_dubiaquercus | Bauerus_dubiaquercus | NT | Unknown | tropical | nocturnal | 22.9 | 265235.241 | 9.287 |
| mammals | Bdeogale_crassicauda | Bdeogale_crassicauda | LC | Unknown | tropical | nocturnal | 1550 | 1497508.737 | 9.873 |
| mammals | Bdeogale_jacksoni | Bdeogale_jacksoni | NT | Decreasing | tropical | nocturnal | 2500 | 63953.373 | 17.736 |
| mammals | Bdeogale_nigripes | Bdeogale_nigripes | LC | Decreasing | tropical | nocturnal | 2500 | 2110590.779 | 5.832 |
| mammals | Bdeogale_omnivora | Bdeogale_omnivora | VU | Decreasing | tropical | nocturnal | 1750 | 28492.999 | 11.045 |
| mammals | Beamys_hindei | Beamys_hindei | LC | Decreasing | tropical | nocturnal | 75.8 | 337356.597 | 11.471 |

| | | | | | | | | | |
|---|---|---|---|---|---|---|---|---|---|
| mammals | Belomys_pearsonii | NA | DD | Unknown | NA | NA | NA | NA | NA |
| mammals | Berylmys_berdmorei | Berylmys_berdmorei | LC | Stable | tropical | nocturnal | 235 | 660824.557 | 12.548 |
| mammals | Berylmys_bowersii | Berylmys_bowersi | LC | Decreasing | tropical | nocturnal | 300 | 2558279.084 | 12.332 |
| mammals | Berylmys_manipulus | NA | DD | Unknown | NA | NA | NA | NA | NA |
| mammals | Bettongia_gaimardi | Bettongia_gaimardi | NT | Unknown | temperate | nocturnal | 1660 | 29089.488 | 8.068 |
| mammals | Bettongia_lesueur | Bettongia_lesueur | NT | Increasing | temperate | nocturnal | 1500 | 979.816 | 7.563 |
| mammals | Bettongia_penicillata | Bettongia_penicillata | CR | Decreasing | temperate | nocturnal | 1300 | 4058.273 | 7.775 |
| mammals | Bettongia_tropica | Bettongia_tropica | EN | Decreasing | tropical | nocturnal | 1200 | 396.783 | 4.255 |
| mammals | Bibimys_chacoensis | Bibimys_chacoensis | LC | Stable | temperate | nocturnal | 28 | 27945.906 | 11.932 |
| mammals | Bibimys_labiosus | Bibimys_labiosus | LC | Decreasing | temperate | nocturnal | 29 | 361277.465 | 13.77 |
| mammals | Bibimys_torresi | Bibimys_torresi | VU | Decreasing | temperate | nocturnal | 28 | 12372.235 | 12.292 |
| mammals | Blarina_brevicauda | Blarina_brevicauda | LC | Stable | temperate | nocturnal | 28 | 4106919.939 | 14.527 |
| mammals | Blarina_carolinensis | Blarina_carolinensis | LC | Stable | temperate | nocturnal | 13.5 | 866209.842 | 16.071 |
| mammals | Blarinella_quadraticauda | Blarinella_quadraticauda | NT | Unknown | temperate | nocturnal | 12.5 | 17615.231 | 6.858 |
| mammals | Blarinella_wardi | Blarinella_wardi | LC | Unknown | temperate | nocturnal | 7.7 | 114891.806 | 10.047 |
| mammals | Blastocerus_dichotomus | Blastocerus_dichotomus | VU | Decreasing | tropical | nocturnal | 86666.2 | 4234152.83 | 8.219 |
| mammals | Boneia_bidens | Rousettus_bidens | VU | Decreasing | tropical | nocturnal | 171 | 154150.117 | 8.013 |
| mammals | Bos_bison | Bison_bison | NT | Stable | temperate | cathemeral | 579255.3 | 140673.076 | 1.48 |
| mammals | Bos_bonasus | Bison_bonasus | NT | Increasing | temperate | cathemeral | 500000 | 24303.162 | 10.505 |
| mammals | Bos_gaurus | Bos_gaurus | VU | Decreasing | tropical | cathemeral | 825000 | 1370086.781 | 10.188 |
| mammals | Bos_javanicus | Bos_javanicus | EN | Decreasing | tropical | cathemeral | 700000 | 183622.811 | 7.208 |
| mammals | Bos_mutus | Bos_mutus | VU | Decreasing | temperate | cathemeral | 546250 | 469488.494 | 0.522 |
| mammals | Brachiones_przewalskii | Brachiones_przewalskii | LC | Unknown | temperate | nocturnal | 22 | 862411.507 | 2.602 |
| mammals | Brachyphylla_cavernarum | Brachyphylla_cavernarum | LC | Unknown | tropical | nocturnal | 46.2 | 14907.704 | 25.517 |
| mammals | Brachyphylla_nana | Brachyphylla_nana | LC | Stable | tropical | nocturnal | 35.8 | 195038.675 | 16.234 |
| mammals | Brachytarsomys_albicauda | Brachytarsomys_albicauda | LC | Unknown | tropical | nocturnal | 200 | 64739.799 | 8.146 |
| mammals | Brachytarsomys_villosus | Brachytarsomys_villosa | VU | Unknown | tropical | nocturnal | 220 | 10127.468 | 4.818 |
| mammals | Brachyteles_arachnoides | Brachyteles_arachnoides | CR | Decreasing | temperate | diurnal | 13499.9 | 73233.964 | 14.488 |
| mammals | Brachyteles_hypoxanthus | Brachyteles_hypoxanthus | CR | Decreasing | tropical | diurnal | 8865 | 96658.147 | 12.506 |
| mammals | Brachyuromys_betsileoensis | Brachyuromys_betsileoensis | LC | Unknown | tropical | cathemeral | 92.5 | 42770.178 | 9.386 |
| mammals | Brachyuromys_ramirohitra | Brachyuromys_ramirohitra | LC | Unknown | tropical | cathemeral | 90 | 68444.56 | 7.964 |
| mammals | Bradypus_torquatus | Bradypus_torquatus | VU | Decreasing | tropical | cathemeral | 3900 | 80086.158 | 11.099 |
| mammals | Bradypus_tridactylus | Bradypus_tridactylus | LC | Decreasing | tropical | cathemeral | 3850 | 1043271.316 | 1.421 |
| mammals | Bradypus_variegatus | Bradypus_variegatus | LC | Decreasing | tropical | cathemeral | 4335 | 10100677.31 | 5.7 |

| | | | | | | | | | |
|---|---|---|---|---|---|---|---|---|---|
| mammals | Brassomys_albidens | NA | DD | Unknown | NA | NA | NA | NA | NA |
| mammals | Brucepattersonius_griserufescens | NA | DD | Unknown | NA | NA | NA | NA | NA |
| mammals | Brucepattersonius_iheringi | Brucepattersonius_iheringi | LC | Decreasing | temperate | nocturnal | 43 | 187534.26 | 12.007 |
| mammals | Bubalus_arnee | Bubalus_arnee | EN | Decreasing | temperate | cathemeral | 725000 | 13690.352 | 6.653 |
| mammals | Bubalus_depressicornis | Bubalus_depressicornis | EN | Decreasing | tropical | cathemeral | 88000 | 107246.617 | 6.764 |
| mammals | Bubalus_mindorensis | Bubalus_mindorensis | CR | Decreasing | tropical | diurnal | 224000 | 98.749 | 7.531 |
| mammals | Bubalus_quarlesi | Bubalus_quarlesi | EN | Decreasing | tropical | cathemeral | 76000 | 70025.754 | 6.147 |
| mammals | Budorcas_taxicolor | Budorcas_taxicolor | VU | Decreasing | temperate | diurnal | 302000 | 440420.955 | 7.193 |
| mammals | Bullimus_bagobus | Bullimus_bagobus | LC | Stable | tropical | nocturnal | 432.5 | 118864.049 | 12.099 |
| mammals | Bullimus_gamay | Bullimus_gamay | VU | Decreasing | tropical | nocturnal | 368 | 252.918 | 14.437 |
| mammals | Bullimus_luzonicus | Bullimus_luzonicus | LC | Unknown | tropical | cathemeral | 445 | 38308.973 | 10.654 |
| mammals | Bunolagus_monticularis | Bunolagus_monticularis | CR | Decreasing | temperate | nocturnal | 1750 | 45473.925 | 7.18 |
| mammals | Bunomys_andrewsi | Bunomys_andrewsi | LC | Unknown | tropical | nocturnal | 125 | 117011.328 | 9.813 |
| mammals | Bunomys_chrysocomus | Bunomys_chrysocomus | LC | Stable | tropical | nocturnal | 116 | 169337.754 | 9.276 |
| mammals | Bunomys_coelestis | Bunomys_coelestis | EN | Decreasing | tropical | nocturnal | 120 | 133.477 | 8.414 |
| mammals | Bunomys_penitus | Bunomys_penitus | LC | Unknown | tropical | nocturnal | 170 | 4215.326 | 5.483 |
| mammals | Bunomys_prolatus | Bunomys_prolatus | EN | Decreasing | tropical | nocturnal | 131 | 782.224 | 3.203 |
| mammals | Burramys_parvus | Burramys_parvus | CR | Decreasing | temperate | nocturnal | 41.5 | 1197.791 | 5.571 |
| mammals | Cabassous_centralis | NA | DD | Unknown | NA | NA | NA | NA | NA |
| mammals | Cabassous_chacoensis | Cabassous_chacoensis | NT | Unknown | temperate | nocturnal | 1490 | 478652.233 | 4.903 |
| mammals | Cabassous_tatouay | Cabassous_tatouay | LC | Unknown | temperate | nocturnal | 5350 | 2538377.787 | 11.785 |
| mammals | Cabassous_unicinctus | Cabassous_unicinctus | LC | Unknown | tropical | nocturnal | 4800 | 9761562.721 | 4.42 |
| mammals | Cabreramops_aequatorianus | Molossops_aequatorianus | EN | Unknown | tropical | nocturnal | 5 | 3326.776 | 21.949 |
| mammals | Cacajao_calvus | Cacajao_calvus | VU | Decreasing | tropical | diurnal | 5796 | 210952.494 | 1.199 |
| mammals | Cacajao_hosomi | Cacajao_hosomi | VU | Decreasing | tropical | diurnal | 3100 | 69745.449 | 0.468 |
| mammals | Caenolestes_caniventer | Caenolestes_caniventer | NT | Decreasing | tropical | nocturnal | 40 | 19664.021 | 11.674 |
| mammals | Caenolestes_condorensis | Caenolestes_condorensis | VU | Unknown | tropical | nocturnal | 48 | 78.683 | 1.692 |
| mammals | Caenolestes_convelatus | Caenolestes_convelatus | VU | Decreasing | tropical | nocturnal | 40 | 22780.355 | 8.629 |
| mammals | Caenolestes_fuliginosus | Caenolestes_fuliginosus | LC | Unknown | tropical | nocturnal | 27.8 | 180934.655 | 11.824 |
| mammals | Caenolestes_sangay | Caenolestes_sangay | VU | Unknown | tropical | nocturnal | 41.5 | 879.699 | 4.973 |
| mammals | Callicebus_barbarabrownae | Callicebus_barbarabrownae | CR | Decreasing | tropical | diurnal | 1228.7 | 129698.416 | 9.316 |
| mammals | Callicebus_coimbrai | Callicebus_coimbrai | EN | Decreasing | tropical | diurnal | 1181.7 | 38585.35 | 13.632 |
| mammals | Callicebus_melanochir | Callicebus_melanochir | VU | Decreasing | tropical | diurnal | 1322.4 | 99867.074 | 8.847 |
| mammals | Callicebus_nigrifrons | Callicebus_nigrifrons | NT | Decreasing | tropical | diurnal | 1215.1 | 490584.706 | 13.87 |

| | | | | | | | | | |
|---|---|---|---|---|---|---|---|---|---|
| mammals | Callicebus_personatus | Callicebus_personatus | VU | Decreasing | tropical | diurnal | 1350 | 142975.24 | 11.066 |
| mammals | Callimico_goeldii | Callimico_goeldii | VU | Decreasing | tropical | diurnal | 480 | 602352.683 | 1.554 |
| mammals | Callistomys_pictus | Callistomys_pictus | EN | Decreasing | tropical | nocturnal | 519 | 27156.594 | 9.846 |
| mammals | Callithrix_aurita | Callithrix_aurita | EN | Decreasing | tropical | diurnal | 342 | 159614.292 | 14.504 |
| mammals | Callithrix_flaviceps | Callithrix_flaviceps | CR | Decreasing | tropical | diurnal | 342 | 24732.63 | 12.032 |
| mammals | Callithrix_geoffroyi | Callithrix_geoffroyi | LC | Decreasing | tropical | diurnal | 342 | 124967.934 | 9.032 |
| mammals | Callithrix_jacchus | Callithrix_jacchus | LC | Decreasing | tropical | diurnal | 292 | 943369.79 | 7.725 |
| mammals | Callithrix_kuhlii | Callithrix_kuhlii | VU | Decreasing | tropical | diurnal | 342 | 45700.724 | 8.679 |
| mammals | Callithrix_penicillata | Callithrix_penicillata | LC | Decreasing | tropical | diurnal | 342 | 1309808.489 | 10.509 |
| mammals | Callosciurus_adamsi | Callosciurus_adamsi | NT | Decreasing | tropical | diurnal | 134.5 | 97681.773 | 6.517 |
| mammals | Callosciurus_baluensis | Callosciurus_baluensis | LC | Decreasing | tropical | diurnal | 356 | 51137.125 | 4.061 |
| mammals | Callosciurus_caniceps | Callosciurus_caniceps | LC | Stable | tropical | diurnal | 257 | 650673.882 | 14.811 |
| mammals | Callosciurus_erythraeus | Callosciurus_erythraeus | LC | Stable | tropical | diurnal | 280 | 4042787.099 | 12.907 |
| mammals | Callosciurus_finlaysonii | Callosciurus_finlaysonii | LC | Stable | tropical | diurnal | 278 | 730177.832 | 14.877 |
| mammals | Callosciurus_inornatus | Callosciurus_inornatus | LC | Stable | tropical | diurnal | 325 | 309660.485 | 9.624 |
| mammals | Callosciurus_melanogaster | Callosciurus_melanogaster | VU | Decreasing | tropical | cathemeral | 278 | 6004.449 | 6.223 |
| mammals | Callosciurus_nigrovittatus | Callosciurus_nigrovittatus | LC | Stable | tropical | diurnal | 202 | 708578.846 | 12.726 |
| mammals | Callosciurus_notatus | Callosciurus_notatus | LC | Increasing | tropical | diurnal | 197.5 | 1496326.218 | 9.309 |
| mammals | Callosciurus_orestes | Callosciurus_orestes | LC | Stable | tropical | diurnal | 278 | 50506.521 | 3.185 |
| mammals | Callosciurus_phayrei | Callosciurus_phayrei | LC | Unknown | tropical | diurnal | 269 | 153735.302 | 9.89 |
| mammals | Callosciurus_prevostii | Callosciurus_prevostii | LC | Decreasing | tropical | diurnal | 361.5 | 1334755.191 | 7.799 |
| mammals | Callosciurus_pygerythrus | Callosciurus_pygerythrus | LC | Stable | tropical | diurnal | 252 | 587628.608 | 11.055 |
| mammals | Callosciurus_quinquestriatus | Callosciurus_quinquestriatus | LC | Unknown | temperate | diurnal | 318.6 | 40860.997 | 8.246 |
| mammals | Callospermophilus_lateralis | Callospermophilus_lateralis | LC | Stable | temperate | diurnal | 191 | 1704582.976 | 8.486 |
| mammals | Callospermophilus_madrensis | Callospermophilus_madrensis | NT | Unknown | temperate | diurnal | 275 | 60013.579 | 4.326 |
| mammals | Callospermophilus_saturatus | Callospermophilus_saturatus | LC | Stable | temperate | diurnal | 220 | 74904.169 | 8.186 |
| mammals | Calomys_boliviae | Calomys_boliviae | LC | Stable | tropical | diurnal | 27 | 129638.124 | 7.014 |
| mammals | Calomys_callidus | Calomys_callidus | LC | Unknown | temperate | nocturnal | 27 | 55966.07 | 9.276 |
| mammals | Calomys_callosus | Calomys_callosus | LC | Stable | tropical | nocturnal | 48 | 3145757.288 | 7.966 |
| mammals | Calomys_expulsus | Calomys_expulsus | LC | Increasing | tropical | nocturnal | 28.2 | 1360452.919 | 8.675 |
| mammals | Calomys_hummelincki | Calomys_hummelincki | LC | Decreasing | tropical | nocturnal | 27 | 312459.193 | 7.958 |
| mammals | Calomys_laucha | Calomys_laucha | LC | Unknown | temperate | nocturnal | 14 | 1437875.389 | 10.391 |
| mammals | Calomys_lepidus | Calomys_lepidus | LC | Stable | tropical | nocturnal | 26.6 | 584231.18 | 7.215 |
| mammals | Calomys_musculinus | Calomys_musculinus | LC | Stable | temperate | nocturnal | 20.1 | 2024366.277 | 7.325 |

| | | | | | | | | | |
|---|---|---|---|---|---|---|---|---|---|
| mammals | Calomys_sorella | Calomys_sorellus | LC | Stable | tropical | nocturnal | 20 | 244104.406 | 8.707 |
| mammals | Calomys_tener | Calomys_tener | LC | Increasing | tropical | nocturnal | 13.8 | 1710353.169 | 10.825 |
| mammals | Calomys_tocantinsi | Calomys_tocantinsi | LC | Unknown | tropical | nocturnal | 25.3 | 132795.309 | 5.218 |
| mammals | Calomyscus_bailwardi | Calomyscus_bailwardi | LC | Unknown | temperate | nocturnal | 20.4 | 185798.115 | 14.831 |
| mammals | Calomyscus_baluchi | Calomyscus_baluchi | LC | Unknown | temperate | nocturnal | 20.1 | 426505.707 | 10.08 |
| mammals | Calomyscus_elburzensis | Calomyscus_elburzensis | LC | Unknown | temperate | nocturnal | 17 | 231907.31 | 9.87 |
| mammals | Calomyscus_hotsoni | Calomyscus_hotsoni | LC | Unknown | temperate | nocturnal | 16.3 | 141022.256 | 5.012 |
| mammals | Calomyscus_mystax | Calomyscus_mystax | LC | Stable | temperate | nocturnal | 17 | 208687.975 | 13.187 |
| mammals | Calomyscus_tsolovi | NA | DD | Unknown | NA | NA | NA | NA | NA |
| mammals | Calomyscus_urartensis | Calomyscus_urartensis | LC | Unknown | temperate | nocturnal | 16.5 | 4812.865 | 13.964 |
| mammals | Caluromys_derbianus | Caluromys_derbianus | LC | Decreasing | tropical | nocturnal | 297 | 792215.256 | 10.531 |
| mammals | Caluromys_lanatus | Caluromys_lanatus | LC | Decreasing | tropical | nocturnal | 325 | 5110003.562 | 5.911 |
| mammals | Caluromys_philander | Caluromys_philander | LC | Decreasing | tropical | nocturnal | 229.3 | 3711568.69 | 4.955 |
| mammals | Caluromysiops_irruptus | Caluromysiops_irrupta | LC | Decreasing | tropical | nocturnal | 250 | 182315.307 | 2.592 |
| mammals | Camelus_ferus | Camelus_ferus | CR | Decreasing | temperate | diurnal | 690000 | 202139.125 | 0.501 |
| mammals | Canis_aureus | Canis_aureus | LC | Increasing | temperate | cathemeral | 10345.2 | 10746781.913 | 13.505 |
| mammals | Canis_latrans | Canis_latrans | LC | Increasing | temperate | cathemeral | 13406.3 | 14921849.182 | 10.195 |
| mammals | Canis_lupus | Canis_lupus | LC | Stable | temperate | cathemeral | 32183.3 | 56006225.875 | 6.855 |
| mammals | Canis_rufus | Canis_rufus | CR | Decreasing | temperate | nocturnal | 22983.2 | 3120.731 | 9.296 |
| mammals | Canis_simensis | Canis_simensis | EN | Decreasing | tropical | diurnal | 10000 | 6913.953 | 11.756 |
| mammals | Cannomys_badius | Cannomys_badius | LC | Unknown | tropical | nocturnal | 275 | 1043848.447 | 11.279 |
| mammals | Cansumys_canus | Cansumys_canus | LC | Unknown | temperate | nocturnal | 90.5 | 161030.746 | 14.115 |
| mammals | Capra_aegagrus | Capra_aegagrus | NT | Stable | temperate | diurnal | 33500 | 1215040.107 | 11.321 |
| mammals | Capra_caucasica | Capra_caucasica | EN | Decreasing | temperate | diurnal | 55000 | 7868.956 | 5.371 |
| mammals | Capra_cylindricornis | Capra_cylindricornis | NT | Stable | temperate | diurnal | 50000 | 22805.654 | 6.236 |
| mammals | Capra_falconeri | Capra_falconeri | NT | Increasing | temperate | diurnal | 41000 | 25328.49 | 8.12 |
| mammals | Capra_ibex | Capra_ibex | LC | Stable | temperate | diurnal | 85166.5 | 19233.354 | 8.056 |
| mammals | Capra_nubiana | Capra_nubiana | VU | Decreasing | temperate | diurnal | 62624.8 | 342497.976 | 4.075 |
| mammals | Capra_pyrenaica | Capra_pyrenaica | LC | Increasing | temperate | diurnal | 50000 | 92676.018 | 13.885 |
| mammals | Capra_sibirica | Capra_sibirica | NT | Decreasing | temperate | diurnal | 130000 | 1562755.821 | 4.446 |
| mammals | Capra_walie | Capra_walie | VU | Increasing | tropical | diurnal | 100000 | 844.924 | 11.712 |
| mammals | Capreolus_capreolus | Capreolus_capreolus | LC | Increasing | temperate | cathemeral | 22500 | 6468697.037 | 17.217 |
| mammals | Capreolus_pygargus | Capreolus_pygargus | LC | Decreasing | temperate | cathemeral | 43752.2 | 11998106.577 | 7.589 |
| mammals | Capricornis_crispus | Capricornis_crispus | LC | Stable | temperate | diurnal | 37160 | 160080.995 | 18.167 |

| | | | | | | | | | |
|---|---|---|---|---|---|---|---|---|---|
| mammals | Capricornis_rubidus | Capricornis_rubidus | VU | Decreasing | temperate | cathemeral | 87500 | 86336.309 | 5.861 |
| mammals | Capricornis_sumatraensis | Capricornis_sumatraensis | VU | Decreasing | tropical | cathemeral | 87500 | 3511835.455 | 10.349 |
| mammals | Capricornis_swinhoei | Capricornis_swinhoei | LC | Unknown | temperate | diurnal | 21000 | 14411.896 | 6.923 |
| mammals | Capromys_pilorides | Capromys_pilorides | LC | Stable | tropical | nocturnal | 3090 | 111094.565 | 16.064 |
| mammals | Caracal_aurata | Caracal_aurata | VU | Decreasing | tropical | cathemeral | 10650 | 4129643.023 | 7.542 |
| mammals | Caracal_caracal | Caracal_caracal | LC | Unknown | tropical | nocturnal | 13749.9 | 17205653.2 | 8.912 |
| mammals | Cardiocranius_paradoxus | NA | DD | Unknown | NA | NA | NA | NA | NA |
| mammals | Cardioderma_cor | Cardioderma_cor | LC | Unknown | tropical | nocturnal | 27.8 | 2246034.166 | 9.276 |
| mammals | Carlito_syrichta | Tarsius_syrichta | NT | Decreasing | tropical | nocturnal | 113.5 | 120136.856 | 12.09 |
| mammals | Carollia_brevicauda | Carollia_brevicauda | LC | Stable | tropical | nocturnal | 14.4 | 10749190.806 | 4.6 |
| mammals | Carollia_castanea | Carollia_castanea | LC | Stable | tropical | nocturnal | 13.3 | 2501247.335 | 7.292 |
| mammals | Carollia_manu | Carollia_manu | LC | Stable | tropical | nocturnal | 18.7 | 31962.745 | 6.366 |
| mammals | Carollia_perspicillata | Carollia_perspicillata | LC | Stable | tropical | nocturnal | 18 | 13796363.062 | 5.975 |
| mammals | Carollia_sowelli | Carollia_sowelli | LC | Stable | tropical | nocturnal | 16.5 | 790178.17 | 10.622 |
| mammals | Carollia_subrufa | Carollia_subrufa | LC | Stable | tropical | nocturnal | 15.6 | 212261.468 | 14.328 |
| mammals | Carpomys_melanurus | NA | DD | Unknown | NA | NA | NA | NA | NA |
| mammals | Carpomys_phaeurus | Carpomys_phaeurus | LC | Unknown | tropical | nocturnal | 123 | 4215.026 | 8.968 |
| mammals | Carterodon_sulcidens | NA | DD | Unknown | NA | NA | NA | NA | NA |
| mammals | Casinycteris_argynnis | Casinycteris_argynnis | LC | Unknown | tropical | nocturnal | 29.9 | 1707684.632 | 5.454 |
| mammals | Casinycteris_ophiodon | NA | not_available | not_available | NA | NA | NA | NA | NA |
| mammals | Cassistrellus_dimissus | NA | DD | Unknown | NA | NA | NA | NA | NA |
| mammals | Castor_canadensis | Castor_canadensis | LC | Stable | temperate | nocturnal | 21820 | 15707994.783 | 8.375 |
| mammals | Castor_fiber | Castor_fiber | LC | Increasing | temperate | nocturnal | 19000 | 3573387.443 | 11.753 |
| mammals | Castoria_angustidens | Akodon_serrensis | LC | Unknown | temperate | cathemeral | 28.3 | 275871.625 | 12.617 |
| mammals | Catopuma_badia | Catopuma_badia | EN | Decreasing | tropical | diurnal | 2500 | 268234.069 | 2.693 |
| mammals | Catopuma_temminckii | Catopuma_temminckii | NT | Decreasing | temperate | nocturnal | 11500 | 337441.822 | 5.597 |
| mammals | Cavia_aperea | Cavia_aperea | LC | Stable | tropical | diurnal | 549 | 6132281.514 | 8.877 |
| mammals | Cavia_fulgida | Cavia_fulgida | LC | Stable | tropical | diurnal | 282.5 | 684399.493 | 11.883 |
| mammals | Cavia_magna | Cavia_magna | LC | Stable | temperate | diurnal | 460 | 104005.766 | 11.974 |
| mammals | Cavia_patzelti | NA | DD | Unknown | NA | NA | NA | NA | NA |
| mammals | Cavia_tschudii | Cavia_tschudii | LC | Stable | tropical | nocturnal | 1000 | 810000.646 | 7.95 |
| mammals | Cebuella_pygmaea | Cebuella_pygmaea | VU | Decreasing | tropical | diurnal | 70 | 497010.926 | 1.863 |
| mammals | Cebus_albifrons | Cebus_albifrons | LC | Decreasing | tropical | diurnal | 2629 | 2701375.069 | 2.966 |
| mammals | Cebus_kaapori | Cebus_kaapori | CR | Decreasing | tropical | diurnal | 2600 | 190775.708 | 4.599 |

| | | | | | | | | | |
|---|---|---|---|---|---|---|---|---|---|
| mammals | Centronycteris_centralis | Centronycteris_centralis | LC | Unknown | tropical | nocturnal | 5.5 | 2377722.801 | 7.08 |
| mammals | Centronycteris_maximiliani | Centronycteris_maximiliani | LC | Unknown | tropical | nocturnal | 5 | 5647629.783 | 3.175 |
| mammals | Centurio_senex | Centurio_senex | LC | Stable | tropical | nocturnal | 21.2 | 2084372.663 | 10.361 |
| mammals | Cephalopachus_bancanus | Tarsius_bancanus | VU | Decreasing | tropical | nocturnal | 94.3 | 847615.953 | 6.392 |
| mammals | Cephalophorus_callipygus | Cephalophus_callipygus | LC | Decreasing | tropical | diurnal | 18249.9 | 745835.021 | 5.675 |
| mammals | Cephalophorus_leucogaster | Cephalophus_leucogaster | NT | Decreasing | tropical | diurnal | 12699.9 | 1031856.958 | 5.606 |
| mammals | Cephalophorus_natalensis | Cephalophus_natalensis | LC | Decreasing | tropical | diurnal | 12000 | 1245390.438 | 10.19 |
| mammals | Cephalophorus_niger | Cephalophus_niger | LC | Decreasing | tropical | cathemeral | 20400 | 761614.686 | 15.029 |
| mammals | Cephalophorus_nigrifrons | Cephalophus_nigrifrons | LC | Decreasing | tropical | diurnal | 13899.8 | 3193058.952 | 6.3 |
| mammals | Cephalophorus_ogilbyi | Cephalophus_ogilbyi | LC | Decreasing | tropical | diurnal | 20000 | 504344.051 | 9.601 |
| mammals | Cephalophorus_rufilatus | Cephalophus_rufilatus | LC | Decreasing | tropical | diurnal | 12000 | 2928978.081 | 10.226 |
| mammals | Cephalophorus_weynsi | Cephalophus_weynsi | LC | Decreasing | tropical | diurnal | 17000 | 1484366.45 | 6.309 |
| mammals | Cephalophula_zebra | Cephalophus_zebra | VU | Decreasing | tropical | diurnal | 17500.1 | 59057.427 | 9.547 |
| mammals | Cephalophus_dorsalis | Cephalophus_dorsalis | NT | Decreasing | tropical | diurnal | 20000 | 3572525.785 | 7.623 |
| mammals | Cephalophus_jentinki | Cephalophus_jentinki | EN | Decreasing | tropical | nocturnal | 70000.3 | 59693.104 | 9.566 |
| mammals | Cephalophus_silvicultor | Cephalophus_silvicultor | NT | Decreasing | tropical | diurnal | 72500.3 | 6162602.941 | 7.671 |
| mammals | Cephalophus_spadix | Cephalophus_spadix | EN | Decreasing | tropical | nocturnal | 56000.3 | 6572.389 | 8.478 |
| mammals | Ceratotherium_simum | Ceratotherium_simum | NT | Decreasing | tropical | cathemeral | 2949986.3 | 12627012.023 | 7.471 |
| mammals | Cercartetus_caudatus | Cercartetus_caudatus | LC | Stable | tropical | nocturnal | 25 | 308509.796 | 4.311 |
| mammals | Cercartetus_concinnus | Cercartetus_concinnus | LC | Stable | temperate | nocturnal | 13 | 581319.927 | 8.162 |
| mammals | Cercartetus_lepidus | Cercartetus_lepidus | LC | Stable | temperate | nocturnal | 7 | 103584.915 | 6.55 |
| mammals | Cercartetus_nana | Cercartetus_nanus | LC | Decreasing | temperate | nocturnal | 24 | 386605.427 | 8.488 |
| mammals | Cercocebus_agilis | Cercocebus_agilis | LC | Decreasing | tropical | diurnal | 8023.4 | 990299.295 | 4.828 |
| mammals | Cercocebus_atys | Cercocebus_atys | VU | Decreasing | tropical | diurnal | 7643.4 | 316365.525 | 13.814 |
| mammals | Cercocebus_chrysogaster | Cercocebus_chrysogaster | EN | Decreasing | tropical | diurnal | 9475.6 | 78980.884 | 4.811 |
| mammals | Cercocebus_galeritus | Cercocebus_galeritus | CR | Decreasing | tropical | diurnal | 7383 | 214.534 | 14.616 |
| mammals | Cercocebus_lunulatus | Cercocebus_lunulatus | EN | Decreasing | tropical | diurnal | 7275 | 164399.839 | 17.081 |
| mammals | Cercocebus_sanjei | Cercocebus_sanjei | EN | Decreasing | tropical | diurnal | 8023.4 | 188.106 | 8.947 |
| mammals | Cercocebus_torquatus | Cercocebus_torquatus | EN | Decreasing | tropical | diurnal | 8135.4 | 263787.993 | 11.995 |
| mammals | Cercopithecus_ascanius | Cercopithecus_ascanius | LC | Decreasing | tropical | diurnal | 3550 | 2619625.407 | 7.084 |
| mammals | Cercopithecus_campbelli | Cercopithecus_campbelli | NT | Decreasing | tropical | diurnal | 4100 | 313600.257 | 14.035 |
| mammals | Cercopithecus_cephus | Cercopithecus_cephus | LC | Unknown | tropical | diurnal | 3485 | 835775.133 | 5.841 |
| mammals | Cercopithecus_denti | Cercopithecus_denti | LC | Decreasing | tropical | diurnal | 4050 | 408498.007 | 6.378 |
| mammals | Cercopithecus_diana | Cercopithecus_diana | EN | Decreasing | tropical | diurnal | 4175 | 221588.188 | 14.643 |

| | | | | | | | | | |
|---|---|---|---|---|---|---|---|---|---|
| mammals | Cercopithecus_erythrogaster | Cercopithecus_erythrogaster | EN | Decreasing | tropical | diurnal | 2500 | 73797.058 | 17.783 |
| mammals | Cercopithecus_erythrotis | Cercopithecus_erythrotis | VU | Decreasing | tropical | diurnal | 3918.3 | 54822.563 | 9.635 |
| mammals | Cercopithecus_hamlyni | Cercopithecus_hamlyni | VU | Decreasing | tropical | diurnal | 6875 | 225790.335 | 4.461 |
| mammals | Cercopithecus_lowei | Cercopithecus_lowei | VU | Decreasing | tropical | diurnal | 3950 | 264353.546 | 15.991 |
| mammals | Cercopithecus_mitis | Cercopithecus_mitis | LC | Decreasing | tropical | diurnal | 5000 | 2345573.669 | 8.903 |
| mammals | Cercopithecus_mona | Cercopithecus_mona | NT | Decreasing | tropical | diurnal | 4208.3 | 649990.561 | 13.206 |
| mammals | Cercopithecus_neglectus | Cercopithecus_neglectus | LC | Unknown | tropical | diurnal | 5480 | 2289730.955 | 6.123 |
| mammals | Cercopithecus_nictitans | Cercopithecus_nictitans | NT | Decreasing | tropical | diurnal | 5803.3 | 1354989.936 | 7.914 |
| mammals | Cercopithecus_petaurista | Cercopithecus_petaurista | NT | Decreasing | tropical | diurnal | 2775 | 552996.171 | 14.833 |
| mammals | Cercopithecus_roloway | Cercopithecus_roloway | CR | Decreasing | tropical | diurnal | 4550 | 132057.774 | 17.116 |
| mammals | Cercopithecus_sclateri | Cercopithecus_sclateri | EN | Decreasing | tropical | diurnal | 3375 | 34781.605 | 21.331 |
| mammals | Cerdocyon_thous | Cerdocyon_thous | LC | Stable | tropical | cathemeral | 5240 | 8433351.958 | 8.287 |
| mammals | Cerradomys_maracajuensis | Cerradomys_maracajuensis | LC | Unknown | tropical | nocturnal | 91.1 | 963856.023 | 9.093 |
| mammals | Cerradomys_marinhus | Cerradomys_marinhus | LC | Unknown | tropical | nocturnal | 130 | 31994.478 | 6.647 |
| mammals | Cerradomys_scotti | Cerradomys_scotti | LC | Unknown | tropical | nocturnal | 92 | 1644216.947 | 8.361 |
| mammals | Cerradomys_subflavus | Cerradomys_subflavus | LC | Decreasing | tropical | nocturnal | 96.4 | 1461038.816 | 10.324 |
| mammals | Cervus_albirostris | Cervus_albirostris | VU | Unknown | temperate | cathemeral | 125000 | 747388.722 | 5.913 |
| mammals | Cervus_canadensis | Cervus_canadensis | LC | Increasing | temperate | cathemeral | 131250 | 6042820.882 | 6.334 |
| mammals | Cervus_elaphus | Cervus_elaphus | LC | Increasing | temperate | cathemeral | 131250 | 4274919.308 | 18.015 |
| mammals | Chacodelphys_formosa | Chacodelphys_formosa | NT | Decreasing | temperate | nocturnal | 10 | 3712.72 | 9.958 |
| mammals | Chaetocauda_sichuanensis | NA | DD | Unknown | NA | NA | NA | NA | NA |
| mammals | Chaetodipus_arenarius | Chaetodipus_arenarius | LC | Unknown | temperate | nocturnal | 23 | 88085.086 | 3.055 |
| mammals | Chaetodipus_artus | Chaetodipus_artus | LC | Stable | temperate | nocturnal | 20 | 89263.003 | 8.245 |
| mammals | Chaetodipus_baileyi | Chaetodipus_baileyi | LC | Stable | temperate | nocturnal | 26.4 | 283199.1 | 8.035 |
| mammals | Chaetodipus_californicus | Chaetodipus_californicus | LC | Stable | temperate | nocturnal | 22 | 106977.894 | 14.658 |
| mammals | Chaetodipus_eremicus | Chaetodipus_eremicus | LC | Unknown | temperate | nocturnal | 19 | 538896.375 | 7.254 |
| mammals | Chaetodipus_fallax | Chaetodipus_fallax | LC | Unknown | temperate | nocturnal | 18.7 | 46901.826 | 11.335 |
| mammals | Chaetodipus_formosus | Chaetodipus_formosus | LC | Stable | temperate | nocturnal | 19.5 | 322064.93 | 7.522 |
| mammals | Chaetodipus_goldmani | Chaetodipus_goldmani | NT | Decreasing | temperate | nocturnal | 23 | 83540.147 | 7.206 |
| mammals | Chaetodipus_hispidus | Chaetodipus_hispidus | LC | Stable | temperate | nocturnal | 32 | 2300275.399 | 12.297 |
| mammals | Chaetodipus_intermedius | Chaetodipus_intermedius | LC | Stable | temperate | nocturnal | 16.5 | 510455.583 | 8.782 |
| mammals | Chaetodipus_lineatus | NA | DD | Unknown | NA | NA | NA | NA | NA |
| mammals | Chaetodipus_nelsoni | Chaetodipus_nelsoni | LC | Stable | temperate | nocturnal | 15.8 | 564413.41 | 7.133 |
| mammals | Chaetodipus_penicillatus | Chaetodipus_penicillatus | LC | Stable | temperate | nocturnal | 16.9 | 395569.822 | 8.035 |

| | | | | | | | | | |
|---|---|---|---|---|---|---|---|---|---|
| mammals | Chaetodipus_pernix | Chaetodipus_pernix | LC | Stable | temperate | nocturnal | 17 | 99068.669 | 8.459 |
| mammals | Chaetodipus_rudinoris | Chaetodipus_rudinoris | LC | Stable | temperate | nocturnal | 26 | 118040.425 | 4.27 |
| mammals | Chaetodipus_spinatus | Chaetodipus_spinatus | LC | Unknown | temperate | nocturnal | 16.4 | 125751.336 | 4.948 |
| mammals | Chaetomys_subspinosus | Chaetomys_subspinosus | VU | Decreasing | tropical | nocturnal | 1300 | 127236.51 | 10.943 |
| mammals | Chaetophractus_vellerosus | Chaetophractus_vellerosus | LC | Stable | temperate | diurnal | 1030 | 1902395.92 | 6.78 |
| mammals | Chaetophractus_villosus | Chaetophractus_villosus | LC | Stable | temperate | nocturnal | 4540 | 2550820.675 | 7.215 |
| mammals | Chalinolobus_dwyeri | Chalinolobus_dwyeri | VU | Decreasing | temperate | nocturnal | 9 | 540243.075 | 7.823 |
| mammals | Chalinolobus_gouldii | Chalinolobus_gouldii | LC | Stable | temperate | nocturnal | 14.5 | 7369434.572 | 4.311 |
| mammals | Chalinolobus_morio | Chalinolobus_morio | LC | Stable | temperate | nocturnal | 8.8 | 2316599.988 | 7.183 |
| mammals | Chalinolobus_neocaledonicus | Chalinolobus_neocaledonicus | EN | Decreasing | tropical | nocturnal | 11.9 | 16190.554 | 7.435 |
| mammals | Chalinolobus_nigrogriseus | Chalinolobus_nigrogriseus | LC | Stable | tropical | nocturnal | 8.8 | 1476865.098 | 3.314 |
| mammals | Chalinolobus_picatus | Chalinolobus_picatus | NT | Decreasing | temperate | nocturnal | 5.1 | 1154162.955 | 6.244 |
| mammals | Chalinolobus_tuberculatus | Chalinolobus_tuberculatus | CR | Decreasing | temperate | nocturnal | 8.5 | 160397.198 | 8.085 |
| mammals | Cheirogaleus_crossleyi | Cheirogaleus_crossleyi | VU | Decreasing | tropical | nocturnal | 375 | 20521.66 | 8.44 |
| mammals | Cheirogaleus_major | Cheirogaleus_major | VU | Decreasing | tropical | nocturnal | 450 | 6465.229 | 9.239 |
| mammals | Cheirogaleus_medius | Cheirogaleus_medius | VU | Decreasing | tropical | nocturnal | 181 | 8993.79 | 7.479 |
| mammals | Cheirogaleus_minusculus | NA | not_available | not_available | NA | NA | NA | NA | NA |
| mammals | Cheirogaleus_sibreei | Cheirogaleus_sibreei | CR | Decreasing | tropical | nocturnal | 285.5 | 501.851 | 8.197 |
| mammals | Cheiromeles_parvidens | Cheiromeles_parvidens | LC | Unknown | tropical | nocturnal | 114.6 | 443817.371 | 12.073 |
| mammals | Cheiromeles_torquatus | Cheiromeles_torquatus | LC | Unknown | tropical | nocturnal | 163.5 | 1330218.926 | 8.968 |
| mammals | Chelemys_megalonyx | Chelemys_megalonyx | NT | Decreasing | temperate | cathemeral | 50.8 | 64386.716 | 12.167 |
| mammals | Cheracebus_lucifer | Callicebus_lucifer | LC | Unknown | tropical | diurnal | 1329 | 255694.179 | 0.939 |
| mammals | Cheracebus_lugens | Callicebus_lugens | LC | Unknown | tropical | diurnal | 1203.6 | 1160872.175 | 1.444 |
| mammals | Cheracebus_medemi | Callicebus_medemi | VU | Decreasing | tropical | diurnal | 1299.6 | 14310.132 | 3.041 |
| mammals | Cheracebus_regulus | Callicebus_regulus | LC | Unknown | tropical | diurnal | 1273.6 | 221340.265 | 0.52 |
| mammals | Cheracebus_torquatus | Callicebus_torquatus | LC | Stable | tropical | diurnal | 1050 | 189486.257 | 0.468 |
| mammals | Chibchanomys_trichotis | NA | DD | Unknown | NA | NA | NA | NA | NA |
| mammals | Chilomys_instans | Chilomys_instans | LC | Stable | tropical | nocturnal | 19 | 197768.174 | 10.98 |
| mammals | Chilonatalus_micropus | Chilonatalus_micropus | VU | Decreasing | tropical | nocturnal | 3.5 | 14935.119 | 17.69 |
| mammals | Chilonatalus_tumidifrons | Chilonatalus_tumidifrons | NT | Unknown | temperate | nocturnal | 3.3 | 4209.138 | 5.588 |
| mammals | Chimarrogale_hantu | Chimarrogale_hantu | NT | Decreasing | tropical | diurnal | 55 | 12589.799 | 5.461 |
| mammals | Chimarrogale_himalayica | Chimarrogale_himalayica | LC | Unknown | temperate | nocturnal | 34.8 | 3308910.204 | 14.754 |
| mammals | Chimarrogale_phaeura | NA | EN | Decreasing | NA | NA | NA | NA | NA |
| mammals | Chimarrogale_platycephala | Chimarrogale_platycephalus | LC | Decreasing | temperate | cathemeral | 32 | 206390.277 | 18.749 |

| | | | | | | | | | |
|---|---|---|---|---|---|---|---|---|---|
| mammals | Chimarrogale_styani | Chimarrogale_styani | LC | Unknown | temperate | nocturnal | 25.8 | 840671.907 | 9.068 |
| mammals | Chinchilla_chinchilla | Chinchilla_chinchilla | EN | Decreasing | tropical | nocturnal | 500 | 410196.989 | 4.285 |
| mammals | Chinchilla_lanigera | Chinchilla_lanigera | EN | Decreasing | temperate | nocturnal | 417 | 24492.797 | 9.524 |
| mammals | Chinchillula_sahamae | Chinchillula_sahamae | LC | Stable | tropical | nocturnal | 169.8 | 160699.618 | 8.282 |
| mammals | Chionomys_gud | Chionomys_gud | LC | Unknown | temperate | cathemeral | 93.5 | 78030.914 | 8.124 |
| mammals | Chionomys_nivalis | Chionomys_nivalis | LC | Stable | temperate | cathemeral | 54 | 890858.692 | 13.689 |
| mammals | Chionomys_roberti | Chionomys_roberti | LC | Unknown | temperate | nocturnal | 93.5 | 73076.508 | 9.519 |
| mammals | Chiroderma_doriae | Chiroderma_doriae | LC | Unknown | tropical | nocturnal | 19.9 | 2630455.51 | 11.328 |
| mammals | Chiroderma_improvisum | Chiroderma_improvisum | EN | Unknown | tropical | nocturnal | 35 | 1571.087 | 23.619 |
| mammals | Chiroderma_salvini | Chiroderma_salvini | LC | Stable | tropical | nocturnal | 25.5 | 3419518.378 | 8.176 |
| mammals | Chiroderma_trinitatum | Chiroderma_trinitatum | LC | Unknown | tropical | nocturnal | 13.7 | 9386551.23 | 3.947 |
| mammals | Chiroderma_villosum | Chiroderma_villosum | LC | Stable | tropical | nocturnal | 23.1 | 10414722.694 | 4.633 |
| mammals | Chiromyscus_chiropus | Chiromyscus_chiropus | LC | Stable | tropical | nocturnal | 77.8 | 426711.866 | 8.919 |
| mammals | Chiromyscus_langbianis | Niviventer_langbianis | LC | Decreasing | tropical | nocturnal | 69.8 | 632381.023 | 8.757 |
| mammals | Chironax_melanocephalus | Chironax_melanocephalus | LC | Decreasing | tropical | nocturnal | 16.7 | 177749.702 | 11.671 |
| mammals | Chironectes_minimus | Chironectes_minimus | LC | Decreasing | tropical | nocturnal | 946 | 6225494.1 | 6.842 |
| mammals | Chiropodomys_calamianensis | NA | DD | Unknown | NA | NA | NA | NA | NA |
| mammals | Chiropodomys_gliroides | Chiropodomys_gliroides | LC | Stable | tropical | nocturnal | 21.5 | 1252844.381 | 12.249 |
| mammals | Chiropodomys_karlkoopmani | Chiropodomys_karlkoopmani | VU | Decreasing | tropical | nocturnal | 29.1 | 4489.84 | 5.283 |
| mammals | Chiropodomys_major | Chiropodomys_major | LC | Unknown | tropical | nocturnal | 42 | 153559.718 | 6.807 |
| mammals | Chiropodomys_muroides | NA | DD | Unknown | NA | NA | NA | NA | NA |
| mammals | Chiropodomys_pusillus | NA | DD | Unknown | NA | NA | NA | NA | NA |
| mammals | Chiropotes_albinasus | Chiropotes_albinasus | VU | Decreasing | tropical | diurnal | 2900 | 981561.865 | 2.05 |
| mammals | Chiropotes_chiropotes | Chiropotes_chiropotes | LC | Stable | tropical | diurnal | 3174 | 1363871.244 | 1.122 |
| mammals | Chiropotes_satanas | Chiropotes_satanas | EN | Decreasing | tropical | diurnal | 3000 | 273123.321 | 4.866 |
| mammals | Chiropotes_utahicki | Chiropotes_utahickae | VU | Decreasing | tropical | diurnal | 2784.7 | 352115.26 | 3.081 |
| mammals | Chiruromys_forbesi | Chiruromys_forbesi | LC | Stable | tropical | nocturnal | 105.2 | 33438.759 | 3.898 |
| mammals | Chiruromys_lamia | Chiruromys_lamia | LC | Unknown | tropical | nocturnal | 47.2 | 34006.1 | 3.307 |
| mammals | Chiruromys_vates | Chiruromys_vates | LC | Stable | tropical | nocturnal | 45.6 | 100855.758 | 2.343 |
| mammals | Chlorocebus_aethiops | Chlorocebus_aethiops | LC | Decreasing | tropical | diurnal | 3975 | 1136054.635 | 10.511 |
| mammals | Chlorocebus_cynosuros | Chlorocebus_cynosuros | LC | Stable | tropical | diurnal | 3975 | 3083230.357 | 6.84 |
| mammals | Chlorocebus_djamdjamensis | Chlorocebus_djamdjamensis | VU | Decreasing | tropical | diurnal | 3975 | 9379.715 | 13.712 |
| mammals | Chlorocebus_dryas | Cercopithecus_dryas | EN | Unknown | tropical | diurnal | 3075 | 41073.211 | 3.26 |
| mammals | Chlorocebus_pygerythrus | Chlorocebus_pygerythrus | LC | Decreasing | tropical | diurnal | 3975 | 4652959.278 | 10.202 |

| | | | | | | | | | |
|---|---|---|---|---|---|---|---|---|---|
| mammals | Chlorocebus_sabaeus | Chlorocebus_sabaeus | LC | Decreasing | tropical | diurnal | 3975 | 1439329.714 | 12.809 |
| mammals | Chlorocebus_tantalus | Chlorocebus_tantalus | LC | Stable | tropical | diurnal | 4650 | 3943411.206 | 10.183 |
| mammals | Chodsigoa_caovansunga | NA | DD | Unknown | NA | NA | NA | NA | NA |
| mammals | Chodsigoa_hypsibia | Chodsigoa_hypsibia | LC | Unknown | temperate | nocturnal | 15.2 | 429528.198 | 10.143 |
| mammals | Chodsigoa_parca | Chodsigoa_parca | LC | Unknown | temperate | nocturnal | 11.1 | 583110.913 | 12.014 |
| mammals | Chodsigoa_parva | NA | DD | Unknown | NA | NA | NA | NA | NA |
| mammals | Chodsigoa_salenskii | NA | DD | Unknown | NA | NA | NA | NA | NA |
| mammals | Chodsigoa_smithii | Chodsigoa_smithii | NT | Decreasing | temperate | nocturnal | 14.4 | 48110.62 | 7.995 |
| mammals | Chodsigoa_sodalis | NA | DD | Unknown | NA | NA | NA | NA | NA |
| mammals | Choeroniscus_godmani | Choeroniscus_godmani | LC | Unknown | tropical | nocturnal | 7.9 | 2679326.392 | 7.255 |
| mammals | Choeroniscus_minor | Choeroniscus_minor | LC | Unknown | tropical | nocturnal | 8.7 | 5047598.735 | 3.469 |
| mammals | Choeroniscus_periosus | Choeroniscus_periosus | VU | Unknown | tropical | nocturnal | 10.5 | 64960.827 | 5.319 |
| mammals | Choeronycteris_mexicana | Choeronycteris_mexicana | NT | Unknown | tropical | nocturnal | 16.6 | 1777869.418 | 8.639 |
| mammals | Choeropsis_liberiensis | Choeropsis_liberiensis | EN | Decreasing | tropical | nocturnal | 235001.2 | 155982.356 | 13.365 |
| mammals | Choloepus_didactylus | Choloepus_didactylus | LC | Decreasing | tropical | nocturnal | 5160 | 4166311.38 | 1.554 |
| mammals | Choloepus_hoffmanni | Choloepus_hoffmanni | LC | Decreasing | tropical | nocturnal | 4750 | 2348289.731 | 5.629 |
| mammals | Chrotogale_owstoni | Chrotogale_owstoni | EN | Decreasing | tropical | nocturnal | 3250 | 238124.711 | 10.106 |
| mammals | Chrotomys_gonzalesi | Chrotomys_gonzalesi | NT | Decreasing | tropical | diurnal | 140.2 | 64.41 | 7.682 |
| mammals | Chrotomys_mindorensis | Chrotomys_mindorensis | LC | Unknown | tropical | cathemeral | 179 | 35894.727 | 15.556 |
| mammals | Chrotomys_silaceus | Chrotomys_silaceus | LC | Unknown | tropical | diurnal | 84.1 | 4484.591 | 8.963 |
| mammals | Chrotomys_whiteheadi | Chrotomys_whiteheadi | LC | Stable | tropical | cathemeral | 140.7 | 16762.785 | 10.116 |
| mammals | Chrotopterus_auritus | Chrotopterus_auritus | LC | Stable | tropical | nocturnal | 77.9 | 13116792.96 | 5.809 |
| mammals | Chrysocyon_brachyurus | Chrysocyon_brachyurus | NT | Unknown | tropical | cathemeral | 23249.8 | 4335925.956 | 9.066 |
| mammals | Cistugo_lesueuri | Cistugo_lesueuri | LC | Decreasing | temperate | nocturnal | 6.4 | 70747.332 | 11.732 |
| mammals | Cistugo_seabrae | Cistugo_seabrae | LC | Unknown | tropical | nocturnal | 5.4 | 316783.726 | 3.753 |
| mammals | Civettictis_civetta | Civettictis_civetta | LC | Unknown | tropical | nocturnal | 12000 | 15260382.485 | 9.362 |
| mammals | Clethrionomys_californicus | Myodes_californicus | LC | Stable | temperate | nocturnal | 18.3 | 200390.005 | 10.622 |
| mammals | Clethrionomys_centralis | Myodes_centralis | LC | Unknown | temperate | nocturnal | 32 | 182570.354 | 5.94 |
| mammals | Clethrionomys_gapperi | Myodes_gapperi | LC | Stable | temperate | cathemeral | 19 | 8370648.959 | 6.757 |
| mammals | Clethrionomys_glareolus | Myodes_glareolus | LC | Stable | temperate | cathemeral | 21.2 | 8679125.162 | 13.03 |
| mammals | Clethrionomys_rutilus | Myodes_rutilus | LC | Unknown | temperate | cathemeral | 28.2 | 18530154.723 | 2.842 |
| mammals | Cloeotis_percivali | Cloeotis_percivali | LC | Unknown | tropical | nocturnal | 4.1 | 528464.528 | 11.649 |
| mammals | Clyomys_laticeps | Clyomys_laticeps | LC | Unknown | tropical | diurnal | 201 | 1699627.394 | 11.552 |
| mammals | Cnephaeus_bottae | Eptesicus_bottae | LC | Stable | tropical | nocturnal | 14.2 | 1067625.067 | 9.529 |

| | | | | | | | | | |
|---|---|---|---|---|---|---|---|---|---|
| mammals | Cnephaeus_floweri | Eptesicus_floweri | LC | Stable | tropical | nocturnal | 6.7 | 1474708.136 | 6.509 |
| mammals | Cnephaeus_gobiensis | Eptesicus_gobiensis | LC | Unknown | temperate | nocturnal | 10 | 4527879.707 | 5.414 |
| mammals | Cnephaeus_hottentotus | Eptesicus_hottentotus | LC | Unknown | tropical | nocturnal | 30 | 2853241.801 | 8.56 |
| mammals | Cnephaeus_isabellinus | Eptesicus_isabellinus | LC | Unknown | temperate | nocturnal | 21 | 851997.503 | 13.878 |
| mammals | Cnephaeus_japonensis | Eptesicus_japonensis | VU | Decreasing | temperate | nocturnal | 10.1 | 1460.469 | 12.228 |
| mammals | Cnephaeus_nilssonii | Eptesicus_nilssonii | LC | Stable | temperate | nocturnal | 10.4 | 10473774.101 | 9.229 |
| mammals | Cnephaeus_pachyotis | Eptesicus_pachyotis | LC | Unknown | temperate | nocturnal | 8.3 | 1205037.528 | 8.386 |
| mammals | Cnephaeus_platyops | NA | DD | Unknown | NA | NA | NA | NA | NA |
| mammals | Cnephaeus_serotinus | Eptesicus_serotinus | LC | Stable | temperate | nocturnal | 22.9 | 10184906.678 | 13.887 |
| mammals | Cnephaeus_tatei | NA | DD | Unknown | NA | NA | NA | NA | NA |
| mammals | Coccymys_ruemmleri | Coccymys_ruemmleri | LC | Unknown | tropical | nocturnal | 29.7 | 31663.248 | 4.487 |
| mammals | Coelops_frithii | Coelops_frithii | NT | Decreasing | tropical | nocturnal | 7.4 | 2491060.602 | 13.359 |
| mammals | Coelops_robinsoni | Coelops_robinsoni | VU | Decreasing | tropical | nocturnal | 6.5 | 137740.901 | 3.764 |
| mammals | Coendou_bicolor | Coendou_bicolor | LC | Unknown | tropical | nocturnal | 4050 | 1229394.858 | 4.072 |
| mammals | Coendou_ichillus | NA | DD | Unknown | NA | NA | NA | NA | NA |
| mammals | Coendou_insidiosus | Coendou_insidiosus | LC | Unknown | tropical | nocturnal | 1000 | 171746.003 | 10.07 |
| mammals | Coendou_mexicanus | Coendou_mexicanus | LC | Unknown | tropical | nocturnal | 2000 | 707181.017 | 11.022 |
| mammals | Coendou_prehensilis | Coendou_prehensilis | LC | Stable | tropical | nocturnal | 4400 | 12257463.454 | 5.427 |
| mammals | Coendou_quichua | Coendou_quichua | DD | Decreasing | tropical | nocturnal | 2133.1 | 462672.509 | 10.466 |
| mammals | Coendou_roosmalenorum | NA | DD | Unknown | NA | NA | NA | NA | NA |
| mammals | Coendou_spinosus | Coendou_spinosus | LC | Unknown | temperate | nocturnal | 750.8 | 1317961.387 | 12.388 |
| mammals | Coendou_vestitus | Coendou_vestitus | DD | Decreasing | tropical | nocturnal | 736 | 1445.687 | 14.873 |
| mammals | Coleura_afra | Coleura_afra | LC | Unknown | tropical | nocturnal | 10.7 | 2796285.82 | 9.213 |
| mammals | Coleura_seychellensis | NA | CR | Decreasing | NA | NA | NA | NA | NA |
| mammals | Colobus_angolensis | Colobus_angolensis | VU | Decreasing | tropical | diurnal | 14462.2 | 2120633.556 | 6.504 |
| mammals | Colobus_guereza | Colobus_guereza | LC | Decreasing | tropical | diurnal | 16498 | 2594832.204 | 7.383 |
| mammals | Colobus_polykomos | Colobus_polykomos | EN | Decreasing | tropical | diurnal | 9089.6 | 335688.54 | 13.86 |
| mammals | Colobus_satanas | Colobus_satanas | VU | Decreasing | tropical | diurnal | 11993.8 | 327231.882 | 6.318 |
| mammals | Colobus_vellerosus | Colobus_vellerosus | CR | Decreasing | tropical | diurnal | 11505.2 | 470710.482 | 14.365 |
| mammals | Colomys_goslingi | Colomys_goslingi | LC | Stable | tropical | nocturnal | 62.5 | 904776.062 | 8.915 |
| mammals | Condylura_cristata | Condylura_cristata | LC | Stable | temperate | cathemeral | 84 | 3486994.163 | 9.849 |
| mammals | Conepatus_chinga | Conepatus_chinga | LC | Decreasing | temperate | cathemeral | 1917.5 | 5532259.187 | 8.472 |
| mammals | Conepatus_leuconotus | Conepatus_leuconotus | LC | Decreasing | temperate | nocturnal | 3500 | 2498458.411 | 9.914 |
| mammals | Conepatus_semistriatus | Conepatus_semistriatus | LC | Unknown | tropical | nocturnal | 1200 | 2021448.583 | 10.147 |

| | | | | | | | | | |
|---|---|---|---|---|---|---|---|---|---|
| mammals | Congomys_lukolelae | Praomys_lukolelae | LC | Unknown | tropical | nocturnal | 41.6 | 208467.114 | 4.571 |
| mammals | Congomys_verschureni | NA | DD | Unknown | NA | NA | NA | NA | NA |
| mammals | Congosorex_phillipsorum | Congosorex_phillipsorum | CR | Decreasing | tropical | nocturnal | 7.9 | 85.623 | 4.831 |
| mammals | Congosorex_polli | NA | DD | Unknown | NA | NA | NA | NA | NA |
| mammals | Conilurus_penicillatus | Conilurus_penicillatus | VU | Decreasing | tropical | nocturnal | 153.5 | 346343.981 | 2.167 |
| mammals | Connochaetes_gnou | Connochaetes_gnou | LC | Increasing | temperate | cathemeral | 179999 | 403590.451 | 11.073 |
| mammals | Connochaetes_taurinus | Connochaetes_taurinus | LC | Stable | tropical | cathemeral | 179999 | 3103294.401 | 7.173 |
| mammals | Cormura_brevirostris | Cormura_brevirostris | LC | Unknown | tropical | nocturnal | 9.2 | 7826267.289 | 2.87 |
| mammals | Corynorhinus_mexicanus | Corynorhinus_mexicanus | NT | Decreasing | tropical | nocturnal | 7.7 | 514936.647 | 11.062 |
| mammals | Corynorhinus_rafinesquii | Corynorhinus_rafinesquii | LC | Increasing | temperate | nocturnal | 9.1 | 1336202.322 | 17.044 |
| mammals | Corynorhinus_townsendii | Corynorhinus_townsendii | LC | Stable | temperate | nocturnal | 9.9 | 4596886.834 | 9.829 |
| mammals | Craseomys_andersoni | Myodes_andersoni | LC | Stable | temperate | nocturnal | 25.5 | 71603.536 | 13.351 |
| mammals | Craseomys_regulus | Myodes_regulus | LC | Stable | temperate | cathemeral | 38.6 | 149439.314 | 19.594 |
| mammals | Craseomys_rex | Myodes_rex | LC | Stable | temperate | cathemeral | 55.5 | 58397.477 | 7.424 |
| mammals | Craseomys_rufocanus | Myodes_rufocanus | LC | Decreasing | temperate | cathemeral | 37 | 13399426.01 | 3.332 |
| mammals | Craseomys_smithii | Myodes_smithii | LC | Stable | temperate | cathemeral | 35 | 118619.05 | 16.956 |
| mammals | Craseonycteris_thonglongyai | Craseonycteris_thonglongyai | NT | Decreasing | tropical | nocturnal | 2.2 | 7954.014 | 17.354 |
| mammals | Crateromys_heaneyi | Crateromys_heaneyi | EN | Decreasing | tropical | nocturnal | 994 | 2464.518 | 7.93 |
| mammals | Crateromys_paulus | NA | DD | Unknown | NA | NA | NA | NA | NA |
| mammals | Crateromys_schadenbergi | Crateromys_schadenbergi | EN | Decreasing | tropical | nocturnal | 1050 | 2699.229 | 9.704 |
| mammals | Cremnomys_cutchicus | Cremnomys_cutchicus | LC | Stable | temperate | nocturnal | 59.8 | 534108.225 | 18.575 |
| mammals | Cremnomys_elvira | Cremnomys_elvira | CR | Decreasing | tropical | nocturnal | 77.1 | 15.315 | 23.776 |
| mammals | Cricetomys_emini | Cricetomys_emini | LC | Stable | tropical | nocturnal | 1300 | 3732493.145 | 8.245 |
| mammals | Cricetomys_gambianus | Cricetomys_gambianus | LC | Stable | tropical | nocturnal | 1271 | 4141042.439 | 11.158 |
| mammals | Cricetulus_barabensis | Cricetulus_barabensis | LC | Unknown | temperate | nocturnal | 22.9 | 3692081.347 | 10.789 |
| mammals | Cricetulus_longicaudatus | Cricetulus_longicaudatus | LC | Unknown | temperate | nocturnal | 22.1 | 2465565.116 | 7.699 |
| mammals | Cricetulus_sokolovi | Cricetulus_sokolovi | LC | Unknown | temperate | nocturnal | 24 | 358309.598 | 5.034 |
| mammals | Cricetus_cricetus | Cricetus_cricetus | CR | Decreasing | temperate | nocturnal | 510 | 4841407.636 | 11.383 |
| mammals | Crocidura_aleksandrisi | Crocidura_aleksandrisi | LC | Stable | temperate | nocturnal | 5.25 | 42683.872 | 9.023 |
| mammals | Crocidura_allex | Crocidura_allex | LC | Stable | tropical | nocturnal | 7 | 8036.615 | 7.978 |
| mammals | Crocidura_andamanensis | Crocidura_andamanensis | CR | Decreasing | tropical | nocturnal | 22.8 | 46.38 | 3.551 |
| mammals | Crocidura_ansellorum | Crocidura_ansellorum | EN | Decreasing | tropical | nocturnal | 4.6 | 2208.853 | 8.478 |
| mammals | Crocidura_arabica | Crocidura_arabica | LC | Unknown | tropical | nocturnal | 2.8 | 1146.284 | 18.506 |
| mammals | Crocidura_arispa | Crocidura_arispa | LC | Unknown | temperate | nocturnal | 9 | 1050.709 | 11.218 |

| | | | | | | | | | |
|---|---|---|---|---|---|---|---|---|---|
| mammals | Crocidura_armenica | NA | DD | Unknown | NA | NA | NA | NA | NA |
| mammals | Crocidura_attenuata | Crocidura_attenuata | LC | Unknown | tropical | nocturnal | 14.3 | 5024744.531 | 13.417 |
| mammals | Crocidura_attila | Crocidura_attila | LC | Unknown | tropical | nocturnal | 5 | 830715.104 | 6.186 |
| mammals | Crocidura_baileyi | Crocidura_baileyi | LC | Decreasing | tropical | nocturnal | 15.3 | 19271.55 | 13.274 |
| mammals | Crocidura_baluensis | Crocidura_baluensis | LC | Stable | tropical | nocturnal | 15.5 | 2805.714 | 4.479 |
| mammals | Crocidura_beata | Crocidura_beatus | LC | Decreasing | tropical | cathemeral | 10.6 | 57076.38 | 12.168 |
| mammals | Crocidura_beccarii | Crocidura_beccarii | LC | Decreasing | tropical | nocturnal | 7 | 91817.471 | 7.823 |
| mammals | Crocidura_bottegi | NA | DD | Unknown | NA | NA | NA | NA | NA |
| mammals | Crocidura_bottegoides | Crocidura_bottegoides | EN | Decreasing | tropical | nocturnal | 3 | 10830.026 | 14.759 |
| mammals | Crocidura_brunnea | Crocidura_brunnea | LC | Unknown | tropical | nocturnal | 14.9 | 8474.396 | 11.963 |
| mammals | Crocidura_buettikoferi | Crocidura_buettikoferi | LC | Unknown | tropical | nocturnal | 11 | 47433.211 | 17.434 |
| mammals | Crocidura_caliginea | Crocidura_caliginea | LC | Unknown | tropical | nocturnal | 8.3 | 76479.264 | 4.907 |
| mammals | Crocidura_canariensis | Crocidura_canariensis | EN | Decreasing | temperate | nocturnal | 7.5 | 2520.602 | 17.129 |
| mammals | Crocidura_caspica | NA | DD | Unknown | NA | NA | NA | NA | NA |
| mammals | Crocidura_cinderella | Crocidura_cinderella | LC | Unknown | tropical | nocturnal | 6.5 | 839382.093 | 10.178 |
| mammals | Crocidura_congobelgica | Crocidura_congobelgica | LC | Unknown | tropical | nocturnal | 17 | 99831.31 | 4.124 |
| mammals | Crocidura_crenata | Crocidura_crenata | LC | Unknown | tropical | nocturnal | 6.5 | 493947.053 | 5.629 |
| mammals | Crocidura_crossei | Crocidura_crossei | LC | Unknown | tropical | nocturnal | 6.9 | 1109826.838 | 14.107 |
| mammals | Crocidura_cyanea | Crocidura_cyanea | LC | Stable | temperate | nocturnal | 9.4 | 2725072.427 | 9.596 |
| mammals | Crocidura_denti | Crocidura_denti | LC | Unknown | tropical | nocturnal | 7 | 1196040.502 | 7.06 |
| mammals | Crocidura_desperata | Crocidura_desperata | VU | Decreasing | tropical | nocturnal | 21.5 | 16890.688 | 15.311 |
| mammals | Crocidura_dhofarensis | NA | DD | Unknown | NA | NA | NA | NA | NA |
| mammals | Crocidura_dolichura | Crocidura_dolichura | LC | Unknown | tropical | nocturnal | 5.7 | 1488528.975 | 7.651 |
| mammals | Crocidura_douceti | Crocidura_douceti | LC | Unknown | tropical | nocturnal | 4.3 | 101866.794 | 13.975 |
| mammals | Crocidura_dsinezumi | Crocidura_dsinezumi | LC | Stable | temperate | nocturnal | 10.2 | 292835.79 | 21.025 |
| mammals | Crocidura_eisentrauti | Crocidura_eisentrauti | VU | Stable | tropical | cathemeral | 9.1 | 363.731 | 8.928 |
| mammals | Crocidura_elgonius | Crocidura_elgonius | LC | Decreasing | tropical | nocturnal | 6.5 | 83847.677 | 17.087 |
| mammals | Crocidura_elongata | Crocidura_elongata | LC | Decreasing | tropical | nocturnal | 17.8 | 128094.518 | 7.866 |
| mammals | Crocidura_erica | NA | DD | Unknown | NA | NA | NA | NA | NA |
| mammals | Crocidura_fischeri | NA | DD | Unknown | NA | NA | NA | NA | NA |
| mammals | Crocidura_flavescens | Crocidura_flavescens | LC | Unknown | temperate | nocturnal | 28.5 | 292163.803 | 12.704 |
| mammals | Crocidura_floweri | NA | DD | Unknown | NA | NA | NA | NA | NA |
| mammals | Crocidura_foxi | Crocidura_foxi | LC | Unknown | tropical | nocturnal | 19 | 1660078.469 | 10.855 |
| mammals | Crocidura_fuliginosa | Crocidura_fuliginosa | LC | Unknown | tropical | nocturnal | 12 | 2850808.631 | 12.625 |

| | | | | | | | | | |
|---|---|---|---|---|---|---|---|---|---|
| mammals | Crocidura_fulvastra | Crocidura_fulvastra | LC | Stable | tropical | nocturnal | 12.7 | 4336348.884 | 9.386 |
| mammals | Crocidura_fumosa | Crocidura_fumosa | LC | Decreasing | tropical | nocturnal | 14.5 | 18541.186 | 12.078 |
| mammals | Crocidura_fuscomurina | Crocidura_fuscomurina | LC | Unknown | tropical | nocturnal | 3.9 | 9829229.265 | 9.768 |
| mammals | Crocidura_glassi | Crocidura_glassi | NT | Decreasing | tropical | nocturnal | 13.6 | 170233.353 | 12.37 |
| mammals | Crocidura_goliath | Crocidura_goliath | LC | Unknown | tropical | nocturnal | 54.8 | 1015996.101 | 5.983 |
| mammals | Crocidura_gracilipes | NA | DD | Unknown | NA | NA | NA | NA | NA |
| mammals | Crocidura_grandiceps | Crocidura_grandiceps | NT | Unknown | tropical | nocturnal | 23 | 34361.332 | 19.233 |
| mammals | Crocidura_grandis | NA | DD | Unknown | NA | NA | NA | NA | NA |
| mammals | Crocidura_grassei | Crocidura_grassei | LC | Unknown | tropical | nocturnal | 13.5 | 356574.619 | 5.362 |
| mammals | Crocidura_grayi | Crocidura_grayi | LC | Unknown | tropical | cathemeral | 10.3 | 116385.301 | 14.53 |
| mammals | Crocidura_greenwoodi | Crocidura_greenwoodi | LC | Unknown | tropical | nocturnal | 11.5 | 51685.879 | 14.479 |
| mammals | Crocidura_harenna | Crocidura_harenna | CR | Decreasing | tropical | nocturnal | 8.5 | 107.925 | 7.086 |
| mammals | Crocidura_hildegardeae | Crocidura_hildegardeae | LC | Unknown | tropical | nocturnal | 9.5 | 2340544.711 | 9.164 |
| mammals | Crocidura_hilliana | NA | DD | Unknown | NA | NA | NA | NA | NA |
| mammals | Crocidura_hispida | NA | VU | Unknown | NA | NA | NA | NA | NA |
| mammals | Crocidura_horsfieldii | NA | DD | Unknown | NA | NA | NA | NA | NA |
| mammals | Crocidura_hutanis | Crocidura_hutanis | LC | Unknown | tropical | nocturnal | 11 | 101089.698 | 9.883 |
| mammals | Crocidura_indochinensis | Crocidura_indochinensis | LC | Unknown | tropical | nocturnal | 6.3 | 667710.312 | 11.452 |
| mammals | Crocidura_jacksoni | Crocidura_jacksoni | LC | Stable | tropical | nocturnal | 7 | 911505.576 | 13.723 |
| mammals | Crocidura_jenkinsi | Crocidura_jenkinsi | CR | Decreasing | tropical | nocturnal | 12.6 | 76.384 | 5.326 |
| mammals | Crocidura_jouvenetae | Crocidura_jouvenetae | LC | Unknown | tropical | nocturnal | 9.5 | 222302.34 | 14.791 |
| mammals | Crocidura_katinka | NA | DD | Unknown | NA | NA | NA | NA | NA |
| mammals | Crocidura_kegoensis | NA | DD | Unknown | NA | NA | NA | NA | NA |
| mammals | Crocidura_lamottei | Crocidura_lamottei | LC | Unknown | tropical | nocturnal | 20.5 | 1559816.205 | 13.046 |
| mammals | Crocidura_lanosa | Crocidura_lanosa | VU | Decreasing | tropical | nocturnal | 22.6 | 4797.252 | 15.854 |
| mammals | Crocidura_lasiura | Crocidura_lasiura | LC | Unknown | temperate | cathemeral | 14.8 | 1577925.438 | 15.048 |
| mammals | Crocidura_latona | Crocidura_latona | LC | Unknown | tropical | nocturnal | 6.6 | 145294.019 | 4 |
| mammals | Crocidura_lea | Crocidura_lea | LC | Unknown | tropical | nocturnal | 6.6 | 116322.607 | 7.983 |
| mammals | Crocidura_lepidura | Crocidura_lepidura | LC | Stable | tropical | nocturnal | 18.5 | 232052.321 | 9.816 |
| mammals | Crocidura_leucodon | Crocidura_leucodon | LC | Unknown | temperate | cathemeral | 9.8 | 3927700.064 | 18.122 |
| mammals | Crocidura_levicula | Crocidura_levicula | LC | Unknown | tropical | diurnal | 1.8 | 42377.708 | 7.505 |
| mammals | Crocidura_littoralis | Crocidura_littoralis | LC | Unknown | tropical | nocturnal | 16 | 1900103.4 | 6.64 |
| mammals | Crocidura_longipes | NA | DD | Unknown | NA | NA | NA | NA | NA |
| mammals | Crocidura_lucina | Crocidura_lucina | VU | Stable | tropical | nocturnal | 12.2 | 5552.601 | 13.16 |

| | | | | | | | | | |
|---|---|---|---|---|---|---|---|---|---|
| mammals | Crocidura_ludia | Crocidura_ludia | LC | Unknown | tropical | nocturnal | 6.5 | 496220.864 | 5.215 |
| mammals | Crocidura_luna | Crocidura_luna | LC | Unknown | tropical | nocturnal | 14.5 | 1833287.874 | 10.906 |
| mammals | Crocidura_lusitania | Crocidura_lusitania | LC | Unknown | tropical | nocturnal | 2 | 2052701.842 | 6.908 |
| mammals | Crocidura_macarthuri | Crocidura_macarthuri | LC | Unknown | tropical | nocturnal | 13 | 467262.622 | 10.011 |
| mammals | Crocidura_macmillani | Crocidura_macmillani | NT | Unknown | tropical | nocturnal | 13.1 | 22024.396 | 11.236 |
| mammals | Crocidura_macowi | NA | DD | Unknown | NA | NA | NA | NA | NA |
| mammals | Crocidura_malayana | Crocidura_malayana | LC | Unknown | tropical | nocturnal | 12.3 | 198370.391 | 13.537 |
| mammals | Crocidura_manengubae | Crocidura_manengubae | VU | Decreasing | tropical | nocturnal | 14 | 5777.111 | 15.002 |
| mammals | Crocidura_maquassiensis | Crocidura_maquassiensis | LC | Unknown | temperate | nocturnal | 6 | 334700.905 | 13.416 |
| mammals | Crocidura_mariquensis | Crocidura_mariquensis | LC | Unknown | tropical | nocturnal | 11.3 | 2082090.811 | 8.315 |
| mammals | Crocidura_maurisca | Crocidura_maurisca | LC | Unknown | tropical | nocturnal | 14 | 572595.601 | 5.877 |
| mammals | Crocidura_maxi | Crocidura_maxi | LC | Unknown | tropical | nocturnal | 7.5 | 217572.153 | 18.846 |
| mammals | Crocidura_mindorus | NA | DD | Unknown | NA | NA | NA | NA | NA |
| mammals | Crocidura_miya | Crocidura_miya | EN | Decreasing | tropical | nocturnal | 10.2 | 330.459 | 24.022 |
| mammals | Crocidura_monax | NA | DD | Unknown | NA | NA | NA | NA | NA |
| mammals | Crocidura_monticola | Crocidura_monticola | LC | Unknown | tropical | nocturnal | 3 | 1393960.569 | 8.894 |
| mammals | Crocidura_montis | Crocidura_montis | LC | Unknown | tropical | nocturnal | 14.5 | 21776.072 | 13.056 |
| mammals | Crocidura_muricauda | Crocidura_muricauda | LC | Unknown | tropical | nocturnal | 5.5 | 403240.81 | 15.85 |
| mammals | Crocidura_musseri | NA | DD | Unknown | NA | NA | NA | NA | NA |
| mammals | Crocidura_mutesae | NA | DD | Unknown | NA | NA | NA | NA | NA |
| mammals | Crocidura_nana | NA | DD | Unknown | NA | NA | NA | NA | NA |
| mammals | Crocidura_nanilla | Crocidura_nanilla | LC | Stable | tropical | nocturnal | 6.5 | 757818.831 | 10.925 |
| mammals | Crocidura_negrina | Crocidura_negrina | EN | Decreasing | tropical | nocturnal | 13.4 | 461.638 | 10.364 |
| mammals | Crocidura_nicobarica | Crocidura_nicobarica | CR | Decreasing | tropical | nocturnal | 28.9 | 90.208 | 6.404 |
| mammals | Crocidura_nigeriae | Crocidura_nigeriae | LC | Unknown | tropical | nocturnal | 23 | 225668.75 | 16.428 |
| mammals | Crocidura_nigricans | Crocidura_nigricans | LC | Unknown | tropical | nocturnal | 20 | 128601.799 | 10.496 |
| mammals | Crocidura_nigripes | Crocidura_nigripes | LC | Unknown | tropical | nocturnal | 13.2 | 80622.629 | 7.896 |
| mammals | Crocidura_nigrofusca | Crocidura_nigrofusca | LC | Unknown | tropical | nocturnal | 20 | 3815204.408 | 8.475 |
| mammals | Crocidura_nimbae | Crocidura_nimbae | NT | Decreasing | tropical | nocturnal | 15.5 | 45466.8 | 14.247 |
| mammals | Crocidura_niobe | Crocidura_niobe | LC | Unknown | tropical | nocturnal | 16 | 20530.136 | 12.948 |
| mammals | Crocidura_obscurior | Crocidura_obscurior | LC | Unknown | tropical | nocturnal | 3.6 | 372998.111 | 15.866 |
| mammals | Crocidura_olivieri | Crocidura_olivieri | LC | Unknown | tropical | nocturnal | 58 | 15840910.302 | 9.176 |
| mammals | Crocidura_orientalis | Crocidura_orientalis | LC | Unknown | tropical | nocturnal | 11.3 | 664.601 | 16.955 |
| mammals | Crocidura_orii | Crocidura_orii | EN | Decreasing | temperate | nocturnal | 14.4 | 945.177 | 15.343 |

| | | | | | | | | | |
|---|---|---|---|---|---|---|---|---|---|
| mammals | Crocidura_pachyura | Crocidura_pachyura | LC | Unknown | temperate | nocturnal | 14.4 | 64302.221 | 19.205 |
| mammals | Crocidura_palawanensis | Crocidura_palawanensis | LC | Unknown | tropical | nocturnal | 10.6 | 11807.349 | 8.996 |
| mammals | Crocidura_paradoxura | Crocidura_paradoxura | LC | Decreasing | tropical | nocturnal | 8.2 | 105412.678 | 7.684 |
| mammals | Crocidura_parvipes | Crocidura_parvipes | LC | Stable | tropical | nocturnal | 12.6 | 5594890.999 | 7.177 |
| mammals | Crocidura_pasha | Crocidura_pasha | LC | Unknown | tropical | nocturnal | 6.5 | 548549.264 | 9.313 |
| mammals | Crocidura_phaeura | Crocidura_phaeura | EN | Unknown | tropical | nocturnal | 5.7 | 897.11 | 11.747 |
| mammals | Crocidura_phanluongi | NA | LC | Unknown | NA | NA | NA | NA | NA |
| mammals | Crocidura_phuquocensis | NA | DD | Unknown | NA | NA | NA | NA | NA |
| mammals | Crocidura_picea | Crocidura_picea | EN | Decreasing | tropical | nocturnal | 12.5 | 6186.953 | 10.845 |
| mammals | Crocidura_pitmani | NA | DD | Unknown | NA | NA | NA | NA | NA |
| mammals | Crocidura_planiceps | NA | DD | Unknown | NA | NA | NA | NA | NA |
| mammals | Crocidura_poensis | Crocidura_poensis | LC | Unknown | tropical | nocturnal | 16.6 | 1557929.717 | 11.426 |
| mammals | Crocidura_polia | NA | DD | Unknown | NA | NA | NA | NA | NA |
| mammals | Crocidura_pullata | NA | DD | Unknown | NA | NA | NA | NA | NA |
| mammals | Crocidura_raineyi | NA | DD | Unknown | NA | NA | NA | NA | NA |
| mammals | Crocidura_ramona | Crocidura_ramona | LC | Unknown | temperate | nocturnal | 4.5 | 1510.987 | 11.166 |
| mammals | Crocidura_rapax | NA | DD | Unknown | NA | NA | NA | NA | NA |
| mammals | Crocidura_religiosa | NA | DD | Unknown | NA | NA | NA | NA | NA |
| mammals | Crocidura_rhoditis | Crocidura_rhoditis | LC | Stable | tropical | nocturnal | 14 | 97286.89 | 9.623 |
| mammals | Crocidura_roosevelti | Crocidura_roosevelti | LC | Unknown | tropical | nocturnal | 7.6 | 1976021.415 | 7.678 |
| mammals | Crocidura_russula | Crocidura_russula | LC | Stable | temperate | cathemeral | 11.5 | 1926182.44 | 19.581 |
| mammals | Crocidura_selina | NA | DD | Unknown | NA | NA | NA | NA | NA |
| mammals | Crocidura_serezkyensis | Crocidura_serezkyensis | LC | Unknown | temperate | nocturnal | 10.1 | 280073.57 | 11.746 |
| mammals | Crocidura_shantungensis | Crocidura_shantungensis | LC | Stable | temperate | cathemeral | 5.5 | 3049262.93 | 15.757 |
| mammals | Crocidura_sibirica | Crocidura_sibirica | LC | Unknown | temperate | nocturnal | 73 | 6044227.822 | 4.736 |
| mammals | Crocidura_sicula | Crocidura_sicula | LC | Unknown | temperate | cathemeral | 6.9 | 26033.946 | 23.224 |
| mammals | Crocidura_silacea | Crocidura_silacea | LC | Stable | tropical | nocturnal | 8 | 339606.763 | 13.462 |
| mammals | Crocidura_smithii | Crocidura_smithii | LC | Unknown | tropical | nocturnal | 10.8 | 185984.456 | 7.641 |
| mammals | Crocidura_sokolovi | NA | DD | Unknown | NA | NA | NA | NA | NA |
| mammals | Crocidura_somalica | Crocidura_somalica | LC | Unknown | tropical | nocturnal | 11.5 | 175965.751 | 6.437 |
| mammals | Crocidura_stenocephala | Crocidura_stenocephala | EN | Decreasing | tropical | nocturnal | 21 | 1374.891 | 18.6 |
| mammals | Crocidura_suaveolens | Crocidura_suaveolens | LC | Stable | temperate | nocturnal | 5.2 | 10161128.927 | 7.952 |
| mammals | Crocidura_susiana | NA | DD | Unknown | NA | NA | NA | NA | NA |
| mammals | Crocidura_tanakae | Crocidura_tanakae | LC | Unknown | temperate | nocturnal | 10.5 | 82788.976 | 15.82 |

| | | | | | | | | | |
|---|---|---|---|---|---|---|---|---|---|
| mammals | Crocidura_tansaniana | Crocidura_tansaniana | NT | Unknown | tropical | nocturnal | 19 | 4971.377 | 17.484 |
| mammals | Crocidura_tarella | Crocidura_tarella | EN | Decreasing | tropical | nocturnal | 16.4 | 2336.464 | 14.026 |
| mammals | Crocidura_tarfayaensis | Crocidura_tarfayensis | DD | Decreasing | temperate | nocturnal | 6.5 | 267266.721 | 3.245 |
| mammals | Crocidura_telfordi | Crocidura_telfordi | VU | Decreasing | tropical | cathemeral | 18 | 4256.997 | 7.654 |
| mammals | Crocidura_tenuis | NA | DD | Unknown | NA | NA | NA | NA | NA |
| mammals | Crocidura_theresae | Crocidura_theresae | LC | Unknown | tropical | nocturnal | 17.6 | 336307.094 | 15.283 |
| mammals | Crocidura_thomensis | NA | EN | Decreasing | NA | NA | NA | NA | NA |
| mammals | Crocidura_turba | Crocidura_turba | LC | Unknown | tropical | nocturnal | 14.5 | 4878103.987 | 7.904 |
| mammals | Crocidura_ultima | NA | DD | Unknown | NA | NA | NA | NA | NA |
| mammals | Crocidura_usambarae | Crocidura_usambarae | VU | Unknown | tropical | nocturnal | 9.7 | 5471.909 | 17.216 |
| mammals | Crocidura_viaria | Crocidura_viaria | LC | Stable | tropical | nocturnal | 17 | 8205161.015 | 8.49 |
| mammals | Crocidura_virgata | Crocidura_virgata | LC | Unknown | tropical | nocturnal | 7 | 84.379 | 12.73 |
| mammals | Crocidura_voi | Crocidura_voi | LC | Stable | tropical | nocturnal | 13.4 | 3003725.295 | 10.608 |
| mammals | Crocidura_vorax | Crocidura_vorax | LC | Unknown | tropical | nocturnal | 6.7 | 1238060.044 | 12.283 |
| mammals | Crocidura_vosmaeri | NA | DD | Unknown | NA | NA | NA | NA | NA |
| mammals | Crocidura_watasei | Crocidura_watasei | LC | Stable | temperate | nocturnal | 5.5 | 2460.829 | 23.657 |
| mammals | Crocidura_whitakeri | Crocidura_whitakeri | LC | Unknown | temperate | nocturnal | 5.5 | 759432.735 | 13.04 |
| mammals | Crocidura_wimmeri | Crocidura_wimmeri | CR | Unknown | tropical | nocturnal | 23.5 | 110.52 | 33.042 |
| mammals | Crocidura_wuchihensis | NA | DD | Unknown | NA | NA | NA | NA | NA |
| mammals | Crocidura_xantippe | Crocidura_xantippe | LC | Stable | tropical | nocturnal | 15 | 26975.22 | 11.913 |
| mammals | Crocidura_yankariensis | Crocidura_yankariensis | LC | Stable | tropical | nocturnal | 5 | 1389110.096 | 10.573 |
| mammals | Crocidura_zaphiri | NA | DD | Unknown | NA | NA | NA | NA | NA |
| mammals | Crocidura_zarudnyi | Crocidura_zarudnyi | LC | Unknown | temperate | nocturnal | 6.7 | 543621.678 | 6.447 |
| mammals | Crocidura_zimmeri | NA | DD | Unknown | NA | NA | NA | NA | NA |
| mammals | Crocidura_zimmermanni | Crocidura_zimmermanni | EN | Decreasing | temperate | nocturnal | 7.5 | 900.2 | 10.217 |
| mammals | Crocuta_crocuta | Crocuta_crocuta | LC | Decreasing | tropical | nocturnal | 62999.9 | 14474886.614 | 9.673 |
| mammals | Crossarchus_alexandri | Crossarchus_alexandri | LC | Decreasing | tropical | diurnal | 1500 | 1412932.503 | 5.993 |
| mammals | Crossarchus_ansorgei | Crossarchus_ansorgei | LC | Decreasing | tropical | diurnal | 700 | 602126.503 | 4.717 |
| mammals | Crossarchus_obscurus | Crossarchus_obscurus | LC | Unknown | tropical | diurnal | 1250 | 492155.149 | 15.203 |
| mammals | Crossarchus_platycephalus | Crossarchus_platycephalus | LC | Unknown | tropical | diurnal | 1000 | 515163.865 | 9.839 |
| mammals | Crossomys_moncktoni | Crossomys_moncktoni | LC | Unknown | tropical | diurnal | 165 | 138694.412 | 4.768 |
| mammals | Crunomys_celebensis | Crunomys_celebensis | DD | Decreasing | tropical | diurnal | 55 | 10307.78 | 6.375 |
| mammals | Crunomys_fallax | NA | DD | Unknown | NA | NA | NA | NA | NA |
| mammals | Crunomys_melanius | Crunomys_melanius | LC | Unknown | tropical | diurnal | 42.5 | 34392.853 | 13.78 |

| | | | | | | | | | |
|---|---|---|---|---|---|---|---|---|---|
| mammals | Crunomys_suncoides | NA | DD | Unknown | NA | NA | NA | NA | NA |
| mammals | Cryptonanus_agricolai | NA | DD | Unknown | NA | NA | NA | NA | NA |
| mammals | Cryptonanus_chacoensis | Cryptonanus_chacoensis | LC | Stable | temperate | nocturnal | 15.5 | 880492.787 | 7.988 |
| mammals | Cryptonanus_guahybae | NA | DD | Unknown | NA | NA | NA | NA | NA |
| mammals | Cryptonanus_unduaviensis | NA | DD | Unknown | NA | NA | NA | NA | NA |
| mammals | Cryptoprocta_ferox | Cryptoprocta_ferox | VU | Decreasing | tropical | cathemeral | 9500 | 442881.074 | 8.61 |
| mammals | Cryptotis_alticola | NA | DD | Unknown | NA | NA | NA | NA | NA |
| mammals | Cryptotis_endersi | Cryptotis_endersi | EN | Unknown | tropical | cathemeral | 8 | 1049.02 | 7.874 |
| mammals | Cryptotis_equatoris | Cryptotis_equatoris | LC | Stable | tropical | cathemeral | 7.8 | 33341.39 | 11.144 |
| mammals | Cryptotis_goldmani | Cryptotis_goldmani | LC | Stable | tropical | cathemeral | 8 | 47998.509 | 7.07 |
| mammals | Cryptotis_goodwini | Cryptotis_goodwini | LC | Stable | tropical | nocturnal | 14.5 | 39439.36 | 14.206 |
| mammals | Cryptotis_gracilis | Cryptotis_gracilis | LC | Unknown | tropical | nocturnal | 7 | 12411.964 | 10.819 |
| mammals | Cryptotis_griseoventris | Cryptotis_griseoventris | EN | Decreasing | tropical | nocturnal | 9.5 | 156.785 | 7.624 |
| mammals | Cryptotis_magnus | Cryptotis_magna | VU | Decreasing | tropical | nocturnal | 7 | 14384.644 | 6.857 |
| mammals | Cryptotis_mayensis | Cryptotis_mayensis | LC | Decreasing | tropical | nocturnal | 5 | 136563.229 | 6.429 |
| mammals | Cryptotis_medellinius | Cryptotis_medellinia | LC | Stable | tropical | cathemeral | 16.2 | 14038.817 | 14.879 |
| mammals | Cryptotis_meridensis | Cryptotis_meridensis | VU | Stable | tropical | diurnal | 11.6 | 13485.207 | 12.075 |
| mammals | Cryptotis_merriami | Cryptotis_merriami | LC | Stable | tropical | nocturnal | 5.9 | 95424.314 | 11.956 |
| mammals | Cryptotis_merus | Cryptotis_mera | EN | Decreasing | tropical | cathemeral | 7 | 2258.785 | 1.922 |
| mammals | Cryptotis_mexicanus | Cryptotis_mexicana | LC | Decreasing | tropical | nocturnal | 7 | 46362.728 | 9.695 |
| mammals | Cryptotis_montivagus | Cryptotis_montivaga | LC | Stable | tropical | cathemeral | 11.3 | 29768.535 | 11.505 |
| mammals | Cryptotis_orophilus | NA | DD | Unknown | NA | NA | NA | NA | NA |
| mammals | Cryptotis_parvus | Cryptotis_parva | LC | Stable | temperate | cathemeral | 5 | 4122005.688 | 16.415 |
| mammals | Cryptotis_peregrinus | NA | DD | Unknown | NA | NA | NA | NA | NA |
| mammals | Cryptotis_peruviensis | NA | DD | Unknown | NA | NA | NA | NA | NA |
| mammals | Cryptotis_phillipsii | Cryptotis_phillipsii | VU | Decreasing | tropical | nocturnal | 8 | 5969.886 | 6.619 |
| mammals | Cryptotis_squamipes | Cryptotis_squamipes | LC | Unknown | tropical | cathemeral | 11 | 38079.812 | 10.246 |
| mammals | Cryptotis_tamensis | Cryptotis_tamensis | LC | Unknown | tropical | cathemeral | 13.9 | 5984.295 | 8.94 |
| mammals | Cryptotis_thomasi | Cryptotis_thomasi | LC | Stable | tropical | cathemeral | 10 | 23699.206 | 8.971 |
| mammals | Cryptotis_tropicalis | NA | DD | Unknown | NA | NA | NA | NA | NA |
| mammals | Ctenodactylus_gundi | Ctenodactylus_gundi | LC | Unknown | temperate | diurnal | 289 | 132731.286 | 7.052 |
| mammals | Ctenodactylus_vali | NA | DD | Unknown | NA | NA | NA | NA | NA |
| mammals | Ctenomys_peruanus | Ctenomys_peruanus | LC | Stable | tropical | diurnal | 490 | 5464.143 | 7.626 |
| mammals | Cuniculus_paca | Cuniculus_paca | LC | Stable | tropical | nocturnal | 8000 | 13436656.961 | 5.969 |

| | | | | | | | | | |
|---|---|---|---|---|---|---|---|---|---|
| mammals | Cuniculus_taczanowskii | Cuniculus_taczanowskii | NT | Decreasing | tropical | nocturnal | 8999.9 | 406861.256 | 7.732 |
| mammals | Cuon_alpinus | Cuon_alpinus | EN | Decreasing | temperate | cathemeral | 12760 | 3444014.514 | 11.34 |
| mammals | Cuscomys_ashaninka | NA | DD | Unknown | NA | NA | NA | NA | NA |
| mammals | Cuscomys_oblativus | NA | DD | Unknown | NA | NA | NA | NA | NA |
| mammals | Cyclopes_didactylus | Cyclopes_didactylus | LC | Unknown | tropical | nocturnal | 329.5 | 7892585.181 | 3.509 |
| mammals | Cynictis_penicillata | Cynictis_penicillata | LC | Stable | temperate | diurnal | 836 | 2252374.388 | 7.15 |
| mammals | Cynocephalus_volans | Cynocephalus_volans | LC | Stable | tropical | nocturnal | 1250 | 120524.268 | 12.096 |
| mammals | Cynogale_bennettii | Cynogale_bennettii | EN | Decreasing | tropical | nocturnal | 4500 | 1090109.434 | 8.889 |
| mammals | Cynomops_abrasus | NA | DD | Unknown | NA | NA | NA | NA | NA |
| mammals | Cynomops_greenhalli | Cynomops_greenhalli | LC | Unknown | tropical | nocturnal | 15 | 4472130.313 | 4.806 |
| mammals | Cynomops_mexicanus | Cynomops_mexicanus | LC | Unknown | tropical | nocturnal | 20 | 236547.604 | 9.025 |
| mammals | Cynomops_milleri | Cynomops_milleri | LC | Unknown | tropical | nocturnal | 16 | 2203941.825 | 2.199 |
| mammals | Cynomops_planirostris | Cynomops_planirostris | LC | Unknown | tropical | nocturnal | 12.4 | 11852682.307 | 5.097 |
| mammals | Cynomys_gunnisoni | Cynomys_gunnisoni | LC | Decreasing | temperate | diurnal | 925 | 309504.09 | 9.208 |
| mammals | Cynomys_leucurus | Cynomys_leucurus | LC | Decreasing | temperate | diurnal | 908.5 | 269601.971 | 9.732 |
| mammals | Cynomys_ludovicianus | Cynomys_ludovicianus | LC | Decreasing | temperate | diurnal | 1364 | 1820824.736 | 12.111 |
| mammals | Cynomys_mexicanus | Cynomys_mexicanus | EN | Decreasing | temperate | diurnal | 900 | 13330.503 | 5.684 |
| mammals | Cynomys_parvidens | Cynomys_parvidens | EN | Decreasing | temperate | diurnal | 900 | 31805.77 | 8.408 |
| mammals | Cynopterus_brachyotis | Cynopterus_brachyotis | LC | Unknown | tropical | nocturnal | 31.2 | 2699824.262 | 12.335 |
| mammals | Cynopterus_horsfieldii | Cynopterus_horsfieldii | LC | Unknown | tropical | nocturnal | 55.1 | 1485603.334 | 9.257 |
| mammals | Cynopterus_luzoniensis | Cynopterus_luzoniensis | LC | Stable | tropical | nocturnal | 27.9 | 455804.74 | 12.042 |
| mammals | Cynopterus_minutus | Cynopterus_minutus | LC | Decreasing | tropical | nocturnal | 25.8 | 1461294.071 | 8.66 |
| mammals | Cynopterus_nusatenggara | Cynopterus_nusatenggara | LC | Stable | tropical | nocturnal | 36.4 | 85669.017 | 13.966 |
| mammals | Cynopterus_sphinx | Cynopterus_sphinx | LC | Increasing | tropical | nocturnal | 43.7 | 6456517.373 | 14.525 |
| mammals | Cynopterus_titthaecheilus | Cynopterus_titthaecheilus | LC | Stable | tropical | nocturnal | 56.3 | 598087.135 | 12.741 |
| mammals | Cyttarops_alecto | Cyttarops_alecto | LC | Unknown | tropical | nocturnal | 5.3 | 687747.273 | 2.284 |
| mammals | Dacnomys_millardi | NA | DD | Unknown | NA | NA | NA | NA | NA |
| mammals | Dactylomys_boliviensis | Dactylomys_boliviensis | LC | Stable | tropical | nocturnal | 728.8 | 336905.94 | 2.376 |
| mammals | Dactylomys_dactylinus | Dactylomys_dactylinus | LC | Stable | tropical | nocturnal | 650 | 3306574.863 | 1.91 |
| mammals | Dactylomys_peruanus | NA | DD | Unknown | NA | NA | NA | NA | NA |
| mammals | Dactylopsila_megalura | Dactylopsila_megalura | LC | Unknown | tropical | nocturnal | 387.5 | 62974.324 | 4.115 |
| mammals | Dactylopsila_palpator | Dactylonax_palpator | LC | Unknown | tropical | nocturnal | 398 | 213289.372 | 4.461 |
| mammals | Dactylopsila_tatei | Dactylopsila_tatei | EN | Unknown | tropical | nocturnal | 252 | 144.867 | 4.59 |
| mammals | Dactylopsila_trivirgata | Dactylopsila_trivirgata | LC | Decreasing | tropical | nocturnal | 413.5 | 765397.192 | 3.367 |

| | | | | | | | | | |
|---|---|---|---|---|---|---|---|---|---|
| mammals | Dama_dama | Dama_dama | LC | Unknown | temperate | cathemeral | 56250 | 4018821.402 | 20.075 |
| mammals | Dama_mesopotamica | Dama_mesopotamica | EN | Increasing | temperate | nocturnal | 72500 | 614.206 | 12.192 |
| mammals | Damaliscus_lunatus | Damaliscus_lunatus | LC | Decreasing | tropical | diurnal | 136000.3 | 1840650.421 | 7.07 |
| mammals | Damaliscus_pygargus | Damaliscus_pygargus | LC | Increasing | temperate | cathemeral | 102000 | 395523.616 | 12.347 |
| mammals | Daptomys_mussoi | Neusticomys_mussoi | VU | Decreasing | tropical | nocturnal | 40 | 19038.667 | 15.045 |
| mammals | Daptomys_oyapocki | NA | DD | Unknown | NA | NA | NA | NA | NA |
| mammals | Daptomys_venezuelae | Neusticomys_venezuelae | VU | Decreasing | tropical | nocturnal | 47.1 | 665777.524 | 3.2 |
| mammals | Dasycercus_blythi | Dasycercus_blythi | LC | Unknown | temperate | nocturnal | 83.8 | 1471984.401 | 2.025 |
| mammals | Dasycercus_cristicauda | Dasycercus_cristicauda | NT | Stable | temperate | nocturnal | 100 | 114850.908 | 1.575 |
| mammals | Dasykaluta_rosamondae | Dasykaluta_rosamondae | LC | Unknown | tropical | diurnal | 28.8 | 386598.868 | 3.282 |
| mammals | Dasymys_foxi | NA | DD | Unknown | NA | NA | NA | NA | NA |
| mammals | Dasymys_incomtus | Dasymys_incomtus | LC | Unknown | tropical | diurnal | 90.2 | 6435516.366 | 8.862 |
| mammals | Dasymys_montanus | Dasymys_montanus | EN | Decreasing | tropical | cathemeral | 82 | 1879.354 | 9.833 |
| mammals | Dasymys_nudipes | NA | DD | Unknown | NA | NA | NA | NA | NA |
| mammals | Dasymys_rufulus | Dasymys_rufulus | LC | Unknown | tropical | nocturnal | 85 | 1666893.487 | 13.89 |
| mammals | Dasyprocta_azarae | Dasyprocta_azarae | DD | Decreasing | tropical | diurnal | 2310 | 1667306.947 | 10.67 |
| mammals | Dasyprocta_coibae | Dasyprocta_coibae | NT | Unknown | tropical | diurnal | 2195.5 | 497.243 | 1.061 |
| mammals | Dasyprocta_fuliginosa | Dasyprocta_fuliginosa | LC | Stable | tropical | diurnal | 3500 | 2612229.73 | 1.889 |
| mammals | Dasyprocta_guamara | Dasyprocta_guamara | NT | Decreasing | tropical | diurnal | 2650 | 7993.764 | 2.313 |
| mammals | Dasyprocta_kalinowskii | NA | DD | Unknown | NA | NA | NA | NA | NA |
| mammals | Dasyprocta_leporina | Dasyprocta_leporina | LC | Stable | tropical | diurnal | 3020 | 4557304.869 | 5.261 |
| mammals | Dasyprocta_mexicana | Dasyprocta_mexicana | CR | Decreasing | tropical | diurnal | 5000 | 107846.18 | 15.109 |
| mammals | Dasyprocta_prymnolopha | Dasyprocta_prymnolopha | LC | Unknown | tropical | diurnal | 2900 | 1929753.313 | 7.596 |
| mammals | Dasyprocta_punctata | Dasyprocta_punctata | LC | Stable | tropical | diurnal | 2675 | 1306837.669 | 10.985 |
| mammals | Dasyprocta_ruatanica | Dasyprocta_ruatanica | EN | Decreasing | tropical | diurnal | 1659.9 | 156.605 | 20.035 |
| mammals | Dasypus_kappleri | Dasypus_kappleri | LC | Unknown | tropical | nocturnal | 9500 | 6028983.441 | 2.095 |
| mammals | Dasypus_mazzai | NA | DD | Unknown | NA | NA | NA | NA | NA |
| mammals | Dasypus_novemcinctus | Dasypus_novemcinctus | LC | Stable | tropical | cathemeral | 4203.8 | 18207313.673 | 7.421 |
| mammals | Dasypus_pilosus | NA | DD | Unknown | NA | NA | NA | NA | NA |
| mammals | Dasypus_sabanicola | Dasypus_sabanicola | NT | Decreasing | tropical | diurnal | 1150 | 441453.933 | 6.596 |
| mammals | Dasypus_septemcinctus | Dasypus_septemcinctus | LC | Unknown | tropical | cathemeral | 1526.7 | 6904376.956 | 8.407 |
| mammals | Dasyuroides_byrnei | Dasyuroides_byrnei | VU | Decreasing | temperate | nocturnal | 103.8 | 71511.287 | 3.081 |
| mammals | Dasyurus_albopunctatus | Dasyurus_albopunctatus | NT | Decreasing | tropical | diurnal | 637 | 580310.276 | 3.801 |
| mammals | Dasyurus_geoffroii | Dasyurus_geoffroii | NT | Stable | temperate | nocturnal | 1100 | 188679.587 | 9.97 |

| | | | | | | | | | |
|---|---|---|---|---|---|---|---|---|---|
| mammals | Dasyurus_hallucatus | Dasyurus_hallucatus | EN | Decreasing | tropical | nocturnal | 525 | 331682.507 | 4.013 |
| mammals | Dasyurus_maculatus | Dasyurus_maculatus | NT | Decreasing | temperate | cathemeral | 5500 | 464968.774 | 8.256 |
| mammals | Dasyurus_spartacus | Dasyurus_spartacus | NT | Unknown | tropical | nocturnal | 1212.6 | 26551.56 | 3.912 |
| mammals | Dasyurus_viverrinus | Dasyurus_viverrinus | EN | Decreasing | temperate | nocturnal | 1090 | 63883.697 | 5.844 |
| mammals | Daubentonia_madagascariensis | Daubentonia_madagascariensis | EN | Decreasing | tropical | nocturnal | 2500 | 124673.999 | 7.451 |
| mammals | Delanymys_brooksi | Delanymys_brooksi | VU | Decreasing | tropical | nocturnal | 5 | 3946.317 | 12.963 |
| mammals | Delomys_dorsalis | Delomys_dorsalis | LC | Decreasing | temperate | nocturnal | 67.5 | 349472.703 | 12.139 |
| mammals | Delomys_sublineatus | Delomys_sublineatus | LC | Unknown | tropical | nocturnal | 90 | 150814.591 | 12.988 |
| mammals | Deltamys_kempi | Deltamys_kempi | LC | Unknown | temperate | nocturnal | 26.4 | 141581.878 | 11.154 |
| mammals | Dendrogale_melanura | Dendrogale_melanura | DD | Decreasing | tropical | diurnal | 60 | 33542.556 | 4.351 |
| mammals | Dendrogale_murina | Dendrogale_murina | LC | Decreasing | tropical | diurnal | 50 | 204367.54 | 12.2 |
| mammals | Dendrohyrax_arboreus | Dendrohyrax_arboreus | LC | Decreasing | tropical | cathemeral | 2950 | 2446114.615 | 10.031 |
| mammals | Dendrohyrax_validus | Dendrohyrax_validus | NT | Decreasing | tropical | nocturnal | 2430 | 37809.884 | 13.089 |
| mammals | Dendrolagus_bennettianus | Dendrolagus_bennettianus | NT | Stable | tropical | nocturnal | 10950 | 3562.458 | 2.36 |
| mammals | Dendrolagus_dorianus | Dendrolagus_dorianus | VU | Decreasing | tropical | diurnal | 10068.3 | 39847.247 | 3.162 |
| mammals | Dendrolagus_goodfellowi | Dendrolagus_goodfellowi | EN | Decreasing | tropical | diurnal | 8025 | 114894.765 | 4.95 |
| mammals | Dendrolagus_lumholtzi | Dendrolagus_lumholtzi | NT | Unknown | tropical | cathemeral | 6750 | 7383.082 | 4.212 |
| mammals | Dendrolagus_matschiei | Dendrolagus_matschiei | EN | Decreasing | tropical | diurnal | 6960 | 13584.63 | 4.064 |
| mammals | Dendrolagus_mbaiso | Dendrolagus_mbaiso | EN | Decreasing | tropical | diurnal | 8750 | 5558.339 | 4.992 |
| mammals | Dendrolagus_notatus | Dendrolagus_notatus | EN | Decreasing | tropical | diurnal | 6250 | 84949.684 | 5.864 |
| mammals | Dendrolagus_pulcherrimus | Dendrolagus_pulcherrimus | CR | Stable | tropical | diurnal | 6536.8 | 2990.486 | 4.219 |
| mammals | Dendrolagus_scottae | Dendrolagus_scottae | CR | Stable | tropical | diurnal | 10375 | 1246.358 | 3.783 |
| mammals | Dendrolagus_spadix | Dendrolagus_spadix | VU | Decreasing | tropical | cathemeral | 9100 | 49872.442 | 2.083 |
| mammals | Dendrolagus_stellarum | Dendrolagus_stellarum | VU | Decreasing | tropical | diurnal | 8666.7 | 46918.516 | 3.974 |
| mammals | Dendrolagus_ursinus | Dendrolagus_ursinus | VU | Decreasing | tropical | diurnal | 13250 | 28295.883 | 2.826 |
| mammals | Dendromus_insignis | Dendromus_insignis | LC | Stable | tropical | nocturnal | 11.1 | 26896.865 | 16.05 |
| mammals | Dendromus_kahuziensis | Dendromus_kahuziensis | CR | Decreasing | tropical | nocturnal | 12 | 75.006 | 9.134 |
| mammals | Dendromus_lovati | Dendromus_lovati | LC | Decreasing | tropical | nocturnal | 12 | 38666.694 | 15.11 |
| mammals | Dendromus_mesomelas | Dendromus_mesomelas | LC | Stable | tropical | nocturnal | 11.1 | 588635.037 | 9.637 |
| mammals | Dendromus_messorius | Dendromus_messorius | LC | Stable | tropical | nocturnal | 14.6 | 60941.424 | 9.94 |
| mammals | Dendromus_mystacalis | Dendromus_mystacalis | LC | Stable | tropical | nocturnal | 7.7 | 2937067.579 | 10.443 |
| mammals | Dendromus_nyasae | Dendromus_kivu | LC | Stable | tropical | nocturnal | 11 | 27532.342 | 16.948 |
| mammals | Dendromus_oreas | Dendromus_oreas | LC | Unknown | tropical | nocturnal | 10.7 | 1370.2 | 11.162 |
| mammals | Dendromus_vernayi | NA | DD | Unknown | NA | NA | NA | NA | NA |

| | | | | | | | | | |
|---|---|---|---|---|---|---|---|---|---|
| mammals | Dendroprionomys_rousseloti | NA | DD | Unknown | NA | NA | NA | NA | NA |
| mammals | Deomys_ferrugineus | Deomys_ferrugineus | LC | Stable | tropical | nocturnal | 62 | 1293572.477 | 6.941 |
| mammals | Dephomys_defua | Dephomys_defua | LC | Unknown | tropical | nocturnal | 43.1 | 426036.662 | 15.803 |
| mammals | Dermanura_anderseni | Dermanura_anderseni | LC | Unknown | tropical | nocturnal | 138 | 4072157.249 | 2.307 |
| mammals | Dermanura_azteca | Dermanura_azteca | LC | Unknown | tropical | nocturnal | 20.4 | 606576.244 | 11.146 |
| mammals | Dermanura_cinerea | Dermanura_cinerea | LC | Unknown | tropical | nocturnal | 12.6 | 4322309.531 | 6.238 |
| mammals | Dermanura_glauca | Dermanura_glauca | LC | Unknown | tropical | nocturnal | 11.1 | 2660226.905 | 4.295 |
| mammals | Dermanura_gnoma | Dermanura_gnoma | LC | Unknown | tropical | nocturnal | 10.5 | 10008664.031 | 4.177 |
| mammals | Dermanura_phaeotis | Dermanura_phaeotis | LC | Stable | tropical | nocturnal | 11.6 | 3737367.004 | 6.642 |
| mammals | Dermanura_rosenbergi | NA | DD | Unknown | NA | NA | NA | NA | NA |
| mammals | Dermanura_tolteca | Dermanura_tolteca | LC | Unknown | tropical | nocturnal | 15.4 | 849394.914 | 10.594 |
| mammals | Dermanura_watsoni | Dermanura_watsoni | LC | Stable | tropical | nocturnal | 12 | 554677.136 | 9.029 |
| mammals | Desmalopex_leucoptera | Pteropus_leucopterus | VU | Decreasing | tropical | nocturnal | 340 | 52582.636 | 13.038 |
| mammals | Desmana_moschata | Desmana_moschata | CR | Decreasing | temperate | nocturnal | 383 | 123164.035 | 11.504 |
| mammals | Desmodilliscus_braueri | Desmodilliscus_braueri | LC | Stable | tropical | nocturnal | 10 | 3380204.826 | 5.838 |
| mammals | Desmodillus_auricularis | Desmodillus_auricularis | LC | Stable | temperate | nocturnal | 53.3 | 1955073.28 | 6.53 |
| mammals | Desmodus_rotundus | Desmodus_rotundus | LC | Stable | tropical | nocturnal | 31.2 | 17726517.724 | 6.374 |
| mammals | Desmomys_harringtoni | Desmomys_harringtoni | LC | Unknown | tropical | nocturnal | 90 | 265251.583 | 12.775 |
| mammals | Desmomys_yaldeni | Desmomys_yaldeni | VU | Unknown | tropical | nocturnal | 47 | 9679.527 | 11.943 |
| mammals | Diaemus_youngii | Diaemus_youngi | LC | Unknown | tropical | nocturnal | 36.3 | 13059176.179 | 5.735 |
| mammals | Dicerorhinus_sumatrensis | Dicerorhinus_sumatrensis | CR | Decreasing | tropical | cathemeral | 1266667 | 960082.542 | 7.258 |
| mammals | Diceros_bicornis | Diceros_bicornis | CR | Increasing | tropical | cathemeral | 1180510.9 | 19274959.561 | 7.979 |
| mammals | Diclidurus_albus | Diclidurus_albus | LC | Unknown | tropical | nocturnal | 15.6 | 12498370.932 | 5.541 |
| mammals | Diclidurus_ingens | NA | DD | Unknown | NA | NA | NA | NA | NA |
| mammals | Diclidurus_isabella | Diclidurus_isabella | LC | Unknown | tropical | nocturnal | 14 | 1061836.667 | 1.071 |
| mammals | Diclidurus_scutatus | Diclidurus_scutatus | LC | Unknown | tropical | nocturnal | 13.3 | 1540656.031 | 2.073 |
| mammals | Dicotyles_tajacu | Pecari_tajacu | LC | Stable | tropical | cathemeral | 21266.7 | 15938720.066 | 6.476 |
| mammals | Dicrostonyx_hudsonius | Dicrostonyx_hudsonius | LC | Stable | temperate | cathemeral | 57 | 492354.102 | 0.212 |
| mammals | Dicrostonyx_nelsoni | NA | LC | Stable | NA | NA | NA | NA | NA |
| mammals | Dicrostonyx_nunatakensis | Dicrostonyx_nunatakensis | LC | Unknown | temperate | cathemeral | 58.5 | 54570.076 | 0.317 |
| mammals | Dicrostonyx_unalascensis | NA | DD | Unknown | NA | NA | NA | NA | NA |
| mammals | Didelphis_albiventris | Didelphis_albiventris | LC | Stable | tropical | nocturnal | 904 | 6795565.816 | 8.883 |
| mammals | Didelphis_aurita | Didelphis_aurita | LC | Stable | tropical | nocturnal | 1164 | 1446051.047 | 12.92 |
| mammals | Didelphis_imperfecta | Didelphis_imperfecta | LC | Stable | tropical | nocturnal | 873.7 | 530382.959 | 1.068 |

| | | | | | | | | | |
|---|---|---|---|---|---|---|---|---|---|
| mammals | Didelphis_marsupialis | Didelphis_marsupialis | LC | Stable | tropical | nocturnal | 1091.2 | 9324398.927 | 4.481 |
| mammals | Didelphis_pernigra | Didelphis_pernigra | LC | Stable | tropical | nocturnal | 1138.5 | 1033727.886 | 9.09 |
| mammals | Didelphis_virginiana | Didelphis_virginiana | LC | Increasing | temperate | nocturnal | 2195.5 | 6033229.978 | 14.603 |
| mammals | Dinaromys_bogdanovi | Dinaromys_bogdanovi | VU | Decreasing | temperate | nocturnal | 56 | 79580.559 | 13.871 |
| mammals | Dinomys_branickii | Dinomys_branickii | LC | Unknown | tropical | nocturnal | 12500 | 1390156.7 | 4.734 |
| mammals | Diomys_crumpi | NA | DD | Unknown | NA | NA | NA | NA | NA |
| mammals | Diphylla_ecaudata | Diphylla_ecaudata | LC | Stable | tropical | nocturnal | 29.3 | 8092435.593 | 8.034 |
| mammals | Diplogale_hosei | Diplogale_hosei | VU | Decreasing | tropical | nocturnal | 1300 | 100926.11 | 2.421 |
| mammals | Diplomys_caniceps | NA | DD | Unknown | NA | NA | NA | NA | NA |
| mammals | Diplomys_labilis | Diplomys_labilis | LC | Stable | tropical | nocturnal | 227.5 | 110966.521 | 5.343 |
| mammals | Diplothrix_legata | Diplothrix_legata | EN | Decreasing | temperate | nocturnal | 505.9 | 799.969 | 13.773 |
| mammals | Dipodomys_agilis | Dipodomys_agilis | LC | Decreasing | temperate | nocturnal | 57.9 | 38595.524 | 14.584 |
| mammals | Dipodomys_californicus | Dipodomys_californicus | LC | Stable | temperate | nocturnal | 85 | 86209.649 | 11.693 |
| mammals | Dipodomys_compactus | Dipodomys_compactus | LC | Stable | temperate | nocturnal | 49.3 | 49440.903 | 15.792 |
| mammals | Dipodomys_deserti | Dipodomys_deserti | LC | Stable | temperate | nocturnal | 104.5 | 307685.283 | 7.62 |
| mammals | Dipodomys_elator | Dipodomys_elator | VU | Decreasing | temperate | nocturnal | 77.5 | 10767.664 | 14.485 |
| mammals | Dipodomys_gravipes | Dipodomys_gravipes | CR | Unknown | temperate | nocturnal | 84 | 5681.977 | 5.144 |
| mammals | Dipodomys_heermanni | Dipodomys_heermanni | LC | Unknown | temperate | nocturnal | 72 | 70208.591 | 18.169 |
| mammals | Dipodomys_ingens | Dipodomys_ingens | EN | Decreasing | temperate | nocturnal | 133.9 | 6784.268 | 14.98 |
| mammals | Dipodomys_microps | Dipodomys_microps | LC | Stable | temperate | cathemeral | 54.6 | 392308.36 | 7.568 |
| mammals | Dipodomys_nelsoni | Dipodomys_nelsoni | LC | Stable | temperate | nocturnal | 88.6 | 192588.892 | 5.538 |
| mammals | Dipodomys_nitratoides | Dipodomys_nitratoides | VU | Decreasing | temperate | nocturnal | 43.9 | 11331.603 | 19.066 |
| mammals | Dipodomys_ordii | Dipodomys_ordii | LC | Stable | temperate | nocturnal | 60.4 | 3419731.414 | 10.307 |
| mammals | Dipodomys_panamintinus | Dipodomys_panamintinus | LC | Stable | temperate | nocturnal | 74.7 | 60988.137 | 9.496 |
| mammals | Dipodomys_phillipsii | Dipodomys_phillipsii | LC | Stable | tropical | nocturnal | 41 | 62377.861 | 19.312 |
| mammals | Dipodomys_simulans | Dipodomys_simulans | LC | Unknown | temperate | nocturnal | 57.5 | 97814.574 | 10.445 |
| mammals | Dipodomys_spectabilis | Dipodomys_spectabilis | NT | Decreasing | temperate | cathemeral | 135.9 | 523883.186 | 9.359 |
| mammals | Dipodomys_stephensi | Dipodomys_stephensi | VU | Decreasing | temperate | nocturnal | 69.8 | 9357.276 | 21.502 |
| mammals | Dipodomys_venustus | Dipodomys_venustus | LC | Decreasing | temperate | nocturnal | 72 | 16356.394 | 17.569 |
| mammals | Dipus_sagitta | Dipus_sagitta | LC | Stable | temperate | nocturnal | 85.7 | 3156284.933 | 4.513 |
| mammals | Distoechurus_pennatus | Distoechurus_pennatus | LC | Stable | tropical | nocturnal | 44.1 | 536111.971 | 3.761 |
| mammals | Dobsonia_anderseni | Dobsonia_anderseni | LC | Stable | tropical | nocturnal | 204.1 | 46526.996 | 4.523 |
| mammals | Dobsonia_beauforti | Dobsonia_beauforti | LC | Unknown | tropical | nocturnal | 159 | 10000.152 | 4.82 |
| mammals | Dobsonia_chapmani | Dobsonia_chapmani | CR | Decreasing | tropical | nocturnal | 134 | 29.113 | 22.865 |

| | | | | | | | | | |
|---|---|---|---|---|---|---|---|---|---|
| mammals | Dobsonia_crenulata | Dobsonia_crenulata | LC | Stable | tropical | nocturnal | 272 | 209291.771 | 8.952 |
| mammals | Dobsonia_emersa | Dobsonia_emersa | VU | Unknown | tropical | nocturnal | 195.5 | 2533.405 | 8.96 |
| mammals | Dobsonia_exoleta | Dobsonia_exoleta | LC | Unknown | tropical | nocturnal | 249 | 179910.944 | 9.293 |
| mammals | Dobsonia_inermis | Dobsonia_inermis | LC | Stable | tropical | nocturnal | 149.5 | 34750.34 | 4.78 |
| mammals | Dobsonia_minor | Dobsonia_minor | LC | Unknown | tropical | nocturnal | 78.2 | 614847.582 | 3.085 |
| mammals | Dobsonia_moluccensis | Dobsonia_moluccensis | LC | Stable | tropical | nocturnal | 381.4 | 39785.126 | 6.446 |
| mammals | Dobsonia_pannietensis | Dobsonia_pannietensis | NT | Decreasing | tropical | nocturnal | 267.7 | 5506.824 | 5.2 |
| mammals | Dobsonia_peronii | Dobsonia_peronii | LC | Stable | tropical | nocturnal | 216.9 | 87783.692 | 13.997 |
| mammals | Dobsonia_praedatrix | Dobsonia_praedatrix | LC | Unknown | tropical | nocturnal | 188.7 | 44746.121 | 4.427 |
| mammals | Dobsonia_viridis | Dobsonia_viridis | LC | Stable | tropical | nocturnal | 238 | 27753.009 | 7.258 |
| mammals | Dolichotis_patagonum | Dolichotis_patagonum | NT | Decreasing | temperate | diurnal | 8000 | 1242816.997 | 6.823 |
| mammals | Dolichotis_salinicola | Dolichotis_salinicola | LC | Stable | temperate | diurnal | 1600 | 735968.049 | 5.41 |
| mammals | Dologale_dybowskii | NA | DD | Unknown | NA | NA | NA | NA | NA |
| mammals | Dorcatragus_megalotis | Dorcatragus_megalotis | VU | Decreasing | tropical | diurnal | 10249.9 | 51752.99 | 8.372 |
| mammals | Dorcopsis_muelleri | Dorcopsis_muelleri | LC | Stable | tropical | nocturnal | 5500 | 125655.341 | 3.177 |
| mammals | Dorcopsulus_macleayi | Dorcopsulus_macleayi | LC | Stable | tropical | nocturnal | 2988 | 25885.916 | 2.679 |
| mammals | Dorcopsulus_vanheurni | Dorcopsulus_vanheurni | NT | Decreasing | tropical | diurnal | 1785.8 | 179392.504 | 4.668 |
| mammals | Doryrhina_camerunensis | NA | DD | Unknown | NA | NA | NA | NA | NA |
| mammals | Doryrhina_cyclops | Hipposideros_cyclops | LC | Decreasing | tropical | nocturnal | 33.5 | 2331176.89 | 10.312 |
| mammals | Dremomys_gularis | NA | DD | Unknown | NA | NA | NA | NA | NA |
| mammals | Dremomys_lokriah | Dremomys_lokriah | LC | Decreasing | temperate | diurnal | 227 | 361878.673 | 6.421 |
| mammals | Dremomys_pernyi | Dremomys_pernyi | LC | Stable | temperate | diurnal | 198.6 | 1752951.561 | 12.694 |
| mammals | Dremomys_pyrrhomerus | Dremomys_pyrrhomerus | LC | Unknown | temperate | diurnal | 245 | 273521.526 | 14.882 |
| mammals | Dremomys_rufigenis | Dremomys_rufigenis | LC | Stable | tropical | diurnal | 200 | 1600228.668 | 11.09 |
| mammals | Dromiciops_gliroides | Dromiciops_gliroides | NT | Decreasing | temperate | nocturnal | 22.3 | 139169.355 | 9.054 |
| mammals | Dryomys_laniger | NA | DD | Unknown | NA | NA | NA | NA | NA |
| mammals | Dryomys_nitedula | Dryomys_nitedula | LC | Unknown | temperate | nocturnal | 28.9 | 5228746.46 | 13.35 |
| mammals | Dyacopterus_brooksi | Dyacopterus_brooksi | VU | Decreasing | tropical | nocturnal | 88.5 | 429470.115 | 9.673 |
| mammals | Dyacopterus_spadiceus | Dyacopterus_spadiceus | NT | Decreasing | tropical | nocturnal | 84.2 | 445030.607 | 9.137 |
| mammals | Dymecodon_pilirostris | Dymecodon_pilirostris | LC | Stable | temperate | cathemeral | 16.2 | 38974.785 | 12.004 |
| mammals | Echimys_chrysurus | Echimys_chrysurus | LC | Stable | tropical | nocturnal | 652.5 | 1583343.242 | 2.044 |
| mammals | Echimys_saturnus | NA | DD | Unknown | NA | NA | NA | NA | NA |
| mammals | Echimys_vieirai | NA | DD | Unknown | NA | NA | NA | NA | NA |
| mammals | Echinops_telfairi | Echinops_telfairi | LC | Stable | tropical | nocturnal | 180 | 111004.162 | 9.569 |

| | | | | | | | | | |
|---|---|---|---|---|---|---|---|---|---|
| mammals | Echinosorex_gymnura | Echinosorex_gymnura | LC | Unknown | tropical | nocturnal | 957 | 1373927.428 | 8.026 |
| mammals | Echiothrix_centrosa | Echiothrix_centrosa | NT | Decreasing | tropical | nocturnal | 260.9 | 34850.353 | 7.691 |
| mammals | Echiothrix_leucura | Echiothrix_leucura | EN | Decreasing | tropical | nocturnal | 267.3 | 7189.865 | 12.488 |
| mammals | Echymipera_clara | Echymipera_clara | LC | Unknown | tropical | nocturnal | 1236.5 | 175468.598 | 3.157 |
| mammals | Echymipera_davidi | Echymipera_davidi | EN | Unknown | tropical | nocturnal | 950 | 278.313 | 9.271 |
| mammals | Echymipera_echinista | NA | DD | Unknown | NA | NA | NA | NA | NA |
| mammals | Echymipera_kalubu | Echymipera_kalubu | LC | Stable | tropical | nocturnal | 782.2 | 840230.18 | 3.49 |
| mammals | Echymipera_rufescens | Echymipera_rufescens | LC | Decreasing | tropical | nocturnal | 1100 | 859314.675 | 3.383 |
| mammals | Ectophylla_alba | Ectophylla_alba | NT | Decreasing | tropical | nocturnal | 5.8 | 124361.573 | 7.426 |
| mammals | Eidolon_dupreanum | Eidolon_dupreanum | VU | Decreasing | tropical | nocturnal | 283.8 | 460027.211 | 8.95 |
| mammals | Eidolon_helvum | Eidolon_helvum | NT | Decreasing | tropical | nocturnal | 253.3 | 11789001.27 | 9.871 |
| mammals | Eira_barbara | Eira_barbara | LC | Decreasing | tropical | diurnal | 3910 | 13375343.987 | 5.981 |
| mammals | Elephantulus_fuscipes | NA | DD | Unknown | NA | NA | NA | NA | NA |
| mammals | Elephantulus_fuscus | NA | DD | Unknown | NA | NA | NA | NA | NA |
| mammals | Elephantulus_myurus | Elephantulus_myurus | LC | Stable | temperate | cathemeral | 52.4 | 968379.599 | 11.592 |
| mammals | Elephas_maximus | Elephas_maximus | EN | Decreasing | temperate | cathemeral | 3160000 | 894367.58 | 9.325 |
| mammals | Eligmodontia_moreni | Eligmodontia_moreni | LC | Stable | temperate | nocturnal | 18 | 46842.775 | 3.542 |
| mammals | Eligmodontia_morgani | Eligmodontia_morgani | LC | Stable | temperate | nocturnal | 16.5 | 371796.23 | 5.26 |
| mammals | Eligmodontia_puerulus | Eligmodontia_puerulus | LC | Stable | tropical | nocturnal | 28.5 | 447158.566 | 5.798 |
| mammals | Eligmodontia_typus | Eligmodontia_typus | LC | Stable | temperate | nocturnal | 17.3 | 922783.357 | 5.674 |
| mammals | Eliomys_melanurus | Eliomys_melanurus | LC | Unknown | temperate | nocturnal | 51.8 | 148691.204 | 9.779 |
| mammals | Eliomys_munbyanus | Eliomys_munbyanus | LC | Stable | temperate | nocturnal | 52 | 873740.078 | 11.975 |
| mammals | Eliomys_quercinus | Eliomys_quercinus | NT | Decreasing | temperate | nocturnal | 115 | 2362160.86 | 18.671 |
| mammals | Eliurus_antsingy | NA | DD | Unknown | NA | NA | NA | NA | NA |
| mammals | Eliurus_carletoni | Eliurus_carletoni | LC | Unknown | tropical | nocturnal | 88.5 | 2411.252 | 7.428 |
| mammals | Eliurus_danieli | Eliurus_danieli | LC | Unknown | tropical | nocturnal | 95.5 | 866.351 | 6.807 |
| mammals | Eliurus_grandidieri | Eliurus_grandidieri | LC | Unknown | tropical | nocturnal | 51 | 60596.301 | 7.167 |
| mammals | Eliurus_majori | Eliurus_majori | LC | Unknown | tropical | nocturnal | 100 | 88713.442 | 7.311 |
| mammals | Eliurus_minor | Eliurus_minor | LC | Unknown | tropical | nocturnal | 35 | 114231.274 | 7.889 |
| mammals | Eliurus_myoxinus | Eliurus_myoxinus | LC | Unknown | tropical | nocturnal | 62.5 | 279015.095 | 8.458 |
| mammals | Eliurus_penicillatus | Eliurus_penicillatus | EN | Unknown | tropical | nocturnal | 100 | 1756.7 | 6.652 |
| mammals | Eliurus_petteri | Eliurus_petteri | EN | Unknown | tropical | nocturnal | 75 | 1491.975 | 8.683 |
| mammals | Eliurus_tanala | Eliurus_tanala | LC | Unknown | tropical | nocturnal | 82.5 | 120856.311 | 7.78 |
| mammals | Eliurus_webbi | Eliurus_webbi | LC | Unknown | tropical | nocturnal | 72.5 | 133408.493 | 7.99 |

| | | | | | | | | | |
|---|---|---|---|---|---|---|---|---|---|
| mammals | Emballonura_alecto | Emballonura_alecto | LC | Decreasing | tropical | diurnal | 5.3 | 1212058.67 | 8.01 |
| mammals | Emballonura_beccarii | Emballonura_beccarii | LC | Unknown | tropical | diurnal | 4.2 | 91119.17 | 4.146 |
| mammals | Emballonura_dianae | Emballonura_dianae | LC | Unknown | tropical | diurnal | 8 | 90456.793 | 3.918 |
| mammals | Emballonura_furax | Emballonura_furax | LC | Stable | tropical | diurnal | 11.2 | 67865.15 | 3.776 |
| mammals | Emballonura_monticola | Emballonura_monticola | LC | Stable | tropical | diurnal | 5.7 | 1690479.34 | 9.224 |
| mammals | Emballonura_raffrayana | Emballonura_raffrayana | LC | Unknown | tropical | diurnal | 5.6 | 216455.311 | 4.732 |
| mammals | Emballonura_semicaudata | Emballonura_semicaudata | EN | Decreasing | tropical | diurnal | 6.3 | 20505.941 | 9.137 |
| mammals | Emballonura_serii | Emballonura_serii | VU | Decreasing | tropical | diurnal | 8.8 | 11497.34 | 5.656 |
| mammals | Enchisthenes_hartii | Enchisthenes_hartii | LC | Unknown | tropical | nocturnal | 16 | 3104106.007 | 8.455 |
| mammals | Eoglaucomys_fimbriatus | Eoglaucomys_fimbriatus | LC | Unknown | temperate | diurnal | 712 | 188936.352 | 9.616 |
| mammals | Eolagurus_przewalskii | Eolagurus_przewalskii | LC | Unknown | temperate | nocturnal | 16 | 903281.892 | 3.145 |
| mammals | Eonycteris_major | Eonycteris_major | NT | Unknown | tropical | nocturnal | 72.8 | 732628.361 | 5.565 |
| mammals | Eonycteris_robusta | Eonycteris_robusta | VU | Decreasing | tropical | nocturnal | 77.5 | 114506.378 | 13.661 |
| mammals | Eonycteris_spelaea | Eonycteris_spelaea | LC | Decreasing | tropical | nocturnal | 58.1 | 3525628.505 | 11.864 |
| mammals | Eothenomys_eleusis | NA | LC/LC | Unknown/Unknown | NA | NA | NA | NA | NA |
| mammals | Eothenomys_melanogaster | Eothenomys_melanogaster | LC | Stable | temperate | cathemeral | 27 | 1722986.44 | 13.437 |
| mammals | Eozapus_setchuanus | Eozapus_setchuanus | LC | Unknown | temperate | nocturnal | 32.6 | 545018.875 | 8.502 |
| mammals | Episoriculus_caudatus | Episoriculus_caudatus | LC | Unknown | temperate | nocturnal | 9.1 | 315602.528 | 9.283 |
| mammals | Episoriculus_leucops | Episoriculus_leucops | LC | Unknown | temperate | nocturnal | 8.2 | 236569.088 | 6.052 |
| mammals | Episoriculus_macrurus | Episoriculus_macrurus | LC | Unknown | temperate | nocturnal | 13.6 | 375510.083 | 10.436 |
| mammals | Epixerus_ebii | Epixerus_ebii | LC | Unknown | tropical | diurnal | 592 | 540525.79 | 11.98 |
| mammals | Epomophorus_angolensis | Epomophorus_angolensis | NT | Decreasing | tropical | nocturnal | 131.7 | 383579.711 | 9.032 |
| mammals | Epomophorus_anselli | NA | DD | Unknown | NA | NA | NA | NA | NA |
| mammals | Epomophorus_crypturus | Epomophorus_crypturus | LC | Unknown | tropical | nocturnal | 111.1 | 2084376.206 | 9.696 |
| mammals | Epomophorus_dobsonii | Epomops_dobsonii | LC | Stable | tropical | nocturnal | 120 | 1367022.747 | 7.286 |
| mammals | Epomophorus_gambianus | Epomophorus_gambianus | LC | Unknown | tropical | nocturnal | 128.9 | 3791163.452 | 10.99 |
| mammals | Epomophorus_grandis | NA | DD | Unknown | NA | NA | NA | NA | NA |
| mammals | Epomophorus_intermedius | NA | DD | Unknown | NA | NA | NA | NA | NA |
| mammals | Epomophorus_labiatus | Epomophorus_labiatus | LC | Stable | tropical | nocturnal | 63 | 1765029.783 | 11.411 |
| mammals | Epomophorus_minor | Epomophorus_minor | LC | Stable | tropical | nocturnal | 50 | 2256487.091 | 10.765 |
| mammals | Epomophorus_pusillus | Micropteropus_pusillus | LC | Stable | tropical | nocturnal | 26.6 | 5380916.58 | 9.519 |
| mammals | Epomophorus_wahlbergi | Epomophorus_wahlbergi | LC | Stable | tropical | nocturnal | 91.6 | 4999921.477 | 9.667 |
| mammals | Epomops_buettikoferi | Epomops_buettikoferi | LC | Decreasing | tropical | nocturnal | 133.8 | 1079556.029 | 13.957 |
| mammals | Epomops_franqueti | Epomops_franqueti | LC | Stable | tropical | nocturnal | 118.9 | 4509721.408 | 7.241 |

| | | | | | | | | | |
|---|---|---|---|---|---|---|---|---|---|
| mammals | Eptesicus_fuscus | Eptesicus_fuscus | LC | Increasing | temperate | nocturnal | 17.5 | 12798347.84 | 11.925 |
| mammals | Equus_africanus | Equus_africanus | CR | Decreasing | tropical | cathemeral | 325000 | 84849.872 | 5.965 |
| mammals | Equus_ferus | Equus_ferus | EN | Increasing | temperate | cathemeral | 250000 | 26271.984 | 5.676 |
| mammals | Equus_grevyi | Equus_grevyi | EN | Stable | tropical | cathemeral | 407999.8 | 90832.523 | 5.936 |
| mammals | Equus_kiang | Equus_kiang | LC | Stable | temperate | cathemeral | 275000 | 1785558.324 | 2.797 |
| mammals | Equus_quagga | Equus_quagga | NT | Decreasing | tropical | cathemeral | 276000 | 2314209.977 | 8.93 |
| mammals | Equus_zebra | Equus_zebra | VU | Increasing | tropical | diurnal | 286998.7 | 321622.17 | 5.209 |
| mammals | Eremitalpa_granti | Eremitalpa_granti | LC | Unknown | temperate | nocturnal | 26.1 | 50294.062 | 5.861 |
| mammals | Eremodipus_lichtensteini | Eremodipus_lichtensteini | LC | Unknown | temperate | nocturnal | 51 | 559492.755 | 5.79 |
| mammals | Erethizon_dorsatum | Erethizon_dorsatum | LC | Stable | temperate | nocturnal | 7085.3 | 12063228.063 | 6.151 |
| mammals | Erinaceus_amurensis | Erinaceus_amurensis | LC | Stable | temperate | nocturnal | 800 | 4254779.309 | 14.115 |
| mammals | Erinaceus_concolor | Erinaceus_concolor | LC | Unknown | temperate | nocturnal | 719 | 1346308.199 | 15.472 |
| mammals | Erinaceus_europaeus | Erinaceus_europaeus | LC | Stable | temperate | nocturnal | 771 | 3809035.097 | 17.674 |
| mammals | Erinaceus_roumanicus | Erinaceus_roumanicus | LC | Stable | temperate | nocturnal | 745 | 6029711.838 | 13.904 |
| mammals | Eropeplus_canus | NA | VU | Decreasing | NA | NA | NA | NA | NA |
| mammals | Erophylla_bombifrons | Erophylla_bombifrons | LC | Stable | tropical | nocturnal | 14.3 | 82891.949 | 16.985 |
| mammals | Erophylla_sezekorni | Erophylla_sezekorni | LC | Unknown | tropical | nocturnal | 15.8 | 122945.384 | 16.428 |
| mammals | Erythrocebus_patas | Erythrocebus_patas | NT | Decreasing | tropical | diurnal | 8090 | 6604128.329 | 9.544 |
| mammals | Euchoreutes_naso | Euchoreutes_naso | LC | Unknown | temperate | nocturnal | 24.1 | 1674821.34 | 3.177 |
| mammals | Euderma_maculatum | Euderma_maculatum | LC | Stable | temperate | nocturnal | 15.7 | 2104986.074 | 8.524 |
| mammals | Eudiscopus_denticulus | Eudiscopus_denticulus | LC | Unknown | tropical | nocturnal | 5 | 5126.251 | 7.403 |
| mammals | Eudorcas_albonotata | Eudorcas_albonotata | LC | Stable | tropical | diurnal | 25000 | 128521.801 | 7.434 |
| mammals | Eudorcas_thomsonii | Eudorcas_thomsonii | LC | Decreasing | tropical | cathemeral | 20499.8 | 157281.782 | 11.18 |
| mammals | Eulemur_albifrons | Eulemur_albifrons | VU | Decreasing | tropical | cathemeral | 2300 | 19643.187 | 6.248 |
| mammals | Eulemur_cinereiceps | Eulemur_cinereiceps | CR | Decreasing | tropical | cathemeral | 1920 | 1817.018 | 5.682 |
| mammals | Eulemur_collaris | Eulemur_collaris | EN | Decreasing | tropical | cathemeral | 2375 | 8156.849 | 5.541 |
| mammals | Eulemur_coronatus | Eulemur_coronatus | EN | Decreasing | tropical | cathemeral | 1650 | 6521.702 | 8.232 |
| mammals | Eulemur_flavifrons | Eulemur_flavifrons | CR | Decreasing | tropical | cathemeral | 1850 | 2450.843 | 6.721 |
| mammals | Eulemur_fulvus | Eulemur_fulvus | VU | Decreasing | tropical | cathemeral | 2500 | 48944.481 | 7.628 |
| mammals | Eulemur_macaco | Eulemur_macaco | EN | Decreasing | tropical | cathemeral | 1900 | 7963.782 | 6.521 |
| mammals | Eulemur_mongoz | Eulemur_mongoz | CR | Decreasing | tropical | cathemeral | 1350 | 8051.596 | 10.602 |
| mammals | Eulemur_rubriventer | Eulemur_rubriventer | VU | Decreasing | tropical | cathemeral | 2000 | 51229.156 | 6.231 |
| mammals | Eulemur_rufifrons | Eulemur_rufifrons | VU | Decreasing | tropical | cathemeral | 1820 | 32696.596 | 7.397 |
| mammals | Eulemur_rufus | Eulemur_rufus | VU | Decreasing | tropical | cathemeral | 2375 | 37372.745 | 7.686 |

| | | | | | | | | | |
|---|---|---|---|---|---|---|---|---|---|
| mammals | Eulemur_sanfordi | Eulemur_sanfordi | EN | Decreasing | tropical | cathemeral | 2250 | 3830.387 | 9.674 |
| mammals | Eumops_auripendulus | Eumops_auripendulus | LC | Unknown | tropical | nocturnal | 28.5 | 14106769.899 | 5.952 |
| mammals | Eumops_bonariensis | Eumops_bonariensis | LC | Unknown | temperate | nocturnal | 11.9 | 1161090.374 | 10.214 |
| mammals | Eumops_dabbenei | Eumops_dabbenei | LC | Unknown | tropical | nocturnal | 104.4 | 2407709.823 | 6.887 |
| mammals | Eumops_floridanus | Eumops_floridanus | VU | Decreasing | temperate | nocturnal | 38.4 | 999.179 | 28.213 |
| mammals | Eumops_glaucinus | Eumops_glaucinus | LC | Unknown | tropical | nocturnal | 35.4 | 8971227.623 | 5.386 |
| mammals | Eumops_hansae | Eumops_hansae | LC | Unknown | tropical | nocturnal | 15.1 | 6416223.235 | 2.989 |
| mammals | Eumops_maurus | Eumops_maurus | DD | Decreasing | tropical | nocturnal | 36.2 | 1223598.05 | 2.045 |
| mammals | Eumops_patagonicus | Eumops_patagonicus | LC | Stable | temperate | nocturnal | 10 | 1535787.62 | 8.681 |
| mammals | Eumops_perotis | Eumops_perotis | LC | Decreasing | tropical | nocturnal | 51.1 | 11695739.961 | 6.098 |
| mammals | Eumops_trumbulli | Eumops_trumbulli | LC | Unknown | tropical | nocturnal | 36.4 | 4744456.772 | 1.475 |
| mammals | Eumops_underwoodi | Eumops_underwoodi | LC | Unknown | tropical | nocturnal | 54.7 | 955823.887 | 10.16 |
| mammals | Eumops_wilsoni | NA | DD | Unknown | NA | NA | NA | NA | NA |
| mammals | Euneomys_chinchilloides | NA | DD | Unknown | NA | NA | NA | NA | NA |
| mammals | Euoticus_elegantulus | Euoticus_elegantulus | LC | Unknown | tropical | nocturnal | 300 | 782197.192 | 5.786 |
| mammals | Euoticus_pallidus | Euoticus_pallidus | NT | Decreasing | tropical | nocturnal | 230 | 95102.992 | 13.759 |
| mammals | Eupetaurus_cinereus | Eupetaurus_cinereus | EN | Decreasing | temperate | nocturnal | 2500 | 36743.241 | 5.678 |
| mammals | Euphractus_sexcinctus | Euphractus_sexcinctus | LC | Stable | tropical | diurnal | 4782.9 | 7716332.833 | 7.819 |
| mammals | Eupleres_goudotii | Eupleres_goudotii | VU | Decreasing | tropical | cathemeral | 1850 | 105598.782 | 7.284 |
| mammals | Euryoryzomys_macconnelli | Euryoryzomys_macconnelli | LC | Unknown | tropical | nocturnal | 58 | 4294868.378 | 1.854 |
| mammals | Euryoryzomys_nitidus | Euryoryzomys_nitidus | LC | Stable | tropical | nocturnal | 55.2 | 1303446.66 | 3.445 |
| mammals | Euryoryzomys_russatus | Euryoryzomys_russatus | LC | Stable | tropical | nocturnal | 60.5 | 1009408.82 | 13.525 |
| mammals | Euryzygomatomys_spinosus | Euryzygomatomys_spinosus | LC | Unknown | temperate | nocturnal | 187.5 | 807543.661 | 13.198 |
| mammals | Eutamias_sibiricus | Eutamias_sibiricus | LC | Stable | temperate | diurnal | 85 | 14527269.109 | 5.244 |
| mammals | Euxerus_erythropus | Xerus_erythropus | LC | Stable | tropical | diurnal | 502.4 | 8816274.077 | 9.799 |
| mammals | Exilisciurus_concinnus | Exilisciurus_concinnus | LC | Stable | tropical | diurnal | 30.9 | 120457.566 | 12.097 |
| mammals | Exilisciurus_exilis | NA | DD | Unknown | NA | NA | NA | NA | NA |
| mammals | Exilisciurus_whiteheadi | Exilisciurus_whiteheadi | LC | Stable | tropical | diurnal | 20 | 155450.153 | 4.083 |
| mammals | Falsistrellus_mackenziei | Falsistrellus_mackenziei | NT | Decreasing | temperate | nocturnal | 21 | 24586.801 | 9.616 |
| mammals | Falsistrellus_tasmaniensis | Falsistrellus_tasmaniensis | VU | Decreasing | temperate | nocturnal | 22.7 | 386354.107 | 9.227 |
| mammals | Felis_bieti | Felis_bieti | VU | Decreasing | temperate | nocturnal | 7750 | 940432.421 | 8.474 |
| mammals | Felis_chaus | Felis_chaus | LC | Decreasing | temperate | nocturnal | 7393 | 7405493.524 | 14.592 |
| mammals | Felis_margarita | Felis_margarita | LC | Unknown | temperate | nocturnal | 2525 | 3701188.812 | 3.414 |
| mammals | Felis_nigripes | Felis_nigripes | VU | Decreasing | temperate | nocturnal | 1300 | 1948137.93 | 7.161 |

| | | | | | | | | | |
|---|---|---|---|---|---|---|---|---|---|
| mammals | Felis_silvestris | Felis_silvestris | LC | Unknown | temperate | nocturnal | 5500 | 1931656.661 | 17.261 |
| mammals | Felovia_vae | Felovia_vae | LC | Unknown | tropical | diurnal | 205 | 183984.78 | 4.655 |
| mammals | Feroculus_feroculus | Feroculus_feroculus | EN | Decreasing | tropical | nocturnal | 36 | 2647.266 | 17.229 |
| mammals | Fossa_fossana | Fossa_fossana | VU | Decreasing | tropical | nocturnal | 1500 | 91103.591 | 6.961 |
| mammals | Frateromys_fratrorum | Bunomys_fratrorum | VU | Decreasing | tropical | nocturnal | 135 | 7086.729 | 12.25 |
| mammals | Funambulus_layardi | Funambulus_layardi | VU | Decreasing | tropical | diurnal | 168 | 1276.826 | 9.02 |
| mammals | Funambulus_palmarum | Funambulus_palmarum | LC | Increasing | tropical | diurnal | 108.4 | 1263136.904 | 18.959 |
| mammals | Funambulus_pennantii | Funambulus_pennantii | LC | Unknown | temperate | diurnal | 99.1 | 2908289.198 | 16.838 |
| mammals | Funambulus_sublineatus | Funambulus_sublineatus | VU | Decreasing | tropical | diurnal | 70.9 | 21379.114 | 11.824 |
| mammals | Funambulus_tristriatus | Funambulus_tristriatus | LC | Decreasing | tropical | diurnal | 139 | 79432.106 | 17.041 |
| mammals | Funisciurus_anerythrus | Funisciurus_anerythrus | LC | Unknown | tropical | diurnal | 218 | 2279777.188 | 6.639 |
| mammals | Funisciurus_bayonii | NA | DD | Unknown | NA | NA | NA | NA | NA |
| mammals | Funisciurus_carruthersi | Funisciurus_carruthersi | LC | Unknown | tropical | diurnal | 286 | 93369.08 | 15.502 |
| mammals | Funisciurus_congicus | Funisciurus_congicus | LC | Stable | tropical | diurnal | 112 | 1735003.199 | 6.912 |
| mammals | Funisciurus_duchaillui | NA | DD | Unknown | NA | NA | NA | NA | NA |
| mammals | Funisciurus_isabella | Funisciurus_isabella | LC | Unknown | tropical | diurnal | 109 | 570175.352 | 6.167 |
| mammals | Funisciurus_lemniscatus | Funisciurus_lemniscatus | LC | Unknown | tropical | diurnal | 141 | 610756.521 | 6.45 |
| mammals | Funisciurus_leucogenys | Funisciurus_leucogenys | LC | Unknown | tropical | diurnal | 250 | 411996.776 | 12.86 |
| mammals | Funisciurus_pyrropus | Funisciurus_pyrropus | LC | Stable | tropical | diurnal | 297 | 2153133.029 | 9.746 |
| mammals | Funisciurus_substriatus | NA | DD | Unknown | NA | NA | NA | NA | NA |
| mammals | Furipterus_horrens | Furipterus_horrens | LC | Unknown | tropical | nocturnal | 3.2 | 9816319.149 | 5.154 |
| mammals | Galago_gallarum | Galago_gallarum | LC | Stable | tropical | nocturnal | 250 | 307921.617 | 10.364 |
| mammals | Galago_matschiei | Galago_matschiei | LC | Decreasing | tropical | nocturnal | 210 | 86725.873 | 14.669 |
| mammals | Galago_moholi | Galago_moholi | LC | Stable | tropical | nocturnal | 160 | 3985248.481 | 7.592 |
| mammals | Galago_senegalensis | Galago_senegalensis | LC | Decreasing | tropical | nocturnal | 227.7 | 7758268.689 | 10.881 |
| mammals | Galagoides_demidoff | Galagoides_demidoff | LC | Stable | tropical | nocturnal | 60.1 | 4348190.695 | 7.956 |
| mammals | Galagoides_thomasi | Galagoides_thomasi | LC | Stable | tropical | nocturnal | 99.7 | 4407656.997 | 8.449 |
| mammals | Galea_flavidens | Galea_flavidens | LC | Unknown | tropical | diurnal | 450 | 32759.674 | 7.559 |
| mammals | Galea_leucoblephara | Galea_leucoblephara | LC | Unknown | temperate | diurnal | 230 | 1914449.5 | 6.805 |
| mammals | Galea_musteloides | NA | DD | Unknown | NA | NA | NA | NA | NA |
| mammals | Galegeeska_revoilii | NA | DD | Unknown | NA | NA | NA | NA | NA |
| mammals | Galemys_pyrenaicus | Galemys_pyrenaicus | EN | Decreasing | temperate | nocturnal | 60.2 | 130939.775 | 15.506 |
| mammals | Galeopterus_variegatus | Galeopterus_variegatus | LC | Decreasing | tropical | nocturnal | 1012.5 | 1410929.24 | 8.64 |
| mammals | Galictis_cuja | Galictis_cuja | LC | Unknown | tropical | diurnal | 1000 | 7672218.09 | 8.862 |

| | | | | | | | | | |
|---|---|---|---|---|---|---|---|---|---|
| mammals | Galictis_vittata | Galictis_vittata | LC | Stable | tropical | diurnal | 3200 | 10014922.31 | 4.303 |
| mammals | Galidia_elegans | Galidia_elegans | LC | Decreasing | tropical | diurnal | 800 | 92492.351 | 6.855 |
| mammals | Galidictis_fasciata | Galidictis_fasciata | VU | Decreasing | tropical | nocturnal | 550 | 83601.291 | 7.105 |
| mammals | Gardnerycteris_crenulata | Mimon_crenulatum | LC | Stable | tropical | nocturnal | 13.8 | 11134091.342 | 4.872 |
| mammals | Gardnerycteris_koepckeae | NA | DD | Unknown | NA | NA | NA | NA | NA |
| mammals | Gazella_bennettii | Gazella_bennettii | LC | Decreasing | temperate | nocturnal | 18916.7 | 2764528.242 | 12.272 |
| mammals | Gazella_cuvieri | Gazella_cuvieri | VU | Decreasing | temperate | nocturnal | 25000 | 152444.46 | 8.581 |
| mammals | Gazella_dorcas | Gazella_dorcas | VU | Decreasing | tropical | nocturnal | 23000.1 | 9905633.094 | 2.487 |
| mammals | Gazella_gazella | Gazella_gazella | EN | Decreasing | temperate | diurnal | 22749.9 | 14845.713 | 28.244 |
| mammals | Gazella_subgutturosa | Gazella_subgutturosa | VU | Decreasing | temperate | diurnal | 28500 | 7107547.319 | 6.341 |
| mammals | Genetta_abyssinica | NA | DD | Unknown | NA | NA | NA | NA | NA |
| mammals | Genetta_angolensis | Genetta_angolensis | LC | Unknown | tropical | nocturnal | 1650 | 1399075.843 | 7.506 |
| mammals | Genetta_bourloni | Genetta_bourloni | VU | Decreasing | tropical | nocturnal | 1750 | 146864.614 | 13.79 |
| mammals | Genetta_fieldiana | Genetta_maculata | LC | Unknown | tropical | nocturnal | 2225 | 12856912.482 | 8.593 |
| mammals | Genetta_genetta | Genetta_genetta | LC | Stable | tropical | nocturnal | 1800 | 14894292.42 | 9.658 |
| mammals | Genetta_johnstoni | Genetta_johnstoni | NT | Decreasing | tropical | nocturnal | 2225 | 307033.322 | 14.578 |
| mammals | Genetta_pardina | Genetta_pardina | LC | Unknown | tropical | nocturnal | 3100 | 1435327.914 | 13.015 |
| mammals | Genetta_piscivora | Genetta_piscivora | NT | Decreasing | tropical | nocturnal | 3000 | 223601.457 | 4.548 |
| mammals | Genetta_poensis | NA | DD | Unknown | NA | NA | NA | NA | NA |
| mammals | Genetta_servalina | Genetta_servalina | LC | Unknown | tropical | nocturnal | 1055 | 3171148.458 | 6.975 |
| mammals | Genetta_thierryi | Genetta_thierryi | LC | Unknown | tropical | nocturnal | 1400 | 2584305.828 | 13.042 |
| mammals | Genetta_tigrina | Genetta_tigrina | LC | Stable | temperate | nocturnal | 2225 | 212859.61 | 11.782 |
| mammals | Genetta_victoriae | Genetta_victoriae | LC | Unknown | tropical | nocturnal | 3000 | 542907.302 | 6.281 |
| mammals | Geocapromys_brownii | Geocapromys_brownii | EN | Decreasing | tropical | nocturnal | 1500 | 987.877 | 14.162 |
| mammals | Geogale_aurita | Geogale_aurita | LC | Unknown | temperate | nocturnal | 7 | 10985.161 | 10.585 |
| mammals | Geosciurus_inauris | Xerus_inauris | LC | Stable | temperate | diurnal | 529 | 1515178.636 | 6.462 |
| mammals | Geosciurus_princeps | Xerus_princeps | LC | Stable | tropical | diurnal | 636 | 238199.934 | 5.613 |
| mammals | Geoxus_valdivianus | Geoxus_valdivianus | LC | Stable | temperate | cathemeral | 31.5 | 232674.621 | 7.351 |
| mammals | Gerbilliscus_afer | Gerbilliscus_afra | LC | Unknown | temperate | nocturnal | 97 | 100211.974 | 10.18 |
| mammals | Gerbilliscus_boehmi | Gerbilliscus_boehmi | LC | Stable | tropical | nocturnal | 146 | 1160649 | 9.449 |
| mammals | Gerbilliscus_brantsii | Gerbilliscus_brantsii | LC | Unknown | temperate | nocturnal | 114.4 | 2084250.523 | 7.364 |
| mammals | Gerbilliscus_gambianus | Gerbilliscus_gambiana | LC | Stable | tropical | nocturnal | 114.5 | 281829.323 | 12.205 |
| mammals | Gerbilliscus_guineae | Gerbilliscus_guineae | LC | Stable | tropical | nocturnal | 102.5 | 892568.329 | 12.111 |
| mammals | Gerbilliscus_inclusus | Gerbilliscus_inclusus | LC | Unknown | tropical | nocturnal | 118.5 | 218326.242 | 9.602 |

| | | | | | | | | | |
|---|---|---|---|---|---|---|---|---|---|
| mammals | Gerbilliscus_kempii | Gerbilliscus_kempi | LC | Unknown | tropical | nocturnal | 101 | 4610375.984 | 10.039 |
| mammals | Gerbilliscus_leucogaster | Gerbilliscus_leucogaster | LC | Stable | tropical | nocturnal | 72.8 | 5765221.891 | 7.869 |
| mammals | Gerbilliscus_nigricaudus | Gerbilliscus_nigricaudus | LC | Stable | tropical | nocturnal | 123 | 610487.087 | 8.57 |
| mammals | Gerbilliscus_paeba | Gerbillurus_paeba | LC | Stable | temperate | nocturnal | 23.3 | 2191068.628 | 6.207 |
| mammals | Gerbilliscus_phillipsi | Gerbilliscus_phillipsi | LC | Unknown | tropical | nocturnal | 62.9 | 71840.695 | 8.543 |
| mammals | Gerbilliscus_robustus | Gerbilliscus_robustus | LC | Stable | tropical | nocturnal | 96 | 5165821.444 | 9.569 |
| mammals | Gerbilliscus_setzeri | Gerbillurus_setzeri | LC | Stable | tropical | nocturnal | 38.3 | 48792.268 | 2.768 |
| mammals | Gerbilliscus_tytonis | Gerbillurus_tytonis | LC | Stable | temperate | nocturnal | 31 | 56156.998 | 2.488 |
| mammals | Gerbilliscus_validus | Gerbilliscus_validus | LC | Stable | tropical | nocturnal | 107.7 | 4626297.077 | 8.55 |
| mammals | Gerbilliscus_vallinus | Gerbillurus_vallinus | LC | Stable | temperate | nocturnal | 35.3 | 286817.658 | 4.811 |
| mammals | Gerbillus_acticola | NA | DD | Unknown | NA | NA | NA | NA | NA |
| mammals | Gerbillus_agag | NA | DD | Unknown | NA | NA | NA | NA | NA |
| mammals | Gerbillus_amoenus | Gerbillus_amoenus | LC | Unknown | temperate | nocturnal | 25 | 855071.287 | 2.269 |
| mammals | Gerbillus_andersoni | Gerbillus_andersoni | LC | Unknown | temperate | nocturnal | 24.5 | 117593.891 | 11.762 |
| mammals | Gerbillus_aquilus | Gerbillus_aquilus | LC | Unknown | temperate | nocturnal | 43.8 | 559283.138 | 3.76 |
| mammals | Gerbillus_bottai | NA | DD | Unknown | NA | NA | NA | NA | NA |
| mammals | Gerbillus_brockmani | NA | DD | Unknown | NA | NA | NA | NA | NA |
| mammals | Gerbillus_campestris | Gerbillus_campestris | LC | Unknown | temperate | nocturnal | 28.4 | 1163165.522 | 9.724 |
| mammals | Gerbillus_cheesmani | Gerbillus_cheesmani | LC | Stable | temperate | nocturnal | 28 | 2285825.369 | 4.457 |
| mammals | Gerbillus_dasyurus | Gerbillus_dasyurus | LC | Stable | temperate | nocturnal | 23.5 | 1542832.661 | 7.011 |
| mammals | Gerbillus_dunni | Gerbillus_dunni | LC | Unknown | tropical | nocturnal | 24.2 | 359416.913 | 8.756 |
| mammals | Gerbillus_famulus | Gerbillus_famulus | LC | Unknown | tropical | nocturnal | 37 | 52564.295 | 12.569 |
| mammals | Gerbillus_floweri | Gerbillus_floweri | LC | Stable | temperate | nocturnal | 56 | 103318.614 | 7.868 |
| mammals | Gerbillus_gerbillus | Gerbillus_gerbillus | LC | Stable | temperate | nocturnal | 25 | 9376064.671 | 2.216 |
| mammals | Gerbillus_gleadowi | Gerbillus_gleadowi | LC | Decreasing | temperate | nocturnal | 24 | 183769.51 | 12.569 |
| mammals | Gerbillus_grobbeni | NA | DD | Unknown | NA | NA | NA | NA | NA |
| mammals | Gerbillus_harwoodi | Gerbillus_harwoodi | LC | Stable | tropical | nocturnal | 11.1 | 205253.703 | 13.853 |
| mammals | Gerbillus_henleyi | Gerbillus_henleyi | LC | Unknown | tropical | nocturnal | 8 | 3846929.152 | 5.932 |
| mammals | Gerbillus_hesperinus | Gerbillus_hesperinus | EN | Decreasing | temperate | nocturnal | 30.2 | 724.542 | 17.697 |
| mammals | Gerbillus_hoogstraali | Gerbillus_hoogstraali | VU | Unknown | temperate | nocturnal | 24.2 | 9461.752 | 18.309 |
| mammals | Gerbillus_jamesi | NA | DD | Unknown | NA | NA | NA | NA | NA |
| mammals | Gerbillus_latastei | NA | DD | Unknown | NA | NA | NA | NA | NA |
| mammals | Gerbillus_lowei | NA | DD | Unknown | NA | NA | NA | NA | NA |
| mammals | Gerbillus_mackilligini | Gerbillus_mackillingini | LC | Stable | tropical | nocturnal | 17.2 | 200370.274 | 2.579 |

| | | | | | | | | | |
|---|---|---|---|---|---|---|---|---|---|
| mammals | Gerbillus_maghrebi | Gerbillus_maghrebi | LC | Unknown | temperate | nocturnal | 46 | 55225.667 | 17.791 |
| mammals | Gerbillus_mesopotamiae | Gerbillus_mesopotamiae | LC | Decreasing | temperate | nocturnal | 20 | 219864.081 | 14.697 |
| mammals | Gerbillus_muriculus | NA | DD | Unknown | NA | NA | NA | NA | NA |
| mammals | Gerbillus_nancillus | NA | DD | Unknown | NA | NA | NA | NA | NA |
| mammals | Gerbillus_nanus | Gerbillus_nanus | LC | Stable | temperate | nocturnal | 21.5 | 10213065.717 | 4.671 |
| mammals | Gerbillus_nigeriae | Gerbillus_nigeriae | LC | Stable | tropical | nocturnal | 26.8 | 71424.327 | 2.96 |
| mammals | Gerbillus_occiduus | NA | DD | Unknown | NA | NA | NA | NA | NA |
| mammals | Gerbillus_poecilops | Gerbillus_poecilops | LC | Unknown | tropical | nocturnal | 31.5 | 102195.847 | 12.581 |
| mammals | Gerbillus_principulus | NA | DD | Unknown | NA | NA | NA | NA | NA |
| mammals | Gerbillus_pulvinatus | NA | LC | Unknown | NA | NA | NA | NA | NA |
| mammals | Gerbillus_pusillus | Gerbillus_pusillus | LC | Unknown | tropical | nocturnal | 17.4 | 570316.355 | 9.622 |
| mammals | Gerbillus_pyramidum | Gerbillus_pyramidum | LC | Stable | tropical | nocturnal | 52 | 3344944.486 | 3.313 |
| mammals | Gerbillus_rosalinda | Gerbillus_rosalinda | LC | Unknown | tropical | nocturnal | 27.6 | 166259.447 | 14.94 |
| mammals | Gerbillus_rupicola | Gerbillus_rupicola | LC | Unknown | tropical | nocturnal | 37 | 42517.337 | 9.533 |
| mammals | Gerbillus_simoni | Gerbillus_simoni | LC | Unknown | temperate | nocturnal | 17.4 | 259962.225 | 11.522 |
| mammals | Gerbillus_somalicus | NA | DD | Unknown | NA | NA | NA | NA | NA |
| mammals | Gerbillus_stigmonyx | NA | DD | Unknown | NA | NA | NA | NA | NA |
| mammals | Gerbillus_tarabuli | Gerbillus_tarabuli | LC | Stable | temperate | nocturnal | 42.5 | 6332711.384 | 2.226 |
| mammals | Gerbillus_watersi | Gerbillus_watersi | LC | Unknown | tropical | nocturnal | 10.4 | 191177.362 | 8.581 |
| mammals | Giraffa_camelopardalis | Giraffa_camelopardalis | VU | Decreasing | tropical | diurnal | 899994.8 | 1794018.082 | 7.309 |
| mammals | Glaucomys_sabrinus | Glaucomys_sabrinus | LC | Stable | temperate | nocturnal | 166 | 7852908.626 | 4.249 |
| mammals | Glaucomys_volans | Glaucomys_volans | LC | Stable | temperate | nocturnal | 63.9 | 3848743.428 | 17.061 |
| mammals | Glauconycteris_alboguttata | Glauconycteris_alboguttata | LC | Unknown | tropical | nocturnal | 8.9 | 632684.443 | 5.184 |
| mammals | Glauconycteris_argentata | Glauconycteris_argentata | LC | Unknown | tropical | nocturnal | 9.1 | 3996259.903 | 8.029 |
| mammals | Glauconycteris_beatrix | Glauconycteris_beatrix | LC | Unknown | tropical | nocturnal | 6.5 | 1964722.551 | 6.701 |
| mammals | Glauconycteris_curryae | NA | DD | Unknown | NA | NA | NA | NA | NA |
| mammals | Glauconycteris_egeria | NA | DD | Unknown | NA | NA | NA | NA | NA |
| mammals | Glauconycteris_gleni | NA | DD | Unknown | NA | NA | NA | NA | NA |
| mammals | Glauconycteris_humeralis | NA | DD | Unknown | NA | NA | NA | NA | NA |
| mammals | Glauconycteris_kenyacola | NA | DD | Unknown | NA | NA | NA | NA | NA |
| mammals | Glauconycteris_machadoi | NA | DD | Unknown | NA | NA | NA | NA | NA |
| mammals | Glauconycteris_poensis | Glauconycteris_poensis | LC | Unknown | tropical | nocturnal | 7.1 | 1530190.039 | 10.262 |
| mammals | Glauconycteris_superba | Glauconycteris_superba | LC | Unknown | tropical | nocturnal | 11.9 | 933.919 | 18.998 |
| mammals | Glauconycteris_variegata | Glauconycteris_variegata | LC | Unknown | tropical | nocturnal | 11.1 | 8695501.174 | 9.105 |

| | | | | | | | | | |
|---|---|---|---|---|---|---|---|---|---|
| mammals | Glironia_venusta | Glironia_venusta | LC | Unknown | tropical | nocturnal | 114 | 2883884.404 | 2.11 |
| mammals | Glirulus_japonicus | Glirulus_japonicus | LC | Unknown | temperate | nocturnal | 27 | 211072.726 | 18.567 |
| mammals | Glis_glis | Glis_glis | LC | Unknown | temperate | nocturnal | 125 | 3718880.114 | 18.988 |
| mammals | Glischropus_javanus | NA | DD | Unknown | NA | NA | NA | NA | NA |
| mammals | Glischropus_tylopus | Glischropus_tylopus | LC | Stable | tropical | nocturnal | 4.4 | 1373589.237 | 8.018 |
| mammals | Glossophaga_commissarisi | Glossophaga_commissarisi | LC | Stable | tropical | nocturnal | 8.9 | 3716729.942 | 4.947 |
| mammals | Glossophaga_leachii | Glossophaga_leachii | LC | Stable | tropical | nocturnal | 10.1 | 480889.971 | 12.248 |
| mammals | Glossophaga_longirostris | Glossophaga_longirostris | LC | Unknown | tropical | nocturnal | 11.9 | 1533778.451 | 6.161 |
| mammals | Glossophaga_morenoi | Glossophaga_morenoi | LC | Stable | tropical | nocturnal | 8.4 | 327072.966 | 10.607 |
| mammals | Glossophaga_soricina | Glossophaga_soricina | LC | Stable | tropical | nocturnal | 9.4 | 14858417.39 | 6.235 |
| mammals | Glyphonycteris_behnii | NA | DD | Unknown | NA | NA | NA | NA | NA |
| mammals | Glyphonycteris_daviesi | Glyphonycteris_daviesi | LC | Unknown | tropical | nocturnal | 18.3 | 4708611.59 | 1.744 |
| mammals | Glyphonycteris_sylvestris | Glyphonycteris_sylvestris | LC | Unknown | tropical | nocturnal | 8.8 | 6011996.714 | 3.412 |
| mammals | Glyphotes_simus | NA | DD | Unknown | NA | NA | NA | NA | NA |
| mammals | Golunda_ellioti | Golunda_ellioti | LC | Stable | tropical | cathemeral | 60.9 | 2864575.376 | 18.214 |
| mammals | Gorilla_beringei | Gorilla_beringei | CR | Decreasing | tropical | diurnal | 130000 | 48938.488 | 5.619 |
| mammals | Gorilla_gorilla | Gorilla_gorilla | CR | Decreasing | tropical | diurnal | 120950 | 696516.85 | 5.635 |
| mammals | Gracilinanus_aceramarcae | Gracilinanus_aceramarcae | LC | Unknown | tropical | nocturnal | 20.5 | 27595.481 | 5.658 |
| mammals | Gracilinanus_agilis | Gracilinanus_agilis | LC | Unknown | tropical | nocturnal | 22 | 3609141.759 | 8.206 |
| mammals | Gracilinanus_dryas | Gracilinanus_dryas | LC | Decreasing | tropical | nocturnal | 18 | 30098.197 | 13.406 |
| mammals | Gracilinanus_emiliae | NA | DD | Unknown | NA | NA | NA | NA | NA |
| mammals | Gracilinanus_marica | Gracilinanus_marica | LC | Decreasing | tropical | nocturnal | 23.8 | 329096.247 | 12.613 |
| mammals | Gracilinanus_microtarsus | Gracilinanus_microtarsus | LC | Unknown | temperate | nocturnal | 31 | 631663.976 | 12.835 |
| mammals | Grammomys_aridulus | NA | DD | Unknown | NA | NA | NA | NA | NA |
| mammals | Grammomys_buntingi | NA | DD | Unknown | NA | NA | NA | NA | NA |
| mammals | Grammomys_caniceps | NA | DD | Unknown | NA | NA | NA | NA | NA |
| mammals | Grammomys_cometes | Grammomys_cometes | LC | Unknown | temperate | nocturnal | 48.5 | 417998.035 | 9.514 |
| mammals | Grammomys_dolichurus | Grammomys_dolichurus | LC | Unknown | tropical | nocturnal | 41.2 | 3257251.677 | 10.235 |
| mammals | Grammomys_dryas | Grammomys_dryas | LC | Decreasing | tropical | nocturnal | 39.1 | 16649.423 | 17.443 |
| mammals | Grammomys_gigas | Grammomys_gigas | EN | Unknown | tropical | nocturnal | 39.1 | 1912.136 | 4.469 |
| mammals | Grammomys_ibeanus | Grammomys_ibeanus | LC | Unknown | tropical | nocturnal | 41.2 | 215705.079 | 12.942 |
| mammals | Grammomys_macmillani | Grammomys_macmillani | LC | Unknown | tropical | nocturnal | 36 | 243998.47 | 8.733 |
| mammals | Grammomys_minnae | Grammomys_minnae | VU | Decreasing | tropical | nocturnal | 36 | 10922.728 | 10.862 |
| mammals | Grammomys_selousi | NA | LC | Unknown | NA | NA | NA | NA | NA |

| | | | | | | | | | |
|---|---|---|---|---|---|---|---|---|---|
| mammals | Graomys_chacoensis | NA | DD | Unknown | NA | NA | NA | NA | NA |
| mammals | Graomys_domorum | Graomys_domorum | LC | Stable | tropical | nocturnal | 102 | 101529.242 | 7.196 |
| mammals | Graomys_griseoflavus | Graomys_griseoflavus | LC | Stable | temperate | nocturnal | 67.5 | 2012497.833 | 5.924 |
| mammals | Graphiurus_christyi | Graphiurus_christyi | LC | Unknown | tropical | nocturnal | 22.7 | 394070.482 | 6.637 |
| mammals | Graphiurus_crassicaudatus | NA | DD | Unknown | NA | NA | NA | NA | NA |
| mammals | Graphiurus_johnstoni | NA | DD | Unknown | NA | NA | NA | NA | NA |
| mammals | Graphiurus_kelleni | Graphiurus_kelleni | LC | Unknown | tropical | nocturnal | 20.1 | 2773770.976 | 11.736 |
| mammals | Graphiurus_lorraineus | Graphiurus_lorraineus | LC | Unknown | tropical | nocturnal | 16.8 | 3472143.755 | 7.799 |
| mammals | Graphiurus_microtis | Graphiurus_microtis | LC | Unknown | tropical | nocturnal | 15.5 | 3300456.429 | 9.313 |
| mammals | Graphiurus_monardi | NA | DD | Unknown | NA | NA | NA | NA | NA |
| mammals | Graphiurus_murinus | Graphiurus_murinus | LC | Stable | temperate | nocturnal | 30 | 966155.445 | 12.755 |
| mammals | Graphiurus_nagtglasii | Graphiurus_nagtglasii | LC | Unknown | tropical | nocturnal | 45 | 882351.203 | 13.33 |
| mammals | Graphiurus_ocularis | Graphiurus_ocularis | LC | Unknown | temperate | nocturnal | 30.3 | 323125.648 | 8.903 |
| mammals | Graphiurus_platyops | Graphiurus_platyops | LC | Unknown | tropical | nocturnal | 46.1 | 340551.544 | 11.468 |
| mammals | Graphiurus_rupicola | Graphiurus_rupicola | LC | Unknown | temperate | nocturnal | 53.6 | 54377.698 | 5.075 |
| mammals | Graphiurus_surdus | NA | DD | Unknown | NA | NA | NA | NA | NA |
| mammals | Gulo_gulo | Gulo_gulo | LC | Decreasing | temperate | cathemeral | 17012.6 | 24394335.156 | 2.257 |
| mammals | Gyldenstolpia_fronto | Kunsia_fronto | CR | Unknown | temperate | nocturnal | 168 | 37194.696 | 9.452 |
| mammals | Gymnobelideus_leadbeateri | Gymnobelideus_leadbeateri | CR | Decreasing | temperate | nocturnal | 127 | 3023.861 | 7.925 |
| mammals | Gymnuromys_roberti | Gymnuromys_roberti | LC | Unknown | tropical | nocturnal | 97.5 | 77428.377 | 7.559 |
| mammals | Habromys_chinanteco | Habromys_chinanteco | CR | Decreasing | tropical | nocturnal | 40 | 83.037 | 3.339 |
| mammals | Habromys_delicatulus | Habromys_delicatulus | EN | Decreasing | tropical | nocturnal | 10.4 | 820.417 | 21.727 |
| mammals | Habromys_ixtlani | Habromys_ixtlani | CR | Decreasing | tropical | nocturnal | 93.4 | 92.151 | 5.306 |
| mammals | Habromys_lepturus | Habromys_lepturus | CR | Decreasing | tropical | nocturnal | 85 | 86.004 | 5.637 |
| mammals | Habromys_lophurus | Habromys_lophurus | NT | Decreasing | tropical | nocturnal | 40 | 25199.025 | 15.204 |
| mammals | Habromys_schmidlyi | Habromys_schmidlyi | CR | Decreasing | tropical | nocturnal | 13.5 | 80.947 | 11.214 |
| mammals | Habromys_simulatus | Habromys_simulatus | CR | Decreasing | tropical | nocturnal | 40 | 18416.036 | 17.59 |
| mammals | Hadromys_humei | Hadromys_humei | EN | Decreasing | temperate | nocturnal | 59 | 2051.048 | 15.764 |
| mammals | Hadromys_yunnanensis | NA | DD | Unknown | NA | NA | NA | NA | NA |
| mammals | Haeromys_margarettae | NA | DD | Unknown | NA | NA | NA | NA | NA |
| mammals | Haeromys_minahassae | Haeromys_minahassae | NT | Decreasing | tropical | nocturnal | 14.7 | 43679.089 | 8.853 |
| mammals | Haeromys_pusillus | Haeromys_pusillus | VU | Decreasing | tropical | nocturnal | 10.4 | 51651.595 | 5.321 |
| mammals | Handleyomys_alfaroi | Handleyomys_alfaroi | LC | Stable | tropical | nocturnal | 33.3 | 838400.353 | 11.46 |
| mammals | Handleyomys_chapmani | Handleyomys_chapmani | VU | Unknown | tropical | nocturnal | 50 | 39012.723 | 11.631 |

| | | | | | | | | | |
|---|---|---|---|---|---|---|---|---|---|
| mammals | Handleyomys_fuscatus | Handleyomys_fuscatus | LC | Decreasing | tropical | nocturnal | 49.5 | 79253.018 | 10.254 |
| mammals | Handleyomys_intectus | Handleyomys_intectus | LC | Decreasing | tropical | nocturnal | 60.5 | 9776.98 | 16.117 |
| mammals | Handleyomys_melanotis | Handleyomys_melanotis | LC | Unknown | tropical | nocturnal | 50 | 138450.878 | 8.499 |
| mammals | Handleyomys_rhabdops | Handleyomys_rhabdops | EN | Decreasing | tropical | nocturnal | 33 | 3686.426 | 15.702 |
| mammals | Handleyomys_rostratus | Handleyomys_rostratus | LC | Stable | tropical | nocturnal | 43.5 | 548403.744 | 10.743 |
| mammals | Hapalemur_alaotrensis | Hapalemur_alaotrensis | CR | Decreasing | tropical | cathemeral | 1240 | 216.328 | 7.33 |
| mammals | Hapalemur_aureus | Hapalemur_aureus | CR | Decreasing | tropical | cathemeral | 1450 | 2392.795 | 6.422 |
| mammals | Hapalemur_griseus | Hapalemur_griseus | VU | Decreasing | tropical | cathemeral | 900 | 90255.607 | 7.504 |
| mammals | Hapalemur_meridionalis | Hapalemur_meridionalis | VU | Decreasing | tropical | cathemeral | 900 | 10993.968 | 5.741 |
| mammals | Hapalemur_occidentalis | Hapalemur_occidentalis | VU | Decreasing | tropical | cathemeral | 850 | 26197.054 | 6.14 |
| mammals | Hapalomys_delacouri | Hapalomys_delacouri | NT | Decreasing | tropical | nocturnal | 64.3 | 392004.203 | 11.978 |
| mammals | Hapalomys_longicaudatus | Hapalomys_longicaudatus | EN | Decreasing | tropical | nocturnal | 70 | 2090.617 | 15.332 |
| mammals | Haplonycteris_fischeri | Haplonycteris_fischeri | LC | Stable | tropical | nocturnal | 18 | 264779.284 | 13.851 |
| mammals | Harpiocephalus_harpia | Harpiocephalus_harpia | LC | Decreasing | tropical | nocturnal | 16.3 | 995261.952 | 13.255 |
| mammals | Harpiola_grisea | NA | DD | Unknown | NA | NA | NA | NA | NA |
| mammals | Harpiola_isodon | Harpiola_isodon | LC | Unknown | temperate | nocturnal | 5.7 | 12490.233 | 5.487 |
| mammals | Harpyionycteris_celebensis | Harpyionycteris_celebensis | NT | Decreasing | tropical | nocturnal | 104 | 169411.15 | 9.274 |
| mammals | Harpyionycteris_whiteheadi | Harpyionycteris_whiteheadi | LC | Stable | tropical | nocturnal | 110.5 | 171865.466 | 13.283 |
| mammals | Heimyscus_fumosus | Heimyscus_fumosus | LC | Stable | tropical | nocturnal | 14.3 | 504708.993 | 5.234 |
| mammals | Helarctos_malayanus | Helarctos_malayanus | VU | Decreasing | tropical | cathemeral | 46500 | 3561925.137 | 10.35 |
| mammals | Heliosciurus_gambianus | Heliosciurus_gambianus | LC | Unknown | tropical | diurnal | 295 | 4714373.114 | 9.45 |
| mammals | Heliosciurus_mutabilis | Heliosciurus_mutabilis | LC | Unknown | tropical | diurnal | 390 | 1063837.253 | 10.019 |
| mammals | Heliosciurus_punctatus | NA | DD | Unknown | NA | NA | NA | NA | NA |
| mammals | Heliosciurus_rufobrachium | Heliosciurus_rufobrachium | LC | Unknown | tropical | diurnal | 371.5 | 2935776.368 | 10.729 |
| mammals | Heliosciurus_ruwenzorii | Heliosciurus_ruwenzorii | LC | Unknown | tropical | diurnal | 291 | 140580.948 | 16.037 |
| mammals | Heliosciurus_undulatus | NA | DD | Unknown | NA | NA | NA | NA | NA |
| mammals | Helogale_hirtula | Helogale_hirtula | LC | Unknown | tropical | diurnal | 289 | 912821.589 | 8.172 |
| mammals | Helogale_parvula | Helogale_parvula | LC | Stable | tropical | diurnal | 300 | 6918503.914 | 8.811 |
| mammals | Hemibelideus_lemuroides | Hemibelideus_lemuroides | NT | Stable | tropical | nocturnal | 952.5 | 2913.768 | 3.875 |
| mammals | Hemicentetes_nigriceps | Hemicentetes_nigriceps | LC | Unknown | tropical | nocturnal | 113 | 39236.488 | 10.916 |
| mammals | Hemicentetes_semispinosus | Hemicentetes_semispinosus | LC | Unknown | tropical | cathemeral | 137.8 | 122554.358 | 8.633 |
| mammals | Hemiechinus_auritus | Hemiechinus_auritus | LC | Unknown | temperate | nocturnal | 342 | 8465510.752 | 7.475 |
| mammals | Hemiechinus_collaris | Hemiechinus_collaris | LC | Unknown | temperate | nocturnal | 141 | 272308.566 | 14.337 |
| mammals | Hemigalus_derbyanus | Hemigalus_derbyanus | NT | Decreasing | tropical | nocturnal | 2322.5 | 1354858.335 | 7.868 |

| | | | | | | | | | |
|---|---|---|---|---|---|---|---|---|---|
| mammals | Hemitragus_jemlahicus | Hemitragus_jemlahicus | NT | Decreasing | temperate | diurnal | 35200 | 55836.041 | 7.598 |
| mammals | Herpailurus_yagouaroundi | Herpailurus_yagouaroundi | LC | Decreasing | tropical | diurnal | 7874.9 | 14850896.612 | 6.054 |
| mammals | Herpestes_flavescens | Herpestes_flavescens | LC | Unknown | tropical | diurnal | 750 | 218032.509 | 6.765 |
| mammals | Herpestes_ichneumon | Herpestes_ichneumon | LC | Stable | tropical | nocturnal | 5175 | 14478898.03 | 10.004 |
| mammals | Herpestes_ochraceus | Herpestes_ochraceus | LC | Unknown | tropical | diurnal | 498 | 506949.093 | 8.655 |
| mammals | Herpestes_pulverulentus | Herpestes_pulverulentus | LC | Stable | temperate | diurnal | 797 | 770354.788 | 8.534 |
| mammals | Herpestes_sanguineus | Herpestes_sanguineus | LC | Stable | tropical | diurnal | 550 | 16793634.037 | 9.362 |
| mammals | Hesperoptenus_blanfordi | Hesperoptenus_blanfordi | LC | Unknown | tropical | nocturnal | 6.5 | 460430.298 | 12.191 |
| mammals | Hesperoptenus_doriae | NA | DD | Unknown | NA | NA | NA | NA | NA |
| mammals | Hesperoptenus_tickelli | Hesperoptenus_tickelli | LC | Unknown | tropical | nocturnal | 18 | 1051792.798 | 14.441 |
| mammals | Hesperoptenus_tomesi | Hesperoptenus_tomesi | VU | Decreasing | tropical | nocturnal | 31 | 44815.716 | 9.655 |
| mammals | Heterogeomys_cavator | Orthogeomys_cavator | LC | Increasing | tropical | cathemeral | 650 | 5662.47 | 8.159 |
| mammals | Heterogeomys_cherriei | Heterogeomys_cherriei | LC | Stable | tropical | cathemeral | 225 | 10525.051 | 14.646 |
| mammals | Heterogeomys_dariensis | Heterogeomys_dariensis | LC | Stable | tropical | cathemeral | 438 | 13438.011 | 4.206 |
| mammals | Heterogeomys_heterodus | Orthogeomys_heterodus | LC | Increasing | tropical | cathemeral | 615 | 1191.9 | 25.61 |
| mammals | Heterogeomys_hispidus | Orthogeomys_hispidus | LC | Stable | tropical | cathemeral | 500 | 397264.662 | 10.59 |
| mammals | Heterogeomys_lanius | Orthogeomys_lanius | CR | Decreasing | tropical | cathemeral | 500 | 87.686 | 16.851 |
| mammals | Heterogeomys_underwoodi | Orthogeomys_underwoodi | LC | Increasing | tropical | cathemeral | 250 | 6276.292 | 14.107 |
| mammals | Heterohyrax_brucei | Heterohyrax_brucei | LC | Unknown | tropical | diurnal | 2625 | 5937564.871 | 9.715 |
| mammals | Heteromys_adspersus | Heteromys_adspersus | LC | Stable | tropical | nocturnal | 65 | 22429.24 | 14.086 |
| mammals | Heteromys_anomalus | Heteromys_anomalus | LC | Unknown | tropical | nocturnal | 70 | 449459.767 | 12.553 |
| mammals | Heteromys_australis | Heteromys_australis | LC | Stable | tropical | nocturnal | 57 | 226886.784 | 9.274 |
| mammals | Heteromys_desmarestianus | Heteromys_desmarestianus | LC | Stable | tropical | nocturnal | 72.5 | 476861.294 | 10.347 |
| mammals | Heteromys_gaumeri | Heteromys_gaumeri | LC | Stable | tropical | nocturnal | 63.6 | 167156.989 | 6.534 |
| mammals | Heteromys_irroratus | Heteromys_irroratus | LC | Stable | tropical | nocturnal | 50 | 606822.969 | 11.484 |
| mammals | Heteromys_nelsoni | Heteromys_nelsoni | EN | Decreasing | tropical | nocturnal | 54.2 | 4649.722 | 10.935 |
| mammals | Heteromys_oasicus | Heteromys_oasicus | EN | Decreasing | tropical | nocturnal | 43.8 | 2513.722 | 16.811 |
| mammals | Heteromys_oresterus | Heteromys_oresterus | LC | Stable | tropical | nocturnal | 74.8 | 1327.429 | 9.026 |
| mammals | Heteromys_pictus | Heteromys_pictus | LC | Decreasing | tropical | nocturnal | 40 | 439292.261 | 8.779 |
| mammals | Heteromys_salvini | Heteromys_salvini | LC | Decreasing | tropical | nocturnal | 42 | 169498.086 | 15.094 |
| mammals | Heteromys_spectabilis | Heteromys_spectabilis | EN | Decreasing | tropical | nocturnal | 65 | 4803.801 | 7.402 |
| mammals | Heteromys_teleus | Heteromys_teleus | VU | Decreasing | tropical | nocturnal | 88 | 39648.506 | 14.122 |
| mammals | Hippocamelus_antisensis | Hippocamelus_antisensis | VU | Decreasing | tropical | diurnal | 68599.4 | 424715.684 | 7.861 |
| mammals | Hippopotamus_amphibius | Hippopotamus_amphibius | VU | Stable | tropical | cathemeral | 1417490 | 1233077.107 | 7.337 |

| | | | | | | | | | |
|---|---|---|---|---|---|---|---|---|---|
| mammals | Hipposideros_abae | Hipposideros_abae | LC | Unknown | tropical | nocturnal | 25.5 | 2324228.535 | 11.736 |
| mammals | Hipposideros_armiger | Hipposideros_armiger | LC | Unknown | tropical | nocturnal | 53 | 4499367.472 | 13.415 |
| mammals | Hipposideros_ater | Hipposideros_ater | LC | Unknown | tropical | nocturnal | 5.8 | 3676234.681 | 9.064 |
| mammals | Hipposideros_beatus | Hipposideros_beatus | LC | Decreasing | tropical | nocturnal | 6.7 | 1636043.796 | 8.145 |
| mammals | Hipposideros_bicolor | Hipposideros_bicolor | LC | Stable | tropical | nocturnal | 7.6 | 1591830.812 | 9.489 |
| mammals | Hipposideros_boeadii | NA | DD | Unknown | NA | NA | NA | NA | NA |
| mammals | Hipposideros_breviceps | NA | DD | Unknown | NA | NA | NA | NA | NA |
| mammals | Hipposideros_caffer | Hipposideros_caffer | LC | Decreasing | tropical | nocturnal | 9.4 | 1264715.775 | 11.117 |
| mammals | Hipposideros_calcaratus | Hipposideros_calcaratus | LC | Stable | tropical | nocturnal | 11.4 | 217484.075 | 4.54 |
| mammals | Hipposideros_cervinus | Hipposideros_cervinus | LC | Decreasing | tropical | nocturnal | 9.3 | 1964927.006 | 7.107 |
| mammals | Hipposideros_cineraceus | Hipposideros_cineraceus | LC | Unknown | tropical | nocturnal | 4.7 | 2205964.575 | 12.019 |
| mammals | Hipposideros_coronatus | NA | DD | Unknown | NA | NA | NA | NA | NA |
| mammals | Hipposideros_corynophyllus | Hipposideros_corynophyllus | LC | Unknown | tropical | nocturnal | 14.5 | 1988.09 | 2.175 |
| mammals | Hipposideros_coxi | Hipposideros_coxi | EN | Decreasing | tropical | nocturnal | 9.3 | 2684.567 | 15.554 |
| mammals | Hipposideros_crumeniferus | NA | DD | Unknown | NA | NA | NA | NA | NA |
| mammals | Hipposideros_curtus | Hipposideros_curtus | EN | Decreasing | tropical | nocturnal | 7.5 | 136020.662 | 9.668 |
| mammals | Hipposideros_demissus | Hipposideros_demissus | EN | Unknown | tropical | nocturnal | 24.9 | 3048.08 | 4.456 |
| mammals | Hipposideros_diadema | Hipposideros_diadema | LC | Decreasing | tropical | nocturnal | 43.8 | 3318837.887 | 9.547 |
| mammals | Hipposideros_dinops | Hipposideros_dinops | VU | Decreasing | tropical | nocturnal | 71.9 | 27145.986 | 4.736 |
| mammals | Hipposideros_doriae | Hipposideros_doriae | NT | Decreasing | tropical | nocturnal | 5 | 1251661.751 | 7.506 |
| mammals | Hipposideros_durgadasi | Hipposideros_durgadasi | VU | Decreasing | tropical | nocturnal | 5.6 | 19888.856 | 16.148 |
| mammals | Hipposideros_dyacorum | Hipposideros_dyacorum | LC | Unknown | tropical | nocturnal | 6.1 | 764978.901 | 5.879 |
| mammals | Hipposideros_edwardshilli | Hipposideros_edwardshilli | VU | Unknown | tropical | nocturnal | 11.3 | 1985.792 | 1.969 |
| mammals | Hipposideros_fuliginosus | Hipposideros_fuliginosus | LC | Decreasing | tropical | nocturnal | 16.6 | 1334587.136 | 7.392 |
| mammals | Hipposideros_fulvus | Hipposideros_fulvus | LC | Stable | tropical | nocturnal | 8.5 | 2455054.706 | 18.271 |
| mammals | Hipposideros_galeritus | Hipposideros_galeritus | LC | Unknown | tropical | nocturnal | 8.2 | 2775105.768 | 12.906 |
| mammals | Hipposideros_grandis | Hipposideros_grandis | LC | Unknown | tropical | nocturnal | 16.3 | 185987.165 | 10.322 |
| mammals | Hipposideros_halophyllus | Hipposideros_halophyllus | VU | Decreasing | tropical | nocturnal | 4 | 23215.653 | 14.614 |
| mammals | Hipposideros_hypophyllus | NA | CR | Decreasing | NA | NA | NA | NA | NA |
| mammals | Hipposideros_inexpectatus | NA | DD | Unknown | NA | NA | NA | NA | NA |
| mammals | Hipposideros_inornatus | Hipposideros_inornatus | VU | Decreasing | tropical | nocturnal | 28.5 | 6625.21 | 2.418 |
| mammals | Hipposideros_jonesi | Hipposideros_jonesi | NT | Decreasing | tropical | nocturnal | 5.5 | 1271130.066 | 13.283 |
| mammals | Hipposideros_khaokhouayensis | Hipposideros_khaokhouayensis | VU | Decreasing | tropical | nocturnal | 8.7 | 17926.372 | 7.112 |
| mammals | Hipposideros_lamottei | Hipposideros_lamottei | CR | Decreasing | tropical | nocturnal | 13.1 | 77.517 | 11.326 |

| | | | | | | | | | |
|---|---|---|---|---|---|---|---|---|---|
| mammals | Hipposideros_lankadiva | Hipposideros_lankadiva | LC | Unknown | temperate | nocturnal | 52.4 | 1547925.258 | 17.426 |
| mammals | Hipposideros_larvatus | Hipposideros_larvatus | LC | Unknown | tropical | nocturnal | 19.9 | 3295221.612 | 11.906 |
| mammals | Hipposideros_lekaguli | Hipposideros_lekaguli | NT | Decreasing | tropical | nocturnal | 31.1 | 96007.541 | 12.652 |
| mammals | Hipposideros_lylei | Hipposideros_lylei | LC | Decreasing | tropical | nocturnal | 40 | 1013744.148 | 11.119 |
| mammals | Hipposideros_macrobullatus | NA | DD | Unknown | NA | NA | NA | NA | NA |
| mammals | Hipposideros_madurae | Hipposideros_madurae | NT | Decreasing | tropical | nocturnal | 13.9 | 66492.708 | 23.775 |
| mammals | Hipposideros_maggietaylorae | Hipposideros_maggietaylorae | LC | Stable | tropical | nocturnal | 18.5 | 263505.955 | 3.732 |
| mammals | Hipposideros_marisae | Hipposideros_marisae | VU | Decreasing | tropical | nocturnal | 6.1 | 49715.658 | 14.192 |
| mammals | Hipposideros_megalotis | Hipposideros_megalotis | LC | Unknown | tropical | nocturnal | 5.6 | 943711.899 | 9.877 |
| mammals | Hipposideros_muscinus | Hipposideros_muscinus | LC | Unknown | tropical | nocturnal | 8.6 | 116471.973 | 4.533 |
| mammals | Hipposideros_nequam | NA | DD | Unknown | NA | NA | NA | NA | NA |
| mammals | Hipposideros_obscurus | Hipposideros_obscurus | LC | Unknown | tropical | nocturnal | 9.5 | 276180.193 | 13.918 |
| mammals | Hipposideros_orbiculus | Hipposideros_orbiculus | VU | Decreasing | tropical | nocturnal | 9.8 | 1922.508 | 19.646 |
| mammals | Hipposideros_papua | Hipposideros_papua | LC | Stable | tropical | nocturnal | 10.1 | 50777.06 | 4.302 |
| mammals | Hipposideros_pelingensis | Hipposideros_pelingensis | NT | Decreasing | tropical | nocturnal | 56.2 | 177328.161 | 9.276 |
| mammals | Hipposideros_pomona | Hipposideros_pomona | EN | Decreasing | tropical | nocturnal | 6.2 | 26967.906 | 14.079 |
| mammals | Hipposideros_pygmaeus | Hipposideros_pygmaeus | LC | Unknown | tropical | nocturnal | 3.5 | 90619.523 | 14.671 |
| mammals | Hipposideros_ridleyi | Hipposideros_ridleyi | VU | Decreasing | tropical | nocturnal | 8.8 | 218978.281 | 10.142 |
| mammals | Hipposideros_rotalis | Hipposideros_rotalis | LC | Unknown | tropical | nocturnal | 9.3 | 29363.541 | 6.416 |
| mammals | Hipposideros_ruber | Hipposideros_ruber | LC | Unknown | tropical | cathemeral | 10.1 | 9062130.348 | 9.038 |
| mammals | Hipposideros_scutinares | Hipposideros_scutinares | VU | Decreasing | tropical | nocturnal | 40 | 34112.177 | 7.246 |
| mammals | Hipposideros_semoni | Hipposideros_semoni | LC | Unknown | tropical | nocturnal | 7.5 | 76340.616 | 2.616 |
| mammals | Hipposideros_sorenseni | Hipposideros_sorenseni | EN | Unknown | tropical | nocturnal | 20.2 | 408.881 | 15.705 |
| mammals | Hipposideros_speoris | Hipposideros_speoris | LC | Stable | tropical | nocturnal | 10.4 | 1221761.433 | 19.644 |
| mammals | Hipposideros_stenotis | Hipposideros_stenotis | VU | Unknown | tropical | nocturnal | 12 | 489413.65 | 1.809 |
| mammals | Hipposideros_sumbae | Hipposideros_sumbae | LC | Decreasing | tropical | nocturnal | 13.3 | 43581.579 | 13.429 |
| mammals | Hipposideros_swinhoii | Hipposideros_pratti | LC | Unknown | temperate | nocturnal | 56 | 2242954.876 | 14.85 |
| mammals | Hipposideros_turpis | Hipposideros_turpis | EN | Decreasing | temperate | nocturnal | 33.4 | 553.867 | 13.789 |
| mammals | Hipposideros_wollastoni | Hipposideros_wollastoni | LC | Unknown | tropical | nocturnal | 6.8 | 33302.603 | 3.224 |
| mammals | Hippotragus_equinus | Hippotragus_equinus | LC | Decreasing | tropical | diurnal | 269997.7 | 5571099.891 | 8.951 |
| mammals | Hippotragus_niger | Hippotragus_niger | LC | Stable | tropical | cathemeral | 227499.3 | 3013917.711 | 9.148 |
| mammals | Histiotus_alienus | NA | DD | Unknown | NA | NA | NA | NA | NA |
| mammals | Histiotus_humboldti | NA | DD | Unknown | NA | NA | NA | NA | NA |
| mammals | Histiotus_laephotis | Histiotus_laephotis | LC | Decreasing | tropical | nocturnal | 11.3 | 823944.646 | 6.911 |

| | | | | | | | | | |
|---|---|---|---|---|---|---|---|---|---|
| mammals | Histiotus_macrotus | Histiotus_macrotus | LC | Unknown | temperate | nocturnal | 11 | 462171.758 | 7.366 |
| mammals | Histiotus_magellanicus | Histiotus_magellanicus | LC | Unknown | temperate | nocturnal | 14 | 330089.678 | 4.228 |
| mammals | Histiotus_montanus | Histiotus_montanus | LC | Unknown | tropical | nocturnal | 9.1 | 3290996.152 | 6.844 |
| mammals | Histiotus_velatus | NA | DD | Unknown | NA | NA | NA | NA | NA |
| mammals | Hodomys_alleni | Hodomys_alleni | LC | Stable | tropical | nocturnal | 367.6 | 214221.357 | 10.081 |
| mammals | Holochilus_brasiliensis | Holochilus_brasiliensis | LC | Stable | temperate | nocturnal | 155 | 2878494.863 | 11.432 |
| mammals | Holochilus_chacarius | Holochilus_chacarius | LC | Decreasing | temperate | nocturnal | 204 | 766923.361 | 7.301 |
| mammals | Holochilus_sciureus | Holochilus_sciureus | LC | Stable | tropical | nocturnal | 171 | 9320092.416 | 3.766 |
| mammals | Hoolock_hoolock | Hoolock_hoolock | EN | Decreasing | tropical | diurnal | 6500 | 319902.675 | 10.344 |
| mammals | Hoolock_leuconedys | Hoolock_leuconedys | VU | Decreasing | temperate | diurnal | 7200 | 112318.947 | 7.912 |
| mammals | Hoplomys_gymnurus | Hoplomys_gymnurus | LC | Stable | tropical | nocturnal | 240 | 332781.883 | 7.701 |
| mammals | Hsunycteris_thomasi | Lonchophylla_thomasi | LC | Unknown | tropical | nocturnal | 6.9 | 8383354.136 | 3.566 |
| mammals | Hyaena_hyaena | Hyaena_hyaena | NT | Decreasing | tropical | nocturnal | 41705.1 | 23460484.545 | 7.692 |
| mammals | Hybomys_basilii | Hybomys_basilii | EN | Unknown | tropical | nocturnal | 53.4 | 588.929 | 9.936 |
| mammals | Hybomys_lunaris | Hybomys_lunaris | VU | Stable | tropical | nocturnal | 53.4 | 2216.953 | 10.741 |
| mammals | Hybomys_rufocanus | NA | EN/EN | Decreasing/Unknown | NA | NA | NA | NA | NA |
| mammals | Hybomys_univittatus | Hybomys_univittatus | LC | Stable | tropical | cathemeral | 47.6 | 2240781.542 | 6.193 |
| mammals | Hydrictis_maculicollis | Hydrictis_maculicollis | NT | Decreasing | tropical | diurnal | 4000 | 9962749.182 | 8.726 |
| mammals | Hydrochoerus_hydrochaeris | Hydrochoerus_hydrochaeris | LC | Stable | tropical | cathemeral | 62449.6 | 13395889.234 | 5.602 |
| mammals | Hydrochoerus_isthmius | NA | DD | Unknown | NA | NA | NA | NA | NA |
| mammals | Hydromys_chrysogaster | Hydromys_chrysogaster | LC | Unknown | temperate | nocturnal | 479.8 | 3705391.857 | 5.628 |
| mammals | Hydromys_hussoni | NA | DD | Unknown | NA | NA | NA | NA | NA |
| mammals | Hydromys_neobrittanicus | NA | DD | Unknown | NA | NA | NA | NA | NA |
| mammals | Hydromys_ziegleri | NA | DD | Unknown | NA | NA | NA | NA | NA |
| mammals | Hydropotes_inermis | Hydropotes_inermis | VU | Decreasing | temperate | cathemeral | 14000 | 181922.922 | 23.731 |
| mammals | Hyemoschus_aquaticus | Hyemoschus_aquaticus | LC | Decreasing | tropical | nocturnal | 10850 | 2450418.943 | 6.996 |
| mammals | Hyladelphys_kalinowskii | Hyladelphys_kalinowskii | LC | Unknown | tropical | nocturnal | 16.3 | 2405718.44 | 1.154 |
| mammals | Hylaeamys_megacephalus | Hylaeamys_megacephalus | LC | Unknown | tropical | nocturnal | 57.8 | 5971593.793 | 4.789 |
| mammals | Hylaeamys_perenensis | Hylaeamys_perenensis | LC | Stable | tropical | nocturnal | 60.6 | 1991968.514 | 1.112 |
| mammals | Hylaeamys_tatei | NA | DD | Unknown | NA | NA | NA | NA | NA |
| mammals | Hylaeamys_yunganus | Hylaeamys_yunganus | LC | Stable | tropical | nocturnal | 60.5 | 5299586.675 | 2.057 |
| mammals | Hylobates_agilis | Hylobates_agilis | EN | Decreasing | tropical | diurnal | 5650 | 385727.495 | 10.076 |
| mammals | Hylobates_albibarbis | Hylobates_albibarbis | EN | Decreasing | tropical | diurnal | 5650 | 199695.864 | 4.957 |
| mammals | Hylobates_klossii | Hylobates_klossii | EN | Decreasing | tropical | diurnal | 5800 | 6004.48 | 6.223 |

| | | | | | | | | | |
|---|---|---|---|---|---|---|---|---|---|
| mammals | Hylobates_lar | Hylobates_lar | EN | Decreasing | tropical | diurnal | 4892 | 638118.633 | 12.694 |
| mammals | Hylobates_moloch | Hylobates_moloch | EN | Decreasing | tropical | diurnal | 6500 | 39231.971 | 18.199 |
| mammals | Hylobates_muelleri | Hylobates_muelleri | EN | Decreasing | tropical | diurnal | 6000 | 103190.456 | 7.354 |
| mammals | Hylobates_pileatus | Hylobates_pileatus | EN | Decreasing | tropical | diurnal | 5440 | 121563.528 | 11.809 |
| mammals | Hylochoerus_meinertzhageni | Hylochoerus_meinertzhageni | LC | Decreasing | tropical | diurnal | 188500 | 2149889.59 | 8.602 |
| mammals | Hylomys_parvus | Hylomys_parvus | VU | Decreasing | tropical | cathemeral | 38.3 | 9948.264 | 6.397 |
| mammals | Hylomys_suillus | Hylomys_suillus | LC | Stable | tropical | diurnal | 65.8 | 1634250.044 | 11.527 |
| mammals | Hylomyscus_aeta | Hylomyscus_aeta | LC | Stable | tropical | nocturnal | 25.1 | 2618038.49 | 6.304 |
| mammals | Hylomyscus_alleni | Hylomyscus_alleni | LC | Stable | tropical | nocturnal | 18.7 | 1510866.935 | 9.552 |
| mammals | Hylomyscus_baeri | Hylomyscus_baeri | EN | Decreasing | tropical | nocturnal | 31.8 | 53952.098 | 16.773 |
| mammals | Hylomyscus_carillus | Hylomyscus_carillus | LC | Unknown | tropical | nocturnal | 26.3 | 280113.319 | 7.378 |
| mammals | Hylomyscus_denniae | Hylomyscus_denniae | LC | Unknown | tropical | nocturnal | 30 | 17471.524 | 14.311 |
| mammals | Hylomyscus_grandis | Hylomyscus_grandis | EN | Decreasing | tropical | nocturnal | 33 | 422.335 | 12.547 |
| mammals | Hylomyscus_pamfi | NA | DD | Unknown | NA | NA | NA | NA | NA |
| mammals | Hylomyscus_parvus | Hylomyscus_parvus | LC | Stable | tropical | nocturnal | 12.7 | 491928.867 | 5.318 |
| mammals | Hylomyscus_stella | Hylomyscus_stella | LC | Stable | tropical | nocturnal | 20.2 | 89591.255 | 11.517 |
| mammals | Hylomyscus_walterverheyeni | Hylomyscus_walterverheyeni | LC | Unknown | tropical | nocturnal | 20 | 253573.047 | 4.734 |
| mammals | Hylonycteris_underwoodi | Hylonycteris_underwoodi | LC | Stable | tropical | nocturnal | 6.7 | 201148.613 | 11.212 |
| mammals | Hylopetes_alboniger | Hylopetes_alboniger | LC | Decreasing | tropical | nocturnal | 255.4 | 1399086.79 | 10.439 |
| mammals | Hylopetes_bartelsi | NA | DD | Unknown | NA | NA | NA | NA | NA |
| mammals | Hylopetes_nigripes | Hylopetes_nigripes | NT | Decreasing | tropical | nocturnal | 534 | 11504.737 | 9.021 |
| mammals | Hylopetes_phayrei | Hylopetes_phayrei | LC | Stable | tropical | nocturnal | 227 | 1105366.586 | 10.698 |
| mammals | Hylopetes_platyurus | NA | DD | Unknown | NA | NA | NA | NA | NA |
| mammals | Hylopetes_sagitta | NA | DD | Unknown | NA | NA | NA | NA | NA |
| mammals | Hylopetes_sipora | Hylopetes_sipora | EN | Decreasing | tropical | nocturnal | 89.2 | 616.075 | 9.935 |
| mammals | Hylopetes_spadiceus | Hylopetes_spadiceus | LC | Unknown | tropical | nocturnal | 84.3 | 1145175.088 | 10.154 |
| mammals | Hylopetes_winstoni | NA | DD | Unknown | NA | NA | NA | NA | NA |
| mammals | Hyomys_dammermani | NA | DD | Unknown | NA | NA | NA | NA | NA |
| mammals | Hyomys_goliath | Hyomys_goliath | LC | Unknown | tropical | nocturnal | 817.5 | 110796.727 | 5.121 |
| mammals | Hyosciurus_heinrichi | Hyosciurus_heinrichi | LC | Unknown | tropical | diurnal | 296 | 32128.583 | 6.434 |
| mammals | Hyosciurus_ileile | Hyosciurus_ileile | VU | Decreasing | tropical | diurnal | 315 | 9499.848 | 4.842 |
| mammals | Hypogeomys_antimena | Hypogeomys_antimena | CR | Decreasing | tropical | nocturnal | 1250 | 193.69 | 4.718 |
| mammals | Hypsignathus_monstrosus | Hypsignathus_monstrosus | LC | Unknown | tropical | nocturnal | 335.1 | 2860674.044 | 8.469 |
| mammals | Hypsiprymnodon_moschatus | Hypsiprymnodon_moschatus | LC | Stable | tropical | diurnal | 520 | 10503.392 | 4.12 |

| | | | | | | | | | |
|---|---|---|---|---|---|---|---|---|---|
| mammals | Hypsugo_affinis | Falsistrellus_affinis | LC | Unknown | temperate | nocturnal | 4.9 | 351917.821 | 9.718 |
| mammals | Hypsugo_alaschanicus | Pipistrellus_alaschanicus | LC | Unknown | temperate | nocturnal | 8.2 | 2990994.154 | 15.461 |
| mammals | Hypsugo_arabicus | NA | DD | Unknown | NA | NA | NA | NA | NA |
| mammals | Hypsugo_ariel | NA | DD | Unknown | NA | NA | NA | NA | NA |
| mammals | Hypsugo_cadornae | Pipistrellus_cadornae | LC | Unknown | tropical | nocturnal | 6.7 | 513196.505 | 11.034 |
| mammals | Hypsugo_imbricatus | Pipistrellus_imbricatus | LC | Decreasing | tropical | nocturnal | 5.7 | 219247.733 | 17.943 |
| mammals | Hypsugo_kitcheneri | NA | DD | Unknown | NA | NA | NA | NA | NA |
| mammals | Hypsugo_macrotis | NA | DD | Unknown | NA | NA | NA | NA | NA |
| mammals | Hypsugo_mordax | NA | DD | Unknown | NA | NA | NA | NA | NA |
| mammals | Hypsugo_musciculus | NA | DD | Unknown | NA | NA | NA | NA | NA |
| mammals | Hypsugo_petersi | NA | DD | Unknown | NA | NA | NA | NA | NA |
| mammals | Hypsugo_pulveratus | Pipistrellus_pulveratus | LC | Unknown | temperate | nocturnal | 5.5 | 2653163.505 | 13.354 |
| mammals | Hypsugo_savii | Hypsugo_savii | LC | Stable | temperate | nocturnal | 6.4 | 4335433.908 | 13.977 |
| mammals | Hypsugo_vordermanni | NA | DD | Unknown | NA | NA | NA | NA | NA |
| mammals | Hystrix_africaeaustralis | Hystrix_africaeaustralis | LC | Stable | tropical | nocturnal | 14936 | 7623760.023 | 8.396 |
| mammals | Hystrix_brachyura | Hystrix_brachyura | LC | Decreasing | tropical | nocturnal | 8450 | 5786235.96 | 12.374 |
| mammals | Hystrix_crassispinis | Hystrix_crassispinis | LC | Stable | tropical | nocturnal | 5445 | 732627.518 | 5.565 |
| mammals | Hystrix_cristata | Hystrix_cristata | LC | Unknown | tropical | nocturnal | 16250 | 5343513.373 | 11.642 |
| mammals | Hystrix_indica | Hystrix_indica | LC | Stable | temperate | nocturnal | 12435.8 | 7063667.05 | 13.26 |
| mammals | Hystrix_javanica | Hystrix_javanica | LC | Stable | tropical | nocturnal | 23014.4 | 168922.881 | 20.782 |
| mammals | Hystrix_pumila | Hystrix_pumila | VU | Decreasing | tropical | nocturnal | 2740 | 12398.137 | 9.288 |
| mammals | Hystrix_sumatrae | Hystrix_sumatrae | LC | Stable | tropical | nocturnal | 3260 | 429467.753 | 9.673 |
| mammals | Ia_io | Ia_io | NT | Decreasing | temperate | nocturnal | 49.2 | 1506489.992 | 11.552 |
| mammals | Ichneumia_albicauda | Ichneumia_albicauda | LC | Stable | tropical | nocturnal | 3500 | 14289624.18 | 9.83 |
| mammals | Ichthyomys_hydrobates | Ichthyomys_hydrobates | LC | Decreasing | tropical | cathemeral | 66.4 | 160034.844 | 12.748 |
| mammals | Ichthyomys_pittieri | Ichthyomys_pittieri | NT | Decreasing | tropical | nocturnal | 69.1 | 28379.941 | 15.158 |
| mammals | Ichthyomys_stolzmanni | NA | DD | Unknown | NA | NA | NA | NA | NA |
| mammals | Ichthyomys_tweedii | NA | DD | Unknown | NA | NA | NA | NA | NA |
| mammals | Ictidomys_mexicanus | Ictidomys_mexicanus | LC | Stable | temperate | diurnal | 189.6 | 715736.914 | 12.807 |
| mammals | Ictidomys_tridecemlineatus | Ictidomys_tridecemlineatus | LC | Stable | temperate | diurnal | 131.7 | 3679544.743 | 15.032 |
| mammals | Ictonyx_striatus | Ictonyx_striatus | LC | Stable | tropical | nocturnal | 1300 | 15345019.386 | 9.354 |
| mammals | Idionycteris_phyllotis | Idionycteris_phyllotis | LC | Stable | temperate | nocturnal | 12 | 1390365.361 | 8.89 |
| mammals | Idiurus_macrotis | Idiurus_macrotis | LC | Unknown | tropical | nocturnal | 30 | 619575.855 | 10.298 |
| mammals | Idiurus_zenkeri | Idiurus_zenkeri | LC | Unknown | tropical | nocturnal | 100 | 277784.739 | 7.286 |

| | | | | | | | | | |
|---|---|---|---|---|---|---|---|---|---|
| mammals | Indri_indri | Indri_indri | CR | Decreasing | tropical | nocturnal | 8000 | 30054.878 | 6.269 |
| mammals | Iomys_horsfieldii | Iomys_horsfieldii | LC | Stable | tropical | nocturnal | 147.5 | 806688.737 | 10.279 |
| mammals | Iomys_sipora | Iomys_sipora | EN | Decreasing | tropical | diurnal | 152.3 | 1242.491 | 8.588 |
| mammals | Irenomys_tarsalis | Irenomys_tarsalis | LC | Stable | temperate | nocturnal | 43.2 | 145039.907 | 5.458 |
| mammals | Isoodon_auratus | Isoodon_auratus | VU | Decreasing | tropical | nocturnal | 425 | 20997.783 | 0.952 |
| mammals | Isoodon_macrourus | Isoodon_macrourus | LC | Stable | tropical | nocturnal | 1600 | 1220483.405 | 4.669 |
| mammals | Isoodon_obesulus | Isoodon_obesulus | LC | Decreasing | temperate | nocturnal | 775 | 203133.139 | 8.191 |
| mammals | Isothrix_barbarabrownae | NA | DD | Unknown | NA | NA | NA | NA | NA |
| mammals | Isothrix_bistriata | Isothrix_bistriata | LC | Unknown | tropical | nocturnal | 445 | 1821069.619 | 2.731 |
| mammals | Isothrix_negrensis | Isothrix_negrensis | LC | Unknown | tropical | nocturnal | 467.7 | 294631.838 | 0.611 |
| mammals | Isothrix_pagurus | Isothrix_pagurus | LC | Unknown | tropical | nocturnal | 210 | 351407.388 | 1.627 |
| mammals | Isothrix_sinnamariensis | Isothrix_sinnamariensis | LC | Unknown | tropical | nocturnal | 260 | 4835.345 | 0.968 |
| mammals | Isthmomys_pirrensis | Isthmomys_pirrensis | LC | Unknown | tropical | nocturnal | 138 | 5206.223 | 1.975 |
| mammals | Jaculus_blanfordi | Jaculus_blanfordi | LC | Decreasing | temperate | nocturnal | 54.7 | 1305249.837 | 6.169 |
| mammals | Jaculus_jaculus | Jaculus_jaculus | LC | Unknown | temperate | nocturnal | 56.6 | 8111416.601 | 4.356 |
| mammals | Jaculus_orientalis | Jaculus_orientalis | LC | Stable | temperate | nocturnal | 138 | 434055.502 | 14.944 |
| mammals | Juliomys_pictipes | Juliomys_pictipes | LC | Unknown | temperate | nocturnal | 22.9 | 437708.817 | 13.602 |
| mammals | Juliomys_rimofrons | Juliomys_rimofrons | NT | Unknown | tropical | nocturnal | 19.8 | 10671.223 | 10.014 |
| mammals | Juscelinomys_huanchacae | NA | DD | Unknown | NA | NA | NA | NA | NA |
| mammals | Kadarsanomys_sodyi | Kadarsanomys_sodyi | EN | Unknown | tropical | nocturnal | 199.5 | 265.889 | 12.113 |
| mammals | Kannabateomys_amblyonyx | Kannabateomys_amblyonyx | LC | Unknown | temperate | nocturnal | 600 | 916239.9 | 13.716 |
| mammals | Kerivoula_africana | Kerivoula_africana | EN | Decreasing | tropical | nocturnal | 15.2 | 31520.502 | 10.231 |
| mammals | Kerivoula_agnella | Kerivoula_agnella | EN | Decreasing | tropical | nocturnal | 11.3 | 3220.914 | 4.39 |
| mammals | Kerivoula_argentata | Kerivoula_argentata | LC | Unknown | tropical | nocturnal | 10 | 2952181.027 | 9.487 |
| mammals | Kerivoula_cuprosa | NA | DD | Unknown | NA | NA | NA | NA | NA |
| mammals | Kerivoula_eriophora | NA | DD | Unknown | NA | NA | NA | NA | NA |
| mammals | Kerivoula_flora | Kerivoula_flora | VU | Decreasing | tropical | nocturnal | 6 | 49016.829 | 14.805 |
| mammals | Kerivoula_hardwickii | Kerivoula_hardwickii | LC | Stable | tropical | nocturnal | 4.6 | 3435503.961 | 10.932 |
| mammals | Kerivoula_intermedia | Kerivoula_intermedia | NT | Decreasing | tropical | nocturnal | 3.6 | 867950.159 | 6.699 |
| mammals | Kerivoula_kachinensis | Kerivoula_kachinensis | LC | Stable | tropical | nocturnal | 4.8 | 643276.889 | 10.001 |
| mammals | Kerivoula_krauensis | Kerivoula_krauensis | NT | Decreasing | tropical | nocturnal | 3 | 2799.993 | 5.127 |
| mammals | Kerivoula_lanosa | Kerivoula_lanosa | LC | Unknown | tropical | nocturnal | 6.5 | 9072400.794 | 8.498 |
| mammals | Kerivoula_lenis | Kerivoula_lenis | LC | Unknown | tropical | nocturnal | 6.8 | 129194.294 | 12.408 |
| mammals | Kerivoula_minuta | Kerivoula_minuta | NT | Decreasing | tropical | nocturnal | 2.1 | 808998.475 | 6.074 |

| | | | | | | | | | |
|---|---|---|---|---|---|---|---|---|---|
| mammals | Kerivoula_muscina | Kerivoula_muscina | LC | Unknown | tropical | nocturnal | 5.2 | 216623.442 | 3.63 |
| mammals | Kerivoula_myrella | Kerivoula_myrella | VU | Decreasing | tropical | nocturnal | 11.3 | 38091.102 | 4.306 |
| mammals | Kerivoula_papillosa | Kerivoula_papillosa | LC | Stable | tropical | nocturnal | 9.8 | 789927.979 | 10.179 |
| mammals | Kerivoula_pellucida | Kerivoula_pellucida | NT | Decreasing | tropical | nocturnal | 4.1 | 1213322.065 | 9.468 |
| mammals | Kerivoula_phalaena | Kerivoula_phalaena | LC | Unknown | tropical | nocturnal | 3 | 1206399.669 | 8.382 |
| mammals | Kerivoula_picta | Kerivoula_picta | NT | Decreasing | tropical | nocturnal | 4.5 | 2925844.479 | 13.234 |
| mammals | Kerivoula_smithii | Kerivoula_smithii | LC | Unknown | tropical | nocturnal | 13.1 | 1100356.074 | 5.642 |
| mammals | Kerivoula_titania | Kerivoula_titania | LC | Unknown | tropical | nocturnal | 4.6 | 966966.943 | 12.961 |
| mammals | Kerivoula_whiteheadi | Kerivoula_whiteheadi | LC | Unknown | tropical | nocturnal | 3.9 | 242485.406 | 11.676 |
| mammals | Kobus_kob | Kobus_kob | LC | Decreasing | tropical | cathemeral | 78500.1 | 3004386.388 | 9.687 |
| mammals | Kobus_leche | Kobus_leche | NT | Decreasing | tropical | diurnal | 110749 | 202159.549 | 5.305 |
| mammals | Kobus_megaceros | Kobus_megaceros | EN | Decreasing | tropical | diurnal | 87500.4 | 110602.966 | 8.065 |
| mammals | Komodomys_rintjanus | Komodomys_rintjanus | VU | Stable | tropical | nocturnal | 115 | 16442.847 | 12.545 |
| mammals | Kunsia_tomentosus | Kunsia_tomentosus | LC | Decreasing | tropical | cathemeral | 115.6 | 405854.328 | 4.171 |
| mammals | Laephotis_angolensis | NA | DD | Unknown | NA | NA | NA | NA | NA |
| mammals | Laephotis_capensis | Neoromicia_capensis | LC | Stable | tropical | nocturnal | 6 | 13760702.434 | 8.522 |
| mammals | Laephotis_malagasyensis | Neoromicia_malagasyensis | VU | Unknown | tropical | nocturnal | 6.5 | 7033.93 | 7.484 |
| mammals | Laephotis_matroka | Neoromicia_matroka | LC | Unknown | tropical | nocturnal | 6.5 | 33499.49 | 10.929 |
| mammals | Laephotis_namibensis | Laephotis_namibensis | LC | Stable | temperate | nocturnal | 7.7 | 27774.566 | 5.455 |
| mammals | Laephotis_wintoni | Laephotis_wintoni | LC | Unknown | tropical | nocturnal | 6 | 1918973.172 | 10.197 |
| mammals | Lagidium_viscacia | Lagidium_viscacia | LC | Stable | temperate | cathemeral | 1540 | 1776083.67 | 5.686 |
| mammals | Lagidium_wolffsohni | NA | DD | Unknown | NA | NA | NA | NA | NA |
| mammals | Lagorchestes_conspicillatus | Lagorchestes_conspicillatus | LC | Decreasing | tropical | nocturnal | 3050 | 1884281.953 | 3.192 |
| mammals | Lagorchestes_hirsutus | Lagorchestes_hirsutus | VU | Stable | temperate | nocturnal | 1265 | 95.234 | 3.466 |
| mammals | Lagostomus_maximus | Lagostomus_maximus | LC | Unknown | temperate | nocturnal | 4647.5 | 1466480.45 | 8.566 |
| mammals | Lagostrophus_fasciatus | Lagostrophus_fasciatus | VU | Stable | temperate | nocturnal | 1700 | 137.024 | 5.152 |
| mammals | Lagothrix_flavicauda | Oreonax_flavicauda | CR | Decreasing | tropical | diurnal | 6800 | 24210.46 | 3.739 |
| mammals | Lagothrix_lagothricha | Lagothrix_lagotricha | VU | Decreasing | tropical | diurnal | 6788.9 | 2568088.693 | 1.447 |
| mammals | Lagurus_lagurus | Lagurus_lagurus | LC | Stable | temperate | cathemeral | 20.3 | 4713486.428 | 8.839 |
| mammals | Lama_guanicoe | Lama_guanicoe | LC | Increasing | temperate | diurnal | 120000 | 1658706.173 | 5.243 |
| mammals | Lama_vicugna | Vicugna_vicugna | LC | Increasing | tropical | diurnal | 47499.6 | 619657.908 | 6.172 |
| mammals | Lampronycteris_brachyotis | Lampronycteris_brachyotis | LC | Stable | tropical | nocturnal | 11 | 4319292.52 | 2.845 |
| mammals | Laonastes_aenigmamus | Laonastes_aenigmamus | LC | Decreasing | tropical | cathemeral | 375 | 6488.204 | 6.416 |
| mammals | Lariscus_hosei | Lariscus_hosei | LC | Decreasing | tropical | diurnal | 170.7 | 52688.425 | 3.704 |

| | | | | | | | | | |
|---|---|---|---|---|---|---|---|---|---|
| mammals | Lariscus_insignis | Lariscus_insignis | LC | Decreasing | tropical | diurnal | 175 | 1276285.786 | 9.754 |
| mammals | Lariscus_niobe | NA | DD | Unknown | NA | NA | NA | NA | NA |
| mammals | Lariscus_obscurus | Lariscus_obscurus | NT | Decreasing | tropical | diurnal | 188.3 | 6004.261 | 6.223 |
| mammals | Lasionycteris_noctivagans | Lasionycteris_noctivagans | LC | Stable | temperate | nocturnal | 10.5 | 10110807.92 | 11.832 |
| mammals | Lasiopodomys_brandtii | Lasiopodomys_brandtii | LC | Stable | temperate | diurnal | 93.8 | 819069.453 | 7.285 |
| mammals | Lasiorhinus_krefftii | Lasiorhinus_krefftii | CR | Stable | tropical | nocturnal | 31450 | 91.215 | 8.813 |
| mammals | Lasiurus_atratus | Lasiurus_atratus | LC | Unknown | tropical | nocturnal | 19.4 | 399779.311 | 1.849 |
| mammals | Lasiurus_blossevillii | Lasiurus_blossevillii | LC | Unknown | tropical | nocturnal | 13 | 18711014.82 | 6.931 |
| mammals | Lasiurus_borealis | Lasiurus_borealis | LC | Stable | temperate | nocturnal | 10.7 | 4881497.799 | 16.616 |
| mammals | Lasiurus_castaneus | NA | DD | Unknown | NA | NA | NA | NA | NA |
| mammals | Lasiurus_cinereus | Lasiurus_cinereus | LC | Unknown | temperate | nocturnal | 25.3 | 21071173.085 | 9.354 |
| mammals | Lasiurus_degelidus | Lasiurus_degelidus | VU | Unknown | tropical | nocturnal | 12.8 | 11021.007 | 19.74 |
| mammals | Lasiurus_ega | Lasiurus_ega | LC | Unknown | tropical | nocturnal | 13.2 | 13205185.72 | 6.473 |
| mammals | Lasiurus_egregius | NA | DD | Unknown | NA | NA | NA | NA | NA |
| mammals | Lasiurus_insularis | Lasiurus_insularis | VU | Unknown | tropical | nocturnal | 31.6 | 106633.252 | 16.283 |
| mammals | Lasiurus_intermedius | Lasiurus_intermedius | LC | Unknown | temperate | nocturnal | 23 | 1748790.816 | 12.809 |
| mammals | Lasiurus_minor | Lasiurus_minor | VU | Decreasing | tropical | nocturnal | 9.4 | 84473.907 | 16.774 |
| mammals | Lasiurus_pfeifferi | Lasiurus_pfeifferi | NT | Unknown | tropical | nocturnal | 11.1 | 96779.232 | 16.415 |
| mammals | Lasiurus_seminolus | Lasiurus_seminolus | LC | Unknown | temperate | nocturnal | 9.9 | 1193528.476 | 16.184 |
| mammals | Lasiurus_varius | Lasiurus_varius | LC | Unknown | temperate | nocturnal | 11.4 | 545210.339 | 6.018 |
| mammals | Lasiurus_xanthinus | Lasiurus_xanthinus | LC | Stable | temperate | nocturnal | 10.7 | 1737395.754 | 8.335 |
| mammals | Latidens_salimalii | Latidens_salimalii | EN | Decreasing | tropical | nocturnal | 55.2 | 18947.746 | 13.667 |
| mammals | Lavia_frons | Lavia_frons | LC | Stable | tropical | diurnal | 23.9 | 7569286.872 | 9.571 |
| mammals | Leggadina_forresti | Leggadina_forresti | LC | Unknown | temperate | nocturnal | 20 | 1981608.763 | 3.019 |
| mammals | Leggadina_lakedownensis | Leggadina_lakedownensis | LC | Decreasing | tropical | nocturnal | 17.5 | 1323526.627 | 2.803 |
| mammals | Leimacomys_buettneri | NA | DD | Unknown | NA | NA | NA | NA | NA |
| mammals | Leiuromys_occasius | NA | DD | Unknown | NA | NA | NA | NA | NA |
| mammals | Lemmus_amurensis | Lemmus_amurensis | LC | Unknown | temperate | cathemeral | 33.6 | 2123204.58 | 1.453 |
| mammals | Lemmus_lemmus | Lemmus_lemmus | LC | Stable | temperate | cathemeral | 47.5 | 758090.79 | 5.444 |
| mammals | Lemniscomys_barbarus | Lemniscomys_barbarus | LC | Stable | temperate | nocturnal | 49 | 249140.731 | 18.896 |
| mammals | Lemniscomys_griselda | Lemniscomys_griselda | LC | Stable | tropical | nocturnal | 58 | 599556.141 | 6.253 |
| mammals | Lemniscomys_hoogstraali | NA | DD | Unknown | NA | NA | NA | NA | NA |
| mammals | Lemniscomys_linulus | Lemniscomys_linulus | LC | Unknown | tropical | nocturnal | 27 | 266618.198 | 10.359 |
| mammals | Lemniscomys_macculus | Lemniscomys_macculus | LC | Stable | tropical | cathemeral | 30.3 | 992343.248 | 10.482 |

| | | | | | | | | | |
|---|---|---|---|---|---|---|---|---|---|
| mammals | Lemniscomys_mittendorfi | Lemniscomys_mittendorfi | LC | Stable | tropical | nocturnal | 28 | 404.854 | 12.419 |
| mammals | Lemniscomys_rosalia | Lemniscomys_rosalia | LC | Stable | tropical | nocturnal | 54.2 | 2429390.548 | 9.345 |
| mammals | Lemniscomys_roseveari | NA | DD | Unknown | NA | NA | NA | NA | NA |
| mammals | Lemur_catta | Lemur_catta | EN | Decreasing | tropical | cathemeral | 2900 | 99812.555 | 9.598 |
| mammals | Lenomys_meyeri | Lenomys_meyeri | LC | Decreasing | tropical | nocturnal | 480 | 116927.085 | 10.147 |
| mammals | Lenothrix_cana | Lenothrix_canus | LC | Unknown | tropical | nocturnal | 135 | 238775.298 | 9.36 |
| mammals | Leontopithecus_caissara | Leontopithecus_caissara | EN | Decreasing | temperate | diurnal | 572 | 205.644 | 2.153 |
| mammals | Leontopithecus_chrysomelas | Leontopithecus_chrysomelas | EN | Decreasing | tropical | diurnal | 700 | 20187.301 | 8.255 |
| mammals | Leontopithecus_chrysopygus | Leontopithecus_chrysopygus | EN | Decreasing | tropical | diurnal | 700 | 63670.785 | 13.846 |
| mammals | Leontopithecus_rosalia | Leontopithecus_rosalia | EN | Decreasing | tropical | diurnal | 535.5 | 3990.552 | 18.849 |
| mammals | Leopardus_colocola | Leopardus_colocolo | NT | Decreasing | temperate | cathemeral | 3935 | 4786752.179 | 7.092 |
| mammals | Leopardus_geoffroyi | Leopardus_geoffroyi | LC | Stable | temperate | cathemeral | 2230 | 3529650.057 | 7.511 |
| mammals | Leopardus_guigna | Leopardus_guigna | VU | Decreasing | temperate | nocturnal | 3590 | 317797.041 | 7.654 |
| mammals | Leopardus_guttulus | Leopardus_guttulus | VU | Decreasing | tropical | cathemeral | 2500 | 2146517.132 | 11.524 |
| mammals | Leopardus_jacobita | Leopardus_jacobita | EN | Decreasing | tropical | diurnal | 9170 | 612427.861 | 6.222 |
| mammals | Leopardus_pardalis | Leopardus_pardalis | LC | Decreasing | tropical | cathemeral | 11900.1 | 14546411.352 | 5.962 |
| mammals | Leopardus_tigrinus | Leopardus_tigrinus | VU | Decreasing | tropical | cathemeral | 2250 | 9389963.336 | 4.161 |
| mammals | Leopardus_wiedii | Leopardus_wiedii | NT | Decreasing | tropical | cathemeral | 3250 | 12288755.81 | 5.719 |
| mammals | Leopoldamys_ciliatus | Leopoldamys_ciliatus | LC | Decreasing | tropical | nocturnal | 362.5 | 199687.854 | 7.851 |
| mammals | Leopoldamys_edwardsi | Leopoldamys_edwardsi | LC | Unknown | temperate | nocturnal | 300 | 2061787.685 | 12.887 |
| mammals | Leopoldamys_milleti | Leopoldamys_milleti | LC | Stable | tropical | nocturnal | 410.8 | 5654.281 | 9.138 |
| mammals | Leopoldamys_neilli | Leopoldamys_neilli | LC | Unknown | tropical | nocturnal | 219 | 109202.32 | 16.241 |
| mammals | Leopoldamys_sabanus | Leopoldamys_sabanus | LC | Stable | tropical | nocturnal | 338.3 | 2538761.32 | 10.482 |
| mammals | Leopoldamys_siporanus | Leopoldamys_siporanus | VU | Decreasing | tropical | nocturnal | 576 | 6004.261 | 6.223 |
| mammals | Lepilemur_aeeclis | Lepilemur_aeeclis | EN | Decreasing | tropical | nocturnal | 867.5 | 363.582 | 9.156 |
| mammals | Lepilemur_ahmansoni | Lepilemur_ahmansonorum | CR | Decreasing | tropical | nocturnal | 610 | 281.282 | 5.122 |
| mammals | Lepilemur_ankaranensis | Lepilemur_ankaranensis | EN | Decreasing | tropical | nocturnal | 750 | 1576.597 | 6.979 |
| mammals | Lepilemur_betsileo | Lepilemur_betsileo | EN | Decreasing | tropical | nocturnal | 1150 | 2186.317 | 6.884 |
| mammals | Lepilemur_dorsalis | Lepilemur_dorsalis | EN | Decreasing | tropical | nocturnal | 855 | 4754.324 | 5.465 |
| mammals | Lepilemur_fleuretae | Lepilemur_fleuretae | EN | Decreasing | temperate | nocturnal | 980 | 1496.081 | 5.973 |
| mammals | Lepilemur_grewcocki | Lepilemur_grewcockorum | CR | Decreasing | tropical | nocturnal | 780 | 139.293 | 10.868 |
| mammals | Lepilemur_hollandorum | Lepilemur_hollandorum | CR | Decreasing | tropical | nocturnal | 1000 | 282.147 | 4.846 |
| mammals | Lepilemur_hubbardi | Lepilemur_hubbardorum | EN | Decreasing | tropical | nocturnal | 990 | 212.991 | 7.853 |
| mammals | Lepilemur_jamesi | Lepilemur_jamesorum | CR | Decreasing | tropical | nocturnal | 780 | 64.437 | 9.714 |

| | | | | | | | | | |
|---|---|---|---|---|---|---|---|---|---|
| mammals | Lepilemur_leucopus | Lepilemur_leucopus | EN | Decreasing | temperate | nocturnal | 525 | 1953.795 | 9.985 |
| mammals | Lepilemur_microdon | Lepilemur_microdon | EN | Decreasing | tropical | nocturnal | 900 | 2505.77 | 5.99 |
| mammals | Lepilemur_milanoii | Lepilemur_milanoii | EN | Decreasing | tropical | nocturnal | 720 | 2566.824 | 7.578 |
| mammals | Lepilemur_mustelinus | Lepilemur_mustelinus | VU | Decreasing | tropical | nocturnal | 1000 | 12959.451 | 6.941 |
| mammals | Lepilemur_otto | Lepilemur_otto | EN | Decreasing | tropical | nocturnal | 938 | 3486.437 | 9.618 |
| mammals | Lepilemur_petteri | Lepilemur_petteri | EN | Decreasing | temperate | nocturnal | 630 | 13303.914 | 9.355 |
| mammals | Lepilemur_randrianasoloi | Lepilemur_randrianasoloi | EN | Decreasing | tropical | nocturnal | 780 | 1884.65 | 3.729 |
| mammals | Lepilemur_ruficaudatus | Lepilemur_ruficaudatus | CR | Decreasing | tropical | nocturnal | 750 | 15786.758 | 8.04 |
| mammals | Lepilemur_sahamalaza | Lepilemur_sahamalazensis | CR | Decreasing | tropical | nocturnal | 789.5 | 436.881 | 7.853 |
| mammals | Lepilemur_scottorum | Lepilemur_scottorum | EN | Decreasing | tropical | nocturnal | 880 | 1992.434 | 3.648 |
| mammals | Lepilemur_seali | Lepilemur_seali | VU | Decreasing | tropical | nocturnal | 950 | 7397.992 | 6.622 |
| mammals | Lepilemur_septentrionalis | Lepilemur_septentrionalis | CR | Decreasing | tropical | nocturnal | 750 | 139.307 | 12.617 |
| mammals | Lepilemur_tymerlachsoni | Lepilemur_tymerlachsoni | CR | Decreasing | tropical | nocturnal | 880 | 52.926 | 16.703 |
| mammals | Lepilemur_wrighti | Lepilemur_wrightae | EN | Decreasing | tropical | nocturnal | 950 | 584.741 | 4.142 |
| mammals | Leporillus_conditor | Leporillus_conditor | NT | Stable | temperate | nocturnal | 350 | 705.737 | 6.113 |
| mammals | Leptomys_elegans | Leptomys_elegans | LC | Unknown | tropical | nocturnal | 84.8 | 73683.009 | 4.054 |
| mammals | Leptomys_ernstmayri | Leptomys_ernstmayri | LC | Unknown | tropical | nocturnal | 42.9 | 49834.37 | 4.718 |
| mammals | Leptomys_signatus | Leptomys_signatus | LC | Unknown | tropical | nocturnal | 54.4 | 764.468 | 2.289 |
| mammals | Leptonycteris_curasoae | Leptonycteris_curasoae | VU | Decreasing | tropical | nocturnal | 23.4 | 840356.495 | 11.097 |
| mammals | Leptonycteris_nivalis | Leptonycteris_nivalis | EN | Decreasing | tropical | nocturnal | 25.6 | 949993.755 | 10.069 |
| mammals | Leptonycteris_yerbabuenae | Leptonycteris_yerbabuenae | NT | Decreasing | temperate | nocturnal | 20 | 1233797.183 | 9.761 |
| mammals | Lepus_alleni | Lepus_alleni | LC | Unknown | temperate | nocturnal | 3685 | 233589.802 | 8.244 |
| mammals | Lepus_americanus | Lepus_americanus | LC | Stable | temperate | nocturnal | 1710 | 9597254.589 | 4.51 |
| mammals | Lepus_arcticus | Lepus_arcticus | LC | Unknown | temperate | nocturnal | 4405 | 2556757.847 | 0.104 |
| mammals | Lepus_brachyurus | Lepus_brachyurus | LC | Stable | temperate | nocturnal | 2350 | 286560.637 | 21.086 |
| mammals | Lepus_californicus | Lepus_californicus | LC | Decreasing | temperate | nocturnal | 2422.5 | 4392320.271 | 10.868 |
| mammals | Lepus_callotis | Lepus_callotis | VU | Decreasing | tropical | nocturnal | 2500 | 378241.513 | 12.446 |
| mammals | Lepus_capensis | Lepus_capensis | LC | Decreasing | tropical | nocturnal | 3261.7 | 11230966.342 | 7.735 |
| mammals | Lepus_castroviejoi | Lepus_castroviejoi | VU | Decreasing | temperate | nocturnal | 2750 | 4980.046 | 10.189 |
| mammals | Lepus_comus | Lepus_comus | LC | Decreasing | temperate | nocturnal | 1460 | 418232.269 | 12.761 |
| mammals | Lepus_coreanus | Lepus_coreanus | LC | Decreasing | temperate | nocturnal | 1700 | 217852.409 | 17.353 |
| mammals | Lepus_corsicanus | Lepus_corsicanus | VU | Decreasing | temperate | nocturnal | 3000 | 114558.639 | 22.309 |
| mammals | Lepus_europaeus | Lepus_europaeus | LC | Decreasing | temperate | cathemeral | 3740 | 14444573.951 | 12.762 |
| mammals | Lepus_fagani | Lepus_fagani | LC | Unknown | tropical | nocturnal | 3500 | 195235.017 | 12.744 |

| | | | | | | | | | |
|---|---|---|---|---|---|---|---|---|---|
| mammals | Lepus_flavigularis | Lepus_flavigularis | EN | Decreasing | tropical | nocturnal | 3000 | 100.001 | 10.382 |
| mammals | Lepus_granatensis | Lepus_granatensis | LC | Stable | temperate | nocturnal | 4010 | 533085.017 | 17.131 |
| mammals | Lepus_habessinicus | Lepus_habessinicus | LC | Unknown | tropical | nocturnal | 1962.1 | 1258924.318 | 8.534 |
| mammals | Lepus_hainanus | Lepus_hainanus | EN | Decreasing | tropical | nocturnal | 2000 | 586.555 | 24.69 |
| mammals | Lepus_mandshuricus | Lepus_mandshuricus | LC | Unknown | temperate | nocturnal | 1800 | 1091802.072 | 9.474 |
| mammals | Lepus_microtis | Lepus_victoriae | LC | Stable | tropical | nocturnal | 1750 | 11320167.709 | 9.787 |
| mammals | Lepus_nigricollis | Lepus_nigricollis | LC | Unknown | tropical | nocturnal | 2298.5 | 3430852.143 | 18.35 |
| mammals | Lepus_oiostolus | Lepus_oiostolus | LC | Unknown | temperate | nocturnal | 3000 | 2759213.207 | 4.286 |
| mammals | Lepus_othus | Lepus_othus | LC | Unknown | temperate | nocturnal | 4806 | 203837.722 | 0.897 |
| mammals | Lepus_peguensis | Lepus_peguensis | LC | Stable | tropical | nocturnal | 2105 | 1243043.051 | 12.126 |
| mammals | Lepus_saxatilis | Lepus_saxatilis | LC | Decreasing | temperate | nocturnal | 2363.3 | 193852.402 | 8.223 |
| mammals | Lepus_sinensis | Lepus_sinensis | LC | Unknown | temperate | nocturnal | 1510 | 957212.127 | 15.825 |
| mammals | Lepus_starcki | Lepus_starcki | LC | Unknown | tropical | nocturnal | 2750 | 60357.605 | 15.244 |
| mammals | Lepus_timidus | Lepus_timidus | LC | Stable | temperate | nocturnal | 3048 | 20131533.222 | 4.593 |
| mammals | Lepus_tolai | Lepus_tolai | LC | Unknown | temperate | nocturnal | 1589 | 9351540.867 | 9.4 |
| mammals | Lepus_townsendii | Lepus_townsendii | LC | Decreasing | temperate | nocturnal | 1555 | 3453856.922 | 12.119 |
| mammals | Lepus_yarkandensis | Lepus_yarkandensis | NT | Decreasing | temperate | nocturnal | 1500 | 581120.533 | 2.924 |
| mammals | Lestodelphys_halli | Lestodelphys_halli | LC | Decreasing | temperate | nocturnal | 76 | 309930.903 | 5.739 |
| mammals | Lestoros_inca | Lestoros_inca | LC | Unknown | tropical | nocturnal | 21.1 | 62779.888 | 3.712 |
| mammals | Leucocephalophus_adersi | Cephalophus_adersi | VU | Decreasing | tropical | diurnal | 9250 | 5035.199 | 8.01 |
| mammals | Liberiictis_kuhni | Liberiictis_kuhni | VU | Decreasing | tropical | diurnal | 2150 | 77730.786 | 12.176 |
| mammals | Lichonycteris_obscura | Lichonycteris_obscura | LC | Unknown | tropical | nocturnal | 5.6 | 696682.6 | 9.883 |
| mammals | Limnomys_bryophilus | Limnomys_bryophilus | LC | Stable | tropical | nocturnal | 64.1 | 1479.534 | 6.242 |
| mammals | Limnomys_sibuanus | Limnomys_sibuanus | LC | Stable | tropical | nocturnal | 60 | 1999.965 | 8.094 |
| mammals | Lionycteris_spurrelli | Lionycteris_spurrelli | LC | Stable | tropical | nocturnal | 8.4 | 7274395.187 | 3.287 |
| mammals | Litocranius_walleri | Litocranius_walleri | NT | Decreasing | tropical | diurnal | 37999.7 | 1418356.417 | 8.1 |
| mammals | Lonchophylla_bokermanni | Lonchophylla_bokermanni | EN | Unknown | tropical | nocturnal | 9.6 | 938.831 | 10.797 |
| mammals | Lonchophylla_chocoana | NA | DD | Unknown | NA | NA | NA | NA | NA |
| mammals | Lonchophylla_concava | Lonchophylla_concava | LC | Unknown | tropical | nocturnal | 7.4 | 206491.241 | 11.287 |
| mammals | Lonchophylla_dekeyseri | Lonchophylla_dekeyseri | EN | Decreasing | tropical | nocturnal | 11.2 | 28619.179 | 8.947 |
| mammals | Lonchophylla_handleyi | Lonchophylla_handleyi | LC | Unknown | tropical | nocturnal | 18.5 | 91306.654 | 3.313 |
| mammals | Lonchophylla_hesperia | Lonchophylla_hesperia | NT | Decreasing | tropical | nocturnal | 9.8 | 275064.216 | 6.078 |
| mammals | Lonchophylla_mordax | Lonchophylla_mordax | NT | Unknown | tropical | nocturnal | 9 | 20483.631 | 13.407 |
| mammals | Lonchophylla_orcesi | NA | DD | Unknown | NA | NA | NA | NA | NA |

| | | | | | | | | | |
|---|---|---|---|---|---|---|---|---|---|
| mammals | Lonchophylla_robusta | Lonchophylla_robusta | LC | Unknown | tropical | nocturnal | 17.2 | 1116127.265 | 10.656 |
| mammals | Lonchorhina_aurita | Lonchorhina_aurita | LC | Stable | tropical | nocturnal | 15.7 | 11577000.206 | 5.49 |
| mammals | Lonchorhina_fernandezi | Lonchorhina_fernandezi | EN | Decreasing | tropical | nocturnal | 8.7 | 105607.867 | 0.667 |
| mammals | Lonchorhina_inusitata | NA | DD | Unknown | NA | NA | NA | NA | NA |
| mammals | Lonchorhina_marinkellei | Lonchorhina_marinkellei | VU | Unknown | tropical | nocturnal | 17 | 161297.63 | 0.934 |
| mammals | Lonchorhina_orinocensis | Lonchorhina_orinocensis | VU | Decreasing | tropical | nocturnal | 8.8 | 386541.587 | 2.944 |
| mammals | Lonchothrix_emiliae | Lonchothrix_emiliae | LC | Stable | tropical | nocturnal | 138.2 | 378168.057 | 2.016 |
| mammals | Lontra_canadensis | Lontra_canadensis | LC | Stable | temperate | nocturnal | 8087.4 | 19086449.764 | 7.095 |
| mammals | Lontra_longicaudis | Lontra_longicaudis | NT | Decreasing | tropical | diurnal | 6555 | 15024030.121 | 6.262 |
| mammals | Lontra_provocax | Lontra_provocax | EN | Decreasing | temperate | nocturnal | 7500 | 125850.092 | 3.252 |
| mammals | Lophiomys_imhausi | Lophiomys_imhausi | LC | Stable | tropical | nocturnal | 755 | 424237.651 | 10.771 |
| mammals | Lophocebus_albigena | Lophocebus_albigena | VU | Decreasing | tropical | diurnal | 6984.8 | 1508231.453 | 6.787 |
| mammals | Lophocebus_aterrimus | Lophocebus_aterrimus | VU | Decreasing | tropical | diurnal | 6984.8 | 706964.316 | 5.475 |
| mammals | Lophostoma_brasiliense | Lophostoma_brasiliense | LC | Stable | tropical | nocturnal | 9.4 | 8095483.469 | 3.834 |
| mammals | Lophostoma_carrikeri | Lophostoma_carrikeri | LC | Decreasing | tropical | nocturnal | 20.9 | 3390162.363 | 2.049 |
| mammals | Lophostoma_evotis | Lophostoma_evotis | LC | Stable | tropical | nocturnal | 19.8 | 176592.762 | 9.456 |
| mammals | Lophostoma_kalkoae | NA | DD | Unknown | NA | NA | NA | NA | NA |
| mammals | Lophostoma_occidentale | Lophostoma_occidentalis | NT | Decreasing | tropical | nocturnal | 25.6 | 201548.718 | 10.713 |
| mammals | Lophostoma_schulzi | Lophostoma_schulzi | LC | Unknown | tropical | nocturnal | 18.2 | 564364.994 | 1.04 |
| mammals | Lophostoma_silvicola | Lophostoma_silvicolum | LC | Unknown | tropical | nocturnal | 29.7 | 7691508.732 | 3.735 |
| mammals | Lophuromys_brevicaudus | Lophuromys_brevicaudus | NT | Decreasing | tropical | cathemeral | 43.1 | 13838.889 | 15.053 |
| mammals | Lophuromys_chrysopus | Lophuromys_chrysopus | LC | Unknown | tropical | cathemeral | 43.1 | 112209.387 | 9.918 |
| mammals | Lophuromys_dieterleni | Lophuromys_dieterleni | EN | Decreasing | tropical | nocturnal | 63.6 | 400.743 | 12.426 |
| mammals | Lophuromys_eisentrauti | Lophuromys_eisentrauti | CR | Decreasing | tropical | nocturnal | 30 | 55.885 | 14.31 |
| mammals | Lophuromys_flavopunctatus | Lophuromys_flavopunctatus | LC | Unknown | tropical | nocturnal | 60 | 1820860.792 | 7.334 |
| mammals | Lophuromys_huttereri | Lophuromys_huttereri | LC | Unknown | tropical | nocturnal | 39.9 | 44203.65 | 5.471 |
| mammals | Lophuromys_luteogaster | Lophuromys_luteogaster | LC | Unknown | tropical | nocturnal | 34 | 63598.452 | 4.508 |
| mammals | Lophuromys_medicaudatus | Lophuromys_medicaudatus | VU | Decreasing | tropical | nocturnal | 35 | 7510.562 | 16.498 |
| mammals | Lophuromys_melanonyx | Lophuromys_melanonyx | VU | Unknown | tropical | diurnal | 102.1 | 10810.375 | 13.22 |
| mammals | Lophuromys_nudicaudus | Lophuromys_nudicaudus | LC | Unknown | tropical | diurnal | 39.5 | 612364.736 | 5.87 |
| mammals | Lophuromys_rahmi | Lophuromys_rahmi | NT | Unknown | tropical | nocturnal | 38 | 4293.325 | 14.695 |
| mammals | Lophuromys_roseveari | Lophuromys_roseveari | LC | Unknown | tropical | nocturnal | 63.5 | 771.072 | 10.531 |
| mammals | Lophuromys_sikapusi | Lophuromys_sikapusi | LC | Unknown | tropical | nocturnal | 74.1 | 3320549.701 | 9.486 |
| mammals | Lophuromys_woosnami | Lophuromys_woosnami | LC | Unknown | tropical | nocturnal | 42.2 | 52330.476 | 18.481 |

| | | | | | | | | | |
|---|---|---|---|---|---|---|---|---|---|
| mammals | Lorentzimys_nouhuysii | Lorentzimys_nouhuysi | LC | Stable | tropical | cathemeral | 14.2 | 251852.19 | 4.216 |
| mammals | Loris_lydekkerianus | Loris_lydekkerianus | NT | Decreasing | tropical | nocturnal | 235 | 493888.944 | 17.752 |
| mammals | Loris_tardigradus | Loris_tardigradus | EN | Decreasing | tropical | nocturnal | 227.7 | 6904.001 | 23.541 |
| mammals | Loxodonta_africana | Loxodonta_africana | EN | Decreasing | tropical | cathemeral | 5300000 | 3213906.059 | 6.603 |
| mammals | Loxodontomys_micropus | Loxodontomys_micropus | LC | Stable | temperate | nocturnal | 72.7 | 452856.897 | 6.404 |
| mammals | Lundomys_molitor | Lundomys_molitor | LC | Decreasing | temperate | nocturnal | 238.5 | 295487.687 | 10.024 |
| mammals | Lupulella_adustus | Canis_adustus | LC | Stable | tropical | nocturnal | 10249.9 | 10714383.053 | 9.644 |
| mammals | Lupulella_mesomelas | Canis_mesomelas | LC | Stable | temperate | cathemeral | 8500 | 6453409.895 | 8.754 |
| mammals | Lutra_capensis | Aonyx_capensis | NT | Decreasing | tropical | nocturnal | 18999.8 | 11817070.922 | 10.263 |
| mammals | Lutra_cinerea | Aonyx_cinereus | VU | Decreasing | tropical | diurnal | 1747.6 | 4793188.51 | 12.104 |
| mammals | Lutra_congica | Aonyx_congicus | NT | Decreasing | tropical | nocturnal | 23800.2 | 2924429.887 | 6.434 |
| mammals | Lutra_lutra | Lutra_lutra | NT | Decreasing | temperate | nocturnal | 8785.1 | 27067759.291 | 9.499 |
| mammals | Lutra_perspicillata | Lutrogale_perspicillata | VU | Decreasing | tropical | diurnal | 9483.3 | 4875750.854 | 15.066 |
| mammals | Lutra_sumatrana | Lutra_sumatrana | EN | Decreasing | tropical | cathemeral | 5500 | 865485.002 | 7.719 |
| mammals | Lutreolina_crassicaudata | Lutreolina_crassicaudata | LC | Unknown | tropical | nocturnal | 537.3 | 3049187.09 | 8.834 |
| mammals | Lutreolina_massoia | Lutreolina_massoia | LC | Unknown | tropical | nocturnal | 284 | 96708.518 | 6.829 |
| mammals | Lycalopex_culpaeus | Lycalopex_culpaeus | LC | Stable | temperate | cathemeral | 9832.4 | 2667800.128 | 5.81 |
| mammals | Lycalopex_fulvipes | Lycalopex_fulvipes | EN | Decreasing | temperate | diurnal | 2837.5 | 15381.477 | 6.198 |
| mammals | Lycalopex_grisea | Lycalopex_griseus | LC | Stable | temperate | cathemeral | 8280 | 1618133.157 | 6.1 |
| mammals | Lycalopex_gymnocerca | Lycalopex_gymnocercus | LC | Stable | temperate | cathemeral | 4690 | 2776265.541 | 8.146 |
| mammals | Lycalopex_sechurae | Lycalopex_sechurae | NT | Unknown | tropical | nocturnal | 4000 | 73116.044 | 8.627 |
| mammals | Lycalopex_vetula | Lycalopex_vetulus | NT | Decreasing | tropical | nocturnal | 5350 | 2052494.179 | 9.074 |
| mammals | Lycaon_pictus | Lycaon_pictus | EN | Decreasing | tropical | cathemeral | 22050.1 | 1272642.801 | 5.189 |
| mammals | Lyncodon_patagonicus | Lyncodon_patagonicus | LC | Unknown | temperate | nocturnal | 225 | 1386425.691 | 6.513 |
| mammals | Lynx_canadensis | Lynx_canadensis | LC | Stable | temperate | cathemeral | 9373.2 | 8328044.88 | 2.372 |
| mammals | Lynx_lynx | Lynx_lynx | LC | Stable | temperate | cathemeral | 17950 | 20817635.888 | 4.936 |
| mammals | Lynx_pardinus | Lynx_pardinus | EN | Increasing | temperate | nocturnal | 9400 | 1191.692 | 12.707 |
| mammals | Lynx_rufus | Lynx_rufus | LC | Stable | temperate | cathemeral | 8904.1 | 9879513.025 | 12.076 |
| mammals | Lyroderma_lyra | Megaderma_lyra | LC | Unknown | tropical | nocturnal | 45.5 | 6060994.625 | 15.525 |
| mammals | Macaca_arctoides | Macaca_arctoides | VU | Decreasing | tropical | diurnal | 5000 | 1712459.601 | 11.133 |
| mammals | Macaca_assamensis | Macaca_assamensis | NT | Decreasing | tropical | diurnal | 12000 | 1808926.188 | 9.443 |
| mammals | Macaca_cyclopis | Macaca_cyclopis | LC | Stable | temperate | diurnal | 6000 | 26772.6 | 19.19 |
| mammals | Macaca_fascicularis | Macaca_fascicularis | EN | Decreasing | tropical | diurnal | 3991.5 | 2431069.105 | 10.872 |
| mammals | Macaca_fuscata | Macaca_fuscata | LC | Stable | temperate | diurnal | 10400 | 157042.032 | 18.538 |

| | | | | | | | | | |
|---|---|---|---|---|---|---|---|---|---|
| mammals | Macaca_hecki | Macaca_hecki | VU | Decreasing | tropical | diurnal | 9000 | 23994.724 | 7.813 |
| mammals | Macaca_leonina | Macaca_leonina | VU | Decreasing | tropical | diurnal | 6500 | 1548715.299 | 10.461 |
| mammals | Macaca_maura | Macaca_maura | EN | Decreasing | tropical | diurnal | 7300 | 19910.243 | 19.137 |
| mammals | Macaca_mulatta | Macaca_mulatta | LC | Unknown | temperate | diurnal | 3713 | 7472315.402 | 13.745 |
| mammals | Macaca_nemestrina | Macaca_nemestrina | EN | Decreasing | tropical | diurnal | 6296.5 | 1343992.195 | 7.925 |
| mammals | Macaca_nigra | Macaca_nigra | CR | Decreasing | tropical | diurnal | 6400 | 7665.756 | 11.495 |
| mammals | Macaca_nigrescens | Macaca_nigrescens | VU | Decreasing | tropical | diurnal | 9500 | 9771.557 | 8.822 |
| mammals | Macaca_ochreata | Macaca_ochreata | VU | Decreasing | tropical | diurnal | 2600 | 35625.648 | 7.746 |
| mammals | Macaca_pagensis | Macaca_pagensis | CR | Decreasing | tropical | diurnal | 6280 | 2140.853 | 8.542 |
| mammals | Macaca_radiata | Macaca_radiata | VU | Decreasing | tropical | diurnal | 3690 | 620918.135 | 21.25 |
| mammals | Macaca_siberu | Macaca_siberu | EN | Decreasing | tropical | diurnal | 4526.7 | 3863.628 | 4.963 |
| mammals | Macaca_silenus | Macaca_silenus | EN | Decreasing | tropical | diurnal | 4750 | 27905.779 | 13.22 |
| mammals | Macaca_sinica | Macaca_sinica | EN | Decreasing | tropical | diurnal | 3590 | 46294.772 | 16.578 |
| mammals | Macaca_sylvanus | Macaca_sylvanus | EN | Decreasing | temperate | diurnal | 12200 | 25901.573 | 10.401 |
| mammals | Macaca_thibetana | Macaca_thibetana | NT | Decreasing | temperate | diurnal | 12000 | 1327321.499 | 13.052 |
| mammals | Macaca_tonkeana | Macaca_tonkeana | VU | Decreasing | tropical | diurnal | 9000 | 81339.184 | 7.565 |
| mammals | Macroderma_gigas | Macroderma_gigas | VU | Decreasing | tropical | nocturnal | 126.4 | 1156786.383 | 3.52 |
| mammals | Macrogalidia_musschenbroekii | Macrogalidia_musschenbroekii | VU | Decreasing | tropical | nocturnal | 4000 | 34097.052 | 8.874 |
| mammals | Macroglossus_minimus | Macroglossus_minimus | LC | Stable | tropical | nocturnal | 16.4 | 3590246.002 | 7.716 |
| mammals | Macroglossus_sobrinus | Macroglossus_sobrinus | LC | Stable | tropical | nocturnal | 22.1 | 2435640.09 | 12.703 |
| mammals | Macronycteris_commersonii | Hipposideros_commersoni | NT | Decreasing | tropical | nocturnal | 99.1 | 366161.233 | 8.351 |
| mammals | Macronycteris_gigas | Hipposideros_gigas | LC | Unknown | tropical | nocturnal | 166.5 | 3577138.019 | 7.834 |
| mammals | Macronycteris_thomensis | Hipposideros_thomensis | LC | Stable | tropical | nocturnal | 67 | 856.808 | 12.812 |
| mammals | Macronycteris_vittata | Hipposideros_vittatus | NT | Decreasing | tropical | nocturnal | 127.9 | 4006579.669 | 9.499 |
| mammals | Macrophyllum_macrophyllum | Macrophyllum_macrophyllum | LC | Unknown | tropical | nocturnal | 8.1 | 13066617.649 | 5.783 |
| mammals | Macropus_fuliginosus | Macropus_fuliginosus | LC | Increasing | temperate | nocturnal | 22000 | 1607008.423 | 6.653 |
| mammals | Macropus_giganteus | Macropus_giganteus | LC | Stable | temperate | cathemeral | 25875 | 2231237.803 | 6.959 |
| mammals | Macroscelides_micus | Macroscelides_micus | LC | Unknown | tropical | nocturnal | 27 | 947.119 | 5.007 |
| mammals | Macrotarsomys_bastardi | Macrotarsomys_bastardi | LC | Stable | tropical | nocturnal | 29.5 | 152421.055 | 9.158 |
| mammals | Macrotarsomys_ingens | Macrotarsomys_ingens | EN | Decreasing | tropical | nocturnal | 63.5 | 2998.651 | 10.851 |
| mammals | Macrotarsomys_petteri | NA | DD | Unknown | NA | NA | NA | NA | NA |
| mammals | Macrotis_lagotis | Macrotis_lagotis | VU | Decreasing | tropical | nocturnal | 1350 | 1079998.511 | 1.42 |
| mammals | Macrotus_californicus | Macrotus_californicus | LC | Stable | temperate | nocturnal | 11.6 | 630012.554 | 7.077 |
| mammals | Macrotus_waterhousii | Macrotus_waterhousii | LC | Stable | tropical | nocturnal | 12.8 | 817691.315 | 11.92 |

| | | | | | | | | | |
|---|---|---|---|---|---|---|---|---|---|
| mammals | Macruromys_major | Macruromys_major | LC | Stable | tropical | nocturnal | 350 | 164573.027 | 4.581 |
| mammals | Madoqua_guentheri | Madoqua_guentheri | LC | Stable | tropical | diurnal | 7500 | 1270987.216 | 8.013 |
| mammals | Madoqua_kirkii | Madoqua_kirkii | LC | Stable | tropical | cathemeral | 5300.1 | 1336497.003 | 9.15 |
| mammals | Madoqua_piacentinii | Madoqua_piacentinii | DD | Decreasing | tropical | cathemeral | 2500 | 158878.14 | 7.65 |
| mammals | Madoqua_saltiana | Madoqua_saltiana | LC | Stable | tropical | nocturnal | 3250 | 1098767.218 | 8.14 |
| mammals | Madromys_blanfordi | Madromys_blanfordi | LC | Unknown | tropical | nocturnal | 141 | 990642.407 | 19.156 |
| mammals | Makalata_didelphoides | Makalata_didelphoides | LC | Unknown | tropical | nocturnal | 400 | 3605587.128 | 2.861 |
| mammals | Makalata_macrura | Makalata_macrura | LC | Unknown | tropical | nocturnal | 439 | 2457872.708 | 1.34 |
| mammals | Malacomys_cansdalei | Malacomys_cansdalei | LC | Unknown | tropical | nocturnal | 71 | 240684.253 | 15.849 |
| mammals | Malacomys_edwardsi | Malacomys_edwardsi | LC | Unknown | tropical | nocturnal | 65 | 482419.363 | 16.149 |
| mammals | Malacomys_longipes | Malacomys_longipes | LC | Unknown | tropical | nocturnal | 95.6 | 1218174.721 | 7.114 |
| mammals | Malacothrix_typica | Malacothrix_typica | LC | Stable | temperate | nocturnal | 13.4 | 1387360.957 | 6.672 |
| mammals | Mallomys_aroaensis | Mallomys_aroaensis | LC | Unknown | tropical | nocturnal | 1664 | 136592.077 | 4.722 |
| mammals | Mallomys_gunung | Mallomys_gunung | EN | Decreasing | tropical | nocturnal | 2000 | 928 | 4.676 |
| mammals | Mallomys_istapantap | Mallomys_istapantap | LC | Unknown | tropical | nocturnal | 1950 | 75500.662 | 5.752 |
| mammals | Mallomys_rothschildi | Mallomys_rothschildi | LC | Stable | tropical | nocturnal | 1157.5 | 190242.218 | 4.644 |
| mammals | Mammelomys_lanosus | Mammelomys_lanosus | LC | Unknown | tropical | nocturnal | 117.2 | 86029.861 | 4.837 |
| mammals | Mammelomys_rattoides | Mammelomys_rattoides | LC | Unknown | tropical | nocturnal | 209.5 | 125631.707 | 2.711 |
| mammals | Mandrillus_leucophaeus | Mandrillus_leucophaeus | EN | Decreasing | tropical | diurnal | 15000 | 17586.985 | 5.102 |
| mammals | Mandrillus_sphinx | Mandrillus_sphinx | VU | Decreasing | tropical | diurnal | 18249.9 | 333380.84 | 6.481 |
| mammals | Manis_crassicaudata | Manis_crassicaudata | EN | Decreasing | tropical | nocturnal | 8457.3 | 2908388.908 | 18.601 |
| mammals | Manis_culionensis | Manis_culionensis | CR | Decreasing | tropical | nocturnal | 5250 | 13297.088 | 9.379 |
| mammals | Manis_javanica | Manis_javanica | CR | Decreasing | tropical | nocturnal | 5150 | 2319150.581 | 9.605 |
| mammals | Manis_pentadactyla | Manis_pentadactyla | CR | Decreasing | temperate | nocturnal | 4675 | 2757986.814 | 12.462 |
| mammals | Margaretamys_beccarii | Margaretamys_beccarii | DD | Decreasing | tropical | nocturnal | 70.4 | 109189.618 | 8.833 |
| mammals | Marmosa_alstoni | Marmosa_alstoni | LC | Stable | tropical | nocturnal | 132.3 | 213003.015 | 8.531 |
| mammals | Marmosa_andersoni | NA | DD | Unknown | NA | NA | NA | NA | NA |
| mammals | Marmosa_demerarae | Marmosa_demerarae | LC | Stable | tropical | nocturnal | 75.8 | 7826251.562 | 4.012 |
| mammals | Marmosa_lepida | Marmosa_lepida | LC | Stable | tropical | nocturnal | 14 | 4163942.393 | 1.718 |
| mammals | Marmosa_mexicana | Marmosa_mexicana | LC | Stable | tropical | nocturnal | 49.5 | 927634.962 | 10.568 |
| mammals | Marmosa_murina | Marmosa_murina | LC | Stable | tropical | nocturnal | 26 | 6450138.447 | 3.125 |
| mammals | Marmosa_paraguayana | Marmosa_paraguayanus | LC | Stable | tropical | nocturnal | 74.4 | 962990.454 | 12.881 |
| mammals | Marmosa_phaea | Marmosa_phaea | VU | Decreasing | tropical | nocturnal | 80.3 | 27306.97 | 8.356 |
| mammals | Marmosa_robinsoni | Marmosa_robinsoni | LC | Stable | tropical | nocturnal | 56.7 | 588222.595 | 11.584 |

| | | | | | | | | | |
|---|---|---|---|---|---|---|---|---|---|
| mammals | Marmosa_rubra | NA | DD | Unknown | NA | NA | NA | NA | NA |
| mammals | Marmosa_tyleriana | NA | DD | Unknown | NA | NA | NA | NA | NA |
| mammals | Marmosa_xerophila | Marmosa_xerophila | VU | Decreasing | tropical | nocturnal | 46.2 | 20304.005 | 12.006 |
| mammals | Marmosops_caucae | Marmosops_neblina | LC | Stable | tropical | nocturnal | 44 | 1675827.061 | 3.509 |
| mammals | Marmosops_creightoni | NA | DD | Unknown | NA | NA | NA | NA | NA |
| mammals | Marmosops_fuscatus | Marmosops_fuscatus | DD | Decreasing | tropical | nocturnal | 60 | 79790.459 | 13.797 |
| mammals | Marmosops_handleyi | Marmosops_handleyi | CR | Decreasing | tropical | nocturnal | 30.7 | 68.388 | 12.431 |
| mammals | Marmosops_incanus | Marmosops_incanus | LC | Unknown | tropical | nocturnal | 62.3 | 605182.477 | 10.717 |
| mammals | Marmosops_invictus | Marmosops_invictus | LC | Stable | tropical | nocturnal | 29.2 | 20699.918 | 8.414 |
| mammals | Marmosops_noctivagus | Marmosops_noctivagus | LC | Stable | tropical | nocturnal | 45 | 2087824.428 | 1.749 |
| mammals | Marmosops_ocellatus | Marmosops_ocellatus | LC | Stable | tropical | nocturnal | 37 | 189468.073 | 6.029 |
| mammals | Marmosops_parvidens | Marmosops_parvidens | LC | Unknown | tropical | nocturnal | 15 | 1459355.694 | 1.258 |
| mammals | Marmosops_paulensis | Marmosops_paulensis | LC | Unknown | tropical | nocturnal | 40 | 74073.183 | 13.83 |
| mammals | Marmosops_pinheiroi | Marmosops_pinheiroi | LC | Unknown | tropical | nocturnal | 26 | 586486.631 | 2.076 |
| mammals | Marmota_baibacina | Marmota_baibacina | LC | Unknown | temperate | diurnal | 7500 | 606964.307 | 6.962 |
| mammals | Marmota_bobak | Marmota_bobak | LC | Stable | temperate | diurnal | 5500 | 790385.686 | 9.221 |
| mammals | Marmota_broweri | Marmota_broweri | LC | Stable | temperate | diurnal | 3405 | 218588.161 | 0.736 |
| mammals | Marmota_caligata | Marmota_caligata | LC | Stable | temperate | diurnal | 7230 | 2006099.66 | 1.881 |
| mammals | Marmota_camtschatica | Marmota_camtschatica | LC | Unknown | temperate | diurnal | 3500 | 1023357.208 | 1.026 |
| mammals | Marmota_caudata | Marmota_caudata | LC | Unknown | temperate | diurnal | 4350 | 547910.243 | 7.764 |
| mammals | Marmota_flaviventris | Marmota_flaviventris | LC | Stable | temperate | diurnal | 3350 | 1718256.719 | 9.684 |
| mammals | Marmota_himalayana | Marmota_himalayana | LC | Unknown | temperate | diurnal | 5000 | 2387800.78 | 5.373 |
| mammals | Marmota_marmota | Marmota_marmota | LC | Stable | temperate | diurnal | 2010 | 213044.429 | 16.497 |
| mammals | Marmota_menzbieri | Marmota_menzbieri | VU | Decreasing | temperate | diurnal | 3250 | 2230.594 | 7.543 |
| mammals | Marmota_monax | Marmota_monax | LC | Stable | temperate | diurnal | 3801.7 | 7611091.829 | 8.656 |
| mammals | Marmota_olympus | Marmota_olympus | LC | Decreasing | temperate | diurnal | 6300 | 5368.557 | 3.095 |
| mammals | Marmota_sibirica | Marmota_sibirica | EN | Decreasing | temperate | diurnal | 8000 | 1534491.666 | 5.661 |
| mammals | Marmota_vancouverensis | Marmota_vancouverensis | CR | Decreasing | temperate | diurnal | 6300 | 2075.021 | 3.94 |
| mammals | Martes_americana | Martes_americana | LC | Decreasing | temperate | cathemeral | 1250 | 7637867.169 | 2.386 |
| mammals | Martes_flavigula | Martes_flavigula | LC | Decreasing | tropical | cathemeral | 1842.5 | 6697917.893 | 11.305 |
| mammals | Martes_foina | Martes_foina | LC | Stable | temperate | cathemeral | 1540.8 | 12454427.458 | 12.016 |
| mammals | Martes_gwatkinsii | Martes_gwatkinsii | VU | Stable | tropical | diurnal | 2043 | 30613.022 | 13.184 |
| mammals | Martes_martes | Martes_martes | LC | Stable | temperate | cathemeral | 1300 | 9929653.827 | 13.772 |
| mammals | Martes_melampus | Martes_melampus | LC | Stable | temperate | cathemeral | 1000 | 279683.216 | 19.759 |

| | | | | | | | | | |
|---|---|---|---|---|---|---|---|---|---|
| mammals | Martes_zibellina | Martes_zibellina | LC | Increasing | temperate | cathemeral | 1130 | 10081922.237 | 2.339 |
| mammals | Massoutiera_mzabi | Massoutiera_mzabi | LC | Unknown | tropical | diurnal | 194 | 652446.502 | 1.935 |
| mammals | Mastacomys_fuscus | Mastacomys_fuscus | NT | Decreasing | temperate | nocturnal | 122 | 65318.33 | 5.652 |
| mammals | Mastomys_angolensis | NA | LC | Stable | NA | NA | NA | NA | NA |
| mammals | Mastomys_awashensis | Mastomys_awashensis | LC | Unknown | tropical | nocturnal | 46 | 29412.027 | 12.903 |
| mammals | Mastomys_coucha | Mastomys_coucha | LC | Stable | temperate | nocturnal | 31 | 1271248.171 | 9.152 |
| mammals | Mastomys_erythroleucus | Mastomys_erythroleucus | LC | Stable | tropical | nocturnal | 46.5 | 7098448.734 | 10.852 |
| mammals | Mastomys_huberti | Mastomys_huberti | LC | Increasing | tropical | nocturnal | 55.3 | 578164.785 | 11.48 |
| mammals | Mastomys_kollmannspergeri | Mastomys_kollmannspergeri | LC | Stable | tropical | nocturnal | 94.7 | 1619743.207 | 8.852 |
| mammals | Mastomys_natalensis | Mastomys_natalensis | LC | Stable | tropical | nocturnal | 36.4 | 16655440.98 | 9.218 |
| mammals | Mastomys_shortridgei | Mastomys_shortridgei | LC | Stable | tropical | nocturnal | 46 | 96016.156 | 4.327 |
| mammals | Maxomys_alticola | Maxomys_alticola | LC | Stable | tropical | nocturnal | 90 | 3957.806 | 6.761 |
| mammals | Maxomys_baeodon | NA | DD | Unknown | NA | NA | NA | NA | NA |
| mammals | Maxomys_bartelsii | Maxomys_bartelsii | LC | Stable | tropical | nocturnal | 102 | 7430.612 | 15.97 |
| mammals | Maxomys_dollmani | NA | DD | Unknown | NA | NA | NA | NA | NA |
| mammals | Maxomys_hellwaldii | Maxomys_hellwaldii | LC | Decreasing | tropical | nocturnal | 203 | 144666.76 | 10.009 |
| mammals | Maxomys_hylomyoides | NA | DD | Unknown | NA | NA | NA | NA | NA |
| mammals | Maxomys_inas | Maxomys_inas | LC | Stable | tropical | nocturnal | 90 | 12645.857 | 6.19 |
| mammals | Maxomys_inflatus | Maxomys_inflatus | VU | Decreasing | tropical | nocturnal | 152 | 77602.801 | 8.46 |
| mammals | Maxomys_moi | Maxomys_moi | LC | Decreasing | tropical | nocturnal | 147 | 38181.71 | 7.737 |
| mammals | Maxomys_musschenbroekii | Maxomys_musschenbroekii | LC | Stable | tropical | nocturnal | 84.8 | 169337.754 | 9.276 |
| mammals | Maxomys_ochraceiventer | NA | DD | Unknown | NA | NA | NA | NA | NA |
| mammals | Maxomys_pagensis | Maxomys_pagensis | VU | Decreasing | tropical | nocturnal | 196 | 6004.261 | 6.223 |
| mammals | Maxomys_panglima | Maxomys_panglima | LC | Stable | tropical | nocturnal | 189 | 13144.027 | 9.302 |
| mammals | Maxomys_rajah | Maxomys_rajah | VU | Decreasing | tropical | nocturnal | 150.3 | 1317194.645 | 7.827 |
| mammals | Maxomys_surifer | Maxomys_surifer | LC | Decreasing | tropical | nocturnal | 150.4 | 2819980.274 | 11.098 |
| mammals | Maxomys_tajuddinii | Maxomys_tajuddinii | LC | Unknown | tropical | nocturnal | 60 | 463605.443 | 7.984 |
| mammals | Maxomys_wattsi | Maxomys_wattsi | EN | Decreasing | tropical | nocturnal | 147 | 782.224 | 3.203 |
| mammals | Maxomys_whiteheadi | Maxomys_whiteheadi | VU | Decreasing | tropical | nocturnal | 56 | 1321686.457 | 7.855 |
| mammals | Mazama_americana | NA | DD | Unknown | NA | NA | NA | NA | NA |
| mammals | Mazama_jucunda | Mazama_bororo | VU | Decreasing | temperate | nocturnal | 25000 | 237946.403 | 13.697 |
| mammals | Mazama_nanus | Mazama_nana | VU | Unknown | temperate | cathemeral | 16499.9 | 447853.259 | 12.372 |
| mammals | Mazama_rufina | Mazama_rufina | VU | Decreasing | tropical | cathemeral | 25999.8 | 85280.751 | 6.217 |
| mammals | Mazama_temama | Mazama_temama | DD | Decreasing | tropical | cathemeral | 25000 | 854551.716 | 10.246 |

| | | | | | | | | | |
|---|---|---|---|---|---|---|---|---|---|
| mammals | Megadendromus_nikolausi | Megadendromus_nikolausi | VU | Unknown | tropical | nocturnal | 52 | 6974.965 | 12.352 |
| mammals | Megaderma_spasma | Megaderma_spasma | LC | Unknown | tropical | nocturnal | 25.4 | 4056887.364 | 11.522 |
| mammals | Megadontomys_cryophilus | Megadontomys_cryophilus | EN | Decreasing | tropical | nocturnal | 32 | 390.375 | 4.475 |
| mammals | Megadontomys_nelsoni | Megadontomys_nelsoni | EN | Decreasing | tropical | nocturnal | 80.1 | 19798.124 | 14.689 |
| mammals | Megadontomys_thomasi | Megadontomys_thomasi | EN | Stable | tropical | nocturnal | 111 | 5103.337 | 5.715 |
| mammals | Megaerops_ecaudatus | Megaerops_ecaudatus | LC | Stable | tropical | nocturnal | 24.7 | 1274119.398 | 7.845 |
| mammals | Megaerops_kusnotoi | Megaerops_kusnotoi | VU | Decreasing | tropical | nocturnal | 20.1 | 16180.297 | 20.104 |
| mammals | Megaerops_niphanae | Megaerops_niphanae | LC | Stable | tropical | nocturnal | 32.6 | 1321362.379 | 12.932 |
| mammals | Megaloglossus_azagnyi | Megaloglossus_azagnyi | LC | Unknown | tropical | nocturnal | 11.5 | 310126.151 | 15.619 |
| mammals | Megaloglossus_woermanni | Megaloglossus_woermanni | LC | Stable | tropical | nocturnal | 15.7 | 3120610.924 | 6.831 |
| mammals | Megasorex_gigas | Megasorex_gigas | LC | Unknown | tropical | cathemeral | 20 | 150206.002 | 8.866 |
| mammals | Melanomys_caliginosus | Melanomys_caliginosus | LC | Stable | tropical | diurnal | 41 | 748135.409 | 10.563 |
| mammals | Melanomys_robustulus | Melanomys_robustulus | LC | Stable | tropical | nocturnal | 53.5 | 22782.28 | 5.392 |
| mammals | Melanomys_zunigae | Melanomys_zunigae | CR | Decreasing | tropical | nocturnal | 53.5 | 228.847 | 12.093 |
| mammals | Melasmothrix_naso | NA | DD | Unknown | NA | NA | NA | NA | NA |
| mammals | Meles_anakuma | Meles_anakuma | LC | Decreasing | temperate | nocturnal | 11500 | 203248.428 | 19.002 |
| mammals | Meles_leucurus | Meles_leucurus | LC | Unknown | temperate | nocturnal | 6250 | 15959053.38 | 8.203 |
| mammals | Meles_meles | Meles_meles | LC | Stable | temperate | nocturnal | 13000 | 9801554.241 | 15.608 |
| mammals | Mellivora_capensis | Mellivora_capensis | LC | Decreasing | tropical | nocturnal | 8500 | 29722162.158 | 8.713 |
| mammals | Melogale_everetti | Melogale_everetti | EN | Decreasing | tropical | nocturnal | 1500 | 19.936 | 41.517 |
| mammals | Melogale_moschata | Melogale_moschata | LC | Stable | tropical | nocturnal | 805.3 | 2960755.418 | 13.775 |
| mammals | Melogale_orientalis | Melogale_orientalis | LC | Unknown | tropical | nocturnal | 2000 | 131392.637 | 22.341 |
| mammals | Melogale_personata | Melogale_personata | LC | Unknown | tropical | nocturnal | 1702.5 | 1971107.055 | 11.596 |
| mammals | Melomys_aerosus | Melomys_aerosus | EN | Decreasing | tropical | nocturnal | 97.3 | 1314.218 | 6.602 |
| mammals | Melomys_arcium | NA | DD | Unknown | NA | NA | NA | NA | NA |
| mammals | Melomys_bannisteri | Melomys_bannisteri | EN | Decreasing | tropical | nocturnal | 52.8 | 566.769 | 10.245 |
| mammals | Melomys_bougainville | NA | DD | Unknown | NA | NA | NA | NA | NA |
| mammals | Melomys_burtoni | Melomys_burtoni | LC | Stable | tropical | nocturnal | 68.5 | 492110.478 | 3.836 |
| mammals | Melomys_capensis | Melomys_capensis | LC | Stable | tropical | nocturnal | 70 | 31410.625 | 2.337 |
| mammals | Melomys_caurinus | Melomys_caurinus | EN | Decreasing | tropical | nocturnal | 148.1 | 929.157 | 9.342 |
| mammals | Melomys_cervinipes | Melomys_cervinipes | LC | Stable | temperate | nocturnal | 78 | 217518.123 | 7.101 |
| mammals | Melomys_cooperae | NA | DD | Unknown | NA | NA | NA | NA | NA |
| mammals | Melomys_dollmani | Melomys_dollmani | LC | Stable | tropical | nocturnal | 77.7 | 47118.681 | 6.601 |
| mammals | Melomys_fraterculus | Melomys_fraterculus | EN | Decreasing | tropical | nocturnal | 47.8 | 155.743 | 6.306 |

| | | | | | | | | | |
|---|---|---|---|---|---|---|---|---|---|
| mammals | Melomys_frigicola | Melomys_frigicola | LC | Unknown | tropical | nocturnal | 71 | 2691.304 | 7.367 |
| mammals | Melomys_fulgens | NA | DD | Unknown | NA | NA | NA | NA | NA |
| mammals | Melomys_howi | NA | DD | Unknown | NA | NA | NA | NA | NA |
| mammals | Melomys_leucogaster | Melomys_leucogaster | LC | Stable | tropical | nocturnal | 108.5 | 243984.082 | 2.449 |
| mammals | Melomys_lutillus | Melomys_lutillus | LC | Unknown | tropical | nocturnal | 37.4 | 340703.58 | 2.847 |
| mammals | Melomys_matambuai | Melomys_matambuai | EN | Decreasing | tropical | nocturnal | 145 | 1830.07 | 6.315 |
| mammals | Melomys_obiensis | Melomys_obiensis | LC | Unknown | tropical | nocturnal | 71 | 2708.401 | 4.762 |
| mammals | Melomys_paveli | NA | DD | Unknown | NA | NA | NA | NA | NA |
| mammals | Melomys_rufescens | Melomys_rufescens | LC | Stable | tropical | nocturnal | 60.1 | 769118.529 | 3.371 |
| mammals | Melomys_talaudium | Melomys_talaudium | EN | Decreasing | tropical | nocturnal | 143.1 | 929.157 | 9.342 |
| mammals | Melonycteris_melanops | Melonycteris_melanops | LC | Stable | tropical | nocturnal | 49 | 45239.909 | 4.413 |
| mammals | Melursus_ursinus | Melursus_ursinus | VU | Decreasing | tropical | cathemeral | 93130 | 1080994.278 | 15.675 |
| mammals | Menetes_berdmorei | Menetes_berdmorei | LC | Stable | tropical | diurnal | 190 | 1232123.007 | 13.502 |
| mammals | Mephitis_macroura | Mephitis_macroura | LC | Increasing | temperate | nocturnal | 801.3 | 1783431.713 | 9.261 |
| mammals | Mephitis_mephitis | Mephitis_mephitis | LC | Stable | temperate | nocturnal | 2085 | 12271610.665 | 10.772 |
| mammals | Meriones_arimalius | Meriones_arimalius | LC | Unknown | tropical | diurnal | 87.6 | 257097.806 | 2.138 |
| mammals | Meriones_crassus | Meriones_crassus | LC | Unknown | temperate | nocturnal | 80 | 7571087.959 | 4.228 |
| mammals | Meriones_grandis | Meriones_grandis | LC | Stable | temperate | nocturnal | 205 | 708604.474 | 13.553 |
| mammals | Meriones_hurrianae | Meriones_hurrianae | LC | Unknown | temperate | diurnal | 70 | 480817.194 | 13.902 |
| mammals | Meriones_libycus | Meriones_libycus | LC | Stable | temperate | nocturnal | 91.3 | 7831992.136 | 7.13 |
| mammals | Meriones_meridianus | Meriones_meridianus | LC | Unknown | temperate | nocturnal | 41 | 6163493.31 | 6.523 |
| mammals | Meriones_persicus | Meriones_persicus | LC | Unknown | temperate | nocturnal | 100 | 2222403.477 | 8.829 |
| mammals | Meriones_psammophilus | Meriones_chengi | LC | Unknown | temperate | nocturnal | 91.3 | 43171.296 | 3.795 |
| mammals | Meriones_rex | Meriones_rex | LC | Unknown | tropical | nocturnal | 125.3 | 205685.994 | 9.857 |
| mammals | Meriones_sacramenti | Meriones_sacramenti | VU | Decreasing | temperate | nocturnal | 200 | 8212.933 | 23.579 |
| mammals | Meriones_shawii | Meriones_shawi | LC | Stable | temperate | nocturnal | 90.6 | 620432.298 | 11.999 |
| mammals | Meriones_tamariscinus | Meriones_tamariscinus | LC | Unknown | temperate | nocturnal | 120 | 2568252.741 | 6.437 |
| mammals | Meriones_tristrami | Meriones_tristrami | LC | Stable | temperate | nocturnal | 105 | 901513.449 | 15.936 |
| mammals | Meriones_unguiculatus | Meriones_unguiculatus | LC | Unknown | temperate | nocturnal | 53.3 | 3276858.639 | 6.792 |
| mammals | Meriones_vinogradovi | Meriones_vinogradovi | LC | Unknown | temperate | cathemeral | 150 | 304025.531 | 16.395 |
| mammals | Meriones_zarudnyi | NA | DD | Unknown | NA | NA | NA | NA | NA |
| mammals | Mesechinus_dauuricus | Mesechinus_dauuricus | LC | Stable | temperate | nocturnal | 342 | 5038641.964 | 7.382 |
| mammals | Mesechinus_hughi | Mesechinus_hughi | LC | Decreasing | temperate | nocturnal | 199.3 | 243269.234 | 14.69 |
| mammals | Mesembriomys_gouldii | Mesembriomys_gouldii | VU | Decreasing | tropical | nocturnal | 716.5 | 237211.175 | 3.196 |

| | | | | | | | | | |
|---|---|---|---|---|---|---|---|---|---|
| mammals | Mesembriomys_macrurus | Mesembriomys_macrurus | NT | Decreasing | tropical | nocturnal | 267 | 34779.613 | 1.051 |
| mammals | Mesocapromys_auritus | Mesocapromys_auritus | EN | Unknown | tropical | nocturnal | 820 | 90.212 | 2.466 |
| mammals | Mesocapromys_melanurus | Mysateles_melanurus | VU | Decreasing | tropical | nocturnal | 1218 | 31192.616 | 16.525 |
| mammals | Mesocricetus_auratus | Mesocricetus_auratus | EN | Decreasing | temperate | nocturnal | 105 | 4704.206 | 23.225 |
| mammals | Mesocricetus_brandti | Mesocricetus_brandti | NT | Decreasing | temperate | nocturnal | 198 | 732898.237 | 14.504 |
| mammals | Mesocricetus_newtoni | Mesocricetus_newtoni | NT | Decreasing | temperate | nocturnal | 97.5 | 38236.103 | 18.434 |
| mammals | Mesocricetus_raddei | Mesocricetus_raddei | LC | Stable | temperate | nocturnal | 260 | 141380.031 | 15.045 |
| mammals | Mesomys_hispidus | Mesomys_hispidus | LC | Unknown | tropical | nocturnal | 175 | 4948976.376 | 1.626 |
| mammals | Mesomys_leniceps | NA | DD | Unknown | NA | NA | NA | NA | NA |
| mammals | Mesomys_occultus | Mesomys_occultus | LC | Unknown | tropical | nocturnal | 146.7 | 53277.641 | 0.482 |
| mammals | Mesomys_stimulax | Mesomys_stimulax | LC | Unknown | tropical | nocturnal | 139 | 437362.908 | 2.745 |
| mammals | Mesophylla_macconnelli | Mesophylla_macconnelli | LC | Unknown | tropical | nocturnal | 6.9 | 8817651.771 | 3.702 |
| mammals | Metachirus_nudicaudatus | Metachirus_nudicaudatus | LC | Stable | tropical | nocturnal | 375 | 10341442.399 | 4.977 |
| mammals | Micaelamys_granti | Aethomys_granti | LC | Unknown | temperate | nocturnal | 40 | 197352.611 | 7.697 |
| mammals | Micaelamys_namaquensis | Aethomys_namaquensis | LC | Stable | temperate | nocturnal | 50.6 | 3338532.919 | 8.281 |
| mammals | Mico_acariensis | Mico_acariensis | LC | Unknown | tropical | diurnal | 472 | 22441.861 | 0.843 |
| mammals | Mico_argentatus | Mico_argentatus | LC | Decreasing | tropical | diurnal | 440 | 137206.256 | 2.782 |
| mammals | Mico_chrysoleucos | Mico_chrysoleucos | LC | Decreasing | tropical | diurnal | 539.3 | 36452.862 | 1.658 |
| mammals | Mico_emiliae | Mico_emiliae | LC | Unknown | tropical | diurnal | 335 | 151986.899 | 3.372 |
| mammals | Mico_humeralifer | Mico_humeralifer | NT | Decreasing | tropical | diurnal | 350 | 63580.261 | 1.245 |
| mammals | Mico_humilis | Mico_humilis | LC | Unknown | tropical | diurnal | 167.5 | 25728.974 | 0.975 |
| mammals | Mico_intermedius | Mico_intermedius | LC | Decreasing | tropical | diurnal | 425 | 62624.114 | 1.04 |
| mammals | Mico_leucippe | Mico_leucippe | LC | Decreasing | tropical | diurnal | 440 | 70830.614 | 2.227 |
| mammals | Mico_marcai | Mico_marcai | VU | Decreasing | tropical | diurnal | 295 | 7000.76 | 2.395 |
| mammals | Mico_mauesi | Mico_mauesi | LC | Unknown | tropical | diurnal | 360 | 29585.793 | 0.562 |
| mammals | Mico_melanurus | Mico_melanurus | NT | Decreasing | tropical | diurnal | 302.3 | 769939.423 | 5.016 |
| mammals | Mico_nigriceps | Mico_nigriceps | NT | Decreasing | tropical | diurnal | 370 | 31646.701 | 0.936 |
| mammals | Mico_rondoni | Mico_rondoni | VU | Decreasing | tropical | diurnal | 329.6 | 70576.071 | 4.494 |
| mammals | Mico_saterei | Mico_saterei | LC | Unknown | tropical | diurnal | 431.6 | 19280.586 | 0.409 |
| mammals | Microcavia_australis | Microcavia_australis | LC | Unknown | temperate | diurnal | 286.1 | 1741369.307 | 6.429 |
| mammals | Microcavia_niata | Microcavia_niata | LC | Stable | tropical | diurnal | 255.2 | 155322.589 | 6.801 |
| mammals | Microcavia_shiptoni | Microcavia_shiptoni | LC | Unknown | temperate | diurnal | 185 | 39515.225 | 2.949 |
| mammals | Microcebus_arnholdi | Microcebus_arnholdi | VU | Decreasing | tropical | nocturnal | 45 | 771.772 | 6.037 |
| mammals | Microcebus_berthae | Microcebus_berthae | CR | Decreasing | tropical | nocturnal | 30.5 | 546.445 | 5.291 |

| | | | | | | | | | |
|---|---|---|---|---|---|---|---|---|---|
| mammals | Microcebus_bongolavensis | Microcebus_bongolavensis | EN | Decreasing | tropical | nocturnal | 54 | 788.893 | 8.695 |
| mammals | Microcebus_danfossi | Microcebus_danfossi | VU | Decreasing | tropical | nocturnal | 63 | 938.204 | 7.01 |
| mammals | Microcebus_gerpi | Microcebus_gerpi | CR | Decreasing | tropical | nocturnal | 68 | 20.684 | 6.39 |
| mammals | Microcebus_griseorufus | Microcebus_griseorufus | LC | Decreasing | temperate | nocturnal | 62.5 | 35820.437 | 11.973 |
| mammals | Microcebus_jollyae | Microcebus_jollyae | EN | Decreasing | tropical | nocturnal | 61 | 156.831 | 10.423 |
| mammals | Microcebus_lehilahytsara | Microcebus_lehilahytsara | NT | Decreasing | tropical | nocturnal | 46.5 | 58151.4 | 10.088 |
| mammals | Microcebus_macarthurii | Microcebus_macarthurii | EN | Decreasing | tropical | nocturnal | 53 | 677.162 | 4.251 |
| mammals | Microcebus_mamiratra | Microcebus_mamiratra | EN | Decreasing | tropical | nocturnal | 61 | 135.398 | 12.963 |
| mammals | Microcebus_margotmarshae | Microcebus_margotmarshae | EN | Decreasing | tropical | nocturnal | 49 | 903.114 | 5.982 |
| mammals | Microcebus_marohita | Microcebus_marohita | CR | Decreasing | tropical | nocturnal | 85 | 28.416 | 11.022 |
| mammals | Microcebus_murinus | Microcebus_murinus | LC | Decreasing | temperate | nocturnal | 60 | 90390.117 | 8.149 |
| mammals | Microcebus_myoxinus | Microcebus_myoxinus | VU | Decreasing | tropical | nocturnal | 47.5 | 31213.797 | 7.679 |
| mammals | Microcebus_ravelobensis | Microcebus_ravelobensis | VU | Decreasing | tropical | nocturnal | 72 | 3336.832 | 10.108 |
| mammals | Microcebus_rufus | Microcebus_rufus | VU | Decreasing | tropical | nocturnal | 42.5 | 2607.395 | 6.38 |
| mammals | Microcebus_sambiranensis | Microcebus_sambiranensis | EN | Decreasing | tropical | nocturnal | 44 | 705.407 | 4.214 |
| mammals | Microcebus_simmonsi | Microcebus_simmonsi | EN | Decreasing | tropical | nocturnal | 65 | 737.654 | 6.093 |
| mammals | Microcebus_tavaratra | Microcebus_tavaratra | VU | Decreasing | tropical | nocturnal | 61 | 4231.199 | 8.021 |
| mammals | Microdillus_peeli | NA | DD | Unknown | NA | NA | NA | NA | NA |
| mammals | Microdipodops_megacephalus | Microdipodops_megacephalus | LC | Decreasing | temperate | nocturnal | 10.5 | 226040.896 | 7.794 |
| mammals | Microdipodops_pallidus | Microdipodops_pallidus | LC | Decreasing | temperate | nocturnal | 12.5 | 31230.105 | 6.665 |
| mammals | Microgale_cowani | Microgale_cowani | LC | Decreasing | tropical | nocturnal | 13.8 | 93169.763 | 7.334 |
| mammals | Microgale_drouhardi | Microgale_drouhardi | LC | Decreasing | tropical | nocturnal | 10.5 | 103819.858 | 7.678 |
| mammals | Microgale_dryas | Microgale_dryas | VU | Decreasing | tropical | nocturnal | 40 | 9465.399 | 5.43 |
| mammals | Microgale_fotsifotsy | Microgale_fotsifotsy | LC | Decreasing | tropical | nocturnal | 7.7 | 111846.773 | 7.527 |
| mammals | Microgale_gracilis | Microgale_gracilis | LC | Decreasing | tropical | nocturnal | 23.3 | 77835.525 | 7.923 |
| mammals | Microgale_gymnorhyncha | Microgale_gymnorhyncha | LC | Decreasing | tropical | nocturnal | 18.2 | 77232.217 | 7.97 |
| mammals | Microgale_jenkinsae | Microgale_jenkinsae | EN | Decreasing | tropical | nocturnal | 5.1 | 1383.993 | 8.356 |
| mammals | Microgale_jobihely | Microgale_jobihely | EN | Unknown | tropical | nocturnal | 9 | 1096.542 | 7.655 |
| mammals | Microgale_longicaudata | Microgale_longicaudata | LC | Decreasing | tropical | nocturnal | 7.3 | 1048.64 | 8.624 |
| mammals | Microgale_majori | Microgale_majori | LC | Decreasing | tropical | nocturnal | 5.8 | 107492.216 | 7.684 |
| mammals | Microgale_mergulus | Limnogale_mergulus | VU | Decreasing | tropical | nocturnal | 78.6 | 56589.478 | 11.194 |
| mammals | Microgale_monticola | Microgale_monticola | VU | Stable | tropical | nocturnal | 13.9 | 759.564 | 5.609 |
| mammals | Microgale_nasoloi | Microgale_nasoloi | VU | Decreasing | tropical | nocturnal | 10 | 1193.561 | 8.277 |
| mammals | Microgale_parvula | Microgale_parvula | LC | Decreasing | tropical | nocturnal | 3.4 | 92421.46 | 7.361 |

| | | | | | | | | | |
|---|---|---|---|---|---|---|---|---|---|
| mammals | Microgale_principula | Microgale_principula | LC | Decreasing | tropical | nocturnal | 10.2 | 89104.731 | 8.032 |
| mammals | Microgale_pusilla | Microgale_pusilla | LC | Stable | tropical | nocturnal | 3.4 | 60647.463 | 8.946 |
| mammals | Microgale_soricoides | Microgale_soricoides | LC | Decreasing | tropical | nocturnal | 18.7 | 102314.277 | 7.614 |
| mammals | Microgale_taiva | Microgale_taiva | LC | Decreasing | tropical | nocturnal | 12.4 | 43037.755 | 6.489 |
| mammals | Microgale_thomasi | Microgale_thomasi | LC | Decreasing | tropical | nocturnal | 22.9 | 22816.325 | 6.811 |
| mammals | Microhydromys_argenteus | NA | DD | Unknown | NA | NA | NA | NA | NA |
| mammals | Microhydromys_richardsoni | NA | DD | Unknown | NA | NA | NA | NA | NA |
| mammals | Micromys_minutus | Micromys_minutus | LC | Stable | temperate | nocturnal | 6 | 18265246.854 | 12.407 |
| mammals | Micronomus_norfolkensis | Mormopterus_norfolkensis | NT | Decreasing | temperate | nocturnal | 8 | 74501.367 | 11.133 |
| mammals | Micronycteris_brosseti | NA | DD | Unknown | NA | NA | NA | NA | NA |
| mammals | Micronycteris_buriri | NA | DD | Unknown | NA | NA | NA | NA | NA |
| mammals | Micronycteris_giovanniae | NA | DD | Unknown | NA | NA | NA | NA | NA |
| mammals | Micronycteris_hirsuta | Micronycteris_hirsuta | LC | Unknown | tropical | nocturnal | 12 | 6515654.571 | 4.019 |
| mammals | Micronycteris_matses | NA | DD | Unknown | NA | NA | NA | NA | NA |
| mammals | Micronycteris_megalotis | Micronycteris_megalotis | LC | Unknown | tropical | nocturnal | 6.1 | 12095131.872 | 5.166 |
| mammals | Micronycteris_microtis | Micronycteris_microtis | LC | Stable | tropical | nocturnal | 5.6 | 4395480.999 | 3.592 |
| mammals | Micronycteris_minuta | Micronycteris_minuta | LC | Unknown | tropical | nocturnal | 6.5 | 12093270.586 | 5.42 |
| mammals | Micronycteris_sanborni | Micronycteris_sanborni | LC | Unknown | tropical | nocturnal | 6.3 | 137406.736 | 7.192 |
| mammals | Micronycteris_schmidtorum | Micronycteris_schmidtorum | LC | Stable | tropical | nocturnal | 7.5 | 6504803.42 | 4.716 |
| mammals | Microperoryctes_aplini | Microperoryctes_aplini | VU | Unknown | tropical | nocturnal | 79.3 | 76.019 | 6.285 |
| mammals | Microperoryctes_longicauda | Microperoryctes_longicauda | LC | Unknown | tropical | nocturnal | 538 | 232311.305 | 4.323 |
| mammals | Microperoryctes_murinus | Microperoryctes_murina | VU | Unknown | tropical | nocturnal | 99.8 | 607.82 | 3.74 |
| mammals | Microperoryctes_papuensis | Microperoryctes_papuensis | LC | Unknown | tropical | nocturnal | 158 | 19193.605 | 2.802 |
| mammals | Micropotamogale_lamottei | Micropotamogale_lamottei | VU | Decreasing | tropical | nocturnal | 69.6 | 2100.006 | 16.304 |
| mammals | Micropotamogale_ruwenzorii | Micropotamogale_ruwenzorii | LC | Unknown | tropical | nocturnal | 98.5 | 52418.555 | 14.905 |
| mammals | Microryzomys_minutus | Microryzomys_minutus | LC | Unknown | tropical | nocturnal | 13.5 | 638218.249 | 8.152 |
| mammals | Microsciurus_alfari | Microsciurus_alfari | LC | Stable | tropical | diurnal | 87.5 | 117452.515 | 10.146 |
| mammals | Microsciurus_flaviventer | Microsciurus_flaviventer | LC | Unknown | tropical | diurnal | 92 | 2575239.408 | 1.641 |
| mammals | Microsciurus_mimulus | Microsciurus_mimulus | LC | Stable | tropical | diurnal | 120 | 160232.779 | 7.42 |
| mammals | Microsciurus_santanderensis | NA | DD | Unknown | NA | NA | NA | NA | NA |
| mammals | Microtus_abbreviatus | Microtus_abbreviatus | LC | Stable | temperate | cathemeral | 62 | 1432006.914 | 0.682 |
| mammals | Microtus_afghanus | Blanfordimys_afghanus | LC | Unknown | temperate | diurnal | 30.8 | 409814.267 | 9.939 |
| mammals | Microtus_agrestis | Microtus_agrestis | LC | Stable | temperate | cathemeral | 42.5 | 10818739.679 | 9.775 |
| mammals | Microtus_anatolicus | NA | DD | Unknown | NA | NA | NA | NA | NA |

| | | | | | | | | | |
|---|---|---|---|---|---|---|---|---|---|
| mammals | Microtus_arvalis | Microtus_arvalis | LC | Stable | temperate | cathemeral | 28 | 6887242.436 | 15.393 |
| mammals | Microtus_brachycercus | Microtus_brachycercus | LC | Stable | temperate | nocturnal | 19 | 70234.695 | 23.328 |
| mammals | Microtus_bucharensis | Blanfordimys_bucharensis | LC | Unknown | temperate | cathemeral | 45 | 116187.808 | 12.438 |
| mammals | Microtus_cabrerae | Microtus_cabrerae | NT | Decreasing | temperate | diurnal | 52.5 | 119784.07 | 16.148 |
| mammals | Microtus_canicaudus | Microtus_canicaudus | LC | Stable | temperate | cathemeral | 28.5 | 8888.221 | 28.583 |
| mammals | Microtus_chrotorrhinus | Microtus_chrotorrhinus | LC | Decreasing | temperate | diurnal | 39 | 1384481.213 | 6.764 |
| mammals | Microtus_felteni | Microtus_felteni | LC | Unknown | temperate | nocturnal | 21 | 35699.48 | 13.064 |
| mammals | Microtus_guatemalensis | Microtus_guatemalensis | NT | Decreasing | tropical | diurnal | 42 | 26501.098 | 14.674 |
| mammals | Microtus_irani | Microtus_irani | VU | Unknown | temperate | cathemeral | 37.1 | 1637.835 | 29.05 |
| mammals | Microtus_longicaudus | Microtus_longicaudus | LC | Stable | temperate | cathemeral | 46.7 | 4021651.477 | 6.431 |
| mammals | Microtus_majori | Microtus_majori | LC | Stable | temperate | nocturnal | 24.5 | 142325.882 | 12.884 |
| mammals | Microtus_mexicanus | Microtus_mexicanus | LC | Unknown | temperate | diurnal | 35 | 854502.221 | 9.261 |
| mammals | Microtus_montanus | Microtus_montanus | LC | Stable | temperate | cathemeral | 36.3 | 1453057.655 | 9.425 |
| mammals | Microtus_oaxacensis | Microtus_oaxacensis | EN | Decreasing | tropical | cathemeral | 39.2 | 510.441 | 3.642 |
| mammals | Microtus_ochrogaster | Microtus_ochrogaster | LC | Stable | temperate | cathemeral | 38 | 3243052.721 | 15.341 |
| mammals | Microtus_oregoni | Microtus_oregoni | LC | Stable | temperate | nocturnal | 20.4 | 206842.167 | 10.958 |
| mammals | Microtus_pennsylvanicus | Microtus_pennsylvanicus | LC | Stable | temperate | cathemeral | 36.8 | 11962526.598 | 6.624 |
| mammals | Microtus_quasiater | Microtus_quasiater | NT | Decreasing | tropical | cathemeral | 40 | 19611.134 | 15.135 |
| mammals | Microtus_richardsoni | Microtus_richardsoni | LC | Stable | temperate | nocturnal | 85 | 859823.369 | 8.074 |
| mammals | Microtus_rossiaemeridionalis | NA | not_available | not_available | NA | NA | NA | NA | NA |
| mammals | Microtus_schelkovnikovi | Microtus_schelkovnikovi | LC | Decreasing | temperate | cathemeral | 26 | 21598.458 | 12.414 |
| mammals | Microtus_townsendii | Microtus_townsendii | LC | Stable | temperate | cathemeral | 64.8 | 213940.737 | 10.531 |
| mammals | Microtus_transcaspicus | Microtus_transcaspicus | LC | Unknown | temperate | cathemeral | 38.8 | 11320.19 | 10.374 |
| mammals | Microtus_xanthognathus | Microtus_xanthognathus | LC | Stable | temperate | cathemeral | 125.8 | 2263668.69 | 1.171 |
| mammals | Microtus_yuldaschi | Neodon_juldaschi | LC | Unknown | temperate | cathemeral | 44.6 | 311957.47 | 7.81 |
| mammals | Mictomys_borealis | Synaptomys_borealis | LC | Unknown | temperate | cathemeral | 21.3 | 6774122.508 | 2.178 |
| mammals | Mimetillus_moloneyi | Mimetillus_moloneyi | LC | Unknown | tropical | nocturnal | 8.8 | 3621345.779 | 10.069 |
| mammals | Mimon_bennettii | Mimon_bennettii | LC | Stable | tropical | nocturnal | 25.2 | 5698708.121 | 7.767 |
| mammals | Mimon_cozumelae | Mimon_cozumelae | LC | Stable | tropical | nocturnal | 12.4 | 803519.036 | 10.078 |
| mammals | Mindomys_hammondi | Mindomys_hammondi | EN | Decreasing | tropical | nocturnal | 60.5 | 16230.497 | 13.326 |
| mammals | Miniopterus_africanus | NA | DD | Unknown | NA | NA | NA | NA | NA |
| mammals | Miniopterus_australis | Miniopterus_australis | LC | Stable | tropical | nocturnal | 6.2 | 2201727.946 | 7.783 |
| mammals | Miniopterus_fraterculus | Miniopterus_fraterculus | LC | Unknown | tropical | nocturnal | 7.3 | 1668579.509 | 9.834 |
| mammals | Miniopterus_fuscus | Miniopterus_fuscus | EN | Decreasing | temperate | nocturnal | 10.7 | 2987.771 | 22.088 |

| | | | | | | | | | |
|---|---|---|---|---|---|---|---|---|---|
| mammals | Miniopterus_gleni | Miniopterus_gleni | LC | Unknown | tropical | nocturnal | 14 | 417528.436 | 8.386 |
| mammals | Miniopterus_griveaudi | NA | DD | Unknown | NA | NA | NA | NA | NA |
| mammals | Miniopterus_inflatus | Miniopterus_inflatus | LC | Unknown | tropical | nocturnal | 14.2 | 2350003.003 | 6.697 |
| mammals | Miniopterus_macrocneme | Miniopterus_macrocneme | LC | Unknown | tropical | nocturnal | 9.2 | 470913.204 | 4.405 |
| mammals | Miniopterus_magnater | Miniopterus_magnater | LC | Unknown | tropical | nocturnal | 14.1 | 3161072.067 | 10.015 |
| mammals | Miniopterus_majori | Miniopterus_majori | LC | Unknown | tropical | nocturnal | 8 | 249561.262 | 8.336 |
| mammals | Miniopterus_manavi | Miniopterus_manavi | LC | Unknown | tropical | nocturnal | 5 | 511543.161 | 8.485 |
| mammals | Miniopterus_medius | Miniopterus_medius | LC | Unknown | tropical | nocturnal | 9 | 1256827.57 | 8.224 |
| mammals | Miniopterus_minor | NA | DD | Unknown | NA | NA | NA | NA | NA |
| mammals | Miniopterus_natalensis | Miniopterus_natalensis | LC | Unknown | tropical | nocturnal | 12.5 | 3787497.563 | 9.305 |
| mammals | Miniopterus_newtoni | NA | DD | Unknown | NA | NA | NA | NA | NA |
| mammals | Miniopterus_paululus | Miniopterus_paululus | LC | Unknown | tropical | nocturnal | 6 | 278632.689 | 13.739 |
| mammals | Miniopterus_petersoni | NA | DD | Unknown | NA | NA | NA | NA | NA |
| mammals | Miniopterus_pusillus | Miniopterus_pusillus | LC | Unknown | tropical | nocturnal | 8.6 | 1106938.271 | 12.833 |
| mammals | Miniopterus_robustior | Miniopterus_robustior | EN | Decreasing | tropical | nocturnal | 8.1 | 1081.819 | 8.429 |
| mammals | Miniopterus_schreibersii | Miniopterus_schreibersii | VU | Decreasing | temperate | nocturnal | 11.1 | 2898979.404 | 17.762 |
| mammals | Miniopterus_shortridgei | NA | DD | Unknown | NA | NA | NA | NA | NA |
| mammals | Miniopterus_sororculus | Miniopterus_sororculus | LC | Unknown | tropical | nocturnal | 8 | 91220.222 | 9.984 |
| mammals | Miniopterus_tristis | Miniopterus_tristis | LC | Unknown | tropical | nocturnal | 15.8 | 732372.808 | 8.89 |
| mammals | Miopithecus_ogouensis | Miopithecus_ogouensis | NT | Decreasing | tropical | diurnal | 1278.3 | 486827.109 | 6.113 |
| mammals | Miopithecus_talapoin | Miopithecus_talapoin | VU | Decreasing | tropical | diurnal | 1499.2 | 399163.104 | 7.933 |
| mammals | Mirimiri_acrodonta | NA | CR | Decreasing | NA | NA | NA | NA | NA |
| mammals | Mirostrellus_joffrei | NA | DD | Unknown | NA | NA | NA | NA | NA |
| mammals | Mirza_coquereli | Mirza_coquereli | EN | Decreasing | tropical | nocturnal | 308 | 16761.47 | 7.163 |
| mammals | Mirza_zaza | Mirza_zaza | VU | Decreasing | tropical | nocturnal | 290 | 9851.881 | 7.321 |
| mammals | Molossops_neglectus | NA | DD | Unknown | NA | NA | NA | NA | NA |
| mammals | Molossops_temminckii | Molossops_temminckii | LC | Unknown | tropical | nocturnal | 5.9 | 11605510.25 | 5.837 |
| mammals | Molossus_aztecus | Molossus_aztecus | LC | Decreasing | tropical | nocturnal | 14.3 | 20680.329 | 10.628 |
| mammals | Molossus_coibensis | Molossus_coibensis | LC | Unknown | tropical | nocturnal | 13 | 6800728.085 | 3.354 |
| mammals | Molossus_currentium | Molossus_currentium | LC | Unknown | tropical | nocturnal | 17.8 | 1057644.541 | 5.432 |
| mammals | Molossus_molossus | Molossus_molossus | LC | Unknown | tropical | nocturnal | 13.6 | 15767723.285 | 6.361 |
| mammals | Molossus_pretiosus | Molossus_pretiosus | LC | Unknown | tropical | nocturnal | 27 | 2034434.035 | 7.486 |
| mammals | Molossus_rufus | Molossus_rufus | LC | Stable | tropical | nocturnal | 31.7 | 14800275.008 | 6.081 |
| mammals | Molossus_sinaloae | Molossus_sinaloae | LC | Stable | tropical | nocturnal | 22.6 | 2024567.249 | 9.608 |

| | | | | | | | | | |
|---|---|---|---|---|---|---|---|---|---|
| mammals | Monodelphis_adusta | Monodelphis_adusta | LC | Stable | tropical | nocturnal | 35 | 1083732.498 | 5.485 |
| mammals | Monodelphis_americana | Monodelphis_americana | LC | Decreasing | tropical | diurnal | 19.5 | 1595848.058 | 11.5 |
| mammals | Monodelphis_brevicaudata | Monodelphis_brevicaudata | LC | Unknown | tropical | nocturnal | 69.6 | 1828323.549 | 1.319 |
| mammals | Monodelphis_dimidiata | Monodelphis_dimidiata | LC | Stable | temperate | diurnal | 58 | 1931861.771 | 12.375 |
| mammals | Monodelphis_domestica | Monodelphis_domestica | LC | Stable | tropical | nocturnal | 71.4 | 4111650.305 | 8.009 |
| mammals | Monodelphis_emiliae | Monodelphis_emiliae | LC | Unknown | tropical | nocturnal | 40 | 1139235.367 | 1.514 |
| mammals | Monodelphis_iheringi | Monodelphis_iheringi | DD | Decreasing | tropical | nocturnal | 112 | 677583.586 | 12.192 |
| mammals | Monodelphis_kunsi | Monodelphis_kunsi | LC | Stable | tropical | diurnal | 11.3 | 1203676.446 | 7.495 |
| mammals | Monodelphis_osgoodi | Monodelphis_osgoodi | LC | Unknown | tropical | nocturnal | 112 | 98261.37 | 5.345 |
| mammals | Monodelphis_reigi | Monodelphis_reigi | VU | Unknown | tropical | nocturnal | 19 | 107.296 | 4.088 |
| mammals | Monodelphis_ronaldi | Monodelphis_ronaldi | LC | Stable | tropical | nocturnal | 52.9 | 574.8 | 0.724 |
| mammals | Monodelphis_scalops | Monodelphis_scalops | LC | Decreasing | tropical | diurnal | 741 | 352992.862 | 13.573 |
| mammals | Monophyllus_plethodon | Monophyllus_plethodon | LC | Unknown | tropical | nocturnal | 14.8 | 5400.192 | 22.602 |
| mammals | Monophyllus_redmani | Monophyllus_redmani | LC | Stable | tropical | nocturnal | 8.7 | 202239.21 | 16.713 |
| mammals | Montemys_delectorum | Praomys_delectorum | LC | Stable | tropical | nocturnal | 36 | 56002.783 | 13.096 |
| mammals | Monticolomys_koopmani | Monticolomys_koopmani | LC | Unknown | tropical | nocturnal | 25 | 20443.331 | 7.908 |
| mammals | Mops_aloysiisabaudiae | Chaerephon_aloysiisabaudiae | LC | Decreasing | tropical | nocturnal | 25.7 | 1117207.365 | 7.016 |
| mammals | Mops_ansorgei | Chaerephon_ansorgei | LC | Stable | tropical | nocturnal | 16.1 | 5633242.365 | 9.326 |
| mammals | Mops_bemmeleni | Chaerephon_bemmeleni | LC | Unknown | tropical | nocturnal | 19 | 671185.433 | 14.561 |
| mammals | Mops_bivittatus | Chaerephon_bivittatus | LC | Stable | tropical | nocturnal | 15 | 1892463.639 | 10.191 |
| mammals | Mops_brachypterus | Mops_brachypterus | LC | Unknown | tropical | nocturnal | 16 | 1924932.172 | 9.02 |
| mammals | Mops_bregullae | Chaerephon_bregullae | EN | Decreasing | tropical | nocturnal | 19.9 | 20613.225 | 8.715 |
| mammals | Mops_chapini | Chaerephon_chapini | LC | Unknown | tropical | nocturnal | 10 | 4301386.964 | 8.194 |
| mammals | Mops_condylurus | Mops_condylurus | LC | Unknown | tropical | nocturnal | 28.4 | 9108820.051 | 9.84 |
| mammals | Mops_congicus | Mops_congicus | LC | Unknown | tropical | nocturnal | 43 | 528771.157 | 6.288 |
| mammals | Mops_demonstrator | Mops_demonstrator | LC | Decreasing | tropical | nocturnal | 23.5 | 1886714.305 | 9.085 |
| mammals | Mops_gallagheri | NA | DD | Unknown | NA | NA | NA | NA | NA |
| mammals | Mops_jobensis | Chaerephon_jobensis | LC | Stable | tropical | nocturnal | 16.9 | 2340174.94 | 3 |
| mammals | Mops_jobimena | Chaerephon_jobimena | LC | Unknown | tropical | nocturnal | 14.3 | 186331.766 | 7.804 |
| mammals | Mops_johorensis | Chaerephon_johorensis | VU | Decreasing | tropical | nocturnal | 20 | 91529.835 | 10.973 |
| mammals | Mops_leucostigma | Mops_leucostigma | LC | Unknown | tropical | nocturnal | 17.5 | 214048.942 | 7.988 |
| mammals | Mops_major | Chaerephon_major | LC | Stable | tropical | nocturnal | 14.8 | 1759227.746 | 12.599 |
| mammals | Mops_midas | Mops_midas | LC | Decreasing | tropical | nocturnal | 45.2 | 415330.64 | 10.123 |
| mammals | Mops_mops | Mops_mops | NT | Decreasing | tropical | nocturnal | 31 | 46635.737 | 7.634 |

| | | | | | | | | | |
|---|---|---|---|---|---|---|---|---|---|
| mammals | Mops_nanulus | Mops_nanulus | LC | Unknown | tropical | nocturnal | 9.2 | 2775168.768 | 7.574 |
| mammals | Mops_niangarae | NA | DD | Unknown | NA | NA | NA | NA | NA |
| mammals | Mops_nigeriae | Chaerephon_nigeriae | LC | Unknown | tropical | nocturnal | 19.8 | 4801793.706 | 8.465 |
| mammals | Mops_niveiventer | Mops_niveiventer | LC | Unknown | tropical | nocturnal | 21.7 | 1867655.713 | 6.947 |
| mammals | Mops_petersoni | Mops_petersoni | NT | Decreasing | tropical | nocturnal | 12.4 | 41534.636 | 11.204 |
| mammals | Mops_plicatus | Chaerephon_plicatus | LC | Unknown | tropical | nocturnal | 17.1 | 2462336.036 | 11.426 |
| mammals | Mops_pumilus | Chaerephon_pumilus | LC | Unknown | tropical | nocturnal | 11.3 | 6943070.971 | 11.602 |
| mammals | Mops_russatus | NA | DD | Unknown | NA | NA | NA | NA | NA |
| mammals | Mops_sarasinorum | NA | DD | Unknown | NA | NA | NA | NA | NA |
| mammals | Mops_solomonis | Chaerephon_solomonis | EN | Decreasing | tropical | nocturnal | 12.3 | 7070.414 | 2.438 |
| mammals | Mops_spurrelli | Mops_spurrelli | LC | Unknown | tropical | nocturnal | 8.9 | 1162141.901 | 8.555 |
| mammals | Mops_thersites | Mops_thersites | LC | Stable | tropical | nocturnal | 22 | 2248550.682 | 8.115 |
| mammals | Mops_tomensis | Chaerephon_tomensis | EN | Decreasing | tropical | nocturnal | 7.1 | 656.644 | 14.135 |
| mammals | Mops_trevori | Mops_trevori | DD | Decreasing | tropical | nocturnal | 28.9 | 426472.231 | 15.28 |
| mammals | Mormoops_blainvillei | Mormoops_blainvillei | LC | Unknown | tropical | nocturnal | 8.6 | 200604.528 | 16.76 |
| mammals | Mormoops_megalophylla | Mormoops_megalophylla | LC | Decreasing | tropical | nocturnal | 15.8 | 3738685.972 | 9.319 |
| mammals | Mormopterus_acetabulosus | Mormopterus_acetabulosus | EN | Decreasing | tropical | nocturnal | 14.1 | 1872.583 | 25.918 |
| mammals | Mormopterus_doriae | NA | DD | Unknown | NA | NA | NA | NA | NA |
| mammals | Mormopterus_jugularis | Mormopterus_jugularis | LC | Unknown | tropical | nocturnal | 11.5 | 235404.565 | 9.445 |
| mammals | Mormopterus_kalinowskii | Mormopterus_kalinowskii | LC | Unknown | tropical | nocturnal | 17.6 | 390233.296 | 6.18 |
| mammals | Mormopterus_minutus | Mormopterus_minutus | VU | Decreasing | tropical | nocturnal | 6.7 | 45074.174 | 15.817 |
| mammals | Mormopterus_phrudus | Mormopterus_phrudus | VU | Unknown | tropical | nocturnal | 9.3 | 1298.379 | 6.511 |
| mammals | Moschiola_indica | Moschiola_indica | LC | Unknown | tropical | nocturnal | 3000 | 2015179.407 | 18.683 |
| mammals | Moschiola_meminna | Moschiola_meminna | LC | Unknown | tropical | nocturnal | 2445 | 53308.917 | 17.32 |
| mammals | Moschus_anhuiensis | Moschus_anhuiensis | EN | Decreasing | temperate | nocturnal | 8400 | 12999.742 | 10.526 |
| mammals | Moschus_berezovskii | Moschus_berezovskii | EN | Decreasing | temperate | nocturnal | 9000 | 1605153.638 | 12.39 |
| mammals | Moschus_chrysogaster | Moschus_chrysogaster | EN | Decreasing | temperate | nocturnal | 14500 | 1421239.733 | 6.757 |
| mammals | Moschus_cupreus | Moschus_cupreus | EN | Decreasing | temperate | nocturnal | 14500 | 127719.941 | 5.297 |
| mammals | Moschus_fuscus | Moschus_fuscus | EN | Decreasing | temperate | nocturnal | 12500 | 213233.22 | 4.913 |
| mammals | Moschus_leucogaster | Moschus_leucogaster | EN | Decreasing | temperate | nocturnal | 15500 | 225843.572 | 7.356 |
| mammals | Mosia_nigrescens | Mosia_nigrescens | LC | Stable | tropical | nocturnal | 3.3 | 1090668.426 | 4.604 |
| mammals | Mungos_gambianus | Mungos_gambianus | LC | Stable | tropical | diurnal | 1500 | 1634057.426 | 13.876 |
| mammals | Mungos_mungo | Mungos_mungo | LC | Stable | tropical | diurnal | 1925 | 10331332.005 | 8.605 |
| mammals | Mungotictis_decemlineata | Mungotictis_decemlineata | EN | Decreasing | tropical | diurnal | 650 | 13303.642 | 7.542 |

| | | | | | | | | | |
|---|---|---|---|---|---|---|---|---|---|
| mammals | Muntiacus_atherodes | Muntiacus_atherodes | NT | Decreasing | tropical | diurnal | 15600 | 732628.361 | 5.565 |
| mammals | Muntiacus_crinifrons | Muntiacus_crinifrons | VU | Decreasing | temperate | cathemeral | 36141 | 113889.541 | 13.78 |
| mammals | Muntiacus_feae | NA | DD | Unknown | NA | NA | NA | NA | NA |
| mammals | Muntiacus_muntjak | Muntiacus_muntjak | LC | Decreasing | tropical | cathemeral | 15925.8 | 1494965.818 | 9.312 |
| mammals | Muntiacus_putaoensis | Muntiacus_putaoensis | DD | Decreasing | temperate | cathemeral | 12000 | 108084.896 | 5.92 |
| mammals | Muntiacus_reevesi | Muntiacus_reevesi | LC | Decreasing | temperate | cathemeral | 13500 | 2322121.553 | 16.437 |
| mammals | Muntiacus_rooseveltorum | Muntiacus_rooseveltorum | DD | Decreasing | tropical | cathemeral | 15000 | 422808.603 | 9.845 |
| mammals | Muntiacus_truongsonensis | Muntiacus_truongsonensis | DD | Decreasing | tropical | cathemeral | 15000 | 288756.893 | 8.325 |
| mammals | Muntiacus_vaginalis | Muntiacus_vaginalis | LC | Decreasing | tropical | cathemeral | 24000 | 5872023.275 | 15.506 |
| mammals | Muntiacus_vuquangensis | Muntiacus_vuquangensis | CR | Decreasing | tropical | diurnal | 34000 | 110001.596 | 7.812 |
| mammals | Murexia_habbema | Murexia_habbema | LC | Stable | tropical | nocturnal | 34.8 | 143836.687 | 4.888 |
| mammals | Murexia_longicaudata | Murexia_longicaudata | LC | Unknown | tropical | cathemeral | 335.8 | 515141.485 | 3.881 |
| mammals | Murexia_melanura | Murexia_melanurus | LC | Stable | tropical | nocturnal | 47.8 | 330753.341 | 3.974 |
| mammals | Murexia_rothschildi | Murexia_rothschildi | NT | Unknown | tropical | nocturnal | 43.8 | 14840.559 | 2.773 |
| mammals | Murina_aenea | Murina_aenea | VU | Decreasing | tropical | nocturnal | 7.4 | 200318.274 | 6.883 |
| mammals | Murina_aurata | NA | DD | Unknown | NA | NA | NA | NA | NA |
| mammals | Murina_cyclotis | Murina_cyclotis | LC | Unknown | temperate | nocturnal | 9.3 | 1100512.772 | 11.454 |
| mammals | Murina_florium | Murina_florium | LC | Unknown | tropical | nocturnal | 4.3 | 159006.847 | 6.396 |
| mammals | Murina_fusca | NA | DD | Unknown | NA | NA | NA | NA | NA |
| mammals | Murina_harrisoni | Murina_harrisoni | LC | Unknown | tropical | nocturnal | 4.3 | 5103.032 | 9.675 |
| mammals | Murina_hilgendorfi | Murina_hilgendorfi | LC | Unknown | temperate | nocturnal | 8.1 | 3719628.22 | 5.57 |
| mammals | Murina_huttonii | Murina_huttoni | LC | Unknown | temperate | nocturnal | 7.6 | 530181.174 | 11.38 |
| mammals | Murina_leucogaster | Murina_leucogaster | LC | Unknown | temperate | nocturnal | 7.6 | 1806033.933 | 15.847 |
| mammals | Murina_puta | Murina_puta | LC | Stable | temperate | nocturnal | 8.3 | 35902.514 | 20.066 |
| mammals | Murina_rozendaali | Murina_rozendaali | VU | Decreasing | tropical | nocturnal | 4.6 | 736664.337 | 5.591 |
| mammals | Murina_ryukyuana | Murina_ryukyuana | EN | Decreasing | temperate | nocturnal | 4.5 | 1084.572 | 14.64 |
| mammals | Murina_suilla | Murina_suilla | LC | Unknown | tropical | nocturnal | 3.6 | 1442016.705 | 9.088 |
| mammals | Murina_tubinaris | NA | DD | Unknown | NA | NA | NA | NA | NA |
| mammals | Murina_ussuriensis | Murina_ussuriensis | LC | Unknown | temperate | nocturnal | 5.3 | 773194.69 | 12.666 |
| mammals | Mus_baoulei | Mus_baoulei | LC | Stable | tropical | nocturnal | 7.8 | 105999.238 | 15.33 |
| mammals | Mus_booduga | Mus_booduga | LC | Stable | tropical | nocturnal | 11.6 | 3186564.993 | 18.258 |
| mammals | Mus_bufo | Mus_bufo | LC | Stable | tropical | nocturnal | 10.4 | 69837.537 | 18.057 |
| mammals | Mus_callewaerti | NA | DD | Unknown | NA | NA | NA | NA | NA |
| mammals | Mus_caroli | Mus_caroli | LC | Stable | tropical | nocturnal | 11.3 | 1841648.169 | 14.312 |

| | | | | | | | | | |
|---|---|---|---|---|---|---|---|---|---|
| mammals | Mus_cervicolor | Mus_cervicolor | LC | Stable | tropical | nocturnal | 17.7 | 251716.075 | 15.595 |
| mammals | Mus_cookii | Mus_cookii | LC | Stable | temperate | nocturnal | 20.5 | 863039.307 | 10.109 |
| mammals | Mus_crociduroides | NA | DD | Unknown | NA | NA | NA | NA | NA |
| mammals | Mus_cypriacus | Mus_cypriacus | LC | Unknown | temperate | nocturnal | 50.6 | 9272.655 | 19.898 |
| mammals | Mus_famulus | Mus_famulus | EN | Unknown | tropical | nocturnal | 22.9 | 605.668 | 10.698 |
| mammals | Mus_fernandoni | NA | EN | Unknown | NA | NA | NA | NA | NA |
| mammals | Mus_fragilicauda | Mus_fragilicauda | LC | Unknown | tropical | nocturnal | 12 | 4661.405 | 15.703 |
| mammals | Mus_goundae | NA | DD | Unknown | NA | NA | NA | NA | NA |
| mammals | Mus_haussa | Mus_haussa | LC | Stable | tropical | nocturnal | 4.2 | 2515879.354 | 11.029 |
| mammals | Mus_imberbis | Muriculus_imberbis | LC | Decreasing | tropical | nocturnal | 18.5 | 35840.014 | 16.159 |
| mammals | Mus_indutus | Mus_indutus | LC | Stable | tropical | nocturnal | 6.2 | 1481732.921 | 5.668 |
| mammals | Mus_macedonicus | Mus_macedonicus | LC | Stable | temperate | nocturnal | 15 | 1386803.652 | 15.447 |
| mammals | Mus_mahomet | Mus_mahomet | LC | Stable | tropical | nocturnal | 10 | 418704.621 | 13.13 |
| mammals | Mus_mattheyi | Mus_mattheyi | LC | Stable | tropical | nocturnal | 4.5 | 333722.409 | 11.986 |
| mammals | Mus_mayori | Mus_mayori | VU | Unknown | tropical | nocturnal | 29.4 | 1122.551 | 18.375 |
| mammals | Mus_minutoides | Mus_minutoides | LC | Stable | tropical | nocturnal | 6.8 | 4215726.736 | 9.884 |
| mammals | Mus_musculoides | Mus_musculoides | LC | Unknown | tropical | nocturnal | 5.6 | 5014869.289 | 10.812 |
| mammals | Mus_musculus | Mus_musculus | LC | Stable | temperate | nocturnal | 16.1 | 84352137.994 | 9.092 |
| mammals | Mus_neavei | NA | DD | Unknown | NA | NA | NA | NA | NA |
| mammals | Mus_oubanguii | NA | DD | Unknown | NA | NA | NA | NA | NA |
| mammals | Mus_pahari | Mus_pahari | LC | Stable | tropical | nocturnal | 25.7 | 1683214.503 | 10.693 |
| mammals | Mus_platythrix | Mus_platythrix | LC | Stable | tropical | nocturnal | 25.2 | 511752.214 | 19.785 |
| mammals | Mus_setulosus | Mus_setulosus | LC | Stable | tropical | nocturnal | 11.1 | 1024297.29 | 12.26 |
| mammals | Mus_setzeri | Mus_setzeri | LC | Stable | tropical | nocturnal | 6.7 | 157445.805 | 4.313 |
| mammals | Mus_shortridgei | Mus_shortridgei | LC | Stable | tropical | nocturnal | 34.6 | 268868.68 | 8.502 |
| mammals | Mus_sorella | Mus_sorella | LC | Stable | tropical | nocturnal | 13.7 | 52516.104 | 7.458 |
| mammals | Mus_spicilegus | Mus_spicilegus | LC | Decreasing | temperate | nocturnal | 14 | 828198.668 | 18.766 |
| mammals | Mus_spretus | Mus_spretus | LC | Stable | temperate | nocturnal | 12.8 | 963695.111 | 17.161 |
| mammals | Mus_tenellus | Mus_tenellus | LC | Stable | tropical | nocturnal | 11 | 491701.51 | 9.717 |
| mammals | Mus_terricolor | Mus_terricolor | LC | Stable | tropical | nocturnal | 9 | 2890511.676 | 18.509 |
| mammals | Mus_triton | Mus_triton | LC | Stable | tropical | cathemeral | 12.1 | 1251170.509 | 11.175 |
| mammals | Mus_vulcani | Mus_vulcani | LC | Stable | tropical | nocturnal | 20.2 | 184.939 | 11.839 |
| mammals | Muscardinus_avellanarius | Muscardinus_avellanarius | LC | Unknown | temperate | nocturnal | 27.5 | 3613069.831 | 19.676 |
| mammals | Musonycteris_harrisoni | Musonycteris_harrisoni | VU | Decreasing | tropical | nocturnal | 11.8 | 54476.966 | 8.999 |

| | | | | | | | | | |
|---|---|---|---|---|---|---|---|---|---|
| mammals | Mustela_altaica | Mustela_altaica | NT | Decreasing | temperate | nocturnal | 171 | 5661382.613 | 6.267 |
| mammals | Mustela_erminea | Mustela_erminea | LC | Stable | temperate | cathemeral | 119.4 | 41482667.378 | 5.898 |
| mammals | Mustela_eversmanii | Mustela_eversmanii | LC | Decreasing | temperate | nocturnal | 1350 | 13299450.446 | 9.273 |
| mammals | Mustela_itatsi | Mustela_itatsi | NT | Decreasing | temperate | nocturnal | 565 | 361776.184 | 19.826 |
| mammals | Mustela_kathiah | Mustela_kathiah | LC | Stable | tropical | cathemeral | 208.1 | 2773336.236 | 12.688 |
| mammals | Mustela_lutreola | Mustela_lutreola | CR | Decreasing | temperate | nocturnal | 440 | 7371841.887 | 14.043 |
| mammals | Mustela_lutreolina | Mustela_lutreolina | LC | Stable | tropical | nocturnal | 706.6 | 84498.729 | 9.895 |
| mammals | Mustela_nigripes | Mustela_nigripes | EN | Decreasing | temperate | nocturnal | 850 | 2515859.228 | 11.254 |
| mammals | Mustela_nivalis | Mustela_nivalis | LC | Stable | temperate | cathemeral | 137.5 | 43300266.986 | 7.14 |
| mammals | Mustela_nudipes | Mustela_nudipes | LC | Decreasing | tropical | cathemeral | 500 | 1346098.668 | 7.896 |
| mammals | Mustela_putorius | Mustela_putorius | LC | Decreasing | temperate | cathemeral | 915.4 | 6011048.913 | 16.459 |
| mammals | Mustela_sibirica | Mustela_sibirica | LC | Stable | temperate | nocturnal | 405 | 13304653.934 | 8.289 |
| mammals | Mustela_strigidorsa | Mustela_strigidorsa | LC | Stable | tropical | cathemeral | 1500 | 1377923.661 | 10.064 |
| mammals | Mydaus_javanensis | Mydaus_javanensis | LC | Stable | tropical | nocturnal | 2500 | 105559.279 | 5.702 |
| mammals | Mydaus_marchei | Mydaus_marchei | LC | Stable | tropical | nocturnal | 1414.2 | 12447.239 | 9.283 |
| mammals | Mylomys_dybowskii | Mylomys_dybowskii | LC | Unknown | tropical | diurnal | 114.4 | 1279787.083 | 9.182 |
| mammals | Mylomys_rex | NA | DD | Unknown | NA | NA | NA | NA | NA |
| mammals | Myocastor_coypus | Myocastor_coypus | LC | Decreasing | temperate | nocturnal | 6937.5 | 3902051.503 | 8.213 |
| mammals | Myoictis_leucura | NA | DD | Unknown | NA | NA | NA | NA | NA |
| mammals | Myoictis_melas | Myoictis_melas | LC | Unknown | tropical | diurnal | 230 | 398695.852 | 3.364 |
| mammals | Myoictis_wallacii | Myoictis_wallacei | LC | Unknown | tropical | diurnal | 230 | 56148.796 | 3.159 |
| mammals | Myoictis_wavica | NA | DD | Unknown | NA | NA | NA | NA | NA |
| mammals | Myomimus_personatus | NA | DD | Unknown | NA | NA | NA | NA | NA |
| mammals | Myomimus_roachi | Myomimus_roachi | VU | Decreasing | temperate | nocturnal | 45.5 | 21889.242 | 17.341 |
| mammals | Myomimus_setzeri | NA | DD | Unknown | NA | NA | NA | NA | NA |
| mammals | Myomyscus_verreauxi | Myomyscus_verreauxii | LC | Stable | temperate | nocturnal | 41 | 97856.177 | 10.481 |
| mammals | Myonycteris_angolensis | Lissonycteris_angolensis | LC | Decreasing | tropical | nocturnal | 74.6 | 9083671.621 | 8.504 |
| mammals | Myonycteris_brachycephala | Myonycteris_brachycephala | EN | Decreasing | tropical | nocturnal | 35.8 | 690.852 | 10.678 |
| mammals | Myonycteris_leptodon | Myonycteris_leptodon | LC | Unknown | tropical | nocturnal | 42 | 835056.633 | 14.202 |
| mammals | Myonycteris_relicta | Myonycteris_relicta | LC | Decreasing | tropical | nocturnal | 52 | 142204.808 | 10.939 |
| mammals | Myonycteris_torquata | Myonycteris_torquata | LC | Stable | tropical | nocturnal | 42.2 | 3651482.596 | 6.019 |
| mammals | Myoprocta_acouchy | Myoprocta_acouchy | LC | Unknown | tropical | diurnal | 600 | 993244.796 | 1.354 |
| mammals | Myoprocta_pratti | Myoprocta_pratti | LC | Stable | tropical | diurnal | 908 | 2337165.235 | 1.674 |
| mammals | Myopterus_daubentonii | NA | DD | Unknown | NA | NA | NA | NA | NA |

| | | | | | | | | | |
|---|---|---|---|---|---|---|---|---|---|
| mammals | Myopterus_whitleyi | Myopterus_whitleyi | LC | Decreasing | tropical | nocturnal | 11.5 | 1914285.176 | 5.966 |
| mammals | Myopus_schisticolor | Myopus_schisticolor | LC | Unknown | temperate | cathemeral | 25 | 10209782.765 | 2.504 |
| mammals | Myosciurus_pumilio | Myosciurus_pumilio | LC | Unknown | tropical | diurnal | 165 | 346127.668 | 7.154 |
| mammals | Myosorex_babaulti | Myosorex_babaulti | LC | Decreasing | tropical | cathemeral | 17 | 24423.894 | 19.193 |
| mammals | Myosorex_blarina | Myosorex_blarina | EN | Decreasing | tropical | nocturnal | 14.2 | 2272.006 | 11.109 |
| mammals | Myosorex_cafer | Myosorex_cafer | VU | Decreasing | temperate | nocturnal | 13.4 | 65136.024 | 15.272 |
| mammals | Myosorex_eisentrauti | Myosorex_eisentrauti | CR | Unknown | tropical | cathemeral | 13.8 | 49.407 | 12.798 |
| mammals | Myosorex_geata | Myosorex_geata | EN | Decreasing | tropical | cathemeral | 9.4 | 463.331 | 12.061 |
| mammals | Myosorex_kihaulei | Myosorex_kihaulei | EN | Decreasing | tropical | nocturnal | 9.5 | 1615.626 | 14.883 |
| mammals | Myosorex_longicaudatus | Myosorex_longicaudatus | EN | Decreasing | temperate | cathemeral | 13 | 7729.014 | 10.298 |
| mammals | Myosorex_okuensis | Myosorex_okuensis | VU | Decreasing | tropical | cathemeral | 11.8 | 4126.801 | 13.541 |
| mammals | Myosorex_rumpii | Myosorex_rumpii | EN | Decreasing | tropical | cathemeral | 15 | 254.13 | 8.79 |
| mammals | Myosorex_schalleri | NA | DD | Unknown | NA | NA | NA | NA | NA |
| mammals | Myosorex_sclateri | Myosorex_sclateri | VU | Decreasing | temperate | cathemeral | 20.9 | 20830.845 | 15.913 |
| mammals | Myosorex_tenuis | Myosorex_tenuis | EN | Decreasing | tropical | cathemeral | 9.4 | 4371.799 | 10.679 |
| mammals | Myosorex_varius | Myosorex_varius | LC | Stable | temperate | cathemeral | 13.2 | 424218.627 | 12.175 |
| mammals | Myotis_adversus | Myotis_adversus | LC | Stable | tropical | nocturnal | 10.6 | 55894.179 | 18.584 |
| mammals | Myotis_albescens | Myotis_albescens | LC | Stable | tropical | nocturnal | 5.3 | 14595013.005 | 6.002 |
| mammals | Myotis_alcathoe | NA | DD | Unknown | NA | NA | NA | NA | NA |
| mammals | Myotis_altarium | Myotis_altarium | LC | Unknown | temperate | nocturnal | 11 | 1646300.535 | 15.029 |
| mammals | Myotis_anjouanensis | NA | DD | Unknown | NA | NA | NA | NA | NA |
| mammals | Myotis_annectans | Myotis_annectans | LC | Unknown | tropical | nocturnal | 9.8 | 296170.997 | 8.666 |
| mammals | Myotis_atacamensis | Myotis_atacamensis | EN | Unknown | tropical | nocturnal | 7.1 | 253619.58 | 5.439 |
| mammals | Myotis_ater | Myotis_ater | LC | Unknown | tropical | nocturnal | 6 | 127895.393 | 7.376 |
| mammals | Myotis_auriculus | Myotis_auriculus | LC | Unknown | temperate | nocturnal | 5 | 617481.09 | 7.302 |
| mammals | Myotis_australis | NA | DD | Unknown | NA | NA | NA | NA | NA |
| mammals | Myotis_austroriparius | Myotis_austroriparius | LC | Stable | temperate | nocturnal | 6.8 | 832901.782 | 15.941 |
| mammals | Myotis_bechsteinii | Myotis_bechsteinii | NT | Decreasing | temperate | nocturnal | 8.5 | 2676244.149 | 20.933 |
| mammals | Myotis_blythii | Myotis_blythii | LC | Decreasing | temperate | nocturnal | 22.5 | 6194956.012 | 14.268 |
| mammals | Myotis_bocagii | Myotis_bocagii | LC | Unknown | tropical | nocturnal | 7.8 | 1819217.096 | 9.983 |
| mammals | Myotis_bombinus | Myotis_bombinus | NT | Decreasing | temperate | nocturnal | 6.9 | 1653772.778 | 5.008 |
| mammals | Myotis_brandtii | Myotis_brandtii | LC | Stable | temperate | nocturnal | 5.3 | 5471819.619 | 14.456 |
| mammals | Myotis_bucharensis | NA | DD | Unknown | NA | NA | NA | NA | NA |
| mammals | Myotis_californicus | Myotis_californicus | LC | Unknown | temperate | nocturnal | 4.6 | 3985147.134 | 8.659 |

| | | | | | | | | | |
|---|---|---|---|---|---|---|---|---|---|
| mammals | Myotis_capaccinii | Myotis_capaccinii | VU | Decreasing | temperate | nocturnal | 7.4 | 1229394.764 | 18.323 |
| mammals | Myotis_chiloensis | Myotis_chiloensis | LC | Unknown | temperate | nocturnal | 6.8 | 502327.681 | 6.391 |
| mammals | Myotis_chinensis | Myotis_chinensis | LC | Unknown | temperate | nocturnal | 42 | 2369550.789 | 13.67 |
| mammals | Myotis_ciliolabrum | Myotis_ciliolabrum | LC | Stable | temperate | nocturnal | 4.9 | 4652123.122 | 9.753 |
| mammals | Myotis_cobanensis | NA | DD | Unknown | NA | NA | NA | NA | NA |
| mammals | Myotis_dasycneme | Myotis_dasycneme | NT | Decreasing | temperate | nocturnal | 14.1 | 5448983.292 | 11.927 |
| mammals | Myotis_daubentonii | Myotis_daubentonii | LC | Stable | temperate | nocturnal | 7.4 | 7012608.325 | 16.315 |
| mammals | Myotis_davidii | Myotis_davidii | LC | Unknown | temperate | nocturnal | 5.3 | 6721248.697 | 13.293 |
| mammals | Myotis_dieteri | NA | DD | Unknown | NA | NA | NA | NA | NA |
| mammals | Myotis_dinellii | Myotis_dinellii | LC | Stable | temperate | nocturnal | 8.4 | 1666009.123 | 7.295 |
| mammals | Myotis_dominicensis | Myotis_dominicensis | VU | Unknown | tropical | nocturnal | 3.2 | 2234.993 | 20.596 |
| mammals | Myotis_elegans | Myotis_elegans | LC | Unknown | tropical | nocturnal | 4.1 | 587280.012 | 10.621 |
| mammals | Myotis_emarginatus | Myotis_emarginatus | LC | Stable | temperate | nocturnal | 7.7 | 4646898.861 | 16.547 |
| mammals | Myotis_evotis | Myotis_evotis | LC | Stable | temperate | nocturnal | 6.7 | 3163297.003 | 9.234 |
| mammals | Myotis_fimbriatus | Myotis_fimbriatus | LC | Unknown | temperate | nocturnal | 7.6 | 1861445.652 | 15.373 |
| mammals | Myotis_findleyi | Myotis_findleyi | EN | Decreasing | tropical | nocturnal | 4.2 | 396.768 | 3.705 |
| mammals | Myotis_formosus | Myotis_formosus | NT | Decreasing | temperate | nocturnal | 7 | 621913.003 | 15.198 |
| mammals | Myotis_fortidens | Myotis_fortidens | LC | Unknown | tropical | nocturnal | 4.4 | 474091.812 | 9.969 |
| mammals | Myotis_frater | Myotis_frater | LC | Stable | temperate | nocturnal | 7.6 | 1270255.424 | 14.374 |
| mammals | Myotis_gomantongensis | Myotis_gomantongensis | LC | Unknown | tropical | nocturnal | 9.4 | 712150.583 | 5.524 |
| mammals | Myotis_goudotii | Myotis_goudoti | LC | Unknown | tropical | nocturnal | 7 | 504612.718 | 8.432 |
| mammals | Myotis_grisescens | Myotis_grisescens | VU | Increasing | temperate | nocturnal | 10.5 | 853015.925 | 17.273 |
| mammals | Myotis_hasseltii | Myotis_hasseltii | LC | Unknown | tropical | nocturnal | 8.8 | 1330531.462 | 9.484 |
| mammals | Myotis_hermani | NA | DD | Unknown | NA | NA | NA | NA | NA |
| mammals | Myotis_horsfieldii | Myotis_horsfieldii | LC | Stable | tropical | nocturnal | 6.3 | 3208239.96 | 12.273 |
| mammals | Myotis_ikonnikovi | Myotis_ikonnikovi | LC | Decreasing | temperate | nocturnal | 3.7 | 5170695.326 | 5.971 |
| mammals | Myotis_insularum | NA | DD | Unknown | NA | NA | NA | NA | NA |
| mammals | Myotis_keaysi | Myotis_keaysi | LC | Unknown | tropical | nocturnal | 5.3 | 2736294.005 | 9.419 |
| mammals | Myotis_keenii | Myotis_keenii | LC | Stable | temperate | nocturnal | 6.3 | 109436.681 | 4.934 |
| mammals | Myotis_laniger | Myotis_laniger | LC | Unknown | temperate | nocturnal | 3.7 | 5482276.069 | 9.349 |
| mammals | Myotis_leibii | Myotis_leibii | EN | Decreasing | temperate | nocturnal | 5.1 | 1423165.264 | 17.598 |
| mammals | Myotis_levis | Myotis_levis | LC | Unknown | temperate | nocturnal | 5.5 | 1393858.707 | 12.177 |
| mammals | Myotis_longipes | NA | DD | Unknown | NA | NA | NA | NA | NA |
| mammals | Myotis_lucifugus | Myotis_lucifugus | EN | Decreasing | temperate | nocturnal | 7.6 | 12626918.361 | 8.308 |

| | | | | | | | | | |
|---|---|---|---|---|---|---|---|---|---|
| mammals | Myotis_macrodactylus | Myotis_macrodactylus | LC | Unknown | temperate | nocturnal | 7.2 | 682552.489 | 16.669 |
| mammals | Myotis_macropus | Myotis_macropus | LC | Unknown | temperate | nocturnal | 10 | 1326845.823 | 5.276 |
| mammals | Myotis_macrotarsus | Myotis_macrotarsus | LC | Stable | tropical | nocturnal | 12.5 | 282020.113 | 13.686 |
| mammals | Myotis_martiniquensis | Myotis_martiniquensis | NT | Unknown | tropical | nocturnal | 5 | 3812.793 | 23.596 |
| mammals | Myotis_moluccarum | Myotis_moluccarum | LC | Unknown | tropical | nocturnal | 9.2 | 825396.208 | 4.962 |
| mammals | Myotis_montivagus | NA | DD | Unknown | NA | NA | NA | NA | NA |
| mammals | Myotis_morrisi | NA | DD | Unknown | NA | NA | NA | NA | NA |
| mammals | Myotis_muricola | Myotis_muricola | LC | Stable | tropical | nocturnal | 4.1 | 4387144.012 | 10.989 |
| mammals | Myotis_myotis | Myotis_myotis | LC | Stable | temperate | nocturnal | 34 | 3874644.449 | 19.762 |
| mammals | Myotis_mystacinus | Myotis_mystacinus | LC | Unknown | temperate | nocturnal | 6 | 4973961.412 | 16.966 |
| mammals | Myotis_nattereri | Myotis_nattereri | LC | Increasing | temperate | nocturnal | 7.1 | 4383587.36 | 18.875 |
| mammals | Myotis_nesopolus | Myotis_nesopolus | LC | Unknown | tropical | nocturnal | 3.5 | 54642.105 | 13.297 |
| mammals | Myotis_nigricans | Myotis_nigricans | LC | Stable | tropical | nocturnal | 4.2 | 14375820.501 | 6.005 |
| mammals | Myotis_nipalensis | Myotis_nipalensis | LC | Unknown | temperate | nocturnal | 6.3 | 4946026.259 | 7.972 |
| mammals | Myotis_nyctor | Myotis_nyctor | VU | Unknown | tropical | nocturnal | 5 | 757.991 | 27.094 |
| mammals | Myotis_occultus | Myotis_occultus | LC | Stable | temperate | nocturnal | 8 | 996794.931 | 9.066 |
| mammals | Myotis_oxyotus | Myotis_oxyotus | LC | Unknown | tropical | nocturnal | 5.5 | 2601562.792 | 7.572 |
| mammals | Myotis_pequinius | Myotis_pequinius | LC | Unknown | temperate | nocturnal | 17.1 | 765279.791 | 21.858 |
| mammals | Myotis_pilosus | Myotis_pilosus | VU | Decreasing | temperate | nocturnal | 15.6 | 1162073.811 | 16.803 |
| mammals | Myotis_planiceps | Myotis_planiceps | EN | Decreasing | temperate | nocturnal | 3.1 | 12295.177 | 5.149 |
| mammals | Myotis_pruinosus | Myotis_pruinosus | EN | Unknown | temperate | nocturnal | 2.7 | 67885.316 | 15.555 |
| mammals | Myotis_punicus | NA | DD | Unknown | NA | NA | NA | NA | NA |
| mammals | Myotis_ridleyi | Myotis_ridleyi | NT | Decreasing | tropical | nocturnal | 4.5 | 864276.584 | 6.629 |
| mammals | Myotis_riparius | Myotis_riparius | LC | Stable | tropical | nocturnal | 5.6 | 14110136.661 | 5.854 |
| mammals | Myotis_rosseti | Myotis_rosseti | LC | Unknown | tropical | nocturnal | 3.3 | 467848.387 | 15.87 |
| mammals | Myotis_ruber | Myotis_ruber | NT | Unknown | temperate | nocturnal | 5 | 1469392.942 | 12.509 |
| mammals | Myotis_rufopictus | NA | DD | Unknown | NA | NA | NA | NA | NA |
| mammals | Myotis_schaubi | NA | DD | Unknown | NA | NA | NA | NA | NA |
| mammals | Myotis_scotti | Myotis_scotti | VU | Unknown | tropical | nocturnal | 8.7 | 185917.728 | 12.88 |
| mammals | Myotis_septentrionalis | Myotis_septentrionalis | NT | Decreasing | temperate | nocturnal | 7.5 | 4946390.383 | 11.575 |
| mammals | Myotis_sicarius | Myotis_sicarius | VU | Unknown | temperate | nocturnal | 13.1 | 66155.333 | 15.404 |
| mammals | Myotis_siligorensis | Myotis_siligorensis | LC | Unknown | tropical | nocturnal | 2.7 | 1309758.19 | 11.302 |
| mammals | Myotis_simus | NA | DD | Unknown | NA | NA | NA | NA | NA |
| mammals | Myotis_sodalis | Myotis_sodalis | NT | Stable | temperate | nocturnal | 6.9 | 1564971.581 | 18.076 |

| | | | | | | | | | |
|---|---|---|---|---|---|---|---|---|---|
| mammals | Myotis_stalkeri | Myotis_stalkeri | LC | Unknown | tropical | nocturnal | 17 | 4156.526 | 3.684 |
| mammals | Myotis_thysanodes | Myotis_thysanodes | LC | Stable | temperate | nocturnal | 8.1 | 3472250.621 | 9.612 |
| mammals | Myotis_tricolor | Myotis_tricolor | LC | Unknown | tropical | nocturnal | 13.9 | 4268392.074 | 10.05 |
| mammals | Myotis_velifer | Myotis_velifer | LC | Stable | temperate | nocturnal | 10.1 | 1956978.828 | 10.67 |
| mammals | Myotis_vivesi | Myotis_vivesi | VU | Decreasing | temperate | nocturnal | 25 | 6940.564 | 4.895 |
| mammals | Myotis_volans | Myotis_volans | LC | Stable | temperate | nocturnal | 8.5 | 4711968.978 | 8.716 |
| mammals | Myotis_welwitschii | Myotis_welwitschii | LC | Unknown | tropical | nocturnal | 15.5 | 4311880.238 | 10.235 |
| mammals | Myotis_yanbarensis | Myotis_yanbarensis | CR | Decreasing | temperate | nocturnal | 4.3 | 31.567 | 11.262 |
| mammals | Myotis_yumanensis | Myotis_yumanensis | LC | Stable | temperate | nocturnal | 5.4 | 4063522.234 | 8.683 |
| mammals | Myrmecobius_fasciatus | Myrmecobius_fasciatus | EN | Decreasing | temperate | diurnal | 471.5 | 2502.5 | 8.36 |
| mammals | Myrmecophaga_tridactyla | Myrmecophaga_tridactyla | VU | Decreasing | tropical | cathemeral | 22333.2 | 12924626.053 | 5.527 |
| mammals | Mysateles_prehensilis | Mysateles_prehensilis | NT | Unknown | tropical | nocturnal | 1800 | 106633.596 | 16.283 |
| mammals | Mystacina_tuberculata | Mystacina_tuberculata | VU | Decreasing | temperate | nocturnal | 12.6 | 13655.018 | 5.986 |
| mammals | Mystromys_albicaudatus | Mystromys_albicaudatus | VU | Decreasing | temperate | nocturnal | 86.9 | 501156.54 | 12.174 |
| mammals | Myzopoda_aurita | Myzopoda_aurita | LC | Unknown | tropical | nocturnal | 9.3 | 107284.175 | 8.182 |
| mammals | Myzopoda_schliemanni | Myzopoda_schliemanni | LC | Unknown | tropical | nocturnal | 9.1 | 36340 | 8.871 |
| mammals | Naemorhedus_baileyi | Naemorhedus_baileyi | VU | Decreasing | temperate | diurnal | 27487.5 | 124540.262 | 3.023 |
| mammals | Naemorhedus_caudatus | Naemorhedus_caudatus | VU | Decreasing | temperate | nocturnal | 27000 | 184715.634 | 9.686 |
| mammals | Nandinia_binotata | Nandinia_binotata | LC | Unknown | tropical | nocturnal | 2000 | 6021747.817 | 8.898 |
| mammals | Nanger_dama | Nanger_dama | CR | Decreasing | tropical | diurnal | 72999.5 | 230062.58 | 2.273 |
| mammals | Nanger_granti | Nanger_granti | LC | Decreasing | tropical | diurnal | 54999.7 | 785168.609 | 8.98 |
| mammals | Nanger_soemmerringii | Nanger_soemmerringii | VU | Decreasing | tropical | diurnal | 40000 | 291391.028 | 7.325 |
| mammals | Nannosciurus_melanotis | Nannosciurus_melanotis | LC | Decreasing | tropical | diurnal | 50 | 1074970.762 | 7.903 |
| mammals | Nanonycteris_veldkampii | Nanonycteris_veldkampii | LC | Unknown | tropical | nocturnal | 21.4 | 1711765.058 | 12.514 |
| mammals | Napaeozapus_insignis | Napaeozapus_insignis | LC | Stable | temperate | nocturnal | 22.3 | 2010052.08 | 9.074 |
| mammals | Nasalis_larvatus | Nasalis_larvatus | EN | Decreasing | tropical | diurnal | 9000 | 538836.412 | 5.731 |
| mammals | Nasua_narica | Nasua_narica | LC | Decreasing | tropical | diurnal | 4030 | 2698807.164 | 9.363 |
| mammals | Nasua_nasua | Nasua_nasua | LC | Decreasing | tropical | diurnal | 3793.8 | 10629424.442 | 4.757 |
| mammals | Nasua_olivacea | Nasuella_olivacea | NT | Decreasing | tropical | diurnal | 1340 | 266801.183 | 10.857 |
| mammals | Natalus_jamaicensis | Natalus_jamaicensis | CR | Decreasing | tropical | nocturnal | 6.7 | 11024.67 | 19.739 |
| mammals | Natalus_macrourus | Natalus_espiritosantensis | NT | Unknown | tropical | nocturnal | 6 | 2483320.535 | 6.937 |
| mammals | Natalus_major | Natalus_major | NT | Decreasing | tropical | nocturnal | 7.1 | 74327.944 | 15.805 |
| mammals | Natalus_mexicanus | Natalus_mexicanus | LC | Stable | tropical | nocturnal | 5.3 | 2430998.384 | 8.924 |
| mammals | Natalus_primus | Natalus_primus | VU | Decreasing | tropical | nocturnal | 9.1 | 3130.365 | 10.21 |

| | | | | | | | | | |
|---|---|---|---|---|---|---|---|---|---|
| mammals | Natalus_stramineus | Natalus_stramineus | LC | Unknown | tropical | nocturnal | 5 | 4059.92 | 22.269 |
| mammals | Natalus_tumidirostris | Natalus_tumidirostris | LC | Unknown | tropical | nocturnal | 6.4 | 1390815.57 | 6.373 |
| mammals | Neacomys_guianae | Neacomys_guianae | LC | Stable | tropical | nocturnal | 14.2 | 289438.839 | 1.55 |
| mammals | Neacomys_minutus | Neacomys_minutus | LC | Stable | tropical | nocturnal | 10.4 | 183731.317 | 0.478 |
| mammals | Neacomys_musseri | Neacomys_musseri | LC | Stable | tropical | nocturnal | 12.1 | 140347.467 | 0.873 |
| mammals | Neacomys_pictus | NA | DD | Unknown | NA | NA | NA | NA | NA |
| mammals | Neacomys_spinosus | Neacomys_spinosus | LC | Stable | tropical | nocturnal | 19 | 3102258.027 | 3.431 |
| mammals | Neacomys_tenuipes | Neacomys_tenuipes | LC | Stable | tropical | nocturnal | 19 | 224158.499 | 11.397 |
| mammals | Necromys_amoenus | Necromys_amoenus | LC | Stable | tropical | diurnal | 29 | 164451.1 | 8.978 |
| mammals | Necromys_lactens | Necromys_lactens | LC | Decreasing | tropical | cathemeral | 32.9 | 163777.507 | 7.275 |
| mammals | Necromys_lasiurus | Necromys_lasiurus | LC | Unknown | tropical | diurnal | 39.9 | 6100713.853 | 8.73 |
| mammals | Necromys_lenguarum | Necromys_lenguarum | LC | Stable | tropical | diurnal | 29.9 | 592293.391 | 5.24 |
| mammals | Necromys_obscurus | Necromys_obscurus | LC | Decreasing | temperate | diurnal | 40.7 | 45303.396 | 12.817 |
| mammals | Necromys_urichi | Necromys_urichi | LC | Unknown | tropical | cathemeral | 34 | 339590.636 | 4.118 |
| mammals | Nectogale_elegans | Nectogale_elegans | LC | Unknown | temperate | cathemeral | 32.5 | 555414.401 | 9.107 |
| mammals | Nectomys_apicalis | Nectomys_apicalis | LC | Stable | tropical | nocturnal | 297.4 | 1077402.36 | 1.724 |
| mammals | Nectomys_magdalenae | NA | DD | Unknown | NA | NA | NA | NA | NA |
| mammals | Nectomys_palmipes | Nectomys_palmipes | LC | Unknown | tropical | nocturnal | 197 | 165524.74 | 3.751 |
| mammals | Nectomys_rattus | Nectomys_rattus | LC | Stable | tropical | nocturnal | 248.8 | 7115323.825 | 3.525 |
| mammals | Nectomys_squamipes | Nectomys_squamipes | LC | Unknown | tropical | nocturnal | 190.7 | 2344287.567 | 11.838 |
| mammals | Nelsonia_goldmani | Nelsonia_goldmani | EN | Decreasing | tropical | nocturnal | 28.5 | 4224.241 | 11.727 |
| mammals | Nelsonia_neotomodon | Nelsonia_neotomodon | LC | Decreasing | temperate | nocturnal | 80 | 77275.933 | 6.043 |
| mammals | Neodon_clarkei | Microtus_clarkei | LC | Stable | temperate | cathemeral | 55.3 | 209251.528 | 9.996 |
| mammals | Neodon_fuscus | Lasiopodomys_fuscus | LC | Unknown | temperate | diurnal | 42.5 | 48850.659 | 4.591 |
| mammals | Neodon_irene | Neodon_irene | LC | Unknown | temperate | nocturnal | 44.6 | 635022.839 | 8.149 |
| mammals | Neodon_leucurus | Phaiomys_leucurus | LC | Unknown | temperate | diurnal | 38 | 2268403.081 | 2.856 |
| mammals | Neoeptesicus_andinus | Eptesicus_andinus | LC | Unknown | tropical | nocturnal | 9.3 | 811200.274 | 9.975 |
| mammals | Neoeptesicus_brasiliensis | Eptesicus_brasiliensis | LC | Unknown | tropical | nocturnal | 9.5 | 13117424.915 | 5.789 |
| mammals | Neoeptesicus_chiriquinus | Eptesicus_chiriquinus | LC | Unknown | tropical | nocturnal | 11.3 | 274035.826 | 8.856 |
| mammals | Neoeptesicus_diminutus | Eptesicus_diminutus | LC | Unknown | tropical | nocturnal | 6 | 2385956.805 | 11.358 |
| mammals | Neoeptesicus_furinalis | Eptesicus_furinalis | LC | Unknown | tropical | nocturnal | 7.7 | 16292708.034 | 6.353 |
| mammals | Neoeptesicus_innoxius | Eptesicus_innoxius | NT | Unknown | tropical | nocturnal | 8.4 | 167224.006 | 10.355 |
| mammals | Neofelis_diardi | Neofelis_diardi | VU | Decreasing | tropical | cathemeral | 20500 | 510006.384 | 3.858 |
| mammals | Neofelis_nebulosa | Neofelis_nebulosa | VU | Decreasing | tropical | nocturnal | 20500 | 5183470.916 | 12.537 |

| | | | | | | | | | |
|---|---|---|---|---|---|---|---|---|---|
| mammals | Neofiber_alleni | Neofiber_alleni | LC | Decreasing | temperate | nocturnal | 266 | 119935.196 | 16.92 |
| mammals | Neogale_africana | Mustela_africana | LC | Unknown | tropical | diurnal | 537 | 5209972.535 | 1.642 |
| mammals | Neogale_felipei | Mustela_felipei | VU | Decreasing | tropical | cathemeral | 211.3 | 57459.898 | 9.912 |
| mammals | Neogale_frenata | Mustela_frenata | LC | Stable | tropical | cathemeral | 147 | 12272775.386 | 12.256 |
| mammals | Neogale_vison | Neovison_vison | LC | Stable | temperate | cathemeral | 945 | 14573783.894 | 7.809 |
| mammals | Neomicroxus_bogotensis | Neomicroxus_bogotensis | LC | Stable | tropical | nocturnal | 13 | 60125.082 | 11.898 |
| mammals | Neomicroxus_latebricola | Neomicroxus_latebricola | EN | Decreasing | tropical | nocturnal | 39 | 29917.191 | 10.554 |
| mammals | Neomys_anomalus | Neomys_anomalus | LC | Decreasing | temperate | cathemeral | 16 | 3282589.819 | 19.017 |
| mammals | Neomys_fodiens | Neomys_fodiens | LC | Stable | temperate | cathemeral | 14.3 | 13089290.242 | 11.091 |
| mammals | Neomys_teres | Neomys_teres | LC | Unknown | temperate | nocturnal | 12.9 | 252150.28 | 12.666 |
| mammals | Neonycteris_pusilla | Neonycteris_pusilla | DD | Decreasing | tropical | nocturnal | 7.3 | 9047.871 | 0.98 |
| mammals | Neophascogale_lorentzii | Neophascogale_lorentzii | LC | Unknown | tropical | diurnal | 212 | 120447.436 | 4.384 |
| mammals | Neoplatymops_mattogrossensis | Molossops_mattogrossensis | LC | Unknown | tropical | nocturnal | 6.8 | 6688972.688 | 4.012 |
| mammals | Neopteryx_frosti | Neopteryx_frosti | EN | Decreasing | tropical | nocturnal | 225 | 6744.001 | 7.135 |
| mammals | Neoromicia_anchietae | Pipistrellus_anchietae | LC | Unknown | tropical | nocturnal | 4.6 | 1457004.258 | 8.521 |
| mammals | Neoromicia_guineensis | Neoromicia_guineensis | LC | Unknown | tropical | nocturnal | 3.1 | 3582874.947 | 10.394 |
| mammals | Neoromicia_somalica | Neoromicia_somalica | LC | Unknown | tropical | nocturnal | 3.8 | 6070953.476 | 8.784 |
| mammals | Neoromicia_zuluensis | Neoromicia_zuluensis | LC | Unknown | tropical | nocturnal | 3.9 | 3479467.786 | 8.757 |
| mammals | Neotamias_alpinus | Neotamias_alpinus | LC | Stable | temperate | diurnal | 36.6 | 8864.498 | 2.2 |
| mammals | Neotamias_amoenus | Neotamias_amoenus | LC | Stable | temperate | diurnal | 50.5 | 1102204.161 | 8.39 |
| mammals | Neotamias_bulleri | Neotamias_bulleri | VU | Decreasing | tropical | diurnal | 100 | 15555.606 | 5.556 |
| mammals | Neotamias_canipes | Neotamias_canipes | LC | Stable | temperate | diurnal | 70.4 | 18918.833 | 9.035 |
| mammals | Neotamias_cinereicollis | Neotamias_cinereicollis | LC | Stable | temperate | diurnal | 61.7 | 74133.928 | 7.945 |
| mammals | Neotamias_dorsalis | Neotamias_dorsalis | LC | Stable | temperate | diurnal | 71.1 | 632158.149 | 8.031 |
| mammals | Neotamias_durangae | Neotamias_durangae | LC | Unknown | temperate | diurnal | 85 | 47076.706 | 5.006 |
| mammals | Neotamias_merriami | Neotamias_merriami | LC | Stable | temperate | diurnal | 74.8 | 72203.811 | 10.375 |
| mammals | Neotamias_minimus | Neotamias_minimus | LC | Stable | temperate | diurnal | 135.3 | 5243481.993 | 5.959 |
| mammals | Neotamias_obscurus | Neotamias_obscurus | LC | Stable | temperate | diurnal | 73 | 13472.167 | 5.394 |
| mammals | Neotamias_ochrogenys | Neotamias_ochrogenys | LC | Stable | temperate | diurnal | 91.7 | 9534.328 | 10.532 |
| mammals | Neotamias_palmeri | Neotamias_palmeri | EN | Decreasing | temperate | diurnal | 69.4 | 303.954 | 6.279 |
| mammals | Neotamias_panamintinus | Neotamias_panamintinus | LC | Stable | temperate | diurnal | 52.3 | 37380.508 | 5.504 |
| mammals | Neotamias_quadrimaculatus | Neotamias_quadrimaculatus | LC | Stable | temperate | diurnal | 85.2 | 25543.021 | 7.841 |
| mammals | Neotamias_quadrivittatus | Neotamias_quadrivittatus | LC | Stable | temperate | diurnal | 62.2 | 296948.736 | 9.553 |
| mammals | Neotamias_ruficaudus | Neotamias_ruficaudus | LC | Stable | temperate | diurnal | 68 | 147850.628 | 7.659 |

| | | | | | | | | | |
|---|---|---|---|---|---|---|---|---|---|
| mammals | Neotamias_rufus | Neotamias_rufus | LC | Stable | temperate | diurnal | 53.6 | 141470.692 | 8.688 |
| mammals | Neotamias_senex | Neotamias_senex | LC | Stable | temperate | diurnal | 89.3 | 99353.863 | 8.066 |
| mammals | Neotamias_siskiyou | Neotamias_siskiyou | LC | Stable | temperate | diurnal | 75 | 22528.817 | 8.969 |
| mammals | Neotamias_sonomae | Neotamias_sonomae | LC | Stable | temperate | diurnal | 75 | 45779.921 | 10.783 |
| mammals | Neotamias_speciosus | Neotamias_speciosus | LC | Stable | temperate | diurnal | 62 | 67800.855 | 8.139 |
| mammals | Neotamias_townsendii | Neotamias_townsendii | LC | Stable | temperate | diurnal | 74.8 | 151287.627 | 11.76 |
| mammals | Neotamias_umbrinus | Neotamias_umbrinus | LC | Stable | temperate | diurnal | 63 | 295485.183 | 7.548 |
| mammals | Neotoma_albigula | Neotoma_albigula | LC | Stable | temperate | nocturnal | 206 | 834340.746 | 7.849 |
| mammals | Neotoma_angustapalata | Neotoma_angustapalata | NT | Unknown | tropical | nocturnal | 198 | 3326.832 | 4.149 |
| mammals | Neotoma_bryanti | Neotoma_bryanti | LC | Decreasing | temperate | nocturnal | 182 | 271446.006 | 8.86 |
| mammals | Neotoma_chrysomelas | Neotoma_chrysomelas | LC | Stable | tropical | nocturnal | 169.5 | 30203.597 | 12.428 |
| mammals | Neotoma_cinerea | Neotoma_cinerea | LC | Stable | temperate | nocturnal | 299.2 | 3608407.828 | 7.271 |
| mammals | Neotoma_devia | Neotoma_devia | LC | Stable | temperate | nocturnal | 200 | 73683.608 | 9.181 |
| mammals | Neotoma_floridana | Neotoma_floridana | LC | Decreasing | temperate | nocturnal | 244.7 | 1871086.898 | 16.065 |
| mammals | Neotoma_fuscipes | Neotoma_fuscipes | LC | Stable | temperate | nocturnal | 229.8 | 171574.867 | 12.742 |
| mammals | Neotoma_goldmani | Neotoma_goldmani | LC | Decreasing | temperate | cathemeral | 198 | 170757.259 | 6.327 |
| mammals | Neotoma_lepida | Neotoma_lepida | LC | Stable | temperate | nocturnal | 163.8 | 675858.655 | 7.893 |
| mammals | Neotoma_leucodon | Neotoma_leucodon | LC | Stable | temperate | nocturnal | 251.1 | 1076957.591 | 9.622 |
| mammals | Neotoma_macrotis | Neotoma_macrotis | LC | Unknown | temperate | nocturnal | 198 | 160394.345 | 10.576 |
| mammals | Neotoma_magister | Neotoma_magister | NT | Decreasing | temperate | nocturnal | 412.5 | 313436.475 | 17.349 |
| mammals | Neotoma_mexicana | Neotoma_mexicana | LC | Stable | temperate | nocturnal | 203 | 1784429.133 | 9.249 |
| mammals | Neotoma_micropus | Neotoma_micropus | LC | Stable | temperate | nocturnal | 237.5 | 1063795.714 | 11.516 |
| mammals | Neotoma_nelsoni | Neotoma_nelsoni | CR | Decreasing | tropical | nocturnal | 198 | 45.828 | 23.729 |
| mammals | Neotoma_palatina | Neotoma_palatina | VU | Decreasing | tropical | nocturnal | 198 | 16059.981 | 5.981 |
| mammals | Neotoma_phenax | Neotoma_phenax | LC | Decreasing | temperate | nocturnal | 227.5 | 42633.375 | 10.822 |
| mammals | Neotoma_stephensi | Neotoma_stephensi | LC | Stable | temperate | nocturnal | 152.5 | 226739.185 | 8.796 |
| mammals | Neotomodon_alstoni | Neotomodon_alstoni | LC | Stable | tropical | nocturnal | 40 | 27485.604 | 18.455 |
| mammals | Neotomys_ebriosus | Neotomys_ebriosus | LC | Stable | tropical | cathemeral | 64.5 | 688578.677 | 6.593 |
| mammals | Neotragus_pygmaeus | Neotragus_pygmaeus | LC | Decreasing | tropical | nocturnal | 4000 | 455193.269 | 15.487 |
| mammals | Nephelomys_albigularis | Nephelomys_albigularis | LC | Decreasing | tropical | nocturnal | 60.5 | 398224.724 | 9.068 |
| mammals | Nephelomys_auriventer | Nephelomys_auriventer | LC | Unknown | tropical | cathemeral | 117.5 | 14415.635 | 5.558 |
| mammals | Nephelomys_caracolus | Nephelomys_caracolus | LC | Unknown | tropical | nocturnal | 93 | 4076.968 | 18.259 |
| mammals | Nephelomys_devius | Nephelomys_devius | LC | Stable | tropical | nocturnal | 150 | 9590.998 | 10.005 |
| mammals | Nephelomys_keaysi | Nephelomys_keaysi | LC | Decreasing | tropical | nocturnal | 163.2 | 67023.197 | 5.96 |

| | | | | | | | | | |
|---|---|---|---|---|---|---|---|---|---|
| mammals | Nephelomys_levipes | Nephelomys_levipes | LC | Decreasing | tropical | cathemeral | 112.8 | 157127.896 | 6.957 |
| mammals | Nephelomys_meridensis | Nephelomys_meridensis | LC | Stable | tropical | nocturnal | 99.3 | 14351.082 | 11.699 |
| mammals | Nesogale_dobsoni | Microgale_dobsoni | LC | Decreasing | tropical | nocturnal | 38 | 148370.959 | 8.305 |
| mammals | Nesogale_talazaci | Microgale_talazaci | LC | Decreasing | tropical | nocturnal | 48 | 92303.033 | 7.355 |
| mammals | Nesolagus_netscheri | NA | DD | Unknown | NA | NA | NA | NA | NA |
| mammals | Nesolagus_timminsi | Nesolagus_timminsi | EN | Decreasing | tropical | nocturnal | 2107.4 | 24821.829 | 6.556 |
| mammals | Nesomys_audeberti | Nesomys_audeberti | LC | Decreasing | tropical | diurnal | 212.5 | 90723.076 | 7.75 |
| mammals | Nesomys_lambertoni | Nesomys_lambertoni | EN | Unknown | tropical | diurnal | 155 | 2220.494 | 4.911 |
| mammals | Nesomys_rufus | Nesomys_rufus | LC | Unknown | tropical | diurnal | 162.5 | 106783.166 | 7.466 |
| mammals | Nesonycteris_fardoulisi | Melonycteris_fardoulisi | NT | Decreasing | tropical | nocturnal | 48.4 | 16829.601 | 4.752 |
| mammals | Nesonycteris_woodfordi | Melonycteris_woodfordi | LC | Unknown | tropical | nocturnal | 29.2 | 16782.922 | 4.849 |
| mammals | Nesoryzomys_fernandinae | NA | VU | Unknown | NA | NA | NA | NA | NA |
| mammals | Nesoryzomys_narboroughi | NA | VU | Decreasing | NA | NA | NA | NA | NA |
| mammals | Nesoryzomys_swarthi | NA | VU | Unknown | NA | NA | NA | NA | NA |
| mammals | Nesotragus_batesi | Neotragus_batesi | LC | Unknown | tropical | cathemeral | 3000 | 802403.341 | 5.932 |
| mammals | Nesotragus_moschatus | Nesotragus_moschatus | LC | Stable | tropical | diurnal | 6500 | 1370859 | 9.893 |
| mammals | Neurotrichus_gibbsii | Neurotrichus_gibbsii | LC | Stable | temperate | cathemeral | 8.9 | 221364.522 | 11.401 |
| mammals | Neusticomys_monticolus | Neusticomys_monticolus | LC | Decreasing | tropical | nocturnal | 39.5 | 65828.271 | 11.761 |
| mammals | Neusticomys_orcesi | NA | DD | Unknown | NA | NA | NA | NA | NA |
| mammals | Nilgiritragus_hylocrius | Nilgiritragus_hylocrius | EN | Decreasing | tropical | diurnal | 75000 | 2096.837 | 11.356 |
| mammals | Ningaui_ridei | Ningaui_ridei | LC | Stable | temperate | nocturnal | 8.5 | 2252519.904 | 2.131 |
| mammals | Ningaui_timealeyi | Ningaui_timealeyi | LC | Stable | tropical | nocturnal | 5.7 | 250377.148 | 4.398 |
| mammals | Ningaui_yvonneae | Ningaui_yvonneae | LC | Stable | temperate | nocturnal | 7 | 308584.026 | 4.112 |
| mammals | Niviventer_andersoni | Niviventer_andersoni | LC | Unknown | temperate | nocturnal | 147 | 188299.344 | 9.984 |
| mammals | Niviventer_brahma | Niviventer_brahma | LC | Stable | temperate | nocturnal | 97.3 | 32329.196 | 3.73 |
| mammals | Niviventer_cameroni | Niviventer_cameroni | VU | Decreasing | tropical | nocturnal | 86.8 | 9974.68 | 5.201 |
| mammals | Niviventer_confucianus | Niviventer_confucianus | LC | Stable | temperate | nocturnal | 65 | 3971381.007 | 15.273 |
| mammals | Niviventer_coninga | Niviventer_coninga | LC | Unknown | temperate | nocturnal | 142 | 28365.687 | 24.345 |
| mammals | Niviventer_cremoriventer | Niviventer_cremoriventer | LC | Unknown | tropical | nocturnal | 69.7 | 1248372.601 | 9.39 |
| mammals | Niviventer_culturatus | Niviventer_culturatus | LC | Unknown | tropical | nocturnal | 90.5 | 3736.757 | 4.241 |
| mammals | Niviventer_eha | Niviventer_eha | LC | Stable | temperate | nocturnal | 53.6 | 183562.824 | 6.192 |
| mammals | Niviventer_excelsior | Niviventer_excelsior | LC | Unknown | temperate | nocturnal | 102 | 150624.337 | 8.402 |
| mammals | Niviventer_fraternus | Niviventer_fraternus | LC | Unknown | tropical | nocturnal | 86.8 | 111954.436 | 7.703 |
| mammals | Niviventer_fulvescens | Niviventer_fulvescens | LC | Decreasing | tropical | nocturnal | 82.5 | 4720379.168 | 12.938 |

| | | | | | | | | | |
|---|---|---|---|---|---|---|---|---|---|
| mammals | Niviventer_hinpoon | Niviventer_hinpoon | EN | Unknown | tropical | nocturnal | 61 | 2929.085 | 10.229 |
| mammals | Niviventer_lepturus | Niviventer_lepturus | LC | Stable | tropical | nocturnal | 84.8 | 7576.82 | 16.122 |
| mammals | Niviventer_niviventer | Niviventer_niviventer | LC | Unknown | temperate | nocturnal | 100 | 252993.531 | 11.743 |
| mammals | Niviventer_rapit | Niviventer_rapit | LC | Unknown | tropical | nocturnal | 86.8 | 250763.404 | 2.938 |
| mammals | Niviventer_tenaster | Niviventer_tenaster | LC | Unknown | tropical | nocturnal | 103 | 266057.263 | 9.384 |
| mammals | Noctilio_albiventris | Noctilio_albiventris | LC | Stable | tropical | nocturnal | 30.8 | 12855674.662 | 5.627 |
| mammals | Noctilio_leporinus | Noctilio_leporinus | LC | Unknown | tropical | nocturnal | 58.3 | 14625648.252 | 6.139 |
| mammals | Nomascus_concolor | Nomascus_concolor | CR | Decreasing | tropical | diurnal | 7500 | 26326.314 | 8.011 |
| mammals | Nomascus_gabriellae | Nomascus_gabriellae | EN | Decreasing | tropical | diurnal | 8000 | 39684.779 | 10.288 |
| mammals | Nomascus_hainanus | Nomascus_hainanus | CR | Stable | tropical | diurnal | 7500 | 164.972 | 14.722 |
| mammals | Nomascus_leucogenys | Nomascus_leucogenys | CR | Decreasing | tropical | diurnal | 8000 | 51340.524 | 7.724 |
| mammals | Nomascus_nasutus | Nomascus_nasutus | CR | Decreasing | tropical | diurnal | 8000 | 8311.896 | 13.653 |
| mammals | Nomascus_siki | Nomascus_siki | CR | Decreasing | tropical | diurnal | 8000 | 26549.037 | 5.129 |
| mammals | Notamacropus_agilis | Macropus_agilis | LC | Decreasing | tropical | nocturnal | 15000 | 1325773.815 | 2.662 |
| mammals | Notamacropus_dorsalis | Macropus_dorsalis | LC | Decreasing | temperate | nocturnal | 11250 | 570362.344 | 6.721 |
| mammals | Notamacropus_eugenii | Macropus_eugenii | LC | Stable | temperate | nocturnal | 6500 | 16450.349 | 8.03 |
| mammals | Notamacropus_irma | Macropus_irma | LC | Stable | temperate | diurnal | 8000 | 152879.451 | 10.605 |
| mammals | Notamacropus_parma | Macropus_parma | NT | Unknown | temperate | nocturnal | 4500 | 50622.903 | 7.032 |
| mammals | Notamacropus_parryi | Macropus_parryi | LC | Stable | tropical | nocturnal | 13500 | 362773.542 | 6.785 |
| mammals | Notamacropus_rufogriseus | Macropus_rufogriseus | LC | Stable | temperate | nocturnal | 16850 | 652547.91 | 7.822 |
| mammals | Nothocricetulus_migratorius | Cricetulus_migratorius | LC | Unknown | temperate | nocturnal | 38.5 | 7677961.276 | 10.41 |
| mammals | Notiomys_edwardsii | Notiomys_edwardsii | LC | Stable | temperate | nocturnal | 21.3 | 267121.196 | 5.348 |
| mammals | Notiosorex_cockrumi | Notiosorex_cockrumi | LC | Stable | temperate | nocturnal | 4 | 66204.036 | 7.952 |
| mammals | Notiosorex_crawfordi | Notiosorex_crawfordi | LC | Stable | temperate | cathemeral | 4.4 | 2165720.112 | 9.727 |
| mammals | Notiosorex_evotis | Notiosorex_evotis | LC | Unknown | tropical | nocturnal | 5.4 | 89137.276 | 10.767 |
| mammals | Notocitellus_adocetus | Notocitellus_adocetus | LC | Stable | tropical | diurnal | 125 | 31693.547 | 9.562 |
| mammals | Notocitellus_annulatus | Notocitellus_annulatus | LC | Unknown | tropical | diurnal | 500 | 44731.304 | 9.449 |
| mammals | Notomys_alexis | Notomys_alexis | LC | Stable | temperate | nocturnal | 35 | 2927774.013 | 2.309 |
| mammals | Notomys_aquilo | Notomys_aquilo | EN | Decreasing | tropical | nocturnal | 39 | 2410.569 | 3.315 |
| mammals | Notomys_cervinus | Notomys_cervinus | NT | Stable | temperate | nocturnal | 35 | 83793.887 | 2.526 |
| mammals | Notomys_fuscus | Notomys_fuscus | VU | Unknown | temperate | nocturnal | 35 | 105460.258 | 2.395 |
| mammals | Notomys_mitchellii | Notomys_mitchellii | LC | Stable | temperate | nocturnal | 52 | 646728.271 | 5.477 |
| mammals | Notopteris_macdonaldii | Notopteris_macdonaldi | VU | Decreasing | tropical | nocturnal | 65.1 | 19268.554 | 9.061 |
| mammals | Notopteris_neocaledonicus | Notopteris_neocaledonica | EN | Decreasing | tropical | nocturnal | 45.1 | 1254.139 | 6.508 |

| | | | | | | | | | |
|---|---|---|---|---|---|---|---|---|---|
| mammals | Nyctalus_aviator | Nyctalus_aviator | NT | Decreasing | temperate | nocturnal | 38.3 | 846242.206 | 18.153 |
| mammals | Nyctalus_azoreum | Nyctalus_azoreum | VU | Decreasing | temperate | nocturnal | 7.1 | 2149.993 | 19.105 |
| mammals | Nyctalus_furvus | Nyctalus_furvus | EN | Decreasing | temperate | nocturnal | 27.6 | 11560.156 | 14.521 |
| mammals | Nyctalus_lasiopterus | Nyctalus_lasiopterus | VU | Decreasing | temperate | nocturnal | 49.7 | 2917259.325 | 16.517 |
| mammals | Nyctalus_leisleri | Nyctalus_leisleri | LC | Unknown | temperate | nocturnal | 13.3 | 5170891.767 | 18.605 |
| mammals | Nyctalus_montanus | Nyctalus_montanus | LC | Decreasing | temperate | nocturnal | 14.2 | 123577.43 | 13.648 |
| mammals | Nyctalus_noctula | Nyctalus_noctula | LC | Unknown | temperate | nocturnal | 28 | 7210092.693 | 16.451 |
| mammals | Nyctalus_plancyi | Nyctalus_plancyi | LC | Unknown | temperate | nocturnal | 24.1 | 2017209.653 | 16.458 |
| mammals | Nyctereutes_procyonoides | Nyctereutes_procyonoides | LC | Stable | temperate | cathemeral | 4040 | 11549507.057 | 13.759 |
| mammals | Nycteris_arge | Nycteris_arge | LC | Stable | tropical | nocturnal | 12 | 3980021.529 | 8.78 |
| mammals | Nycteris_aurita | Nycteris_aurita | LC | Unknown | tropical | nocturnal | 10 | 1789679.26 | 9.696 |
| mammals | Nycteris_gambiensis | Nycteris_gambiensis | LC | Unknown | tropical | nocturnal | 7 | 1861005.066 | 13.253 |
| mammals | Nycteris_grandis | Nycteris_grandis | LC | Decreasing | tropical | nocturnal | 30.2 | 4404698.714 | 8.882 |
| mammals | Nycteris_hispida | Nycteris_hispida | LC | Stable | tropical | nocturnal | 8.6 | 11511751.466 | 9.131 |
| mammals | Nycteris_intermedia | Nycteris_intermedia | LC | Decreasing | tropical | nocturnal | 10.7 | 2619716.274 | 7.386 |
| mammals | Nycteris_javanica | Nycteris_javanica | VU | Decreasing | tropical | nocturnal | 17.7 | 145230.845 | 21.988 |
| mammals | Nycteris_macrotis | Nycteris_macrotis | LC | Unknown | tropical | nocturnal | 14.5 | 12141283.975 | 9.61 |
| mammals | Nycteris_major | NA | DD | Unknown | NA | NA | NA | NA | NA |
| mammals | Nycteris_nana | Nycteris_nana | LC | Unknown | tropical | nocturnal | 7 | 2938142.019 | 7.238 |
| mammals | Nycteris_parisii | NA | DD | Unknown | NA | NA | NA | NA | NA |
| mammals | Nycteris_thebaica | Nycteris_thebaica | LC | Unknown | tropical | nocturnal | 9.8 | 14428246.95 | 9.645 |
| mammals | Nycteris_tragata | Nycteris_tragata | NT | Decreasing | tropical | nocturnal | 14.4 | 1394635.244 | 8.048 |
| mammals | Nycteris_woodi | Nycteris_woodi | LC | Decreasing | tropical | nocturnal | 7.6 | 620419.004 | 11.713 |
| mammals | Nycticebus_bengalensis | Nycticebus_bengalensis | EN | Decreasing | tropical | nocturnal | 1500 | 2049989.224 | 11.8 |
| mammals | Nycticebus_coucang | Nycticebus_coucang | EN | Decreasing | tropical | nocturnal | 803.5 | 441468.371 | 10.483 |
| mammals | Nycticebus_menagensis | Nycticebus_menagensis | VU | Decreasing | tropical | nocturnal | 803.5 | 727974.521 | 5.546 |
| mammals | Nycticeinops_crassulus | Pipistrellus_crassulus | LC | Unknown | tropical | nocturnal | 6.1 | 2601004.407 | 6.601 |
| mammals | Nycticeinops_eisentrauti | NA | DD | Unknown | NA | NA | NA | NA | NA |
| mammals | Nycticeinops_grandidieri | NA | DD | Unknown | NA | NA | NA | NA | NA |
| mammals | Nycticeinops_schlieffenii | Nycticeinops_schlieffeni | LC | Unknown | tropical | nocturnal | 5 | 5499366.939 | 10.14 |
| mammals | Nycticeius_cubanus | Nycticeius_cubanus | NT | Unknown | tropical | nocturnal | 5.5 | 29183.543 | 16.53 |
| mammals | Nycticeius_humeralis | Nycticeius_humeralis | LC | Stable | temperate | nocturnal | 9 | 2883260.061 | 17.353 |
| mammals | Nyctiellus_lepidus | Nyctiellus_lepidus | LC | Unknown | tropical | nocturnal | 2.4 | 110184.099 | 16.146 |
| mammals | Nyctimene_aello | Nyctimene_aello | LC | Stable | tropical | nocturnal | 84.3 | 621332.74 | 3.038 |

| | | | | | | | | | |
|---|---|---|---|---|---|---|---|---|---|
| mammals | Nyctimene_albiventer | Nyctimene_albiventer | LC | Stable | tropical | nocturnal | 29.9 | 748029.694 | 3.303 |
| mammals | Nyctimene_cephalotes | Nyctimene_cephalotes | LC | Unknown | tropical | nocturnal | 44.5 | 239455.094 | 9.558 |
| mammals | Nyctimene_certans | Nyctimene_certans | LC | Stable | tropical | nocturnal | 42.1 | 212379.588 | 4.385 |
| mammals | Nyctimene_cyclotis | NA | DD | Unknown | NA | NA | NA | NA | NA |
| mammals | Nyctimene_draconilla | NA | DD | Unknown | NA | NA | NA | NA | NA |
| mammals | Nyctimene_keasti | Nyctimene_keasti | NT | Decreasing | tropical | nocturnal | 44.6 | 47545.665 | 12.783 |
| mammals | Nyctimene_major | Nyctimene_major | LC | Unknown | tropical | nocturnal | 104.2 | 70315.077 | 4.725 |
| mammals | Nyctimene_malaitensis | Nyctimene_malaitensis | LC | Unknown | tropical | nocturnal | 75.4 | 7178.487 | 5.12 |
| mammals | Nyctimene_masalai | NA | DD | Unknown | NA | NA | NA | NA | NA |
| mammals | Nyctimene_rabori | Nyctimene_rabori | EN | Decreasing | tropical | nocturnal | 67.5 | 30770.514 | 17.947 |
| mammals | Nyctimene_robinsoni | Nyctimene_robinsoni | LC | Unknown | tropical | nocturnal | 49.5 | 351781.04 | 5.292 |
| mammals | Nyctimene_sanctacrucis | NA | DD | Unknown | NA | NA | NA | NA | NA |
| mammals | Nyctimene_varia | Nyctimene_minutus | VU | Decreasing | tropical | nocturnal | 33.1 | 1746.8 | 5.708 |
| mammals | Nyctimene_vizcaccia | Nyctimene_vizcaccia | LC | Unknown | tropical | nocturnal | 40 | 70176.73 | 4.566 |
| mammals | Nyctinomops_aurispinosus | Nyctinomops_aurispinosus | LC | Unknown | tropical | nocturnal | 18 | 3566231.211 | 10.748 |
| mammals | Nyctinomops_femorosaccus | Nyctinomops_femorosaccus | LC | Stable | temperate | nocturnal | 14.6 | 1251911.46 | 7.155 |
| mammals | Nyctinomops_laticaudatus | Nyctinomops_laticaudatus | LC | Unknown | tropical | nocturnal | 12.2 | 13101230.905 | 6.271 |
| mammals | Nyctinomops_macrotis | Nyctinomops_macrotis | LC | Unknown | tropical | nocturnal | 18.4 | 15743813.979 | 6.313 |
| mammals | Nyctomys_sumichrasti | Nyctomys_sumichrasti | LC | Stable | tropical | nocturnal | 49.4 | 673131.854 | 11.258 |
| mammals | Nyctophilus_arnhemensis | Nyctophilus_arnhemensis | LC | Stable | tropical | nocturnal | 6.7 | 704005.75 | 2.141 |
| mammals | Nyctophilus_bifax | Nyctophilus_bifax | LC | Decreasing | tropical | nocturnal | 9.8 | 400008.966 | 5.033 |
| mammals | Nyctophilus_geoffroyi | Nyctophilus_geoffroyi | LC | Stable | temperate | nocturnal | 7.3 | 7272506.251 | 4.276 |
| mammals | Nyctophilus_gouldi | Nyctophilus_gouldi | LC | Decreasing | temperate | nocturnal | 10 | 1499403.091 | 7.91 |
| mammals | Nyctophilus_heran | NA | DD | Unknown | NA | NA | NA | NA | NA |
| mammals | Nyctophilus_microdon | Nyctophilus_microdon | LC | Unknown | tropical | nocturnal | 8 | 32941.357 | 6.146 |
| mammals | Nyctophilus_microtis | Nyctophilus_microtis | LC | Stable | tropical | nocturnal | 7.1 | 166521.905 | 4.402 |
| mammals | Nyctophilus_sherrini | Nyctophilus_sherrini | VU | Decreasing | temperate | nocturnal | 15.5 | 48998.26 | 7.109 |
| mammals | Nyctophilus_walkeri | Nyctophilus_walkeri | LC | Stable | tropical | nocturnal | 4.3 | 397742.811 | 1.754 |
| mammals | Ochotona_alpina | Ochotona_alpina | LC | Stable | temperate | diurnal | 263 | 575019.438 | 3.765 |
| mammals | Ochotona_cansus | Ochotona_cansus | LC | Unknown | temperate | diurnal | 74.5 | 700340.63 | 10.024 |
| mammals | Ochotona_collaris | Ochotona_collaris | LC | Unknown | temperate | diurnal | 129 | 846284.432 | 0.708 |
| mammals | Ochotona_coreana | NA | DD | Unknown | NA | NA | NA | NA | NA |
| mammals | Ochotona_curzoniae | Ochotona_curzoniae | LC | Decreasing | temperate | diurnal | 162.5 | 2250961.787 | 3.982 |
| mammals | Ochotona_dauurica | Ochotona_dauurica | LC | Unknown | temperate | diurnal | 130 | 2054450.612 | 7.774 |

| | | | | | | | | | |
|---|---|---|---|---|---|---|---|---|---|
| mammals | Ochotona_erythrotis | Ochotona_erythrotis | LC | Unknown | temperate | diurnal | 268 | 961201.752 | 6.351 |
| mammals | Ochotona_forresti | Ochotona_forresti | LC | Unknown | temperate | cathemeral | 129 | 38532.763 | 4.526 |
| mammals | Ochotona_hoffmanni | Ochotona_hoffmanni | EN | Unknown | temperate | diurnal | 285 | 1241.7 | 5.503 |
| mammals | Ochotona_hyperborea | Ochotona_hyperborea | LC | Stable | temperate | diurnal | 156 | 8990041.403 | 1.686 |
| mammals | Ochotona_iliensis | Ochotona_iliensis | EN | Decreasing | temperate | cathemeral | 233.5 | 12780.675 | 3.285 |
| mammals | Ochotona_koslowi | Ochotona_koslowi | EN | Decreasing | temperate | diurnal | 165 | 44741.585 | 0.25 |
| mammals | Ochotona_ladacensis | Ochotona_ladacensis | LC | Unknown | temperate | diurnal | 239 | 786588.389 | 1.255 |
| mammals | Ochotona_macrotis | Ochotona_macrotis | LC | Unknown | temperate | diurnal | 166 | 2931184.447 | 4.117 |
| mammals | Ochotona_mantchurica | Ochotona_mantchurica | LC | Unknown | temperate | diurnal | 160 | 336661.148 | 4.656 |
| mammals | Ochotona_nubrica | Ochotona_nubrica | LC | Unknown | temperate | diurnal | 115.5 | 491702.49 | 4.591 |
| mammals | Ochotona_opaca | Ochotona_opaca | LC | Stable | temperate | cathemeral | 136.5 | 179949.759 | 5.915 |
| mammals | Ochotona_pallasii | Ochotona_pallasii | LC | Stable | temperate | diurnal | 214 | 941438.89 | 4.654 |
| mammals | Ochotona_princeps | Ochotona_princeps | LC | Decreasing | temperate | diurnal | 157.6 | 894668.897 | 6.575 |
| mammals | Ochotona_pusilla | Ochotona_pusilla | LC | Decreasing | temperate | nocturnal | 142.5 | 1530161.497 | 8.963 |
| mammals | Ochotona_roylii | Ochotona_roylei | LC | Stable | temperate | diurnal | 260 | 493849.813 | 5.056 |
| mammals | Ochotona_rufescens | Ochotona_rufescens | LC | Stable | temperate | diurnal | 250 | 946121.476 | 10.559 |
| mammals | Ochotona_rutila | Ochotona_rutila | LC | Stable | temperate | diurnal | 270 | 206939.191 | 6.965 |
| mammals | Ochotona_thibetana | Ochotona_thibetana | LC | Unknown | temperate | diurnal | 104 | 729354.295 | 7.763 |
| mammals | Ochotona_thomasi | Ochotona_thomasi | LC | Unknown | temperate | diurnal | 77.5 | 232122.673 | 6.734 |
| mammals | Ochotona_turuchanensis | Ochotona_turuchanensis | LC | Unknown | temperate | diurnal | 148.2 | 1642659.489 | 1.209 |
| mammals | Ochromyscus_brockmani | Myomyscus_brockmani | LC | Stable | tropical | nocturnal | 29.5 | 1713270.655 | 7.962 |
| mammals | Ochromyscus_yemeni | NA | DD | Unknown | NA | NA | NA | NA | NA |
| mammals | Ochrotomys_nuttalli | Ochrotomys_nuttalli | LC | Stable | temperate | nocturnal | 22.4 | 1444556.94 | 16.222 |
| mammals | Octodon_bridgesii | Octodon_bridgesi | VU | Decreasing | temperate | nocturnal | 150 | 120967.758 | 9.067 |
| mammals | Octodon_degus | Octodon_degus | LC | Unknown | temperate | diurnal | 210 | 63501.347 | 12.65 |
| mammals | Octodon_lunatus | Octodon_lunatus | NT | Decreasing | temperate | cathemeral | 233 | 25576.987 | 13.116 |
| mammals | Octodon_pacificus | Octodon_pacificus | CR | Decreasing | temperate | diurnal | 290 | 53.041 | 12.811 |
| mammals | Octodontomys_gliroides | Octodontomys_gliroides | LC | Stable | tropical | nocturnal | 150 | 487312.915 | 5.986 |
| mammals | Octomys_mimax | Octomys_mimax | LC | Unknown | temperate | nocturnal | 131 | 118944.764 | 3.208 |
| mammals | Oecomys_bicolor | Oecomys_bicolor | LC | Unknown | tropical | nocturnal | 34 | 8594483.012 | 3.466 |
| mammals | Oecomys_flavicans | Oecomys_flavicans | LC | Stable | tropical | nocturnal | 73.4 | 42577.437 | 14.817 |
| mammals | Oecomys_mamorae | Oecomys_mamorae | LC | Stable | tropical | nocturnal | 62.5 | 651947.359 | 5.577 |
| mammals | Oecomys_phaeotis | Oecomys_phaeotis | LC | Stable | tropical | nocturnal | 73.4 | 117475.218 | 3.27 |
| mammals | Oecomys_speciosus | Oecomys_speciosus | LC | Stable | tropical | nocturnal | 73.4 | 380157.556 | 7.495 |

| | | | | | | | | | |
|---|---|---|---|---|---|---|---|---|---|
| mammals | Oecomys_superans | Oecomys_superans | LC | Stable | tropical | nocturnal | 73.4 | 1547918.826 | 1.647 |
| mammals | Oecomys_trinitatis | Oecomys_trinitatis | LC | Stable | tropical | nocturnal | 73.4 | 4830193.884 | 5.968 |
| mammals | Oenomys_hypoxanthus | Oenomys_hypoxanthus | LC | Stable | tropical | cathemeral | 90 | 4172195.574 | 7.772 |
| mammals | Oenomys_ornatus | NA | DD | Unknown | NA | NA | NA | NA | NA |
| mammals | Okapia_johnstoni | Okapia_johnstoni | EN | Decreasing | tropical | diurnal | 230001.1 | 264220.215 | 4.099 |
| mammals | Olallamys_albicauda | NA | DD | Unknown | NA | NA | NA | NA | NA |
| mammals | Olallamys_edax | NA | DD | Unknown | NA | NA | NA | NA | NA |
| mammals | Oligoryzomys_andinus | Oligoryzomys_andinus | LC | Stable | tropical | nocturnal | 25.2 | 260991.295 | 8.204 |
| mammals | Oligoryzomys_arenalis | Oligoryzomys_arenalis | LC | Decreasing | tropical | nocturnal | 25.2 | 96928.401 | 6.561 |
| mammals | Oligoryzomys_brendae | NA | DD | Unknown | NA | NA | NA | NA | NA |
| mammals | Oligoryzomys_chacoensis | Oligoryzomys_chacoensis | LC | Stable | tropical | nocturnal | 23 | 760937.185 | 5.162 |
| mammals | Oligoryzomys_destructor | Oligoryzomys_destructor | LC | Stable | tropical | nocturnal | 25.2 | 661051.395 | 7.21 |
| mammals | Oligoryzomys_flavescens | Oligoryzomys_flavescens | LC | Stable | tropical | nocturnal | 21.3 | 4603205.002 | 10.504 |
| mammals | Oligoryzomys_fulvescens | Oligoryzomys_fulvescens | LC | Stable | tropical | nocturnal | 25 | 2895296.895 | 6.762 |
| mammals | Oligoryzomys_griseolus | Oligoryzomys_griseolus | LC | Decreasing | tropical | nocturnal | 25.2 | 52670.17 | 11.552 |
| mammals | Oligoryzomys_longicaudatus | Oligoryzomys_longicaudatus | LC | Stable | temperate | nocturnal | 27 | 745244.388 | 5.856 |
| mammals | Oligoryzomys_microtis | Oligoryzomys_microtis | LC | Stable | tropical | nocturnal | 22.5 | 2722586.198 | 2.645 |
| mammals | Oligoryzomys_nigripes | Oligoryzomys_nigripes | LC | Stable | tropical | nocturnal | 20.5 | 3221464.717 | 11.308 |
| mammals | Oligoryzomys_stramineus | Oligoryzomys_stramineus | LC | Unknown | tropical | nocturnal | 25 | 530369.854 | 8.655 |
| mammals | Oligoryzomys_vegetus | Oligoryzomys_vegetus | LC | Stable | tropical | nocturnal | 15 | 9901.691 | 9.13 |
| mammals | Ondatra_zibethicus | Ondatra_zibethicus | LC | Stable | temperate | cathemeral | 1065.7 | 35093842.141 | 7.933 |
| mammals | Onychogalea_frenata | Onychogalea_fraenata | VU | Stable | temperate | nocturnal | 5500 | 4205.866 | 4.84 |
| mammals | Onychogalea_unguifera | Onychogalea_unguifera | LC | Unknown | tropical | nocturnal | 6650 | 951054.222 | 2.348 |
| mammals | Onychomys_arenicola | Onychomys_arenicola | LC | Stable | temperate | nocturnal | 30 | 686486.362 | 7.556 |
| mammals | Onychomys_leucogaster | Onychomys_leucogaster | LC | Stable | temperate | nocturnal | 27.9 | 3841451.168 | 11.942 |
| mammals | Onychomys_torridus | Onychomys_torridus | LC | Stable | temperate | nocturnal | 25 | 767649.884 | 8.304 |
| mammals | Oreamnos_americanus | Oreamnos_americanus | LC | Stable | temperate | cathemeral | 72500.3 | 838023.448 | 2.261 |
| mammals | Oreoryzomys_balneator | NA | DD | Unknown | NA | NA | NA | NA | NA |
| mammals | Oreoscaptor_mizura | Euroscaptor_mizura | LC | Decreasing | temperate | cathemeral | 35.5 | 15898.037 | 11.891 |
| mammals | Oreotragus_oreotragus | Oreotragus_oreotragus | LC | Stable | tropical | diurnal | 12999.9 | 5220692.402 | 9.633 |
| mammals | Orientallactaga_balikunica | Allactaga_balikunica | LC | Unknown | temperate | nocturnal | 72.5 | 234834.395 | 1.947 |
| mammals | Orientallactaga_bullata | Allactaga_bullata | LC | Unknown | temperate | nocturnal | 87 | 1563011.218 | 3.974 |
| mammals | Orientallactaga_sibirica | Allactaga_sibirica | LC | Unknown | temperate | nocturnal | 97.5 | 5412810.797 | 6.18 |
| mammals | Ornithorhynchus_anatinus | Ornithorhynchus_anatinus | NT | Decreasing | temperate | cathemeral | 1484.3 | 901638.005 | 8.319 |

| | | | | | | | | | |
|---|---|---|---|---|---|---|---|---|---|
| mammals | Orthogeomys_grandis | Orthogeomys_cuniculus | LC | Stable | tropical | cathemeral | 500 | 140934.705 | 11.102 |
| mammals | Orycteropus_afer | Orycteropus_afer | LC | Unknown | tropical | nocturnal | 52350.4 | 19273002.491 | 8.966 |
| mammals | Oryctolagus_cuniculus | Oryctolagus_cuniculus | EN | Decreasing | temperate | cathemeral | 1958.3 | 634229.133 | 17.457 |
| mammals | Oryx_beisa | Oryx_beisa | EN | Decreasing | tropical | cathemeral | 170000 | 975973.872 | 6.89 |
| mammals | Oryx_gazella | Oryx_gazella | LC | Stable | temperate | cathemeral | 169001.3 | 1766027.324 | 5.677 |
| mammals | Oryx_leucoryx | Oryx_leucoryx | VU | Stable | tropical | cathemeral | 87700 | 20909.658 | 2.404 |
| mammals | Oryzomys_couesi | Oryzomys_couesi | LC | Unknown | tropical | nocturnal | 69.3 | 1491857.118 | 10.959 |
| mammals | Oryzomys_dimidiatus | NA | DD | Unknown | NA | NA | NA | NA | NA |
| mammals | Oryzomys_gorgasi | Oryzomys_gorgasi | EN | Decreasing | tropical | cathemeral | 60.5 | 161378.005 | 14.156 |
| mammals | Oryzomys_palustris | Oryzomys_palustris | LC | Stable | temperate | nocturnal | 53.9 | 1508984.729 | 16.636 |
| mammals | Oryzorictes_hova | Oryzorictes_hova | LC | Decreasing | tropical | nocturnal | 36.5 | 124191.677 | 7.889 |
| mammals | Oryzorictes_tetradactylus | NA | DD | Unknown | NA | NA | NA | NA | NA |
| mammals | Osgoodomys_banderanus | Osgoodomys_banderanus | LC | Unknown | tropical | nocturnal | 50 | 179803.687 | 9.791 |
| mammals | Osphranter_antilopinus | Macropus_antilopinus | LC | Decreasing | tropical | nocturnal | 27250 | 653127.964 | 2.004 |
| mammals | Osphranter_bernardus | Macropus_bernardus | NT | Unknown | tropical | nocturnal | 17000 | 30111.041 | 0.611 |
| mammals | Osphranter_robustus | Macropus_robustus | LC | Stable | temperate | nocturnal | 21262.5 | 6349650.737 | 3.876 |
| mammals | Osphranter_rufus | Macropus_rufus | LC | Stable | temperate | nocturnal | 46250 | 5360863.501 | 3.425 |
| mammals | Otocolobus_manul | Otocolobus_manul | LC | Decreasing | temperate | cathemeral | 3500 | 8387217.253 | 6.394 |
| mammals | Otocyon_megalotis | Otocyon_megalotis | LC | Stable | temperate | nocturnal | 4150 | 4957043.238 | 8.155 |
| mammals | Otohylomys_megalotis | NA | DD | Unknown | NA | NA | NA | NA | NA |
| mammals | Otolemur_crassicaudatus | Otolemur_crassicaudatus | LC | Stable | tropical | nocturnal | 1500 | 4845511.434 | 8.86 |
| mammals | Otolemur_garnettii | Otolemur_garnettii | LC | Decreasing | tropical | nocturnal | 760 | 568222.413 | 10.645 |
| mammals | Otomops_formosus | NA | DD | Unknown | NA | NA | NA | NA | NA |
| mammals | Otomops_johnstonei | NA | DD | Unknown | NA | NA | NA | NA | NA |
| mammals | Otomops_madagascariensis | Otomops_madagascariensis | LC | Unknown | tropical | nocturnal | 25.5 | 3539.5 | 5.235 |
| mammals | Otomops_martiensseni | Otomops_martiensseni | NT | Decreasing | tropical | nocturnal | 34.3 | 5186383.984 | 8.632 |
| mammals | Otomops_papuensis | NA | DD | Unknown | NA | NA | NA | NA | NA |
| mammals | Otomops_wroughtoni | NA | DD | Unknown | NA | NA | NA | NA | NA |
| mammals | Otomys_anchietae | Otomys_anchietae | LC | Unknown | tropical | diurnal | 206.5 | 71608.939 | 8.069 |
| mammals | Otomys_angoniensis | Otomys_angoniensis | LC | Stable | tropical | cathemeral | 121.9 | 595014.774 | 10.911 |
| mammals | Otomys_barbouri | Otomys_barbouri | EN | Decreasing | tropical | diurnal | 107.4 | 398.1 | 7.187 |
| mammals | Otomys_burtoni | Otomys_burtoni | EN | Decreasing | tropical | diurnal | 90 | 383.856 | 8.809 |
| mammals | Otomys_cuanzensis | Otomys_cuanzensis | LC | Stable | tropical | diurnal | 90.2 | 115191.666 | 7.955 |
| mammals | Otomys_denti | Otomys_denti | LC | Decreasing | tropical | diurnal | 120 | 80971.542 | 16.929 |

| | | | | | | | | | |
|---|---|---|---|---|---|---|---|---|---|
| mammals | Otomys_irroratus | Otomys_irroratus | LC | Decreasing | temperate | diurnal | 118 | 183846.805 | 9.821 |
| mammals | Otomys_karoensis | Otomys_saundersiae | LC | Stable | temperate | diurnal | 103 | 254423.212 | 10.139 |
| mammals | Otomys_lacustris | Otomys_lacustris | LC | Decreasing | tropical | diurnal | 92.6 | 34134.569 | 13.41 |
| mammals | Otomys_laminatus | Otomys_laminatus | NT | Decreasing | temperate | diurnal | 145 | 86794.597 | 15.003 |
| mammals | Otomys_occidentalis | Otomys_occidentalis | VU | Decreasing | tropical | diurnal | 72.8 | 10690.744 | 8.422 |
| mammals | Otomys_sloggetti | Otomys_sloggetti | LC | Increasing | temperate | diurnal | 106.3 | 156904.327 | 10.736 |
| mammals | Otomys_tropicalis | Otomys_tropicalis | LC | Stable | tropical | diurnal | 108.8 | 143756.978 | 16.486 |
| mammals | Otomys_typus | Otomys_typus | LC | Unknown | tropical | cathemeral | 133.4 | 96218.784 | 12.916 |
| mammals | Otomys_unisulcatus | Otomys_unisulcatus | LC | Stable | temperate | diurnal | 124.5 | 374301.339 | 7.867 |
| mammals | Otonycteris_hemprichii | Otonycteris_hemprichii | LC | Unknown | temperate | nocturnal | 23.8 | 1283936.359 | 8.811 |
| mammals | Otonyctomys_hatti | Otonyctomys_hatti | LC | Unknown | tropical | nocturnal | 36.2 | 127970.466 | 6.523 |
| mammals | Otopteropus_cartilagonodus | Otopteropus_cartilagonodus | LC | Decreasing | tropical | nocturnal | 16.8 | 105071.322 | 14.89 |
| mammals | Otospermophilus_beecheyi | Otospermophilus_beecheyi | LC | Stable | temperate | diurnal | 578.5 | 467246.645 | 12.01 |
| mammals | Otospermophilus_variegatus | Otospermophilus_variegatus | LC | Stable | temperate | diurnal | 691.6 | 2304576.892 | 9.114 |
| mammals | Ototylomys_phyllotis | Ototylomys_phyllotis | LC | Stable | tropical | nocturnal | 120 | 491821.325 | 10.062 |
| mammals | Ourebia_ourebi | Ourebia_ourebi | LC | Decreasing | tropical | cathemeral | 17250 | 7497157.678 | 10.006 |
| mammals | Ovibos_moschatus | Ovibos_moschatus | LC | Decreasing | temperate | cathemeral | 340501.1 | 3760945.123 | 0.232 |
| mammals | Ovis_dalli | Ovis_dalli | LC | Stable | temperate | diurnal | 55650.6 | 816225.071 | 0.486 |
| mammals | Ovis_nivicola | Ovis_nivicola | LC | Unknown | temperate | diurnal | 90000 | 1369336.66 | 0.758 |
| mammals | Oxymycterus_amazonicus | Oxymycterus_amazonicus | LC | Unknown | tropical | nocturnal | 25.2 | 775764.675 | 1.916 |
| mammals | Oxymycterus_caparaoe | Oxymycterus_caparoae | LC | Unknown | tropical | diurnal | 66.4 | 63202.747 | 13.404 |
| mammals | Oxymycterus_dasytrichus | Oxymycterus_dasytrichus | LC | Stable | tropical | cathemeral | 72.5 | 1718385.511 | 11.102 |
| mammals | Oxymycterus_delator | Oxymycterus_delator | LC | Stable | temperate | diurnal | 81.5 | 82738.639 | 13.39 |
| mammals | Oxymycterus_hiska | Oxymycterus_hiska | LC | Stable | tropical | cathemeral | 12.3 | 34290.912 | 5.68 |
| mammals | Oxymycterus_hucucha | Oxymycterus_hucucha | EN | Decreasing | tropical | cathemeral | 68 | 3965.437 | 7.951 |
| mammals | Oxymycterus_inca | Oxymycterus_inca | LC | Stable | tropical | diurnal | 35 | 360850.754 | 5.419 |
| mammals | Oxymycterus_josei | Oxymycterus_josei | NT | Decreasing | temperate | diurnal | 66.4 | 22829.997 | 13.783 |
| mammals | Oxymycterus_nasutus | Oxymycterus_nasutus | LC | Unknown | temperate | diurnal | 68 | 558988.936 | 12.223 |
| mammals | Oxymycterus_paramensis | Oxymycterus_paramensis | LC | Stable | tropical | cathemeral | 42 | 286552.677 | 7.651 |
| mammals | Oxymycterus_quaestor | Oxymycterus_quaestor | LC | Unknown | temperate | diurnal | 75.5 | 784234.915 | 13.616 |
| mammals | Oxymycterus_rufus | Oxymycterus_rufus | LC | Decreasing | temperate | cathemeral | 75.4 | 280710.357 | 10.892 |
| mammals | Ozimops_beccarii | Mormopterus_beccarii | LC | Unknown | tropical | nocturnal | 14.9 | 90003.553 | 6.252 |
| mammals | Ozimops_loriae | Mormopterus_loriae | DD | Stable | tropical | nocturnal | 8.9 | 3195.173 | 10.127 |
| mammals | Ozimops_planiceps | Mormopterus_planiceps | LC | Decreasing | temperate | nocturnal | 8.8 | 3437648.755 | 5.782 |

| | | | | | | | | | |
|---|---|---|---|---|---|---|---|---|---|
| mammals | Ozotoceros_bezoarticus | Ozotoceros_bezoarticus | NT | Decreasing | tropical | cathemeral | 40000 | 468419.422 | 5.809 |
| mammals | Pachyuromys_duprasi | Pachyuromys_duprasi | LC | Stable | temperate | nocturnal | 47.5 | 1305337.932 | 8.232 |
| mammals | Paguma_larvata | Paguma_larvata | LC | Decreasing | tropical | nocturnal | 4300 | 6472276.596 | 11.776 |
| mammals | Palawanomys_furvus | Palawanomys_furvus | LC | Unknown | tropical | nocturnal | 102.5 | 878.13 | 4.474 |
| mammals | Palawanosorex_ater | NA | DD | Unknown | NA | NA | NA | NA | NA |
| mammals | Pan_paniscus | Pan_paniscus | EN | Decreasing | tropical | diurnal | 34000.1 | 415766.365 | 4.37 |
| mammals | Pan_troglodytes | Pan_troglodytes | EN | Decreasing | tropical | diurnal | 42500 | 2314271.999 | 7.299 |
| mammals | Panthera_leo | Panthera_leo | VU | Decreasing | tropical | cathemeral | 161499.1 | 26107440.051 | 8.889 |
| mammals | Panthera_onca | Panthera_onca | NT | Decreasing | tropical | cathemeral | 100000 | 9025908.691 | 3.24 |
| mammals | Panthera_pardus | Panthera_pardus | VU | Decreasing | tropical | cathemeral | 54999.7 | 34544707.146 | 11.194 |
| mammals | Panthera_tigris | Panthera_tigris | EN | Decreasing | temperate | cathemeral | 162564 | 11805900.961 | 14.823 |
| mammals | Panthera_uncia | Panthera_uncia | VU | Decreasing | temperate | cathemeral | 44167 | 3496431.337 | 4.384 |
| mammals | Papagomys_armandvillei | Papagomys_armandvillei | NT | Decreasing | tropical | nocturnal | 1048.1 | 13936.036 | 12.484 |
| mammals | Papio_anubis | Papio_anubis | LC | Stable | tropical | diurnal | 28329.2 | 7814659.709 | 10.218 |
| mammals | Papio_cynocephalus | Papio_cynocephalus | LC | Stable | tropical | diurnal | 19484 | 1667455.195 | 10.329 |
| mammals | Papio_hamadryas | Papio_hamadryas | LC | Increasing | tropical | diurnal | 14972.2 | 610339.022 | 9.501 |
| mammals | Papio_kindae | Papio_kindae | LC | Stable | tropical | diurnal | 13000 | 2214120.948 | 6.974 |
| mammals | Papio_papio | Papio_papio | NT | Decreasing | tropical | diurnal | 15500.2 | 494995.574 | 10.686 |
| mammals | Papio_ursinus | Papio_ursinus | LC | Decreasing | temperate | diurnal | 27939.1 | 3348995.376 | 8.532 |
| mammals | Parablarinella_griselda | Blarinella_griselda | LC | Unknown | temperate | nocturnal | 8 | 439053.96 | 13.18 |
| mammals | Parachoerus_wagneri | Catagonus_wagneri | EN | Decreasing | tropical | diurnal | 35566.4 | 304464.107 | 4.349 |
| mammals | Paracrocidura_graueri | NA | DD | Unknown | NA | NA | NA | NA | NA |
| mammals | Paracrocidura_maxima | Paracrocidura_maxima | LC | Decreasing | tropical | nocturnal | 16 | 23382.417 | 14.38 |
| mammals | Paracrocidura_schoutedeni | Paracrocidura_schoutedeni | LC | Unknown | tropical | nocturnal | 13 | 671227.208 | 5.972 |
| mammals | Paracynictis_selousi | Paracynictis_selousi | LC | Unknown | tropical | nocturnal | 1640 | 1796820.965 | 8.361 |
| mammals | Paradipus_ctenodactylus | Paradipus_ctenodactylus | LC | Unknown | temperate | nocturnal | 109 | 386135.515 | 5.589 |
| mammals | Paradoxurus_hermaphroditus | Paradoxurus_hermaphroditus | LC | Decreasing | tropical | nocturnal | 3156.7 | 7641014.956 | 13.869 |
| mammals | Paradoxurus_jerdoni | Paradoxurus_jerdoni | LC | Stable | tropical | nocturnal | 2780.8 | 66739.957 | 13.215 |
| mammals | Paradoxurus_zeylonensis | Paradoxurus_zeylonensis | LC | Unknown | tropical | nocturnal | 2780.8 | 18391.662 | 19.866 |
| mammals | Paraechinus_aethiopicus | Paraechinus_aethiopicus | LC | Stable | temperate | nocturnal | 188 | 3874413.08 | 5.611 |
| mammals | Paraechinus_hypomelas | Paraechinus_hypomelas | LC | Unknown | temperate | nocturnal | 213 | 2536454.331 | 8.162 |
| mammals | Paraechinus_micropus | Paraechinus_micropus | LC | Decreasing | temperate | nocturnal | 171 | 414921.362 | 13.536 |
| mammals | Paraechinus_nudiventris | Paraechinus_nudiventris | LC | Unknown | tropical | nocturnal | 175.3 | 9534.749 | 21.767 |
| mammals | Paragalago_cocos | Galagoides_cocos | LC | Decreasing | tropical | nocturnal | 144 | 9853.783 | 12.647 |

| | | | | | | | | | |
|---|---|---|---|---|---|---|---|---|---|
| mammals | Paragalago_granti | Galagoides_granti | LC | Decreasing | tropical | nocturnal | 158.7 | 367704.733 | 9.029 |
| mammals | Paragalago_orinus | Galagoides_orinus | VU | Decreasing | tropical | nocturnal | 97 | 18913.305 | 13.337 |
| mammals | Paragalago_rondoensis | Galagoides_rondoensis | EN | Decreasing | tropical | nocturnal | 84.2 | 126.804 | 12.617 |
| mammals | Paragalago_zanzibaricus | Galagoides_zanzibaricus | NT | Decreasing | tropical | nocturnal | 100.5 | 7077.55 | 15.586 |
| mammals | Parahyaena_brunnea | Parahyaena_brunnea | NT | Stable | temperate | nocturnal | 32200.3 | 2290722.985 | 7.182 |
| mammals | Paraleptomys_rufilatus | Paraleptomys_rufilatus | EN | Unknown | tropical | nocturnal | 55 | 586.356 | 3.764 |
| mammals | Paraleptomys_wilhelmina | NA | DD | Unknown | NA | NA | NA | NA | NA |
| mammals | Paramelomys_levipes | Paramelomys_levipes | LC | Unknown | tropical | nocturnal | 86.7 | 27271.383 | 3.393 |
| mammals | Paramelomys_lorentzii | Paramelomys_lorentzii | LC | Unknown | tropical | nocturnal | 150 | 175784.197 | 2.243 |
| mammals | Paramelomys_mollis | Paramelomys_mollis | LC | Stable | tropical | nocturnal | 90.1 | 225112.951 | 4.371 |
| mammals | Paramelomys_moncktoni | Paramelomys_moncktoni | LC | Unknown | tropical | nocturnal | 95.4 | 79468.891 | 4.911 |
| mammals | Paramelomys_naso | Paramelomys_naso | LC | Unknown | tropical | nocturnal | 129.7 | 115422.817 | 2.456 |
| mammals | Paramelomys_platyops | Paramelomys_platyops | LC | Stable | tropical | nocturnal | 104.2 | 496373.431 | 3.336 |
| mammals | Paramelomys_rubex | Paramelomys_rubex | LC | Stable | tropical | nocturnal | 46.6 | 234666.787 | 4.422 |
| mammals | Paramelomys_steini | NA | DD | Unknown | NA | NA | NA | NA | NA |
| mammals | Parantechinus_apicalis | Parantechinus_apicalis | EN | Decreasing | temperate | nocturnal | 68.8 | 2838.735 | 4.487 |
| mammals | Paranyctimene_raptor | Paranyctimene_raptor | LC | Unknown | tropical | nocturnal | 24.3 | 356130.327 | 3.205 |
| mammals | Parastrellus_hesperus | Pipistrellus_hesperus | LC | Stable | temperate | nocturnal | 3.7 | 2686280.323 | 8.857 |
| mammals | Paratriaenops_auritus | Paratriaenops_auritus | VU | Unknown | tropical | nocturnal | 5 | 10567.577 | 9.042 |
| mammals | Paratriaenops_furcula | Paratriaenops_furculus | LC | Unknown | tropical | nocturnal | 5.5 | 57158.769 | 8.05 |
| mammals | Paraxerus_alexandri | Paraxerus_alexandri | LC | Unknown | tropical | diurnal | 52 | 259068.847 | 7.448 |
| mammals | Paraxerus_boehmi | Paraxerus_boehmi | LC | Unknown | tropical | diurnal | 70 | 956025.61 | 8.631 |
| mammals | Paraxerus_cepapi | Paraxerus_cepapi | LC | Stable | tropical | diurnal | 180 | 2692246.213 | 8.533 |
| mammals | Paraxerus_cooperi | NA | DD | Unknown | NA | NA | NA | NA | NA |
| mammals | Paraxerus_flavovittis | Paraxerus_flavovittis | LC | Unknown | tropical | diurnal | 160 | 666787.981 | 9.55 |
| mammals | Paraxerus_lucifer | NA | DD | Unknown | NA | NA | NA | NA | NA |
| mammals | Paraxerus_ochraceus | Paraxerus_ochraceus | LC | Stable | tropical | diurnal | 104.3 | 547064.702 | 11.455 |
| mammals | Paraxerus_palliatus | Paraxerus_palliatus | LC | Unknown | tropical | diurnal | 309.3 | 1045560.147 | 11.16 |
| mammals | Paraxerus_poensis | Paraxerus_poensis | LC | Unknown | tropical | diurnal | 124.5 | 1570579.529 | 9.318 |
| mammals | Paraxerus_vexillarius | Paraxerus_vexillarius | LC | Unknown | tropical | diurnal | 675 | 39824.991 | 13.001 |
| mammals | Paraxerus_vincenti | Paraxerus_vincenti | EN | Decreasing | tropical | diurnal | 337.6 | 2218.844 | 12.338 |
| mammals | Pardofelis_marmorata | Pardofelis_marmorata | NT | Decreasing | temperate | cathemeral | 2854.3 | 1388973.702 | 6.905 |
| mammals | Paremballonura_atrata | Paremballonura_atrata | LC | Unknown | tropical | diurnal | 4.3 | 105682.264 | 8.127 |
| mammals | Paremballonura_tiavato | Paremballonura_tiavato | LC | Unknown | tropical | nocturnal | 3.3 | 149432.764 | 7.939 |

| | | | | | | | | | |
|---|---|---|---|---|---|---|---|---|---|
| mammals | Parotomys_brantsii | Parotomys_brantsii | LC | Stable | temperate | diurnal | 129.5 | 679427.622 | 6.588 |
| mammals | Parotomys_littledalei | Parotomys_littledalei | LC | Stable | temperate | diurnal | 127 | 597770.682 | 5.304 |
| mammals | Passalites_nemorivagus | Mazama_nemorivaga | LC | Decreasing | tropical | cathemeral | 15000 | 7647916.969 | 3.022 |
| mammals | Pattonomys_carrikeri | NA | DD | Unknown | NA | NA | NA | NA | NA |
| mammals | Pattonomys_flavidus | NA | DD | Unknown | NA | NA | NA | NA | NA |
| mammals | Pattonomys_semivillosus | Pattonomys_semivillosus | LC | Unknown | tropical | nocturnal | 200 | 70580.132 | 14.652 |
| mammals | Paucidentomys_vermidax | NA | DD | Unknown | NA | NA | NA | NA | NA |
| mammals | Paulamys_naso | Paulamys_naso | EN | Decreasing | tropical | nocturnal | 123 | 748.498 | 10.588 |
| mammals | Paynomys_macronyx | Chelemys_macronyx | LC | Unknown | temperate | nocturnal | 73.3 | 322022.594 | 6.421 |
| mammals | Pectinator_spekei | Pectinator_spekei | LC | Unknown | tropical | diurnal | 160 | 419154.665 | 8.043 |
| mammals | Pedetes_capensis | Pedetes_capensis | LC | Unknown | tropical | nocturnal | 3155 | 3817904.34 | 7.323 |
| mammals | Pedetes_surdaster | Pedetes_surdaster | LC | Unknown | tropical | nocturnal | 3500 | 457191.762 | 11.722 |
| mammals | Pekania_pennanti | Martes_pennanti | LC | Unknown | temperate | cathemeral | 4000 | 3721001.426 | 5.608 |
| mammals | Pelea_capreolus | Pelea_capreolus | NT | Decreasing | temperate | cathemeral | 20000 | 557077.483 | 10.647 |
| mammals | Pelomys_campanae | Pelomys_campanae | LC | Unknown | tropical | nocturnal | 79.9 | 656702.215 | 8.345 |
| mammals | Pelomys_fallax | Pelomys_fallax | LC | Unknown | tropical | cathemeral | 130 | 3495837.145 | 8.837 |
| mammals | Pelomys_hopkinsi | NA | DD | Unknown | NA | NA | NA | NA | NA |
| mammals | Pelomys_isseli | Pelomys_isseli | NT | Unknown | tropical | nocturnal | 60 | 162.933 | 28.156 |
| mammals | Pelomys_minor | Pelomys_minor | LC | Unknown | tropical | nocturnal | 50 | 309824.591 | 5.197 |
| mammals | Pentalagus_furnessi | Pentalagus_furnessi | EN | Decreasing | temperate | nocturnal | 2396.3 | 370.201 | 13.193 |
| mammals | Penthetor_lucasii | Penthetor_lucasi | LC | Decreasing | tropical | nocturnal | 38.2 | 1288201.064 | 7.633 |
| mammals | Perameles_bougainville | Perameles_bougainville | VU | Stable | temperate | nocturnal | 226 | 235.895 | 5.385 |
| mammals | Perameles_gunnii | Perameles_gunnii | VU | Decreasing | temperate | nocturnal | 870 | 31989.71 | 8.397 |
| mammals | Perameles_nasuta | Perameles_nasuta | LC | Unknown | temperate | nocturnal | 975 | 411183.136 | 8.153 |
| mammals | Perimyotis_subflavus | Pipistrellus_subflavus | VU | Decreasing | temperate | nocturnal | 5.8 | 4264005.661 | 16.532 |
| mammals | Perodicticus_edwardsi | Perodicticus_edwardsi | LC | Stable | tropical | nocturnal | 1210 | 2432405.685 | 6.638 |
| mammals | Perodicticus_ibeanus | Perodicticus_ibeanus | LC | Decreasing | tropical | nocturnal | 1014 | 861196.586 | 8.278 |
| mammals | Perodicticus_potto | Perodicticus_potto | NT | Decreasing | tropical | nocturnal | 1100 | 680452.649 | 15.38 |
| mammals | Perognathus_alticola | Perognathus_alticola | VU | Decreasing | temperate | nocturnal | 24 | 11069.462 | 16.827 |
| mammals | Perognathus_amplus | Perognathus_amplus | LC | Stable | temperate | nocturnal | 11.7 | 122119.519 | 9.392 |
| mammals | Perognathus_fasciatus | Perognathus_fasciatus | LC | Stable | temperate | nocturnal | 11.6 | 1039860.874 | 12.479 |
| mammals | Perognathus_flavescens | Perognathus_flavescens | LC | Stable | temperate | nocturnal | 8.8 | 1542733.953 | 13.681 |
| mammals | Perognathus_flavus | Perognathus_flavus | LC | Stable | temperate | nocturnal | 7.7 | 1540956.831 | 9.837 |
| mammals | Perognathus_inornatus | Perognathus_inornatus | LC | Unknown | temperate | nocturnal | 10.9 | 73759.511 | 18.344 |

| | | | | | | | | | |
|---|---|---|---|---|---|---|---|---|---|
| mammals | Perognathus_longimembris | Perognathus_longimembris | LC | Stable | temperate | nocturnal | 7.6 | 520454.392 | 9.152 |
| mammals | Perognathus_merriami | Perognathus_merriami | LC | Stable | temperate | nocturnal | 6.9 | 693402.514 | 12.308 |
| mammals | Perognathus_parvus | Perognathus_parvus | LC | Stable | temperate | nocturnal | 21.8 | 921305.427 | 9.214 |
| mammals | Peromyscus_attwateri | Peromyscus_attwateri | LC | Stable | temperate | nocturnal | 27.9 | 452891.853 | 14.955 |
| mammals | Peromyscus_aztecus | Peromyscus_aztecus | LC | Stable | tropical | nocturnal | 40 | 218569.364 | 11.713 |
| mammals | Peromyscus_beatae | Peromyscus_beatae | LC | Unknown | tropical | nocturnal | 20.7 | 145828.321 | 12.208 |
| mammals | Peromyscus_boylii | Peromyscus_boylii | LC | Stable | temperate | nocturnal | 21.4 | 2530151.143 | 9.876 |
| mammals | Peromyscus_bullatus | Peromyscus_bullatus | CR | Decreasing | tropical | nocturnal | 40 | 65.19 | 23.802 |
| mammals | Peromyscus_californicus | Peromyscus_californicus | LC | Stable | temperate | nocturnal | 43.5 | 132623.266 | 12.861 |
| mammals | Peromyscus_caniceps | Peromyscus_caniceps | CR | Decreasing | temperate | nocturnal | 30.2 | 18.1 | 0.891 |
| mammals | Peromyscus_crinitus | Peromyscus_crinitus | LC | Stable | temperate | nocturnal | 18 | 913321.966 | 8.274 |
| mammals | Peromyscus_difficilis | Peromyscus_difficilis | LC | Stable | temperate | nocturnal | 28 | 546382.943 | 9.054 |
| mammals | Peromyscus_eremicus | Peromyscus_eremicus | LC | Stable | temperate | nocturnal | 23.7 | 1217908.661 | 7.647 |
| mammals | Peromyscus_eva | Peromyscus_eva | LC | Stable | temperate | nocturnal | 22 | 39058.947 | 5.005 |
| mammals | Peromyscus_fraterculus | Peromyscus_fraterculus | LC | Unknown | temperate | nocturnal | 24.3 | 144188.192 | 6.632 |
| mammals | Peromyscus_furvus | NA | DD | Unknown | NA | NA | NA | NA | NA |
| mammals | Peromyscus_gossypinus | Peromyscus_gossypinus | LC | Stable | temperate | nocturnal | 29.4 | 1148138.973 | 16.045 |
| mammals | Peromyscus_grandis | Peromyscus_grandis | NT | Stable | tropical | nocturnal | 71 | 3981.898 | 12.961 |
| mammals | Peromyscus_gratus | Peromyscus_gratus | LC | Unknown | temperate | nocturnal | 27.4 | 598242.302 | 9.966 |
| mammals | Peromyscus_guatemalensis | Peromyscus_guatemalensis | LC | Stable | tropical | nocturnal | 40 | 17875.067 | 13.938 |
| mammals | Peromyscus_gymnotis | Peromyscus_gymnotis | LC | Decreasing | tropical | nocturnal | 40 | 42202.778 | 18.094 |
| mammals | Peromyscus_hooperi | Peromyscus_hooperi | LC | Stable | temperate | nocturnal | 36 | 19804.777 | 4.522 |
| mammals | Peromyscus_hylocetes | Peromyscus_hylocetes | LC | Stable | tropical | nocturnal | 42.6 | 37378.531 | 16.932 |
| mammals | Peromyscus_keeni | Peromyscus_keeni | LC | Stable | temperate | nocturnal | 21.3 | 212135.71 | 2.78 |
| mammals | Peromyscus_leucopus | Peromyscus_leucopus | LC | Stable | temperate | nocturnal | 21.2 | 5831351.766 | 15.192 |
| mammals | Peromyscus_levipes | Peromyscus_levipes | LC | Unknown | tropical | nocturnal | 23 | 160994.64 | 15.348 |
| mammals | Peromyscus_madrensis | Peromyscus_madrensis | EN | Decreasing | tropical | nocturnal | 57 | 415.335 | 3.633 |
| mammals | Peromyscus_maniculatus | Peromyscus_maniculatus | LC | Stable | temperate | nocturnal | 21.3 | 13030881.644 | 9.358 |
| mammals | Peromyscus_mayensis | Peromyscus_mayensis | CR | Decreasing | tropical | nocturnal | 32 | 89.083 | 14.509 |
| mammals | Peromyscus_megalops | Peromyscus_megalops | LC | Unknown | tropical | nocturnal | 66.2 | 44159.056 | 7.857 |
| mammals | Peromyscus_mekisturus | Peromyscus_mekisturus | CR | Unknown | tropical | nocturnal | 60 | 7008.654 | 16.752 |
| mammals | Peromyscus_melanocarpus | Peromyscus_melanocarpus | EN | Unknown | tropical | nocturnal | 59 | 3955.827 | 4.634 |
| mammals | Peromyscus_melanophrys | Peromyscus_melanophrys | LC | Stable | tropical | nocturnal | 40 | 512462.172 | 10.456 |
| mammals | Peromyscus_melanotis | Peromyscus_melanotis | LC | Unknown | temperate | nocturnal | 39.6 | 494800.924 | 10.378 |

| | | | | | | | | | |
|---|---|---|---|---|---|---|---|---|---|
| mammals | Peromyscus_melanurus | Peromyscus_melanurus | EN | Stable | tropical | nocturnal | 40 | 3783.265 | 6.001 |
| mammals | Peromyscus_merriami | Peromyscus_merriami | LC | Unknown | temperate | nocturnal | 40 | 197642.763 | 7.616 |
| mammals | Peromyscus_mexicanus | Peromyscus_mexicanus | LC | Stable | tropical | nocturnal | 32.6 | 367383.466 | 12.027 |
| mammals | Peromyscus_nasutus | Peromyscus_nasutus | LC | Unknown | temperate | nocturnal | 28 | 576501.021 | 8.985 |
| mammals | Peromyscus_ochraventer | Peromyscus_ochraventer | EN | Unknown | tropical | nocturnal | 40 | 4659.359 | 6.661 |
| mammals | Peromyscus_pectoralis | Peromyscus_pectoralis | LC | Stable | temperate | nocturnal | 39 | 832035.554 | 9.444 |
| mammals | Peromyscus_perfulvus | Peromyscus_perfulvus | LC | Stable | tropical | nocturnal | 40 | 66536.097 | 9.145 |
| mammals | Peromyscus_polionotus | Peromyscus_polionotus | LC | Unknown | temperate | nocturnal | 13 | 376320.252 | 17.365 |
| mammals | Peromyscus_polius | Peromyscus_polius | NT | Decreasing | temperate | nocturnal | 40 | 14189.417 | 6.77 |
| mammals | Peromyscus_sagax | NA | DD | Unknown | NA | NA | NA | NA | NA |
| mammals | Peromyscus_schmidlyi | Peromyscus_schmidlyi | LC | Unknown | temperate | nocturnal | 21.4 | 19073.515 | 4.386 |
| mammals | Peromyscus_sejugis | Peromyscus_sejugis | EN | Decreasing | temperate | nocturnal | 21.5 | 13.589 | 1.352 |
| mammals | Peromyscus_simulus | Peromyscus_simulus | VU | Decreasing | tropical | nocturnal | 40 | 11840.649 | 12.282 |
| mammals | Peromyscus_slevini | Peromyscus_slevini | CR | Decreasing | temperate | nocturnal | 53.6 | 39.511 | 1.229 |
| mammals | Peromyscus_spicilegus | Peromyscus_spicilegus | LC | Decreasing | tropical | nocturnal | 36 | 125505.294 | 6.698 |
| mammals | Peromyscus_stephani | Peromyscus_stephani | CR | Decreasing | temperate | nocturnal | 26.1 | 41.202 | 1.574 |
| mammals | Peromyscus_stirtoni | Peromyscus_stirtoni | LC | Stable | tropical | nocturnal | 29.2 | 26277.367 | 13.582 |
| mammals | Peromyscus_truei | Peromyscus_truei | LC | Stable | temperate | nocturnal | 27.4 | 1452308.013 | 10.288 |
| mammals | Peromyscus_winkelmanni | Peromyscus_winkelmanni | EN | Decreasing | tropical | nocturnal | 53.8 | 301.035 | 4.904 |
| mammals | Peromyscus_yucatanicus | Peromyscus_yucatanicus | LC | Stable | tropical | nocturnal | 26.3 | 137965.811 | 6.128 |
| mammals | Peromyscus_zarhynchus | Peromyscus_zarhynchus | VU | Decreasing | tropical | nocturnal | 40 | 10773.67 | 10.555 |
| mammals | Peropteryx_kappleri | Peropteryx_kappleri | LC | Unknown | tropical | nocturnal | 9.4 | 7287612.735 | 4.589 |
| mammals | Peropteryx_leucoptera | Peropteryx_leucoptera | LC | Unknown | tropical | nocturnal | 6 | 5089787.566 | 2.238 |
| mammals | Peropteryx_macrotis | Peropteryx_macrotis | LC | Stable | tropical | nocturnal | 5.4 | 12697128.888 | 5.585 |
| mammals | Peropteryx_pallidoptera | NA | DD | Unknown | NA | NA | NA | NA | NA |
| mammals | Peropteryx_trinitatis | NA | DD | Unknown | NA | NA | NA | NA | NA |
| mammals | Peroryctes_broadbenti | Peroryctes_broadbenti | EN | Decreasing | tropical | nocturnal | 4800 | 31530.635 | 4.913 |
| mammals | Peroryctes_raffrayanus | Peroryctes_raffrayana | LC | Unknown | tropical | nocturnal | 838.5 | 298144.48 | 4.478 |
| mammals | Petaurillus_emiliae | NA | DD | Unknown | NA | NA | NA | NA | NA |
| mammals | Petaurillus_hosei | NA | DD | Unknown | NA | NA | NA | NA | NA |
| mammals | Petaurillus_kinlochii | NA | DD | Unknown | NA | NA | NA | NA | NA |
| mammals | Petaurista_alborufus | Petaurista_alborufus | LC | Unknown | temperate | nocturnal | 4290 | 943217.74 | 13.544 |
| mammals | Petaurista_elegans | Petaurista_elegans | LC | Stable | tropical | nocturnal | 997 | 2490770.446 | 10.99 |
| mammals | Petaurista_leucogenys | Petaurista_leucogenys | LC | Unknown | temperate | nocturnal | 1178 | 251379.478 | 20.151 |

| | | | | | | | | | |
|---|---|---|---|---|---|---|---|---|---|
| mammals | Petaurista_magnificus | Petaurista_magnificus | LC | Decreasing | temperate | nocturnal | 1475.5 | 142043.462 | 6.946 |
| mammals | Petaurista_mechukaensis | NA | DD | Unknown | NA | NA | NA | NA | NA |
| mammals | Petaurista_mishmiensis | NA | NT | Unknown | NA | NA | NA | NA | NA |
| mammals | Petaurista_nobilis | Petaurista_nobilis | NT | Decreasing | temperate | nocturnal | 2710 | 44806.376 | 7.54 |
| mammals | Petaurista_petaurista | Petaurista_petaurista | LC | Decreasing | tropical | nocturnal | 1660.1 | 3195973.778 | 10.412 |
| mammals | Petaurista_philippensis | Petaurista_philippensis | LC | Decreasing | tropical | nocturnal | 1676 | 3398886.324 | 12.893 |
| mammals | Petaurista_xanthotis | Petaurista_xanthotis | LC | Unknown | temperate | nocturnal | 1900 | 730211.785 | 9.086 |
| mammals | Petauroides_volans | Petauroides_volans | VU | Decreasing | temperate | nocturnal | 1300 | 633491.265 | 7.129 |
| mammals | Petaurus_abidi | Petaurus_abidi | CR | Decreasing | tropical | nocturnal | 278 | 92.806 | 3.976 |
| mammals | Petaurus_australis | Petaurus_australis | NT | Decreasing | temperate | nocturnal | 575 | 231711.033 | 7.355 |
| mammals | Petaurus_biacensis | Petaurus_biacensis | LC | Unknown | tropical | nocturnal | 88.4 | 2533.472 | 8.959 |
| mammals | Petaurus_breviceps | Petaurus_breviceps | LC | Stable | tropical | nocturnal | 106.3 | 2761627.499 | 5.325 |
| mammals | Petaurus_gracilis | Petaurus_gracilis | EN | Decreasing | tropical | nocturnal | 353 | 3309.374 | 6.104 |
| mammals | Petaurus_norfolcensis | Petaurus_norfolcensis | LC | Decreasing | temperate | nocturnal | 230 | 788590.476 | 7.147 |
| mammals | Petinomys_crinitus | Petinomys_crinitus | LC | Stable | tropical | nocturnal | 113 | 33685.249 | 11.452 |
| mammals | Petinomys_fuscocapillus | Petinomys_fuscocapillus | LC | Decreasing | tropical | nocturnal | 794.5 | 56586.731 | 17.419 |
| mammals | Petinomys_genibarbis | Petinomys_genibarbis | VU | Decreasing | tropical | nocturnal | 110 | 844696.57 | 8.929 |
| mammals | Petinomys_hageni | Petinomys_hageni | DD | Decreasing | tropical | nocturnal | 388 | 187.196 | 15.534 |
| mammals | Petinomys_lugens | Petinomys_lugens | VU | Decreasing | tropical | nocturnal | 433 | 5106.487 | 5.825 |
| mammals | Petinomys_mindanensis | Petinomys_mindanensis | LC | Unknown | tropical | nocturnal | 285 | 32438.599 | 11.454 |
| mammals | Petinomys_setosus | Petinomys_setosus | VU | Decreasing | tropical | nocturnal | 38.4 | 715360.468 | 9.119 |
| mammals | Petinomys_vordermanni | Petinomys_vordermanni | VU | Decreasing | tropical | nocturnal | 40.4 | 114244.837 | 7.229 |
| mammals | Petrogale_assimilis | Petrogale_assimilis | LC | Stable | tropical | nocturnal | 4500 | 208649.694 | 5.022 |
| mammals | Petrogale_brachyotis | Petrogale_brachyotis | LC | Unknown | tropical | nocturnal | 4050 | 221775.599 | 1.86 |
| mammals | Petrogale_burbidgei | Petrogale_burbidgei | NT | Unknown | tropical | nocturnal | 1258 | 17334.32 | 1.129 |
| mammals | Petrogale_coenensis | Petrogale_coenensis | EN | Decreasing | tropical | nocturnal | 4375 | 10779.954 | 1.763 |
| mammals | Petrogale_concinna | Petrogale_concinna | EN | Decreasing | tropical | nocturnal | 1400 | 82698.688 | 1.02 |
| mammals | Petrogale_godmani | Petrogale_godmani | NT | Decreasing | tropical | nocturnal | 4750 | 36293.794 | 2.201 |
| mammals | Petrogale_herberti | Petrogale_herberti | LC | Unknown | temperate | nocturnal | 5150 | 281939.808 | 6.243 |
| mammals | Petrogale_inornata | Petrogale_inornata | LC | Unknown | tropical | nocturnal | 4600 | 85125.443 | 6.328 |
| mammals | Petrogale_lateralis | Petrogale_lateralis | VU | Decreasing | tropical | nocturnal | 4625 | 113472.063 | 2.657 |
| mammals | Petrogale_mareeba | Petrogale_mareeba | NT | Stable | tropical | nocturnal | 4150 | 19944.213 | 3.929 |
| mammals | Petrogale_penicillata | Petrogale_penicillata | VU | Decreasing | temperate | nocturnal | 7100 | 101650.442 | 6.799 |
| mammals | Petrogale_persephone | Petrogale_persephone | EN | Decreasing | tropical | nocturnal | 6200 | 4210.626 | 5.698 |

| | | | | | | | | | |
|---|---|---|---|---|---|---|---|---|---|
| mammals | Petrogale_purpureicollis | Petrogale_purpureicollis | NT | Decreasing | tropical | nocturnal | 4500 | 80361.155 | 4.836 |
| mammals | Petrogale_rothschildi | Petrogale_rothschildi | LC | Unknown | tropical | nocturnal | 3400 | 142489.183 | 4.103 |
| mammals | Petrogale_sharmani | Petrogale_sharmani | VU | Unknown | tropical | nocturnal | 4250 | 16921.712 | 3.568 |
| mammals | Petrogale_xanthopus | Petrogale_xanthopus | NT | Unknown | temperate | nocturnal | 8500 | 24922.636 | 5.714 |
| mammals | Petromus_typicus | Petromus_typicus | LC | Stable | tropical | diurnal | 224 | 372746.367 | 5.247 |
| mammals | Petromyscus_barbouri | Petromyscus_barbouri | LC | Stable | temperate | nocturnal | 20.1 | 38176.271 | 7.309 |
| mammals | Petromyscus_collinus | Petromyscus_collinus | LC | Stable | temperate | nocturnal | 20.1 | 497480.985 | 5.329 |
| mammals | Petromyscus_monticularis | Petromyscus_monticularis | LC | Stable | temperate | nocturnal | 20.7 | 186365.108 | 4.673 |
| mammals | Petromyscus_shortridgei | Petromyscus_shortridgei | LC | Stable | tropical | nocturnal | 46.5 | 123751.783 | 5.695 |
| mammals | Phacochoerus_aethiopicus | Phacochoerus_aethiopicus | LC | Decreasing | tropical | diurnal | 71000.3 | 318268.664 | 8.786 |
| mammals | Phacochoerus_africanus | Phacochoerus_africanus | LC | Decreasing | tropical | cathemeral | 82500 | 9957729.535 | 9.572 |
| mammals | Phaenomys_ferrugineus | Phaenomys_ferrugineus | EN | Decreasing | tropical | nocturnal | 93.8 | 36149.629 | 15.141 |
| mammals | Phalanger_carmelitae | Phalanger_carmelitae | LC | Stable | tropical | nocturnal | 2047.5 | 198474.773 | 4.578 |
| mammals | Phalanger_gymnotis | Phalanger_gymnotis | LC | Decreasing | tropical | nocturnal | 2705 | 616455.781 | 3.759 |
| mammals | Phalanger_intercastellanus | Phalanger_intercastellanus | LC | Decreasing | tropical | nocturnal | 1747.5 | 63331.478 | 4.807 |
| mammals | Phalanger_lullulae | Phalanger_lullulae | EN | Stable | tropical | nocturnal | 1632.5 | 886.299 | 3.983 |
| mammals | Phalanger_matabiru | NA | VU | Stable | NA | NA | NA | NA | NA |
| mammals | Phalanger_matanim | Phalanger_matanim | CR | Unknown | tropical | nocturnal | 1550 | 153.145 | 3.823 |
| mammals | Phalanger_mimicus | Phalanger_mimicus | LC | Decreasing | tropical | nocturnal | 1850 | 233040.721 | 2.589 |
| mammals | Phalanger_orientalis | Phalanger_orientalis | LC | Stable | tropical | nocturnal | 2168.8 | 456940.116 | 4.763 |
| mammals | Phalanger_ornatus | Phalanger_ornatus | LC | Stable | tropical | nocturnal | 1787.5 | 22280.653 | 6.413 |
| mammals | Phalanger_pelengensis | Strigocuscus_pelengensis | LC | Stable | tropical | nocturnal | 1103 | 6411.265 | 6.651 |
| mammals | Phalanger_rothschildi | Phalanger_rothschildi | LC | Stable | tropical | nocturnal | 1365 | 2774.761 | 4.822 |
| mammals | Phalanger_sericeus | Phalanger_sericeus | LC | Unknown | tropical | nocturnal | 2003 | 197226.979 | 4.582 |
| mammals | Phalanger_vestitus | Phalanger_vestitus | LC | Decreasing | tropical | nocturnal | 1850 | 88965.087 | 5.03 |
| mammals | Phaner_electromontis | Phaner_electromontis | EN | Decreasing | tropical | nocturnal | 383.6 | 1662.997 | 6.451 |
| mammals | Phaner_furcifer | Phaner_furcifer | EN | Decreasing | tropical | nocturnal | 400 | 20954.538 | 5.712 |
| mammals | Phaner_pallescens | Phaner_pallescens | EN | Decreasing | tropical | nocturnal | 327 | 47678.92 | 7.503 |
| mammals | Phaner_parienti | Phaner_parienti | EN | Decreasing | tropical | nocturnal | 363.4 | 1912.238 | 5.784 |
| mammals | Pharotis_imogene | Pharotis_imogene | CR | Stable | tropical | nocturnal | 10 | 537.257 | 8.474 |
| mammals | Phascogale_calura | Phascogale_calura | NT | Stable | temperate | nocturnal | 51.5 | 46849.833 | 12.291 |
| mammals | Phascogale_pirata | Phascogale_pirata | VU | Unknown | tropical | nocturnal | 181.5 | 40381.031 | 2.125 |
| mammals | Phascogale_tapoatafa | Phascogale_tapoatafa | NT | Decreasing | temperate | nocturnal | 193.5 | 272010.805 | 8.233 |
| mammals | Phascolarctos_cinereus | Phascolarctos_cinereus | VU | Decreasing | temperate | nocturnal | 10250 | 1102194.403 | 7.513 |

| | | | | | | | | | |
|---|---|---|---|---|---|---|---|---|---|
| mammals | Phascolosorex_doriae | Phascolosorex_doriae | LC | Unknown | tropical | diurnal | 168.3 | 82246.536 | 4.077 |
| mammals | Phascolosorex_dorsalis | Phascolosorex_dorsalis | LC | Stable | tropical | diurnal | 138.5 | 174794.952 | 4.599 |
| mammals | Phataginus_tetradactylus | Phataginus_tetradactyla | VU | Decreasing | tropical | diurnal | 2750 | 2929072.012 | 7.859 |
| mammals | Phataginus_tricuspis | Phataginus_tricuspis | EN | Decreasing | tropical | nocturnal | 2066.7 | 5954123.196 | 8.314 |
| mammals | Phenacomys_intermedius | Phenacomys_intermedius | LC | Stable | temperate | nocturnal | 25.2 | 524347.744 | 7.595 |
| mammals | Phenacomys_ungava | Phenacomys_ungava | LC | Stable | temperate | nocturnal | 32.5 | 5082619.312 | 2.066 |
| mammals | Philander_andersoni | Philander_andersoni | LC | Stable | tropical | nocturnal | 325 | 1346166.853 | 1.285 |
| mammals | Philander_mcilhennyi | Philander_mcilhennyi | LC | Unknown | tropical | cathemeral | 513.9 | 443217.679 | 0.789 |
| mammals | Philander_opossum | Philander_opossum | LC | Stable | tropical | nocturnal | 585.5 | 5893336.84 | 4.357 |
| mammals | Philander_quica | Philander_frenatus | LC | Unknown | tropical | nocturnal | 167.7 | 921860.966 | 12.398 |
| mammals | Philantomba_maxwellii | Philantomba_maxwellii | LC | Decreasing | tropical | diurnal | 12000 | 872152.03 | 13.423 |
| mammals | Philantomba_monticola | Philantomba_monticola | LC | Decreasing | tropical | diurnal | 6250 | 6383795.312 | 7.375 |
| mammals | Philantomba_walteri | NA | DD | Unknown | NA | NA | NA | NA | NA |
| mammals | Philetor_brachypterus | Philetor_brachypterus | LC | Stable | tropical | nocturnal | 10.9 | 1292109.23 | 5.79 |
| mammals | Phloeomys_cumingi | Phloeomys_cumingi | LC | Decreasing | tropical | nocturnal | 1946.3 | 19097.58 | 15.184 |
| mammals | Phloeomys_pallidus | Phloeomys_pallidus | LC | Stable | tropical | nocturnal | 2250 | 43063.295 | 8.453 |
| mammals | Phodopus_campbelli | Phodopus_campbelli | LC | Stable | temperate | nocturnal | 23.4 | 2049854.902 | 6.958 |
| mammals | Phodopus_roborovskii | Phodopus_roborovskii | LC | Stable | temperate | nocturnal | 18.8 | 3141864.39 | 5.217 |
| mammals | Phodopus_sungorus | Phodopus_sungorus | LC | Unknown | temperate | nocturnal | 33.8 | 1181966.721 | 9.79 |
| mammals | Phoniscus_atrox | Phoniscus_atrox | NT | Decreasing | tropical | nocturnal | 4.3 | 198710.727 | 12.847 |
| mammals | Phoniscus_jagorii | Phoniscus_jagorii | LC | Unknown | tropical | nocturnal | 4.7 | 347840.557 | 9.75 |
| mammals | Phoniscus_papuensis | Phoniscus_papuensis | VU | Unknown | temperate | nocturnal | 6.2 | 474747.756 | 5.911 |
| mammals | Phylloderma_stenops | Phylloderma_stenops | LC | Stable | tropical | nocturnal | 53.7 | 10972630.636 | 4.988 |
| mammals | Phyllomys_blainvilii | Phyllomys_blainvillii | LC | Unknown | tropical | nocturnal | 243.3 | 332344.105 | 9.305 |
| mammals | Phyllomys_brasiliensis | Phyllomys_brasiliensis | EN | Decreasing | tropical | nocturnal | 298.6 | 2858.165 | 10.47 |
| mammals | Phyllomys_dasythrix | Phyllomys_dasythrix | LC | Unknown | temperate | nocturnal | 260 | 276267.04 | 11.828 |
| mammals | Phyllomys_kerri | NA | DD | Unknown | NA | NA | NA | NA | NA |
| mammals | Phyllomys_lamarum | NA | DD | Unknown | NA | NA | NA | NA | NA |
| mammals | Phyllomys_lundi | Phyllomys_lundi | EN | Decreasing | tropical | nocturnal | 173.5 | 16102.604 | 15.268 |
| mammals | Phyllomys_mantiqueirensis | Phyllomys_mantiqueirensis | CR | Decreasing | tropical | nocturnal | 207 | 16165.799 | 13.462 |
| mammals | Phyllomys_medius | Phyllomys_medius | LC | Unknown | temperate | nocturnal | 229.9 | 501981.637 | 13.817 |
| mammals | Phyllomys_nigrispinus | Phyllomys_nigrispinus | LC | Unknown | temperate | nocturnal | 224.3 | 174482.499 | 14.954 |
| mammals | Phyllomys_pattoni | Phyllomys_pattoni | LC | Unknown | tropical | nocturnal | 229.9 | 302917.215 | 11.346 |
| mammals | Phyllomys_sulinus | NA | DD | Unknown | NA | NA | NA | NA | NA |

| | | | | | | | | | |
|---|---|---|---|---|---|---|---|---|---|
| mammals | Phyllomys_thomasi | Phyllomys_thomasi | EN | Decreasing | temperate | nocturnal | 560.2 | 341.49 | 6.283 |
| mammals | Phyllomys_unicolor | Phyllomys_unicolor | CR | Decreasing | tropical | nocturnal | 439 | 4485.124 | 6.527 |
| mammals | Phyllonycteris_aphylla | Phyllonycteris_aphylla | CR | Decreasing | tropical | nocturnal | 24.5 | 215.186 | 25.067 |
| mammals | Phyllonycteris_poeyi | Phyllonycteris_poeyi | LC | Unknown | tropical | nocturnal | 22 | 182992.967 | 16.028 |
| mammals | Phyllops_falcatus | Phyllops_falcatus | LC | Stable | tropical | nocturnal | 19.5 | 181362.688 | 16.077 |
| mammals | Phyllostomus_discolor | Phyllostomus_discolor | LC | Stable | tropical | nocturnal | 34.1 | 12498386.825 | 5.569 |
| mammals | Phyllostomus_elongatus | Phyllostomus_elongatus | LC | Unknown | tropical | nocturnal | 40.4 | 10601026.638 | 4.623 |
| mammals | Phyllostomus_hastatus | Phyllostomus_hastatus | LC | Stable | tropical | nocturnal | 87.2 | 12629829.36 | 5.499 |
| mammals | Phyllostomus_latifolius | Phyllostomus_latifolius | LC | Unknown | tropical | nocturnal | 134 | 1553256.895 | 1.093 |
| mammals | Phyllotis_amicus | Phyllotis_amicus | LC | Stable | tropical | nocturnal | 20.2 | 71787.376 | 6.674 |
| mammals | Phyllotis_andium | Phyllotis_andium | LC | Stable | tropical | nocturnal | 53 | 124063.11 | 8.18 |
| mammals | Phyllotis_anitae | NA | DD | Unknown | NA | NA | NA | NA | NA |
| mammals | Phyllotis_bonariensis | Phyllotis_bonariensis | NT | Decreasing | temperate | nocturnal | 42.5 | 27439.825 | 13.485 |
| mammals | Phyllotis_caprinus | Phyllotis_caprinus | LC | Stable | tropical | cathemeral | 50.8 | 42361.2 | 8.424 |
| mammals | Phyllotis_darwinii | Phyllotis_darwini | LC | Stable | temperate | nocturnal | 50.8 | 130478.736 | 10.513 |
| mammals | Phyllotis_definitus | Phyllotis_definitus | EN | Decreasing | tropical | cathemeral | 89 | 20652.777 | 8.066 |
| mammals | Phyllotis_gerbillus | Phyllotis_gerbillus | LC | Stable | tropical | nocturnal | 17.4 | 33728.372 | 7.507 |
| mammals | Phyllotis_haggardi | Phyllotis_haggardi | LC | Stable | tropical | nocturnal | 42.5 | 34812.809 | 12.712 |
| mammals | Phyllotis_limatus | Phyllotis_limatus | LC | Stable | tropical | nocturnal | 36.9 | 137878.403 | 5.598 |
| mammals | Phyllotis_magister | Phyllotis_magister | LC | Stable | tropical | cathemeral | 68.5 | 58908.139 | 5.401 |
| mammals | Phyllotis_osgoodi | NA | DD | Unknown | NA | NA | NA | NA | NA |
| mammals | Phyllotis_xanthopygus | Phyllotis_xanthopygus | LC | Stable | temperate | nocturnal | 56.3 | 1499341.73 | 5.676 |
| mammals | Piliocolobus_badius | Piliocolobus_badius | EN | Decreasing | tropical | diurnal | 7750.1 | 342829.966 | 14.533 |
| mammals | Piliocolobus_bouvieri | NA | EN | Decreasing | NA | NA | NA | NA | NA |
| mammals | Piliocolobus_epieni | NA | CR | Decreasing | NA | NA | NA | NA | NA |
| mammals | Piliocolobus_gordonorum | Piliocolobus_gordonorum | VU | Decreasing | tropical | diurnal | 8023.4 | 1709.899 | 5.133 |
| mammals | Piliocolobus_kirkii | Piliocolobus_kirkii | EN | Decreasing | tropical | diurnal | 5630 | 1581.63 | 20.564 |
| mammals | Piliocolobus_oustaleti | Piliocolobus_oustaleti | VU | Decreasing | tropical | diurnal | 10350 | 424140.16 | 5.64 |
| mammals | Piliocolobus_pennantii | Piliocolobus_pennantii | CR | Decreasing | tropical | diurnal | 10896 | 161.411 | 6.728 |
| mammals | Piliocolobus_preussi | Piliocolobus_preussi | CR | Decreasing | tropical | diurnal | 9750 | 4263.463 | 4.434 |
| mammals | Piliocolobus_rufomitratus | Piliocolobus_rufomitratus | CR | Decreasing | tropical | diurnal | 8000 | 201.248 | 15.562 |
| mammals | Piliocolobus_tephrosceles | Piliocolobus_tephrosceles | EN | Decreasing | tropical | diurnal | 9700 | 3893.263 | 8.641 |
| mammals | Piliocolobus_tholloni | NA | VU | Decreasing | NA | NA | NA | NA | NA |
| mammals | Pilonycteris_celebensis | Rousettus_celebensis | LC | Decreasing | tropical | nocturnal | 56.3 | 181192.803 | 9.25 |

| | | | | | | | | | |
|---|---|---|---|---|---|---|---|---|---|
| mammals | Pipanacoctomys_aureus | Tympanoctomys_aureus | CR | Decreasing | temperate | nocturnal | 139.4 | 16.885 | 1.123 |
| mammals | Pipistrellus_abramus | Pipistrellus_abramus | LC | Stable | temperate | nocturnal | 5.6 | 3910979.374 | 16.78 |
| mammals | Pipistrellus_adamsi | Pipistrellus_adamsi | LC | Stable | tropical | nocturnal | 4.5 | 277615.884 | 1.809 |
| mammals | Pipistrellus_aero | NA | DD | Unknown | NA | NA | NA | NA | NA |
| mammals | Pipistrellus_angulatus | Pipistrellus_angulatus | LC | Stable | tropical | nocturnal | 3.3 | 344529.747 | 4.277 |
| mammals | Pipistrellus_ceylonicus | Pipistrellus_ceylonicus | LC | Stable | tropical | nocturnal | 8.4 | 2718309.287 | 17.308 |
| mammals | Pipistrellus_collinus | Pipistrellus_collinus | LC | Stable | tropical | nocturnal | 6 | 201503.033 | 4.434 |
| mammals | Pipistrellus_coromandra | Pipistrellus_coromandra | LC | Unknown | tropical | nocturnal | 4.6 | 3655673.731 | 14.39 |
| mammals | Pipistrellus_endoi | Pipistrellus_endoi | NT | Decreasing | temperate | nocturnal | 4.2 | 22538.229 | 18.48 |
| mammals | Pipistrellus_hanaki | Pipistrellus_hanaki | VU | Decreasing | temperate | nocturnal | 6 | 48435.963 | 10.801 |
| mammals | Pipistrellus_hesperidus | Pipistrellus_hesperidus | LC | Unknown | tropical | nocturnal | 5.7 | 1600945.63 | 12.563 |
| mammals | Pipistrellus_inexspectatus | NA | DD | Unknown | NA | NA | NA | NA | NA |
| mammals | Pipistrellus_javanicus | Pipistrellus_javanicus | LC | Stable | tropical | nocturnal | 4.8 | 3666505.455 | 12.342 |
| mammals | Pipistrellus_kuhlii | Pipistrellus_kuhlii | LC | Unknown | temperate | nocturnal | 5.9 | 10905819.431 | 11.542 |
| mammals | Pipistrellus_maderensis | Pipistrellus_maderensis | EN | Decreasing | temperate | nocturnal | 7 | 5017.464 | 18.701 |
| mammals | Pipistrellus_minahassae | NA | DD | Unknown | NA | NA | NA | NA | NA |
| mammals | Pipistrellus_nanulus | Pipistrellus_nanulus | LC | Unknown | tropical | nocturnal | 2.5 | 1565043.123 | 13.06 |
| mammals | Pipistrellus_nathusii | Pipistrellus_nathusii | LC | Unknown | temperate | nocturnal | 7.3 | 5890730.923 | 18.278 |
| mammals | Pipistrellus_papuanus | Pipistrellus_papuanus | LC | Stable | tropical | nocturnal | 4.7 | 406795.995 | 3.822 |
| mammals | Pipistrellus_paterculus | Pipistrellus_paterculus | LC | Unknown | tropical | nocturnal | 5.3 | 782192.183 | 10.458 |
| mammals | Pipistrellus_permixtus | NA | DD | Unknown | NA | NA | NA | NA | NA |
| mammals | Pipistrellus_pipistrellus | Pipistrellus_pipistrellus | LC | Stable | temperate | nocturnal | 5.7 | 7619643.117 | 15.844 |
| mammals | Pipistrellus_pygmaeus | Pipistrellus_pygmaeus | LC | Unknown | temperate | nocturnal | 5.7 | 1960533.948 | 19.302 |
| mammals | Pipistrellus_raceyi | NA | DD | Unknown | NA | NA | NA | NA | NA |
| mammals | Pipistrellus_rusticus | Pipistrellus_rusticus | LC | Unknown | tropical | nocturnal | 4.6 | 3459632.36 | 10.643 |
| mammals | Pipistrellus_stenopterus | Pipistrellus_stenopterus | LC | Unknown | tropical | nocturnal | 15.2 | 641151.188 | 9.733 |
| mammals | Pipistrellus_tenuis | NA | LC | Stable | NA | NA | NA | NA | NA |
| mammals | Pipistrellus_wattsi | Pipistrellus_wattsi | LC | Unknown | tropical | nocturnal | 3.3 | 14693.099 | 6.198 |
| mammals | Pipistrellus_westralis | Pipistrellus_westralis | LC | Stable | tropical | nocturnal | 3.1 | 189808.93 | 2.246 |
| mammals | Pithecheir_melanurus | Pithecheir_melanurus | VU | Decreasing | tropical | nocturnal | 108 | 132.57 | 25.708 |
| mammals | Pithecheir_parvus | NA | DD | Unknown | NA | NA | NA | NA | NA |
| mammals | Pithecia_aequatorialis | Pithecia_aequatorialis | LC | Decreasing | tropical | diurnal | 2250 | 45738.292 | 1.474 |
| mammals | Pithecia_albicans | Pithecia_albicans | LC | Decreasing | tropical | diurnal | 2800 | 63173.782 | 0.63 |
| mammals | Pithecia_chrysocephala | Pithecia_chrysocephala | LC | Decreasing | tropical | diurnal | 1900 | 114364.756 | 1.611 |

| | | | | | | | | | |
|---|---|---|---|---|---|---|---|---|---|
| mammals | Pithecia_milleri | Pithecia_milleri | VU | Decreasing | tropical | diurnal | 1537.5 | 55168.456 | 6.172 |
| mammals | Pithecia_pithecia | Pithecia_pithecia | LC | Decreasing | tropical | diurnal | 1375.5 | 886392.045 | 1.566 |
| mammals | Pithecia_vanzolinii | Pithecia_vanzolinii | DD | Decreasing | tropical | diurnal | 2241 | 49143.579 | 1.014 |
| mammals | Plagiodontia_aedium | Plagiodontia_aedium | EN | Decreasing | tropical | nocturnal | 2630 | 10576.599 | 9.742 |
| mammals | Planigale_ingrami | Planigale_ingrami | LC | Stable | tropical | nocturnal | 4.3 | 617152.388 | 2.97 |
| mammals | Planigale_maculata | Planigale_maculata | LC | Stable | tropical | nocturnal | 11 | 1233506.988 | 4.128 |
| mammals | Planigale_novaeguineae | Planigale_novaeguineae | LC | Unknown | tropical | nocturnal | 14.9 | 30188.899 | 4.21 |
| mammals | Planigale_tenuirostris | Planigale_tenuirostris | LC | Stable | temperate | nocturnal | 6.1 | 1617101.253 | 5.285 |
| mammals | Platacanthomys_lasiurus | Platacanthomys_lasiurus | VU | Decreasing | tropical | nocturnal | 75 | 28208.761 | 14.646 |
| mammals | Platalina_genovensium | Platalina_genovensium | NT | Unknown | tropical | nocturnal | 47 | 300140.5 | 7.832 |
| mammals | Platymops_setiger | Platymops_setiger | LC | Unknown | tropical | nocturnal | 13 | 229068.205 | 10.57 |
| mammals | Platyrrhinus_albericoi | Platyrrhinus_albericoi | LC | Unknown | tropical | nocturnal | 55 | 608162.993 | 4.527 |
| mammals | Platyrrhinus_aquilus | NA | DD | Unknown | NA | NA | NA | NA | NA |
| mammals | Platyrrhinus_aurarius | Platyrrhinus_aurarius | LC | Stable | tropical | nocturnal | 34.2 | 349895.004 | 0.91 |
| mammals | Platyrrhinus_brachycephalus | Platyrrhinus_brachycephalus | LC | Stable | tropical | nocturnal | 14.1 | 6159441.37 | 2.29 |
| mammals | Platyrrhinus_dorsalis | Platyrrhinus_dorsalis | LC | Unknown | tropical | nocturnal | 26 | 767245.605 | 11.949 |
| mammals | Platyrrhinus_helleri | Platyrrhinus_helleri | LC | Stable | tropical | nocturnal | 13.5 | 10016789.877 | 4.657 |
| mammals | Platyrrhinus_infuscus | Platyrrhinus_infuscus | LC | Unknown | tropical | nocturnal | 50.2 | 1466979.139 | 5.128 |
| mammals | Platyrrhinus_ismaeli | Platyrrhinus_ismaeli | NT | Decreasing | tropical | nocturnal | 37.1 | 129434.932 | 6.1 |
| mammals | Platyrrhinus_lineatus | Platyrrhinus_lineatus | LC | Stable | tropical | nocturnal | 24.4 | 5807566.654 | 9.385 |
| mammals | Platyrrhinus_masu | Platyrrhinus_masu | LC | Unknown | tropical | nocturnal | 27 | 172371.142 | 2.884 |
| mammals | Platyrrhinus_matapalensis | Platyrrhinus_matapalensis | NT | Decreasing | tropical | nocturnal | 18 | 80808.01 | 13.553 |
| mammals | Platyrrhinus_nitelinea | NA | DD | Unknown | NA | NA | NA | NA | NA |
| mammals | Platyrrhinus_recifinus | Platyrrhinus_recifinus | LC | Unknown | tropical | nocturnal | 17.7 | 1009873.95 | 12.4 |
| mammals | Platyrrhinus_umbratus | NA | DD | Unknown | NA | NA | NA | NA | NA |
| mammals | Platyrrhinus_vittatus | Platyrrhinus_vittatus | LC | Unknown | tropical | nocturnal | 41.4 | 769716.345 | 11.329 |
| mammals | Plecotus_auritus | Plecotus_auritus | LC | Stable | temperate | nocturnal | 7.8 | 6103621.632 | 17.697 |
| mammals | Plecotus_austriacus | Plecotus_austriacus | NT | Decreasing | temperate | nocturnal | 7.3 | 3363262.952 | 20.425 |
| mammals | Plecotus_balensis | NA | DD | Unknown | NA | NA | NA | NA | NA |
| mammals | Plecotus_christii | NA | DD | Unknown | NA | NA | NA | NA | NA |
| mammals | Plecotus_kolombatovici | Plecotus_kolombatovici | LC | Decreasing | temperate | nocturnal | 7 | 201013.868 | 15.276 |
| mammals | Plecotus_macrobullaris | Plecotus_macrobullaris | LC | Decreasing | temperate | nocturnal | 8.5 | 1010971.419 | 14.918 |
| mammals | Plecotus_ognevi | Plecotus_ognevi | LC | Unknown | temperate | nocturnal | 8.4 | 5958715.331 | 5.641 |
| mammals | Plecotus_sacrimontis | Plecotus_sacrimontis | LC | Stable | temperate | nocturnal | 9.3 | 206693.021 | 16.4 |

| | | | | | | | | | |
|---|---|---|---|---|---|---|---|---|---|
| mammals | Plecotus_sardus | Plecotus_sardus | CR | Decreasing | temperate | nocturnal | 9.8 | 5346.977 | 15.914 |
| mammals | Plecotus_taivanus | Plecotus_taivanus | NT | Decreasing | temperate | nocturnal | 6.5 | 17199.913 | 11.739 |
| mammals | Plecotus_teneriffae | Plecotus_teneriffae | CR | Decreasing | temperate | nocturnal | 8 | 2808.504 | 18.747 |
| mammals | Plecturocebus_aureipalatii | Callicebus_aureipalatii | LC | Stable | tropical | diurnal | 950 | 58371.593 | 2.692 |
| mammals | Plecturocebus_baptista | Callicebus_baptista | LC | Stable | tropical | diurnal | 956.5 | 14740.991 | 2.12 |
| mammals | Plecturocebus_bernhardi | Callicebus_bernhardi | LC | Stable | tropical | diurnal | 950 | 119557.518 | 1.936 |
| mammals | Plecturocebus_brunneus | Callicebus_brunneus | VU | Decreasing | tropical | diurnal | 992.4 | 184934.529 | 3.517 |
| mammals | Plecturocebus_caligatus | Callicebus_caligatus | LC | Unknown | tropical | diurnal | 992.4 | 195914.355 | 1.944 |
| mammals | Plecturocebus_caquetensis | NA | CR | Decreasing | NA | NA | NA | NA | NA |
| mammals | Plecturocebus_cinerascens | Callicebus_cinerascens | LC | Unknown | tropical | diurnal | 992.4 | 210384.504 | 1.242 |
| mammals | Plecturocebus_cupreus | Callicebus_cupreus | LC | Unknown | tropical | diurnal | 915 | 776067.491 | 0.757 |
| mammals | Plecturocebus_discolor | Callicebus_discolor | LC | Unknown | tropical | diurnal | 917.7 | 356789.184 | 2.396 |
| mammals | Plecturocebus_donacophilus | Callicebus_donacophilus | LC | Unknown | tropical | diurnal | 795 | 163767.86 | 4.967 |
| mammals | Plecturocebus_hoffmannsi | Callicebus_hoffmannsi | LC | Unknown | tropical | diurnal | 992.4 | 92127.776 | 1.328 |
| mammals | Plecturocebus_modestus | Callicebus_modestus | EN | Decreasing | tropical | diurnal | 992.4 | 6539.926 | 6.614 |
| mammals | Plecturocebus_moloch | Callicebus_moloch | LC | Unknown | tropical | diurnal | 854.7 | 944033.501 | 2.677 |
| mammals | Plecturocebus_oenanthe | Callicebus_oenanthe | CR | Decreasing | tropical | diurnal | 992.4 | 9106.744 | 7.842 |
| mammals | Plecturocebus_olallae | Callicebus_olallae | CR | Decreasing | tropical | diurnal | 992.4 | 677.034 | 7.771 |
| mammals | Plecturocebus_ornatus | Callicebus_ornatus | VU | Decreasing | tropical | diurnal | 832.6 | 23835.554 | 7.739 |
| mammals | Plecturocebus_pallescens | Callicebus_pallescens | LC | Decreasing | tropical | diurnal | 832.1 | 323682.676 | 4.438 |
| mammals | Plecturocebus_stephennashi | NA | DD | Unknown | NA | NA | NA | NA | NA |
| mammals | Plerotes_anchietae | Plerotes_anchietae | LC | Unknown | tropical | nocturnal | 38 | 796433.155 | 6.658 |
| mammals | Podogymnura_aureospinula | Podogymnura_aureospinula | EN | Decreasing | tropical | nocturnal | 247 | 1377.622 | 12.035 |
| mammals | Podogymnura_truei | Podogymnura_truei | LC | Stable | tropical | nocturnal | 114 | 5291.514 | 8.389 |
| mammals | Podomys_floridanus | Podomys_floridanus | NT | Decreasing | temperate | nocturnal | 30.8 | 47852.638 | 22.46 |
| mammals | Podoxymys_roraimae | Podoxymys_roraimae | VU | Unknown | tropical | nocturnal | 33.8 | 16615.078 | 2.476 |
| mammals | Poecilictis_libyca | Ictonyx_libycus | LC | Unknown | temperate | nocturnal | 2062.5 | 5082352.293 | 8.176 |
| mammals | Poecilogale_albinucha | Poecilogale_albinucha | LC | Unknown | tropical | nocturnal | 340 | 5663520.452 | 9.316 |
| mammals | Poelagus_marjorita | Poelagus_marjorita | LC | Decreasing | tropical | nocturnal | 2500 | 123506.317 | 5.997 |
| mammals | Poemys_melanotis | Dendromus_melanotis | LC | Stable | tropical | nocturnal | 9.1 | 4305594.488 | 8.946 |
| mammals | Poemys_nyikae | Dendromus_nyikae | LC | Unknown | tropical | nocturnal | 15 | 2375792.291 | 8.112 |
| mammals | Pogonomelomys_bruijnii | Pogonomelomys_bruijnii | LC | Decreasing | tropical | nocturnal | 178.8 | 26888.31 | 4.411 |
| mammals | Pogonomelomys_mayeri | Pogonomelomys_mayeri | LC | Unknown | tropical | nocturnal | 109.7 | 102264.114 | 3.942 |
| mammals | Pogonomys_championi | Pogonomys_championi | DD | Stable | tropical | nocturnal | 50.5 | 220.674 | 1.782 |

| | | | | | | | | | |
|---|---|---|---|---|---|---|---|---|---|
| mammals | Pogonomys_fergussoniensis | Pogonomys_fergussoniensis | EN | Decreasing | tropical | nocturnal | 128.5 | 2050.819 | 5.055 |
| mammals | Pogonomys_loriae | Pogonomys_loriae | LC | Unknown | tropical | nocturnal | 95.1 | 151035.631 | 4.566 |
| mammals | Pogonomys_macrourus | Pogonomys_macrourus | LC | Unknown | tropical | nocturnal | 54 | 317142.642 | 4.207 |
| mammals | Pogonomys_sylvestris | Pogonomys_sylvestris | LC | Unknown | tropical | nocturnal | 43.4 | 117161.339 | 4.975 |
| mammals | Poiana_leightoni | Poiana_leightoni | VU | Decreasing | tropical | nocturnal | 600 | 49336.175 | 10.684 |
| mammals | Poiana_richardsonii | Poiana_richardsonii | LC | Unknown | tropical | nocturnal | 500 | 2017237.125 | 5.746 |
| mammals | Poliocitellus_franklinii | Poliocitellus_franklinii | LC | Decreasing | temperate | diurnal | 363 | 1657975.706 | 16.272 |
| mammals | Pongo_abelii | Pongo_abelii | CR | Decreasing | tropical | diurnal | 56750 | 33185.922 | 5.95 |
| mammals | Pongo_pygmaeus | Pongo_pygmaeus | CR | Decreasing | tropical | diurnal | 57150 | 249610.106 | 4.34 |
| mammals | Porcula_salvania | Porcula_salvania | EN | Unknown | temperate | diurnal | 7718 | 881.679 | 11.93 |
| mammals | Potamochoerus_larvatus | Potamochoerus_larvatus | LC | Stable | tropical | nocturnal | 97500 | 5881488.572 | 9.888 |
| mammals | Potamochoerus_porcus | Potamochoerus_porcus | LC | Decreasing | tropical | nocturnal | 70000.3 | 4799557.048 | 8.006 |
| mammals | Potamogale_velox | Potamogale_velox | LC | Decreasing | tropical | nocturnal | 660 | 3803856.976 | 6.537 |
| mammals | Potorous_longipes | Potorous_longipes | VU | Decreasing | temperate | nocturnal | 1900 | 4229.452 | 6.408 |
| mammals | Potorous_tridactylus | Potorous_tridactylus | NT | Decreasing | temperate | nocturnal | 1100 | 125038.539 | 7.9 |
| mammals | Potos_flavus | Potos_flavus | LC | Decreasing | tropical | nocturnal | 3000 | 12691116.841 | 5.577 |
| mammals | Praomys_daltoni | Praomys_daltoni | LC | Stable | tropical | nocturnal | 38.5 | 3198976.106 | 12.409 |
| mammals | Praomys_degraaffi | Praomys_degraaffi | LC | Decreasing | tropical | nocturnal | 45.5 | 12854.583 | 11.871 |
| mammals | Praomys_hartwigi | Praomys_hartwigi | VU | Decreasing | tropical | nocturnal | 57.2 | 7809.654 | 12.81 |
| mammals | Praomys_jacksoni | Praomys_jacksoni | LC | Stable | tropical | nocturnal | 42 | 3744052.126 | 7.064 |
| mammals | Praomys_minor | Praomys_minor | LC | Unknown | tropical | nocturnal | 42.8 | 8768.524 | 8.405 |
| mammals | Praomys_misonnei | Praomys_misonnei | LC | Stable | tropical | nocturnal | 39.1 | 144160.072 | 3.997 |
| mammals | Praomys_morio | Praomys_morio | EN | Decreasing | tropical | nocturnal | 30 | 1343.909 | 10.185 |
| mammals | Praomys_mutoni | Praomys_mutoni | NT | Unknown | tropical | nocturnal | 52 | 14876.308 | 9.722 |
| mammals | Praomys_obscurus | Praomys_obscurus | EN | Decreasing | tropical | nocturnal | 47.2 | 3123.226 | 9.526 |
| mammals | Praomys_petteri | Praomys_petteri | LC | Stable | tropical | nocturnal | 49.3 | 218482.97 | 7.051 |
| mammals | Praomys_rostratus | Praomys_rostratus | LC | Unknown | tropical | nocturnal | 52 | 612377.039 | 14.857 |
| mammals | Praomys_tullbergi | Praomys_tullbergi | LC | Stable | tropical | nocturnal | 35.8 | 932845.126 | 14.559 |
| mammals | Presbytis_chrysomelas | Presbytis_chrysomelas | CR | Decreasing | tropical | diurnal | 7000 | 39860.909 | 5.767 |
| mammals | Presbytis_comata | Presbytis_comata | VU | Decreasing | tropical | diurnal | 6250 | 41162.76 | 18.553 |
| mammals | Presbytis_femoralis | Presbytis_femoralis | CR | Decreasing | tropical | diurnal | 7000 | 11279.71 | 18.197 |
| mammals | Presbytis_frontata | Presbytis_frontata | VU | Decreasing | tropical | diurnal | 6050 | 313412.235 | 4.167 |
| mammals | Presbytis_hosei | Presbytis_hosei | VU | Decreasing | tropical | diurnal | 6500 | 125048.689 | 4.555 |
| mammals | Presbytis_melalophos | Presbytis_melalophos | EN | Decreasing | tropical | diurnal | 6300 | 94639.092 | 8.837 |

| | | | | | | | | | |
|---|---|---|---|---|---|---|---|---|---|
| mammals | Presbytis_natunae | Presbytis_natunae | VU | Decreasing | tropical | diurnal | 4920 | 1643.704 | 9.354 |
| mammals | Presbytis_potenziani | Presbytis_potenziani | CR | Decreasing | tropical | diurnal | 6450 | 2140.853 | 8.542 |
| mammals | Presbytis_rubicunda | Presbytis_rubicunda | VU | Decreasing | tropical | diurnal | 6300 | 503494.897 | 4.772 |
| mammals | Presbytis_siamensis | Presbytis_siamensis | NT | Decreasing | tropical | diurnal | 7000 | 160647.489 | 10.569 |
| mammals | Presbytis_thomasi | Presbytis_thomasi | VU | Decreasing | tropical | diurnal | 6690 | 60508.915 | 8.542 |
| mammals | Priodontes_maximus | Priodontes_maximus | VU | Decreasing | tropical | nocturnal | 45359.7 | 9614613.558 | 3.841 |
| mammals | Prionailurus_bengalensis | Prionailurus_bengalensis | LC | Stable | temperate | cathemeral | 3300 | 8627600.165 | 13.481 |
| mammals | Prionailurus_planiceps | Prionailurus_planiceps | EN | Decreasing | tropical | nocturnal | 6750 | 148872.288 | 4.242 |
| mammals | Prionailurus_rubiginosus | Prionailurus_rubiginosus | NT | Decreasing | tropical | nocturnal | 1384.4 | 2416123.723 | 18.708 |
| mammals | Prionailurus_viverrinus | Prionailurus_viverrinus | VU | Decreasing | tropical | nocturnal | 9140.3 | 235697.123 | 16.784 |
| mammals | Prionodon_linsang | Prionodon_linsang | LC | Decreasing | tropical | nocturnal | 700 | 1447867.954 | 8.029 |
| mammals | Prionodon_pardicolor | Prionodon_pardicolor | LC | Decreasing | tropical | nocturnal | 512 | 2342407.952 | 10.755 |
| mammals | Prionomys_batesi | NA | DD | Unknown | NA | NA | NA | NA | NA |
| mammals | Procapra_gutturosa | Procapra_gutturosa | LC | Stable | temperate | diurnal | 27750 | 715708.203 | 4.74 |
| mammals | Procapra_przewalskii | Procapra_przewalskii | EN | Increasing | temperate | diurnal | 27500 | 1076.301 | 9.604 |
| mammals | Procavia_capensis | Procavia_capensis | LC | Stable | tropical | diurnal | 3030 | 15715542.142 | 8.523 |
| mammals | Procolobus_verus | Procolobus_verus | VU | Decreasing | tropical | diurnal | 4000 | 412576.702 | 15.462 |
| mammals | Procyon_cancrivorus | Procyon_cancrivorus | LC | Decreasing | tropical | nocturnal | 6949.9 | 12754113.762 | 5.762 |
| mammals | Procyon_lotor | Procyon_lotor | LC | Increasing | temperate | cathemeral | 6550 | 11684834.857 | 12.18 |
| mammals | Procyon_pygmaeus | Procyon_pygmaeus | CR | Decreasing | tropical | nocturnal | 2500 | 488.354 | 10.206 |
| mammals | Proechimys_brevicauda | Proechimys_brevicauda | LC | Unknown | tropical | nocturnal | 285 | 1460844.711 | 2.322 |
| mammals | Proechimys_canicollis | Proechimys_canicollis | LC | Stable | tropical | nocturnal | 285 | 32608.098 | 16.288 |
| mammals | Proechimys_chrysaeolus | NA | DD | Unknown | NA | NA | NA | NA | NA |
| mammals | Proechimys_cuvieri | Proechimys_cuvieri | LC | Unknown | tropical | nocturnal | 330 | 3500125.381 | 1.397 |
| mammals | Proechimys_decumanus | Proechimys_decumanus | NT | Decreasing | tropical | nocturnal | 285 | 65536.943 | 13.563 |
| mammals | Proechimys_echinothrix | Proechimys_echinothrix | LC | Unknown | tropical | nocturnal | 309.6 | 277424.044 | 0.472 |
| mammals | Proechimys_gardneri | NA | DD | Unknown | NA | NA | NA | NA | NA |
| mammals | Proechimys_goeldii | Proechimys_goeldii | LC | Stable | tropical | nocturnal | 285 | 1029773.277 | 2.175 |
| mammals | Proechimys_guairae | Proechimys_guairae | LC | Unknown | tropical | nocturnal | 400 | 329800.189 | 11.837 |
| mammals | Proechimys_guyannensis | Proechimys_guyannensis | LC | Unknown | tropical | nocturnal | 316 | 1497170.028 | 1.431 |
| mammals | Proechimys_hoplomyoides | NA | DD | Unknown | NA | NA | NA | NA | NA |
| mammals | Proechimys_kulinae | NA | DD | Unknown | NA | NA | NA | NA | NA |
| mammals | Proechimys_longicaudatus | Proechimys_longicaudatus | LC | Unknown | tropical | nocturnal | 205 | 1349084.798 | 7.188 |
| mammals | Proechimys_mincae | NA | DD | Unknown | NA | NA | NA | NA | NA |

| | | | | | | | | | |
|---|---|---|---|---|---|---|---|---|---|
| mammals | Proechimys_oconnelli | NA | DD | Unknown | NA | NA | NA | NA | NA |
| mammals | Proechimys_pattoni | Proechimys_pattoni | LC | Unknown | tropical | nocturnal | 178.2 | 157595.734 | 1.078 |
| mammals | Proechimys_quadruplicatus | Proechimys_quadruplicatus | LC | Stable | tropical | nocturnal | 285 | 1498656.176 | 1.236 |
| mammals | Proechimys_roberti | Proechimys_roberti | LC | Stable | tropical | nocturnal | 285 | 1703345.297 | 4.878 |
| mammals | Proechimys_semispinosus | Proechimys_semispinosus | LC | Stable | tropical | nocturnal | 285 | 348118.048 | 9.005 |
| mammals | Proechimys_simonsi | Proechimys_simonsi | LC | Unknown | tropical | nocturnal | 285 | 1935012.331 | 2.083 |
| mammals | Proechimys_steerei | Proechimys_steerei | LC | Unknown | tropical | nocturnal | 323 | 1677776.522 | 1.261 |
| mammals | Proechimys_trinitatis | NA | DD | Unknown | NA | NA | NA | NA | NA |
| mammals | Proedromys_bedfordi | NA | VU | Decreasing | NA | NA | NA | NA | NA |
| mammals | Prolemur_simus | Prolemur_simus | CR | Decreasing | tropical | cathemeral | 2350 | 1709.042 | 6.392 |
| mammals | Promops_centralis | Promops_centralis | LC | Unknown | tropical | nocturnal | 26.4 | 4729320.399 | 7.072 |
| mammals | Promops_davisoni | NA | DD | Unknown | NA | NA | NA | NA | NA |
| mammals | Promops_nasutus | Promops_nasutus | LC | Unknown | tropical | nocturnal | 15.2 | 8029235.887 | 4.812 |
| mammals | Pronolagus_crassicaudatus | Pronolagus_crassicaudatus | LC | Decreasing | temperate | nocturnal | 2089 | 198444.314 | 13.694 |
| mammals | Pronolagus_randensis | Pronolagus_randensis | LC | Decreasing | tropical | nocturnal | 2335 | 675441.128 | 9.902 |
| mammals | Pronolagus_rupestris | Pronolagus_rupestris | LC | Unknown | temperate | nocturnal | 1754 | 461340.914 | 9.707 |
| mammals | Propithecus_candidus | Propithecus_candidus | CR | Decreasing | tropical | diurnal | 5750 | 1360.656 | 5.64 |
| mammals | Propithecus_coquereli | Propithecus_coquereli | CR | Decreasing | tropical | diurnal | 3900 | 7389.955 | 8.866 |
| mammals | Propithecus_coronatus | Propithecus_coronatus | CR | Decreasing | tropical | diurnal | 3900 | 45223.885 | 8.501 |
| mammals | Propithecus_deckenii | Propithecus_deckenii | CR | Decreasing | tropical | diurnal | 4000 | 25611.858 | 7.219 |
| mammals | Propithecus_diadema | Propithecus_diadema | CR | Decreasing | tropical | diurnal | 5250 | 16421.244 | 6.494 |
| mammals | Propithecus_edwardsi | Propithecus_edwardsi | EN | Decreasing | tropical | diurnal | 5750 | 908.696 | 6.343 |
| mammals | Propithecus_perrieri | Propithecus_perrieri | CR | Decreasing | tropical | diurnal | 4350 | 62.3 | 5.795 |
| mammals | Propithecus_tattersalli | Propithecus_tattersalli | CR | Decreasing | tropical | diurnal | 3500 | 1772.872 | 7.646 |
| mammals | Propithecus_verreauxi | Propithecus_verreauxi | CR | Decreasing | tropical | diurnal | 3250 | 30758.376 | 8.316 |
| mammals | Prosciurillus_abstrusus | NA | DD | Unknown | NA | NA | NA | NA | NA |
| mammals | Prosciurillus_leucomus | Prosciurillus_leucomus | LC | Unknown | tropical | diurnal | 128 | 37690.891 | 9.023 |
| mammals | Prosciurillus_murinus | Prosciurillus_murinus | LC | Unknown | tropical | diurnal | 72.9 | 67498.16 | 9.564 |
| mammals | Prosciurillus_rosenbergii | Prosciurillus_rosenbergii | LC | Stable | tropical | diurnal | 222.6 | 555.948 | 11.586 |
| mammals | Prosciurillus_weberi | Prosciurillus_weberi | EN | Unknown | tropical | diurnal | 132.9 | 1824.084 | 15.163 |
| mammals | Proteles_cristatus | Proteles_cristata | LC | Stable | temperate | nocturnal | 10000 | 5997315.148 | 8.381 |
| mammals | Protoxerus_aubinnii | Protoxerus_aubinnii | NT | Unknown | tropical | diurnal | 350 | 134472.882 | 13.047 |
| mammals | Protoxerus_stangeri | Protoxerus_stangeri | LC | Unknown | tropical | diurnal | 621.5 | 3388918.949 | 8.713 |
| mammals | Psammomys_obesus | Psammomys_obesus | LC | Stable | temperate | cathemeral | 369.6 | 2313613.016 | 5.24 |

| | | | | | | | | | |
|---|---|---|---|---|---|---|---|---|---|
| mammals | Psammomys_vexillaris | NA | DD | Unknown | NA | NA | NA | NA | NA |
| mammals | Pseudantechinus_bilarni | Pseudantechinus_bilarni | LC | Decreasing | tropical | nocturnal | 27.5 | 134277.934 | 1.495 |
| mammals | Pseudantechinus_macdonnellensis | Pseudantechinus_macdonnellensis | LC | Unknown | temperate | nocturnal | 32.5 | 1267010.727 | 1.642 |
| mammals | Pseudantechinus_mimulus | Pseudantechinus_mimulus | NT | Decreasing | tropical | nocturnal | 17 | 3092.032 | 5.259 |
| mammals | Pseudantechinus_ningbing | Pseudantechinus_ningbing | LC | Unknown | tropical | nocturnal | 20 | 208494.502 | 2.215 |
| mammals | Pseudantechinus_woolleyae | Pseudantechinus_woolleyae | LC | Unknown | temperate | nocturnal | 40 | 645661.157 | 4.25 |
| mammals | Pseudocheirus_occidentalis | Pseudocheirus_occidentalis | CR | Decreasing | temperate | nocturnal | 987.5 | 18007.813 | 9.231 |
| mammals | Pseudocheirus_peregrinus | Pseudocheirus_peregrinus | LC | Stable | temperate | nocturnal | 900 | 916198.631 | 8.059 |
| mammals | Pseudochirops_albertisii | Pseudochirops_albertisii | NT | Decreasing | tropical | nocturnal | 784.9 | 25738.523 | 3.533 |
| mammals | Pseudochirops_archeri | Pseudochirops_archeri | NT | Decreasing | tropical | nocturnal | 1091.5 | 5210.844 | 3.823 |
| mammals | Pseudochirops_corinnae | Pseudochirops_corinnae | NT | Decreasing | tropical | nocturnal | 1123 | 185300.979 | 4.674 |
| mammals | Pseudochirops_coronatus | Pseudochirops_coronatus | VU | Decreasing | tropical | nocturnal | 1500 | 7454.749 | 2.901 |
| mammals | Pseudochirops_cupreus | Pseudochirops_cupreus | LC | Stable | tropical | nocturnal | 1724.5 | 197734.762 | 4.57 |
| mammals | Pseudochirops_dahlii | Petropseudes_dahli | LC | Unknown | tropical | nocturnal | 1640 | 198708.378 | 1.912 |
| mammals | Pseudochirulus_canescens | Pseudochirulus_canescens | LC | Decreasing | tropical | nocturnal | 300 | 459129.801 | 3.264 |
| mammals | Pseudochirulus_caroli | Pseudochirulus_caroli | LC | Unknown | tropical | nocturnal | 440 | 37928.033 | 4.885 |
| mammals | Pseudochirulus_cinereus | Pseudochirulus_cinereus | NT | Decreasing | tropical | nocturnal | 1000 | 895.814 | 1.056 |
| mammals | Pseudochirulus_forbesi | Pseudochirulus_forbesi | LC | Unknown | tropical | nocturnal | 649 | 47168.026 | 3.338 |
| mammals | Pseudochirulus_herbertensis | Pseudochirulus_herbertensis | LC | Stable | tropical | nocturnal | 1110.5 | 3432.756 | 3.709 |
| mammals | Pseudochirulus_larvatus | Pseudochirulus_larvatus | LC | Decreasing | tropical | nocturnal | 948.8 | 121062.006 | 5.305 |
| mammals | Pseudochirulus_mayeri | Pseudochirulus_mayeri | LC | Decreasing | tropical | nocturnal | 151.8 | 84923.68 | 4.656 |
| mammals | Pseudochirulus_schlegelii | Pseudochirulus_schlegeli | VU | Decreasing | tropical | nocturnal | 274.5 | 10303.009 | 2.729 |
| mammals | Pseudohydromys_berniceae | NA | LC | Unknown | NA | NA | NA | NA | NA |
| mammals | Pseudohydromys_ellermani | Pseudohydromys_ellermani | LC | Unknown | tropical | nocturnal | 22.5 | 109570.264 | 5.323 |
| mammals | Pseudohydromys_fuscus | Pseudohydromys_fuscus | LC | Stable | tropical | nocturnal | 18.8 | 66904.571 | 6.332 |
| mammals | Pseudois_nayaur | Pseudois_nayaur | LC | Unknown | temperate | diurnal | 45000 | 2858794.559 | 4.714 |
| mammals | Pseudomys_albocinereus | Pseudomys_albocinereus | LC | Stable | temperate | nocturnal | 30.5 | 221496.578 | 9.427 |
| mammals | Pseudomys_apodemoides | Pseudomys_apodemoides | LC | Stable | temperate | nocturnal | 20 | 35937.018 | 8.565 |
| mammals | Pseudomys_australis | Pseudomys_australis | VU | Decreasing | temperate | nocturnal | 65 | 199918.689 | 3.253 |
| mammals | Pseudomys_bolami | Pseudomys_bolami | LC | Stable | temperate | nocturnal | 15.5 | 746188.585 | 3.39 |
| mammals | Pseudomys_calabyi | Pseudomys_calabyi | VU | Decreasing | tropical | nocturnal | 16 | 20211.852 | 2.535 |
| mammals | Pseudomys_chapmani | Pseudomys_chapmani | LC | Stable | tropical | nocturnal | 12 | 299581.297 | 3.941 |
| mammals | Pseudomys_delicatulus | Pseudomys_delicatulus | LC | Unknown | tropical | nocturnal | 11 | 1842508.198 | 3.741 |
| mammals | Pseudomys_desertor | Pseudomys_desertor | LC | Decreasing | temperate | diurnal | 25 | 5158688.13 | 3.175 |

| | | | | | | | | | |
|---|---|---|---|---|---|---|---|---|---|
| mammals | Pseudomys_fumeus | Pseudomys_fumeus | VU | Decreasing | temperate | nocturnal | 70 | 16696.309 | 6.315 |
| mammals | Pseudomys_gouldii | NA | EX | not_available | NA | NA | NA | NA | NA |
| mammals | Pseudomys_gracilicaudatus | Pseudomys_gracilicaudatus | LC | Decreasing | tropical | nocturnal | 74.8 | 337741.326 | 6.795 |
| mammals | Pseudomys_hermannsburgensis | Pseudomys_hermannsburgensis | LC | Stable | temperate | nocturnal | 12 | 3846942.822 | 2.729 |
| mammals | Pseudomys_higginsi | Pseudomys_higginsi | LC | Stable | temperate | nocturnal | 67 | 64548.424 | 5.864 |
| mammals | Pseudomys_johnsoni | Pseudomys_johnsoni | LC | Stable | tropical | nocturnal | 12 | 766530.121 | 2.091 |
| mammals | Pseudomys_nanus | Pseudomys_nanus | LC | Decreasing | tropical | nocturnal | 34 | 1006949.822 | 1.913 |
| mammals | Pseudomys_novaehollandiae | Pseudomys_novaehollandiae | VU | Decreasing | temperate | nocturnal | 20 | 89948.949 | 6.854 |
| mammals | Pseudomys_occidentalis | Pseudomys_occidentalis | NT | Decreasing | temperate | nocturnal | 34 | 51491.415 | 9.51 |
| mammals | Pseudomys_oralis | Pseudomys_oralis | VU | Decreasing | temperate | nocturnal | 95 | 26516.215 | 5.194 |
| mammals | Pseudomys_patrius | Pseudomys_patrius | LC | Unknown | tropical | nocturnal | 14.5 | 404417.544 | 5.591 |
| mammals | Pseudomys_shortridgei | Pseudomys_shortridgei | NT | Decreasing | temperate | diurnal | 70 | 32075.836 | 8.749 |
| mammals | Pseudoromicia_brunnea | Neoromicia_brunnea | NT | Decreasing | tropical | nocturnal | 9.9 | 757189.018 | 11.229 |
| mammals | Pseudoromicia_rendalli | Neoromicia_rendalli | LC | Unknown | tropical | nocturnal | 6.3 | 8376297.098 | 10.203 |
| mammals | Pseudoromicia_tenuipinnis | Neoromicia_tenuipinnis | LC | Unknown | tropical | nocturnal | 5.3 | 4247435.612 | 9.078 |
| mammals | Pseudoryx_nghetinhensis | Pseudoryx_nghetinhensis | CR | Decreasing | tropical | diurnal | 85000 | 13994.994 | 5.296 |
| mammals | Pseudoryzomys_simplex | Pseudoryzomys_simplex | LC | Unknown | tropical | nocturnal | 51.3 | 3116984.095 | 8.21 |
| mammals | Pseudosoriculus_fumidus | Episoriculus_fumidus | LC | Unknown | temperate | cathemeral | 6.7 | 12808.648 | 8.235 |
| mammals | Ptenochirus_jagorii | Ptenochirus_jagori | LC | Stable | tropical | nocturnal | 77.3 | 271017.144 | 13.899 |
| mammals | Ptenochirus_minor | Ptenochirus_minor | LC | Stable | tropical | nocturnal | 36.3 | 118793.761 | 12.099 |
| mammals | Ptenochirus_wetmorei | Megaerops_wetmorei | VU | Decreasing | tropical | nocturnal | 18.5 | 165916.685 | 8.322 |
| mammals | Pteralopex_anceps | Pteralopex_anceps | EN | Decreasing | tropical | nocturnal | 560.4 | 8736.504 | 6.301 |
| mammals | Pteralopex_atrata | Pteralopex_atrata | EN | Decreasing | tropical | nocturnal | 472 | 5153.554 | 5.854 |
| mammals | Pteralopex_flanneryi | Pteralopex_flanneryi | CR | Decreasing | tropical | nocturnal | 670.2 | 14052.402 | 4.859 |
| mammals | Pteralopex_taki | Pteralopex_taki | VU | Decreasing | tropical | nocturnal | 286.7 | 3316.801 | 2.66 |
| mammals | Pteromys_momonga | Pteromys_momonga | LC | Unknown | temperate | nocturnal | 151.8 | 168642.957 | 17.72 |
| mammals | Pteromys_volans | Pteromys_volans | LC | Decreasing | temperate | nocturnal | 143.8 | 13737442.366 | 3.883 |
| mammals | Pteromyscus_pulverulentus | Pteromyscus_pulverulentus | EN | Decreasing | tropical | nocturnal | 334.3 | 393205.985 | 8.837 |
| mammals | Pteronotus_davyi | Pteronotus_davyi | LC | Stable | tropical | nocturnal | 9.2 | 3461633.974 | 7.669 |
| mammals | Pteronotus_gymnonotus | Pteronotus_gymnonotus | LC | Stable | tropical | nocturnal | 14.4 | 3202565.123 | 6.108 |
| mammals | Pteronotus_macleayii | Pteronotus_macleayii | LC | Stable | tropical | nocturnal | 5.4 | 119896.668 | 16.516 |
| mammals | Pteronotus_mesoamericanus | Pteronotus_mesoamericanus | LC | Unknown | tropical | nocturnal | 17.75 | 539712.782 | 10.81 |
| mammals | Pteronotus_paraguanensis | Pteronotus_paraguanensis | EN | Decreasing | tropical | nocturnal | 13.9 | 142.806 | 18.868 |
| mammals | Pteronotus_parnellii | Pteronotus_parnellii | LC | Stable | tropical | nocturnal | 19.7 | 200546.183 | 16.762 |

| | | | | | | | | | |
|---|---|---|---|---|---|---|---|---|---|
| mammals | Pteronotus_personatus | Pteronotus_personatus | LC | Stable | tropical | nocturnal | 8.7 | 4748166.222 | 7.244 |
| mammals | Pteronotus_quadridens | Pteronotus_quadridens | LC | Stable | tropical | nocturnal | 5.2 | 200546.183 | 16.762 |
| mammals | Pteronotus_rubiginosus | Pteronotus_rubiginosus | LC | Unknown | tropical | nocturnal | 23.5 | 4505960.104 | 2.18 |
| mammals | Pteronura_brasiliensis | Pteronura_brasiliensis | EN | Decreasing | tropical | diurnal | 23999.9 | 6631572.763 | 2.467 |
| mammals | Pteropus_admiralitatum | Pteropus_admiralitatum | LC | Stable | tropical | nocturnal | 299.9 | 75834.564 | 4.57 |
| mammals | Pteropus_aldabrensis | NA | EN | Stable | NA | NA | NA | NA | NA |
| mammals | Pteropus_alecto | Pteropus_alecto | LC | Unknown | tropical | nocturnal | 643.6 | 1427125.684 | 4.506 |
| mammals | Pteropus_anetianus | Pteropus_anetianus | VU | Decreasing | tropical | nocturnal | 426 | 11890.491 | 5.905 |
| mammals | Pteropus_aruensis | Pteropus_aruensis | CR | Unknown | tropical | nocturnal | 743.2 | 7030.018 | 4.214 |
| mammals | Pteropus_caniceps | Pteropus_caniceps | VU | Decreasing | tropical | nocturnal | 505 | 22506.982 | 6.522 |
| mammals | Pteropus_capistratus | Pteropus_capistratus | VU | Decreasing | tropical | nocturnal | 223 | 36299.768 | 4.207 |
| mammals | Pteropus_chrysoproctus | Pteropus_chrysoproctus | VU | Decreasing | tropical | nocturnal | 710 | 29919.416 | 6.883 |
| mammals | Pteropus_cognatus | Pteropus_cognatus | VU | Decreasing | tropical | nocturnal | 302.5 | 3106.404 | 4.63 |
| mammals | Pteropus_conspicillatus | Pteropus_conspicillatus | EN | Decreasing | tropical | nocturnal | 775.7 | 219525.84 | 4.504 |
| mammals | Pteropus_dasymallus | Pteropus_dasymallus | VU | Decreasing | temperate | nocturnal | 435 | 2890.665 | 19.91 |
| mammals | Pteropus_faunulus | Pteropus_faunulus | EN | Decreasing | tropical | nocturnal | 288 | 602.969 | 10.194 |
| mammals | Pteropus_fundatus | Pteropus_fundatus | EN | Unknown | tropical | nocturnal | 205.7 | 379.794 | 2.578 |
| mammals | Pteropus_gilliardi | Pteropus_gilliardorum | VU | Unknown | tropical | nocturnal | 400 | 3297.525 | 2.833 |
| mammals | Pteropus_griseus | Pteropus_griseus | VU | Decreasing | tropical | nocturnal | 269 | 222132.823 | 10.038 |
| mammals | Pteropus_hypomelanus | Pteropus_hypomelanus | NT | Decreasing | tropical | nocturnal | 428.1 | 524209.43 | 11.128 |
| mammals | Pteropus_intermedius | NA | DD | Unknown | NA | NA | NA | NA | NA |
| mammals | Pteropus_keyensis | NA | DD | Unknown | NA | NA | NA | NA | NA |
| mammals | Pteropus_livingstonii | Pteropus_livingstonii | CR | Decreasing | tropical | nocturnal | 588.5 | 265.682 | 13.08 |
| mammals | Pteropus_lombocensis | NA | DD | Unknown | NA | NA | NA | NA | NA |
| mammals | Pteropus_lylei | Pteropus_lylei | VU | Decreasing | temperate | nocturnal | 316.7 | 106826.975 | 20.183 |
| mammals | Pteropus_macrotis | Pteropus_macrotis | LC | Stable | tropical | nocturnal | 357 | 619521.294 | 3.075 |
| mammals | Pteropus_mahaganus | Pteropus_mahaganus | LC | Decreasing | tropical | nocturnal | 251.4 | 12629.986 | 4.854 |
| mammals | Pteropus_mariannus | Pteropus_mariannus | EN | Decreasing | temperate | nocturnal | 453.5 | 2221.338 | 24.201 |
| mammals | Pteropus_melanopogon | Pteropus_melanopogon | EN | Decreasing | tropical | nocturnal | 860 | 31588.241 | 6.639 |
| mammals | Pteropus_melanotus | Pteropus_melanotus | VU | Decreasing | tropical | diurnal | 375 | 399.667 | 8.693 |
| mammals | Pteropus_molossinus | Pteropus_molossinus | EN | Stable | tropical | diurnal | 178 | 352.295 | 12.251 |
| mammals | Pteropus_neohibernicus | Pteropus_neohibernicus | LC | Stable | tropical | nocturnal | 966.8 | 672585.197 | 3.128 |
| mammals | Pteropus_niger | Pteropus_niger | EN | Decreasing | tropical | nocturnal | 456.3 | 4413.639 | 22.868 |
| mammals | Pteropus_nitendiensis | Pteropus_nitendiensis | EN | Decreasing | tropical | nocturnal | 265 | 563.676 | 4.364 |

| | | | | | | | | | |
|---|---|---|---|---|---|---|---|---|---|
| mammals | Pteropus_ocularis | Pteropus_ocularis | VU | Decreasing | tropical | nocturnal | 220 | 26749.185 | 7.104 |
| mammals | Pteropus_ornatus | Pteropus_ornatus | VU | Decreasing | tropical | nocturnal | 330 | 17983.231 | 7.482 |
| mammals | Pteropus_personatus | Pteropus_personatus | LC | Stable | tropical | nocturnal | 127.5 | 25277.05 | 6.337 |
| mammals | Pteropus_pohlei | Pteropus_pohlei | VU | Decreasing | tropical | nocturnal | 343 | 2597.564 | 6.372 |
| mammals | Pteropus_poliocephalus | Pteropus_poliocephalus | VU | Decreasing | temperate | nocturnal | 749.9 | 454359.021 | 9.893 |
| mammals | Pteropus_pselaphon | NA | EN | Stable | NA | NA | NA | NA | NA |
| mammals | Pteropus_pumilus | Pteropus_pumilus | NT | Decreasing | tropical | nocturnal | 183.4 | 57993.959 | 15.815 |
| mammals | Pteropus_rayneri | Pteropus_rayneri | NT | Decreasing | tropical | nocturnal | 724.5 | 30622.524 | 4.754 |
| mammals | Pteropus_rennelli | Pteropus_rennelli | EN | Decreasing | tropical | nocturnal | 296.4 | 544.102 | 3.368 |
| mammals | Pteropus_rodricensis | NA | EN | Increasing | NA | NA | NA | NA | NA |
| mammals | Pteropus_rufus | Pteropus_rufus | VU | Decreasing | tropical | nocturnal | 361 | 185368.315 | 9.09 |
| mammals | Pteropus_samoensis | Pteropus_samoensis | NT | Decreasing | tropical | cathemeral | 310 | 20144.58 | 9.392 |
| mammals | Pteropus_scapulatus | Pteropus_scapulatus | LC | Unknown | temperate | nocturnal | 447.5 | 3033575.196 | 5.002 |
| mammals | Pteropus_seychellensis | Pteropus_seychellensis | LC | Stable | tropical | nocturnal | 457 | 2730.91 | 17.982 |
| mammals | Pteropus_speciosus | NA | DD | Unknown | NA | NA | NA | NA | NA |
| mammals | Pteropus_temminckii | Pteropus_temminckii | VU | Decreasing | tropical | nocturnal | 250 | 26749.128 | 7.104 |
| mammals | Pteropus_tonganus | Pteropus_tonganus | LC | Decreasing | tropical | nocturnal | 418.3 | 57838.601 | 7.366 |
| mammals | Pteropus_tuberculatus | Pteropus_tuberculatus | EN | Decreasing | tropical | nocturnal | 267.1 | 281.253 | 2.077 |
| mammals | Pteropus_ualanus | Pteropus_ualanus | EN | Stable | tropical | nocturnal | 348.9 | 109.223 | 10.433 |
| mammals | Pteropus_vampyrus | Pteropus_vampyrus | EN | Decreasing | tropical | nocturnal | 935.9 | 2043987.347 | 10.79 |
| mammals | Pteropus_vetulus | Pteropus_vetulus | NT | Decreasing | tropical | nocturnal | 148 | 16532.637 | 7.541 |
| mammals | Pteropus_voeltzkowi | Pteropus_voeltzkowi | VU | Increasing | tropical | nocturnal | 527.5 | 908.531 | 17.669 |
| mammals | Pteropus_woodfordi | Pteropus_woodfordi | LC | Decreasing | tropical | nocturnal | 119.3 | 13702.957 | 4.815 |
| mammals | Ptilocercus_lowii | Ptilocercus_lowii | LC | Decreasing | tropical | nocturnal | 41.9 | 356708.955 | 9.202 |
| mammals | Pudu_mephistophiles | Pudu_mephistophiles | DD | Decreasing | tropical | nocturnal | 9600 | 130792.064 | 9.115 |
| mammals | Puma_concolor | Puma_concolor | LC | Decreasing | tropical | cathemeral | 51600 | 22659138.002 | 6.632 |
| mammals | Punomys_kofordi | Punomys_kofordi | NT | Unknown | tropical | cathemeral | 82 | 1443.91 | 9.307 |
| mammals | Punomys_lemminus | Punomys_lemminus | VU | Unknown | tropical | diurnal | 84.8 | 47849.817 | 7.826 |
| mammals | Pygathrix_cinerea | Pygathrix_cinerea | CR | Decreasing | tropical | diurnal | 8310 | 24681.135 | 11.034 |
| mammals | Pygathrix_nemaeus | Pygathrix_nemaeus | CR | Decreasing | tropical | diurnal | 8310 | 91649.514 | 7.54 |
| mammals | Pygathrix_nigripes | Pygathrix_nigripes | CR | Decreasing | tropical | diurnal | 8310 | 95056.802 | 11.072 |
| mammals | Pygeretmus_platyurus | Pygeretmus_platyurus | LC | Unknown | temperate | nocturnal | 26.4 | 926949.308 | 5.672 |
| mammals | Pygeretmus_pumilio | Pygeretmus_pumilio | LC | Unknown | temperate | nocturnal | 52 | 3430883.498 | 6.446 |
| mammals | Pygeretmus_shitkovi | Pygeretmus_zhitkovi | LC | Unknown | temperate | nocturnal | 56.5 | 48808.719 | 9.296 |

| | | | | | | | | | |
|---|---|---|---|---|---|---|---|---|---|
| mammals | Pygoderma_bilabiatum | Pygoderma_bilabiatum | LC | Unknown | tropical | nocturnal | 18.5 | 2957165.72 | 10.974 |
| mammals | Rangifer_tarandus | Rangifer_tarandus | VU | Decreasing | temperate | cathemeral | 86034 | 18784282.673 | 1.252 |
| mammals | Raphicerus_campestris | Raphicerus_campestris | LC | Stable | tropical | cathemeral | 10500 | 4047877.073 | 8.069 |
| mammals | Raphicerus_melanotis | Raphicerus_melanotis | LC | Stable | temperate | nocturnal | 10249.9 | 160973.791 | 10.308 |
| mammals | Raphicerus_sharpei | Raphicerus_sharpei | LC | Stable | tropical | nocturnal | 9499.9 | 2462567.949 | 9.34 |
| mammals | Rattus_adustus | Rattus_adustus | DD | Decreasing | tropical | nocturnal | 158 | 399.965 | 8.692 |
| mammals | Rattus_andamanensis | Rattus_andamanensis | LC | Stable | tropical | nocturnal | 129 | 1747879.157 | 10.491 |
| mammals | Rattus_arfakiensis | NA | DD | Unknown | NA | NA | NA | NA | NA |
| mammals | Rattus_argentiventer | Rattus_argentiventer | LC | Stable | tropical | nocturnal | 133.1 | 2183638.249 | 10.795 |
| mammals | Rattus_arrogans | NA | LC | Stable | NA | NA | NA | NA | NA |
| mammals | Rattus_baluensis | Rattus_baluensis | LC | Stable | tropical | nocturnal | 108.8 | 709.869 | 8.639 |
| mammals | Rattus_blangorum | NA | DD | Unknown | NA | NA | NA | NA | NA |
| mammals | Rattus_bontanus | NA | DD | Unknown | NA | NA | NA | NA | NA |
| mammals | Rattus_burrus | NA | EN | Decreasing | NA | NA | NA | NA | NA |
| mammals | Rattus_ceramicus | Nesoromys_ceramicus | EN | Decreasing | tropical | nocturnal | 65.5 | 443.241 | 5.886 |
| mammals | Rattus_colletti | Rattus_colletti | LC | Unknown | tropical | nocturnal | 61 | 86055.59 | 2.201 |
| mammals | Rattus_elaphinus | Rattus_elaphinus | NT | Decreasing | tropical | nocturnal | 131.4 | 4062.508 | 5.228 |
| mammals | Rattus_enganus | Rattus_enganus | DD | Decreasing | tropical | nocturnal | 305 | 399.965 | 8.692 |
| mammals | Rattus_everetti | Rattus_everetti | LC | Stable | tropical | nocturnal | 255 | 241325.765 | 13.251 |
| mammals | Rattus_exulans | Rattus_exulans | LC | Stable | tropical | nocturnal | 47.3 | 4966659.436 | 9.529 |
| mammals | Rattus_feliceus | Rattus_feliceus | NT | Decreasing | tropical | nocturnal | 306 | 17400.278 | 7.056 |
| mammals | Rattus_fuscipes | Rattus_fuscipes | LC | Stable | temperate | nocturnal | 133 | 486689.526 | 8.545 |
| mammals | Rattus_giluwensis | Rattus_giluwensis | LC | Unknown | tropical | nocturnal | 116.7 | 3283.059 | 6.839 |
| mammals | Rattus_hainaldi | Rattus_hainaldi | EN | Decreasing | tropical | nocturnal | 81 | 748.498 | 10.588 |
| mammals | Rattus_hoffmanni | Rattus_hoffmanni | LC | Stable | tropical | nocturnal | 147 | 168758.239 | 9.259 |
| mammals | Rattus_hoogerwerfi | Rattus_hoogerwerfi | VU | Decreasing | tropical | nocturnal | 165 | 650.572 | 3.055 |
| mammals | Rattus_jobiensis | Rattus_jobiensis | LC | Unknown | tropical | nocturnal | 380 | 5155.67 | 7.553 |
| mammals | Rattus_koopmani | NA | DD | Unknown | NA | NA | NA | NA | NA |
| mammals | Rattus_korinchi | NA | DD | Unknown | NA | NA | NA | NA | NA |
| mammals | Rattus_leucopus | Rattus_leucopus | LC | Stable | tropical | nocturnal | 171.7 | 365095.78 | 2.848 |
| mammals | Rattus_losea | Rattus_losea | LC | Increasing | temperate | nocturnal | 77 | 2038643.177 | 15.673 |
| mammals | Rattus_lugens | Rattus_lugens | VU | Decreasing | tropical | nocturnal | 224 | 6004.261 | 6.223 |
| mammals | Rattus_lutreola | Rattus_lutreolus | LC | Decreasing | temperate | cathemeral | 106 | 260763.051 | 8.742 |
| mammals | Rattus_marmosurus | Rattus_marmosurus | LC | Decreasing | tropical | nocturnal | 217 | 93655.514 | 8.045 |

| | | | | | | | | | |
|---|---|---|---|---|---|---|---|---|---|
| mammals | Rattus_mindorensis | Rattus_mindorensis | LC | Unknown | tropical | nocturnal | 136 | 5016.035 | 6.753 |
| mammals | Rattus_mollicomulus | NA | LC | Stable | NA | NA | NA | NA | NA |
| mammals | Rattus_montanus | Rattus_montanus | EN | Unknown | tropical | diurnal | 136 | 315.95 | 10.3 |
| mammals | Rattus_mordax | Rattus_mordax | LC | Stable | tropical | nocturnal | 230 | 51171.652 | 5.292 |
| mammals | Rattus_morotaiensis | Rattus_morotaiensis | LC | Unknown | tropical | nocturnal | 148.3 | 22281.508 | 6.413 |
| mammals | Rattus_niobe | Rattus_niobe | LC | Stable | tropical | nocturnal | 46.1 | 110059.911 | 5.195 |
| mammals | Rattus_nitidus | Rattus_nitidus | LC | Stable | temperate | cathemeral | 133.5 | 3665292.809 | 12.681 |
| mammals | Rattus_norvegicus | Rattus_norvegicus | LC | Stable | temperate | nocturnal | 356.7 | 19191093.605 | 11.222 |
| mammals | Rattus_novaeguineae | Rattus_novaeguineae | LC | Unknown | tropical | nocturnal | 133 | 34947.82 | 4.787 |
| mammals | Rattus_omichlodes | NA | DD | Unknown | NA | NA | NA | NA | NA |
| mammals | Rattus_osgoodi | Rattus_osgoodi | LC | Stable | tropical | nocturnal | 93 | 6032.165 | 9.251 |
| mammals | Rattus_palmarum | Rattus_palmarum | VU | Unknown | tropical | nocturnal | 491 | 1092.556 | 5.441 |
| mammals | Rattus_pelurus | NA | DD | Unknown | NA | NA | NA | NA | NA |
| mammals | Rattus_pococki | NA | LC | Unknown | NA | NA | NA | NA | NA |
| mammals | Rattus_praetor | Rattus_praetor | LC | Stable | tropical | nocturnal | 204.1 | 310739.949 | 3.328 |
| mammals | Rattus_pyctoris | Rattus_pyctoris | LC | Unknown | temperate | nocturnal | 136 | 1957430.215 | 8.623 |
| mammals | Rattus_rattus | Rattus_rattus | LC | Stable | tropical | nocturnal | 178.3 | 28316841.32 | 13.223 |
| mammals | Rattus_richardsoni | Rattus_richardsoni | VU | Decreasing | tropical | nocturnal | 64.3 | 6974.601 | 5.465 |
| mammals | Rattus_salocco | NA | DD | Unknown | NA | NA | NA | NA | NA |
| mammals | Rattus_satarae | Rattus_satarae | VU | Decreasing | tropical | nocturnal | 180 | 3126.118 | 15.225 |
| mammals | Rattus_simalurensis | NA | EN | Decreasing | NA | NA | NA | NA | NA |
| mammals | Rattus_sordidus | Rattus_sordidus | LC | Stable | tropical | nocturnal | 157.5 | 258376.798 | 3.503 |
| mammals | Rattus_steini | Rattus_steini | LC | Stable | tropical | nocturnal | 155 | 234808.036 | 4.323 |
| mammals | Rattus_stoicus | Rattus_stoicus | VU | Stable | tropical | nocturnal | 323 | 261.766 | 8.693 |
| mammals | Rattus_tanezumi | Rattus_tanezumi | LC | Increasing | tropical | nocturnal | 134 | 9717429.762 | 13.475 |
| mammals | Rattus_tawitawiensis | NA | DD | Unknown | NA | NA | NA | NA | NA |
| mammals | Rattus_timorensis | NA | DD | Unknown | NA | NA | NA | NA | NA |
| mammals | Rattus_tiomanicus | Rattus_tiomanicus | LC | Increasing | tropical | nocturnal | 130 | 1495112.483 | 9.258 |
| mammals | Rattus_tunneyi | Rattus_tunneyi | LC | Unknown | tropical | nocturnal | 124 | 762536.611 | 4.068 |
| mammals | Rattus_vandeuseni | Rattus_vandeuseni | EN | Decreasing | tropical | nocturnal | 67.3 | 2014.707 | 2.702 |
| mammals | Rattus_verecundus | Rattus_verecundus | LC | Stable | tropical | nocturnal | 108.5 | 152812.926 | 4.615 |
| mammals | Rattus_villosissimus | Rattus_villosissimus | LC | Unknown | tropical | nocturnal | 134 | 2155750.144 | 2.787 |
| mammals | Rattus_xanthurus | Rattus_xanthurus | NT | Decreasing | tropical | nocturnal | 303 | 15866.834 | 10.4 |
| mammals | Ratufa_affinis | Ratufa_affinis | NT | Decreasing | tropical | diurnal | 1156.3 | 1335699.565 | 7.943 |

| | | | | | | | | | |
|---|---|---|---|---|---|---|---|---|---|
| mammals | Ratufa_bicolor | Ratufa_bicolor | NT | Decreasing | tropical | diurnal | 2050 | 2557024.665 | 11.387 |
| mammals | Ratufa_indica | Ratufa_indica | LC | Decreasing | tropical | diurnal | 1060 | 355292.043 | 15.555 |
| mammals | Ratufa_macroura | Ratufa_macroura | NT | Decreasing | tropical | diurnal | 1327.1 | 29502.646 | 17.245 |
| mammals | Redunca_arundinum | Redunca_arundinum | LC | Stable | tropical | cathemeral | 58000.3 | 4764551.682 | 8.191 |
| mammals | Redunca_fulvorufula | Redunca_fulvorufula | EN | Decreasing | temperate | cathemeral | 29499.9 | 1044828.846 | 11.009 |
| mammals | Redunca_redunca | Redunca_redunca | LC | Decreasing | tropical | cathemeral | 44050.4 | 5901393.909 | 10.652 |
| mammals | Reithrodon_auritus | Reithrodon_auritus | LC | Stable | temperate | nocturnal | 70.9 | 1191439.988 | 7.394 |
| mammals | Reithrodon_typicus | Reithrodon_typicus | LC | Stable | temperate | nocturnal | 65.8 | 309299.71 | 10.259 |
| mammals | Reithrodontomys_brevirostris | Reithrodontomys_brevirostris | LC | Stable | tropical | nocturnal | 12.9 | 20841.435 | 15.069 |
| mammals | Reithrodontomys_burti | NA | DD | Unknown | NA | NA | NA | NA | NA |
| mammals | Reithrodontomys_chrysopsis | Reithrodontomys_chrysopsis | LC | Unknown | tropical | nocturnal | 19 | 53802.407 | 16.869 |
| mammals | Reithrodontomys_creper | Reithrodontomys_creper | LC | Stable | tropical | nocturnal | 22.8 | 12286.38 | 9.448 |
| mammals | Reithrodontomys_darienensis | Reithrodontomys_darienensis | LC | Stable | tropical | nocturnal | 13 | 43587.413 | 8.412 |
| mammals | Reithrodontomys_fulvescens | Reithrodontomys_fulvescens | LC | Stable | temperate | nocturnal | 11.4 | 2632629.574 | 11.619 |
| mammals | Reithrodontomys_gracilis | Reithrodontomys_gracilis | LC | Stable | tropical | nocturnal | 12.3 | 325050.229 | 10.48 |
| mammals | Reithrodontomys_hirsutus | Reithrodontomys_hirsutus | VU | Decreasing | tropical | nocturnal | 20 | 9699.533 | 8.442 |
| mammals | Reithrodontomys_humulis | Reithrodontomys_humulis | LC | Stable | temperate | nocturnal | 8.3 | 1296370.654 | 16.602 |
| mammals | Reithrodontomys_megalotis | Reithrodontomys_megalotis | LC | Stable | temperate | nocturnal | 9.4 | 5116788.756 | 11.439 |
| mammals | Reithrodontomys_mexicanus | Reithrodontomys_mexicanus | LC | Stable | tropical | nocturnal | 16 | 540022.718 | 11.065 |
| mammals | Reithrodontomys_microdon | Reithrodontomys_microdon | LC | Decreasing | tropical | nocturnal | 20 | 77650.03 | 16.694 |
| mammals | Reithrodontomys_montanus | Reithrodontomys_montanus | LC | Stable | temperate | nocturnal | 10.9 | 1846897.399 | 12.574 |
| mammals | Reithrodontomys_paradoxus | NA | DD | Unknown | NA | NA | NA | NA | NA |
| mammals | Reithrodontomys_raviventris | Reithrodontomys_raviventris | EN | Decreasing | temperate | nocturnal | 11.1 | 971.418 | 23.739 |
| mammals | Reithrodontomys_rodriguezi | Reithrodontomys_rodriguezi | LC | Stable | tropical | nocturnal | 31.2 | 1719.804 | 16.598 |
| mammals | Reithrodontomys_spectabilis | Reithrodontomys_spectabilis | CR | Decreasing | tropical | nocturnal | 20.2 | 488.275 | 10.21 |
| mammals | Reithrodontomys_sumichrasti | Reithrodontomys_sumichrasti | LC | Stable | tropical | nocturnal | 19 | 328552.487 | 11.838 |
| mammals | Reithrodontomys_tenuirostris | Reithrodontomys_tenuirostris | EN | Decreasing | tropical | nocturnal | 20 | 14140.467 | 17.126 |
| mammals | Reithrodontomys_zacatecae | Reithrodontomys_zacatecae | LC | Unknown | temperate | nocturnal | 9.7 | 211192.084 | 6.835 |
| mammals | Rhabdomys_pumilio | Rhabdomys_pumilio | LC | Stable | temperate | diurnal | 41.1 | 134988.05 | 10.247 |
| mammals | Rhagomys_longilingua | Rhagomys_longilingua | LC | Unknown | tropical | nocturnal | 30 | 2294.231 | 5.466 |
| mammals | Rheithrosciurus_macrotis | Rheithrosciurus_macrotis | VU | Decreasing | tropical | diurnal | 1225 | 732598.545 | 5.564 |
| mammals | Rheomys_mexicanus | Rheomys_mexicanus | EN | Unknown | tropical | cathemeral | 40 | 4237.406 | 7.442 |
| mammals | Rheomys_raptor | Rheomys_raptor | LC | Stable | tropical | cathemeral | 38 | 12331.439 | 8.275 |
| mammals | Rheomys_thomasi | Rheomys_thomasi | NT | Unknown | tropical | cathemeral | 40 | 27299.261 | 14.194 |

| | | | | | | | | | |
|---|---|---|---|---|---|---|---|---|---|
| mammals | Rheomys_underwoodi | Rheomys_underwoodi | LC | Stable | tropical | cathemeral | 66.2 | 5344.815 | 6.22 |
| mammals | Rhinoceros_sondaicus | Rhinoceros_sondaicus | CR | Stable | tropical | cathemeral | 1750000 | 125954.911 | 22.364 |
| mammals | Rhinoceros_unicornis | Rhinoceros_unicornis | VU | Increasing | temperate | cathemeral | 1602333 | 551653.911 | 16.983 |
| mammals | Rhinolophus_acuminatus | Rhinolophus_acuminatus | LC | Unknown | tropical | nocturnal | 12.1 | 1135409.323 | 14.379 |
| mammals | Rhinolophus_adami | NA | DD | Unknown | NA | NA | NA | NA | NA |
| mammals | Rhinolophus_affinis | Rhinolophus_affinis | LC | Stable | tropical | nocturnal | 13.6 | 5505773.099 | 12.687 |
| mammals | Rhinolophus_alcyone | Rhinolophus_alcyone | LC | Unknown | tropical | nocturnal | 18.8 | 2859335.955 | 7.772 |
| mammals | Rhinolophus_arcuatus | Rhinolophus_arcuatus | DD | Stable | tropical | nocturnal | 8.8 | 291355.825 | 13.581 |
| mammals | Rhinolophus_beddomei | Rhinolophus_beddomei | LC | Stable | tropical | nocturnal | 14.5 | 186306.709 | 17.969 |
| mammals | Rhinolophus_blasii | Rhinolophus_blasii | LC | Decreasing | temperate | nocturnal | 9.7 | 3410468.096 | 11.653 |
| mammals | Rhinolophus_bocharicus | Rhinolophus_bocharicus | LC | Stable | temperate | nocturnal | 15.2 | 924906.686 | 9.193 |
| mammals | Rhinolophus_borneensis | Rhinolophus_borneensis | LC | Unknown | tropical | nocturnal | 7.9 | 1426753.691 | 8.986 |
| mammals | Rhinolophus_canuti | Rhinolophus_canuti | VU | Decreasing | tropical | nocturnal | 12.7 | 10574.157 | 23.243 |
| mammals | Rhinolophus_capensis | Rhinolophus_capensis | LC | Stable | temperate | nocturnal | 12.8 | 242578.759 | 9.793 |
| mammals | Rhinolophus_celebensis | Rhinolophus_celebensis | LC | Unknown | tropical | nocturnal | 6.7 | 308233.828 | 14.888 |
| mammals | Rhinolophus_clivosus | Rhinolophus_clivosus | LC | Unknown | tropical | nocturnal | 15.7 | 3659664.552 | 9.533 |
| mammals | Rhinolophus_coelophyllus | Rhinolophus_coelophyllus | LC | Stable | tropical | nocturnal | 7.1 | 426055.556 | 14.647 |
| mammals | Rhinolophus_cognatus | Rhinolophus_cognatus | EN | Decreasing | tropical | nocturnal | 5.4 | 3077.32 | 7.717 |
| mammals | Rhinolophus_convexus | NA | DD | Unknown | NA | NA | NA | NA | NA |
| mammals | Rhinolophus_creaghi | Rhinolophus_creaghi | LC | Decreasing | tropical | nocturnal | 12 | 161673.082 | 7.959 |
| mammals | Rhinolophus_darlingi | Rhinolophus_darlingi | LC | Unknown | tropical | nocturnal | 9.5 | 1360113.629 | 11.279 |
| mammals | Rhinolophus_deckenii | Rhinolophus_deckenii | NT | Decreasing | tropical | nocturnal | 16.8 | 187337.721 | 9.328 |
| mammals | Rhinolophus_denti | Rhinolophus_denti | LC | Unknown | tropical | nocturnal | 6.2 | 1704313.392 | 7.306 |
| mammals | Rhinolophus_eloquens | Rhinolophus_eloquens | LC | Unknown | tropical | nocturnal | 19.1 | 711922.767 | 12.813 |
| mammals | Rhinolophus_euryale | Rhinolophus_euryale | NT | Decreasing | temperate | nocturnal | 8.2 | 2991976.979 | 17.282 |
| mammals | Rhinolophus_euryotis | Rhinolophus_euryotis | LC | Unknown | tropical | nocturnal | 14.3 | 878387.639 | 4.956 |
| mammals | Rhinolophus_ferrumequinum | Rhinolophus_ferrumequinum | LC | Decreasing | temperate | nocturnal | 21.1 | 9751019.671 | 14.889 |
| mammals | Rhinolophus_formosae | Rhinolophus_formosae | LC | Decreasing | temperate | nocturnal | 20 | 28759.051 | 15.802 |
| mammals | Rhinolophus_fumigatus | Rhinolophus_fumigatus | LC | Unknown | tropical | nocturnal | 12.9 | 8364570.957 | 9.325 |
| mammals | Rhinolophus_guineensis | Rhinolophus_guineensis | EN | Decreasing | tropical | nocturnal | 10.4 | 213322.918 | 13.351 |
| mammals | Rhinolophus_hildebrandtii | Rhinolophus_hildebrandtii | LC | Unknown | tropical | nocturnal | 25.8 | 3184495.863 | 10.693 |
| mammals | Rhinolophus_hilli | Rhinolophus_hilli | CR | Decreasing | tropical | nocturnal | 16.5 | 122.033 | 10.49 |
| mammals | Rhinolophus_hillorum | Rhinolophus_hillorum | VU | Decreasing | tropical | nocturnal | 14.8 | 28211.393 | 13.815 |
| mammals | Rhinolophus_hipposideros | Rhinolophus_hipposideros | LC | Decreasing | temperate | nocturnal | 4.7 | 6256257.88 | 16.135 |

| | | | | | | | | | |
|---|---|---|---|---|---|---|---|---|---|
| mammals | Rhinolophus_inops | Rhinolophus_inops | LC | Stable | tropical | nocturnal | 13.5 | 226653.042 | 13.874 |
| mammals | Rhinolophus_keyensis | NA | DD | Unknown | NA | NA | NA | NA | NA |
| mammals | Rhinolophus_landeri | Rhinolophus_landeri | LC | Unknown | tropical | nocturnal | 8.9 | 12125111.266 | 9.5 |
| mammals | Rhinolophus_lepidus | Rhinolophus_lepidus | LC | Unknown | tropical | nocturnal | 5.7 | 3509104.608 | 16.377 |
| mammals | Rhinolophus_luctus | Rhinolophus_luctus | LC | Unknown | tropical | nocturnal | 33.5 | 5655752.199 | 11.289 |
| mammals | Rhinolophus_maclaudi | Rhinolophus_maclaudi | EN | Decreasing | tropical | nocturnal | 13.1 | 8188.631 | 14.397 |
| mammals | Rhinolophus_macrotis | Rhinolophus_macrotis | LC | Stable | temperate | nocturnal | 6.2 | 2405475.714 | 12.175 |
| mammals | Rhinolophus_madurensis | Rhinolophus_madurensis | VU | Decreasing | tropical | nocturnal | 6.7 | 4927.496 | 26.317 |
| mammals | Rhinolophus_maendeleo | NA | DD | Unknown | NA | NA | NA | NA | NA |
| mammals | Rhinolophus_malayanus | Rhinolophus_malayanus | LC | Stable | tropical | nocturnal | 6.7 | 1922229.023 | 12.612 |
| mammals | Rhinolophus_marshalli | Rhinolophus_marshalli | LC | Unknown | tropical | nocturnal | 5 | 714382.374 | 12.38 |
| mammals | Rhinolophus_megaphyllus | Rhinolophus_megaphyllus | LC | Unknown | tropical | nocturnal | 10.1 | 876558.898 | 5.879 |
| mammals | Rhinolophus_mehelyi | Rhinolophus_mehelyi | VU | Decreasing | temperate | nocturnal | 12.3 | 1976290.665 | 15.3 |
| mammals | Rhinolophus_mitratus | NA | DD | Unknown | NA | NA | NA | NA | NA |
| mammals | Rhinolophus_montanus | Rhinolophus_montanus | EN | Unknown | tropical | nocturnal | 6.7 | 71.282 | 11.244 |
| mammals | Rhinolophus_nereis | NA | DD | Unknown | NA | NA | NA | NA | NA |
| mammals | Rhinolophus_osgoodi | Rhinolophus_osgoodi | LC | Stable | temperate | nocturnal | 7.3 | 769589.593 | 13.44 |
| mammals | Rhinolophus_pearsonii | Rhinolophus_pearsonii | LC | Unknown | tropical | nocturnal | 11.8 | 3176331.023 | 13.394 |
| mammals | Rhinolophus_philippinensis | Rhinolophus_philippinensis | LC | Unknown | tropical | nocturnal | 10.3 | 247505.548 | 6.106 |
| mammals | Rhinolophus_pusillus | Rhinolophus_pusillus | LC | Stable | tropical | nocturnal | 7.2 | 3582812.621 | 12.855 |
| mammals | Rhinolophus_rex | Rhinolophus_rex | EN | Decreasing | temperate | nocturnal | 11.3 | 2144413.653 | 13.343 |
| mammals | Rhinolophus_robinsoni | Rhinolophus_robinsoni | NT | Decreasing | tropical | nocturnal | 8.5 | 47297.552 | 12.464 |
| mammals | Rhinolophus_rouxii | Rhinolophus_rouxii | LC | Unknown | temperate | nocturnal | 11.8 | 821256.077 | 18.219 |
| mammals | Rhinolophus_rufus | Rhinolophus_rufus | NT | Decreasing | tropical | nocturnal | 30 | 34730.176 | 14.456 |
| mammals | Rhinolophus_ruwenzorii | Rhinolophus_ruwenzorii | EN | Decreasing | tropical | nocturnal | 17.6 | 27330.971 | 14.965 |
| mammals | Rhinolophus_sakejiensis | NA | DD | Unknown | NA | NA | NA | NA | NA |
| mammals | Rhinolophus_sedulus | Rhinolophus_sedulus | NT | Decreasing | tropical | nocturnal | 8.8 | 805437.73 | 6.263 |
| mammals | Rhinolophus_shameli | Rhinolophus_shameli | LC | Stable | tropical | nocturnal | 9.6 | 580977.065 | 13.727 |
| mammals | Rhinolophus_shortridgei | NA | DD | Unknown | NA | NA | NA | NA | NA |
| mammals | Rhinolophus_siamensis | Rhinolophus_siamensis | LC | Unknown | tropical | nocturnal | 5.1 | 1047159.94 | 12.694 |
| mammals | Rhinolophus_silvestris | Rhinolophus_silvestris | DD | Decreasing | tropical | nocturnal | 16.8 | 136275.249 | 5.958 |
| mammals | Rhinolophus_simulator | Rhinolophus_simulator | LC | Decreasing | tropical | nocturnal | 8.1 | 1911252.193 | 10.742 |
| mammals | Rhinolophus_sinicus | Rhinolophus_sinicus | LC | Unknown | temperate | nocturnal | 11.6 | 2168584.379 | 14.292 |
| mammals | Rhinolophus_stheno | Rhinolophus_stheno | LC | Stable | tropical | nocturnal | 7.9 | 852346.828 | 12.615 |

| | | | | | | | | | |
|---|---|---|---|---|---|---|---|---|---|
| mammals | Rhinolophus_subbadius | Rhinolophus_subbadius | LC | Decreasing | temperate | nocturnal | 6.6 | 76243.223 | 13.762 |
| mammals | Rhinolophus_subrufus | NA | DD | Unknown | NA | NA | NA | NA | NA |
| mammals | Rhinolophus_swinnyi | Rhinolophus_swinnyi | LC | Unknown | tropical | nocturnal | 7.1 | 1347873.844 | 10.4 |
| mammals | Rhinolophus_thomasi | Rhinolophus_thomasi | LC | Unknown | tropical | nocturnal | 8.3 | 1408310.583 | 12.635 |
| mammals | Rhinolophus_trifoliatus | Rhinolophus_trifoliatus | NT | Decreasing | tropical | nocturnal | 14.5 | 1344638.629 | 8.176 |
| mammals | Rhinolophus_virgo | Rhinolophus_virgo | LC | Stable | tropical | nocturnal | 6 | 269684.683 | 13.905 |
| mammals | Rhinolophus_yunanensis | Rhinolophus_yunanensis | LC | Unknown | tropical | nocturnal | 19.7 | 401250.629 | 8.748 |
| mammals | Rhinolophus_ziama | Rhinolophus_ziama | EN | Decreasing | tropical | nocturnal | 22.3 | 2057.051 | 11.991 |
| mammals | Rhinonicteris_aurantia | Rhinonicteris_aurantia | LC | Unknown | tropical | nocturnal | 8.8 | 747496.387 | 2.441 |
| mammals | Rhinophylla_alethina | Rhinophylla_alethina | NT | Unknown | tropical | nocturnal | 14 | 125700.979 | 9.327 |
| mammals | Rhinophylla_fischerae | Rhinophylla_fischerae | LC | Stable | tropical | nocturnal | 10.6 | 4564653.785 | 2.342 |
| mammals | Rhinophylla_pumilio | Rhinophylla_pumilio | LC | Unknown | tropical | nocturnal | 9.6 | 8334315.242 | 3.305 |
| mammals | Rhinopithecus_avunculus | Rhinopithecus_avunculus | CR | Decreasing | tropical | diurnal | 8000 | 10209.638 | 11.075 |
| mammals | Rhinopithecus_bieti | Rhinopithecus_bieti | EN | Stable | temperate | diurnal | 9960 | 29673.802 | 6.906 |
| mammals | Rhinopithecus_brelichi | Rhinopithecus_brelichi | CR | Decreasing | temperate | diurnal | 11500 | 67.48 | 10.288 |
| mammals | Rhinopithecus_roxellana | Rhinopithecus_roxellana | EN | Decreasing | temperate | diurnal | 12125 | 250586.154 | 11.451 |
| mammals | Rhinopithecus_strykeri | Rhinopithecus_strykeri | CR | Decreasing | temperate | diurnal | 11250 | 203.435 | 1.249 |
| mammals | Rhinopoma_hardwickii | Rhinopoma_hardwickii | LC | Stable | tropical | nocturnal | 14.2 | 16931124.568 | 6.81 |
| mammals | Rhinopoma_macinnesi | NA | DD | Unknown | NA | NA | NA | NA | NA |
| mammals | Rhinopoma_microphyllum | Rhinopoma_microphyllum | LC | Stable | temperate | nocturnal | 27.9 | 7151525.959 | 9.473 |
| mammals | Rhinopoma_muscatellum | Rhinopoma_muscatellum | LC | Unknown | temperate | nocturnal | 9.1 | 580313.148 | 9.509 |
| mammals | Rhinosciurus_laticaudatus | Rhinosciurus_laticaudatus | NT | Decreasing | tropical | diurnal | 228.3 | 911388.272 | 9.109 |
| mammals | Rhipidomys_austrinus | Rhipidomys_austrinus | LC | Stable | tropical | nocturnal | 89 | 138614.187 | 6.283 |
| mammals | Rhipidomys_caucensis | NA | DD | Unknown | NA | NA | NA | NA | NA |
| mammals | Rhipidomys_couesi | Rhipidomys_couesi | LC | Stable | tropical | nocturnal | 89 | 144541.233 | 12.327 |
| mammals | Rhipidomys_emiliae | Rhipidomys_emiliae | LC | Unknown | tropical | nocturnal | 96.5 | 698505.619 | 3.792 |
| mammals | Rhipidomys_fulviventer | Rhipidomys_fulviventer | LC | Unknown | tropical | nocturnal | 89 | 52004.661 | 12.459 |
| mammals | Rhipidomys_gardneri | Rhipidomys_gardneri | LC | Unknown | tropical | nocturnal | 71.1 | 152165.93 | 1.009 |
| mammals | Rhipidomys_latimanus | Rhipidomys_latimanus | LC | Decreasing | tropical | nocturnal | 57.5 | 157706.169 | 11.224 |
| mammals | Rhipidomys_leucodactylus | Rhipidomys_leucodactylus | LC | Unknown | tropical | nocturnal | 80 | 4402230.204 | 1.863 |
| mammals | Rhipidomys_macconnelli | Rhipidomys_macconnelli | LC | Stable | tropical | nocturnal | 41.6 | 189757.686 | 0.977 |
| mammals | Rhipidomys_macrurus | Rhipidomys_macrurus | LC | Unknown | tropical | nocturnal | 79.9 | 1565931.641 | 9.041 |
| mammals | Rhipidomys_mastacalis | Rhipidomys_mastacalis | LC | Unknown | tropical | nocturnal | 77.5 | 633285.532 | 12.778 |
| mammals | Rhipidomys_modicus | NA | LC | Decreasing | NA | NA | NA | NA | NA |

| | | | | | | | | | |
|---|---|---|---|---|---|---|---|---|---|
| mammals | Rhipidomys_nitela | Rhipidomys_nitela | LC | Stable | tropical | nocturnal | 56 | 1447250.059 | 1.244 |
| mammals | Rhipidomys_ochrogaster | NA | DD | Unknown | NA | NA | NA | NA | NA |
| mammals | Rhipidomys_venezuelae | Rhipidomys_venezuelae | LC | Stable | tropical | nocturnal | 90 | 172678.318 | 12.699 |
| mammals | Rhipidomys_venustus | Rhipidomys_venustus | LC | Unknown | tropical | nocturnal | 89 | 8414.418 | 13.564 |
| mammals | Rhipidomys_wetzeli | Rhipidomys_wetzeli | LC | Stable | tropical | nocturnal | 89 | 138914.108 | 0.898 |
| mammals | Rhizomys_sinensis | Rhizomys_sinensis | LC | Unknown | temperate | nocturnal | 865 | 2057627.575 | 13.982 |
| mammals | Rhizomys_sumatrensis | Rhizomys_sumatrensis | LC | Unknown | tropical | nocturnal | 1625 | 1372330.121 | 10.011 |
| mammals | Rhogeessa_aenea | Rhogeessa_aeneus | LC | Stable | tropical | nocturnal | 3.4 | 144231.54 | 6.275 |
| mammals | Rhogeessa_genowaysi | Rhogeessa_genowaysi | EN | Decreasing | tropical | nocturnal | 5 | 913.219 | 8.252 |
| mammals | Rhogeessa_hussoni | NA | DD | Unknown | NA | NA | NA | NA | NA |
| mammals | Rhogeessa_io | Rhogeessa_io | LC | Unknown | tropical | nocturnal | 5 | 4609642.847 | 5.689 |
| mammals | Rhogeessa_minutilla | Rhogeessa_minutilla | VU | Decreasing | tropical | nocturnal | 3.7 | 663829.689 | 10.656 |
| mammals | Rhogeessa_mira | Rhogeessa_mira | VU | Stable | tropical | nocturnal | 3.1 | 3345.677 | 9.322 |
| mammals | Rhogeessa_parvula | Rhogeessa_parvula | LC | Stable | tropical | nocturnal | 4.4 | 448909.734 | 9.139 |
| mammals | Rhogeessa_tumida | Rhogeessa_tumida | LC | Stable | tropical | nocturnal | 4.5 | 462224.299 | 12.612 |
| mammals | Rhombomys_opimus | Rhombomys_opimus | LC | Stable | temperate | diurnal | 144 | 4585921.877 | 5.523 |
| mammals | Rhynchocyon_chrysopygus | Rhynchocyon_chrysopygus | EN | Decreasing | tropical | diurnal | 540 | 3574.33 | 16.27 |
| mammals | Rhynchocyon_cirnei | Rhynchocyon_cirnei | LC | Decreasing | tropical | diurnal | 490 | 1130811.53 | 8.323 |
| mammals | Rhynchocyon_petersi | Rhynchocyon_petersi | LC | Decreasing | tropical | diurnal | 424 | 31086.283 | 12.867 |
| mammals | Rhynchocyon_udzungwensis | Rhynchocyon_udzungwensis | VU | Unknown | tropical | diurnal | 711 | 320.402 | 3.096 |
| mammals | Rhynchogale_melleri | Rhynchogale_melleri | LC | Unknown | tropical | nocturnal | 2500 | 2215668.763 | 9.285 |
| mammals | Rhyncholestes_raphanurus | Rhyncholestes_raphanurus | NT | Decreasing | temperate | nocturnal | 21 | 40999.524 | 9.081 |
| mammals | Rhynchomys_banahao | Rhynchomys_banahao | LC | Unknown | tropical | nocturnal | 155 | 61.734 | 8.535 |
| mammals | Rhynchomys_isarogensis | Rhynchomys_isarogensis | VU | Stable | tropical | nocturnal | 136 | 46.195 | 7.305 |
| mammals | Rhynchomys_soricoides | Rhynchomys_soricoides | NT | Decreasing | tropical | cathemeral | 156 | 4215.026 | 8.968 |
| mammals | Rhynchomys_tapulao | NA | DD | Unknown | NA | NA | NA | NA | NA |
| mammals | Rhynchonycteris_naso | Rhynchonycteris_naso | LC | Unknown | tropical | nocturnal | 3.8 | 11508826.658 | 4.945 |
| mammals | Rhyneptesicus_nasutus | Eptesicus_nasutus | LC | Unknown | temperate | nocturnal | 7.9 | 1866190.539 | 6.744 |
| mammals | Rousettus_aegyptiacus | Rousettus_aegyptiacus | LC | Stable | tropical | nocturnal | 132.2 | 4071413.311 | 11.287 |
| mammals | Rousettus_amplexicaudatus | Rousettus_amplexicaudatus | LC | Unknown | tropical | nocturnal | 82.5 | 4281498.886 | 9.551 |
| mammals | Rousettus_leschenaultii | Rousettus_leschenaultii | NT | Decreasing | tropical | nocturnal | 97.1 | 6873172.217 | 15.398 |
| mammals | Rousettus_linduensis | NA | DD | Unknown | NA | NA | NA | NA | NA |
| mammals | Rousettus_madagascariensis | Rousettus_madagascariensis | VU | Decreasing | tropical | nocturnal | 62.1 | 292740.755 | 8.684 |
| mammals | Rousettus_obliviosus | Rousettus_obliviosus | VU | Unknown | tropical | nocturnal | 44.1 | 1667.125 | 16.549 |

| | | | | | | | | | |
|---|---|---|---|---|---|---|---|---|---|
| mammals | Rousettus_spinalatus | Rousettus_spinalatus | VU | Decreasing | tropical | nocturnal | 90.8 | 84003.773 | 6.023 |
| mammals | Rubrisciurus_rubriventer | Rubrisciurus_rubriventer | LC | Unknown | tropical | diurnal | 688.3 | 169893.702 | 9.283 |
| mammals | Rucervus_duvaucelii | Rucervus_duvaucelii | VU | Decreasing | temperate | diurnal | 143000 | 34612.901 | 15.994 |
| mammals | Rungwecebus_kipunji | Rungwecebus_kipunji | EN | Decreasing | tropical | diurnal | 13000 | 71.061 | 5.865 |
| mammals | Rupicapra_pyrenaica | Rupicapra_pyrenaica | LC | Increasing | temperate | diurnal | 30000 | 15275.945 | 9.643 |
| mammals | Rupicapra_rupicapra | Rupicapra_rupicapra | LC | Stable | temperate | cathemeral | 26100 | 209047.997 | 13.13 |
| mammals | Rusa_alfredi | Rusa_alfredi | EN | Decreasing | tropical | nocturnal | 47500 | 1689.09 | 7.011 |
| mammals | Rusa_marianna | Rusa_marianna | VU | Decreasing | tropical | nocturnal | 50000 | 112256.431 | 9.462 |
| mammals | Rusa_timorensis | Rusa_timorensis | VU | Decreasing | tropical | diurnal | 63000 | 11319.645 | 12.885 |
| mammals | Rusa_unicolor | Rusa_unicolor | VU | Decreasing | tropical | cathemeral | 180344.4 | 7570390.032 | 13.628 |
| mammals | Ruwenzorisorex_suncoides | Ruwenzorisorex_suncoides | VU | Unknown | tropical | nocturnal | 18.2 | 22305.789 | 18.465 |
| mammals | Saccolaimus_flaviventris | Saccolaimus_flaviventris | LC | Decreasing | temperate | nocturnal | 42.1 | 6052125.983 | 4.201 |
| mammals | Saccolaimus_mixtus | Saccolaimus_mixtus | NT | Decreasing | tropical | nocturnal | 23 | 53569.26 | 2.739 |
| mammals | Saccolaimus_peli | Saccolaimus_peli | LC | Unknown | tropical | nocturnal | 100 | 1959912.673 | 7.614 |
| mammals | Saccolaimus_saccolaimus | Saccolaimus_saccolaimus | LC | Stable | tropical | nocturnal | 41.8 | 1930053.007 | 12.656 |
| mammals | Saccopteryx_antioquensis | Saccopteryx_antioquensis | EN | Unknown | tropical | nocturnal | 5 | 3092.046 | 10.243 |
| mammals | Saccopteryx_bilineata | Saccopteryx_bilineata | LC | Unknown | tropical | nocturnal | 7.8 | 12632095.593 | 5.661 |
| mammals | Saccopteryx_canescens | Saccopteryx_canescens | LC | Unknown | tropical | nocturnal | 3.4 | 6384043.266 | 3.176 |
| mammals | Saccopteryx_gymnura | NA | DD | Unknown | NA | NA | NA | NA | NA |
| mammals | Saccopteryx_leptura | Saccopteryx_leptura | LC | Unknown | tropical | nocturnal | 4.7 | 12289438.468 | 5.536 |
| mammals | Saccostomus_campestris | Saccostomus_campestris | LC | Stable | tropical | nocturnal | 51.2 | 5347600.615 | 7.94 |
| mammals | Saccostomus_mearnsi | Saccostomus_mearnsi | LC | Stable | tropical | nocturnal | 45.5 | 667846.57 | 9.846 |
| mammals | Saguinus_bicolor | Saguinus_bicolor | CR | Decreasing | tropical | diurnal | 465 | 10128.756 | 5.555 |
| mammals | Saguinus_fuscicollis | Saguinus_fuscicollis | LC | Decreasing | tropical | diurnal | 387 | 434351.007 | 0.769 |
| mammals | Saguinus_geoffroyi | Saguinus_geoffroyi | NT | Decreasing | tropical | diurnal | 486.5 | 75592.617 | 5.044 |
| mammals | Saguinus_imperator | Saguinus_imperator | LC | Decreasing | tropical | diurnal | 400 | 234318.648 | 1.37 |
| mammals | Saguinus_inustus | Saguinus_inustus | LC | Stable | tropical | diurnal | 410.5 | 380406.835 | 0.953 |
| mammals | Saguinus_labiatus | Saguinus_labiatus | LC | Decreasing | tropical | diurnal | 575 | 286447.289 | 1.646 |
| mammals | Saguinus_leucopus | Saguinus_leucopus | VU | Decreasing | tropical | diurnal | 440 | 56861.638 | 12.082 |
| mammals | Saguinus_martinsi | Saguinus_martinsi | NT | Decreasing | tropical | diurnal | 603.6 | 42109.163 | 1.358 |
| mammals | Saguinus_midas | Saguinus_midas | LC | Stable | tropical | diurnal | 540 | 863250.692 | 1.217 |
| mammals | Saguinus_mystax | Saguinus_mystax | LC | Decreasing | tropical | diurnal | 618 | 589295.469 | 0.753 |
| mammals | Saguinus_niger | Saguinus_niger | VU | Decreasing | tropical | diurnal | 591.2 | 281250.693 | 2.961 |
| mammals | Saguinus_nigricollis | Saguinus_nigricollis | LC | Decreasing | tropical | diurnal | 462 | 211767.622 | 3.132 |

| | | | | | | | | | |
|---|---|---|---|---|---|---|---|---|---|
| mammals | Saguinus_oedipus | Saguinus_oedipus | CR | Decreasing | tropical | diurnal | 430 | 51980.63 | 15.538 |
| mammals | Saguinus_tripartitus | Saguinus_tripartitus | NT | Decreasing | tropical | diurnal | 393.5 | 25699.951 | 1.427 |
| mammals | Saguinus_weddelli | Saguinus_melanoleucus | LC | Unknown | tropical | diurnal | 491 | 597636.46 | 1.917 |
| mammals | Saiga_tatarica | Saiga_tatarica | NT | Increasing | temperate | diurnal | 29000 | 749562.587 | 5.719 |
| mammals | Saimiri_boliviensis | Saimiri_boliviensis | LC | Decreasing | tropical | diurnal | 615 | 1373872.354 | 2.951 |
| mammals | Saimiri_oerstedii | Saimiri_oerstedii | EN | Decreasing | tropical | diurnal | 278.5 | 10599.788 | 14.037 |
| mammals | Saimiri_sciureus | Saimiri_sciureus | LC | Decreasing | tropical | diurnal | 743.2 | 956591.723 | 1.28 |
| mammals | Saimiri_ustus | Saimiri_ustus | NT | Decreasing | tropical | diurnal | 1000 | 876710.21 | 2.121 |
| mammals | Saimiri_vanzolinii | Saimiri_vanzolinii | EN | Decreasing | tropical | diurnal | 769.6 | 2930.59 | 1.03 |
| mammals | Salanoia_concolor | Salanoia_concolor | VU | Decreasing | tropical | diurnal | 650 | 18642.912 | 7.557 |
| mammals | Salinomys_delicatus | NA | DD | Unknown | NA | NA | NA | NA | NA |
| mammals | Salpingotus_crassicauda | Salpingotus_crassicauda | LC | Unknown | temperate | nocturnal | 12 | 1359343.506 | 3.812 |
| mammals | Salpingotus_heptneri | NA | DD | Unknown | NA | NA | NA | NA | NA |
| mammals | Salpingotus_kozlovi | Salpingotus_kozlovi | LC | Unknown | temperate | nocturnal | 9.5 | 1541055.817 | 4 |
| mammals | Salpingotus_michaelis | NA | DD | Unknown | NA | NA | NA | NA | NA |
| mammals | Salpingotus_pallidus | NA | DD | Unknown | NA | NA | NA | NA | NA |
| mammals | Santamartamys_rufodorsalis | Santamartamys_rufodorsalis | CR | Unknown | tropical | nocturnal | 144.8 | 89.523 | 11.158 |
| mammals | Sapajus_apella | Sapajus_apella | LC | Decreasing | tropical | diurnal | 2500 | 5970729.543 | 2.187 |
| mammals | Sapajus_flavius | Sapajus_flavius | EN | Decreasing | tropical | diurnal | 2550 | 21664.316 | 16.305 |
| mammals | Sapajus_libidinosus | Sapajus_libidinosus | NT | Decreasing | tropical | diurnal | 3200 | 2612578.687 | 7.853 |
| mammals | Sapajus_xanthosternos | Sapajus_xanthosternos | CR | Decreasing | tropical | diurnal | 3200 | 466598.875 | 9.276 |
| mammals | Sarcophilus_harrisii | Sarcophilus_harrisii | EN | Decreasing | temperate | nocturnal | 11000 | 64284.135 | 5.859 |
| mammals | Sauromys_petrophilus | Sauromys_petrophilus | LC | Stable | temperate | nocturnal | 13.9 | 728212.456 | 9.317 |
| mammals | Saxatilomys_paulinae | NA | DD | Unknown | NA | NA | NA | NA | NA |
| mammals | Scapteromys_aquaticus | Scapteromys_aquaticus | LC | Stable | temperate | nocturnal | 116.7 | 260142.726 | 9.92 |
| mammals | Scapteromys_tumidus | Scapteromys_tumidus | LC | Stable | temperate | nocturnal | 146 | 258565.206 | 10.137 |
| mammals | Scarturus_elater | Allactaga_elater | LC | Decreasing | temperate | nocturnal | 59 | 3246015.698 | 7.287 |
| mammals | Scarturus_euphraticus | Allactaga_euphratica | LC | Decreasing | temperate | nocturnal | 70 | 758072.397 | 8.695 |
| mammals | Scarturus_hotsoni | Allactaga_hotsoni | LC | Unknown | temperate | nocturnal | 76.5 | 498712.568 | 4.412 |
| mammals | Scarturus_tetradactylus | NA | DD | Unknown | NA | NA | NA | NA | NA |
| mammals | Scarturus_vinogradovi | Allactaga_vinogradovi | LC | Decreasing | temperate | nocturnal | 75.2 | 52562.374 | 12.515 |
| mammals | Scarturus_williamsi | Allactaga_williamsi | LC | Decreasing | temperate | nocturnal | 100 | 705703.699 | 14.759 |
| mammals | Sciurillus_pusillus | Sciurillus_pusillus | LC | Unknown | tropical | diurnal | 39 | 1209632.3 | 1.978 |
| mammals | Sciurocheirus_alleni | Sciurocheirus_alleni | NT | Decreasing | tropical | nocturnal | 332.5 | 94727.984 | 13.82 |

| | | | | | | | | | |
|---|---|---|---|---|---|---|---|---|---|
| mammals | Sciurocheirus_gabonensis | Sciurocheirus_gabonensis | LC | Unknown | tropical | nocturnal | 271.6 | 530152.238 | 5.408 |
| mammals | Sciurotamias_davidianus | Sciurotamias_davidianus | LC | Unknown | temperate | diurnal | 433.5 | 1330267.362 | 15.286 |
| mammals | Sciurotamias_forresti | Sciurotamias_forresti | LC | Unknown | temperate | diurnal | 433.5 | 312768.841 | 11.881 |
| mammals | Sciurus_aberti | Sciurus_aberti | LC | Stable | temperate | diurnal | 624 | 270389.017 | 7.968 |
| mammals | Sciurus_aestuans | Sciurus_aestuans | LC | Unknown | tropical | diurnal | 185 | 6983651.392 | 6.321 |
| mammals | Sciurus_alleni | Sciurus_alleni | LC | Decreasing | temperate | diurnal | 434.5 | 48935.032 | 8.867 |
| mammals | Sciurus_anomalus | Sciurus_anomalus | LC | Decreasing | temperate | diurnal | 256 | 614332.404 | 14.818 |
| mammals | Sciurus_arizonensis | NA | DD | Unknown | NA | NA | NA | NA | NA |
| mammals | Sciurus_aureogaster | Sciurus_aureogaster | LC | Stable | tropical | diurnal | 595 | 620614.53 | 11.923 |
| mammals | Sciurus_carolinensis | Sciurus_carolinensis | LC | Increasing | temperate | diurnal | 506.5 | 4083873.203 | 17.26 |
| mammals | Sciurus_colliaei | Sciurus_colliaei | LC | Unknown | temperate | diurnal | 498 | 81302.042 | 7.236 |
| mammals | Sciurus_deppei | Sciurus_deppei | LC | Stable | tropical | diurnal | 190 | 694996.561 | 10.788 |
| mammals | Sciurus_granatensis | Sciurus_granatensis | LC | Stable | tropical | diurnal | 250 | 988528.232 | 11.994 |
| mammals | Sciurus_griseus | Sciurus_griseus | LC | Unknown | temperate | diurnal | 731 | 376963.854 | 11.951 |
| mammals | Sciurus_ignitus | Sciurus_ignitus | LC | Unknown | tropical | diurnal | 190 | 2092301.555 | 3.836 |
| mammals | Sciurus_igniventris | Sciurus_igniventris | LC | Unknown | tropical | diurnal | 700 | 2756262.842 | 1.698 |
| mammals | Sciurus_lis | Sciurus_lis | LC | Stable | temperate | diurnal | 264.3 | 214979.467 | 20.198 |
| mammals | Sciurus_nayaritensis | Sciurus_nayaritensis | LC | Unknown | temperate | diurnal | 697 | 249137.653 | 6.431 |
| mammals | Sciurus_niger | Sciurus_niger | LC | Stable | temperate | diurnal | 761.9 | 4360630.241 | 16.663 |
| mammals | Sciurus_oculatus | Sciurus_oculatus | LC | Unknown | tropical | diurnal | 650 | 101820.663 | 16.781 |
| mammals | Sciurus_pucheranii | NA | DD | Unknown | NA | NA | NA | NA | NA |
| mammals | Sciurus_pyrrhinus | NA | DD | Unknown | NA | NA | NA | NA | NA |
| mammals | Sciurus_richmondi | Sciurus_richmondi | NT | Unknown | tropical | diurnal | 205 | 22521.101 | 9.877 |
| mammals | Sciurus_spadiceus | Sciurus_spadiceus | LC | Unknown | tropical | diurnal | 403.3 | 3247882.946 | 2.473 |
| mammals | Sciurus_stramineus | Sciurus_stramineus | LC | Unknown | tropical | diurnal | 433.3 | 156433.128 | 10.394 |
| mammals | Sciurus_variegatoides | Sciurus_variegatoides | LC | Stable | tropical | diurnal | 485 | 392638.554 | 11.707 |
| mammals | Sciurus_vulgaris | Sciurus_vulgaris | LC | Unknown | temperate | diurnal | 333 | 27053169.45 | 7.598 |
| mammals | Sciurus_yucatanensis | Sciurus_yucatanensis | LC | Stable | tropical | diurnal | 225 | 194232.524 | 6.915 |
| mammals | Scleronycteris_ega | Scleronycteris_ega | DD | Decreasing | tropical | nocturnal | 49 | 974464.32 | 0.812 |
| mammals | Scolomys_melanops | Scolomys_melanops | LC | Unknown | tropical | nocturnal | 26.5 | 100718.941 | 2.513 |
| mammals | Scolomys_ucayalensis | Scolomys_ucayalensis | LC | Unknown | tropical | nocturnal | 26.5 | 197626.023 | 0.93 |
| mammals | Scoteanax_rueppellii | Scoteanax_rueppellii | LC | Stable | temperate | nocturnal | 26.5 | 301736.576 | 7.307 |
| mammals | Scotinomys_teguina | Scotinomys_teguina | LC | Stable | tropical | diurnal | 11.3 | 129224.408 | 11.664 |
| mammals | Scotinomys_xerampelinus | Scotinomys_xerampelinus | LC | Stable | tropical | diurnal | 15 | 5119.333 | 7.653 |

| | | | | | | | | | |
|---|---|---|---|---|---|---|---|---|---|
| mammals | Scotoecus_albofuscus | NA | DD | Unknown | NA | NA | NA | NA | NA |
| mammals | Scotoecus_hirundo | Scotoecus_hirundo | LC | Unknown | tropical | nocturnal | 9.3 | 4917152.73 | 10.017 |
| mammals | Scotoecus_pallidus | Scotoecus_pallidus | LC | Unknown | temperate | nocturnal | 11.65 | 187893.76 | 19.513 |
| mammals | Scotomanes_ornatus | Scotomanes_ornatus | LC | Unknown | tropical | nocturnal | 25 | 2690534.096 | 13.252 |
| mammals | Scotonycteris_zenkeri | Scotonycteris_zenkeri | NT | Unknown | tropical | nocturnal | 20.8 | 114273.669 | 13.389 |
| mammals | Scotophilus_collinus | Scotophilus_collinus | LC | Unknown | tropical | nocturnal | 23.4 | 210159.195 | 13.63 |
| mammals | Scotophilus_dinganii | Scotophilus_dinganii | LC | Unknown | tropical | nocturnal | 25.1 | 10952348.342 | 9.239 |
| mammals | Scotophilus_heathii | Scotophilus_heathii | LC | Stable | tropical | nocturnal | 34.6 | 5950831.342 | 15.529 |
| mammals | Scotophilus_kuhlii | Scotophilus_kuhlii | LC | Stable | tropical | nocturnal | 20.7 | 5208270.37 | 14.534 |
| mammals | Scotophilus_leucogaster | Scotophilus_leucogaster | LC | Unknown | tropical | nocturnal | 20.2 | 5430385.925 | 10.338 |
| mammals | Scotophilus_marovaza | Scotophilus_marovaza | LC | Unknown | tropical | nocturnal | 14.2 | 75618.293 | 7.896 |
| mammals | Scotophilus_nigrita | Scotophilus_nigrita | LC | Decreasing | tropical | nocturnal | 101 | 2196022.142 | 10.961 |
| mammals | Scotophilus_nucella | NA | DD | Unknown | NA | NA | NA | NA | NA |
| mammals | Scotophilus_nux | Scotophilus_nux | LC | Unknown | tropical | nocturnal | 30 | 951762.195 | 9.6 |
| mammals | Scotophilus_robustus | Scotophilus_robustus | LC | Unknown | tropical | nocturnal | 44.8 | 398693.89 | 8.513 |
| mammals | Scotophilus_tandrefana | NA | DD | Unknown | NA | NA | NA | NA | NA |
| mammals | Scotophilus_viridis | Scotophilus_viridis | LC | Unknown | tropical | nocturnal | 19.6 | 4650910.92 | 10.818 |
| mammals | Scotorepens_balstoni | Scotorepens_balstoni | LC | Decreasing | temperate | nocturnal | 11.7 | 4980250.106 | 4.156 |
| mammals | Scotorepens_greyii | Scotorepens_greyii | LC | Stable | tropical | nocturnal | 10 | 4064788.846 | 4.052 |
| mammals | Scotorepens_orion | Scotorepens_orion | LC | Unknown | tropical | nocturnal | 11 | 265076.922 | 8.405 |
| mammals | Scotorepens_sanborni | Scotorepens_sanborni | LC | Stable | tropical | nocturnal | 8 | 702866.884 | 4.331 |
| mammals | Scotozous_dormeri | Scotozous_dormeri | LC | Stable | temperate | nocturnal | 6.9 | 2183092.096 | 18.836 |
| mammals | Scutisorex_somereni | Scutisorex_somereni | LC | Unknown | tropical | cathemeral | 91.5 | 434028.617 | 7.611 |
| mammals | Sekeetamys_calurus | Sekeetamys_calurus | LC | Stable | temperate | cathemeral | 55.3 | 200686.833 | 4.385 |
| mammals | Selevinia_betpakdalaensis | NA | DD | Unknown | NA | NA | NA | NA | NA |
| mammals | Semnopithecus_ajax | Semnopithecus_ajax | EN | Decreasing | temperate | diurnal | 20150 | 8509.043 | 11.04 |
| mammals | Semnopithecus_entellus | Semnopithecus_entellus | LC | Decreasing | tropical | diurnal | 14500 | 558928.443 | 16.482 |
| mammals | Semnopithecus_hector | Semnopithecus_hector | NT | Decreasing | temperate | diurnal | 17350 | 101204.736 | 14.395 |
| mammals | Semnopithecus_hypoleucos | Semnopithecus_hypoleucos | LC | Decreasing | tropical | diurnal | 8950 | 407292.34 | 21.078 |
| mammals | Semnopithecus_johnii | Trachypithecus_johnii | VU | Stable | tropical | diurnal | 11921 | 24067.171 | 16.856 |
| mammals | Semnopithecus_priam | Semnopithecus_priam | NT | Decreasing | tropical | diurnal | 12450 | 190726.316 | 20.234 |
| mammals | Semnopithecus_schistaceus | Semnopithecus_schistaceus | LC | Decreasing | temperate | diurnal | 17350 | 311696.722 | 9.69 |
| mammals | Semnopithecus_vetulus | Trachypithecus_vetulus | EN | Decreasing | tropical | diurnal | 8830 | 36899.918 | 18.115 |
| mammals | Serengetimys_pernanus | NA | DD | Unknown | NA | NA | NA | NA | NA |

| | | | | | | | | | |
|---|---|---|---|---|---|---|---|---|---|
| mammals | Setifer_setosus | Setifer_setosus | LC | Stable | tropical | nocturnal | 294 | 590831.629 | 8.745 |
| mammals | Setonix_brachyurus | Setonix_brachyurus | VU | Decreasing | temperate | nocturnal | 3250 | 18067.243 | 8.246 |
| mammals | Sicista_betulina | Sicista_betulina | LC | Unknown | temperate | nocturnal | 8 | 6815198.931 | 7.531 |
| mammals | Sicista_caucasica | Sicista_caucasica | NT | Unknown | temperate | nocturnal | 10 | 6718.827 | 5.547 |
| mammals | Sicista_caudata | NA | DD | Unknown | NA | NA | NA | NA | NA |
| mammals | Sicista_concolor | Sicista_concolor | LC | Unknown | temperate | nocturnal | 6.5 | 527987.061 | 10.108 |
| mammals | Sicista_kazbegica | Sicista_kazbegica | EN | Unknown | temperate | nocturnal | 7 | 3375.933 | 7.979 |
| mammals | Sicista_kluchorica | Sicista_kluchorica | LC | Unknown | temperate | nocturnal | 7.5 | 7368.936 | 6.308 |
| mammals | Sicista_napaea | Sicista_napaea | LC | Unknown | temperate | nocturnal | 10 | 117911.871 | 7.145 |
| mammals | Sicista_pseudonapaea | NA | DD | Unknown | NA | NA | NA | NA | NA |
| mammals | Sicista_severtzovi | Sicista_severtzovi | LC | Stable | temperate | nocturnal | 11.2 | 350083.033 | 17.334 |
| mammals | Sicista_strandi | Sicista_strandi | LC | Stable | temperate | nocturnal | 10.3 | 652702.095 | 15.976 |
| mammals | Sicista_subtilis | Sicista_subtilis | LC | Decreasing | temperate | nocturnal | 11.8 | 2576602.47 | 9.536 |
| mammals | Sicista_tianschanica | Sicista_tianshanica | LC | Unknown | temperate | nocturnal | 11.5 | 358985.782 | 6.168 |
| mammals | Sigmodon_alleni | Sigmodon_alleni | VU | Decreasing | tropical | cathemeral | 179 | 174869.567 | 8.796 |
| mammals | Sigmodon_alstoni | Sigmodon_alstoni | LC | Unknown | tropical | cathemeral | 55.7 | 840163.265 | 5.067 |
| mammals | Sigmodon_arizonae | Sigmodon_arizonae | LC | Unknown | temperate | cathemeral | 198 | 345280.161 | 7.538 |
| mammals | Sigmodon_fulviventer | Sigmodon_fulviventer | LC | Stable | temperate | cathemeral | 214 | 464112.148 | 8.668 |
| mammals | Sigmodon_hirsutus | Sigmodon_hirsutus | LC | Increasing | tropical | cathemeral | 171.5 | 761233.046 | 12.467 |
| mammals | Sigmodon_hispidus | Sigmodon_hispidus | LC | Increasing | temperate | cathemeral | 92.4 | 2615707.968 | 15.061 |
| mammals | Sigmodon_inopinatus | Sigmodon_inopinatus | VU | Decreasing | tropical | cathemeral | 140.5 | 13693.252 | 14.15 |
| mammals | Sigmodon_leucotis | Sigmodon_leucotis | LC | Unknown | tropical | diurnal | 135.5 | 282532.309 | 11.473 |
| mammals | Sigmodon_mascotensis | Sigmodon_mascotensis | LC | Stable | tropical | cathemeral | 120 | 252559.649 | 9.411 |
| mammals | Sigmodon_ochrognathus | Sigmodon_ochrognathus | LC | Unknown | temperate | diurnal | 122 | 464511.072 | 7.092 |
| mammals | Sigmodon_peruanus | Sigmodon_peruanus | LC | Stable | tropical | cathemeral | 88.7 | 98145.692 | 12.268 |
| mammals | Sigmodon_planifrons | NA | not_available | not_available | NA | NA | NA | NA | NA |
| mammals | Sigmodon_toltecus | Sigmodon_toltecus | LC | Unknown | tropical | diurnal | 50 | 416823.485 | 9.911 |
| mammals | Sigmodontomys_alfari | Sigmodontomys_alfari | LC | Stable | tropical | nocturnal | 50 | 383737.193 | 9.714 |
| mammals | Simias_concolor | Simias_concolor | CR | Decreasing | tropical | diurnal | 7980 | 6004.48 | 6.223 |
| mammals | Sminthopsis_archeri | NA | DD | Unknown | NA | NA | NA | NA | NA |
| mammals | Sminthopsis_bindi | Sminthopsis_bindi | NT | Decreasing | tropical | nocturnal | 16 | 45581.567 | 1.962 |
| mammals | Sminthopsis_butleri | Sminthopsis_butleri | VU | Decreasing | tropical | nocturnal | 25 | 7067.236 | 2.458 |
| mammals | Sminthopsis_crassicaudata | Sminthopsis_crassicaudata | LC | Stable | temperate | nocturnal | 15 | 4137398.352 | 4.933 |
| mammals | Sminthopsis_dolichura | Sminthopsis_dolichura | LC | Stable | temperate | nocturnal | 13.6 | 551815.736 | 5.712 |

| | | | | | | | | | |
|---|---|---|---|---|---|---|---|---|---|
| mammals | Sminthopsis_douglasi | Sminthopsis_douglasi | NT | Decreasing | tropical | nocturnal | 55 | 47129.402 | 5.265 |
| mammals | Sminthopsis_fuliginosa | Sminthopsis_griseoventer | LC | Unknown | temperate | nocturnal | 17.5 | 144138.804 | 8.535 |
| mammals | Sminthopsis_gilberti | Sminthopsis_gilberti | LC | Stable | temperate | nocturnal | 19.5 | 76628.817 | 10.67 |
| mammals | Sminthopsis_granulipes | Sminthopsis_granulipes | LC | Unknown | temperate | nocturnal | 25 | 102873.959 | 11.265 |
| mammals | Sminthopsis_hirtipes | Sminthopsis_hirtipes | LC | Unknown | temperate | nocturnal | 15 | 1035105.776 | 1.441 |
| mammals | Sminthopsis_leucopus | Sminthopsis_leucopus | LC | Stable | temperate | nocturnal | 26 | 96976.298 | 7.853 |
| mammals | Sminthopsis_macroura | Sminthopsis_macroura | LC | Unknown | temperate | nocturnal | 20 | 4300281.356 | 3.235 |
| mammals | Sminthopsis_murina | Sminthopsis_murina | LC | Decreasing | temperate | nocturnal | 17 | 1210760.962 | 7.845 |
| mammals | Sminthopsis_ooldea | Sminthopsis_ooldea | LC | Unknown | temperate | nocturnal | 11 | 1133609.414 | 1.656 |
| mammals | Sminthopsis_psammophila | Sminthopsis_psammophila | VU | Decreasing | temperate | nocturnal | 33.5 | 7619.609 | 2.388 |
| mammals | Sminthopsis_virginiae | Sminthopsis_virginiae | LC | Stable | tropical | nocturnal | 35.3 | 362600.722 | 2.424 |
| mammals | Sminthopsis_youngsoni | Sminthopsis_youngsoni | LC | Stable | tropical | nocturnal | 10.1 | 1373345.181 | 1.492 |
| mammals | Smutsia_gigantea | Smutsia_gigantea | EN | Decreasing | tropical | nocturnal | 32999.7 | 3497994.29 | 7.927 |
| mammals | Smutsia_temminckii | Smutsia_temminckii | VU | Decreasing | tropical | cathemeral | 7179.9 | 7124374.963 | 8.979 |
| mammals | Solenodon_paradoxus | Solenodon_paradoxus | LC | Unknown | tropical | nocturnal | 900 | 13314.145 | 10.175 |
| mammals | Solisorex_pearsoni | Solisorex_pearsoni | EN | Decreasing | tropical | nocturnal | 66.6 | 481.979 | 19.185 |
| mammals | Solomys_ponceleti | Solomys_ponceleti | CR | Decreasing | tropical | nocturnal | 1000 | 11785.138 | 5.197 |
| mammals | Solomys_salamonis | NA | DD | Unknown | NA | NA | NA | NA | NA |
| mammals | Solomys_salebrosus | Solomys_salebrosus | VU | Unknown | tropical | nocturnal | 397 | 9993.959 | 4.954 |
| mammals | Solomys_sapientis | Solomys_sapientis | EN | Decreasing | tropical | nocturnal | 225.5 | 3824.601 | 2.927 |
| mammals | Sommeromys_macrorhinos | NA | DD | Unknown | NA | NA | NA | NA | NA |
| mammals | Sooretamys_angouya | Sooretamys_angouya | LC | Unknown | temperate | nocturnal | 100.3 | 1389336.811 | 11.898 |
| mammals | Sorex_alpinus | Sorex_alpinus | NT | Decreasing | temperate | cathemeral | 7.8 | 422525.149 | 16.382 |
| mammals | Sorex_antinorii | Sorex_antinorii | LC | Unknown | temperate | cathemeral | 11.5 | 189627.142 | 21.554 |
| mammals | Sorex_araneus | Sorex_araneus | LC | Stable | temperate | cathemeral | 10 | 13236507.592 | 9.478 |
| mammals | Sorex_arcticus | Sorex_arcticus | LC | Stable | temperate | cathemeral | 8.2 | 4533841.872 | 4.907 |
| mammals | Sorex_arizonae | Sorex_arizonae | LC | Unknown | temperate | cathemeral | 2.4 | 62195.043 | 8.197 |
| mammals | Sorex_asper | Sorex_asper | LC | Unknown | temperate | cathemeral | 7.9 | 379195.761 | 7.849 |
| mammals | Sorex_bedfordiae | Sorex_bedfordiae | LC | Unknown | temperate | nocturnal | 6.6 | 585011.117 | 10.1 |
| mammals | Sorex_bendirii | Sorex_bendirii | LC | Unknown | temperate | diurnal | 16.1 | 207238.101 | 11.614 |
| mammals | Sorex_buchariensis | NA | DD | Unknown | NA | NA | NA | NA | NA |
| mammals | Sorex_caecutiens | Sorex_caecutiens | LC | Stable | temperate | cathemeral | 5.4 | 17639588.441 | 4.295 |
| mammals | Sorex_camtschaticus | Sorex_camtschatica | LC | Unknown | temperate | cathemeral | 4.6 | 447577.752 | 1.516 |
| mammals | Sorex_cansulus | NA | DD | Unknown | NA | NA | NA | NA | NA |

| | | | | | | | | | |
|---|---|---|---|---|---|---|---|---|---|
| mammals | Sorex_coronatus | Sorex_coronatus | LC | Stable | temperate | cathemeral | 9.6 | 884132.192 | 23.087 |
| mammals | Sorex_cylindricauda | Sorex_cylindricauda | LC | Unknown | temperate | cathemeral | 6.1 | 327292.253 | 9.241 |
| mammals | Sorex_daphaenodon | Sorex_daphaenodon | LC | Stable | temperate | cathemeral | 6.8 | 8192141.479 | 3.819 |
| mammals | Sorex_emarginatus | Sorex_emarginatus | LC | Decreasing | tropical | cathemeral | 7 | 30452.151 | 7.113 |
| mammals | Sorex_excelsus | Sorex_excelsus | LC | Unknown | temperate | nocturnal | 7.5 | 200703.772 | 7.785 |
| mammals | Sorex_fumeus | Sorex_fumeus | LC | Stable | temperate | cathemeral | 7.7 | 1508616.008 | 14.354 |
| mammals | Sorex_gracillimus | Sorex_gracillimus | LC | Stable | temperate | nocturnal | 4.5 | 1136758.485 | 4.33 |
| mammals | Sorex_granarius | Sorex_granarius | LC | Unknown | temperate | cathemeral | 6.3 | 69946.848 | 19.533 |
| mammals | Sorex_haydeni | Sorex_haydeni | LC | Stable | temperate | diurnal | 3.5 | 1160249.702 | 15.106 |
| mammals | Sorex_hosonoi | Sorex_hosonoi | LC | Unknown | temperate | cathemeral | 3.3 | 10546.77 | 10.229 |
| mammals | Sorex_isodon | Sorex_isodon | LC | Unknown | temperate | cathemeral | 9.4 | 12216462.267 | 4.141 |
| mammals | Sorex_ixtlanensis | NA | DD | Unknown | NA | NA | NA | NA | NA |
| mammals | Sorex_jacksoni | Sorex_jacksoni | LC | Unknown | temperate | cathemeral | 3.5 | 4929.275 | 0.583 |
| mammals | Sorex_kozlovi | NA | DD | Unknown | NA | NA | NA | NA | NA |
| mammals | Sorex_leucogaster | NA | DD | Unknown | NA | NA | NA | NA | NA |
| mammals | Sorex_longirostris | Sorex_longirostris | LC | Stable | temperate | cathemeral | 3.6 | 1643808.719 | 16.969 |
| mammals | Sorex_lyelli | Sorex_lyelli | LC | Stable | temperate | cathemeral | 5 | 3090.756 | 4.067 |
| mammals | Sorex_maritimensis | Sorex_maritimensis | LC | Decreasing | temperate | cathemeral | 7.3 | 61267.676 | 10.556 |
| mammals | Sorex_mediopua | Sorex_mediopua | LC | Unknown | tropical | cathemeral | 8 | 56066.925 | 11.906 |
| mammals | Sorex_merriami | Sorex_merriami | LC | Unknown | temperate | cathemeral | 5.9 | 1800350.218 | 9.688 |
| mammals | Sorex_milleri | Sorex_milleri | VU | Decreasing | temperate | cathemeral | 7 | 11692.588 | 9.933 |
| mammals | Sorex_minutissimus | Sorex_minutissimus | LC | Unknown | temperate | cathemeral | 2.5 | 17514810.101 | 4.247 |
| mammals | Sorex_monticolus | Sorex_monticolus | LC | Stable | temperate | cathemeral | 5.3 | 4535520.827 | 3.664 |
| mammals | Sorex_nanus | Sorex_nanus | LC | Unknown | temperate | cathemeral | 2.4 | 997639.341 | 9.697 |
| mammals | Sorex_navigator | NA | DD | Unknown | NA | NA | NA | NA | NA |
| mammals | Sorex_oreopolus | Sorex_oreopolus | LC | Stable | tropical | cathemeral | 7 | 14769.791 | 26.581 |
| mammals | Sorex_orizabae | Sorex_orizabae | LC | Stable | tropical | cathemeral | 4.8 | 56542.846 | 18.131 |
| mammals | Sorex_pacificus | Sorex_pacificus | LC | Stable | temperate | cathemeral | 6.7 | 93854.054 | 11.153 |
| mammals | Sorex_palustris | Sorex_palustris | LC | Stable | temperate | cathemeral | 13.5 | 6242380.887 | 6.398 |
| mammals | Sorex_planiceps | Sorex_planiceps | LC | Unknown | temperate | cathemeral | 10.5 | 27733.294 | 7.413 |
| mammals | Sorex_portenkoi | NA | DD | Unknown | NA | NA | NA | NA | NA |
| mammals | Sorex_preblei | Sorex_preblei | LC | Unknown | temperate | cathemeral | 3.1 | 688410.089 | 9.21 |
| mammals | Sorex_pribilofensis | NA | EN | Unknown | NA | NA | NA | NA | NA |
| mammals | Sorex_raddei | Sorex_raddei | LC | Unknown | temperate | nocturnal | 10 | 245664.418 | 12.348 |

| | | | | | | | | | |
|---|---|---|---|---|---|---|---|---|---|
| mammals | Sorex_roboratus | Sorex_roboratus | LC | Unknown | temperate | cathemeral | 9.1 | 9303837.797 | 2.139 |
| mammals | Sorex_rohweri | Sorex_rohweri | LC | Unknown | temperate | cathemeral | 5.1 | 24366.372 | 10.608 |
| mammals | Sorex_samniticus | Sorex_samniticus | LC | Unknown | temperate | cathemeral | 8.3 | 157801.637 | 24.085 |
| mammals | Sorex_satunini | Sorex_satunini | LC | Unknown | temperate | nocturnal | 7.5 | 412610.262 | 13.446 |
| mammals | Sorex_saussurei | Sorex_saussurei | LC | Stable | tropical | cathemeral | 4.7 | 87917.992 | 15.001 |
| mammals | Sorex_sclateri | Sorex_sclateri | CR | Decreasing | tropical | cathemeral | 7 | 33.506 | 13.393 |
| mammals | Sorex_shinto | Sorex_shinto | LC | Stable | temperate | nocturnal | 5.9 | 27674.38 | 12.271 |
| mammals | Sorex_sinalis | NA | DD | Unknown | NA | NA | NA | NA | NA |
| mammals | Sorex_sonomae | Sorex_sonomae | LC | Stable | temperate | nocturnal | 8.3 | 143305.118 | 11.045 |
| mammals | Sorex_tenellus | Sorex_tenellus | LC | Stable | temperate | cathemeral | 3.8 | 65722.259 | 5.293 |
| mammals | Sorex_thibetanus | NA | DD | Unknown | NA | NA | NA | NA | NA |
| mammals | Sorex_trowbridgii | Sorex_trowbridgii | LC | Stable | temperate | cathemeral | 3.8 | 377575.969 | 11.139 |
| mammals | Sorex_tundrensis | Sorex_tundrensis | LC | Stable | temperate | cathemeral | 7.7 | 15993094.218 | 3.349 |
| mammals | Sorex_ugyunak | Sorex_ugyunak | LC | Stable | temperate | cathemeral | 3.6 | 1412342.544 | 0.218 |
| mammals | Sorex_unguiculatus | Sorex_unguiculatus | LC | Stable | temperate | cathemeral | 9.9 | 1194028.932 | 5.231 |
| mammals | Sorex_vagrans | Sorex_vagrans | LC | Stable | temperate | cathemeral | 5.9 | 973482.077 | 9.602 |
| mammals | Sorex_ventralis | Sorex_ventralis | LC | Stable | tropical | cathemeral | 7 | 63024.279 | 14.503 |
| mammals | Sorex_veraecrucis | Sorex_veraecrucis | LC | Stable | tropical | cathemeral | 7.6 | 417038.618 | 11.088 |
| mammals | Sorex_veraepacis | Sorex_veraepacis | LC | Stable | tropical | cathemeral | 7.5 | 90682.591 | 9.678 |
| mammals | Sorex_volnuchini | Sorex_volnuchini | LC | Unknown | temperate | cathemeral | 5.5 | 434037.433 | 13.675 |
| mammals | Soricomys_kalinga | Soricomys_kalinga | LC | Unknown | tropical | diurnal | 25.5 | 83.15 | 2.196 |
| mammals | Soricomys_musseri | Soricomys_musseri | LC | Unknown | tropical | diurnal | 26.5 | 3298.95 | 6.825 |
| mammals | Soriculus_nigrescens | Soriculus_nigrescens | LC | Unknown | temperate | nocturnal | 14 | 136320.638 | 5.639 |
| mammals | Spalacopus_cyanus | Spalacopus_cyanus | LC | Unknown | temperate | diurnal | 127.5 | 76144.182 | 11.541 |
| mammals | Speothos_venaticus | Speothos_venaticus | NT | Decreasing | tropical | diurnal | 6000 | 11389107.508 | 5.377 |
| mammals | Spermophilopsis_leptodactyla | Spermophilopsis_leptodactylus | LC | Unknown | temperate | diurnal | 600 | 1268483.868 | 7.594 |
| mammals | Spermophilus_alaschanicus | Spermophilus_alashanicus | LC | Decreasing | temperate | diurnal | 208 | 487834.925 | 12.464 |
| mammals | Spermophilus_brevicauda | Spermophilus_brevicauda | LC | Unknown | temperate | diurnal | 289.5 | 564431.589 | 5.853 |
| mammals | Spermophilus_citellus | Spermophilus_citellus | EN | Decreasing | temperate | diurnal | 290 | 432578.76 | 20.532 |
| mammals | Spermophilus_dauricus | Spermophilus_dauricus | LC | Unknown | temperate | diurnal | 161.3 | 1379395.545 | 14.072 |
| mammals | Spermophilus_erythrogenys | Spermophilus_erythrogenys | LC | Stable | temperate | diurnal | 355 | 1727062.329 | 7.656 |
| mammals | Spermophilus_fulvus | Spermophilus_fulvus | LC | Unknown | temperate | diurnal | 596 | 1851642.095 | 7.508 |
| mammals | Spermophilus_major | Spermophilus_major | LC | Unknown | temperate | diurnal | 786 | 629933.297 | 12.961 |
| mammals | Spermophilus_musicus | Spermophilus_musicus | NT | Unknown | temperate | diurnal | 316.9 | 1036.685 | 11.434 |

| | | | | | | | | | |
|---|---|---|---|---|---|---|---|---|---|
| mammals | Spermophilus_nilkaensis | Spermophilus_ralli | LC | Unknown | temperate | diurnal | 347.5 | 39704.858 | 8.846 |
| mammals | Spermophilus_pallidicauda | Spermophilus_pallidicauda | LC | Unknown | temperate | diurnal | 439.5 | 699451.663 | 5.271 |
| mammals | Spermophilus_pygmaeus | Spermophilus_pygmaeus | LC | Decreasing | temperate | diurnal | 136 | 1810870.846 | 9.29 |
| mammals | Spermophilus_relictus | Spermophilus_relictus | LC | Unknown | temperate | diurnal | 600 | 188222.908 | 9.601 |
| mammals | Spermophilus_suslicus | Spermophilus_suslicus | NT | Decreasing | temperate | diurnal | 252 | 804158.503 | 16.536 |
| mammals | Spermophilus_taurensis | Spermophilus_taurensis | LC | Unknown | temperate | diurnal | 200.7 | 17713.103 | 11.304 |
| mammals | Spermophilus_xanthoprymnus | Spermophilus_xanthoprymnus | NT | Decreasing | temperate | diurnal | 289.7 | 475379.397 | 13.831 |
| mammals | Sphaerias_blanfordi | Sphaerias_blanfordi | LC | Stable | tropical | nocturnal | 28.2 | 481371.884 | 10.059 |
| mammals | Sphaeronycteris_toxophyllum | Sphaeronycteris_toxophyllum | LC | Unknown | tropical | nocturnal | 16.2 | 3805535.483 | 3.145 |
| mammals | Spilocuscus_kraemeri | Spilocuscus_kraemeri | NT | Decreasing | tropical | nocturnal | 2300 | 1892.779 | 6.243 |
| mammals | Spilocuscus_maculatus | Spilocuscus_maculatus | LC | Stable | tropical | nocturnal | 5106.8 | 886129.18 | 3.531 |
| mammals | Spilocuscus_papuensis | Spilocuscus_papuensis | VU | Unknown | tropical | cathemeral | 5735.5 | 3142.734 | 2.642 |
| mammals | Spilocuscus_rufoniger | Spilocuscus_rufoniger | CR | Decreasing | tropical | cathemeral | 6000 | 178086.303 | 3.588 |
| mammals | Spilocuscus_wilsoni | Spilocuscus_wilsoni | CR | Decreasing | tropical | cathemeral | 3262.3 | 2524.159 | 8.959 |
| mammals | Spilogale_angustifrons | Spilogale_angustifrons | LC | Stable | tropical | nocturnal | 386.5 | 777447.141 | 11.762 |
| mammals | Spilogale_gracilis | Spilogale_gracilis | LC | Decreasing | temperate | nocturnal | 521.8 | 3825446.678 | 9.03 |
| mammals | Spilogale_putorius | Spilogale_putorius | VU | Decreasing | temperate | nocturnal | 341 | 2979584.787 | 15.329 |
| mammals | Spilogale_pygmaea | Spilogale_pygmaea | VU | Decreasing | tropical | nocturnal | 235 | 95518.339 | 8.951 |
| mammals | Srilankamys_ohiensis | Srilankamys_ohiensis | VU | Decreasing | tropical | nocturnal | 125.5 | 1707.502 | 11.213 |
| mammals | Steatomys_caurinus | Steatomys_caurinus | LC | Stable | tropical | nocturnal | 44.8 | 1058295.65 | 13.701 |
| mammals | Steatomys_cuppedius | Steatomys_cuppedius | LC | Stable | tropical | nocturnal | 16.2 | 1075014.2 | 13.427 |
| mammals | Steatomys_jacksoni | NA | DD | Unknown | NA | NA | NA | NA | NA |
| mammals | Steatomys_krebsii | Steatomys_krebsii | LC | Stable | temperate | nocturnal | 20.7 | 979358.6 | 7.786 |
| mammals | Steatomys_opimus | Steatomys_opimus | LC | Unknown | tropical | nocturnal | 42.7 | 721011.85 | 5.139 |
| mammals | Steatomys_parvus | Steatomys_parvus | LC | Stable | tropical | nocturnal | 8.9 | 1128038.118 | 9.408 |
| mammals | Steatomys_pratensis | Steatomys_pratensis | LC | Stable | tropical | nocturnal | 48 | 3477980.207 | 7.835 |
| mammals | Stenocephalemys_albipes | Stenocephalemys_albipes | LC | Stable | tropical | nocturnal | 59.5 | 360891.896 | 13.024 |
| mammals | Stenocephalemys_albocaudatus | Stenocephalemys_albocaudata | LC | Stable | tropical | nocturnal | 144 | 12140.109 | 13.757 |
| mammals | Stenocephalemys_griseicauda | Stenocephalemys_griseicauda | LC | Decreasing | tropical | nocturnal | 85.7 | 38987.75 | 13.525 |
| mammals | Stenocephalemys_ruppi | NA | DD | Unknown | NA | NA | NA | NA | NA |
| mammals | Stenocranius_gregalis | Microtus_gregalis | LC | Stable | temperate | cathemeral | 47.5 | 4537963.905 | 7.159 |
| mammals | Stenoderma_rufum | Stenoderma_rufum | NT | Unknown | tropical | nocturnal | 20.7 | 9281.028 | 27.106 |
| mammals | Stenonycteris_lanosa | Rousettus_lanosus | LC | Decreasing | tropical | nocturnal | 103 | 209037.618 | 14.356 |
| mammals | Stochomys_longicaudatus | Stochomys_longicaudatus | LC | Unknown | tropical | nocturnal | 71 | 1866744.352 | 6.66 |

| | | | | | | | | | |
|---|---|---|---|---|---|---|---|---|---|
| mammals | Strigocuscus_celebensis | Strigocuscus_celebensis | NT | Decreasing | tropical | nocturnal | 500 | 172973.193 | 9.355 |
| mammals | Sturnira_aratathomasi | Sturnira_aratathomasi | LC | Unknown | tropical | nocturnal | 48.6 | 72657.454 | 11.67 |
| mammals | Sturnira_bidens | Sturnira_bidens | LC | Stable | tropical | nocturnal | 17.6 | 705384.621 | 9.912 |
| mammals | Sturnira_bogotensis | Sturnira_bogotensis | LC | Stable | tropical | nocturnal | 20.8 | 501247.49 | 11.061 |
| mammals | Sturnira_erythromos | Sturnira_erythromos | LC | Stable | tropical | nocturnal | 15.1 | 2534366.142 | 8.257 |
| mammals | Sturnira_koopmanhilli | NA | DD | Unknown | NA | NA | NA | NA | NA |
| mammals | Sturnira_lilium | Sturnira_lilium | LC | Stable | tropical | nocturnal | 19.1 | 4859393.426 | 9.777 |
| mammals | Sturnira_ludovici | Sturnira_ludovici | LC | Unknown | tropical | nocturnal | 21 | 1216315.862 | 10.025 |
| mammals | Sturnira_luisi | Sturnira_luisi | LC | Unknown | tropical | nocturnal | 21 | 340279.766 | 9.298 |
| mammals | Sturnira_magna | Sturnira_magna | LC | Stable | tropical | nocturnal | 48.6 | 1567758.189 | 3.953 |
| mammals | Sturnira_mistratensis | NA | DD | Unknown | NA | NA | NA | NA | NA |
| mammals | Sturnira_mordax | Sturnira_mordax | LC | Stable | tropical | nocturnal | 25.1 | 28315.491 | 11.697 |
| mammals | Sturnira_nana | Sturnira_nana | EN | Unknown | tropical | nocturnal | 10.9 | 2319.44 | 3.69 |
| mammals | Sturnira_oporaphilum | Sturnira_oporaphilum | LC | Unknown | tropical | nocturnal | 26.5 | 1015753.634 | 6.257 |
| mammals | Sturnira_perla | NA | DD | Unknown | NA | NA | NA | NA | NA |
| mammals | Sturnira_sorianoi | NA | DD | Unknown | NA | NA | NA | NA | NA |
| mammals | Sturnira_tildae | Sturnira_tildae | LC | Stable | tropical | nocturnal | 24 | 10661362.097 | 4.764 |
| mammals | Styloctenium_mindorense | Styloctenium_mindorensis | EN | Decreasing | tropical | nocturnal | 212 | 485.35 | 10.653 |
| mammals | Styloctenium_wallacei | Styloctenium_wallacei | NT | Decreasing | tropical | nocturnal | 173 | 170647.356 | 9.284 |
| mammals | Stylodipus_andrewsi | Stylodipus_andrewsi | LC | Unknown | temperate | nocturnal | 60 | 940743.685 | 5.433 |
| mammals | Stylodipus_sungorus | Stylodipus_sungorus | LC | Decreasing | temperate | nocturnal | 76.9 | 61392.538 | 3.652 |
| mammals | Stylodipus_telum | Stylodipus_telum | LC | Decreasing | temperate | nocturnal | 80 | 1340954.309 | 6.254 |
| mammals | Subulo_gouazoubira | Mazama_gouazoubira | LC | Decreasing | tropical | cathemeral | 16300.1 | 5746702.264 | 9.341 |
| mammals | Suncus_aequatorius | Suncus_aequatorius | EN | Unknown | tropical | nocturnal | 8.8 | 491.261 | 18.951 |
| mammals | Suncus_dayi | Suncus_dayi | EN | Decreasing | tropical | nocturnal | 9.4 | 209.341 | 10.319 |
| mammals | Suncus_etruscus | Suncus_etruscus | LC | Unknown | temperate | nocturnal | 2.1 | 3765817.869 | 14.362 |
| mammals | Suncus_fellowesgordoni | NA | EN | Decreasing | NA | NA | NA | NA | NA |
| mammals | Suncus_hosei | NA | DD | Unknown | NA | NA | NA | NA | NA |
| mammals | Suncus_hututsi | NA | DD | Unknown | NA | NA | NA | NA | NA |
| mammals | Suncus_infinitesimus | Suncus_infinitesimus | LC | Unknown | temperate | cathemeral | 3 | 131972.951 | 14.199 |
| mammals | Suncus_lixa | Suncus_lixus | LC | Unknown | tropical | nocturnal | 4.9 | 1437689.013 | 8.01 |
| mammals | Suncus_malayanus | Suncus_malayanus | LC | Unknown | tropical | nocturnal | 5 | 171596.116 | 13.629 |
| mammals | Suncus_megalurus | Suncus_megalura | LC | Unknown | tropical | nocturnal | 4.7 | 4240345.548 | 10.934 |
| mammals | Suncus_mertensi | Suncus_mertensi | EN | Decreasing | tropical | nocturnal | 8 | 28.326 | 18.257 |

| | | | | | | | | | |
|---|---|---|---|---|---|---|---|---|---|
| mammals | Suncus_montanus | Suncus_montanus | VU | Decreasing | tropical | nocturnal | 18 | 5369.616 | 17.249 |
| mammals | Suncus_murinus | Suncus_murinus | LC | Stable | tropical | nocturnal | 61.5 | 9425536.555 | 13.954 |
| mammals | Suncus_remyi | Suncus_remyi | LC | Unknown | tropical | nocturnal | 1.8 | 150166.811 | 3.156 |
| mammals | Suncus_stoliczkanus | Suncus_stoliczkanus | LC | Unknown | tropical | nocturnal | 9.4 | 1280662.879 | 17.34 |
| mammals | Suncus_varilla | Suncus_varilla | LC | Unknown | tropical | nocturnal | 5 | 2013476.897 | 9.87 |
| mammals | Suncus_zeylanicus | Suncus_zeylanicus | EN | Decreasing | tropical | nocturnal | 29.8 | 456.492 | 12.476 |
| mammals | Sundamys_annandalei | Rattus_annandalei | LC | Decreasing | tropical | nocturnal | 216.1 | 124978.866 | 12.607 |
| mammals | Sundamys_infraluteus | Sundamys_infraluteus | LC | Unknown | tropical | nocturnal | 418.5 | 102571.949 | 4.782 |
| mammals | Sundamys_maxi | Sundamys_maxi | VU | Decreasing | tropical | nocturnal | 361 | 5127.641 | 15.989 |
| mammals | Sundamys_muelleri | Sundamys_muelleri | LC | Decreasing | tropical | nocturnal | 275.7 | 1408490.368 | 8.104 |
| mammals | Sundasciurus_brookei | Sundasciurus_brookei | LC | Unknown | tropical | diurnal | 115.5 | 259522.753 | 3.11 |
| mammals | Sundasciurus_davensis | NA | DD | Unknown | NA | NA | NA | NA | NA |
| mammals | Sundasciurus_everetti | Dremomys_everetti | LC | Stable | tropical | diurnal | 147.5 | 65727.42 | 3.342 |
| mammals | Sundasciurus_fraterculus | Sundasciurus_fraterculus | VU | Decreasing | tropical | diurnal | 58.6 | 6004.261 | 6.223 |
| mammals | Sundasciurus_hippurus | Sundasciurus_hippurus | NT | Decreasing | tropical | diurnal | 406.3 | 1314932.177 | 7.817 |
| mammals | Sundasciurus_hoogstraali | Sundasciurus_hoogstraali | LC | Unknown | tropical | diurnal | 266.3 | 943.552 | 12.539 |
| mammals | Sundasciurus_jentinki | Sundasciurus_jentinki | LC | Unknown | tropical | diurnal | 76 | 50301.98 | 3.128 |
| mammals | Sundasciurus_juvencus | Sundasciurus_juvencus | LC | Stable | tropical | diurnal | 259 | 7784.606 | 8.925 |
| mammals | Sundasciurus_lowii | Sundasciurus_lowii | LC | Stable | tropical | diurnal | 87.5 | 1335061.879 | 7.925 |
| mammals | Sundasciurus_mindanensis | Sundasciurus_mindanensis | LC | Unknown | tropical | diurnal | 285 | 94133.375 | 12.032 |
| mammals | Sundasciurus_moellendorffi | Sundasciurus_moellendorffi | NT | Decreasing | tropical | diurnal | 252 | 527.188 | 10.299 |
| mammals | Sundasciurus_philippinensis | Sundasciurus_philippinensis | LC | Stable | tropical | diurnal | 244 | 120457.566 | 12.097 |
| mammals | Sundasciurus_rabori | NA | DD | Unknown | NA | NA | NA | NA | NA |
| mammals | Sundasciurus_samarensis | Sundasciurus_samarensis | LC | Unknown | tropical | diurnal | 225 | 19561.713 | 11.315 |
| mammals | Sundasciurus_steerii | Sundasciurus_steerii | LC | Stable | tropical | diurnal | 252 | 3901.168 | 8.999 |
| mammals | Sundasciurus_tenuis | Sundasciurus_tenuis | LC | Decreasing | tropical | diurnal | 73.8 | 1210112.927 | 8.028 |
| mammals | Surdisorex_norae | Surdisorex_norae | LC | Stable | tropical | cathemeral | 9 | 3057.853 | 13.405 |
| mammals | Surdisorex_polulus | Surdisorex_polulus | DD | Stable | tropical | cathemeral | 9 | 962.089 | 3.08 |
| mammals | Suricata_suricatta | Suricata_suricatta | LC | Stable | temperate | diurnal | 725.5 | 1758699.399 | 6.77 |
| mammals | Sus_ahoenobarbus | Sus_ahoenobarbus | NT | Decreasing | tropical | diurnal | 56749.7 | 10580.518 | 8.457 |
| mammals | Sus_barbatus | Sus_barbatus | VU | Decreasing | tropical | diurnal | 70500 | 813194.655 | 5.382 |
| mammals | Sus_cebifrons | Sus_cebifrons | CR | Decreasing | tropical | nocturnal | 32500 | 2381.998 | 8.648 |
| mammals | Sus_celebensis | Sus_celebensis | NT | Decreasing | tropical | diurnal | 55000 | 257141.822 | 9.843 |
| mammals | Sus_oliveri | Sus_oliveri | VU | Decreasing | tropical | nocturnal | 44801.2 | 2450.555 | 7.291 |

| | | | | | | | | | |
|---|---|---|---|---|---|---|---|---|---|
| mammals | Sus_philippensis | Sus_philippensis | VU | Decreasing | tropical | nocturnal | 44253.5 | 64212.435 | 9.352 |
| mammals | Sus_scrofa | Sus_scrofa | LC | Unknown | temperate | cathemeral | 101052.1 | 28026537.211 | 12.987 |
| mammals | Sus_verrucosus | Sus_verrucosus | EN | Decreasing | tropical | nocturnal | 92500 | 4537.91 | 16.973 |
| mammals | Syconycteris_australis | Syconycteris_australis | LC | Stable | tropical | nocturnal | 17.1 | 1053996.418 | 3.871 |
| mammals | Syconycteris_carolinae | Syconycteris_carolinae | NT | Decreasing | tropical | nocturnal | 39 | 19996.954 | 6.533 |
| mammals | Syconycteris_hobbit | Syconycteris_hobbit | LC | Decreasing | tropical | nocturnal | 18.3 | 4176.044 | 4.156 |
| mammals | Sylvicapra_grimmia | Sylvicapra_grimmia | LC | Decreasing | tropical | diurnal | 19499.8 | 13576010.775 | 9.37 |
| mammals | Sylvilagus_aquaticus | Sylvilagus_aquaticus | LC | Decreasing | temperate | nocturnal | 2135.8 | 909647.525 | 15.83 |
| mammals | Sylvilagus_brasiliensis | Sylvilagus_brasiliensis | EN | Decreasing | tropical | nocturnal | 950 | 7611.783 | 15.372 |
| mammals | Sylvilagus_dicei | Sylvilagus_dicei | VU | Decreasing | tropical | nocturnal | 1442.9 | 5963.594 | 18.618 |
| mammals | Sylvilagus_graysoni | Sylvilagus_graysoni | EN | Decreasing | tropical | nocturnal | 964 | 415.252 | 3.632 |
| mammals | Sylvilagus_holzneri | NA | EN/VU | Unknown/Decreasing | NA | NA | NA | NA | NA |
| mammals | Sylvilagus_holzneri | NA | EN/VU | Unknown/Decreasing | NA | NA | NA | NA | NA |
| mammals | Sylvilagus_insonus | NA | DD | Unknown | NA | NA | NA | NA | NA |
| mammals | Sylvilagus_obscurus | Sylvilagus_obscurus | NT | Decreasing | temperate | diurnal | 653.1 | 94346.012 | 13.447 |
| mammals | Sylvilagus_palustris | Sylvilagus_palustris | LC | Unknown | temperate | nocturnal | 1500 | 361966.322 | 17.273 |
| mammals | Sylvisorex_camerunensis | Sylvisorex_camerunensis | VU | Decreasing | tropical | cathemeral | 5 | 6332.792 | 12.785 |
| mammals | Sylvisorex_granti | Sylvisorex_granti | LC | Unknown | tropical | cathemeral | 4 | 29983.029 | 14.215 |
| mammals | Sylvisorex_howelli | Sylvisorex_howelli | LC | Decreasing | tropical | nocturnal | 4.3 | 6764.488 | 14.089 |
| mammals | Sylvisorex_isabellae | Sylvisorex_isabellae | VU | Decreasing | tropical | cathemeral | 6.8 | 677.19 | 12.306 |
| mammals | Sylvisorex_johnstoni | Sylvisorex_johnstoni | LC | Unknown | tropical | cathemeral | 3.2 | 970551.988 | 8.891 |
| mammals | Sylvisorex_konganensis | NA | DD | Unknown | NA | NA | NA | NA | NA |
| mammals | Sylvisorex_lunaris | Sylvisorex_lunaris | NT | Decreasing | tropical | cathemeral | 18.5 | 13561.079 | 16.842 |
| mammals | Sylvisorex_morio | Sylvisorex_morio | EN | Decreasing | tropical | cathemeral | 8 | 444.444 | 8.98 |
| mammals | Sylvisorex_ollula | Sylvisorex_ollula | LC | Unknown | tropical | cathemeral | 13.2 | 1407399.493 | 5.991 |
| mammals | Sylvisorex_oriundus | NA | DD | Unknown | NA | NA | NA | NA | NA |
| mammals | Sylvisorex_pluvialis | NA | DD | Unknown | NA | NA | NA | NA | NA |
| mammals | Sylvisorex_vulcanorum | Sylvisorex_vulcanorum | LC | Decreasing | tropical | cathemeral | 3.5 | 16604.761 | 12.351 |
| mammals | Symphalangus_syndactylus | Symphalangus_syndactylus | EN | Decreasing | tropical | diurnal | 10000 | 340358.941 | 9.104 |
| mammals | Synaptomys_cooperi | Synaptomys_cooperi | LC | Unknown | temperate | cathemeral | 31.9 | 3223190.972 | 14.11 |
| mammals | Syncerus_caffer | Syncerus_caffer | NT | Decreasing | tropical | cathemeral | 580002.7 | 8443088.906 | 8.467 |
| mammals | Syntheosciurus_brochus | Syntheosciurus_brochus | DD | Decreasing | tropical | diurnal | 132.8 | 12107.716 | 10.822 |
| mammals | Tachyglossus_aculeatus | Tachyglossus_aculeatus | LC | Stable | temperate | nocturnal | 3025 | 7766679.886 | 4.25 |
| mammals | Tadarida_aegyptiaca | Tadarida_aegyptiaca | LC | Unknown | temperate | nocturnal | 16.8 | 4174304.731 | 12.019 |

| | | | | | | | | | |
|---|---|---|---|---|---|---|---|---|---|
| mammals | Tadarida_brasiliensis | Tadarida_brasiliensis | LC | Stable | temperate | nocturnal | 12.5 | 13833395.317 | 10.092 |
| mammals | Tadarida_fulminans | Tadarida_fulminans | LC | Stable | tropical | nocturnal | 35.3 | 295187 | 12.003 |
| mammals | Tadarida_insignis | NA | DD | Unknown | NA | NA | NA | NA | NA |
| mammals | Tadarida_latouchei | Tadarida_latouchei | EN | Decreasing | temperate | nocturnal | 22 | 186758.506 | 16.555 |
| mammals | Tadarida_lobata | Tadarida_lobata | LC | Unknown | tropical | nocturnal | 26 | 186860.184 | 14.193 |
| mammals | Tadarida_teniotis | Tadarida_teniotis | LC | Unknown | temperate | nocturnal | 27 | 3893776.109 | 14.51 |
| mammals | Tadarida_ventralis | NA | DD | Unknown | NA | NA | NA | NA | NA |
| mammals | Taeromys_callitrichus | NA | DD | Unknown | NA | NA | NA | NA | NA |
| mammals | Taeromys_celebensis | Taeromys_celebensis | LC | Decreasing | tropical | nocturnal | 323 | 86669.866 | 8.13 |
| mammals | Taeromys_dominator | Paruromys_dominator | LC | Stable | tropical | nocturnal | 342 | 169337.754 | 9.276 |
| mammals | Taeromys_hamatus | NA | DD | Unknown | NA | NA | NA | NA | NA |
| mammals | Taeromys_microbullatus | NA | DD | Unknown | NA | NA | NA | NA | NA |
| mammals | Taeromys_punicans | NA | DD | Unknown | NA | NA | NA | NA | NA |
| mammals | Taeromys_taerae | Taeromys_taerae | VU | Unknown | tropical | nocturnal | 247 | 4355.207 | 8.079 |
| mammals | Tamandua_mexicana | Tamandua_mexicana | LC | Unknown | tropical | cathemeral | 4210 | 1495856.05 | 10.76 |
| mammals | Tamandua_tetradactyla | Tamandua_tetradactyla | LC | Unknown | tropical | diurnal | 5515.1 | 12732832.025 | 5.378 |
| mammals | Tamias_striatus | Tamias_striatus | LC | Stable | temperate | diurnal | 111.9 | 4197616.711 | 14.632 |
| mammals | Tamiasciurus_douglasii | Tamiasciurus_douglasii | LC | Stable | temperate | diurnal | 225 | 511249.763 | 9.623 |
| mammals | Tamiasciurus_hudsonicus | Tamiasciurus_hudsonicus | LC | Stable | temperate | diurnal | 201.2 | 10363663.519 | 6.11 |
| mammals | Tamiops_maritimus | Tamiops_maritimus | LC | Stable | tropical | diurnal | 70.3 | 1718636.154 | 13.896 |
| mammals | Tamiops_mcclellandii | Tamiops_macclellandii | LC | Stable | tropical | diurnal | 55.5 | 1464987.459 | 11.002 |
| mammals | Tamiops_rodolphii | Tamiops_rodolphii | LC | Stable | tropical | diurnal | 56 | 590763.668 | 13.976 |
| mammals | Tamiops_swinhoei | Tamiops_swinhoei | LC | Stable | temperate | diurnal | 85 | 869533.077 | 10.731 |
| mammals | Tapecomys_primus | Tapecomys_primus | LC | Decreasing | tropical | nocturnal | 71.5 | 2330.952 | 6.846 |
| mammals | Tapecomys_wolffsohni | Phyllotis_wolffsohni | LC | Stable | tropical | cathemeral | 42.5 | 83243.936 | 9.25 |
| mammals | Taphozous_achates | NA | DD | Unknown | NA | NA | NA | NA | NA |
| mammals | Taphozous_australis | Taphozous_australis | NT | Decreasing | tropical | nocturnal | 22.9 | 38340.92 | 5.761 |
| mammals | Taphozous_hamiltoni | NA | DD | Unknown | NA | NA | NA | NA | NA |
| mammals | Taphozous_hildegardeae | Taphozous_hildegardeae | EN | Decreasing | tropical | nocturnal | 29.6 | 44478.917 | 14.493 |
| mammals | Taphozous_hilli | Taphozous_hilli | LC | Stable | tropical | nocturnal | 22 | 1411632.65 | 1.913 |
| mammals | Taphozous_kapalgensis | Taphozous_kapalgensis | LC | Unknown | tropical | nocturnal | 26 | 97559.252 | 2.192 |
| mammals | Taphozous_longimanus | Taphozous_longimanus | LC | Stable | tropical | nocturnal | 26.9 | 4903146.147 | 14.689 |
| mammals | Taphozous_mauritianus | Taphozous_mauritianus | LC | Unknown | tropical | nocturnal | 27.3 | 10153657.375 | 8.259 |
| mammals | Taphozous_melanopogon | Taphozous_melanopogon | LC | Stable | tropical | nocturnal | 25.9 | 5670373.5 | 13.296 |

| | | | | | | | | | |
|---|---|---|---|---|---|---|---|---|---|
| mammals | Taphozous_nudiventris | Taphozous_nudiventris | LC | Stable | tropical | nocturnal | 30.1 | 1206353.709 | 16.025 |
| mammals | Taphozous_perforatus | Taphozous_perforatus | LC | Stable | temperate | nocturnal | 27.3 | 2114774.55 | 10.704 |
| mammals | Taphozous_theobaldi | Taphozous_theobaldi | LC | Unknown | tropical | nocturnal | 39.8 | 1042380.039 | 13.631 |
| mammals | Taphozous_troughtoni | Taphozous_troughtoni | LC | Decreasing | tropical | nocturnal | 32.5 | 741835.35 | 5.239 |
| mammals | Tapirus_bairdii | Tapirus_bairdii | EN | Decreasing | tropical | nocturnal | 299999.1 | 673656.443 | 8.483 |
| mammals | Tapirus_indicus | Tapirus_indicus | EN | Decreasing | tropical | nocturnal | 296250 | 89491.877 | 5.794 |
| mammals | Tapirus_pinchaque | Tapirus_pinchaque | EN | Decreasing | tropical | cathemeral | 148949.8 | 49536.771 | 5.969 |
| mammals | Tapirus_terrestris | Tapirus_terrestris | VU | Decreasing | tropical | cathemeral | 207500.9 | 13052361.742 | 5.58 |
| mammals | Tarsipes_rostratus | Tarsipes_rostratus | LC | Stable | temperate | nocturnal | 9 | 236675.497 | 9.918 |
| mammals | Tarsius_dentatus | Tarsius_dentatus | VU | Decreasing | tropical | nocturnal | 107 | 35043.151 | 6.489 |
| mammals | Tarsius_lariang | Tarsius_lariang | DD | Decreasing | tropical | nocturnal | 110 | 40261.175 | 9.543 |
| mammals | Tarsius_pelengensis | Tarsius_pelengensis | EN | Decreasing | tropical | nocturnal | 170 | 2615.018 | 9.222 |
| mammals | Tarsius_pumilus | Tarsius_pumilus | EN | Decreasing | tropical | nocturnal | 27.5 | 395.745 | 5.927 |
| mammals | Tarsius_sangirensis | Tarsius_sangirensis | EN | Decreasing | tropical | nocturnal | 171.3 | 556.374 | 11.586 |
| mammals | Tarsius_tarsier | Tarsius_tarsier | VU | Decreasing | tropical | nocturnal | 200 | 37585.801 | 8.626 |
| mammals | Tarsius_tumpara | Tarsius_tumpara | CR | Decreasing | tropical | nocturnal | 104 | 115.183 | 16.337 |
| mammals | Tarsius_wallacei | Tarsius_wallacei | VU | Decreasing | tropical | nocturnal | 106 | 4159.037 | 7.494 |
| mammals | Tarsomys_apoensis | Tarsomys_apoensis | LC | Stable | tropical | cathemeral | 75 | 4440.286 | 7.543 |
| mammals | Tarsomys_echinatus | NA | VU | Decreasing | NA | NA | NA | NA | NA |
| mammals | Tateomys_macrocercus | NA | DD | Unknown | NA | NA | NA | NA | NA |
| mammals | Tateomys_rhinogradoides | Tateomys_rhinogradoides | DD | Stable | tropical | nocturnal | 82.8 | 2159.53 | 4.917 |
| mammals | Tatera_indica | Tatera_indica | LC | Unknown | tropical | nocturnal | 166.3 | 4786124.727 | 15.069 |
| mammals | Taterillus_arenarius | Taterillus_arenarius | LC | Unknown | tropical | nocturnal | 47 | 402966.823 | 3.724 |
| mammals | Taterillus_congicus | Taterillus_congicus | LC | Unknown | tropical | nocturnal | 54.5 | 1317164.847 | 5.389 |
| mammals | Taterillus_emini | Taterillus_emini | LC | Unknown | tropical | nocturnal | 52 | 3860531.19 | 9.715 |
| mammals | Taterillus_gracilis | Taterillus_gracilis | LC | Stable | tropical | nocturnal | 48 | 2210732.886 | 13.396 |
| mammals | Taterillus_lacustris | Taterillus_lacustris | LC | Unknown | tropical | nocturnal | 41.5 | 39538.931 | 15.4 |
| mammals | Taterillus_petteri | Taterillus_petteri | LC | Stable | tropical | nocturnal | 50.8 | 178975.98 | 9.359 |
| mammals | Taterillus_pygargus | Taterillus_pygargus | LC | Stable | tropical | nocturnal | 60.2 | 365303.217 | 12.306 |
| mammals | Taterillus_tranieri | Taterillus_tranieri | LC | Unknown | tropical | nocturnal | 46.8 | 93503.586 | 8.066 |
| mammals | Taxidea_taxus | Taxidea_taxus | LC | Decreasing | temperate | nocturnal | 7107.6 | 8851818.096 | 11.076 |
| mammals | Tayassu_pecari | Tayassu_pecari | VU | Decreasing | tropical | cathemeral | 32233.7 | 13750306.265 | 5.676 |
| mammals | Tenrec_ecaudatus | Tenrec_ecaudatus | LC | Stable | tropical | nocturnal | 868 | 590834.3 | 8.745 |
| mammals | Thainycteris_aureocollaris | Arielulus_aureocollaris | LC | Unknown | tropical | nocturnal | 14.5 | 302997.639 | 11.584 |

| | | | | | | | | | |
|---|---|---|---|---|---|---|---|---|---|
| mammals | Thainycteris_torquata | Arielulus_torquatus | LC | Stable | temperate | nocturnal | 6.3 | 23909.893 | 18.192 |
| mammals | Thallomys_loringi | Thallomys_loringi | LC | Unknown | tropical | nocturnal | 80 | 56457.551 | 10.656 |
| mammals | Thallomys_nigricauda | Thallomys_nigricauda | LC | Unknown | tropical | nocturnal | 124.7 | 1214564.264 | 5.129 |
| mammals | Thallomys_paedulcus | Thallomys_paedulcus | LC | Unknown | tropical | nocturnal | 80.5 | 5096542.573 | 9.022 |
| mammals | Thallomys_shortridgei | NA | DD | Unknown | NA | NA | NA | NA | NA |
| mammals | Thalpomys_cerradensis | Thalpomys_cerradensis | LC | Unknown | tropical | nocturnal | 24 | 488719.397 | 8.709 |
| mammals | Thalpomys_lasiotis | Thalpomys_lasiotis | LC | Unknown | tropical | nocturnal | 24 | 555211.185 | 9.244 |
| mammals | Thamnomys_kempi | Thamnomys_kempi | VU | Unknown | tropical | nocturnal | 75.1 | 5110.786 | 12.992 |
| mammals | Thamnomys_kuru | Grammomys_kuru | LC | Unknown | tropical | nocturnal | 52 | 2284482.444 | 8.704 |
| mammals | Thamnomys_schoutedeni | NA | DD | Unknown | NA | NA | NA | NA | NA |
| mammals | Thamnomys_venustus | Thamnomys_venustus | LC | Decreasing | tropical | nocturnal | 61.9 | 10760.923 | 16.559 |
| mammals | Thaptomys_nigrita | Thaptomys_nigrita | LC | Unknown | tropical | diurnal | 19.9 | 1057629.59 | 12.309 |
| mammals | Theropithecus_gelada | Theropithecus_gelada | LC | Decreasing | tropical | diurnal | 17049.8 | 104365.324 | 13.739 |
| mammals | Thomasomys_aureus | Thomasomys_aureus | LC | Decreasing | tropical | nocturnal | 88 | 557724.619 | 9.322 |
| mammals | Thomasomys_baeops | Thomasomys_baeops | LC | Decreasing | tropical | nocturnal | 36 | 86495.215 | 11.295 |
| mammals | Thomasomys_bombycinus | Thomasomys_bombycinus | VU | Unknown | tropical | nocturnal | 114.5 | 17506.045 | 10.257 |
| mammals | Thomasomys_caudivarius | Thomasomys_caudivarius | LC | Decreasing | tropical | nocturnal | 95.9 | 34453.218 | 9.972 |
| mammals | Thomasomys_cinereus | Thomasomys_cinereus | LC | Stable | tropical | nocturnal | 77 | 26409.049 | 9.395 |
| mammals | Thomasomys_cinnameus | Thomasomys_cinnameus | LC | Unknown | tropical | nocturnal | 17 | 27426.453 | 11.315 |
| mammals | Thomasomys_erro | Thomasomys_erro | LC | Unknown | tropical | nocturnal | 46.3 | 13995.296 | 12.71 |
| mammals | Thomasomys_hudsoni | Thomasomys_hudsoni | VU | Unknown | tropical | nocturnal | 23.1 | 12772.012 | 11.634 |
| mammals | Thomasomys_incanus | Thomasomys_incanus | LC | Decreasing | tropical | nocturnal | 77 | 19613.627 | 6.228 |
| mammals | Thomasomys_kalinowskii | Thomasomys_kalinowskii | LC | Decreasing | tropical | nocturnal | 77 | 41986.575 | 6.077 |
| mammals | Thomasomys_paramorum | Thomasomys_paramorum | LC | Unknown | tropical | nocturnal | 21.5 | 29904.442 | 8.802 |
| mammals | Thomasomys_praetor | Thomasomys_praetor | NT | Unknown | tropical | nocturnal | 107.8 | 55615.888 | 8.536 |
| mammals | Thomasomys_pyrrhonotus | Thomasomys_pyrrhonotus | DD | Decreasing | tropical | nocturnal | 77 | 49532.683 | 8.63 |
| mammals | Thomasomys_taczanowskii | Thomasomys_taczanowskii | LC | Unknown | tropical | nocturnal | 77 | 151302.468 | 9.312 |
| mammals | Thomasomys_ucucha | Thomasomys_ucucha | VU | Decreasing | tropical | nocturnal | 36 | 16490.792 | 11.789 |
| mammals | Thomasomys_vestitus | Thomasomys_vestitus | LC | Unknown | tropical | nocturnal | 76.5 | 6459.223 | 11.605 |
| mammals | Thomasomys_vulcani | Thomasomys_vulcani | LC | Stable | tropical | nocturnal | 39.1 | 16199.502 | 11.001 |
| mammals | Thoopterus_nigrescens | Thoopterus_nigrescens | LC | Unknown | tropical | nocturnal | 60.4 | 182866.823 | 9.13 |
| mammals | Thrichomys_apereoides | Thrichomys_apereoides | LC | Unknown | tropical | nocturnal | 339 | 1168838.93 | 8.487 |
| mammals | Thryonomys_gregorianus | Thryonomys_gregorianus | LC | Unknown | tropical | nocturnal | 2707.1 | 2375922.351 | 11.362 |
| mammals | Thryonomys_swinderianus | Thryonomys_swinderianus | LC | Unknown | tropical | nocturnal | 3180 | 7541449.629 | 9.883 |

| | | | | | | | | | |
|---|---|---|---|---|---|---|---|---|---|
| mammals | Thylamys_elegans | Thylamys_elegans | LC | Stable | temperate | nocturnal | 28.9 | 142691.222 | 11.713 |
| mammals | Thylamys_karimii | Thylamys_karimii | VU | Decreasing | tropical | nocturnal | 28 | 1431448.592 | 7.346 |
| mammals | Thylamys_macrurus | Thylamys_macrurus | NT | Decreasing | tropical | nocturnal | 20.7 | 88390.177 | 9.544 |
| mammals | Thylamys_pallidior | Thylamys_pallidior | LC | Unknown | temperate | nocturnal | 14.9 | 1493250.669 | 5.371 |
| mammals | Thylamys_pusillus | Thylamys_pusillus | LC | Decreasing | temperate | nocturnal | 41 | 699825.628 | 6.458 |
| mammals | Thylamys_tatei | NA | DD | Unknown | NA | NA | NA | NA | NA |
| mammals | Thylamys_velutinus | Thylamys_velutinus | NT | Decreasing | tropical | nocturnal | 20.7 | 565210.768 | 12.757 |
| mammals | Thylamys_venustus | NA | DD | Unknown | NA | NA | NA | NA | NA |
| mammals | Thylogale_billardierii | Thylogale_billardierii | LC | Stable | temperate | nocturnal | 5450 | 66448.237 | 5.95 |
| mammals | Thylogale_brownii | Thylogale_browni | VU | Decreasing | tropical | nocturnal | 5478 | 269510.344 | 4.715 |
| mammals | Thylogale_brunii | Thylogale_brunii | VU | Decreasing | tropical | nocturnal | 3620 | 108853.519 | 3.115 |
| mammals | Thylogale_calabyi | Thylogale_calabyi | EN | Decreasing | tropical | nocturnal | 4326.1 | 1002.372 | 5.724 |
| mammals | Thylogale_stigmatica | Thylogale_stigmatica | LC | Decreasing | tropical | cathemeral | 4600 | 221199.779 | 6.176 |
| mammals | Thylogale_thetis | Thylogale_thetis | LC | Stable | temperate | cathemeral | 5400 | 73431.485 | 6.955 |
| mammals | Thyroptera_devivoi | NA | DD | Unknown | NA | NA | NA | NA | NA |
| mammals | Thyroptera_discifera | Thyroptera_discifera | LC | Unknown | tropical | nocturnal | 3.1 | 5711490.832 | 3.48 |
| mammals | Thyroptera_lavali | NA | DD | Unknown | NA | NA | NA | NA | NA |
| mammals | Thyroptera_tricolor | Thyroptera_tricolor | LC | Unknown | tropical | nocturnal | 4.4 | 8555467.311 | 4.45 |
| mammals | Thyroptera_wynneae | NA | DD | Unknown | NA | NA | NA | NA | NA |
| mammals | Tlacuatzin_canescens | Tlacuatzin_canescens | LC | Stable | tropical | nocturnal | 60 | 479508.388 | 8.258 |
| mammals | Tokudaia_muenninki | Tokudaia_muenninki | CR | Decreasing | temperate | nocturnal | 177.4 | 240.568 | 13.775 |
| mammals | Tokudaia_tokunoshimensis | NA | EN | Decreasing | NA | NA | NA | NA | NA |
| mammals | Tolypeutes_matacus | Tolypeutes_matacus | NT | Decreasing | temperate | nocturnal | 1200 | 1198272.043 | 6.167 |
| mammals | Tolypeutes_tricinctus | Tolypeutes_tricinctus | VU | Decreasing | tropical | nocturnal | 1487 | 932819.599 | 7.643 |
| mammals | Tomopeas_ravum | Tomopeas_ravus | EN | Decreasing | tropical | nocturnal | 5.6 | 112800.735 | 8.6 |
| mammals | Tonatia_bidens | NA | DD | Unknown | NA | NA | NA | NA | NA |
| mammals | Tonatia_saurophila | Tonatia_saurophila | LC | Stable | tropical | nocturnal | 30 | 6509470.684 | 3.522 |
| mammals | Tonkinomys_daovantieni | NA | DD | Unknown | NA | NA | NA | NA | NA |
| mammals | Toromys_grandis | Toromys_grandis | LC | Unknown | tropical | nocturnal | 584 | 176731.299 | 3.079 |
| mammals | Trachops_cirrhosus | Trachops_cirrhosus | LC | Stable | tropical | nocturnal | 37.7 | 12642871.515 | 5.613 |
| mammals | Trachypithecus_auratus | Trachypithecus_auratus | VU | Decreasing | tropical | diurnal | 7100 | 120357.236 | 22.208 |
| mammals | Trachypithecus_barbei | Trachypithecus_barbei | VU | Decreasing | tropical | diurnal | 7500 | 3817.011 | 4.124 |
| mammals | Trachypithecus_cristatus | Trachypithecus_cristatus | VU | Decreasing | tropical | diurnal | 5250 | 1213231.284 | 7.561 |
| mammals | Trachypithecus_delacouri | Trachypithecus_delacouri | CR | Decreasing | tropical | diurnal | 8899.9 | 6008.398 | 17.338 |

| | | | | | | | | | |
|---|---|---|---|---|---|---|---|---|---|
| mammals | Trachypithecus_francoisi | Trachypithecus_francoisi | EN | Decreasing | temperate | diurnal | 7325 | 248808.276 | 15.348 |
| mammals | Trachypithecus_geei | Trachypithecus_geei | EN | Decreasing | temperate | diurnal | 8350 | 7413.458 | 11.163 |
| mammals | Trachypithecus_germaini | Trachypithecus_germaini | EN | Decreasing | tropical | diurnal | 7179 | 261220.837 | 13.178 |
| mammals | Trachypithecus_hatinhensis | Trachypithecus_hatinhensis | EN | Decreasing | tropical | diurnal | 6323.1 | 18902.385 | 8.504 |
| mammals | Trachypithecus_laotum | Trachypithecus_laotum | EN | Decreasing | tropical | diurnal | 5848.6 | 2849.698 | 6.305 |
| mammals | Trachypithecus_obscurus | Trachypithecus_obscurus | EN | Decreasing | tropical | diurnal | 7000 | 259473.527 | 12.367 |
| mammals | Trachypithecus_phayrei | Trachypithecus_phayrei | EN | Decreasing | tropical | diurnal | 8400 | 512798.887 | 11.678 |
| mammals | Trachypithecus_pileatus | Trachypithecus_pileatus | VU | Decreasing | temperate | diurnal | 11450 | 305059.049 | 11.487 |
| mammals | Trachypithecus_poliocephalus | NA | CR | Increasing | NA | NA | NA | NA | NA |
| mammals | Trachypithecus_shortridgei | Trachypithecus_shortridgei | EN | Decreasing | temperate | diurnal | 28500 | 10597.022 | 2.898 |
| mammals | Tragelaphus_angasii | Tragelaphus_angasii | LC | Stable | temperate | diurnal | 86600.4 | 197902.695 | 7.967 |
| mammals | Tragelaphus_buxtoni | Tragelaphus_buxtoni | EN | Decreasing | tropical | nocturnal | 215000.8 | 6875.038 | 10.108 |
| mammals | Tragelaphus_eurycerus | Tragelaphus_eurycerus | NT | Decreasing | tropical | nocturnal | 329003.1 | 2217282.34 | 7.209 |
| mammals | Tragelaphus_imberbis | Tragelaphus_imberbis | NT | Decreasing | tropical | nocturnal | 81575.5 | 1488947.506 | 8.492 |
| mammals | Tragelaphus_oryx | Tragelaphus_oryx | LC | Stable | tropical | nocturnal | 569993.6 | 6776785.864 | 8.459 |
| mammals | Tragelaphus_scriptus | Tragelaphus_scriptus | LC | Stable | tropical | nocturnal | 43250.4 | 12064054.459 | 9.948 |
| mammals | Tragelaphus_spekii | Tragelaphus_spekii | LC | Decreasing | tropical | diurnal | 77999.2 | 4392202.517 | 6.883 |
| mammals | Tragelaphus_strepsiceros | Tragelaphus_strepsiceros | LC | Stable | tropical | cathemeral | 213501 | 6085607.88 | 8.46 |
| mammals | Tragulus_javanicus | NA | DD | Unknown | NA | NA | NA | NA | NA |
| mammals | Tragulus_kanchil | Tragulus_kanchil | LC | Unknown | tropical | cathemeral | 2000 | 2303447.463 | 9.487 |
| mammals | Tragulus_napu | Tragulus_napu | LC | Decreasing | tropical | cathemeral | 4000 | 1402060.128 | 7.771 |
| mammals | Tragulus_nigricans | Tragulus_nigricans | EN | Decreasing | tropical | nocturnal | 2473.8 | 506.71 | 7.507 |
| mammals | Tragulus_versicolor | Tragulus_versicolor | DD | Decreasing | tropical | nocturnal | 1700 | 1479.888 | 12.941 |
| mammals | Tragulus_williamsoni | Tragulus_williamsoni | DD | Decreasing | tropical | nocturnal | 1935.5 | 316.191 | 9.962 |
| mammals | Transandinomys_bolivaris | Transandinomys_bolivaris | LC | Stable | tropical | nocturnal | 60.5 | 271356.278 | 7.454 |
| mammals | Transandinomys_talamancae | Transandinomys_talamancae | LC | Stable | tropical | nocturnal | 55 | 510964.247 | 11.437 |
| mammals | Tremarctos_ornatus | Tremarctos_ornatus | VU | Decreasing | tropical | diurnal | 140000.6 | 218405.24 | 4.672 |
| mammals | Triaenops_menamena | Triaenops_rufus | LC | Unknown | tropical | nocturnal | 10.5 | 284167.072 | 7.875 |
| mammals | Triaenops_persicus | Triaenops_persicus | LC | Unknown | temperate | nocturnal | 12.6 | 1163737.954 | 11.118 |
| mammals | Trichosurus_caninus | Trichosurus_caninus | LC | Stable | temperate | nocturnal | 3500 | 148314.195 | 7.785 |
| mammals | Trichosurus_cunninghami | Trichosurus_cunninghami | LC | Stable | temperate | nocturnal | 3270 | 119403.447 | 8.269 |
| mammals | Trichosurus_vulpecula | Trichosurus_vulpecula | LC | Decreasing | temperate | nocturnal | 2650 | 2704232.993 | 6.469 |
| mammals | Trichys_fasciculata | Trichys_fasciculata | LC | Stable | tropical | nocturnal | 1655 | 1189879.282 | 7.397 |
| mammals | Trinomys_albispinus | Trinomys_albispinus | LC | Unknown | tropical | nocturnal | 285 | 70209.403 | 11.409 |

| | | | | | | | | | |
|---|---|---|---|---|---|---|---|---|---|
| mammals | Trinomys_dimidiatus | Trinomys_dimidiatus | LC | Unknown | tropical | nocturnal | 167.6 | 63989.208 | 14.772 |
| mammals | Trinomys_iheringi | Trinomys_iheringi | LC | Stable | tropical | nocturnal | 203.5 | 702152.55 | 12.163 |
| mammals | Trinomys_mirapitanga | NA | DD | Unknown | NA | NA | NA | NA | NA |
| mammals | Trinomys_moojeni | Trinomys_moojeni | EN | Decreasing | tropical | nocturnal | 207 | 4672.074 | 9.46 |
| mammals | Trinomys_paratus | NA | DD | Unknown | NA | NA | NA | NA | NA |
| mammals | Trinomys_setosus | Trinomys_setosus | LC | Unknown | tropical | nocturnal | 285 | 414903.904 | 11.038 |
| mammals | Trinomys_yonenagae | Trinomys_yonenagae | EN | Decreasing | tropical | nocturnal | 200.2 | 4982.01 | 3.971 |
| mammals | Trinycteris_nicefori | Trinycteris_nicefori | LC | Unknown | tropical | nocturnal | 8.2 | 4253378.914 | 4.271 |
| mammals | Trogopterus_xanthipes | Trogopterus_xanthipes | NT | Decreasing | temperate | nocturnal | 744.9 | 1062333.593 | 13.211 |
| mammals | Tryphomys_adustus | NA | DD | Unknown | NA | NA | NA | NA | NA |
| mammals | Tscherskia_triton | Tscherskia_triton | LC | Unknown | temperate | nocturnal | 166.5 | 3140943.025 | 15.386 |
| mammals | Tupaia_belangeri | Tupaia_belangeri | LC | Stable | tropical | diurnal | 200 | 2888122.744 | 12.218 |
| mammals | Tupaia_chrysogaster | Tupaia_chrysogaster | VU | Unknown | tropical | diurnal | 177.7 | 6002.771 | 6.223 |
| mammals | Tupaia_dorsalis | Tupaia_dorsalis | DD | Decreasing | tropical | diurnal | 265 | 449710.803 | 4.895 |
| mammals | Tupaia_everetti | Urogale_everetti | LC | Stable | tropical | diurnal | 350 | 95385.362 | 12.031 |
| mammals | Tupaia_glis | Tupaia_glis | LC | Decreasing | tropical | diurnal | 169.8 | 190363.117 | 13.734 |
| mammals | Tupaia_gracilis | Tupaia_gracilis | LC | Decreasing | tropical | diurnal | 74.6 | 643734.653 | 5.12 |
| mammals | Tupaia_javanica | Tupaia_javanica | LC | Decreasing | tropical | diurnal | 105 | 248043.794 | 12.851 |
| mammals | Tupaia_longipes | Tupaia_longipes | LC | Decreasing | tropical | diurnal | 196 | 161229.925 | 5.564 |
| mammals | Tupaia_minor | Tupaia_minor | LC | Decreasing | tropical | diurnal | 64.3 | 1264126.169 | 7.823 |
| mammals | Tupaia_montana | Tupaia_montana | LC | Unknown | tropical | diurnal | 199 | 179713.652 | 3.121 |
| mammals | Tupaia_nicobarica | Tupaia_nicobarica | EN | Decreasing | tropical | diurnal | 170.3 | 1108.483 | 4.111 |
| mammals | Tupaia_palawanensis | Tupaia_palawanensis | LC | Stable | tropical | diurnal | 218 | 13167.701 | 9.329 |
| mammals | Tupaia_picta | Tupaia_picta | LC | Decreasing | tropical | diurnal | 147.5 | 154832.286 | 5.502 |
| mammals | Tupaia_splendidula | Tupaia_splendidula | LC | Decreasing | tropical | diurnal | 160 | 411411.087 | 6.001 |
| mammals | Tupaia_tana | Tupaia_tana | LC | Decreasing | tropical | diurnal | 197.8 | 999970.616 | 7.104 |
| mammals | Tylomys_bullaris | Tylomys_bullaris | CR | Decreasing | tropical | nocturnal | 280 | 713.639 | 6.133 |
| mammals | Tylomys_mirae | Tylomys_mirae | LC | Unknown | tropical | nocturnal | 183.6 | 125758.518 | 6.991 |
| mammals | Tylomys_nudicaudus | Tylomys_nudicaudus | LC | Stable | tropical | nocturnal | 100 | 489186.845 | 11.884 |
| mammals | Tylomys_panamensis | NA | DD | Unknown | NA | NA | NA | NA | NA |
| mammals | Tylonycteris_pachypus | Tylonycteris_pachypus | LC | Unknown | tropical | nocturnal | 3.6 | 4182645.855 | 11.794 |
| mammals | Tylonycteris_robustula | Tylonycteris_robustula | LC | Unknown | tropical | nocturnal | 7.6 | 2721209.15 | 12.019 |
| mammals | Tympanoctomys_barrerae | Tympanoctomys_barrerae | NT | Unknown | temperate | nocturnal | 70 | 280711.692 | 4.432 |
| mammals | Tympanoctomys_kirchnerorum | NA | DD | Unknown | NA | NA | NA | NA | NA |

| | | | | | | | | | |
|---|---|---|---|---|---|---|---|---|---|
| mammals | Typhlomys_cinereus | Typhlomys_cinereus | LC | Decreasing | temperate | nocturnal | 24.5 | 1030421.04 | 13.431 |
| mammals | Typomys_planifrons | Hybomys_planifrons | LC | Stable | tropical | diurnal | 49.6 | 239879.554 | 13.97 |
| mammals | Typomys_trivirgatus | Hybomys_trivirgatus | LC | Stable | tropical | cathemeral | 56.9 | 387198.511 | 16.039 |
| mammals | Uranomys_ruddi | Uranomys_ruddi | LC | Decreasing | tropical | nocturnal | 40.4 | 4057097.01 | 10.584 |
| mammals | Urocitellus_armatus | Urocitellus_armatus | LC | Unknown | temperate | diurnal | 313 | 176257.798 | 9.208 |
| mammals | Urocitellus_beldingi | Urocitellus_beldingi | LC | Stable | temperate | diurnal | 280.5 | 337683.745 | 9.237 |
| mammals | Urocitellus_brunneus | Urocitellus_brunneus | EN | Increasing | temperate | diurnal | 300 | 6763.229 | 12.631 |
| mammals | Urocitellus_canus | Urocitellus_canus | LC | Unknown | temperate | diurnal | 154 | 217761.205 | 9.638 |
| mammals | Urocitellus_columbianus | Urocitellus_columbianus | LC | Stable | temperate | diurnal | 493 | 532364.643 | 6.753 |
| mammals | Urocitellus_elegans | Urocitellus_elegans | LC | Unknown | temperate | diurnal | 453.6 | 282690.693 | 10.165 |
| mammals | Urocitellus_mollis | Urocitellus_mollis | LC | Stable | temperate | diurnal | 165.4 | 416260.768 | 8.331 |
| mammals | Urocitellus_parryii | Urocitellus_parryii | LC | Unknown | temperate | diurnal | 760 | 5346615.243 | 0.654 |
| mammals | Urocitellus_richardsonii | Urocitellus_richardsonii | LC | Stable | temperate | diurnal | 406 | 900811.965 | 14.993 |
| mammals | Urocitellus_townsendii | Urocitellus_townsendii | VU | Decreasing | temperate | diurnal | 213 | 13910.872 | 15.062 |
| mammals | Urocitellus_undulatus | Urocitellus_undulatus | LC | Stable | temperate | diurnal | 840 | 2310791.437 | 4.603 |
| mammals | Urocitellus_washingtoni | Urocitellus_washingtoni | NT | Decreasing | temperate | diurnal | 207.5 | 17294.85 | 14.326 |
| mammals | Urocricetus_alticola | Cricetulus_alticola | LC | Unknown | temperate | nocturnal | 35 | 140657.571 | 2.451 |
| mammals | Urocricetus_kamensis | Cricetulus_kamensis | LC | Unknown | temperate | cathemeral | 30 | 163548.694 | 7.5 |
| mammals | Urocyon_cinereoargenteus | Urocyon_cinereoargenteus | LC | Stable | temperate | nocturnal | 3833.7 | 9037640.209 | 13.189 |
| mammals | Urocyon_littoralis | Urocyon_littoralis | NT | Increasing | temperate | nocturnal | 1896 | 911.163 | 8.564 |
| mammals | Uroderma_bilobatum | Uroderma_bilobatum | LC | Stable | tropical | nocturnal | 16.7 | 10739162.989 | 4.407 |
| mammals | Uroderma_magnirostrum | Uroderma_magnirostrum | LC | Stable | tropical | nocturnal | 16.9 | 10831975.702 | 4.534 |
| mammals | Uromys_anak | Uromys_anak | LC | Unknown | tropical | nocturnal | 664.5 | 207596.3 | 4.511 |
| mammals | Uromys_caudimaculatus | Uromys_caudimaculatus | LC | Stable | tropical | nocturnal | 606.8 | 880888.113 | 3.406 |
| mammals | Uromys_hadrourus | Uromys_hadrourus | NT | Stable | tropical | nocturnal | 194 | 635.188 | 1.011 |
| mammals | Uromys_neobrittanicus | Uromys_neobritannicus | NT | Decreasing | tropical | nocturnal | 612.5 | 35333.443 | 4.192 |
| mammals | Uromys_rex | Uromys_rex | EN | Decreasing | tropical | nocturnal | 420 | 5461.348 | 5.671 |
| mammals | Uropsilus_andersoni | NA | DD | Unknown | NA | NA | NA | NA | NA |
| mammals | Uropsilus_gracilis | Uropsilus_gracilis | LC | Unknown | temperate | cathemeral | 18.4 | 765423.281 | 12.826 |
| mammals | Uropsilus_investigator | NA | DD | Unknown | NA | NA | NA | NA | NA |
| mammals | Uropsilus_soricipes | Uropsilus_soricipes | LC | Unknown | temperate | cathemeral | 16 | 148166.635 | 9.98 |
| mammals | Urotrichus_talpoides | Urotrichus_talpoides | LC | Stable | temperate | cathemeral | 45 | 286809.075 | 21.094 |
| mammals | Ursus_americanus | Ursus_americanus | LC | Increasing | temperate | cathemeral | 99949.4 | 16113743.925 | 8.264 |
| mammals | Ursus_arctos | Ursus_arctos | LC | Stable | temperate | cathemeral | 180520.4 | 36100052.355 | 6.046 |

| | | | | | | | | | |
|---|---|---|---|---|---|---|---|---|---|
| mammals | Ursus_maritimus | Ursus_maritimus | VU | Unknown | temperate | cathemeral | 388750.4 | 22605413.562 | 0.35 |
| mammals | Ursus_thibetanus | Ursus_thibetanus | VU | Decreasing | temperate | cathemeral | 77500 | 8529242.114 | 12.865 |
| mammals | Urva_auropunctata | Herpestes_auropunctatus | LC | Unknown | temperate | diurnal | 483.5 | 4171313.507 | 13.854 |
| mammals | Urva_brachyura | Herpestes_brachyurus | NT | Decreasing | tropical | diurnal | 1853.5 | 1271946.618 | 7.464 |
| mammals | Urva_edwardsii | Herpestes_edwardsii | LC | Stable | tropical | diurnal | 1324.2 | 4314164.448 | 15.762 |
| mammals | Urva_fusca | Herpestes_fuscus | LC | Stable | tropical | nocturnal | 2700 | 42989.008 | 15.744 |
| mammals | Urva_javanica | Herpestes_javanicus | LC | Unknown | tropical | nocturnal | 750 | 2125157.912 | 14.314 |
| mammals | Urva_semitorquata | Herpestes_semitorquatus | NT | Decreasing | tropical | diurnal | 3500 | 1174536.962 | 7.105 |
| mammals | Urva_smithii | Herpestes_smithii | LC | Unknown | tropical | diurnal | 1782 | 1895779.204 | 18.991 |
| mammals | Urva_urva | Herpestes_urva | LC | Decreasing | tropical | diurnal | 1863.2 | 4060874.84 | 13.08 |
| mammals | Urva_vitticollis | Herpestes_vitticollis | LC | Stable | tropical | diurnal | 2575.6 | 152078.876 | 15.301 |
| mammals | Vampyressa_melissa | Vampyressa_melissa | VU | Stable | tropical | nocturnal | 16.3 | 445446.343 | 5.732 |
| mammals | Vampyressa_pusilla | NA | DD | Unknown | NA | NA | NA | NA | NA |
| mammals | Vampyressa_thyone | Vampyressa_thyone | LC | Unknown | tropical | nocturnal | 8.5 | 4543755.924 | 5.532 |
| mammals | Vampyriscus_bidens | Vampyriscus_bidens | LC | Unknown | tropical | nocturnal | 11.6 | 6731906.155 | 2.476 |
| mammals | Vampyriscus_brocki | Vampyriscus_brocki | LC | Unknown | tropical | nocturnal | 10 | 927195.884 | 1.2 |
| mammals | Vampyriscus_nymphaeus | Vampyriscus_nymphaea | LC | Unknown | tropical | nocturnal | 11.8 | 248097.109 | 8.131 |
| mammals | Vampyrodes_caraccioli | Vampyrodes_caraccioli | LC | Unknown | tropical | nocturnal | 34.7 | 7326122.458 | 2.679 |
| mammals | Vampyrodes_major | Vampyrodes_major | LC | Unknown | tropical | nocturnal | 38.5 | 880555.254 | 10.276 |
| mammals | Vampyrum_spectrum | Vampyrum_spectrum | NT | Decreasing | tropical | nocturnal | 167.7 | 5277255.153 | 5.826 |
| mammals | Vandeleuria_nilagiricus | NA | EN | Decreasing | NA | NA | NA | NA | NA |
| mammals | Vandeleuria_nolthenii | Vandeleuria_nolthenii | EN | Decreasing | tropical | nocturnal | 56.6 | 316.664 | 13.014 |
| mammals | Vandeleuria_oleraceus | Vandeleuria_oleracea | LC | Stable | tropical | nocturnal | 10 | 3789829.902 | 15.788 |
| mammals | Vansonia_rueppellii | Pipistrellus_rueppellii | LC | Unknown | tropical | nocturnal | 6.9 | 2970000.576 | 10.322 |
| mammals | Varecia_rubra | Varecia_rubra | CR | Decreasing | tropical | diurnal | 3450 | 5271.839 | 5.271 |
| mammals | Varecia_variegata | Varecia_variegata | CR | Decreasing | tropical | diurnal | 3350 | 20717.203 | 5.779 |
| mammals | Vernaya_fulva | Vernaya_fulva | LC | Unknown | temperate | nocturnal | 24.9 | 339732.288 | 10.415 |
| mammals | Vespadelus_baverstocki | Vespadelus_baverstocki | LC | Stable | temperate | nocturnal | 5 | 2261700.397 | 4.063 |
| mammals | Vespadelus_caurinus | Vespadelus_caurinus | LC | Stable | tropical | nocturnal | 4 | 606796.834 | 2.178 |
| mammals | Vespadelus_darlingtoni | Vespadelus_darlingtoni | LC | Stable | temperate | nocturnal | 6 | 503722.133 | 9.601 |
| mammals | Vespadelus_douglasorum | Vespadelus_douglasorum | LC | Unknown | tropical | nocturnal | 5 | 132951.747 | 2.699 |
| mammals | Vespadelus_finlaysoni | Vespadelus_finlaysoni | LC | Stable | tropical | nocturnal | 5 | 3435392.144 | 2.626 |
| mammals | Vespadelus_pumilus | Vespadelus_pumilus | LC | Stable | tropical | nocturnal | 4.4 | 148605.622 | 7.987 |
| mammals | Vespadelus_regulus | Vespadelus_regulus | LC | Stable | temperate | nocturnal | 5.4 | 1241504.74 | 8.897 |

| | | | | | | | | | |
|---|---|---|---|---|---|---|---|---|---|
| mammals | Vespadelus_troughtoni | Vespadelus_troughtoni | LC | Stable | tropical | nocturnal | 5.5 | 895634.146 | 5.762 |
| mammals | Vespadelus_vulturnus | Vespadelus_vulturnus | LC | Decreasing | temperate | nocturnal | 4.1 | 1401978.479 | 8.475 |
| mammals | Vespertilio_murinus | Vespertilio_murinus | LC | Stable | temperate | nocturnal | 18.7 | 15761265.224 | 9.863 |
| mammals | Vespertilio_sinensis | Vespertilio_sinensis | LC | Unknown | temperate | nocturnal | 24.5 | 6003456.597 | 13.747 |
| mammals | Viverra_megaspila | Viverra_megaspila | EN | Decreasing | tropical | nocturnal | 9250 | 1341800.272 | 11.899 |
| mammals | Viverra_tangalunga | Viverra_tangalunga | LC | Stable | tropical | nocturnal | 6885 | 1602751.768 | 8.792 |
| mammals | Viverra_zibetha | Viverra_zibetha | LC | Decreasing | tropical | nocturnal | 9500 | 5069770.834 | 13.185 |
| mammals | Viverricula_indica | Viverricula_indica | LC | Stable | tropical | nocturnal | 2908.4 | 8473961.846 | 15.27 |
| mammals | Voalavo_antsahabensis | Voalavo_antsahabensis | EN | Unknown | tropical | nocturnal | 21.8 | 299.494 | 7.88 |
| mammals | Voalavo_gymnocaudus | Voalavo_gymnocaudus | LC | Stable | tropical | nocturnal | 21.8 | 1144.09 | 6.327 |
| mammals | Volemys_millicens | NA | NT | Decreasing | NA | NA | NA | NA | NA |
| mammals | Volemys_musseri | NA | DD | Unknown | NA | NA | NA | NA | NA |
| mammals | Vombatus_ursinus | Vombatus_ursinus | LC | Stable | temperate | nocturnal | 26000 | 310428.62 | 8.107 |
| mammals | Vormela_peregusna | Vormela_peregusna | VU | Decreasing | temperate | nocturnal | 450.4 | 7929465.091 | 8.884 |
| mammals | Vulpes_bengalensis | Vulpes_bengalensis | LC | Decreasing | tropical | nocturnal | 2726 | 3129366.449 | 17.549 |
| mammals | Vulpes_cana | Vulpes_cana | LC | Stable | temperate | nocturnal | 936.9 | 3142580.11 | 7.116 |
| mammals | Vulpes_chama | Vulpes_chama | LC | Stable | temperate | nocturnal | 2955 | 2126479.341 | 7.335 |
| mammals | Vulpes_corsac | Vulpes_corsac | LC | Unknown | temperate | nocturnal | 2400 | 8976804.646 | 6.888 |
| mammals | Vulpes_ferrilata | Vulpes_ferrilata | LC | Unknown | temperate | nocturnal | 5000 | 2231673.465 | 4.16 |
| mammals | Vulpes_lagopus | Vulpes_lagopus | LC | Stable | temperate | nocturnal | 4867.7 | 10726428.996 | 0.583 |
| mammals | Vulpes_macrotis | Vulpes_macrotis | LC | Decreasing | temperate | nocturnal | 2050 | 1611407.338 | 8.035 |
| mammals | Vulpes_pallida | Vulpes_pallida | LC | Unknown | tropical | nocturnal | 2800 | 4152381.022 | 9.091 |
| mammals | Vulpes_rueppellii | Vulpes_rueppellii | LC | Stable | tropical | nocturnal | 2865 | 12646617.267 | 2.665 |
| mammals | Vulpes_velox | Vulpes_velox | LC | Stable | temperate | nocturnal | 2197.5 | 1550926.193 | 13.175 |
| mammals | Vulpes_vulpes | Vulpes_vulpes | LC | Stable | temperate | cathemeral | 5318.2 | 72402469.182 | 7.879 |
| mammals | Vulpes_zerda | Vulpes_zerda | LC | Stable | temperate | nocturnal | 1100 | 8675207.948 | 1.935 |
| mammals | Wallabia_bicolor | Wallabia_bicolor | LC | Increasing | temperate | nocturnal | 15000 | 1212921.515 | 8.17 |
| mammals | Wiedomys_cerradensis | NA | DD | Unknown | NA | NA | NA | NA | NA |
| mammals | Wiedomys_pyrrhorinos | Wiedomys_pyrrhorhinos | LC | Unknown | tropical | nocturnal | 46.7 | 694703.086 | 9.32 |
| mammals | Wilfredomys_oenax | Wilfredomys_oenax | EN | Unknown | temperate | nocturnal | 46.8 | 512619.206 | 11.341 |
| mammals | Wyulda_squamicaudata | Wyulda_squamicaudata | NT | Decreasing | tropical | nocturnal | 1675 | 82875.635 | 2.012 |
| mammals | Xanthonycticebus_pygmaeus | Nycticebus_pygmaeus | EN | Decreasing | tropical | nocturnal | 900 | 481890.254 | 11.108 |
| mammals | Xenogale_naso | Herpestes_naso | LC | Decreasing | tropical | diurnal | 3000 | 2656378.519 | 6.078 |
| mammals | Xenomys_nelsoni | Xenomys_nelsoni | EN | Decreasing | tropical | nocturnal | 130 | 9111.056 | 11.6 |

| | | | | | | | | | |
|---|---|---|---|---|---|---|---|---|---|
| mammals | Xenuromys_barbatus | Xenuromys_barbatus | LC | Unknown | tropical | cathemeral | 1000 | 591368.224 | 3.766 |
| mammals | Xeromys_myoides | Xeromys_myoides | VU | Decreasing | tropical | nocturnal | 41 | 12793.233 | 6.292 |
| mammals | Xeronycteris_vieirai | NA | DD | Unknown | NA | NA | NA | NA | NA |
| mammals | Xerospermophilus_mohavensis | Xerospermophilus_mohavensis | NT | Decreasing | temperate | diurnal | 190 | 22711.458 | 8.58 |
| mammals | Xerospermophilus_perotensis | Xerospermophilus_perotensis | EN | Decreasing | tropical | diurnal | 140 | 2655.847 | 17.253 |
| mammals | Xerospermophilus_spilosoma | Xerospermophilus_spilosoma | LC | Stable | temperate | diurnal | 89 | 1712244.655 | 10.216 |
| mammals | Xerospermophilus_tereticaudus | Xerospermophilus_tereticaudus | LC | Stable | temperate | diurnal | 156.5 | 309353.217 | 8.164 |
| mammals | Xerus_rutilus | Xerus_rutilus | LC | Stable | tropical | diurnal | 317.5 | 2086690.867 | 9.032 |
| mammals | Zaedyus_pichiy | Zaedyus_pichiy | NT | Decreasing | temperate | diurnal | 1380 | 1362776.269 | 5.676 |
| mammals | Zaglossus_attenboroughi | Zaglossus_attenboroughi | CR | Decreasing | tropical | nocturnal | 4332.1 | 15.36 | 5.927 |
| mammals | Zaglossus_bartoni | Zaglossus_bartoni | VU | Decreasing | tropical | nocturnal | 9413.8 | 174289.765 | 4.773 |
| mammals | Zaglossus_bruijnii | Zaglossus_bruijnii | CR | Decreasing | tropical | nocturnal | 7800 | 91613.586 | 3.234 |
| mammals | Zapus_hudsonius | Zapus_hudsonius | LC | Stable | temperate | nocturnal | 17.1 | 8737824.371 | 8.583 |
| mammals | Zapus_princeps | Zapus_princeps | LC | Stable | temperate | nocturnal | 29 | 3278014.042 | 8.642 |
| mammals | Zapus_trinotatus | Zapus_trinotatus | LC | Unknown | temperate | nocturnal | 27.5 | 162624.786 | 11.613 |
| mammals | Zelotomys_hildegardeae | Zelotomys_hildegardeae | LC | Stable | tropical | nocturnal | 60 | 3051583.274 | 9.005 |
| mammals | Zelotomys_woosnami | Zelotomys_woosnami | LC | Stable | tropical | nocturnal | 54.1 | 674859.921 | 5.119 |
| mammals | Zenkerella_insignis | Zenkerella_insignis | LC | Unknown | tropical | cathemeral | 200 | 280311.123 | 5.413 |
| mammals | Zygodontomys_brevicauda | Zygodontomys_brevicauda | LC | Stable | tropical | nocturnal | 52.2 | 1694720.069 | 5.726 |
| mammals | Zygodontomys_brunneus | Zygodontomys_brunneus | LC | Stable | tropical | nocturnal | 75.6 | 75737.704 | 15.944 |
| mammals | Zyzomys_argurus | Zyzomys_argurus | LC | Stable | tropical | nocturnal | 36 | 1188571.347 | 3.297 |
| mammals | Zyzomys_maini | Zyzomys_maini | VU | Decreasing | tropical | nocturnal | 94 | 21158.545 | 1.073 |
| mammals | Zyzomys_palatalis | Zyzomys_palatalis | CR | Unknown | tropical | nocturnal | 123 | 2582.43 | 1.927 |
| mammals | Zyzomys_pedunculatus | Zyzomys_pedunculatus | CR | Decreasing | temperate | nocturnal | 100 | 7110.096 | 2.888 |
| mammals | Zyzomys_woodwardi | Zyzomys_woodwardi | LC | Stable | tropical | nocturnal | 136 | 35599.281 | 1.279 |
